# TDP-43 pathology links innate and adaptive immunity in amyotrophic lateral sclerosis

**DOI:** 10.1101/2024.01.07.574541

**Authors:** Baggio A. Evangelista, Joey V. Ragusa, Kyle Pellegrino, Yija Wu, Ivana Yoseli Quiroga-Barber, Shannon R. Cahalan, Omeed K. Arooji, Jillann A. Madren, Sally Schroeter, Joe Cozzarin, Ling Xie, Xian Chen, Kristen K. White, J. Ashley Ezzell, Marie A. Iannone, Sarah Cohen, Rebecca E. Traub, Xiaoyan Li, Richard Bedlack, Douglas H. Phanstiel, Rick Meeker, Natalie Stanley, Todd J. Cohen

**Author notes:** Co-corresponding authors: Todd J. Cohen, Ph.D., Baggio A. Evangelista, Ph.D.

## Abstract

Amyotrophic lateral sclerosis is the most common fatal motor neuron disease. Approximately 90% of ALS patients exhibit pathology of the master RNA regulator, Transactive Response DNA Binding protein (TDP-43). Despite the prevalence TDP-43 pathology in ALS motor neurons, recent findings suggest immune dysfunction is a determinant of disease progression in patients. Whether TDP-43 pathology elicits disease-modifying immune responses in ALS remains underexplored. In this study, we demonstrate that TDP-43 pathology is internalized by antigen presenting cells, causes vesicle rupture, and leads to innate and adaptive immune cell activation. Using a multiplex imaging platform, we observed interactions between innate and adaptive immune cells near TDP-43 pathological lesions in ALS brain. We used a mass cytometry-based whole-blood stimulation assay to provide evidence that ALS patient peripheral immune cells exhibit responses to TDP-43 aggregates. Taken together, this study provides a novel link between TDP-43 pathology and ALS immune dysfunction, and further highlights the translational and diagnostic implications of monitoring and manipulating the ALS immune response.

## Introduction

ALS is the most common, fatal motor neuron disease worldwide. The disease is primarily characterized by the progressive degeneration of motor neurons in the motor cortex and spinal cord (1). The average post-diagnosis survival interval is approximately 2-5 years (2), however a subset of patients can survive >10 years post-diagnosis (3). Recent evidence suggests systemic immune perturbations modify ALS presentation and progression (4–6). Indeed, disparities in the levels of nearly every immune population including innate and adaptive immune cells have been documented in ALS (7–11). Additionally, loss-of-function mutations in several ALS-associated risk genes, such as C9ORF72 (chromosome 9 open reading frame 72) and TBK1 (TANK-binding kinase1) were shown to promote autoimmune-like inflammatory responses (12, 13). However, C9ORF72 and TBK1 mutations only constitute 5-7% and 1%, respectively, of all ALS cases (14, 15). In 2006, Neumann and colleagues identified pathological inclusions of TDP-43 as an underlying feature to motor neuron degeneration in the vast majority of nearly all ALS cases, including > 90% of those with sporadic ALS (16, 17).

TDP-43 regulates the expression or stability of approximately 6,000 transcripts across diverse biological processes in diverse cell types such as neuroglia and peripheral immune cells (18–22). In disease, TDP-43 mislocalizes to the cytoplasm and forms post-translationally modified aggregates (23–25). We recently demonstrated that TDP-43 aggregates generated in human cells and then isolated by biochemical methods contained nearly 2,000 sequestered proteins. These include classically defined damage associated molecular patterns (DAMPs) such as heat shock proteins, heterogeneous nuclear ribonucleoproteins, and nuclear pore proteins. We also demonstrated that monocyte-derived macrophages and microglia internalize TDP-43 aggregates (26). Other studies have shown that TDP-43 dysfunction leads to dysregulation of STING antiviral immune responses in neurons (27). Here, we aimed to address the following question: do pathological TDP-43 species serve as antigens that stimulate disease-modifying immune cells that are relevant to ALS?

## Results

### TDP-43 aggregates are phagocytosed, trafficked to autophagolysosomes, and promote acute activation of primary monocyte-derived macrophages

Our prior study showed that immunopurified, human TDP-43 aggregates (referred to as **TDP-43a**) are readily internalized by both primary murine microglia and primary human monocyte-derived macrophages (hMDM)(26). Additionally, we noted transcriptional similarity between human microglia and our hMDM cultures (28). For this reason, we used hMDM for studying interactions with TDP-43a. We first assessed the dynamics of TDP-43a internalization using biochemical methods. By immunoblot, we observed significant dose-dependent aggregate internalization by hMDM following stimulation with TDP-43a at 0.025% (**p = 0.0051) and 0.1% (****p < 0.0001) (Figure 1A). Laser-scanning confocal microscopy confirmed internalization via active, actin-dependent transport (Figure 1B). These results suggest that actin-dependent phagocytosis is the dominant mode of TDP-43a internalization. However, several internalization pathways could employ actin-polymerization. Thus, to conclusively assess phagocytosis as the dominant route by which TDP-43a is internalized, we used a panel of pharmacological inhibitors to target different internalization routes. These included Dynasore to inhibit dynamin-mediated endocytosis and 5-(N-ethyl-N-isopropyl)-Amiloride (EIPA) to inhibit micropinocytosis. Following inhibition and stimulation with TDP-43a, we counterstained the hMDM cell membrane with wheat-germ agglutinin (WGA) and quantified complete internalization using laser-scanning confocal microscopy (Figure 1C-D). With respect to vehicle control (27.13% ± 3.528%), a partial inhibitory effect was observed with Dynasore (15.75% ± 2.84%; *p = 0.018) and EIPA (15.13% ± 2.88%; *p = 0.011); however, cytochalasin D treatment consistently inhibited internalization with 100% efficacy (****p < 0.0001). This led us to assess the functional response of hMDM to TDP-43a internalization using a systems-based proteomics approach.

**Figure 1.**
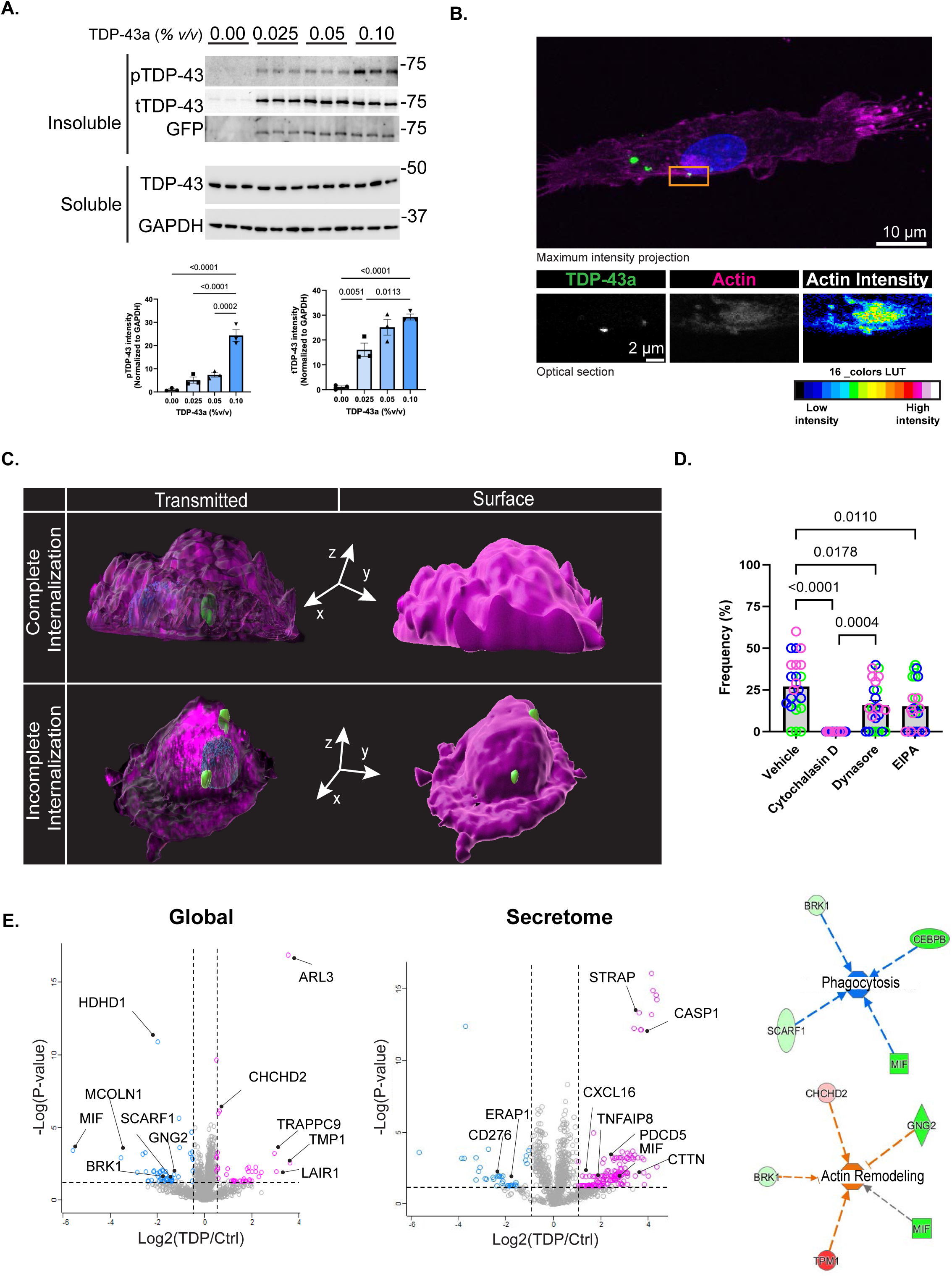
TDP-43 aggregates are phagocytosed, trafficked to autophagolysosomes, and promote acute activation of primary human monocyte-derived macrophages. **(A)** Immunoblot depicts the dose-dependent uptake of insoluble, hyper-phosphorylated, GFP-tagged TDP-43a in primary hMDM**. (B)** Laser-scanning confocal micrograph of early internalization of TDP-43a in primary hMDM counter-stained with phalloidin. Pseudo-colored phalloidin stain depicts relative actin intensity. **(C)** Representative 3-dimensional renderings of complete and incomplete TDP-43a internalization with respect to hMDM plasma membrane. Green denotes TDP-43a, magenta denotes wheat-germ agglutinin. **(D)** Quantification of complete TDP-43a internalization in the presence of phagocytosis inhibitor Cytochalasin D, endocytosis inhibitor Dynasore, and micropinocytosis inhibitor EIPA. Data points depict the frequency of cells with at least one completely internalized TDP-43a particle relative to the total number of cells in a randomized field of view. Color depicts data point with respect to donor genotype. N = 3 independent experiments from 3 different genotypes. *P < 0.05, **P < 0.01, ***P < 0.001, ****P < 0.0001. Ordinary one-Way ANOVA with Tukey’s multiple comparison’s test. Scale-bar 10-µm. **(E)** Volcano plots depict differentially expressed intracellular (left) and secreted (middle) proteomes from primary hMDM cultures stimulated with TDP-43a. Network analyses depict differentially expressed proteins related to phagocytosis (upper right) and actin remodeling (lower right) pathways. Performed in triplicate (n = 3); FDR < 0.05; student’s *t-*test.

We performed combinatorial secretome and proteomics analysis on hMDM cultures stimulated with TDP-43a for 16 hours and identified several features of macrophage activation in the global intracellular proteome and secretome. These included increased secretion of macrophage migration inhibitory factor (MIF) (Log_2_fold = 3.84; FDR < 0.05) and IL-1 converting enzyme or caspase 1 (Log_2_fold = 3.67; FDR < 0.05), and reduced secretion of immunosuppressive immune checkpoint molecule CD276 (Log_2_fold = -2.14; FDR < 0.05), among others (Supplementary Tables S1A-B). By merging these data into a network analysis, we identified upregulated signaling pathways including actin remodeling, phagocytosis, and autophagy (Figure 1E). These findings confirm that extracellular TDP-43a augments innate immune signaling and scavenging pathways, supporting the ‘DAMP-like’ properties of TDP-43a.

### TDP-43a compromises autophagy and promotes vesicle rupture

Our proteomics analysis highlighted autophagy as a significant response following TDP-43a stimulation. Given the intimate link between TDP-43 dysfunction and autophagic dysregulation (29), and the fact that autophagic dysregulation can drive maladaptive immune responses (30), we next interrogated TDP-43a trafficking and autophagic integrity in hMDM. We first performed immunofluorescence and Airyscan confocal imaging where we observed co-localization between TDP-43a and markers of early endosomes (Rab5) (Figure 2A). We then validated lysosomal targeting by performing live-cell imaging using the vesicle acidification probe, LysoTracker-Red. Indeed, we observed co-localization between TDP-43a and LysoTracker-positive vesicles (Figure 2B). Furthermore, Airyscan confocal imaging of hMDM stimulated with TDP-43a for 16 hours revealed the presence of LC3-positive autophagosomes co-localizing with galectin-3 and TDP-43a, suggesting TDP-43a burden promotes vesicle rupture and downstream vesicle turnover (Figure 2C)(31, 32). We further characterized the ultrastructural integrity of vesicles by transmission electron microscopy (TEM) of hMDM stimulated with TDP-43a for 1 hour and 16 hours to simulate acute and chronic exposures, respectively (Figure 2D). By 16 hours we observed greater incidences of irregular, enlarged vesicles. We also identified features of vesicle degradation and turnover, indicated by increased presence of multivesicular bodies (MVBs). These ultrastructural features were not present in unstimulated or 1-hour stimulated hMDM. To validate whether autophagic processing is abnormal in the presence of pathological TDP-43a, hMDM were stimulated with TDP-43a and lysates were collected at 0, 0.5, 4, 8, 16, and 24 hours. Lysates were then probed for markers of autophagic flux, LC3-I and LC3-II, to assess autophagy induction (Figure 2E). We noted a distinctive loss of both LC3B-I and LC3B-II, rather than ratiometric loss of LC3B-I and gain of LC3B-II, which is indicative of dysregulated autophagy (33). Additional readouts of autophagic dysfunction include increased expression of the autophagy adaptor p62 (34). Indeed, increased p62 levels correlated with LC3B-I and LC3B-II between 8 and 24 hours (Spearman’s r > 0.4) (Figure 2E). Together, these results suggest that TDP-43 pathology induces autophagic dysfunction and vesicular damage characterized by the mobilization of the inflammatory lectin galectin-3.

**Figure 2.**
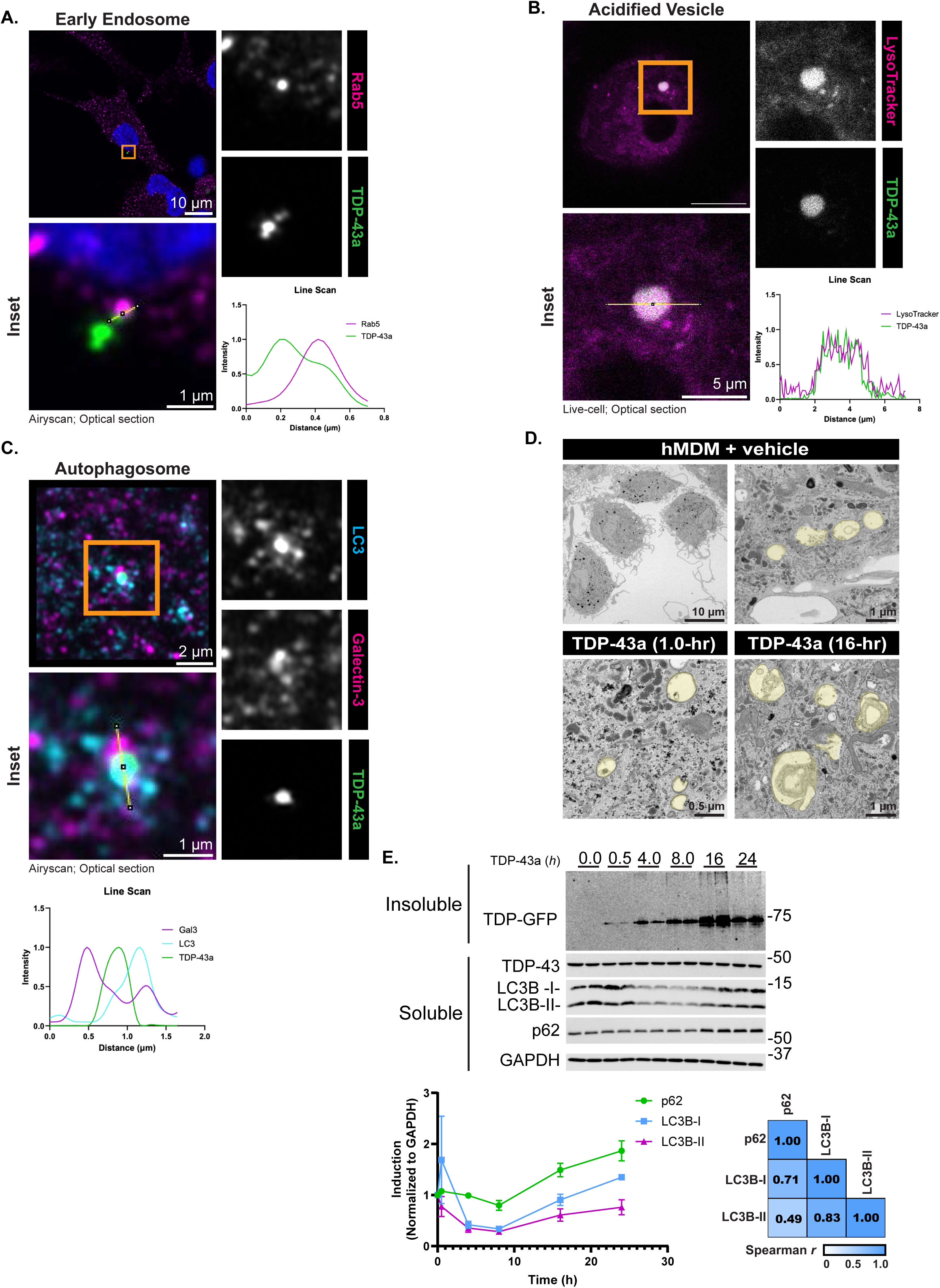
TDP-43 aggregates compromise autophagy and promote vesicle rupture. Airyscan and confocal micrographs (630X) illustrate the extent of TDP-43a co-localization with early endosome marker Rab5a **(A)**, acidified vesicle marker LysoTracker **(B)**, and ruptured autophagosome markers LC3B and Galectin-3 **(C)**. **(D)** Transmission electron micrographs depict whole-cell hMDM (top left; scale-bar 10-µm, hMDM vesicles following vehicle treatment (upper right; scale-bar 1-µm), and vesicles following TDP-43a treatment for 1-hour (lower left; scale-bar 0.5-µm) and 16-hours (lower right; scale-bar 1-µm). Representative vesicles are shaded yellow. **(E)** Representative immunoblots of LC3-I, LC3-II, and p62 kinetics in hMDM stimulated with TDP-43a. N = 2 independent experiments. Spearman’s *r* correlation analysis of LC3B-I, LC3B-II, and p62 changes after incubation with TDP-43a.

### TDP-43a drives a reactive transcriptome in primary monocyte-derived macrophages

To determine if TDP-43a elicited an innate immune response at the transcriptional level, we conducted RNA-sequencing on hMDM treated with either PBS or TDP-43a for 12 hours. We identified 262 differentially expressed genes (Figure 3A; DESeq2, p < 0.01, absolute fold change > 1.15), including those with known roles in innate immune response including galectin-3, TLR4, CCL22, SIGLEC1, and complement receptor 1 (CR1). Additional differentially expressed genes included signaling regulators JAK1 and STAT5B, suggesting broad immune modulation following TDP-43a stimulation. We note that these changes were more targeted and specific (both in number and magnitude) compared to a 6-day, ‘chronic’ amyloid beta oligomer (_o_Aβ_1-42_) stimulation (712 differentially expressed genes) and a more potent 12-hour lipopolysaccharide (LPS) stimulation (5940 differentially expressed genes) (Figure 3B; Supplementary Table S3A-C). TDP-43a gene expression differences were distinct compared to _o_Aβ_1-42_, as only 48 genes showed a common stimulation-dependent regulation, and among these, the magnitude of differential expression was unique between TDP-43a and _o_Aβ_1-42_ (Figure 3C). Gene ontology (GO) clustering indicated that TDP-43a preferentially regulated innate immune responses (cytokine production involved in immune response), blood cell differentiation (myeloid cell differentiation and regulation of hematopoiesis) and interleukin-2 signaling (Figure 3D). Importantly, interleukin-2 signaling is a major orchestrator of innate-to-adaptive immune activation, predominantly through T-cell activation and maintenance. These data suggest that internalized TDP-43a can activate innate immune responses and thereby influence the crosstalk between innate and adaptive immune cells.

**Figure 3.**
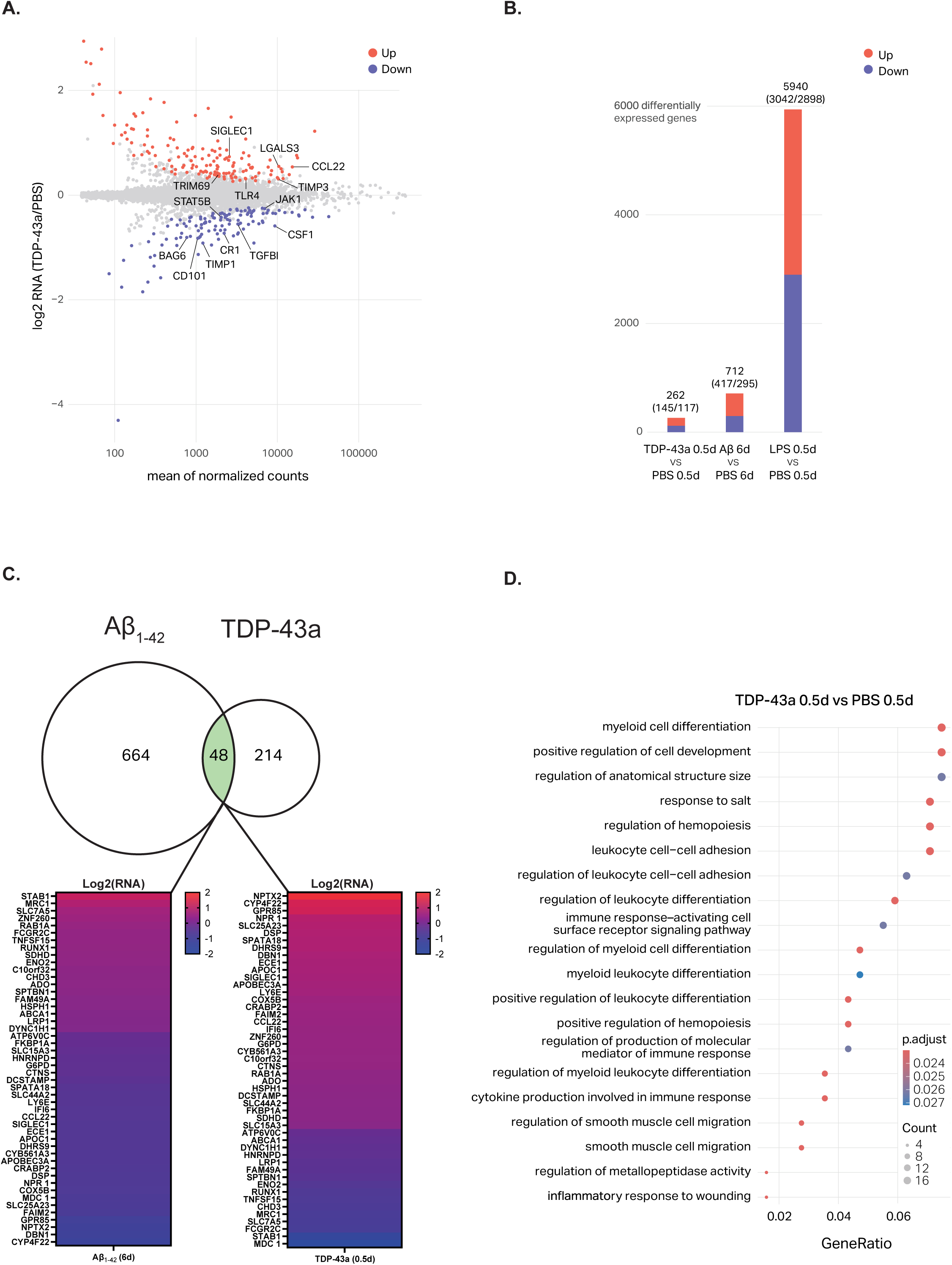
Distinct and specific gene expression changes in primary human monocyte-derived macrophages challenged with TDP43 aggregates. **(A)** An MA plot depicts differential analysis of RNA-seq data of cells treated with either TDP-43a or PBS for 12 hours. Differential genes are depicted in blue or red. **(B)** A bar plot depicts the number of differential genes detected when comparing TDP-43a, AB, or LPS to cells treated with PBS for the equivalent amount of time. Red represents upregulated genes and blue represents down-regulated genes. **(C)** Gene Ontology enrichment analysis depicts functional gene clusters that were enriched in those genes upregulated in response to 12 hours of TDP43a treatment. **(D)** Venn diagram depicting unique and conserved genes that differentially expressed following TDP-43a or _o_Aβ_1-42_ treatment. Heat maps depict the magnitude of expression change of each of the 48 conserved genes between TDP-43a and _o_Aβ_1-42_ treatments. Performed in duplicate, n = 2.

### TDP-43a promotes global immunophenotypic changes

Given that there are numerous immune cell types with the potential to phagocytose and react to TDP-43a, we asked how broadly TDP-43a impacts the human immune landscape using *ex vivo* human immune cell cultures. Based on our mechanistic studies of TDP-43a internalization in hMDM, we reasoned that any professional phagocyte may internalize TDP-43a. To address this question, we optimized a novel mass cytometry-based aggregate internalization assay, that we termed *Aggre-*Gate, in bulk PBMCs that allowed for simultaneous global immunophenotyping and cell population-wide localization of TDP-43a (Figure 4A schematic). To monitor localization of TDP-43a, aggregates were first coupled to a Tellurium-Maleimide adduct (130Te) and used to stimulate bulk primary human PBMC cultures. We confirmed that our assay follows linear kinetics of internalization in phagocytes such as classical monocytes by traditional Boolean gating (Figure 4B-C and Supplementary Figure S4A-B). We next used Spanning-Tree Progression Analysis of Density-Normalized Events (SPADE) to stratify PBMCs and overlay relative 130Te intensity to identify TDP-43a internalization on a cell cluster-specific basis. We observed TDP-43a internalization by all available phagocytic and antigen presenting cells in PBMC preparations, including classical monocytes (77.13% ± 3.19%; ****p < 0.0001) and dendritic cells (65.28% ± 8.91%; ****p < 0.0001) by SPADE and manual analyses. Strikingly, this analysis platform identified TDP-43a internalization in B-cells, an unconventional phagocyte implicated in both antigen-presentation and antibody production (8.683% ± 3.63%; *p = 0.0263). Finally, as expected, classically described non-phagocytic cells such as CD8 T-cells (^ns^p > 0.999) and natural killer (NK) cells (^ns^p = 0.9978) did not significantly internalize TDP-43a (Figure 4E-F).

**Figure 4.**
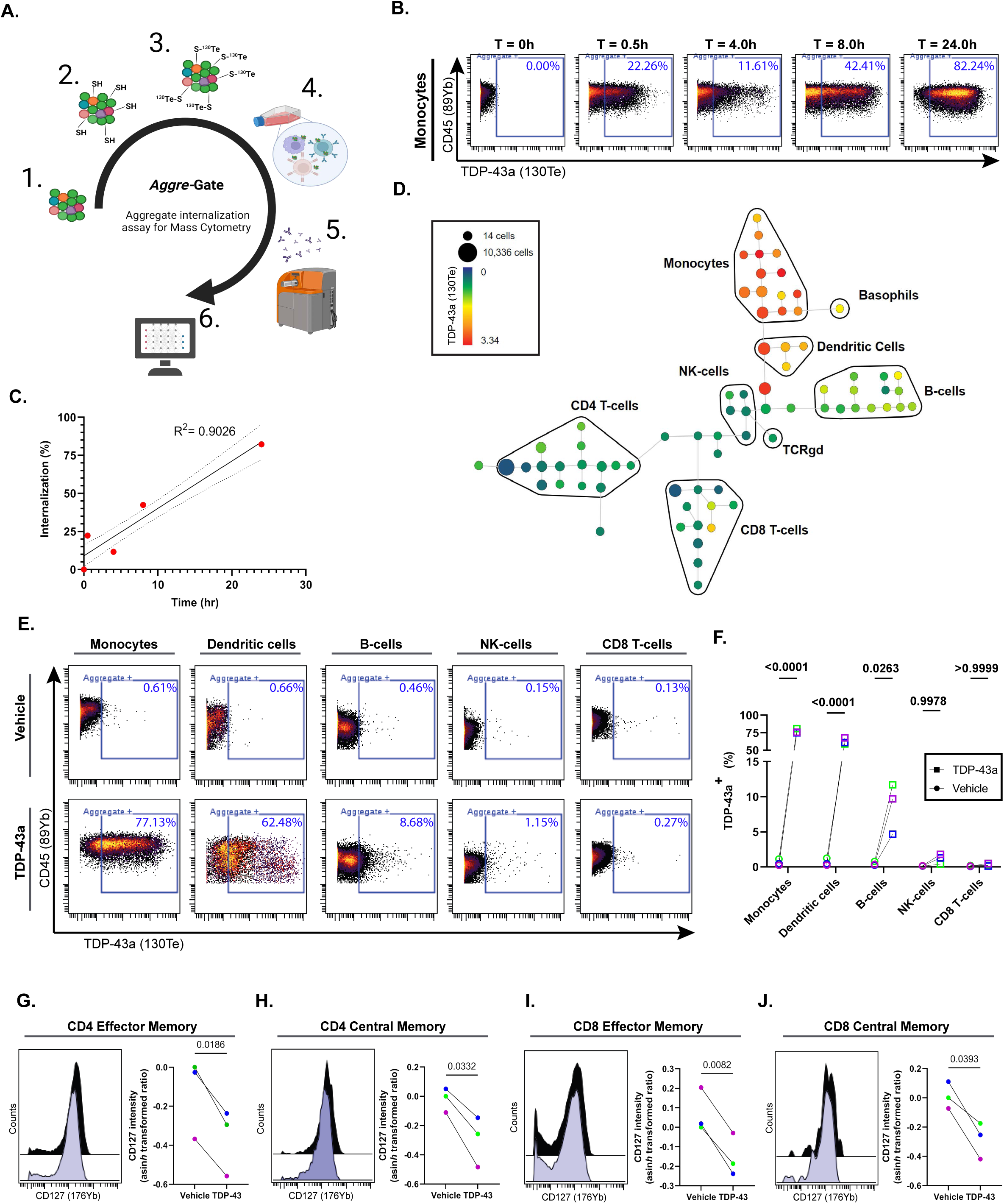
Global immunophenotypic changes in response to TDP43 aggregates. **(A)** Schematic of mass cytometry-based TDP-43a internalization assay, *Aggre-*Gate. Purified TDP-43a is partially reduced with DTT and coupled to Tellurium maleimide (^130^TeMal) via thiol-reactive chemistry. Labeled TDP-43a are added to cell cultures and internalization is determined by Boolean gating and/or Spanning-tree Progression of Density-normalized Events (SPADE) following mass cytometry analysis. **(B)** Concatenated dot-plots depict kinetics of TDP-43a internalization in classical monocytes from N = 3 independent cultures of a single genotype. **(C)** Linear regression of the percent classical monocytes positive for TDP-43a with increasing incubation time with TDP-43a. Gating scheme for identification of total monocytes is illustrated in Supplementary Figure 4B. **(E)** Concatenated dot-plots of monocytes, dendritic cells, B-cells, natural killer (NK) cells, and CD8 T-cells positive for TDP-43a. **(F)** Quantification of the percent of cells positive for TDP-43a following a 24-hour stimulation of bulk PBMC from N = 3 genotypically unique donors (magenta, blue, and green datapoints). Analyzed by Two-way ANOVA with Sidak’s multiple comparison’s test. Gating scheme for identification of immune cell subsets is illustrated in Supplementary Figure 4B. **(G-J)** Quantification of relative CD127 expression levels as a readout of T-cell effector activity following TDP-43a stimulation from N = 3 genetically unique donors. Arcsin(h) transformed ratios of CD127 intensity across CD4 effector **(G)**, CD4 central **(H)**, CD8 effector **(I)**, and CD8 central **(J)** memory T-cells. *P < 0.05, **P < 0.01, ***P < 0.001, ****P < 0.0001. Parametric, paired student’s *t*-test.

In parallel, our TDP-43a mass cytometry assay also allowed us to profile adaptive immune cell activation following acute, 24-hour stimulation with TDP-43a. Our phenotyping strategy included markers of proliferation, early activation, tissue residency/homing, and immune checkpoints (Table 4). Unless otherwise specified, CD4 and CD8 T-cell subsets refer to the TCRαβ subtype, as this is the predominant TCR responsible for antigen recognition. Both CD4 and CD8 T-cell subsets exhibited significant reduction in surface expression of CD127 levels following TDP-43a treatment, which is associated with immune activation, increased effector activity, and senescence (35–37). CD127 downregulation was evident in CD4 effector T-cells (-0.232 ± 0.06 transformed intensity units; *p = 0.0186) (Figure 4G), CD4 central memory T-cells (-0.276 ± 0.09 transformed intensity units; *p = 0.0332) (Figure 4H), CD8 effector T-cells (-0.225 ± 0.04 transformed intensity units; **p = 0.0082) (Figure 4I), and CD8 central memory T-cells (-0.261 ± 0.08 transformed intensity units; *p = 0.0393) (Figure 4J).

### TDP-43a stimulation promotes antigen presentation and activation of naïve T-cells

Classically, protein-driven T-cell immunity requires an essential intercellular signaling event between antigen presenting cells and naïve CD4 and/or CD8 T-cells before memory responses are established. This activation can be monitored via transient intracellular calcium release within T-cells, which is the downstream result of T-cell receptor-mediated PKCγ1 signal transduction (38). To address antigen presentation capabilities of TDP-43a-primed innate immune cells, we first adopted an established co-culture assay using Raji Burkitt lymphoma and Jurkat T-cell lines (39). This model was chosen as Raji and Jurkat cell lines have compatible MHC and TCRs, a necessity when assessing classical antigen presentation (40). Raji B-cells were labeled with CellTracker DeepRed and stimulated with immunopurified TDP-43a or isotype control product for 16-hours. Jurkat T-cells were labeled with CellTracker green and added to Raji cultures for 45 minutes and labeled with phalloidin to detect actin-dense immune synapses that form at junctions between APC and T-cells. The frequency of actin-dense intercellular contacts was quantified using laser-scanning confocal microscopy (Figure 5A-B). Phytohemagglutinin (PHA) was used as a positive control to non-specifically activate Raji cells and encourage immune synapse formation. TDP-43a (3.507% ± 0.17%; ****p < 0.0001) and PHA (2.533% ± 0.27%; ****p < 0.0001) stimulated Raji cells exhibited significantly higher frequencies of actin-polarized cell clusters with Jurkat T-cells relative to isotype control (0.513% ± 0.25%). We note that TDP-43a-stimulated Raji cells showed an even higher level of synapse formation compared to PHA (**p = 0.0049). However, since PHA is not a peptide antigen, we cannot directly compare the antigenic potency between TDP-43a and PHA.

**Figure 5.**
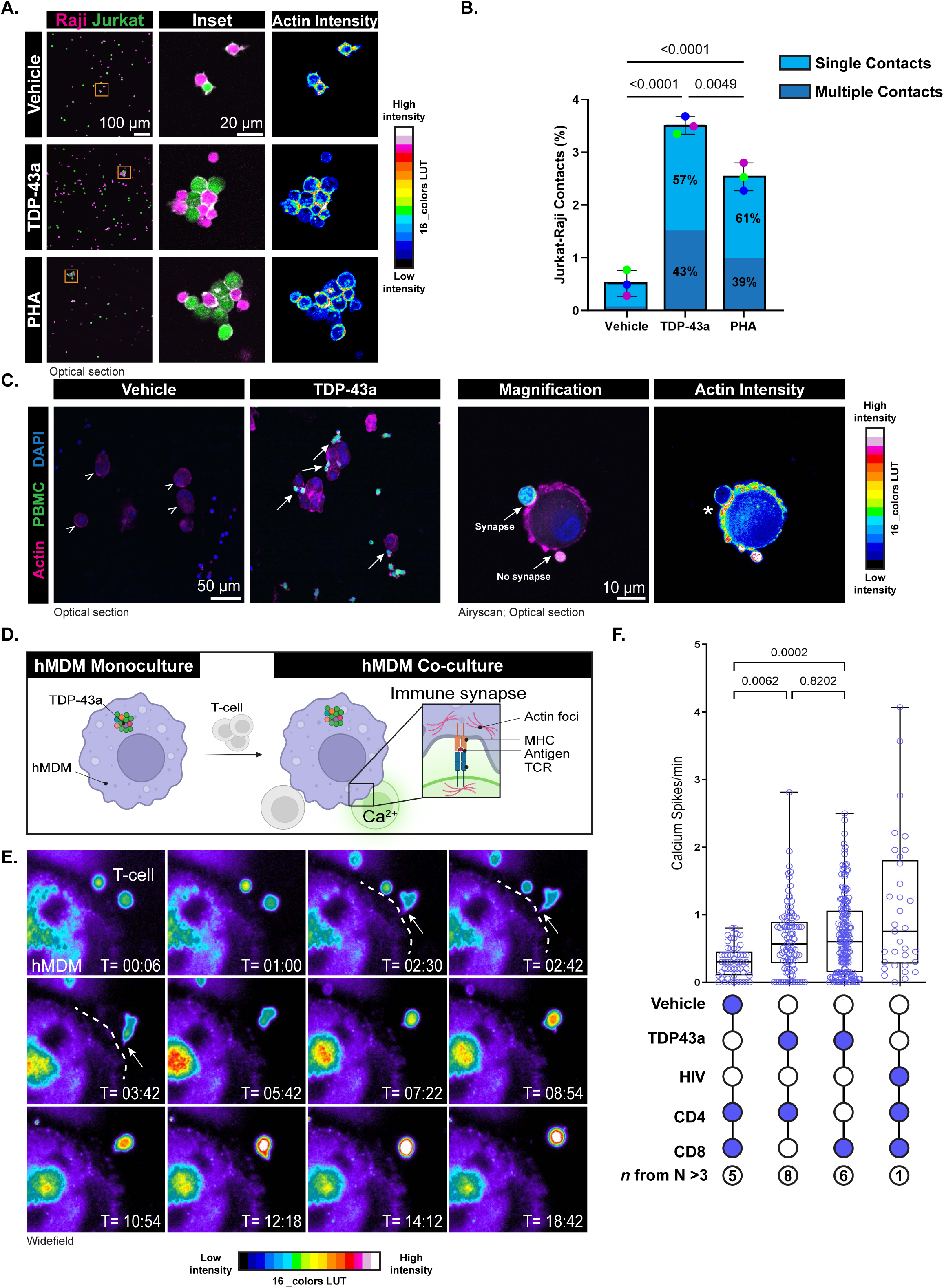
TDP-43 aggregates stimulate antigen presentation and activation of naïve T-cells. **(A)** Representative confocal micrographs (200X) of Raji-Jurkat synapses in the presence of isotype control (vehicle) or TDP-43a purification products, or PHA. Pseudo-colored actin intensity images illustrate intercellular contacts characterized by actin polarization. **(B)** Quantification of **A** by One-way ANOVA with Tukey’s multiple comparisons test. Graph depicts the frequency of cells forming contacts relative to the total number of cells in a randomized field of view. Data bars represent relative percentages of single and multiple cell contacts per condition, relative to the number of contacts in an image. Data points represent N = 3 independent passages. **(C)** Representative confocal micrographs (left; 200X) of primary hMDM immune synapses in the presence of isotype control (vehicle) or TDP-43a purification product. Arrowheads indicate hMDM (ActiStain-555; magenta) and arrows indicate regions of intercellular contact between hMDM and syngeneic PBMCs (CellTracker Green). High magnification Airyscan micrograph (right; 630X) illustrates actin-polarized contact between hMDM and PBMC (asterisk). **(D)** Schematic of live-cell calcium imaging co-culture assay to monitor naïve T-cell activation following TDP-43a stimulation of hMDM. **(E)** Time-series widefield fluorescence micrographs of intercellular contact between hMDM and naïve CD8 T-cell with intracellular calcium release. Dashed line denotes hMDM plasma membrane, and arrowheads denote site of intercellular contact. **(F)** Quantification of T-cell calcium signaling in both naïve CD4 and CD8 T-cell co-cultures with hMDM. Each data point represents single-cell calcium signaling events from a 40-minute imaging period within a single field of view. One field of view per coverslip. Inscribed values denote the total number of coverslips (n) analyzed across all genotypes (N). All data points originated from N = 3-5 unique genotypes. *P < 0.05, **P < 0.01, ***P < 0.001, ****P < 0.0001. Ordinary one-Way ANOVA with Dunnett’s multiple comparisons test.

To further validate this finding in a primary human system, we used syngeneic macrophage-mononuclear cell co-cultures to ensure HLA haplotype compatibility to appropriately assess TCR signaling. In this paradigm, hMDM were stimulated with TDP-43a or isotype control material for 16-hours. In parallel, frozen, syngeneic peripheral blood mononuclear cells (PBMCs) were thawed and allowed to recover overnight. At the time of assay, PBMCs were labeled with CellTracker Green (CTG), co-cultured with macrophages for 45 minutes, and subsequently fixed and labeled with phalloidin prior to analysis by Airyscan confocal microscopy. Strikingly, we noticed the presence of CTG-positive macrophage synapses with TDP-43a treatment (Figure 5C), but a complete absence of CTG labeled cells in the isotype control treatment. The lack of synapses in the isotype control condition is likely due to the inability to elicit hMDM activation leading to reduced chemotactic cues that recruit CTG-positive PBMCs. Supporting this possibility, our secretome analysis revealed significant increase in a notable T-cell chemotactic factor, CXCL16 (Log2fold change 2.836, FDR < 0.05, *p = 0.034) following TDP-43a stimulation. We then performed a separate live-cell imaging analysis where mixed hMDM-PBMC cultures were imaged immediately following the addition of TDP-43a or isotype control product. Compared to control, TDP-43a stimulated cultures exhibited robust migration of PBMCS to hMDM that were actively interacting with aggregates visible by phase-contrast microscopy (Supplementary Video SV1). Taken together, these data suggest hMDM stimulated with TDP-43a exhibit heightened chemo-attractive properties that induce recruitment and engagement of adaptive immune cells.

Our data so far suggests that TDP-43 aggregates can drive immune synapses between antigen presenting cells and adaptive immune cells in several co-culture models of TDP-43a stimulation. However, it is still unclear whether immune synapse formation in response to TDP-43a leads to cellular activation. Since sustained memory T-cell responses indicate prior activation of naïve T-cells, we asked whether TDP-43a drives antigen presentation and activation of naïve T-cell populations.

To address this question, we turned to a sensitive live-cell calcium imaging and co-culture assay (Figure 5D schematic). A functional readout of T-cell activation during antigen presentation is the immediate release of intracellular calcium following TCR engagement (38). Thus, we stimulated hMDM for 16 hours with TDP-43a or isotype control product. In parallel, syngeneic naïve CD4 and CD8 T-cells were enriched by negative magnetic activated cell sorting (MACS). Here, naïve T-cells are defined as CD45RO-, CD45RA+, CCR7+, and CD27+ (Supplementary Figure S5A). As a positive control, we included co-cultures where hMDM were stimulated with HIV virions, a previously established method to induce hMDM-T-cell antigen presentation and calcium signaling (41). Next, hMDM and either naïve CD4 or CD8 T-cells were co-cultured and immediately imaged for 45 minutes to monitor migration, cellular contact, and calcium activity. Strikingly, we observed recruitment of T-cells to the hMDM plasma membrane (Figure 5E). Following recruitment, T-cells formed transient immune synapses, and elicited rapid intracellular calcium responses. Quantification of calcium spiking events revealed significant increases in both naïve CD4 (0.634 spikes/minute; **p = 0.0062) and CD8 T-cells (0.698 spikes/minute; ***p = 0.0002) following co-culture with TDP-43a-stimulated hMDM relative to isotype control stimulations (0.308 spikes/minute) (Figure 5F). These findings suggest TDP-43a can drive naïve T-cell activation via contact-mediated antigen presentation, and thus satisfy early requirements for formation of conventional memory T-cell responses.

### Imaging mass cytometry reveals antigen presenting microenvironments in ALS brain

To support our *in vitro* and *ex vivo* assays, we determined whether the molecular and cellular features of innate-to-adaptive immune activation are present in the ALS brain. Using formalin-fixed paraffin embedded ALS and control brain sections, we performed a high parameter imaging mass cytometry assay to enable the simultaneous labeling of TDP-43 pathology (Figure 6A) as well as a panel of heterogenous brain cell types including brain-resident immune cells (Figure 6B) using a 17-plex antibody panel that included functional markers of antigen presentation and memory T-cells (Table 5). We observed *in vivo* features of innate and adaptive immune activation in ALS tissue that recapitulated features identified *in vitro*. These included microenvironments containing activated microglia (CD68+) associated with pathological TDP-43 (phosphor-serine 409/410) (Figure 6B, arrowheads). Additionally, we identified a subset of myeloid cells that were positive for CD163, a marker associated with infiltrating macrophages (42), galectin-3, and MHC-II (HLA-DR). We also observed intercellular contacts between galectin-3+, HLA-DR+, CD163+ myeloid cells and CD8 T-cells (Figure 6B, white arrows). Interestingly, these CD8 T-cells were also positive for the memory T-cell marker CD45RO but were granzyme B-negative, although granzyme B-positive lymphocytes negative for T-cell markers were identified in the gray matter (Supplementary Figure S6).

**Figure 6.**
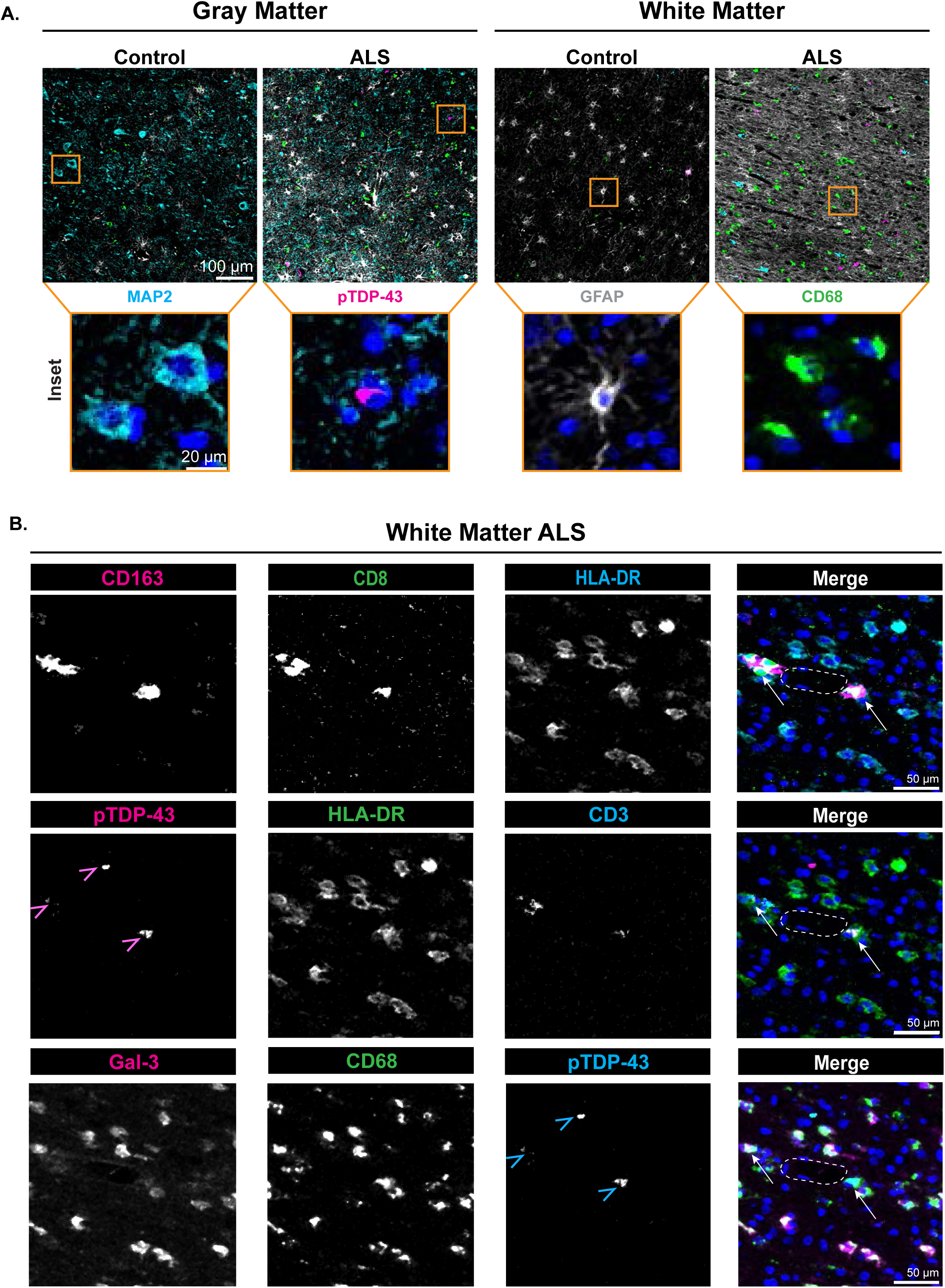
Imaging mass cytometry (IMC) profile of immune microenvironments near sites of TDP-43 pathology in ALS motor cortex. **(A)** Qualitative, representative overlays of the gray (left) and white (right) matter of motor cortex from a single control and ALS motor cortex tissue section. Insets represent magnified views of single feature markers for neurons (MAP2), TDP-43 pathology (pTDP-43), astrocytes (GFAP), and microglia (CD68). Nuclei counter-labeled blue. **(B)** Overlays and corresponding single-channel images of activated myeloid cells/microglia expressing antigen presentation machinery (Gal3, CD68, HLA-DR, CD163) and interacting with CD3+CD8+ T-cells in the parenchyma of ALS white matter. Nuclei counter-labeled blue. Vasculature marked with dashed line. Arrowheads denote pTDP-43 pathology, arrows denote interactions between antigen presenting cells, CD8 T-cells, and pTDP-43. Images were despeckled and adjusted similarly between control and ALS for brightness/contrast using FIJI.

### CD8 T-cell subpopulations show divergent responses to TDP-43a in ALS patients

Our data thus far suggest TDP-43 pathology alters immunophenotypes both *in vitro* and *ex vivo*. However, it remains unknown whether immune responses to TDP-43 pathology are clinically relevant. Therefore, we interrogated the relationship between immune populations, TDP-43a reactivity, and clinical progression of ALS. First, we employed a whole-blood immune profiling assay (Figure 7A), which allows us to analyze the entirety of a patient’s immune profile without fractionation and cryopreservation, which minimizes frequency artifacts in sensitive cell types such as monocytes and NK cells (43). Furthermore, we were able to characterize granulocytes that are typically absent during PBMC profiling protocols (44).

**Figure 7.**
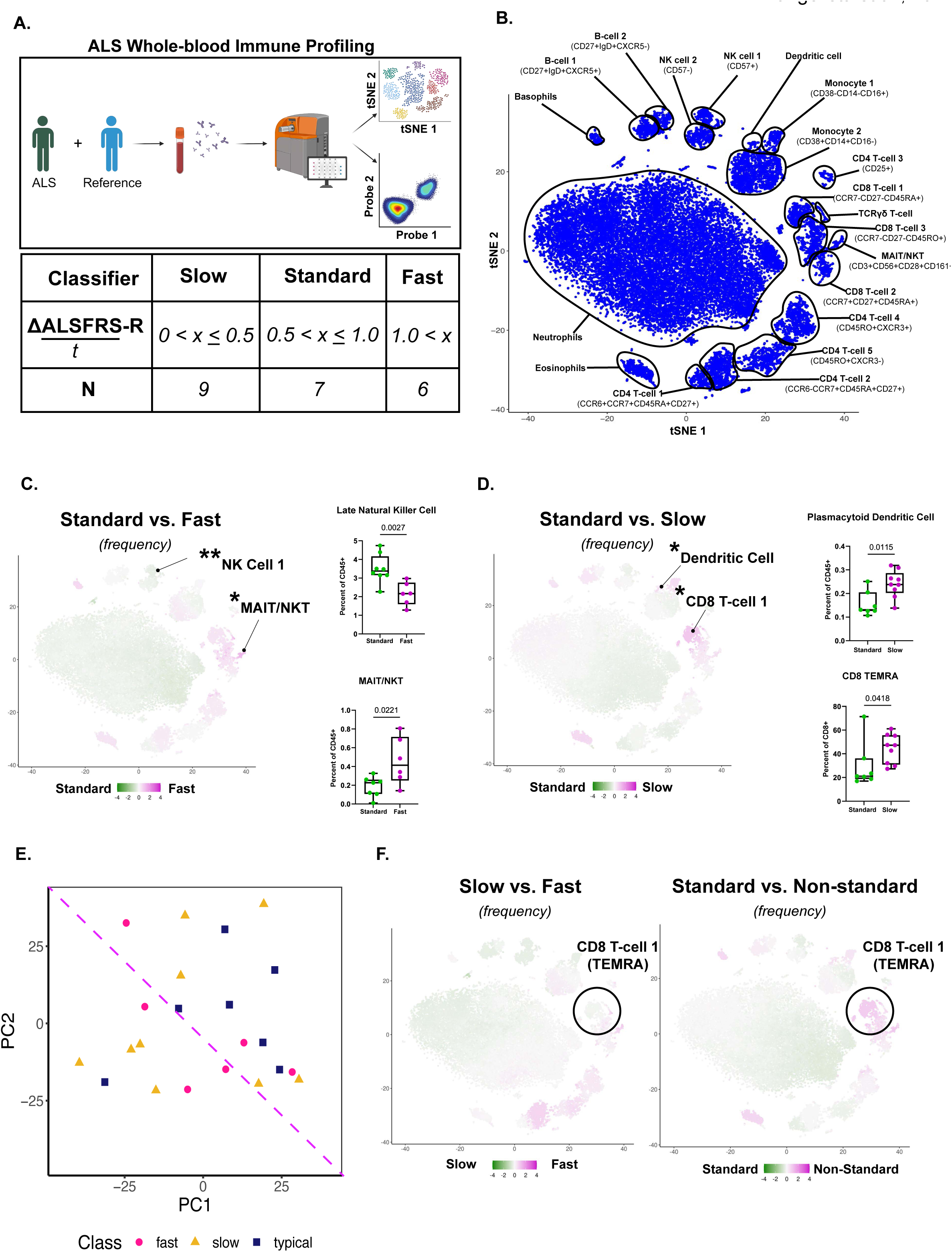
Whole-blood immune profiling reveals niche immune populations that correlate with ALS progression-rate. **(A)** Schematic depicts whole-blood profiling strategy using computational and manual analyses (top). Table summarizes sample sizes and progression rate classifiers determined using the rate of change of the revised ALS functional rating scale (ALSFRS-R). **(B)** Annotated tSNE plot denotes immune sub-populations and associated phenotyping markers of whole-blood leukocytes. **(C)** VoPo analyses and resulting per-cluster enrichment scores between standard- and fast-progressing ALS patients. Quantification of manual gating for NK cell 1 cluster and NKT cell cluster depicts reduced late natural killer cell frequencies (top) and increased natural killer T-cells (bottom) in fast-progressing patients relative to standard-progressing patients. Analyzed by Mann-Whitney U test. *P < 0.05, **P < 0.01, ***P < 0.001, ****P < 0.0001. **(D)** VoPo analysis and cluster-specific enrichment scores between standard- and slow-progressing ALS patients. Quantification of manual gating for dendritic cell cluster and CD8 T-cell cluster 1 depicts increased plasmacytoid dendritic cell (top) and CD8 TEMRA cell (bottom) frequencies in slow-progressing patients relative to standard-progressing patients. Analyzed by Mann-Whitney U test. *P < 0.05, **P < 0.01, ***P < 0.001, ****P < 0.0001. **(E)** Principal component analysis of ALS whole-blood profiles depicts discrete clustering of standard- and non-standard (fast + slow) patient whole-blood profiles. **(F)** VoPo analyses and resulting per-cluster enrichment scores between slow- and fast-progressing patients (left) and standard- and non-standard-progressing patients (right). Circle illustrates penetrance of CD8 TEMRA enrichment score in non-standard ALS patients.

ALS patients were categorized as fast-progressing (ΔALSFRS/*t* > 1.0; n=6), standard-progressing (0.5 < ΔALSFRS/*t* ≤ 1.0; n=8), and-slow progressing (ΔALSFRS/*t* ≤ 0.5; n=9) based on the rate of change to their revised ALS functional rating scale (ALSFRS-R) calculated between clinical visits (Table 6). Per-patient immune profiles assayed through mass cytometry were then analyzed with VoPo, a machine learning pipeline for identifying immunological correlates of particular clinical outcomes (Stanley et al., 2020). Briefly, Vopo first integrates cells across all profiled samples to identify a common set of cell-populations in an agnostic manner. Frequencies and functional marker readouts within each of these uncovered populations can then be systematically compared to identify distinct cell subtypes that may be protective against fast progression (45). Our analyses of the patient cohorts included several patient group comparisons to highlight the most relevant immune populations in each of the fast, standard, or slow progression categories (Figure 7B and Supplementary Figure S7).

Using VoPo, we first compared whole-blood profiles between fast- and standard-progressing ALS patients. Candidates were validated by manual gating and statistical analysis. We identified a statistically significant increased frequency of the mucosal associated invariant T-cell/natural killer T-cell (MAIT/NKT) niche in fast-progressing patients compared to standard-progressing patients (0.10% vs. 0.23% of CD45+ leukocytes, respectively; *p = 0.0221, Mann-Whitney U test). Additionally, we observed a significant reduction in senescent NK cells in fast-progressing patients (3.48% vs 2.16% of CD45+ leukocytes, respectively; **p = 0.0027, Mann-Whitney U test) (Figure 7C). We next compared whole-blood profiles between standard- and slow-progressing patients. We identified a significant increase in plasmacytoid dendritic cells (0.13% vs 0.24% of CD45+ leukocytes, respectively; *p = 0.0115, Mann-Whitney U test) and terminally differentiated CD8 T-cells re-expressing CD45RA (21.07% vs 47.21% of CD8+ T-cells, respectively; *p = 0.0418, Mann-Whitney U test) in slow-progressing patients relative to standard-progressing (Figure 7D). We observed an increasing trend (though not significant) in myeloid dendritic cells (^ns^p = 0.0549) and eosinophils (^ns^p = 0.175, Mann-Whitney U test) in slow-progressing patients relative to standard-progressing patients.

We next wanted to assess whether immune population differences can be used to segregate slow-, standard-, and fast-progressing patients as a proof-of-principle for predictive modeling of ALS progression. To do so, we performed principal component analysis on whole blood profiles. At this sample size, we identified relative segregation of standard-progressing patients, while slow- and fast-progressing patients appeared to cluster together (Figure 7E). In further support of this, VoPo analysis revealed a lack of differentiating signal when comparing slow-against fast-progressing patients in several clusters, including the CD8 TEMRA cluster (CD8 T-cell 1). However, signal in this cluster returned when we compared standard to non-standard profiles (aggregated slow- and fast-progressing profiles), with enrichment in the non-standard progressing group (Figure 7F). Although slow- and fast-progressing patients had similar CD8 TEMRA profiles based on VoPo, we suspected that this particular sub-population may be functionally distinct and/or uniquely regulated in slow-versus fast-progressing subtypes. Indeed, CD8 TEMRA have been characterized as functionally dichotomous, with cytotoxic and regulatory roles governed by the IL-7/CD127 (IL-7R) axis (45). We reasoned that *bona fide* functional readouts of CD8 TEMRA activity would provide additional information for distinguishing between fast- and slow-progressing immune profiles.

To test this, we performed a proof-of-concept clinical case assessment. We implemented a blinded whole-blood stimulation and mass cytometry assay with the goal of capturing functional differences, rather than population differences, of ALS CD8 TEMRA following stimulation with TDP-43a. We performed the assay on 3 different ALS donors, one each of a fast-, standard-, and slow-progressing patient, and 3 same-visit control samples taken from either a caretaker or cohabitating donor that is considered environmentally matched to the respective patient. This design allowed us to reference ALS responses against natural immunological variability that might arise from different genotypes and environments. Patient and control whole blood were stimulated for 16-hours with TDP-43a or dimethyl sulfoxide (DMSO) to establish baseline. We first opted to assess markers of CD8 TEMRA senescence, an inherent property of cytotoxic T-cells with regulatory and pathological implications (45). Specifically, we calculated the ratio of proliferative (CD127+CD57-) to senescent-like (CD127-CD57+) CD8 TEMRA. Each ratio was normalized to the respective DMSO control. Notably, the slow-progressing patient TEMRA exhibited a 2.74-fold increase (non-inferential) in conversion of proliferative CD127+CD57-cells to CD127-CD57+ senescent-like TEMRA following stimulation with TDP-43a. This conversion was absent in the matched control, as well as fast- and standard-progressing patients and respective controls (Figure 8A). Next, we assessed expression changes of CD103 as a readout of tissue-residency, a property that can minimize or exacerbate chronic tissue damage (46). Remarkably, the slow-progressing patient TEMRA population exhibited the largest loss of CD103+ TEMRA (-2.75%; non-inferential), compared to controls, standard-progressing (-0.76%; non-inferential) and fast-progressing (+0.14%; non-inferential) patients (Figure 8B). While this data indicates proof-of-concept functional differences across all three disease progression states, we emphasize that larger sample sizes for whole-blood stimulation and profiling will be required for future quantitative ALS biomarker studies. These data suggest that ALS patients exhibit dichotomous immune responses to human TDP-43 pathology that may delineate slow-, standard-, and fast-progressing ALS patients. We note that, since TDP-43a contains numerous co-aggregating factors that are targets of innate and adaptive immunity, the identity of the specific antigen(s) triggering this response is not yet known. Regardless of the exact self-antigens, these data warrant further investigation into TDP-43a reactivity and the potential for immune monitoring as a readout for ALS phenotypes.

**Figure 8.**
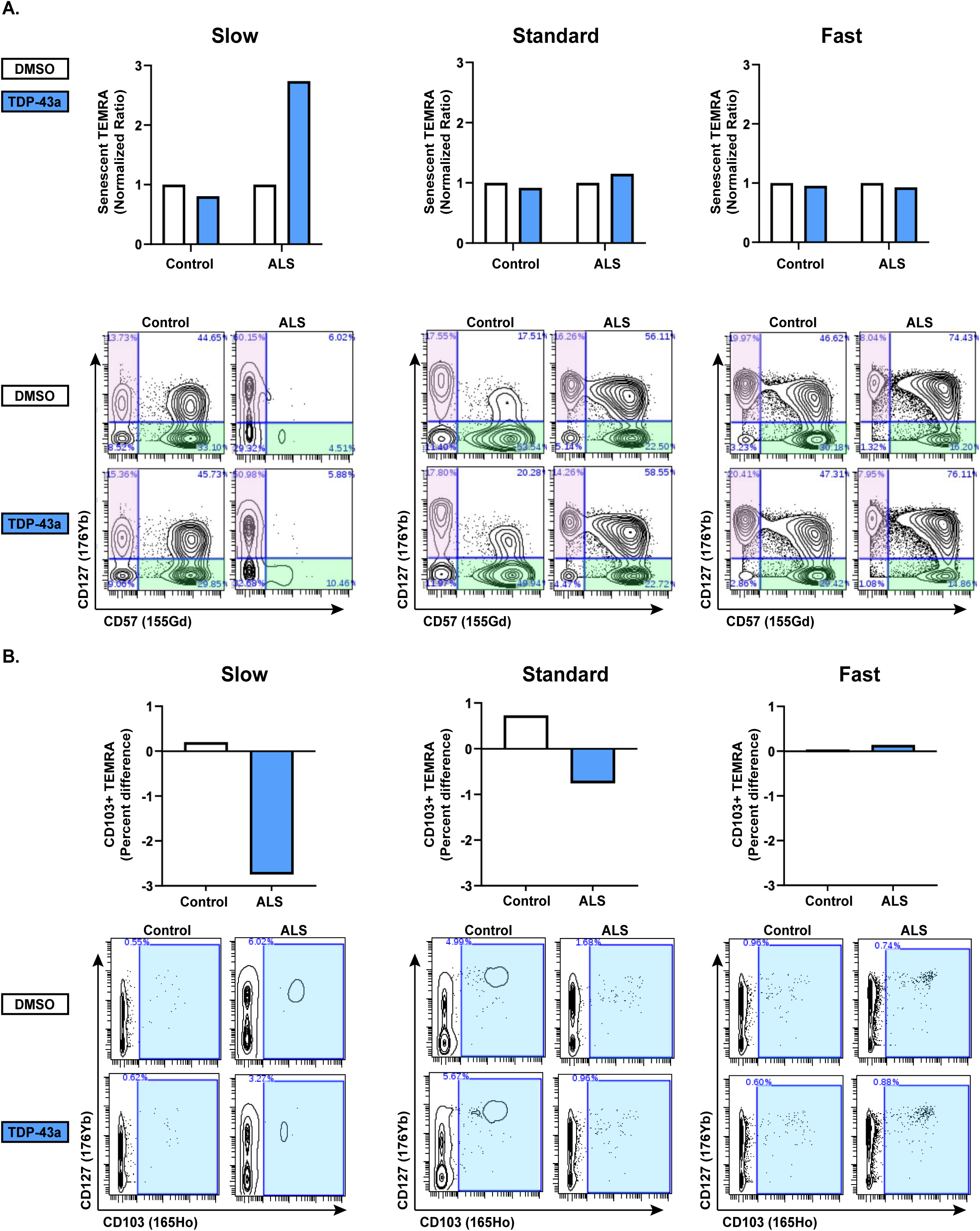
Clinical case assessment: TDP-43a whole-blood stimulation induces functional responses that are dependent on disease progression rate. **(A)** Normalized ratios of senescent (CD127-CD57+) to proliferative (CD127+CD57-) CD8 TEMRA T-cells following TDP-43a stimulation of slow-, standard-, and fast-progressing patient whole-blood cultures (top). Control samples are environmentally matched to each ALS patient. Mass cytometry contour plots depict frequency changes of proliferative (magenta) and senescent (green) CD8 TEMRA cells. Non-inferential, N = 1 patient and control pair, per progression-rate category. **(B)** Percent difference of CD103+ CD8 TEMRA cells as a readout of tissue-homing potential following TDP-43a stimulation of slow-, standard-, and fast-progressing patient whole-blood cultures (top). Control samples are environmentally matched to each ALS patient. Mass cytometry contour plots depict CD103+ CD8 TEMRA cells (cyan). Non-inferential, N = 1 patient and control pair, per progression-rate category.

## Discussion

We discovered a novel link between innate and adaptive immune activation in response to pathological TDP-43a. It was recently demonstrated that ALS adaptive immune responses are restricted to self-proteins (8) and not pure TDP-43 protein (47). Using physiologically relevant TDP-43 aggregates containing accessory co-aggregating DAMPs that we isolated from human cells, we provide evidence that this pathological TDP-43 species acts like a classically defined antigen with potential to induce innate and adaptive immune activation. We assessed the ability of TDP-43 aggregates to 1) serve as a substrate for innate immune cell scavenging, 2) drive innate immune cell activation, 3) augment antigen presentation to naïve CD4 and CD8 T-cells, and 4) elicit memory T-cell responses in ALS immune cells.

When TDP-43a was used to stimulate hMDM, we observed phagocytosis-dependent internalization that correlated with general immune reactivity (i.e., release of pro-inflammatory factors such as caspase 1, MIF, and CXCL16), vesicle rupture, and autophagic dysfunction. It remains to be explored whether this autophagic dysfunction is a key event in the activation of *both* CD4 and CD8 T-cells described herein. Classically, cargo destined for degradation via autophagy is loaded onto MHC-II for antigen presentation to CD4 T-cells. Conversely, intracellular antigenic cargo is degraded via proteasome and loaded onto MHC-I via TAP for antigen presentation to CD8 T-cells. In certain host-defense states, non-conventional presentation routes of internalized cargo can lead to both class I and class II presentation. Such events are termed *cross-antigen presentation* and are characterized by downstream activation of both CD4 and CD8 T-cells (48). Our observations that TDP-43a promotes autophagic dysfunction and vesicle rupture as evidenced by galectin-3 imaging, TEM, proteomics analyses, and biochemical assays, leads us to speculate that a vesicle escape mechanism may drive cross-presentation naïve CD4 and CD8 T-cell activation and T-cell memory responses. While it has been shown that TDP-43 pathology can compromise both autophagy and proteasomal processing in ALS-affected neurons, further experiments in immune cells are warranted to assess this hypothesis.

Our RNA-sequencing data further validated the galectin-3 feed-forward pathway of innate immune activation following vesicle rupture, including increased expression of *LGALS3* and *TLR4* transcript in hMDM following TDP-43a stimulation. We also gleaned information regarding ALS immunophenotypes including reduced *TGFβ* and *STAT5* transcript levels, factors necessary for maintenance of regulatory T-cells (49) that are reduced in ALS patients and inversely correlate with autoimmune-type Th17 T-cells (50–52). Moreover, we observed increased expression of *CCL22*, a homing cytokine for CCR4-expressing NKT cells (53–55), which our VoPo analysis highlighted in fast-progressing ALS patients. Finally, GO Term analysis highlighted the IL-2 signaling, a major T-cell regulatory pathway, following TDP-43a stimulation. These data further support intercellular, innate-to-adaptive immune signaling as a link between early innate responses to TDP-43a and clinically relevant adaptive immune signatures in ALS patients.

Using multiple primary and cell culture models of antigen presentation, we demonstrated that TDP-43a stimulation of antigen presenting cells formed immune synapses with T-cells (Figure 5). With respect to naïve T-cells, we observed intracellular activation via calcium signaling, a hallmark event for TCR-mediated T-cell activation and adaptive immunity. Our high parameter neuroimmune imaging platform highlighted intercellular contacts between innate and adaptive immune cells surrounding TDP-43 pathology *in the ALS brain*. We emphasize that the exact peptide(s) presented on MHC in these experiments are unidentified. At the moment, we cannot 1) conclude what protein within the TDP-43 aggregate is responsible for eliciting this reaction, nor 2) assume it is necessarily the *same* peptide sequence or protein source across human donors. Additional variables could include the highly polymorphic nature of MHC and its impact on peptide-selection and presentation, or natural variation of thymic selection against anti-self T-cells that may exist on an individualistic, disease-specific, or disease subtype-specific basis. Nevertheless, these are certainly topics for future global immunomic studies.

Finally, we implemented machine learning to identify peripheral immune signatures among ALS patients that were associated with ALS progression rates. Surprisingly, ALS subtype-specific immune populations exhibited functional responses when stimulated with TDP-43 pathology. We were surprised to observe that, relative to standard-progressing patients, both slow and fast-progressors exhibited increased representation of various innate and adaptive immune populations, one of these being CD8 TEMRA cells. We note an overall higher representation of CD8 TEMRA cells in ALS, which is in line with previous reports (8). However, we found that CD8 TEMRA cells—typically regarded as terminally differentiated with high cytolytic potential—were overrepresented in non-standard-progressing ALS patients. One consideration is that TEMRA cells could have dichotomous, regulatory properties in slow progressors but cytotoxic properties in fast-progressors. For example, if CD8 TEMRA cells can adopt senescent phenotypes (CD127-CD57+) and regress from sites of tissue inflammation (CD103-), a patient may be protected from advanced ALS phenotypes. However, if CD8 TEMRA maintain proliferative capacity (CD127+CD57-) and reside in the CNS (CD103+), then off-target tissue damage may expedite disease progression. We therefore suggest implementation of functional assays, rather than solely population-based assays, that may provide greater clinical utility when combined with standard profiling approaches. By analyzing whole-blood, but not PBMCs, within 4-hours of a patient visit, we were able to identify changes to NKT cells in fast-progressing ALS. Though larger sample sizes in the future are necessary to extend these findings, we suspect that critical cellular and functional information may be lacking in cryopreserved PBMC preparations (8, 56, 57). Very few studies have been able to evaluate NKT cells, which are significantly impacted by cryopreservation damage following PBMC isolation and storage (43) (58). In summary, this study sheds light on how TDP-43 proteinopathy may be intimately linked to immune dysfunction in ALS and further highlights whole-blood immune assays as a potential precision-medicine approach for diagnosing and predicting clinical features of ALS progression.

## Materials and Methods

### Reagents, cell lines, and cell culture

Raji Burkitt lymphoma cell line (ATCC, CCL-86) and Jurkat T-cell clone E6-1 (ATCC, TIB-152) were procured from UNC Tissue Culture Facility (UNC-TCF). Cells were cultured in vented T-75 flasks in Roswell Park Memorial Institute (RPMI) 1640 media (Thermo Scientific, A1049101) supplemented with 10% fetal bovine serum, 2.0 mM L-glutamine, and 1X penicillin/streptomycin. Cells were maintained between 1.0-2.0x^6^ cells/mL. To passage, cells were collected by centrifugation at 300X RCF for 5 minutes. The cell pellet was resuspended in 1.0 mL of calcium/magnesium-free PBS and incubated at 37°C for 5 minutes. Cell suspension was strained through a 40-micron nylon strainer, counted as above, and re-seeded to a fresh T-75 flask. For a complete list of key resources, see Table 1.

**Table 1.**
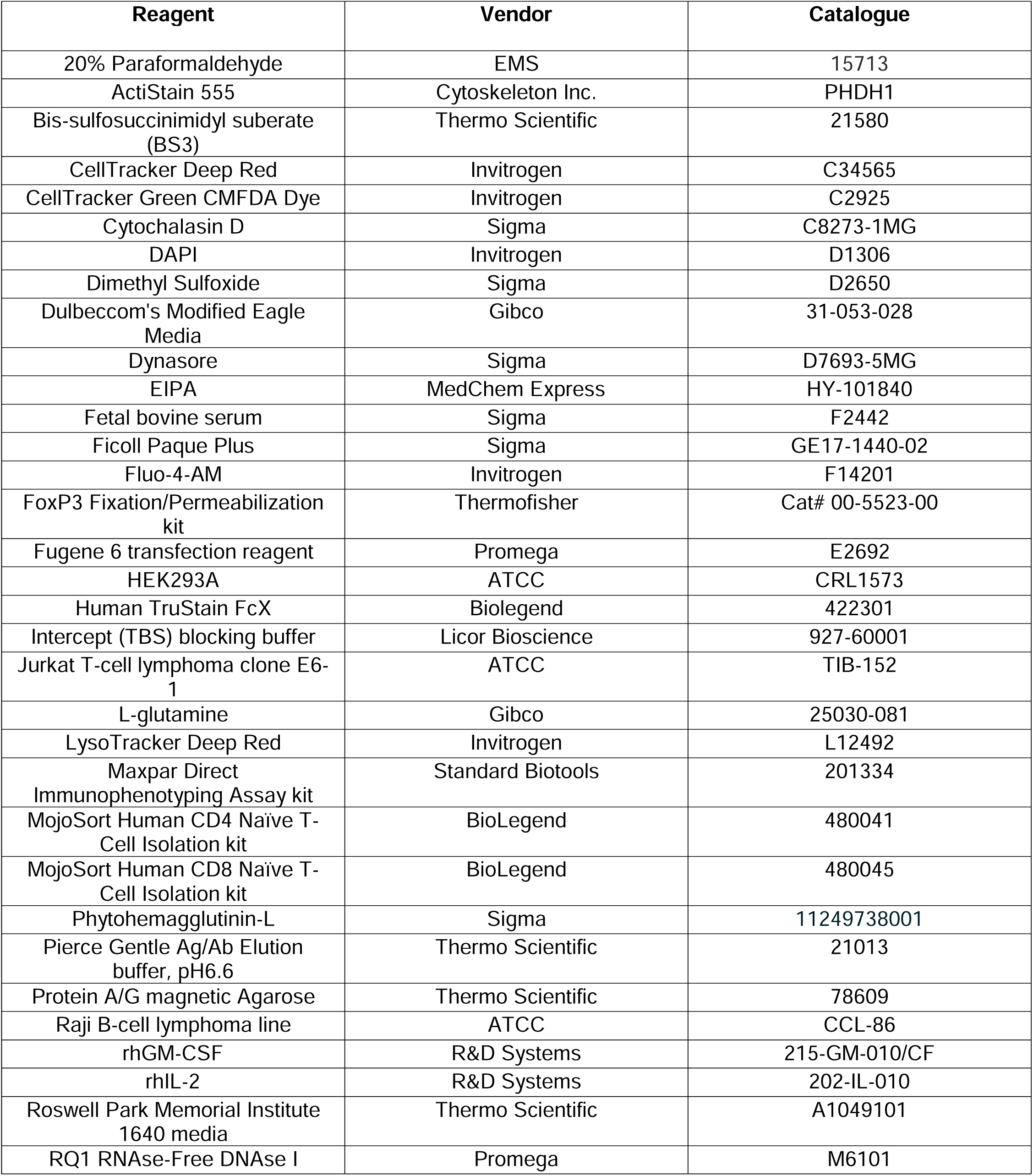
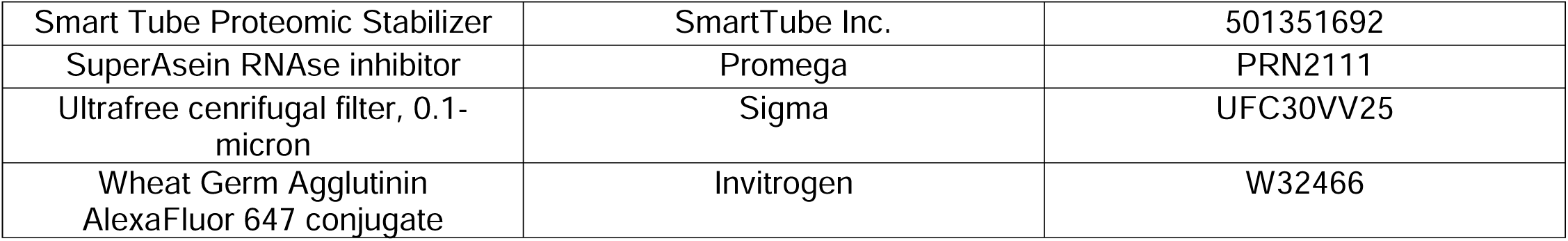
Key Resource Table.

| Reagent | Vendor | Catalogue |
| --- | --- | --- |
| 20% Paraformaldehyde | EMS | 15713 |
| ActiStain 555 | Cytoskeleton Inc. | PHDH1 |
| Bis-sulfosuccinimidyl suberate (BS3) | Thermo Scientific | 21580 |
| CellTracker Deep Red | Invitrogen | C34565 |
| CellTracker Green CMFDA Dye | Invitrogen | C2925 |
| Cytochalasin D | Sigma | C8273-1MG |
| DAPI | Invitrogen | D1306 |
| Dimethyl Sulfoxide | Sigma | D2650 |
| Dulbeccom's Modified Eagle Media | Gibco | 31-053-028 |
| Dynasore | Sigma | D7693-5MG |
| EIPA | MedChem Express | HY-101840 |
| Fetal bovine serum | Sigma | F2442 |
| Ficoll Paque Plus | Sigma | GE17-1440-02 |
| Fluo-4-AM | Invitrogen | F14201 |
| FoxP3 Fixation/Permeabilization kit | Thermofisher | Cat# 00-5523-00 |
| Fugene 6 transfection reagent | Promega | E2692 |
| HEK293A | ATCC | CRL1573 |
| Human TruStain FcX | Biolegend | 422301 |
| Intercept (TBS) blocking buffer | Licor Bioscience | 927-60001 |
| Jurkat T-cell lymphoma clone E6-1 | ATCC | TIB-152 |
| L-glutamine | Gibco | 25030-081 |
| LysoTracker Deep Red | Invitrogen | L12492 |
| Maxpar Direct Immunophenotyping Assay kit | Standard Biotools | 201334 |
| MojoSort Human CD4 Naïve T-Cell Isolation kit | BioLegend | 480041 |
| MojoSort Human CD8 Naïve T-Cell Isolation kit | BioLegend | 480045 |
| Phytohemagglutinin-L | Sigma | 11249738001 |
| Pierce Gentle Ag/Ab Elution buffer, pH6.6 | Thermo Scientific | 21013 |
| Protein A/G magnetic Agarose | Thermo Scientific | 78609 |
| Raji B-cell lymphoma line | ATCC | CCL-86 |
| rhGM-CSF | R&D Systems | 215-GM-010/CF |
| rhIL-2 | R&D Systems | 202-IL-010 |
| Roswell Park Memorial Institute 1640 media | Thermo Scientific | A1049101 |
| RQ1 RNase-Free DNase I | Promega | M6101 |
| Smart Tube Proteomic Stabilizer | SmartTube Inc. | 501351692 |
| SuperAsein RNase inhibitor | Promega | PRN2111 |
| Ultrafree cenrifugal filter, 0.1-micron | Sigma | UFC30VV25 |
| Wheat Germ Agglutinin AlexaFluor 647 conjugate | Invitrogen | W32466 |

### Peripheral Blood Mononuclear Cell (PBMC) Isolation

PBMCs were isolated from de-identified buffy-coated leukocytes obtained from New York Blood Center (Long Island City, NY) following screening against bacteremia and viremia. Briefly, anticoagulated blood (heparin or ACD citrate as indicated) was diluted 1:1 with 1X sterile phosphate-buffered saline (PBS). Diluted blood was overlaid atop Ficoll-Hypaque density gradient (Sigma, GE17-1440-02) and centrifuged at 120X RCF for 20 minutes with zero brake. Buffy coat was isolated, washed in 1X PBS, centrifuged as above twice, and cleared with 1X erythrocyte (RBC) lysis buffer for 10 minutes with gentle agitation. Cleaned PBMCs were then washed with 1X PBS, counted using Trypan Blue exclusion hemocytometry, and plated according to assay or differentiation protocol. For storage of PBMCs, cells were cryopreserved in 10% DMSO in antibiotic-free media containing 10% FBS (Sigma, F2442), and 2 mM L-Glutamine (Gibco, 25030-081) in Dulbecco’s Modified Eagle Media (DMEM; Gibco, 31-053-028).

### Primary human monocyte-derived macrophage (hMDM) culture

Freshly isolated PBMCs were plated at a density of 1.0x^6^ cells/mL into ultralow adhesion plastic tissue culture dishes (Corning CLS3473), glass bottom slides (Cellvis, C8-1.5H-N), or glass-bottom culture dishes (Nunc, 150680) in complete DMEM consisting of 10% FBS, 2.0 mM L-glutamine, 1X Penicillin/Streptamycin (Gibco, 15140122). Monocytes were allowed to adhere to substrate for 5 days at 37°C, 5% CO_2_. At day 5 in culture, media was exchanged with complete media plus 15 ng/mL carrier-free recombinant human granulocyte-macrophage colony stimulating factor (rhGM-CSF; R&D, 215-GM-010/CF). Thereafter, differentiated macrophages were maintained with half-volume media exchanges performed every other day.

### TDP-43 aggregate purification

The following is an abridged protocol based on Evangelista et al., 2023. All buffers and plastics were sterile and procedures performed in a laminar flow hood. Briefly, HEK293A (ATCC, CRL1573) cells (ATCC, CRL1573) were seeded to 15-cm tissue culture plates such that they reached 80% confluence at day of transfection. Cells were transfected with 8 µg of pcDNA3.1-GFP-mNLS-K145Q using Fugene 6 Transfection reagent (Promega, E2692) per manufacturer protocol. Cells were harvested 48-hours post-transfection in sterile 1X RIPA buffer supplemented with protease, phosphatase, and RNAse inhibitors (Proega, PRN2111). Cells were passed through 18-, 21-, and 25-gage syringes, vortexed at top speed for 30 seconds, and then treated with RQ1 RNAse-free DNAse (Promega, M6101) for 30 minutes at 37°C. Sarkosyl was added to a final concentration of 0.5%. Lysates were sheared as above, then centrifuged at 100,000X RCF for one hour at 4°C. The supernatants were removed. Crude pellets were stored at -80 indefinitely until immunoprecipitation. To prepare immunoprecipitation matrix, 30 µL of Protein A/G magnetic agarose (Thermo Scientific, 78609) was incubated with 15 µg of TDP-43 antibody (Proteintech, 10782-2-AP) for one hour at 4°C. Residual antibody was removed, beads were washed twice with 1X RIPA, and twice with 1X PBS. BS3 cross-linker (Thermo Scientific, 21580) was diluted to 5 mM with 1X sterile PBS. Antibody-bead complexes were crosslinked for 30 minutes at room temperature on an orbital shaker. Crosslinking was quenched with a 1:10 ratio of 1.0 M Tris pH 7.5. Non-crosslinked antibody was removed with 2 washes of Pierce Gentle Antibody elution buffer (Thermo Scientific, 21013), and the beads washed twice more with 1X RIPA. Beads were then blocked overnight in non-protein block buffer (Licor Bioscience, 927-60001) with 1mM DTT and 1.0% Tween-20 at 4°C with orbital rotation. For control samples, Rabbit isotype control antibody was used in place of 10782-2-AP. To immunoprecipitated TDP-43 aggregates, pellets were retrieved from -80°C. and sonicated in 500 µL of extraction buffer (1X RIPA supplemented with 0.5% Sarkosyl, 1.0% Tween-20, 1.0 mM DTT, protease, phosphatase, and RNAse inhbitors). Bead slurry and lysate were combined and allowed to bind overnight at 4°C with constant rotation. The next day, supernatants were removed, beads were washed 3X in extraction buffer, and aggregates were eluted with a 3-series wash with 5.0 M NaCl warmed to 55°C. Aggregates were pelleted at 100,000x RCF for 1 hour, and sonicated into a 1.0 mL solution (15 second pulses, 30 second intermission on ice, for 4 cycles) using sterile 1X PBS unless otherwise specified.

### Immunofluorescence analysis

Coverslips were gently rinsed in room temperature PBS and then immediately fixed in fresh, 4.0% paraformaldehyde (PFA; Electron Microscopy Services, 15713S) for 10 minutes at room temperature. Cells were rinsed 3X in PBS, then permeabilized using 0.2% Triton X-100 in PBS for 8 minutes at room temperature. Cells were blocked in 2.0% normal goat serum (Sigma, NS02L-1ML) + 0.2% Triton X-100 (PBSTN) for one hour at room temperature. Cells were incubated with primary antibodies in 2.0% PBSTN overnight at 4°C. Cells were rinsed in PBST and subsequently stained with fluorophore conjugated secondary antibodies in PBSTN for one hour at room temperature protected from light. Cells were preserved in ProLong Diamond anti-fade mounting medium (Thermo Scientific, P36965).

### hMDM aggregate uptake assay for confocal microscopy

hMDM were cultured in glass bottom dishes. Cells were pre-charged with 10 µM Cytochalasin D (Sigma, C8273), 80 µM Dynasore (Sigma, D7693), 10 µM EIPA (MedChem Express, HY-101840), or 0.05% DMSO control for 30 minutes at 37°C. Cells were then stimulated with 0.025% (v/v) TDP-43a and returned to incubator for 4 hours. Cells were rinsed 1X in PBS, then immediately fixed for 10 minutes at room temperature with 4.0% PFA. Cells were rinsed 2X in PBS and stained with 10 µg/mL Wheat-Germ Agglutinin (WGA) AlexaFluor 647 conjugate (Invitrogen, W32466) for 10 minutes at room temperature. Cells were rinsed 2X in PBS and mounted with ProLong Diamond antifade mounting media.

### Confocal microscopy and live-cell imaging

Images were obtained on an inverted Zeiss 800/Airyscan laser-scanning confocal microscope fitted with 405, 488, 561, and 640 nm diode lasers and gallium arsenide phosphide (GaAsP) detectors. Live cells were incubated in a 37°C heated stage in a humidified 5% CO_2_ chamber. Image analyses were performed in FIJI (version 1.53t) (59). Brightness and contrast were similarly adjusted between treatments.

### Solubility fractionation and immunoblotting

6-well tissue culture plates were removed of media and rinsed 1X in 1.0 mL PBS. Cells were collected in 250 µL of 1X radioimmunoprecipitation assay buffer (RIPA) by scraping over ice. In all cases, RIPA buffer was supplemented with protease and phosphatase inhibitors. Lysates were sonicated 20-times at 25% amplitude with a QSonica hand sonicator probe over ice. Samples were centrifuged at 21,000X RCF for 30 minutes at 4°C. Supernatants were transferred to chilled microcentrifuge tubes and stored at -80°C as the soluble fraction. Pellets containing insoluble fraction were resuspended in Urea plus protease and phosphatase inhibitors, and sonicated for 10 pulses as above. Samples were centrifuged as above, and the supernatants stored as the insoluble fraction. To prepare samples for sodium dodecyl sulfate polyacrylamide gel electrophoresis (SDS-PAGE) analysis, protein lysate concentrations were normalized by bicinchoninic assay. 5.0-10.0 µg of protein lysate were denatured in 1X reducing Laemmli buffer, boiled at 98°C for 5 minutes, and electrophoresed on 12% SDS-PAGE gel. Proteins were transferred to nitrocellulose membrane at 4°C, 100V, for 75 minutes. Membranes were blocked in Tris buffered saline-0.2% Tween (TBS-T) with 2.0% skim-milk for 30 minutes at room temperature with rocking. Membranes were incubated with primary antibodies (Table 2) at 4°C overnight in 0.2% Milk-TBS. Membranes were washed 3X in TBS-T, incubated with respective secondary antibodies (Table 3) for 60 minutes at room temperature with rocking and washed as above. Membranes were imaged by enhanced chemiluminescence and quantified using ImageStudo Lite.

**Table 2.** Primary antibodies.

| Antibody | Vendor | Dilution | Catalogue | RRID |
| --- | --- | --- | --- | --- |
| TDP-43 pS409/410 | Proteintech | IB: 1:1000 | 80007-1-RR | AB_2882937 |
| Total TDP-43 | Proteintech | IB: 1:2000<br>TDiP: 40 µg | 10782-2-AP | AB_615042 |
| GFP | Aves | IB: 1:1000 | GFP1202 | AB_2734732 |
| GAPDH | Millipore | IB: 1:1000 | ABS16 | AB_10806772 |
| LC3B | CST | IB: 1:1000<br>IF: 1:100 | 2775S | AB_915950 |
| p62 | CST | IB: 1:1000 | 5114S | AB_10624872 |
| Galectin-3 | BioLegend | IF: 1:100 | 126701 | AB_1134255 |
| Rab5a | Santa Cruz | IF: 1:100 | SC46692 | AB_628191 |

**Table 3.** Secondary antibodies.

| Antibody | Vendor | Dilution | Catalogue |
| --- | --- | --- | --- |
| HRP Goat anti-Rabbit | Invitrogen | IB: 1: 3000 | A31458 |
| HRP Goat anti-Chicken | Invitrogen | IB:1: 3000 | A16054 |
| HRP Goat anti-Mouse | Invitrogen | IB:1:1000 | A31430 |
| AlexaFluor 594 conjugated goat anti-Rabbit | Invitrogen | IF: 1:500 | A11012 |
| AlexaFluor 594 conjugated goat anti-Mouse | Invitrogen | IF: 1:500 | A32742 |
| AlexaFluor 647 conjugated goat anti-Rabbit | Invitrogen | IF: 1:500 | A32733 |

**Table 4.** Suspension mass cytometry probes.

| Assembly | Isotope | Antibody | Clone | Dilution |
| --- | --- | --- | --- | --- |
| MDIPA Kit | 89Y | CD45 | HI30 |  |
|  | 141Pr | CD196/CCR6 | G034E3 |  |
|  | 143Nd | CD123 | 6H6 |  |
|  | 144Nd | CD196/CCR6 | HIB19 |  |
|  | 145Nd | CD4 | RPA-T4 |  |
|  | 146Nd | CD8a | RPA-T8 |  |
|  | 147Sm | CD11c | Bu15 |  |
|  | 148Nd | CD16 | 3G8 |  |
|  | 149Sm | CD45RO | UCHL1 |  |
|  | 150Nd | CD45RA | HI100 |  |
|  | 151Eu | CD161 | HP-3G10 |  |
|  | 152Sm | CD194/CCR4 | L291H4 |  |
|  | 153Eu | CD25 | BC96 |  |
|  | 154Sm | CD27 | O323 |  |
|  | 155Gd | CD57 | HNK-1 |  |
|  | 156Gd | CD183/CXCR3 | G025H7 |  |
|  | 158Gd | CD185/CXCR5 | J252D4 |  |
|  | 160Gd | CD28 | CD28.2 |  |
|  | 161Dy | CD38 | HB-7 |  |
|  | 163Dy | CD56/NCACM | NCAM16.2 |  |
|  | 164Dy | TCRgd | B1 |  |
|  | 166Er | CD294 | BM16 |  |
|  | 167Er | CD197/CCR7 | G043H7 |  |
|  | 168Er | CD14 | 63D3 |  |
|  | 170Er | CD3 | UCHT1 |  |
|  | 171Yb | CD20 | 2H7 |  |
|  | 172Yb | CD66b | G10F5 |  |
|  | 173Yb | HLA-DR | LN3 |  |
|  | 174Yb | IgD | IA6-2 |  |
|  | 176Yb | CD127 | A019D5 |  |
|  | 103Rh | Cell-ID Intercalator |  |  |
| **Expansion Panel** | 169Tm | CD69 | FN50 | 0.125 µL/test |
|  | 165Ho | CD103 | Ber-ACT8 | 0.125 µL/test |
|  | 159Tb | CD366 | F38-2E2 | 2.0 µL/test |
|  | 162Dy | CD95 | DX2 | 0.125 µL/test |
|  | 142Nd | CD134 | ACT35 | 1.0 µL/test |
|  | 209Bi | CD137 | 4B4-1 | 1.0 µL/test |
|  | 175Lu | CD279 | EH12.2H7 | 0.25 µL/test |
|  | 198Pt | Ki67 | Ki-67 | 0.25 µL/test |
|  | 159Tb | HLA-ABC | W6/32 | 0.25 µL/test |
| **Barcodes** | 76Se | 76SeMal | In-house | Cell: 98 µM |
|  | 77Se | 77SeMal | In-house | Cell: 98 µM |
|  | 78Se | 78SeMal | In-house | Cell: 98 µM |
|  | 124Te | 124TeMal | In-house | Cell: 2 µM |
|  | 126Te | 126TeMal | In-house | Cell: 2 µM |
|  | 128Te | 128TeMal | In-house | Cell: 2 µM |
|  | 130Te | 130TeMal | In-house | Cell: 2 µM<br>TDiP: 100 µM |

**Table 5.** Imaging mass cytometry probes.

| Assembly | Isotope | Antibody | Clone | Dilution | Conjugation |
| --- | --- | --- | --- | --- | --- |
| Neural Landscape | 150Nd | Total TDP-43 | 6H6E12 | 1:250 | In-house |
|  | 159Tb | CD68 | KP1 | 1:200 | SBT |
|  | 161Dy | pS409/410 (pTDP-43) | 1D3 | 1:100 | In-house |
|  | 164Dy | GFAP | GA-5 | 1:700 | In-house |
|  | 169Tm | Myelin Basic Protein | EPR21188 | 1:800 | In-house |
|  | 194Pt | MAP2 | MT-08 | 1:150 | In-house |
| Immune Landscape | 143Nd | CD45RA | HI100 | 1:200 | In-house |
|  | 144Nd | CD31 | EPR3094 | 1:50 | In-house |
|  | 147Sm | CD163 | EDHu-1 | 1:200 | SBT |
|  | 153Eu | Mac2 (Galectin-3) | M3/38 | 1:50 | SBT |
|  | 156Gd | CD4 | EPR6855 | 1:200 | SBT |
|  | 162Dy | CD8a | C8/144B | 1:200 | SBT |
|  | 167Er | Granzyme B | EPR20129-217 | 1:750 | SBT |
|  | 170Er | CD3 | Polyclonal | 1:200 | SBT |
|  | 173Yb | CD45RO | UCHL1 | 1:200 | SBT |
|  | 174Yb | HLA-DR | LN3 | 1:50 | SBT |
|  | 191/193Ir | DNA | N/A | 1:800 | SBT |

**Table 6.** Patient demographics for whole-blood analysis.

| Identifier | Age of onset | Sex | ALSFRS-R/time* | Classifier | HIV | Riluzole* | Edaravone* | Sodium phenylbutyrate and taurursodiol* |
| --- | --- | --- | --- | --- | --- | --- | --- | --- |
| Duke-812 | 78 | F | 0.60 | Standard | no risks | no | no | no |
| Duke-813 | 78 | F | 0.15 | Slow | no risks | yes | no | no |
| Duke-814 | 52 | F | 1.00 | Standard | negative | no | no | no |
| Duke-815 | 61 | F | 0.70 | Standard | no risks | yes | no | no |
| Duke-816 | 74 | F | 0.07 | Slow | negative | no | no | no |
| Duke-817 | 60 | M | 0.42 | Slow | no risks | no | no | no |
| Duke-818 | 79 | M | 0.37 | Slow | no risks | yes | no | no |
| Duke-819 | 44 | F | 4.50 | Fast | no risks | yes | no | no |
| Duke-824 | 65 | M | 0.38 | Slow | negative | no | no | no |
| Duke-825 | 56 | M | 0.20 | Slow | negative | no | no | no |
| Duke-826 | 73 | M | 1.40 | Fast | negative | yes | no | no |
| Duke-827 | 61 | M | 1.40 | Fast | no risks | yes | no | no |
| Duke-828 | 65 | M | 1.00 | Standard | no risks | yes | no | no |
| Duke-829 | 55 | F | 0.40 | Slow | no risks | no | no | no |
| Duke-830 | 68 | M | 0.74 | Standard | negative | no | no | no |
| Duke-831 | 61 | F | 1.20 | Fast | negative | no | no | no |
| Duke-832 | 59 | F | 2.20 | Fast | negative | yes | no | no |
| Duke-835 | 75 | M | 0.50 | Standard | negative | yes | no | no |
| **Duke-836** | 67 | M | 0.60 | Standard | negative | yes | yes | yes |
| **Duke-838** | 74 | F | 0.35 | Slow | no risks | no | no | no |
| **UNC-003** | 72 | M | 0.00 | Slow | no risks | yes | no | no |
| **UNC-004** | 50 | M | 4.00 | Fast | No risks | yes | yes | no |

*\* At time of sample acquisition*

### Secretome and proteome analysis

Differentiated macrophages in 10-cm dishes underwent complete media change to serum-free DMEM. Cells were then stimulated with 0.025% (v/v) aggregate-containing media or equal volume of isotype control material and allowed to incubate for 16-hours. Cell culture media was removed and centrifuged at 300 RCF for 5 minutes at 4°C to remove cell debris, followed by filtration through a 0.2 µm syringe filter. Cells were harvested by gently scraping in ice-cold PBS, centrifuged as above and supernatant was removed. All components were then stored at -80°C until a full set of replicates was obtained. For media analysis, proteins were precipitated with deoxycholate and trichloroacetic acid. Next, protein precipitates and cell pellets were solubilized in 8M urea. Proteins were then reduced, alkylated, and digested with Trypsin at room temperature overnight. Digested peptides were then acidified with trifluoroacetic acid, desalted, and reconstituted in 0.1% formic acid. Samples were analyzed by reverse phase liquid chromatography (LC)-MS/MS using Velos Orbitrap mass spectrometer (Thermo). Experiments were calibrated to a mass accuracy of < 0.05.

### Electron Microscopy

Adherent cells were fixed in 2% paraformaldehyde/2.5% glutaraldehyde in 0.15 M sodium phosphate buffer, pH 7.4 for 1 hour at room temperature, then stored in fixative at 4°C until processed for TEM. Fixative was aspirated and cells were washed three times in 0.15 M sodium phosphate buffer. After washing, reduced osmium tetroxide (1% osmium tetroxide with 1.25% potassium ferrocyanide in 0.15 M sodium phosphate buffer) was added to cover the cellular monolayer and incubated at room temperature for 1 hour. Cells were washed three times with deionized water and dehydrated through an ascending series of ethanol (30%, 50%, 75%, 90%, 100%, 100% 100%). They were then infiltrated with three exchanges of Polybed 812 epoxy resin (Polysciences, Inc., Warrington, PA) before being embedded in fresh 100% Polybed 812 epoxy resin and allowed to cure at 60°C until hardened. Resin blocks were sectioned at 80 nm using a diamond ultra knife on a Leica UCT7 ultramicrotome and mounted on 200 mesh copper grids. Grids were stained with 4% aqueous uranyl acetate for 12 minutes followed by Reynold’s lead citrate for 8 minutes, based on Reynolds et al., 1963 (60). Samples were viewed using a JEOL JEM-1230 transmission electron microscope operating at 80 kV (JEOL USA, Inc., Peabody, MA) and images were obtained using a Gatan Orius SC1000 CCD Digital Camera and Gatan Microscopy Suite 3.0 software (Gatan, Inc., Pleasanton, CA).

### RNA Sequencing

#### RNA-seq library preparation

Total RNA was extracted using RNeasy Mini Kit (QIAGEN) guided by the manufacturer’s instructions. Each sample’s RNA concentration and RNA integrity number (RIN) were measured using Qubit and Agilent Tapestation 4150 system. The depletion of ribosomal RNAs (rRNAs) and the generation of stranded RNA-seq libraries were performed using KAPA RNA HyperPrep with RiboErase kit with 500 ng of isolated RNA as input and following the guidelines. Final libraries were quantified and normalized by Qubit and DNA Tapestation and then combined to a total 4 nM library. The combined library was subsequently subjected to paired-end 75bp read sequencing in the Illumina Nextseq550 platform.

#### RNA-seq data analysis

The quality of the reads was assessed by FastQC (version 0.11.9). Low-quality reads and adapters were trimmed by using Trim Galore (version 0.6.7). The remaining reads were mapped to the hg19 transcriptome (GENCODE release19) and quantified by Salmon (v. 1.10.0). The estimated counts for each sample were summarized by using tximport (v. 1.28.0). Differential gene analysis was applied to the counts matrix with DESeq2 (v. 1.40.2) using a design adjusting for technical bias when calculating differences between treatment groups (∼Replicates+Condition). The method “apeglm” was used for effect size shrinkage (LFC estimates). Genes with an FDR-adjusted p-value below 0.05 (Wald test) and an absolute fold change greater than 1.15 were selected as the differential genes when comparing the treated samples to their corresponding controls. The gene symbols and Entrez IDs were annotated to the data by using the package AnnotationHub (v. 3.8.0) and annotables (v. 0.2.0). The universal gene list and differential gene lists were then prepared for generating functional enrichment results using over-representation analysis from the GO database by clusterProfiler (v. 4.8.3), along with its complementary packages. The GO terms were identified with both FDR-adjusted p-value and multiple testing corrected p-value lower than 0.05. The number of overlap genes between different conditions was conducted by the package eulerr (v. 7.0.0) displaying the proportion of area related to the numbers.

### Aggre-Gate: TDP-43a internalization assay by mass cytometry

#### ^130^Tellurium maleimide (130TeMal) coupling

TDP-43a were purified as above. After the final spin, the supernatant was removed and replaced with 500 µL of sterile Maxpar PBS, vortexed for 5 seconds, and collected by centrifugation for 30 minutes at 100,000x RCF. The supernatant was removed and 100 mL of fresh sterile Maxpar PBS was added. The pellet was sonicated for two cycles of 20, 1-second pulses at 25% amplitude with a 30 second incubation on ice between cycles. Next, 1M DTT was added to a final concentration of 0.5 mM and the sample partially reduced for 10 minutes at room temperature. The reaction was quenched with 900 µL of sterile Maxpar Cell Staining Buffer (CSB), followed by centrifugation at 100,000 RCF for 30 minutes. The supernatant was removed, replaced with 1.0 mL of sterile Maxpar PBS, vortexed, and centrifuged as above. The partially reduced TDP-43a pellet was sonicated in 100 µL Maxpar PBS as above, and 1.1 µL of 10 mM 130TeMal was added for a final concentration of 100 µM. Coupling occurred for 15 minutes at room temperature with constant mixing by vortex at low speed. The reaction was quenched and centrifuged as above. 1.0 mL of sterile PBS was added to the TDP-43a pellet and sonicated for two, 20-pulse cycles on ice as above. Aggregate preparations were then tested for microbial and endotoxin contamination by culturing a sample in nutrient agar followed by OD600 measurement (<0.001 arbitrary absorbance units) and LAL assay (<0.05 EU/mL), respectively.

#### Cell stimulations

On Day 0, PBMCs were thawed in pre-warmed complete RPMI1640 supplemented with 50 units/mL of benzonase endonuclease. Cells were pelleted at 300x RCF for 3 minutes and the supernatant discarded. Cells were resuspended in 5.0 mL of complete RPMI1640 plus benzonase and transferred to a vented T-25 flask, allowed to recover overnight. The next morning, cells were collected by centrifugation. The cells were resuspended in complete media to count viable cells by trypan blue exclusion. 2.0e6 cells were added to a 24-well round-bottom tissue culture plate in complete RPMI. As indicated, TDP-43a was added to a final concentration of 0.025% (v/v) and returned to 5% CO_2_ incubation. Stimulations were performed in reverse chronological order (i.e. starting with longest time-points) then collected at once for cytometry staining to normalize phenotypic drift in bulk immune cultures along incubation.

#### Mass cytometry staining and acquisition

Following stimulation, cells were harvested in Maxpar PBS and centrifuged at 300x RCF for 5 minutes. As needed, cells were barcoded using 1.0 µM TeMal and Selenium maleimide (SeMal) barcodes in Maxpar PBS for 15 minutes at room temperature. TeMal isotopes were synthesized by Dr.Youngran Seo at the University of North Carolina Department of Chemistry according to the published protocol of *Willis et al* (61). Reactions were quenched with 3 volumes of CSB and centrifuged at 300x RCF for 5 minutes. Cells were pooled in 100 µL volumes of CSB. Fc receptors were blocked with 10 µL TruStain FCX receptor block (BioLegend, 422301) for 10 minutes at room temperature. Blocked cells were added to 110 µL of CSB and transferred to a Maxpar Direct Immunophenotyping Assay (MDIPA; Standard BioTools, 201334) tube along with 50 µL of custom checkpoint antibody cocktail (For complete suspension mass cytometry antibody panels see Table 4). Cells were stained for 30 minutes at room temperature. Cells were washed twice in 3.0 with mL of CSB followed by centrifugation. Intracellular Ki67 staining was performed using the FoxP3 transcription factor staining kit per manufacturer protocol (ThermoFisher, 00-5523-00). Cells were fixed in 2% paraformaldehyde for 1 hour at 4°C, pelleted at 800X RCF for 7 minutes, and resuspended in 1.0 mL of Maxpar Fix/Perm buffer containing 1:3000 Cell ID-Iridium DNA intercalator for overnight incubation at 4°C. The next day, cells were pelleted at 800 RCF for 7 minutes, resuspended in 200 µL of Fix/Perm buffer, and stored at -80°C until batch analysis. On day of analysis, cells were washed once with CSB, once with Cell Acquisition Solution (CAS; Standard BioTools), then filtered through a 40-micron filter and diluted in CAS containing 10% EQ Calibration Beads (Standard BioTools) at 0.5 million cells per mL and acquired on a mass cytometer (Helios). Mass cytometry data were normalized using Fluigidm CyTOF software (v7.0). Sample files were pre-processed to remove beads, debris, dead cells, and doublets for further downstream analysis. For SPADE analysis, down-sampling was set to 10%, with a node setting of 75.

### Raji-Jurkat immune synapse assay

Raji cells were stained with 1.0 µM Celltracker Deep Red (Invitrogen, C34565) for 30 minutes in supplement-free RPMI1640 at 37°C. 2.0e6 cells were added to vented flow cytometry tubes in complete RPMI 1640. As indicated 0.025% (v/v) TDP-43a, isovolumetric equivalent of isotype control material, or (0.5) µg/mL phytohemagglutinin (PHA) were added to tubes for a 16-hour simulation period at 37°C. The next day, Jurkat T-cells were incubated with 1.0 µM CellTracker Green (Invitrogen, C2925) for 30 minutes at 37°C. 2.0e6 Jurkat cells were added to flow cytometry tubes containing Raji cells and incubated at 37°C for 45 minutes. Cells were gently pipetted and strained through a 70 µm strainer directly into 1 volume equivalent of 8% PFA (for a final concentration of 4%). Cells were fixed for 10 minutes at room temperature in the dark. Cells were pelleted at 800 RCF for 5 minutes, then resuspended in PHEM buffer (5 mM HEPES, 60 mM PIPES, 10 mM EGTA, 2 mM MgCl_2,_ pH 7.0 with KOH) plus 0.1% (w/v) digitonin. To label immunological synapses, ActiStain-555 (phalloidin) was added to a final concentration of 14 µM and incubated for 30 minutes at room temperature in the dark. Cells were then rinsed in 1X PBS and incubated with 1ug/mL DAPI in PBS prior to mounting on microscope slides with Prolong Diamond Antifade mounting media. Synapse frequencies were quantified as at least 1 Raji and 1 Jurkat T-cell containing a polarized actin contact site from randomized fields of view obtained on a 20X air objective. Descriptively, synapses were then categorized as pair (one Raji per one Jurkat) or clusters.

### Macrophage-PBMC co-culture assay

hMDM were sub-cultured to glass coverslips. hMDM were stimulated with 0.025% (v/v) immunopurified TDP-43a and volumetric equivalent of isotype control product for 16 hours. In parallel, cryopreserved syngeneic PBMCs were thawed and allowed to recover overnight in complete RPMI 1640 plus benzonase as above. The next day, 2e6 PBMCs per condition were labeled with 1 µM CellTracker green for 30 minutes at 37°C in supplement-free RPMI 1640. The reaction was quenched and washed with complete media. 1e6 labeled PBMCs were added drop wise to stimulated hMDM and were returned to 37°C, 5% CO_2_ for 45 minutes. Without disturbing the cells, a 1-well volume equivalent of 8% PFA was added per well (final concentration 4%) and allowed to incubate for 10 minutes at room temperature. Media was removed and fresh fixative was added for an additional 10 minutes. Coverslips were rinsed and permeabilized in PHEM buffer. Actin was stained using Acti-Stain 555 as above, nuclei were counter-labeled with DAPI, and immune synapses were imaged on an LSM 800 confocal microscope at 20X and 63X magnification with Airyscan.

### hMDM-T-cell calcium imaging assay

Macrophages were stimulated for 16-hours with 0.025% (v/v) TDiP aggregates or isovolumetric isotype control material in complete media in Nunc glass-bottom microscope dishes. In parallel, naïve CD4 and CD8 T-cells were isolated from syngeneic cryopreserved PBMC aliquots using the MojoSort magnetic negative selection kits (BioLegend, 480041 and 480045). Naïve T-cells were incubated in complete medium supplemented with 10 IU/mL recombinant human IL-2 (R&D Systems, 202-IL-050/CF). As a positive control, macrophages were treated with 10 pg/mL neutralized, CCR5 tropic, HIV virions. Following macrophage stimulations, macrophages and T-cells were separately loaded with calcium indicator dye using a modified protocol. Here, cells were washed in HEPES-buffered artificial cerebrospinal fluid (137 mM NaCl, 5 mM KCl, 2.3 mM CaCl_2_, 1.3 mM MgCl_2_, 20 mM glucose, and 10 mM HEPES, pH 7.4) and pre-charged with 25% (v/v) Fluo-4 acetoxymethyl (Invitrogen, # F14201) for 30 minutes. T-cells were added to macrophages at a ratio of 1 (macrophage) : 2 (T-cell) and imaged on an Olympus IX71 inverted microscope. Time-lapsed images were measured every 6 seconds for 6 minutes for 20 minutes using Metamorph (Molecular Devices). Change to fluorescence intensity was calculated using baseline calcium levels per cell.

### Qualitative Imaging Mass Cytometry

#### Antibody Conjugation

Antibody panel configuration was performed by Standard BioTools to ensure optimal isotope assignment based on antigen abundance. Approximately 100 µg of each antibody (carrier-free) were coupled to lanthanide metals using previously described protocols with the Maxpar X8 labeling kit (Standard BioTools, 201169B). Briefly, antibodies were partially reduced with TCEP buffer (SBT, 77720) at 37°C. Following 112 reduction, antibodies were then incubated with an excess of metal-loaded MaxPar X8 polymer for 90 min at 37°C. Labeled antibodies were purified using a 50 kDa size exclusion centrifugal filter unit. Final antibody concentration was determined by A_280_. For complete list of IMC probes, refer to Table 5.

#### Tissue staining

10 µM formalin-fixed paraffin embedded (FFPE) spinal cord sections from ALS and control spinal patients were obtained from the Veterans Affairs ALS Brain Biorepository. FFPE sections were de-paraffinized in xylene followed by progressive rehydration in a series of graded alcohols. Heat-induced epitope retrieval with pH 9.0 buffer was performed prior to blocking in PBS + 3% BSA for 45 minutes. Tissues were then incubated with antibodies overnight at 4°C in PBS + 0.5% BSA. Tissues were washed in PBS + 0.2% Triton X-100 prior to nuclear staining with Iridium intercalator (1:800) for 30 minutes. Finally, tissues were washed with water and dehydrated at room temperature. Regions of interest (ROI) were previously determined on an immediate serial section using chromogenic immunohistochemistry, focusing on areas positive for pS409/410 (1D3). 1 mm x 1 mm ROIs were placed around 2 pS409/410-positive lesions. Synchronized laser ablation was performed at 200 Hz over a 2-hour period, generating regions with 1 µm^2^ resolution. Data were exported to MCD Viewer (Standard BioTools) where they were converted to 32-bit TIFF files. Each channel was despeckled and composite images were created using FIJI (version 1.53t). Brightness and contrast for each marker was adjusted similarly across each ROI.

### Whole blood immune profiling and stimulation

#### General immunophenotyping

ALS patient blood was collected at Duke University and UNC Chapel Hill Neurology clinics under approved Institutional Review. Blood was collected in 10 mL ACD citrate collection tubes and processed within 4 hours of collection. 300 µL of anticoagulated blood were then added to one tube of the Maxpar Direct Immunophenotyping Assay and incubated for 30 minutes at room temperature. Blood was briefly agitated by tapping every 10 minutes to ensure even staining. Stained blood was then transferred to cryovials containing 420 µL of SmartTube Proteomic Stabilizing agent (SmartTube Inc, 501351692), mixed, and allowed to stabilize for 10 minutes at room temperature. Samples were then stored at -80°C until batch processing and mass cytometry analysis. One day prior to analysis, stabilized blood was thawed at room temperature and transferred to 10 mL of 1X Thaw/Lyse buffer (SmartTube Inc.) and allowed to incubate at room temperature for 10 minutes with orbital rotation. Lysed material was pelleted by centrifugation at 500 RCF for 5 minutes. Lysis was repeated one time. Cells were then washed in 5 mL of CSB and resuspended in 1mL of 1X Maxpar PBS. Cells were strained through 70 µM strainer caps into flow cytometry tubes containing 1 mL of 4% fresh PFA (final concentration 2%). Cells were fixed for 1 hour at 4°C. Cells were pelleted at 800 RCF for 7 minutes, then resuspended in 1 mL of Maxpar Fix/Perm buffer containing 1:3000 Cell-ID Iridium Intercalator. All samples contained 500,000 universal donor PBMCs barcoded with 130TeMal to aid in normalization. A minimum of 300,000 events were collected in all conditions.

#### Whole-blood TDP-43a stimulations

For whole-blood stimulation assays, ALS and control patient blood was collected in 5 mL Heparin tubes. We opted for heparin tubes as these appeared to be preferred for whole-blood stimulation assays, although a comparative analysis suggests no significant difference in lymphocyte-antigen response between ACD citrate and heparin (62). 300µL of blood was added to sterile vented flow cytometry tubes and received either 3.0% (v/v) TDP-43a, 0.5 µg/mL PHA, or isovolumetric DMSO control that never exceeded 0.05% in culture. Samples were incubated for 16-hours in 5% CO_2_ at 37°C. Blood was transferred to MDIPA reaction tubes with 50 µL of immune checkpoint antibody cocktail. A metal-minus-one control for CD103 gating was established by staining whole blood with MDIPA, excluding the CD103 probe. Following staining, blood was stabilized, cleaned, and prepared for batch analysis as above.

### Statistical analysis and data reporting

Where appropriate, data were assessed for normality by Kruskall-Wallis test. Non-parametric data were analyzed by non-parametric two-tailed t-test, or multivariable Mann-Whitney U-test. Parametric data were analyzed by two-tailed t-test, one-way ANOVA with Bonferroni correction or Tukey’s multiple comparison test, or Two-way ANOVA with Sidak’s multiple comparison test. For correlation analysis, Spearman’s *r* was used. For primary PBMC stimulation data, paired t-tests were used to match genotypes. Arcsin(*h*) transformed data of surface marker expression changes were analyzed via paired t-test. Statistical analysis was performed using GraphPad Prism Version 9 for Windows (San Diego, CA). Cut-off for statistical significance were as follows: *p < 0.05, **p < 0.01, ***p < 0.001, ****p < 0.0001.

## Contributions

B.A.E. conceptualized study, designed, and performed experimentation. J.V.R. and S.C. assisted in confocal microscopy experimentation and analysis. K.J.P and S.R.C. assisted in mass cytometry analysis and biochemical assays. O.K.A assisted in biochemical assays. J.M. and K.W. performed electron microscopy sample preparation and image acquisition. S.S., J.C., imaging mass cytometry panel design and image acquisition. L.X. and X.C. mass spectrometry analysis. J.A.E., histology. M.A.I, suspension mass cytometry panel design and sample acquisition. R.T., X.L., and R.B., patient care for study participants, managed clinical data, and sample acquisition. N.S., VoPo analysis of mass cytometry datasets. R.M. assisted with live-cell calcium imaging and analysis, and primary cell culture. T.J.C. supervised generation, acquisition, and analyses of all research data. B.A.E. wrote the manuscript, which was reviewed and edited by all co-authors.

## Study Approval

The study was approved by the Institutional Review Pro00109640 of Duke University and 22-2638 of University of North Carolina at Chapel Hill. All participants were studied after providing written informed consent.

## Acknowledgments

The Microscopy Services Laboratory, Department of Pathology and Laboratory Medicine, is supported in part by P30 CA016086 Cancer Center Core Support Grant to the UNC Lineberger Comprehensive Cancer Center. UNC Mass Cytometry Core University Cancer Research Fund (UCRF) UNC Cancer Center Core Support Grant P30CA016086. J.V.R. and S.C. were supported by the National Institute of General Medical Sciences of the National Institutes of Health under award number R35 GM133460. T.J.C. and was supported in part by National Institute of Neurological Disorders and Stroke R01 NS105981. D.H.P, I.Y.Q., Y.W., and T.J.C were supported in part by the National Institute of Aging R01 AG066871. Clinical work was supported in part by the North Carolina Translational and Clinical Sciences (TraCS) pilot award 550KR282107 awarded to R.T. and X.L. B.A.E was supported in part by the National Institute of Neurological Disorders and Stroke F31 NS122242. S.R.C was supported by National Institutes of Aging T35 AG038047. ALS tissue specimens were provided by the Department of Veterans Affairs Biorepository, VA Merit review BX002466. This work was supported by the Standard BioTools Discovery Lab at Standard BioTools Inc. Assay development such as staining, titrations, and data acquisition were performed as paid services. We’d like to thank N. Lane, K. Mottershead, R.J. Perna, A. Sidders, E. Swanson, for strategic discussion. We’d like to thank C. Simmons, M. Chopra, M. Ward, and H. Zampa for patient recruitment, enrollment, and sample acquisition. Finally, we sincerely thank the ALS patients and family members for their selfless contribution to this body of work.

---

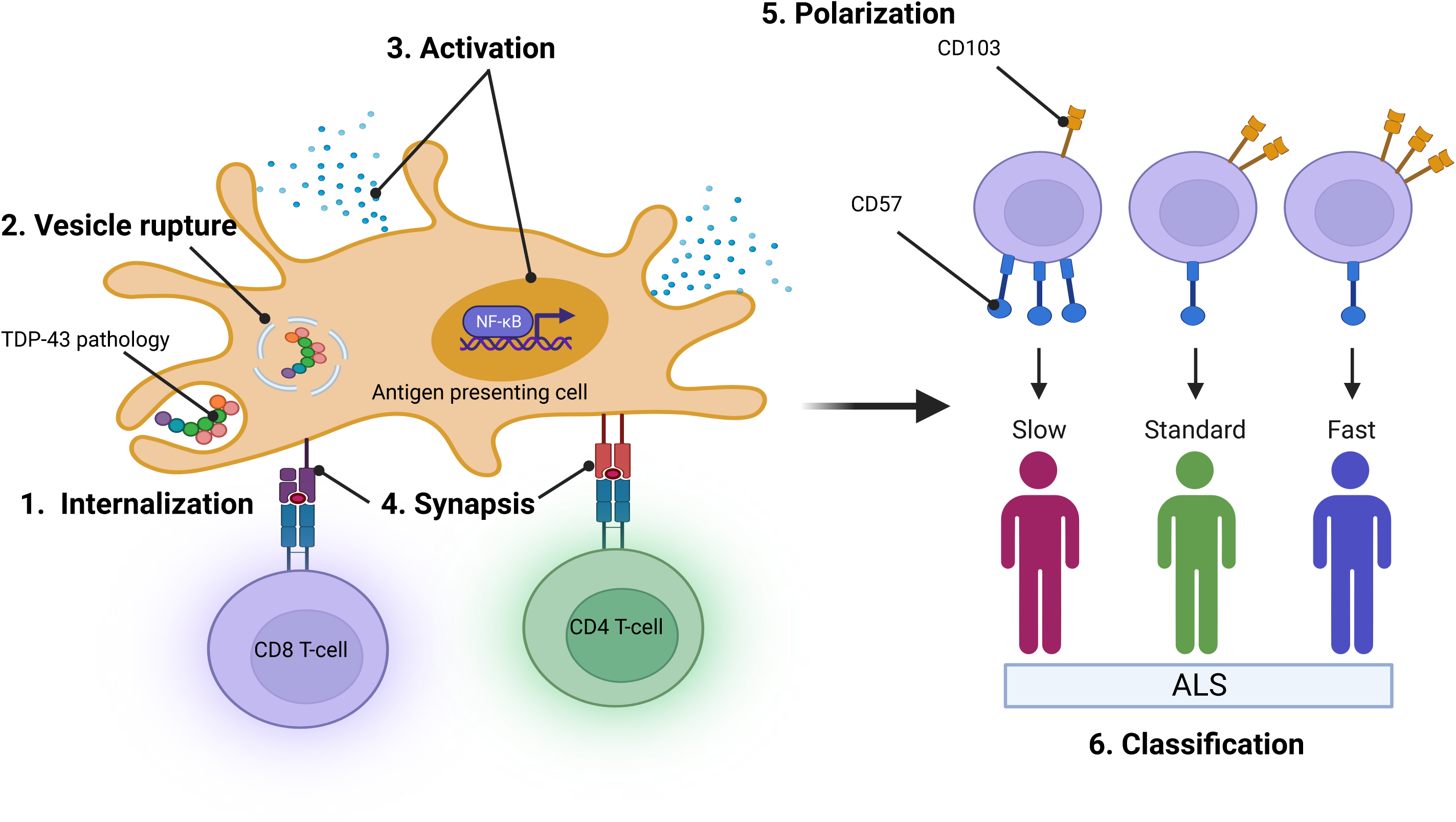

