## Supplemental Figures 4,5,6,7 for "TDP-43 pathology links innate and adaptive immunity in amyotrophic lateral sclerosis"

**Supplementary Figure S4A. Pre-processing and de-barcoding of suspension CyTOF datasets**

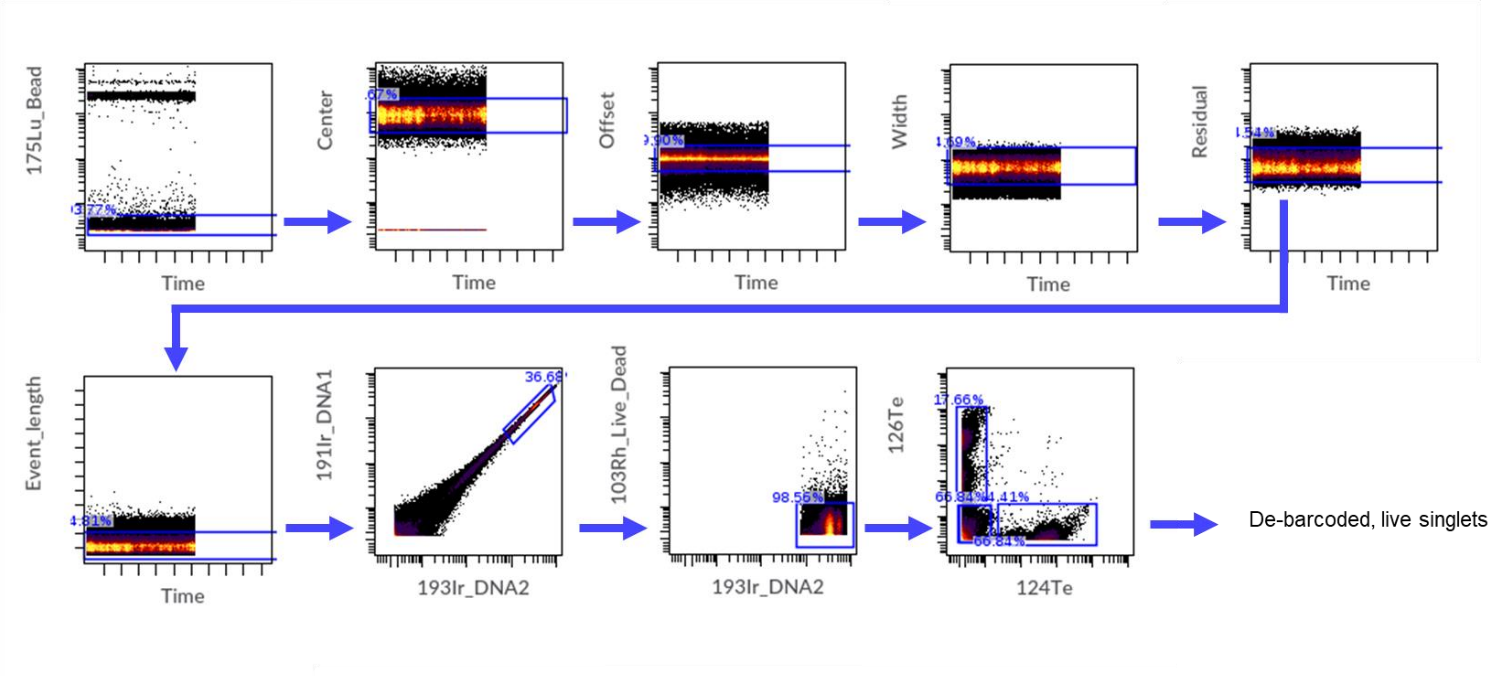

#### Supplementary Figure S4B. Gating strategy for CyTOF datasets of bulk PBMC cultures

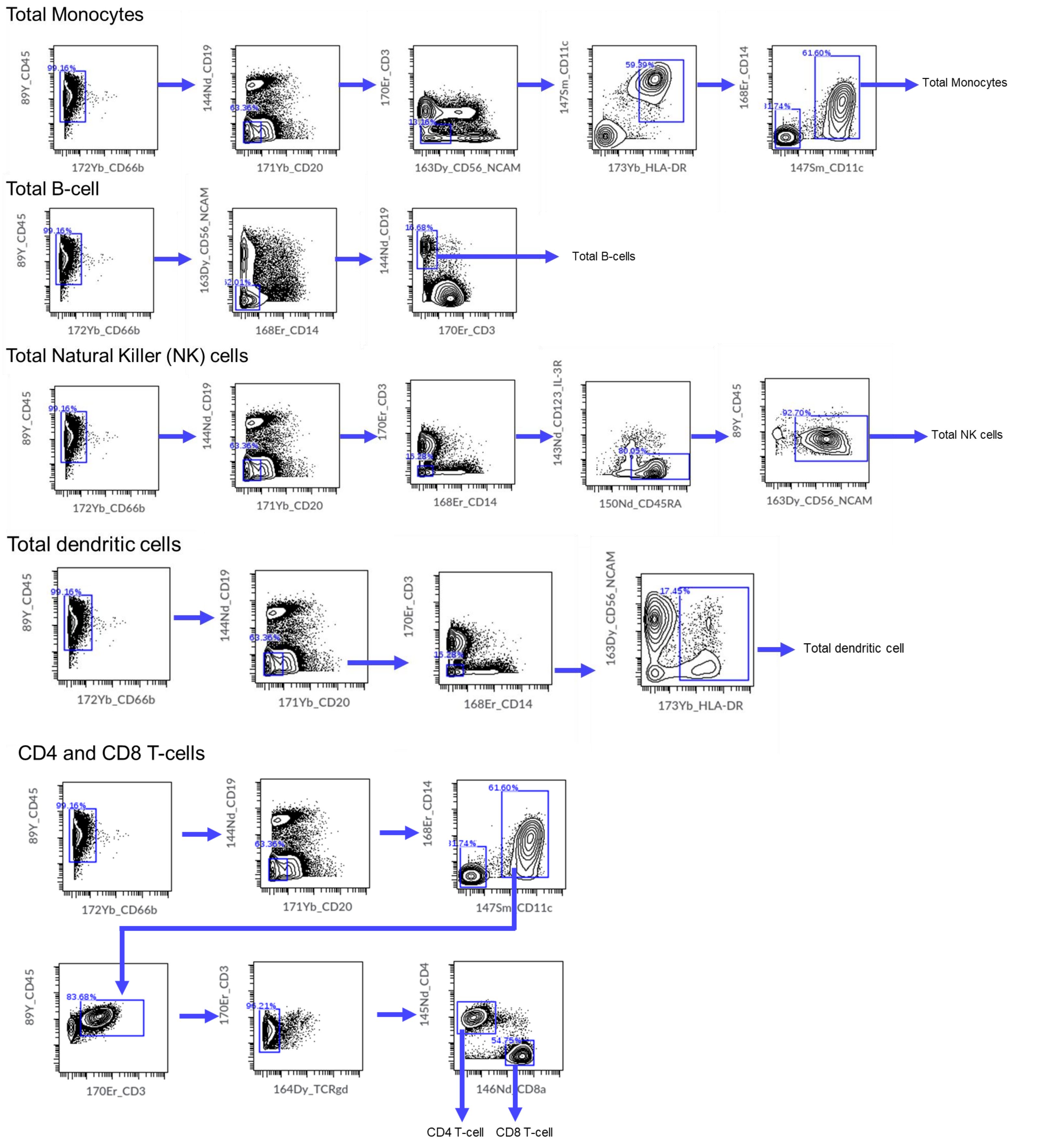

**Supplementary Figure S4C. Gating strategy for CyTOF datasets of T-cell subsets**

T-cell Subsets

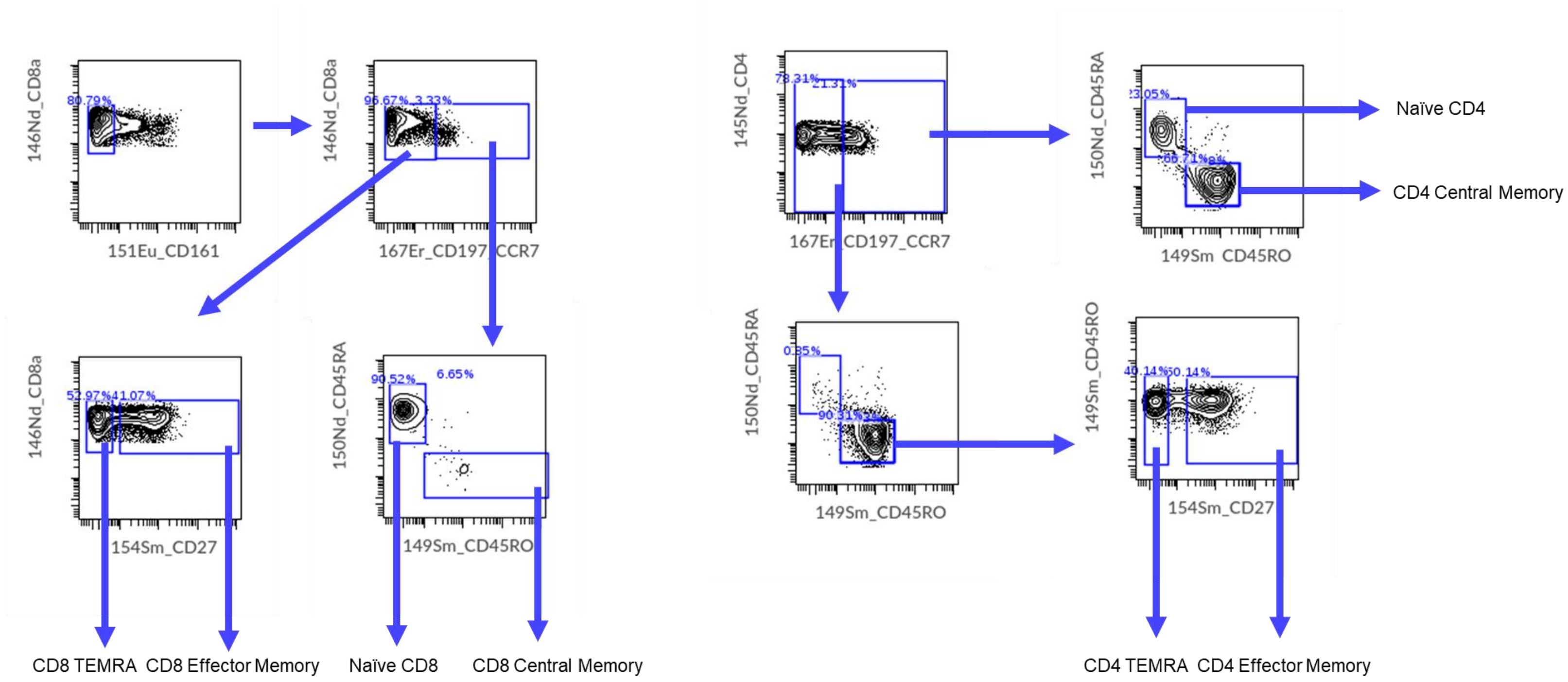

### Supplementary Figure S5A. Enrichment of naïve CD4 and CD8 T-cells by magnetic activated cell sorting

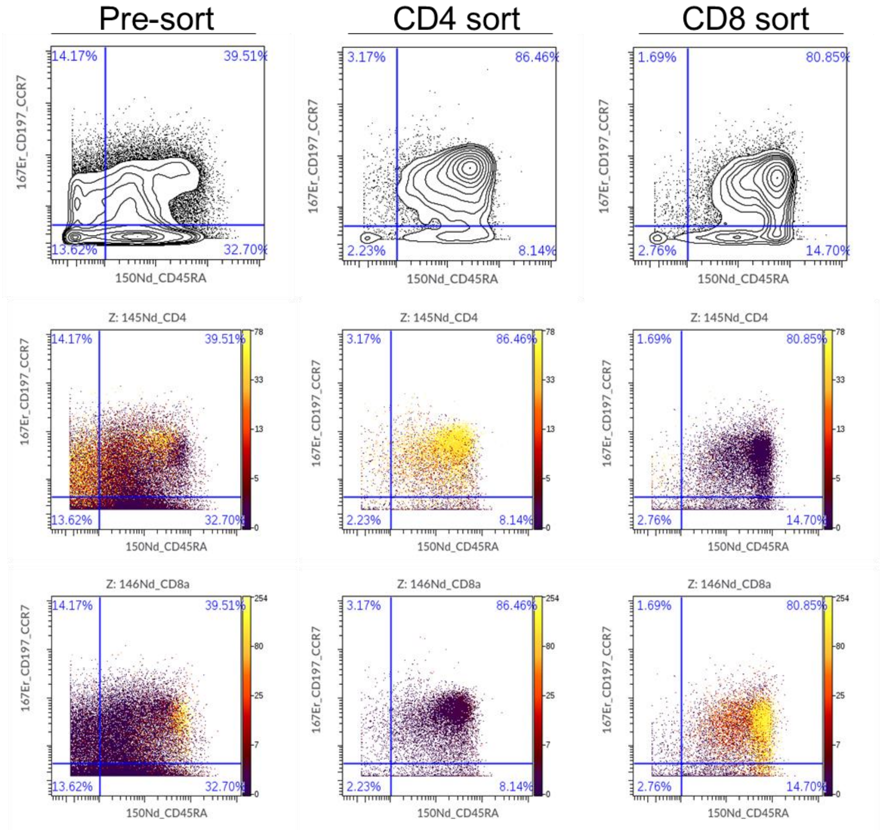

### Supplementary Figure S6. Imaging mass cytometry channels (1 of 3)

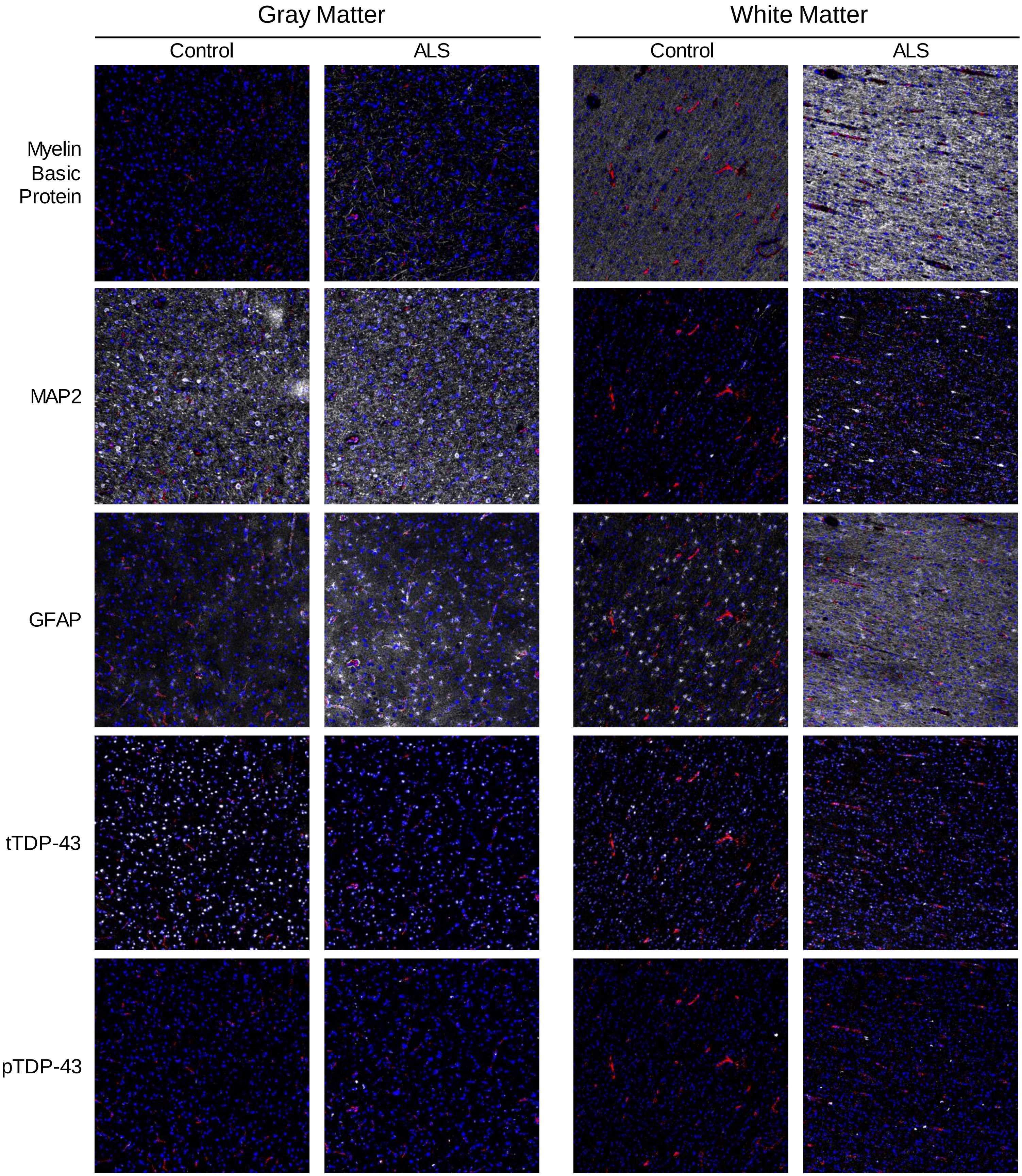


Nuclei (blue); vasculature (CD31; red); target (white)

### Supplementary Figure S6. Imaging mass cytometry channels (2 of 3)

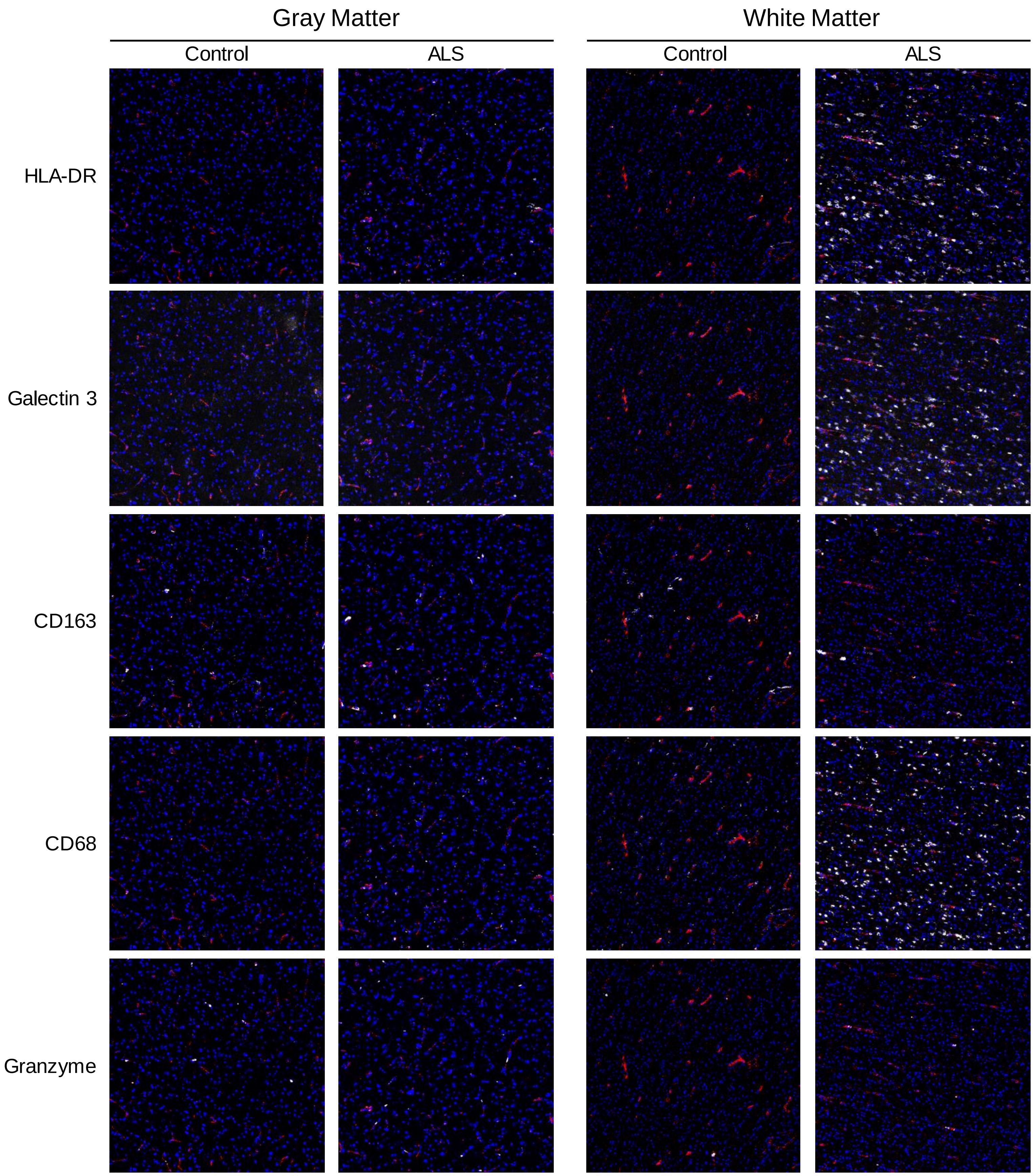


Nuclei (blue); vasculature (CD31; red); target (white)

### Supplementary Figure S6. Imaging mass cytometry channels (3 of 3)

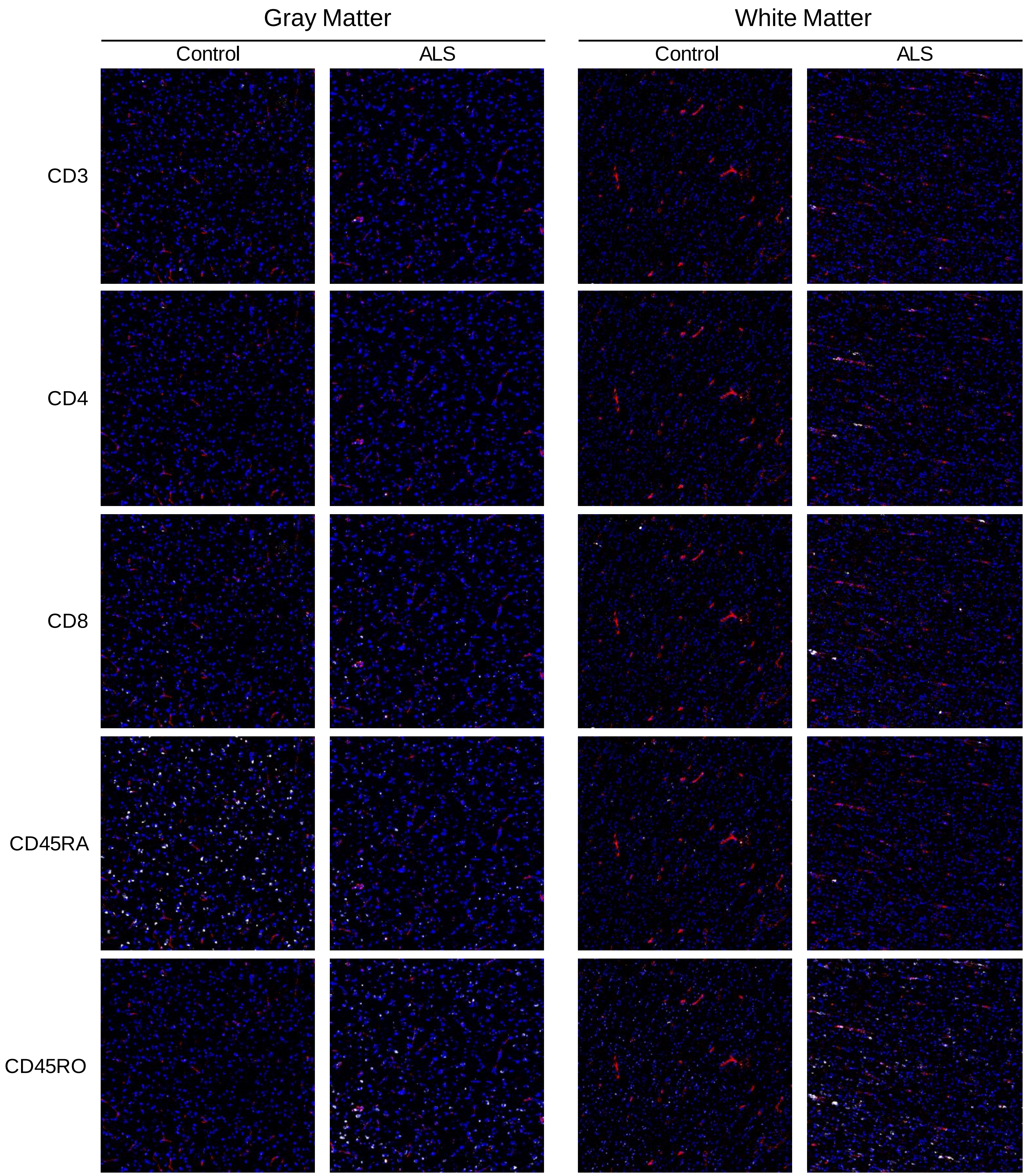


Nuclei (blue); vasculature (CD31; red); target (white)

#### Supplementary Figure S7. Gating strategy for CyTOF datasets of ALS whole blood

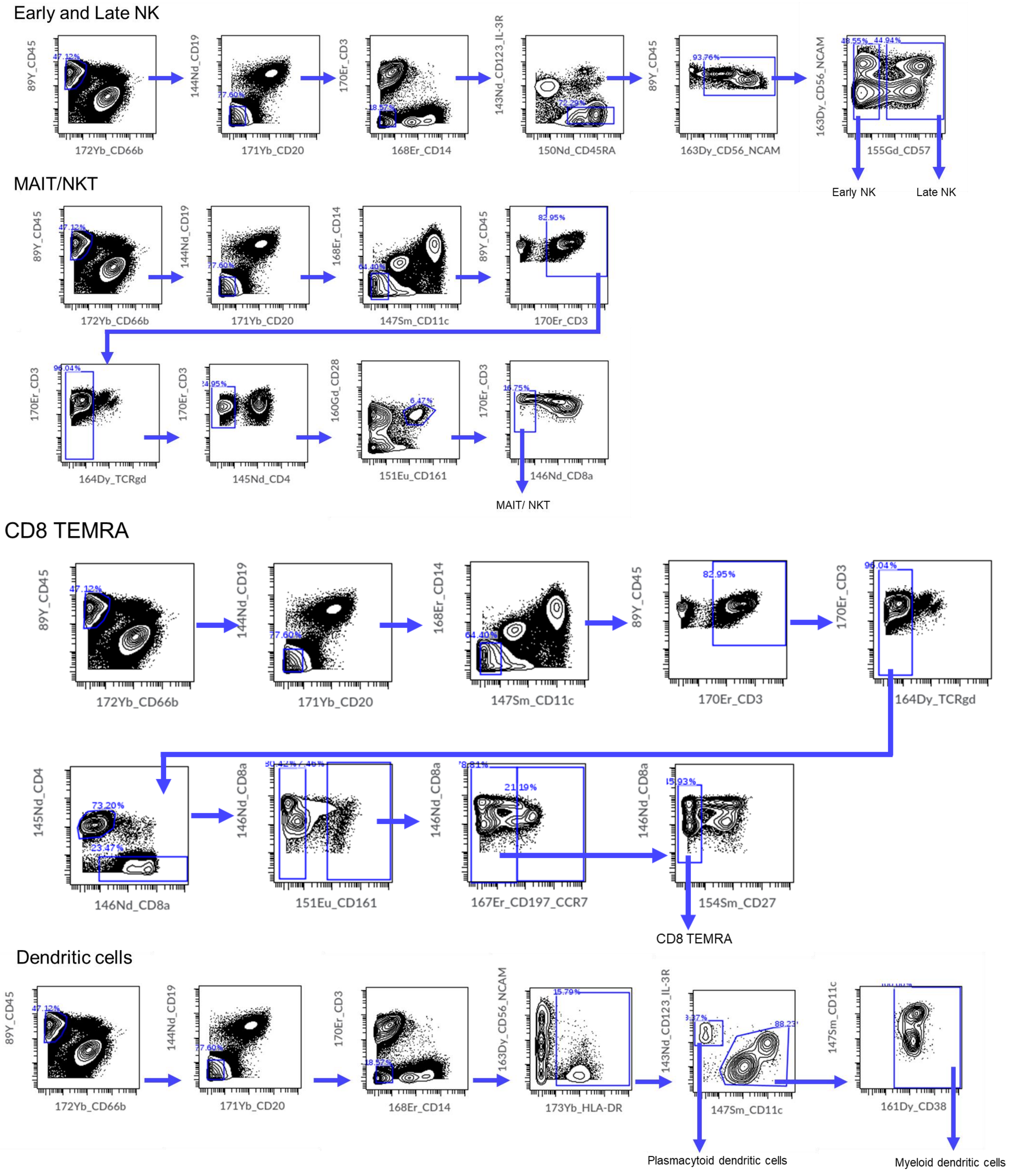
