## Supplementary Table S1A for "TDP-43 pathology links innate and adaptive immunity in amyotrophic lateral sclerosis"

| Gene name | -LOG(P-value) | P-value | log2(TDP/crl) | TDP/crl_imp cons | Protein IDs | Majority protein IDs | Protein ID | Protein names |
|---|---|---|---|---|---|---|---|---|
| TPM1 | 2.566966 | 0.00271 | 3.61789 | up3 | F5H7S3;H7 | F5H7S3;H7 | F5H7S3 | Tropomyos |
| ARL3 | 16.88511 | 1.3E-17 | 3.559849 | up3 | P36405 | P36405 | P36405 | ADP-ribosy |
| LAIR1 | 1.980202 | 0.010466 | 3.053663 | up3 | A0A0G2JNI | A0A0G2JNI | A0A0G2JNI | Leukocyte- |
| TRAPPC9 | 3.24099 | 0.000574 | 2.975781 | up2 | Q96Q05;H( | Q96Q05;H( | Q96Q05 | Trafficking |
| ATP11B | 1.637126 | 0.023061 | 2.309852 | up2 | Q9Y2G3;H7 | Q9Y2G3;H7 | Q9Y2G3 | Probable pl |
| GYS1 | 1.981418 | 0.010437 | 2.174451 | up2 | P13807;M( | P13807 | P13807 | Glycogen [s |
| BLOC1S6 | 1.897603 | 0.012659 | 2.414231 | up2 | H3BST1;Q9 | H3BST1;Q9 | H3BST1 | Biogenesis |
| TLN2 | 1.30675 | 0.049346 | 2.017599 | up2 | Q9Y4G6;AC | Q9Y4G6;AC | Q9Y4G6 | Talin-2 |
| HIPK4 | 2.254268 | 0.005568 | 2.183624 | up2 | Q8NE63 | Q8NE63 | Q8NE63 | Homeodom |
| DCAF7 | 1.989775 | 0.010238 | 2.004544 | up2 | A0A087WV | A0A087WV | A0A087WV | DDB1- and |
| FASTKD1 | 1.303707 | 0.049693 | 1.243008 | up1 | Q53R41;E7 | Q53R41;E7 | Q53R41 | FAST kinase |
| GPAA1 | 1.392701 | 0.040485 | 1.897217 | up1 | O43292;E9 | O43292;E9 | O43292 | Glycosylph |
| ALG6 | 1.306482 | 0.049376 | 1.743188 | up1 | Q9Y672;A0 | Q9Y672 | Q9Y672 | Dolichyl py |
| FAM208A | 1.44513 | 0.035881 | 1.673333 | up1 | A0A087X0I | A0A087X0I | A0A087X0I | Protein FAM |
| CDC40 | 1.306261 | 0.049401 | 1.653601 | up1 | Q5SRN1;O( | Q5SRN1;O( | Q5SRN1 | Pre-mRNA- |
| DGUOK | 1.303271 | 0.049743 | 1.93545 | up1 | Q16854 | Q16854 | Q16854 | Deoxyguan |
| NUDT16L1 | 1.303816 | 0.04968 | 1.604408 | up1 | Q9BRJ7;K7 | Q9BRJ7;K7 | Q9BRJ7 | Protein syn |
| CAMLG | 1.306834 | 0.049336 | 1.512568 | up1 | P49069;D6 | P49069 | P49069 | Calcium sig |
| EPS8L2 | 1.300628 | 0.050046 | 1.315781 | up1 | E9PLH1;A0 | E9PLH1;A0 | E9PLH1 | Epidermal g |
| DHODH | 1.388897 | 0.040842 | 1.209878 | up1 | Q02127;I3I | Q02127;I3I | Q02127 | Dihydroorc |
| EXOSC3 | 2.172954 | 0.006715 | 1.798467 | up1 | Q9NQT5 | Q9NQT5 | Q9NQT5 | Exosome co |
| FAR1 | 2.252573 | 0.00559 | 1.88962 | up1 | E9PNW8;Q | E9PNW8;Q | E9PNW8 | Fatty acyl-C |
| TMEM222 | 1.987577 | 0.01029 | 1.866145 | up1 | H0Y8D0;H( | H0Y8D0;H( | H0Y8D0 | Transmemb |
| ULK3 | 1.300243 | 0.050091 | 1.733959 | up1 | Q6PHR2 | Q6PHR2 | Q6PHR2 | Serine/thre |
| PHLDA3 | 1.304671 | 0.049583 | 1.258878 | up1 | Q9Y5J5 | Q9Y5J5 | Q9Y5J5 | Pleckstrin h |
| PPWD1 | 1.304137 | 0.049644 | 1.223155 | up1 | F5H7P7;Q9 | F5H7P7;Q9 | F5H7P7 | Peptidylpro |
| QPCT | 1.300465 | 0.050065 | 1.151202 | up1 | B5MCZ9;Q1 | B5MCZ9;Q1 | B5MCZ9 | Glutaminyl |
| PLIN2 | 9.671006 | 2.13E-10 | 0.502661 | up0.5 | Q99541;Q5 | Q99541 | Q99541 | Perilipin-2 |
| THOC7 | 2.918743 | 0.001206 | 0.541811 | up0.5 | Q6I9Y2;A0 | Q6I9Y2;A0 | Q6I9Y2 | THO compl |
| PPAP2B | 6.110166 | 7.76E-07 | 0.636199 | up0.5 | O14495;AC | O14495 | O14495 | Lipid phosp |
| NDUFB11 | 2.012305 | 0.009721 | 0.513199 | up0.5 | Q9NX14 | Q9NX14 | Q9NX14 | NADH dehy |
| DDX41 | 2.989797 | 0.001024 | 0.503068 | up0.5 | Q9UJV9;A0 | Q9UJV9 | Q9UJV9 | Probable AT |
| POLR2H | 2.188667 | 0.006476 | 0.969492 | up0.5 | C9JLU1;P5 | C9JLU1;P5 | C9JLU1 | DNA-direct |
| TMEM106E | 2.292274 | 0.005102 | 0.542791 | up0.5 | Q9NUM4;F | Q9NUM4;F | Q9NUM4 | Transmemb |
| CHCHD2 | 5.963961 | 1.09E-06 | 0.582704 | up0.5 | Q9Y6H1;Q5 | Q9Y6H1;Q5 | Q9Y6H1 | Coiled-coil- |
| AKAP2 | 3.291474 | 0.000511 | 0.543036 | up0.5 | Q9Y2D5;C9 | Q9Y2D5;C9 | Q9Y2D5 | A-kinase an |
| ACTN1 | 1.746839 | 0.017913 | 0.931628 | up0.5 | H9KV75;H7 | H9KV75 | H9KV75 | |
| HIST2H3PS | 1.53431 | 0.029221 | 0.570691 | up0.5 | Q5TEC6;A0 | Q5TEC6 | Q5TEC6 | Histone H3 |
| C14orf159 | 2.128396 | 0.007441 | 0.563838 | up0.5 | H0YB09;A0 | H0YB09;A0 | H0YB09 | |
| MIF | 3.400478 | 0.000398 | -5.57777 | down5 | P14174 | P14174 | P14174 | Macrophag |
| NDUFV3 | 2.903837 | 0.001248 | -3.55998 | down3 | P56181 | P56181 | P56181 | NADH dehy |
| DNAJC16 | 1.993415 | 0.010153 | -2.15464 | down2 | Q9Y2G8;Q5 | Q9Y2G8;Q5 | Q9Y2G8 | DnaJ homo |
| DFNA5 | 3.254265 | 0.000557 | -2.5073 | down2 | C9JSR4;O6 | C9JSR4;O6 | C9JSR4 | Non-syndro |
| HRSP12 | 1.931009 | 0.011722 | -2.85974 | down2 | H0YBX3;HC | H0YBX3;HC | H0YBX3 | Ribonuclea |
| DNAJB6 | 1.812547 | 0.015398 | -2.02566 | down2 | A0A0J9YX6 | A0A0J9YX6 | A0A0J9YX6 | DnaJ homo |

| | | | | | | | | |
|---|---|---|---|---|---|---|---|---|
| SRSF5 | 1.304638 | 0.049586 | -2.39018 | down2 | Q13243;B4 | Q13243;B4 | Q13243 | Serine/argi |
| MRPS21 | 1.343541 | 0.045338 | -2.2602 | down2 | P82921 | P82921 | P82921 | 28S ribosor |
| CEBPB | 1.306245 | 0.049403 | -2.41667 | down2 | P17676 | P17676 | P17676 | CCAAT/enh |
| NOL6 | 3.175066 | 0.000668 | -2.61785 | down2 | A0A0A0MR | A0A0A0MR | A0A0A0MR | Nucleolar p |
| PRDM2 | 1.981513 | 0.010435 | -2.1766 | down2 | Q13029 | Q13029 | Q13029 | PR domain |
| MCOLN1 | 1.990728 | 0.010216 | -2.48098 | down2 | Q9GZU1;M | Q9GZU1;M | Q9GZU1 | Mucolipin- |
| HPGD | 3.638013 | 0.00023 | -1.06765 | down1 | P15428;E9 | P15428;E9 | P15428 | 15-hydroxy |
| GAS7 | 1.478367 | 0.033238 | -1.58732 | down1 | O60861;AC | O60861;AC | O60861 | Growth arr |
| PLEKHO1 | 1.401378 | 0.039685 | -1.47082 | down1 | Q53GL0;Q5 | Q53GL0 | Q53GL0 | Pleckstrin h |
| RRP9 | 1.393123 | 0.040446 | -1.63531 | down1 | O43818 | O43818 | O43818 | U3 small nu |
| CRYZL1 | 1.326014 | 0.047205 | -1.92377 | down1 | A6NMA8;O | A6NMA8;O | A6NMA8 | Quinone ox |
| EMB | 1.305131 | 0.04953 | -1.46872 | down1 | D6RDX7;Q( | D6RDX7;Q( | D6RDX7 | Embigin |
| TIPRL | 1.430152 | 0.037141 | -1.84545 | down1 | O75663 | O75663 | O75663 | TIP41-like p |
| STX17 | 1.616528 | 0.024181 | -1.12245 | down1 | P56962;E9 | P56962;E9 | P56962 | Syntaxin-1 |
| COQ10B | 1.391289 | 0.040617 | -1.36674 | down1 | B8ZZV9;Q9 | B8ZZV9;Q9 | B8ZZV9 | Coenzyme ( |
| ITM2B | 1.349056 | 0.044766 | -1.40033 | down1 | Q9Y287;A0 | Q9Y287;A0 | Q9Y287 | Integral me |
| TRAPPC1 | 1.320526 | 0.047805 | -1.40809 | down1 | Q9Y5R8;I3I | Q9Y5R8 | Q9Y5R8 | Trafficking p |
| AFTPH | 1.549122 | 0.028241 | -1.5014 | down1 | Q6ULP2;B8 | Q6ULP2;B8 | Q6ULP2 | Aftiphilin |
| CD37 | 1.748613 | 0.01784 | -1.63995 | down1 | M0QZP6;B7 | M0QZP6;B7 | M0QZP6 | Tetraspanin |
| TOM1L2 | 1.304613 | 0.049589 | -1.7283 | down1 | Q6ZVM7;B7 | Q6ZVM7;B7 | Q6ZVM7 | TOM1-like p |
| GPR107 | 1.565348 | 0.027205 | -1.78858 | down1 | Q5VW38;G | Q5VW38;G | Q5VW38 | Protein GP |
| SPATS2L | 1.991891 | 0.010188 | -1.91259 | down1 | Q9NUQ6;F8 | Q9NUQ6;F8 | Q9NUQ6 | SPATS2-like |
| SMDT1 | 1.350403 | 0.044627 | -1.61491 | down1 | Q9H4I9 | Q9H4I9 | Q9H4I9 | Essential M |
| LRP10 | 5.633204 | 2.33E-06 | -1.07403 | down1 | H0YHW7;Q | H0YHW7;Q | H0YHW7 | Low-densit |
| NACC1 | 1.760111 | 0.017374 | -1.30566 | down1 | Q96RE7;Q9 | Q96RE7;Q9 | Q96RE7 | Nucleus acc |
| DOK2 | 1.557563 | 0.027697 | -1.37737 | down1 | G3V114;E5 | G3V114;E5 | G3V114 | Docking pro |
| MRPL33 | 1.375395 | 0.042131 | -1.38768 | down1 | O75394 | O75394 | O75394 | 39S ribosor |
| CTSO | 1.991776 | 0.010191 | -1.65516 | down1 | P43234 | P43234 | P43234 | Cathepsin ( |
| LYRM2 | 1.319962 | 0.047867 | -1.6913 | down1 | Q9NU23 | Q9NU23 | Q9NU23 | LYR motif-c |
| COA5 | 1.993428 | 0.010152 | -1.76899 | down1 | B9A057;Q8 | B9A057;Q8 | B9A057 | Cytochrom |
| RRAGD | 2.0572 | 0.008766 | -1.79307 | down1 | Q9NQL2 | Q9NQL2 | Q9NQL2 | Ras-related |
| UBL5 | 1.929003 | 0.011776 | -1.87077 | down1 | Q9BZL1 | Q9BZL1 | Q9BZL1 | Ubiquitin-l |
| GNG2 | 1.4823 | 0.032938 | -1.883 | down1 | P59768;G3 | P59768;G3 | P59768 | Guanine nu |
| HDHD1 | 10.89699 | 1.27E-11 | -1.97883 | down1 | Q08623 | Q08623 | Q08623 | Pseudouric |
| CHMP2A | 1.684339 | 0.020685 | -1.99084 | down1 | M0R1L7;M | M0R1L7;M | M0R1L7 | Charged mu |
| AMPD2 | 2.817728 | 0.001521 | -0.49937 | down0.5 | A0A5F9UK9 | A0A5F9UK9 | A0A5F9UK9 | AMP deami |
| BRK1 | 2.03478 | 0.00923 | -0.50956 | down0.5 | Q8WUW1 | Q8WUW1 | Q8WUW1 | Protein BRI |
| SELT | 2.904377 | 0.001246 | -0.53492 | down0.5 | A0A087WV | A0A087WV | A0A087WV | Selenoprot |
| RPL29 | 3.331994 | 0.000466 | -0.51803 | down0.5 | A0A3B3ITT | A0A3B3ITT | A0A3B3ITT | 60S ribosor |
| MOB1B | 4.670082 | 2.14E-05 | -0.5621 | down0.5 | Q7L9L4;Q9 | Q7L9L4;Q9 | Q7L9L4 | MOB kinase |
| NEDD8 | 3.238013 | 0.000578 | -0.61596 | down0.5 | Q15843;F8 | Q15843;F8 | Q15843 | NEDD8 |
| SCARF1 | 1.306316 | 0.049395 | -0.53592 | down0.5 | Q14162;AC | Q14162;AC | Q14162 | Scavenger r |
| DYNC1H1 | 0.183599 | 0.655241 | 0.019303 | | Q14204;AC | Q14204 | Q14204 | Cytoplasmi |
| AHNAK | 0.060764 | 0.869432 | 0.012887 | | Q09666;E9 | Q09666 | Q09666 | Neuroblast |
| PRKDC | 0.562931 | 0.27357 | 0.052275 | | P78527;F5 | P78527 | P78527 | DNA-depen |
| RNF213 | 0.396049 | 0.401745 | 0.089957 | | A0A0A0MT | A0A0A0MT | A0A0A0MT | E3 ubiquiti |
| FLNA | 0.595323 | 0.253909 | 0.117796 | | P21333;Q6 | P21333;Q6 | P21333 | Filamin-A |
| PLEC | 1.34634 | 0.045046 | 0.091132 | | Q15149;E9 | Q15149 | Q15149 | Plectin |

| | | | | | | | |
|---|---|---|---|---|---|---|---|
| LRP1 | 0.290172 | 0.512658 | 0.040362 | Q07954;Q6 | Q07954 | Q07954 | Prolow-der |
| VPS13C | 0.920447 | 0.120103 | 0.094898 | Q709C8;A0 | Q709C8 | Q709C8 | Vacuolar pr |
| TLN1 | 0.71293 | 0.193674 | -0.05834 | Q9Y490 | Q9Y490 | Q9Y490 | Talin-1 |
| MYH9 | 0.379955 | 0.416912 | 0.048793 | P35579;Q5 | P35579 | P35579 | Myosin-9 |
| GOLGB1 | 1.718221 | 0.019133 | 0.152884 | Q14789;E7 | Q14789;E7 | Q14789 | Golgin subf |
| GCN1L1 | 0.15518 | 0.699552 | -0.04241 | Q92616 | Q92616 | Q92616 | Translation |
| IQGAP1 | 0.506319 | 0.31166 | 0.051931 | P46940;A0 | P46940;A0 | P46940 | Ras GTPase- |
| UTRN | 1.545536 | 0.028475 | 0.096352 | P46939;Q5 | P46939 | P46939 | Utrophin |
| SNRNP200 | 0.842079 | 0.143854 | 0.085114 | O75643;A0 | O75643;A0 | O75643 | U5 small nu |
| PRPF8 | 0.009095 | 0.979276 | 0.001307 | Q6P2Q9;I3 | Q6P2Q9 | Q6P2Q9 | Pre-mRNA-p |
| KTN1 | 2.869971 | 0.001349 | 0.119807 | Q86UP2;G3 | Q86UP2 | Q86UP2 | Kinectin |
| IQGAP2 | 0.102901 | 0.78904 | 0.016506 | Q13576;F5 | Q13576;F5 | Q13576 | Ras GTPase- |
| FASN | 0.416101 | 0.383618 | -0.07121 | P49327;A0 | P49327;A0 | P49327 | Fatty acid s |
| RRBP1 | 0.080451 | 0.8309 | 0.008546 | Q9P2E9;A0 | Q9P2E9;A0 | Q9P2E9 | Ribosome-b |
| CLTC | 0.080123 | 0.831529 | 0.004928 | A0A087WV | A0A087WV | A0A087WV | Clathrin he |
| IGF2R | 0.10306 | 0.788752 | -0.01332 | P11717;S4 | P11717 | P11717 | Cation-inde |
| LRPPRC | 0.395167 | 0.402562 | 0.052655 | P42704;A0 | P42704 | P42704 | Leucine-ric |
| EPRS | 0.216283 | 0.607739 | 0.032133 | P07814;V9 | P07814;V9 | P07814 | Bifunctiona |
| EEA1 | 1.098592 | 0.079691 | 0.195546 | Q15075 | Q15075 | Q15075 | Early endos |
| SPTAN1 | 1.01594 | 0.096396 | 0.140382 | Q13813;A0 | Q13813;A0 | Q13813 | Spectrin alp |
| COPA | 0.325609 | 0.472488 | 0.067988 | P53621;A0 | P53621;A0 | P53621 | Coatomer s |
| VWA8 | 0.058419 | 0.87414 | 0.010631 | A3KMH1 | A3KMH1 | A3KMH1 | von Willebr |
| DNAJC13 | 0.807245 | 0.155867 | -0.06212 | O75165;A0 | O75165 | O75165 | DnaJ homo |
| FLNB | 0.875856 | 0.13309 | 0.109181 | O75369;E7 | O75369;E7 | O75369 | Filamin-B |
| UGGT1 | 0.01648 | 0.962763 | 0.00184 | Q9NYU2;H7 | Q9NYU2 | Q9NYU2 | UDP-glucos |
| OGDH | 2.633008 | 0.002328 | 0.157963 | Q02218;E9 | Q02218;E9 | Q02218 | 2-oxoglutar |
| SND1 | 2.274549 | 0.005314 | 0.09075 | Q7KZF4;H7 | Q7KZF4 | Q7KZF4 | Staphyloco |
| DCTN1 | 0.916184 | 0.121288 | 0.069009 | E7EX90;Q1 | E7EX90;Q1 | E7EX90 | Dynactin su |
| MYOF | 0.070379 | 0.850395 | -0.00946 | Q9NZM1 | Q9NZM1 | Q9NZM1 | Myoferlin |
| VCL | 0.035558 | 0.921388 | -0.0102 | P18206;A0 | P18206 | P18206 | Vinculin |
| MRC1 | 2.925365 | 0.001188 | 0.305828 | P22897 | P22897 | P22897 | Macrophag |
| WDFY4 | 0.320151 | 0.478463 | 0.05817 | Q6ZS81 | Q6ZS81 | Q6ZS81 | WD repeat- |
| EIF3A | 2.146863 | 0.007131 | 0.109173 | Q14152 | Q14152 | Q14152 | Eukaryotic |
| NUMA1 | 0.092161 | 0.808796 | -0.02096 | Q14980;A0 | Q14980;A0 | Q14980 | Nuclear mit |
| VCP | 1.045426 | 0.090069 | 0.075724 | P55072 | P55072 | P55072 | Transitiona |
| VIM | 0.147057 | 0.71276 | 0.024487 | P08670;B0 | P08670;B0 | P08670 | Vimentin |
| TPR | 1.505104 | 0.031253 | 0.150095 | P12270;Q5 | P12270 | P12270 | Nucleoprot |
| ITPR2 | 0.506848 | 0.31128 | 0.076022 | Q14571;H7 | Q14571 | Q14571 | Inositol 1,4 |
| PTPRC | 0.408788 | 0.390133 | 0.039683 | P08575;M3 | P08575;M3 | P08575 | Receptor-ty |
| APOBR | 1.245996 | 0.056755 | 0.117796 | A0A4V7I44 | A0A4V7I44 | A0A4V7I44 | Apolipoprc |
| ITPR1 | 0.824267 | 0.149876 | 0.121243 | Q14643;A0 | Q14643;A0 | Q14643 | Inositol 1,4 |
| MYO1C | 0.372308 | 0.424318 | 0.025656 | F5H6E2;O0 | F5H6E2;O0 | F5H6E2 | Unconvent |
| DHX9 | 0.097658 | 0.798623 | 0.015507 | Q08211 | Q08211 | Q08211 | ATP-depend |
| DMXL2 | 0.030852 | 0.931425 | -0.00428 | Q8TDJ6;H0 | Q8TDJ6 | Q8TDJ6 | DmX-like pr |
| MVP | 1.753498 | 0.01764 | 0.117408 | Q14764;H3 | Q14764 | Q14764 | Major vault |
| LCP1 | 0.742056 | 0.181111 | -0.14008 | P13796;Q5 | P13796 | P13796 | Plastin-2 |
| DMXL1 | 0.553058 | 0.279861 | 0.109015 | Q9Y485;F5 | Q9Y485;F5 | Q9Y485 | DmX-like pr |
| ACLY | 0.417426 | 0.382449 | -0.05658 | P53396;K7 | P53396 | P53396 | ATP-citrate |

| | | | | | | | |
|---|---|---|---|---|---|---|---|
| ITGAX | 0.370528 | 0.426061 | 0.056851 | P20702;H3 | P20702;H3 | P20702 | Integrin alp |
| EEF2 | 0.179699 | 0.661152 | 0.03256 | P13639;M( | P13639 | P13639 | Elongation |
| DOCK8 | 0.402146 | 0.396145 | -0.03247 | Q8NF50;A2 | Q8NF50;A2 | Q8NF50 | Dedicator c |
| HDLBP | 0.307528 | 0.492575 | 0.036773 | A0A024R4I | A0A024R4I | A0A024R4I | Vigilin |
| ESYT1 | 0.282332 | 0.521997 | 0.060513 | Q9BSJ8;F8 | Q9BSJ8 | Q9BSJ8 | Extended sy |
| NSF | 0.029927 | 0.93341 | 0.004974 | I3L0N3;P4 | I3L0N3;P4 | I3L0N3 | Vesicle-fusi |
| PI4KA | 1.58691 | 0.025887 | 0.088647 | P42356;A8 | P42356 | P42356 | Phosphatic |
| ATP13A1 | 0.948529 | 0.112583 | 0.090932 | Q9HD20;H | Q9HD20 | Q9HD20 | Manganese |
| COPG1 | 0.106608 | 0.782333 | 0.026225 | Q9Y678;H0 | Q9Y678 | Q9Y678 | Coatomer s |
| PCK2 | 1.062396 | 0.086617 | 0.078743 | Q16822;H( | Q16822;H( | Q16822 | Phosphoen |
| MTHFD1 | 0.553226 | 0.279752 | -0.10837 | P11586;V9 | P11586;V9 | P11586 | C-1-tetrahy |
| LARS | 0.35479 | 0.441784 | -0.05435 | A0A6I8PIV | A0A6I8PIV | A0A6I8PIV | Leucine--tR |
| HADHA | 0.423653 | 0.377005 | 0.058358 | H0YFD6;P4 | H0YFD6;P4 | H0YFD6 | Trifunction |
| GPD2 | 0.154554 | 0.700562 | 0.022722 | P43304;F5 | P43304 | P43304 | Glycerol-3- |
| PITRM1 | 1.540255 | 0.028823 | 0.171969 | Q5JRX3;A0 | Q5JRX3;A0 | Q5JRX3 | Presequenc |
| GLG1 | 0.365996 | 0.430531 | 0.042584 | Q92896;H3 | Q92896;H3 | Q92896 | Golgi appar |
| SAMHD1 | 0.320765 | 0.477788 | -0.03775 | Q9Y3Z3;A0 | Q9Y3Z3;A0 | Q9Y3Z3 | Deoxynucle |
| HK1 | 0.281812 | 0.522622 | -0.02691 | P19367;B1 | P19367 | P19367 | Hexokinase |
| H6PD | 0.062009 | 0.866945 | 0.009893 | O95479 | O95479 | O95479 | GDH/6PGL |
| HSD17B4 | 0.052516 | 0.886102 | 0.007361 | P51659;A0 | P51659;A0 | P51659 | Peroxisoma |
| ITGAM | 0.008635 | 0.980313 | 0.001115 | P11215;H3 | P11215 | P11215 | Integrin alp |
| HSPA4 | 0.00667 | 0.984759 | 0.001382 | P34932;A0 | P34932;A0 | P34932 | Heat shock |
| UBR4 | 0.722741 | 0.189347 | 0.138207 | Q5T4S7;A0 | Q5T4S7 | Q5T4S7 | E3 ubiquiti |
| PSMD2 | 0.525584 | 0.298137 | 0.055881 | Q13200;H7 | Q13200 | Q13200 | 26S protea |
| NBAS | 0.501835 | 0.314894 | 0.054983 | A2RRP1;H0 | A2RRP1;H0 | A2RRP1 | Neuroblast |
| SF3B1 | 0.041893 | 0.908045 | -0.00698 | O75533;H7 | O75533 | O75533 | Splicing fac |
| RAD50 | 1.749722 | 0.017794 | 0.129337 | Q92878;A0 | Q92878;A0 | Q92878 | DNA repair |
| PSMD1 | 1.347368 | 0.04494 | 0.090967 | Q99460;A0 | Q99460;A0 | Q99460 | 26S protea |
| COPB1 | 0.729677 | 0.186347 | 0.145808 | P53618;E9 | P53618;E9 | P53618 | Coatomer s |
| ANPEP | 0.693326 | 0.202616 | 0.07552 | P15144;H0 | P15144 | P15144 | Aminopept |
| ACO2 | 0.693204 | 0.202673 | 0.075242 | Q99798;A2 | Q99798;A2 | Q99798 | Aconitate h |
| UPF1 | 0.594837 | 0.254193 | 0.05252 | Q92900 | Q92900 | Q92900 | Regulator c |
| HSP90B1 | 0.203902 | 0.625314 | 0.032894 | P14625;Q9 | P14625 | P14625 | Endoplasm |
| CAND1 | 0.169001 | 0.677639 | -0.03283 | Q86VP6;A0 | Q86VP6;A0 | Q86VP6 | Cullin-asso |
| SEPTIN9 | 1.356389 | 0.044016 | 0.132727 | Q9UHD8;K | Q9UHD8 | Q9UHD8 | Septin-9 |
| HUWE1 | 1.081217 | 0.082944 | 0.111622 | Q7Z6Z7;H0 | Q7Z6Z7 | Q7Z6Z7 | E3 ubiquiti |
| PLXNB2 | 0.673419 | 0.21212 | 0.05534 | O15031;E2 | O15031 | O15031 | Plexin-B2 |
| EIF4G1 | 0.478358 | 0.332385 | -0.04297 | Q04637;E7 | Q04637;E7 | Q04637 | Eukaryotic |
| LONP1 | 0.473273 | 0.3363 | 0.048353 | K7EKE6;P3 | K7EKE6;P3 | K7EKE6 | Lon protea |
| NES | 0.027781 | 0.938035 | 0.006857 | P48681 | P48681 | P48681 | Nestin |
| PDCD6IP | 0.022387 | 0.949758 | -0.00312 | Q8WUM4; | Q8WUM4 | Q8WUM4 | Programme |
| RPN1 | 1.609649 | 0.024567 | 0.11567 | P04843;B7 | P04843;B7 | P04843 | Dolichyl-di |
| ATP1A1 | 1.371414 | 0.042519 | -0.06833 | P05023;M( | P05023 | P05023 | Sodium/po |
| MYO1F | 1.141237 | 0.072238 | -0.08955 | O00160;M | O00160 | O00160 | Unconvent |
| HSPA9 | 0.672543 | 0.212548 | 0.064212 | P38646;A0 | P38646 | P38646 | Stress-70 p |
| HSPD1 | 0.485829 | 0.326717 | 0.069306 | P10809;E7 | P10809 | P10809 | 60 kDa hea |
| ERAP1 | 0.473984 | 0.33575 | -0.04538 | Q9NZ08;H( | Q9NZ08 | Q9NZ08 | Endoplasm |
| CALR | 0.211695 | 0.614192 | -0.0187 | P27797;K7 | P27797 | P27797 | Calreticulir |

| | | | | | | | |
|---|---|---|---|---|---|---|---|
| UBA1 | 0.203586 | 0.625769 | -0.0351 | P22314;Q5 | P22314 | P22314 | Ubiquitin-l |
| IARS2 | 0.885406 | 0.130195 | 0.095464 | Q9NSE4 | Q9NSE4 | Q9NSE4 | Isoleucine-- |
| TRIP11 | 0.797947 | 0.15924 | 0.107446 | Q15643;H0 | Q15643;H0 | Q15643 | Thyroid rec |
| INF2 | 0.795987 | 0.159961 | 0.070136 | Q27J81;A0 | Q27J81;A0 | Q27J81 | Inverted fo |
| HK3 | 0.764827 | 0.171859 | -0.18458 | P52790;H0 | P52790 | P52790 | Hexokinase |
| ATP6V1A | 0.72278 | 0.18933 | -0.05096 | P38606;C9 | P38606 | P38606 | V-type prot |
| P4HB | 0.435995 | 0.366442 | 0.051396 | P07237;H7 | P07237;H7 | P07237 | Protein dis |
| FLII | 0.018497 | 0.958303 | 0.003073 | Q13045;J3 | Q13045 | Q13045 | Protein flig |
| TEX2 | 1.743973 | 0.018031 | 0.149878 | Q8IWB9;A0 | Q8IWB9;A0 | Q8IWB9 | Testis-expre |
| FAM120A | 2.315641 | 0.004835 | 0.144866 | Q9NZB2;A0 | Q9NZB2 | Q9NZB2 | Constitutiv |
| FKBP15 | 1.108039 | 0.077976 | 0.141315 | Q5T1M5 | Q5T1M5 | Q5T1M5 | FK506-binc |
| ACADVL | 0.951818 | 0.111733 | 0.07116 | P49748;G3 | P49748 | P49748 | Very long-c |
| PLXND1 | 0.766557 | 0.171176 | 0.060507 | Q9Y4D7;H0 | Q9Y4D7 | Q9Y4D7 | Plexin-D1 |
| XRCC5 | 0.58802 | 0.258214 | 0.026958 | P13010;C9 | P13010 | P13010 | X-ray repair |
| USO1 | 0.505575 | 0.312194 | 0.052151 | O60763;RE | O60763 | O60763 | General ves |
| TGM2 | 0.466131 | 0.341876 | -0.11908 | P21980;A2 | P21980 | P21980 | Protein-glu |
| ACTN4 | 0.424888 | 0.375934 | 0.047937 | O43707;F5 | O43707;F5 | O43707 | Alpha-actir |
| VARS | 0.338507 | 0.458662 | 0.048304 | P26640;A0 | P26640;A0 | P26640 | Valine--tRN |
| PML | 0.30483 | 0.495644 | 0.064862 | P29590;H3 | P29590;H3 | P29590 | Protein PM |
| PTPN6 | 0.247293 | 0.565858 | -0.04802 | P29350;F5 | P29350 | P29350 | Tyrosine-pr |
| MAP4 | 0.230622 | 0.588001 | 0.042166 | E7EVA0;P2 | E7EVA0;P2 | E7EVA0 | Microtubul |
| ROCK1 | 0.190677 | 0.644648 | 0.04229 | Q13464;A0 | Q13464;A0 | Q13464 | Rho-associa |
| UFL1 | 0.157776 | 0.695384 | -0.0238 | O94874 | O94874 | O94874 | E3 UFM1-p |
| MYO6 | 0.118329 | 0.761503 | 0.022607 | A0A1Y0BRI | A0A1Y0BRI | A0A1Y0BRI | Unconvent |
| ITGB2 | 0.062193 | 0.866577 | 0.007789 | A0A494C0 | A0A494C0 | A0A494C0 | Integrin be |
| MYO9B | 0.039426 | 0.913216 | -0.0031 | M0R300;M | M0R300;M | M0R300 | Unconvent |
| MICAL1 | 3.222854 | 0.000599 | 0.183038 | Q8TDZ2;H0 | Q8TDZ2 | Q8TDZ2 | Protein-me |
| SF3B2 | 1.64176 | 0.022816 | 0.136287 | Q13435;E9 | Q13435;E9 | Q13435 | Splicing fac |
| NDUFS1 | 1.242883 | 0.057163 | 0.13448 | P28331;B4 | P28331;B4 | P28331 | NADH-ubiq |
| SEC31A | 0.84544 | 0.142745 | 0.105652 | O94979;D6 | O94979;D6 | O94979 | Protein tra |
| CANX | 0.801954 | 0.157778 | 0.062084 | P27824;H0 | P27824 | P27824 | Calnexin |
| HYOU1 | 0.687589 | 0.205311 | 0.071063 | A0A494C0 | A0A494C0 | A0A494C0 | Hypoxia up |
| PC | 0.639254 | 0.229481 | 0.079415 | P11498;A0 | P11498;A0 | P11498 | Pyruvate ca |
| HK2 | 0.607408 | 0.24694 | 0.115737 | P52789;E9 | P52789;E9 | P52789 | Hexokinase |
| ABCC1 | 0.5562 | 0.277843 | -0.03607 | P33527;I3I | P33527;I3I | P33527 | Multidrug r |
| COPB2 | 0.310865 | 0.488805 | 0.055755 | P35606;D6 | P35606 | P35606 | Coatomer s |
| ERAP2 | 0.284 | 0.519996 | -0.04611 | Q6P179;D6 | Q6P179;D6 | Q6P179 | Endoplasm |
| KIAA0196 | 0.209256 | 0.617653 | 0.046338 | Q12768;E7 | Q12768;E7 | Q12768 | WASH com |
| GOLGA3 | 0.568962 | 0.269797 | 0.052872 | Q08378;A0 | Q08378 | Q08378 | Golgin subf |
| PLCB2 | 0.525164 | 0.298425 | 0.101327 | Q00722;Q9 | Q00722 | Q00722 | 1-phosphat |
| STAT1 | 0.37325 | 0.4234 | -0.0581 | A0A669KB6 | A0A669KB6 | A0A669KB6 | Signal trans |
| PNPLA6 | 0.199485 | 0.631706 | 0.020304 | A0A384DV | A0A384DV | A0A384DV | Neuropathy |
| HTT | 0.152046 | 0.704618 | 0.044901 | P42858;A0 | P42858 | P42858 | Huntingtin |
| LMNB1 | 0.071563 | 0.84808 | 0.013096 | P20700;E9 | P20700;E9 | P20700 | Lamin-B1 |
| POR | 1.308771 | 0.049117 | 0.067759 | P16435;H0 | P16435;H0 | P16435 | NADPH--cy |
| PARP1 | 0.962238 | 0.109084 | 0.122983 | P09874;Q5 | P09874 | P09874 | Poly [ADP-r |
| ITGAV | 0.559915 | 0.275477 | -0.05555 | P06756;H7 | P06756 | P06756 | Integrin alp |
| FERMT3 | 0.310153 | 0.489607 | -0.0639 | Q86UX7;F5 | Q86UX7;F5 | Q86UX7 | Fermitin fa |

| | | | | | | | |
|---|---|---|---|---|---|---|---|
| VPS35 | 0.298637 | 0.502763 | 0.08267 | Q96QK1;I3 | Q96QK1 | Q96QK1 | Vacuolar pr |
| USP15 | 0.190785 | 0.644489 | -0.03534 | Q9Y4E8;H0 | Q9Y4E8 | Q9Y4E8 | Ubiquitin c |
| DOCK2 | 0.17852 | 0.662949 | -0.02928 | Q92608;E5 | Q92608;E5 | Q92608 | Dedicator c |
| EFTUD2 | 0.064943 | 0.861106 | 0.007744 | Q15029;K7 | Q15029 | Q15029 | 116 kDa U5 |
| MYO18A | 0.00922 | 0.978993 | -0.0014 | A0A0D9SFK | A0A0D9SFK | A0A0D9SFK | Unconvent |
| TFRC | 1.694638 | 0.0202 | 0.205599 | P02786;G3 | P02786;G3 | P02786 | Transferrin |
| MIA3 | 2.016997 | 0.009616 | 0.13952 | Q5JRA6;A0 | Q5JRA6;A0 | Q5JRA6 | Melanoma |
| SPG11 | 0.993803 | 0.101437 | 0.080534 | Q96JI7;C4E | Q96JI7;C4E | Q96JI7 | Spatacsin |
| SF3B3 | 0.964451 | 0.10853 | 0.102854 | Q15393;J3 | Q15393 | Q15393 | Splicing fac |
| USP9X | 0.594213 | 0.254558 | 0.067274 | Q93008;O( | Q93008 | Q93008 | Probable ul |
| DARS | 0.452614 | 0.352684 | -0.05502 | P14868;H7 | P14868 | P14868 | Aspartate-- |
| KIF5B | 0.352537 | 0.444081 | -0.06569 | P33176;O6 | P33176 | P33176 | Kinesin-1 h |
| ACO1 | 0.307941 | 0.492107 | -0.09315 | P21399;F5 | P21399 | P21399 | Cytoplasmi |
| TOP2B | 0.288663 | 0.514443 | -0.04065 | Q02880;E9 | Q02880;E9 | Q02880 | DNA topois |
| MSN | 0.261891 | 0.547153 | -0.04049 | P26038;V9 | P26038 | P26038 | Moesin |
| TKT | 0.236203 | 0.580494 | -0.07368 | P29401;A0 | P29401;A0 | P29401 | Transketola |
| AP2A1 | 2.005805 | 0.009867 | 0.071013 | O95782 | O95782 | O95782 | AP-2 compl |
| ACAD9 | 0.777061 | 0.167086 | 0.060992 | Q9H845;H( | Q9H845;H( | Q9H845 | Acyl-CoA de |
| KIAA1033 | 0.711099 | 0.194491 | 0.091369 | A0A087X2! | A0A087X2! | A0A087X2! | WASH com |
| KPNB1 | 0.528332 | 0.296256 | 0.055987 | Q14974;J3 | Q14974;J3 | Q14974 | Importin su |
| MYO1D | 0.517075 | 0.304036 | 0.080359 | O94832;J3 | O94832;J3 | O94832 | Unconvent |
| MYO1G | 0.508879 | 0.309829 | -0.12364 | B0I1T2;A0/ | B0I1T2;A0/ | B0I1T2 | Unconvent |
| HSPA5 | 0.401289 | 0.396927 | 0.058879 | P11021 | P11021 | P11021 | 78 kDa gluc |
| NUP210 | 0.138737 | 0.726545 | 0.025327 | Q8TEM1 | Q8TEM1 | Q8TEM1 | Nuclear por |
| SQRDL | 0.082478 | 0.827031 | 0.009481 | Q9Y6N5;H3 | Q9Y6N5;H3 | Q9Y6N5 | Sulfide:quir |
| DDX1 | 2.311154 | 0.004885 | 0.137978 | A0A087X2( | A0A087X2( | A0A087X2( | ATP-depenc |
| TRAP1 | 2.311016 | 0.004886 | 0.133875 | Q12931;I3I | Q12931;I3I | Q12931 | Heat shock |
| ACSL1 | 1.937729 | 0.011542 | 0.075567 | P33121;E7 | P33121;E7 | P33121 | Long-chain |
| ANXA2 | 1.153117 | 0.070288 | -0.17741 | P07355;H0 | P07355;H0 | P07355 | Annexin A2 |
| VPS11 | 0.859514 | 0.138193 | 0.085564 | Q9H270;A( | Q9H270;A( | Q9H270 | Vacuolar pr |
| ILF3 | 0.727593 | 0.187244 | 0.057771 | Q12906;K7 | Q12906 | Q12906 | Interleukin |
| MMP9 | 0.69929 | 0.199853 | -0.14064 | P14780 | P14780 | P14780 | Matrix met |
| EPB41L2 | 0.58163 | 0.262042 | 0.08038 | E9PHY5;O4 | E9PHY5;O4 | E9PHY5 | Band 4.1-lil |
| TCIRG1 | 0.579179 | 0.263524 | 0.054069 | Q13488;E9 | Q13488;E9 | Q13488 | V-type prot |
| COLGALT1 | 0.578872 | 0.263711 | 0.072097 | Q8NBJ5;M( | Q8NBJ5 | Q8NBJ5 | Procollager |
| PARP14 | 0.474324 | 0.335487 | -0.08966 | Q460N5;H7 | Q460N5;H7 | Q460N5 | Poly [ADP-r |
| ZC3HAV1 | 0.376946 | 0.419811 | 0.044444 | Q7Z2W4;C! | Q7Z2W4;C! | Q7Z2W4 | Zinc finger ( |
| CPT2 | 0.321463 | 0.477021 | 0.026489 | P23786;A0 | P23786;A0 | P23786 | Carnitine O |
| CCT5 | 0.304542 | 0.495973 | 0.035777 | P48643;E7 | P48643;E7 | P48643 | T-complex p |
| AP1G1 | 0.279887 | 0.524944 | -0.04713 | O43747;H3 | O43747 | O43747 | AP-1 compl |
| EML4 | 0.241746 | 0.573131 | 0.040137 | Q9HC35;B5 | Q9HC35;B5 | Q9HC35 | Echinoderm |
| LTA4H | 0.198899 | 0.632558 | -0.08698 | P09960;B4 | P09960 | P09960 | Leukotriene |
| STIP1 | 0.176572 | 0.665929 | -0.05184 | P31948;F5 | P31948 | P31948 | Stress-indu |
| EMC1 | 0.041188 | 0.909519 | -0.00806 | Q8N766;A( | Q8N766 | Q8N766 | ER membra |
| DDB1 | 0.040641 | 0.910665 | 0.003716 | Q16531;F5 | Q16531;F5 | Q16531 | DNA damag |
| AP3B1 | 0.030322 | 0.932564 | 0.009304 | O00203;A( | O00203 | O00203 | AP-3 compl |
| ARAP1 | 0.018131 | 0.959111 | -0.00538 | Q96P48;E7 | Q96P48;E7 | Q96P48 | Arf-GAP wit |
| USP7 | 1.351343 | 0.04453 | 0.102606 | Q93009;A( | Q93009;A( | Q93009 | Ubiquitin c |

| | | | | | | | |
|---|---|---|---|---|---|---|---|
| PARP4 | 1.188197 | 0.064834 | 0.14749 | Q9UKK3 | Q9UKK3 | Q9UKK3 | Poly [ADP-r |
| LAP3 | 1.153769 | 0.070183 | -0.1942 | P28838;HC | P28838 | P28838 | Cytosol am |
| WARS | 0.739274 | 0.182275 | -0.17735 | P23381;G3 | P23381;G3 | P23381 | Tryptophar |
| HNRNPUL2 | 0.689492 | 0.204413 | -0.06245 | Q1KMD3;H | Q1KMD3;H | Q1KMD3 | Heterogene |
| ATP6V1B2 | 0.365497 | 0.431026 | -0.0419 | P21281;HC | P21281 | P21281 | V-type prot |
| CYP27A1 | 0.312333 | 0.487155 | 0.061633 | Q02318;C9 | Q02318 | Q02318 | Sterol 26-h |
| XRCC6 | 0.202869 | 0.626802 | 0.019265 | P12956;B1 | P12956;B1 | P12956 | X-ray repair |
| XPO1 | 0.20236 | 0.627538 | 0.030483 | O14980;C9 | O14980 | O14980 | Exportin-1 |
| GTF2I | 0.035182 | 0.922186 | -0.0052 | A0A494C1I | A0A494C1I | A0A494C1I | General tra |
| ALDH4A1 | 1.54199 | 0.028708 | 0.167655 | P30038;Q5 | P30038 | P30038 | Delta-1-pyr |
| GLUD1 | 1.295898 | 0.050594 | 0.099505 | P00367;P4 | P00367;P4 | P00367 | Glutamate |
| VPS39 | 1.280153 | 0.052462 | 0.091732 | Q96JC1;H3 | Q96JC1 | Q96JC1 | Vam6/Vps3 |
| TNS3 | 0.961587 | 0.109248 | -0.08972 | Q68CZ2;E9 | Q68CZ2;E9 | Q68CZ2 | Tensin-3 |
| C16orf62 | 0.961134 | 0.109362 | 0.119632 | Q7Z3J2;E7I | Q7Z3J2;E7I | Q7Z3J2 | UPF0505 p |
| ATP2A2 | 0.816336 | 0.152638 | 0.089067 | P16615;H7 | P16615;H7 | P16615 | Sarcoplasm |
| TCP1 | 0.780029 | 0.165948 | 0.080277 | P17987;E7 | P17987;E7 | P17987 | T-complex |
| TRIM28 | 0.709351 | 0.195276 | 0.082817 | Q13263;M | Q13263 | Q13263 | Transcripti |
| CCT8 | 0.681278 | 0.208316 | 0.079062 | P50990;H7 | P50990;H7 | P50990 | T-complex |
| CHD4 | 0.588463 | 0.257951 | -0.09085 | A0A2R8Y4 | A0A2R8Y4 | A0A2R8Y4 | Chromodor |
| WDR7 | 0.564214 | 0.272763 | -0.08879 | Q9Y4E6;A2 | Q9Y4E6;A2 | Q9Y4E6 | WD repeat- |
| PLXNC1 | 0.335237 | 0.462129 | 0.03957 | O60486;B4 | O60486;B4 | O60486 | Plexin-C1 |
| SACM1L | 0.30106 | 0.499965 | 0.037773 | A0A5F9ZHI | A0A5F9ZHI | A0A5F9ZHI | Phosphatic |
| PRPF6 | 0.277103 | 0.52832 | -0.03918 | O94906 | O94906 | O94906 | Pre-mRNA- |
| SHMT2 | 0.264756 | 0.543556 | 0.037941 | P34897;HC | P34897;HC | P34897 | Serine hydr |
| CAT | 2.399204 | 0.003988 | 0.151389 | P04040 | P04040 | P04040 | Catalase |
| ZFYVE26 | 1.427687 | 0.037352 | 0.123199 | Q68DK2;G | Q68DK2;G | Q68DK2 | Zinc finger |
| NCKAP1L | 0.770503 | 0.169628 | 0.116776 | P55160;F8 | P55160 | P55160 | Nck-associa |
| NPEPPS | 0.744399 | 0.180136 | -0.16077 | E9PLK3;P5 | E9PLK3;P5 | E9PLK3 | Puromycin |
| TOMM70A | 0.611402 | 0.244679 | 0.072181 | O94826 | O94826 | O94826 | Mitochond |
| DDX46 | 0.513216 | 0.306749 | 0.048806 | Q7L014;A0 | Q7L014;A0 | Q7L014 | Probable A |
| GCC2 | 0.478919 | 0.331957 | -0.06177 | Q8IWJ2;B8 | Q8IWJ2;B8 | Q8IWJ2 | GRIP and c |
| TRPV2 | 0.395461 | 0.40229 | 0.019209 | Q9Y5S1;H7 | Q9Y5S1 | Q9Y5S1 | Transient re |
| WDR1 | 0.30238 | 0.498449 | -0.06186 | O75083;D6 | O75083 | O75083 | WD repeat- |
| HNRNPU | 0.232955 | 0.58485 | 0.037704 | Q00839;AC | Q00839;AC | Q00839 | Heterogene |
| MAN2A1 | 0.199052 | 0.632337 | -0.04053 | Q16706;H | Q16706 | Q16706 | Alpha-manı |
| OSBPL8 | 0.132018 | 0.737873 | -0.02665 | Q9BZF1;F8 | Q9BZF1;F8 | Q9BZF1 | Oxysterol-b |
| DARS2 | 0.109423 | 0.777279 | 0.014564 | Q6PI48;A0 | Q6PI48;A0 | Q6PI48 | Aspartate-- |
| DNM1L | 0.103865 | 0.78729 | 0.040061 | O00429;F8 | O00429 | O00429 | Dynamin-1 |
| PFKL | 0.02601 | 0.941868 | 0.006389 | P17858;F8 | P17858 | P17858 | ATP-depend |
| OSBPL1A | 2.436967 | 0.003656 | 0.226397 | Q9BXW6;E | Q9BXW6 | Q9BXW6 | Oxysterol-b |
| TUFM | 2.062144 | 0.008667 | 0.189988 | P49411;H3 | P49411 | P49411 | Elongation |
| EIF3C | 2.91763 | 0.001209 | 0.184411 | Q99613;B5 | Q99613;B5 | Q99613 | Eukaryotic |
| CCDC93 | 1.443005 | 0.036057 | 0.18292 | Q567U6;F8 | Q567U6;F8 | Q567U6 | Coiled-coil |
| AGPS | 1.440536 | 0.036263 | 0.15535 | O00116;AC | O00116;AC | O00116 | Alkyldihydr |
| RANBP2 | 2.238318 | 0.005777 | 0.148364 | P49792;Q9 | P49792 | P49792 | E3 SUMO-p |
| ACSL3 | 1.48753 | 0.032544 | 0.126707 | O95573;F5 | O95573 | O95573 | Long-chain |
| LACTB | 1.430373 | 0.037122 | 0.107444 | P83111;HC | P83111 | P83111 | Serine beta |
| IPO5 | 2.103658 | 0.007877 | -0.11447 | O00410;H | O00410;H | O00410 | Importin-5 |

| | | | | | | | |
|---|---|---|---|---|---|---|---|
| MAN2B1 | 1.030365 | 0.093247 | 0.156309 | O00754;M( | O00754 | O00754 | Lysosomal a |
| TYMP | 0.937759 | 0.115409 | -0.1796 | P19971;C9 | P19971;C9 | P19971 | Thymidine |
| CCT7 | 0.901073 | 0.125582 | 0.086763 | Q99832;F8 | Q99832 | Q99832 | T-complex p |
| SMC3 | 0.706576 | 0.196528 | 0.097303 | Q9UQE7 | Q9UQE7 | Q9UQE7 | Structural r |
| SFPQ | 0.671212 | 0.2132 | -0.05501 | P23246;H0 | P23246 | P23246 | Splicing fac |
| CKAP4 | 0.580668 | 0.262623 | 0.072469 | Q07065 | Q07065 | Q07065 | Cytoskeleto |
| MATR3 | 0.563551 | 0.27318 | 0.069669 | P43243;A8 | P43243;A8 | P43243 | Matrin-3 |
| PDXDC1 | 0.535869 | 0.29116 | 0.218626 | Q6P996;H3 | Q6P996;H3 | Q6P996 | Pyridoxal-d |
| ALDH18A1 | 0.525449 | 0.298229 | 0.078718 | P54886 | P54886 | P54886 | Delta-1-pyr |
| PSMD3 | 0.513506 | 0.306545 | 0.058479 | O43242;H0 | O43242 | O43242 | 26S proteas |
| KARS | 0.454656 | 0.35103 | -0.05732 | Q15046;H3 | Q15046 | Q15046 | Lysine--tRN |
| LNPEP | 0.447333 | 0.356999 | 0.074529 | Q9UIQ6 | Q9UIQ6 | Q9UIQ6 | Leucyl-cyst |
| MCCC1 | 0.386408 | 0.410763 | 0.054887 | Q96RQ3;E9 | Q96RQ3;E9 | Q96RQ3 | Methylcrot |
| ITGA6 | 0.216235 | 0.607806 | 0.019385 | P23229;C9 | P23229 | P23229 | Integrin alp |
| MTOR | 0.208319 | 0.618986 | 0.032419 | P42345;B1 | P42345 | P42345 | Serine/thre |
| DHX15 | 0.206864 | 0.621064 | 0.023764 | O43143 | O43143 | O43143 | Pre-mRNA-s |
| WDR11 | 0.187229 | 0.649786 | 0.040741 | Q9BZH6;S4 | Q9BZH6 | Q9BZH6 | WD repeat- |
| VPS18 | 0.181157 | 0.658935 | 0.024234 | Q9P253;H0 | Q9P253 | Q9P253 | Vacuolar pr |
| RARS | 0.133222 | 0.735831 | -0.01671 | P54136;E5 | P54136 | P54136 | Arginine--tR |
| CCT2 | 0.084025 | 0.824091 | 0.011867 | P78371;F5 | P78371;F5 | P78371 | T-complex p |
| GARS | 0.083734 | 0.824643 | -0.01549 | P41250;H7 | P41250 | P41250 | Glycine--tR |
| GAA | 0.052471 | 0.886195 | 0.01562 | P10253;I3I | P10253 | P10253 | Lysosomal a |
| SEC63 | 0.048759 | 0.893801 | -0.00935 | Q9UGP8;A6 | Q9UGP8 | Q9UGP8 | Translocati |
| TAP2 | 0.021945 | 0.950725 | 0.006091 | X5CMH5;A0 | X5CMH5;A0 | X5CMH5 | Antigen pep |
| WDFY1 | 2.541059 | 0.002877 | 0.149955 | Q8IWB7;C9 | Q8IWB7 | Q8IWB7 | WD repeat |
| PECAM1 | 1.540659 | 0.028797 | 0.087204 | P16284;A0 | P16284 | P16284 | Platelet end |
| CAPN1 | 1.825675 | 0.014939 | -0.09746 | P07384;E9 | P07384 | P07384 | Calpain-1 c |
| FAM129A | 0.93833 | 0.115258 | 0.128166 | Q9BZQ8;H0 | Q9BZQ8 | Q9BZQ8 | Protein Nib |
| PCCA | 0.90873 | 0.123387 | 0.115747 | P05165;A0 | P05165;A0 | P05165 | Propionyl-C |
| SF3A1 | 0.836645 | 0.145665 | 0.102991 | Q15459;H7 | Q15459 | Q15459 | Splicing fac |
| ALDH1A1 | 0.675013 | 0.211343 | -0.17527 | P00352;Q5 | P00352 | P00352 | Retinal deh |
| SDHA | 0.493396 | 0.321073 | 0.06926 | P31040;D6 | P31040;D6 | P31040 | Succinate d |
| ACSL4 | 0.462254 | 0.344942 | -0.02812 | O60488;D6 | O60488 | O60488 | Long-chain |
| CNDP2 | 0.454092 | 0.351486 | -0.12143 | Q96KP4;J3 | Q96KP4 | Q96KP4 | Cytosolic n |
| ANXA5 | 0.378894 | 0.417933 | -0.06338 | P08758;D6 | P08758;D6 | P08758 | Annexin A5 |
| EPHX1 | 0.372154 | 0.424469 | 0.043171 | P07099;B1 | P07099 | P07099 | Epoxide hy |
| USP5 | 0.336694 | 0.460581 | -0.08708 | P45974;F5 | P45974 | P45974 | Ubiquitin c |
| EIF3B | 0.332013 | 0.465572 | 0.029533 | P55884;C9 | P55884 | P55884 | Eukaryotic |
| ATP5B | 0.240335 | 0.574996 | 0.037853 | P06576;H0 | P06576;H0 | P06576 | ATP synthas |
| IFI16 | 0.216256 | 0.607777 | 0.043736 | Q16666;A0 | Q16666 | Q16666 | Gamma-int |
| ACOX3 | 0.190767 | 0.644515 | 0.033888 | O15254;D6 | O15254 | O15254 | Peroxisoma |
| AFG3L2 | 0.147349 | 0.71228 | 0.024113 | Q9Y4W6;K7 | Q9Y4W6 | Q9Y4W6 | AFG3-like p |
| SCP2 | 0.126094 | 0.748008 | 0.02195 | P22307;E9 | P22307 | P22307 | Non-specifi |
| ARCN1 | 0.051405 | 0.888373 | 0.014513 | P48444;B0 | P48444;B0 | P48444 | Coatomer s |
| AMPD3 | 0.028379 | 0.936745 | 0.007362 | Q01432;E9 | Q01432;E9 | Q01432 | AMP deami |
| ECM29 | 0.026628 | 0.940528 | -0.00469 | Q5VYK3;J3I | Q5VYK3;J3I | Q5VYK3 | Proteasome |
| ACOX1 | 0.02486 | 0.944366 | -0.00478 | Q15067;K7 | Q15067 | Q15067 | Peroxisoma |
| NPC1 | 2.277992 | 0.005272 | 0.177895 | O15118;K7 | O15118;K7 | O15118 | Niemann-P |

| | | | | | | | |
|---|---|---|---|---|---|---|---|
| GLS | 1.573611 | 0.026692 | 0.150319 | O94925;B7 | O94925 | O94925 | Glutaminas |
| LETM1 | 1.962876 | 0.010892 | 0.148356 | O95202 | O95202 | O95202 | LETM1 and |
| ITGB1 | 2.250427 | 0.005618 | 0.134992 | P05556;E7 | P05556 | P05556 | Integrin be |
| MARS | 1.389019 | 0.04083 | 0.112631 | P56192;H0 | P56192 | P56192 | Methionine |
| TNPO1 | 1.502258 | 0.031459 | 0.10108 | Q92973;O1 | Q92973 | Q92973 | Transportir |
| PRKCD | 1.369941 | 0.042664 | 0.061755 | Q05655;A0 | Q05655 | Q05655 | Protein kin |
| ANXA4 | 2.01119 | 0.009746 | -0.19738 | P09525;Q6 | P09525;Q6 | P09525 | Annexin A4 |
| AIFM1 | 1.200357 | 0.063044 | 0.111827 | O95831;E9 | O95831;E9 | O95831 | Apoptosis-i |
| VPS16 | 1.199881 | 0.063113 | 0.151441 | Q9H269;Q5 | Q9H269 | Q9H269 | Vacuolar pr |
| LRRFIP1 | 1.174228 | 0.066953 | 0.113338 | Q32MZ4;C9 | Q32MZ4 | Q32MZ4 | Leucine-ric |
| ATP6V1C1 | 1.13952 | 0.072524 | -0.07199 | P21283;E7 | P21283;E7 | P21283 | V-type prot |
| LMNB2 | 0.950412 | 0.112095 | 0.11072 | Q03252 | Q03252 | Q03252 | Lamin-B2 |
| ATIC | 0.927783 | 0.118091 | -0.16693 | P31939;H7 | P31939 | P31939 | Bifunctiona |
| PYGL | 0.875949 | 0.133061 | -0.21735 | P06737;E9 | P06737;E9 | P06737 | Glycogen p |
| RECQL | 0.698033 | 0.200432 | 0.122651 | P46063;F8 | P46063 | P46063 | ATP-depend |
| DDX5 | 0.650498 | 0.223616 | 0.067711 | J3KTA4;P17 | J3KTA4;P17 | J3KTA4 | Probable AT |
| PTPN1 | 0.570573 | 0.268798 | 0.09086 | P18031;B4 | P18031;B4 | P18031 | Tyrosine-pr |
| CORO1C | 0.547834 | 0.283247 | 0.098742 | Q9ULV4;B4 | Q9ULV4;B4 | Q9ULV4 | Coronin-1C |
| PGD | 0.436003 | 0.366435 | -0.10026 | P52209;K7 | P52209 | P52209 | 6-phosphog |
| MFN2 | 0.395686 | 0.402081 | 0.039376 | O95140;Q5 | O95140 | O95140 | Mitofusin-2 |
| TBXAS1 | 0.379827 | 0.417036 | -0.02907 | Q53F23;P2 | Q53F23;P2 | Q53F23 | Thromboxa |
| CPT1A | 0.301216 | 0.499785 | 0.039156 | P50416;H3 | P50416 | P50416 | Carnitine O |
| PSMC2 | 0.283855 | 0.52017 | 0.029192 | P35998;C9 | P35998 | P35998 | 26S proteas |
| YARS | 0.194909 | 0.638397 | -0.03413 | P54577;A0 | P54577;A0 | P54577 | Tyrosine--tR |
| PLOD3 | 0.16538 | 0.683314 | 0.035405 | O60568;H7 | O60568 | O60568 | Procollagen |
| PDIA3 | 0.165143 | 0.683686 | 0.024457 | P30101 | P30101 | P30101 | Protein disu |
| NUCB1 | 0.158644 | 0.693994 | -0.0179 | Q02818;H7 | Q02818;H7 | Q02818 | Nucleobind |
| ABHD16A | 0.044368 | 0.902883 | -0.00791 | A0A0G2JJD | A0A0G2JJD | A0A0G2JJD | Abhydrolas |
| SAMD9L | 0.029039 | 0.935322 | 0.00613 | Q8IVG5;A0 | Q8IVG5 | Q8IVG5 | Sterile alph |
| CCT4 | 0.014143 | 0.96796 | -0.00179 | P50991 | P50991 | P50991 | T-complex p |
| GFPT1 | 0.010266 | 0.976639 | -0.00344 | Q06210;A0 | Q06210;A0 | Q06210 | Glutamine- |
| AP2A2 | 1.725984 | 0.018794 | 0.102314 | O94973;A0 | O94973;A0 | O94973 | AP-2 compl |
| SEPTIN7 | 1.470973 | 0.033809 | 0.094201 | Q16181;E7 | Q16181;E7 | Q16181 | Septin-7 |
| DIAPH1 | 1.143665 | 0.071835 | -0.10553 | H9KV28;A0 | H9KV28;A0 | H9KV28 | Protein dia |
| LSS | 1.114683 | 0.076792 | 0.089023 | P48449;A0 | P48449;A0 | P48449 | Lanosterol |
| ALDH2 | 1.01246 | 0.097172 | 0.163323 | P05091;F8 | P05091 | P05091 | Aldehyde d |
| ACIN1 | 0.922747 | 0.119468 | 0.117676 | S4R3H4;E7 | S4R3H4;E7 | S4R3H4 | Apoptotic c |
| MUT | 0.866389 | 0.136023 | 0.082297 | P22033 | P22033 | P22033 | Methylmalo |
| PI4K2A | 0.826294 | 0.149179 | 0.105407 | Q9BTU6;E9 | Q9BTU6;E9 | Q9BTU6 | Phosphatid |
| CYFIP1 | 0.814417 | 0.153314 | -0.10977 | Q7L576;A0 | Q7L576;A0 | Q7L576 | Cytoplasmi |
| TRIM25 | 0.777676 | 0.166849 | 0.135984 | Q14258;A0 | Q14258;A0 | Q14258 | E3 ubiquiti |
| SMC1A | 0.701392 | 0.198888 | 0.10446 | G8JLG1;Q1 | G8JLG1;Q1 | G8JLG1 | Structural m |
| RDX | 0.598297 | 0.252176 | 0.117478 | P35241;A0 | P35241;A0 | P35241 | Radixin |
| SUPT16H | 0.528477 | 0.296158 | 0.111763 | Q9Y5B9 | Q9Y5B9 | Q9Y5B9 | FACT comp |
| EXOC4 | 0.499097 | 0.316886 | 0.07983 | Q96A65 | Q96A65 | Q96A65 | Exocyst cor |
| RRP12 | 0.479165 | 0.331768 | 0.053925 | Q5JTH9;A0 | Q5JTH9 | Q5JTH9 | RRP12-like |
| ARPC2 | 0.406571 | 0.392129 | -0.05706 | O15144;H7 | O15144 | O15144 | Actin-relate |
| TAP1 | 0.403908 | 0.394541 | -0.0919 | Q03518;A0 | Q03518;A0 | Q03518 | Antigen pep |

| | | | | | | | |
|---|---|---|---|---|---|---|---|
| CDC42BPB | 0.390369 | 0.407034 | 0.064219 | Q9Y5S2;H0 | Q9Y5S2;H0 | Q9Y5S2 | Serine/thre |
| UGP2 | 0.365934 | 0.430592 | -0.06348 | Q16851;A0 | Q16851;A0 | Q16851 | UTP--gluco: |
| CAPN2 | 0.311663 | 0.487907 | -0.05229 | P17655;C9 | P17655 | P17655 | Calpain-2 c |
| STK10 | 0.267767 | 0.539801 | 0.046 | O94804;H0 | O94804 | O94804 | Serine/thre |
| TPP2 | 0.254308 | 0.55679 | 0.028306 | P29144;Q5 | P29144;Q5 | P29144 | Tripeptidyl |
| AIM1 | 0.253907 | 0.557304 | 0.024329 | A0A0J9YWI | A0A0J9YWI | A0A0J9YWI | Absent in m |
| CCT3 | 0.251237 | 0.560742 | 0.028686 | P49368;B4 | P49368;B4 | P49368 | T-complex p |
| ABCD1 | 0.220923 | 0.60128 | 0.034916 | P33897;A6 | P33897 | P33897 | ATP-bindin |
| HSP90AA1 | 0.155023 | 0.699805 | -0.03994 | P07900;G3 | P07900 | P07900 | Heat shock |
| LIMA1 | 0.125191 | 0.749564 | 0.043539 | Q9UHB6;F8 | Q9UHB6;F8 | Q9UHB6 | LIM domair |
| CORO7-PAM | 0.099833 | 0.794635 | -0.01964 | A0A0A6YYL | A0A0A6YYL | A0A0A6YYL | Coronin;Co |
| SCFD1 | 0.097016 | 0.799805 | -0.01634 | Q8WVM8;J | Q8WVM8 | Q8WVM8 | Sec1 family |
| EEF1A1 | 0.094602 | 0.804262 | 0.01574 | P68104;Q5 | P68104;Q5 | P68104 | Elongation |
| SRPR | 0.094148 | 0.805103 | 0.018681 | P08240 | P08240 | P08240 | Signal recog |
| PFKP | 0.089072 | 0.81457 | 0.018718 | Q01813;B1 | Q01813;B1 | Q01813 | ATP-depend |
| DENND4C | 0.052924 | 0.88527 | -0.01295 | Q5VZ89;X6 | Q5VZ89 | Q5VZ89 | DENN doma |
| PYGB | 0.018934 | 0.95734 | -0.00588 | P11216;H0 | P11216 | P11216 | Glycogen p |
| GOLGA2 | 1.879304 | 0.013204 | 0.177231 | Q08379;A0 | Q08379 | Q08379 | Golgin subf |
| CHIT1 | 1.596816 | 0.025304 | 0.170961 | Q13231;D6 | Q13231;D6 | Q13231 | Chitotriosi |
| AP1B1 | 1.95315 | 0.011139 | 0.15535 | Q10567;C9 | Q10567;C9 | Q10567 | AP-1 compl |
| NCF2 | 1.15112 | 0.070612 | -0.15651 | P19878;B1 | P19878 | P19878 | Neutrophil |
| SART3 | 0.981089 | 0.104451 | 0.119454 | A0A499FI3 | A0A499FI3 | A0A499FI3 | Squamous c |
| NUP93 | 0.975905 | 0.105705 | 0.078793 | Q8N1F7;H3 | Q8N1F7;H3 | Q8N1F7 | Nuclear por |
| GFM1 | 0.96036 | 0.109557 | 0.100923 | Q96RP9;C9 | Q96RP9;C9 | Q96RP9 | Elongation |
| VPS41 | 0.892303 | 0.128144 | 0.089844 | P49754;C9 | P49754 | P49754 | Vacuolar pr |
| RNPEP | 0.889034 | 0.129112 | -0.17693 | Q9H4A4;A6 | Q9H4A4;A6 | Q9H4A4 | Aminopept |
| VAC14 | 0.842676 | 0.143656 | 0.082924 | Q08AM6;H | Q08AM6 | Q08AM6 | Protein VAC |
| SRP68 | 0.737001 | 0.183231 | 0.071526 | Q9UHB9 | Q9UHB9 | Q9UHB9 | Signal recog |
| NUP155 | 0.688948 | 0.204669 | 0.120359 | O75694;E9 | O75694;E9 | O75694 | Nuclear por |
| EIF4G2 | 0.646883 | 0.225485 | 0.064795 | P78344;H0 | P78344;H0 | P78344 | Eukaryotic |
| ATG7 | 0.645812 | 0.226041 | -0.13821 | O95352;C9 | O95352 | O95352 | Ubiquitin-l |
| ASCC3 | 0.623331 | 0.238051 | 0.077382 | Q8N3C0;E5 | Q8N3C0 | Q8N3C0 | Activating s |
| PGK1 | 0.529085 | 0.295744 | -0.11745 | P00558;P0 | P00558 | P00558 | Phosphogly |
| GBP5 | 0.279776 | 0.525078 | -0.16383 | Q96PP8;E7 | Q96PP8;E7 | Q96PP8 | Guanylate-l |
| ATP2B4 | 0.227695 | 0.591977 | 0.018921 | P23634;H0 | P23634 | P23634 | Plasma mer |
| ARHGEF1 | 0.20725 | 0.620512 | 0.045232 | M0QZR4;Q | M0QZR4;Q | M0QZR4 | Rho guanin |
| NONO | 0.191482 | 0.643455 | 0.027542 | Q15233;H7 | Q15233;H7 | Q15233 | Non-POU d |
| EZR | 0.173456 | 0.670724 | 0.034738 | E7EQR4;P1 | E7EQR4;P1 | E7EQR4 | Ezrin |
| RMDN3 | 0.173119 | 0.671245 | 0.025247 | Q96TC7;H0 | Q96TC7;H0 | Q96TC7 | Regulator c |
| DOPEY2 | 0.163837 | 0.685745 | 0.032555 | Q9Y3R5;F8 | Q9Y3R5 | Q9Y3R5 | Protein dop |
| ASPH | 0.134094 | 0.734355 | -0.01494 | Q12797;E5 | Q12797 | Q12797 | Aspartyl/as |
| GALNT7 | 0.122844 | 0.753625 | -0.01789 | Q86SF2;H0 | Q86SF2;H0 | Q86SF2 | N-acetylgal |
| CKAP5 | 0.075962 | 0.839533 | -0.01211 | Q14008;H0 | Q14008 | Q14008 | Cytoskeleto |
| SMARCC2 | 0.059894 | 0.871176 | -0.01165 | F8VXC8;Q8 | F8VXC8;Q8 | F8VXC8 | SWI/SNF co |
| SLC25A24 | 0.004878 | 0.98883 | -0.00068 | Q6NUK1;A0 | Q6NUK1 | Q6NUK1 | Calcium-bi |
| GOT2 | 3.570138 | 0.000269 | 0.196115 | P00505 | P00505 | P00505 | Aspartate a |
| HNRNPA2B | 2.654993 | 0.002213 | 0.149007 | P22626;A0 | P22626;A0 | P22626 | Heterogene |
| SRRM2 | 1.478024 | 0.033264 | 0.143086 | Q9UQ35;I3 | Q9UQ35 | Q9UQ35 | Serine/argi |

| | | | | | | | |
|---|---|---|---|---|---|---|---|
| NDUFV1 | 2.43221 | 0.003696 | 0.10605 | G3V0I5;P4 | G3V0I5;P4 | G3V0I5 | NADH dehy |
| EIF3L | 1.437192 | 0.036543 | 0.055278 | Q9Y262;B0 | Q9Y262;B0 | Q9Y262 | Eukaryotic |
| TBL2 | 2.315991 | 0.004831 | -0.16794 | E9PF19;Q9 | E9PF19;Q9 | E9PF19 | Transducin |
| CTTN | 1.281948 | 0.052246 | 0.114128 | Q14247;H | Q14247 | Q14247 | Src substra |
| CES1 | 1.055 | 0.088105 | 0.335689 | P23141;H3 | P23141 | P23141 | Liver carbo |
| FGR | 0.929304 | 0.117678 | 0.058948 | P09769;Q5 | P09769;Q5 | P09769 | Tyrosine-pr |
| CDK5RAP3 | 0.884781 | 0.130382 | -0.0748 | Q96JB5;J3 | Q96JB5;J3 | Q96JB5 | CDK5 regul |
| CAD | 0.814565 | 0.153262 | 0.126266 | F8VPD4;P2 | F8VPD4;P2 | F8VPD4 | CAD protei |
| A2M | 0.719889 | 0.190595 | 0.050215 | P01023;F8 | P01023 | P01023 | Alpha-2-ma |
| ASAH1 | 0.63319 | 0.232707 | -0.09188 | Q13510;A0 | Q13510;A0 | Q13510 | Acid ceram |
| NCL | 0.59224 | 0.255717 | 0.070717 | P19338;H7 | P19338;H7 | P19338 | Nucleolin |
| ZYX | 0.581483 | 0.26213 | 0.07529 | Q15942;H | Q15942;H | Q15942 | Zyxin |
| HADHB | 0.36401 | 0.432504 | 0.031274 | P55084;F5 | P55084;F5 | P55084 | Trifunction |
| STIM1 | 0.357083 | 0.439458 | 0.055217 | Q13586;G | Q13586;G | Q13586 | Stromal int |
| RPL4 | 0.346961 | 0.44982 | 0.023801 | P36578;H3 | P36578;H3 | P36578 | 60S ribosor |
| ANKFY1 | 0.337044 | 0.46021 | 0.088326 | Q9P2R3;I3 | Q9P2R3 | Q9P2R3 | Rabankyrin |
| UQCRC1 | 0.302631 | 0.498161 | -0.03645 | P31930 | P31930 | P31930 | Cytochrom |
| FH | 0.291173 | 0.511479 | 0.047384 | P07954 | P07954 | P07954 | Fumarate h |
| PLCG2 | 0.275978 | 0.52969 | -0.05578 | A0A0A0MR | A0A0A0MR | A0A0A0MR | Phosphoin |
| ACTR3 | 0.260225 | 0.549256 | 0.040828 | P61158;B4 | P61158;B4 | P61158 | Actin-relate |
| OAS3 | 0.244865 | 0.569029 | 0.049282 | Q9Y6K5;F8 | Q9Y6K5 | Q9Y6K5 | 2-5-oligoac |
| RAB11FIP1 | 0.24293 | 0.571571 | 0.037711 | Q6WKZ4;E | Q6WKZ4;E | Q6WKZ4 | Rab11 fami |
| STXBP2 | 0.177538 | 0.664449 | -0.01364 | Q15833;M | Q15833;M | Q15833 | Syntaxin-bi |
| CUL2 | 0.147219 | 0.712494 | 0.042463 | A0A0A0MT | A0A0A0MT | A0A0A0MT | Cullin-2 |
| FAM129B | 0.090543 | 0.811814 | -0.02788 | Q96TA1 | Q96TA1 | Q96TA1 | Niban-like |
| ACADM | 0.077973 | 0.835654 | -0.01386 | Q5T4U5;P1 | Q5T4U5;P1 | Q5T4U5 | Medium-ch |
| XRN2 | 0.063519 | 0.863935 | -0.00979 | Q9H0D6 | Q9H0D6 | Q9H0D6 | 5-3 exoribo |
| ABCF1 | 2.782534 | 0.00165 | 0.171975 | Q8NE71;H | Q8NE71;H | Q8NE71 | ATP-bindin |
| PDHA1 | 1.603512 | 0.024917 | 0.162017 | P08559;Q5 | P08559 | P08559 | Pyruvate de |
| MTDH | 1.9032 | 0.012497 | 0.152524 | Q86UE4;E5 | Q86UE4;E5 | Q86UE4 | Protein LYR |
| ADAM10 | 1.331151 | 0.04665 | 0.080346 | O14672;H | O14672 | O14672 | Disintegrin |
| GCLC | 1.570335 | 0.026895 | -0.30266 | P48506;A0 | P48506;A0 | P48506 | Glutamate- |
| LY75 | 1.134435 | 0.073378 | 0.171508 | O60449 | O60449 | O60449 | Lymphocyt |
| PA2G4 | 1.067744 | 0.085557 | 0.103884 | Q9UQ80;F | Q9UQ80;F | Q9UQ80 | Proliferatio |
| CCDC22 | 1.013274 | 0.09699 | 0.126686 | O60826 | O60826 | O60826 | Coiled-coil |
| ATP6V1H | 1.00354 | 0.099188 | -0.10303 | Q9UI12;G3 | Q9UI12;G3 | Q9UI12 | V-type prot |
| AP2B1 | 0.782171 | 0.165131 | 0.072531 | P63010;A0 | P63010;A0 | P63010 | AP-2 compl |
| GSR | 0.767705 | 0.170724 | 0.045746 | P00390;A0 | P00390 | P00390 | Glutathion |
| SRP72 | 0.679621 | 0.209112 | 0.109309 | O76094;D | O76094;D | O76094 | Signal reco |
| GBF1 | 0.598357 | 0.252141 | -0.06328 | A0A669KB | A0A669KB | A0A669KB | Golgi-speci |
| PPP2R1A | 0.55524 | 0.278458 | -0.08825 | P30153;B3 | P30153;B3 | P30153 | Serine/thre |
| NLN | 0.544886 | 0.285177 | 0.081632 | Q9BYT8;E9 | Q9BYT8;E9 | Q9BYT8 | Neurolysin, |
| ARHGEF2 | 0.542353 | 0.286845 | 0.074331 | V9GYM8;A | V9GYM8;A | V9GYM8 | Rho guanin |
| OAS2 | 0.526255 | 0.297677 | 0.084468 | A0A087X0 | A0A087X0 | A0A087X0 | 2-5-oligoac |
| CCAR2 | 0.498465 | 0.317347 | -0.09719 | Q8N163;H | Q8N163;H | Q8N163 | Cell cycle a |
| PMPCA | 0.482628 | 0.329133 | 0.084423 | Q10713;Q5 | Q10713 | Q10713 | Mitochond |
| SEPTIN2 | 0.473987 | 0.335748 | 0.04786 | Q15019;B5 | Q15019;B5 | Q15019 | Septin-2 |
| TLR8 | 0.438111 | 0.364661 | 0.079947 | Q9NR97 | Q9NR97 | Q9NR97 | Toll-like rec |

| | | | | | | | |
|---|---|---|---|---|---|---|---|
| ESYT2 | 0.341058 | 0.455976 | -0.03588 | A0A499FIX | A0A499FIX | A0A499FIX | Extended sy |
| CYP1B1 | 0.259191 | 0.550565 | -0.04024 | Q16678;A0 | Q16678 | Q16678 | Cytochrom |
| UBA6 | 0.228686 | 0.590628 | -0.05251 | A0AVT1;H0 | A0AVT1 | A0AVT1 | Ubiquitin-l |
| CCDC47 | 0.163025 | 0.687028 | 0.025312 | Q96A33;A0 | Q96A33;A0 | Q96A33 | Coiled-coil |
| KHSRP | 0.15618 | 0.697943 | 0.020347 | Q92945;A0 | Q92945;A0 | Q92945 | Far upstrea |
| HSP90AB1 | 0.134258 | 0.734078 | -0.03189 | P08238;Q5 | P08238 | P08238 | Heat shock |
| PADI2 | 0.122037 | 0.755028 | -0.04504 | Q9Y2J8 | Q9Y2J8 | Q9Y2J8 | Protein-arg |
| TBC1D9B | 0.08245 | 0.827086 | 0.026139 | Q66K14;G3 | Q66K14 | Q66K14 | TBC1 doma |
| MCCC2 | 0.077858 | 0.835876 | 0.012801 | Q9HCC0;D6 | Q9HCC0;D6 | Q9HCC0 | Methylcrot |
| ALDH3A2 | 0.063887 | 0.863203 | 0.010607 | P51648;A0 | P51648;A0 | P51648 | Fatty aldeh |
| ANXA1 | 0.027442 | 0.938767 | 0.00432 | P04083;Q5 | P04083;Q5 | P04083 | Annexin A1 |
| FECH | 3.107828 | 0.00078 | 0.243197 | P22830;A0 | P22830;A0 | P22830 | Ferrochelat |
| SEC24C | 2.000195 | 0.009996 | 0.241735 | P53992;G5 | P53992;G5 | P53992 | Protein trar |
| PSAP | 2.080606 | 0.008306 | 0.172106 | P07602;C9 | P07602;C9 | P07602 | Prosaposin |
| UQCRC2 | 2.172912 | 0.006716 | 0.138695 | P22695;H3 | P22695;H3 | P22695 | Cytochrom |
| ACAA2 | 1.505662 | 0.031213 | 0.126357 | P42765;A0 | P42765;A0 | P42765 | 3-ketoacyl- |
| SEC23A | 2.337569 | 0.004597 | 0.112336 | Q15436;F5 | Q15436;F5 | Q15436 | Protein trar |
| PSMC4 | 1.31867 | 0.04801 | 0.100289 | P43686 | P43686 | P43686 | 26S proteas |
| ANXA11 | 1.400959 | 0.039723 | -0.06987 | P50995;H0 | P50995 | P50995 | Annexin A1 |
| ME1 | 1.310663 | 0.048903 | -0.23134 | P48163 | P48163 | P48163 | NADP-depe |
| CARS | 1.092722 | 0.080775 | 0.129862 | P49589;B4 | P49589;B4 | P49589 | Cysteine--tl |
| MOGS | 1.073707 | 0.08439 | 0.125511 | Q13724;A0 | Q13724;A0 | Q13724 | Mannosyl-o |
| ALDH5A1 | 1.069154 | 0.08528 | 0.105837 | P51649;C9 | P51649;C9 | P51649 | Succinate-s |
| ATP11A | 1.064826 | 0.086134 | 0.084646 | H0Y547;E9 | H0Y547;E9 | H0Y547 | Phospholip |
| ACSF2 | 1.05981 | 0.087134 | 0.151277 | Q96CM8;E9 | Q96CM8;E9 | Q96CM8 | Acyl-CoA sy |
| EPS15 | 0.68719 | 0.205499 | 0.093796 | P42566;B1 | P42566;B1 | P42566 | Epidermal g |
| NSDHL | 0.650753 | 0.223484 | 0.071229 | Q15738;C9 | Q15738;C9 | Q15738 | Sterol-4-alp |
| PNPT1 | 0.585732 | 0.259578 | 0.071986 | Q8TCS8;H7 | Q8TCS8 | Q8TCS8 | Polyribonu |
| GNPAT | 0.548869 | 0.282573 | 0.094613 | O15228;Q5 | O15228;Q5 | O15228 | Dihydroxya |
| NSUN2 | 0.519338 | 0.302456 | 0.055077 | Q08J23;A0 | Q08J23 | Q08J23 | tRNA (cytos |
| GDI2 | 0.467773 | 0.340586 | -0.12355 | P50395;Q5 | P50395;Q5 | P50395 | Rab GDP di |
| ACE | 0.438339 | 0.364469 | 0.126298 | P12821;A0 | P12821;A0 | P12821 | Angiotensir |
| PHB | 0.410997 | 0.388153 | -0.06036 | P35232;C9 | P35232;C9 | P35232 | Prohibitin |
| ACSF3 | 0.400149 | 0.39797 | -0.05269 | Q4G176;A0 | Q4G176;A0 | Q4G176 | Acyl-CoA sy |
| IDH1 | 0.374669 | 0.422018 | -0.11419 | O75874;C9 | O75874 | O75874 | Isocitrate d |
| RAB3GAP2 | 0.245975 | 0.567577 | 0.030572 | Q9H2M9 | Q9H2M9 | Q9H2M9 | Rab3 GTPas |
| KYNU | 0.237775 | 0.578396 | -0.0548 | Q16719;A0 | Q16719;A0 | Q16719 | Kynurenina |
| IDH2 | 0.234957 | 0.582161 | 0.030997 | P48735;H0 | P48735 | P48735 | Isocitrate d |
| SEC23B | 0.228766 | 0.590519 | 0.041841 | A0A2R8YFH | A0A2R8YFH | A0A2R8YFH | Protein trar |
| DHCR24 | 0.205441 | 0.623102 | 0.023878 | A0A0A0MT | A0A0A0MT | A0A0A0MT | Delta(24)-st |
| SRRT | 0.17557 | 0.667468 | -0.02079 | Q9BXP5;H7 | Q9BXP5;H7 | Q9BXP5 | Serrate RNA |
| VAT1 | 0.160444 | 0.691123 | -0.04071 | Q99536;K7 | Q99536 | Q99536 | Synaptic ve |
| ITGB5 | 0.13793 | 0.727897 | 0.01925 | P18084;H7 | P18084;H7 | P18084 | Integrin be |
| ACOT9 | 0.094163 | 0.805076 | -0.01465 | Q9Y305;H7 | Q9Y305;H7 | Q9Y305 | Acyl-coenzy |
| FMNL2 | 0.083538 | 0.825016 | -0.00938 | Q96PY5;C9 | Q96PY5;C9 | Q96PY5 | Formin-like |
| CCDC88A | 0.067821 | 0.855419 | -0.01503 | Q3V6T2;A0 | Q3V6T2;A0 | Q3V6T2 | Girdin |
| PPP1R12A | 0.053644 | 0.883803 | 0.012182 | O14974;F8 | O14974;F8 | O14974 | Protein pho |
| DPYSL2 | 0.014155 | 0.967932 | -0.00307 | A0A1C7CY) | A0A1C7CY) | A0A1C7CY) | Dihydropyr |

| | | | | | | | |
|---|---|---|---|---|---|---|---|
| WFS1 | 1.67577 | 0.021097 | 0.24337 | A0A669KA) | A0A669KA) | A0A669KA) | Wolframin |
| DDX17 | 1.882701 | 0.013101 | 0.202135 | A0A1X7SBZ | A0A1X7SBZ | A0A1X7SBZ | Probable A |
| ITGA5 | 2.895898 | 0.001271 | 0.182427 | P08648;H0 | P08648 | P08648 | Integrin alp |
| EIF5B | 3.071027 | 0.000849 | 0.16676 | A0A087WU | A0A087WU | A0A087WU | Eukaryotic |
| CRAT | 1.531793 | 0.029391 | 0.10421 | P43155;H0 | P43155 | P43155 | Carnitine O |
| SLC27A3 | 1.316184 | 0.048285 | -0.10153 | X6R3N0;Q5 | X6R3N0;Q5 | X6R3N0 | Long-chain |
| MYO1E | 4.986561 | 1.03E-05 | -0.39912 | Q12965;H0 | Q12965 | Q12965 | Unconvent |
| MOSPD2 | 1.224855 | 0.059586 | 0.115716 | Q8NHP6;R4 | Q8NHP6;R4 | Q8NHP6 | Motile sper |
| PLAA | 1.119676 | 0.075914 | 0.183079 | Q9Y263;E5 | Q9Y263;E5 | Q9Y263 | Phospholip |
| SNTB1 | 1.060953 | 0.086905 | 0.083474 | Q13884;A0 | Q13884;A0 | Q13884 | Beta-1-synt |
| PRPF19 | 0.968044 | 0.107636 | 0.116343 | Q9UMS4;F5 | Q9UMS4 | Q9UMS4 | Pre-mRNA-p |
| PSMC3 | 0.884606 | 0.130435 | 0.07211 | R4GNH3;P1 | R4GNH3;P1 | R4GNH3 | 26S proteas |
| PGM2 | 0.788803 | 0.162629 | -0.17214 | Q96G03;E7 | Q96G03 | Q96G03 | Phosphoglu |
| CLPX | 0.780462 | 0.165782 | 0.090248 | O76031;H0 | O76031 | O76031 | ATP-depend |
| SPECC1 | 0.719075 | 0.190952 | 0.092456 | Q5M775;J3 | Q5M775;J3 | Q5M775 | Cytospin-B |
| CPOX | 0.649879 | 0.223934 | 0.09749 | P36551;H0 | P36551 | P36551 | Oxygen-dep |
| HEXB | 0.631301 | 0.233721 | 0.149914 | P07686;Q5 | P07686;Q5 | P07686 | Beta-hexosa |
| AARS | 0.603861 | 0.248965 | 0.11561 | P49588;H3 | P49588 | P49588 | Alanine--tR |
| ALDH9A1 | 0.582325 | 0.261622 | -0.08586 | P49189 | P49189 | P49189 | 4-trimethyl |
| FKBP5 | 0.561943 | 0.274193 | 0.087406 | Q13451 | Q13451 | Q13451 | Peptidyl-pr |
| STAT3 | 0.535309 | 0.291535 | 0.097707 | P40763;G8 | P40763;G8 | P40763 | Signal trans |
| CSDE1 | 0.480845 | 0.330488 | -0.04955 | O75534;E9 | O75534;E9 | O75534 | Cold shock |
| GAPDH | 0.433171 | 0.368832 | -0.06654 | P04406;E7 | P04406;E7 | P04406 | Glyceraldeh |
| MIPEP | 0.393097 | 0.404485 | 0.053637 | Q99797 | Q99797 | Q99797 | Mitochond |
| TNFAIP2 | 0.366452 | 0.430079 | 0.062527 | Q03169;H0 | Q03169 | Q03169 | Tumor necr |
| ACAT1 | 0.344665 | 0.452205 | 0.05537 | P24752;A0 | P24752;A0 | P24752 | Acetyl-CoA |
| GPD1 | 0.303715 | 0.496918 | -0.13585 | P21695;Q8 | P21695 | P21695 | Glycerol-3- |
| PSMD11 | 0.26105 | 0.548214 | -0.02182 | O00231;J3 | O00231 | O00231 | 26S proteas |
| CAST | 0.25282 | 0.558702 | 0.050941 | A0A0C4DG | A0A0C4DG | A0A0C4DG | Calpastatin |
| STRN | 0.224762 | 0.595989 | 0.024308 | O43815;G3 | O43815 | O43815 | Striatin |
| SUCLG2 | 0.212442 | 0.613138 | -0.02098 | Q96I99;E9I | Q96I99;E9I | Q96I99 | Succinyl-Co |
| ERMP1 | 0.19288 | 0.641387 | -0.02005 | Q7Z2K6;E7 | Q7Z2K6;E7 | Q7Z2K6 | Endoplasm |
| FBP1 | 0.182887 | 0.656317 | -0.07188 | P09467;A0 | P09467;A0 | P09467 | Fructose-1, |
| CTSD | 0.128309 | 0.744202 | 0.023706 | A0A1B0GVI | A0A1B0GVI | A0A1B0GVI | Cathepsin D |
| DLAT | 0.119584 | 0.759304 | -0.02479 | P10515;H0 | P10515;H0 | P10515 | Dihydrolipo |
| EHHADH | 0.105481 | 0.784367 | -0.02059 | Q08426;C9 | Q08426 | Q08426 | Peroxisoma |
| EXOC1 | 0.098323 | 0.797401 | 0.023698 | Q9NV70 | Q9NV70 | Q9NV70 | Exocyst cor |
| ERP44 | 0.080982 | 0.829884 | 0.01426 | Q9BS26 | Q9BS26 | Q9BS26 | Endoplasm |
| NUP205 | 0.076196 | 0.839081 | -0.01038 | Q92621;U3 | Q92621 | Q92621 | Nuclear por |
| ALOX5 | 0.073659 | 0.843998 | 0.011606 | P09917;A0 | P09917 | P09917 | Arachidona |
| OPTN | 0.057908 | 0.875169 | -0.01307 | Q96CV9;H7 | Q96CV9 | Q96CV9 | Optineurin |
| SEC23IP | 0.046573 | 0.898311 | 0.00747 | Q9Y6Y8;A0 | Q9Y6Y8 | Q9Y6Y8 | SEC23-inter |
| HNRNPM | 0.026608 | 0.940572 | 0.006232 | P52272;A0 | P52272;A0 | P52272 | Heterogene |
| SLC25A13 | 0.022981 | 0.94846 | -0.0054 | Q9UJS0;R4 | Q9UJS0 | Q9UJS0 | Calcium-bir |
| UBE4A | 0.005859 | 0.986599 | -0.00086 | Q14139;B7 | Q14139 | Q14139 | Ubiquitin c |
| TRMT10C | 5.086544 | 8.19E-06 | 0.403534 | Q7L0Y3;C9 | Q7L0Y3;C9 | Q7L0Y3 | Mitochond |
| CYP51A1 | 3.286818 | 0.000517 | 0.275459 | A0A0C4DFI | A0A0C4DFI | A0A0C4DFI | Lanosterol |
| DHX30 | 2.106968 | 0.007817 | 0.225843 | H7BXY3;Q7 | H7BXY3;Q7 | H7BXY3 | Putative AT |

| | | | | | | | |
|---|---|---|---|---|---|---|---|
| OGT | 1.871571 | 0.013441 | 0.190307 | O15294;C9 | O15294 | O15294 | UDP-N-acet |
| AARS2 | 2.15657 | 0.006973 | 0.154774 | Q5JTZ9 | Q5JTZ9 | Q5JTZ9 | Alanine--tR |
| VPS33A | 1.349341 | 0.044736 | 0.135208 | Q96AX1;A0 | Q96AX1;A0 | Q96AX1 | Vacuolar pr |
| RTCB | 1.788439 | 0.016276 | 0.123143 | Q9Y3I0 | Q9Y3I0 | Q9Y3I0 | tRNA-splici |
| VASP | 2.942263 | 0.001142 | -0.18906 | P50552;K7 | P50552 | P50552 | Vasodilator |
| SUCLA2 | 1.256476 | 0.055402 | 0.099841 | A0A2R8YD( | A0A2R8YD( | A0A2R8YD( | Succinyl-Co |
| SARS2 | 1.127143 | 0.07462 | 0.147249 | M0QWZ7;( | M0QWZ7;( | M0QWZ7 | Serine--tRN |
| PRAM1 | 1.104852 | 0.07855 | 0.095136 | Q96QH2;M | Q96QH2 | Q96QH2 | PML-RARA- |
| SERBP1 | 1.07633 | 0.083882 | 0.084065 | Q8NC51 | Q8NC51 | Q8NC51 | Plasminoge |
| SLC25A12 | 0.844672 | 0.142997 | 0.090773 | O75746;F8 | O75746 | O75746 | Calcium-bir |
| ABCD3 | 0.833628 | 0.14668 | 0.087608 | A0A3B3ITW | A0A3B3ITW | A0A3B3ITW | ATP-bindin |
| TXNRD2 | 0.668916 | 0.214331 | 0.090816 | A0A182DW | A0A182DW | A0A182DW | Thioredoxir |
| PLOD1 | 0.621253 | 0.239192 | 0.096421 | Q02809;Q5 | Q02809 | Q02809 | Procollager |
| CNP | 0.606672 | 0.247359 | -0.05011 | P09543;K7 | P09543 | P09543 | 2,3-cyclic-r |
| VDAC1 | 0.553895 | 0.279322 | 0.070359 | P21796;C9 | P21796;C9 | P21796 | Voltage-dep |
| DLST | 0.496518 | 0.318773 | 0.081373 | P36957;Q8 | P36957;Q8 | P36957 | Dihydrolipo |
| KIAA1598 | 0.438022 | 0.364736 | -0.11864 | A0MZ66 | A0MZ66 | A0MZ66 | Shootin-1 |
| MROH1 | 0.424874 | 0.375946 | 0.100583 | E9PHY8;Q8 | E9PHY8;Q8 | E9PHY8 | Maestro he |
| PSMD12 | 0.376217 | 0.420516 | -0.04899 | O00232;J3 | O00232 | O00232 | 26S proteas |
| NCEH1 | 0.358117 | 0.438412 | -0.03992 | Q6PIU2;A0 | Q6PIU2;A0 | Q6PIU2 | Neutral cho |
| SARS | 0.354116 | 0.44247 | -0.05851 | Q5T5C7;P4 | Q5T5C7;P4 | Q5T5C7 | Serine--tRN |
| TARS2 | 0.314838 | 0.484353 | 0.0449 | Q9BW92;U | Q9BW92;U | Q9BW92 | Threonine-- |
| MOV10 | 0.294968 | 0.507028 | 0.039202 | Q9HCE1;Q5 | Q9HCE1;Q5 | Q9HCE1 | Putative he |
| NOP56 | 0.290838 | 0.511873 | 0.047725 | O00567;A0 | O00567;A0 | O00567 | Nucleolar p |
| ALDH7A1 | 0.28147 | 0.523034 | 0.045089 | A0A1B0GT( | A0A1B0GT( | A0A1B0GT( | Alpha-amir |
| ALCAM | 0.257997 | 0.552081 | -0.06939 | Q13740;F5 | Q13740;F5 | Q13740 | CD166 anti |
| PDIA6 | 0.252695 | 0.558862 | 0.03605 | Q15084 | Q15084 | Q15084 | Protein disu |
| RPL3 | 0.230235 | 0.588525 | -0.01645 | P39023;G5 | P39023;G5 | P39023 | 60S ribosor |
| ABCE1 | 0.178582 | 0.662855 | -0.04352 | P61221;D6 | P61221;D6 | P61221 | ATP-bindin |
| ALDOA | 0.14491 | 0.716291 | 0.023584 | P04075;J3 | P04075;J3 | P04075 | Fructose-bi |
| PSMD13 | 0.104992 | 0.785251 | -0.01168 | Q9UNM6;A | Q9UNM6;A | Q9UNM6 | 26S proteas |
| PLBD1 | 0.089792 | 0.813219 | 0.017563 | Q6P4A8;F5 | Q6P4A8 | Q6P4A8 | Phospholip |
| DBT | 0.081521 | 0.828855 | -0.01344 | P11182;Q5 | P11182;Q5 | P11182 | Lipoamide |
| SNX5 | 0.076906 | 0.837711 | -0.02052 | Q9Y5X3;Q5 | Q9Y5X3;Q5 | Q9Y5X3 | Sorting nex |
| DLD | 0.048223 | 0.894905 | 0.006729 | P09622;E9 | P09622;E9 | P09622 | Dihydrolipo |
| ILF2 | 0.042153 | 0.907501 | 0.00724 | Q12905;B4 | Q12905;B4 | Q12905 | Interleukin |
| RBM14 | 0.032128 | 0.928693 | 0.007066 | Q96PK6;A0 | Q96PK6 | Q96PK6 | RNA-bindir |
| LRPAP1 | 2.830878 | 0.001476 | 0.297955 | P30533;A0 | P30533;A0 | P30533 | Alpha-2-ma |
| OAT | 4.360942 | 4.36E-05 | 0.264093 | P04181 | P04181 | P04181 | Ornithine a |
| TBRG4 | 1.856823 | 0.013905 | 0.226758 | Q969Z0;C9 | Q969Z0;C9 | Q969Z0 | Protein TBF |
| HNRNPA3 | 2.705178 | 0.001972 | 0.184363 | P51991;H7 | P51991 | P51991 | Heterogene |
| SDHB | 2.950967 | 0.00112 | 0.179525 | P21912;A0 | P21912;A0 | P21912 | Succinate c |
| EIF2S1 | 1.934326 | 0.011633 | 0.157622 | P05198;G3 | P05198;G3 | P05198 | Eukaryotic |
| EIF3D | 2.95748 | 0.001103 | 0.152935 | O15371;B0 | O15371 | O15371 | Eukaryotic |
| NDUFS2 | 2.080833 | 0.008302 | 0.152223 | O75306 | O75306 | O75306 | NADH dehy |
| KIF2A | 1.440514 | 0.036265 | 0.142347 | O00139;D6 | O00139 | O00139 | Kinesin-like |
| RPL6 | 2.321851 | 0.004766 | 0.112177 | Q02878;F8 | Q02878 | Q02878 | 60S ribosor |
| IVD | 1.709274 | 0.019531 | 0.093791 | A0A0A0MT | A0A0A0MT | A0A0A0MT | Isovaleryl-C |

| | | | | | | | |
|---|---|---|---|---|---|---|---|
| GPC4 | 2.611791 | 0.002445 | -0.16447 | O75487;Q9 | O75487 | O75487 | Glypican-4, |
| THRAP3 | 1.225867 | 0.059447 | 0.137335 | A0A3B3ITZ9 | A0A3B3ITZ9 | A0A3B3ITZ9 | Thyroid hor |
| LARS2 | 1.21312 | 0.061218 | 0.121587 | Q15031;E9 | Q15031;E9 | Q15031 | Probable le |
| EIF2S2 | 0.995339 | 0.101079 | 0.100126 | P20042 | P20042 | P20042 | Eukaryotic |
| TOP1 | 0.979757 | 0.104771 | 0.146797 | P11387;E5 | P11387 | P11387 | DNA topois |
| NDUFA9 | 0.960468 | 0.10953 | 0.08731 | Q16795;A0 | Q16795;A0 | Q16795 | NADH dehy |
| DOCK7 | 0.958633 | 0.109994 | -0.10579 | Q96N67;A0 | Q96N67;A0 | Q96N67 | Dedicator o |
| ATL1 | 0.913968 | 0.121908 | 0.096622 | Q8WXF7;G | Q8WXF7 | Q8WXF7 | Atlastin-1 |
| RANGAP1 | 0.873853 | 0.133705 | 0.090009 | P46060;H0 | P46060 | P46060 | Ran GTPase |
| ARHGAP1 | 0.850345 | 0.141142 | 0.119352 | Q07960;H0 | Q07960;H0 | Q07960 | Rho GTPase |
| PCCB | 0.75177 | 0.177105 | 0.087502 | E7EUY3;E9I | E7EUY3;E9I | E7EUY3 | Propionyl-C |
| RPS3 | 0.743794 | 0.180387 | 0.067449 | P23396;E9 | P23396;E9 | P23396 | 40S ribosor |
| CDC5L | 0.710351 | 0.194827 | 0.125069 | Q99459 | Q99459 | Q99459 | Cell divisior |
| HNRNPUL1 | 0.697433 | 0.200709 | 0.120657 | Q9BUJ2;A0 | Q9BUJ2;A0 | Q9BUJ2 | Heterogene |
| STT3B | 0.644689 | 0.226627 | 0.072773 | Q8TCJ2 | Q8TCJ2 | Q8TCJ2 | Dolichyl-di |
| GBA | 0.635344 | 0.231556 | -0.08655 | P04062;A0 | P04062;A0 | P04062 | Glucosylcer |
| GALNT12 | 0.58224 | 0.261674 | -0.07504 | Q8IXK2 | Q8IXK2 | Q8IXK2 | Polypeptid |
| RUVBL2 | 0.576108 | 0.265395 | 0.063634 | Q9Y230;M0 | Q9Y230;M0 | Q9Y230 | RuvB-like 2 |
| NUP98 | 0.565878 | 0.27172 | 0.093836 | P52948;A0 | P52948;A0 | P52948 | Nuclear por |
| MLYCD | 0.5406 | 0.288005 | 0.064606 | O95822;V9 | O95822 | O95822 | Malonyl-Co |
| VPS45 | 0.53823 | 0.289581 | -0.07116 | Q9NRW7;A | Q9NRW7;A | Q9NRW7 | Vacuolar pr |
| AKAP13 | 0.528505 | 0.296138 | 0.070842 | Q12802;A0 | Q12802;A0 | Q12802 | A-kinase an |
| PTCD3 | 0.4896 | 0.323892 | -0.06844 | Q96EY7;B8 | Q96EY7 | Q96EY7 | Pentatricop |
| CYP2S1 | 0.453469 | 0.351991 | 0.06898 | Q96SQ9;M0 | Q96SQ9 | Q96SQ9 | Cytochrom |
| MRE11A | 0.435342 | 0.366993 | -0.07384 | P49959;F8 | P49959;F8 | P49959 | Double-stra |
| AKR1B1 | 0.392368 | 0.405165 | -0.08965 | P15121;E9 | P15121;E9 | P15121 | Aldose redu |
| ENO1 | 0.382736 | 0.414251 | -0.09606 | P06733;A0 | P06733;A0 | P06733 | Alpha-enola |
| DTX3L | 0.368777 | 0.427783 | 0.050118 | Q8TDB6 | Q8TDB6 | Q8TDB6 | E3 ubiquiti |
| STXBP3 | 0.356257 | 0.440294 | 0.030201 | O00186 | O00186 | O00186 | Syntaxin-bi |
| NCF1 | 0.348864 | 0.447853 | -0.05228 | P14598;A8 | P14598;A8 | P14598 | Neutrophil |
| PCYOX1 | 0.345333 | 0.451509 | -0.04554 | Q9UHG3;F8 | Q9UHG3 | Q9UHG3 | Prenylcyste |
| ITGAL | 0.341263 | 0.455761 | 0.061574 | P20701;H3 | P20701 | P20701 | Integrin alp |
| EVL | 0.316427 | 0.482584 | 0.059573 | Q9UI08;H0 | Q9UI08;H0 | Q9UI08 | Ena/VASP-li |
| ATP6V0A1 | 0.264385 | 0.54402 | 0.021729 | Q93050;B7 | Q93050;B7 | Q93050 | V-type prot |
| HNRNPR | 0.243424 | 0.570921 | 0.024035 | O43390;B4 | O43390;B4 | O43390 | Heterogene |
| NARS | 0.231047 | 0.587426 | 0.060128 | O43776;K7 | O43776 | O43776 | Asparagine- |
| FSCN1 | 0.226872 | 0.5931 | -0.07142 | Q16658;C9 | Q16658 | Q16658 | Fascin |
| MFN1 | 0.213088 | 0.612226 | 0.047036 | Q8IWA4;H7 | Q8IWA4;H7 | Q8IWA4 | Mitofusin-1 |
| TWF2 | 0.18397 | 0.654682 | 0.033732 | Q6IBS0;D6I | Q6IBS0;D6I | Q6IBS0 | Twinfilin-2 |
| SLC4A2 | 0.171985 | 0.673 | -0.02126 | P04920;C9 | P04920 | P04920 | Anion exch |
| PHB2 | 0.159482 | 0.692657 | 0.015472 | Q99623;J3 | Q99623;J3 | Q99623 | Prohibitin- |
| ITGA3 | 0.152894 | 0.703244 | 0.021479 | P26006;K7 | P26006 | P26006 | Integrin alp |
| NCBP1 | 0.152145 | 0.704458 | -0.03947 | Q09161;F2 | Q09161 | Q09161 | Nuclear cap |
| HEXA | 0.14793 | 0.711328 | 0.042213 | H3BP20;P0 | H3BP20;P0 | H3BP20 | Beta-hexos |
| CALU | 0.1333 | 0.735698 | 0.027101 | O43852;H0 | O43852;H0 | O43852 | Calumenin |
| RPN2 | 0.111707 | 0.773202 | 0.012749 | P04844;Q5 | P04844 | P04844 | Dolichyl-di |
| SF3A3 | 0.080733 | 0.830361 | 0.012778 | Q12874 | Q12874 | Q12874 | Splicing fac |
| SERPINH1 | 0.058413 | 0.874151 | -0.00888 | P50454;E9 | P50454;E9 | P50454 | Serpin H1 |

| | | | | | | | |
|---|---|---|---|---|---|---|---|
| CPNE3 | 0.045027 | 0.901515 | -0.00586 | O75131;A0 | O75131;A0 | O75131 | Copine-3 |
| EHD4 | 0.002276 | 0.994772 | 0.000586 | Q9H223 | Q9H223 | Q9H223 | EH domain- |
| GRN | 2.842727 | 0.001436 | 0.239532 | P28799;K7 | P28799;K7 | P28799 | Granulins;A |
| TOM1 | 3.920497 | 0.00012 | 0.198785 | O60784;B0 | O60784;B0 | O60784 | Target of M |
| MTHFD1L | 1.745288 | 0.017977 | 0.177561 | A0A087WV | A0A087WV | A0A087WV | Monofunct |
| IDH3A | 1.821258 | 0.015092 | 0.15853 | P50213;H0 | P50213;H0 | P50213 | Isocitrate d |
| EIF4A3 | 1.433306 | 0.036872 | 0.145185 | P38919;I3I | P38919;I3I | P38919 | Eukaryotic |
| FARSB | 1.886522 | 0.012986 | 0.125018 | Q9NSD9 | Q9NSD9 | Q9NSD9 | Phenylalan |
| PPA2 | 1.434279 | 0.036789 | 0.077857 | Q9H2U2;H | Q9H2U2;H | Q9H2U2 | Inorganic p |
| USP14 | 1.448964 | 0.035566 | -0.10711 | P54578;A6 | P54578;A6 | P54578 | Ubiquitin c |
| RPA1 | 1.29391 | 0.050827 | 0.154414 | P27694;I3I | P27694 | P27694 | Replication |
| PSME2 | 1.236319 | 0.058034 | -0.18992 | A0A087X1 | A0A087X1 | A0A087X1 | Proteasom |
| GBE1 | 1.124437 | 0.075087 | -0.22034 | Q04446;E9 | Q04446;E9 | Q04446 | 1,4-alpha-g |
| RAB14 | 1.051786 | 0.088759 | 0.106248 | P61106;X6 | P61106;X6 | P61106 | Ras-related |
| LRRC47 | 1.028331 | 0.093685 | 0.125503 | Q8N1G4 | Q8N1G4 | Q8N1G4 | Leucine-ric |
| PSMC5 | 1.011745 | 0.097332 | 0.116607 | P62195;J3 | P62195;J3 | P62195 | 26S protea |
| SYNCRIP | 0.977759 | 0.105255 | 0.071909 | O60506;B7 | O60506;B7 | O60506 | Heterogene |
| APMAP | 0.944321 | 0.113679 | 0.14332 | Q9HDC9;H | Q9HDC9;H | Q9HDC9 | Adipocyte p |
| BCLAF1 | 0.909147 | 0.123269 | 0.125088 | Q9NYF8;E9 | Q9NYF8;E9 | Q9NYF8 | Bcl-2-assoc |
| ILK | 0.900191 | 0.125837 | -0.13508 | A0A0A0MT | A0A0A0MT | A0A0A0MT | Integrin-lin |
| GAK | 0.846959 | 0.142246 | 0.132028 | O14976;A0 | O14976 | O14976 | Cyclin-G-as |
| SNX27 | 0.812422 | 0.15402 | 0.171096 | Q96L92;A0 | Q96L92;A0 | Q96L92 | Sorting nex |
| TIMM44 | 0.784834 | 0.164122 | 0.073506 | O43615;M | O43615;M | O43615 | Mitochond |
| PDPR | 0.778261 | 0.166625 | 0.092336 | Q8NCN5;A | Q8NCN5;A | Q8NCN5 | Pyruvate de |
| UGDH | 0.757937 | 0.174608 | -0.18376 | O60701;E7 | O60701 | O60701 | UDP-glucos |
| DNPEP | 0.664277 | 0.216632 | 0.113384 | E5RIA4;E7E | E5RIA4;E7E | E5RIA4 | Aspartyl am |
| CYP20A1 | 0.663465 | 0.217038 | 0.042189 | Q6UW02;E | Q6UW02;E | Q6UW02 | Cytochrom |
| DDX6 | 0.655012 | 0.221303 | 0.067239 | P26196;Q8 | P26196 | P26196 | Probable A |
| CYP4F22 | 0.650465 | 0.223632 | 0.07023 | Q6NT55;F8 | Q6NT55 | Q6NT55 | Cytochrom |
| PLD1 | 0.628626 | 0.235166 | -0.11188 | Q13393;C9 | Q13393 | Q13393 | Phospholip |
| PEPD | 0.601373 | 0.250396 | -0.1637 | P12955;A0 | P12955;A0 | P12955 | Xaa-Pro dip |
| RFTN1 | 0.548727 | 0.282666 | 0.080308 | Q14699;G3 | Q14699;G3 | Q14699 | Raftlin |
| ILVBL | 0.540205 | 0.288267 | 0.066985 | A1L0T0;M0 | A1L0T0;M0 | A1L0T0 | Acetolactat |
| CS | 0.514316 | 0.305974 | -0.06149 | O75390;B4 | O75390;B4 | O75390 | Citrate synt |
| FARSA | 0.50992 | 0.309086 | 0.036724 | Q9Y285;K7 | Q9Y285;K7 | Q9Y285 | Phenylalan |
| SMARCA5 | 0.476979 | 0.333443 | -0.0757 | O60264 | O60264 | O60264 | SWI/SNF-re |
| SCARB2 | 0.474374 | 0.335449 | -0.05786 | A0A1W2PS | A0A1W2PS | A0A1W2PS | Lysosome m |
| SYK | 0.464948 | 0.342809 | -0.07558 | P43405 | P43405 | P43405 | Tyrosine-pr |
| TCERG1 | 0.428629 | 0.37271 | -0.06008 | O14776;G3 | O14776;G3 | O14776 | Transcripti |
| RBM25 | 0.400172 | 0.397949 | -0.04867 | P49756;E9 | P49756 | P49756 | RNA-bindir |
| EEF1G | 0.353116 | 0.443491 | 0.053662 | P26641 | P26641 | P26641 | Elongation |
| LMAN1 | 0.342821 | 0.454128 | 0.041526 | P49257 | P49257 | P49257 | Protein ERG |
| GPI | 0.306196 | 0.494088 | -0.08817 | P06744;A0 | P06744;A0 | P06744 | Glucose-6-p |
| PSMA1 | 0.304322 | 0.496224 | 0.053271 | P25786;F5 | P25786;F5 | P25786 | Proteasom |
| UFSP2 | 0.2927 | 0.509683 | 0.043461 | Q9NUQ7;D | Q9NUQ7;D | Q9NUQ7 | Ufm1-speci |
| PRKCSH | 0.288677 | 0.514426 | 0.043601 | K7ELL7;P1 | K7ELL7;P1 | K7ELL7 | Glucosidas |
| CMAS | 0.259908 | 0.549657 | -0.03132 | Q8NFW8;F | Q8NFW8 | Q8NFW8 | N-acylneur |
| ZFYVE16 | 0.244497 | 0.569512 | 0.073816 | Q7Z3T8;D6 | Q7Z3T8 | Q7Z3T8 | Zinc finger |

| | | | | | | | |
|---|---|---|---|---|---|---|---|
| RETSAT | 0.244204 | 0.569897 | -0.02774 | Q6NUM9;H | Q6NUM9;H | Q6NUM9 | All-trans-re |
| SNX1 | 0.242194 | 0.572541 | -0.04735 | Q13596;H( | Q13596;H( | Q13596 | Sorting nex |
| GNPTAB | 0.237629 | 0.57859 | -0.02785 | Q3T906;H0 | Q3T906 | Q3T906 | N-acetylglu |
| NBEAL2 | 0.216236 | 0.607805 | -0.0472 | A0A494C1\ | A0A494C1\ | A0A494C1\ | Neurobeacl |
| PGM1 | 0.215851 | 0.608344 | -0.0763 | P36871;A0 | P36871;A0 | P36871 | Phosphoglu |
| SLC25A11 | 0.185203 | 0.652825 | 0.02872 | Q02978;I3I | Q02978;I3I | Q02978 | Mitochond |
| TMEM43 | 0.181491 | 0.65843 | 0.031379 | Q9BTV4;V9 | Q9BTV4 | Q9BTV4 | Transmemb |
| ZW10 | 0.174269 | 0.66947 | -0.04575 | O43264;F5 | O43264 | O43264 | Centromer |
| GNA13 | 0.172526 | 0.672162 | -0.01389 | Q14344 | Q14344 | Q14344 | Guanine nu |
| PTPN12 | 0.164435 | 0.684802 | 0.049037 | Q05209;H( | Q05209 | Q05209 | Tyrosine-pr |
| CCAR1 | 0.159104 | 0.69326 | -0.02701 | Q8IX12;A0, | Q8IX12;A0, | Q8IX12 | Cell divisior |
| CORO1A | 0.156151 | 0.69799 | 0.061807 | P31146;H3 | P31146;H3 | P31146 | Coronin-1A |
| KCNMA1 | 0.131087 | 0.739456 | 0.016932 | A0A1W2PS | A0A1W2PS | A0A1W2PS | Calcium-ac |
| LPCAT2 | 0.118817 | 0.760646 | -0.01819 | Q7L5N7;H3 | Q7L5N7 | Q7L5N7 | Lysophospl |
| CYBB | 0.113325 | 0.770327 | 0.016586 | P04839;Q5 | P04839 | P04839 | Cytochrom |
| PSMC6 | 0.112063 | 0.772569 | 0.014407 | A0A087X2I | A0A087X2I | A0A087X2I | 26S proteas |
| AP1M1 | 0.095564 | 0.802483 | -0.01637 | Q9BXS5;E7 | Q9BXS5;E7 | Q9BXS5 | AP-1 compl |
| TBC1D2 | 0.07945 | 0.832817 | 0.013059 | Q9BYX2 | Q9BYX2 | Q9BYX2 | TBC1 doma |
| GART | 0.069749 | 0.851631 | -0.01406 | P22102;F8 | P22102 | P22102 | Trifunction |
| CORO1B | 0.048885 | 0.893541 | 0.010531 | Q9BR76;A( | Q9BR76 | Q9BR76 | Coronin-1E |
| PABPC1 | 0.008708 | 0.980149 | -0.00096 | P11940;A0 | P11940;A0 | P11940 | Polyadenyl |
| ERO1L | 0.005895 | 0.986519 | 0.000992 | Q96HE7;G3 | Q96HE7 | Q96HE7 | ERO1-like p |
| NRP2 | 4.048005 | 8.95E-05 | 0.217647 | O60462;C9 | O60462 | O60462 | Neuropilin- |
| ANKRD44 | 1.652515 | 0.022258 | 0.185066 | A0A2R8Y7Y | A0A2R8Y7Y | A0A2R8Y7Y | Serine/thre |
| TRAPPC10 | 1.623311 | 0.023806 | 0.169981 | P48553;A0 | P48553 | P48553 | Trafficking |
| BCAP31 | 2.25607 | 0.005545 | 0.132573 | P51572;A0 | P51572;A0 | P51572 | B-cell recep |
| AP2M1 | 2.470756 | 0.003383 | 0.132495 | A0A087WY | A0A087WY | A0A087WY | AP-2 compl |
| ETFDH | 2.01005 | 0.009771 | 0.122899 | Q16134;D6 | Q16134 | Q16134 | Electron tra |
| ABHD12 | 1.293144 | 0.050916 | 0.135692 | Q8N2K0;A( | Q8N2K0;A( | Q8N2K0 | Monoacylg |
| PARP9 | 1.177216 | 0.066494 | -0.13946 | Q8IXQ6;G5 | Q8IXQ6;G5 | Q8IXQ6 | Poly [ADP-r |
| CYB5R1 | 1.098238 | 0.079756 | 0.129878 | Q9UHQ9;H | Q9UHQ9;H | Q9UHQ9 | NADH-cyto |
| SFXN3 | 0.971707 | 0.106732 | 0.091304 | Q9BWM7;A | Q9BWM7;A | Q9BWM7 | Sideroflexir |
| ALDH6A1 | 0.829421 | 0.148108 | 0.158143 | Q02252;G3 | Q02252 | Q02252 | Methylmal |
| BCL2L13 | 0.828221 | 0.148518 | 0.12565 | Q9BXK5;A( | Q9BXK5;A( | Q9BXK5 | Bcl-2-like p |
| PMPCB | 0.746232 | 0.179377 | 0.109507 | O75439;G3 | O75439;G3 | O75439 | Mitochond |
| ACBD3 | 0.699448 | 0.19978 | 0.108192 | Q9H3P7 | Q9H3P7 | Q9H3P7 | Golgi reside |
| DYNC1I2 | 0.693837 | 0.202378 | 0.091784 | Q13409;E7 | Q13409 | Q13409 | Cytoplasmi |
| CLCN7 | 0.685446 | 0.206326 | 0.091485 | P51798;H( | P51798;H( | P51798 | H(+)/Cl(-) ex |
| CTSB | 0.639736 | 0.229226 | 0.099341 | P07858;E9 | P07858 | P07858 | Cathepsin E |
| RPS3A | 0.561011 | 0.274783 | 0.070564 | P61247;D6 | P61247;D6 | P61247 | 40S ribosor |
| APOE | 0.560748 | 0.274949 | -0.10737 | P02649;H( | P02649;H( | P02649 | Apolipoprc |
| EXOC2 | 0.532602 | 0.293358 | -0.05885 | Q96KP1;Q2 | Q96KP1 | Q96KP1 | Exocyst cor |
| DBNL | 0.502517 | 0.3144 | -0.0655 | Q9UJU6;B4 | Q9UJU6;B4 | Q9UJU6 | Drebrin-like |
| MNDA | 0.489705 | 0.323814 | -0.07301 | P41218;H( | P41218 | P41218 | Myeloid cel |
| EIF2A | 0.445471 | 0.358532 | 0.067551 | Q9BY44;F8 | Q9BY44;F8 | Q9BY44 | Eukaryotic |
| EIF3E | 0.440787 | 0.36242 | 0.054451 | P60228;E5 | P60228;E5 | P60228 | Eukaryotic |
| NAMPT | 0.435732 | 0.366664 | -0.07573 | P43490;A0 | P43490;A0 | P43490 | Nicotinami |
| SLC27A4 | 0.384604 | 0.412473 | 0.04968 | Q6P1M0 | Q6P1M0 | Q6P1M0 | Long-chain |

| | | | | | | | | |
|---|---|---|---|---|---|---|---|---|
| PSMC1 | 0.374975 | 0.421721 | 0.037765 | | P62191;G3 | P62191 | P62191 | 26S proteas |
| PTBP1 | 0.345677 | 0.451152 | 0.02142 | | P26599;A0 | P26599;A0 | P26599 | Polypyrimi |
| MAP1S | 0.28577 | 0.517881 | 0.055432 | | Q66K74;M | Q66K74 | Q66K74 | Microtubul |
| LRRC8A | 0.26631 | 0.541614 | -0.03148 | | Q8IWT6;M | Q8IWT6 | Q8IWT6 | Volume-reg |
| SLC25A20 | 0.253677 | 0.557601 | 0.050875 | | O43772;C9 | O43772;C9 | O43772 | Mitochond |
| HCFC1 | 0.242583 | 0.572028 | -0.05208 | | P51610;A6 | P51610;A6 | P51610 | Host cell fa |
| MICU1 | 0.228359 | 0.591073 | 0.043877 | | F6XEV2;A0 | F6XEV2;A0 | F6XEV2 | Calcium up |
| ABR | 0.216763 | 0.607067 | -0.03582 | | Q12979;A0 | Q12979;A0 | Q12979 | Active brea |
| SSB | 0.214738 | 0.609905 | 0.039622 | | P05455;E7 | P05455 | P05455 | Lupus La pr |
| APEX1 | 0.213453 | 0.611712 | -0.07444 | | P27695;G3 | P27695;G3 | P27695 | DNA-(apuri |
| GZMA | 0.209482 | 0.617331 | 0.097215 | | P12544 | P12544 | P12544 | Granzyme A |
| TARS | 0.172084 | 0.672846 | 0.029044 | | P26639;D6 | P26639 | P26639 | Threonine-- |
| EIF3I | 0.170656 | 0.675063 | 0.021764 | | Q13347;Q5 | Q13347 | Q13347 | Eukaryotic |
| CAPG | 0.145218 | 0.715784 | -0.05236 | | P40121;E7 | P40121;E7 | P40121 | Macrophag |
| NPLOC4 | 0.122361 | 0.754464 | 0.027532 | | Q8TAT6;I3L | Q8TAT6 | Q8TAT6 | Nuclear pro |
| PPP1R18 | 0.120794 | 0.757192 | -0.02685 | | Q6NYC8;A0 | Q6NYC8;A0 | Q6NYC8 | Phostensin |
| FUCA1 | 0.115669 | 0.766181 | -0.0237 | | P04066 | P04066 | P04066 | Tissue alpha |
| DDOST | 0.092445 | 0.808268 | 0.028555 | | P39656;A0 | P39656;A0 | P39656 | Dolichyl-di |
| DCTN2 | 0.089327 | 0.814092 | -0.01147 | | Q13561;F8 | Q13561;F8 | Q13561 | Dynactin su |
| NDUFA10 | 0.088015 | 0.816555 | -0.01396 | | A0A087WX | A0A087WX | A0A087WX | NADH dehy |
| AK4 | 0.087331 | 0.817841 | 0.007351 | | P27144;F8 | P27144;F8 | P27144 | Adenylate k |
| ANXA7 | 0.086731 | 0.818972 | 0.016163 | | P20073 | P20073 | P20073 | Annexin A7 |
| MCU | 0.07759 | 0.836392 | 0.008378 | | Q8NE86;S4 | Q8NE86;S4 | Q8NE86 | Calcium un |
| ERBB2IP | 0.057674 | 0.875642 | 0.008764 | | Q96RT1;B4 | Q96RT1 | Q96RT1 | Protein LAP |
| AGL | 0.054893 | 0.881266 | -0.01149 | | P35573;A0 | P35573 | P35573 | Glycogen d |
| KMO | 0.023248 | 0.947876 | 0.006598 | | O15229;H0 | O15229 | O15229 | Kynurenine |
| POLR2A | 0.020737 | 0.953373 | -0.0032 | | P24928 | P24928 | P24928 | DNA-direct |
| HSDL2 | 0.01891 | 0.957392 | 0.003569 | | Q6YN16 | Q6YN16 | Q6YN16 | Hydroxyste |
| DNM2 | 0.011303 | 0.97431 | 0.002708 | | P50570;E5 | P50570 | P50570 | Dynamin-2 |
| ACSL5 | 0.010833 | 0.975364 | -0.00203 | | Q9ULC5;A0 | Q9ULC5 | Q9ULC5 | Long-chain |
| PLEKHO2 | 0.00112 | 0.997425 | -0.00033 | | Q8TD55;C9 | Q8TD55 | Q8TD55 | Pleckstrin h |
| CARS2 | 3.126128 | 0.000748 | 0.246883 | | Q9HA77;H0 | Q9HA77 | Q9HA77 | Probable cy |
| CLCC1 | 2.099705 | 0.007949 | 0.196744 | | Q96S66;Q5 | Q96S66 | Q96S66 | Chloride ch |
| ELAVL1 | 4.117247 | 7.63E-05 | 0.183794 | | Q15717;M | Q15717 | Q15717 | ELAV-like pr |
| DNAJC11 | 2.114331 | 0.007685 | 0.180651 | | Q9NVH1;B1 | Q9NVH1 | Q9NVH1 | DnaJ homo |
| LDHD | 2.210719 | 0.006156 | 0.179156 | | Q86WU2 | Q86WU2 | Q86WU2 | Probable D |
| FDXR | 1.909723 | 0.012311 | 0.156626 | | P22570;A0 | P22570;A0 | P22570 | NADPH:adr |
| AK2 | 2.102755 | 0.007893 | 0.142344 | | F8W1A4;P5 | F8W1A4;P5 | F8W1A4 | Adenylate k |
| DNAJC10 | 1.878088 | 0.013241 | 0.141147 | | Q8IXB1;A0 | Q8IXB1 | Q8IXB1 | DnaJ homo |
| TMEM63A | 2.588551 | 0.002579 | 0.131818 | | O94886;Q2 | O94886 | O94886 | CSC1-like p |
| MTA2 | 1.83822 | 0.014514 | 0.130992 | | O94776;E9 | O94776 | O94776 | Metastasis- |
| AK3 | 1.668027 | 0.021477 | 0.125713 | | Q9UIJ7 | Q9UIJ7 | Q9UIJ7 | GTP:AMP p |
| PLD3 | 2.09062 | 0.008117 | -0.21631 | | Q8IV08;M0 | Q8IV08 | Q8IV08 | Phospholip |
| NOP58 | 1.270204 | 0.053678 | 0.200701 | | Q9Y2X3;H7 | Q9Y2X3 | Q9Y2X3 | Nucleolar p |
| TTLL12 | 1.207024 | 0.062083 | 0.18353 | | Q14166;V9 | Q14166 | Q14166 | Tubulin--ty |
| PGAM1 | 1.198021 | 0.063384 | -0.20996 | | P18669;P1 | P18669 | P18669 | Phosphogly |
| SWAP70 | 1.17987 | 0.066089 | 0.158098 | | Q9UH65;E7 | Q9UH65;E7 | Q9UH65 | Switch-asso |
| NAGK | 1.108256 | 0.077937 | -0.12676 | | Q9UJ70;H7 | Q9UJ70;H7 | Q9UJ70 | N-acetyl-D- |

| | | | | | | | |
|---|---|---|---|---|---|---|---|
| SRSF1 | 1.077394 | 0.083677 | 0.10079 | J3KTL2;Q0 | J3KTL2;Q0 | J3KTL2 | Serine/argi |
| EPS15L1 | 1.029621 | 0.093407 | 0.104172 | M0R165;Q | M0R165;Q | M0R165 | Epidermal |
| VDAC2 | 0.995648 | 0.101007 | 0.062223 | P45880;A0 | P45880;A0 | P45880 | Voltage-de |
| ACAA1 | 0.991188 | 0.10205 | 0.098949 | P09110;C9 | P09110;C9 | P09110 | 3-ketoacyl- |
| ATP6V0D2 | 0.98447 | 0.103641 | 0.077832 | Q8N8Y2;E5 | Q8N8Y2 | Q8N8Y2 | V-type prot |
| GALNT2 | 0.914152 | 0.121856 | -0.13167 | Q10471 | Q10471 | Q10471 | Polypeptid |
| UNC13D | 0.841423 | 0.144071 | 0.122266 | Q70J99;K7 | Q70J99 | Q70J99 | Protein un |
| SGPL1 | 0.836361 | 0.14576 | 0.054414 | O95470;H | O95470 | O95470 | Sphingosin |
| MDH2 | 0.783386 | 0.16467 | 0.05164 | P40926;G3 | P40926;G3 | P40926 | Malate deh |
| LIMS1 | 0.750863 | 0.177475 | -0.15126 | P48059;A0 | P48059;A0 | P48059 | LIM and ser |
| GALNS | 0.740262 | 0.18186 | 0.087955 | P34059;H3 | P34059 | P34059 | N-acetylgal |
| UBA2 | 0.71211 | 0.194039 | 0.13188 | Q9UBT2;K7 | Q9UBT2;K7 | Q9UBT2 | SUMO-acti |
| POLR2B | 0.685594 | 0.206256 | -0.0947 | C9J4M6;C | C9J4M6;C | C9J4M6 | DNA-direct |
| GIMAP4 | 0.671019 | 0.213295 | -0.12474 | Q9NUV9;G | Q9NUV9;G | Q9NUV9 | GTPase IM |
| SLC27A1 | 0.655096 | 0.22126 | 0.074375 | Q6PCB7;M | Q6PCB7 | Q6PCB7 | Long-chain |
| CYB5R3 | 0.627034 | 0.236029 | 0.104151 | P00387;B1 | P00387;B1 | P00387 | NADH-cyto |
| ATP5A1 | 0.621637 | 0.238981 | 0.056194 | P25705;K7 | P25705 | P25705 | ATP syntha |
| CSE1L | 0.527048 | 0.297134 | 0.124485 | P55060 | P55060 | P55060 | Exportin-2 |
| TPI1 | 0.523967 | 0.299249 | -0.08216 | P60174;U3 | P60174;U3 | P60174 | Triosephos |
| GOLGA5 | 0.494099 | 0.320554 | -0.10084 | Q8TBA6;H | Q8TBA6 | Q8TBA6 | Golgin sub |
| RAB7A | 0.488108 | 0.325006 | 0.0631 | P51149;C9 | P51149;C9 | P51149 | Ras-related |
| ACBD5 | 0.466153 | 0.341859 | -0.06299 | B7Z2R7;Q5 | B7Z2R7;Q5 | B7Z2R7 | Acyl-CoA-bi |
| DNAJC3 | 0.45168 | 0.353444 | 0.064328 | Q13217;X | Q13217;X | Q13217 | DnaJ homo |
| ICAM1 | 0.443742 | 0.359963 | 0.107572 | P05362;K7 | P05362 | P05362 | Intercellula |
| PUF60 | 0.43026 | 0.371313 | -0.0873 | A0A0J9YYL | A0A0J9YYL | A0A0J9YYL | Poly(U)-bin |
| CCT6A | 0.38802 | 0.409242 | -0.04974 | P40227;J3 | P40227 | P40227 | T-complex |
| PSME1 | 0.381472 | 0.415459 | -0.06723 | Q06323;H | Q06323 | Q06323 | Proteasom |
| DDX23 | 0.374609 | 0.422076 | -0.07203 | Q9BUQ8;H | Q9BUQ8 | Q9BUQ8 | Probable A |
| OXR1 | 0.315239 | 0.483906 | 0.099574 | Q8N573;H | Q8N573;H | Q8N573 | Oxidation r |
| PRDX6 | 0.299159 | 0.502158 | -0.07644 | P30041 | P30041 | P30041 | Peroxiredo |
| RTN1 | 0.291929 | 0.510589 | 0.064508 | Q16799;Q | Q16799;Q | Q16799 | Reticulon-1 |
| DPP3 | 0.274381 | 0.531642 | -0.07571 | Q9NY33;G | Q9NY33;G | Q9NY33 | Dipeptidyl |
| DDX42 | 0.267974 | 0.539543 | 0.037669 | Q86XP3;A | Q86XP3;A | Q86XP3 | ATP-depen |
| ETFB | 0.240798 | 0.574384 | 0.040663 | P38117;M | P38117;M | P38117 | Electron tr |
| RBM47 | 0.232745 | 0.585133 | 0.024677 | A0AV96;D6 | A0AV96;D6 | A0AV96 | RNA-bindi |
| SAMM50 | 0.199095 | 0.632274 | 0.032032 | Q9Y512 | Q9Y512 | Q9Y512 | Sorting an |
| SNX2 | 0.191422 | 0.643544 | 0.05188 | O60749;D | O60749 | O60749 | Sorting nex |
| ABCB7 | 0.1895 | 0.646398 | 0.027573 | A0A5F9ZA | A0A5F9ZA | A0A5F9ZA | ATP-bindin |
| TOR4A | 0.151208 | 0.70598 | -0.02668 | Q9NXH8 | Q9NXH8 | Q9NXH8 | Torsin-4A |
| CAP1 | 0.147776 | 0.711581 | 0.03998 | Q01518;Q | Q01518 | Q01518 | Adenylyl cy |
| STT3A | 0.145853 | 0.714739 | -0.02782 | P46977;A0 | P46977 | P46977 | Dolichyl-di |
| NUP107 | 0.135792 | 0.73149 | 0.023588 | P57740;H | P57740 | P57740 | Nuclear po |
| RAB3GAP1 | 0.0988 | 0.796525 | 0.025002 | Q15042 | Q15042 | Q15042 | Rab3 GTPa |
| H2AFY | 0.094229 | 0.804953 | 0.018579 | O75367;B4 | O75367;B4 | O75367 | Core histor |
| UBA7 | 0.084739 | 0.822736 | -0.02238 | P41226 | P41226 | P41226 | Ubiquitin- |
| CSTF3 | 0.075932 | 0.839591 | 0.017955 | Q12996;E9 | Q12996;E9 | Q12996 | Cleavage st |
| FLOT2 | 0.068217 | 0.85464 | 0.013317 | J3QLD9;E7 | J3QLD9;E7 | J3QLD9 | Flotillin-2 |
| ATP13A3 | 0.066873 | 0.857288 | -0.00728 | A0A2R8Y6 | A0A2R8Y6 | A0A2R8Y6 | Probable c |

| | | | | | | | |
|---|---|---|---|---|---|---|---|
| NCKAP1 | 0.042676 | 0.906408 | -0.0108 | Q9Y2A7 | Q9Y2A7 | Q9Y2A7 | Nck-associa |
| RMDN2 | 0.026192 | 0.941473 | -0.00521 | Q96LZ7;A0 | Q96LZ7;A0 | Q96LZ7 | Regulator c |
| RBM39 | 0.01251 | 0.971606 | 0.002611 | Q14498;G3 | Q14498;G3 | Q14498 | RNA-bindir |
| COG4 | 0.004093 | 0.99062 | 0.001619 | A0A6I8PIQ | A0A6I8PIQ | A0A6I8PIQ | Conserved |
| EIF3H | 3.412955 | 0.000386 | 0.230066 | B3KS98;O1 | B3KS98;O1 | B3KS98 | Eukaryotic |
| PDS5A | 1.777464 | 0.016693 | 0.221757 | Q29RF7;H0 | Q29RF7 | Q29RF7 | Sister chror |
| BASP1 | 1.600292 | 0.025102 | 0.195097 | P80723;U3 | P80723 | P80723 | Brain acid s |
| RDH11 | 2.065448 | 0.008601 | 0.181606 | Q8TC12;H0 | Q8TC12;H0 | Q8TC12 | Retinol deh |
| PBXIP1 | 2.123505 | 0.007525 | 0.166232 | Q96AQ6;Q | Q96AQ6 | Q96AQ6 | Pre-B-cell le |
| ECHS1 | 1.407421 | 0.039136 | 0.125994 | P30084 | P30084 | P30084 | Enoyl-CoA h |
| PICALM | 3.069492 | 0.000852 | 0.115098 | Q13492;H0 | Q13492 | Q13492 | Phosphatic |
| EIF2S3 | 1.941074 | 0.011453 | 0.094034 | P41091;Q2 | P41091;Q2 | P41091 | Eukaryotic |
| ERLIN2 | 1.835332 | 0.014611 | -0.12658 | O94905;E5 | O94905;E5 | O94905 | Erlin-2 |
| DYSF | 2.261818 | 0.005472 | -0.17153 | O75923 | O75923 | O75923 | Dysferlin |
| COG7 | 1.273108 | 0.05332 | -0.20505 | P83436 | P83436 | P83436 | Conserved |
| SRC | 1.222173 | 0.059955 | 0.068175 | P12931;H2 | P12931 | P12931 | Proto-onco |
| ACAP2 | 1.088866 | 0.081496 | -0.17458 | A0A087X1I | A0A087X1I | A0A087X1I | Arf-GAP wit |
| COMT | 1.046341 | 0.089879 | 0.133864 | P21964;E7 | P21964;E7 | P21964 | Catechol O- |
| NDUFS3 | 1.038932 | 0.091426 | 0.099531 | O75489;E9 | O75489 | O75489 | NADH dehy |
| MPP1 | 1.033581 | 0.092559 | 0.055584 | Q00013;C9 | Q00013;C9 | Q00013 | 55 kDa eryt |
| GNS | 1.021547 | 0.09516 | 0.107968 | P15586;H7 | P15586;H7 | P15586 | N-acetylglu |
| HSD17B10 | 0.983327 | 0.103914 | 0.079727 | Q99714;Q5 | Q99714;Q5 | Q99714 | 3-hydroxya |
| PSMB2 | 0.939194 | 0.115029 | -0.13589 | P49721;A0 | P49721;A0 | P49721 | Proteasom |
| AHCY | 0.919017 | 0.120499 | -0.17091 | P23526 | P23526 | P23526 | Adenosylho |
| BCKDK | 0.90036 | 0.125788 | 0.116749 | O14874;H3 | O14874;H3 | O14874 | [3-methyl-2 |
| QRSL1 | 0.81919 | 0.151639 | 0.171503 | Q9H0R6;X6 | Q9H0R6;X6 | Q9H0R6 | Glutamyl-t |
| PTGES2 | 0.800398 | 0.158344 | 0.117118 | Q9H7Z7;A6 | Q9H7Z7;A6 | Q9H7Z7 | Prostagland |
| MYBBP1A | 0.788365 | 0.162793 | -0.10263 | Q9BQG0;I3 | Q9BQG0;I3 | Q9BQG0 | Myb-bindir |
| HCLS1 | 0.769412 | 0.170054 | 0.074304 | E7EVW7;P1 | E7EVW7;P1 | E7EVW7 | Hematopoi |
| CNOT1 | 0.734827 | 0.184151 | -0.09201 | A5YKK6;H3 | A5YKK6 | A5YKK6 | CCR4-NOT t |
| PDXK | 0.711763 | 0.194195 | -0.23048 | O00764;F2 | O00764;F2 | O00764 | Pyridoxal k |
| SUPV3L1 | 0.697501 | 0.200678 | 0.139899 | Q8IYB8;B1 | Q8IYB8 | Q8IYB8 | ATP-depend |
| CPSF1 | 0.687639 | 0.205287 | 0.120202 | Q10570;A0 | Q10570 | Q10570 | Cleavage ar |
| U2AF2 | 0.654624 | 0.221501 | 0.096409 | P26368;K7 | P26368;K7 | P26368 | Splicing fac |
| ABHD10 | 0.64478 | 0.226579 | 0.099023 | Q9NUJ1;F8 | Q9NUJ1 | Q9NUJ1 | Mycopheno |
| SPG7 | 0.636637 | 0.230868 | 0.085116 | Q9UQ90;A0 | Q9UQ90;A0 | Q9UQ90 | Paraplegin |
| RHOT2 | 0.600277 | 0.251029 | 0.08379 | Q8IXI1;H3E | Q8IXI1 | Q8IXI1 | Mitochond |
| SEC24B | 0.59746 | 0.252662 | 0.077472 | O95487 | O95487 | O95487 | Protein trai |
| SH3BP1 | 0.549234 | 0.282336 | 0.089934 | Q9Y3L3;Q6 | Q9Y3L3;Q6 | Q9Y3L3 | SH3 domai |
| G6PD | 0.533478 | 0.292767 | -0.0735 | P11413;E9 | P11413;E9 | P11413 | Glucose-6- |
| SEC24D | 0.518818 | 0.302818 | 0.132449 | O94855;E9 | O94855;E9 | O94855 | Protein trai |
| ALDH16A1 | 0.514232 | 0.306033 | -0.10767 | Q8IZ83;F5H | Q8IZ83;F5H | Q8IZ83 | Aldehyde d |
| ALDH1B1 | 0.487145 | 0.325728 | 0.06997 | P30837;A0 | P30837 | P30837 | Aldehyde d |
| HARS2 | 0.486866 | 0.325937 | -0.04576 | A0A2R8Y5F | A0A2R8Y5F | A0A2R8Y5F | Probable hi |
| GPNMB | 0.440713 | 0.362483 | -0.08438 | Q14956;A0 | Q14956;A0 | Q14956 | Transmemb |
| TROVE2 | 0.438473 | 0.364357 | 0.080426 | P10155;G5 | P10155 | P10155 | 60 kDa SS-A |
| PREP | 0.427838 | 0.373389 | -0.09958 | P48147;A0 | P48147;A0 | P48147 | Prolyl endo |
| RUVBL1 | 0.378839 | 0.417985 | 0.04999 | Q9Y265;E7 | Q9Y265;E7 | Q9Y265 | RuvB-like 1 |

| | | | | | | | |
|---|---|---|---|---|---|---|---|
| ACADS | 0.377815 | 0.418972 | 0.065866 | P16219;E9 | P16219;E9 | P16219 | Short-chair |
| SNX6 | 0.355356 | 0.441209 | 0.09463 | Q9UNH7;A( | Q9UNH7;A( | Q9UNH7 | Sorting nex |
| PLBD2 | 0.349811 | 0.446878 | 0.076478 | Q8NHP8 | Q8NHP8 | Q8NHP8 | Putative ph |
| SON | 0.313455 | 0.485898 | 0.073815 | P18583;H7 | P18583;H7 | P18583 | Protein SOI |
| HNRNPL | 0.308167 | 0.49185 | 0.030835 | P14866;M( | P14866;M( | P14866 | Heterogene |
| PSTPIP1 | 0.288884 | 0.514181 | 0.042325 | J3KPG6;O4 | J3KPG6;O4 | J3KPG6 | Proline-seri |
| ACADSB | 0.266468 | 0.541417 | 0.051985 | P45954 | P45954 | P45954 | Short/bran |
| CCDC6 | 0.253005 | 0.558464 | -0.05738 | Q16204 | Q16204 | Q16204 | Coiled-coil |
| ATP6V1E1 | 0.236928 | 0.579525 | -0.02327 | P36543;C9 | P36543;C9 | P36543 | V-type prot |
| PLIN3 | 0.235437 | 0.581518 | -0.06768 | O60664;K7 | O60664;K7 | O60664 | Perilipin-3 |
| FLOT1 | 0.235048 | 0.582039 | 0.027923 | O75955;AC | O75955;AC | O75955 | Flotillin-1 |
| PRKAR2A | 0.214134 | 0.610753 | -0.01441 | P13861;H7 | P13861 | P13861 | cAMP-depe |
| G3BP2 | 0.212434 | 0.613149 | 0.020816 | Q9UN86;D( | Q9UN86;D( | Q9UN86 | Ras GTPase- |
| VWA5A | 0.205115 | 0.62357 | 0.03011 | O00534;B4 | O00534 | O00534 | von Willebr |
| ARHGAP17 | 0.199528 | 0.631643 | -0.0435 | Q68EM7;C9 | Q68EM7 | Q68EM7 | Rho GTPase |
| SLTM | 0.198805 | 0.632696 | 0.040575 | Q9NWH9;H | Q9NWH9;H | Q9NWH9 | SAFB-like tr |
| SERPINB6 | 0.198281 | 0.63346 | 0.043404 | A0A024QZ) | A0A024QZ) | A0A024QZ) | Serpin B6 |
| OAS1 | 0.164384 | 0.684882 | 0.030348 | P00973;F8 | P00973;F8 | P00973 | 2-5-oligoac |
| LMAN2 | 0.150757 | 0.706712 | 0.01721 | Q12907;D6 | Q12907;D6 | Q12907 | Vesicular ir |
| ACTR2 | 0.15065 | 0.706887 | -0.02607 | P61160;F5 | P61160;F5 | P61160 | Actin-relate |
| CAPRIN1 | 0.142987 | 0.719471 | -0.02246 | Q14444;G3 | Q14444;G3 | Q14444 | Caprin-1 |
| PLCD1 | 0.114163 | 0.768842 | -0.0157 | P51178 | P51178 | P51178 | 1-phosphat |
| PSMD6 | 0.098893 | 0.796355 | -0.01311 | Q15008;C9 | Q15008 | Q15008 | 26S proteas |
| LASP1 | 0.0903 | 0.812269 | -0.00934 | Q14847;C9 | Q14847;C9 | Q14847 | LIM and SH |
| FUBP1 | 0.088776 | 0.815125 | -0.0203 | Q96AE4;E9 | Q96AE4;E9 | Q96AE4 | Far upstrea |
| STX11 | 0.084746 | 0.822724 | 0.022232 | O75558 | O75558 | O75558 | Syntaxin-1: |
| HNRNPA1 | 0.072255 | 0.84673 | 0.009643 | F8W6I7;P0 | F8W6I7;P0 | F8W6I7 | Heterogene |
| TMEM214 | 0.047001 | 0.897426 | -0.01356 | Q6NUQ4;H | Q6NUQ4;H | Q6NUQ4 | Transmemb |
| HTATSF1 | 0.038656 | 0.914838 | -0.00896 | O43719;Q5 | O43719 | O43719 | HIV Tat-spe |
| DYNC1LI1 | 0.026233 | 0.941384 | 0.003249 | Q9Y6G9;E9 | Q9Y6G9;E9 | Q9Y6G9 | Cytoplasmi |
| GLA | 0.013454 | 0.969496 | -0.00409 | P06280;A0 | P06280;A0 | P06280 | Alpha-galac |
| IDH3B | 0.013299 | 0.969842 | -0.00229 | A0A087WZ | A0A087WZ | A0A087WZ | Isocitrate d |
| PDHB | 0.001929 | 0.995569 | 0.000453 | P11177;C9 | P11177;C9 | P11177 | Pyruvate de |
| SQSTM1 | 4.338729 | 4.58E-05 | 0.414563 | Q13501;E7 | Q13501;E7 | Q13501 | Sequestoso |
| EIF4B | 2.421225 | 0.003791 | 0.333862 | E7EX17;P2: | E7EX17;P2: | E7EX17 | Eukaryotic |
| HNRNPF | 2.05989 | 0.008712 | 0.236663 | P52597;A0 | P52597 | P52597 | Heterogene |
| AIFM2 | 2.082798 | 0.008264 | 0.219388 | Q9BRQ8 | Q9BRQ8 | Q9BRQ8 | Apoptosis-i |
| YARS2 | 3.188693 | 0.000648 | 0.209491 | Q9Y2Z4;H0 | Q9Y2Z4;H0 | Q9Y2Z4 | Tyrosine--tl |
| APLP2 | 2.854574 | 0.001398 | 0.205633 | Q06481;E9 | Q06481 | Q06481 | Amyloid-lik |
| C21orf33 | 1.928167 | 0.011799 | 0.201359 | P0DPI2;A0, | P0DPI2;A0, | P0DPI2 | |
| MRPS5 | 1.980222 | 0.010466 | 0.172161 | P82675 | P82675 | P82675 | 28S ribosor |
| STAU1 | 1.758022 | 0.017457 | 0.122806 | A0A087X1/ | A0A087X1/ | A0A087X1/ | Double-stra |
| PLA2G4A | 1.475415 | 0.033465 | -0.12993 | P47712 | P47712 | P47712 | Cytosolic p |
| SERPINB1 | 1.728189 | 0.018699 | -0.23875 | P30740;C9 | P30740 | P30740 | Leukocyte e |
| PIK3R4 | 1.221601 | 0.060034 | 0.332353 | Q99570;D6 | Q99570 | Q99570 | Phosphoin( |
| TALDO1 | 1.16508 | 0.068379 | -0.2538 | P37837;F2 | P37837;F2 | P37837 | Transaldola |
| COPE | 1.133902 | 0.073468 | 0.114627 | O14579;M( | O14579;M( | O14579 | Coatomer s |
| ATM | 1.0949 | 0.080371 | 0.152705 | Q13315;E9 | Q13315;E9 | Q13315 | Serine-prot |

| | | | | | | | | |
|---|---|---|---|---|---|---|---|---|
| CLYBL | 1.029393 | 0.093456 | 0.089141 | | Q8N0X4;Q5 | Q8N0X4 | Q8N0X4 | Citrate lyas |
| SRP54 | 0.996459 | 0.100819 | 0.128546 | | P61011;G3 | P61011;G3 | P61011 | Signal recog |
| LYN | 0.989661 | 0.102409 | 0.088867 | | P07948;E5 | P07948 | P07948 | Tyrosine-pr |
| RPS9 | 0.988015 | 0.102798 | -0.08318 | | P46781;B5 | P46781;B5 | P46781 | 40S ribosor |
| EXOC7 | 0.937318 | 0.115527 | 0.126077 | | A0A0A0MS | A0A0A0MS | A0A0A0MS | Exocyst cor |
| NLRX1 | 0.935525 | 0.116005 | 0.164625 | | Q86UT6;C9 | Q86UT6 | Q86UT6 | NLR family |
| HP1BP3 | 0.926452 | 0.118453 | 0.149406 | | Q5SSJ5;B0( | Q5SSJ5;B0( | Q5SSJ5 | Heterochro |
| EIF3M | 0.922899 | 0.119427 | 0.097118 | | Q7L2H7;J3 | Q7L2H7;J3 | Q7L2H7 | Eukaryotic |
| RCN1 | 0.891208 | 0.128467 | 0.093864 | | Q15293;E9 | Q15293 | Q15293 | Reticulocal |
| PLEK | 0.834771 | 0.146295 | -0.19465 | | P08567 | P08567 | P08567 | Pleckstrin |
| GBP2 | 0.768099 | 0.170569 | -0.15186 | | P32456;F6 | P32456 | P32456 | Interferon-i |
| ARMCX3 | 0.754264 | 0.17609 | -0.06088 | | Q9UH62;H: | Q9UH62 | Q9UH62 | Armadillo r |
| ADPGK | 0.750459 | 0.17764 | 0.086124 | | Q9BRR6;H3 | Q9BRR6 | Q9BRR6 | ADP-depen |
| APEH | 0.696094 | 0.201329 | -0.1441 | | C9JIF9;P13 | C9JIF9;P13 | C9JIF9 | Acylamino- |
| RPL5 | 0.683944 | 0.207041 | 0.061241 | | P46777;A0 | P46777;A0 | P46777 | 60S ribosor |
| PAFAH1B1 | 0.682701 | 0.207634 | -0.09195 | | P43034;I3I | P43034 | P43034 | Platelet-act |
| COQ5 | 0.652012 | 0.222837 | -0.07735 | | Q5HYK3;F8 | Q5HYK3;F8 | Q5HYK3 | 2-methoxy- |
| PPP1R7 | 0.63794 | 0.230176 | -0.11823 | | Q15435;H7 | Q15435;H7 | Q15435 | Protein pho |
| CTSS | 0.631903 | 0.233398 | 0.077518 | | P25774;U3 | P25774 | P25774 | Cathepsin S |
| EIF5 | 0.616691 | 0.241718 | 0.100113 | | P55010;H0 | P55010;H0 | P55010 | Eukaryotic |
| DES | 0.578198 | 0.26412 | 0.252949 | | P17661 | P17661 | P17661 | Desmin |
| DHRS9 | 0.573036 | 0.267278 | -0.16304 | | Q9BPW9;C | Q9BPW9 | Q9BPW9 | Dehydroger |
| STK11IP | 0.570786 | 0.268667 | 0.070656 | | Q8N1F8;H7 | Q8N1F8 | Q8N1F8 | Serine/thre |
| CLU | 0.558557 | 0.27634 | 0.147762 | | P10909;H0 | P10909;H0 | P10909 | Clusterin;C |
| VAPA | 0.545295 | 0.284908 | 0.06504 | | Q9P0L0;J3 | Q9P0L0 | Q9P0L0 | Vesicle-asso |
| TOR1AIP2 | 0.544342 | 0.285534 | 0.058111 | | Q8NFQ8 | Q8NFQ8 | Q8NFQ8 | Torsin-1A-ii |
| TSFM | 0.537699 | 0.289935 | 0.072735 | | P43897;C9 | P43897;C9 | P43897 | Elongation |
| TRIM54 | 0.522511 | 0.300254 | -0.06317 | | Q9BYV2;Q9 | Q9BYV2 | Q9BYV2 | Tripartite m |
| HMOX2 | 0.485235 | 0.327163 | 0.096531 | | A0A087WT | A0A087WT | A0A087WT | Heme oxyge |
| PNN | 0.481157 | 0.33025 | 0.066472 | | Q9H307;G3 | Q9H307 | Q9H307 | Pinin |
| ATP6V1D | 0.471065 | 0.338014 | 0.064174 | | Q9Y5K8;G3 | Q9Y5K8;G3 | Q9Y5K8 | V-type prot |
| PNP | 0.464091 | 0.343486 | -0.10042 | | P00491;G3 | P00491;G3 | P00491 | Purine nucl |
| GSTO1 | 0.439407 | 0.363574 | -0.12575 | | P78417;Q5 | P78417;Q5 | P78417 | Glutathion |
| TAGLN2 | 0.435491 | 0.366867 | -0.09025 | | P37802;X6 | P37802;X6 | P37802 | Transgelin- |
| DHRS7 | 0.372103 | 0.424519 | -0.02842 | | Q9Y394;H0 | Q9Y394;H0 | Q9Y394 | Dehydroger |
| HGS | 0.365878 | 0.430648 | 0.045464 | | O14964;I3I | O14964;I3I | O14964 | Hepatocyte |
| INPP5D | 0.352232 | 0.444394 | 0.065625 | | Q92835;H( | Q92835;H( | Q92835 | Phosphatic |
| ACSS1 | 0.345535 | 0.4513 | 0.067922 | | Q9NUB1;Q | Q9NUB1 | Q9NUB1 | Acetyl-coer |
| NUP133 | 0.324275 | 0.473942 | -0.04009 | | Q8WUM0 | Q8WUM0 | Q8WUM0 | Nuclear por |
| MMAA | 0.321214 | 0.477294 | -0.0344 | | A0A3B3ITP | A0A3B3ITP | A0A3B3ITP | Methylmal |
| SART1 | 0.3052 | 0.495222 | -0.05718 | | O43290;E9 | O43290 | O43290 | U4/U6.U5 t |
| PDLIM5 | 0.303821 | 0.496797 | 0.057714 | | Q96HC4;H( | Q96HC4;H( | Q96HC4 | PDZ and LIN |
| OSBP | 0.298333 | 0.503115 | 0.06224 | | P22059;H0 | P22059 | P22059 | Oxysterol-b |
| THOC2 | 0.284631 | 0.519241 | 0.053811 | | Q8NI27;A0 | Q8NI27;A0 | Q8NI27 | THO compl |
| DERA | 0.264092 | 0.544387 | -0.07285 | | Q9Y315;G3 | Q9Y315;G3 | Q9Y315 | Deoxyribos |
| PAK2 | 0.245621 | 0.56804 | 0.031851 | | Q13177;H7 | Q13177 | Q13177 | Serine/thre |
| CPSF3 | 0.229589 | 0.589401 | -0.0523 | | Q9UKF6;G5 | Q9UKF6;G5 | Q9UKF6 | Cleavage ar |
| MGLL | 0.213117 | 0.612185 | 0.035077 | | Q99685;A0 | Q99685;A0 | Q99685 | Monoglyce |

| | | | | | | | |
|---|---|---|---|---|---|---|---|
| TXNDC5 | 0.212539 | 0.613 | -0.04296 | Q8NBS9 | Q8NBS9 | Q8NBS9 | Thioredoxi |
| CDC37 | 0.211478 | 0.6145 | 0.063791 | Q16543;K7 | Q16543;K7 | Q16543 | Hsp90 co-c |
| ATP6V0D1 | 0.20978 | 0.616907 | -0.01849 | F5GYQ1;P6 | F5GYQ1;P6 | F5GYQ1 | V-type prot |
| ACAT2 | 0.17045 | 0.675383 | -0.06441 | Q9BWD1 | Q9BWD1 | Q9BWD1 | Acetyl-CoA |
| CCBL2 | 0.163733 | 0.68591 | 0.031966 | Q6YP21;A0 | Q6YP21;A0 | Q6YP21 | Kynurenine |
| ARSB | 0.158667 | 0.693957 | 0.059264 | P15848;A0 | P15848;A0 | P15848 | Arylsulfatas |
| PSMD5 | 0.15256 | 0.703785 | 0.039982 | Q16401;F2 | Q16401 | Q16401 | 26S proteas |
| ABCC10 | 0.146709 | 0.713331 | 0.03606 | Q5T3U5;H( | Q5T3U5 | Q5T3U5 | Multidrug r |
| ELMO1 | 0.136009 | 0.731123 | 0.023594 | Q92556;H7 | Q92556 | Q92556 | Engulfment |
| ARHGDIB | 0.134351 | 0.733921 | -0.02672 | P52566;H0 | P52566;H0 | P52566 | Rho GDP-di |
| FCHO2 | 0.125776 | 0.748555 | -0.02123 | Q0JRZ9;H7 | Q0JRZ9;H7 | Q0JRZ9 | F-BAR doma |
| CLIC4 | 0.123054 | 0.753261 | 0.022804 | Q9Y696 | Q9Y696 | Q9Y696 | Chloride in |
| SCCPDH | 0.119012 | 0.760305 | 0.032165 | Q8NBX0 | Q8NBX0 | Q8NBX0 | Saccharopi |
| DPP7 | 0.11579 | 0.765967 | 0.02758 | Q9UHL4;R4 | Q9UHL4;R4 | Q9UHL4 | Dipeptidyl |
| SOD2 | 0.110429 | 0.77548 | 0.022679 | P04179;F5 | P04179;F5 | P04179 | Superoxide |
| ZMPSTE24 | 0.100506 | 0.793403 | 0.023777 | O75844 | O75844 | O75844 | CAAX preny |
| IPO7 | 0.074013 | 0.84331 | -0.01374 | O95373;E9 | O95373 | O95373 | Importin-7 |
| SLFN5 | 0.068489 | 0.854104 | -0.02039 | Q08AF3;B4 | Q08AF3;B4 | Q08AF3 | Schlafen fa |
| YWHAE | 0.067314 | 0.856418 | 0.018892 | P62258;B4 | P62258 | P62258 | 14-3-3 prot |
| NFS1 | 0.067191 | 0.85666 | 0.014231 | Q9Y697;F2 | Q9Y697;F2 | Q9Y697 | Cysteine de |
| SCPEP1 | 0.065406 | 0.860188 | 0.018672 | Q9HB40;I3 | Q9HB40 | Q9HB40 | Retinoid-in |
| EXOC5 | 0.063269 | 0.864433 | -0.01553 | A0A0A0MS | A0A0A0MS | A0A0A0MS | Exocyst cor |
| P4HA1 | 0.056431 | 0.878151 | 0.015908 | P13674 | P13674 | P13674 | Prolyl 4-hy |
| TMX3 | 0.041876 | 0.908079 | 0.006731 | Q96JJ7;B4I | Q96JJ7 | Q96JJ7 | Protein dis |
| ATP2C1 | 0.025673 | 0.942599 | -0.00469 | B4E2Q0;P9 | B4E2Q0;P9 | B4E2Q0 | Calcium-tra |
| DKC1 | 0.020954 | 0.952898 | -0.00515 | O60832;C9 | O60832;C9 | O60832 | H/ACA ribo |
| CPD | 3.39222 | 0.000405 | 0.391669 | O75976;J3 | O75976 | O75976 | Carboxypep |
| GRSF1 | 2.992097 | 0.001018 | 0.321143 | H0Y8R1;F5 | H0Y8R1;F5 | H0Y8R1 | G-rich sequ |
| SNRNP70 | 2.741519 | 0.001813 | 0.303739 | P08621;M( | P08621 | P08621 | U1 small nu |
| ACAD8 | 1.771879 | 0.016909 | 0.233691 | Q9UKU7;E9 | Q9UKU7 | Q9UKU7 | Isobutyryl- |
| MANBA | 1.557145 | 0.027724 | 0.229944 | O00462;A0 | O00462;A0 | O00462 | Beta-mann |
| POLDIP2 | 2.073692 | 0.008439 | 0.209 | Q9Y2S7;B4 | Q9Y2S7;B4 | Q9Y2S7 | Polymerase |
| HACL1 | 2.165707 | 0.006828 | 0.199128 | Q9UJ83;B3 | Q9UJ83;B3 | Q9UJ83 | 2-hydroxya |
| SKIV2L2 | 1.408815 | 0.039011 | 0.1789 | P42285;H0 | P42285 | P42285 | Superkiller |
| SUCLG1 | 2.363574 | 0.004329 | 0.175327 | P53597;A0 | P53597;A0 | P53597 | Succinyl-Co |
| STRAP | 2.189376 | 0.006466 | 0.170759 | Q9Y3F4;H0 | Q9Y3F4 | Q9Y3F4 | Serine-thre |
| SUPT5H | 2.781584 | 0.001654 | 0.154699 | O00267 | O00267 | O00267 | Transcripti |
| TM9SF4 | 1.349336 | 0.044737 | 0.115118 | A0A0C4DFI | A0A0C4DFI | A0A0C4DFI | Transmemb |
| GNAI2 | 1.601925 | 0.025008 | 0.090251 | P04899;P6 | P04899 | P04899 | Guanine nu |
| SLC25A35 | 1.305845 | 0.049449 | 0.069002 | Q3KQZ1 | Q3KQZ1 | Q3KQZ1 | Solute carri |
| PSMA7 | 2.147005 | 0.007128 | -0.10755 | O14818;H( | O14818;H( | O14818 | Proteasom |
| EFHD2 | 1.968631 | 0.010749 | -0.24078 | Q96C19;H( | Q96C19;H( | Q96C19 | EF-hand do |
| FAM49B | 2.179849 | 0.006609 | -0.24112 | Q9NUQ9;E5 | Q9NUQ9;E5 | Q9NUQ9 | Protein FAM |
| LDHA | 1.571794 | 0.026804 | -0.24125 | P00338;F5 | P00338 | P00338 | L-lactate de |
| GBP1 | 2.600223 | 0.002511 | -0.32363 | P32455 | P32455 | P32455 | Interferon-i |
| FAF2 | 1.225109 | 0.059551 | 0.127003 | Q96CS3;D6 | Q96CS3 | Q96CS3 | FAS-associa |
| DAP3 | 1.196372 | 0.063625 | 0.130417 | P51398;V9 | P51398;V9 | P51398 | 28S ribosor |
| CECR1 | 1.138787 | 0.072646 | 0.192133 | B4E3Q4;Q9 | B4E3Q4;Q9 | B4E3Q4 | Adenosine |

| | | | | | | | | |
|---|---|---|---|---|---|---|---|---|
| IDE | 1.120261 | 0.075812 | 0.095402 | | P14735;A0 | P14735;A0 | P14735 | Insulin-deg |
| CSNK2A2 | 1.086771 | 0.08189 | -0.10294 | | P19784;H3 | P19784;H3 | P19784 | Casein kina |
| RPL8 | 1.021626 | 0.095142 | 0.087396 | | P62917;E9 | P62917;E9 | P62917 | 60S riboso |
| SAFB | 0.969098 | 0.107375 | 0.126771 | | Q15424;K7 | Q15424 | Q15424 | Scaffold att |
| CLIC1 | 0.958724 | 0.109971 | -0.16934 | | O00299 | O00299 | O00299 | Chloride in |
| RPS8 | 0.924628 | 0.118952 | 0.061713 | | P62241;Q5 | P62241;Q5 | P62241 | 40S riboso |
| SEC13 | 0.909083 | 0.123287 | 0.080471 | | P55735;A8 | P55735;A8 | P55735 | Protein SE |
| CBR1 | 0.89224 | 0.128162 | -0.18193 | | P16152;E9 | P16152;E9 | P16152 | Carbonyl re |
| FASTKD5 | 0.856138 | 0.139272 | -0.14635 | | Q7L8L6 | Q7L8L6 | Q7L8L6 | FAST kinase |
| CHCHD3 | 0.850634 | 0.141048 | 0.100068 | | C9JRZ6;Q9 | C9JRZ6;Q9 | C9JRZ6 | MICOS com |
| PRKAA1 | 0.832744 | 0.146979 | 0.086005 | | Q13131;Q9 | Q13131 | Q13131 | 5-AMP-acti |
| PRMT1 | 0.794349 | 0.160565 | 0.15504 | | Q99873;E9 | Q99873;E9 | Q99873 | Protein arg |
| PRPF40A | 0.713441 | 0.193446 | 0.102719 | | O75400;F5 | O75400 | O75400 | Pre-mRNA- |
| IL4I1 | 0.684337 | 0.206853 | 0.088401 | | Q96RQ9;M | Q96RQ9 | Q96RQ9 | L-amino-ac |
| BTK | 0.637113 | 0.230615 | 0.098532 | | Q06187;U3 | Q06187;U3 | Q06187 | Tyrosine-pr |
| CHID1 | 0.618186 | 0.240887 | 0.095971 | | Q9BWS9;E9 | Q9BWS9;E9 | Q9BWS9 | Chitinase d |
| MSR1 | 0.615864 | 0.242179 | 0.101206 | | P21757;B4 | P21757;B4 | P21757 | Macrophag |
| STAT5A | 0.594687 | 0.25428 | 0.136659 | | K7EK35;P4 | K7EK35;P4 | K7EK35 | Signal trans |
| ACP2 | 0.588455 | 0.257956 | 0.079217 | | P11117;E9 | P11117;E9 | P11117 | Lysosomal |
| HSPA1B | 0.553273 | 0.279722 | 0.06258 | | A0A0G2JIW | A0A0G2JIW | A0A0G2JIW | Heat shock |
| COG2 | 0.550443 | 0.281551 | -0.10247 | | Q14746;B7 | Q14746;B7 | Q14746 | Conserved |
| PTGS1 | 0.548171 | 0.283028 | -0.07789 | | P23219;A0 | P23219;A0 | P23219 | Prostagland |
| MRPS22 | 0.53653 | 0.290717 | 0.06964 | | G5E9W7;G | G5E9W7;G | G5E9W7 | 28S riboso |
| TAOK3 | 0.520968 | 0.301322 | 0.131045 | | Q9H2K8;G3 | Q9H2K8;G3 | Q9H2K8 | Serine/thre |
| ATP2A3 | 0.520204 | 0.301853 | -0.11033 | | Q93084 | Q93084 | Q93084 | Sarcoplasm |
| GRHPR | 0.508795 | 0.309888 | 0.079299 | | Q9UBQ7;U | Q9UBQ7;U | Q9UBQ7 | Glyoxylate |
| GSTK1 | 0.497557 | 0.318012 | 0.057262 | | Q9Y2Q3;E9 | Q9Y2Q3;E9 | Q9Y2Q3 | Glutathion |
| LEMD2 | 0.448333 | 0.356178 | 0.05321 | | Q8NC56;H0 | Q8NC56 | Q8NC56 | LEM domai |
| ADAM17 | 0.445938 | 0.358147 | -0.07268 | | P78536;A6 | P78536 | P78536 | Disintegrin |
| PGAM5 | 0.42277 | 0.377772 | -0.04279 | | Q96HS1;F5 | Q96HS1 | Q96HS1 | Serine/thre |
| ATP9B | 0.415738 | 0.383939 | 0.068787 | | O43861;F6 | O43861 | O43861 | Probable p |
| CSK | 0.395348 | 0.402394 | -0.10322 | | P41240;H3 | P41240 | P41240 | Tyrosine-pr |
| ADK | 0.38294 | 0.414057 | -0.09381 | | P55263;A0 | P55263;A0 | P55263 | Adenosine |
| PREB | 0.371759 | 0.424855 | 0.047356 | | Q9HCU5;B5 | Q9HCU5;B5 | Q9HCU5 | Prolactin re |
| SEL1L | 0.361279 | 0.435233 | 0.068149 | | Q9UBV2;G3 | Q9UBV2 | Q9UBV2 | Protein sel- |
| RPS2 | 0.345951 | 0.450867 | -0.04341 | | P15880;H0 | P15880;H0 | P15880 | 40S riboso |
| PSPC1 | 0.31653 | 0.482469 | -0.0476 | | Q8WXF1;X0 | Q8WXF1 | Q8WXF1 | Paraspeckle |
| MDH1 | 0.304696 | 0.495797 | -0.10765 | | P40925;A0 | P40925;A0 | P40925 | Malate deh |
| KIAA0930 | 0.245044 | 0.568795 | -0.03528 | | Q6ICG6;B0 | Q6ICG6;B0 | Q6ICG6 | Uncharacte |
| RPS5 | 0.216386 | 0.607594 | -0.03014 | | M0R0F0;P4 | M0R0F0;P4 | M0R0F0 | 40S riboso |
| GDI1 | 0.215967 | 0.608181 | -0.06044 | | P31150;G5 | P31150 | P31150 | Rab GDP di |
| OCIAD1 | 0.207243 | 0.620522 | 0.03362 | | Q9NX40;D6 | Q9NX40;D6 | Q9NX40 | OCIA doma |
| LDHB | 0.201262 | 0.629127 | 0.064629 | | A0A5F9ZHN | A0A5F9ZHN | A0A5F9ZHN | L-lactate de |
| RDH14 | 0.198612 | 0.632978 | 0.032293 | | Q9HBH5;C9 | Q9HBH5 | Q9HBH5 | Retinol deh |
| OSBPL11 | 0.188103 | 0.648481 | 0.056437 | | Q9BXB4 | Q9BXB4 | Q9BXB4 | Oxysterol-b |
| EMC2 | 0.177094 | 0.665129 | 0.036005 | | Q15006;H0 | Q15006 | Q15006 | ER membra |
| CLPTM1 | 0.168665 | 0.678164 | -0.02514 | | O96005;K7 | O96005 | O96005 | Cleft lip an |
| MAT2A | 0.149337 | 0.709027 | -0.02828 | | P31153;Q0 | P31153 | P31153 | S-adenosyl |

| | | | | | | | | |
|---|---|---|---|---|---|---|---|---|
| CMPK1 | 0.148102 | 0.711047 | -0.04311 | | P30085;Q5 | P30085;Q5 | P30085 | UMP-CMP k |
| SUN2 | 0.146582 | 0.71354 | 0.030745 | | Q9UH99;B( | Q9UH99 | Q9UH99 | SUN domai |
| COPS4 | 0.145111 | 0.71596 | 0.028593 | | Q9BT78;D6 | Q9BT78;D6 | Q9BT78 | COP9 signa |
| MRPS27 | 0.128169 | 0.744442 | -0.02225 | | Q92552;G5 | Q92552;G5 | Q92552 | 28S ribosor |
| TPP1 | 0.124886 | 0.750091 | 0.034478 | | O14773;A0 | O14773;A0 | O14773 | Tripeptidyl |
| CD97 | 0.111373 | 0.773798 | -0.00938 | | P48960;K7 | P48960 | P48960 | CD97 antig |
| ANXA6 | 0.103123 | 0.788637 | -0.04068 | | P08133;E7 | P08133;E7 | P08133 | Annexin A6 |
| PIEZO1 | 0.097511 | 0.798894 | 0.028555 | | Q92508;E7 | Q92508 | Q92508 | Piezo-type r |
| AKR1A1 | 0.090655 | 0.811605 | 0.026352 | | P14550;Q5 | P14550 | P14550 | Alcohol deh |
| GRB2 | 0.089134 | 0.814453 | -0.01318 | | P62993;J3 | P62993 | P62993 | Growth fac |
| ALDH3B1 | 0.054738 | 0.88158 | -0.00635 | | P43353;Q9 | P43353;Q9 | P43353 | Aldehyde d |
| ATP5F1 | 0.030937 | 0.931244 | 0.006761 | | P24539;Q5 | P24539;Q5 | P24539 | ATP syntha |
| PRCP | 0.024743 | 0.944619 | -0.00703 | | P42785;E9 | P42785;E9 | P42785 | Lysosomal |
| TRIM21 | 0.012437 | 0.971768 | -0.00231 | | P19474;H0 | P19474 | P19474 | E3 ubiquiti |
| CECR5 | 0.011132 | 0.974694 | 0.00217 | | Q9BXW7;A | Q9BXW7 | Q9BXW7 | Cat eye syn |
| CPNE1 | 0.011024 | 0.974936 | 0.00237 | | Q99829;F2 | Q99829;F2 | Q99829 | Copine-1 |
| EXOC8 | 0.007381 | 0.983148 | 0.002306 | | Q8IYI6 | Q8IYI6 | Q8IYI6 | Exocyst cor |
| CCZ1B | 0.006218 | 0.985785 | 0.000959 | | P86790;P8 | P86790;P8 | P86790 | Vacuolar fu |
| DPM1 | 0.002646 | 0.993925 | -0.00054 | | Q5QPK2;H( | Q5QPK2;H( | Q5QPK2 | Dolichol-ph |
| CD74 | 1.766055 | 0.017137 | 0.201296 | | P04233;H0 | P04233;H0 | P04233 | HLA class II |
| PSMD7 | 1.733864 | 0.018456 | 0.186674 | | P51665;H3 | P51665;H3 | P51665 | 26S protea |
| DCXR | 1.922341 | 0.011958 | 0.186467 | | Q7Z4W1;J3 | Q7Z4W1;J3 | Q7Z4W1 | L-xylulose r |
| CPM | 2.261196 | 0.00548 | 0.176538 | | P14384;F8 | P14384;F8 | P14384 | Carboxypep |
| AP5Z1 | 1.595305 | 0.025392 | 0.159937 | | O43299;A0 | O43299;A0 | O43299 | AP-5 compl |
| PPIB | 3.709603 | 0.000195 | 0.157299 | | P23284 | P23284 | P23284 | Peptidyl-pr |
| TPD52L2 | 1.511803 | 0.030775 | 0.153309 | | O43399;A0 | O43399;A0 | O43399 | Tumor prot |
| EIF3G | 2.011454 | 0.00974 | 0.146704 | | O75821;K7 | O75821;K7 | O75821 | Eukaryotic |
| LACTB2 | 1.870584 | 0.013472 | 0.111839 | | Q53H82 | Q53H82 | Q53H82 | Beta-lactam |
| PSMA6 | 1.986536 | 0.010315 | 0.109465 | | P60900;G3 | P60900;G3 | P60900 | Proteasom |
| NMT1 | 1.511758 | 0.030778 | 0.098653 | | P30419;B7 | P30419 | P30419 | Glycylpepti |
| C14orf166 | 1.530303 | 0.029492 | 0.060635 | | Q9Y224;G3 | Q9Y224;G3 | Q9Y224 | UPF0568 p |
| SPG21 | 1.733606 | 0.018467 | -0.07709 | | Q9NZD8;H( | Q9NZD8;H( | Q9NZD8 | Maspardin |
| ANO6 | 2.456071 | 0.003499 | -0.1632 | | Q4KMQ2 | Q4KMQ2 | Q4KMQ2 | Anoctamin |
| TOR1A | 1.409802 | 0.038922 | -0.22857 | | O14656;A0 | O14656;A0 | O14656 | Torsin-1A |
| DHRS4 | 1.293519 | 0.050872 | 0.134599 | | Q9BTZ2;F5 | Q9BTZ2;F5 | Q9BTZ2 | Dehydroger |
| APBB1IP | 1.250477 | 0.056172 | -0.16919 | | Q7Z5R6;C9 | Q7Z5R6 | Q7Z5R6 | Amyloid be |
| ETFA | 1.218064 | 0.060525 | 0.129891 | | P13804;H0 | P13804;H0 | P13804 | Electron tra |
| TXNL1 | 1.202998 | 0.062662 | -0.16054 | | O43396;K7 | O43396;K7 | O43396 | Thioredoxi |
| DNAJB12 | 1.05546 | 0.088012 | -0.09643 | | J3KPS0;Q9 | J3KPS0;Q9 | J3KPS0 | DnaJ homo |
| OXCT1 | 0.987633 | 0.102889 | 0.133749 | | P55809;E9 | P55809;E9 | P55809 | Succinyl-Co |
| FARP1 | 0.983593 | 0.10385 | 0.254588 | | C9JME2;Q9 | C9JME2;Q9 | C9JME2 | FERM, Rho( |
| SKAP2 | 0.969871 | 0.107184 | 0.117278 | | O75563;C9 | O75563 | O75563 | Src kinase-a |
| CFL1 | 0.959958 | 0.109658 | -0.2153 | | P23528;G3 | P23528;G3 | P23528 | Cofilin-1 |
| HDHD3 | 0.956604 | 0.110509 | 0.197552 | | Q9BSH5 | Q9BSH5 | Q9BSH5 | Haloacid de |
| C17orf59 | 0.937491 | 0.115481 | 0.089426 | | Q96GS4 | Q96GS4 | Q96GS4 | Uncharacte |
| ATP6AP1 | 0.930986 | 0.117223 | -0.08783 | | Q15904;A0 | Q15904;A0 | Q15904 | V-type prot |
| SLC3A2 | 0.903926 | 0.12476 | -0.09686 | | F5GZS6;J3K | F5GZS6;J3K | F5GZS6 | 4F2 cell-sur |
| FDFT1 | 0.836264 | 0.145793 | 0.138187 | | P37268;E9 | P37268;E9 | P37268 | Squalene sy |

| | | | | | | | |
|---|---|---|---|---|---|---|---|
| EHD1 | 0.825075 | 0.149598 | 0.125566 | A0A024R57 | A0A024R57 | A0A024R57 | EH domain- |
| SPTBN2 | 0.81981 | 0.151422 | 0.220848 | O15020;A4 | O15020 | O15020 | Spectrin be |
| BCKDHA | 0.798461 | 0.159052 | 0.099886 | F5H5P2;P1 | F5H5P2;P1 | F5H5P2 | 2-oxoisoval |
| DNAJB11 | 0.781565 | 0.165362 | 0.102106 | Q9UBS4;H7 | Q9UBS4;H7 | Q9UBS4 | DnaJ homo |
| DNAJA1 | 0.778747 | 0.166438 | 0.11201 | P31689 | P31689 | P31689 | DnaJ homo |
| GLOD4 | 0.769204 | 0.170136 | -0.08812 | F6TLX2;Q9 | F6TLX2;Q9 | F6TLX2 | Glyoxalase |
| CA2 | 0.752103 | 0.176969 | -0.24091 | P00918;E5 | P00918 | P00918 | Carbonic ar |
| NIPSNAP1 | 0.747377 | 0.178905 | 0.078592 | Q9BPW8;H | Q9BPW8;H | Q9BPW8 | Protein Nip |
| TMOD3 | 0.733302 | 0.184798 | 0.10096 | Q9NYL9;H0 | Q9NYL9;H0 | Q9NYL9 | Tropomodu |
| PPP2CA | 0.731388 | 0.185614 | -0.09035 | P67775;P6 | P67775;P6 | P67775 | Serine/thre |
| DNAJA3 | 0.719876 | 0.1906 | 0.095757 | Q96EY1;I3L | Q96EY1 | Q96EY1 | DnaJ homo |
| MAVS | 0.695707 | 0.201508 | 0.08221 | Q7Z434 | Q7Z434 | Q7Z434 | Mitochond |
| ECI2 | 0.69016 | 0.204099 | 0.082439 | A0A0C4DG | A0A0C4DG | A0A0C4DG | Enoyl-CoA d |
| ATP2B1 | 0.685098 | 0.206492 | 0.161006 | P20020;E7 | P20020;E7 | P20020 | Plasma mer |
| EIF3F | 0.682892 | 0.207543 | -0.09543 | O00303;H0 | O00303 | O00303 | Eukaryotic |
| RPS6KA3 | 0.660519 | 0.218515 | 0.092658 | B4DG22;B1 | B4DG22;B1 | B4DG22 | Ribosomal |
| GIMAP1-GI | 0.636264 | 0.231066 | 0.103722 | A0A087WT | A0A087WT | A0A087WT | GTPase IMA |
| CRYZ | 0.592769 | 0.255406 | 0.070608 | Q08257;A6 | Q08257;A6 | Q08257 | Quinone ox |
| ADH5 | 0.591313 | 0.256264 | -0.11953 | P11766;H0 | P11766;H0 | P11766 | Alcohol del |
| PDHX | 0.569337 | 0.269565 | 0.067608 | O00330;H0 | O00330;H0 | O00330 | Pyruvate de |
| HIST1H4A | 0.546846 | 0.283893 | 0.082502 | P62805 | P62805 | P62805 | Histone H4 |
| TRABD | 0.544067 | 0.285715 | 0.055264 | J3KPT4;Q9 | J3KPT4;Q9 | J3KPT4 | TraB domai |
| FAM114A2 | 0.525139 | 0.298443 | 0.056689 | Q9NRY5;E7 | Q9NRY5;E7 | Q9NRY5 | Protein FAN |
| PTPRJ | 0.515705 | 0.304996 | 0.082544 | A0A087WT | A0A087WT | A0A087WT | Protein-tyr |
| HPCAL1 | 0.50019 | 0.316089 | 0.071243 | P37235;E9 | P37235;E9 | P37235 | Hippocalcin |
| KIAA2013 | 0.495672 | 0.319395 | 0.05532 | Q8IYS2 | Q8IYS2 | Q8IYS2 | Uncharacte |
| PPA1 | 0.49079 | 0.323005 | -0.07733 | Q15181;Q5 | Q15181;Q5 | Q15181 | Inorganic p |
| ODR4 | 0.455475 | 0.350369 | 0.067411 | Q5SWX8 | Q5SWX8 | Q5SWX8 | Protein odr |
| RAB18 | 0.449119 | 0.355534 | -0.05763 | Q9NP72;A0 | Q9NP72;A0 | Q9NP72 | Ras-related |
| KLC1 | 0.440888 | 0.362337 | -0.07314 | Q07866;G3 | Q07866;G3 | Q07866 | Kinesin ligh |
| SSRP1 | 0.415124 | 0.384482 | 0.079335 | Q08945;E9 | Q08945 | Q08945 | FACT comp |
| PCMT1 | 0.388077 | 0.409188 | -0.0692 | A0A0A0MR | A0A0A0MR | A0A0A0MR | Protein-L-is |
| AGK | 0.371299 | 0.425305 | 0.060899 | A0A3B3ISZ | A0A3B3ISZ | A0A3B3ISZ | Acylglycero |
| NANS | 0.37097 | 0.425628 | -0.12684 | Q9NR45;Q5 | Q9NR45 | Q9NR45 | Sialic acid s |
| GHDC | 0.334655 | 0.462748 | -0.0243 | Q8N2G8;K7 | Q8N2G8;K7 | Q8N2G8 | GH3 domai |
| HSD17B12 | 0.332924 | 0.464596 | 0.043184 | Q53GQ0;A0 | Q53GQ0;A0 | Q53GQ0 | Very-long-c |
| AIMP1 | 0.326233 | 0.471809 | 0.053758 | Q12904;A0 | Q12904;A0 | Q12904 | Aminoacyl |
| DPP9 | 0.323246 | 0.475066 | 0.067728 | Q86TI2;M0 | Q86TI2;M0 | Q86TI2 | Dipeptidyl |
| TTC37 | 0.319428 | 0.479261 | 0.077958 | Q6PGP7;D6 | Q6PGP7 | Q6PGP7 | Tetratricop |
| PPP1R21 | 0.314451 | 0.484785 | 0.077726 | Q6ZMI0;F8 | Q6ZMI0 | Q6ZMI0 | Protein pho |
| GIT2 | 0.314376 | 0.484869 | 0.057305 | Q14161;R4 | Q14161;R4 | Q14161 | ARF GTPase |
| DDRGK1 | 0.311148 | 0.488486 | -0.08625 | Q96HY6;A0 | Q96HY6;A0 | Q96HY6 | DDRGK dor |
| HMGCL | 0.296414 | 0.505343 | 0.035165 | P35914;H0 | P35914 | P35914 | Hydroxyme |
| LGALS3 | 0.294933 | 0.507069 | -0.05641 | P17931;G3 | P17931;G3 | P17931 | Galectin-3; |
| NCF4 | 0.272199 | 0.53432 | -0.05213 | Q15080;A0 | Q15080;A0 | Q15080 | Neutrophil |
| PLA2G7 | 0.269075 | 0.538177 | 0.033015 | Q13093 | Q13093 | Q13093 | Platelet-act |
| RPL7 | 0.252671 | 0.558893 | 0.04576 | P18124;A8 | P18124;A8 | P18124 | 60S ribosor |
| FAM91A1 | 0.216211 | 0.60784 | -0.04059 | Q658Y4;E7 | Q658Y4;E7 | Q658Y4 | Protein FAN |

| | | | | | | | |
|---|---|---|---|---|---|---|---|
| PRKAR1A | 0.211019 | 0.61515 | -0.04645 | P10644;K7 | P10644;K7 | P10644 | cAMP-depe |
| RPS11 | 0.207088 | 0.620744 | -0.02347 | P62280;M | P62280;M | P62280 | 40S riboso |
| DNAJC1 | 0.206022 | 0.622269 | 0.045604 | Q96KC8;Q | Q96KC8 | Q96KC8 | DnaJ homo |
| HADH | 0.202967 | 0.626661 | 0.031793 | A0A0A0MS | A0A0A0MS | A0A0A0MS | Hydroxyac |
| ABCB10 | 0.172575 | 0.672086 | -0.03306 | Q9NRK6 | Q9NRK6 | Q9NRK6 | ATP-bindin |
| PTPRA | 0.169483 | 0.676889 | -0.02081 | P18433;Q5 | P18433 | P18433 | Receptor-t |
| WASF2 | 0.168619 | 0.678236 | 0.026609 | Q9Y6W5;A | Q9Y6W5 | Q9Y6W5 | Wiskott-Al |
| FNDC3A | 0.15929 | 0.692962 | 0.045802 | Q9Y2H6;G | Q9Y2H6;G | Q9Y2H6 | Fibronectir |
| SLC25A19 | 0.158729 | 0.693859 | 0.027005 | Q9HC21;J3 | Q9HC21 | Q9HC21 | Mitochond |
| EPB41L1 | 0.158516 | 0.694199 | -0.02638 | A0A0C4DH | A0A0C4DH | A0A0C4DH | Band 4.1-li |
| NRP1 | 0.130999 | 0.739607 | -0.02525 | O14786;E9 | O14786;E9 | O14786 | Neuropilin |
| TRIP12 | 0.112028 | 0.77263 | 0.028656 | Q14669;C | Q14669 | Q14669 | E3 ubiquiti |
| ALG11 | 0.108771 | 0.778446 | 0.024263 | Q2TAA5;A0 | Q2TAA5 | Q2TAA5 | GDP-Man: |
| TAPBP | 0.10784 | 0.780118 | -0.02241 | A0A0G2JK | A0A0G2JK | A0A0G2JK | Tapasin |
| TUBG1 | 0.095849 | 0.801957 | 0.012983 | P23258;Q | P23258;Q | P23258 | Tubulin ga |
| COG3 | 0.088088 | 0.816417 | 0.015632 | Q96JB2;A0 | Q96JB2 | Q96JB2 | Conserved |
| GLB1 | 0.082343 | 0.827289 | -0.02212 | P16278;E7 | P16278;E7 | P16278 | Beta-galact |
| G3BP1 | 0.078587 | 0.834475 | -0.00984 | Q13283;E5 | Q13283;E5 | Q13283 | Ras GTPase |
| RPS6KA1 | 0.078424 | 0.834787 | -0.02056 | E9PGT3;Q1 | E9PGT3;Q1 | E9PGT3 | Ribosomal |
| INPP5K | 0.06706 | 0.85692 | 0.011673 | Q9BT40;C9 | Q9BT40;C9 | Q9BT40 | Inositol po |
| LGALS3BP | 0.060557 | 0.869847 | -0.00758 | Q08380;K7 | Q08380;K7 | Q08380 | Galectin-3- |
| DTYMK | 0.060418 | 0.870126 | 0.011162 | P23919;H7 | P23919;H7 | P23919 | Thymidylat |
| ERLEC1 | 0.045197 | 0.901162 | 0.009507 | Q96DZ1 | Q96DZ1 | Q96DZ1 | Endoplasm |
| TM9SF3 | 0.041595 | 0.908667 | 0.005286 | Q9HD45;Q | Q9HD45;Q | Q9HD45 | Transmemb |
| LRRC59 | 0.036439 | 0.919519 | -0.00594 | Q96AG4;I3 | Q96AG4 | Q96AG4 | Leucine-ric |
| ALAS1 | 0.03351 | 0.925741 | -0.01401 | P13196;H7 | P13196 | P13196 | 5-aminolev |
| UPP1 | 0.032155 | 0.928635 | 0.011779 | Q16831;C | Q16831;C | Q16831 | Uridine ph |
| TOMM40 | 0.030946 | 0.931223 | -0.00609 | O96008;K7 | O96008 | O96008 | Mitochond |
| PIGS | 0.027418 | 0.938819 | -0.00693 | Q96S52;K7 | Q96S52 | Q96S52 | GPI transan |
| HSDL1 | 0.023045 | 0.94832 | -0.0044 | Q3SXM5;H | Q3SXM5 | Q3SXM5 | Inactive hy |
| ABI1 | 0.021786 | 0.951072 | -0.00505 | A0A0A0MR | A0A0A0MR | A0A0A0MR | Abl interac |
| ALG2 | 0.02071 | 0.953432 | -0.00225 | Q9H553;A | Q9H553 | Q9H553 | Alpha-1,3/ |
| PRDX1 | 0.010749 | 0.975552 | -0.00223 | Q06830;A | Q06830;A | Q06830 | Peroxiredo |
| TOLLIP | 2.406306 | 0.003924 | 0.350258 | Q9H0E2;F2 | Q9H0E2;F2 | Q9H0E2 | Toll-intera |
| PLAUR | 2.24831 | 0.005645 | 0.277812 | Q03405;M | Q03405;M | Q03405 | Urokinase |
| HMOX1 | 2.979889 | 0.001047 | 0.270814 | P09601;B1 | P09601;B1 | P09601 | Heme oxyg |
| SFXN1 | 1.490181 | 0.032346 | 0.220414 | Q9H9B4;D | Q9H9B4 | Q9H9B4 | Sideroflexi |
| RPL15 | 1.726834 | 0.018757 | 0.205902 | P61313;E7 | P61313;E7 | P61313 | 60S riboso |
| SEC22B | 2.745748 | 0.001796 | 0.193615 | O75396;A | O75396 | O75396 | Vesicle-tra |
| ITSN2 | 1.570041 | 0.026913 | 0.185367 | A0A087W | A0A087W | A0A087W | Intersectin |
| ADCK3 | 1.401453 | 0.039678 | 0.168573 | Q8NI60 | Q8NI60 | Q8NI60 | Atypical ki |
| RPL10A | 1.631254 | 0.023375 | 0.155327 | P62906 | P62906 | P62906 | 60S riboso |
| PSMB1 | 2.823621 | 0.001501 | 0.146473 | P20618 | P20618 | P20618 | Proteasom |
| CYC1 | 1.703105 | 0.019811 | 0.143658 | P08574 | P08574 | P08574 | Cytochrom |
| ETHE1 | 2.018224 | 0.009589 | 0.138442 | O95571;M | O95571;M | O95571 | Persulfide |
| SOAT1 | 1.700343 | 0.019937 | 0.136304 | P35610;B1 | P35610;B1 | P35610 | Sterol O-ac |
| EIF2B5 | 2.178031 | 0.006637 | 0.134598 | Q13144;A | Q13144;A | Q13144 | Translation |
| GPX1 | 1.584439 | 0.026035 | 0.129869 | P07203;A0 | P07203;A0 | P07203 | Glutathion |

| | | | | | | | |
|---|---|---|---|---|---|---|---|
| TECR | 1.299143 | 0.050218 | 0.122982 | | Q9NZ01;M | Q9NZ01;M | Q9NZ01 | Very-long-c |
| RAB21 | 1.866972 | 0.013584 | 0.117744 | | Q9UL25 | Q9UL25 | Q9UL25 | Ras-related |
| LPXN | 1.634356 | 0.023208 | 0.107186 | | O60711;B7 | O60711;B7 | O60711 | Leupaxin |
| ATP9A | 2.265419 | 0.005427 | 0.099026 | | O75110;A0 | O75110;A0 | O75110 | Probable pl |
| CD44 | 1.679941 | 0.020896 | 0.098229 | | H0YD13;P1 | H0YD13;P1 | H0YD13 | CD44 antig |
| STX4 | 1.897189 | 0.012671 | 0.080684 | | Q12846;C9 | Q12846 | Q12846 | Syntaxin-4 |
| CHI3L1 | 1.735695 | 0.018378 | -0.13525 | | P36222 | P36222 | P36222 | Chitinase-3 |
| BLVRA | 1.535437 | 0.029145 | -0.24252 | | P53004;C9 | P53004 | P53004 | Biliverdin r |
| PYCARD | 1.794181 | 0.016063 | -0.27354 | | Q9ULZ3;H3 | Q9ULZ3 | Q9ULZ3 | Apoptosis-a |
| BPNT1 | 2.041464 | 0.009089 | -0.27783 | | O95861;A6 | O95861;A6 | O95861 | 3(2),5-bispl |
| FUBP3 | 1.293574 | 0.050866 | 0.168264 | | Q96I24;A0 | Q96I24 | Q96I24 | Far upstrea |
| PACSIN2 | 1.265919 | 0.05421 | -0.22911 | | Q9UNF0;A0 | Q9UNF0;A0 | Q9UNF0 | Protein kin |
| CAB39 | 1.229499 | 0.058952 | -0.22868 | | Q9Y376;A0 | Q9Y376;A0 | Q9Y376 | Calcium-bi |
| CTSK | 1.197874 | 0.063405 | 0.185299 | | P43235;Q5 | P43235 | P43235 | Cathepsin K |
| NEK9 | 1.184046 | 0.065457 | 0.259041 | | Q8TD19;G3 | Q8TD19 | Q8TD19 | Serine/thre |
| STX12 | 1.14081 | 0.072309 | 0.123708 | | Q86Y82;B1 | Q86Y82;B1 | Q86Y82 | Syntaxin-1 |
| PDK3 | 1.12026 | 0.075812 | 0.123248 | | Q15120 | Q15120 | Q15120 | [Pyruvate d |
| ATAD1 | 1.102936 | 0.078898 | 0.125872 | | Q8NBU5 | Q8NBU5 | Q8NBU5 | ATPase fam |
| SPATA20 | 1.092686 | 0.080782 | 0.094831 | | A0A0U1RR | A0A0U1RR | A0A0U1RR | Spermatog |
| MRPL44 | 1.035473 | 0.092157 | 0.164696 | | Q9H9J2 | Q9H9J2 | Q9H9J2 | 39S riboson |
| MMP14 | 1.019179 | 0.09568 | -0.107 | | P50281;F8 | P50281 | P50281 | Matrix met |
| MTCH1 | 1.016478 | 0.096277 | 0.169039 | | Q9NZJ7;H0 | Q9NZJ7;H0 | Q9NZJ7 | Mitochond |
| HSD17B11 | 1.010128 | 0.097695 | 0.102342 | | Q8NBQ5;D | Q8NBQ5;D | Q8NBQ5 | Estradiol 1 |
| NPM1 | 1.001262 | 0.09971 | -0.13504 | | P06748;E5 | P06748 | P06748 | Nucleopho |
| RPL13 | 0.963896 | 0.108669 | 0.139132 | | P26373;J3 | P26373;J3 | P26373 | 60S riboson |
| MREG | 0.917018 | 0.121055 | 0.091234 | | Q8N565;C9 | Q8N565;C9 | Q8N565 | Melanoregu |
| TM9SF2 | 0.916225 | 0.121276 | 0.115562 | | Q99805 | Q99805 | Q99805 | Transmemb |
| DNAJA2 | 0.910659 | 0.12284 | -0.10719 | | O60884;A0 | O60884 | O60884 | DnaJ homo |
| SRI | 0.907317 | 0.123789 | 0.120609 | | C9J0K6;P3 | C9J0K6;P3 | C9J0K6 | Sorcin |
| NDRG1 | 0.905719 | 0.124246 | 0.102982 | | Q92597;E7 | Q92597 | Q92597 | Protein ND |
| PRPF4 | 0.879929 | 0.131847 | 0.124098 | | O43172 | O43172 | O43172 | U4/U6 sma |
| RPS16 | 0.819094 | 0.151672 | 0.101865 | | P62249;M | P62249;M | P62249 | 40S riboson |
| ARFGEF2 | 0.733956 | 0.18452 | 0.135381 | | Q9Y6D5 | Q9Y6D5 | Q9Y6D5 | Brefeldin A- |
| HTATIP2 | 0.733187 | 0.184847 | -0.05048 | | Q9BUP3;E9 | Q9BUP3;E9 | Q9BUP3 | Oxidoreduc |
| VDAC3 | 0.716685 | 0.192006 | 0.113917 | | Q9Y277;E5 | Q9Y277;E5 | Q9Y277 | Voltage-dep |
| MTHFD2 | 0.713261 | 0.193526 | 0.115535 | | P13995;B8 | P13995;B8 | P13995 | Bifunctiona |
| XPNPEP1 | 0.712736 | 0.19376 | -0.15609 | | Q9NQW7;C | Q9NQW7;C | Q9NQW7 | Xaa-Pro am |
| RPSA | 0.700182 | 0.199443 | 0.07655 | | C9J9K3;A0 | C9J9K3;A0 | C9J9K3 | 40S riboson |
| MRPL39 | 0.699141 | 0.199921 | 0.083985 | | Q9NYK5;C9 | Q9NYK5;C9 | Q9NYK5 | 39S riboson |
| GGT1 | 0.684869 | 0.2066 | -0.08767 | | P19440;E7 | P19440;E7 | P19440 | Gamma-glu |
| RENBP | 0.66793 | 0.214818 | -0.10765 | | P51606;A6 | P51606;A6 | P51606 | N-acylgluco |
| PDCD4 | 0.652865 | 0.2224 | 0.105699 | | Q53EL6;Q5 | Q53EL6 | Q53EL6 | Programme |
| SEC61A1 | 0.630684 | 0.234054 | 0.077209 | | B4DR61;P6 | B4DR61;P6 | B4DR61 | Protein tran |
| TST | 0.630236 | 0.234295 | 0.080366 | | Q16762;B1 | Q16762 | Q16762 | Thiosulfate |
| PLRG1 | 0.613842 | 0.243309 | 0.089952 | | O43660;H0 | O43660 | O43660 | Pleiotropic |
| CTSC | 0.567136 | 0.270934 | 0.098462 | | P53634;H0 | P53634 | P53634 | Dipeptidyl |
| GCDH | 0.567003 | 0.271017 | 0.084716 | | Q92947;K7 | Q92947 | Q92947 | Glutaryl-Co |
| ITPRIP | 0.555612 | 0.27822 | -0.07462 | | Q8IWB1;RE | Q8IWB1 | Q8IWB1 | Inositol 1,4 |

| | | | | | | | | |
|---|---|---|---|---|---|---|---|---|
| COL4A3BP | 0.547546 | 0.283435 | 0.129447 | | A0A2R8Y7( | A0A2R8Y7( | A0A2R8Y7( | Collagen ty |
| MAN1A1 | 0.482061 | 0.329564 | -0.08212 | | P33908 | P33908 | P33908 | Mannosyl-c |
| UQCRFS1 | 0.481625 | 0.329894 | 0.060124 | | P47985;P0 | P47985;P0 | P47985 | Cytochrom |
| YWHAG | 0.479521 | 0.331496 | -0.10217 | | P61981 | P61981 | P61981 | 14-3-3 prot |
| BZW1 | 0.466518 | 0.341571 | 0.102949 | | Q7L1Q6;C9 | Q7L1Q6;C9 | Q7L1Q6 | Basic leucir |
| SLC12A7 | 0.458983 | 0.34755 | 0.06932 | | A0A0G2JN\ | A0A0G2JN\ | A0A0G2JN\ | Solute carri |
| UCHL5 | 0.448911 | 0.355705 | -0.07217 | | Q5LJA5;Q5 | Q5LJA5;Q5 | Q5LJA5 | Ubiquitin c |
| CTSA | 0.437293 | 0.365348 | 0.065943 | | X6R5C5;X6 | X6R5C5;X6 | X6R5C5 | Carboxypep |
| TOR3A | 0.436334 | 0.366156 | -0.09826 | | Q9H497;Q5 | Q9H497;Q5 | Q9H497 | Torsin-3A |
| RABEP1 | 0.431735 | 0.370054 | -0.17774 | | Q15276;K7 | Q15276 | Q15276 | Rab GTPase |
| PALLD | 0.427103 | 0.374022 | 0.097345 | | Q8WX93;H | Q8WX93;H | Q8WX93 | Palladin |
| TRIM4 | 0.420572 | 0.379689 | 0.166104 | | Q9C037;H7 | Q9C037 | Q9C037 | E3 ubiquiti |
| C2CD2L | 0.39656 | 0.401273 | -0.07498 | | O14523;E9 | O14523;E9 | O14523 | C2 domain- |
| ARHGEF7 | 0.376683 | 0.420065 | 0.058073 | | B1ALK7;A0 | B1ALK7;A0 | B1ALK7 | Rho guanin |
| F11R | 0.371297 | 0.425308 | -0.04994 | | Q9Y624;A0 | Q9Y624;A0 | Q9Y624 | Junctional |
| SRPRB | 0.367426 | 0.429116 | -0.04823 | | Q9Y5M8;H | Q9Y5M8;H | Q9Y5M8 | Signal recog |
| PSMA4 | 0.356573 | 0.439974 | -0.05765 | | P25789;H0 | P25789;H0 | P25789 | Proteasom |
| MICU2 | 0.348689 | 0.448034 | 0.032747 | | Q8IYU8;A0 | Q8IYU8 | Q8IYU8 | Calcium up |
| SEC24A | 0.3446 | 0.452273 | 0.071111 | | O95486;H0 | O95486 | O95486 | Protein trai |
| RPLP0 | 0.342909 | 0.454037 | 0.053705 | | P05388;F8 | P05388;F8 | P05388 | 60S acidic r |
| FKBP4 | 0.323035 | 0.475296 | 0.07739 | | Q02790;H0 | Q02790 | Q02790 | Peptidyl-pr |
| PIGT | 0.29968 | 0.501556 | 0.051121 | | Q969N2;A0 | Q969N2;A0 | Q969N2 | GPI transan |
| GOT1 | 0.28155 | 0.522937 | -0.09895 | | P17174 | P17174 | P17174 | Aspartate a |
| GSS | 0.28046 | 0.524252 | -0.11058 | | A0A2R8Y43 | A0A2R8Y43 | A0A2R8Y43 | Glutathion |
| P2RX7 | 0.277953 | 0.527287 | -0.05056 | | Q99572;F5 | Q99572 | Q99572 | P2X purino |
| HDGF | 0.271091 | 0.535685 | -0.06029 | | P51858 | P51858 | P51858 | Hepatoma- |
| HIBADH | 0.267138 | 0.540582 | 0.063244 | | P31937;H7 | P31937 | P31937 | 3-hydroxyi: |
| HARS | 0.261977 | 0.547045 | -0.07964 | | P12081;B3 | P12081;B3 | P12081 | Histidine--t |
| PRDX4 | 0.261344 | 0.547843 | 0.056178 | | Q13162;H7 | Q13162;H7 | Q13162 | Peroxiredo |
| RALY | 0.247782 | 0.56522 | -0.04165 | | Q5QPL9;Q9 | Q5QPL9;Q9 | Q5QPL9 | RNA-bindir |
| CYP4V2 | 0.247401 | 0.565717 | 0.053838 | | Q6ZWL3;H | Q6ZWL3 | Q6ZWL3 | Cytochrom |
| ETF1 | 0.244092 | 0.570043 | -0.04242 | | P62495;B7 | P62495;B7 | P62495 | Eukaryotic |
| GUF1 | 0.240856 | 0.574307 | 0.059168 | | Q8N442 | Q8N442 | Q8N442 | Translation |
| IDH3G | 0.228286 | 0.591173 | 0.041968 | | P51553;G5 | P51553;G5 | P51553 | Isocitrate d |
| YWHAZ | 0.226112 | 0.594139 | 0.068507 | | P63104;E7 | P63104;E7 | P63104 | 14-3-3 prot |
| TAPBPL | 0.223986 | 0.597054 | 0.050475 | | Q9BX59;G3 | Q9BX59 | Q9BX59 | Tapasin-rel |
| MX2 | 0.219547 | 0.603188 | 0.041587 | | P20592;C9 | P20592 | P20592 | Interferon-i |
| WDR26 | 0.209774 | 0.616916 | 0.029903 | | A0A499FIZ( | A0A499FIZ( | A0A499FIZ( | WD repeat- |
| RPL11 | 0.194344 | 0.639229 | 0.036097 | | Q5VVC8;P6 | Q5VVC8;P6 | Q5VVC8 | 60S ribosor |
| CC2D1B | 0.186918 | 0.650252 | 0.037145 | | Q5T0F9;H7 | Q5T0F9;H7 | Q5T0F9 | Coiled-coil |
| ATP5C1 | 0.183789 | 0.654955 | 0.024521 | | P36542 | P36542 | P36542 | ATP syntha: |
| PSMD14 | 0.179548 | 0.661381 | -0.03868 | | O00487 | O00487 | O00487 | 26S protea: |
| DEK | 0.168692 | 0.678122 | 0.030879 | | H0Y993;P3 | H0Y993;P3 | H0Y993 | Protein DEI |
| AHCYL1 | 0.163195 | 0.686759 | -0.02691 | | O43865 | O43865 | O43865 | Putative ad |
| SCIN | 0.161504 | 0.68944 | -0.05918 | | Q9Y6U3;E5 | Q9Y6U3 | Q9Y6U3 | Adseverin |
| FUCA2 | 0.132312 | 0.737375 | -0.01584 | | Q9BTY2;Q7 | Q9BTY2 | Q9BTY2 | Plasma alpl |
| MGAT1 | 0.111514 | 0.773546 | 0.030769 | | P26572;D6 | P26572 | P26572 | Alpha-1,3-r |
| MAN1B1 | 0.110285 | 0.775738 | 0.021962 | | Q9UKM7;H | Q9UKM7;H | Q9UKM7 | Endoplasm |

| | | | | | | | |
|---|---|---|---|---|---|---|---|
| ATP1B1 | 0.108439 | 0.779042 | -0.01679 | P05026;V9 | P05026;V9 | P05026 | Sodium/po |
| ADAMDEC1 | 0.108316 | 0.779263 | 0.022672 | O15204 | O15204 | O15204 | ADAM DEC1 |
| PCBP1 | 0.10488 | 0.785453 | 0.015341 | Q15365;H3 | Q15365 | Q15365 | Poly(rC)-bir |
| LBR | 0.104852 | 0.785503 | -0.04655 | Q14739;A0 | Q14739;A0 | Q14739 | Lamin-B rec |
| VAMP7 | 0.093997 | 0.805384 | -0.01166 | P51809;E5 | P51809 | P51809 | Vesicle-asso |
| SDSL | 0.089139 | 0.814444 | -0.03291 | Q96GA7;F8 | Q96GA7 | Q96GA7 | Serine dehy |
| CPVL | 0.071576 | 0.848055 | 0.0188 | Q9H3G5;H7 | Q9H3G5;H7 | Q9H3G5 | Probable se |
| PAFAH2 | 0.070458 | 0.85024 | -0.01454 | Q99487;Q5 | Q99487 | Q99487 | Platelet-act |
| ATG9A | 0.065858 | 0.859294 | -0.00944 | Q7Z3C6;H7 | Q7Z3C6 | Q7Z3C6 | Autophagy- |
| METAP1 | 0.064875 | 0.861241 | -0.02327 | P53582;H0 | P53582 | P53582 | Methionine |
| MAT2B | 0.064165 | 0.862651 | -0.01018 | Q9NZL9;H7 | Q9NZL9 | Q9NZL9 | Methionine |
| BCAT2 | 0.062717 | 0.865532 | 0.015625 | M0QXF9;O | M0QXF9;O | M0QXF9 | Branched-c |
| PSM8 | 0.033756 | 0.925217 | 0.005149 | X5D2R7;P2 | X5D2R7;P2 | X5D2R7 | Proteasome |
| PFN1 | 0.031251 | 0.93057 | -0.00854 | P07737;K7 | P07737;K7 | P07737 | Profilin-1 |
| ALDH1A2 | 0.029135 | 0.935114 | -0.0118 | O94788;H0 | O94788;H0 | O94788 | Retinal deh |
| CHMP4B | 0.024164 | 0.945881 | -0.00554 | Q9H444;P5 | Q9H444 | Q9H444 | Charged mu |
| MLEC | 0.021748 | 0.951156 | -0.00372 | Q14165;F5 | Q14165;F5 | Q14165 | Malectin |
| PPM1G | 0.01609 | 0.96363 | -0.00623 | O15355 | O15355 | O15355 | Protein pho |
| HNRNPLL | 0.009889 | 0.977487 | 0.001772 | B7WPG3;C | B7WPG3;C | B7WPG3 | Heterogene |
| GBAS | 0.003091 | 0.992908 | -0.00095 | O75323;H7 | O75323 | O75323 | Protein Nip |
| ACOT7 | 0.001588 | 0.99635 | -0.00041 | O00154;K7 | O00154;K7 | O00154 | Cytosolic ac |
| SDCBP | 1.487862 | 0.032519 | 0.281169 | O00560;G5 | O00560;G5 | O00560 | Syntenin-1 |
| NUMB | 2.773599 | 0.001684 | 0.254665 | P49757;G3 | P49757;G3 | P49757 | Protein nur |
| NEMF | 2.185618 | 0.006522 | 0.245134 | G3V5V3;O6 | G3V5V3;O6 | G3V5V3 | Nuclear exp |
| SCAMP1 | 1.593263 | 0.025512 | 0.242909 | O15126;A0 | O15126;A0 | O15126 | Secretory c |
| ATP7A | 1.723616 | 0.018897 | 0.23442 | Q04656 | Q04656 | Q04656 | Copper-trar |
| CLINT1 | 1.601779 | 0.025016 | 0.231781 | Q14677;H0 | Q14677 | Q14677 | Clathrin int |
| GPRIN3 | 1.802833 | 0.015746 | 0.223397 | Q6ZVF9 | Q6ZVF9 | Q6ZVF9 | G protein-r |
| ACP5 | 1.525393 | 0.029827 | 0.2214 | P13686;K7 | P13686 | P13686 | Tartrate-res |
| RHOG | 2.660765 | 0.002184 | 0.221017 | P84095 | P84095 | P84095 | Rho-related |
| PTPN9 | 2.078518 | 0.008346 | 0.170288 | P43378;A0 | P43378;A0 | P43378 | Tyrosine-pr |
| TRIM33 | 1.732754 | 0.018503 | 0.144913 | Q9UPN9;H0 | Q9UPN9;H0 | Q9UPN9 | E3 ubiquiti |
| CHP1 | 1.570681 | 0.026873 | 0.139068 | Q99653;H0 | Q99653;H0 | Q99653 | Calcineurin |
| PPP1R9B | 1.881305 | 0.013143 | 0.128819 | Q96SB3;Q9 | Q96SB3 | Q96SB3 | Neurabin-2 |
| ATP5O | 2.11549 | 0.007665 | 0.128181 | P48047;A0 | P48047;A0 | P48047 | ATP synthas |
| CAPZA2 | 1.4217 | 0.03787 | 0.106068 | P47755;F8 | P47755;F8 | P47755 | F-actin-cap |
| TM9SF1 | 1.299869 | 0.050134 | 0.098613 | E9PSI1;E9P | E9PSI1;E9P | E9PSI1 | Transmemb |
| RRAGA | 2.491201 | 0.003227 | -0.12585 | Q7L523;Q5 | Q7L523;Q5 | Q7L523 | Ras-related |
| BCAP29 | 1.436521 | 0.0366 | -0.13016 | A0A669KBI | A0A669KBI | A0A669KBI | B-cell recep |
| HM13 | 1.76257 | 0.017275 | -0.1314 | A0A3B3IUE | A0A3B3IUE | A0A3B3IUE | Minor histo |
| PSME3 | 1.600581 | 0.025085 | -0.1315 | K9J957;P6 | K9J957;P6 | K9J957 | Proteasome |
| DAB2 | 1.890701 | 0.012862 | -0.15132 | P98082;D6 | P98082 | P98082 | Disabled ho |
| PRPF4B | 1.219974 | 0.06026 | 0.126371 | Q13523;H0 | Q13523;H0 | Q13523 | Serine/thre |
| BDH1 | 1.191373 | 0.064362 | 0.167975 | Q02338;E9 | Q02338;E9 | Q02338 | D-beta-hyd |
| SCFD2 | 1.189127 | 0.064695 | -0.15388 | Q8WU76;A | Q8WU76;A | Q8WU76 | Sec1 family |
| EDC4 | 1.187893 | 0.064879 | 0.138311 | Q6P2E9;I3I | Q6P2E9 | Q6P2E9 | Enhancer o |
| DDX24 | 1.179183 | 0.066194 | 0.160295 | G3V529;Q9 | G3V529;Q9 | G3V529 | ATP-depenc |
| SNRNP40 | 1.162025 | 0.068861 | 0.104292 | Q96DI7;Q9 | Q96DI7;Q9 | Q96DI7 | U5 small nu |

| | | | | | | | |
|---|---|---|---|---|---|---|---|
| GRPEL1 | 1.15259 | 0.070374 | 0.207472 | Q9HAV7 | Q9HAV7 | Q9HAV7 | GrpE protei |
| CAPNS1 | 1.085186 | 0.082189 | -0.14455 | A0A0C4DG | A0A0C4DG | A0A0C4DG | Calpain sma |
| FAM213A | 1.074242 | 0.084287 | -0.10153 | Q9BRX8 | Q9BRX8 | Q9BRX8 | Redox-regu |
| EIF2AK2 | 0.966405 | 0.108043 | 0.066085 | P19525;F8 | P19525 | P19525 | Interferon-i |
| HNRNPA0 | 0.934418 | 0.116301 | 0.141909 | Q13151 | Q13151 | Q13151 | Heterogene |
| GTPBP4 | 0.92834 | 0.11794 | -0.27333 | Q9BZE4;Q5 | Q9BZE4;Q5 | Q9BZE4 | Nucleolar G |
| ST14 | 0.911563 | 0.122585 | 0.130744 | Q9Y5Y6 | Q9Y5Y6 | Q9Y5Y6 | Suppressor |
| RPS10 | 0.874504 | 0.133505 | -0.08608 | P46783;F6 | P46783;F6 | P46783 | 40S riboson |
| ACOT8 | 0.859705 | 0.138132 | 0.14114 | O14734;E9 | O14734;E9 | O14734 | Acyl-coenzy |
| SLC26A11 | 0.858147 | 0.138629 | 0.117242 | Q86WA9;I3 | Q86WA9 | Q86WA9 | Sodium-inc |
| ENG | 0.855394 | 0.13951 | -0.07531 | P17813;F5 | P17813;F5 | P17813 | Endoglin |
| RAB20 | 0.851104 | 0.140895 | 0.132607 | Q9NX57 | Q9NX57 | Q9NX57 | Ras-related |
| DDAH2 | 0.806499 | 0.156135 | 0.127914 | O95865;A0 | O95865;A0 | O95865 | N(G),N(G)-c |
| DAGLB | 0.806358 | 0.156186 | -0.09389 | Q8NCG7;E7 | Q8NCG7;E7 | Q8NCG7 | Sn1-specific |
| LUC7L3 | 0.788829 | 0.162619 | 0.162166 | J3KPP4;O9 | J3KPP4;O9 | J3KPP4 | Luc7-like pr |
| PPID | 0.78163 | 0.165337 | -0.17926 | Q08752 | Q08752 | Q08752 | Peptidyl-pr |
| LNP | 0.779489 | 0.166154 | 0.137666 | Q9C0E8;C9 | Q9C0E8 | Q9C0E8 | Protein lun |
| RCN2 | 0.763804 | 0.172265 | 0.129077 | Q14257;H0 | Q14257;H0 | Q14257 | Reticulocal |
| EEF1D | 0.755735 | 0.175495 | 0.106359 | P29692;E9 | P29692;E9 | P29692 | Elongation |
| SF1 | 0.742169 | 0.181063 | 0.08401 | Q15637;C9 | Q15637 | Q15637 | Splicing fac |
| ANO10 | 0.73142 | 0.185601 | 0.059403 | Q9NW15;C | Q9NW15;C | Q9NW15 | Anoctamin |
| RAN | 0.700989 | 0.199072 | 0.097947 | B5MDF5;P6 | B5MDF5;P6 | B5MDF5 | GTP-bindin |
| EIF2B3 | 0.69934 | 0.19983 | 0.083074 | Q9NR50;A0 | Q9NR50 | Q9NR50 | Translation |
| STOML2 | 0.681117 | 0.208393 | 0.08758 | Q9UJZ1;A0 | Q9UJZ1;A0 | Q9UJZ1 | Stomatin-li |
| ADCY7 | 0.677574 | 0.2101 | -0.08561 | P51828;H3 | P51828;H3 | P51828 | Adenylate c |
| DDO | 0.673666 | 0.211999 | 0.068718 | Q99489;F2 | Q99489 | Q99489 | D-aspartate |
| NIT2 | 0.649639 | 0.224058 | -0.16898 | Q9NQR4;H7 | Q9NQR4;H7 | Q9NQR4 | Omega-ami |
| SEPTIN10 | 0.649639 | 0.224058 | 0.187849 | E7EW69;E7 | E7EW69;E7 | E7EW69 | Septin-10 |
| HSPB1 | 0.627359 | 0.235853 | -0.14033 | P04792;F8 | P04792;F8 | P04792 | Heat shock |
| PEBP1 | 0.625213 | 0.237021 | -0.15571 | P30086 | P30086 | P30086 | Phosphatic |
| CTNNA1 | 0.621908 | 0.238832 | 0.133657 | P35221;G3 | P35221;G3 | P35221 | Catenin alp |
| EXOC6 | 0.620609 | 0.239547 | 0.157609 | E7EW84;Q8 | E7EW84;Q8 | E7EW84 | Exocyst cor |
| ALDH1L2 | 0.610453 | 0.245215 | 0.847165 | A0A494C1I | A0A494C1I | A0A494C1I | Mitochond |
| GRIPAP1 | 0.603142 | 0.249378 | 0.158908 | Q4V328;A0 | Q4V328;A0 | Q4V328 | GRIP1-asso |
| VAV1 | 0.573762 | 0.266832 | 0.133299 | F5H5P4;Q9 | F5H5P4;Q9 | F5H5P4 | Proto-onco |
| AP5B1 | 0.570744 | 0.268693 | 0.089029 | Q2VPB7 | Q2VPB7 | Q2VPB7 | AP-5 compl |
| MRPS9 | 0.564744 | 0.27243 | 0.11165 | P82933;H7 | P82933 | P82933 | 28S riboson |
| STX5 | 0.5545 | 0.278933 | 0.085821 | Q13190;E9 | Q13190;E9 | Q13190 | Syntaxin-5 |
| WDR82 | 0.541826 | 0.287193 | 0.074226 | Q6UXN9;C9 | Q6UXN9 | Q6UXN9 | WD repeat- |
| MAOA | 0.521292 | 0.301098 | 0.10371 | P21397 | P21397 | P21397 | Amine oxid |
| MRPL37 | 0.518235 | 0.303225 | 0.07995 | Q9BZE1;S4I | Q9BZE1;S4I | Q9BZE1 | 39S riboson |
| STAT6 | 0.517308 | 0.303873 | -0.09431 | A0A494C0T | A0A494C0T | A0A494C0T | Signal trans |
| KPNA4 | 0.511078 | 0.308263 | 0.032615 | O00629;H7 | O00629 | O00629 | Importin su |
| SORT1 | 0.504089 | 0.313264 | -0.08289 | Q99523;A0 | Q99523 | Q99523 | Sortilin |
| BUB3 | 0.502883 | 0.314136 | 0.061473 | O43684;J3 | O43684;J3 | O43684 | Mitotic che |
| WDR37 | 0.497466 | 0.318079 | -0.1608 | A0A3B3IT4 | A0A3B3IT4 | A0A3B3IT4 | WD repeat- |
| LGALS1 | 0.487651 | 0.325348 | 0.138312 | P09382;F8 | P09382 | P09382 | Galectin-1 |
| API5 | 0.487386 | 0.325547 | 0.060028 | G3V1C3;Q9 | G3V1C3;Q9 | G3V1C3 | Apoptosis i |

| | | | | | | | |
|---|---|---|---|---|---|---|---|
| MYH10 | 0.441593 | 0.361748 | 0.166692 | P35580 | P35580 | P35580 | Myosin-10 |
| TRAPPC8 | 0.435292 | 0.367035 | 0.06522 | Q9Y2L5;J3( | Q9Y2L5;J3( | Q9Y2L5 | Trafficking |
| RABGGTA | 0.431508 | 0.370248 | -0.10473 | Q92696;H( | Q92696;H( | Q92696 | Geranylger |
| RPS7 | 0.419016 | 0.381052 | -0.12553 | P62081;A0 | P62081;A0 | P62081 | 40S ribosor |
| U2SURP | 0.397948 | 0.399993 | -0.05058 | O15042;E7 | O15042;E7 | O15042 | U2 snRNP-a |
| AGPAT9 | 0.396179 | 0.401625 | -0.04047 | Q53EU6 | Q53EU6 | Q53EU6 | Glycerol-3- |
| GOLGA1 | 0.390731 | 0.406695 | 0.08779 | Q92805;Q5 | Q92805 | Q92805 | Golgin subf |
| MX1 | 0.385381 | 0.411737 | -0.18382 | P20591;F8 | P20591;F8 | P20591 | Interferon-i |
| CUL4A | 0.384963 | 0.412132 | -0.07099 | Q13619;A( | Q13619;A( | Q13619 | Cullin-4A |
| IL16 | 0.361581 | 0.434929 | 0.133095 | H0YLH9;Q1 | H0YLH9;Q1 | H0YLH9 | Pro-interle |
| MAPK13 | 0.359969 | 0.436547 | -0.09037 | O15264;Q5 | O15264 | O15264 | Mitogen-ac |
| SKIV2L | 0.354322 | 0.44226 | 0.075542 | Q15477;H7 | Q15477 | Q15477 | Helicase SK |
| ERP29 | 0.353957 | 0.442633 | 0.042946 | P30040;F8 | P30040;F8 | P30040 | Endoplasm |
| SPR | 0.346676 | 0.450116 | 0.066225 | P35270;RE | P35270 | P35270 | Sepiapterir |
| DBN1 | 0.322288 | 0.476115 | -0.0825 | Q16643;D6 | Q16643;D6 | Q16643 | Drebrin |
| XPC | 0.30999 | 0.48979 | 0.085324 | Q01831;E7 | Q01831 | Q01831 | DNA repair |
| RHOF | 0.302273 | 0.498571 | 0.063117 | Q9HBH0;V9 | Q9HBH0;V9 | Q9HBH0 | Rho-relatec |
| ARHGEF6 | 0.299824 | 0.501391 | 0.068066 | Q15052 | Q15052 | Q15052 | Rho guanin |
| TES | 0.289721 | 0.513191 | 0.061962 | Q9UGI8;F8 | Q9UGI8 | Q9UGI8 | Testin |
| PARK7 | 0.284936 | 0.518876 | 0.048985 | Q99497;K7 | Q99497;K7 | Q99497 | Protein deg |
| SDF2L1 | 0.272829 | 0.533545 | 0.045189 | Q9HCN8 | Q9HCN8 | Q9HCN8 | Stromal cel |
| BIN2 | 0.262635 | 0.546217 | -0.05612 | Q9UBW5;S | Q9UBW5;S | Q9UBW5 | Bridging in |
| RSU1 | 0.251048 | 0.560986 | -0.07837 | Q15404 | Q15404 | Q15404 | Ras suppres |
| TBC1D5 | 0.250508 | 0.561684 | -0.07088 | Q92609;C9 | Q92609;C9 | Q92609 | TBC1 doma |
| ECHDC1 | 0.246108 | 0.567403 | 0.035563 | Q9NTX5;H( | Q9NTX5;H( | Q9NTX5 | Ethylmalor |
| HTRA2 | 0.231314 | 0.587064 | 0.056613 | O43464;A( | O43464;A( | O43464 | Serine prot |
| PON2 | 0.226378 | 0.593776 | 0.050191 | A0A0J9YXF | A0A0J9YXF | A0A0J9YXF | Serum para |
| WAS | 0.207401 | 0.620295 | 0.032286 | P42768;C9 | P42768;C9 | P42768 | Wiskott-Alc |
| HAGH | 0.202659 | 0.627106 | -0.05821 | Q16775;H3 | Q16775;H3 | Q16775 | Hydroxyacy |
| PUS1 | 0.202349 | 0.627555 | 0.037866 | F5H1S9;G8 | F5H1S9;G8 | F5H1S9 | tRNA pseuc |
| DYNC1LI2 | 0.177745 | 0.664133 | 0.039607 | O43237;B4 | O43237;B4 | O43237 | Cytoplasmi |
| EXOC3 | 0.176394 | 0.666202 | -0.03507 | O60645;D6 | O60645;D6 | O60645 | Exocyst cor |
| MESDC2 | 0.172709 | 0.671878 | 0.035101 | Q14696;H( | Q14696 | Q14696 | LDLR chape |
| NAA10 | 0.172463 | 0.67226 | -0.05595 | P41227;F8 | P41227;F8 | P41227 | N-alpha-ace |
| MRPL15 | 0.167826 | 0.679476 | 0.03034 | Q9P015;E5 | Q9P015;E5 | Q9P015 | 39S ribosor |
| RASAL3 | 0.164413 | 0.684837 | 0.072285 | Q86YV0;M( | Q86YV0 | Q86YV0 | RAS proteir |
| MAPK1 | 0.162968 | 0.68712 | -0.04014 | P28482 | P28482 | P28482 | Mitogen-ac |
| PPT1 | 0.154594 | 0.700496 | 0.042696 | A0A2C9F2F | A0A2C9F2F | A0A2C9F2F | Palmitoyl-p |
| PEX14 | 0.143254 | 0.719029 | 0.035625 | O75381;K7 | O75381 | O75381 | Peroxisoma |
| PXN | 0.126115 | 0.747971 | -0.02381 | F5GZ78;A0 | F5GZ78;A0 | F5GZ78 | Paxillin |
| HMHA1 | 0.106988 | 0.78165 | 0.035539 | F5H1R4;K7 | F5H1R4;K7 | F5H1R4 | Minor histc |
| ARFGEF1 | 0.09642 | 0.800903 | 0.039111 | Q9Y6D6;E5 | Q9Y6D6;E5 | Q9Y6D6 | Brefeldin A- |
| SERPINB9 | 0.093528 | 0.806255 | 0.035238 | P50453 | P50453 | P50453 | Serpin B9 |
| ESD | 0.091468 | 0.810088 | -0.03455 | P10768;U3 | P10768;U3 | P10768 | S-formylglu |
| STOM | 0.085277 | 0.821718 | 0.018001 | P27105;F8 | P27105 | P27105 | Erythrocyte |
| ITGB3 | 0.062842 | 0.865282 | 0.010835 | P05106;H3 | P05106;H3 | P05106 | Integrin be |
| GMPS | 0.059774 | 0.871416 | -0.02538 | P49915 | P49915 | P49915 | GMP synth |
| RNH1 | 0.05628 | 0.878456 | -0.0171 | P13489;E9 | P13489 | P13489 | Ribonuclea |

| | | | | | | | |
|---|---|---|---|---|---|---|---|
| RAP1GDS1 | 0.045714 | 0.900089 | -0.00906 | P52306;H( | P52306 | P52306 | Rap1 GTPa |
| SPTLC2 | 0.042845 | 0.906055 | 0.013423 | O15270;H( | O15270;H( | O15270 | Serine palm |
| EOGT | 0.036297 | 0.91982 | -0.01024 | Q5NDL2;F5 | Q5NDL2 | Q5NDL2 | EGF domai |
| GGH | 0.035955 | 0.920545 | -0.01089 | Q92820 | Q92820 | Q92820 | Gamma-glu |
| ARRB2 | 0.030176 | 0.932877 | 0.005286 | P32121;K7 | P32121;K7 | P32121 | Beta-arresti |
| GK | 0.027918 | 0.93774 | 0.006053 | P32189;Q1 | P32189;Q1 | P32189 | Glycerol ki |
| GSTP1 | 0.012642 | 0.971311 | 0.003139 | P09211;A8 | P09211;A8 | P09211 | Glutathion |
| STAT2 | 0.00835 | 0.980956 | 0.002366 | A0A494C1( | A0A494C1( | A0A494C1( | Signal trans |
| DHX29 | 3.462514 | 0.000345 | 0.328989 | A0A087WY | A0A087WY | A0A087WY | ATP-depen |
| ABHD6 | 1.320627 | 0.047794 | 0.303509 | Q9BV23;C9 | Q9BV23 | Q9BV23 | Monoacylg |
| TNKS1BP1 | 2.032939 | 0.00927 | 0.278539 | Q9C0C2;A0 | Q9C0C2 | Q9C0C2 | 182 kDa ta |
| FAM136A | 5.172786 | 6.72E-06 | 0.276407 | Q96C01;E7 | Q96C01;E7 | Q96C01 | Protein FA |
| PRPF31 | 3.528648 | 0.000296 | 0.266134 | E7EVX8;Q8 | E7EVX8;Q8 | E7EVX8 | U4/U6 sma |
| P2RX4 | 2.927936 | 0.00118 | 0.250882 | Q99571;F6 | Q99571;F6 | Q99571 | P2X purino |
| CYP27B1 | 2.307368 | 0.004928 | 0.243465 | O15528;V9 | O15528 | O15528 | 25-hydroxy |
| SEPTIN11 | 1.835669 | 0.014599 | 0.232201 | D6RGI3;D6 | D6RGI3;D6 | D6RGI3 | Septin-11 |
| GLYR1 | 1.88697 | 0.012973 | 0.224561 | K7EMM8;Q | K7EMM8;Q | K7EMM8 | Putative ox |
| ECH1 | 1.768999 | 0.017022 | 0.156399 | Q13011;M( | Q13011;M( | Q13011 | Delta(3,5)-I |
| DCTN4 | 1.74989 | 0.017787 | 0.150857 | Q9UJW0;E5 | Q9UJW0 | Q9UJW0 | Dynactin su |
| VAPB | 1.829356 | 0.014813 | 0.145974 | O95292;E5 | O95292 | O95292 | Vesicle-ass |
| RHOT1 | 1.420332 | 0.03799 | 0.145384 | H7BXZ6;Q8 | H7BXZ6;Q8 | H7BXZ6 | Mitochond |
| SNAP23 | 1.543101 | 0.028635 | 0.135955 | O00161;H3 | O00161;H3 | O00161 | Synaptoso |
| NADK2 | 1.510523 | 0.030866 | 0.132648 | Q4G0N4;B7 | Q4G0N4;B7 | Q4G0N4 | NAD kinase |
| DDX39B | 1.406374 | 0.039231 | 0.131715 | Q13838;Q5 | Q13838;Q5 | Q13838 | Spliceosom |
| GNAQ | 1.485369 | 0.032706 | -0.0908 | P50148;B1 | P50148 | P50148 | Guanine nu |
| RMDN1 | 1.401363 | 0.039686 | -0.12639 | Q96DB5;E5 | Q96DB5;E5 | Q96DB5 | Regulator o |
| TRA2B | 1.588656 | 0.025784 | -0.13906 | P62995;H7 | P62995;H7 | P62995 | Transforme |
| RAB10 | 2.240748 | 0.005744 | -0.15642 | P61026 | P61026 | P61026 | Ras-related |
| PDCD6 | 1.510981 | 0.030833 | -0.17072 | O75340;A0 | O75340;A0 | O75340 | Programme |
| PGLS | 1.546467 | 0.028414 | -0.27701 | O95336;M( | O95336;M( | O95336 | 6-phospho |
| SLC9A3R1 | 1.279708 | 0.052516 | 0.221651 | O14745;J3 | O14745;J3 | O14745 | Na(+)/H(+) e |
| IMPDH2 | 1.241737 | 0.057314 | 0.146475 | P12268;H( | P12268;H( | P12268 | Inosine-5-m |
| TRIM38 | 1.215678 | 0.060859 | -0.21382 | O00635 | O00635 | O00635 | E3 ubiquiti |
| RAB7B | 1.211975 | 0.06138 | -0.11506 | Q96AH8;A( | Q96AH8;A( | Q96AH8 | Ras-related |
| MAP7D1 | 1.202806 | 0.062689 | 0.188623 | Q3KQU3;H( | Q3KQU3 | Q3KQU3 | MAP7 dom |
| ATP1B3 | 1.201939 | 0.062815 | 0.089559 | P54709;C9 | P54709;C9 | P54709 | Sodium/po |
| POFUT1 | 1.197547 | 0.063453 | 0.117038 | Q9H488 | Q9H488 | Q9H488 | GDP-fucose |
| NUP160 | 1.185061 | 0.065304 | 0.159985 | Q12769;G3 | Q12769;G3 | Q12769 | Nuclear po |
| CD109 | 1.181763 | 0.065802 | 0.155898 | Q6YHK3 | Q6YHK3 | Q6YHK3 | CD109 anti |
| CACYBP | 1.145573 | 0.07152 | 0.097391 | Q9HB71 | Q9HB71 | Q9HB71 | Calcyclin-b |
| VPS26B | 1.128642 | 0.074363 | 0.142824 | Q4G0F5;A0 | Q4G0F5 | Q4G0F5 | Vacuolar p |
| DDI2 | 1.113414 | 0.077017 | 0.094572 | Q5TDH0;H( | Q5TDH0 | Q5TDH0 | Protein DD |
| DAZAP1 | 1.08556 | 0.082118 | 0.079489 | K7EQ55;K7 | K7EQ55;K7 | K7EQ55 | DAZ-associ |
| NDUFA12 | 1.05076 | 0.088969 | 0.099349 | Q9UI09;F8 | Q9UI09 | Q9UI09 | NADH dehy |
| GLUL | 1.049629 | 0.089201 | -0.20263 | P15104;A0 | P15104;A0 | P15104 | Glutamine |
| TIMM50 | 1.019145 | 0.095688 | 0.077643 | Q3ZCQ8;M( | Q3ZCQ8;M( | Q3ZCQ8 | Mitochond |
| HOOK3 | 1.01001 | 0.097721 | 0.191536 | Q86VS8;H( | Q86VS8 | Q86VS8 | Protein Ho |
| PRDX2 | 1.006297 | 0.098561 | -0.11733 | P32119;A6 | P32119;A6 | P32119 | Peroxiredo |

| | | | | | | | |
|---|---|---|---|---|---|---|---|
| COPG2 | 1.002963 | 0.09932 | 0.206901 | Q9UBF2 | Q9UBF2 | Q9UBF2 | Coatomer s |
| TARDBP | 0.996308 | 0.100854 | 0.264254 | Q13148;A0 | Q13148;A0 | Q13148 | TAR DNA-bi |
| HLA-DRA | 0.990493 | 0.102213 | 0.149014 | A0A0G2JM | A0A0G2JM | A0A0G2JM | HLA class II |
| ERGIC1 | 0.942776 | 0.114084 | 0.076603 | Q969X5;F2 | Q969X5 | Q969X5 | Endoplasm |
| TACO1 | 0.939223 | 0.115021 | 0.050716 | Q9BSH4 | Q9BSH4 | Q9BSH4 | Translation |
| COG5 | 0.917386 | 0.120952 | -0.18972 | Q9UP83;U3 | Q9UP83 | Q9UP83 | Conserved |
| OTUB1 | 0.892819 | 0.127992 | -0.08697 | J3KR44;Q9 | J3KR44;Q9 | J3KR44 | Ubiquitin t |
| QKI | 0.890548 | 0.128662 | -0.08508 | Q96PU8;F5 | Q96PU8 | Q96PU8 | Protein qua |
| CARM1 | 0.885133 | 0.130277 | -0.07289 | Q86X55;K7 | Q86X55;K7 | Q86X55 | Histone-arg |
| BRCC3 | 0.837831 | 0.145268 | -0.13426 | P46736;H9 | P46736;H9 | P46736 | Lys-63-spec |
| CAPZA1 | 0.796832 | 0.15965 | 0.055311 | P52907 | P52907 | P52907 | F-actin-cap |
| IGF2BP2 | 0.780419 | 0.165799 | 0.140753 | F8W930;Q | F8W930;Q | F8W930 | Insulin-like |
| CDC42 | 0.760071 | 0.173752 | 0.061017 | P60953;A0 | P60953;A0 | P60953 | Cell division |
| WDR81 | 0.75493 | 0.175821 | 0.159893 | Q562E7;A0 | Q562E7;A0 | Q562E7 | WD repeat- |
| RUFY1 | 0.743803 | 0.180384 | 0.172944 | Q96T51;H0 | Q96T51 | Q96T51 | RUN and FY |
| RBBP4 | 0.729803 | 0.186293 | -0.06835 | Q09028;H0 | Q09028;H0 | Q09028 | Histone-bir |
| NCLN | 0.724816 | 0.188445 | 0.062056 | Q969V3;K7 | Q969V3;K7 | Q969V3 | Nicalin |
| MTPAP | 0.721658 | 0.18982 | 0.205606 | Q9NVV4;Q5 | Q9NVV4 | Q9NVV4 | Poly(A) RNA |
| ERLIN1 | 0.68484 | 0.206614 | -0.07706 | O75477;B0 | O75477;B0 | O75477 | Erlin-1 |
| PSMA5 | 0.683599 | 0.207205 | 0.04094 | P28066 | P28066 | P28066 | Proteasom |
| HNRNPH1 | 0.683502 | 0.207252 | 0.060312 | P31943;G8 | P31943;G8 | P31943 | Heterogene |
| PPIA | 0.665108 | 0.216218 | -0.16955 | P62937;F8 | P62937;F8 | P62937 | Peptidyl-pr |
| CLIC2 | 0.654891 | 0.221365 | 0.119943 | O15247;A6 | O15247;A6 | O15247 | Chloride in |
| APP | 0.609503 | 0.245752 | 0.186983 | P05067;A0 | P05067;A0 | P05067 | Amyloid be |
| DSTN | 0.609251 | 0.245895 | -0.12736 | P60981;F6 | P60981;F6 | P60981 | Destrin |
| GLRX3 | 0.600548 | 0.250872 | -0.12986 | O76003 | O76003 | O76003 | Glutaredox |
| PTPN11 | 0.59712 | 0.25286 | -0.95172 | Q06124;A0 | Q06124 | Q06124 | Tyrosine-pr |
| COG1 | 0.557085 | 0.277278 | -0.10228 | A0A087WV | A0A087WV | A0A087WV | Conserved |
| MRPS25 | 0.553172 | 0.279787 | 0.072477 | P82663;E7 | P82663;E7 | P82663 | 28S ribosor |
| RAD21 | 0.551118 | 0.281113 | 0.111416 | O60216;E5 | O60216 | O60216 | Double-stra |
| RBM4 | 0.544324 | 0.285546 | -0.10299 | Q9BWF3;E9 | Q9BWF3;E9 | Q9BWF3 | RNA-bindir |
| RSL1D1 | 0.537552 | 0.290033 | 0.101177 | O76021;J3 | O76021;J3 | O76021 | Ribosomal |
| PRPF3 | 0.535315 | 0.291531 | 0.164659 | O43395 | O43395 | O43395 | U4/U6 sma |
| TMEM173 | 0.493182 | 0.321231 | 0.105701 | Q86WV6;A | Q86WV6;A | Q86WV6 | Stimulator |
| VPS4B | 0.486345 | 0.326328 | 0.080274 | O75351;K7 | O75351 | O75351 | Vacuolar pr |
| RPS6 | 0.46933 | 0.339367 | 0.058741 | P62753;A2 | P62753;A2 | P62753 | 40S ribosor |
| VPS53 | 0.46122 | 0.345764 | 0.118243 | Q5VIR6;F6 | Q5VIR6;F6 | Q5VIR6 | Vacuolar pr |
| SUMF2 | 0.456396 | 0.349626 | 0.044273 | J3KQJ1;Q8 | J3KQJ1;Q8 | J3KQJ1 | Sulfatase-m |
| PCYT1A | 0.456027 | 0.349923 | -0.11608 | C9JEJ2;P49 | C9JEJ2;P49 | C9JEJ2 | Choline-ph |
| ARPC3 | 0.451709 | 0.35342 | -0.0905 | O15145;C9 | O15145 | O15145 | Actin-relate |
| IKBKB | 0.449382 | 0.355319 | -0.12856 | A0A499FJS | A0A499FJS | A0A499FJS | Inhibitor o |
| RDH13 | 0.44082 | 0.362393 | 0.123374 | Q8NBN7;G | Q8NBN7;G | Q8NBN7 | Retinol deh |
| UFD1L | 0.429622 | 0.371859 | 0.054094 | Q92890;C9 | Q92890 | Q92890 | Ubiquitin f |
| NEU1 | 0.423579 | 0.377069 | 0.045129 | Q99519;E9 | Q99519;E9 | Q99519 | Sialidase-1 |
| CTSZ | 0.418095 | 0.381861 | 0.061387 | Q9UBR2 | Q9UBR2 | Q9UBR2 | Cathepsin Z |
| WDR61 | 0.416207 | 0.383525 | 0.060674 | Q9GZS3;H0 | Q9GZS3;H0 | Q9GZS3 | WD repeat- |
| ZFR | 0.415654 | 0.384013 | 0.106353 | Q96KR1;H0 | Q96KR1 | Q96KR1 | Zinc finger |
| NDUFB10 | 0.412926 | 0.386433 | -0.04333 | O96000;H3 | O96000;H3 | O96000 | NADH dehy |

| | | | | | | | |
|---|---|---|---|---|---|---|---|
| ATP6V0A2 | 0.410033 | 0.389015 | -0.06129 | Q9Y487;Q8 | Q9Y487;Q8 | Q9Y487 | V-type prot |
| CTBP1 | 0.408774 | 0.390145 | -0.05376 | Q13363;D6 | Q13363;D6 | Q13363 | C-terminal- |
| PRDX3 | 0.403702 | 0.394728 | 0.074778 | P30048 | P30048 | P30048 | Thioredoxi |
| HEBP1 | 0.396228 | 0.40158 | 0.127391 | Q9NRV9;A0 | Q9NRV9;A0 | Q9NRV9 | Heme-bind |
| PSMA3 | 0.364968 | 0.431551 | -0.0443 | P25788;G3 | P25788 | P25788 | Proteasome |
| PDLIM4 | 0.363343 | 0.433169 | 0.093712 | P50479;C9 | P50479 | P50479 | PDZ and LIM |
| CDKN2AIP | 0.363011 | 0.4335 | 0.090444 | Q9NXV6;J3 | Q9NXV6 | Q9NXV6 | CDKN2A-in |
| AKR7A2 | 0.35499 | 0.441581 | -0.03835 | O43488;H3 | O43488;H3 | O43488 | Aflatoxin B |
| TIMMDC1 | 0.351086 | 0.445568 | 0.050365 | Q9NPL8;C9 | Q9NPL8;C9 | Q9NPL8 | Complex I a |
| STX7 | 0.332173 | 0.4654 | 0.05663 | O15400 | O15400 | O15400 | Syntaxin-7 |
| PNKD | 0.332155 | 0.46542 | -0.03322 | Q8N490 | Q8N490 | Q8N490 | Probable hy |
| SPRYD4 | 0.325773 | 0.47231 | 0.045835 | Q8WW59 | Q8WW59 | Q8WW59 | SPRY doma |
| SPTLC1 | 0.321466 | 0.477017 | -0.08096 | O15269;A0 | O15269 | O15269 | Serine palm |
| ASAP1 | 0.321303 | 0.477196 | 0.092394 | A0A0A0MR | A0A0A0MR | A0A0A0MR | Arf-GAP wit |
| UBTF | 0.320981 | 0.477551 | 0.052092 | E9PKP7;P1 | E9PKP7;P1 | E9PKP7 | Nucleolar t |
| COPS2 | 0.307423 | 0.492693 | -0.04447 | P61201;B4 | P61201;B4 | P61201 | COP9 signa |
| LPCAT1 | 0.306055 | 0.494248 | 0.04378 | Q8NF37;A0 | Q8NF37;A0 | Q8NF37 | Lysophosph |
| YBX1 | 0.291685 | 0.510876 | -0.02415 | P67809;H0 | P67809;H0 | P67809 | Nuclease-se |
| OLA1 | 0.291253 | 0.511384 | -0.06765 | J3KQ32;Q9 | J3KQ32;Q9 | J3KQ32 | Obg-like AT |
| APOOL | 0.28347 | 0.520631 | 0.048667 | Q6UXV4;A0 | Q6UXV4;A0 | Q6UXV4 | MICOS com |
| PSTPIP2 | 0.270608 | 0.53628 | -0.06818 | Q9H939 | Q9H939 | Q9H939 | Proline-seri |
| MTX1 | 0.269518 | 0.537628 | 0.035161 | Q13505 | Q13505 | Q13505 | Metaxin-1 |
| PURB | 0.26444 | 0.543951 | 0.04911 | Q96QR8 | Q96QR8 | Q96QR8 | Transcripti |
| PPP2R2A | 0.263645 | 0.544948 | -0.04426 | P63151;A0 | P63151;A0 | P63151 | Serine/thre |
| AP3D1 | 0.251092 | 0.560929 | 0.075562 | O14617;A0 | O14617 | O14617 | AP-3 compl |
| HACD3 | 0.250747 | 0.561375 | 0.040628 | Q9P035;H3 | Q9P035;H3 | Q9P035 | Very-long-c |
| RPL24 | 0.246181 | 0.567308 | 0.037982 | C9JXB8;C9 | C9JXB8;C9 | C9JXB8 | 60S riboson |
| DECR1 | 0.228983 | 0.590224 | 0.037817 | Q16698;E5 | Q16698;E5 | Q16698 | 2,4-dienoyl |
| USE1 | 0.227477 | 0.592274 | 0.047678 | Q9NZ43;G8 | Q9NZ43;G8 | Q9NZ43 | Vesicle tran |
| NAPA | 0.219913 | 0.60268 | -0.04607 | P54920;M0 | P54920;M0 | P54920 | Alpha-solub |
| RAB32 | 0.216157 | 0.607915 | 0.042978 | Q13637 | Q13637 | Q13637 | Ras-related |
| TACC1 | 0.215585 | 0.608716 | -0.05859 | O75410;E7 | O75410;E7 | O75410 | Transformi |
| CUL4B | 0.210454 | 0.615951 | 0.042309 | A0A669KA | A0A669KA | A0A669KA | Cullin-4B |
| EMD | 0.204574 | 0.624348 | 0.04494 | P50402;Q5 | P50402;Q5 | P50402 | Emerin |
| PGRMC1 | 0.19749 | 0.634614 | 0.025912 | O00264 | O00264 | O00264 | Membrane- |
| BAG2 | 0.191953 | 0.642758 | 0.044623 | O95816 | O95816 | O95816 | BAG family |
| RRAS | 0.188245 | 0.648268 | -0.02125 | P10301 | P10301 | P10301 | Ras-related |
| PIGK | 0.185206 | 0.65282 | 0.015547 | Q92643;A6 | Q92643;A6 | Q92643 | GPI-anchor |
| HSD3B7 | 0.171784 | 0.673311 | -0.02016 | Q9H2F3;I3 | Q9H2F3;I3 | Q9H2F3 | 3 beta-hydr |
| RP2 | 0.163075 | 0.686949 | 0.019488 | O75695 | O75695 | O75695 | Protein XRP |
| DHRS7B | 0.137198 | 0.729124 | -0.01806 | A0A0C4DG | A0A0C4DG | A0A0C4DG | Dehydroger |
| PRDX5 | 0.134288 | 0.734027 | 0.019343 | P30044 | P30044 | P30044 | Peroxiredo |
| SULT1C2 | 0.130872 | 0.739824 | -0.04925 | O00338;B8 | O00338;B8 | O00338 | Sulfotransf |
| ENDOD1 | 0.114435 | 0.768361 | -0.0291 | O94919 | O94919 | O94919 | Endonuclea |
| L2HGDH | 0.098017 | 0.797964 | 0.033319 | Q9H9P8;C9 | Q9H9P8;C9 | Q9H9P8 | L-2-hydroxy |
| EIF2B2 | 0.094751 | 0.803987 | -0.02591 | P49770;H0 | P49770 | P49770 | Translation |
| SLK | 0.088599 | 0.815456 | -0.01847 | Q9H2G2 | Q9H2G2 | Q9H2G2 | STE20-like s |
| MRPS26 | 0.087163 | 0.818157 | -0.02213 | Q9BYN8 | Q9BYN8 | Q9BYN8 | 28S ribosor |

| | | | | | | | |
|---|---|---|---|---|---|---|---|
| AGPAT4 | 0.085417 | 0.821453 | -0.02585 | Q9NRZ5;H( | Q9NRZ5 | Q9NRZ5 | 1-acyl-sn-gl |
| MRPL19 | 0.077649 | 0.836278 | 0.017176 | P49406;S4 | P49406;S4 | P49406 | 39S ribosor |
| PDE12 | 0.075878 | 0.839697 | -0.01739 | Q6L8Q7;F6 | Q6L8Q7;F6 | Q6L8Q7 | 2,5-phosph |
| GOLGA4 | 0.063359 | 0.864252 | 0.023447 | H0Y6I0;Q1 | H0Y6I0;Q1 | H0Y6I0 | Golgin subf |
| GOLPH3 | 0.057768 | 0.875451 | -0.01622 | Q9H4A6;D( | Q9H4A6 | Q9H4A6 | Golgi phos |
| COPS5 | 0.052566 | 0.886002 | -0.01045 | Q92905;E5 | Q92905 | Q92905 | COP9 signa |
| AIP | 0.052283 | 0.886579 | -0.01797 | O00170;E9 | O00170;E9 | O00170 | AH recepto |
| FBL | 0.051495 | 0.888189 | -0.01932 | P22087;M( | P22087;M( | P22087 | rRNA 2-O-m |
| RPL17 | 0.049271 | 0.892749 | -0.01205 | A0A087WX | A0A087WX | A0A087WX | 60S ribosor |
| ATAD3A | 0.046481 | 0.898501 | -0.00668 | Q9NVI7;Q5 | Q9NVI7 | Q9NVI7 | ATPase fam |
| STX18 | 0.039534 | 0.912989 | 0.006946 | Q9P2W9;D | Q9P2W9;D | Q9P2W9 | Syntaxin-18 |
| LMAN2L | 0.037154 | 0.918008 | -0.01295 | Q9H0V9;F8 | Q9H0V9 | Q9H0V9 | VIP36-like |
| KIF13B | 0.03645 | 0.919496 | -0.01097 | Q9NQT8;E7 | Q9NQT8;E7 | Q9NQT8 | Kinesin-like |
| ELMOD2 | 0.035428 | 0.921663 | 0.006591 | Q8IZ81;D6 | Q8IZ81;D6 | Q8IZ81 | ELMO doma |
| H1FX | 0.033739 | 0.925255 | -0.00793 | Q92522 | Q92522 | Q92522 | Histone H1 |
| STK4 | 0.022109 | 0.950366 | -0.00782 | Q13043;F5 | Q13043;F5 | Q13043 | Serine/thre |
| SNX9 | 0.016348 | 0.963056 | -0.00327 | Q9Y5X1 | Q9Y5X1 | Q9Y5X1 | Sorting nex |
| TRMT1L | 0.012733 | 0.971106 | 0.003364 | Q7Z2T5;X6 | Q7Z2T5;X6 | Q7Z2T5 | TRMT1-like |
| NT5C2 | 0.005391 | 0.987664 | -0.00163 | P49902;Q5 | P49902;Q5 | P49902 | Cytosolic p |
| MRPS18B | 0.003124 | 0.992833 | 0.000623 | Q9Y676;A0 | Q9Y676;A0 | Q9Y676 | 28S ribosor |
| COL6A1 | 1.501288 | 0.031529 | 0.484405 | A0A087X0S | A0A087X0S | A0A087X0S | Collagen al |
| MAP7 | 1.318016 | 0.048082 | 0.373243 | Q14244;A0 | Q14244;A0 | Q14244 | Ensconsin |
| EIF2B4 | 3.875135 | 0.000133 | 0.35571 | A0A087WT | A0A087WT | A0A087WT | Translation |
| SLC38A10 | 2.017958 | 0.009595 | 0.319928 | Q9HBR0;I3 | Q9HBR0 | Q9HBR0 | Putative so |
| EIF3J | 3.892733 | 0.000128 | 0.314307 | O75822;H( | O75822 | O75822 | Eukaryotic |
| CPSF7 | 2.505249 | 0.003124 | 0.266047 | Q8N684;F5 | Q8N684;F5 | Q8N684 | Cleavage ar |
| HDGFRP2 | 1.822674 | 0.015043 | 0.265605 | Q7Z4V5;M( | Q7Z4V5 | Q7Z4V5 | Hepatoma- |
| SEPTIN6 | 2.735485 | 0.001839 | 0.254726 | B1AMS2;Q | B1AMS2;Q | B1AMS2 | Septin-6 |
| PLA2G15 | 1.876075 | 0.013302 | 0.249251 | Q8NCC3;H3 | Q8NCC3;H3 | Q8NCC3 | Group XV p |
| DGKZ | 1.315815 | 0.048326 | 0.239156 | E9PNL8;Q1 | E9PNL8;Q1 | E9PNL8 | Diacylglyce |
| RAB6A | 2.364955 | 0.004316 | 0.233097 | P20340;Q9 | P20340 | P20340 | Ras-related |
| DSCR3 | 1.954858 | 0.011095 | 0.204406 | O14972;A8 | O14972;A8 | O14972 | Down synd |
| SLIRP | 2.100815 | 0.007928 | 0.200154 | Q9GZT3;A0 | Q9GZT3;A0 | Q9GZT3 | SRA stem-lo |
| WBSCR16 | 3.001078 | 0.000998 | 0.190463 | Q96I51;A0 | Q96I51;A0 | Q96I51 | Williams-B |
| SCAMP3 | 1.913582 | 0.012202 | 0.188465 | O14828 | O14828 | O14828 | Secretory c |
| PTBP3 | 1.891448 | 0.01284 | 0.177093 | O95758;X6 | O95758 | O95758 | Polypyrimi |
| ABHD11 | 4.868882 | 1.35E-05 | 0.176086 | Q8NFV4;H7 | Q8NFV4 | Q8NFV4 | Alpha/beta |
| YWHAH | 1.540742 | 0.028791 | 0.169974 | Q04917;A2 | Q04917 | Q04917 | 14-3-3 prot |
| LSP1 | 2.99257 | 0.001017 | 0.166136 | P33241;E7 | P33241;E7 | P33241 | Lymphocyt |
| FXR1 | 1.698399 | 0.020026 | 0.147636 | P51114;B4 | P51114;B4 | P51114 | Fragile X m |
| ARF6 | 1.4039 | 0.039455 | 0.136511 | P62330 | P62330 | P62330 | ADP-ribosy |
| EIF3K | 1.340033 | 0.045705 | 0.119905 | Q9UBQ5;K7 | Q9UBQ5;K7 | Q9UBQ5 | Eukaryotic |
| STX3 | 1.586575 | 0.025907 | 0.104651 | A0A0C4DG | A0A0C4DG | A0A0C4DG | Syntaxin-3 |
| AGPAT3 | 1.408801 | 0.039012 | -0.15542 | Q9NRZ7;C9 | Q9NRZ7 | Q9NRZ7 | 1-acyl-sn-gl |
| THEM6 | 1.553703 | 0.027945 | -0.17933 | Q8WUY1;H | Q8WUY1 | Q8WUY1 | Protein THI |
| CDK5 | 1.385611 | 0.041152 | -0.18854 | Q00535;F8 | Q00535 | Q00535 | Cyclin-depe |
| MRPS2 | 1.914742 | 0.012169 | -0.1981 | Q9Y399;Q5 | Q9Y399;Q5 | Q9Y399 | 28S ribosor |
| MAPRE1 | 2.004394 | 0.009899 | -0.20149 | Q15691 | Q15691 | Q15691 | Microtubul |

| | | | | | | | |
|---|---|---|---|---|---|---|---|
| PSMB3 | 1.65885 | 0.021936 | -0.22893 | P49720;A0 | P49720;A0 | P49720 | Proteasom |
| NIPSNAP3A | 2.029116 | 0.009352 | -0.291 | Q9UFN0;Q9 | Q9UFN0 | Q9UFN0 | Protein Nip |
| CYP4X1 | 1.819237 | 0.015162 | -0.30424 | Q8N118 | Q8N118 | Q8N118 | Cytochrom |
| TIGAR | 2.257105 | 0.005532 | -0.4249 | Q9NQ88;A0 | Q9NQ88;A0 | Q9NQ88 | Fructose-2, |
| DDX3X | 1.267169 | 0.054054 | 0.128321 | A0A2U3TZJ | A0A2U3TZJ | A0A2U3TZJ | ATP-depend |
| UAP1L1 | 1.262774 | 0.054604 | -0.25057 | Q3KQV9;A0 | Q3KQV9 | Q3KQV9 | UDP-N-acet |
| CTBS | 1.247509 | 0.056558 | 0.261407 | Q01459 | Q01459 | Q01459 | Di-N-acetyl |
| MRPL1 | 1.243173 | 0.057125 | 0.174537 | Q9BYD6;H0 | Q9BYD6;H0 | Q9BYD6 | 39S riboson |
| TLDC1 | 1.19105 | 0.06441 | 0.134133 | Q6P9B6;H3 | Q6P9B6;H3 | Q6P9B6 | TLD domain |
| HLA-DMB | 1.139227 | 0.072573 | 0.138506 | P28068;A0 | P28068;A0 | P28068 | HLA class II |
| SPTBN1 | 1.083695 | 0.082472 | 0.329005 | Q01082;A0 | Q01082;A0 | Q01082 | Spectrin be |
| EIF4E | 1.075397 | 0.084063 | -0.11693 | D6RBW1;P | D6RBW1;P | D6RBW1 | Eukaryotic |
| SCARB1 | 1.07335 | 0.08446 | 0.275047 | B7ZKQ9;F5 | B7ZKQ9;F5 | B7ZKQ9 | Scavenger r |
| TK2 | 1.015661 | 0.096458 | 0.090714 | O00142;H3 | O00142;H3 | O00142 | Thymidine |
| SH3GLB1 | 1.006971 | 0.098408 | -0.14795 | Q9Y371;A0 | Q9Y371;A0 | Q9Y371 | Endophilin |
| COMMD10 | 0.986278 | 0.10321 | 0.111843 | Q9Y6G5;D6 | Q9Y6G5;D6 | Q9Y6G5 | COMM dom |
| FAHD1 | 0.964152 | 0.108605 | 0.132576 | Q6P587 | Q6P587 | Q6P587 | Acylpyruva |
| CTSL | 0.95209 | 0.111663 | 0.086376 | P07711;Q5 | P07711;Q5 | P07711 | Cathepsin L |
| ECM1 | 0.906487 | 0.124026 | 0.16261 | Q16610 | Q16610 | Q16610 | Extracellula |
| CSNK2A1 | 0.896483 | 0.126916 | 0.051671 | P68400;Q8 | P68400;Q8 | P68400 | Casein kina |
| LARP4B | 0.868784 | 0.135274 | 0.146649 | Q92615;H0 | Q92615;H0 | Q92615 | La-related p |
| NDUFB9 | 0.85773 | 0.138762 | 0.085416 | Q9Y6M9;E9 | Q9Y6M9;E9 | Q9Y6M9 | NADH dehy |
| OSTF1 | 0.850182 | 0.141194 | -0.11018 | Q92882 | Q92882 | Q92882 | Osteoclast- |
| BRIX1 | 0.842464 | 0.143726 | 0.083735 | Q8TDN6 | Q8TDN6 | Q8TDN6 | Ribosome b |
| NDUFA8 | 0.832702 | 0.146993 | -0.0946 | P51970 | P51970 | P51970 | NADH dehy |
| MBNL1 | 0.821304 | 0.150902 | 0.187142 | C9JP00;A0 | C9JP00;A0 | C9JP00 | Musclebin |
| CCDC90B | 0.798198 | 0.159148 | 0.106595 | Q9GZT6;H0 | Q9GZT6;H0 | Q9GZT6 | Coiled-coil |
| RCC1 | 0.776248 | 0.167399 | 0.152993 | P18754;C9 | P18754;C9 | P18754 | Regulator c |
| CIAO1 | 0.772454 | 0.168868 | -0.0717 | O76071 | O76071 | O76071 | Probable cy |
| MRPL46 | 0.767289 | 0.170888 | 0.103566 | Q9H2W6;A | Q9H2W6 | Q9H2W6 | 39S riboson |
| B3GALTL | 0.761327 | 0.17325 | -0.13769 | Q6Y288 | Q6Y288 | Q6Y288 | Beta-1,3-gl |
| MRPS23 | 0.753472 | 0.176412 | -0.12361 | Q9Y3D9;J3 | Q9Y3D9;J3 | Q9Y3D9 | 28S riboson |
| FDX1 | 0.743812 | 0.18038 | 0.065392 | P10109 | P10109 | P10109 | Adrenodox |
| RINT1 | 0.739783 | 0.182061 | 0.147362 | Q6NUQ1 | Q6NUQ1 | Q6NUQ1 | RAD50-inte |
| THNSL1 | 0.735051 | 0.184055 | -0.11415 | Q8IYQ7 | Q8IYQ7 | Q8IYQ7 | Threonine s |
| COMMD9 | 0.720198 | 0.190459 | 0.132027 | Q9P000;E9 | Q9P000 | Q9P000 | COMM dom |
| SETD3 | 0.715945 | 0.192333 | 0.154046 | Q86TU7;Q6 | Q86TU7;Q6 | Q86TU7 | Histone-lys |
| AP3M1 | 0.700884 | 0.199121 | 0.123545 | Q9Y2T2;E5 | Q9Y2T2 | Q9Y2T2 | AP-3 compl |
| NUDT19 | 0.699925 | 0.199561 | -0.12337 | A8MXV4 | A8MXV4 | A8MXV4 | Nucleoside |
| RPS19 | 0.675527 | 0.211093 | 0.060087 | P39019;A0 | P39019;A0 | P39019 | 40S riboson |
| CASP1 | 0.664414 | 0.216564 | -0.10778 | P29466;G3 | P29466;G3 | P29466 | Caspase-1;C |
| DHX36 | 0.663355 | 0.217092 | 0.163012 | E7EWK3;Q9 | E7EWK3;Q9 | E7EWK3 | ATP-depend |
| NAA15 | 0.653179 | 0.222239 | 0.149016 | Q9BXJ9;A0 | Q9BXJ9;A0 | Q9BXJ9 | N-alpha-ace |
| EXD2 | 0.64935 | 0.224207 | 0.132143 | Q9NVH0;C9 | Q9NVH0 | Q9NVH0 | Exonucleas |
| RPL12 | 0.648355 | 0.224722 | 0.088716 | P30050 | P30050 | P30050 | 60S riboson |
| AHSA1 | 0.622058 | 0.238749 | 0.126659 | O95433;H0 | O95433;H0 | O95433 | Activator o |
| RAB9A | 0.587318 | 0.258632 | 0.084348 | P51151;Q9 | P51151 | P51151 | Ras-related |
| MRPL45 | 0.571635 | 0.268142 | 0.102924 | A0A087X2I | A0A087X2I | A0A087X2I | 39S riboson |

| | | | | | | | |
|---|---|---|---|---|---|---|---|
| CELF2 | 0.526358 | 0.297606 | 0.059545 | V9GYD9;Q5 | V9GYD9;Q5 | V9GYD9 | CUGBP Elav |
| FKBP3 | 0.510414 | 0.308735 | 0.077217 | Q00688;G3 | Q00688 | Q00688 | Peptidyl-pr |
| NIT1 | 0.506792 | 0.311321 | 0.082151 | Q86X76 | Q86X76 | Q86X76 | Nitrilase hc |
| SUPT6H | 0.493562 | 0.32095 | -1.5932 | Q7KZ85 | Q7KZ85 | Q7KZ85 | Transcripti |
| PSMD8 | 0.491775 | 0.322274 | -0.05151 | R4GMR5;P | R4GMR5;P | R4GMR5 | 26S proteas |
| GOSR1 | 0.480448 | 0.33079 | 0.074306 | K7EJC8;E9F | K7EJC8;E9F | K7EJC8 | Golgi SNAP |
| SDF4 | 0.475915 | 0.33426 | 0.114215 | A0A5F9UP | A0A5F9UP | A0A5F9UP | 45 kDa calc |
| TMEM33 | 0.471292 | 0.337838 | 0.084854 | P57088;D6 | P57088;D6 | P57088 | Transmemb |
| MVD | 0.461986 | 0.345155 | -0.12989 | P53602;H3 | P53602;H3 | P53602 | Diphospho |
| APPL1 | 0.452478 | 0.352795 | 0.09944 | Q9UKG1;C | Q9UKG1 | Q9UKG1 | DCC-intera |
| FABP3 | 0.436452 | 0.366056 | -0.21463 | P05413;S4 | P05413;S4 | P05413 | Fatty acid-b |
| ELAC2 | 0.431609 | 0.370162 | -0.05557 | Q9BQ52;G5 | Q9BQ52;G5 | Q9BQ52 | Zinc phospl |
| NUP54 | 0.412988 | 0.386377 | 0.064265 | Q7Z3B4 | Q7Z3B4 | Q7Z3B4 | Nucleopori |
| SYMPK | 0.392314 | 0.405216 | 0.107581 | Q92797;A0 | Q92797;A0 | Q92797 | Symplekin |
| PGP | 0.382844 | 0.414149 | -0.16816 | A6NDG6;H | A6NDG6 | A6NDG6 | Phosphogly |
| MT-CO2 | 0.37378 | 0.422883 | -0.05224 | P00403 | P00403 | P00403 | Cytochrom |
| EMC8 | 0.362668 | 0.433843 | -0.05402 | O43402;M | O43402;M | O43402 | ER membra |
| LMF2 | 0.3554 | 0.441164 | -0.07886 | Q9BU23 | Q9BU23 | Q9BU23 | Lipase matu |
| DDX18 | 0.354171 | 0.442414 | 0.078729 | Q9NVP1;H | Q9NVP1 | Q9NVP1 | ATP-depend |
| ABCB8 | 0.352116 | 0.444512 | 0.07254 | Q9NUT2;C | Q9NUT2 | Q9NUT2 | ATP-bindin |
| ATP5H | 0.34413 | 0.452762 | -0.04561 | O75947;F5 | O75947 | O75947 | ATP syntha |
| IFIT3 | 0.34403 | 0.452866 | -0.1015 | O14879 | O14879 | O14879 | Interferon-i |
| MRPS31 | 0.325346 | 0.472775 | 0.072457 | Q92665 | Q92665 | Q92665 | 28S ribosor |
| LPP | 0.31919 | 0.479524 | -0.09894 | Q93052;A0 | Q93052;A0 | Q93052 | Lipoma-pre |
| SMU1 | 0.317297 | 0.481618 | 0.076614 | Q2TAY7 | Q2TAY7 | Q2TAY7 | WD40 repe |
| NPL | 0.299851 | 0.501359 | -0.14614 | Q9BXD5;A | Q9BXD5;A | Q9BXD5 | N-acetylneu |
| HNRNPDL | 0.280337 | 0.5244 | -0.09954 | A0A087WU | A0A087WU | A0A087WU | Heterogene |
| GALE | 0.279868 | 0.524967 | -0.07908 | Q14376;Q5 | Q14376;Q5 | Q14376 | UDP-glucos |
| DRG1 | 0.278407 | 0.526736 | -0.07664 | Q9Y295;H0 | Q9Y295 | Q9Y295 | Developme |
| HS2ST1 | 0.276935 | 0.528525 | -0.05922 | Q7LGA3;K7 | Q7LGA3 | Q7LGA3 | Heparan su |
| ACTR1A | 0.276313 | 0.529282 | -0.06009 | P61163;R4 | P61163;R4 | P61163 | Alpha-centr |
| PET112 | 0.2763 | 0.529297 | 0.099396 | D6REA0;O7 | D6REA0;O7 | D6REA0 | Glutamyl-t |
| TRAPPC11 | 0.275396 | 0.530401 | 0.073314 | Q7Z392;D6 | Q7Z392;D6 | Q7Z392 | Trafficking |
| UBAP2L | 0.272361 | 0.534121 | 0.030229 | F8W726;Q | F8W726;Q | F8W726 | Ubiquitin-a |
| RAB12 | 0.27051 | 0.536401 | 0.04391 | A0A3B3ITT | A0A3B3ITT | A0A3B3ITT | Ras-related |
| PITPNA | 0.260551 | 0.548844 | -0.08845 | F5GWE5;Q | F5GWE5;Q | F5GWE5 | Phosphatic |
| HDAC1 | 0.245898 | 0.567678 | 0.034692 | Q13547;Q5 | Q13547;Q5 | Q13547 | Histone dea |
| ARMC10 | 0.240024 | 0.575408 | 0.054884 | Q8N2F6;A0 | Q8N2F6;A0 | Q8N2F6 | Armadillo r |
| RTCA | 0.232721 | 0.585166 | -0.06551 | O00442;A6 | O00442 | O00442 | RNA 3-term |
| BAG6 | 0.232505 | 0.585457 | 0.039185 | P46379;A0 | P46379;A0 | P46379 | Large prolir |
| THUMPD1 | 0.230072 | 0.588746 | -0.07301 | Q9NXG2;H | Q9NXG2 | Q9NXG2 | THUMP dor |
| PRRC1 | 0.22653 | 0.593567 | -0.0555 | Q96M27 | Q96M27 | Q96M27 | Protein PRI |
| DIABLO | 0.221284 | 0.600781 | -0.04196 | Q9NR28;H | Q9NR28 | Q9NR28 | Diablo hom |
| PAICS | 0.221105 | 0.601028 | -0.06294 | P22234;E9 | P22234;E9 | P22234 | Multifuncti |
| RPL32 | 0.218633 | 0.60446 | 0.039643 | D3YTB1;F8 | D3YTB1;F8 | D3YTB1 | 60S ribosor |
| PSMD4 | 0.218198 | 0.605065 | -0.0191 | P55036;Q5 | P55036;Q5 | P55036 | 26S proteas |
| SAE1 | 0.213482 | 0.611672 | 0.064562 | Q9UBE0;M | Q9UBE0;M | Q9UBE0 | SUMO-activ |
| ASNA1 | 0.212577 | 0.612946 | 0.042617 | O43681 | O43681 | O43681 | ATPase ASN |

| | | | | | | | |
|---|---|---|---|---|---|---|---|
| PSMA2 | 0.194692 | 0.638716 | -0.02713 | A0A024RA! | A0A024RA! | A0A024RA! | Proteasom |
| GALM | 0.190046 | 0.645585 | -0.09583 | Q96C23;H7 | Q96C23;H7 | Q96C23 | Aldose 1-ep |
| SIAE | 0.187272 | 0.649722 | 0.04667 | Q9HAT2 | Q9HAT2 | Q9HAT2 | Sialate O-ac |
| NAGA | 0.184566 | 0.653783 | 0.048389 | P17050 | P17050 | P17050 | Alpha-N-ac |
| PELP1 | 0.183135 | 0.655941 | 0.049273 | Q8IZL8;C9J | Q8IZL8;C9J | Q8IZL8 | Proline-, gl |
| SET | 0.181825 | 0.657923 | -0.04925 | A0A0C4DF\ | A0A0C4DF\ | A0A0C4DF\ | Protein SET |
| DECR2 | 0.181437 | 0.65851 | 0.052997 | Q9NUI1;A0 | Q9NUI1;A0 | Q9NUI1 | Peroxisoma |
| BAZ1B | 0.174511 | 0.669097 | 0.090193 | Q9UIG0 | Q9UIG0 | Q9UIG0 | Tyrosine-pr |
| NAT10 | 0.17127 | 0.674108 | -0.05521 | Q9H0A0;A( | Q9H0A0;A( | Q9H0A0 | N-acetyltra |
| LRCH1 | 0.168964 | 0.677697 | -0.03459 | Q9Y2L9;C9 | Q9Y2L9 | Q9Y2L9 | Leucine-ric |
| RPL9 | 0.16581 | 0.682637 | -0.03071 | P32969;A0 | P32969;A0 | P32969 | 60S ribosor |
| TP53I3 | 0.162384 | 0.688044 | -0.07963 | Q53FA7;H7 | Q53FA7 | Q53FA7 | Quinone ox |
| ECI1 | 0.161947 | 0.688736 | -0.02615 | P42126;Q9 | P42126;Q9 | P42126 | Enoyl-CoA ( |
| COX4I1 | 0.134072 | 0.734393 | -0.01894 | P13073;H3 | P13073;H3 | P13073 | Cytochrom |
| SPCS2 | 0.131565 | 0.738643 | -0.0206 | A0A087WL | A0A087WL | A0A087WL | Signal pept |
| ARSA | 0.129677 | 0.741862 | -0.0245 | A0A0C4DF/ | A0A0C4DF/ | A0A0C4DF/ | Arylsulfatas |
| EDF1 | 0.126705 | 0.746955 | -0.02209 | O60869 | O60869 | O60869 | Endothelial |
| TUBB6 | 0.126137 | 0.747933 | 0.022041 | Q9BUF5;K7 | Q9BUF5;K7 | Q9BUF5 | Tubulin bet |
| LAMP1 | 0.122648 | 0.753966 | -0.02444 | P11279 | P11279 | P11279 | Lysosome-a |
| STK24 | 0.112297 | 0.772153 | -0.03014 | B4DR80;Q9 | B4DR80;Q9 | B4DR80 | Serine/thre |
| METTL7B | 0.111453 | 0.773654 | 0.029205 | A0A087WZ | A0A087WZ | A0A087WZ | Methyltran |
| STX8 | 0.109577 | 0.777004 | 0.022073 | Q9UNK0;I3 | Q9UNK0;I3 | Q9UNK0 | Syntaxin-8 |
| TMF1 | 0.104399 | 0.786323 | -0.03726 | P82094;A0 | P82094;A0 | P82094 | TATA eleme |
| NPEPL1 | 0.100291 | 0.793796 | 0.037202 | Q8NDH3;H | Q8NDH3 | Q8NDH3 | Probable ar |
| RTN4IP1 | 0.09352 | 0.806269 | 0.025009 | Q8WWV3;( | Q8WWV3;( | Q8WWV3 | Reticulon-4 |
| YWHAQ | 0.082455 | 0.827075 | 0.025598 | P27348;E9 | P27348;E9 | P27348 | 14-3-3 prot |
| COX5A | 0.080926 | 0.829992 | 0.019787 | P20674;H3 | P20674;H3 | P20674 | Cytochrom |
| CELF1 | 0.08005 | 0.831668 | -0.02243 | G5EA30;Q9 | G5EA30;Q9 | G5EA30 | CUGBP Elav |
| HIBCH | 0.078266 | 0.835091 | -0.01297 | Q6NVY1;B8 | Q6NVY1 | Q6NVY1 | 3-hydroxyis |
| SF3A2 | 0.076724 | 0.838061 | -0.01624 | Q15428;K7 | Q15428;K7 | Q15428 | Splicing fac |
| RPS17 | 0.07413 | 0.843082 | 0.015818 | H3BNC9;P( | H3BNC9;P( | H3BNC9 | 40S ribosor |
| TWF1 | 0.074085 | 0.843169 | -0.03051 | Q12792;F8 | Q12792;F8 | Q12792 | Twinfilin-1 |
| M6PR | 0.07401 | 0.843316 | 0.010694 | P20645;F5 | P20645;F5 | P20645 | Cation-dep |
| LIPA | 0.067797 | 0.855467 | 0.018421 | A0A0A0MT | A0A0A0MT | A0A0A0MT | Lysosomal a |
| COQ6 | 0.059253 | 0.872462 | 0.011967 | Q9Y2Z9;A0 | Q9Y2Z9;A0 | Q9Y2Z9 | Ubiquinone |
| TEX264 | 0.054649 | 0.88176 | -0.01218 | Q9Y6I9;C9J | Q9Y6I9;C9J | Q9Y6I9 | Testis-expre |
| TMED10 | 0.050218 | 0.890804 | -0.01217 | P49755;G3 | P49755 | P49755 | Transmemb |
| NAPG | 0.046381 | 0.898708 | 0.015031 | Q99747;J3 | Q99747 | Q99747 | Gamma-sol |
| MRPS30 | 0.042458 | 0.906863 | -0.01867 | Q9NP92 | Q9NP92 | Q9NP92 | 28S ribosor |
| OXSR1 | 0.036863 | 0.918621 | 0.008272 | O95747;C9 | O95747;C9 | O95747 | Serine/thre |
| LYZ | 0.030656 | 0.931845 | 0.004885 | P61626;F8 | P61626;F8 | P61626 | Lysozyme C |
| ARMC8 | 0.024568 | 0.945001 | 0.005415 | Q8IUR7;B7 | Q8IUR7;B7 | Q8IUR7 | Armadillo r |
| PRMT5 | 0.023475 | 0.947381 | -0.00804 | O14744;G3 | O14744 | O14744 | Protein arg |
| GNAI3 | 0.013846 | 0.968622 | -0.00398 | P08754;A0 | P08754 | P08754 | Guanine nu |
| LEMD3 | 0.011437 | 0.97401 | 0.003442 | Q9Y2U8 | Q9Y2U8 | Q9Y2U8 | Inner nucle |
| GORASP2 | 0.009398 | 0.978592 | -0.00173 | Q9H8Y8;F8 | Q9H8Y8 | Q9H8Y8 | Golgi reasse |
| PLXDC2 | 0.007379 | 0.983153 | 0.001657 | Q6UX71 | Q6UX71 | Q6UX71 | Plexin dom |
| PLEKHM2 | 0.003322 | 0.992381 | 0.000989 | A0A2R8Y57 | A0A2R8Y57 | A0A2R8Y57 | Pleckstrin h |

| | | | | | | | |
|---|---|---|---|---|---|---|---|
| GIT1 | 0.001471 | 0.996618 | 0.000633 | J3QRU8;A0 | J3QRU8;A0 | J3QRU8 | ARF GTPase |
| NAGLU | 1.504205 | 0.031318 | 0.390259 | P54802;K7 | P54802 | P54802 | Alpha-N-ac |
| EIF4A2 | 4.360604 | 4.36E-05 | 0.327392 | Q14240;E7 | Q14240;E7 | Q14240 | Eukaryotic |
| FAM162A | 2.877468 | 0.001326 | 0.285454 | Q96A26;F8 | Q96A26;F8 | Q96A26 | Protein FAI |
| SCO1 | 2.331879 | 0.004657 | 0.284974 | O75880;J3 | O75880;J3 | O75880 | Protein SC |
| MANF | 4.151743 | 7.05E-05 | 0.264402 | P55145;H7 | P55145 | P55145 | Mesenceph |
| GOPC | 2.036218 | 0.0092 | 0.263743 | Q9HD26;F5 | Q9HD26;F5 | Q9HD26 | Golgi-assoc |
| TGOLN2 | 1.804173 | 0.015697 | 0.260541 | A0A5H1ZRI | A0A5H1ZRI | A0A5H1ZRI | Trans-Golgi |
| RNASET2 | 1.578365 | 0.026402 | 0.25957 | D6RHI9;A0 | D6RHI9;A0 | D6RHI9 | Ribonuclea |
| DHTKD1 | 3.33721 | 0.00046 | 0.256292 | Q96HY7;H7 | Q96HY7 | Q96HY7 | Probable 2- |
| SEC16A | 1.601154 | 0.025052 | 0.253467 | A0A3F2YN› | A0A3F2YN› | A0A3F2YN› | Protein tra |
| PRKAG1 | 1.712554 | 0.019384 | 0.24526 | F8VYY9;P5 | F8VYY9;P5 | F8VYY9 | 5-AMP-acti |
| ITFG1 | 3.626821 | 0.000236 | 0.240601 | Q8TB96;F5 | Q8TB96;F5 | Q8TB96 | T-cell immu |
| MMS19 | 1.749169 | 0.017817 | 0.233912 | Q96T76;H0 | Q96T76 | Q96T76 | MMS19 nu |
| MARCKS | 1.459219 | 0.034736 | 0.217647 | P29966 | P29966 | P29966 | Myristoylat |
| SLC15A4 | 2.665841 | 0.002159 | 0.216075 | Q8N697;B6 | Q8N697 | Q8N697 | Solute carri |
| PNPLA8 | 1.452731 | 0.035259 | 0.199064 | A0A0C4DG | A0A0C4DG | A0A0C4DG | Calcium-in |
| SAP18 | 1.36455 | 0.043197 | 0.198333 | O00422;H7 | O00422;H7 | O00422 | Histone dea |
| IGSF6 | 1.50169 | 0.0315 | 0.196484 | O95976;H3 | O95976 | O95976 | Immunoglo |
| SLC30A9 | 1.754484 | 0.0176 | 0.174714 | Q6PML9;D | Q6PML9 | Q6PML9 | Zinc transp |
| RPL28 | 2.451727 | 0.003534 | 0.156943 | H0YKD8;P4 | H0YKD8;P4 | H0YKD8 | 60S riboso |
| MSRA | 2.113329 | 0.007703 | 0.148173 | Q9UJ68;E5 | Q9UJ68;E5 | Q9UJ68 | Mitochond |
| LOH12CR1 | 1.910262 | 0.012295 | 0.141821 | G3V1P3;Q | G3V1P3;Q | G3V1P3 | Loss of hete |
| PDLIM7 | 1.325213 | 0.047292 | 0.136878 | Q9NR12;D6 | Q9NR12 | Q9NR12 | PDZ and LI |
| CD36 | 1.798714 | 0.015896 | 0.133409 | E7EU05;P1 | E7EU05;P1 | E7EU05 | Platelet gly |
| EEF1E1 | 1.579805 | 0.026314 | 0.133224 | O43324;D6 | O43324;D6 | O43324 | Eukaryotic |
| EVI2B | 1.38562 | 0.041151 | 0.086024 | P34910 | P34910 | P34910 | Protein EVI |
| PEF1 | 1.312445 | 0.048703 | -0.14337 | Q9UBV8 | Q9UBV8 | Q9UBV8 | Peflin |
| C11orf54 | 1.67369 | 0.021199 | -0.37352 | Q9H0W9;E | Q9H0W9;E | Q9H0W9 | Ester hydro |
| ADPRHL2 | 2.367147 | 0.004294 | -0.3798 | Q9NX46 | Q9NX46 | Q9NX46 | Poly(ADP-ri |
| PBDC1 | 1.283541 | 0.052055 | -0.19782 | Q9BVG4;A6 | Q9BVG4 | Q9BVG4 | Protein PB |
| LPGAT1 | 1.267383 | 0.054028 | -0.2022 | Q92604 | Q92604 | Q92604 | Acyl-CoA:ly |
| RPL19 | 1.233084 | 0.058468 | -0.08422 | J3QR09;J3I | J3QR09;J3I | J3QR09 | Ribosomal |
| OTUD4 | 1.222823 | 0.059865 | 0.225309 | Q01804;D6 | Q01804 | Q01804 | OTU domai |
| THOC6 | 1.206674 | 0.062134 | 0.219212 | Q86W42 | Q86W42 | Q86W42 | THO compl |
| COX6B1 | 1.189474 | 0.064644 | 0.093224 | P14854;A0 | P14854;A0 | P14854 | Cytochrom |
| CSTF1 | 1.148721 | 0.071003 | 0.202523 | Q05048;A0 | Q05048;A0 | Q05048 | Cleavage sti |
| LCP2 | 1.133904 | 0.073468 | -0.12932 | Q13094;A0 | Q13094;A0 | Q13094 | Lymphocyt |
| PPIF | 1.110139 | 0.0776 | 0.167761 | P30405;R4 | P30405 | P30405 | Peptidyl-pr |
| TCF25 | 1.073461 | 0.084438 | -0.23934 | Q9BQ70;H3 | Q9BQ70;H3 | Q9BQ70 | Transcripti |
| PEX11B | 1.040411 | 0.091115 | -0.07903 | O96011;H7 | O96011 | O96011 | Peroxisoma |
| TPMT | 1.022412 | 0.09497 | -0.2734 | P51580 | P51580 | P51580 | Thiopurine |
| ACTL6A | 1.019393 | 0.095633 | -0.16485 | O96019;H7 | O96019;H7 | O96019 | Actin-like p |
| HIP1 | 1.007232 | 0.098348 | 1.565696 | O00291 | O00291 | O00291 | Huntingtin |
| FAM98B | 1.005852 | 0.098662 | 0.098558 | Q52LJ0;H0 | Q52LJ0 | Q52LJ0 | Protein FAI |
| PLEKHF2 | 0.993609 | 0.101482 | 0.188319 | Q9H8W4 | Q9H8W4 | Q9H8W4 | Pleckstrin h |
| RAB13 | 0.98409 | 0.103731 | -0.10668 | P51153;A0 | P51153 | P51153 | Ras-related |
| HNRNPD | 0.977044 | 0.105428 | 0.077896 | Q14103;H0 | Q14103;H0 | Q14103 | Heterogene |

| | | | | | | | |
|---|---|---|---|---|---|---|---|
| SH3BGRL | 0.962165 | 0.109103 | -0.23994 | O75368 | O75368 | O75368 | SH3 domai |
| PDCD11 | 0.96149 | 0.109272 | 0.148142 | A0A3B3IUD | A0A3B3IUD | A0A3B3IUD | Protein RRP |
| MTX2 | 0.94557 | 0.113352 | -0.13434 | O75431;C9 | O75431;C9 | O75431 | Metaxin-2 |
| TTN | 0.939412 | 0.114971 | 0.622825 | A0A0A0MT | A0A0A0MT | A0A0A0MT | Titin |
| MRPS35 | 0.920433 | 0.120107 | 0.133356 | P82673;H0 | P82673 | P82673 | 28S riboso |
| RAB43 | 0.903875 | 0.124774 | -0.13911 | Q86YS6;C9 | Q86YS6 | Q86YS6 | Ras-related |
| GALK1 | 0.885821 | 0.13007 | 0.158923 | P51570;K7 | P51570 | P51570 | Galactokin |
| NAP1L1 | 0.879266 | 0.132049 | -0.05983 | P55209;F8 | P55209;F8 | P55209 | Nucleosom |
| HCCS | 0.869751 | 0.134974 | 0.110387 | P53701 | P53701 | P53701 | Cytochrom |
| PTPMT1 | 0.861313 | 0.137622 | 0.146346 | Q8WUK0;E | Q8WUK0 | Q8WUK0 | Phosphatic |
| GANAB | 0.859046 | 0.138342 | 0.077384 | Q14697;F5 | Q14697;F5 | Q14697 | Neutral alp |
| PPP2R4 | 0.809539 | 0.155046 | -0.20648 | Q15257;F6 | Q15257;F6 | Q15257 | Serine/thre |
| NOP2 | 0.804323 | 0.156919 | 0.132267 | P46087;F5 | P46087 | P46087 | Probable 28 |
| PLA2G4C | 0.803846 | 0.157092 | 0.154156 | Q9UP65;M | Q9UP65 | Q9UP65 | Cytosolic p |
| AGPAT2 | 0.792048 | 0.161418 | -0.12497 | O15120 | O15120 | O15120 | 1-acyl-sn-gl |
| TMEM30A | 0.784681 | 0.164179 | 0.093097 | Q9NV96;A0 | Q9NV96;A0 | Q9NV96 | Cell cycle c |
| NENF | 0.780164 | 0.165896 | 0.094052 | Q9UMX5 | Q9UMX5 | Q9UMX5 | Neudesin |
| BCAS2 | 0.772074 | 0.169015 | -0.10692 | O75934 | O75934 | O75934 | Pre-mRNA- |
| UQCRB | 0.758586 | 0.174347 | -0.06785 | P14927;B7 | P14927;B7 | P14927 | Cytochrom |
| NDUFA13 | 0.745126 | 0.179835 | -0.09414 | Q9P0J0;B4 | Q9P0J0;B4 | Q9P0J0 | NADH dehy |
| SPINT1 | 0.738155 | 0.182745 | -0.09369 | O43278;H3 | O43278;H3 | O43278 | Kunitz-type |
| CPSF6 | 0.738115 | 0.182762 | 0.117268 | F8WJN3;Q1 | F8WJN3;Q1 | F8WJN3 | Cleavage ar |
| RARS2 | 0.736716 | 0.183352 | -0.18904 | Q5T160;H0 | Q5T160 | Q5T160 | Probable ar |
| SUGT1 | 0.730762 | 0.185882 | 0.083287 | Q9Y2Z0 | Q9Y2Z0 | Q9Y2Z0 | Suppressor |
| SLC29A3 | 0.728874 | 0.186692 | 0.075389 | Q9BZD2;A0 | Q9BZD2;A0 | Q9BZD2 | Equilibrativ |
| BSG | 0.724328 | 0.188657 | -0.07128 | P35613;A0 | P35613;A0 | P35613 | Basigin |
| OXSM | 0.718587 | 0.191167 | 0.114517 | Q9NWU1;C | Q9NWU1 | Q9NWU1 | 3-oxoacyl-[ |
| MTAP | 0.696409 | 0.201183 | -0.17223 | B4DUC8;Q1 | B4DUC8;Q1 | B4DUC8 | S-methyl-5- |
| TMED7 | 0.686399 | 0.205874 | 0.078654 | Q9Y3B3;A0 | Q9Y3B3;A0 | Q9Y3B3 | Transmemb |
| SNW1 | 0.684516 | 0.206768 | 0.141376 | Q13573;G3 | Q13573;G3 | Q13573 | SNW doma |
| NCSTN | 0.682902 | 0.207538 | 0.092754 | Q92542;H0 | Q92542;H0 | Q92542 | Nicastrin |
| GNB2L1 | 0.677464 | 0.210153 | 0.064053 | P63244;J3 | P63244;J3 | P63244 | Guanine nu |
| BCAT1 | 0.676795 | 0.210477 | -0.19534 | P54687;F5 | P54687;F5 | P54687 | Branched-c |
| RPS18 | 0.673327 | 0.212164 | 0.097102 | P62269;A0 | P62269 | P62269 | 40S riboso |
| NME3 | 0.672627 | 0.212507 | 0.123608 | Q13232;H3 | Q13232;H3 | Q13232 | Nucleoside |
| FCGR1A | 0.668984 | 0.214297 | 0.09933 | P12314;A0 | P12314;A0 | P12314 | High affinit |
| ABHD3 | 0.666945 | 0.215306 | 0.134004 | Q8WU67;J1 | Q8WU67;J1 | Q8WU67 | Phospholip |
| NNT | 0.657427 | 0.220076 | 0.068018 | Q13423;A0 | Q13423;A0 | Q13423 | NAD(P) tran |
| IDO1 | 0.653908 | 0.221867 | -0.83371 | P14902;E5 | P14902 | P14902 | Indoleamir |
| MRAS | 0.651114 | 0.223298 | -0.10768 | O14807;C9 | O14807 | O14807 | Ras-related |
| GDAP1 | 0.630161 | 0.234336 | 0.051794 | Q8TB36;E5 | Q8TB36 | Q8TB36 | Ganglioside |
| GDAP2 | 0.607361 | 0.246967 | 0.077038 | Q9NXN4 | Q9NXN4 | Q9NXN4 | Ganglioside |
| TPT1 | 0.587072 | 0.258778 | -0.1087 | Q5W0H4;A | Q5W0H4;A | Q5W0H4 | Translation |
| GNB4 | 0.582963 | 0.261238 | -0.07589 | Q9HAV0;C9 | Q9HAV0 | Q9HAV0 | Guanine nu |
| RPL31 | 0.574748 | 0.266227 | -0.04802 | P62899;H7 | P62899;H7 | P62899 | 60S riboso |
| NFKB1 | 0.573179 | 0.26719 | 0.138291 | A0A494C1I | A0A494C1I | A0A494C1I | Nuclear fac |
| SPP1 | 0.571501 | 0.268225 | 0.171291 | P10451;D6 | P10451 | P10451 | Osteoponti |
| MAPK3 | 0.564561 | 0.272545 | -0.14463 | P27361;E9 | P27361;E9 | P27361 | Mitogen-ac |

| | | | | | | | |
|---|---|---|---|---|---|---|---|
| RAB5C | 0.560408 | 0.275165 | -0.05618 | P51148;K7 | P51148;K7 | P51148 | Ras-related |
| CSTB | 0.557205 | 0.277201 | -0.15178 | P04080;A0 | P04080;A0 | P04080 | Cystatin-B |
| BICD2 | 0.548839 | 0.282593 | 0.171345 | Q8TD16;A( | Q8TD16 | Q8TD16 | Protein bic |
| PTDSS1 | 0.548781 | 0.282631 | 0.092568 | P48651 | P48651 | P48651 | Phosphatic |
| NUP62 | 0.53532 | 0.291528 | -0.08612 | P37198;M( | P37198;M( | P37198 | Nuclear po |
| RAB31 | 0.53113 | 0.294354 | 0.062852 | Q13636;J3 | Q13636 | Q13636 | Ras-related |
| SLC30A1 | 0.526483 | 0.297521 | -0.08036 | Q9Y6M5 | Q9Y6M5 | Q9Y6M5 | Zinc transp |
| GSTZ1 | 0.525921 | 0.297906 | 0.097161 | A0A0C4DFI | A0A0C4DFI | A0A0C4DFI | Maleylacet |
| PTPRE | 0.513135 | 0.306807 | -0.10688 | P23469;S4 | P23469;S4 | P23469 | Receptor-ty |
| RPS12 | 0.491356 | 0.322585 | 0.069005 | P25398 | P25398 | P25398 | 40S riboso |
| OS9 | 0.490087 | 0.323529 | 0.131097 | Q13438;B4 | Q13438;B4 | Q13438 | Protein OS- |
| VPS29 | 0.484948 | 0.32738 | 0.114799 | Q9UBQ0;F8 | Q9UBQ0;F8 | Q9UBQ0 | Vacuolar pr |
| KCTD12 | 0.471267 | 0.337857 | -0.15726 | Q96CX2;Q6 | Q96CX2 | Q96CX2 | BTB/POZ do |
| COMMD8 | 0.466165 | 0.341849 | 0.096879 | Q9NX08 | Q9NX08 | Q9NX08 | COMM dom |
| CCDC51 | 0.443476 | 0.360183 | 0.126867 | Q96ER9;C9 | Q96ER9 | Q96ER9 | Coiled-coil |
| RPL21 | 0.442604 | 0.360908 | -0.08304 | P46778;G3 | P46778;G3 | P46778 | 60S riboso |
| AIMP2 | 0.439941 | 0.363127 | 0.091221 | Q13155;A8 | Q13155;A8 | Q13155 | Aminoacyl |
| PPCS | 0.437739 | 0.364973 | -0.10839 | Q9HAB8;Q5 | Q9HAB8 | Q9HAB8 | Phosphopa |
| LDAH | 0.432225 | 0.369636 | 0.142752 | B5MDU6;Q | B5MDU6;Q | B5MDU6 | UPF0554 p |
| GNPDA1 | 0.412613 | 0.386712 | -0.10262 | P46926;D6 | P46926;D6 | P46926 | Glucosamir |
| PPP3CA | 0.411988 | 0.387268 | -0.10475 | Q08209;E7 | Q08209;E7 | Q08209 | Serine/thre |
| RAB1A | 0.395664 | 0.402102 | -0.06704 | P62820;E7 | P62820;E7 | P62820 | Ras-related |
| UBE2J1 | 0.392852 | 0.404714 | -0.04277 | Q9Y385 | Q9Y385 | Q9Y385 | Ubiquitin-c |
| NDUFV2 | 0.392846 | 0.40472 | 0.039921 | E7EPT4;P1 | E7EPT4;P1 | E7EPT4 | NADH dehy |
| CSRP1 | 0.373731 | 0.42293 | 0.091818 | P21291;E9 | P21291;E9 | P21291 | Cysteine an |
| COTL1 | 0.371445 | 0.425163 | -0.08102 | Q14019;H3 | Q14019;H3 | Q14019 | Coactosin-l |
| PRKCB | 0.37032 | 0.426265 | -0.06256 | P05771;I3I | P05771 | P05771 | Protein kin |
| SLC16A3 | 0.369363 | 0.427205 | 0.059486 | O15427;J3 | O15427;J3 | O15427 | Monocarbo |
| SPN | 0.360467 | 0.436046 | 0.038555 | P16150;C9 | P16150;C9 | P16150 | Leukosialin |
| COMMD3 | 0.355772 | 0.440786 | 0.0679 | Q9UBI1;R4 | Q9UBI1;R4 | Q9UBI1 | COMM dom |
| SARNP | 0.355704 | 0.440856 | 0.046132 | P82979;H0 | P82979;H0 | P82979 | SAP domair |
| SLC25A22 | 0.350316 | 0.446359 | 0.084647 | Q9H936;E9 | Q9H936;E9 | Q9H936 | Mitochond |
| NDUFS8 | 0.349543 | 0.447153 | 0.042131 | O00217;E9 | O00217;E9 | O00217 | NADH dehy |
| ERGIC3 | 0.342802 | 0.454148 | 0.066084 | H0Y5K5;Q9 | H0Y5K5;Q9 | H0Y5K5 | Endoplasm |
| TCOF1 | 0.340964 | 0.456075 | -0.51373 | Q13428;A0 | Q13428;A0 | Q13428 | Treacle pro |
| ACTR10 | 0.319986 | 0.478645 | -0.06785 | Q9NZ32;V9 | Q9NZ32;V9 | Q9NZ32 | Actin-relate |
| EXOSC10 | 0.319613 | 0.479057 | 0.499558 | Q01780;K7 | Q01780 | Q01780 | Exosome co |
| PPP1R10 | 0.311142 | 0.488492 | 0.119832 | Q96QC0 | Q96QC0 | Q96QC0 | Serine/thre |
| PARS2 | 0.300907 | 0.500142 | -0.06448 | Q7L3T8 | Q7L3T8 | Q7L3T8 | Probable pr |
| BLVRB | 0.285231 | 0.518524 | 0.069923 | P30043;A0 | P30043;A0 | P30043 | Flavin redu |
| ENO2 | 0.285104 | 0.518676 | -0.08184 | P09104;F5 | P09104;F5 | P09104 | Gamma-en |
| SIL1 | 0.275703 | 0.530025 | -0.05826 | D6REA1;Q9 | D6REA1;Q9 | D6REA1 | Nucleotide |
| RAD23B | 0.270923 | 0.535892 | -0.04083 | P54727;Q5 | P54727 | P54727 | UV excisior |
| DDX21 | 0.270351 | 0.536598 | 0.097696 | Q9NR30 | Q9NR30 | Q9NR30 | Nucleolar R |
| PEX3 | 0.267592 | 0.540018 | -0.03504 | P56589;Q7 | P56589 | P56589 | Peroxisoma |
| U2AF1 | 0.263914 | 0.54461 | 0.054242 | P0DN76;Q( | P0DN76;Q( | P0DN76 | Splicing fac |
| GALNT6 | 0.261274 | 0.547931 | 0.087218 | Q8NCL4;S4 | Q8NCL4;S4 | Q8NCL4 | Polypeptid |
| CYCS | 0.255901 | 0.554752 | -0.06419 | C9JFR7;P9 | C9JFR7;P9 | C9JFR7 | Cytochrom |

| | | | | | | | |
|---|---|---|---|---|---|---|---|
| CBR3 | 0.250432 | 0.561782 | -0.07286 | O75828 | O75828 | O75828 | Carbonyl re |
| HNRNPH3 | 0.24404 | 0.570111 | -0.03137 | P31942 | P31942 | P31942 | Heterogene |
| MBOAT7 | 0.237913 | 0.578212 | -0.04708 | Q96N66;A9 | Q96N66 | Q96N66 | Lysophosph |
| TXLNA | 0.235736 | 0.581118 | -0.07332 | P40222 | P40222 | P40222 | Alpha-taxili |
| MRPL13 | 0.231535 | 0.586766 | -0.03199 | Q9BYD1;E5 | Q9BYD1;E5 | Q9BYD1 | 39S riboso |
| LGALS9 | 0.229797 | 0.589119 | -0.03487 | O00182;J3 | O00182 | O00182 | Galectin-9 |
| GYG1 | 0.22961 | 0.589372 | 0.046798 | P46976;C9 | P46976;C9 | P46976 | Glycogenin |
| SQLE | 0.223175 | 0.59817 | -0.04947 | E7EVQ6;Q1 | E7EVQ6;Q1 | E7EVQ6 | Squalene m |
| MRPS10 | 0.216854 | 0.60694 | -0.04932 | P82664 | P82664 | P82664 | 28S riboso |
| COQ9 | 0.216786 | 0.607036 | 0.030412 | O75208;H3 | O75208;H3 | O75208 | Ubiquinon |
| LEPRE1 | 0.212259 | 0.613396 | 0.057984 | Q32P28;E2 | Q32P28 | Q32P28 | Prolyl 3-hy |
| GRK6 | 0.208127 | 0.61926 | 0.074629 | D6RHX8;F8 | D6RHX8;F8 | D6RHX8 | G protein-c |
| GLIPR2 | 0.206591 | 0.621453 | -0.03878 | Q9H4G4;Q | Q9H4G4;Q | Q9H4G4 | Golgi-assoc |
| PSMD9 | 0.204985 | 0.623756 | 0.049189 | O00233;J3 | O00233;J3 | O00233 | 26S protea |
| FTH1 | 0.203152 | 0.626394 | -0.06416 | P02794;G3 | P02794;G3 | P02794 | Ferritin hea |
| RPL36 | 0.200253 | 0.63059 | 0.056499 | Q9Y3U8;J3 | Q9Y3U8 | Q9Y3U8 | 60S riboso |
| C8orf82 | 0.196157 | 0.636565 | 0.051816 | Q6P1X6;H0 | Q6P1X6;H0 | Q6P1X6 | UPF0598 p |
| ACTR1B | 0.193096 | 0.641068 | 0.046309 | P42025 | P42025 | P42025 | Beta-centra |
| EIF5A | 0.192783 | 0.64153 | -0.04226 | P63241;I3I | P63241;I3I | P63241 | Eukaryotic |
| APRT | 0.189385 | 0.646569 | 0.083114 | P07741;H3 | P07741;H3 | P07741 | Adenine ph |
| PCYT2 | 0.185496 | 0.652384 | -0.05902 | I3L1R7;Q9 | I3L1R7;Q9 | I3L1R7 | Ethanolami |
| PARVG | 0.180812 | 0.65946 | -0.0503 | Q9HBI0;B0 | Q9HBI0 | Q9HBI0 | Gamma-par |
| BRE | 0.163745 | 0.685891 | 0.080463 | Q9NXR7;F8 | Q9NXR7;F8 | Q9NXR7 | BRCA1-A co |
| RPL23A | 0.163326 | 0.686553 | 0.042759 | P62750;H7 | P62750;H7 | P62750 | 60S riboso |
| KHDRBS1 | 0.163228 | 0.686708 | 0.022123 | Q07666;Q5 | Q07666 | Q07666 | KH domain |
| ADRM1 | 0.146345 | 0.71393 | -0.02684 | Q16186;A0 | Q16186;A0 | Q16186 | Proteasom |
| DLG1 | 0.138812 | 0.72642 | -0.03101 | Q12959;A0 | Q12959;A0 | Q12959 | Disks large |
| BNIP1 | 0.124932 | 0.750011 | -0.02667 | Q12981 | Q12981 | Q12981 | Vesicle tra |
| GM2A | 0.121262 | 0.756376 | 0.030608 | P17900;H0 | P17900 | P17900 | Ganglioside |
| SLC25A29 | 0.113549 | 0.769929 | 0.042494 | Q8N8R3;G3 | Q8N8R3 | Q8N8R3 | Mitochond |
| KDSR | 0.104544 | 0.78606 | 0.037238 | Q06136;K7 | Q06136;K7 | Q06136 | 3-ketodihy |
| UQCC1 | 0.096858 | 0.800096 | -0.01357 | Q9NVA1;B1 | Q9NVA1;B1 | Q9NVA1 | Ubiquinol- |
| RAB5B | 0.094198 | 0.805012 | 0.022328 | P61020;F8 | P61020;F8 | P61020 | Ras-related |
| CLPP | 0.092539 | 0.808093 | 0.026783 | Q16740;M | Q16740;M | Q16740 | ATP-depen |
| PRKRA | 0.081468 | 0.828958 | -0.01065 | O75569;G5 | O75569;G5 | O75569 | Interferon- |
| HEATR1 | 0.078705 | 0.834247 | -0.03365 | Q5T3Q7;Q9 | Q5T3Q7;Q9 | Q5T3Q7 | HEAT repea |
| MAP1A | 0.078 | 0.835603 | 0.017184 | E9PGC8;P7 | E9PGC8;P7 | E9PGC8 | Microtubul |
| CLPB | 0.0755 | 0.840426 | 0.013073 | Q9H078;A0 | Q9H078;A0 | Q9H078 | Caseinolyti |
| RHOC | 0.07435 | 0.842655 | 0.008644 | Q5JR08;P0 | Q5JR08;P0 | Q5JR08 | Rho-relate |
| MRPL47 | 0.069086 | 0.852932 | 0.02001 | Q9HD33 | Q9HD33 | Q9HD33 | 39S riboso |
| NMES1 | 0.058885 | 0.873203 | 0.009395 | Q9C002;H0 | Q9C002;H0 | Q9C002 | Normal mu |
| ISOC2 | 0.057805 | 0.875378 | 0.014178 | Q96AB3;K7 | Q96AB3;K7 | Q96AB3 | Isochorism |
| GRAMD4 | 0.05374 | 0.883608 | 0.007586 | Q6IC98;H0 | Q6IC98 | Q6IC98 | GRAM dom |
| HDAC2 | 0.047835 | 0.895705 | 0.011624 | Q92769;H3 | Q92769 | Q92769 | Histone de |
| NUDT5 | 0.047045 | 0.897336 | -0.01579 | A6NFX8;Q9 | A6NFX8;Q9 | A6NFX8 | ADP-sugar |
| ROCK2 | 0.040722 | 0.910496 | 0.010607 | O75116;E9 | O75116;E9 | O75116 | Rho-associ |
| TMLHE | 0.038039 | 0.916137 | -0.01517 | Q9NVH6 | Q9NVH6 | Q9NVH6 | Trimethylly |
| ARHGDIA | 0.036688 | 0.918992 | 0.013861 | P52565;J3 | P52565;J3 | P52565 | Rho GDP-di |

| | | | | | | | |
|---|---|---|---|---|---|---|---|
| COG6 | 0.031435 | 0.930176 | -0.0111 | Q9Y2V7 | Q9Y2V7 | Q9Y2V7 | Conserved ( |
| IFIT1 | 0.028105 | 0.937336 | -0.00747 | P09914;Q5 | P09914 | P09914 | Interferon-i |
| UXS1 | 0.024833 | 0.944425 | -0.0076 | Q8NBZ7;C9 | Q8NBZ7 | Q8NBZ7 | UDP-glucur |
| THOC1 | 0.020613 | 0.953646 | -0.00668 | A0A087WV | A0A087WV | A0A087WV | THO compl |
| TMX1 | 0.018068 | 0.95925 | -0.00458 | Q9H3N1;G | Q9H3N1 | Q9H3N1 | Thioredoxin |
| RPL35A | 0.015755 | 0.964373 | -0.00738 | P18077;C9 | P18077;C9 | P18077 | 60S ribosor |
| TMED9 | 0.01574 | 0.964407 | -0.00527 | Q9BVK6 | Q9BVK6 | Q9BVK6 | Transmemb |
| SNX3 | 0.01397 | 0.968345 | -0.00615 | O60493 | O60493 | O60493 | Sorting nex |
| PMM2 | 0.01394 | 0.968412 | -0.00554 | O15305;H3 | O15305;H3 | O15305 | Phosphoma |
| MTIF2 | 0.008682 | 0.980208 | 0.002337 | P46199;H7 | P46199 | P46199 | Translation |
| CNBP | 1.964331 | 0.010856 | 0.422546 | P62633;D6 | P62633 | P62633 | Cellular nu |
| HLA-E | 1.858232 | 0.01386 | 0.422214 | P13747 | P13747 | P13747 | HLA class I h |
| SNX8 | 1.451317 | 0.035374 | 0.406583 | Q9Y5X2;C9 | Q9Y5X2;C9 | Q9Y5X2 | Sorting nex |
| PIK3C3 | 2.470874 | 0.003382 | 0.405206 | A0A1W2PS | A0A1W2PS | A0A1W2PS | Phosphatic |
| HLA-F | 3.407003 | 0.000392 | 0.385497 | A0A0G2JH6 | A0A0G2JH6 | A0A0G2JH6 | HLA class I h |
| ADAM9 | 2.218911 | 0.006041 | 0.291874 | Q13443;A0 | Q13443;A0 | Q13443 | Disintegrin |
| RPL18 | 3.445777 | 0.000358 | 0.255627 | G3V203;H0 | G3V203;H0 | G3V203 | 60S ribosor |
| SEC62 | 1.547756 | 0.02833 | 0.253987 | Q99442;F8 | Q99442;F8 | Q99442 | Translocati |
| NQO1 | 1.447638 | 0.035675 | 0.240943 | B4DLR8;P1 | B4DLR8;P1 | B4DLR8 | NAD(P)H de |
| GHITM | 2.168808 | 0.006779 | 0.227231 | Q9H3K2 | Q9H3K2 | Q9H3K2 | Growth hor |
| SNRPA1 | 1.60665 | 0.024737 | 0.217607 | P09661;H0 | P09661;H0 | P09661 | U2 small nu |
| ZADH2 | 1.534262 | 0.029224 | 0.206165 | Q8N4Q0;J3 | Q8N4Q0 | Q8N4Q0 | Zinc-bindir |
| MRPS7 | 1.64606 | 0.022591 | 0.198209 | J3QLS3;Q9 | J3QLS3;Q9 | J3QLS3 | 28S ribosor |
| WDR48 | 1.376231 | 0.04205 | 0.187329 | Q8TAF3;F8 | Q8TAF3 | Q8TAF3 | WD repeat- |
| NDUFA7 | 1.657204 | 0.022019 | 0.173311 | O95182;M0 | O95182;M0 | O95182 | NADH dehy |
| NUDT21 | 1.404545 | 0.039396 | 0.17156 | O43809;H3 | O43809;H3 | O43809 | Cleavage ar |
| DHRS1 | 1.489901 | 0.032367 | 0.163634 | Q96LJ7;H0 | Q96LJ7;H0 | Q96LJ7 | Dehydroger |
| SLC25A5 | 1.615487 | 0.024239 | 0.153315 | P05141;Q9 | P05141 | P05141 | ADP/ATP tr |
| EEF1B2 | 1.389236 | 0.04081 | 0.138593 | P24534;C9 | P24534 | P24534 | Elongation |
| NUP88 | 1.315067 | 0.04841 | 0.133598 | Q99567;J3 | Q99567;J3 | Q99567 | Nuclear por |
| RAB1B | 1.834431 | 0.014641 | 0.125136 | Q9H0U4;E9 | Q9H0U4;E9 | Q9H0U4 | Ras-related |
| SRSF2 | 1.437284 | 0.036536 | 0.114864 | J3KP15;Q0 | J3KP15;Q0 | J3KP15 | Serine/argi |
| RHOA | 1.784659 | 0.016419 | 0.110506 | P61586;C9 | P61586;C9 | P61586 | Transformi |
| HLA-B | 1.531356 | 0.02942 | 0.091604 | P01889;A0 | P01889;A0 | P01889 | HLA class I h |
| BAIAP2 | 2.075655 | 0.008401 | -0.11927 | I3L4C2;Q9 | I3L4C2;Q9 | I3L4C2 | Brain-speci |
| IFI35 | 1.41203 | 0.038723 | -0.17117 | P80217 | P80217 | P80217 | Interferon-i |
| IDI1 | 1.510188 | 0.03089 | -0.18005 | Q13907;C9 | Q13907 | Q13907 | Isopenteny |
| CD84 | 1.740206 | 0.018188 | -0.19472 | Q9UIB8 | Q9UIB8 | Q9UIB8 | SLAM famil |
| LAMP2 | 2.51016 | 0.003089 | -0.19567 | P13473;H0 | P13473 | P13473 | Lysosome-a |
| AGPAT5 | 3.656839 | 0.00022 | -0.2884 | Q9NUQ2;H | Q9NUQ2 | Q9NUQ2 | 1-acyl-sn-gl |
| LPL | 3.741455 | 0.000181 | -0.4269 | P06858;E7 | P06858 | P06858 | Lipoprotein |
| ARF4 | 1.29341 | 0.050885 | -0.16782 | P18085;C9 | P18085;C9 | P18085 | ADP-ribosy |
| DNAJC7 | 1.278832 | 0.052622 | -0.2045 | Q99615;A0 | Q99615;A0 | Q99615 | DnaJ homo |
| CNPY3 | 1.229146 | 0.059 | 0.167841 | Q9BT09;A0 | Q9BT09 | Q9BT09 | Protein car |
| NRBP1 | 1.215139 | 0.060934 | 0.225368 | F8W6G1;Q | F8W6G1;Q | F8W6G1 | Nuclear rec |
| MAGT1 | 1.21089 | 0.061533 | -0.14317 | A0A087WU | A0A087WU | A0A087WU | Magnesium |
| ADSS | 1.183184 | 0.065587 | -0.21034 | P30520 | P30520 | P30520 | Adenylosuc |
| SAR1A | 1.182414 | 0.065703 | -0.21375 | Q9NR31;H0 | Q9NR31;H0 | Q9NR31 | GTP-bindin |

| | | | | | | | |
|---|---|---|---|---|---|---|---|
| ELMO2 | 1.138654 | 0.072668 | -0.1475 | Q5JVZ5;Q9 | Q5JVZ5;Q9 | Q5JVZ5 | Engulfment |
| RNF13 | 1.135418 | 0.073212 | 0.133959 | O43567;C9 | O43567;C9 | O43567 | E3 ubiquiti |
| RRAGC | 1.089563 | 0.081365 | -0.09333 | Q9HB90 | Q9HB90 | Q9HB90 | Ras-related |
| RAB2B | 1.077093 | 0.083735 | 0.12161 | Q8WUD1;A | Q8WUD1;A | Q8WUD1 | Ras-related |
| BMP2K | 1.074205 | 0.084294 | -0.17202 | Q9NSY1;H0 | Q9NSY1;H0 | Q9NSY1 | BMP-2-indu |
| SLC1A4 | 1.073056 | 0.084517 | -0.08748 | P43007 | P43007 | P43007 | Neutral am |
| APOBEC3G | 1.063714 | 0.086355 | -0.2359 | Q9HC16 | Q9HC16 | Q9HC16 | DNA dC->dU |
| RBBP7 | 1.004222 | 0.099032 | 0.122702 | Q16576;E9 | Q16576;E9 | Q16576 | Histone-bir |
| RAB35 | 0.943867 | 0.113798 | 0.161554 | Q15286;F5 | Q15286;F5 | Q15286 | Ras-related |
| HLA-DPA1 | 0.934499 | 0.116279 | 0.117963 | P20036;J3I | P20036;J3I | P20036 | HLA class II |
| STRIP1 | 0.927308 | 0.11822 | -2.413 | Q5VSL9;Q9 | Q5VSL9 | Q5VSL9 | Striatin-int |
| FLAD1 | 0.926654 | 0.118398 | 0.161657 | Q8NFF5;Q5 | Q8NFF5;Q5 | Q8NFF5 | FAD syntha |
| EMC7 | 0.925152 | 0.118809 | 0.066688 | Q9NPA0;H( | Q9NPA0;H( | Q9NPA0 | ER membra |
| SLC33A1 | 0.903892 | 0.12477 | -0.16776 | O00400;H7 | O00400;H7 | O00400 | Acetyl-coer |
| CALM2 | 0.874806 | 0.133412 | -0.16381 | P0DP25;P0 | P0DP25;P0 | P0DP25 | |
| TGFB1 | 0.870952 | 0.134601 | -0.22744 | P01137;A0 | P01137;A0 | P01137 | Transformi |
| MAGOH | 0.852313 | 0.140503 | -0.09105 | P61326;Q9 | P61326;Q9 | P61326 | Protein ma |
| COMMD2 | 0.847564 | 0.142048 | -0.13051 | Q86X83;F8 | Q86X83;F8 | Q86X83 | COMM don |
| RAB42 | 0.838638 | 0.144998 | 0.079262 | Q8N4Z0 | Q8N4Z0 | Q8N4Z0 | Putative Ra |
| EML3 | 0.834827 | 0.146276 | 0.183312 | G3V195;B7 | G3V195;B7 | G3V195 | Echinoderm |
| EEFSEC | 0.828877 | 0.148294 | 0.275421 | P57772;C9 | P57772;C9 | P57772 | Selenocyste |
| CSNK2B | 0.827457 | 0.148779 | -0.05638 | Q5SRQ6;P6 | Q5SRQ6;P6 | Q5SRQ6 | Casein kina |
| ZNF326 | 0.816351 | 0.152633 | -0.18799 | Q5BKZ1;A0 | Q5BKZ1;A0 | Q5BKZ1 | DBIRD com |
| NUP214 | 0.790763 | 0.161896 | 0.172641 | A0A494C1I | A0A494C1I | A0A494C1I | Nuclear po |
| YKT6 | 0.777316 | 0.166987 | -0.17509 | O15498;H7 | O15498 | O15498 | Synaptobre |
| PIP4K2C | 0.760816 | 0.173454 | -0.1191 | Q8TBX8;F8 | Q8TBX8;F8 | Q8TBX8 | Phosphatic |
| GPX4 | 0.747039 | 0.179045 | -0.11933 | A0A087X2I | A0A087X2I | A0A087X2I | Glutathion |
| GIMAP8 | 0.746258 | 0.179367 | -0.18837 | Q8ND71 | Q8ND71 | Q8ND71 | GTPase IMA |
| SNAP29 | 0.744872 | 0.17994 | -0.10448 | O95721;C9 | O95721;C9 | O95721 | Synaptosor |
| CRABP2 | 0.732186 | 0.185274 | -0.24839 | P29373;Q5 | P29373;Q5 | P29373 | Cellular ret |
| CD276 | 0.731424 | 0.185599 | 0.064139 | A0A0C4DG | A0A0C4DG | A0A0C4DG | CD276 anti |
| ENPP4 | 0.718371 | 0.191262 | 0.101274 | Q9Y6X5 | Q9Y6X5 | Q9Y6X5 | Bis(5-adenc |
| CTR9 | 0.716454 | 0.192108 | 0.152548 | Q6PD62;H( | Q6PD62 | Q6PD62 | RNA polym |
| PARVB | 0.699615 | 0.199703 | 0.139893 | Q9HBI1;A0 | Q9HBI1;A0 | Q9HBI1 | Beta-parvir |
| PPP6C | 0.685452 | 0.206323 | -0.12275 | O00743;Q5 | O00743 | O00743 | Serine/thre |
| RAB5A | 0.684147 | 0.206944 | -0.11514 | P20339;F8 | P20339 | P20339 | Ras-related |
| ATP5J | 0.683595 | 0.207207 | 0.116626 | P18859;A8 | P18859;A8 | P18859 | ATP synthas |
| CDC73 | 0.67606 | 0.210834 | -0.12344 | A0A3B3IRP | A0A3B3IRP | A0A3B3IRP | Parafibrom |
| PRRC2C | 0.666245 | 0.215653 | 0.115219 | E7EPN9;Q9 | E7EPN9;Q9 | E7EPN9 | Protein PRI |
| MRPS34 | 0.655904 | 0.220849 | 0.070537 | C9JJ19;P82 | C9JJ19;P82 | C9JJ19 | 28S riboson |
| IKBIP | 0.654412 | 0.221609 | 0.221015 | Q70UQ0 | Q70UQ0 | Q70UQ0 | Inhibitor of |
| C18orf8 | 0.647097 | 0.225374 | 0.229783 | K7ENL9;Q9 | K7ENL9;Q9 | K7ENL9 | Uncharacte |
| COMMD1 | 0.630531 | 0.234136 | 0.066498 | Q8N668;E9 | Q8N668 | Q8N668 | COMM don |
| IMPA1 | 0.615587 | 0.242333 | -0.17505 | P29218;H0 | P29218;H0 | P29218 | Inositol mc |
| EARS2 | 0.609649 | 0.245669 | 0.119835 | H3BTB7;Q5 | H3BTB7;Q5 | H3BTB7 | Probable gl |
| GLIPR1 | 0.603217 | 0.249335 | 0.087767 | P48060;B4 | P48060 | P48060 | Glioma pat |
| LAMTOR2 | 0.596969 | 0.252948 | -0.06293 | Q9Y2Q5 | Q9Y2Q5 | Q9Y2Q5 | Ragulator c |
| SMARCE1 | 0.591298 | 0.256272 | -0.07831 | A0A2R8Y76 | A0A2R8Y76 | A0A2R8Y76 | SWI/SNF-re |

| | | | | | | | |
|---|---|---|---|---|---|---|---|
| NDRG3 | 0.584052 | 0.260584 | -0.15107 | Q9UGV2;Q | Q9UGV2;Q | Q9UGV2 | Protein ND |
| PTPN23 | 0.574089 | 0.266631 | 0.129702 | Q9H3S7;C9 | Q9H3S7 | Q9H3S7 | Tyrosine-pr |
| PDP1 | 0.5664 | 0.271394 | -0.08671 | Q9P0J1;E5 | Q9P0J1 | Q9P0J1 | [Pyruvate d |
| RBM15 | 0.560161 | 0.275321 | 0.099444 | A0A494C0( | A0A494C0( | A0A494C0( | Putative RN |
| TBCB | 0.55156 | 0.280828 | 0.09722 | Q99426;K7 | Q99426;K7 | Q99426 | Tubulin-fol |
| SPARC | 0.55103 | 0.28117 | -0.1192 | P09486;F5 | P09486;F5 | P09486 | SPARC |
| MRPS15 | 0.548496 | 0.282816 | 0.078769 | P82914 | P82914 | P82914 | 28S ribosor |
| RPS4X | 0.548223 | 0.282994 | 0.030031 | P62701 | P62701 | P62701 | 40S ribosor |
| TPM3 | 0.539877 | 0.288485 | 0.073399 | A0A087WV | A0A087WV | A0A087WWU8 | |
| SRSF7 | 0.538998 | 0.289069 | -0.11254 | A0A0B4J1Z | A0A0B4J1Z | A0A0B4J1Z | Serine/argi |
| TMED4 | 0.534542 | 0.292051 | 0.104272 | Q7Z7H5;F8 | Q7Z7H5;F8 | Q7Z7H5 | Transmemb |
| DHRS11 | 0.533455 | 0.292783 | -0.19159 | Q6UWP2;A | Q6UWP2;A | Q6UWP2 | Dehydroger |
| TBL1XR1 | 0.526922 | 0.29722 | 0.091831 | Q9BZK7;A0 | Q9BZK7;A0 | Q9BZK7 | F-box-like/\ |
| SLC4A1AP | 0.526164 | 0.297739 | 0.106958 | A0A087X0I | A0A087X0I | A0A087X0I | Kanadaptir |
| SMEK1 | 0.520738 | 0.301482 | 0.843005 | G3V5Z3;Q6 | G3V5Z3;Q6 | G3V5Z3 | Serine/thre |
| CPSF2 | 0.517891 | 0.303465 | 0.159145 | Q9P2I0;H0 | Q9P2I0 | Q9P2I0 | Cleavage ar |
| GLRX | 0.517221 | 0.303934 | -0.12846 | P35754 | P35754 | P35754 | Glutaredox |
| ATG5 | 0.516357 | 0.304539 | 0.068137 | Q9H1Y0;L7 | Q9H1Y0;L7 | Q9H1Y0 | Autophagy |
| UBXN4 | 0.499638 | 0.316491 | 0.08118 | Q92575;F8 | Q92575 | Q92575 | UBX domai |
| FDPS | 0.483787 | 0.328256 | -0.05947 | P14324;A0 | P14324;A0 | P14324 | Farnesyl py |
| MRPL38 | 0.473888 | 0.335824 | 0.083329 | Q96DV4 | Q96DV4 | Q96DV4 | 39S ribosor |
| SRSF4 | 0.465528 | 0.342352 | 0.069912 | Q08170;A0 | Q08170;A0 | Q08170 | Serine/argi |
| DCUN1D1 | 0.456928 | 0.349198 | -0.11097 | Q96GG9;C9 | Q96GG9;C9 | Q96GG9 | DCN1-like p |
| USP39 | 0.455897 | 0.350028 | -0.06426 | A0A087X1I | A0A087X1I | A0A087X1I | U4/U6.U5 t |
| ATPAF2 | 0.453423 | 0.352028 | 0.105426 | Q8N5M1;C | Q8N5M1;C | Q8N5M1 | ATP synthas |
| SNRPD2 | 0.446976 | 0.357293 | -0.05113 | P62316;K7 | P62316;K7 | P62316 | Small nucle |
| CD40 | 0.441068 | 0.362187 | 0.118082 | P25942;A0 | P25942;A0 | P25942 | Tumor necr |
| STUB1 | 0.438261 | 0.364535 | 0.189914 | Q9UNE7;H3 | Q9UNE7;H3 | Q9UNE7 | E3 ubiquiti |
| BIN1 | 0.428182 | 0.373094 | -0.21363 | O00499 | O00499 | O00499 | Myc box-de |
| POGLUT1 | 0.421834 | 0.378587 | -0.12025 | Q8NBL1;H7 | Q8NBL1;H7 | Q8NBL1 | Protein O-g |
| CD2AP | 0.421759 | 0.378653 | 0.101378 | Q9Y5K6 | Q9Y5K6 | Q9Y5K6 | CD2-associ |
| TRIOBP | 0.421031 | 0.379288 | 0.107388 | Q9H2D6;H( | Q9H2D6 | Q9H2D6 | TRIO and F- |
| SRSF9 | 0.402512 | 0.395811 | -0.04465 | Q13242;H( | Q13242;H( | Q13242 | Serine/argi |
| LCK | 0.399326 | 0.398726 | -0.48867 | P06239;Q5 | P06239;Q5 | P06239 | Tyrosine-pr |
| HPRT1 | 0.396616 | 0.401221 | -0.08344 | P00492 | P00492 | P00492 | Hypoxanth |
| RPL23 | 0.378145 | 0.418654 | 0.050825 | P62829;J3 | P62829;J3 | P62829 | 60S ribosor |
| VTI1B | 0.369494 | 0.427077 | 0.055282 | Q9UEU0;H( | Q9UEU0 | Q9UEU0 | Vesicle trar |
| EMR2 | 0.366922 | 0.429614 | 0.053538 | Q9UHX3;M | Q9UHX3;M | Q9UHX3 | EGF-like mc |
| ARPC5 | 0.366581 | 0.429951 | 0.067029 | O15511;B1 | O15511;B1 | O15511 | Actin-relate |
| NDUFB7 | 0.353283 | 0.443319 | 0.057044 | P17568;M( | P17568 | P17568 | NADH dehy |
| SEPHS2 | 0.352947 | 0.443663 | 0.06064 | Q99611 | Q99611 | Q99611 | Selenide, w |
| MAN2B2 | 0.350886 | 0.445773 | 0.117428 | Q9Y2E5;E9 | Q9Y2E5;E9 | Q9Y2E5 | Epididymis |
| PDCD10 | 0.349646 | 0.447048 | -0.04402 | Q9BUL8;C9 | Q9BUL8;C9 | Q9BUL8 | Programme |
| DICER1 | 0.345183 | 0.451665 | -0.55655 | Q9UPY3 | Q9UPY3 | Q9UPY3 | Endoribonu |
| SEH1L | 0.338087 | 0.459106 | 0.184131 | K7ELV2;Q9 | K7ELV2;Q9 | K7ELV2 | Nucleopori |
| ATG3 | 0.331928 | 0.465663 | -0.23534 | Q9NT62;F8 | Q9NT62 | Q9NT62 | Ubiquitin-l |
| HNRNPH2 | 0.327333 | 0.470616 | -0.04931 | P55795 | P55795 | P55795 | Heterogene |
| LRMP | 0.317273 | 0.481645 | 0.10038 | F5H006;A0 | F5H006;A0 | F5H006 | Lymphoid-r |

| | | | | | | | |
|---|---|---|---|---|---|---|---|
| OSGEP | 0.31653 | 0.482469 | -0.12912 | Q9NPF4;G3 | Q9NPF4 | Q9NPF4 | Probable tF |
| ICAM3 | 0.316269 | 0.482759 | 0.101648 | K7ERN2;P3 | K7ERN2;P3 | K7ERN2 | Intercellula |
| CRTAP | 0.312173 | 0.487335 | -0.07383 | C9JP16;O7 | C9JP16;O7 | C9JP16 | Cartilage-as |
| PKN1 | 0.312118 | 0.487396 | 0.040299 | Q16512;K7 | Q16512 | Q16512 | Serine/thre |
| IFI30 | 0.299562 | 0.501693 | 0.084431 | P13284;M( | P13284 | P13284 | Gamma-int |
| COPS3 | 0.295071 | 0.506908 | 0.049139 | Q9UNS2;H7 | Q9UNS2;H7 | Q9UNS2 | COP9 signa |
| MFF | 0.292137 | 0.510344 | 0.070422 | Q9GZY8;A0 | Q9GZY8;A0 | Q9GZY8 | Mitochond |
| CLPTM1L | 0.288368 | 0.514792 | 0.057378 | Q96KA5;G5 | Q96KA5;G5 | Q96KA5 | Cleft lip and |
| SCYL1 | 0.287829 | 0.515432 | -0.07104 | E9PS17;E9I | E9PS17;E9I | E9PS17 | N-terminal |
| PSMB9 | 0.279937 | 0.524883 | 0.028597 | A2ACR1;A0 | A2ACR1;A0 | A2ACR1 | Proteasome |
| LAMTOR1 | 0.277274 | 0.528112 | -0.05043 | Q6IAA8;F5( | Q6IAA8;F5( | Q6IAA8 | Ragulator c |
| MCTS1 | 0.273608 | 0.532589 | 0.085062 | Q9ULC4;A0 | Q9ULC4;A0 | Q9ULC4 | Malignant T |
| SPON2 | 0.268862 | 0.538441 | 0.057007 | Q9BUD6;D( | Q9BUD6;D( | Q9BUD6 | Spondin-2 |
| TRAPPC3 | 0.268482 | 0.538912 | -0.06681 | O43617;A0 | O43617;A0 | O43617 | Trafficking p |
| PGRMC2 | 0.264139 | 0.544328 | 0.052568 | O15173;U3 | O15173 | O15173 | Membrane- |
| DOCK10 | 0.261983 | 0.547037 | 0.04427 | A0A2R8YD8 | A0A2R8YD8 | A0A2R8YD8 | Dedicator c |
| GIMAP2 | 0.26008 | 0.54944 | 0.09142 | Q9UG22 | Q9UG22 | Q9UG22 | GTPase IMA |
| PARP12 | 0.256766 | 0.553648 | -0.08333 | Q9H0J9;G5 | Q9H0J9 | Q9H0J9 | Poly [ADP-r |
| LPCAT3 | 0.254136 | 0.557011 | -0.03781 | Q6P1A2;F5 | Q6P1A2;F5 | Q6P1A2 | Lysophosph |
| ADSL | 0.246394 | 0.56703 | -0.04203 | P30566;A0 | P30566;A0 | P30566 | Adenylosuc |
| MIA2 | 0.245816 | 0.567785 | 0.043456 | G3V599;Q9 | G3V599;Q9 | G3V599 | Melanoma |
| HNRNPAB | 0.245797 | 0.56781 | -0.04014 | D6R9P3;D6 | D6R9P3;D6 | D6R9P3 | Heterogene |
| PSMB4 | 0.235316 | 0.58168 | 0.04859 | P28070 | P28070 | P28070 | Proteasome |
| RAB27A | 0.230864 | 0.587673 | -0.06329 | P51159;H3 | P51159;H3 | P51159 | Ras-related |
| ARL1 | 0.227163 | 0.592703 | -0.03185 | P40616;F8 | P40616;F8 | P40616 | ADP-ribosy |
| CUL5 | 0.221851 | 0.599997 | -0.34791 | Q93034;H( | Q93034 | Q93034 | Cullin-5 |
| CTNND1 | 0.219098 | 0.603813 | 0.047093 | A0A669KB6 | A0A669KB6 | A0A669KB6 | Catenin del |
| DCTN3 | 0.216712 | 0.607139 | -0.05751 | O75935;X6 | O75935;X6 | O75935 | Dynactin su |
| MRPL4 | 0.210634 | 0.615696 | 0.041758 | Q9BYD3;K7 | Q9BYD3;K7 | Q9BYD3 | 39S ribosor |
| COX7A2L | 0.209563 | 0.617215 | 0.031567 | O14548;E5 | O14548;E5 | O14548 | Cytochrom |
| DAPK1 | 0.208032 | 0.619395 | -0.04434 | P53355;F8 | P53355 | P53355 | Death-asso |
| GIMAP1 | 0.199287 | 0.631994 | -0.03679 | Q8WWP7 | Q8WWP7 | Q8WWP7 | GTPase IMA |
| YME1L1 | 0.196766 | 0.635674 | -0.06106 | Q96I63;Q9 | Q96I63;Q9 | Q96I63 | ATP-depend |
| DUSP3 | 0.195069 | 0.638162 | -0.08747 | P51452;K7 | P51452;K7 | P51452 | Dual specifi |
| ALDOC | 0.186366 | 0.65108 | -0.06966 | P09972;A8 | P09972;A8 | P09972 | Fructose-bi |
| GLO1 | 0.186316 | 0.651154 | -0.06488 | Q04760 | Q04760 | Q04760 | Lactoylglut |
| RPL7A | 0.184898 | 0.653284 | 0.020154 | P62424;Q5 | P62424;Q5 | P62424 | 60S ribosor |
| IST1 | 0.183138 | 0.655936 | 0.037788 | P53990;H3 | P53990;H3 | P53990 | IST1 homol |
| CUL1 | 0.180821 | 0.659445 | 0.082755 | A0A590UJF | A0A590UJF | A0A590UJF | Cullin-1 |
| METAP2 | 0.172736 | 0.671837 | 0.061275 | P50579;F8 | P50579;F8 | P50579 | Methionine |
| REEP5 | 0.171098 | 0.674377 | 0.022691 | Q00765;E2 | Q00765;E2 | Q00765 | Receptor ex |
| MMAB | 0.15769 | 0.69552 | 0.043206 | Q96EY8;S4I | Q96EY8;S4I | Q96EY8 | Cob(I)yrinic |
| WDR91 | 0.154046 | 0.701381 | 0.033212 | C9J1X0;A4I | C9J1X0;A4I | C9J1X0 | WD repeat- |
| SNRPB2 | 0.153136 | 0.702852 | 0.028749 | P08579 | P08579 | P08579 | U2 small nu |
| WDR18 | 0.151243 | 0.705923 | 0.028495 | U3KQC1;Q9 | U3KQC1;Q9 | U3KQC1 | WD repeat- |
| SASH3 | 0.151004 | 0.706311 | -0.05002 | O75995 | O75995 | O75995 | SAM and SH |
| ARFIP1 | 0.150246 | 0.707544 | 0.054504 | P53367 | P53367 | P53367 | Arfaptin-1 |
| HSPH1 | 0.145158 | 0.715883 | 0.021767 | Q92598;R4 | Q92598 | Q92598 | Heat shock |

| | | | | | | | |
|---|---|---|---|---|---|---|---|
| PANK4 | 0.128365 | 0.744107 | -0.02976 | Q9NVE7;E9 | Q9NVE7;E9 | Q9NVE7 | Pantothena |
| TBK1 | 0.125893 | 0.748354 | -0.03286 | A0A494C14 | A0A494C14 | A0A494C14 | Serine/thre |
| SLC2A5 | 0.114904 | 0.76753 | 0.022858 | P22732;K7 | P22732 | P22732 | Solute carri |
| UBQLN1 | 0.097847 | 0.798275 | 0.018594 | Q9UMX0;Q | Q9UMX0;Q | Q9UMX0 | Ubiquilin-1 |
| SEPHS1 | 0.095442 | 0.802708 | -0.0283 | P49903;Q5 | P49903 | P49903 | Selenide, w |
| NMNAT1 | 0.095109 | 0.803325 | 0.031871 | Q9HAN9;B1 | Q9HAN9;B1 | Q9HAN9 | Nicotinami |
| TMEM126A | 0.087765 | 0.817025 | 0.016601 | Q9H061;E9 | Q9H061;E9 | Q9H061 | Transmemb |
| BCKDHB | 0.083407 | 0.825265 | -0.01743 | P21953 | P21953 | P21953 | 2-oxoisoval |
| ATXN2L | 0.078834 | 0.834001 | 0.015807 | H3BUF6;Q8 | H3BUF6;Q8 | H3BUF6 | Ataxin-2-lik |
| C16orf58 | 0.078525 | 0.834593 | -0.01551 | H3BR29;H3 | H3BR29;H3 | H3BR29 | RUS1 family |
| HS1BP3 | 0.076459 | 0.838573 | -0.03185 | Q53T59;F6 | Q53T59;F6 | Q53T59 | HCLS1-binc |
| TCEA1 | 0.073495 | 0.844315 | 0.01795 | A0A1W2PF | A0A1W2PF | A0A1W2PF | Transcripti |
| TMED2 | 0.072625 | 0.84601 | 0.015027 | Q15363;F5 | Q15363;F5 | Q15363 | Transmemb |
| CDS2 | 0.071752 | 0.847712 | 0.012893 | O95674 | O95674 | O95674 | Phosphatic |
| SFXN2 | 0.066846 | 0.857342 | -0.01879 | Q96NB2;R4 | Q96NB2;R4 | Q96NB2 | Sideroflexir |
| ENOSF1 | 0.065598 | 0.859809 | -0.02119 | A0A3F2YNX | A0A3F2YNX | A0A3F2YNX | Mitochond |
| GIGYF2 | 0.064148 | 0.862684 | 0.031237 | I1E4Y6;Q6Y | I1E4Y6;Q6Y | I1E4Y6 | PERQ amin |
| DOK3 | 0.060632 | 0.869697 | -0.01831 | D6RAM3;Q | D6RAM3;Q | D6RAM3 | Docking pr |
| TSN | 0.056851 | 0.877302 | -0.01833 | Q15631;E9 | Q15631;E9 | Q15631 | Translin |
| SLC25A6 | 0.056061 | 0.8789 | -0.00735 | P12236 | P12236 | P12236 | ADP/ATP tr |
| THOC5 | 0.055265 | 0.880512 | 0.021071 | Q13769;F8 | Q13769;F8 | Q13769 | THO compl |
| FAM175B | 0.053919 | 0.883245 | 0.011416 | Q15018 | Q15018 | Q15018 | BRISC comp |
| S100A11 | 0.05374 | 0.883609 | -0.01544 | P31949 | P31949 | P31949 | Protein S10 |
| NUDC | 0.041413 | 0.909047 | 0.007192 | Q9Y266;A0 | Q9Y266 | Q9Y266 | Nuclear mi |
| UBE2M | 0.037793 | 0.916656 | -0.01012 | P61081;M0 | P61081 | P61081 | NEDD8-cor |
| CLTA | 0.034631 | 0.923356 | 0.006039 | P09496;F8 | P09496;F8 | P09496 | Clathrin lig |
| LIMK1 | 0.03107 | 0.930958 | -0.009 | E9PC47;P5 | E9PC47;P5 | E9PC47 | LIM domai |
| KIF1B | 0.030834 | 0.931464 | -0.01033 | A0A087WV | A0A087WV | A0A087WV | Kinesin-like |
| COX5B | 0.02487 | 0.944343 | -0.00485 | P10606 | P10606 | P10606 | Cytochrom |
| DHX38 | 0.024622 | 0.944882 | -0.0151 | Q92620;J3 | Q92620 | Q92620 | Pre-mRNA- |
| RAE1 | 0.024141 | 0.94593 | 0.006404 | P78406;E9 | P78406;E9 | P78406 | mRNA expo |
| PAPSS2 | 0.013492 | 0.96941 | 0.006214 | O95340 | O95340 | O95340 | Bifunctiona |
| SSBP1 | 0.011265 | 0.974396 | 0.003413 | Q04837;A0 | Q04837;A0 | Q04837 | Single-strar |
| TMX2 | 0.010622 | 0.975839 | -0.00192 | Q9Y320;A0 | Q9Y320 | Q9Y320 | Thioredoxi |
| RPS13 | 0.004334 | 0.99007 | -0.00088 | P62277;J3 | P62277;J3 | P62277 | 40S riboso |
| SCRIB | 3.537726 | 0.00029 | 0.48037 | A0A0G2JM | A0A0G2JM | A0A0G2JM | Protein scri |
| ME3 | 1.750202 | 0.017775 | 0.457758 | E9PMB9;Q1 | E9PMB9;Q1 | E9PMB9 | Malic enzy |
| SYVN1 | 1.671824 | 0.02129 | 0.442601 | Q86TM6;H0 | Q86TM6 | Q86TM6 | E3 ubiquiti |
| C17orf62 | 1.423217 | 0.037738 | 0.399353 | Q9BQA9;J3 | Q9BQA9;J3 | Q9BQA9 | Uncharacte |
| NDUFA6 | 4.127737 | 7.45E-05 | 0.398644 | P56556;A0 | P56556;A0 | P56556 | NADH dehy |
| PDLIM2 | 1.966804 | 0.010794 | 0.377183 | A0A087WV | A0A087WV | A0A087WV | PDZ and LIM |
| FIG4 | 1.994821 | 0.01012 | 0.366421 | Q92562;Q5 | Q92562 | Q92562 | Polyphosph |
| SH3KBP1 | 1.464347 | 0.034328 | 0.337794 | Q5JPT2;Q9 | Q5JPT2;Q9 | Q5JPT2 | SH3 domai |
| EWSR1 | 2.822194 | 0.001506 | 0.326494 | B0QYK0;Q0 | B0QYK0;Q0 | B0QYK0 | RNA-bindir |
| CPQ | 1.418479 | 0.038152 | 0.304928 | Q9Y646;E5 | Q9Y646 | Q9Y646 | Carboxype |
| CCDC109B | 1.885602 | 0.013014 | 0.30127 | Q9NWR8 | Q9NWR8 | Q9NWR8 | Calcium un |
| WARS2 | 1.782112 | 0.016515 | 0.28165 | Q9UGM6 | Q9UGM6 | Q9UGM6 | Tryptophar |
| DAG1 | 1.46663 | 0.034148 | 0.281412 | Q14118 | Q14118 | Q14118 | Dystroglyca |

| | | | | | | | |
|---|---|---|---|---|---|---|---|
| APOA1BP | 1.736624 | 0.018339 | 0.270389 | Q8NCW5;C | Q8NCW5 | Q8NCW5 | NAD(P)H-hy |
| SUMF1 | 1.415267 | 0.038436 | 0.260103 | Q8NBK3;F5 | Q8NBK3;F5 | Q8NBK3 | Sulfatase-m |
| AGFG1 | 3.068061 | 0.000855 | 0.259479 | P52594;B8 | P52594;B8 | P52594 | Arf-GAP dor |
| GPR108 | 2.441459 | 0.003619 | 0.25216 | Q9NPR9;M | Q9NPR9;M | Q9NPR9 | Protein GP |
| ARFGAP1 | 1.386554 | 0.041063 | 0.24694 | F8W1U7;F8 | F8W1U7;F8 | F8W1U7 | ADP-ribosy |
| CYB5A | 2.734926 | 0.001841 | 0.24677 | P00167;J3 | P00167 | P00167 | Cytochrom |
| BAK1 | 3.197257 | 0.000635 | 0.243233 | Q16611;A0 | Q16611;A0 | Q16611 | Bcl-2 homc |
| SCO2 | 1.908193 | 0.012354 | 0.242593 | O43819;A0 | O43819 | O43819 | Protein SC |
| MRPS36 | 3.099202 | 0.000796 | 0.235536 | P82909;D6 | P82909 | P82909 | 28S ribosor |
| MRPL43 | 1.342291 | 0.045468 | 0.211969 | H0Y6Y8;B1 | H0Y6Y8;B1 | H0Y6Y8 | 39S ribosor |
| FKBP8 | 1.777523 | 0.016691 | 0.203923 | Q14318;U3 | Q14318;U3 | Q14318 | Peptidyl-pr |
| RNMT | 2.015376 | 0.009652 | 0.197052 | O43148;K7 | O43148;K7 | O43148 | mRNA cap g |
| FTSJ3 | 2.208632 | 0.006185 | 0.193823 | Q8IY81 | Q8IY81 | Q8IY81 | pre-rRNA p |
| TIAL1 | 2.458491 | 0.003479 | 0.189256 | Q01085;E7 | Q01085;E7 | Q01085 | Nucleolysir |
| UNC45A | 1.716748 | 0.019198 | 0.181976 | A0A1W2PN | A0A1W2PN | A0A1W2PN | Protein unc |
| SKP1 | 1.727404 | 0.018733 | 0.181051 | E5RJR5;P63 | E5RJR5;P63 | E5RJR5 | S-phase kin |
| CCDC53 | 1.782066 | 0.016517 | 0.177238 | F5GZ97;Q9 | F5GZ97;Q9 | F5GZ97 | WASH com |
| VAMP8 | 1.718842 | 0.019105 | 0.157557 | Q9BV40;B8 | Q9BV40;B8 | Q9BV40 | Vesicle-assc |
| HMGB1 | 2.347407 | 0.004494 | 0.157453 | Q5T7C4;P0 | Q5T7C4;P0 | Q5T7C4 | High mobil |
| RPS14 | 2.094984 | 0.008036 | 0.15515 | P62263;A0 | P62263;A0 | P62263 | 40S ribosor |
| FAHD2A | 1.814375 | 0.015333 | 0.150368 | Q96GK7;Q6 | Q96GK7;Q6 | Q96GK7 | Fumarylace |
| MTHFS | 1.387094 | 0.041012 | 0.139897 | P49914;A0 | P49914;A0 | P49914 | 5-formyltet |
| SPNS1 | 1.435681 | 0.036671 | 0.130825 | Q9H2V7;H3 | Q9H2V7;H3 | Q9H2V7 | Protein spi |
| HVCN1 | 1.323578 | 0.04747 | 0.125509 | Q96D96;F8 | Q96D96 | Q96D96 | Voltage-gat |
| GMFG | 1.769266 | 0.017011 | -0.10873 | O60234;M | O60234;M | O60234 | Glia matura |
| ARL6IP5 | 1.864869 | 0.01365 | -0.16631 | O75915;F8 | O75915 | O75915 | PRA1 family |
| TRAPPC4 | 1.515872 | 0.030488 | -0.17572 | E9PQE8;Q9 | E9PQE8;Q9 | E9PQE8 | Trafficking |
| UQCRQ | 1.763468 | 0.01724 | -0.18806 | O14949 | O14949 | O14949 | Cytochrom |
| VBP1 | 1.399803 | 0.039829 | -0.22456 | P61758 | P61758 | P61758 | Prefoldin su |
| SRSF3 | 3.138572 | 0.000727 | -0.24482 | P84103;A0 | P84103;A0 | P84103 | Serine/argi |
| SRP14 | 3.274711 | 0.000531 | -0.26072 | P37108;H0 | P37108;H0 | P37108 | Signal reco |
| TRAPPC5 | 1.486327 | 0.032634 | -0.26249 | Q8IUR0;M0 | Q8IUR0;M0 | Q8IUR0 | Trafficking |
| RBP1 | 1.32729 | 0.047066 | -0.36036 | P09455;A0 | P09455;A0 | P09455 | Retinol-bin |
| VTA1 | 2.152021 | 0.007047 | -0.41263 | Q9NP79;A0 | Q9NP79;A0 | Q9NP79 | Vacuolar pr |
| UBE2L3 | 1.735056 | 0.018405 | -0.42501 | P68036;A0 | P68036 | P68036 | Ubiquitin-c |
| CTSH | 1.266439 | 0.054145 | 0.285776 | A0A087X0I | A0A087X0I | A0A087X0I | Pro-cathep |
| PDK2 | 1.265429 | 0.054271 | 0.154739 | Q15119;D6 | Q15119;D6 | Q15119 | [Pyruvate d |
| ARF5 | 1.257455 | 0.055277 | 0.195098 | P84085;C9 | P84085;C9 | P84085 | ADP-ribosy |
| IKBKG | 1.240418 | 0.057489 | 0.212962 | A0A087WU | A0A087WU | A0A087WU | NF-kappa-B |
| TAF15 | 1.214523 | 0.061021 | 0.367204 | Q92804;A0 | Q92804;A0 | Q92804 | TATA-bindi |
| SNRPN | 1.208567 | 0.061863 | 0.144435 | J3QLE5;P63 | J3QLE5;P63 | J3QLE5 | Small nucle |
| LYPLA1 | 1.199698 | 0.06314 | -0.21484 | O75608;A0 | O75608;A0 | O75608 | Acyl-protei |
| MRPL28 | 1.19845 | 0.063321 | 0.432542 | Q13084;A2 | Q13084;A2 | Q13084 | 39S ribosor |
| MBOAT1 | 1.191445 | 0.064351 | -0.24044 | Q6ZNC8 | Q6ZNC8 | Q6ZNC8 | Lysophospl |
| ZFPL1 | 1.187644 | 0.064917 | -0.1634 | O95159;E9 | O95159;E9 | O95159 | Zinc finger |
| DLGAP4 | 1.183771 | 0.065498 | -0.20947 | F8WF49;A0 | F8WF49;A0 | F8WF49 | Disks large- |
| APOO | 1.175788 | 0.066713 | 0.142684 | H7C1U8;Q9 | H7C1U8;Q9 | H7C1U8 | Apolipoprc |
| RANBP1 | 1.175322 | 0.066785 | 0.210537 | P43487;F6 | P43487;F6 | P43487 | Ran-specifi |

| | | | | | | | | |
|---|---|---|---|---|---|---|---|---|
| SIRPA | 1.17204 | 0.067292 | 0.11516 | | P78324;Q9 | P78324 | P78324 | Tyrosine-pr |
| TXN | 1.170995 | 0.067454 | -0.19228 | | P10599 | P10599 | P10599 | Thioredoxi |
| CBX3 | 1.141424 | 0.072206 | 0.107464 | | Q13185;S4 | Q13185 | Q13185 | Chromobo: |
| TXNDC12 | 1.137494 | 0.072863 | 0.180687 | | O95881;V9 | O95881 | O95881 | Thioredoxi |
| HLA-DRB3 | 1.133129 | 0.073599 | 0.127829 | | A0A182DW | A0A182DW | A0A182DW | HLA class II |
| TAPT1 | 1.13162 | 0.073855 | 0.262824 | | Q6NXT6;H0 | Q6NXT6 | Q6NXT6 | Transmemb |
| GADD45GII | 1.122969 | 0.075341 | 0.16149 | | Q8TAE8 | Q8TAE8 | Q8TAE8 | Growth arr |
| PAFAH1B2 | 1.085423 | 0.082144 | -0.22807 | | P68402;J3I | P68402 | P68402 | Platelet-act |
| GNPTG | 1.060474 | 0.087001 | 0.150492 | | Q9UJJ9;H0 | Q9UJJ9 | Q9UJJ9 | N-acetylglu |
| PDS5B | 1.047822 | 0.089573 | 0.163519 | | Q9NTI5;A9I | Q9NTI5 | Q9NTI5 | Sister chror |
| RAB8A | 1.047453 | 0.089649 | -0.1321 | | P61006;H0 | P61006 | P61006 | Ras-related |
| GCA | 1.042522 | 0.090673 | -0.13671 | | H7C2Z6;H7 | H7C2Z6;H7 | H7C2Z6 | Grancalcin |
| RANBP3 | 1.038059 | 0.09161 | 0.214141 | | B7Z7F3;Q9 | B7Z7F3;Q9 | B7Z7F3 | Ran-bindin |
| RPL34 | 1.037489 | 0.09173 | -0.14217 | | P49207 | P49207 | P49207 | 60S riboso |
| COQ3 | 1.036216 | 0.091999 | 0.234768 | | Q9NZJ6;F6 | Q9NZJ6 | Q9NZJ6 | Ubiquinon |
| CNN2 | 1.029059 | 0.093528 | 0.278685 | | B4DDF4;B4 | B4DDF4;B4 | B4DDF4 | Calponin;C |
| HSD17B7 | 0.989983 | 0.102333 | 0.194067 | | P56937 | P56937 | P56937 | 3-keto-ster |
| RAB22A | 0.9819 | 0.104256 | 0.132408 | | Q9UL26 | Q9UL26 | Q9UL26 | Ras-related |
| H2AFY2 | 0.963108 | 0.108866 | 0.296298 | | Q9P0M6;Q | Q9P0M6;Q | Q9P0M6 | Core histor |
| SIDT2 | 0.951375 | 0.111847 | 0.170243 | | C9JBG5;Q8 | C9JBG5;Q8 | C9JBG5 | SID1 transn |
| LRRFIP2 | 0.946523 | 0.113104 | 1.657579 | | Q9Y608;C9 | Q9Y608;C9 | Q9Y608 | Leucine-ric |
| TRAM1 | 0.935608 | 0.115982 | 0.100644 | | G3XAN4;Q | G3XAN4;Q | G3XAN4 | Translocati |
| MRPS6 | 0.91282 | 0.122231 | -0.17703 | | P82932 | P82932 | P82932 | 28S riboso |
| NSFL1C | 0.91058 | 0.122863 | 0.121776 | | Q9UNZ2;F2 | Q9UNZ2;F2 | Q9UNZ2 | NSFL1 cofa |
| RPA2 | 0.89933 | 0.126087 | 0.18578 | | P15927;Q5 | P15927;Q5 | P15927 | Replicatior |
| PSAT1 | 0.898562 | 0.12631 | -0.32224 | | Q9Y617 | Q9Y617 | Q9Y617 | Phosphose |
| PHF5A | 0.895212 | 0.127288 | 0.124007 | | Q7RTV0 | Q7RTV0 | Q7RTV0 | PHD finger- |
| CPPED1 | 0.889346 | 0.129019 | -0.35127 | | Q9BRF8 | Q9BRF8 | Q9BRF8 | Serine/thre |
| SRSF10 | 0.872783 | 0.134035 | 0.178333 | | Q5JRI1;O7 | Q5JRI1;O7 | Q5JRI1 | Serine/argi |
| DAK | 0.867559 | 0.135657 | -0.97648 | | Q3LXA3;H0 | Q3LXA3;H0 | Q3LXA3 | Bifunctiona |
| RPRD1B | 0.865261 | 0.136376 | 0.132352 | | Q9NQG5;E | Q9NQG5 | Q9NQG5 | Regulation |
| MPC2 | 0.843606 | 0.143349 | -0.1135 | | Q5R3B4;O | Q5R3B4;O | Q5R3B4 | Mitochond |
| ZC3H15 | 0.80986 | 0.154932 | 0.147587 | | Q8WU90;H | Q8WU90 | Q8WU90 | Zinc finger |
| ARFGAP2 | 0.802107 | 0.157722 | -1.14881 | | A0A0D9SF7 | A0A0D9SF7 | A0A0D9SF7 | ADP-ribosy |
| UBE2N | 0.794558 | 0.160488 | -0.2282 | | P61088;F8 | P61088;F8 | P61088 | Ubiquitin-c |
| QDPR | 0.788989 | 0.162559 | -0.20279 | | P09417;B7 | P09417;B7 | P09417 | Dihydropte |
| PISD | 0.774616 | 0.168029 | 0.196083 | | Q9UG56;H | Q9UG56;H | Q9UG56 | Phosphatic |
| SIGLEC7 | 0.744113 | 0.180255 | 0.15187 | | Q9Y286;M | Q9Y286 | Q9Y286 | Sialic acid-l |
| DCAKD | 0.742252 | 0.181029 | 0.215009 | | K7ESP4;Q8 | K7ESP4;Q8 | K7ESP4 | Dephosphc |
| RALB | 0.733099 | 0.184885 | -0.09415 | | P11234;C9 | P11234;C9 | P11234 | Ras-related |
| FAM134C | 0.709414 | 0.195248 | -0.13995 | | Q86VR2;K7 | Q86VR2;K7 | Q86VR2 | Protein FAI |
| HMGB2 | 0.704534 | 0.197454 | 0.227891 | | P26583;D6 | P26583;D6 | P26583 | High mobil |
| POLR2E | 0.697703 | 0.200585 | -0.06793 | | P19388;A0 | P19388;A0 | P19388 | DNA-direct |
| IK | 0.692571 | 0.202969 | -0.1606 | | Q13123;D6 | Q13123;D6 | Q13123 | Protein Rec |
| EPB41 | 0.684342 | 0.206851 | 0.107607 | | P11171;A0 | P11171;A0 | P11171 | Protein 4.1 |
| MYCBP | 0.684243 | 0.206898 | 0.122004 | | Q99417;A0 | Q99417;A0 | Q99417 | C-Myc-binc |
| THOC3 | 0.68326 | 0.207367 | 0.104432 | | Q96J01;D6 | Q96J01 | Q96J01 | THO compl |
| SCD | 0.678567 | 0.20962 | 0.088084 | | O00767 | O00767 | O00767 | Acyl-CoA de |

| | | | | | | | | |
|---|---|---|---|---|---|---|---|---|
| SMCHD1 | 0.652952 | 0.222355 | 0.738666 | | A6NHR9;J3 | A6NHR9 | A6NHR9 | Structural r |
| PPP1R8 | 0.638054 | 0.230115 | 0.094345 | | Q12972;A0 | Q12972;A0 | Q12972 | Nuclear inh |
| CLN5 | 0.620524 | 0.239594 | -0.12791 | | O75503;A0 | O75503;A0 | O75503 | Ceroid-lipo |
| Nucb2 | 0.614905 | 0.242714 | 0.714308 | | Q2L696;P8 | Q2L696;P8 | Q2L696 | Nucleobinc |
| FAM134A | 0.60867 | 0.246224 | 0.12834 | | Q8NC44;C9 | Q8NC44 | Q8NC44 | Protein FAM |
| VPS51 | 0.608129 | 0.246531 | 0.206938 | | Q9UID3;E9 | Q9UID3 | Q9UID3 | Vacuolar pr |
| CTNNBL1 | 0.581584 | 0.262069 | -0.13038 | | Q8WYA6;A( | Q8WYA6;A( | Q8WYA6 | Beta-cateni |
| ITCH | 0.579487 | 0.263337 | -0.87703 | | A0A590UK | A0A590UK | A0A590UK | E3 ubiquiti |
| METTL7A | 0.572501 | 0.267608 | -0.07899 | | Q9H8H3;H( | Q9H8H3;H( | Q9H8H3 | Methyltran |
| DPYSL3 | 0.57092 | 0.268584 | 1.225525 | | Q14195;H0 | Q14195;H0 | Q14195 | Dihydropyr |
| RPL27A | 0.568193 | 0.270276 | 0.113579 | | P46776;E9 | P46776;E9 | P46776 | 60S ribosor |
| PCNA | 0.564811 | 0.272389 | 0.127207 | | P12004 | P12004 | P12004 | Proliferatir |
| NDUFS7 | 0.561853 | 0.27425 | 0.156604 | | O75251;F5 | O75251;F5 | O75251 | NADH dehy |
| COMTD1 | 0.5616 | 0.27441 | -0.08381 | | Q86VU5;R4 | Q86VU5;R4 | Q86VU5 | Catechol O- |
| UCHL3 | 0.551032 | 0.281169 | -0.14361 | | P15374;A0 | P15374;A0 | P15374 | Ubiquitin c |
| CLCN3 | 0.542162 | 0.286971 | 0.071188 | | P51790;A0 | P51790;A0 | P51790 | H(+)/Cl(-) ex |
| SGTA | 0.531826 | 0.293883 | -1.3507 | | O43765;K7 | O43765;K7 | O43765 | Small gluta |
| MGST3 | 0.527311 | 0.296954 | 0.108609 | | O14880;Q5 | O14880;Q5 | O14880 | Microsoma |
| VTI1A | 0.51789 | 0.303466 | -0.7845 | | Q96AJ9 | Q96AJ9 | Q96AJ9 | Vesicle trar |
| LILRA2 | 0.512942 | 0.306943 | 0.131464 | | A8MZH0;A( | A8MZH0;A( | A8MZH0 | Leukocyte i |
| RAB4B | 0.511569 | 0.307915 | 0.063725 | | P61018;M( | P61018;M( | P61018 | Ras-related |
| GMIP | 0.506167 | 0.311769 | -0.14398 | | Q9P107 | Q9P107 | Q9P107 | GEM-intera |
| ISOC1 | 0.504211 | 0.313176 | 0.099435 | | Q96CN7;D6 | Q96CN7;D6 | Q96CN7 | Isochorism |
| PURA | 0.503053 | 0.314012 | 0.083487 | | Q00577;A0 | Q00577 | Q00577 | Transcripti |
| AKAP8L | 0.502754 | 0.314229 | -0.1121 | | Q9ULX6 | Q9ULX6 | Q9ULX6 | A-kinase an |
| RAB23 | 0.497449 | 0.31809 | -0.10267 | | Q9ULC3 | Q9ULC3 | Q9ULC3 | Ras-related |
| GPS1 | 0.494796 | 0.32004 | -0.68398 | | C9JFE4;Q1 | C9JFE4;Q1 | C9JFE4 | COP9 signa |
| RHEB | 0.492886 | 0.32145 | 0.090482 | | Q15382;C9 | Q15382;C9 | Q15382 | GTP-bindin |
| TRIM56 | 0.490945 | 0.322891 | 0.089256 | | Q9BRZ2;C9 | Q9BRZ2;C9 | Q9BRZ2 | E3 ubiquiti |
| YWHAB | 0.484591 | 0.327649 | -0.10102 | | P31946;Q4 | P31946 | P31946 | 14-3-3 prot |
| RBM12 | 0.470265 | 0.338637 | -0.64505 | | Q9NTZ6 | Q9NTZ6 | Q9NTZ6 | RNA-bindir |
| RALA | 0.470075 | 0.338786 | 0.046192 | | P11233;H7 | P11233;H7 | P11233 | Ras-related |
| SRP9 | 0.469732 | 0.339053 | -0.09072 | | P49458;Q6 | P49458;Q6 | P49458 | Signal recog |
| PPIL4 | 0.457105 | 0.349056 | -0.60991 | | Q8WUA2 | Q8WUA2 | Q8WUA2 | Peptidyl-pr |
| RCC2 | 0.450875 | 0.354099 | 0.09359 | | Q9P258 | Q9P258 | Q9P258 | Protein RC( |
| DUT | 0.438692 | 0.364173 | 0.062491 | | P33316;H0 | P33316;H0 | P33316 | Deoxyuridi |
| SRSF6 | 0.428433 | 0.372878 | -0.07454 | | A0A590UJK | A0A590UJK | A0A590UJK | Serine/argi |
| FNTA | 0.426144 | 0.374849 | -0.07135 | | B3KVN2;P4 | B3KVN2;P4 | B3KVN2 | Protein farr |
| FNDC3B | 0.425667 | 0.375261 | 0.082328 | | Q53EP0 | Q53EP0 | Q53EP0 | Fibronectir |
| RAB24 | 0.422286 | 0.378194 | 0.042102 | | Q969Q5;F8 | Q969Q5;F8 | Q969Q5 | Ras-related |
| ACOT13 | 0.42138 | 0.378983 | 0.076051 | | Q9NPJ3 | Q9NPJ3 | Q9NPJ3 | Acyl-coenzy |
| ANKLE2 | 0.410213 | 0.388854 | -0.19338 | | Q86XL3;F5 | Q86XL3 | Q86XL3 | Ankyrin rep |
| PPP5C | 0.388884 | 0.408429 | 0.123903 | | P53041;H0 | P53041;H0 | P53041 | Serine/thre |
| ATPIF1 | 0.388208 | 0.409065 | 0.126305 | | Q9UII2;A0A | Q9UII2;A0A | Q9UII2 | ATPase inhi |
| ANP32A | 0.385562 | 0.411565 | -0.07475 | | H0YN26;P3 | H0YN26;P3 | H0YN26 | Acidic leuci |
| LAMTOR3 | 0.380997 | 0.415913 | 0.072917 | | Q9UHA4 | Q9UHA4 | Q9UHA4 | Ragulator c |
| ZNF207 | 0.38011 | 0.416763 | 0.069892 | | J3QRS9;O4 | J3QRS9;O4 | J3QRS9 | BUB3-inter |
| GAPVD1 | 0.376306 | 0.42043 | -0.1114 | | F8W9S7;Q1 | F8W9S7;Q1 | F8W9S7 | GTPase-acti |

| | | | | | | | |
|---|---|---|---|---|---|---|---|
| DDX60 | 0.369874 | 0.426703 | 0.487948 | Q8IY21;H0Y | Q8IY21 | Q8IY21 | Probable AT |
| HMGA1 | 0.36442 | 0.432096 | -0.06953 | P17096;A0 | P17096 | P17096 | High mobil |
| NAA50 | 0.359725 | 0.436793 | 0.080277 | E7EQ69;Q9 | E7EQ69;Q9 | E7EQ69 | N-alpha-ace |
| RMND1 | 0.35114 | 0.445512 | 0.05783 | A0A087WX | A0A087WX | A0A087WX | Required fo |
| NIF3L1 | 0.344341 | 0.452542 | -0.05738 | Q9GZT8;E7 | Q9GZT8;E7 | Q9GZT8 | NIF3-like pr |
| RPL14 | 0.34417 | 0.452721 | -0.0744 | E7EPB3;P5 | E7EPB3;P5 | E7EPB3 | 60S ribosor |
| RPS23 | 0.34262 | 0.454339 | -0.06711 | P62266;D6 | P62266;D6 | P62266 | 40S ribosor |
| BTF3L4 | 0.341709 | 0.455293 | 0.074328 | Q96K17;E9 | Q96K17;E9 | Q96K17 | Transcripti |
| TEX10 | 0.337646 | 0.459572 | -0.07477 | Q9NXF1;A0 | Q9NXF1 | Q9NXF1 | Testis-expre |
| COMMD4 | 0.332581 | 0.464963 | 0.056194 | A0A0B4J28 | A0A0B4J28 | A0A0B4J28 | COMM dom |
| SPG20 | 0.330875 | 0.466794 | -0.07224 | Q8N0X7 | Q8N0X7 | Q8N0X7 | Spartin |
| HERC4 | 0.317074 | 0.481865 | -0.10393 | Q5GLZ8;H0 | Q5GLZ8 | Q5GLZ8 | Probable E3 |
| DHCR7 | 0.316656 | 0.48233 | -0.08397 | Q9UBM7;E9 | Q9UBM7;E9 | Q9UBM7 | 7-dehydroc |
| FMNL1 | 0.300602 | 0.500493 | 0.038719 | O95466;A0 | O95466;A0 | O95466 | Formin-like |
| TSSC1 | 0.29869 | 0.502701 | 0.077812 | A8MUM1;C | A8MUM1;C | A8MUM1 | Protein TSS |
| HEATR5B | 0.290947 | 0.511744 | -0.08681 | Q9P2D3 | Q9P2D3 | Q9P2D3 | HEAT repea |
| RAB2A | 0.290631 | 0.512117 | -0.07466 | P61019;E9 | P61019;E9 | P61019 | Ras-related |
| GNA11 | 0.286947 | 0.516479 | -0.03861 | P29992;K7 | P29992;K7 | P29992 | Guanine nu |
| TUBGCP2 | 0.285944 | 0.517674 | -0.74293 | Q9BSJ2;F2 | Q9BSJ2;F2 | Q9BSJ2 | Gamma-tul |
| EIF4A1 | 0.281036 | 0.523557 | 0.054624 | P60842;J3 | P60842;J3 | P60842 | Eukaryotic |
| MRPL2 | 0.2785 | 0.526623 | 0.050546 | A0A669KA | A0A669KA | A0A669KA | 39S ribosor |
| SLC1A5 | 0.278468 | 0.526662 | 0.040737 | Q15758;M | Q15758;M | Q15758 | Neutral am |
| XRN1 | 0.27781 | 0.52746 | 0.59785 | Q8IZH2;H7 | Q8IZH2;H7 | Q8IZH2 | 5-3 exoribo |
| INTS3 | 0.276922 | 0.52854 | -0.90694 | Q68E01;A0 | Q68E01 | Q68E01 | Integrator |
| GALNT1 | 0.268344 | 0.539083 | -0.06487 | Q10472;K7 | Q10472 | Q10472 | Polypeptid |
| TPRG1L | 0.266527 | 0.541343 | -0.05942 | Q5T0D9 | Q5T0D9 | Q5T0D9 | Tumor prot |
| NOL9 | 0.25594 | 0.554702 | 0.065875 | Q5SY16 | Q5SY16 | Q5SY16 | Polynucleo |
| NAP1L4 | 0.245208 | 0.568581 | -0.06656 | Q99733;C9 | Q99733;C9 | Q99733 | Nucleosom |
| IRF5 | 0.24157 | 0.573364 | -0.07677 | Q13568;C9 | Q13568 | Q13568 | Interferon |
| ST13 | 0.235224 | 0.581803 | -0.04237 | P50502;Q3 | P50502;Q3 | P50502 | Hsc70-inter |
| SRRM1 | 0.223201 | 0.598135 | -0.07238 | A9Z1X7;Q8 | A9Z1X7;Q8 | A9Z1X7 | Serine/argi |
| PSMB10 | 0.216878 | 0.606907 | 0.029591 | P40306;J3 | P40306 | P40306 | Proteasom |
| VKORC1L1 | 0.209144 | 0.617812 | -0.05867 | A0A3B3ISV | A0A3B3ISV | A0A3B3ISV | Vitamin K e |
| MOSPD1 | 0.207061 | 0.620782 | -0.07023 | Q9UJG1 | Q9UJG1 | Q9UJG1 | Motile sper |
| COQ7 | 0.201139 | 0.629304 | -0.06039 | H3BP28;Q9 | H3BP28;Q9 | H3BP28 | 5-demetho |
| LYPLAL1 | 0.198088 | 0.633741 | -0.04182 | Q5VWZ2 | Q5VWZ2 | Q5VWZ2 | Lysophosph |
| NASP | 0.196225 | 0.636466 | -0.35775 | P49321;E9 | P49321;E9 | P49321 | Nuclear aut |
| ALG1 | 0.192304 | 0.642238 | -0.06051 | Q9BT22;K7 | Q9BT22 | Q9BT22 | Chitobiosyl |
| FN1 | 0.191149 | 0.643949 | -0.43043 | P02751;H0 | P02751 | P02751 | Fibronectin |
| EIF2B1 | 0.189606 | 0.64624 | -0.03755 | Q14232;F5 | Q14232;F5 | Q14232 | Translation |
| MRPL10 | 0.187263 | 0.649736 | 0.044655 | Q7Z7H8;F2 | Q7Z7H8 | Q7Z7H8 | 39S ribosor |
| RPA3 | 0.185336 | 0.652625 | 0.034833 | P35244;B5 | P35244;B5 | P35244 | Replication |
| OPLAH | 0.184122 | 0.654453 | -0.42631 | O14841 | O14841 | O14841 | 5-oxoprolir |
| ARHGAP30 | 0.183709 | 0.655075 | 0.037795 | Q7Z6I6;A0 | Q7Z6I6;A0 | Q7Z6I6 | Rho GTPase |
| BANF1 | 0.18336 | 0.655602 | -0.03727 | O75531;E9 | O75531 | O75531 | Barrier-to-a |
| WDR5 | 0.177981 | 0.663773 | 0.047276 | P61964;V9 | P61964 | P61964 | WD repeat- |
| APOBEC3C | 0.177067 | 0.66517 | 0.029436 | Q9NRW3;C | Q9NRW3 | Q9NRW3 | DNA dC->dU |
| PPOX | 0.167475 | 0.680025 | -0.05335 | A0A494C14 | A0A494C14 | A0A494C14 | Protoporph |

| | | | | | | | |
|---|---|---|---|---|---|---|---|
| CDV3 | 0.166244 | 0.681955 | 0.046426 | Q9UKY7;H( | Q9UKY7;H( | Q9UKY7 | Protein CD' |
| DDX27 | 0.164432 | 0.684807 | 0.030909 | B7Z6D5;Q9 | B7Z6D5;Q9 | B7Z6D5 | Probable A' |
| USP10 | 0.153492 | 0.702276 | 0.040828 | Q14694;H3 | Q14694 | Q14694 | Ubiquitin c |
| SF3B4 | 0.151765 | 0.705075 | -0.0271 | Q15427;Q5 | Q15427;Q5 | Q15427 | Splicing fac |
| YLPM1 | 0.147876 | 0.711416 | 0.03573 | F8VU51;P4 | F8VU51;P4 | F8VU51 | YLP motif-c |
| SAR1B | 0.141539 | 0.721873 | 0.032794 | Q9Y6B6;Q9 | Q9Y6B6;Q9 | Q9Y6B6 | GTP-bindin |
| MRPL3 | 0.13816 | 0.727512 | 0.030744 | H0Y9G6;E7 | H0Y9G6;E7 | H0Y9G6 | 39S riboso |
| TIMM13 | 0.132988 | 0.736227 | 0.024021 | Q9Y5L4;K7 | Q9Y5L4;K7 | Q9Y5L4 | Mitochond |
| TPCN1 | 0.120608 | 0.757516 | -0.02413 | B7Z3R2;Q9 | B7Z3R2;Q9 | B7Z3R2 | Two pore c |
| PSMB7 | 0.11411 | 0.768935 | 0.033584 | Q99436;Q5 | Q99436 | Q99436 | Proteasom |
| PTGES3 | 0.111309 | 0.77391 | 0.027724 | Q15185;A0 | Q15185;A0 | Q15185 | Prostaglan |
| SLAMF7 | 0.105071 | 0.785108 | 0.030807 | Q9NQ25;B4 | Q9NQ25;B4 | Q9NQ25 | SLAM famil |
| GMPPA | 0.102741 | 0.78933 | -0.0423 | Q96IJ6;A0A | Q96IJ6;A0A | Q96IJ6 | Mannose-1 |
| PROSC | 0.101893 | 0.790873 | -0.0328 | O94903;E5 | O94903;E5 | O94903 | Proline syn |
| NPC2 | 0.100751 | 0.792955 | 0.022783 | J3KMY5;G3 | J3KMY5;G3 | J3KMY5 | Epididymal |
| FYB | 0.093581 | 0.806157 | 0.024418 | O15117;D6 | O15117 | O15117 | FYN-bindin |
| TSNAX | 0.092483 | 0.808197 | -0.04014 | Q99598;C4 | Q99598;C4 | Q99598 | Translin-as |
| LAS1L | 0.090685 | 0.81155 | 0.022895 | Q9Y4W2 | Q9Y4W2 | Q9Y4W2 | Ribosomal |
| LARP7 | 0.089509 | 0.813751 | 0.046142 | Q4G0J3;D6 | Q4G0J3;D6 | Q4G0J3 | La-related p |
| NUDT9 | 0.088209 | 0.81619 | -0.01063 | Q9BW91;D | Q9BW91;D | Q9BW91 | ADP-ribose |
| BTN3A3 | 0.080592 | 0.830631 | -0.01581 | O00478;C9 | O00478;C9 | O00478 | Butyrophili |
| SLC25A40 | 0.078588 | 0.834472 | -0.03562 | Q8TBP6 | Q8TBP6 | Q8TBP6 | Solute carri |
| HINT1 | 0.074512 | 0.842341 | -0.02067 | P49773;D6 | P49773;D6 | P49773 | Histidine tr |
| STX6 | 0.074368 | 0.842621 | 0.020411 | O43752 | O43752 | O43752 | Syntaxin-6 |
| EMC4 | 0.063044 | 0.86488 | -0.01882 | Q5J8M3;H( | Q5J8M3;H( | Q5J8M3 | ER membra |
| AP3S1 | 0.062755 | 0.865455 | -0.01586 | Q92572;F5 | Q92572;F5 | Q92572 | AP-3 compl |
| C2orf47 | 0.062062 | 0.866838 | 0.020235 | Q8WWC4;I | Q8WWC4 | Q8WWC4 | Uncharacte |
| EMC10 | 0.053386 | 0.884329 | 0.01114 | M0R2A0;Q | M0R2A0;Q | M0R2A0 | ER membra |
| ARL8B | 0.050108 | 0.89103 | 0.012307 | Q9NVJ2 | Q9NVJ2 | Q9NVJ2 | ADP-ribosy |
| FKBP2 | 0.049441 | 0.892398 | -0.00861 | A0A494C1I | A0A494C1I | A0A494C1I | Peptidyl-pr |
| SORD | 0.045538 | 0.900455 | -0.01812 | A0A6I8PIS1 | A0A6I8PIS1 | A0A6I8PIS1 | Sorbitol de |
| STEAP3 | 0.042246 | 0.907307 | -0.01394 | B8ZZX6;Q6 | B8ZZX6;Q6 | B8ZZX6 | Metallored |
| FEN1 | 0.033499 | 0.925766 | -0.01302 | P39748;I3I | P39748 | P39748 | Flap endon |
| EIF6 | 0.029031 | 0.93534 | -0.00674 | P56537;B7 | P56537 | P56537 | Eukaryotic |
| RPL30 | 0.024706 | 0.9447 | 0.003711 | P62888;E5 | P62888;E5 | P62888 | 60S riboso |
| RHBDF2 | 0.023374 | 0.947603 | 0.006531 | Q6PJF5 | Q6PJF5 | Q6PJF5 | Inactive rh |
| FNBP1 | 0.020972 | 0.952859 | 0.006875 | B7ZL14;Q9 | B7ZL14;Q9 | B7ZL14 | Formin-bin |
| GOSR2 | 0.018505 | 0.958286 | 0.004255 | O14653;I3I | O14653;I3I | O14653 | Golgi SNAP |
| SELENOF | 0.018289 | 0.958763 | 0.004353 | A0A087X1( | A0A087X1( | A0A087X1( | 15 kDa sele |
| RAB38 | 0.004736 | 0.989154 | -0.00124 | P57729;H0 | P57729;H0 | P57729 | Ras-related |
| LYPLA2 | 0.002939 | 0.993255 | -0.00094 | O95372;Q5 | O95372;Q5 | O95372 | Acyl-protei |
| DKFZp566H | 0.001311 | 0.996986 | 0.000292 | Q9UFM8;Q | Q9UFM8;Q | Q9UFM8 | Neuroplast |
| CCRL2 | 2.998863 | 0.001003 | 0.465751 | O00421;C9 | O00421 | O00421 | C-C chemol |
| COA3 | 4.247887 | 5.65E-05 | 0.420593 | Q9Y2R0;K7 | Q9Y2R0;K7 | Q9Y2R0 | Cytochrom |
| NDUFB4 | 3.843377 | 0.000143 | 0.381474 | O95168;F2 | O95168;F2 | O95168 | NADH dehy |
| RCSD1 | 1.762088 | 0.017295 | 0.326447 | B7ZKW8;Q | B7ZKW8;Q | B7ZKW8 | CapZ-intera |
| PPP4C | 2.45886 | 0.003476 | 0.305501 | H3BTA2;P6 | H3BTA2;P6 | H3BTA2 | Serine/thre |
| RPS28 | 2.465039 | 0.003427 | 0.277745 | P62857 | P62857 | P62857 | 40S riboso |

| | | | | | | | |
|---|---|---|---|---|---|---|---|
| FAM3C | 1.617519 | 0.024126 | 0.273031 | Q92520;C9 | Q92520 | Q92520 | Protein FAI |
| EMC3 | 2.013569 | 0.009692 | 0.272381 | Q9P0I2;S4I | Q9P0I2 | Q9P0I2 | ER membra |
| SF3B5 | 2.310102 | 0.004897 | 0.263816 | Q9BWJ5 | Q9BWJ5 | Q9BWJ5 | Splicing fac |
| NDUFS4 | 1.899559 | 0.012602 | 0.263353 | O43181;D6 | O43181 | O43181 | NADH dehy |
| ABAT | 1.561745 | 0.027432 | 0.252892 | H3BRN4;H3 | H3BRN4;H3 | H3BRN4 | 4-aminobu |
| TRIAP1 | 3.004868 | 0.000989 | 0.224902 | O43715 | O43715 | O43715 | TP53-regul |
| ATP6V1F | 1.333106 | 0.04644 | 0.220511 | Q16864 | Q16864 | Q16864 | V-type prot |
| BLOC1S1 | 2.292833 | 0.005095 | 0.208337 | P78537;G8 | P78537;G8 | P78537 | Biogenesis |
| PIP4K2A | 2.843622 | 0.001433 | 0.202161 | P48426;H7 | P48426;H7 | P48426 | Phosphatic |
| UBE2I | 1.826995 | 0.014894 | 0.200797 | P63279;H3 | P63279;H3 | P63279 | SUMO-conj |
| RPS15 | 1.804779 | 0.015675 | 0.195318 | K7ELC2;K7 | K7ELC2;K7 | K7ELC2 | 40S ribosor |
| CRLF3 | 1.363763 | 0.043275 | 0.192885 | Q8IUI8;J3C | Q8IUI8 | Q8IUI8 | Cytokine re |
| TIMM9 | 1.366551 | 0.042998 | 0.185865 | Q9Y5J7;A0 | Q9Y5J7;A0 | Q9Y5J7 | Mitochond |
| RAB29 | 2.058285 | 0.008744 | 0.181422 | O14966;E9 | O14966 | O14966 | Ras-related |
| HNRNPK | 3.385556 | 0.000412 | 0.179164 | P61978;S4 | P61978 | P61978 | Heterogene |
| GOLIM4 | 1.357894 | 0.043864 | 0.178116 | F8W785;O | F8W785;O | F8W785 | Golgi integ |
| UQCC2 | 1.62389 | 0.023774 | 0.169275 | Q9BRT2;V5 | Q9BRT2;V5 | Q9BRT2 | Ubiquinol- |
| FAM21A | 1.319517 | 0.047916 | 0.168938 | Q641Q2;F8 | Q641Q2;F8 | Q641Q2 | WASH com |
| NDUFS5 | 2.056321 | 0.008784 | 0.163804 | O43920 | O43920 | O43920 | NADH dehy |
| WDR43 | 1.439775 | 0.036327 | 0.161389 | Q15061;C9 | Q15061 | Q15061 | WD repeat- |
| CYBA | 1.867924 | 0.013554 | 0.152105 | P13498;H3 | P13498 | P13498 | Cytochrom |
| MTCH2 | 1.597139 | 0.025285 | 0.148747 | Q9Y6C9 | Q9Y6C9 | Q9Y6C9 | Mitochond |
| LMBRD2 | 1.33543 | 0.046192 | 0.146238 | Q68DH5 | Q68DH5 | Q68DH5 | LMBR1 dor |
| RBM8A | 1.325525 | 0.047258 | -0.17637 | Q9Y5S9;A0 | Q9Y5S9 | Q9Y5S9 | RNA-bindir |
| SLC12A9 | 1.544027 | 0.028574 | -0.19385 | Q9BXP2 | Q9BXP2 | Q9BXP2 | Solute carri |
| KPNA1 | 1.807281 | 0.015585 | -0.23099 | P52294;C9 | P52294 | P52294 | Importin su |
| UBLCP1 | 1.506225 | 0.031173 | -0.24024 | Q8WVY7 | Q8WVY7 | Q8WVY7 | Ubiquitin-l |
| UBE2Z | 1.346279 | 0.045053 | -0.26134 | Q9H832;I3 | Q9H832 | Q9H832 | Ubiquitin-c |
| PEA15 | 1.620767 | 0.023946 | -0.28998 | Q15121;B1 | Q15121;B1 | Q15121 | Astrocytic |
| MRTO4 | 2.112276 | 0.007722 | -0.30908 | Q9UKD2 | Q9UKD2 | Q9UKD2 | mRNA turn |
| OSBPL9 | 1.278446 | 0.052669 | -0.2811 | Q96SU4;A0 | Q96SU4 | Q96SU4 | Oxysterol-b |
| PNPO | 1.277007 | 0.052844 | -0.20974 | J3QQZ9;Q9 | J3QQZ9;Q9 | J3QQZ9 | Pyridoxine- |
| VPS25 | 1.270854 | 0.053598 | -0.23527 | Q9BRG1;K7 | Q9BRG1;K7 | Q9BRG1 | Vacuolar pr |
| FAM134B | 1.259567 | 0.055009 | 0.273893 | Q9H6L5;H0 | Q9H6L5 | Q9H6L5 | Protein FAI |
| NFU1 | 1.253422 | 0.055793 | 0.149804 | Q9UMS0;C | Q9UMS0;C | Q9UMS0 | NFU1 iron- |
| SPCS3 | 1.25312 | 0.055832 | -0.14184 | P61009 | P61009 | P61009 | Signal pept |
| PYCR2 | 1.250433 | 0.056178 | 0.188107 | A0A087WT | A0A087WT | A0A087WT | Pyrroline-5 |
| CERS2 | 1.247376 | 0.056575 | -0.21679 | Q96G23;H0 | Q96G23;H0 | Q96G23 | Ceramide s |
| HINT2 | 1.234426 | 0.058287 | 0.128936 | Q9BX68 | Q9BX68 | Q9BX68 | Histidine tr |
| IL18 | 1.231223 | 0.058719 | -0.3132 | Q14116 | Q14116 | Q14116 | Interleukin |
| HSPE1-MOI | 1.206448 | 0.062166 | 0.077216 | S4R3N1;Q9 | S4R3N1;Q9 | S4R3N1 | MOB-like pr |
| TECPR1 | 1.203858 | 0.062538 | 1.751343 | Q7Z6L1;C9 | Q7Z6L1 | Q7Z6L1 | Tectonin be |
| NMI | 1.197579 | 0.063448 | -0.18843 | Q13287;C9 | Q13287 | Q13287 | N-myc-inte |
| PIK3AP1 | 1.176154 | 0.066657 | 0.165463 | Q6ZUJ8 | Q6ZUJ8 | Q6ZUJ8 | Phosphoin |
| ATL2 | 1.150004 | 0.070794 | -0.14835 | Q8NHH9;B | Q8NHH9;B | Q8NHH9 | Atlastin-2 |
| DDT | 1.144947 | 0.071623 | -0.27593 | P30046;J3 | P30046;J3 | P30046 | D-dopachro |
| ATP6V1G1 | 1.105512 | 0.078431 | -0.1471 | O75348;F2 | O75348 | O75348 | V-type prot |
| NUP153 | 1.08966 | 0.081347 | 0.329591 | P49790 | P49790 | P49790 | Nuclear po |

| | | | | | | | |
|---|---|---|---|---|---|---|---|
| HINT3 | 1.071536 | 0.084813 | 0.121798 | Q9NQE9 | Q9NQE9 | Q9NQE9 | Histidine tr |
| ARHGAP4 | 1.071077 | 0.084903 | -0.25817 | E9PCM6;E7 | E9PCM6;E7 | E9PCM6 | Rho GTPase |
| PCBD2 | 1.069174 | 0.085276 | -0.14141 | H0YA52;Q9 | H0YA52;Q9 | H0YA52 | Pterin-4-alp |
| TNS1 | 1.064174 | 0.086263 | -0.15444 | A0A494C0( | A0A494C0( | A0A494C0( | Tensin-1 |
| ALYREF | 1.060824 | 0.086931 | 0.135023 | E9PB61;Q8 | E9PB61;Q8 | E9PB61 | THO compl |
| CHCHD6 | 1.047806 | 0.089576 | 0.19306 | Q9BRQ6;H( | Q9BRQ6;H( | Q9BRQ6 | MICOS com |
| OXNAD1 | 1.041214 | 0.090947 | -1.62164 | U3KQU8;C9 | U3KQU8;C9 | U3KQU8 | Oxidoreduc |
| IKBKE | 1.029541 | 0.093424 | -1.45508 | A0A075B7I | A0A075B7I | A0A075B7I | Inhibitor of |
| RPF2 | 1.024934 | 0.09442 | 0.197988 | Q9H7B2;Q5 | Q9H7B2;Q5 | Q9H7B2 | Ribosome p |
| TREX1 | 0.999502 | 0.100115 | 0.141773 | Q9NSU2;C9 | Q9NSU2;C9 | Q9NSU2 | Three-prim |
| UMPS | 0.996519 | 0.100805 | 0.151035 | P11172;E9 | P11172;E9 | P11172 | Uridine 5-m |
| PTMA | 0.971702 | 0.106733 | -0.41397 | B8ZZQ6;P0 | B8ZZQ6;P0 | B8ZZQ6 | Prothymos |
| NUPL1 | 0.961002 | 0.109395 | 0.122994 | Q5JRG1;H7 | Q5JRG1;H7 | Q5JRG1 | Nucleopori |
| RAB4A | 0.934595 | 0.116253 | -0.19569 | P20338;A0 | P20338;A0 | P20338 | Ras-related |
| HNMT | 0.915729 | 0.121415 | -0.21148 | P50135 | P50135 | P50135 | Histamine I |
| SLC30A5 | 0.914456 | 0.121771 | 0.142115 | Q8TAD4;Q9 | Q8TAD4;Q9 | Q8TAD4 | Zinc transp |
| OXA1L | 0.913957 | 0.121911 | 0.210481 | A0A087X0I | A0A087X0I | A0A087X0I | Mitochond |
| PHYH | 0.907502 | 0.123737 | 0.179593 | B1ALH6;O1 | B1ALH6;O1 | B1ALH6 | Phytanoyl- |
| OCRL | 0.902722 | 0.125106 | 1.173714 | Q01968;A0 | Q01968 | Q01968 | Inositol pol |
| TFG | 0.880979 | 0.131529 | 0.122609 | Q92734;C9 | Q92734;C9 | Q92734 | Protein TFG |
| CUL3 | 0.865347 | 0.136349 | 0.15549 | Q13618;A0 | Q13618;A0 | Q13618 | Cullin-3 |
| FIS1 | 0.860319 | 0.137937 | -0.19451 | Q9Y3D6 | Q9Y3D6 | Q9Y3D6 | Mitochond |
| SYNE1 | 0.850927 | 0.140952 | 0.187349 | E7ENN3;A0 | E7ENN3;A0 | E7ENN3 | Nesprin-1 |
| PTER | 0.836193 | 0.145817 | -0.26901 | Q96BW5;A | Q96BW5;A | Q96BW5 | Phosphotri |
| PTGDS | 0.834236 | 0.146475 | -0.11 | P41222;H0 | P41222;H0 | P41222 | Prostagland |
| SCAMP2 | 0.828008 | 0.148591 | 0.132428 | O15127;H3 | O15127;H3 | O15127 | Secretory c |
| GATM | 0.783331 | 0.164691 | 0.120198 | P50440;H0 | P50440 | P50440 | Glycine am |
| LANCL2 | 0.781048 | 0.165559 | 0.136462 | Q9NS86 | Q9NS86 | Q9NS86 | LanC-like pr |
| SEPN1 | 0.780104 | 0.165919 | -1.09786 | Q9NZV5;H9 | Q9NZV5;H9 | Q9NZV5 | Selenoprot |
| AQR | 0.754091 | 0.176161 | -0.10801 | O60306 | O60306 | O60306 | Intron-binc |
| INPPL1 | 0.737982 | 0.182818 | -0.16047 | A0A0A0MT | A0A0A0MT | A0A0A0MT | Phosphatic |
| TRAPPC12 | 0.733521 | 0.184705 | -1.07696 | Q8WVT3;A( | Q8WVT3;A( | Q8WVT3 | Trafficking |
| COPS6 | 0.731666 | 0.185496 | 0.101954 | Q7L5N1;E7 | Q7L5N1;E7 | Q7L5N1 | COP9 signa |
| ALG9 | 0.730663 | 0.185925 | 0.102191 | A0A087WZ | A0A087WZ | A0A087WZ | Alpha-1,2-r |
| BAX | 0.729744 | 0.186318 | 0.096691 | Q07812;K4 | Q07812;K4 | Q07812 | Apoptosis r |
| FARS2 | 0.720383 | 0.190378 | -0.14866 | O95363;A0 | O95363;A0 | O95363 | Phenylalan |
| CHERP | 0.714759 | 0.19286 | -0.13567 | J3QK89;Q8 | J3QK89;Q8 | J3QK89 | Calcium ho |
| FAM45A | 0.702697 | 0.198291 | 0.240617 | Q8TCE6;A0 | Q8TCE6;A0 | Q8TCE6 | Protein FAM |
| UBE2L6 | 0.696382 | 0.201196 | -0.56909 | O14933;E9 | O14933;E9 | O14933 | Ubiquitin/I |
| GSPT1 | 0.688695 | 0.204788 | -0.10897 | P15170;H3 | P15170;H3 | P15170 | Eukaryotic |
| XPO7 | 0.688687 | 0.204792 | -0.12692 | Q9UIA9;E7 | Q9UIA9;E7 | Q9UIA9 | Exportin-7 |
| PTGR1 | 0.688547 | 0.204858 | -0.2984 | Q14914;F2 | Q14914;F2 | Q14914 | Prostagland |
| RRP1 | 0.678733 | 0.20954 | 0.103652 | P56182;D6 | P56182 | P56182 | Ribosomal |
| RABL6 | 0.671848 | 0.212888 | -0.19108 | F2Z2T0;Q3 | F2Z2T0;Q3 | F2Z2T0 | Rab-like pro |
| PPP6R1 | 0.670615 | 0.213494 | -0.11737 | Q9UPN7;K7 | Q9UPN7 | Q9UPN7 | Serine/thre |
| RPS26 | 0.663098 | 0.217221 | 0.103757 | P62854;Q5 | P62854;Q5 | P62854 | 40S riboson |
| TOMM22 | 0.657528 | 0.220025 | 0.109024 | Q9NS69 | Q9NS69 | Q9NS69 | Mitochond |
| TBCA | 0.651795 | 0.222949 | -0.14563 | E5RIW3;E5 | E5RIW3;E5 | E5RIW3 | Tubulin-spe |

| | | | | | | | |
|---|---|---|---|---|---|---|---|
| SLC25A16 | 0.650015 | 0.223865 | 1.098965 | P16260;V9 | P16260 | P16260 | Graves dise |
| PUM1 | 0.648534 | 0.224629 | 0.092067 | E9PR38;H0 | E9PR38;H0 | E9PR38 | Pumilio ho |
| MRPS17 | 0.645973 | 0.225958 | -0.06894 | Q9Y2R5;I3I | Q9Y2R5;I3I | Q9Y2R5 | 28S riboson |
| TCEB1 | 0.638335 | 0.229967 | -0.10102 | Q15369;E5 | Q15369;E5 | Q15369 | Transcripti |
| PAPSS1 | 0.623446 | 0.237987 | -0.15509 | O43252 | O43252 | O43252 | Bifunctiona |
| GMPR2 | 0.614224 | 0.243095 | -0.81572 | Q9P2T1;H0 | Q9P2T1;H0 | Q9P2T1 | GMP reduc |
| SGPP1 | 0.596434 | 0.25326 | 0.092303 | Q9BX95 | Q9BX95 | Q9BX95 | Sphingosin |
| POLDIP3 | 0.592637 | 0.255484 | -0.18519 | F6VRR5;Q9 | F6VRR5;Q9 | F6VRR5 | Polymerase |
| RBM17 | 0.58809 | 0.258173 | -0.12605 | Q96I25;Q5 | Q96I25;Q5 | Q96I25 | Splicing fac |
| BLMH | 0.585445 | 0.25975 | -0.94733 | Q13867;K7 | Q13867;K7 | Q13867 | Bleomycin |
| VPS8 | 0.577897 | 0.264304 | 0.605179 | Q8N3P4;C9 | Q8N3P4;C9 | Q8N3P4 | Vacuolar pr |
| TRNT1 | 0.576209 | 0.265333 | 0.132928 | Q96Q11;F8 | Q96Q11 | Q96Q11 | CCA tRNA n |
| PIR | 0.57414 | 0.2666 | -0.18526 | O00625 | O00625 | O00625 | Pirin |
| MRRF | 0.572938 | 0.267339 | 0.12651 | Q96E11;X6 | Q96E11;X6 | Q96E11 | Ribosome-r |
| RAI14 | 0.564189 | 0.272779 | 0.193797 | Q9P0K7;D6 | Q9P0K7 | Q9P0K7 | Ankycorbin |
| EIF4H | 0.562435 | 0.273883 | -0.09378 | Q15056 | Q15056 | Q15056 | Eukaryotic |
| KXD1 | 0.561044 | 0.274762 | -0.11356 | M0R0B4;M | M0R0B4;M | M0R0B4 | KxDL motif |
| UBE3A | 0.552614 | 0.280147 | -1.03046 | A0A1B0GVI | A0A1B0GVI | A0A1B0GVI | Ubiquitin-p |
| NCS1 | 0.55089 | 0.281261 | 1.232017 | P62166 | P62166 | P62166 | Neuronal ca |
| RPLP2 | 0.541621 | 0.287328 | 0.053507 | P05387;H0 | P05387 | P05387 | 60S acidic r |
| MON1B | 0.54061 | 0.287999 | 1.27308 | Q7L1V2;E7 | Q7L1V2;E7 | Q7L1V2 | Vacuolar fu |
| CNOT10 | 0.540221 | 0.288256 | 0.082643 | Q9H9A5;E9 | Q9H9A5;E9 | Q9H9A5 | CCR4-NOT t |
| DOCK5 | 0.528288 | 0.296287 | 2.417077 | Q9H7D0;H0 | Q9H7D0;H0 | Q9H7D0 | Dedicator o |
| LARP1 | 0.524776 | 0.298692 | -1.33528 | Q6PKG0;E5 | Q6PKG0;E5 | Q6PKG0 | La-related p |
| FRG1 | 0.517694 | 0.303603 | 0.70769 | Q14331;E9 | Q14331;E9 | Q14331 | Protein FRG |
| NDUFB6 | 0.50843 | 0.310149 | -0.169 | O95139;A0 | O95139;A0 | O95139 | NADH dehy |
| ACOT4 | 0.505278 | 0.312408 | -0.7633 | Q8N9L9 | Q8N9L9 | Q8N9L9 | Acyl-coenzy |
| TRAPPC6B | 0.499475 | 0.31661 | 0.220127 | Q86SZ2;G3 | Q86SZ2;G3 | Q86SZ2 | Trafficking p |
| RPE | 0.499309 | 0.316731 | -0.13454 | Q96AT9;C9 | Q96AT9;C9 | Q96AT9 | Ribulose-ph |
| COX20 | 0.493648 | 0.320887 | 0.092225 | Q5RI15 | Q5RI15 | Q5RI15 | Cytochrom |
| BUD31 | 0.491279 | 0.322642 | 0.093514 | P41223;C9 | P41223;C9 | P41223 | Protein BU |
| MECP2 | 0.488547 | 0.324678 | 0.104351 | P51608;A0 | P51608;A0 | P51608 | Methyl-CpG |
| CRK | 0.48274 | 0.329048 | -0.10947 | P46108;I3I | P46108;I3I | P46108 | Adapter mo |
| ACP1 | 0.477322 | 0.333179 | -0.12073 | P24666;G5 | P24666;G5 | P24666 | Low molecu |
| DNAJB1 | 0.474171 | 0.335606 | -0.10065 | P25685;M0 | P25685;M0 | P25685 | DnaJ homo |
| DNASE2 | 0.474076 | 0.335679 | -0.0734 | O00115;K7 | O00115;K7 | O00115 | Deoxyribon |
| MFSD12 | 0.467731 | 0.340619 | 0.100801 | Q6NUT3;K7 | Q6NUT3;K7 | Q6NUT3 | Major facili |
| PAIP1 | 0.463688 | 0.343805 | -0.98811 | D6REB4;Q9 | D6REB4;Q9 | D6REB4 | Polyadenyl |
| MAPKAPK3 | 0.454497 | 0.351158 | 0.116065 | C9J8E1;Q1 | C9J8E1;Q1 | C9J8E1 | MAP kinase |
| RBMX | 0.445441 | 0.358558 | 0.099157 | P38159;H3 | P38159;H3 | P38159 | RNA-bindin |
| IMPAD1 | 0.442037 | 0.361379 | 0.087136 | Q9NX62;H0 | Q9NX62 | Q9NX62 | Inositol mo |
| PPP4R2 | 0.440096 | 0.362997 | 0.134274 | Q9NY27;C9 | Q9NY27;C9 | Q9NY27 | Serine/thre |
| KPNA3 | 0.431277 | 0.370444 | 0.051379 | O00505;H0 | O00505 | O00505 | Importin su |
| GCLM | 0.415791 | 0.383892 | -0.11073 | P48507 | P48507 | P48507 | Glutamate- |
| ZZEF1 | 0.41496 | 0.384627 | 0.58779 | O43149 | O43149 | O43149 | Zinc finger |
| H1F0 | 0.41487 | 0.384707 | 0.058677 | P07305 | P07305 | P07305 | Histone H1 |
| NDUFB8 | 0.411438 | 0.387759 | 0.081502 | O95169;E9 | O95169;E9 | O95169 | NADH dehy |
| AGPAT6 | 0.410577 | 0.388529 | 0.08335 | Q86UL3;E5 | Q86UL3 | Q86UL3 | Glycerol-3- |

| | | | | | | | |
|---|---|---|---|---|---|---|---|
| NTPCR | 0.409178 | 0.389782 | 0.119784 | Q9BSD7;Q5 | Q9BSD7;Q5 | Q9BSD7 | Cancer-rela |
| LSM2 | 0.408229 | 0.390635 | 0.050311 | Q9Y333 | Q9Y333 | Q9Y333 | U6 snRNA-a |
| ARHGAP10 | 0.398437 | 0.399542 | -0.55953 | A1A4S6;H0 | A1A4S6 | A1A4S6 | Rho GTPase |
| WDR77 | 0.391833 | 0.405665 | -0.71334 | Q9BQA1;H( | Q9BQA1;H( | Q9BQA1 | Methylosor |
| NDUFB1 | 0.391271 | 0.40619 | 0.039187 | O75438 | O75438 | O75438 | NADH dehy |
| FRMD8 | 0.391093 | 0.406356 | -0.66631 | Q9BZ67;E9 | Q9BZ67;E9 | Q9BZ67 | FERM doma |
| MAPK14 | 0.389943 | 0.407434 | -0.17942 | B4E0K5;Q1 | B4E0K5;Q1 | B4E0K5 | Mitogen-ac |
| GGA1 | 0.387445 | 0.409784 | -0.61413 | Q9UJY5;B0 | Q9UJY5;B0 | Q9UJY5 | ADP-ribosy |
| TMEM192 | 0.379289 | 0.417552 | 0.076418 | Q8IY95;D6I | Q8IY95 | Q8IY95 | Transmemb |
| ARMC1 | 0.377812 | 0.418975 | 0.079617 | Q9NVT9;E5 | Q9NVT9;E5 | Q9NVT9 | Armadillo r |
| TBCE | 0.375398 | 0.42131 | -0.50164 | Q15813;A0 | Q15813;A0 | Q15813 | Tubulin-spe |
| SLC35A4 | 0.373602 | 0.423056 | 0.065739 | L0R6Q1 | L0R6Q1 | L0R6Q1 | |
| TRIM22 | 0.371176 | 0.425426 | 0.09695 | C9JWC5;Q8 | C9JWC5;Q8 | C9JWC5 | E3 ubiquiti |
| WIBG | 0.36891 | 0.427651 | 0.058322 | Q9BRP8 | Q9BRP8 | Q9BRP8 | Partner of Y |
| RABGGTB | 0.366893 | 0.429642 | -0.17638 | P53611;Q5 | P53611;Q5 | P53611 | Geranylgera |
| BTF3 | 0.36516 | 0.43136 | -0.08765 | P20290;D6 | P20290;D6 | P20290 | Transcriptio |
| UNG | 0.361581 | 0.434929 | 0.114798 | P13051;F5 | P13051;F5 | P13051 | Uracil-DNA |
| SDF2 | 0.359682 | 0.436836 | 0.044626 | Q99470;K7 | Q99470;K7 | Q99470 | Stromal cel |
| FBXW11 | 0.350094 | 0.446587 | -0.07272 | Q9UKB1;B7 | Q9UKB1;B7 | Q9UKB1 | F-box/WD r |
| ARL8A | 0.349838 | 0.446851 | -0.06543 | Q96BM9;A( | Q96BM9 | Q96BM9 | ADP-ribosy |
| TCEB2 | 0.340891 | 0.456152 | 0.048976 | Q15370;B8 | Q15370;B8 | Q15370 | Transcriptio |
| GUSB | 0.32743 | 0.470511 | 0.080342 | P08236;F8 | P08236 | P08236 | Beta-glucur |
| ATP5S | 0.323727 | 0.47454 | 0.068388 | A0A5K1VW | A0A5K1VW | A0A5K1VW | ATP synthas |
| BLOC1S2 | 0.321109 | 0.477409 | 0.062692 | Q6QNY1;J3 | Q6QNY1 | Q6QNY1 | Biogenesis o |
| DOCK11 | 0.31426 | 0.484999 | -0.40967 | A6NIW2;Q5 | A6NIW2;Q5 | A6NIW2 | Dedicator o |
| GDE1 | 0.313066 | 0.486334 | -0.04694 | Q9NZC3;H3 | Q9NZC3 | Q9NZC3 | Glyceropho |
| GSTT1 | 0.310996 | 0.488657 | -0.19926 | P30711;A0 | P30711;A0 | P30711 | Glutathion |
| DBI | 0.309586 | 0.490246 | -0.1064 | P07108;A0 | P07108;A0 | P07108 | Acyl-CoA-bi |
| CMPK2 | 0.303232 | 0.497471 | 0.824277 | Q5EBM0 | Q5EBM0 | Q5EBM0 | UMP-CMP k |
| ALG5 | 0.302843 | 0.497917 | 0.060039 | Q9Y673 | Q9Y673 | Q9Y673 | Dolichyl-ph |
| MPST | 0.297939 | 0.503571 | -0.05067 | P25325;B1 | P25325;B1 | P25325 | 3-mercapto |
| MFSD10 | 0.29768 | 0.503871 | 0.069847 | D6RE79;Q1 | D6RE79;Q1 | D6RE79 | Major facili |
| BABAM1 | 0.29621 | 0.50558 | -0.0629 | Q9NWV8;N | Q9NWV8;N | Q9NWV8 | BRISC and B |
| NDUFA5 | 0.294726 | 0.507311 | 0.034639 | Q16718;A0 | Q16718;A0 | Q16718 | NADH dehy |
| RPL27 | 0.283052 | 0.521133 | 0.038989 | P61353;K7 | P61353;K7 | P61353 | 60S ribosor |
| EML2 | 0.281952 | 0.522454 | -0.62958 | A0A0C4DG | A0A0C4DG | A0A0C4DG | Echinoderm |
| NDUFAF2 | 0.281497 | 0.523002 | 0.06664 | Q8N183;D6 | Q8N183;D6 | Q8N183 | Mimitin, m |
| NUFIP2 | 0.280153 | 0.524623 | 0.093993 | Q7Z417 | Q7Z417 | Q7Z417 | Nuclear frag |
| RPS21 | 0.278255 | 0.526921 | -0.02435 | Q8WVC2;P | Q8WVC2;P | Q8WVC2 | 40S ribosor |
| TMCO3 | 0.272145 | 0.534386 | 0.055088 | Q6UWJ1;A( | Q6UWJ1;A( | Q6UWJ1 | Transmemb |
| XAB2 | 0.269399 | 0.537776 | -0.31901 | Q9HCS7 | Q9HCS7 | Q9HCS7 | Pre-mRNA-s |
| SURF1 | 0.26332 | 0.545356 | 0.050296 | Q15526;A0 | Q15526;A0 | Q15526 | Surfeit locu |
| ABCF2 | 0.263311 | 0.545368 | -0.07 | Q9UG63;C9 | Q9UG63;C9 | Q9UG63 | ATP-bindin |
| COASY | 0.25445 | 0.556609 | -0.05291 | Q13057;K7 | Q13057 | Q13057 | Bifunctiona |
| DYNLRB1 | 0.254028 | 0.55715 | 0.080319 | Q9NP97;B1 | Q9NP97;B1 | Q9NP97 | Dynein ligh |
| GTF2F2 | 0.242976 | 0.57151 | 0.080417 | P13984 | P13984 | P13984 | General tra |
| PIP5K1C | 0.227247 | 0.592588 | -0.04563 | O60331 | O60331 | O60331 | Phosphatic |
| SPPL2A | 0.227114 | 0.59277 | 0.052346 | Q8TCT8;H0 | Q8TCT8;H0 | Q8TCT8 | Signal pept |

| | | | | | | | |
|---|---|---|---|---|---|---|---|
| RPS4Y1 | 0.205985 | 0.622322 | -0.02902 | P22090;C9 | P22090;C9 | P22090 | 40S ribosor |
| SSR4 | 0.19727 | 0.634936 | -0.04303 | P51571;A6 | P51571;A6 | P51571 | Translocon |
| MTPN | 0.195418 | 0.637649 | -0.07067 | P58546;C9 | P58546 | P58546 | Myotrophi |
| COA7 | 0.193599 | 0.640325 | -0.04176 | Q96BR5 | Q96BR5 | Q96BR5 | Cytochrom |
| PLGRKT | 0.189653 | 0.64617 | -0.03242 | Q9HBL7 | Q9HBL7 | Q9HBL7 | Plasminoge |
| B3GAT3 | 0.188568 | 0.647786 | 0.036114 | O94766;G3 | O94766;G3 | O94766 | Galactosylg |
| CDK11A | 0.186454 | 0.650948 | 0.053544 | Q5QPR4;A0 | Q5QPR4;A0 | Q5QPR4 | Cyclin-depe |
| PDCD5 | 0.183407 | 0.655531 | -0.04002 | O14737;K7 | O14737;K7 | O14737 | Programme |
| ITPA | 0.181027 | 0.659133 | -0.04153 | Q9BY32 | Q9BY32 | Q9BY32 | Inosine trip |
| TBC1D15 | 0.167504 | 0.679979 | -0.22684 | Q8TC07;C9 | Q8TC07;C9 | Q8TC07 | TBC1 doma |
| MPO | 0.166443 | 0.681643 | -0.10238 | P05164;J3 | P05164 | P05164 | Myeloperox |
| HSD11B1 | 0.16634 | 0.681805 | 0.052021 | P28845;A0 | P28845;A0 | P28845 | Corticoster |
| BPHL | 0.165862 | 0.682556 | -0.05561 | Q86WA6;Q | Q86WA6;Q | Q86WA6 | Valacyclovi |
| EFR3A | 0.165152 | 0.683673 | -0.02944 | Q14156;A0 | Q14156;A0 | Q14156 | Protein EFF |
| PPIG | 0.163455 | 0.686348 | -0.05392 | Q13427;C9 | Q13427;C9 | Q13427 | Peptidyl-pr |
| VPS52 | 0.163252 | 0.68667 | 0.047602 | Q8N1B4;A0 | Q8N1B4 | Q8N1B4 | Vacuolar pr |
| STBD1 | 0.161584 | 0.689313 | -0.03114 | O95210 | O95210 | O95210 | Starch-binc |
| CD9 | 0.158978 | 0.693461 | -0.03187 | A6NNI4;A0 | A6NNI4;A0 | A6NNI4 | Tetraspanir |
| MIOS | 0.1563 | 0.69775 | -0.04249 | Q9NXC5;C9 | Q9NXC5 | Q9NXC5 | WD repeat- |
| PSMD10 | 0.155382 | 0.699226 | -0.04381 | O75832;B1 | O75832;B1 | O75832 | 26S proteas |
| FAM63A | 0.150904 | 0.706474 | -0.04069 | Q8N5J2;Q8 | Q8N5J2 | Q8N5J2 | Protein FAM |
| SOD1 | 0.150211 | 0.707602 | 0.042954 | P00441;H7 | P00441;H7 | P00441 | Superoxide |
| TUBA4A | 0.131622 | 0.738547 | -0.04182 | P68366;C9 | P68366 | P68366 | Tubulin alp |
| LETMD1 | 0.126428 | 0.747433 | -0.04263 | F8VZA2;Q6 | F8VZA2;Q6 | F8VZA2 | LETM1 dom |
| DNAJC8 | 0.125801 | 0.748512 | -0.03066 | O75937;S4 | O75937 | O75937 | DnaJ homo |
| REL | 0.125103 | 0.749717 | 0.033907 | Q04864;A0 | Q04864 | Q04864 | Proto-onco |
| CYB5B | 0.124261 | 0.751172 | 0.027881 | H3BUX2;O4 | H3BUX2;O4 | H3BUX2 | Cytochrom |
| MYL12A | 0.119646 | 0.759196 | -0.01971 | J3QRS3;P19 | J3QRS3;P19 | J3QRS3 | Myosin regi |
| MRPL16 | 0.111795 | 0.773046 | -0.04057 | Q9NX20;E9 | Q9NX20;E9 | Q9NX20 | 39S ribosor |
| PTK2B | 0.110019 | 0.776213 | 0.02154 | Q14289;H0 | Q14289 | Q14289 | Protein-tyr |
| RAB34 | 0.093422 | 0.806451 | -0.03517 | C9JBG0;Q9 | C9JBG0;Q9 | C9JBG0 | Ras-related |
| MRPS28 | 0.08566 | 0.820994 | -0.02022 | Q9Y2Q9;H7 | Q9Y2Q9;H7 | Q9Y2Q9 | 28S ribosor |
| UCHL1 | 0.082015 | 0.827914 | -0.03794 | P09936;D6 | P09936;D6 | P09936 | Ubiquitin c |
| PIK3R1 | 0.078126 | 0.83536 | 0.01769 | P27986;H0 | P27986 | P27986 | Phosphatic |
| CRELD1 | 0.075715 | 0.840011 | -0.02985 | Q96HD1;A0 | Q96HD1;A0 | Q96HD1 | Cysteine-ric |
| CASP3 | 0.07279 | 0.845687 | -0.0236 | P42574;C9 | P42574;C9 | P42574 | Caspase-3;C |
| TMEM199 | 0.072488 | 0.846276 | 0.011482 | Q8N511;J3 | Q8N511;J3 | Q8N511 | Transmemb |
| DNAJC5 | 0.067524 | 0.856004 | -0.0111 | Q9H3Z4 | Q9H3Z4 | Q9H3Z4 | DnaJ homo |
| TMEM38B | 0.066594 | 0.857839 | -0.02106 | Q9NVV0;A0 | Q9NVV0;A0 | Q9NVV0 | Trimeric in |
| SLC1A3 | 0.055495 | 0.880045 | 0.016156 | A0A087X0U | A0A087X0U | A0A087X0U | Amino acid |
| COX17 | 0.054797 | 0.881461 | 0.0154 | Q14061;H7 | Q14061;H7 | Q14061 | Cytochrom |
| RPP30 | 0.054226 | 0.882621 | -0.01924 | P78346;Q5 | P78346;Q5 | P78346 | Ribonuclea |
| PHF6 | 0.044855 | 0.901873 | 0.008787 | Q5JRC6;A0 | Q5JRC6;A0 | Q5JRC6 | PHD finger |
| NDUFA11 | 0.043632 | 0.904416 | -0.00866 | Q86Y39;K7 | Q86Y39;K7 | Q86Y39 | NADH dehy |
| NME1-NME | 0.033713 | 0.92531 | -0.00913 | Q32Q12;J3 | Q32Q12;J3 | Q32Q12 | Nucleoside |
| FKBP1A | 0.031877 | 0.929229 | 0.009242 | P62942;Q5 | P62942;Q5 | P62942 | Peptidyl-pr |
| SMS | 0.029968 | 0.933324 | 0.012862 | P52788;H7 | P52788;H7 | P52788 | Spermine sy |
| MRPL53 | 0.029605 | 0.934105 | 0.004683 | Q96EL3;B9 | Q96EL3 | Q96EL3 | 39S ribosor |

| | | | | | | | |
|---|---|---|---|---|---|---|---|
| OPA3 | 0.029039 | 0.935322 | 0.005696 | Q9H6K4;B4 | Q9H6K4;B4 | Q9H6K4 | Optic atrop |
| NUP85 | 0.027647 | 0.938324 | -0.00753 | J3QL54;Q9 | J3QL54;Q9 | J3QL54 | Nuclear por |
| RER1 | 0.02275 | 0.948965 | 0.003386 | O15258;Q5 | O15258;Q5 | O15258 | Protein REF |
| SLC17A5 | 0.022176 | 0.950219 | -0.00627 | Q9NRA2 | Q9NRA2 | Q9NRA2 | Sialin |
| YBX3 | 0.021563 | 0.951562 | -0.00488 | P16989;A0 | P16989 | P16989 | Y-box-bindi |
| TMED5 | 0.018428 | 0.958456 | 0.005819 | Q9Y3A6;B1 | Q9Y3A6;B1 | Q9Y3A6 | Transmemb |
| PPP1CB | 0.018326 | 0.958681 | 0.003116 | P62140;E7 | P62140;E7 | P62140 | Serine/thre |
| CCDC127 | 0.014765 | 0.966574 | -0.00553 | Q96BQ5 | Q96BQ5 | Q96BQ5 | Coiled-coil |
| RBMS1 | 0.010563 | 0.975972 | -0.00247 | E7ETU5;F6 | E7ETU5;F6 | E7ETU5 | RNA-bindir |
| SCAF11 | 0.006786 | 0.984496 | 0.002894 | A0A0A0MT | A0A0A0MT | A0A0A0MT | Protein SCA |
| CLTB | 0.002696 | 0.993812 | 0.000871 | P09497;HC | P09497 | P09497 | Clathrin lig |
| ISCU | 1.663035 | 0.021725 | 0.491895 | Q9H1K1;F5 | Q9H1K1;F5 | Q9H1K1 | Iron-sulfur |
| SRP19 | 2.533251 | 0.002929 | 0.431288 | P09132;A0 | P09132;A0 | P09132 | Signal recog |
| FAM213B | 2.317523 | 0.004814 | 0.391524 | A0A2P0CTE | A0A2P0CTE | A0A2P0CTE | Prostamide |
| C10orf11 | 1.754778 | 0.017588 | 0.360718 | A0A087WV | A0A087WV | A0A087WV | Leucine-ric |
| WDR12 | 3.526407 | 0.000298 | 0.352635 | Q9GZL7 | Q9GZL7 | Q9GZL7 | Ribosome b |
| RPS20 | 3.683891 | 0.000207 | 0.34232 | P60866;G3 | P60866;G3 | P60866 | 40S ribosor |
| MRPS16 | 2.80556 | 0.001565 | 0.332153 | Q9Y3D3;A6 | Q9Y3D3;A6 | Q9Y3D3 | 28S ribosor |
| ENTPD1 | 1.811208 | 0.015445 | 0.297995 | P49961;A0 | P49961 | P49961 | Ectonucleo |
| SPRYD7 | 1.651068 | 0.022332 | 0.288375 | Q5W111 | Q5W111 | Q5W111 | SPRY doma |
| FAM98A | 2.825769 | 0.001494 | 0.283275 | Q8NCA5;E9 | Q8NCA5;E9 | Q8NCA5 | Protein FAN |
| DDX39A | 2.443982 | 0.003598 | 0.271187 | O00148;K7 | O00148;K7 | O00148 | ATP-depend |
| SLC25A4 | 3.290227 | 0.000513 | 0.25851 | P12235;V9 | P12235;V9 | P12235 | ADP/ATP tr |
| SYNJ2BP-C | 1.896989 | 0.012677 | 0.232248 | A0A087X1I | A0A087X1I | A0A087X1I | Synaptojan |
| PTRH2 | 1.627522 | 0.023576 | 0.231041 | J3KQ48;Q9 | J3KQ48;Q9 | J3KQ48 | Peptidyl-tR |
| PAF1 | 2.369822 | 0.004268 | 0.228093 | Q8N7H5;B4 | Q8N7H5 | Q8N7H5 | RNA polym |
| PAM | 1.302396 | 0.049843 | 0.211346 | P19021;H7 | P19021 | P19021 | Peptidyl-gl |
| IBA57 | 1.493671 | 0.032087 | 0.205449 | Q5T440 | Q5T440 | Q5T440 | Putative tra |
| SLC16A1 | 1.601454 | 0.025035 | 0.203581 | P53985;Q5 | P53985;Q5 | P53985 | Monocarbo |
| TIMM8B | 1.355063 | 0.044151 | 0.186474 | Q9Y5J9;G3 | Q9Y5J9;G3 | Q9Y5J9 | Mitochond |
| CHCHD4 | 1.516127 | 0.03047 | 0.16854 | Q8N4Q1 | Q8N4Q1 | Q8N4Q1 | Mitochond |
| RAP2B | 1.325616 | 0.047248 | 0.137094 | P61225 | P61225 | P61225 | Ras-related |
| PABPN1 | 1.579617 | 0.026326 | 0.114248 | Q86U42;B4 | Q86U42;B4 | Q86U42 | Polyadenyl |
| RBM3 | 1.612272 | 0.024419 | -0.13772 | P98179 | P98179 | P98179 | RNA-bindir |
| YIPF3 | 1.456755 | 0.034934 | -0.19854 | D6RFI3;E7E | D6RFI3;E7E | D6RFI3 | Protein YIP |
| TIMP2 | 1.331872 | 0.046572 | -0.21402 | P16035;B4 | P16035;B4 | P16035 | Metallopro |
| CDIPT | 1.968748 | 0.010746 | -0.21838 | O14735;B3 | O14735;B3 | O14735 | CDP-diacylg |
| SLC9A7 | 2.921402 | 0.001198 | -0.24895 | A0A087WX | A0A087WX | A0A087WX | Sodium/hy |
| TMEM245 | 1.535187 | 0.029162 | -0.29192 | Q9H330;H7 | Q9H330;H7 | Q9H330 | Transmemb |
| CISD1 | 3.813666 | 0.000154 | -0.39389 | Q9NZ45 | Q9NZ45 | Q9NZ45 | CDGSH iror |
| S100A8 | 2.075045 | 0.008413 | -0.40154 | P05109 | P05109 | P05109 | Protein S10 |
| TMEM165 | 1.288173 | 0.051502 | 0.122336 | Q9HC07;V9 | Q9HC07;V9 | Q9HC07 | Transmemb |
| ARHGAP18 | 1.282168 | 0.052219 | -1.11835 | Q8N392 | Q8N392 | Q8N392 | Rho GTPase |
| WASH3P | 1.241762 | 0.057311 | 0.202516 | C4AMC7;Q | C4AMC7;Q | C4AMC7 | Putative W |
| IAH1 | 1.216886 | 0.06069 | -0.44727 | H7C5G1;Q | H7C5G1;Q | H7C5G1 | Isoamyl ace |
| ARPC5L | 1.209339 | 0.061753 | -0.17855 | Q9BPX5 | Q9BPX5 | Q9BPX5 | Actin-relate |
| TIMM23 | 1.195491 | 0.063754 | 0.228562 | O14925;Q5 | O14925;Q5 | O14925 | Mitochond |
| ABCF3 | 1.165685 | 0.068283 | -0.30997 | Q9NUQ8 | Q9NUQ8 | Q9NUQ8 | ATP-bindin |

| | | | | | | | |
|---|---|---|---|---|---|---|---|
| AGO1 | 1.165179 | 0.068363 | 0.073537 | A0A6I8PTZ | A0A6I8PTZ | A0A6I8PTZ | Protein arg |
| UBXN1 | 1.160765 | 0.069061 | -1.63175 | E9PRQ7;Q0 | E9PRQ7;Q0 | E9PRQ7 | UBX domai |
| CHORDC1 | 1.154611 | 0.070047 | 0.182281 | Q9UHD1;E | Q9UHD1;E | Q9UHD1 | Cysteine an |
| ARIH1 | 1.147429 | 0.071215 | 0.149899 | Q9Y4X5;A0 | Q9Y4X5 | Q9Y4X5 | E3 ubiquiti |
| ATP5D | 1.098967 | 0.079622 | 0.121169 | P30049 | P30049 | P30049 | ATP syntha |
| HLA-DQB1 | 1.096943 | 0.079994 | 0.208861 | A2AAZ0;Q5 | A2AAZ0;Q5 | A2AAZ0 | |
| THEMIS2 | 1.080273 | 0.083124 | -0.10839 | Q5TEJ8;H7 | Q5TEJ8;H7 | Q5TEJ8 | Protein THI |
| C10orf32-A | 1.052846 | 0.088543 | 0.130309 | A0A0B4J1F | A0A0B4J1F | A0A0B4J1F | UPF0693 p |
| PFDN5 | 1.052448 | 0.088624 | 0.129849 | Q99471;H3 | Q99471;H3 | Q99471 | Prefoldin su |
| TP53RK | 1.039724 | 0.091259 | 0.151076 | Q96S44;Q5 | Q96S44 | Q96S44 | TP53-regul |
| ERH | 1.035054 | 0.092246 | 0.207887 | P84090;G3 | P84090;G3 | P84090 | Enhancer o |
| SIRT2 | 1.033679 | 0.092538 | 1.959311 | A0A0A0MR | A0A0A0MR | A0A0A0MR | NAD-depen |
| LHFPL2 | 1.021317 | 0.09521 | -0.13196 | Q6ZUX7 | Q6ZUX7 | Q6ZUX7 | Lipoma HM |
| S100A6 | 1.017935 | 0.095954 | -0.1403 | R4GN98;P | R4GN98;P | R4GN98 | Protein S1 |
| RQCD1 | 0.983096 | 0.103969 | -1.08578 | Q92600 | Q92600 | Q92600 | Cell differe |
| LACE1 | 0.977738 | 0.10526 | -0.13225 | Q8WV93;H | Q8WV93;H | Q8WV93 | Lactation e |
| RPL13A | 0.969761 | 0.107211 | 0.138693 | P40429;M | P40429;M | P40429 | 60S riboso |
| CLN6 | 0.965458 | 0.108278 | -0.18477 | Q9NWW5; | Q9NWW5; | Q9NWW5 | Ceroid-lipo |
| ARHGEF11 | 0.961022 | 0.10939 | 0.20276 | O15085 | O15085 | O15085 | Rho guanin |
| SLC30A7 | 0.95376 | 0.111235 | 0.180582 | Q8NEW0;H | Q8NEW0 | Q8NEW0 | Zinc transp |
| PCYOX1L | 0.94888 | 0.112492 | 0.160379 | Q8NBM8;E | Q8NBM8;E | Q8NBM8 | Prenylcyste |
| EIF4E2 | 0.938563 | 0.115196 | 0.245996 | C9JEL3;B9 | C9JEL3;B9 | C9JEL3 | Eukaryotic |
| ZAP70 | 0.921213 | 0.119891 | -1.79922 | P43403 | P43403 | P43403 | Tyrosine-pr |
| PRPF38A | 0.921187 | 0.119898 | 1.183425 | Q8NAV1 | Q8NAV1 | Q8NAV1 | Pre-mRNA- |
| POLR2C | 0.918514 | 0.120639 | -0.1214 | P19387;H3 | P19387 | P19387 | DNA-direct |
| C9orf64 | 0.914285 | 0.121819 | -0.2148 | Q5T6V5;Q5 | Q5T6V5;Q5 | Q5T6V5 | UPF0553 p |
| CMC4 | 0.903798 | 0.124796 | 0.151136 | P56277 | P56277 | P56277 | Cx9C motif |
| STARD3NL | 0.88589 | 0.13005 | 1.372536 | C9JPX5;C9 | C9JPX5;C9 | C9JPX5 | MLN64 N-t |
| TMEM87B | 0.876258 | 0.132967 | -0.15044 | A0A494BZ | A0A494BZ | A0A494BZ | Transmemb |
| NDUFAF3 | 0.8496 | 0.141384 | 0.170876 | Q9BU61 | Q9BU61 | Q9BU61 | NADH dehy |
| CCDC115 | 0.838079 | 0.145185 | -0.13424 | Q96NT0;B8 | Q96NT0;B8 | Q96NT0 | Coiled-coil |
| THEM4 | 0.837286 | 0.14545 | -0.16832 | Q5T1C6;F6 | Q5T1C6 | Q5T1C6 | Acyl-coenz |
| CIRBP | 0.833209 | 0.146822 | -0.0963 | Q14011;K7 | Q14011;K7 | Q14011 | Cold-induc |
| PDDC1 | 0.830083 | 0.147883 | 0.219216 | Q8NB37;E9 | Q8NB37;E9 | Q8NB37 | Parkinson c |
| WBP11 | 0.825102 | 0.149588 | 0.198078 | Q9Y2W2;F | Q9Y2W2 | Q9Y2W2 | WW domai |
| FAM210A | 0.823676 | 0.150081 | 0.162705 | K7EK00;Q9 | K7EK00;Q9 | K7EK00 | Protein FA |
| FUNDC2 | 0.816576 | 0.152554 | -0.1767 | Q9BWH2;H | Q9BWH2 | Q9BWH2 | FUN14 don |
| ARPC4-TTLI | 0.812709 | 0.153919 | -0.10487 | F8WCF6;P5 | F8WCF6;P5 | F8WCF6 | Actin-relat |
| ENOPH1 | 0.800999 | 0.158125 | -0.41379 | Q9UHY7;A0 | Q9UHY7;A0 | Q9UHY7 | Enolase-ph |
| IFIT2 | 0.795622 | 0.160095 | -0.36857 | A0A087X2 | A0A087X2 | A0A087X2 | Interferon- |
| TRA2A | 0.776398 | 0.167341 | -0.18071 | Q13595 | Q13595 | Q13595 | Transforme |
| EFTUD1 | 0.757 | 0.174985 | -0.84939 | Q7Z2Z2;A0 | Q7Z2Z2;A0 | Q7Z2Z2 | Elongation |
| XPNPEP3 | 0.7557 | 0.175509 | 0.976086 | Q9NQH7 | Q9NQH7 | Q9NQH7 | Probable X |
| NARS2 | 0.703834 | 0.197772 | 0.140717 | Q96I59;E9 | Q96I59;E9 | Q96I59 | Probable as |
| UACA | 0.703698 | 0.197835 | 0.775248 | F5H2B9;H0 | F5H2B9;H0 | F5H2B9 | Uveal autoa |
| PPIL1 | 0.696818 | 0.200994 | 0.111364 | Q9Y3C6 | Q9Y3C6 | Q9Y3C6 | Peptidyl-pr |
| FUS | 0.694802 | 0.201929 | 0.099781 | P35637;H3 | P35637;H3 | P35637 | RNA-bindir |
| NECAP2 | 0.677948 | 0.209919 | -1.10217 | Q9NVZ3;F6 | Q9NVZ3;F6 | Q9NVZ3 | Adaptin ea |

| | | | | | | | |
|---|---|---|---|---|---|---|---|
| FASTKD2 | 0.674556 | 0.211565 | 0.246022 | Q9NYY8 | Q9NYY8 | Q9NYY8 | FAST kinase |
| IVNS1ABP | 0.665341 | 0.216102 | -0.14313 | Q9Y6Y0;X6 | Q9Y6Y0 | Q9Y6Y0 | Influenza vi |
| SEPTIN8 | 0.661672 | 0.217935 | -0.71836 | A6NMH6;Q | A6NMH6;Q | A6NMH6 | Septin-8 |
| ABHD2 | 0.65771 | 0.219933 | -1.30497 | P08910 | P08910 | P08910 | Abhydrolas |
| FAM50A | 0.651498 | 0.223101 | -0.17467 | Q14320;B0 | Q14320;B0 | Q14320 | Protein FAM |
| CCPG1 | 0.651313 | 0.223196 | -0.61547 | Q9ULG6;H3 | Q9ULG6 | Q9ULG6 | Cell cycle p |
| PRKACA | 0.639857 | 0.229162 | 0.065151 | P17612;Q1 | P17612;Q1 | P17612 | cAMP-depe |
| TSG101 | 0.638592 | 0.229831 | 0.11896 | F5H442;Q9 | F5H442;Q9 | F5H442 | Tumor susc |
| MT-ND4 | 0.636423 | 0.230981 | -0.13958 | P03905 | P03905 | P03905 | NADH-ubiq |
| OGFR | 0.62887 | 0.235034 | -0.7322 | Q9NZT2;A0 | Q9NZT2;A0 | Q9NZT2 | Opioid grov |
| TMX4 | 0.62873 | 0.235109 | 0.098666 | Q9H1E5;E9 | Q9H1E5 | Q9H1E5 | Thioredoxi |
| WNK1 | 0.627412 | 0.235824 | 0.176017 | Q9H4A3;F5 | Q9H4A3;F5 | Q9H4A3 | Serine/thre |
| ADD1 | 0.621117 | 0.239267 | -0.7623 | E7EV99;E7I | E7EV99;E7I | E7EV99 | Alpha-addu |
| MTARC1 | 0.616955 | 0.241571 | -1.33892 | Q5VT66;H7 | Q5VT66;H7 | Q5VT66 | Mitochond |
| LRCH4 | 0.614215 | 0.2431 | 0.159035 | O75427 | O75427 | O75427 | Leucine-ric |
| RBM10 | 0.612966 | 0.2438 | 0.759946 | P98175 | P98175 | P98175 | RNA-bindir |
| GGCT | 0.595107 | 0.254035 | -0.12827 | O75223;M | O75223;M | O75223 | Gamma-glu |
| S100A9 | 0.587834 | 0.258325 | -0.16254 | P06702 | P06702 | P06702 | Protein S10 |
| CHMP5 | 0.585029 | 0.259999 | -0.17755 | Q9NZZ3 | Q9NZZ3 | Q9NZZ3 | Charged mu |
| GZMB | 0.579445 | 0.263363 | -0.66776 | J3KQ52;P1 | J3KQ52;P1 | J3KQ52 | Granzyme E |
| PRPSAP2 | 0.575435 | 0.265806 | -0.60922 | O60256;E7 | O60256;E7 | O60256 | Phosphorib |
| SLC22A18 | 0.574386 | 0.266449 | -0.11245 | E9PRM7;A0 | E9PRM7;A0 | E9PRM7 | Solute carri |
| PTPN7 | 0.573953 | 0.266715 | -0.80537 | P35236;J3I | P35236;J3I | P35236 | Tyrosine-pr |
| SNX4 | 0.572855 | 0.26739 | 1.177238 | O95219 | O95219 | O95219 | Sorting nex |
| ABHD14B | 0.568964 | 0.269796 | -0.14291 | Q96IU4;F8 | Q96IU4;F8 | Q96IU4 | Alpha/beta |
| MT-ND5 | 0.551906 | 0.280604 | 0.126096 | P03915 | P03915 | P03915 | NADH-ubiq |
| TXNDC17 | 0.550106 | 0.281769 | -0.17359 | Q9BRA2;I3I | Q9BRA2;I3I | Q9BRA2 | Thioredoxi |
| ARHGAP9 | 0.54208 | 0.287025 | 1.173585 | Q9BRR9;R4 | Q9BRR9;R4 | Q9BRR9 | Rho GTPase |
| NUBP2 | 0.529591 | 0.295399 | 0.491095 | Q9Y5Y2;H3 | Q9Y5Y2;H3 | Q9Y5Y2 | Cytosolic F |
| CD46 | 0.515806 | 0.304926 | 0.107472 | P15529 | P15529 | P15529 | Membrane |
| MRPS33 | 0.511715 | 0.307811 | -0.10903 | Q9Y291;C9 | Q9Y291;C9 | Q9Y291 | 28S riboson |
| RAC1 | 0.505892 | 0.311967 | -0.09823 | P63000;P6 | P63000;P6 | P63000 | Ras-related |
| MRPL14 | 0.505132 | 0.312513 | -0.09266 | Q6P1L8 | Q6P1L8 | Q6P1L8 | 39S riboson |
| BLOC1S5 | 0.498427 | 0.317375 | -0.50607 | Q8TDH9;G5 | Q8TDH9 | Q8TDH9 | Biogenesis |
| STAM | 0.497384 | 0.318138 | -0.10001 | Q92783;A6 | Q92783 | Q92783 | Signal trans |
| DOCK4 | 0.496403 | 0.318857 | -1.19467 | H0Y599;Q8 | H0Y599;Q8 | H0Y599 | Dedicator c |
| FAM105A | 0.494753 | 0.320071 | -0.64903 | Q9NUU6 | Q9NUU6 | Q9NUU6 | Inactive ub |
| MSMO1 | 0.492868 | 0.321464 | -0.25887 | Q15800;D6 | Q15800;D6 | Q15800 | Methylster |
| ALAD | 0.490054 | 0.323553 | 0.658133 | P13716;B7 | P13716 | P13716 | Delta-amin |
| LSM8 | 0.487452 | 0.325498 | -0.09197 | F2Z2Y6;O9 | F2Z2Y6;O9 | F2Z2Y6 | U6 snRNA-a |
| RPL22 | 0.48324 | 0.32867 | -0.06822 | P35268;K7 | P35268;K7 | P35268 | 60S riboson |
| TTYH3 | 0.47429 | 0.335513 | -0.79702 | Q9C0H2;H7 | Q9C0H2 | Q9C0H2 | Protein twe |
| HECTD3 | 0.460648 | 0.34622 | -0.1121 | Q5T447 | Q5T447 | Q5T447 | E3 ubiquiti |
| XRCC4 | 0.459817 | 0.346883 | -0.54449 | Q13426 | Q13426 | Q13426 | DNA repair |
| SIPA1 | 0.450967 | 0.354024 | 0.783888 | F6RY50;Q9 | F6RY50;Q9 | F6RY50 | Signal-indu |
| ALG12 | 0.447006 | 0.357268 | 0.139748 | Q9BV10;H7 | Q9BV10 | Q9BV10 | Dol-P-Man: |
| MRPL24 | 0.446896 | 0.357358 | 0.075486 | Q96A35;X6 | Q96A35 | Q96A35 | 39S riboson |
| MPV17 | 0.44449 | 0.359344 | 0.226252 | B5MC53;B5 | B5MC53;B5 | B5MC53 | Protein Mp |

| | | | | | | | |
|---|---|---|---|---|---|---|---|
| C19orf66 | 0.443657 | 0.360033 | 0.100322 | Q9NUL5;K7 | Q9NUL5 | Q9NUL5 | UPF0515 p |
| MLKL | 0.439922 | 0.363144 | 0.511395 | Q8NB16;I3 | Q8NB16;I3 | Q8NB16 | Mixed linea |
| NACA | 0.431555 | 0.370207 | 0.062537 | F8VZJ2;F8V | F8VZJ2;F8V | F8VZJ2 | Nascent po |
| GPR89B | 0.428221 | 0.37306 | -0.20495 | P0CG08;B7 | P0CG08;B7 | P0CG08 | Golgi pH re |
| ARFRP1 | 0.419083 | 0.380993 | 0.121438 | Q13795 | Q13795 | Q13795 | ADP-ribosy |
| ZRANB2 | 0.418355 | 0.381632 | 0.110706 | O95218 | O95218 | O95218 | Zinc finger |
| ANP32B | 0.416065 | 0.38365 | -0.08985 | Q92688 | Q92688 | Q92688 | Acidic leuci |
| DNAJB2 | 0.412659 | 0.38667 | -0.06042 | P25686;C9 | P25686;C9 | P25686 | DnaJ homo |
| PKM | 0.40266 | 0.395676 | -0.08775 | P14618;B4 | P14618;B4 | P14618 | Pyruvate ki |
| SLC44A1 | 0.400961 | 0.397227 | -0.0912 | Q8WWI5 | Q8WWI5 | Q8WWI5 | Choline tra |
| QIL1 | 0.394092 | 0.40356 | 0.136935 | A0A140TA8 | A0A140TA8 | A0A140TA8 | Protein QIL |
| RABL3 | 0.389813 | 0.407556 | 0.106475 | C9JXM3;Q5 | C9JXM3;Q5 | C9JXM3 | Rab-like pro |
| MCAT | 0.376514 | 0.420229 | -0.40503 | Q8IVS2 | Q8IVS2 | Q8IVS2 | Malonyl-Co |
| RAP2C | 0.37112 | 0.425481 | -0.06052 | Q9Y3L5;A0 | Q9Y3L5;A0 | Q9Y3L5 | Ras-related |
| C5AR1 | 0.369341 | 0.427227 | -0.07626 | P21730 | P21730 | P21730 | C5a anaphy |
| PFDN6 | 0.363049 | 0.433462 | 0.055962 | O15212;A2 | O15212;A2 | O15212 | Prefoldin su |
| UNC93B1 | 0.362876 | 0.433635 | 0.062064 | Q9H1C4;E9 | Q9H1C4 | Q9H1C4 | Protein unc |
| CARKD | 0.349306 | 0.447398 | -0.09465 | Q8IW45 | Q8IW45 | Q8IW45 | ATP-depend |
| PARN | 0.346181 | 0.450629 | -0.53082 | A0A494BZU | A0A494BZU | A0A494BZU | Poly(A)-spe |
| PEX16 | 0.343887 | 0.453015 | -0.07745 | Q9Y5Y5;E9 | Q9Y5Y5;E9 | Q9Y5Y5 | Peroxisoma |
| UFC1 | 0.331337 | 0.466298 | -0.13369 | Q9Y3C8 | Q9Y3C8 | Q9Y3C8 | Ubiquitin-f |
| ROGDI | 0.32206 | 0.476366 | -0.47079 | Q9GZN7;K7 | Q9GZN7;K7 | Q9GZN7 | Protein rog |
| VPS33B | 0.318822 | 0.479931 | -0.10022 | F5H008;Q9 | F5H008;Q9 | F5H008 | Vacuolar pr |
| RRS1 | 0.318453 | 0.480338 | -0.60091 | Q15050 | Q15050 | Q15050 | Ribosome b |
| SH3GL1 | 0.316207 | 0.482829 | 0.089801 | Q99961;M0 | Q99961;M0 | Q99961 | Endophilin |
| TLE3 | 0.309091 | 0.490805 | -0.37416 | A0A0D9SES | A0A0D9SES | A0A0D9SES | Transducin |
| MYDGF | 0.308122 | 0.491901 | -0.06622 | Q969H8;M | Q969H8;M | Q969H8 | Myeloid-de |
| SIRT5 | 0.304814 | 0.495662 | -0.07141 | Q9NXA8 | Q9NXA8 | Q9NXA8 | NAD-depen |
| EPN1 | 0.294149 | 0.507985 | -0.47774 | Q9Y6I3;D6 | Q9Y6I3 | Q9Y6I3 | Epsin-1 |
| SNAPIN | 0.289204 | 0.513802 | 0.093027 | O95295 | O95295 | O95295 | SNARE-asso |
| MRPL22 | 0.275825 | 0.529877 | -0.59571 | J3KQY1;E7I | J3KQY1;E7I | J3KQY1 | 39S riboson |
| TFAM | 0.274979 | 0.53091 | -0.06119 | Q00059;H7 | Q00059 | Q00059 | Transcripti |
| SMPD2 | 0.272392 | 0.534082 | -0.1043 | O60906;H0 | O60906 | O60906 | Sphingomy |
| RTN2 | 0.271155 | 0.535605 | -0.04782 | K7EMR7;O7 | K7EMR7;O7 | K7EMR7 | Reticulon;F |
| DHRS3 | 0.270348 | 0.536602 | 0.616648 | O75911;Q5 | O75911;Q5 | O75911 | Short-chair |
| SEPTIN1 | 0.263507 | 0.545121 | 0.64972 | J3KNL2;Q8 | J3KNL2;Q8 | J3KNL2 | Septin-1 |
| NUP37 | 0.261943 | 0.547088 | -0.3899 | Q8NFH4;H0 | Q8NFH4 | Q8NFH4 | Nucleopori |
| FAS | 0.250185 | 0.562102 | -0.06658 | P25445;K9 | P25445 | P25445 | Tumor necr |
| TRMT5 | 0.243593 | 0.570699 | -0.48841 | Q32P41;G3 | Q32P41 | Q32P41 | tRNA (guan |
| USP30 | 0.241474 | 0.57349 | 0.261632 | B3KUS5;Q7 | B3KUS5;Q7 | B3KUS5 | Ubiquitin c |
| GAR1 | 0.239309 | 0.576356 | 0.052781 | Q9NY12 | Q9NY12 | Q9NY12 | H/ACA ribo |
| ARRB1 | 0.231577 | 0.586709 | -0.07353 | P49407;E9 | P49407;E9 | P49407 | Beta-arresti |
| LGALS8 | 0.229018 | 0.590177 | 0.078108 | F6V2D4;O0 | F6V2D4;O0 | F6V2D4 | Galectin;Ga |
| MEF2BNB | 0.227949 | 0.591631 | 0.057335 | Q96FH0;K7 | Q96FH0;K7 | Q96FH0 | Protein ME |
| REEP3 | 0.227668 | 0.592014 | 0.036351 | Q6NUK4;A0 | Q6NUK4;A0 | Q6NUK4 | Receptor ex |
| LYAR | 0.224484 | 0.596371 | 0.338627 | Q9NX58;D6 | Q9NX58 | Q9NX58 | Cell growth |
| TMEM205 | 0.210289 | 0.616185 | 0.065926 | K7ELQ9;Q6 | K7ELQ9;Q6 | K7ELQ9 | Transmemb |
| TMUB2 | 0.208754 | 0.618366 | -0.03127 | Q71RG4;K7 | Q71RG4;K7 | Q71RG4 | Transmemb |

| | | | | | | | |
|---|---|---|---|---|---|---|---|
| AP2S1 | 0.20315 | 0.626398 | -0.03709 | M0R0N4;M | M0R0N4;M | M0R0N4 | AP-2 compl |
| GMFB | 0.20263 | 0.627147 | -0.11728 | P60983;G3 | P60983;G3 | P60983 | Glia matura |
| TRIM47 | 0.196274 | 0.636394 | 0.536946 | Q96LD4;A0 | Q96LD4 | Q96LD4 | Tripartite m |
| HIST1H1B | 0.194613 | 0.638833 | 0.075039 | P16401 | P16401 | P16401 | Histone H1 |
| GCHFR | 0.190687 | 0.644634 | -0.09361 | P30047;H0 | P30047;H0 | P30047 | GTP cycloh |
| ADRBK1 | 0.188114 | 0.648464 | -0.32451 | P25098;E9 | P25098 | P25098 | Beta-adren |
| TMEM55B | 0.186348 | 0.651106 | 0.039288 | Q86T03;G3 | Q86T03;G3 | Q86T03 | Type 1 pho |
| CYGB | 0.182061 | 0.657565 | -0.07189 | Q8WWM9; | Q8WWM9; | Q8WWM9 | Cytoglobin |
| SSR1 | 0.176492 | 0.666052 | -0.03639 | A0A3B3IRT | A0A3B3IRT | A0A3B3IRT | Translocon |
| FAH | 0.175801 | 0.667113 | 0.269927 | P16930;H0 | P16930;H0 | P16930 | Fumarylace |
| OSTM1 | 0.174159 | 0.669639 | 0.036752 | Q86WC4;A | Q86WC4 | Q86WC4 | Osteopetro |
| GRAMD1A | 0.173764 | 0.670248 | -0.7977 | M0QZ12;Q | M0QZ12;Q | M0QZ12 | GRAM dom |
| C19orf52 | 0.172572 | 0.672091 | 0.03807 | Q9BSF4;K7 | Q9BSF4 | Q9BSF4 | Uncharacte |
| SFXN4 | 0.164838 | 0.684166 | -0.42164 | Q6P4A7;B1 | Q6P4A7 | Q6P4A7 | Sideroflexi |
| IRF2BPL | 0.162137 | 0.688436 | 0.277283 | Q9H1B7 | Q9H1B7 | Q9H1B7 | Interferon |
| MAP2K1 | 0.16069 | 0.690732 | -0.05673 | Q02750;H3 | Q02750 | Q02750 | Dual specifi |
| IARS | 0.160333 | 0.691301 | 0.027 | P41252;A0 | P41252;A0 | P41252 | Isoleucine- |
| FAM96A | 0.158169 | 0.694754 | -0.04913 | Q9H5X1;H0 | Q9H5X1;H0 | Q9H5X1 | MIP18 fami |
| TRIM14 | 0.149173 | 0.709295 | -0.35654 | Q14142;F2 | Q14142 | Q14142 | Tripartite m |
| SSFA2 | 0.1404 | 0.723769 | 0.044044 | P28290;E9 | P28290;E9 | P28290 | Sperm-spec |
| LGMN | 0.139295 | 0.725612 | -0.03564 | Q99538;G3 | Q99538;G3 | Q99538 | Legumain |
| TMEM11 | 0.136034 | 0.731081 | 0.04203 | P17152 | P17152 | P17152 | Transmemb |
| CRYL1 | 0.127785 | 0.745101 | 0.062413 | Q9Y2S2;A0 | Q9Y2S2;A0 | Q9Y2S2 | Lambda-cry |
| GNG5 | 0.127602 | 0.745414 | 0.0232 | P63218 | P63218 | P63218 | Guanine nu |
| RBM26 | 0.112654 | 0.771518 | 0.280279 | Q5T8P6;A0 | Q5T8P6;A0 | Q5T8P6 | RNA-bindir |
| VPS28 | 0.110838 | 0.77475 | 0.022749 | Q9UK41;E9 | Q9UK41;E9 | Q9UK41 | Vacuolar pr |
| ACTN1 | 0.109238 | 0.777609 | 0.025151 | P12814;G3 | P12814 | P12814 | Alpha-actin |
| RNF40 | 0.106791 | 0.782004 | -0.01735 | H3BP71;O7 | H3BP71;O7 | H3BP71 | E3 ubiquiti |
| SF3B6 | 0.103366 | 0.788195 | 0.027983 | Q9Y3B4 | Q9Y3B4 | Q9Y3B4 | Splicing fac |
| SLC12A6 | 0.100937 | 0.792616 | 0.029718 | Q9UHW9;E | Q9UHW9;E | Q9UHW9 | Solute carri |
| TIMM8A | 0.09739 | 0.799115 | -0.02778 | O60220 | O60220 | O60220 | Mitochond |
| MPDU1 | 0.094027 | 0.805328 | 0.027193 | J3QS48;I3L | J3QS48;I3L | J3QS48 | Mannose-P |
| NADK | 0.089867 | 0.813079 | 0.027977 | O95544;A0 | O95544;A0 | O95544 | NAD kinase |
| C9orf89 | 0.082603 | 0.826793 | 0.022741 | Q96LW7 | Q96LW7 | Q96LW7 | Bcl10-inter |
| FHIT | 0.081614 | 0.828678 | -0.03105 | P49789;E9 | P49789 | P49789 | Bis(5-aden |
| CD274 | 0.081562 | 0.828778 | -0.03778 | Q9NZQ7 | Q9NZQ7 | Q9NZQ7 | Programme |
| TTC1 | 0.073751 | 0.843818 | 0.157616 | Q99614;H0 | Q99614;H0 | Q99614 | Tetratricop |
| SH3BGRL3 | 0.073338 | 0.844622 | -0.01845 | Q9H299;Q5 | Q9H299;Q5 | Q9H299 | SH3 domai |
| CTDNEP1 | 0.069494 | 0.852129 | 0.214681 | I3L1D9;I3L | I3L1D9;I3L | I3L1D9 | CTD nuclea |
| RAC2 | 0.068667 | 0.853755 | -0.01194 | P15153;B1 | P15153;B1 | P15153 | Ras-related |
| F8A1 | 0.064608 | 0.861771 | -0.14562 | P23610 | P23610 | P23610 | Factor VIII i |
| MICALL1 | 0.063214 | 0.864541 | -0.03432 | Q8N3F8;B0 | Q8N3F8 | Q8N3F8 | MICAL-like |
| ZC3H14 | 0.061789 | 0.867383 | -0.02143 | G3V256;Q6 | G3V256;Q6 | G3V256 | Zinc finger |
| SERPING1 | 0.059972 | 0.87102 | 0.157909 | E9PGN7;P0 | E9PGN7;P0 | E9PGN7 | Plasma pro |
| TIMM21 | 0.059683 | 0.871599 | -0.02559 | Q9BVV7;J3 | Q9BVV7 | Q9BVV7 | Mitochond |
| CD48 | 0.0581 | 0.874782 | 0.021045 | A0A087X1S | A0A087X1S | A0A087X1S | CD48 antig |
| COPZ1 | 0.057845 | 0.875295 | -0.01361 | F8VVA7;P6 | F8VVA7;P6 | F8VVA7 | Coatomer s |
| SP100 | 0.055512 | 0.880011 | -0.02004 | P23497;E9 | P23497;E9 | P23497 | Nuclear aut |

| | | | | | | | |
|---|---|---|---|---|---|---|---|
| S100A10 | 0.051873 | 0.887416 | 0.013947 | P60903 | P60903 | P60903 | Protein S10 |
| WIPF1 | 0.051532 | 0.888113 | 0.011241 | O43516;E9 | O43516;E9 | O43516 | WAS/WASL |
| NDUFA2 | 0.051515 | 0.888148 | -0.00814 | O43678 | O43678 | O43678 | NADH dehy |
| LRRC57 | 0.050731 | 0.889753 | 0.015698 | H3BSW0;A( | H3BSW0;A( | H3BSW0 | Leucine-ric |
| NCK1 | 0.049447 | 0.892386 | 0.01059 | P16333 | P16333 | P16333 | Cytoplasmi |
| TIMM10B | 0.049385 | 0.892514 | 0.013653 | Q9Y5J6;F2; | Q9Y5J6 | Q9Y5J6 | Mitochond |
| ISG15 | 0.0413 | 0.909284 | -0.01074 | P05161;A0 | P05161;A0 | P05161 | Ubiquitin-l |
| SAFB2 | 0.038575 | 0.915008 | 0.016287 | Q14151;K7 | Q14151 | Q14151 | Scaffold att |
| TMEM87A | 0.033701 | 0.925335 | 0.013446 | Q8NBN3 | Q8NBN3 | Q8NBN3 | Transmemb |
| NHP2L1 | 0.031003 | 0.931102 | 0.005101 | B1AHD1;A( | B1AHD1;A( | B1AHD1 | NHP2-like p |
| TMPPE | 0.03097 | 0.931173 | 0.010199 | Q6ZT21 | Q6ZT21 | Q6ZT21 | Transmemb |
| TXNRD1 | 0.029388 | 0.934569 | -0.00807 | A0A087WS | A0A087WS | A0A087WS | Thioredoxi |
| TMPO | 0.027641 | 0.938338 | -0.00682 | P42167;G5 | P42167;G5 | P42167 | Lamina-asso |
| COMMD7 | 0.019542 | 0.956 | 0.00425 | A0A2R8Y62 | A0A2R8Y62 | A0A2R8Y62 | COMM don |
| RPS24 | 0.016021 | 0.963782 | -0.00276 | E7ETK0;A0/ | E7ETK0;A0/ | E7ETK0 | 40S riboson |
| MGAT2 | 0.013754 | 0.968826 | -0.00361 | Q10469 | Q10469 | Q10469 | Alpha-1,6-r |
| AKAP5 | 0.012415 | 0.971817 | 0.029941 | P24588 | P24588 | P24588 | A-kinase an |
| CCDC58 | 0.00924 | 0.978948 | -0.00223 | C9JQ41;H7 | C9JQ41;H7 | C9JQ41 | Coiled-coil |
| SNRPD3 | 0.006325 | 0.985543 | -0.00169 | P62318;H3 | P62318;H3 | P62318 | Small nucle |
| GABARAPL: | 0.002699 | 0.993805 | 0.000869 | P60520;H3 | P60520;H3 | P60520 | Gamma-am |
| MGST1 | 0.00162 | 0.996278 | 0.000569 | P10620;F5 | P10620;F5 | P10620 | Microsoma |
| CISD2 | 0.000785 | 0.998194 | -0.00024 | Q8N5K1;D( | Q8N5K1;D( | Q8N5K1 | CDGSH iron |
| COG8 | 0.000365 | 0.999161 | 0.001056 | B4DYU2;Q9 | B4DYU2;Q9 | B4DYU2 | Conserved |
| PPP6R3 | 0.479846 | 0.331249 | -0.75661 | H7BXH2;E9 | H7BXH2;E9 | H7BXH2 | Serine/thre |
| LUZP1 | 0.365248 | 0.431273 | -0.44187 | Q86V48;E5 | Q86V48;E5 | Q86V48 | Leucine zip |
| TMEM63B | 3.686027 | 0.000206 | 0.45259 | H0YCP6;H3 | H0YCP6;H3 | H0YCP6 | CSC1-like p |
| NUP35 | 2.328022 | 0.004699 | 0.446953 | Q8NFH5;C9 | Q8NFH5;C9 | Q8NFH5 | Nucleopori |
| ZC3H7A | 3.338396 | 0.000459 | 0.435555 | Q8IWR0;I3 | Q8IWR0;I3 | Q8IWR0 | Zinc finger |
| CEPT1 | 2.336828 | 0.004604 | 0.422301 | Q9Y6K0;A0 | Q9Y6K0;A0 | Q9Y6K0 | Choline/eth |
| NME6 | 1.899663 | 0.012599 | 0.40004 | C9J9V6;A0/ | C9J9V6;A0/ | C9J9V6 | Nucleoside |
| MRPL11 | 2.205436 | 0.006231 | 0.390944 | Q9Y3B7;H0 | Q9Y3B7 | Q9Y3B7 | 39S riboson |
| EXOG | 3.386309 | 0.000411 | 0.349311 | Q9Y2C4;H7 | Q9Y2C4;H7 | Q9Y2C4 | Nuclease EX |
| BID | 2.146477 | 0.007137 | 0.338075 | P55957;B2 | P55957 | P55957 | BH3-intera |
| SLC39A11 | 3.472214 | 0.000337 | 0.326364 | J3QLB2;Q8 | J3QLB2;Q8 | J3QLB2 | Zinc transp |
| SUB1 | 2.177553 | 0.006644 | 0.320835 | P53999 | P53999 | P53999 | Activated R |
| BOLA3 | 1.613716 | 0.024338 | 0.31158 | Q53S33 | Q53S33 | Q53S33 | BolA-like pr |
| CD99 | 1.66049 | 0.021853 | 0.288257 | P14209;A0 | P14209;A0 | P14209 | CD99 antig |
| DIS3 | 1.819531 | 0.015152 | 0.274699 | Q9Y2L1;G3 | Q9Y2L1;G3 | Q9Y2L1 | Exosome co |
| MTIF3 | 2.469866 | 0.003389 | 0.272606 | Q9H2K0 | Q9H2K0 | Q9H2K0 | Translation |
| UQCRH | 2.071278 | 0.008486 | 0.263673 | P07919;A0 | P07919 | P07919 | Cytochrom |
| SNRPA | 2.576142 | 0.002654 | 0.257826 | P09012;M( | P09012;M( | P09012 | U1 small nu |
| VAMP3 | 2.78406 | 0.001644 | 0.251617 | Q15836;K7 | Q15836 | Q15836 | Vesicle-asso |
| HLA-DRB1 | 3.531815 | 0.000294 | 0.242267 | Q5Y7D1;A( | Q5Y7D1;A( | Q5Y7D1 | |
| AUP1 | 1.90598 | 0.012417 | 0.237411 | Q9Y679 | Q9Y679 | Q9Y679 | Ancient ubi |
| GMPPB | 1.634062 | 0.023224 | 0.226777 | Q9Y5P6 | Q9Y5P6 | Q9Y5P6 | Mannose-1 |
| NDUFB5 | 2.131406 | 0.007389 | 0.225116 | E7EWP0;H( | E7EWP0;H( | E7EWP0 | NADH dehy |
| MMGT1 | 2.10052 | 0.007934 | 0.221468 | Q8N4V1 | Q8N4V1 | Q8N4V1 | Membrane |
| ATP5I | 2.474626 | 0.003353 | 0.211272 | P56385 | P56385 | P56385 | ATP syntha |

| | | | | | | | |
|---|---|---|---|---|---|---|---|
| SYAP1 | 1.392272 | 0.040525 | 0.188884 | Q96A49 | Q96A49 | Q96A49 | Synapse-ass |
| TBC1D10B | 1.730452 | 0.018601 | 0.188832 | Q4KMP7;H | Q4KMP7 | Q4KMP7 | TBC1 doma |
| NDUFC2 | 1.47854 | 0.033225 | 0.173895 | O95298;E9 | O95298 | O95298 | NADH dehy |
| LSM4 | 1.934854 | 0.011618 | 0.166854 | V9GZ56;Q9 | V9GZ56;Q9 | V9GZ56 | U6 snRNA-a |
| | 1.595599 | 0.025375 | 0.140007 | A0A182DW | A0A182DW | A0A182DWE9 | |
| LMNA | 1.377066 | 0.04197 | 0.073008 | P02545;H0 | P02545 | P02545 | Prelamin-A |
| COX6C | 3.089191 | 0.000814 | -0.15811 | P09669 | P09669 | P09669 | Cytochrom |
| APOC1 | 1.622265 | 0.023864 | -0.20521 | K7ERI9;P02 | K7ERI9;P02 | K7ERI9 | Apolipopro |
| CMTM6 | 1.620816 | 0.023943 | -0.21413 | Q9NX76 | Q9NX76 | Q9NX76 | CKLF-like M |
| SLC5A3 | 3.846247 | 0.000142 | -0.43813 | P53794 | P53794 | P53794 | Sodium/my |
| GATC | 1.786246 | 0.016359 | -0.45723 | H0YIV9;O43 | H0YIV9;O43 | H0YIV9 | Glutamyl-tI |
| IFIT5 | 1.288626 | 0.051449 | -0.37798 | Q13325 | Q13325 | Q13325 | Interferon-i |
| SEC11A | 1.284451 | 0.051946 | -0.14171 | H0YNG3;P6 | H0YNG3;P6 | H0YNG3 | Signal pept |
| CLUH | 1.278265 | 0.052691 | 1.341603 | K7EIG1;I3L | K7EIG1;I3L | K7EIG1 | Clustered m |
| DCTN5 | 1.275557 | 0.05302 | -0.12903 | H3BR94;Q9 | H3BR94;Q9 | H3BR94 | Dynactin su |
| CHMP6 | 1.262813 | 0.054599 | 0.216246 | I3L4G8;I3L | I3L4G8;I3L | I3L4G8 | Charged mu |
| PES1 | 1.247208 | 0.056597 | 0.236355 | B3KXD6;B5 | B3KXD6;B5 | B3KXD6 | Pescadillo h |
| YTHDC1 | 1.184605 | 0.065373 | -0.29497 | Q96MU7;J3 | Q96MU7;J3 | Q96MU7 | YTH domain |
| LSM12 | 1.179018 | 0.066219 | -2.03051 | Q3MHD2;K | Q3MHD2;K | Q3MHD2 | Protein LSM |
| RBM42 | 1.159432 | 0.069274 | 1.277994 | K7EP90;Q9 | K7EP90;Q9 | K7EP90 | RNA-bindin |
| TOMM6 | 1.134804 | 0.073315 | -0.21226 | Q96B49 | Q96B49 | Q96B49 | Mitochond |
| QPCTL | 1.13118 | 0.07393 | 0.247895 | Q9NXS2;K7 | Q9NXS2 | Q9NXS2 | Glutaminyl |
| CHCHD5 | 1.104715 | 0.078575 | 0.262637 | Q9BSY4;F8 | Q9BSY4 | Q9BSY4 | Coiled-coil- |
| TBC1D2B | 1.087319 | 0.081786 | -1.27347 | Q9UPU7 | Q9UPU7 | Q9UPU7 | TBC1 doma |
| RILP | 1.071873 | 0.084748 | 1.343062 | Q96NA2;I3 | Q96NA2;I3 | Q96NA2 | Rab-interac |
| PSIP1 | 1.048795 | 0.089373 | 1.556519 | O75475 | O75475 | O75475 | PC4 and SF |
| COX15 | 1.045865 | 0.089978 | 0.221577 | Q7KZN9;A0 | Q7KZN9;A0 | Q7KZN9 | Cytochrom |
| MRPL9 | 1.042123 | 0.090756 | 0.251525 | Q9BYD2;Q5 | Q9BYD2;Q5 | Q9BYD2 | 39S ribosom |
| SLC9A1 | 1.028638 | 0.093619 | -0.24353 | P19634;B1 | P19634 | P19634 | Sodium/hy |
| GATAD2A | 1.022489 | 0.094954 | -1.3209 | Q86YP4;C9 | Q86YP4;C9 | Q86YP4 | Transcripti |
| PGM3 | 1.01978 | 0.095548 | -0.34321 | A0A494C0 | A0A494C0 | A0A494C0 | Phosphoac |
| TM6SF1 | 1.014626 | 0.096688 | -1.47035 | Q9BZW5;H | Q9BZW5;H | Q9BZW5 | Transmemb |
| TSPO | 1.004973 | 0.098861 | 0.293612 | B1AH87;P3 | B1AH87;P3 | B1AH87 | Translocato |
| GXYLT1 | 0.97953 | 0.104826 | -1.061 | Q4G148 | Q4G148 | Q4G148 | Glucoside x |
| TMED3 | 0.969692 | 0.107228 | 0.240459 | Q9Y3Q3;G3 | Q9Y3Q3;G3 | Q9Y3Q3 | Transmemb |
| RIPK2 | 0.954454 | 0.111057 | 1.608255 | O43353 | O43353 | O43353 | Receptor-in |
| POM121C | 0.9491 | 0.112435 | -1.39809 | A8CG34;Q9 | A8CG34;Q9 | A8CG34 | Nuclear env |
| ATPAF1 | 0.94065 | 0.114644 | 0.142844 | H0YEW4;A8 | H0YEW4;A8 | H0YEW4 | ATP syntha |
| SNRPF | 0.93017 | 0.117444 | 0.199598 | P62306;A0 | P62306;A0 | P62306 | Small nucle |
| TMCO1 | 0.902045 | 0.125301 | 0.148309 | Q9UM00;J3 | Q9UM00;J3 | Q9UM00 | Transmemb |
| WBP2 | 0.898557 | 0.126311 | 2.004763 | K7EIJ0;K7E | K7EIJ0;K7E | K7EIJ0 | WW domai |
| EPDR1 | 0.895375 | 0.12724 | 0.938825 | Q9UM22;D | Q9UM22;D | Q9UM22 | Mammalian |
| SYNGR2 | 0.891349 | 0.128425 | -0.15392 | O43760;K7 | O43760;K7 | O43760 | Synaptogyr |
| AGAP3 | 0.890779 | 0.128594 | 1.079329 | Q96P47;H0 | Q96P47;H0 | Q96P47 | Arf-GAP wit |
| PSD3 | 0.887563 | 0.12955 | 1.155251 | E5RJ29;Q9 | E5RJ29;Q9 | E5RJ29 | PH and SEC |
| PYURF | 0.876715 | 0.132826 | -0.22684 | Q96I23 | Q96I23 | Q96I23 | Protein pre |
| HNRNPC | 0.87137 | 0.134471 | 0.104843 | G3V4C1;B2 | G3V4C1;B2 | G3V4C1 | |
| TRAPPC2B | 0.859399 | 0.13823 | -0.1928 | P0DI82;P0 | P0DI82;P0 | P0DI82 | Trafficking |

| | | | | | | | |
|---|---|---|---|---|---|---|---|
| ERGIC2 | 0.854402 | 0.139829 | -0.10593 | A0A087WU | A0A087WU | A0A087WU | Endoplasm |
| RAVER1 | 0.853741 | 0.140042 | -0.26471 | A0A087WZ | A0A087WZ | A0A087WZ | Ribonucleo |
| RIN3 | 0.850277 | 0.141164 | -0.16653 | Q8TB24;A0 | Q8TB24;A0 | Q8TB24 | Ras and Rab |
| NFAM1 | 0.838724 | 0.144969 | 0.204894 | Q8NET5;F8 | Q8NET5;F8 | Q8NET5 | NFAT activa |
| TBC1D1 | 0.835742 | 0.145968 | -0.80658 | Q86TI0;H0 | Q86TI0;H0 | Q86TI0 | TBC1 doma |
| HACD2 | 0.835542 | 0.146035 | 0.324299 | Q6Y1H2 | Q6Y1H2 | Q6Y1H2 | Very-long-c |
| ASRGL1 | 0.818277 | 0.151958 | -0.13848 | Q7L266;A0 | Q7L266;A0 | Q7L266 | Isoaspartyl |
| RAB8B | 0.818123 | 0.152012 | 0.109022 | Q92930;H0 | Q92930;H0 | Q92930 | Ras-related |
| SNRPC | 0.811837 | 0.154228 | 0.174738 | P09234;A0 | P09234;A0 | P09234 | U1 small nu |
| RNF170 | 0.803564 | 0.157194 | -0.092 | Q96K19;E9 | Q96K19 | Q96K19 | E3 ubiquiti |
| TYROBP | 0.792457 | 0.161266 | 0.148982 | O43914;X6 | O43914;X6 | O43914 | TYRO prote |
| MRPS11 | 0.791108 | 0.161768 | 0.116083 | P82912;H0 | P82912;H0 | P82912 | 28S ribosor |
| YIF1B | 0.790999 | 0.161808 | -0.166 | Q5BJH7;K7 | Q5BJH7;K7 | Q5BJH7 | Protein YIF |
| SEPTIN3 | 0.784043 | 0.164421 | -0.15846 | B1AHR1;A0 | B1AHR1;A0 | B1AHR1 | Neuronal-s |
| LIN7C | 0.782958 | 0.164832 | 0.88469 | G3V1D4;Q | G3V1D4;Q | G3V1D4 | Protein lin- |
| H2AFV | 0.781183 | 0.165507 | 0.145251 | Q71UI9;P0 | Q71UI9;P0 | Q71UI9 | Histone H2 |
| CD59 | 0.781103 | 0.165538 | 0.125596 | E9PNW4;A | E9PNW4;A | E9PNW4 | CD59 glyco |
| CD33 | 0.772278 | 0.168936 | 0.194736 | P20138;B4 | P20138;B4 | P20138 | Myeloid cel |
| TMEM70 | 0.76989 | 0.169867 | -0.07813 | Q9BUB7 | Q9BUB7 | Q9BUB7 | Transmemb |
| TOMM40L | 0.766221 | 0.171308 | -1.09022 | Q969M1 | Q969M1 | Q969M1 | Mitochond |
| HIST1H1C | 0.76535 | 0.171652 | 0.17838 | P16403 | P16403 | P16403 | Histone H1 |
| SARAF | 0.759141 | 0.174124 | 0.951157 | E5RFJ9;E5R | E5RFJ9;E5R | E5RFJ9 | Store-opera |
| NCOA5 | 0.754166 | 0.17613 | -0.11899 | Q9HCD5;Q | Q9HCD5 | Q9HCD5 | Nuclear rec |
| SLC7A8 | 0.75055 | 0.177603 | -1.63973 | Q9UHI5;E9 | Q9UHI5;E9 | Q9UHI5 | Large neutr |
| SMARCD2 | 0.748956 | 0.178256 | 0.237666 | J3KMX2;B9 | J3KMX2;B9 | J3KMX2 | SWI/SNF-re |
| TMED1 | 0.725339 | 0.188218 | -0.13047 | Q13445;K7 | Q13445;K7 | Q13445 | Transmemb |
| PAK1 | 0.713255 | 0.193529 | -1.194 | Q13153;B3 | Q13153;B3 | Q13153 | Serine/thre |
| MT-CO1 | 0.695168 | 0.201759 | -0.14559 | P00395 | P00395 | P00395 | Cytochrom |
| PPP3R1 | 0.69201 | 0.203231 | -0.10862 | F6U1T9;D3 | F6U1T9;D3 | F6U1T9 | Calcineurin |
| TRAPPC2L | 0.686298 | 0.205922 | -1.10135 | H3BUK6;A0 | H3BUK6;A0 | H3BUK6 | Trafficking |
| SLC25A25 | 0.678268 | 0.209764 | -1.37909 | Q6KCM7;M | Q6KCM7;M | Q6KCM7 | Calcium-bi |
| GGCX | 0.667491 | 0.215035 | 0.109855 | P38435 | P38435 | P38435 | Vitamin K-c |
| NDUFS6 | 0.659806 | 0.218874 | 0.099108 | O75380;D6 | O75380;D6 | O75380 | NADH dehy |
| UBE2G2 | 0.659722 | 0.218916 | 0.126643 | P60604 | P60604 | P60604 | Ubiquitin-c |
| TRIP6 | 0.650137 | 0.223802 | -0.57605 | H7BZE2;Q1 | H7BZE2;Q1 | H7BZE2 | Thyroid rec |
| TMEM115 | 0.646384 | 0.225744 | 1.48071 | Q12893 | Q12893 | Q12893 | Transmemb |
| MYL9 | 0.637864 | 0.230217 | -0.17175 | P24844 | P24844 | P24844 | Myosin regu |
| CRELD2 | 0.635502 | 0.231472 | 0.117703 | A6PWM2;C | A6PWM2;C | A6PWM2 | Cysteine-ric |
| MYD88 | 0.630883 | 0.233947 | -1.11909 | H0Y4G9;Q | H0Y4G9;Q | H0Y4G9 | Myeloid dif |
| SLC38A9 | 0.629212 | 0.234849 | 0.080708 | E7ESU6;D6 | E7ESU6;D6 | E7ESU6 | Sodium-cou |
| CCDC124 | 0.628896 | 0.23502 | 0.675521 | Q96CT7;M0 | Q96CT7 | Q96CT7 | Coiled-coil |
| JAK1 | 0.625184 | 0.237037 | 0.862574 | A0A5F9ZH0 | A0A5F9ZH0 | A0A5F9ZH0 | Tyrosine-pr |
| CRKL | 0.620408 | 0.239658 | -0.75359 | P46109 | P46109 | P46109 | Crk-like pro |
| IGHM | 0.618957 | 0.24046 | 0.632967 | P01871 | P01871 | P01871 | Ig mu chain |
| PITPNB | 0.615814 | 0.242206 | -0.13483 | P48739;A0 | P48739;A0 | P48739 | Phosphatid |
| AIF1 | 0.615079 | 0.242617 | -0.82065 | Q5STX8;P5 | Q5STX8;P5 | Q5STX8 | Allograft in |
| SLC6A12 | 0.589059 | 0.257597 | 0.252778 | P48065;F5 | P48065 | P48065 | Sodium- an |
| MRPL18 | 0.587347 | 0.258615 | 0.187374 | Q9H0U6 | Q9H0U6 | Q9H0U6 | 39S ribosor |

| | | | | | | | |
|---|---|---|---|---|---|---|---|
| PELO | 0.576315 | 0.265268 | 0.795244 | Q9BRX2 | Q9BRX2 | Q9BRX2 | Protein pel |
| STAG2 | 0.574518 | 0.266368 | 0.586978 | Q8N3U4;B1 | Q8N3U4 | Q8N3U4 | Cohesin sul |
| TSTA3 | 0.57173 | 0.268084 | -1.16598 | E9PLH9;E9I | E9PLH9;E9I | E9PLH9 | GDP-L-fuco |
| CASP7 | 0.56013 | 0.27534 | 0.203827 | A0A5F9ZHZ | A0A5F9ZHZ | A0A5F9ZHZ | Caspase-7;C |
| RHBDD1 | 0.557475 | 0.277029 | 0.711066 | Q8TEB9;B9 | Q8TEB9;B9 | Q8TEB9 | Rhomboid- |
| MARCHF5 | 0.555596 | 0.27823 | 0.095459 | Q9NX47 | Q9NX47 | Q9NX47 | E3 ubiquiti |
| SERPINB8 | 0.555247 | 0.278454 | -0.95866 | P50452;H7 | P50452;H7 | P50452 | Serpin B8 |
| STMN1 | 0.55148 | 0.280879 | -0.85532 | A2A2D0;P1 | A2A2D0;P1 | A2A2D0 | Stathmin;S |
| NAAA | 0.551262 | 0.281021 | -1.07446 | Q02083;R4 | Q02083;R4 | Q02083 | N-acylethar |
| NDUFAF5 | 0.547739 | 0.28331 | 0.175488 | Q5TEU4;B3 | Q5TEU4;B3 | Q5TEU4 | NADH dehy |
| LANCL1 | 0.543647 | 0.285992 | -1.09439 | O43813;E9 | O43813;E9 | O43813 | LanC-like pr |
| HMGN1 | 0.539066 | 0.289024 | -0.13193 | A6NL93;P0 | A6NL93;P0 | A6NL93 | Non-histon |
| PACS1 | 0.534841 | 0.29185 | -1.26048 | Q6VY07;B4 | Q6VY07;B4 | Q6VY07 | Phosphofur |
| HECTD1 | 0.533841 | 0.292523 | 0.634846 | A0A087X2I | A0A087X2I | A0A087X2I | E3 ubiquiti |
| CMC1 | 0.533459 | 0.29278 | 0.057516 | Q7Z7K0;C9 | Q7Z7K0;C9 | Q7Z7K0 | COX assemb |
| ISCA2 | 0.529371 | 0.295549 | -0.11377 | Q86U28;J3 | Q86U28;J3 | Q86U28 | Iron-sulfur |
| NT5C3A | 0.529238 | 0.295639 | -0.60492 | X6RM59;Q | X6RM59;Q | X6RM59 | 5-nucleotic |
| SERINC1 | 0.526641 | 0.297412 | 0.121424 | Q9NRX5 | Q9NRX5 | Q9NRX5 | Serine inco |
| TWSG1 | 0.514255 | 0.306017 | -0.10607 | Q9GZX9;J3 | Q9GZX9;J3 | Q9GZX9 | Twisted gas |
| ANKRD13A | 0.503249 | 0.313871 | 0.549152 | Q8IZ07;H0 | Q8IZ07;H0 | Q8IZ07 | Ankyrin rep |
| C1orf123 | 0.502338 | 0.31453 | -0.63887 | Q9NWV4 | Q9NWV4 | Q9NWV4 | UPF0587 p |
| MAPRE2 | 0.500264 | 0.316035 | -0.10951 | Q15555;K7 | Q15555;K7 | Q15555 | Microtubul |
| PPIE | 0.495128 | 0.319795 | -0.65321 | Q9UNP9;E9 | Q9UNP9;E9 | Q9UNP9 | Peptidyl-pr |
| ZNF622 | 0.493661 | 0.320878 | 0.801893 | Q969S3 | Q969S3 | Q969S3 | Zinc finger |
| ADO | 0.488755 | 0.324523 | 0.706607 | Q96SZ5 | Q96SZ5 | Q96SZ5 | 2-aminoeth |
| DNPH1 | 0.487595 | 0.325391 | -0.15837 | H0Y8X4;O4 | H0Y8X4;O4 | H0Y8X4 | 2-deoxynuc |
| CCDC132 | 0.484492 | 0.327724 | 0.988942 | Q96JG6;H7 | Q96JG6;H7 | Q96JG6 | Coiled-coil |
| ARMT1 | 0.4804 | 0.330826 | -0.10469 | Q9H993;F5 | Q9H993;F5 | Q9H993 | Protein-glu |
| GLS | 0.479517 | 0.3315 | 0.104556 | B8ZZA8;B8 | B8ZZA8;B8 | B8ZZA8 | |
| NAE1 | 0.478569 | 0.332224 | 0.587213 | Q13564;H3 | Q13564;H3 | Q13564 | NEDD8-acti |
| BOLA2 | 0.474106 | 0.335656 | -0.14862 | Q9H3K6;A0 | Q9H3K6;A0 | Q9H3K6 | BolA-like pr |
| SMARCA2 | 0.473172 | 0.336378 | -0.13 | F6VDE0;P5 | F6VDE0;P5 | F6VDE0 | Probable gl |
| NOP10 | 0.471459 | 0.337708 | -0.17962 | Q9NPE3;H0 | Q9NPE3;H0 | Q9NPE3 | H/ACA ribo |
| C6orf203 | 0.468074 | 0.34035 | 0.785102 | A0A0D9SEI | A0A0D9SEI | A0A0D9SEI | Uncharacte |
| GORASP1 | 0.464508 | 0.343156 | 0.708264 | Q9BQQ3;H7 | Q9BQQ3;H7 | Q9BQQ3 | Golgi reasse |
| CBX5 | 0.463784 | 0.343729 | -0.63581 | P45973;F8 | P45973 | P45973 | Chromobox |
| RNPS1 | 0.460638 | 0.346228 | 0.874598 | H3BV80;H3 | H3BV80;H3 | H3BV80 | RNA-bindir |
| MVK | 0.460427 | 0.346396 | -0.10739 | F5H8H2;Q0 | F5H8H2;Q0 | F5H8H2 | Mevalonate |
| NUTF2 | 0.45821 | 0.348169 | 0.077122 | P61970;H3 | P61970;H3 | P61970 | Nuclear tra |
| TIMM10 | 0.455535 | 0.35032 | 0.083176 | P62072 | P62072 | P62072 | Mitochond |
| FMR1 | 0.450955 | 0.354034 | 0.091375 | A8MQB8;G | A8MQB8;G | A8MQB8 | Fragile X m |
| COMMD6 | 0.442342 | 0.361125 | -0.64065 | Q7Z4G1;B0 | Q7Z4G1;B0 | Q7Z4G1 | COMM dom |
| COA6 | 0.439881 | 0.363177 | 0.072907 | Q5JTJ3 | Q5JTJ3 | Q5JTJ3 | Cytochrom |
| FYTTD1 | 0.439264 | 0.363694 | 0.23437 | Q96QD9;H9 | Q96QD9;H9 | Q96QD9 | UAP56-inte |
| PAP1 | 0.432022 | 0.369809 | -0.4948 | Q8TD49;Q9 | Q8TD49;Q9 | Q8TD49 | Methyltran |
| FHL1 | 0.42565 | 0.375275 | -0.64123 | Q5JXI2;A0A | Q5JXI2;A0A | Q5JXI2 | Four and a |
| MINOS1 | 0.42218 | 0.378286 | 0.189288 | A0A087WU | A0A087WU | A0A087WU | MICOS com |
| KDELC2 | 0.416979 | 0.382843 | -0.56108 | Q7Z4H8 | Q7Z4H8 | Q7Z4H8 | KDEL motif |

| | | | | | | | |
|---|---|---|---|---|---|---|---|
| CSTA | 0.409852 | 0.389178 | -0.12062 | P01040;C9 | P01040;C9 | P01040 | Cystatin-A; |
| RRM2B | 0.409127 | 0.389828 | 0.107377 | Q7LG56;H( | Q7LG56;H( | Q7LG56 | Ribonucleo |
| CD53 | 0.408847 | 0.39008 | -0.82886 | P19397 | P19397 | P19397 | Leukocyte s |
| ENDOG | 0.407602 | 0.3912 | -0.55849 | Q14249 | Q14249 | Q14249 | Endonuclea |
| LSM3 | 0.405713 | 0.392904 | -0.69238 | P62310 | P62310 | P62310 | U6 snRNA-a |
| NDUFAF4 | 0.396167 | 0.401637 | -0.10591 | Q9P032 | Q9P032 | Q9P032 | NADH dehy |
| ASCC1 | 0.384933 | 0.412161 | -0.67087 | A0A5F9ZH7 | A0A5F9ZH7 | A0A5F9ZH7 | Activating s |
| PRPSAP1 | 0.384753 | 0.412332 | 0.461959 | Q14558;C9 | Q14558;C9 | Q14558 | Phosphorib |
| RNF114 | 0.376593 | 0.420152 | -0.51602 | A0A096LN\ | A0A096LN\ | A0A096LN\ | E3 ubiquiti |
| BNIP2 | 0.372157 | 0.424466 | 0.18329 | J3KN59;Q1 | J3KN59;Q1 | J3KN59 | BCL2/aden |
| HSPE1 | 0.370816 | 0.425779 | -0.03599 | P61604;B8 | P61604;B8 | P61604 | 10 kDa hea |
| DERL1 | 0.368098 | 0.428452 | 0.078054 | Q9BUN8;B4 | Q9BUN8;B4 | Q9BUN8 | Derlin-1 |
| ATP5L | 0.365132 | 0.431388 | 0.046609 | E9PN17;A0 | E9PN17;A0 | E9PN17 | ATP synthas |
| DYNLT3 | 0.361495 | 0.435016 | 0.099408 | A6NGJ0;P5 | A6NGJ0;P5 | A6NGJ0 | Dynein ligh |
| APOC4-APC | 0.35088 | 0.445779 | -0.06066 | K7ER74;P0 | K7ER74;P0 | K7ER74 | Apolipopro |
| RPL37A | 0.34959 | 0.447105 | 0.073202 | C9J4Z3;P6: | C9J4Z3;P6: | C9J4Z3 | 60S riboson |
| ERC1 | 0.34768 | 0.449076 | 0.322522 | G8JLD3;X6 | G8JLD3;X6 | G8JLD3 | ELKS/Rab6- |
| PILRA | 0.345485 | 0.451352 | -0.51166 | Q9UKJ1;C9 | Q9UKJ1;C9 | Q9UKJ1 | Paired imm |
| DPAGT1 | 0.34081 | 0.456237 | -0.15577 | A0A1W2PF | A0A1W2PF | A0A1W2PF | UDP-N-acet |
| C10orf76 | 0.338487 | 0.458684 | 0.62504 | Q5T2E6;Q5 | Q5T2E6 | Q5T2E6 | UPF0668 p |
| HYPK | 0.334364 | 0.463059 | -0.53043 | Q9NX55;J3 | Q9NX55;J3 | Q9NX55 | Huntingtin |
| TRAFD1 | 0.327053 | 0.470919 | -0.36902 | O14545;F8 | O14545;F8 | O14545 | TRAF-type z |
| MPC1 | 0.321454 | 0.47703 | 0.153911 | Q9Y5U8 | Q9Y5U8 | Q9Y5U8 | Mitochond |
| COMMD5 | 0.313803 | 0.485509 | -0.05362 | Q9GZQ3;E9 | Q9GZQ3;E9 | Q9GZQ3 | COMM dom |
| RPS25 | 0.302061 | 0.498814 | -0.07097 | P62851 | P62851 | P62851 | 40S riboson |
| ARPC1B | 0.300144 | 0.501022 | 0.043131 | O15143;C9 | O15143;C9 | O15143 | Actin-relate |
| CETN2 | 0.29476 | 0.507271 | -0.04995 | P41208;Q1 | P41208 | P41208 | Centrin-2 |
| NUP43 | 0.293917 | 0.508256 | -0.07278 | Q8NFH3;B4 | Q8NFH3 | Q8NFH3 | Nucleopori |
| HLA-A | 0.292899 | 0.509449 | -0.05198 | Q53Z42;A0 | Q53Z42;A0 | Q53Z42 | |
| DYNLT1 | 0.287643 | 0.515652 | 0.054458 | P63172 | P63172 | P63172 | Dynein ligh |
| RPL35 | 0.28274 | 0.521506 | -0.0635 | P42766;F2 | P42766;F2 | P42766 | 60S riboson |
| POP1 | 0.281172 | 0.523393 | -0.35669 | Q99575 | Q99575 | Q99575 | Ribonuclea |
| FAM160B1 | 0.2807 | 0.523962 | -0.7135 | Q5W0V3;Q | Q5W0V3;Q | Q5W0V3 | Protein FAN |
| TXN2 | 0.280477 | 0.524231 | 0.043923 | Q99757;M( | Q99757;M( | Q99757 | Thioredoxi |
| SLC25A1 | 0.278905 | 0.526132 | -0.0328 | P53007 | P53007 | P53007 | Tricarboxyl |
| CD82 | 0.278232 | 0.526948 | 0.092151 | P27701;E9 | P27701 | P27701 | CD82 antig |
| PIN1 | 0.277095 | 0.528329 | -0.68679 | Q13526;K7 | Q13526;K7 | Q13526 | Peptidyl-pr |
| MRPL23 | 0.274649 | 0.531313 | 0.834911 | Q16540;A6 | Q16540;A6 | Q16540 | 39S riboson |
| CMC2 | 0.258044 | 0.552022 | -0.1305 | Q9NRP2;H3 | Q9NRP2;H3 | Q9NRP2 | COX assemb |
| CD300LF | 0.257225 | 0.553063 | 0.02993 | J3KS52;Q8 | J3KS52;Q8 | J3KS52 | CMRF35-lik |
| SLC11A1 | 0.255416 | 0.555372 | 0.120993 | P49279;Q9 | P49279;Q9 | P49279 | Natural resi |
| TNFAIP8 | 0.247086 | 0.566127 | -0.07899 | O95379;D6 | O95379;D6 | O95379 | Tumor necr |
| FAM107B | 0.241377 | 0.573619 | 0.059217 | C9J6N5;C9 | C9J6N5;C9 | C9J6N5 | Protein FAN |
| PRKAB1 | 0.239155 | 0.57656 | -0.03271 | Q9Y478;F5 | Q9Y478;F5 | Q9Y478 | 5-AMP-acti |
| GATAD2B | 0.237387 | 0.578913 | 0.055523 | A0A0U1RRI | A0A0U1RRI | A0A0U1RRI | Transcripti |
| KCNN4 | 0.23571 | 0.581153 | 0.384045 | M0R1J0;D1 | M0R1J0;D1 | M0R1J0 | Intermedia |
| TRMT2A | 0.227573 | 0.592144 | 0.478529 | F2Z2W7;Q8 | F2Z2W7;Q8 | F2Z2W7 | tRNA (uraci |
| MFHAS1 | 0.223786 | 0.59733 | -0.7613 | Q9Y4C4 | Q9Y4C4 | Q9Y4C4 | Malignant f |

| | | | | | | | |
|---|---|---|---|---|---|---|---|
| ANP32E | 0.222347 | 0.599312 | 0.1518 | Q9BTT0;E9I | Q9BTT0;E9I | Q9BTT0 | Acidic leuci |
| FABP5 | 0.218886 | 0.604107 | -0.07612 | Q01469 | Q01469 | Q01469 | Fatty acid-b |
| COX7C | 0.216413 | 0.607557 | -0.07719 | P15954;D6 | P15954;D6 | P15954 | Cytochrom |
| KIAA0391 | 0.21378 | 0.611252 | 0.612225 | O15091;A0 | O15091 | O15091 | Mitochond |
| NUBPL | 0.212875 | 0.612527 | 0.086142 | Q8TB37;F8 | Q8TB37;F8 | Q8TB37 | Iron-sulfur |
| GID8 | 0.208753 | 0.618368 | -0.05908 | Q9NWU2 | Q9NWU2 | Q9NWU2 | Glucose-inc |
| BST1 | 0.20503 | 0.623692 | -0.47697 | H0Y984;A6 | H0Y984;A6 | H0Y984 | ADP-ribosy |
| TTYH2 | 0.204596 | 0.624315 | 0.447289 | B4DKD1;Q9 | B4DKD1;Q9 | B4DKD1 | Protein twe |
| TMEM104 | 0.198673 | 0.632888 | 0.060169 | Q8NE00;J3 | Q8NE00;J3 | Q8NE00 | Transmemb |
| SFXN5 | 0.192178 | 0.642424 | 0.05257 | Q8TD22;C9 | Q8TD22;C9 | Q8TD22 | Sideroflexir |
| SUMO1 | 0.189213 | 0.646826 | -0.03005 | B8ZZN6;P6 | B8ZZN6;P6 | B8ZZN6 | Small ubiqu |
| CREG1 | 0.188136 | 0.648431 | 0.045161 | A0A3B3IRL | A0A3B3IRL | A0A3B3IRL | Protein CRI |
| PSEN2 | 0.183535 | 0.655337 | 0.041949 | B1AP22;P4 | B1AP22;P4 | B1AP22 | Presenilin;I |
| ACY1 | 0.181425 | 0.658529 | -0.39392 | A0A1B0GU | A0A1B0GU | A0A1B0GU | Aminoacyla |
| TAOK2 | 0.181295 | 0.658726 | -0.18916 | Q9UL54 | Q9UL54 | Q9UL54 | Serine/thre |
| LAMTOR4 | 0.176514 | 0.666019 | -0.03854 | Q0VGL1;A0 | Q0VGL1;A0 | Q0VGL1 | Ragulator c |
| ARHGAP25 | 0.175161 | 0.668096 | 0.032421 | P42331;H7 | P42331;H7 | P42331 | Rho GTPase |
| ADAR | 0.170684 | 0.675019 | 0.030154 | P55265 | P55265 | P55265 | Double-stra |
| NUP50 | 0.16894 | 0.677735 | -0.04963 | Q9UKX7 | Q9UKX7 | Q9UKX7 | Nuclear por |
| YY1 | 0.163019 | 0.687038 | 0.072189 | P25490;H0 | P25490;H0 | P25490 | Transcripti |
| ZCCHC8 | 0.158262 | 0.694605 | -0.36401 | Q6NZY4;F5 | Q6NZY4;F5 | Q6NZY4 | Zinc finger |
| PPM1A | 0.153105 | 0.702902 | -0.05161 | P35813 | P35813 | P35813 | Protein pho |
| MRPS12 | 0.151992 | 0.704707 | 0.037013 | O15235 | O15235 | O15235 | 28S riboson |
| MRPL48 | 0.149053 | 0.709491 | -0.04085 | F5H702;Q9 | F5H702;Q9 | F5H702 | 39S riboson |
| TOMM5 | 0.143929 | 0.717911 | -0.03244 | F8W8Z9;Q8 | F8W8Z9;Q8 | F8W8Z9 | Mitochond |
| YIF1A | 0.143555 | 0.71853 | 0.039118 | A6NGW1;C | A6NGW1;C | A6NGW1 | Protein YIF |
| CD86 | 0.142617 | 0.720084 | 0.042794 | P42081;C9 | P42081;C9 | P42081 | T-lymphocy |
| MCFD2 | 0.139839 | 0.724705 | -0.04226 | Q8NI22;H7 | Q8NI22;H7 | Q8NI22 | Multiple co |
| MCOLN3 | 0.137653 | 0.728362 | 0.041095 | Q8TDD5;B1 | Q8TDD5;B1 | Q8TDD5 | Mucolipin- |
| GPSM3 | 0.13478 | 0.733195 | -0.03622 | E9PIP3;Q9 | E9PIP3;Q9 | E9PIP3 | G-protein-s |
| VPS26A | 0.1281 | 0.744561 | 0.039187 | O75436;S4 | O75436 | O75436 | Vacuolar pr |
| TNFAIP8L2 | 0.127997 | 0.744737 | 0.075674 | Q6P589 | Q6P589 | Q6P589 | Tumor necr |
| CSTF2 | 0.119489 | 0.759471 | -0.01938 | E7EWR4;P3 | E7EWR4;P3 | E7EWR4 | Cleavage sti |
| MRPS18A | 0.118815 | 0.76065 | -0.03241 | Q9NVS2;Q5 | Q9NVS2;Q5 | Q9NVS2 | 28S riboson |
| AUH | 0.11776 | 0.7625 | -0.03616 | Q13825 | Q13825 | Q13825 | Methylglut |
| EBNA1BP2 | 0.107966 | 0.779891 | 0.05251 | H7C2Q8;Q9 | H7C2Q8;Q9 | H7C2Q8 | Probable rF |
| MCEE | 0.103364 | 0.788199 | -0.06097 | Q96PE7;F5 | Q96PE7;F5 | Q96PE7 | Methylmal |
| BDH2 | 0.101678 | 0.791264 | 0.041663 | Q9BUT1;D6 | Q9BUT1;D6 | Q9BUT1 | 3-hydroxyb |
| PDAP1 | 0.100399 | 0.793598 | 0.198912 | Q13442;F8 | Q13442 | Q13442 | 28 kDa hea |
| SNRPG | 0.088964 | 0.814772 | -0.01187 | Q49AN9;F5 | Q49AN9;F5 | Q49AN9 | Small nucle |
| SYNE2 | 0.088028 | 0.816529 | 0.233569 | G3V5X4;A0 | G3V5X4;A0 | G3V5X4 | Nesprin-2 |
| PPP2R5C | 0.084262 | 0.82364 | -0.01984 | H0YJ75;Q1 | H0YJ75;Q1 | H0YJ75 | Serine/thre |
| MT-CO3 | 0.083352 | 0.825369 | 0.02676 | P00414 | P00414 | P00414 | Cytochrom |
| ALG8 | 0.083227 | 0.825607 | -0.01557 | Q9BVK2;E9 | Q9BVK2;E9 | Q9BVK2 | Probable de |
| LSM6 | 0.082832 | 0.826357 | -0.02295 | P62312 | P62312 | P62312 | U6 snRNA-a |
| RFT1 | 0.081804 | 0.828317 | 0.022275 | Q96AA3;B5 | Q96AA3;B5 | Q96AA3 | Protein RFT |
| STIM2 | 0.080298 | 0.831193 | 0.12905 | H7C5C6;H7 | H7C5C6;H7 | H7C5C6 | Stromal int |
| RBMXL1 | 0.078367 | 0.834897 | -0.02572 | Q96E39;A0 | Q96E39 | Q96E39 | RNA bindin |

| | | | | | | | |
|---|---|---|---|---|---|---|---|
| DAD1 | 0.077337 | 0.836879 | 0.012174 | F5GXX5;P6 | F5GXX5;P6 | F5GXX5 | Dolichyl-di |
| FAM96B | 0.073729 | 0.843861 | 0.125312 | Q9Y3D0;J3 | Q9Y3D0;J3 | Q9Y3D0 | Mitotic spi |
| GPR180 | 0.067979 | 0.855108 | 0.01965 | Q86V85 | Q86V85 | Q86V85 | Integral me |
| POLR2G | 0.062812 | 0.865343 | 0.016733 | P62487;E9 | P62487;E9 | P62487 | DNA-direct |
| RBM22 | 0.058317 | 0.874346 | 0.019156 | Q9NW64;E | Q9NW64;E | Q9NW64 | Pre-mRNA- |
| CCDC134 | 0.055689 | 0.879653 | -0.02047 | Q9H6E4;B0 | Q9H6E4;B0 | Q9H6E4 | Coiled-coil |
| CREB1 | 0.052029 | 0.887096 | 0.015583 | P16220;E9 | P16220;E9 | P16220 | Cyclic AMP |
| NOMO2 | 0.050673 | 0.889871 | 0.009129 | Q5JPE7;J3 | Q5JPE7;J3 | Q5JPE7 | Nodal mod |
| PWP1 | 0.049442 | 0.892397 | 0.137154 | Q13610;B4 | Q13610;B4 | Q13610 | Periodic try |
| MECR | 0.049115 | 0.893068 | 0.00998 | Q9BV79;H3 | Q9BV79 | Q9BV79 | Trans-2-enc |
| DCK | 0.048791 | 0.893736 | 0.01514 | P27707;D6 | P27707;D6 | P27707 | Deoxycytid |
| TMEM55A | 0.048519 | 0.894294 | -0.09629 | E5RIY0;E5R | E5RIY0;E5R | E5RIY0 | Type 2 pho |
| SLC43A2 | 0.04615 | 0.899187 | 0.023999 | Q8N370 | Q8N370 | Q8N370 | Large neutr |
| PPP1R12C | 0.044985 | 0.901602 | -0.01313 | K7EL81;Q9 | K7EL81;Q9 | K7EL81 | Protein pho |
| SDHC | 0.04389 | 0.903878 | 0.00905 | Q99643 | Q99643 | Q99643 | Succinate c |
| PPIL3 | 0.03639 | 0.919623 | 0.09732 | Q9H2H8;C9 | Q9H2H8;C9 | Q9H2H8 | Peptidyl-pr |
| PPP1R2 | 0.03381 | 0.925103 | 0.078346 | E7EMN6;P4 | E7EMN6;P4 | E7EMN6 | Protein pho |
| MS4A1 | 0.033386 | 0.926007 | 0.020465 | E9PKH8;P1 | E9PKH8;P1 | E9PKH8 | B-lymphoc |
| COX11 | 0.030029 | 0.933191 | -0.11023 | Q9Y6N1;B4 | Q9Y6N1;B4 | Q9Y6N1 | Cytochrom |
| PHPT1 | 0.029884 | 0.933503 | -0.01083 | Q9NRX4 | Q9NRX4 | Q9NRX4 | 14 kDa pho |
| FTL | 0.027434 | 0.938785 | 0.01522 | P02792 | P02792 | P02792 | Ferritin ligh |
| C14orf1 | 0.021797 | 0.95105 | -0.00322 | Q9UKR5 | Q9UKR5 | Q9UKR5 | Probable er |
| GFM2 | 0.020258 | 0.954426 | 0.044964 | Q969S9;D6 | Q969S9;D6 | Q969S9 | Ribosome- |
| S100A4 | 0.015265 | 0.965461 | 0.004205 | P26447 | P26447 | P26447 | Protein S10 |
| CCDC88B | 0.015221 | 0.96556 | 0.027725 | A6NC98;H7 | A6NC98;H7 | A6NC98 | Coiled-coil |
| MYADM | 0.011507 | 0.973853 | 0.003463 | Q96S97;C9 | Q96S97;C9 | Q96S97 | Myeloid-as |
| MRPS18C | 0.010738 | 0.975578 | 0.03708 | Q9Y3D5;H0 | Q9Y3D5;H0 | Q9Y3D5 | 28S riboso |
| PNPLA7 | 0.010505 | 0.976102 | -0.03516 | Q6ZV29 | Q6ZV29 | Q6ZV29 | Patatin-like |
| KIAA0020 | 0.010151 | 0.976898 | 0.020319 | Q15397 | Q15397 | Q15397 | Pumilio do |
| DNAJB14 | 0.009555 | 0.97824 | -0.01632 | Q8TBM8;H | Q8TBM8 | Q8TBM8 | DnaJ homo |
| FGFR1OP2 | 0.008919 | 0.979673 | 0.017866 | Q9NVK5;F5 | Q9NVK5 | Q9NVK5 | FGFR1 onc |
| TUBB | 0.006268 | 0.985671 | 0.001307 | Q5JP53;P0 | Q5JP53;P0 | Q5JP53 | Tubulin bet |
| HIGD2A | 0.006234 | 0.985747 | 0.002236 | Q9BW72 | Q9BW72 | Q9BW72 | HIG1 doma |
| KPNA2 | 0.005504 | 0.987406 | -0.01481 | P52292;J3 | P52292;J3 | P52292 | Importin su |
| MRPL41 | 0.002472 | 0.994325 | 0.010483 | Q8IXM3 | Q8IXM3 | Q8IXM3 | 39S riboso |
| NUP188 | 0.002333 | 0.994643 | 0.0061 | Q5SRE5;H7 | Q5SRE5 | Q5SRE5 | Nucleopori |
| ACSS2 | 1.002936 | 0.099326 | -1.91857 | Q9NR19;F8 | Q9NR19 | Q9NR19 | Acetyl-coe |
| MRPL40 | 0.907698 | 0.123681 | 1.649262 | Q9NQ50 | Q9NQ50 | Q9NQ50 | 39S riboso |
| SNRPD1 | 3.063552 | 0.000864 | 0.465629 | P62314;J3 | P62314 | P62314 | Small nucle |
| MRPS24 | 1.344493 | 0.045238 | 0.377323 | Q96EL2;F8 | Q96EL2 | Q96EL2 | 28S riboso |
| RPL26L1 | 2.961534 | 0.001093 | 0.361341 | Q9UNX3;E5 | Q9UNX3;E5 | Q9UNX3 | 60S riboso |
| KCMF1 | 1.798592 | 0.0159 | 0.327814 | Q9P0J7 | Q9P0J7 | Q9P0J7 | E3 ubiquiti |
| FXR2 | 1.420514 | 0.037974 | 0.32416 | P51116 | P51116 | P51116 | Fragile X m |
| ABHD17B | 2.72132 | 0.0019 | 0.284837 | Q5VST6 | Q5VST6 | Q5VST6 | Alpha/beta |
| DPY19L1 | 1.988259 | 0.010274 | 0.284197 | Q2PZI1;A0 | Q2PZI1;A0 | Q2PZI1 | Probable C |
| IGKC | 1.317607 | 0.048127 | 0.262728 | A0A5H1ZR | A0A5H1ZR | A0A5H1ZR | Ig kappa ch |
| ATP6V0C | 1.444422 | 0.03594 | 0.253524 | P27449;H3 | P27449 | P27449 | V-type prot |
| ARL15 | 2.415379 | 0.003843 | 0.247997 | A0A087WU | A0A087WU | A0A087WU | ADP-ribosy |

| | | | | | | | |
|---|---|---|---|---|---|---|---|
| DHRSX | 1.765574 | 0.017156 | 0.235348 | Q8N5I4;C9 | Q8N5I4 | Q8N5I4 | Dehydroger |
| CCNY | 1.640918 | 0.02286 | 0.231318 | Q8ND76;R4 | Q8ND76 | Q8ND76 | Cyclin-Y |
| LYRM7 | 2.044738 | 0.009021 | 0.207782 | Q5U5X0;D6 | Q5U5X0 | Q5U5X0 | Complex III |
| SYNGR1 | 2.798054 | 0.001592 | 0.204073 | B5MCD7;O | B5MCD7;O | B5MCD7 | Synaptogyr |
| USMG5 | 2.981972 | 0.001042 | 0.197276 | Q96IX5 | Q96IX5 | Q96IX5 | Up-regulate |
| MBP | 1.297655 | 0.05039 | 0.19656 | J3KS94;J3C | J3KS94;J3C | J3KS94 | Myelin basi |
| RPL26 | 1.541607 | 0.028734 | 0.188655 | P61254;J3 | P61254;J3 | P61254 | 60S ribosor |
| TIA1 | 1.844541 | 0.014304 | 0.184594 | F8W8I6;P3 | F8W8I6;P3 | F8W8I6 | Nucleolysir |
| CD47 | 2.199244 | 0.006321 | -0.16893 | A0A2R8Y48 | A0A2R8Y48 | A0A2R8Y48 | Leukocyte s |
| RPLP1 | 2.039833 | 0.009124 | -0.19102 | P05386 | P05386 | P05386 | 60S acidic r |
| RTN3 | 1.603236 | 0.024932 | -0.24473 | O95197;F5 | O95197;F5 | O95197 | Reticulon-3 |
| AP1S1 | 1.644897 | 0.022652 | -0.26616 | A0A2R8YGI | A0A2R8YGI | A0A2R8YGI | AP-1 compl |
| RP13-360B | 1.827208 | 0.014886 | -0.34239 | Q9H617;Q5 | Q9H617;Q5 | Q9H617 | Transmemb |
| NDUFA4 | 1.823469 | 0.015015 | -0.37283 | O00483 | O00483 | O00483 | Cytochrom |
| FCER1G | 2.418529 | 0.003815 | -0.37671 | P30273;A6 | P30273 | P30273 | High affinit |
| RPS29 | 2.995143 | 0.001011 | -0.43739 | A0A2R8Y85 | A0A2R8Y85 | A0A2R8Y85 | 40S ribosor |
| YIPF6 | 1.255678 | 0.055504 | 0.151113 | Q96EC8;B1 | Q96EC8 | Q96EC8 | Protein YIP |
| AGA | 1.251617 | 0.056025 | 0.281941 | P20933;HC | P20933;HC | P20933 | N(4)-(beta-I |
| CD300A | 1.235036 | 0.058206 | -0.26035 | Q9UGN4;J3 | Q9UGN4 | Q9UGN4 | CMRF35-lik |
| DGAT1 | 1.225744 | 0.059464 | 0.423904 | O75907;AC | O75907;AC | O75907 | Diacylglyce |
| PLXNA1 | 1.213707 | 0.061135 | -1.71415 | Q9UIW2 | Q9UIW2 | Q9UIW2 | Plexin-A1 |
| MRPL27 | 1.210052 | 0.061652 | -1.48292 | H7C5U8;Q9 | H7C5U8;Q9 | H7C5U8 | 39S ribosor |
| GLRX5 | 1.208694 | 0.061845 | 0.174712 | Q86SX6 | Q86SX6 | Q86SX6 | Glutaredox |
| CD63 | 1.195879 | 0.063697 | -0.97273 | F8VV56;F8 | F8VV56;F8 | F8VV56 | Tetraspanir |
| TSPAN15 | 1.193745 | 0.064011 | -1.86591 | H7C285;O9 | H7C285;O9 | H7C285 | Tetraspanir |
| CAB39L | 1.185283 | 0.065271 | -0.30459 | Q5TAW7;Q | Q5TAW7;Q | Q5TAW7 | Calcium-bir |
| RAPGEF1 | 1.171614 | 0.067357 | 0.631179 | Q13905;HC | Q13905;HC | Q13905 | Rap guanin |
| ASCC2 | 1.170781 | 0.067487 | -1.61183 | Q9H1I8;B1 | Q9H1I8;B1 | Q9H1I8 | Activating s |
| RAP2A | 1.170642 | 0.067508 | -1.75547 | P10114;F6 | P10114;F6 | P10114 | Ras-related |
| TMEM9B | 1.170572 | 0.067519 | -1.619 | Q9NQ34 | Q9NQ34 | Q9NQ34 | Transmemb |
| SSU72 | 1.128872 | 0.074324 | 1.485429 | Q9NP77 | Q9NP77 | Q9NP77 | RNA polym |
| RBX1 | 1.105945 | 0.078353 | 1.791169 | P62877 | P62877 | P62877 | E3 ubiquiti |
| HLA-DQA1 | 1.098656 | 0.079679 | 0.249335 | Q5Y7D2;PC | Q5Y7D2;PC | Q5Y7D2 | HLA class II |
| DERL2 | 1.097273 | 0.079933 | -2.2136 | Q9GZP9;I3 | Q9GZP9;I3 | Q9GZP9 | Derlin-2 |
| OVCA2 | 1.09064 | 0.081163 | -1.64975 | Q8WZ82 | Q8WZ82 | Q8WZ82 | Ovarian car |
| PTPN18 | 1.059134 | 0.08727 | -0.41433 | Q99952;E7 | Q99952;E7 | Q99952 | Tyrosine-pr |
| RPS27A | 1.050094 | 0.089106 | 0.121578 | P62979;A0 | P62979;A0 | P62979 | Ubiquitin-4 |
| CIRH1A | 1.048456 | 0.089442 | -0.10829 | H3BSH7;Q9 | H3BSH7;Q9 | H3BSH7 | Cirhin |
| SNRPE | 1.044653 | 0.090229 | -0.17585 | P62304;A6 | P62304;A6 | P62304 | Small nucle |
| TOR1B | 1.037642 | 0.091698 | 0.27269 | O14657;HC | O14657;HC | O14657 | Torsin-1B |
| SMAD2 | 1.028939 | 0.093554 | 0.211391 | B7Z5N5;Q1 | B7Z5N5;Q1 | B7Z5N5 | Mothers ag |
| CLEC12A | 1.028679 | 0.09361 | 0.318554 | Q5QGZ9 | Q5QGZ9 | Q5QGZ9 | C-type lecti |
| SLC35E1 | 1.028578 | 0.093631 | 0.179024 | Q96K37;H7 | Q96K37;H7 | Q96K37 | Solute carri |
| LY96 | 1.021695 | 0.095127 | 1.087345 | Q9Y6Y9 | Q9Y6Y9 | Q9Y6Y9 | Lymphocyt |
| PVRL2 | 1.015005 | 0.096604 | -0.148 | K7EKE8;Q9 | K7EKE8;Q9 | K7EKE8 | Nectin-2 |
| MRPL32 | 1.011957 | 0.097284 | 1.866271 | Q9BYC8 | Q9BYC8 | Q9BYC8 | 39S ribosor |
| EMILIN2 | 1.003851 | 0.099117 | -1.18233 | Q9BXX0 | Q9BXX0 | Q9BXX0 | EMILIN-2 |
| ICT1 | 0.982289 | 0.104162 | 1.273495 | J3KS15;Q1 | J3KS15;Q1 | J3KS15 | Peptidyl-tR |

| | | | | | | | |
|---|---|---|---|---|---|---|---|
| COA4 | 0.967637 | 0.107736 | -0.96325 | Q9NYJ1 | Q9NYJ1 | Q9NYJ1 | Cytochrom |
| SLC29A1 | 0.958104 | 0.110128 | 1.488957 | Q99808 | Q99808 | Q99808 | Equilibrativ |
| TEFM | 0.9465 | 0.11311 | 0.171657 | Q96QE5;J3 | Q96QE5;J3 | Q96QE5 | Transcriptio |
| ABRACL | 0.938724 | 0.115153 | -0.31625 | Q9P1F3;Q5 | Q9P1F3 | Q9P1F3 | Costars fam |
| LILRB1 | 0.928164 | 0.117987 | 0.186323 | Q8NHL6;A0 | Q8NHL6;A0 | Q8NHL6 | Leukocyte i |
| DENR | 0.926253 | 0.118508 | -0.16698 | O43583;F8 | O43583;F8 | O43583 | Density-reg |
| LXN | 0.921998 | 0.119675 | -0.26885 | Q9BS40;H7 | Q9BS40 | Q9BS40 | Latexin |
| PDIA5 | 0.860917 | 0.137747 | 0.137306 | Q14554;H7 | Q14554 | Q14554 | Protein disu |
| TSC2 | 0.849006 | 0.141577 | 0.208514 | A0A2R8Y6( | A0A2R8Y6( | A0A2R8Y6( | Tuberin |
| CHMP1A | 0.841439 | 0.144066 | -0.7061 | F8VUA2;Q9 | F8VUA2;Q9 | F8VUA2 | Charged mu |
| TM7SF3 | 0.840226 | 0.144469 | -1.11922 | H0YFW7;Q9 | H0YFW7;Q9 | H0YFW7 | Transmemb |
| SNX18 | 0.822406 | 0.15052 | 0.244214 | Q96RF0 | Q96RF0 | Q96RF0 | Sorting nex |
| IPO9 | 0.814743 | 0.153199 | -0.18856 | Q96P70 | Q96P70 | Q96P70 | Importin-9 |
| UBA5 | 0.812545 | 0.153977 | -0.23725 | E7EWE1;E7 | E7EWE1;E7 | E7EWE1 | Ubiquitin-l |
| PSMB5 | 0.806752 | 0.156044 | 0.912053 | P28074;H0 | P28074;H0 | P28074 | Proteasome |
| USP33 | 0.802857 | 0.15745 | 0.210943 | Q8TEY7;H0 | Q8TEY7 | Q8TEY7 | Ubiquitin c |
| STAM2 | 0.802265 | 0.157665 | -1.09669 | O75886 | O75886 | O75886 | Signal trans |
| HDHD2 | 0.792893 | 0.161104 | -0.11345 | K7ER15;Q9 | K7ER15;Q9 | K7ER15 | Haloacid de |
| C14orf159 | 0.791446 | 0.161642 | 0.109179 | Q7Z3D6 | Q7Z3D6 | Q7Z3D6 | UPF0317 p |
| GCSH | 0.785156 | 0.164 | 0.189858 | A0A1W2P( | A0A1W2P( | A0A1W2P( | Glycine clea |
| | 0.767562 | 0.17078 | 0.132321 | B4DLN1;Q9 | B4DLN1 | B4DLN1 | |
| DGKA | 0.754246 | 0.176098 | 0.755246 | P23743;G3 | P23743;G3 | P23743 | Diacylglyce |
| PCNP | 0.749962 | 0.177844 | 0.185146 | Q8WW12 | Q8WW12 | Q8WW12 | PEST protec |
| AK1 | 0.745127 | 0.179834 | 1.2759 | P00568;Q5 | P00568;Q5 | P00568 | Adenylate k |
| PPIH | 0.743688 | 0.180432 | -0.1021 | C9JQD4;O4 | C9JQD4;O4 | C9JQD4 | Peptidyl-pr |
| GTF2F1 | 0.740923 | 0.181584 | 0.232817 | M0R0Z3;P3 | M0R0Z3;P3 | M0R0Z3 | General tra |
| GSTM1 | 0.733683 | 0.184636 | -0.16178 | P09488;E7 | P09488;E7 | P09488 | Glutathion |
| BMS1 | 0.731711 | 0.185477 | -1.25922 | Q14692 | Q14692 | Q14692 | Ribosome b |
| COPS7A | 0.72819 | 0.186986 | 0.652939 | F5H7C6;F5 | F5H7C6;F5 | F5H7C6 | COP9 signa |
| YIPF4 | 0.727057 | 0.187475 | 0.881776 | Q9BSR8;H7 | Q9BSR8;H7 | Q9BSR8 | Protein YIP |
| AP5S1 | 0.722433 | 0.189481 | -0.82505 | Q9NUS5;Q5 | Q9NUS5;Q5 | Q9NUS5 | AP-5 compl |
| PIGU | 0.71921 | 0.190893 | 1.289275 | Q9H490 | Q9H490 | Q9H490 | Phosphatic |
| CNNM4 | 0.717577 | 0.191612 | 0.141607 | Q6P4Q7;Q9 | Q6P4Q7 | Q6P4Q7 | Metal trans |
| FAU | 0.683707 | 0.207154 | -1.77981 | P62861;E9 | P62861;E9 | P62861 | 40S ribosor |
| TBL3 | 0.682846 | 0.207565 | -1.06962 | Q12788;J3 | Q12788;J3 | Q12788 | Transducin |
| MRPL30 | 0.68127 | 0.20832 | 0.188335 | B8ZZV5;Q8 | B8ZZV5;Q8 | B8ZZV5 | 39S ribosor |
| FAM210B | 0.67404 | 0.211816 | -0.69871 | Q96KR6 | Q96KR6 | Q96KR6 | Protein FAM |
| FXN | 0.659816 | 0.218869 | 0.105427 | C9JAX1;Q1 | C9JAX1;Q1 | C9JAX1 | Frataxin, m |
| CARD9 | 0.649393 | 0.224185 | -0.90551 | Q9H257 | Q9H257 | Q9H257 | Caspase rec |
| HMGN3 | 0.648486 | 0.224654 | 1.386321 | A0A087WZ | A0A087WZ | A0A087WZ | High mobil |
| EPT1 | 0.648253 | 0.224775 | 0.100679 | Q9C0D9;A0 | Q9C0D9;A0 | Q9C0D9 | Ethanolami |
| UBTD1 | 0.63242 | 0.23312 | -0.67642 | Q9HAC8 | Q9HAC8 | Q9HAC8 | Ubiquitin c |
| RNMTL1 | 0.607398 | 0.246946 | 1.048086 | Q9HC36;I3 | Q9HC36;I3 | Q9HC36 | rRNA methy |
| EXOSC2 | 0.605781 | 0.247867 | 0.693173 | Q13868;A3 | Q13868;A3 | Q13868 | Exosome co |
| WDR33 | 0.602339 | 0.249839 | -0.53281 | Q9C0J8;C9 | Q9C0J8;C9 | Q9C0J8 | pre-mRNA 3 |
| LAPTM5 | 0.594963 | 0.254119 | 0.787131 | Q13571 | Q13571 | Q13571 | Lysosomal- |
| PRKACB | 0.594099 | 0.254625 | 0.091303 | A0A0A0MS | A0A0A0MS | A0A0A0MS | cAMP-depe |
| KIAA0100 | 0.593543 | 0.254951 | 0.841751 | K7EQ86;Q0 | K7EQ86;Q0 | K7EQ86 | Protein KIA |

| | | | | | | | |
|---|---|---|---|---|---|---|---|
| WRB | 0.593258 | 0.255119 | 0.909224 | H7C384;A0 | H7C384;A0 | H7C384 | Tail-anchor |
| RBM27 | 0.5925 | 0.255564 | -1.20287 | U3KPZ7;Q9 | U3KPZ7;Q9 | U3KPZ7 | RNA-bindir |
| CLN8 | 0.587204 | 0.2587 | -1.61567 | Q9UBY8;A0 | Q9UBY8;A0 | Q9UBY8 | Protein CLN |
| SIRPB1 | 0.56541 | 0.272013 | -0.71872 | Q5TFQ8;H3 | Q5TFQ8 | Q5TFQ8 | Signal-regu |
| GLRX2 | 0.558402 | 0.276438 | -0.12867 | Q9NS18 | Q9NS18 | Q9NS18 | Glutaredox |
| MFAP1 | 0.552366 | 0.280307 | -0.10063 | P55081 | P55081 | P55081 | Microfibril |
| IMMT | 0.549529 | 0.282144 | 0.030637 | Q16891;B9 | Q16891;B9 | Q16891 | MICOS com |
| CST3 | 0.546568 | 0.284074 | 0.158511 | P01034 | P01034 | P01034 | Cystatin-C |
| SLC2A6 | 0.542351 | 0.286846 | 0.087303 | Q9UGQ3;F2 | Q9UGQ3 | Q9UGQ3 | Solute carri |
| HPS6 | 0.539408 | 0.288796 | 0.906282 | Q86YV9 | Q86YV9 | Q86YV9 | Hermansky |
| TCEAL5 | 0.539275 | 0.288885 | 0.1543 | Q5H9L2;Q9 | Q5H9L2;Q9 | Q5H9L2 | Transcripti |
| C18orf32 | 0.536135 | 0.290981 | 0.218462 | Q8TCD1 | Q8TCD1 | Q8TCD1 | UPF0729 p |
| HSPA13 | 0.535598 | 0.291341 | 0.118381 | P48723 | P48723 | P48723 | Heat shock |
| GBP4 | 0.534475 | 0.292096 | -0.13465 | Q96PP9;A0 | Q96PP9 | Q96PP9 | Guanylate-l |
| EXOSC7 | 0.531751 | 0.293934 | -0.65985 | Q15024 | Q15024 | Q15024 | Exosome co |
| CUX1 | 0.527077 | 0.297114 | 0.097227 | Q13948 | Q13948 | Q13948 | Protein CAS |
| EBP | 0.522999 | 0.299917 | 0.091899 | Q15125;C9 | Q15125;C9 | Q15125 | 3-beta-hydr |
| WDR44 | 0.518712 | 0.302892 | 0.393973 | H7BY83;Q5 | H7BY83;Q5 | H7BY83 | WD repeat- |
| PIP4K2B | 0.518384 | 0.303121 | 0.609503 | P78356 | P78356 | P78356 | Phosphatic |
| EIF4G3 | 0.515105 | 0.305418 | 0.399792 | A0A0A0MS | A0A0A0MS | A0A0A0MS | Eukaryotic |
| CHMP2B | 0.514523 | 0.305828 | 0.634639 | Q9UQN3;A( | Q9UQN3;A( | Q9UQN3 | Charged mu |
| UQCR10 | 0.507212 | 0.31102 | 0.98922 | Q9UDW1 | Q9UDW1 | Q9UDW1 | Cytochrom |
| CBFB | 0.496493 | 0.318792 | -0.58107 | Q13951;A0 | Q13951;A0 | Q13951 | Core-bindir |
| TFB1M | 0.495718 | 0.319361 | 0.113679 | Q8WVM0 | Q8WVM0 | Q8WVM0 | Dimethylac |
| CASC4 | 0.481104 | 0.33029 | -0.46891 | Q6P4E1;H0 | Q6P4E1;H0 | Q6P4E1 | Protein CAS |
| BCAR1 | 0.477109 | 0.333343 | 0.480876 | A0A2R8Y5E | A0A2R8Y5E | A0A2R8Y5E | Breast canc |
| PCID2 | 0.476765 | 0.333607 | -0.0668 | Q5JVF3 | Q5JVF3 | Q5JVF3 | PCI domair |
| SMNDC1 | 0.474294 | 0.33551 | 0.308898 | O75940 | O75940 | O75940 | Survival of r |
| PTDSS2 | 0.4705 | 0.338454 | 0.445011 | Q9BVG9;A0 | Q9BVG9;A0 | Q9BVG9 | Phosphatic |
| MAP2K3 | 0.469507 | 0.339229 | -0.1802 | P46734;P5 | P46734;P5 | P46734 | Dual specifi |
| YTHDF2 | 0.461105 | 0.345856 | 0.110581 | Q9Y5A9 | Q9Y5A9 | Q9Y5A9 | YTH domair |
| HLA-B | 0.454233 | 0.351372 | 0.083633 | Q2L6G2;D5 | Q2L6G2;D5 | Q2L6G2 | |
| CDA | 0.449841 | 0.354943 | 0.569981 | P32320 | P32320 | P32320 | Cytidine de |
| HEBP2 | 0.43707 | 0.365536 | 0.060613 | Q9Y5Z4;Q5 | Q9Y5Z4;Q5 | Q9Y5Z4 | Heme-bind |
| RAP1A | 0.420321 | 0.379908 | -0.06739 | P62834;A0 | P62834;A0 | P62834 | Ras-related |
| TOMM20 | 0.419494 | 0.380633 | -0.78851 | Q15388 | Q15388 | Q15388 | Mitochond |
| SLMAP | 0.415041 | 0.384556 | -0.56259 | A0A5F9VB9 | A0A5F9VB9 | A0A5F9VB9 | Sarcolemm |
| CHMP1B | 0.413564 | 0.385865 | 0.96204 | Q7LBR1 | Q7LBR1 | Q7LBR1 | Charged mu |
| RTN4 | 0.397803 | 0.400126 | 0.078264 | F8W914;H7 | F8W914 | F8W914 | Reticulon |
| NHLRC3 | 0.393534 | 0.404079 | 0.11504 | Q5JS37;C9 | Q5JS37;C9 | Q5JS37 | NHL repeat |
| GOLGA7 | 0.383168 | 0.41384 | -0.12436 | Q7Z5G4;M( | Q7Z5G4 | Q7Z5G4 | Golgin subf |
| VAMP5 | 0.376387 | 0.420352 | -0.52172 | O95183 | O95183 | O95183 | Vesicle-asso |
| TMEM256 | 0.375682 | 0.421035 | -0.61939 | K7ERE1;Q8 | K7ERE1;Q8 | K7ERE1 | Transmemb |
| TMBIM6 | 0.37164 | 0.424972 | -0.06932 | F8VQW0;F8 | F8VQW0;F8 | F8VQW0 | Bax inhibit |
| TREM2 | 0.370316 | 0.426269 | -0.11448 | Q9NZC2 | Q9NZC2 | Q9NZC2 | Triggering r |
| VMP1 | 0.366238 | 0.430291 | -0.1031 | Q96GC9;K7 | Q96GC9;K7 | Q96GC9 | Vacuole me |
| PQLC1 | 0.364411 | 0.432104 | -0.73546 | Q8N2U9;K7 | Q8N2U9 | Q8N2U9 | PQ-loop rep |
| ATXN2 | 0.354788 | 0.441786 | -0.2768 | A0A5F9ZH8 | A0A5F9ZH8 | A0A5F9ZH8 | Ataxin-2 |

| | | | | | | | |
|---|---|---|---|---|---|---|---|
| RAB3D | 0.351573 | 0.445068 | 0.396585 | O95716;M( | O95716 | O95716 | Ras-related |
| CD5 | 0.350926 | 0.445732 | -0.4334 | P06127 | P06127 | P06127 | T-cell surfac |
| SNX12 | 0.336899 | 0.460364 | -0.39129 | A0A087X0I | A0A087X0I | A0A087X0I | Sorting nex |
| POLR1C | 0.334581 | 0.462827 | -0.68283 | A0A2R8YEY | A0A2R8YEY | A0A2R8YEY | DNA-direct |
| CPSF4 | 0.318158 | 0.480665 | 0.844359 | C9JEV9;O9! | C9JEV9;O9! | C9JEV9 | Cleavage ar |
| OSBPL3 | 0.318084 | 0.480746 | -0.70193 | Q9H4L5 | Q9H4L5 | Q9H4L5 | Oxysterol-b |
| MT-ND1 | 0.308986 | 0.490924 | -0.17677 | P03886 | P03886 | P03886 | NADH-ubiq |
| HMGN4 | 0.307313 | 0.492819 | 0.076982 | O00479 | O00479 | O00479 | High mobil |
| CIAPIN1 | 0.306746 | 0.493462 | -0.53038 | H3BQ23;H3 | H3BQ23;H3 | H3BQ23 | Anamorsin |
| UROD | 0.293001 | 0.50933 | -0.37174 | A0A494BZY | A0A494BZY | A0A494BZY | Uroporphy |
| MRPL50 | 0.292587 | 0.509816 | 0.133388 | Q8N5N7 | Q8N5N7 | Q8N5N7 | 39S riboson |
| KCNAB2 | 0.292129 | 0.510353 | -0.05712 | Q13303;A0 | Q13303;A0 | Q13303 | Voltage-gat |
| CD68 | 0.291618 | 0.510954 | 0.184577 | P34810;J3I | P34810 | P34810 | Macrosialin |
| HLA-DPB1 | 0.291117 | 0.511544 | -0.10359 | P04440;H0 | P04440;H0 | P04440 | HLA class II |
| ATOX1 | 0.288377 | 0.514782 | -0.08096 | E5RGN3;E5 | E5RGN3;E5 | E5RGN3 | Copper tran |
| C6orf120 | 0.286096 | 0.517492 | 0.541795 | Q7Z4R8 | Q7Z4R8 | Q7Z4R8 | UPF0669 p |
| CDK9 | 0.285514 | 0.518186 | 0.288937 | P50750;X6 | P50750 | P50750 | Cyclin-depe |
| GNB2 | 0.28278 | 0.521459 | -0.09571 | P62879;C9 | P62879;C9 | P62879 | Guanine nu |
| NDUFAF1 | 0.282068 | 0.522314 | -0.40546 | H0YL22;Q9 | H0YL22;Q9 | H0YL22 | Complex I i |
| IGBP1 | 0.280543 | 0.524151 | 0.352848 | P78318 | P78318 | P78318 | Immunoglo |
| CD6 | 0.273216 | 0.533069 | 0.363119 | G5E973;P3 | G5E973;P3 | G5E973 | T-cell differ |
| RNASE6 | 0.268909 | 0.538383 | 0.751525 | Q93091 | Q93091 | Q93091 | Ribonuclea |
| ARFIP2 | 0.258409 | 0.551558 | -0.58811 | P53365;A0 | P53365;A0 | P53365 | Arfaptin-2 |
| AKT1 | 0.253055 | 0.5584 | 0.487122 | P31749;G3 | P31749 | P31749 | RAC-alpha s |
| ATP1B4 | 0.248253 | 0.564608 | -0.43928 | B7ZKW0;Q | B7ZKW0;Q | B7ZKW0 | Protein ATF |
| RGS19 | 0.23755 | 0.578696 | 0.048267 | P49795 | P49795 | P49795 | Regulator o |
| COX7A2 | 0.23526 | 0.581754 | -0.03861 | H0UI06;P1 | H0UI06;P1 | H0UI06 | Cytochrom |
| MIEF1 | 0.232027 | 0.586101 | -0.50369 | B0QY95;Q9 | B0QY95;Q9 | B0QY95 | Mitochond |
| MFSD1 | 0.230193 | 0.588582 | 0.049486 | A0A494C1( | A0A494C1( | A0A494C1( | Major facili |
| PTPRCAP | 0.228298 | 0.591156 | -0.43544 | Q14761 | Q14761 | Q14761 | Protein tyr |
| RPS15A | 0.226138 | 0.594103 | -0.03706 | P62244;I3I | P62244;I3I | P62244 | 40S riboson |
| CWC15 | 0.225687 | 0.59472 | -0.29574 | Q9P013 | Q9P013 | Q9P013 | Spliceosom |
| DDX58 | 0.221649 | 0.600276 | -0.79693 | A2A376;O9 | A2A376;O9 | A2A376 | Probable A |
| NUB1 | 0.221018 | 0.601149 | -0.51326 | H3BM14;H | H3BM14;H | H3BM14 | NEDD8 ulti |
| MRPL21 | 0.217831 | 0.605577 | -0.29763 | Q7Z2W9;F! | Q7Z2W9 | Q7Z2W9 | 39S riboson |
| ACOT1 | 0.217103 | 0.606592 | 0.036178 | Q86TX2;G3 | Q86TX2;G3 | Q86TX2 | Acyl-coenzy |
| SLC2A1 | 0.216714 | 0.607136 | -0.04171 | P11166 | P11166 | P11166 | Solute carri |
| CTCF | 0.215854 | 0.60834 | -0.07025 | A0A2R8Y59 | A0A2R8Y59 | A0A2R8Y59 | Transcriptio |
| RBM7 | 0.210254 | 0.616234 | -0.34207 | J3KPD3;Q9 | J3KPD3;Q9 | J3KPD3 | RNA-bindin |
| FCGRT | 0.205951 | 0.62237 | 0.481673 | P55899;M( | P55899;M( | P55899 | IgG recepto |
| PFDN1 | 0.200547 | 0.630163 | -0.44287 | E5RGS4;O6 | E5RGS4;O6 | E5RGS4 | Prefoldin su |
| CSF1 | 0.193596 | 0.64033 | 0.060813 | E9PJA2;P0! | E9PJA2;P0! | E9PJA2 | Macrophag |
| IMPDH1 | 0.190192 | 0.645369 | 0.301616 | A0A3B3IRL | A0A3B3IRL | A0A3B3IRL | Inosine-5-m |
| RGS10 | 0.189499 | 0.6464 | -0.04244 | O43665 | O43665 | O43665 | Regulator o |
| PPIC | 0.185996 | 0.651635 | 0.037232 | P45877 | P45877 | P45877 | Peptidyl-pr |
| PFDN2 | 0.185466 | 0.652431 | -0.03283 | Q9UHV9 | Q9UHV9 | Q9UHV9 | Prefoldin su |
| CD2 | 0.180287 | 0.660256 | -0.20874 | P06729;Q5 | P06729 | P06729 | T-cell surfac |
| MYLK | 0.179472 | 0.661498 | 0.051588 | Q15746 | Q15746 | Q15746 | Myosin ligh |

| | | | | | | | |
|---|---|---|---|---|---|---|---|
| SLC2A3 | 0.170328 | 0.675573 | 0.0341 | P11169;Q8 | P11169;Q8 | P11169 | Solute carri |
| YTHDF3 | 0.167548 | 0.679911 | 0.032724 | A0A024R7\ | A0A024R7\ | A0A024R7\ | YTH domair |
| CD55 | 0.166072 | 0.682225 | -0.33319 | H3BLV0;B1 | H3BLV0;B1 | H3BLV0 | Complemer |
| PPIP5K2 | 0.157562 | 0.695726 | -0.42467 | O43314;AC | O43314;AC | O43314 | Inositol hex |
| RUFY3 | 0.155772 | 0.698599 | 0.222088 | Q7L099;D6 | Q7L099 | Q7L099 | Protein RU |
| RPL38 | 0.150476 | 0.70717 | -0.03924 | P63173;J3 | P63173;J3 | P63173 | 60S ribosor |
| TNPO3 | 0.146288 | 0.714023 | -0.15101 | E9PFH4;C9 | E9PFH4;C9 | E9PFH4 | Transportir |
| CD81 | 0.144529 | 0.71692 | -0.05028 | E9PJK1;E9F | E9PJK1;E9F | E9PJK1 | Tetraspanir |
| MP68 | 0.141555 | 0.721847 | 0.021955 | P56378;G3 | P56378;G3 | P56378 | 6.8 kDa mit |
| UFM1 | 0.141388 | 0.722124 | 0.065824 | P61960;HC | P61960;HC | P61960 | Ubiquitin-f |
| SURF4 | 0.135257 | 0.732391 | -0.04262 | Q5T8U5;O1 | Q5T8U5;O1 | Q5T8U5 | Surfeit locu |
| SLC15A3 | 0.134256 | 0.734081 | 0.026928 | Q8IY34;H0 | Q8IY34 | Q8IY34 | Solute carri |
| NRBF2 | 0.128556 | 0.74378 | -0.02343 | Q96F24 | Q96F24 | Q96F24 | Nuclear rec |
| UBE2D2 | 0.119051 | 0.760237 | -0.03256 | D6RFM0;D | D6RFM0;D | D6RFM0 | Ubiquitin-c |
| MRPS14 | 0.115169 | 0.767062 | 0.035 | O60783 | O60783 | O60783 | 28S ribosor |
| GNB1 | 0.109663 | 0.77685 | 0.022067 | P62873;F6 | P62873;F6 | P62873 | Guanine nu |
| KDELR1 | 0.109114 | 0.777832 | -0.04481 | P24390;M( | P24390;M( | P24390 | ER lumen p |
| MAN1A2 | 0.097714 | 0.79852 | -0.1768 | O60476;H( | O60476 | O60476 | Mannosyl-c |
| PODXL | 0.095415 | 0.802759 | -0.03491 | O00592 | O00592 | O00592 | Podocalyxi |
| PPP1CA | 0.079213 | 0.833273 | 0.016798 | P62136;E9 | P62136;E9 | P62136 | Serine/thre |
| SDR39U1 | 0.073984 | 0.843367 | 0.137085 | Q9NRG7 | Q9NRG7 | Q9NRG7 | Epimerase f |
| RPS15A | 0.072324 | 0.846595 | -0.01205 | H3BN98;I3 | H3BN98;I3 | H3BN98 | |
| FGD5 | 0.072315 | 0.846613 | -0.0857 | Q6ZNL6;B7 | Q6ZNL6;B7 | Q6ZNL6 | FYVE, RhoG |
| STARD3 | 0.070609 | 0.849946 | -0.17922 | Q14849;J3 | Q14849;J3 | Q14849 | StAR-relate |
| LRRCC1 | 0.070494 | 0.85017 | 0.241522 | Q9C099 | Q9C099 | Q9C099 | Leucine-ric |
| SMARCB1 | 0.070171 | 0.850803 | 0.014802 | G5E975;Q1 | G5E975;Q1 | G5E975 | SWI/SNF-re |
| HFE | 0.068739 | 0.853613 | 0.111869 | F8W7W8;C | F8W7W8;C | F8W7W8 | Hereditary |
| HPS3 | 0.062723 | 0.86552 | 0.124978 | Q969F9;G5 | Q969F9;G5 | Q969F9 | Hermansky |
| PRPS1 | 0.059896 | 0.871172 | 0.017186 | B1ALA9;P6 | B1ALA9;P6 | B1ALA9 | Ribose-pho |
| BZW2 | 0.05978 | 0.871404 | -0.15485 | E7ETZ4;Q9 | E7ETZ4;Q9 | E7ETZ4 | Basic leucir |
| BAD | 0.059687 | 0.871592 | 0.107326 | Q92934;F5 | Q92934;F5 | Q92934 | Bcl2-associ |
| GOLT1B | 0.053958 | 0.883165 | 0.015471 | F5H6U7;Q9 | F5H6U7;Q9 | F5H6U7 | Vesicle trar |
| SEC22A | 0.049141 | 0.893016 | -0.01259 | C9JDJ8;C9J | C9JDJ8;C9J | C9JDJ8 | Vesicle-traf |
| TXNL4A | 0.048977 | 0.893353 | -0.00945 | K7EPA6;K7 | K7EPA6;K7 | K7EPA6 | Thioredoxi |
| IL1RN | 0.04392 | 0.903816 | -0.01961 | P18510 | P18510 | P18510 | Interleukin |
| ACP6 | 0.043904 | 0.90385 | 0.139383 | Q9NPH0;A( | Q9NPH0;A( | Q9NPH0 | Lysophospl |
| DEFA3 | 0.040871 | 0.910183 | 0.030398 | P59666;P5 | P59666;P5 | P59666 | Neutrophil |
| MORC3 | 0.036835 | 0.918681 | 0.068493 | Q14149 | Q14149 | Q14149 | MORC fami |
| CHCHD7 | 0.035704 | 0.921076 | -0.01686 | J3KPV1;Q9 | J3KPV1;Q9 | J3KPV1 | Coiled-coil- |
| MRPL17 | 0.032748 | 0.927369 | 0.009078 | E9PKV2;Q9 | E9PKV2;Q9 | E9PKV2 | 39S ribosor |
| SLC25A3 | 0.03028 | 0.932654 | -0.00597 | Q00325 | Q00325 | Q00325 | Phosphate |
| ENSA | 0.028547 | 0.936383 | 0.008259 | A6NMQ3;Q | A6NMQ3;Q | A6NMQ3 | Alpha-endc |
| SRD5A3 | 0.026329 | 0.941177 | -0.00954 | Q9H8P0;H( | Q9H8P0 | Q9H8P0 | Polyprenol |
| GAL | 0.026149 | 0.941567 | 0.02044 | P22466 | P22466 | P22466 | Galanin pe |
| TMEM109 | 0.024468 | 0.945219 | -0.00677 | Q9BVC6 | Q9BVC6 | Q9BVC6 | Transmemb |
| CD151 | 0.023558 | 0.9472 | 0.004996 | E9PRJ3;K4I | E9PRJ3;K4I | E9PRJ3 | Tetraspanir |
| HLA-DMA | 0.01715 | 0.96128 | 0.005678 | A0A140T98 | A0A140T98 | A0A140T98 | HLA class II |
| TIMM17A | 0.016367 | 0.963014 | 0.052181 | Q99595 | Q99595 | Q99595 | Mitochond |

| | | | | | | | |
|---|---|---|---|---|---|---|---|
| ARL2 | 0.015665 | 0.964573 | 0.052228 | P36404;V9 | P36404 | P36404 | ADP-ribosy |
| CWF19L1 | 0.008131 | 0.981451 | 0.028587 | Q69YN2;A0 | Q69YN2;A0 | Q69YN2 | CWF19-like |
| REEP4 | 0.007571 | 0.982717 | 0.001643 | E5RGS2;Q9 | E5RGS2;Q9 | E5RGS2 | Receptor ex |
| NOC2L | 0.004388 | 0.989947 | 0.001962 | Q9Y3T9 | Q9Y3T9 | Q9Y3T9 | Nucleolar c |
| EDEM3 | 0.004353 | 0.990027 | -0.01491 | Q9BZQ6 | Q9BZQ6 | Q9BZQ6 | ER degrada |
| RGCC | 0.003798 | 0.991293 | -0.00097 | Q9H4X1 | Q9H4X1 | Q9H4X1 | Regulator c |
| TMEM168 | 1.10305 | 0.078877 | -1.61249 | H7C268;Q9 | H7C268;Q9 | H7C268 | Transmemb |
| PDPK1 | 1.01346 | 0.096948 | -1.38804 | E9PER6;Q6 | E9PER6;Q6 | E9PER6 | Putative 3- |
| PPP1CC | 0.940767 | 0.114613 | -1.14615 | F8VYE8;P3 | F8VYE8;P3 | F8VYE8 | Serine/thre |
| PSMB6 | 0.900949 | 0.125618 | -1.48075 | P28072;A0 | P28072;A0 | P28072 | Proteasom |
| AMFR | 1.29083 | 0.051188 | 1.00861 | Q9UKV5;H3 | Q9UKV5;H3 | Q9UKV5 | E3 ubiquiti |
| DCSTAMP | 0.587301 | 0.258642 | 0.800973 | Q9H295 | Q9H295 | Q9H295 | Dendritic c |
| NOLC1 | 0.367034 | 0.429503 | -0.56542 | A0A0A0MR | A0A0A0MR | A0A0A0MR | Nucleolar a |
| SHC1 | 0.172979 | 0.671462 | 0.358453 | H0Y539;Q5 | H0Y539;Q5 | H0Y539 | SHC-transfo |
| CST7 | 0.035543 | 0.921419 | 0.090169 | O76096 | O76096 | O76096 | Cystatin-F |
| CRIP1 | 1.466381 | 0.034168 | 0.450609 | P50238;K4 | P50238;K4 | P50238 | Cysteine-ric |
| CDC42SE2 | 2.428184 | 0.003731 | 0.369044 | Q9NRR3;D6 | Q9NRR3;D6 | Q9NRR3 | CDC42 sma |
| PTP4A2 | 2.81402 | 0.001535 | 0.307093 | E9PMY3;E9 | E9PMY3;E9 | E9PMY3 | Protein tyr |
| SLC35F6 | 3.593165 | 0.000255 | 0.284017 | Q8N357;F8 | Q8N357;F8 | Q8N357 | Solute carri |
| MT-ATP6 | 1.380457 | 0.041643 | 0.198434 | P00846 | P00846 | P00846 | ATP syntha |
| TOR1AIP1 | 1.480538 | 0.033072 | 0.149624 | A0A0A0MS | A0A0A0MS | A0A0A0MS | Torsin-1A-i |
| RAB11B | 1.446598 | 0.03576 | 0.069108 | Q15907;M | Q15907 | Q15907 | Ras-related |
| CLN3 | 1.426923 | 0.037418 | -0.15362 | Q13286;O9 | Q13286;O9 | Q13286 | Battenin |
| EIF1 | 1.321076 | 0.047745 | -0.15958 | P41567;K7 | P41567;K7 | P41567 | Eukaryotic |
| CD58 | 1.525692 | 0.029806 | -0.21979 | P19256;B1 | P19256;B1 | P19256 | Lymphocyt |
| TMEM189- | 1.72362 | 0.018896 | -0.26359 | I3L0A0;Q1 | I3L0A0;Q1 | I3L0A0 | Ubiquitin-c |
| SEC61G | 1.851858 | 0.014065 | -0.29333 | P60059 | P60059 | P60059 | Protein tra |
| BST2 | 1.36336 | 0.043315 | -0.31177 | Q10589 | Q10589 | Q10589 | Bone marro |
| ATP5J2-PT | 1.788524 | 0.016273 | -0.39239 | G3V325;C9 | G3V325;C9 | G3V325 | |
| HAT1 | 1.294847 | 0.050717 | 1.478616 | O14929 | O14929 | O14929 | Histone ace |
| AFAP1L1 | 1.29381 | 0.050838 | -1.21201 | Q8TED9 | Q8TED9 | Q8TED9 | Actin filam |
| COX6A1 | 1.284815 | 0.051902 | 0.131895 | P12074 | P12074 | P12074 | Cytochrom |
| RCOR1 | 1.269004 | 0.053827 | -1.72455 | Q9UKL0 | Q9UKL0 | Q9UKL0 | REST corep |
| YPEL5 | 1.250699 | 0.056144 | -1.22802 | P62699 | P62699 | P62699 | Protein yip |
| NDUFA3 | 1.247358 | 0.056577 | 0.16232 | O95167;H7 | O95167;H7 | O95167 | NADH dehy |
| PFDN4 | 1.22608 | 0.059418 | -1.70342 | E9PQY2;Q9 | E9PQY2;Q9 | E9PQY2 | Prefoldin s |
| MAP2K2 | 1.219822 | 0.060281 | -0.27362 | P36507;G5 | P36507;G5 | P36507 | Dual specif |
| ARFGAP3 | 1.184911 | 0.065326 | 0.869603 | Q9NP61 | Q9NP61 | Q9NP61 | ADP-ribosy |
| CEBPA | 1.177212 | 0.066495 | 1.266753 | P49715 | P49715 | P49715 | CCAAT/enh |
| DAP | 1.152366 | 0.07041 | 1.184457 | P51397 | P51397 | P51397 | Death-asso |
| CHI3L1 | 1.11961 | 0.075926 | -1.67684 | H0Y3U8 | H0Y3U8 | H0Y3U8 | |
| CPEB2 | 1.10953 | 0.077709 | 0.785602 | H0Y9D9;A0 | H0Y9D9;A0 | H0Y9D9 | Cytoplasmi |
| SLC37A2 | 1.092839 | 0.080753 | 1.931205 | Q8TED4 | Q8TED4 | Q8TED4 | Sugar phos |
| PDLIM1 | 1.079715 | 0.083231 | -1.88032 | O00151 | O00151 | O00151 | PDZ and LI |
| HIGD1A | 1.077824 | 0.083594 | -2.18482 | C9JAW5;C9 | C9JAW5;C9 | C9JAW5 | HIG1 doma |
| SPATA13 | 1.064893 | 0.086121 | -1.26317 | E9PFR9;J3 | E9PFR9;J3 | E9PFR9 | Spermatog |
| BRI3 | 1.057547 | 0.08759 | 1.400683 | O95415 | O95415 | O95415 | Brain prote |
| ACYP2 | 1.045985 | 0.089953 | -0.83915 | U3KPX1;E9 | U3KPX1;E9 | U3KPX1 | Acylphosph |

| | | | | | | | |
|---|---|---|---|---|---|---|---|
| NDUFB3 | 1.013445 | 0.096952 | -0.24444 | O43676;C9 | O43676;C9 | O43676 | NADH dehy |
| SYPL1 | 1.001773 | 0.099593 | -2.09115 | C9JYN0;H7 | C9JYN0;H7 | C9JYN0 | Synaptophy |
| PEX13 | 0.985742 | 0.103338 | -1.0266 | Q92968 | Q92968 | Q92968 | Peroxisoma |
| SSR3 | 0.959671 | 0.109731 | 0.075176 | Q9UNL2;C9 | Q9UNL2;C9 | Q9UNL2 | Translocon |
| TMEM176E | 0.934606 | 0.11625 | -1.62606 | Q3YBM2 | Q3YBM2 | Q3YBM2 | Transmemb |
| PPAN | 0.933648 | 0.116507 | -0.27899 | H7C446;A8 | H7C446;A8 | H7C446 | Suppressor |
| HLA-Cw | 0.905264 | 0.124376 | -2.01512 | Q5RIP0;A0 | Q5RIP0;A0 | Q5RIP0 | |
| GZMM | 0.8955 | 0.127204 | -0.75573 | U3KQV5;P5 | U3KQV5;P5 | U3KQV5 | Granzyme M |
| PARL | 0.857904 | 0.138706 | -0.43959 | F8WCQ4;A0 | F8WCQ4;A0 | F8WCQ4 | Presenilins- |
| QARS | 0.834088 | 0.146525 | -0.10147 | P47897;A0 | P47897 | P47897 | Glutamine- |
| MBD2 | 0.832268 | 0.14714 | 0.866366 | Q9UBB5 | Q9UBB5 | Q9UBB5 | Methyl-CpG |
| CYSTM1 | 0.806251 | 0.156225 | 0.145962 | Q9H1C7 | Q9H1C7 | Q9H1C7 | Cysteine-ric |
| ARF1 | 0.799551 | 0.158653 | -0.09939 | P84077 | P84077 | P84077 | ADP-ribosy |
| ORMDL2 | 0.782156 | 0.165137 | -1.77005 | Q53FV1;F8 | Q53FV1;F8 | Q53FV1 | ORM1-like p |
| MTFMT | 0.77232 | 0.168919 | -0.82777 | Q96DP5;H3 | Q96DP5;H3 | Q96DP5 | Methionyl- |
| PQLC3 | 0.768493 | 0.170414 | -0.74623 | G5E9P4;C9 | G5E9P4;C9 | G5E9P4 | PQ-loop rep |
| GSN | 0.758915 | 0.174215 | -0.16863 | Q5T0I0 | Q5T0I0 | Q5T0I0 | |
| ATP5EP2 | 0.751712 | 0.177128 | -0.18343 | Q5VTU8;P5 | Q5VTU8;P5 | Q5VTU8 | ATP syntha |
| SLC43A3 | 0.751699 | 0.177134 | -0.76809 | Q8NBI5;E9 | Q8NBI5;E9 | Q8NBI5 | Solute carri |
| SGTB | 0.748874 | 0.178289 | -0.69168 | D6RFW1;Q | D6RFW1;Q | D6RFW1 | Small gluta |
| LUC7L2 | 0.722927 | 0.189266 | -0.05261 | A0A0A6YYJ | A0A0A6YYJ | A0A0A6YYJ8 | |
| NDUFAB1 | 0.718116 | 0.191375 | -0.25066 | O14561;H3 | O14561;H3 | O14561 | Acyl carrier |
| AGPAT1 | 0.716318 | 0.192168 | 1.107317 | Q99943 | Q99943 | Q99943 | 1-acyl-sn-gl |
| PTMS | 0.71366 | 0.193348 | -0.81456 | P20962;F5 | P20962;F5 | P20962 | Parathymo |
| ITGB6 | 0.700562 | 0.199268 | -1.66843 | A0A087WX | A0A087WX | A0A087WX | Integrin be |
| SRXN1 | 0.698236 | 0.200338 | 0.124133 | Q9BYN0 | Q9BYN0 | Q9BYN0 | Sulfiredoxi |
| HSD17B8 | 0.698015 | 0.20044 | 0.10117 | Q92506 | Q92506 | Q92506 | Estradiol 17 |
| ALOX5AP | 0.687253 | 0.205469 | -0.11431 | A0A087WV | A0A087WV | A0A087WV | Arachidona |
| RIPK1 | 0.681409 | 0.208253 | -1.0272 | Q13546 | Q13546 | Q13546 | Receptor-in |
| DEGS1 | 0.680065 | 0.208898 | -0.21823 | O15121 | O15121 | O15121 | Sphingolipi |
| LHPP | 0.676531 | 0.210605 | -0.2582 | Q9H008;Q5 | Q9H008;Q5 | Q9H008 | Phospholys |
| UBL3 | 0.673353 | 0.212152 | -0.55226 | O95164 | O95164 | O95164 | Ubiquitin-l |
| HBA2 | 0.667767 | 0.214898 | -1.45446 | G3V1N2;P6 | G3V1N2;P6 | G3V1N2 | Hemoglobi |
| TMEM167A | 0.666103 | 0.215723 | -1.90126 | Q8TBQ9 | Q8TBQ9 | Q8TBQ9 | Protein kish |
| SYP | 0.658859 | 0.219352 | 0.952417 | P08247;H7 | P08247;H7 | P08247 | Synaptophy |
| SREK1 | 0.65705 | 0.220267 | -0.97767 | Q8WXA9 | Q8WXA9 | Q8WXA9 | Splicing reg |
| RALGAPB | 0.653637 | 0.222005 | 1.174311 | Q86X10;A2 | Q86X10;A2 | Q86X10 | Ral GTPase- |
| SFT2D3 | 0.648449 | 0.224673 | 1.198922 | Q587I9 | Q587I9 | Q587I9 | Vesicle tran |
| CHMP3 | 0.638848 | 0.229695 | -0.8254 | Q9Y3E7;A0 | Q9Y3E7;A0 | Q9Y3E7 | Charged mu |
| FAM46C | 0.631655 | 0.233531 | -0.24969 | Q5VWP2 | Q5VWP2 | Q5VWP2 | Protein FAM |
| STX16-NPEI | 0.629474 | 0.234707 | -0.05169 | H3BU86;O1 | H3BU86;O1 | H3BU86 | Syntaxin-16 |
| DOLPP1 | 0.618412 | 0.240762 | -0.5494 | Q86YN1 | Q86YN1 | Q86YN1 | Dolichyldip |
| SPCS1 | 0.61511 | 0.2426 | 0.073238 | A0A5F9YFS | A0A5F9YFS | A0A5F9YFS | Signal pept |
| RASA4 | 0.612594 | 0.244009 | 1.145154 | H0YB35;E7 | H0YB35;E7 | H0YB35 | Ras GTPase- |
| RNF121 | 0.609775 | 0.245598 | -0.7442 | Q9H920;E9 | Q9H920;E9 | Q9H920 | RING finger |
| SMAP1 | 0.607669 | 0.246792 | -1.54455 | A0A087X1) | A0A087X1) | A0A087X1) | Stromal me |
| APH1A | 0.607205 | 0.247056 | -0.94367 | Q96BI3;Q5 | Q96BI3;Q5 | Q96BI3 | Gamma-sec |
| GPATCH8 | 0.599744 | 0.251337 | -0.79176 | Q9UKJ3 | Q9UKJ3 | Q9UKJ3 | G patch dor |

| | | | | | | | |
|---|---|---|---|---|---|---|---|
| TMEM50A | 0.596442 | 0.253255 | -0.6196 | O95807 | O95807 | O95807 | Transmemb |
| HIST1H1E | 0.596018 | 0.253502 | 1.24277 | P10412;Q0 | P10412 | P10412 | Histone H1 |
| PANX1 | 0.591292 | 0.256276 | 1.419879 | Q96RD7 | Q96RD7 | Q96RD7 | Pannexin-1 |
| C20orf24 | 0.580201 | 0.262905 | 0.099012 | Q9BUV8 | Q9BUV8 | Q9BUV8 | Uncharacte |
| IGLL5 | 0.572898 | 0.267363 | 1.061575 | P0DOY3;P0 | P0DOY3;P0 | P0DOY3 | Ig lambda-1 |
| MR1 | 0.567764 | 0.270543 | -0.81889 | Q95460 | Q95460 | Q95460 | Major histo |
| SDHAF2 | 0.561943 | 0.274193 | 0.143972 | F5GYJ5;Q9 | F5GYJ5;Q9 | F5GYJ5 | Succinate c |
| LYRM4 | 0.551994 | 0.280547 | 0.113837 | Q9HD34;C9 | Q9HD34;C9 | Q9HD34 | LYR motif-c |
| WDR75 | 0.545925 | 0.284495 | 0.73635 | Q8IWA0 | Q8IWA0 | Q8IWA0 | WD repeat- |
| FRYL | 0.538099 | 0.289668 | -0.44665 | O94915;A0 | O94915;A0 | O94915 | Protein furr |
| HLA-DQA1 | 0.536957 | 0.290431 | 0.139396 | E9PMV2 | E9PMV2 | E9PMV2 | |
| NCBP2 | 0.529693 | 0.295329 | -0.16346 | P52298;C9 | P52298;C9 | P52298 | Nuclear cap |
| ACTG1 | 0.528003 | 0.296481 | -0.04338 | P63261;I3I | P63261;I3I | P63261 | Actin, cytop |
| SRM | 0.525611 | 0.298118 | -0.97931 | K7ESL0;P19 | K7ESL0;P19 | K7ESL0 | Spermidine |
| SLC28A3 | 0.518221 | 0.303234 | -1.12293 | Q9HAS3 | Q9HAS3 | Q9HAS3 | Solute carri |
| FDX1L | 0.517424 | 0.303791 | -0.80695 | Q6P4F2;A0 | Q6P4F2;A0 | Q6P4F2 | Adrenodox |
| ELOVL1 | 0.510235 | 0.308863 | -0.23492 | Q9BW60 | Q9BW60 | Q9BW60 | Elongation |
| VKORC1 | 0.508976 | 0.309759 | -1.05629 | I3L3B4;F8V | I3L3B4;F8V | I3L3B4 | Vitamin K e |
| RPL10 | 0.508385 | 0.310181 | 0.047017 | X1WI28;P2 | X1WI28;P2 | X1WI28 | 60S riboson |
| PTTG1IP | 0.500882 | 0.315586 | 0.571929 | P53801;A8 | P53801;A8 | P53801 | Pituitary tu |
| TOMM7 | 0.500278 | 0.316025 | -0.94557 | Q9P0U1;B5 | Q9P0U1;B5 | Q9P0U1 | Mitochond |
| LIMD2 | 0.49935 | 0.316702 | -0.65364 | J3QQM5;Q | J3QQM5;Q | J3QQM5 | LIM domair |
| CMTM3 | 0.498266 | 0.317493 | 0.129057 | H3BUD5;Q | H3BUD5;Q | H3BUD5 | CKLF-like M |
| CYB561A3 | 0.496864 | 0.31852 | -1.29921 | Q8NBI2 | Q8NBI2 | Q8NBI2 | Cytochrom |
| DTNBP1 | 0.496603 | 0.318711 | 0.597417 | D6RJC6;A0 | D6RJC6;A0 | D6RJC6 | Dysbindin |
| RNASE3 | 0.495264 | 0.319695 | 0.128952 | P12724 | P12724 | P12724 | Eosinophil |
| PRPS2 | 0.49153 | 0.322456 | -0.10623 | P11908;H7 | P11908;H7 | P11908 | Ribose-pho |
| ZC3H4 | 0.48073 | 0.330575 | 0.442774 | M0QY97;Q | M0QY97;Q | M0QY97 | Zinc finger |
| TMEM14C | 0.479059 | 0.33185 | 0.184783 | Q9P0S9 | Q9P0S9 | Q9P0S9 | Transmemb |
| TAX1BP3 | 0.470198 | 0.33869 | -0.13641 | O14907 | O14907 | O14907 | Tax1-bindir |
| GK2 | 0.458255 | 0.348133 | 0.093496 | Q14410 | Q14410 | Q14410 | Glycerol kir |
| HSPA8 | 0.453864 | 0.351671 | -0.02957 | P11142;E9 | P11142;E9 | P11142 | Heat shock |
| TMBIM1 | 0.453684 | 0.351816 | 0.666146 | C9JEN3;C9 | C9JEN3;C9 | C9JEN3 | Protein life |
| GEMIN5 | 0.446268 | 0.357875 | -1.04704 | Q8TEQ6 | Q8TEQ6 | Q8TEQ6 | Gem-associ |
| RTN4 | 0.444093 | 0.359672 | 0.072669 | Q9NQC3 | Q9NQC3 | Q9NQC3 | Reticulon-4 |
| MARK2 | 0.442329 | 0.361136 | -0.66469 | E7ETY4;E9F | E7ETY4;E9F | E7ETY4 | Serine/thre |
| GFER | 0.44211 | 0.361318 | 0.754026 | H3BRW3;H | H3BRW3;H | H3BRW3 | Sulfhydryl c |
| IER3IP1 | 0.422805 | 0.377742 | -0.91269 | A0A1W2PN | A0A1W2PN | A0A1W2PN | Immediate |
| C6orf47 | 0.411876 | 0.387369 | 0.66902 | O95873 | O95873 | O95873 | Uncharacte |
| C15orf38-A | 0.409262 | 0.389707 | -0.75718 | A0A0A6YYH | A0A0A6YYH | A0A0A6YYH | Arpin |
| TMSB4X | 0.390269 | 0.407128 | 1.459531 | P62328;CC | P62328 | P62328 | Thymosin b |
| GABARAPL1 | 0.378317 | 0.418488 | -0.59388 | F5GZY7;Q9 | F5GZY7;Q9 | F5GZY7 | Gamma-am |
| OSCAR | 0.376201 | 0.420532 | -0.40603 | A0A0A0MR | A0A0A0MR | A0A0A0MR | Osteoclast- |
| ARGLU1 | 0.363139 | 0.433372 | -0.59869 | Q9NWB6 | Q9NWB6 | Q9NWB6 | Arginine an |
| GNG10 | 0.359014 | 0.437508 | 0.070791 | P50151 | P50151 | P50151 | Guanine nu |
| hCG_2004! | 0.358702 | 0.437823 | -0.71939 | U3KQA5;A0 | U3KQA5;A0 | U3KQA5 | Casein kina |
| HLA-DRB5 | 0.353844 | 0.442748 | 1.124801 | Q30154 | Q30154 | Q30154 | HLA class II |
| SARDH | 0.351802 | 0.444834 | -0.31522 | A0A0A0MT | A0A0A0MT | A0A0A0MT | Sarcosine d |

| | | | | | | | | |
|---|---|---|---|---|---|---|---|---|
| APH1B | 0.348655 | 0.448069 | 0.353831 | | Q8WW43;I | Q8WW43;I | Q8WW43 | Gamma-sec |
| CIR1 | 0.337846 | 0.45936 | 0.143954 | | Q86X95 | Q86X95 | Q86X95 | Corepresso |
| MCUR1 | 0.335214 | 0.462154 | -0.44125 | | Q96AQ8;U3 | Q96AQ8;U3 | Q96AQ8 | Mitochond |
| IMPACT | 0.32208 | 0.476343 | -0.59067 | | J3KT25;J3C | J3KT25;J3C | J3KT25 | Protein IMI |
| DNAJB4 | 0.302661 | 0.498126 | 1.159509 | | Q9UDY4;C9 | Q9UDY4 | Q9UDY4 | DnaJ homo |
| NCK1 | 0.295875 | 0.50597 | 0.110624 | | C9JAB9;H7 | C9JAB9 | C9JAB9 | |
| LSM5 | 0.273949 | 0.53217 | 0.803579 | | Q9Y4Y9;B8 | Q9Y4Y9;B8 | Q9Y4Y9 | U6 snRNA-a |
| SMIM20 | 0.271692 | 0.534943 | 0.420054 | | Q8N5G0;E5 | Q8N5G0;E5 | Q8N5G0 | Small integ |
| CHCHD1 | 0.263027 | 0.545724 | -0.55364 | | Q96BP2;A6 | Q96BP2;A6 | Q96BP2 | Coiled-coil- |
| IMP3 | 0.262619 | 0.546237 | 0.455943 | | Q9NV31 | Q9NV31 | Q9NV31 | U3 small nu |
| DNTTIP2 | 0.254603 | 0.556412 | 0.448426 | | Q5QJE6;J3I | Q5QJE6;J3I | Q5QJE6 | Deoxynucle |
| CLTCL1 | 0.254222 | 0.556901 | -1.01757 | | P53675;A0 | P53675;A0 | P53675 | Clathrin he |
| PRKCA | 0.252678 | 0.558884 | 0.464178 | | P17252;J3I | P17252 | P17252 | Protein kin |
| TPRKB | 0.250517 | 0.561672 | 0.634123 | | Q9Y3C4 | Q9Y3C4 | Q9Y3C4 | EKC/KEOPS |
| C4orf32 | 0.25049 | 0.561707 | 0.285704 | | Q8N8J7 | Q8N8J7 | Q8N8J7 | Uncharacte |
| MT-ATP8 | 0.247379 | 0.565745 | 0.943884 | | P03928 | P03928 | P03928 | ATP synthas |
| WASL | 0.246715 | 0.566611 | 0.290671 | | O00401 | O00401 | O00401 | Neural Wisl |
| COPS7B | 0.243808 | 0.570417 | 0.447701 | | A0A087X1I | A0A087X1I | A0A087X1I | COP9 signa |
| MRPL57 | 0.241244 | 0.573794 | 0.097107 | | Q9BQC6 | Q9BQC6 | Q9BQC6 | Ribosomal |
| NDUFC1 | 0.236198 | 0.580499 | 0.053566 | | O43677 | O43677 | O43677 | NADH dehy |
| DPY30 | 0.229758 | 0.589172 | -0.59443 | | Q9C005 | Q9C005 | Q9C005 | Protein dpy |
| G6PC3 | 0.229286 | 0.589812 | 0.704631 | | Q9BUM1;K | Q9BUM1;K | Q9BUM1 | Glucose-6-p |
| TMEM127 | 0.225793 | 0.594575 | 0.715709 | | O75204 | O75204 | O75204 | Transmemb |
| ABHD5 | 0.225207 | 0.595378 | 0.521796 | | Q8WTS1;F8 | Q8WTS1;F8 | Q8WTS1 | 1-acylglyce |
| TP53I11 | 0.222845 | 0.598625 | -0.06943 | | E9PN66;E9 | E9PN66;E9 | E9PN66 | Tumor prot |
| DPM3 | 0.217417 | 0.606154 | 0.100929 | | Q9P2X0 | Q9P2X0 | Q9P2X0 | Dolichol-ph |
| SDHD | 0.216923 | 0.606844 | -0.56155 | | O14521 | O14521 | O14521 | Succinate d |
| SLC44A2 | 0.214502 | 0.610236 | -0.0862 | | Q8IWA5 | Q8IWA5 | Q8IWA5 | Choline tra |
| LSM7 | 0.214399 | 0.610381 | -0.77772 | | Q9UK45;A0 | Q9UK45;A0 | Q9UK45 | U6 snRNA-a |
| MAP4K4 | 0.211885 | 0.613925 | 0.035557 | | E7ENQ1;G3 | E7ENQ1;G3 | E7ENQ1 | Mitogen-ac |
| PXMP4 | 0.210265 | 0.616219 | 0.332514 | | Q9Y6I8;B4I | Q9Y6I8;B4I | Q9Y6I8 | Peroxisoma |
| MRPL20 | 0.192084 | 0.642564 | -0.35241 | | Q9BYC9;I3I | Q9BYC9;I3I | Q9BYC9 | 39S riboson |
| PSEN1 | 0.19172 | 0.643102 | 0.068418 | | P49768;E7 | P49768;E7 | P49768 | Presenilin-1 |
| PRNP | 0.187273 | 0.649721 | -0.29945 | | X6RKS3;A2 | X6RKS3;A2 | X6RKS3 | Major prior |
| CAPZB | 0.186712 | 0.650561 | 0.017497 | | B1AK88;B1 | B1AK88;B1 | B1AK88 | |
| DAGLA | 0.178752 | 0.662595 | -0.39579 | | Q9Y4D2 | Q9Y4D2 | Q9Y4D2 | Sn1-specific |
| TAMM41 | 0.176933 | 0.665376 | 0.381232 | | Q96BW9;H | Q96BW9;H | Q96BW9 | Phosphatic |
| BCL2L1 | 0.175975 | 0.666846 | -0.46457 | | Q5QP56;Q9 | Q5QP56;Q9 | Q5QP56 | Bcl-2-like p |
| ZFYVE21 | 0.172705 | 0.671885 | -0.34908 | | Q9BQ24;G3 | Q9BQ24;G3 | Q9BQ24 | Zinc finger I |
| TMEM258 | 0.170444 | 0.675391 | 0.731081 | | F5GWH5;P | F5GWH5;P | F5GWH5 | Transmemb |
| NT5DC1 | 0.167519 | 0.679956 | -0.25541 | | Q5TFE4;H0 | Q5TFE4;H0 | Q5TFE4 | 5-nucleotic |
| CHTOP | 0.156389 | 0.697607 | -0.05432 | | A0A087X1I | A0A087X1I | A0A087X1I | Chromatin |
| SRSF11 | 0.14951 | 0.708745 | 0.04791 | | Q5T760;Q0 | Q5T760;Q0 | Q5T760 | Serine/argi |
| APOL2 | 0.146834 | 0.713125 | 0.034748 | | J3KQL8 | J3KQL8 | J3KQL8 | |
| JAGN1 | 0.133257 | 0.735771 | -0.06197 | | Q8N5M9;A | Q8N5M9;A | Q8N5M9 | Protein jagu |
| TPM4 | 0.111916 | 0.77283 | 0.022648 | | P67936;A0 | P67936;A0 | P67936 | Tropomyos |
| MRPL51 | 0.10783 | 0.780135 | 0.244333 | | Q4U2R6;M | Q4U2R6;M | Q4U2R6 | 39S riboson |
| ME2 | 0.099902 | 0.794508 | 0.012492 | | P23368;A0 | P23368;A0 | P23368 | NAD-depen |

| | | | | | | | |
|---|---|---|---|---|---|---|---|
| HMGN2 | 0.098751 | 0.796616 | -0.13538 | P05204 | P05204 | P05204 | Non-histon |
| S1PR2 | 0.094141 | 0.805118 | 0.175393 | O95136 | O95136 | O95136 | Sphingosin |
| GLMP | 0.082755 | 0.826505 | 0.326713 | A0A087WV | A0A087WV | A0A087WV | Glycosylate |
| AAK1 | 0.075763 | 0.839918 | -0.12132 | E9PG46;Q2 | E9PG46;Q2 | E9PG46 | AP2-associa |
| MAP2K4 | 0.074543 | 0.842281 | -0.15402 | P45985 | P45985 | P45985 | Dual specifi |
| MPHOSPH1 | 0.071015 | 0.849151 | 0.166118 | O00566;U3 | O00566;U3 | O00566 | U3 small nu |
| EXOSC4 | 0.066768 | 0.857496 | -0.16987 | Q9NPD3;E9 | Q9NPD3;E9 | Q9NPD3 | Exosome co |
| SLC35B2 | 0.065681 | 0.859646 | -0.02035 | Q8TB61 | Q8TB61 | Q8TB61 | Adenosine 3 |
| STX2 | 0.063129 | 0.864712 | -0.11229 | P32856 | P32856 | P32856 | Syntaxin-2 |
| C17orf49 | 0.059044 | 0.872883 | 0.182196 | F8W038;H( | F8W038;H( | F8W038 | Chromatin |
| ATP6AP2 | 0.058508 | 0.873962 | 0.019581 | A0A1B0GV' | A0A1B0GV' | A0A1B0GV' | Renin recep |
| LMBRD1 | 0.056988 | 0.877025 | 0.028011 | A0A3B3IT3 | A0A3B3IT3 | A0A3B3IT3 | Probable ly |
| CCL3 | 0.056817 | 0.87737 | -0.08027 | P10147;A0 | P10147;A0 | P10147 | C-C motif cl |
| ANAPC7 | 0.052463 | 0.886211 | 0.095571 | Q9UJX3;H0 | Q9UJX3;H0 | Q9UJX3 | Anaphase-p |
| CYTH1 | 0.05031 | 0.890615 | 0.090967 | Q15438;K7 | Q15438;K7 | Q15438 | Cytohesin-1 |
| SLC37A4 | 0.049148 | 0.893 | -0.01353 | U3KQS2;U3 | U3KQS2;U3 | U3KQS2 | Glucose-6-p |
| SPAG9 | 0.045035 | 0.901499 | 0.012082 | O60271;D6 | O60271 | O60271 | C-Jun-amin |
| DR1 | 0.044762 | 0.902065 | 0.018556 | Q01658 | Q01658 | Q01658 | Protein Dr1 |
| C14orf142 | 0.043749 | 0.904172 | 0.029758 | Q9BXV9 | Q9BXV9 | Q9BXV9 | Uncharacte |
| SCAF4 | 0.03588 | 0.920705 | 0.090068 | O95104 | O95104 | O95104 | Splicing fac |
| PLEKHA2 | 0.031577 | 0.929871 | 0.097056 | E5RHB5;A0 | E5RHB5;A0 | E5RHB5 | Pleckstrin h |
| TMEM208 | 0.029467 | 0.9344 | 0.095154 | Q9BTX3;H3 | Q9BTX3;H3 | Q9BTX3 | Transmemb |
| PRPF38B | 0.027569 | 0.938493 | 0.048124 | Q5VTL8 | Q5VTL8 | Q5VTL8 | Pre-mRNA-s |
| OSGEPL1 | 0.025518 | 0.942936 | 0.025409 | Q9H4B0;E5 | Q9H4B0;E5 | Q9H4B0 | Probable tR |
| SLC31A1 | 0.025415 | 0.94316 | 0.004757 | O15431 | O15431 | O15431 | High affinit |
| NDUFA1 | 0.024987 | 0.944089 | -0.04905 | O15239 | O15239 | O15239 | NADH dehy |
| NTMT1 | 0.020368 | 0.954184 | -0.02586 | Q9BV86 | Q9BV86 | Q9BV86 | N-terminal |
| CLIP1 | 0.017738 | 0.95998 | 0.001891 | P30622;A0 | P30622;A0 | P30622 | CAP-Gly do |
| NUDT2 | 0.017525 | 0.960451 | -0.03828 | P50583 | P50583 | P50583 | Bis(5-nucle |
| SREK1 | 0.017503 | 0.960499 | -0.03833 | A0A087WY | A0A087WY | A0A087WY03 | |
| PPTC7 | 0.016247 | 0.963281 | 0.027417 | Q8NI37 | Q8NI37 | Q8NI37 | Protein pho |
| | 0.015685 | 0.964527 | -0.01431 | A0A2R8Y61 | A0A2R8Y61 | A0A2R8Y611 | |
| GTF3C3 | 0.004951 | 0.988665 | -0.01543 | A0A494C19 | A0A494C19 | A0A494C19 | General tra |
| NINJ1 | 0.004152 | 0.990486 | 0.001634 | Q92982 | Q92982 | Q92982 | Ninjurin-1 |
| CASK | 0.003288 | 0.992459 | -0.00627 | A0A2R8Y3E | A0A2R8Y3E | A0A2R8Y3E | Peripheral p |
| NRAS | 0.000979 | 0.997749 | -0.00025 | P01111;G3 | P01111 | P01111 | GTPase NRa |

| Gene names | id | Peptides | Razor + unique peptides | Unique peptides | Sequence coverage [%] | Unique + razor sequence coverage [%] | Unique sequence coverage [%] | Mol. weight [kDa] |
|---|---|---|---|---|---|---|---|---|
| TPM1 | 417 | 9 | 4 | 4 | 27.3 | 18.4 | 18.4 | 28.524 |
| ARL3 | 2235 | 3 | 3 | 3 | 23.1 | 23.1 | 23.1 | 20.455 |
| LAIR1 | 249 | 3 | 3 | 3 | 14.9 | 14.9 | 14.9 | 29.692 |
| TRAPPC9 | 1054 | 5 | 5 | 5 | 5.3 | 5.3 | 5.3 | 128.53 |
| ATP11B | 4556 | 6 | 5 | 5 | 7 | 6.3 | 6.3 | 134.19 |
| GYS1 | 1919 | 4 | 4 | 4 | 8.4 | 8.4 | 8.4 | 83.785 |
| BLOC1S6 | 1103 | 3 | 3 | 3 | 32.8 | 32.8 | 32.8 | 20.265 |
| TLN2 | 4619 | 20 | 3 | 3 | 6.4 | 1 | 1 | 271.61 |
| HIPK4 | 3563 | 2 | 2 | 2 | 2.9 | 2.9 | 2.9 | 69.424 |
| DCAF7 | 56 | 2 | 2 | 2 | 10.7 | 10.7 | 10.7 | 23.431 |
| FASTKD1 | 3129 | 4 | 4 | 4 | 5.3 | 5.3 | 5.3 | 97.41 |
| GPAA1 | 1432 | 4 | 4 | 4 | 7.1 | 7.1 | 7.1 | 67.622 |
| ALG6 | 4659 | 3 | 3 | 3 | 5.7 | 5.7 | 5.7 | 58.12 |
| FAM208A | 88 | 3 | 3 | 3 | 3.5 | 3.5 | 3.5 | 117.31 |
| CDC40 | 1496 | 3 | 3 | 3 | 8.2 | 8.2 | 8.2 | 60.83 |
| DGUOK | 3076 | 3 | 3 | 3 | 23.8 | 23.8 | 23.8 | 32.055 |
| NUDT16L1 | 3955 | 3 | 3 | 3 | 14.2 | 14.2 | 14.2 | 23.338 |
| CAMLG | 2360 | 3 | 3 | 3 | 22 | 22 | 22 | 32.952 |
| EPS8L2 | 392 | 3 | 3 | 3 | 52.4 | 52.4 | 52.4 | 11.703 |
| DHODH | 2743 | 3 | 3 | 3 | 10.6 | 10.6 | 10.6 | 42.867 |
| EXOSC3 | 4236 | 2 | 2 | 2 | 10.5 | 10.5 | 10.5 | 29.572 |
| FAR1 | 874 | 2 | 2 | 2 | 7.5 | 7.5 | 7.5 | 37.481 |
| TMEM222 | 1042 | 2 | 2 | 2 | 38.8 | 38.8 | 38.8 | 8.8913 |
| ULK3 | 3281 | 2 | 2 | 2 | 4.9 | 4.9 | 4.9 | 53.444 |
| PHLDA3 | 4634 | 2 | 2 | 2 | 14.2 | 14.2 | 14.2 | 13.891 |
| PPWD1 | 934 | 2 | 2 | 2 | 4.2 | 4.2 | 4.2 | 70.276 |
| QPCT | 603 | 2 | 2 | 2 | 10.2 | 10.2 | 10.2 | 32.107 |
| PLIN2 | 3903 | 22 | 22 | 22 | 67.7 | 67.7 | 67.7 | 48.075 |
| THOC7 | 3240 | 6 | 6 | 6 | 37.7 | 37.7 | 37.7 | 23.743 |
| PPAP2B | 1335 | 5 | 5 | 5 | 23.5 | 23.5 | 23.5 | 35.116 |
| NDUFB11 | 4322 | 5 | 5 | 5 | 40.5 | 40.5 | 40.5 | 17.316 |
| DDX41 | 4471 | 5 | 5 | 5 | 9.2 | 9.2 | 9.2 | 69.837 |
| POLR2H | 682 | 4 | 4 | 4 | 33.8 | 33.8 | 33.8 | 16.925 |
| TMEM106E | 4278 | 4 | 4 | 4 | 17.5 | 17.5 | 17.5 | 31.127 |
| CHCHD2;CI | 4673 | 3 | 3 | 3 | 33.8 | 33.8 | 33.8 | 15.512 |
| AKAP2 | 4555 | 3 | 3 | 3 | 2.8 | 2.8 | 2.8 | 94.659 |
| ACTN1 | 1139 | 57 | 2 | 2 | 76 | 1.3 | 1.3 | 94.825 |
| HIST2H3PS | 3203 | 6 | 2 | 2 | 41.2 | 22.1 | 22.1 | 15.43 |
| C14orf159 | 1052 | 25 | 2 | 2 | 61.3 | 6.6 | 6.6 | 60.523 |
| MIF | 1923 | 2 | 2 | 2 | 17.4 | 17.4 | 17.4 | 12.476 |
| NDUFV3 | 2520 | 2 | 2 | 2 | 34.3 | 34.3 | 34.3 | 11.94 |
| DNAJC16 | 4557 | 7 | 7 | 7 | 10.6 | 10.6 | 10.6 | 90.59 |
| DFNA5 | 691 | 5 | 5 | 5 | 19.1 | 19.1 | 19.1 | 39.685 |
| HRSP12 | 1053 | 3 | 3 | 3 | 31.5 | 31.5 | 31.5 | 13.715 |
| DNAJB6;DN | 267 | 4 | 3 | 3 | 14.1 | 11.1 | 11.1 | 36.657 |

| | | | | | | | | |
|---|---|---|---|---|---|---|---|---|
| SRSF5 | 2855 | 4 | 3 | 3 | 19.9 | 16.5 | 16.5 | 31.263 |
| MRPS21 | 2699 | 2 | 2 | 2 | 29.9 | 29.9 | 29.9 | 10.688 |
| CEBPB | 1981 | 2 | 2 | 2 | 6.7 | 6.7 | 6.7 | 36.105 |
| NOL6 | 139 | 2 | 2 | 2 | 3.7 | 3.7 | 3.7 | 112.16 |
| PRDM2 | 2832 | 2 | 2 | 2 | 1.3 | 1.3 | 1.3 | 188.91 |
| MCOLN1 | 4082 | 2 | 2 | 2 | 7.4 | 7.4 | 7.4 | 65.022 |
| HPGD | 1944 | 5 | 5 | 5 | 24.4 | 24.4 | 24.4 | 28.977 |
| GAS7 | 1514 | 5 | 5 | 5 | 10.1 | 10.1 | 10.1 | 54.371 |
| PLEKHO1 | 3125 | 4 | 4 | 4 | 12.2 | 12.2 | 12.2 | 46.236 |
| RRP9 | 1470 | 4 | 4 | 4 | 17.1 | 17.1 | 17.1 | 51.84 |
| CRYZL1 | 530 | 4 | 4 | 4 | 24.9 | 24.9 | 24.9 | 37.061 |
| EMB | 742 | 3 | 3 | 3 | 13.6 | 13.6 | 13.6 | 30.874 |
| TIPRL | 1569 | 3 | 3 | 3 | 17.6 | 17.6 | 17.6 | 31.444 |
| STX17 | 2532 | 3 | 3 | 3 | 12.3 | 12.3 | 12.3 | 33.403 |
| COQ10B | 635 | 3 | 3 | 3 | 13.3 | 13.3 | 13.3 | 24.344 |
| ITM2B | 4549 | 3 | 3 | 3 | 16.9 | 16.9 | 16.9 | 30.338 |
| TRAPPC1 | 4643 | 3 | 3 | 3 | 23.4 | 23.4 | 23.4 | 16.831 |
| AFTPH | 631 | 3 | 3 | 3 | 3.8 | 3.8 | 3.8 | 102.11 |
| CD37 | 620 | 3 | 3 | 3 | 16.5 | 16.5 | 16.5 | 27.592 |
| TOM1L2 | 3317 | 3 | 3 | 3 | 9.1 | 9.1 | 9.1 | 55.556 |
| GPR107 | 1020 | 3 | 3 | 3 | 5.3 | 5.3 | 5.3 | 66.99 |
| SPATS2L | 4281 | 3 | 3 | 3 | 7.3 | 7.3 | 7.3 | 61.728 |
| SMDT1 | 4143 | 2 | 2 | 2 | 26.2 | 26.2 | 26.2 | 11.441 |
| LRP10 | 1067 | 2 | 2 | 2 | 4.4 | 4.4 | 4.4 | 49.326 |
| NACC1;NA( | 3869 | 2 | 2 | 2 | 4.7 | 4.7 | 4.7 | 57.258 |
| DOK2 | 771 | 2 | 2 | 2 | 21.1 | 21.1 | 21.1 | 13.564 |
| MRPL33 | 1543 | 2 | 2 | 2 | 38.5 | 38.5 | 38.5 | 7.6191 |
| CTSO | 2299 | 2 | 2 | 2 | 10.9 | 10.9 | 10.9 | 35.957 |
| LYRM2 | 4272 | 2 | 2 | 2 | 22.7 | 22.7 | 22.7 | 10.449 |
| COA5 | 637 | 2 | 2 | 2 | 31.5 | 31.5 | 31.5 | 8.0572 |
| RRAGD | 4234 | 11 | 2 | 2 | 29.5 | 5.5 | 5.5 | 45.587 |
| UBL5 | 4061 | 2 | 2 | 2 | 27.4 | 27.4 | 27.4 | 8.5468 |
| GNG2 | 2542 | 2 | 2 | 2 | 36.6 | 36.6 | 36.6 | 7.8501 |
| HDHD1 | 2794 | 2 | 2 | 2 | 11 | 11 | 11 | 25.249 |
| CHMP2A | 1271 | 2 | 2 | 2 | 11.3 | 11.3 | 11.3 | 18.739 |
| AMPD2 | 456 | 10 | 10 | 10 | 14.1 | 14.1 | 14.1 | 94.889 |
| BRK1 | 3643 | 6 | 6 | 6 | 66.7 | 66.7 | 66.7 | 8.7448 |
| SELT | 42 | 4 | 4 | 4 | 21.5 | 21.5 | 21.5 | 22.344 |
| RPL29 | 403 | 3 | 3 | 3 | 18 | 18 | 18 | 18.512 |
| MOB1B;MC | 3347 | 3 | 3 | 3 | 16.2 | 16.2 | 16.2 | 25.091 |
| NEDD8;NEI | 3042 | 2 | 2 | 2 | 30.9 | 30.9 | 30.9 | 9.0714 |
| SCARF1 | 2933 | 2 | 2 | 2 | 3 | 3 | 3 | 87.386 |
| DYNC1H1 | 2937 | 241 | 241 | 241 | 62 | 62 | 62 | 532.4 |
| AHNAK | 2802 | 227 | 227 | 226 | 66.8 | 66.8 | 66.7 | 629.09 |
| PRKDC | 2685 | 177 | 177 | 177 | 50.2 | 50.2 | 50.2 | 469.08 |
| RNF213 | 159 | 148 | 148 | 148 | 36.5 | 36.5 | 36.5 | 591.4 |
| FLNA | 2037 | 158 | 158 | 142 | 73.8 | 73.8 | 69.3 | 280.74 |
| PLEC | 3002 | 377 | 377 | 132 | 74.9 | 74.9 | 21 | 531.78 |

| | | | | | | | | |
|---|---|---|---|---|---|---|---|---|
| LRP1 | 2784 | 127 | 127 | 126 | 37.5 | 37.5 | 37.4 | 504.6 |
| VPS13C | 3320 | 122 | 122 | 122 | 44.2 | 44.2 | 44.2 | 422.39 |
| TLN1 | 4613 | 133 | 133 | 116 | 69.2 | 69.2 | 64.1 | 269.76 |
| MYH9 | 2223 | 139 | 139 | 110 | 66.5 | 66.5 | 54.5 | 226.53 |
| GOLGB1 | 2972 | 102 | 102 | 102 | 39.7 | 39.7 | 39.7 | 376.01 |
| GCN1L1 | 3686 | 102 | 102 | 102 | 48.7 | 48.7 | 48.7 | 292.75 |
| IQGAP1 | 2331 | 100 | 100 | 98 | 71.5 | 71.5 | 70.5 | 189.25 |
| UTRN | 2330 | 96 | 96 | 94 | 38.1 | 38.1 | 37.6 | 394.46 |
| SNRNP200 | 1568 | 87 | 87 | 87 | 57.6 | 57.6 | 57.6 | 244.5 |
| PRPF8 | 3267 | 87 | 87 | 87 | 48.7 | 48.7 | 48.7 | 273.6 |
| KTN1 | 3403 | 86 | 86 | 86 | 62.4 | 62.4 | 62.4 | 156.27 |
| IQGAP2 | 2896 | 88 | 86 | 86 | 66.5 | 65.5 | 65.5 | 180.58 |
| FASN | 2366 | 85 | 85 | 85 | 50.9 | 50.9 | 50.9 | 273.42 |
| RRBP1 | 4392 | 81 | 81 | 80 | 67.3 | 67.3 | 66.8 | 152.45 |
| CLTC | 49 | 97 | 97 | 78 | 69.9 | 69.9 | 58.4 | 192.06 |
| IGF2R | 1884 | 77 | 77 | 77 | 41.3 | 41.3 | 41.3 | 274.37 |
| LRPPRC | 2290 | 76 | 76 | 75 | 67.6 | 67.6 | 67.1 | 157.9 |
| EPRS | 1790 | 75 | 75 | 74 | 62.9 | 62.9 | 62.4 | 170.59 |
| EEA1 | 2995 | 72 | 72 | 72 | 55.3 | 55.3 | 55.3 | 162.46 |
| SPTAN1 | 2909 | 72 | 72 | 72 | 39.5 | 39.5 | 39.5 | 284.54 |
| COPA | 2481 | 69 | 69 | 69 | 65.3 | 65.3 | 65.3 | 138.34 |
| VWA8 | 503 | 69 | 69 | 69 | 48 | 48 | 48 | 214.82 |
| DNAJC13 | 1526 | 67 | 67 | 67 | 43.3 | 43.3 | 43.3 | 254.41 |
| FLNB | 1538 | 79 | 67 | 65 | 40.9 | 37.2 | 35.9 | 278.16 |
| UGGT1 | 4350 | 64 | 64 | 64 | 58.5 | 58.5 | 58.5 | 177.19 |
| OGDH | 2744 | 62 | 62 | 62 | 65.8 | 65.8 | 65.8 | 115.93 |
| SND1 | 3328 | 62 | 62 | 62 | 76 | 76 | 76 | 102 |
| DCTN1;DKF | 817 | 59 | 59 | 59 | 61.5 | 61.5 | 61.5 | 139.09 |
| MYOF | 4364 | 142 | 142 | 59 | 73.8 | 73.8 | 29.4 | 234.71 |
| VCL | 1992 | 59 | 59 | 59 | 56.9 | 56.9 | 56.9 | 123.8 |
| MRC1 | 2063 | 58 | 58 | 58 | 49.4 | 49.4 | 49.4 | 166.01 |
| WDFY4 | 3311 | 58 | 58 | 58 | 25.6 | 25.6 | 25.6 | 353.61 |
| EIF3A | 2931 | 57 | 57 | 57 | 44.5 | 44.5 | 44.5 | 166.57 |
| NUMA1 | 2978 | 57 | 57 | 57 | 35.8 | 35.8 | 35.8 | 238.26 |
| VCP | 2505 | 68 | 68 | 55 | 76.3 | 76.3 | 61.9 | 89.321 |
| VIM | 1810 | 62 | 62 | 55 | 89.7 | 89.7 | 83.5 | 53.651 |
| TPR | 1894 | 54 | 54 | 54 | 26.9 | 26.9 | 26.9 | 267.29 |
| ITPR2 | 2958 | 67 | 67 | 54 | 31.8 | 31.8 | 27.7 | 308.06 |
| PTPRC | 1806 | 54 | 54 | 54 | 45.2 | 45.2 | 45.2 | 147.48 |
| APOBR | 445 | 53 | 53 | 53 | 51.4 | 51.4 | 51.4 | 114.87 |
| ITPR1 | 404 | 65 | 53 | 53 | 30.4 | 26.7 | 26.7 | 313.93 |
| MYO1C | 930 | 53 | 53 | 53 | 58.3 | 58.3 | 58.3 | 118.99 |
| DHX9 | 2788 | 53 | 53 | 53 | 53.4 | 53.4 | 53.4 | 140.96 |
| DMXL2 | 3619 | 55 | 55 | 53 | 26.9 | 26.9 | 26.4 | 339.64 |
| MVP | 2971 | 52 | 52 | 52 | 73 | 73 | 73 | 99.326 |
| LCP1 | 1917 | 52 | 52 | 52 | 84.7 | 84.7 | 84.7 | 70.288 |
| DMXL1 | 918 | 54 | 52 | 52 | 27.8 | 27.2 | 27.2 | 337.83 |
| ACLY | 2475 | 52 | 52 | 52 | 56.3 | 56.3 | 56.3 | 120.84 |

| | | | | | | | | |
|---|---|---|---|---|---|---|---|---|
| ITGAX | 2030 | 54 | 54 | 52 | 61.7 | 61.7 | 59.5 | 127.83 |
| EEF2 | 1911 | 53 | 53 | 52 | 69.3 | 69.3 | 68.2 | 95.337 |
| DOCK8 | 3570 | 54 | 54 | 51 | 32.4 | 32.4 | 31.2 | 238.53 |
| HDLBP | 2 | 51 | 51 | 51 | 47.7 | 47.7 | 47.7 | 141.44 |
| ESYT1 | 3972 | 51 | 51 | 51 | 57.9 | 57.9 | 57.9 | 122.85 |
| NSF | 1143 | 52 | 52 | 51 | 74.3 | 74.3 | 73.3 | 82.091 |
| PI4KA | 2286 | 50 | 50 | 50 | 33.1 | 33.1 | 33.1 | 236.83 |
| ATP13A1 | 4204 | 50 | 50 | 50 | 52 | 52 | 52 | 132.95 |
| COPG1 | 4662 | 55 | 55 | 50 | 78.8 | 78.8 | 72.3 | 97.717 |
| PCK2 | 3073 | 49 | 49 | 49 | 75.6 | 75.6 | 75.6 | 70.698 |
| MTHFD1 | 1883 | 50 | 50 | 49 | 71.3 | 71.3 | 70.3 | 101.56 |
| LARS | 485 | 49 | 49 | 49 | 52 | 52 | 52 | 134.34 |
| HADHA | 1064 | 48 | 48 | 48 | 68.6 | 68.6 | 68.6 | 86.371 |
| GPD2 | 2301 | 48 | 48 | 48 | 75.4 | 75.4 | 75.4 | 80.852 |
| PITRM1 | 3154 | 47 | 47 | 47 | 64.4 | 64.4 | 64.4 | 117.41 |
| GLG1 | 3704 | 47 | 47 | 47 | 47.5 | 47.5 | 47.5 | 134.55 |
| SAMHD1 | 4610 | 47 | 47 | 47 | 78.3 | 78.3 | 78.3 | 72.2 |
| HK1 | 2003 | 56 | 56 | 47 | 59.3 | 59.3 | 51.8 | 102.48 |
| H6PD | 1653 | 47 | 47 | 47 | 68.1 | 68.1 | 68.1 | 88.892 |
| HSD17B4 | 2442 | 47 | 47 | 47 | 70.5 | 70.5 | 70.5 | 79.685 |
| ITGAM | 1874 | 49 | 47 | 47 | 44.3 | 42 | 42 | 127.18 |
| HSPA4 | 2214 | 49 | 49 | 47 | 67.4 | 67.4 | 64.5 | 94.33 |
| UBR4 | 3194 | 46 | 46 | 46 | 12.9 | 12.9 | 12.9 | 573.83 |
| PSMD2 | 2850 | 46 | 46 | 46 | 65 | 65 | 65 | 100.2 |
| NBAS | 502 | 46 | 46 | 46 | 27.8 | 27.8 | 27.8 | 268.57 |
| SF3B1 | 1560 | 46 | 46 | 46 | 48.5 | 48.5 | 48.5 | 145.83 |
| RAD50 | 3701 | 45 | 45 | 45 | 38 | 38 | 38 | 153.89 |
| PSMD1 | 3893 | 45 | 45 | 45 | 66.4 | 66.4 | 66.4 | 105.84 |
| COPB1 | 2480 | 45 | 45 | 45 | 59.4 | 59.4 | 59.4 | 107.14 |
| ANPEP | 1939 | 45 | 45 | 45 | 52.2 | 52.2 | 52.2 | 109.54 |
| ACO2 | 3924 | 45 | 45 | 45 | 71.9 | 71.9 | 71.9 | 85.424 |
| UPF1 | 3705 | 45 | 45 | 45 | 51.4 | 51.4 | 51.4 | 124.34 |
| HSP90B1 | 1930 | 46 | 46 | 45 | 62.5 | 62.5 | 61 | 92.468 |
| CAND1 | 3407 | 45 | 45 | 45 | 49.1 | 49.1 | 49.1 | 136.37 |
| SEPTIN9 | 4439 | 44 | 44 | 44 | 76.3 | 76.3 | 76.3 | 65.401 |
| HUWE1 | 3380 | 44 | 44 | 44 | 14.3 | 14.3 | 14.3 | 481.89 |
| PLXNB2 | 1376 | 45 | 45 | 44 | 33.1 | 33.1 | 32.7 | 205.12 |
| EIF4G1 | 2760 | 50 | 50 | 44 | 35.1 | 35.1 | 31.5 | 175.49 |
| LONP1 | 1217 | 44 | 44 | 44 | 67.5 | 67.5 | 67.5 | 95.179 |
| NES | 2354 | 44 | 44 | 44 | 34.4 | 34.4 | 34.4 | 177.44 |
| PDCD6IP | 3642 | 44 | 44 | 44 | 57.3 | 57.3 | 57.3 | 96.022 |
| RPN1 | 1739 | 43 | 43 | 43 | 72.2 | 72.2 | 72.2 | 68.569 |
| ATP1A1 | 1743 | 47 | 47 | 43 | 50.8 | 50.8 | 47.2 | 112.89 |
| MYO1F | 1284 | 49 | 49 | 43 | 52.8 | 52.8 | 46.6 | 124.84 |
| HSPA9 | 2254 | 44 | 44 | 43 | 73.8 | 73.8 | 72.8 | 73.68 |
| HSPD1 | 1864 | 43 | 43 | 43 | 76.8 | 76.8 | 76.8 | 61.054 |
| ERAP1 | 4353 | 43 | 43 | 43 | 52.6 | 52.6 | 52.6 | 107.23 |
| CALR | 2128 | 44 | 44 | 43 | 85.4 | 85.4 | 85.4 | 48.141 |

| | | | | | | | | |
|---|---|---|---|---|---|---|---|---|
| UBA1 | 2055 | 43 | 43 | 43 | 61.8 | 61.8 | 61.8 | 117.85 |
| IARS2 | 4265 | 42 | 42 | 42 | 51.1 | 51.1 | 51.1 | 113.79 |
| TRIP11 | 3031 | 42 | 42 | 42 | 27.6 | 27.6 | 27.6 | 227.58 |
| INF2 | 3080 | 42 | 42 | 42 | 49.8 | 49.8 | 49.8 | 135.62 |
| HK3 | 2465 | 45 | 43 | 42 | 78.4 | 76.4 | 76.4 | 99.024 |
| ATP6V1A | 2253 | 42 | 42 | 42 | 86.4 | 86.4 | 86.4 | 68.303 |
| P4HB | 1779 | 42 | 42 | 42 | 80.3 | 80.3 | 80.3 | 57.116 |
| FLII | 2834 | 42 | 42 | 42 | 44.2 | 44.2 | 44.2 | 144.75 |
| TEX2 | 3449 | 41 | 41 | 41 | 44 | 44 | 44 | 125.3 |
| FAM120A | 4357 | 41 | 41 | 41 | 50.4 | 50.4 | 50.4 | 121.89 |
| FKBP15 | 3186 | 41 | 41 | 41 | 44.9 | 44.9 | 44.9 | 133.63 |
| ACADVL | 2379 | 41 | 41 | 41 | 74.8 | 74.8 | 74.8 | 70.389 |
| PLXND1 | 4616 | 42 | 41 | 41 | 33.1 | 32.8 | 32.8 | 212 |
| XRCC5 | 1903 | 41 | 41 | 41 | 74.6 | 74.6 | 74.6 | 82.704 |
| USO1 | 1508 | 42 | 42 | 41 | 50.6 | 50.6 | 49.9 | 107.89 |
| TGM2 | 2047 | 42 | 42 | 41 | 71.3 | 71.3 | 71.3 | 77.328 |
| ACTN4 | 1458 | 66 | 66 | 41 | 74.6 | 74.6 | 53.1 | 104.85 |
| VARS | 2116 | 41 | 41 | 41 | 47.8 | 47.8 | 47.8 | 140.47 |
| PML | 2149 | 41 | 41 | 41 | 58 | 58 | 58 | 97.55 |
| PTPN6 | 2145 | 41 | 41 | 41 | 77.5 | 77.5 | 77.5 | 67.56 |
| MAP4 | 805 | 41 | 41 | 41 | 23.7 | 23.7 | 23.7 | 245.44 |
| ROCK1 | 2877 | 43 | 43 | 41 | 34.4 | 34.4 | 32.9 | 158.17 |
| UFL1 | 1612 | 41 | 41 | 41 | 55.3 | 55.3 | 55.3 | 89.594 |
| MYO6 | 334 | 41 | 41 | 41 | 44.4 | 44.4 | 44.4 | 148.6 |
| ITGB2 | 426 | 41 | 41 | 41 | 62.9 | 62.9 | 62.9 | 84.79 |
| MYO9B | 1266 | 41 | 41 | 41 | 29.2 | 29.2 | 29.2 | 234.07 |
| MICAL1 | 3622 | 41 | 41 | 40 | 54.9 | 54.9 | 53.9 | 117.87 |
| SF3B2 | 2872 | 40 | 40 | 40 | 56.3 | 56.3 | 56.3 | 100.23 |
| NDUFS1 | 2137 | 40 | 40 | 40 | 72.5 | 72.5 | 72.5 | 79.467 |
| SEC31A | 749 | 40 | 40 | 40 | 44 | 44 | 44 | 133.01 |
| CANX | 2129 | 41 | 40 | 40 | 57.4 | 57.4 | 57.4 | 67.567 |
| HYOU1 | 420 | 40 | 40 | 40 | 51.6 | 51.6 | 51.6 | 111.41 |
| PC | 1882 | 40 | 40 | 40 | 50.7 | 50.7 | 50.7 | 129.63 |
| HK2 | 2464 | 45 | 40 | 40 | 60 | 54.3 | 54.3 | 102.38 |
| ABCC1 | 2206 | 40 | 40 | 40 | 34.1 | 34.1 | 34.1 | 171.59 |
| COPB2 | 2225 | 40 | 40 | 40 | 63.6 | 63.6 | 63.6 | 102.49 |
| ERAP2 | 3260 | 40 | 40 | 40 | 50.5 | 50.5 | 50.5 | 110.46 |
| KIAA0196 | 2813 | 40 | 40 | 40 | 47.1 | 47.1 | 47.1 | 134.28 |
| GOLGA3 | 2790 | 39 | 39 | 39 | 34.7 | 34.7 | 34.7 | 167.35 |
| PLCB2 | 2727 | 39 | 39 | 39 | 43.5 | 43.5 | 43.5 | 134.02 |
| STAT1 | 482 | 40 | 40 | 39 | 66.7 | 66.7 | 64.7 | 87.03 |
| PNPLA6 | 380 | 42 | 42 | 39 | 39.6 | 39.6 | 38.4 | 149.92 |
| HTT | 2295 | 39 | 39 | 39 | 19.7 | 19.7 | 19.7 | 347.6 |
| LMNB1 | 2028 | 41 | 40 | 39 | 63 | 61.6 | 60.1 | 66.408 |
| POR | 1966 | 38 | 38 | 38 | 65.9 | 65.9 | 65.9 | 76.689 |
| PARP1 | 1835 | 38 | 38 | 38 | 45.1 | 45.1 | 45.1 | 113.08 |
| ITGAV | 1774 | 38 | 38 | 38 | 41.4 | 41.4 | 41.4 | 116.04 |
| FERMT3 | 3405 | 42 | 42 | 38 | 70.8 | 70.8 | 66.9 | 75.952 |

| | | | | | | | | |
|---|---|---|---|---|---|---|---|---|
| VPS35 | 3866 | 38 | 38 | 38 | 56.9 | 56.9 | 56.9 | 91.706 |
| USP15 | 4618 | 40 | 40 | 38 | 52.7 | 52.7 | 50.8 | 112.42 |
| DOCK2 | 3683 | 38 | 38 | 38 | 28.9 | 28.9 | 28.9 | 211.95 |
| EFTUD2 | 2985 | 39 | 38 | 38 | 56.4 | 55.3 | 55.3 | 109.43 |
| MYO18A | 231 | 38 | 38 | 38 | 26.3 | 26.3 | 26.3 | 231.1 |
| TFRC | 1714 | 37 | 37 | 37 | 53.2 | 53.2 | 53.2 | 84.87 |
| MIA3 | 3153 | 37 | 37 | 37 | 22.9 | 22.9 | 22.9 | 213.7 |
| SPG11 | 3826 | 37 | 37 | 37 | 22 | 22 | 22 | 278.86 |
| SF3B3 | 3017 | 37 | 37 | 37 | 41.3 | 41.3 | 41.3 | 135.58 |
| USP9X | 3716 | 37 | 37 | 37 | 20.5 | 20.5 | 20.5 | 292.28 |
| DARS | 1934 | 37 | 37 | 37 | 71.7 | 71.7 | 71.7 | 57.136 |
| KIF5B | 2203 | 48 | 48 | 37 | 59.7 | 59.7 | 47.2 | 109.68 |
| ACO1 | 2039 | 38 | 38 | 37 | 51.9 | 51.9 | 51.1 | 98.398 |
| TOP2B | 2753 | 37 | 37 | 37 | 29.9 | 29.9 | 29.9 | 183.26 |
| MSN | 2106 | 51 | 51 | 37 | 75.6 | 75.6 | 57.2 | 67.819 |
| TKT | 2147 | 37 | 37 | 37 | 84.1 | 84.1 | 84.1 | 67.877 |
| AP2A1 | 1669 | 46 | 46 | 36 | 59.3 | 59.3 | 47.5 | 107.54 |
| ACAD9 | 4160 | 36 | 36 | 36 | 70 | 70 | 70 | 68.76 |
| KIAA1033 | 109 | 36 | 36 | 36 | 41.7 | 41.7 | 41.7 | 136.5 |
| KPNB1 | 2977 | 36 | 36 | 36 | 50.7 | 50.7 | 50.7 | 97.169 |
| MYO1D | 1609 | 37 | 36 | 36 | 44.2 | 43.3 | 43.3 | 116.2 |
| MYO1G | 552 | 37 | 37 | 36 | 50.1 | 50.1 | 49.2 | 116.44 |
| HSPA5 | 1866 | 37 | 37 | 36 | 59.8 | 59.8 | 57.3 | 72.332 |
| NUP210 | 3628 | 36 | 36 | 36 | 31.6 | 31.6 | 31.6 | 205.11 |
| SQRDL | 4682 | 36 | 36 | 36 | 76.4 | 76.4 | 76.4 | 49.96 |
| DDX1 | 113 | 35 | 35 | 35 | 66.8 | 66.8 | 66.8 | 73.975 |
| TRAP1 | 2826 | 35 | 35 | 35 | 51.8 | 51.8 | 51.8 | 80.109 |
| ACSL1 | 2202 | 38 | 38 | 35 | 59.9 | 59.9 | 56 | 77.942 |
| ANXA2;AN) | 1781 | 35 | 35 | 35 | 82.3 | 82.3 | 82.3 | 38.604 |
| VPS11 | 4110 | 35 | 35 | 35 | 49.1 | 49.1 | 49.1 | 107.84 |
| ILF3 | 2824 | 39 | 39 | 35 | 51.8 | 51.8 | 48 | 95.337 |
| MMP9 | 1931 | 35 | 35 | 35 | 57.6 | 57.6 | 57.6 | 78.457 |
| EPB41L2 | 847 | 39 | 39 | 35 | 49.3 | 49.3 | 46.7 | 104.36 |
| TCIRG1 | 2878 | 35 | 35 | 35 | 50.5 | 50.5 | 50.5 | 92.967 |
| COLGALT1 | 3535 | 35 | 35 | 35 | 60.6 | 60.6 | 60.6 | 71.635 |
| PARP14 | 3106 | 35 | 35 | 35 | 26.8 | 26.8 | 26.8 | 202.8 |
| ZC3HAV1 | 3355 | 36 | 36 | 35 | 49.9 | 49.9 | 49 | 101.43 |
| CPT2 | 2079 | 35 | 35 | 35 | 63.1 | 63.1 | 63.1 | 73.776 |
| CCT5 | 2352 | 35 | 35 | 35 | 80 | 80 | 80 | 59.67 |
| AP1G1 | 1461 | 35 | 35 | 35 | 49.9 | 49.9 | 49.9 | 91.35 |
| EML4 | 4194 | 35 | 35 | 35 | 46.5 | 46.5 | 46.5 | 108.91 |
| LTA4H | 1840 | 35 | 35 | 35 | 68.1 | 68.1 | 68.1 | 69.284 |
| STIP1 | 2191 | 35 | 35 | 35 | 62.6 | 62.6 | 62.6 | 62.639 |
| EMC1 | 3525 | 36 | 36 | 35 | 51.3 | 51.3 | 49.2 | 111.76 |
| DDB1 | 3052 | 35 | 35 | 35 | 41.8 | 41.8 | 41.8 | 126.97 |
| AP3B1 | 1290 | 41 | 41 | 35 | 42.2 | 42.2 | 36.2 | 121.32 |
| ARAP1 | 3851 | 35 | 35 | 35 | 34.3 | 34.3 | 34.3 | 162.19 |
| USP7 | 3717 | 34 | 34 | 34 | 44.2 | 44.2 | 44.2 | 128.3 |

| | | | | | | | | |
|---|---|---|---|---|---|---|---|---|
| PARP4 | 4485 | 34 | 34 | 34 | 26.5 | 26.5 | 26.5 | 192.59 |
| LAP3 | 2140 | 34 | 34 | 34 | 82.1 | 82.1 | 82.1 | 56.166 |
| WARS | 2071 | 34 | 34 | 34 | 81.5 | 81.5 | 81.5 | 53.165 |
| HNRNPUL2 | 3079 | 34 | 34 | 34 | 53 | 53 | 53 | 85.104 |
| ATP6V1B2 | 2034 | 34 | 34 | 34 | 84.7 | 84.7 | 84.7 | 56.5 |
| CYP27A1 | 2746 | 34 | 34 | 34 | 71.6 | 71.6 | 71.6 | 60.234 |
| XRCC6 | 1902 | 34 | 34 | 34 | 57.6 | 57.6 | 57.6 | 69.842 |
| XPO1 | 1374 | 34 | 34 | 34 | 42.4 | 42.4 | 42.4 | 123.38 |
| GTF2I | 434 | 34 | 34 | 34 | 32.5 | 32.5 | 32.5 | 131.05 |
| ALDH4A1 | 2153 | 33 | 33 | 33 | 71 | 71 | 71 | 61.719 |
| GLUD1;GLU | 1687 | 33 | 33 | 33 | 62.2 | 62.2 | 62.2 | 61.397 |
| VPS39 | 3824 | 33 | 33 | 33 | 51.2 | 51.2 | 51.2 | 101.81 |
| TNS3 | 3231 | 35 | 35 | 33 | 39 | 39 | 38.1 | 155.26 |
| C16orf62 | 3362 | 33 | 33 | 33 | 45 | 45 | 45 | 109.56 |
| ATP2A2 | 1967 | 51 | 51 | 33 | 53.5 | 53.5 | 34.5 | 114.76 |
| TCP1 | 1986 | 33 | 33 | 33 | 70.7 | 70.7 | 70.7 | 60.343 |
| TRIM28 | 2856 | 33 | 33 | 33 | 63.8 | 63.8 | 63.8 | 88.549 |
| CCT8 | 2419 | 33 | 33 | 33 | 69.7 | 69.7 | 69.7 | 59.62 |
| CHD4 | 211 | 44 | 44 | 33 | 30.4 | 30.4 | 22.8 | 215.49 |
| WDR7 | 4617 | 33 | 33 | 33 | 33.7 | 33.7 | 33.7 | 163.81 |
| PLXNC1 | 1492 | 34 | 33 | 33 | 29 | 28.5 | 28.5 | 175.74 |
| SACM1L | 465 | 33 | 33 | 33 | 61.6 | 61.6 | 61.6 | 69.359 |
| PRPF6 | 1615 | 33 | 33 | 33 | 41 | 41 | 41 | 106.92 |
| SHMT2 | 2212 | 33 | 33 | 33 | 73.2 | 73.2 | 73.2 | 55.992 |
| CAT | 1724 | 32 | 32 | 32 | 70.6 | 70.6 | 70.6 | 59.755 |
| ZFYVE26 | 3233 | 32 | 32 | 32 | 16.2 | 16.2 | 16.2 | 284.57 |
| NCKAP1L | 2509 | 32 | 32 | 32 | 43.2 | 43.2 | 43.2 | 128.15 |
| NPEPPS | 865 | 32 | 32 | 32 | 42.6 | 42.6 | 42.6 | 102.99 |
| TOMM70A | 1608 | 32 | 32 | 32 | 69.9 | 69.9 | 69.9 | 67.454 |
| DDX46 | 202 | 33 | 33 | 32 | 39.5 | 39.5 | 38.8 | 117.36 |
| GCC2 | 3450 | 32 | 32 | 32 | 21.1 | 21.1 | 21.1 | 195.91 |
| TRPV2 | 4644 | 32 | 32 | 32 | 47.6 | 47.6 | 47.6 | 85.98 |
| WDR1 | 1522 | 32 | 32 | 32 | 81 | 81 | 81 | 66.193 |
| HNRNPU | 2729 | 32 | 32 | 32 | 47.3 | 47.3 | 47.3 | 90.583 |
| MAN2A1 | 3064 | 32 | 32 | 32 | 39.6 | 39.6 | 39.6 | 131.14 |
| OSBPL8 | 4058 | 32 | 32 | 32 | 46.8 | 46.8 | 46.8 | 101.19 |
| DARS2 | 3282 | 32 | 32 | 32 | 62.2 | 62.2 | 62.2 | 73.562 |
| DNM1L | 1312 | 32 | 32 | 32 | 60.6 | 60.6 | 60.6 | 81.876 |
| PFKL | 1983 | 36 | 36 | 32 | 58.7 | 58.7 | 52.9 | 85.018 |
| OSBPL1A | 4036 | 31 | 31 | 31 | 46.9 | 46.9 | 46.9 | 108.47 |
| TUFM | 2370 | 31 | 31 | 31 | 74.6 | 74.6 | 74.6 | 49.541 |
| EIF3C;EIF3C | 3911 | 31 | 31 | 31 | 41.1 | 41.1 | 41.1 | 105.34 |
| CCDC93 | 982 | 31 | 31 | 31 | 54.5 | 54.5 | 54.5 | 73.197 |
| AGPS | 1278 | 31 | 31 | 31 | 63.5 | 63.5 | 63.5 | 72.911 |
| RANBP2 | 2390 | 47 | 47 | 31 | 20.2 | 20.2 | 13.5 | 358.2 |
| ACSL3 | 1661 | 34 | 34 | 31 | 59.3 | 59.3 | 57.8 | 80.419 |
| LACTB | 2705 | 31 | 31 | 31 | 57.4 | 57.4 | 57.4 | 60.693 |
| IPO5 | 1307 | 38 | 38 | 31 | 52.8 | 52.8 | 43.6 | 123.63 |

| | | | | | | | | |
|---|---|---|---|---|---|---|---|---|
| MAN2B1 | 1331 | 31 | 31 | 31 | 37.7 | 37.7 | 37.7 | 113.74 |
| TYMP | 2013 | 31 | 31 | 31 | 80.1 | 80.1 | 80.1 | 49.955 |
| CCT7 | 3928 | 31 | 31 | 31 | 73.8 | 73.8 | 73.8 | 59.366 |
| SMC3 | 4535 | 31 | 31 | 31 | 35.2 | 35.2 | 35.2 | 141.54 |
| SFPQ | 2067 | 32 | 32 | 31 | 46.3 | 46.3 | 45.1 | 76.149 |
| CKAP4 | 2780 | 31 | 31 | 31 | 58.3 | 58.3 | 58.3 | 66.022 |
| MATR3 | 272 | 31 | 31 | 31 | 52.9 | 52.9 | 52.9 | 94.622 |
| PDXDC1 | 3275 | 31 | 31 | 31 | 58.2 | 58.2 | 58.2 | 86.706 |
| ALDH18A1 | 2499 | 31 | 31 | 31 | 51.3 | 51.3 | 51.3 | 87.301 |
| PSMD3 | 1427 | 31 | 31 | 31 | 59.4 | 59.4 | 59.4 | 60.977 |
| KARS | 2989 | 31 | 31 | 31 | 58.5 | 58.5 | 58.5 | 68.047 |
| LNPEP | 4458 | 31 | 31 | 31 | 43.3 | 43.3 | 43.3 | 117.35 |
| MCCC1 | 3872 | 31 | 31 | 31 | 63.2 | 63.2 | 63.2 | 80.472 |
| ITGA6 | 2066 | 31 | 31 | 31 | 37.4 | 37.4 | 37.4 | 126.6 |
| MTOR | 2285 | 31 | 31 | 31 | 19.4 | 19.4 | 19.4 | 288.89 |
| DHX15 | 1421 | 32 | 32 | 31 | 50.1 | 50.1 | 48.9 | 90.932 |
| WDR11 | 4059 | 31 | 31 | 31 | 38.3 | 38.3 | 38.3 | 136.68 |
| VPS18 | 4385 | 31 | 31 | 31 | 44.7 | 44.7 | 44.7 | 110.18 |
| RARS | 2490 | 31 | 31 | 31 | 54.2 | 54.2 | 54.2 | 75.378 |
| CCT2 | 2682 | 31 | 31 | 31 | 76.4 | 76.4 | 76.4 | 57.488 |
| GARS | 2276 | 31 | 31 | 31 | 51.4 | 51.4 | 51.4 | 83.165 |
| GAA | 1853 | 31 | 31 | 31 | 42.2 | 42.2 | 42.2 | 105.32 |
| SEC63 | 4428 | 31 | 31 | 31 | 50.3 | 50.3 | 50.3 | 87.996 |
| TAP2 | 4744 | 33 | 33 | 31 | 59.7 | 59.7 | 58.6 | 77.694 |
| WDFY1 | 3448 | 30 | 30 | 30 | 79 | 79 | 79 | 46.323 |
| PECAM1 | 1961 | 30 | 30 | 30 | 51.5 | 51.5 | 51.5 | 82.521 |
| CAPN1 | 1782 | 30 | 30 | 30 | 50.8 | 50.8 | 50.8 | 81.889 |
| FAM129A | 4063 | 30 | 30 | 30 | 38.4 | 38.4 | 38.4 | 103.13 |
| PCCA | 1750 | 30 | 30 | 30 | 52.9 | 52.9 | 52.9 | 80.058 |
| SF3A1 | 3026 | 30 | 30 | 30 | 45.3 | 45.3 | 45.3 | 88.885 |
| ALDH1A1 | 1686 | 34 | 34 | 30 | 79.6 | 79.6 | 76 | 54.861 |
| SDHA | 2177 | 30 | 30 | 30 | 62.7 | 62.7 | 62.7 | 72.691 |
| ACSL4 | 1493 | 33 | 30 | 30 | 56 | 54.4 | 54.4 | 79.187 |
| CNDP2 | 3834 | 30 | 30 | 30 | 77.3 | 77.3 | 77.3 | 52.878 |
| ANXA5 | 1812 | 30 | 30 | 30 | 80.9 | 80.9 | 80.9 | 35.936 |
| EPHX1 | 1777 | 30 | 30 | 30 | 73.6 | 73.6 | 73.6 | 52.948 |
| USP5 | 2313 | 30 | 30 | 30 | 47.9 | 47.9 | 47.9 | 95.785 |
| EIF3B | 2516 | 30 | 30 | 30 | 47.7 | 47.7 | 47.7 | 92.48 |
| ATP5B | 1766 | 30 | 30 | 30 | 78.6 | 78.6 | 78.6 | 56.559 |
| IFI16 | 3061 | 33 | 33 | 30 | 48.4 | 48.4 | 44.7 | 88.255 |
| ACOX3 | 1400 | 30 | 30 | 30 | 61.6 | 61.6 | 61.6 | 77.628 |
| AFG3L2 | 4623 | 31 | 31 | 30 | 43.8 | 43.8 | 42.8 | 88.583 |
| SCP2 | 2054 | 30 | 30 | 30 | 67.6 | 67.6 | 67.6 | 58.993 |
| ARCN1 | 2346 | 30 | 30 | 30 | 56.6 | 56.6 | 56.6 | 57.21 |
| AMPD3 | 2731 | 30 | 30 | 30 | 54.1 | 54.1 | 54.1 | 88.811 |
| ECM29;KIA | 3219 | 30 | 30 | 30 | 21.2 | 21.2 | 21.2 | 204.29 |
| ACOX1 | 2994 | 30 | 30 | 30 | 65.2 | 65.2 | 65.2 | 74.423 |
| NPC1 | 1383 | 29 | 29 | 29 | 28.7 | 28.7 | 28.7 | 142.17 |

| | | | | | | | | |
|---|---|---|---|---|---|---|---|---|
| GLS | 1618 | 33 | 33 | 29 | 63.8 | 63.8 | 51.6 | 73.46 |
| LETM1 | 1632 | 29 | 29 | 29 | 49.9 | 49.9 | 49.9 | 83.353 |
| ITGB1 | 1760 | 29 | 29 | 29 | 40.2 | 40.2 | 40.2 | 88.414 |
| MARS | 2522 | 29 | 29 | 29 | 46.2 | 46.2 | 46.2 | 101.11 |
| TNPO1 | 3714 | 29 | 29 | 29 | 48.6 | 48.6 | 48.6 | 102.35 |
| PRKCD | 2771 | 29 | 29 | 29 | 56.7 | 56.7 | 56.7 | 77.504 |
| ANXA4 | 1828 | 29 | 29 | 29 | 81.8 | 81.8 | 81.8 | 35.882 |
| AIFM1 | 1673 | 29 | 29 | 29 | 59.1 | 59.1 | 59.1 | 66.9 |
| VPS16 | 4109 | 29 | 29 | 29 | 54.1 | 54.1 | 54.1 | 94.693 |
| LRRFIP1 | 3090 | 30 | 30 | 29 | 43.4 | 43.4 | 42 | 89.252 |
| ATP6V1C1 | 2035 | 29 | 29 | 29 | 75.1 | 75.1 | 75.1 | 43.941 |
| LMNB2 | 2756 | 31 | 29 | 29 | 46.5 | 43.7 | 43.7 | 69.948 |
| ATIC | 2186 | 29 | 29 | 29 | 61.8 | 61.8 | 61.8 | 64.615 |
| PYGL | 1771 | 36 | 30 | 29 | 55.7 | 47.1 | 45.8 | 97.147 |
| RECQL | 2316 | 29 | 29 | 29 | 59.9 | 59.9 | 59.9 | 73.457 |
| DDX5 | 1190 | 38 | 38 | 29 | 59.6 | 59.6 | 47.7 | 69.086 |
| PTPN1 | 1987 | 29 | 29 | 29 | 67.6 | 67.6 | 67.6 | 49.966 |
| CORO1C | 4501 | 29 | 29 | 29 | 56.3 | 56.3 | 56.3 | 53.248 |
| PGD | 2452 | 29 | 29 | 29 | 75.8 | 75.8 | 75.8 | 53.139 |
| MFN2 | 1623 | 29 | 29 | 29 | 47.8 | 47.8 | 47.8 | 86.401 |
| TBXAS1 | 2085 | 29 | 29 | 29 | 60.3 | 60.3 | 60.3 | 60.649 |
| CPT1A | 2407 | 29 | 29 | 29 | 45.4 | 45.4 | 45.4 | 88.367 |
| PSMC2 | 2232 | 29 | 29 | 29 | 70.9 | 70.9 | 70.9 | 48.633 |
| YARS | 2491 | 29 | 29 | 29 | 61.2 | 61.2 | 61.2 | 59.143 |
| PLOD3 | 1498 | 29 | 29 | 29 | 57.7 | 57.7 | 57.7 | 84.784 |
| PDIA3 | 2166 | 37 | 36 | 29 | 72.5 | 69.5 | 55.2 | 56.782 |
| NUCB1 | 2751 | 29 | 29 | 29 | 60.3 | 60.3 | 60.3 | 53.879 |
| ABHD16A | 246 | 29 | 29 | 29 | 49.3 | 49.3 | 49.3 | 67.518 |
| SAMD9L | 3440 | 31 | 31 | 29 | 26 | 26 | 24.8 | 184.53 |
| CCT4 | 2420 | 29 | 29 | 29 | 72.4 | 72.4 | 72.4 | 57.924 |
| GFPT1 | 2775 | 29 | 29 | 29 | 53.2 | 53.2 | 53.2 | 78.806 |
| AP2A2 | 1619 | 38 | 28 | 28 | 53.2 | 41 | 41 | 103.96 |
| SEPTIN7 | 3047 | 29 | 29 | 28 | 65.9 | 65.9 | 64.3 | 50.679 |
| DIAPH1 | 240 | 28 | 28 | 28 | 30.3 | 30.3 | 30.3 | 136.85 |
| LSS | 2347 | 28 | 28 | 28 | 49.5 | 49.5 | 49.5 | 83.308 |
| ALDH2 | 1746 | 32 | 28 | 28 | 77 | 73.9 | 73.9 | 56.381 |
| ACIN1 | 4741 | 28 | 28 | 28 | 26.6 | 26.6 | 26.6 | 145.44 |
| MUT | 2048 | 28 | 28 | 28 | 52.3 | 52.3 | 52.3 | 83.134 |
| PI4K2A | 3981 | 28 | 28 | 28 | 63.7 | 63.7 | 63.7 | 54.022 |
| CYFIP1 | 3340 | 49 | 49 | 28 | 45.9 | 45.9 | 27.2 | 145.18 |
| TRIM25 | 2944 | 28 | 28 | 28 | 63 | 63 | 63 | 70.973 |
| SMC1A | 1028 | 28 | 28 | 28 | 27.7 | 27.7 | 27.7 | 140.86 |
| RDX | 2217 | 43 | 29 | 28 | 62.4 | 44.3 | 43.1 | 68.563 |
| SUPT16H | 4632 | 28 | 28 | 28 | 32.9 | 32.9 | 32.9 | 119.91 |
| EXOC4 | 3740 | 28 | 28 | 28 | 38.8 | 38.8 | 38.8 | 110.5 |
| RRP12 | 3156 | 28 | 28 | 28 | 30.5 | 30.5 | 30.5 | 143.7 |
| ARPC2 | 1389 | 28 | 28 | 28 | 78.7 | 78.7 | 78.7 | 34.333 |
| TAP1 | 287 | 29 | 28 | 28 | 50.5 | 48.6 | 48.6 | 87.217 |

| | | | | | | | | |
|---|---|---|---|---|---|---|---|---|
| CDC42BPB | 4645 | 28 | 28 | 28 | 22 | 22 | 22 | 194.31 |
| UGP2 | 3075 | 28 | 28 | 28 | 64.2 | 64.2 | 64.2 | 56.94 |
| CAPN2 | 1979 | 28 | 28 | 28 | 54.9 | 54.9 | 54.9 | 79.994 |
| STK10 | 1606 | 29 | 29 | 28 | 36.9 | 36.9 | 35.7 | 112.13 |
| TPP2 | 2143 | 28 | 28 | 28 | 30.3 | 30.3 | 30.3 | 138.35 |
| AIM1 | 265 | 28 | 28 | 28 | 22 | 22 | 22 | 231.71 |
| CCT3 | 2367 | 28 | 28 | 28 | 66.2 | 66.2 | 66.2 | 60.533 |
| ABCD1 | 2207 | 28 | 28 | 28 | 48.6 | 48.6 | 48.6 | 82.936 |
| HSP90AA1 | 1792 | 42 | 42 | 28 | 62.8 | 62.8 | 45.4 | 84.659 |
| LIMA1 | 4436 | 28 | 28 | 28 | 43.7 | 43.7 | 43.7 | 85.225 |
| CORO7-PAN | 174 | 28 | 28 | 28 | 48.6 | 48.6 | 48.6 | 114.16 |
| SCFD1 | 3647 | 28 | 28 | 28 | 66 | 66 | 66 | 72.379 |
| EEF1A1;EEF | 2668 | 28 | 28 | 28 | 69 | 69 | 69 | 50.14 |
| SRPR | 1801 | 28 | 28 | 28 | 53.1 | 53.1 | 53.1 | 69.81 |
| PFKP | 2738 | 31 | 28 | 28 | 53.2 | 48.6 | 48.6 | 85.595 |
| DENND4C | 3222 | 28 | 28 | 28 | 21.4 | 21.4 | 21.4 | 212.71 |
| PYGB | 1875 | 38 | 38 | 28 | 55.6 | 55.6 | 41.9 | 96.695 |
| GOLGA2 | 2791 | 27 | 27 | 27 | 35.6 | 35.6 | 35.6 | 113.08 |
| CHIT1 | 2852 | 27 | 27 | 27 | 73.8 | 73.8 | 73.8 | 51.681 |
| AP1B1 | 2808 | 48 | 48 | 27 | 60.5 | 60.5 | 40.3 | 104.64 |
| NCF2 | 2012 | 27 | 27 | 27 | 58.6 | 58.6 | 58.6 | 59.761 |
| SART3 | 439 | 27 | 27 | 27 | 37.2 | 37.2 | 37.2 | 111.96 |
| NUP93 | 3488 | 27 | 27 | 27 | 37.2 | 37.2 | 37.2 | 93.487 |
| GFM1 | 3871 | 27 | 27 | 27 | 51.5 | 51.5 | 51.5 | 83.471 |
| VPS41 | 2380 | 27 | 27 | 27 | 41.3 | 41.3 | 41.3 | 98.565 |
| RNPEP | 4138 | 27 | 27 | 27 | 53.7 | 53.7 | 53.7 | 72.595 |
| VAC14 | 2798 | 27 | 27 | 27 | 45 | 45 | 45 | 87.972 |
| SRP68 | 4437 | 27 | 27 | 27 | 49 | 49 | 49 | 70.729 |
| NUP155 | 1570 | 27 | 27 | 27 | 32.5 | 32.5 | 32.5 | 155.2 |
| EIF4G2 | 2678 | 27 | 27 | 27 | 33.3 | 33.3 | 33.3 | 102.36 |
| ATG7 | 1642 | 27 | 27 | 27 | 56.6 | 56.6 | 56.6 | 77.959 |
| ASCC3 | 3500 | 27 | 27 | 27 | 17.6 | 17.6 | 17.6 | 251.46 |
| PGK1 | 1697 | 27 | 27 | 27 | 78.2 | 78.2 | 78.2 | 44.614 |
| GBP5 | 3855 | 29 | 29 | 27 | 60.9 | 60.9 | 58.9 | 66.616 |
| ATP2B4 | 2077 | 40 | 40 | 27 | 37.1 | 37.1 | 24.8 | 137.92 |
| ARHGEF1 | 1263 | 27 | 27 | 27 | 33.8 | 33.8 | 33.8 | 108.32 |
| NONO | 3005 | 30 | 29 | 27 | 56.3 | 54.6 | 51.6 | 54.231 |
| EZR | 794 | 38 | 27 | 27 | 59 | 43.9 | 43.9 | 69.371 |
| RMDN3 | 3888 | 27 | 27 | 27 | 55.7 | 55.7 | 55.7 | 52.118 |
| DOPEY2 | 4607 | 27 | 27 | 27 | 16.7 | 16.7 | 16.7 | 258.23 |
| ASPH | 2816 | 27 | 27 | 27 | 43.4 | 43.4 | 43.4 | 85.862 |
| GALNT7 | 3387 | 27 | 27 | 27 | 52.2 | 52.2 | 52.2 | 75.388 |
| CKAP5 | 2919 | 27 | 27 | 27 | 17.9 | 17.9 | 17.9 | 225.49 |
| SMARCC2 | 958 | 27 | 27 | 27 | 27.3 | 27.3 | 27.3 | 136.18 |
| SLC25A24 | 3251 | 27 | 27 | 27 | 69.8 | 69.8 | 69.8 | 53.354 |
| GOT2 | 1696 | 26 | 26 | 26 | 70.5 | 70.5 | 70.5 | 47.517 |
| HNRNPA2B | 2059 | 27 | 27 | 26 | 71.7 | 71.7 | 71.7 | 37.429 |
| SRRM2 | 4533 | 26 | 26 | 26 | 14.1 | 14.1 | 14.1 | 299.61 |

| | | | | | | | | |
|---|---|---|---|---|---|---|---|---|
| NDUFV1 | 991 | 26 | 26 | 26 | 68.3 | 68.3 | 68.3 | 50.054 |
| EIF3L | 4541 | 26 | 26 | 26 | 56.6 | 56.6 | 56.6 | 66.726 |
| TBL2 | 833 | 26 | 26 | 26 | 66.7 | 66.7 | 66.7 | 45.935 |
| CTTN | 2941 | 26 | 26 | 26 | 51.8 | 51.8 | 51.8 | 61.585 |
| CES1 | 2064 | 26 | 26 | 26 | 63.1 | 63.1 | 63.1 | 62.52 |
| FGR | 1834 | 27 | 27 | 26 | 66.2 | 66.2 | 64.7 | 59.478 |
| CDK5RAP3 | 3823 | 26 | 26 | 26 | 50.4 | 50.4 | 50.4 | 56.92 |
| CAD | 946 | 26 | 26 | 26 | 19.6 | 19.6 | 19.6 | 236.02 |
| A2M | 1702 | 33 | 33 | 26 | 33.2 | 33.2 | 27.8 | 163.29 |
| ASAH1 | 298 | 26 | 26 | 26 | 58 | 58 | 58 | 44.659 |
| NCL | 2002 | 26 | 26 | 26 | 33.7 | 33.7 | 33.7 | 76.613 |
| ZYX | 3045 | 26 | 26 | 26 | 57.7 | 57.7 | 57.7 | 61.277 |
| HADHB | 2507 | 26 | 26 | 26 | 66.7 | 66.7 | 66.7 | 51.294 |
| STIM1 | 2897 | 27 | 27 | 26 | 52.7 | 52.7 | 50.4 | 77.422 |
| RPL4 | 2241 | 26 | 26 | 26 | 47.8 | 47.8 | 47.8 | 47.697 |
| ANKFY1 | 4396 | 26 | 26 | 26 | 33 | 33 | 33 | 128.4 |
| UQCRC1 | 2184 | 27 | 27 | 26 | 67.3 | 67.3 | 65.8 | 52.645 |
| FH | 1796 | 26 | 26 | 26 | 73.9 | 73.9 | 73.9 | 54.636 |
| PLCG2 | 131 | 26 | 26 | 26 | 27.5 | 27.5 | 27.5 | 146.06 |
| ACTR3 | 2568 | 26 | 26 | 26 | 67.5 | 67.5 | 67.5 | 47.371 |
| OAS3 | 4678 | 26 | 26 | 26 | 33.9 | 33.9 | 33.9 | 121.17 |
| RAB11FIP1 | 3302 | 26 | 26 | 26 | 28.2 | 28.2 | 28.2 | 137.17 |
| STXBP2 | 3040 | 33 | 33 | 26 | 67.3 | 67.3 | 55.5 | 66.452 |
| CUL2 | 163 | 26 | 26 | 26 | 44.3 | 44.3 | 44.3 | 88.505 |
| FAM129B | 3887 | 26 | 26 | 26 | 44 | 44 | 44 | 84.137 |
| ACADM | 1879 | 26 | 26 | 26 | 57.3 | 57.3 | 57.3 | 50.27 |
| XRN2 | 4090 | 26 | 26 | 26 | 45.5 | 45.5 | 45.5 | 108.58 |
| ABCF1 | 3564 | 25 | 25 | 25 | 37.6 | 37.6 | 37.6 | 95.925 |
| PDHA1 | 1803 | 25 | 25 | 25 | 65.1 | 65.1 | 65.1 | 43.295 |
| MTDH | 3401 | 25 | 25 | 25 | 50.9 | 50.9 | 50.9 | 63.836 |
| ADAM10 | 1348 | 25 | 25 | 25 | 46 | 46 | 46 | 84.141 |
| GCLC | 2348 | 25 | 25 | 25 | 51.5 | 51.5 | 51.5 | 72.765 |
| LY75 | 1489 | 25 | 25 | 25 | 18.4 | 18.4 | 18.4 | 198.31 |
| PA2G4 | 4534 | 25 | 25 | 25 | 73.9 | 73.9 | 73.9 | 43.786 |
| CCDC22 | 1512 | 25 | 25 | 25 | 50.7 | 50.7 | 50.7 | 70.755 |
| ATP6V1H | 4450 | 25 | 25 | 25 | 64.8 | 64.8 | 64.8 | 55.882 |
| AP2B1 | 2648 | 48 | 27 | 25 | 65.2 | 44.7 | 41.1 | 104.55 |
| GSR | 1689 | 25 | 25 | 25 | 71.3 | 71.3 | 71.3 | 56.256 |
| SRP72 | 1601 | 25 | 25 | 25 | 56 | 56 | 56 | 74.605 |
| GBF1 | 479 | 25 | 25 | 25 | 20.1 | 20.1 | 20.1 | 210.17 |
| PPP2R1A | 2167 | 25 | 25 | 25 | 50.9 | 50.9 | 50.9 | 65.308 |
| NLN | 4051 | 25 | 25 | 25 | 43.8 | 43.8 | 43.8 | 80.651 |
| ARHGEF2 | 469 | 25 | 25 | 25 | 36.8 | 36.8 | 36.8 | 116.07 |
| OAS2 | 95 | 25 | 25 | 25 | 43.1 | 43.1 | 43.1 | 83.234 |
| CCAR2 | 3485 | 25 | 25 | 25 | 34.9 | 34.9 | 34.9 | 102.9 |
| PMPCA | 2811 | 25 | 25 | 25 | 58.1 | 58.1 | 58.1 | 58.252 |
| SEPTIN2 | 2982 | 25 | 25 | 25 | 75.6 | 75.6 | 75.6 | 41.487 |
| TLR8 | 4246 | 25 | 25 | 25 | 28.9 | 28.9 | 28.9 | 119.83 |

| | | | | | | | | |
|---|---|---|---|---|---|---|---|---|
| ESYT2 | 441 | 25 | 25 | 25 | 40.6 | 40.6 | 40.6 | 93.977 |
| CYP1B1 | 3062 | 25 | 25 | 25 | 53.4 | 53.4 | 53.4 | 60.845 |
| UBA6 | 490 | 25 | 25 | 25 | 34.9 | 34.9 | 34.9 | 117.97 |
| CCDC47 | 3737 | 25 | 25 | 25 | 48.2 | 48.2 | 48.2 | 55.873 |
| KHSRP | 3711 | 26 | 26 | 25 | 44.6 | 44.6 | 43.5 | 73.114 |
| HSP90AB1 | 1800 | 42 | 28 | 25 | 62.7 | 44.3 | 40.1 | 83.263 |
| PADI2 | 4559 | 25 | 25 | 25 | 49.3 | 49.3 | 49.3 | 75.563 |
| TBC1D9B | 3229 | 25 | 25 | 25 | 27.3 | 27.3 | 27.3 | 140.52 |
| MCCC2 | 4196 | 25 | 25 | 25 | 59.1 | 59.1 | 59.1 | 61.332 |
| ALDH3A2 | 466 | 26 | 26 | 25 | 56.7 | 56.7 | 54.2 | 54.847 |
| ANXA1 | 1729 | 25 | 25 | 25 | 69.1 | 69.1 | 69.1 | 38.714 |
| FECH | 2062 | 24 | 24 | 24 | 70 | 70 | 70 | 47.862 |
| SEC24C | 2488 | 25 | 25 | 24 | 33.7 | 33.7 | 32.4 | 118.32 |
| PSAP | 1784 | 32 | 32 | 24 | 69.7 | 69.7 | 55 | 58.112 |
| UQCRC2 | 2060 | 24 | 24 | 24 | 74.4 | 74.4 | 74.4 | 48.442 |
| ACAA2 | 2291 | 24 | 24 | 24 | 79.1 | 79.1 | 79.1 | 41.924 |
| SEC23A | 3024 | 27 | 27 | 24 | 55.2 | 55.2 | 52.3 | 86.16 |
| PSMC4 | 2307 | 24 | 24 | 24 | 69.4 | 69.4 | 69.4 | 47.366 |
| ANXA11 | 2421 | 25 | 25 | 24 | 42.2 | 42.2 | 40.4 | 54.389 |
| ME1 | 2344 | 24 | 24 | 24 | 69.2 | 69.2 | 69.2 | 64.149 |
| CARS | 2373 | 24 | 24 | 24 | 40.4 | 40.4 | 40.4 | 85.472 |
| MOGS | 2905 | 28 | 28 | 24 | 52.3 | 52.3 | 45.4 | 91.916 |
| ALDH5A1 | 2441 | 24 | 24 | 24 | 59.4 | 59.4 | 59.4 | 57.214 |
| ATP11A | 831 | 25 | 25 | 24 | 31.4 | 31.4 | 30.6 | 106.51 |
| ACSF2 | 3766 | 24 | 24 | 24 | 55.1 | 55.1 | 55.1 | 68.124 |
| EPS15 | 2287 | 24 | 24 | 24 | 37.7 | 37.7 | 37.7 | 98.655 |
| NSDHL | 3034 | 24 | 24 | 24 | 76.9 | 76.9 | 76.9 | 41.9 |
| PNPT1 | 3609 | 24 | 24 | 24 | 36.7 | 36.7 | 36.7 | 85.95 |
| GNPAT | 1395 | 24 | 24 | 24 | 49.7 | 49.7 | 49.7 | 77.187 |
| NSUN2 | 2799 | 24 | 24 | 24 | 40.7 | 40.7 | 40.7 | 86.47 |
| GDI2 | 2405 | 34 | 34 | 24 | 84 | 84 | 62.2 | 50.663 |
| ACE | 1899 | 24 | 24 | 24 | 27.5 | 27.5 | 27.5 | 149.71 |
| PHB | 2215 | 24 | 24 | 24 | 86.4 | 86.4 | 86.4 | 29.804 |
| ACSF3 | 3113 | 24 | 24 | 24 | 51.6 | 51.6 | 51.6 | 64.129 |
| IDH1 | 1581 | 25 | 25 | 24 | 69.6 | 69.6 | 67.4 | 46.659 |
| RAB3GAP2 | 4119 | 24 | 24 | 24 | 22.9 | 22.9 | 22.9 | 155.98 |
| KYNU | 3066 | 24 | 24 | 24 | 68.4 | 68.4 | 68.4 | 52.351 |
| IDH2 | 2356 | 25 | 24 | 24 | 59.3 | 59.3 | 59.3 | 50.909 |
| SEC23B | 374 | 27 | 24 | 24 | 55.1 | 52.2 | 52.2 | 84.472 |
| DHCR24 | 162 | 24 | 24 | 24 | 50.8 | 50.8 | 50.8 | 61.446 |
| SRRT | 4033 | 24 | 24 | 24 | 34.4 | 34.4 | 34.4 | 100.67 |
| VAT1 | 3901 | 24 | 24 | 24 | 68.7 | 68.7 | 68.7 | 41.92 |
| ITGB5 | 1989 | 24 | 24 | 24 | 42.3 | 42.3 | 42.3 | 88.053 |
| ACOT9 | 4577 | 24 | 24 | 24 | 66.3 | 66.3 | 66.3 | 49.901 |
| FMNL2 | 3858 | 28 | 28 | 24 | 27.8 | 27.8 | 24.4 | 123.32 |
| CCDC88A | 378 | 24 | 24 | 24 | 14.2 | 14.2 | 14.2 | 216.04 |
| PPP1R12A | 1372 | 24 | 24 | 24 | 28.5 | 28.5 | 28.5 | 115.28 |
| DPYSL2 | 312 | 28 | 28 | 24 | 61.2 | 61.2 | 52.7 | 73.502 |

| | | | | | | | | |
|---|---|---|---|---|---|---|---|---|
| WFS1 | 475 | 23 | 23 | 23 | 32.3 | 32.3 | 32.3 | 101.54 |
| DDX17 | 333 | 32 | 23 | 23 | 52.4 | 42.4 | 42.4 | 80.253 |
| ITGA5 | 1809 | 23 | 23 | 23 | 32.6 | 32.6 | 32.6 | 114.54 |
| EIF5B | 34 | 23 | 23 | 23 | 24.6 | 24.6 | 24.6 | 138.68 |
| CRAT | 2298 | 23 | 23 | 23 | 44.1 | 44.1 | 44.1 | 70.857 |
| SLC27A3 | 3163 | 26 | 26 | 23 | 39 | 39 | 34.5 | 87.323 |
| MYO1E | 2827 | 29 | 23 | 23 | 32.9 | 26.8 | 26.8 | 127.06 |
| MOSPD2 | 3583 | 23 | 23 | 23 | 52.9 | 52.9 | 52.9 | 59.745 |
| PLAA | 4542 | 23 | 23 | 23 | 37.7 | 37.7 | 37.7 | 87.156 |
| SNTB1 | 2914 | 23 | 23 | 23 | 59.3 | 59.3 | 59.3 | 58.06 |
| PRPF19 | 4508 | 23 | 23 | 23 | 81 | 81 | 81 | 55.18 |
| PSMC3 | 4691 | 23 | 23 | 23 | 71.9 | 71.9 | 71.9 | 47.352 |
| PGM2 | 3795 | 23 | 23 | 23 | 47.2 | 47.2 | 47.2 | 68.283 |
| CLPX | 1598 | 23 | 23 | 23 | 47.1 | 47.1 | 47.1 | 69.223 |
| SPECC1 | 3165 | 23 | 23 | 23 | 26.4 | 26.4 | 26.4 | 118.58 |
| CPOX | 2240 | 23 | 23 | 23 | 63.4 | 63.4 | 63.4 | 50.151 |
| HEXB | 1785 | 23 | 23 | 23 | 46.8 | 46.8 | 46.8 | 63.111 |
| AARS | 2372 | 23 | 23 | 23 | 32.2 | 32.2 | 32.2 | 106.81 |
| ALDH9A1 | 2361 | 23 | 23 | 23 | 57.3 | 57.3 | 57.3 | 53.801 |
| FKBP5 | 2876 | 23 | 23 | 23 | 58.2 | 58.2 | 58.2 | 51.212 |
| STAT3 | 2265 | 23 | 23 | 23 | 43.6 | 43.6 | 43.6 | 88.067 |
| CSDE1 | 1561 | 23 | 23 | 23 | 36.3 | 36.3 | 36.3 | 88.884 |
| GAPDH | 1735 | 23 | 23 | 23 | 82.1 | 82.1 | 82.1 | 36.053 |
| MIPEP | 3923 | 23 | 23 | 23 | 40.1 | 40.1 | 40.1 | 80.64 |
| TNFAIP2 | 2755 | 23 | 23 | 23 | 44.6 | 44.6 | 44.6 | 72.66 |
| ACAT1 | 2088 | 23 | 23 | 23 | 78.5 | 78.5 | 78.5 | 45.199 |
| GPD1 | 2040 | 23 | 23 | 23 | 87.4 | 87.4 | 87.4 | 37.567 |
| PSMD11 | 1293 | 23 | 23 | 23 | 61.6 | 61.6 | 61.6 | 47.463 |
| CAST | 207 | 23 | 23 | 23 | 42.2 | 42.2 | 42.2 | 80.998 |
| STRN | 1469 | 24 | 24 | 23 | 42.7 | 42.7 | 41.8 | 86.131 |
| SUCLG2 | 3818 | 23 | 23 | 23 | 65.5 | 65.5 | 65.5 | 46.51 |
| ERMP1 | 3353 | 23 | 23 | 23 | 34.5 | 34.5 | 34.5 | 100.23 |
| FBP1 | 1823 | 26 | 26 | 23 | 76.9 | 76.9 | 71.3 | 36.842 |
| CTSD | 306 | 24 | 24 | 23 | 71.4 | 71.4 | 69.2 | 44.237 |
| DLAT | 1857 | 23 | 23 | 23 | 48.8 | 48.8 | 48.8 | 68.996 |
| EHHADH | 2793 | 23 | 23 | 23 | 40.2 | 40.2 | 40.2 | 79.494 |
| EXOC1 | 4291 | 23 | 23 | 23 | 37.6 | 37.6 | 37.6 | 101.98 |
| ERP44 | 3965 | 23 | 23 | 23 | 68 | 68 | 68 | 46.971 |
| NUP205 | 3688 | 23 | 23 | 23 | 14.2 | 14.2 | 14.2 | 227.92 |
| ALOX5 | 1837 | 23 | 23 | 23 | 43.3 | 43.3 | 43.3 | 77.982 |
| OPTN | 3771 | 23 | 23 | 23 | 44.7 | 44.7 | 44.7 | 65.921 |
| SEC23IP | 4688 | 23 | 23 | 23 | 33.3 | 33.3 | 33.3 | 111.08 |
| HNRNPM | 2453 | 43 | 43 | 23 | 71.6 | 71.6 | 36 | 77.515 |
| SLC25A13 | 4469 | 30 | 30 | 23 | 56.7 | 56.7 | 48.7 | 74.175 |
| UBE4A | 2927 | 23 | 23 | 23 | 33.5 | 33.5 | 33.5 | 122.56 |
| TRMT10C | 3331 | 22 | 22 | 22 | 63.5 | 63.5 | 63.5 | 47.346 |
| CYP51A1 | 188 | 22 | 22 | 22 | 56.2 | 56.2 | 56.2 | 57.278 |
| DHX30 | 1116 | 22 | 22 | 22 | 23.8 | 23.8 | 23.8 | 130.55 |

| | | | | | | | | |
|---|---|---|---|---|---|---|---|---|
| OGT | 1406 | 22 | 22 | 22 | 29.7 | 29.7 | 29.7 | 116.92 |
| AARS2 | 3158 | 22 | 22 | 22 | 34 | 34 | 34 | 107.34 |
| VPS33A | 3750 | 22 | 22 | 22 | 47.8 | 47.8 | 47.8 | 67.61 |
| RTCB | 4602 | 22 | 22 | 22 | 55.2 | 55.2 | 55.2 | 55.21 |
| VASP | 2414 | 22 | 22 | 22 | 51.3 | 51.3 | 51.3 | 39.829 |
| SUCLA2 | 373 | 22 | 22 | 22 | 58 | 58 | 58 | 43.841 |
| SARS2 | 1255 | 22 | 22 | 22 | 55.6 | 55.6 | 55.6 | 58.182 |
| PRAM1 | 3865 | 22 | 22 | 22 | 47.2 | 47.2 | 47.2 | 73.968 |
| SERBP1 | 3547 | 22 | 22 | 22 | 54.9 | 54.9 | 54.9 | 44.965 |
| SLC25A12 | 1572 | 29 | 22 | 22 | 56.6 | 48.7 | 48.7 | 74.761 |
| ABCD3 | 405 | 22 | 22 | 22 | 41.4 | 41.4 | 41.4 | 75.421 |
| TXNRD2 | 293 | 23 | 22 | 22 | 72.3 | 69.2 | 69.2 | 53.567 |
| PLOD1 | 2750 | 22 | 22 | 22 | 37 | 37 | 37 | 83.549 |
| CNP | 1829 | 22 | 22 | 22 | 57 | 57 | 57 | 47.578 |
| VDAC1 | 2043 | 22 | 22 | 22 | 84.5 | 84.5 | 84.5 | 30.772 |
| DLST | 2243 | 22 | 22 | 22 | 50.1 | 50.1 | 50.1 | 48.755 |
| KIAA1598 | 491 | 22 | 22 | 22 | 43.4 | 43.4 | 43.4 | 71.639 |
| MROH1 | 848 | 22 | 22 | 22 | 16.3 | 16.3 | 16.3 | 180.28 |
| PSMD12 | 1294 | 22 | 22 | 22 | 56.8 | 56.8 | 56.8 | 52.904 |
| NCEH1 | 3283 | 22 | 22 | 22 | 68.9 | 68.9 | 68.9 | 45.807 |
| SARS | 2374 | 22 | 22 | 22 | 48.7 | 48.7 | 48.7 | 61.312 |
| TARS2 | 4017 | 22 | 22 | 22 | 39.7 | 39.7 | 39.7 | 81.035 |
| MOV10 | 4199 | 22 | 22 | 22 | 31.2 | 31.2 | 31.2 | 113.67 |
| NOP56 | 1324 | 22 | 22 | 22 | 52.5 | 52.5 | 52.5 | 66.049 |
| ALDH7A1 | 295 | 22 | 22 | 22 | 51.1 | 51.1 | 51.1 | 58.139 |
| ALCAM | 2906 | 22 | 22 | 22 | 50.4 | 50.4 | 50.4 | 65.102 |
| PDIA6 | 2997 | 22 | 22 | 22 | 62.3 | 62.3 | 62.3 | 48.121 |
| RPL3 | 2257 | 22 | 22 | 22 | 56.1 | 56.1 | 56.1 | 46.108 |
| ABCE1 | 2573 | 22 | 22 | 22 | 45.9 | 45.9 | 45.9 | 67.314 |
| ALDOA | 1727 | 27 | 27 | 22 | 84.6 | 84.6 | 70.3 | 39.42 |
| PSMD13 | 4516 | 22 | 22 | 22 | 74.5 | 74.5 | 74.5 | 42.945 |
| PLBD1 | 3269 | 22 | 22 | 22 | 47.9 | 47.9 | 47.9 | 63.254 |
| DBT | 1873 | 22 | 22 | 22 | 55.2 | 55.2 | 55.2 | 53.486 |
| SNX5 | 4650 | 22 | 22 | 22 | 63.4 | 63.4 | 63.4 | 46.816 |
| DLD | 1831 | 22 | 22 | 22 | 67.4 | 67.4 | 67.4 | 54.177 |
| ILF2 | 2823 | 22 | 22 | 22 | 73.6 | 73.6 | 73.6 | 43.062 |
| RBM14 | 3854 | 22 | 22 | 22 | 39.3 | 39.3 | 39.3 | 69.491 |
| LRPAP1 | 2172 | 21 | 21 | 21 | 52.9 | 52.9 | 52.9 | 41.465 |
| OAT | 1731 | 21 | 21 | 21 | 73.6 | 73.6 | 73.6 | 48.534 |
| TBRG4 | 3736 | 21 | 21 | 21 | 53.2 | 53.2 | 53.2 | 70.737 |
| HNRNPA3 | 2451 | 23 | 23 | 21 | 51.3 | 51.3 | 51.3 | 39.594 |
| SDHB | 2044 | 21 | 21 | 21 | 65 | 65 | 65 | 31.629 |
| EIF2S1 | 1751 | 21 | 21 | 21 | 62.5 | 62.5 | 62.5 | 36.112 |
| EIF3D | 1409 | 21 | 21 | 21 | 54.6 | 54.6 | 54.6 | 63.972 |
| NDUFS2 | 1531 | 21 | 21 | 21 | 62.9 | 62.9 | 62.9 | 52.545 |
| KIF2A | 1279 | 21 | 21 | 21 | 35.4 | 35.4 | 35.4 | 79.954 |
| RPL6 | 2752 | 21 | 21 | 21 | 51 | 51 | 51 | 32.728 |
| IVD | 157 | 21 | 21 | 21 | 63.6 | 63.6 | 63.6 | 46.319 |

| | | | | | | | | |
|---|---|---|---|---|---|---|---|---|
| GPC4 | 1554 | 21 | 21 | 21 | 52.9 | 52.9 | 52.9 | 62.411 |
| THRAP3 | 407 | 22 | 22 | 21 | 27.5 | 27.5 | 27.5 | 103.52 |
| LARS2 | 2986 | 21 | 21 | 21 | 36.3 | 36.3 | 36.3 | 101.97 |
| EIF2S2 | 2015 | 21 | 21 | 21 | 55 | 55 | 55 | 38.388 |
| TOP1 | 1880 | 21 | 21 | 21 | 32.3 | 32.3 | 32.3 | 90.725 |
| NDUFA9 | 3071 | 21 | 21 | 21 | 59.9 | 59.9 | 59.9 | 42.509 |
| DOCK7 | 3845 | 25 | 23 | 21 | 16 | 15.1 | 14 | 242.56 |
| ATL1 | 3660 | 22 | 22 | 21 | 45.2 | 45.2 | 43.2 | 63.543 |
| RANGAP1 | 2315 | 21 | 21 | 21 | 48.2 | 48.2 | 48.2 | 63.541 |
| ARHGAP1 | 2785 | 21 | 21 | 21 | 74.7 | 74.7 | 74.7 | 50.435 |
| PCCB | 804 | 21 | 21 | 21 | 58.5 | 58.5 | 58.5 | 55.708 |
| RPS3 | 2072 | 22 | 22 | 21 | 84.8 | 84.8 | 81.1 | 26.688 |
| CDC5L | 3892 | 21 | 21 | 21 | 33.5 | 33.5 | 33.5 | 92.25 |
| HNRNPUL1 | 3991 | 21 | 21 | 21 | 29.9 | 29.9 | 29.9 | 95.737 |
| STT3B | 3608 | 22 | 22 | 21 | 24.2 | 24.2 | 23.4 | 93.673 |
| GBA | 1725 | 21 | 21 | 21 | 55.6 | 55.6 | 55.6 | 59.716 |
| GALNT12 | 3460 | 22 | 22 | 21 | 49.9 | 49.9 | 47.7 | 66.938 |
| RUVBL2 | 4540 | 21 | 21 | 21 | 52.1 | 52.1 | 52.1 | 51.156 |
| NUP98 | 2469 | 21 | 21 | 21 | 15.9 | 15.9 | 15.9 | 197.58 |
| MLYCD | 1672 | 22 | 22 | 21 | 53.8 | 53.8 | 52.3 | 55.003 |
| VPS45 | 4255 | 21 | 21 | 21 | 43 | 43 | 43 | 65.076 |
| AKAP13 | 2818 | 21 | 21 | 21 | 9.6 | 9.6 | 9.6 | 307.55 |
| PTCD3 | 3788 | 21 | 21 | 21 | 47 | 47 | 47 | 78.549 |
| CYP2S1 | 3881 | 21 | 21 | 21 | 61.1 | 61.1 | 61.1 | 55.816 |
| MRE11A | 2397 | 21 | 21 | 21 | 34.2 | 34.2 | 34.2 | 80.592 |
| AKR1B1 | 1938 | 21 | 21 | 21 | 67.1 | 67.1 | 67.1 | 35.853 |
| ENO1 | 1770 | 24 | 24 | 21 | 69.4 | 69.4 | 60.8 | 47.168 |
| DTX3L | 3616 | 21 | 21 | 21 | 40.8 | 40.8 | 40.8 | 83.553 |
| STXBP3 | 1288 | 21 | 21 | 21 | 42.2 | 42.2 | 42.2 | 67.764 |
| NCF1;NCF1 | 1928 | 21 | 21 | 21 | 66.4 | 66.4 | 66.4 | 44.681 |
| PCYOX1 | 4440 | 21 | 21 | 21 | 57.2 | 57.2 | 57.2 | 56.639 |
| ITGAL | 2029 | 21 | 21 | 21 | 25.6 | 25.6 | 25.6 | 128.77 |
| EVL | 4448 | 29 | 29 | 21 | 66.3 | 66.3 | 48.6 | 44.619 |
| ATP6V0A1 | 3719 | 30 | 30 | 21 | 42.2 | 42.2 | 27.2 | 96.412 |
| HNRNPR | 1436 | 30 | 30 | 21 | 53.1 | 53.1 | 42.2 | 70.942 |
| NARS | 1466 | 22 | 22 | 21 | 38.5 | 38.5 | 37 | 62.942 |
| FSCN1 | 3060 | 21 | 21 | 21 | 52.5 | 52.5 | 52.5 | 54.529 |
| MFN1 | 3445 | 21 | 21 | 21 | 35.6 | 35.6 | 35.6 | 84.159 |
| TWF2 | 3243 | 24 | 24 | 21 | 75.4 | 75.4 | 69.3 | 39.548 |
| SLC4A2 | 1742 | 22 | 22 | 21 | 25.4 | 25.4 | 24.5 | 137.01 |
| PHB2 | 3913 | 21 | 21 | 21 | 77.6 | 77.6 | 77.6 | 33.296 |
| ITGA3 | 2105 | 21 | 21 | 21 | 21.4 | 21.4 | 21.4 | 116.61 |
| NCBP1 | 2801 | 21 | 21 | 21 | 39.6 | 39.6 | 39.6 | 91.838 |
| HEXA | 1093 | 21 | 21 | 21 | 42.6 | 42.6 | 42.6 | 61.999 |
| CALU | 1474 | 21 | 21 | 21 | 67.3 | 67.3 | 67.3 | 37.106 |
| RPN2 | 1740 | 26 | 26 | 21 | 62 | 62 | 52.1 | 69.283 |
| SF3A3 | 2820 | 21 | 21 | 21 | 52.7 | 52.7 | 52.7 | 58.848 |
| SERPINH1 | 2411 | 21 | 21 | 21 | 60 | 60 | 60 | 46.44 |

| | | | | | | | | |
|---|---|---|---|---|---|---|---|---|
| CPNE3 | 1525 | 22 | 22 | 21 | 50.1 | 50.1 | 50.1 | 60.13 |
| EHD4 | 4107 | 25 | 25 | 21 | 60.4 | 60.4 | 52.9 | 61.174 |
| GRN | 2139 | 20 | 20 | 20 | 45.4 | 45.4 | 45.4 | 63.544 |
| TOM1 | 1510 | 20 | 20 | 20 | 55.5 | 55.5 | 55.5 | 53.818 |
| MTHFD1L | 47 | 21 | 20 | 20 | 32.4 | 31.3 | 31.3 | 99.31 |
| IDH3A | 2402 | 20 | 20 | 20 | 51.6 | 51.6 | 51.6 | 39.591 |
| EIF4A3 | 2255 | 25 | 20 | 20 | 59.4 | 53.3 | 53.3 | 46.871 |
| FARSB | 4264 | 20 | 20 | 20 | 43.5 | 43.5 | 43.5 | 66.115 |
| PPA2 | 4120 | 22 | 22 | 20 | 72.8 | 72.8 | 65.3 | 37.92 |
| USP14 | 2492 | 21 | 21 | 20 | 56.1 | 56.1 | 56.1 | 56.068 |
| RPA1 | 2125 | 20 | 20 | 20 | 46.1 | 46.1 | 46.1 | 68.137 |
| PSME2 | 108 | 20 | 20 | 20 | 71.3 | 71.3 | 71.3 | 29.126 |
| GBE1 | 2759 | 20 | 20 | 20 | 44.6 | 44.6 | 44.6 | 80.473 |
| RAB14 | 2567 | 20 | 20 | 20 | 86 | 86 | 86 | 23.897 |
| LRRC47 | 3491 | 20 | 20 | 20 | 45.5 | 45.5 | 45.5 | 63.472 |
| PSMC5 | 2598 | 20 | 20 | 20 | 58.4 | 58.4 | 58.4 | 45.626 |
| SYNCRIP | 1495 | 29 | 20 | 20 | 55.2 | 44.1 | 44.1 | 69.602 |
| APMAP | 4209 | 20 | 20 | 20 | 57.2 | 57.2 | 57.2 | 46.48 |
| BCLAF1 | 4345 | 21 | 20 | 20 | 25.2 | 24 | 24 | 106.12 |
| ILK | 161 | 20 | 20 | 20 | 50.9 | 50.9 | 50.9 | 54.611 |
| GAK | 1373 | 20 | 20 | 20 | 19.7 | 19.7 | 19.7 | 143.19 |
| SNX27 | 3837 | 20 | 20 | 20 | 47.9 | 47.9 | 47.9 | 61.264 |
| TIMM44 | 1449 | 20 | 20 | 20 | 47.8 | 47.8 | 47.8 | 51.355 |
| PDPR | 3553 | 20 | 20 | 20 | 28.4 | 28.4 | 28.4 | 99.363 |
| UGDH | 1502 | 20 | 20 | 20 | 58.7 | 58.7 | 58.7 | 55.023 |
| DNPEP | 769 | 20 | 20 | 20 | 60.1 | 60.1 | 60.1 | 51.998 |
| CYP20A1 | 3292 | 20 | 20 | 20 | 59.7 | 59.7 | 59.7 | 52.432 |
| DDX6 | 2107 | 20 | 20 | 20 | 59.4 | 59.4 | 59.4 | 54.416 |
| CYP4F22 | 3250 | 20 | 20 | 20 | 43.1 | 43.1 | 43.1 | 61.958 |
| PLD1 | 2865 | 20 | 20 | 20 | 23.9 | 23.9 | 23.9 | 124.18 |
| PEPD | 1901 | 20 | 20 | 20 | 53.5 | 53.5 | 53.5 | 54.548 |
| RFTN1 | 2967 | 20 | 20 | 20 | 59.2 | 59.2 | 59.2 | 63.145 |
| ILVBL | 495 | 20 | 20 | 20 | 42.6 | 42.6 | 42.6 | 67.867 |
| CS | 580 | 22 | 22 | 20 | 56.2 | 56.2 | 50.9 | 51.712 |
| FARSA | 4547 | 20 | 20 | 20 | 50.8 | 50.8 | 50.8 | 57.563 |
| SMARCA5 | 1483 | 28 | 28 | 20 | 32.3 | 32.3 | 24.5 | 121.9 |
| SCARB2 | 331 | 20 | 20 | 20 | 52.6 | 52.6 | 52.6 | 53.891 |
| SYK | 2304 | 20 | 20 | 20 | 44.6 | 44.6 | 44.6 | 72.065 |
| TCERG1 | 1354 | 20 | 20 | 20 | 20.5 | 20.5 | 20.5 | 123.9 |
| RBM25 | 2382 | 20 | 20 | 20 | 27 | 27 | 27 | 100.18 |
| EEF1G | 2117 | 20 | 20 | 20 | 65 | 65 | 65 | 50.118 |
| LMAN1 | 2363 | 20 | 20 | 20 | 56.1 | 56.1 | 56.1 | 57.548 |
| GPI | 1772 | 25 | 25 | 20 | 57.3 | 57.3 | 45.2 | 63.146 |
| PSMA1 | 2101 | 20 | 20 | 20 | 88.6 | 88.6 | 88.6 | 29.555 |
| UFSP2 | 4282 | 20 | 20 | 20 | 62.7 | 62.7 | 62.7 | 53.261 |
| PRKCSH | 1224 | 27 | 27 | 20 | 60 | 60 | 39.6 | 60.192 |
| CMAS | 3579 | 20 | 20 | 20 | 66.8 | 66.8 | 66.8 | 48.379 |
| ZFYVE16 | 3363 | 20 | 20 | 20 | 20 | 20 | 20 | 168.9 |

| | | | | | | | | |
|---|---|---|---|---|---|---|---|---|
| RETSAT | 3253 | 20 | 20 | 20 | 46.7 | 46.7 | 46.7 | 66.819 |
| SNX1 | 2899 | 22 | 22 | 20 | 46 | 46 | 42 | 59.069 |
| GNPTAB | 3101 | 20 | 20 | 20 | 20.6 | 20.6 | 20.6 | 143.62 |
| NBEAL2 | 438 | 20 | 20 | 20 | 11.8 | 11.8 | 11.8 | 298.4 |
| PGM1 | 2242 | 20 | 20 | 20 | 49.5 | 49.5 | 49.5 | 61.448 |
| SLC25A11 | 2754 | 20 | 20 | 20 | 71.7 | 71.7 | 71.7 | 34.061 |
| TMEM43 | 3982 | 20 | 20 | 20 | 56 | 56 | 56 | 44.875 |
| ZW10 | 1429 | 20 | 20 | 20 | 38.5 | 38.5 | 38.5 | 88.828 |
| GNA13 | 2949 | 21 | 21 | 20 | 59.4 | 59.4 | 57.3 | 44.049 |
| PTPN12 | 2768 | 20 | 20 | 20 | 34.7 | 34.7 | 34.7 | 88.105 |
| CCAR1 | 3457 | 20 | 20 | 20 | 23.2 | 23.2 | 23.2 | 132.82 |
| CORO1A | 2178 | 26 | 25 | 20 | 54.9 | 54.9 | 42.7 | 51.026 |
| KCNMA1 | 325 | 20 | 20 | 20 | 20.1 | 20.1 | 20.1 | 131.03 |
| LPCAT2 | 3342 | 20 | 20 | 20 | 45.2 | 45.2 | 45.2 | 60.207 |
| CYBB | 1738 | 20 | 20 | 20 | 42.6 | 42.6 | 42.6 | 65.335 |
| PSMC6 | 116 | 20 | 20 | 20 | 59.3 | 59.3 | 59.3 | 45.796 |
| AP1M1 | 4034 | 20 | 20 | 20 | 53.2 | 53.2 | 53.2 | 48.586 |
| TBC1D2 | 4053 | 20 | 20 | 20 | 25.8 | 25.8 | 25.8 | 105.41 |
| GART | 2052 | 20 | 20 | 20 | 26.4 | 26.4 | 26.4 | 107.77 |
| CORO1B | 3951 | 20 | 20 | 20 | 46.6 | 46.6 | 46.6 | 54.234 |
| PABPC1;PA | 1887 | 32 | 32 | 20 | 53.1 | 53.1 | 33 | 70.67 |
| ERO1L | 3807 | 26 | 26 | 20 | 69 | 69 | 56.4 | 54.392 |
| NRP2 | 1490 | 19 | 19 | 19 | 29.1 | 29.1 | 29.1 | 104.83 |
| ANKRD44 | 365 | 21 | 21 | 19 | 31.1 | 31.1 | 28.5 | 114.3 |
| TRAPPC10 | 2350 | 19 | 19 | 19 | 20.6 | 20.6 | 20.6 | 142.19 |
| BCAP31 | 2437 | 19 | 19 | 19 | 58.9 | 58.9 | 58.9 | 27.991 |
| AP2M1 | 69 | 19 | 19 | 19 | 48.4 | 48.4 | 48.4 | 49.526 |
| ETFDH | 3046 | 19 | 19 | 19 | 47.5 | 47.5 | 47.5 | 68.495 |
| ABHD12 | 3495 | 19 | 19 | 19 | 58.3 | 58.3 | 58.3 | 45.096 |
| PARP9 | 3463 | 19 | 19 | 19 | 29.3 | 29.3 | 29.3 | 96.342 |
| CYB5R1 | 4443 | 19 | 19 | 19 | 66.6 | 66.6 | 66.6 | 34.094 |
| SFXN3 | 4022 | 20 | 20 | 19 | 76 | 76 | 70.1 | 35.503 |
| ALDH6A1 | 2745 | 19 | 19 | 19 | 52.3 | 52.3 | 52.3 | 57.839 |
| BCL2L13 | 4031 | 19 | 19 | 19 | 60.4 | 60.4 | 60.4 | 52.723 |
| PMPCB | 1551 | 20 | 19 | 19 | 47.9 | 46.4 | 46.4 | 54.366 |
| ACBD3 | 4130 | 19 | 19 | 19 | 48.7 | 48.7 | 48.7 | 60.593 |
| DYNC1I2 | 2867 | 21 | 21 | 19 | 54.4 | 54.4 | 49.8 | 71.456 |
| CLCN7 | 2446 | 19 | 19 | 19 | 30.6 | 30.6 | 30.6 | 88.678 |
| CTSB | 1791 | 19 | 19 | 19 | 57.8 | 57.8 | 57.8 | 37.821 |
| RPS3A | 2576 | 19 | 19 | 19 | 61 | 61 | 61 | 29.945 |
| APOE | 1712 | 23 | 23 | 19 | 73.2 | 73.2 | 61.5 | 36.154 |
| EXOC2 | 3833 | 19 | 19 | 19 | 23.2 | 23.2 | 23.2 | 104.07 |
| DBNL | 4470 | 19 | 19 | 19 | 54.4 | 54.4 | 54.4 | 48.207 |
| MNDA | 2270 | 19 | 19 | 19 | 46.2 | 46.2 | 46.2 | 45.836 |
| EIF2A | 4040 | 19 | 19 | 19 | 52.1 | 52.1 | 52.1 | 64.989 |
| EIF3E | 2545 | 19 | 19 | 19 | 50.1 | 50.1 | 50.1 | 52.22 |
| NAMPT | 2306 | 19 | 19 | 19 | 51.9 | 51.9 | 51.9 | 55.52 |
| SLC27A4 | 3264 | 19 | 19 | 19 | 42.1 | 42.1 | 42.1 | 72.063 |

| | | | | | | | | |
|---|---|---|---|---|---|---|---|---|
| PSMC1 | 2597 | 19 | 19 | 19 | 62.3 | 62.3 | 62.3 | 49.184 |
| PTBP1 | 2114 | 22 | 22 | 19 | 72.7 | 72.7 | 68 | 57.221 |
| MAP1S | 3230 | 19 | 19 | 19 | 27.6 | 27.6 | 27.6 | 112.21 |
| LRRC8A | 3453 | 20 | 20 | 19 | 36.5 | 36.5 | 35.4 | 94.198 |
| SLC25A20 | 1465 | 19 | 19 | 19 | 75.1 | 75.1 | 75.1 | 32.943 |
| HCFC1 | 514 | 19 | 19 | 19 | 12.4 | 12.4 | 12.4 | 208.73 |
| MICU1 | 276 | 19 | 19 | 19 | 45 | 45 | 45 | 54.578 |
| ABR | 263 | 22 | 22 | 19 | 33.9 | 33.9 | 30.6 | 97.597 |
| SSB | 1759 | 19 | 19 | 19 | 47.8 | 47.8 | 47.8 | 46.836 |
| APEX1 | 2126 | 19 | 19 | 19 | 71.4 | 71.4 | 71.4 | 35.554 |
| GZMA | 1896 | 19 | 19 | 19 | 71.8 | 71.8 | 71.8 | 28.999 |
| TARS | 2115 | 23 | 23 | 19 | 38.2 | 38.2 | 33.5 | 83.434 |
| EIF3I | 2862 | 19 | 19 | 19 | 76.3 | 76.3 | 76.3 | 36.501 |
| CAPG | 2259 | 19 | 19 | 19 | 58.6 | 58.6 | 58.6 | 38.498 |
| NPLOC4 | 3593 | 19 | 19 | 19 | 47.9 | 47.9 | 47.9 | 68.119 |
| PPP1R18 | 241 | 19 | 19 | 19 | 40 | 40 | 40 | 67.942 |
| FUCA1 | 1726 | 19 | 19 | 19 | 50 | 50 | 50 | 53.688 |
| DDOST | 218 | 19 | 19 | 19 | 58.3 | 58.3 | 58.3 | 50.8 |
| DCTN2 | 2891 | 19 | 19 | 19 | 66.1 | 66.1 | 66.1 | 44.23 |
| NDUFA10 | 62 | 19 | 19 | 19 | 64.5 | 64.5 | 64.5 | 40.836 |
| AK4 | 2119 | 19 | 19 | 19 | 85.7 | 85.7 | 85.7 | 25.268 |
| ANXA7 | 2016 | 19 | 19 | 19 | 38.1 | 38.1 | 38.1 | 52.739 |
| MCU | 3565 | 19 | 19 | 19 | 47.3 | 47.3 | 47.3 | 39.866 |
| ERBB2IP | 3874 | 19 | 19 | 19 | 17.1 | 17.1 | 17.1 | 158.3 |
| AGL | 2222 | 19 | 19 | 19 | 17.7 | 17.7 | 17.7 | 174.76 |
| KMO | 1396 | 19 | 19 | 19 | 58.6 | 58.6 | 58.6 | 55.809 |
| POLR2A | 2090 | 19 | 19 | 19 | 13.3 | 13.3 | 13.3 | 217.17 |
| HSDL2 | 3306 | 19 | 19 | 19 | 58.6 | 58.6 | 58.6 | 45.394 |
| DNM2 | 2415 | 34 | 34 | 19 | 47.9 | 47.9 | 28.7 | 98.063 |
| ACSL5 | 4499 | 22 | 20 | 19 | 44.9 | 42.2 | 40.8 | 75.99 |
| PLEKHO2 | 3615 | 19 | 19 | 19 | 53.1 | 53.1 | 53.1 | 53.349 |
| CARS2 | 4174 | 18 | 18 | 18 | 38.1 | 38.1 | 38.1 | 62.223 |
| CLCC1 | 3878 | 18 | 18 | 18 | 44.1 | 44.1 | 44.1 | 62.022 |
| ELAVL1 | 3033 | 20 | 20 | 18 | 69.6 | 69.6 | 64.1 | 36.091 |
| DNAJC11 | 4297 | 18 | 18 | 18 | 41.1 | 41.1 | 41.1 | 63.277 |
| LDHD | 3415 | 18 | 18 | 18 | 61.9 | 61.9 | 61.9 | 54.87 |
| FDXR | 2057 | 18 | 18 | 18 | 52.5 | 52.5 | 52.5 | 53.836 |
| AK2 | 965 | 18 | 18 | 18 | 76.3 | 76.3 | 76.3 | 25.63 |
| DNAJC10 | 3458 | 18 | 18 | 18 | 27.6 | 27.6 | 27.6 | 91.079 |
| TMEM63A | 1613 | 18 | 18 | 18 | 23.2 | 23.2 | 23.2 | 92.125 |
| MTA2 | 1604 | 22 | 22 | 18 | 43.4 | 43.4 | 36.5 | 75.022 |
| AK3 | 4457 | 18 | 18 | 18 | 74.9 | 74.9 | 74.9 | 25.565 |
| PLD3 | 3437 | 18 | 18 | 18 | 49.4 | 49.4 | 49.4 | 54.705 |
| NOP58 | 4573 | 19 | 19 | 18 | 42.3 | 42.3 | 42.3 | 59.578 |
| TTLL12 | 2935 | 18 | 18 | 18 | 46.3 | 46.3 | 46.3 | 74.403 |
| PGAM1 | 1997 | 18 | 18 | 18 | 79.1 | 79.1 | 79.1 | 28.804 |
| SWAP70 | 4433 | 18 | 18 | 18 | 28.7 | 28.7 | 28.7 | 68.997 |
| NAGK | 4462 | 18 | 18 | 18 | 70.9 | 70.9 | 70.9 | 37.375 |

| | | | | | | | | |
|---|---|---|---|---|---|---|---|---|
| SRSF1 | 1192 | 18 | 18 | 18 | 54.9 | 54.9 | 54.9 | 28.329 |
| EPS15L1 | 1269 | 18 | 18 | 18 | 31.1 | 31.1 | 31.1 | 83.474 |
| VDAC2 | 2310 | 18 | 18 | 18 | 73.1 | 73.1 | 73.1 | 31.566 |
| ACAA1 | 1816 | 18 | 18 | 18 | 70.3 | 70.3 | 70.3 | 44.292 |
| ATP6V0D2 | 3529 | 18 | 18 | 18 | 64.3 | 64.3 | 64.3 | 40.426 |
| GALNT2 | 2806 | 19 | 19 | 18 | 44.1 | 44.1 | 42.4 | 64.732 |
| UNC13D | 3321 | 18 | 18 | 18 | 23.9 | 23.9 | 23.9 | 123.28 |
| SGPL1 | 1651 | 18 | 18 | 18 | 43.8 | 43.8 | 43.8 | 63.523 |
| MDH2 | 2267 | 18 | 18 | 18 | 65.7 | 65.7 | 65.7 | 35.503 |
| LIMS1 | 2340 | 18 | 18 | 18 | 66.2 | 66.2 | 66.2 | 37.251 |
| GALNS | 2210 | 18 | 18 | 18 | 46.9 | 46.9 | 46.9 | 58.025 |
| UBA2 | 4410 | 18 | 18 | 18 | 37.7 | 37.7 | 37.7 | 71.223 |
| POLR2B | 642 | 18 | 18 | 18 | 22.5 | 22.5 | 22.5 | 121.37 |
| GIMAP4 | 4288 | 18 | 18 | 18 | 58.4 | 58.4 | 58.4 | 37.533 |
| SLC27A1 | 3277 | 18 | 18 | 18 | 38.4 | 38.4 | 38.4 | 71.107 |
| CYB5R3 | 1688 | 18 | 18 | 18 | 71.8 | 71.8 | 71.8 | 34.234 |
| ATP5A1 | 2099 | 33 | 33 | 18 | 64.4 | 64.4 | 37.3 | 59.75 |
| CSE1L | 2504 | 18 | 18 | 18 | 25.3 | 25.3 | 25.3 | 110.42 |
| TPI1 | 2544 | 18 | 18 | 18 | 77.3 | 77.3 | 77.3 | 30.791 |
| GOLGA5 | 3598 | 18 | 18 | 18 | 31.2 | 31.2 | 31.2 | 83.023 |
| RAB7A | 2426 | 18 | 18 | 18 | 77.3 | 77.3 | 77.3 | 23.489 |
| ACBD5 | 610 | 18 | 18 | 18 | 36.5 | 36.5 | 36.5 | 58.792 |
| DNAJC3 | 2851 | 18 | 18 | 18 | 51 | 51 | 51 | 57.579 |
| ICAM1 | 1753 | 18 | 18 | 18 | 36.7 | 36.7 | 36.7 | 57.825 |
| PUF60 | 264 | 18 | 18 | 18 | 47.9 | 47.9 | 47.9 | 54.626 |
| CCT6A | 2261 | 24 | 24 | 18 | 55.7 | 55.7 | 42.7 | 58.024 |
| PSME1 | 2776 | 18 | 18 | 18 | 73.5 | 73.5 | 73.5 | 28.723 |
| DDX23 | 3995 | 18 | 18 | 18 | 28.9 | 28.9 | 28.9 | 95.581 |
| OXR1 | 3513 | 18 | 18 | 18 | 30.1 | 30.1 | 30.1 | 97.969 |
| PRDX6 | 2155 | 18 | 18 | 18 | 76.8 | 76.8 | 76.8 | 25.035 |
| RTN1 | 3072 | 18 | 18 | 18 | 34.5 | 34.5 | 34.5 | 83.617 |
| DPP3 | 4344 | 18 | 18 | 18 | 44.4 | 44.4 | 44.4 | 82.588 |
| DDX42 | 3424 | 18 | 18 | 18 | 28.1 | 28.1 | 28.1 | 102.97 |
| ETFB | 2250 | 18 | 18 | 18 | 72.5 | 72.5 | 72.5 | 27.843 |
| RBM47 | 488 | 18 | 18 | 18 | 43.2 | 43.2 | 43.2 | 64.098 |
| SAMM50 | 4627 | 18 | 18 | 18 | 44.3 | 44.3 | 44.3 | 51.976 |
| SNX2 | 1506 | 20 | 18 | 18 | 42.8 | 38.7 | 38.7 | 58.47 |
| ABCB7 | 460 | 18 | 18 | 18 | 34.3 | 34.3 | 34.3 | 83.27 |
| TOR4A | 4336 | 18 | 18 | 18 | 56.3 | 56.3 | 56.3 | 46.914 |
| CAP1 | 2734 | 30 | 30 | 18 | 65.1 | 65.1 | 35.6 | 51.901 |
| STT3A | 2333 | 19 | 18 | 18 | 24.5 | 23.5 | 23.5 | 80.529 |
| NUP107 | 2535 | 18 | 18 | 18 | 24.5 | 24.5 | 24.5 | 106.37 |
| RAB3GAP1 | 2988 | 18 | 18 | 18 | 24.7 | 24.7 | 24.7 | 110.52 |
| H2AFY | 1536 | 20 | 20 | 18 | 60.5 | 60.5 | 53.5 | 39.617 |
| UBA7 | 2273 | 18 | 18 | 18 | 32.8 | 32.8 | 32.8 | 111.69 |
| CSTF3 | 2830 | 18 | 18 | 18 | 33.3 | 33.3 | 33.3 | 82.921 |
| FLOT2 | 780 | 18 | 18 | 18 | 49.5 | 49.5 | 49.5 | 47.142 |
| ATP13A3 | 360 | 18 | 18 | 18 | 23 | 23 | 23 | 130.81 |

| | | | | | | | | |
|---|---|---|---|---|---|---|---|---|
| NCKAP1 | 4552 | 18 | 18 | 18 | 25.5 | 25.5 | 25.5 | 128.79 |
| RMDN2 | 3841 | 18 | 18 | 18 | 51.7 | 51.7 | 51.7 | 47.399 |
| RBM39 | 2954 | 18 | 18 | 18 | 47 | 47 | 47 | 59.379 |
| COG4 | 142 | 18 | 18 | 18 | 30.9 | 30.9 | 30.9 | 86.765 |
| EIF3H | 571 | 17 | 17 | 17 | 59.8 | 59.8 | 59.8 | 41.581 |
| PDS5A | 3081 | 18 | 18 | 17 | 20.4 | 20.4 | 19.4 | 150.83 |
| BASP1 | 2690 | 17 | 17 | 17 | 81.1 | 81.1 | 81.1 | 22.693 |
| RDH11 | 3605 | 17 | 17 | 17 | 70.1 | 70.1 | 70.1 | 35.386 |
| PBXIP1 | 3747 | 17 | 17 | 17 | 28.6 | 28.6 | 28.6 | 80.642 |
| ECHS1 | 2163 | 17 | 17 | 17 | 69.3 | 69.3 | 69.3 | 31.387 |
| PICALM | 2879 | 23 | 23 | 17 | 42.9 | 42.9 | 34.2 | 70.754 |
| EIF2S3;EIF2 | 2268 | 17 | 17 | 17 | 49.2 | 49.2 | 49.2 | 51.109 |
| ERLIN2 | 768 | 19 | 19 | 17 | 77 | 77 | 69.9 | 37.839 |
| DYSF | 1587 | 20 | 17 | 17 | 14.8 | 13.4 | 13.4 | 237.29 |
| COG7 | 2706 | 17 | 17 | 17 | 33.2 | 33.2 | 33.2 | 86.343 |
| SRC | 1900 | 24 | 22 | 17 | 51.3 | 48.5 | 40.5 | 59.834 |
| ACAP2 | 104 | 17 | 17 | 17 | 27.4 | 27.4 | 27.4 | 88.02 |
| COMT | 2046 | 17 | 17 | 17 | 71.2 | 71.2 | 71.2 | 30.037 |
| NDUFS3 | 1555 | 17 | 17 | 17 | 69.3 | 69.3 | 69.3 | 30.241 |
| MPP1 | 2720 | 17 | 17 | 17 | 56.7 | 56.7 | 56.7 | 52.296 |
| GNS | 1947 | 17 | 17 | 17 | 36.4 | 36.4 | 36.4 | 62.081 |
| HSD17B10 | 3917 | 17 | 17 | 17 | 90.4 | 90.4 | 90.4 | 26.923 |
| PSMB2 | 2378 | 17 | 17 | 17 | 87.1 | 87.1 | 87.1 | 22.836 |
| AHCY | 2074 | 17 | 17 | 17 | 52.1 | 52.1 | 52.1 | 47.716 |
| BCKDK | 1360 | 17 | 17 | 17 | 65.5 | 65.5 | 65.5 | 46.36 |
| QRSL1 | 4094 | 17 | 17 | 17 | 47 | 47 | 47 | 57.46 |
| PTGES2 | 4157 | 17 | 17 | 17 | 56.2 | 56.2 | 56.2 | 41.943 |
| MYBBP1A | 3947 | 17 | 17 | 17 | 15.2 | 15.2 | 15.2 | 148.85 |
| HCLS1 | 810 | 17 | 17 | 17 | 45.4 | 45.4 | 45.4 | 49.704 |
| CNOT1 | 508 | 17 | 17 | 17 | 9.6 | 9.6 | 9.6 | 266.94 |
| PDXK | 1333 | 17 | 17 | 17 | 75 | 75 | 75 | 35.102 |
| SUPV3L1 | 3468 | 17 | 17 | 17 | 28.6 | 28.6 | 28.6 | 87.99 |
| CPSF1 | 2809 | 17 | 17 | 17 | 15 | 15 | 15 | 160.88 |
| U2AF2 | 2109 | 17 | 17 | 17 | 50.9 | 50.9 | 50.9 | 53.5 |
| ABHD10 | 4275 | 17 | 17 | 17 | 68.3 | 68.3 | 68.3 | 33.932 |
| SPG7 | 347 | 17 | 17 | 17 | 33 | 33 | 33 | 88.234 |
| RHOT2 | 3459 | 18 | 18 | 17 | 48.1 | 48.1 | 46.3 | 68.117 |
| SEC24B | 1655 | 19 | 19 | 17 | 21.1 | 21.1 | 19.9 | 137.42 |
| SH3BP1 | 4603 | 17 | 17 | 17 | 35.5 | 35.5 | 35.5 | 75.712 |
| G6PD | 1881 | 47 | 47 | 17 | 87.8 | 87.8 | 31.7 | 59.256 |
| SEC24D | 1611 | 18 | 17 | 17 | 26.8 | 25.4 | 25.4 | 113.01 |
| ALDH16A1 | 3476 | 17 | 17 | 17 | 31 | 31 | 31 | 85.126 |
| ALDH1B1 | 2176 | 19 | 17 | 17 | 50.3 | 48.5 | 48.5 | 57.206 |
| HARS2 | 354 | 23 | 23 | 17 | 51.2 | 51.2 | 39.8 | 57.531 |
| GPNMB | 2975 | 17 | 17 | 17 | 36 | 36 | 36 | 63.922 |
| TROVE2 | 1851 | 17 | 17 | 17 | 37.7 | 37.7 | 37.7 | 60.67 |
| PREP | 2343 | 17 | 17 | 17 | 35.4 | 35.4 | 35.4 | 80.699 |
| RUVBL1 | 4543 | 17 | 17 | 17 | 53.7 | 53.7 | 53.7 | 50.227 |

| | | | | | | | | |
|---|---|---|---|---|---|---|---|---|
| ACADS | 1957 | 17 | 17 | 17 | 58.5 | 58.5 | 58.5 | 44.297 |
| SNX6 | 4514 | 17 | 17 | 17 | 51.2 | 51.2 | 51.2 | 46.648 |
| PLBD2 | 3584 | 17 | 17 | 17 | 34.8 | 34.8 | 34.8 | 65.471 |
| SON | 1996 | 17 | 17 | 17 | 9.9 | 9.9 | 9.9 | 263.83 |
| HNRNPL | 1933 | 27 | 27 | 17 | 77.8 | 77.8 | 54 | 64.132 |
| PSTPIP1 | 1167 | 17 | 17 | 17 | 40.4 | 40.4 | 40.4 | 47.332 |
| ACADSB | 2311 | 17 | 17 | 17 | 57.6 | 57.6 | 57.6 | 47.485 |
| CCDC6 | 3049 | 17 | 17 | 17 | 38 | 38 | 38 | 53.29 |
| ATP6V1E1 | 2239 | 17 | 17 | 17 | 73.9 | 73.9 | 73.9 | 26.145 |
| PLIN3 | 1500 | 17 | 17 | 17 | 54.6 | 54.6 | 54.6 | 47.074 |
| FLOT1 | 1593 | 17 | 17 | 17 | 50.8 | 50.8 | 50.8 | 47.355 |
| PRKAR2A | 1920 | 20 | 20 | 17 | 58.7 | 58.7 | 51.5 | 45.518 |
| G3BP2 | 4511 | 18 | 18 | 17 | 45 | 45 | 43.4 | 54.12 |
| VWA5A | 1321 | 17 | 17 | 17 | 29.5 | 29.5 | 29.5 | 86.488 |
| ARHGAP17 | 3235 | 17 | 17 | 17 | 27.6 | 27.6 | 27.6 | 95.436 |
| SLTM | 4314 | 17 | 17 | 17 | 21.3 | 21.3 | 21.3 | 117.15 |
| SERPINB6 | 0 | 18 | 18 | 17 | 61.6 | 61.6 | 59.5 | 43.024 |
| OAS1 | 1701 | 17 | 17 | 17 | 49.5 | 49.5 | 49.5 | 46.028 |
| LMAN2 | 2825 | 17 | 17 | 17 | 57.3 | 57.3 | 57.3 | 40.228 |
| ACTR2 | 2569 | 17 | 17 | 17 | 49 | 49 | 49 | 44.76 |
| CAPRIN1 | 2952 | 17 | 17 | 17 | 23.4 | 23.4 | 23.4 | 78.365 |
| PLCD1 | 2430 | 17 | 17 | 17 | 36.1 | 36.1 | 36.1 | 85.664 |
| PSMD6 | 2980 | 17 | 17 | 17 | 50.6 | 50.6 | 50.6 | 45.531 |
| LASP1 | 2973 | 17 | 17 | 17 | 64.8 | 64.8 | 64.8 | 29.717 |
| FUBP1 | 3742 | 19 | 18 | 17 | 37.3 | 36 | 34.6 | 67.56 |
| STX11 | 1562 | 17 | 17 | 17 | 70.4 | 70.4 | 70.4 | 33.195 |
| HNRNPA1;I | 968 | 20 | 17 | 17 | 56 | 56 | 56 | 33.155 |
| TMEM214 | 3255 | 17 | 17 | 17 | 29.8 | 29.8 | 29.8 | 77.15 |
| HTATSF1 | 1460 | 17 | 17 | 17 | 27.8 | 27.8 | 27.8 | 85.852 |
| DYNC1LI1 | 4672 | 19 | 19 | 17 | 52 | 52 | 46.7 | 56.578 |
| GLA | 1765 | 17 | 17 | 17 | 50.6 | 50.6 | 50.6 | 48.766 |
| IDH3B | 83 | 17 | 17 | 17 | 54.5 | 54.5 | 54.5 | 42.41 |
| PDHB | 1872 | 17 | 17 | 17 | 69.1 | 69.1 | 69.1 | 39.233 |
| SQSTM1 | 2881 | 16 | 16 | 16 | 63.6 | 63.6 | 63.6 | 47.687 |
| EIF4B | 816 | 16 | 16 | 16 | 29.1 | 29.1 | 29.1 | 69.697 |
| HNRNPF | 2461 | 18 | 16 | 16 | 67.7 | 61.2 | 61.2 | 45.671 |
| AIFM2 | 3958 | 16 | 16 | 16 | 64.9 | 64.9 | 64.9 | 40.526 |
| YARS2 | 4575 | 16 | 16 | 16 | 54.5 | 54.5 | 54.5 | 53.198 |
| APLP2 | 2777 | 17 | 17 | 16 | 27.7 | 27.7 | 26.6 | 86.955 |
| C21orf33 | 186 | 16 | 16 | 16 | 71.6 | 71.6 | 71.6 | 28.17 |
| MRPS5 | 2695 | 16 | 16 | 16 | 37.2 | 37.2 | 37.2 | 48.006 |
| STAU1 | 99 | 16 | 16 | 16 | 41.6 | 41.6 | 41.6 | 54.945 |
| PLA2G4A | 2334 | 16 | 16 | 16 | 33.4 | 33.4 | 33.4 | 85.238 |
| SERPINB1 | 2175 | 17 | 17 | 16 | 57.3 | 57.3 | 54.6 | 42.741 |
| PIK3R4 | 3904 | 16 | 16 | 16 | 15.9 | 15.9 | 15.9 | 153.1 |
| TALDO1 | 2249 | 16 | 16 | 16 | 53.7 | 53.7 | 53.7 | 37.54 |
| COPE | 1343 | 16 | 16 | 16 | 65.3 | 65.3 | 65.3 | 34.482 |
| ATM | 2860 | 16 | 16 | 16 | 6.4 | 6.4 | 6.4 | 350.68 |

| | | | | | | | | |
|---|---|---|---|---|---|---|---|---|
| CLYBL | 3480 | 16 | 16 | 16 | 59.1 | 59.1 | 59.1 | 37.359 |
| SRP54 | 2560 | 16 | 16 | 16 | 42.9 | 42.9 | 42.9 | 55.704 |
| LYN | 1794 | 19 | 16 | 16 | 50.8 | 45.9 | 45.9 | 58.573 |
| RPS9 | 2327 | 16 | 16 | 16 | 52.1 | 52.1 | 52.1 | 22.591 |
| EXOC7 | 146 | 16 | 16 | 16 | 30.2 | 30.2 | 30.2 | 78.808 |
| NLRX1 | 3404 | 16 | 16 | 16 | 25.2 | 25.2 | 25.2 | 107.61 |
| HP1BP3 | 3176 | 16 | 16 | 16 | 31.3 | 31.3 | 31.3 | 61.206 |
| EIF3M | 3336 | 16 | 16 | 16 | 53.5 | 53.5 | 53.5 | 42.502 |
| RCN1 | 3010 | 16 | 16 | 16 | 59.2 | 59.2 | 59.2 | 38.89 |
| PLEK | 1804 | 16 | 16 | 16 | 54.9 | 54.9 | 54.9 | 40.124 |
| GBP2 | 2199 | 23 | 18 | 16 | 42.3 | 38.4 | 34 | 67.208 |
| ARMCX3 | 4432 | 16 | 16 | 16 | 55.1 | 55.1 | 55.1 | 42.5 |
| ADPGK | 3959 | 16 | 16 | 16 | 55.9 | 55.9 | 55.9 | 54.088 |
| APEH | 675 | 16 | 16 | 16 | 32.8 | 32.8 | 32.8 | 81.673 |
| RPL5 | 2325 | 16 | 16 | 16 | 49.2 | 49.2 | 49.2 | 34.362 |
| PAFAH1B1 | 2297 | 16 | 16 | 16 | 56.8 | 56.8 | 56.8 | 46.637 |
| COQ5 | 3149 | 16 | 16 | 16 | 62.1 | 62.1 | 62.1 | 37.14 |
| PPP1R7 | 3023 | 16 | 16 | 16 | 63.9 | 63.9 | 63.9 | 41.564 |
| CTSS | 2100 | 16 | 16 | 16 | 69.5 | 69.5 | 69.5 | 37.495 |
| EIF5 | 2502 | 16 | 16 | 16 | 37.8 | 37.8 | 37.8 | 49.222 |
| DES | 1980 | 21 | 16 | 16 | 49.6 | 43.4 | 43.4 | 53.535 |
| DHRS9 | 3935 | 16 | 16 | 16 | 60.2 | 60.2 | 60.2 | 35.227 |
| STK11IP | 3489 | 16 | 16 | 16 | 20.9 | 20.9 | 20.9 | 120.26 |
| CLU | 1865 | 16 | 16 | 16 | 36.3 | 36.3 | 36.3 | 52.494 |
| VAPA | 4377 | 17 | 17 | 16 | 65.5 | 65.5 | 60.6 | 27.893 |
| TOR1AIP2 | 3576 | 17 | 16 | 16 | 55.5 | 53.4 | 53.4 | 51.263 |
| TSFM | 2308 | 16 | 16 | 16 | 67.4 | 67.4 | 67.4 | 35.39 |
| TRIM54 | 4052 | 16 | 16 | 16 | 58.4 | 58.4 | 58.4 | 40.3 |
| HMOX2 | 17 | 16 | 16 | 16 | 51.6 | 51.6 | 51.6 | 41.669 |
| PNN | 4123 | 16 | 16 | 16 | 27.9 | 27.9 | 27.9 | 81.627 |
| ATP6V1D | 4639 | 16 | 16 | 16 | 57.5 | 57.5 | 57.5 | 28.262 |
| PNP | 1694 | 16 | 16 | 16 | 79.6 | 79.6 | 79.6 | 32.118 |
| GSTO1 | 2684 | 16 | 16 | 16 | 59.3 | 59.3 | 59.3 | 27.566 |
| TAGLN2 | 2248 | 16 | 16 | 16 | 89.9 | 89.9 | 89.9 | 22.391 |
| DHRS7 | 4586 | 16 | 16 | 16 | 56.9 | 56.9 | 56.9 | 38.298 |
| HGS | 1369 | 16 | 16 | 16 | 22.1 | 22.1 | 22.1 | 86.191 |
| INPP5D | 3700 | 17 | 17 | 16 | 22.9 | 22.9 | 22 | 133.29 |
| ACSS1 | 4273 | 16 | 16 | 16 | 36 | 36 | 36 | 74.856 |
| NUP133 | 3641 | 16 | 16 | 16 | 23.8 | 23.8 | 23.8 | 128.98 |
| MMAA | 400 | 16 | 16 | 16 | 46.7 | 46.7 | 46.7 | 44.22 |
| SART1 | 1431 | 16 | 16 | 16 | 27 | 27 | 27 | 90.254 |
| PDLIM5 | 3805 | 16 | 16 | 16 | 32.4 | 32.4 | 32.4 | 63.944 |
| OSBP | 2049 | 16 | 16 | 16 | 26.1 | 26.1 | 26.1 | 89.42 |
| THOC2 | 3587 | 16 | 16 | 16 | 12.6 | 12.6 | 12.6 | 182.77 |
| DERA | 4578 | 16 | 16 | 16 | 65.4 | 65.4 | 65.4 | 35.23 |
| PAK2 | 2847 | 25 | 25 | 16 | 56.9 | 56.9 | 41.4 | 58.042 |
| CPSF3 | 4481 | 16 | 16 | 16 | 36.8 | 36.8 | 36.8 | 77.485 |
| MGLL | 3916 | 16 | 16 | 16 | 69.6 | 69.6 | 69.6 | 33.261 |

| | | | | | | | | |
|---|---|---|---|---|---|---|---|---|
| TXNDC5 | 3542 | 16 | 16 | 16 | 50 | 50 | 50 | 47.628 |
| CDC37 | 3055 | 16 | 16 | 16 | 37 | 37 | 37 | 44.468 |
| ATP6V0D1 | 903 | 16 | 16 | 16 | 46.4 | 46.4 | 46.4 | 44.659 |
| ACAT2 | 4018 | 16 | 16 | 16 | 59.4 | 59.4 | 59.4 | 41.35 |
| CCBL2 | 3307 | 16 | 16 | 16 | 46.5 | 46.5 | 46.5 | 51.4 |
| ARSB | 1948 | 16 | 16 | 16 | 37 | 37 | 37 | 59.687 |
| PSMD5 | 3050 | 16 | 16 | 16 | 43.7 | 43.7 | 43.7 | 56.195 |
| ABCC10 | 3190 | 16 | 16 | 16 | 14.5 | 14.5 | 14.5 | 161.63 |
| ELMO1 | 3674 | 20 | 20 | 16 | 37.4 | 37.4 | 33 | 83.829 |
| ARHGDIB | 2459 | 16 | 16 | 16 | 74.6 | 74.6 | 74.6 | 22.988 |
| FCHO2 | 2803 | 16 | 16 | 16 | 27.2 | 27.2 | 27.2 | 88.923 |
| CLIC4 | 4664 | 17 | 17 | 16 | 76.7 | 76.7 | 73.5 | 28.772 |
| SCCPDH | 3544 | 16 | 16 | 16 | 61.1 | 61.1 | 61.1 | 47.151 |
| DPP7 | 4442 | 16 | 16 | 16 | 43.5 | 43.5 | 43.5 | 54.341 |
| SOD2 | 1730 | 16 | 16 | 16 | 86.9 | 86.9 | 86.9 | 24.75 |
| ZMPSTE24 | 1579 | 16 | 16 | 16 | 39.6 | 39.6 | 39.6 | 54.812 |
| IPO7 | 1644 | 16 | 16 | 16 | 22.2 | 22.2 | 22.2 | 119.52 |
| SLFN5 | 2797 | 16 | 16 | 16 | 25.7 | 25.7 | 25.7 | 101.05 |
| YWHAE | 2602 | 19 | 19 | 16 | 69 | 69 | 62 | 29.174 |
| NFS1 | 4665 | 16 | 16 | 16 | 44 | 44 | 44 | 50.195 |
| SCPEP1 | 4182 | 16 | 16 | 16 | 38.9 | 38.9 | 38.9 | 50.83 |
| EXOC5 | 149 | 16 | 16 | 16 | 28 | 28 | 28 | 82.118 |
| P4HA1 | 1913 | 16 | 16 | 16 | 38.2 | 38.2 | 38.2 | 61.049 |
| TMX3 | 3827 | 16 | 16 | 16 | 38.5 | 38.5 | 38.5 | 51.871 |
| ATP2C1 | 595 | 16 | 16 | 16 | 23.9 | 23.9 | 23.9 | 104.71 |
| DKC1 | 1513 | 16 | 16 | 16 | 39.5 | 39.5 | 39.5 | 57.673 |
| CPD | 1594 | 15 | 15 | 15 | 14.6 | 14.6 | 14.6 | 152.93 |
| GRSF1 | 925 | 15 | 15 | 15 | 51.1 | 51.1 | 51.1 | 47.27 |
| SNRNP70 | 1808 | 15 | 15 | 15 | 37.8 | 37.8 | 37.8 | 51.556 |
| ACAD8 | 4489 | 15 | 15 | 15 | 47.5 | 47.5 | 47.5 | 45.069 |
| MANBA | 1314 | 16 | 16 | 15 | 20.4 | 20.4 | 19.6 | 100.89 |
| POLDIP2 | 4568 | 15 | 15 | 15 | 58.2 | 58.2 | 58.2 | 42.033 |
| HACL1 | 4464 | 15 | 15 | 15 | 37.9 | 37.9 | 37.9 | 63.728 |
| SKIV2L2 | 2283 | 15 | 15 | 15 | 19.1 | 19.1 | 19.1 | 117.8 |
| SUCLG1 | 2477 | 15 | 15 | 15 | 41.6 | 41.6 | 41.6 | 36.249 |
| STRAP | 4601 | 15 | 15 | 15 | 56.3 | 56.3 | 56.3 | 38.438 |
| SUPT5H | 1298 | 15 | 15 | 15 | 19.3 | 19.3 | 19.3 | 121 |
| TM9SF4 | 190 | 15 | 15 | 15 | 28.2 | 28.2 | 28.2 | 72.54 |
| GNAI2 | 1741 | 22 | 22 | 15 | 67.9 | 67.9 | 46.5 | 40.45 |
| SLC25A35 | 3097 | 15 | 15 | 15 | 74.7 | 74.7 | 74.7 | 32.437 |
| PSMA7 | 1357 | 15 | 15 | 15 | 65.3 | 65.3 | 65.3 | 27.887 |
| EFHD2 | 3763 | 15 | 15 | 15 | 45.8 | 45.8 | 45.8 | 26.697 |
| FAM49B | 4284 | 17 | 17 | 15 | 68.5 | 68.5 | 61.7 | 36.748 |
| LDHA | 1685 | 16 | 16 | 15 | 60.2 | 60.2 | 56.6 | 36.688 |
| GBP1 | 2198 | 23 | 23 | 15 | 47.6 | 47.6 | 30.2 | 67.93 |
| FAF2 | 3769 | 15 | 15 | 15 | 43.4 | 43.4 | 43.4 | 52.623 |
| DAP3 | 2432 | 15 | 15 | 15 | 49 | 49 | 49 | 45.566 |
| CECR1 | 596 | 15 | 15 | 15 | 40.7 | 40.7 | 40.7 | 54.36 |

| | | | | | | | | |
|---|---|---|---|---|---|---|---|---|
| IDE | 391 | 15 | 15 | 15 | 20.5 | 20.5 | 20.5 | 117.97 |
| CSNK2A2 | 2011 | 15 | 15 | 15 | 50 | 50 | 50 | 41.213 |
| RPL8 | 2640 | 15 | 15 | 15 | 49 | 49 | 49 | 28.024 |
| SAFB | 3020 | 23 | 23 | 15 | 30.7 | 30.7 | 19 | 102.64 |
| CLIC1 | 1300 | 15 | 15 | 15 | 80.9 | 80.9 | 80.9 | 26.922 |
| RPS8 | 2599 | 15 | 15 | 15 | 59.1 | 59.1 | 59.1 | 24.205 |
| SEC13 | 2513 | 15 | 15 | 15 | 62.4 | 62.4 | 62.4 | 35.54 |
| CBR1 | 1956 | 17 | 17 | 15 | 75.1 | 75.1 | 64.6 | 30.375 |
| FASTKD5 | 3345 | 15 | 15 | 15 | 28.9 | 28.9 | 28.9 | 86.573 |
| CHCHD3 | 689 | 15 | 15 | 15 | 55.6 | 55.6 | 55.6 | 26.753 |
| PRKAA1 | 2840 | 15 | 15 | 15 | 34 | 34 | 34 | 64.009 |
| PRMT1 | 3929 | 15 | 15 | 15 | 48.8 | 48.8 | 48.8 | 42.461 |
| PRPF40A | 1545 | 15 | 15 | 15 | 24.6 | 24.6 | 24.6 | 108.8 |
| IL4I1 | 3873 | 15 | 15 | 15 | 31.6 | 31.6 | 31.6 | 62.88 |
| BTK | 2774 | 15 | 15 | 15 | 29.3 | 29.3 | 29.3 | 76.28 |
| CHID1 | 4023 | 15 | 15 | 15 | 55.2 | 55.2 | 55.2 | 44.94 |
| MSR1 | 2042 | 15 | 15 | 15 | 44.6 | 44.6 | 44.6 | 49.762 |
| STAT5A;STA | 1214 | 15 | 15 | 15 | 23.5 | 23.5 | 23.5 | 87.361 |
| ACP2 | 1867 | 15 | 15 | 15 | 38.5 | 38.5 | 38.5 | 48.344 |
| HSPA1B;HS | 244 | 32 | 29 | 15 | 68.4 | 64.2 | 40.3 | 70.108 |
| COG2 | 2969 | 15 | 15 | 15 | 29.1 | 29.1 | 29.1 | 83.207 |
| PTGS1 | 2065 | 15 | 15 | 15 | 37.9 | 37.9 | 37.9 | 68.686 |
| MRPS22 | 1022 | 15 | 15 | 15 | 54.9 | 54.9 | 54.9 | 36.805 |
| TAOK3 | 4118 | 18 | 18 | 15 | 27.3 | 27.3 | 24.8 | 105.4 |
| ATP2A3 | 3722 | 25 | 17 | 15 | 29.4 | 21.2 | 19.6 | 113.98 |
| GRHPR | 4407 | 15 | 15 | 15 | 53 | 53 | 53 | 35.668 |
| GSTK1 | 4563 | 15 | 15 | 15 | 77 | 77 | 77 | 25.497 |
| LEMD2 | 3548 | 15 | 15 | 15 | 30.8 | 30.8 | 30.8 | 56.974 |
| ADAM17 | 2686 | 15 | 15 | 15 | 27.9 | 27.9 | 27.9 | 93.02 |
| PGAM5 | 3808 | 15 | 15 | 15 | 63 | 63 | 63 | 32.004 |
| ATP9B | 1475 | 19 | 19 | 15 | 25 | 25 | 20.6 | 129.3 |
| CSK | 2275 | 16 | 16 | 15 | 46.2 | 46.2 | 44 | 50.704 |
| ADK | 2510 | 15 | 15 | 15 | 53.3 | 53.3 | 53.3 | 40.545 |
| PREB | 4203 | 15 | 15 | 15 | 61.6 | 61.6 | 61.6 | 45.468 |
| SEL1L | 4411 | 15 | 15 | 15 | 32.4 | 32.4 | 32.4 | 88.754 |
| RPS2 | 1949 | 15 | 15 | 15 | 49.1 | 49.1 | 49.1 | 31.324 |
| PSPC1 | 3659 | 17 | 15 | 15 | 41.1 | 38.4 | 38.4 | 58.743 |
| MDH1 | 2266 | 15 | 15 | 15 | 59.3 | 59.3 | 59.3 | 36.426 |
| KIAA0930 | 3245 | 15 | 15 | 15 | 44.8 | 44.8 | 44.8 | 45.794 |
| RPS5 | 1265 | 15 | 15 | 15 | 72.5 | 72.5 | 72.5 | 22.391 |
| GDI1 | 2179 | 25 | 15 | 15 | 71.6 | 49.9 | 49.9 | 50.582 |
| OCIAD1 | 4324 | 15 | 15 | 15 | 55.1 | 55.1 | 55.1 | 27.626 |
| LDHB | 464 | 16 | 15 | 15 | 51.9 | 51.9 | 51.9 | 37.294 |
| RDH14 | 4186 | 15 | 15 | 15 | 57.4 | 57.4 | 57.4 | 36.864 |
| OSBPL11 | 4028 | 16 | 16 | 15 | 32.8 | 32.8 | 31.5 | 83.642 |
| EMC2 | 2979 | 15 | 15 | 15 | 60.3 | 60.3 | 60.3 | 34.833 |
| CLPTM1 | 1680 | 15 | 15 | 15 | 36.8 | 36.8 | 36.8 | 76.096 |
| MAT2A | 2180 | 15 | 15 | 15 | 44.3 | 44.3 | 44.3 | 43.66 |

| | | | | | | | | |
|---|---|---|---|---|---|---|---|---|
| CMPK1 | 2164 | 15 | 15 | 15 | 67.9 | 67.9 | 67.9 | 22.222 |
| SUN2 | 4434 | 15 | 15 | 15 | 32.9 | 32.9 | 32.9 | 80.31 |
| COPS4 | 3978 | 15 | 15 | 15 | 50.2 | 50.2 | 50.2 | 46.268 |
| MRPS27 | 3673 | 15 | 15 | 15 | 43.7 | 43.7 | 43.7 | 47.611 |
| TPP1 | 1353 | 15 | 15 | 15 | 49 | 49 | 49 | 61.247 |
| CD97 | 2358 | 18 | 18 | 15 | 33.2 | 33.2 | 28.3 | 91.868 |
| ANXA6 | 1797 | 41 | 41 | 15 | 61.7 | 61.7 | 29.4 | 75.872 |
| PIEZO1 | 3668 | 15 | 15 | 15 | 7.3 | 7.3 | 7.3 | 286.79 |
| AKR1A1 | 1927 | 15 | 15 | 15 | 63.7 | 63.7 | 63.7 | 36.573 |
| GRB2 | 2645 | 15 | 15 | 15 | 71.4 | 71.4 | 71.4 | 25.206 |
| ALDH3B1 | 2302 | 16 | 15 | 15 | 50.2 | 47.6 | 47.6 | 51.839 |
| ATP5F1 | 2084 | 15 | 15 | 15 | 53.9 | 53.9 | 53.9 | 28.908 |
| PRCP | 2294 | 15 | 15 | 15 | 40.9 | 40.9 | 40.9 | 55.799 |
| TRIM21 | 2008 | 15 | 15 | 15 | 39.4 | 39.4 | 39.4 | 54.169 |
| CECR5 | 4037 | 15 | 15 | 15 | 49.9 | 49.9 | 49.9 | 46.321 |
| CPNE1 | 888 | 15 | 15 | 15 | 34.3 | 34.3 | 34.3 | 59.058 |
| EXOC8 | 3470 | 15 | 15 | 15 | 24.6 | 24.6 | 24.6 | 81.798 |
| CCZ1B;CCZ | 2716 | 15 | 15 | 15 | 39.2 | 39.2 | 39.2 | 55.866 |
| DPM1 | 1031 | 15 | 15 | 15 | 62 | 62 | 62 | 32.426 |
| CD74 | 1732 | 14 | 14 | 14 | 44.9 | 44.9 | 44.9 | 33.515 |
| PSMD7 | 2443 | 14 | 14 | 14 | 59.9 | 59.9 | 59.9 | 37.025 |
| DCXR | 3373 | 14 | 14 | 14 | 73.4 | 73.4 | 73.4 | 25.913 |
| CPM | 1926 | 14 | 14 | 14 | 34.3 | 34.3 | 34.3 | 50.513 |
| AP5Z1 | 1433 | 14 | 14 | 14 | 25.8 | 25.8 | 25.8 | 88.604 |
| PPIB | 2069 | 14 | 14 | 14 | 57.4 | 57.4 | 57.4 | 23.742 |
| TPD52L2 | 1439 | 14 | 14 | 14 | 78.2 | 78.2 | 78.2 | 22.237 |
| EIF3G | 1574 | 14 | 14 | 14 | 43.8 | 43.8 | 43.8 | 35.611 |
| LACTB2 | 3127 | 14 | 14 | 14 | 58.3 | 58.3 | 58.3 | 32.805 |
| PSMA6 | 2553 | 14 | 14 | 14 | 71.1 | 71.1 | 71.1 | 27.399 |
| NMT1 | 2170 | 16 | 16 | 14 | 41.3 | 41.3 | 36.3 | 56.806 |
| C14orf166 | 4539 | 14 | 14 | 14 | 62.3 | 62.3 | 62.3 | 28.068 |
| SPG21 | 4360 | 14 | 14 | 14 | 53.6 | 53.6 | 53.6 | 34.96 |
| ANO6 | 3115 | 14 | 14 | 14 | 21.5 | 21.5 | 21.5 | 106.16 |
| TOR1A | 1346 | 14 | 14 | 14 | 47.6 | 47.6 | 47.6 | 37.808 |
| DHRS4 | 3985 | 14 | 14 | 14 | 62.6 | 62.6 | 62.6 | 29.537 |
| APBB1IP | 3377 | 15 | 15 | 14 | 29.6 | 29.6 | 28.2 | 73.182 |
| ETFA | 1918 | 14 | 14 | 14 | 64 | 64 | 64 | 35.079 |
| TXNL1 | 1438 | 14 | 14 | 14 | 65.1 | 65.1 | 65.1 | 32.251 |
| DNAJB12 | 1170 | 14 | 14 | 14 | 40.1 | 40.1 | 40.1 | 45.489 |
| OXCT1 | 2515 | 14 | 14 | 14 | 45.8 | 45.8 | 45.8 | 56.157 |
| FARP1 | 683 | 14 | 14 | 14 | 20 | 20 | 20 | 122.09 |
| SKAP2 | 1563 | 14 | 14 | 14 | 53.2 | 53.2 | 53.2 | 41.216 |
| CFL1 | 2075 | 15 | 15 | 14 | 77.7 | 77.7 | 73.5 | 18.502 |
| HDHD3 | 3970 | 14 | 14 | 14 | 73.3 | 73.3 | 73.3 | 28 |
| C17orf59 | 3802 | 14 | 14 | 14 | 59.7 | 59.7 | 59.7 | 37.226 |
| ATP6AP1 | 3043 | 14 | 14 | 14 | 32.8 | 32.8 | 32.8 | 52.025 |
| SLC3A2 | 908 | 22 | 22 | 14 | 46.1 | 46.1 | 31.4 | 64.872 |
| FDFT1 | 2247 | 14 | 14 | 14 | 40.8 | 40.8 | 40.8 | 48.115 |

| | | | | | | | | |
|---|---|---|---|---|---|---|---|---|
| EHD1 | 4 | 23 | 21 | 14 | 63 | 59.3 | 40.9 | 61.927 |
| SPTBN2 | 1375 | 15 | 15 | 14 | 11.3 | 11.3 | 10.9 | 271.32 |
| BCKDHA | 926 | 14 | 14 | 14 | 38.4 | 38.4 | 38.4 | 54.202 |
| DNAJB11 | 4409 | 14 | 14 | 14 | 47.5 | 47.5 | 47.5 | 40.513 |
| DNAJA1 | 2182 | 14 | 14 | 14 | 51.4 | 51.4 | 51.4 | 44.868 |
| GLOD4 | 939 | 14 | 14 | 14 | 31.9 | 31.9 | 31.9 | 54.719 |
| CA2 | 1700 | 14 | 14 | 14 | 69.6 | 69.6 | 69.6 | 29.246 |
| NIPSNAP1 | 3934 | 16 | 16 | 14 | 62 | 62 | 59.2 | 33.31 |
| TMOD3 | 4348 | 14 | 14 | 14 | 55.7 | 55.7 | 55.7 | 39.594 |
| PPP2CA;PP | 2662 | 16 | 16 | 14 | 66.7 | 66.7 | 61.8 | 35.594 |
| DNAJA3 | 3787 | 14 | 14 | 14 | 31.9 | 31.9 | 31.9 | 52.488 |
| MAVS | 3366 | 14 | 14 | 14 | 51.3 | 51.3 | 51.3 | 56.527 |
| ECI2 | 204 | 14 | 14 | 14 | 56.6 | 56.6 | 56.6 | 40.183 |
| ATP2B1 | 2014 | 26 | 14 | 14 | 26.4 | 15.2 | 15.2 | 134.68 |
| EIF3F | 1301 | 14 | 14 | 14 | 46.5 | 46.5 | 46.5 | 37.563 |
| RPS6KA3 | 567 | 21 | 21 | 14 | 34.7 | 34.7 | 23.9 | 80.583 |
| GIMAP1-GI | 20 | 20 | 20 | 14 | 52.3 | 52.3 | 35.2 | 57.051 |
| CRYZ | 2789 | 14 | 14 | 14 | 56.2 | 56.2 | 56.2 | 35.206 |
| ADH5 | 1885 | 14 | 14 | 14 | 59.9 | 59.9 | 59.9 | 39.724 |
| PDHX | 1302 | 14 | 14 | 14 | 34.3 | 34.3 | 34.3 | 54.122 |
| HIST1H4A | 2625 | 14 | 14 | 14 | 67 | 67 | 67 | 11.367 |
| TRABD | 1171 | 14 | 14 | 14 | 50.7 | 50.7 | 50.7 | 42.692 |
| FAM114A2 | 4258 | 14 | 14 | 14 | 40 | 40 | 40 | 55.468 |
| PTPRJ | 21 | 14 | 14 | 14 | 14.9 | 14.9 | 14.9 | 146.07 |
| HPCAL1;HF | 2246 | 14 | 14 | 14 | 67.4 | 67.4 | 67.4 | 22.313 |
| KIAA2013 | 3472 | 14 | 14 | 14 | 33 | 33 | 33 | 69.156 |
| PPA1 | 3003 | 14 | 14 | 14 | 63 | 63 | 63 | 32.66 |
| ODR4 | 3178 | 14 | 14 | 14 | 37.2 | 37.2 | 37.2 | 51.103 |
| RAB18 | 4211 | 14 | 14 | 14 | 75.7 | 75.7 | 75.7 | 22.977 |
| KLC1 | 807 | 19 | 19 | 14 | 41.5 | 41.5 | 29.8 | 65.309 |
| SSRP1 | 2796 | 14 | 14 | 14 | 30 | 30 | 30 | 81.074 |
| PCMT1 | 133 | 14 | 14 | 14 | 61.8 | 61.8 | 61.8 | 30.315 |
| AGK | 395 | 14 | 14 | 14 | 54 | 54 | 54 | 39.032 |
| NANS | 4244 | 14 | 14 | 14 | 58.8 | 58.8 | 58.8 | 40.307 |
| GHDC | 3494 | 14 | 14 | 14 | 36.2 | 36.2 | 36.2 | 57.522 |
| HSD17B12 | 3126 | 14 | 14 | 14 | 53.8 | 53.8 | 53.8 | 34.324 |
| AIMP1 | 2822 | 14 | 14 | 14 | 65.1 | 65.1 | 65.1 | 34.352 |
| DPP9 | 3392 | 14 | 14 | 14 | 24.7 | 24.7 | 24.7 | 98.262 |
| TTC37 | 3280 | 14 | 14 | 14 | 13.9 | 13.9 | 13.9 | 175.48 |
| PPP1R21 | 3308 | 14 | 14 | 14 | 25.5 | 25.5 | 25.5 | 88.313 |
| GIT2 | 2932 | 17 | 17 | 14 | 29.4 | 29.4 | 25.2 | 84.542 |
| DDRGK1 | 3810 | 14 | 14 | 14 | 41.7 | 41.7 | 41.7 | 35.61 |
| HMGCL | 2231 | 14 | 14 | 14 | 47.4 | 47.4 | 47.4 | 34.36 |
| LGALS3 | 1985 | 14 | 14 | 14 | 47.6 | 47.6 | 47.6 | 26.152 |
| NCF4 | 2996 | 14 | 14 | 14 | 49.3 | 49.3 | 49.3 | 39.031 |
| PLA2G7 | 2838 | 14 | 14 | 14 | 45.6 | 45.6 | 45.6 | 50.077 |
| RPL7 | 1991 | 14 | 14 | 14 | 52 | 52 | 52 | 29.225 |
| FAM91A1 | 3228 | 14 | 14 | 14 | 26 | 26 | 26 | 93.908 |

| | | | | | | | | |
|---|---|---|---|---|---|---|---|---|
| PRKAR1A | 1862 | 14 | 14 | 14 | 42 | 42 | 42 | 42.981 |
| RPS11 | 2606 | 14 | 14 | 14 | 62 | 62 | 62 | 18.431 |
| DNAJC1 | 3832 | 14 | 14 | 14 | 27.6 | 27.6 | 27.6 | 63.882 |
| HADH | 147 | 14 | 14 | 14 | 69.7 | 69.7 | 69.7 | 34.594 |
| ABCB10 | 4249 | 15 | 14 | 14 | 27.2 | 26.2 | 26.2 | 79.147 |
| PTPRA | 1994 | 17 | 17 | 14 | 31.3 | 31.3 | 26.9 | 90.718 |
| WASF2 | 4685 | 14 | 14 | 14 | 36.3 | 36.3 | 36.3 | 54.283 |
| FNDC3A | 4558 | 14 | 14 | 14 | 16.4 | 16.4 | 16.4 | 131.85 |
| SLC25A19 | 4193 | 14 | 14 | 14 | 51.9 | 51.9 | 51.9 | 35.511 |
| EPB41L1 | 222 | 23 | 22 | 14 | 37 | 35.7 | 21.6 | 98.374 |
| NRP1 | 1355 | 14 | 14 | 14 | 26.7 | 26.7 | 26.7 | 103.13 |
| TRIP12 | 2960 | 14 | 14 | 14 | 9.2 | 9.2 | 9.2 | 220.43 |
| ALG11 | 3084 | 14 | 14 | 14 | 42.7 | 42.7 | 42.7 | 55.65 |
| TAPBP | 247 | 14 | 14 | 14 | 58.5 | 58.5 | 58.5 | 47.606 |
| TUBG1;TUE | 2068 | 14 | 14 | 14 | 53.7 | 53.7 | 53.7 | 51.169 |
| COG3 | 3822 | 14 | 14 | 14 | 24.2 | 24.2 | 24.2 | 94.095 |
| GLB1 | 1960 | 14 | 14 | 14 | 28.1 | 28.1 | 28.1 | 76.074 |
| G3BP1 | 2857 | 15 | 14 | 14 | 39.5 | 37.8 | 37.8 | 52.164 |
| RPS6KA1 | 842 | 20 | 14 | 14 | 33.4 | 24.4 | 24.4 | 81.471 |
| INPP5K | 3977 | 14 | 14 | 14 | 39.7 | 39.7 | 39.7 | 51.09 |
| LGALS3BP | 2792 | 14 | 14 | 14 | 34.7 | 34.7 | 34.7 | 65.33 |
| DTYMK | 2080 | 14 | 14 | 14 | 61.8 | 61.8 | 61.8 | 23.819 |
| ERLEC1 | 3779 | 14 | 14 | 14 | 36.4 | 36.4 | 36.4 | 54.858 |
| TM9SF3 | 4208 | 14 | 14 | 14 | 25 | 25 | 25 | 67.887 |
| LRRC59 | 3744 | 14 | 14 | 14 | 45.9 | 45.9 | 45.9 | 34.93 |
| ALAS1 | 1906 | 14 | 14 | 14 | 31.4 | 31.4 | 31.4 | 70.58 |
| UPP1 | 3074 | 14 | 14 | 14 | 58.4 | 58.4 | 58.4 | 33.934 |
| TOMM40 | 1681 | 14 | 14 | 14 | 59.3 | 59.3 | 59.3 | 37.893 |
| PIGS | 3876 | 14 | 14 | 14 | 38.6 | 38.6 | 38.6 | 61.655 |
| HSDL1 | 3100 | 14 | 14 | 14 | 47.3 | 47.3 | 47.3 | 37.001 |
| ABI1 | 138 | 14 | 14 | 14 | 33.5 | 33.5 | 33.5 | 51.769 |
| ALG2 | 4146 | 14 | 14 | 14 | 42.3 | 42.3 | 42.3 | 47.091 |
| PRDX1 | 2778 | 18 | 18 | 14 | 73.9 | 73.9 | 60.8 | 22.11 |
| TOLLIP | 4091 | 13 | 13 | 13 | 59.1 | 59.1 | 59.1 | 30.281 |
| PLAUR | 2757 | 13 | 13 | 13 | 52.8 | 52.8 | 52.8 | 36.978 |
| HMOX1 | 1830 | 13 | 13 | 13 | 55.9 | 55.9 | 55.9 | 32.818 |
| SFXN1 | 4170 | 14 | 13 | 13 | 58.4 | 52.5 | 52.5 | 35.619 |
| RPL15 | 2578 | 13 | 13 | 13 | 49.5 | 49.5 | 49.5 | 24.146 |
| SEC22B | 1544 | 13 | 13 | 13 | 51.6 | 51.6 | 51.6 | 24.593 |
| ITSN2 | 43 | 13 | 13 | 13 | 9.4 | 9.4 | 9.4 | 191.6 |
| ADCK3 | 3589 | 13 | 13 | 13 | 29.8 | 29.8 | 29.8 | 71.949 |
| RPL10A | 2638 | 13 | 13 | 13 | 50.2 | 50.2 | 50.2 | 24.831 |
| PSMB1 | 2025 | 13 | 13 | 13 | 58.9 | 58.9 | 58.9 | 26.489 |
| CYC1 | 1805 | 13 | 13 | 13 | 56 | 56 | 56 | 35.422 |
| ETHE1 | 1660 | 13 | 13 | 13 | 75.2 | 75.2 | 75.2 | 27.873 |
| SOAT1 | 2226 | 13 | 13 | 13 | 26.2 | 26.2 | 26.2 | 64.734 |
| EIF2B5 | 410 | 13 | 13 | 13 | 29.8 | 29.8 | 29.8 | 80.379 |
| GPX1 | 33 | 13 | 13 | 13 | 80.3 | 80.3 | 80.3 | 22.088 |

| | | | | | | | | |
|---|---|---|---|---|---|---|---|---|
| TECR | 4352 | 13 | 13 | 13 | 36.4 | 36.4 | 36.4 | 36.034 |
| RAB21 | 4494 | 13 | 13 | 13 | 58.2 | 58.2 | 58.2 | 24.347 |
| LPXN | 1503 | 13 | 13 | 13 | 49.2 | 49.2 | 49.2 | 43.332 |
| ATP9A | 1523 | 17 | 13 | 13 | 20.2 | 15.3 | 15.3 | 118.58 |
| CD44 | 1058 | 13 | 13 | 13 | 47.6 | 47.6 | 47.6 | 22.683 |
| STX4 | 2819 | 13 | 13 | 13 | 52.2 | 52.2 | 52.2 | 34.18 |
| CHI3L1 | 2233 | 22 | 22 | 13 | 66.6 | 66.6 | 39.9 | 42.625 |
| BLVRA | 2470 | 13 | 13 | 13 | 44.9 | 44.9 | 44.9 | 33.428 |
| PYCARD | 4504 | 13 | 13 | 13 | 82.6 | 82.6 | 82.6 | 21.627 |
| BPNT1 | 1674 | 13 | 13 | 13 | 53.9 | 53.9 | 53.9 | 33.392 |
| FUBP3 | 3813 | 15 | 13 | 13 | 35.3 | 32.3 | 32.3 | 61.64 |
| PACSIN2 | 4513 | 13 | 13 | 13 | 35.8 | 35.8 | 35.8 | 55.738 |
| CAB39 | 4584 | 16 | 16 | 13 | 48.1 | 48.1 | 40.5 | 39.869 |
| CTSK | 2300 | 14 | 14 | 13 | 68.1 | 68.1 | 66 | 36.966 |
| NEK9 | 3612 | 13 | 13 | 13 | 17 | 17 | 17 | 107.17 |
| STX12 | 3426 | 13 | 13 | 13 | 68.5 | 68.5 | 68.5 | 31.642 |
| PDK3 | 2999 | 13 | 13 | 13 | 47 | 47 | 47 | 46.938 |
| ATAD1 | 3543 | 13 | 13 | 13 | 51.8 | 51.8 | 51.8 | 40.744 |
| SPATA20 | 278 | 13 | 13 | 13 | 17.3 | 17.3 | 17.3 | 88.051 |
| MRPL44 | 4172 | 13 | 13 | 13 | 44.3 | 44.3 | 44.3 | 37.535 |
| MMP14 | 2404 | 13 | 13 | 13 | 29 | 29 | 29 | 65.893 |
| MTCH1 | 4362 | 13 | 13 | 13 | 35.7 | 35.7 | 35.7 | 41.544 |
| HSD17B11 | 3541 | 13 | 13 | 13 | 62 | 62 | 62 | 32.935 |
| NPM1 | 1773 | 13 | 13 | 13 | 55.1 | 55.1 | 55.1 | 32.575 |
| RPL13 | 2110 | 13 | 13 | 13 | 45.5 | 45.5 | 45.5 | 24.261 |
| MREG | 3512 | 13 | 13 | 13 | 62.1 | 62.1 | 62.1 | 24.927 |
| TM9SF2 | 3925 | 13 | 13 | 13 | 28.8 | 28.8 | 28.8 | 75.775 |
| DNAJA2 | 1517 | 13 | 13 | 13 | 51.7 | 51.7 | 51.7 | 45.745 |
| SRI | 641 | 13 | 13 | 13 | 70.3 | 70.3 | 70.3 | 17.605 |
| NDRG1 | 3679 | 13 | 13 | 13 | 62.2 | 62.2 | 62.2 | 42.835 |
| PRPF4 | 1424 | 13 | 13 | 13 | 37.2 | 37.2 | 37.2 | 58.449 |
| RPS16 | 2601 | 13 | 13 | 13 | 65.1 | 65.1 | 65.1 | 16.445 |
| ARFGEF2 | 4668 | 22 | 22 | 13 | 17.4 | 17.4 | 10.7 | 202.04 |
| HTATIP2 | 3994 | 13 | 13 | 13 | 50.4 | 50.4 | 50.4 | 27.049 |
| VDAC3 | 4546 | 13 | 13 | 13 | 58.7 | 58.7 | 58.7 | 30.658 |
| MTHFD2 | 1922 | 13 | 13 | 13 | 55.7 | 55.7 | 55.7 | 37.895 |
| XPNPEP1 | 4238 | 14 | 13 | 13 | 26.8 | 25.7 | 25.7 | 69.917 |
| RPSA | 198 | 13 | 13 | 13 | 68.4 | 68.4 | 68.4 | 29.404 |
| MRPL39 | 4347 | 13 | 13 | 13 | 43.8 | 43.8 | 43.8 | 38.711 |
| GGT1;GGT2 | 2007 | 13 | 13 | 13 | 35.3 | 35.3 | 35.3 | 61.409 |
| RENBP | 2439 | 13 | 13 | 13 | 37.2 | 37.2 | 37.2 | 48.83 |
| PDCD4 | 3119 | 13 | 13 | 13 | 43.9 | 43.9 | 43.9 | 51.735 |
| SEC61A1 | 587 | 13 | 13 | 13 | 24.7 | 24.7 | 24.7 | 52.949 |
| TST | 3067 | 13 | 13 | 13 | 65.3 | 65.3 | 65.3 | 33.429 |
| PLRG1 | 1451 | 13 | 13 | 13 | 35.8 | 35.8 | 35.8 | 57.193 |
| CTSC | 2482 | 13 | 13 | 13 | 41.7 | 41.7 | 41.7 | 51.853 |
| GCDH | 3712 | 13 | 13 | 13 | 45.9 | 45.9 | 45.9 | 48.127 |
| ITPRIP | 3447 | 13 | 13 | 13 | 31.1 | 31.1 | 31.1 | 62.059 |

| | | | | | | | | |
|---|---|---|---|---|---|---|---|---|
| COL4A3BP | 362 | 13 | 13 | 13 | 33.6 | 33.6 | 33.6 | 64.357 |
| MAN1A1 | 2208 | 13 | 13 | 13 | 35.5 | 35.5 | 35.5 | 72.968 |
| UQCRFS1;U | 2338 | 13 | 13 | 13 | 55.8 | 55.8 | 55.8 | 29.668 |
| YWHAG | 2591 | 18 | 15 | 13 | 69.6 | 62.3 | 55.9 | 28.302 |
| BZW1 | 3332 | 14 | 14 | 13 | 39.1 | 39.1 | 37.5 | 48.043 |
| SLC12A7 | 258 | 14 | 14 | 13 | 15.3 | 15.3 | 14 | 119.13 |
| UCHL5 | 3164 | 13 | 13 | 13 | 52.7 | 52.7 | 52.7 | 40.423 |
| CTSA | 1860 | 13 | 13 | 13 | 33.1 | 33.1 | 33.1 | 54.255 |
| TOR3A | 4137 | 13 | 13 | 13 | 47.1 | 47.1 | 47.1 | 46.198 |
| RABEP1 | 3007 | 13 | 13 | 13 | 18.2 | 18.2 | 18.2 | 99.289 |
| PALLD | 3657 | 13 | 13 | 13 | 13.4 | 13.4 | 13.4 | 150.56 |
| TRIM4 | 4067 | 13 | 13 | 13 | 34 | 34 | 34 | 57.46 |
| C2CD2L | 1337 | 13 | 13 | 13 | 27.6 | 27.6 | 27.6 | 76.18 |
| ARHGEF7 | 375 | 17 | 17 | 13 | 32.8 | 32.8 | 26.1 | 79.831 |
| F11R | 4657 | 13 | 13 | 13 | 47.8 | 47.8 | 47.8 | 32.583 |
| SRPRB | 4641 | 13 | 13 | 13 | 48 | 48 | 48 | 29.702 |
| PSMA4 | 2103 | 13 | 13 | 13 | 67.8 | 67.8 | 67.8 | 29.483 |
| MICU2 | 3473 | 13 | 13 | 13 | 47 | 47 | 47 | 49.666 |
| SEC24A | 1654 | 15 | 13 | 13 | 17.4 | 15.9 | 15.9 | 119.75 |
| RPLP0;RPLI | 1756 | 13 | 13 | 13 | 56.2 | 56.2 | 56.2 | 34.273 |
| FKBP4 | 2749 | 13 | 13 | 13 | 33.8 | 33.8 | 33.8 | 51.804 |
| PIGT | 314 | 13 | 13 | 13 | 26.8 | 26.8 | 26.8 | 65.699 |
| GOT1 | 1975 | 13 | 13 | 13 | 53.8 | 53.8 | 53.8 | 46.247 |
| GSS | 339 | 13 | 13 | 13 | 36.7 | 36.7 | 36.7 | 49.976 |
| P2RX7 | 3906 | 13 | 13 | 13 | 28.7 | 28.7 | 28.7 | 68.584 |
| HDGF | 2449 | 13 | 13 | 13 | 65.8 | 65.8 | 65.8 | 26.788 |
| HIBADH | 2185 | 13 | 13 | 13 | 64.3 | 64.3 | 64.3 | 35.329 |
| HARS | 1890 | 19 | 13 | 13 | 45 | 33.6 | 33.6 | 57.41 |
| PRDX4 | 2846 | 16 | 13 | 13 | 62.4 | 56.8 | 56.8 | 30.54 |
| RALY | 3170 | 13 | 13 | 13 | 51.5 | 51.5 | 51.5 | 24.665 |
| CYP4V2 | 3318 | 13 | 13 | 13 | 27.4 | 27.4 | 27.4 | 60.723 |
| ETF1 | 2618 | 13 | 13 | 13 | 41.9 | 41.9 | 41.9 | 49.03 |
| GUF1 | 3504 | 13 | 13 | 13 | 26.2 | 26.2 | 26.2 | 74.327 |
| IDH3G | 2434 | 13 | 13 | 13 | 50.9 | 50.9 | 50.9 | 42.794 |
| YWHAZ | 2650 | 17 | 14 | 13 | 59.2 | 51.8 | 48.2 | 27.745 |
| TAPBPL | 4024 | 13 | 13 | 13 | 34.2 | 34.2 | 34.2 | 50.182 |
| MX2 | 2024 | 15 | 15 | 13 | 29 | 29 | 24.1 | 82.088 |
| WDR26 | 442 | 13 | 13 | 13 | 22.2 | 22.2 | 22.2 | 81.327 |
| RPL11 | 2639 | 13 | 13 | 13 | 71.3 | 71.3 | 71.3 | 19.024 |
| CC2D1B | 1127 | 13 | 13 | 13 | 19.3 | 19.3 | 19.3 | 94.223 |
| ATP5C1 | 2238 | 13 | 13 | 13 | 49.7 | 49.7 | 49.7 | 32.996 |
| PSMD14 | 1318 | 13 | 13 | 13 | 55.5 | 55.5 | 55.5 | 34.577 |
| DEK | 1048 | 13 | 13 | 13 | 33.4 | 33.4 | 33.4 | 43.247 |
| AHCYL1 | 1476 | 20 | 20 | 13 | 51.3 | 51.3 | 36.4 | 58.951 |
| SCIN | 4683 | 13 | 13 | 13 | 32.9 | 32.9 | 32.9 | 80.488 |
| FUCA2 | 3984 | 13 | 13 | 13 | 30.2 | 30.2 | 30.2 | 54.066 |
| MGAT1 | 2112 | 13 | 13 | 13 | 38.7 | 38.7 | 38.7 | 50.878 |
| MAN1B1 | 4487 | 13 | 13 | 13 | 27.5 | 27.5 | 27.5 | 79.579 |

| | | | | | | | | |
|---|---|---|---|---|---|---|---|---|
| ATP1B1 | 1744 | 13 | 13 | 13 | 40.6 | 40.6 | 40.6 | 35.061 |
| ADAMDEC1 | 1392 | 13 | 13 | 13 | 46.6 | 46.6 | 46.6 | 52.775 |
| PCBP1 | 3012 | 18 | 18 | 13 | 84.8 | 84.8 | 69.4 | 37.497 |
| LBR | 2968 | 13 | 13 | 13 | 22.3 | 22.3 | 22.3 | 70.702 |
| VAMP7 | 2447 | 13 | 13 | 13 | 63.6 | 63.6 | 63.6 | 24.935 |
| SDSL | 3797 | 13 | 13 | 13 | 58.7 | 58.7 | 58.7 | 34.674 |
| CPVL | 4125 | 13 | 13 | 13 | 30.3 | 30.3 | 30.3 | 54.163 |
| PAFAH2 | 3896 | 13 | 13 | 13 | 45.9 | 45.9 | 45.9 | 44.035 |
| ATG9A | 3360 | 13 | 13 | 13 | 22.6 | 22.6 | 22.6 | 94.446 |
| METAP1 | 2476 | 13 | 13 | 13 | 49.2 | 49.2 | 49.2 | 43.215 |
| MAT2B | 4363 | 13 | 13 | 13 | 56.3 | 56.3 | 56.3 | 37.551 |
| BCAT2 | 1257 | 13 | 13 | 13 | 31.9 | 31.9 | 31.9 | 49.967 |
| PSM8;PSMI | 2131 | 13 | 13 | 13 | 56.9 | 56.9 | 56.9 | 30.354 |
| PFN1 | 1787 | 13 | 13 | 13 | 93.6 | 93.6 | 93.6 | 15.054 |
| ALDH1A2 | 1605 | 15 | 13 | 13 | 35.7 | 34 | 34 | 56.723 |
| CHMP4B | 4134 | 13 | 13 | 13 | 76.3 | 76.3 | 76.3 | 24.95 |
| MLEC | 2934 | 13 | 13 | 13 | 46.2 | 46.2 | 46.2 | 32.233 |
| PPM1G | 1408 | 13 | 13 | 13 | 31.3 | 31.3 | 31.3 | 59.271 |
| HNRNPLL | 609 | 14 | 13 | 13 | 36.2 | 34.8 | 34.8 | 56.45 |
| GBAS | 1532 | 15 | 13 | 13 | 58.7 | 55.9 | 55.9 | 33.742 |
| ACOT7 | 1283 | 13 | 13 | 13 | 50.3 | 50.3 | 50.3 | 41.796 |
| SDCBP | 1322 | 12 | 12 | 12 | 62.4 | 62.4 | 62.4 | 32.444 |
| NUMB | 2383 | 14 | 14 | 12 | 26.3 | 26.3 | 22.9 | 70.803 |
| NEMF | 1014 | 12 | 12 | 12 | 16.5 | 16.5 | 16.5 | 113.95 |
| SCAMP1 | 1386 | 12 | 12 | 12 | 42.6 | 42.6 | 42.6 | 37.92 |
| ATP7A | 2761 | 12 | 12 | 12 | 12.1 | 12.1 | 12.1 | 163.37 |
| CLINT1 | 2961 | 12 | 12 | 12 | 22.6 | 22.6 | 22.6 | 68.259 |
| GPRIN3 | 3316 | 12 | 12 | 12 | 19.6 | 19.6 | 19.6 | 82.438 |
| ACP5 | 1914 | 12 | 12 | 12 | 45.5 | 45.5 | 45.5 | 36.598 |
| RHOG | 2710 | 13 | 13 | 12 | 82.7 | 82.7 | 77 | 21.308 |
| PTPN9 | 125 | 12 | 12 | 12 | 27.2 | 27.2 | 27.2 | 68.019 |
| TRIM33 | 4527 | 12 | 12 | 12 | 12.8 | 12.8 | 12.8 | 122.53 |
| CHP1 | 3915 | 12 | 12 | 12 | 67.7 | 67.7 | 67.7 | 22.456 |
| PPP1R9B | 3880 | 12 | 12 | 12 | 20.2 | 20.2 | 20.2 | 89.333 |
| ATP5O | 2339 | 12 | 12 | 12 | 62.4 | 62.4 | 62.4 | 23.277 |
| CAPZA2 | 2335 | 14 | 14 | 12 | 74.1 | 74.1 | 64.7 | 32.949 |
| TM9SF1 | 884 | 12 | 12 | 12 | 18.2 | 18.2 | 18.2 | 92.386 |
| RRAGA;RRA | 3339 | 12 | 12 | 12 | 44.1 | 44.1 | 44.1 | 36.566 |
| BCAP29 | 480 | 12 | 12 | 12 | 22.8 | 22.8 | 22.8 | 41.682 |
| HM13 | 411 | 12 | 12 | 12 | 28.6 | 28.6 | 28.6 | 46.313 |
| PSME3 | 1253 | 12 | 12 | 12 | 61.9 | 61.9 | 61.9 | 26.867 |
| DAB2 | 2717 | 12 | 12 | 12 | 21.8 | 21.8 | 21.8 | 82.447 |
| PRPF4B | 2883 | 12 | 12 | 12 | 17 | 17 | 17 | 116.99 |
| BDH1 | 2747 | 12 | 12 | 12 | 49 | 49 | 49 | 38.157 |
| SCFD2 | 3635 | 12 | 12 | 12 | 22.8 | 22.8 | 22.8 | 75.126 |
| EDC4 | 3266 | 12 | 12 | 12 | 12.9 | 12.9 | 12.9 | 151.66 |
| DDX24 | 1010 | 12 | 12 | 12 | 18.1 | 18.1 | 18.1 | 91.48 |
| SNRNP40;D | 3776 | 12 | 12 | 12 | 50.1 | 50.1 | 50.1 | 39.31 |

| | | | | | | | | |
|---|---|---|---|---|---|---|---|---|
| GRPEL1 | 4181 | 12 | 12 | 12 | 53.9 | 53.9 | 53.9 | 24.279 |
| CAPNS1 | 215 | 12 | 12 | 12 | 51.9 | 51.9 | 51.9 | 33.786 |
| FAM213A | 3963 | 12 | 12 | 12 | 45.4 | 45.4 | 45.4 | 25.764 |
| EIF2AK2 | 2009 | 12 | 12 | 12 | 24.9 | 24.9 | 24.9 | 62.094 |
| HNRNPA0 | 2842 | 12 | 12 | 12 | 54.8 | 54.8 | 54.8 | 30.84 |
| GTPBP4 | 4057 | 12 | 12 | 12 | 20.3 | 20.3 | 20.3 | 73.964 |
| ST14 | 4653 | 12 | 12 | 12 | 20.1 | 20.1 | 20.1 | 94.769 |
| RPS10;RPS1 | 2328 | 12 | 12 | 12 | 60 | 60 | 60 | 18.898 |
| ACOT8 | 1349 | 12 | 12 | 12 | 48 | 48 | 48 | 35.914 |
| SLC26A11 | 3413 | 12 | 12 | 12 | 25.6 | 25.6 | 25.6 | 65.298 |
| ENG | 1982 | 12 | 12 | 12 | 28 | 28 | 28 | 70.577 |
| RAB20 | 4328 | 12 | 12 | 12 | 65.4 | 65.4 | 65.4 | 26.277 |
| DDAH2 | 1675 | 12 | 12 | 12 | 66.3 | 66.3 | 66.3 | 29.644 |
| DAGLB | 3551 | 12 | 12 | 12 | 24.6 | 24.6 | 24.6 | 73.731 |
| LUC7L3 | 1169 | 12 | 12 | 12 | 29 | 29 | 29 | 58.22 |
| PPID | 2795 | 13 | 12 | 12 | 42.4 | 40.5 | 40.5 | 40.763 |
| LNP | 4071 | 12 | 12 | 12 | 36 | 36 | 36 | 47.739 |
| RCN2 | 2943 | 12 | 12 | 12 | 51.1 | 51.1 | 51.1 | 36.876 |
| EEF1D | 2150 | 23 | 23 | 12 | 72.2 | 72.2 | 38.8 | 31.121 |
| SF1 | 3030 | 12 | 12 | 12 | 25.2 | 25.2 | 25.2 | 68.329 |
| ANO10 | 4311 | 12 | 12 | 12 | 24.1 | 24.1 | 24.1 | 76.328 |
| RAN | 605 | 12 | 12 | 12 | 43.8 | 43.8 | 43.8 | 26.224 |
| EIF2B3 | 4245 | 12 | 12 | 12 | 32.5 | 32.5 | 32.5 | 50.24 |
| STOML2 | 4476 | 12 | 12 | 12 | 47.8 | 47.8 | 47.8 | 38.534 |
| ADCY7 | 2448 | 12 | 12 | 12 | 15.3 | 15.3 | 15.3 | 120.31 |
| DDO | 3897 | 12 | 12 | 12 | 48.4 | 48.4 | 48.4 | 37.534 |
| NIT2 | 4235 | 12 | 12 | 12 | 65.6 | 65.6 | 65.6 | 30.608 |
| SEPTIN10 | 607 | 14 | 12 | 12 | 44.4 | 39.8 | 39.8 | 52.145 |
| HSPB1 | 1737 | 12 | 12 | 12 | 78.5 | 78.5 | 78.5 | 22.782 |
| PEBP1 | 2165 | 12 | 12 | 12 | 80.7 | 80.7 | 80.7 | 21.057 |
| CTNNA1 | 1016 | 14 | 14 | 12 | 23.7 | 23.7 | 21 | 100.07 |
| EXOC6 | 812 | 12 | 12 | 12 | 21.5 | 21.5 | 21.5 | 81.851 |
| ALDH1L2 | 435 | 12 | 12 | 12 | 17.4 | 17.4 | 17.4 | 102.85 |
| GRIPAP1 | 3117 | 12 | 12 | 12 | 20.2 | 20.2 | 20.2 | 96.004 |
| VAV1 | 927 | 12 | 12 | 12 | 18 | 18 | 18 | 91.729 |
| AP5B1 | 3087 | 12 | 12 | 12 | 21.4 | 21.4 | 21.4 | 93.948 |
| MRPS9 | 2701 | 12 | 12 | 12 | 43.2 | 43.2 | 43.2 | 45.834 |
| STX5 | 2849 | 12 | 12 | 12 | 39.7 | 39.7 | 39.7 | 39.672 |
| WDR82 | 3300 | 12 | 12 | 12 | 44.4 | 44.4 | 44.4 | 35.079 |
| MAOA | 2038 | 13 | 13 | 12 | 35.5 | 35.5 | 33.4 | 59.681 |
| MRPL37 | 4056 | 12 | 12 | 12 | 42.8 | 42.8 | 42.8 | 48.117 |
| STAT6 | 317 | 12 | 12 | 12 | 24.6 | 24.6 | 24.6 | 80.305 |
| KPNA4 | 1328 | 19 | 19 | 12 | 56.8 | 56.8 | 43.4 | 57.886 |
| SORT1 | 3900 | 12 | 12 | 12 | 22.7 | 22.7 | 22.7 | 92.067 |
| BUB3 | 1457 | 12 | 12 | 12 | 56.1 | 56.1 | 56.1 | 37.154 |
| WDR37 | 397 | 12 | 12 | 12 | 29.3 | 29.3 | 29.3 | 63.833 |
| LGALS1 | 1819 | 12 | 12 | 12 | 94.1 | 94.1 | 94.1 | 14.716 |
| API5 | 995 | 12 | 12 | 12 | 30.8 | 30.8 | 30.8 | 57.56 |

| | | | | | | | | |
|---|---|---|---|---|---|---|---|---|
| MYH10 | 2224 | 30 | 12 | 12 | 16.9 | 7.9 | 7.9 | 229 |
| TRAPPC8 | 4561 | 12 | 12 | 12 | 10.9 | 10.9 | 10.9 | 161 |
| RABGGTA | 3692 | 12 | 12 | 12 | 28.6 | 28.6 | 28.6 | 65.071 |
| RPS7 | 2593 | 12 | 12 | 12 | 60.8 | 60.8 | 60.8 | 22.127 |
| U2SURP | 1377 | 12 | 12 | 12 | 12 | 12 | 12 | 118.29 |
| AGPAT9 | 3121 | 14 | 14 | 12 | 46.1 | 46.1 | 44 | 48.705 |
| GOLGA1 | 3698 | 12 | 12 | 12 | 20.6 | 20.6 | 20.6 | 88.183 |
| MX1 | 2023 | 14 | 12 | 12 | 32.5 | 27.2 | 27.2 | 75.519 |
| CUL4A | 2902 | 19 | 19 | 12 | 33.1 | 33.1 | 23.2 | 87.679 |
| IL16 | 1079 | 12 | 12 | 12 | 26.7 | 26.7 | 26.7 | 64.547 |
| MAPK13 | 1403 | 12 | 12 | 12 | 49.9 | 49.9 | 49.9 | 42.089 |
| SKIV2L | 3027 | 12 | 12 | 12 | 16.1 | 16.1 | 16.1 | 137.75 |
| ERP29 | 2154 | 12 | 12 | 12 | 56.3 | 56.3 | 56.3 | 28.993 |
| SPR | 2220 | 12 | 12 | 12 | 52.9 | 52.9 | 52.9 | 28.048 |
| DBN1 | 3059 | 12 | 12 | 12 | 23.9 | 23.9 | 23.9 | 71.428 |
| XPC | 2739 | 12 | 12 | 12 | 19.9 | 19.9 | 19.9 | 105.95 |
| RHOF | 4185 | 12 | 12 | 12 | 80.1 | 80.1 | 80.1 | 23.625 |
| ARHGEF6 | 2991 | 16 | 12 | 12 | 26.8 | 20.7 | 20.7 | 87.498 |
| TES | 4425 | 12 | 12 | 12 | 38.5 | 38.5 | 38.5 | 47.996 |
| PARK7 | 3898 | 12 | 12 | 12 | 67.7 | 67.7 | 67.7 | 19.891 |
| SDF2L1 | 4201 | 12 | 12 | 12 | 78.3 | 78.3 | 78.3 | 23.598 |
| BIN2 | 4413 | 12 | 12 | 12 | 27.6 | 27.6 | 27.6 | 61.874 |
| RSU1 | 3019 | 12 | 12 | 12 | 62.5 | 62.5 | 62.5 | 31.54 |
| TBC1D5 | 3684 | 12 | 12 | 12 | 24.7 | 24.7 | 24.7 | 89.003 |
| ECHDC1 | 4270 | 12 | 12 | 12 | 55 | 55 | 55 | 33.698 |
| HTRA2 | 1441 | 12 | 12 | 12 | 51.3 | 51.3 | 51.3 | 48.84 |
| PON2 | 270 | 12 | 12 | 12 | 63.7 | 63.7 | 63.7 | 41.527 |
| WAS | 2293 | 12 | 12 | 12 | 40.8 | 40.8 | 40.8 | 52.912 |
| HAGH | 3069 | 12 | 12 | 12 | 57.1 | 57.1 | 57.1 | 33.805 |
| PUS1 | 915 | 12 | 12 | 12 | 39.6 | 39.6 | 39.6 | 41.729 |
| DYNC1LI2 | 1426 | 14 | 12 | 12 | 42.7 | 37 | 37 | 54.099 |
| EXOC3 | 1499 | 12 | 12 | 12 | 20.7 | 20.7 | 20.7 | 85.566 |
| MESDC2 | 2965 | 12 | 12 | 12 | 54.7 | 54.7 | 54.7 | 26.076 |
| NAA10 | 2274 | 12 | 12 | 12 | 66.8 | 66.8 | 66.8 | 26.458 |
| MRPL15 | 4369 | 12 | 12 | 12 | 36.1 | 36.1 | 36.1 | 33.419 |
| RASAL3 | 3430 | 13 | 13 | 12 | 15.7 | 15.7 | 15 | 111.9 |
| MAPK1 | 2138 | 16 | 16 | 12 | 65.3 | 65.3 | 52.5 | 41.389 |
| PPT1 | 337 | 12 | 12 | 12 | 46.2 | 46.2 | 46.2 | 34.136 |
| PEX14 | 1542 | 12 | 12 | 12 | 46.2 | 46.2 | 46.2 | 41.236 |
| PXN | 300 | 12 | 12 | 12 | 27.2 | 27.2 | 27.2 | 64.232 |
| HMHA1 | 914 | 12 | 12 | 12 | 13.4 | 13.4 | 13.4 | 112.47 |
| ARFGEF1 | 4669 | 21 | 12 | 12 | 13.1 | 6.7 | 6.7 | 208.76 |
| SERPINB9 | 2410 | 14 | 13 | 12 | 46.3 | 43.6 | 40.4 | 42.403 |
| ESD | 1863 | 12 | 12 | 12 | 70.9 | 70.9 | 70.9 | 31.462 |
| STOM | 2118 | 12 | 12 | 12 | 61.1 | 61.1 | 61.1 | 31.73 |
| ITGB3 | 1084 | 12 | 12 | 12 | 19.2 | 19.2 | 19.2 | 87.057 |
| GMPS | 2395 | 12 | 12 | 12 | 24.8 | 24.8 | 24.8 | 76.715 |
| RNH1 | 1909 | 24 | 24 | 12 | 79.6 | 79.6 | 39.9 | 49.973 |

| | | | | | | | | |
|---|---|---|---|---|---|---|---|---|
| RAP1GDS1 | 2457 | 13 | 13 | 12 | 34.6 | 34.6 | 30.3 | 66.316 |
| SPTLC2 | 1405 | 12 | 12 | 12 | 29.5 | 29.5 | 29.5 | 62.924 |
| EOGT | 3166 | 12 | 12 | 12 | 29 | 29 | 29 | 62.01 |
| GGH | 3699 | 12 | 12 | 12 | 36.2 | 36.2 | 36.2 | 35.964 |
| ARRB2 | 2194 | 13 | 13 | 12 | 41.6 | 41.6 | 39.9 | 46.105 |
| GK;GK3P | 2195 | 15 | 15 | 12 | 37.4 | 37.4 | 29 | 61.244 |
| GSTP1 | 1818 | 12 | 12 | 12 | 61 | 61 | 61 | 23.356 |
| STAT2 | 430 | 12 | 12 | 12 | 20.5 | 20.5 | 20.5 | 94.234 |
| DHX29 | 74 | 11 | 11 | 11 | 10.5 | 10.5 | 10.5 | 155.29 |
| ABHD6 | 3999 | 11 | 11 | 11 | 46.6 | 46.6 | 46.6 | 38.33 |
| TNKS1BP1 | 4070 | 11 | 11 | 11 | 9.7 | 9.7 | 9.7 | 181.79 |
| FAM136A | 3762 | 11 | 11 | 11 | 81.2 | 81.2 | 81.2 | 15.641 |
| PRPF31 | 811 | 11 | 11 | 11 | 34.9 | 34.9 | 34.9 | 54.766 |
| P2RX4 | 3905 | 11 | 11 | 11 | 40.5 | 40.5 | 40.5 | 43.369 |
| CYP27B1 | 1419 | 11 | 11 | 11 | 34.8 | 34.8 | 34.8 | 56.503 |
| SEPTIN11 | 748 | 20 | 20 | 11 | 52.5 | 52.5 | 34.8 | 49.005 |
| GLYR1 | 1231 | 11 | 11 | 11 | 29.8 | 29.8 | 29.8 | 57.215 |
| ECH1 | 2831 | 16 | 16 | 11 | 71 | 71 | 48.2 | 35.816 |
| DCTN4 | 4472 | 11 | 11 | 11 | 36.3 | 36.3 | 36.3 | 52.337 |
| VAPB | 1636 | 12 | 11 | 11 | 50.6 | 50.6 | 50.6 | 27.228 |
| RHOT1 | 1117 | 12 | 11 | 11 | 27 | 25.1 | 25.1 | 68.449 |
| SNAP23 | 1285 | 11 | 11 | 11 | 70.6 | 70.6 | 70.6 | 23.354 |
| NADK2 | 3111 | 11 | 11 | 11 | 35.7 | 35.7 | 35.7 | 49.432 |
| DDX39B | 2911 | 24 | 24 | 11 | 67.3 | 67.3 | 38.6 | 48.991 |
| GNAQ | 2400 | 18 | 17 | 11 | 59.6 | 57.4 | 34.8 | 42.142 |
| RMDN1 | 3775 | 11 | 11 | 11 | 37.3 | 37.3 | 37.3 | 35.808 |
| TRA2B | 2646 | 13 | 13 | 11 | 41.7 | 41.7 | 41.7 | 33.665 |
| RAB10 | 2564 | 13 | 12 | 11 | 66.5 | 61 | 57.5 | 22.541 |
| PDCD6 | 1533 | 11 | 11 | 11 | 67.5 | 67.5 | 67.5 | 21.868 |
| PGLS | 1640 | 11 | 11 | 11 | 57.4 | 57.4 | 57.4 | 27.547 |
| SLC9A3R1 | 1352 | 11 | 11 | 11 | 44.4 | 44.4 | 44.4 | 38.868 |
| IMPDH2 | 1893 | 12 | 12 | 11 | 34 | 34 | 32.5 | 55.804 |
| TRIM38 | 1329 | 11 | 11 | 11 | 25.8 | 25.8 | 25.8 | 53.415 |
| RAB7B | 3745 | 11 | 11 | 11 | 66.8 | 66.8 | 66.8 | 22.511 |
| MAP7D1 | 3095 | 11 | 11 | 11 | 13.9 | 13.9 | 13.9 | 92.819 |
| ATP1B3 | 2494 | 11 | 11 | 11 | 43 | 43 | 43 | 31.512 |
| POFUT1 | 4135 | 11 | 11 | 11 | 46.4 | 46.4 | 46.4 | 43.955 |
| NUP160 | 993 | 11 | 11 | 11 | 10.7 | 10.7 | 10.7 | 162.12 |
| CD109 | 3305 | 11 | 11 | 11 | 9.9 | 9.9 | 9.9 | 161.69 |
| CACYBP | 4183 | 11 | 11 | 11 | 58.3 | 58.3 | 58.3 | 26.21 |
| VPS26B | 3108 | 13 | 11 | 11 | 48.2 | 46.1 | 46.1 | 39.154 |
| DDI2 | 3202 | 11 | 11 | 11 | 32.6 | 32.6 | 32.6 | 44.522 |
| DAZAP1 | 1213 | 11 | 11 | 11 | 37.2 | 37.2 | 37.2 | 43.312 |
| NDUFA12 | 4449 | 11 | 11 | 11 | 74.5 | 74.5 | 74.5 | 17.114 |
| GLUL | 1937 | 11 | 11 | 11 | 37.8 | 37.8 | 37.8 | 42.064 |
| TIMM50 | 3105 | 11 | 11 | 11 | 34.3 | 34.3 | 34.3 | 39.646 |
| HOOK3 | 3409 | 11 | 11 | 11 | 18 | 18 | 18 | 83.125 |
| PRDX2 | 2193 | 12 | 11 | 11 | 57.1 | 51.5 | 51.5 | 21.892 |

| | | | | | | | | |
|---|---|---|---|---|---|---|---|---|
| COPG2 | 4400 | 16 | 11 | 11 | 26.6 | 20.1 | 20.1 | 97.621 |
| TARDBP;TD | 2841 | 11 | 11 | 11 | 41.3 | 41.3 | 41.3 | 44.739 |
| HLA-DRA | 253 | 12 | 12 | 11 | 35 | 35 | 35 | 28.621 |
| ERGIC1 | 3735 | 11 | 11 | 11 | 46.2 | 46.2 | 46.2 | 32.592 |
| TACO1 | 3969 | 11 | 11 | 11 | 53.5 | 53.5 | 53.5 | 32.477 |
| COG5 | 4523 | 11 | 11 | 11 | 16.2 | 16.2 | 16.2 | 92.742 |
| OTUB1 | 1180 | 11 | 11 | 11 | 48.9 | 48.9 | 48.9 | 31.44 |
| QKI | 3857 | 11 | 11 | 11 | 43.4 | 43.4 | 43.4 | 37.67 |
| CARM1 | 3419 | 11 | 11 | 11 | 27.5 | 27.5 | 27.5 | 65.853 |
| BRCC3 | 2323 | 11 | 11 | 11 | 49.1 | 49.1 | 49.1 | 36.072 |
| CAPZA1 | 2468 | 13 | 11 | 11 | 73.1 | 63.6 | 63.6 | 32.922 |
| IGF2BP2 | 978 | 12 | 12 | 11 | 25.3 | 25.3 | 23.1 | 66.785 |
| CDC42 | 2555 | 12 | 11 | 11 | 63.4 | 57.6 | 57.6 | 21.258 |
| WDR81 | 271 | 11 | 11 | 11 | 10.2 | 10.2 | 10.2 | 211.69 |
| RUFY1 | 3885 | 27 | 27 | 11 | 48 | 48 | 24.3 | 79.817 |
| RBBP4 | 2800 | 16 | 16 | 11 | 58.8 | 58.8 | 48.2 | 47.655 |
| NCLN | 3733 | 25 | 25 | 11 | 47.8 | 47.8 | 23.6 | 62.974 |
| MTPAP | 4308 | 11 | 11 | 11 | 29.9 | 29.9 | 29.9 | 66.171 |
| ERLIN1 | 1553 | 13 | 11 | 11 | 50.6 | 47.1 | 47.1 | 39.171 |
| PSMA5 | 2132 | 11 | 11 | 11 | 60.2 | 60.2 | 60.2 | 26.411 |
| HNRNPH1 | 1026 | 21 | 21 | 11 | 56.3 | 56.3 | 34.3 | 49.229 |
| PPIA | 2641 | 11 | 11 | 11 | 69.1 | 69.1 | 69.1 | 18.012 |
| CLIC2 | 1399 | 12 | 11 | 11 | 67.2 | 64 | 64 | 28.356 |
| APP | 1745 | 12 | 11 | 11 | 17 | 16 | 16 | 86.942 |
| DSTN | 2556 | 12 | 11 | 11 | 68.5 | 64.2 | 64.2 | 18.506 |
| GLRX3 | 1596 | 11 | 11 | 11 | 47.5 | 47.5 | 47.5 | 37.432 |
| PTPN11 | 2773 | 11 | 11 | 11 | 22.3 | 22.3 | 22.3 | 68.01 |
| COG1 | 44 | 11 | 11 | 11 | 15.7 | 15.7 | 15.7 | 104.79 |
| MRPS25 | 2692 | 11 | 11 | 11 | 65.9 | 65.9 | 65.9 | 20.116 |
| RAD21 | 1479 | 11 | 11 | 11 | 19.2 | 19.2 | 19.2 | 71.689 |
| RBM4;RBM | 4019 | 11 | 11 | 11 | 37.9 | 37.9 | 37.9 | 40.313 |
| RSL1D1 | 1597 | 11 | 11 | 11 | 28.4 | 28.4 | 28.4 | 54.972 |
| PRPF3 | 1437 | 11 | 11 | 11 | 22.4 | 22.4 | 22.4 | 77.528 |
| TMEM173 | 3416 | 11 | 11 | 11 | 45.6 | 45.6 | 45.6 | 42.192 |
| VPS4B | 1535 | 13 | 13 | 11 | 38.1 | 38.1 | 33.6 | 49.301 |
| RPS6 | 2624 | 11 | 11 | 11 | 36.9 | 36.9 | 36.9 | 28.68 |
| VPS53 | 3211 | 11 | 11 | 11 | 21.5 | 21.5 | 21.5 | 94.404 |
| SUMF2 | 1177 | 11 | 11 | 11 | 34.7 | 34.7 | 34.7 | 35.922 |
| PCYT1A | 669 | 11 | 11 | 11 | 34.7 | 34.7 | 34.7 | 43.263 |
| ARPC3 | 1390 | 11 | 11 | 11 | 66.3 | 66.3 | 66.3 | 20.546 |
| IKBKB | 444 | 11 | 11 | 11 | 21.4 | 21.4 | 21.4 | 79.104 |
| RDH13 | 3540 | 11 | 11 | 11 | 39 | 39 | 39 | 35.932 |
| UFD1L | 3703 | 11 | 11 | 11 | 47.2 | 47.2 | 47.2 | 34.5 |
| NEU1 | 3899 | 11 | 11 | 11 | 30.8 | 30.8 | 30.8 | 45.467 |
| CTSZ | 4408 | 11 | 11 | 11 | 48.8 | 48.8 | 48.8 | 33.868 |
| WDR61 | 4078 | 11 | 11 | 11 | 52.1 | 52.1 | 52.1 | 33.58 |
| ZFR | 3835 | 11 | 11 | 11 | 14.4 | 14.4 | 14.4 | 117.01 |
| NDUFB10 | 1679 | 11 | 11 | 11 | 54.7 | 54.7 | 54.7 | 20.776 |

| | | | | | | | | |
|---|---|---|---|---|---|---|---|---|
| ATP6V0A2 | 4612 | 12 | 11 | 11 | 18.6 | 17.6 | 17.6 | 98.081 |
| CTBP1 | 2863 | 13 | 13 | 11 | 46.6 | 46.6 | 42.7 | 47.535 |
| PRDX3 | 2160 | 11 | 11 | 11 | 55.1 | 55.1 | 55.1 | 27.692 |
| HEBP1 | 4253 | 11 | 11 | 11 | 67.7 | 67.7 | 67.7 | 21.097 |
| PSMA3 | 2102 | 13 | 13 | 11 | 51.8 | 51.8 | 40 | 28.433 |
| PDLIM4 | 2412 | 11 | 11 | 11 | 49.1 | 49.1 | 49.1 | 35.398 |
| CDKN2AIP | 4341 | 11 | 11 | 11 | 24.1 | 24.1 | 24.1 | 61.124 |
| AKR7A2 | 1442 | 11 | 11 | 11 | 41.5 | 41.5 | 41.5 | 39.589 |
| TIMMDC1 | 4222 | 11 | 11 | 11 | 40 | 40 | 40 | 32.177 |
| STX7 | 1411 | 11 | 11 | 11 | 56.3 | 56.3 | 56.3 | 29.815 |
| PNKD | 3505 | 11 | 11 | 11 | 42.1 | 42.1 | 42.1 | 42.875 |
| SPRYD4 | 3652 | 11 | 11 | 11 | 68.1 | 68.1 | 68.1 | 23.128 |
| SPTLC1 | 1404 | 11 | 11 | 11 | 30 | 30 | 30 | 52.743 |
| ASAP1 | 129 | 11 | 11 | 11 | 14.1 | 14.1 | 14.1 | 124.77 |
| UBTF | 861 | 11 | 11 | 11 | 20.1 | 20.1 | 20.1 | 87.435 |
| COPS2 | 2571 | 11 | 11 | 11 | 31.6 | 31.6 | 31.6 | 51.596 |
| LPCAT1 | 3569 | 11 | 11 | 11 | 29.4 | 29.4 | 29.4 | 59.151 |
| YBX1 | 2663 | 16 | 16 | 11 | 66.7 | 66.7 | 54 | 35.924 |
| OLA1 | 1173 | 11 | 11 | 11 | 38.5 | 38.5 | 38.5 | 46.937 |
| APOOL | 71 | 11 | 11 | 11 | 62.3 | 62.3 | 62.3 | 29.159 |
| PSTPIP2 | 4167 | 11 | 11 | 11 | 39.2 | 39.2 | 39.2 | 38.858 |
| MTX1 | 2882 | 11 | 11 | 11 | 27.5 | 27.5 | 27.5 | 51.462 |
| PURB | 3867 | 11 | 11 | 11 | 56.4 | 56.4 | 56.4 | 33.24 |
| PPP2R2A | 2651 | 11 | 11 | 11 | 40 | 40 | 40 | 51.691 |
| AP3D1 | 1345 | 42 | 42 | 11 | 50.8 | 50.8 | 12.7 | 130.16 |
| HACD3 | 4371 | 11 | 11 | 11 | 30.7 | 30.7 | 30.7 | 43.159 |
| RPL24 | 684 | 11 | 11 | 11 | 54.5 | 54.5 | 54.5 | 14.369 |
| DECR1 | 3063 | 21 | 21 | 11 | 75.5 | 75.5 | 40.9 | 36.067 |
| USE1 | 4355 | 11 | 11 | 11 | 37.1 | 37.1 | 37.1 | 29.371 |
| NAPA | 2500 | 19 | 19 | 11 | 79.3 | 79.3 | 51.5 | 33.232 |
| RAB32 | 2904 | 13 | 13 | 11 | 67.6 | 67.6 | 58.2 | 24.997 |
| TACC1 | 1546 | 13 | 13 | 11 | 19.4 | 19.4 | 17 | 87.793 |
| CUL4B | 476 | 18 | 11 | 11 | 24 | 15.6 | 15.6 | 102.53 |
| EMD | 2406 | 11 | 11 | 11 | 44.5 | 44.5 | 44.5 | 28.994 |
| PGRMC1 | 1297 | 12 | 12 | 11 | 59 | 59 | 54.4 | 21.671 |
| BAG2 | 1671 | 11 | 11 | 11 | 64.5 | 64.5 | 64.5 | 23.772 |
| RRAS | 1854 | 12 | 12 | 11 | 58.7 | 58.7 | 58.7 | 23.48 |
| PIGK | 3689 | 11 | 11 | 11 | 41.3 | 41.3 | 41.3 | 45.251 |
| HSD3B7 | 4114 | 11 | 11 | 11 | 48.5 | 48.5 | 48.5 | 41.016 |
| RP2 | 1571 | 11 | 11 | 11 | 32 | 32 | 32 | 39.641 |
| DHRS7B | 217 | 11 | 11 | 11 | 43.2 | 43.2 | 43.2 | 33.524 |
| PRDX5 | 2157 | 11 | 11 | 11 | 65.9 | 65.9 | 65.9 | 22.086 |
| SULT1C2 | 1303 | 11 | 11 | 11 | 53.4 | 53.4 | 53.4 | 34.88 |
| ENDOD1 | 1617 | 11 | 11 | 11 | 40.8 | 40.8 | 40.8 | 55.016 |
| L2HGDH | 4173 | 11 | 11 | 11 | 38.9 | 38.9 | 38.9 | 50.315 |
| EIF2B2 | 2386 | 11 | 11 | 11 | 41.3 | 41.3 | 41.3 | 38.989 |
| SLK | 4115 | 12 | 11 | 11 | 12.5 | 11.6 | 11.6 | 142.69 |
| MRPS26 | 4050 | 11 | 11 | 11 | 51.2 | 51.2 | 51.2 | 24.211 |

| | | | | | | | | |
|---|---|---|---|---|---|---|---|---|
| AGPAT4 | 4259 | 11 | 11 | 11 | 29.6 | 29.6 | 29.6 | 44.021 |
| MRPL19 | 2368 | 11 | 11 | 11 | 37.7 | 37.7 | 37.7 | 33.535 |
| PDE12 | 3248 | 11 | 11 | 11 | 25.9 | 25.9 | 25.9 | 67.351 |
| GOLGA4 | 1040 | 11 | 11 | 11 | 5.9 | 5.9 | 5.9 | 246.6 |
| GOLPH3 | 4139 | 11 | 11 | 11 | 44.6 | 44.6 | 44.6 | 33.81 |
| COPS5 | 3707 | 11 | 11 | 11 | 37.7 | 37.7 | 37.7 | 37.578 |
| AIP | 1286 | 11 | 11 | 11 | 47.9 | 47.9 | 47.9 | 37.636 |
| FBL | 2050 | 14 | 14 | 11 | 53.3 | 53.3 | 49.8 | 33.784 |
| RPL17;RPL1 | 64 | 11 | 11 | 11 | 56.8 | 56.8 | 56.8 | 19.586 |
| ATAD3A | 4299 | 29 | 29 | 11 | 42.6 | 42.6 | 16.2 | 71.368 |
| STX18 | 750 | 11 | 11 | 11 | 46 | 46 | 46 | 38.673 |
| LMAN2L | 4097 | 11 | 11 | 11 | 46.8 | 46.8 | 46.8 | 39.71 |
| KIF13B | 4237 | 12 | 12 | 11 | 7.1 | 7.1 | 6.6 | 202.79 |
| ELMOD2 | 3475 | 11 | 11 | 11 | 35.8 | 35.8 | 35.8 | 34.96 |
| H1FX | 3670 | 11 | 11 | 11 | 49.3 | 49.3 | 49.3 | 22.487 |
| STK4 | 2833 | 11 | 11 | 11 | 28.7 | 28.7 | 28.7 | 55.63 |
| SNX9 | 4648 | 11 | 11 | 11 | 25 | 25 | 25 | 66.591 |
| TRMT1L | 3354 | 11 | 11 | 11 | 24.4 | 24.4 | 24.4 | 81.746 |
| NT5C2 | 2392 | 11 | 11 | 11 | 29.6 | 29.6 | 29.6 | 64.969 |
| MRPS18B | 4661 | 11 | 11 | 11 | 59.7 | 59.7 | 59.7 | 29.395 |
| COL6A1 | 94 | 10 | 10 | 10 | 14.6 | 14.6 | 14.6 | 108.34 |
| MAP7 | 2940 | 10 | 10 | 10 | 12 | 12 | 12 | 84.051 |
| EIF2B4 | 19 | 10 | 10 | 10 | 33.8 | 33.8 | 33.8 | 57.495 |
| SLC38A10 | 4190 | 10 | 10 | 10 | 10.9 | 10.9 | 10.9 | 119.76 |
| EIF3J | 1575 | 10 | 10 | 10 | 50.8 | 50.8 | 50.8 | 29.062 |
| CPSF7 | 3522 | 10 | 10 | 10 | 26.3 | 26.3 | 26.3 | 52.049 |
| HDGFRP2 | 3372 | 11 | 11 | 10 | 21.5 | 21.5 | 18.9 | 74.316 |
| SEPTIN6 | 560 | 17 | 10 | 10 | 50.6 | 38.1 | 38.1 | 49.303 |
| PLA2G15 | 3550 | 10 | 10 | 10 | 28.6 | 28.6 | 28.6 | 46.657 |
| DGKZ | 873 | 10 | 10 | 10 | 19.9 | 19.9 | 19.9 | 78.927 |
| RAB6A | 2022 | 14 | 14 | 10 | 56.2 | 56.2 | 46.6 | 23.593 |
| DSCR3 | 1371 | 10 | 10 | 10 | 45.1 | 45.1 | 45.1 | 33.01 |
| SLIRP | 4079 | 10 | 10 | 10 | 77.1 | 77.1 | 77.1 | 12.349 |
| WBSCR16 | 3815 | 10 | 10 | 10 | 36.2 | 36.2 | 36.2 | 49.996 |
| SCAMP3 | 1358 | 10 | 10 | 10 | 42.9 | 42.9 | 42.9 | 38.287 |
| PTBP3 | 1668 | 13 | 10 | 10 | 37 | 32.4 | 32.4 | 59.689 |
| ABHD11 | 3578 | 10 | 10 | 10 | 59.7 | 59.7 | 59.7 | 34.69 |
| YWHAH | 2765 | 13 | 10 | 10 | 54.1 | 50 | 50 | 28.218 |
| LSP1 | 2204 | 20 | 20 | 10 | 70.5 | 70.5 | 32.4 | 37.191 |
| FXR1 | 2422 | 13 | 13 | 10 | 31.4 | 31.4 | 26.7 | 69.72 |
| ARF6 | 2615 | 10 | 10 | 10 | 62.9 | 62.9 | 62.9 | 20.082 |
| EIF3K | 4406 | 10 | 10 | 10 | 66.5 | 66.5 | 66.5 | 25.059 |
| STX3 | 209 | 10 | 10 | 10 | 48.5 | 48.5 | 48.5 | 23.243 |
| AGPAT3 | 4260 | 10 | 10 | 10 | 25.3 | 25.3 | 25.3 | 43.38 |
| THEM6 | 3644 | 10 | 10 | 10 | 54.8 | 54.8 | 54.8 | 23.865 |
| CDK5 | 2723 | 11 | 11 | 10 | 48.3 | 48.3 | 45.5 | 33.304 |
| MRPS2 | 4587 | 10 | 10 | 10 | 32.4 | 32.4 | 32.4 | 33.249 |
| MAPRE1 | 3032 | 12 | 12 | 10 | 63.8 | 63.8 | 57.5 | 29.999 |

| | | | | | | | | |
|---|---|---|---|---|---|---|---|---|
| PSMB3 | 2377 | 10 | 10 | 10 | 54.6 | 54.6 | 54.6 | 22.949 |
| NIPSNAP3A | 4421 | 10 | 10 | 10 | 54.7 | 54.7 | 54.7 | 28.466 |
| CYP4X1 | 3482 | 10 | 10 | 10 | 24.8 | 24.8 | 24.8 | 58.875 |
| TIGAR | 4228 | 10 | 10 | 10 | 57 | 57 | 57 | 30.062 |
| DDX3X | 228 | 36 | 35 | 10 | 57.9 | 57.9 | 16.3 | 73.214 |
| UAP1L1 | 3096 | 10 | 10 | 10 | 27.2 | 27.2 | 27.2 | 57.03 |
| CTBS | 2732 | 10 | 10 | 10 | 39 | 39 | 39 | 43.759 |
| MRPL1 | 4046 | 10 | 10 | 10 | 39.1 | 39.1 | 39.1 | 36.908 |
| TLDC1 | 3276 | 10 | 10 | 10 | 35.7 | 35.7 | 35.7 | 50.993 |
| HLA-DMB | 2133 | 10 | 10 | 10 | 64.3 | 64.3 | 64.3 | 28.943 |
| SPTBN1 | 37 | 11 | 10 | 10 | 5.7 | 5.2 | 5.2 | 274.61 |
| EIF4E | 740 | 10 | 10 | 10 | 56.7 | 56.7 | 56.7 | 28.512 |
| SCARB1 | 623 | 10 | 10 | 10 | 26.2 | 26.2 | 26.2 | 53.847 |
| TK2 | 1280 | 10 | 10 | 10 | 43.4 | 43.4 | 43.4 | 31.004 |
| SH3GLB1 | 4583 | 11 | 11 | 10 | 40.3 | 40.3 | 38.1 | 40.796 |
| COMMD10 | 4671 | 10 | 10 | 10 | 54 | 54 | 54 | 22.966 |
| FAHD1 | 3273 | 10 | 10 | 10 | 60.7 | 60.7 | 60.7 | 24.843 |
| CTSL | 1786 | 11 | 10 | 10 | 49.8 | 47.7 | 47.7 | 37.564 |
| ECM1 | 3057 | 10 | 10 | 10 | 23 | 23 | 23 | 60.673 |
| CSNK2A1;C | 351 | 10 | 10 | 10 | 31.5 | 31.5 | 31.5 | 45.143 |
| LARP4B | 3685 | 10 | 10 | 10 | 19.9 | 19.9 | 19.9 | 80.551 |
| NDUFB9 | 4680 | 10 | 10 | 10 | 65.9 | 65.9 | 65.9 | 21.831 |
| OSTF1 | 3702 | 10 | 10 | 10 | 57.9 | 57.9 | 57.9 | 23.787 |
| BRIX1 | 3620 | 10 | 10 | 10 | 35.7 | 35.7 | 35.7 | 41.401 |
| NDUFA8 | 2450 | 10 | 10 | 10 | 65.7 | 65.7 | 65.7 | 20.105 |
| MBNL1 | 122 | 10 | 10 | 10 | 30.7 | 30.7 | 30.7 | 37.898 |
| CCDC90B | 4080 | 10 | 10 | 10 | 39.4 | 39.4 | 39.4 | 29.505 |
| RCC1 | 1998 | 10 | 10 | 10 | 37.8 | 37.8 | 37.8 | 44.969 |
| CIAO1 | 1600 | 10 | 10 | 10 | 51.3 | 51.3 | 51.3 | 37.84 |
| MRPL46 | 4122 | 10 | 10 | 10 | 45.2 | 45.2 | 45.2 | 31.705 |
| B3GALTL | 3304 | 10 | 10 | 10 | 38.6 | 38.6 | 38.6 | 56.563 |
| MRPS23 | 4599 | 10 | 10 | 10 | 60.5 | 60.5 | 60.5 | 21.77 |
| FDX1 | 1847 | 10 | 10 | 10 | 39.1 | 39.1 | 39.1 | 19.393 |
| RINT1 | 3254 | 10 | 10 | 10 | 17.8 | 17.8 | 17.8 | 90.631 |
| THNSL1 | 3471 | 10 | 10 | 10 | 17.9 | 17.9 | 17.9 | 83.069 |
| COMMD9 | 4367 | 10 | 10 | 10 | 71.2 | 71.2 | 71.2 | 21.819 |
| SETD3 | 3394 | 10 | 10 | 10 | 18.5 | 18.5 | 18.5 | 67.256 |
| AP3M1 | 4569 | 10 | 10 | 10 | 42.1 | 42.1 | 42.1 | 46.939 |
| NUDT19 | 549 | 10 | 10 | 10 | 36.8 | 36.8 | 36.8 | 42.233 |
| RPS19 | 2256 | 10 | 10 | 10 | 58.6 | 58.6 | 58.6 | 16.06 |
| CASP1 | 2148 | 12 | 12 | 10 | 38.9 | 38.9 | 34.4 | 45.158 |
| DHX36 | 813 | 10 | 10 | 10 | 15.4 | 15.4 | 15.4 | 91.43 |
| NAA15 | 4030 | 14 | 14 | 10 | 16.6 | 16.6 | 12.2 | 101.27 |
| EXD2 | 4296 | 10 | 10 | 10 | 21.9 | 21.9 | 21.9 | 70.352 |
| RPL12 | 2162 | 10 | 10 | 10 | 63.6 | 63.6 | 63.6 | 17.818 |
| AHSA1 | 1649 | 10 | 10 | 10 | 45.6 | 45.6 | 45.6 | 38.274 |
| RAB9A | 2427 | 10 | 10 | 10 | 59.7 | 59.7 | 59.7 | 22.837 |
| MRPL45 | 111 | 10 | 10 | 10 | 42.8 | 42.8 | 42.8 | 35.29 |

| | | | | | | | | |
|---|---|---|---|---|---|---|---|---|
| CELF2 | 268 | 11 | 11 | 10 | 30 | 30 | 27.7 | 46.621 |
| FKBP3 | 2726 | 10 | 10 | 10 | 50.4 | 50.4 | 50.4 | 25.177 |
| NIT1 | 3420 | 10 | 10 | 10 | 41.9 | 41.9 | 41.9 | 35.896 |
| SUPT6H | 3327 | 10 | 10 | 10 | 7.6 | 7.6 | 7.6 | 199.07 |
| PSMD8 | 2351 | 10 | 10 | 10 | 38 | 38 | 38 | 32.551 |
| GOSR1 | 826 | 10 | 10 | 10 | 46.2 | 46.2 | 46.2 | 26.989 |
| SDF4 | 457 | 10 | 10 | 10 | 47 | 47 | 47 | 40.936 |
| TMEM33 | 2533 | 10 | 10 | 10 | 39.3 | 39.3 | 39.3 | 27.978 |
| MVD | 2478 | 10 | 10 | 10 | 47.2 | 47.2 | 47.2 | 43.404 |
| APPL1 | 4482 | 10 | 10 | 10 | 20.9 | 20.9 | 20.9 | 79.663 |
| FABP3 | 1758 | 11 | 11 | 10 | 71.4 | 71.4 | 66.2 | 14.858 |
| ELAC2 | 3938 | 10 | 10 | 10 | 17.8 | 17.8 | 17.8 | 92.218 |
| NUP54 | 3359 | 10 | 10 | 10 | 24.7 | 24.7 | 24.7 | 55.435 |
| SYMPK | 3696 | 10 | 10 | 10 | 13 | 13 | 13 | 141.15 |
| PGP | 512 | 10 | 10 | 10 | 45.2 | 45.2 | 45.2 | 34.006 |
| MT-CO2 | 1691 | 10 | 10 | 10 | 48 | 48 | 48 | 25.565 |
| EMC8 | 1440 | 10 | 10 | 10 | 56.2 | 56.2 | 56.2 | 23.773 |
| LMF2 | 3986 | 10 | 10 | 10 | 22.9 | 22.9 | 22.9 | 79.697 |
| DDX18 | 4303 | 10 | 10 | 10 | 19.7 | 19.7 | 19.7 | 75.406 |
| ABCB8 | 4286 | 11 | 10 | 10 | 18.6 | 17.6 | 17.6 | 79.988 |
| ATP5H | 1592 | 10 | 10 | 10 | 69.6 | 69.6 | 69.6 | 18.491 |
| IFIT3 | 1361 | 10 | 10 | 10 | 29.4 | 29.4 | 29.4 | 55.984 |
| MRPS31 | 3690 | 10 | 10 | 10 | 37.2 | 37.2 | 37.2 | 45.318 |
| LPP | 3720 | 10 | 10 | 10 | 26.1 | 26.1 | 26.1 | 65.746 |
| SMU1 | 3086 | 10 | 10 | 10 | 24.8 | 24.8 | 24.8 | 57.543 |
| NPL | 4029 | 10 | 10 | 10 | 39.1 | 39.1 | 39.1 | 35.162 |
| HNRNPDL | 32 | 12 | 10 | 10 | 26.7 | 24 | 24 | 40.04 |
| GALE | 2950 | 10 | 10 | 10 | 40.8 | 40.8 | 40.8 | 38.281 |
| DRG1 | 4551 | 10 | 10 | 10 | 34.6 | 34.6 | 34.6 | 40.542 |
| HS2ST1 | 3350 | 10 | 10 | 10 | 38.8 | 38.8 | 38.8 | 41.881 |
| ACTR1A | 2570 | 19 | 19 | 10 | 66.5 | 66.5 | 40.7 | 42.613 |
| PET112;GA | 744 | 10 | 10 | 10 | 34.5 | 34.5 | 34.5 | 57.653 |
| TRAPPC11 | 3358 | 10 | 10 | 10 | 14.9 | 14.9 | 14.9 | 128.88 |
| UBAP2L | 969 | 10 | 10 | 10 | 14.3 | 14.3 | 14.3 | 113.63 |
| RAB12 | 402 | 10 | 10 | 10 | 32.1 | 32.1 | 32.1 | 36.315 |
| PITPNA | 891 | 11 | 11 | 10 | 47.8 | 47.8 | 44.4 | 31.802 |
| HDAC1 | 2889 | 16 | 16 | 10 | 42.1 | 42.1 | 30.9 | 55.102 |
| ARMC10 | 3493 | 10 | 10 | 10 | 44.9 | 44.9 | 44.9 | 37.54 |
| RTCA | 1313 | 10 | 10 | 10 | 41.8 | 41.8 | 41.8 | 39.336 |
| BAG6;BAT3 | 8 | 10 | 10 | 10 | 13.1 | 13.1 | 13.1 | 119.41 |
| THUMPD1 | 4335 | 10 | 10 | 10 | 34.6 | 34.6 | 34.6 | 39.315 |
| PRRC1 | 3842 | 10 | 10 | 10 | 31.2 | 31.2 | 31.2 | 46.701 |
| DIABLO | 4241 | 10 | 10 | 10 | 51 | 51 | 51 | 27.131 |
| PAICS | 2053 | 10 | 10 | 10 | 33.9 | 33.9 | 33.9 | 47.079 |
| RPL32 | 724 | 10 | 10 | 10 | 57.1 | 57.1 | 57.1 | 15.616 |
| PSMD4 | 2503 | 13 | 13 | 10 | 54.1 | 54.1 | 38.2 | 40.736 |
| SAE1 | 4399 | 10 | 10 | 10 | 41.3 | 41.3 | 41.3 | 38.449 |
| ASNA1 | 1456 | 10 | 10 | 10 | 35.6 | 35.6 | 35.6 | 38.792 |

| | | | | | | | | |
|---|---|---|---|---|---|---|---|---|
| PSMA2 | 6 | 10 | 10 | 10 | 59.4 | 59.4 | 59.4 | 25.898 |
| GALM | 3764 | 10 | 10 | 10 | 47.4 | 47.4 | 47.4 | 37.765 |
| SIAE | 4179 | 10 | 10 | 10 | 30.4 | 30.4 | 30.4 | 58.314 |
| NAGA | 1970 | 10 | 10 | 10 | 32.6 | 32.6 | 32.6 | 46.564 |
| PELP1 | 3478 | 10 | 10 | 10 | 15 | 15 | 15 | 119.7 |
| SET;SETSIP | 194 | 10 | 10 | 10 | 52.3 | 52.3 | 52.3 | 31.124 |
| DECR2 | 4274 | 10 | 10 | 10 | 42.1 | 42.1 | 42.1 | 30.777 |
| BAZ1B | 4455 | 10 | 10 | 10 | 8.7 | 8.7 | 8.7 | 170.9 |
| NAT10 | 4088 | 10 | 10 | 10 | 12 | 12 | 12 | 115.73 |
| LRCH1 | 4562 | 11 | 11 | 10 | 21.8 | 21.8 | 20.3 | 80.874 |
| RPL9 | 2201 | 10 | 10 | 10 | 65.1 | 65.1 | 65.1 | 21.863 |
| TP53I3 | 3123 | 10 | 10 | 10 | 43.7 | 43.7 | 43.7 | 35.536 |
| ECI1;DCI | 2280 | 10 | 10 | 10 | 47.7 | 47.7 | 47.7 | 32.816 |
| COX4I1 | 1905 | 10 | 10 | 10 | 47.3 | 47.3 | 47.3 | 19.576 |
| SPCS2 | 29 | 10 | 10 | 10 | 46.7 | 46.7 | 46.7 | 25.09 |
| ARSA | 196 | 10 | 10 | 10 | 32.6 | 32.6 | 32.6 | 53.806 |
| EDF1 | 1515 | 10 | 10 | 10 | 54.1 | 54.1 | 54.1 | 16.368 |
| TUBB6 | 3990 | 22 | 12 | 10 | 70.4 | 45.1 | 40.4 | 49.857 |
| LAMP1 | 1878 | 10 | 10 | 10 | 18.7 | 18.7 | 18.7 | 44.882 |
| STK24 | 588 | 14 | 14 | 10 | 43 | 43 | 32.8 | 45.835 |
| METTL7B | 84 | 10 | 10 | 10 | 44.9 | 44.9 | 44.9 | 31.288 |
| STX8 | 4515 | 10 | 10 | 10 | 61.4 | 61.4 | 61.4 | 26.906 |
| TMF1 | 2691 | 10 | 10 | 10 | 10.8 | 10.8 | 10.8 | 122.84 |
| NPEPL1 | 3561 | 10 | 10 | 10 | 36.5 | 36.5 | 36.5 | 55.86 |
| RTN4IP1 | 3656 | 10 | 10 | 10 | 37.4 | 37.4 | 37.4 | 43.589 |
| YWHAQ | 2121 | 14 | 10 | 10 | 53.5 | 42.4 | 42.4 | 27.764 |
| COX5A | 2027 | 10 | 10 | 10 | 52.7 | 52.7 | 52.7 | 16.762 |
| CELF1 | 1023 | 11 | 10 | 10 | 22.4 | 20.4 | 20.4 | 55.142 |
| HIBCH | 3257 | 18 | 18 | 10 | 51 | 51 | 30.3 | 43.482 |
| SF3A2 | 3022 | 10 | 10 | 10 | 24.6 | 24.6 | 24.6 | 49.255 |
| RPS17 | 1090 | 10 | 10 | 10 | 14.6 | 14.6 | 14.6 | 64.532 |
| TWF1 | 2815 | 13 | 10 | 10 | 43.7 | 37.7 | 37.7 | 40.282 |
| M6PR | 2026 | 10 | 10 | 10 | 41.9 | 41.9 | 41.9 | 30.993 |
| LIPA | 153 | 10 | 10 | 10 | 54.4 | 54.4 | 54.4 | 32.57 |
| COQ6 | 4576 | 12 | 12 | 10 | 39.3 | 39.3 | 35.7 | 50.869 |
| TEX264 | 4676 | 10 | 10 | 10 | 43.5 | 43.5 | 43.5 | 34.188 |
| TMED10 | 2381 | 10 | 10 | 10 | 42.9 | 42.9 | 42.9 | 24.976 |
| NAPG | 3921 | 10 | 10 | 10 | 36.5 | 36.5 | 36.5 | 34.746 |
| MRPS30 | 4213 | 10 | 10 | 10 | 29.2 | 29.2 | 29.2 | 50.364 |
| OXSR1 | 1667 | 12 | 12 | 10 | 32.6 | 32.6 | 27.3 | 58.022 |
| LYZ | 2583 | 10 | 10 | 10 | 68.9 | 68.9 | 68.9 | 16.537 |
| ARMC8 | 3436 | 10 | 10 | 10 | 16 | 16 | 16 | 75.509 |
| PRMT5 | 1351 | 10 | 10 | 10 | 28.4 | 28.4 | 28.4 | 72.683 |
| GNAI3 | 1811 | 17 | 10 | 10 | 63.8 | 42.4 | 42.4 | 40.532 |
| LEMD3 | 4570 | 10 | 10 | 10 | 13.1 | 13.1 | 13.1 | 99.996 |
| GORASP2 | 4164 | 10 | 10 | 10 | 32.3 | 32.3 | 32.3 | 47.145 |
| PLXDC2 | 3299 | 10 | 10 | 10 | 23.3 | 23.3 | 23.3 | 59.582 |
| PLEKHM2 | 349 | 10 | 10 | 10 | 15.4 | 15.4 | 15.4 | 108.6 |

| | | | | | | | | |
|---|---|---|---|---|---|---|---|---|
| GIT1 | 214 | 13 | 10 | 10 | 24.4 | 19.7 | 19.7 | 76.805 |
| NAGLU | 2498 | 9 | 9 | 9 | 21.3 | 21.3 | 21.3 | 82.265 |
| EIF4A2 | 2939 | 22 | 9 | 9 | 63.1 | 36.4 | 36.4 | 46.402 |
| FAM162A | 971 | 9 | 9 | 9 | 42.2 | 42.2 | 42.2 | 17.342 |
| SCO1 | 1196 | 9 | 9 | 9 | 26.9 | 26.9 | 26.9 | 33.814 |
| MANF | 2508 | 9 | 9 | 9 | 46.2 | 46.2 | 46.2 | 20.7 |
| GOPC | 4205 | 9 | 9 | 9 | 26.6 | 26.6 | 26.6 | 50.519 |
| TGOLN2 | 458 | 9 | 9 | 9 | 22.3 | 22.3 | 22.3 | 51.018 |
| RNASET2 | 82 | 9 | 9 | 9 | 37.2 | 37.2 | 37.2 | 29.114 |
| DHTKD1 | 3811 | 9 | 9 | 9 | 18.3 | 18.3 | 18.3 | 103.08 |
| SEC16A | 413 | 9 | 9 | 9 | 6.7 | 6.7 | 6.7 | 228.87 |
| PRKAG1 | 961 | 9 | 9 | 9 | 48.2 | 48.2 | 48.2 | 31.905 |
| ITFG1 | 3597 | 9 | 9 | 9 | 23 | 23 | 23 | 68.107 |
| MMS19 | 3886 | 9 | 9 | 9 | 13.5 | 13.5 | 13.5 | 113.29 |
| MARCKS | 2151 | 9 | 9 | 9 | 46.7 | 46.7 | 46.7 | 31.554 |
| SLC15A4 | 3523 | 9 | 9 | 9 | 16.6 | 16.6 | 16.6 | 62.033 |
| PNPLA8 | 201 | 10 | 9 | 9 | 25.4 | 24.3 | 24.3 | 71.877 |
| SAP18 | 1310 | 9 | 9 | 9 | 65.4 | 65.4 | 65.4 | 17.561 |
| IGSF6 | 1678 | 9 | 9 | 9 | 36.5 | 36.5 | 36.5 | 27.013 |
| SLC30A9 | 3287 | 9 | 9 | 9 | 17.3 | 17.3 | 17.3 | 63.514 |
| RPL28 | 1074 | 9 | 9 | 9 | 40.6 | 40.6 | 40.6 | 19.072 |
| MSRA | 4461 | 9 | 9 | 9 | 54.5 | 54.5 | 54.5 | 26.132 |
| LOH12CR1 | 998 | 9 | 9 | 9 | 70.6 | 70.6 | 70.6 | 20.247 |
| PDLIM7 | 4239 | 9 | 9 | 9 | 27.6 | 27.6 | 27.6 | 49.844 |
| CD36 | 802 | 9 | 9 | 9 | 27 | 27 | 27 | 52.062 |
| EEF1E1;EEF | 1434 | 9 | 9 | 9 | 60.9 | 60.9 | 60.9 | 19.81 |
| EVI2B | 2213 | 9 | 9 | 9 | 19.6 | 19.6 | 19.6 | 48.666 |
| PEF1 | 4412 | 9 | 9 | 9 | 44 | 44 | 44 | 30.381 |
| C11orf54 | 4098 | 9 | 9 | 9 | 42.5 | 42.5 | 42.5 | 35.117 |
| ADPRHL2 | 4325 | 9 | 9 | 9 | 38.6 | 38.6 | 38.6 | 38.946 |
| PBDC1 | 4007 | 9 | 9 | 9 | 43.8 | 43.8 | 43.8 | 26.056 |
| LPGAT1 | 3682 | 9 | 9 | 9 | 25.4 | 25.4 | 25.4 | 43.089 |
| RPL19 | 1191 | 9 | 9 | 9 | 31.1 | 31.1 | 31.1 | 23.134 |
| OTUD4 | 2737 | 9 | 9 | 9 | 10.9 | 10.9 | 10.9 | 124.04 |
| THOC6 | 3411 | 9 | 9 | 9 | 36.1 | 36.1 | 36.1 | 37.535 |
| COX6B1 | 1932 | 9 | 9 | 9 | 83.7 | 83.7 | 83.7 | 10.192 |
| CSTF1 | 2767 | 9 | 9 | 9 | 33.4 | 33.4 | 33.4 | 48.357 |
| LCP2 | 235 | 9 | 9 | 9 | 20.1 | 20.1 | 20.1 | 60.187 |
| PPIF | 2169 | 11 | 10 | 9 | 65.2 | 61.8 | 58.5 | 22.04 |
| TCF25 | 3941 | 9 | 9 | 9 | 23.1 | 23.1 | 23.1 | 76.666 |
| PEX11B | 1682 | 9 | 9 | 9 | 32.4 | 32.4 | 32.4 | 28.431 |
| TPMT | 2438 | 9 | 9 | 9 | 35.5 | 35.5 | 35.5 | 28.18 |
| ACTL6A | 1683 | 9 | 9 | 9 | 33.1 | 33.1 | 33.1 | 47.46 |
| HIP1 | 1299 | 9 | 9 | 9 | 14.5 | 14.5 | 14.5 | 116.22 |
| FAM98B | 3118 | 12 | 12 | 9 | 40.4 | 40.4 | 31.2 | 45.547 |
| PLEKHF2 | 4163 | 9 | 9 | 9 | 41 | 41 | 41 | 27.797 |
| RAB13 | 2428 | 12 | 9 | 9 | 64.5 | 51.7 | 51.7 | 22.774 |
| HNRNPD | 2923 | 16 | 16 | 9 | 35.2 | 35.2 | 18.3 | 38.434 |

| | | | | | | | | |
|---|---|---|---|---|---|---|---|---|
| SH3BGRL | 1537 | 9 | 9 | 9 | 85.1 | 85.1 | 85.1 | 12.774 |
| PDCD11 | 412 | 9 | 9 | 9 | 6.9 | 6.9 | 6.9 | 208.83 |
| MTX2 | 1548 | 9 | 9 | 9 | 46.8 | 46.8 | 46.8 | 29.763 |
| TTN | 167 | 9 | 9 | 9 | 0.3 | 0.3 | 0.3 | 3994.6 |
| MRPS35 | 2694 | 9 | 9 | 9 | 41.2 | 41.2 | 41.2 | 36.844 |
| RAB43 | 3429 | 9 | 9 | 9 | 50 | 50 | 50 | 23.339 |
| GALK1 | 2435 | 9 | 9 | 9 | 32.9 | 32.9 | 32.9 | 42.272 |
| NAP1L1 | 924 | 11 | 11 | 9 | 45.8 | 45.8 | 43 | 45.374 |
| HCCS | 2484 | 9 | 9 | 9 | 45.9 | 45.9 | 45.9 | 30.601 |
| PTPMT1 | 3640 | 9 | 9 | 9 | 53.7 | 53.7 | 53.7 | 22.843 |
| GANAB | 2966 | 61 | 61 | 9 | 70.1 | 70.1 | 10.5 | 106.87 |
| PPP2R4 | 3006 | 9 | 9 | 9 | 37.4 | 37.4 | 37.4 | 40.667 |
| NOP2 | 2317 | 9 | 9 | 9 | 13.4 | 13.4 | 13.4 | 89.301 |
| PLA2G4C | 4522 | 9 | 9 | 9 | 25 | 25 | 25 | 60.938 |
| AGPAT2 | 1384 | 9 | 9 | 9 | 43.9 | 43.9 | 43.9 | 30.914 |
| TMEM30A | 4292 | 9 | 9 | 9 | 28.5 | 28.5 | 28.5 | 40.683 |
| NENF | 4510 | 9 | 9 | 9 | 57.6 | 57.6 | 57.6 | 18.856 |
| BCAS2 | 1588 | 9 | 9 | 9 | 65.3 | 65.3 | 65.3 | 26.131 |
| UQCRB | 1936 | 9 | 9 | 9 | 58.6 | 58.6 | 58.6 | 13.53 |
| NDUFA13 | 4373 | 9 | 9 | 9 | 66 | 66 | 66 | 16.698 |
| SPINT1 | 1430 | 9 | 9 | 9 | 20.6 | 20.6 | 20.6 | 58.397 |
| CPSF6 | 990 | 9 | 9 | 9 | 25.9 | 25.9 | 25.9 | 52.269 |
| RARS2 | 3184 | 9 | 9 | 9 | 21.6 | 21.6 | 21.6 | 65.505 |
| SUGT1 | 4574 | 9 | 9 | 9 | 36.7 | 36.7 | 36.7 | 41.024 |
| SLC29A3 | 4055 | 9 | 9 | 9 | 24.4 | 24.4 | 24.4 | 51.814 |
| BSG | 2227 | 9 | 9 | 9 | 34 | 34 | 34 | 42.2 |
| OXSM | 4316 | 9 | 9 | 9 | 40.3 | 40.3 | 40.3 | 48.842 |
| MTAP | 589 | 9 | 9 | 9 | 43 | 43 | 43 | 33.186 |
| TMED7;TMI | 170 | 9 | 9 | 9 | 50 | 50 | 50 | 25.171 |
| SNW1 | 2895 | 9 | 9 | 9 | 22.8 | 22.8 | 22.8 | 61.494 |
| NCSTN | 3672 | 9 | 9 | 9 | 18.6 | 18.6 | 18.6 | 78.41 |
| GNB2L1 | 2658 | 22 | 22 | 9 | 87.7 | 87.7 | 38.8 | 35.076 |
| BCAT1 | 2493 | 9 | 9 | 9 | 37.3 | 37.3 | 37.3 | 42.966 |
| RPS18 | 2604 | 9 | 9 | 9 | 48.7 | 48.7 | 48.7 | 17.718 |
| NME3 | 2853 | 9 | 9 | 9 | 56.2 | 56.2 | 56.2 | 19.015 |
| FCGR1A;FC | 1895 | 9 | 9 | 9 | 23.3 | 23.3 | 23.3 | 42.631 |
| ABHD3 | 3634 | 9 | 9 | 9 | 34.2 | 34.2 | 34.2 | 46.008 |
| NNT | 2868 | 50 | 50 | 9 | 47.4 | 47.4 | 7.9 | 113.89 |
| IDO1 | 1935 | 9 | 9 | 9 | 34.5 | 34.5 | 34.5 | 45.326 |
| MRAS | 1356 | 9 | 9 | 9 | 55.8 | 55.8 | 55.8 | 23.846 |
| GDAP1 | 3594 | 9 | 9 | 9 | 39.1 | 39.1 | 39.1 | 41.345 |
| GDAP2 | 4338 | 9 | 9 | 9 | 27.2 | 27.2 | 27.2 | 56.224 |
| TPT1 | 185 | 9 | 9 | 9 | 41.5 | 41.5 | 41.5 | 21.525 |
| GNB4 | 4180 | 17 | 17 | 9 | 71.2 | 71.2 | 45 | 37.567 |
| RPL31 | 2637 | 9 | 9 | 9 | 53.6 | 53.6 | 53.6 | 14.463 |
| NFKB1 | 431 | 9 | 9 | 9 | 24.7 | 24.7 | 24.7 | 56.051 |
| SPP1 | 1856 | 9 | 9 | 9 | 42.4 | 42.4 | 42.4 | 35.422 |
| MAPK3 | 2122 | 13 | 9 | 9 | 48.8 | 36.7 | 36.7 | 43.135 |

| | | | | | | | | |
|---|---|---|---|---|---|---|---|---|
| RAB5C | 2425 | 13 | 13 | 9 | 78.7 | 78.7 | 63.9 | 23.482 |
| CSTB | 1728 | 9 | 9 | 9 | 79.6 | 79.6 | 79.6 | 11.139 |
| BICD2 | 3611 | 9 | 9 | 9 | 13.3 | 13.3 | 13.3 | 93.532 |
| PTDSS1 | 2353 | 9 | 9 | 9 | 23.5 | 23.5 | 23.5 | 55.527 |
| NUP62 | 2245 | 9 | 9 | 9 | 25.3 | 25.3 | 25.3 | 53.254 |
| RAB31 | 2903 | 11 | 11 | 9 | 72.7 | 72.7 | 61.3 | 21.569 |
| SLC30A1 | 4679 | 9 | 9 | 9 | 31.2 | 31.2 | 31.2 | 55.299 |
| GSTZ1 | 189 | 9 | 9 | 9 | 53.7 | 53.7 | 53.7 | 24.114 |
| PTPRE | 2073 | 12 | 9 | 9 | 21.6 | 16.6 | 16.6 | 80.641 |
| RPS12 | 2094 | 9 | 9 | 9 | 71.2 | 71.2 | 71.2 | 14.515 |
| OS9 | 2873 | 9 | 9 | 9 | 23.5 | 23.5 | 23.5 | 75.561 |
| VPS29 | 4405 | 9 | 9 | 9 | 57.1 | 57.1 | 57.1 | 20.505 |
| KCTD12 | 3772 | 9 | 9 | 9 | 38.2 | 38.2 | 38.2 | 35.7 |
| COMMD8 | 4321 | 9 | 9 | 9 | 72.7 | 72.7 | 72.7 | 21.09 |
| CCDC51 | 3785 | 9 | 9 | 9 | 36.3 | 36.3 | 36.3 | 45.811 |
| RPL21 | 2326 | 9 | 9 | 9 | 46.9 | 46.9 | 46.9 | 18.565 |
| AIMP2 | 2844 | 9 | 9 | 9 | 46.6 | 46.6 | 46.6 | 35.348 |
| PPCS | 4175 | 9 | 9 | 9 | 28.3 | 28.3 | 28.3 | 34.005 |
| LDAH;C2or | 606 | 9 | 9 | 9 | 34.6 | 34.6 | 34.6 | 42.668 |
| GNPDA1 | 2329 | 13 | 13 | 9 | 67.8 | 67.8 | 47.1 | 32.668 |
| PPP3CA;PP | 2787 | 9 | 9 | 9 | 22.8 | 22.8 | 22.8 | 58.687 |
| RAB1A | 2626 | 17 | 17 | 9 | 76.6 | 76.6 | 39 | 22.677 |
| UBE2J1 | 4585 | 9 | 9 | 9 | 40.6 | 40.6 | 40.6 | 35.198 |
| NDUFV2 | 788 | 9 | 9 | 9 | 46.4 | 46.4 | 46.4 | 27.907 |
| CSRP1 | 2036 | 9 | 9 | 9 | 60.6 | 60.6 | 60.6 | 20.567 |
| COTL1 | 2921 | 9 | 9 | 9 | 74.6 | 74.6 | 74.6 | 15.945 |
| PRKCB | 1761 | 16 | 16 | 9 | 35.6 | 35.6 | 20.6 | 76.868 |
| SLC16A3 | 1412 | 9 | 9 | 9 | 22.2 | 22.2 | 22.2 | 49.469 |
| SPN | 1955 | 9 | 9 | 9 | 30.5 | 30.5 | 30.5 | 40.321 |
| COMMD3;( | 4402 | 9 | 9 | 9 | 57.4 | 57.4 | 57.4 | 22.151 |
| SARNP | 2703 | 9 | 9 | 9 | 56.7 | 56.7 | 56.7 | 23.671 |
| SLC25A22 | 4166 | 9 | 9 | 9 | 44.6 | 44.6 | 44.6 | 34.47 |
| NDUFS8 | 1291 | 9 | 9 | 9 | 48.6 | 48.6 | 48.6 | 23.705 |
| ERGIC3 | 1038 | 9 | 9 | 9 | 35 | 35 | 35 | 44.568 |
| TCOF1 | 2871 | 9 | 9 | 9 | 8.5 | 8.5 | 8.5 | 152.1 |
| ACTR10 | 4354 | 9 | 9 | 9 | 30.2 | 30.2 | 30.2 | 46.306 |
| EXOSC10 | 2736 | 9 | 9 | 9 | 13.1 | 13.1 | 13.1 | 100.83 |
| PPP1R10 | 3860 | 9 | 9 | 9 | 12.7 | 12.7 | 12.7 | 99.057 |
| PARS2 | 3338 | 9 | 9 | 9 | 32 | 32 | 32 | 53.262 |
| BLVRB | 2156 | 9 | 9 | 9 | 65.5 | 65.5 | 65.5 | 22.119 |
| ENO2 | 1815 | 12 | 9 | 9 | 47.5 | 34.8 | 34.8 | 47.268 |
| SIL1 | 745 | 9 | 9 | 9 | 27.1 | 27.1 | 27.1 | 52.748 |
| RAD23B | 2497 | 11 | 11 | 9 | 32.5 | 32.5 | 28.4 | 43.171 |
| DDX21 | 4242 | 9 | 9 | 9 | 15.1 | 15.1 | 15.1 | 87.343 |
| PEX3 | 2530 | 9 | 9 | 9 | 41 | 41 | 41 | 42.139 |
| U2AF1 | 1844 | 9 | 9 | 9 | 47.1 | 47.1 | 47.1 | 27.872 |
| GALNT6 | 3552 | 9 | 9 | 9 | 26.4 | 26.4 | 26.4 | 71.158 |
| CYCS | 674 | 9 | 9 | 9 | 67.3 | 67.3 | 67.3 | 11.333 |

| | | | | | | | | |
|---|---|---|---|---|---|---|---|---|
| CBR3 | 1576 | 11 | 9 | 9 | 50.9 | 40.4 | 40.4 | 30.85 |
| HNRNPH3 | 2188 | 11 | 9 | 9 | 45.7 | 42.8 | 42.8 | 36.926 |
| MBOAT7 | 3844 | 9 | 9 | 9 | 25.6 | 25.6 | 25.6 | 52.764 |
| TXLNA | 2260 | 9 | 9 | 9 | 18.3 | 18.3 | 18.3 | 61.89 |
| MRPL13 | 4043 | 9 | 9 | 9 | 63.5 | 63.5 | 63.5 | 20.692 |
| LGALS9 | 1287 | 15 | 15 | 9 | 51.8 | 51.8 | 29.3 | 39.518 |
| GYG1 | 2332 | 9 | 9 | 9 | 23.4 | 23.4 | 23.4 | 39.383 |
| SQLE | 809 | 9 | 9 | 9 | 30.7 | 30.7 | 30.7 | 53.067 |
| MRPS10 | 2693 | 9 | 9 | 9 | 53.2 | 53.2 | 53.2 | 22.999 |
| COQ9 | 1528 | 9 | 9 | 9 | 51.3 | 51.3 | 51.3 | 35.509 |
| LEPRE1 | 3091 | 9 | 9 | 9 | 18.2 | 18.2 | 18.2 | 83.393 |
| GRK6 | 754 | 9 | 9 | 9 | 26.5 | 26.5 | 26.5 | 54.483 |
| GLIPR2 | 4142 | 9 | 9 | 9 | 73.4 | 73.4 | 73.4 | 17.218 |
| PSMD9 | 1295 | 9 | 9 | 9 | 44.4 | 44.4 | 44.4 | 24.682 |
| FTH1 | 1716 | 9 | 9 | 9 | 56.8 | 56.8 | 56.8 | 21.225 |
| RPL36 | 4609 | 9 | 9 | 9 | 35.2 | 35.2 | 35.2 | 12.254 |
| C8orf82 | 3265 | 9 | 9 | 9 | 55.1 | 55.1 | 55.1 | 23.889 |
| ACTR1B | 2278 | 18 | 9 | 9 | 62.8 | 37 | 37 | 42.293 |
| EIF5A;EIF5/ | 2657 | 9 | 9 | 9 | 80.5 | 80.5 | 80.5 | 16.832 |
| APRT | 1789 | 9 | 9 | 9 | 56.1 | 56.1 | 56.1 | 19.608 |
| PCYT2 | 1146 | 9 | 9 | 9 | 36 | 36 | 36 | 41.395 |
| PARVG | 4187 | 9 | 9 | 9 | 33.8 | 33.8 | 33.8 | 37.485 |
| BRE | 4339 | 9 | 9 | 9 | 32.1 | 32.1 | 32.1 | 43.551 |
| RPL23A | 2623 | 9 | 9 | 9 | 47.4 | 47.4 | 47.4 | 17.695 |
| KHDRBS1 | 2781 | 9 | 9 | 9 | 26.2 | 26.2 | 26.2 | 48.227 |
| ADRM1 | 3048 | 9 | 9 | 9 | 27 | 27 | 27 | 42.153 |
| DLG1 | 192 | 9 | 9 | 9 | 14 | 14 | 14 | 100.45 |
| BNIP1 | 2829 | 9 | 9 | 9 | 42.1 | 42.1 | 42.1 | 26.132 |
| GM2A | 1984 | 9 | 9 | 9 | 62.7 | 62.7 | 62.7 | 20.838 |
| SLC25A29 | 3528 | 9 | 9 | 9 | 48.5 | 48.5 | 48.5 | 32.062 |
| KDSR | 1247 | 9 | 9 | 9 | 33.7 | 33.7 | 33.7 | 36.187 |
| UQCC1 | 4293 | 9 | 9 | 9 | 32.4 | 32.4 | 32.4 | 34.6 |
| RAB5B | 2563 | 13 | 9 | 9 | 79.5 | 64.7 | 64.7 | 23.707 |
| CLPP | 1272 | 9 | 9 | 9 | 37.5 | 37.5 | 37.5 | 30.18 |
| PRKRA | 1564 | 9 | 9 | 9 | 43.5 | 43.5 | 43.5 | 34.404 |
| HEATR1 | 3189 | 9 | 9 | 9 | 6 | 6 | 6 | 233.27 |
| MAP1A | 838 | 9 | 9 | 9 | 3.9 | 3.9 | 3.9 | 331.25 |
| CLPB | 4087 | 25 | 25 | 9 | 46.4 | 46.4 | 12.9 | 78.728 |
| RHOC | 1798 | 17 | 17 | 9 | 84 | 84 | 39.9 | 21.523 |
| MRPL47 | 4206 | 9 | 9 | 9 | 31.6 | 31.6 | 31.6 | 29.45 |
| NMES1;C15 | 4065 | 9 | 9 | 9 | 62.7 | 62.7 | 62.7 | 9.6172 |
| ISOC2 | 3741 | 9 | 9 | 9 | 75.6 | 75.6 | 75.6 | 22.337 |
| GRAMD4 | 3244 | 9 | 9 | 9 | 20.9 | 20.9 | 20.9 | 66.407 |
| HDAC2 | 3694 | 15 | 9 | 9 | 38.1 | 27 | 27 | 55.364 |
| NUDT5 | 517 | 9 | 9 | 9 | 51.7 | 51.7 | 51.7 | 25.895 |
| ROCK2 | 1524 | 11 | 9 | 9 | 9.9 | 8.5 | 8.5 | 160.9 |
| TMLHE | 4298 | 9 | 9 | 9 | 30.2 | 30.2 | 30.2 | 49.517 |
| ARHGDIA | 2458 | 9 | 9 | 9 | 67.2 | 67.2 | 67.2 | 23.207 |

| | | | | | | | | |
|---|---|---|---|---|---|---|---|---|
| COG6 | 4571 | 9 | 9 | 9 | 25.9 | 25.9 | 25.9 | 73.278 |
| IFIT1 | 1836 | 9 | 9 | 9 | 25.5 | 25.5 | 25.5 | 55.36 |
| UXS1 | 3545 | 9 | 9 | 9 | 36.7 | 36.7 | 36.7 | 47.576 |
| THOC1 | 59 | 9 | 9 | 9 | 19.5 | 19.5 | 19.5 | 75.651 |
| TMX1 | 4129 | 9 | 9 | 9 | 26.4 | 26.4 | 26.4 | 31.791 |
| RPL35A | 1988 | 9 | 9 | 9 | 53.6 | 53.6 | 53.6 | 12.538 |
| TMED9 | 4010 | 10 | 10 | 9 | 43.4 | 43.4 | 36.2 | 27.277 |
| SNX3 | 1494 | 11 | 11 | 9 | 56.8 | 56.8 | 46.9 | 18.762 |
| PMM2 | 1407 | 9 | 9 | 9 | 48.8 | 48.8 | 48.8 | 28.082 |
| MTIF2 | 2321 | 9 | 9 | 9 | 18.4 | 18.4 | 18.4 | 81.316 |
| CNBP | 2619 | 8 | 8 | 8 | 50.3 | 50.3 | 50.3 | 19.463 |
| HLA-E | 1916 | 13 | 8 | 8 | 42.2 | 27.9 | 27.9 | 40.057 |
| SNX8 | 4649 | 8 | 8 | 8 | 22.6 | 22.6 | 22.6 | 52.569 |
| PIK3C3 | 330 | 11 | 11 | 8 | 14.3 | 14.3 | 10.6 | 101.7 |
| HLA-F | 239 | 16 | 10 | 8 | 43 | 31.2 | 21.9 | 50.428 |
| ADAM9 | 489 | 8 | 8 | 8 | 12.2 | 12.2 | 12.2 | 90.555 |
| RPL18 | 999 | 8 | 8 | 8 | 42.7 | 42.7 | 42.7 | 18.756 |
| SEC62 | 3891 | 8 | 8 | 8 | 21.3 | 21.3 | 21.3 | 45.861 |
| NQO1 | 585 | 8 | 8 | 8 | 43.1 | 43.1 | 43.1 | 22.793 |
| GHITM | 4126 | 8 | 8 | 8 | 18.6 | 18.6 | 18.6 | 37.205 |
| SNRPA1 | 1832 | 8 | 8 | 8 | 41.2 | 41.2 | 41.2 | 28.415 |
| ZADH2 | 3507 | 8 | 8 | 8 | 26.3 | 26.3 | 26.3 | 40.14 |
| MRPS7 | 1201 | 8 | 8 | 8 | 35.4 | 35.4 | 35.4 | 31.704 |
| WDR48 | 3592 | 8 | 8 | 8 | 15.7 | 15.7 | 15.7 | 76.21 |
| NDUFA7 | 1629 | 8 | 8 | 8 | 64.6 | 64.6 | 64.6 | 12.551 |
| NUDT21 | 1467 | 8 | 8 | 8 | 49.8 | 49.8 | 49.8 | 26.227 |
| DHRS1 | 3839 | 8 | 8 | 8 | 39.6 | 39.6 | 39.6 | 33.909 |
| SLC25A5 | 1748 | 24 | 9 | 8 | 51.3 | 25.2 | 22.5 | 32.852 |
| EEF1B2 | 2083 | 11 | 8 | 8 | 58.2 | 49.8 | 49.8 | 24.763 |
| NUP88 | 1156 | 8 | 8 | 8 | 11.5 | 11.5 | 11.5 | 83.541 |
| RAB1B | 4095 | 16 | 8 | 8 | 74.6 | 36.3 | 36.3 | 22.171 |
| SRSF2 | 1164 | 8 | 8 | 8 | 53.4 | 53.4 | 53.4 | 15.371 |
| RHOA | 2581 | 16 | 8 | 8 | 81.9 | 38.9 | 38.9 | 21.768 |
| HLA-B | 1709 | 27 | 22 | 8 | 56.9 | 43.6 | 16.3 | 40.46 |
| BAIAP2 | 1150 | 8 | 8 | 8 | 19.9 | 19.9 | 19.9 | 61.381 |
| IFI35 | 2688 | 8 | 8 | 8 | 33.6 | 33.6 | 33.6 | 31.546 |
| IDI1 | 2917 | 8 | 8 | 8 | 47.6 | 47.6 | 47.6 | 26.319 |
| CD84 | 4452 | 8 | 8 | 8 | 33 | 33 | 33 | 38.782 |
| LAMP2 | 1908 | 8 | 8 | 8 | 17.6 | 17.6 | 17.6 | 44.96 |
| AGPAT5 | 4280 | 8 | 8 | 8 | 29.9 | 29.9 | 29.9 | 42.072 |
| LPL | 1775 | 8 | 8 | 8 | 32.2 | 32.2 | 32.2 | 53.162 |
| ARF4 | 1990 | 12 | 8 | 8 | 80 | 51.1 | 51.1 | 20.511 |
| DNAJC7 | 483 | 8 | 8 | 8 | 18.6 | 18.6 | 18.6 | 56.44 |
| CNPY3 | 3975 | 8 | 8 | 8 | 32 | 32 | 32 | 30.748 |
| NRBP1 | 967 | 8 | 8 | 8 | 25.8 | 25.8 | 25.8 | 60.829 |
| MAGT1 | 27 | 8 | 8 | 8 | 19.9 | 19.9 | 19.9 | 41.531 |
| ADSS | 2171 | 8 | 8 | 8 | 25.7 | 25.7 | 25.7 | 50.097 |
| SAR1A | 4243 | 12 | 12 | 8 | 73.7 | 73.7 | 59.6 | 22.367 |

| | | | | | | | | |
|---|---|---|---|---|---|---|---|---|
| ELMO2 | 3161 | 12 | 8 | 8 | 21.7 | 17.3 | 17.3 | 82.364 |
| RNF13 | 1446 | 8 | 8 | 8 | 24.1 | 24.1 | 24.1 | 42.813 |
| RRAGC | 4184 | 17 | 17 | 8 | 54.9 | 54.9 | 30.8 | 44.223 |
| RAB2B | 3638 | 17 | 17 | 8 | 81.9 | 81.9 | 46.3 | 24.214 |
| BMP2K | 4267 | 9 | 9 | 8 | 12.3 | 12.3 | 11.3 | 129.17 |
| SLC1A4 | 2296 | 8 | 8 | 8 | 22.6 | 22.6 | 22.6 | 55.722 |
| APOBEC3G | 4192 | 9 | 9 | 8 | 32.8 | 32.8 | 28.9 | 46.407 |
| RBBP7 | 3056 | 13 | 8 | 8 | 51.3 | 40.7 | 40.7 | 47.82 |
| RAB35 | 3008 | 8 | 8 | 8 | 49.8 | 49.8 | 49.8 | 23.025 |
| HLA-DPA1 | 282 | 9 | 9 | 8 | 48.8 | 48.8 | 48.8 | 29.38 |
| STRIP1 | 3212 | 8 | 8 | 8 | 13.6 | 13.6 | 13.6 | 95.575 |
| FLAD1 | 3571 | 8 | 8 | 8 | 28.1 | 28.1 | 28.1 | 65.265 |
| EMC7 | 4215 | 8 | 8 | 8 | 41.3 | 41.3 | 41.3 | 26.47 |
| SLC33A1 | 1305 | 8 | 8 | 8 | 17.7 | 17.7 | 17.7 | 60.908 |
| CALM2;CAL | 1845 | 10 | 10 | 8 | 82.6 | 82.6 | 72.5 | 16.837 |
| TGFB1 | 443 | 8 | 8 | 8 | 33.1 | 33.1 | 33.1 | 44.341 |
| MAGOH;M/ | 2579 | 8 | 8 | 8 | 59.6 | 59.6 | 59.6 | 17.163 |
| COMMD2 | 3421 | 8 | 8 | 8 | 55.3 | 55.3 | 55.3 | 22.745 |
| RAB42 | 3510 | 8 | 8 | 8 | 57.3 | 57.3 | 57.3 | 24.133 |
| EML3 | 608 | 9 | 9 | 8 | 19.9 | 19.9 | 17.5 | 73.281 |
| EEFSEC | 2537 | 8 | 8 | 8 | 17.6 | 17.6 | 17.6 | 65.304 |
| CSNK2B;CS | 2664 | 8 | 8 | 8 | 53.4 | 53.4 | 53.4 | 26.925 |
| ZNF326 | 3144 | 8 | 8 | 8 | 20.4 | 20.4 | 20.4 | 65.653 |
| NUP214 | 432 | 8 | 8 | 8 | 5.5 | 5.5 | 5.5 | 212.48 |
| YKT6 | 1416 | 8 | 8 | 8 | 41.9 | 41.9 | 41.9 | 22.417 |
| PIP4K2C | 3603 | 8 | 8 | 8 | 27.3 | 27.3 | 27.3 | 47.299 |
| GPX4 | 16 | 8 | 8 | 8 | 45.4 | 45.4 | 45.4 | 22.024 |
| GIMAP8 | 3556 | 8 | 8 | 8 | 15.6 | 15.6 | 15.6 | 74.889 |
| SNAP29 | 1666 | 8 | 8 | 8 | 39.1 | 39.1 | 39.1 | 28.97 |
| CRABP2 | 2146 | 8 | 8 | 8 | 58 | 58 | 58 | 15.693 |
| CD276 | 212 | 8 | 8 | 8 | 34.5 | 34.5 | 34.5 | 41.767 |
| ENPP4 | 4686 | 8 | 8 | 8 | 29.8 | 29.8 | 29.8 | 51.641 |
| CTR9 | 3279 | 8 | 8 | 8 | 10.1 | 10.1 | 10.1 | 133.5 |
| PARVB | 4188 | 8 | 8 | 8 | 26.4 | 26.4 | 26.4 | 41.714 |
| PPP6C | 1330 | 8 | 8 | 8 | 27.9 | 27.9 | 27.9 | 35.144 |
| RAB5A | 2021 | 12 | 8 | 8 | 78.6 | 63.7 | 63.7 | 23.658 |
| ATP5J | 1999 | 8 | 8 | 8 | 55.6 | 55.6 | 55.6 | 12.587 |
| CDC73 | 384 | 8 | 8 | 8 | 23.2 | 23.2 | 23.2 | 54.155 |
| PRRC2C | 787 | 8 | 8 | 8 | 3.5 | 3.5 | 3.5 | 308.77 |
| MRPS34 | 678 | 8 | 8 | 8 | 48.4 | 48.4 | 48.4 | 26.332 |
| IKBIP | 3322 | 8 | 8 | 8 | 25.7 | 25.7 | 25.7 | 39.309 |
| C18orf8 | 1236 | 8 | 8 | 8 | 16.6 | 16.6 | 16.6 | 69.389 |
| COMMD1 | 3521 | 8 | 8 | 8 | 64.7 | 64.7 | 64.7 | 21.178 |
| IMPA1 | 2144 | 8 | 8 | 8 | 31.4 | 31.4 | 31.4 | 30.188 |
| EARS2 | 1105 | 8 | 8 | 8 | 31.2 | 31.2 | 31.2 | 56.825 |
| GLIPR1 | 2341 | 8 | 8 | 8 | 43.2 | 43.2 | 43.2 | 30.366 |
| LAMTOR2 | 4564 | 8 | 8 | 8 | 90.4 | 90.4 | 90.4 | 13.507 |
| SMARCE1 | 346 | 9 | 8 | 8 | 37.9 | 35.4 | 35.4 | 32.319 |

| | | | | | | | | |
|---|---|---|---|---|---|---|---|---|
| NDRG3 | 4431 | 8 | 8 | 8 | 38.1 | 38.1 | 38.1 | 41.408 |
| PTPN23 | 4131 | 8 | 8 | 8 | 7.5 | 7.5 | 7.5 | 178.97 |
| PDP1 | 4374 | 8 | 8 | 8 | 21.6 | 21.6 | 21.6 | 61.053 |
| RBM15 | 58 | 8 | 8 | 8 | 14.1 | 14.1 | 14.1 | 98.013 |
| TBCB | 3889 | 8 | 8 | 8 | 34.8 | 34.8 | 34.8 | 27.325 |
| SPARC | 1824 | 8 | 8 | 8 | 42.9 | 42.9 | 42.9 | 34.632 |
| MRPS15 | 2698 | 8 | 8 | 8 | 28.4 | 28.4 | 28.4 | 29.842 |
| RPS4X | 2621 | 20 | 20 | 8 | 64.3 | 64.3 | 31.9 | 29.597 |
| TPM3 | 60 | 29 | 29 | 8 | 73.6 | 73.6 | 21.6 | 26.42 |
| SRSF7 | 177 | 9 | 9 | 8 | 62.8 | 62.8 | 62.8 | 15.763 |
| TMED4 | 3383 | 9 | 8 | 8 | 41.9 | 41.9 | 41.9 | 25.943 |
| DHRS11 | 3297 | 8 | 8 | 8 | 45.8 | 45.8 | 45.8 | 28.308 |
| TBL1XR1 | 4060 | 8 | 8 | 8 | 24.7 | 24.7 | 24.7 | 55.594 |
| SLC4A1AP | 91 | 8 | 8 | 8 | 19.1 | 19.1 | 19.1 | 82.89 |
| SMEK1 | 1015 | 8 | 8 | 8 | 17.4 | 17.4 | 17.4 | 82.03 |
| CPSF2 | 4394 | 8 | 8 | 8 | 16.2 | 16.2 | 16.2 | 88.486 |
| GLRX | 2229 | 8 | 8 | 8 | 78.3 | 78.3 | 78.3 | 11.776 |
| ATG5 | 4106 | 8 | 8 | 8 | 33.5 | 33.5 | 33.5 | 32.447 |
| UBXN4 | 3677 | 8 | 8 | 8 | 17.1 | 17.1 | 17.1 | 56.777 |
| FDPS | 1925 | 8 | 8 | 8 | 20.5 | 20.5 | 20.5 | 48.275 |
| MRPL38 | 3778 | 8 | 8 | 8 | 30 | 30 | 30 | 44.596 |
| SRSF4 | 2786 | 12 | 12 | 8 | 20.9 | 20.9 | 15.2 | 56.678 |
| DCUN1D1 | 3800 | 8 | 8 | 8 | 34 | 34 | 34 | 30.124 |
| USP39 | 100 | 8 | 8 | 8 | 25.5 | 25.5 | 25.5 | 56.399 |
| ATPAF2 | 3518 | 8 | 8 | 8 | 46.4 | 46.4 | 46.4 | 32.772 |
| SNRPD2 | 2612 | 8 | 8 | 8 | 62.7 | 62.7 | 62.7 | 13.527 |
| CD40 | 2104 | 8 | 8 | 8 | 53.4 | 53.4 | 53.4 | 30.619 |
| STUB1 | 4512 | 8 | 8 | 8 | 28.7 | 28.7 | 28.7 | 34.856 |
| BIN1 | 1319 | 8 | 8 | 8 | 22.9 | 22.9 | 22.9 | 64.699 |
| POGLUT1 | 3537 | 8 | 8 | 8 | 25.5 | 25.5 | 25.5 | 46.189 |
| CD2AP | 4638 | 8 | 8 | 8 | 18.6 | 18.6 | 18.6 | 71.45 |
| TRIOBP | 4113 | 8 | 8 | 8 | 4 | 4 | 4 | 261.37 |
| SRSF9 | 2854 | 8 | 8 | 8 | 32.1 | 32.1 | 32.1 | 25.542 |
| LCK | 1764 | 8 | 8 | 8 | 22.2 | 22.2 | 22.2 | 58 |
| HPRT1 | 1695 | 8 | 8 | 8 | 53.2 | 53.2 | 53.2 | 24.579 |
| RPL23 | 2627 | 8 | 8 | 8 | 60 | 60 | 60 | 14.865 |
| VTI1B | 4417 | 8 | 8 | 8 | 40.9 | 40.9 | 40.9 | 26.688 |
| EMR2;ADGI | 4446 | 11 | 8 | 8 | 22.4 | 17.4 | 17.4 | 90.471 |
| ARPC5 | 1417 | 9 | 9 | 8 | 66.9 | 66.9 | 61.6 | 16.32 |
| NDUFB7 | 1977 | 8 | 8 | 8 | 64.2 | 64.2 | 64.2 | 16.402 |
| SEPHS2 | 3910 | 8 | 8 | 8 | 29.2 | 29.2 | 29.2 | 47.305 |
| MAN2B2 | 825 | 8 | 8 | 8 | 11.6 | 11.6 | 11.6 | 113.98 |
| PDCD10 | 3992 | 8 | 8 | 8 | 43.9 | 43.9 | 43.9 | 24.701 |
| DICER1 | 4530 | 8 | 8 | 8 | 6.1 | 6.1 | 6.1 | 218.68 |
| SEH1L | 1226 | 8 | 8 | 8 | 45.6 | 45.6 | 45.6 | 29.541 |
| ATG3 | 4268 | 8 | 8 | 8 | 33.8 | 33.8 | 33.8 | 35.864 |
| HNRNPH2 | 2514 | 17 | 8 | 8 | 48.3 | 30.1 | 30.1 | 49.263 |
| LRMP | 302 | 8 | 8 | 8 | 23.1 | 23.1 | 23.1 | 50.424 |

| | | | | | | | | |
|---|---|---|---|---|---|---|---|---|
| OSGEP | 4218 | 8 | 8 | 8 | 40 | 40 | 40 | 36.426 |
| ICAM3 | 1250 | 8 | 8 | 8 | 21.5 | 21.5 | 21.5 | 51.377 |
| CRTAP | 685 | 8 | 8 | 8 | 23.5 | 23.5 | 23.5 | 41.432 |
| PKN1 | 3051 | 8 | 8 | 8 | 10.2 | 10.2 | 10.2 | 103.93 |
| IFI30 | 1907 | 8 | 8 | 8 | 36.4 | 36.4 | 36.4 | 27.963 |
| COPS3 | 4518 | 8 | 8 | 8 | 31.7 | 31.7 | 31.7 | 47.873 |
| MFF | 4084 | 8 | 8 | 8 | 41.2 | 41.2 | 41.2 | 38.464 |
| CLPTM1L | 3831 | 8 | 8 | 8 | 22.1 | 22.1 | 22.1 | 62.228 |
| SCYL1 | 856 | 8 | 8 | 8 | 15.9 | 15.9 | 15.9 | 85.718 |
| PSMB9 | 245 | 8 | 8 | 8 | 59.2 | 59.2 | 59.2 | 20.96 |
| LAMTOR1 | 3241 | 8 | 8 | 8 | 74.5 | 74.5 | 74.5 | 17.745 |
| MCTS1 | 4498 | 8 | 8 | 8 | 63 | 63 | 63 | 20.555 |
| SPON2 | 3989 | 8 | 8 | 8 | 28.7 | 28.7 | 28.7 | 35.846 |
| TRAPPC3 | 1450 | 8 | 8 | 8 | 46.7 | 46.7 | 46.7 | 20.274 |
| PGRMC2 | 1391 | 9 | 8 | 8 | 40.4 | 36.3 | 36.3 | 23.818 |
| DOCK10 | 371 | 9 | 9 | 8 | 5.7 | 5.7 | 5.2 | 251.31 |
| GIMAP2 | 4422 | 8 | 8 | 8 | 28.5 | 28.5 | 28.5 | 38.017 |
| PARP12 | 4092 | 9 | 8 | 8 | 14.7 | 13.6 | 13.6 | 79.063 |
| LPCAT3 | 3261 | 8 | 8 | 8 | 17.2 | 17.2 | 17.2 | 56.034 |
| ADSL | 169 | 8 | 8 | 8 | 26.7 | 26.7 | 26.7 | 54.889 |
| MIA2 | 1011 | 23 | 23 | 8 | 18.7 | 18.7 | 7.5 | 151.72 |
| HNRNPAB | 730 | 8 | 8 | 8 | 27.9 | 27.9 | 27.9 | 30.302 |
| PSMB4 | 2134 | 8 | 8 | 8 | 48.5 | 48.5 | 48.5 | 29.204 |
| RAB27A | 2429 | 8 | 8 | 8 | 43.4 | 43.4 | 43.4 | 24.868 |
| ARL1 | 2264 | 8 | 8 | 8 | 72.9 | 72.9 | 72.9 | 20.417 |
| CUL5 | 3718 | 8 | 8 | 8 | 11.8 | 11.8 | 11.8 | 90.954 |
| CTNND1 | 478 | 8 | 8 | 8 | 13 | 13 | 13 | 99.077 |
| DCTN3 | 1589 | 8 | 8 | 8 | 51.1 | 51.1 | 51.1 | 21.119 |
| MRPL4 | 4045 | 8 | 8 | 8 | 48.2 | 48.2 | 48.2 | 34.919 |
| COX7A2L | 1339 | 8 | 8 | 8 | 63.2 | 63.2 | 63.2 | 12.615 |
| DAPK1 | 2472 | 8 | 8 | 8 | 7.6 | 7.6 | 7.6 | 160.04 |
| GIMAP1 | 3655 | 14 | 8 | 8 | 57.5 | 29.1 | 29.1 | 34.369 |
| YME1L1 | 3817 | 8 | 8 | 8 | 19.5 | 19.5 | 19.5 | 82.665 |
| DUSP3 | 2433 | 8 | 8 | 8 | 57.8 | 57.8 | 57.8 | 20.478 |
| ALDOC | 1841 | 13 | 8 | 8 | 44.2 | 32.7 | 32.7 | 39.455 |
| GLO1 | 2762 | 8 | 8 | 8 | 59.2 | 59.2 | 59.2 | 20.777 |
| RPL7A | 2616 | 16 | 16 | 8 | 51.1 | 51.1 | 26.7 | 29.995 |
| IST1 | 2487 | 8 | 8 | 8 | 25.3 | 25.3 | 25.3 | 39.75 |
| CUL1 | 221 | 8 | 8 | 8 | 13.5 | 13.5 | 13.5 | 84.571 |
| METAP2 | 2416 | 8 | 8 | 8 | 23.2 | 23.2 | 23.2 | 52.891 |
| REEP5 | 2728 | 8 | 8 | 8 | 43.4 | 43.4 | 43.4 | 21.493 |
| MMAB | 3789 | 8 | 8 | 8 | 38.8 | 38.8 | 38.8 | 27.388 |
| WDR91 | 505 | 8 | 8 | 8 | 16.2 | 16.2 | 16.2 | 79.239 |
| SNRPB2 | 1807 | 11 | 11 | 8 | 48 | 48 | 40.4 | 25.486 |
| WDR18 | 4000 | 8 | 8 | 8 | 30.2 | 30.2 | 30.2 | 43.237 |
| SASH3 | 1595 | 8 | 8 | 8 | 35.5 | 35.5 | 35.5 | 41.595 |
| ARFIP1 | 2473 | 8 | 8 | 8 | 34 | 34 | 34 | 41.738 |
| HSPH1 | 3680 | 38 | 37 | 8 | 58.3 | 57.3 | 11.4 | 96.864 |

| | | | | | | | | |
|---|---|---|---|---|---|---|---|---|
| PANK4 | 4295 | 8 | 8 | 8 | 14.4 | 14.4 | 14.4 | 85.99 |
| TBK1 | 421 | 8 | 8 | 8 | 16.6 | 16.6 | 16.6 | 80.791 |
| SLC2A5 | 2061 | 8 | 8 | 8 | 11.4 | 11.4 | 11.4 | 54.973 |
| UBQLN1;UI | 4509 | 8 | 8 | 8 | 20.5 | 20.5 | 20.5 | 62.518 |
| SEPHS1 | 2393 | 8 | 8 | 8 | 36.5 | 36.5 | 36.5 | 42.91 |
| NMNAT1 | 4177 | 8 | 8 | 8 | 39.4 | 39.4 | 39.4 | 31.932 |
| TMEM126A | 4086 | 8 | 8 | 8 | 45.1 | 45.1 | 45.1 | 21.527 |
| BCKDHB | 2045 | 8 | 8 | 8 | 44.4 | 44.4 | 44.4 | 43.122 |
| ATXN2L | 1109 | 8 | 8 | 8 | 10 | 10 | 10 | 112.71 |
| C16orf58 | 1099 | 8 | 8 | 8 | 23.4 | 23.4 | 23.4 | 46.342 |
| HS1BP3 | 3132 | 8 | 8 | 8 | 29.3 | 29.3 | 29.3 | 42.78 |
| TCEA1 | 326 | 8 | 8 | 8 | 31.6 | 31.6 | 31.6 | 33.935 |
| TMED2 | 3011 | 8 | 8 | 8 | 38.8 | 38.8 | 38.8 | 22.761 |
| CDS2 | 1664 | 9 | 9 | 8 | 25.6 | 25.6 | 22.2 | 51.417 |
| SFXN2 | 3847 | 8 | 8 | 8 | 34.5 | 34.5 | 34.5 | 36.231 |
| ENOSF1 | 414 | 8 | 8 | 8 | 20.5 | 20.5 | 20.5 | 53.063 |
| GIGYF2 | 1140 | 8 | 8 | 8 | 7.2 | 7.2 | 7.2 | 152.53 |
| DOK3 | 734 | 8 | 8 | 8 | 26.6 | 26.6 | 26.6 | 47.601 |
| TSN | 841 | 8 | 8 | 8 | 40.4 | 40.4 | 40.4 | 26.183 |
| SLC25A6 | 1892 | 28 | 28 | 8 | 75.5 | 75.5 | 28.2 | 32.866 |
| THOC5 | 2907 | 8 | 8 | 8 | 19.5 | 19.5 | 19.5 | 78.507 |
| FAM175B | 2981 | 8 | 8 | 8 | 19.8 | 19.8 | 19.8 | 46.9 |
| S100A11 | 2192 | 8 | 8 | 8 | 83.8 | 83.8 | 83.8 | 11.74 |
| NUDC | 4544 | 8 | 8 | 8 | 26.6 | 26.6 | 26.6 | 38.242 |
| UBE2M | 2565 | 8 | 8 | 8 | 51.9 | 51.9 | 51.9 | 20.9 |
| CLTA | 1826 | 8 | 8 | 8 | 23 | 23 | 23 | 27.076 |
| LIMK1 | 824 | 8 | 8 | 8 | 14.8 | 14.8 | 14.8 | 75.526 |
| KIF1B;KIF1I | 54 | 9 | 9 | 8 | 5.9 | 5.9 | 5.2 | 202.01 |
| COX5B | 1859 | 8 | 8 | 8 | 65.9 | 65.9 | 65.9 | 13.696 |
| DHX38 | 3687 | 8 | 8 | 8 | 10.2 | 10.2 | 10.2 | 140.5 |
| RAE1 | 2683 | 8 | 8 | 8 | 37 | 37 | 37 | 40.968 |
| PAPSS2 | 1641 | 9 | 9 | 8 | 23 | 23 | 20.7 | 69.5 |
| SSBP1 | 2763 | 8 | 8 | 8 | 52.7 | 52.7 | 52.7 | 17.259 |
| TMX2 | 4580 | 8 | 8 | 8 | 32.1 | 32.1 | 32.1 | 34.037 |
| RPS13 | 2605 | 8 | 8 | 8 | 48.3 | 48.3 | 48.3 | 17.222 |
| SCRIB | 256 | 7 | 7 | 7 | 7.1 | 7.1 | 7.1 | 165.27 |
| ME3 | 869 | 8 | 7 | 7 | 20.9 | 19.3 | 19.3 | 64.277 |
| SYVN1 | 3393 | 7 | 7 | 7 | 10.5 | 10.5 | 10.5 | 67.684 |
| C17orf62 | 3943 | 7 | 7 | 7 | 33.7 | 33.7 | 33.7 | 20.774 |
| NDUFA6 | 379 | 7 | 7 | 7 | 50 | 50 | 50 | 15.136 |
| PDLIM2 | 45 | 7 | 7 | 7 | 34.2 | 34.2 | 34.2 | 34.767 |
| FIG4 | 3675 | 7 | 7 | 7 | 10.7 | 10.7 | 10.7 | 103.63 |
| SH3KBP1 | 3152 | 7 | 7 | 7 | 14.6 | 14.6 | 14.6 | 70.443 |
| EWSR1 | 554 | 7 | 7 | 7 | 15.2 | 15.2 | 15.2 | 64.929 |
| CPQ | 4658 | 7 | 7 | 7 | 24.2 | 24.2 | 24.2 | 51.887 |
| CCDC109B | 4315 | 7 | 7 | 7 | 25.3 | 25.3 | 25.3 | 39.082 |
| WARS2 | 4426 | 7 | 7 | 7 | 32.8 | 32.8 | 32.8 | 40.146 |
| DAG1 | 2925 | 7 | 7 | 7 | 13.1 | 13.1 | 13.1 | 97.44 |

| | | | | | | | | |
|---|---|---|---|---|---|---|---|---|
| APOA1BP | 3555 | 7 | 7 | 7 | 33 | 33 | 33 | 31.674 |
| SUMF1 | 3536 | 7 | 7 | 7 | 31.8 | 31.8 | 31.8 | 40.556 |
| AGFG1 | 2460 | 7 | 7 | 7 | 19 | 19 | 19 | 58.259 |
| GPR108 | 4224 | 7 | 7 | 7 | 13.8 | 13.8 | 13.8 | 60.632 |
| ARFGAP1 | 957 | 7 | 7 | 7 | 36.9 | 36.9 | 36.9 | 29.571 |
| CYB5A | 1684 | 7 | 7 | 7 | 56 | 56 | 56 | 15.33 |
| BAK1 | 3058 | 7 | 7 | 7 | 50.2 | 50.2 | 50.2 | 23.408 |
| SCO2 | 1471 | 7 | 7 | 7 | 38.7 | 38.7 | 38.7 | 29.81 |
| MRPS36 | 2696 | 7 | 7 | 7 | 73.8 | 73.8 | 73.8 | 11.466 |
| MRPL43 | 557 | 7 | 7 | 7 | 49.1 | 49.1 | 49.1 | 18.657 |
| FKBP8 | 2946 | 13 | 13 | 7 | 35.2 | 35.2 | 17.7 | 44.561 |
| RNMT | 1422 | 7 | 7 | 7 | 21.4 | 21.4 | 21.4 | 54.843 |
| FTSJ3 | 3466 | 7 | 7 | 7 | 10.2 | 10.2 | 10.2 | 96.557 |
| TIAL1 | 2730 | 10 | 10 | 7 | 42.7 | 42.7 | 30.1 | 41.59 |
| UNC45A | 318 | 7 | 7 | 7 | 9 | 9 | 9 | 118.39 |
| SKP1 | 775 | 7 | 7 | 7 | 39.3 | 39.3 | 39.3 | 18.72 |
| CCDC53 | 905 | 7 | 7 | 7 | 47.7 | 47.7 | 47.7 | 21.044 |
| VAMP8 | 4001 | 7 | 7 | 7 | 55 | 55 | 55 | 11.438 |
| HMGB1;HN | 1821 | 9 | 9 | 7 | 47.5 | 47.5 | 37.3 | 18.311 |
| RPS14 | 366 | 7 | 7 | 7 | 65.6 | 65.6 | 65.6 | 16.273 |
| FAHD2A;FA | 3801 | 7 | 7 | 7 | 38.2 | 38.2 | 38.2 | 34.596 |
| MTHFS;ST2 | 2394 | 7 | 7 | 7 | 31.5 | 31.5 | 31.5 | 23.255 |
| SPNS1 | 4121 | 7 | 7 | 7 | 20.5 | 20.5 | 20.5 | 56.629 |
| HVCN1 | 3774 | 7 | 7 | 7 | 35.2 | 35.2 | 35.2 | 31.683 |
| GMFG | 1481 | 9 | 9 | 7 | 78.2 | 78.2 | 52.8 | 16.801 |
| ARL6IP5 | 1586 | 7 | 7 | 7 | 28.7 | 28.7 | 28.7 | 21.614 |
| TRAPPC4 | 876 | 7 | 7 | 7 | 44.9 | 44.9 | 44.9 | 19.483 |
| UQCRQ | 1367 | 7 | 7 | 7 | 57.3 | 57.3 | 57.3 | 9.9062 |
| VBP1 | 2584 | 7 | 7 | 7 | 46.7 | 46.7 | 46.7 | 22.626 |
| SRSF3 | 2712 | 8 | 7 | 7 | 42.7 | 42.7 | 42.7 | 19.329 |
| SRP14 | 2244 | 7 | 7 | 7 | 45.6 | 45.6 | 45.6 | 14.57 |
| TRAPPC5 | 3435 | 7 | 7 | 7 | 47.9 | 47.9 | 47.9 | 20.783 |
| RBP1 | 1822 | 7 | 7 | 7 | 60.7 | 60.7 | 60.7 | 15.85 |
| VTA1 | 68 | 7 | 7 | 7 | 26.7 | 26.7 | 26.7 | 33.879 |
| UBE2L3 | 2667 | 7 | 7 | 7 | 65.6 | 65.6 | 65.6 | 17.861 |
| CTSH | 87 | 7 | 7 | 7 | 30.7 | 30.7 | 30.7 | 36.269 |
| PDK2 | 2998 | 7 | 7 | 7 | 26.8 | 26.8 | 26.8 | 46.153 |
| ARF5 | 2708 | 12 | 7 | 7 | 80 | 51.1 | 51.1 | 20.529 |
| IKBKG | 35 | 7 | 7 | 7 | 18.7 | 18.7 | 18.7 | 47.384 |
| TAF15 | 3697 | 9 | 9 | 7 | 19.6 | 19.6 | 15.7 | 61.829 |
| SNRPN;SNF | 1198 | 7 | 7 | 7 | 35.5 | 35.5 | 35.5 | 17.546 |
| LYPLA1 | 1566 | 7 | 7 | 7 | 63 | 63 | 63 | 24.669 |
| MRPL28 | 2837 | 7 | 7 | 7 | 35.5 | 35.5 | 35.5 | 30.156 |
| MBOAT1 | 3309 | 7 | 7 | 7 | 14.9 | 14.9 | 14.9 | 56.557 |
| ZFPL1 | 1624 | 7 | 7 | 7 | 30.3 | 30.3 | 30.3 | 34.114 |
| DLGAP4 | 184 | 7 | 7 | 7 | 36.9 | 36.9 | 36.9 | 26.941 |
| APOO | 1128 | 7 | 7 | 7 | 65.2 | 65.2 | 65.2 | 20.08 |
| RANBP1 | 2305 | 7 | 7 | 7 | 58.2 | 58.2 | 58.2 | 23.31 |

| | | | | | | | | |
|---|---|---|---|---|---|---|---|---|
| SIRPA | 2676 | 18 | 18 | 7 | 52.6 | 52.6 | 29.2 | 54.966 |
| TXN | 1858 | 7 | 7 | 7 | 62.9 | 62.9 | 62.9 | 11.737 |
| CBX3 | 2848 | 9 | 9 | 7 | 58.5 | 58.5 | 43.2 | 20.811 |
| TXNDC12 | 1677 | 7 | 7 | 7 | 58.7 | 58.7 | 58.7 | 19.206 |
| HLA-DRB3;I | 294 | 14 | 14 | 7 | 62 | 62 | 26.3 | 29.941 |
| TAPT1 | 3258 | 7 | 7 | 7 | 16.8 | 16.8 | 16.8 | 64.259 |
| GADD45GII | 3591 | 7 | 7 | 7 | 38.3 | 38.3 | 38.3 | 25.384 |
| PAFAH1B2 | 2672 | 7 | 7 | 7 | 53.3 | 53.3 | 53.3 | 25.569 |
| GNPTG | 4468 | 7 | 7 | 7 | 46.2 | 46.2 | 46.2 | 33.973 |
| PDS5B | 4269 | 8 | 7 | 7 | 8.9 | 7.9 | 7.9 | 164.67 |
| RAB8A | 2558 | 12 | 10 | 7 | 53.6 | 44.9 | 30.4 | 23.668 |
| GCA | 1115 | 7 | 7 | 7 | 54.8 | 54.8 | 54.8 | 16.712 |
| RANBP3 | 617 | 9 | 9 | 7 | 22.1 | 22.1 | 16.4 | 52.894 |
| RPL34 | 2362 | 7 | 7 | 7 | 37.6 | 37.6 | 37.6 | 13.293 |
| COQ3 | 4361 | 7 | 7 | 7 | 26.3 | 26.3 | 26.3 | 41.054 |
| CNN2 | 577 | 7 | 7 | 7 | 27.5 | 27.5 | 27.5 | 32.616 |
| HSD17B7 | 2531 | 7 | 7 | 7 | 29 | 29 | 29 | 38.206 |
| RAB22A | 4495 | 9 | 7 | 7 | 58.2 | 46.9 | 46.9 | 21.855 |
| H2AFY2 | 4378 | 9 | 7 | 7 | 34.4 | 27.4 | 27.4 | 40.058 |
| SIDT2 | 663 | 7 | 7 | 7 | 9.7 | 9.7 | 9.7 | 96.728 |
| LRRFIP2 | 4655 | 8 | 7 | 7 | 15 | 13.3 | 13.3 | 82.17 |
| TRAM1 | 1017 | 7 | 7 | 7 | 18.4 | 18.4 | 18.4 | 33.435 |
| MRPS6 | 2700 | 7 | 7 | 7 | 48 | 48 | 48 | 14.226 |
| NSFL1C | 4521 | 7 | 7 | 7 | 23.5 | 23.5 | 23.5 | 40.572 |
| RPA2 | 1951 | 7 | 7 | 7 | 48.9 | 48.9 | 48.9 | 29.247 |
| PSAT1 | 4656 | 7 | 7 | 7 | 20.3 | 20.3 | 20.3 | 40.422 |
| PHF5A | 3352 | 7 | 7 | 7 | 62.7 | 62.7 | 62.7 | 12.405 |
| CPPED1 | 3953 | 7 | 7 | 7 | 31.5 | 31.5 | 31.5 | 35.548 |
| SRSF10 | 1556 | 7 | 7 | 7 | 27.9 | 27.9 | 27.9 | 20.913 |
| DAK;TKFC | 3098 | 7 | 7 | 7 | 17.9 | 17.9 | 17.9 | 58.946 |
| RPRD1B | 4232 | 7 | 7 | 7 | 32.5 | 32.5 | 32.5 | 36.899 |
| MPC2 | 1659 | 7 | 7 | 7 | 69.5 | 69.5 | 69.5 | 11.681 |
| ZC3H15 | 3636 | 7 | 7 | 7 | 19 | 19 | 19 | 48.602 |
| ARFGAP2 | 229 | 7 | 7 | 7 | 23.9 | 23.9 | 23.9 | 42.153 |
| UBE2N;UBE | 2566 | 7 | 7 | 7 | 60.5 | 60.5 | 60.5 | 17.138 |
| QDPR | 1820 | 7 | 7 | 7 | 40.2 | 40.2 | 40.2 | 25.789 |
| PISD | 4423 | 7 | 7 | 7 | 22 | 22 | 22 | 46.671 |
| SIGLEC7 | 4548 | 7 | 7 | 7 | 19.5 | 19.5 | 19.5 | 51.142 |
| DCAKD | 1252 | 7 | 7 | 7 | 45 | 45 | 45 | 24.177 |
| RALB | 1877 | 14 | 7 | 7 | 68 | 34.5 | 34.5 | 23.408 |
| FAM134C | 3408 | 7 | 7 | 7 | 25.5 | 25.5 | 25.5 | 51.396 |
| HMGB2 | 2113 | 9 | 7 | 7 | 37.8 | 30.1 | 30.1 | 24.033 |
| POLR2E | 2005 | 7 | 7 | 7 | 54.8 | 54.8 | 54.8 | 24.551 |
| IK | 2839 | 7 | 7 | 7 | 12.9 | 12.9 | 12.9 | 65.601 |
| EPB41 | 364 | 12 | 9 | 7 | 16.1 | 12.8 | 10.2 | 97.016 |
| MYCBP | 38 | 7 | 7 | 7 | 65 | 65 | 65 | 11.967 |
| THOC3 | 3821 | 7 | 7 | 7 | 29.3 | 29.3 | 29.3 | 38.771 |
| SCD | 1334 | 7 | 7 | 7 | 21.4 | 21.4 | 21.4 | 41.522 |

| | | | | | | | | |
|---|---|---|---|---|---|---|---|---|
| SMCHD1 | 524 | 7 | 7 | 7 | 4.3 | 4.3 | 4.3 | 226.37 |
| PPP1R8 | 2828 | 7 | 7 | 7 | 39.6 | 39.6 | 39.6 | 38.478 |
| CLN5 | 1557 | 7 | 7 | 7 | 19.8 | 19.8 | 19.8 | 41.496 |
| Nucb2;NU( | 2689 | 7 | 7 | 7 | 21.3 | 21.3 | 21.3 | 46.629 |
| FAM134A | 3546 | 7 | 7 | 7 | 20.4 | 20.4 | 20.4 | 57.829 |
| VPS51 | 4453 | 7 | 7 | 7 | 13.6 | 13.6 | 13.6 | 86.041 |
| CTNNBL1 | 28 | 7 | 7 | 7 | 18.5 | 18.5 | 18.5 | 65.173 |
| ITCH | 453 | 7 | 7 | 7 | 11.3 | 11.3 | 11.3 | 88.805 |
| METTL7A | 1068 | 7 | 7 | 7 | 44.7 | 44.7 | 44.7 | 28.319 |
| DPYSL3 | 2936 | 11 | 8 | 7 | 29.5 | 21.8 | 20.5 | 61.963 |
| RPL27A | 2324 | 7 | 7 | 7 | 42.6 | 42.6 | 42.6 | 16.561 |
| PCNA | 1888 | 7 | 7 | 7 | 36.4 | 36.4 | 36.4 | 28.768 |
| NDUFS7 | 1530 | 7 | 7 | 7 | 46 | 46 | 46 | 23.563 |
| COMTD1 | 3410 | 7 | 7 | 7 | 39.3 | 39.3 | 39.3 | 28.808 |
| UCHL3 | 1943 | 7 | 7 | 7 | 49.1 | 49.1 | 49.1 | 26.182 |
| CLCN3 | 2444 | 7 | 7 | 7 | 7.9 | 7.9 | 7.9 | 90.965 |
| SGTA | 1464 | 7 | 7 | 7 | 28.8 | 28.8 | 28.8 | 34.063 |
| MGST3 | 1362 | 7 | 7 | 7 | 73.7 | 73.7 | 73.7 | 16.516 |
| VTI1A | 3746 | 7 | 7 | 7 | 39.2 | 39.2 | 39.2 | 25.217 |
| LILRA2 | 252 | 7 | 7 | 7 | 24.7 | 24.7 | 24.7 | 49.956 |
| RAB4B | 2561 | 12 | 12 | 7 | 65.3 | 65.3 | 44.1 | 23.586 |
| GMIP | 4383 | 12 | 12 | 7 | 15.3 | 15.3 | 10.1 | 106.68 |
| ISOC1 | 3767 | 7 | 7 | 7 | 27.5 | 27.5 | 27.5 | 32.236 |
| PURA | 2724 | 7 | 7 | 7 | 32.9 | 32.9 | 32.9 | 34.91 |
| AKAP8L | 4502 | 7 | 7 | 7 | 16.4 | 16.4 | 16.4 | 71.639 |
| RAB23 | 4497 | 7 | 7 | 7 | 30.8 | 30.8 | 30.8 | 26.659 |
| GPS1 | 672 | 7 | 7 | 7 | 19.3 | 19.3 | 19.3 | 53.371 |
| RHEB | 3015 | 7 | 7 | 7 | 33.2 | 33.2 | 33.2 | 20.497 |
| TRIM56 | 3964 | 7 | 7 | 7 | 14.7 | 14.7 | 14.7 | 81.487 |
| YWHAB | 2189 | 13 | 7 | 7 | 55.7 | 38.2 | 38.2 | 28.082 |
| RBM12 | 4271 | 7 | 7 | 7 | 10.1 | 10.1 | 10.1 | 97.394 |
| RALA | 1876 | 14 | 14 | 7 | 57.3 | 57.3 | 23.8 | 23.567 |
| SRP9;DKFZ | 2371 | 7 | 7 | 7 | 57 | 57 | 57 | 10.112 |
| PPIL4 | 3637 | 7 | 7 | 7 | 21.1 | 21.1 | 21.1 | 57.224 |
| RCC2 | 4386 | 7 | 7 | 7 | 19 | 19 | 19 | 56.084 |
| DUT | 2205 | 7 | 7 | 7 | 38.1 | 38.1 | 38.1 | 26.563 |
| SRSF6 | 452 | 11 | 7 | 7 | 38.1 | 25.7 | 25.7 | 26.306 |
| FNTA | 574 | 7 | 7 | 7 | 32.5 | 32.5 | 32.5 | 27.443 |
| FNDC3B | 3120 | 7 | 7 | 7 | 6.8 | 6.8 | 6.8 | 132.89 |
| RAB24 | 3730 | 7 | 7 | 7 | 39.9 | 39.9 | 39.9 | 23.124 |
| ACOT13 | 4221 | 7 | 7 | 7 | 68.6 | 68.6 | 68.6 | 14.96 |
| ANKLE2 | 3423 | 7 | 7 | 7 | 11.3 | 11.3 | 11.3 | 104.11 |
| PPP5C | 1060 | 7 | 7 | 7 | 21.2 | 21.2 | 21.2 | 56.878 |
| ATPIF1 | 4456 | 7 | 7 | 7 | 29.2 | 29.2 | 29.2 | 12.249 |
| ANP32A | 1081 | 9 | 9 | 7 | 48.6 | 48.6 | 48.6 | 19.997 |
| LAMTOR3 | 4435 | 7 | 7 | 7 | 80.6 | 80.6 | 80.6 | 13.623 |
| ZNF207 | 1206 | 7 | 7 | 7 | 12.6 | 12.6 | 12.6 | 52.541 |
| GAPVD1 | 980 | 11 | 11 | 7 | 9.3 | 9.3 | 6.7 | 157.46 |

| | | | | | | | | |
|---|---|---|---|---|---|---|---|---|
| DDX60 | 3464 | 8 | 8 | 7 | 5.4 | 5.4 | 4.6 | 197.85 |
| HMGA1 | 1973 | 7 | 7 | 7 | 54.2 | 54.2 | 54.2 | 11.676 |
| NAA50 | 791 | 7 | 7 | 7 | 48.8 | 48.8 | 48.8 | 19.311 |
| RMND1 | 66 | 7 | 7 | 7 | 27.6 | 27.6 | 27.6 | 32.26 |
| NIF3L1 | 4081 | 7 | 7 | 7 | 25.5 | 25.5 | 25.5 | 41.968 |
| RPL14 | 786 | 7 | 7 | 7 | 50 | 50 | 50 | 14.558 |
| RPS23 | 2603 | 7 | 7 | 7 | 49 | 49 | 49 | 15.807 |
| BTF3L4 | 3828 | 7 | 7 | 7 | 65.2 | 65.2 | 65.2 | 17.27 |
| TEX10 | 4334 | 7 | 7 | 7 | 10.8 | 10.8 | 10.8 | 105.67 |
| COMMD4 | 181 | 7 | 7 | 7 | 64.4 | 64.4 | 64.4 | 19.382 |
| SPG20 | 3481 | 7 | 7 | 7 | 17.7 | 17.7 | 17.7 | 72.832 |
| HERC4 | 3147 | 7 | 7 | 7 | 11.9 | 11.9 | 11.9 | 118.56 |
| DHCR7 | 4404 | 7 | 7 | 7 | 16.8 | 16.8 | 16.8 | 54.489 |
| FMNL1 | 1650 | 22 | 18 | 7 | 24.2 | 20.8 | 8.5 | 121.85 |
| TSSC1 | 546 | 7 | 7 | 7 | 25.8 | 25.8 | 25.8 | 46.319 |
| HEATR5B | 4390 | 7 | 7 | 7 | 5.2 | 5.2 | 5.2 | 224.3 |
| RAB2A | 2562 | 16 | 7 | 7 | 71.7 | 35.8 | 35.8 | 23.545 |
| GNA11 | 2152 | 16 | 9 | 7 | 54.9 | 30.1 | 24.2 | 42.123 |
| TUBGCP2 | 3971 | 7 | 7 | 7 | 9.9 | 9.9 | 9.9 | 102.53 |
| EIF4A1 | 2551 | 25 | 25 | 7 | 61.3 | 61.3 | 23.9 | 46.153 |
| MRPL2 | 477 | 7 | 7 | 7 | 42.9 | 42.9 | 42.9 | 26.04 |
| SLC1A5 | 3036 | 7 | 7 | 7 | 15.7 | 15.7 | 15.7 | 56.598 |
| XRN1 | 3477 | 7 | 7 | 7 | 5.3 | 5.3 | 5.3 | 194.1 |
| INTS3 | 3234 | 7 | 7 | 7 | 12.4 | 12.4 | 12.4 | 118.07 |
| GALNT1 | 2807 | 7 | 7 | 7 | 17.5 | 17.5 | 17.5 | 64.218 |
| TPRG1L | 3181 | 7 | 7 | 7 | 48.9 | 48.9 | 48.9 | 30.211 |
| NOL9 | 3179 | 7 | 7 | 7 | 16.2 | 16.2 | 16.2 | 79.322 |
| NAP1L4 | 3919 | 9 | 7 | 7 | 25.3 | 22.4 | 22.4 | 42.823 |
| IRF5 | 2893 | 7 | 7 | 7 | 17.5 | 17.5 | 17.5 | 56.043 |
| ST13;ST13F | 2413 | 13 | 13 | 7 | 39.8 | 39.8 | 20.6 | 41.331 |
| SRRM1 | 551 | 7 | 7 | 7 | 9.2 | 9.2 | 9.2 | 103.39 |
| PSMB10 | 2262 | 7 | 7 | 7 | 34.1 | 34.1 | 34.1 | 28.936 |
| VKORC1L1 | 393 | 7 | 7 | 7 | 34.1 | 34.1 | 34.1 | 23.499 |
| MOSPD1 | 4467 | 7 | 7 | 7 | 42.3 | 42.3 | 42.3 | 24.086 |
| COQ7 | 1094 | 7 | 7 | 7 | 52.1 | 52.1 | 52.1 | 21.871 |
| LYPLAL1 | 3218 | 7 | 7 | 7 | 45.6 | 45.6 | 45.6 | 26.316 |
| NASP | 2365 | 7 | 7 | 7 | 13.1 | 13.1 | 13.1 | 85.237 |
| ALG1 | 3976 | 7 | 7 | 7 | 22.4 | 22.4 | 22.4 | 52.518 |
| FN1 | 1713 | 7 | 7 | 7 | 4 | 4 | 4 | 272.32 |
| EIF2B1 | 2938 | 7 | 7 | 7 | 27.9 | 27.9 | 27.9 | 33.712 |
| MRPL10 | 3384 | 7 | 7 | 7 | 37.5 | 37.5 | 37.5 | 29.282 |
| RPA3 | 2218 | 7 | 7 | 7 | 67.8 | 67.8 | 67.8 | 13.569 |
| OPLAH | 1359 | 7 | 7 | 7 | 9 | 9 | 9 | 137.46 |
| ARHGAP30 | 3378 | 8 | 8 | 7 | 8.5 | 8.5 | 7.9 | 118.58 |
| BANF1 | 1559 | 7 | 7 | 7 | 62.9 | 62.9 | 62.9 | 10.058 |
| WDR5 | 2588 | 7 | 7 | 7 | 34.7 | 34.7 | 34.7 | 36.588 |
| APOBEC3C | 4254 | 8 | 8 | 7 | 58.9 | 58.9 | 52.6 | 22.826 |
| PPOX | 429 | 7 | 7 | 7 | 24.1 | 24.1 | 24.1 | 46.457 |

| | | | | | | | | |
|---|---|---|---|---|---|---|---|---|
| CDV3 | 4492 | 7 | 7 | 7 | 57.8 | 57.8 | 57.8 | 27.335 |
| DDX27 | 616 | 7 | 7 | 7 | 10.6 | 10.6 | 10.6 | 86.604 |
| USP10 | 2964 | 7 | 7 | 7 | 14.8 | 14.8 | 14.8 | 87.133 |
| SF3B4 | 3021 | 7 | 7 | 7 | 28.1 | 28.1 | 28.1 | 44.385 |
| YLPM1 | 951 | 7 | 7 | 7 | 4.9 | 4.9 | 4.9 | 160.74 |
| SAR1B;DKF | 4666 | 11 | 7 | 7 | 73.7 | 59.6 | 59.6 | 22.41 |
| MRPL3 | 799 | 7 | 7 | 7 | 27.3 | 27.3 | 27.3 | 40.47 |
| TIMM13 | 4640 | 7 | 7 | 7 | 77.9 | 77.9 | 77.9 | 10.5 |
| TPCN1 | 611 | 7 | 7 | 7 | 12.2 | 12.2 | 12.2 | 87.317 |
| PSMB7 | 3890 | 7 | 7 | 7 | 37.2 | 37.2 | 37.2 | 29.965 |
| PTGES3 | 76 | 7 | 7 | 7 | 41.9 | 41.9 | 41.9 | 18.697 |
| SLAMF7 | 4225 | 7 | 7 | 7 | 31 | 31 | 31 | 37.421 |
| GMPPA | 3819 | 7 | 7 | 7 | 21 | 21 | 21 | 46.291 |
| PROSC | 1614 | 7 | 7 | 7 | 33.1 | 33.1 | 33.1 | 30.344 |
| NPC2 | 782 | 7 | 7 | 7 | 56.1 | 56.1 | 56.1 | 16.23 |
| FYB | 1382 | 7 | 7 | 7 | 12.3 | 12.3 | 12.3 | 85.386 |
| TSNAX;DISC | 3909 | 7 | 7 | 7 | 35.5 | 35.5 | 35.5 | 33.112 |
| LAS1L | 4622 | 7 | 7 | 7 | 10.8 | 10.8 | 10.8 | 83.064 |
| LARP7 | 3110 | 7 | 7 | 7 | 16.2 | 16.2 | 16.2 | 66.898 |
| NUDT9 | 4016 | 7 | 7 | 7 | 27.7 | 27.7 | 27.7 | 39.125 |
| BTN3A3 | 1315 | 11 | 11 | 7 | 24 | 24 | 15.9 | 65.001 |
| SLC25A40 | 3601 | 7 | 7 | 7 | 24.6 | 24.6 | 24.6 | 38.124 |
| HINT1 | 2387 | 7 | 7 | 7 | 77.8 | 77.8 | 77.8 | 13.802 |
| STX6 | 1462 | 7 | 7 | 7 | 41.2 | 41.2 | 41.2 | 29.176 |
| EMC4 | 3150 | 7 | 7 | 7 | 50.3 | 50.3 | 50.3 | 20.086 |
| AP3S1 | 3676 | 7 | 7 | 7 | 47.7 | 47.7 | 47.7 | 21.732 |
| C2orf47 | 3653 | 7 | 7 | 7 | 26.8 | 26.8 | 26.8 | 32.544 |
| EMC10 | 1273 | 7 | 7 | 7 | 19.7 | 19.7 | 19.7 | 38.975 |
| ARL8B | 4300 | 17 | 17 | 7 | 68.8 | 68.8 | 42.5 | 21.539 |
| FKBP2 | 436 | 7 | 7 | 7 | 51.5 | 51.5 | 51.5 | 18.562 |
| SORD | 484 | 7 | 7 | 7 | 34.5 | 34.5 | 34.5 | 27.076 |
| STEAP3 | 636 | 7 | 7 | 7 | 22.1 | 22.1 | 22.1 | 50.628 |
| FEN1 | 2258 | 7 | 7 | 7 | 28.9 | 28.9 | 28.9 | 42.592 |
| EIF6 | 2528 | 7 | 7 | 7 | 55.9 | 55.9 | 55.9 | 26.599 |
| RPL30 | 2635 | 7 | 7 | 7 | 66.1 | 66.1 | 66.1 | 12.784 |
| RHBDF2 | 3285 | 7 | 7 | 7 | 11.7 | 11.7 | 11.7 | 96.685 |
| FNBP1 | 626 | 7 | 7 | 7 | 12.3 | 12.3 | 12.3 | 64.071 |
| GOSR2 | 323 | 7 | 7 | 7 | 53.3 | 53.3 | 53.3 | 24.775 |
| SELENOF | 103 | 7 | 7 | 7 | 50.6 | 50.6 | 50.6 | 17.942 |
| RAB38 | 2534 | 9 | 7 | 7 | 55 | 45 | 45 | 23.712 |
| LYPLA2 | 1643 | 7 | 7 | 7 | 47.6 | 47.6 | 47.6 | 24.737 |
| DKFZp566H | 4420 | 7 | 7 | 7 | 22.7 | 22.7 | 22.7 | 34.815 |
| CCRL2 | 1309 | 6 | 6 | 6 | 17.7 | 17.7 | 17.7 | 39.513 |
| COA3 | 4566 | 6 | 6 | 6 | 47.2 | 47.2 | 47.2 | 11.731 |
| NDUFB4 | 1627 | 6 | 6 | 6 | 42.6 | 42.6 | 42.6 | 15.208 |
| RCSD1 | 625 | 6 | 6 | 6 | 19.7 | 19.7 | 19.7 | 41.261 |
| PPP4C | 1104 | 8 | 6 | 6 | 34.5 | 28.8 | 28.8 | 30.561 |
| RPS28 | 2631 | 6 | 6 | 6 | 56.5 | 56.5 | 56.5 | 7.8409 |

| | | | | | | | | |
|---|---|---|---|---|---|---|---|---|
| FAM3C | 3669 | 6 | 6 | 6 | 30.8 | 30.8 | 30.8 | 24.68 |
| EMC3 | 4372 | 8 | 8 | 6 | 44.1 | 44.1 | 32.6 | 29.952 |
| SF3B5 | 4021 | 6 | 6 | 6 | 87.2 | 87.2 | 87.2 | 10.135 |
| NDUFS4 | 1425 | 6 | 6 | 6 | 46.3 | 46.3 | 46.3 | 20.108 |
| ABAT | 1092 | 6 | 6 | 6 | 18.6 | 18.6 | 18.6 | 58.14 |
| TRIAP1 | 1459 | 6 | 6 | 6 | 73.7 | 73.7 | 73.7 | 8.7858 |
| ATP6V1F | 3077 | 6 | 6 | 6 | 81.5 | 81.5 | 81.5 | 13.37 |
| BLOC1S1 | 2687 | 6 | 6 | 6 | 55.6 | 55.6 | 55.6 | 17.262 |
| PIP4K2A | 2345 | 9 | 9 | 6 | 25.4 | 25.4 | 17 | 46.224 |
| UBE2I | 2660 | 6 | 6 | 6 | 60.8 | 60.8 | 60.8 | 18.007 |
| RPS15 | 1223 | 6 | 6 | 6 | 53.3 | 53.3 | 53.3 | 17.723 |
| CRLF3 | 3434 | 6 | 6 | 6 | 19.9 | 19.9 | 19.9 | 49.765 |
| TIMM9 | 4636 | 6 | 6 | 6 | 77.5 | 77.5 | 77.5 | 10.378 |
| RAB29 | 1370 | 7 | 6 | 6 | 46.3 | 40.9 | 40.9 | 23.155 |
| HNRNPK | 2590 | 27 | 27 | 6 | 67.4 | 67.4 | 17.5 | 50.976 |
| GOLIM4 | 970 | 6 | 6 | 6 | 10.6 | 10.6 | 10.6 | 78.759 |
| UQCC2 | 3961 | 6 | 6 | 6 | 46 | 46 | 46 | 14.875 |
| FAM21A | 3227 | 29 | 29 | 6 | 35.2 | 35.2 | 6.6 | 147.18 |
| NDUFS5 | 1478 | 6 | 6 | 6 | 57.5 | 57.5 | 57.5 | 12.517 |
| WDR43 | 2993 | 6 | 6 | 6 | 12.1 | 12.1 | 12.1 | 74.89 |
| CYBA | 1910 | 6 | 6 | 6 | 37.4 | 37.4 | 37.4 | 21.012 |
| MTCH2 | 4667 | 16 | 16 | 6 | 51.2 | 51.2 | 11.9 | 33.331 |
| LMBRD2 | 3232 | 6 | 6 | 6 | 11.7 | 11.7 | 11.7 | 81.171 |
| RBM8A | 4646 | 6 | 6 | 6 | 56.3 | 56.3 | 56.3 | 19.889 |
| SLC12A9 | 4032 | 6 | 6 | 6 | 10.2 | 10.2 | 10.2 | 96.109 |
| KPNA1 | 2455 | 12 | 12 | 6 | 29 | 29 | 14.7 | 60.221 |
| UBLCP1 | 3649 | 6 | 6 | 6 | 20.1 | 20.1 | 20.1 | 36.804 |
| UBE2Z | 4159 | 6 | 6 | 6 | 19.8 | 19.8 | 19.8 | 38.21 |
| PEA15 | 3000 | 6 | 6 | 6 | 57.7 | 57.7 | 57.7 | 15.04 |
| MRTO4 | 4480 | 6 | 6 | 6 | 31.8 | 31.8 | 31.8 | 27.56 |
| OSBPL9 | 3883 | 6 | 6 | 6 | 10.7 | 10.7 | 10.7 | 83.184 |
| PNPO | 1203 | 6 | 6 | 6 | 37.4 | 37.4 | 37.4 | 27.312 |
| VPS25 | 3954 | 6 | 6 | 6 | 35.2 | 35.2 | 35.2 | 20.747 |
| FAM134B | 4151 | 6 | 6 | 6 | 16.3 | 16.3 | 16.3 | 54.68 |
| NFU1 | 4507 | 6 | 6 | 6 | 30.3 | 30.3 | 30.3 | 28.462 |
| SPCS3 | 2559 | 6 | 6 | 6 | 33.9 | 33.9 | 33.9 | 20.313 |
| PYCR2 | 23 | 6 | 6 | 6 | 37.4 | 37.4 | 37.4 | 25.868 |
| CERS2 | 3796 | 6 | 6 | 6 | 16.6 | 16.6 | 16.6 | 44.876 |
| HINT2 | 4025 | 6 | 6 | 6 | 52.1 | 52.1 | 52.1 | 17.162 |
| IL18 | 2924 | 6 | 6 | 6 | 34.7 | 34.7 | 34.7 | 22.326 |
| HSPE1-MOI | 4742 | 11 | 11 | 6 | 52.1 | 52.1 | 35.2 | 29.736 |
| TECPR1 | 3379 | 6 | 6 | 6 | 6.9 | 6.9 | 6.9 | 129.69 |
| NMI | 2858 | 6 | 6 | 6 | 22.5 | 22.5 | 22.5 | 35.056 |
| PIK3AP1 | 3313 | 6 | 6 | 6 | 10.2 | 10.2 | 10.2 | 90.397 |
| ATL2 | 3581 | 7 | 6 | 6 | 14.9 | 13 | 13 | 66.228 |
| DDT;DDTL | 2158 | 6 | 6 | 6 | 56.8 | 56.8 | 56.8 | 12.712 |
| ATP6V1G1 | 1534 | 7 | 7 | 6 | 66.1 | 66.1 | 53.4 | 13.757 |
| NUP153 | 2389 | 6 | 6 | 6 | 4.9 | 4.9 | 4.9 | 153.94 |

| | | | | | | | | |
|---|---|---|---|---|---|---|---|---|
| HINT3 | 4231 | 6 | 6 | 6 | 50 | 50 | 50 | 20.361 |
| ARHGAP4 | 793 | 6 | 6 | 6 | 10.3 | 10.3 | 10.3 | 85.38 |
| PCBD2 | 1051 | 6 | 6 | 6 | 58 | 58 | 58 | 13.409 |
| TNS1 | 345 | 8 | 6 | 6 | 6.2 | 5.5 | 5.5 | 195.47 |
| ALYREF | 820 | 6 | 6 | 6 | 26.9 | 26.9 | 26.9 | 27.557 |
| CHCHD6 | 3957 | 6 | 6 | 6 | 36.2 | 36.2 | 36.2 | 26.457 |
| OXNAD1 | 681 | 6 | 6 | 6 | 20.9 | 20.9 | 20.9 | 34.725 |
| IKBKE | 12 | 6 | 6 | 6 | 9.4 | 9.4 | 9.4 | 73.75 |
| RPF2 | 4155 | 6 | 6 | 6 | 24.2 | 24.2 | 24.2 | 35.582 |
| TREX1 | 4266 | 6 | 6 | 6 | 34.7 | 34.7 | 34.7 | 33.212 |
| UMPS | 1871 | 6 | 6 | 6 | 20.8 | 20.8 | 20.8 | 52.221 |
| PTMA | 633 | 6 | 6 | 6 | 36.4 | 36.4 | 36.4 | 11.758 |
| NUPL1 | 1119 | 6 | 6 | 6 | 20.3 | 20.3 | 20.3 | 49.095 |
| RAB4A | 2020 | 8 | 6 | 6 | 46.8 | 40.4 | 40.4 | 24.389 |
| HNMT | 2399 | 6 | 6 | 6 | 32.9 | 32.9 | 32.9 | 33.295 |
| SLC30A5 | 3590 | 6 | 6 | 6 | 11.9 | 11.9 | 11.9 | 84.046 |
| OXA1L | 90 | 6 | 6 | 6 | 14.9 | 14.9 | 14.9 | 48.519 |
| PHYH | 559 | 6 | 6 | 6 | 21.5 | 21.5 | 21.5 | 36.965 |
| OCRL | 2740 | 6 | 6 | 6 | 8.1 | 8.1 | 8.1 | 104.2 |
| TFG | 3693 | 6 | 6 | 6 | 22.2 | 22.2 | 22.2 | 43.447 |
| CUL3 | 2901 | 6 | 6 | 6 | 8.3 | 8.3 | 8.3 | 88.929 |
| FIS1 | 4598 | 6 | 6 | 6 | 44.7 | 44.7 | 44.7 | 16.937 |
| SYNE1 | 200 | 6 | 6 | 6 | 1 | 1 | 1 | 964.83 |
| PTER | 3759 | 6 | 6 | 6 | 26.1 | 26.1 | 26.1 | 39.017 |
| PTGDS | 2271 | 6 | 6 | 6 | 47.4 | 47.4 | 47.4 | 21.029 |
| SCAMP2 | 1387 | 6 | 6 | 6 | 17 | 17 | 17 | 36.648 |
| GATM | 2408 | 6 | 6 | 6 | 25.8 | 25.8 | 25.8 | 48.455 |
| LANCL2 | 4263 | 6 | 6 | 6 | 18.2 | 18.2 | 18.2 | 50.854 |
| SEPN1 | 1138 | 6 | 6 | 6 | 11.5 | 11.5 | 11.5 | 65.812 |
| AQR | 1485 | 6 | 6 | 6 | 6.1 | 6.1 | 6.1 | 171.29 |
| INPPL1 | 165 | 7 | 6 | 6 | 7.3 | 6.5 | 6.5 | 126.52 |
| TRAPPC12; | 3648 | 6 | 6 | 6 | 13.9 | 13.9 | 13.9 | 79.374 |
| COPS6 | 778 | 6 | 6 | 6 | 32.1 | 32.1 | 32.1 | 36.163 |
| ALG9 | 80 | 6 | 6 | 6 | 10.7 | 10.7 | 10.7 | 95.921 |
| BAX | 2782 | 7 | 7 | 6 | 41.1 | 41.1 | 35.4 | 21.184 |
| FARS2 | 401 | 6 | 6 | 6 | 17.5 | 17.5 | 17.5 | 52.356 |
| CHERP | 1194 | 6 | 6 | 6 | 10.4 | 10.4 | 10.4 | 104.93 |
| FAM45A;FA | 3607 | 6 | 6 | 6 | 26.3 | 26.3 | 26.3 | 40.513 |
| UBE2L6 | 1366 | 6 | 6 | 6 | 62.1 | 62.1 | 62.1 | 17.769 |
| GSPT1 | 1941 | 14 | 14 | 6 | 33.5 | 33.5 | 18.8 | 55.755 |
| XPO7 | 4451 | 6 | 6 | 6 | 6.8 | 6.8 | 6.8 | 123.91 |
| PTGR1 | 2974 | 6 | 6 | 6 | 27.4 | 27.4 | 27.4 | 35.869 |
| RRP1 | 2521 | 6 | 6 | 6 | 16.7 | 16.7 | 16.7 | 52.839 |
| RABL6 | 886 | 6 | 6 | 6 | 17.8 | 17.8 | 17.8 | 53.859 |
| PPP6R1 | 4526 | 6 | 6 | 6 | 9.5 | 9.5 | 9.5 | 96.723 |
| RPS26;RPS: | 2630 | 6 | 6 | 6 | 50.4 | 50.4 | 50.4 | 13.015 |
| TOMM22 | 4262 | 6 | 6 | 6 | 66.2 | 66.2 | 66.2 | 15.521 |
| TBCA | 770 | 6 | 6 | 6 | 58.3 | 58.3 | 58.3 | 10.08 |

| | | | | | | | | |
|---|---|---|---|---|---|---|---|---|
| SLC25A16 | 1959 | 6 | 6 | 6 | 18.1 | 18.1 | 18.1 | 36.223 |
| PUM1 | 879 | 6 | 6 | 6 | 9 | 9 | 9 | 100.5 |
| MRPS17;h( | 1142 | 6 | 6 | 6 | 60 | 60 | 60 | 14.502 |
| TCEB1 | 767 | 6 | 6 | 6 | 66.1 | 66.1 | 66.1 | 12.473 |
| PAPSS1 | 1428 | 7 | 6 | 6 | 16.3 | 14.1 | 14.1 | 70.832 |
| GMPR2 | 57 | 6 | 6 | 6 | 31.6 | 31.6 | 31.6 | 37.874 |
| SGPP1 | 4026 | 6 | 6 | 6 | 11.6 | 11.6 | 11.6 | 49.107 |
| POLDIP3 | 942 | 6 | 6 | 6 | 18.5 | 18.5 | 18.5 | 48.103 |
| RBM17 | 3814 | 6 | 6 | 6 | 19 | 19 | 19 | 44.961 |
| BLMH | 2912 | 6 | 6 | 6 | 20.4 | 20.4 | 20.4 | 52.562 |
| VPS8 | 679 | 6 | 6 | 6 | 6.3 | 6.3 | 6.3 | 161.75 |
| TRNT1 | 3859 | 6 | 6 | 6 | 17.5 | 17.5 | 17.5 | 50.127 |
| PIR | 1326 | 6 | 6 | 6 | 35.9 | 35.9 | 35.9 | 32.113 |
| MRRF | 3780 | 6 | 6 | 6 | 36.3 | 36.3 | 36.3 | 29.277 |
| RAI14 | 4376 | 6 | 6 | 6 | 7.1 | 7.1 | 7.1 | 110.04 |
| EIF4H | 2992 | 6 | 6 | 6 | 41.1 | 41.1 | 41.1 | 27.385 |
| KXD1 | 1256 | 6 | 6 | 6 | 67.1 | 67.1 | 67.1 | 9.8313 |
| UBE3A | 307 | 6 | 6 | 6 | 10.5 | 10.5 | 10.5 | 93.543 |
| NCS1 | 2596 | 6 | 6 | 6 | 40.5 | 40.5 | 40.5 | 21.878 |
| RPLP2 | 1755 | 7 | 7 | 6 | 91.3 | 91.3 | 76.5 | 11.665 |
| MON1B | 3333 | 6 | 6 | 6 | 18.5 | 18.5 | 18.5 | 59.217 |
| CNOT10 | 4169 | 6 | 6 | 6 | 9.1 | 9.1 | 9.1 | 82.309 |
| DOCK5 | 4156 | 6 | 6 | 6 | 5.4 | 5.4 | 5.4 | 215.31 |
| LARP1 | 3286 | 6 | 6 | 6 | 7.4 | 7.4 | 7.4 | 123.51 |
| FRG1 | 2948 | 6 | 6 | 6 | 28.7 | 28.7 | 28.7 | 29.172 |
| NDUFB6 | 1622 | 6 | 6 | 6 | 53.1 | 53.1 | 53.1 | 15.489 |
| ACOT4 | 3530 | 7 | 6 | 6 | 29.7 | 24 | 24 | 46.326 |
| TRAPPC6B | 3389 | 6 | 6 | 6 | 42.4 | 42.4 | 42.4 | 17.983 |
| RPE | 3749 | 6 | 6 | 6 | 40.8 | 40.8 | 40.8 | 24.927 |
| COX20 | 3172 | 6 | 6 | 6 | 53.4 | 53.4 | 53.4 | 13.291 |
| BUD31 | 2272 | 6 | 6 | 6 | 47.9 | 47.9 | 47.9 | 17 |
| MECP2 | 2440 | 11 | 11 | 6 | 29.6 | 29.6 | 16 | 52.44 |
| CRK | 2319 | 6 | 6 | 6 | 25.7 | 25.7 | 25.7 | 33.83 |
| ACP1 | 2087 | 6 | 6 | 6 | 40.5 | 40.5 | 40.5 | 18.042 |
| DNAJB1 | 2097 | 7 | 7 | 6 | 24.7 | 24.7 | 21.2 | 38.044 |
| DNASE2 | 1277 | 6 | 6 | 6 | 15.8 | 15.8 | 15.8 | 39.58 |
| MFSD12 | 3256 | 6 | 6 | 6 | 21 | 21 | 21 | 52.075 |
| PAIP1 | 747 | 6 | 6 | 6 | 18.5 | 18.5 | 18.5 | 39.908 |
| MAPKAPK3 | 653 | 7 | 7 | 6 | 29.5 | 29.5 | 27.1 | 37.389 |
| RBMX | 2251 | 21 | 21 | 6 | 49.4 | 49.4 | 14.6 | 42.331 |
| IMPAD1 | 4330 | 6 | 6 | 6 | 17.3 | 17.3 | 17.3 | 38.681 |
| PPP4R2 | 4343 | 6 | 6 | 6 | 20.1 | 20.1 | 20.1 | 46.898 |
| KPNA3 | 1320 | 13 | 6 | 6 | 32.8 | 19.4 | 19.4 | 57.81 |
| GCLM | 2349 | 6 | 6 | 6 | 28.1 | 28.1 | 28.1 | 30.727 |
| ZZEF1 | 1423 | 6 | 6 | 6 | 3.2 | 3.2 | 3.2 | 331.07 |
| H1F0 | 1780 | 6 | 6 | 6 | 27.3 | 27.3 | 27.3 | 20.863 |
| NDUFB8 | 1628 | 6 | 6 | 6 | 38.7 | 38.7 | 38.7 | 21.766 |
| AGPAT6 | 3402 | 8 | 6 | 6 | 22.4 | 15.8 | 15.8 | 52.07 |

| | | | | | | | | |
|---|---|---|---|---|---|---|---|---|
| NTPCR | 3967 | 6 | 6 | 6 | 46.8 | 46.8 | 46.8 | 20.713 |
| LSM2 | 4581 | 6 | 6 | 6 | 61.1 | 61.1 | 61.1 | 10.834 |
| ARHGAP10 | 493 | 7 | 7 | 6 | 9 | 9 | 8.1 | 89.374 |
| WDR77 | 3942 | 6 | 6 | 6 | 32.5 | 32.5 | 32.5 | 36.724 |
| NDUFB1 | 1550 | 6 | 6 | 6 | 72.4 | 72.4 | 72.4 | 6.9611 |
| FRMD8 | 4054 | 6 | 6 | 6 | 26.1 | 26.1 | 26.1 | 51.217 |
| MAPK14 | 592 | 6 | 6 | 6 | 41 | 41 | 41 | 32.357 |
| GGA1 | 4475 | 6 | 6 | 6 | 20.2 | 20.2 | 20.2 | 70.383 |
| TMEM192 | 3467 | 6 | 6 | 6 | 36.5 | 36.5 | 36.5 | 30.922 |
| ARMC1 | 4306 | 6 | 6 | 6 | 34.4 | 34.4 | 34.4 | 31.28 |
| TBCE | 356 | 6 | 6 | 6 | 15.9 | 15.9 | 15.9 | 59.345 |
| SLC35A4 | 1254 | 6 | 6 | 6 | 42.7 | 42.7 | 42.7 | 11.133 |
| TRIM22 | 693 | 6 | 6 | 6 | 22 | 22 | 22 | 34.866 |
| WIBG | 3956 | 6 | 6 | 6 | 33.8 | 33.8 | 33.8 | 22.655 |
| RABGGTB | 2479 | 6 | 6 | 6 | 23.6 | 23.6 | 23.6 | 36.924 |
| BTF3 | 2019 | 6 | 6 | 6 | 50 | 50 | 50 | 22.168 |
| UNG;DKFZp | 1904 | 6 | 6 | 6 | 38 | 38 | 38 | 34.645 |
| SDF2 | 3894 | 6 | 6 | 6 | 40.3 | 40.3 | 40.3 | 23.026 |
| FBXW11;B | 4479 | 6 | 6 | 6 | 12.7 | 12.7 | 12.7 | 62.09 |
| ARL8A | 3754 | 16 | 6 | 6 | 74.7 | 36 | 36 | 21.416 |
| TCEB2 | 3014 | 6 | 6 | 6 | 48.3 | 48.3 | 48.3 | 13.133 |
| GUSB | 1799 | 6 | 6 | 6 | 9.4 | 9.4 | 9.4 | 74.731 |
| ATP5S | 474 | 6 | 6 | 6 | 39 | 39 | 39 | 23.226 |
| BLOC1S2 | 3289 | 6 | 6 | 6 | 46.5 | 46.5 | 46.5 | 15.961 |
| DOCK11 | 525 | 7 | 6 | 6 | 5 | 4.5 | 4.5 | 238.1 |
| GDE1 | 4359 | 6 | 6 | 6 | 24.5 | 24.5 | 24.5 | 37.718 |
| GSTT1 | 2174 | 6 | 6 | 6 | 40 | 40 | 40 | 27.335 |
| DBI | 1778 | 6 | 6 | 6 | 60.9 | 60.9 | 60.9 | 10.044 |
| CMPK2 | 3145 | 6 | 6 | 6 | 18.5 | 18.5 | 18.5 | 49.447 |
| ALG5 | 4660 | 6 | 6 | 6 | 23.8 | 23.8 | 23.8 | 36.946 |
| MPST | 2093 | 6 | 6 | 6 | 32.7 | 32.7 | 32.7 | 33.178 |
| MFSD10 | 743 | 6 | 6 | 6 | 17.9 | 17.9 | 17.9 | 50.431 |
| BABAM1 | 4319 | 6 | 6 | 6 | 29.5 | 29.5 | 29.5 | 36.56 |
| NDUFA5 | 3065 | 6 | 6 | 6 | 66.4 | 66.4 | 66.4 | 13.459 |
| RPL27 | 2580 | 6 | 6 | 6 | 44.9 | 44.9 | 44.9 | 15.798 |
| EML2 | 216 | 6 | 6 | 6 | 20.1 | 20.1 | 20.1 | 46.644 |
| NDUFAF2 | 3486 | 6 | 6 | 6 | 36.7 | 36.7 | 36.7 | 19.856 |
| NUFIP2 | 3365 | 6 | 6 | 6 | 10.1 | 10.1 | 10.1 | 76.12 |
| RPS21 | 2656 | 6 | 6 | 6 | 61.7 | 61.7 | 61.7 | 8.85 |
| TMCO3 | 3296 | 6 | 6 | 6 | 10 | 10 | 10 | 75.597 |
| XAB2 | 4202 | 6 | 6 | 6 | 9.1 | 9.1 | 9.1 | 100.01 |
| SURF1 | 3028 | 6 | 6 | 6 | 24.3 | 24.3 | 24.3 | 33.331 |
| ABCF2 | 4424 | 6 | 6 | 6 | 12.5 | 12.5 | 12.5 | 71.289 |
| COASY | 2835 | 6 | 6 | 6 | 14.7 | 14.7 | 14.7 | 62.328 |
| DYNLRB1 | 4214 | 6 | 6 | 6 | 69.8 | 69.8 | 69.8 | 10.921 |
| GTF2F2 | 1921 | 6 | 6 | 6 | 31.7 | 31.7 | 31.7 | 28.38 |
| PIP5K1C | 1486 | 9 | 9 | 6 | 19.2 | 19.2 | 14.8 | 73.259 |
| SPPL2A | 3610 | 6 | 6 | 6 | 15.6 | 15.6 | 15.6 | 58.143 |

| | | | | | | | | |
|---|---|---|---|---|---|---|---|---|
| RPS4Y1 | 2051 | 19 | 10 | 6 | 60.5 | 38 | 28.1 | 29.455 |
| SSR4 | 2436 | 6 | 6 | 6 | 37 | 37 | 37 | 18.998 |
| MTPN | 2540 | 6 | 6 | 6 | 61.9 | 61.9 | 61.9 | 12.895 |
| COA7 | 3758 | 6 | 6 | 6 | 38.5 | 38.5 | 38.5 | 25.709 |
| PLGRKT | 4189 | 6 | 6 | 6 | 34 | 34 | 34 | 17.201 |
| B3GAT3 | 992 | 6 | 6 | 6 | 31.9 | 31.9 | 31.9 | 37.121 |
| CDK11A;CD | 225 | 6 | 6 | 6 | 8.2 | 8.2 | 8.2 | 86.823 |
| PDCD5 | 1350 | 6 | 6 | 6 | 55.2 | 55.2 | 55.2 | 14.285 |
| ITPA | 4039 | 6 | 6 | 6 | 41.8 | 41.8 | 41.8 | 21.445 |
| TBC1D15 | 3604 | 6 | 6 | 6 | 8.1 | 8.1 | 8.1 | 79.49 |
| MPO | 1749 | 6 | 6 | 6 | 10.9 | 10.9 | 10.9 | 83.868 |
| HSD11B1 | 2141 | 6 | 6 | 6 | 37.3 | 37.3 | 37.3 | 32.401 |
| BPHL | 3412 | 6 | 6 | 6 | 33 | 33 | 33 | 32.542 |
| EFR3A | 305 | 6 | 6 | 6 | 11.6 | 11.6 | 11.6 | 92.923 |
| PPIG | 2870 | 6 | 6 | 6 | 8.9 | 8.9 | 8.9 | 88.616 |
| VPS52 | 3487 | 6 | 6 | 6 | 11.2 | 11.2 | 11.2 | 82.22 |
| STBD1 | 1633 | 6 | 6 | 6 | 20.4 | 20.4 | 20.4 | 39.007 |
| CD9 | 341 | 6 | 6 | 6 | 33.3 | 33.3 | 33.3 | 17.764 |
| MIOS | 4333 | 6 | 6 | 6 | 9.9 | 9.9 | 9.9 | 98.583 |
| PSMD10 | 1577 | 6 | 6 | 6 | 42.5 | 42.5 | 42.5 | 24.428 |
| FAM63A | 3516 | 6 | 6 | 6 | 17.5 | 17.5 | 17.5 | 51.777 |
| SOD1 | 1693 | 6 | 6 | 6 | 61 | 61 | 61 | 15.936 |
| TUBA4A | 2670 | 28 | 6 | 6 | 78.3 | 28.1 | 28.1 | 49.924 |
| LETMD1 | 962 | 6 | 6 | 6 | 25.5 | 25.5 | 25.5 | 34.046 |
| DNAJC8 | 1590 | 6 | 6 | 6 | 30 | 30 | 30 | 29.841 |
| REL | 2764 | 6 | 6 | 6 | 12.3 | 12.3 | 12.3 | 68.519 |
| CYB5B | 1111 | 6 | 6 | 6 | 66 | 66 | 66 | 16.079 |
| MYL12A;MY | 1205 | 12 | 12 | 6 | 64.4 | 64.4 | 32.2 | 20.457 |
| MRPL16 | 4323 | 6 | 6 | 6 | 24.7 | 24.7 | 24.7 | 28.449 |
| PTK2B | 2945 | 6 | 6 | 6 | 8.1 | 8.1 | 8.1 | 115.87 |
| RAB34 | 662 | 6 | 6 | 6 | 33.3 | 33.3 | 33.3 | 23.74 |
| MRPS28 | 4565 | 9 | 9 | 6 | 62 | 62 | 42.2 | 20.843 |
| UCHL1 | 1838 | 6 | 6 | 6 | 43 | 43 | 43 | 24.824 |
| PIK3R1 | 2130 | 6 | 6 | 6 | 12 | 12 | 12 | 83.597 |
| CRELD1 | 3806 | 6 | 6 | 6 | 18.1 | 18.1 | 18.1 | 45.439 |
| CASP3 | 2288 | 6 | 6 | 6 | 27.8 | 27.8 | 27.8 | 31.608 |
| TMEM199 | 3511 | 6 | 6 | 6 | 38.5 | 38.5 | 38.5 | 23.13 |
| DNAJC5 | 4133 | 6 | 6 | 6 | 42.4 | 42.4 | 42.4 | 22.149 |
| TMEM38B | 4307 | 6 | 6 | 6 | 26.8 | 26.8 | 26.8 | 32.509 |
| SLC1A3 | 18 | 6 | 6 | 6 | 18.8 | 18.8 | 18.8 | 47.099 |
| COX17 | 2922 | 6 | 6 | 6 | 81 | 81 | 81 | 6.9151 |
| RPP30 | 2679 | 6 | 6 | 6 | 29.9 | 29.9 | 29.9 | 29.321 |
| PHF6 | 234 | 6 | 6 | 6 | 20.4 | 20.4 | 20.4 | 36.43 |
| NDUFA11 | 3425 | 6 | 6 | 6 | 56.7 | 56.7 | 56.7 | 14.852 |
| NME1-NME | 3093 | 15 | 15 | 6 | 71.2 | 71.2 | 24 | 32.642 |
| FKBP1A;FKI | 2642 | 6 | 6 | 6 | 75.9 | 75.9 | 75.9 | 11.951 |
| SMS | 2463 | 6 | 6 | 6 | 18.9 | 18.9 | 18.9 | 41.268 |
| MRPL53 | 3784 | 6 | 6 | 6 | 72.3 | 72.3 | 72.3 | 12.107 |

| | | | | | | | | |
|---|---|---|---|---|---|---|---|---|
| OPA3 | 4150 | 6 | 6 | 6 | 29.1 | 29.1 | 29.1 | 19.996 |
| NUP85 | 1195 | 6 | 6 | 6 | 16.5 | 16.5 | 16.5 | 62.101 |
| RER1 | 1401 | 6 | 6 | 6 | 33.7 | 33.7 | 33.7 | 22.958 |
| SLC17A5 | 4247 | 6 | 6 | 6 | 14.9 | 14.9 | 14.9 | 54.639 |
| YBX3 | 1969 | 11 | 6 | 6 | 31.2 | 20.2 | 20.2 | 40.089 |
| TMED5 | 4588 | 6 | 6 | 6 | 39.3 | 39.3 | 39.3 | 26.005 |
| PPP1CB | 2595 | 18 | 18 | 6 | 64.2 | 64.2 | 26.9 | 37.186 |
| CCDC127 | 3757 | 6 | 6 | 6 | 33.1 | 33.1 | 33.1 | 30.834 |
| RBMS1 | 798 | 6 | 6 | 6 | 23.8 | 23.8 | 23.8 | 40.147 |
| SCAF11 | 166 | 6 | 6 | 6 | 6.7 | 6.7 | 6.7 | 106.01 |
| CLTB | 1827 | 6 | 6 | 6 | 22.7 | 22.7 | 22.7 | 25.19 |
| ISCU | 4105 | 5 | 5 | 5 | 28.1 | 28.1 | 28.1 | 17.999 |
| SRP19 | 1817 | 5 | 5 | 5 | 46.5 | 46.5 | 46.5 | 16.156 |
| FAM213B | 154 | 5 | 5 | 5 | 27.6 | 27.6 | 27.6 | 24.278 |
| C10orf11 | 55 | 5 | 5 | 5 | 29.2 | 29.2 | 29.2 | 25.689 |
| WDR12 | 4074 | 5 | 5 | 5 | 17 | 17 | 17 | 47.707 |
| RPS20 | 2552 | 5 | 5 | 5 | 32.8 | 32.8 | 32.8 | 13.373 |
| MRPS16 | 4596 | 5 | 5 | 5 | 51.1 | 51.1 | 51.1 | 15.345 |
| ENTPD1 | 2398 | 5 | 5 | 5 | 12 | 12 | 12 | 57.964 |
| SPRYD7 | 3225 | 5 | 5 | 5 | 31.6 | 31.6 | 31.6 | 21.666 |
| FAM98A | 3549 | 8 | 5 | 5 | 22.2 | 14.5 | 14.5 | 55.272 |
| DDX39A | 1281 | 21 | 8 | 5 | 58.8 | 30 | 22.5 | 49.129 |
| SLC25A4 | 1891 | 22 | 5 | 5 | 58.4 | 17.1 | 17.1 | 33.064 |
| SYNJ2BP-C( | 77 | 5 | 5 | 5 | 48.7 | 48.7 | 48.7 | 17.188 |
| PTRH2 | 1175 | 5 | 5 | 5 | 47.2 | 47.2 | 47.2 | 19.325 |
| PAF1 | 3526 | 5 | 5 | 5 | 12.1 | 12.1 | 12.1 | 59.975 |
| PAM | 2001 | 5 | 5 | 5 | 8.1 | 8.1 | 8.1 | 108.33 |
| IBA57 | 3191 | 5 | 5 | 5 | 26.1 | 26.1 | 26.1 | 38.155 |
| SLC16A1 | 2486 | 5 | 5 | 5 | 9.4 | 9.4 | 9.4 | 53.944 |
| TIMM8B | 4637 | 5 | 5 | 5 | 60.2 | 60.2 | 60.2 | 9.3435 |
| CHCHD4 | 3508 | 5 | 5 | 5 | 47.2 | 47.2 | 47.2 | 15.996 |
| RAP2B | 2575 | 10 | 5 | 5 | 74.3 | 36.6 | 36.6 | 20.504 |
| PABPN1 | 3398 | 5 | 5 | 5 | 22.9 | 22.9 | 22.9 | 32.749 |
| RBM3 | 2719 | 5 | 5 | 5 | 55.4 | 55.4 | 55.4 | 17.17 |
| YIPF3 | 752 | 5 | 5 | 5 | 30.9 | 30.9 | 30.9 | 28.797 |
| TIMP2 | 1953 | 5 | 5 | 5 | 40.5 | 40.5 | 40.5 | 24.399 |
| CDIPT | 576 | 5 | 5 | 5 | 27.2 | 27.2 | 27.2 | 23.539 |
| SLC9A7 | 63 | 5 | 5 | 5 | 7.9 | 7.9 | 7.9 | 80.215 |
| TMEM245 | 4124 | 5 | 5 | 5 | 6.7 | 6.7 | 6.7 | 97.356 |
| CISD1 | 4356 | 5 | 5 | 5 | 52.8 | 52.8 | 52.8 | 12.199 |
| S100A8 | 1747 | 5 | 5 | 5 | 49.5 | 49.5 | 49.5 | 10.834 |
| TMEM165 | 4191 | 5 | 5 | 5 | 21.9 | 21.9 | 21.9 | 34.905 |
| ARHGAP18 | 3499 | 5 | 5 | 5 | 7.8 | 7.8 | 7.8 | 74.976 |
| WASH3P;W | 639 | 7 | 7 | 5 | 21.8 | 21.8 | 15.1 | 49.994 |
| IAH1 | 1136 | 5 | 5 | 5 | 28.9 | 28.9 | 28.9 | 25.085 |
| ARPC5L | 3936 | 6 | 5 | 5 | 47.7 | 42.5 | 42.5 | 16.941 |
| TIMM23;TI | 1364 | 5 | 5 | 5 | 53.1 | 53.1 | 53.1 | 21.943 |
| ABCF3 | 4283 | 5 | 5 | 5 | 8.7 | 8.7 | 8.7 | 79.744 |

| | | | | | | | | |
|---|---|---|---|---|---|---|---|---|
| AGO1;AGO4 | 487 | 12 | 12 | 5 | 17.6 | 17.6 | 7.5 | 101.07 |
| UBXN1 | 881 | 5 | 5 | 5 | 35.8 | 35.8 | 35.8 | 32.913 |
| CHORDC1 | 4438 | 5 | 5 | 5 | 23.2 | 23.2 | 23.2 | 37.489 |
| ARIH1 | 4624 | 5 | 5 | 5 | 12 | 12 | 12 | 64.117 |
| ATP5D | 2161 | 5 | 5 | 5 | 61.9 | 61.9 | 61.9 | 17.49 |
| HLA-DQB1 | 500 | 13 | 13 | 5 | 60.3 | 60.3 | 33.5 | 25.961 |
| THEMIS2 | 3204 | 5 | 5 | 5 | 11.5 | 11.5 | 11.5 | 72.048 |
| C10orf32-A | 176 | 5 | 5 | 5 | 70.8 | 70.8 | 70.8 | 11.695 |
| PFDN5 | 3895 | 5 | 5 | 5 | 49.4 | 49.4 | 49.4 | 17.328 |
| TP53RK | 3875 | 5 | 5 | 5 | 27.3 | 27.3 | 27.3 | 28.16 |
| ERH | 2709 | 5 | 5 | 5 | 51 | 51 | 51 | 12.259 |
| SIRT2 | 73 | 5 | 5 | 5 | 28.6 | 28.6 | 28.6 | 26.712 |
| LHFPL2 | 3314 | 5 | 5 | 5 | 21.1 | 21.1 | 21.1 | 24.486 |
| S100A6 | 1768 | 5 | 5 | 5 | 48.2 | 48.2 | 48.2 | 9.681 |
| RQCD1 | 3681 | 5 | 5 | 5 | 23.7 | 23.7 | 23.7 | 33.631 |
| LACE1 | 3645 | 5 | 5 | 5 | 16.4 | 16.4 | 16.4 | 54.844 |
| RPL13A | 2263 | 13 | 13 | 5 | 44.8 | 44.8 | 21.7 | 23.577 |
| CLN6 | 4320 | 5 | 5 | 5 | 21.5 | 21.5 | 21.5 | 35.919 |
| ARHGEF11 | 1378 | 5 | 5 | 5 | 4.9 | 4.9 | 4.9 | 167.7 |
| SLC30A7 | 3567 | 5 | 5 | 5 | 14.9 | 14.9 | 14.9 | 41.625 |
| PCYOX1L | 3538 | 5 | 5 | 5 | 17.6 | 17.6 | 17.6 | 54.646 |
| EIF4E2 | 628 | 5 | 5 | 5 | 26.8 | 26.8 | 26.8 | 24.602 |
| ZAP70 | 2303 | 5 | 5 | 5 | 13.9 | 13.9 | 13.9 | 69.872 |
| PRPF38A | 3531 | 5 | 5 | 5 | 18.6 | 18.6 | 18.6 | 37.476 |
| POLR2C | 2004 | 5 | 5 | 5 | 27.6 | 27.6 | 27.6 | 31.441 |
| C9orf64 | 3195 | 5 | 5 | 5 | 17.3 | 17.3 | 17.3 | 39.028 |
| CMC4 | 2523 | 5 | 5 | 5 | 57.4 | 57.4 | 57.4 | 7.7469 |
| STARD3NL | 680 | 5 | 5 | 5 | 31.1 | 31.1 | 31.1 | 23.505 |
| TMEM87B | 418 | 5 | 5 | 5 | 10.6 | 10.6 | 10.6 | 62.592 |
| NDUFAF3 | 3987 | 5 | 5 | 5 | 34.2 | 34.2 | 34.2 | 20.35 |
| CCDC115 | 3848 | 5 | 5 | 5 | 50.6 | 50.6 | 50.6 | 19.76 |
| THEM4 | 3185 | 5 | 5 | 5 | 23.3 | 23.3 | 23.3 | 27.129 |
| CIRBP | 2920 | 5 | 5 | 5 | 38.4 | 38.4 | 38.4 | 18.648 |
| PDDC1 | 3533 | 5 | 5 | 5 | 39.5 | 39.5 | 39.5 | 23.297 |
| WBP11 | 4572 | 5 | 5 | 5 | 8.6 | 8.6 | 8.6 | 69.997 |
| FAM210A | 1211 | 5 | 5 | 5 | 21.3 | 21.3 | 21.3 | 30.292 |
| FUNDC2 | 4020 | 5 | 5 | 5 | 33.9 | 33.9 | 33.9 | 20.675 |
| ARPC4-TTLL | 985 | 13 | 13 | 5 | 67.4 | 67.4 | 24.9 | 21.058 |
| ENOPH1 | 4447 | 5 | 5 | 5 | 27.2 | 27.2 | 27.2 | 28.932 |
| IFIT2 | 110 | 5 | 5 | 5 | 13.3 | 13.3 | 13.3 | 55.984 |
| TRA2A | 2898 | 7 | 5 | 5 | 26.2 | 22.7 | 22.7 | 32.688 |
| EFTUD1 | 3357 | 5 | 5 | 5 | 5.9 | 5.9 | 5.9 | 125.43 |
| XPNPEP3 | 4233 | 5 | 5 | 5 | 20.9 | 20.9 | 20.9 | 57.033 |
| NARS2 | 3816 | 5 | 5 | 5 | 11.5 | 11.5 | 11.5 | 54.089 |
| UACA | 919 | 5 | 5 | 5 | 4.1 | 4.1 | 4.1 | 150.53 |
| PPIL1 | 4593 | 5 | 5 | 5 | 51.8 | 51.8 | 51.8 | 18.237 |
| FUS | 2228 | 7 | 5 | 5 | 20.7 | 16.3 | 16.3 | 53.425 |
| NECAP2 | 4309 | 5 | 5 | 5 | 29.7 | 29.7 | 29.7 | 28.338 |

| | | | | | | | | |
|---|---|---|---|---|---|---|---|---|
| FASTKD2 | 4351 | 5 | 5 | 5 | 11.7 | 11.7 | 11.7 | 81.462 |
| IVNS1ABP | 4687 | 5 | 5 | 5 | 12 | 12 | 12 | 71.729 |
| SEPTIN8 | 531 | 9 | 5 | 5 | 28.6 | 20.2 | 20.2 | 49.915 |
| ABHD2 | 1813 | 5 | 5 | 5 | 23.1 | 23.1 | 23.1 | 48.314 |
| FAM50A | 2947 | 5 | 5 | 5 | 19.2 | 19.2 | 19.2 | 40.241 |
| CCPG1 | 4500 | 5 | 5 | 5 | 7.4 | 7.4 | 7.4 | 87.339 |
| PRKACA;KII | 1978 | 14 | 14 | 5 | 42.5 | 42.5 | 13.7 | 40.589 |
| TSG101 | 922 | 5 | 5 | 5 | 14 | 14 | 14 | 40.917 |
| MT-ND4 | 1720 | 5 | 5 | 5 | 14.4 | 14.4 | 14.4 | 51.58 |
| OGFR | 136 | 5 | 5 | 5 | 9 | 9 | 9 | 73.324 |
| TMX4 | 4103 | 5 | 5 | 5 | 15.2 | 15.2 | 15.2 | 38.952 |
| WNK1 | 894 | 5 | 5 | 5 | 2.8 | 2.8 | 2.8 | 250.79 |
| ADD1 | 784 | 5 | 5 | 5 | 11.7 | 11.7 | 11.7 | 70.055 |
| MTARC1 | 3214 | 5 | 5 | 5 | 19.9 | 19.9 | 19.9 | 37.499 |
| LRCH4 | 1547 | 7 | 7 | 5 | 19.8 | 19.8 | 13.9 | 73.449 |
| RBM10 | 2718 | 5 | 5 | 5 | 8 | 8 | 8 | 103.53 |
| GGCT | 1529 | 5 | 5 | 5 | 30.9 | 30.9 | 30.9 | 21.007 |
| S100A9 | 1767 | 5 | 5 | 5 | 60.5 | 60.5 | 60.5 | 13.242 |
| CHMP5 | 4366 | 5 | 5 | 5 | 26.5 | 26.5 | 26.5 | 24.57 |
| GZMB | 1176 | 6 | 6 | 5 | 28.9 | 28.9 | 24.3 | 26.417 |
| PRPSAP2 | 1482 | 5 | 5 | 5 | 17.9 | 17.9 | 17.9 | 40.925 |
| SLC22A18 | 7 | 5 | 5 | 5 | 18.4 | 18.4 | 18.4 | 34.952 |
| PTPN7 | 1181 | 5 | 5 | 5 | 22.8 | 22.8 | 22.8 | 40.529 |
| SNX4 | 1635 | 5 | 5 | 5 | 13.6 | 13.6 | 13.6 | 51.908 |
| ABHD14B | 981 | 5 | 5 | 5 | 35.7 | 35.7 | 35.7 | 22.345 |
| MT-ND5 | 1721 | 5 | 5 | 5 | 14.9 | 14.9 | 14.9 | 67.026 |
| TXNDC17 | 3952 | 5 | 5 | 5 | 51.2 | 51.2 | 51.2 | 13.941 |
| ARHGAP9 | 3960 | 5 | 5 | 5 | 10 | 10 | 10 | 83.259 |
| NUBP2 | 4651 | 5 | 5 | 5 | 27.7 | 27.7 | 27.7 | 28.825 |
| CD46 | 1945 | 5 | 5 | 5 | 11.2 | 11.2 | 11.2 | 43.747 |
| MRPS33 | 4550 | 5 | 5 | 5 | 63.2 | 63.2 | 63.2 | 12.629 |
| RAC1;RAC3 | 2647 | 9 | 5 | 5 | 58.3 | 34.4 | 34.4 | 21.45 |
| MRPL14 | 3263 | 5 | 5 | 5 | 44.1 | 44.1 | 44.1 | 15.947 |
| BLOC1S5 | 3618 | 5 | 5 | 5 | 31 | 31 | 31 | 21.609 |
| STAM | 3695 | 7 | 7 | 5 | 18.9 | 18.9 | 15.4 | 59.179 |
| DOCK4 | 1037 | 5 | 5 | 5 | 3.7 | 3.7 | 3.7 | 228.52 |
| FAM105A | 4287 | 5 | 5 | 5 | 13.2 | 13.2 | 13.2 | 42.195 |
| MSMO1 | 3038 | 5 | 5 | 5 | 20.8 | 20.8 | 20.8 | 35.215 |
| ALAD | 1915 | 5 | 5 | 5 | 25.5 | 25.5 | 25.5 | 36.294 |
| LSM8 | 890 | 5 | 5 | 5 | 92.5 | 92.5 | 92.5 | 7.3713 |
| RPL22 | 2219 | 5 | 5 | 5 | 57 | 57 | 57 | 14.787 |
| TTYH3 | 4072 | 5 | 5 | 5 | 12 | 12 | 12 | 57.544 |
| HECTD3 | 3192 | 5 | 5 | 5 | 9.2 | 9.2 | 9.2 | 97.112 |
| XRCC4 | 2869 | 5 | 5 | 5 | 19 | 19 | 19 | 38.286 |
| SIPA1 | 938 | 5 | 5 | 5 | 5.4 | 5.4 | 5.4 | 101.82 |
| ALG12 | 3998 | 5 | 5 | 5 | 24.2 | 24.2 | 24.2 | 54.654 |
| MRPL24 | 3738 | 5 | 5 | 5 | 26.4 | 26.4 | 26.4 | 24.915 |
| MPV17 | 599 | 5 | 5 | 5 | 34.7 | 34.7 | 34.7 | 18.628 |

| | | | | | | | | |
|---|---|---|---|---|---|---|---|---|
| C19orf66 | 4276 | 5 | 5 | 5 | 19.6 | 19.6 | 19.6 | 33.11 |
| MLKL | 3532 | 5 | 5 | 5 | 11.7 | 11.7 | 11.7 | 54.478 |
| NACA | 819 | 5 | 5 | 5 | 45.6 | 45.6 | 45.6 | 15.016 |
| GPR89B;GF | 621 | 5 | 5 | 5 | 15.8 | 15.8 | 15.8 | 52.916 |
| ARFRP1 | 2908 | 5 | 5 | 5 | 35.8 | 35.8 | 35.8 | 22.614 |
| ZRANB2 | 1634 | 5 | 5 | 5 | 16.7 | 16.7 | 16.7 | 37.404 |
| ANP32B | 3691 | 7 | 5 | 5 | 29.5 | 24.3 | 24.3 | 28.787 |
| DNAJB2 | 2098 | 6 | 6 | 5 | 21.6 | 21.6 | 18.5 | 35.58 |
| PKM | 1929 | 48 | 48 | 5 | 80.2 | 80.2 | 9.4 | 57.936 |
| SLC44A1 | 3654 | 5 | 5 | 5 | 9.4 | 9.4 | 9.4 | 73.301 |
| QIL1 | 290 | 5 | 5 | 5 | 62.1 | 62.1 | 62.1 | 15.436 |
| RABL3 | 694 | 5 | 5 | 5 | 34.4 | 34.4 | 34.4 | 23.78 |
| MCAT | 3442 | 5 | 5 | 5 | 17.7 | 17.7 | 17.7 | 42.961 |
| RAP2C | 4604 | 11 | 11 | 5 | 79.8 | 79.8 | 33.9 | 20.745 |
| C5AR1 | 2041 | 5 | 5 | 5 | 13.7 | 13.7 | 13.7 | 39.335 |
| PFDN6 | 1393 | 5 | 5 | 5 | 41.9 | 41.9 | 41.9 | 14.582 |
| UNC93B1 | 4100 | 5 | 5 | 5 | 10.7 | 10.7 | 10.7 | 66.63 |
| CARKD | 3443 | 5 | 5 | 5 | 21.9 | 21.9 | 21.9 | 36.576 |
| PARN | 415 | 5 | 5 | 5 | 16.6 | 16.6 | 16.6 | 56.046 |
| PEX16 | 4652 | 5 | 5 | 5 | 22.9 | 22.9 | 22.9 | 38.628 |
| UFC1 | 4594 | 5 | 5 | 5 | 25.1 | 25.1 | 25.1 | 19.458 |
| ROGDI | 4075 | 5 | 5 | 5 | 24.4 | 24.4 | 24.4 | 32.254 |
| VPS33B | 910 | 5 | 5 | 5 | 12.9 | 12.9 | 12.9 | 67.605 |
| RRS1 | 2990 | 5 | 5 | 5 | 18.1 | 18.1 | 18.1 | 41.193 |
| SH3GL1 | 3933 | 5 | 5 | 5 | 25.8 | 25.8 | 25.8 | 41.489 |
| TLE3 | 226 | 5 | 5 | 5 | 11 | 11 | 11 | 83.183 |
| MYDGF | 3726 | 5 | 5 | 5 | 32.9 | 32.9 | 32.9 | 18.795 |
| SIRT5 | 4332 | 5 | 5 | 5 | 23.9 | 23.9 | 23.9 | 33.881 |
| EPN1 | 4674 | 6 | 6 | 5 | 16.3 | 16.3 | 14.2 | 60.293 |
| SNAPIN | 1637 | 5 | 5 | 5 | 61 | 61 | 61 | 14.874 |
| MRPL22 | 797 | 5 | 5 | 5 | 22.8 | 22.8 | 22.8 | 26.475 |
| TFAM | 2721 | 5 | 5 | 5 | 17.5 | 17.5 | 17.5 | 29.096 |
| SMPD2 | 1520 | 5 | 5 | 5 | 22.9 | 22.9 | 22.9 | 47.645 |
| RTN2 | 1233 | 5 | 5 | 5 | 18.5 | 18.5 | 18.5 | 29.519 |
| DHRS3 | 1585 | 5 | 5 | 5 | 22.5 | 22.5 | 22.5 | 33.548 |
| SEPTIN1 | 1161 | 5 | 5 | 5 | 12.6 | 12.6 | 12.6 | 46.918 |
| NUP37 | 3574 | 5 | 5 | 5 | 29.4 | 29.4 | 29.4 | 36.707 |
| FAS | 2095 | 5 | 5 | 5 | 22.1 | 22.1 | 22.1 | 37.732 |
| TRMT5 | 3092 | 5 | 5 | 5 | 10.4 | 10.4 | 10.4 | 58.246 |
| USP30 | 572 | 5 | 5 | 5 | 13.2 | 13.2 | 13.2 | 55.108 |
| GAR1 | 4342 | 5 | 5 | 5 | 25.8 | 25.8 | 25.8 | 22.348 |
| ARRB1 | 2369 | 12 | 11 | 5 | 41.6 | 40 | 18.4 | 47.065 |
| LGALS8 | 940 | 5 | 5 | 5 | 18 | 18 | 18 | 33.985 |
| MEF2BNB | 3792 | 5 | 5 | 5 | 54.6 | 54.6 | 54.6 | 13.403 |
| REEP3 | 3252 | 5 | 5 | 5 | 15.7 | 15.7 | 15.7 | 29.264 |
| LYAR | 4329 | 5 | 5 | 5 | 18.5 | 18.5 | 18.5 | 43.614 |
| TMEM205 | 1225 | 5 | 5 | 5 | 31.2 | 31.2 | 31.2 | 19.27 |
| TMUB2 | 3325 | 5 | 5 | 5 | 22.7 | 22.7 | 22.7 | 33.788 |

| | | | | | | | | |
|---|---|---|---|---|---|---|---|---|
| AP2S1 | 1260 | 5 | 5 | 5 | 31.9 | 31.9 | 31.9 | 17.303 |
| GMFB | 2557 | 7 | 5 | 5 | 62.7 | 37.3 | 37.3 | 16.713 |
| TRIM47 | 3838 | 5 | 5 | 5 | 9.9 | 9.9 | 9.9 | 69.531 |
| HIST1H1B | 1963 | 7 | 5 | 5 | 29.2 | 22.1 | 22.1 | 22.58 |
| GCHFR | 2159 | 5 | 5 | 5 | 82.1 | 82.1 | 82.1 | 9.6981 |
| ADRBK1 | 2091 | 5 | 5 | 5 | 9.4 | 9.4 | 9.4 | 79.573 |
| TMEM55B | 3390 | 6 | 6 | 5 | 22.4 | 22.4 | 18.8 | 29.469 |
| CYGB | 1230 | 5 | 5 | 5 | 29.5 | 29.5 | 29.5 | 21.404 |
| SSR1 | 386 | 5 | 5 | 5 | 30.6 | 30.6 | 30.6 | 32.449 |
| FAH | 1968 | 5 | 5 | 5 | 17.2 | 17.2 | 17.2 | 46.374 |
| OSTM1 | 3414 | 5 | 5 | 5 | 15.9 | 15.9 | 15.9 | 37.256 |
| GRAMD1A | 1261 | 5 | 5 | 5 | 9.2 | 9.2 | 9.2 | 89.734 |
| C19orf52 | 3968 | 5 | 5 | 5 | 28.5 | 28.5 | 28.5 | 29.233 |
| SFXN4 | 3268 | 5 | 5 | 5 | 19 | 19 | 19 | 37.998 |
| IRF2BPL | 4099 | 5 | 5 | 5 | 7.3 | 7.3 | 7.3 | 82.658 |
| MAP2K1 | 2748 | 8 | 8 | 5 | 29.5 | 29.5 | 19.3 | 43.439 |
| IARS | 2277 | 51 | 51 | 5 | 51.4 | 51.4 | 6.8 | 144.5 |
| FAM96A | 4147 | 5 | 5 | 5 | 44.4 | 44.4 | 44.4 | 18.355 |
| TRIM14 | 2928 | 5 | 5 | 5 | 12 | 12 | 12 | 49.772 |
| SSFA2 | 845 | 5 | 5 | 5 | 4.4 | 4.4 | 4.4 | 138.38 |
| LGMN | 3902 | 5 | 5 | 5 | 20.1 | 20.1 | 20.1 | 49.411 |
| TMEM11 | 1974 | 5 | 5 | 5 | 30.7 | 30.7 | 30.7 | 21.541 |
| CRYL1 | 4567 | 5 | 5 | 5 | 18.2 | 18.2 | 18.2 | 35.419 |
| GNG5 | 2655 | 5 | 5 | 5 | 79.4 | 79.4 | 79.4 | 7.3184 |
| RBM26 | 89 | 6 | 6 | 5 | 7.6 | 7.6 | 6.5 | 113.6 |
| VPS28 | 4477 | 5 | 5 | 5 | 41.2 | 41.2 | 41.2 | 25.425 |
| ACTN1 | 1898 | 65 | 41 | 5 | 81.6 | 60.9 | 10.1 | 103.06 |
| RNF40 | 1095 | 6 | 6 | 5 | 7.3 | 7.3 | 5.8 | 109.05 |
| SF3B6 | 4589 | 5 | 5 | 5 | 47.2 | 47.2 | 47.2 | 14.585 |
| SLC12A6 | 4445 | 11 | 11 | 5 | 9.8 | 9.8 | 5.1 | 127.62 |
| TIMM8A | 1480 | 5 | 5 | 5 | 55.7 | 55.7 | 55.7 | 10.998 |
| MPDU1 | 1147 | 5 | 5 | 5 | 51.5 | 51.5 | 51.5 | 10.978 |
| NADK | 1656 | 5 | 5 | 5 | 19.7 | 19.7 | 19.7 | 49.228 |
| C9orf89 | 3840 | 5 | 5 | 5 | 21.9 | 21.9 | 21.9 | 25.589 |
| FHIT | 2388 | 5 | 5 | 5 | 42.9 | 42.9 | 42.9 | 16.858 |
| CD274 | 4365 | 5 | 5 | 5 | 21.7 | 21.7 | 21.7 | 33.275 |
| TTC1 | 3912 | 5 | 5 | 5 | 20.2 | 20.2 | 20.2 | 33.526 |
| SH3BGRL3 | 4112 | 5 | 5 | 5 | 76.3 | 76.3 | 76.3 | 10.438 |
| CTDNEP1 | 1145 | 5 | 5 | 5 | 44.6 | 44.6 | 44.6 | 23.03 |
| RAC2 | 1940 | 9 | 8 | 5 | 58.3 | 52.6 | 34.4 | 21.429 |
| F8A1 | 2076 | 5 | 5 | 5 | 20.5 | 20.5 | 20.5 | 39.103 |
| MICALL1 | 3502 | 5 | 5 | 5 | 8.6 | 8.6 | 8.6 | 93.44 |
| ZC3H14 | 1000 | 5 | 5 | 5 | 11.4 | 11.4 | 11.4 | 68.349 |
| SERPING1 | 839 | 5 | 5 | 5 | 10.5 | 10.5 | 10.5 | 59.491 |
| TIMM21 | 4013 | 5 | 5 | 5 | 22.6 | 22.6 | 22.6 | 28.202 |
| CD48 | 106 | 5 | 5 | 5 | 22.2 | 22.2 | 22.2 | 28.855 |
| COPZ1 | 955 | 5 | 5 | 5 | 31.3 | 31.3 | 31.3 | 22.34 |
| SP100 | 846 | 6 | 6 | 5 | 9.6 | 9.6 | 8.3 | 100.42 |

| | | | | | | | | |
|---|---|---|---|---|---|---|---|---|
| S100A10 | 2554 | 5 | 5 | 5 | 45.4 | 45.4 | 45.4 | 11.203 |
| WIPF1 | 1445 | 5 | 5 | 5 | 16.5 | 16.5 | 16.5 | 51.258 |
| NDUFA2 | 1455 | 5 | 5 | 5 | 61.6 | 61.6 | 61.6 | 10.921 |
| LRRC57 | 394 | 5 | 5 | 5 | 35.8 | 35.8 | 35.8 | 16.324 |
| NCK1 | 1962 | 13 | 13 | 5 | 58.6 | 58.6 | 34 | 42.864 |
| TIMM10B | 4635 | 5 | 5 | 5 | 57.3 | 57.3 | 57.3 | 11.586 |
| ISG15 | 120 | 5 | 5 | 5 | 45.5 | 45.5 | 45.5 | 17.887 |
| SAFB2 | 2930 | 13 | 5 | 5 | 17.7 | 6.5 | 6.5 | 107.47 |
| TMEM87A | 3539 | 5 | 5 | 5 | 9.7 | 9.7 | 9.7 | 63.429 |
| NHP2L1 | 409 | 5 | 5 | 5 | 44.7 | 44.7 | 44.7 | 14.627 |
| TMPPE | 3312 | 5 | 5 | 5 | 19.6 | 19.6 | 19.6 | 49.452 |
| TXNRD1 | 15 | 23 | 23 | 5 | 55.6 | 55.6 | 12.8 | 70.755 |
| TMPO | 2282 | 14 | 14 | 5 | 44.5 | 44.5 | 14.1 | 50.67 |
| COMMD7 | 340 | 5 | 5 | 5 | 36.8 | 36.8 | 36.8 | 20.151 |
| RPS24 | 367 | 5 | 5 | 5 | 40.5 | 40.5 | 40.5 | 15.197 |
| MGAT2 | 2805 | 5 | 5 | 5 | 17.2 | 17.2 | 17.2 | 51.55 |
| AKAP5 | 2086 | 5 | 5 | 5 | 14.8 | 14.8 | 14.8 | 47.088 |
| CCDC58 | 686 | 5 | 5 | 5 | 44.6 | 44.6 | 44.6 | 15.296 |
| SNRPD3 | 2613 | 5 | 5 | 5 | 53.2 | 53.2 | 53.2 | 13.916 |
| GABARAPL: | 2547 | 6 | 6 | 5 | 59.8 | 59.8 | 50.4 | 13.667 |
| MGST1 | 1861 | 5 | 5 | 5 | 45.2 | 45.2 | 45.2 | 17.598 |
| CISD2 | 3517 | 5 | 5 | 5 | 45.2 | 45.2 | 45.2 | 15.278 |
| COG8 | 591 | 5 | 5 | 5 | 9.9 | 9.9 | 9.9 | 71.208 |
| PPP6R3 | 859 | 4 | 4 | 4 | 5.2 | 5.2 | 5.2 | 92.407 |
| LUZP1 | 759 | 4 | 4 | 4 | 4.7 | 4.7 | 4.7 | 120.27 |
| TMEM63B | 1056 | 4 | 4 | 4 | 9.4 | 9.4 | 9.4 | 60.414 |
| NUP35 | 3575 | 4 | 4 | 4 | 19 | 19 | 19 | 34.773 |
| ZC3H7A | 3452 | 5 | 5 | 4 | 7.6 | 7.6 | 6 | 110.54 |
| CEPT1 | 4677 | 4 | 4 | 4 | 11.3 | 11.3 | 11.3 | 46.553 |
| NME6 | 203 | 4 | 4 | 4 | 38.5 | 38.5 | 38.5 | 20.061 |
| MRPL11 | 4590 | 4 | 4 | 4 | 27.6 | 27.6 | 27.6 | 20.683 |
| EXOG | 4554 | 4 | 4 | 4 | 14.1 | 14.1 | 14.1 | 41.084 |
| BID | 2518 | 4 | 4 | 4 | 33.3 | 33.3 | 33.3 | 21.994 |
| SLC39A11 | 1197 | 4 | 4 | 4 | 27.6 | 27.6 | 27.6 | 19.293 |
| SUB1 | 2489 | 4 | 4 | 4 | 34.6 | 34.6 | 34.6 | 14.395 |
| BOLA3 | 3131 | 4 | 4 | 4 | 26.2 | 26.2 | 26.2 | 12.114 |
| CD99 | 1924 | 4 | 4 | 4 | 48.1 | 48.1 | 48.1 | 18.848 |
| DIS3 | 4560 | 4 | 4 | 4 | 5.3 | 5.3 | 5.3 | 109 |
| MTIF3 | 4117 | 4 | 4 | 4 | 16.2 | 16.2 | 16.2 | 31.725 |
| UQCRH | 1793 | 4 | 4 | 4 | 38.5 | 38.5 | 38.5 | 10.739 |
| SNRPA | 1814 | 11 | 8 | 4 | 44 | 37.9 | 17.7 | 31.279 |
| VAMP3 | 3041 | 7 | 7 | 4 | 53 | 53 | 29 | 11.309 |
| HLA-DRB1 | 3226 | 11 | 8 | 4 | 55.6 | 42.9 | 20.3 | 30.12 |
| AUP1 | 4663 | 4 | 4 | 4 | 9.8 | 9.8 | 9.8 | 45.786 |
| GMPPB | 4642 | 4 | 4 | 4 | 19.7 | 19.7 | 19.7 | 39.834 |
| NDUFB5 | 814 | 4 | 4 | 4 | 22.6 | 22.6 | 22.6 | 20.038 |
| MMGT1 | 3509 | 4 | 4 | 4 | 49.6 | 49.6 | 49.6 | 14.686 |
| ATP5I | 2527 | 4 | 4 | 4 | 62.3 | 62.3 | 62.3 | 7.9331 |

| | | | | | | | | |
|---|---|---|---|---|---|---|---|---|
| SYAP1 | 3739 | 4 | 4 | 4 | 15.9 | 15.9 | 15.9 | 39.933 |
| TBC1D10B | 3114 | 4 | 4 | 4 | 6.6 | 6.6 | 6.6 | 87.198 |
| NDUFC2 | 1638 | 4 | 4 | 4 | 24.4 | 24.4 | 24.4 | 14.187 |
| LSM4 | 4626 | 4 | 4 | 4 | 19.3 | 19.3 | 19.3 | 25.733 |
| | 292 | 12 | 12 | 4 | 46.5 | 46.5 | 19.4 | 29.357 |
| LMNA | 1711 | 59 | 59 | 4 | 76.7 | 76.7 | 14.2 | 74.139 |
| COX6C | 1833 | 4 | 4 | 4 | 30.7 | 30.7 | 30.7 | 8.7813 |
| APOC1 | 1249 | 4 | 4 | 4 | 35.1 | 35.1 | 35.1 | 8.647 |
| CMTM6 | 4331 | 4 | 4 | 4 | 25.7 | 25.7 | 25.7 | 20.419 |
| SLC5A3 | 2485 | 4 | 4 | 4 | 8.5 | 8.5 | 8.5 | 79.693 |
| GATC | 1069 | 6 | 6 | 4 | 57.1 | 57.1 | 33.9 | 18.606 |
| IFIT5 | 2861 | 4 | 4 | 4 | 11.6 | 11.6 | 11.6 | 55.846 |
| SEC11A | 1082 | 9 | 9 | 4 | 47.9 | 47.9 | 17.8 | 18.651 |
| CLUH | 424 | 4 | 4 | 4 | 3.3 | 3.3 | 3.3 | 140.49 |
| DCTN5 | 1100 | 4 | 4 | 4 | 29.7 | 29.7 | 29.7 | 16.91 |
| CHMP6 | 1149 | 4 | 4 | 4 | 41 | 41 | 41 | 20.167 |
| PES1 | 575 | 4 | 4 | 4 | 10.2 | 10.2 | 10.2 | 51.464 |
| YTHDC1 | 1204 | 4 | 4 | 4 | 5.5 | 5.5 | 5.5 | 84.699 |
| LSM12 | 3099 | 4 | 4 | 4 | 28.7 | 28.7 | 28.7 | 21.701 |
| RBM42 | 1239 | 4 | 4 | 4 | 9.8 | 9.8 | 9.8 | 48.024 |
| TOMM6 | 3752 | 4 | 4 | 4 | 70.3 | 70.3 | 70.3 | 8.0019 |
| QPCTL | 4340 | 4 | 4 | 4 | 19.6 | 19.6 | 19.6 | 42.924 |
| CHCHD5 | 3974 | 4 | 4 | 4 | 54.5 | 54.5 | 54.5 | 12.395 |
| TBC1D2B | 4529 | 4 | 4 | 4 | 4.6 | 4.6 | 4.6 | 109.88 |
| RILP | 3846 | 4 | 4 | 4 | 12.2 | 12.2 | 12.2 | 44.2 |
| PSIP1 | 1552 | 4 | 4 | 4 | 9.8 | 9.8 | 9.8 | 60.103 |
| COX15 | 3329 | 4 | 4 | 4 | 14.1 | 14.1 | 14.1 | 46.03 |
| MRPL9 | 4044 | 4 | 4 | 4 | 15.4 | 15.4 | 15.4 | 30.243 |
| SLC9A1 | 2010 | 4 | 4 | 4 | 6.7 | 6.7 | 6.7 | 90.762 |
| GATAD2A | 3428 | 6 | 4 | 4 | 11.1 | 7.3 | 7.3 | 68.062 |
| PGM3 | 423 | 13 | 13 | 4 | 37.7 | 37.7 | 19 | 55.386 |
| TM6SF1 | 4064 | 4 | 4 | 4 | 10.8 | 10.8 | 10.8 | 41.636 |
| TSPO | 555 | 4 | 4 | 4 | 43 | 43 | 43 | 11.896 |
| GXYLT1 | 3112 | 4 | 4 | 4 | 10.2 | 10.2 | 10.2 | 50.566 |
| TMED3 | 4606 | 4 | 4 | 4 | 26.7 | 26.7 | 26.7 | 24.777 |
| RIPK2 | 1435 | 4 | 4 | 4 | 10.9 | 10.9 | 10.9 | 61.194 |
| POM121C;I | 538 | 4 | 4 | 4 | 4.9 | 4.9 | 4.9 | 125.09 |
| ATPAF1 | 542 | 4 | 4 | 4 | 26.4 | 26.4 | 26.4 | 19.91 |
| SNRPF | 2608 | 4 | 4 | 4 | 48.8 | 48.8 | 48.8 | 9.7251 |
| TMCO1 | 4505 | 4 | 4 | 4 | 19.2 | 19.2 | 19.2 | 27.079 |
| WBP2 | 518 | 4 | 4 | 4 | 25.7 | 25.7 | 25.7 | 18.224 |
| EPDR1 | 4506 | 4 | 4 | 4 | 19.2 | 19.2 | 19.2 | 25.437 |
| SYNGR2 | 1463 | 5 | 5 | 4 | 20.5 | 20.5 | 16.1 | 24.81 |
| AGAP3 | 3850 | 5 | 5 | 4 | 8.2 | 8.2 | 6.4 | 95.043 |
| PSD3 | 772 | 4 | 4 | 4 | 6.8 | 6.8 | 6.8 | 108.99 |
| PYURF | 3812 | 4 | 4 | 4 | 55.3 | 55.3 | 55.3 | 12.654 |
| HNRNPC | 568 | 22 | 22 | 4 | 61.6 | 61.6 | 8.2 | 32.238 |
| TRAPPC2B; | 1843 | 4 | 4 | 4 | 49.3 | 49.3 | 49.3 | 16.445 |

| | | | | | | | | |
|---|---|---|---|---|---|---|---|---|
| ERGIC2 | 24 | 4 | 4 | 4 | 13.5 | 13.5 | 13.5 | 43.432 |
| RAVER1 | 78 | 4 | 4 | 4 | 10.8 | 10.8 | 10.8 | 77.843 |
| RIN3 | 61 | 4 | 4 | 4 | 6.4 | 6.4 | 6.4 | 107.85 |
| NFAM1 | 3566 | 4 | 4 | 4 | 17.4 | 17.4 | 17.4 | 29.685 |
| TBC1D1 | 3391 | 4 | 4 | 4 | 4.3 | 4.3 | 4.3 | 133.08 |
| HACD2 | 3303 | 4 | 4 | 4 | 16.1 | 16.1 | 16.1 | 28.368 |
| ASRGL1 | 3335 | 4 | 4 | 4 | 19.2 | 19.2 | 19.2 | 32.054 |
| RAB8B | 3709 | 9 | 4 | 4 | 45.9 | 22.7 | 22.7 | 23.584 |
| SNRPC | 137 | 4 | 4 | 4 | 25.2 | 25.2 | 25.2 | 17.394 |
| RNF170 | 3829 | 4 | 4 | 4 | 15.1 | 15.1 | 15.1 | 29.814 |
| TYROBP | 1477 | 4 | 4 | 4 | 36.3 | 36.3 | 36.3 | 12.179 |
| MRPS11 | 2697 | 4 | 4 | 4 | 34.5 | 34.5 | 34.5 | 20.616 |
| YIF1B | 3142 | 7 | 7 | 4 | 41.4 | 41.4 | 21 | 34.435 |
| SEPTIN3 | 344 | 4 | 4 | 4 | 15.6 | 15.6 | 15.6 | 33.601 |
| LIN7C | 996 | 4 | 4 | 4 | 24.9 | 24.9 | 24.9 | 19.186 |
| H2AFV;H2A | 1842 | 6 | 4 | 4 | 53.9 | 46.9 | 46.9 | 13.509 |
| CD59 | 377 | 4 | 4 | 4 | 29.6 | 29.6 | 29.6 | 11.985 |
| CD33 | 2017 | 4 | 4 | 4 | 14 | 14 | 14 | 39.825 |
| TMEM70 | 3988 | 4 | 4 | 4 | 24.6 | 24.6 | 24.6 | 28.969 |
| TOMM40L | 3727 | 4 | 4 | 4 | 16.6 | 16.6 | 16.6 | 33.916 |
| HIST1H1C | 1965 | 11 | 11 | 4 | 36.2 | 36.2 | 12.2 | 21.364 |
| SARAF | 758 | 4 | 4 | 4 | 35.4 | 35.4 | 35.4 | 16.085 |
| NCOA5 | 4197 | 4 | 4 | 4 | 10.7 | 10.7 | 10.7 | 65.536 |
| SLC7A8 | 4441 | 4 | 4 | 4 | 9.3 | 9.3 | 9.3 | 58.381 |
| SMARCD2 | 638 | 4 | 4 | 4 | 15.1 | 15.1 | 15.1 | 52.238 |
| TMED1 | 2875 | 4 | 4 | 4 | 20.7 | 20.7 | 20.7 | 25.206 |
| PAK1 | 2843 | 10 | 4 | 4 | 25.7 | 14.5 | 14.5 | 60.646 |
| MT-CO1 | 1690 | 4 | 4 | 4 | 13.6 | 13.6 | 13.6 | 57.041 |
| PPP3R1 | 723 | 5 | 5 | 4 | 43.1 | 43.1 | 34.4 | 18.208 |
| TRAPPC2L | 182 | 4 | 4 | 4 | 50.5 | 50.5 | 50.5 | 12.561 |
| SLC25A25;S | 3247 | 4 | 4 | 4 | 7.9 | 7.9 | 7.9 | 52.662 |
| GGCX | 2252 | 4 | 4 | 4 | 6.5 | 6.5 | 6.5 | 87.56 |
| NDUFS6 | 1541 | 4 | 4 | 4 | 48.4 | 48.4 | 48.4 | 13.711 |
| UBE2G2 | 2549 | 4 | 4 | 4 | 42.4 | 42.4 | 42.4 | 18.566 |
| TRIP6 | 1123 | 4 | 4 | 4 | 27 | 27 | 27 | 23.04 |
| TMEM115 | 2821 | 4 | 4 | 4 | 17.4 | 17.4 | 17.4 | 38.197 |
| MYL9 | 2089 | 10 | 4 | 4 | 62.2 | 29.1 | 29.1 | 19.827 |
| CRELD2 | 536 | 4 | 4 | 4 | 18.3 | 18.3 | 18.3 | 25.129 |
| MYD88 | 1034 | 4 | 4 | 4 | 18 | 18 | 18 | 35.444 |
| SLC38A9 | 731 | 4 | 4 | 4 | 61.8 | 61.8 | 61.8 | 14.859 |
| CCDC124 | 3770 | 4 | 4 | 4 | 18.8 | 18.8 | 18.8 | 25.835 |
| JAK1 | 462 | 4 | 4 | 4 | 4.4 | 4.4 | 4.4 | 114.07 |
| CRKL | 2320 | 4 | 4 | 4 | 21.5 | 21.5 | 21.5 | 33.777 |
| IGHM | 1708 | 4 | 4 | 4 | 12.4 | 12.4 | 12.4 | 49.439 |
| PITPNB | 2357 | 11 | 10 | 4 | 53.1 | 49.8 | 16.6 | 31.54 |
| AIF1 | 2501 | 4 | 4 | 4 | 31.1 | 31.1 | 31.1 | 18.038 |
| SLC6A12 | 2342 | 4 | 4 | 4 | 6.7 | 6.7 | 6.7 | 69.368 |
| MRPL18 | 4096 | 4 | 4 | 4 | 28.9 | 28.9 | 28.9 | 20.576 |

| | | | | | | | | |
|---|---|---|---|---|---|---|---|---|
| PELO | 3962 | 4 | 4 | 4 | 12.2 | 12.2 | 12.2 | 43.359 |
| STAG2 | 3503 | 5 | 5 | 4 | 5.2 | 5.2 | 4.5 | 141.32 |
| TSTA3 | 266 | 4 | 4 | 4 | 50.8 | 50.8 | 50.8 | 12.788 |
| CASP7 | 134 | 4 | 4 | 4 | 20.2 | 20.2 | 20.2 | 34.413 |
| RHBDD1 | 3625 | 4 | 4 | 4 | 21.6 | 21.6 | 21.6 | 35.823 |
| MARCHF5 | 4326 | 4 | 4 | 4 | 17.3 | 17.3 | 17.3 | 31.231 |
| SERPINB8 | 2409 | 7 | 4 | 4 | 21.9 | 13.9 | 13.9 | 42.766 |
| STMN1;STN | 496 | 4 | 4 | 4 | 43.5 | 43.5 | 43.5 | 9.7931 |
| NAAA | 2742 | 4 | 4 | 4 | 12.5 | 12.5 | 12.5 | 40.066 |
| NDUFAF5 | 3205 | 4 | 4 | 4 | 16.2 | 16.2 | 16.2 | 38.918 |
| LANCL1 | 1468 | 4 | 4 | 4 | 17.5 | 17.5 | 17.5 | 45.283 |
| HMGN1 | 528 | 4 | 4 | 4 | 47.8 | 47.8 | 47.8 | 9.5355 |
| PACS1 | 3301 | 4 | 4 | 4 | 6.4 | 6.4 | 6.4 | 104.9 |
| HECTD1 | 115 | 4 | 4 | 4 | 1.9 | 1.9 | 1.9 | 289.63 |
| CMC1 | 3385 | 4 | 4 | 4 | 43.4 | 43.4 | 43.4 | 12.49 |
| ISCA2 | 3397 | 4 | 4 | 4 | 33.8 | 33.8 | 33.8 | 16.476 |
| NT5C3A | 4093 | 6 | 6 | 4 | 21.1 | 21.1 | 14.5 | 37.405 |
| SERINC1 | 4257 | 4 | 4 | 4 | 10.8 | 10.8 | 10.8 | 50.494 |
| TWSG1 | 4083 | 4 | 4 | 4 | 24.2 | 24.2 | 24.2 | 25.017 |
| ANKRD13A | 3474 | 4 | 4 | 4 | 7.6 | 7.6 | 7.6 | 67.618 |
| C1orf123 | 4318 | 4 | 4 | 4 | 33.8 | 33.8 | 33.8 | 18.048 |
| MAPRE2 | 3029 | 6 | 5 | 4 | 21.1 | 18.7 | 15.6 | 37.031 |
| PPIE | 4517 | 5 | 4 | 4 | 30.2 | 27.9 | 27.9 | 33.43 |
| ZNF622 | 3731 | 4 | 4 | 4 | 12.6 | 12.6 | 12.6 | 54.271 |
| ADO | 3884 | 4 | 4 | 4 | 16.7 | 16.7 | 16.7 | 29.751 |
| DNPH1 | 1046 | 4 | 4 | 4 | 28.4 | 28.4 | 28.4 | 25.868 |
| CCDC132 | 3825 | 4 | 4 | 4 | 6.3 | 6.3 | 6.3 | 111.17 |
| ARMT1 | 4168 | 4 | 4 | 4 | 8.8 | 8.8 | 8.8 | 51.172 |
| GLS | 629 | 8 | 4 | 4 | 73.4 | 24.9 | 24.9 | 18.634 |
| NAE1 | 2892 | 4 | 4 | 4 | 10.1 | 10.1 | 10.1 | 60.246 |
| BOLA2;BOL | 4127 | 4 | 4 | 4 | 61.6 | 61.6 | 61.6 | 10.116 |
| SMARCA2 | 941 | 17 | 17 | 4 | 12.4 | 12.4 | 3.4 | 172.59 |
| NOP10 | 4217 | 4 | 4 | 4 | 70.3 | 70.3 | 70.3 | 7.7059 |
| C6orf203 | 224 | 4 | 4 | 4 | 14.7 | 14.7 | 14.7 | 28.457 |
| GORASP1 | 3949 | 4 | 4 | 4 | 15.2 | 15.2 | 15.2 | 46.482 |
| CBX5 | 2312 | 5 | 4 | 4 | 31.4 | 25.1 | 25.1 | 22.225 |
| RNPS1 | 1086 | 4 | 4 | 4 | 29.4 | 29.4 | 29.4 | 24.561 |
| MVK | 935 | 4 | 4 | 4 | 20.9 | 20.9 | 20.9 | 37.171 |
| NUTF2 | 2589 | 4 | 4 | 4 | 62.2 | 62.2 | 62.2 | 14.478 |
| TIMM10 | 2592 | 4 | 4 | 4 | 32.2 | 32.2 | 32.2 | 10.333 |
| FMR1 | 541 | 7 | 5 | 4 | 13.4 | 10.7 | 7.4 | 65.821 |
| COMMD6 | 3367 | 4 | 4 | 4 | 55.3 | 55.3 | 55.3 | 9.638 |
| COA6 | 3157 | 4 | 4 | 4 | 37.6 | 37.6 | 37.6 | 14.116 |
| FYTTD1 | 3861 | 4 | 4 | 4 | 16 | 16 | 16 | 35.818 |
| PAP1;METT | 3614 | 4 | 4 | 4 | 17 | 17 | 17 | 32.884 |
| FHL1 | 3162 | 4 | 4 | 4 | 18.1 | 18.1 | 18.1 | 23.701 |
| MINOS1;MI | 26 | 4 | 4 | 4 | 77.4 | 77.4 | 77.4 | 7.036 |
| KDELC2 | 3369 | 4 | 4 | 4 | 13.4 | 13.4 | 13.4 | 58.572 |

| | | | | | | | | |
|---|---|---|---|---|---|---|---|---|
| CSTA | 1704 | 4 | 4 | 4 | 50 | 50 | 50 | 11.006 |
| RRM2B | 3349 | 4 | 4 | 4 | 14 | 14 | 14 | 40.736 |
| CD53 | 2006 | 4 | 4 | 4 | 11.9 | 11.9 | 11.9 | 24.341 |
| ENDOG | 2942 | 4 | 4 | 4 | 18.9 | 18.9 | 18.9 | 32.62 |
| LSM3 | 2609 | 4 | 4 | 4 | 41.2 | 41.2 | 41.2 | 11.845 |
| NDUFAF4 | 4370 | 4 | 4 | 4 | 23.4 | 23.4 | 23.4 | 20.266 |
| ASCC1 | 463 | 4 | 4 | 4 | 14.2 | 14.2 | 14.2 | 36.876 |
| PRPSAP1 | 2956 | 4 | 4 | 4 | 15.4 | 15.4 | 15.4 | 39.393 |
| RNF114 | 118 | 4 | 4 | 4 | 24.5 | 24.5 | 24.5 | 20.885 |
| BNIP2 | 1158 | 4 | 4 | 4 | 10.3 | 10.3 | 10.3 | 48.844 |
| HSPE1 | 2582 | 9 | 4 | 4 | 75.5 | 32.4 | 32.4 | 10.932 |
| DERL1 | 593 | 4 | 4 | 4 | 16.3 | 16.3 | 16.3 | 28.8 |
| ATP5L | 406 | 4 | 4 | 4 | 60.5 | 60.5 | 60.5 | 8.4518 |
| DYNLT3 | 519 | 4 | 4 | 4 | 55.9 | 55.9 | 55.9 | 16.979 |
| APOC4-APC | 1246 | 4 | 4 | 4 | 28.1 | 28.1 | 28.1 | 20.049 |
| RPL37A | 648 | 4 | 4 | 4 | 57.4 | 57.4 | 57.4 | 7.624 |
| ERC1 | 1027 | 4 | 4 | 4 | 4.1 | 4.1 | 4.1 | 124.82 |
| PILRA | 4483 | 4 | 4 | 4 | 19.8 | 19.8 | 19.8 | 34.004 |
| DPAGT1 | 322 | 4 | 4 | 4 | 8.8 | 8.8 | 8.8 | 42.587 |
| C10orf76 | 3188 | 4 | 4 | 4 | 10.2 | 10.2 | 10.2 | 78.71 |
| HYPK | 4327 | 4 | 4 | 4 | 39.5 | 39.5 | 39.5 | 14.665 |
| TRAFD1 | 1338 | 4 | 4 | 4 | 8.9 | 8.9 | 8.9 | 64.84 |
| MPC1 | 4647 | 8 | 8 | 4 | 73.4 | 73.4 | 27.5 | 12.347 |
| COMMD5 | 4077 | 4 | 4 | 4 | 26.8 | 26.8 | 26.8 | 24.67 |
| RPS25 | 2629 | 4 | 4 | 4 | 29.6 | 29.6 | 29.6 | 13.742 |
| ARPC1B | 1388 | 23 | 23 | 4 | 70.2 | 70.2 | 15.1 | 40.949 |
| CETN2 | 2269 | 4 | 4 | 4 | 30.2 | 30.2 | 30.2 | 19.738 |
| NUP43 | 3573 | 4 | 4 | 4 | 20 | 20 | 20 | 42.15 |
| HLA-A | 281 | 30 | 30 | 4 | 67.9 | 67.9 | 7.1 | 40.921 |
| DYNLT1 | 2653 | 4 | 4 | 4 | 60.2 | 60.2 | 60.2 | 12.452 |
| RPL35 | 2292 | 4 | 4 | 4 | 27.6 | 27.6 | 27.6 | 14.551 |
| POP1 | 3907 | 4 | 4 | 4 | 5.7 | 5.7 | 5.7 | 114.71 |
| FAM160B1 | 3224 | 4 | 4 | 4 | 10.5 | 10.5 | 10.5 | 86.557 |
| TXN2 | 3922 | 4 | 4 | 4 | 36.7 | 36.7 | 36.7 | 18.383 |
| SLC25A1 | 2471 | 16 | 16 | 4 | 55.6 | 55.6 | 13.2 | 34.012 |
| CD82 | 2127 | 4 | 4 | 4 | 16.1 | 16.1 | 16.1 | 29.625 |
| PIN1 | 2884 | 4 | 4 | 4 | 47.2 | 47.2 | 47.2 | 18.243 |
| MRPL23 | 3054 | 4 | 4 | 4 | 46.4 | 46.4 | 46.4 | 17.781 |
| CMC2 | 4251 | 4 | 4 | 4 | 64.6 | 64.6 | 64.6 | 9.4598 |
| CD300LF | 1185 | 4 | 4 | 4 | 13.4 | 13.4 | 13.4 | 33.797 |
| SLC11A1 | 2364 | 4 | 4 | 4 | 13.1 | 13.1 | 13.1 | 59.872 |
| TNFAIP8 | 1645 | 4 | 4 | 4 | 24.2 | 24.2 | 24.2 | 23.003 |
| FAM107B | 311 | 4 | 4 | 4 | 34.7 | 34.7 | 34.7 | 14.844 |
| PRKAB1 | 4611 | 4 | 4 | 4 | 32.2 | 32.2 | 32.2 | 30.382 |
| GATAD2B | 279 | 6 | 6 | 4 | 13.2 | 13.2 | 9 | 63.461 |
| KCNN4 | 721 | 4 | 4 | 4 | 32.1 | 32.1 | 32.1 | 18.367 |
| TRMT2A | 889 | 4 | 4 | 4 | 8.7 | 8.7 | 8.7 | 70.834 |
| MFHAS1 | 4614 | 4 | 4 | 4 | 4.8 | 4.8 | 4.8 | 116.95 |

| | | | | | | | | |
|---|---|---|---|---|---|---|---|---|
| ANP32E | 3980 | 4 | 4 | 4 | 27.6 | 27.6 | 27.6 | 30.692 |
| FABP5 | 2733 | 11 | 11 | 4 | 77.8 | 77.8 | 19.3 | 15.164 |
| COX7C | 1952 | 4 | 4 | 4 | 63.5 | 63.5 | 63.5 | 7.2454 |
| KIAA0391 | 1379 | 4 | 4 | 4 | 14.8 | 14.8 | 14.8 | 67.315 |
| NUBPL | 3595 | 4 | 4 | 4 | 20.7 | 20.7 | 20.7 | 34.082 |
| GID8 | 4317 | 4 | 4 | 4 | 30.3 | 30.3 | 30.3 | 26.748 |
| BST1 | 509 | 4 | 4 | 4 | 33.9 | 33.9 | 33.9 | 20.971 |
| TTYH2 | 582 | 4 | 4 | 4 | 6.4 | 6.4 | 6.4 | 56.32 |
| TMEM104 | 3562 | 4 | 4 | 4 | 12.7 | 12.7 | 12.7 | 55.678 |
| SFXN5 | 3613 | 4 | 4 | 4 | 16.5 | 16.5 | 16.5 | 37.124 |
| SUMO1 | 632 | 4 | 4 | 4 | 29.5 | 29.5 | 29.5 | 16.645 |
| CREG1 | 381 | 4 | 4 | 4 | 25.9 | 25.9 | 25.9 | 28.058 |
| PSEN2 | 566 | 6 | 6 | 4 | 21.2 | 21.2 | 16.2 | 53.732 |
| ACY1;ABHD | 303 | 4 | 4 | 4 | 18.2 | 18.2 | 18.2 | 42.179 |
| TAOK2 | 4496 | 6 | 4 | 4 | 6.1 | 5.3 | 5.3 | 138.25 |
| LAMTOR4 | 2804 | 4 | 4 | 4 | 36.4 | 36.4 | 36.4 | 10.741 |
| ARHGAP25 | 2284 | 30 | 30 | 4 | 47.3 | 47.3 | 5.9 | 73.434 |
| ADAR | 2511 | 37 | 37 | 4 | 37.2 | 37.2 | 3.4 | 136.06 |
| NUP50 | 4491 | 4 | 4 | 4 | 12 | 12 | 12 | 50.144 |
| YY1;YY2 | 2096 | 4 | 4 | 4 | 10.6 | 10.6 | 10.6 | 44.712 |
| ZCCHC8 | 3259 | 4 | 4 | 4 | 7.4 | 7.4 | 7.4 | 78.576 |
| PPM1A | 2230 | 10 | 10 | 4 | 34.3 | 34.3 | 13.4 | 42.447 |
| MRPS12 | 1397 | 4 | 4 | 4 | 37.7 | 37.7 | 37.7 | 15.173 |
| MRPL48 | 933 | 4 | 4 | 4 | 38.9 | 38.9 | 38.9 | 12.77 |
| TOMM5 | 976 | 4 | 4 | 4 | 34.7 | 34.7 | 34.7 | 10.976 |
| YIF1A | 521 | 4 | 4 | 4 | 24.1 | 24.1 | 24.1 | 26.377 |
| CD86 | 2279 | 4 | 4 | 4 | 13.1 | 13.1 | 13.1 | 37.682 |
| MCFD2 | 3586 | 4 | 4 | 4 | 68.5 | 68.5 | 68.5 | 16.39 |
| MCOLN3 | 3617 | 4 | 4 | 4 | 7.8 | 7.8 | 7.8 | 64.247 |
| GPSM3 | 851 | 4 | 4 | 4 | 39.5 | 39.5 | 39.5 | 17.639 |
| VPS26A | 1549 | 16 | 16 | 4 | 59 | 59 | 15.3 | 38.169 |
| TNFAIP8L2 | 3274 | 4 | 4 | 4 | 31.5 | 31.5 | 31.5 | 20.555 |
| CSTF2 | 815 | 9 | 9 | 4 | 20.4 | 20.4 | 9 | 62.942 |
| MRPS18A | 4305 | 4 | 4 | 4 | 27.6 | 27.6 | 27.6 | 22.184 |
| AUH | 2910 | 4 | 4 | 4 | 16.5 | 16.5 | 16.5 | 35.608 |
| EBNA1BP2 | 1132 | 4 | 4 | 4 | 11.4 | 11.4 | 11.4 | 40.684 |
| MCEE | 3853 | 4 | 4 | 4 | 26.1 | 26.1 | 26.1 | 18.749 |
| BDH2 | 3996 | 4 | 4 | 4 | 23.7 | 23.7 | 23.7 | 26.724 |
| PDAP1 | 2874 | 4 | 4 | 4 | 19.9 | 19.9 | 19.9 | 20.63 |
| SNRPG;SNF | 911 | 4 | 4 | 4 | 79.7 | 79.7 | 79.7 | 7.1013 |
| SYNE2 | 128 | 4 | 4 | 4 | 0.8 | 0.8 | 0.8 | 787.71 |
| PPP2R5C | 1072 | 8 | 8 | 4 | 19.9 | 19.9 | 10.3 | 64.063 |
| MT-CO3 | 1692 | 4 | 4 | 4 | 13.4 | 13.4 | 13.4 | 29.95 |
| ALG8 | 4009 | 4 | 4 | 4 | 10.6 | 10.6 | 10.6 | 60.087 |
| LSM6 | 2610 | 4 | 4 | 4 | 45 | 45 | 45 | 9.1275 |
| RFT1 | 604 | 4 | 4 | 4 | 6.8 | 6.8 | 6.8 | 60.335 |
| STIM2 | 332 | 5 | 4 | 4 | 10.6 | 7.8 | 7.8 | 66.1 |
| RBMXL1 | 3781 | 19 | 4 | 4 | 46.7 | 9 | 9 | 42.141 |

| | | | | | | | | |
|---|---|---|---|---|---|---|---|---|
| DAD1 | 898 | 4 | 4 | 4 | 47.1 | 47.1 | 47.1 | 9.5541 |
| FAM96B | 4595 | 4 | 4 | 4 | 42.9 | 42.9 | 42.9 | 17.663 |
| GPR180 | 3406 | 4 | 4 | 4 | 12.3 | 12.3 | 12.3 | 49.395 |
| POLR2G | 2617 | 4 | 4 | 4 | 30.2 | 30.2 | 30.2 | 19.294 |
| RBM22 | 4312 | 4 | 4 | 4 | 11 | 11 | 11 | 46.895 |
| CCDC134 | 4149 | 4 | 4 | 4 | 23.6 | 23.6 | 23.6 | 26.56 |
| CREB1;CREI | 1958 | 4 | 4 | 4 | 17.3 | 17.3 | 17.3 | 36.688 |
| NOMO2;NC | 3151 | 57 | 57 | 4 | 66.5 | 66.5 | 6 | 139.44 |
| PWP1 | 581 | 4 | 4 | 4 | 11.4 | 11.4 | 11.4 | 55.827 |
| MECR | 4002 | 4 | 4 | 4 | 13.4 | 13.4 | 13.4 | 40.461 |
| DCK | 751 | 4 | 4 | 4 | 14.6 | 14.6 | 14.6 | 30.518 |
| TMEM55A | 761 | 5 | 4 | 4 | 39.2 | 33.1 | 33.1 | 17.745 |
| SLC43A2 | 3498 | 4 | 4 | 4 | 6.9 | 6.9 | 6.9 | 62.746 |
| PPP1R12C | 1222 | 4 | 4 | 4 | 8.5 | 8.5 | 8.5 | 80.199 |
| SDHC | 3914 | 4 | 4 | 4 | 38.5 | 38.5 | 38.5 | 18.61 |
| PPIL3 | 4116 | 4 | 4 | 4 | 36 | 36 | 36 | 18.154 |
| PPP1R2;PP | 781 | 4 | 4 | 4 | 30 | 30 | 30 | 19.245 |
| MS4A1 | 860 | 4 | 4 | 4 | 18.3 | 18.3 | 18.3 | 31.609 |
| COX11 | 4681 | 4 | 4 | 4 | 21 | 21 | 21 | 31.43 |
| PHPT1 | 4256 | 4 | 4 | 4 | 51.2 | 51.2 | 51.2 | 13.832 |
| FTL | 1715 | 4 | 4 | 4 | 29.1 | 29.1 | 29.1 | 20.019 |
| C14orf1 | 4488 | 4 | 4 | 4 | 23.6 | 23.6 | 23.6 | 15.864 |
| GFM2 | 3732 | 4 | 4 | 4 | 8.1 | 8.1 | 8.1 | 86.6 |
| S100A4 | 2111 | 4 | 4 | 4 | 36.6 | 36.6 | 36.6 | 11.728 |
| CCDC88B | 510 | 4 | 4 | 4 | 3.5 | 3.5 | 3.5 | 164.81 |
| MYADM | 3879 | 4 | 4 | 4 | 18.9 | 18.9 | 18.9 | 35.273 |
| MRPS18C | 4597 | 4 | 4 | 4 | 28.2 | 28.2 | 28.2 | 15.85 |
| PNPLA7 | 3315 | 7 | 4 | 4 | 7.3 | 5.4 | 5.4 | 145.7 |
| KIAA0020 | 3018 | 4 | 4 | 4 | 9.6 | 9.6 | 9.6 | 73.584 |
| DNAJB14 | 3600 | 4 | 4 | 4 | 11.6 | 11.6 | 11.6 | 42.515 |
| FGFR1OP2 | 4301 | 4 | 4 | 4 | 23.3 | 23.3 | 23.3 | 29.426 |
| TUBB | 1783 | 25 | 5 | 4 | 73.9 | 20.2 | 16.7 | 47.766 |
| HIGD2A | 4015 | 4 | 4 | 4 | 72.6 | 72.6 | 72.6 | 11.528 |
| KPNA2 | 2454 | 4 | 4 | 4 | 9.8 | 9.8 | 9.8 | 57.861 |
| MRPL41 | 3462 | 4 | 4 | 4 | 51.1 | 51.1 | 51.1 | 15.383 |
| NUP188 | 3174 | 4 | 4 | 4 | 2.7 | 2.7 | 2.7 | 196.04 |
| ACSS2 | 4240 | 3 | 3 | 3 | 11.4 | 11.4 | 11.4 | 78.579 |
| MRPL40 | 4227 | 3 | 3 | 3 | 20.4 | 20.4 | 20.4 | 24.49 |
| SNRPD1 | 2611 | 3 | 3 | 3 | 37.8 | 37.8 | 37.8 | 13.281 |
| MRPS24 | 3783 | 3 | 3 | 3 | 22.2 | 22.2 | 22.2 | 19.015 |
| RPL26L1 | 4519 | 10 | 3 | 3 | 45.5 | 12.4 | 12.4 | 17.256 |
| KCMF1 | 4375 | 3 | 3 | 3 | 10 | 10 | 10 | 41.945 |
| FXR2 | 2423 | 6 | 3 | 3 | 12.9 | 7.7 | 7.7 | 74.222 |
| ABHD17B | 3213 | 3 | 3 | 3 | 16.3 | 16.3 | 16.3 | 32.214 |
| DPY19L1 | 310 | 3 | 3 | 3 | 6.7 | 6.7 | 6.7 | 77.318 |
| IGKC | 472 | 3 | 3 | 3 | 49.5 | 49.5 | 49.5 | 11.609 |
| ATP6V0C | 2123 | 3 | 3 | 3 | 36.1 | 36.1 | 36.1 | 15.736 |
| ARL15 | 36 | 3 | 3 | 3 | 21.1 | 21.1 | 21.1 | 22.876 |

| | | | | | | | | |
|---|---|---|---|---|---|---|---|---|
| DHRSX | 3515 | 3 | 3 | 3 | 12.1 | 12.1 | 12.1 | 36.443 |
| CCNY | 3557 | 4 | 4 | 3 | 15.2 | 15.2 | 12.9 | 39.336 |
| LYRM7 | 3210 | 3 | 3 | 3 | 39.4 | 39.4 | 39.4 | 11.955 |
| SYNGR1 | 601 | 3 | 3 | 3 | 20.1 | 20.1 | 20.1 | 18.465 |
| USMG5 | 3820 | 3 | 3 | 3 | 44.8 | 44.8 | 44.8 | 6.4575 |
| MBP | 1186 | 3 | 3 | 3 | 27.7 | 27.7 | 27.7 | 17.442 |
| RPL26 | 2577 | 10 | 10 | 3 | 45.5 | 45.5 | 12.4 | 17.258 |
| TIA1 | 975 | 8 | 6 | 3 | 32.5 | 26.2 | 15.3 | 42.835 |
| CD47 | 342 | 3 | 3 | 3 | 15.3 | 15.3 | 15.3 | 20.313 |
| RPLP1 | 1754 | 4 | 3 | 3 | 89.5 | 74.6 | 74.6 | 11.514 |
| RTN3 | 1631 | 4 | 4 | 3 | 5.8 | 5.8 | 2 | 112.61 |
| AP1S1 | 376 | 4 | 4 | 3 | 25.5 | 25.5 | 20.4 | 18.602 |
| RP13-360B | 3208 | 3 | 3 | 3 | 11.2 | 11.2 | 11.2 | 29.231 |
| NDUFA4 | 1317 | 3 | 3 | 3 | 37 | 37 | 37 | 9.3697 |
| FCER1G | 2168 | 3 | 3 | 3 | 32.6 | 32.6 | 32.6 | 9.6674 |
| RPS29 | 368 | 3 | 3 | 3 | 21.8 | 21.8 | 21.8 | 10.256 |
| YIPF6 | 3782 | 3 | 3 | 3 | 9.3 | 9.3 | 9.3 | 26.256 |
| AGA | 2032 | 3 | 3 | 3 | 10.7 | 10.7 | 10.7 | 37.208 |
| CD300A | 4427 | 3 | 3 | 3 | 14.7 | 14.7 | 14.7 | 33.2 |
| DGAT1 | 1584 | 3 | 3 | 3 | 10 | 10 | 10 | 55.278 |
| PLXNA1 | 4460 | 4 | 4 | 3 | 3 | 3 | 2.4 | 211.06 |
| MRPL27 | 1137 | 3 | 3 | 3 | 23.5 | 23.5 | 23.5 | 14.731 |
| GLRX5 | 3388 | 3 | 3 | 3 | 28 | 28 | 28 | 16.628 |
| CD63 | 954 | 3 | 3 | 3 | 27.6 | 27.6 | 27.6 | 16.021 |
| TSPAN15 | 1131 | 3 | 3 | 3 | 26.9 | 26.9 | 26.9 | 18.219 |
| CAB39L | 3199 | 6 | 3 | 3 | 23.6 | 14.3 | 14.3 | 32.423 |
| RAPGEF1 | 2916 | 3 | 3 | 3 | 4.8 | 4.8 | 4.8 | 120.55 |
| ASCC2 | 4104 | 3 | 3 | 3 | 5.7 | 5.7 | 5.7 | 86.359 |
| RAP2A | 1848 | 9 | 3 | 3 | 65.6 | 19.7 | 19.7 | 20.615 |
| TMEM9B | 4226 | 3 | 3 | 3 | 31.3 | 31.3 | 31.3 | 22.531 |
| SSU72 | 4212 | 3 | 3 | 3 | 16.5 | 16.5 | 16.5 | 22.574 |
| RBX1 | 2633 | 3 | 3 | 3 | 30.6 | 30.6 | 30.6 | 12.274 |
| HLA-DQA1 | 1710 | 5 | 4 | 3 | 22 | 19.3 | 13.8 | 27.805 |
| DERL2 | 4076 | 3 | 3 | 3 | 20.5 | 20.5 | 20.5 | 27.567 |
| OVCA2 | 3663 | 3 | 3 | 3 | 28.2 | 28.2 | 28.2 | 24.418 |
| PTPN18 | 3932 | 3 | 3 | 3 | 9.1 | 9.1 | 9.1 | 50.482 |
| RPS27A | 2643 | 7 | 7 | 3 | 48.7 | 48.7 | 21.2 | 17.965 |
| CIRH1A | 1102 | 3 | 3 | 3 | 4.4 | 4.4 | 4.4 | 77.942 |
| SNRPE | 2607 | 3 | 3 | 3 | 52.2 | 52.2 | 52.2 | 10.803 |
| TOR1B | 1347 | 3 | 3 | 3 | 13.7 | 13.7 | 13.7 | 37.978 |
| SMAD2;SM | 615 | 3 | 3 | 3 | 7.7 | 7.7 | 7.7 | 48.669 |
| CLEC12A | 3167 | 3 | 3 | 3 | 13.2 | 13.2 | 13.2 | 30.761 |
| SLC35E1 | 3830 | 3 | 3 | 3 | 11.7 | 11.7 | 11.7 | 44.772 |
| LY96 | 4689 | 3 | 3 | 3 | 22.5 | 22.5 | 22.5 | 18.545 |
| PVRL2 | 1218 | 3 | 3 | 3 | 16.2 | 16.2 | 16.2 | 22.642 |
| MRPL32 | 4041 | 3 | 3 | 3 | 23.9 | 23.9 | 23.9 | 21.405 |
| EMILIN2 | 4038 | 3 | 3 | 3 | 4.3 | 4.3 | 4.3 | 115.69 |
| ICT1 | 1184 | 3 | 3 | 3 | 19.3 | 19.3 | 19.3 | 21.905 |

| | | | | | | | | |
|---|---|---|---|---|---|---|---|---|
| COA4 | 4346 | 3 | 3 | 3 | 42.5 | 42.5 | 42.5 | 10.134 |
| SLC29A1 | 3926 | 3 | 3 | 3 | 8.6 | 8.6 | 8.6 | 50.219 |
| TEFM | 3863 | 3 | 3 | 3 | 11.9 | 11.9 | 11.9 | 41.676 |
| ABRACL | 4384 | 3 | 3 | 3 | 46.9 | 46.9 | 46.9 | 9.0564 |
| LILRB1 | 14 | 11 | 8 | 3 | 21.4 | 17.4 | 6.5 | 70.819 |
| DENR | 1447 | 3 | 3 | 3 | 23.7 | 23.7 | 23.7 | 22.092 |
| LXN | 3966 | 3 | 3 | 3 | 17.6 | 17.6 | 17.6 | 25.75 |
| PDIA5 | 2955 | 3 | 3 | 3 | 6 | 6 | 6 | 59.594 |
| TSC2 | 361 | 3 | 3 | 3 | 2.5 | 2.5 | 2.5 | 149.4 |
| CHMP1A | 952 | 3 | 3 | 3 | 14.4 | 14.4 | 14.4 | 19.531 |
| TM7SF3 | 1065 | 3 | 3 | 3 | 23 | 23 | 23 | 16.609 |
| SNX18 | 3870 | 3 | 3 | 3 | 9.9 | 9.9 | 9.9 | 68.894 |
| IPO9 | 3852 | 3 | 3 | 3 | 2.6 | 2.6 | 2.6 | 115.96 |
| UBA5 | 790 | 3 | 3 | 3 | 19 | 19 | 19 | 38.753 |
| PSMB5 | 2136 | 3 | 3 | 3 | 13.3 | 13.3 | 13.3 | 28.48 |
| USP33 | 3630 | 3 | 3 | 3 | 3.7 | 3.7 | 3.7 | 106.73 |
| STAM2 | 1583 | 5 | 3 | 3 | 11 | 7.4 | 7.4 | 58.164 |
| HDHD2 | 1245 | 3 | 3 | 3 | 24.5 | 24.5 | 24.5 | 22.49 |
| C14orf159 | 3361 | 26 | 26 | 3 | 57.3 | 57.3 | 7.1 | 66.436 |
| GCSH | 327 | 3 | 3 | 3 | 67 | 67 | 67 | 10.999 |
| | 584 | 10 | 10 | 3 | 33.7 | 33.7 | 7.9 | 48.099 |
| DGKA | 2078 | 3 | 3 | 3 | 7.3 | 7.3 | 7.3 | 82.629 |
| PCNP | 3650 | 3 | 3 | 3 | 20.2 | 20.2 | 20.2 | 18.925 |
| AK1 | 1698 | 3 | 3 | 3 | 24.2 | 24.2 | 24.2 | 21.635 |
| PPIH | 687 | 3 | 3 | 3 | 14.6 | 14.6 | 14.6 | 15.8 |
| GTF2F1 | 1267 | 3 | 3 | 3 | 29.9 | 29.9 | 29.9 | 14.685 |
| GSTM1 | 1825 | 12 | 12 | 3 | 56.4 | 56.4 | 12.8 | 25.712 |
| BMS1 | 2963 | 3 | 3 | 3 | 3.3 | 3.3 | 3.3 | 145.81 |
| COPS7A | 899 | 3 | 3 | 3 | 20.8 | 20.8 | 20.8 | 21.306 |
| YIPF4 | 3973 | 3 | 3 | 3 | 12.7 | 12.7 | 12.7 | 27.082 |
| AP5S1 | 4285 | 3 | 3 | 3 | 20.5 | 20.5 | 20.5 | 22.522 |
| PIGU | 4136 | 3 | 3 | 3 | 8.7 | 8.7 | 8.7 | 50.051 |
| CNNM4 | 3272 | 4 | 4 | 3 | 6.5 | 6.5 | 5.4 | 86.606 |
| FAU | 878 | 3 | 3 | 3 | 32.2 | 32.2 | 32.2 | 6.6478 |
| TBL3 | 1162 | 3 | 3 | 3 | 4.5 | 4.5 | 4.5 | 89.034 |
| MRPL30 | 634 | 3 | 3 | 3 | 32.7 | 32.7 | 32.7 | 11.935 |
| FAM210B | 3836 | 3 | 3 | 3 | 20.8 | 20.8 | 20.8 | 20.424 |
| FXN | 659 | 3 | 3 | 3 | 36.3 | 36.3 | 36.3 | 14.91 |
| CARD9 | 4108 | 3 | 3 | 3 | 5.2 | 5.2 | 5.2 | 62.24 |
| HMGN3 | 81 | 4 | 4 | 3 | 23.8 | 23.8 | 23.8 | 13.956 |
| EPT1 | 52 | 3 | 3 | 3 | 13.4 | 13.4 | 13.4 | 45.228 |
| UBTD1 | 4176 | 6 | 6 | 3 | 24.7 | 24.7 | 10.6 | 25.938 |
| RNMTL1 | 4195 | 3 | 3 | 3 | 11.2 | 11.2 | 11.2 | 47.019 |
| EXOSC2 | 2913 | 3 | 3 | 3 | 17.4 | 17.4 | 17.4 | 32.789 |
| WDR33 | 4073 | 3 | 3 | 3 | 2.5 | 2.5 | 2.5 | 145.89 |
| LAPTM5 | 2894 | 3 | 3 | 3 | 12.2 | 12.2 | 12.2 | 29.937 |
| PRKACB | 144 | 12 | 3 | 3 | 43.4 | 15.1 | 15.1 | 41.308 |
| KIAA0100 | 1241 | 3 | 3 | 3 | 2.1 | 2.1 | 2.1 | 237.3 |

| | | | | | | | | |
|---|---|---|---|---|---|---|---|---|
| WRB | 390 | 3 | 3 | 3 | 30.2 | 30.2 | 30.2 | 15.977 |
| RBM27 | 4395 | 4 | 3 | 3 | 6.2 | 5.1 | 5.1 | 114.54 |
| CLN8 | 4414 | 3 | 3 | 3 | 18.2 | 18.2 | 18.2 | 32.787 |
| SIRPB1 | 3207 | 14 | 3 | 3 | 41.7 | 12.1 | 12.1 | 43.359 |
| GLRX2 | 4261 | 3 | 3 | 3 | 27.4 | 27.4 | 27.4 | 18.051 |
| MFAP1 | 2506 | 4 | 4 | 3 | 10.5 | 10.5 | 8.9 | 51.958 |
| IMMT | 3078 | 61 | 61 | 3 | 74.3 | 74.3 | 8.3 | 83.677 |
| CST3 | 1703 | 3 | 3 | 3 | 32.9 | 32.9 | 32.9 | 15.799 |
| SLC2A6 | 4429 | 3 | 3 | 3 | 7.7 | 7.7 | 7.7 | 54.539 |
| HPS6 | 3431 | 3 | 3 | 3 | 5.9 | 5.9 | 5.9 | 82.974 |
| TCEAL5;TCE | 3148 | 3 | 3 | 3 | 14.6 | 14.6 | 14.6 | 23.307 |
| C18orf32 | 3606 | 3 | 3 | 3 | 36.8 | 36.8 | 36.8 | 8.6693 |
| HSPA13 | 2355 | 3 | 3 | 3 | 9.1 | 9.1 | 9.1 | 51.927 |
| GBP4 | 3856 | 3 | 3 | 3 | 5.5 | 5.5 | 5.5 | 73.164 |
| EXOSC7 | 2983 | 3 | 3 | 3 | 15.8 | 15.8 | 15.8 | 31.821 |
| CUX1 | 2918 | 10 | 10 | 3 | 15 | 15 | 4.1 | 77.454 |
| EBP | 3001 | 3 | 3 | 3 | 16.5 | 16.5 | 16.5 | 26.352 |
| WDR44 | 1118 | 3 | 3 | 3 | 4.1 | 4.1 | 4.1 | 89.847 |
| PIP4K2B | 2680 | 6 | 3 | 3 | 16.1 | 7.9 | 7.9 | 47.377 |
| EIF4G3 | 145 | 9 | 3 | 3 | 6.2 | 3 | 3 | 180.5 |
| CHMP2B | 4536 | 3 | 3 | 3 | 13.6 | 13.6 | 13.6 | 23.906 |
| UQCR10 | 4415 | 3 | 3 | 3 | 50.8 | 50.8 | 50.8 | 7.3084 |
| CBFB | 422 | 3 | 3 | 3 | 17.6 | 17.6 | 17.6 | 21.508 |
| TFB1M | 3646 | 3 | 3 | 3 | 17.1 | 17.1 | 17.1 | 39.542 |
| CASC4 | 3270 | 3 | 3 | 3 | 7.6 | 7.6 | 7.6 | 49.497 |
| BCAR1 | 343 | 3 | 3 | 3 | 5.4 | 5.4 | 5.4 | 77.76 |
| PCID2 | 3160 | 3 | 3 | 3 | 10 | 10 | 10 | 46.029 |
| SMNDC1 | 1591 | 3 | 3 | 3 | 12.6 | 12.6 | 12.6 | 26.711 |
| PTDSS2 | 4008 | 3 | 3 | 3 | 7.8 | 7.8 | 7.8 | 56.252 |
| MAP2K3;M | 2322 | 4 | 4 | 3 | 14.4 | 14.4 | 12.4 | 39.318 |
| YTHDF2 | 4630 | 6 | 3 | 3 | 11.1 | 6.7 | 6.7 | 62.333 |
| HLA-B | 726 | 15 | 5 | 3 | 43.4 | 14.1 | 6.6 | 40.479 |
| CDA | 2196 | 3 | 3 | 3 | 39.7 | 39.7 | 39.7 | 16.185 |
| HEBP2 | 4654 | 3 | 3 | 3 | 19.5 | 19.5 | 19.5 | 22.875 |
| RAP1A | 2628 | 14 | 14 | 3 | 83.7 | 83.7 | 28.3 | 20.987 |
| TOMM20 | 3016 | 3 | 3 | 3 | 27.6 | 27.6 | 27.6 | 16.298 |
| SLMAP | 451 | 3 | 3 | 3 | 8.8 | 8.8 | 8.8 | 85.923 |
| CHMP1B | 3348 | 3 | 3 | 3 | 12.1 | 12.1 | 12.1 | 22.109 |
| RTN4 | 977 | 14 | 14 | 3 | 58.6 | 58.6 | 4.6 | 37.144 |
| NHLRC3 | 3155 | 3 | 3 | 3 | 11 | 11 | 11 | 38.282 |
| GOLGA7 | 3374 | 4 | 4 | 3 | 32.1 | 32.1 | 26.3 | 15.824 |
| VAMP5 | 1630 | 3 | 3 | 3 | 28.4 | 28.4 | 28.4 | 12.805 |
| TMEM256 | 1248 | 3 | 3 | 3 | 77.9 | 77.9 | 77.9 | 7.1301 |
| TMBIM6 | 947 | 3 | 3 | 3 | 68.3 | 68.3 | 68.3 | 4.7756 |
| TREM2 | 4358 | 3 | 3 | 3 | 27.4 | 27.4 | 27.4 | 25.447 |
| VMP1 | 3798 | 3 | 3 | 3 | 13.5 | 13.5 | 13.5 | 46.237 |
| PQLC1 | 3496 | 4 | 4 | 3 | 15.5 | 15.5 | 12.5 | 30.477 |
| ATXN2 | 352 | 3 | 3 | 3 | 9.8 | 9.8 | 9.8 | 38.545 |

| | | | | | | | | |
|---|---|---|---|---|---|---|---|---|
| RAB3D | 1665 | 5 | 3 | 3 | 26 | 19.2 | 19.2 | 24.267 |
| CD5 | 1763 | 3 | 3 | 3 | 6.5 | 6.5 | 6.5 | 54.577 |
| SNX12 | 93 | 5 | 3 | 3 | 20.9 | 16.9 | 16.9 | 19.825 |
| POLR1C | 353 | 3 | 3 | 3 | 14.1 | 14.1 | 14.1 | 32.917 |
| CPSF4 | 670 | 3 | 3 | 3 | 23.7 | 23.7 | 23.7 | 23.653 |
| OSBPL3 | 4144 | 3 | 3 | 3 | 4.4 | 4.4 | 4.4 | 101.22 |
| MT-ND1 | 1718 | 3 | 3 | 3 | 8.8 | 8.8 | 8.8 | 35.66 |
| HMGN4 | 1316 | 3 | 3 | 3 | 24.4 | 24.4 | 24.4 | 9.5388 |
| CIAPIN1 | 1098 | 3 | 3 | 3 | 40.3 | 40.3 | 40.3 | 13.633 |
| UROD | 309 | 3 | 3 | 3 | 22.7 | 22.7 | 22.7 | 31.778 |
| MRPL50 | 3520 | 3 | 3 | 3 | 34.2 | 34.2 | 34.2 | 18.325 |
| KCNAB2 | 448 | 15 | 15 | 3 | 57.5 | 57.5 | 13.4 | 41 |
| CD68 | 2211 | 3 | 3 | 3 | 22 | 22 | 22 | 37.408 |
| HLA-DPB1 | 1736 | 10 | 3 | 3 | 43 | 10.5 | 10.5 | 29.159 |
| ATOX1 | 763 | 3 | 3 | 3 | 44.1 | 44.1 | 44.1 | 6.3304 |
| C6orf120 | 3371 | 3 | 3 | 3 | 45 | 45 | 45 | 20.772 |
| CDK9 | 2418 | 4 | 3 | 3 | 11.8 | 9.7 | 9.7 | 42.777 |
| GNB2 | 2634 | 16 | 6 | 3 | 69.1 | 40 | 21.8 | 37.331 |
| NDUFAF1 | 1076 | 3 | 3 | 3 | 14.6 | 14.6 | 14.6 | 30.042 |
| IGBP1 | 2675 | 3 | 3 | 3 | 12.1 | 12.1 | 12.1 | 39.221 |
| CD6 | 1018 | 3 | 3 | 3 | 7.9 | 7.9 | 7.9 | 49.8 |
| RNASE6 | 3723 | 3 | 3 | 3 | 30.7 | 30.7 | 30.7 | 17.196 |
| ARFIP2 | 102 | 3 | 3 | 3 | 13.8 | 13.8 | 13.8 | 37.855 |
| AKT1 | 2183 | 3 | 3 | 3 | 8.3 | 8.3 | 8.3 | 55.686 |
| ATP1B4 | 624 | 3 | 3 | 3 | 10.2 | 10.2 | 10.2 | 36.735 |
| RGS19 | 2391 | 3 | 3 | 3 | 21.2 | 21.2 | 21.2 | 24.635 |
| COX7A2 | 1029 | 3 | 3 | 3 | 40.9 | 40.9 | 40.9 | 12.844 |
| MIEF1 | 553 | 3 | 3 | 3 | 16.3 | 16.3 | 16.3 | 53.331 |
| MFSD1 | 428 | 3 | 3 | 3 | 7.5 | 7.5 | 7.5 | 46.618 |
| PTPRCAP | 2970 | 3 | 3 | 3 | 15.5 | 15.5 | 15.5 | 21.196 |
| RPS15A | 2600 | 9 | 9 | 3 | 70 | 70 | 27.7 | 14.839 |
| CWC15 | 4368 | 3 | 3 | 3 | 14 | 14 | 14 | 26.624 |
| DDX58 | 498 | 3 | 3 | 3 | 4.8 | 4.8 | 4.8 | 82.612 |
| NUB1 | 1083 | 3 | 3 | 3 | 6.6 | 6.6 | 6.6 | 71.921 |
| MRPL21 | 3356 | 3 | 3 | 3 | 15.1 | 15.1 | 15.1 | 22.814 |
| ACOT1 | 3396 | 26 | 26 | 3 | 82.4 | 82.4 | 13.5 | 46.277 |
| SLC2A1 | 1869 | 3 | 3 | 3 | 7.7 | 7.7 | 7.7 | 54.083 |
| CTCF | 350 | 3 | 3 | 3 | 4.1 | 4.1 | 4.1 | 72.999 |
| RBM7 | 1165 | 3 | 3 | 3 | 16.9 | 16.9 | 16.9 | 30.59 |
| FCGRT | 2517 | 3 | 3 | 3 | 14.2 | 14.2 | 14.2 | 39.743 |
| PFDN1 | 765 | 3 | 3 | 3 | 38.5 | 38.5 | 38.5 | 13.466 |
| CSF1 | 852 | 3 | 3 | 3 | 14.4 | 14.4 | 14.4 | 31.171 |
| IMPDH1 | 382 | 4 | 3 | 3 | 7.9 | 6.3 | 6.3 | 54.272 |
| RGS10 | 1452 | 3 | 3 | 3 | 24.9 | 24.9 | 24.9 | 21.21 |
| PPIC | 2309 | 3 | 3 | 3 | 14.6 | 14.6 | 14.6 | 22.763 |
| PFDN2 | 4444 | 3 | 3 | 3 | 29.2 | 29.2 | 29.2 | 16.648 |
| CD2 | 1769 | 3 | 3 | 3 | 10.3 | 10.3 | 10.3 | 39.448 |
| MYLK | 3035 | 3 | 3 | 3 | 2.8 | 2.8 | 2.8 | 210.71 |

| | | | | | | | | |
|---|---|---|---|---|---|---|---|---|
| SLC2A3;SLC | 1870 | 3 | 3 | 3 | 7.9 | 7.9 | 7.9 | 53.924 |
| YTHDF3 | 5 | 9 | 9 | 3 | 22.5 | 22.5 | 10.5 | 58.311 |
| CD55 | 565 | 3 | 3 | 3 | 12.6 | 12.6 | 12.6 | 35.695 |
| PPIP5K2 | 85 | 6 | 6 | 3 | 8 | 8 | 4.4 | 140.41 |
| RUFY3 | 3330 | 4 | 3 | 3 | 9.2 | 7.5 | 7.5 | 52.964 |
| RPL38 | 2654 | 3 | 3 | 3 | 38.6 | 38.6 | 38.6 | 8.2178 |
| TNPO3 | 652 | 3 | 3 | 3 | 4.9 | 4.9 | 4.9 | 96.669 |
| CD81 | 532 | 3 | 3 | 3 | 35.8 | 35.8 | 35.8 | 17.963 |
| MP68;C14c | 2525 | 3 | 3 | 3 | 39.7 | 39.7 | 39.7 | 6.662 |
| UFM1 | 2587 | 3 | 3 | 3 | 60 | 60 | 60 | 9.1175 |
| SURF4 | 1402 | 6 | 6 | 3 | 39.8 | 39.8 | 21 | 21.127 |
| SLC15A3 | 3465 | 9 | 9 | 3 | 22.4 | 22.4 | 3.1 | 63.559 |
| NRBF2 | 3790 | 3 | 3 | 3 | 10.5 | 10.5 | 10.5 | 32.377 |
| UBE2D2;UE | 70 | 3 | 3 | 3 | 54.1 | 54.1 | 54.1 | 11.065 |
| MRPS14 | 1509 | 3 | 3 | 3 | 28.9 | 28.9 | 28.9 | 15.139 |
| GNB1 | 2632 | 16 | 12 | 3 | 69.1 | 50.9 | 9.7 | 37.377 |
| KDELR1 | 2082 | 3 | 3 | 3 | 20.8 | 20.8 | 20.8 | 24.542 |
| MAN1A2 | 1491 | 3 | 3 | 3 | 6.4 | 6.4 | 6.4 | 73.003 |
| PODXL | 1325 | 3 | 3 | 3 | 6.5 | 6.5 | 6.5 | 58.635 |
| PPP1CA | 2594 | 17 | 5 | 3 | 62.1 | 22.7 | 11.5 | 37.512 |
| SDR39U1 | 4248 | 3 | 3 | 3 | 13.3 | 13.3 | 13.3 | 31.076 |
| RPS15A | 1088 | 9 | 3 | 3 | 42.6 | 19.4 | 19.4 | 27.168 |
| FGD5 | 627 | 3 | 3 | 3 | 2.7 | 2.7 | 2.7 | 159.89 |
| STARD3 | 1199 | 3 | 3 | 3 | 12.4 | 12.4 | 12.4 | 50.501 |
| LRRCC1 | 4068 | 3 | 3 | 3 | 2.9 | 2.9 | 2.9 | 119.59 |
| SMARCB1 | 1019 | 3 | 3 | 3 | 11.2 | 11.2 | 11.2 | 45.05 |
| HFE | 973 | 3 | 3 | 3 | 13.1 | 13.1 | 13.1 | 38.871 |
| HPS3 | 1021 | 3 | 3 | 3 | 4.9 | 4.9 | 4.9 | 113.73 |
| PRPS1 | 558 | 11 | 11 | 3 | 53.7 | 53.7 | 15.1 | 31.392 |
| BZW2 | 801 | 4 | 3 | 3 | 11 | 9.3 | 9.3 | 46.913 |
| BAD | 3710 | 3 | 3 | 3 | 20.8 | 20.8 | 20.8 | 18.392 |
| GOLT1B | 932 | 3 | 3 | 3 | 18.8 | 18.8 | 18.8 | 14.307 |
| SEC22A | 666 | 3 | 3 | 3 | 28.3 | 28.3 | 28.3 | 16.089 |
| TXNL4A | 1210 | 3 | 3 | 3 | 92.6 | 92.6 | 92.6 | 6.3042 |
| IL1RN | 1995 | 3 | 3 | 3 | 30.5 | 30.5 | 30.5 | 20.055 |
| ACP6 | 4219 | 3 | 3 | 3 | 11.9 | 11.9 | 11.9 | 48.886 |
| DEFA3;DEF/ | 2541 | 3 | 3 | 3 | 20.2 | 20.2 | 20.2 | 10.245 |
| MORC3 | 2929 | 3 | 3 | 3 | 5.4 | 5.4 | 5.4 | 107.11 |
| CHCHD7 | 1172 | 3 | 3 | 3 | 40.4 | 40.4 | 40.4 | 11.137 |
| MRPL17 | 863 | 3 | 3 | 3 | 16.2 | 16.2 | 16.2 | 16.371 |
| SLC25A3 | 2722 | 16 | 16 | 3 | 51.7 | 51.7 | 13.5 | 40.094 |
| ENSA | 533 | 4 | 4 | 3 | 30.7 | 30.7 | 22.9 | 15.634 |
| SRD5A3 | 4162 | 3 | 3 | 3 | 9.7 | 9.7 | 9.7 | 36.521 |
| GAL | 2056 | 3 | 3 | 3 | 39.8 | 39.8 | 39.8 | 13.302 |
| TMEM109 | 4006 | 3 | 3 | 3 | 9.1 | 9.1 | 9.1 | 26.21 |
| CD151 | 870 | 3 | 3 | 3 | 14 | 14 | 14 | 19.792 |
| HLA-DMA | 284 | 3 | 3 | 3 | 26.6 | 26.6 | 26.6 | 19.15 |
| TIMM17A | 3908 | 3 | 3 | 3 | 31.6 | 31.6 | 31.6 | 18.023 |

| | | | | | | | | |
|---|---|---|---|---|---|---|---|---|
| ARL2 | 2234 | 3 | 3 | 3 | 21.7 | 21.7 | 21.7 | 20.878 |
| CWF19L1 | 3236 | 3 | 3 | 3 | 11.2 | 11.2 | 11.2 | 60.618 |
| REEP4 | 764 | 3 | 3 | 3 | 17.1 | 17.1 | 17.1 | 28.214 |
| NOC2L | 4608 | 3 | 3 | 3 | 4 | 4 | 4 | 84.918 |
| EDEM3 | 4062 | 3 | 3 | 3 | 4.8 | 4.8 | 4.8 | 104.66 |
| RGCC | 4145 | 3 | 3 | 3 | 21.9 | 21.9 | 21.9 | 14.559 |
| TMEM168 | 1130 | 2 | 2 | 2 | 11.6 | 11.6 | 11.6 | 29.105 |
| PDPK1;PDF | 832 | 2 | 2 | 2 | 4.9 | 4.9 | 4.9 | 60.328 |
| PPP1CC | 960 | 15 | 2 | 2 | 57.9 | 6.2 | 6.2 | 34.95 |
| PSMB6 | 2135 | 2 | 2 | 2 | 20.9 | 20.9 | 20.9 | 25.357 |
| AMFR | 4490 | 2 | 2 | 2 | 3.3 | 3.3 | 3.3 | 72.995 |
| DCSTAMP | 4111 | 2 | 2 | 2 | 4.9 | 4.9 | 4.9 | 53.392 |
| NOLC1 | 135 | 3 | 3 | 2 | 4 | 4 | 2.5 | 74.543 |
| SHC1 | 1035 | 2 | 2 | 2 | 8.1 | 8.1 | 8.1 | 32.507 |
| CST7 | 1603 | 2 | 2 | 2 | 24.1 | 24.1 | 24.1 | 16.454 |
| CRIP1 | 2403 | 2 | 2 | 2 | 27.3 | 27.3 | 27.3 | 8.5328 |
| CDC42SE2 | 4252 | 2 | 2 | 2 | 66.7 | 66.7 | 66.7 | 9.2234 |
| PTP4A2;PTI | 854 | 2 | 2 | 2 | 46 | 46 | 46 | 7.2082 |
| SLC35F6 | 3497 | 2 | 2 | 2 | 7 | 7 | 7 | 40.214 |
| MT-ATP6 | 1699 | 2 | 2 | 2 | 8.4 | 8.4 | 8.4 | 24.817 |
| TOR1AIP1 | 152 | 21 | 21 | 2 | 51.3 | 51.3 | 4.8 | 52.406 |
| RAB11B | 3044 | 16 | 16 | 2 | 67.4 | 67.4 | 13.3 | 24.488 |
| CLN3 | 230 | 2 | 2 | 2 | 7.3 | 7.3 | 7.3 | 47.622 |
| EIF1 | 1229 | 6 | 6 | 2 | 75.2 | 75.2 | 26.5 | 12.732 |
| CD58 | 562 | 2 | 2 | 2 | 10 | 10 | 10 | 28.147 |
| TMEM189-I | 1141 | 10 | 10 | 2 | 32.2 | 32.2 | 6.8 | 42.208 |
| SEC61G | 2543 | 2 | 2 | 2 | 36.8 | 36.8 | 36.8 | 7.7412 |
| BST2 | 2810 | 2 | 2 | 2 | 13.9 | 13.9 | 13.9 | 19.769 |
| ATP5J2-PTC | 1003 | 4 | 4 | 2 | 6.4 | 6.4 | 3.2 | 84.109 |
| HAT1 | 1365 | 2 | 2 | 2 | 9.1 | 9.1 | 9.1 | 49.512 |
| AFAP1L1 | 3627 | 2 | 2 | 2 | 4.3 | 4.3 | 4.3 | 86.431 |
| COX6A1 | 1889 | 4 | 2 | 2 | 62.4 | 26.6 | 26.6 | 12.155 |
| RCOR1 | 4486 | 2 | 2 | 2 | 5.4 | 5.4 | 5.4 | 53.327 |
| YPEL5 | 2620 | 2 | 2 | 2 | 21.5 | 21.5 | 21.5 | 13.841 |
| NDUFA3 | 1626 | 2 | 2 | 2 | 50 | 50 | 50 | 9.2787 |
| PFDN4 | 877 | 2 | 2 | 2 | 19.9 | 19.9 | 19.9 | 15.598 |
| MAP2K2 | 2237 | 6 | 3 | 2 | 18.8 | 8.8 | 5.8 | 44.424 |
| ARFGAP3 | 4210 | 2 | 2 | 2 | 5 | 5 | 5 | 56.928 |
| CEBPA | 2376 | 2 | 2 | 2 | 6.1 | 6.1 | 6.1 | 37.561 |
| DAP | 2431 | 2 | 2 | 2 | 31.4 | 31.4 | 31.4 | 11.164 |
| CHI3L1 | 1033 | 11 | 2 | 2 | 79.6 | 12.5 | 12.5 | 16.714 |
| CPEB2;CPEI | 450 | 2 | 2 | 2 | 17.4 | 17.4 | 17.4 | 18.644 |
| SLC37A2 | 3626 | 2 | 2 | 2 | 5.4 | 5.4 | 5.4 | 54.436 |
| PDLIM1 | 1282 | 2 | 2 | 2 | 6.4 | 6.4 | 6.4 | 36.071 |
| HIGD1A;HI( | 658 | 2 | 2 | 2 | 32.5 | 32.5 | 32.5 | 8.8131 |
| SPATA13 | 140 | 2 | 2 | 2 | 6.8 | 6.8 | 6.8 | 58.977 |
| BRI3 | 1648 | 2 | 2 | 2 | 16 | 16 | 16 | 13.645 |
| ACYP2 | 834 | 2 | 2 | 2 | 33.3 | 33.3 | 33.3 | 7.2321 |

| | | | | | | | | |
|---|---|---|---|---|---|---|---|---|
| NDUFB3 | 1453 | 2 | 2 | 2 | 21.4 | 21.4 | 21.4 | 11.402 |
| SYPL1 | 695 | 2 | 2 | 2 | 11.6 | 11.6 | 11.6 | 24.784 |
| PEX13 | 3713 | 2 | 2 | 2 | 7.7 | 7.7 | 7.7 | 44.129 |
| SSR3 | 656 | 2 | 2 | 2 | 11.9 | 11.9 | 11.9 | 21.08 |
| TMEM176E | 3102 | 2 | 2 | 2 | 8.1 | 8.1 | 8.1 | 29.056 |
| PPAN;PPAN | 172 | 2 | 2 | 2 | 11.5 | 11.5 | 11.5 | 23.108 |
| HLA-Cw;HL | 238 | 19 | 3 | 2 | 49.2 | 6.6 | 3 | 40.912 |
| GZMM | 2424 | 2 | 2 | 2 | 9.6 | 9.6 | 9.6 | 23.546 |
| PARL | 320 | 2 | 2 | 2 | 13.3 | 13.3 | 13.3 | 19.605 |
| QARS | 2337 | 35 | 34 | 2 | 57.3 | 56.3 | 4.4 | 87.798 |
| MBD2 | 4398 | 2 | 2 | 2 | 6.6 | 6.6 | 6.6 | 43.254 |
| CYSTM1 | 4101 | 2 | 2 | 2 | 11.3 | 11.3 | 11.3 | 10.631 |
| ARF1 | 2707 | 13 | 13 | 2 | 80.1 | 80.1 | 16 | 20.697 |
| ORMDL2 | 3124 | 3 | 3 | 2 | 26.1 | 26.1 | 16.3 | 17.363 |
| MTFMT | 3777 | 2 | 2 | 2 | 5.7 | 5.7 | 5.7 | 43.832 |
| PQLC3 | 646 | 2 | 2 | 2 | 11.8 | 11.8 | 11.8 | 18.608 |
| GSN | 3182 | 12 | 2 | 2 | 56.2 | 3.8 | 3.8 | 28.953 |
| ATP5EP2;A | 2526 | 2 | 2 | 2 | 29.4 | 29.4 | 29.4 | 5.8068 |
| SLC43A3 | 882 | 2 | 2 | 2 | 4.1 | 4.1 | 4.1 | 54.528 |
| SGTB | 753 | 2 | 2 | 2 | 13.3 | 13.3 | 13.3 | 21.83 |
| LUC7L2 | 173 | 16 | 16 | 2 | 40.6 | 40.6 | 4.8 | 54.223 |
| NDUFAB1 | 1340 | 3 | 3 | 2 | 15.4 | 15.4 | 9.6 | 17.417 |
| AGPAT1 | 3931 | 2 | 2 | 2 | 10.6 | 10.6 | 10.6 | 31.716 |
| PTMS | 2033 | 2 | 2 | 2 | 22.5 | 22.5 | 22.5 | 11.53 |
| ITGB6 | 65 | 2 | 2 | 2 | 2.6 | 2.6 | 2.6 | 75.849 |
| SRXN1 | 4049 | 2 | 2 | 2 | 24.8 | 24.8 | 24.8 | 14.259 |
| HSD17B8 | 3667 | 2 | 2 | 2 | 10.7 | 10.7 | 10.7 | 26.973 |
| ALOX5AP | 51 | 2 | 2 | 2 | 9.6 | 9.6 | 9.6 | 24.208 |
| RIPK1 | 2888 | 2 | 2 | 2 | 4 | 4 | 4 | 75.93 |
| DEGS1 | 1385 | 2 | 2 | 2 | 10.8 | 10.8 | 10.8 | 37.866 |
| LHPP | 4085 | 2 | 2 | 2 | 10.7 | 10.7 | 10.7 | 29.165 |
| UBL3 | 1625 | 2 | 2 | 2 | 23.1 | 23.1 | 23.1 | 13.157 |
| HBA2;HBA1 | 997 | 6 | 2 | 2 | 41.8 | 18.2 | 18.2 | 11.948 |
| TMEM167A | 3602 | 2 | 2 | 2 | 25 | 25 | 25 | 8.0598 |
| SYP | 1802 | 2 | 2 | 2 | 7 | 7 | 7 | 33.845 |
| SREK1 | 3658 | 2 | 2 | 2 | 4.7 | 4.7 | 4.7 | 59.38 |
| RALGAPB | 3417 | 2 | 2 | 2 | 3.3 | 3.3 | 3.3 | 166.8 |
| SFT2D3 | 3136 | 2 | 2 | 2 | 14.4 | 14.4 | 14.4 | 21.789 |
| CHMP3 | 4600 | 2 | 2 | 2 | 7.7 | 7.7 | 7.7 | 25.073 |
| FAM46C | 3217 | 2 | 2 | 2 | 5.4 | 5.4 | 5.4 | 44.944 |
| STX16-NPE | 1107 | 9 | 9 | 2 | 35.6 | 35.6 | 6.5 | 43.229 |
| DOLPP1 | 3427 | 2 | 2 | 2 | 13.4 | 13.4 | 13.4 | 27.03 |
| SPCS1 | 459 | 2 | 2 | 2 | 26.5 | 26.5 | 26.5 | 11.805 |
| RASA4;RAS | 795 | 2 | 2 | 2 | 9.5 | 9.5 | 9.5 | 30.476 |
| RNF121 | 4165 | 2 | 2 | 2 | 9.2 | 9.2 | 9.2 | 37.882 |
| SMAP1 | 107 | 2 | 2 | 2 | 2.8 | 2.8 | 2.8 | 49.219 |
| APH1A | 3753 | 2 | 2 | 2 | 5.3 | 5.3 | 5.3 | 28.996 |
| GPATCH8 | 4484 | 2 | 2 | 2 | 2 | 2 | 2 | 164.2 |

| | | | | | | | | |
|---|---|---|---|---|---|---|---|---|
| TMEM50A | 1670 | 2 | 2 | 2 | 11.5 | 11.5 | 11.5 | 17.4 |
| HIST1H1E | 1855 | 9 | 2 | 2 | 34.2 | 11 | 11 | 21.865 |
| PANX1 | 3868 | 2 | 2 | 2 | 10.6 | 10.6 | 10.6 | 48.05 |
| C20orf24 | 3997 | 2 | 2 | 2 | 11.7 | 11.7 | 11.7 | 15.487 |
| IGLL5;IGLC | 179 | 2 | 2 | 2 | 28.3 | 28.3 | 28.3 | 11.265 |
| MR1 | 3724 | 2 | 2 | 2 | 8.2 | 8.2 | 8.2 | 39.366 |
| SDHAF2 | 901 | 2 | 2 | 2 | 15.6 | 15.6 | 15.6 | 18.388 |
| LYRM4 | 4207 | 2 | 2 | 2 | 25.3 | 25.3 | 25.3 | 10.758 |
| WDR75 | 3444 | 2 | 2 | 2 | 4.7 | 4.7 | 4.7 | 94.498 |
| FRYL | 1616 | 2 | 2 | 2 | 1.2 | 1.2 | 1.2 | 339.59 |
| HLA-DQA1 | 871 | 5 | 4 | 2 | 25.7 | 20.1 | 8.9 | 23.673 |
| NCBP2 | 2456 | 2 | 2 | 2 | 16.7 | 16.7 | 16.7 | 18.001 |
| ACTG1 | 2659 | 28 | 28 | 2 | 86.1 | 86.1 | 4.8 | 41.792 |
| SRM | 1251 | 2 | 2 | 2 | 28.6 | 28.6 | 28.6 | 12.982 |
| SLC28A3 | 4178 | 2 | 2 | 2 | 4.2 | 4.2 | 4.2 | 76.929 |
| FDX1L | 3271 | 2 | 2 | 2 | 12.4 | 12.4 | 12.4 | 19.888 |
| ELOVL1 | 4014 | 2 | 2 | 2 | 9 | 9 | 9 | 32.662 |
| VKORC1 | 867 | 2 | 2 | 2 | 29.8 | 29.8 | 29.8 | 11.082 |
| RPL10 | 2124 | 11 | 11 | 2 | 50.5 | 50.5 | 14.5 | 22.975 |
| PTTG1IP | 548 | 2 | 2 | 2 | 14.4 | 14.4 | 14.4 | 20.324 |
| TOMM7 | 4381 | 2 | 2 | 2 | 47.3 | 47.3 | 47.3 | 6.2484 |
| LIMD2 | 1202 | 2 | 2 | 2 | 25.7 | 25.7 | 25.7 | 12.013 |
| CMTM3 | 1108 | 2 | 2 | 2 | 26.7 | 26.7 | 26.7 | 16.423 |
| CYB561A3 | 3534 | 2 | 2 | 2 | 7.4 | 7.4 | 7.4 | 27.214 |
| DTNBP1 | 75 | 2 | 2 | 2 | 13.8 | 13.8 | 13.8 | 21.437 |
| RNASE3 | 1897 | 2 | 2 | 2 | 15 | 15 | 15 | 18.385 |
| PRPS2 | 1886 | 9 | 3 | 2 | 40.6 | 15.7 | 11.6 | 34.769 |
| ZC3H4 | 1258 | 2 | 2 | 2 | 3.3 | 3.3 | 3.3 | 95.425 |
| TMEM14C | 4380 | 2 | 2 | 2 | 24.1 | 24.1 | 24.1 | 11.564 |
| TAX1BP3 | 1363 | 2 | 2 | 2 | 28.2 | 28.2 | 28.2 | 13.735 |
| GK2 | 2951 | 5 | 2 | 2 | 13.6 | 5.1 | 5.1 | 60.593 |
| HSPA8 | 1868 | 35 | 33 | 2 | 68.7 | 66.3 | 5.7 | 70.897 |
| TMBIM1 | 590 | 2 | 2 | 2 | 14.2 | 14.2 | 14.2 | 11.535 |
| GEMIN5 | 3629 | 2 | 2 | 2 | 1.9 | 1.9 | 1.9 | 168.59 |
| RTN4 | 4230 | 13 | 2 | 2 | 18 | 4.8 | 4.8 | 129.93 |
| MARK2 | 800 | 3 | 3 | 2 | 4.7 | 4.7 | 3.4 | 78.452 |
| GFER | 1101 | 2 | 2 | 2 | 33.9 | 33.9 | 33.9 | 11.66 |
| IER3IP1 | 315 | 2 | 2 | 2 | 40.6 | 40.6 | 40.6 | 7.6079 |
| C6orf47 | 1676 | 2 | 2 | 2 | 7.5 | 7.5 | 7.5 | 31.71 |
| C15orf38-A | 171 | 2 | 2 | 2 | 9.4 | 9.4 | 9.4 | 43.881 |
| TMSB4X | 2614 | 3 | 3 | 2 | 40.9 | 40.9 | 25 | 5.0526 |
| GABARAPL1 | 909 | 5 | 4 | 2 | 54.3 | 54.3 | 12.9 | 8.577 |
| OSCAR | 39 | 2 | 2 | 2 | 15.1 | 15.1 | 15.1 | 27.602 |
| ARGLU1 | 4313 | 2 | 2 | 2 | 6.2 | 6.2 | 6.2 | 33.216 |
| GNG10 | 2401 | 5 | 5 | 2 | 79.4 | 79.4 | 26.5 | 7.2053 |
| hCG_2004 | 275 | 2 | 2 | 2 | 5.7 | 5.7 | 5.7 | 48.586 |
| HLA-DRB5 | 3089 | 9 | 3 | 2 | 39.5 | 5.6 | 5.6 | 30.056 |
| SARDH | 155 | 2 | 2 | 2 | 4.3 | 4.3 | 4.3 | 67.216 |

| | | | | | | | | |
|---|---|---|---|---|---|---|---|---|
| APH1B | 3651 | 2 | 2 | 2 | 7.8 | 7.8 | 7.8 | 28.46 |
| CIR1 | 3422 | 2 | 2 | 2 | 5.1 | 5.1 | 5.1 | 52.313 |
| MCUR1 | 3748 | 2 | 2 | 2 | 8.4 | 8.4 | 8.4 | 39.694 |
| IMPACT | 399 | 2 | 2 | 2 | 11.1 | 11.1 | 11.1 | 20.506 |
| DNAJB4 | 4416 | 3 | 2 | 2 | 10.1 | 6.5 | 6.5 | 37.806 |
| NCK1 | 657 | 10 | 2 | 2 | 62.1 | 8.6 | 8.6 | 20.068 |
| LSM5 | 4625 | 2 | 2 | 2 | 46.2 | 46.2 | 46.2 | 9.9374 |
| SMIM20 | 3514 | 2 | 2 | 2 | 34.3 | 34.3 | 34.3 | 7.7019 |
| CHCHD1 | 3756 | 2 | 2 | 2 | 22.9 | 22.9 | 22.9 | 13.475 |
| IMP3 | 4290 | 2 | 2 | 2 | 17.9 | 17.9 | 17.9 | 21.85 |
| DNTTIP2 | 3168 | 2 | 2 | 2 | 3.6 | 3.6 | 3.6 | 84.468 |
| CLTCL1 | 2483 | 21 | 2 | 2 | 13.5 | 1.7 | 1.7 | 187.03 |
| PRKCA | 1976 | 9 | 2 | 2 | 18.6 | 3.6 | 3.6 | 76.749 |
| TPRKB | 4592 | 2 | 2 | 2 | 21.1 | 21.1 | 21.1 | 19.661 |
| C4orf32 | 3527 | 2 | 2 | 2 | 19.7 | 19.7 | 19.7 | 14.652 |
| MT-ATP8 | 1723 | 2 | 2 | 2 | 35.3 | 35.3 | 35.3 | 7.9916 |
| WASL | 1306 | 2 | 2 | 2 | 3.8 | 3.8 | 3.8 | 54.826 |
| COPS7B | 105 | 2 | 2 | 2 | 12.9 | 12.9 | 12.9 | 23.556 |
| MRPL57 | 3944 | 2 | 2 | 2 | 31.4 | 31.4 | 31.4 | 12.266 |
| NDUFC1 | 1454 | 2 | 2 | 2 | 32.9 | 32.9 | 32.9 | 8.7341 |
| DPY30 | 4066 | 2 | 2 | 2 | 36.4 | 36.4 | 36.4 | 11.25 |
| G6PC3 | 3993 | 2 | 2 | 2 | 8.7 | 8.7 | 8.7 | 38.734 |
| TMEM127 | 1527 | 2 | 2 | 2 | 21.8 | 21.8 | 21.8 | 25.841 |
| ABHD5 | 3631 | 2 | 2 | 2 | 12 | 12 | 12 | 39.095 |
| TP53I11 | 850 | 2 | 2 | 2 | 40.3 | 40.3 | 40.3 | 6.8649 |
| DPM3 | 4397 | 2 | 2 | 2 | 23.9 | 23.9 | 23.9 | 10.094 |
| SDHD | 1336 | 2 | 2 | 2 | 20.8 | 20.8 | 20.8 | 17.043 |
| SLC44A2 | 3446 | 5 | 5 | 2 | 10.8 | 10.8 | 5.4 | 80.123 |
| LSM7 | 4478 | 2 | 2 | 2 | 49.5 | 49.5 | 49.5 | 11.602 |
| MAP4K4 | 227 | 13 | 13 | 2 | 12.6 | 12.6 | 2.7 | 132.16 |
| PXMP4 | 4675 | 2 | 2 | 2 | 8.5 | 8.5 | 8.5 | 24.264 |
| MRPL20 | 4042 | 2 | 2 | 2 | 14.8 | 14.8 | 14.8 | 17.442 |
| PSEN1 | 2385 | 4 | 2 | 2 | 13.1 | 7.9 | 7.9 | 52.667 |
| PRNP | 497 | 2 | 2 | 2 | 9.2 | 9.2 | 9.2 | 23.554 |
| CAPZB | 556 | 17 | 17 | 2 | 58.1 | 58.1 | 5.3 | 33.781 |
| DAGLA | 4615 | 2 | 2 | 2 | 4.7 | 4.7 | 4.7 | 114.95 |
| TAMM41 | 3760 | 2 | 2 | 2 | 8.4 | 8.4 | 8.4 | 51.066 |
| BCL2L1 | 2783 | 2 | 2 | 2 | 10.5 | 10.5 | 10.5 | 20.244 |
| ZFYVE21 | 3937 | 2 | 2 | 2 | 12 | 12 | 12 | 26.505 |
| TMEM258 | 892 | 2 | 2 | 2 | 16.3 | 16.3 | 16.3 | 5.7065 |
| NT5DC1 | 3206 | 2 | 2 | 2 | 6.4 | 6.4 | 6.4 | 51.844 |
| CHTOP | 101 | 2 | 2 | 2 | 38.2 | 38.2 | 38.2 | 8.5199 |
| SRSF11 | 2770 | 4 | 4 | 2 | 15.4 | 15.4 | 8 | 42.316 |
| APOL2 | 1178 | 22 | 22 | 2 | 46.5 | 46.5 | 4.2 | 48.915 |
| JAGN1 | 3519 | 2 | 2 | 2 | 13.1 | 13.1 | 13.1 | 21.125 |
| TPM4 | 2665 | 25 | 17 | 2 | 64.9 | 50.4 | 11.3 | 28.521 |
| MRPL51 | 3116 | 2 | 2 | 2 | 17.2 | 17.2 | 17.2 | 15.095 |
| ME2 | 2070 | 35 | 35 | 2 | 82.9 | 82.9 | 2.7 | 65.443 |

| | | | | | | | | |
|---|---|---|---|---|---|---|---|---|
| HMGN2 | 1752 | 3 | 2 | 2 | 25.6 | 25.6 | 25.6 | 9.3926 |
| S1PR2 | 1621 | 2 | 2 | 2 | 7.9 | 7.9 | 7.9 | 38.867 |
| GLMP | 41 | 2 | 2 | 2 | 14.4 | 14.4 | 14.4 | 34.576 |
| AAK1 | 837 | 3 | 2 | 2 | 5.2 | 3.4 | 3.4 | 73.769 |
| MAP2K4 | 2314 | 3 | 2 | 2 | 8.3 | 6.5 | 6.5 | 44.287 |
| MPHOSPH1 | 1323 | 2 | 2 | 2 | 5.4 | 5.4 | 5.4 | 78.863 |
| EXOSC4 | 4216 | 2 | 2 | 2 | 8.2 | 8.2 | 8.2 | 26.383 |
| SLC35B2 | 3596 | 2 | 2 | 2 | 5.3 | 5.3 | 5.3 | 47.514 |
| STX2 | 2200 | 2 | 2 | 2 | 9.7 | 9.7 | 9.7 | 33.341 |
| C17orf49;R | 964 | 2 | 2 | 2 | 21.7 | 21.7 | 21.7 | 16.29 |
| ATP6AP2 | 308 | 13 | 13 | 2 | 44.3 | 44.3 | 8.7 | 38.386 |
| LMBRD1 | 396 | 2 | 2 | 2 | 5.9 | 5.9 | 5.9 | 63.817 |
| CCL3;CCL1 | 1850 | 2 | 2 | 2 | 22.8 | 22.8 | 22.8 | 10.085 |
| ANAPC7 | 4474 | 2 | 2 | 2 | 4.2 | 4.2 | 4.2 | 66.855 |
| CYTH1;CYTH | 3025 | 3 | 3 | 2 | 7.5 | 7.5 | 4.5 | 46.412 |
| SLC37A4 | 1473 | 2 | 2 | 2 | 6.7 | 6.7 | 6.7 | 48.922 |
| SPAG9 | 1484 | 46 | 46 | 2 | 49.4 | 49.4 | 2 | 146.2 |
| DR1 | 2735 | 2 | 2 | 2 | 27.3 | 27.3 | 27.3 | 19.443 |
| C14orf142 | 4035 | 2 | 2 | 2 | 34 | 34 | 34 | 10.859 |
| SCAF4 | 1620 | 4 | 4 | 2 | 4.5 | 4.5 | 2.9 | 125.87 |
| PLEKHA2 | 79 | 2 | 2 | 2 | 9.2 | 9.2 | 9.2 | 27.837 |
| TMEM208 | 1087 | 2 | 2 | 2 | 22 | 22 | 22 | 19.642 |
| PRPF38B | 3215 | 2 | 2 | 2 | 5.1 | 5.1 | 5.1 | 64.467 |
| OSGEPL1 | 4140 | 2 | 2 | 2 | 6.8 | 6.8 | 6.8 | 45.122 |
| SLC31A1 | 1413 | 2 | 2 | 2 | 15.8 | 15.8 | 15.8 | 21.09 |
| NDUFA1 | 1398 | 2 | 2 | 2 | 20 | 20 | 20 | 8.0723 |
| NTMT1 | 4004 | 2 | 2 | 2 | 10.3 | 10.3 | 10.3 | 25.387 |
| CLIP1 | 2173 | 45 | 45 | 2 | 36 | 36 | 1.3 | 162.24 |
| NUDT2 | 2417 | 2 | 2 | 2 | 23.1 | 23.1 | 23.1 | 16.829 |
| SREK1 | 67 | 2 | 2 | 2 | 34 | 34 | 34 | 10.713 |
| PPTC7 | 3588 | 2 | 2 | 2 | 9.2 | 9.2 | 9.2 | 32.645 |
| | 359 | 33 | 2 | 2 | 48.4 | 1.9 | 1.9 | 106.19 |
| GTF3C3 | 437 | 2 | 2 | 2 | 4.3 | 4.3 | 4.3 | 71.458 |
| NINJ1 | 3715 | 2 | 2 | 2 | 21.1 | 21.1 | 21.1 | 16.345 |
| CASK | 338 | 2 | 2 | 2 | 5.4 | 5.4 | 5.4 | 63.534 |
| NRAS | 1705 | 10 | 10 | 2 | 69.3 | 69.3 | 12.7 | 21.229 |

| Q-value | Score | Intensity | MS/MS count | ctrl_1_1 | ctrl_1_2 | ctrl_2_1 | ctrl_2_2 | ctrl_3_1 |
|---|---|---|---|---|---|---|---|---|
| 0 | 5.7635 | 6.4E+08 | 15 | NaN | NaN | NaN | NaN | NaN |
| 0 | 4.7833 | 3.63E+08 | 15 | NaN | NaN | NaN | NaN | NaN |
| 0 | 5.7863 | 1.41E+09 | 8 | NaN | NaN | NaN | NaN | NaN |
| 0 | 6.1332 | 3.64E+08 | 12 | NaN | NaN | NaN | NaN | NaN |
| 0 | 14.259 | 3.99E+08 | 19 | NaN | NaN | 25.369 | NaN | NaN |
| 0 | 9.1941 | 2.37E+08 | 7 | NaN | NaN | NaN | NaN | NaN |
| 0 | 22.832 | 4.12E+08 | 7 | 25.4357 | NaN | NaN | NaN | NaN |
| 0.003156 | 1.9517 | 5.13E+08 | 4 | NaN | NaN | NaN | NaN | NaN |
| 0.000698 | 2.4801 | 1.19E+09 | 34 | 24.737 | 24.8426 | 25.3559 | NaN | 25.3865 |
| 0 | 3.0513 | 2.24E+08 | 10 | NaN | NaN | NaN | NaN | NaN |
| 0 | 5.3107 | 1.43E+08 | 5 | NaN | NaN | NaN | NaN | NaN |
| 0 | 7.2203 | 4.8E+08 | 16 | NaN | 25.0816 | 25.2814 | NaN | 25.2442 |
| 0 | 3.6302 | 3.89E+08 | 12 | NaN | NaN | NaN | NaN | NaN |
| 0 | 3.7151 | 3.31E+08 | 11 | 24.6906 | 24.722 | NaN | NaN | 24.6073 |
| 0 | 7.3912 | 1.75E+08 | 4 | NaN | NaN | NaN | NaN | NaN |
| 0 | 13.796 | 3.21E+08 | 7 | NaN | NaN | NaN | NaN | NaN |
| 0 | 3.7639 | 2.41E+08 | 3 | NaN | NaN | NaN | NaN | NaN |
| 0 | 10.757 | 2.26E+08 | 3 | NaN | NaN | NaN | NaN | NaN |
| 0 | 3.9033 | 1.66E+08 | 10 | NaN | NaN | NaN | NaN | NaN |
| 0 | 12.588 | 2.56E+08 | 11 | 23.9177 | NaN | 24.1054 | 23.9456 | 24.0296 |
| 0 | 5.7706 | 2.03E+08 | 8 | NaN | 24.1667 | NaN | NaN | 23.9998 |
| 0 | 3.4614 | 2.09E+08 | 7 | NaN | NaN | 24.2421 | NaN | 24.0135 |
| 0 | 4.9855 | 1.71E+08 | 4 | NaN | NaN | NaN | NaN | NaN |
| 0 | 2.987 | 2.64E+08 | 9 | NaN | NaN | NaN | NaN | NaN |
| 0.000471 | 2.6899 | 1.56E+08 | 11 | NaN | NaN | NaN | NaN | NaN |
| 0.003599 | 1.9101 | 1.64E+08 | 4 | NaN | NaN | NaN | NaN | NaN |
| 0 | 5.1936 | 2.06E+08 | 3 | NaN | NaN | NaN | NaN | NaN |
| 0 | 318.89 | 1.62E+10 | 252 | 29.8214 | 29.8695 | 29.9539 | 29.9087 | 29.8853 |
| 0 | 15.971 | 7.42E+08 | 28 | 25.5677 | 25.2965 | 25.52 | 25.1435 | 25.3405 |
| 0 | 23.666 | 6.09E+08 | 21 | 24.9335 | 25.0555 | 24.8924 | 25.2453 | 25.1484 |
| 0 | 28.451 | 1.87E+09 | 30 | 26.8142 | 26.868 | 26.7831 | 26.8009 | 26.7173 |
| 0 | 3.9522 | 3.87E+08 | 19 | 24.3434 | 24.3624 | 24.3473 | 24.3288 | 24.7394 |
| 0 | 6.3041 | 7.46E+08 | 13 | 25.7133 | 24.5536 | 25.0983 | 25.1507 | 25.6495 |
| 0 | 16.817 | 8.51E+08 | 33 | 25.4555 | 25.3981 | 25.7252 | 25.2895 | 25.6731 |
| 0 | 32.297 | 4.9E+09 | 52 | 28.1391 | 28.1402 | 28.1771 | 27.9794 | 28.1451 |
| 0 | 4.0989 | 6.94E+08 | 19 | 25.1139 | 25.1295 | 25.3794 | 25.3491 | 25.3959 |
| 0 | 7.1075 | 1.73E+09 | 21 | 26.3098 | 26.0071 | 26.1505 | 26.342 | 26.5988 |
| 0 | 8.9271 | 1.65E+10 | 42 | 29.4848 | 29.4744 | 30.2122 | 30.2944 | 29.8257 |
| 0 | 9.1794 | 3.27E+09 | 56 | 27.1002 | 27.1608 | 27.8915 | 27.5626 | 27.7236 |
| 0 | 14.558 | 3.34E+09 | 36 | 28.5637 | 28.523 | 28.3321 | 28.0863 | 27.9568 |
| 0 | 4.1106 | 6.79E+08 | 16 | 25.7309 | 25.9992 | 26.0061 | 26.0517 | 25.9897 |
| 0 | 6.7777 | 3.6E+08 | 10 | 24.9113 | 24.871 | 24.7398 | NaN | NaN |
| 0.000699 | 2.5235 | 1.78E+08 | 4 | 24.5941 | 24.8324 | NaN | 24.4992 | 24.5029 |
| 0 | 28.105 | 6.95E+08 | 10 | 25.952 | 26.0218 | 25.2968 | 26.4202 | 25.9209 |
| 0 | 6.5961 | 2.1E+08 | 16 | 24.5407 | NaN | 24.5637 | 24.6123 | 24.7239 |

| | | | | | | | | |
|---|---|---|---|---|---|---|---|---|
| 0.000471 | 2.6949 | 1.09E+09 | 15 | 26.3716 | 26.4758 | 26.3578 | 26.235 | 26.4591 |
| 0 | 32.406 | 7.15E+08 | 18 | 26.0542 | 25.85 | 26.2942 | 26.2258 | 25.869 |
| 0 | 3.185 | 5.74E+08 | 8 | 26.5546 | 26.4204 | NaN | NaN | NaN |
| 0 | 3.5425 | 2.97E+08 | 9 | 24.6222 | 24.7129 | 24.9581 | 24.8095 | 24.7457 |
| 0.008863 | 1.4513 | 3.24E+08 | 7 | 25.0147 | 24.9205 | 25.0041 | NaN | 24.5203 |
| 0 | 15.01 | 2.77E+08 | 7 | 25.1294 | NaN | 25.2827 | 25.4278 | NaN |
| 0 | 11.651 | 5.22E+08 | 16 | 25.7329 | 26.0351 | 25.1881 | 25.2336 | 25.7638 |
| 0 | 2.985 | 2.76E+08 | 12 | 24.6275 | 24.424 | 24.7196 | 24.6423 | 24.6673 |
| 0 | 5.0493 | 2.35E+08 | 9 | 24.5371 | 24.2405 | 24.5033 | 24.2151 | 24.6526 |
| 0 | 27.517 | 3.26E+08 | 8 | 24.7166 | 24.8853 | 24.9384 | 24.552 | 24.8641 |
| 0 | 20.846 | 5.06E+08 | 13 | 25.3851 | 25.5884 | 25.2975 | 25.2602 | 25.4231 |
| 0 | 3.2907 | 1.77E+08 | 5 | NaN | NaN | 24.5916 | 24.4032 | NaN |
| 0 | 5.5733 | 4.14E+08 | 18 | 24.9718 | 25.2415 | 25.0307 | 24.9158 | 25.2043 |
| 0 | 3.1601 | 1.3E+08 | 13 | 23.6312 | 23.2908 | 22.8635 | 23.356 | 23.7485 |
| 0 | 5.9475 | 1.45E+08 | 11 | NaN | 24.0428 | 24.1666 | 24.1092 | 23.8878 |
| 0 | 4.6867 | 2.46E+08 | 10 | 24.3042 | 24.1758 | 24.5501 | 24.2122 | 24.6969 |
| 0 | 9.8489 | 2.61E+08 | 23 | 24.4316 | 24.1483 | 24.1953 | 24.3621 | 24.5445 |
| 0.000924 | 2.3584 | 1.74E+08 | 8 | 23.7861 | 24.2609 | 23.637 | 23.9246 | NaN |
| 0 | 5.7588 | 2.89E+08 | 12 | 24.0231 | 24.0848 | 24.0522 | 23.9818 | 24.7104 |
| 0 | 4.6205 | 3.81E+08 | 5 | NaN | 24.9592 | NaN | 25.2375 | NaN |
| 0 | 12.708 | 3.46E+08 | 27 | 25.107 | 24.9396 | 25.0114 | 24.8698 | 24.8883 |
| 0 | 4.8936 | 2.33E+08 | 2 | NaN | 24.5669 | 24.3441 | NaN | 24.4488 |
| 0 | 85.656 | 4.84E+08 | 20 | 25.3137 | 25.5416 | 25.4531 | 25.4753 | 25.2841 |
| 0.003158 | 1.9577 | 64002000 | 7 | 22.1963 | 22.6726 | 22.766 | 22.5763 | 22.9952 |
| 0 | 3.4954 | 1.17E+08 | 12 | 23.4907 | 23.4952 | NaN | 23.5617 | 23.6503 |
| 0.000922 | 2.3305 | 1.4E+08 | 12 | NaN | 23.5778 | 23.74 | 23.884 | 23.8186 |
| 0 | 4.5715 | 1.33E+08 | 8 | 24.0075 | NaN | 24.4308 | 24.2711 | 23.9075 |
| 0 | 7.5281 | 1.64E+08 | 4 | NaN | NaN | 24.0797 | 24.0874 | 23.9781 |
| 0 | 5.819 | 3.88E+08 | 15 | 24.9615 | 24.8346 | 24.9362 | 25.3284 | 24.7972 |
| 0.004907 | 1.7961 | 1.66E+08 | 8 | 24.221 | NaN | 24.3138 | 24.2957 | 24.1836 |
| 0 | 3.2821 | 2.35E+08 | 8 | 24.1362 | 24.3051 | 24.2603 | 24.1516 | 24.2006 |
| 0 | 3.8982 | 1.99E+08 | 11 | 24.4505 | 24.6535 | 24.4123 | 24.332 | 24.3988 |
| 0 | 5.7399 | 4.03E+08 | 18 | 25.0096 | 25.0707 | 25.0732 | 25.15 | 25.2188 |
| 0 | 3.7904 | 1.03E+08 | 4 | 23.5784 | 23.8635 | 23.5089 | 23.5366 | 23.5169 |
| 0 | 3.9697 | 2.84E+08 | 9 | 24.6344 | NaN | 24.704 | 24.5202 | 24.6974 |
| 0 | 10.92 | 6.93E+08 | 28 | 25.7519 | 25.7533 | 26.0736 | 25.441 | 26.0795 |
| 0 | 14.035 | 1.63E+09 | 29 | 27.2501 | 27.1137 | 27.0646 | 26.9404 | 27.0186 |
| 0 | 6.4368 | 7.37E+08 | 29 | 26.0068 | 25.9911 | 26.493 | 25.9127 | 25.9387 |
| 0 | 5.129 | 5.01E+09 | 41 | 28.7313 | 28.6125 | 28.897 | 28.7242 | 28.741 |
| 0 | 6.7254 | 3.34E+08 | 24 | 24.8907 | 24.9322 | 24.9302 | 24.7233 | 24.9683 |
| 0 | 15.594 | 8.88E+08 | 37 | 26.4488 | 26.2921 | 26.4432 | 25.9744 | 26.1734 |
| 0 | 5.1129 | 83378000 | 5 | 22.494 | 22.6811 | 22.7052 | 22.5888 | 22.8025 |
| 0 | 323.31 | 1.37E+11 | 2466 | 33.2351 | 33.2306 | 33.1983 | 33.1914 | 33.3403 |
| 0 | 323.31 | 4.09E+11 | 3470 | 34.8909 | 34.8737 | 34.9925 | 34.9698 | 34.5923 |
| 0 | 323.31 | 7.68E+10 | 1581 | 32.3866 | 32.3601 | 32.4954 | 32.5041 | 32.3342 |
| 0 | 323.31 | 2.31E+10 | 846 | 30.4863 | 30.4801 | 30.5387 | 30.5145 | 30.9045 |
| 0 | 323.31 | 4.93E+11 | 3081 | 35.1027 | 35.0739 | 35.089 | 35.0878 | 34.9432 |
| 0 | 323.31 | 4.47E+11 | 4762 | 34.9293 | 34.9633 | 34.9974 | 35.0196 | 34.8109 |

| | | | | | | | | |
|---|---|---|---|---|---|---|---|---|
| 0 | 323.31 | 4.02E+10 | 1113 | 31.4986 | 31.5293 | 31.6175 | 31.5976 | 31.3335 |
| 0 | 323.31 | 2.5E+10 | 779 | 30.7078 | 30.6894 | 30.7246 | 30.6931 | 30.8281 |
| 0 | 323.31 | 1.63E+11 | 1777 | 33.5352 | 33.5332 | 33.4595 | 33.4519 | 33.6396 |
| 0 | 323.31 | 3.5E+11 | 2162 | 34.6182 | 34.6657 | 34.4234 | 34.4447 | 34.6582 |
| 0 | 323.31 | 1.42E+10 | 588 | 29.8352 | 29.9027 | 30.0237 | 30.0003 | 29.9237 |
| 0 | 323.31 | 1.97E+10 | 638 | 30.443 | 30.3806 | 30.3511 | 30.3367 | 30.6063 |
| 0 | 323.31 | 1.98E+11 | 1431 | 33.7969 | 33.7827 | 33.7165 | 33.7 | 33.751 |
| 0 | 323.31 | 1.82E+10 | 625 | 30.3115 | 30.3459 | 30.3021 | 30.3489 | 30.2389 |
| 0 | 323.31 | 2.93E+10 | 684 | 31.015 | 30.9821 | 31.076 | 30.9968 | 30.9352 |
| 0 | 323.31 | 2.37E+10 | 709 | 30.7725 | 30.7138 | 30.8192 | 30.7713 | 30.6712 |
| 0 | 323.31 | 4.56E+10 | 783 | 31.6288 | 31.5878 | 31.6053 | 31.6059 | 31.6344 |
| 0 | 323.31 | 7.24E+10 | 1038 | 32.2983 | 32.2724 | 32.2728 | 32.2505 | 32.3933 |
| 0 | 323.31 | 2.9E+10 | 640 | 31.0861 | 31.0734 | 30.8507 | 30.8152 | 31.2171 |
| 0 | 323.31 | 9.23E+10 | 1036 | 32.6508 | 32.6611 | 32.7135 | 32.7192 | 32.702 |
| 0 | 323.31 | 3.54E+11 | 1898 | 34.6494 | 34.6199 | 34.5821 | 34.6077 | 34.6512 |
| 0 | 323.31 | 2.45E+10 | 692 | 30.7461 | 30.7722 | 30.8105 | 30.8736 | 30.7341 |
| 0 | 323.31 | 8.9E+10 | 965 | 32.6541 | 32.6185 | 32.687 | 32.6837 | 32.5382 |
| 0 | 323.31 | 3.32E+10 | 673 | 31.2239 | 31.1782 | 31.084 | 31.0866 | 31.2744 |
| 0 | 323.31 | 3.99E+10 | 736 | 31.2787 | 31.2866 | 31.1984 | 31.2484 | 31.5662 |
| 0 | 286.97 | 9.32E+09 | 323 | 29.4058 | 29.3775 | 29.4146 | 29.3443 | 29.2212 |
| 0 | 323.31 | 4.16E+10 | 711 | 31.4421 | 31.4229 | 31.3981 | 31.3327 | 31.6482 |
| 0 | 307.57 | 1.77E+10 | 459 | 30.3162 | 30.2927 | 30.3599 | 30.3637 | 30.2699 |
| 0 | 323.31 | 1.41E+10 | 453 | 30.051 | 29.9982 | 29.9802 | 30.05 | 29.9449 |
| 0 | 280.28 | 9.11E+09 | 375 | 29.3589 | 29.3221 | 29.3267 | 29.294 | 29.2561 |
| 0 | 323.31 | 6.74E+10 | 861 | 32.2537 | 32.2428 | 32.3038 | 32.2871 | 32.2107 |
| 0 | 323.31 | 1.33E+11 | 1098 | 33.1217 | 33.0974 | 33.2015 | 33.1982 | 33.1266 |
| 0 | 323.31 | 8.96E+10 | 942 | 32.594 | 32.6341 | 32.6285 | 32.6697 | 32.5468 |
| 0 | 323.31 | 2.11E+10 | 511 | 30.4828 | 30.4837 | 30.5241 | 30.4529 | 30.5951 |
| 0 | 323.31 | 3.65E+11 | 2505 | 34.6497 | 34.6476 | 34.6958 | 34.6732 | 34.7072 |
| 0 | 314.1 | 1.96E+10 | 475 | 30.6773 | 30.6054 | 30.3288 | 30.2931 | 30.374 |
| 0 | 323.31 | 4.91E+10 | 656 | 31.6737 | 31.6386 | 31.7144 | 31.6952 | 31.4357 |
| 0 | 323.31 | 1.44E+10 | 424 | 30.0557 | 30.0219 | 29.9745 | 29.9061 | 29.9964 |
| 0 | 244.43 | 2.08E+10 | 540 | 30.4275 | 30.4621 | 30.491 | 30.4617 | 30.5293 |
| 0 | 323.31 | 1.12E+10 | 409 | 29.6111 | 29.6238 | 29.7783 | 29.6904 | 29.6325 |
| 0 | 323.31 | 1.43E+11 | 1116 | 33.3243 | 33.2728 | 33.2445 | 33.2271 | 33.2939 |
| 0 | 323.31 | 2.65E+12 | 5423 | 37.4785 | 37.4915 | 37.5908 | 37.5126 | 37.3782 |
| 0 | 266.77 | 1.17E+10 | 338 | 29.5965 | 29.5763 | 29.6961 | 29.7004 | 29.671 |
| 0 | 323.31 | 1.4E+10 | 479 | 29.9936 | 30.0193 | 30.0846 | 29.9882 | 29.8638 |
| 0 | 323.31 | 1.06E+11 | 943 | 32.841 | 32.8811 | 32.8989 | 32.8527 | 32.8581 |
| 0 | 323.31 | 5.91E+10 | 801 | 31.8757 | 31.8764 | 32.0618 | 32.0345 | 31.9792 |
| 0 | 292.31 | 1.16E+10 | 361 | 29.7236 | 29.6671 | 29.8004 | 29.8036 | 29.3744 |
| 0 | 323.31 | 2.59E+10 | 523 | 30.8724 | 30.915 | 30.8278 | 30.8297 | 30.7918 |
| 0 | 323.31 | 4.36E+10 | 641 | 31.6021 | 31.5938 | 31.6385 | 31.6695 | 31.5653 |
| 0 | 182.54 | 7.53E+09 | 321 | 28.9947 | 29.0334 | 29.0325 | 29.0701 | 29.1849 |
| 0 | 323.31 | 9.55E+10 | 865 | 32.7016 | 32.687 | 32.5803 | 32.5622 | 32.7613 |
| 0 | 323.31 | 3.88E+11 | 1180 | 35.0473 | 35.0519 | 34.6781 | 34.5886 | 34.6865 |
| 0 | 323.31 | 8.16E+09 | 244 | 28.8929 | 28.9765 | 29.0517 | 29.02 | 29.3891 |
| 0 | 231.57 | 2.12E+10 | 431 | 30.595 | 30.5374 | 30.4653 | 30.414 | 30.6972 |

| | | | | | | | | |
|---|---|---|---|---|---|---|---|---|
| 0 | 323.31 | 1.26E+11 | 891 | 33.1122 | 33.1231 | 33.2406 | 33.247 | 32.9562 |
| 0 | 323.31 | 7.84E+10 | 748 | 32.6511 | 32.5957 | 32.3112 | 32.319 | 32.3466 |
| 0 | 315.97 | 9.02E+09 | 286 | 29.3255 | 29.3147 | 29.3497 | 29.3458 | 29.4306 |
| 0 | 323.31 | 2.3E+10 | 433 | 30.6196 | 30.614 | 30.74 | 30.7533 | 30.6194 |
| 0 | 323.31 | 8.75E+10 | 719 | 32.6777 | 32.6518 | 32.7217 | 32.7851 | 32.3575 |
| 0 | 323.31 | 5.22E+10 | 681 | 31.92 | 31.913 | 31.8907 | 31.9049 | 31.7846 |
| 0 | 149.73 | 7.1E+09 | 185 | 28.9321 | 28.9386 | 28.9062 | 28.8922 | 29.0055 |
| 0 | 323.31 | 3.27E+10 | 568 | 31.0821 | 31.1219 | 31.2375 | 31.1587 | 31.1513 |
| 0 | 323.31 | 3.24E+10 | 607 | 31.0201 | 31.039 | 31.0739 | 31.0162 | 31.3919 |
| 0 | 323.31 | 1.43E+11 | 899 | 33.2857 | 33.2873 | 33.4105 | 33.3602 | 33.1762 |
| 0 | 323.31 | 3.89E+10 | 491 | 31.6852 | 31.6579 | 31.2348 | 31.2688 | 31.5408 |
| 0 | 248.54 | 1.29E+10 | 322 | 29.9119 | 29.8847 | 29.7921 | 29.7934 | 29.9252 |
| 0 | 323.31 | 3.96E+11 | 1335 | 34.7949 | 34.7698 | 34.8134 | 34.8054 | 34.6423 |
| 0 | 323.31 | 1.03E+11 | 743 | 32.8123 | 32.8116 | 32.8761 | 32.9242 | 32.7907 |
| 0 | 323.31 | 2.32E+10 | 557 | 30.6488 | 30.674 | 30.7469 | 30.7566 | 30.4397 |
| 0 | 323.31 | 1.86E+10 | 408 | 30.4052 | 30.3866 | 30.4406 | 30.5164 | 30.2525 |
| 0 | 323.31 | 9.81E+10 | 750 | 32.7108 | 32.6737 | 32.8219 | 32.7513 | 32.9095 |
| 0 | 323.31 | 9.44E+10 | 708 | 32.71 | 32.6804 | 32.8708 | 32.8049 | 32.7171 |
| 0 | 323.31 | 5.4E+10 | 714 | 31.866 | 31.9052 | 31.774 | 31.8556 | 31.9811 |
| 0 | 323.31 | 2.24E+11 | 1132 | 33.9138 | 33.9157 | 34.024 | 33.9676 | 33.92 |
| 0 | 323.31 | 8.02E+10 | 680 | 32.4828 | 32.4322 | 32.6234 | 32.5947 | 32.4011 |
| 0 | 323.31 | 2.35E+10 | 426 | 30.838 | 30.8274 | 30.5551 | 30.5459 | 30.7933 |
| 0 | 121.4 | 3.88E+09 | 129 | 27.8457 | 27.8857 | 27.8838 | 27.96 | 28.2641 |
| 0 | 310.18 | 2.24E+10 | 446 | 30.5647 | 30.5585 | 30.5917 | 30.5523 | 30.7367 |
| 0 | 154.8 | 6.64E+09 | 256 | 28.9061 | 28.7552 | 28.9319 | 28.9795 | 28.803 |
| 0 | 323.31 | 1.7E+10 | 419 | 30.234 | 30.2262 | 30.3414 | 30.3471 | 30.2235 |
| 0 | 198.65 | 7.41E+09 | 293 | 28.9524 | 28.8975 | 28.9543 | 28.9838 | 29.0056 |
| 0 | 323.31 | 2.52E+10 | 491 | 30.7654 | 30.7355 | 30.7213 | 30.671 | 30.8772 |
| 0 | 323.31 | 2.42E+10 | 426 | 30.5998 | 30.5621 | 30.4989 | 30.486 | 30.8729 |
| 0 | 323.31 | 6.29E+10 | 667 | 32.1293 | 32.0994 | 32.0969 | 32.1166 | 32.095 |
| 0 | 323.31 | 2.8E+11 | 1049 | 34.2644 | 34.2388 | 34.2761 | 34.3411 | 34.2306 |
| 0 | 216.18 | 9.85E+09 | 311 | 29.4238 | 29.4053 | 29.3983 | 29.4258 | 29.4877 |
| 0 | 323.31 | 2.35E+11 | 1136 | 34.0096 | 34.0265 | 34.0509 | 33.9998 | 33.9947 |
| 0 | 252.28 | 1.47E+10 | 368 | 30.2195 | 30.1269 | 29.9516 | 29.8944 | 30.0142 |
| 0 | 323.31 | 5.27E+10 | 514 | 31.7981 | 31.8196 | 31.7356 | 31.7856 | 31.8646 |
| 0 | 98.703 | 5.07E+09 | 135 | 28.4301 | 28.2862 | 28.4241 | 28.3984 | 28.4773 |
| 0 | 296.79 | 1.02E+10 | 340 | 29.4746 | 29.4768 | 29.5423 | 29.6038 | 29.3872 |
| 0 | 323.31 | 2.4E+10 | 429 | 30.6642 | 30.7633 | 30.6779 | 30.7283 | 30.8569 |
| 0 | 323.31 | 1.4E+11 | 797 | 33.2743 | 33.2467 | 33.3165 | 33.3344 | 33.2191 |
| 0 | 273.41 | 8.25E+09 | 283 | 29.0221 | 29.0308 | 29.3296 | 29.3819 | 29.2168 |
| 0 | 323.31 | 2.02E+10 | 408 | 30.5748 | 30.5452 | 30.3697 | 30.3921 | 30.5184 |
| 0 | 323.31 | 1.56E+11 | 762 | 33.3616 | 33.327 | 33.442 | 33.3752 | 33.3557 |
| 0 | 323.31 | 1.33E+11 | 722 | 33.2255 | 33.2386 | 33.2909 | 33.2739 | 33.2507 |
| 0 | 323.31 | 4.02E+10 | 569 | 31.5821 | 31.5567 | 31.5549 | 31.532 | 31.517 |
| 0 | 323.31 | 3.73E+11 | 1477 | 34.6803 | 34.6849 | 34.649 | 34.7176 | 34.617 |
| 0 | 323.31 | 3.16E+11 | 2000 | 34.4389 | 34.4091 | 34.4425 | 34.5031 | 34.3441 |
| 0 | 323.31 | 9.99E+10 | 695 | 32.8484 | 32.8583 | 32.7887 | 32.8164 | 32.8524 |
| 0 | 323.31 | 5.12E+11 | 1716 | 35.1073 | 35.1217 | 35.1687 | 35.1846 | 35.2096 |

| | | | | | | | | |
|---|---|---|---|---|---|---|---|---|
| 0 | 323.31 | 2.98E+10 | 428 | 31.1845 | 31.1739 | 30.945 | 31.0069 | 31.0609 |
| 0 | 323.31 | 4.68E+10 | 612 | 31.7433 | 31.6828 | 31.6981 | 31.7866 | 31.5095 |
| 0 | 107.49 | 3.78E+09 | 155 | 27.8573 | 27.7708 | 27.972 | 27.9992 | 28.2067 |
| 0 | 323.31 | 1.86E+10 | 413 | 30.3489 | 30.3946 | 30.401 | 30.3761 | 30.2784 |
| 0 | 323.31 | 1.02E+11 | 613 | 33.1591 | 33.148 | 32.7375 | 32.706 | 32.827 |
| 0 | 323.31 | 1.87E+11 | 927 | 33.7676 | 33.7235 | 33.6669 | 33.6666 | 33.7645 |
| 0 | 323.31 | 3.88E+11 | 1216 | 34.6748 | 34.7227 | 34.7876 | 34.7161 | 34.7049 |
| 0 | 296.54 | 1.33E+10 | 380 | 29.8602 | 29.8472 | 29.7745 | 29.741 | 29.9862 |
| 0 | 258.6 | 1.07E+10 | 284 | 29.5006 | 29.5354 | 29.6 | 29.5109 | 29.4255 |
| 0 | 323.31 | 1.25E+10 | 370 | 29.6594 | 29.6487 | 29.7022 | 29.7053 | 29.7827 |
| 0 | 323.31 | 1.99E+10 | 401 | 30.3161 | 30.3705 | 30.2949 | 30.3922 | 30.4898 |
| 0 | 323.31 | 6.72E+10 | 680 | 32.166 | 32.1141 | 32.2118 | 32.2418 | 32.2105 |
| 0 | 174.97 | 6.44E+09 | 207 | 28.7926 | 28.7491 | 28.9042 | 28.9699 | 28.7574 |
| 0 | 323.31 | 4.36E+10 | 508 | 31.5482 | 31.584 | 31.6905 | 31.6071 | 31.6465 |
| 0 | 323.31 | 3.36E+10 | 493 | 31.1215 | 31.0709 | 31.1721 | 31.234 | 31.2892 |
| 0 | 323.31 | 2.38E+11 | 876 | 34.342 | 34.3545 | 33.7511 | 33.7512 | 34.2264 |
| 0 | 323.31 | 1.79E+11 | 1216 | 33.7199 | 33.6889 | 33.4716 | 33.522 | 33.6466 |
| 0 | 302.01 | 1.29E+10 | 318 | 29.7989 | 29.8159 | 29.7196 | 29.8359 | 29.8545 |
| 0 | 323.31 | 2.84E+10 | 435 | 30.7996 | 30.8124 | 30.9982 | 31.0153 | 31.0287 |
| 0 | 323.31 | 5.31E+10 | 542 | 32.0583 | 32.106 | 31.7861 | 31.7094 | 31.9367 |
| 0 | 214.98 | 1.9E+10 | 397 | 30.3866 | 30.3758 | 30.3458 | 30.3921 | 30.3752 |
| 0 | 100.26 | 4.89E+09 | 176 | 28.2185 | 28.4225 | 28.2502 | 28.1892 | 28.482 |
| 0 | 286.77 | 2.34E+10 | 458 | 30.7155 | 30.707 | 30.7996 | 30.7613 | 30.678 |
| 0 | 170.06 | 7.37E+09 | 238 | 29.1169 | 29.1509 | 29.1619 | 29.1684 | 28.7873 |
| 0 | 323.31 | 1.62E+11 | 980 | 33.483 | 33.5152 | 33.5763 | 33.6075 | 33.3815 |
| 0 | 153.5 | 5.74E+09 | 189 | 28.6875 | 28.71 | 28.6711 | 28.7254 | 28.6472 |
| 0 | 318.38 | 1.79E+10 | 381 | 30.2664 | 30.286 | 30.1858 | 30.1736 | 30.1766 |
| 0 | 323.31 | 8E+09 | 264 | 29.1046 | 29.0641 | 29.1428 | 29.198 | 29.0346 |
| 0 | 323.31 | 4.75E+10 | 509 | 31.6887 | 31.6658 | 31.6866 | 31.7476 | 31.5997 |
| 0 | 262.4 | 1.44E+10 | 396 | 29.9195 | 29.8471 | 29.9273 | 29.9136 | 30.016 |
| 0 | 323.31 | 1.45E+11 | 932 | 33.2928 | 33.2811 | 33.2685 | 33.3624 | 33.3178 |
| 0 | 323.31 | 4.27E+10 | 491 | 31.549 | 31.5664 | 31.5849 | 31.5345 | 31.5721 |
| 0 | 250.35 | 1.47E+10 | 383 | 30.0177 | 30.0602 | 30.1378 | 30.1146 | 29.8572 |
| 0 | 323.31 | 4.41E+10 | 515 | 31.3215 | 31.293 | 31.6292 | 31.5786 | 31.8077 |
| 0 | 203.87 | 1.06E+10 | 304 | 29.5375 | 29.5326 | 29.6535 | 29.6542 | 29.5966 |
| 0 | 323.31 | 2.45E+10 | 360 | 30.6986 | 30.625 | 30.6425 | 30.6444 | 30.94 |
| 0 | 323.31 | 4.35E+10 | 515 | 31.5765 | 31.5854 | 31.5896 | 31.65 | 31.706 |
| 0 | 146.75 | 1.71E+10 | 260 | 30.2163 | 30.2175 | 30.1961 | 30.2517 | 30.3708 |
| 0 | 199.11 | 4.38E+09 | 198 | 28.2148 | 28.2224 | 28.3619 | 28.4172 | 28.2661 |
| 0 | 136.52 | 7.45E+09 | 250 | 29.0375 | 29.0802 | 28.8334 | 28.8175 | 29.0689 |
| 0 | 323.31 | 7.77E+10 | 736 | 32.5445 | 32.5481 | 32.2845 | 32.2485 | 32.6061 |
| 0 | 231.75 | 1.34E+10 | 291 | 29.8988 | 29.9449 | 29.8944 | 29.9982 | 29.862 |
| 0 | 172.35 | 5.01E+09 | 147 | 28.4493 | 28.4661 | 28.3347 | 28.265 | 28.6323 |
| 0 | 323.31 | 3.5E+10 | 478 | 31.2855 | 31.2584 | 31.2545 | 31.3466 | 31.2583 |
| 0 | 323.31 | 3.29E+10 | 505 | 31.1157 | 31.1219 | 31.1599 | 31.2036 | 31.1597 |
| 0 | 188.75 | 9.84E+09 | 288 | 29.3309 | 29.3964 | 29.3839 | 29.361 | 29.433 |
| 0 | 137.14 | 1.01E+10 | 292 | 29.4099 | 29.4467 | 29.6212 | 29.6374 | 29.4867 |
| 0 | 323.31 | 6.87E+10 | 667 | 32.4635 | 32.4124 | 32.1622 | 32.0779 | 32.2435 |

| | | | | | | | | |
|---|---|---|---|---|---|---|---|---|
| 0 | 323.31 | 3.99E+10 | 489 | 31.3585 | 31.3587 | 31.3197 | 31.3138 | 31.5997 |
| 0 | 222.4 | 7.8E+09 | 237 | 29.2374 | 29.2147 | 28.9836 | 29.0061 | 29.201 |
| 0 | 116.68 | 5.26E+09 | 188 | 28.6467 | 28.563 | 28.538 | 28.4479 | 28.6254 |
| 0 | 323.31 | 1.39E+10 | 346 | 29.9702 | 29.9606 | 29.9701 | 30.0659 | 29.8939 |
| 0 | 113.88 | 4.42E+09 | 148 | 28.3192 | 28.241 | 28.1533 | 28.2379 | 28.4345 |
| 0 | 323.31 | 1.53E+11 | 701 | 33.1966 | 33.1893 | 33.3185 | 33.2616 | 33.4233 |
| 0 | 76.107 | 9.79E+09 | 209 | 29.4063 | 29.4289 | 29.4315 | 29.4516 | 29.4785 |
| 0 | 87.144 | 3.93E+09 | 180 | 28.0122 | 28.0261 | 27.9908 | 28.2197 | 28.1431 |
| 0 | 312.27 | 1.46E+10 | 353 | 30.0206 | 29.9951 | 29.991 | 30.0552 | 29.9231 |
| 0 | 140.32 | 4.65E+09 | 209 | 28.2822 | 28.3088 | 28.2283 | 28.2004 | 28.4189 |
| 0 | 140.39 | 1.51E+10 | 343 | 30.101 | 30.1084 | 29.9666 | 29.9489 | 30.2099 |
| 0 | 323.31 | 1.01E+10 | 362 | 29.5405 | 29.5255 | 29.3626 | 29.3296 | 29.7115 |
| 0 | 323.31 | 2.34E+10 | 325 | 31.039 | 31.0282 | 30.6522 | 30.5442 | 30.6785 |
| 0 | 128.34 | 5.71E+09 | 180 | 28.7568 | 28.7062 | 28.6909 | 28.7406 | 28.5865 |
| 0 | 323.31 | 1.37E+11 | 834 | 33.4638 | 33.4351 | 33.1169 | 33.1407 | 33.2199 |
| 0 | 323.31 | 9.78E+10 | 677 | 32.9439 | 32.9146 | 32.7713 | 32.7327 | 32.6863 |
| 0 | 253.63 | 3.53E+10 | 459 | 31.2535 | 31.2497 | 31.2775 | 31.2887 | 31.2647 |
| 0 | 267.93 | 2.34E+10 | 396 | 30.6682 | 30.6971 | 30.7029 | 30.7303 | 30.6474 |
| 0 | 188.56 | 8.46E+09 | 229 | 29.2445 | 29.196 | 29.1611 | 29.1672 | 29.231 |
| 0 | 323.31 | 2.78E+10 | 384 | 31.0252 | 31.0582 | 30.8622 | 30.9094 | 30.9229 |
| 0 | 129.45 | 7.11E+09 | 262 | 29.0468 | 29.073 | 29.0505 | 29.0769 | 28.765 |
| 0 | 299.04 | 7.53E+09 | 243 | 29.0138 | 29.0654 | 29.0555 | 29.1036 | 29.2063 |
| 0 | 323.31 | 2.73E+11 | 1395 | 34.2651 | 34.2191 | 34.222 | 34.2238 | 34.1776 |
| 0 | 300.19 | 9.88E+09 | 267 | 29.4922 | 29.5292 | 29.4963 | 29.4622 | 29.434 |
| 0 | 323.31 | 2.68E+11 | 1134 | 34.196 | 34.198 | 34.2319 | 34.335 | 34.1561 |
| 0 | 323.31 | 1.53E+10 | 339 | 30.0048 | 29.9542 | 29.9947 | 30.0454 | 30.1269 |
| 0 | 323.31 | 3.54E+10 | 378 | 31.2418 | 31.2607 | 31.2543 | 31.2511 | 31.2114 |
| 0 | 304.19 | 1.21E+11 | 650 | 33.0097 | 32.9921 | 33.0506 | 33.0441 | 33.0492 |
| 0 | 323.31 | 1.28E+12 | 1468 | 36.6819 | 36.6636 | 36.6958 | 36.6458 | 36.3866 |
| 0 | 175.52 | 8.33E+09 | 236 | 29.1397 | 29.0634 | 29.1203 | 29.1246 | 29.273 |
| 0 | 323.31 | 3.05E+10 | 582 | 31.0067 | 31.0425 | 31.1271 | 31.1516 | 31.0167 |
| 0 | 194.61 | 3.47E+10 | 424 | 31.4644 | 31.4225 | 31.5997 | 31.4815 | 31.0656 |
| 0 | 200.57 | 9.84E+09 | 362 | 29.3685 | 29.3374 | 29.3207 | 29.4114 | 29.4468 |
| 0 | 323.31 | 1.39E+11 | 845 | 33.2077 | 33.1124 | 33.2953 | 33.2635 | 33.3003 |
| 0 | 314.37 | 3.29E+10 | 414 | 31.0891 | 31.1006 | 31.3302 | 31.3231 | 31.1102 |
| 0 | 113.03 | 4.61E+09 | 164 | 28.6257 | 28.5762 | 28.1857 | 28.1164 | 28.4381 |
| 0 | 171.54 | 7.76E+09 | 238 | 29.0286 | 29.029 | 29.0594 | 29.1685 | 29.1476 |
| 0 | 323.31 | 4.22E+10 | 522 | 31.5852 | 31.5581 | 31.5875 | 31.6225 | 31.476 |
| 0 | 323.31 | 2.73E+10 | 432 | 30.89 | 30.9389 | 30.7172 | 30.7668 | 30.9571 |
| 0 | 200.29 | 1.38E+10 | 282 | 29.8807 | 29.8163 | 29.8683 | 29.8951 | 30.1347 |
| 0 | 149.26 | 1.31E+10 | 303 | 29.7657 | 29.8041 | 29.7721 | 29.816 | 29.8489 |
| 0 | 323.31 | 6.35E+10 | 619 | 32.5072 | 32.4574 | 32.1988 | 32.1639 | 31.8764 |
| 0 | 323.31 | 5.44E+10 | 391 | 32.0704 | 32.0429 | 31.7165 | 31.8072 | 31.9882 |
| 0 | 291.24 | 1.86E+10 | 385 | 30.3549 | 30.3037 | 30.5539 | 30.4672 | 30.3606 |
| 0 | 213.28 | 8.85E+09 | 277 | 29.2505 | 29.3104 | 29.3352 | 29.3966 | 29.2914 |
| 0 | 199.29 | 1.29E+10 | 345 | 29.694 | 29.7178 | 29.713 | 29.67 | 30.0428 |
| 0 | 137.29 | 6.87E+09 | 139 | 28.9107 | 28.9889 | 28.8859 | 28.8032 | 29.0002 |
| 0 | 194.73 | 7.06E+09 | 236 | 28.8859 | 28.8631 | 29.0125 | 28.972 | 28.9875 |

| | | | | | | | | |
|---|---|---|---|---|---|---|---|---|
| 0 | 143.53 | 8.8E+09 | 281 | 29.2009 | 29.0651 | 29.1557 | 29.0462 | 29.392 |
| 0 | 323.31 | 6.73E+10 | 522 | 32.4833 | 32.4993 | 32.1607 | 32.2081 | 32.2856 |
| 0 | 323.31 | 3.41E+11 | 1072 | 34.8408 | 34.7928 | 34.3709 | 34.3923 | 34.7453 |
| 0 | 323.31 | 3.01E+10 | 467 | 31.0347 | 31.0847 | 31.1291 | 31.1161 | 31.1424 |
| 0 | 323.31 | 1.4E+11 | 899 | 33.3435 | 33.3404 | 33.2784 | 33.2219 | 33.341 |
| 0 | 323.31 | 1.25E+11 | 868 | 33.1747 | 33.1811 | 33.1662 | 33.1897 | 32.9164 |
| 0 | 266.93 | 4.02E+10 | 441 | 31.4563 | 31.4181 | 31.5518 | 31.5755 | 31.4594 |
| 0 | 169.68 | 9.22E+09 | 252 | 29.3589 | 29.3436 | 29.2089 | 29.264 | 29.48 |
| 0 | 155.11 | 7.8E+09 | 266 | 29.0127 | 29.1273 | 29.3427 | 29.1932 | 29.1049 |
| 0 | 323.31 | 6.51E+10 | 627 | 32.1312 | 32.0843 | 32.1685 | 32.1749 | 32.0346 |
| 0 | 323.31 | 1.31E+11 | 807 | 33.1619 | 33.107 | 33.1943 | 33.2023 | 33.0937 |
| 0 | 96.144 | 6.79E+09 | 246 | 28.8317 | 28.8208 | 28.8232 | 28.8161 | 28.8908 |
| 0 | 243.27 | 8.8E+09 | 218 | 29.3291 | 29.3379 | 29.3489 | 29.2594 | 29.4122 |
| 0 | 169.46 | 8.31E+09 | 213 | 29.0921 | 29.0717 | 29.1566 | 29.0229 | 29.1913 |
| 0 | 323.31 | 1.5E+11 | 982 | 33.2988 | 33.305 | 33.2936 | 33.3445 | 33.3459 |
| 0 | 323.31 | 2.36E+10 | 357 | 30.697 | 30.6501 | 30.6091 | 30.5425 | 30.7819 |
| 0 | 306.63 | 1.41E+10 | 314 | 29.8647 | 29.7776 | 29.9975 | 29.9881 | 29.9335 |
| 0 | 221.48 | 2.32E+10 | 338 | 30.6791 | 30.6531 | 30.5137 | 30.5211 | 30.7727 |
| 0 | 114.06 | 7.89E+09 | 271 | 29.0273 | 29.1132 | 29.2146 | 29.3287 | 29.2763 |
| 0 | 141.41 | 7.88E+09 | 258 | 29.0254 | 28.9901 | 29.1466 | 29.1167 | 29.2073 |
| 0 | 136.58 | 5.83E+09 | 207 | 28.7199 | 28.7032 | 28.7468 | 28.7618 | 28.6391 |
| 0 | 323.31 | 2.78E+10 | 341 | 30.9364 | 30.9356 | 30.9635 | 30.9714 | 30.9285 |
| 0 | 130.34 | 3.34E+09 | 166 | 27.9579 | 27.8741 | 27.8765 | 28.0037 | 27.897 |
| 0 | 323.31 | 4.62E+10 | 445 | 31.7272 | 31.7013 | 31.6703 | 31.6314 | 31.6491 |
| 0 | 323.31 | 4.77E+10 | 430 | 31.755 | 31.7216 | 31.6842 | 31.7399 | 31.5233 |
| 0 | 57.904 | 3.99E+09 | 142 | 28.0494 | 27.9482 | 28.1037 | 28.2287 | 28.2348 |
| 0 | 323.31 | 1.01E+10 | 274 | 29.3967 | 29.3668 | 29.2431 | 29.3095 | 29.5648 |
| 0 | 213.56 | 9.99E+09 | 251 | 29.8209 | 29.7768 | 29.4584 | 29.4232 | 29.4728 |
| 0 | 323.31 | 2.38E+10 | 324 | 30.7636 | 30.647 | 30.8005 | 30.8092 | 30.6064 |
| 0 | 195.41 | 6.26E+09 | 227 | 28.7903 | 28.6791 | 28.8466 | 28.838 | 28.7629 |
| 0 | 52.726 | 2.25E+09 | 120 | 27.3042 | 27.1986 | 27.2628 | 27.439 | 27.4656 |
| 0 | 323.31 | 6.38E+10 | 572 | 32.1762 | 32.1829 | 32.1877 | 32.1697 | 32.1135 |
| 0 | 323.31 | 5.25E+10 | 509 | 32.1505 | 32.123 | 31.8009 | 31.7897 | 31.8115 |
| 0 | 323.31 | 7.94E+10 | 528 | 32.4693 | 32.4935 | 32.4694 | 32.5466 | 32.4424 |
| 0 | 181.87 | 7.91E+09 | 241 | 29.1447 | 29.0919 | 29.2577 | 29.0912 | 29.2019 |
| 0 | 183.63 | 1.02E+10 | 260 | 29.4287 | 29.4646 | 29.739 | 29.6266 | 29.4855 |
| 0 | 323.31 | 1.2E+10 | 291 | 29.829 | 29.7337 | 29.8756 | 29.7752 | 29.6203 |
| 0 | 323.31 | 9.33E+09 | 284 | 29.3826 | 29.3938 | 29.2223 | 29.1152 | 29.5404 |
| 0 | 323.31 | 2.99E+10 | 411 | 31.1422 | 31.064 | 30.9209 | 30.9285 | 31.155 |
| 0 | 179.61 | 1.08E+10 | 192 | 29.502 | 29.5609 | 29.4397 | 29.4949 | 29.5972 |
| 0 | 323.31 | 1.33E+11 | 638 | 33.131 | 33.0919 | 33.1975 | 33.2386 | 33.0403 |
| 0 | 168.61 | 1.78E+10 | 336 | 30.0547 | 30.1747 | 30.2289 | 30.2165 | 30.2438 |
| 0 | 79.774 | 4.28E+09 | 175 | 28.1186 | 28.1961 | 27.9974 | 28.1005 | 28.2024 |
| 0 | 323.31 | 2.55E+10 | 388 | 30.6935 | 30.7427 | 30.8887 | 30.853 | 30.7196 |
| 0 | 175.73 | 5.15E+09 | 219 | 28.3921 | 28.3744 | 28.5838 | 28.5616 | 28.4485 |
| 0 | 323.31 | 2.02E+10 | 359 | 30.379 | 30.3545 | 30.587 | 30.4814 | 30.3822 |
| 0 | 323.31 | 1.11E+11 | 620 | 32.8863 | 32.8485 | 33.0207 | 32.9387 | 32.853 |
| 0 | 313.25 | 9.85E+09 | 306 | 29.5679 | 29.4819 | 29.5994 | 29.5245 | 29.4549 |

| | | | | | | | | |
|---|---|---|---|---|---|---|---|---|
| 0 | 323.31 | 2.67E+10 | 378 | 30.897 | 30.8364 | 30.9452 | 30.9595 | 30.692 |
| 0 | 323.31 | 4.08E+10 | 374 | 31.7445 | 31.8066 | 31.5244 | 31.5025 | 31.5334 |
| 0 | 298.92 | 1.96E+10 | 286 | 30.4215 | 30.4267 | 30.2864 | 30.2449 | 30.4683 |
| 0 | 112.43 | 4.5E+09 | 108 | 28.124 | 28.1381 | 28.2321 | 28.281 | 28.1426 |
| 0 | 323.31 | 4.33E+10 | 347 | 31.5817 | 31.6116 | 31.682 | 31.6995 | 31.5923 |
| 0 | 323.31 | 1.73E+10 | 343 | 30.241 | 30.223 | 30.359 | 30.3625 | 30.1343 |
| 0 | 323.31 | 2.31E+10 | 414 | 30.6087 | 30.6194 | 30.7404 | 30.7199 | 30.6706 |
| 0 | 294.24 | 8.27E+09 | 222 | 28.877 | 28.9153 | 28.6907 | 28.8619 | 29.3868 |
| 0 | 202.17 | 6.91E+09 | 242 | 28.8967 | 28.7701 | 29.0323 | 29.015 | 28.9258 |
| 0 | 168.21 | 1.34E+10 | 294 | 29.7905 | 29.8191 | 29.8131 | 29.7853 | 29.9378 |
| 0 | 223.43 | 1.54E+10 | 311 | 30.0436 | 30.1159 | 30.0045 | 30.0271 | 30.2765 |
| 0 | 323.31 | 9.95E+09 | 321 | 29.5189 | 29.4525 | 29.5422 | 29.5039 | 29.2932 |
| 0 | 323.31 | 1.64E+10 | 309 | 30.2301 | 30.1931 | 30.253 | 30.2103 | 30.0668 |
| 0 | 245.86 | 6.12E+09 | 202 | 28.8287 | 28.6937 | 28.839 | 28.817 | 28.7264 |
| 0 | 101.78 | 3.97E+09 | 114 | 28.029 | 28.0799 | 28.186 | 28.192 | 28.1866 |
| 0 | 148.72 | 1.55E+10 | 351 | 30.0285 | 30.0036 | 30.0739 | 30.101 | 30.226 |
| 0 | 135.3 | 5.54E+09 | 153 | 28.4651 | 28.4251 | 28.5361 | 28.6318 | 28.7917 |
| 0 | 143.31 | 1.05E+10 | 269 | 29.5914 | 29.5504 | 29.5245 | 29.4893 | 29.6288 |
| 0 | 265.38 | 1.21E+10 | 330 | 29.7007 | 29.7134 | 29.6879 | 29.6714 | 29.8864 |
| 0 | 323.31 | 2.8E+10 | 361 | 30.9053 | 30.9539 | 30.8753 | 30.8616 | 31.0518 |
| 0 | 323.31 | 3.01E+10 | 365 | 31.2414 | 31.1849 | 31.0086 | 30.9942 | 31.0354 |
| 0 | 323.31 | 4.09E+10 | 482 | 31.6129 | 31.5354 | 31.7 | 31.6869 | 31.2465 |
| 0 | 255.2 | 1.27E+10 | 320 | 29.843 | 29.8387 | 29.9449 | 29.9168 | 29.7901 |
| 0 | 323.31 | 1.61E+11 | 545 | 33.4274 | 33.387 | 33.397 | 33.4633 | 33.6211 |
| 0 | 323.31 | 1.99E+10 | 342 | 30.3835 | 30.4217 | 30.4883 | 30.4977 | 30.3007 |
| 0 | 323.31 | 1.73E+10 | 295 | 30.2787 | 30.2544 | 30.3204 | 30.2278 | 30.1601 |
| 0 | 223.56 | 2.17E+10 | 314 | 30.6515 | 30.6399 | 30.6081 | 30.5763 | 30.6359 |
| 0 | 149.51 | 1.56E+10 | 280 | 30.07 | 30.026 | 29.9293 | 29.8455 | 30.2366 |
| 0 | 125.94 | 1.3E+10 | 234 | 29.9467 | 29.9246 | 29.9964 | 29.9025 | 29.6354 |
| 0 | 143.72 | 8.95E+09 | 209 | 29.2982 | 29.2957 | 29.2999 | 29.2964 | 29.2171 |
| 0 | 323.31 | 9.46E+10 | 670 | 33.0927 | 33.0189 | 32.7593 | 32.7002 | 32.6184 |
| 0 | 323.31 | 1.1E+11 | 695 | 32.8806 | 32.9102 | 33.0157 | 32.9697 | 32.8184 |
| 0 | 273.55 | 2.37E+10 | 363 | 30.6971 | 30.7407 | 30.8126 | 30.711 | 30.7653 |
| 0 | 316.96 | 3.53E+10 | 398 | 31.626 | 31.5951 | 31.2921 | 31.2208 | 31.1713 |
| 0 | 323.31 | 4.23E+11 | 785 | 34.921 | 34.9451 | 35.0455 | 34.9986 | 34.7846 |
| 0 | 323.31 | 7.41E+10 | 654 | 32.3853 | 32.3705 | 32.4135 | 32.4499 | 32.2215 |
| 0 | 168.77 | 5.27E+09 | 174 | 28.8364 | 28.8031 | 28.5038 | 28.4226 | 28.4415 |
| 0 | 308.74 | 1.11E+10 | 319 | 29.6136 | 29.6394 | 29.5675 | 29.5182 | 29.679 |
| 0 | 323.31 | 7.62E+11 | 1772 | 35.7077 | 35.6795 | 35.7289 | 35.7292 | 35.616 |
| 0 | 157.16 | 1.47E+10 | 295 | 29.9775 | 29.9271 | 30.049 | 29.9255 | 30.0251 |
| 0 | 320.37 | 1.2E+10 | 247 | 29.6922 | 29.6955 | 29.8155 | 29.6315 | 29.7577 |
| 0 | 323.31 | 2.33E+10 | 328 | 30.735 | 30.686 | 30.7679 | 30.8008 | 30.577 |
| 0 | 323.31 | 1.07E+11 | 454 | 32.8892 | 32.9322 | 32.9919 | 32.9862 | 32.7232 |
| 0 | 110.57 | 1.5E+10 | 306 | 30.0426 | 30.0032 | 29.8315 | 29.9096 | 30.213 |
| 0 | 130.55 | 8.17E+09 | 270 | 29.2657 | 29.2425 | 28.9544 | 29.0001 | 29.3322 |
| 0 | 152.14 | 4.6E+09 | 164 | 28.4252 | 28.3547 | 28.3064 | 28.1749 | 28.4201 |
| 0 | 323.31 | 2.59E+10 | 391 | 30.9282 | 30.8777 | 30.9594 | 30.9534 | 30.7758 |
| 0 | 292.47 | 2.96E+10 | 339 | 31.0091 | 30.9771 | 30.9732 | 30.9549 | 30.9467 |

| | | | | | | | | |
|---|---|---|---|---|---|---|---|---|
| 0 | 323.31 | 1.26E+11 | 688 | 33.0414 | 33.0167 | 33.1511 | 33.17 | 33.0181 |
| 0 | 323.31 | 4.49E+10 | 415 | 31.6291 | 31.5507 | 31.6369 | 31.6045 | 31.5056 |
| 0 | 323.31 | 3.4E+10 | 381 | 31.1814 | 31.1372 | 31.2766 | 31.1974 | 31.1172 |
| 0 | 183.44 | 6.97E+09 | 227 | 28.836 | 28.8184 | 28.8238 | 28.8247 | 28.9973 |
| 0 | 323.31 | 1.24E+10 | 225 | 29.6873 | 29.751 | 29.6683 | 29.685 | 29.7937 |
| 0 | 264.46 | 1.03E+10 | 228 | 29.499 | 29.4576 | 29.4083 | 29.4679 | 29.528 |
| 0 | 323.31 | 1.03E+11 | 443 | 32.936 | 32.95 | 32.9773 | 32.9185 | 32.9037 |
| 0 | 323.31 | 5.25E+10 | 446 | 31.8245 | 31.8073 | 31.946 | 31.8717 | 31.7797 |
| 0 | 274.75 | 1.01E+10 | 189 | 29.3953 | 29.4001 | 29.2795 | 29.4111 | 29.5021 |
| 0 | 236.2 | 1.19E+10 | 270 | 29.5841 | 29.6458 | 29.6455 | 29.6087 | 29.7505 |
| 0 | 323.31 | 3.82E+10 | 287 | 31.463 | 31.445 | 31.4695 | 31.392 | 31.4421 |
| 0 | 133.54 | 6.36E+09 | 179 | 28.7984 | 28.7803 | 28.7621 | 28.8343 | 28.6879 |
| 0 | 130.34 | 1.01E+10 | 247 | 29.8634 | 29.8466 | 29.4381 | 29.4627 | 29.4676 |
| 0 | 230.8 | 9.06E+09 | 246 | 29.5936 | 29.6066 | 29.2837 | 29.2822 | 29.4181 |
| 0 | 323.31 | 1.81E+10 | 367 | 30.0012 | 30.1029 | 30.2961 | 30.2556 | 30.4407 |
| 0 | 323.31 | 2.86E+10 | 501 | 30.8574 | 30.8451 | 31.0821 | 31.0548 | 31.0093 |
| 0 | 323.31 | 3.12E+10 | 412 | 30.9951 | 30.9514 | 30.9787 | 31.1173 | 31.1351 |
| 0 | 323.31 | 7.99E+10 | 409 | 32.6317 | 32.631 | 32.4415 | 32.405 | 32.2273 |
| 0 | 323.31 | 1.62E+11 | 556 | 33.7747 | 33.7476 | 33.3665 | 33.3768 | 33.5775 |
| 0 | 214.98 | 7.62E+09 | 233 | 29.1246 | 29.0947 | 29.1206 | 29.0709 | 28.9958 |
| 0 | 244.43 | 2.92E+10 | 349 | 31.0785 | 31.1081 | 31.0651 | 31.1041 | 30.977 |
| 0 | 136.48 | 9.96E+09 | 253 | 29.3522 | 29.4303 | 29.5206 | 29.5799 | 29.4959 |
| 0 | 185.71 | 1.3E+10 | 233 | 29.8354 | 29.7763 | 29.7528 | 29.7151 | 29.904 |
| 0 | 60.34 | 6.22E+09 | 211 | 28.8712 | 28.9517 | 28.6685 | 28.6518 | 28.8503 |
| 0 | 214.3 | 8.61E+09 | 193 | 29.2231 | 29.3004 | 29.3674 | 29.4208 | 29.1376 |
| 0 | 323.31 | 3.56E+11 | 988 | 34.5994 | 34.6039 | 34.6475 | 34.6156 | 34.6579 |
| 0 | 323.31 | 2.8E+10 | 404 | 30.9626 | 30.9133 | 31.0386 | 31.0566 | 30.9847 |
| 0 | 219.29 | 1.77E+10 | 302 | 30.2792 | 30.2155 | 30.3705 | 30.3268 | 30.3212 |
| 0 | 121.44 | 4.12E+09 | 177 | 28.121 | 28.0981 | 28.0809 | 27.9349 | 28.3344 |
| 0 | 323.31 | 1.9E+10 | 328 | 30.4111 | 30.3904 | 30.3386 | 30.3289 | 30.5244 |
| 0 | 137.3 | 6.48E+09 | 200 | 28.8844 | 28.8661 | 28.6541 | 28.5918 | 29.1973 |
| 0 | 316.51 | 1.34E+10 | 303 | 29.8719 | 29.8017 | 29.9548 | 29.8893 | 29.7585 |
| 0 | 323.31 | 3.52E+10 | 415 | 31.073 | 31.1245 | 31.2008 | 31.1485 | 31.2497 |
| 0 | 192.33 | 4.19E+09 | 150 | 28.3276 | 28.2835 | 28.1798 | 28.2615 | 28.3263 |
| 0 | 98.866 | 1.09E+10 | 259 | 29.5951 | 29.5545 | 29.58 | 29.576 | 29.5587 |
| 0 | 323.31 | 1.51E+11 | 675 | 33.3823 | 33.3797 | 33.4821 | 33.4459 | 33.0974 |
| 0 | 50.846 | 4.65E+09 | 177 | 28.2184 | 28.3564 | 28.4345 | 28.4347 | 28.3634 |
| 0 | 282.03 | 1.44E+10 | 259 | 29.9753 | 29.9354 | 29.9858 | 29.9845 | 29.9215 |
| 0 | 252.91 | 2.93E+10 | 374 | 31.0178 | 30.989 | 31.057 | 31.1094 | 30.896 |
| 0 | 185.25 | 3.8E+10 | 497 | 31.4596 | 31.4542 | 31.3516 | 31.3533 | 31.5677 |
| 0 | 323.31 | 1.21E+10 | 323 | 29.7096 | 29.6691 | 29.4893 | 29.5535 | 29.8227 |
| 0 | 71.74 | 3.55E+09 | 109 | 27.8278 | 27.9113 | 27.877 | 28.007 | 27.9929 |
| 0 | 205.55 | 2.25E+10 | 322 | 30.5176 | 30.5306 | 30.3897 | 30.4209 | 30.7582 |
| 0 | 57.753 | 2.78E+09 | 122 | 27.395 | 27.2118 | 27.6246 | 27.5996 | 27.7705 |
| 0 | 132.89 | 4.2E+09 | 164 | 28.0742 | 28.0077 | 28.1504 | 28.0741 | 28.2728 |
| 0 | 126.77 | 3.48E+09 | 143 | 27.9324 | 27.8923 | 28.0487 | 28.0156 | 27.9009 |
| 0 | 323.31 | 5.26E+10 | 540 | 32.0552 | 32.0504 | 31.776 | 31.7738 | 31.8965 |
| 0 | 323.31 | 1.52E+11 | 650 | 33.3342 | 33.4035 | 33.4474 | 33.398 | 33.5805 |

| | | | | | | | | |
|---|---|---|---|---|---|---|---|---|
| 0 | 56.556 | 2.85E+09 | 70 | 27.6348 | 27.6339 | 27.494 | 27.4834 | 27.7927 |
| 0 | 323.31 | 2.11E+10 | 303 | 30.6884 | 30.6758 | 30.4058 | 30.4173 | 30.6919 |
| 0 | 235.59 | 1.05E+10 | 262 | 29.7503 | 29.6989 | 29.52 | 29.486 | 29.5907 |
| 0 | 251.86 | 6.79E+09 | 242 | 28.9636 | 28.9664 | 28.6817 | 28.7212 | 29.0048 |
| 0 | 130.69 | 6.54E+09 | 200 | 28.8982 | 28.9498 | 28.9002 | 28.8025 | 28.8634 |
| 0 | 323.31 | 6.29E+09 | 164 | 28.7674 | 28.7642 | 28.805 | 28.6849 | 28.9373 |
| 0 | 310.46 | 2.74E+10 | 367 | 30.9559 | 30.92 | 30.8093 | 30.843 | 31.0229 |
| 0 | 195.56 | 3.17E+10 | 376 | 31.1758 | 31.0945 | 31.2258 | 31.1585 | 31.0796 |
| 0 | 323.31 | 1.73E+11 | 953 | 33.6709 | 33.6881 | 33.3982 | 33.46 | 33.6853 |
| 0 | 87.725 | 5.92E+09 | 187 | 28.5805 | 28.6732 | 28.4524 | 28.4787 | 28.952 |
| 0 | 149.56 | 8.62E+09 | 234 | 29.3592 | 29.425 | 29.1159 | 29.1621 | 29.3168 |
| 0 | 323.31 | 1.66E+10 | 294 | 30.3051 | 30.2218 | 30.2728 | 30.2667 | 30.1645 |
| 0 | 323.31 | 2.31E+11 | 1034 | 34.0368 | 33.9504 | 33.8639 | 33.8303 | 34.071 |
| 0 | 165.36 | 8.31E+09 | 202 | 29.2423 | 29.1204 | 29.2659 | 29.2399 | 29.2389 |
| 0 | 323.31 | 1.58E+10 | 258 | 30.1133 | 30.1831 | 30.0589 | 29.8894 | 30.2177 |
| 0 | 111.46 | 3.67E+09 | 131 | 28.0787 | 28.0968 | 27.9442 | 27.7938 | 28.1371 |
| 0 | 194.43 | 7.85E+09 | 245 | 29.1949 | 29.2021 | 29.0732 | 28.9819 | 29.2072 |
| 0 | 138.06 | 4.74E+09 | 169 | 28.2073 | 28.158 | 28.3902 | 28.3275 | 28.2902 |
| 0 | 323.31 | 1.57E+11 | 623 | 33.3892 | 33.3603 | 33.4996 | 33.5028 | 33.2336 |
| 0 | 323.31 | 4.59E+10 | 566 | 31.5609 | 31.5438 | 31.5779 | 31.4802 | 31.6928 |
| 0 | 108.27 | 1.51E+10 | 246 | 30.1647 | 30.1887 | 29.9382 | 29.9683 | 30.3527 |
| 0 | 237.9 | 6.2E+09 | 194 | 28.5559 | 28.597 | 28.7885 | 28.8735 | 28.7645 |
| 0 | 140.84 | 3.98E+09 | 165 | 28.1774 | 28.1123 | 28.1635 | 28.1775 | 28.0505 |
| 0 | 147.25 | 9.19E+09 | 242 | 29.3392 | 29.4073 | 29.3689 | 29.3382 | 29.215 |
| 0 | 63.917 | 5.96E+09 | 156 | 28.6613 | 28.6152 | 28.6059 | 28.6548 | 28.7567 |
| 0 | 264.1 | 6.95E+09 | 230 | 29.3309 | 29.3039 | 28.9381 | 28.9515 | 28.8856 |
| 0 | 123.72 | 7.36E+09 | 181 | 28.9725 | 29.0325 | 28.9794 | 28.982 | 29.0225 |
| 0 | 143.86 | 5.38E+09 | 149 | 28.5559 | 28.4828 | 28.513 | 28.4204 | 28.6728 |
| 0 | 130.82 | 4.32E+09 | 112 | 28.1864 | 28.1748 | 28.3546 | 28.3481 | 28.1146 |
| 0 | 191.13 | 4.86E+09 | 194 | 28.384 | 28.4845 | 28.2889 | 28.2976 | 28.5074 |
| 0 | 156.68 | 7.21E+09 | 187 | 29.2057 | 29.1816 | 29.0669 | 28.9987 | 29.066 |
| 0 | 45.864 | 2.06E+09 | 96 | 27.2522 | 27.0848 | 27.0885 | 27.0702 | 27.2652 |
| 0 | 323.31 | 1.01E+11 | 619 | 33.169 | 33.1369 | 32.8007 | 32.764 | 32.6713 |
| 0 | 323.31 | 3.08E+10 | 403 | 30.7816 | 30.9166 | 30.7709 | 30.835 | 31.7334 |
| 0 | 323.31 | 2.09E+10 | 407 | 30.5476 | 30.5402 | 30.6111 | 30.5555 | 30.5003 |
| 0 | 163.25 | 6.33E+09 | 191 | 28.8656 | 28.7519 | 28.5884 | 28.5582 | 28.9504 |
| 0 | 323.31 | 2.43E+10 | 334 | 30.7206 | 30.7158 | 30.7674 | 30.8225 | 30.7441 |
| 0 | 155.62 | 1.63E+10 | 290 | 30.2302 | 30.287 | 30.0704 | 30.0948 | 30.1567 |
| 0 | 323.31 | 5.65E+10 | 385 | 31.8805 | 31.9047 | 32.0439 | 32.0515 | 31.9806 |
| 0 | 80.399 | 3.56E+09 | 114 | 27.9372 | 28.0174 | 27.8727 | 27.8599 | 28.0695 |
| 0 | 323.31 | 2.16E+10 | 328 | 30.586 | 30.5321 | 30.664 | 30.658 | 30.6502 |
| 0 | 149.38 | 9.23E+09 | 286 | 29.2873 | 29.3894 | 29.3397 | 29.3778 | 29.4041 |
| 0 | 40.117 | 2.33E+09 | 72 | 27.4877 | 27.5017 | 27.2644 | 27.447 | 27.386 |
| 0 | 64.212 | 3.64E+09 | 148 | 27.9768 | 27.9459 | 27.9439 | 28.2291 | 28.0736 |
| 0 | 323.31 | 5.05E+10 | 338 | 31.8536 | 31.7904 | 31.889 | 31.9314 | 31.7179 |
| 0 | 323.31 | 7.09E+10 | 514 | 32.1141 | 32.1309 | 32.3228 | 32.2717 | 32.2178 |
| 0 | 323.31 | 1.7E+11 | 675 | 33.4144 | 33.4736 | 33.4933 | 33.5559 | 33.4945 |
| 0 | 129.77 | 5.43E+09 | 167 | 28.5411 | 28.5101 | 28.6501 | 28.5757 | 28.3955 |

| | | | | | | | | |
|---|---|---|---|---|---|---|---|---|
| 0 | 231.48 | 2.29E+10 | 267 | 30.6494 | 30.6456 | 30.6546 | 30.6678 | 30.5329 |
| 0 | 323.31 | 1.13E+10 | 270 | 29.6265 | 29.655 | 29.5988 | 29.5938 | 29.5794 |
| 0 | 136.28 | 1.2E+10 | 254 | 29.7737 | 29.8027 | 29.8159 | 29.7615 | 29.8783 |
| 0 | 159.36 | 8.78E+09 | 256 | 29.258 | 29.2678 | 29.2717 | 29.3145 | 29.1012 |
| 0 | 323.31 | 3.37E+10 | 361 | 30.9879 | 31.0427 | 31.3201 | 31.2137 | 30.6551 |
| 0 | 323.31 | 2.7E+10 | 321 | 30.8463 | 30.8927 | 30.9375 | 30.9517 | 30.8831 |
| 0 | 133.68 | 1.23E+10 | 228 | 29.817 | 29.7757 | 29.8261 | 29.9114 | 29.7905 |
| 0 | 99.269 | 2.67E+09 | 91 | 27.4764 | 27.489 | 27.4792 | 27.2083 | 27.6682 |
| 0 | 77.261 | 5.33E+09 | 181 | 28.5363 | 28.5403 | 28.6031 | 28.6428 | 28.5115 |
| 0 | 323.31 | 1.76E+11 | 592 | 33.6957 | 33.6703 | 33.7637 | 33.8431 | 33.4803 |
| 0 | 223.92 | 3.92E+10 | 294 | 31.3943 | 31.4296 | 31.4783 | 31.5175 | 31.3697 |
| 0 | 323.31 | 2.25E+10 | 362 | 30.5451 | 30.6095 | 30.608 | 30.6303 | 30.6428 |
| 0 | 323.31 | 1.92E+11 | 736 | 33.7989 | 33.7622 | 33.8142 | 33.7189 | 33.6308 |
| 0 | 308.32 | 8.2E+09 | 266 | 29.1121 | 29.068 | 29.2274 | 29.1718 | 29.185 |
| 0 | 298.67 | 5.42E+10 | 457 | 31.9363 | 31.9087 | 31.9739 | 31.9726 | 31.8368 |
| 0 | 109.44 | 4.54E+09 | 156 | 28.2645 | 28.3129 | 28.0803 | 28.1402 | 28.4141 |
| 0 | 323.31 | 1.17E+11 | 707 | 32.9861 | 32.9847 | 33.1353 | 33.0947 | 33.0388 |
| 0 | 323.31 | 1.1E+11 | 540 | 32.8986 | 32.8772 | 33.0159 | 33.023 | 32.855 |
| 0 | 84.907 | 4.4E+09 | 124 | 28.4339 | 28.3838 | 28.1198 | 28.0763 | 28.4883 |
| 0 | 323.31 | 7.12E+10 | 477 | 32.4164 | 32.3968 | 32.2235 | 32.1987 | 32.2614 |
| 0 | 65.892 | 4.3E+09 | 170 | 28.1065 | 28.1491 | 28.1275 | 28.0988 | 28.4402 |
| 0 | 119.59 | 1.52E+10 | 180 | 29.9916 | 29.9939 | 30.1606 | 30.0241 | 30.0099 |
| 0 | 323.31 | 4.08E+10 | 517 | 31.4919 | 31.5389 | 31.4732 | 31.5248 | 31.5696 |
| 0 | 207.44 | 7.86E+09 | 240 | 29.1703 | 29.0964 | 28.9062 | 28.9339 | 29.2019 |
| 0 | 323.31 | 1.51E+10 | 243 | 30.1416 | 30.1 | 30.0074 | 29.9828 | 30.0913 |
| 0 | 323.31 | 9.36E+10 | 472 | 32.6679 | 32.7108 | 32.819 | 32.7383 | 32.6822 |
| 0 | 112.37 | 5.27E+09 | 179 | 28.475 | 28.4217 | 28.6984 | 28.5913 | 28.5271 |
| 0 | 254.69 | 6.2E+09 | 215 | 28.6617 | 28.6545 | 28.7732 | 28.7061 | 28.743 |
| 0 | 323.31 | 4.12E+10 | 295 | 31.4382 | 31.4691 | 31.5351 | 31.6115 | 31.3353 |
| 0 | 215.33 | 1.7E+10 | 297 | 30.0549 | 30.0574 | 30.246 | 30.2087 | 30.1881 |
| 0 | 290.89 | 7.82E+09 | 221 | 29.096 | 29.1056 | 29.1518 | 29.1534 | 29.004 |
| 0 | 166.74 | 6.29E+09 | 193 | 29.1949 | 29.2086 | 28.7256 | 28.7431 | 28.8754 |
| 0 | 44.631 | 4.54E+09 | 118 | 28.2041 | 28.1816 | 28.1364 | 28.1324 | 28.3567 |
| 0 | 323.31 | 1.37E+10 | 327 | 30.0099 | 30.0125 | 29.8735 | 29.8989 | 29.7878 |
| 0 | 139.37 | 7.16E+09 | 233 | 28.884 | 28.8526 | 28.8801 | 28.8716 | 29.0522 |
| 0 | 323.31 | 6.62E+10 | 365 | 32.2899 | 32.2608 | 32.221 | 32.1539 | 32.3474 |
| 0 | 323.31 | 1.55E+10 | 343 | 30.1276 | 30.1365 | 30.1649 | 30.0966 | 30.0073 |
| 0 | 323.31 | 3.37E+10 | 318 | 31.2825 | 31.2031 | 31.1621 | 31.1986 | 31.2305 |
| 0 | 282.38 | 7.54E+09 | 201 | 28.8846 | 28.908 | 28.8849 | 28.9157 | 29.0701 |
| 0 | 73.173 | 3.12E+09 | 105 | 27.8826 | 27.7313 | 27.8888 | 27.8111 | 27.8921 |
| 0 | 172.62 | 2.11E+10 | 313 | 30.7559 | 30.7648 | 30.4915 | 30.5081 | 30.6331 |
| 0 | 158.2 | 7.76E+09 | 198 | 29.0684 | 29.0533 | 29.2931 | 29.1312 | 28.9424 |
| 0 | 196.17 | 4.39E+09 | 131 | 28.2138 | 28.3394 | 28.1454 | 28.1466 | 28.318 |
| 0 | 179.92 | 1.02E+10 | 228 | 29.4111 | 29.293 | 29.3424 | 29.4307 | 29.6336 |
| 0 | 122.28 | 6.92E+09 | 164 | 28.953 | 28.9436 | 29.1228 | 29.0328 | 29.0643 |
| 0 | 198.87 | 1.57E+10 | 241 | 30.1164 | 30.1269 | 30.2266 | 30.1852 | 29.9349 |
| 0 | 323.31 | 3.85E+10 | 288 | 31.3613 | 31.3525 | 31.4263 | 31.3667 | 31.4868 |
| 0 | 69.585 | 4.51E+09 | 176 | 28.4944 | 28.4956 | 28.358 | 28.3991 | 28.0598 |

| | | | | | | | | |
|---|---|---|---|---|---|---|---|---|
| 0 | 300.01 | 7.03E+09 | 229 | 29.0065 | 28.9079 | 29.0605 | 29.0696 | 28.9938 |
| 0 | 207.63 | 2.26E+10 | 307 | 30.6655 | 30.6614 | 30.8012 | 30.7837 | 30.5442 |
| 0 | 72.673 | 4.27E+09 | 156 | 28.5083 | 28.466 | 27.9854 | 28.0362 | 28.3986 |
| 0 | 260.21 | 1.17E+10 | 275 | 29.7421 | 29.7047 | 29.6996 | 29.7314 | 29.6168 |
| 0 | 153.02 | 1.28E+10 | 252 | 29.7419 | 29.6661 | 29.8779 | 29.9229 | 29.8787 |
| 0 | 323.31 | 7.53E+10 | 490 | 32.3693 | 32.3876 | 32.1441 | 32.0392 | 32.403 |
| 0 | 201.6 | 9.13E+09 | 183 | 29.6182 | 29.7054 | 29.4511 | 29.3787 | 28.9908 |
| 0 | 68.136 | 2.98E+09 | 92 | 27.5926 | 27.7766 | 27.516 | 27.6015 | 27.8606 |
| 0 | 160.89 | 1.03E+10 | 234 | 29.5687 | 29.5794 | 29.5962 | 29.6176 | 29.425 |
| 0 | 246.91 | 3.04E+10 | 313 | 31.0641 | 31.0241 | 31.1657 | 31.1494 | 31.0467 |
| 0 | 323.31 | 3.28E+11 | 595 | 34.5318 | 34.4885 | 34.5965 | 34.5901 | 34.46 |
| 0 | 319.01 | 1.24E+10 | 296 | 29.6918 | 29.7326 | 29.6362 | 29.6241 | 29.6232 |
| 0 | 109.25 | 4.73E+09 | 131 | 28.3378 | 28.2452 | 28.1749 | 28.1162 | 28.3713 |
| 0 | 323.31 | 2.72E+11 | 813 | 33.8328 | 33.8159 | 33.8037 | 33.7982 | 33.6086 |
| 0 | 323.31 | 1.48E+11 | 487 | 33.2799 | 33.2835 | 33.3832 | 33.3955 | 33.2825 |
| 0 | 323.31 | 1.65E+11 | 633 | 33.4689 | 33.4913 | 33.4992 | 33.4651 | 33.4158 |
| 0 | 239.61 | 1.1E+10 | 214 | 29.6033 | 29.5549 | 29.5274 | 29.5461 | 29.6137 |
| 0 | 323.31 | 1.32E+10 | 253 | 29.7596 | 29.7359 | 29.8287 | 29.7403 | 29.9056 |
| 0 | 270.86 | 7.22E+10 | 408 | 32.3594 | 32.3532 | 32.335 | 32.4107 | 32.3494 |
| 0 | 196.84 | 6.79E+09 | 193 | 29.1966 | 29.1225 | 29.0392 | 28.9594 | 28.9902 |
| 0 | 188.72 | 3.84E+09 | 126 | 28.0734 | 28.0267 | 27.9902 | 27.9366 | 28.1221 |
| 0 | 306.41 | 1.55E+10 | 293 | 30.0349 | 30.0554 | 30.187 | 30.2075 | 29.9289 |
| 0 | 322.53 | 1.54E+10 | 212 | 30.1334 | 30.1064 | 30.0785 | 30.1177 | 29.9881 |
| 0 | 68.157 | 4.73E+09 | 144 | 28.4057 | 28.2519 | 28.388 | 28.4176 | 28.3833 |
| 0 | 323.31 | 2.53E+10 | 317 | 30.8108 | 30.7782 | 30.8814 | 30.8405 | 30.6045 |
| 0 | 129.77 | 3.5E+09 | 126 | 27.9916 | 27.8628 | 27.7341 | 27.8276 | 28.0467 |
| 0 | 277.51 | 2.21E+10 | 234 | 30.7204 | 30.6386 | 30.6777 | 30.6395 | 30.4822 |
| 0 | 118.43 | 4.99E+09 | 163 | 28.4201 | 28.5152 | 28.4737 | 28.5051 | 28.4473 |
| 0 | 106.02 | 4.11E+09 | 109 | 28.0096 | 28.0281 | 28.2242 | 28.294 | 28.243 |
| 0 | 78.253 | 3.8E+09 | 124 | 28.0134 | 28.0315 | 28.1506 | 28.0505 | 28.1092 |
| 0 | 323.31 | 6.63E+10 | 493 | 32.6211 | 32.5521 | 32.0658 | 32.0787 | 32.1694 |
| 0 | 63.241 | 4.65E+09 | 138 | 28.4486 | 28.3086 | 28.4217 | 28.5414 | 28.1282 |
| 0 | 323.31 | 1.03E+11 | 564 | 32.8802 | 32.8616 | 32.9834 | 32.9581 | 32.7843 |
| 0 | 68.881 | 8.58E+09 | 231 | 29.2918 | 29.2522 | 29.3991 | 29.2958 | 29.1645 |
| 0 | 323.31 | 7.65E+10 | 407 | 32.7486 | 32.8195 | 32.2828 | 32.3238 | 32.2847 |
| 0 | 102.93 | 3.15E+09 | 153 | 27.8038 | 27.8437 | 27.6248 | 27.6996 | 27.9131 |
| 0 | 149.96 | 2.43E+10 | 390 | 30.9369 | 30.938 | 30.6925 | 30.5961 | 30.7338 |
| 0 | 204.47 | 8.17E+10 | 472 | 32.5374 | 32.4841 | 32.5917 | 32.517 | 32.3838 |
| 0 | 152.1 | 5.64E+09 | 167 | 28.6002 | 28.5601 | 28.4725 | 28.5199 | 28.8104 |
| 0 | 323.31 | 5.36E+10 | 389 | 31.8094 | 31.8541 | 31.9828 | 32.0148 | 31.8 |
| 0 | 73.671 | 5.75E+09 | 117 | 28.6754 | 28.6411 | 28.7663 | 28.6388 | 28.7676 |
| 0 | 323.31 | 1.22E+11 | 713 | 33.2803 | 33.2229 | 33.0052 | 33.0527 | 32.9681 |
| 0 | 314.25 | 1.29E+10 | 280 | 29.7069 | 29.7061 | 29.9101 | 29.9185 | 29.7557 |
| 0 | 323.31 | 5.39E+10 | 419 | 31.8742 | 31.8458 | 31.9587 | 31.9395 | 31.9685 |
| 0 | 66.14 | 4.21E+09 | 167 | 28.1052 | 28.1706 | 28.2766 | 28.1979 | 28.2527 |
| 0 | 96.621 | 3.45E+09 | 126 | 27.8572 | 27.9524 | 27.9844 | 27.8843 | 28.078 |
| 0 | 242.34 | 5.43E+09 | 156 | 28.5659 | 28.5857 | 28.4608 | 28.5582 | 28.6557 |
| 0 | 323.31 | 4.61E+10 | 504 | 31.7006 | 31.7077 | 31.6028 | 31.5515 | 31.78 |

| | | | | | | | | |
|---|---|---|---|---|---|---|---|---|
| 0 | 140.82 | 6.39E+09 | 170 | 28.6926 | 28.7136 | 28.7349 | 28.6291 | 28.679 |
| 0 | 251.61 | 1.57E+10 | 279 | 29.9126 | 29.9237 | 30.1658 | 29.9969 | 30.0904 |
| 0 | 306.04 | 1.1E+10 | 201 | 29.521 | 29.4731 | 29.5251 | 29.572 | 29.508 |
| 0 | 77.066 | 3.91E+09 | 152 | 28.0251 | 27.9986 | 28.1435 | 28.0373 | 28.0492 |
| 0 | 196.16 | 7.66E+09 | 242 | 29.0152 | 29.0113 | 29.1038 | 29.0861 | 29.0383 |
| 0 | 323.31 | 3.41E+10 | 420 | 31.3181 | 31.3398 | 31.4058 | 31.3131 | 31.2638 |
| 0 | 51.198 | 4.68E+09 | 109 | 28.5513 | 28.6094 | 28.6177 | 28.5641 | 28.6048 |
| 0 | 144.79 | 8.76E+09 | 172 | 29.16 | 29.2133 | 29.3238 | 29.245 | 29.2187 |
| 0 | 105.33 | 2.68E+09 | 137 | 27.4503 | 27.5319 | 27.2841 | 27.259 | 27.6715 |
| 0 | 160.38 | 7.79E+09 | 229 | 29.0823 | 29.1333 | 29.051 | 29.1142 | 29.0387 |
| 0 | 323.31 | 1.56E+10 | 268 | 30.0497 | 29.9595 | 30.0602 | 30.1523 | 30.0967 |
| 0 | 314.89 | 1.24E+10 | 239 | 29.7474 | 29.8247 | 29.6467 | 29.6498 | 29.7953 |
| 0 | 120.34 | 7.94E+09 | 196 | 29.4741 | 29.3355 | 29.1655 | 29.0866 | 29.0582 |
| 0 | 82.362 | 4.3E+09 | 163 | 28.0739 | 28.2207 | 28.2287 | 28.3675 | 28.1774 |
| 0 | 81.338 | 3.51E+09 | 147 | 27.8176 | 27.7572 | 27.8799 | 27.8431 | 28.046 |
| 0 | 246.56 | 2.28E+10 | 328 | 30.7026 | 30.6563 | 30.7238 | 30.719 | 30.5247 |
| 0 | 323.31 | 1.64E+11 | 482 | 33.5625 | 33.4824 | 33.5615 | 33.6362 | 33.1491 |
| 0 | 83.686 | 2.78E+09 | 113 | 27.8424 | 27.8501 | 27.3898 | 27.3421 | 27.6058 |
| 0 | 323.31 | 2.36E+10 | 246 | 30.8435 | 30.8029 | 30.9044 | 30.856 | 30.606 |
| 0 | 115.19 | 5.64E+09 | 147 | 28.5261 | 28.4041 | 28.6038 | 28.5803 | 28.8197 |
| 0 | 197.83 | 4.61E+09 | 162 | 28.3807 | 28.1387 | 28.1641 | 28.1962 | 28.5205 |
| 0 | 153.82 | 4.66E+09 | 161 | 28.3446 | 28.3811 | 28.2508 | 28.354 | 28.4563 |
| 0 | 323.31 | 2.9E+11 | 938 | 34.5343 | 34.5238 | 34.2997 | 34.2885 | 34.2709 |
| 0 | 59.585 | 3.39E+09 | 116 | 27.9644 | 27.7877 | 28.0444 | 27.9409 | 27.7972 |
| 0 | 88.348 | 4.65E+09 | 150 | 28.2503 | 28.2819 | 28.1831 | 28.2718 | 28.4626 |
| 0 | 323.31 | 8.97E+10 | 372 | 32.6294 | 32.6822 | 32.7035 | 32.7392 | 32.521 |
| 0 | 144.65 | 1.56E+10 | 232 | 30.6504 | 30.6367 | 29.9258 | 29.9337 | 29.934 |
| 0 | 140.24 | 1.42E+10 | 255 | 30.0166 | 30.0214 | 29.9454 | 29.9937 | 30.0141 |
| 0 | 221.74 | 6.88E+09 | 156 | 29.0322 | 28.9438 | 28.8651 | 28.7755 | 28.9213 |
| 0 | 102.84 | 3.93E+09 | 148 | 28.1124 | 28.1475 | 28.1461 | 28.0533 | 28.094 |
| 0 | 179.76 | 2.85E+10 | 252 | 31.0377 | 31.0057 | 31.1163 | 31.0314 | 30.9628 |
| 0 | 217.93 | 1.59E+10 | 199 | 30.1616 | 30.2035 | 30.2062 | 30.1383 | 30.1488 |
| 0 | 323.31 | 1.51E+11 | 533 | 33.6702 | 33.6284 | 33.3615 | 33.359 | 33.1627 |
| 0 | 323.31 | 4.6E+11 | 1047 | 34.9996 | 34.9852 | 35.0779 | 35.0866 | 34.8157 |
| 0 | 264.61 | 2.71E+10 | 284 | 31.0088 | 30.8658 | 30.9794 | 31.0056 | 30.9109 |
| 0 | 46.461 | 3.64E+09 | 120 | 28.0375 | 28.0626 | 28.2028 | 28.1789 | 27.8483 |
| 0 | 52.562 | 3.66E+09 | 120 | 28.0732 | 27.9617 | 27.9882 | 27.8266 | 28.1187 |
| 0 | 323.31 | 5.43E+10 | 392 | 31.9068 | 31.8807 | 31.9895 | 31.9635 | 31.903 |
| 0 | 56.431 | 2.17E+09 | 102 | 27.1597 | 27.1068 | 27.2401 | 27.1523 | 27.3089 |
| 0 | 102.16 | 6.98E+09 | 188 | 29.0523 | 28.9373 | 29.1148 | 29.0731 | 28.7934 |
| 0 | 115.6 | 5.1E+09 | 193 | 28.4592 | 28.66 | 28.3867 | 28.302 | 28.6866 |
| 0 | 122.97 | 4.86E+09 | 157 | 28.5508 | 28.3659 | 28.3382 | 28.3617 | 28.5 |
| 0 | 323.31 | 4.67E+10 | 571 | 31.4927 | 31.5398 | 31.7828 | 31.7653 | 31.7927 |
| 0 | 184.57 | 2.03E+10 | 365 | 30.5311 | 30.4954 | 30.6089 | 30.5707 | 30.4345 |
| 0 | 61.009 | 3.21E+09 | 88 | 27.7596 | 27.7445 | 27.9224 | 27.8853 | 27.9623 |
| 0 | 71.596 | 5.15E+09 | 137 | 28.3446 | 28.3244 | 28.3316 | 28.2916 | 28.2406 |
| 0 | 245.82 | 1.47E+10 | 219 | 29.9448 | 29.9268 | 29.9498 | 29.8898 | 29.8427 |
| 0 | 82.399 | 3.46E+09 | 142 | 27.9428 | 27.8489 | 27.7227 | 27.901 | 27.8179 |

| | | | | | | | | |
|---|---|---|---|---|---|---|---|---|
| 0 | 52.202 | 2.26E+09 | 86 | 27.3293 | 27.0323 | 27.2853 | 27.3405 | 27.2455 |
| 0 | 86.21 | 2.5E+09 | 124 | 27.4228 | 27.4342 | 27.4558 | 27.4755 | 27.3526 |
| 0 | 112.97 | 5.85E+09 | 158 | 28.4699 | 28.598 | 28.6647 | 28.5936 | 28.84 |
| 0 | 170.86 | 1.46E+10 | 245 | 29.8762 | 29.9068 | 29.9428 | 30.017 | 30.0142 |
| 0 | 127.33 | 1.24E+10 | 239 | 29.9008 | 29.8365 | 29.975 | 29.8259 | 29.9503 |
| 0 | 143.26 | 2.1E+10 | 232 | 30.5724 | 30.5605 | 30.4916 | 30.5118 | 30.4626 |
| 0 | 94.729 | 4.81E+09 | 131 | 28.3596 | 28.3393 | 28.5114 | 28.2998 | 28.2373 |
| 0 | 34.284 | 2.95E+09 | 172 | 27.6515 | 27.6584 | 27.6016 | 27.701 | 27.6069 |
| 0 | 323.31 | 2.3E+10 | 282 | 30.6485 | 30.6324 | 30.6895 | 30.6375 | 30.5656 |
| 0 | 168.11 | 1.07E+10 | 216 | 29.5123 | 29.4981 | 29.614 | 29.5761 | 29.4693 |
| 0 | 267.95 | 1.02E+10 | 254 | 29.4477 | 29.4404 | 29.5311 | 29.5345 | 29.4638 |
| 0 | 302.83 | 1.45E+10 | 275 | 29.9824 | 29.9523 | 30.068 | 30.1488 | 29.8619 |
| 0 | 113.18 | 6.1E+09 | 161 | 28.7606 | 28.7777 | 28.7742 | 28.8873 | 28.5682 |
| 0 | 240.61 | 8.46E+09 | 188 | 29.3336 | 29.2824 | 29.2801 | 29.3664 | 29.1524 |
| 0 | 323.31 | 1.98E+11 | 1229 | 33.6562 | 33.6731 | 33.7589 | 33.7817 | 33.5189 |
| 0 | 312.63 | 5.28E+10 | 409 | 31.8583 | 31.8428 | 31.8602 | 31.8968 | 31.8117 |
| 0 | 133.11 | 3.07E+09 | 109 | 28.0615 | 28.0245 | 27.5792 | 27.7236 | 27.7419 |
| 0 | 42.28 | 2.75E+09 | 113 | 27.6926 | 27.6145 | 27.4279 | 27.4007 | 27.5733 |
| 0 | 172.83 | 9.68E+09 | 184 | 29.4297 | 29.4467 | 29.3476 | 29.3268 | 29.6371 |
| 0 | 323.31 | 1.34E+11 | 564 | 33.2608 | 33.267 | 33.3145 | 33.2351 | 33.2408 |
| 0 | 84.287 | 4.02E+09 | 138 | 28.3189 | 28.36 | 28.1595 | 27.9594 | 28.2159 |
| 0 | 55.38 | 3.64E+09 | 108 | 28.0141 | 28.1815 | 28.0101 | 28.0394 | 28.0171 |
| 0 | 52.765 | 2.55E+09 | 88 | 27.4788 | 27.4652 | 27.3934 | 27.3516 | 27.5831 |
| 0 | 129.18 | 6.46E+09 | 220 | 28.9416 | 28.8618 | 28.9502 | 28.8393 | 28.6901 |
| 0 | 278.02 | 1.32E+10 | 219 | 29.9491 | 29.9068 | 29.9467 | 29.9276 | 29.6679 |
| 0 | 105.49 | 7.04E+09 | 140 | 28.7536 | 28.7904 | 28.9969 | 29.0613 | 29.1013 |
| 0 | 323.31 | 1.07E+11 | 543 | 32.8969 | 32.8641 | 32.9755 | 32.9573 | 32.8367 |
| 0 | 313.76 | 7.14E+10 | 355 | 32.3309 | 32.3822 | 32.3436 | 32.3613 | 32.2672 |
| 0 | 176.69 | 5.92E+09 | 152 | 28.9874 | 28.908 | 28.7019 | 28.5293 | 28.7024 |
| 0 | 323.31 | 1.11E+11 | 619 | 33 | 33.0518 | 32.7519 | 32.7993 | 32.9931 |
| 0 | 189.38 | 1.42E+10 | 196 | 30.0028 | 29.99 | 29.9282 | 29.9337 | 30.1046 |
| 0 | 290.44 | 1.8E+10 | 228 | 30.3427 | 30.3868 | 30.4836 | 30.5007 | 30.1453 |
| 0 | 87.986 | 7.56E+09 | 187 | 29.1097 | 29.0946 | 29.1516 | 29.2347 | 28.9548 |
| 0 | 121.39 | 7.51E+09 | 190 | 29.1059 | 29.1647 | 29.0385 | 28.9252 | 29.0985 |
| 0 | 323.31 | 1.11E+11 | 836 | 32.9394 | 32.9254 | 32.995 | 32.9362 | 32.8405 |
| 0 | 245.12 | 2.35E+10 | 256 | 30.7085 | 30.6993 | 30.7637 | 30.8224 | 30.5651 |
| 0 | 92.479 | 8.95E+09 | 188 | 29.2173 | 29.2621 | 29.4535 | 29.4031 | 29.3206 |
| 0 | 133.48 | 1.49E+10 | 198 | 29.8487 | 29.8536 | 30.0247 | 29.9372 | 29.8354 |
| 0 | 243.04 | 1.87E+10 | 233 | 30.3316 | 30.2784 | 30.2945 | 30.2584 | 30.1803 |
| 0 | 265.82 | 6.83E+09 | 151 | 28.7795 | 28.7068 | 28.8818 | 28.92 | 28.7835 |
| 0 | 323.31 | 7.06E+10 | 384 | 32.0949 | 32.1729 | 32.3344 | 32.2059 | 32.2188 |
| 0 | 182.55 | 4.72E+10 | 338 | 31.6669 | 31.6459 | 31.7014 | 31.7196 | 31.5402 |
| 0 | 173.11 | 1.49E+10 | 228 | 29.9349 | 29.9168 | 29.9625 | 29.9229 | 30.0578 |
| 0 | 323.31 | 1.07E+10 | 231 | 29.519 | 29.503 | 29.4738 | 29.4967 | 29.5036 |
| 0 | 323.31 | 1.98E+10 | 327 | 30.4673 | 30.4909 | 30.4016 | 30.3934 | 30.2921 |
| 0 | 123.04 | 2.88E+09 | 132 | 27.6832 | 27.5623 | 27.5219 | 27.5214 | 27.6184 |
| 0 | 208.69 | 5.16E+10 | 378 | 31.808 | 31.8655 | 31.7729 | 31.8849 | 31.7204 |
| 0 | 323.31 | 2.67E+10 | 292 | 30.8176 | 30.8924 | 30.9325 | 30.9436 | 30.7832 |

| | | | | | | | | |
|---|---|---|---|---|---|---|---|---|
| 0 | 323.31 | 1.13E+10 | 239 | 29.6794 | 29.6751 | 29.7696 | 29.8925 | 29.7422 |
| 0 | 93.786 | 7.54E+09 | 192 | 28.8646 | 28.9177 | 29.1093 | 29.0213 | 28.9813 |
| 0 | 52.639 | 2.87E+09 | 115 | 27.6318 | 27.5651 | 27.7915 | 27.6375 | 27.6423 |
| 0 | 274.99 | 9.55E+09 | 247 | 29.2688 | 29.3138 | 29.3634 | 29.2206 | 29.4432 |
| 0 | 118.94 | 4.06E+09 | 175 | 28.1018 | 28.0515 | 28.1349 | 28.06 | 28.0484 |
| 0 | 107.18 | 1.79E+10 | 292 | 30.3383 | 30.2732 | 30.2834 | 30.3436 | 30.2328 |
| 0 | 51.98 | 2.51E+09 | 88 | 27.5928 | 27.6345 | 27.6279 | 27.5248 | 27.5125 |
| 0 | 128.1 | 5.77E+09 | 153 | 28.6227 | 28.5306 | 28.7147 | 28.5897 | 28.6164 |
| 0 | 101.06 | 6.87E+09 | 127 | 28.9814 | 28.8837 | 28.712 | 28.8639 | 28.9126 |
| 0 | 304.12 | 1.43E+10 | 254 | 29.9235 | 29.9014 | 29.8027 | 29.7454 | 30.038 |
| 0 | 190.37 | 7.37E+09 | 175 | 29.009 | 29.0065 | 29.1117 | 29.1542 | 28.893 |
| 0 | 303.11 | 5.58E+10 | 399 | 31.9931 | 31.962 | 31.8869 | 31.9337 | 31.8623 |
| 0 | 54.301 | 2.73E+09 | 144 | 27.5635 | 27.5666 | 27.5965 | 27.6427 | 27.3925 |
| 0 | 86.025 | 6.74E+09 | 147 | 28.7004 | 28.6845 | 28.9487 | 28.9297 | 28.9108 |
| 0 | 136.69 | 1.66E+10 | 258 | 30.0092 | 30.0868 | 30.1002 | 30.1744 | 30.1788 |
| 0 | 323.31 | 2.24E+10 | 311 | 30.6449 | 30.6392 | 30.8882 | 30.8164 | 30.5577 |
| 0 | 167.84 | 1.55E+10 | 257 | 30.0835 | 30.1303 | 30.1507 | 30.1394 | 30.2112 |
| 0 | 38.774 | 4.42E+09 | 124 | 28.3147 | 28.3185 | 28.2123 | 28.2371 | 28.2389 |
| 0 | 48.104 | 2.4E+09 | 65 | 27.2321 | 27.4725 | 27.373 | 27.5965 | 27.295 |
| 0 | 128.52 | 7.17E+09 | 167 | 28.9018 | 28.9305 | 29.0678 | 29.0698 | 28.8711 |
| 0 | 111.9 | 3.59E+09 | 100 | 27.9857 | 28.0388 | 27.985 | 28.0001 | 28.0325 |
| 0 | 51.599 | 2.2E+09 | 102 | 27.1266 | 27.1345 | 27.2537 | 27.3265 | 27.3449 |
| 0 | 86.809 | 4.67E+09 | 147 | 28.5245 | 28.3024 | 28.3817 | 28.5295 | 28.3541 |
| 0 | 155.19 | 6.75E+09 | 164 | 28.8424 | 28.8163 | 29.0164 | 29.1035 | 28.8098 |
| 0 | 55.371 | 2.81E+09 | 111 | 27.7049 | 27.5199 | 27.6369 | 27.7332 | 27.6793 |
| 0 | 323.31 | 2.66E+10 | 260 | 31.1059 | 31.2036 | 30.8722 | 30.8461 | 30.8315 |
| 0 | 323.31 | 1.74E+11 | 838 | 33.8733 | 33.8976 | 33.4905 | 33.4808 | 33.3859 |
| 0 | 161.06 | 3.77E+09 | 141 | 28.1668 | 28.2208 | 28.0212 | 28.0161 | 27.9495 |
| 0 | 170.16 | 1.13E+10 | 213 | 29.737 | 29.7417 | 29.6652 | 29.8114 | 29.606 |
| 0 | 111.74 | 1.05E+10 | 184 | 29.6122 | 29.5964 | 29.4062 | 29.45 | 29.7456 |
| 0 | 234.14 | 1.68E+10 | 263 | 30.2813 | 30.2587 | 30.3306 | 30.3028 | 30.2014 |
| 0 | 323.31 | 4.72E+09 | 153 | 28.3259 | 28.2968 | 28.3775 | 28.3168 | 28.398 |
| 0 | 323.31 | 2.15E+10 | 360 | 30.5889 | 30.6472 | 30.4536 | 30.4161 | 30.6007 |
| 0 | 323.31 | 4.46E+10 | 478 | 31.6025 | 31.6117 | 31.7141 | 31.6565 | 31.5934 |
| 0 | 323.31 | 2.4E+10 | 384 | 30.7044 | 30.7096 | 30.7582 | 30.7299 | 30.7196 |
| 0 | 177.96 | 1.01E+10 | 202 | 29.6635 | 29.5704 | 29.3668 | 29.3114 | 29.3501 |
| 0 | 245.54 | 8.52E+09 | 176 | 29.5899 | 29.6373 | 29.0884 | 29.1947 | 29.0664 |
| 0 | 72.67 | 3.35E+09 | 112 | 27.8368 | 27.8451 | 27.8054 | 27.9484 | 27.8513 |
| 0 | 323.31 | 1.8E+10 | 294 | 30.4577 | 30.3436 | 30.1985 | 30.2199 | 30.3068 |
| 0 | 106.05 | 3E+09 | 91 | 27.708 | 27.7691 | 27.8147 | 27.8473 | 27.7001 |
| 0 | 323.31 | 6.4E+10 | 350 | 32.218 | 32.2231 | 32.1337 | 32.2052 | 32.072 |
| 0 | 167.25 | 1.24E+10 | 193 | 29.7565 | 29.7541 | 29.9649 | 29.9204 | 29.6305 |
| 0 | 53.449 | 3.57E+09 | 122 | 27.8512 | 27.8216 | 28.1411 | 28.1055 | 28.0773 |
| 0 | 316.82 | 3.42E+10 | 283 | 31.2976 | 31.2262 | 31.3653 | 31.3448 | 30.9466 |
| 0 | 323.31 | 4.13E+10 | 325 | 31.5061 | 31.4823 | 31.5675 | 31.673 | 31.4622 |
| 0 | 323.31 | 1.38E+11 | 602 | 33.3043 | 33.2883 | 33.3186 | 33.3217 | 33.1734 |
| 0 | 222.11 | 6.7E+09 | 183 | 28.8779 | 28.8046 | 28.9627 | 28.9353 | 28.8744 |
| 0 | 166.45 | 1.8E+10 | 286 | 30.2907 | 30.2486 | 30.4403 | 30.3434 | 30.3253 |

| | | | | | | | | |
|---|---|---|---|---|---|---|---|---|
| 0 | 304.26 | 1.36E+10 | 219 | 29.8812 | 29.9214 | 30.0283 | 30.0381 | 29.9255 |
| 0 | 163.72 | 7.86E+09 | 203 | 29.2563 | 29.2644 | 28.9773 | 28.9257 | 29.2079 |
| 0 | 217.3 | 1.9E+10 | 279 | 30.1868 | 30.1489 | 30.309 | 30.3299 | 30.3592 |
| 0 | 284.97 | 1.87E+10 | 215 | 30.2028 | 30.2064 | 30.3008 | 30.3387 | 30.2855 |
| 0 | 39.075 | 2.69E+09 | 101 | 27.568 | 27.5453 | 27.5359 | 27.6452 | 27.3669 |
| 0 | 323.31 | 4.09E+10 | 308 | 31.3692 | 31.3426 | 31.5305 | 31.469 | 31.4751 |
| 0 | 119.11 | 9.78E+09 | 255 | 29.2649 | 29.3082 | 29.4814 | 29.4512 | 29.3966 |
| 0 | 96.739 | 4.86E+09 | 172 | 28.4972 | 28.376 | 28.3648 | 28.4508 | 28.4149 |
| 0 | 323.31 | 3.79E+10 | 309 | 31.4716 | 31.4099 | 31.4079 | 31.4048 | 31.3077 |
| 0 | 198.93 | 4.62E+09 | 166 | 28.4245 | 28.6146 | 28.373 | 28.4019 | 28.4228 |
| 0 | 176.79 | 3.4E+09 | 113 | 27.7188 | 27.8021 | 27.7066 | 27.8701 | 27.9468 |
| 0 | 323.31 | 4.75E+10 | 288 | 31.8684 | 31.8481 | 31.6944 | 31.7138 | 31.9609 |
| 0 | 87.384 | 4.35E+09 | 140 | 28.6187 | 28.6108 | 28.2861 | 28.2304 | 28.2486 |
| 0 | 323.31 | 4.38E+10 | 278 | 31.5576 | 31.5138 | 31.6123 | 31.6269 | 31.5209 |
| 0 | 80.955 | 2.54E+09 | 123 | 27.4745 | 27.4442 | 27.2287 | 27.2849 | 27.5961 |
| 0 | 306.75 | 1.16E+10 | 163 | 29.5605 | 29.5848 | 29.5389 | 29.4432 | 29.7117 |
| 0 | 323.31 | 3.06E+10 | 297 | 31.118 | 31.0769 | 31.1126 | 31.1241 | 30.9828 |
| 0 | 190.45 | 2.7E+10 | 253 | 30.9347 | 30.8388 | 31.0164 | 30.9335 | 30.7137 |
| 0 | 76.82 | 6.31E+09 | 150 | 28.602 | 28.6544 | 28.775 | 28.9419 | 28.7663 |
| 0 | 138.47 | 7.11E+09 | 180 | 29.1578 | 29.2175 | 28.9794 | 28.8706 | 29.0477 |
| 0 | 60.065 | 2.3E+09 | 107 | 27.0567 | 27.1747 | 27.1163 | 27.2519 | 27.45 |
| 0 | 72.788 | 4.34E+09 | 124 | 28.0198 | 28.0116 | 28.0446 | 27.9702 | 28.2915 |
| 0 | 59.226 | 9.45E+09 | 175 | 29.3799 | 29.3363 | 29.3381 | 29.4939 | 29.3262 |
| 0 | 41.953 | 3.08E+09 | 104 | 27.6984 | 27.655 | 27.8399 | 27.8267 | 27.7409 |
| 0 | 87.211 | 3.18E+09 | 95 | 28.1562 | 27.9401 | 27.7124 | 27.6483 | 27.9859 |
| 0 | 82.301 | 6.02E+09 | 153 | 28.5231 | 28.6558 | 28.437 | 28.614 | 28.8247 |
| 0 | 196.39 | 6.32E+09 | 154 | 28.7823 | 28.7723 | 28.8417 | 28.8538 | 28.811 |
| 0 | 162.43 | 6.93E+09 | 159 | 28.7693 | 28.8429 | 28.983 | 28.8878 | 29.0349 |
| 0 | 77.442 | 8.32E+09 | 180 | 29.1281 | 29.1746 | 29.1908 | 29.2522 | 29.1354 |
| 0 | 45.039 | 2.57E+09 | 73 | 27.5976 | 27.6046 | 27.7614 | 27.651 | 27.4222 |
| 0 | 246.14 | 5.58E+09 | 168 | 29.0264 | 29.0229 | 28.6649 | 28.6108 | 28.6259 |
| 0 | 248.4 | 5.69E+09 | 151 | 28.5232 | 28.6167 | 28.5359 | 28.5558 | 28.791 |
| 0 | 323.31 | 1.52E+10 | 264 | 30.0475 | 30.0342 | 30.1975 | 30.2041 | 29.9373 |
| 0 | 323.31 | 1.55E+11 | 507 | 33.487 | 33.4407 | 33.5292 | 33.4833 | 33.413 |
| 0 | 208.39 | 7.58E+09 | 218 | 29.0683 | 29.0782 | 29.0794 | 29.0594 | 28.9505 |
| 0 | 148.21 | 3.26E+09 | 146 | 27.926 | 27.853 | 27.9328 | 27.9639 | 27.8038 |
| 0 | 323.31 | 7.41E+10 | 292 | 32.4815 | 32.4127 | 32.4789 | 32.4274 | 32.3905 |
| 0 | 176.18 | 5.96E+09 | 152 | 28.7767 | 28.808 | 28.8291 | 28.6746 | 28.7634 |
| 0 | 31.356 | 2.23E+09 | 88 | 27.3659 | 27.3609 | 27.2665 | 27.3466 | 27.3844 |
| 0 | 60.297 | 5.41E+09 | 124 | 28.6376 | 28.5085 | 28.6213 | 28.5899 | 28.6151 |
| 0 | 278.64 | 4.9E+10 | 329 | 31.686 | 31.6233 | 31.6414 | 31.5815 | 31.8702 |
| 0 | 323.31 | 2.9E+10 | 254 | 30.9548 | 30.9112 | 31.0152 | 31.0434 | 31.0733 |
| 0 | 323.31 | 4.67E+10 | 399 | 32.0166 | 31.9671 | 31.7187 | 31.6315 | 31.5963 |
| 0 | 323.31 | 2.45E+10 | 269 | 30.7323 | 30.7505 | 30.6855 | 30.6721 | 30.7764 |
| 0 | 135.59 | 4.98E+09 | 157 | 28.4053 | 28.4274 | 28.5614 | 28.4445 | 28.4607 |
| 0 | 323.31 | 5.5E+10 | 405 | 31.9072 | 31.8861 | 31.9475 | 31.9713 | 31.9372 |
| 0 | 133.4 | 5.54E+09 | 163 | 28.511 | 28.5102 | 28.7129 | 28.7309 | 28.6896 |
| 0 | 51.773 | 1.98E+09 | 69 | 27.0422 | 26.9749 | 26.9174 | 26.9585 | 27.2059 |

| | | | | | | | | |
|---|---|---|---|---|---|---|---|---|
| 0 | 147.79 | 5.14E+09 | 138 | 28.5031 | 28.4842 | 28.4165 | 28.6535 | 28.5735 |
| 0 | 83.218 | 8.67E+09 | 189 | 29.3844 | 29.3568 | 29.0725 | 29.1404 | 29.4138 |
| 0 | 111.26 | 7.3E+09 | 164 | 28.9393 | 29.0268 | 29.1373 | 29.0567 | 29.1449 |
| 0 | 68.673 | 2.19E+09 | 72 | 27.3358 | 27.2972 | 27.073 | 27.0279 | 27.4743 |
| 0 | 40.184 | 2.5E+09 | 122 | 27.8768 | 27.7605 | 27.6034 | 27.4488 | 27.2263 |
| 0 | 323.31 | 4.5E+10 | 302 | 31.5607 | 31.564 | 31.7735 | 31.754 | 31.5824 |
| 0 | 323.31 | 3.07E+10 | 359 | 31.0416 | 31.0007 | 31.2505 | 31.1471 | 31.0234 |
| 0 | 171.36 | 4.15E+09 | 140 | 28.2195 | 28.211 | 28.2396 | 28.1994 | 28.2916 |
| 0 | 175.65 | 1.9E+10 | 274 | 30.4421 | 30.4648 | 30.4787 | 30.3484 | 30.3303 |
| 0 | 144.75 | 3.03E+09 | 117 | 27.7732 | 27.6792 | 27.4957 | 27.5452 | 28.0132 |
| 0 | 61.273 | 3.88E+09 | 154 | 28.035 | 28.1733 | 28.2166 | 28.2875 | 28.0454 |
| 0 | 199.5 | 2.78E+10 | 256 | 30.9935 | 30.9994 | 30.6945 | 30.7866 | 31.0169 |
| 0 | 45.333 | 2.9E+09 | 122 | 27.7029 | 27.695 | 27.8398 | 27.7811 | 27.5516 |
| 0 | 190.81 | 1.63E+10 | 260 | 30.1585 | 30.1361 | 30.2144 | 30.1719 | 30.2211 |
| 0 | 323.31 | 1.3E+11 | 387 | 33.1374 | 33.1118 | 33.2175 | 33.3161 | 33.1042 |
| 0 | 291.24 | 1.55E+10 | 230 | 30.0625 | 30.0328 | 30.0029 | 30.0196 | 30.187 |
| 0 | 67.549 | 8.7E+09 | 170 | 29.1828 | 29.2253 | 29.1986 | 29.1674 | 29.4283 |
| 0 | 78.867 | 3.9E+09 | 117 | 28.1639 | 28.1255 | 28.005 | 27.9932 | 28.1729 |
| 0 | 75.964 | 2.91E+09 | 88 | 27.8607 | 27.8399 | 27.6273 | 27.493 | 27.6662 |
| 0 | 150.14 | 1.98E+10 | 174 | 30.47 | 30.4752 | 30.3279 | 30.2406 | 30.5778 |
| 0 | 323.31 | 4.21E+10 | 421 | 31.555 | 31.5743 | 31.5596 | 31.563 | 31.5982 |
| 0 | 323.31 | 5.13E+10 | 368 | 31.7932 | 31.7875 | 31.9004 | 31.9085 | 31.8392 |
| 0 | 89.25 | 2.47E+09 | 110 | 27.3399 | 27.2552 | 27.3549 | 27.2895 | 27.2386 |
| 0 | 72.34 | 2.77E+09 | 101 | 27.5006 | 27.5143 | 27.3645 | 27.3344 | 27.6505 |
| 0 | 81.762 | 1.54E+09 | 63 | 26.7295 | 26.5417 | 26.5936 | 26.7049 | 26.8035 |
| 0 | 94.484 | 1.11E+11 | 287 | 32.8197 | 32.8556 | 32.8995 | 32.9312 | 32.837 |
| 0 | 230.94 | 1.74E+10 | 176 | 30.2226 | 30.1781 | 30.1443 | 30.2671 | 30.1231 |
| 0 | 249.78 | 6.07E+09 | 154 | 28.6414 | 28.6903 | 28.7885 | 28.6881 | 28.6695 |
| 0 | 286.23 | 2.6E+10 | 291 | 30.7677 | 30.7187 | 30.7941 | 30.8083 | 30.853 |
| 0 | 54.918 | 3.98E+09 | 104 | 28.3779 | 28.1948 | 28.2609 | 28.1125 | 28.1768 |
| 0 | 201.94 | 3.86E+10 | 327 | 31.3201 | 31.267 | 31.4941 | 31.5823 | 31.2705 |
| 0 | 323.31 | 7.47E+10 | 388 | 32.3536 | 32.2753 | 32.3786 | 32.4054 | 32.2711 |
| 0 | 68.554 | 4.91E+09 | 137 | 28.4189 | 28.4271 | 28.5113 | 28.5465 | 28.2947 |
| 0 | 303.62 | 1.72E+10 | 285 | 30.1621 | 30.0456 | 30.3689 | 30.3428 | 30.184 |
| 0 | 231.23 | 9.35E+09 | 165 | 29.353 | 29.3269 | 29.5484 | 29.4589 | 29.174 |
| 0 | 244.28 | 7.84E+09 | 194 | 28.8495 | 28.9045 | 29.2025 | 29.0667 | 29.2502 |
| 0 | 218.41 | 1.14E+10 | 213 | 29.6367 | 29.6717 | 29.4843 | 29.5374 | 29.7264 |
| 0 | 267.2 | 1.42E+10 | 196 | 29.8984 | 29.8713 | 30.0634 | 29.9901 | 29.9286 |
| 0 | 323.31 | 3.76E+11 | 1184 | 34.629 | 34.5353 | 34.6497 | 34.706 | 34.4135 |
| 0 | 307.31 | 5.14E+10 | 317 | 31.7194 | 31.7888 | 31.8809 | 31.8721 | 31.7499 |
| 0 | 323.31 | 8.18E+10 | 290 | 32.3773 | 32.377 | 32.6102 | 32.5946 | 32.7259 |
| 0 | 43.61 | 2.28E+09 | 95 | 27.5077 | 27.2984 | 27.2518 | 27.2473 | 27.4742 |
| 0 | 323.31 | 9.71E+09 | 210 | 29.5526 | 29.505 | 29.4026 | 29.4167 | 29.5095 |
| 0 | 227.42 | 1.79E+10 | 228 | 30.4185 | 30.3739 | 30.4735 | 30.5303 | 30.2103 |
| 0 | 51.44 | 2.55E+09 | 57 | 27.5199 | 27.516 | 27.3787 | 27.3194 | 27.4693 |
| 0 | 151.07 | 7.72E+09 | 204 | 29.1842 | 29.102 | 29.0772 | 29.0656 | 29.1603 |
| 0 | 128.89 | 4.32E+09 | 107 | 28.515 | 28.4246 | 28.1355 | 28.097 | 28.3456 |
| 0 | 124.16 | 3.19E+09 | 135 | 27.8013 | 27.8195 | 27.8773 | 27.885 | 27.7238 |

| | | | | | | | | |
|---|---|---|---|---|---|---|---|---|
| 0 | 284 | 1.15E+10 | 183 | 29.7363 | 29.7369 | 29.6208 | 29.6128 | 29.6377 |
| 0 | 323.31 | 6.92E+10 | 427 | 32.2303 | 32.2329 | 32.328 | 32.3011 | 32.2856 |
| 0 | 109.21 | 4.69E+09 | 131 | 28.2675 | 28.3001 | 28.3263 | 28.3204 | 28.475 |
| 0 | 108.83 | 3.32E+09 | 142 | 28.0037 | 27.9234 | 27.9874 | 27.8339 | 27.9147 |
| 0 | 77.579 | 1.47E+10 | 185 | 29.9716 | 29.9498 | 30.0033 | 29.9721 | 30.1342 |
| 0 | 32.074 | 1.48E+09 | 74 | 26.5986 | 26.6532 | 26.7457 | 26.7985 | 26.7932 |
| 0 | 99.135 | 5.38E+09 | 153 | 28.3179 | 28.4867 | 28.4854 | 28.6567 | 28.6593 |
| 0 | 72.187 | 4.64E+09 | 177 | 28.4199 | 28.4868 | 28.3364 | 28.3741 | 28.311 |
| 0 | 74.655 | 8.33E+09 | 186 | 29.1432 | 29.0568 | 29.089 | 29.1057 | 29.3349 |
| 0 | 285.6 | 1.37E+10 | 185 | 29.9393 | 29.8519 | 29.891 | 29.8513 | 30.0821 |
| 0 | 145.29 | 8.89E+09 | 231 | 29.2751 | 29.2425 | 29.0404 | 29.0958 | 29.3007 |
| 0 | 54.553 | 3.92E+09 | 167 | 28.2645 | 28.2795 | 28.0293 | 28.0016 | 28.1521 |
| 0 | 158.88 | 1.16E+10 | 197 | 29.6421 | 29.6417 | 29.6679 | 29.6703 | 29.7661 |
| 0 | 323.31 | 1.62E+11 | 532 | 33.7928 | 33.746 | 33.4059 | 33.3939 | 33.3618 |
| 0 | 242.81 | 5.93E+09 | 132 | 28.7473 | 28.694 | 28.5883 | 28.5276 | 28.7986 |
| 0 | 53.491 | 4.1E+09 | 106 | 28.1083 | 28.1915 | 28.0742 | 28.0905 | 28.2736 |
| 0 | 132.89 | 1.81E+10 | 274 | 30.4077 | 30.3882 | 30.4971 | 30.4923 | 30.2058 |
| 0 | 323.31 | 6.15E+10 | 380 | 32.0523 | 32.0379 | 32.1857 | 32.1253 | 32.1226 |
| 0 | 259.6 | 1.77E+10 | 215 | 30.3648 | 30.3285 | 30.2929 | 30.2261 | 30.2709 |
| 0 | 307.17 | 1.63E+10 | 193 | 30.2799 | 30.3386 | 30.1871 | 30.2471 | 30.0884 |
| 0 | 191.33 | 2.02E+10 | 303 | 30.4565 | 30.4778 | 30.6001 | 30.5225 | 30.4798 |
| 0 | 249.14 | 1.9E+10 | 216 | 30.3417 | 30.3404 | 30.458 | 30.4782 | 30.4152 |
| 0 | 323.31 | 2.78E+10 | 304 | 30.9195 | 30.9168 | 30.9966 | 30.9984 | 30.9254 |
| 0 | 118.62 | 2.74E+09 | 142 | 27.6563 | 27.653 | 27.6469 | 27.7729 | 27.6471 |
| 0 | 32.358 | 1.87E+09 | 59 | 27.1171 | 27.1364 | 26.9189 | 27.0764 | 26.9775 |
| 0 | 69.11 | 4.6E+09 | 117 | 28.4348 | 28.4536 | 28.3984 | 28.4929 | 28.2053 |
| 0 | 33.548 | 1.8E+09 | 56 | 27.0184 | 27.1369 | 27.051 | 26.9926 | 26.9621 |
| 0 | 90.947 | 6.04E+09 | 127 | 28.6372 | 28.7968 | 28.9373 | 28.8457 | 28.6731 |
| 0 | 323.31 | 1.87E+10 | 406 | 30.4057 | 30.3284 | 30.26 | 30.1831 | 30.54 |
| 0 | 108.75 | 6.19E+09 | 159 | 28.756 | 28.7977 | 28.7419 | 28.786 | 28.8283 |
| 0 | 249.24 | 1.36E+10 | 178 | 30.0169 | 30.0064 | 29.7898 | 29.9221 | 29.921 |
| 0 | 44.424 | 4.64E+09 | 130 | 28.2982 | 28.3133 | 28.2836 | 28.2884 | 28.0645 |
| 0 | 144.23 | 4.06E+09 | 117 | 28.0141 | 27.9357 | 28.2207 | 28.1533 | 27.9951 |
| 0 | 207.26 | 1.06E+10 | 183 | 29.5002 | 29.5067 | 29.5184 | 29.4982 | 29.4492 |
| 0 | 81.442 | 6.18E+09 | 142 | 28.6752 | 28.684 | 28.7562 | 28.7541 | 28.66 |
| 0 | 323.31 | 1.13E+10 | 177 | 29.6838 | 29.6842 | 29.6288 | 29.5175 | 29.5104 |
| 0 | 143.25 | 9.87E+09 | 177 | 29.356 | 29.2749 | 29.5332 | 29.4939 | 29.3778 |
| 0 | 323.31 | 2.84E+10 | 354 | 30.9568 | 30.9912 | 30.9408 | 30.9945 | 30.7963 |
| 0 | 89.061 | 2.98E+09 | 107 | 27.6141 | 27.5706 | 27.8048 | 27.7866 | 27.5784 |
| 0 | 115.94 | 1.06E+10 | 158 | 29.4901 | 29.5099 | 29.4941 | 29.5698 | 29.5321 |
| 0 | 52.217 | 3.56E+09 | 122 | 27.9242 | 27.9293 | 27.9905 | 27.9532 | 28.001 |
| 0 | 207.43 | 2.22E+10 | 231 | 30.6556 | 30.6265 | 30.6224 | 30.6516 | 30.451 |
| 0 | 323.31 | 2.46E+10 | 321 | 30.9056 | 30.9172 | 31.0534 | 31.057 | 30.7179 |
| 0 | 102.9 | 4.7E+09 | 160 | 28.2999 | 28.3838 | 28.4082 | 28.4731 | 28.2403 |
| 0 | 58.646 | 3.51E+09 | 115 | 28.0862 | 27.9531 | 27.8195 | 27.7362 | 27.7869 |
| 0 | 323.31 | 5.61E+10 | 355 | 32.3495 | 32.3263 | 31.9617 | 31.9664 | 31.9647 |
| 0 | 34.059 | 2.38E+09 | 104 | 27.3583 | 27.3742 | 27.3323 | 27.1183 | 27.2871 |
| 0 | 156.5 | 1.46E+10 | 228 | 30.2912 | 30.224 | 30.0139 | 30.0952 | 30.0032 |

| | | | | | | | | |
|---|---|---|---|---|---|---|---|---|
| 0 | 79.138 | 1.33E+10 | 291 | 29.7167 | 29.7984 | 29.9602 | 29.8873 | 29.8506 |
| 0 | 194.37 | 3.35E+09 | 114 | 27.9264 | 27.8184 | 27.7886 | 27.6557 | 27.9884 |
| 0 | 323.31 | 1.06E+11 | 564 | 32.8556 | 32.8884 | 32.841 | 32.8399 | 32.7582 |
| 0 | 323.31 | 7.19E+10 | 452 | 32.2432 | 32.2361 | 32.2892 | 32.2162 | 32.3424 |
| 0 | 323.31 | 2.79E+10 | 297 | 30.8402 | 30.775 | 30.9138 | 30.8993 | 30.7792 |
| 0 | 109.21 | 5.64E+09 | 139 | 28.7249 | 28.7861 | 28.8482 | 28.8088 | 28.6183 |
| 0 | 49.828 | 2.81E+09 | 75 | 27.3319 | 27.4336 | 27.5904 | 27.6722 | 27.7069 |
| 0 | 130.44 | 6.48E+09 | 182 | 28.8119 | 28.7968 | 28.8203 | 28.9419 | 28.7901 |
| 0 | 323.31 | 2E+11 | 530 | 33.7581 | 33.7434 | 33.8312 | 33.8302 | 33.6952 |
| 0 | 95.406 | 8.57E+09 | 195 | 29.367 | 29.3542 | 29.2157 | 29.259 | 29.4393 |
| 0 | 78.874 | 8.77E+09 | 161 | 29.1396 | 29.2262 | 29.4095 | 29.4405 | 29.1611 |
| 0 | 74.61 | 4.39E+09 | 99 | 28.2731 | 28.2374 | 28.2229 | 28.0989 | 28.344 |
| 0 | 42.538 | 1.43E+09 | 52 | 26.7674 | 26.4937 | 26.6898 | 26.7153 | 26.7974 |
| 0 | 206.44 | 7.86E+09 | 167 | 29.236 | 29.2497 | 29.0213 | 28.9832 | 29.3006 |
| 0 | 33.528 | 3.13E+09 | 96 | 27.9146 | 27.8219 | 27.8954 | 27.6842 | 27.5542 |
| 0 | 323.31 | 5.68E+10 | 452 | 31.856 | 31.8533 | 32.087 | 31.9599 | 31.9333 |
| 0 | 323.31 | 5.67E+11 | 1472 | 35.2259 | 35.2931 | 35.2779 | 35.3071 | 35.2166 |
| 0 | 107.54 | 4.2E+09 | 79 | 28.3982 | 28.3647 | 27.9729 | 27.9655 | 28.3073 |
| 0 | 305.3 | 7.1E+10 | 398 | 32.5703 | 32.5192 | 32.3912 | 32.2618 | 32.208 |
| 0 | 83.151 | 2.22E+09 | 97 | 27.2994 | 27.4016 | 27.2495 | 27.3536 | 27.4982 |
| 0 | 323.31 | 1.48E+11 | 446 | 33.4008 | 33.3453 | 33.4189 | 33.3774 | 33.2415 |
| 0 | 91.713 | 3.94E+09 | 108 | 28.1608 | 28.2111 | 28.1208 | 28.0928 | 28.203 |
| 0 | 247.34 | 9.43E+09 | 167 | 29.3733 | 29.3405 | 29.4975 | 29.3663 | 29.3271 |
| 0 | 110.42 | 1.2E+10 | 225 | 29.563 | 29.4776 | 29.6055 | 29.5684 | 29.9585 |
| 0 | 107.6 | 5.29E+09 | 151 | 28.4749 | 28.4872 | 28.6503 | 28.6792 | 28.5536 |
| 0 | 275.9 | 2.13E+10 | 254 | 30.6132 | 30.5997 | 30.536 | 30.5046 | 30.6613 |
| 0 | 177.94 | 2.78E+10 | 329 | 30.9318 | 30.985 | 30.9189 | 30.7908 | 31.1146 |
| 0 | 30.541 | 2.17E+09 | 65 | 27.2917 | 27.3787 | 27.423 | 27.3608 | 27.143 |
| 0 | 43.441 | 1.92E+09 | 84 | 27.0771 | 27.0446 | 26.8599 | 27.1421 | 26.9752 |
| 0 | 176.08 | 2.48E+10 | 232 | 31.1258 | 31.1214 | 30.6488 | 30.6215 | 30.6958 |
| 0 | 117.21 | 4.82E+09 | 137 | 28.4002 | 28.4384 | 28.666 | 28.6082 | 28.1528 |
| 0 | 199.15 | 4.02E+09 | 110 | 28.4479 | 28.5601 | 28.033 | 28.1109 | 27.9846 |
| 0 | 32.619 | 2.23E+09 | 90 | 27.1341 | 27.1119 | 27.3017 | 27.1296 | 27.2584 |
| 0 | 277.1 | 2.3E+10 | 267 | 30.6559 | 30.6332 | 30.7989 | 30.6688 | 30.6223 |
| 0 | 108.01 | 5.46E+09 | 140 | 28.6237 | 28.6055 | 28.6965 | 28.5586 | 28.6082 |
| 0 | 286.05 | 2.18E+10 | 244 | 30.623 | 30.603 | 30.708 | 30.6778 | 30.486 |
| 0 | 132.37 | 7.5E+09 | 185 | 29.0645 | 29.0928 | 28.8887 | 28.9343 | 29.0409 |
| 0 | 143.76 | 5.15E+09 | 152 | 28.4751 | 28.3245 | 28.5331 | 28.6442 | 28.435 |
| 0 | 64.445 | 5.13E+09 | 155 | 28.6148 | 28.4218 | 28.6324 | 28.6759 | 28.542 |
| 0 | 323.31 | 9.43E+10 | 457 | 32.8236 | 32.8433 | 32.4987 | 32.5475 | 32.709 |
| 0 | 97.171 | 2.1E+10 | 207 | 30.5193 | 30.5598 | 30.6545 | 30.7186 | 30.4912 |
| 0 | 44.994 | 2.16E+09 | 66 | 27.1806 | 27.4057 | 27.3832 | 27.3254 | 27.2247 |
| 0 | 53.386 | 3.25E+09 | 121 | 27.7699 | 27.7534 | 27.8339 | 27.7899 | 27.935 |
| 0 | 308.83 | 4.35E+10 | 493 | 31.6304 | 31.6214 | 31.6279 | 31.691 | 31.4999 |
| 0 | 76.454 | 2.9E+09 | 101 | 27.8817 | 27.856 | 27.5045 | 27.5391 | 27.6783 |
| 0 | 63.599 | 3.05E+09 | 109 | 27.8788 | 27.7484 | 27.753 | 27.7853 | 27.737 |
| 0 | 62.191 | 3.86E+09 | 119 | 27.9914 | 28.0689 | 28.1666 | 28.2575 | 28.1393 |
| 0 | 95.058 | 3.01E+09 | 82 | 27.745 | 27.7417 | 27.6362 | 27.8051 | 27.804 |

| | | | | | | | | |
|---|---|---|---|---|---|---|---|---|
| 0 | 75.641 | 2.59E+09 | 91 | 27.5044 | 27.3331 | 27.4508 | 27.3632 | 27.7935 |
| 0 | 152 | 6.07E+09 | 142 | 28.688 | 28.7756 | 28.7545 | 28.7635 | 28.7773 |
| 0 | 194.52 | 6.79E+09 | 172 | 28.8842 | 28.9217 | 29.0838 | 28.9319 | 28.8931 |
| 0 | 41.018 | 1.59E+09 | 66 | 26.7216 | 26.7366 | 26.8293 | 26.8316 | 26.8479 |
| 0 | 96.378 | 8.88E+09 | 149 | 29.1484 | 29.185 | 29.0556 | 29.1343 | 29.2099 |
| 0 | 60.276 | 1.61E+09 | 79 | 26.5678 | 26.722 | 26.9258 | 26.5795 | 26.857 |
| 0 | 207.27 | 1.77E+10 | 316 | 30.1031 | 30.1935 | 30.3791 | 30.3333 | 30.0612 |
| 0 | 301.35 | 3.31E+10 | 359 | 31.1236 | 31.0589 | 31.0732 | 31.1126 | 31.1338 |
| 0 | 35.884 | 2.89E+09 | 89 | 27.6506 | 27.6356 | 27.6705 | 27.7145 | 27.5287 |
| 0 | 323.31 | 1.15E+11 | 416 | 32.9873 | 33.0098 | 32.9634 | 33.0378 | 32.8238 |
| 0 | 323.31 | 1.87E+10 | 250 | 30.363 | 30.2955 | 30.3628 | 30.2686 | 30.3111 |
| 0 | 257.03 | 1.64E+10 | 209 | 30.173 | 30.1793 | 30.141 | 30.0807 | 30.2219 |
| 0 | 188.5 | 1.54E+10 | 172 | 30.1408 | 30.1277 | 30.1718 | 30.2207 | 30.2206 |
| 0 | 44.8 | 1.93E+09 | 66 | 27.2582 | 27.3571 | 27.1902 | 27.1009 | 27.2235 |
| 0 | 24.221 | 1.61E+09 | 58 | 26.9213 | 27.2043 | 26.6794 | 27.0787 | 26.8874 |
| 0 | 303.05 | 1.25E+10 | 209 | 29.725 | 29.7164 | 29.7475 | 29.7289 | 29.8557 |
| 0 | 22.083 | 1.17E+09 | 57 | 26.4325 | 26.5055 | 26.3878 | 26.2448 | 26.521 |
| 0 | 323.31 | 6.36E+10 | 360 | 32.1469 | 32.0621 | 32.2014 | 32.2261 | 31.9076 |
| 0 | 141.27 | 1.55E+10 | 260 | 30.1452 | 30.1676 | 30.0461 | 30.0911 | 29.9981 |
| 0 | 313.69 | 6.24E+09 | 180 | 28.7448 | 28.7587 | 28.7502 | 28.7486 | 28.852 |
| 0 | 301.41 | 4.94E+10 | 292 | 31.7242 | 31.7483 | 31.8183 | 31.744 | 31.5573 |
| 0 | 323.31 | 3.42E+10 | 290 | 31.2241 | 31.2059 | 31.2639 | 31.2704 | 31.2245 |
| 0 | 143.03 | 1.47E+10 | 168 | 30.0907 | 30.0285 | 30.055 | 30.1163 | 30.1409 |
| 0 | 159.25 | 9.42E+09 | 174 | 29.6319 | 29.5892 | 29.3706 | 29.3753 | 29.4527 |
| 0 | 228.82 | 2.34E+10 | 234 | 30.6549 | 30.5398 | 30.5928 | 30.797 | 30.676 |
| 0 | 101.91 | 2.16E+09 | 78 | 27.1388 | 27.1732 | 27.3402 | 27.2042 | 27.1563 |
| 0 | 129.98 | 1.2E+10 | 183 | 29.7028 | 29.775 | 29.8008 | 29.7672 | 29.5077 |
| 0 | 18.736 | 1.25E+09 | 54 | 26.3522 | 26.5162 | 26.5374 | 26.792 | 26.559 |
| 0 | 150.89 | 1.03E+10 | 219 | 29.5454 | 29.4997 | 29.4788 | 29.4003 | 29.4756 |
| 0 | 40.339 | 1.25E+09 | 44 | 26.5271 | 26.6694 | 26.3657 | 26.5777 | 26.5457 |
| 0 | 185.44 | 1.29E+10 | 193 | 30.2373 | 30.3286 | 29.8363 | 29.8632 | 29.7564 |
| 0 | 34.59 | 2.08E+09 | 72 | 27.2747 | 27.2409 | 27.2157 | 27.3179 | 27.0067 |
| 0 | 23.815 | 1.32E+09 | 39 | 26.4724 | 26.1843 | 26.5645 | 26.3285 | 26.4888 |
| 0 | 150.97 | 9.3E+09 | 196 | 29.2512 | 29.2463 | 29.3249 | 29.416 | 29.3264 |
| 0 | 227.82 | 3.73E+10 | 280 | 31.3303 | 31.3146 | 31.4299 | 31.4251 | 31.2728 |
| 0 | 53.482 | 3.72E+09 | 90 | 28.1278 | 27.9543 | 28.186 | 28.0537 | 27.8504 |
| 0 | 217.35 | 6.08E+09 | 159 | 28.7821 | 28.6879 | 28.7893 | 28.8234 | 28.7157 |
| 0 | 90.613 | 2.23E+09 | 108 | 27.0525 | 27.1819 | 27.3204 | 27.1409 | 27.4107 |
| 0 | 138.8 | 4.29E+09 | 148 | 28.2799 | 28.2752 | 28.0213 | 28.024 | 28.3253 |
| 0 | 323.31 | 1.55E+11 | 690 | 33.545 | 33.5638 | 33.312 | 33.2239 | 33.5958 |
| 0 | 53.552 | 2.44E+09 | 72 | 27.4574 | 27.3369 | 27.1968 | 27.0003 | 27.5968 |
| 0 | 142.59 | 5.93E+09 | 158 | 28.9477 | 28.8997 | 28.6914 | 28.6377 | 28.7639 |
| 0 | 95.921 | 4.14E+09 | 124 | 28.222 | 28.0923 | 28.2481 | 28.3731 | 28.0577 |
| 0 | 50.588 | 4.53E+09 | 118 | 28.4611 | 28.4814 | 28.3299 | 28.2853 | 28.3842 |
| 0 | 323.31 | 1.62E+11 | 299 | 33.5143 | 33.5325 | 33.6545 | 33.6959 | 33.2915 |
| 0 | 62.563 | 3.25E+09 | 121 | 27.7982 | 27.7059 | 27.7188 | 27.714 | 27.8349 |
| 0 | 73.717 | 3.27E+09 | 134 | 28.19 | 28.2419 | 27.7141 | 27.643 | 27.9911 |
| 0 | 139.96 | 5E+09 | 127 | 28.4098 | 28.4747 | 28.5418 | 28.4767 | 28.4215 |

| | | | | | | | | |
|---|---|---|---|---|---|---|---|---|
| 0 | 117.2 | 1.51E+10 | 204 | 30.1618 | 30.1266 | 30.1986 | 30.0999 | 29.939 |
| 0 | 115.38 | 7.36E+09 | 155 | 29.1949 | 29.0539 | 28.7029 | 28.8561 | 29.0411 |
| 0 | 199.39 | 1.52E+10 | 153 | 30.02 | 30.0625 | 30.243 | 30.2072 | 29.7718 |
| 0 | 42.076 | 1.84E+09 | 81 | 26.8753 | 26.8908 | 27.1391 | 27.0951 | 26.9618 |
| 0 | 323.31 | 4.42E+10 | 394 | 31.6089 | 31.5871 | 31.6281 | 31.6314 | 31.5896 |
| 0 | 183.52 | 4.64E+09 | 145 | 28.4007 | 28.468 | 28.1994 | 28.2145 | 28.3781 |
| 0 | 228.8 | 1.16E+10 | 170 | 29.6391 | 29.6012 | 29.7677 | 29.7481 | 29.6533 |
| 0 | 83.039 | 2.78E+09 | 87 | 27.7304 | 27.734 | 27.4442 | 27.5191 | 27.7532 |
| 0 | 242.29 | 6.12E+10 | 243 | 32.1227 | 32.1603 | 32.0387 | 32.0124 | 32.1592 |
| 0 | 230.49 | 7.18E+09 | 134 | 29.1939 | 29.0722 | 29.2171 | 29.1325 | 28.8647 |
| 0 | 86.062 | 3.72E+09 | 142 | 27.9998 | 27.974 | 28.0593 | 28.1133 | 28.1345 |
| 0 | 235.42 | 1.04E+10 | 187 | 29.6033 | 29.616 | 29.528 | 29.5411 | 29.5145 |
| 0 | 104.73 | 9.55E+09 | 204 | 29.4308 | 29.3813 | 29.4614 | 29.3293 | 29.4939 |
| 0 | 62.494 | 2.07E+09 | 88 | 27.2565 | 27.2012 | 27.1809 | 27.0077 | 27.3597 |
| 0 | 72.434 | 3.78E+09 | 113 | 28.0338 | 28.0049 | 27.914 | 28.0558 | 28.1923 |
| 0 | 84.582 | 1.92E+09 | 85 | 26.8476 | 27.07 | 27.1106 | 27.0017 | 27.2788 |
| 0 | 167.46 | 1E+10 | 174 | 29.4447 | 29.4674 | 29.4372 | 29.52 | 29.4604 |
| 0 | 106.77 | 7.35E+09 | 140 | 28.962 | 29.0644 | 29.0207 | 28.9183 | 29.2149 |
| 0 | 323.31 | 3.22E+10 | 251 | 31.1383 | 31.1147 | 31.1905 | 31.1649 | 31.2161 |
| 0 | 226.54 | 5.49E+10 | 295 | 32.0716 | 32.0524 | 31.883 | 31.8505 | 31.9309 |
| 0 | 98.628 | 1.04E+10 | 165 | 29.6207 | 29.5974 | 29.5735 | 29.5235 | 29.5362 |
| 0 | 168.87 | 3.37E+09 | 105 | 27.9031 | 27.8437 | 27.8566 | 27.8115 | 27.9986 |
| 0 | 88.585 | 6.27E+09 | 203 | 28.8871 | 28.7847 | 28.6757 | 28.7555 | 28.8456 |
| 0 | 86.25 | 2.95E+10 | 238 | 31.0485 | 31.0684 | 31.0537 | 31.0154 | 31.0639 |
| 0 | 105.15 | 3.65E+09 | 143 | 27.951 | 27.8771 | 27.8716 | 27.8805 | 28.1811 |
| 0 | 138.38 | 1.07E+10 | 212 | 29.4921 | 29.4009 | 29.5054 | 29.5277 | 29.6858 |
| 0 | 323.31 | 7.89E+10 | 375 | 32.3864 | 32.4027 | 32.4379 | 32.5053 | 32.4347 |
| 0 | 48.421 | 2.07E+09 | 81 | 27.1484 | 26.9983 | 27.4338 | 27.3272 | 27.2291 |
| 0 | 108.29 | 1.86E+09 | 80 | 26.8605 | 27.0062 | 27.064 | 27.0632 | 27.0098 |
| 0 | 116.72 | 5.81E+09 | 179 | 28.6739 | 28.7028 | 28.6697 | 28.6408 | 28.7757 |
| 0 | 310.59 | 1.86E+10 | 267 | 30.3909 | 30.435 | 30.611 | 30.5528 | 30.1747 |
| 0 | 151 | 1.97E+10 | 231 | 30.4349 | 30.4271 | 30.516 | 30.5681 | 30.4763 |
| 0 | 265.34 | 4.77E+10 | 372 | 31.7944 | 31.7643 | 31.7594 | 31.8091 | 31.6723 |
| 0 | 323.31 | 7.13E+09 | 133 | 28.6484 | 28.6781 | 28.7449 | 28.654 | 28.8872 |
| 0 | 92.574 | 2.08E+09 | 89 | 26.7995 | 26.9243 | 26.9418 | 26.9508 | 27.2619 |
| 0 | 321.08 | 1.98E+10 | 177 | 30.2413 | 30.2175 | 30.384 | 30.2489 | 30.4648 |
| 0 | 170.63 | 6.67E+09 | 148 | 28.6859 | 28.7002 | 28.6981 | 28.6251 | 28.8529 |
| 0 | 76.403 | 4.12E+09 | 110 | 28.0456 | 28.0806 | 28.1903 | 28.2375 | 28.0637 |
| 0 | 77.505 | 3.26E+09 | 122 | 27.7589 | 27.715 | 27.7977 | 27.8446 | 27.7314 |
| 0 | 287.75 | 4.77E+10 | 427 | 31.5572 | 31.5856 | 31.5981 | 31.6944 | 31.6327 |
| 0 | 36.705 | 2.92E+09 | 110 | 27.4807 | 27.7352 | 27.6564 | 27.6631 | 27.504 |
| 0 | 97.978 | 2.71E+09 | 116 | 27.5515 | 27.5253 | 27.5513 | 27.5333 | 27.5243 |
| 0 | 57.516 | 3.04E+09 | 111 | 27.8245 | 27.8455 | 27.7657 | 27.7934 | 27.9132 |
| 0 | 122.64 | 1.7E+10 | 220 | 30.6146 | 30.6024 | 30.3427 | 30.1871 | 30.2856 |
| 0 | 33.504 | 4.58E+09 | 50 | 28.0316 | 28.2223 | 27.472 | 28.3827 | 28.3295 |
| 0 | 84.256 | 4.48E+10 | 214 | 31.8201 | 31.7766 | 31.647 | 31.7791 | 31.7797 |
| 0 | 259.03 | 1.33E+10 | 213 | 29.8118 | 29.7299 | 29.7832 | 29.7448 | 29.9359 |
| 0 | 15.439 | 2.19E+09 | 24 | 27.1941 | 27.0468 | 27.3008 | 27.0692 | 27.232 |

| | | | | | | | | |
|---|---|---|---|---|---|---|---|---|
| 0 | 146.39 | 1.11E+10 | 143 | 29.6226 | 29.713 | 29.6495 | 29.6711 | 29.4519 |
| 0 | 154.68 | 4E+09 | 120 | 28.0819 | 28.0142 | 27.9736 | 27.915 | 28.2509 |
| 0 | 203.56 | 6.62E+09 | 131 | 28.822 | 28.8321 | 28.893 | 28.7713 | 28.9439 |
| 0 | 29.858 | 2.92E+10 | 163 | 31.0911 | 31.1527 | 31.1594 | 31.1248 | 31.005 |
| 0 | 78.82 | 1.88E+09 | 69 | 26.9027 | 26.9618 | 26.9681 | 26.9435 | 26.9983 |
| 0 | 40.247 | 2.48E+09 | 76 | 27.6017 | 27.4219 | 27.6024 | 27.4521 | 27.1352 |
| 0 | 37.794 | 7.49E+09 | 114 | 28.9268 | 28.9759 | 29.1479 | 29.0178 | 28.9495 |
| 0 | 165.26 | 7.92E+09 | 142 | 29.0873 | 28.9624 | 29.0402 | 29.0356 | 29.1736 |
| 0 | 323.31 | 2.63E+10 | 244 | 30.7164 | 30.6988 | 30.792 | 30.9664 | 30.8895 |
| 0 | 323.31 | 3.82E+10 | 244 | 31.7029 | 31.7072 | 31.3084 | 31.2776 | 31.5448 |
| 0 | 165.58 | 5.68E+09 | 179 | 28.8872 | 28.8977 | 28.4838 | 28.4408 | 28.8262 |
| 0 | 190.48 | 7.12E+09 | 159 | 29.0417 | 29.0564 | 29.1099 | 29.1014 | 28.9391 |
| 0 | 323.31 | 3.41E+10 | 247 | 31.2103 | 31.1269 | 31.3279 | 31.2543 | 31.191 |
| 0 | 56.174 | 4.26E+09 | 120 | 28.5391 | 28.5048 | 28.2655 | 28.1407 | 28.2952 |
| 0 | 105.28 | 5.41E+10 | 281 | 31.8932 | 31.8906 | 31.9542 | 31.9703 | 31.776 |
| 0 | 120.79 | 9.6E+09 | 198 | 29.5997 | 29.5221 | 29.4495 | 29.3797 | 29.4235 |
| 0 | 92.685 | 8.24E+09 | 148 | 29.2278 | 29.2154 | 29.3157 | 29.3109 | 29.2884 |
| 0 | 323.31 | 5.5E+09 | 138 | 28.8863 | 28.8435 | 28.5119 | 28.4657 | 28.7101 |
| 0 | 323.31 | 1.36E+11 | 362 | 33.2814 | 33.2771 | 33.3031 | 33.2176 | 33.0676 |
| 0 | 45.571 | 3.79E+09 | 159 | 27.9566 | 28.1337 | 27.8454 | 27.8347 | 28.2072 |
| 0 | 138.02 | 4.51E+09 | 93 | 27.9519 | 27.9589 | 27.726 | 27.7901 | 28.7507 |
| 0 | 155.79 | 2.44E+10 | 215 | 30.655 | 30.6031 | 30.7635 | 30.6704 | 31.158 |
| 0 | 36.857 | 2.74E+09 | 92 | 27.5867 | 27.6209 | 27.6277 | 27.6602 | 27.4556 |
| 0 | 108.97 | 4.07E+09 | 90 | 27.9175 | 27.8793 | 28.0564 | 28.021 | 28.3706 |
| 0 | 174.53 | 3.71E+10 | 361 | 31.2993 | 31.3081 | 31.429 | 31.4217 | 31.3163 |
| 0 | 213.56 | 7.44E+09 | 199 | 28.9676 | 28.907 | 29.1449 | 29.153 | 29.042 |
| 0 | 83.407 | 1.19E+10 | 163 | 29.6509 | 29.6508 | 29.7422 | 29.7506 | 29.7572 |
| 0 | 50.586 | 2.2E+09 | 71 | 27.2685 | 27.4839 | 27.237 | 27.372 | 27.2425 |
| 0 | 323.31 | 1.09E+10 | 215 | 29.4548 | 29.4752 | 29.7383 | 29.634 | 29.6123 |
| 0 | 37.124 | 2.46E+09 | 117 | 27.2744 | 27.3761 | 27.4055 | 27.5837 | 27.3394 |
| 0 | 103.93 | 1.6E+10 | 211 | 30.227 | 30.1985 | 30.0196 | 29.9943 | 30.0271 |
| 0 | 242.39 | 9.84E+09 | 203 | 29.8262 | 29.7703 | 29.4375 | 29.3811 | 29.383 |
| 0 | 144.38 | 1.94E+10 | 276 | 30.7748 | 30.8471 | 30.4117 | 30.3053 | 30.337 |
| 0 | 267.44 | 4.28E+10 | 249 | 31.8941 | 31.8501 | 31.5069 | 31.4739 | 31.5825 |
| 0 | 112.37 | 4.99E+10 | 273 | 31.907 | 31.7871 | 31.8404 | 31.8382 | 31.7303 |
| 0 | 38.516 | 3.34E+09 | 100 | 27.8145 | 27.852 | 27.6802 | 27.9071 | 27.9796 |
| 0 | 100.75 | 2.69E+09 | 79 | 27.6053 | 27.4917 | 27.3876 | 27.3857 | 27.64 |
| 0 | 66.102 | 4.84E+09 | 148 | 28.3342 | 28.459 | 28.4705 | 28.5096 | 28.3512 |
| 0 | 61.596 | 2.24E+09 | 78 | 27.3703 | 27.3313 | 27.4686 | 27.3998 | 27.1697 |
| 0 | 63.418 | 4.63E+09 | 134 | 28.402 | 28.4565 | 28.3345 | 28.5141 | 28.3076 |
| 0 | 34.256 | 1.37E+09 | 62 | 26.5532 | 26.4285 | 26.8099 | 26.6239 | 26.8323 |
| 0 | 38.985 | 2.8E+09 | 100 | 27.519 | 27.6158 | 27.6148 | 27.5582 | 27.7372 |
| 0 | 40.829 | 2.18E+09 | 68 | 27.2993 | 27.27 | 27.1496 | 26.9711 | 27.3222 |
| 0 | 27.677 | 2.22E+09 | 78 | 27.3194 | 27.3166 | 27.3558 | 27.1316 | 27.3083 |
| 0 | 154.19 | 3.52E+09 | 105 | 28.0984 | 28.1113 | 27.8859 | 27.9295 | 27.8617 |
| 0 | 146.93 | 9.72E+09 | 186 | 29.498 | 29.479 | 29.3202 | 29.3101 | 29.4484 |
| 0 | 41.994 | 1.82E+09 | 61 | 27.07 | 27.2359 | 27.174 | 26.8699 | 27.0368 |
| 0 | 218.57 | 2.01E+10 | 249 | 30.5989 | 30.6239 | 30.4277 | 30.4517 | 30.3419 |

| | | | | | | | | |
|---|---|---|---|---|---|---|---|---|
| 0 | 215.39 | 1.41E+10 | 197 | 30.0246 | 29.9881 | 30.0878 | 30.0423 | 30.0073 |
| 0 | 84.478 | 5.62E+09 | 120 | 28.5566 | 28.5568 | 28.4346 | 28.4816 | 28.714 |
| 0 | 323.31 | 1.11E+11 | 253 | 32.9274 | 32.9465 | 33.0379 | 33.043 | 32.8998 |
| 0 | 323.31 | 4.51E+10 | 241 | 31.8717 | 31.8515 | 31.4568 | 31.5078 | 31.6416 |
| 0 | 57.34 | 3.89E+09 | 99 | 28.145 | 28.3076 | 28.2296 | 28.095 | 28.002 |
| 0 | 33.912 | 2.92E+09 | 105 | 27.7392 | 27.7607 | 27.9663 | 27.8506 | 27.4398 |
| 0 | 89.834 | 2.48E+09 | 96 | 27.6942 | 27.6769 | 27.2109 | 27.1479 | 27.3881 |
| 0 | 70.46 | 1.54E+09 | 50 | 26.869 | 26.8087 | 26.8038 | 26.7391 | 26.7374 |
| 0 | 90.505 | 4.52E+09 | 135 | 28.3634 | 28.3987 | 28.0909 | 28.3277 | 28.3486 |
| 0 | 199.41 | 2.49E+10 | 185 | 31.049 | 31.0614 | 30.7433 | 30.6291 | 30.7337 |
| 0 | 106.64 | 4.11E+09 | 134 | 28.2991 | 28.2468 | 28.2106 | 28.332 | 28.1517 |
| 0 | 211.61 | 8.24E+09 | 186 | 29.3632 | 29.3327 | 29.0879 | 29.0255 | 29.1489 |
| 0 | 323.31 | 1.76E+10 | 207 | 30.2936 | 30.2812 | 30.4178 | 30.4222 | 30.1334 |
| 0 | 190.27 | 7.89E+09 | 166 | 29.2396 | 29.2853 | 29.248 | 29.238 | 28.8691 |
| 0 | 323.31 | 2.87E+11 | 951 | 34.1877 | 34.1289 | 34.377 | 34.3104 | 34.3342 |
| 0 | 151.61 | 1.9E+10 | 234 | 30.3951 | 30.3775 | 30.4721 | 30.4256 | 30.435 |
| 0 | 52.068 | 3.24E+09 | 98 | 27.9057 | 27.9299 | 27.7064 | 27.7526 | 27.9375 |
| 0 | 31.829 | 2.23E+09 | 60 | 27.1655 | 27.2766 | 27.1703 | 27.1207 | 27.4494 |
| 0 | 185.44 | 7.08E+10 | 282 | 32.4218 | 32.4479 | 32.1142 | 32.1759 | 32.2688 |
| 0 | 89.403 | 5.72E+09 | 144 | 28.6296 | 28.7422 | 28.7891 | 28.8364 | 28.4685 |
| 0 | 88.172 | 4.45E+10 | 264 | 31.7002 | 31.6254 | 31.7812 | 31.7698 | 31.4514 |
| 0 | 32.829 | 2.11E+09 | 82 | 27.2537 | 27.1085 | 27.0923 | 27.23 | 27.4521 |
| 0 | 72.857 | 3.39E+09 | 134 | 27.9295 | 27.8748 | 28.0958 | 28.1214 | 27.7733 |
| 0 | 129.3 | 1.07E+10 | 173 | 29.4956 | 29.5058 | 29.7299 | 29.6551 | 29.5368 |
| 0 | 79.545 | 3.06E+09 | 99 | 27.718 | 27.8102 | 27.7429 | 27.8283 | 27.7974 |
| 0 | 51.875 | 3.71E+09 | 102 | 28.0997 | 28.0251 | 28.1586 | 28.0524 | 27.9725 |
| 0 | 32.018 | 9.12E+09 | 47 | 28.8588 | 29.1886 | 29.1652 | 29.2763 | 29.0299 |
| 0 | 174.3 | 4.06E+09 | 139 | 27.9803 | 28.0617 | 27.9729 | 27.8542 | 28.102 |
| 0 | 83.717 | 6.63E+09 | 147 | 28.6212 | 28.6989 | 28.7891 | 28.8722 | 28.6715 |
| 0 | 100.4 | 2.47E+09 | 106 | 27.3395 | 27.3976 | 27.5069 | 27.4423 | 27.1264 |
| 0 | 22.949 | 1.56E+09 | 57 | 26.5462 | 26.729 | 26.7512 | 26.9507 | 26.4403 |
| 0 | 38.764 | 2.87E+09 | 66 | 27.6982 | 27.5455 | 27.599 | 27.5324 | 27.5027 |
| 0 | 136.98 | 1.5E+09 | 63 | 26.6048 | 26.6013 | 26.6944 | 26.6287 | 26.4947 |
| 0 | 28.083 | 1.8E+09 | 72 | 26.8501 | 27.1157 | 26.9519 | 27.0328 | 26.9159 |
| 0 | 236.08 | 2.86E+10 | 282 | 30.8911 | 30.8226 | 31.0565 | 30.9647 | 30.8764 |
| 0 | 129.34 | 5.38E+09 | 125 | 28.5759 | 28.4206 | 28.4003 | 28.3918 | 28.5261 |
| 0 | 52.036 | 2.37E+09 | 77 | 27.2932 | 27.2562 | 27.3147 | 27.3504 | 27.426 |
| 0 | 162.41 | 8.07E+09 | 170 | 29.0831 | 29.0997 | 29.207 | 29.1521 | 29.0976 |
| 0 | 323.31 | 1.27E+11 | 705 | 33.0551 | 33.0569 | 33.0753 | 33.164 | 32.9894 |
| 0 | 191.62 | 6.33E+09 | 163 | 28.8285 | 28.735 | 28.8803 | 28.7371 | 28.8291 |
| 0 | 227.87 | 1.69E+10 | 178 | 30.3045 | 30.2694 | 30.2924 | 30.2651 | 30.3515 |
| 0 | 81.263 | 4.72E+09 | 127 | 28.7266 | 28.7202 | 28.4345 | 28.409 | 28.42 |
| 0 | 146.22 | 1.04E+10 | 227 | 29.8727 | 29.8587 | 29.4994 | 29.5014 | 29.5798 |
| 0 | 231.01 | 4.61E+10 | 266 | 32.0849 | 32.0231 | 31.8423 | 31.8232 | 31.6196 |
| 0 | 289.91 | 1.16E+10 | 307 | 29.9525 | 29.8919 | 29.706 | 29.5468 | 30.0309 |
| 0 | 152.14 | 6.51E+09 | 157 | 28.8155 | 28.6917 | 28.9458 | 28.8782 | 28.7226 |
| 0 | 97.466 | 1.05E+10 | 174 | 29.3578 | 29.4273 | 29.5723 | 29.5412 | 29.4642 |
| 0 | 51.077 | 3.84E+09 | 121 | 28.0019 | 27.9955 | 28.3041 | 28.2121 | 27.7984 |

| | | | | | | | | |
|---|---|---|---|---|---|---|---|---|
| 0 | 38.408 | 2.33E+09 | 93 | 27.3921 | 27.3939 | 27.1933 | 27.3179 | 27.2928 |
| 0 | 51.949 | 2.83E+09 | 106 | 27.5915 | 27.5507 | 27.5953 | 27.5881 | 27.7806 |
| 0 | 176.8 | 2.5E+10 | 188 | 30.7988 | 30.8018 | 30.8095 | 30.7923 | 30.6543 |
| 0 | 208.83 | 5.76E+09 | 165 | 28.5941 | 28.6028 | 28.6021 | 28.7347 | 28.5672 |
| 0 | 264.15 | 2.98E+10 | 256 | 31.4472 | 31.3836 | 30.9925 | 30.9826 | 31.1062 |
| 0 | 186.81 | 5.84E+10 | 283 | 32.0365 | 32.0322 | 32.0579 | 32.0196 | 31.9372 |
| 0 | 204.2 | 1.09E+10 | 158 | 29.5622 | 29.575 | 29.5939 | 29.4371 | 29.5758 |
| 0 | 173.39 | 1.42E+10 | 188 | 30.3908 | 30.3055 | 29.975 | 29.9344 | 29.929 |
| 0 | 58.327 | 1.94E+09 | 64 | 27.2228 | 27.0077 | 27.4802 | 27.0839 | 27.2197 |
| 0 | 141.27 | 2.9E+10 | 200 | 30.9706 | 30.9615 | 31.0722 | 30.9819 | 30.8831 |
| 0 | 81.937 | 1.67E+09 | 59 | 26.8533 | 26.7782 | 26.6885 | 26.8792 | 26.9536 |
| 0 | 62.055 | 3.29E+09 | 84 | 27.5392 | 27.5218 | 27.8647 | 27.8083 | 28.0205 |
| 0 | 58.053 | 3.44E+09 | 97 | 27.853 | 27.7327 | 28.1011 | 27.9777 | 27.9019 |
| 0 | 37.468 | 2.62E+09 | 116 | 27.5026 | 27.4594 | 27.7075 | 27.5776 | 27.3348 |
| 0 | 33.592 | 2.59E+09 | 96 | 27.6031 | 27.582 | 27.3002 | 27.3581 | 27.5619 |
| 0 | 261.21 | 7.2E+09 | 152 | 29.0049 | 29.0723 | 29.008 | 29.0428 | 28.9113 |
| 0 | 282.95 | 8.17E+09 | 167 | 29.1049 | 29.1407 | 29.345 | 29.2588 | 29.0744 |
| 0 | 34.743 | 1.44E+09 | 61 | 26.7767 | 26.5676 | 26.1907 | 26.4244 | 26.7596 |
| 0 | 306.82 | 2.95E+10 | 382 | 31.0469 | 31.0133 | 31.1733 | 31.1188 | 30.8208 |
| 0 | 323.31 | 4.52E+10 | 391 | 31.7198 | 31.664 | 31.5874 | 31.5582 | 31.6057 |
| 0 | 37.161 | 3.22E+09 | 50 | 27.6579 | 27.7068 | 27.7631 | 27.7129 | 28.0527 |
| 0 | 148.92 | 6.58E+09 | 154 | 28.9883 | 28.9615 | 28.8192 | 28.7824 | 28.9627 |
| 0 | 113.88 | 5.66E+09 | 136 | 28.6902 | 28.5982 | 28.7096 | 28.6158 | 28.548 |
| 0 | 53.434 | 2.28E+09 | 86 | 27.3019 | 27.2473 | 27.0492 | 27.0877 | 27.3678 |
| 0 | 66.918 | 4.4E+09 | 140 | 28.3712 | 28.3164 | 28.3349 | 28.235 | 28.4281 |
| 0 | 93.238 | 8.12E+09 | 118 | 29.2388 | 29.3106 | 29.178 | 29.1185 | 29.0052 |
| 0 | 323.31 | 1.06E+11 | 405 | 32.8485 | 32.8351 | 32.8809 | 32.8644 | 32.8114 |
| 0 | 59.882 | 3.98E+09 | 125 | 28.0901 | 28.0496 | 28.1936 | 28.1766 | 28.0861 |
| 0 | 59.792 | 2E+09 | 61 | 27.196 | 27.1605 | 27.4326 | 27.2464 | 27.026 |
| 0 | 36.87 | 4.31E+09 | 134 | 28.302 | 28.3112 | 28.2383 | 28.3622 | 28.2397 |
| 0 | 63.203 | 1.75E+09 | 69 | 27.0488 | 26.8219 | 26.959 | 27.0422 | 26.9033 |
| 0 | 35.036 | 3.54E+09 | 103 | 28.3492 | 28.2701 | 27.8506 | 27.7205 | 27.9777 |
| 0 | 148.75 | 2.66E+09 | 97 | 27.857 | 27.8833 | 27.5043 | 27.3175 | 27.6095 |
| 0 | 153.87 | 6.78E+09 | 128 | 28.8787 | 28.8589 | 28.9524 | 28.9717 | 28.8549 |
| 0 | 221.32 | 5.75E+09 | 146 | 28.6755 | 28.6119 | 28.6379 | 28.7399 | 28.6171 |
| 0 | 55.847 | 3.98E+10 | 205 | 31.5752 | 31.531 | 31.4545 | 31.556 | 31.4683 |
| 0 | 221.3 | 3.94E+09 | 84 | 28.1575 | 28.0886 | 28.2714 | 28.3435 | 28.046 |
| 0 | 210.21 | 4.54E+10 | 286 | 32.0609 | 32.0559 | 31.5534 | 31.5651 | 31.4877 |
| 0 | 25.736 | 2.6E+09 | 95 | 27.464 | 27.6003 | 27.5857 | 27.5462 | 27.4262 |
| 0 | 189.85 | 3.73E+10 | 193 | 31.3371 | 31.4047 | 31.434 | 31.4593 | 31.4211 |
| 0 | 153.71 | 4.59E+09 | 119 | 28.7377 | 28.5899 | 28.1726 | 28.1936 | 28.384 |
| 0 | 81.925 | 1.47E+10 | 244 | 29.9351 | 29.907 | 30.2077 | 30.0172 | 30.0821 |
| 0 | 224.31 | 3.7E+10 | 195 | 31.7005 | 31.6735 | 31.2045 | 31.1734 | 31.0958 |
| 0 | 323.31 | 4.2E+09 | 161 | 28.2274 | 28.2581 | 28.328 | 28.2832 | 28.1471 |
| 0 | 64.756 | 2.01E+09 | 79 | 27.0423 | 26.9728 | 27.0455 | 26.9982 | 27.3336 |
| 0 | 225.78 | 6.82E+09 | 141 | 28.8674 | 28.9529 | 29.0445 | 28.9426 | 28.9517 |
| 0 | 323.31 | 8.8E+09 | 129 | 29.3625 | 29.2494 | 29.417 | 29.3462 | 29.2024 |
| 0 | 38.109 | 4.11E+09 | 104 | 28.265 | 28.2303 | 27.9911 | 28.0961 | 28.3385 |

| | | | | | | | | |
|---|---|---|---|---|---|---|---|---|
| 0 | 97.312 | 3.51E+09 | 136 | 28.2373 | 28.3964 | 27.9235 | 27.8168 | 27.8338 |
| 0 | 91.59 | 4.05E+09 | 117 | 28.2324 | 28.076 | 28.0522 | 28.2842 | 28.2289 |
| 0 | 61.199 | 2.31E+09 | 74 | 27.4391 | 27.4365 | 27.2536 | 27.1669 | 27.4387 |
| 0 | 174.76 | 3.7E+09 | 125 | 28.062 | 28.0383 | 28.211 | 28.1727 | 27.9574 |
| 0 | 227.85 | 3.84E+10 | 254 | 31.4368 | 31.438 | 31.6375 | 31.5911 | 31.1763 |
| 0 | 323.31 | 4.12E+10 | 335 | 31.5265 | 31.464 | 31.6073 | 31.5756 | 31.5503 |
| 0 | 323.31 | 2.78E+10 | 470 | 30.8767 | 30.8935 | 31.0003 | 30.958 | 31.0808 |
| 0 | 23.792 | 1.36E+09 | 36 | 26.6565 | 26.6721 | 26.5369 | 26.6628 | 26.4577 |
| 0 | 216.37 | 1.86E+10 | 242 | 30.5505 | 30.5978 | 30.3314 | 30.2699 | 30.1583 |
| 0 | 65.364 | 1E+10 | 190 | 29.5415 | 29.6043 | 29.4088 | 29.3277 | 29.4995 |
| 0 | 173.72 | 5.6E+09 | 139 | 28.6693 | 28.664 | 28.7276 | 28.6568 | 28.5841 |
| 0 | 323.31 | 6.53E+10 | 378 | 32.0993 | 32.0841 | 32.3476 | 32.2685 | 32.1844 |
| 0 | 323.31 | 2.14E+10 | 228 | 30.6174 | 30.6197 | 30.7722 | 30.7142 | 30.4074 |
| 0 | 64.696 | 3.37E+09 | 87 | 27.9115 | 27.9674 | 27.8255 | 27.7858 | 27.9904 |
| 0 | 115.68 | 5.28E+09 | 85 | 28.6216 | 28.6425 | 28.5395 | 28.6197 | 28.5469 |
| 0 | 166.53 | 6.24E+09 | 145 | 28.7725 | 28.7723 | 28.9151 | 28.847 | 28.8216 |
| 0 | 22.964 | 5.91E+09 | 69 | 28.8412 | 28.7135 | 28.5373 | 28.723 | 28.6626 |
| 0 | 26.204 | 1.6E+09 | 58 | 26.757 | 26.8836 | 26.8985 | 26.9131 | 26.914 |
| 0 | 82.04 | 1.28E+10 | 144 | 29.7947 | 29.8057 | 29.9205 | 29.8571 | 29.8297 |
| 0 | 251.66 | 5.85E+10 | 384 | 31.8657 | 31.8766 | 32.0197 | 31.88 | 31.8515 |
| 0 | 117.47 | 9.39E+09 | 189 | 29.2572 | 29.2681 | 29.1652 | 29.339 | 29.4242 |
| 0 | 70.849 | 5.34E+09 | 124 | 28.4892 | 28.5424 | 28.5343 | 28.5088 | 28.4448 |
| 0 | 206.73 | 1.23E+10 | 198 | 29.8837 | 29.7022 | 29.6967 | 29.7314 | 29.6167 |
| 0 | 45.178 | 2.71E+09 | 63 | 27.5797 | 27.6212 | 27.4456 | 27.5546 | 27.4982 |
| 0 | 166.6 | 9.38E+10 | 337 | 32.6653 | 32.6611 | 32.6315 | 32.6367 | 32.5775 |
| 0 | 273.18 | 1.09E+10 | 165 | 29.5749 | 29.5596 | 29.5024 | 29.3869 | 29.5331 |
| 0 | 229.4 | 3.73E+09 | 79 | 27.8284 | 27.9231 | 28.1058 | 27.9988 | 28.0624 |
| 0 | 87.468 | 5.04E+09 | 134 | 28.4617 | 28.4153 | 28.5434 | 28.4021 | 28.3433 |
| 0 | 192.77 | 2.29E+10 | 177 | 30.621 | 30.6227 | 30.6021 | 30.6154 | 30.6561 |
| 0 | 57.868 | 7.24E+09 | 129 | 29.0049 | 29.0457 | 28.9845 | 28.9428 | 28.9324 |
| 0 | 140.26 | 9.11E+09 | 129 | 29.28 | 29.2571 | 29.3437 | 29.2967 | 29.3363 |
| 0 | 92.682 | 4.84E+09 | 118 | 28.4934 | 28.5484 | 28.5024 | 28.391 | 28.5061 |
| 0 | 47.439 | 3.61E+09 | 110 | 28.1396 | 28.0687 | 28.1577 | 28.1842 | 28.0696 |
| 0 | 37.646 | 2.33E+09 | 85 | 27.4706 | 27.3866 | 27.7534 | 27.5916 | 27.3229 |
| 0 | 204.64 | 1.16E+10 | 128 | 29.5529 | 29.6243 | 29.739 | 29.7633 | 29.5717 |
| 0 | 104.4 | 4.07E+09 | 116 | 28.1962 | 28.2179 | 28.2078 | 28.1534 | 28.3593 |
| 0 | 131.66 | 4.09E+10 | 236 | 31.4964 | 31.5088 | 31.4785 | 31.5198 | 31.4151 |
| 0 | 68.208 | 3.89E+09 | 110 | 28.2945 | 28.3119 | 28.0664 | 28.1668 | 28.1383 |
| 0 | 33.818 | 5.25E+09 | 103 | 28.5598 | 28.6877 | 28.6768 | 28.5655 | 28.5359 |
| 0 | 121.45 | 2.45E+09 | 93 | 27.4358 | 27.44 | 27.297 | 27.4133 | 27.3315 |
| 0 | 81.828 | 2.48E+09 | 74 | 27.4053 | 27.4458 | 27.5134 | 27.5551 | 26.9539 |
| 0 | 68.785 | 4.78E+09 | 94 | 28.3541 | 28.3189 | 28.2711 | 28.2396 | 28.304 |
| 0 | 201.87 | 1.23E+11 | 385 | 33.5433 | 33.4973 | 33.0964 | 33.0919 | 33.0488 |
| 0 | 31 | 2.52E+09 | 87 | 27.5425 | 27.4743 | 27.4907 | 27.5389 | 27.1051 |
| 0 | 88.54 | 3.24E+09 | 123 | 27.6682 | 27.7549 | 27.8721 | 27.9344 | 27.8019 |
| 0 | 103.48 | 1.53E+10 | 158 | 30.0581 | 30.1807 | 30.2613 | 30.2526 | 30.1125 |
| 0 | 297.9 | 2.82E+10 | 315 | 30.9611 | 30.9664 | 31.0933 | 31.0368 | 31.1047 |
| 0 | 36.504 | 2.39E+09 | 55 | 27.3719 | 27.2618 | 27.3751 | 27.2192 | 27.3836 |

| | | | | | | | | |
|---|---|---|---|---|---|---|---|---|
| 0 | 230.2 | 9.3E+09 | 210 | 29.3126 | 29.2273 | 29.2107 | 29.0802 | 29.4783 |
| 0 | 51.245 | 1.49E+09 | 42 | 27.0215 | 26.7057 | 26.8504 | 26.7064 | 26.2983 |
| 0 | 33.868 | 2.46E+09 | 85 | 27.355 | 27.3366 | 27.4225 | 27.6193 | 27.4174 |
| 0 | 188.76 | 1.69E+10 | 171 | 30.2134 | 30.1853 | 30.16 | 30.2193 | 30.2413 |
| 0 | 113.46 | 8.19E+09 | 162 | 28.9989 | 29.0409 | 29.1541 | 29.0279 | 29.2343 |
| 0 | 45.238 | 3.19E+09 | 115 | 28.001 | 28.0163 | 27.9444 | 27.8697 | 27.7001 |
| 0 | 188.25 | 1.03E+10 | 145 | 30.0849 | 30.094 | 29.3981 | 29.4113 | 29.4498 |
| 0 | 290.96 | 1.02E+10 | 152 | 29.4061 | 29.3009 | 29.5759 | 29.5522 | 29.4532 |
| 0 | 311.2 | 5.56E+09 | 136 | 28.7023 | 28.6352 | 28.3575 | 28.504 | 28.6923 |
| 0 | 266.04 | 1.1E+10 | 175 | 29.7325 | 29.7004 | 29.5254 | 29.6058 | 29.7242 |
| 0 | 57.003 | 3.26E+09 | 100 | 27.7457 | 27.7372 | 27.7972 | 27.7547 | 27.9735 |
| 0 | 323.31 | 6.1E+09 | 154 | 28.7565 | 28.6613 | 28.8969 | 28.7807 | 28.6917 |
| 0 | 204.97 | 4.69E+09 | 133 | 28.4691 | 28.3767 | 28.428 | 28.3654 | 28.267 |
| 0 | 27.483 | 1.44E+09 | 56 | 26.4924 | 26.5928 | 26.8622 | 26.7017 | 26.4996 |
| 0 | 103.37 | 7.33E+09 | 116 | 29.0405 | 28.9352 | 29.1618 | 29.0283 | 29.1517 |
| 0 | 71.04 | 3.32E+09 | 89 | 27.9339 | 27.8557 | 27.6707 | 27.7095 | 27.9678 |
| 0 | 323.31 | 8.14E+09 | 183 | 29.0329 | 28.9938 | 29.2431 | 29.2843 | 29.1652 |
| 0 | 61.63 | 6.52E+09 | 133 | 28.8857 | 28.8244 | 28.7585 | 28.8751 | 28.8946 |
| 0 | 167.66 | 3.99E+09 | 95 | 28.4954 | 28.4939 | 28.1613 | 28.1263 | 28.0689 |
| 0 | 52.653 | 4.14E+09 | 124 | 28.1116 | 28.1975 | 28.123 | 28.2875 | 28.2206 |
| 0 | 79.52 | 2.16E+11 | 350 | 33.9501 | 33.941 | 33.9031 | 33.9053 | 33.6964 |
| 0 | 57.707 | 3.13E+09 | 88 | 27.6292 | 27.8132 | 27.7916 | 27.9084 | 27.6802 |
| 0 | 42.257 | 2.24E+09 | 106 | 27.232 | 27.2278 | 27.2399 | 27.2966 | 27.4122 |
| 0 | 34.088 | 1.26E+09 | 59 | 26.5849 | 26.5003 | 26.1851 | 26.4589 | 26.5188 |
| 0 | 142.39 | 1.18E+10 | 154 | 29.7153 | 29.6519 | 29.683 | 29.6995 | 29.6971 |
| 0 | 59.898 | 2.37E+09 | 81 | 27.4278 | 27.3388 | 27.5111 | 27.3597 | 27.2702 |
| 0 | 93.05 | 3.53E+09 | 108 | 28.1704 | 28.2035 | 27.8633 | 27.8237 | 28.0046 |
| 0 | 79.388 | 4.53E+09 | 125 | 28.2887 | 28.3134 | 28.3268 | 28.4416 | 28.2874 |
| 0 | 185.46 | 2.75E+10 | 203 | 30.9309 | 30.9649 | 30.9174 | 30.9074 | 31.0745 |
| 0 | 110.79 | 4.11E+09 | 150 | 28.2245 | 28.2655 | 28.1112 | 28.0978 | 28.3904 |
| 0 | 69.857 | 2.89E+09 | 108 | 27.3961 | 27.4855 | 27.8003 | 27.557 | 27.8063 |
| 0 | 182.2 | 1.05E+10 | 168 | 29.705 | 29.764 | 29.4653 | 29.5627 | 29.4885 |
| 0 | 109.71 | 5.99E+09 | 107 | 28.6881 | 28.7985 | 28.7777 | 28.7573 | 28.6125 |
| 0 | 112.24 | 8.21E+09 | 142 | 29.4647 | 29.3084 | 29.0142 | 29.0956 | 29.236 |
| 0 | 118.98 | 2.66E+09 | 77 | 27.6207 | 27.5712 | 27.5156 | 27.5949 | 27.596 |
| 0 | 233.42 | 8.74E+10 | 196 | 32.5561 | 32.5567 | 32.6401 | 32.6235 | 32.5922 |
| 0 | 152.95 | 5.97E+09 | 130 | 28.6454 | 28.6964 | 28.7058 | 28.5452 | 28.7595 |
| 0 | 94.136 | 1.29E+09 | 48 | 26.6407 | 26.5516 | 26.5458 | 26.244 | 26.4774 |
| 0 | 31.959 | 1.23E+09 | 48 | 26.1964 | 26.5274 | 26.3341 | 26.226 | 26.7157 |
| 0 | 35.056 | 1.25E+09 | 41 | 26.382 | 26.2644 | 26.3666 | 26.1996 | 26.6701 |
| 0 | 103.26 | 2.88E+09 | 96 | 27.5431 | 27.6083 | 27.6114 | 27.597 | 27.8081 |
| 0 | 174.68 | 5.86E+09 | 115 | 28.723 | 28.5763 | 28.7373 | 28.7698 | 28.7822 |
| 0 | 211.04 | 3.27E+10 | 229 | 31.1904 | 31.1915 | 31.2826 | 31.2584 | 31.1191 |
| 0 | 121.49 | 1.43E+11 | 458 | 33.4806 | 33.5142 | 33.2143 | 33.4248 | 33.194 |
| 0 | 23.326 | 1.95E+09 | 85 | 27.2347 | 27.184 | 27.0134 | 26.8532 | 27.3139 |
| 0 | 151.66 | 3.17E+09 | 100 | 27.8007 | 27.7932 | 27.9454 | 27.9076 | 27.7503 |
| 0 | 127.86 | 4.42E+10 | 221 | 31.6416 | 31.7477 | 31.651 | 31.7099 | 31.4963 |
| 0 | 102.25 | 1.66E+09 | 69 | 26.6871 | 26.8697 | 26.9329 | 26.9032 | 26.9808 |

| | | | | | | | | |
|---|---|---|---|---|---|---|---|---|
| 0 | 78.444 | 8.79E+09 | 180 | 29.4152 | 29.3885 | 29.2211 | 29.1125 | 29.3847 |
| 0 | 83.601 | 3.73E+10 | 168 | 31.4086 | 31.4878 | 31.3348 | 31.3983 | 31.3749 |
| 0 | 33.968 | 2.33E+09 | 94 | 27.2644 | 27.3297 | 27.3939 | 27.4132 | 27.3716 |
| 0 | 323.31 | 3.6E+10 | 249 | 31.3641 | 31.3587 | 31.4336 | 31.4007 | 31.205 |
| 0 | 52.216 | 1.59E+09 | 80 | 26.8457 | 26.8089 | 26.9248 | 26.8153 | 26.7659 |
| 0 | 45.123 | 2.44E+09 | 110 | 27.3774 | 27.4815 | 27.511 | 27.3448 | 27.5025 |
| 0 | 149.89 | 3.79E+09 | 117 | 28.2245 | 28.069 | 28.0164 | 27.9125 | 28.1583 |
| 0 | 43.771 | 1.94E+09 | 62 | 26.8345 | 26.9565 | 27.2804 | 27.2104 | 27.1103 |
| 0 | 179.93 | 4.75E+09 | 98 | 28.3039 | 28.4097 | 28.5121 | 28.5037 | 28.2651 |
| 0 | 189.99 | 9.74E+09 | 190 | 29.4487 | 29.5288 | 29.2315 | 29.4363 | 29.4715 |
| 0 | 60.668 | 2.94E+09 | 87 | 27.7491 | 27.6417 | 27.8388 | 27.8313 | 27.6858 |
| 0 | 37.469 | 1.42E+09 | 47 | 26.4934 | 26.5963 | 26.9492 | 26.5622 | 26.5841 |
| 0 | 61.932 | 2.99E+09 | 77 | 27.7176 | 27.6899 | 27.7197 | 27.7831 | 27.8367 |
| 0 | 202.41 | 9.21E+10 | 607 | 32.4889 | 32.6179 | 32.6264 | 32.6478 | 32.7922 |
| 0 | 41.747 | 2.06E+09 | 49 | 27.1491 | 27.2022 | 27.1979 | 27.1536 | 27.1682 |
| 0 | 87.059 | 1.79E+09 | 60 | 26.9325 | 26.8253 | 26.9173 | 27.012 | 27.2083 |
| 0 | 76.628 | 1.16E+10 | 143 | 29.7579 | 29.7383 | 29.8729 | 29.8418 | 29.4729 |
| 0 | 168.4 | 1.21E+10 | 199 | 29.7365 | 29.6816 | 29.835 | 29.764 | 29.7677 |
| 0 | 92.797 | 2.58E+09 | 99 | 27.6985 | 27.6818 | 27.3091 | 27.4272 | 27.5008 |
| 0 | 39.06 | 2.82E+09 | 105 | 27.5874 | 27.5802 | 27.5625 | 27.681 | 27.6785 |
| 0 | 323.31 | 6.35E+09 | 103 | 28.7211 | 28.7368 | 28.8578 | 28.8326 | 28.9541 |
| 0 | 19.164 | 2.07E+09 | 78 | 27.2755 | 27.3456 | 27.2298 | 27.2483 | 27.0503 |
| 0 | 29.081 | 1.64E+09 | 81 | 26.7977 | 26.7487 | 26.9132 | 27.0232 | 26.9382 |
| 0 | 197.22 | 1.41E+10 | 176 | 30.0125 | 29.9516 | 30.0664 | 30.0044 | 29.9371 |
| 0 | 240.05 | 2.99E+10 | 191 | 31.1267 | 31.1031 | 31.0572 | 31.0486 | 31.0299 |
| 0 | 27.54 | 2.56E+09 | 60 | 27.3838 | 27.3499 | 27.3408 | 27.2468 | 27.8144 |
| 0 | 152.64 | 5.83E+09 | 143 | 28.6811 | 28.667 | 28.5825 | 28.5727 | 28.7958 |
| 0 | 323.31 | 2.85E+10 | 249 | 31.0187 | 30.8545 | 31.1181 | 31.0981 | 30.9647 |
| 0 | 204.72 | 3.32E+09 | 108 | 27.7881 | 27.9148 | 28.0585 | 27.9811 | 27.8085 |
| 0 | 47.522 | 2.45E+09 | 82 | 27.4107 | 27.3689 | 27.4693 | 27.4934 | 27.5295 |
| 0 | 110.87 | 5.1E+09 | 126 | 28.561 | 28.611 | 28.4344 | 28.4028 | 28.5852 |
| 0 | 110.81 | 8.27E+09 | 153 | 29.1742 | 29.1934 | 29.1931 | 29.3228 | 29.1834 |
| 0 | 188.01 | 1.88E+11 | 471 | 33.8011 | 33.8278 | 33.5417 | 33.6276 | 33.7171 |
| 0 | 54.074 | 8.86E+09 | 118 | 29.2385 | 29.1072 | 28.9804 | 29.1247 | 29.071 |
| 0 | 302.8 | 8.57E+09 | 155 | 28.9769 | 28.9193 | 29.1911 | 29.1617 | 29.1789 |
| 0 | 167.97 | 2.25E+10 | 236 | 30.3916 | 30.4821 | 30.4516 | 30.4771 | 30.6158 |
| 0 | 38.957 | 8.03E+09 | 118 | 29.0867 | 29.0237 | 29.1619 | 29.1476 | 28.9335 |
| 0 | 44.25 | 3.54E+10 | 167 | 31.217 | 31.1787 | 31.242 | 31.2978 | 31.0578 |
| 0 | 198.87 | 4.94E+10 | 286 | 31.6694 | 31.653 | 31.7915 | 31.7401 | 31.6313 |
| 0 | 55.645 | 1.23E+09 | 61 | 26.37 | 26.5457 | 26.1315 | 26.4147 | 26.299 |
| 0 | 39.638 | 2.74E+09 | 104 | 27.5413 | 27.6574 | 27.6361 | 27.5519 | 27.4245 |
| 0 | 132.2 | 3.3E+10 | 267 | 31.1543 | 31.0596 | 31.0459 | 31.1861 | 31.0459 |
| 0 | 151.67 | 1.58E+10 | 92 | 30.0289 | 30.0483 | 30.0714 | 30.0288 | 30.1743 |
| 0 | 323.31 | 3.59E+10 | 299 | 31.0907 | 31.1877 | 31.335 | 31.4348 | 31.2677 |
| 0 | 243.93 | 3.27E+10 | 227 | 31.0924 | 31.0282 | 31.2494 | 31.1759 | 31.1449 |
| 0 | 242.76 | 1.25E+10 | 196 | 29.6695 | 29.7312 | 29.8484 | 29.7809 | 29.706 |
| 0 | 51.908 | 2.21E+09 | 64 | 27.2857 | 27.1726 | 27.2099 | 27.333 | 27.2772 |
| 0 | 176.12 | 2.25E+10 | 260 | 30.6836 | 30.6174 | 30.657 | 30.6142 | 30.4715 |

| | | | | | | | | |
|---|---|---|---|---|---|---|---|---|
| 0 | 42.96 | 2.13E+10 | 236 | 30.4394 | 30.4118 | 30.5755 | 30.6345 | 30.5578 |
| 0 | 209.2 | 1.1E+10 | 209 | 29.5325 | 29.5205 | 29.5938 | 29.6122 | 29.5786 |
| 0 | 52.896 | 9.45E+09 | 181 | 29.2845 | 29.2806 | 29.2624 | 29.3638 | 29.5032 |
| 0 | 39.881 | 1.76E+09 | 54 | 26.8383 | 26.8869 | 26.9301 | 26.9347 | 27.023 |
| 0 | 76.497 | 4.22E+10 | 227 | 31.5377 | 31.5726 | 31.5505 | 31.5223 | 31.4628 |
| 0 | 177.71 | 7.83E+09 | 143 | 29.0635 | 29.139 | 29.1598 | 29.1338 | 29.0208 |
| 0 | 323.31 | 2.02E+10 | 319 | 30.6005 | 30.6228 | 30.5661 | 30.687 | 30.4543 |
| 0 | 30.962 | 5.42E+09 | 97 | 28.8468 | 29.0142 | 28.5401 | 28.4193 | 28.6717 |
| 0 | 89.685 | 7.04E+09 | 118 | 29.3474 | 29.3196 | 29.0442 | 28.9761 | 29.0706 |
| 0 | 21.865 | 2.51E+09 | 67 | 27.876 | 27.9148 | 27.512 | 27.6 | 27.4741 |
| 0 | 50.994 | 4.93E+09 | 83 | 28.0983 | 27.9814 | 28.3619 | 28.2875 | 28.1605 |
| 0 | 134.35 | 4.61E+09 | 129 | 28.4935 | 28.4616 | 28.3164 | 28.325 | 28.6111 |
| 0 | 37.68 | 3.44E+09 | 126 | 28.1007 | 28.2807 | 27.6299 | 27.8394 | 27.8126 |
| 0 | 323.31 | 2.9E+10 | 240 | 30.9586 | 30.8384 | 31.0841 | 31.0497 | 30.816 |
| 0 | 50.578 | 1.09E+09 | 47 | 26.0616 | 25.9777 | 26.0729 | 25.7442 | 26.3229 |
| 0 | 323.31 | 1.84E+10 | 173 | 30.2886 | 30.3595 | 30.3069 | 30.2714 | 30.2667 |
| 0 | 125.16 | 3.34E+09 | 114 | 27.8811 | 27.8914 | 27.9079 | 27.9308 | 27.8711 |
| 0 | 192.77 | 3.32E+09 | 116 | 27.8553 | 27.7995 | 27.8409 | 27.9557 | 27.8243 |
| 0 | 10.047 | 1.05E+09 | 43 | 26.1408 | 26.0818 | 26.1392 | 26.3264 | 26.1383 |
| 0 | 31.457 | 3.7E+09 | 108 | 28.0128 | 28.0347 | 28.0904 | 28.0734 | 27.8253 |
| 0 | 251.42 | 6.61E+09 | 120 | 28.9267 | 28.9818 | 29.0054 | 28.9574 | 28.9852 |
| 0 | 41.782 | 5.74E+09 | 114 | 28.2719 | 28.3649 | 28.7136 | 28.6442 | 28.5768 |
| 0 | 111.45 | 7.87E+09 | 141 | 29.1572 | 29.0927 | 29.153 | 29.2443 | 29.0586 |
| 0 | 323.31 | 5.34E+10 | 189 | 32.0321 | 32.0173 | 31.9935 | 32.0005 | 31.805 |
| 0 | 36.342 | 2.83E+10 | 126 | 30.9261 | 30.9321 | 30.9786 | 31.0784 | 30.7958 |
| 0 | 133.84 | 1.24E+10 | 158 | 29.8385 | 29.7702 | 29.8401 | 29.7856 | 29.6289 |
| 0 | 183.28 | 8.36E+09 | 212 | 29.1765 | 29.0112 | 29.2726 | 29.2695 | 29.2069 |
| 0 | 172.06 | 5.73E+09 | 137 | 28.6874 | 28.6614 | 28.7588 | 28.732 | 28.8158 |
| 0 | 59.265 | 1.21E+10 | 125 | 29.6724 | 29.667 | 29.6465 | 29.6649 | 29.7057 |
| 0 | 94.422 | 7.81E+09 | 162 | 29.1123 | 29.0314 | 29.0729 | 29.0488 | 29.1237 |
| 0 | 39.223 | 2.07E+09 | 92 | 27.1929 | 27.1779 | 27.1957 | 27.1449 | 27.0648 |
| 0 | 29.361 | 2.23E+10 | 195 | 30.5948 | 30.5434 | 30.67 | 30.6771 | 30.4988 |
| 0 | 78.38 | 2.91E+09 | 107 | 27.392 | 27.5741 | 27.4783 | 27.5728 | 27.8096 |
| 0 | 110.52 | 8.47E+09 | 137 | 29.1701 | 29.2665 | 29.23 | 29.2358 | 29.3382 |
| 0 | 323.31 | 3.65E+10 | 274 | 31.2499 | 31.2086 | 31.3529 | 31.3772 | 31.2001 |
| 0 | 166.68 | 6.58E+09 | 139 | 28.7247 | 28.6912 | 28.9327 | 28.9496 | 28.9185 |
| 0 | 31.799 | 1.34E+09 | 50 | 26.7434 | 27.0342 | 26.6196 | 26.4844 | 26.5158 |
| 0 | 323.31 | 8.08E+10 | 251 | 32.536 | 32.466 | 32.523 | 32.4397 | 32.4241 |
| 0 | 27.976 | 2.14E+09 | 80 | 27.2313 | 27.34 | 27.2272 | 27.3838 | 26.9973 |
| 0 | 78.339 | 3.31E+09 | 86 | 27.8754 | 27.8849 | 27.9557 | 27.8985 | 27.9823 |
| 0 | 45.743 | 5.98E+09 | 99 | 28.9996 | 28.9325 | 28.5937 | 28.7788 | 28.8192 |
| 0 | 117.2 | 3.01E+09 | 92 | 27.6488 | 27.6862 | 27.6772 | 27.604 | 27.7648 |
| 0 | 110.06 | 1.66E+10 | 201 | 30.1646 | 30.18 | 30.2572 | 30.1251 | 30.2185 |
| 0 | 130.08 | 1.12E+10 | 151 | 29.5919 | 29.6553 | 29.6416 | 29.6892 | 29.5537 |
| 0 | 19.89 | 1.11E+09 | 53 | 26.354 | 26.2321 | 26.2149 | 26.3746 | 26.2201 |
| 0 | 323.31 | 1.95E+10 | 301 | 30.479 | 30.443 | 30.5541 | 30.5724 | 30.1728 |
| 0 | 225.57 | 6.32E+09 | 137 | 28.8837 | 28.8051 | 28.8916 | 28.8713 | 28.7501 |
| 0 | 21.627 | 1.53E+09 | 48 | 26.6929 | 26.6921 | 26.8445 | 26.8362 | 26.9599 |

| | | | | | | | | |
|---|---|---|---|---|---|---|---|---|
| 0 | 57.149 | 1.9E+09 | 92 | 27.1587 | 27.1242 | 26.8941 | 26.9041 | 26.9994 |
| 0 | 50.015 | 3.05E+09 | 64 | 27.7626 | 27.5937 | 27.7959 | 27.7705 | 27.8108 |
| 0 | 273.81 | 4.2E+10 | 361 | 31.5649 | 31.4695 | 31.6254 | 31.4684 | 31.4976 |
| 0 | 244.88 | 4.59E+10 | 184 | 31.7976 | 31.84 | 31.6959 | 31.6723 | 31.7024 |
| 0 | 49.924 | 5.23E+09 | 98 | 28.5934 | 28.6053 | 28.3077 | 28.2425 | 28.5905 |
| 0 | 32.636 | 1.4E+09 | 76 | 26.4649 | 26.5357 | 26.564 | 26.6451 | 26.7643 |
| 0 | 52.624 | 1.81E+09 | 71 | 27.1544 | 27.1411 | 27.0115 | 27.1679 | 27.0986 |
| 0 | 219.48 | 2.82E+10 | 182 | 31.0056 | 31.1061 | 30.9089 | 30.9084 | 30.8439 |
| 0 | 66.486 | 1.46E+09 | 49 | 26.6951 | 26.8024 | 26.7685 | 26.7167 | 26.6353 |
| 0 | 72.224 | 1.66E+09 | 61 | 26.9814 | 26.8368 | 26.8542 | 27.0253 | 26.9956 |
| 0 | 50.759 | 3.18E+09 | 75 | 27.6705 | 27.3279 | 27.4341 | 27.6075 | 27.8078 |
| 0 | 26.639 | 5.52E+09 | 44 | 28.5994 | 28.5499 | 28.6851 | 28.7635 | 28.2647 |
| 0 | 41.079 | 1.89E+09 | 85 | 27.0622 | 27.0649 | 27.3085 | 27.1132 | 27.1987 |
| 0 | 106.54 | 3.5E+09 | 121 | 27.9458 | 27.8998 | 27.7785 | 27.9427 | 28.0972 |
| 0 | 128.32 | 3.42E+09 | 112 | 27.8584 | 27.9301 | 28.0961 | 28.0641 | 27.8287 |
| 0 | 51.141 | 8.58E+09 | 99 | 29.2858 | 29.316 | 29.337 | 29.3672 | 29.2911 |
| 0 | 323.31 | 2.03E+10 | 163 | 30.4821 | 30.5114 | 30.5572 | 30.5148 | 30.5861 |
| 0 | 323.31 | 8.44E+09 | 167 | 29.2493 | 29.2736 | 29.2544 | 29.249 | 29.1595 |
| 0 | 36.093 | 1.47E+09 | 30 | 26.6118 | 26.6878 | 26.5601 | 26.447 | 26.8858 |
| 0 | 296.15 | 6.21E+10 | 306 | 32.1657 | 32.1268 | 32.1163 | 32.0998 | 32.0362 |
| 0 | 31.06 | 1.86E+09 | 82 | 27.0583 | 26.9446 | 26.869 | 26.7615 | 27.1417 |
| 0 | 27.203 | 3.28E+09 | 100 | 27.8069 | 27.6362 | 27.9176 | 27.977 | 28.0009 |
| 0 | 52.403 | 1.57E+09 | 44 | 27.1829 | 27.2898 | 26.7192 | 26.6966 | 26.6488 |
| 0 | 34.343 | 4.57E+09 | 68 | 28.7398 | 28.7053 | 28.1177 | 28.3394 | 28.0258 |
| 0 | 44.665 | 1.67E+09 | 60 | 26.8534 | 26.7567 | 26.9823 | 27.1311 | 26.9389 |
| 0 | 165.99 | 4.71E+09 | 111 | 28.3767 | 28.3666 | 28.3205 | 28.4515 | 28.577 |
| 0 | 323.31 | 3.38E+10 | 241 | 31.3021 | 31.2751 | 31.2862 | 31.2775 | 31.0643 |
| 0 | 26.891 | 1.83E+09 | 48 | 27.3615 | 27.303 | 26.8013 | 26.8942 | 27.1611 |
| 0 | 144.67 | 1.81E+10 | 268 | 30.2703 | 30.1358 | 30.3822 | 30.3644 | 30.3578 |
| 0 | 31.95 | 9.43E+09 | 110 | 29.3904 | 29.437 | 29.4849 | 29.3208 | 29.4387 |
| 0 | 144.63 | 2.28E+09 | 93 | 27.4072 | 27.2415 | 27.4314 | 27.2128 | 27.4263 |
| 0 | 89.494 | 4.19E+09 | 122 | 28.3163 | 28.4034 | 28.159 | 28.1187 | 28.2332 |
| 0 | 34.574 | 1.62E+09 | 72 | 26.8751 | 26.8827 | 26.8234 | 26.7772 | 26.7874 |
| 0 | 323.31 | 1.83E+10 | 225 | 30.2975 | 30.2465 | 30.4701 | 30.2958 | 30.465 |
| 0 | 323.31 | 1.11E+11 | 287 | 33.0889 | 33.1097 | 32.768 | 32.7315 | 32.8688 |
| 0 | 122.56 | 9.6E+09 | 150 | 29.3167 | 29.2477 | 29.5029 | 29.4507 | 29.4096 |
| 0 | 85.082 | 1.79E+09 | 83 | 27.0842 | 27.0076 | 27.0937 | 26.8844 | 27.0584 |
| 0 | 33.922 | 1.58E+09 | 49 | 26.7506 | 26.8341 | 26.6656 | 26.7336 | 26.898 |
| 0 | 97.401 | 2.17E+10 | 218 | 30.601 | 30.7738 | 30.5175 | 30.599 | 30.3758 |
| 0 | 21.768 | 2.03E+09 | 40 | 27.2104 | 27.1718 | 26.8466 | 27.2024 | 27.2692 |
| 0 | 150.92 | 7.03E+10 | 315 | 32.3274 | 32.2969 | 32.2842 | 32.4216 | 32.1969 |
| 0 | 140.46 | 7.07E+09 | 123 | 28.9525 | 28.9325 | 28.9191 | 28.7233 | 29.1036 |
| 0 | 42.49 | 3.35E+09 | 94 | 27.975 | 27.817 | 27.8879 | 27.9071 | 27.7593 |
| 0 | 133.14 | 9.73E+09 | 169 | 29.5939 | 29.4415 | 29.3538 | 29.3714 | 29.5344 |
| 0 | 86.879 | 2.22E+09 | 66 | 27.5777 | 27.638 | 27.1018 | 26.8359 | 27.3936 |
| 0 | 82.274 | 4.25E+09 | 92 | 28.266 | 28.2489 | 28.4045 | 28.3142 | 28.1887 |
| 0 | 127.48 | 6.66E+09 | 137 | 28.7785 | 28.7995 | 28.9868 | 28.9884 | 28.9567 |
| 0 | 62.905 | 3.06E+09 | 103 | 27.8289 | 27.7555 | 27.8718 | 27.8263 | 27.727 |

| | | | | | | | | |
|---|---|---|---|---|---|---|---|---|
| 0 | 56.246 | 1.23E+10 | 161 | 29.6936 | 29.7458 | 29.7357 | 29.8438 | 29.8677 |
| 0 | 202.6 | 3.13E+09 | 106 | 27.6624 | 27.6587 | 27.7471 | 27.7117 | 28.0049 |
| 0 | 323.31 | 2.74E+10 | 262 | 30.894 | 30.9561 | 30.9744 | 30.8631 | 30.9657 |
| 0 | 74.755 | 3.17E+09 | 67 | 27.6736 | 27.877 | 27.8492 | 27.8813 | 27.8647 |
| 0 | 70.03 | 7.55E+09 | 142 | 29.1473 | 29.1464 | 29.1655 | 29.1132 | 29.0197 |
| 0 | 68.655 | 5.15E+09 | 107 | 28.8413 | 28.8959 | 28.4113 | 28.251 | 28.4908 |
| 0 | 71.125 | 5.12E+09 | 158 | 28.5715 | 28.6059 | 28.6872 | 28.6653 | 28.2569 |
| 0 | 22.793 | 2.08E+09 | 79 | 27.2148 | 27.2913 | 27.2237 | 27.201 | 27.2415 |
| 0 | 42.075 | 1.54E+09 | 56 | 26.7049 | 26.827 | 26.9261 | 26.8358 | 26.7208 |
| 0 | 45.536 | 1.64E+09 | 73 | 26.8182 | 26.7213 | 26.6276 | 26.9662 | 27.0436 |
| 0 | 47.525 | 3.93E+09 | 125 | 28.1805 | 28.1839 | 28.0523 | 27.9964 | 28.2065 |
| 0 | 36.2 | 1.95E+09 | 76 | 27.2624 | 27.1948 | 27.1632 | 27.1523 | 26.9882 |
| 0 | 226.64 | 2.07E+10 | 167 | 30.597 | 30.5892 | 30.3748 | 30.4292 | 30.5876 |
| 0 | 204.05 | 1.21E+11 | 342 | 33.3723 | 33.3465 | 32.9776 | 32.9136 | 32.9907 |
| 0 | 129.91 | 2.97E+09 | 88 | 27.9463 | 27.9219 | 27.3608 | 27.3656 | 27.9018 |
| 0 | 130.19 | 5.96E+09 | 97 | 28.6583 | 28.8191 | 28.5729 | 28.5754 | 28.785 |
| 0 | 315.47 | 1.49E+10 | 212 | 30.0107 | 30.0621 | 30.1097 | 30.109 | 29.9898 |
| 0 | 31.476 | 1.64E+09 | 85 | 26.5217 | 26.9201 | 26.8246 | 26.6851 | 27.1631 |
| 0 | 26.275 | 1.72E+09 | 60 | 26.8297 | 26.9255 | 26.9978 | 26.9125 | 27.043 |
| 0 | 174.22 | 1.42E+10 | 179 | 30.0018 | 30.0894 | 30.1344 | 30.1681 | 29.8109 |
| 0 | 114.73 | 1.03E+10 | 147 | 29.7205 | 29.747 | 29.3514 | 29.3391 | 29.4783 |
| 0 | 323.31 | 5.23E+09 | 132 | 28.1075 | 28.213 | 28.3401 | 28.3251 | 28.5905 |
| 0 | 44.052 | 3.08E+09 | 95 | 27.4781 | 27.5583 | 27.7051 | 27.715 | 27.751 |
| 0 | 27.204 | 1.39E+09 | 52 | 26.4183 | 26.4472 | 26.4188 | 26.5925 | 26.5582 |
| 0 | 67.057 | 2.56E+09 | 71 | 27.2321 | 27.3063 | 27.5157 | 27.3976 | 27.3813 |
| 0 | 20.704 | 1E+09 | 35 | 26.152 | 26.0304 | 26.0526 | 25.9275 | 26.1196 |
| 0 | 52.969 | 4.66E+09 | 121 | 28.2393 | 28.1764 | 28.2496 | 28.3515 | 28.2734 |
| 0 | 55.072 | 1E+09 | 46 | 25.7566 | 26.122 | 26.221 | 25.9691 | 26.0512 |
| 0 | 173.96 | 2.79E+10 | 291 | 30.8134 | 30.7968 | 31.0548 | 30.9653 | 30.7368 |
| 0 | 323.31 | 2.56E+10 | 227 | 30.651 | 30.6637 | 30.8424 | 30.6949 | 30.6505 |
| 0 | 45.973 | 1.59E+09 | 67 | 26.6308 | 26.8473 | 26.7393 | 26.6591 | 26.7064 |
| 0 | 26.661 | 1.62E+09 | 31 | 26.7187 | 26.7651 | 26.888 | 26.7724 | 26.8507 |
| 0 | 138.39 | 8.77E+09 | 180 | 29.29 | 29.2154 | 29.2044 | 29.2464 | 29.1126 |
| 0 | 33.882 | 1.44E+09 | 68 | 26.6067 | 26.7027 | 26.5418 | 26.6457 | 26.5599 |
| 0 | 323.31 | 9.21E+10 | 300 | 32.6276 | 32.6249 | 32.6666 | 32.6818 | 32.5416 |
| 0 | 323.31 | 3.17E+10 | 289 | 31.1952 | 31.1872 | 31.0272 | 31.0368 | 31.0797 |
| 0 | 36.682 | 2.42E+09 | 86 | 27.3835 | 27.4558 | 27.4209 | 27.3235 | 27.3876 |
| 0 | 300.82 | 7.74E+09 | 139 | 29.2198 | 29.222 | 29.2121 | 29.1789 | 29.1031 |
| 0 | 19.272 | 3.83E+09 | 90 | 28.0783 | 28.062 | 28.1543 | 28.1965 | 28.2837 |
| 0 | 207.33 | 3.98E+10 | 295 | 31.5724 | 31.5185 | 31.4816 | 31.6052 | 31.5299 |
| 0 | 75.034 | 3.71E+09 | 111 | 28.0443 | 27.9721 | 28.1467 | 28.1486 | 28.2441 |
| 0 | 126.26 | 3.53E+09 | 98 | 28.1016 | 28.1185 | 28.0945 | 28.086 | 28.0315 |
| 0 | 49.719 | 1.74E+09 | 68 | 26.8869 | 26.8666 | 27.0161 | 26.9297 | 26.8301 |
| 0 | 79.916 | 3.61E+09 | 106 | 28.0811 | 28.0466 | 27.9265 | 27.9773 | 27.6839 |
| 0 | 22.193 | 1.86E+09 | 69 | 27.104 | 27.089 | 27.0026 | 27.3144 | 27.1846 |
| 0 | 89.971 | 1.75E+09 | 65 | 26.9668 | 26.8855 | 26.9303 | 26.8089 | 26.9394 |
| 0 | 18.976 | 7.17E+08 | 41 | 25.4565 | 25.6002 | 25.4633 | 25.6962 | 25.5994 |
| 0 | 56.545 | 3.04E+09 | 89 | 27.7418 | 27.6964 | 27.7052 | 27.7415 | 27.6685 |

| | | | | | | | | |
|---|---|---|---|---|---|---|---|---|
| 0 | 171.6 | 4.31E+09 | 118 | 27.9728 | 27.9504 | 28.2705 | 28.3373 | 28.171 |
| 0 | 323.31 | 1.76E+10 | 166 | 30.5699 | 30.5246 | 30.1829 | 30.2666 | 30.3483 |
| 0 | 46.566 | 4.55E+10 | 118 | 31.6465 | 31.6688 | 31.8839 | 31.829 | 31.6404 |
| 0 | 23.88 | 1.93E+09 | 56 | 27.2188 | 27.0759 | 27.1613 | 27.0985 | 26.9747 |
| 0 | 125.39 | 1.43E+10 | 137 | 29.4321 | 29.3339 | 29.6906 | 29.6352 | 29.5814 |
| 0 | 17.253 | 1.07E+09 | 39 | 26.0918 | 26.3134 | 26.3091 | 26.3041 | 26.2871 |
| 0 | 26.573 | 1.49E+09 | 74 | 26.7248 | 26.834 | 26.6029 | 26.771 | 26.6174 |
| 0 | 62.775 | 1.29E+10 | 211 | 29.915 | 29.8829 | 29.955 | 29.9106 | 29.8015 |
| 0 | 160.33 | 2.39E+09 | 84 | 27.3043 | 27.338 | 27.36 | 27.4242 | 27.3388 |
| 0 | 58.56 | 6.05E+09 | 95 | 28.7823 | 28.6901 | 28.8783 | 28.763 | 28.5702 |
| 0 | 108.26 | 2.41E+09 | 86 | 27.3864 | 27.3709 | 27.4818 | 27.5597 | 27.4954 |
| 0 | 24.912 | 1.74E+09 | 61 | 26.8557 | 27.2248 | 26.8303 | 26.8014 | 26.6542 |
| 0 | 102.79 | 2.86E+09 | 103 | 27.6409 | 27.7841 | 27.4686 | 27.548 | 27.6404 |
| 0 | 52.454 | 1.55E+09 | 71 | 27.0524 | 26.8049 | 26.8696 | 26.9276 | 26.8453 |
| 0 | 79.779 | 2.6E+09 | 64 | 27.2845 | 27.2241 | 27.5651 | 27.464 | 27.5298 |
| 0 | 11.752 | 1.04E+09 | 49 | 26.4168 | 26.4573 | 25.8815 | 26.0307 | 26.4799 |
| 0 | 56.221 | 3.38E+09 | 158 | 27.683 | 27.6229 | 28.0373 | 27.942 | 27.7077 |
| 0 | 164.22 | 6.08E+09 | 88 | 28.722 | 28.7855 | 28.7951 | 28.7478 | 28.5846 |
| 0 | 323.31 | 3.98E+10 | 275 | 31.4509 | 31.3618 | 31.3698 | 31.3862 | 31.4804 |
| 0 | 65.513 | 3.85E+09 | 152 | 27.9221 | 27.9142 | 28.0957 | 28.1109 | 28.1019 |
| 0 | 197.66 | 6.11E+09 | 114 | 28.7497 | 28.7812 | 28.7686 | 28.6794 | 28.6273 |
| 0 | 35.406 | 1.42E+10 | 198 | 30.1202 | 30.1184 | 29.7936 | 29.7636 | 29.8087 |
| 0 | 38.013 | 1.64E+09 | 81 | 26.8633 | 26.7948 | 26.8341 | 26.9225 | 26.6759 |
| 0 | 312.87 | 1.16E+10 | 195 | 29.7279 | 29.6754 | 29.7602 | 29.7416 | 29.5863 |
| 0 | 13.907 | 9.76E+08 | 18 | 26.12 | 26.1195 | 26.3162 | 26.3013 | 26.0794 |
| 0 | 27.924 | 4.29E+09 | 92 | 28.3195 | 28.1632 | 28.1699 | 28.2402 | 28.2826 |
| 0 | 92.412 | 4.67E+09 | 81 | 28.7547 | 28.6523 | 28.4088 | 28.4487 | 28.3422 |
| 0 | 31.265 | 1.1E+09 | 30 | 26.5002 | 26.4 | 26.1437 | 26.1666 | 25.969 |
| 0 | 305.34 | 1.75E+10 | 161 | 30.4992 | 30.5115 | 30.3636 | 30.2581 | 30.2644 |
| 0 | 242.9 | 1.79E+10 | 184 | 30.7469 | 30.689 | 30.3266 | 30.3546 | 30.1159 |
| 0 | 56.604 | 1.42E+09 | 42 | 26.7598 | 26.8486 | 26.4553 | 26.2352 | 26.7107 |
| 0 | 29.806 | 1.26E+09 | 37 | 26.2961 | 26.5327 | 25.9841 | 26.3133 | 26.6779 |
| 0 | 8.5871 | 7.36E+08 | 21 | 25.7922 | 25.5911 | 25.9163 | 25.7407 | 25.2911 |
| 0 | 64.551 | 2.31E+09 | 70 | 27.0637 | 26.9807 | 26.9241 | 26.9331 | 27.467 |
| 0 | 19.068 | 8.3E+08 | 38 | 26.005 | 25.6479 | 25.4069 | 25.7767 | 26.0523 |
| 0 | 59.443 | 1.52E+09 | 35 | 26.7718 | 26.7997 | 26.9037 | 26.6734 | 26.7028 |
| 0 | 74.062 | 2.75E+09 | 95 | 27.5505 | 27.6425 | 27.7432 | 27.6296 | 27.4706 |
| 0 | 25.556 | 2.33E+09 | 55 | 27.3327 | 27.2923 | 27.3758 | 27.3434 | 27.2353 |
| 0 | 59.394 | 1.49E+09 | 68 | 26.6118 | 26.6653 | 26.5873 | 26.7336 | 26.8078 |
| 0 | 61.456 | 6.08E+09 | 143 | 28.7384 | 28.6962 | 28.5491 | 28.6994 | 28.7628 |
| 0 | 69.894 | 2.9E+09 | 82 | 27.7354 | 27.6688 | 27.7508 | 27.8277 | 27.4852 |
| 0 | 40.312 | 2.49E+09 | 86 | 27.5825 | 27.5033 | 27.2916 | 27.371 | 27.6956 |
| 0 | 240.4 | 9.04E+09 | 171 | 29.3633 | 29.3819 | 29.3245 | 29.2764 | 29.4252 |
| 0 | 26.429 | 1.44E+09 | 49 | 26.7155 | 26.6429 | 26.8809 | 26.8346 | 26.6968 |
| 0 | 78.683 | 5.56E+09 | 127 | 28.5292 | 28.4856 | 28.6262 | 28.5921 | 28.6496 |
| 0 | 43.385 | 9.23E+08 | 47 | 26.0329 | 25.7313 | 26.076 | 26.2649 | 26.2353 |
| 0 | 320.27 | 1.88E+11 | 649 | 33.8786 | 33.7845 | 33.6647 | 33.7413 | 33.2605 |
| 0 | 184.8 | 4.58E+09 | 109 | 28.1736 | 28.2054 | 28.3609 | 28.3929 | 28.4837 |

| | | | | | | | | |
|---|---|---|---|---|---|---|---|---|
| 0 | 33.817 | 6.82E+08 | 31 | 25.3176 | 25.4254 | 25.3999 | 25.1686 | 25.7773 |
| 0 | 22.523 | 1.37E+09 | 29 | 26.4003 | 26.5741 | 26.7285 | 26.3669 | 26.6466 |
| 0 | 49.517 | 1.13E+09 | 49 | 26.5943 | 26.548 | 26.3027 | 26.2306 | 26.2942 |
| 0 | 257.97 | 4.98E+10 | 138 | 31.9199 | 31.8588 | 31.9072 | 31.8617 | 31.7785 |
| 0 | 14.451 | 8.35E+08 | 31 | 25.9802 | 25.8982 | 26.1635 | 25.9637 | 26.0537 |
| 0 | 80.462 | 5.43E+09 | 124 | 28.6457 | 28.6718 | 28.6392 | 28.6762 | 28.6482 |
| 0 | 27.631 | 9.52E+08 | 36 | 25.8317 | 25.8253 | 26.1619 | 26.0062 | 26.1983 |
| 0 | 92.256 | 2.05E+09 | 56 | 27.3052 | 27.1852 | 26.9319 | 26.551 | 27.7343 |
| 0 | 48.806 | 2.28E+09 | 90 | 27.3493 | 27.3713 | 27.2945 | 27.3848 | 27.4624 |
| 0 | 53.23 | 1.73E+09 | 58 | 26.8959 | 26.9884 | 26.7415 | 26.7191 | 26.887 |
| 0 | 45.292 | 1.92E+09 | 59 | 27.3016 | 27.2365 | 26.9608 | 27.0502 | 27.0696 |
| 0 | 31.615 | 1.24E+09 | 25 | 26.4638 | 26.3883 | 26.211 | 26.4747 | 26.5988 |
| 0 | 266.68 | 5.24E+10 | 310 | 31.8261 | 31.8423 | 31.8523 | 31.8376 | 31.7124 |
| 0 | 109.9 | 6.05E+09 | 120 | 28.965 | 28.8921 | 28.5823 | 28.6274 | 28.5846 |
| 0 | 76.785 | 2.34E+09 | 76 | 27.2516 | 27.4787 | 27.1816 | 27.3736 | 27.6589 |
| 0 | 20.872 | 1.14E+09 | 51 | 26.0669 | 26.0748 | 26.6151 | 26.4498 | 26.1677 |
| 0 | 93.852 | 2.37E+09 | 85 | 27.3579 | 27.3647 | 27.448 | 27.1868 | 27.5347 |
| 0 | 61.271 | 1.17E+09 | 43 | 26.3003 | 26.1679 | 26.1679 | 26.346 | 26.5659 |
| 0 | 22.057 | 1.67E+09 | 91 | 27.0335 | 26.963 | 26.679 | 26.6148 | 26.9785 |
| 0 | 66.697 | 1.17E+10 | 168 | 29.9205 | 29.8829 | 29.5909 | 29.5923 | 29.6026 |
| 0 | 154.82 | 5.03E+09 | 111 | 28.487 | 28.3829 | 28.4672 | 28.49 | 28.5356 |
| 0 | 80.285 | 2.38E+09 | 90 | 27.4792 | 27.5297 | 27.1972 | 27.2733 | 27.5167 |
| 0 | 99.085 | 8.22E+09 | 140 | 29.4084 | 29.432 | 29.0938 | 29.2455 | 29.2401 |
| 0 | 14.62 | 1.15E+09 | 22 | 26.1013 | 26.5833 | 26.6161 | 26.2443 | 26.5785 |
| 0 | 237.97 | 3.97E+09 | 70 | 28.1941 | 28.1154 | 28.249 | 28.1219 | 28.0228 |
| 0 | 128.55 | 3.56E+09 | 84 | 27.9233 | 27.9006 | 28.04 | 28.2345 | 27.9463 |
| 0 | 174.51 | 8.17E+09 | 179 | 29.3479 | 29.3095 | 29.2818 | 29.2134 | 29.0107 |
| 0 | 73.574 | 2.94E+09 | 78 | 27.8209 | 27.7836 | 27.6574 | 27.4541 | 27.631 |
| 0 | 67.897 | 4.13E+09 | 109 | 28.4739 | 28.398 | 28.1195 | 28.2174 | 28.1193 |
| 0 | 17.581 | 1.26E+09 | 52 | 26.2582 | 26.3928 | 26.5379 | 26.5582 | 26.5833 |
| 0 | 89.144 | 2.21E+09 | 78 | 27.2644 | 27.2456 | 27.1441 | 27.2737 | 27.1984 |
| 0 | 25.606 | 2.03E+09 | 64 | 27.1939 | 27.0506 | 27.0595 | 27.2104 | 27.4016 |
| 0 | 221.63 | 2.06E+10 | 200 | 30.4531 | 30.3087 | 30.6317 | 30.5995 | 30.563 |
| 0 | 37.578 | 2.15E+09 | 90 | 27.1954 | 27.4328 | 27.16 | 27.1385 | 27.3198 |
| 0 | 27.25 | 2.7E+09 | 84 | 27.6806 | 27.644 | 27.7117 | 27.6424 | 27.3754 |
| 0 | 21.649 | 1.01E+09 | 40 | 25.8895 | 26.2291 | 25.8325 | 26.0989 | 26.2506 |
| 0 | 270.8 | 9.76E+09 | 176 | 29.6681 | 29.6032 | 29.4566 | 29.3621 | 29.3501 |
| 0 | 310.42 | 4.47E+10 | 231 | 31.693 | 31.6765 | 31.762 | 31.7783 | 31.4014 |
| 0 | 87.59 | 3.37E+09 | 77 | 27.6246 | 27.6484 | 27.9148 | 28.0397 | 28.0197 |
| 0 | 35.719 | 2.57E+09 | 107 | 27.4829 | 27.4886 | 27.504 | 27.5619 | 27.6882 |
| 0 | 20.945 | 8.75E+08 | 43 | 25.856 | 25.9409 | 25.9014 | 25.6265 | 26.0668 |
| 0 | 24.18 | 1.04E+09 | 49 | 26.2331 | 25.8526 | 26.1057 | 25.8697 | 26.552 |
| 0 | 59.7 | 2.18E+09 | 78 | 27.3845 | 27.3471 | 27.1565 | 27.2011 | 27.2074 |
| 0 | 193.73 | 7.7E+09 | 140 | 29.4748 | 29.4975 | 28.8954 | 28.9312 | 29.0053 |
| 0 | 323.31 | 2.65E+10 | 169 | 30.837 | 30.7727 | 30.7597 | 30.7793 | 31.0197 |
| 0 | 52.39 | 1.78E+09 | 94 | 26.862 | 26.9209 | 27.0555 | 27.0577 | 27.0625 |
| 0 | 37.48 | 1.16E+09 | 54 | 26.1526 | 26.2528 | 26.2736 | 26.1109 | 26.5757 |
| 0 | 323.31 | 3.4E+10 | 295 | 31.5208 | 31.4901 | 31.2175 | 31.1453 | 31.14 |

| | | | | | | | | |
|---|---|---|---|---|---|---|---|---|
| 0 | 50.786 | 1.71E+09 | 54 | 27.0799 | 26.9596 | 26.9029 | 26.804 | 26.9094 |
| 0 | 42.993 | 1.3E+09 | 54 | 26.5295 | 26.3149 | 26.6535 | 26.6461 | 26.6602 |
| 0 | 28.208 | 2.07E+09 | 79 | 26.9496 | 27.124 | 27.1224 | 27.2792 | 27.3743 |
| 0 | 56.181 | 4.21E+09 | 115 | 28.1233 | 28.1447 | 28.5284 | 28.4798 | 28.1762 |
| 0 | 64.884 | 1.56E+09 | 76 | 26.8533 | 26.7448 | 26.6937 | 26.5535 | 26.8971 |
| 0 | 40.464 | 3.66E+09 | 124 | 28.0693 | 28.031 | 27.9306 | 27.9604 | 28.1113 |
| 0 | 323.31 | 1.38E+10 | 144 | 30.1799 | 30.157 | 29.9506 | 29.8652 | 29.796 |
| 0 | 25.503 | 8.98E+08 | 35 | 26.1 | 26.0537 | 25.5683 | 25.8346 | 26.0721 |
| 0 | 12.375 | 8.85E+08 | 22 | 25.7929 | 25.9514 | 25.8521 | 25.8831 | 25.7228 |
| 0 | 26.024 | 1.01E+09 | 36 | 25.9791 | 25.8855 | 26.0488 | 26.0798 | 26.0103 |
| 0 | 34.66 | 1.22E+09 | 43 | 26.0488 | 26.0894 | 26.3783 | 26.3342 | 26.382 |
| 0 | 31.251 | 3.92E+09 | 88 | 27.9403 | 28.0915 | 27.9503 | 27.9972 | 27.8775 |
| 0 | 23.56 | 1.9E+09 | 60 | 26.9841 | 26.9515 | 26.8655 | 27.0846 | 26.9392 |
| 0 | 217.53 | 5.43E+09 | 136 | 28.3989 | 28.4441 | 28.6105 | 28.6128 | 28.3523 |
| 0 | 46.95 | 1.43E+09 | 55 | 26.5352 | 26.6061 | 26.396 | 26.5863 | 26.5675 |
| 0 | 323.31 | 2.79E+10 | 287 | 30.8311 | 30.8535 | 30.7022 | 30.795 | 30.8625 |
| 0 | 66.017 | 1.75E+09 | 74 | 26.8795 | 26.8954 | 26.8839 | 26.9329 | 26.8353 |
| 0 | 255.18 | 3.04E+10 | 329 | 31.0028 | 31.0126 | 31.0341 | 30.9813 | 30.895 |
| 0 | 92.515 | 3.21E+09 | 119 | 27.8191 | 27.8002 | 27.8297 | 27.6019 | 27.799 |
| 0 | 54.972 | 9.49E+09 | 151 | 29.1859 | 29.278 | 29.4279 | 29.4224 | 29.3151 |
| 0 | 25.003 | 1.16E+09 | 50 | 26.3936 | 26.4245 | 26.3734 | 26.2698 | 26.2436 |
| 0 | 76.59 | 9.22E+09 | 142 | 29.3658 | 29.4133 | 29.3298 | 29.2879 | 29.1014 |
| 0 | 43.305 | 1.38E+09 | 37 | 26.4165 | 26.5879 | 26.6999 | 26.5557 | 26.4828 |
| 0 | 323.31 | 2.99E+10 | 319 | 30.9329 | 30.782 | 31.016 | 31.0029 | 31.1204 |
| 0 | 78.448 | 8.73E+09 | 194 | 29.3937 | 29.3596 | 29.3896 | 29.3971 | 29.3253 |
| 0 | 75.708 | 6.26E+09 | 132 | 28.8829 | 28.8547 | 28.8898 | 28.95 | 28.8673 |
| 0 | 154.46 | 7.06E+09 | 136 | 29.078 | 29.0614 | 28.9929 | 29.192 | 29.0793 |
| 0 | 108.83 | 3.88E+10 | 161 | 31.4352 | 31.5122 | 31.4716 | 31.5408 | 31.621 |
| 0 | 152.78 | 6.87E+09 | 125 | 29.1823 | 29.112 | 28.9436 | 28.9452 | 28.9259 |
| 0 | 21.749 | 2.08E+09 | 52 | 27.4698 | 27.6686 | 27.5802 | 27.3403 | 27.1485 |
| 0 | 27.144 | 3.1E+09 | 91 | 27.7149 | 27.741 | 27.3698 | 27.4009 | 27.7112 |
| 0 | 36.598 | 1.71E+09 | 60 | 26.9302 | 26.7738 | 26.7427 | 26.7633 | 26.9263 |
| 0 | 19.432 | 8.6E+08 | 43 | 25.9944 | 25.7959 | 25.9368 | 26.1281 | 26.0856 |
| 0 | 68.699 | 5.19E+09 | 98 | 28.6902 | 28.5765 | 28.6609 | 28.5496 | 28.7004 |
| 0 | 20.712 | 1.02E+09 | 50 | 25.9024 | 26.0018 | 26.1311 | 25.8871 | 26.2603 |
| 0 | 81.918 | 1.96E+10 | 171 | 30.4114 | 30.3533 | 30.4887 | 30.4032 | 30.423 |
| 0 | 42.984 | 4.01E+09 | 115 | 28.1704 | 28.0061 | 28.1166 | 28.0545 | 28.0327 |
| 0 | 23.255 | 8.47E+08 | 46 | 25.8093 | 25.7808 | 26.0709 | 25.6989 | 25.7291 |
| 0 | 27.085 | 1.11E+09 | 65 | 26.1326 | 26.006 | 26.4427 | 26.0297 | 26.2817 |
| 0 | 25.629 | 3.28E+09 | 92 | 27.9126 | 27.8614 | 27.7973 | 27.8061 | 27.7574 |
| 0 | 57.57 | 2.16E+09 | 98 | 26.9945 | 27.2082 | 27.2376 | 27.1451 | 27.2081 |
| 0 | 20.406 | 1.58E+09 | 59 | 26.721 | 26.7859 | 26.8465 | 26.6031 | 26.7035 |
| 0 | 60.697 | 7.81E+09 | 124 | 29.0501 | 28.9409 | 28.9808 | 29.0594 | 29.1363 |
| 0 | 63.825 | 6.74E+09 | 142 | 28.9703 | 28.9425 | 28.8763 | 28.768 | 28.7425 |
| 0 | 67.859 | 2.08E+09 | 54 | 27.4703 | 27.3245 | 27.3041 | 27.1297 | 27.1865 |
| 0 | 94.091 | 6.16E+09 | 118 | 28.7595 | 28.8309 | 28.8351 | 28.7908 | 28.6674 |
| 0 | 13.493 | 5.73E+08 | 23 | 25.0529 | 25.2694 | 25.037 | 25.2198 | 25.3663 |
| 0 | 52.109 | 5.24E+09 | 103 | 28.6603 | 28.6091 | 28.6213 | 28.4271 | 28.629 |

| | | | | | | | | |
|---|---|---|---|---|---|---|---|---|
| 0 | 52.99 | 1.43E+09 | 44 | 26.3391 | 26.3496 | 26.5075 | 26.3726 | 26.7832 |
| 0 | 158.73 | 6.73E+09 | 90 | 28.6043 | 28.5689 | 28.7907 | 28.7847 | 28.7523 |
| 0 | 155.46 | 1.19E+11 | 351 | 32.9929 | 32.9046 | 32.9162 | 32.9972 | 33.0578 |
| 0 | 211.18 | 1.11E+10 | 166 | 29.5889 | 29.5931 | 29.6787 | 29.6922 | 29.5177 |
| 0 | 91.986 | 6.88E+09 | 125 | 28.9564 | 28.9187 | 28.9531 | 28.9498 | 28.9227 |
| 0 | 11.562 | 1.07E+09 | 35 | 26.5497 | 26.3588 | 26.1824 | 26.3784 | 26.519 |
| 0 | 53.526 | 3.42E+09 | 105 | 28.0274 | 27.9568 | 27.8703 | 27.9337 | 28.0368 |
| 0 | 16.146 | 2.27E+09 | 56 | 27.2848 | 27.3489 | 27.4562 | 27.3482 | 27.4375 |
| 0 | 71.051 | 2.27E+09 | 63 | 27.2783 | 27.3843 | 27.3552 | 27.327 | 27.3963 |
| 0 | 38.601 | 2.04E+09 | 74 | 27.1697 | 27.2066 | 27.2086 | 27.1253 | 27.4147 |
| 0 | 323.31 | 4.25E+10 | 294 | 31.5975 | 31.6236 | 31.4717 | 31.4779 | 31.5456 |
| 0 | 31.535 | 1.03E+09 | 37 | 26.2021 | 26.1837 | 26.0296 | 26.1359 | 25.9448 |
| 0 | 177.85 | 3.32E+10 | 263 | 31.1399 | 31.1723 | 31.2277 | 31.0534 | 31.1099 |
| 0 | 28.052 | 1.04E+09 | 27 | 26.2993 | 26.1781 | 26.1678 | 25.8526 | 26.0888 |
| 0 | 200.74 | 6.48E+09 | 214 | 28.5465 | 28.5085 | 28.6896 | 28.6142 | 28.8034 |
| 0 | 212.87 | 1.06E+10 | 160 | 29.5779 | 29.5875 | 29.6387 | 29.6171 | 29.6055 |
| 0 | 323.31 | 1.46E+10 | 251 | 29.9624 | 29.9736 | 30.0288 | 29.9286 | 30.0607 |
| 0 | 17.826 | 8.34E+08 | 30 | 25.9445 | 25.8592 | 26.0011 | 25.5442 | 25.838 |
| 0 | 71.035 | 6.21E+09 | 131 | 28.9304 | 28.9377 | 28.7877 | 28.8355 | 28.87 |
| 0 | 323.31 | 2.18E+10 | 137 | 30.6708 | 30.5957 | 30.5589 | 30.6235 | 30.5407 |
| 0 | 302.99 | 4.42E+10 | 280 | 31.4909 | 31.4868 | 31.6506 | 31.6924 | 31.599 |
| 0 | 108.2 | 1.07E+11 | 242 | 33.3601 | 33.3067 | 32.8183 | 32.8195 | 32.7322 |
| 0 | 156.77 | 4.27E+09 | 143 | 28.2878 | 28.292 | 27.9672 | 28.0755 | 28.1718 |
| 0 | 13.143 | 9.13E+08 | 28 | 25.8268 | 25.588 | 26.0376 | 25.9922 | 25.8594 |
| 0 | 24.797 | 2.65E+09 | 100 | 27.9041 | 27.9098 | 27.6099 | 27.5734 | 27.4739 |
| 0 | 60.087 | 2.46E+09 | 69 | 27.8339 | 27.6559 | 27.1967 | 27.524 | 27.3533 |
| 0 | 26.696 | 1.06E+09 | 30 | 26.5305 | 26.6352 | 26.6577 | 26.0389 | 26.4084 |
| 0 | 15.974 | 1.12E+09 | 40 | 26.217 | 26.3001 | 26.4314 | 26.4227 | 26.4506 |
| 0 | 110.75 | 3.27E+09 | 105 | 27.8455 | 27.7211 | 27.8086 | 27.7676 | 27.7042 |
| 0 | 17.426 | 9.76E+08 | 49 | 26.1213 | 25.9342 | 26.1984 | 26.0553 | 25.9248 |
| 0 | 61.6 | 8.99E+08 | 46 | 26.1281 | 25.9817 | 25.9578 | 26.1064 | 25.9555 |
| 0 | 53.208 | 7.88E+09 | 126 | 29.1323 | 29.0643 | 29.1624 | 29.3506 | 28.8574 |
| 0 | 17.468 | 1.06E+09 | 38 | 25.8528 | 26.2859 | 26.4079 | 26.0033 | 25.8702 |
| 0 | 143.77 | 3.45E+09 | 93 | 27.7819 | 27.7871 | 27.8515 | 27.8196 | 27.8948 |
| 0 | 36.774 | 3.1E+09 | 109 | 27.8001 | 27.9358 | 27.5447 | 27.5887 | 27.8159 |
| 0 | 85.185 | 2.34E+10 | 127 | 30.6732 | 30.6579 | 30.7463 | 30.6188 | 30.6773 |
| 0 | 25.331 | 7.47E+08 | 23 | 25.7755 | 25.5534 | 25.4236 | 25.8483 | 25.5288 |
| 0 | 89.546 | 1.55E+10 | 167 | 30.0605 | 30.135 | 30.1094 | 30.1505 | 30.0623 |
| 0 | 26.14 | 2.35E+09 | 82 | 27.4511 | 27.4484 | 27.2977 | 27.4441 | 27.4735 |
| 0 | 74.848 | 1.54E+10 | 159 | 30.3312 | 30.3246 | 30.039 | 30.1334 | 30.1422 |
| 0 | 20.256 | 6.41E+08 | 19 | 25.8457 | 25.7701 | 25.1042 | 25.4235 | 25.6383 |
| 0 | 15.406 | 1.31E+09 | 37 | 26.6581 | 26.5185 | 26.5892 | 26.7169 | 26.496 |
| 0 | 51.24 | 1.98E+09 | 71 | 27.0998 | 27.1478 | 27.0602 | 27.0336 | 26.9579 |
| 0 | 23.882 | 2.14E+09 | 79 | 27.2283 | 27.1476 | 27.1647 | 27.2866 | 27.2421 |
| 0 | 162.22 | 8.55E+10 | 262 | 32.6363 | 32.6132 | 32.5637 | 32.6003 | 32.4343 |
| 0 | 87.438 | 3.14E+09 | 102 | 27.9446 | 27.7911 | 27.7019 | 27.6064 | 27.8754 |
| 0 | 43.963 | 1.4E+09 | 61 | 26.4214 | 26.3832 | 26.8164 | 26.6647 | 26.5777 |
| 0 | 100.59 | 1.29E+10 | 132 | 29.9459 | 29.8946 | 29.8726 | 29.9307 | 29.7901 |

| | | | | | | | | |
|---|---|---|---|---|---|---|---|---|
| 0 | 49.953 | 2.68E+09 | 84 | 27.6675 | 27.567 | 27.5702 | 27.5378 | 27.6113 |
| 0 | 114.22 | 2.89E+09 | 80 | 27.5776 | 27.5438 | 27.7514 | 27.8743 | 27.6615 |
| 0 | 323.31 | 1.72E+11 | 579 | 33.4648 | 33.4791 | 33.6468 | 33.6974 | 33.4925 |
| 0 | 141.72 | 7.94E+09 | 101 | 29.2435 | 29.3152 | 29.0873 | 29.1988 | 28.7464 |
| 0 | 105.99 | 1.26E+10 | 160 | 29.8536 | 29.8078 | 29.8443 | 29.7069 | 29.8858 |
| 0 | 119.4 | 3.19E+09 | 98 | 27.904 | 27.9663 | 27.6971 | 27.6994 | 27.6219 |
| 0 | 61.259 | 1.02E+09 | 60 | 25.9506 | 25.8699 | 26.3806 | 26.0151 | 26.2292 |
| 0 | 103.57 | 1.14E+10 | 181 | 29.8469 | 29.7978 | 29.613 | 29.6488 | 29.6348 |
| 0 | 103 | 3.74E+09 | 97 | 27.9817 | 28.0263 | 28.0845 | 28.1304 | 28.0328 |
| 0 | 323.31 | 2.56E+10 | 182 | 30.8525 | 30.8319 | 30.9067 | 30.8303 | 30.7785 |
| 0 | 91.777 | 4.32E+09 | 112 | 28.3067 | 28.1885 | 28.2132 | 28.2768 | 28.4367 |
| 0 | 47.534 | 8.21E+09 | 114 | 29.1388 | 29.0967 | 29.1987 | 29.2067 | 29.1425 |
| 0 | 55.146 | 2.83E+09 | 56 | 27.7582 | 27.798 | 27.6391 | 27.6442 | 27.5588 |
| 0 | 21.622 | 8.34E+08 | 25 | 25.9363 | 26.1568 | 25.6367 | 25.831 | 25.6961 |
| 0 | 20.783 | 1.9E+09 | 37 | 27.0235 | 27.2843 | 27.0071 | 26.9866 | 26.9554 |
| 0 | 19.651 | 1.12E+09 | 46 | 26.2678 | 26.2269 | 26.341 | 26.2748 | 26.3832 |
| 0 | 106.7 | 2.28E+09 | 80 | 27.3876 | 27.3519 | 27.2742 | 27.4858 | 27.3351 |
| 0 | 323.31 | 1.91E+10 | 241 | 30.3965 | 30.3531 | 30.5258 | 30.5562 | 30.3893 |
| 0 | 41.329 | 3.88E+09 | 96 | 28.2911 | 28.3381 | 27.9715 | 27.9205 | 28.2123 |
| 0 | 110.41 | 8.97E+09 | 159 | 29.3224 | 29.2965 | 29.3977 | 29.4103 | 29.2624 |
| 0 | 25.73 | 1.67E+09 | 65 | 26.9891 | 26.8083 | 26.6331 | 26.7033 | 27.1971 |
| 0 | 72.225 | 1.09E+10 | 110 | 29.5099 | 29.4953 | 29.6906 | 29.6131 | 29.6247 |
| 0 | 35.914 | 3.2E+09 | 69 | 27.81 | 27.7119 | 27.8946 | 27.8163 | 27.929 |
| 0 | 79.002 | 2.88E+09 | 106 | 27.8043 | 27.9246 | 27.6502 | 27.6047 | 27.6968 |
| 0 | 323.31 | 1.23E+10 | 333 | 29.6518 | 29.5721 | 29.6247 | 29.5705 | 29.9521 |
| 0 | 126.21 | 1.21E+10 | 148 | 29.7348 | 29.6832 | 29.8052 | 29.8562 | 29.6839 |
| 0 | 23.106 | 1.52E+10 | 113 | 30.0926 | 30.1663 | 30.0303 | 30.1719 | 29.9214 |
| 0 | 323.31 | 1.28E+11 | 506 | 33.1071 | 33.0762 | 33.2601 | 33.1428 | 33.118 |
| 0 | 16.521 | 8.41E+08 | 55 | 25.9202 | 25.7998 | 25.7848 | 26.0212 | 25.6863 |
| 0 | 323.31 | 2.73E+10 | 313 | 31.0753 | 31.113 | 30.9633 | 30.936 | 30.8711 |
| 0 | 121.72 | 5.94E+09 | 158 | 28.6333 | 28.7046 | 28.8926 | 28.8605 | 28.6402 |
| 0 | 28.15 | 1.24E+09 | 45 | 26.202 | 26.3212 | 26.8497 | 26.4656 | 26.5448 |
| 0 | 50.019 | 7.88E+08 | 43 | 25.7205 | 25.5576 | 25.6456 | 25.9098 | 25.8161 |
| 0 | 114.52 | 2.86E+09 | 81 | 27.4627 | 27.5259 | 27.817 | 27.8241 | 27.7446 |
| 0 | 69.422 | 8.05E+09 | 100 | 29.2842 | 29.2729 | 29.0423 | 29.2268 | 29.091 |
| 0 | 62.681 | 1.46E+09 | 52 | 26.6573 | 26.5871 | 26.8187 | 26.6203 | 26.5539 |
| 0 | 244.63 | 6.51E+09 | 99 | 28.8357 | 28.7735 | 28.9791 | 28.9737 | 28.7815 |
| 0 | 72.246 | 3.84E+09 | 110 | 28.1476 | 28.0704 | 28.1704 | 28.0819 | 28.0968 |
| 0 | 92.869 | 4.51E+09 | 148 | 28.3099 | 28.3192 | 28.4574 | 28.3674 | 28.3146 |
| 0 | 92.16 | 7.66E+09 | 127 | 29.0321 | 29.0847 | 29.0871 | 29.0186 | 29.0917 |
| 0 | 39.793 | 4.35E+09 | 110 | 28.3346 | 28.2344 | 28.3221 | 28.3393 | 28.3427 |
| 0 | 293.46 | 3.6E+10 | 293 | 31.3738 | 31.2973 | 31.4512 | 31.3031 | 31.2391 |
| 0 | 71.988 | 2.3E+09 | 74 | 27.6277 | 27.6994 | 27.2465 | 27.234 | 27.3168 |
| 0 | 77.295 | 1.82E+09 | 50 | 26.9902 | 27.0408 | 27.0484 | 26.9054 | 27.0926 |
| 0 | 29.402 | 1.32E+09 | 44 | 26.4232 | 26.8957 | 26.5792 | 26.6502 | 26.2946 |
| 0 | 19.973 | 2.1E+09 | 80 | 27.138 | 27.094 | 27.2785 | 27.1014 | 27.579 |
| 0 | 12.527 | 6.36E+08 | 27 | 25.6138 | 25.705 | 25.2549 | 25.4666 | 25.5952 |
| 0 | 19.885 | 1.49E+09 | 54 | 26.6573 | 26.7955 | 26.8772 | 26.9165 | 26.6721 |

| | | | | | | | | |
|---|---|---|---|---|---|---|---|---|
| 0 | 26.391 | 2.26E+09 | 101 | 27.4692 | 27.4885 | 27.4235 | 27.4371 | 27.1444 |
| 0 | 10.819 | 1.28E+09 | 39 | 26.5431 | 26.5452 | 26.7017 | 26.4659 | 26.2829 |
| 0 | 20.744 | 1.78E+09 | 61 | 26.9841 | 27.0653 | 26.9895 | 27.1097 | 26.7806 |
| 0 | 8.9354 | 5.82E+08 | 24 | 25.7717 | 24.9867 | 25.2832 | 25.5809 | 25.3891 |
| 0 | 24.562 | 2.33E+09 | 64 | 27.3365 | 27.2484 | 27.3654 | 27.321 | 27.5494 |
| 0 | 21.758 | 1.22E+09 | 72 | 26.4432 | 26.4446 | 26.3782 | 26.3236 | 26.5587 |
| 0 | 60.96 | 1.91E+09 | 61 | 27.1415 | 27.0242 | 26.9769 | 26.9071 | 27.1786 |
| 0 | 79.226 | 7.24E+09 | 167 | 29.0695 | 28.9164 | 29.2486 | 29.0592 | 28.977 |
| 0 | 80.195 | 2.25E+10 | 239 | 30.6419 | 30.6261 | 30.6996 | 30.794 | 30.5343 |
| 0 | 218.52 | 9.96E+09 | 226 | 29.4634 | 29.4266 | 29.5869 | 29.4906 | 29.4515 |
| 0 | 94.249 | 3.15E+09 | 96 | 27.8734 | 27.7736 | 27.9033 | 27.8644 | 27.7195 |
| 0 | 56.669 | 2.63E+09 | 72 | 27.6639 | 27.5248 | 27.6535 | 27.4899 | 27.5751 |
| 0 | 21.513 | 8.58E+08 | 47 | 25.8627 | 25.8836 | 25.7422 | 25.7333 | 26.0212 |
| 0 | 27.138 | 4.2E+09 | 85 | 28.2271 | 28.1302 | 28.335 | 28.1665 | 28.295 |
| 0 | 42.948 | 4.44E+09 | 115 | 28.3816 | 28.4434 | 28.3675 | 28.4643 | 28.047 |
| 0 | 21.708 | 1.94E+09 | 54 | 27.2987 | 27.0087 | 26.9866 | 27.1281 | 27.0889 |
| 0 | 105.58 | 2.19E+09 | 69 | 27.131 | 27.2361 | 27.2223 | 27.3634 | 27.4916 |
| 0 | 58.776 | 1.93E+09 | 45 | 27.1714 | 27.1001 | 27.0418 | 27.1722 | 27.1521 |
| 0 | 39.824 | 2.1E+09 | 76 | 27.3604 | 27.4047 | 26.9852 | 26.9216 | 27.3572 |
| 0 | 125.73 | 3.07E+09 | 66 | 27.7607 | 27.7977 | 28.0209 | 27.8561 | 27.6405 |
| 0 | 32.82 | 1.31E+09 | 31 | 26.3821 | 26.2227 | 26.6919 | 26.565 | 25.985 |
| 0 | 10.114 | 6.83E+08 | 43 | 25.083 | 25.2059 | 25.3608 | 25.3365 | 25.7521 |
| 0 | 81.112 | 1.56E+09 | 49 | 26.6547 | 26.4352 | 26.5282 | 26.6671 | 26.5503 |
| 0 | 21.263 | 5.2E+08 | 33 | 25.1923 | 25.1206 | 25.1553 | 24.884 | 25.0068 |
| 0 | 51.53 | 2.01E+09 | 58 | 26.9432 | 27.0087 | 26.9292 | 27.0742 | 27.0336 |
| 0 | 44.49 | 1.24E+09 | 70 | 26.2622 | 26.175 | 26.4633 | 26.3294 | 26.3941 |
| 0 | 37.453 | 1.78E+09 | 79 | 26.6211 | 26.6651 | 26.9039 | 26.9576 | 27.009 |
| 0 | 262.23 | 3.64E+09 | 91 | 27.8904 | 27.8569 | 27.6841 | 27.8946 | 27.9642 |
| 0 | 71.207 | 8.9E+09 | 97 | 29.3852 | 29.2605 | 29.2995 | 29.4305 | 29.0355 |
| 0 | 21.876 | 1.27E+09 | 33 | 26.3075 | 26.5333 | 26.1997 | 26.0908 | 26.5802 |
| 0 | 62.676 | 2.64E+10 | 166 | 30.6423 | 30.6824 | 30.7753 | 30.5968 | 30.7161 |
| 0 | 50.575 | 2.14E+09 | 54 | 27.1472 | 26.9601 | 27.2087 | 27.1609 | 27.1754 |
| 0 | 68.181 | 7.72E+09 | 128 | 28.9298 | 28.9596 | 29.0701 | 29.1674 | 28.9426 |
| 0 | 57.703 | 2.31E+09 | 66 | 27.2622 | 27.3084 | 27.2965 | 27.2969 | 27.2408 |
| 0 | 204.55 | 7.94E+09 | 128 | 29.0416 | 29.0189 | 29.161 | 29.1339 | 28.9786 |
| 0 | 188.81 | 4.43E+09 | 108 | 28.2811 | 28.2239 | 28.1616 | 28.2664 | 28.2261 |
| 0 | 66.919 | 2.62E+09 | 78 | 27.4788 | 27.4974 | 27.4056 | 27.4094 | 27.4591 |
| 0 | 323.31 | 8.1E+09 | 80 | 29.1982 | 29.1931 | 29.0471 | 29.0266 | 29.0555 |
| 0 | 323.31 | 5.47E+10 | 394 | 31.8053 | 31.864 | 31.9182 | 31.8091 | 31.8412 |
| 0 | 32.642 | 2.39E+09 | 73 | 27.3434 | 27.3334 | 27.3685 | 27.5609 | 27.18 |
| 0 | 211.45 | 1.33E+10 | 136 | 29.6915 | 29.7136 | 29.8757 | 29.7902 | 29.8166 |
| 0 | 46.061 | 5E+09 | 99 | 28.4537 | 28.4168 | 28.3843 | 28.3662 | 28.4426 |
| 0 | 27.024 | 3.53E+09 | 84 | 27.9085 | 27.9278 | 27.8624 | 27.9237 | 27.9192 |
| 0 | 16.886 | 2.96E+09 | 60 | 27.8506 | 27.7618 | 27.8269 | 27.7303 | 27.7756 |
| 0 | 27.842 | 3.35E+09 | 107 | 27.8671 | 27.906 | 27.9833 | 28.0969 | 28.1305 |
| 0 | 25.801 | 3.98E+09 | 75 | 28.3626 | 28.1869 | 28.0423 | 27.9324 | 27.895 |
| 0 | 9.9495 | 1.52E+09 | 66 | 26.685 | 26.7892 | 26.9625 | 27.0197 | 26.9441 |
| 0 | 323.31 | 7.36E+09 | 152 | 29.303 | 29.2274 | 29.1015 | 28.9408 | 29.1401 |

| | | | | | | | | |
|---|---|---|---|---|---|---|---|---|
| 0 | 116.76 | 1.07E+10 | 140 | 29.8282 | 29.7095 | 29.5944 | 29.5715 | 29.6937 |
| 0 | 123.7 | 4.83E+09 | 103 | 28.4927 | 28.4955 | 28.6556 | 28.7027 | 28.5353 |
| 0 | 20.726 | 1.2E+09 | 51 | 26.4315 | 26.5042 | 26.5828 | 26.5003 | 26.6874 |
| 0 | 48.552 | 3.18E+09 | 66 | 28.3381 | 28.421 | 27.8027 | 27.8788 | 27.7911 |
| 0 | 201.66 | 1.9E+10 | 358 | 30.2704 | 30.2985 | 30.2372 | 30.2296 | 30.502 |
| 0 | 30.562 | 1.4E+09 | 66 | 27.0149 | 26.9529 | 26.6442 | 26.7249 | 26.5911 |
| 0 | 32.194 | 2.03E+09 | 66 | 27.1583 | 27.1171 | 27.355 | 27.1693 | 26.8474 |
| 0 | 157.05 | 2.93E+09 | 92 | 27.6583 | 27.6954 | 27.686 | 27.7548 | 27.4969 |
| 0 | 25.936 | 1.82E+09 | 47 | 27.0151 | 27.0206 | 27.0519 | 26.9772 | 26.8809 |
| 0 | 251.05 | 3.6E+10 | 169 | 31.419 | 31.3127 | 31.2324 | 31.2419 | 31.3423 |
| 0 | 13.232 | 1.27E+09 | 31 | 26.2998 | 26.528 | 26.2025 | 26.403 | 26.2067 |
| 0 | 29.471 | 3.66E+09 | 70 | 27.9896 | 28.1565 | 28.13 | 27.9372 | 28.0739 |
| 0 | 37.338 | 1.64E+09 | 49 | 26.774 | 26.9897 | 26.9235 | 26.9725 | 26.6082 |
| 0 | 24.434 | 5.14E+09 | 98 | 28.3912 | 28.4856 | 28.563 | 28.6565 | 28.3774 |
| 0 | 31.349 | 2.99E+09 | 112 | 27.9241 | 27.8805 | 27.6339 | 27.7307 | 27.888 |
| 0 | 93.801 | 1.78E+09 | 76 | 26.9626 | 26.7627 | 26.8656 | 27.059 | 26.8801 |
| 0 | 117.36 | 6.77E+09 | 139 | 28.855 | 28.8942 | 28.9276 | 28.984 | 28.7555 |
| 0 | 303.13 | 2.86E+10 | 251 | 30.8404 | 30.7944 | 30.8865 | 30.8505 | 30.7637 |
| 0 | 24.482 | 1.07E+09 | 57 | 26.0764 | 26.2706 | 26.2754 | 26.1764 | 26.3177 |
| 0 | 73.591 | 3.84E+09 | 107 | 28.0984 | 27.9961 | 28.1659 | 28.1306 | 28.0765 |
| 0 | 34.931 | 1.36E+09 | 29 | 26.5851 | 26.6112 | 26.6214 | 26.5055 | 26.3687 |
| 0 | 107.53 | 1.71E+10 | 170 | 30.217 | 30.0947 | 30.227 | 30.1957 | 30.0152 |
| 0 | 204.56 | 1.14E+10 | 135 | 29.8934 | 29.8045 | 29.6996 | 29.5417 | 29.7216 |
| 0 | 16.575 | 1.32E+09 | 42 | 26.6321 | 26.5004 | 26.6162 | 26.4233 | 26.6079 |
| 0 | 53.022 | 9.73E+09 | 209 | 29.4803 | 29.4411 | 29.5648 | 29.5427 | 29.4755 |
| 0 | 22.62 | 2.56E+09 | 63 | 27.1999 | 27.1499 | 27.2932 | 27.5356 | 27.5951 |
| 0 | 58.841 | 4.28E+09 | 91 | 28.1575 | 28.0797 | 28.3638 | 28.3871 | 28.1484 |
| 0 | 39.154 | 1.51E+09 | 70 | 26.7326 | 26.5004 | 26.9144 | 26.8615 | 26.5305 |
| 0 | 122.68 | 3.3E+09 | 100 | 27.9046 | 27.9701 | 27.8948 | 27.8538 | 27.9072 |
| 0 | 65.089 | 2.65E+09 | 58 | 27.5462 | 27.3948 | 27.5988 | 27.5704 | 27.4965 |
| 0 | 49.105 | 2.73E+09 | 56 | 27.8599 | 27.8416 | 27.7731 | 27.7033 | 27.3825 |
| 0 | 32.252 | 1.28E+09 | 52 | 26.6484 | 26.6353 | 26.5859 | 26.5446 | 26.312 |
| 0 | 214.18 | 4.06E+10 | 106 | 31.462 | 31.5258 | 31.4867 | 31.6268 | 31.3922 |
| 0 | 21.771 | 7.66E+08 | 32 | 25.7655 | 25.5486 | 25.5493 | 25.6769 | 25.9059 |
| 0 | 32.002 | 1.18E+09 | 40 | 26.5551 | 26.6182 | 26.5284 | 26.5648 | 26.3162 |
| 0 | 167.25 | 5.67E+09 | 99 | 28.5146 | 28.362 | 28.5556 | 28.5483 | 28.566 |
| 0 | 24.11 | 1.04E+09 | 47 | 26.0904 | 25.7767 | 26.3465 | 26.2825 | 26.2593 |
| 0 | 111.2 | 4.33E+09 | 129 | 28.2468 | 28.1068 | 28.0768 | 28.0515 | 28.3606 |
| 0 | 25.989 | 2.07E+09 | 59 | 27.3156 | 27.2775 | 27.4114 | 27.291 | 27.1588 |
| 0 | 28.138 | 2.72E+10 | 105 | 30.9519 | 30.9929 | 30.9453 | 30.9546 | 30.7672 |
| 0 | 55.11 | 2.45E+09 | 105 | 27.572 | 27.6296 | 27.2526 | 27.4245 | 27.6175 |
| 0 | 13.57 | 8.61E+08 | 27 | 25.7538 | 25.7078 | 25.9412 | 25.9757 | 26.0043 |
| 0 | 38.188 | 1.18E+09 | 73 | 26.2463 | 26.4864 | 26.0048 | 26.0852 | 26.4417 |
| 0 | 23.56 | 1.15E+09 | 44 | 26.5999 | 26.033 | 26.3352 | 26.3498 | 26.3567 |
| 0 | 204.88 | 3.61E+10 | 179 | 31.3694 | 31.2796 | 31.3838 | 31.3446 | 31.2226 |
| 0 | 37.046 | 1.41E+09 | 47 | 26.7811 | 26.7895 | 26.2626 | 26.3646 | 26.7031 |
| 0 | 94.757 | 4.7E+09 | 91 | 28.4676 | 28.2988 | 28.3338 | 28.5141 | 28.2575 |
| 0 | 23.383 | 2.34E+09 | 79 | 27.3858 | 27.3847 | 27.4229 | 27.44 | 27.1335 |

| | | | | | | | | |
|---|---|---|---|---|---|---|---|---|
| 0 | 62.595 | 3.76E+09 | 100 | 28.0171 | 27.9909 | 28.1018 | 28.2575 | 27.9078 |
| 0 | 129.15 | 6.01E+09 | 119 | 28.7029 | 28.8431 | 28.6999 | 28.6927 | 28.6054 |
| 0 | 40.295 | 3.06E+09 | 97 | 27.7034 | 27.8123 | 27.598 | 27.8134 | 27.7048 |
| 0 | 12.224 | 2.29E+10 | 12 | 30.9293 | 30.878 | 31.0443 | 30.7414 | 30.6511 |
| 0 | 42.543 | 4.11E+09 | 74 | 28.2321 | 28.1721 | 28.1464 | 28.1145 | 28.3862 |
| 0 | 24.805 | 3.81E+09 | 83 | 27.9827 | 28.0559 | 28.0717 | 28.0943 | 28.0606 |
| 0 | 137.85 | 1.47E+09 | 65 | 26.6857 | 26.3699 | 26.8082 | 26.9683 | 26.6711 |
| 0 | 51.615 | 1.48E+10 | 135 | 29.8947 | 29.9252 | 30.0352 | 29.9874 | 29.9773 |
| 0 | 93.457 | 2.1E+09 | 72 | 27.4831 | 27.4771 | 26.8796 | 27.1091 | 27.3606 |
| 0 | 24.778 | 1.11E+09 | 28 | 26.2717 | 26.0872 | 26.3575 | 26.4867 | 26.3101 |
| 0 | 50.74 | 9.66E+09 | 107 | 29.9994 | 29.9728 | 29.501 | 29.361 | 28.9739 |
| 0 | 15.674 | 1.18E+09 | 39 | 26.354 | 26.3423 | 26.4056 | 26.5923 | 26.3057 |
| 0 | 65.917 | 1.42E+09 | 47 | 26.5813 | 26.6499 | 26.637 | 26.7019 | 26.6807 |
| 0 | 17.569 | 1.37E+09 | 28 | 26.5541 | 26.5477 | 26.7818 | 26.4468 | 26.5946 |
| 0 | 31.078 | 1.38E+09 | 64 | 27.281 | 26.9599 | 26.5679 | 26.4696 | 26.4194 |
| 0 | 283.96 | 1E+11 | 167 | 32.9464 | 32.872 | 32.8602 | 32.9476 | 32.7346 |
| 0 | 59.013 | 2.68E+09 | 68 | 27.6593 | 27.5266 | 27.6282 | 27.6446 | 27.587 |
| 0 | 129.22 | 2.04E+09 | 44 | 27.0784 | 27.3376 | 27.4372 | 27.2755 | 27.1083 |
| 0 | 18.12 | 9.77E+08 | 38 | 26.0086 | 26.2317 | 26.2737 | 25.9525 | 26.0516 |
| 0 | 16.34 | 9.53E+08 | 45 | 25.9702 | 26.1884 | 25.9779 | 25.9398 | 26.2532 |
| 0 | 113.63 | 4.89E+10 | 278 | 31.7832 | 31.7373 | 31.887 | 31.884 | 31.758 |
| 0 | 32.786 | 1.11E+09 | 42 | 26.2625 | 26.3423 | 25.9945 | 26.1477 | 26.7626 |
| 0 | 80.496 | 3.36E+09 | 112 | 27.9385 | 27.8941 | 28.1056 | 27.936 | 27.8386 |
| 0 | 63.767 | 2.06E+09 | 47 | 27.3947 | 27.0926 | 27.1174 | 27.2535 | 27.2447 |
| 0 | 52.683 | 2.31E+09 | 95 | 27.0188 | 27.0564 | 27.3679 | 27.3977 | 27.4193 |
| 0 | 36.11 | 1.53E+09 | 52 | 27.1981 | 27.2116 | 26.6067 | 26.7955 | 26.5091 |
| 0 | 241.4 | 9.77E+09 | 95 | 29.3538 | 29.4615 | 29.5486 | 29.539 | 29.4006 |
| 0 | 139.74 | 2.08E+09 | 66 | 27.4281 | 27.4622 | 27.104 | 26.9324 | 27.4342 |
| 0 | 47.98 | 9.54E+08 | 29 | 26.2474 | 26.2229 | 26.1219 | 26.1368 | 25.7314 |
| 0 | 64.726 | 2.92E+09 | 79 | 27.7974 | 27.8649 | 27.8538 | 27.789 | 27.648 |
| 0 | 323.31 | 1.67E+10 | 239 | 30.3216 | 30.3065 | 30.2682 | 30.2739 | 30.1819 |
| 0 | 37.999 | 1.03E+09 | 26 | 26.0882 | 26.2848 | 26.2878 | 26.3022 | 25.9961 |
| 0 | 13.251 | 9.32E+08 | 33 | 26.1065 | 26.0705 | 25.9051 | 26.0128 | 26.0182 |
| 0 | 135.82 | 3.05E+09 | 71 | 27.74 | 27.8108 | 27.8784 | 27.7664 | 27.7365 |
| 0 | 28.671 | 1.76E+09 | 68 | 26.9096 | 26.9788 | 26.9884 | 26.8202 | 27.0622 |
| 0 | 172.68 | 1.73E+09 | 58 | 27.3655 | 27.2482 | 26.6224 | 26.7054 | 26.7516 |
| 0 | 42.077 | 3.71E+09 | 91 | 27.921 | 28.056 | 28.0811 | 28.1295 | 27.99 |
| 0 | 159.01 | 3.82E+09 | 81 | 28.1265 | 28.143 | 28.1613 | 28.1836 | 27.9937 |
| 0 | 39.242 | 1.91E+09 | 78 | 27.2929 | 27.3539 | 26.8415 | 26.8076 | 27.1267 |
| 0 | 36.269 | 1.65E+09 | 51 | 26.7671 | 26.9127 | 26.7447 | 26.8401 | 27.1135 |
| 0 | 53.699 | 1.54E+09 | 68 | 26.7064 | 26.6451 | 26.7781 | 26.4451 | 27.2097 |
| 0 | 102.17 | 3.72E+09 | 102 | 28.0365 | 28.0138 | 28.1383 | 27.9817 | 28.0873 |
| 0 | 221.52 | 6.29E+09 | 133 | 28.8429 | 28.7763 | 29.0161 | 29.0319 | 28.7045 |
| 0 | 76.206 | 1.34E+09 | 41 | 26.5657 | 26.6863 | 26.2213 | 26.6451 | 26.6398 |
| 0 | 50.869 | 2.64E+10 | 150 | 30.942 | 30.8443 | 30.9511 | 30.9503 | 30.7695 |
| 0 | 155.7 | 5.8E+09 | 111 | 28.7182 | 28.6413 | 28.7252 | 28.6957 | 28.7283 |
| 0 | 16.841 | 1.48E+09 | 61 | 26.5466 | 26.5772 | 26.6414 | 26.5514 | 26.85 |
| 0 | 44.834 | 3.7E+09 | 86 | 28.2283 | 28.0746 | 27.9282 | 27.9411 | 27.8921 |

| | | | | | | | | |
|---|---|---|---|---|---|---|---|---|
| 0 | 160.25 | 1.76E+10 | 143 | 30.2875 | 30.2944 | 30.3045 | 30.1501 | 30.4124 |
| 0 | 76.961 | 4.63E+09 | 87 | 28.9289 | 28.7029 | 27.9154 | 28.0543 | 28.3205 |
| 0 | 41.729 | 2.71E+09 | 53 | 27.6464 | 27.471 | 27.8107 | 27.6378 | 27.3372 |
| 0 | 111.17 | 4.04E+09 | 113 | 28.2763 | 28.1465 | 28.2316 | 28.3767 | 27.9928 |
| 0 | 25.019 | 1.09E+09 | 27 | 26.3756 | 26.1116 | 26.2333 | 26.2586 | 26.1 |
| 0 | 323.31 | 2.6E+10 | 154 | 30.792 | 30.8087 | 30.8025 | 30.8696 | 31.0007 |
| 0 | 16.878 | 1.88E+09 | 74 | 26.796 | 26.9817 | 27.1553 | 27.2277 | 27.1056 |
| 0 | 13.488 | 5.16E+08 | 21 | 25.4892 | 25.6908 | 24.8921 | 25.1261 | 24.6954 |
| 0 | 11.681 | 7.32E+08 | 29 | 26.0664 | 25.7025 | 25.7296 | 25.5802 | 25.6087 |
| 0 | 63.63 | 1.51E+09 | 61 | 26.7632 | 26.9676 | 26.8176 | 26.8142 | 26.9002 |
| 0 | 262.36 | 3.29E+10 | 232 | 31.309 | 31.3346 | 31.1366 | 31.266 | 31.134 |
| 0 | 82.986 | 2.27E+09 | 68 | 27.7201 | 27.7844 | 27.2095 | 27.1893 | 27.0905 |
| 0 | 86.467 | 1.7E+10 | 144 | 30.2802 | 30.2269 | 30.3261 | 30.3802 | 30.2553 |
| 0 | 86.226 | 5.78E+10 | 201 | 32.1022 | 32.0613 | 32.1551 | 31.9804 | 32.0062 |
| 0 | 323.31 | 9.31E+09 | 122 | 29.3357 | 29.4147 | 29.5174 | 29.5169 | 29.3653 |
| 0 | 59.419 | 1.93E+09 | 65 | 27.0857 | 27.0848 | 27.2837 | 27.251 | 26.9884 |
| 0 | 19.505 | 2.87E+09 | 68 | 27.4913 | 27.6391 | 27.7839 | 27.6653 | 27.691 |
| 0 | 125.47 | 8.07E+09 | 107 | 29.1249 | 29.0073 | 28.8848 | 28.9126 | 29.0594 |
| 0 | 201.04 | 9.25E+10 | 150 | 32.7148 | 32.6249 | 32.7758 | 32.7711 | 32.6725 |
| 0 | 54.548 | 2.38E+09 | 85 | 27.5569 | 27.5522 | 27.2206 | 27.3642 | 27.4114 |
| 0 | 50.163 | 9.33E+09 | 97 | 29.1477 | 29.2436 | 29.451 | 29.4242 | 29.5459 |
| 0 | 69.02 | 3.11E+09 | 85 | 27.84 | 27.859 | 27.9461 | 27.9066 | 27.6121 |
| 0 | 18.052 | 1.99E+09 | 33 | 27.4451 | 27.3268 | 27.3989 | 27.1576 | 27.0215 |
| 0 | 144.76 | 2.01E+09 | 57 | 27.0626 | 27.028 | 27.2395 | 27.0548 | 27.2162 |
| 0 | 45.51 | 1.06E+09 | 36 | 26.1867 | 25.9446 | 26.3804 | 26.6182 | 26.215 |
| 0 | 284.44 | 1.93E+10 | 137 | 30.5415 | 30.5081 | 30.2245 | 30.3401 | 30.4646 |
| 0 | 167.26 | 8.54E+10 | 261 | 32.5448 | 32.5385 | 32.6788 | 32.6796 | 32.5277 |
| 0 | 28.539 | 1.96E+09 | 74 | 26.9558 | 26.8934 | 27.1872 | 27.1303 | 27.2042 |
| 0 | 278.3 | 1.5E+10 | 229 | 30.0786 | 30.0846 | 30.2372 | 30.1962 | 29.9702 |
| 0 | 34.078 | 3.76E+09 | 119 | 27.9677 | 27.8419 | 28.2038 | 28.1573 | 28.1693 |
| 0 | 323.31 | 2.38E+10 | 152 | 30.7169 | 30.6163 | 30.8213 | 30.9186 | 30.6226 |
| 0 | 54.51 | 1.75E+09 | 56 | 27.1419 | 27.1412 | 26.6929 | 26.8967 | 26.9542 |
| 0 | 177.27 | 9.93E+09 | 132 | 29.3666 | 29.4531 | 29.512 | 29.6375 | 29.5023 |
| 0 | 323.31 | 6.85E+10 | 315 | 32.2151 | 32.1695 | 32.3132 | 32.3644 | 32.1626 |
| 0 | 45.91 | 3.01E+09 | 97 | 27.8145 | 27.711 | 27.8562 | 27.7491 | 27.6587 |
| 0 | 323.31 | 5.02E+09 | 116 | 28.4423 | 28.5229 | 28.4096 | 28.413 | 28.5458 |
| 0 | 220.31 | 3.82E+10 | 205 | 31.3736 | 31.4192 | 31.4572 | 31.5042 | 31.3354 |
| 0 | 106.51 | 1.25E+09 | 51 | 26.6161 | 26.395 | 26.7249 | 26.7056 | 26.2673 |
| 0 | 34.734 | 1.65E+09 | 62 | 27.0473 | 27.0287 | 26.8427 | 27.1038 | 26.8007 |
| 0 | 31.76 | 1.87E+09 | 99 | 27.0316 | 27.1766 | 26.8474 | 26.9131 | 27.099 |
| 0 | 68.622 | 6.17E+10 | 127 | 32.1191 | 32.0745 | 32.1545 | 32.2531 | 32.0078 |
| 0 | 11.142 | 9.35E+08 | 47 | 26.0953 | 26.1082 | 25.9085 | 25.921 | 26.0866 |
| 0 | 33.259 | 1.37E+09 | 35 | 26.7794 | 26.6 | 26.4057 | 26.3412 | 26.6659 |
| 0 | 323.31 | 1.03E+10 | 125 | 29.4092 | 29.4075 | 29.7185 | 29.603 | 29.5591 |
| 0 | 12.95 | 7.02E+08 | 31 | 25.5239 | 25.5814 | 25.7813 | 25.8607 | 25.4175 |
| 0 | 103.53 | 7.43E+09 | 99 | 28.9377 | 28.9551 | 29.1029 | 29.0149 | 29.105 |
| 0 | 15.639 | 9.35E+08 | 44 | 25.9483 | 26.0022 | 26.1816 | 26.182 | 26.1417 |
| 0 | 47.906 | 1.46E+09 | 41 | 26.832 | 26.4234 | 26.7301 | 26.5803 | 26.8028 |

| | | | | | | | | |
|---|---|---|---|---|---|---|---|---|
| 0 | 23.769 | 9.65E+08 | 34 | 25.8585 | 26.0796 | 26.0392 | 25.7207 | 26.2787 |
| 0 | 27.193 | 9.56E+08 | 27 | 26.1345 | 25.8089 | 25.8864 | 26.3285 | 25.7009 |
| 0 | 157.4 | 2.72E+09 | 78 | 27.5533 | 27.5179 | 27.2911 | 27.342 | 27.4678 |
| 0 | 180.95 | 1.18E+10 | 152 | 29.5401 | 29.6365 | 29.7283 | 29.5237 | 29.5121 |
| 0 | 19.759 | 5.33E+09 | 100 | 28.3011 | 28.3078 | 28.545 | 28.475 | 28.5483 |
| 0 | 30.174 | 1.1E+10 | 143 | 29.5231 | 29.5452 | 29.4257 | 29.4955 | 29.4229 |
| 0 | 22.773 | 1.12E+09 | 43 | 25.976 | 25.9666 | 26.2045 | 25.9835 | 26.3107 |
| 0 | 77.209 | 1.02E+09 | 65 | 25.816 | 25.7914 | 26.244 | 26.1935 | 26.0627 |
| 0 | 157.04 | 5.18E+09 | 90 | 28.4219 | 28.4456 | 28.6169 | 28.4315 | 28.2634 |
| 0 | 22.017 | 1.15E+09 | 32 | 26.2741 | 26.1037 | 26.142 | 26.3398 | 26.2058 |
| 0 | 30.772 | 7.49E+08 | 40 | 25.4037 | 25.5494 | 25.6855 | 25.4734 | 25.8498 |
| 0 | 34.854 | 9.7E+08 | 37 | 26.0808 | 26.0283 | 25.823 | 25.9839 | 26.1638 |
| 0 | 39.107 | 2.6E+09 | 67 | 27.4324 | 27.3865 | 27.3924 | 27.3471 | 27.2972 |
| 0 | 21.636 | 8.67E+08 | 33 | 25.7924 | 25.8055 | 25.796 | 25.8154 | 26.0365 |
| 0 | 67.984 | 2.58E+09 | 110 | 27.324 | 27.3551 | 27.4113 | 27.5281 | 27.4558 |
| 0 | 26.985 | 7.79E+08 | 41 | 25.5608 | 25.621 | 25.7041 | 25.6257 | 25.7769 |
| 0 | 36.008 | 1.11E+09 | 24 | 26.3618 | 26.073 | 26.3914 | 26.5016 | 26.3089 |
| 0 | 50.029 | 2.06E+09 | 58 | 26.8035 | 26.8616 | 27.0601 | 27.1734 | 26.9996 |
| 0 | 189.41 | 2.78E+09 | 83 | 27.3318 | 27.4118 | 27.5443 | 27.5693 | 27.6256 |
| 0 | 37.688 | 1.1E+09 | 39 | 26.2645 | 26.2933 | 26.1751 | 26.281 | 26.0668 |
| 0 | 26.731 | 2.72E+10 | 65 | 30.8044 | 30.9719 | 30.8757 | 30.8466 | 30.8091 |
| 0 | 41.472 | 2.72E+09 | 72 | 27.5214 | 27.5455 | 27.4715 | 27.5495 | 27.5345 |
| 0 | 44.443 | 2.39E+09 | 101 | 27.2687 | 27.3057 | 27.4771 | 27.4215 | 27.2911 |
| 0 | 35.625 | 1.74E+09 | 72 | 26.8404 | 26.6995 | 26.9389 | 26.9819 | 26.8046 |
| 0 | 80.265 | 2.87E+09 | 58 | 27.6444 | 27.5851 | 27.567 | 27.568 | 27.7082 |
| 0 | 24.275 | 2.51E+09 | 66 | 27.3558 | 27.307 | 27.3303 | 27.3449 | 27.4622 |
| 0 | 196.58 | 9.21E+09 | 115 | 29.2982 | 29.2489 | 29.3472 | 29.411 | 29.2477 |
| 0 | 158.83 | 3.96E+09 | 86 | 28.2804 | 28.1293 | 28.2384 | 28.3712 | 28.2819 |
| 0 | 21.351 | 1.24E+09 | 52 | 26.8955 | 27.0443 | 26.6014 | 26.429 | 26.4784 |
| 0 | 58.705 | 1.09E+09 | 50 | 26.5529 | 26.4969 | 26.5401 | 26.4862 | 26.3742 |
| 0 | 99.008 | 2.84E+09 | 73 | 27.9623 | 27.96 | 27.6534 | 27.6007 | 27.8038 |
| 0 | 34.573 | 1.2E+09 | 39 | 26.6102 | 26.5184 | 26.5191 | 26.5285 | 26.4995 |
| 0 | 79.33 | 2.55E+10 | 104 | 30.9202 | 30.9461 | 30.8327 | 30.9371 | 30.8022 |
| 0 | 26.656 | 5.23E+08 | 28 | 25.0489 | 24.9137 | 24.9693 | 25.1355 | 25.2208 |
| 0 | 21.554 | 9.04E+08 | 32 | 25.7081 | 25.954 | 25.7538 | 25.9807 | 25.7702 |
| 0 | 161.63 | 4.08E+10 | 243 | 31.49 | 31.4571 | 31.4139 | 31.5765 | 31.3558 |
| 0 | 61.091 | 1.94E+09 | 64 | 26.998 | 26.9189 | 27.1913 | 26.8443 | 27.1068 |
| 0 | 23.659 | 1.94E+09 | 67 | 27.2363 | 27.1147 | 27.1406 | 27.1178 | 27.2602 |
| 0 | 309.73 | 2.31E+10 | 128 | 30.4485 | 30.6421 | 30.7599 | 30.7404 | 30.6001 |
| 0 | 26.677 | 6.17E+08 | 20 | 25.5426 | 25.6877 | 25.827 | 25.6558 | 25.7038 |
| 0 | 17.202 | 2.74E+09 | 52 | 27.6405 | 27.6415 | 27.5333 | 27.8072 | 27.6266 |
| 0 | 13.354 | 9.86E+08 | 60 | 26.5594 | 26.7362 | 26.0938 | 26.1325 | 26.066 |
| 0 | 50.685 | 1.9E+09 | 46 | 26.9734 | 27.0748 | 27.2889 | 27.1733 | 27.237 |
| 0 | 19.882 | 7.31E+08 | 13 | NaN | 25.4521 | 25.7034 | NaN | 25.9076 |
| 0 | 112.29 | 2.27E+09 | 77 | 27.1822 | 27.2041 | 27.2926 | 27.2968 | 27.4225 |
| 0 | 37.616 | 9.94E+08 | 53 | 26.1969 | 26.1157 | 25.8722 | 25.8032 | 26.099 |
| 0 | 120.73 | 6.4E+09 | 94 | 28.8369 | 28.7452 | 29.0743 | 28.966 | 28.7863 |
| 0 | 222.79 | 3.37E+10 | 240 | 31.002 | 31.0271 | 31.0576 | 31.067 | 31.0114 |

| | | | | | | | | |
|---|---|---|---|---|---|---|---|---|
| 0 | 27.901 | 9.68E+09 | 69 | 29.8157 | 29.8192 | 29.454 | 29.404 | 29.3014 |
| 0 | 10.185 | 1.62E+09 | 27 | 26.9715 | 26.7665 | 26.858 | 26.8321 | 26.6678 |
| 0 | 283.04 | 8.22E+09 | 113 | 29.3441 | 29.2488 | 29.5018 | 29.3326 | 29.1986 |
| 0.002508 | 2.0911 | 1.99E+09 | 13 | 26.6553 | 26.6521 | 26.6798 | 25.2951 | 27.4812 |
| 0 | 119.64 | 3.08E+09 | 58 | 27.8288 | 27.5555 | 27.7329 | 27.9197 | 27.6464 |
| 0 | 40.563 | 4.9E+09 | 41 | 28.3755 | 28.4822 | 28.3796 | 28.2257 | 28.3904 |
| 0 | 32.923 | 9.21E+08 | 49 | 26.1173 | 26.0797 | 25.7752 | 25.6951 | 25.8938 |
| 0 | 307.2 | 1.03E+10 | 154 | 29.6003 | 29.592 | 29.5445 | 29.4398 | 29.5609 |
| 0 | 33.747 | 2.58E+09 | 78 | 27.3775 | 27.4398 | 27.6031 | 27.4122 | 27.5588 |
| 0 | 37.598 | 2.66E+09 | 84 | 27.4454 | 27.4099 | 27.8026 | 27.5944 | 27.3383 |
| 0 | 323.31 | 1.04E+11 | 967 | 32.7434 | 32.7512 | 32.9412 | 32.9462 | 32.7889 |
| 0 | 49.165 | 2.09E+09 | 70 | 27.515 | 27.5394 | 27.0636 | 27.1288 | 27.2225 |
| 0 | 15.898 | 1.68E+09 | 47 | 26.8404 | 27.0521 | 26.9442 | 26.7735 | 26.7085 |
| 0 | 18.789 | 1.16E+09 | 53 | 26.5065 | 26.3291 | 26.2898 | 26.2406 | 25.9481 |
| 0 | 69.564 | 1.94E+09 | 43 | 27.0706 | 26.9957 | 27.4436 | 27.2833 | 27.2074 |
| 0 | 51.896 | 1.86E+09 | 47 | 26.9556 | 26.9898 | 26.9454 | 27.2759 | 27.0462 |
| 0 | 79.553 | 4.3E+09 | 74 | 28.3611 | 28.1942 | 28.2192 | 28.2559 | 28.0649 |
| 0 | 98.067 | 1.31E+09 | 40 | 26.6422 | 26.484 | 26.6738 | 26.7757 | 26.6683 |
| 0 | 51.427 | 2.82E+10 | 149 | 31.0364 | 30.9913 | 31.0437 | 31.1067 | 30.9981 |
| 0 | 80.306 | 7.66E+09 | 138 | 29.1959 | 29.168 | 29.166 | 29.2616 | 29.0785 |
| 0 | 45.063 | 7.92E+08 | 29 | 25.6806 | 25.9108 | 25.8841 | 26.1175 | 25.9431 |
| 0 | 50.468 | 2.42E+09 | 75 | 27.3154 | 27.3209 | 27.4921 | 27.3582 | 27.363 |
| 0 | 26.68 | 9.77E+08 | 32 | 26.3822 | 26.1672 | 26.1302 | 26.5544 | 26.0644 |
| 0 | 32.584 | 1.45E+09 | 68 | 26.7331 | 26.6294 | 26.6195 | 26.5302 | 26.5914 |
| 0 | 76.196 | 7.42E+09 | 117 | 29.0028 | 28.9848 | 29.1528 | 29.0008 | 29.0396 |
| 0 | 186.03 | 2.28E+10 | 96 | 30.6422 | 30.6568 | 30.8082 | 30.7414 | 30.647 |
| 0 | 74.031 | 1.33E+09 | 53 | 26.6974 | 26.4619 | 26.426 | 26.4771 | 26.4789 |
| 0 | 46.101 | 2.13E+09 | 73 | 27.5353 | 27.6038 | 27.2897 | 27.0585 | 27.2233 |
| 0 | 97.117 | 1.45E+10 | 120 | 29.8583 | 29.8289 | 29.982 | 29.915 | 30.0362 |
| 0 | 19.501 | 1.14E+09 | 34 | 26.0286 | 26.2573 | 26.2826 | 26.3838 | 26.5446 |
| 0 | 75.179 | 2.64E+09 | 74 | 27.5452 | 27.4913 | 27.65 | 27.6724 | 27.3896 |
| 0 | 323.31 | 8.25E+10 | 371 | 32.519 | 32.4863 | 32.5544 | 32.582 | 32.4639 |
| 0 | 182.18 | 7.36E+09 | 121 | 29.4571 | 29.4456 | 29.0968 | 29.1002 | 28.8261 |
| 0 | 37.514 | 2.77E+10 | 112 | 30.9388 | 30.969 | 30.9733 | 30.9838 | 30.7627 |
| 0 | 20.577 | 1.13E+09 | 57 | 26.2929 | 26.2608 | 26.4196 | 26.3229 | 26.1207 |
| 0 | 8.7221 | 9.11E+08 | 38 | 25.9414 | 25.9091 | 26.0046 | 25.7848 | 26.038 |
| 0 | 21.007 | 2.27E+09 | 56 | 27.0054 | 27.1775 | 27.2975 | 27.1132 | 27.4425 |
| 0 | 323.31 | 7.12E+10 | 684 | 32.3037 | 32.332 | 32.3263 | 32.3257 | 32.1838 |
| 0 | 36.061 | 5.08E+08 | 18 | 25.1305 | 25.2217 | 24.5467 | 24.555 | 25.875 |
| 0 | 23.129 | 2.21E+09 | 62 | 27.4599 | 27.3179 | 27.3844 | 27.2924 | 27.3096 |
| 0 | 57.686 | 2.71E+09 | 82 | 27.5596 | 27.5517 | 27.5305 | 27.5891 | 27.5342 |
| 0 | 35.331 | 1.02E+09 | 34 | 26.1019 | 26.061 | 26.0815 | 26.1544 | 26.1634 |
| 0 | 88.603 | 6.7E+09 | 94 | 29.1434 | 29.1528 | 28.7911 | 28.7759 | 28.9783 |
| 0 | 323.31 | 5.26E+10 | 281 | 31.911 | 31.9562 | 32.0511 | 32.0111 | 31.7331 |
| 0 | 20.146 | 1.94E+10 | 77 | 30.4571 | 30.4435 | 30.6453 | 30.4621 | 30.4371 |
| 0 | 16.192 | 7.73E+08 | 39 | 25.7736 | 25.6287 | 25.3027 | 25.6799 | 25.8172 |
| 0 | 136.2 | 3.29E+09 | 91 | 27.4909 | 27.4028 | 27.6636 | 27.7734 | 27.7864 |
| 0 | 15.763 | 1.24E+09 | 44 | 26.8207 | 26.6759 | 26.1913 | 26.3276 | 26.6558 |

| | | | | | | | | |
|---|---|---|---|---|---|---|---|---|
| 0 | 235.07 | 4.28E+10 | 292 | 31.5698 | 31.5961 | 31.6938 | 31.6218 | 31.5804 |
| 0 | 261.92 | 8.76E+10 | 277 | 33.0515 | 32.9185 | 32.7422 | 32.7535 | 32.308 |
| 0 | 13.691 | 5.36E+08 | 26 | 25.0521 | 25.0138 | 25.0972 | 25.16 | 25.2226 |
| 0 | 37.941 | 2.82E+09 | 73 | 27.5709 | 27.4809 | 27.6816 | 27.6989 | 27.6384 |
| 0 | 39.807 | 1.64E+09 | 56 | 27.0872 | 26.9428 | 26.9753 | 26.8178 | 26.8143 |
| 0 | 63.641 | 5.18E+09 | 137 | 28.4127 | 28.3946 | 28.586 | 28.5386 | 28.5627 |
| 0 | 62.511 | 3.93E+09 | 88 | 28.0669 | 28.0819 | 28.3025 | 28.059 | 28.218 |
| 0 | 28.711 | 4.32E+09 | 85 | 28.3942 | 28.2239 | 28.2917 | 28.3585 | 28.2099 |
| 0 | 11.266 | 8.04E+08 | 37 | 26.0114 | 25.8825 | 26.0907 | 26.071 | 25.8863 |
| 0 | 323.31 | 3.89E+10 | 288 | 31.4545 | 31.4531 | 31.4084 | 31.5155 | 31.3502 |
| 0 | 34.618 | 1.06E+09 | 37 | 26.0427 | 26.1988 | 26.2473 | 26.3066 | 25.9436 |
| 0 | 154.8 | 1.61E+10 | 163 | 30.1845 | 30.1024 | 29.8988 | 30.1586 | 30.1346 |
| 0 | 42.922 | 2.27E+09 | 67 | 27.5657 | 27.476 | 27.4573 | 27.2343 | 27.335 |
| 0 | 80.05 | 4.22E+09 | 66 | 28.0939 | 27.8879 | 28.1901 | 28.0837 | 28.172 |
| 0 | 48.193 | 1.03E+09 | 39 | 26.2318 | 26.4232 | 26.2382 | 26.1415 | 26.1114 |
| 0 | 41.277 | 1.95E+10 | 133 | 30.5894 | 30.5647 | 30.7177 | 30.5206 | 30.26 |
| 0 | 107.85 | 3.21E+09 | 77 | 27.7314 | 27.7119 | 27.581 | 27.6306 | 27.973 |
| 0 | 15.064 | 9.33E+08 | 51 | 26.3734 | 26.4604 | 25.8801 | 25.9354 | 26.1295 |
| 0 | 35.38 | 1.44E+09 | 40 | 26.6316 | 26.6366 | 26.6143 | 26.7919 | 26.3547 |
| 0 | 103.38 | 6.74E+09 | 138 | 29.1731 | 29.1551 | 28.7204 | 28.8454 | 28.8847 |
| 0 | 16.746 | 1.04E+09 | 45 | 26.0849 | 26.4174 | 26.3025 | 26.2883 | 26.3399 |
| 0 | 323.31 | 1.2E+11 | 456 | 32.9487 | 32.9769 | 33.1323 | 33.08 | 33.1241 |
| 0 | 48.962 | 3.58E+09 | 82 | 27.9906 | 27.9129 | 28.1351 | 28.0045 | 28.0618 |
| 0 | 61.428 | 7.88E+09 | 131 | 29.1202 | 29.134 | 29.1931 | 29.0896 | 28.9993 |
| 0 | 204.61 | 7.5E+09 | 120 | 29.1389 | 29.1123 | 28.8978 | 28.9669 | 28.9356 |
| 0 | 65.205 | 1.32E+10 | 113 | 30.1705 | 30.1261 | 29.9513 | 29.8386 | 29.802 |
| 0 | 122.73 | 8.1E+09 | 164 | 29.1764 | 29.1731 | 29.3058 | 29.25 | 29.2432 |
| 0 | 79.218 | 6.22E+09 | 115 | 28.6832 | 28.757 | 28.9015 | 28.8613 | 28.8137 |
| 0 | 30.019 | 8.22E+09 | 131 | 29.1573 | 29.2089 | 29.2107 | 29.2441 | 29.1063 |
| 0 | 62.88 | 3.79E+09 | 91 | 27.9576 | 27.9106 | 28.1054 | 28.0182 | 28.1282 |
| 0 | 57.973 | 3.68E+09 | 68 | 27.8878 | 27.84 | 28.0661 | 28.0767 | 28.1281 |
| 0 | 59.599 | 2.34E+09 | 69 | 27.295 | 27.1119 | 27.2705 | 27.3245 | 27.5014 |
| 0 | 60.881 | 7.18E+09 | 77 | 29.0551 | 29.1124 | 29.0773 | 28.9186 | 28.8843 |
| 0 | 26.874 | 1.3E+09 | 38 | 26.3663 | 26.3723 | 26.543 | 26.461 | 26.5553 |
| 0 | 16.164 | 5.39E+08 | 24 | 25.3585 | 25.4202 | 25.5222 | 25.6862 | 25.1434 |
| 0 | 39.563 | 1.73E+09 | 42 | 27.0663 | 27.102 | 26.8335 | 26.9259 | 27.0636 |
| 0 | 4.7859 | 6.12E+08 | 13 | 25.5953 | 25.5982 | NaN | 25.559 | 25.7629 |
| 0 | 10.147 | 4.18E+08 | 24 | 24.4182 | 24.5663 | 24.9797 | 24.7115 | 24.8986 |
| 0 | 30.501 | 1.26E+09 | 27 | 26.5361 | 26.5323 | 26.7862 | 26.5786 | 26.3296 |
| 0 | 45.76 | 4.46E+09 | 139 | 28.5713 | 28.5828 | 28.1098 | 28.1674 | 28.1484 |
| 0 | 58.642 | 1.99E+09 | 57 | 27.5948 | 27.3623 | 26.9268 | 26.9895 | 27.0588 |
| 0 | 25.73 | 1.77E+09 | 58 | 27.024 | 27.1106 | 26.9122 | 27.2641 | 26.9165 |
| 0 | 140.2 | 7.35E+09 | 97 | 29.0606 | 29.1098 | 28.9553 | 28.9787 | 29.0942 |
| 0 | 17.719 | 9.09E+08 | 35 | 25.9575 | 25.698 | 25.8956 | 26.15 | 25.9383 |
| 0 | 82.708 | 1.8E+09 | 59 | 26.996 | 27.0285 | 27.0071 | 27.1313 | 27.0298 |
| 0 | 101.33 | 6.1E+09 | 113 | 28.5617 | 28.5705 | 28.7437 | 28.8586 | 28.8977 |
| 0 | 120.67 | 2.61E+09 | 68 | 27.4092 | 27.6019 | 27.6551 | 27.7208 | 27.4468 |
| 0 | 148.68 | 9.84E+10 | 212 | 32.8376 | 32.8813 | 32.8736 | 32.8071 | 32.654 |

| | | | | | | | | |
|---|---|---|---|---|---|---|---|---|
| 0 | 33.634 | 2.28E+09 | 62 | 27.7009 | 27.7106 | 27.3 | 27.126 | 27.1909 |
| 0 | 207.86 | 8.73E+09 | 143 | 29.1552 | 29.1915 | 29.3184 | 29.4487 | 29.2547 |
| 0 | 84.694 | 9.83E+09 | 116 | 29.4846 | 29.3258 | 29.4767 | 29.6176 | 29.5554 |
| 0 | 13.915 | 7.85E+08 | 34 | 25.8455 | 25.4637 | 25.9842 | 25.8447 | 25.9154 |
| 0 | 23.526 | 2E+09 | 62 | 27.3054 | 27.2399 | 27.2482 | 27.1912 | 27.0181 |
| 0 | 256.66 | 4.71E+10 | 276 | 31.8141 | 31.7724 | 31.7466 | 31.6781 | 31.6767 |
| 0 | 33.028 | 2.48E+09 | 59 | 27.6787 | 27.6231 | 27.2495 | 27.4015 | 27.3516 |
| 0 | 29.958 | 2.26E+09 | 59 | 27.3371 | 27.4679 | 27.4738 | 27.3758 | 27.3367 |
| 0 | 21.826 | 2.14E+09 | 59 | 27.0963 | 27.1276 | 27.4919 | 27.3795 | 27.263 |
| 0 | 206.34 | 7.27E+09 | 117 | 29.0405 | 28.9918 | 28.9862 | 29.1649 | 28.914 |
| 0 | 37.213 | 9.57E+08 | 47 | 25.9522 | 26.026 | 26.1901 | 25.9322 | 26.2301 |
| 0 | 7.7921 | 6.51E+08 | 9 | 25.5015 | 25.3737 | 25.6202 | 25.4332 | 25.4469 |
| 0 | 145.57 | 2.7E+10 | 198 | 30.976 | 31.0387 | 30.9683 | 30.9224 | 30.8393 |
| 0 | 29.732 | 1.25E+09 | 68 | 26.7173 | 26.6672 | 26.1785 | 26.3711 | 26.4044 |
| 0 | 14.185 | 1.62E+09 | 70 | 26.4767 | 26.6617 | 26.8704 | 27.0674 | 27.1138 |
| 0 | 26.994 | 1.36E+10 | 117 | 29.8883 | 29.9676 | 29.9268 | 29.9485 | 29.7509 |
| 0 | 50.959 | 2.23E+09 | 60 | 27.2169 | 27.147 | 27.6019 | 27.5547 | 27.0755 |
| 0 | 72.741 | 5.19E+09 | 112 | 28.4338 | 28.258 | 28.385 | 28.3608 | 28.4904 |
| 0 | 74.341 | 1.29E+10 | 139 | 30.1168 | 30.1007 | 29.6337 | 29.7909 | 29.7191 |
| 0 | 152.33 | 4.98E+09 | 94 | 28.8384 | 28.8609 | 28.233 | 28.1685 | 28.352 |
| 0 | 23.87 | 1.37E+09 | 31 | 26.6838 | 26.8428 | 26.4214 | 26.3421 | 26.8894 |
| 0 | 44.905 | 3.5E+09 | 88 | 27.9891 | 28.16 | 27.7311 | 27.8717 | 28.0612 |
| 0 | 22.768 | 1.32E+09 | 39 | 26.7581 | 26.4097 | 26.0848 | 26.3166 | 26.4426 |
| 0 | 82.936 | 1.98E+10 | 169 | 30.3671 | 30.4542 | 30.5692 | 30.5185 | 30.3469 |
| 0 | 305.73 | 1.04E+10 | 117 | 29.4784 | 29.4447 | 29.6044 | 29.6042 | 29.5465 |
| 0 | 51.631 | 1.5E+09 | 71 | 26.7083 | 26.7045 | 26.8202 | 26.7579 | 26.8215 |
| 0 | 16.464 | 5.56E+08 | 21 | 25.3425 | 25.2871 | 25.2843 | 25.2837 | 25.4067 |
| 0 | 13.69 | 1.25E+09 | 51 | 26.2783 | 26.462 | 26.5997 | 26.3631 | 26.5486 |
| 0 | 252.42 | 9.24E+10 | 276 | 32.7758 | 32.7446 | 32.693 | 32.7018 | 32.4115 |
| 0 | 51.267 | 1.38E+09 | 62 | 26.6299 | 26.6814 | 26.7576 | 26.765 | 26.1919 |
| 0 | 26.562 | 2.55E+09 | 52 | 27.4984 | 27.3412 | 27.6126 | 27.6252 | 27.4191 |
| 0 | 79.046 | 2.45E+09 | 68 | 27.4248 | 27.4903 | 27.5324 | 27.5949 | 27.4071 |
| 0 | 94.813 | 4.54E+09 | 126 | 28.233 | 28.2962 | 28.3661 | 28.397 | 28.3618 |
| 0 | 49.401 | 2.87E+09 | 73 | 27.6656 | 27.4936 | 27.8667 | 27.7593 | 27.6553 |
| 0 | 29.015 | 1.79E+09 | 68 | 26.8934 | 26.9044 | 27.0602 | 27.0809 | 26.8906 |
| 0 | 17.082 | 9.21E+08 | 23 | 26.1258 | 26.0294 | 26.1963 | 26.102 | 25.7324 |
| 0 | 12.45 | 5.79E+08 | 22 | 25.5194 | 25.1557 | 25.4125 | 25.2989 | 25.4239 |
| 0 | 162.88 | 8.4E+09 | 229 | 29.1631 | 29.2091 | 29.3364 | 29.2995 | 29.191 |
| 0 | 261.58 | 3.88E+10 | 310 | 31.4927 | 31.4846 | 31.3203 | 31.4014 | 31.4458 |
| 0 | 12.929 | 1.07E+09 | 60 | 26.3822 | 26.3239 | 26.4317 | 26.4571 | 26.0194 |
| 0 | 234.26 | 6.91E+10 | 210 | 32.2906 | 32.2335 | 32.2901 | 32.2533 | 32.1724 |
| 0 | 131.76 | 1.34E+10 | 158 | 29.8482 | 29.8449 | 30.1498 | 30.073 | 29.6849 |
| 0 | 23.393 | 9.22E+08 | 36 | 26.1119 | 25.9989 | 26.1126 | 26.1575 | 25.9624 |
| 0 | 45.048 | 1.88E+09 | 61 | 26.8126 | 26.9369 | 27.2911 | 27.1997 | 27.1685 |
| 0 | 21.902 | 2.71E+09 | 72 | 27.7657 | 27.7593 | 27.546 | 27.5062 | 27.6487 |
| 0 | 23.989 | 5.32E+08 | 23 | 25.2485 | 25.3896 | 25.0968 | 24.957 | 25.5074 |
| 0 | 18.316 | 1.42E+09 | 40 | 26.6822 | 26.6465 | 26.7837 | 26.8937 | 26.6036 |
| 0 | 132.06 | 9.38E+09 | 160 | 29.7629 | 29.6783 | 29.1257 | 29.2438 | 29.1242 |

| | | | | | | | | |
|---|---|---|---|---|---|---|---|---|
| 0 | 51.188 | 1.88E+09 | 37 | 26.8471 | 26.9357 | 26.9991 | 26.9886 | 27.2046 |
| 0 | 17.186 | 6.41E+08 | 26 | 25.4759 | 25.2285 | 25.4515 | 25.4738 | 25.7555 |
| 0 | 21.617 | 8.86E+08 | 15 | 26.0167 | 25.7764 | 25.986 | 25.9327 | 26.0254 |
| 0 | 65.01 | 1.09E+09 | 33 | 26.2371 | 25.9529 | 26.5084 | 26.519 | 26.2687 |
| 0 | 18.302 | 1.06E+10 | 109 | 29.5901 | 29.5917 | 29.5787 | 29.5394 | 29.5616 |
| 0 | 10.364 | 9.19E+09 | 73 | 29.1329 | 29.3374 | 29.4019 | 29.2656 | 29.369 |
| 0 | 217.72 | 2.54E+10 | 151 | 30.8469 | 30.864 | 30.9356 | 30.7389 | 30.8275 |
| 0 | 49.287 | 9.88E+09 | 112 | 29.5433 | 29.5265 | 29.2788 | 29.2508 | 29.5204 |
| 0 | 19.571 | 1.14E+09 | 28 | 26.6521 | 26.5105 | 26.3664 | 26.171 | 26.1957 |
| 0 | 31.961 | 4.49E+09 | 67 | 28.4396 | 28.4643 | 28.5822 | 28.3966 | 28.1536 |
| 0 | 23.048 | 1.91E+09 | 72 | 26.4742 | 26.66 | 26.7718 | 26.7461 | 27.2207 |
| 0 | 66.125 | 8.38E+08 | 41 | 25.4018 | 25.5484 | 25.5484 | 25.4782 | 25.8334 |
| 0 | 33.773 | 8.8E+08 | 15 | 25.823 | 25.6152 | 25.5855 | 25.8467 | 25.6964 |
| 0 | 13.575 | 2.13E+09 | 25 | 26.8053 | 27.2265 | 27.1524 | 26.6931 | 27.1876 |
| 0 | 95.71 | 4.94E+09 | 137 | 28.263 | 28.1547 | 28.2152 | 28.3055 | 28.2748 |
| 0 | 17.008 | 6.39E+08 | 35 | 25.279 | 25.326 | 25.315 | 25.5868 | 25.4532 |
| 0 | 26.031 | 1.79E+10 | 103 | 30.2977 | 30.2591 | 30.1947 | 30.298 | 30.0946 |
| 0 | 16.06 | 2.6E+09 | 62 | 27.2692 | 27.5047 | 27.3588 | 27.7247 | 27.3298 |
| 0 | 64.291 | 1.18E+09 | 59 | 26.2288 | 26.4843 | 26.2526 | 26.2043 | 25.9892 |
| 0 | 18.923 | 3.93E+09 | 68 | 27.949 | 27.9987 | 28.0146 | 28.2091 | 27.9995 |
| 0 | 62.474 | 2.69E+09 | 86 | 27.3866 | 27.519 | 27.6005 | 27.5091 | 27.3845 |
| 0 | 24.954 | 1.48E+09 | 67 | 26.6074 | 26.7264 | 26.7148 | 26.6055 | 26.5225 |
| 0 | 38.787 | 2.59E+09 | 75 | 27.6127 | 27.5373 | 27.5707 | 27.4159 | 27.31 |
| 0 | 29.1 | 1.11E+09 | 42 | 25.9701 | 26.2153 | 26.2356 | 26.3338 | 26.1987 |
| 0 | 40.665 | 3.63E+09 | 79 | 27.8145 | 27.9774 | 27.8897 | 27.8007 | 28.0691 |
| 0 | 35.848 | 4.68E+09 | 61 | 28.2082 | 28.1111 | 28.4342 | 28.3365 | 28.4037 |
| 0 | 53.017 | 1.59E+09 | 59 | 26.8088 | 26.7274 | 26.8123 | 26.9122 | 26.5948 |
| 0 | 149.94 | 5.8E+10 | 206 | 31.9592 | 31.9585 | 32.0282 | 31.8901 | 31.8501 |
| 0 | 147.44 | 1.73E+10 | 112 | 30.0738 | 30.0343 | 30.0531 | 29.9496 | 30.2383 |
| 0 | 11.224 | 6.79E+08 | 23 | 25.483 | 25.6549 | 25.4693 | 25.5897 | 25.5413 |
| 0 | 147.01 | 2.33E+10 | 190 | 30.6036 | 30.6035 | 30.6923 | 30.6688 | 30.5998 |
| 0 | 114.49 | 6.54E+09 | 62 | 28.6488 | 28.7546 | 28.9879 | 28.8242 | 28.8757 |
| 0 | 51.446 | 9.42E+09 | 120 | 29.3383 | 29.3866 | 29.2034 | 29.3662 | 29.3793 |
| 0 | 323.31 | 8.17E+10 | 479 | 32.4016 | 32.4081 | 32.5151 | 32.556 | 32.4809 |
| 0 | 20.222 | 2.42E+09 | 31 | 27.3983 | 27.5131 | 27.6108 | 27.4692 | 27.5699 |
| 0 | 14.427 | 1.24E+09 | 57 | 26.6475 | 26.6085 | 26.3938 | 26.4228 | 26.6951 |
| 0 | 58.446 | 3.31E+09 | 82 | 28.1605 | 28.0997 | 27.791 | 27.8901 | 28.026 |
| 0 | 46.565 | 6.7E+09 | 102 | 28.9868 | 28.9639 | 29.0809 | 29.1383 | 28.9529 |
| 0 | 82.93 | 3.23E+10 | 204 | 31.2511 | 31.1609 | 31.2258 | 31.2734 | 31.191 |
| 0 | 54.388 | 1.75E+09 | 38 | 27.1619 | 27.0228 | 27.2345 | 27.0935 | 27.2159 |
| 0 | 28.772 | 2.19E+09 | 41 | 27.5234 | 27.4707 | 27.5353 | 27.6065 | 27.4805 |
| 0 | 35.807 | 9.78E+09 | 78 | 29.5783 | 29.6106 | 29.4068 | 29.4118 | 29.632 |
| 0 | 10.461 | 6.66E+08 | 23 | 25.6464 | 25.7794 | 25.2836 | 25.6162 | 25.8456 |
| 0 | 78.056 | 7.46E+09 | 68 | 28.9851 | 28.9628 | 29.0313 | 29.0528 | 28.8191 |
| 0 | 48.411 | 1.08E+09 | 32 | 26.0418 | 26.1971 | 25.987 | 25.9129 | 26.5377 |
| 0 | 145.89 | 6.39E+09 | 86 | 28.9165 | 28.9767 | 28.9607 | 28.9728 | 28.761 |
| 0 | 35.074 | 1.52E+09 | 60 | 27.135 | 27.1736 | 26.6651 | 26.6389 | 26.8993 |
| 0 | 143.2 | 1.54E+10 | 192 | 30.1923 | 30.2649 | 30.0334 | 30.1694 | 30.3704 |

| | | | | | | | | |
|---|---|---|---|---|---|---|---|---|
| 0 | 32.954 | 7.28E+08 | 26 | 25.7877 | 25.7673 | 25.6762 | 25.9252 | 25.7182 |
| 0 | 21.996 | 3.13E+09 | 45 | 27.7882 | 27.794 | 27.835 | 27.86 | 27.7253 |
| 0 | 103.54 | 8.41E+09 | 207 | 29.2615 | 29.1469 | 29.3326 | 29.3325 | 29.3131 |
| 0 | 129.12 | 2.17E+10 | 219 | 30.5002 | 30.5135 | 30.5838 | 30.625 | 30.5477 |
| 0 | 31.856 | 1.47E+09 | 47 | 26.8202 | 27.1095 | 26.7393 | 26.9991 | 26.5958 |
| 0 | 169.45 | 5.81E+09 | 116 | 28.6538 | 28.6869 | 28.7466 | 28.8583 | 28.7348 |
| 0 | 19.967 | 1.28E+09 | 54 | 26.5728 | 26.4851 | 26.8007 | 26.7381 | 26.5895 |
| 0 | 100.92 | 2.9E+09 | 89 | 27.7578 | 27.518 | 27.5512 | 27.5933 | 27.6971 |
| 0 | 44.59 | 6.41E+09 | 89 | 28.5203 | 28.5145 | 28.8592 | 28.8449 | 28.8039 |
| 0 | 214.46 | 5.27E+10 | 209 | 31.7466 | 31.6116 | 31.9117 | 31.8424 | 31.9073 |
| 0 | 8.0267 | 9.51E+08 | 14 | 26.7807 | 25.909 | 26.7035 | 26.6512 | 27.1024 |
| 0 | 14.718 | 1.25E+09 | 23 | 26.602 | 26.4413 | 26.2819 | 26.1151 | 26.4297 |
| 0 | 53.949 | 4.05E+09 | 93 | 28.1846 | 28.1502 | 28.1286 | 28.2009 | 28.0273 |
| 0 | 22.794 | 1.53E+09 | 53 | 26.6787 | 26.697 | 26.9402 | 26.9686 | 26.8928 |
| 0 | 323.31 | 7.61E+10 | 174 | 32.6646 | 32.6212 | 32.4624 | 32.4781 | 32.3221 |
| 0 | 44.632 | 9.36E+08 | 53 | 25.9692 | 26.0934 | 26.3008 | 26.2848 | 26.2554 |
| 0 | 39.937 | 4E+09 | 91 | 28.1905 | 28.268 | 28.2403 | 28.3152 | 28.1665 |
| 0 | 37.316 | 2.16E+09 | 51 | 27.3558 | 27.2009 | 27.2564 | 27.3698 | 27.3349 |
| 0 | 98.789 | 2.7E+09 | 69 | 27.6083 | 27.5806 | 27.62 | 27.5738 | 27.4663 |
| 0 | 25.67 | 1.05E+09 | 40 | 26.2391 | 26.2138 | 25.8951 | 25.9905 | 26.2528 |
| 0 | 9.3157 | 6.36E+08 | 31 | 25.0952 | 25.0444 | 25.3072 | 25.3798 | 25.5901 |
| 0 | 72.755 | 3.09E+09 | 70 | 27.7842 | 27.7571 | 27.7407 | 27.7745 | 27.8456 |
| 0 | 64.715 | 1.05E+09 | 41 | 26.1829 | 26.2367 | 26.487 | 26.4501 | 26.4261 |
| 0 | 7.5409 | 5.37E+08 | 15 | 25.0957 | 25.0747 | 25.1034 | 25.2635 | 25.0158 |
| 0 | 15.144 | 8.79E+08 | 29 | 26.3379 | 26.202 | 25.8374 | 25.8037 | 25.9781 |
| 0 | 15.513 | 1E+09 | 26 | 26.3293 | 26.1184 | 26.0395 | 26.1252 | 26.2568 |
| 0 | 17.636 | 3.74E+09 | 76 | 28.2217 | 28.1403 | 28.1566 | 28.1907 | 28.1017 |
| 0 | 11.847 | 3.72E+08 | 17 | 24.7119 | 24.804 | 24.5209 | 24.7903 | 25.2247 |
| 0 | 14.383 | 8.44E+08 | 51 | 25.9629 | 25.889 | 26.0812 | 26.1628 | 25.8506 |
| 0 | 268.31 | 1.43E+10 | 83 | 30.5786 | 30.6215 | 29.8419 | 29.8557 | 29.8702 |
| 0 | 323.31 | 9.2E+09 | 116 | 29.3707 | 29.3445 | 29.3811 | 29.418 | 29.1849 |
| 0 | 22.873 | 1.59E+09 | 62 | 26.8246 | 26.7356 | 26.8471 | 26.8151 | 26.892 |
| 0 | 23.511 | 7.4E+08 | 18 | 25.6005 | 25.5254 | 25.4135 | 25.7946 | 25.7363 |
| 0 | 38.455 | 1.38E+09 | 59 | 26.6919 | 26.5802 | 26.4073 | 26.4258 | 26.4701 |
| 0 | 36.442 | 1.1E+09 | 62 | 26.3554 | 26.4032 | 26.2515 | 26.3087 | 26.368 |
| 0 | 68.656 | 4.51E+09 | 77 | 28.2787 | 28.3507 | 28.3691 | 28.3984 | 28.4945 |
| 0 | 323.31 | 3.27E+10 | 162 | 31.2079 | 31.1578 | 31.225 | 31.1851 | 31.087 |
| 0 | 17.494 | 5.4E+08 | 22 | 25.3114 | 25.5196 | 25.0904 | 25.3655 | 25.5574 |
| 0 | 20.996 | 7.23E+08 | 40 | 25.4745 | 25.5161 | 25.6657 | 25.6521 | 25.7984 |
| 0 | 70.894 | 1.77E+09 | 68 | 27.0652 | 26.9841 | 27.1008 | 26.9365 | 26.8566 |
| 0 | 104.03 | 9.79E+08 | 44 | 26.0226 | 26.0639 | 26.5024 | 26.1145 | 25.8406 |
| 0 | 8.1539 | 1.11E+09 | 26 | 26.3347 | 26.1453 | 26.406 | 26.0257 | 25.8541 |
| 0 | 231.05 | 3.4E+09 | 49 | 27.8492 | 27.9895 | 27.7617 | 27.8734 | 27.9675 |
| 0 | 17.468 | 1.04E+09 | 50 | 26.7133 | 26.5539 | 26.0881 | 26.1275 | 26.2366 |
| 0 | 64.43 | 1.46E+09 | 35 | 26.2274 | 26.7878 | 26.7756 | 26.6214 | 26.7814 |
| 0 | 29.987 | 1.28E+09 | 43 | 26.34 | 26.4549 | 26.4045 | 26.6315 | 26.5006 |
| 0 | 149.86 | 1.28E+10 | 95 | 29.8859 | 29.9503 | 29.8156 | 29.9008 | 29.75 |
| 0 | 34.175 | 7.61E+08 | 37 | 25.8257 | 25.9173 | 25.8711 | 25.8743 | 25.7027 |

| | | | | | | | | |
|---|---|---|---|---|---|---|---|---|
| 0 | 17.19 | 1.61E+09 | 36 | 27.0339 | 27.0495 | 26.8663 | 26.8554 | 26.8137 |
| 0 | 25.12 | 5.44E+08 | 15 | 25.2923 | 25.3585 | 25.3246 | 24.9836 | 25.1071 |
| 0 | 20.623 | 1.08E+09 | 33 | 26.1301 | 26.3655 | 26.2014 | 26.3241 | 26.4496 |
| 0 | 13.282 | 8.2E+08 | 23 | 25.9074 | 25.8352 | 25.9405 | 25.911 | 25.8573 |
| 0 | 16.419 | 4.3E+08 | 32 | 24.7846 | 25.0017 | 24.9599 | 24.7582 | 25.0143 |
| 0 | 20.821 | 2.14E+09 | 47 | 27.2305 | 27.3294 | 27.5027 | 27.461 | 27.0738 |
| 0 | 4.9676 | 9.16E+08 | 20 | 26.1075 | 25.935 | 25.9661 | 26.1227 | 25.7959 |
| 0 | 143.22 | 4.86E+10 | 345 | 31.7345 | 31.7749 | 31.8054 | 31.7253 | 31.7095 |
| 0 | 295.59 | 1.06E+11 | 488 | 32.9156 | 32.9517 | 32.7234 | 32.8171 | 32.8315 |
| 0 | 49.157 | 8.2E+09 | 98 | 29.0594 | 29.1839 | 29.4575 | 29.3138 | 29.3099 |
| 0 | 63.629 | 4.64E+09 | 78 | 28.469 | 28.163 | 28.3791 | 28.2204 | 28.4414 |
| 0 | 145.37 | 9.99E+08 | 33 | 26.6308 | 26.5299 | 25.9654 | 25.9639 | 26.0576 |
| 0 | 25.672 | 6.37E+08 | 24 | 25.4865 | 25.4027 | 25.5103 | 25.4701 | 25.6301 |
| 0 | 13.735 | 8.61E+08 | 26 | 25.6379 | 25.6892 | 25.9272 | 25.8599 | 26.1254 |
| 0 | 48.81 | 1E+09 | 23 | 26.1229 | 26.1687 | 26.1617 | 26.2377 | 26.2268 |
| 0 | 26.528 | 8.59E+08 | 44 | 25.9717 | 25.8193 | 25.9021 | 25.9776 | 25.7573 |
| 0 | 281.61 | 1.63E+10 | 124 | 30.6185 | 30.5038 | 30.1846 | 30.1207 | 30.0937 |
| 0 | 44.965 | 1.24E+09 | 51 | 26.4852 | 26.4293 | 26.4843 | 26.3114 | 26.3879 |
| 0 | 139.42 | 4.55E+09 | 94 | 28.3557 | 28.2105 | 28.3053 | 28.2864 | 28.2666 |
| 0 | 110.25 | 4.29E+09 | 71 | 28.4507 | 28.4009 | 28.1016 | 28.2221 | 28.2229 |
| 0 | 23.906 | 1.16E+09 | 42 | 26.4071 | 26.3567 | 26.2997 | 26.5067 | 26.2148 |
| 0 | 72.869 | 4.66E+09 | 114 | 28.0878 | 28.3095 | 28.431 | 28.3792 | 28.339 |
| 0 | 119.8 | 1.51E+09 | 35 | 26.9243 | 26.9473 | 26.5721 | 26.6471 | 26.8679 |
| 0 | 26.458 | 1.18E+09 | 47 | 26.4266 | 26.4991 | 26.4268 | 26.3649 | 26.3074 |
| 0 | 53.21 | 1.14E+09 | 39 | 26.3722 | 26.2701 | 26.1705 | 26.3882 | 26.3501 |
| 0 | 63.419 | 6.91E+09 | 112 | 28.9598 | 29.0386 | 29.0761 | 28.9692 | 28.894 |
| 0 | 167.77 | 9.31E+09 | 109 | 29.2206 | 29.1255 | 29.2393 | 29.2265 | 29.5604 |
| 0 | 11.599 | 1.39E+09 | 26 | 26.0909 | 26.1829 | 26.1292 | 26.5167 | 26.5613 |
| 0 | 33.22 | 1.48E+09 | 31 | 27.0196 | 26.7003 | 26.7367 | 26.655 | 26.8027 |
| 0 | 13.183 | 8.73E+08 | 23 | 26.1705 | 26.1749 | 26.1412 | 26.1043 | 25.9784 |
| 0 | 23.477 | 1.92E+09 | 37 | 26.9056 | 26.877 | 26.6962 | 26.5759 | 27.1645 |
| 0 | 9.4978 | 2.09E+09 | 25 | 26.9168 | 27.1581 | 27.4788 | 27.1115 | 27.0907 |
| 0 | 10.127 | 2.16E+09 | 51 | 27.2577 | 27.3382 | 27.4201 | 27.2417 | 27.2848 |
| 0 | 15.722 | 3.88E+08 | 14 | 24.8034 | 24.9172 | 24.7562 | 24.6377 | 25.0505 |
| 0 | 61.352 | 8.05E+08 | 40 | 26.136 | 26.0604 | 25.6754 | 25.7739 | 25.9208 |
| 0 | 53.204 | 1.75E+10 | 206 | 30.3802 | 30.3002 | 30.2684 | 30.3128 | 30.1452 |
| 0 | 163.45 | 3.91E+09 | 72 | 28.2157 | 28.2066 | 28.1508 | 28.1477 | 27.9265 |
| 0 | 169.86 | 3.05E+09 | 102 | 27.821 | 27.6966 | 27.9473 | 27.7701 | 27.567 |
| 0 | 288 | 1.78E+10 | 130 | 30.4073 | 30.4909 | 30.1771 | 30.1439 | 30.1975 |
| 0 | 26.324 | 3.58E+09 | 71 | 27.815 | 28.0036 | 28.0161 | 27.7073 | 27.9705 |
| 0 | 35.363 | 1.2E+09 | 34 | 26.5266 | 26.3917 | 26.1597 | 26.3539 | 26.5323 |
| 0 | 16.693 | 8.71E+08 | 30 | 26.3146 | 25.8313 | 26.1924 | 25.7862 | 25.3789 |
| 0 | 50.984 | 3E+09 | 72 | 27.8741 | 27.7206 | 27.6268 | 27.6572 | 27.8448 |
| 0 | 10.518 | 7.26E+08 | 14 | 25.5402 | 25.7974 | 25.8462 | 25.7161 | 25.7838 |
| 0 | 16.793 | 1.75E+09 | 32 | 26.0878 | 26.4033 | 26.9631 | 27.3753 | 26.8174 |
| 0 | 31.709 | 1.45E+09 | 43 | 26.9157 | 26.8299 | 26.3605 | 26.6306 | 26.8993 |
| 0 | 174.29 | 8.4E+09 | 108 | 29.1953 | 29.2887 | 29.3394 | 29.2276 | 29.2392 |
| 0 | 30.53 | 8.86E+08 | 40 | 25.9522 | 25.6727 | 26.002 | 26.1853 | 25.974 |

| | | | | | | | | |
|---|---|---|---|---|---|---|---|---|
| 0 | 22.502 | 7.72E+08 | 31 | 26.0097 | 26.0949 | 25.7586 | 25.7057 | 25.8147 |
| 0 | 9.4775 | 1.04E+09 | 28 | 25.934 | 26.1109 | 25.9488 | 26.1436 | 26.2439 |
| 0 | 10.971 | 8.14E+08 | 33 | 25.9584 | 25.8318 | 25.8677 | 26.0445 | 25.5786 |
| 0 | 8.9317 | 7.94E+08 | 20 | 25.925 | 25.7452 | 25.678 | 25.7481 | 25.8155 |
| 0 | 323.31 | 2.2E+11 | 355 | 33.7384 | 33.8711 | 34.0485 | 34.0277 | 33.7014 |
| 0 | 164.89 | 1.6E+09 | 41 | 26.7712 | 26.7744 | 26.8464 | 26.6846 | 26.9488 |
| 0 | 28.319 | 1.79E+09 | 40 | 26.8937 | 26.7565 | 27.0299 | 27.1404 | 27.1163 |
| 0 | 41.398 | 2.49E+09 | 62 | 27.2408 | 27.3897 | 27.5512 | 27.4798 | 27.5103 |
| 0 | 47.331 | 1.12E+09 | 48 | 26.2236 | 26.2049 | 26.5734 | 26.07 | 26.5373 |
| 0 | 72.537 | 1.03E+10 | 85 | 29.5319 | 29.4448 | 29.5246 | 29.3991 | 29.6196 |
| 0 | 215.94 | 1.75E+10 | 153 | 30.2546 | 30.2638 | 30.5553 | 30.3841 | 30.3241 |
| 0 | 33.285 | 1.72E+09 | 48 | 27.2703 | 27.0434 | 26.8631 | 26.4836 | 26.8154 |
| 0 | 26.065 | 3.93E+09 | 51 | 28.0684 | 27.9844 | 28.2493 | 28.1124 | 28.1103 |
| 0 | 36.559 | 1.71E+09 | 55 | 26.8893 | 27.0967 | 27.0111 | 27.0508 | 26.8452 |
| 0 | 323.31 | 4.26E+09 | 85 | 28.3697 | 28.1663 | 28.2133 | 28.3444 | 28.1342 |
| 0 | 7.4244 | 7.31E+08 | 15 | 25.6503 | 25.713 | 25.7218 | 25.6933 | 25.5689 |
| 0 | 20.238 | 9.4E+08 | 41 | 25.6401 | 25.8414 | 25.8959 | 26.171 | 26.2034 |
| 0 | 9.3948 | 4.1E+08 | 13 | 24.6654 | 25.0641 | 24.7229 | 24.8272 | 25.1639 |
| 0 | 25.562 | 2.6E+09 | 77 | 27.4402 | 27.6036 | 27.5531 | 27.5901 | 27.6217 |
| 0 | 13.969 | 9.87E+08 | 18 | 26.2647 | 26.1707 | 26.0539 | 26.2143 | 26.3342 |
| 0 | 181.29 | 5.08E+09 | 182 | 28.5085 | 28.479 | 28.4828 | 28.5515 | 28.4392 |
| 0 | 29.077 | 8.03E+09 | 74 | 29.2139 | 29.1361 | 29.1934 | 29.1948 | 29.2079 |
| 0 | 323.31 | 1.09E+10 | 130 | 29.676 | 29.6983 | 29.5002 | 29.6358 | 29.5711 |
| 0 | 17.799 | 2.24E+09 | 60 | 27.4391 | 27.2814 | 27.3526 | 27.2894 | 27.4744 |
| 0 | 161.55 | 7.31E+09 | 100 | 29.0524 | 28.974 | 29.0403 | 29.0603 | 29.0503 |
| 0 | 7.1097 | 5.16E+08 | 14 | 25.1905 | 25.1984 | 25.0575 | 25.246 | 24.9516 |
| 0 | 12.958 | 6.69E+08 | 21 | 25.6346 | 25.4642 | 25.5846 | 25.4716 | 25.6978 |
| 0 | 59.634 | 1.86E+09 | 39 | 27.0751 | 27.2537 | 27.0408 | 26.9166 | 26.9811 |
| 0 | 33.481 | 2.52E+09 | 69 | 27.5137 | 27.4845 | 27.5554 | 27.5319 | 27.3381 |
| 0 | 123.79 | 8.53E+09 | 81 | 29.2391 | 29.1787 | 29.367 | 29.3062 | 29.2475 |
| 0 | 10.61 | 5.41E+08 | 22 | 25.137 | 25.3377 | 25.3632 | 25.3086 | 25.5413 |
| 0 | 305.92 | 6.39E+09 | 84 | 28.8537 | 28.7193 | 28.848 | 29.0063 | 28.9088 |
| 0 | 90.185 | 1.23E+09 | 39 | 26.6087 | 26.0736 | 26.5935 | 26.6277 | 26.6438 |
| 0 | 29.112 | 1.64E+09 | 74 | 27.3944 | 27.3388 | 26.9402 | 26.6618 | 26.5275 |
| 0 | 61.086 | 3.46E+09 | 33 | 27.9582 | 27.8591 | 27.8482 | 27.5169 | 27.8638 |
| 0 | 30.835 | 1.57E+09 | 33 | 27.2757 | 27.0696 | 26.5752 | 26.8852 | 26.7236 |
| 0 | 140.53 | 4.78E+10 | 260 | 31.785 | 31.778 | 31.8547 | 31.7635 | 31.658 |
| 0 | 43.706 | 1.68E+09 | 53 | 26.9801 | 27.0327 | 26.8091 | 26.6698 | 26.9876 |
| 0 | 12.907 | 6.81E+08 | 13 | 25.5621 | 25.0301 | 25.5949 | 25.4805 | 25.8069 |
| 0 | 22.323 | 1.13E+09 | 41 | 26.2823 | 26.0529 | 26.3532 | 26.2996 | 26.3507 |
| 0 | 47.072 | 2.69E+10 | 93 | 30.7294 | 30.8116 | 30.9917 | 31.043 | 30.9484 |
| 0 | 102.53 | 1.51E+09 | 58 | 26.6838 | 26.4373 | 26.8948 | 26.9556 | 26.6786 |
| 0 | 18.979 | 6.18E+08 | 44 | 25.5939 | 25.4112 | 25.3698 | 25.4991 | 25.4138 |
| 0 | 45.682 | 3.72E+09 | 109 | 27.9668 | 27.9785 | 28.1689 | 28.1372 | 28.0215 |
| 0 | 32.226 | 1.14E+09 | 33 | 26.3433 | 26.2628 | 26.4771 | 26.3729 | 26.4208 |
| 0 | 16.187 | 1.94E+09 | 30 | 27.3693 | 27.3238 | 26.9908 | 26.8888 | 26.7892 |
| 0 | 31.277 | 1.51E+09 | 50 | 26.5566 | 26.5776 | 26.5835 | 26.9905 | 26.6455 |
| 0 | 323.31 | 1.6E+10 | 347 | 30.2107 | 30.2542 | 29.958 | 30.0484 | 30.2123 |

| | | | | | | | | |
|---|---|---|---|---|---|---|---|---|
| 0 | 9.4399 | 8.59E+08 | 18 | 26.0454 | 25.7685 | 25.6833 | 26.0287 | 25.943 |
| 0 | 20.925 | 8.15E+08 | 32 | 26.0617 | 25.699 | 25.8553 | 25.9623 | 25.8482 |
| 0 | 39.129 | 3.64E+09 | 95 | 28.0571 | 28.1171 | 28.206 | 28.1494 | 27.8424 |
| 0 | 54.981 | 2.43E+09 | 83 | 27.4613 | 27.4675 | 27.3658 | 27.2361 | 27.4983 |
| 0 | 65.887 | 1.95E+09 | 54 | 27.159 | 27.176 | 26.9687 | 26.9394 | 27.2174 |
| 0 | 27.455 | 1.11E+09 | 29 | 26.2508 | 25.8977 | 26.478 | 26.4103 | 26.2466 |
| 0 | 38.404 | 3.63E+09 | 96 | 28.0083 | 28.1215 | 28.0973 | 27.9866 | 28.0074 |
| 0 | 56.267 | 2.68E+09 | 70 | 27.7808 | 27.6086 | 27.6786 | 27.6516 | 27.481 |
| 0 | 33.54 | 1.38E+09 | 63 | 26.6103 | 26.5886 | 26.4868 | 26.5832 | 26.7264 |
| 0 | 18.876 | 1.33E+09 | 28 | 26.471 | 26.5134 | 26.7864 | 26.7345 | 26.4551 |
| 0 | 65.317 | 1.24E+09 | 35 | 26.6031 | 26.5325 | 26.3369 | 26.3005 | 26.4799 |
| 0 | 18.163 | 7.75E+08 | 45 | 25.8452 | 25.7495 | 25.6125 | 25.7763 | 25.7432 |
| 0 | 90.081 | 2.55E+10 | 134 | 30.7978 | 30.8701 | 30.8176 | 30.9213 | 30.8345 |
| 0 | 142.42 | 2.79E+10 | 168 | 31.0274 | 30.9251 | 30.9352 | 31.0877 | 30.9265 |
| 0 | 14.539 | 8.81E+08 | 33 | 25.9747 | 25.9344 | 26.1509 | 26.1941 | 25.6724 |
| 0 | 27.276 | 7.2E+08 | 34 | 25.9066 | 25.8486 | 25.941 | 25.5407 | 25.5437 |
| 0 | 11.803 | 5.94E+08 | 30 | 25.0759 | 25.0821 | 25.1723 | 25.5873 | 25.633 |
| 0 | 17.114 | 6.49E+08 | 18 | 25.8179 | 25.4232 | 25.6455 | 25.4874 | 25.4492 |
| 0 | 45.276 | 2.55E+09 | 45 | 27.5397 | 27.5637 | 27.5796 | 27.4296 | 27.4147 |
| 0 | 323.31 | 2.21E+11 | 727 | 33.874 | 33.8763 | 33.983 | 34.0026 | 33.8411 |
| 0 | 27.82 | 1.72E+09 | 54 | 26.9039 | 27.1377 | 26.9775 | 27.0518 | 26.8174 |
| 0 | 18.216 | 1.33E+09 | 63 | 26.5256 | 26.6577 | 26.5316 | 26.3872 | 26.5919 |
| 0 | 323.31 | 3.05E+11 | 792 | 34.4788 | 34.4855 | 34.5851 | 34.4945 | 34.342 |
| 0 | 13.907 | 1.54E+09 | 38 | 26.597 | 26.693 | 26.8586 | 26.8868 | 26.8655 |
| 0 | 28.172 | 1.6E+09 | 56 | 27.0038 | 26.9546 | 26.7247 | 26.7773 | 26.8042 |
| 0 | 10.81 | 1.21E+10 | 89 | 29.7012 | 29.6698 | 29.6244 | 29.526 | 29.768 |
| 0 | 8.0157 | 5.27E+08 | 15 | 25.4519 | 25.3 | 24.8934 | 25.1257 | 25.4258 |
| 0 | 21.813 | 6.7E+08 | 43 | 25.3661 | 25.5776 | 25.3771 | 25.5487 | 25.6841 |
| 0 | 117.05 | 5.33E+10 | 354 | 31.7875 | 31.8625 | 31.9383 | 31.801 | 31.6798 |
| 0 | 10.536 | 7.42E+08 | 20 | 25.4186 | 25.7068 | 25.7946 | 25.5587 | 25.6463 |
| 0 | 25.785 | 2.23E+09 | 53 | 27.3998 | 27.3221 | 27.319 | 27.4243 | 27.2462 |
| 0 | 18.686 | 8.89E+08 | 41 | 26.1713 | 26.1924 | 25.66 | 25.9232 | 26.2485 |
| 0 | 178.23 | 1.57E+10 | 116 | 30.0324 | 30.0158 | 30.3316 | 30.2151 | 30.1044 |
| 0 | 19.245 | 2.7E+09 | 72 | 27.6582 | 27.5082 | 27.6623 | 27.6936 | 27.7212 |
| 0 | 19.093 | 2.19E+10 | 129 | 30.662 | 30.6308 | 30.6522 | 30.6382 | 30.5046 |
| 0 | 16.035 | 3.4E+09 | 27 | 27.2148 | 27.3907 | 27.4387 | 27.3274 | 27.2001 |
| 0 | 70.735 | 1.37E+09 | 38 | 26.6242 | 26.5849 | 26.3277 | 26.5247 | 26.1852 |
| 0 | 6.4978 | 9.8E+08 | 31 | 25.3926 | 25.5122 | 26.0127 | 26.0405 | 25.9723 |
| 0 | 18.878 | 7.41E+09 | 80 | 28.4341 | 28.4234 | 28.9555 | 28.8239 | 28.9675 |
| 0 | 64.04 | 3.21E+09 | 78 | 27.6064 | 27.7833 | 27.7434 | 27.7001 | 27.4361 |
| 0 | 18.302 | 6.97E+08 | 25 | 25.5997 | 25.4765 | 25.4609 | 25.1497 | 25.3043 |
| 0 | 14.176 | 1.24E+09 | 23 | 25.9613 | 26.3466 | 26.1527 | 26.2865 | 26.3922 |
| 0 | 40.01 | 6.57E+08 | 29 | 25.4908 | 25.2521 | 25.3053 | 25.1991 | 25.4126 |
| 0 | 72.058 | 4.45E+09 | 105 | 27.8801 | 27.9773 | 28.2761 | 28.2537 | 28.215 |
| 0 | 56.034 | 1.6E+09 | 49 | 26.5846 | 26.5308 | 26.7525 | 26.9084 | 26.241 |
| 0 | 89.635 | 1.15E+09 | 36 | 26.1397 | 26.0854 | 26.3169 | 26.1548 | 26.3422 |
| 0 | 18.983 | 8.95E+08 | 16 | 25.9573 | 25.9549 | 25.974 | 25.9727 | 25.6778 |
| 0 | 28.52 | 1.06E+09 | 44 | 26.0573 | 25.9209 | 26.2146 | 25.9114 | 26.2027 |

| | | | | | | | | |
|---|---|---|---|---|---|---|---|---|
| 0 | 23.422 | 1.45E+09 | 39 | 26.7946 | 26.6258 | 26.7061 | 26.5764 | 26.454 |
| 0 | 28.239 | 1.64E+09 | 54 | 26.4201 | 26.6667 | 26.9397 | 26.8165 | 26.7831 |
| 0 | 71.676 | 1.31E+09 | 46 | 26.3217 | 26.2523 | 26.4892 | 26.4865 | 26.4471 |
| 0 | 20.293 | 8.64E+08 | 41 | 25.7714 | 25.6957 | 25.8873 | 26.0054 | 25.7254 |
| 0 | 23.78 | 7.09E+08 | 37 | 25.5589 | 25.4489 | 25.56 | 25.6626 | 25.5601 |
| 0 | 208.74 | 2.02E+10 | 180 | 30.3388 | 30.3702 | 30.5447 | 30.4887 | 30.2202 |
| 0 | 138.23 | 3.98E+09 | 68 | 27.9636 | 27.8888 | 27.9655 | 27.9954 | 27.9904 |
| 0 | 23.567 | 1.09E+09 | 55 | 26.1034 | 26.2425 | 26.3311 | 26.3307 | 25.8543 |
| 0 | 84.529 | 6.53E+09 | 116 | 28.7914 | 28.7816 | 28.6678 | 28.7505 | 28.7128 |
| 0 | 17.292 | 3.44E+09 | 68 | 27.8287 | 27.8315 | 27.8931 | 27.7706 | 27.6418 |
| 0 | 126.7 | 4.26E+09 | 111 | 28.0916 | 28.1953 | 28.2502 | 28.0934 | 28.2012 |
| 0 | 25.74 | 1.02E+09 | 38 | 26.1459 | 25.8542 | 26.0992 | 26.103 | 26.1128 |
| 0 | 37.51 | 4.59E+08 | 11 | 24.9568 | 24.9435 | 24.8419 | 24.8033 | 25.042 |
| 0 | 117 | 2.58E+09 | 92 | 27.4998 | 27.4364 | 27.4158 | 27.414 | 27.3365 |
| 0 | 23.068 | 7.26E+08 | 14 | 25.542 | 25.7027 | 25.5508 | 25.4189 | 25.5447 |
| 0 | 68.755 | 3.55E+09 | 93 | 27.8489 | 27.8129 | 27.9006 | 27.9054 | 27.9925 |
| 0 | 34.063 | 1.49E+09 | 52 | 26.5638 | 26.623 | 26.5015 | 26.5443 | 26.6558 |
| 0 | 37.596 | 1.44E+10 | 83 | 29.8746 | 29.9035 | 30.0452 | 29.9474 | 29.9324 |
| 0 | 38.207 | 1.04E+10 | 163 | 29.4611 | 29.3582 | 29.3971 | 29.435 | 29.5853 |
| 0 | 323.31 | 3.59E+10 | 177 | 31.2955 | 31.2886 | 31.2917 | 31.3141 | 31.1769 |
| 0 | 71.176 | 2.31E+09 | 68 | 27.3421 | 27.3585 | 27.4186 | 27.1885 | 27.2179 |
| 0 | 34.81 | 1.57E+09 | 77 | 26.7838 | 26.7769 | 26.5676 | 26.8569 | 26.7553 |
| 0 | 92.383 | 3.13E+09 | 85 | 27.7933 | 27.7103 | 27.8157 | 27.8645 | 27.6195 |
| 0 | 92.093 | 1.4E+10 | 117 | 29.8639 | 29.8523 | 29.9974 | 30.0065 | 29.8439 |
| 0 | 272.35 | 6.22E+09 | 79 | 28.8612 | 28.9705 | 28.783 | 28.9013 | 28.7376 |
| 0 | 101.65 | 4.58E+10 | 156 | 31.7165 | 31.6878 | 31.7038 | 31.7746 | 31.5596 |
| 0 | 11.946 | 1.14E+09 | 44 | 26.3325 | 26.4839 | 26.2779 | 26.4475 | 26.3781 |
| 0 | 76.658 | 1.18E+10 | 101 | 29.6804 | 29.7534 | 29.7925 | 29.9635 | 29.7845 |
| 0 | 15.133 | 2.38E+09 | 38 | 27.6523 | 27.5737 | 27.5111 | 27.3493 | 27.6839 |
| 0 | 92.596 | 1.35E+10 | 102 | 30.0109 | 30.072 | 30.1617 | 30.1179 | 29.8596 |
| 0 | 61.175 | 4.84E+09 | 50 | 28.6236 | 28.4775 | 28.65 | 28.5138 | 28.6211 |
| 0 | 22.155 | 9.26E+08 | 38 | 26.2298 | 26.2071 | 25.8512 | 26.3821 | 26.2473 |
| 0 | 11.626 | 8.85E+08 | 35 | 26.533 | 26.6098 | 25.7595 | 25.8349 | 26.3009 |
| 0 | 17.357 | 1.18E+09 | 31 | 26.6218 | 26.8845 | 26.4537 | 26.3217 | 26.7132 |
| 0 | 50.78 | 4.35E+09 | 67 | 28.8883 | 28.819 | 28.3375 | 28.4691 | 28.1231 |
| 0 | 35.348 | 4E+09 | 69 | 28.0403 | 27.9482 | 28.2754 | 28.2341 | 27.7357 |
| 0 | 67.032 | 1.01E+09 | 36 | 25.9645 | 26.0257 | 26.0263 | 26.1496 | 25.9044 |
| 0 | 88.111 | 3.3E+09 | 58 | 27.831 | 27.7398 | 27.5267 | 27.672 | 27.9091 |
| 0 | 25.052 | 5.51E+08 | 39 | 25.339 | 25.0969 | 25.1172 | 25.1106 | 25.2818 |
| 0 | 38.008 | 6.54E+09 | 80 | 28.2602 | 28.2924 | 28.7244 | 28.8241 | 28.8924 |
| 0 | 31.902 | 9.09E+09 | 117 | 29.2584 | 29.2286 | 29.3207 | 29.2993 | 29.2717 |
| 0 | 104.93 | 6.11E+09 | 88 | 29.0507 | 29.054 | 28.8829 | 28.8415 | 28.758 |
| 0 | 10.872 | 1.18E+09 | 25 | 26.3442 | 26.4093 | 26.3863 | 26.2629 | 26.3381 |
| 0 | 42.562 | 1.12E+09 | 46 | 26.4553 | 26.3569 | 26.6502 | 26.463 | 26.3942 |
| 0 | 16.114 | 1.79E+09 | 69 | 27.0097 | 27.0093 | 27.1931 | 27.2121 | 27.0876 |
| 0 | 14.289 | 8.71E+08 | 37 | 26.1252 | 26.1533 | 26.0698 | 25.9347 | 26.0426 |
| 0 | 36.222 | 2.62E+09 | 73 | 27.5134 | 27.4722 | 27.5171 | 27.5704 | 27.4467 |
| 0 | 27.295 | 2.34E+09 | 64 | 27.4845 | 27.4413 | 27.16 | 27.1475 | 27.0704 |

| | | | | | | | | |
|---|---|---|---|---|---|---|---|---|
| 0 | 232.14 | 1.3E+10 | 220 | 29.6976 | 29.7137 | 29.92 | 29.7518 | 29.8032 |
| 0 | 39.298 | 4.73E+10 | 234 | 31.9928 | 32.0447 | 31.7066 | 31.6829 | 31.6775 |
| 0 | 154.27 | 5.68E+09 | 101 | 28.5554 | 28.5773 | 28.5621 | 28.5463 | 28.6631 |
| 0 | 101.27 | 5.06E+09 | 114 | 28.332 | 28.4595 | 28.5494 | 28.491 | 28.2805 |
| 0 | 259.15 | 9.5E+10 | 447 | 32.5749 | 32.5785 | 32.6739 | 32.5534 | 32.7487 |
| 0 | 12.435 | 1.09E+09 | 30 | 26.0515 | 26.1025 | 25.9788 | 25.8174 | 26.1687 |
| 0 | 17.954 | 9.68E+08 | 30 | 26.0722 | 26.1388 | 26.2342 | 26.1465 | 25.8034 |
| 0 | 40.602 | 2.98E+09 | 63 | 28.263 | 28.0474 | 27.6881 | 27.8243 | 27.6232 |
| 0 | 140.24 | 3.28E+09 | 62 | 27.6538 | 27.6064 | 27.9976 | 27.7818 | 27.7676 |
| 0 | 11.943 | 6.83E+08 | 22 | 25.683 | 25.4897 | 25.3308 | 25.5905 | 25.511 |
| 0 | 116.86 | 1.16E+10 | 115 | 29.6822 | 29.6987 | 29.8016 | 29.7585 | 29.7414 |
| 0 | 27.715 | 2.89E+09 | 102 | 27.8236 | 27.6631 | 27.7775 | 27.7271 | 27.8036 |
| 0 | 35.585 | 9.74E+08 | 55 | 25.8365 | 25.8393 | 26.1017 | 26.0951 | 25.9172 |
| 0 | 11.268 | 1.31E+10 | 94 | 29.9061 | 29.9507 | 30.0054 | 29.9613 | 29.874 |
| 0 | 39.46 | 9.59E+08 | 31 | 25.7482 | 26.2647 | 26.0224 | 26.0364 | 25.718 |
| 0 | 68.766 | 2.59E+09 | 48 | 27.5764 | 27.3466 | 27.1799 | 27.0969 | 27.4206 |
| 0 | 33.054 | 9.11E+08 | 32 | 26.0907 | 25.9438 | 26.0099 | 25.7779 | 26.1435 |
| 0 | 88.981 | 1.73E+09 | 56 | 26.8591 | 26.8949 | 27.0648 | 27.0807 | 26.695 |
| 0 | 70.233 | 2.03E+09 | 54 | 27.1938 | 27.1705 | 27.099 | 27.2294 | 26.1736 |
| 0 | 18.556 | 1.67E+09 | 71 | 26.9099 | 26.9185 | 26.984 | 26.9673 | 26.6726 |
| 0 | 11.589 | 3.85E+08 | 10 | NaN | NaN | NaN | NaN | NaN |
| 0 | 105.97 | 2.17E+09 | 83 | 27.3576 | 27.1746 | 27.1739 | 27.2922 | 27.2559 |
| 0 | 7.5399 | 1.12E+09 | 35 | 26.4897 | 26.5369 | 26.4982 | 26.4479 | 26.3964 |
| 0 | 29.986 | 1.35E+09 | 54 | 26.6298 | 26.5679 | 26.5672 | 26.331 | 26.4166 |
| 0 | 42.666 | 2.92E+09 | 57 | 27.6539 | 27.6373 | 27.2994 | 27.5181 | 27.7129 |
| 0 | 4.6842 | 4.56E+08 | 17 | 25.74 | 25.5249 | 24.8457 | 24.5626 | 25.1567 |
| 0 | 42.172 | 2.24E+09 | 55 | 27.2893 | 27.1543 | 27.3826 | 27.2949 | 27.3362 |
| 0 | 14.08 | 7.92E+08 | 24 | 26.1471 | 26.2989 | 25.859 | 26.0341 | 25.8572 |
| 0 | 20.758 | 2.47E+09 | 60 | 27.302 | 27.2373 | 27.51 | 27.4783 | 27.3943 |
| 0 | 7.4934 | 2.93E+08 | 12 | 24.5659 | 24.512 | 24.2802 | 24.3091 | 24.6039 |
| 0 | 56.817 | 2.43E+09 | 70 | 27.1475 | 27.1182 | 27.4185 | 27.3344 | 27.4598 |
| 0 | 30.256 | 1.55E+10 | 108 | 30.1647 | 30.1394 | 30.3313 | 30.2671 | 30.1388 |
| 0 | 9.8998 | 4.15E+08 | 27 | 24.7476 | 24.9792 | 24.6786 | 24.9261 | 24.9496 |
| 0 | 7.3751 | 4.41E+08 | 12 | 25.075 | 25.0828 | 24.894 | 24.9537 | 25.2962 |
| 0 | 22.456 | 3.68E+09 | 60 | 28.4701 | 28.6408 | 27.9252 | 28.0851 | 27.9143 |
| 0 | 72.691 | 1.84E+09 | 41 | 27.3127 | 27.0936 | 27.3595 | 27.3734 | 26.9742 |
| 0 | 6.8787 | 7.05E+08 | 18 | 25.6812 | 25.6222 | 25.5752 | 25.4445 | 25.7159 |
| 0 | 51.127 | 8.07E+08 | 43 | 25.6232 | 25.6386 | 25.7276 | 25.9721 | 25.7577 |
| 0 | 10.844 | 5.03E+08 | 20 | 25.2939 | 24.8993 | 25.2192 | 25.0224 | 24.9268 |
| 0 | 35.201 | 1.94E+10 | 149 | 30.3413 | 30.4252 | 30.6371 | 30.4526 | 30.4788 |
| 0 | 37.233 | 2.8E+09 | 63 | 27.7281 | 27.5372 | 27.8921 | 27.7866 | 27.7347 |
| 0 | 22.091 | 4.07E+09 | 86 | 28.1928 | 28.1056 | 27.8013 | 27.8831 | 28.1421 |
| 0 | 46.462 | 1.42E+09 | 49 | 26.6409 | 26.7685 | 26.8515 | 26.6338 | 26.6749 |
| 0 | 10.143 | 3.98E+08 | 22 | 24.6617 | 24.9907 | 25.0646 | 24.974 | 24.7354 |
| 0 | 16.136 | 8.13E+08 | 29 | 25.6048 | 25.8746 | 25.8414 | 25.8934 | 25.8675 |
| 0 | 16.422 | 1.48E+09 | 29 | 26.6553 | 26.5257 | 26.8881 | 26.3748 | 26.8237 |
| 0 | 15.978 | 6.52E+08 | 39 | 25.6425 | 25.3734 | 25.3282 | 25.5961 | 25.4986 |
| 0 | 36.114 | 2.44E+09 | 62 | 27.5038 | 27.4737 | 27.3758 | 27.4268 | 27.3163 |

| | | | | | | | | |
|---|---|---|---|---|---|---|---|---|
| 0 | 10.662 | 4.58E+08 | 17 | 25.1768 | 24.7917 | 25.093 | NaN | 24.9772 |
| 0 | 28.953 | 1.18E+09 | 32 | 26.2766 | 26.1058 | 26.424 | 26.4379 | 26.4472 |
| 0 | 126.95 | 1.67E+09 | 61 | 27.1089 | 26.9544 | 27.2517 | 26.9429 | 26.7367 |
| 0 | 7.5735 | 4.57E+08 | 14 | 25.1264 | NaN | 25.1692 | 24.8653 | 24.9014 |
| 0 | 24.272 | 8.24E+08 | 32 | 25.6667 | 25.7041 | 25.9303 | 26.0306 | 25.8355 |
| 0 | 17.184 | 5.5E+08 | 21 | 25.0241 | 25.0009 | 25.0515 | 24.9243 | 25.4972 |
| 0 | 55.305 | 9.57E+08 | 39 | 26.0246 | 26.0341 | 26.0545 | 26.2698 | 26.047 |
| 0 | 6.968 | 3.81E+08 | 14 | 24.8992 | 24.4751 | 24.9941 | 24.7967 | 24.7746 |
| 0 | 98.64 | 6.64E+09 | 85 | 28.9154 | 28.8146 | 29.1284 | 29.0511 | 28.8269 |
| 0 | 12.915 | 4.88E+08 | 12 | NaN | 25.3574 | 25.2651 | NaN | NaN |
| 0 | 35.6 | 1.63E+10 | 100 | 30.141 | 30.2058 | 30.0305 | 30.1715 | 30.0629 |
| 0 | 144.14 | 1.83E+09 | 66 | 27.0888 | 26.9556 | 26.931 | 26.8185 | 27.0979 |
| 0 | 89.58 | 2.47E+09 | 53 | 27.2764 | 27.3551 | 27.5515 | 27.346 | 27.3027 |
| 0 | 20.359 | 1.07E+09 | 28 | 26.2789 | 26.1085 | 26.4738 | 26.3947 | 26.3586 |
| 0 | 37.463 | 1.93E+09 | 41 | 27.5094 | 27.4095 | 27.1632 | 26.7863 | 27.1253 |
| 0 | 10.949 | 8.4E+08 | 61 | 25.7524 | 25.8948 | 25.978 | 25.7882 | 25.9187 |
| 0 | 9.0648 | 6.87E+08 | 16 | NaN | 26.083 | 25.8037 | 25.8073 | 25.8296 |
| 0 | 323.31 | 3.54E+10 | 200 | 31.2527 | 31.1927 | 31.4684 | 31.3616 | 31.1159 |
| 0 | 49.045 | 7.88E+08 | 18 | 25.7751 | 25.8739 | 26.1379 | 26.258 | 25.7964 |
| 0 | 30.42 | 5.94E+08 | 20 | 24.9493 | 25.3494 | 25.1867 | 25.3865 | 25.5029 |
| 0 | 80.075 | 5.93E+09 | 129 | 28.6841 | 28.6537 | 28.6709 | 28.8002 | 28.6399 |
| 0 | 26.369 | 1.36E+09 | 56 | 26.5923 | 26.6359 | 26.4107 | 26.6262 | 26.8927 |
| 0 | 24.834 | 1.33E+09 | 47 | 26.8237 | 26.734 | 26.264 | 26.5265 | 26.4191 |
| 0 | 104.5 | 2.16E+09 | 67 | 27.4066 | 27.314 | 27.2375 | 27.2391 | 27.1568 |
| 0 | 10.57 | 1.42E+09 | 37 | 26.6169 | 26.6671 | 26.6939 | 26.7457 | 26.7423 |
| 0 | 10.106 | 8.78E+08 | 30 | 25.9877 | 26.1814 | 26.105 | 26.0354 | 25.9394 |
| 0 | 9.7296 | 6.05E+08 | 15 | 25.477 | 25.6135 | 25.4759 | 25.5071 | 25.399 |
| 0 | 29.344 | 2.4E+09 | 58 | 27.3003 | 27.3187 | 27.5325 | 27.3205 | 27.2981 |
| 0 | 102.44 | 8.33E+08 | 43 | 26.0939 | 25.8583 | 25.7008 | 25.6877 | 26.1286 |
| 0 | 323.31 | 2.52E+10 | 104 | 31.0649 | 30.989 | 30.6939 | 30.7137 | 30.8419 |
| 0 | 10.158 | 5.48E+08 | 27 | 25.1216 | 25.2372 | 25.4946 | 25.6442 | 25.4691 |
| 0 | 323.31 | 3.5E+10 | 232 | 31.2375 | 31.3156 | 31.3437 | 31.2887 | 31.2043 |
| 0 | 14.034 | 3.86E+09 | 40 | 28.0865 | 28.174 | 28.1493 | 27.9644 | 28.1461 |
| 0 | 12.223 | 5.11E+08 | 13 | 25.4271 | 25.5401 | 25.1401 | 25.2279 | 25.0217 |
| 0 | 13.323 | 8.72E+08 | 32 | 25.922 | 25.9906 | 25.9496 | 25.6802 | 26.022 |
| 0 | 51.419 | 1.56E+09 | 64 | 26.8651 | 26.7212 | 26.8274 | 26.8344 | 26.7074 |
| 0 | 17.485 | 2.22E+09 | 58 | 26.9408 | 27.0061 | 27.178 | 27.1518 | 27.1483 |
| 0 | 9.9983 | 6.21E+08 | 40 | 25.5439 | 25.6878 | 25.4341 | 25.2602 | 25.6752 |
| 0 | 18.147 | 5.72E+08 | 35 | 25.2664 | 25.379 | 25.4848 | 25.407 | 24.964 |
| 0 | 43.313 | 1.9E+09 | 60 | 27.1137 | 27.116 | 27.0128 | 27.063 | 27.0868 |
| 0 | 238.18 | 3.12E+10 | 209 | 30.9977 | 31.017 | 31.1877 | 31.1036 | 30.8752 |
| 0 | 25.786 | 6.18E+08 | 32 | 25.3697 | 25.4942 | 25.4502 | 25.5304 | 25.5626 |
| 0 | 34.944 | 4.21E+08 | 13 | 25.0296 | 25.204 | 24.5628 | 24.6646 | 24.7493 |
| 0 | 30.253 | 6.11E+09 | 88 | 28.3207 | 28.3804 | 28.7889 | 28.7675 | 28.8272 |
| 0 | 323.31 | 1.19E+10 | 120 | 29.777 | 29.6781 | 29.7385 | 29.61 | 29.7597 |
| 0 | 176.14 | 1.59E+10 | 193 | 30.0786 | 30.1095 | 30.2151 | 30.1299 | 29.9751 |
| 0 | 14.945 | 1.91E+09 | 41 | 26.8859 | 26.8498 | 27.0578 | 27.0569 | 27.254 |
| 0 | 16.059 | 8.18E+08 | 61 | 26.0088 | 25.8227 | 25.7912 | 25.8177 | 26.0275 |

| | | | | | | | | |
|---|---|---|---|---|---|---|---|---|
| 0 | 11.192 | 4.22E+08 | 16 | 24.9174 | 24.7919 | 25.123 | NaN | 25.3636 |
| 0 | 16.711 | 3.21E+09 | 69 | 27.7749 | 27.7126 | 28.0679 | 27.7754 | 27.8195 |
| 0 | 8.9447 | 5.7E+08 | 33 | 25.2845 | 25.2177 | 25.2333 | 25.0911 | 25.4919 |
| 0 | 15.929 | 1.12E+09 | 32 | 26.2939 | 26.228 | 26.3093 | 26.1684 | 26.2653 |
| 0 | 13.844 | 7.84E+08 | 27 | 25.9633 | 25.8684 | 25.8909 | 25.7193 | 25.8619 |
| 0 | 22.312 | 1.74E+10 | 93 | 30.4263 | 30.5034 | 30.2062 | 30.3773 | 30.1696 |
| 0 | 58.516 | 9.01E+09 | 107 | 29.3955 | 29.3531 | 29.3982 | 29.348 | 29.2751 |
| 0 | 54.859 | 5.13E+09 | 85 | 28.7197 | 28.5393 | 28.5832 | 28.55 | 28.2412 |
| 0 | 19.892 | 7.34E+08 | 29 | 25.6471 | 25.8041 | 25.9412 | 25.7381 | 25.7705 |
| 0 | 79.264 | 2.58E+09 | 87 | 27.3876 | 27.272 | 27.501 | 27.406 | 27.6867 |
| 0 | 42.177 | 9.52E+08 | 33 | 26.4187 | 26.0915 | 26.0669 | 25.8805 | 26.0179 |
| 0 | 8.854 | 1E+09 | 13 | 26.2859 | 26.3952 | 25.91 | 26.0932 | 26.0994 |
| 0 | 36.266 | 2.32E+09 | 63 | 27.3471 | 27.3493 | 27.4113 | 27.3755 | 27.4319 |
| 0 | 61.343 | 3.71E+09 | 129 | 27.9553 | 27.9907 | 27.9373 | 27.9568 | 28.0535 |
| 0 | 26.64 | 9.96E+08 | 21 | 26.1323 | 26.217 | 25.8951 | 26.117 | 26.2065 |
| 0 | 31.812 | 6.61E+08 | 24 | 25.8077 | 25.5746 | 25.4522 | 25.4284 | 25.8777 |
| 0 | 176.39 | 1.84E+10 | 103 | 30.49 | 30.385 | 30.4849 | 30.4694 | 30.4017 |
| 0 | 33.531 | 2.63E+09 | 78 | 27.6081 | 27.4908 | 27.6789 | 27.7248 | 27.5686 |
| 0 | 9.5792 | 4.41E+08 | 6 | NaN | NaN | 25.3484 | NaN | 25.1762 |
| 0 | 323.31 | 4.42E+10 | 307 | 31.674 | 31.6483 | 31.4561 | 31.4798 | 31.6435 |
| 0 | 29.703 | 1.13E+09 | 34 | 26.4623 | 26.26 | 26.4545 | 26.406 | 26.1576 |
| 0 | 30.383 | 3.72E+09 | 60 | 27.9338 | 28.0336 | 28.2349 | 28.2076 | 28.054 |
| 0 | 7.6478 | 3.26E+08 | 8 | 24.4342 | NaN | NaN | NaN | 24.4344 |
| 0 | 25.14 | 1.55E+09 | 12 | 27.1683 | 26.71 | 26.8208 | NaN | 26.6375 |
| 0 | 40.325 | 1.13E+09 | 43 | 26.5123 | 26.1687 | 26.4887 | 26.4978 | 26.3796 |
| 0 | 54.398 | 1.89E+09 | 52 | 27.1727 | 27.2869 | 27.336 | 27.1058 | 26.8693 |
| 0 | 17.483 | 8.22E+08 | 25 | 25.7588 | 25.9876 | 26.0155 | 25.7881 | 25.7055 |
| 0 | 80.221 | 1.01E+09 | 48 | 26.1644 | 26.269 | 26.0957 | 25.9763 | 26.3115 |
| 0 | 15.11 | 8.47E+08 | 31 | 26.2245 | 26.044 | 25.5858 | 25.7437 | 25.8516 |
| 0 | 215.04 | 1.88E+10 | 187 | 30.558 | 30.508 | 30.2182 | 30.239 | 30.4614 |
| 0 | 52.517 | 1.44E+09 | 49 | 26.6439 | 26.8209 | 26.6859 | 26.8679 | 26.4338 |
| 0 | 60.667 | 1.08E+10 | 123 | 29.5744 | 29.536 | 29.5455 | 29.5006 | 29.555 |
| 0 | 17.053 | 1.68E+09 | 52 | 26.6806 | 26.7052 | 26.9939 | 27.1081 | 26.9903 |
| 0 | 16.269 | 6.91E+08 | 39 | 25.7342 | 25.7155 | 25.5024 | 25.7674 | 25.5414 |
| 0 | 10.726 | 1.15E+09 | 61 | 26.1365 | 26.2684 | 26.5612 | 26.4358 | 26.4584 |
| 0 | 38.547 | 1.57E+09 | 51 | 26.7904 | 26.7812 | 26.8299 | 26.8734 | 26.6019 |
| 0 | 13.222 | 7.2E+08 | 16 | 25.1496 | 25.5607 | 26.1339 | 25.2667 | 25.5227 |
| 0 | 26.134 | 1.53E+09 | 34 | 26.458 | 26.7253 | 26.6999 | 26.7816 | 27.018 |
| 0 | 6.5442 | 3.25E+08 | 10 | NaN | NaN | 24.36 | 24.4094 | 24.4487 |
| 0 | 11.028 | 8.44E+08 | 39 | 26.1244 | 26.0309 | 25.783 | 25.8955 | 25.9044 |
| 0 | 20.765 | 1.36E+09 | 59 | 26.5968 | 26.4996 | 26.6619 | 26.9069 | 26.5073 |
| 0 | 17.504 | 1.71E+09 | 43 | 27.1067 | 26.885 | 26.73 | 26.7098 | 26.9293 |
| 0 | 8.1135 | 3.51E+08 | 12 | 24.7199 | 24.555 | 24.4524 | 24.407 | NaN |
| 0 | 10.27 | 5.28E+08 | 37 | 25.1987 | 25.3517 | 24.9103 | 25.3414 | 25.2701 |
| 0 | 24.706 | 7.74E+09 | 75 | 29.2191 | 29.2731 | 29.2002 | 29.1107 | 28.974 |
| 0 | 16.573 | 9.41E+08 | 42 | 25.9945 | 25.9009 | 26.0626 | 26.1355 | 25.9713 |
| 0 | 36.32 | 2.3E+09 | 69 | 27.4327 | 27.3997 | 27.2746 | 27.4692 | 27.3081 |
| 0 | 11.936 | 6.37E+08 | 22 | 25.6809 | 25.5344 | 25.5847 | 25.6463 | 25.408 |

| | | | | | | | | |
|---|---|---|---|---|---|---|---|---|
| 0 | 82.592 | 1.5E+09 | 53 | 26.897 | 26.8709 | 26.4868 | 26.7403 | 26.7327 |
| 0 | 8.5362 | 3.95E+08 | 28 | 24.8215 | 24.7841 | 24.8416 | 24.8649 | 24.7927 |
| 0 | 15.355 | 9.68E+08 | 31 | 25.928 | 26.0166 | 26.4169 | 26.0524 | 26.0194 |
| 0 | 74.223 | 3.7E+09 | 75 | 28.0559 | 28.0853 | 28.132 | 28.112 | 28.0394 |
| 0 | 7.629 | 4.27E+08 | 19 | 24.7166 | 24.5832 | 24.7912 | 24.7587 | 24.8542 |
| 0 | 117.23 | 4.53E+09 | 93 | 28.351 | 28.2652 | 28.2368 | 28.3241 | 28.4204 |
| 0 | 12.705 | 1.34E+09 | 55 | 26.5697 | 26.5896 | 26.8006 | 26.7762 | 26.4627 |
| 0 | 80.151 | 4.75E+09 | 64 | 28.5059 | 28.4877 | 28.3821 | 28.5617 | 28.2059 |
| 0 | 15.404 | 9.83E+08 | 34 | 26.1709 | 26.0641 | 26.2685 | 26.2839 | 26.1557 |
| 0 | 19.221 | 2.09E+09 | 65 | 27.2502 | 27.1502 | 27.2086 | 27.281 | 27.1857 |
| 0 | 76.639 | 8.88E+09 | 72 | 29.3902 | 29.545 | 29.0555 | 29.1308 | 29.2645 |
| 0 | 15.813 | 1.73E+09 | 50 | 26.9484 | 26.7653 | 26.8784 | 26.8805 | 26.9913 |
| 0 | 9.6252 | 5.42E+08 | 34 | 25.246 | 25.3963 | 24.7108 | 25.2178 | 25.3586 |
| 0 | 19.26 | 1.07E+09 | 37 | 26.4592 | 26.1946 | 26.2721 | 26.2931 | 26.3012 |
| 0 | 71.435 | 3.2E+10 | 148 | 30.8104 | 30.9547 | 31.2029 | 31.1908 | 31.0663 |
| 0 | 15.211 | 7E+08 | 38 | 25.5593 | 25.5867 | 25.2391 | 25.7604 | 25.6988 |
| 0 | 20.278 | 9.88E+08 | 31 | 26.2106 | 26.4875 | 26.1343 | 25.756 | 26.2614 |
| 0 | 21.29 | 5.83E+08 | 28 | 25.292 | 25.5029 | 25.1825 | 25.3723 | 25.3545 |
| 0 | 16.993 | 4.54E+08 | 28 | 24.7328 | 24.2787 | 25.2098 | 25.048 | 25.1696 |
| 0 | 28.796 | 1.12E+09 | 47 | 26.4266 | 26.3187 | 26.3496 | 26.3205 | 26.324 |
| 0 | 24.545 | 1.46E+09 | 64 | 26.7257 | 26.805 | 26.6335 | 26.8588 | 26.7214 |
| 0 | 9.384 | 7.78E+08 | 20 | 25.3415 | 26.017 | 26.1997 | 25.9758 | 25.4635 |
| 0 | 99.396 | 1.25E+10 | 121 | 30.0549 | 30.0116 | 29.7158 | 29.7378 | 29.6828 |
| 0 | 59.578 | 1.42E+09 | 36 | 26.6865 | 26.5836 | 26.5595 | 26.4419 | 26.7803 |
| 0 | 18.268 | 1.13E+09 | 45 | 26.6207 | 26.2587 | 26.4169 | 26.4438 | 26.1701 |
| 0 | 45.206 | 1.85E+09 | 58 | 26.9303 | 26.865 | 26.9315 | 27.0276 | 27.1206 |
| 0 | 17.359 | 2.67E+09 | 55 | 27.2822 | 27.3872 | 27.6788 | 27.7181 | 27.598 |
| 0 | 65.593 | 1.3E+09 | 47 | 26.5634 | 26.6787 | 26.5494 | 26.6373 | 26.3636 |
| 0 | 323.31 | 7.27E+10 | 401 | 32.1287 | 32.1649 | 32.5226 | 32.4437 | 32.3456 |
| 0 | 48.556 | 1.75E+10 | 70 | 30.2471 | 30.3368 | 30.447 | 30.2746 | 30.1769 |
| 0 | 14.036 | 6.94E+08 | 43 | 25.764 | 25.806 | 25.6326 | 25.4703 | 25.5387 |
| 0 | 125.56 | 2.56E+09 | 68 | 27.7443 | 27.4871 | 27.6944 | 27.6611 | 27.2686 |
| 0 | 46.144 | 1.49E+09 | 27 | 26.4921 | 26.5043 | 26.7891 | 26.6037 | 27.0811 |
| 0 | 196.86 | 5.26E+09 | 77 | 28.7955 | 28.6663 | 28.5842 | 28.4311 | 28.5439 |
| 0 | 153.71 | 2.83E+10 | 165 | 31.0592 | 31.0022 | 30.9116 | 31.0831 | 30.9644 |
| 0 | 18.222 | 8.91E+08 | 43 | 25.9733 | 25.8808 | 26.1284 | 26.1285 | 26.1261 |
| 0 | 17.638 | 9.06E+08 | 33 | 26.0313 | 26.1798 | 26.0253 | 25.6097 | 26.078 |
| 0 | 110.19 | 5.32E+09 | 111 | 28.45 | 28.4281 | 28.5984 | 28.5425 | 28.7198 |
| 0 | 50.902 | 5.08E+09 | 56 | 28.4795 | 28.4766 | 28.6826 | 28.6376 | 28.427 |
| 0 | 16.784 | 9.15E+08 | 44 | 26.154 | 26.0934 | 26.0713 | 26.2299 | 26.0224 |
| 0 | 40.963 | 2.9E+09 | 67 | 27.972 | 27.9966 | 27.5894 | 27.4694 | 27.6649 |
| 0 | 24.681 | 4.57E+09 | 71 | 28.3242 | 28.2724 | 28.3211 | 28.4066 | 28.4288 |
| 0 | 6.8171 | 8.2E+08 | 25 | 25.4339 | 25.4552 | 25.6495 | 25.6937 | 25.5227 |
| 0 | 11.029 | 2.58E+09 | 52 | 27.3397 | 27.3348 | 27.2232 | 27.1693 | 27.2266 |
| 0 | 53.484 | 6.41E+09 | 89 | 28.7693 | 28.7203 | 28.655 | 28.6263 | 28.4235 |
| 0 | 7.0554 | 3.77E+08 | 13 | 24.483 | 24.679 | 24.4203 | 24.3907 | 24.6533 |
| 0 | 28.532 | 8.74E+08 | 44 | 25.6294 | 25.7332 | 25.5867 | 25.8032 | 25.9021 |
| 0 | 68.061 | 1.21E+10 | 94 | 29.6654 | 29.6302 | 29.6319 | 29.6226 | 29.4797 |

| | | | | | | | | |
|---|---|---|---|---|---|---|---|---|
| 0 | 32.51 | 1.25E+09 | 29 | 26.0489 | 26.1942 | 26.1713 | 26.1899 | 26.0453 |
| 0 | 197.54 | 3.79E+09 | 94 | 28.0919 | 27.9143 | 28.0452 | 28.1081 | 27.7437 |
| 0 | 23.924 | 2.88E+09 | 78 | 27.4014 | 27.4253 | 27.6964 | 27.4584 | 27.4296 |
| 0 | 26.251 | 9.24E+09 | 72 | 29.2447 | 29.284 | 29.4148 | 29.1917 | 29.2297 |
| 0 | 7.213 | 8.79E+08 | 32 | 25.6506 | 26.0263 | 26.0116 | 26.0071 | 25.9364 |
| 0 | 12.778 | 8.12E+08 | 50 | 25.684 | 25.878 | 25.801 | 25.7403 | 25.7063 |
| 0 | 176.16 | 1.75E+10 | 126 | 30.3108 | 30.2644 | 30.169 | 30.0784 | 30.2425 |
| 0 | 60.438 | 3.76E+09 | 57 | 28.0217 | 27.9906 | 27.9173 | 28.0451 | 27.8642 |
| 0 | 92.617 | 2.6E+09 | 88 | 27.5196 | 27.4858 | 27.3157 | 27.3616 | 27.4631 |
| 0 | 104.73 | 4.61E+09 | 54 | 28.4392 | 28.3794 | 28.2115 | 28.1147 | 28.1683 |
| 0 | 76.78 | 2.31E+10 | 136 | 30.6732 | 30.6624 | 30.4993 | 30.7026 | 30.4239 |
| 0 | 20.447 | 5.06E+08 | 14 | 25.1033 | 25.205 | 25.2314 | 25.0843 | 25.2195 |
| 0 | 30.111 | 3.57E+09 | 52 | 27.8391 | 27.8037 | 28.0216 | 28.0993 | 27.8168 |
| 0 | 169.43 | 1.59E+09 | 43 | 26.7402 | 26.8247 | 26.5627 | 26.7033 | 26.7281 |
| 0 | 323.31 | 7.3E+10 | 571 | 32.1813 | 32.1561 | 32.2569 | 32.2985 | 32.3524 |
| 0 | 15.337 | 5.56E+08 | 39 | 25.0903 | 25.0091 | 25.2028 | 25.3659 | 25.4844 |
| 0 | 12.494 | 1.51E+09 | 44 | 26.6267 | 26.7283 | 26.6812 | 26.7313 | 26.7082 |
| 0 | 323.31 | 4.59E+09 | 146 | 28.294 | 28.1974 | 28.1582 | 28.2833 | 28.3049 |
| 0 | 23.652 | 6.29E+09 | 79 | 28.7731 | 28.8484 | 28.7253 | 28.7044 | 28.6114 |
| 0 | 16.524 | 7.56E+08 | 41 | 25.7686 | 25.656 | 25.7883 | 25.7901 | 25.6744 |
| 0 | 19.947 | 1.56E+10 | 134 | 30.015 | 29.8826 | 29.9621 | 30.0464 | 29.9877 |
| 0 | 263.55 | 4.72E+10 | 391 | 31.6497 | 31.5672 | 31.6742 | 31.7559 | 31.6121 |
| 0 | 14.011 | 6.97E+08 | 33 | 25.5664 | 25.622 | 25.6366 | 25.7456 | 25.4383 |
| 0 | 59.231 | 3.99E+09 | 76 | 28.2039 | 28.2131 | 28.3134 | 28.3025 | 28.2594 |
| 0 | 66.517 | 1.71E+09 | 49 | 27.1056 | 26.9062 | 27.3024 | 27.0047 | 26.9539 |
| 0 | 52.463 | 2.33E+09 | 61 | 27.4158 | 27.4626 | 27.4625 | 27.5638 | 27.6304 |
| 0 | 12.679 | 7.6E+08 | 44 | 26.0267 | 26.0134 | 25.8116 | 25.7463 | 25.8489 |
| 0 | 13.29 | 7.74E+08 | 37 | 25.919 | 26.1525 | 25.5444 | 25.9474 | 26.0599 |
| 0 | 80.657 | 4.11E+09 | 72 | 28.5851 | 28.3769 | 28.3151 | 28.1305 | 28.2311 |
| 0 | 6.7634 | 6.2E+08 | 18 | 25.7428 | 25.5236 | 25.4465 | 25.4735 | 25.5347 |
| 0 | 10.397 | 4.71E+08 | 19 | 25.4778 | 25.5419 | 24.9649 | 25.0608 | 25.3244 |
| 0 | 15.316 | 1.4E+09 | 40 | 26.8916 | 26.9523 | 26.646 | 26.7351 | 26.6237 |
| 0 | 10.458 | 4.38E+08 | 18 | 25.287 | 25.1966 | 24.9261 | 25.017 | 24.9417 |
| 0 | 35.973 | 8.83E+08 | 36 | 25.7952 | 25.8091 | 26.0725 | 25.7046 | 25.8815 |
| 0 | 62.587 | 2.72E+09 | 60 | 27.5883 | 27.5559 | 27.4727 | 27.6078 | 27.4122 |
| 0 | 13.814 | 5.61E+09 | 54 | 28.6917 | 28.6815 | 28.7612 | 28.8407 | 28.7001 |
| 0 | 22.608 | 1.19E+09 | 44 | 26.4445 | 26.3977 | 26.4098 | 26.3008 | 26.25 |
| 0 | 23.48 | 1.74E+09 | 47 | 26.9663 | 27.1224 | 27.2028 | 27.162 | 26.9519 |
| 0 | 270 | 8.58E+09 | 78 | 29.1664 | 29.1282 | 29.2364 | 29.202 | 29.2191 |
| 0 | 6.4523 | 4.51E+08 | 23 | 25.5108 | 25.1495 | 24.9157 | 25.4552 | 24.9845 |
| 0 | 51.427 | 4.61E+10 | 180 | 31.6515 | 31.6177 | 31.6433 | 31.6685 | 31.6439 |
| 0 | 11.486 | 2.95E+08 | 10 | NaN | NaN | NaN | NaN | 24.3237 |
| 0 | 11.983 | 6.83E+08 | 36 | 25.696 | 25.6633 | 25.6411 | 25.5589 | 25.8005 |
| 0 | 19.178 | 6.69E+08 | 35 | 25.3287 | 25.3302 | 25.5338 | 25.5986 | 25.4438 |
| 0 | 12.533 | 1.19E+09 | 35 | 26.512 | 26.4278 | 26.5375 | 26.5075 | 26.4934 |
| 0 | 13.058 | 3.58E+09 | 52 | 28.5412 | 28.4262 | 28.0955 | 27.9184 | 27.8764 |
| 0 | 166.28 | 4.64E+10 | 131 | 31.7707 | 31.9967 | 31.7007 | 31.804 | 31.7296 |
| 0 | 6.4051 | 2.96E+08 | 23 | 23.8387 | 24.1204 | 23.884 | 24.3685 | 24.3873 |

| | | | | | | | | |
|---|---|---|---|---|---|---|---|---|
| 0 | 16.295 | 2.23E+09 | 41 | 27.2643 | 27.342 | 27.183 | 27.2512 | 27.259 |
| 0 | 24.247 | 4.65E+08 | 24 | 25.1798 | 25.3124 | 24.9317 | 24.9953 | 25.2511 |
| 0 | 20.934 | 2.32E+09 | 44 | 27.6164 | 27.4791 | 27.4362 | 27.4597 | 27.2312 |
| 0 | 7.8779 | 4.6E+08 | 12 | 24.9776 | 25.2162 | 25.1 | 25.4336 | 25.1692 |
| 0 | 276.07 | 6.62E+09 | 88 | 28.8545 | 28.7696 | 28.7733 | 28.9417 | 28.7842 |
| 0 | 8.8283 | 1.68E+09 | 26 | 26.8065 | 26.7182 | 27.0019 | 26.9478 | 26.8778 |
| 0 | 5.3487 | 3.1E+08 | 14 | 24.7301 | 24.7224 | 25.074 | 24.8039 | NaN |
| 0 | 3.7249 | 6.31E+08 | 9 | 25.3521 | 25.5117 | 25.2523 | 25.3465 | 25.7361 |
| 0 | 10.1 | 6.56E+08 | 17 | 25.6655 | 25.494 | 25.4996 | 25.678 | 25.1662 |
| 0 | 18 | 8.69E+08 | 26 | 25.7218 | 25.9098 | 26.1293 | 25.9662 | 25.8636 |
| 0 | 8.5065 | 7.6E+08 | 21 | 25.7235 | 25.8804 | 25.6434 | 25.6194 | 25.7133 |
| 0 | 77.85 | 5.2E+09 | 64 | 28.5497 | 28.5731 | 28.4651 | 28.5506 | 28.8859 |
| 0 | 16.452 | 8.65E+08 | 33 | 25.7508 | 25.8658 | 25.9968 | 25.8178 | 25.9877 |
| 0 | 25.04 | 1.18E+09 | 48 | 26.6449 | 26.4939 | 26.5393 | 26.4894 | 26.5135 |
| 0 | 12.665 | 6.12E+08 | 26 | 25.695 | 25.9863 | 25.3573 | 25.6638 | 25.368 |
| 0 | 15.919 | 8.02E+08 | 29 | 25.6525 | 25.8544 | 25.7895 | 25.7799 | 25.8099 |
| 0 | 16.556 | 1.16E+09 | 54 | 26.194 | 26.034 | 26.6984 | 26.2961 | 26.168 |
| 0 | 5.5191 | 6.11E+08 | 30 | 25.5612 | 25.2907 | 25.6 | 25.368 | 25.0583 |
| 0 | 5.6081 | 4.1E+08 | 7 | NaN | NaN | 24.7753 | 24.8774 | 24.9887 |
| 0 | 153.12 | 2.87E+09 | 60 | 27.8209 | 27.6519 | 27.6454 | 27.3144 | 27.7197 |
| 0 | 6.7459 | 2.61E+08 | 16 | 24.3665 | 23.867 | 24.0838 | 24.1858 | 24.0609 |
| 0 | 89.258 | 7.73E+09 | 86 | 29.2668 | 29.365 | 29.3011 | 29.3366 | 29.0688 |
| 0 | 3.2767 | 3.74E+09 | 24 | 27.8395 | 28.1161 | 28.129 | 27.9035 | 27.9693 |
| 0 | 43.62 | 5.48E+08 | 33 | 25.8729 | 25.8474 | 25.0747 | 25.1467 | 25.3662 |
| 0 | 110.75 | 7.51E+09 | 68 | 29.1708 | 29.1693 | 29.0492 | 29.2556 | 29.0466 |
| 0 | 183.22 | 1.12E+10 | 81 | 29.4542 | 29.5839 | 29.7288 | 29.6239 | 29.5327 |
| 0 | 81.009 | 1.19E+09 | 34 | 26.2354 | 26.278 | 26.6264 | 26.3304 | 26.3141 |
| 0 | 17.676 | 6.05E+08 | 24 | 25.2479 | 25.6918 | 25.4838 | 25.3499 | 25.3985 |
| 0 | 5.8587 | 4.57E+08 | 12 | 25.103 | 24.8241 | 24.8759 | 25.2214 | 24.926 |
| 0 | 21.35 | 4.85E+08 | 12 | 25.1904 | 25.1009 | 25.1246 | 25.0587 | 25.1463 |
| 0 | 29.716 | 5.78E+08 | 23 | 25.1661 | 25.3461 | 25.3784 | 25.7389 | 25.674 |
| 0 | 20.363 | 4.27E+08 | 25 | 24.8777 | 25.0235 | 25.1433 | 25.1383 | 25.056 |
| 0 | 21.886 | 1.26E+09 | 40 | 26.4167 | 26.501 | 26.3285 | 26.3107 | 26.5204 |
| 0 | 35.52 | 8.12E+08 | 37 | 25.7208 | 25.8784 | 25.7443 | 25.83 | 25.9063 |
| 0 | 41.136 | 1.57E+09 | 66 | 26.7411 | 26.6477 | 26.7659 | 26.8103 | 26.7882 |
| 0 | 9.1046 | 5.07E+08 | 17 | 25.0135 | 25.1654 | 25.52 | 25.3149 | 25.295 |
| 0 | 22.739 | 7.21E+08 | 22 | 25.9876 | 25.9292 | 25.7914 | 25.7759 | 25.7035 |
| 0 | 24.993 | 8.91E+08 | 41 | 26.166 | 26.0451 | 25.6415 | 25.7169 | 25.7462 |
| 0 | 23.849 | 2.16E+09 | 34 | 27.7555 | 27.6932 | 27.1865 | 27.1145 | 27.3813 |
| 0 | 45.158 | 2.51E+09 | 85 | 27.5185 | 27.4904 | 27.3813 | 27.3767 | 27.6701 |
| 0 | 30.077 | 5.75E+08 | 20 | 25.6822 | 25.6555 | 25.4564 | 25.2494 | 25.2399 |
| 0 | 56.48 | 1.1E+09 | 36 | 26.9206 | 26.7938 | 26.4824 | 26.2951 | 26.0614 |
| 0 | 20.179 | 5.44E+08 | 29 | 24.9908 | 25.3076 | 25.2228 | 25.4151 | 25.2694 |
| 0 | 11.715 | 5.89E+08 | 27 | 25.3837 | 25.3513 | 25.5463 | 25.3293 | 25.5565 |
| 0 | 11.333 | 4.95E+08 | 21 | 25.1914 | 25.2175 | 25.0462 | 25.0835 | 25.4186 |
| 0 | 25.289 | 1.64E+10 | 81 | 30.0691 | 30.1072 | 30.3838 | 30.218 | 30.0264 |
| 0 | 167.4 | 5.09E+09 | 107 | 28.5916 | 28.5568 | 28.57 | 28.5001 | 28.211 |
| 0 | 10.7 | 1.24E+09 | 39 | 26.6792 | 26.8709 | 26.4198 | 26.4488 | 26.2956 |

| | | | | | | | | |
|---|---|---|---|---|---|---|---|---|
| 0 | 7.5632 | 3.25E+08 | 15 | 24.548 | 24.4526 | 24.7418 | NaN | NaN |
| 0 | 10.313 | 5.15E+08 | 18 | 25.1006 | 25.3554 | 25.0672 | 25.2284 | 25.1826 |
| 0 | 20.926 | 1.42E+09 | 45 | 26.7685 | 26.6931 | 26.8018 | 26.7301 | 26.6369 |
| 0 | 107.76 | 3.79E+09 | 67 | 28.2171 | 28.2028 | 28.0664 | 27.9814 | 28.1306 |
| 0 | 8.9677 | 5.23E+08 | 16 | 25.0527 | 24.9511 | 25.1035 | 25.5119 | 25.3398 |
| 0 | 22.687 | 6.51E+08 | 10 | 25.8769 | 25.7907 | 25.7164 | 25.5755 | 25.564 |
| 0 | 6.6396 | 5.6E+08 | 30 | 25.1911 | 25.2678 | 25.4954 | 25.3326 | 25.5036 |
| 0 | 7.9308 | 4.9E+08 | 27 | 25.0306 | 24.8295 | 25.3711 | 25.2383 | 25.268 |
| 0 | 7.8379 | 5.46E+08 | 17 | 25.3025 | 25.2358 | 25.4464 | 25.238 | 25.409 |
| 0 | 32.866 | 1.09E+09 | 29 | 26.6064 | 26.6565 | 26.501 | 26.5385 | 26.2458 |
| 0 | 9.2666 | 3.49E+08 | 12 | 24.5335 | 24.4762 | 24.435 | NaN | 24.7907 |
| 0 | 13.388 | 1.05E+09 | 33 | 26.1223 | 26.076 | 26.2249 | 26.2905 | 25.9612 |
| 0 | 17.937 | 1.08E+09 | 50 | 26.7829 | 26.862 | 26.3003 | 26.0339 | 26.1117 |
| 0 | 144.04 | 2.48E+09 | 65 | 27.4572 | 27.4549 | 27.4826 | 27.4867 | 27.4651 |
| 0 | 8.4702 | 2.56E+08 | 11 | 24.0481 | 23.7475 | 23.968 | 23.8939 | 24.3833 |
| 0 | 23.989 | 3.48E+09 | 61 | 28.1337 | 28.1897 | 27.8433 | 28.0157 | 27.9651 |
| 0 | 99.489 | 1.03E+09 | 47 | 26.2956 | 26.2757 | 26.3818 | 26.1818 | 26.2443 |
| 0 | 9.1617 | 5.09E+08 | 8 | 24.866 | NaN | 24.6461 | NaN | NaN |
| 0 | 13.044 | 1.14E+09 | 12 | 26.4777 | NaN | 27.1162 | 26.6553 | NaN |
| 0 | 183.56 | 6.53E+10 | 153 | 32.2316 | 32.1557 | 32.2835 | 32.1519 | 32.129 |
| 0 | 9.9555 | 5.38E+08 | 11 | 25.4592 | NaN | NaN | NaN | 25.3099 |
| 0 | 8.7413 | 6.52E+08 | 14 | 25.447 | 25.3992 | 25.4796 | 25.5239 | 25.4767 |
| 0 | 3.6238 | 5.98E+09 | 10 | NaN | NaN | 28.8057 | NaN | 29.1718 |
| 0 | 9.112 | 6.32E+08 | 16 | NaN | 25.6238 | 25.5448 | 26.0517 | NaN |
| 0 | 8.3288 | 6E+08 | 23 | 25.3171 | NaN | 25.5512 | 25.4702 | 25.4902 |
| 0 | 14.842 | 1.72E+09 | 52 | 27.1108 | 26.7681 | 27.0225 | 27.1342 | 27.0517 |
| 0 | 13.418 | 8.03E+08 | 19 | 25.8564 | 25.87 | 25.9725 | 25.9235 | 26.0438 |
| 0 | 62.219 | 9.45E+08 | 45 | 26.1695 | 25.8843 | 26.0028 | 26.061 | 25.8265 |
| 0 | 9.7195 | 7.29E+08 | 27 | 25.9883 | 26.2225 | 25.718 | 25.4429 | 25.8289 |
| 0 | 41.697 | 2.08E+09 | 45 | 27.2103 | 27.3301 | 27.1484 | 27.3082 | 27.1249 |
| 0 | 23.975 | 8.56E+08 | 53 | 25.7303 | 25.785 | 25.9376 | 25.9422 | 25.9711 |
| 0 | 42.182 | 1.49E+09 | 78 | 26.7777 | 26.8978 | 26.7423 | 26.72 | 26.5285 |
| 0 | 22.034 | 1E+09 | 42 | 26.4431 | 26.4768 | 26.1642 | 26.1136 | 26.051 |
| 0 | 68.126 | 1.1E+09 | 36 | 26.6162 | 26.4955 | 26.4731 | 26.1004 | 26.1178 |
| 0 | 30.942 | 8.62E+08 | 25 | 26.3245 | 26.0005 | 25.62 | 25.9567 | 26.0903 |
| 0 | 13.395 | 3.59E+09 | 77 | 28.1712 | 28.0907 | 28.1481 | 28.1551 | 27.8576 |
| 0 | 86.749 | 5.53E+09 | 124 | 28.3074 | 28.4184 | 28.653 | 28.6743 | 28.6396 |
| 0 | 24.086 | 3.38E+08 | 14 | 25.5382 | 25.3387 | NaN | NaN | 24.6875 |
| 0 | 11.398 | 9.77E+08 | 31 | 25.97 | 26.098 | 25.8678 | 25.9363 | 26.1756 |
| 0 | 166 | 1.45E+10 | 244 | 29.7537 | 29.8441 | 29.9904 | 30.0435 | 30.0628 |
| 0 | 7.4475 | 4.16E+08 | 27 | 24.7729 | 24.7425 | 24.9198 | 24.9148 | 24.903 |
| 0 | 15.066 | 6.1E+08 | 32 | 25.0738 | 25.0514 | 25.7825 | 25.3065 | 25.5292 |
| 0 | 17.986 | 1.28E+09 | 37 | 26.6084 | 26.5542 | 26.4176 | 26.4267 | 26.507 |
| 0 | 9.3128 | 6.38E+08 | 35 | 25.9152 | 25.8836 | 25.3495 | 25.3943 | 25.467 |
| 0 | 12.714 | 5.45E+08 | 14 | 25.2058 | NaN | 25.1812 | 25.4998 | 25.6531 |
| 0 | 11.089 | 7.85E+09 | 50 | 29.072 | 29.0537 | 28.8985 | 29.2177 | 29.1566 |
| 0 | 35.238 | 3.35E+09 | 67 | 27.8611 | 27.8826 | 27.714 | 27.871 | 27.7554 |
| 0 | 28.131 | 9.48E+08 | 43 | 25.8271 | 26.0272 | 26.0527 | 26.1287 | 26.035 |

| | | | | | | | | |
|---|---|---|---|---|---|---|---|---|
| 0 | 12.512 | 7.6E+08 | 21 | 25.4266 | 25.5123 | 25.8563 | 25.9131 | 25.7104 |
| 0 | 67.607 | 1.97E+09 | 48 | 27.1356 | 27.1394 | 27.0971 | 27.3028 | 27.1157 |
| 0 | 7.9581 | 5.41E+08 | 40 | 25.2862 | 25.3457 | 25.2957 | 25.1115 | 25.3701 |
| 0 | 18.883 | 1.12E+09 | 27 | 26.2701 | 26.2877 | 26.3042 | 26.1238 | 26.7148 |
| 0 | 17.515 | 3.2E+09 | 71 | 27.6887 | 27.6255 | 27.9197 | 27.7585 | 27.6994 |
| 0 | 7.5523 | 8.67E+08 | 11 | 26.0595 | 26.1955 | 25.8706 | 26.1657 | 26.1382 |
| 0 | 63.594 | 1.39E+09 | 54 | 27.1453 | 26.9964 | 26.5693 | 26.5707 | 26.6776 |
| 0 | 14.481 | 7.06E+08 | 21 | 25.6463 | 25.9275 | 25.6607 | 25.4675 | 25.7066 |
| 0 | 19.585 | 4.1E+09 | 67 | 28.089 | 28.1138 | 28.2774 | 28.1619 | 28.2148 |
| 0 | 15.574 | 1.15E+09 | 28 | 26.5495 | 26.4092 | 26.4373 | 26.3487 | 26.1887 |
| 0 | 16.177 | 4.51E+08 | 11 | 24.8761 | 25.1387 | 25.0683 | 25.1469 | 25.2856 |
| 0 | 37.264 | 1.56E+09 | 53 | 26.75 | 26.8394 | 26.6579 | 26.5686 | 26.87 |
| 0 | 16.128 | 6.79E+08 | 29 | 25.5685 | 25.4843 | 25.467 | 25.1499 | 25.6835 |
| 0 | 66.301 | 1.56E+09 | 42 | 26.7318 | 26.5888 | 26.9284 | 26.7594 | 26.7339 |
| 0 | 28.376 | 6.59E+08 | 17 | 25.8553 | 26.2154 | 25.6768 | 25.2074 | 25.7323 |
| 0 | 304.57 | 8.27E+09 | 89 | 29.2794 | 29.315 | 29.2839 | 29.2348 | 29.3559 |
| 0 | 19.238 | 8.46E+08 | 38 | 25.6768 | 25.5119 | 25.988 | 25.7856 | 26.016 |
| 0 | 214.01 | 2.09E+09 | 55 | 26.9985 | 27.2185 | 27.1978 | 27.3103 | 27.2546 |
| 0 | 7.3046 | 2.99E+08 | 14 | 24.4376 | 24.3097 | 24.3727 | 24.3223 | 24.6745 |
| 0 | 22.494 | 7.14E+09 | 113 | 28.9435 | 28.8718 | 28.7897 | 28.9914 | 28.7891 |
| 0 | 29.23 | 3.28E+09 | 57 | 27.8673 | 27.8277 | 27.945 | 27.8499 | 27.811 |
| 0 | 8.0493 | 4.83E+08 | 32 | 25.0896 | 25.0105 | 25.3811 | 25.1899 | 24.8573 |
| 0 | 11.726 | 1E+09 | 32 | 25.9068 | 26.0951 | 26.1841 | 26.2173 | 26.2567 |
| 0 | 59.165 | 1.6E+09 | 53 | 26.8255 | 26.7729 | 26.9546 | 26.6092 | 26.8571 |
| 0 | 6.1317 | 4.06E+08 | 6 | 24.6677 | 24.6853 | 24.7804 | 24.7218 | 25.0431 |
| 0 | 46.996 | 2.35E+09 | 43 | 27.3906 | 27.3449 | 27.5089 | 27.3755 | 27.3187 |
| 0 | 25.634 | 9.89E+08 | 34 | 26.8121 | 26.6811 | 26.1859 | 26.2465 | 25.6785 |
| 0 | 323.31 | 3.69E+10 | 97 | 31.701 | 31.7889 | 31.4155 | 31.4401 | 31.0164 |
| 0 | 10.466 | 8.24E+08 | 11 | 25.9289 | 25.9328 | 25.8287 | NaN | 25.9026 |
| 0 | 47.653 | 5.01E+09 | 55 | 28.308 | 28.3436 | 28.5285 | 28.4597 | 28.4622 |
| 0 | 21.257 | 1.72E+09 | 55 | 27.0032 | 27.0884 | 27.0719 | 26.9002 | 26.8399 |
| 0 | 14.429 | 1.28E+09 | 24 | 26.4713 | 26.6091 | 26.5088 | 26.6968 | 26.3932 |
| 0 | 28.41 | 5.39E+08 | 31 | 25.3841 | 25.2862 | 25.1702 | 25.2487 | 25.4027 |
| 0 | 131.85 | 1.57E+10 | 88 | 30.2129 | 30.1278 | 30.2512 | 30.0764 | 30.0738 |
| 0 | 53.702 | 1.96E+10 | 180 | 30.4947 | 30.4622 | 30.5534 | 30.5628 | 30.2692 |
| 0 | 8.2122 | 4.48E+08 | 8 | 25.2278 | 25.3575 | NaN | 24.9687 | 25.1043 |
| 0 | 28.162 | 3.9E+09 | 32 | 28.1079 | 28.4263 | 27.9408 | 28.1898 | 28.1863 |
| 0 | 18.441 | 5.29E+08 | 30 | 24.9387 | 25.2494 | 25.5688 | 25.1104 | 25.1699 |
| 0 | 173.45 | 1.68E+10 | 103 | 30.2889 | 30.2723 | 30.289 | 30.2519 | 30.1939 |
| 0 | 14.461 | 1.01E+09 | 51 | 26.1882 | 26.029 | 26.2171 | 26.3736 | 26.0492 |
| 0 | 4.7019 | 2.92E+08 | 13 | 24.231 | 24.3955 | 24.0217 | 24.5539 | 24.5097 |
| 0 | 9.9694 | 1.06E+09 | 33 | 26.3673 | 26.2169 | 26.4035 | 26.329 | 26.1181 |
| 0 | 4.3445 | 5.01E+08 | 15 | 25.2772 | 25.2972 | 24.9988 | 25.4667 | 25.2522 |
| 0 | 10.993 | 9.25E+08 | 44 | 26.0997 | 25.9871 | 25.7585 | 26.0986 | 26.3018 |
| 0 | 21.145 | 3.16E+09 | 34 | 27.8254 | 27.7094 | 27.979 | 27.5955 | 27.656 |
| 0 | 12.071 | 9.47E+08 | 33 | 26.0145 | 25.7364 | 25.9163 | 25.7486 | 26.3918 |
| 0 | 46.796 | 1.43E+09 | 61 | 26.6003 | 26.6798 | 26.7274 | 26.7575 | 26.5599 |
| 0 | 15.606 | 2.74E+09 | 34 | 27.567 | 27.7075 | 27.6115 | 27.5717 | 27.49 |

| | | | | | | | | |
|---|---|---|---|---|---|---|---|---|
| 0 | 102.44 | 3.31E+09 | 79 | 28.0243 | 28.0083 | 27.9825 | 27.8723 | 27.7389 |
| 0 | 62.765 | 1.93E+10 | 105 | 30.3705 | 30.3977 | 30.6431 | 30.3474 | 30.5369 |
| 0 | 96.599 | 8.65E+09 | 96 | 29.6447 | 29.6039 | 29.2283 | 29.1897 | 29.0525 |
| 0 | 16.47 | 7.49E+08 | 53 | 25.8209 | 25.8201 | 25.8418 | 25.8276 | 25.8322 |
| 0 | 18.904 | 7.91E+09 | 64 | 29.1836 | 29.0927 | 29.3046 | 29.1794 | 29.1012 |
| 0 | 21.779 | 9.86E+08 | 40 | 26.0606 | 26.1987 | 26.1217 | 26.0441 | 26.073 |
| 0 | 8.0176 | 6.21E+08 | 42 | 25.2833 | 25.3435 | 25.6878 | 25.3702 | 25.5775 |
| 0 | 22.218 | 4.95E+08 | 46 | 25.3521 | 25.1844 | 25.0413 | 25.1489 | 25.2281 |
| 0 | 50.285 | 1.49E+09 | 56 | 27.0912 | 26.8451 | 26.6276 | 26.6197 | 26.6469 |
| 0 | 4.8402 | 3.25E+08 | 24 | 24.4378 | 24.6775 | 24.6073 | 24.462 | 24.1013 |
| 0 | 7.4311 | 5.01E+08 | 27 | 25.1426 | 24.9422 | 24.7941 | 25.1189 | 25.3953 |
| 0 | 142.36 | 3.02E+09 | 66 | 27.578 | 27.6091 | 27.7762 | 27.9873 | 27.7462 |
| 0 | 35.207 | 1.46E+09 | 37 | 26.6598 | 26.808 | 26.9478 | 26.8327 | 26.6683 |
| 0 | 8.1368 | 8.03E+08 | 21 | 25.7713 | 25.8214 | 25.8332 | 25.9101 | 25.8318 |
| 0 | 17.09 | 5.81E+08 | 31 | 25.4267 | 25.5152 | 25.3888 | 25.6834 | 25.3314 |
| 0 | 19.504 | 4.96E+08 | 32 | 25.0978 | 25.2685 | 25.1555 | 25.354 | 25.2045 |
| 0 | 9.761 | 9.62E+08 | 47 | 26.165 | 26.1642 | 25.964 | 26.1822 | 26.0196 |
| 0 | 57.86 | 3.16E+10 | 137 | 31.2277 | 31.234 | 31.2009 | 31.2657 | 31.0405 |
| 0 | 34.38 | 6.05E+08 | 16 | 25.602 | 25.5009 | 25.727 | 25.5547 | 25.1675 |
| 0 | 24.732 | 1.85E+09 | 56 | 27.3157 | 27.3259 | 26.7515 | 26.9192 | 26.9812 |
| 0 | 14.511 | 5.14E+08 | 42 | 25.1073 | 24.8199 | 25.239 | 25.218 | 25.4235 |
| 0 | 275.15 | 8.12E+09 | 102 | 29.3663 | 29.3674 | 28.9973 | 29.0304 | 29.1235 |
| 0 | 82.332 | 3.59E+09 | 55 | 28.0844 | 28.014 | 27.7879 | 27.8657 | 28.229 |
| 0 | 10.313 | 7.05E+08 | 30 | 25.7139 | 25.7996 | 25.768 | 25.7109 | 25.7757 |
| 0 | 28.412 | 1.91E+09 | 48 | 27.0043 | 27.1864 | 27.1414 | 26.8775 | 27.1256 |
| 0 | 13.021 | 3.3E+08 | 16 | 24.6375 | 24.3809 | 24.3847 | 24.4111 | 24.6707 |
| 0 | 136.11 | 1.81E+10 | 125 | 30.2463 | 30.24 | 30.3628 | 30.4101 | 30.4085 |
| 0 | 323.31 | 2.84E+10 | 204 | 31.1474 | 31.1269 | 30.9244 | 30.9033 | 30.9898 |
| 0 | 12.327 | 1.02E+09 | 54 | 26.2474 | 26.409 | 26.3367 | 26.4565 | 26.1523 |
| 0 | 67.511 | 8.03E+08 | 18 | 25.8237 | 25.8433 | 25.7225 | 25.7328 | 25.9032 |
| 0 | 10.823 | 6.47E+08 | 41 | 25.171 | 25.3582 | 25.6456 | 25.6334 | 25.5737 |
| 0 | 21.795 | 2.13E+09 | 62 | 27.485 | 27.2848 | 27.3798 | 27.361 | 27.0833 |
| 0 | 59.057 | 2.64E+09 | 46 | 27.9976 | 27.965 | 27.5795 | 27.5431 | 27.0897 |
| 0 | 19.714 | 6.99E+08 | 20 | 25.6532 | 25.7155 | 25.4848 | 25.6426 | 25.9245 |
| 0 | 83.581 | 1.43E+09 | 58 | 26.6176 | 26.6831 | 26.7888 | 26.7123 | 26.7234 |
| 0 | 9.3139 | 6.05E+08 | 16 | 25.6519 | 25.6635 | 25.3334 | 25.366 | 25.5209 |
| 0 | 16.719 | 2.2E+09 | 41 | 27.2965 | 27.2341 | 27.2974 | 27.3163 | 27.4036 |
| 0 | 175.88 | 5.81E+09 | 71 | 28.6142 | 28.6177 | 28.6977 | 28.8389 | 28.7117 |
| 0 | 20.316 | 2.24E+09 | 48 | 27.1603 | 27.3048 | 27.1994 | 27.3233 | 27.5466 |
| 0 | 30.267 | 2.7E+09 | 77 | 27.6257 | 27.3707 | 27.8514 | 27.485 | 27.1762 |
| 0 | 78.493 | 3.76E+09 | 68 | 27.751 | 27.902 | 28.0902 | 28.1502 | 28.219 |
| 0 | 34.294 | 8.91E+08 | 23 | 25.9144 | 25.9199 | 26.009 | 25.9548 | 25.9446 |
| 0 | 11.401 | 5.01E+08 | 37 | 25.1649 | 25.0571 | 25.3308 | 25.1673 | 25.0611 |
| 0 | 19.126 | 2.32E+09 | 63 | 27.2938 | 27.3661 | 27.4941 | 27.3441 | 27.2849 |
| 0 | 108.26 | 2.99E+10 | 278 | 31.2668 | 31.3066 | 31.0569 | 30.9824 | 30.9207 |
| 0 | 134.4 | 1.5E+10 | 128 | 30.3448 | 30.3687 | 29.919 | 29.8672 | 29.8607 |
| 0 | 22.3 | 9.39E+08 | 31 | 26.3204 | 26.2232 | 25.6957 | 26.3313 | 25.826 |
| 0 | 35.986 | 1.54E+09 | 66 | 26.7046 | 26.7563 | 26.889 | 26.7905 | 26.8443 |

| | | | | | | | | |
|---|---|---|---|---|---|---|---|---|
| 0 | 5.4311 | 1.13E+09 | 37 | 26.3084 | 26.2625 | 26.3258 | 26.4595 | 26.3065 |
| 0 | 20.649 | 6.39E+08 | 28 | 25.655 | 25.4493 | 25.7148 | 25.5561 | 25.3905 |
| 0 | 38.078 | 6.26E+09 | 86 | 28.8034 | 28.735 | 28.7953 | 28.8916 | 28.7613 |
| 0 | 20.34 | 2.48E+09 | 40 | 27.5088 | 27.2962 | 27.5209 | 27.4796 | 27.5067 |
| 0 | 55.661 | 7.34E+08 | 48 | 25.7914 | 25.8165 | 25.7195 | 25.7064 | 25.6019 |
| 0 | 107.04 | 9.84E+09 | 50 | 29.5885 | 29.6303 | 29.4906 | 29.4608 | 29.4044 |
| 0 | 307.53 | 3.01E+10 | 279 | 31.0415 | 30.991 | 30.9956 | 31.0776 | 31.1714 |
| 0 | 135.28 | 8.61E+08 | 44 | 26.0879 | 25.9087 | 26.0434 | 25.9458 | 26.1061 |
| 0 | 33.305 | 2.56E+09 | 52 | 27.4812 | 27.513 | 27.4431 | 27.4579 | 27.5269 |
| 0 | 13.075 | 2.79E+08 | 30 | 23.8502 | 24.2701 | 24.2502 | 24.4895 | 24.5749 |
| 0 | 5.5791 | 2.82E+09 | 44 | 27.4886 | 27.5026 | 27.7818 | 27.5795 | 27.6623 |
| 0 | 7.9253 | 9.9E+08 | 24 | 25.2613 | 26.0867 | 25.9697 | 25.6789 | 25.6636 |
| 0 | 21.47 | 7.76E+08 | 22 | 25.2107 | 25.3359 | 25.6301 | 25.6965 | 25.5598 |
| 0 | 22.391 | 1.81E+09 | 52 | 27.0148 | 27.0995 | 26.8412 | 26.8001 | 26.5686 |
| 0 | 8.6346 | 3.96E+08 | 28 | 24.7256 | 24.5592 | 24.8443 | 24.4994 | 24.4111 |
| 0 | 15.962 | 6.33E+08 | 28 | 25.4147 | 25.3659 | 25.3587 | 25.2769 | 25.1488 |
| 0 | 21.823 | 1.25E+10 | 122 | 29.6527 | 29.656 | 29.701 | 29.7016 | 29.5523 |
| 0 | 10.46 | 1.25E+09 | 29 | 26.0108 | 26.1875 | 26.4037 | 26.4238 | 26.4656 |
| 0 | 25.215 | 8.83E+08 | 27 | 25.9879 | 25.8092 | 25.7812 | 25.8573 | 25.7505 |
| 0 | 20.266 | 9.43E+08 | 40 | 25.7421 | 25.662 | 26.0574 | 26.1931 | 25.9074 |
| 0 | 20.366 | 1.23E+09 | 52 | 26.2589 | 26.3846 | 26.2935 | 26.1262 | 26.4208 |
| 0 | 59.027 | 5.13E+09 | 90 | 28.2721 | 28.1148 | 28.3957 | 28.4218 | 28.5231 |
| 0 | 26.056 | 3.75E+09 | 69 | 27.9146 | 28.0045 | 27.869 | 27.8223 | 27.9819 |
| 0 | 40.164 | 1.61E+09 | 62 | 26.7019 | 26.8333 | 26.8503 | 26.732 | 26.5346 |
| 0 | 323.31 | 4.42E+09 | 103 | 28.3509 | 28.2172 | 27.9584 | 28.2103 | 28.1862 |
| 0 | 7.8677 | 3.25E+08 | 8 | 24.3471 | 24.3118 | 24.5288 | 24.2736 | 24.4804 |
| 0 | 5.5963 | 6.71E+08 | 16 | 25.4631 | 25.4805 | 25.4852 | 25.5281 | 25.4153 |
| 0 | 22.305 | 8.34E+08 | 41 | 25.7929 | 25.8534 | 25.9061 | 25.8203 | 25.7669 |
| 0 | 24.442 | 2.19E+09 | 53 | 27.2032 | 27.0996 | 27.1602 | 27.0461 | 27.2989 |
| 0 | 9.6286 | 7.6E+08 | 32 | 25.6663 | 25.6266 | 25.612 | 25.8014 | 25.4359 |
| 0 | 204.54 | 2.47E+09 | 67 | 27.4024 | 27.2955 | 27.2559 | 27.4677 | 27.263 |
| 0 | 89.342 | 1.31E+10 | 81 | 29.7602 | 29.7372 | 29.9771 | 29.9378 | 29.6302 |
| 0 | 25.42 | 2.27E+09 | 62 | 27.1652 | 27.3025 | 27.2553 | 27.3612 | 27.2323 |
| 0 | 123.54 | 3.88E+09 | 77 | 28.2105 | 28.2632 | 28.1512 | 28.105 | 28.1789 |
| 0 | 19.677 | 1.08E+09 | 26 | 26.4161 | 26.2415 | 26.4553 | 26.4743 | 26.2589 |
| 0 | 62.082 | 1.68E+09 | 31 | 26.8597 | 26.9869 | 27.072 | 26.9372 | 27.1744 |
| 0 | 16.452 | 6.22E+09 | 67 | 29.0542 | 28.8757 | 29.0145 | 28.8953 | 28.8614 |
| 0 | 8.2139 | 1.02E+09 | 45 | 26.1411 | 26.1833 | 26.2271 | 26.2954 | 26.1949 |
| 0 | 55.377 | 9.18E+08 | 36 | 26.1285 | 26.0831 | 26.267 | 26.4324 | 26.1404 |
| 0 | 79.885 | 6.98E+09 | 88 | 29.1373 | 29.1232 | 29.1478 | 29.2966 | 29.1493 |
| 0 | 18.039 | 4.57E+08 | 28 | 25.1281 | 25.4399 | 24.9127 | 25.2646 | 25.1725 |
| 0 | 78.273 | 2.42E+09 | 45 | 27.4892 | 27.5123 | 27.4522 | 27.3934 | 27.3183 |
| 0 | 4.8741 | 1.57E+08 | 7 | 24.101 | 23.6745 | 23.8273 | 23.8877 | 23.8884 |
| 0 | 29.913 | 1.19E+09 | 40 | 26.3444 | 26.342 | 26.059 | 26.1728 | 26.4417 |
| 0 | 11.079 | 6.13E+08 | 12 | 26.0439 | 25.8002 | 25.4447 | 25.4847 | 25.7 |
| 0 | 15.464 | 2.14E+09 | 49 | 27.4727 | 27.4199 | 27.3594 | 27.1485 | 27.33 |
| 0 | 46.11 | 1.67E+09 | 31 | 26.8271 | 26.8127 | 27.0873 | 26.8923 | 26.8173 |
| 0 | 6.3376 | 2.52E+08 | 6 | 24.3189 | 24.0047 | 23.9506 | 24.035 | 24.5521 |

| | | | | | | | | |
|---|---|---|---|---|---|---|---|---|
| 0 | 23.038 | 1.35E+09 | 69 | 26.4434 | 26.6087 | 26.5585 | 26.5976 | 26.544 |
| 0 | 41.719 | 4.2E+08 | 10 | 24.9767 | 24.8765 | 24.9758 | 24.7809 | 25.3576 |
| 0 | 22.383 | 5.29E+08 | 15 | 25.054 | 25.043 | 25.1006 | 25.1318 | 25.0781 |
| 0 | 18.648 | 4.21E+08 | 8 | 24.8879 | 24.7943 | 24.7426 | 24.7183 | 24.7027 |
| 0 | 123.67 | 4.42E+10 | 217 | 31.5884 | 31.5793 | 31.5936 | 31.5563 | 31.4446 |
| 0 | 263.42 | 2.44E+10 | 221 | 30.5777 | 30.5769 | 30.6104 | 30.5114 | 30.8574 |
| 0 | 34.412 | 6.84E+08 | 34 | 25.7315 | 25.819 | 25.5629 | 25.553 | 25.7725 |
| 0 | 44.789 | 2.9E+09 | 57 | 27.6617 | 27.6053 | 27.7866 | 27.6566 | 27.5936 |
| 0 | 32.872 | 1.13E+09 | 36 | 26.3626 | 26.0909 | 26.4552 | 26.2668 | 26.2699 |
| 0 | 22.501 | 6.3E+08 | 15 | 25.3337 | 25.399 | 25.2607 | 25.4085 | 25.5678 |
| 0 | 145.15 | 8.23E+09 | 69 | 29.0566 | 28.9638 | 29.251 | 29.1094 | 29.2025 |
| 0 | 5.862 | 4.09E+08 | 8 | NaN | NaN | NaN | NaN | NaN |
| 0 | 17.577 | 1.25E+09 | 55 | 26.3049 | 26.4946 | 26.5487 | 26.6214 | 26.7348 |
| 0 | 49.194 | 2.04E+11 | 104 | 33.9712 | 33.8524 | 33.9445 | 33.9023 | 33.9623 |
| 0 | 8.4099 | 2.68E+08 | 15 | 23.7663 | 23.7919 | 23.7833 | NaN | NaN |
| 0 | 36.003 | 6.63E+08 | 30 | 25.6885 | 25.7436 | 25.7345 | 25.688 | 25.5078 |
| 0 | 43.288 | 2.15E+10 | 189 | 30.5876 | 30.5537 | 30.3307 | 30.468 | 30.3828 |
| 0 | 42.819 | 2.07E+09 | 73 | 27.1294 | 27.1939 | 27.5254 | 27.5945 | 27.0736 |
| 0 | 10.149 | 4.14E+08 | 18 | 24.8445 | 24.8311 | 24.4161 | 24.7268 | 25.041 |
| 0 | 135.24 | 3.54E+09 | 46 | 27.8086 | 27.6913 | 27.9543 | 27.7554 | 27.9943 |
| 0 | 18.879 | 1.01E+09 | 35 | 26.1792 | 26.2194 | 26.0059 | 26.1822 | 25.9032 |
| 0 | 7.3543 | 5.33E+08 | 18 | 24.8834 | 24.9572 | 25.1186 | 25.3672 | 25.1407 |
| 0 | 20.459 | 4.3E+08 | 10 | 25.1415 | 25.2975 | NaN | 24.9408 | 25.7308 |
| 0 | 11.307 | 3.72E+08 | 17 | NaN | 25.1715 | 24.9996 | 24.8672 | NaN |
| 0 | 17.543 | 8.61E+08 | 24 | 25.985 | 25.9321 | 25.9349 | 25.9691 | 25.8486 |
| 0 | 7.9448 | 3.41E+08 | 18 | 24.948 | 25.0115 | 24.4203 | 24.6786 | 24.4215 |
| 0 | 10.125 | 9.11E+08 | 59 | 25.8138 | 25.6526 | 26.0493 | 25.9138 | 26.1217 |
| 0 | 9.2311 | 6.3E+08 | 21 | 25.7149 | 25.5562 | 25.2504 | 25.5566 | NaN |
| 0 | 10.956 | 1.24E+09 | 43 | 26.5015 | 26.4461 | 26.6319 | 26.695 | 26.4037 |
| 0 | 44.726 | 9.82E+08 | 36 | 26.1121 | 26.3505 | 26.118 | 25.8729 | 26.0942 |
| 0 | 30.468 | 1.33E+09 | 37 | 26.6092 | 26.7344 | 26.6931 | 26.7543 | 26.633 |
| 0 | 31.088 | 1.1E+09 | 43 | 26.4041 | 26.4835 | 26.3551 | 26.329 | 26.4864 |
| 0 | 89.874 | 2.6E+09 | 53 | 27.532 | 27.5709 | 27.6246 | 27.6392 | 27.4697 |
| 0 | 32.242 | 1.26E+09 | 45 | 26.4244 | 26.2024 | 26.3445 | 26.1024 | 26.4717 |
| 0 | 7.3265 | 3.09E+08 | 26 | 24.2879 | 24.5339 | 24.2628 | 24.3776 | 24.3685 |
| 0 | 10.695 | 4.48E+08 | 24 | 25.0196 | 24.5645 | 25.1308 | 25.0601 | 25.0022 |
| 0 | 26.834 | 4.43E+09 | 36 | 28.1466 | 28.1333 | 28.848 | 28.3711 | 28.3885 |
| 0 | 218.93 | 3.22E+10 | 230 | 31.2661 | 31.3146 | 31.1089 | 31.2372 | 31.2098 |
| 0 | 33.128 | 1.64E+09 | 31 | 27.1882 | 27.3072 | 27.2419 | 26.7878 | 26.8131 |
| 0 | 6.8991 | 1.78E+08 | 10 | 23.3163 | 23.5445 | 23.2818 | 23.3745 | 24.3367 |
| 0 | 51.107 | 1.66E+09 | 56 | 27.0897 | 27.0437 | 27.133 | 27.0989 | 26.838 |
| 0 | 6.3777 | 5.05E+08 | 9 | 25.4068 | 25.3406 | 25.5925 | 25.0214 | 24.894 |
| 0 | 9.3052 | 7.73E+08 | 17 | 25.6966 | NaN | 25.5429 | 25.5258 | 25.4312 |
| 0 | 4.8191 | 4.39E+08 | 12 | 24.8126 | 25.1383 | 25.069 | 24.8602 | 24.7592 |
| 0 | 6.9687 | 2.1E+08 | 19 | NaN | 23.9513 | 23.6965 | 24.1342 | NaN |
| 0 | 19.372 | 1.3E+09 | 31 | 26.614 | 26.5092 | 26.5133 | 26.5021 | 26.4249 |
| 0 | 107.92 | 3.66E+09 | 65 | 27.8181 | 27.7684 | 28.114 | 27.8892 | 28.0418 |
| 0 | 17.319 | 1.42E+09 | 25 | 26.7861 | 26.9091 | 26.7618 | 26.8376 | 26.6271 |

| | | | | | | | | |
|---|---|---|---|---|---|---|---|---|
| 0 | 17.227 | 6.99E+08 | 17 | 25.4791 | 25.4643 | 25.4788 | 25.6372 | 25.4104 |
| 0 | 14.241 | 4.53E+08 | 12 | 25.1346 | 24.8799 | 24.7877 | 25.0692 | 25.3895 |
| 0 | 18.557 | 4.09E+08 | 16 | 24.96 | 24.8749 | 24.6657 | 24.7713 | 25.2752 |
| 0 | 13.748 | 3.74E+08 | 10 | NaN | 24.9907 | 25.1679 | NaN | 25.6701 |
| 0 | 8.1021 | 5.48E+08 | 26 | 25.3723 | 25.07 | 25.2715 | 25.2559 | 25.4995 |
| 0 | 4.1138 | 2.9E+08 | 14 | 24.5793 | 24.5682 | 24.5851 | 24.7524 | 24.5324 |
| 0 | 55.885 | 6.08E+09 | 166 | 28.8137 | 28.781 | 28.6221 | 28.7066 | 28.744 |
| 0 | 27.528 | 9.36E+08 | 27 | 25.9805 | 25.9402 | 25.7379 | 25.9279 | 26.1126 |
| 0 | 23.038 | 3.1E+09 | 60 | 27.727 | 27.8859 | 28.1065 | 28.0255 | 27.674 |
| 0 | 4.8582 | 2.94E+08 | 7 | 23.7364 | 24.0651 | 24.0943 | 24.1704 | 24.3611 |
| 0 | 52.63 | 2.12E+09 | 52 | 27.1648 | 27.1698 | 27.3011 | 27.4276 | 27.1446 |
| 0 | 9.4034 | 2.87E+08 | 24 | 24.1088 | 24.3106 | 23.938 | 24.3996 | 24.3852 |
| 0 | 15.118 | 5.25E+08 | 14 | 25.4917 | 25.2979 | 25.2208 | 25.1972 | 25.4626 |
| 0 | 7.6123 | 5.67E+08 | 14 | 25.5949 | 25.6747 | 25.6168 | 25.4882 | 25.0926 |
| 0 | 25.185 | 7.18E+08 | 13 | 25.7384 | 25.5329 | 25.55 | 25.5858 | 25.5187 |
| 0 | 10.242 | 5.19E+08 | 13 | 25.1999 | NaN | 25.0508 | 25.3065 | 24.9881 |
| 0 | 12.212 | 9.59E+08 | 33 | 26.3254 | 26.4662 | 26.1455 | 26.0361 | 26.0037 |
| 0 | 51.852 | 1.59E+09 | 29 | 26.8048 | 26.9578 | 26.5154 | 26.8545 | 27.0683 |
| 0 | 89.083 | 7.87E+08 | 26 | 25.8015 | 25.6831 | 25.9739 | 25.6871 | 25.873 |
| 0 | 15.482 | 3.89E+08 | 25 | 24.7949 | 24.8107 | 24.5709 | 24.6578 | 25.3135 |
| 0 | 4.8405 | 3.72E+08 | 5 | 24.7725 | 24.85 | 24.769 | 24.7811 | 24.7251 |
| 0 | 106.15 | 1.8E+09 | 47 | 27.0538 | 26.9361 | 27.0868 | 27.1331 | 27.1167 |
| 0 | 11.087 | 6.49E+08 | 23 | 25.9177 | 25.6121 | 25.7311 | 25.748 | 25.8542 |
| 0 | 35.319 | 3.59E+08 | 8 | 24.9742 | NaN | NaN | NaN | 24.7171 |
| 0 | 25.385 | 1.76E+09 | 55 | 27.383 | 27.4109 | 26.81 | 26.906 | 26.916 |
| 0 | 142.53 | 2.18E+09 | 58 | 27.0667 | 27.0447 | 27.5465 | 27.2996 | 27.2127 |
| 0 | 8.82 | 1.87E+09 | 44 | 27.6058 | 27.5737 | 26.8994 | 26.8165 | 26.9024 |
| 0 | 9.7512 | 3.24E+08 | 15 | 24.902 | NaN | 25.2505 | NaN | NaN |
| 0 | 8.1814 | 2.69E+08 | 14 | 24.2864 | 24.4697 | 24.0859 | NaN | 24.0113 |
| 0 | 5.1286 | 6.2E+08 | 29 | 25.6184 | 25.5514 | 25.2373 | 25.3752 | 25.2912 |
| 0 | 8.1717 | 1.01E+09 | 34 | 26.3324 | 26.2681 | 26.3449 | 26.209 | 26.2284 |
| 0 | 15.488 | 3.98E+09 | 73 | 28.281 | 28.1922 | 28.2449 | 28.3213 | 28.1559 |
| 0 | 32.722 | 8.03E+08 | 41 | 25.9659 | 25.878 | 26.0396 | 25.9469 | 25.9181 |
| 0 | 5.9776 | 2.82E+08 | 16 | 24.2326 | 24.2515 | 24.2894 | 24.407 | 24.7575 |
| 0 | 47.111 | 1.47E+09 | 33 | 26.7461 | 26.8196 | 26.5866 | 26.5871 | 26.8421 |
| 0 | 9.3101 | 6.34E+08 | 8 | 25.5791 | 25.5319 | NaN | 25.3534 | 25.4522 |
| 0 | 4.2872 | 5.39E+08 | 20 | 25.3444 | 25.5165 | 25.3922 | 25.4511 | 25.1905 |
| 0 | 66.852 | 1.47E+09 | 42 | 26.4241 | 26.3598 | 26.8675 | 26.7787 | 26.9251 |
| 0 | 7.5829 | 5.27E+08 | 13 | 25.4035 | 25.4401 | NaN | 25.356 | 25.2331 |
| 0 | 19.049 | 1.64E+09 | 36 | 26.7665 | 26.817 | 27.015 | 26.8096 | 27.1216 |
| 0 | 45.722 | 2.95E+10 | 76 | 31.1733 | 31.1784 | 31.2453 | 31.099 | 30.9273 |
| 0 | 37.873 | 1.05E+09 | 24 | 26.3453 | 26.4308 | 26.3545 | 26.4169 | 26.4191 |
| 0 | 11.241 | 3.98E+08 | 17 | 24.9201 | 24.9027 | 24.9619 | 24.6458 | 25.0395 |
| 0 | 10.322 | 3.64E+08 | 14 | 24.3587 | 24.6693 | 24.8598 | 24.7634 | 24.7946 |
| 0.000695 | 2.4158 | 2.76E+08 | 7 | NaN | 24.7175 | 24.3455 | NaN | NaN |
| 0 | 35.032 | 1.14E+09 | 28 | 26.2379 | 25.9016 | 26.4412 | 26.6665 | 26.2898 |
| 0 | 9.117 | 1.05E+09 | 39 | 26.266 | 26.1454 | 26.1191 | 26.4171 | 26.0646 |
| 0 | 323.31 | 9.08E+09 | 55 | 29.3819 | 29.1384 | 29.6357 | 29.4691 | 29.2441 |

| | | | | | | | | |
|---|---|---|---|---|---|---|---|---|
| 0 | 6.4164 | 2.96E+08 | 17 | 24.282 | 24.4782 | 24.065 | 24.7108 | 24.2744 |
| 0 | 7.4863 | 3.83E+08 | 16 | 24.9888 | 24.9344 | NaN | 24.6826 | 24.6738 |
| 0 | 108.8 | 9.57E+09 | 90 | 29.4819 | 29.4317 | 29.5186 | 29.3525 | 29.3702 |
| 0 | 12.036 | 8.91E+08 | 29 | 26.1052 | 26.2188 | 25.95 | 26.2175 | 26.1757 |
| 0 | 31.857 | 8.02E+08 | 35 | 25.9731 | 25.9084 | 25.7543 | 26.0486 | 25.8534 |
| 0 | 45.393 | 4.61E+08 | 30 | 25.0311 | 25.004 | 24.7712 | 24.7647 | 25.1214 |
| 0 | 15.709 | 2.99E+09 | 49 | 27.9124 | 27.8069 | 27.5225 | 27.7703 | 27.8061 |
| 0 | 13.127 | 6.19E+08 | 28 | 25.5678 | 25.4432 | 25.3864 | 25.562 | 25.6342 |
| 0 | 323.31 | 5.24E+11 | 1826 | 35.4328 | 35.4301 | 35.0631 | 35.0655 | 35.1038 |
| 0 | 15.38 | 7.75E+08 | 20 | 25.7265 | 25.7905 | 25.7921 | 25.6608 | 25.5905 |
| 0 | 136.17 | 6.64E+09 | 92 | 28.8802 | 28.7985 | 28.9696 | 29.0316 | 28.759 |
| 0 | 12.523 | 1.25E+09 | 25 | 26.4755 | 26.4174 | 26.4319 | 26.4152 | 26.4576 |
| 0 | 4.4611 | 2.9E+08 | 18 | 24.4288 | 24.256 | 24.4923 | 24.4609 | 24.3489 |
| 0 | 206.05 | 1.11E+10 | 148 | 29.72 | 29.7053 | 29.6687 | 29.6921 | 29.6223 |
| 0 | 41.3 | 4.52E+09 | 74 | 28.2672 | 28.1955 | 28.6573 | 28.6332 | 28.2967 |
| 0 | 22.274 | 1.02E+09 | 38 | 26.0609 | 26.1232 | 26.1572 | 25.9871 | 26.2041 |
| 0 | 40.144 | 1.82E+09 | 42 | 26.8791 | 26.8934 | 27.0812 | 27.1484 | 27.0804 |
| 0 | 10.495 | 5.79E+08 | 22 | 25.51 | 25.5623 | 25.2199 | 25.3015 | 25.4986 |
| 0 | 18.9 | 6.05E+08 | 24 | 25.4443 | 25.3045 | 25.5851 | 25.7822 | 25.5038 |
| 0 | 15.889 | 6.58E+08 | 20 | 25.4604 | 25.345 | 25.6312 | 25.9136 | 25.7351 |
| 0 | 4.0635 | 9.26E+08 | 22 | 26.5308 | 26.3043 | 25.9695 | 25.724 | 25.9712 |
| 0 | 15.372 | 5.37E+08 | 15 | 25.4819 | 25.2206 | 25.0808 | 25.0023 | 25.3575 |
| 0 | 15.396 | 6.05E+08 | 22 | 25.4681 | 25.5757 | 25.2151 | 25.2793 | 25.7407 |
| 0 | 8.7577 | 2.93E+08 | 8 | NaN | 24.7519 | 24.6731 | 24.9414 | 24.557 |
| 0 | 82.395 | 8.21E+08 | 31 | 25.9729 | 25.9949 | 25.604 | 25.6903 | 25.9588 |
| 0 | 7.1275 | 3.42E+08 | 12 | 24.358 | 24.4332 | 24.5409 | 24.5443 | 24.5357 |
| 0 | 11.292 | 1.65E+10 | 80 | 30.1971 | 30.2441 | 30.4277 | 30.1886 | 30.1908 |
| 0 | 33.21 | 8.78E+08 | 34 | 26.1242 | 26.0305 | 26.0903 | 26.0985 | 25.8037 |
| 0 | 30.618 | 6.93E+08 | 34 | 25.6623 | 25.4748 | 25.4452 | 25.5498 | 25.5933 |
| 0 | 107.51 | 3.65E+09 | 72 | 28.1545 | 28.1299 | 27.9059 | 28.0474 | 27.7835 |
| 0 | 14.14 | 3.94E+08 | 28 | 25.1225 | NaN | 24.9778 | 24.9704 | 25.0559 |
| 0 | 10.212 | 1.79E+09 | 30 | 27.1125 | 27.2209 | 27.1913 | 27.1307 | 26.8284 |
| 0 | 8.8714 | 4.08E+08 | 20 | 24.8538 | 24.736 | 24.6981 | 25.4218 | 24.5658 |
| 0 | 8.9285 | 6.42E+08 | 35 | 25.4125 | 25.3411 | 25.5539 | 25.7143 | 25.5896 |
| 0 | 5.2705 | 4.83E+08 | 7 | 25.0739 | 24.954 | 25.3798 | NaN | NaN |
| 0 | 4.7932 | 2.94E+08 | 11 | 24.9484 | NaN | NaN | NaN | 24.7707 |
| 0 | 16.691 | 4.91E+08 | 26 | 25.3072 | 25.0544 | 25.2019 | 25.1347 | 24.9062 |
| 0 | 14.967 | 5.54E+08 | 27 | 25.0416 | 25.2844 | 25.6261 | 25.3361 | 25.2351 |
| 0 | 5.0679 | 1.95E+08 | 6 | NaN | 24.3705 | NaN | NaN | 24.3361 |
| 0 | 4.7439 | 2.41E+08 | 13 | 24.0641 | NaN | 24.3067 | 24.5699 | 24.1465 |
| 0 | 13.07 | 1.1E+09 | 29 | 26.2253 | 26.117 | 26.3414 | 26.5518 | 25.9995 |
| 0 | 190.03 | 4.36E+09 | 124 | 28.5187 | 28.5053 | 28.1899 | 28.0617 | 28.2822 |
| 0 | 5.6744 | 3.85E+08 | 23 | 24.964 | 24.7491 | 24.7224 | 24.8606 | 24.5074 |
| 0 | 9.2198 | 7.82E+08 | 36 | 25.8851 | 25.7109 | 25.9632 | 25.7133 | 25.8863 |
| 0 | 12.931 | 1.34E+09 | 64 | 26.531 | 26.6878 | 26.6249 | 26.5956 | 26.4458 |
| 0 | 8.3315 | 1.94E+08 | 10 | NaN | NaN | 23.532 | 23.6585 | NaN |
| 0 | 27.983 | 3.53E+09 | 47 | 27.7919 | 27.9337 | 28.2152 | 28.1471 | 27.8127 |
| 0 | 12.45 | 1.62E+09 | 27 | 26.8637 | 26.911 | 26.7935 | 26.8342 | 26.9709 |

| | | | | | | | | |
|---|---|---|---|---|---|---|---|---|
| 0 | 20.785 | 3.67E+09 | 65 | 28.105 | 28.0428 | 28.042 | 28.182 | 27.962 |
| 0 | 32.469 | 1.28E+09 | 27 | 26.981 | 26.817 | 26.2014 | 26.4145 | 26.4753 |
| 0 | 10.328 | 3.91E+08 | 6 | 25.1931 | 25.324 | NaN | NaN | NaN |
| 0 | 15.302 | 1.96E+10 | 127 | 30.4188 | 30.4267 | 30.4107 | 30.405 | 30.2881 |
| 0 | 116.46 | 7.06E+09 | 90 | 29.3048 | 29.2897 | 29.0653 | 28.9656 | 28.6961 |
| 0 | 10.45 | 2.09E+08 | 13 | 23.9032 | 24.0144 | NaN | 24.1137 | 24.0963 |
| 0 | 36.774 | 2.63E+09 | 62 | 27.5156 | 27.5458 | 27.5774 | 27.5597 | 27.6213 |
| 0 | 10.246 | 8.14E+08 | 36 | 26.3408 | 26.2497 | 25.5863 | 25.5388 | 25.7283 |
| 0 | 37.567 | 8.46E+09 | 63 | 29.3062 | 29.3232 | 29.346 | 29.3241 | 29.0872 |
| 0 | 11.881 | 4.1E+08 | 21 | 25.7822 | 24.8859 | 24.9136 | NaN | 25.079 |
| 0 | 21.54 | 4.2E+09 | 47 | 28.1203 | 28.0943 | 28.16 | 28.1486 | 28.2912 |
| 0 | 6.21 | 2.56E+09 | 9 | 27.2923 | 27.3632 | 27.1763 | 27.5136 | NaN |
| 0 | 16.797 | 1.09E+09 | 25 | 26.2083 | 26.2748 | 26.2436 | 26.2483 | 26.234 |
| 0 | 15.78 | 3.31E+08 | 10 | 24.6881 | 24.4191 | 24.8715 | 24.9944 | NaN |
| 0 | 6.9749 | 2.05E+08 | 19 | 24.235 | NaN | NaN | 24.1922 | 24.0093 |
| 0 | 67.154 | 2.8E+09 | 67 | 27.981 | 28.0015 | 27.4968 | 27.5583 | 27.4455 |
| 0 | 323.31 | 2.21E+10 | 510 | 30.6008 | 30.5952 | 30.5558 | 30.4446 | 30.7057 |
| 0 | 11.319 | 6.85E+08 | 38 | 25.6743 | 25.7046 | 25.3842 | 25.2237 | 25.8008 |
| 0.000922 | 2.3322 | 2.13E+08 | 5 | 24.6256 | 24.4338 | NaN | 24.286 | NaN |
| 0 | 6.5786 | 3.09E+08 | 20 | 24.418 | 24.2656 | 24.5678 | 24.6102 | 24.5758 |
| 0 | 29.887 | 4.94E+09 | 65 | 28.3598 | 28.2808 | 28.5401 | 28.6397 | 28.4446 |
| 0 | 11.677 | 2.36E+09 | 58 | 27.0129 | 27.1739 | 27.4999 | 27.7366 | 27.3831 |
| 0 | 10.868 | 3.48E+08 | 14 | 25.1445 | 24.8851 | 24.3483 | 24.3087 | 24.2163 |
| 0 | 114.06 | 1.61E+10 | 88 | 30.1835 | 30.1182 | 30.2866 | 30.2724 | 30.0009 |
| 0 | 9.9305 | 3.05E+08 | 18 | NaN | NaN | 24.5532 | 24.5731 | 24.4713 |
| 0 | 75.853 | 1.15E+09 | 33 | 26.3006 | 26.1858 | 26.3944 | 26.1229 | 26.5531 |
| 0 | 323.31 | 2.06E+11 | 974 | 33.9661 | 33.9552 | 33.7379 | 33.7676 | 33.7198 |
| 0 | 3.3279 | 3.11E+08 | 17 | 24.3867 | 24.4865 | 24.5847 | 24.3979 | 24.4768 |
| 0 | 25.033 | 1.49E+09 | 54 | 26.5109 | 26.567 | 26.8771 | 26.6948 | 26.892 |
| 0 | 41.442 | 9.43E+08 | 68 | 26.2495 | 26.0074 | 26.1153 | 26.2293 | 25.7956 |
| 0 | 88.985 | 4.88E+09 | 51 | 28.5325 | 28.5262 | 28.3651 | 28.6865 | 28.3333 |
| 0 | 64.131 | 6.86E+09 | 66 | 28.9465 | 28.985 | 28.8872 | 28.9805 | 28.7573 |
| 0 | 16.97 | 6.88E+08 | 29 | 25.5197 | 25.6877 | 25.4583 | 25.2776 | 25.5836 |
| 0 | 9.7326 | 1.75E+09 | 58 | 27.0786 | 27.2156 | 26.8324 | 26.5646 | 27.0227 |
| 0 | 6.8641 | 5.02E+08 | 18 | 25.1852 | 25.1812 | 25.2945 | 25.1562 | 24.9668 |
| 0 | 18.729 | 8.86E+08 | 36 | 26.0244 | 25.9315 | 25.6994 | 25.8871 | 26.3781 |
| 0 | 6.8184 | 3.96E+08 | 17 | NaN | 25.1778 | 24.5143 | 24.4743 | 24.7981 |
| 0 | 96.952 | 3.1E+10 | 97 | 31.2355 | 31.28 | 30.9007 | 30.9789 | 30.731 |
| 0 | 7.9536 | 4.23E+08 | 8 | 24.8414 | 24.9319 | NaN | NaN | 24.9397 |
| 0 | 133.65 | 4.1E+10 | 182 | 31.6268 | 31.6161 | 31.4483 | 31.6373 | 31.4283 |
| 0 | 5.8526 | 4.44E+08 | 17 | 25.1069 | NaN | 25.0166 | 25.0045 | 25.1295 |
| 0 | 10.905 | 7.23E+08 | 16 | 26.0389 | 26.4767 | 25.2946 | 25.3219 | 25.5116 |
| 0 | 3.617 | 8.96E+08 | 24 | 25.6991 | 26.0355 | 26.2295 | 26.2655 | 26.083 |
| 0 | 5.9566 | 5.51E+08 | 19 | 25.3291 | NaN | 25.2921 | 25.4518 | 25.5753 |
| 0 | 5.388 | 6.17E+08 | 27 | 25.7279 | 25.7056 | 25.5078 | 25.3564 | 25.1767 |
| 0 | 10.338 | 9.69E+08 | 31 | 26.0674 | 25.9324 | 26.2726 | 25.9426 | 26.2183 |
| 0 | 15.416 | 3.33E+09 | 57 | 27.8768 | 27.9489 | 27.7895 | 27.8266 | 27.9078 |
| 0 | 26.617 | 1.37E+09 | 45 | 26.7178 | 26.4109 | 26.6446 | 26.6327 | 26.5489 |

| | | | | | | | | |
|---|---|---|---|---|---|---|---|---|
| 0 | 110.39 | 8.1E+10 | 342 | 32.4088 | 32.4163 | 32.7215 | 32.4723 | 32.3264 |
| 0 | 33.371 | 7.9E+08 | 31 | 25.8523 | 25.5606 | 25.6998 | 25.5768 | 25.8781 |
| 0 | 88.208 | 3.08E+09 | 65 | 27.7829 | 27.7549 | 27.9404 | 27.7856 | 27.7287 |
| 0 | 10.195 | 5.63E+08 | 31 | 25.5577 | 25.3594 | 25.1737 | 25.1918 | 25.2753 |
| 0 | 126.12 | 3.46E+09 | 149 | 27.9777 | 27.9507 | 28.0091 | 28.0691 | 27.8038 |
| 0 | 13.208 | 5.56E+08 | 21 | 25.1946 | 25.221 | 25.1548 | 25.5518 | 25.2672 |
| 0 | 23.265 | 1.18E+09 | 32 | 26.5544 | 26.5309 | 26.3274 | 26.3539 | 26.3665 |
| 0 | 6.2891 | 4.59E+08 | 21 | 24.6953 | 24.7673 | 25.1931 | 24.8939 | 24.9632 |
| 0 | 10.419 | 6E+08 | 26 | 25.4018 | 25.5917 | 25.509 | 25.3465 | 25.3909 |
| 0 | 38.403 | 3.41E+09 | 49 | 27.9276 | 27.9936 | 27.97 | 27.9281 | 28.005 |
| 0 | 14.883 | 1.07E+09 | 37 | 26.1231 | 26.0599 | 26.5583 | 26.3377 | 26.0531 |
| 0 | 323.31 | 2.59E+10 | 261 | 31.0576 | 31.0129 | 30.6825 | 30.652 | 30.8457 |
| 0 | 119.73 | 6.05E+09 | 123 | 28.7147 | 28.7217 | 28.8748 | 28.7437 | 28.7656 |
| 0 | 104.92 | 1.36E+09 | 25 | 26.3804 | 26.552 | 26.6498 | 26.5772 | 26.7138 |
| 0 | 10.933 | 1.57E+10 | 73 | 30.2027 | 30.2156 | 30.1716 | 30.1947 | 29.9974 |
| 0 | 8.0162 | 4.12E+08 | 14 | 24.7046 | 24.7408 | 24.8245 | 24.9355 | 25.0356 |
| 0 | 45.437 | 3E+08 | 12 | NaN | NaN | NaN | NaN | 24.7768 |
| 0 | 37.068 | 2.47E+09 | 64 | 27.4609 | 27.4262 | 27.4758 | 27.5848 | 27.3534 |
| 0 | 223.16 | 9.52E+09 | 66 | 29.5564 | 29.4214 | 29.3509 | 29.471 | 29.2737 |
| 0 | 12.507 | 8.14E+08 | 42 | 25.9231 | 26.1172 | 25.7304 | 25.6495 | 25.8495 |
| 0 | 58.764 | 4.82E+10 | 124 | 31.3934 | 31.5383 | 31.9184 | 32.0095 | 31.7294 |
| 0 | 89.194 | 3.29E+09 | 75 | 28.088 | 27.7484 | 27.7465 | 27.9174 | 27.8207 |
| 0 | 5.0005 | 2.33E+08 | 12 | NaN | NaN | NaN | 24.6835 | 24.6598 |
| 0 | 23.588 | 2.56E+08 | 15 | 24.4727 | 24.9564 | 24.5325 | NaN | 24.212 |
| 0 | 5.0519 | 2.11E+08 | 16 | 23.7533 | 23.7185 | 23.7807 | 23.8128 | 23.8276 |
| 0 | 6.7031 | 1.53E+10 | 44 | 29.8799 | 29.8468 | 29.8683 | 29.7335 | 29.881 |
| 0 | 6.6402 | 3.66E+08 | 17 | 24.5252 | 24.3401 | 24.4215 | 24.1088 | 24.4086 |
| 0 | 8.6323 | 4.26E+08 | 21 | 24.6678 | 24.5452 | 24.7262 | 24.6328 | 24.6352 |
| 0.000697 | 2.4592 | 2.17E+09 | 27 | 26.8541 | 27.0098 | 27.1245 | 27.1936 | 27.2009 |
| 0 | 12.296 | 5.47E+08 | 17 | 24.8448 | 25.0446 | 25.2729 | 24.7736 | 24.9823 |
| 0 | 24.684 | 8.13E+08 | 34 | 25.4694 | 25.6851 | 25.8371 | 25.8336 | 25.4874 |
| 0 | 6.4659 | 4.6E+08 | 28 | 24.7529 | 24.8418 | 24.8872 | 25.0291 | 24.8102 |
| 0 | 15.22 | 6.29E+08 | 35 | 25.4152 | 25.3083 | 25.3689 | 25.096 | 25.3656 |
| 0 | 11.048 | 1.93E+09 | 32 | 27.0491 | 26.9664 | 26.8722 | 26.8203 | 26.8093 |
| 0 | 8.2747 | 2.1E+09 | 49 | 27.1147 | 26.9953 | 27.0355 | 26.975 | 27.0764 |
| 0 | 11.78 | 1.22E+09 | 36 | 25.9747 | 26.0268 | 26.2092 | 25.9835 | 26.4507 |
| 0 | 75.566 | 1.96E+09 | 45 | 26.9715 | 26.9343 | 27.0919 | 27.0598 | 26.9349 |
| 0 | 11.064 | 4.12E+08 | 12 | 24.728 | 24.5957 | 24.5462 | 24.776 | 25.022 |
| 0.000476 | 2.8785 | 4.71E+08 | 17 | 24.8695 | 24.8233 | 24.9299 | 24.9908 | 24.9315 |
| 0 | 323.31 | 2.39E+10 | 83 | 30.6597 | 30.7448 | 30.487 | 30.5729 | 30.4297 |
| 0 | 63.554 | 6.02E+09 | 85 | 28.5011 | 28.4479 | 28.6633 | 28.6811 | 28.6219 |
| 0 | 211.27 | 1.3E+10 | 104 | 29.5871 | 29.7117 | 29.8744 | 29.796 | 29.7037 |
| 0 | 323.31 | 5.31E+10 | 366 | 31.6662 | 31.6705 | 31.704 | 31.7367 | 31.7922 |
| 0 | 6.9871 | 5.46E+08 | 23 | 25.1485 | 25.2405 | 25.3241 | 25.1407 | 25.3035 |
| 0 | 7.9287 | 6.95E+08 | 19 | 25.6619 | 25.6331 | 25.353 | 25.4245 | 25.7263 |
| 0 | 7.7923 | 2.87E+09 | 56 | 27.5304 | 27.5142 | 27.6203 | 27.5793 | 27.5568 |
| 0 | 33.173 | 2.07E+09 | 35 | 27.074 | 27.0331 | 27.1225 | 27.1369 | 27.0022 |
| 0 | 16.202 | 1.12E+10 | 104 | 29.5741 | 29.5755 | 29.5476 | 29.6144 | 29.4944 |

| | | | | | | | | |
|---|---|---|---|---|---|---|---|---|
| 0 | 73.558 | 4.27E+08 | 16 | 24.8931 | 25.0469 | 24.6661 | 24.9825 | 24.7408 |
| 0 | 13.289 | 3.65E+08 | 22 | 24.623 | 24.5488 | 24.5895 | 24.4135 | 24.67 |
| 0 | 5.3181 | 2.33E+09 | 36 | 27.285 | 27.396 | 27.3815 | 27.3059 | 27.1315 |
| 0 | 12.858 | 1.14E+09 | 35 | 26.3582 | 26.175 | 26.4631 | 26.3396 | 26.2508 |
| 0 | 323.31 | 6.05E+10 | 335 | 31.9248 | 31.9137 | 31.9279 | 32.0919 | 32.0787 |
| 0 | 323.31 | 1.59E+11 | 996 | 33.4131 | 33.4046 | 33.529 | 33.4992 | 33.4156 |
| 0.000701 | 2.5525 | 1.82E+10 | 36 | 30.4887 | 30.5455 | 30.4074 | 30.4637 | 30.3528 |
| 0 | 10.425 | 1.14E+10 | 60 | 29.6379 | 29.842 | 29.8781 | 29.8338 | 29.7288 |
| 0 | 23.471 | 1.77E+09 | 31 | 27.0913 | 27.036 | 27.1079 | 27.1436 | 27.1894 |
| 0 | 15.396 | 1.01E+09 | 29 | 26.5405 | 26.5002 | 26.3798 | 26.441 | 26.2972 |
| 0 | 30.477 | 7.58E+09 | 72 | 29.4081 | 29.4174 | 28.9788 | 29.3018 | 29.5938 |
| 0 | 4.9491 | 3.2E+08 | 19 | 24.5426 | 24.8246 | 24.4367 | 24.1419 | 25.1259 |
| 0 | 85.898 | 5.83E+09 | 108 | 28.6908 | 28.712 | 28.715 | 28.6913 | 28.6587 |
| 0 | 4.3467 | 1.59E+08 | 7 | NaN | NaN | NaN | NaN | NaN |
| 0 | 79.2 | 1.12E+09 | 35 | 26.4088 | 26.5162 | 26.3752 | 26.3712 | 26.303 |
| 0 | 148.79 | 1.8E+09 | 50 | 27.0137 | 26.9399 | 27.0441 | 27.0198 | 26.6946 |
| 0 | 7.2417 | 4.12E+08 | 31 | 24.7084 | 24.7282 | 25.0004 | 24.6225 | 24.89 |
| 0 | 3.2333 | 3.39E+08 | 15 | 24.6001 | 24.7229 | 24.9287 | 24.6382 | 24.7831 |
| 0 | 16.655 | 6.78E+08 | 23 | 25.7927 | 25.822 | 25.8584 | 25.9913 | 25.7917 |
| 0 | 12.789 | 1.46E+08 | 9 | NaN | NaN | 23.6804 | NaN | NaN |
| 0 | 16.158 | 3.8E+09 | 34 | 28.3331 | 28.3181 | 28.201 | 28.4968 | 27.9777 |
| 0 | 22.026 | 6.59E+08 | 19 | 25.1714 | 25.3437 | 25.129 | 25.8318 | 25.4121 |
| 0 | 20.725 | 9.15E+08 | 33 | 25.7613 | 25.8346 | 25.9344 | 25.9553 | 25.8531 |
| 0 | 3.8741 | 2.03E+08 | 8 | 24.1043 | 24.0035 | 24.0893 | NaN | 24.1023 |
| 0 | 7.2671 | 4.32E+08 | 8 | 24.805 | 24.8359 | NaN | NaN | 25.0258 |
| 0 | 8.7567 | 3.38E+08 | 7 | NaN | NaN | NaN | NaN | NaN |
| 0 | 15.178 | 1.12E+09 | 30 | 26.5074 | 26.2262 | 26.2818 | 26.2907 | 26.1012 |
| 0 | 8.5312 | 5.87E+08 | 29 | 25.558 | 25.6688 | 25.2385 | 25.4001 | 24.903 |
| 0 | 49.247 | 7.9E+08 | 31 | 25.93 | 25.978 | 26.1593 | 25.9373 | 25.6911 |
| 0 | 4.3878 | 2.18E+08 | 5 | 24.2066 | 24.2097 | 24.1478 | 24.3855 | NaN |
| 0 | 68.637 | 1.7E+09 | 43 | 27.6695 | 27.3753 | 26.737 | 26.6978 | 26.9611 |
| 0 | 7.9814 | 5.89E+08 | 21 | 25.421 | 25.4363 | 25.7923 | 25.7623 | 25.3499 |
| 0 | 37.696 | 5.75E+10 | 85 | 31.4886 | 31.6251 | 32.1637 | 32.3393 | 31.8816 |
| 0 | 5.3599 | 2.9E+08 | 5 | 24.4363 | 24.7361 | 24.554 | 24.5913 | 24.5594 |
| 0 | 20.745 | 6.11E+08 | 23 | 25.0828 | 25.3449 | 25.5153 | 25.4569 | 25.3717 |
| 0 | 7.4512 | 3.51E+08 | 7 | NaN | 24.9446 | NaN | NaN | 25.2163 |
| 0 | 7.0463 | 2.66E+08 | 14 | 24.491 | 24.3578 | NaN | 24.5247 | 24.5103 |
| 0 | 6.6084 | 6.03E+08 | 27 | 25.3924 | 25.4556 | 25.3602 | 25.5308 | 25.0813 |
| 0 | 42.469 | 6.35E+09 | 64 | 28.7371 | 28.6715 | 28.827 | 28.7186 | 28.5635 |
| 0 | 12.684 | 4.82E+09 | 57 | 28.0932 | 28.1784 | 28.6077 | 28.3667 | 28.3928 |
| 0 | 5.0394 | 6.46E+08 | 16 | 26.1537 | 26.2014 | NaN | NaN | NaN |
| 0 | 7.5808 | 2.51E+08 | 19 | 24.189 | 24.4177 | NaN | 24.3407 | NaN |
| 0 | 5.6833 | 3.39E+09 | 40 | 27.9224 | 28.0268 | 28.1225 | 28.1498 | 27.9017 |
| 0 | 11.701 | 3.49E+08 | 16 | 24.3712 | NaN | NaN | 24.641 | 24.6022 |
| 0 | 3.755 | 3.82E+08 | 11 | 25.1611 | 25.0154 | 24.4609 | NaN | NaN |
| 0 | 28.553 | 9.99E+08 | 29 | 25.8677 | 26.4373 | 26.4061 | 26.309 | 26.2523 |
| 0 | 305.72 | 4.94E+10 | 340 | 31.6738 | 31.6694 | 31.7612 | 31.8415 | 31.6883 |
| 0 | 22.515 | 7.1E+08 | 25 | 26.031 | 25.8929 | 25.7686 | 25.7183 | 25.6881 |

| | | | | | | | | |
|---|---|---|---|---|---|---|---|---|
| 0 | 5.7314 | 6.71E+08 | 28 | 25.7539 | 25.6089 | 25.6776 | 25.5419 | 25.7569 |
| 0 | 89.978 | 5.63E+08 | 25 | 25.4609 | 25.1925 | 25.4019 | 25.8138 | 25.2659 |
| 0 | 17.824 | 6.09E+08 | 25 | 25.5548 | 25.603 | 25.546 | 25.5649 | 25.5197 |
| 0 | 9.5146 | 1.15E+09 | 35 | 26.2965 | 26.52 | 26.1895 | 26.0735 | 26.2013 |
| 0 | 3.897 | 1.86E+08 | 11 | 23.5783 | 23.6648 | 23.4809 | 23.6468 | 24.4213 |
| 0 | 8.8674 | 1.02E+09 | 33 | 25.6549 | 25.688 | 26.0632 | 26.0837 | 26.0529 |
| 0 | 20.454 | 3.9E+08 | 23 | 25.0567 | 24.9973 | 24.593 | 24.8843 | 24.8242 |
| 0 | 25.984 | 3.6E+09 | 43 | 27.8252 | 27.9544 | 27.8434 | 27.9757 | 27.9932 |
| 0 | 10.25 | 2.11E+09 | 21 | 27.0497 | 27.0083 | 27.1464 | 27.1427 | 27.1249 |
| 0 | 27.218 | 1.31E+09 | 36 | 26.7066 | 26.6225 | 26.5869 | 26.5388 | 26.6057 |
| 0 | 170.98 | 2.53E+09 | 47 | 27.3864 | 27.3465 | 27.5992 | 27.6645 | 27.3145 |
| 0 | 19.641 | 1.04E+09 | 41 | 26.0766 | 26.0681 | 26.033 | 26.2929 | 26.1699 |
| 0 | 109.56 | 3.48E+09 | 59 | 27.9021 | 27.9463 | 28.2511 | 28.1805 | 28.0702 |
| 0.000476 | 2.8762 | 2.46E+09 | 24 | 27.5744 | 27.7127 | 27.298 | 27.5499 | 27.7049 |
| 0 | 6.2331 | 5.44E+08 | 13 | 25.0476 | 25.1873 | 25.2341 | 25.1129 | 25.0383 |
| 0 | 61.766 | 5.16E+09 | 84 | 28.407 | 28.3956 | 28.6574 | 28.4054 | 28.396 |
| 0 | 39.208 | 1.14E+10 | 61 | 29.599 | 29.6981 | 29.5006 | 29.6056 | 29.5735 |
| 0 | 40.227 | 8.85E+08 | 26 | 25.834 | 26.0848 | 25.8668 | 26.0129 | 25.6406 |
| 0 | 24.295 | 2.76E+09 | 55 | 27.6701 | 27.6954 | 27.5769 | 27.6748 | 27.7606 |
| 0 | 5.098 | 3.97E+08 | 21 | 25.0177 | 24.9088 | 24.83 | 25.0994 | 25.0817 |
| 0 | 15.899 | 3.34E+10 | 151 | 31.0841 | 31.1449 | 31.1571 | 31.2429 | 31.0095 |
| 0 | 35.825 | 3.16E+08 | 24 | 24.5897 | 24.8459 | 24.6317 | NaN | NaN |
| 0 | 7.7192 | 6.87E+08 | 18 | 25.6879 | 25.6463 | 25.6586 | 25.6036 | 25.8327 |
| 0 | 5.5417 | 5.66E+08 | 19 | NaN | 25.7151 | 25.6817 | 25.9171 | 26.0682 |
| 0 | 26.649 | 4.43E+08 | 26 | 24.6774 | 24.7739 | 25.1516 | 24.6361 | 25.0823 |
| 0 | 9.6478 | 1.33E+09 | 41 | 26.7479 | 26.6455 | 26.542 | 26.6342 | 26.6203 |
| 0 | 6.498 | 6.2E+08 | 10 | 25.539 | 25.5373 | 25.7503 | 25.1748 | 25.5363 |
| 0 | 66.777 | 1.71E+10 | 81 | 30.4266 | 30.2869 | 30.3041 | 30.276 | 30.0981 |
| 0 | 15.104 | 1.25E+09 | 44 | 26.7169 | 26.5785 | 26.459 | 26.4647 | 26.5505 |
| 0 | 7.3765 | 4.87E+08 | 29 | 25.3593 | 25.2351 | 24.9898 | 25.2098 | 25.4393 |
| 0 | 3.258 | 9.47E+08 | 14 | 26.3603 | 26.3027 | 26.289 | 26.2211 | 25.9656 |
| 0 | 5.0968 | 5.02E+08 | 14 | 25.1627 | 25.1259 | 25.2781 | 25.0563 | 24.9175 |
| 0 | 212.98 | 5.97E+09 | 88 | 28.5789 | 28.6326 | 28.8729 | 28.658 | 28.6351 |
| 0 | 6.0568 | 1.06E+09 | 20 | 26.2309 | 25.9934 | 26.343 | 26.2904 | 26.1713 |
| 0 | 5.4714 | 2.76E+08 | 16 | 24.3366 | 24.2729 | 24.4887 | 24.2574 | 24.3746 |
| 0 | 5.8201 | 6.06E+08 | 12 | NaN | NaN | 25.7043 | 25.8217 | 25.4256 |
| 0 | 42.071 | 9.89E+08 | 46 | 26.2737 | 26.4324 | 26.1567 | 25.9608 | 26.1315 |
| 0 | 7.7415 | 6.45E+08 | 27 | 25.4227 | 25.5076 | 25.553 | 25.504 | 25.5945 |
| 0 | 7.1217 | 3.5E+08 | 9 | 24.7818 | 24.8812 | NaN | 25.0221 | 24.5812 |
| 0 | 9.116 | 7.23E+08 | 30 | 25.6673 | 25.4485 | 25.6558 | 25.7082 | 25.7431 |
| 0 | 6.9249 | 3.87E+08 | 15 | NaN | 24.869 | 24.7661 | 24.7882 | 24.7254 |
| 0 | 4.8217 | 1.45E+08 | 4 | NaN | 23.5181 | NaN | 23.8434 | NaN |
| 0 | 7.079 | 5.09E+08 | 23 | 25.6066 | 25.3368 | 25.3821 | 25.1023 | 25.2757 |
| 0 | 34.572 | 3.02E+08 | 9 | 24.2804 | NaN | 24.0527 | 24.2404 | 24.7054 |
| 0 | 34.199 | 1.86E+09 | 78 | 27.3951 | 27.3761 | 26.9175 | 26.9781 | 27.0064 |
| 0 | 31.44 | 6.57E+08 | 25 | 25.6731 | 25.7692 | 25.8881 | 25.8334 | 25.4441 |
| 0 | 6.5047 | 4.38E+08 | 30 | 24.5955 | 24.8233 | 24.679 | 24.4723 | 25.3998 |
| 0 | 6.1911 | 7.77E+08 | 22 | 25.9474 | 25.4563 | 25.4495 | 25.8042 | 25.8094 |

| | | | | | | | | |
|---|---|---|---|---|---|---|---|---|
| 0 | 5.0829 | 1.56E+08 | 9 | NaN | NaN | NaN | 24.0385 | NaN |
| 0 | 5.3808 | 3.53E+08 | 14 | 24.7463 | NaN | 24.59 | 24.6248 | 24.5109 |
| 0 | 8.9766 | 3.52E+08 | 13 | 25.1926 | 25.0789 | 25.25 | NaN | 25.0898 |
| 0 | 8.7199 | 1.02E+09 | 16 | 26.2915 | 26.3977 | 25.6558 | 26.3195 | 26.1686 |
| 0 | 80.694 | 5.25E+08 | 13 | 25.0576 | 25.0307 | 25.2101 | 25.7327 | NaN |
| 0 | 10.822 | 9.23E+08 | 32 | 26.0103 | 25.8666 | 26.0837 | 26.0144 | 26.0516 |
| 0 | 3.015 | 3.02E+08 | 5 | NaN | 24.5045 | NaN | 24.518 | NaN |
| 0 | 4.1168 | 3.68E+08 | 16 | 24.7261 | 24.9072 | 24.58 | 24.766 | 24.8621 |
| 0 | 3.4372 | 2.97E+08 | 9 | 24.9367 | 24.8614 | 24.9451 | NaN | NaN |
| 0 | 17.614 | 4.92E+08 | 25 | 25.227 | 25.3644 | 25.1173 | 25.1908 | 24.9211 |
| 0 | 8.5296 | 3.76E+08 | 9 | NaN | 24.995 | 24.9619 | 24.954 | 25.0265 |
| 0 | 15.865 | 1.55E+09 | 42 | 26.5855 | 26.7778 | 27.016 | 26.9539 | 26.8129 |
| 0 | 9.086 | 6.33E+08 | 16 | 25.4962 | 25.7476 | NaN | 25.2512 | 25.675 |
| 0 | 3.4861 | 3.99E+08 | 9 | 24.9528 | 24.9064 | NaN | 24.8559 | 25.0321 |
| 0 | 56.257 | 4.34E+09 | 43 | 28.1996 | 28.2192 | 28.4311 | 28.3028 | 28.2501 |
| 0 | 8.9268 | 4.78E+08 | 25 | 25.3393 | 25.0175 | 25.2269 | 25.0811 | 24.8118 |
| 0 | 7.8867 | 3.64E+08 | 22 | 25.0047 | 25.1894 | 24.4944 | 24.5884 | 24.994 |
| 0 | 38.41 | 7E+08 | 37 | 25.584 | 25.4063 | 25.3641 | 25.7517 | 25.5989 |
| 0 | 6.7778 | 4.66E+08 | 17 | 24.9173 | 25.1845 | 25.1187 | 25.2216 | 25.1686 |
| 0 | 5.9694 | 3.13E+08 | 16 | 24.2406 | NaN | 24.1392 | 24.1592 | 24.7732 |
| 0 | 9.0529 | 4.68E+08 | 26 | 25.1965 | 25.235 | 25.1933 | 25.401 | 25.116 |
| 0 | 13.307 | 7.78E+08 | 23 | 25.7449 | 26.009 | 25.4828 | 26.0488 | 25.9907 |
| 0 | 17.967 | 4.93E+08 | 18 | 25.0144 | 25.2779 | 25.4108 | 25.303 | 25.3244 |
| 0 | 7.6521 | 2.91E+08 | 4 | NaN | NaN | NaN | NaN | NaN |
| 0 | 3.9339 | 2.92E+08 | 14 | NaN | 23.7131 | 23.7629 | 23.976 | 24.0901 |
| 0 | 29.722 | 7.5E+08 | 27 | 26.2953 | 26.1849 | 25.6304 | 25.7522 | 25.7897 |
| 0 | 6.7974 | 3.48E+08 | 13 | NaN | NaN | NaN | 25.0124 | 25.1987 |
| 0 | 9.8772 | 4.12E+08 | 30 | 25.132 | 25.3058 | 24.7917 | 24.7024 | 25.0289 |
| 0 | 191.09 | 4.73E+09 | 44 | 28.3296 | 28.1807 | 28.5947 | 28.2632 | 28.2223 |
| 0.005977 | 1.7185 | 4.25E+08 | 13 | 24.8342 | 24.8269 | 24.83 | 24.806 | NaN |
| 0 | 22.837 | 6.23E+08 | 23 | 25.9855 | 25.7516 | 25.3152 | 25.4423 | 25.3771 |
| 0 | 32.526 | 1.97E+09 | 72 | 27.2192 | 27.1705 | 27.2663 | 27.1627 | 27.1516 |
| 0 | 13.834 | 6.24E+08 | 24 | 25.3848 | 25.6001 | 25.9339 | 25.7292 | 25.4502 |
| 0 | 5.6781 | 1.09E+09 | 8 | 26.2706 | 26.2871 | 26.2529 | 26.502 | NaN |
| 0 | 31.919 | 7.67E+08 | 23 | 25.8882 | NaN | 25.9082 | 26.044 | 25.5457 |
| 0 | 4.0216 | 5.27E+08 | 10 | 24.9763 | 25.2813 | 25.2555 | 25.6003 | 25.702 |
| 0 | 11.513 | 1.56E+09 | 33 | 26.9618 | 26.9908 | 26.887 | NaN | 26.7344 |
| 0 | 7.0557 | 6.49E+08 | 21 | 25.7667 | 25.7747 | 25.3691 | 25.3769 | 25.5893 |
| 0 | 66.057 | 3.63E+09 | 55 | 28.2509 | 28.1192 | 27.8483 | 27.7999 | 27.9148 |
| 0 | 74.246 | 2.38E+09 | 44 | 27.2621 | 27.2504 | 27.4668 | 27.2954 | 27.3204 |
| 0 | 7.6701 | 5.73E+08 | 13 | 25.374 | 25.2698 | 25.3932 | 25.2197 | 25.376 |
| 0 | 22.43 | 5.9E+08 | 22 | 25.4359 | 25.3993 | 25.4497 | 25.5719 | 25.4344 |
| 0 | 106.58 | 6.09E+09 | 44 | 28.7969 | 28.8093 | 28.8057 | 28.7339 | 28.7468 |
| 0 | 17.533 | 5.48E+08 | 12 | 25.091 | 25.1947 | 25.1137 | 25.1653 | 25.1441 |
| 0 | 5.9161 | 3.38E+08 | 10 | 24.6727 | 24.5554 | 24.5225 | 25.0356 | 24.6105 |
| 0 | 3.9127 | 6.8E+08 | 16 | 25.7387 | 25.7992 | 25.5984 | 25.7853 | 25.4988 |
| 0 | 68.173 | 1.09E+10 | 42 | 29.0415 | 28.7773 | 29.8266 | 29.8515 | 29.6857 |
| 0 | 8.8467 | 4.74E+08 | 16 | 25.0001 | 25.1196 | 25.416 | 25.1674 | 25.1908 |

| | | | | | | | | |
|---|---|---|---|---|---|---|---|---|
| 0 | 18.963 | 2.89E+09 | 56 | 28.1565 | 28.1433 | 27.5544 | 27.6129 | 27.4647 |
| 0 | 19.526 | 4.31E+08 | 20 | 24.8471 | 24.8869 | 24.8542 | 25.1061 | 24.6894 |
| 0 | 3.1175 | 1.69E+09 | 20 | 26.8355 | 26.8342 | 26.8523 | 27.0147 | 27.1427 |
| 0 | 8.2506 | 4.87E+08 | 20 | 25.3139 | 25.2235 | 25.1457 | 25.4912 | 25.0287 |
| 0 | 103.41 | 9.7E+08 | 19 | 26.2246 | 26.1568 | 26.0278 | 26.0641 | 26.1238 |
| 0 | 14.157 | 1.66E+09 | 28 | 27.0543 | 26.9898 | 26.9273 | 26.9063 | 27.0354 |
| 0 | 6.1133 | 3.31E+08 | 9 | 24.4526 | NaN | 24.7894 | 24.8238 | 24.1981 |
| 0 | 9.9637 | 3.99E+08 | 12 | 24.7891 | 24.559 | 24.843 | NaN | 24.9294 |
| 0 | 8.8988 | 4.94E+08 | 23 | 25.1675 | 25.2015 | 24.97 | 25.1551 | 25.2458 |
| 0 | 8.1184 | 7.1E+08 | 33 | 25.3493 | 25.2832 | 25.737 | 25.3687 | 25.6026 |
| 0 | 6.0599 | 2.36E+10 | 42 | 30.7977 | 30.7736 | 30.7611 | 30.8203 | 30.7112 |
| 0 | 15.623 | 2.66E+09 | 50 | 27.6106 | 27.593 | 27.5336 | 27.4088 | 27.511 |
| 0 | 56.078 | 2.16E+10 | 129 | 30.644 | 30.6435 | 30.622 | 30.6918 | 30.4283 |
| 0 | 31.009 | 2.2E+09 | 33 | 26.9212 | 27.1254 | 27.3315 | 27.4152 | 27.3577 |
| 0 | 14.854 | 1.32E+09 | 41 | 26.6845 | 26.6638 | 26.5093 | 26.669 | 26.5922 |
| 0 | 51.572 | 1.07E+10 | 77 | 29.5003 | 29.5297 | 29.5765 | 29.5478 | 29.5809 |
| 0 | 3.5554 | 2.1E+08 | 12 | NaN | 23.9691 | 23.9258 | 23.9815 | 24.1275 |
| 0 | 3.907 | 5.58E+08 | 12 | 25.3025 | 25.2441 | 25.2481 | 25.0007 | 25.2671 |
| 0 | 4.5231 | 1.15E+09 | 24 | 26.3859 | 26.5484 | 26.3137 | 26.3926 | 26.3163 |
| 0 | 4.8923 | 2.96E+08 | 9 | 24.5097 | 24.4911 | 24.3429 | 24.8122 | NaN |
| 0 | 13.455 | 6.73E+08 | 25 | 25.5789 | 25.7263 | 25.2719 | 25.5732 | 25.5645 |
| 0 | 3.5694 | 1.01E+08 | 11 | 23.0197 | 22.7807 | NaN | NaN | 23.8116 |
| 0 | 323.31 | 1.15E+10 | 99 | 29.7631 | 29.6421 | 29.919 | 29.8914 | 29.5218 |
| 0 | 20.76 | 1.3E+09 | 30 | 26.5253 | 26.5213 | 26.6648 | 26.531 | 26.5285 |
| 0 | 9.0009 | 8.41E+09 | 70 | 29.3939 | 29.253 | 29.3562 | 29.2334 | 29.1041 |
| 0 | 323.31 | 6.37E+10 | 372 | 32.2597 | 32.2654 | 32.069 | 31.9962 | 32.1082 |
| 0 | 6.3491 | 6.22E+08 | 17 | 25.8497 | 25.5374 | 25.4704 | 25.4631 | 25.3734 |
| 0 | 29.391 | 8.25E+08 | 32 | 25.7684 | 25.7921 | 25.9443 | 26.2067 | 26.0831 |
| 0 | 323.31 | 2.07E+11 | 821 | 33.7893 | 33.7682 | 33.9751 | 33.8377 | 33.8948 |
| 0 | 129.77 | 2.77E+09 | 56 | 27.5531 | 27.6589 | 27.5681 | 27.6332 | 27.6488 |
| 0 | 10.337 | 9.69E+09 | 46 | 29.4138 | 29.3675 | 29.6682 | 29.6067 | 29.3081 |
| 0 | 19.582 | 3.32E+08 | 7 | 24.5886 | 24.5793 | 24.6302 | 24.6146 | 24.6714 |
| 0 | 8.5526 | 3.68E+08 | 7 | 25.1666 | NaN | NaN | NaN | 25.2797 |
| 0 | 178.58 | 1.04E+10 | 99 | 29.5327 | 29.4848 | 29.5652 | 29.586 | 29.5038 |
| 0 | 174.21 | 2.61E+10 | 221 | 30.781 | 30.8413 | 31.0105 | 30.9002 | 30.9251 |
| 0 | 9.8389 | 2.32E+09 | 46 | 27.3217 | 27.2944 | 27.0077 | 27.1723 | 27.7079 |
| 0 | 34.974 | 5.78E+08 | 18 | 25.6721 | 25.885 | NaN | 25.2691 | 25.3559 |
| 0 | 8.4948 | 9.73E+08 | 25 | 26.3562 | 26.5753 | NaN | 26.3298 | NaN |
| 0 | 5.1487 | 1.41E+09 | 26 | 26.4445 | 26.8713 | 26.9345 | 26.2732 | 26.2914 |
| 0 | 8.7867 | 1.43E+09 | 49 | 26.6555 | 26.6834 | 26.7507 | 26.5995 | 26.7071 |
| 0 | 11.914 | 6.53E+08 | 21 | 25.369 | 25.5879 | 25.7321 | 25.5839 | 25.5905 |
| 0 | 77.305 | 1.05E+09 | 46 | 26.4306 | 26.3501 | 26.3734 | 26.3879 | 26.2967 |
| 0 | 24.9 | 5.19E+08 | 19 | 25.3981 | 25.3976 | 24.9803 | 25.05 | 25.183 |
| 0 | 17.475 | 6.36E+08 | 18 | 25.6554 | 25.5353 | 25.4612 | 25.3436 | 25.5681 |
| 0 | 16.721 | 1.61E+09 | 26 | 26.3754 | 26.1501 | 26.3372 | 26.3773 | 26.4972 |
| 0 | 5.5164 | 5.5E+08 | 15 | 25.7204 | 25.5253 | 25.8306 | 25.4628 | 25.2688 |
| 0 | 7.3965 | 2.8E+08 | 6 | 24.3488 | NaN | NaN | NaN | NaN |
| 0.009454 | 1.3883 | 7.26E+08 | 7 | NaN | 26.1709 | NaN | NaN | 26.1523 |

| | | | | | | | | |
|---|---|---|---|---|---|---|---|---|
| 0 | 17.885 | 1.18E+09 | 18 | 26.1247 | 26.0729 | 26.015 | 26.1055 | 25.9407 |
| 0 | 234.4 | 5.78E+10 | 237 | 32.452 | 32.4599 | 31.8282 | 31.8058 | 31.817 |
| 0 | 27.352 | 1.25E+10 | 90 | 29.9835 | 29.8912 | 30.0328 | 30.0177 | 29.7933 |
| 0 | 6.4816 | 5.07E+08 | 6 | NaN | 25.3624 | NaN | 25.4783 | NaN |
| 0 | 22.489 | 9.5E+08 | 26 | 26.2526 | 26.3898 | 26.0933 | 26.0542 | 25.673 |
| 0 | 62.317 | 1.17E+09 | 30 | 26.445 | 26.4214 | 26.525 | 26.2447 | 26.5032 |
| 0 | 10.168 | 4.21E+08 | 19 | 25.0561 | 25.0975 | 24.8033 | 24.9344 | NaN |
| 0.007002 | 1.5687 | 3.44E+08 | 8 | 25.0066 | 24.7649 | 25.0346 | NaN | 24.9441 |
| 0 | 23.015 | 9.36E+08 | 26 | 25.9459 | 26.0079 | 26.139 | 26.0658 | 26.0306 |
| 0 | 12.13 | 6.18E+08 | 18 | 25.4718 | 25.2768 | 25.5113 | 25.5221 | 25.346 |
| 0 | 17.104 | 9.4E+08 | 29 | 26.1175 | 26.1994 | 26.1029 | 26.2872 | 26.1593 |
| 0 | 173.71 | 6.76E+09 | 96 | 28.8016 | 28.7731 | 28.9166 | 28.8522 | 28.6989 |
| 0 | 71.657 | 1.78E+09 | 46 | 26.7773 | 26.9102 | 27.0872 | 26.9014 | 27.0605 |
| 0 | 8.8347 | 2.17E+08 | 10 | 24.2588 | 24.5143 | NaN | NaN | 24.1337 |
| 0 | 12.245 | 2.24E+08 | 8 | 23.7739 | 23.5793 | 23.6159 | 23.9535 | 23.8518 |
| 0 | 35.198 | 4.7E+09 | 65 | 28.3498 | 28.5557 | 28.5376 | 28.3833 | 28.2861 |
| 0 | 167.32 | 1.29E+10 | 252 | 29.809 | 29.8418 | 29.7174 | 29.6424 | 29.9492 |
| 0 | 277.43 | 8.08E+09 | 261 | 29.0334 | 29.1242 | 29.1102 | 29.1835 | 29.1438 |
| 0 | 18.969 | 2.93E+08 | 24 | 24.4428 | 24.4263 | 24.1195 | 24.3312 | 24.4472 |
| 0 | 3.2999 | 2.41E+08 | 18 | 23.5553 | 23.7126 | 23.9952 | 24.1143 | 24.3799 |
| 0 | 5.2215 | 2.26E+08 | 9 | NaN | 24.4402 | NaN | 24.3754 | 24.5116 |
| 0 | 24.813 | 1.13E+09 | 60 | 26.6088 | 26.5253 | 26.214 | 25.9777 | 26.4281 |
| 0 | 5.7036 | 8.29E+08 | 23 | 25.6382 | 25.8089 | 25.8901 | 26.2085 | 25.9198 |
| 0 | 14.364 | 4.54E+08 | 33 | 24.9198 | 24.8761 | 25.2392 | 25.1768 | 25.0388 |
| 0 | 4.352 | 2.73E+09 | 34 | 27.6944 | 27.6127 | 27.7108 | 27.7542 | 27.5154 |
| 0 | 7.138 | 4.28E+08 | 16 | 24.6785 | 24.7039 | 24.8925 | 25.2684 | 24.9603 |
| 0 | 7.286 | 1.1E+09 | 34 | 26.0474 | 26.4163 | 26.1521 | 25.9954 | 26.466 |
| 0 | 162.75 | 1.26E+10 | 93 | 29.7666 | 29.5635 | 29.9417 | 29.9011 | 29.8946 |
| 0 | 7.1076 | 7.49E+08 | 28 | 25.5 | 25.6854 | 26.0722 | 25.9122 | 25.5425 |
| 0 | 9.1621 | 5.34E+08 | 21 | 24.7816 | 25.2995 | 25.296 | 25.2063 | 25.439 |
| 0 | 316.47 | 1.29E+10 | 179 | 29.7667 | 29.726 | 29.7008 | 29.7143 | 29.9238 |
| 0 | 6.353 | 2.61E+08 | 16 | 23.8506 | 23.9619 | 23.8446 | 24.226 | 24.4074 |
| 0 | 83.067 | 1.69E+09 | 61 | 26.8853 | 26.9768 | 27.0938 | 27.0075 | 26.969 |
| 0 | 5.2488 | 9.33E+08 | 21 | 26.1971 | 26.0658 | 26.1354 | 26.1654 | 26.1164 |
| 0 | 29.825 | 7.93E+08 | 23 | 25.9239 | 25.7515 | 26.2228 | 25.7529 | 25.683 |
| 0 | 4.9 | 3.74E+08 | 31 | 24.5482 | 24.5429 | 24.9592 | 24.8521 | 24.8325 |
| 0 | 4.4052 | 5.87E+08 | 20 | 25.4333 | 24.6017 | 25.7091 | 25.8294 | 25.2092 |
| 0 | 19.46 | 1.18E+09 | 37 | 26.8386 | 26.6443 | 26.0578 | 26.2143 | 26.2316 |
| 0 | 13.392 | 3.45E+08 | 14 | 24.6074 | 24.8313 | 24.635 | 24.4778 | 24.4223 |
| 0 | 18.578 | 5.1E+09 | 34 | 28.5746 | 28.6038 | 28.4777 | 28.6158 | 28.4385 |
| 0.005559 | 1.7649 | 6.94E+08 | 12 | 25.7311 | 25.7472 | 25.6877 | 24.828 | 25.7933 |
| 0 | 11.999 | 8.83E+08 | 34 | 26.1376 | 26.1559 | 25.9082 | 25.8369 | 25.9232 |
| 0 | 21.066 | 4.04E+09 | 60 | 27.8174 | 27.9173 | 28.3173 | 28.1734 | 28.2271 |
| 0 | 9.8274 | 4.94E+08 | 22 | 25.3365 | 25.0265 | 25.1491 | 25.1569 | 25.1691 |
| 0 | 6.7418 | 1.31E+09 | 33 | 26.4271 | 26.4693 | 26.442 | 26.4712 | 26.5759 |
| 0 | 3.3556 | 6.35E+08 | 25 | 25.3323 | 25.4548 | 25.6384 | 25.641 | 25.3504 |
| 0 | 6.7863 | 3.78E+08 | 16 | 25.0069 | NaN | 25.176 | 25.0914 | 24.917 |
| 0 | 3.7965 | 4.3E+08 | 28 | 24.8878 | 24.8975 | 24.9349 | 24.9781 | 25.0658 |

| | | | | | | | | |
|---|---|---|---|---|---|---|---|---|
| 0 | 21.938 | 7.33E+09 | 37 | 29.0839 | 29.0233 | 28.998 | 29.089 | 29.0506 |
| 0 | 4.9747 | 2.37E+08 | 13 | 24.096 | 24.0883 | NaN | 24.0602 | 24.2492 |
| 0 | 13.273 | 3.18E+08 | 21 | 24.4113 | 24.4547 | 24.7096 | 24.6192 | 24.3651 |
| 0 | 7.7547 | 3.86E+08 | 15 | 24.9099 | 24.9522 | 24.5144 | 24.9955 | 24.8565 |
| 0 | 4.7009 | 1.37E+09 | 21 | 26.1016 | 26.4841 | 26.5045 | 26.6232 | 26.5422 |
| 0 | 20.559 | 5.75E+08 | 15 | 25.3513 | 25.0866 | 25.6009 | 25.369 | 25.3859 |
| 0 | 5.527 | 4.7E+08 | 21 | 24.9649 | 25.2605 | 24.9243 | 25.1278 | 25.1468 |
| 0 | 323.31 | 3.26E+10 | 666 | 31.1804 | 31.1319 | 31.2391 | 31.2502 | 31.1365 |
| 0 | 3.3745 | 4.01E+08 | 6 | 24.8394 | NaN | NaN | 24.8846 | 24.7982 |
| 0 | 3.7773 | 3.7E+08 | 15 | 24.7783 | 24.743 | 24.8935 | 24.621 | 24.7128 |
| 0 | 18.258 | 2.99E+08 | 26 | 24.7812 | 24.4339 | 24.0417 | 24.2762 | 24.3593 |
| 0 | 8.0893 | 6.24E+08 | 16 | 25.0809 | 25.1529 | 25.3813 | 25.3802 | 25.2322 |
| 0 | 5.133 | 1.68E+08 | 20 | 23.2874 | 23.4743 | 23.9277 | 23.5755 | 23.314 |
| 0 | 7.5343 | 6.61E+08 | 24 | 25.3485 | 25.1806 | 25.2394 | 25.2438 | 25.3614 |
| 0 | 30.805 | 5.4E+09 | 68 | 28.4993 | 28.5757 | 28.7047 | 28.5535 | 28.4052 |
| 0 | 16.841 | 6.3E+08 | 21 | 25.4749 | NaN | 25.7311 | 25.4325 | 25.4658 |
| 0 | 7.785 | 4.08E+08 | 19 | 25.0177 | 25.1029 | NaN | 25.016 | 24.7622 |
| 0 | 23.213 | 4.48E+08 | 20 | 24.8293 | 24.9719 | 24.7392 | 24.8363 | 25.1002 |
| 0 | 12.338 | 5.94E+08 | 21 | NaN | NaN | 26.2208 | 26.1864 | 25.6048 |
| 0 | 26.68 | 2.11E+09 | 71 | 27.5399 | 27.6452 | 27.1757 | 27.1835 | 26.9554 |
| 0 | 10.48 | 1.53E+09 | 43 | 26.2694 | 26.304 | 27.2319 | 26.8769 | 26.9617 |
| 0 | 12.595 | 1.6E+09 | 40 | 26.8283 | 26.8072 | 26.8371 | 26.8923 | 26.7615 |
| 0 | 7.8506 | 3.7E+08 | 16 | 24.5762 | 24.9095 | 24.9685 | NaN | 24.625 |
| 0 | 8.39 | 2.77E+10 | 63 | 31.0737 | 31.0682 | 31.037 | 30.8718 | 30.8021 |
| 0 | 3.5438 | 2.48E+08 | 12 | NaN | 24.1737 | 24.1081 | 23.7681 | 24.4973 |
| 0 | 78.449 | 1.78E+09 | 24 | 26.7975 | 26.8401 | 27.0394 | 27.0316 | 27.1061 |
| 0 | 84.367 | 5.06E+08 | 15 | NaN | NaN | 25.4529 | 25.8071 | 25.3975 |
| 0.008478 | 1.5069 | 4.42E+08 | 6 | 25.0982 | 25.3806 | 25.2779 | NaN | NaN |
| 0 | 5.6715 | 3.2E+08 | 7 | NaN | 24.3839 | 24.47 | 24.5374 | 24.7257 |
| 0 | 3.8609 | 1.29E+08 | 10 | NaN | 23.2939 | 23.4768 | 23.5664 | NaN |
| 0 | 8.9983 | 2.92E+08 | 19 | 24.218 | 24.6091 | 24.5753 | NaN | 24.54 |
| 0 | 108.05 | 3.95E+10 | 184 | 31.6084 | 31.4441 | 31.4811 | 31.231 | 31.418 |
| 0 | 13.843 | 2.57E+09 | 23 | 27.4176 | 27.4101 | 27.8106 | 27.7804 | 27.7042 |
| 0 | 5.2693 | 2.71E+08 | 6 | 24.5268 | 24.4298 | NaN | NaN | 24.5487 |
| 0 | 20.004 | 9.45E+08 | 14 | 26.5691 | 26.2007 | NaN | 26.5492 | 26.2849 |
| 0 | 6.3057 | 2.81E+08 | 9 | 24.4375 | NaN | NaN | NaN | 24.3152 |
| 0 | 15.091 | 4.77E+08 | 15 | 25.6193 | 25.4378 | 25.3452 | NaN | 25.5297 |
| 0 | 7.1668 | 4.24E+08 | 12 | 24.8826 | 24.7746 | NaN | NaN | NaN |
| 0 | 20.022 | 5.86E+09 | 69 | 28.484 | 28.3514 | 28.4102 | 28.6109 | 28.431 |
| 0 | 9.0612 | 4.96E+08 | 27 | 24.7549 | 24.98 | 24.9928 | 25.0588 | 24.8701 |
| 0.000473 | 2.779 | 7.91E+08 | 21 | 25.6034 | 25.6152 | 25.6873 | 25.7643 | 25.4564 |
| 0 | 7.6353 | 3.06E+08 | 17 | 24.3222 | 24.1474 | 23.95 | 24.4445 | 24.4623 |
| 0 | 44.935 | 4.05E+08 | 12 | 24.7594 | 24.7592 | 24.555 | 24.6108 | 24.6025 |
| 0.000697 | 2.4551 | 3.55E+08 | 10 | 24.6524 | 24.437 | 24.5759 | 24.299 | 24.5571 |
| 0 | 5.4327 | 4.31E+08 | 24 | 24.9154 | 24.9185 | 24.8277 | 24.7727 | 24.7329 |
| 0 | 38.63 | 3.78E+08 | 14 | 24.6593 | 24.4181 | 24.4809 | 24.7183 | 24.7633 |
| 0 | 148.47 | 1.55E+10 | 147 | 29.874 | 29.6479 | 30.0096 | 29.7275 | 30.1435 |
| 0 | 19.632 | 8.27E+08 | 30 | 25.8435 | 25.8502 | 25.8681 | 25.8197 | 25.6117 |

| | | | | | | | | |
|---|---|---|---|---|---|---|---|---|
| 0 | 7.35 | 4.53E+08 | 28 | 25.0226 | 24.893 | 24.9468 | 24.9795 | 24.9483 |
| 0 | 23.588 | 1.07E+09 | 32 | 26.3187 | 26.1501 | 26.1788 | 26.0847 | 26.1099 |
| 0 | 29.056 | 1.61E+09 | 34 | 26.8019 | 26.7718 | 26.8065 | 26.7497 | 26.717 |
| 0 | 19.82 | 8.12E+09 | 60 | 28.9956 | 29.1478 | 29.0911 | 29.1296 | 29.0163 |
| 0 | 109.39 | 1.24E+10 | 33 | 29.6125 | 29.5917 | 29.7736 | 29.6787 | 29.6871 |
| 0 | 6.2099 | 1.11E+09 | 53 | 26.3118 | 26.3845 | 25.9748 | 26.1483 | 26.2465 |
| 0 | 19.123 | 1.83E+10 | 148 | 30.2154 | 30.3157 | 30.2225 | 30.2842 | 30.1799 |
| 0 | 8.5459 | 5.89E+08 | 34 | 25.2358 | 25.3645 | 25.3876 | 25.016 | 25.3145 |
| 0 | 3.4589 | 9.76E+09 | 27 | 29.6254 | 29.6595 | 29.5419 | 29.5115 | 29.5345 |
| 0 | 81.716 | 1.92E+10 | 126 | 30.5665 | 30.4419 | 30.6074 | 30.5627 | 30.521 |
| 0 | 71.561 | 9.11E+09 | 45 | 29.5176 | 29.4675 | 29.5427 | 29.6438 | 29.4423 |
| 0 | 82.133 | 8.26E+08 | 30 | 25.9397 | 25.9449 | 25.88 | 25.9225 | 26.1117 |
| 0 | 12.037 | 5.73E+08 | 25 | 25.5013 | 25.4701 | 25.2324 | 25.393 | 25.624 |
| 0 | 5.9705 | 1.4E+09 | 26 | 26.9842 | 27.058 | 26.7218 | 26.9586 | 26.6917 |
| 0 | 4.1868 | 4.13E+09 | 23 | 28.3162 | 28.3743 | 28.2372 | 28.4498 | 28.4635 |
| 0 | 19.087 | 8.31E+09 | 66 | 29.6434 | 29.5014 | 29.506 | 29.3819 | 29.2667 |
| 0 | 11.944 | 6.8E+08 | 26 | 25.4862 | 25.4772 | 25.6944 | 25.4947 | 25.4955 |
| 0 | 4.9133 | 3.09E+08 | 19 | 24.252 | 24.2041 | 24.5172 | 24.3079 | 24.145 |
| 0 | 14.736 | 4.46E+08 | 11 | 24.9198 | 25.0739 | 25.1953 | 24.9367 | 25.3698 |
| 0 | 30.276 | 9.96E+08 | 39 | 25.2814 | 25.8204 | 25.9402 | 26.2045 | 25.9328 |
| 0 | 6.7422 | 4.3E+08 | 14 | 25.0711 | 25.0381 | 25.1348 | 24.9972 | 25.1489 |
| 0 | 5.2102 | 3.24E+08 | 15 | 24.6042 | 24.8568 | 24.5953 | 24.498 | 24.4659 |
| 0 | 80.196 | 3.26E+09 | 46 | 27.7554 | 27.7795 | 27.9625 | 27.761 | 27.6609 |
| 0 | 23.366 | 7.11E+09 | 49 | 29.5273 | 29.5045 | 29.1643 | 29.5953 | 29.5886 |
| 0 | 5.9615 | 4.57E+08 | 22 | 25.2838 | 25.5554 | 25.5316 | 25.6419 | 25.4333 |
| 0 | 6.8339 | 5.05E+08 | 15 | 25.63 | 25.6192 | 25.0343 | 25.0274 | 25.2541 |
| 0 | 6.5168 | 85889000 | 7 | NaN | NaN | NaN | NaN | NaN |
| 0 | 19.582 | 4.34E+08 | 7 | 24.9378 | 24.8383 | 25.0071 | 25.0671 | 25.1094 |
| 0.002719 | 2.0333 | 3.54E+08 | 8 | 24.8734 | NaN | 25.0496 | 25.0195 | 24.8597 |
| 0 | 14.658 | 4.07E+08 | 16 | 24.944 | 25.057 | 24.6797 | 25.0438 | 25.1348 |
| 0 | 3.2811 | 1.69E+08 | 10 | NaN | NaN | NaN | 24.1355 | NaN |
| 0 | 42.818 | 9.31E+08 | 19 | NaN | NaN | 25.7905 | 25.9311 | 25.9138 |
| 0 | 18.166 | 2.37E+09 | 56 | 27.2211 | 27.2524 | 27.2932 | 27.4192 | 27.4473 |
| 0 | 41.829 | 7.1E+08 | 15 | 25.5912 | 25.779 | 26.2664 | NaN | 26.0684 |
| 0 | 4.2671 | 4.18E+08 | 21 | 25.0305 | 24.8527 | 25.1007 | 25.3803 | 25.1924 |
| 0 | 5.2436 | 4.37E+08 | 15 | 25.084 | 24.9592 | 24.7995 | 25.1345 | 25.3738 |
| 0 | 160.49 | 1.28E+11 | 416 | 33.1039 | 33.043 | 33.1993 | 33.0696 | 33.0417 |
| 0 | 5.7607 | 2.17E+08 | 11 | 24.0529 | 23.9271 | 24.1278 | 24.223 | 23.8597 |
| 0 | 35.594 | 4.29E+09 | 29 | 28.2719 | 28.4448 | 28.3981 | 28.5342 | 28.276 |
| 0 | 8.5152 | 2.64E+08 | 10 | 24.0092 | 24.0821 | 24.4054 | 23.9062 | 24.0865 |
| 0.0007 | 2.5251 | 9.56E+08 | 11 | 25.8604 | 26.166 | 26.0696 | 25.8928 | 26.0186 |
| 0 | 13.233 | 3.98E+08 | 23 | 24.4176 | 24.3512 | 25.2271 | 24.9715 | 24.3686 |
| 0 | 37.01 | 5.86E+08 | 27 | 25.53 | 25.2334 | 25.0356 | 25.3975 | 25.309 |
| 0 | 11.287 | 2.79E+08 | 13 | NaN | NaN | 24.6187 | 24.1345 | 24.4616 |
| 0 | 5.2778 | 2.31E+08 | 21 | 23.979 | 23.9919 | 24.2703 | 24.2439 | 24.2173 |
| 0 | 11.385 | 3.95E+08 | 22 | NaN | NaN | NaN | 25.4447 | NaN |
| 0 | 10.209 | 3.11E+08 | 7 | 24.8212 | 24.7226 | 25.0078 | 25.2358 | 24.5104 |
| 0 | 5.6527 | 4.28E+08 | 17 | 24.9498 | 24.9137 | NaN | 25.0117 | NaN |

| | | | | | | | | |
|---|---|---|---|---|---|---|---|---|
| 0 | 4.4412 | 1.8E+08 | 12 | 23.492 | 23.7221 | 23.7018 | 23.7567 | 23.9356 |
| 0 | 8.4911 | 2.83E+08 | 15 | NaN | NaN | 24.8469 | NaN | 24.7743 |
| 0 | 12.159 | 4.92E+08 | 21 | 25.1269 | 24.7382 | 25.2954 | 25.0794 | 24.8491 |
| 0 | 28.449 | 7.09E+09 | 41 | 29.5058 | 29.4444 | 28.8502 | 28.9594 | 28.9921 |
| 0 | 26.214 | 1.09E+09 | 49 | 25.8469 | 26.0803 | 26.3072 | 26.1563 | 26.2861 |
| 0 | 102.53 | 3.64E+08 | 24 | 24.8783 | 24.9281 | 24.8655 | 24.6404 | 24.7157 |
| 0 | 16.057 | 4.95E+08 | 18 | 25.5993 | 25.8447 | 24.9954 | 25.1057 | 24.9292 |
| 0 | 4.6823 | 2.83E+08 | 22 | 24.3395 | 24.2641 | 24.3162 | 24.1021 | 24.1774 |
| 0.003817 | 1.8859 | 4.77E+08 | 13 | 24.6561 | 24.9495 | 25.0439 | 24.9026 | 25.3471 |
| 0 | 3.4564 | 2.81E+08 | 12 | 24.4256 | 24.5457 | 24.1965 | 24.6048 | 24.3672 |
| 0 | 3.8005 | 3.66E+08 | 9 | 24.7432 | 24.9552 | 24.8962 | 25.2311 | 24.8349 |
| 0 | 19.463 | 4.79E+08 | 25 | 25.0401 | 24.9464 | 25.2308 | 24.9591 | 25.1058 |
| 0 | 3.2686 | 2.14E+08 | 20 | 24.077 | 23.8609 | 23.9069 | 23.9664 | 23.9989 |
| 0 | 37.277 | 5.83E+08 | 16 | 25.688 | 25.6827 | 25.2997 | 25.24 | 25.5095 |
| 0 | 3.7938 | 1.72E+08 | 10 | NaN | 23.4616 | 23.6798 | 23.6358 | NaN |
| 0 | 4.2158 | 1.53E+08 | 9 | 23.1378 | 23.486 | 23.4906 | 23.3617 | 23.335 |
| 0 | 34.937 | 4.1E+08 | 15 | 24.9137 | 25.0309 | 24.8016 | 24.8152 | 25.0096 |
| 0 | 5.3817 | 3.78E+08 | 14 | 24.9493 | 24.8964 | 24.9865 | 24.8078 | 24.6482 |
| 0 | 321.48 | 2.45E+10 | 321 | 30.7617 | 30.7189 | 30.8758 | 30.8015 | 30.5736 |
| 0 | 104.09 | 3.68E+09 | 49 | 27.9736 | 28.0107 | 28.1548 | 28.0254 | 27.7972 |
| 0 | 43.241 | 6.89E+09 | 67 | 28.9702 | 28.9223 | 29.1337 | 28.9731 | 28.6427 |
| 0 | 6.1196 | 3.18E+08 | 8 | 24.4144 | 24.7551 | 24.0636 | NaN | 24.5759 |
| 0.000469 | 2.6202 | 2.83E+08 | 4 | 24.164 | 24.0837 | 24.0791 | 24.3498 | 24.4216 |
| 0 | 19.349 | 6.37E+08 | 22 | 25.852 | 26.1461 | NaN | 25.4612 | NaN |
| 0 | 4.0431 | 3.79E+08 | 21 | 25.028 | 24.7529 | 24.847 | 24.6742 | 24.676 |
| 0 | 8.8075 | 4.34E+08 | 14 | 24.6222 | 24.5421 | 24.6006 | 24.8087 | 25.2246 |
| 0 | 65.427 | 2.19E+09 | 79 | 27.6193 | 27.7028 | 27.3057 | 27.0834 | 27.2134 |
| 0 | 11.622 | 2.64E+08 | 5 | NaN | 24.4653 | 24.4548 | 24.5357 | 24.4663 |
| 0 | 12.354 | 2.81E+08 | 9 | 24.4212 | NaN | 24.3239 | 24.1595 | 24.3301 |
| 0 | 3.3215 | 5.37E+08 | 18 | NaN | 25.1488 | 25.3147 | 25.2185 | 24.9175 |
| 0 | 6.4662 | 5.01E+08 | 26 | 25.3864 | 24.9436 | 25.4963 | 25.2122 | 25.4431 |
| 0 | 19.692 | 7.48E+08 | 23 | 26.1255 | 25.8068 | NaN | 25.9488 | NaN |
| 0 | 15.369 | 3.55E+08 | 15 | 24.7398 | 24.3994 | 24.4054 | 24.6183 | 24.5092 |
| 0.002509 | 2.0931 | 8.64E+08 | 14 | NaN | 26.556 | 26.4524 | 26.7176 | 26.402 |
| 0.004031 | 1.8537 | 2.77E+08 | 7 | 24.7451 | 24.462 | 24.5229 | 24.5899 | NaN |
| 0 | 5.4536 | 2.12E+08 | 21 | 23.5249 | 23.6092 | 23.6474 | 23.9965 | 24.0459 |
| 0 | 4.185 | 3.89E+08 | 5 | 24.8179 | 25.0373 | 24.8674 | 24.7659 | 24.962 |
| 0 | 18.937 | 1.32E+09 | 31 | 26.5615 | 26.5042 | 26.5015 | 26.5901 | 26.535 |
| 0.002283 | 2.1419 | 2.67E+08 | 8 | 24.1096 | NaN | 23.9332 | NaN | 24.5065 |
| 0 | 4.7878 | 1.13E+09 | 16 | 26.0206 | 26.3568 | NaN | 26.5838 | NaN |
| 0 | 12.961 | 1.48E+09 | 29 | 26.7767 | 26.6647 | 26.7529 | 26.6016 | 26.6716 |
| 0 | 5.1892 | 3.69E+08 | 21 | 24.8643 | 24.9312 | 24.9619 | 25.0465 | 24.6358 |
| 0 | 6.5525 | 2.71E+08 | 8 | NaN | NaN | 24.5886 | NaN | NaN |
| 0 | 12.105 | 4.21E+08 | 21 | 24.9912 | 24.8463 | 25.0868 | 25.0554 | 24.6721 |
| 0 | 4.8335 | 2.43E+08 | 20 | 24.037 | 24.3642 | 24.442 | 24.2883 | 24.193 |
| 0.000476 | 2.8891 | 6.11E+08 | 11 | 25.2343 | 25.2463 | NaN | 25.5445 | 25.5463 |
| 0 | 12.26 | 1.15E+09 | 31 | 26.3427 | 26.399 | 26.3299 | 26.3234 | 26.3052 |
| 0 | 6.2801 | 2.32E+08 | 4 | NaN | NaN | NaN | 24.0676 | NaN |

| | | | | | | | | |
|---|---|---|---|---|---|---|---|---|
| 0 | 5.9882 | 8.9E+08 | 16 | 25.9456 | 25.8046 | NaN | 26.1473 | 26.0683 |
| 0.000695 | 2.4066 | 3.85E+08 | 5 | 24.6324 | NaN | NaN | 24.9398 | 25.6828 |
| 0 | 11.078 | 1.01E+09 | 14 | 26.4082 | NaN | 26.6798 | NaN | 26.2285 |
| 0 | 13.964 | 5.7E+08 | 18 | 25.316 | 25.3485 | 25.3335 | 25.3897 | 25.5836 |
| 0 | 17.361 | 4.18E+08 | 16 | 25.2718 | 24.7012 | 24.7889 | 25.0544 | 24.883 |
| 0 | 5.2445 | 3.62E+08 | 20 | 24.6023 | 24.8628 | 24.886 | 24.8545 | 24.6287 |
| 0 | 323.31 | 1.15E+11 | 964 | 32.938 | 32.981 | 33.0692 | 33.0464 | 32.978 |
| 0 | 35.913 | 1.95E+09 | 23 | 27.1724 | 27.0336 | 27.3893 | 27.3091 | 26.5983 |
| 0 | 11.966 | 2.09E+09 | 46 | 27.1198 | 27.1017 | 27.1441 | 27.291 | 27.2474 |
| 0 | 3.7686 | 1.74E+08 | 5 | NaN | NaN | NaN | NaN | NaN |
| 0 | 3.9703 | 2.92E+08 | 23 | 24.1261 | 24.0935 | 24.1514 | 24.3352 | 24.4223 |
| 0 | 6.5332 | 7.35E+08 | 22 | 25.3049 | 25.4885 | 25.6406 | 25.7197 | 25.5088 |
| 0 | 5.3048 | 5.39E+08 | 21 | 25.116 | 25.1405 | 25.2847 | 25.2642 | 25.082 |
| 0 | 5.409 | 2.11E+08 | 11 | 23.9099 | 23.9152 | 23.8068 | 23.773 | 24.4638 |
| 0.000474 | 2.8134 | 1.66E+08 | 5 | 23.7622 | 23.6254 | 23.7353 | 23.7962 | NaN |
| 0 | 29.944 | 6.75E+08 | 60 | 25.4907 | 25.7268 | 25.5584 | 25.4776 | 25.4193 |
| 0 | 26.133 | 1.39E+10 | 66 | 29.7955 | 29.8857 | 29.8473 | 29.9948 | 29.9323 |
| 0 | 3.9115 | 1.73E+08 | 17 | 23.5022 | 23.5603 | 23.9395 | 23.5139 | NaN |
| 0 | 4.9708 | 4.31E+08 | 10 | 24.8495 | 25.0133 | 24.8117 | NaN | 25.1537 |
| 0 | 4.3837 | 1.76E+08 | 4 | 23.8683 | 23.6479 | 23.6455 | NaN | 23.6235 |
| 0.005567 | 1.7772 | 1.67E+08 | 8 | NaN | 23.7122 | NaN | NaN | NaN |
| 0 | 43.628 | 1.95E+09 | 34 | NaN | 27.2365 | 27.1145 | 27.3208 | 27.0268 |
| 0 | 8.5052 | 1.8E+08 | 7 | 23.7939 | 23.8431 | 23.4253 | 24.0215 | 24.1956 |
| 0 | 11.453 | 4.05E+08 | 6 | 24.7607 | 24.8497 | 24.9573 | 24.7223 | 24.8549 |
| 0 | 3.7567 | 2.57E+08 | 12 | 24.2296 | 24.2248 | 24.4301 | 24.3144 | 24.314 |
| 0.001609 | 2.262 | 1.3E+08 | 8 | 23.0087 | 23.1748 | 22.9142 | NaN | NaN |
| 0 | 3.4985 | 2.32E+08 | 10 | 23.8876 | 24.1427 | 24.1486 | 24.135 | 24.1212 |
| 0 | 5.2356 | 1.34E+08 | 18 | 23.0701 | 22.9227 | 23.2402 | NaN | 23.5291 |
| 0 | 3.1993 | 2.39E+08 | 14 | NaN | 24.186 | 24.2132 | 24.199 | 24.2827 |
| 0 | 9.5933 | 9.33E+08 | 27 | 26.2462 | 26.0786 | 25.7471 | 26.0221 | 26.3502 |
| 0 | 6.3167 | 2.86E+08 | 17 | 24.1347 | 24.5388 | 24.1339 | 24.3376 | 24.3242 |
| 0 | 77.146 | 1.78E+10 | 96 | 30.0246 | 30.0452 | 30.3424 | 30.1962 | 30.3066 |
| 0 | 36.613 | 4.5E+08 | 20 | 25.0282 | 25.0881 | 25.0291 | 25.1424 | 25.0745 |
| 0 | 8.9508 | 2.6E+08 | 11 | 24.2446 | 24.0854 | 24.1717 | 24.2439 | 24.0552 |
| 0 | 323.31 | 1.04E+11 | 419 | 32.8931 | 32.9175 | 32.9752 | 32.9694 | 32.8089 |
| 0 | 27.5 | 1.2E+09 | 30 | 26.8045 | 26.7928 | 26.5754 | 26.8569 | 26.3689 |
| 0 | 15.209 | 4.94E+08 | 16 | 25.1976 | 25.3953 | 25.2427 | 24.9406 | 25.4446 |
| 0.00047 | 2.6686 | 1.76E+09 | 16 | 25.3657 | 25.2905 | NaN | NaN | NaN |
| 0 | 323.31 | 6.34E+10 | 511 | 31.9792 | 31.9305 | 32.1782 | 32.1067 | 32.1133 |
| 0 | 7.6505 | 6.67E+08 | 30 | 25.5939 | 25.65 | 25.637 | 25.4382 | 25.3873 |
| 0 | 10.008 | 8.66E+08 | 31 | 25.9232 | 25.8395 | 25.8853 | 25.8407 | 26.4272 |
| 0 | 27.685 | 4.83E+08 | 15 | 25.1973 | 24.939 | 24.884 | 25.219 | 25.3421 |
| 0 | 12.719 | 8.46E+08 | 24 | 25.9635 | 26.0338 | 26.0135 | 25.8932 | 25.6784 |
| 0 | 4.7089 | 1.3E+09 | 34 | 26.672 | 26.5825 | 26.6129 | 26.5661 | 26.6125 |
| 0 | 22.712 | 1.08E+09 | 32 | 26.3093 | 26.352 | 26.3747 | 26.0254 | 26.513 |
| 0 | 80.791 | 1.78E+09 | 26 | 27.0966 | 27.0828 | 26.7305 | 27.1137 | 27.0361 |
| 0 | 7.5943 | 3.13E+08 | 17 | 24.6879 | 25.0494 | 24.7958 | 25.0407 | 25.6081 |
| 0.007422 | 1.5502 | 1.67E+08 | 9 | 23.3185 | 23.6539 | 23.7296 | 23.5609 | 23.425 |

| | | | | | | | | |
|---|---|---|---|---|---|---|---|---|
| 0 | 6.0543 | 3.01E+08 | 14 | NaN | 24.6293 | 24.5614 | 24.862 | 24.2351 |
| 0 | 5.2503 | 1.77E+08 | 18 | 23.6731 | 23.503 | 23.7038 | 23.3123 | 23.8766 |
| 0.00598 | 1.7225 | 2.9E+08 | 11 | 24.2736 | 24.823 | 24.2898 | 24.2691 | 24.2862 |
| 0.000469 | 2.6324 | 2.27E+08 | 6 | NaN | NaN | 24.6742 | 24.651 | 24.7203 |
| 0 | 8.5604 | 4.4E+08 | 13 | NaN | NaN | NaN | 25.7365 | 25.373 |
| 0 | 3.4457 | 4.72E+08 | 15 | 25.31 | 25.3778 | 25.2419 | 25.2793 | 24.9946 |
| 0 | 36.768 | 1.31E+09 | 24 | 26.5454 | 26.6362 | 26.6764 | 26.5722 | 26.6523 |
| 0 | 7.2927 | 8.01E+08 | 23 | 25.6969 | 25.9248 | 25.5553 | 26.1038 | 25.8044 |
| 0 | 11.8 | 2.57E+08 | 10 | 24.9579 | 24.5053 | 24.2374 | 24.1781 | 24.0755 |
| 0 | 9.3667 | 3.29E+08 | 11 | 24.8859 | 24.6753 | 24.7717 | 24.5012 | 24.3675 |
| 0 | 56.519 | 1.41E+09 | 27 | 26.7308 | 26.8011 | 26.3526 | 26.6238 | 26.3507 |
| 0 | 239.97 | 1.27E+10 | 208 | 29.8678 | 29.9037 | 29.7249 | 29.6667 | 29.933 |
| 0 | 196.59 | 6.29E+10 | 130 | 31.2601 | 31.1653 | 32.4032 | 32.4637 | 32.0231 |
| 0 | 126.6 | 9.34E+09 | 64 | 29.4648 | 29.3608 | 29.345 | 29.3091 | 29.5957 |
| 0 | 34.418 | 2.26E+09 | 42 | 27.7188 | 27.5357 | 27.2981 | 27.2308 | 27.2101 |
| 0 | 24.759 | 9.81E+08 | 22 | 26.3336 | 26.2071 | NaN | 26.358 | 26.605 |
| 0 | 16.59 | 91402000 | 11 | NaN | NaN | 22.9329 | NaN | 22.9093 |
| 0 | 323.31 | 3.56E+10 | 142 | 31.2943 | 31.1785 | 31.4642 | 31.4752 | 31.3056 |
| 0 | 5.8556 | 1.78E+08 | 10 | 23.6842 | 23.8746 | NaN | 23.8326 | 23.8669 |
| 0 | 4.3257 | 97277000 | 7 | NaN | 23.1859 | NaN | NaN | 23.3115 |
| 0 | 3.0996 | 93449000 | 6 | NaN | NaN | 23.2754 | NaN | 23.4142 |
| 0 | 8.1213 | 5.48E+08 | 18 | 25.3536 | NaN | NaN | NaN | 25.4538 |
| 0 | 8.921 | 4.32E+08 | 5 | 25.0188 | 25.1545 | 25.0581 | NaN | 25.2815 |
| 0 | 3.1668 | 3.04E+08 | 4 | 24.5301 | 24.6669 | 24.5008 | NaN | NaN |
| 0 | 4.1943 | 1.92E+08 | 5 | 23.8224 | NaN | NaN | 23.8964 | 24.3881 |
| 0 | 8.4663 | 5.5E+08 | 26 | 25.3872 | 25.2006 | 25.3212 | 25.2479 | 25.2169 |
| 0 | 22.406 | 1.92E+10 | 61 | 30.4455 | 30.3546 | 30.5726 | 30.5342 | 30.3925 |
| 0 | 6.9252 | 2.56E+08 | 9 | 24.4772 | 24.651 | 24.3332 | NaN | 24.6181 |
| 0 | 6.3051 | 6.3E+08 | 30 | 25.4565 | 25.3627 | 25.6857 | 25.6768 | 25.4095 |
| 0 | 2.9633 | 1.47E+08 | 8 | 24.2295 | NaN | 24.0532 | NaN | 24.193 |
| 0 | 322.35 | 3.14E+10 | 156 | 31.1192 | 31.1277 | 31.0777 | 31.2325 | 31.1414 |
| 0 | 3.656 | 3.26E+08 | 18 | 24.1765 | 24.0762 | 24.6637 | 24.5398 | 24.5349 |
| 0 | 4.1563 | 1.14E+09 | 4 | NaN | 26.3665 | NaN | 26.3986 | 26.404 |
| 0 | 7.1559 | 3.03E+08 | 13 | 24.6107 | NaN | NaN | 24.3003 | 24.8215 |
| 0 | 3.2127 | 3.68E+08 | 15 | 24.503 | 24.6094 | 24.7137 | 24.8505 | 24.4871 |
| 0 | 323.31 | 3.25E+10 | 437 | 31.3096 | 31.2532 | 31.1809 | 31.2279 | 31.0109 |
| 0 | 33.981 | 5.1E+08 | 26 | 25.1892 | 25.1749 | 25.2182 | 25.0956 | 25.3636 |
| 0.003157 | 1.9556 | 47820000 | 6 | NaN | 21.838 | 21.882 | 22.0912 | 22.0314 |
| 0 | 3.262 | 2.12E+08 | 9 | 23.9102 | 23.8607 | 24.0004 | 24.1971 | NaN |
| 0 | 7.3646 | 4.41E+08 | 8 | NaN | NaN | 25.5391 | 25.2052 | 24.9165 |
| 0 | 10.17 | 3.93E+08 | 9 | 24.7758 | 25.0761 | 24.6094 | NaN | 24.5795 |
| 0 | 7.3167 | 4.74E+08 | 8 | 25.0387 | 25.1587 | 25.2043 | 25.0959 | 25.2755 |
| 0.000238 | 2.9328 | 3.06E+08 | 14 | 23.6411 | 24.2073 | NaN | 23.546 | 24.0939 |
| 0 | 8.9879 | 4.58E+08 | 15 | 25.0827 | 24.8962 | 24.829 | 25.2259 | 25.131 |
| 0 | 3.8828 | 4.63E+08 | 23 | 25.0025 | 25.0058 | 25.2101 | 25.1837 | 24.9807 |
| 0 | 30.484 | 7.55E+08 | 25 | 25.831 | 25.8825 | 25.6034 | 25.8027 | 25.6787 |
| 0 | 4.9944 | 2.09E+08 | 21 | 23.6259 | 23.8855 | 23.1349 | 23.9975 | 24.3908 |
| 0.005558 | 1.7616 | 8.04E+08 | 14 | 25.9164 | 25.9226 | 25.4194 | 26.0118 | 25.9223 |

| | | | | | | | | |
|---|---|---|---|---|---|---|---|---|
| 0 | 59.059 | 1.19E+09 | 43 | 26.3242 | 26.3372 | 26.4241 | 26.3965 | 26.541 |
| 0 | 29.771 | 1.41E+09 | 70 | 26.6468 | 26.6539 | 26.4369 | 26.6447 | 26.6288 |
| 0.000469 | 2.6243 | 1.89E+08 | 11 | 23.6629 | 24.1256 | NaN | NaN | 24.3516 |
| 0 | 23.853 | 3.76E+08 | 5 | 24.8092 | 24.9835 | NaN | 24.7806 | NaN |
| 0 | 16.093 | 1.95E+08 | 23 | 23.7063 | 23.5367 | 23.7572 | NaN | 23.9493 |
| 0 | 4.6786 | 6.53E+09 | 67 | 28.8334 | 28.7884 | 28.982 | 28.8753 | 28.803 |
| 0 | 6.5203 | 1.94E+08 | 9 | 23.7764 | 23.6786 | 23.7377 | 23.7416 | 23.9027 |
| 0 | 323.31 | 1.44E+10 | 54 | 30.0969 | 29.9304 | 30.1261 | 30.2006 | 30.0157 |
| 0.009653 | 1.3694 | 7.65E+09 | 24 | 29.1708 | 29.1548 | 29.1267 | 29.1035 | 28.9561 |
| 0 | 7.0895 | 2.12E+09 | 21 | 27.5664 | 27.4261 | 26.9223 | 27.0246 | 27.1924 |
| 0 | 140.03 | 1.36E+10 | 82 | 30.0623 | 30.0765 | 30.005 | 30.1747 | 29.9146 |
| 0 | 143.51 | 5.68E+09 | 112 | 28.8036 | 28.5788 | 28.7032 | 28.5794 | 28.6758 |
| 0.000471 | 2.6975 | 1.75E+08 | 18 | 23.5537 | 23.6888 | 23.5094 | 23.7316 | 23.7762 |
| 0 | 19.327 | 2.44E+09 | 41 | 27.533 | 27.7762 | 27.4594 | 27.3533 | 27.3207 |
| 0 | 6.8289 | 7.45E+08 | 28 | 25.5139 | 25.6449 | 26.0166 | 25.8543 | 25.4249 |
| 0 | 323.31 | 6.69E+10 | 342 | 32.2016 | 32.1773 | 32.3677 | 32.3199 | 32.0397 |
| 0 | 33.048 | 8E+08 | 18 | 25.3431 | 25.8331 | 26.0838 | 25.7923 | 25.8932 |
| 0 | 4.8577 | 2.19E+08 | 5 | 23.7337 | NaN | 24.0102 | 23.8337 | 23.9384 |
| 0 | 17.726 | 3.68E+08 | 24 | 25.0156 | 24.781 | 24.708 | 24.8287 | 24.6952 |
| 0 | 175.99 | 1.01E+10 | 69 | 29.452 | 29.5776 | 29.5459 | 29.3122 | 29.4793 |
| 0 | 20.904 | 2.78E+08 | 10 | 24.2044 | 24.2342 | 24.3996 | 24.4556 | 23.8881 |
| 0 | 8.9308 | 1.25E+10 | 32 | 29.7474 | 29.85 | 29.8785 | 29.9971 | 29.7957 |
| 0 | 10.203 | 2.01E+08 | 10 | 23.816 | 23.7483 | 23.469 | 23.7681 | 23.5607 |
| 0 | 62.962 | 6.03E+08 | 22 | 25.8524 | 25.7321 | 25.5196 | NaN | 25.5283 |
| 0.007417 | 1.5432 | 4.66E+08 | 5 | NaN | 25.2973 | 25.2037 | NaN | NaN |
| 0 | 7.4406 | 4.57E+08 | 8 | 24.9846 | 25.1056 | 25.0901 | 24.7803 | 25.1287 |
| 0 | 5.6568 | 2.15E+08 | 10 | NaN | 24.0053 | 24.2209 | 24.0631 | 23.8784 |
| 0.00316 | 1.9613 | 1.98E+08 | 7 | NaN | 23.9582 | NaN | 23.8442 | 23.8677 |
| 0 | 86.679 | 3.66E+09 | 99 | 28.1986 | 28.2418 | 27.8304 | 27.7288 | 28.0075 |
| 0 | 7.7494 | 3.86E+08 | 14 | 24.6089 | 24.6728 | 24.545 | NaN | 24.8739 |
| 0 | 4.0027 | 1.3E+08 | 16 | NaN | NaN | 23.3652 | 23.5289 | NaN |
| 0 | 13.208 | 1.94E+09 | 40 | 27.2278 | 27.0173 | 27.0911 | 27.2837 | 26.9694 |
| 0 | 4.3802 | 5.64E+08 | 16 | 25.4112 | 25.4232 | 25.3883 | 25.3103 | 25.2839 |
| 0 | 18.808 | 5.38E+08 | 19 | 25.4084 | 25.3823 | 25.2978 | 25.1256 | 25.1769 |
| 0 | 108.27 | 2.44E+09 | 54 | 27.7776 | 27.814 | 27.0856 | 27.1527 | 27.5228 |
| 0 | 5.6118 | 3.21E+08 | 6 | NaN | 25.0197 | 25.0124 | 24.9946 | NaN |
| 0 | 7.383 | 2.12E+09 | 41 | 26.9857 | 27.1086 | 26.7421 | 26.8569 | 27.7144 |
| 0 | 3.2064 | 2.68E+08 | 11 | NaN | 23.9672 | 24.6182 | 24.1405 | 24.2701 |
| 0 | 5.9848 | 3.88E+08 | 19 | 24.712 | 24.7297 | 24.8541 | 24.6578 | 25.055 |
| 0 | 3.2712 | 3.06E+08 | 21 | 24.5273 | 24.6476 | 24.2912 | 24.6764 | 24.2584 |
| 0 | 155.86 | 1.06E+11 | 437 | 32.8604 | 32.8773 | 32.9357 | 32.996 | 32.8382 |
| 0 | 9.1743 | 7.09E+08 | 24 | 25.7368 | 25.7561 | 25.654 | 25.4998 | 25.7269 |
| 0 | 5.554 | 8.12E+08 | 42 | 25.5809 | 25.7132 | 25.8742 | 25.9075 | 26.1474 |
| 0 | 8.8442 | 5.87E+08 | 24 | 24.9737 | 25.6218 | 25.6445 | 25.5993 | 25.5893 |
| 0 | 9.9832 | 4.27E+09 | 35 | 28.1805 | 28.2791 | 28.2754 | 28.3679 | 28.3036 |
| 0.000694 | 2.4004 | 3.09E+09 | 20 | 27.651 | 27.7123 | 27.8358 | 27.9729 | 27.7586 |
| 0 | 142.01 | 1.18E+10 | 53 | 29.849 | 29.9015 | 29.7003 | 29.4838 | 29.6635 |
| 0 | 4.8525 | 4.21E+08 | 8 | 25.1958 | 25.1757 | 25.0975 | 25.4751 | NaN |

| | | | | | | | | |
|---|---|---|---|---|---|---|---|---|
| 0 | 11.226 | 3.53E+08 | 8 | NaN | 25.348 | 25.3968 | 24.7208 | NaN |
| 0 | 5.7477 | 1.32E+09 | 8 | 26.6231 | NaN | 26.4694 | 26.4815 | 26.7972 |
| 0 | 5.7791 | 4.76E+08 | 21 | 25.0771 | 25.0581 | 25.2408 | 24.9842 | 25.0557 |
| 0 | 6.2762 | 2.64E+08 | 14 | 24.0641 | 24.4324 | 24.0715 | 24.4791 | 24.1225 |
| 0 | 4.3698 | 5.2E+08 | 11 | NaN | 25.5043 | NaN | 25.4345 | 25.5082 |
| 0 | 6.302 | 4.57E+08 | 21 | 24.933 | 25.1266 | 25.0467 | 25.0258 | 25.0772 |
| 0 | 3.0802 | 2.67E+08 | 8 | 24.7262 | 24.6676 | 24.7218 | 24.8064 | NaN |
| 0 | 4.2821 | 2.2E+08 | 3 | NaN | 24.3741 | 24.3418 | NaN | 24.473 |
| 0 | 6.9629 | 7.67E+08 | 18 | 26.1965 | 26.0055 | 26.0907 | 25.9236 | 25.8893 |
| 0 | 3.6352 | 3.2E+08 | 7 | 24.5908 | 24.7088 | NaN | NaN | 24.7596 |
| 0 | 3.2942 | 1.99E+08 | 8 | 23.5711 | 23.7621 | NaN | 23.781 | NaN |
| 0.000473 | 2.7543 | 2.14E+08 | 4 | NaN | NaN | NaN | NaN | NaN |
| 0.000701 | 2.5469 | 1.55E+09 | 17 | NaN | 23.5392 | 23.5831 | 23.8532 | 23.7873 |
| 0 | 4.9308 | 2.4E+08 | 5 | 23.9064 | 23.9958 | 24.1104 | 24.1372 | NaN |
| 0.000476 | 2.8717 | 2.11E+08 | 6 | 24.3414 | NaN | NaN | NaN | 24.2862 |
| 0 | 5.0122 | 1.33E+09 | 39 | 26.1812 | 26.34 | 25.8671 | 26.2966 | 26.4404 |
| 0 | 31.77 | 9.52E+08 | 21 | 25.9657 | 25.8383 | 25.9187 | 26.0699 | 25.6767 |
| 0 | 24.12 | 4.88E+08 | 18 | 25.2572 | 24.9916 | 24.978 | 24.8036 | 24.8507 |
| 0 | 8.6444 | 3.11E+09 | 33 | 27.7165 | 27.6671 | 27.5483 | 27.7528 | 27.5157 |
| 0 | 6.9497 | 1.04E+10 | 34 | 29.1861 | 29.4698 | 29.3353 | 29.7087 | 29.4741 |
| 0 | 204.06 | 1.06E+10 | 245 | 29.3702 | 29.4545 | 29.5549 | 29.626 | 29.4877 |
| 0 | 323.31 | 5.81E+10 | 252 | 31.9791 | 31.9513 | 32.0024 | 32.0025 | 32.0015 |
| 0 | 5.7681 | 4.54E+08 | 14 | 24.8447 | 25.0488 | 25.2253 | 25.0959 | 25.2284 |
| 0 | 76.366 | 3.51E+09 | 40 | 28.204 | 28.1683 | 27.8038 | 27.9873 | 28.0525 |
| 0 | 35.141 | 9.58E+08 | 28 | 26.0635 | 26.4264 | 26.2032 | 26.4097 | 26.1313 |
| 0 | 22.898 | 2.14E+09 | 55 | 27.6527 | 27.6653 | 27.2283 | 27.3566 | 27.2762 |
| 0 | 12.481 | 2.57E+09 | 32 | 27.8016 | 27.6669 | 27.7286 | 27.7457 | 27.6595 |
| 0 | 7.7772 | 7.17E+08 | 23 | 25.7005 | 25.6641 | 25.8068 | 25.7607 | 26.1326 |
| 0 | 9.7462 | 2.68E+10 | 39 | 31.1311 | 31.1058 | 31.1492 | 31.1123 | 30.9459 |
| 0 | 4.4947 | 2.97E+08 | 5 | NaN | NaN | NaN | NaN | NaN |
| 0 | 3.4653 | 1.63E+08 | 7 | 24.3252 | 23.9256 | 23.8212 | NaN | NaN |
| 0 | 68.788 | 1.8E+10 | 128 | 30.4274 | 30.2493 | 30.2401 | 30.2127 | 30.1724 |
| 0 | 3.9991 | 3.23E+08 | 6 | 24.7393 | 24.8223 | NaN | 24.8349 | 24.8073 |
| 0 | 4.3071 | 2.07E+08 | 10 | 24.0804 | 24.0038 | 24.0855 | 24.0152 | 24.2128 |
| 0 | 17.594 | 2.7E+09 | 22 | 27.5641 | 27.4466 | 27.6267 | 27.4006 | 27.5324 |
| 0 | 8.7152 | 3.88E+08 | 14 | 25.1282 | 25.0408 | 25.044 | 24.9083 | 25.1579 |
| 0 | 5.8235 | 3.12E+08 | 21 | 24.6346 | 24.949 | 24.6021 | 24.3752 | 24.4916 |
| 0.008664 | 1.4699 | 1.26E+08 | 5 | NaN | NaN | NaN | NaN | NaN |
| 0.000698 | 2.4959 | 1.38E+08 | 3 | 23.5973 | NaN | NaN | NaN | NaN |
| 0.000468 | 2.6009 | 2.76E+08 | 11 | 24.2027 | NaN | 24.4517 | NaN | 24.2816 |
| 0 | 7.4753 | 3.19E+08 | 12 | 24.883 | 25.0045 | 24.7708 | 24.7364 | 24.827 |
| 0 | 6.1501 | 2.38E+08 | 13 | 24.0574 | 23.9457 | 23.6749 | NaN | 23.8348 |
| 0 | 11.725 | 7.4E+08 | 15 | NaN | NaN | 25.9777 | 25.9894 | 26.1436 |
| 0 | 3.0014 | 3.35E+08 | 12 | 24.9992 | NaN | NaN | 25.202 | 25.3384 |
| 0 | 32.521 | 7.88E+08 | 26 | 25.7452 | 25.6979 | NaN | 26.0816 | 26.0189 |
| 0 | 4.8459 | 1.69E+08 | 7 | 24.2904 | 23.9797 | 24.1041 | 24.0144 | NaN |
| 0 | 17.16 | 4.79E+08 | 14 | 24.7431 | 24.7194 | NaN | 25.5388 | NaN |
| 0.000474 | 2.8001 | 1.02E+08 | 14 | 23.6856 | 23.7586 | 22.9955 | 23.2988 | NaN |

| | | | | | | | | |
|---|---|---|---|---|---|---|---|---|
| 0 | 7.7184 | 2.23E+09 | 30 | 27.5535 | 27.424 | 27.4892 | 27.4615 | 27.4721 |
| 0 | 5.975 | 1.12E+09 | 22 | 25.882 | 25.9392 | NaN | 25.6563 | 25.8411 |
| 0 | 9.4448 | 1.97E+08 | 9 | 23.8486 | 23.7971 | 23.8287 | 23.8822 | 23.9405 |
| 0 | 8.9373 | 4.12E+09 | 32 | 28.2325 | 28.1126 | 28.2194 | 28.2303 | 28.0917 |
| 0 | 3.0801 | 4.6E+08 | 14 | 24.6555 | 24.8198 | NaN | 24.8886 | 25.356 |
| 0 | 3.3953 | 29190000 | 10 | 22.029 | 22.5351 | 22.1411 | NaN | NaN |
| 0 | 3.9743 | 5.97E+08 | 18 | 25.7252 | 25.6987 | 26.148 | 25.8173 | 25.8761 |
| 0.002939 | 1.9956 | 1.6E+08 | 5 | 23.7935 | 23.7033 | 23.6495 | 24.1142 | 23.5942 |
| 0.009261 | 1.4113 | 1.24E+08 | 5 | 23.1636 | 23.167 | 23.7408 | 23.1656 | 23.2432 |
| 0 | 323.31 | 1.11E+10 | 299 | 29.6644 | 29.6431 | 29.6427 | 29.6031 | 29.7302 |
| 0 | 10.82 | 2.62E+08 | 11 | NaN | NaN | 24.4435 | 24.1068 | 23.9589 |
| 0 | 24.666 | 1.31E+09 | 34 | 26.4546 | 26.4624 | 26.7137 | 26.582 | 26.3327 |
| 0 | 249.5 | 6.3E+10 | 294 | 32.22 | 32.2353 | 32.0171 | 32.0546 | 32.1615 |
| 0 | 85.982 | 1.8E+09 | 18 | 26.9035 | 27.0288 | 27.2546 | 26.9726 | 27.1756 |
| 0.000472 | 2.7069 | 1.4E+08 | 11 | NaN | 23.8731 | 23.7276 | 23.8413 | 23.7461 |
| 0.000468 | 2.6095 | 3.8E+08 | 11 | 24.8744 | 24.708 | 24.786 | 24.8318 | 24.7776 |
| 0 | 4.2758 | 3.74E+09 | 68 | 28.07 | 27.9857 | 28.3841 | 28.4666 | 27.8889 |
| 0.002728 | 2.0636 | 5.19E+09 | 27 | 28.3935 | 28.4793 | 28.6407 | 28.7499 | 28.63 |
| 0.000698 | 2.469 | 2.19E+08 | 17 | 23.7874 | 24.0161 | 23.8022 | 24.2367 | 24.1296 |
| 0 | 3.4984 | 3.07E+08 | 14 | 24.9301 | 24.512 | 24.2898 | 24.5025 | 24.8083 |
| 0 | 79.928 | 4.72E+09 | 149 | 28.3619 | 28.409 | 28.4371 | 28.4934 | 28.409 |
| 0 | 15.842 | 8.38E+09 | 79 | 29.401 | 29.4349 | 29.3552 | 29.2135 | 29.1423 |
| 0 | 8.0938 | 2.07E+08 | 14 | NaN | NaN | NaN | 23.9613 | NaN |
| 0.000475 | 2.8259 | 4.49E+08 | 21 | 25.1482 | 25.3945 | 24.7133 | 25.2622 | 25.3794 |
| 0.004035 | 1.8714 | 7.38E+08 | 15 | 26.1665 | 26.0981 | 26.292 | 26.0529 | 26.1014 |
| 0 | 8.4274 | 5.35E+08 | 14 | 25.3413 | 25.1438 | 25.1347 | 25.3574 | 25.1287 |
| 0 | 4.4092 | 2.76E+08 | 19 | 24.1595 | 24.1418 | 24.3717 | 24.1608 | 24.4113 |
| 0 | 6.7379 | 2.22E+09 | 13 | 27.5029 | 27.5016 | 27.5176 | 27.4358 | 27.3319 |
| 0 | 3.4777 | 2.37E+08 | 6 | 24.2714 | NaN | 23.8253 | NaN | 24.6868 |
| 0 | 5.039 | 7.26E+08 | 32 | 25.8527 | 25.6678 | 25.8649 | 26.0043 | 25.5803 |
| 0 | 3.4722 | 2.94E+08 | 13 | 24.987 | 24.9295 | 24.4239 | 24.3713 | 24.2972 |
| 0 | 5.2132 | 8.07E+08 | 22 | 25.7256 | 25.5785 | 27.8668 | 25.6756 | 25.193 |
| 0 | 3.0137 | 5.27E+08 | 19 | NaN | 24.2874 | 24.2678 | 24.3884 | 25.8952 |
| 0 | 3.2378 | 1.35E+09 | 21 | 26.2901 | NaN | 27.1437 | 27.0799 | 26.9398 |
| 0.001834 | 2.2441 | 3.59E+08 | 18 | 24.7449 | 24.7271 | 25.0382 | 25.2668 | NaN |
| 0 | 3.8495 | 1.41E+08 | 10 | NaN | NaN | 24.5283 | 24.257 | NaN |
| 0 | 6.5637 | 2.55E+08 | 2 | NaN | NaN | NaN | NaN | NaN |
| 0.000696 | 2.4349 | 7.48E+08 | 18 | 25.997 | NaN | 26.044 | NaN | 25.9998 |
| 0.002734 | 2.0844 | 1.79E+08 | 15 | 23.829 | 23.9602 | 23.9634 | 23.8431 | 24.0552 |
| 0.007835 | 1.5148 | 1.66E+09 | 7 | 27.4077 | 27.4001 | 26.3357 | 26.6536 | 26.9832 |
| 0 | 82.261 | 2.8E+09 | 69 | 27.654 | 27.5715 | 27.8009 | 27.5535 | 27.7506 |
| 0.000701 | 2.5463 | 2.45E+08 | 9 | 24.3914 | 24.4504 | 24.2343 | 24.5659 | 24.1815 |
| 0 | 5.9639 | 4.64E+09 | 35 | 28.308 | 28.4751 | 28.4985 | 28.3348 | 28.2143 |
| 0 | 10.569 | 2.56E+08 | 9 | NaN | 24.8265 | NaN | NaN | NaN |
| 0 | 44.224 | 4.83E+08 | 21 | 25.1217 | 25.159 | 25.4618 | 25.2951 | 25.174 |
| 0 | 3.6469 | 4.22E+08 | 16 | 26.2407 | 26.227 | NaN | 25.9988 | NaN |
| 0.000698 | 2.4674 | 1.84E+08 | 4 | NaN | NaN | 24.3631 | 24.2643 | NaN |
| 0 | 5.0822 | 1.56E+08 | 5 | NaN | 23.8542 | NaN | 24.0089 | 23.86 |

| | | | | | | | | |
|---|---|---|---|---|---|---|---|---|
| 0.002715 | 2.0129 | 91829000 | 12 | 23.1667 | 23.1973 | 23.621 | 23.4171 | NaN |
| 0 | 4.426 | 1.94E+09 | 23 | NaN | 26.3636 | 26.5754 | 28.3912 | 26.1663 |
| 0 | 4.0958 | 8.09E+08 | 4 | NaN | 25.8036 | NaN | NaN | NaN |
| 0 | 5.8635 | 1E+09 | 15 | 25.9684 | 25.9314 | 26.1367 | 26.0405 | 26.3736 |
| 0 | 5.2098 | 2.25E+08 | 8 | NaN | NaN | NaN | NaN | 24.8815 |
| 0 | 6.5213 | 3.45E+08 | 18 | 24.7317 | 24.7287 | 24.7133 | 24.5063 | 24.455 |
| 0 | 24.258 | 6.03E+08 | 29 | 25.5342 | 25.4788 | 25.3363 | 25.5591 | 25.3697 |
| 0 | 8.6938 | 9.17E+08 | 23 | 25.9781 | 26.0494 | 26.1444 | 26.0504 | 25.8678 |
| 0.000697 | 2.4508 | 6.03E+08 | 18 | NaN | 25.6059 | 25.6547 | 25.5382 | 25.2453 |
| 0 | 7.205 | 1.93E+08 | 2 | 23.957 | 23.8742 | 23.9194 | 23.8819 | 24.3181 |
| 0 | 19.022 | 5.66E+09 | 68 | 28.4183 | 28.3755 | 28.627 | 28.5689 | 28.8427 |
| 0 | 5.7982 | 2.47E+08 | 12 | 23.8178 | 23.9675 | 24.3124 | 24.1198 | 24.603 |
| 0 | 323.31 | 1.32E+12 | 3061 | 36.5419 | 36.5709 | 36.4019 | 36.3714 | 36.3899 |
| 0 | 4.1907 | 2.16E+08 | 7 | 24.7578 | 24.9135 | 24.1486 | 24.4182 | NaN |
| 0 | 4.3924 | 4.01E+08 | 6 | 25.3369 | 25.0081 | NaN | 25.2001 | 25.2866 |
| 0.002285 | 2.1507 | 2.02E+08 | 11 | 24.279 | 24.1526 | NaN | 24.1608 | 24.1154 |
| 0 | 8.4151 | 3.21E+09 | 24 | 28.1008 | 27.9776 | 28.0661 | 28.0729 | 27.9668 |
| 0 | 10.991 | 2.37E+09 | 16 | 27.3464 | 27.3602 | 27.6487 | 28.1793 | 27.3789 |
| 0 | 134.25 | 3.82E+10 | 162 | 31.3966 | 31.426 | 31.5054 | 31.4097 | 31.3158 |
| 0 | 6.1663 | 4.37E+08 | 22 | 24.5082 | 24.4689 | 24.8974 | 24.909 | NaN |
| 0 | 5.7079 | 1.75E+09 | 18 | 26.9995 | 27.0339 | 27.0693 | 27.222 | 27.0774 |
| 0 | 57.091 | 5.38E+08 | 23 | 25.2377 | 25.2254 | 25.1142 | 25.3623 | 25.4898 |
| 0 | 6.6148 | 5.13E+08 | 16 | 24.9345 | 25.0708 | 25.2137 | 25.2522 | 25.0655 |
| 0.003823 | 1.9039 | 6.83E+08 | 7 | 25.7376 | NaN | 25.9307 | 25.7057 | 25.8773 |
| 0 | 4.2223 | 1.68E+08 | 7 | 23.652 | NaN | 23.203 | NaN | 23.5703 |
| 0 | 3.8701 | 2.92E+08 | 12 | 24.3697 | 24.2867 | 24.1743 | 24.0746 | 24.356 |
| 0 | 49.379 | 6.41E+08 | 33 | 25.4318 | 25.5461 | 25.4516 | 25.3375 | 25.8268 |
| 0 | 6.3414 | 2.43E+08 | 12 | 24.103 | 24.1711 | 24.1858 | 24.0828 | 24.2445 |
| 0 | 6.6735 | 5.77E+09 | 37 | 28.8471 | 28.8119 | 28.7739 | 28.8293 | 28.5001 |
| 0 | 17.062 | 8.03E+08 | 23 | 26.1879 | 26.1012 | 25.629 | 25.7794 | 25.8586 |
| 0 | 14.672 | 6.64E+08 | 26 | 25.5642 | 25.6891 | 25.4448 | 25.6997 | 25.4527 |
| 0 | 323.31 | 2.78E+11 | 1091 | 34.3138 | 34.3053 | 34.2833 | 34.2784 | 34.2815 |
| 0 | 4.7326 | 6.82E+08 | 22 | 25.6862 | 25.5757 | 25.6882 | NaN | 25.6618 |
| 0.008671 | 1.4787 | 9.81E+08 | 11 | 25.9498 | 25.7766 | 25.7186 | 26.1726 | 26.3865 |
| 0 | 201.19 | 9.05E+09 | 74 | 29.4052 | 29.3213 | 29.34 | 29.3739 | 29.1818 |
| 0 | 3.8584 | 1.64E+08 | 12 | 23.6087 | 23.6875 | NaN | 23.4866 | 24.429 |
| 0 | 6.5673 | 2.38E+08 | 16 | 24.5759 | 24.67 | NaN | 24.2215 | NaN |
| 0 | 56.992 | 2.35E+09 | 31 | 27.6246 | 27.3355 | 27.4721 | 27.5271 | 27.224 |
| 0 | 4.3537 | 2.45E+08 | 7 | NaN | 24.1641 | 24.1884 | NaN | 24.1796 |
| 0 | 14.959 | 1.83E+08 | 3 | NaN | 24.3608 | 24.0668 | 24.1189 | 24.3598 |
| 0 | 6.9871 | 4.25E+10 | 52 | 31.7965 | 31.8303 | 31.53 | 31.6583 | NaN |
| 0 | 6.4856 | 6.8E+08 | 17 | 25.8782 | 25.9937 | 25.4983 | 25.5714 | 25.709 |
| 0 | 5.0642 | 2.71E+08 | 5 | 24.3482 | 24.3441 | 24.4977 | 24.6052 | 24.2093 |
| 0.003155 | 1.9467 | 1.21E+08 | 11 | NaN | 23.9576 | 24.6128 | 23.9941 | NaN |
| 0 | 114.85 | 1.14E+10 | 93 | 29.7451 | 29.801 | 29.6179 | 29.6392 | 29.5101 |
| 0.000475 | 2.8579 | 2.4E+08 | 8 | NaN | NaN | 24.7029 | 24.7209 | 25.0411 |
| 0 | 3.1007 | 1.28E+10 | 21 | NaN | 30.0037 | 29.805 | 29.9651 | 30.0693 |
| 0.000696 | 2.4381 | 1.86E+08 | 6 | 23.763 | 23.9871 | 24.016 | 23.8604 | 23.6417 |

| | | | | | | | | |
|---|---|---|---|---|---|---|---|---|
| 0.005552 | 1.7518 | 2.37E+08 | 8 | 23.9316 | 24.0554 | 24.6402 | 24.1969 | 24.115 |
| 0 | 4.608 | 1.56E+08 | 21 | 23.4246 | 23.5017 | 23.3898 | 23.7424 | 23.023 |
| 0 | 14.663 | 4.19E+08 | 18 | 24.7636 | 24.7931 | 25.2636 | 24.9027 | 24.7934 |
| 0.007414 | 1.5392 | 2.59E+08 | 5 | 24.0612 | 24.1142 | 24.1204 | 24.0488 | 24.251 |
| 0.000693 | 2.3873 | 2.02E+09 | 15 | NaN | 26.6645 | NaN | 26.5154 | NaN |
| 0.006621 | 1.6833 | 2.88E+08 | 17 | 24.6317 | 24.5637 | 24.491 | 24.2819 | 23.9538 |
| 0 | 27.33 | 9.06E+08 | 10 | 26.4012 | 25.7086 | 26.1485 | 26.1766 | NaN |
| 0 | 6.5263 | 5.41E+08 | 35 | 25.4779 | 25.6066 | NaN | 25.5199 | 25.3649 |
| 0 | 11.069 | 3.68E+08 | 18 | 24.7328 | 24.8872 | 24.9827 | 24.9476 | 24.8691 |
| 0 | 4.158 | 2.22E+08 | 5 | 24.2337 | 24.3467 | NaN | 24.3458 | NaN |
| 0 | 3.9221 | 1.92E+08 | 3 | NaN | 23.8752 | NaN | 24.0899 | 23.8629 |
| 0 | 15.741 | 1.26E+09 | 29 | 27.2643 | NaN | 27.2216 | 27.0154 | NaN |
| 0 | 5.1194 | 1.62E+08 | 9 | NaN | NaN | 24.1545 | NaN | 23.8911 |
| 0 | 8.3113 | 4.14E+08 | 7 | 25.0397 | NaN | NaN | NaN | 24.8521 |
| 0 | 5.3336 | 2.63E+08 | 15 | 24.3539 | 24.2107 | NaN | 24.5514 | 24.4724 |
| 0 | 8.3517 | 5E+09 | 174 | 27.9308 | NaN | 28.0168 | 28.1974 | 26.8502 |
| 0.007625 | 1.5305 | 99770000 | 13 | 23.2181 | 23.106 | NaN | NaN | NaN |
| 0 | 43.641 | 2.24E+08 | 7 | 24.0919 | 23.9821 | NaN | NaN | NaN |
| 0 | 5.2217 | 8.56E+08 | 30 | 25.6476 | 25.7932 | 25.9888 | 26.2969 | 25.8051 |
| 0 | 10.353 | 1.11E+09 | 44 | 26.3403 | 26.156 | 26.2909 | 26.4243 | 26.3802 |
| 0 | 7.4163 | 5.27E+08 | 3 | 25.3458 | 25.3763 | 25.5718 | 25.548 | 25.6738 |
| 0 | 4.3415 | 8.26E+08 | 8 | NaN | 26.2612 | NaN | 26.5149 | 26.2909 |
| 0 | 25.117 | 6.37E+08 | 10 | 25.7387 | NaN | NaN | 26.0887 | NaN |
| 0 | 4.6745 | 2.58E+08 | 6 | NaN | 24.6588 | 24.7166 | NaN | NaN |
| 0 | 6.3222 | 1.44E+09 | 27 | 26.632 | 26.7187 | 27.0013 | 26.9889 | 26.3931 |
| 0 | 4.3701 | 1.67E+09 | 28 | 26.4449 | 26.5498 | 27.005 | 26.767 | 27.1357 |
| 0 | 14.919 | 2.33E+09 | 28 | 27.1477 | 27.2566 | 27.9342 | 27.4319 | 25.3429 |
| 0 | 9.2892 | 6.42E+08 | 34 | 25.5078 | 25.658 | 25.4859 | 25.2068 | 25.5464 |
| 0 | 15.204 | 1.13E+09 | 13 | 26.436 | NaN | 26.5013 | 26.4615 | NaN |
| 0 | 21.411 | 1.89E+09 | 111 | 26.9518 | 27.2259 | 27.0238 | 26.9951 | 26.9915 |
| 0.004032 | 1.8566 | 1.14E+08 | 8 | NaN | NaN | NaN | NaN | 23.8764 |
| 0 | 3.4616 | 2.66E+08 | 24 | 24.2114 | 24.2897 | 23.8964 | NaN | 24.3233 |
| 0 | 6.4882 | 4.34E+08 | 22 | 24.7455 | 24.6 | 24.9289 | 24.9822 | 24.7716 |
| 0.007011 | 1.5865 | 92514000 | 6 | 23.6435 | 23.4969 | NaN | NaN | NaN |
| 0 | 323.31 | 5.55E+10 | 399 | 31.9724 | 31.9393 | 31.9066 | 31.8474 | 32.0201 |
| 0 | 4.6919 | 2.27E+08 | 5 | 24.4034 | NaN | NaN | 24.6301 | 24.3301 |
| 0 | 9.1227 | 3.22E+08 | 5 | NaN | NaN | NaN | 24.5012 | 24.4614 |
| 0 | 3.3835 | 5.36E+08 | 19 | NaN | 25.4634 | 25.332 | 25.1235 | 25.3591 |
| 0 | 18.377 | 3.06E+08 | 10 | 24.6263 | 24.2139 | NaN | 24.4902 | 24.3327 |
| 0.008047 | 1.511 | 1.87E+09 | 14 | 27.3284 | 27.3434 | NaN | NaN | NaN |
| 0.009458 | 1.3908 | 99969000 | 5 | 23.532 | 23.5349 | NaN | NaN | 23.3495 |
| 0 | 24.919 | 1.34E+09 | 26 | 26.6453 | 26.6312 | 26.7321 | 26.6221 | 26.5619 |
| 0 | 29.098 | 1.41E+09 | 50 | 26.4418 | 26.56 | 26.8367 | 26.672 | 26.5381 |
| 0 | 189.65 | 6.69E+10 | 335 | 32.0583 | 32.0143 | 32.0919 | 32.2669 | 32.3631 |
| 0 | 9.2917 | 2.08E+09 | 38 | 27.07 | 27.0367 | 27.6369 | 27.0737 | 27.2405 |
| 0 | 201.3 | 1.86E+10 | 171 | 30.3986 | 30.4723 | 30.2984 | 30.3204 | 30.338 |
| 0.004472 | 1.8279 | 1.96E+08 | 11 | NaN | 24.1359 | 23.9971 | NaN | 23.9457 |
| 0 | 323.31 | 8.04E+10 | 520 | 32.4932 | 32.534 | 32.5228 | 32.4868 | 32.4516 |

| | | | | | | | | |
|---|---|---|---|---|---|---|---|---|
| 0 | 4.1345 | 2.84E+08 | 27 | 24.0199 | 24.3943 | 23.8651 | 24.2321 | 24.1566 |
| 0 | 12.978 | 2.89E+08 | 16 | 24.7466 | 24.6353 | 24.0674 | NaN | 24.2235 |
| 0 | 10.714 | 2.71E+09 | 28 | 27.6765 | 27.4051 | NaN | 27.7856 | 27.3484 |
| 0.002717 | 2.0191 | 2.11E+08 | 13 | 24.1056 | 24.0947 | NaN | 24.0629 | 24.1229 |
| 0 | 4.0911 | 1.86E+08 | 5 | 24.1521 | 24.3023 | 24.1087 | NaN | 24.3793 |
| 0 | 5.8854 | 2.55E+08 | 10 | 24.2506 | 24.3378 | NaN | 24.3212 | NaN |
| 0 | 3.2394 | 1.74E+08 | 16 | NaN | 24.4095 | 24.877 | NaN | 24.6239 |
| 0 | 4.2494 | 2.76E+08 | 18 | 24.2037 | 24.2275 | 24.3108 | 24.2196 | 24.4378 |
| 0 | 5.3985 | 1.47E+08 | 3 | NaN | 23.8159 | 24.0153 | 23.6434 | NaN |
| 0 | 4.4494 | 4.62E+08 | 17 | NaN | 25.1554 | 25.2062 | 25.0169 | 25.2303 |
| 0 | 221.8 | 4.99E+09 | 130 | 28.3752 | 28.3567 | 28.6792 | 28.5801 | 28.4352 |
| 0 | 132.28 | 2.35E+09 | 27 | 27.4999 | 27.2494 | 27.3942 | 27.2989 | 27.5319 |
| 0 | 3.5346 | 2.67E+08 | 11 | 24.3118 | 23.8186 | 24.503 | 24.2661 | 24.0518 |
| 0 | 3.0341 | 1.2E+08 | 9 | NaN | 23.3859 | 23.4466 | NaN | 23.7839 |
| 0 | 3.5226 | 2.73E+08 | 20 | 24.5945 | 24.6021 | 23.7527 | NaN | 23.9356 |
| 0 | 10.76 | 3.83E+08 | 13 | 24.7399 | 24.6775 | 24.9912 | 24.8819 | 24.6591 |
| 0 | 323.31 | 8.79E+09 | 281 | 29.2924 | 29.2378 | 29.0925 | 29.1655 | 29.4849 |
| 0 | 19.755 | 4.35E+08 | 12 | 24.9793 | 24.9053 | 24.88 | 25.2218 | 24.8693 |
| 0 | 14.381 | 6.27E+08 | 12 | 25.3723 | 25.2096 | 25.536 | 25.41 | 25.6511 |
| 0 | 5.1364 | 2.82E+08 | 23 | NaN | NaN | 24.5329 | 24.3888 | 24.7246 |
| 0.000701 | 2.5464 | 9.02E+08 | 17 | 26.3015 | 26.3215 | 25.8275 | 26.0157 | 25.9561 |
| 0 | 29.82 | 4.68E+08 | 13 | NaN | NaN | 25.0348 | NaN | 24.9753 |
| 0 | 10.989 | 2.36E+08 | 10 | 24.0378 | 24.0498 | 24.1835 | 24.3182 | 24.1004 |
| 0 | 4.0781 | 1.5E+08 | 5 | 23.2364 | 23.1476 | 23.2445 | 23.2022 | 23.2893 |
| 0 | 16.433 | 9.77E+09 | 51 | 29.4001 | 29.4609 | 29.4777 | 29.4655 | 29.4931 |
| 0 | 3.4141 | 4.94E+08 | 16 | 25.7326 | 25.2737 | NaN | 25.082 | 25.7291 |
| 0.00047 | 2.6365 | 1.38E+08 | 4 | 23.458 | 23.4315 | 23.4876 | NaN | 23.4941 |
| 0 | 214.49 | 9.12E+09 | 266 | 29.2826 | 29.2608 | 29.267 | 29.2049 | 29.3177 |
| 0.00403 | 1.8522 | 2.03E+08 | 2 | NaN | 24.5749 | NaN | 23.96 | 23.9225 |
| 0 | 12.432 | 1.81E+08 | 9 | 24.1497 | NaN | 24.2266 | 24.2017 | 24.0088 |
| 0 | 8.1986 | 2.07E+08 | 12 | 23.7889 | NaN | 23.8628 | 24.5252 | 23.8858 |
| 0 | 76.561 | 1.65E+08 | 17 | 23.3355 | 23.0844 | 23.6755 | 23.1782 | 23.5497 |
| 0.000474 | 2.7963 | 3.04E+08 | 8 | 24.8203 | NaN | NaN | 25.1572 | 24.9073 |
| 0 | 62.357 | 9.62E+08 | 32 | 26.0717 | 26.4168 | 26.1193 | 26.1909 | 26.2007 |
| 0 | 6.3073 | 2.65E+08 | 12 | 24.4108 | 24.2161 | NaN | 24.4265 | 24.3209 |
| 0 | 214.8 | 7.95E+09 | 141 | 29.2184 | 29.1144 | 29.2064 | 29.0765 | 29.1274 |

| ctrl_3_2 | TDP_1_1 | TDP_1_2 | TDP_2_1 | TDP_2_2 | TDP_3_1 | TDP_3_2 |
|---|---|---|---|---|---|---|
| NaN | NaN | 28.4116 | 25.0438 | 25.2062 | 26.4185 | 24.6272 |
| NaN | 25.1855 | 25.1358 | 25.2577 | 25.1203 | 25.1888 | 25.0709 |
| NaN | NaN | NaN | 26.5396 | 26.1026 | 26.3395 | 25.7403 |
| NaN | NaN | 25.1574 | 24.8806 | 25.1402 | 25.4991 | 25.1773 |
| NaN | 24.9608 | 25.2812 | 25.1814 | 25.2025 | 25.0023 | NaN |
| NaN | NaN | 24.8139 | 24.8469 | 25.1758 | 24.6101 | NaN |
| 25.3015 | 25.3346 | 25.1241 | 25.2476 | 25.3778 | 25.1084 | 25.4301 |
| NaN | NaN | NaN | 25.6798 | 25.596 | 25.6298 | NaN |
| 25.3911 | 27.0729 | 27.1162 | 26.2807 | 26.3977 | 27.0015 | 26.5458 |
| NaN | NaN | 24.7831 | 24.5592 | 24.4689 | 24.6161 | NaN |
| NaN | NaN | NaN | NaN | 24.238 | 24.0331 | 23.9869 |
| NaN | 25.3705 | 25.305 | 25.3887 | 25.277 | 25.1881 | 25.2613 |
| NaN | NaN | NaN | 25.0108 | 25.124 | 25.1244 | NaN |
| NaN | 25.0476 | 24.7843 | 24.6783 | 24.6557 | 24.7182 | 24.9757 |
| NaN | NaN | NaN | 24.8175 | 24.9573 | 24.9469 | NaN |
| NaN | 25.5656 | 25.6298 | 25.2173 | NaN | NaN | NaN |
| NaN | NaN | NaN | NaN | 24.7332 | 24.6904 | 25.0029 |
| NaN | NaN | NaN | 24.6194 | 24.6116 | 24.6444 | NaN |
| NaN | NaN | NaN | 24.0097 | NaN | 24.2909 | 24.3941 |
| NaN | 24.6503 | 24.6067 | 24.3919 | 24.522 | 24.2456 | 24.041 |
| NaN | 24.3731 | 24.4578 | 24.0632 | 24.1744 | 24.0554 | 24.2334 |
| NaN | 24.5621 | 24.408 | 24.3225 | 24.084 | 24.2562 | 24.3605 |
| NaN | NaN | NaN | 24.4807 | 24.5635 | 24.3033 | 24.2493 |
| NaN | NaN | NaN | NaN | 25.3292 | 24.7895 | 25.0851 |
| NaN | NaN | NaN | 24.0696 | NaN | 24.0371 | 24.2466 |
| NaN | 24.1801 | NaN | 23.9406 | NaN | 24.0182 | NaN |
| NaN | NaN | 24.0719 | NaN | 23.9155 | 23.7198 | NaN |
| 29.8936 | 30.4117 | 30.414 | 30.3788 | 30.3573 | 30.394 | 30.3927 |
| 25.5103 | 25.8791 | 26.1971 | 26.037 | 26.1296 | 25.8804 | 25.5061 |
| 25.0914 | 25.6382 | 25.6738 | 25.7294 | 25.7127 | 25.7881 | 25.6415 |
| 26.8182 | 27.2046 | 27.3084 | 27.7658 | 27.7464 | 27.1077 | 26.7479 |
| 24.8009 | 24.8833 | 25.0346 | 24.9829 | 24.7676 | 25.2221 | 25.05 |
| 24.6696 | 26.3219 | 26.9289 | 25.9086 | 25.7594 | 26.2193 | 25.5139 |
| 25.5484 | 25.9841 | 25.8126 | 26.6765 | 26.1615 | 25.9418 | 25.7701 |
| 28.2261 | 28.8992 | 28.7724 | 28.7098 | 28.6196 | 28.6018 | 28.7002 |
| 25.4812 | 25.6927 | 25.7899 | 25.9131 | 26.2491 | 25.8183 | 25.644 |
| 26.6137 | 27.9143 | 27.7673 | 26.303 | 26.2631 | 27.5943 | 27.7697 |
| 29.7342 | 30.7626 | 30.4552 | 30.912 | 30.4741 | 30.061 | 29.785 |
| 27.6562 | 27.6716 | 27.8273 | 28.1715 | 28.2911 | 28.239 | 28.2775 |
| 27.9996 | 27.9949 | NaN | NaN | NaN | NaN | NaN |
| 25.9812 | 26.3989 | NaN | NaN | NaN | NaN | NaN |
| 24.8057 | NaN | NaN | NaN | NaN | NaN | NaN |
| 24.6152 | NaN | NaN | NaN | NaN | NaN | NaN |
| 25.8548 | NaN | 25.9708 | NaN | NaN | 25.9373 | NaN |
| 24.5946 | 24.4814 | NaN | NaN | NaN | NaN | NaN |

| | | | | | | |
|---|---|---|---|---|---|---|
| 26.3891 | 26.2989 | 26.3367 | NaN | NaN | 26.5117 | NaN |
| 26.0288 | 26.2796 | 26.1346 | NaN | 25.5465 | NaN | NaN |
| 26.3251 | NaN | NaN | NaN | NaN | NaN | NaN |
| 24.6823 | NaN | 24.8237 | NaN | NaN | NaN | NaN |
| NaN | NaN | NaN | NaN | NaN | NaN | NaN |
| 25.446 | NaN | NaN | NaN | NaN | NaN | NaN |
| 25.9139 | 24.4126 | 24.6219 | 24.5439 | 25.1577 | 24.4328 | 24.2925 |
| 24.6439 | 24.3541 | 24.606 | NaN | NaN | NaN | 24.4406 |
| 24.5582 | NaN | 24.4727 | NaN | 24.2811 | NaN | 24.3281 |
| 24.7354 | NaN | 24.6919 | 24.832 | NaN | 24.5561 | NaN |
| 25.5724 | 25.338 | 25.4173 | NaN | NaN | NaN | 25.4289 |
| 24.6175 | NaN | NaN | NaN | NaN | NaN | NaN |
| 25.4248 | 25.4214 | 25.0238 | NaN | 24.471 | NaN | NaN |
| 24.2232 | 22.6323 | NaN | NaN | 23.3657 | 23.5804 | NaN |
| 23.9346 | 23.5522 | 23.5884 | NaN | NaN | NaN | NaN |
| 24.1644 | NaN | NaN | 24.2868 | 24.3002 | 24.3147 | NaN |
| 24.6718 | 24.3204 | 24.3742 | NaN | NaN | 24.4103 | NaN |
| 23.9905 | NaN | NaN | NaN | NaN | NaN | 24.1907 |
| 23.9406 | NaN | 24.5058 | NaN | NaN | 24.0474 | NaN |
| 24.9731 | NaN | NaN | NaN | NaN | NaN | NaN |
| 24.7609 | NaN | NaN | 24.5979 | 24.8631 | NaN | 24.5843 |
| 24.5157 | NaN | NaN | NaN | NaN | NaN | NaN |
| 25.5118 | NaN | NaN | 25.1152 | 24.9961 | 24.7894 | 24.7893 |
| 22.8378 | NaN | NaN | NaN | NaN | NaN | NaN |
| 23.5701 | NaN | NaN | 23.5341 | NaN | NaN | NaN |
| 23.6298 | NaN | NaN | NaN | NaN | 23.986 | NaN |
| 23.6389 | NaN | 23.8012 | NaN | NaN | NaN | 23.3286 |
| 24.1858 | NaN | NaN | NaN | NaN | NaN | NaN |
| 24.8511 | NaN | NaN | 25.0436 | 25.1267 | 24.5908 | NaN |
| NaN | NaN | NaN | NaN | NaN | NaN | NaN |
| 24.5134 | NaN | 24.2697 | NaN | NaN | NaN | 24.139 |
| 24.3293 | 24.5703 | 24.3814 | NaN | NaN | NaN | NaN |
| 25.3909 | 25.3541 | 25.132 | NaN | NaN | NaN | 24.3292 |
| 23.4686 | NaN | NaN | NaN | NaN | NaN | NaN |
| 24.5219 | NaN | NaN | NaN | 24.7329 | NaN | NaN |
| 25.9271 | 25.3071 | 25.3459 | 25.3906 | 25.0924 | 25.3479 | 25.5462 |
| 27.2168 | 26.13 | 26.2199 | 26.9039 | 27.0584 | 26.5297 | 26.7048 |
| 25.7692 | 25.474 | 25.7919 | 25.4789 | 25.3279 | 25.4134 | 25.4159 |
| 28.6372 | 28.1705 | 28.1049 | 28.5392 | 28.4129 | 27.9251 | 28.0824 |
| 24.7543 | 24.4875 | 24.3821 | 24.1368 | 24.392 | 24.1004 | 24.3278 |
| 26.2417 | 25.5258 | 25.9387 | 25.2996 | 25.8905 | 25.5053 | 25.7179 |
| 23.3789 | 22.5032 | 22.5595 | NaN | 22.6132 | NaN | 22.5591 |
| 33.3493 | 33.296 | 33.2503 | 33.2046 | 33.1911 | 33.3356 | 33.3832 |
| 34.6678 | 34.9858 | 34.9209 | 34.773 | 34.754 | 34.8144 | 34.8163 |
| 32.3545 | 32.5571 | 32.5441 | 32.4511 | 32.4546 | 32.3891 | 32.3526 |
| 30.9022 | 30.7505 | 30.6719 | 30.9268 | 30.855 | 30.5733 | 30.5885 |
| 34.9742 | 35.1767 | 35.0985 | 34.9069 | 34.9391 | 35.4295 | 35.427 |
| 34.8401 | 35.0748 | 35.052 | 34.9787 | 34.9472 | 35.0432 | 35.0113 |

| | | | | | | |
|---|---|---|---|---|---|---|
| 31.3377 | 31.6618 | 31.5652 | 31.4526 | 31.4871 | 31.5048 | 31.485 |
| 30.8876 | 30.8301 | 30.7711 | 30.799 | 30.7338 | 31.0081 | 30.958 |
| 33.6559 | 33.473 | 33.3934 | 33.4939 | 33.4803 | 33.5412 | 33.5434 |
| 34.695 | 34.6054 | 34.5512 | 34.5813 | 34.6063 | 34.7382 | 34.7156 |
| 29.8461 | 30.2163 | 30.1712 | 30.084 | 30.0705 | 29.9775 | 29.9296 |
| 30.6045 | 30.3173 | 30.1117 | 30.3516 | 30.3145 | 30.6442 | 30.7284 |
| 33.7942 | 33.7489 | 33.6905 | 33.7587 | 33.7554 | 33.9462 | 33.9532 |
| 30.2374 | 30.4313 | 30.2824 | 30.3735 | 30.3317 | 30.4712 | 30.4728 |
| 30.962 | 31.2462 | 31.1826 | 31.0546 | 31.1004 | 30.9436 | 30.9505 |
| 30.6587 | 30.8262 | 30.7885 | 30.8002 | 30.7882 | 30.6191 | 30.5924 |
| 31.651 | 31.7664 | 31.8 | 31.7844 | 31.7609 | 31.6497 | 31.6709 |
| 32.3925 | 32.2797 | 32.1962 | 32.252 | 32.257 | 32.5115 | 32.4825 |
| 31.2013 | 30.9572 | 30.9038 | 30.9116 | 30.893 | 31.0764 | 31.0747 |
| 32.7308 | 32.7449 | 32.6247 | 32.7795 | 32.8121 | 32.6838 | 32.5837 |
| 34.6481 | 34.7062 | 34.6629 | 34.5881 | 34.5888 | 34.6008 | 34.6411 |
| 30.7947 | 30.941 | 30.8792 | 30.733 | 30.7096 | 30.691 | 30.6975 |
| 32.532 | 32.86 | 32.7847 | 32.6266 | 32.6768 | 32.5331 | 32.5482 |
| 31.3539 | 31.2015 | 31.1668 | 31.1253 | 31.1766 | 31.3761 | 31.3475 |
| 31.5536 | 31.4127 | 31.3246 | 31.5494 | 31.4799 | 31.7569 | 31.7818 |
| 29.1323 | 29.5236 | 29.6002 | 29.2234 | 29.3208 | 29.5237 | 29.5464 |
| 31.7649 | 31.39 | 31.4126 | 31.585 | 31.5679 | 31.7276 | 31.7336 |
| 30.3364 | 30.5493 | 30.5082 | 30.2177 | 30.281 | 30.1797 | 30.2667 |
| 30.0326 | 30.0774 | 29.9258 | 29.8727 | 29.8319 | 30.004 | 29.9724 |
| 29.2752 | 29.4401 | 29.2916 | 29.2767 | 29.2733 | 29.6225 | 29.5841 |
| 32.2306 | 32.3847 | 32.305 | 32.2462 | 32.282 | 32.1748 | 32.147 |
| 33.1414 | 33.4304 | 33.3763 | 33.2901 | 33.3036 | 33.2141 | 33.22 |
| 32.5388 | 32.7563 | 32.7036 | 32.66 | 32.6915 | 32.6627 | 32.6823 |
| 30.6399 | 30.5714 | 30.5208 | 30.6034 | 30.5464 | 30.7056 | 30.645 |
| 34.7387 | 34.8089 | 34.7183 | 34.7163 | 34.7427 | 34.5359 | 34.5335 |
| 30.3953 | 30.4401 | 30.3936 | 30.2327 | 30.2361 | 30.6861 | 30.6241 |
| 31.4979 | 32.0741 | 32.0498 | 31.8037 | 31.8476 | 31.9174 | 31.7979 |
| 29.9969 | 30.079 | 29.9816 | 29.8294 | 29.8836 | 30.2907 | 30.2364 |
| 30.581 | 30.6925 | 30.6329 | 30.5779 | 30.5235 | 30.5953 | 30.5857 |
| 29.7602 | 29.8228 | 29.8784 | 29.7171 | 29.6972 | 29.4178 | 29.4373 |
| 33.2781 | 33.3258 | 33.2797 | 33.3033 | 33.2584 | 33.4844 | 33.4433 |
| 37.4331 | 37.6652 | 37.6578 | 37.339 | 37.3668 | 37.4588 | 37.5439 |
| 29.6508 | 29.9566 | 29.9238 | 29.83 | 29.8058 | 29.6147 | 29.6608 |
| 29.8171 | 30.2832 | 30.1376 | 29.9731 | 29.9586 | 29.9278 | 29.9425 |
| 32.8666 | 33.0516 | 32.9281 | 32.938 | 32.9563 | 32.7798 | 32.7827 |
| 32.0095 | 32.0418 | 32.0055 | 32.22 | 32.2198 | 32.0881 | 31.9687 |
| 29.4814 | 29.7851 | 29.8341 | 29.6611 | 29.6929 | 29.829 | 29.7758 |
| 30.8627 | 30.9624 | 30.9121 | 30.8608 | 30.9021 | 30.7949 | 30.821 |
| 31.596 | 31.7839 | 31.711 | 31.6615 | 31.6958 | 31.447 | 31.459 |
| 29.1454 | 28.9706 | 28.9887 | 29.0246 | 29.0919 | 29.1935 | 29.166 |
| 32.7673 | 32.838 | 32.8046 | 32.7288 | 32.7577 | 32.7718 | 32.8631 |
| 34.7093 | 34.7335 | 34.5881 | 34.5327 | 34.5059 | 34.7956 | 34.7655 |
| 29.3785 | 29.133 | 29.1299 | 29.1873 | 29.2413 | 29.3767 | 29.2946 |
| 30.7959 | 30.5856 | 30.4731 | 30.5157 | 30.4722 | 30.575 | 30.5438 |

| | | | | | | |
|---|---|---|---|---|---|---|
| 32.9692 | 33.3012 | 33.2383 | 32.9914 | 33.0863 | 33.1757 | 33.1966 |
| 32.3427 | 32.4408 | 32.4271 | 32.3778 | 32.3798 | 32.551 | 32.5852 |
| 29.4431 | 29.4118 | 29.3883 | 29.3389 | 29.377 | 29.2507 | 29.2479 |
| 30.7258 | 30.8698 | 30.7997 | 30.7089 | 30.7123 | 30.6076 | 30.5945 |
| 32.3897 | 32.7931 | 32.8162 | 32.5305 | 32.4905 | 32.6343 | 32.682 |
| 31.8018 | 32.0462 | 31.9809 | 31.8871 | 31.8653 | 31.7639 | 31.7016 |
| 28.9592 | 29.1182 | 29.0298 | 29.0569 | 29.0714 | 28.9735 | 28.9159 |
| 31.1815 | 31.4048 | 31.3201 | 31.252 | 31.2769 | 31.1306 | 31.0942 |
| 31.3905 | 31.0939 | 31.0289 | 31.1343 | 31.147 | 31.3204 | 31.3643 |
| 33.2146 | 33.437 | 33.4313 | 33.3288 | 33.3284 | 33.3468 | 33.3346 |
| 31.565 | 31.3693 | 31.2974 | 31.2543 | 31.304 | 31.574 | 31.5032 |
| 29.9609 | 29.7253 | 29.7173 | 29.7419 | 29.7249 | 30.0456 | 29.987 |
| 34.6893 | 35.0317 | 34.9306 | 34.7449 | 34.7536 | 34.6866 | 34.718 |
| 32.8185 | 33.0437 | 33.0172 | 32.7724 | 32.7692 | 32.783 | 32.7842 |
| 30.4942 | 30.9484 | 30.8902 | 30.6933 | 30.7325 | 30.7865 | 30.741 |
| 30.3115 | 30.5049 | 30.5559 | 30.3362 | 30.3556 | 30.4155 | 30.4001 |
| 32.931 | 32.8071 | 32.714 | 32.6672 | 32.7306 | 32.8194 | 32.8334 |
| 32.6978 | 32.8242 | 32.7822 | 32.6791 | 32.676 | 32.6748 | 32.6833 |
| 32.0277 | 31.8791 | 31.975 | 32.0123 | 32.0233 | 31.7857 | 31.7937 |
| 34.0063 | 34.1085 | 34.0921 | 33.912 | 33.9569 | 33.829 | 33.8929 |
| 32.4233 | 32.579 | 32.5169 | 32.4407 | 32.4439 | 32.4739 | 32.5097 |
| 30.7994 | 30.6782 | 30.6089 | 30.6757 | 30.6917 | 30.881 | 30.8318 |
| 28.3323 | 28.3649 | 28.1788 | 28.0729 | 28.112 | 28.2017 | 28.0705 |
| 30.7704 | 30.696 | 30.6122 | 30.6197 | 30.6292 | 30.7898 | 30.7626 |
| 28.849 | 29.0941 | 28.9255 | 28.944 | 28.9186 | 28.8134 | 28.859 |
| 30.2601 | 30.4123 | 30.3758 | 30.2808 | 30.3108 | 30.1172 | 30.0936 |
| 29.0663 | 29.0415 | 29.0099 | 29.2383 | 29.207 | 29.1051 | 29.0341 |
| 30.8834 | 30.8692 | 30.8498 | 30.7959 | 30.8582 | 30.9093 | 30.9173 |
| 30.928 | 30.6028 | 30.6041 | 30.9673 | 30.8281 | 30.9106 | 30.9098 |
| 32.0992 | 32.3535 | 32.2427 | 32.254 | 32.1993 | 32.0256 | 32.0145 |
| 34.2342 | 34.5372 | 34.4523 | 34.2871 | 34.3158 | 34.2209 | 34.2234 |
| 29.4873 | 29.6088 | 29.5338 | 29.4935 | 29.5568 | 29.3453 | 29.4051 |
| 34.0661 | 34.1386 | 34.1918 | 34.1453 | 34.1576 | 33.8579 | 33.8537 |
| 30.0284 | 30.0436 | 29.9011 | 29.881 | 29.8627 | 30.1501 | 30.1995 |
| 31.8812 | 31.9314 | 31.8882 | 31.8128 | 31.835 | 32.1068 | 32.1068 |
| 28.6114 | 28.652 | 28.5451 | 28.401 | 28.5094 | 28.5401 | 28.6496 |
| 29.4175 | 29.6012 | 29.567 | 29.4971 | 29.447 | 29.6068 | 29.5153 |
| 30.916 | 30.7373 | 30.7189 | 30.6854 | 30.7351 | 30.7136 | 30.7585 |
| 33.2453 | 33.488 | 33.4038 | 33.3208 | 33.2716 | 33.1925 | 33.25 |
| 29.2734 | 29.3962 | 29.2894 | 29.2372 | 29.2922 | 29.0198 | 29.0611 |
| 30.6154 | 30.5511 | 30.445 | 30.4482 | 30.4464 | 30.5124 | 30.5939 |
| 33.3685 | 33.6394 | 33.5825 | 33.4513 | 33.4501 | 33.4082 | 33.3925 |
| 33.317 | 33.2947 | 33.1183 | 33.1994 | 33.2432 | 33.1566 | 33.1745 |
| 31.5514 | 31.6079 | 31.5656 | 31.4313 | 31.442 | 31.3757 | 31.3343 |
| 34.6518 | 34.8932 | 34.8374 | 34.715 | 34.6889 | 34.5959 | 34.6554 |
| 34.3382 | 34.7181 | 34.6273 | 34.4041 | 34.3757 | 34.3539 | 34.4125 |
| 32.8755 | 32.8849 | 32.8831 | 32.8435 | 32.8316 | 32.6795 | 32.6449 |
| 35.2507 | 35.2803 | 35.1795 | 35.0886 | 35.1562 | 35.1161 | 35.1097 |

| | | | | | | |
|---|---|---|---|---|---|---|
| 31.1036 | 31.0615 | 30.9498 | 30.9435 | 30.8852 | 31.192 | 31.2321 |
| 31.553 | 31.908 | 31.8298 | 31.7194 | 31.699 | 31.7174 | 31.6725 |
| 28.1526 | 28.1187 | 28.1135 | 28.1497 | 28.1305 | 28.0309 | 28.06 |
| 30.2887 | 30.5882 | 30.395 | 30.4529 | 30.4274 | 30.3434 | 30.3016 |
| 32.8898 | 32.7796 | 32.6402 | 32.4682 | 32.4883 | 32.9952 | 32.9885 |
| 33.8076 | 33.6867 | 33.6303 | 33.6455 | 33.6099 | 33.7321 | 33.7865 |
| 34.7215 | 34.9684 | 34.8597 | 34.7487 | 34.7731 | 34.6716 | 34.6145 |
| 29.9909 | 29.8494 | 29.7345 | 29.8786 | 29.8088 | 29.9807 | 29.9664 |
| 29.4072 | 29.7739 | 29.6495 | 29.7665 | 29.618 | 29.5736 | 29.4973 |
| 29.8202 | 29.8695 | 29.8325 | 29.9731 | 29.9177 | 29.8151 | 29.7799 |
| 30.5163 | 30.4187 | 30.3441 | 30.5058 | 30.5334 | 30.6926 | 30.7331 |
| 32.2189 | 32.4012 | 32.2843 | 32.2562 | 32.3031 | 32.2021 | 32.1431 |
| 28.7892 | 28.915 | 28.8837 | 28.815 | 28.8787 | 28.8788 | 28.9542 |
| 31.5825 | 31.6615 | 31.6355 | 31.6535 | 31.6044 | 31.6406 | 31.625 |
| 31.3383 | 31.2308 | 31.19 | 31.2143 | 31.3313 | 31.2312 | 31.3414 |
| 34.2544 | 34.118 | 33.9955 | 33.9517 | 33.9744 | 33.9287 | 33.997 |
| 33.6769 | 33.6354 | 33.5961 | 33.6344 | 33.6179 | 33.7387 | 33.791 |
| 29.8447 | 29.8674 | 29.6566 | 29.8433 | 29.7506 | 30.035 | 30.0062 |
| 31.1081 | 31.162 | 31.1106 | 31.1391 | 31.1673 | 30.7987 | 30.7737 |
| 31.9125 | 31.9088 | 31.8555 | 31.7451 | 31.7057 | 32.0152 | 31.9905 |
| 30.4775 | 30.3464 | 30.3236 | 30.3184 | 30.2892 | 30.6686 | 30.6598 |
| 28.448 | 28.3133 | 28.14 | 28.3809 | 28.2717 | 28.5899 | 28.5684 |
| 30.7067 | 30.8814 | 30.8393 | 30.7091 | 30.6863 | 30.5482 | 30.561 |
| 28.9903 | 29.2122 | 29.1891 | 29.0752 | 28.9902 | 29.0018 | 29.0429 |
| 33.4127 | 33.5494 | 33.4991 | 33.4448 | 33.4064 | 33.5629 | 33.5606 |
| 28.6644 | 28.6541 | 28.7362 | 28.6115 | 28.6328 | 28.7692 | 28.6831 |
| 30.2175 | 30.3695 | 30.3595 | 30.3475 | 30.3315 | 30.5132 | 30.4829 |
| 29.0551 | 29.2905 | 29.2963 | 29.3507 | 29.278 | 29.1053 | 29.0963 |
| 31.6247 | 32.0644 | 31.862 | 31.7419 | 31.7723 | 31.6608 | 31.7185 |
| 30.0632 | 29.9885 | 29.8449 | 30.0437 | 30.0131 | 30.2093 | 30.2211 |
| 33.3502 | 33.4264 | 33.4355 | 33.4129 | 33.4553 | 33.2398 | 33.2756 |
| 31.5565 | 31.7918 | 31.7159 | 31.6557 | 31.6662 | 31.4964 | 31.4638 |
| 29.8416 | 30.1341 | 30.1999 | 29.9998 | 29.9866 | 30.1193 | 30.066 |
| 31.758 | 31.6462 | 31.533 | 31.7088 | 31.7167 | 31.747 | 31.7306 |
| 29.5911 | 29.6387 | 29.5919 | 29.5151 | 29.5867 | 29.5194 | 29.4973 |
| 30.9536 | 30.6935 | 30.7182 | 30.7683 | 30.7686 | 30.9338 | 30.9561 |
| 31.7194 | 31.7328 | 31.6501 | 31.6927 | 31.6891 | 31.3698 | 31.4157 |
| 30.4364 | 30.126 | 30.1025 | 30.3337 | 30.2979 | 30.5586 | 30.5481 |
| 28.3165 | 28.4338 | 28.4417 | 28.2448 | 28.3359 | 28.3592 | 28.3007 |
| 29.1542 | 29.0418 | 28.9844 | 28.9136 | 29.0139 | 29.3112 | 29.3347 |
| 32.5953 | 32.3968 | 32.3103 | 32.4555 | 32.4088 | 32.4485 | 32.4586 |
| 29.8376 | 30.0704 | 29.8765 | 29.9734 | 29.8523 | 29.9097 | 29.8754 |
| 28.4804 | 28.3126 | 28.3214 | 28.312 | 28.3385 | 28.8379 | 28.7748 |
| 31.2973 | 31.3396 | 31.3147 | 31.469 | 31.4427 | 31.1325 | 31.0806 |
| 31.2109 | 31.3172 | 31.2995 | 31.1918 | 31.2106 | 31.1687 | 31.1904 |
| 29.3812 | 29.6189 | 29.6064 | 29.6036 | 29.6124 | 29.3421 | 29.2409 |
| 29.5361 | 29.4146 | 29.4713 | 29.5844 | 29.5085 | 29.4489 | 29.3769 |
| 32.2809 | 32.1915 | 32.1751 | 32.0254 | 32.0623 | 32.4162 | 32.3865 |

| | | | | | | |
|---|---|---|---|---|---|---|
| 31.5419 | 31.2643 | 31.2331 | 31.4165 | 31.4276 | 31.8305 | 31.8163 |
| 29.1827 | 28.9918 | 28.9868 | 29.0441 | 29.0254 | 29.2467 | 29.3187 |
| 28.7037 | 28.647 | 28.44 | 28.6963 | 28.6692 | 28.5157 | 28.3808 |
| 29.9555 | 30.1046 | 30.0525 | 29.9537 | 29.9844 | 29.9005 | 29.8669 |
| 28.4664 | 28.3086 | 28.2896 | 28.3482 | 28.3478 | 28.2578 | 28.2919 |
| 33.4515 | 33.6477 | 33.6234 | 33.5983 | 33.5403 | 33.3188 | 33.3457 |
| 29.516 | 29.6034 | 29.5288 | 29.7606 | 29.6399 | 29.4897 | 29.5274 |
| 28.1418 | 28.0572 | 28.1561 | 28.1978 | 28.1981 | 28.2236 | 28.184 |
| 29.9586 | 30.2571 | 30.226 | 30.0903 | 30.0992 | 29.9084 | 29.9798 |
| 28.4848 | 28.3829 | 28.2936 | 28.3662 | 28.3235 | 28.5035 | 28.4572 |
| 30.194 | 30.0037 | 29.8799 | 30.0562 | 30.0673 | 30.0741 | 30.1176 |
| 29.7129 | 29.4377 | 29.3988 | 29.3675 | 29.3597 | 29.6083 | 29.6164 |
| 30.6524 | 30.6667 | 30.6489 | 30.3899 | 30.438 | 30.9267 | 30.9654 |
| 28.5268 | 28.7281 | 28.7418 | 28.6357 | 28.6704 | 28.4392 | 28.5485 |
| 33.3145 | 33.2565 | 33.3251 | 33.2122 | 33.1469 | 33.2734 | 33.2339 |
| 32.7136 | 32.6605 | 32.5471 | 32.4352 | 32.4983 | 33.0521 | 33.1273 |
| 31.2841 | 31.4298 | 31.3443 | 31.3317 | 31.3584 | 31.2802 | 31.2999 |
| 30.6739 | 30.8336 | 30.7413 | 30.8228 | 30.8225 | 30.6124 | 30.653 |
| 29.1382 | 29.2017 | 29.1106 | 29.2106 | 29.2128 | 29.4891 | 29.4614 |
| 30.8734 | 31.0704 | 30.9183 | 30.8941 | 30.9256 | 31.0807 | 31.0982 |
| 28.7643 | 29.1611 | 29.0762 | 28.9331 | 28.9203 | 29.061 | 29.107 |
| 29.3234 | 29.166 | 29.0824 | 29.2087 | 29.2159 | 28.6811 | 28.6722 |
| 34.2363 | 34.5274 | 34.3646 | 34.2908 | 34.2967 | 34.0813 | 34.1366 |
| 29.4384 | 29.7224 | 29.7072 | 29.4235 | 29.4186 | 29.3909 | 29.3417 |
| 34.2155 | 34.3582 | 34.2992 | 34.1805 | 34.2415 | 34.1593 | 34.1507 |
| 30.0483 | 30.2127 | 30.0242 | 30.2059 | 30.1725 | 30.1694 | 30.2175 |
| 31.2593 | 31.5141 | 31.4503 | 31.3518 | 31.3805 | 31.2713 | 31.3139 |
| 33.0491 | 33.1621 | 33.1453 | 33.1366 | 33.1246 | 33.0324 | 33.0472 |
| 36.4454 | 36.6816 | 36.5405 | 36.2868 | 36.3107 | 36.2594 | 36.3755 |
| 29.253 | 29.1091 | 29.1733 | 29.285 | 29.213 | 29.3322 | 29.3748 |
| 31.028 | 31.1937 | 31.1079 | 31.2127 | 31.145 | 31.0335 | 31.0264 |
| 31.1321 | 31.4369 | 31.3316 | 31.1285 | 31.0969 | 31.1619 | 31.1661 |
| 29.5564 | 29.3383 | 29.2959 | 29.4953 | 29.5589 | 29.6452 | 29.59 |
| 33.3002 | 33.4588 | 33.3145 | 33.2891 | 33.2656 | 33.246 | 33.2299 |
| 31.1266 | 31.365 | 31.3845 | 31.2041 | 31.1995 | 31.176 | 31.1832 |
| 28.5432 | 28.293 | 28.2807 | 28.3141 | 28.3728 | 28.3277 | 28.3591 |
| 29.2091 | 29.2576 | 29.1574 | 29.2491 | 29.1899 | 29.0378 | 29.0169 |
| 31.4883 | 31.6732 | 31.6439 | 31.5743 | 31.5476 | 31.5163 | 31.5212 |
| 30.9523 | 30.9019 | 30.8479 | 30.8281 | 30.8844 | 30.9771 | 30.9978 |
| 30.1899 | 29.8917 | 29.7756 | 29.9843 | 29.9887 | 29.9324 | 29.9295 |
| 29.9809 | 29.7471 | 29.6461 | 29.8933 | 29.9064 | 30.0268 | 30.0088 |
| 31.9285 | 32.1771 | 31.9878 | 31.7242 | 31.7412 | 32.5146 | 32.4654 |
| 32.0223 | 31.6673 | 31.6394 | 31.8006 | 31.8489 | 32.17 | 32.2105 |
| 30.3916 | 30.5764 | 30.5782 | 30.2463 | 30.3144 | 30.3365 | 30.3317 |
| 29.338 | 29.3844 | 29.2277 | 29.2823 | 29.3098 | 29.3711 | 29.3692 |
| 30.1332 | 29.7415 | 29.6147 | 29.7633 | 29.8519 | 30.0419 | 30.0135 |
| 28.9828 | 28.8669 | 28.9235 | 28.7745 | 28.5947 | 29.259 | 29.1207 |
| 28.8951 | 29.1697 | 29.13 | 28.9876 | 29.0177 | 28.9823 | 28.9445 |

| | | | | | | |
|---|---|---|---|---|---|---|
| 29.4459 | 29.281 | 29.328 | 29.3853 | 29.3939 | 29.4081 | 29.3943 |
| 32.3466 | 32.0618 | 32.0542 | 31.9544 | 31.9993 | 32.3741 | 32.3747 |
| 34.7051 | 34.3925 | 34.2429 | 34.3336 | 34.3264 | 34.7481 | 34.7395 |
| 31.1591 | 31.1739 | 31.0792 | 31.0705 | 31.1151 | 30.8957 | 30.9571 |
| 33.3527 | 33.3919 | 33.2657 | 33.1382 | 33.1356 | 33.3131 | 33.3819 |
| 32.952 | 33.3878 | 33.2702 | 32.9793 | 32.9563 | 33.1763 | 33.1801 |
| 31.4991 | 31.5851 | 31.4005 | 31.4507 | 31.5685 | 31.5396 | 31.5314 |
| 29.4099 | 29.3041 | 29.2315 | 29.3363 | 29.4332 | 29.5565 | 29.3868 |
| 29.1322 | 29.2292 | 29.1825 | 29.0746 | 29.0672 | 29.1642 | 29.1642 |
| 32.0541 | 32.5019 | 32.3774 | 32.2154 | 32.3065 | 32.1033 | 32.1491 |
| 33.1048 | 33.3974 | 33.3176 | 33.2179 | 33.2295 | 33.169 | 33.1296 |
| 28.9637 | 28.9937 | 28.9037 | 28.8854 | 28.8415 | 29.0313 | 29.041 |
| 29.3388 | 29.1235 | 29.2467 | 29.1707 | 29.1719 | 29.3883 | 29.3869 |
| 29.2181 | 29.0873 | 29.1117 | 29.2202 | 29.2001 | 29.4302 | 29.4211 |
| 33.402 | 33.5657 | 33.5218 | 33.4364 | 33.4892 | 33.2501 | 33.2611 |
| 30.8324 | 30.8227 | 30.7397 | 30.7028 | 30.6816 | 30.7677 | 30.8801 |
| 29.9541 | 30.0618 | 30.012 | 30.0931 | 30.1274 | 29.8208 | 29.8974 |
| 30.74 | 30.636 | 30.5798 | 30.7706 | 30.8083 | 30.7956 | 30.7638 |
| 29.2281 | 29.3058 | 29.2571 | 29.1039 | 29.0537 | 28.9167 | 29.006 |
| 29.4264 | 29.1112 | 28.9357 | 29.0599 | 28.9634 | 29.0898 | 29.2197 |
| 28.6941 | 28.8779 | 28.8899 | 28.6776 | 28.5787 | 28.7317 | 28.7464 |
| 30.9624 | 31.15 | 31.1316 | 30.9518 | 30.9885 | 30.8613 | 30.8413 |
| 27.889 | 27.9408 | 27.9518 | 27.9881 | 27.9457 | 27.6262 | 27.8107 |
| 31.708 | 31.9314 | 31.8603 | 31.654 | 31.6828 | 31.5873 | 31.5991 |
| 31.5649 | 31.841 | 31.8054 | 31.7991 | 31.7907 | 31.8422 | 31.8188 |
| 28.1329 | 28.2264 | 28.2049 | 28.2082 | 28.2004 | 28.2335 | 28.3634 |
| 29.5868 | 29.3775 | 29.365 | 29.5551 | 29.5265 | 29.6747 | 29.6697 |
| 29.5207 | 29.6005 | 29.4629 | 29.2057 | 29.1079 | 29.5325 | 29.5987 |
| 30.6611 | 30.9395 | 30.921 | 30.7468 | 30.7095 | 30.6781 | 30.7259 |
| 28.7842 | 28.8974 | 28.834 | 28.8799 | 28.941 | 28.7277 | 28.7139 |
| 27.4404 | 27.2442 | 27.3658 | 27.3772 | 27.3782 | 27.1443 | 27.2302 |
| 32.1327 | 32.2408 | 32.2189 | 32.1203 | 32.1483 | 32.1688 | 32.1807 |
| 31.8365 | 31.9473 | 31.7985 | 31.6832 | 31.7488 | 31.9367 | 32.0263 |
| 32.4199 | 32.723 | 32.6076 | 32.5274 | 32.5533 | 32.3065 | 32.3496 |
| 29.2136 | 29.2972 | 29.2411 | 29.2229 | 29.2269 | 28.9002 | 28.8696 |
| 29.5134 | 29.7524 | 29.6254 | 29.4563 | 29.51 | 29.399 | 29.355 |
| 29.631 | 29.8778 | 29.786 | 29.7184 | 29.7136 | 29.7039 | 29.7525 |
| 29.4225 | 29.2684 | 29.2271 | 29.1882 | 29.0566 | 29.7788 | 29.7982 |
| 31.113 | 31.0362 | 30.9604 | 30.89 | 30.9101 | 31.2427 | 31.3223 |
| 29.5891 | 29.8229 | 29.6309 | 29.6391 | 29.6594 | 29.9553 | 29.8345 |
| 33.0291 | 33.4754 | 33.4263 | 33.2808 | 33.2814 | 33.2233 | 33.181 |
| 30.2955 | 30.4793 | 30.3416 | 30.3895 | 30.4282 | 30.361 | 30.321 |
| 28.2089 | 28.1669 | 28.133 | 28.3006 | 28.2883 | 28.532 | 28.5004 |
| 30.7189 | 31.0597 | 31.0652 | 30.9811 | 30.8839 | 30.7238 | 30.8349 |
| 28.4119 | 28.6376 | 28.6395 | 28.6054 | 28.6726 | 28.5862 | 28.521 |
| 30.4549 | 30.6808 | 30.6718 | 30.5544 | 30.5105 | 30.5184 | 30.4633 |
| 32.8909 | 33.1514 | 33.096 | 32.9869 | 32.9502 | 32.94 | 32.9582 |
| 29.5069 | 29.5157 | 29.451 | 29.3378 | 29.4062 | 29.3682 | 29.3698 |

| | | | | | | |
|---|---|---|---|---|---|---|
| 30.6636 | 31.19 | 31.114 | 30.7732 | 30.8272 | 30.9843 | 31.0428 |
| 31.5181 | 31.4177 | 31.3449 | 31.2332 | 31.186 | 31.6776 | 31.6925 |
| 30.5242 | 30.4574 | 30.4318 | 30.4736 | 30.4043 | 30.5777 | 30.5478 |
| 28.3254 | 28.2998 | 28.2729 | 28.4754 | 28.4849 | 28.1532 | 28.1409 |
| 31.6297 | 31.6694 | 31.546 | 31.6727 | 31.6059 | 31.4426 | 31.5299 |
| 30.1368 | 30.4479 | 30.4387 | 30.3017 | 30.3028 | 30.2216 | 30.1788 |
| 30.6718 | 30.8801 | 30.8372 | 30.8125 | 30.7811 | 30.5741 | 30.5638 |
| 29.4227 | 28.7612 | 28.8352 | 29.2666 | 29.3721 | 29.6136 | 29.6174 |
| 28.8619 | 29.1996 | 29.0178 | 29.0373 | 29.0655 | 28.8074 | 28.8464 |
| 30.0427 | 29.7986 | 29.8671 | 29.9018 | 30.0256 | 29.9828 | 29.9632 |
| 30.3112 | 30.0674 | 29.9754 | 30.0746 | 30.1296 | 30.1169 | 30.071 |
| 29.3754 | 29.8218 | 29.6017 | 29.4351 | 29.4388 | 29.4106 | 29.4253 |
| 30.1105 | 30.4631 | 30.3386 | 30.1092 | 30.1422 | 30.1546 | 30.1852 |
| 28.7926 | 28.8877 | 28.7409 | 28.8607 | 28.8332 | 28.7276 | 28.7636 |
| 28.1026 | 27.9307 | 28.2843 | 28.1753 | 28.068 | 28.235 | 28.2774 |
| 30.2138 | 30.209 | 30.1189 | 30.1903 | 30.1569 | 30.0688 | 30.0456 |
| 28.7906 | 28.8455 | 28.5864 | 28.7225 | 28.7316 | 28.5227 | 28.476 |
| 29.6285 | 29.465 | 29.4208 | 29.6687 | 29.6558 | 29.6792 | 29.6687 |
| 29.9028 | 29.6822 | 29.7022 | 29.7459 | 29.7188 | 29.8212 | 29.7921 |
| 31.0913 | 30.9877 | 31.0196 | 30.9284 | 30.8203 | 31.0604 | 30.9941 |
| 31.0742 | 31.1357 | 31.1207 | 30.9502 | 30.8689 | 31.1834 | 31.1868 |
| 31.2632 | 31.7204 | 31.5926 | 31.3419 | 31.3119 | 31.5578 | 31.6139 |
| 29.8249 | 30.0756 | 30.0152 | 29.7353 | 29.8149 | 29.7041 | 29.7573 |
| 33.6036 | 33.5803 | 33.558 | 33.6912 | 33.6577 | 33.2016 | 33.2473 |
| 30.3398 | 30.5466 | 30.479 | 30.5511 | 30.5716 | 30.6349 | 30.5481 |
| 30.157 | 30.3619 | 30.2885 | 30.2837 | 30.3343 | 30.3952 | 30.2582 |
| 30.756 | 30.5901 | 30.5448 | 30.481 | 30.4945 | 30.6216 | 30.5509 |
| 30.2457 | 30.1753 | 30.0598 | 30.2023 | 30.1322 | 30.2671 | 30.2854 |
| 29.713 | 30.0342 | 30.0816 | 29.8531 | 29.8831 | 29.987 | 29.9743 |
| 29.3295 | 29.5525 | 29.4144 | 29.4756 | 29.5094 | 29.2251 | 29.178 |
| 32.6464 | 32.7638 | 32.6659 | 32.2926 | 32.3537 | 32.8028 | 32.9055 |
| 32.8257 | 33.1983 | 33.0542 | 32.9932 | 32.9324 | 32.8414 | 32.8165 |
| 30.7809 | 30.8105 | 30.762 | 30.7244 | 30.6851 | 30.6731 | 30.6837 |
| 31.2156 | 31.1276 | 31.2003 | 31.0806 | 30.9682 | 31.5193 | 31.4965 |
| 34.7778 | 35.0782 | 34.9696 | 34.7743 | 34.765 | 34.6827 | 34.8224 |
| 32.2455 | 32.3167 | 32.3416 | 32.4906 | 32.5123 | 32.3326 | 32.3514 |
| 28.5306 | 28.4116 | 28.273 | 28.4149 | 28.3873 | 28.769 | 28.7597 |
| 29.7146 | 29.6969 | 29.6736 | 29.5392 | 29.6399 | 29.6437 | 29.7161 |
| 35.6403 | 35.9724 | 35.8434 | 35.6603 | 35.6608 | 35.5925 | 35.5994 |
| 30.1017 | 30.2271 | 30.1358 | 30.2001 | 30.0889 | 29.8015 | 29.8148 |
| 29.8438 | 29.9275 | 29.7879 | 29.8284 | 29.9101 | 29.6581 | 29.5275 |
| 30.6592 | 30.8826 | 30.8971 | 30.6837 | 30.686 | 30.5633 | 30.6579 |
| 32.8155 | 33.0417 | 33.0955 | 32.8629 | 32.8402 | 32.7952 | 32.8343 |
| 30.3258 | 29.8773 | 29.8597 | 30.1407 | 30.097 | 30.2413 | 30.1968 |
| 29.2948 | 28.9468 | 29.1386 | 29.1891 | 29.1584 | 29.3647 | 29.3364 |
| 28.4867 | 28.3637 | 28.2237 | 28.3009 | 28.3053 | 28.4516 | 28.4946 |
| 30.7573 | 31.0768 | 31.0036 | 30.7758 | 30.7636 | 30.7568 | 30.8466 |
| 30.8914 | 31.3144 | 31.2251 | 31.1353 | 31.1019 | 31.036 | 31.007 |

| | | | | | | |
|---|---|---|---|---|---|---|
| 32.9762 | 33.4165 | 33.2894 | 33.1542 | 33.1343 | 33.1707 | 33.1103 |
| 31.5751 | 31.8368 | 31.8783 | 31.6931 | 31.7094 | 31.6754 | 31.599 |
| 31.1359 | 31.397 | 31.3862 | 31.2865 | 31.3264 | 31.2226 | 31.2369 |
| 29.0499 | 28.9708 | 28.9432 | 28.9923 | 28.9755 | 29.0594 | 29.0847 |
| 29.7881 | 29.8555 | 29.7117 | 29.8162 | 29.778 | 29.9507 | 29.8677 |
| 29.5377 | 29.5061 | 29.507 | 29.5185 | 29.5412 | 29.6078 | 29.5884 |
| 32.9784 | 32.8429 | 32.8229 | 32.8702 | 32.8322 | 32.5591 | 32.5524 |
| 31.7449 | 32.114 | 32.0426 | 31.8577 | 31.861 | 31.8699 | 31.8999 |
| 29.5173 | 29.3941 | 29.4863 | 29.5279 | 29.49 | 29.8013 | 29.7144 |
| 29.8301 | 29.7562 | 29.6469 | 29.7221 | 29.8718 | 29.8646 | 29.8832 |
| 31.5197 | 31.2376 | 31.3614 | 31.4221 | 31.4415 | 31.4387 | 31.3981 |
| 28.7674 | 28.9597 | 28.9581 | 28.9545 | 29.0277 | 28.7347 | 28.6599 |
| 29.4828 | 29.2985 | 29.4736 | 29.3286 | 29.3238 | 29.5434 | 29.5917 |
| 29.4482 | 29.0769 | 28.9156 | 29.0735 | 29.0809 | 29.5976 | 29.5838 |
| 30.4923 | 30.3006 | 30.2374 | 30.3699 | 30.4194 | 30.5466 | 30.4506 |
| 30.9611 | 31.0811 | 31.0528 | 31.0727 | 31.1322 | 30.9475 | 30.9297 |
| 31.2017 | 31.2718 | 31.1472 | 31.3127 | 31.2681 | 31.0036 | 30.9209 |
| 32.2094 | 32.4679 | 32.5339 | 32.4319 | 32.4087 | 32.6597 | 32.6363 |
| 33.528 | 33.537 | 33.4127 | 33.2487 | 33.2427 | 33.6708 | 33.6574 |
| 29.0396 | 29.15 | 29.1312 | 29.2211 | 29.1876 | 29.0286 | 28.9639 |
| 30.9962 | 30.9084 | 31.0525 | 31.0658 | 31.0678 | 31.0621 | 30.998 |
| 29.4861 | 29.6768 | 29.6074 | 29.4832 | 29.5346 | 29.4069 | 29.391 |
| 29.99 | 29.8563 | 29.8429 | 29.8115 | 29.8762 | 29.863 | 29.8987 |
| 28.8749 | 28.841 | 28.5562 | 28.7485 | 28.87 | 28.8848 | 28.7632 |
| 29.139 | 29.5311 | 29.4754 | 29.1011 | 29.1782 | 29.2344 | 29.2806 |
| 34.7004 | 34.8083 | 34.7018 | 34.7329 | 34.741 | 34.452 | 34.5353 |
| 30.9541 | 31.0639 | 31.0966 | 30.9704 | 30.878 | 30.8946 | 30.8991 |
| 30.4156 | 30.4345 | 30.3936 | 30.3767 | 30.401 | 30.1593 | 30.1162 |
| 28.3878 | 28.2048 | 28.105 | 28.1333 | 28.269 | 28.1379 | 28.1438 |
| 30.4991 | 30.4843 | 30.4092 | 30.3511 | 30.3154 | 30.4806 | 30.4412 |
| 29.1373 | 29.0231 | 28.9631 | 28.9621 | 28.8968 | 28.645 | 28.8202 |
| 29.8026 | 29.9628 | 29.859 | 29.9104 | 29.9791 | 29.9939 | 29.9874 |
| 31.2865 | 31.2858 | 31.2609 | 31.2295 | 31.2147 | 31.3367 | 31.3208 |
| 28.3092 | 28.1937 | 27.9457 | 28.2613 | 28.2424 | 28.2034 | 28.2082 |
| 29.5856 | 29.816 | 29.7764 | 29.6047 | 29.6528 | 29.5383 | 29.5956 |
| 33.148 | 33.6756 | 33.6216 | 33.3392 | 33.3008 | 33.482 | 33.4962 |
| 28.3047 | 28.6204 | 28.6009 | 28.5324 | 28.4821 | 28.262 | 28.3204 |
| 29.9624 | 30.2488 | 30.1019 | 30.0225 | 30.0263 | 29.9023 | 29.957 |
| 30.8959 | 31.301 | 31.2469 | 31.0424 | 31.0475 | 30.9449 | 31.0147 |
| 31.5844 | 31.2539 | 31.2421 | 31.2645 | 31.2821 | 31.5329 | 31.5369 |
| 29.7548 | 29.7058 | 29.5043 | 29.8265 | 29.7927 | 29.9927 | 29.9929 |
| 27.9881 | 28.1389 | 27.8922 | 28.2111 | 28.1802 | 28.0241 | 27.7843 |
| 30.8265 | 30.4897 | 30.5085 | 30.7909 | 30.7101 | 30.8108 | 30.8382 |
| 27.7676 | 27.5504 | 27.7514 | 27.8494 | 27.7054 | 27.6012 | 27.582 |
| 28.3783 | 28.2628 | 28.0542 | 28.1304 | 28.2876 | 28.3473 | 28.3542 |
| 27.8604 | 28.1942 | 28.0016 | 28.0038 | 27.92 | 27.8998 | 27.9545 |
| 31.9201 | 31.9446 | 31.8801 | 31.7144 | 31.7618 | 31.9287 | 31.9 |
| 33.5935 | 33.6456 | 33.5329 | 33.4178 | 33.4279 | 33.0722 | 33.1093 |

| | | | | | | |
|---|---|---|---|---|---|---|
| 27.7914 | 27.799 | 27.5333 | 27.7574 | 27.5683 | 27.7438 | 27.8137 |
| 30.69 | 30.4761 | 30.4681 | 30.4108 | 30.453 | 30.6881 | 30.6921 |
| 29.5913 | 29.7244 | 29.6839 | 29.3695 | 29.4047 | 29.5272 | 29.6136 |
| 29.0458 | 28.9132 | 28.7918 | 28.9635 | 28.944 | 29.0312 | 29.0157 |
| 28.9524 | 28.9764 | 28.9274 | 28.8103 | 28.7959 | 28.9893 | 29.0372 |
| 28.8813 | 28.8758 | 28.777 | 28.8392 | 28.8513 | 28.7915 | 28.8513 |
| 31.0424 | 30.9573 | 30.9218 | 30.903 | 30.8974 | 31.0713 | 31.015 |
| 31.1201 | 31.3779 | 31.3188 | 31.1236 | 31.1852 | 30.988 | 31.0702 |
| 33.7021 | 33.4353 | 33.3569 | 33.4786 | 33.451 | 33.8424 | 33.8009 |
| 28.8954 | 28.7167 | 28.4981 | 29.0009 | 29.0274 | 28.4647 | 28.5855 |
| 29.361 | 29.2458 | 29.1704 | 29.1928 | 29.1445 | 29.4238 | 29.4449 |
| 30.1893 | 30.4362 | 30.3672 | 30.1428 | 30.1627 | 30.114 | 30.0992 |
| 34.1106 | 34.0899 | 34.0302 | 33.8578 | 33.884 | 34.0122 | 34.0832 |
| 29.2789 | 29.4702 | 29.406 | 29.2508 | 29.2175 | 28.9956 | 29.1583 |
| 30.3871 | 30.2017 | 30.1054 | 30.0788 | 30.0559 | 30.2665 | 30.2534 |
| 28.1557 | 27.9633 | 27.9817 | 27.9654 | 27.7985 | 28.2533 | 28.1663 |
| 29.1571 | 28.921 | 28.9127 | 29.0405 | 29.0273 | 29.4085 | 29.4712 |
| 28.4252 | 28.4682 | 28.4753 | 28.5985 | 28.579 | 28.3326 | 28.4082 |
| 33.2178 | 33.7084 | 33.5602 | 33.4355 | 33.4364 | 33.5452 | 33.5434 |
| 31.7941 | 31.8041 | 31.7599 | 31.7435 | 31.7086 | 31.8267 | 31.7391 |
| 30.3435 | 30.0464 | 29.9201 | 29.9471 | 29.9757 | 30.0329 | 30.0948 |
| 28.7901 | 28.7292 | 28.7067 | 28.9522 | 28.9183 | 28.9349 | 28.8448 |
| 28.1232 | 28.3364 | 28.2066 | 28.2653 | 28.2424 | 28.0504 | 28.1761 |
| 29.2798 | 29.6037 | 29.4352 | 29.4132 | 29.5158 | 29.3099 | 29.2761 |
| 28.7122 | 28.5619 | 28.7143 | 28.7208 | 28.845 | 28.804 | 28.8991 |
| 28.9288 | 28.8099 | 28.7577 | 28.7873 | 28.7422 | 29.1072 | 29.0728 |
| 29.0649 | 29.0769 | 28.9196 | 29.023 | 29.0747 | 29.2664 | 29.1906 |
| 28.6144 | 28.6369 | 28.4927 | 28.5634 | 28.6105 | 28.733 | 28.652 |
| 28.1775 | 28.6049 | 28.4702 | 28.3711 | 28.3878 | 28.1059 | 28.1382 |
| 28.48 | 28.5207 | 28.3522 | 28.4827 | 28.518 | 28.5441 | 28.4135 |
| 29.0656 | 29.0947 | 29.0178 | 28.5837 | 28.7226 | 29.2261 | 29.1103 |
| 27.2753 | 27.2981 | 27.2144 | 27.3142 | 27.3958 | 27.2087 | 27.0693 |
| 32.7216 | 32.8391 | 32.7418 | 32.5711 | 32.6118 | 32.9014 | 32.8936 |
| 31.7098 | 30.8281 | 30.6881 | 31.4531 | 31.4711 | 30.6474 | 30.6764 |
| 30.5013 | 30.6406 | 30.5231 | 30.6528 | 30.5692 | 30.5123 | 30.4715 |
| 29.0412 | 28.7803 | 28.7357 | 28.9376 | 28.8831 | 28.9426 | 28.7478 |
| 30.8143 | 30.8995 | 30.8861 | 30.8272 | 30.8923 | 30.6315 | 30.6134 |
| 30.2209 | 30.3789 | 30.4628 | 30.1813 | 30.0814 | 30.0084 | 30.1557 |
| 31.9786 | 32.1783 | 32.0995 | 31.9821 | 31.9804 | 31.8428 | 31.9083 |
| 28.0587 | 28.0064 | 28.0059 | 27.8564 | 27.7746 | 28.2112 | 28.1562 |
| 30.5643 | 30.5128 | 30.6614 | 30.6881 | 30.6703 | 30.5432 | 30.489 |
| 29.5043 | 29.3885 | 29.3317 | 29.4763 | 29.5128 | 29.255 | 29.2309 |
| 27.2395 | 27.4453 | 27.2718 | 27.2621 | 27.4572 | 27.4285 | 27.3885 |
| 28.0727 | 28.098 | 28.1574 | 28.0907 | 28.0868 | 27.807 | 27.9321 |
| 31.7576 | 31.7985 | 31.9007 | 31.9001 | 31.8776 | 31.7228 | 31.7361 |
| 32.2239 | 32.447 | 32.3725 | 32.3735 | 32.4529 | 32.4172 | 32.3948 |
| 33.5148 | 33.6445 | 33.6844 | 33.726 | 33.6893 | 33.5621 | 33.5344 |
| 28.5537 | 28.7932 | 28.7983 | 28.6727 | 28.7298 | 28.5426 | 28.5482 |

| | | | | | | |
|---|---|---|---|---|---|---|
| 30.6523 | 30.7462 | 30.7874 | 30.7753 | 30.7564 | 30.6564 | 30.7171 |
| 29.5733 | 29.7002 | 29.5771 | 29.681 | 29.6317 | 29.69 | 29.6787 |
| 29.9427 | 29.6383 | 29.6417 | 29.7378 | 29.7884 | 29.5284 | 29.6326 |
| 29.1754 | 29.2085 | 29.236 | 29.4579 | 29.3906 | 29.4058 | 29.3745 |
| 30.6347 | 31.6876 | 31.6237 | 30.9533 | 30.8914 | 31.3536 | 31.3587 |
| 30.869 | 31.072 | 30.8762 | 30.9567 | 31.0104 | 30.9197 | 30.8989 |
| 29.8508 | 29.8112 | 29.9211 | 29.7371 | 29.7364 | 29.6615 | 29.6554 |
| 27.6502 | 27.5476 | 27.6765 | 27.4263 | 27.7125 | 27.704 | 27.6619 |
| 28.5396 | 28.7204 | 28.6568 | 28.5059 | 28.5938 | 28.6111 | 28.5869 |
| 33.5085 | 33.6555 | 33.628 | 33.4379 | 33.4452 | 33.5629 | 33.6808 |
| 31.4433 | 31.685 | 31.5712 | 31.5714 | 31.5214 | 31.3254 | 31.3826 |
| 30.7099 | 30.8101 | 30.7474 | 30.5097 | 30.5185 | 30.8019 | 30.8096 |
| 33.6436 | 33.7571 | 33.8619 | 33.7377 | 33.7435 | 33.7146 | 33.7413 |
| 29.2666 | 29.2189 | 29.2306 | 29.398 | 29.3967 | 29.0578 | 29.0604 |
| 31.8385 | 31.9777 | 31.932 | 31.9793 | 31.9423 | 31.878 | 31.9002 |
| 28.5618 | 28.1125 | 28.2571 | 28.3324 | 28.3028 | 28.6428 | 28.6562 |
| 33.0308 | 33.1181 | 33.1718 | 32.9747 | 32.9731 | 32.9301 | 32.884 |
| 32.8469 | 33.1779 | 33.144 | 32.876 | 32.8666 | 32.8655 | 32.8709 |
| 28.457 | 28.2503 | 28.334 | 28.1735 | 28.1258 | 28.3378 | 28.4031 |
| 32.27 | 32.3947 | 32.2275 | 32.2098 | 32.2145 | 32.4762 | 32.489 |
| 28.5065 | 28.2008 | 28.1555 | 28.3869 | 28.3072 | 28.3241 | 28.3497 |
| 30.0957 | 30.2874 | 30.2342 | 30.0135 | 30.0473 | 29.9431 | 29.9766 |
| 31.5225 | 31.582 | 31.5084 | 31.5534 | 31.5378 | 31.4049 | 31.4526 |
| 29.209 | 28.768 | 28.9942 | 29.1571 | 29.0955 | 29.449 | 29.3086 |
| 30.1158 | 29.9614 | 29.8333 | 29.9148 | 29.7982 | 30.4634 | 30.3005 |
| 32.731 | 32.9475 | 32.8408 | 32.6527 | 32.6554 | 32.613 | 32.5566 |
| 28.6647 | 28.7077 | 28.5261 | 28.5417 | 28.5566 | 28.5304 | 28.4569 |
| 28.7287 | 28.933 | 29.0074 | 28.7904 | 28.9292 | 28.8437 | 28.7953 |
| 31.3335 | 31.8121 | 31.6359 | 31.569 | 31.5252 | 31.59 | 31.5624 |
| 30.179 | 30.4191 | 30.3861 | 30.3504 | 30.2839 | 30.2288 | 30.181 |
| 29.0527 | 29.219 | 29.0624 | 29.1745 | 29.2297 | 29.2195 | 29.1405 |
| 28.8971 | 28.5798 | 28.5952 | 28.4361 | 28.4755 | 28.8848 | 28.8573 |
| 28.5237 | 28.5497 | 28.4833 | 28.5282 | 28.4999 | 28.1987 | 28.3042 |
| 29.7517 | 30.0828 | 29.916 | 29.9532 | 29.9067 | 30.0372 | 30.0616 |
| 29.0492 | 29.0738 | 28.9879 | 28.8886 | 28.9518 | 29.2357 | 29.212 |
| 32.2968 | 32.0038 | 32.134 | 32.1162 | 32.0889 | 32.3002 | 32.3085 |
| 30.0028 | 30.2381 | 30.2333 | 30.073 | 30.0116 | 30.1834 | 30.2315 |
| 31.2959 | 31.3823 | 31.2519 | 31.2394 | 31.2569 | 31.2829 | 31.2338 |
| 29.1047 | 28.8972 | 28.9038 | 29.0437 | 29.0186 | 29.2981 | 29.2624 |
| 27.8705 | 27.6666 | 27.7445 | 27.7743 | 27.6847 | 27.9474 | 27.8793 |
| 30.5531 | 30.5166 | 30.3649 | 30.4742 | 30.4131 | 30.7017 | 30.7064 |
| 28.9575 | 29.3023 | 29.3109 | 29.108 | 29.0255 | 29.1251 | 29.0638 |
| 28.2811 | 28.1824 | 28.1513 | 28.2893 | 28.3218 | 28.466 | 28.4795 |
| 29.6515 | 29.7154 | 29.6499 | 29.4951 | 29.5326 | 29.4413 | 29.4348 |
| 29.0911 | 29.2837 | 29.1173 | 28.8422 | 28.8432 | 28.8023 | 28.736 |
| 30.0029 | 30.4075 | 30.3328 | 30.0583 | 29.9731 | 30.0967 | 30.2312 |
| 31.4634 | 31.5 | 31.4103 | 31.3433 | 31.3638 | 31.5287 | 31.5979 |
| 28.1024 | 28.509 | 28.4668 | 28.3968 | 28.3413 | 28.3603 | 28.3148 |

| | | | | | | |
|---|---|---|---|---|---|---|
| 29.0312 | 29.1399 | 28.9967 | 28.9574 | 28.9807 | 28.937 | 28.8426 |
| 30.5948 | 30.8502 | 30.6871 | 30.6376 | 30.506 | 30.5672 | 30.5612 |
| 28.2948 | 28.3035 | 28.161 | 28.1489 | 28.2151 | 28.2768 | 28.269 |
| 29.662 | 29.9295 | 29.7409 | 29.778 | 29.7387 | 29.5281 | 29.5933 |
| 29.8709 | 29.7832 | 29.7446 | 29.9537 | 29.8494 | 29.8954 | 29.8542 |
| 32.3884 | 32.2086 | 32.1483 | 32.1552 | 32.1163 | 32.4107 | 32.5012 |
| 29.0709 | 29.5647 | 29.4642 | 29.116 | 29.1084 | 29.2772 | 29.4143 |
| 27.8029 | 27.6446 | 27.5691 | 27.4865 | 27.5424 | 28.0583 | 28.006 |
| 29.4284 | 29.7319 | 29.6355 | 29.5452 | 29.5237 | 29.446 | 29.4098 |
| 31.094 | 31.0003 | 31.1658 | 31.2628 | 31.2311 | 31.0062 | 30.9416 |
| 34.4785 | 34.7225 | 34.6154 | 34.4922 | 34.4402 | 34.3928 | 34.5081 |
| 29.6082 | 29.9399 | 29.9559 | 29.9544 | 30.0128 | 29.8072 | 29.7054 |
| 28.3868 | 28.5122 | 28.2276 | 28.5785 | 28.5087 | 28.6454 | 28.6103 |
| 33.6055 | 34.0501 | 33.8984 | 33.8804 | 33.8548 | 33.9395 | 33.8742 |
| 33.22 | 33.5763 | 33.4805 | 33.3834 | 33.4259 | 33.4262 | 33.3843 |
| 33.4537 | 33.7705 | 33.7002 | 33.583 | 33.5414 | 33.4783 | 33.4787 |
| 29.5801 | 29.6665 | 29.5945 | 29.6792 | 29.6724 | 29.6825 | 29.8044 |
| 29.9688 | 29.8546 | 29.9527 | 29.8892 | 29.8948 | 29.981 | 29.9683 |
| 32.3939 | 32.3162 | 32.2711 | 32.3951 | 32.3347 | 32.204 | 32.2613 |
| 28.967 | 28.7595 | 28.6631 | 28.6867 | 28.5694 | 29.0596 | 29.1488 |
| 28.0186 | 28.0077 | 28.059 | 28.2176 | 28.0125 | 28.3403 | 28.3096 |
| 29.9564 | 30.2907 | 30.3536 | 30.1408 | 30.1515 | 30.1311 | 30.0554 |
| 29.8714 | 30.3024 | 30.0443 | 30.2078 | 30.0948 | 30.133 | 30.1483 |
| 28.3393 | 28.576 | 28.4252 | 28.5127 | 28.4694 | 28.325 | 28.3854 |
| 30.6101 | 31.155 | 31.0339 | 30.856 | 30.7621 | 30.8402 | 30.786 |
| 28.1047 | 27.9221 | 28.0084 | 27.9569 | 27.9797 | 28.1663 | 28.0969 |
| 30.4604 | 30.7513 | 30.7912 | 30.6154 | 30.6769 | 30.5832 | 30.6283 |
| 28.4155 | 28.7771 | 28.5818 | 28.4918 | 28.5585 | 28.3935 | 28.4061 |
| 28.0818 | 28.4822 | 28.3657 | 28.0519 | 28.2735 | 28.1044 | 28.1707 |
| 28.1773 | 28.0982 | 28.2999 | 28.2354 | 28.1369 | 28.0291 | 28.0635 |
| 32.1579 | 32.1861 | 32.0722 | 31.9994 | 31.9436 | 32.3551 | 32.3472 |
| 28.0703 | 28.7473 | 28.6844 | 28.1035 | 28.1803 | 28.3879 | 28.5731 |
| 32.825 | 32.851 | 33.0825 | 32.7767 | 32.8237 | 32.6558 | 32.7408 |
| 29.1855 | 29.2206 | 29.317 | 29.3565 | 29.2263 | 29.1034 | 29.0489 |
| 32.3362 | 32.2882 | 32.2576 | 32.1328 | 32.17 | 32.6311 | 32.6308 |
| 27.9344 | 27.8121 | 27.7682 | 27.8519 | 27.8552 | 27.8553 | 27.8602 |
| 30.7622 | 30.6848 | 30.6244 | 30.5098 | 30.6057 | 30.9942 | 30.9119 |
| 32.4486 | 32.6669 | 32.6366 | 32.5143 | 32.5263 | 32.4011 | 32.4033 |
| 28.8073 | 28.5826 | 28.5873 | 28.6676 | 28.595 | 28.8727 | 28.7161 |
| 31.9019 | 32.0075 | 31.9998 | 31.878 | 31.8911 | 31.9084 | 31.8216 |
| 28.774 | 28.5757 | 28.5907 | 28.7855 | 28.8016 | 28.6862 | 28.6988 |
| 33.0043 | 33.1341 | 33.038 | 32.7889 | 32.8202 | 33.2318 | 33.2763 |
| 29.6691 | 29.8474 | 29.8883 | 29.71 | 29.7091 | 29.8016 | 29.8257 |
| 31.9612 | 32.0553 | 32.0422 | 31.9261 | 31.9254 | 31.757 | 31.7541 |
| 28.2812 | 28.2986 | 28.1804 | 28.1181 | 28.1278 | 28.259 | 28.244 |
| 28.1048 | 27.8794 | 27.9135 | 27.9281 | 27.7318 | 28.1961 | 28.1219 |
| 28.7717 | 28.5107 | 28.3471 | 28.6577 | 28.817 | 28.7464 | 28.5921 |
| 31.8357 | 31.7778 | 31.7108 | 31.4969 | 31.5231 | 31.8226 | 31.8285 |

| | | | | | | |
|---|---|---|---|---|---|---|
| 28.7949 | 29.1563 | 29.0567 | 29.0828 | 29.042 | 28.689 | 28.6776 |
| 30.0519 | 30.2178 | 30.2746 | 30.3024 | 30.4048 | 30.1106 | 30.0439 |
| 29.5688 | 29.7631 | 29.6363 | 29.7962 | 29.7826 | 29.7247 | 29.5596 |
| 28.0622 | 28.3341 | 28.2167 | 28.1841 | 28.2721 | 28.1569 | 28.1523 |
| 29.0798 | 29.2405 | 29.2877 | 29.1675 | 29.0838 | 29.1367 | 29.0434 |
| 31.3068 | 31.3396 | 31.2578 | 31.3154 | 31.2089 | 31.0962 | 31.1204 |
| 28.7554 | 28.3717 | 28.2332 | 28.2673 | 28.189 | 28.1019 | 28.1448 |
| 29.2778 | 29.4915 | 29.3842 | 29.4834 | 29.3233 | 29.2596 | 29.191 |
| 27.6351 | 27.4549 | 27.5951 | 27.6383 | 27.5944 | 27.803 | 27.8446 |
| 29.1043 | 29.2096 | 29.1386 | 29.056 | 29.076 | 29.2176 | 29.3267 |
| 30.16 | 30.3458 | 30.2995 | 30.2641 | 30.2297 | 30.0338 | 30.0036 |
| 29.8289 | 29.7557 | 29.7585 | 29.7688 | 29.8612 | 29.8727 | 29.9087 |
| 29.0975 | 29.1078 | 28.8613 | 28.7905 | 28.8544 | 29.281 | 29.2896 |
| 28.2552 | 28.4615 | 28.3685 | 28.3527 | 28.2471 | 28.1344 | 28.3006 |
| 28.0173 | 27.8803 | 27.9023 | 28.1415 | 28.1163 | 27.9555 | 27.9199 |
| 30.5553 | 31.0167 | 30.8647 | 30.5887 | 30.6516 | 30.6636 | 30.6813 |
| 33.2086 | 33.7689 | 33.7188 | 33.3134 | 33.3287 | 33.7017 | 33.6684 |
| 27.4885 | 27.7222 | 27.5877 | 27.7123 | 27.7574 | 27.7761 | 27.6565 |
| 30.6609 | 30.8703 | 30.8439 | 30.5592 | 30.6197 | 30.6076 | 30.6579 |
| 28.7541 | 28.5867 | 28.5529 | 28.773 | 28.7225 | 28.7777 | 28.7997 |
| 28.5894 | 28.5695 | 28.5015 | 28.4363 | 28.3697 | 28.3781 | 28.321 |
| 28.5883 | 28.3265 | 28.3135 | 28.3336 | 28.3743 | 28.3432 | 28.3868 |
| 34.3255 | 34.4239 | 34.3535 | 34.14 | 34.17 | 34.3366 | 34.4194 |
| 27.9356 | 28.1608 | 27.9553 | 27.934 | 28.0144 | 27.8381 | 27.8895 |
| 28.5792 | 28.2971 | 28.2451 | 28.4095 | 28.4314 | 28.4946 | 28.5263 |
| 32.488 | 32.8783 | 32.7861 | 32.5356 | 32.5322 | 32.6316 | 32.7318 |
| 29.9422 | 29.8047 | 29.8367 | 29.8608 | 29.8724 | 30.3837 | 30.4493 |
| 30.0969 | 30.0793 | 30.0462 | 29.9404 | 29.9801 | 29.8887 | 30.0226 |
| 28.9313 | 28.886 | 28.7892 | 28.8383 | 28.8747 | 29.2654 | 29.1213 |
| 28.2756 | 28.1823 | 28.2613 | 28.1122 | 28.0416 | 28.217 | 28.1602 |
| 31.0154 | 31.1211 | 31.0596 | 31.0514 | 30.9668 | 30.9536 | 30.8909 |
| 30.1354 | 30.1854 | 30.3049 | 30.1503 | 30.1121 | 30.0121 | 30.1087 |
| 33.2073 | 33.1821 | 33.2178 | 33.0678 | 33.0292 | 33.7518 | 33.7091 |
| 34.8219 | 35.1098 | 35.0961 | 34.8613 | 34.8124 | 34.9946 | 35.055 |
| 30.8897 | 31.1963 | 31.1057 | 30.8133 | 30.7978 | 30.7895 | 30.8089 |
| 27.9912 | 28.2455 | 28.1163 | 27.9864 | 27.9823 | 27.9284 | 27.939 |
| 28.2459 | 28.1133 | 27.9264 | 27.9903 | 27.8516 | 28.2947 | 28.1801 |
| 31.9107 | 32.1727 | 32.0617 | 31.8809 | 31.9195 | 31.747 | 31.8582 |
| 27.2132 | 27.2385 | 27.2383 | 27.2495 | 27.0256 | 27.2641 | 27.1027 |
| 28.8143 | 28.9847 | 28.962 | 29.0196 | 28.9878 | 28.9308 | 28.97 |
| 28.6903 | 28.5278 | 28.5705 | 28.3512 | 28.444 | 28.5839 | 28.6288 |
| 28.5522 | 28.5576 | 28.3283 | 28.4466 | 28.3505 | 28.4812 | 28.5494 |
| 31.8242 | 31.6858 | 31.8221 | 31.8211 | 31.8234 | 31.5554 | 31.5271 |
| 30.3956 | 30.7927 | 30.6438 | 30.4241 | 30.4649 | 30.3145 | 30.3638 |
| 27.8615 | 27.8917 | 27.8334 | 27.8409 | 27.7186 | 27.9833 | 27.8625 |
| 28.2989 | 28.8142 | 28.8454 | 28.7496 | 28.6694 | 28.6048 | 28.5693 |
| 29.8085 | 30.3502 | 30.281 | 30.1652 | 30.0825 | 30.0775 | 30.0588 |
| 27.76 | 28.3161 | 28.0538 | 28.0636 | 28.0262 | 28.0187 | 27.8699 |

| | | | | | | |
|---|---|---|---|---|---|---|
| 27.2118 | 27.4537 | 27.2715 | 27.3881 | 27.3837 | 27.568 | 27.5215 |
| 27.3493 | 27.6927 | 27.6499 | 27.6229 | 27.4653 | 27.4613 | 27.5269 |
| 28.7361 | 28.7518 | 28.6858 | 28.7783 | 28.7798 | 28.8706 | 28.8472 |
| 30.0652 | 30.0018 | 29.996 | 30.1565 | 30.1157 | 30.1541 | 30.137 |
| 29.9188 | 29.7596 | 29.7926 | 29.5815 | 29.6587 | 29.6921 | 29.7886 |
| 30.4348 | 30.7014 | 30.6681 | 30.669 | 30.6325 | 30.4891 | 30.4726 |
| 28.304 | 28.702 | 28.5999 | 28.4855 | 28.5215 | 28.3002 | 28.3257 |
| 27.5333 | 27.7673 | 27.7823 | 27.5881 | 27.6185 | 27.8609 | 27.7065 |
| 30.6664 | 30.8076 | 30.8796 | 30.6716 | 30.625 | 30.704 | 30.6563 |
| 29.5323 | 29.7851 | 29.7113 | 29.5441 | 29.7193 | 29.5107 | 29.4762 |
| 29.4803 | 29.7458 | 29.7152 | 29.5606 | 29.4884 | 29.4667 | 29.4468 |
| 29.9521 | 30.2627 | 30.218 | 29.9137 | 29.9856 | 30.0436 | 30.0868 |
| 28.6037 | 28.9528 | 29.0603 | 28.7157 | 28.7403 | 28.7187 | 28.7626 |
| 29.2329 | 29.2372 | 29.1137 | 29.3142 | 29.2372 | 29.2054 | 29.2394 |
| 33.6157 | 33.8886 | 33.8502 | 33.7514 | 33.7035 | 33.5937 | 33.6394 |
| 31.8463 | 32.2083 | 32.1284 | 31.8399 | 31.8238 | 31.7399 | 31.864 |
| 27.8845 | 27.5974 | 27.6658 | 27.5678 | 27.4442 | 28.0446 | 27.9838 |
| 27.7487 | 27.4367 | 27.3897 | 27.7246 | 27.6965 | 27.846 | 27.9676 |
| 29.5967 | 29.4184 | 29.4043 | 29.3296 | 29.3963 | 29.5284 | 29.4135 |
| 33.2107 | 33.3569 | 33.3193 | 33.1825 | 33.2243 | 33.0327 | 33.1735 |
| 28.3005 | 28.2519 | 28.2234 | 28.0117 | 28.0519 | 28.2398 | 28.1844 |
| 27.8837 | 28.2668 | 28.1561 | 28.0397 | 27.9777 | 28.0022 | 27.9728 |
| 27.6693 | 27.5936 | 27.4837 | 27.5798 | 27.605 | 27.4221 | 27.4924 |
| 28.707 | 28.9949 | 28.8924 | 29.0086 | 28.9409 | 28.7133 | 28.7262 |
| 29.7296 | 29.9948 | 29.9578 | 29.716 | 29.8362 | 30.0125 | 29.8809 |
| 29.3344 | 29.0678 | 29.0315 | 29.0626 | 29.0295 | 28.6803 | 28.7499 |
| 32.8696 | 33.1206 | 33.0879 | 32.8904 | 32.8746 | 32.7885 | 32.8545 |
| 32.272 | 32.2642 | 32.3138 | 32.3736 | 32.3688 | 32.3002 | 32.238 |
| 28.6985 | 28.698 | 28.7362 | 28.486 | 28.5721 | 28.9635 | 28.8107 |
| 33.0121 | 32.9106 | 32.8344 | 32.9245 | 32.9573 | 33.0695 | 33.0536 |
| 30.0541 | 29.867 | 30.0162 | 29.9428 | 29.9975 | 30.0389 | 30.0808 |
| 30.2169 | 30.4845 | 30.4123 | 30.2468 | 30.2208 | 30.3897 | 30.4271 |
| 29.0844 | 29.21 | 29.2664 | 29.0405 | 29.0392 | 28.9951 | 28.998 |
| 29.1516 | 28.9256 | 28.9685 | 28.8179 | 28.976 | 29.3079 | 29.3654 |
| 32.897 | 33.0329 | 33.0988 | 32.8211 | 32.857 | 32.8839 | 32.8801 |
| 30.6733 | 30.8962 | 30.8191 | 30.7032 | 30.6236 | 30.5897 | 30.6439 |
| 29.3293 | 29.4451 | 29.3868 | 29.489 | 29.4437 | 29.0727 | 29.1909 |
| 29.9077 | 30.0583 | 30.1344 | 30.416 | 30.3463 | 30.1926 | 30.0475 |
| 30.2681 | 30.6596 | 30.586 | 30.5275 | 30.5046 | 30.4828 | 30.4354 |
| 28.6753 | 29.1862 | 29.2169 | 29.0395 | 28.969 | 28.8497 | 28.846 |
| 32.2904 | 32.5012 | 32.4158 | 32.4454 | 32.326 | 32.4061 | 32.3291 |
| 31.6066 | 31.9375 | 31.8875 | 31.7753 | 31.748 | 31.802 | 31.8075 |
| 29.9928 | 30.0415 | 29.9337 | 30.1712 | 30.2441 | 30.1707 | 30.1724 |
| 29.6149 | 29.7617 | 29.637 | 29.596 | 29.6118 | 29.6975 | 29.7245 |
| 30.3781 | 30.6836 | 30.5481 | 30.6144 | 30.5724 | 30.4814 | 30.4369 |
| 27.7568 | 27.832 | 27.6817 | 27.5686 | 27.7648 | 27.842 | 27.8291 |
| 31.7431 | 31.8576 | 31.9068 | 31.937 | 31.887 | 31.9656 | 31.9138 |
| 30.808 | 31.0495 | 30.9316 | 30.9489 | 30.9437 | 30.9345 | 30.9321 |

| | | | | | | |
|---|---|---|---|---|---|---|
| 29.8231 | 29.6818 | 29.6212 | 29.6286 | 29.5724 | 29.5405 | 29.5506 |
| 29.1204 | 29.3253 | 29.2343 | 29.0955 | 29.1338 | 29.0486 | 29.0011 |
| 27.5669 | 27.8053 | 27.9521 | 27.7528 | 27.664 | 27.7628 | 27.6275 |
| 29.511 | 29.3159 | 29.4291 | 29.4876 | 29.529 | 29.5303 | 29.4296 |
| 28.1768 | 28.2531 | 28.2328 | 28.4901 | 28.4269 | 28.0264 | 28.025 |
| 30.2043 | 30.4451 | 30.3954 | 30.4646 | 30.4221 | 30.2859 | 30.1864 |
| 27.4502 | 27.3813 | 27.4535 | 27.3444 | 27.3449 | 27.5048 | 27.6791 |
| 28.7697 | 28.8643 | 28.7472 | 28.8019 | 28.7928 | 28.6517 | 28.5655 |
| 28.8674 | 29.0245 | 28.8643 | 28.8364 | 29.0571 | 29.0671 | 28.9117 |
| 30.1296 | 30.0215 | 29.8879 | 29.9742 | 30.0217 | 30.1724 | 30.179 |
| 28.9054 | 29.1715 | 29.2301 | 28.944 | 29.0217 | 29.1218 | 29.1158 |
| 31.8682 | 32.0949 | 32.0404 | 32.0068 | 32.0462 | 31.8338 | 31.8891 |
| 27.5441 | 27.9286 | 27.7708 | 27.7754 | 27.6987 | 27.5242 | 27.3586 |
| 28.8961 | 29.0672 | 28.9202 | 29.1424 | 29.1291 | 28.8468 | 28.6886 |
| 30.1649 | 30.2523 | 30.2016 | 30.3196 | 30.2605 | 30.1397 | 29.9773 |
| 30.6118 | 30.7006 | 30.722 | 30.4679 | 30.4958 | 30.6088 | 30.6438 |
| 30.2346 | 30.0014 | 30.1359 | 30.2333 | 30.2543 | 29.9442 | 29.9303 |
| 28.3661 | 28.3776 | 28.4238 | 28.4627 | 28.4107 | 28.1983 | 28.1963 |
| 27.4718 | 27.5131 | 27.5957 | 27.6979 | 27.5321 | 27.3546 | 27.3107 |
| 28.9095 | 29.114 | 29.1535 | 29.032 | 29.0385 | 28.849 | 28.9512 |
| 28.1831 | 27.9714 | 27.9785 | 28.1224 | 28.1065 | 27.8011 | 27.8184 |
| 27.525 | 27.283 | 27.3093 | 27.3884 | 27.4047 | 27.3584 | 27.3925 |
| 28.4127 | 28.5756 | 28.3713 | 28.3541 | 28.282 | 28.3398 | 28.1715 |
| 28.7644 | 29.1195 | 28.9011 | 29.0406 | 28.9941 | 28.8504 | 28.8611 |
| 27.7781 | 27.697 | 27.8121 | 27.6568 | 27.6184 | 27.4978 | 27.3273 |
| 30.8506 | 31.0273 | 31.0661 | 30.6189 | 30.6258 | 30.9151 | 30.9186 |
| 33.4492 | 33.5646 | 33.4668 | 33.3441 | 33.2993 | 33.6483 | 33.6779 |
| 27.9598 | 28.1614 | 28.128 | 28.1383 | 28.2258 | 27.9598 | 28.0215 |
| 29.601 | 29.6815 | 29.7738 | 29.7639 | 29.7188 | 29.6885 | 29.717 |
| 29.7548 | 29.4646 | 29.5667 | 29.5395 | 29.4768 | 29.5383 | 29.6658 |
| 30.163 | 30.3713 | 30.319 | 30.2154 | 30.2316 | 30.0571 | 30.0702 |
| 28.4392 | 28.4572 | 28.4809 | 28.6011 | 28.5996 | 28.1851 | 28.1997 |
| 30.6492 | 30.3674 | 30.4719 | 30.7634 | 30.621 | 30.8246 | 30.6647 |
| 31.641 | 31.7736 | 31.7185 | 31.5864 | 31.6281 | 31.6293 | 31.6137 |
| 30.7274 | 30.8439 | 30.8469 | 30.7188 | 30.8098 | 30.6492 | 30.6245 |
| 29.4825 | 29.483 | 29.4394 | 29.2666 | 29.3407 | 29.8036 | 29.7722 |
| 29.1137 | 29.0978 | 29.0368 | 29.1392 | 29.113 | 29.4576 | 29.4175 |
| 28.0752 | 28.0909 | 28.0209 | 28.0798 | 28.0673 | 27.6486 | 27.7367 |
| 30.3672 | 30.2565 | 30.2044 | 30.2625 | 30.2908 | 30.5457 | 30.5361 |
| 27.751 | 27.7761 | 27.7801 | 27.6039 | 27.7693 | 27.8916 | 27.6418 |
| 32.1212 | 32.2656 | 32.2118 | 32.1899 | 32.1948 | 32.1424 | 32.0615 |
| 29.7067 | 29.8288 | 29.8159 | 29.7413 | 29.7926 | 29.8531 | 29.8303 |
| 28.2449 | 28.2805 | 27.992 | 27.9475 | 28.1405 | 27.7995 | 27.8448 |
| 30.9338 | 31.3793 | 31.3192 | 30.9507 | 31.0277 | 31.362 | 31.3289 |
| 31.5032 | 31.8333 | 31.7251 | 31.45 | 31.4319 | 31.429 | 31.4875 |
| 33.211 | 33.4159 | 33.3611 | 33.2486 | 33.2323 | 33.2303 | 33.2057 |
| 28.915 | 28.9136 | 29.0979 | 28.9885 | 28.9059 | 28.8222 | 28.7186 |
| 30.343 | 30.4548 | 30.3475 | 30.3777 | 30.4007 | 30.1693 | 30.1879 |

| | | | | | | |
|---|---|---|---|---|---|---|
| 29.8418 | 29.9303 | 29.9317 | 30.0152 | 30.0349 | 29.8362 | 29.8529 |
| 29.195 | 28.9907 | 28.9202 | 29.1952 | 29.1424 | 29.2963 | 29.2853 |
| 30.4398 | 30.5867 | 30.6021 | 30.5564 | 30.584 | 30.4829 | 30.3988 |
| 30.3114 | 30.5333 | 30.4662 | 30.4586 | 30.3903 | 30.5418 | 30.4483 |
| 27.321 | 27.7761 | 27.7794 | 27.5507 | 27.6527 | 27.6632 | 27.6255 |
| 31.4976 | 31.6222 | 31.6546 | 31.6642 | 31.7369 | 31.5263 | 31.4309 |
| 29.3484 | 29.6274 | 29.5644 | 29.6075 | 29.5664 | 29.4509 | 29.3052 |
| 28.3173 | 28.5953 | 28.638 | 28.4235 | 28.4915 | 28.4862 | 28.5366 |
| 31.2892 | 31.4803 | 31.4767 | 31.514 | 31.4466 | 31.4323 | 31.4085 |
| 28.4178 | 28.3833 | 28.4249 | 28.2361 | 28.3098 | 28.3346 | 28.3234 |
| 27.9807 | 27.8579 | 27.8348 | 27.991 | 28.0698 | 28.1612 | 28.0367 |
| 31.8235 | 31.6275 | 31.3069 | 31.6305 | 31.5578 | 31.7927 | 31.8541 |
| 28.3946 | 28.1964 | 28.0028 | 28.0062 | 28.0099 | 28.4103 | 28.4416 |
| 31.5815 | 31.8525 | 31.7849 | 31.6643 | 31.6816 | 31.5516 | 31.5155 |
| 27.5609 | 27.6795 | 27.4445 | 27.5267 | 27.5505 | 27.5598 | 27.5813 |
| 29.729 | 29.8861 | 29.6999 | 29.6564 | 29.5458 | 29.748 | 29.7314 |
| 31.0307 | 31.239 | 31.2161 | 31.1069 | 31.1866 | 31.0999 | 31.028 |
| 30.7123 | 31.2637 | 31.0565 | 31.0139 | 31.0008 | 30.8354 | 30.839 |
| 28.8348 | 29.0535 | 28.8475 | 28.9946 | 28.9515 | 28.7687 | 28.7091 |
| 29.1489 | 28.7694 | 28.8942 | 28.8463 | 28.8613 | 29.0959 | 29.1443 |
| 27.4101 | 27.3077 | 27.147 | 27.422 | 27.4474 | 27.4682 | 27.4596 |
| 28.3477 | 28.0816 | 28.0052 | 28.2303 | 28.355 | 28.5326 | 28.5073 |
| 29.4105 | 29.5799 | 29.5738 | 29.4178 | 29.4451 | 29.3614 | 29.3479 |
| 27.7325 | 27.9194 | 28.0413 | 27.6689 | 27.8848 | 27.7696 | 27.7635 |
| 28.0087 | 27.5317 | 27.5808 | 27.6175 | 27.5622 | 27.9595 | 28.0972 |
| 28.813 | 28.5409 | 28.6365 | 28.8488 | 28.7545 | 28.8719 | 28.8954 |
| 28.8493 | 28.9154 | 28.9117 | 28.9261 | 28.7767 | 28.86 | 28.7736 |
| 29.0421 | 29.0694 | 28.9434 | 29.0356 | 28.9779 | 28.9107 | 29.0262 |
| 29.1978 | 29.3314 | 29.1969 | 29.4146 | 29.3081 | 29.1802 | 29.069 |
| 27.5077 | 27.7563 | 27.5953 | 27.3773 | 27.2418 | 27.5162 | 27.3863 |
| 28.4942 | 28.7108 | 28.5844 | 28.2886 | 28.2903 | 28.7472 | 28.8414 |
| 28.8103 | 28.8119 | 28.7402 | 28.8026 | 28.7741 | 28.6804 | 28.5056 |
| 29.9746 | 30.2384 | 30.2142 | 30.0761 | 29.987 | 30.1724 | 30.1091 |
| 33.4259 | 33.4814 | 33.6129 | 33.4156 | 33.3531 | 33.2474 | 33.2997 |
| 29.0288 | 29.068 | 29.1618 | 29.0666 | 28.9825 | 29.1551 | 29.0507 |
| 27.9998 | 27.9248 | 28.0457 | 27.9663 | 27.7762 | 27.6956 | 27.6166 |
| 32.3044 | 32.5479 | 32.4388 | 32.3559 | 32.3376 | 32.2716 | 32.1964 |
| 28.822 | 28.7719 | 28.7339 | 28.5256 | 28.4543 | 28.826 | 28.9087 |
| 27.3648 | 27.3381 | 27.3815 | 27.3161 | 27.462 | 27.0325 | 27.1984 |
| 28.4817 | 28.7286 | 28.5677 | 28.436 | 28.519 | 28.3876 | 28.523 |
| 31.8939 | 31.7283 | 31.6571 | 31.7168 | 31.7376 | 31.8763 | 31.9021 |
| 31.0854 | 31.1803 | 31.1484 | 31.0519 | 31.1043 | 30.9385 | 30.9091 |
| 31.5542 | 31.7457 | 31.5001 | 31.4951 | 31.3856 | 31.8761 | 31.9528 |
| 30.8625 | 30.5307 | 30.7033 | 30.8033 | 30.8017 | 30.9923 | 30.9675 |
| 28.496 | 28.7603 | 28.5265 | 28.5166 | 28.5239 | 28.3659 | 28.3629 |
| 31.934 | 32.1955 | 32.1235 | 31.9285 | 31.9521 | 31.8064 | 31.8389 |
| 28.7469 | 28.5584 | 28.7128 | 28.6265 | 28.5583 | 28.6247 | 28.6328 |
| 27.217 | 26.8214 | 26.8667 | 27.0175 | 27.1627 | 27.368 | 27.5224 |

| | | | | | | |
|---|---|---|---|---|---|---|
| 28.6538 | 28.5104 | 28.5932 | 28.5838 | 28.5371 | 28.4472 | 28.4462 |
| 29.4167 | 29.146 | 29.1974 | 29.156 | 29.1715 | 29.4327 | 29.3968 |
| 29.1199 | 29.046 | 29.1988 | 29.0199 | 29.0608 | 28.9356 | 28.9965 |
| 27.4322 | 27.0986 | 27.2156 | 27.1682 | 27.1485 | 27.3168 | 27.4095 |
| 27.2937 | 27.6479 | 27.486 | 27.1347 | 27.1971 | 27.7233 | 27.5627 |
| 31.5604 | 31.7401 | 31.683 | 31.7553 | 31.7435 | 31.5564 | 31.4891 |
| 31.1064 | 31.232 | 31.2163 | 31.1236 | 31.2691 | 30.9232 | 30.9939 |
| 28.3616 | 28.559 | 28.486 | 28.1136 | 28.0208 | 28.0146 | 28.0543 |
| 30.4561 | 30.4066 | 30.4077 | 30.3981 | 30.488 | 30.3541 | 30.3827 |
| 27.883 | 27.7153 | 27.6931 | 27.612 | 27.6102 | 28.1393 | 27.9139 |
| 28.1075 | 28.1667 | 28.1153 | 28.2943 | 28.1881 | 27.9874 | 27.9515 |
| 30.9235 | 31.1165 | 30.997 | 31.2659 | 31.3396 | 30.4961 | 30.5703 |
| 27.5939 | 27.7655 | 27.7084 | 27.7813 | 27.6854 | 27.6393 | 27.6861 |
| 30.2719 | 30.3489 | 30.282 | 30.2425 | 30.1226 | 30.0189 | 30.0499 |
| 33.1491 | 33.2854 | 33.3616 | 33.12 | 33.1533 | 33.091 | 33.1243 |
| 30.2886 | 30.0795 | 30.0659 | 30.1266 | 30.1262 | 30.1554 | 30.1261 |
| 29.5251 | 29.1943 | 29.2686 | 29.3074 | 29.2945 | 29.2974 | 29.267 |
| 28.2072 | 28.0978 | 28.0753 | 28.0555 | 27.992 | 28.218 | 28.3077 |
| 27.7726 | 27.8174 | 27.8562 | 27.6425 | 27.6095 | 27.6052 | 27.6445 |
| 30.5964 | 30.5094 | 30.435 | 30.2566 | 30.3876 | 30.5775 | 30.5849 |
| 31.6058 | 31.6664 | 31.6165 | 31.6383 | 31.6048 | 31.4571 | 31.4669 |
| 31.8556 | 31.9813 | 31.9474 | 31.8674 | 31.908 | 31.6402 | 31.7461 |
| 27.4389 | 27.5048 | 27.5231 | 27.6134 | 27.553 | 27.515 | 27.5137 |
| 27.6925 | 27.5878 | 27.6512 | 27.7723 | 27.7208 | 27.6322 | 27.8029 |
| 26.9208 | 26.9595 | 26.8622 | 26.7686 | 26.9623 | 26.8541 | 26.9072 |
| 32.8411 | 32.9324 | 32.8766 | 33.0991 | 33.0526 | 33.0318 | 32.987 |
| 30.2429 | 30.4066 | 30.2184 | 30.3538 | 30.3484 | 30.3066 | 30.3393 |
| 28.6974 | 28.9085 | 28.859 | 28.8872 | 28.8063 | 28.7436 | 28.7082 |
| 30.8787 | 31.1195 | 31.0399 | 30.9999 | 30.9062 | 30.7983 | 30.7708 |
| 28.1836 | 28.3093 | 28.0599 | 28.1705 | 28.0173 | 27.9556 | 27.9572 |
| 31.2971 | 31.6262 | 31.5921 | 31.4268 | 31.3991 | 31.5269 | 31.4393 |
| 32.3026 | 32.5787 | 32.55 | 32.3833 | 32.3724 | 32.3304 | 32.3195 |
| 28.1731 | 28.8309 | 28.7783 | 28.3454 | 28.4027 | 28.5272 | 28.4359 |
| 30.1503 | 30.509 | 30.4677 | 30.338 | 30.3867 | 30.1519 | 30.1541 |
| 29.2107 | 29.6537 | 29.5489 | 29.394 | 29.3919 | 29.3799 | 29.3607 |
| 29.2724 | 29.2901 | 29.2382 | 29.1436 | 29.226 | 29.1084 | 29.1886 |
| 29.6721 | 29.6656 | 29.6313 | 29.5991 | 29.609 | 29.8962 | 29.8781 |
| 30.0046 | 30.2372 | 30.1689 | 30.0268 | 30.1082 | 29.8524 | 29.9119 |
| 34.4365 | 34.8469 | 34.7915 | 34.4504 | 34.5471 | 34.6572 | 34.6729 |
| 31.7934 | 31.7591 | 31.7983 | 32.0308 | 32.0552 | 31.8177 | 31.7668 |
| 32.7569 | 32.5122 | 32.5803 | 32.573 | 32.5868 | 32.2112 | 32.3343 |
| 27.411 | 27.2472 | 27.3446 | 27.2109 | 27.3287 | 27.3531 | 27.3529 |
| 29.5602 | 29.5749 | 29.6047 | 29.2982 | 29.3382 | 29.3073 | 29.4301 |
| 30.2865 | 30.4735 | 30.4326 | 30.147 | 30.2993 | 30.2143 | 30.2881 |
| 27.5457 | 27.6611 | 27.5185 | 27.3833 | 27.3446 | 27.5321 | 27.7149 |
| 28.9421 | 29.2305 | 28.926 | 29.1696 | 29.1915 | 29.2 | 29.1406 |
| 28.364 | 28.3899 | 28.272 | 28.0727 | 28.1816 | 28.2871 | 28.2241 |
| 27.8032 | 28.0547 | 27.9295 | 27.6939 | 27.8815 | 27.8982 | 27.7505 |

| | | | | | | |
|---|---|---|---|---|---|---|
| 29.7122 | 29.6811 | 29.679 | 29.6605 | 29.6048 | 29.8169 | 29.8409 |
| 32.1772 | 32.2777 | 32.2088 | 32.3131 | 32.2948 | 32.2864 | 32.3028 |
| 28.5388 | 28.3388 | 28.2104 | 28.3611 | 28.3896 | 28.5933 | 28.6675 |
| 27.8377 | 28.0083 | 27.9431 | 27.9432 | 27.872 | 27.7987 | 27.7466 |
| 30.1132 | 30.3682 | 30.2619 | 29.9355 | 29.9544 | 29.9528 | 29.9766 |
| 26.8663 | 26.8126 | 26.7525 | 26.8076 | 26.7378 | 26.3004 | 26.7322 |
| 28.712 | 28.3996 | 28.5671 | 28.7663 | 28.6796 | 28.5958 | 28.573 |
| 28.4239 | 28.4471 | 28.2846 | 28.2912 | 28.1344 | 28.4086 | 28.5713 |
| 29.3957 | 29.0609 | 29.1243 | 29.3124 | 29.2038 | 29.3605 | 29.3012 |
| 30.0839 | 29.4163 | 29.7562 | 29.823 | 29.7365 | 30.2173 | 30.3035 |
| 29.3474 | 29.5012 | 29.3564 | 29.708 | 29.775 | 28.75 | 28.7946 |
| 27.9768 | 28.1098 | 28.2172 | 28.1207 | 28.0117 | 28.1364 | 28.2822 |
| 29.7763 | 29.5731 | 29.7744 | 29.6919 | 29.7483 | 29.6326 | 29.8747 |
| 33.3641 | 33.5816 | 33.488 | 33.1087 | 33.1312 | 33.7082 | 33.7326 |
| 28.9103 | 28.6525 | 28.6435 | 28.5822 | 28.681 | 28.9297 | 28.9424 |
| 28.4049 | 27.9596 | 28.1293 | 28.267 | 28.4245 | 28.073 | 28.1286 |
| 30.1496 | 30.4957 | 30.2931 | 30.2277 | 30.1871 | 30.4441 | 30.3509 |
| 32.0954 | 32.5092 | 32.3642 | 32.1248 | 32.1336 | 31.8307 | 31.8281 |
| 30.4176 | 30.3345 | 30.3825 | 30.1948 | 30.1776 | 30.3639 | 30.3787 |
| 30.1499 | 30.1436 | 30.2326 | 30.242 | 30.3884 | 30.1059 | 30.0947 |
| 30.522 | 30.5525 | 30.5202 | 30.5359 | 30.5753 | 30.4142 | 30.5046 |
| 30.3998 | 30.5528 | 30.3762 | 30.5488 | 30.568 | 30.2884 | 30.1961 |
| 31.0277 | 31.0345 | 31.063 | 30.9548 | 30.9375 | 31.0126 | 30.8322 |
| 27.4186 | 27.6197 | 27.6991 | 27.5533 | 27.7114 | 27.6861 | 27.5777 |
| 27.1519 | 26.8417 | 26.984 | 27.1594 | 27.0352 | 27.2967 | 26.9924 |
| 28.2153 | 28.6449 | 28.5094 | 28.114 | 28.1588 | 28.3572 | 28.4556 |
| 26.9218 | 27.1885 | 26.9306 | 26.8846 | 27.0605 | 26.983 | 27.0165 |
| 28.6712 | 28.7019 | 28.9214 | 28.6296 | 28.7203 | 28.7559 | 28.8536 |
| 30.541 | 30.3172 | 30.2644 | 30.3714 | 30.233 | 30.5229 | 30.5656 |
| 28.897 | 28.964 | 28.9401 | 28.8213 | 28.8024 | 28.5828 | 28.6839 |
| 29.9632 | 29.8769 | 29.8315 | 29.7242 | 29.7426 | 30.2589 | 30.1833 |
| 28.0977 | 28.5524 | 28.3712 | 28.4525 | 28.4907 | 28.4766 | 28.4835 |
| 28.0587 | 28.4125 | 28.3002 | 28.2816 | 28.2645 | 28.1359 | 28.1634 |
| 29.4178 | 29.7416 | 29.6685 | 29.7102 | 29.6551 | 29.6447 | 29.5732 |
| 28.6753 | 29.0104 | 28.9844 | 28.921 | 28.8783 | 28.6618 | 28.8329 |
| 29.5142 | 29.9156 | 29.7427 | 29.7542 | 29.6836 | 29.8432 | 29.6747 |
| 29.2964 | 29.6467 | 29.5871 | 29.558 | 29.5403 | 29.4935 | 29.4465 |
| 30.7843 | 31.1122 | 30.9955 | 31.0756 | 31.0686 | 31.0609 | 31.0052 |
| 27.5854 | 27.7436 | 27.8458 | 27.8038 | 27.8303 | 27.7884 | 27.7747 |
| 29.481 | 29.7797 | 29.6814 | 29.605 | 29.609 | 29.6067 | 29.586 |
| 27.9115 | 28.1032 | 28.0379 | 28.0284 | 28.2649 | 27.9661 | 28.095 |
| 30.4603 | 30.6763 | 30.7454 | 30.7371 | 30.7774 | 30.6129 | 30.6725 |
| 30.7403 | 30.6683 | 30.6586 | 30.5924 | 30.6529 | 30.7744 | 30.7469 |
| 28.0154 | 28.7077 | 28.4709 | 28.6367 | 28.5379 | 28.3651 | 28.3065 |
| 27.8483 | 28.126 | 28.0473 | 27.8464 | 27.8682 | 28.2873 | 28.1561 |
| 31.8937 | 31.9678 | 31.8223 | 31.6711 | 31.7551 | 31.978 | 32.0083 |
| 27.3179 | 27.2013 | 27.4184 | 27.4541 | 27.4522 | 27.71 | 27.5007 |
| 30.0657 | 29.8624 | 29.9338 | 30.0052 | 29.9107 | 30.1388 | 30.0817 |

| | | | | | | |
|---|---|---|---|---|---|---|
| 29.7954 | 30.0569 | 29.908 | 30.0364 | 29.9124 | 29.7943 | 29.9053 |
| 27.8315 | 27.953 | 27.8483 | 27.8449 | 27.9704 | 28.0205 | 27.9968 |
| 32.8339 | 33.0202 | 32.9154 | 32.8958 | 32.9104 | 32.8305 | 32.8179 |
| 32.3784 | 32.5462 | 32.4721 | 32.3599 | 32.3859 | 32.3309 | 32.2043 |
| 30.7988 | 31.059 | 30.9623 | 30.886 | 30.9007 | 30.8521 | 30.8131 |
| 28.7129 | 28.8646 | 28.7372 | 28.6585 | 28.5802 | 28.3973 | 28.4714 |
| 27.7411 | 27.7629 | 27.6067 | 27.7228 | 27.5672 | 27.8126 | 27.7375 |
| 28.9151 | 28.8916 | 28.9587 | 28.9475 | 28.9106 | 28.8092 | 28.885 |
| 33.6961 | 33.878 | 33.8195 | 33.8502 | 33.8329 | 33.7209 | 33.7626 |
| 29.412 | 29.3461 | 29.2714 | 28.8262 | 28.9663 | 29.2818 | 29.4477 |
| 29.2564 | 29.4676 | 29.4127 | 29.2423 | 29.2926 | 29.3639 | 29.382 |
| 28.3847 | 28.2511 | 28.2553 | 28.207 | 28.3278 | 28.6177 | 28.6933 |
| 26.7882 | 26.4399 | 26.6176 | 26.7491 | 26.6682 | 26.7295 | 26.4792 |
| 29.3265 | 28.8913 | 28.9644 | 29.2202 | 29.3409 | 28.9866 | 28.9655 |
| 27.7151 | 27.8419 | 27.8461 | 27.8146 | 27.8225 | 27.8457 | 27.8608 |
| 31.9589 | 32.3174 | 32.2286 | 31.9585 | 31.9899 | 31.9115 | 31.8673 |
| 35.2041 | 35.4609 | 35.3979 | 35.2239 | 35.3148 | 35.2123 | 35.2522 |
| 28.2403 | 28.3664 | 28.0164 | 28.1933 | 28.4284 | 28.4069 | 28.5842 |
| 32.2182 | 32.3366 | 32.248 | 32.2253 | 32.1323 | 32.3936 | 32.34 |
| 27.434 | 27.5707 | 27.4109 | 27.2896 | 27.3184 | 27.0421 | 26.9997 |
| 33.299 | 33.6267 | 33.4986 | 33.3753 | 33.3832 | 33.2503 | 33.3275 |
| 28.3132 | 28.3101 | 28.2159 | 28.0948 | 28.1518 | 27.9855 | 27.9657 |
| 29.3893 | 29.6079 | 29.6265 | 29.3559 | 29.4474 | 29.2359 | 29.4062 |
| 29.9141 | 29.972 | 29.7639 | 29.9825 | 29.8653 | 29.6105 | 29.5384 |
| 28.6031 | 28.6521 | 28.6544 | 28.5647 | 28.6203 | 28.2171 | 28.2159 |
| 30.6571 | 30.5093 | 30.3758 | 30.5193 | 30.4895 | 30.7354 | 30.6444 |
| 31.1029 | 30.8074 | 30.6937 | 30.9855 | 30.8821 | 31.1296 | 30.9424 |
| 27.2654 | 27.4187 | 27.2982 | 27.3639 | 27.2889 | 26.9061 | 27.1547 |
| 27.0992 | 26.9327 | 26.7768 | 26.9493 | 27.0873 | 27.5509 | 27.4986 |
| 30.7646 | 30.6456 | 30.6798 | 30.6945 | 30.652 | 30.9545 | 30.8928 |
| 28.1617 | 28.5062 | 28.5903 | 28.4174 | 28.3488 | 28.5037 | 28.4479 |
| 27.9871 | 28.2269 | 28.1219 | 27.9342 | 27.9538 | 28.23 | 28.2024 |
| 27.3358 | 27.2602 | 27.189 | 27.1404 | 27.4349 | 27.2943 | 27.1786 |
| 30.6957 | 30.9778 | 30.8572 | 30.6551 | 30.6239 | 30.6118 | 30.5931 |
| 28.5482 | 28.8022 | 28.65 | 28.626 | 28.5909 | 28.529 | 28.5907 |
| 30.5271 | 30.8294 | 30.7792 | 30.5798 | 30.576 | 30.5505 | 30.5022 |
| 29.1414 | 28.9867 | 29.028 | 28.8873 | 28.8065 | 29.3561 | 29.4093 |
| 28.6148 | 28.5967 | 28.5388 | 28.5923 | 28.5962 | 28.4234 | 28.4449 |
| 28.3546 | 28.7036 | 28.4217 | 28.448 | 28.5897 | 28.4664 | 28.4521 |
| 32.6816 | 32.5492 | 32.5435 | 32.6187 | 32.6389 | 32.9596 | 33.0336 |
| 30.499 | 30.718 | 30.6267 | 30.6418 | 30.5826 | 30.3451 | 30.3613 |
| 27.1218 | 27.274 | 27.5171 | 27.2165 | 27.2889 | 27.1861 | 27.3004 |
| 27.9306 | 27.5388 | 27.6703 | 27.95 | 27.9671 | 28.1335 | 27.9029 |
| 31.5073 | 31.8918 | 31.721 | 31.4726 | 31.5725 | 31.4809 | 31.5505 |
| 27.773 | 27.7593 | 27.5464 | 27.5716 | 27.4694 | 27.8677 | 27.8839 |
| 27.7367 | 28.104 | 27.9375 | 27.7786 | 27.758 | 27.5457 | 27.623 |
| 28.1404 | 28.4201 | 28.1164 | 28.0497 | 28.1787 | 28.0615 | 28.0174 |
| 27.8486 | 27.8127 | 27.742 | 27.7609 | 27.8346 | 27.6651 | 27.7216 |

| | | | | | | |
|---|---|---|---|---|---|---|
| 27.7897 | 27.4999 | 27.4617 | 27.4782 | 27.4854 | 27.6634 | 27.5814 |
| 28.7787 | 28.9652 | 28.8505 | 28.7216 | 28.8401 | 28.6051 | 28.5238 |
| 28.8923 | 29.1191 | 28.9788 | 29.0126 | 29.0278 | 28.7554 | 28.729 |
| 26.7918 | 26.3388 | 26.5283 | 27.2591 | 26.7991 | 26.9402 | 26.903 |
| 29.2703 | 29.4949 | 29.4122 | 29.4357 | 29.3748 | 29.4091 | 29.2573 |
| 26.765 | 27.022 | 26.9308 | 27.0917 | 27.0591 | 26.7733 | 26.8706 |
| 30.1713 | 30.2935 | 30.3486 | 30.3482 | 30.3002 | 30.6221 | 30.4996 |
| 31.14 | 31.1952 | 31.2736 | 31.5011 | 31.3929 | 31.2064 | 31.1625 |
| 27.4909 | 27.8561 | 27.773 | 27.7584 | 27.895 | 27.7551 | 27.6505 |
| 32.8206 | 33.2415 | 33.066 | 33.0022 | 33.0413 | 33.042 | 33.0058 |
| 30.3455 | 30.3976 | 30.4711 | 30.4158 | 30.4741 | 30.3843 | 30.494 |
| 30.1678 | 30.3509 | 30.2526 | 30.1918 | 30.2541 | 30.2005 | 30.2781 |
| 30.1439 | 29.9631 | 30.1493 | 30.1189 | 30.1284 | 29.9513 | 29.9549 |
| 27.1527 | 27.1661 | 27.099 | 26.9771 | 27.0419 | 27.0151 | 26.9542 |
| 26.9983 | 26.893 | 26.5767 | 26.9058 | 26.7575 | 26.5952 | 26.8109 |
| 29.872 | 29.7788 | 29.8246 | 29.8621 | 29.8538 | 29.8758 | 29.8596 |
| 26.6857 | 26.2382 | 26.0701 | 26.1828 | 26.2826 | 26.505 | 26.4511 |
| 31.9878 | 32.3671 | 32.3839 | 32.1284 | 32.1101 | 32.1881 | 32.1575 |
| 29.9986 | 30.2836 | 30.2382 | 30.2254 | 30.2119 | 30.0871 | 29.9975 |
| 28.8583 | 28.8176 | 28.835 | 28.796 | 28.9115 | 28.7956 | 28.8904 |
| 31.5917 | 31.8953 | 31.9548 | 31.703 | 31.7337 | 31.7284 | 31.8163 |
| 31.2266 | 31.4766 | 31.4244 | 31.2633 | 31.2351 | 31.2438 | 31.2505 |
| 30.2614 | 29.7557 | 29.9195 | 29.9176 | 29.9106 | 30.1438 | 30.2303 |
| 29.4682 | 29.3216 | 29.1991 | 29.0385 | 29.1604 | 29.5614 | 29.5814 |
| 30.7022 | 31.0157 | 30.8932 | 30.7372 | 30.6924 | 30.6373 | 30.6875 |
| 27.1795 | 27.7362 | 27.5131 | 27.2993 | 27.4726 | 26.9908 | 27.2093 |
| 29.5829 | 29.9989 | 29.9717 | 29.6581 | 29.7116 | 29.7987 | 29.7002 |
| 26.5148 | 26.6027 | 26.3711 | 26.3873 | 26.4654 | 26.4541 | 26.3752 |
| 29.5728 | 29.6163 | 29.4906 | 29.596 | 29.7468 | 29.449 | 29.5195 |
| 26.5959 | 26.5013 | 26.3881 | 26.6305 | 26.502 | 26.2728 | 26.4348 |
| 29.6729 | 29.7918 | 29.7825 | 29.407 | 29.299 | 29.9761 | 30.0553 |
| 26.9474 | 27.5872 | 27.4825 | 27.3064 | 27.2651 | 27.1174 | 27.0841 |
| 26.3513 | 26.5549 | 26.7491 | 26.5517 | 26.5789 | 26.4324 | 26.2441 |
| 29.469 | 29.4682 | 29.3948 | 29.6916 | 29.5023 | 29.2464 | 29.3089 |
| 31.2727 | 31.7045 | 31.6048 | 31.383 | 31.2761 | 31.2967 | 31.3745 |
| 27.9827 | 28.2848 | 28.1514 | 28.1477 | 28.009 | 27.9904 | 28.0824 |
| 28.6084 | 29.0502 | 28.9219 | 28.8471 | 28.6769 | 28.7001 | 28.7135 |
| 27.3912 | 27.382 | 27.2214 | 27.3411 | 27.3485 | 27.3563 | 27.3131 |
| 28.3704 | 28.2755 | 28.0875 | 28.404 | 28.3181 | 28.3456 | 28.4049 |
| 33.5799 | 33.3872 | 33.42 | 33.3591 | 33.3781 | 33.4302 | 33.4048 |
| 27.6181 | 27.3835 | 27.3916 | 27.3204 | 27.4653 | 27.6894 | 27.7509 |
| 28.7221 | 28.6542 | 28.5165 | 28.4592 | 28.5298 | 28.8878 | 28.969 |
| 28.0778 | 28.3484 | 28.3588 | 28.1394 | 28.3036 | 28.2456 | 28.0948 |
| 28.3015 | 28.3994 | 28.2606 | 28.3545 | 28.404 | 28.2373 | 28.3132 |
| 33.2134 | 33.421 | 33.319 | 33.3181 | 33.3122 | 33.5229 | 33.5026 |
| 27.843 | 27.7135 | 27.5181 | 28.0079 | 27.9755 | 27.8755 | 28.0067 |
| 27.8335 | 27.8889 | 27.9233 | 27.7487 | 27.7154 | 27.8924 | 27.8475 |
| 28.5026 | 28.7108 | 28.6076 | 28.5161 | 28.3819 | 28.3522 | 28.5585 |

| | | | | | | |
|---|---|---|---|---|---|---|
| 29.9168 | 30.3892 | 30.2352 | 30.1375 | 30.0418 | 30.0531 | 29.9811 |
| 29.0734 | 28.8905 | 28.8846 | 28.9096 | 29.084 | 29.3367 | 29.3846 |
| 29.8835 | 30.2357 | 30.2881 | 29.934 | 29.9283 | 30.1027 | 30.158 |
| 27.0225 | 27.3138 | 27.1734 | 27.1781 | 27.1773 | 26.7335 | 26.8513 |
| 31.6132 | 31.7032 | 31.6742 | 31.6945 | 31.7482 | 31.4843 | 31.5389 |
| 28.4297 | 28.3208 | 28.2684 | 28.3293 | 28.4323 | 28.447 | 28.5466 |
| 29.6857 | 30.0026 | 29.928 | 29.728 | 29.5877 | 29.539 | 29.6216 |
| 27.9256 | 27.8309 | 27.6755 | 27.474 | 27.5211 | 27.4914 | 27.7691 |
| 32.2107 | 32.1542 | 32.0815 | 32.0331 | 32.0083 | 32.1549 | 32.1324 |
| 28.8779 | 29.1136 | 29.0716 | 28.6456 | 28.72 | 29.1689 | 29.2325 |
| 27.9642 | 28.0739 | 28.2101 | 28.0878 | 28.1115 | 27.9967 | 27.9327 |
| 29.5224 | 29.6063 | 29.4566 | 29.5448 | 29.5293 | 29.5752 | 29.5266 |
| 29.4502 | 29.5321 | 29.3722 | 29.449 | 29.538 | 29.3556 | 29.425 |
| 27.2039 | 27.3376 | 27.1538 | 27.1101 | 27.2074 | 27.3051 | 27.2764 |
| 28.3211 | 27.9241 | 27.9038 | 27.9628 | 28.0228 | 28.3239 | 28.1235 |
| 27.193 | 27.2976 | 27.1333 | 27.1552 | 27.0529 | 27.2006 | 26.9055 |
| 29.4787 | 29.6307 | 29.5241 | 29.2524 | 29.2731 | 29.5845 | 29.8041 |
| 29.0863 | 29.2773 | 29.0637 | 29.1846 | 29.0797 | 28.9645 | 28.8789 |
| 31.2051 | 31.3325 | 31.2706 | 31.1775 | 31.1668 | 31.143 | 31.0426 |
| 31.9891 | 32.0394 | 31.9785 | 31.7361 | 31.7895 | 32.0547 | 32.0229 |
| 29.5584 | 29.8252 | 29.5603 | 29.4938 | 29.4163 | 29.4933 | 29.4861 |
| 28.0988 | 27.9263 | 27.8204 | 27.8256 | 27.9191 | 27.9578 | 27.9692 |
| 28.97 | 28.849 | 28.8485 | 28.7357 | 28.7771 | 28.7446 | 28.8851 |
| 31.0755 | 30.9682 | 30.9913 | 30.9865 | 31.0031 | 31.1845 | 31.1358 |
| 28.1082 | 28.0172 | 27.9311 | 28.0728 | 28.1665 | 27.7918 | 27.7682 |
| 29.7871 | 29.5572 | 29.536 | 29.8374 | 29.7876 | 29.4707 | 29.3435 |
| 32.4887 | 32.6081 | 32.5564 | 32.4285 | 32.417 | 32.3899 | 32.3138 |
| 27.2518 | 27.5619 | 27.3187 | 27.2216 | 27.0399 | 27.141 | 27.0242 |
| 27.1155 | 26.8714 | 27.0768 | 27.2279 | 27.1625 | 26.9736 | 26.7533 |
| 28.8015 | 28.736 | 28.7105 | 28.744 | 28.5502 | 28.7573 | 28.7859 |
| 30.1802 | 30.6401 | 30.5037 | 30.1496 | 30.2418 | 30.4372 | 30.3476 |
| 30.4833 | 30.6728 | 30.6042 | 30.4678 | 30.4153 | 30.3042 | 30.4277 |
| 31.7092 | 32.0744 | 31.8746 | 31.5879 | 31.6043 | 31.6907 | 31.6795 |
| 28.9266 | 29.1788 | 29.0902 | 29.3006 | 29.214 | 29.1763 | 29.0667 |
| 27.2303 | 27.1917 | 27.2161 | 27.4405 | 27.4229 | 27.429 | 27.4117 |
| 30.5227 | 30.6908 | 30.5821 | 30.6636 | 30.6826 | 30.4969 | 30.3833 |
| 28.9324 | 29.0718 | 29.1458 | 28.8826 | 28.9184 | 28.8581 | 28.9342 |
| 28.0493 | 28.4297 | 28.3286 | 28.3442 | 28.2744 | 28.2361 | 28.3108 |
| 27.7189 | 28.087 | 27.9638 | 28.0007 | 28.0299 | 27.9325 | 27.7864 |
| 31.667 | 32.0104 | 32.0047 | 31.6956 | 31.8056 | 31.7715 | 31.6554 |
| 27.5679 | 27.7581 | 27.7974 | 27.8443 | 27.8286 | 27.8111 | 27.6007 |
| 27.6308 | 27.8238 | 27.7615 | 27.6043 | 27.5802 | 27.6091 | 27.6742 |
| 27.8957 | 27.8638 | 27.7182 | 27.7199 | 27.7663 | 27.5125 | 27.6779 |
| 30.2675 | 30.2839 | 30.172 | 30.0139 | 30.0426 | 30.1372 | 30.2177 |
| 28.182 | 28.2489 | 28.4783 | 28.7825 | 28.3368 | 28.4614 | 28.3064 |
| 31.8418 | 31.4397 | 31.2838 | 31.3038 | 31.3049 | 31.8484 | 31.941 |
| 29.9983 | 29.9131 | 29.8747 | 29.8781 | 29.9035 | 30.0487 | 30.0735 |
| 27.3026 | 27.1646 | 27.4444 | 27.1513 | 27.3513 | 27.543 | 27.407 |

| | | | | | | |
|---|---|---|---|---|---|---|
| 29.5045 | 29.7631 | 29.6672 | 29.6245 | 29.6327 | 29.7516 | 29.7083 |
| 28.2198 | 28.0657 | 28.075 | 28.2941 | 28.205 | 28.3064 | 28.2804 |
| 28.7723 | 29.0792 | 28.9769 | 28.9181 | 28.949 | 28.8531 | 28.7915 |
| 31.0278 | 31.0689 | 30.9609 | 31.0721 | 31.1323 | 30.9227 | 30.9048 |
| 27.1295 | 26.9999 | 26.9928 | 27.078 | 26.9669 | 27.3407 | 27.282 |
| 27.1691 | 27.7858 | 27.4619 | 27.5358 | 27.5388 | 27.5559 | 27.492 |
| 28.9807 | 29.3654 | 29.3464 | 29.1446 | 29.1892 | 28.9967 | 28.8526 |
| 29.2692 | 29.1154 | 29.1985 | 29.3491 | 29.1619 | 29.1199 | 29.2063 |
| 30.9198 | 30.955 | 30.9201 | 31.0321 | 30.9813 | 30.8432 | 30.8144 |
| 31.5293 | 31.2091 | 31.2264 | 31.1366 | 31.0918 | 31.6229 | 31.6155 |
| 28.8983 | 28.6846 | 28.3679 | 28.5704 | 28.5829 | 28.7455 | 28.5715 |
| 28.9798 | 29.098 | 28.9027 | 29.0161 | 29.0057 | 28.9399 | 28.9006 |
| 31.2734 | 31.5101 | 31.4446 | 31.2387 | 31.2628 | 31.2049 | 31.2395 |
| 28.2616 | 28.0827 | 28.0881 | 28.0238 | 28.0484 | 28.5148 | 28.3845 |
| 31.8602 | 32.0424 | 31.9168 | 31.8145 | 32.005 | 31.9127 | 32.0205 |
| 29.4563 | 29.3091 | 29.3101 | 29.2396 | 29.2855 | 29.5519 | 29.5829 |
| 29.2582 | 29.3201 | 29.3473 | 29.2052 | 29.1859 | 29.1319 | 28.962 |
| 28.7173 | 28.8145 | 28.4545 | 28.5651 | 28.3804 | 28.6075 | 28.6035 |
| 33.1197 | 33.3973 | 33.4394 | 33.1565 | 33.1736 | 33.2862 | 33.2786 |
| 28.1778 | 28.0637 | 28.0312 | 28.0642 | 28.0862 | 28.2345 | 28.2764 |
| 28.6938 | 28.4815 | 28.2897 | 28.6755 | 28.6886 | 28.1631 | 28.0907 |
| 31.1396 | 30.7578 | 30.6066 | 30.874 | 30.9458 | 30.4406 | 30.3865 |
| 27.5347 | 27.7201 | 27.8519 | 27.4733 | 27.633 | 27.6449 | 27.5865 |
| 28.4033 | 28.4967 | 28.4697 | 28.2744 | 28.2978 | 28.0092 | 27.9868 |
| 31.3761 | 31.5879 | 31.5625 | 31.3657 | 31.4098 | 31.2551 | 31.3597 |
| 28.9684 | 29.04 | 29.1855 | 29.0274 | 29.0073 | 29.1651 | 29.1063 |
| 29.7859 | 30.0307 | 29.9053 | 29.6935 | 29.7822 | 29.6247 | 29.7377 |
| 27.3242 | 27.2772 | 27.3104 | 27.1057 | 27.237 | 27.2018 | 27.417 |
| 29.5487 | 29.9081 | 29.9131 | 29.661 | 29.6429 | 29.4375 | 29.4799 |
| 27.5486 | 27.504 | 27.5373 | 27.4486 | 27.6573 | 27.3654 | 27.414 |
| 30.148 | 30.1813 | 30.0778 | 30.0855 | 30.0436 | 30.341 | 30.2703 |
| 29.3727 | 29.5524 | 29.4486 | 29.2402 | 29.2973 | 29.5107 | 29.5191 |
| 30.3805 | 30.4735 | 30.4422 | 30.1002 | 30.1194 | 30.5944 | 30.572 |
| 31.5712 | 31.7069 | 31.5586 | 31.3689 | 31.3811 | 31.6346 | 31.6871 |
| 31.7263 | 31.7587 | 31.8236 | 31.7643 | 31.7782 | 31.7069 | 31.8272 |
| 27.9828 | 27.8212 | 27.9041 | 27.977 | 27.984 | 27.9666 | 27.8361 |
| 27.7113 | 27.4866 | 27.626 | 27.6494 | 27.8383 | 27.4049 | 27.6101 |
| 28.4278 | 28.859 | 28.5856 | 28.3468 | 28.3594 | 28.3611 | 28.4479 |
| 27.3948 | 27.4508 | 27.3456 | 27.2903 | 27.305 | 27.191 | 27.3112 |
| 28.3882 | 28.4629 | 28.3249 | 28.3899 | 28.4193 | 28.2201 | 28.3795 |
| 26.6678 | 26.5999 | 26.7516 | 26.6988 | 26.5777 | 26.5492 | 26.3953 |
| 27.7226 | 27.7528 | 27.4269 | 27.52 | 27.6909 | 27.8298 | 27.8937 |
| 27.3127 | 27.1213 | 27.123 | 27.233 | 27.2357 | 27.4631 | 27.5222 |
| 27.2499 | 27.4939 | 27.5793 | 27.3403 | 27.2476 | 27.2621 | 27.0812 |
| 28.007 | 27.8491 | 27.8533 | 27.7104 | 27.5927 | 28.237 | 28.2143 |
| 29.5384 | 29.4857 | 29.4644 | 29.3289 | 29.3915 | 29.5712 | 29.5435 |
| 27.013 | 27.3283 | 27.0964 | 27.0323 | 26.9036 | 26.7696 | 26.9555 |
| 30.4247 | 30.7119 | 30.5064 | 30.4156 | 30.3691 | 30.4977 | 30.5787 |

| | | | | | | |
|---|---|---|---|---|---|---|
| 29.9687 | 30.2125 | 30.2484 | 29.8665 | 29.8555 | 29.8467 | 29.8315 |
| 28.8955 | 28.2989 | 28.6226 | 28.5736 | 28.6367 | 28.9952 | 28.8949 |
| 32.99 | 32.9374 | 33.0638 | 32.9437 | 32.9763 | 32.8647 | 32.9477 |
| 31.6465 | 31.3104 | 31.475 | 31.4305 | 31.3574 | 32.043 | 31.9732 |
| 27.8946 | 28.3519 | 28.083 | 28.0276 | 28.1508 | 28.0888 | 28.1633 |
| 27.4349 | 28.1791 | 27.9763 | 27.3962 | 27.5472 | 27.7615 | 27.6867 |
| 27.4937 | 27.3998 | 27.4914 | 27.4381 | 27.3771 | 27.4908 | 27.6542 |
| 26.6466 | 27.1469 | 26.7436 | 26.8115 | 26.9361 | 26.6761 | 26.5068 |
| 28.5113 | 28.3666 | 28.2844 | 28.3776 | 28.4227 | 28.2469 | 28.484 |
| 30.7316 | 30.7852 | 30.8125 | 30.7715 | 30.7777 | 30.8689 | 30.772 |
| 28.1496 | 28.4126 | 28.259 | 28.0277 | 28.1027 | 28.1706 | 28.2898 |
| 29.1996 | 29.2224 | 29.1634 | 29.203 | 29.0454 | 29.3596 | 29.3008 |
| 30.1858 | 30.5949 | 30.5927 | 30.0969 | 30.121 | 30.2533 | 30.2683 |
| 28.9961 | 29.3537 | 29.2619 | 29.0327 | 28.9845 | 29.2028 | 29.2058 |
| 34.2884 | 34.4442 | 34.1628 | 34.4217 | 34.4606 | 34.1694 | 34.104 |
| 30.4041 | 30.782 | 30.6102 | 30.3924 | 30.3785 | 30.2578 | 30.2311 |
| 27.9039 | 27.7532 | 27.6606 | 27.8157 | 27.8361 | 27.9939 | 27.9941 |
| 27.6643 | 27.4376 | 27.3375 | 27.2286 | 27.4885 | 27.1019 | 27.1303 |
| 32.2637 | 32.2148 | 32.1887 | 32.1231 | 32.1433 | 32.5202 | 32.6156 |
| 28.5696 | 28.6472 | 28.6761 | 28.9075 | 28.7376 | 28.5721 | 28.5803 |
| 31.3828 | 31.914 | 31.8076 | 31.4458 | 31.4485 | 31.6181 | 31.589 |
| 27.416 | 27.3719 | 27.2319 | 27.2285 | 26.9626 | 27.4062 | 27.2584 |
| 27.8203 | 28.2203 | 28.1953 | 27.8772 | 27.8174 | 27.7659 | 27.8345 |
| 29.6177 | 29.7596 | 29.6481 | 29.6165 | 29.5803 | 29.4798 | 29.4968 |
| 27.7801 | 28.0177 | 27.7226 | 27.8989 | 27.631 | 27.6904 | 27.6882 |
| 28.0654 | 28.3951 | 28.1676 | 27.8977 | 27.9216 | 27.8941 | 28.0669 |
| 29.2573 | 29.589 | 29.5424 | 29.384 | 29.5314 | 29.64 | 29.4394 |
| 27.9681 | 28.4676 | 28.214 | 28.3867 | 28.4224 | 28.0687 | 28.3066 |
| 28.6676 | 28.9068 | 28.9832 | 29.2518 | 29.1535 | 28.9912 | 28.8564 |
| 27.126 | 27.753 | 27.6167 | 27.4386 | 27.4486 | 27.5586 | 27.5254 |
| 26.6913 | 27.0509 | 27.0731 | 26.9342 | 26.7464 | 26.8572 | 26.8266 |
| 27.5476 | 27.8654 | 27.8666 | 27.8727 | 27.8741 | 27.5737 | 27.6268 |
| 26.6634 | 26.8156 | 26.7348 | 26.8048 | 26.8553 | 27.0237 | 26.648 |
| 26.6706 | 27.2309 | 27.0293 | 27.0785 | 27.2222 | 26.9899 | 27.0597 |
| 30.9023 | 31.1921 | 31.1951 | 31.105 | 31.0362 | 31.0003 | 31.0368 |
| 28.5777 | 28.7073 | 28.5395 | 28.7522 | 28.6703 | 28.5594 | 28.6882 |
| 27.299 | 27.4391 | 27.4595 | 27.6042 | 27.4344 | 27.4979 | 27.4325 |
| 29.1422 | 29.3801 | 29.3292 | 29.3327 | 29.1504 | 29.1606 | 29.1196 |
| 32.974 | 33.135 | 33.1843 | 33.1427 | 33.2081 | 33.0695 | 33.1165 |
| 28.805 | 28.9236 | 28.8963 | 28.8987 | 28.8684 | 28.7813 | 28.8608 |
| 30.3694 | 30.2253 | 30.1193 | 30.13 | 30.2723 | 30.1967 | 30.2635 |
| 28.4843 | 28.3984 | 28.1621 | 28.4105 | 28.3628 | 28.252 | 28.1641 |
| 29.6519 | 29.3889 | 29.4717 | 29.3873 | 29.3813 | 29.4359 | 29.452 |
| 31.6284 | 31.718 | 31.6397 | 31.4781 | 31.4141 | 31.6686 | 31.6555 |
| 30.0402 | 29.564 | 29.5704 | 29.5222 | 29.5219 | 29.5402 | 29.508 |
| 28.8055 | 29.0829 | 28.9897 | 28.9446 | 28.9931 | 28.822 | 28.789 |
| 29.4839 | 29.8354 | 29.6466 | 29.4592 | 29.529 | 29.5341 | 29.6249 |
| 27.795 | 28.3898 | 28.2742 | 28.126 | 28.0995 | 28.189 | 28.1813 |

| | | | | | | |
|---|---|---|---|---|---|---|
| 27.4088 | 27.4978 | 27.5345 | 27.3199 | 27.3731 | 27.3763 | 27.4697 |
| 27.772 | 27.6734 | 27.522 | 27.4708 | 27.4505 | 27.579 | 27.5649 |
| 30.6482 | 30.749 | 30.7857 | 30.9304 | 30.9627 | 30.7954 | 30.8062 |
| 28.6996 | 28.7919 | 28.8776 | 28.8272 | 28.9329 | 28.6242 | 28.5072 |
| 31.0293 | 31.1705 | 31.0207 | 30.8681 | 30.8666 | 30.9555 | 31.0439 |
| 31.9595 | 32.1988 | 32.1023 | 31.9818 | 32.0357 | 32.0352 | 32.0595 |
| 29.699 | 29.7493 | 29.6434 | 29.5127 | 29.6263 | 29.6924 | 29.7016 |
| 29.9598 | 30.0574 | 29.9613 | 29.6666 | 29.7052 | 29.9904 | 30.0221 |
| 27.2222 | 27.2445 | 26.8969 | 27.2247 | 27.0761 | 27.029 | 26.8872 |
| 30.9713 | 31.1783 | 31.3027 | 31.0402 | 30.9799 | 31.0182 | 30.9218 |
| 26.8901 | 26.9087 | 26.9343 | 26.8629 | 26.8102 | 26.951 | 27.0917 |
| 28.0251 | 27.87 | 27.8984 | 28.0411 | 28.1096 | 27.9855 | 27.8053 |
| 27.9179 | 28.0024 | 27.8659 | 28.1909 | 28.1227 | 28.0484 | 27.8701 |
| 27.5064 | 27.6874 | 27.5848 | 27.6562 | 27.682 | 27.593 | 27.4151 |
| 27.4726 | 27.7645 | 27.4974 | 27.521 | 27.3726 | 27.6488 | 27.6648 |
| 28.7612 | 29.259 | 29.1474 | 28.8453 | 28.9705 | 29.1493 | 29.0049 |
| 29.0757 | 29.4907 | 29.4571 | 29.1085 | 29.205 | 29.2194 | 29.1261 |
| 26.8049 | 26.7883 | 26.8794 | 26.8344 | 26.6411 | 26.5374 | 26.6633 |
| 30.9294 | 31.233 | 31.187 | 31.0054 | 30.9844 | 31.0623 | 31.1056 |
| 31.6353 | 31.615 | 31.505 | 31.692 | 31.6959 | 31.8173 | 31.8206 |
| 28.16 | 27.6877 | 27.7366 | 27.6451 | 27.7424 | 27.8158 | 27.8109 |
| 29.0455 | 28.9616 | 28.8825 | 28.9156 | 28.962 | 28.7301 | 28.6405 |
| 28.5641 | 28.8084 | 28.7881 | 28.7765 | 28.7405 | 28.5026 | 28.5276 |
| 27.4853 | 27.1442 | 27.0465 | 27.4319 | 27.5946 | 27.6527 | 27.4554 |
| 28.4385 | 28.3304 | 28.3508 | 28.4541 | 28.4295 | 27.888 | 28.0092 |
| 29.0693 | 29.3597 | 29.4189 | 29.1676 | 29.0176 | 29.2245 | 29.208 |
| 32.8483 | 33.0828 | 33.0406 | 32.8936 | 32.8518 | 32.8198 | 32.7438 |
| 28.2303 | 28.2797 | 28.2871 | 28.2284 | 28.2572 | 28.0345 | 28.0588 |
| 27.0781 | 27.0047 | 27.2991 | 26.9901 | 27.1271 | 27.0897 | 27.1927 |
| 28.324 | 28.4091 | 28.3331 | 28.2136 | 28.2318 | 28.1174 | 28.2156 |
| 26.9394 | 27.2753 | 26.9116 | 27.0805 | 27.1168 | 26.8628 | 26.8804 |
| 27.997 | 28.0939 | 28.0911 | 27.9845 | 27.8486 | 27.6957 | 27.8319 |
| 27.5636 | 27.5759 | 27.4736 | 27.3254 | 27.3897 | 27.7399 | 27.6679 |
| 28.8761 | 29.1532 | 29.0134 | 28.8772 | 28.8027 | 28.9856 | 28.8446 |
| 28.6512 | 28.9847 | 28.9513 | 28.696 | 28.5425 | 28.5244 | 28.6436 |
| 31.4599 | 31.4647 | 31.6177 | 31.559 | 31.4692 | 31.257 | 31.4169 |
| 28.1806 | 28.233 | 28.2128 | 28.17 | 28.21 | 27.9959 | 27.9802 |
| 31.6416 | 31.8573 | 31.7185 | 31.2178 | 31.3567 | 31.738 | 31.8304 |
| 27.6645 | 27.6916 | 27.572 | 27.4207 | 27.464 | 27.555 | 27.372 |
| 31.3194 | 31.5626 | 31.454 | 31.2287 | 31.2503 | 31.3157 | 31.3836 |
| 28.2957 | 28.4396 | 28.3683 | 28.1374 | 28.1198 | 28.4867 | 28.4592 |
| 30.0363 | 30.1563 | 30.0769 | 30.1847 | 30.1315 | 29.9448 | 29.8929 |
| 31.1072 | 31.4085 | 31.3044 | 31.2547 | 31.2679 | 31.6263 | 31.481 |
| 28.1472 | 28.4029 | 28.3412 | 28.2748 | 28.3774 | 28.1499 | 28.0386 |
| 27.3371 | 26.9641 | 27.1486 | 27.0138 | 26.986 | 27.449 | 27.5066 |
| 28.8555 | 29.2295 | 29.179 | 28.8829 | 28.8482 | 28.7841 | 28.9069 |
| 29.2701 | 29.4381 | 29.3487 | 29.1782 | 29.3612 | 29.1265 | 29.2441 |
| 28.3868 | 28.1851 | 28.2915 | 28.0589 | 28.1076 | 28.1765 | 28.3185 |

| | | | | | | |
|---|---|---|---|---|---|---|
| 27.7941 | 28.0751 | 27.856 | 27.9484 | 27.8125 | 28.0824 | 27.9688 |
| 28.1137 | 28.3284 | 28.3799 | 28.2438 | 28.2624 | 27.9704 | 27.9869 |
| 27.4969 | 27.5454 | 27.2349 | 27.2743 | 27.397 | 27.561 | 27.3908 |
| 28.0331 | 28.3167 | 28.0054 | 28.0076 | 27.9465 | 28.0566 | 28.0081 |
| 31.1843 | 31.7167 | 31.5792 | 31.3118 | 31.2872 | 31.3914 | 31.3845 |
| 31.5529 | 31.6166 | 31.5584 | 31.5489 | 31.554 | 31.507 | 31.4355 |
| 31.0441 | 31.2414 | 31.1402 | 31.1549 | 31.1061 | 30.4753 | 30.4916 |
| 26.4854 | 27.1076 | 26.5131 | 26.4459 | 26.4877 | 26.5819 | 26.5065 |
| 30.1226 | 30.5143 | 30.337 | 30.1119 | 30.2115 | 30.5269 | 30.487 |
| 29.5434 | 29.4074 | 29.3278 | 29.5065 | 29.5481 | 29.5298 | 29.5265 |
| 28.6762 | 28.6254 | 28.6084 | 28.7231 | 28.7823 | 28.6754 | 28.5253 |
| 32.2424 | 32.4958 | 32.2736 | 32.1548 | 32.1579 | 32.1373 | 32.0475 |
| 30.4356 | 30.6625 | 30.8187 | 30.3618 | 30.3416 | 30.648 | 30.6917 |
| 28.0454 | 27.9489 | 27.6958 | 27.8942 | 27.9437 | 28.037 | 27.9925 |
| 28.4069 | 28.7739 | 28.7026 | 28.4794 | 28.4349 | 28.5008 | 28.4984 |
| 28.7145 | 29.1142 | 28.7086 | 28.7862 | 28.8744 | 28.6857 | 28.688 |
| 28.828 | 28.6289 | 28.4108 | 28.6348 | 28.6404 | 28.9891 | 29.0155 |
| 26.7453 | 27.0011 | 26.8489 | 26.7136 | 26.8925 | 26.7689 | 26.8923 |
| 29.8952 | 29.8791 | 30.0209 | 29.8857 | 30.0026 | 29.6526 | 29.6589 |
| 32.0063 | 32.3168 | 32.2415 | 32.167 | 32.1017 | 31.9518 | 31.9289 |
| 29.3255 | 29.4801 | 29.2789 | 29.4488 | 29.4436 | 29.6934 | 29.5546 |
| 28.4767 | 28.8617 | 28.87 | 28.58 | 28.5638 | 28.5715 | 28.6679 |
| 29.5714 | 29.9621 | 29.8961 | 29.9198 | 29.8331 | 29.8356 | 29.8145 |
| 27.4748 | 27.7133 | 27.8377 | 27.5165 | 27.541 | 27.7203 | 27.805 |
| 32.6146 | 32.8705 | 32.851 | 32.773 | 32.7698 | 32.7244 | 32.7418 |
| 29.6138 | 29.6225 | 29.5199 | 29.647 | 29.6318 | 29.8482 | 29.8212 |
| 28.0579 | 28.1412 | 28.2201 | 28.0939 | 28.142 | 28.1544 | 28.105 |
| 28.4759 | 28.654 | 28.5831 | 28.4988 | 28.5429 | 28.5421 | 28.4918 |
| 30.6367 | 30.6772 | 30.6575 | 30.6685 | 30.7367 | 30.8517 | 30.8192 |
| 28.9679 | 29.1665 | 29.1015 | 29.0149 | 28.9356 | 29.138 | 29.1134 |
| 29.3524 | 29.3101 | 29.4005 | 29.322 | 29.4094 | 29.386 | 29.402 |
| 28.4656 | 28.3675 | 28.4793 | 28.3924 | 28.427 | 28.3737 | 28.4045 |
| 28.08 | 28.069 | 28.006 | 27.9724 | 27.9672 | 27.9076 | 27.7985 |
| 27.461 | 27.3446 | 27.5168 | 27.2308 | 27.3765 | 27.1014 | 27.0446 |
| 29.5456 | 29.9439 | 29.8643 | 29.7495 | 29.6535 | 29.712 | 29.6811 |
| 28.538 | 28.2204 | 28.2143 | 27.8848 | 28.0621 | 28.1656 | 28.1104 |
| 31.3765 | 31.7977 | 31.7056 | 31.6139 | 31.5281 | 31.5123 | 31.4168 |
| 28.2965 | 27.9524 | 27.9625 | 27.9226 | 27.9707 | 28.2143 | 28.2887 |
| 28.5333 | 28.6057 | 28.5528 | 28.471 | 28.5399 | 28.3075 | 28.5036 |
| 27.4924 | 27.5728 | 27.6622 | 27.6859 | 27.6218 | 27.4113 | 27.2584 |
| 27.0478 | 27.7889 | 27.7262 | 27.2837 | 27.2629 | 27.6464 | 27.7409 |
| 28.5287 | 28.3153 | 28.3292 | 28.4208 | 28.476 | 28.646 | 28.5328 |
| 32.9703 | 33.1101 | 33.0895 | 32.7261 | 32.8382 | 33.0339 | 33.1583 |
| 27.2819 | 27.8826 | 27.8605 | 27.5202 | 27.3793 | 27.4869 | 27.4891 |
| 27.7632 | 27.8001 | 27.9754 | 27.78 | 27.8656 | 27.9453 | 27.9648 |
| 30.1273 | 30.2211 | 30.1654 | 30.0376 | 30.0633 | 29.9805 | 29.9975 |
| 31.069 | 31.0709 | 31.0744 | 30.922 | 30.993 | 30.7818 | 30.808 |
| 27.5373 | 27.8298 | 27.5418 | 27.3848 | 27.5192 | 27.3435 | 27.3589 |

| | | | | | | |
|---|---|---|---|---|---|---|
| 29.535 | 29.4556 | 29.2416 | 29.4529 | 29.4599 | 29.4887 | 29.4987 |
| 26.2008 | 26.9865 | 26.8625 | 26.7943 | 26.6403 | 26.7998 | 27.0246 |
| 27.3773 | 27.6701 | 27.5884 | 27.4648 | 27.3573 | 27.4239 | 27.623 |
| 30.2201 | 30.5096 | 30.5256 | 30.2257 | 30.2454 | 30.1833 | 30.1624 |
| 29.3841 | 29.3878 | 29.2981 | 29.3411 | 29.1864 | 29.1951 | 29.1036 |
| 27.721 | 27.8647 | 27.7922 | 27.7223 | 27.7527 | 27.7964 | 27.7954 |
| 29.3621 | 29.3623 | 29.3555 | 29.2737 | 29.122 | 29.6628 | 29.5783 |
| 29.5666 | 29.5243 | 29.6671 | 29.5493 | 29.5602 | 29.5886 | 29.437 |
| 28.6643 | 28.6655 | 28.5506 | 28.6626 | 28.6536 | 28.8676 | 28.7613 |
| 29.7867 | 29.7108 | 29.6511 | 29.4668 | 29.4053 | 29.6674 | 29.6314 |
| 27.8469 | 28.0613 | 28.0148 | 27.9002 | 27.9798 | 27.7447 | 27.7289 |
| 28.6668 | 29.0344 | 28.8911 | 28.7704 | 28.75 | 28.7601 | 28.7413 |
| 28.2256 | 28.5098 | 28.5198 | 28.2674 | 28.3178 | 28.471 | 28.5407 |
| 26.54 | 26.9653 | 26.9171 | 26.4265 | 26.7182 | 26.5503 | 27.0774 |
| 29.1704 | 29.1123 | 29.1649 | 28.9696 | 28.9978 | 28.7594 | 28.9113 |
| 27.9239 | 28.0288 | 27.93 | 27.9475 | 28.081 | 27.7366 | 27.8935 |
| 29.241 | 29.4279 | 29.3459 | 29.3481 | 29.3321 | 29.0835 | 29.0452 |
| 28.8661 | 29.0618 | 29.0979 | 28.8558 | 28.7437 | 28.8911 | 28.8776 |
| 28.031 | 28.1744 | 28.0023 | 27.9843 | 28.0229 | 28.2017 | 28.2741 |
| 28.2202 | 28.4606 | 28.3504 | 28.2714 | 28.1603 | 28.1446 | 28.1787 |
| 33.7303 | 33.9993 | 34.014 | 34.0214 | 34.0645 | 33.7278 | 33.7944 |
| 27.8587 | 27.8862 | 27.8767 | 27.8374 | 27.8669 | 27.807 | 27.7388 |
| 27.4005 | 27.4758 | 27.2733 | 27.291 | 27.26 | 27.4217 | 27.4273 |
| 26.4193 | 26.5395 | 26.6238 | 26.5352 | 26.6902 | 26.3488 | 26.425 |
| 29.6433 | 29.4726 | 29.6713 | 29.7872 | 29.7902 | 29.9349 | 29.8612 |
| 27.3557 | 27.5329 | 27.4717 | 27.5334 | 27.4227 | 27.2952 | 27.3393 |
| 27.9773 | 28.0531 | 28.0525 | 27.8651 | 27.8765 | 27.8606 | 27.8707 |
| 28.2648 | 28.5313 | 28.5512 | 28.4097 | 28.4377 | 28.1874 | 28.2099 |
| 31.1072 | 31.0344 | 31.0164 | 30.9929 | 30.9541 | 30.8025 | 30.7561 |
| 28.3856 | 28.2266 | 28.1211 | 28.0153 | 28.031 | 28.3075 | 28.3347 |
| 27.8085 | 27.7508 | 27.6225 | 27.867 | 27.804 | 27.6937 | 27.5916 |
| 29.5622 | 29.6822 | 29.5443 | 29.2976 | 29.3644 | 29.5955 | 29.6483 |
| 28.6909 | 28.9945 | 28.9829 | 28.7294 | 28.7148 | 28.6482 | 28.6206 |
| 29.1769 | 29.0483 | 29.1093 | 28.7509 | 28.6842 | 29.4645 | 29.4775 |
| 27.5988 | 27.5767 | 27.5368 | 27.572 | 27.6775 | 27.5065 | 27.4819 |
| 32.5924 | 32.7147 | 32.6575 | 32.7245 | 32.7779 | 32.4341 | 32.5113 |
| 28.8395 | 28.877 | 28.649 | 28.5689 | 28.6589 | 28.8373 | 28.9233 |
| 26.5085 | 26.4269 | 26.3932 | 26.4231 | 26.5761 | 26.7869 | 26.7681 |
| 26.4208 | 26.5363 | 26.4706 | 26.3631 | 26.2314 | 26.5623 | 26.7244 |
| 26.6443 | 26.3458 | 26.3673 | 26.4718 | 26.3886 | 26.7866 | 26.6331 |
| 27.7779 | 27.7729 | 27.4936 | 27.7602 | 27.5448 | 27.9029 | 27.8154 |
| 28.8812 | 28.1527 | 28.8211 | 28.8835 | 28.8697 | 28.6011 | 28.6241 |
| 31.0653 | 31.333 | 31.3484 | 31.162 | 31.1564 | 31.1621 | 31.1564 |
| 33.167 | 33.1884 | 33.0819 | 33.2998 | 33.3042 | 33.4384 | 33.3439 |
| 27.2854 | 27.1987 | 26.9469 | 27.1014 | 27.1536 | 27.0954 | 27.0758 |
| 27.6668 | 27.8815 | 27.7163 | 27.9267 | 27.891 | 27.8064 | 27.8403 |
| 31.4776 | 31.8448 | 31.8175 | 31.6915 | 31.6423 | 31.4981 | 31.5044 |
| 27.1164 | 26.6576 | 26.9654 | 26.983 | 26.9027 | 26.9355 | 26.8024 |

| | | | | | | |
|---|---|---|---|---|---|---|
| 29.3513 | 29.2199 | 29.1592 | 29.1954 | 29.0472 | 29.5388 | 29.4341 |
| 31.4473 | 31.5158 | 31.4708 | 31.3865 | 31.2873 | 31.2629 | 31.3875 |
| 27.5076 | 27.4104 | 27.6182 | 27.6788 | 27.4548 | 27.2021 | 27.1896 |
| 31.1754 | 31.5808 | 31.2959 | 31.3342 | 31.2799 | 31.3061 | 31.3314 |
| 26.9463 | 27.0404 | 26.942 | 26.7127 | 26.9039 | 26.7288 | 26.5806 |
| 27.544 | 27.5504 | 27.3015 | 27.3833 | 27.4385 | 27.4966 | 27.4662 |
| 28.0798 | 28.1336 | 28.0161 | 28.0893 | 27.9637 | 28.2667 | 28.1508 |
| 27.1008 | 27.2584 | 27.4953 | 27.1487 | 26.8985 | 26.9819 | 26.9848 |
| 28.3796 | 28.6057 | 28.4741 | 28.4356 | 28.4646 | 28.2446 | 28.3116 |
| 29.5535 | 29.3912 | 29.2294 | 29.4125 | 29.418 | 29.5135 | 29.5477 |
| 27.8347 | 27.9633 | 27.8723 | 27.5682 | 27.6072 | 27.7835 | 27.6354 |
| 26.6674 | 26.8998 | 26.7689 | 26.545 | 26.794 | 26.4595 | 26.5573 |
| 27.7103 | 27.9986 | 27.9096 | 27.7661 | 27.8606 | 27.4862 | 27.5818 |
| 32.6907 | 32.652 | 32.5696 | 32.8185 | 32.7959 | 32.4442 | 32.4492 |
| 27.3347 | 27.231 | 27.3462 | 27.1078 | 27.1699 | 27.3176 | 27.1111 |
| 27.0104 | 26.9607 | 26.988 | 26.8356 | 27.0495 | 27.0882 | 27.0776 |
| 29.5443 | 29.9507 | 29.8197 | 29.4751 | 29.5392 | 29.7115 | 29.5993 |
| 29.8465 | 29.933 | 29.7921 | 29.7538 | 29.6761 | 29.6887 | 29.7285 |
| 27.4961 | 27.6212 | 27.5905 | 27.2685 | 27.2614 | 27.6133 | 27.6353 |
| 27.8194 | 27.8196 | 27.5596 | 27.5613 | 27.7021 | 27.7809 | 27.5556 |
| 28.9199 | 28.8731 | 28.8552 | 28.8742 | 28.8526 | 28.7929 | 28.7288 |
| 27.2038 | 27.4249 | 27.2404 | 27.2415 | 27.2317 | 27.2642 | 27.0177 |
| 26.8113 | 27.029 | 26.9021 | 27.0021 | 26.9294 | 26.8088 | 26.6179 |
| 29.9237 | 29.8965 | 30.108 | 30.0278 | 30.071 | 29.8741 | 29.9499 |
| 31.0731 | 31.2143 | 31.0297 | 31.2238 | 31.0895 | 30.9577 | 30.888 |
| 27.8173 | 27.6582 | 27.6933 | 27.6899 | 27.4902 | 27.1202 | 27.2172 |
| 28.8232 | 28.5223 | 28.4503 | 28.5038 | 28.5597 | 29.0848 | 29.0721 |
| 30.9829 | 31.1766 | 31.0821 | 31.1003 | 30.9366 | 30.8525 | 30.8524 |
| 27.6842 | 28.1314 | 27.9999 | 27.8352 | 27.7496 | 27.7052 | 27.7723 |
| 27.4105 | 27.5091 | 27.6697 | 27.275 | 27.4312 | 27.2791 | 27.492 |
| 28.5691 | 28.3914 | 28.462 | 28.3186 | 28.4912 | 28.7001 | 28.7699 |
| 29.2661 | 29.3192 | 29.2151 | 29.2237 | 29.2661 | 29.1123 | 29.1831 |
| 33.724 | 33.7276 | 33.6934 | 33.5441 | 33.5654 | 33.7918 | 33.9036 |
| 29.0954 | 29.2761 | 29.1758 | 29.6287 | 29.754 | 29.4464 | 29.438 |
| 29.2731 | 29.4961 | 29.4686 | 29.5205 | 29.4257 | 29.2832 | 29.1739 |
| 30.6154 | 30.8572 | 30.7437 | 30.902 | 30.8593 | 30.6742 | 30.6221 |
| 28.9714 | 29.4909 | 29.3554 | 29.3802 | 29.362 | 29.0198 | 29.039 |
| 31.1337 | 31.2019 | 31.2467 | 31.5228 | 31.5183 | 31.563 | 31.3098 |
| 31.6869 | 32.0045 | 31.9781 | 31.928 | 31.8339 | 31.8255 | 31.7639 |
| 26.4777 | 26.5585 | 26.3869 | 26.5889 | 26.574 | 26.6874 | 26.555 |
| 27.4436 | 27.7928 | 27.9523 | 27.6843 | 27.5783 | 27.5579 | 27.7007 |
| 31.0764 | 31.2113 | 31.1295 | 31.355 | 31.4593 | 31.1628 | 31.1823 |
| 30.1556 | 30.2608 | 30.1948 | 30.1645 | 30.2078 | 30.2488 | 30.3094 |
| 31.2361 | 31.4631 | 31.3737 | 31.4452 | 31.4051 | 31.3454 | 31.3814 |
| 31.1388 | 31.39 | 31.3487 | 31.2216 | 31.2254 | 31.2648 | 31.2097 |
| 29.7608 | 29.9198 | 30.0065 | 29.9094 | 29.9532 | 29.7965 | 29.7291 |
| 27.2478 | 27.5004 | 27.3738 | 27.3246 | 27.3771 | 27.2979 | 27.4602 |
| 30.5419 | 30.8706 | 30.7575 | 30.7099 | 30.5799 | 30.7137 | 30.7333 |

| | | | | | | |
|---|---|---|---|---|---|---|
| 30.4847 | 30.8236 | 30.6952 | 30.6017 | 30.6196 | 30.5521 | 30.5495 |
| 29.6268 | 29.7466 | 29.6672 | 29.8243 | 29.7205 | 29.6249 | 29.5872 |
| 29.4175 | 29.4139 | 29.461 | 29.4828 | 29.4601 | 29.456 | 29.4813 |
| 26.8978 | 26.9828 | 27.0629 | 27.0013 | 27.0282 | 26.9953 | 27.0344 |
| 31.4816 | 31.7061 | 31.6466 | 31.5613 | 31.494 | 31.6507 | 31.6583 |
| 29.0936 | 29.2521 | 29.1467 | 29.1567 | 29.1618 | 29.1956 | 29.1815 |
| 30.4679 | 30.3735 | 30.3817 | 30.5213 | 30.534 | 30.3936 | 30.3831 |
| 28.6661 | 28.3657 | 28.4873 | 28.3113 | 28.4792 | 28.4802 | 28.5795 |
| 29.0053 | 29.116 | 28.8289 | 28.7196 | 28.6475 | 28.9036 | 28.9064 |
| 27.4957 | 27.4404 | 27.3319 | 27.2452 | 27.37 | 27.4202 | 27.3979 |
| 28.2763 | 28.5068 | 28.3903 | 28.3122 | 28.4939 | 28.246 | 28.2263 |
| 28.5617 | 27.9405 | 27.9684 | 28.2768 | 28.3065 | 28.4494 | 28.453 |
| 27.8415 | 27.721 | 27.7441 | 27.5549 | 27.5333 | 27.8528 | 27.7266 |
| 30.8448 | 31.3938 | 31.2897 | 30.9171 | 30.999 | 31.0592 | 31.0447 |
| 26.4597 | 26.1521 | 26.2661 | 26.308 | 26.3782 | 26.6564 | 26.4324 |
| 30.3188 | 30.5316 | 30.5396 | 30.4551 | 30.5443 | 30.2648 | 30.2188 |
| 27.6757 | 28.0423 | 27.8423 | 28.0748 | 28.1457 | 27.902 | 27.8904 |
| 27.735 | 28.1177 | 28.1371 | 27.9387 | 27.8794 | 27.78 | 27.913 |
| 26.1585 | 26.243 | 26.1323 | 26.2712 | 26.2058 | 26.3372 | 26.3646 |
| 27.7891 | 28.3139 | 28.3759 | 28.0977 | 28.0918 | 27.9768 | 27.9577 |
| 28.8332 | 29.0228 | 28.9538 | 28.7831 | 28.8603 | 28.7219 | 28.7059 |
| 28.6723 | 28.8226 | 28.7965 | 28.7677 | 28.7797 | 28.4761 | 28.6154 |
| 28.9808 | 29.3777 | 29.1662 | 29.1987 | 29.2739 | 29.2105 | 29.0737 |
| 31.8409 | 31.5643 | 31.9973 | 31.8573 | 31.9251 | 31.728 | 31.807 |
| 30.784 | 30.8877 | 30.8545 | 31.1052 | 31.0528 | 31.2524 | 31.1774 |
| 29.6635 | 29.9058 | 29.985 | 29.8773 | 29.8293 | 29.7407 | 29.736 |
| 29.2425 | 29.5271 | 29.426 | 29.2494 | 29.2762 | 29.2168 | 29.1772 |
| 28.8282 | 28.7209 | 28.8707 | 28.6203 | 28.474 | 28.5373 | 28.6173 |
| 29.6974 | 29.7623 | 29.7277 | 30.0227 | 29.9946 | 29.6782 | 29.5921 |
| 29.068 | 29.2741 | 29.0816 | 29.1182 | 28.9685 | 29.3505 | 29.2822 |
| 27.1203 | 27.5571 | 27.2882 | 27.2232 | 27.3048 | 27.2671 | 27.0006 |
| 30.5506 | 30.6867 | 30.5721 | 30.8469 | 30.8861 | 30.5958 | 30.5583 |
| 27.8937 | 27.7551 | 27.568 | 27.6795 | 27.7312 | 27.8865 | 27.9126 |
| 29.3736 | 29.2815 | 29.2308 | 29.2403 | 29.1797 | 29.2192 | 29.1599 |
| 31.3425 | 31.4754 | 31.5691 | 31.4778 | 31.5351 | 31.255 | 31.1022 |
| 28.8557 | 29.1807 | 29.1215 | 29.0231 | 28.833 | 28.8013 | 28.806 |
| 26.5951 | 26.3918 | 26.3488 | 26.4529 | 26.399 | 26.6359 | 26.8276 |
| 32.4848 | 32.763 | 32.6539 | 32.4114 | 32.4949 | 32.5252 | 32.4844 |
| 27.1781 | 27.4021 | 27.2672 | 27.2344 | 27.3036 | 27.2873 | 27.367 |
| 28.1045 | 27.9617 | 28.0596 | 27.7411 | 27.9051 | 27.7441 | 27.7638 |
| 28.9783 | 28.7341 | 28.6727 | 28.6606 | 28.6056 | 28.9463 | 28.8369 |
| 27.8353 | 28.0476 | 28.0161 | 27.6852 | 27.7902 | 27.6721 | 27.6395 |
| 30.2481 | 30.5527 | 30.2077 | 30.2718 | 30.2294 | 30.1616 | 30.2335 |
| 29.544 | 29.8826 | 29.8582 | 29.6757 | 29.6194 | 29.5592 | 29.5628 |
| 26.3777 | 26.497 | 26.4829 | 26.5381 | 26.3947 | 26.1251 | 26.2753 |
| 30.188 | 30.6364 | 30.5978 | 30.4325 | 30.3555 | 30.4556 | 30.5223 |
| 28.5713 | 29.0885 | 28.9936 | 28.8186 | 28.7907 | 28.815 | 28.7748 |
| 26.9192 | 26.7946 | 26.7303 | 26.849 | 26.8693 | 26.6886 | 26.5652 |

| | | | | | | |
|---|---|---|---|---|---|---|
| 26.9058 | 27.0242 | 26.7462 | 27.1241 | 27.0576 | 27.4602 | 27.3506 |
| 28.0571 | 27.6855 | 27.8377 | 27.6275 | 27.8777 | 27.7312 | 27.5382 |
| 31.4418 | 31.809 | 31.5854 | 31.4748 | 31.5737 | 31.5095 | 31.4761 |
| 31.6651 | 31.4259 | 31.573 | 31.5419 | 31.3902 | 31.9492 | 31.88 |
| 28.5705 | 28.6731 | 28.2557 | 28.4825 | 28.6246 | 28.7909 | 28.7009 |
| 26.7295 | 26.8519 | 26.727 | 26.8054 | 26.616 | 26.5739 | 26.5454 |
| 27.1323 | 27.3351 | 27.1803 | 26.9708 | 26.9167 | 26.964 | 26.9058 |
| 30.9868 | 31.2542 | 31.1359 | 30.881 | 30.9131 | 31.0083 | 30.9629 |
| 26.7266 | 26.5933 | 27.0958 | 26.6122 | 26.5696 | 26.5246 | 26.3597 |
| 27.062 | 25.9963 | 26.8545 | 26.6634 | 26.7275 | 27.2645 | 27.1826 |
| 28.026 | 27.7432 | 27.8177 | 27.7273 | 27.6829 | 27.753 | 27.7337 |
| 28.1241 | 28.7985 | 28.1873 | 28.7783 | 28.5177 | 29.2493 | 28.4523 |
| 27.1203 | 27.2541 | 27.3459 | 26.9753 | 26.9964 | 26.8622 | 26.984 |
| 28.0347 | 27.9125 | 27.81 | 28.0428 | 28.027 | 28.0837 | 28.1712 |
| 27.9612 | 27.9535 | 27.7565 | 27.989 | 28.0019 | 27.929 | 27.8093 |
| 29.3013 | 29.4634 | 29.3051 | 29.2571 | 29.3361 | 29.0463 | 29.201 |
| 30.5713 | 30.3519 | 30.2893 | 30.402 | 30.4683 | 30.6335 | 30.732 |
| 29.2222 | 29.4148 | 29.3071 | 29.2988 | 29.2399 | 29.1815 | 29.1622 |
| 26.9447 | 26.6784 | 26.7525 | 26.6362 | 26.9308 | 26.7104 | 26.8555 |
| 32.0396 | 32.4212 | 32.2726 | 32.011 | 32.0955 | 32.0514 | 32.0549 |
| 27.3307 | 27.0401 | 27.0666 | 26.9494 | 26.9494 | 27.3142 | 27.2505 |
| 27.8909 | 27.9708 | 27.9423 | 27.9509 | 28.0881 | 27.7356 | 27.8485 |
| 26.7474 | 26.9463 | 26.8647 | 26.6659 | 26.3555 | 26.9899 | 26.8689 |
| 28.1224 | 27.9187 | 27.8875 | 28.2978 | 28.3223 | 28.5034 | 28.4573 |
| 27.0058 | 26.9146 | 26.6574 | 26.8488 | 26.9069 | 27.0713 | 26.9659 |
| 28.5863 | 28.3078 | 28.2586 | 28.1971 | 28.2867 | 28.559 | 28.7077 |
| 30.9931 | 31.5575 | 31.4461 | 31.1269 | 31.0484 | 31.1119 | 31.2871 |
| 26.9033 | 27.0715 | 26.8595 | 26.7773 | 26.8075 | 27.1544 | 27.2765 |
| 30.3255 | 30.6429 | 30.4177 | 30.4302 | 30.4218 | 30.1776 | 30.0828 |
| 29.4758 | 29.6125 | 29.4052 | 29.4433 | 29.4372 | 29.1714 | 29.228 |
| 27.2875 | 27.6296 | 27.6328 | 27.2592 | 27.3867 | 27.2109 | 27.2108 |
| 28.2986 | 28.3716 | 28.1944 | 28.1328 | 27.9822 | 28.2717 | 28.3221 |
| 26.9125 | 27.2571 | 27.1255 | 26.6417 | 26.7411 | 26.9086 | 26.7393 |
| 30.3978 | 30.6366 | 30.5523 | 30.3517 | 30.3819 | 30.252 | 30.25 |
| 32.9189 | 32.8325 | 32.7883 | 32.8261 | 32.8134 | 33.333 | 33.3036 |
| 29.4853 | 29.5504 | 29.4538 | 29.5946 | 29.6868 | 29.1505 | 29.2797 |
| 26.9533 | 27.2295 | 27.3085 | 26.8995 | 27.007 | 26.9346 | 26.9518 |
| 26.9484 | 26.8116 | 26.9814 | 26.8926 | 26.8409 | 26.7286 | 26.7547 |
| 30.4828 | 30.6446 | 30.7773 | 30.4421 | 30.484 | 30.5597 | 30.6586 |
| 27.3255 | 27.3659 | 27.2054 | 27.2632 | 27.0807 | 27.1494 | 27.1841 |
| 32.1705 | 32.4781 | 32.2795 | 32.2882 | 32.3401 | 32.2337 | 32.2249 |
| 29.1071 | 28.7921 | 28.7022 | 29.0125 | 28.863 | 29.0777 | 29.0586 |
| 27.9372 | 27.8362 | 27.7963 | 28.021 | 28.1857 | 27.8223 | 27.8072 |
| 29.4665 | 29.5916 | 29.4816 | 29.2471 | 29.3259 | 29.4237 | 29.5301 |
| 27.5124 | 27.1792 | 27.0244 | 27.3465 | 27.3861 | 27.4823 | 27.2858 |
| 28.3165 | 28.4235 | 28.2326 | 28.192 | 28.2089 | 28.311 | 28.276 |
| 28.9157 | 29.3077 | 29.0152 | 28.9663 | 28.9516 | 28.7535 | 28.6159 |
| 27.6618 | 28.1054 | 27.8749 | 27.7062 | 27.7564 | 27.6911 | 27.6691 |

| | | | | | | |
|---|---|---|---|---|---|---|
| 29.8645 | 29.5822 | 29.6834 | 29.8259 | 29.8596 | 29.9074 | 29.7917 |
| 27.9369 | 27.7523 | 27.7336 | 27.9505 | 27.9814 | 27.7536 | 27.6863 |
| 30.971 | 31.0744 | 31.1309 | 30.8845 | 30.9564 | 30.8475 | 30.8225 |
| 27.83 | 27.0867 | 27.9471 | 28.1847 | 28.1113 | 27.6906 | 27.6761 |
| 28.9969 | 29.1019 | 29.2018 | 29.0767 | 29.1369 | 29.0597 | 28.942 |
| 28.4319 | 28.6423 | 28.4897 | 28.2736 | 28.2632 | 28.7311 | 28.7249 |
| 28.3352 | 28.6171 | 28.6275 | 28.3355 | 28.3812 | 28.7156 | 28.5579 |
| 27.3342 | 27.4487 | 27.1746 | 27.3463 | 27.314 | 27.1941 | 26.9416 |
| 26.8827 | 26.944 | 26.7335 | 26.7565 | 26.8124 | 26.885 | 26.7094 |
| 27.0268 | 26.3801 | 26.6718 | 26.914 | 27.0098 | 27.0469 | 27.0416 |
| 28.2493 | 28.044 | 28.0413 | 28.1003 | 28.104 | 28.2761 | 28.242 |
| 26.9309 | 27.3977 | 27.2167 | 27.1539 | 27.1326 | 26.8695 | 27.015 |
| 30.6251 | 30.5029 | 30.4432 | 30.5422 | 30.4808 | 30.6275 | 30.6371 |
| 32.924 | 33.219 | 33.0583 | 32.9214 | 33.0561 | 33.1086 | 33.1102 |
| 27.9298 | 27.6366 | 27.5385 | 27.5997 | 27.6556 | 27.9946 | 27.9304 |
| 28.9867 | 28.5547 | 28.6806 | 28.6758 | 28.8354 | 28.813 | 28.8047 |
| 30.1599 | 30.1967 | 30.1663 | 30.1438 | 30.0333 | 30.0345 | 29.8444 |
| 27.0958 | 26.5711 | 26.6077 | 26.9247 | 27.0067 | 27.0771 | 26.9858 |
| 27.0735 | 26.9935 | 26.9425 | 26.9663 | 27.0993 | 27.0404 | 26.7506 |
| 29.8213 | 30.1731 | 30.3231 | 29.81 | 29.8499 | 29.9482 | 29.9158 |
| 29.5596 | 29.64 | 29.5317 | 29.3994 | 29.3581 | 29.6364 | 29.6276 |
| 28.6906 | 28.8042 | 28.7091 | 28.8432 | 28.6791 | 28.4501 | 28.4682 |
| 27.6889 | 27.8442 | 27.9186 | 28.0915 | 27.7989 | 27.8835 | 27.8877 |
| 26.6292 | 26.7914 | 26.8267 | 26.5547 | 26.9864 | 26.7073 | 26.6684 |
| 27.3942 | 27.8675 | 27.7978 | 27.6431 | 27.5422 | 27.5396 | 27.2944 |
| 26.0462 | 26.4739 | 26.453 | 26.3026 | 26.3666 | 25.9787 | 26.1599 |
| 28.1547 | 28.3454 | 28.1711 | 28.7307 | 28.6524 | 28.4946 | 28.4413 |
| 25.9679 | 26.0872 | 26.214 | 26.3113 | 26.3837 | 26.2019 | 26.23 |
| 30.8002 | 31.3834 | 31.1978 | 31.0045 | 31.0083 | 31.0032 | 30.8984 |
| 30.7255 | 30.9998 | 30.9693 | 30.9985 | 31.0113 | 30.8227 | 30.7525 |
| 26.8573 | 26.9902 | 26.8002 | 26.8265 | 26.9769 | 26.9844 | 26.8837 |
| 26.6938 | 26.9729 | 27.0454 | 27.0066 | 26.9001 | 26.8621 | 26.771 |
| 29.2203 | 29.4653 | 29.4011 | 29.4746 | 29.3314 | 29.1686 | 29.2825 |
| 26.746 | 26.8266 | 26.6621 | 26.8028 | 26.8091 | 26.7801 | 26.695 |
| 32.5255 | 32.8302 | 32.7916 | 32.6953 | 32.7566 | 32.7263 | 32.6371 |
| 31.1006 | 31.313 | 31.2547 | 31.1141 | 31.1189 | 31.2013 | 31.2611 |
| 27.3643 | 27.5372 | 27.3991 | 27.5689 | 27.6129 | 27.4369 | 27.3724 |
| 29.0596 | 29.0195 | 28.9795 | 29.0898 | 29.0206 | 29.0766 | 29.0544 |
| 28.2259 | 27.8666 | 28.1026 | 28.0899 | 28.054 | 28.1354 | 27.9714 |
| 31.5752 | 31.4741 | 31.388 | 31.5352 | 31.4979 | 31.3264 | 31.2729 |
| 28.1887 | 27.9731 | 27.9268 | 28.0085 | 27.9602 | 27.958 | 28.1287 |
| 28.038 | 28.0454 | 28.0187 | 27.8156 | 27.7779 | 28.0251 | 27.8797 |
| 26.8688 | 27.1887 | 27.1674 | 26.9783 | 27.029 | 26.9357 | 26.8573 |
| 27.7271 | 28.2165 | 28.1217 | 28.17 | 28.0008 | 27.9608 | 27.9805 |
| 27.1637 | 27.1427 | 27.0631 | 26.9954 | 26.8853 | 26.7497 | 27.0989 |
| 26.9023 | 27.2877 | 27.1258 | 27.0208 | 27.055 | 26.9193 | 26.8545 |
| 25.8337 | 25.8281 | 25.7986 | 25.6386 | 25.9466 | 25.6101 | 25.7892 |
| 27.6842 | 27.9265 | 27.8736 | 27.7926 | 27.9151 | 27.6067 | 27.7489 |

| | | | | | | |
|---|---|---|---|---|---|---|
| 28.2353 | 28.4961 | 28.526 | 28.4681 | 28.4398 | 28.2068 | 28.0453 |
| 30.2931 | 30.2005 | 30.1198 | 30.1972 | 30.1172 | 30.3652 | 30.3183 |
| 31.6961 | 31.6697 | 31.6139 | 31.7078 | 31.6873 | 31.5773 | 31.4996 |
| 27.021 | 27.1702 | 27.1802 | 27.1549 | 27.1713 | 27.125 | 27.1451 |
| 29.5459 | 29.7837 | 29.7688 | 29.7298 | 29.8171 | 29.4597 | 29.5114 |
| 26.4387 | 25.3904 | 26.2282 | 26.3541 | 26.2058 | 26.1984 | 25.7272 |
| 26.5543 | 27.0692 | 26.6736 | 26.9277 | 26.7369 | 26.7966 | 26.6849 |
| 29.7892 | 29.9915 | 29.8197 | 29.73 | 29.7962 | 29.7032 | 29.6972 |
| 27.3982 | 27.7756 | 27.6994 | 27.4372 | 27.4779 | 27.4293 | 27.1911 |
| 28.5557 | 28.9167 | 29.008 | 28.6652 | 28.7209 | 28.8356 | 28.7967 |
| 27.5699 | 27.4829 | 27.4941 | 27.3155 | 27.4295 | 27.367 | 27.3231 |
| 26.8253 | 26.9597 | 26.95 | 27.0157 | 27.1304 | 26.9356 | 26.9958 |
| 27.6337 | 27.6594 | 27.6402 | 27.7564 | 27.5327 | 27.8861 | 28.0083 |
| 26.7453 | 26.9425 | 26.8651 | 26.6717 | 26.7618 | 26.683 | 26.7577 |
| 27.6333 | 27.6788 | 27.783 | 27.526 | 27.8952 | 27.4045 | 27.3862 |
| 26.5173 | 25.9762 | 25.9754 | 26.2199 | 26.2218 | 26.1607 | 26.1539 |
| 27.9965 | 27.9977 | 28.0621 | 27.9829 | 28.1138 | 27.9358 | 27.7231 |
| 28.5515 | 29.1188 | 28.9929 | 28.7158 | 28.7912 | 28.7021 | 28.6402 |
| 31.5209 | 31.4366 | 31.2938 | 31.5389 | 31.4999 | 31.6967 | 31.7423 |
| 28.0428 | 28.1357 | 27.9562 | 28.0223 | 28.2001 | 28.2552 | 28.1221 |
| 28.6578 | 28.7297 | 28.8864 | 28.8503 | 28.7547 | 28.7049 | 28.6944 |
| 29.9094 | 30.017 | 29.9208 | 30.0073 | 29.9962 | 30.1122 | 30.048 |
| 26.8546 | 26.958 | 27.0157 | 26.8694 | 26.6769 | 26.9586 | 26.9649 |
| 29.4862 | 29.8979 | 29.8678 | 29.7772 | 29.6688 | 29.5946 | 29.6968 |
| 26.3511 | 26.0844 | 26.0462 | 26.3043 | 26.1925 | 26.0725 | 26.0738 |
| 28.3036 | 28.4914 | 28.3142 | 28.2688 | 28.3371 | 28.3171 | 28.1626 |
| 28.3464 | 28.5641 | 28.6594 | 27.9811 | 28.0861 | 28.4206 | 28.2278 |
| 25.838 | 25.9843 | 26.2908 | 26.3156 | 26.3765 | 26.4096 | 26.7679 |
| 30.2806 | 30.1592 | 30.1066 | 30.0449 | 29.9676 | 30.5598 | 30.4973 |
| 30.2026 | 30.4701 | 30.3402 | 30.0549 | 30.0718 | 30.2661 | 30.2981 |
| 26.5258 | 26.8466 | 26.5933 | 26.6271 | 26.6706 | 26.683 | 26.9166 |
| 26.5299 | 26.7316 | 26.2343 | 26.4189 | 26.6675 | 26.6494 | 26.578 |
| NaN | 26.0267 | 25.9349 | 25.6847 | 25.8344 | 25.9056 | 25.628 |
| 27.6103 | 27.3424 | 27.1728 | 27.2257 | 27.3071 | 27.4416 | 27.4427 |
| 26.0828 | 25.9701 | 25.8291 | 26.0148 | 25.956 | 26.0388 | 25.9627 |
| 26.5345 | 26.7945 | 26.972 | 26.706 | 26.7107 | 26.7252 | 27.0117 |
| 27.3342 | 27.959 | 27.8065 | 27.5818 | 27.5712 | 27.6827 | 27.4394 |
| 27.4626 | 27.453 | 27.3819 | 27.7253 | 27.4371 | 27.3239 | 27.2358 |
| 26.7154 | 26.6875 | 26.7402 | 27.0008 | 26.6736 | 26.843 | 26.6216 |
| 28.8206 | 28.9976 | 28.869 | 28.963 | 28.97 | 28.524 | 28.565 |
| 27.5234 | 27.8809 | 27.8742 | 27.6166 | 27.7405 | 27.7545 | 27.6042 |
| 27.6263 | 27.2133 | 27.3065 | 27.5888 | 27.4537 | 27.5618 | 27.3804 |
| 29.3292 | 29.4502 | 29.4358 | 29.3331 | 29.3267 | 29.3502 | 29.4003 |
| 26.8085 | 26.8533 | 26.8603 | 26.7688 | 26.5922 | 26.4899 | 26.5175 |
| 28.7105 | 28.5784 | 28.5989 | 28.8267 | 28.7674 | 28.6636 | 28.5271 |
| 26.2974 | 25.4201 | 25.8167 | 25.9883 | 26.0297 | 26.3448 | 26.0735 |
| 33.2785 | 33.9919 | 33.7988 | 33.4667 | 33.5654 | 33.7874 | 33.8277 |
| 28.432 | 28.5148 | 28.4536 | 28.3663 | 28.3482 | 28.3488 | 28.377 |

| | | | | | | |
|---|---|---|---|---|---|---|
| 25.8201 | 25.479 | 25.2154 | 26.0078 | 26.0952 | 25.6391 | 25.4725 |
| 26.6362 | 26.6487 | 26.4781 | 26.6113 | 26.6283 | 26.6262 | 26.7514 |
| 26.3046 | 26.4043 | 25.9905 | 26.3448 | 25.9969 | 26.5377 | 26.3719 |
| 31.7387 | 31.0686 | 31.979 | 31.7347 | 31.8338 | 31.8568 | 31.8387 |
| 25.895 | 26.0977 | 25.855 | 25.9839 | 25.8271 | 25.953 | 25.9342 |
| 28.629 | 28.7188 | 28.7806 | 28.5435 | 28.5824 | 28.5073 | 28.5347 |
| 26.1194 | 26.2993 | 26.2211 | 26.1326 | 26.221 | 25.7934 | 26.002 |
| 27.8224 | 27.0016 | 26.8089 | 27.3341 | 27.2486 | 27.1623 | 26.8715 |
| 27.4799 | 27.1217 | 27.2869 | 27.1739 | 27.223 | 27.5539 | 27.5568 |
| 26.9825 | 27.177 | 27.0076 | 27.3307 | 27.4162 | 26.5565 | 26.525 |
| 27.1048 | 27.0002 | 27.0213 | 27.0752 | 26.5891 | 27.2237 | 27.2719 |
| 26.4352 | 26.4273 | 26.3575 | 26.5555 | 26.2882 | 26.8288 | 26.5677 |
| 31.7208 | 32.0274 | 31.8966 | 31.8404 | 31.8286 | 31.6924 | 31.7637 |
| 28.687 | 28.7069 | 28.8047 | 28.6399 | 28.7192 | 28.908 | 28.9569 |
| 27.6105 | 27.0771 | 27.133 | 27.5138 | 27.4167 | 27.3822 | 27.537 |
| 26.2879 | 26.2422 | 26.5254 | 26.6359 | 26.2107 | 26.1552 | 26.4048 |
| 27.5644 | 27.7507 | 27.5943 | 27.3776 | 27.3516 | 27.4648 | 27.2962 |
| 26.5677 | 26.1838 | 26.4557 | 26.4467 | 26.3714 | 26.6695 | 26.397 |
| 27.0069 | 26.8382 | 26.8706 | 26.9977 | 27.1366 | 27.0108 | 26.7935 |
| 29.5812 | 29.8106 | 29.8219 | 29.7926 | 29.7344 | 29.6858 | 29.619 |
| 28.5065 | 28.6905 | 28.7032 | 28.5069 | 28.5513 | 28.4085 | 28.2799 |
| 27.5779 | 27.1588 | 27.3435 | 27.2579 | 27.3893 | 27.582 | 27.5059 |
| 29.0465 | 29.1447 | 28.9109 | 28.9675 | 28.9561 | 29.5412 | 29.4755 |
| 26.2136 | 26.3295 | 25.9972 | 26.5442 | 26.3615 | 26.3669 | 26.3124 |
| 28.0212 | 28.3551 | 28.1332 | 28.0393 | 28.0496 | 28.1597 | 28.2008 |
| 27.7567 | 28.2495 | 28.2601 | 27.964 | 27.9803 | 27.8561 | 27.8312 |
| 28.942 | 29.4728 | 29.3545 | 29.1465 | 29.178 | 29.0938 | 29.1608 |
| 27.808 | 27.6583 | 27.7003 | 27.6947 | 27.6934 | 27.7771 | 27.825 |
| 28.149 | 28.4137 | 28.3908 | 27.8043 | 27.9957 | 28.2708 | 28.2526 |
| 26.4048 | 26.702 | 26.4777 | 26.4602 | 26.5922 | 26.3148 | 26.4154 |
| 27.3719 | 27.4208 | 27.3686 | 26.9745 | 27.0946 | 27.4527 | 27.4247 |
| 27.3864 | 27.3115 | 27.2327 | 27.0162 | 27.1158 | 27.1121 | 27.3038 |
| 30.5932 | 30.7263 | 30.7203 | 30.5251 | 30.6062 | 30.3396 | 30.4422 |
| 27.3514 | 26.9002 | 26.879 | 27.2786 | 27.1396 | 27.5528 | 27.5121 |
| 27.5446 | 27.791 | 27.7478 | 27.6369 | 27.5945 | 27.4601 | 27.5503 |
| 26.4332 | 26.497 | 26.2835 | 26.3592 | 26.5149 | 25.8593 | 25.6535 |
| 29.4013 | 29.5643 | 29.3814 | 29.2108 | 29.2026 | 29.6347 | 29.6067 |
| 31.5173 | 31.9434 | 31.8615 | 31.4291 | 31.4144 | 31.7103 | 31.726 |
| 28.0422 | 28.1403 | 28.0005 | 27.8394 | 27.7954 | 27.8235 | 27.904 |
| 27.6875 | 27.5584 | 27.3248 | 27.4522 | 27.5346 | 27.7444 | 27.6559 |
| 26.0867 | 25.7704 | 26.2703 | 26.1504 | 26.0708 | 25.831 | 25.5986 |
| 26.3497 | 26.0745 | 25.8197 | 26.1856 | 26.4612 | 26.4797 | 26.1769 |
| 27.3767 | 27.6602 | 27.4215 | 27.4451 | 27.5456 | 26.9258 | 26.8865 |
| 28.9896 | 29.2036 | 29.2172 | 28.79 | 28.894 | 29.2696 | 29.2122 |
| 30.9801 | 30.9372 | 30.945 | 31.0125 | 30.995 | 30.656 | 30.7107 |
| 27.0646 | 27.1505 | 27.1028 | 27.0632 | 27.0482 | 26.8603 | 26.863 |
| 26.6791 | 26.0881 | 25.8754 | 26.6063 | 26.2627 | 26.4744 | 26.5855 |
| 31.0779 | 31.1472 | 31.1734 | 31.1282 | 31.0715 | 31.4916 | 31.4773 |

| | | | | | | |
|---|---|---|---|---|---|---|
| 26.9248 | 27.0233 | 26.7128 | 26.8375 | 26.9221 | 27.1367 | 26.8939 |
| 26.5353 | 26.8991 | 26.8216 | 26.5091 | 26.4967 | 26.3664 | 26.327 |
| 27.1462 | 27.3634 | 27.3781 | 27.0416 | 27.2152 | 26.9037 | 27.0322 |
| 28.1564 | 28.53 | 28.4208 | 28.1165 | 28.0487 | 28.1819 | 28.2456 |
| 26.8196 | 26.6911 | 26.6667 | 26.7438 | 26.7894 | 26.8898 | 26.8129 |
| 28.1025 | 28.2133 | 28.2658 | 27.8816 | 27.8553 | 28.0407 | 27.9848 |
| 29.8441 | 30.1391 | 30.059 | 29.8294 | 29.8263 | 30.0237 | 29.9341 |
| 26.1257 | 26.0332 | 25.7941 | 26.0245 | 25.9129 | 26.0008 | 26.0029 |
| 25.7336 | 26.2942 | 26.3046 | 26.0965 | 26.0357 | 26.1509 | 26.0281 |
| 25.9474 | 26.6923 | 26.6942 | 26.1259 | 26.0724 | 26.2391 | 25.9481 |
| 26.3292 | 26.3771 | 26.4222 | 26.7619 | 26.6321 | 26.4378 | 26.6022 |
| 28.0349 | 28.2456 | 28.2434 | 28.2462 | 28.238 | 28.2818 | 28.2951 |
| 26.8502 | 27.1227 | 27.1601 | 27.2598 | 27.1737 | 27.3586 | 27.197 |
| 28.4504 | 28.7484 | 28.7283 | 28.8537 | 28.6142 | 28.7644 | 28.6653 |
| 26.5876 | 27.0138 | 26.8623 | 26.7374 | 26.8407 | 26.5989 | 26.6865 |
| 30.9029 | 30.91 | 30.8012 | 31.0215 | 31.121 | 31.2505 | 31.2361 |
| 26.8614 | 27.217 | 27.2061 | 27.2122 | 27.2457 | 26.9175 | 26.8373 |
| 31.0034 | 31.3447 | 31.2504 | 31.0782 | 31.1094 | 31.0328 | 31.052 |
| 27.772 | 28.026 | 27.764 | 27.8585 | 27.9068 | 28.007 | 27.9648 |
| 29.3813 | 29.5214 | 29.5397 | 29.4635 | 29.429 | 29.5693 | 29.3634 |
| 26.2223 | 26.6505 | 26.3698 | 26.5622 | 26.3839 | 26.3465 | 26.4867 |
| 29.2883 | 29.5388 | 29.4692 | 29.4661 | 29.4219 | 29.3723 | 29.3339 |
| 26.6497 | 26.7987 | 26.7841 | 26.6224 | 26.6937 | 26.6465 | 26.6431 |
| 31.0926 | 31.0745 | 31.0422 | 31.1763 | 31.1655 | 31.1821 | 31.0965 |
| 29.2718 | 29.3183 | 29.1399 | 29.3379 | 29.3184 | 29.2356 | 29.2423 |
| 28.8705 | 28.9686 | 28.7413 | 28.7918 | 28.7784 | 28.6894 | 28.5872 |
| 29.0319 | 29.1193 | 28.9535 | 28.8609 | 28.9879 | 28.868 | 28.8115 |
| 31.6052 | 31.4766 | 31.4056 | 31.4459 | 31.342 | 31.267 | 31.3103 |
| 28.8725 | 28.6319 | 28.7728 | 28.9613 | 28.8596 | 28.8354 | 28.8963 |
| 27.113 | 26.907 | 27.276 | 27.0231 | 27.0909 | 27.2405 | 27.121 |
| 27.8113 | 27.6247 | 27.7745 | 28.0776 | 27.9725 | 27.8191 | 27.8105 |
| 26.992 | 27.0242 | 27.113 | 27.0409 | 27.0973 | 26.9707 | 26.761 |
| 26.1918 | 25.4803 | 25.8171 | 25.9932 | 26.0319 | 25.6987 | 25.8285 |
| 28.6156 | 28.6008 | 28.694 | 28.4889 | 28.3608 | 28.5241 | 28.4343 |
| 26.2645 | 26.2323 | 26.2163 | 26.2449 | 26.4575 | 26.3774 | 26.0505 |
| 30.4937 | 30.6492 | 30.6028 | 30.4658 | 30.5166 | 30.4557 | 30.4208 |
| 28.0789 | 28.4185 | 28.1369 | 28.2238 | 28.1668 | 28.1641 | 28.0512 |
| 25.7575 | 26.0475 | 26.1704 | 26.0198 | 25.8803 | 25.8584 | 25.8299 |
| 26.3363 | 26.3398 | 26.3852 | 26.3968 | 26.3556 | 26.4234 | 26.2635 |
| 27.9345 | 28.105 | 27.9593 | 27.8964 | 27.8175 | 27.9627 | 27.9128 |
| 27.2811 | 27.1408 | 27.1799 | 27.3376 | 27.3178 | 27.4488 | 27.5067 |
| 26.8016 | 26.874 | 26.7218 | 26.9038 | 26.8397 | 26.7666 | 26.9232 |
| 29.1535 | 29.045 | 29.1857 | 29.1618 | 29.1803 | 29.0855 | 29.1397 |
| 28.6626 | 28.9804 | 28.967 | 28.8774 | 28.9037 | 28.9444 | 28.8854 |
| 27.3148 | 27.2035 | 27.0209 | 26.9098 | 26.7419 | 27.3 | 27.338 |
| 28.6691 | 28.8924 | 28.8612 | 28.7098 | 28.8109 | 28.9041 | 28.8405 |
| 25.4654 | 25.7799 | 25.27 | 25.2829 | 25.3334 | 25.5025 | 25.3913 |
| 28.6875 | 28.6367 | 28.5499 | 28.5113 | 28.4939 | 28.2522 | 28.4862 |

| | | | | | | |
|---|---|---|---|---|---|---|
| 26.8716 | 26.5757 | 26.6221 | 26.8195 | 26.6446 | 26.847 | 26.9562 |
| 28.7817 | 28.3103 | 28.9168 | 29.225 | 29.1839 | 29.1267 | 29.1055 |
| 33.064 | 33.3292 | 33.2487 | 33.2629 | 33.1908 | 32.8904 | 32.9046 |
| 29.6373 | 29.8 | 29.7209 | 29.658 | 29.6854 | 29.7533 | 29.5498 |
| 28.8928 | 29.084 | 29.0184 | 28.9774 | 28.8871 | 28.9923 | 28.9387 |
| 26.2517 | 25.9893 | 25.9552 | 26.0498 | 26.1989 | 26.361 | 26.5476 |
| 28.0984 | 27.9976 | 27.9722 | 27.8699 | 27.9737 | 27.8441 | 27.7442 |
| 27.5117 | 27.2062 | 27.2284 | 27.2547 | 27.4217 | 27.3849 | 27.3809 |
| 27.5271 | 27.3046 | 27.273 | 27.1962 | 27.368 | 27.3834 | 27.3056 |
| 27.4525 | 27.0121 | 27.1547 | 27.0723 | 27.0112 | 27.0906 | 27.4309 |
| 31.5689 | 31.6693 | 31.5047 | 31.567 | 31.5806 | 31.6645 | 31.6311 |
| 26.2564 | 26.0864 | 26.4657 | 26.3714 | 26.4914 | 26.1248 | 26.0572 |
| 31.2324 | 31.2854 | 31.2825 | 31.195 | 31.0931 | 31.2618 | 31.1839 |
| 26.1611 | 26.4659 | 25.9205 | 26.3698 | 26.1488 | 26.526 | 26.2761 |
| 28.8108 | 28.6489 | 28.4637 | 28.8097 | 28.8254 | 29.1033 | 29.1596 |
| 29.6755 | 29.3678 | 29.632 | 29.6289 | 29.6607 | 29.4975 | 29.5053 |
| 30.0537 | 30.2196 | 30.0417 | 30.0372 | 30.1145 | 29.9484 | 30.0189 |
| 25.6921 | 26.3521 | 26.397 | 25.909 | 25.8108 | 26.058 | 25.5857 |
| 28.7704 | 28.7353 | 28.6835 | 28.8853 | 28.9574 | 28.7651 | 28.6429 |
| 30.613 | 30.6272 | 30.6281 | 30.5897 | 30.6021 | 30.65 | 30.7511 |
| 31.6079 | 31.723 | 31.692 | 31.6892 | 31.6627 | 31.5459 | 31.5766 |
| 32.7603 | 32.9032 | 32.8504 | 32.6198 | 32.6464 | 32.9051 | 32.8549 |
| 28.3528 | 28.3783 | 28.2831 | 28.0997 | 28.1317 | 28.4929 | 28.4812 |
| 26.297 | 26.0733 | 26.2927 | 26.2095 | 26.5323 | 25.8397 | 25.7755 |
| 27.4874 | 27.55 | 27.5354 | 27.3224 | 27.3834 | 27.7085 | 27.6944 |
| 27.3921 | 27.4712 | 27.3171 | 27.272 | 27.1913 | 27.5254 | 27.3997 |
| 26.3707 | 26.4036 | 26.1409 | 26.3492 | 26.2686 | 26.1689 | NaN |
| 26.2813 | 26.0651 | 26.007 | 26.2436 | 26.2545 | 26.3904 | 26.5288 |
| 27.7384 | 27.9132 | 28.065 | 27.6248 | 27.8006 | 27.8178 | 27.7989 |
| 26.1718 | 26.2347 | 26.2004 | 26.4363 | 26.3194 | 26.0303 | 25.8532 |
| 25.882 | 25.8668 | 25.8444 | 25.9874 | 26.2417 | 25.8147 | 25.6385 |
| 29.03 | 29.3465 | 29.3827 | 29.0983 | 28.9966 | 29.1328 | 29.2473 |
| 26.4542 | 26.5528 | 26.3826 | 26.3982 | 26.5 | 25.9535 | 26.0752 |
| 27.9275 | 28.2516 | 28.1597 | 27.9964 | 27.9373 | 27.638 | 27.7136 |
| 27.8648 | 28.021 | 27.8561 | 27.8001 | 27.7038 | 27.8783 | 27.7724 |
| 30.7309 | 30.9281 | 30.7901 | 30.7424 | 30.7947 | 30.6783 | 30.5232 |
| 25.9526 | 25.9618 | 25.5689 | 25.6233 | 25.8549 | 26.0933 | 25.6896 |
| 30.1092 | 30.3002 | 30.2595 | 30.0562 | 30.1096 | 30.0987 | 30.0683 |
| 27.4596 | 26.9741 | 26.938 | 27.5547 | 27.3852 | 27.5184 | 27.5076 |
| 29.9563 | 30.0293 | 29.9478 | 29.9258 | 29.945 | 30.1816 | 30.3543 |
| 25.6867 | 25.3966 | 25.2889 | 25.3617 | 25.3294 | 25.7496 | 25.5708 |
| 25.9615 | 26.9385 | 26.5298 | 26.5289 | 26.5401 | 26.6092 | 26.5339 |
| 27.0881 | 27.0442 | 27.0535 | 26.9507 | 27.2034 | 27.2984 | 27.1618 |
| 27.3354 | 27.3114 | 27.2114 | 27.4501 | 27.1772 | 27.2502 | 27.2752 |
| 32.4987 | 32.7817 | 32.7632 | 32.462 | 32.4514 | 32.5777 | 32.6789 |
| 27.8274 | 27.9785 | 27.8972 | 27.7458 | 27.8105 | 27.7221 | 27.9567 |
| 26.6709 | 26.8889 | 26.957 | 26.6757 | 26.8096 | 26.3742 | 26.467 |
| 29.8968 | 29.9654 | 29.9562 | 29.7886 | 29.875 | 29.7722 | 29.7134 |

| | | | | | | |
|---|---|---|---|---|---|---|
| 27.7509 | 27.7101 | 27.7429 | 27.4117 | 27.5898 | 27.425 | 27.4575 |
| 27.7987 | 27.6598 | 27.5508 | 27.6193 | 27.6954 | 27.6122 | 27.7474 |
| 33.505 | 33.9091 | 33.7859 | 33.5272 | 33.5007 | 33.4649 | 33.5463 |
| 28.91 | 29.2441 | 29.3431 | 28.8651 | 28.8645 | 29.4496 | 29.499 |
| 29.8589 | 29.7341 | 29.7387 | 29.6388 | 29.7431 | 29.8739 | 29.9629 |
| 27.7619 | 27.7725 | 27.6527 | 27.7399 | 27.6851 | 28.2303 | 28.1323 |
| 26.2569 | 26.2231 | 26.2203 | 26.3042 | 26.4809 | 26.0522 | 25.9642 |
| 29.6698 | 29.7609 | 29.691 | 29.5822 | 29.5952 | 29.6612 | 29.6903 |
| 28.0322 | 28.2156 | 28.1052 | 28.2709 | 28.1399 | 27.9775 | 27.8809 |
| 30.6784 | 31.1613 | 30.9646 | 30.8257 | 30.7313 | 30.7482 | 30.7869 |
| 28.4062 | 28.3442 | 28.2552 | 28.2568 | 28.2802 | 28.2474 | 28.2452 |
| 29.2031 | 29.3794 | 29.3961 | 29.1736 | 29.13 | 29.1333 | 29.0491 |
| 27.5941 | 27.9513 | 27.8487 | 27.4919 | 27.4576 | 27.3421 | 27.4151 |
| 25.7591 | 25.8405 | 25.8425 | 25.736 | 25.6926 | 26.1948 | 26.2641 |
| 26.8912 | 27.2479 | 27.0205 | 27.1707 | 27.0307 | 27.019 | 26.9718 |
| 26.5979 | 26.2972 | 26.3067 | 26.3998 | 26.1822 | 26.2857 | 26.3531 |
| 27.2986 | 27.5362 | 27.4704 | 27.48 | 27.4394 | 27.2377 | 27.2321 |
| 30.4225 | 30.4217 | 30.3923 | 30.3847 | 30.4523 | 30.4506 | 30.3968 |
| 28.1686 | 28.1066 | 28.1238 | 27.8902 | 27.8582 | 28.2713 | 28.2461 |
| 29.2389 | 29.563 | 29.585 | 29.3305 | 29.2728 | 29.1874 | 29.2816 |
| 27.1274 | 26.7778 | 26.9205 | 26.6514 | 26.813 | 26.8869 | 26.9997 |
| 29.6478 | 29.6936 | 29.8183 | 29.6158 | 29.5916 | 29.504 | 29.569 |
| 27.7975 | 28.1076 | 28.047 | 27.8449 | 27.6355 | 27.8499 | 27.769 |
| 27.6168 | 27.6817 | 27.5736 | 27.5273 | 27.6576 | 27.7253 | 27.8661 |
| 29.9683 | 29.477 | 29.5114 | 29.8951 | 29.871 | 30.0532 | 29.9851 |
| 29.6389 | 29.6684 | 29.8752 | 29.9428 | 29.8787 | 29.5831 | 29.6978 |
| 30.0293 | 29.9714 | 29.9275 | 30.1715 | 30.1822 | 30.2312 | 30.1557 |
| 33.1395 | 33.3453 | 33.3364 | 33.1765 | 33.2117 | 32.9762 | 33.0244 |
| 25.953 | 25.6039 | 25.9009 | 26.0129 | 26.0764 | 26.0074 | 25.8498 |
| 30.8498 | 31.1571 | 31.1062 | 30.8412 | 30.9487 | 30.7517 | 30.7272 |
| 28.644 | 28.9747 | 28.9408 | 28.7151 | 28.7793 | 28.6577 | 28.5655 |
| 26.4912 | 26.6327 | 26.4703 | 26.4451 | 26.4136 | 26.1432 | 26.418 |
| 25.9697 | 25.6698 | 25.683 | 25.7633 | 25.9312 | 25.9448 | 25.881 |
| 27.6858 | 27.7424 | 27.8969 | 27.7915 | 27.8046 | 27.6444 | 27.4499 |
| 29.1466 | 29.2874 | 29.1967 | 29.2645 | 29.2424 | 29.0598 | 29.1682 |
| 26.9197 | 27.0683 | 26.7887 | 26.5743 | 26.7012 | 26.6558 | 26.6364 |
| 28.8935 | 28.9194 | 28.7883 | 28.8949 | 28.8656 | 28.8807 | 28.7607 |
| 28.1142 | 28.2275 | 28.204 | 28.1334 | 28.0813 | 28.0735 | 28.0549 |
| 28.4015 | 28.4341 | 28.4279 | 28.2126 | 28.2298 | 28.3645 | 28.3802 |
| 29.1804 | 29.0022 | 29.0168 | 29.1511 | 29.2646 | 29.047 | 29.1299 |
| 28.294 | 28.4655 | 28.3853 | 28.1701 | 28.2755 | 28.2913 | 28.1712 |
| 31.3226 | 31.3201 | 31.5675 | 31.3654 | 31.3299 | 31.2719 | 31.2485 |
| 27.1562 | 27.5802 | 27.4266 | 26.9577 | 27.0246 | 27.4375 | 27.5586 |
| 27.1919 | 27.1019 | 26.9362 | 27.0255 | 27.3477 | 26.994 | 26.6895 |
| 26.4937 | 26.9702 | 26.7367 | 26.4481 | 26.5942 | 26.3495 | 26.4378 |
| 27.3752 | 27.0068 | 27.172 | 27.1743 | 27.2496 | 27.4748 | 27.3331 |
| 25.4587 | 25.54 | 25.515 | 25.3776 | 25.4537 | 25.6692 | 25.4279 |
| 26.6603 | 26.7343 | 26.646 | 27.0101 | 26.942 | 26.6133 | 26.5005 |

| | | | | | | |
|---|---|---|---|---|---|---|
| 27.3115 | 27.6903 | 27.6046 | 27.2533 | 27.2849 | 27.2088 | 27.0771 |
| 26.5243 | 26.6357 | 26.5699 | 26.6337 | 26.6213 | 26.4208 | 26.2848 |
| 27.1669 | 27.1006 | 27.1464 | 27.1313 | 26.7572 | 26.9523 | 26.9039 |
| 25.2136 | 25.5703 | 25.6063 | 25.3914 | 25.1564 | 25.2594 | 25.3819 |
| 27.4734 | 27.3886 | 26.9229 | 27.4379 | 27.471 | 27.4551 | 27.5212 |
| 26.6278 | 26.5496 | 26.2916 | 26.3927 | 26.3308 | 26.5273 | 26.6214 |
| 27.4412 | 27.4901 | 27.2645 | 26.9193 | 26.9683 | 26.9229 | 26.9967 |
| 28.972 | 29.3878 | 29.3183 | 29.04 | 29.0396 | 28.6585 | 28.6826 |
| 30.6792 | 30.9198 | 30.7959 | 30.6659 | 30.6196 | 30.54 | 30.3616 |
| 29.4572 | 29.598 | 29.5759 | 29.4883 | 29.4927 | 29.3523 | 29.329 |
| 27.8995 | 28.0645 | 27.9335 | 27.7263 | 27.8449 | 27.758 | 27.7482 |
| 27.5505 | 27.9064 | 27.9101 | 27.3559 | 27.3837 | 27.228 | 27.5958 |
| 26.2259 | 25.9162 | 25.6154 | 25.9083 | 25.8113 | 26.149 | 26.0027 |
| 28.2405 | 28.4402 | 28.3226 | 28.1834 | 28.28 | 28.0329 | 28.1748 |
| 28.2465 | 28.3117 | 28.4101 | 28.4922 | 28.3409 | 28.1493 | 28.1985 |
| 27.0904 | 26.9506 | 27.3393 | 27.4515 | 27.1922 | 26.8819 | 26.7391 |
| 27.5262 | 27.3925 | 27.3604 | 27.3319 | 27.2268 | 27.3437 | 27.2956 |
| 27.1316 | 27.2739 | 27.269 | 27.2126 | 27.0961 | 27.2273 | 26.7104 |
| 27.4116 | 27.4096 | 27.2382 | 27.2547 | 27.0462 | 27.2572 | 27.2251 |
| 27.6907 | 27.721 | 27.924 | 27.7782 | 27.8185 | 27.8683 | 27.6603 |
| 26.079 | 27.2753 | 27.1866 | 26.8276 | 26.2888 | 26.4713 | 26.7826 |
| 25.8635 | 25.5957 | 25.5708 | 26.1627 | 26.0422 | 25.9027 | 25.5672 |
| 26.7205 | 27.104 | 26.9063 | 26.892 | 27.0354 | 26.9083 | 26.8442 |
| 24.8686 | 25.0642 | 25.204 | 25.4587 | 25.5454 | 25.5704 | 25.3045 |
| 27.0726 | 27.3056 | 27.1343 | 27.2728 | 27.4013 | 27.4152 | 27.4181 |
| 26.2698 | 26.7515 | 26.5936 | 26.4989 | 26.723 | 26.5085 | 26.4144 |
| 27.0127 | 26.9949 | 26.9869 | 27.0524 | 27.1577 | 27.2684 | 27.3029 |
| 27.9781 | 28.0963 | 28.0377 | 28.2605 | 28.2465 | 28.1446 | 28.0109 |
| 29.116 | 29.4359 | 29.725 | 29.3156 | 29.506 | 29.5135 | 29.5267 |
| 26.5441 | 26.396 | 26.4675 | 26.7159 | 26.7948 | 26.7347 | 26.5816 |
| 30.7425 | 31.1328 | 31.0457 | 30.9098 | 30.8927 | 30.8172 | 30.7557 |
| 27.2632 | 27.3078 | 27.2672 | 27.256 | 27.2804 | 27.5099 | 27.5207 |
| 29.0089 | 29.3989 | 29.2616 | 29.2001 | 29.2114 | 29.0441 | 29.1633 |
| 27.2662 | 27.4786 | 27.4505 | 27.3542 | 27.6274 | 27.3845 | 27.5187 |
| 28.9975 | 29.4947 | 29.2544 | 29.2201 | 29.1969 | 29.1155 | 29.1807 |
| 28.1088 | 28.6226 | 28.3872 | 28.3624 | 28.29 | 28.407 | 28.2612 |
| 27.4809 | 27.6875 | 27.5673 | 27.6332 | 27.6431 | 27.6311 | 27.6255 |
| 29.0427 | 29.2869 | 29.1484 | 29.1901 | 29.0985 | 29.422 | 29.4369 |
| 31.8622 | 31.9477 | 32.0392 | 32.1099 | 32.0862 | 32.0064 | 31.9075 |
| 27.2831 | 27.5252 | 27.5096 | 27.4253 | 27.5247 | 27.4963 | 27.4741 |
| 29.9466 | 29.8814 | 29.874 | 29.9255 | 29.9164 | 29.9073 | 30.1487 |
| 28.4591 | 28.6875 | 28.5222 | 28.3851 | 28.4149 | 28.6495 | 28.5829 |
| 28.0433 | 28.1455 | 27.9568 | 27.9691 | 28.1105 | 28.0373 | 27.9937 |
| 27.85 | 27.75 | 27.8271 | 27.6218 | 27.6723 | 27.382 | 27.6096 |
| 28.0793 | 27.7892 | 28.0678 | 27.776 | 27.9004 | 27.7247 | 27.729 |
| 28.025 | 27.8248 | 27.8726 | 27.8189 | 27.8008 | 27.9665 | 28.0294 |
| 26.895 | 26.5803 | 26.6374 | 26.8073 | 26.7322 | 26.578 | 26.7715 |
| 29.2157 | 28.9469 | 29.0137 | 28.9045 | 28.8381 | 29.0966 | 28.9196 |

| | | | | | | |
|---|---|---|---|---|---|---|
| 29.7553 | 29.1482 | 29.3478 | 29.6044 | 29.4948 | 29.5754 | 29.6084 |
| 28.5542 | 28.2197 | 28.2991 | 28.5367 | 28.4948 | 28.0223 | 28.1174 |
| 26.6509 | 25.9671 | 26.2213 | 26.5444 | 26.5344 | 26.1205 | 26.1437 |
| 27.8398 | 27.6956 | 27.5298 | 27.5329 | 27.5367 | 27.587 | 27.6401 |
| 30.4968 | 30.3802 | 30.395 | 30.4999 | 30.4841 | 30.5635 | 30.4817 |
| 26.6962 | 26.832 | 26.5461 | 26.2823 | 26.2433 | 26.6422 | 26.5748 |
| 26.7976 | 27.5391 | 27.6241 | 27.0666 | 27.2949 | 27.2693 | 27.2192 |
| 27.3703 | 27.9384 | 27.8054 | 27.9288 | 27.609 | 27.7658 | 27.6616 |
| 26.9599 | 27.3897 | 27.1312 | 26.959 | 27.1161 | 27.0487 | 27.0655 |
| 31.2218 | 31.5572 | 31.5189 | 31.5263 | 31.5118 | 31.2027 | 31.2841 |
| 26.2848 | 26.8615 | 26.68 | 26.821 | 27.0712 | 25.9221 | 26.5429 |
| 28.2278 | 27.9397 | 27.8938 | 28.0597 | 28.0858 | 28.0152 | 27.8194 |
| 26.1307 | 27.2012 | 26.933 | 27.0434 | 27.0142 | 27.0339 | 26.8231 |
| 28.3759 | 28.5424 | 28.6065 | 28.547 | 28.5214 | 28.5705 | 28.606 |
| 27.9412 | 27.5585 | 27.7803 | 27.6519 | 27.607 | 27.5566 | 27.9563 |
| 26.9625 | 26.9745 | 26.8984 | 26.9709 | 26.9917 | 27.1225 | 27.2054 |
| 28.7588 | 29.258 | 29.1189 | 28.9492 | 28.8829 | 28.9147 | 28.8467 |
| 30.7544 | 31.0605 | 30.99 | 30.8599 | 30.9136 | 30.8128 | 30.7714 |
| 26.0464 | 26.5948 | 26.642 | 26.2405 | 26.2652 | 26.2586 | 26.1376 |
| 28.086 | 28.2106 | 28.1715 | 28.0948 | 28.1208 | 28.1778 | 28.088 |
| 26.5526 | 27.0236 | 26.7922 | 26.6647 | 26.5491 | 26.4548 | 26.6402 |
| 30.0603 | 30.2464 | 30.3707 | 30.2258 | 30.2118 | 30.0799 | 30.1878 |
| 29.7564 | 29.6501 | 29.6307 | 29.5086 | 29.4751 | 29.8078 | 29.6836 |
| 26.5764 | 26.6579 | 26.8023 | 26.5112 | 26.6051 | 26.6959 | 26.5865 |
| 29.533 | 29.3421 | 29.4593 | 29.536 | 29.5946 | 29.2529 | 29.285 |
| 27.5637 | 27.4391 | 27.3317 | 27.6589 | 27.9519 | 27.4745 | 27.604 |
| 28.2143 | 28.4616 | 28.4057 | 28.3758 | 28.3556 | 28.1314 | 28.2603 |
| 26.5361 | 27.0127 | 27.0786 | 26.7853 | 26.6975 | 26.7701 | 26.6494 |
| 28.0148 | 27.865 | 27.7535 | 27.735 | 27.8406 | 27.9055 | 28.0155 |
| 27.4077 | 27.8529 | 27.6624 | 27.6006 | 27.5302 | 27.5866 | 27.4032 |
| 27.4366 | 27.502 | 27.6082 | 27.4356 | 27.3926 | 27.6337 | 27.5988 |
| 26.7375 | 26.4284 | 26.4992 | 26.5914 | 26.6334 | 26.2822 | 26.2876 |
| 31.3952 | 31.6381 | 31.5659 | 31.5525 | 31.4616 | 31.4745 | 31.5886 |
| 25.7671 | 26.1847 | 25.9335 | 25.7946 | 25.8812 | 25.5574 | 25.7461 |
| 26.3499 | 26.6088 | 26.4332 | 26.3973 | 26.2565 | 26.1705 | 26.3812 |
| 28.5693 | 28.4072 | 28.5893 | 28.6369 | 28.4592 | 28.9347 | 28.8806 |
| 26.1998 | 26.051 | 26.1499 | 26.4148 | 26.338 | 26.4579 | 26.4678 |
| 28.4506 | 28.3806 | 28.2143 | 28.1964 | 28.2317 | 28.472 | 28.5394 |
| 27.2889 | 27.4351 | 27.3724 | 26.8952 | 27.144 | 27.0284 | 27.1278 |
| 30.8307 | 31.043 | 31.0022 | 30.8546 | 30.9914 | 30.9852 | 30.927 |
| 27.6335 | 27.6691 | 27.3509 | 27.3715 | 27.3701 | 27.4058 | 27.3154 |
| 25.5628 | 26.0277 | 26.1735 | 26.1216 | 26.2353 | 25.6495 | 25.7161 |
| 26.5453 | 26.1847 | 26.305 | 26.5861 | 26.4925 | 26.6033 | 26.5324 |
| 26.0971 | 26.5989 | 26.5529 | 26.497 | 26.2484 | 26.2935 | 26.3738 |
| 31.2156 | 31.6555 | 31.4852 | 31.3396 | 31.2833 | 31.3154 | 31.2689 |
| 26.6362 | 26.8311 | 26.8569 | 26.7015 | 26.6923 | 26.6134 | 26.6019 |
| 28.2326 | 28.6447 | 28.5042 | 28.4314 | 28.4196 | 28.3128 | 28.2977 |
| 27.2393 | 27.6041 | 27.6747 | 27.4572 | 27.203 | 27.3371 | 27.3476 |

| | | | | | | |
|---|---|---|---|---|---|---|
| 27.9942 | 28.1229 | 28.0374 | 28.1501 | 28.1743 | 28.0544 | 28.0876 |
| 28.6831 | 28.7374 | 28.5358 | 28.7894 | 28.7339 | 29.0039 | 28.8901 |
| 27.7174 | 27.72 | 27.5291 | 27.8137 | 27.8522 | 28.0329 | 27.8944 |
| 30.629 | 30.878 | 30.6644 | NaN | 30.68 | 30.8032 | 30.6882 |
| 28.2957 | 28.2496 | 28.1485 | 28.2327 | 28.1227 | 28.2003 | 28.0841 |
| 28.1028 | 28.1262 | 28.0053 | 28.4667 | 28.1453 | 27.9999 | 28.0705 |
| 26.5131 | 26.9468 | 26.7163 | 26.7651 | 26.9216 | 26.8802 | 26.4716 |
| 29.9584 | 30.2816 | 30.1402 | 30.1085 | 30.1561 | 29.7932 | 29.8078 |
| 27.4507 | 26.9441 | 27.2433 | 26.8935 | 27.1707 | 27.3253 | 27.4039 |
| 26.0617 | 26.3415 | 26.0651 | 26.3338 | 26.3189 | 26.4752 | 26.637 |
| 29.048 | 29.5037 | 29.4302 | 28.7958 | 28.8578 | 29.4896 | 29.4912 |
| 26.5206 | 26.2884 | 26.222 | 26.4587 | 26.3737 | 26.4248 | 26.4196 |
| 26.6381 | 27.0113 | 26.7515 | 26.6763 | 26.7179 | 26.5976 | 26.52 |
| 26.4911 | 27.0152 | 26.8579 | 26.8728 | 26.5045 | 26.2839 | 26.5274 |
| 26.3971 | 26.602 | 26.6393 | 26.054 | 26.1765 | 26.7652 | 26.849 |
| 32.6403 | 32.9343 | 32.8017 | 32.7745 | 32.7844 | 32.6523 | 32.7404 |
| 27.7542 | 27.7918 | 27.6911 | 27.5139 | 27.4093 | 27.472 | 27.5975 |
| 27.1666 | 27.1929 | 27.4075 | 27.3262 | 27.0345 | 26.8736 | 27.0958 |
| 25.9795 | 26.3014 | 26.4252 | 26.0397 | 26.1908 | 26.1566 | 25.8563 |
| 26.0488 | 26.0651 | 25.8582 | 26.0405 | 26.2106 | 26.3491 | 26.29 |
| 31.759 | 31.6229 | 31.9152 | 31.8627 | 31.8194 | 31.6253 | 31.6892 |
| 26.6918 | 26.2802 | 26.2269 | 26.1266 | 26.4049 | 26.2236 | 26.33 |
| 27.6827 | 28.2146 | 28.1347 | 27.9704 | 28.0095 | 27.7825 | 27.7186 |
| 27.325 | 26.5636 | 27.1169 | 27.3127 | 27.0902 | 27.3908 | 27.36 |
| 27.5491 | 27.4615 | 27.4055 | 27.5246 | 27.4492 | 27.1342 | 27.2939 |
| 26.6095 | 26.1784 | 26.7464 | 26.4854 | 26.3572 | 27.1997 | 27.0866 |
| 29.5099 | 28.6544 | 29.635 | 29.5297 | 29.5207 | 29.5171 | 29.3594 |
| 27.3241 | 27.3966 | 27.3095 | 26.9717 | 26.9837 | 27.4144 | 27.1346 |
| 26.2313 | 26.3489 | 26.128 | 25.9793 | 25.9892 | 25.7105 | 26.0759 |
| 27.7238 | 27.9466 | 27.9854 | 27.6336 | 27.7075 | 27.4647 | 27.5838 |
| 30.1069 | 30.0648 | 30.0312 | 29.9814 | 30.1257 | 30.4427 | 30.4525 |
| 25.517 | 26.512 | 26.0613 | 26.3093 | 25.9522 | 26.2583 | 25.9793 |
| 26.1508 | 26.4215 | 26.2555 | 25.9996 | 25.6587 | 26.1866 | 26.182 |
| 27.6741 | 27.7727 | 27.9768 | 27.7754 | 27.7214 | 27.7935 | 27.7479 |
| 27.1771 | 27.2271 | 27.0872 | 26.9661 | 26.9441 | 26.923 | 27.0523 |
| 26.6701 | 26.7948 | 26.7617 | 26.7 | 26.6885 | 26.8516 | 27.036 |
| 28.0282 | 28.1151 | 28.0388 | 28.1322 | 28.254 | 27.9183 | 27.9555 |
| 27.8908 | 28.4147 | 28.3758 | 27.9955 | 28.029 | 27.9672 | 28.046 |
| 27.2979 | 26.917 | 27.0733 | 26.938 | 26.9409 | 27.1724 | 27.2859 |
| 26.7601 | 26.9163 | 26.8187 | 26.8011 | 26.8469 | 27.0596 | 26.9309 |
| 27.1026 | 26.6092 | 26.5942 | 26.8854 | 26.6607 | 26.9015 | 26.7979 |
| 28.037 | 28.0086 | 27.5571 | 28.0757 | 27.9543 | 28.2773 | 28.0885 |
| 28.7053 | 28.8729 | 28.9819 | 28.7459 | 28.816 | 28.6239 | 28.7847 |
| 26.7125 | 26.7591 | 26.2289 | 26.6527 | 26.7056 | 26.3224 | 26.4243 |
| 30.8105 | 31.1483 | 31.0371 | 30.9374 | 30.9053 | 30.7202 | 30.7572 |
| 28.8155 | 28.8013 | 28.7105 | 28.6847 | 28.611 | 28.657 | 28.745 |
| 26.8558 | 26.6048 | 26.5564 | 26.6832 | 26.4456 | 27.0184 | 27.1013 |
| 28.0218 | 28.2397 | 27.9679 | 27.9375 | 27.9098 | 28.0202 | 28.2668 |

| | | | | | | |
|---|---|---|---|---|---|---|
| 30.4219 | 30.1819 | 30.2866 | 30.2474 | 30.2026 | 30.4256 | 30.364 |
| 28.0882 | 27.7531 | 28.0376 | 28.3324 | 28.5455 | 28.368 | 28.3986 |
| 27.4243 | 27.3946 | 27.4891 | 27.5456 | 27.5616 | 27.7949 | 27.8216 |
| 28.0464 | 28.4898 | 28.4485 | 28.0009 | 28.0038 | 28.162 | 28.2556 |
| 26.35 | 26.6277 | 26.4165 | 26.4117 | 26.16 | 26.108 | 26.0008 |
| 30.9821 | 30.5253 | 30.6701 | 30.7801 | 30.7325 | 31.1253 | 31.1269 |
| 27.1094 | 27.431 | 27.2506 | 27.1102 | 27.1127 | 26.7032 | 27.086 |
| 24.7896 | 25.2028 | 25.0341 | 25.4235 | 25.6771 | 24.8461 | 25.0407 |
| 25.7577 | 25.5176 | 26.1103 | 25.6375 | 25.3564 | 25.6764 | 25.8158 |
| 26.4643 | 26.8548 | 26.6476 | 26.6842 | 26.6745 | 26.8333 | 26.8253 |
| 31.1981 | 31.3962 | 31.3565 | 31.0339 | 31.0331 | 31.1459 | 31.2283 |
| 27.2171 | 27.2817 | 27.2256 | 26.9048 | 26.9035 | 27.7646 | 27.6529 |
| 30.2078 | 30.4767 | 30.3661 | 30.2503 | 30.1419 | 30.1498 | 30.1348 |
| 32.0225 | 32.2432 | 32.0833 | 32.0289 | 31.9273 | 31.9745 | 31.9569 |
| 29.3398 | 29.51 | 29.2702 | 29.5315 | 29.4643 | 29.2769 | 29.3133 |
| 26.9974 | 27.1247 | 27.3237 | 27.0058 | 26.9899 | 27.0158 | 27.084 |
| 27.7044 | 27.5694 | 27.6045 | 27.8023 | 27.7782 | 27.6328 | 27.4552 |
| 29.2124 | 29.1651 | 29.1127 | 28.9192 | 28.9385 | 29.1423 | 29.0558 |
| 32.6267 | 32.3528 | 32.8056 | 32.7394 | 32.8263 | 32.6837 | 32.6313 |
| 27.5192 | 27.2705 | 27.3391 | 27.3369 | 27.1659 | 27.6517 | 27.6797 |
| 29.5051 | 29.6836 | 29.5822 | 29.3782 | 29.3654 | 29.1848 | 29.2985 |
| 27.7098 | 28.0859 | 27.804 | 27.8495 | 27.7711 | 27.6812 | 27.8144 |
| 26.8439 | 26.9902 | 26.9713 | 27.5483 | 27.308 | 27.006 | 27.1464 |
| 27.1172 | 27.0822 | 26.7964 | 26.9895 | 26.9591 | 27.4757 | 27.6387 |
| 26.2412 | 26.4624 | 26.2672 | 26.254 | 26.3 | 26.2818 | 26.1708 |
| 30.3386 | 30.0046 | 30.355 | 30.4413 | 30.4015 | 30.677 | 30.6915 |
| 32.4707 | 32.9164 | 32.706 | 32.4494 | 32.6383 | 32.3617 | 32.487 |
| 27.2447 | 26.96 | 26.906 | 27.2679 | 27.3796 | 27.0954 | 26.8722 |
| 30.0447 | 30.26 | 30.1105 | 30.04 | 30.1564 | 30.0214 | 29.9454 |
| 28.2002 | 27.892 | 28.1129 | 28.2024 | 28.1733 | 28.0681 | 27.994 |
| 30.6687 | 30.9489 | 30.9125 | 30.6712 | 30.6627 | 30.6659 | 30.5982 |
| 27.0531 | 26.8219 | 26.8888 | 26.6054 | 26.6867 | 27.2904 | 27.4037 |
| 29.4482 | 29.5855 | 29.4637 | 29.4817 | 29.6328 | 29.4091 | 29.4111 |
| 32.2572 | 32.47 | 32.6043 | 32.035 | 32.0694 | 32.163 | 32.2509 |
| 27.7418 | 27.7584 | 28.0466 | 27.7935 | 27.7233 | 27.5396 | 27.7418 |
| 28.5241 | 28.4921 | 28.2886 | 28.6372 | 28.6719 | 28.4897 | 28.2052 |
| 31.4293 | 31.6851 | 31.5546 | 31.4065 | 31.4188 | 31.2718 | 31.1091 |
| 26.3959 | 26.8351 | 26.5723 | 26.561 | 26.6209 | 26.2329 | 26.3728 |
| 26.8357 | 27.3819 | 27.2666 | 26.6377 | 26.4779 | 26.7679 | 27.0148 |
| 27.1892 | 27.172 | 27.0303 | 27.0844 | 26.7982 | 27.1434 | 27.0784 |
| 32.0354 | 32.0236 | 31.9626 | 32.2196 | 32.2045 | 32.0979 | 32.1654 |
| 26.2571 | 25.9614 | 26.1771 | 26.1599 | 26.0726 | 26.1833 | 25.8548 |
| 26.7672 | 26.1544 | 26.6061 | 26.6922 | 26.578 | 26.8088 | 26.6717 |
| 29.5541 | 29.7771 | 29.8293 | 29.335 | 29.4262 | 29.3952 | 29.4647 |
| 25.8547 | 25.9707 | 25.714 | 25.7029 | 25.5483 | 25.6102 | 25.4941 |
| 29.2047 | 29.1227 | 28.8609 | 29.121 | 28.9558 | 29.1165 | 29.133 |
| 25.9032 | 26.0715 | 26.1633 | 25.9341 | 25.904 | 26.2753 | 26.0208 |
| 26.8021 | 26.8651 | 26.5155 | 26.468 | 26.6079 | 26.8866 | 26.8334 |

| | | | | | | |
|---|---|---|---|---|---|---|
| 26.4408 | 25.661 | 25.977 | 26.2806 | 26.3154 | 26.1484 | 26.0388 |
| 25.5129 | 26.501 | 26.3748 | 25.8261 | 26.2226 | 26.3905 | 26.3986 |
| 27.3835 | 27.7454 | 27.6737 | 27.822 | 27.7121 | 27.7904 | 27.7762 |
| 29.5455 | 29.916 | 29.9534 | 29.9584 | 29.9716 | 29.7507 | 29.6489 |
| 28.4268 | 28.7219 | 28.8515 | 28.8251 | 28.8372 | 28.6322 | 28.4462 |
| 29.5096 | 29.6343 | 29.7055 | 29.8204 | 29.857 | 29.7954 | 29.6958 |
| 26.2732 | 26.4723 | 26.4156 | 26.4019 | 26.429 | 26.1434 | 26.4345 |
| 26.0505 | 26.1312 | 26.3452 | 26.3362 | 26.1807 | 26.4396 | 26.2884 |
| 28.3461 | 28.9083 | 28.9821 | 28.6162 | 28.6019 | 28.5101 | 28.4641 |
| 26.2914 | 26.5154 | 26.5371 | 26.4042 | 26.5962 | 26.3828 | 26.4587 |
| 25.6815 | 25.875 | 25.6735 | 25.6612 | 25.8624 | 26.0715 | 26.0205 |
| 25.9041 | 26.3306 | 25.9681 | 26.1852 | 26.4449 | 26.386 | 26.1407 |
| 27.5583 | 27.6742 | 27.6671 | 27.6312 | 27.6703 | 27.6822 | 27.5324 |
| 25.8283 | 26.3657 | 25.8314 | 26.0089 | 26.0043 | 26.1457 | 26.1216 |
| 27.4426 | 27.5644 | 27.5025 | 27.4449 | 27.5151 | 27.9044 | 27.8914 |
| 25.7502 | 25.905 | 25.7294 | 25.8721 | 26.0333 | 25.8729 | 25.9224 |
| 26.1866 | 26.3117 | 26.5848 | 26.6479 | 26.5597 | 26.5338 | 26.3798 |
| 27.2935 | 27.2425 | 27.1817 | 27.2839 | 27.3936 | 27.1237 | 27.1564 |
| 27.7195 | 27.6734 | 27.8455 | 27.8253 | 27.8651 | 27.5802 | 27.5916 |
| 26.2221 | 26.4388 | 26.5468 | 26.3272 | 26.3527 | 26.2007 | 26.485 |
| 30.8011 | 31.0915 | 31.0189 | 30.8636 | 31.0068 | 31.0485 | 31.0211 |
| 27.5896 | 27.5793 | 27.586 | 27.8056 | 27.6218 | 27.7932 | 27.7152 |
| 27.3549 | 27.5231 | 27.4832 | 27.4719 | 27.4688 | 27.6318 | 27.391 |
| 27.0213 | 27.0028 | 26.874 | 27.1227 | 27.0022 | 27.0821 | 27.024 |
| 27.6713 | 27.8305 | 27.8367 | 27.7067 | 27.7645 | 27.5895 | 27.8167 |
| 27.5594 | 27.4301 | 27.4644 | 27.5445 | 27.5024 | 27.5673 | 27.6503 |
| 29.3045 | 29.3347 | 29.3659 | 29.4386 | 29.5072 | 29.3552 | 29.372 |
| 28.1306 | 28.133 | 28.2386 | 28.1729 | 28.0245 | 27.8851 | 28.1175 |
| 26.3604 | 26.4382 | 26.3102 | 26.0655 | 25.9993 | 26.4644 | 26.2905 |
| 26.4125 | 26.3308 | 26.1146 | 25.7499 | 25.8722 | 26.1643 | 26.3525 |
| 27.6844 | 27.4229 | 27.5025 | 27.5225 | 27.4842 | 27.7895 | 27.756 |
| 26.3928 | 25.9475 | 26.5308 | 26.4418 | 26.4604 | 26.2021 | 26.2727 |
| 30.8927 | 30.7356 | 30.7492 | 30.7703 | 30.8356 | 30.7916 | 30.9434 |
| 25.3383 | 25.1004 | 25.0938 | 25.2906 | 25.6033 | 25.4542 | 25.4362 |
| 26.1347 | 26.0692 | 25.971 | 26.1837 | 26.4528 | 25.9139 | 26.0262 |
| 31.3682 | 31.531 | 31.4402 | 31.5819 | 31.5136 | 31.6514 | 31.5028 |
| 27.152 | 27.3036 | 27.3018 | 27.1813 | 27.5704 | 27.0695 | 26.9999 |
| 27.3578 | 27.294 | 27.1315 | 26.9926 | 27.079 | 26.9536 | 27.0008 |
| 30.5384 | 31.0441 | 30.9664 | 30.703 | 30.6244 | 30.6939 | 30.7041 |
| 25.3888 | 25.8432 | 25.5507 | 25.3391 | 25.3299 | 25.1025 | 25.2042 |
| 27.7316 | 27.6348 | 27.6093 | 27.579 | 27.5605 | 27.6072 | 27.5157 |
| 26.1088 | 26.2648 | 26.2119 | 25.7963 | 25.7753 | 25.8713 | 26.1367 |
| 27.2435 | 26.9268 | 27.2778 | 27.11 | 27.023 | 26.7379 | 26.9263 |
| 25.504 | 25.9483 | 25.4587 | 25.8705 | 25.8379 | 26.0678 | 25.9782 |
| 27.372 | 27.3859 | 27.2874 | 27.4397 | 27.2775 | 27.4647 | 27.5063 |
| 26.0826 | 26.1717 | 25.9593 | 26.2694 | 26.0443 | 26.5234 | 26.3315 |
| 28.9363 | 28.9051 | 28.8154 | 28.8089 | 28.7721 | 28.6915 | 28.7121 |
| 31.1077 | 31.0831 | 31.0888 | 31.203 | 31.2741 | 30.9927 | 31.0984 |

| | | | | | | |
|---|---|---|---|---|---|---|
| 29.3238 | 29.2192 | 29.2716 | 29.0107 | 29.0629 | 29.5439 | 29.5702 |
| 26.6377 | 27.1023 | 26.7991 | 26.9532 | 26.7243 | 26.9017 | 27.1418 |
| 29.0304 | 29.1556 | 29.005 | 29.17 | 29.0744 | 29.1315 | 29.3138 |
| 26.8547 | 27.6742 | 27.7062 | 27.2817 | 26.4464 | 27.5004 | 26.7461 |
| 27.7385 | 28.0083 | 27.9786 | 27.6853 | 27.6978 | 27.9459 | 27.906 |
| 28.3529 | 27.9302 | 28.4375 | 28.2077 | 28.1198 | 28.2956 | 28.3807 |
| 26.1148 | 26.1235 | 25.9133 | 26.2115 | 26.3219 | 26.0529 | 26.0063 |
| 29.5577 | 29.5427 | 29.5456 | 29.4218 | 29.3855 | 29.5175 | 29.5231 |
| 27.4503 | 27.758 | 27.6355 | 27.648 | 27.6295 | 27.3702 | 27.4627 |
| 27.4394 | 27.7995 | 27.8203 | 27.6177 | 27.4227 | 27.6292 | 27.6188 |
| 32.7964 | 32.9835 | 32.9962 | 32.8889 | 32.9054 | 32.8123 | 32.8453 |
| 27.2049 | 26.8309 | 26.9352 | 26.9977 | 26.8862 | 27.3006 | 27.4848 |
| 26.806 | 27.2345 | 27.1269 | 26.9973 | 26.9419 | 26.8024 | 26.8152 |
| 26.248 | 26.1694 | 26.4259 | 26.4468 | 26.6759 | 26.452 | 26.3171 |
| 27.1959 | 27.1559 | 27.1034 | 27.0056 | 27.0897 | 26.8644 | 27.2278 |
| 26.9555 | 27.0601 | 27.1863 | 27.2006 | 26.9832 | 27.1486 | 27.1481 |
| 28.0468 | 28.2931 | 28.2262 | 28.3494 | 28.4413 | 28.2209 | 28.1756 |
| 26.5322 | 26.3709 | 26.5659 | 26.5151 | 26.7776 | 26.4346 | 26.4708 |
| 30.9758 | 31.0508 | 31.0721 | 30.8846 | 31.0211 | 30.8913 | 30.8249 |
| 29.0596 | 29.2143 | 29.1357 | 29.1291 | 29.1009 | 28.8316 | 28.9532 |
| 25.9508 | 25.8172 | 25.8017 | 25.8953 | 25.9224 | 25.7173 | 25.771 |
| 27.5692 | 27.7221 | 27.6551 | 27.6382 | 27.4131 | 27.3063 | 27.3876 |
| 25.8961 | 25.8804 | 26.1019 | 25.9266 | 25.7118 | 26.3613 | 26.0783 |
| 26.8159 | 26.5536 | 26.7086 | 26.8297 | 26.7388 | 26.7974 | 26.7912 |
| 29.037 | 29.2604 | 29.2253 | 29.0338 | 29.1197 | 29.0729 | 28.9582 |
| 30.749 | 30.8015 | 30.5516 | 30.5082 | 30.6293 | 30.6821 | 30.6442 |
| 26.2936 | 26.7873 | 26.6246 | 26.4526 | 26.6865 | 26.3734 | 26.5976 |
| 27.3345 | 26.9012 | 27.275 | 27.0479 | 26.9553 | 27.474 | 27.3584 |
| 30.0864 | 30.058 | 30.0647 | 30.1499 | 30.0648 | 29.986 | 29.8552 |
| 26.1266 | 26.4986 | 26.4936 | 26.6835 | 26.3432 | 26.2307 | 26.2222 |
| 27.3772 | 27.6005 | 27.6059 | 27.7997 | 27.6553 | 27.5615 | 27.4593 |
| 32.4698 | 32.7492 | 32.6703 | 32.5225 | 32.4979 | 32.5271 | 32.4929 |
| 28.9241 | 29.05 | 29.0171 | 28.6571 | 28.6259 | 29.1627 | 29.165 |
| 30.7609 | 30.8582 | 30.8643 | 31.2235 | 31.0939 | 31.0034 | 30.9278 |
| 26.3164 | 26.6084 | 26.6561 | 26.2016 | 26.2301 | 26.5195 | 26.2593 |
| 26.0549 | 26.0983 | 26.1594 | 26.147 | 26.1185 | 26.0547 | 25.7509 |
| 27.4996 | 27.4811 | 27.3673 | 27.5204 | 27.5572 | 27.1775 | 27.2362 |
| 32.2419 | 32.5236 | 32.4606 | 32.3172 | 32.2901 | 32.2941 | 32.2361 |
| 26.1253 | NaN | 24.8227 | 25.5631 | 25.2099 | 24.5691 | 24.6872 |
| 27.4189 | 27.4338 | 27.4589 | 26.9756 | 27.3603 | 27.1383 | 27.1701 |
| 27.6609 | 27.7295 | 27.7045 | 27.625 | 27.621 | 27.5635 | 27.4934 |
| 26.4142 | 26.3063 | 26.2142 | 26.1413 | 26.2754 | 26.1572 | 26.3443 |
| 28.9553 | 29.0036 | 28.7802 | 28.6077 | 28.824 | 28.9561 | 28.973 |
| 31.9366 | 31.7203 | 32.0569 | 31.815 | 31.863 | 31.8484 | 31.84 |
| 30.4366 | 30.4692 | 30.3331 | 30.4347 | 30.4843 | 30.473 | 30.3993 |
| 26.0713 | 25.8704 | 25.9247 | 25.8495 | 26.0182 | 25.8387 | 25.6016 |
| 27.8988 | 28.1768 | 28.1295 | 27.9218 | 27.8314 | 27.4077 | 27.5764 |
| 26.5537 | 26.3551 | 26.3746 | 26.1405 | 26.2749 | 26.5119 | 26.7001 |

| | | | | | | |
|---|---|---|---|---|---|---|
| 31.6125 | 31.716 | 31.6734 | 31.5248 | 31.4983 | 31.4571 | 31.4679 |
| 32.3747 | 32.7539 | 32.6162 | 32.4619 | 32.368 | 32.5281 | 32.5096 |
| 25.4291 | 25.392 | 25.8467 | 25.1102 | 24.8895 | 25.5422 | 25.2221 |
| 27.6495 | 27.8163 | 27.7672 | 27.7728 | 27.9203 | 27.5862 | 27.4128 |
| 26.9367 | 26.9334 | 26.8784 | 27.0334 | 26.8866 | 26.7465 | 26.5792 |
| 28.4754 | 28.7146 | 28.6307 | 28.5165 | 28.5943 | 28.3879 | 28.5033 |
| 28.4178 | 28.1858 | 28.1629 | 28.1605 | 28.1762 | 27.9326 | 28.0459 |
| 28.0751 | 28.2558 | 28.5007 | 28.0715 | 28.3388 | 28.378 | 28.5915 |
| 25.482 | 25.8853 | 25.8722 | 25.8496 | 25.707 | 25.6628 | 25.8055 |
| 31.3253 | 31.7605 | 31.4864 | 31.3281 | 31.4739 | 31.4121 | 31.46 |
| 26.1232 | 26.2744 | 26.3937 | 26.291 | 26.6996 | 26.1288 | 25.8613 |
| 30.0525 | 30.0242 | 29.9212 | 30.109 | 30.1365 | 30.5008 | 30.5283 |
| 27.3858 | 26.9977 | 27.0779 | 27.0301 | 26.9657 | 27.7004 | 27.7388 |
| 28.3423 | 28.4713 | 28.1464 | 28.0145 | 28.0465 | 28.3716 | 28.3009 |
| 25.8172 | 26.4722 | 26.7091 | 26.0717 | 26.1816 | 26.2287 | 26.0612 |
| 30.3718 | 30.548 | 30.6217 | 30.4092 | 30.346 | 30.2845 | 30.3165 |
| 28.1097 | 27.7066 | 27.8454 | 27.9533 | 27.8421 | 28.0268 | 27.9109 |
| 26.044 | 26.0632 | 25.974 | 25.9415 | 25.8949 | 26.3178 | 25.9811 |
| 26.6225 | 26.8195 | 27.2724 | 26.2729 | 26.6577 | 26.9418 | 26.5438 |
| 29.0239 | 28.9341 | 28.8697 | 28.5663 | 28.6755 | 29.0087 | 29.1326 |
| 26.2575 | 25.964 | 25.7878 | 26.1739 | 26.4276 | 26.451 | 26.2578 |
| 33.1941 | 33.241 | 33.1891 | 32.9143 | 32.9176 | 32.9229 | 32.8689 |
| 28.0797 | 27.9844 | 28.0715 | 28.0494 | 28.0574 | 27.8361 | 27.9293 |
| 29.0555 | 29.1957 | 29.1931 | 29.2343 | 29.1548 | 29.0297 | 29.0236 |
| 29.0395 | 28.8848 | 28.8285 | 29.0208 | 29.0938 | 29.4142 | 29.3997 |
| 29.714 | 30.0574 | 29.9772 | 29.6747 | 29.7667 | 29.7247 | 29.9157 |
| 29.2782 | 29.4227 | 29.2651 | 29.2028 | 29.2109 | 28.9262 | 29.0236 |
| 28.7057 | 28.9625 | 29.0477 | 28.9003 | 28.8098 | 28.6383 | 28.7208 |
| 29.121 | 29.3308 | 29.0425 | 29.2637 | 29.277 | 29.1586 | 29.2071 |
| 28.1544 | 27.8445 | 28.0892 | 28.0084 | 28.1107 | 28.335 | 28.294 |
| 28.0645 | 28.0628 | 28.0903 | 28.085 | 28.16 | 28.0124 | 27.9296 |
| 27.5513 | 27.692 | 27.3325 | 27.5502 | 27.5573 | 27.2904 | 27.1401 |
| 28.9506 | 29.1603 | 29.125 | 28.9489 | 29.0511 | 28.9364 | 29.0294 |
| 26.7681 | 26.8149 | 26.642 | 26.4064 | 26.5163 | 26.605 | 26.4779 |
| 25.185 | 25.6873 | 25.6097 | 25.6176 | NaN | 25.4449 | 25.2736 |
| 26.8585 | 26.8816 | 26.9796 | 26.9718 | 26.5357 | 27.0866 | 26.9875 |
| 25.5573 | 25.4496 | 25.2398 | 25.501 | 25.8644 | 25.5622 | 25.053 |
| 25.028 | 24.6101 | 24.4785 | 25.1666 | 25.3416 | 24.9237 | 24.8008 |
| 26.5333 | 26.7387 | 26.6252 | 26.5175 | 26.3185 | 26.3879 | 26.3216 |
| 28.2418 | 28.4316 | 28.3894 | 28.1831 | 28.2337 | 28.5301 | 28.4731 |
| 27.0586 | 27.0283 | 27.0135 | 26.8905 | 27.0522 | 27.1923 | 27.3228 |
| 26.8134 | 26.9652 | 26.7439 | 27.1973 | 26.9108 | 26.8974 | 26.9765 |
| 29.1293 | 28.8005 | 28.9435 | 28.9588 | 29.0747 | 29.156 | 29.1495 |
| 25.9275 | 25.9913 | 26.3856 | 26.1338 | 26.3237 | 25.8774 | 25.4411 |
| 26.9901 | 26.9467 | 27.2109 | 27.0585 | 26.8521 | 26.9304 | 26.9741 |
| 28.7468 | 28.9202 | 28.485 | 28.7943 | 28.9148 | 28.7862 | 28.804 |
| 27.1912 | 27.9737 | 27.8373 | 27.3707 | 27.2462 | 27.4506 | 27.67 |
| 32.8361 | 33.0106 | 33.0955 | 32.5856 | 32.5371 | 32.5622 | 32.7135 |

| | | | | | | |
|---|---|---|---|---|---|---|
| 27.3711 | 27.5095 | 27.2369 | 27.2013 | 27.1356 | 27.4956 | 27.3836 |
| 29.3707 | 29.2313 | 29.245 | 29.3263 | 29.356 | 29.2206 | 29.1718 |
| 29.5549 | 29.7353 | 29.5655 | 29.3883 | 29.4574 | 29.2571 | 29.3289 |
| 26.097 | 26.1324 | 25.7094 | 25.5908 | 25.9745 | 25.5328 | 25.7707 |
| 27.1486 | 27.2734 | 27.2063 | 26.9873 | 27.1673 | 27.1308 | 27.1943 |
| 31.7325 | 31.9011 | 31.7899 | 31.5765 | 31.5219 | 31.6518 | 31.7699 |
| 27.4601 | 27.5725 | 27.4949 | 27.2933 | 27.4654 | 27.5619 | 27.6572 |
| 27.4291 | 27.7241 | 27.3145 | 27.4209 | 27.3686 | 27.1421 | 27.1533 |
| 27.3386 | 27.4724 | 27.3966 | 27.2231 | 27.1704 | 27.0975 | 27.0411 |
| 29.0274 | 29.2714 | 29.0403 | 29.006 | 29.0173 | 28.9442 | 29.028 |
| 25.9572 | 26.0857 | 26.5632 | 26.0242 | 25.9602 | 25.8852 | 26.1173 |
| 25.6595 | 26.2546 | 25.4091 | 25.5193 | 25.3276 | 25.5271 | 25.4452 |
| 30.9548 | 31.0989 | 31.0302 | 30.6833 | 30.7637 | 30.8303 | 31.0604 |
| 26.5019 | 26.5661 | 26.6429 | 26.6614 | 26.3271 | 26.5125 | 26.4256 |
| 27.1383 | 26.9751 | 26.84 | 26.9681 | 26.8968 | 26.631 | 26.6323 |
| 29.8655 | 30.2593 | 30.2927 | 29.6776 | 29.7598 | 29.7639 | 29.9333 |
| 27.1406 | 27.4781 | 27.2852 | 27.2061 | 27.2042 | 27.4528 | 27.4212 |
| 28.4473 | 28.0995 | 28.4965 | 28.3183 | 28.4282 | 28.7508 | 28.5599 |
| 29.8061 | 29.7888 | 29.7418 | 29.8022 | 29.8927 | 29.7546 | 29.9337 |
| 28.1024 | 28.9797 | 28.6168 | 28.3783 | 28.2019 | 28.4365 | 28.4407 |
| 26.9784 | 26.887 | 26.5701 | 26.5485 | 26.3902 | 26.6847 | 26.7231 |
| 28.1044 | 27.6051 | 28.0144 | 27.941 | 27.7569 | 28.1768 | 28.1215 |
| 26.5133 | 26.19 | 25.8838 | 26.8064 | 26.3788 | 26.8093 | 26.9397 |
| 30.4121 | 30.8174 | 30.7533 | 30.2842 | 30.4272 | 30.3103 | 30.3321 |
| 29.5822 | 29.4537 | 29.7164 | 29.6689 | 29.4384 | 29.5565 | 29.5594 |
| 26.8542 | 26.7052 | 26.9731 | 26.8561 | 26.7367 | 26.4882 | 26.7462 |
| 25.265 | 24.9738 | 25.0906 | 25.4639 | 25.444 | 25.4132 | 25.2977 |
| 26.574 | 26.5118 | 26.4819 | 26.5288 | 26.6283 | 26.2636 | 26.2513 |
| 32.4123 | 32.937 | 32.746 | 32.5551 | 32.4678 | 32.5876 | 32.6292 |
| 26.1877 | 26.8483 | 26.2047 | 26.5446 | 26.6901 | 26.5576 | 26.6231 |
| 27.5946 | 28.0197 | 27.8111 | 27.5094 | 27.3803 | 27.1964 | 27.3977 |
| 27.5415 | 27.4761 | 27.6864 | 27.3829 | 27.4648 | 27.478 | 27.4215 |
| 28.4221 | 28.5437 | 28.6096 | 28.3528 | 28.4371 | 28.1793 | 28.0878 |
| 27.8166 | 28.0577 | 27.8837 | 27.7668 | 27.7296 | 27.5549 | 27.425 |
| 26.9146 | 26.9428 | 26.9486 | 26.9932 | 26.9028 | 26.8324 | 27.0604 |
| 26.0946 | 26.4396 | 26.2792 | 26.0729 | 26.0761 | 25.5328 | 25.6781 |
| 25.4277 | 25.5107 | 25.5946 | 25.1955 | 25.3351 | 25.4362 | 25.2692 |
| 29.1838 | 29.41 | 29.4218 | 29.1511 | 29.125 | 29.2235 | 29.1301 |
| 31.4264 | 31.3682 | 31.3056 | 31.4974 | 31.4945 | 31.5065 | 31.4508 |
| 25.9586 | 26.414 | 26.4241 | 26.2504 | 26.3333 | 26.0587 | 26.2126 |
| 32.2465 | 32.4272 | 32.3216 | 32.3381 | 32.2216 | 32.0495 | 32.1847 |
| 29.7547 | 29.8692 | 30.0002 | 29.8643 | 29.7232 | 30.054 | 29.9296 |
| 25.9835 | 26.1606 | 26.1805 | 25.9827 | 25.9554 | 26.0272 | 26.066 |
| 27.0704 | 27.2073 | 27.1705 | 27.1816 | 27.0103 | 27.0613 | 26.9179 |
| 27.7396 | 28.0028 | 27.8588 | 27.6255 | 27.6881 | 27.2752 | 27.4205 |
| 25.3058 | 25.1426 | 25.2712 | 25.3193 | 25.2019 | 25.4362 | 25.1975 |
| 26.5689 | 27.3129 | 26.6706 | 26.5537 | 26.3965 | 26.5802 | 26.5736 |
| 29.2197 | 29.5955 | 29.5128 | 29.2228 | 29.2217 | 29.2869 | 29.3983 |

| | | | | | | |
|---|---|---|---|---|---|---|
| 27.3784 | 27.2075 | 26.7416 | 26.9251 | 26.8938 | 27.1642 | 27.3546 |
| 25.7432 | 25.6046 | 25.3948 | 25.4454 | 25.413 | 25.6439 | 25.5818 |
| 26.2739 | 25.9696 | 26.2129 | 26.1659 | 26.0655 | 25.6524 | 25.8992 |
| 26.4429 | 26.6158 | 26.262 | 26.1888 | 26.2553 | 26.3997 | 26.1672 |
| 29.6259 | 29.8483 | 29.7035 | 29.559 | 29.6849 | 29.2908 | 29.3736 |
| 29.5284 | 28.9233 | 29.0941 | 29.6097 | 29.9261 | 29.2914 | 29.1463 |
| 30.85 | 31.2215 | 31.1563 | 30.6708 | 30.6963 | 30.5877 | 30.6988 |
| 29.5904 | 29.1741 | 29.0803 | 29.4008 | 29.2972 | 29.8917 | 29.8292 |
| 26.2374 | 26.3932 | 26.5705 | 25.8474 | 26.2481 | 26.4878 | 26.5528 |
| 28.5911 | 28.7082 | 28.5465 | 28.2933 | 28.3707 | 28.3177 | 28.4049 |
| 27.1565 | 27.1214 | 27.0195 | 27.4374 | 27.361 | 27.2916 | 27.3338 |
| 25.9634 | 25.9379 | 26.224 | 26.3457 | 26.262 | 25.888 | 25.6491 |
| 25.7612 | 25.9014 | 25.5705 | 26.3403 | 26.3244 | 26.6794 | 25.9514 |
| 27.1137 | 27.5522 | 27.5483 | 27.2835 | 27.5635 | 27.3116 | 27.3506 |
| 28.306 | 28.6968 | 28.519 | 28.8046 | 28.8559 | 28.5131 | 28.4426 |
| 25.269 | 25.5715 | 25.6376 | 25.9277 | 25.4895 | 25.7901 | 25.564 |
| 30.0488 | 30.4913 | 30.4549 | 30.3891 | 30.4275 | 30.4278 | 30.536 |
| 27.212 | 27.6143 | 27.6768 | 27.9532 | 27.5319 | 27.5802 | 27.5665 |
| 26.2254 | 26.4807 | 26.171 | 26.5635 | 26.5522 | 26.3591 | 26.7038 |
| 27.9245 | 28.4609 | 28.3142 | 28.2306 | 28.1413 | 28.2064 | 28.1053 |
| 27.463 | 27.9014 | 27.8487 | 27.7493 | 27.7098 | 27.5372 | 27.4219 |
| 26.5627 | 26.9699 | 27.0639 | 26.8618 | 26.8197 | 26.6966 | 26.5643 |
| 27.3713 | 27.8698 | 27.761 | 27.565 | 27.6984 | 27.5978 | 27.515 |
| 26.3002 | 26.5134 | 26.5895 | 26.3759 | 26.366 | 26.3805 | 26.1525 |
| 27.9775 | 27.8856 | 28.0581 | 28.2296 | 28.1247 | 28.1221 | 28.1487 |
| 28.3108 | 28.4266 | 28.4444 | 28.4667 | 28.7211 | 28.343 | 28.4321 |
| 26.7537 | 27.0444 | 26.8046 | 26.8975 | 26.7838 | 27.0669 | 26.9937 |
| 31.8741 | 32.2668 | 32.191 | 32.0439 | 32.0574 | 31.9735 | 31.9475 |
| 30.2856 | 30.2174 | 30.1487 | 30.2324 | 30.2312 | 30.3293 | 30.3073 |
| 25.6021 | 25.865 | 25.6536 | 25.5608 | 25.7982 | 25.5519 | 25.7126 |
| 30.6321 | 30.8112 | 30.8777 | 30.7568 | 30.8136 | 30.6225 | 30.6691 |
| 28.8509 | 28.9361 | 28.9618 | 28.9004 | 28.9483 | 28.9353 | 28.9495 |
| 29.3636 | 29.4373 | 29.3507 | 29.4777 | 29.4112 | 29.5302 | 29.4932 |
| 32.4614 | 32.5094 | 32.5735 | 32.6649 | 32.5952 | 32.5454 | 32.4842 |
| 27.5284 | 27.4298 | 27.3507 | 27.3684 | 27.4521 | 27.4337 | 27.3393 |
| 26.6933 | 26.4543 | 26.2958 | 26.531 | 26.5082 | 26.3935 | 26.2513 |
| 27.9306 | 27.8684 | 27.9665 | 27.8241 | 27.7197 | 27.6615 | 27.7774 |
| 29.0553 | 29.0955 | 28.8937 | 28.7152 | 28.8155 | 28.662 | 28.8279 |
| 31.171 | 31.1857 | 31.082 | 30.986 | 31.0193 | 30.8386 | 30.9877 |
| 27.1355 | 26.872 | 26.9888 | 26.9342 | 26.7847 | 26.8334 | 26.7206 |
| 27.5365 | 26.8611 | 27.3647 | 27.1969 | 27.1337 | 27.0447 | 26.9904 |
| 29.6406 | 29.5166 | 29.337 | 29.1996 | 29.2115 | 29.488 | 29.5204 |
| 25.7779 | 25.3878 | 25.3383 | 25.432 | 25.4438 | 25.639 | 25.4812 |
| 29.0046 | 29.2923 | 29.1952 | 29.2476 | 29.2752 | 28.8806 | 28.9719 |
| 26.1455 | 26.4736 | 26.2506 | 26.4124 | 26.5261 | 26.353 | 26.1585 |
| 28.8835 | 28.9199 | 28.6255 | 28.9085 | 28.8671 | 28.6145 | 28.6767 |
| 26.8125 | 26.7114 | 26.6019 | 26.6109 | 26.5759 | 26.7206 | 26.8417 |
| 30.3218 | 29.8818 | 29.7442 | 29.9265 | 29.9498 | 30.3002 | 30.2673 |

| | | | | | | |
|---|---|---|---|---|---|---|
| 25.8644 | 25.7084 | 25.6647 | 25.3555 | 25.6123 | 25.7032 | 25.8097 |
| 27.5365 | 28.0672 | 27.8608 | 27.8455 | 27.8107 | 27.7676 | 27.991 |
| 29.3217 | 29.1906 | 29.1279 | 29.2905 | 29.3111 | 29.1536 | 29.0744 |
| 30.5478 | 30.8812 | 30.8055 | 30.6724 | 30.648 | 30.5477 | 30.4927 |
| 26.7273 | 26.6079 | 26.6633 | 26.5759 | 26.6084 | 26.8782 | 26.6253 |
| 28.8459 | 28.598 | 28.6736 | 28.7212 | 28.7799 | 28.6481 | 28.5806 |
| 26.5735 | 26.6505 | 26.1017 | 26.5345 | 26.7124 | 26.0693 | 26.2759 |
| 27.6682 | 27.9672 | 27.6213 | 27.743 | 27.8428 | 27.7452 | 27.6022 |
| 28.9793 | 28.8232 | 28.8724 | 29.0932 | 29.033 | 28.8959 | 28.7737 |
| 31.9891 | 32.0212 | 32.0248 | 31.9902 | 32.0063 | 31.8988 | 31.7753 |
| NaN | 26.8785 | 26.9902 | NaN | NaN | NaN | NaN |
| 26.5306 | 26.5247 | 26.6948 | 26.6309 | 26.57 | 26.2826 | 26.6676 |
| 28.2769 | 28.2901 | 28.2378 | 28.2674 | 28.2111 | 28.1584 | 28.2039 |
| 26.8771 | 26.3808 | 26.6938 | 27.0113 | 26.7681 | 26.6338 | 26.5601 |
| 32.4312 | 32.1599 | 32.3659 | 32.162 | 32.1431 | 32.5965 | 32.5694 |
| 26.247 | 26.3928 | 26.2932 | 25.9467 | 25.7013 | 25.6166 | 25.8353 |
| 28.1163 | 28.1632 | 28.2856 | 28.2317 | 28.0494 | 27.9993 | 28.0213 |
| 27.4413 | 27.0261 | 26.9718 | 27.4092 | 27.3543 | 27.2925 | 27.122 |
| 27.5267 | 27.7437 | 27.5045 | 27.7096 | 27.7439 | 27.6287 | 27.521 |
| 26.1745 | 25.918 | 26.3897 | 26.4225 | 26.1299 | 26.5928 | 26.4127 |
| 25.7397 | 25.8905 | 25.4736 | 25.5093 | 26.1811 | 25.4748 | 25.2796 |
| 27.9289 | 27.8113 | 27.7558 | 27.7562 | 27.6714 | 27.7034 | 27.7948 |
| 26.3901 | 26.4964 | 26.3966 | 26.1838 | 26.202 | 26.0291 | 25.737 |
| 25.2135 | 25.4848 | 25.711 | 25.1259 | 24.9839 | 25.1758 | 25.321 |
| 26.0284 | 26.065 | 25.6642 | 26.0808 | 25.6057 | 25.9104 | 25.8109 |
| 26.4376 | 26.0552 | 26.2394 | 25.864 | 26.1071 | 26.15 | 26.1766 |
| 28.0731 | 28.2633 | 28.2548 | 27.8353 | 27.8312 | 27.9463 | 28.0373 |
| 25.1725 | 24.8787 | 24.4844 | 24.856 | 24.7061 | 24.5207 | 24.6482 |
| 25.831 | 25.8376 | 25.9313 | 25.7019 | 25.7476 | 25.9312 | 26.0009 |
| 29.9053 | 29.7495 | 29.764 | 29.6698 | 29.7726 | 30.0702 | 30.1567 |
| 29.2389 | 29.4463 | 29.4747 | 29.3809 | 29.3097 | 29.3383 | 29.3731 |
| 26.797 | 27.1273 | 27.0197 | 27.0364 | 26.8826 | 26.7162 | 26.7367 |
| 25.8622 | 25.9686 | 25.8056 | 25.9665 | 25.9862 | 25.6079 | 25.5131 |
| 26.6721 | 26.8816 | 26.5157 | 26.577 | 26.4041 | 26.9507 | 26.7577 |
| 26.5186 | 26.5226 | 26.4616 | 26.0792 | 26.112 | 26.0523 | 26.2411 |
| 28.4184 | 28.3223 | 28.6043 | 28.2474 | 28.048 | 28.293 | 28.1039 |
| 31.0445 | 31.5766 | 31.4435 | 31.0469 | 31.1581 | 31.1832 | 31.1987 |
| 25.3252 | 24.9986 | 25.3433 | 25.2776 | 25.3227 | 25.3809 | 25.1059 |
| 25.7667 | 25.9559 | 25.6873 | 25.7025 | 25.9239 | 25.7966 | 25.4987 |
| 26.7924 | 27.1184 | 26.9626 | 27.0697 | 26.9956 | 27.0245 | 26.988 |
| 25.4452 | 26.4657 | 25.8823 | 26.1399 | 26.3059 | 26.4447 | 26.0768 |
| 26.4695 | 26.7905 | 26.9734 | 26.1403 | 26.4009 | 26.1766 | 26.1323 |
| 27.8113 | 27.8952 | 27.9973 | 27.9356 | 27.7847 | 27.9919 | 28.0467 |
| 26.1721 | 25.7209 | 26.179 | 26.2972 | 26.3883 | 26.1634 | 26.0924 |
| 26.6772 | 26.644 | 26.8298 | 26.8215 | 26.9035 | 26.6826 | 26.7085 |
| 26.5118 | 26.4938 | 26.6134 | 26.7285 | 26.7123 | 26.4352 | 26.3867 |
| 29.9893 | 29.7897 | 30.0017 | 29.7569 | 29.7983 | 29.8101 | 29.7578 |
| 25.8026 | 25.9144 | 25.7532 | 25.642 | 25.7587 | 25.556 | 25.8995 |

| | | | | | | |
|---|---|---|---|---|---|---|
| 26.884 | 26.2032 | 26.9862 | 26.7309 | 26.7943 | 26.8982 | 26.9837 |
| 25.1298 | 25.2194 | 25.0159 | 25.4967 | 25.1723 | 25.5964 | 25.4734 |
| 26.3798 | 26.2097 | 26.3073 | 26.3876 | 26.3041 | 26.083 | 26.0383 |
| 25.6303 | 26.053 | 25.9091 | 26.1174 | 26.0755 | 25.8887 | 25.6348 |
| 24.8368 | 24.8304 | 25.0627 | 25.1943 | 25.0431 | 25.083 | 24.7253 |
| 27.1098 | 27.4331 | 27.3167 | 27.0027 | 26.9542 | 27.1825 | 27.1028 |
| 25.9956 | 26.2604 | 25.9187 | 26.0872 | 26.094 | 26.0742 | 25.961 |
| 31.758 | 31.8125 | 31.7302 | 31.8523 | 31.8218 | 31.748 | 31.723 |
| 32.8848 | 32.8538 | 32.8763 | 32.8021 | 32.8267 | 33.0999 | 33.1057 |
| 29.2193 | 28.9913 | 29.1049 | 29.4379 | 29.3652 | 28.9952 | 28.9741 |
| 28.4057 | 28.6304 | 28.4597 | 28.4648 | 28.686 | 28.2604 | 28.203 |
| 26.413 | 26.394 | 26.0697 | 25.6674 | 25.7683 | 26.3472 | 26.1645 |
| 25.3421 | 25.7888 | 25.7784 | 25.441 | 25.5221 | 25.5096 | 25.3529 |
| 26.002 | 25.8946 | 25.9113 | 25.8708 | 25.9254 | 26.0113 | 26.2699 |
| NaN | 26.5818 | 26.7009 | 26.0879 | 26.1303 | 26.0465 | 26.0284 |
| 25.8995 | 26.1629 | 26.2664 | 25.7244 | 26.5364 | 25.5994 | 25.9927 |
| 30.0589 | 30.3298 | 30.1388 | 30.0048 | 29.8767 | 30.208 | 30.2512 |
| 26.6058 | 26.6684 | 26.3994 | 26.5278 | 26.6223 | 26.373 | 26.5218 |
| 28.2738 | 28.1523 | 28.3872 | 28.5784 | 28.5749 | 28.208 | 28.2846 |
| 28.3156 | 28.1476 | 28.245 | 28.17 | 28.3038 | 28.2645 | 28.226 |
| 26.2608 | 26.6417 | 26.5979 | 26.3065 | 26.2862 | 26.2381 | 26.4755 |
| 28.4208 | 28.2795 | 28.4583 | 28.2832 | 28.577 | 28.4424 | 28.3463 |
| 26.9707 | 26.8744 | 26.7652 | 26.355 | 26.5301 | 26.8816 | 26.8571 |
| 26.1289 | 26.3376 | 26.2255 | 26.4399 | 26.3413 | 26.1913 | 26.2327 |
| 26.2787 | 26.8093 | 26.5549 | 26.0891 | 26.4181 | 26.2374 | 26.3533 |
| 28.9798 | 29.1183 | 29.0188 | 28.8191 | 28.9183 | 28.8844 | 28.8518 |
| 29.534 | 29.5892 | 29.4771 | 29.5621 | 29.6968 | 29.2275 | 29.0621 |
| 26.7312 | 26.0809 | 26.2798 | 27.2185 | 26.7704 | 26.6538 | 26.3484 |
| 26.8747 | 25.5107 | 26.5511 | 26.7773 | 26.8942 | 26.9763 | 26.7975 |
| 25.5334 | 25.8807 | 25.5156 | 26.0168 | 26.0927 | 25.9726 | 25.9027 |
| 27.0738 | 26.8324 | 26.7535 | 27.0526 | 27.145 | 27.0566 | 27.0611 |
| 27.0296 | 26.8702 | 27.1832 | 27.3507 | 27.5148 | 27.2352 | 27.2758 |
| 27.3223 | 27.4123 | 27.1503 | 27.3173 | 27.3294 | 27.1549 | 27.2329 |
| 24.6765 | 24.908 | 24.7296 | 25.2074 | 24.9054 | NaN | 24.5593 |
| 25.7334 | 25.6568 | 25.7684 | 25.7682 | 26.0129 | 25.6749 | 25.9182 |
| 30.1679 | 30.457 | 30.2944 | 30.4236 | 30.3404 | 30.1785 | 30.1857 |
| 28.0464 | 28.3258 | 28.1399 | 28.0154 | 28.1048 | 28.1348 | 28.3047 |
| 27.7186 | 27.9025 | 27.8393 | 27.8591 | 27.6358 | 27.8418 | 27.7634 |
| 30.273 | 30.3353 | 30.2202 | 30.2426 | 30.2426 | 30.5466 | 30.5045 |
| 28.0559 | 27.8606 | 27.9995 | 28.1183 | 28.099 | 27.9631 | 27.8702 |
| 26.5318 | 26.4941 | 26.319 | 26.4206 | 26.4184 | 26.6089 | 26.5989 |
| 25.7941 | 26.0843 | 25.8487 | 25.9684 | 25.8996 | 25.9606 | 26.2406 |
| 27.9468 | 27.7486 | 27.8018 | 27.6675 | 27.7106 | 27.7438 | 27.7338 |
| 25.3964 | 25.8112 | 25.649 | NaN | 25.7185 | 26.039 | 25.9232 |
| 27.275 | 27.2872 | 26.7053 | 27.3031 | 27.1632 | 26.9946 | 26.5732 |
| 26.6751 | 25.0788 | 26.5873 | 26.4376 | 26.7335 | 26.9348 | 27.127 |
| 29.3409 | 29.4288 | 29.1875 | 29.0758 | 29.2469 | 29.0495 | 29.3468 |
| 26.0564 | 26.4699 | 26.3286 | 26.1019 | 26.0253 | 25.7573 | 25.7618 |

| | | | | | | |
|---|---|---|---|---|---|---|
| 25.7265 | 25.7728 | 25.6907 | 25.5424 | 25.0703 | 26.2197 | 26.0394 |
| 26.5202 | 25.935 | 26.1663 | 26.678 | 26.4138 | 26.2487 | 26.0695 |
| 25.6964 | 25.9854 | 25.8131 | 25.7802 | 25.4532 | 25.6484 | 25.8543 |
| 25.7463 | 25.9704 | 25.7505 | 25.8948 | 25.7224 | 25.859 | 25.7028 |
| 33.6219 | 34.0738 | 34.159 | 33.6261 | 33.617 | 33.971 | 34.0686 |
| 26.9317 | 27.0343 | 26.7864 | 26.8778 | 26.6473 | 26.9122 | 26.9938 |
| 26.9632 | 26.975 | 27.2745 | 27.1463 | 27.0922 | 27.1545 | 26.68 |
| 27.5304 | 27.5908 | 27.6327 | 27.7413 | 27.3732 | 27.4075 | 27.301 |
| 26.5918 | 26.3733 | 26.4111 | 26.2402 | 26.1629 | 26.4274 | 26.1598 |
| 29.5929 | 29.5659 | 29.4596 | 29.4987 | 29.526 | 29.6062 | 29.628 |
| 30.2891 | 30.5173 | 30.3775 | 30.2571 | 30.3158 | 30.2359 | 30.0648 |
| 26.8574 | 26.9308 | 26.8087 | 26.8107 | 26.8862 | 27.2736 | 27.1339 |
| 28.1723 | 28.4461 | 28.1951 | 28.2709 | 28.2486 | 27.9421 | 27.9363 |
| 26.8799 | 26.5425 | 26.6725 | 27.0247 | 27.1324 | 27.0623 | 26.9375 |
| 28.073 | 28.2606 | 28.3696 | 28.3596 | 28.4652 | 28.161 | 28.0003 |
| 25.7746 | 25.9208 | 25.9318 | 25.6374 | 25.6374 | 25.708 | 25.5521 |
| 26.3003 | 25.6922 | 26.2652 | 26.312 | 26.0843 | 26.3328 | 25.9142 |
| 25.2056 | 24.8386 | 24.7488 | 25.0078 | 25.237 | 24.546 | 24.771 |
| 27.6302 | 27.4447 | 27.6308 | 27.6391 | 27.6816 | 27.3398 | 27.4759 |
| 25.9871 | 25.9729 | 26.3246 | 26.1661 | 26.1165 | 26.0886 | 26.1041 |
| 28.3775 | 28.4029 | 28.4013 | 28.7921 | 28.6628 | 28.4548 | 28.3854 |
| 29.2651 | 29.3928 | 29.2529 | 29.1863 | 29.1809 | 28.9894 | 28.9681 |
| 29.3429 | 29.5122 | 29.4185 | 29.5618 | 29.6075 | 29.7647 | 29.851 |
| 27.4078 | 27.6212 | 27.3787 | 27.4785 | 27.415 | 26.98 | 26.9916 |
| 29.137 | 29.175 | 29.1135 | 29.0492 | 28.9636 | 28.7979 | 29.024 |
| 25.1644 | 25.3533 | NaN | 25.4455 | 25.259 | 25.7315 | 25.3316 |
| 25.639 | 25.7014 | 25.821 | 25.8438 | 25.3847 | 25.4382 | 25.5852 |
| 27.11 | 27.0588 | 26.9832 | 26.6317 | 26.8605 | 27.2165 | 27.2815 |
| 27.4151 | 27.7222 | 27.7193 | 27.3069 | 27.3628 | 27.4226 | 27.5554 |
| 29.1587 | 29.4564 | 29.3939 | 29.2487 | 29.2979 | 29.1397 | 29.15 |
| 25.2463 | 25.3235 | 25.3018 | 25.3892 | 25.3378 | 25.3661 | 24.9494 |
| 28.8459 | 28.956 | 28.8308 | 28.817 | 29.0143 | 28.7775 | 28.5657 |
| 26.4725 | 26.7869 | 26.5042 | 26.1366 | 26.4797 | 26.3458 | 26.4002 |
| 26.6893 | 26.9752 | 26.714 | 26.5543 | 26.6003 | 27.1198 | 27.0634 |
| 28.0862 | 27.2932 | 27.8699 | 28.102 | 28.0749 | 27.8264 | 27.5481 |
| 26.6729 | 27.1689 | 26.8045 | 26.7352 | 26.7281 | 26.5162 | 26.8601 |
| 31.6647 | 31.8965 | 31.8228 | 31.7213 | 31.7301 | 31.7549 | 31.6993 |
| 26.9685 | 26.955 | 26.9416 | 26.7406 | 27.022 | 26.8491 | 27.1661 |
| 25.7894 | 26.2127 | 25.1705 | 25.5351 | 25.4763 | 25.6229 | 25.7429 |
| 26.4298 | 25.979 | 26.0546 | 26.4159 | 26.2744 | 26.8051 | 26.6072 |
| 30.9766 | 30.8932 | 31.0007 | 30.9413 | 30.9733 | 30.8835 | 30.9449 |
| 26.8795 | 26.9467 | 26.9685 | 26.8427 | 26.8534 | 26.5048 | 26.6726 |
| 25.2979 | 25.5003 | 25.3507 | 25.2204 | 25.3981 | 25.6018 | 25.7137 |
| 28.058 | 28.341 | 28.2141 | 27.9666 | 28.038 | 28.0064 | 27.9373 |
| 26.2202 | 26.3909 | 26.6615 | 26.2373 | 26.3499 | 26.3695 | 26.259 |
| 26.8984 | 27.1185 | 27.1059 | 26.7614 | 26.7167 | 27.0706 | 27.1869 |
| 27.0474 | 26.4545 | 26.4837 | 27.0207 | 26.7727 | 26.9776 | 27.019 |
| 30.2422 | 30.1607 | 30.0543 | 30.1999 | 30.1552 | 30.277 | 30.2093 |

| | | | | | | |
|---|---|---|---|---|---|---|
| 26.1038 | 25.9333 | 25.8115 | 26.093 | 25.7701 | 26.0202 | 25.7658 |
| 25.8826 | 25.8877 | 25.6744 | 25.9768 | 25.5169 | 26.0057 | 26.0505 |
| 27.81 | 28.0894 | 28.1474 | 28.0055 | 27.9166 | 28.0727 | 28.0877 |
| 27.5831 | 27.4054 | 27.3947 | 27.3961 | 27.3205 | 27.5452 | 27.6617 |
| 27.293 | 26.7948 | 26.9195 | 27.0235 | 27.1345 | 27.3244 | 27.3872 |
| 26.4062 | 26.4838 | 26.5849 | 26.2556 | 26.3941 | 25.9689 | 26.1935 |
| 28.0295 | 28.2515 | 28.2464 | 27.9888 | 27.8343 | 28.0185 | 28.0107 |
| 27.4502 | 27.8026 | 27.5898 | 27.5706 | 27.3594 | 27.6133 | 27.6105 |
| 26.5975 | 26.4924 | 26.7054 | 26.4571 | 26.865 | 26.4836 | 26.684 |
| 26.7622 | 26.7543 | 26.6303 | 26.5746 | 26.5415 | 26.6101 | 26.5187 |
| 26.7543 | 26.5926 | 26.5776 | 26.0572 | 26.0619 | 26.6707 | 26.8561 |
| 25.9074 | 25.574 | 25.5746 | 26.078 | 25.8867 | 25.8365 | 25.7919 |
| 30.7142 | 30.9118 | 31.0846 | 30.8905 | 30.8677 | 30.6129 | 30.6783 |
| 30.8908 | 31.1421 | 31.0761 | 30.8486 | 30.7785 | 31.06 | 30.9648 |
| 25.8624 | 26.1806 | 25.9807 | 25.81 | 25.7386 | 26.0402 | 25.9261 |
| 25.4365 | 25.424 | 25.6682 | 25.6986 | 25.5426 | 25.9264 | 25.8302 |
| 25.6142 | 24.8946 | 25.0838 | 25.5422 | 25.7677 | 25.5394 | 25.5247 |
| 25.6778 | 25.8206 | 25.2083 | 25.5154 | 25.4699 | 25.69 | 25.687 |
| 27.5741 | 27.0732 | 27.482 | 27.4137 | 27.4339 | 27.8494 | 27.7393 |
| 33.8979 | 33.9662 | 33.9918 | 33.9466 | 33.9525 | 33.7558 | 33.8179 |
| 26.7739 | 27.5064 | 26.8019 | 27.153 | 26.7524 | 26.7938 | 26.7812 |
| 26.6583 | 26.4626 | 26.3318 | 26.6096 | 26.7351 | 26.7217 | 26.5601 |
| 34.1442 | 34.707 | 34.5544 | 34.2346 | 34.2739 | 34.223 | 34.4446 |
| 26.9066 | 26.9197 | 26.8062 | 26.7112 | 26.8431 | 26.7157 | 26.8547 |
| 26.8647 | 26.7062 | 26.6009 | 26.7178 | 26.9023 | 27.0037 | 27.1376 |
| 29.837 | 29.5974 | 29.6598 | 29.7912 | 29.8516 | 29.6178 | 29.6449 |
| 25.2359 | 25.4058 | 25.3163 | 25.2186 | 25.2859 | 25.039 | 25.1129 |
| 25.7241 | 25.3525 | 25.3977 | 25.3069 | 25.4909 | 25.9184 | 25.7493 |
| 31.8119 | 31.9985 | 31.9834 | 31.7164 | 31.6897 | 31.7317 | 31.7321 |
| 26.1185 | 24.9264 | 26.1558 | 25.5213 | 25.5217 | 25.9452 | 26.0826 |
| 27.3047 | 27.5958 | 27.5614 | 27.2486 | 27.2666 | 27.0171 | 27.3649 |
| 25.6145 | 26.0491 | 25.8531 | 25.9353 | 26.185 | 26.2886 | 25.536 |
| 30.2157 | 30.4782 | 30.3734 | 30.1266 | 30.0537 | 29.9244 | 29.9792 |
| 27.4672 | 27.7797 | 27.4918 | 27.7126 | 27.5717 | 27.6077 | 27.5356 |
| 30.5292 | 30.8419 | 30.6224 | 30.5704 | 30.6926 | 30.4644 | 30.4199 |
| 27.5954 | 28.0559 | 27.6317 | 27.7727 | 27.7503 | 27.85 | 27.9887 |
| 26.0767 | 27.1976 | 27.2679 | 26.495 | 26.5351 | 26.7949 | 26.7794 |
| 26.2701 | 26.3563 | 26.3301 | 26.454 | 26.5561 | 26.19 | 25.9695 |
| 28.9969 | 29.1334 | 28.8532 | 29.595 | 29.4869 | 29.0254 | 28.9035 |
| 27.553 | 28.0275 | 27.9978 | 28.1011 | 28.0469 | 28.129 | 27.912 |
| 25.4249 | 25.4517 | 25.7341 | 26.1859 | 25.7894 | 25.9001 | 25.6179 |
| 26.4755 | 26.4 | 26.7588 | 26.5736 | 26.3951 | 26.9208 | 26.765 |
| 25.4762 | 25.802 | 25.8791 | 26.0113 | 25.8685 | 25.2263 | 25.3757 |
| 28.2389 | 28.5571 | 28.5487 | 28.3777 | 28.4805 | 28.3827 | 28.4534 |
| 26.2834 | 26.9418 | 27.0586 | 26.5605 | 26.7988 | 26.8102 | 26.9604 |
| 26.0665 | 26.5439 | 26.5066 | 26.8127 | 26.5483 | 26.3033 | 26.1983 |
| 25.7415 | 26.4914 | 26.1929 | 26.0993 | 26.1634 | 25.87 | 26.1513 |
| 26.1532 | 26.2816 | 26.0521 | 26.3053 | 26.3304 | 26.8044 | 26.3748 |

| | | | | | | |
|---|---|---|---|---|---|---|
| 26.2695 | 26.8333 | 26.8124 | 26.7935 | 27.1136 | 26.6917 | 26.8044 |
| 26.7484 | 26.637 | 27.0728 | 26.8647 | 27.0945 | 27.1769 | 27.0893 |
| 26.5144 | 26.6369 | 26.5983 | 26.7487 | 26.7309 | 26.5783 | 26.775 |
| 25.8923 | 26.1539 | 26.2167 | 26.0565 | 26.1416 | 25.9038 | 26.0181 |
| 25.5668 | 25.8195 | 25.6248 | 25.9426 | 26.2295 | 25.5709 | 25.6518 |
| 30.3002 | 30.7474 | 30.6578 | 30.5727 | 30.663 | 30.5443 | 30.5583 |
| 28.0014 | 28.143 | 28.215 | 28.343 | 28.3517 | 28.077 | 28.1348 |
| 26.0736 | 26.4387 | 26.4657 | 26.4642 | 26.3624 | 26.3281 | 26.3319 |
| 28.8079 | 29.1327 | 29.0872 | 28.8726 | 28.8867 | 28.9222 | 29.0237 |
| 27.9233 | 28.3033 | 28.1594 | 28.0888 | 27.7864 | 28.0252 | 27.7978 |
| 28.2438 | 28.5814 | 28.5762 | 28.3634 | 28.2776 | 28.2108 | 28.2897 |
| 26.1723 | 26.2288 | 26.2557 | 26.4373 | 26.3555 | 26.1741 | 26.2182 |
| 25.0695 | 25.1837 | 25.0516 | 25.0402 | 25.1327 | 25.278 | 25.1338 |
| 27.515 | 27.7316 | 27.567 | 27.7193 | 27.6457 | 27.6351 | 27.4543 |
| 25.7116 | 25.6357 | 25.8654 | 25.8354 | 25.7064 | 25.884 | 25.6355 |
| 28.1349 | 28.1627 | 28.1524 | 28.0441 | 28.1627 | 28.2284 | 27.9312 |
| 26.7533 | 26.6629 | 26.6782 | 26.8866 | 26.6815 | 26.9063 | 26.8897 |
| 29.9578 | 30.2233 | 30.1972 | 30.1437 | 30.1504 | 29.9776 | 29.9141 |
| 29.4162 | 29.5542 | 29.5253 | 29.6622 | 29.7105 | 29.568 | 29.5774 |
| 31.1947 | 31.5731 | 31.4928 | 31.381 | 31.3126 | 31.3965 | 31.3364 |
| 27.3277 | 27.5751 | 27.5416 | 27.339 | 27.4453 | 27.3825 | 27.472 |
| 26.7538 | 26.6927 | 26.8555 | 27.0179 | 26.9422 | 26.9103 | 26.915 |
| 27.5723 | 27.9123 | 27.8957 | 27.9182 | 27.7589 | 27.7998 | 27.8757 |
| 29.8545 | 30.135 | 30.0986 | 30.0241 | 30.0325 | 30.0659 | 29.8155 |
| 28.8349 | 28.8097 | 28.7323 | 28.7024 | 28.6989 | 28.765 | 28.7278 |
| 31.5752 | 31.6628 | 31.6077 | 31.3896 | 31.4375 | 31.4554 | 31.4666 |
| 26.7049 | 26.4248 | 26.2207 | 26.2331 | 26.2366 | 26.2162 | 26.2392 |
| 29.8093 | 29.7242 | 29.5884 | 29.4562 | 29.7217 | 29.4484 | 29.7164 |
| 27.6416 | 27.5142 | 27.2516 | 27.0479 | 27.265 | 27.397 | 27.5888 |
| 29.9591 | 29.8311 | 29.8028 | 29.7656 | 29.7954 | 29.6794 | 29.8379 |
| 28.6596 | 28.5013 | 28.3912 | 28.314 | 28.2873 | 28.2775 | 28.21 |
| 26.0376 | 25.935 | 25.5687 | 25.9771 | 25.9528 | 25.8472 | 26.0993 |
| 26.0306 | 26.048 | 25.9388 | 25.8167 | 25.7616 | 25.7159 | 25.6256 |
| 26.6217 | 26.2348 | 26.5946 | 26.0342 | 25.9688 | 26.0732 | 26.235 |
| 28.2402 | 27.7331 | 28.0795 | 28.1372 | 27.9371 | 28.1227 | 28.3174 |
| 27.8489 | 28.3138 | 28.525 | 28.0519 | 27.9648 | 28.523 | 28.4188 |
| 25.9151 | 26.1687 | 26.3858 | 26.248 | 26.0593 | 25.9739 | 26.0783 |
| 27.8945 | 27.8999 | 27.7345 | 27.902 | 27.9131 | 28.098 | 28.1962 |
| 25.1485 | 24.9952 | 25.3149 | 25.6011 | 25.5509 | 25.4937 | 25.4162 |
| 29.0259 | 28.7727 | 29.0467 | 29.2933 | 29.4522 | 28.8418 | 28.816 |
| 29.3664 | 29.6201 | 29.4753 | 29.5428 | 29.486 | 29.3003 | 29.1872 |
| 28.7292 | 28.3213 | 28.846 | 28.5941 | 28.6 | 28.8673 | 28.7985 |
| 25.2834 | 27.0581 | 26.5239 | 26.4206 | 26.5244 | 26.7343 | 26.3583 |
| 26.4803 | 26.5595 | 26.3759 | 25.8192 | 26.1555 | 26.0719 | 26.3751 |
| 27.0975 | 26.9653 | 27.0251 | 27.0161 | 27.1553 | 26.6926 | 26.7745 |
| 26.167 | 26.0772 | 25.8893 | 25.6324 | 25.5899 | 26.1734 | 25.8735 |
| 27.364 | 27.7542 | 27.4909 | 27.8166 | 27.7104 | 27.4672 | 27.5007 |
| 27.3966 | 27.6621 | 27.5966 | 27.5334 | 27.6271 | 27.2532 | 27.2911 |

| | | | | | | |
|---|---|---|---|---|---|---|
| 29.9174 | 29.9394 | 29.7482 | 29.932 | 29.9025 | 30.0456 | 29.9269 |
| 31.66 | 31.8682 | 31.6598 | 31.4977 | 31.4544 | 31.552 | 31.5787 |
| 28.7186 | 28.6303 | 28.6969 | 28.6753 | 28.5818 | 28.8085 | 28.8747 |
| 28.3959 | 28.7862 | 28.7136 | 28.5701 | 28.7882 | 28.3554 | 28.3788 |
| 32.7849 | 32.8305 | 32.7746 | 32.9016 | 32.9014 | 32.6586 | 32.6145 |
| 26.4044 | 26.333 | 26.316 | 26.6211 | 26.6462 | 26.216 | 25.9678 |
| 26.0046 | 26.3326 | 26.2034 | 25.9882 | 26.2619 | 26.2293 | 26.3531 |
| 27.7044 | 27.6724 | 27.6576 | 27.4473 | 27.4551 | 27.7533 | 27.7963 |
| 27.8771 | 28.0602 | 27.8255 | 28.1061 | 27.9553 | 27.7881 | 27.852 |
| 25.4483 | 25.7455 | 25.5841 | 25.8218 | 25.8993 | 25.4586 | 25.5253 |
| 29.7818 | 29.3166 | 29.6628 | 29.7662 | 29.729 | 29.6682 | 29.5287 |
| 27.8173 | 27.7915 | 27.4408 | 27.7905 | 27.7645 | 27.5346 | 27.47 |
| 26.1257 | 26.0613 | 25.7764 | 26.3813 | 26.2073 | 26.3813 | 26.3928 |
| 29.9596 | 29.6393 | 29.6923 | 29.9971 | 30.0621 | 29.7311 | 29.6822 |
| 26.1292 | 26.5266 | 26.4246 | 25.9315 | 26.2114 | 26.0652 | 26.1683 |
| 27.579 | 27.6508 | 27.5991 | 28.2316 | 27.599 | 27.3437 | 27.4473 |
| 25.5057 | 26.1614 | 26.2015 | 26.0476 | 26.0538 | 26.2456 | 25.926 |
| 26.8555 | 27.2007 | 26.9334 | 26.9643 | 27.1079 | 26.9441 | 27.0941 |
| 27.0259 | 27.1504 | 27.3753 | 27.3326 | 27.2034 | 27.3465 | 27.2618 |
| 26.5188 | 27.1777 | 27.1423 | 26.9829 | 26.7724 | 26.9709 | 26.9464 |
| 25.3168 | 24.698 | 25.2243 | 24.8306 | NaN | NaN | 25.3094 |
| 27.2363 | 27.4242 | 27.5486 | 27.319 | 27.3454 | 27.1958 | 27.2613 |
| 26.3887 | 26.6682 | 26.5018 | 26.025 | 26.0946 | 26.1783 | 26.2278 |
| 26.5319 | 26.7107 | 26.7307 | 26.4697 | 26.4361 | 26.6988 | 26.7291 |
| 27.8032 | 27.7461 | 27.5574 | 28.1233 | 27.5921 | 27.8502 | 27.8704 |
| 25.1257 | 24.8358 | 24.6373 | 24.638 | 25.1493 | 24.8013 | 24.9604 |
| 27.2288 | 27.4805 | 27.6258 | 27.2172 | 27.2175 | 27.5011 | 27.3879 |
| 25.4 | 25.6854 | 25.6482 | 25.3065 | 25.1181 | 26.3126 | 25.4179 |
| 27.4818 | 27.7826 | 27.5921 | 27.83 | 27.6728 | 27.1828 | 27.4134 |
| 24.5385 | 24.4672 | 24.3303 | 24.5554 | 24.3977 | NaN | NaN |
| 27.5392 | 27.317 | 27.4151 | 27.4488 | 27.5023 | 27.6286 | 27.4999 |
| 29.9873 | 30.1027 | 30.2232 | 30.1292 | 30.084 | 29.8948 | 29.9137 |
| 24.8442 | 25.2021 | 25.0874 | 25.0518 | 25.16 | 24.6965 | 24.813 |
| 25.3676 | NaN | NaN | 25.1762 | 25.065 | 25.014 | 25.3213 |
| 27.8403 | 27.884 | 27.6569 | 27.9169 | 27.8868 | 28.0222 | 28.1398 |
| 26.8192 | 27.1889 | 27.0781 | 26.6621 | 26.6462 | 27.0413 | 27.0992 |
| 25.2998 | 26.0128 | 26.0431 | 25.4228 | 25.9619 | 25.5573 | 25.5174 |
| 25.8922 | 25.7747 | 25.8157 | 25.8103 | 25.868 | 26.3555 | 25.8985 |
| 25.0101 | 25.491 | 25.3376 | 25.2789 | 25.7258 | 25.0353 | 24.7932 |
| 30.5034 | 30.3626 | 30.4269 | 30.5553 | 30.4615 | 30.2492 | 30.218 |
| 27.8388 | 28.0237 | 27.6225 | 27.5202 | 27.6 | 27.4203 | 27.4911 |
| 28.1972 | 28.3776 | 28.3765 | 28.4809 | 28.7635 | 27.8038 | 27.8872 |
| 26.7695 | 26.6357 | 26.6538 | 26.5906 | 26.6629 | 26.8132 | 26.5753 |
| 24.9152 | 24.8318 | 24.6878 | 25.0015 | 24.9523 | 24.4904 | 24.4139 |
| 25.7722 | 26.0375 | 26.0385 | 25.8159 | 26.1054 | 25.7794 | 25.7226 |
| 26.73 | 26.875 | 26.8008 | 26.6019 | 26.72 | 26.9054 | 26.8265 |
| 25.4501 | 25.6584 | 25.5672 | 25.7771 | 25.66 | 25.3718 | 25.481 |
| 27.4683 | 27.4447 | 27.4671 | 27.2982 | 27.6876 | 27.6572 | 27.5384 |

| | | | | | | |
|---|---|---|---|---|---|---|
| 24.6083 | 25.2172 | 24.9468 | 25.0295 | 25.3727 | 24.8348 | 25.278 |
| 26.4135 | 26.5258 | 26.4787 | 26.5131 | 26.4181 | 26.2147 | 26.5206 |
| 26.7281 | 26.6495 | 26.8312 | 26.7453 | 26.7396 | 27.0145 | 26.9751 |
| 25.0518 | 25.3946 | 25.1887 | 25.2271 | 25.0373 | 25.1239 | 25.0283 |
| 25.8345 | 26.1779 | 25.8561 | 26.0346 | 26.2036 | 25.8531 | 25.6464 |
| 25.3077 | 24.8634 | 24.9267 | 25.5616 | 25.581 | 25.5036 | 25.611 |
| 26.4666 | 26.0349 | 25.659 | 26.0225 | 26.2654 | 26.0314 | 26.1011 |
| 24.8503 | NaN | NaN | 25.1467 | 25.2492 | 25.1188 | 24.8131 |
| 28.9749 | 28.9838 | 28.7891 | 28.8041 | 29.0328 | 28.8446 | 28.783 |
| 25.4481 | 25.3135 | 25.2352 | 25.1438 | NaN | 25.5175 | 25.4137 |
| 30.0931 | 29.9979 | 29.9602 | 30.4378 | 30.5021 | 30.3377 | 30.1507 |
| 26.9136 | 27.4082 | 27.0669 | 27.3072 | 27.1184 | 26.7321 | 26.9358 |
| 27.357 | 27.9344 | 27.8479 | 27.495 | 27.398 | 27.0928 | 27.3603 |
| 26.1412 | 26.1976 | 26.3489 | 26.3059 | 26.1803 | 26.1517 | 26.0683 |
| 27.1344 | 27.2611 | 27.1957 | 26.8763 | 26.8777 | 26.907 | 27.1486 |
| 25.943 | 26.1134 | 25.9867 | 26.0455 | 25.9362 | 25.8532 | 25.767 |
| 25.8343 | NaN | 25.8519 | 26.196 | NaN | 26.0059 | NaN |
| 31.0806 | 31.6425 | 31.4947 | 31.1675 | 31.1472 | 31.3137 | 31.3579 |
| 25.8807 | 26.1712 | 26.0628 | 25.8084 | 25.6182 | NaN | 25.7545 |
| 25.6099 | 25.4833 | 25.5356 | 25.5968 | 25.6781 | 25.2214 | 25.2584 |
| 28.6187 | 28.971 | 28.6986 | 28.6291 | 28.8178 | 28.6867 | 28.6466 |
| 26.8795 | 26.2411 | 26.4506 | 26.2383 | 26.55 | 26.8809 | 26.8125 |
| 26.39 | 26.7223 | 26.6623 | 26.6713 | 26.6078 | 26.4886 | 26.6014 |
| 27.1923 | 27.5829 | 27.48 | 27.1235 | 27.2928 | 27.2081 | 27.36 |
| 26.7762 | 26.9754 | 26.46 | 26.6392 | 26.6388 | 26.6469 | 26.2092 |
| 26.0037 | 26.1083 | 25.9658 | 25.6346 | 25.7977 | 25.8725 | 26.2577 |
| 25.758 | NaN | 25.412 | 25.5515 | 25.6201 | 25.4467 | 25.4964 |
| 27.3904 | 27.4307 | 27.1121 | 27.4016 | 27.4985 | 27.6238 | 27.6369 |
| 25.8299 | 26.1008 | 25.964 | 25.8651 | 25.8744 | 26.0232 | 26.0073 |
| 30.8513 | 30.674 | 30.6936 | 30.6111 | 30.575 | 31.0066 | 30.9881 |
| 25.2422 | 25.2581 | NaN | 24.8482 | 25.3101 | 25.6114 | 25.7107 |
| 31.2362 | 31.403 | 31.4516 | 31.335 | 31.249 | 31.1863 | 31.2782 |
| 28.2797 | 28.0221 | 28.1639 | 28.2623 | 27.91 | 27.7341 | 28.1634 |
| 25.3343 | 25.1635 | 25.2498 | 25.3316 | 25.2178 | 25.4691 | NaN |
| 26.006 | 26.105 | 26.2409 | 25.8654 | 25.8378 | 26.2458 | 25.8371 |
| 26.6909 | 26.9916 | 27.0361 | 26.7977 | 26.7502 | 26.6914 | 26.7542 |
| 27.0996 | 27.2006 | 27.1466 | 27.0825 | 27.0279 | 26.7524 | 26.8674 |
| 25.5179 | 25.4339 | 25.5131 | 25.2788 | 25.5753 | 25.4504 | 25.4396 |
| 25.2139 | 25.4492 | 25.294 | 25.5579 | 25.2585 | 25.3157 | 25.3337 |
| 27.0889 | 27.2509 | 27.1213 | 27.2018 | 27.1629 | 26.9972 | 26.9997 |
| 31.03 | 31.3006 | 31.3193 | 30.9029 | 30.9516 | 31.1195 | 31.0735 |
| 25.7122 | 26.1416 | 25.7537 | 25.1621 | 24.8696 | 24.8747 | 25.1573 |
| 24.7952 | 24.6209 | 25.0055 | 24.7898 | 24.9985 | 25.3899 | 24.9444 |
| 28.8496 | 28.2703 | 28.7201 | 28.8542 | 28.9102 | 29.0037 | 28.9336 |
| 29.7271 | 29.6794 | 29.5617 | 29.4486 | 29.4238 | 29.7696 | 29.9586 |
| 30.2116 | 30.4526 | 30.4082 | 30.0831 | 30.1538 | 30.0128 | 30.047 |
| 27.2437 | 27.2591 | 27.2101 | 27.0325 | 27.2023 | 27.0569 | 27.0066 |
| 26.1583 | 26.0002 | 25.8329 | 25.6233 | 25.3963 | 25.8825 | 26.2225 |

| | | | | | | |
|---|---|---|---|---|---|---|
| 25.0073 | 24.9782 | 25.0552 | 25.4209 | 24.7641 | 24.819 | 24.6935 |
| 27.8402 | 27.6887 | 27.4516 | 27.834 | 27.8771 | 27.8663 | 27.8557 |
| 25.6808 | 25.3558 | 25.3598 | 25.3182 | 25.3305 | 25.5462 | 25.5704 |
| 26.4188 | 26.3588 | 26.1492 | 26.5147 | 26.5071 | 26.3284 | 26.1725 |
| 25.9435 | 25.9505 | 25.9413 | 25.5354 | 25.8674 | 25.8714 | 25.7369 |
| 30.2328 | 30.0437 | 30.0132 | 30.4628 | 30.4415 | 30.2447 | 30.2634 |
| 29.3366 | 28.9382 | 29.2779 | 29.4987 | 29.4839 | 29.2015 | 29.3037 |
| 28.242 | 28.3382 | 28.4578 | 28.619 | 28.6524 | 28.5818 | 28.6723 |
| 25.6312 | 25.9371 | 25.7395 | 25.7317 | 25.3874 | 25.8149 | 25.4731 |
| 27.5544 | 27.6723 | 27.3921 | 27.4644 | 27.4374 | 27.5894 | 27.5894 |
| 26.2893 | 26.1297 | 25.9091 | 25.9319 | 25.985 | 26.1531 | 26.2225 |
| 26.2821 | 26.475 | 26.0434 | 25.5825 | 25.9417 | 26.2039 | 26.1956 |
| 27.5628 | 27.4145 | 27.4592 | 27.4237 | 27.6576 | 26.911 | 27.1082 |
| 28.0299 | 27.969 | 27.8885 | 28.2197 | 28.1004 | 28.0712 | 27.907 |
| 26.1387 | 26.295 | 25.7912 | 26.0683 | 26.1612 | 26.4926 | 26.3652 |
| 25.4727 | 25.7371 | 25.1669 | 25.5194 | 25.7527 | 25.6407 | 25.2755 |
| 30.4526 | 30.7037 | 30.7215 | 30.2163 | 30.2474 | 30.147 | 30.1998 |
| 27.4767 | 27.645 | 27.5541 | 27.6144 | 27.6395 | 27.4404 | 27.4229 |
| 25.541 | 24.7267 | NaN | NaN | NaN | 25.2813 | NaN |
| 31.6996 | 31.6256 | 31.5036 | 31.511 | 31.546 | 31.8728 | 31.87 |
| 26.3476 | 26.6389 | 26.5112 | 26.3682 | 26.2816 | 26.2887 | 26.3026 |
| 28.1101 | 28.1334 | 27.9782 | 28.2938 | 28.1129 | 28.1274 | 28.1727 |
| NaN | NaN | NaN | 24.6069 | 24.8276 | NaN | 24.6212 |
| 26.6449 | 26.7397 | 26.8467 | 26.7575 | 26.5961 | NaN | NaN |
| 26.2334 | 26.4138 | 26.4246 | 26.4336 | 26.4998 | 26.0242 | 26.0953 |
| 27.201 | 27.0269 | 27.2169 | 26.925 | 26.9301 | 27.2611 | 27.2552 |
| 25.8013 | 26.2472 | 25.5229 | 25.889 | 25.9398 | 25.898 | 25.9551 |
| 26.393 | 26.2766 | 26.1513 | 26.3296 | 26.35 | 25.8339 | 25.869 |
| 26.0786 | 25.9898 | 25.6355 | 25.5025 | 25.9848 | 26.0236 | 25.9313 |
| 30.5273 | 30.2937 | 30.2868 | 30.407 | 30.2813 | 30.4874 | 30.5014 |
| 26.6628 | 26.7196 | 26.5956 | 26.8879 | 26.918 | 26.2056 | 26.3542 |
| 29.6535 | 29.4998 | 29.4567 | 29.7659 | 29.5105 | 29.5905 | 29.7191 |
| 27.093 | 26.6263 | 26.9181 | 27.0746 | 27.0687 | 26.9172 | 26.6141 |
| 25.6013 | 26.1415 | 25.7416 | 25.2774 | 25.4523 | 25.3766 | 25.4514 |
| 26.2579 | 26.3223 | 26.4544 | 26.2911 | 26.0073 | 26.6596 | 26.0211 |
| 26.7398 | 26.9565 | 26.8763 | 26.6301 | 26.506 | 26.837 | 26.5598 |
| 25.4402 | NaN | 25.4784 | 26.1177 | 26.1149 | 26.0643 | 25.5518 |
| 27.157 | 26.39 | 26.871 | 26.8219 | 26.7351 | 26.7623 | 26.8964 |
| 24.6984 | NaN | NaN | 24.763 | 24.529 | NaN | 24.4419 |
| 26.1485 | 26.0198 | 25.7604 | 25.9747 | 25.9154 | 25.8561 | 26.1351 |
| 26.3888 | 26.858 | 26.5742 | 26.5376 | 26.4744 | 26.6012 | 26.7837 |
| 27.0072 | 26.9987 | 26.9106 | 26.7442 | 26.934 | 27.0157 | 26.9737 |
| NaN | NaN | NaN | NaN | 24.4692 | 24.732 | 24.7752 |
| 25.2668 | 25.2424 | 25.2678 | 25.1918 | 25.2405 | 25.1409 | 25.4825 |
| 29.0882 | 29.1403 | 29.3235 | 29.0799 | 29.1739 | 28.8147 | 29.1095 |
| 26.3644 | 26.3935 | 26.2348 | 26.074 | 25.8395 | 26.2041 | 25.9671 |
| 27.4113 | 27.6207 | 27.5432 | 27.2911 | 27.3015 | 27.4249 | 27.291 |
| 25.385 | 25.9094 | 25.6973 | 25.1607 | 25.414 | 25.5037 | 25.2341 |

| | | | | | | |
|---|---|---|---|---|---|---|
| 26.6863 | 26.8707 | 26.6772 | 26.6291 | 26.4722 | 27.0469 | 26.9966 |
| 24.8002 | 24.9143 | 24.9718 | 24.806 | 25.0666 | 24.5573 | 24.7744 |
| 26.1574 | 26.2385 | 26.2153 | 26.2702 | 26.1583 | 25.7668 | 26.1867 |
| 28.0732 | 28.1522 | 28.3503 | 27.9802 | 27.9057 | 27.9474 | 27.9994 |
| 24.7768 | 24.6834 | 24.8257 | 25.096 | 24.7441 | 24.4648 | 24.8811 |
| 28.3098 | 28.191 | 28.2119 | 28.2319 | 28.2284 | 28.6091 | 28.6317 |
| 26.4412 | 26.7862 | 26.7985 | 26.4875 | 26.693 | 26.5922 | 26.4671 |
| 28.3699 | 28.6064 | 28.4411 | 28.4796 | 28.2585 | 28.4474 | 28.4243 |
| 26.0689 | 26.3657 | 26.2758 | 26.0851 | 25.916 | 26.0613 | 26.1634 |
| 27.0663 | 27.2887 | 27.1112 | 26.8831 | 27.0283 | 27.5124 | 27.5199 |
| 29.2073 | 29.1191 | 29.0994 | 29.4244 | 29.3994 | 29.3739 | 29.3435 |
| 27.0112 | 27.0305 | 27.1105 | 27.0456 | 27.2165 | 26.666 | 26.5908 |
| 25.5898 | 25.0457 | 25.0591 | 24.9914 | 25.1877 | 25.3997 | 25.5818 |
| 26.2549 | 26.6525 | 26.4249 | 26.2725 | 26.3039 | 26.0949 | 25.8296 |
| 31.0009 | 31.0367 | 31.3169 | 30.9717 | 31.0223 | 31.1113 | 30.9037 |
| 25.8229 | 25.5309 | 25.5825 | 25.5816 | 25.5626 | 25.7416 | 25.8145 |
| 25.9728 | 25.7787 | 25.6869 | 26.0786 | 26.2875 | 26.3492 | 26.4009 |
| 25.5648 | 25.2861 | 25.7309 | 25.2068 | 25.3451 | 25.4521 | 25.3854 |
| 25.3726 | 24.9122 | 24.9854 | 25.3506 | 25.227 | 24.9409 | 24.6723 |
| 26.2929 | 26.2515 | 26.1739 | 26.3263 | 26.4302 | 26.3996 | 26.387 |
| 26.7586 | 26.8948 | 26.9303 | 26.6508 | 26.7808 | 26.6082 | 26.5433 |
| 25.5899 | 26.0562 | 25.6441 | 25.7883 | 25.8217 | 25.6441 | 25.4193 |
| 29.7617 | 30.0967 | 29.984 | 29.7186 | 29.7293 | 29.6838 | 29.6281 |
| 26.6899 | 26.2149 | 26.7123 | 26.6987 | 26.7161 | 26.8107 | 26.7116 |
| 26.2622 | 26.5395 | 26.6259 | 26.1102 | 26.2071 | 26.3812 | 26.1956 |
| 27.2561 | 27.0636 | 26.9417 | 26.8424 | 26.8062 | 27.2162 | 27.1658 |
| 27.6873 | 27.7735 | 27.8764 | 27.5834 | 27.5814 | 27.3313 | 27.327 |
| 26.4841 | 26.6886 | 26.6244 | 26.7169 | 26.5337 | 26.3846 | 26.3953 |
| 32.4887 | 32.519 | 32.5164 | 32.2802 | 32.297 | 32.1898 | 32.3655 |
| 30.1774 | 30.0537 | 30.293 | 30.3646 | 30.29 | 30.3446 | 30.2621 |
| 25.6017 | 25.6679 | 25.3309 | 25.6802 | 25.1585 | 26.0237 | 25.8432 |
| 27.3296 | 27.6137 | 27.7538 | 27.2044 | 27.3554 | 27.5198 | 27.6544 |
| 26.9191 | 26.8538 | 26.7387 | 26.8963 | 26.9565 | 26.4141 | 26.4519 |
| 28.472 | 28.6975 | 28.7704 | 28.4537 | 28.5967 | 28.3763 | 28.5579 |
| 30.9508 | 31.2106 | 30.9516 | 30.9601 | 31.0207 | 30.9296 | 30.921 |
| 26.1797 | 25.9717 | 26.1685 | 26.4054 | 25.7995 | 26.1241 | 25.9869 |
| 26.1825 | 26.0091 | 26.2119 | 26.0367 | 26.0935 | 25.6862 | 26.1104 |
| 28.6631 | 28.6822 | 28.7065 | 28.6415 | 28.6529 | 28.3644 | 28.3797 |
| 28.3767 | 28.7289 | 28.6552 | 28.5382 | 28.2901 | 28.4033 | 28.4904 |
| 25.7447 | 26.1855 | 26.2601 | 25.925 | 25.9461 | 26.0375 | 25.954 |
| 27.4775 | 27.729 | 27.8425 | 27.7699 | 27.5292 | 27.7262 | 27.5672 |
| 28.3379 | 28.0501 | 28.274 | 28.5754 | 28.4072 | 28.3995 | 28.3867 |
| 26.0185 | 25.8924 | 26.0421 | 26.1357 | 26.2306 | 26.0755 | 26.1918 |
| 27.2625 | 27.6071 | 27.5722 | 27.703 | 27.8385 | 27.8445 | 27.5144 |
| 28.52 | 28.9265 | 28.9744 | 29.0165 | 29.1515 | 28.8961 | 29.0382 |
| 24.7894 | 24.7983 | 24.9203 | 24.8195 | 25.3123 | 24.6211 | 24.903 |
| 25.8889 | 26.3024 | 25.8881 | 26.0677 | 25.9705 | 26.0012 | 26.1467 |
| 29.492 | 29.719 | 29.6953 | 29.8703 | 29.8567 | 30.1399 | 29.907 |

| | | | | | | |
|---|---|---|---|---|---|---|
| 26.4099 | 26.4692 | 26.6622 | 26.682 | 26.4727 | 26.2757 | 26.1358 |
| 27.8282 | 28.4365 | 28.3918 | 28.1587 | 28.1367 | 28.1262 | 28.1158 |
| 27.6849 | 27.8168 | 27.7428 | 27.8794 | 27.9228 | 27.713 | 27.6041 |
| 29.0004 | 29.345 | 29.2401 | 29.6162 | 29.6236 | 29.6217 | 29.4989 |
| 25.5358 | 26.1353 | 26.0434 | 26.2096 | 26.2422 | 26.1145 | 25.9401 |
| 25.6927 | 25.9837 | 26.0021 | 26.0558 | 26.0515 | 25.956 | 25.8027 |
| 30.075 | 30.3009 | 30.0384 | 30.4312 | 30.5158 | 30.5237 | 30.6531 |
| 27.9237 | 28.353 | 28.2124 | 27.9784 | 28.097 | 28.1898 | 28.182 |
| 27.4943 | 27.5635 | 27.5885 | 27.7572 | 27.6917 | 27.6803 | 27.572 |
| 28.2802 | 28.5044 | 28.2722 | 28.5136 | 28.3998 | 28.5722 | 28.5359 |
| 30.6563 | 31.0142 | 30.769 | 30.7415 | 30.7392 | 30.6768 | 30.8489 |
| 24.8634 | 25.3926 | 25.2828 | 25.2415 | 25.5347 | 25.0892 | 25.3234 |
| 27.91 | 28.3003 | 28.0567 | 27.9315 | 28.1109 | 28.2603 | 27.946 |
| 26.8078 | 26.941 | 26.9094 | 27.0005 | 26.9626 | 26.7161 | 26.9258 |
| 32.3208 | 32.4575 | 32.4068 | 32.4587 | 32.4806 | 32.4393 | 32.3981 |
| 25.3253 | 25.4272 | 25.4389 | 25.3637 | 25.5215 | 25.4455 | 25.3496 |
| 26.5164 | 27.0296 | 26.741 | 26.9484 | 26.7103 | 26.7266 | 26.852 |
| 28.3979 | 28.2852 | 28.2583 | 28.4694 | 28.3912 | 28.5694 | 28.6759 |
| 28.7336 | 28.8887 | 28.9846 | 29.0081 | 28.9041 | 28.8495 | 28.7439 |
| 25.5722 | 25.9788 | 26.0928 | 25.7649 | 25.793 | 25.7452 | 25.843 |
| 30.1274 | 30.151 | 30.0417 | 30.1844 | 30.2014 | 30.2934 | 30.062 |
| 31.5883 | 31.9703 | 31.8275 | 31.8254 | 31.7728 | 31.7401 | 31.6035 |
| 25.4477 | 25.7131 | 25.659 | 25.8713 | 25.7996 | 25.7157 | 25.5753 |
| 28.3743 | 27.8799 | 28.4297 | 28.0694 | 28.0537 | 28.1075 | 28.0681 |
| 26.7997 | 26.8977 | 26.797 | 26.8812 | 26.8204 | 26.7834 | 26.7297 |
| 27.4633 | 27.4356 | 27.4775 | 27.0729 | 27.0576 | 27.3156 | 27.2533 |
| 26.0025 | 25.8686 | 25.5464 | 25.3773 | 25.5626 | 25.7957 | 25.8574 |
| 25.9599 | 25.4504 | 25.5449 | 25.7949 | 25.5395 | 25.7467 | 25.9386 |
| 28.3759 | 27.8585 | 28.3808 | 27.8942 | 27.8364 | 28.13 | 28.1746 |
| 25.8575 | 25.1354 | 25.0881 | 25.3583 | 25.3426 | 25.2833 | 25.5164 |
| 25.2406 | 24.7709 | 24.8094 | 24.9423 | 25.002 | 25.0234 | 25.376 |
| 26.8295 | 26.6854 | 26.6549 | 26.3657 | 26.2992 | 26.6315 | 26.7831 |
| 25.1684 | 25.0911 | 25.0198 | 24.8879 | 24.4785 | 24.8912 | 24.7569 |
| 25.6018 | 25.9716 | 25.6278 | 26.3082 | 26.2117 | 26.3299 | 26.0588 |
| 27.4518 | 27.8862 | 27.8227 | 27.5518 | 27.6024 | 27.5723 | 27.5522 |
| 28.7759 | 28.7482 | 28.6701 | 28.6676 | 28.6746 | 28.4825 | 28.3571 |
| 26.1429 | 26.6843 | 26.6444 | 26.2578 | 26.5333 | 26.3243 | 26.6302 |
| 26.9859 | 27.0265 | 27.1263 | 26.8495 | 26.7149 | 26.5078 | 26.8655 |
| 29.2865 | 29.5529 | 29.3236 | 29.4369 | 29.2784 | 29.2382 | 29.1822 |
| 25.1216 | 24.8106 | 25.0364 | 24.4649 | 24.8042 | 25.2575 | 24.8843 |
| 31.5815 | 31.8386 | 31.7666 | 31.7073 | 31.7167 | 31.6189 | 31.6216 |
| NaN | NaN | NaN | 25.2232 | 24.9702 | 24.7761 | 24.6622 |
| 25.8338 | 25.6741 | 25.6393 | 25.6616 | 25.4396 | 25.4843 | 25.1642 |
| 25.4896 | 25.4507 | 25.4143 | 25.6503 | 25.6164 | 25.8295 | 25.7561 |
| 26.3923 | 26.292 | 26.5869 | 26.3104 | 26.4229 | 26.0682 | 26.3001 |
| 28.0279 | 27.9992 | 27.8623 | 27.597 | 27.6838 | 28.0643 | 28.0233 |
| 31.6722 | 31.4461 | 31.4974 | 31.6907 | 31.6044 | 31.8102 | 31.7427 |
| 24.4618 | 24.5904 | 24.6632 | 24.6957 | 24.8616 | 24.122 | 24.1054 |

| | | | | | | |
|---|---|---|---|---|---|---|
| 27.2704 | 27.1276 | 27.346 | 27.3817 | 27.456 | 27.5719 | 27.4176 |
| 25.3978 | 24.841 | 25.0152 | 24.9806 | 24.4742 | 25.3227 | 24.8853 |
| 27.4788 | 27.1198 | 27.4522 | 27.1744 | 27.3639 | 27.3411 | 27.4015 |
| 25.0837 | 25.1366 | 25.0334 | 25.0241 | 24.7995 | 24.9433 | 25.1169 |
| 28.8348 | 29.0313 | 29.0144 | 29.1853 | 28.9915 | 28.769 | 28.7767 |
| 26.6955 | 27.0786 | 27.1279 | 27.0427 | 27.3225 | 26.9788 | 26.6555 |
| NaN | 24.8005 | NaN | NaN | NaN | NaN | NaN |
| 25.8458 | NaN | NaN | 25.2021 | 25.5019 | 24.9952 | 25.4147 |
| 25.2566 | 25.7141 | 25.7821 | 25.8723 | 25.4603 | 25.5855 | 25.5334 |
| 25.8703 | 26.2218 | 26.122 | 26.031 | 26.1485 | 25.8955 | 25.8929 |
| 25.6446 | 25.6385 | 25.9317 | 25.6455 | 26.0875 | 25.8597 | 25.9681 |
| 29.0948 | 27.2377 | 28.2962 | 28.5337 | 28.5173 | 28.5685 | 28.482 |
| 25.7552 | 25.8871 | 26.0421 | 26.2236 | 25.972 | 25.8755 | 25.9118 |
| 26.3634 | 26.6602 | 26.638 | 26.1834 | 26.0977 | 26.1141 | 26.1769 |
| 25.3938 | 25.3752 | 25.4183 | 25.0818 | 25.2662 | 25.5921 | 25.4617 |
| 25.8775 | 26.0517 | 25.9992 | 25.8543 | 26.2186 | 25.7961 | 25.6965 |
| 26.1669 | 26.6661 | 26.684 | 26.4539 | 26.5412 | 26.2098 | 26.2652 |
| 25.189 | 25.801 | 25.5026 | 25.5068 | 25.4751 | 25.3223 | 25.537 |
| 25.024 | 25.1257 | 25.0673 | 24.8871 | 24.9505 | 24.9973 | 24.8797 |
| 27.6348 | 27.8677 | 27.7611 | 27.7278 | 27.6777 | 27.7809 | 27.7076 |
| 24.4036 | 24.3249 | 24.1704 | 24.2909 | 24.5238 | 24.3516 | 24.2386 |
| 29.0584 | 29.2248 | 29.1157 | 28.6351 | 28.7992 | 29.1468 | 29.3081 |
| 28.0359 | 28.4185 | 28.5786 | 27.919 | 27.9455 | 28.1423 | 28.1135 |
| 25.4306 | 25.2478 | 25.4501 | 24.7984 | 24.9928 | 25.3149 | 25.3206 |
| 29.0854 | 29.2066 | 29.129 | 28.8994 | 28.948 | 28.8256 | 29.1083 |
| 29.5392 | 30.01 | 29.7761 | 29.6699 | 29.7226 | 29.6306 | 29.4482 |
| 26.3103 | 26.4204 | 26.6899 | 26.4938 | 26.5379 | 26.3782 | 26.2957 |
| 25.2474 | 25.7227 | 25.5196 | 25.6549 | 25.3069 | 25.4613 | 25.5727 |
| 25.0988 | NaN | NaN | 25.1143 | 25.1157 | 25.1678 | 24.8641 |
| 25.3665 | 24.9058 | 25.1829 | 24.9553 | 25.2598 | 24.9363 | 25.0993 |
| 25.4132 | 25.4715 | 25.326 | 25.2279 | 25.1988 | 25.486 | 25.0436 |
| 25.1288 | 25.1271 | 25.0457 | 25.2737 | NaN | NaN | 25.2594 |
| 26.5149 | 26.5672 | 26.4203 | 26.3403 | 26.4784 | 26.6909 | 26.7069 |
| 25.9355 | 26.1378 | 26.0889 | 25.8075 | 25.9137 | 25.934 | 25.7466 |
| 26.8724 | 26.9743 | 27.0151 | 26.9247 | 26.6017 | 26.8833 | 26.8066 |
| 25.1162 | 25.2472 | 25.3024 | 25.0376 | 24.8999 | 24.8511 | 25.1947 |
| 25.6781 | 26.0078 | 25.6693 | 25.5386 | 25.7334 | 25.4032 | 25.6992 |
| 25.5878 | 26.0152 | 25.5063 | 26.0166 | 25.9078 | 26.4075 | 26.4938 |
| 27.5873 | 24.8831 | 27.172 | 27.2896 | 27.4065 | 27.2831 | 27.2695 |
| 27.761 | 27.5838 | 27.446 | 27.2843 | 27.3381 | 27.3473 | 27.5447 |
| 25.2717 | 25.2305 | 25.2364 | 25.2391 | 25.3495 | 25.248 | 25.4901 |
| 26.0292 | 26.31 | 26.1069 | 25.5234 | 25.8447 | 26.5175 | 26.4897 |
| 25.3403 | 25.5271 | 25.3863 | 25.1638 | 25.3587 | 25.3144 | 25.4176 |
| 25.5676 | 24.9111 | 24.8776 | 25.2356 | 25.2991 | 25.6971 | 25.5678 |
| 25.4758 | 24.9081 | 25.2443 | 25.2586 | 25.1038 | 25.079 | 25.135 |
| 30.0772 | 30.1953 | 30.3081 | 30.4645 | 30.2541 | 30.2459 | 30.0364 |
| 28.4447 | 28.8216 | 28.7107 | 28.5183 | 28.4291 | 28.528 | 28.5205 |
| 26.3789 | 26.2383 | 26.4897 | 26.5356 | 26.5272 | 26.141 | 26.2877 |

| | | | | | | |
|---|---|---|---|---|---|---|
| NaN | NaN | 24.4988 | 24.9762 | 24.602 | 24.6841 | 24.775 |
| 25.2172 | 25.519 | 25.3083 | 25.2552 | 25.3353 | 25.1481 | 25.138 |
| 26.6129 | 26.5314 | 26.6179 | 26.7074 | 26.7709 | 26.4977 | 26.7044 |
| 28.3097 | 28.252 | 28.0259 | 27.8118 | 27.9702 | 28.1664 | 28.0758 |
| 25.602 | 25.1326 | 24.9373 | 24.9622 | 25.1587 | 25.0976 | 25.342 |
| 25.5728 | NaN | 25.641 | 25.3469 | 25.4992 | 25.6744 | 25.4406 |
| 25.3498 | 25.4202 | 25.1976 | 25.4713 | 25.6048 | 25.4604 | 25.5397 |
| 25.4463 | 25.1993 | 24.832 | 25.0511 | 25.483 | 24.6517 | 24.8556 |
| 25.5071 | 25.4789 | 25.1169 | 25.3928 | 25.396 | 24.8746 | 25.1232 |
| 26.1702 | 26.4097 | 26.3486 | 25.777 | NaN | 26.3349 | 26.5641 |
| 24.8269 | 24.4642 | 24.8138 | 24.6276 | 24.7301 | 24.9911 | 24.6665 |
| 26.257 | 25.8863 | 26.6296 | 26.2686 | 26.203 | 26.3332 | 26.4086 |
| 26.2423 | 26.2818 | 26.0159 | 26.0868 | 26.1385 | 26.4734 | 26.225 |
| 26.9014 | 27.4033 | 27.3944 | 27.681 | 27.6369 | 27.4601 | 27.4313 |
| 24.086 | 24.024 | 23.7001 | 24.2973 | 24.0875 | 24.5844 | 24.5963 |
| 28.0517 | 28.0113 | 27.9278 | 27.7955 | 27.7227 | 28.0526 | 28.1267 |
| 26.2792 | 26.4335 | 26.2796 | 26.2672 | 26.2261 | 25.9646 | 25.8061 |
| 24.8105 | NaN | NaN | NaN | NaN | NaN | 24.9399 |
| 26.5879 | 26.1318 | 26.29 | 26.2201 | 26.152 | 26.209 | 26.4265 |
| 32.1434 | 32.4269 | 32.2492 | 32.2012 | 32.1723 | 32.2137 | 32.1528 |
| NaN | NaN | 25.0967 | 25.2523 | 25.6991 | 25.5594 | NaN |
| 25.7237 | 25.397 | 25.674 | 25.7969 | 25.6327 | 25.503 | 25.5422 |
| NaN | NaN | NaN | 29.0214 | 28.9706 | 28.8915 | 28.7964 |
| 25.3815 | 25.8259 | 25.5642 | NaN | NaN | NaN | NaN |
| 25.572 | 25.7143 | 25.436 | 25.8129 | 25.5719 | 25.3108 | 25.4008 |
| 27.1566 | 27.3832 | 26.3053 | 26.8059 | 26.8232 | 26.8071 | 27.1052 |
| 25.9768 | 25.8886 | 25.8545 | 26.0134 | 26.0003 | NaN | 25.7064 |
| 25.4901 | 26.2181 | 25.3149 | 26.6224 | 25.9495 | 26.2898 | 26.3602 |
| 25.5783 | 25.6223 | 25.6895 | 25.8815 | 25.4835 | 25.7108 | 25.5841 |
| 27.1244 | 27.4214 | 27.6319 | 27.2696 | 27.2173 | 27.1095 | 27.1498 |
| 25.8531 | 25.7044 | 25.7424 | 26.0743 | 26.1989 | 26.0672 | 25.9931 |
| 26.562 | 26.9564 | 27.0318 | 26.9188 | 26.8289 | 26.5491 | 26.5694 |
| 26.1197 | 26.0951 | 26.0245 | 26.0726 | 25.8542 | 26.3388 | 26.3263 |
| 26.3415 | 26.3163 | 26.5277 | 26.1544 | 25.9329 | 26.3132 | 26.1756 |
| 26.0006 | 25.8161 | 25.9092 | 25.9262 | 25.7899 | 26.0307 | 25.9166 |
| 27.8829 | 27.9699 | 27.9311 | 27.8995 | 27.8808 | 28.1701 | 28.0137 |
| 28.7839 | 28.7581 | 28.6029 | 28.8351 | 28.8866 | 28.5173 | 28.4815 |
| 25.228 | NaN | 24.4903 | 24.5579 | NaN | 24.2156 | NaN |
| 26.0977 | 25.8147 | 25.968 | 26.0373 | 26.5795 | 26.1943 | 26.248 |
| 30.1378 | 30.0755 | 30.1958 | 30.2914 | 30.2194 | 29.8656 | 29.7796 |
| 24.9163 | 25.1872 | 25.0493 | 25.0456 | 24.9612 | 24.8637 | 24.5851 |
| 25.406 | 25.6037 | 25.4609 | 25.7772 | 25.5559 | 25.3689 | 25.1885 |
| 26.4572 | 26.7242 | 26.444 | 26.55 | 26.4158 | 26.5512 | 26.5942 |
| 25.5275 | 25.6773 | 25.463 | 25.2348 | 25.5629 | 25.3451 | 25.5895 |
| 25.038 | 25.1885 | 24.7195 | 25.3697 | 25.0418 | 25.8519 | 25.5333 |
| 29.2218 | 29.335 | 29.1327 | 29.203 | 29.135 | 29.0384 | 29.1284 |
| 27.7701 | 27.6851 | 27.7396 | 28.1862 | 27.944 | 28.0727 | 27.7156 |
| 26.1173 | 26.1378 | 26.2376 | 26.1603 | 26.3014 | 26.117 | 25.7341 |

| | | | | | | |
|---|---|---|---|---|---|---|
| 25.665 | 26.2087 | 25.9093 | 25.45 | 25.5845 | 25.7775 | 25.8724 |
| 27.302 | 27.3535 | 27.3255 | 27.1683 | 27.1168 | 27.1504 | 27.2798 |
| 25.5106 | 25.1947 | NaN | 25.4837 | 25.4625 | 25.354 | 25.4677 |
| 26.6458 | 26.5573 | NaN | 26.5646 | 26.3613 | 26.4723 | 26.5108 |
| 27.7799 | 27.8376 | 27.7624 | 27.7749 | 27.7781 | 27.833 | 27.7211 |
| 25.8519 | 26.1664 | 25.6299 | NaN | 26.0205 | 26.3501 | 26.5166 |
| 26.2196 | 26.7208 | 25.974 | 26.4048 | 26.2139 | 26.9796 | 26.8093 |
| 25.6314 | NaN | 25.7375 | 25.3424 | 25.2647 | 26.1731 | 26.2375 |
| 28.1897 | 28.55 | 28.3543 | 28.264 | 28.2993 | 27.9448 | 28.0928 |
| 26.4237 | 26.6484 | 26.5528 | 26.4392 | 26.1392 | 26.6702 | 26.385 |
| 24.7798 | 25.3579 | 25.037 | NaN | 24.9232 | 25.0724 | 25.295 |
| 26.7943 | 26.7152 | 26.7887 | 26.8371 | 26.7419 | 27.1087 | 26.683 |
| 25.96 | 25.7498 | 25.5729 | 25.5502 | 25.7835 | 25.6708 | 25.5677 |
| 26.8451 | 27.0371 | 26.7089 | 26.6951 | 26.7724 | 26.8953 | 26.8286 |
| 24.9526 | 25.4199 | 25.0774 | 25.4207 | 25.2554 | 25.8309 | 25.5772 |
| 29.1777 | 29.5791 | 29.2744 | 29.3216 | 28.8792 | 29.0022 | 29.0642 |
| 25.9952 | 26.2983 | 26.1531 | 25.8208 | 25.9302 | 25.4978 | 25.962 |
| 27.3291 | 27.2908 | 27.3247 | 27.2228 | 27.3141 | 27.2684 | 27.156 |
| 24.4442 | 23.9928 | 24.3905 | 24.4583 | 24.3731 | 24.4554 | 24.4546 |
| 28.7721 | 28.9665 | 29.0425 | 28.6778 | 28.6085 | 28.6444 | 28.8254 |
| 27.9111 | 28.0461 | 27.8817 | 27.8174 | 27.6937 | 28.0644 | 28.0027 |
| 25.0402 | 25.4789 | 25.2546 | 24.9288 | 25.0175 | 25.2199 | 25.1509 |
| 26.1302 | 26.4175 | 26.2127 | 26.2234 | 26.2947 | 26.1981 | 25.8543 |
| 26.8927 | 27.0871 | 26.9679 | 26.7292 | 26.6258 | 26.94 | 26.9383 |
| 24.9815 | 24.9169 | NaN | 25.0001 | 25.0527 | 24.9032 | 24.9488 |
| 27.5337 | 27.1783 | 27.4442 | 27.5125 | 27.472 | 27.2583 | 27.3254 |
| 25.7777 | 26.6947 | 26.5701 | 25.6005 | 25.5321 | 26.0506 | 25.7381 |
| 31.0979 | 31.55 | 31.545 | 31.1874 | 31.1752 | 31.1515 | 31.2122 |
| NaN | NaN | 26.0462 | 26.2172 | 26.2069 | 25.8216 | 25.8468 |
| 28.3551 | 28.675 | 28.73 | 28.2606 | 28.377 | 28.3336 | 28.4411 |
| 26.8357 | 27.0635 | 26.9544 | 26.9586 | 26.6626 | 26.9549 | 26.8413 |
| 26.2964 | 26.903 | 26.7098 | 26.5028 | 26.4736 | 26.4082 | 26.3973 |
| 25.3905 | 25.3972 | 25.2418 | 25.0668 | 24.9536 | 25.4518 | 25.3938 |
| 30.0697 | 30.2694 | 30.0772 | 30.2566 | 30.1262 | 30.2342 | 30.0559 |
| 30.3043 | 30.5696 | 30.4893 | 30.5092 | 30.5153 | 30.3802 | 30.417 |
| 25.1307 | NaN | NaN | 25.1324 | 24.9804 | 25.3189 | 24.9798 |
| 27.8535 | 28.3457 | 28.1192 | 27.969 | 28.2887 | 28.2619 | 28.1198 |
| 25.211 | 25.4227 | 25.5459 | 25.4796 | 25.4526 | 24.8411 | 25.0703 |
| 30.1439 | 30.3228 | 30.2047 | 30.2739 | 30.1772 | 30.1369 | 30.1784 |
| 26.1879 | 26.2989 | 26.4045 | 26.3711 | 26.0392 | 26.0084 | 26.2534 |
| 24.5336 | 24.4762 | 24.5318 | NaN | 24.5037 | 24.4889 | 24.7307 |
| 26.1123 | 26.3235 | 26.3694 | 26.3508 | 26.5134 | 26.0671 | 26.2245 |
| 24.8751 | 25.328 | 25.1563 | 24.8886 | 25.1292 | 24.9776 | 25.2674 |
| 26.0945 | 26.0713 | 25.8535 | 25.8871 | 25.9732 | 26.1217 | 26.1161 |
| 27.6494 | 27.966 | 27.3657 | 27.7717 | 27.6089 | 28.0756 | 28.1088 |
| 26.3825 | 25.9778 | 26.0917 | 26.0267 | 26.0028 | 26.4182 | 26.1553 |
| 26.7096 | 26.6252 | 26.614 | 26.4049 | 26.4473 | 26.7762 | 26.8932 |
| 27.6129 | 27.9926 | 27.6814 | 27.552 | 27.7535 | 27.5555 | 27.3398 |

| | | | | | | |
|---|---|---|---|---|---|---|
| 27.9251 | 27.9936 | 27.9478 | 27.8221 | 27.9137 | 27.9453 | 27.7548 |
| 30.5267 | 30.7169 | 30.5843 | 30.3058 | 30.3795 | 30.289 | 30.2886 |
| 29.0693 | 29.3985 | 29.0648 | 28.9258 | 29.0709 | 29.5522 | 29.3522 |
| 25.7363 | 25.9382 | 25.8835 | 25.4442 | 25.95 | 25.5833 | 25.8291 |
| 29.1824 | 29.3223 | 29.3427 | 29.0703 | 29.0578 | 29.0096 | 29.0466 |
| 26.2008 | 26.2529 | 26.1786 | 26.3028 | 26.2293 | 25.8168 | 26.1354 |
| 25.4987 | 25.753 | 25.5943 | 25.7075 | 25.5787 | 25.2595 | 25.1893 |
| 25.2431 | 25.0001 | 25.035 | 25.1501 | 25.0127 | 25.4628 | 25.2972 |
| 26.7724 | 26.8008 | 26.9046 | 26.5312 | 26.7316 | 26.6381 | 26.7474 |
| 24.4244 | 24.504 | NaN | 24.9062 | 24.7762 | 24.7949 | 24.7678 |
| 25.4138 | 25.3756 | 25.0232 | 25.4913 | 25.4866 | 24.6799 | 24.1359 |
| 27.7507 | 28.1865 | 28.0682 | 27.6263 | 27.6881 | 27.5646 | 27.626 |
| 26.3708 | 27.0691 | 26.8488 | 26.4932 | 26.474 | 26.4209 | 26.6477 |
| 25.8782 | 25.9378 | 25.9165 | 26.0073 | 25.7122 | 25.5869 | 25.7085 |
| 25.4998 | 25.7936 | 25.5598 | 25.3837 | 25.017 | 25.157 | 25.6109 |
| 24.7883 | 25.4048 | 24.9619 | 25.4069 | 25.1028 | 25.2871 | 24.9907 |
| 26.2162 | 25.9989 | 26.2966 | 26.09 | 26.2039 | 25.8528 | 26.0821 |
| 31.1072 | 31.1269 | 31.2041 | 30.92 | 30.9817 | 31.3069 | 31.3451 |
| 25.3248 | 25.5735 | 25.5714 | 25.3713 | 25.5445 | 25.1505 | 25.4109 |
| 27.0439 | 27.2134 | 27.071 | 26.9012 | 26.7941 | 27.0587 | 27.0362 |
| 25.2702 | 24.952 | 24.9264 | 25.252 | 25.2272 | 25.1987 | 25.2776 |
| 29.1435 | 29.3375 | 29.3438 | 29.3975 | 29.3437 | 28.9326 | 28.931 |
| 28.2405 | 28.3727 | 28.1113 | 27.9846 | 27.9887 | 27.7191 | 27.794 |
| 25.6351 | 26.1347 | 25.8602 | 25.3698 | 25.3395 | 25.6089 | 25.8342 |
| 27.3389 | 27.3036 | 27.14 | 26.9728 | 26.8567 | 27.2043 | 27.0127 |
| 24.6769 | 24.4556 | 24.2095 | 24.7313 | 24.5721 | 24.7646 | 24.6322 |
| 30.4068 | 30.5655 | 30.54 | 30.4071 | 30.4522 | 30.1831 | 30.094 |
| 31.0415 | 31.0629 | 30.9973 | 30.848 | 30.8955 | 31.054 | 31.1575 |
| 25.859 | 26.5456 | 26.493 | 25.9656 | 26.1384 | 26.1007 | 25.9743 |
| 26.0234 | 25.8014 | 25.972 | 25.9705 | 25.6096 | 25.8692 | 25.9554 |
| 25.7697 | 25.187 | 25.4318 | 25.7636 | 25.8336 | 25.4712 | 25.2534 |
| 27.1617 | 27.4581 | 27.4648 | 27.2346 | 27.1192 | 27.2185 | 27.1392 |
| 27.2825 | 27.7168 | 27.4733 | 27.2797 | 27.3299 | 27.7685 | 27.6617 |
| 25.5685 | 25.855 | 25.543 | 25.6329 | 25.5879 | 25.8547 | 25.6218 |
| 26.7977 | 27.2317 | 26.9261 | 26.5565 | 26.6359 | 26.2261 | 26.5674 |
| 25.2203 | 25.3723 | 25.0287 | 25.6141 | 25.4946 | 25.4464 | 25.6583 |
| 27.5434 | 27.403 | 27.4597 | 27.2844 | 27.4544 | 27.2783 | 27.2803 |
| 28.7419 | 28.6947 | 28.5386 | 28.6812 | 28.6958 | 28.9042 | 28.641 |
| 27.5452 | 27.4143 | 27.4921 | 27.2916 | 27.5743 | 26.938 | 27.2429 |
| 27.2861 | 27.4213 | 27.4853 | 27.4921 | 27.3938 | 27.4946 | 27.6048 |
| 28.285 | 28.2898 | 28.1694 | 28.0223 | 27.9375 | 28.1335 | 27.9372 |
| 26.2482 | 25.4202 | 26.0647 | 26.0344 | 26.1189 | 26.2242 | 26.013 |
| 25.188 | 25.299 | 25.2618 | 25.231 | 25.225 | 24.9455 | 25.0595 |
| 27.4638 | 27.5417 | 27.5217 | 27.3964 | 27.3235 | 27.1795 | 27.2321 |
| 30.9169 | 31.1202 | 31.0066 | 30.9279 | 30.8691 | 31.2284 | 31.2435 |
| 30.0246 | 30.0704 | 30.1261 | 29.9963 | 30.2085 | 30.0499 | 29.9893 |
| 26.0276 | 25.9906 | 26.1267 | 25.6872 | 25.9954 | 26.3593 | 26.3424 |
| 26.7853 | 26.9817 | 26.9039 | 26.6736 | 26.7304 | 26.7831 | 26.7256 |

| | | | | | | |
|---|---|---|---|---|---|---|
| 26.558 | 26.3305 | 26.175 | 26.467 | 26.53 | 26.3496 | 26.4027 |
| 25.3712 | 25.2645 | 25.5069 | 25.5179 | 25.728 | 25.7098 | 25.3647 |
| 28.8855 | 28.7962 | 28.7642 | 28.9608 | 28.931 | 28.7625 | 28.6777 |
| 27.6347 | 27.6274 | 27.2993 | 27.3264 | 27.8514 | 27.3967 | 27.408 |
| 25.7986 | 25.5732 | 25.6802 | 25.9143 | 25.5059 | 25.9065 | 25.825 |
| 29.2795 | 29.8582 | 29.6819 | 29.3775 | 29.2513 | 29.3527 | 29.3675 |
| 31.1169 | 30.8443 | 31.0093 | 31.1913 | 31.1032 | 31.1313 | 31.1333 |
| 25.5154 | 26.1797 | 25.5284 | 26.0195 | 25.9031 | 25.8719 | 26.0716 |
| 27.7315 | 27.5214 | 27.7654 | 27.5347 | 27.5841 | 27.4758 | 27.2574 |
| 24.6322 | 24.6093 | 24.5119 | 24.1838 | 24.4746 | 24.1158 | 24.1892 |
| 27.8434 | 27.4645 | 27.3738 | 27.6052 | 27.5993 | 27.9438 | 27.8769 |
| 25.7978 | 25.8001 | 25.8626 | 26.3729 | 26.5634 | 26.5737 | 26.2368 |
| 25.8472 | 25.8192 | 25.8644 | 26.1196 | 25.8944 | 26.0741 | 26.0963 |
| 26.4922 | 27.2873 | 27.3597 | 27.1367 | 27.1096 | 27.2225 | 27.0497 |
| 24.5125 | 24.7008 | 24.7064 | 24.7917 | 25.0388 | 25.3668 | 25.112 |
| 25.2844 | 25.4314 | 25.603 | 25.7512 | 25.7881 | 25.679 | 25.7123 |
| 29.5103 | 30.1411 | 30.0941 | 29.9297 | 29.8553 | 29.9725 | 29.835 |
| 26.2423 | 26.6788 | 26.5606 | 26.5976 | 26.5179 | 26.694 | 26.6776 |
| 25.8291 | 26.0948 | 26.0212 | 26.3913 | 26.4576 | 25.9465 | 25.8917 |
| 25.9735 | 26.3117 | 26.2896 | 26.4498 | 25.9811 | 26.1602 | 26.0735 |
| 26.3856 | 26.6551 | 26.7 | 26.5286 | 26.6734 | 26.6228 | 26.3893 |
| 28.5443 | 28.6757 | 28.6687 | 28.7448 | 28.6543 | 28.6128 | 28.5425 |
| 28.0965 | 28.3232 | 28.2792 | 28.1807 | 28.1265 | 28.1542 | 28.176 |
| 26.6207 | 27.0614 | 27.1168 | 26.7764 | 26.8474 | 26.8301 | 27.034 |
| 28.2289 | 28.6884 | 28.5234 | 28.394 | 28.404 | 28.1912 | 28.3373 |
| 24.3979 | 24.5345 | 24.5648 | 24.7989 | 24.7278 | 24.5352 | 24.5471 |
| 25.5303 | 25.6504 | 26.0711 | 25.4236 | 25.5109 | 25.7846 | 25.73 |
| 25.661 | 26.0988 | 26.3014 | 25.7772 | 26.0399 | 25.8864 | 25.9296 |
| 27.2768 | 27.4143 | 27.35 | 27.4424 | 27.6299 | 27.3351 | 27.1347 |
| 25.7252 | 25.834 | 25.8849 | 26.0479 | 25.9037 | 25.7093 | 25.6064 |
| 27.3797 | 27.4951 | 27.4661 | 27.5699 | 27.7595 | 27.4257 | 27.3594 |
| 29.7466 | 29.8925 | 29.8375 | 29.9262 | 29.9356 | 30.0182 | 30.0016 |
| 27.3905 | 27.3192 | 27.3109 | 27.451 | 27.4467 | 27.4566 | 27.4081 |
| 28.3461 | 28.0993 | 28.2405 | 27.9781 | 28.0598 | 27.994 | 28.057 |
| 26.4116 | 26.4096 | 26.282 | 25.9139 | 26.0805 | 26.1444 | 26.2362 |
| 27.1599 | 26.5931 | 27.0964 | 26.7615 | 27.0108 | 26.7013 | 26.7428 |
| 28.7625 | 28.6409 | 28.7152 | 28.8589 | 28.83 | 28.5312 | 28.5772 |
| 26.1921 | 26.0181 | 25.8689 | 26.1127 | 26.0559 | 25.78 | 25.9045 |
| 26.1343 | 26.0808 | 26.1306 | 26.002 | 26.0402 | 25.6239 | 25.5566 |
| 29.1135 | 28.7605 | 28.8152 | 28.9424 | 28.8994 | 28.5794 | 28.6073 |
| 25.2495 | 24.492 | 24.5065 | 25.041 | 24.97 | 24.9764 | 24.7721 |
| 27.2532 | 27.4831 | 27.6806 | 27.56 | 27.5231 | 27.4904 | 27.4155 |
| 23.6841 | 23.8348 | 23.7904 | NaN | NaN | NaN | 23.9277 |
| 26.1934 | 26.2588 | 26.3061 | 26.345 | 26.5361 | 26.6204 | 26.702 |
| 25.287 | 24.3118 | 25.3732 | 25.2851 | 25.3371 | 25.2195 | 25.5502 |
| 27.4484 | 27.4742 | 27.3035 | 27.1219 | 27.0468 | 27.0359 | 27.1253 |
| 26.3578 | 27.2092 | 27.0587 | 26.8496 | 27.0793 | 26.9619 | 27.0072 |
| 24.8582 | 23.8469 | 24.1058 | 23.9665 | 23.9537 | 23.9631 | 24.0237 |

| | | | | | | |
|---|---|---|---|---|---|---|
| 26.5792 | 26.5459 | 26.6751 | 26.7075 | 26.5915 | 26.6732 | 26.5793 |
| 25.095 | NaN | NaN | NaN | 24.9519 | 25.171 | 25.3492 |
| 25.3719 | 25.5809 | 25.358 | 25.4574 | 25.1576 | 25.1831 | 25.1361 |
| 24.8945 | 24.7853 | 24.7936 | 24.9237 | 24.865 | 25.1058 | 25.1663 |
| 31.4334 | 31.903 | 31.6588 | 31.6114 | 31.663 | 31.5379 | 31.5484 |
| 30.8508 | 31.0438 | 30.9632 | 30.9801 | 31.0525 | 30.6114 | 30.5868 |
| 25.5744 | 25.5274 | 25.4574 | 25.5649 | 25.631 | 25.6447 | 25.5376 |
| 27.6477 | 27.9624 | 27.8439 | 27.5929 | 27.5999 | 27.8628 | 27.8714 |
| 26.1598 | 26.5689 | 26.4493 | 26.3524 | 26.421 | 26.2948 | 26.2979 |
| 25.642 | 25.7145 | 25.4046 | 25.6056 | 25.4407 | 25.7299 | 25.6229 |
| 29.1147 | 29.7233 | 29.5291 | 29.0749 | 29.2332 | 29.1052 | 29.2797 |
| 25.5178 | 25.6267 | NaN | NaN | 25.378 | 25.5331 | 25.5358 |
| 26.7166 | 26.3408 | 26.4777 | 26.4125 | 26.4278 | 26.5559 | 26.4146 |
| 33.9452 | 33.8074 | 33.5244 | 33.7188 | 33.7704 | 33.8302 | 34.0849 |
| 23.9034 | 23.9302 | NaN | NaN | NaN | NaN | NaN |
| 25.6524 | 25.4322 | 25.8356 | 25.5736 | 25.4663 | 25.3889 | 25.5247 |
| 30.4403 | 30.4546 | 30.3614 | 30.7405 | 30.7929 | 30.6193 | 30.6266 |
| 27.285 | 26.859 | 27.0457 | 27.2185 | 27.2081 | 27.2088 | 27.1531 |
| 24.8667 | 24.9228 | 24.9063 | 24.832 | 24.8193 | 25.2241 | 25.2383 |
| 28.1307 | 28.2937 | 28.1204 | 28.1036 | 28.2121 | 27.7831 | 27.9051 |
| 26.1725 | 26.563 | 26.4139 | 26.1934 | 26.2669 | 26.1147 | 26.0729 |
| 25.1522 | 25.0239 | 25.1916 | 25.5777 | 25.8435 | 25.1566 | 25.302 |
| 25.6791 | NaN | NaN | 25.755 | 25.4393 | NaN | NaN |
| 23.8934 | 25.3916 | 24.5202 | 24.9538 | 24.65 | 24.7932 | 24.9234 |
| 26.2428 | 25.8623 | 26.0437 | 25.745 | 25.7496 | 25.9227 | 25.8607 |
| 24.798 | 24.4109 | 24.1904 | 24.5301 | 24.6883 | 24.6418 | 24.5277 |
| 26.1175 | 26.223 | 26.0914 | 26.253 | 26.0106 | 26.0266 | 25.9709 |
| NaN | 25.7344 | 25.8441 | 25.4888 | 25.4083 | 25.3875 | 25.6503 |
| 26.5501 | 26.6476 | 26.6109 | 26.3316 | 26.2552 | 26.3133 | 26.1669 |
| 26.0936 | 26.5178 | 26.2082 | 26.42 | 26.3923 | 25.9195 | 26.2088 |
| 26.541 | 26.8749 | 26.6218 | 26.4673 | 26.4145 | 26.3949 | 26.386 |
| 26.4158 | 26.5703 | 26.3356 | 26.2621 | 26.3918 | 26.0316 | 25.8726 |
| 27.7229 | 27.7038 | 27.5629 | 27.4629 | 27.4551 | 27.4408 | 27.3561 |
| 26.6151 | 26.5828 | 26.2029 | 26.3419 | 26.5452 | 26.957 | 26.8459 |
| 24.2509 | 24.4057 | 24.1058 | 24.3782 | 24.8267 | 24.7775 | 24.7762 |
| 24.8864 | 25.328 | 25.2281 | 25.1072 | 25.0978 | 24.973 | 24.9058 |
| 28.3773 | 28.2463 | 28.1233 | 28.2893 | 28.0729 | 28.1331 | 28.3398 |
| 31.2449 | 31.2524 | 31.0337 | 30.96 | 30.9748 | 31.2438 | 31.2874 |
| 26.8301 | 25.4347 | 26.982 | 26.5544 | 26.6119 | 27.015 | 27.0877 |
| 24.6244 | 23.2173 | 23.287 | 23.5309 | 23.3437 | 23.4837 | 23.4042 |
| 26.7149 | 27.1412 | 26.9039 | 26.6812 | 26.9604 | 26.4539 | 26.6934 |
| 25.3321 | NaN | 25.1382 | 24.7978 | 24.5761 | 25.1768 | 25.2021 |
| 25.7649 | 26.1298 | 25.8561 | 25.9821 | 25.8065 | 25.7623 | 25.881 |
| 24.8155 | 25.3013 | 25.048 | 24.9385 | 25.2537 | 24.7909 | 24.9669 |
| 24.7513 | 23.5973 | 24.1119 | 23.9822 | 24.1934 | 24.2617 | 24.2383 |
| 26.4693 | 26.9084 | 26.7615 | 26.5001 | 26.458 | 26.6393 | 26.4337 |
| 28.1593 | 28.1498 | 28.1536 | 27.9804 | 28.0179 | 28.0823 | 28.0053 |
| 26.7966 | NaN | 26.2355 | 26.6242 | 26.7669 | 26.6309 | 26.2478 |

| | | | | | | |
|---|---|---|---|---|---|---|
| 25.4914 | 26.2884 | 26.0634 | 26.0399 | 25.4393 | 25.4107 | 25.1956 |
| 25.3847 | 24.9407 | 25.1385 | 24.889 | 24.9118 | 24.9653 | 24.9416 |
| 25.3012 | NaN | 24.5718 | 24.6077 | 24.9508 | 24.9808 | 24.827 |
| 24.5696 | NaN | 24.9971 | 24.3715 | NaN | NaN | NaN |
| 25.4807 | 25.0179 | 24.7806 | 25.201 | 24.9943 | 25.3002 | 25.6078 |
| 24.3092 | 24.4538 | 24.6037 | 24.3256 | 24.3976 | 24.2533 | NaN |
| 28.7803 | 28.7819 | 28.6525 | 28.8226 | 28.7438 | 28.9245 | 28.9133 |
| 26.0437 | 25.9956 | 25.7298 | 26.1214 | 26.2137 | 26.1753 | 26.2208 |
| 27.8564 | 27.9903 | 27.9192 | 27.7156 | 27.8197 | 27.5023 | 27.4908 |
| 24.3804 | NaN | NaN | 24.188 | 24.2577 | 24.2835 | 24.4853 |
| 27.077 | 27.4822 | 27.4196 | 27.2139 | 27.3163 | 27.356 | 27.089 |
| 24.4709 | 24.2339 | 24.0632 | 24.3721 | 24.5797 | 24.6064 | 24.8139 |
| 25.5411 | 25.3914 | NaN | 24.8701 | 25.1717 | 25.3737 | 25.2307 |
| NaN | NaN | 25.2117 | NaN | NaN | 25.4539 | 25.568 |
| 25.6947 | 25.7286 | 25.487 | 25.4599 | 26.2128 | 26.026 | 25.6606 |
| 25.323 | 25.7854 | 25.3095 | 25.6795 | 24.9951 | 25.2694 | 24.9892 |
| 26.1296 | 26.0023 | 26.063 | 26.3413 | 26.1723 | 25.9742 | 25.7838 |
| 27.4338 | 26.9737 | 26.6772 | 26.8804 | 26.7783 | 26.6245 | 26.7252 |
| 26.1051 | 26.185 | 25.8688 | 25.432 | 25.2861 | 25.5546 | 25.7319 |
| 25.425 | 24.86 | 24.6034 | 24.992 | 25.153 | NaN | 24.3578 |
| 24.7755 | NaN | 24.4901 | 24.6775 | 24.6538 | 24.8932 | 24.7031 |
| 27.159 | 27.2681 | 27.136 | 26.9461 | 26.9952 | 26.6562 | 26.8092 |
| 25.5828 | 25.9828 | 25.9782 | 25.4871 | 25.5298 | NaN | 25.0357 |
| NaN | 24.5518 | NaN | NaN | 24.6431 | 25.6365 | 25.1234 |
| 26.9637 | 26.7879 | 26.9994 | 26.7872 | 26.8987 | 27.1703 | 26.8887 |
| 27.0861 | 27.467 | 27.4824 | 27.3179 | 27.5281 | 27.0698 | 27.1477 |
| 27.0001 | 27.0944 | 27.0832 | 26.8548 | 26.8859 | 26.9509 | 26.8872 |
| NaN | 25.3462 | NaN | 24.7728 | 25.1953 | 25.0798 | NaN |
| 24.2444 | 24.2035 | 24.1251 | 24.1873 | 24.4778 | 24.2689 | 24.3815 |
| 25.2005 | 25.2227 | 25.3174 | 25.6419 | 25.5467 | 25.6017 | 25.5885 |
| 26.0672 | 26.49 | 26.2699 | 25.9992 | 26.1589 | 25.8453 | 26.0325 |
| 28.0434 | 28.2929 | 27.7734 | 28.1855 | 28.2106 | 27.9474 | 28.2394 |
| 25.8173 | 26.0494 | 26.0016 | 25.9717 | 25.7259 | 25.704 | 25.5573 |
| 24.6026 | NaN | 24.0122 | 24.487 | 24.6824 | 24.2479 | 24.4748 |
| 26.5593 | 26.7891 | 26.6451 | 26.601 | 26.4347 | 26.293 | 26.7778 |
| 25.7197 | 25.5804 | NaN | 25.7335 | NaN | NaN | 25.9543 |
| 25.0353 | 25.4684 | 25.1725 | 25.2698 | 25.3256 | 25.1995 | NaN |
| 27.1481 | 25.4983 | 26.5488 | 26.9767 | 26.884 | 26.474 | 26.5682 |
| 25.3424 | 25.6775 | 25.258 | 25.1548 | 25.2295 | 25.3139 | 25.69 |
| 26.9334 | 26.5543 | 26.9311 | 26.7868 | 26.9968 | 26.936 | 26.7062 |
| 30.9551 | 30.8823 | 31.107 | 31.1532 | 31.0605 | 30.9629 | 31.0033 |
| 25.7624 | 26.1718 | 26.544 | 26.2894 | 26.1209 | NaN | 26.2208 |
| 24.9051 | 24.6363 | 24.7433 | 24.4459 | 24.747 | 25.0491 | 25.0809 |
| 25.2153 | NaN | 24.4031 | 25.1873 | 24.8957 | 24.8496 | 24.4583 |
| 24.6354 | NaN | 24.433 | 24.1121 | 24.4475 | 24.2647 | 24.3444 |
| 26.1405 | 26.8407 | 26.5637 | 26.3018 | 26.1995 | 26.27 | 26.3402 |
| 26.1937 | 26.3652 | 26.5224 | 26.1926 | 26.1788 | 26.1384 | 26.2612 |
| 28.1211 | 29.5198 | 29.2144 | 29.3273 | 29.1126 | 29.5894 | 29.5845 |

| | | | | | | |
|---|---|---|---|---|---|---|
| 24.3621 | 24.3805 | 24.2429 | 24.4947 | 24.5681 | 24.6153 | 24.473 |
| 24.6178 | 24.4863 | 24.763 | 24.7797 | 24.8243 | 24.9072 | 24.8053 |
| 29.306 | 29.6304 | 29.6259 | 29.3089 | 29.3262 | 29.4307 | 29.514 |
| 26.1318 | 26.575 | 26.3061 | 26.0315 | 25.0346 | 25.791 | 25.8309 |
| 25.4349 | 26.2998 | 26.1632 | 25.9174 | 25.9018 | 25.7498 | 25.6694 |
| 25.0895 | 24.7475 | 24.9322 | 24.9184 | 25.2246 | 25.4317 | 25.1917 |
| 27.8462 | 27.4844 | 27.5151 | 27.7279 | 27.5853 | 27.7984 | 28.0141 |
| 25.4912 | 25.6918 | 25.4735 | 25.3524 | 25.4818 | 25.4207 | 25.3021 |
| 35.1551 | 35.0843 | 35.0991 | 34.9445 | 34.9631 | 35.2712 | 35.3618 |
| 25.747 | 25.2967 | 25.7112 | 25.9935 | 25.7326 | 25.5056 | 25.5206 |
| 28.3251 | 29.3395 | 29.2321 | 28.9161 | 28.7085 | 28.6463 | 28.7431 |
| 26.6022 | 26.8654 | 26.7376 | 26.7577 | 26.5191 | 26.512 | 26.0468 |
| 24.3671 | NaN | 24.3362 | 24.2792 | 24.6266 | 24.5568 | 24.525 |
| 29.5429 | 29.8618 | 29.752 | 29.5107 | 29.483 | 29.494 | 29.4867 |
| 28.2249 | 28.2203 | 28.3431 | 28.3921 | 28.3994 | 28.2313 | 28.2311 |
| 26.2218 | 25.9568 | 26.0574 | 26.2096 | 26.287 | 26.2968 | 26.2824 |
| 27.0323 | 26.8815 | 27.1511 | 27.3178 | 27.1017 | 26.9703 | 27.065 |
| 25.3892 | 25.3431 | 25.2645 | 24.9416 | 25.1937 | 25.7211 | 25.4495 |
| 25.1876 | 25.7682 | 25.7371 | 25.3608 | 25.4602 | 25.6961 | NaN |
| 25.6804 | 25.5979 | 25.3036 | 25.5147 | 25.5615 | 25.6306 | 25.6926 |
| 26.143 | 25.9671 | 25.5449 | 25.8332 | 25.8444 | 26.1482 | 26.5029 |
| 25.2118 | NaN | 25.3654 | 25.2136 | 25.493 | 25.4544 | 25.4038 |
| 25.7302 | 25.7753 | 25.3512 | 25.3477 | 25.5553 | 25.3523 | 25.0261 |
| 24.641 | NaN | NaN | 24.666 | 24.3534 | 24.6178 | 24.7217 |
| 25.6117 | 25.8115 | 25.8391 | 25.7718 | 26.0792 | 25.6076 | 26.2622 |
| 24.4349 | 24.194 | NaN | 25.0164 | 25.0724 | 24.3665 | 24.3527 |
| 30.2841 | 30.4859 | 30.2875 | 30.1746 | 30.2779 | 29.9541 | 29.9552 |
| 26.0009 | 26.1025 | 26.0786 | 25.6408 | 26.1181 | 25.708 | 26.0714 |
| 25.456 | NaN | 25.4383 | 25.8187 | 25.6114 | 25.8739 | 25.9727 |
| 27.7145 | 28.0017 | 27.5098 | 28.2015 | 28.0819 | 28.2632 | 28.2357 |
| 24.9858 | NaN | 24.9885 | 24.7595 | 25.0138 | 25.1763 | NaN |
| 26.7758 | 26.8525 | 27.196 | 26.9032 | 27.0077 | 26.8855 | 27.0475 |
| 25.03 | 24.8832 | 24.4838 | 25.1559 | 24.5667 | 24.6683 | 24.9217 |
| 25.6784 | 25.5119 | 25.4666 | 25.4611 | 25.7118 | 25.4108 | 25.4408 |
| 25.2379 | 25.1215 | NaN | 25.368 | 25.328 | 25.1615 | 24.9666 |
| NaN | NaN | 25.0229 | 25.1632 | 25.0313 | NaN | NaN |
| 25.0054 | 25.3641 | 25.352 | 25.2755 | NaN | 25.4246 | 25.2543 |
| 25.4487 | 25.0023 | 25.2394 | 25.4439 | 25.254 | 25.4992 | 25.1338 |
| 24.3483 | NaN | NaN | 24.3459 | 24.1785 | NaN | NaN |
| 24.1831 | 24.1506 | 24.2021 | 24.0406 | 23.9604 | 24.0332 | 24.0533 |
| 26.0914 | 26.2317 | 26.159 | 26.4209 | 26.3565 | 26.2818 | 26.1931 |
| 28.2931 | 28.0201 | 28.009 | 28.1049 | 28.1327 | 28.6177 | 28.5253 |
| 24.8157 | 25.3331 | 25.0724 | 24.5641 | 24.7545 | 24.5381 | 24.8257 |
| 25.7574 | 25.8977 | 25.8695 | 26.1675 | 25.9385 | 25.9266 | 25.4603 |
| 26.5573 | 26.6137 | 26.6558 | 26.8113 | 26.6505 | 26.5288 | 26.4004 |
| NaN | 23.5224 | 23.3992 | NaN | 23.9006 | NaN | NaN |
| 27.835 | 28.3808 | 28.2792 | 27.9935 | 27.9041 | 27.7697 | 27.8038 |
| 26.9827 | 27.0742 | 26.9322 | 26.7206 | 26.8137 | 26.866 | 26.7618 |

| | | | | | | |
|---|---|---|---|---|---|---|
| 28.0464 | 28.2541 | 28.1864 | 27.8423 | 27.948 | 27.8894 | 28.0375 |
| 26.2723 | 25.8191 | 26.4137 | 25.9957 | 26.3132 | 26.824 | 27.0921 |
| NaN | NaN | 24.9936 | NaN | NaN | 25.1373 | 25.2078 |
| 30.2807 | 30.7843 | 30.699 | 30.5405 | 30.704 | 29.9393 | 30.0132 |
| 28.7678 | 29.058 | 28.9579 | 28.3797 | 28.4998 | 29.1884 | 29.4439 |
| 24.2043 | 23.8363 | 24.2057 | 24.3214 | 24.4214 | NaN | NaN |
| 27.5854 | 27.8545 | 27.8127 | 27.6121 | 27.5803 | 27.3251 | 27.4562 |
| 25.9178 | 25.8957 | 25.8974 | 25.6119 | 25.589 | 26.0564 | 25.8799 |
| 29.2193 | 29.1334 | 29.5254 | 29.3061 | 29.2334 | 29.0136 | 29.1759 |
| 24.6602 | 24.5284 | 24.5044 | 24.6173 | 24.7073 | 25.2266 | 24.9565 |
| 28.349 | 28.4015 | 28.3065 | 28.3279 | 28.3258 | 27.9486 | 28.0735 |
| NaN | NaN | NaN | 28.0529 | NaN | 27.2472 | 27.659 |
| 26.3079 | 26.4296 | 26.6334 | 26.281 | 26.0461 | 26.1942 | 26.161 |
| NaN | NaN | NaN | 24.6951 | 25.3294 | 24.8187 | NaN |
| 23.8908 | 23.4935 | 23.5497 | 24.2605 | 24.4827 | 23.8046 | NaN |
| 27.5081 | 27.6849 | 27.5493 | 27.3572 | 27.3715 | 27.9162 | 27.7717 |
| 30.7958 | 30.6429 | 30.5482 | 30.5598 | 30.5691 | 30.8103 | 30.7296 |
| 25.8139 | 25.42 | 25.2787 | 25.6078 | 25.5644 | 25.6824 | 25.7536 |
| NaN | NaN | NaN | 24.8583 | NaN | 24.7479 | NaN |
| 24.465 | 24.7032 | 24.6154 | 24.7785 | 24.4548 | 24.031 | 24.5838 |
| 28.5176 | 28.4639 | 28.8098 | 28.3582 | 28.4006 | 28.2224 | 28.3138 |
| 27.535 | 27.5646 | 27.451 | 27.4868 | 27.5296 | 27.3342 | 27.2272 |
| 24.3706 | 24.4517 | 24.4137 | 24.5104 | 24.3937 | 24.9176 | 24.9608 |
| 30.1305 | 30.3554 | 30.3358 | 30.1081 | 30.0324 | 30.1021 | 30.1975 |
| 24.5176 | 24.8894 | 25.1739 | 24.9035 | 24.8301 | NaN | NaN |
| 26.5144 | 26.3163 | 26.3898 | 26.3963 | 26.4127 | 26.4501 | 26.2424 |
| 33.7631 | 33.8402 | 33.6891 | 33.7017 | 33.7026 | 34.0284 | 34.0985 |
| 24.566 | 24.4914 | 24.5763 | 24.6131 | 24.3222 | 24.4763 | 24.3153 |
| 26.7482 | 26.819 | 26.6375 | 27.061 | 26.7984 | 26.6239 | 26.5182 |
| 25.956 | 26.3076 | 26.3167 | 25.9435 | 26.1472 | 25.8018 | 26.0146 |
| 28.4432 | 28.7838 | 28.652 | 28.2383 | 28.309 | 28.2563 | 28.4805 |
| 29.0105 | 29.1936 | 29.2466 | 29.0052 | 28.9358 | 28.6842 | 28.6647 |
| 25.9159 | 25.5947 | 25.5564 | 25.5239 | 25.3515 | 25.8947 | 25.6894 |
| 27.0677 | 27.0123 | 27.0691 | 26.8451 | 27.0574 | 26.909 | 27.0253 |
| 25.0634 | 25.5338 | 25.3378 | 24.5938 | 25.1615 | 24.9695 | 25.0646 |
| 26.4535 | 26.3006 | 26.2889 | 26.2807 | 25.8431 | 25.7421 | 25.6919 |
| 24.9105 | NaN | 24.7561 | 25.0384 | 25.0364 | 24.9625 | 25.0274 |
| 30.8575 | 31.0131 | 31.0463 | 30.8888 | 30.9441 | 30.9804 | 31.0003 |
| NaN | 24.9431 | NaN | 25.8921 | NaN | 25.1659 | NaN |
| 31.4557 | 31.6105 | 31.6582 | 31.5516 | 31.5502 | 31.3441 | 31.4263 |
| 25.3706 | 25.1335 | 24.8188 | 24.7089 | 25.0248 | NaN | 25.0683 |
| 25.5711 | 25.7579 | 25.7076 | 25.5445 | 25.483 | 25.746 | 25.7698 |
| 25.8722 | 25.9578 | 26.3386 | 25.8228 | 26.2108 | 25.9242 | 25.8019 |
| 25.714 | 25.5119 | 25.5364 | 25.8829 | 26.1896 | 25.189 | NaN |
| 25.4204 | 25.804 | 25.631 | 25.2985 | 25.175 | 25.0759 | 25.7569 |
| 26.2486 | 26.191 | 26.5303 | 26.0391 | 26.3585 | 25.8086 | 25.8809 |
| 27.9357 | 27.7074 | 27.5823 | 28.0307 | 27.9167 | 27.8568 | 28.1097 |
| 26.7421 | 26.5568 | 26.3544 | 26.8983 | 27.0173 | 26.4045 | 26.3454 |

| | | | | | | |
|---|---|---|---|---|---|---|
| 32.4057 | 32.6454 | 32.6248 | 32.242 | 32.2543 | 32.4306 | 32.6377 |
| 25.8017 | 25.6365 | 25.7715 | 25.891 | 25.7582 | 25.8433 | 25.5365 |
| 27.8394 | 27.9664 | 27.8022 | 27.8292 | 27.8157 | 27.7604 | 27.6091 |
| 25.3999 | 25.4684 | 25.4123 | 24.9402 | 25.2092 | 25.5586 | 25.4633 |
| 27.842 | 28.0144 | 27.9234 | 27.8335 | 27.7332 | 28.0359 | 28.1756 |
| 25.3123 | 25.5792 | 25.3211 | 25.3053 | 25.3367 | 25.2715 | 24.9697 |
| 26.4901 | 26.6952 | 26.6095 | 26.3139 | 26.3416 | 26.1478 | 26.4507 |
| 25.1502 | 25.0297 | 24.8257 | 25.187 | 25.4058 | 24.7119 | 24.6007 |
| 25.3872 | 25.9776 | 25.5828 | 25.4701 | 25.447 | 25.0071 | 25.2231 |
| 27.8776 | 28.0966 | 28.1517 | 27.8786 | 27.8901 | 27.8521 | 27.8634 |
| 26.3233 | 26.5015 | 26.2506 | 26.4129 | 26.3052 | 25.9821 | 26.0645 |
| 30.8301 | 30.918 | 30.8798 | 30.6591 | 30.6049 | 31.0047 | 30.9658 |
| 28.8336 | 28.9449 | 28.9088 | 28.8065 | 28.891 | 28.6331 | 28.4287 |
| 26.6373 | 26.4558 | 26.465 | 26.5472 | 26.5348 | 26.7353 | 26.7977 |
| 30.0707 | 30.3308 | 30.1857 | 30.1247 | 30.138 | 29.9874 | 30.0696 |
| 24.8274 | 24.6657 | 24.6913 | 25.135 | 24.9871 | 24.7943 | 24.7733 |
| 24.76 | NaN | NaN | NaN | 23.6233 | 23.8383 | 23.8549 |
| 27.4021 | 27.7401 | 27.5626 | 27.3421 | 27.3019 | 27.2503 | 27.4929 |
| 29.5019 | 29.6905 | 29.6059 | 29.4557 | 29.3796 | 29.239 | 29.1944 |
| 25.9389 | 25.8278 | 25.9305 | 25.5816 | 25.6892 | 26.0854 | 26.0995 |
| 31.7378 | 31.8295 | 31.8277 | 31.7684 | 31.9096 | 31.4943 | 31.5007 |
| 27.9727 | 28.1186 | 28.1916 | 27.7829 | 27.6728 | 27.7104 | 27.8159 |
| 24.7478 | NaN | NaN | 24.73 | 24.6969 | NaN | 24.6704 |
| 24.5009 | 24.0304 | 23.6783 | 24.6014 | NaN | 24.2246 | NaN |
| 23.4784 | NaN | 23.9802 | NaN | 24.1892 | 24.3508 | 24 |
| 29.9802 | 30.4229 | 30.2393 | 30.363 | 30.4138 | 30.4748 | 29.9915 |
| 24.7393 | 24.577 | 24.9161 | 25.0141 | 25.196 | 24.797 | 24.725 |
| 24.6762 | 24.9973 | 25.023 | 25.2726 | 25.2253 | 25.2293 | 24.7493 |
| 27.1215 | 27.7869 | 27.8105 | 27.3597 | 27.1586 | 27.4831 | 27.4394 |
| 25.2473 | 25.5818 | 25.2729 | 25.363 | 25.2352 | 25.8654 | 25.2475 |
| 25.5827 | 26.2647 | 26.3455 | 26.048 | 25.945 | 25.796 | 25.8417 |
| 24.7932 | 25.343 | 24.9486 | 25.2009 | 25.2304 | 25.2366 | 25.2508 |
| 25.2718 | 25.849 | 25.3015 | 25.8836 | 25.7088 | 25.5772 | 25.5343 |
| 26.9209 | 27.1126 | 27.16 | 27.2784 | 27.4213 | 27.2909 | 27.1332 |
| 27.1307 | 27.5052 | 27.3286 | 27.6693 | 27.4976 | 27.1409 | 27.1109 |
| 26.5103 | 26.3093 | 26.4051 | 26.3674 | 26.6587 | 26.6475 | 26.6367 |
| 26.9257 | 27.5781 | 27.5042 | 27.2799 | 27.3118 | 27.0112 | 26.9625 |
| 24.7395 | 25.0772 | 24.7759 | 24.9165 | 25.1856 | 25.1578 | 24.9425 |
| 24.9457 | 25.0893 | 25.0416 | 25.3892 | 25.3586 | 25.0091 | 25.2385 |
| 30.7101 | 30.8845 | 30.7869 | 30.9897 | 31.0188 | 30.9044 | 30.602 |
| 28.7702 | 28.9493 | 28.7929 | 28.9617 | 28.9869 | 28.7932 | 28.7484 |
| 29.7689 | 30.1612 | 29.9704 | 29.9413 | 30.058 | 29.9712 | 29.8495 |
| 31.8272 | 32.1085 | 31.9129 | 32.0525 | 31.9554 | 31.9422 | 31.8789 |
| 25.1665 | 25.6165 | 25.6791 | 25.261 | 25.3859 | 25.2886 | 25.5171 |
| 25.3679 | 25.716 | 25.7819 | 25.5214 | 25.7946 | 25.8347 | 25.8788 |
| 27.6401 | 27.8931 | 27.9872 | 27.7135 | 27.9229 | 27.6836 | 27.5915 |
| 27.1767 | 27.4645 | 27.3987 | 27.3314 | 27.4257 | 27.1223 | 27.1315 |
| 29.3819 | 29.8203 | 29.6854 | 29.8356 | 29.8553 | 29.6481 | 29.6109 |

| | | | | | | |
|---|---|---|---|---|---|---|
| 24.6874 | 25.1874 | 24.8974 | 24.929 | 25.13 | 24.9887 | 25.0176 |
| 24.8305 | 24.86 | 24.7522 | 24.6442 | 24.8803 | 24.8075 | 24.8641 |
| 27.2742 | 27.5621 | 27.386 | 27.5838 | 27.5659 | 27.215 | 27.5048 |
| 26.2619 | 26.4243 | 26.3903 | 26.4951 | 26.3987 | 26.6096 | 26.5317 |
| 32.1113 | 32.278 | 32.1949 | 32.1698 | 32.1338 | 32.0988 | 32.0131 |
| 33.4259 | 33.5712 | 33.5336 | 33.5681 | 33.5504 | 33.4575 | 33.4448 |
| 30.3959 | 30.3223 | 30.2604 | 30.2756 | 30.3502 | 30.2502 | 30.2467 |
| 29.752 | 29.496 | 29.4938 | 29.7507 | 29.8173 | 29.4585 | 29.4249 |
| 27.0484 | 26.9737 | 27.1432 | 26.8257 | 27.0187 | 26.6517 | 26.7189 |
| 26.2568 | 26.1383 | 25.7499 | 26.0105 | 26.0511 | 25.8478 | 25.9893 |
| 29.3148 | 29.3786 | 29.0209 | 28.6475 | 29.0211 | 28.4423 | 28.7609 |
| 24.9987 | 24.3848 | 24.1721 | 24.3374 | 24.4698 | 23.9735 | 24.4649 |
| 28.7765 | 28.6798 | 28.6454 | 28.6609 | 28.6643 | 28.409 | 28.3347 |
| NaN | NaN | NaN | 23.8619 | NaN | 24.7381 | 24.2496 |
| 26.3715 | 26.3587 | 26.2532 | 26.2505 | 26.1595 | 26.4461 | 26.1037 |
| 26.7265 | 27.2602 | 27.129 | 27.316 | 27.2408 | 26.9113 | 26.8787 |
| 24.6288 | 24.8455 | 25.1455 | 25.2593 | 25.1025 | 24.9954 | 24.6483 |
| 24.9546 | 24.9045 | 24.4743 | 23.9183 | 24.524 | 24.5422 | 24.4946 |
| 25.7307 | 25.9885 | 26.073 | NaN | NaN | NaN | 25.9421 |
| NaN | 23.8078 | 23.9116 | 23.9145 | 24.5144 | NaN | NaN |
| 27.8955 | 28.052 | 28.1211 | 28.12 | 27.828 | 27.923 | 27.9045 |
| 25.5916 | 25.5595 | 25.4835 | 25.7006 | 25.9081 | 25.7185 | 25.5968 |
| 26.0172 | 26.4662 | 26.5926 | 26.0744 | 26.0711 | 25.9738 | 25.7536 |
| 24.1767 | 24.0234 | NaN | NaN | NaN | NaN | 24.0118 |
| 25.017 | 25.1617 | 25.2123 | 25.0892 | 25.1445 | 25.2886 | 25.0458 |
| 24.625 | NaN | 24.8362 | 24.6716 | 24.8999 | 24.3564 | NaN |
| 26.1072 | 26.7629 | 26.768 | 26.3565 | 26.3925 | 26.4269 | 26.1371 |
| 25.0078 | 25.5091 | 25.3646 | 25.4515 | 25.6083 | 25.7051 | 25.6466 |
| 25.838 | 25.2956 | 25.4142 | 25.8267 | 25.6672 | 26.054 | 25.8148 |
| 24.2928 | NaN | 24.0489 | 24.4682 | NaN | NaN | NaN |
| 26.9474 | 26.7416 | 26.2555 | 26.6768 | 26.8057 | 26.9767 | 26.8724 |
| 25.7062 | 25.6434 | 25.54 | NaN | NaN | 25.0157 | 25.2469 |
| 31.3564 | 32.0565 | 32.1811 | 32.0652 | 31.9958 | 32.152 | 32.1658 |
| 24.3017 | NaN | NaN | 24.45 | 24.5208 | 24.3487 | 24.2933 |
| 25.3807 | 25.9076 | 25.6987 | 25.4066 | 25.9595 | 25.3624 | 25.2603 |
| NaN | 24.9608 | NaN | 24.9029 | 24.8536 | 24.8956 | 24.9976 |
| 24.5908 | NaN | NaN | NaN | 24.6782 | 24.6079 | NaN |
| 25.3593 | 25.3057 | 25.5465 | 25.4552 | 25.5092 | 25.5052 | 25.7149 |
| 28.7118 | 29.0903 | 28.9423 | 28.7802 | 29.3299 | 28.6454 | 28.6391 |
| 28.4446 | 28.6192 | 28.477 | 28.5232 | 28.5528 | 28.5129 | 28.2882 |
| NaN | 26.087 | 25.4096 | NaN | 25.5605 | 26.1719 | 25.9545 |
| 24.1046 | 24.3028 | 24.3228 | 24.2558 | 24.4118 | 24.4153 | 24.1765 |
| 27.9417 | 28.0219 | 27.9669 | 28.03 | 27.902 | 27.5424 | 27.6782 |
| 24.9742 | 24.763 | 24.6673 | 24.6506 | 24.4218 | 24.7815 | 24.9803 |
| 24.8754 | 25.0228 | 25.0408 | 24.7722 | 24.8451 | 24.9042 | 25.0591 |
| 26.3547 | 25.7026 | 25.9127 | 26.3502 | 26.3651 | 26.0598 | 25.8756 |
| 31.7683 | 31.9555 | 31.952 | 31.9001 | 31.9054 | 31.6903 | 31.6284 |
| 25.3586 | 25.5029 | 25.2348 | 25.48 | 25.7363 | 25.6914 | 25.6551 |

| | | | | | | |
|---|---|---|---|---|---|---|
| 25.5067 | 25.3704 | 25.5683 | 25.7386 | 25.5476 | 25.5159 | 25.4695 |
| 25.5724 | 24.7669 | 25.0606 | 25.0344 | 25.1319 | 25.746 | 25.3794 |
| 25.4125 | 25.4279 | 25.0538 | 25.3071 | 25.2143 | 25.4149 | 25.7837 |
| 26.1349 | 26.4913 | 26.4 | 26.6628 | 26.8165 | 26.1197 | 26.1548 |
| 24.5787 | NaN | 23.6228 | 23.81 | NaN | 23.9163 | 23.9822 |
| 26.2195 | 26.3461 | 26.7529 | 26.5197 | 26.6228 | 25.735 | 25.7315 |
| 24.997 | 24.8549 | 24.6582 | 24.7616 | 24.5325 | 24.9124 | 24.8021 |
| 28.1029 | 28.2041 | 28.2121 | 27.9904 | 28.1075 | 27.9612 | 27.8737 |
| 27.2236 | 27.0313 | 27.0862 | 27.6663 | 27.5587 | 27.2958 | 27.1058 |
| 26.6498 | 26.6788 | 26.4695 | 26.5324 | 26.6851 | 26.4543 | 26.3382 |
| 27.2315 | 27.6389 | 27.7004 | 27.7493 | 27.6268 | 27.384 | 27.3371 |
| 26.276 | 26.3381 | 26.2011 | 26.2153 | 26.5442 | 26.1285 | 26.1857 |
| 28.0733 | 28.2708 | 28.1112 | 27.7515 | 27.6817 | 27.7591 | 27.8532 |
| 27.4554 | 27.4339 | 27.2148 | 27.2502 | 27.2117 | 27.507 | 27.7269 |
| NaN | 25.4236 | 25.5816 | 25.2219 | 25.3561 | 25.4788 | 25.4662 |
| 28.3941 | 28.7344 | 28.5557 | 28.7542 | 28.7834 | 28.4621 | 28.2371 |
| 29.6399 | 29.4549 | 29.6738 | 29.8932 | 30.0005 | 29.7228 | 29.6251 |
| 25.748 | 25.925 | 25.765 | 26.2373 | 25.7607 | 26.3885 | 26.2792 |
| 27.6413 | 27.6784 | 27.4354 | 27.7197 | 27.6747 | 27.5128 | 27.5293 |
| 25.1808 | 25.2194 | 24.9567 | NaN | 25.1945 | NaN | 25.0064 |
| 31.0654 | 31.4524 | 31.3664 | 31.4589 | 31.612 | 30.9604 | 30.9242 |
| 24.5252 | 24.9791 | 24.4906 | 24.4791 | 24.6204 | 24.3521 | 24.5782 |
| 25.8817 | 25.8559 | 25.6294 | 25.6801 | 25.5551 | 25.5111 | 25.3652 |
| 26.1323 | 25.547 | NaN | 25.3427 | 25.5861 | NaN | NaN |
| 24.726 | 24.7839 | 25.4132 | 25.3781 | 25.3595 | 24.7041 | 24.8345 |
| 26.6889 | 26.6206 | 26.8356 | 26.2587 | 26.4116 | 26.3432 | 26.6263 |
| 25.406 | 25.6561 | NaN | NaN | 25.5497 | 25.6589 | 25.7149 |
| 30.0808 | 29.7516 | 30.2042 | 29.9841 | 30.148 | 30.0875 | 30.4236 |
| 26.4947 | 26.6726 | 26.6176 | 26.2846 | 26.2752 | 26.3738 | 26.3886 |
| 25.3454 | 25.4706 | 25.2915 | NaN | 25.6454 | NaN | 25.3633 |
| 26.2197 | NaN | 26.5432 | 26.5087 | 26.4942 | 26.3377 | NaN |
| 25.1558 | 25.3701 | 25.3784 | 25.2759 | 25.2711 | 24.9662 | 25.0938 |
| 28.7784 | 28.9115 | 28.9486 | 28.8155 | 28.8548 | 28.6296 | 28.5904 |
| 26.0412 | 26.5931 | 26.4673 | 26.1484 | 26.0874 | 26.2626 | 26.2714 |
| 24.47 | 24.0019 | NaN | 24.1357 | 24.2502 | 24.4401 | 24.316 |
| NaN | 25.9226 | NaN | 25.7826 | 25.827 | 25.7524 | 25.7511 |
| 26.4944 | 26.2672 | 25.9488 | 25.7395 | 25.8814 | 26.1384 | 26.4435 |
| 25.2406 | 25.5989 | 25.7144 | 25.7904 | 25.5855 | 25.2429 | 25.5965 |
| 24.8058 | 24.5461 | NaN | NaN | NaN | 24.9794 | 24.6321 |
| 25.6869 | 25.9188 | 25.5942 | 25.7036 | 25.8134 | 25.6534 | 25.7108 |
| 24.7794 | 25.1018 | 24.8614 | 25.1216 | 24.8149 | 24.7889 | 24.8926 |
| NaN | NaN | NaN | 24.1348 | 24.663 | 23.3828 | 23.5563 |
| 25.3007 | 25.2926 | 24.9121 | 25.2436 | NaN | 25.2977 | 25.1366 |
| 24.7022 | 25.1266 | 24.7477 | 24.8112 | 24.4837 | 24.0131 | 24.1966 |
| 27.0995 | 26.9703 | 27.0772 | 26.7176 | 26.9122 | 27.107 | 27.1794 |
| 25.6077 | NaN | 25.5641 | 25.4545 | 25.4113 | 25.6529 | 25.6089 |
| 25.0041 | 24.9909 | 25.0562 | 25.815 | 25.129 | 24.7567 | 24.743 |
| 25.7977 | 26.3219 | 25.9838 | 25.3524 | 25.9718 | 25.9982 | 25.7607 |

| | | | | | | |
|---|---|---|---|---|---|---|
| NaN | NaN | 23.9585 | 24.0483 | NaN | NaN | 24.0031 |
| 24.359 | 24.6487 | 24.5431 | 24.8282 | 24.6569 | 24.6619 | 24.6139 |
| 25.0708 | 25.3302 | 25.336 | NaN | NaN | 24.8201 | NaN |
| 25.8126 | 26.8454 | 26.4328 | 26.1794 | 25.9436 | 26.1515 | 26.3159 |
| 25.2007 | 25.253 | 25.1638 | 25.4589 | 25.4579 | 25.3367 | 25.428 |
| 25.9502 | 26.3344 | 26.0043 | 26.1786 | 26.2468 | 25.9092 | 25.8765 |
| 24.6191 | NaN | NaN | NaN | NaN | 24.6896 | NaN |
| 24.8013 | 24.4938 | 24.8956 | 25.5835 | 25.3379 | NaN | NaN |
| NaN | NaN | NaN | NaN | NaN | 25.0964 | NaN |
| 24.4723 | 25.5669 | 25.1177 | 25.1752 | 25.2483 | 25.2918 | 24.9461 |
| 25.1537 | NaN | 25.2695 | 24.9371 | NaN | NaN | 25.1182 |
| 26.954 | 26.492 | 26.3628 | 26.7429 | 26.9713 | 26.9061 | 26.8335 |
| 25.681 | NaN | NaN | NaN | 25.5665 | 25.8125 | 25.7091 |
| 24.865 | 25.1489 | 25.0746 | 25.2822 | 24.3633 | 25.2336 | 24.9188 |
| 28.2397 | 28.3921 | 28.404 | 28.3662 | 28.3925 | 28.17 | 28.2628 |
| 25.0106 | 24.9193 | 25.0782 | 25.2256 | 24.9864 | 24.7269 | 24.8683 |
| 24.9971 | 25.0054 | 24.822 | NaN | 24.7913 | 24.7796 | 24.6399 |
| 25.7115 | 25.5153 | 25.4864 | 25.4796 | 25.8101 | 25.9248 | 25.9287 |
| 25.1616 | 25.0454 | 25.2663 | 25.0433 | 25.23 | 24.724 | 24.8269 |
| 24.6842 | 24.1319 | 24.4786 | 25.3467 | 24.6604 | 24.1752 | 24.0985 |
| 25.0643 | 24.733 | NaN | 25.1435 | 25.1367 | 25.5273 | 25.2324 |
| 25.8535 | 25.6364 | 25.6037 | 25.7137 | 25.9797 | 25.7277 | 25.8116 |
| 25.5367 | 25.2965 | 25.119 | 25.3506 | 24.9188 | NaN | 25.6631 |
| 24.5723 | 24.0877 | 24.5703 | 23.9256 | NaN | NaN | NaN |
| 24.0296 | 25.3683 | 25.2164 | NaN | 24.3038 | 24.2641 | 24.6587 |
| 25.4342 | 26.0277 | 25.5017 | 25.6841 | 25.6367 | 25.6114 | 25.6748 |
| 25.1152 | 25.1528 | 24.9623 | 24.7422 | NaN | 24.5987 | 25.0039 |
| 24.9702 | 24.7769 | 24.7271 | 24.9561 | 24.9391 | 24.8588 | 25.045 |
| 28.5313 | 28.7534 | 28.5157 | 28.3624 | 28.5267 | 28.2181 | 28.3728 |
| 25.1498 | 24.3597 | 24.7139 | 25.1673 | 24.6406 | 25.3955 | 25.2931 |
| 25.4122 | 25.6274 | 25.6321 | 25.524 | 25.3881 | 25.1904 | 25.03 |
| 27.2599 | 27.4881 | 27.3806 | 27.068 | 27.0231 | 26.7511 | 26.7393 |
| 25.4137 | 26.0017 | 25.431 | 24.8799 | 25.5631 | 25.4121 | 25.1465 |
| 26.1313 | 26.4006 | 26.2177 | 26.1644 | 26.4117 | 26.3403 | 26.2199 |
| 25.8259 | 25.7519 | 25.8354 | 25.9617 | 25.9957 | 25.8032 | 25.7137 |
| 25.3271 | 25.4516 | 25.3668 | 25.3647 | 25.6069 | 24.9376 | NaN |
| 26.6506 | 27.3439 | 27.071 | 26.7943 | 26.6655 | 26.5035 | 26.694 |
| 25.6623 | 25.3345 | 25.5612 | 25.234 | 25.3882 | 25.633 | 25.7436 |
| 27.897 | 28.0636 | 28.008 | 28.0141 | 28.0553 | 28.1906 | 27.9614 |
| 27.3562 | 27.5628 | 27.6276 | 27.413 | 27.4745 | 27.256 | 27.1164 |
| 25.3941 | 25.7204 | 25.4827 | 25.6429 | 25.2579 | 25.2355 | 25.2357 |
| 25.662 | 25.0632 | 25.1151 | 25.4898 | NaN | 25.8439 | 25.9973 |
| 28.6481 | 29.0787 | 28.9954 | 28.7357 | 28.8064 | 28.5991 | 28.7629 |
| 25.1971 | 25.7794 | 25.8661 | 25.278 | 24.2309 | 25.5309 | 25.6267 |
| 24.718 | 24.8946 | 24.7949 | 24.715 | 24.6691 | 24.4723 | NaN |
| 25.6092 | NaN | 25.4469 | 25.8203 | 25.6129 | 25.8619 | 25.8402 |
| 29.6922 | 29.3179 | 29.9127 | 29.5151 | 29.7826 | 29.7827 | 29.6994 |
| 25.2345 | 25.2548 | 25.5293 | 24.9542 | 25.0129 | NaN | 25.4108 |

| | | | | | | |
|---|---|---|---|---|---|---|
| 27.5198 | 27.6167 | 27.5345 | 27.6505 | 27.749 | 27.5704 | 27.6066 |
| 24.9233 | 25.042 | 25.1266 | 24.8785 | 24.5378 | 25.2932 | 25.0732 |
| 27.3857 | 27.0273 | 27.0275 | NaN | 27.3639 | 27.0984 | 26.9749 |
| 25.1119 | 25.5346 | 25.578 | 25.1113 | 25.0347 | NaN | 25.1053 |
| 26.2211 | 26.549 | NaN | 26.0928 | 26.1518 | 25.878 | 26.3924 |
| 26.6381 | 26.3215 | 26.8323 | 26.9213 | 26.8509 | 26.9641 | 27.0256 |
| 24.8268 | NaN | NaN | 24.8295 | 24.3649 | 24.1681 | 24.1029 |
| 24.9053 | 24.978 | 24.6722 | 24.7356 | 24.7252 | 24.5426 | 24.7439 |
| 25.218 | 25.2468 | 25.2535 | 25.4258 | 25.3829 | 24.9528 | NaN |
| 25.6883 | 25.5747 | 25.0912 | 25.6373 | 25.3151 | 26.4903 | 26.0202 |
| 30.6733 | 30.8476 | 30.692 | 30.7857 | 30.75 | 30.6125 | 30.6335 |
| 27.6255 | 27.855 | 27.7841 | 27.6147 | 27.716 | 27.2511 | 27.53 |
| 30.476 | 30.755 | 30.7276 | 30.5799 | 30.6051 | 30.5238 | 30.5938 |
| 27.2995 | 27.5159 | 27.3109 | 27.2216 | 26.9406 | 27.5386 | 27.5195 |
| 26.5335 | 26.7698 | 26.6273 | 26.642 | 26.4607 | 26.5123 | 26.2763 |
| 29.4048 | 29.3707 | 29.3116 | 29.8617 | 29.6089 | 29.7657 | 29.6604 |
| 24.0776 | 23.7367 | 23.7351 | 24.109 | 23.9961 | 24.0794 | 23.9603 |
| 25.5046 | 24.6718 | NaN | 25.6689 | 25.7495 | 25.5272 | 25.2796 |
| 26.3357 | 25.3867 | 26.4717 | 26.7136 | 26.5885 | 26.0422 | 26.1553 |
| NaN | 24.7125 | NaN | 24.6231 | 24.5144 | 24.7623 | 24.8938 |
| 25.556 | NaN | 25.3754 | 25.4248 | 25.6541 | 25.9433 | 26.0906 |
| 23.2542 | 22.6585 | NaN | 23.2154 | 23.1782 | NaN | NaN |
| 28.7627 | 30.001 | 29.1999 | 29.8357 | 29.7682 | 29.777 | 29.8417 |
| 26.6847 | 26.6451 | 26.5689 | 26.2285 | 26.4324 | 26.6406 | 26.6183 |
| 29.1427 | 29.1643 | 28.9469 | 29.3818 | 29.4919 | 29.0994 | 28.9733 |
| 32.1438 | 32.2552 | 32.1279 | 32.0679 | 32.0692 | 32.2805 | 32.3005 |
| 25.3897 | 25.4232 | 25.4837 | 25.4502 | 25.508 | 25.4831 | 25.4358 |
| 25.9028 | 26.1992 | 25.9656 | 25.821 | 25.7609 | 25.6215 | 25.8925 |
| 33.9146 | 34.0052 | 33.8737 | 33.8738 | 33.9092 | 33.6081 | 33.5978 |
| 27.5112 | 27.3634 | 27.6158 | 27.5825 | 27.6342 | 27.7884 | 27.9159 |
| 29.4339 | 29.5023 | 29.2321 | 29.6569 | 29.5347 | 29.1976 | 29.2936 |
| 24.7244 | 24.8547 | 24.8359 | 25.0559 | NaN | 24.6699 | 24.6521 |
| 25.3286 | NaN | NaN | 24.97 | NaN | 24.9239 | NaN |
| 29.4979 | 29.8304 | 29.6986 | 29.5301 | 29.4479 | 29.4892 | 29.4377 |
| 30.9029 | 30.9627 | 30.9612 | 30.8459 | 30.8788 | 30.7234 | 30.7922 |
| 27.7466 | 27.4071 | 27.3186 | 27.5062 | 27.4637 | 27.3126 | 27.7953 |
| 25.3489 | NaN | NaN | 25.1399 | 25.1573 | 25.6841 | 25.8291 |
| 26.1865 | 26.724 | 26.4096 | NaN | 26.3507 | 26.2934 | 26.2795 |
| 26.8728 | 26.3141 | 25.7287 | 26.8761 | 26.7474 | 26.5686 | 26.6698 |
| 26.6488 | 26.916 | 26.6274 | 26.6606 | 26.6535 | 26.655 | 26.7121 |
| 24.8977 | 25.833 | 25.0025 | 25.5622 | 25.2959 | 25.7337 | 26.0598 |
| 25.8059 | 26.4977 | 25.802 | 26.2619 | 26.2436 | 26.1294 | 26.2361 |
| 25.1236 | 25.3684 | 25.1061 | 25.323 | 25.4904 | 25.0175 | 25.1825 |
| 25.5893 | 25.4151 | 25.4199 | 25.582 | 25.4356 | 25.4941 | 25.6098 |
| 26.3596 | 26.5118 | 26.641 | 26.0323 | 26.3503 | 26.4984 | 26.3962 |
| NaN | 25.6004 | 25.5397 | 25.2162 | 24.8827 | 25.2731 | 25.2001 |
| 24.4447 | NaN | NaN | 24.2615 | 24.5949 | NaN | 24.4083 |
| 26.114 | 26.1767 | 26.0927 | NaN | NaN | NaN | NaN |

| | | | | | | |
|---|---|---|---|---|---|---|
| 26.9102 | 25.8931 | 26.0844 | 27.4545 | 26.5412 | 26.0604 | 26.0461 |
| 31.8933 | 32.0461 | 31.9848 | 31.8004 | 31.8288 | 32.1566 | 31.9827 |
| 29.2871 | 29.9552 | 29.367 | 29.8624 | 29.9279 | 29.6924 | 29.7376 |
| NaN | NaN | NaN | 25.2866 | 25.2783 | 25.5492 | NaN |
| 25.9124 | 26.5105 | 26.5448 | 25.77 | 25.9876 | 25.9526 | 26.1266 |
| 26.5727 | 26.7184 | 26.6929 | 26.0903 | 26.2149 | 26.3283 | 26.3128 |
| 25.0769 | NaN | NaN | 24.939 | 25.5799 | 25.2371 | 24.7504 |
| NaN | 24.9239 | NaN | 24.6597 | 24.8474 | 24.7453 | 24.8577 |
| 25.7741 | 25.6105 | 26.4063 | 26.1268 | 25.9865 | 25.9652 | 26.229 |
| 25.4455 | 25.6082 | 25.1244 | 25.75 | 25.7152 | 25.4043 | 25.2868 |
| 26.0434 | 26.2243 | 26.0827 | 25.9075 | 26.234 | 26.0561 | 26.2246 |
| 28.8679 | 28.6381 | 28.9342 | 28.9631 | 29.1858 | 28.8749 | 28.5853 |
| 27.1238 | 27.0098 | 27.1268 | 27.2169 | 27.1456 | 26.8345 | 26.7786 |
| 24.4351 | NaN | NaN | NaN | 24.2178 | 24.4658 | 24.6948 |
| 23.9785 | 23.8991 | 23.9525 | NaN | 24.2284 | 23.6862 | 24.2518 |
| 28.4241 | 28.5513 | 28.4176 | 28.3726 | 28.6023 | 28.2492 | 28.1123 |
| 30.0428 | 29.8665 | 29.8163 | 29.7313 | 29.8079 | 29.9979 | 29.9772 |
| 29.1804 | 29.3555 | 29.2857 | 29.1895 | 29.1927 | 28.9727 | 28.9603 |
| 24.5179 | 24.1729 | 24.3685 | 24.7697 | 24.4101 | 24.1265 | 24.1394 |
| 24.4147 | 23.7242 | 23.8761 | 24.3312 | 24.2913 | 24.1373 | 24.2451 |
| 24.3221 | 24.7209 | 24.7447 | NaN | NaN | 24.3996 | NaN |
| 26.4462 | 26.1517 | 25.9848 | 26.2387 | 26.5053 | 26.5132 | 26.4968 |
| 25.8059 | 25.7307 | 25.7672 | 26.0208 | 26.0157 | 25.9242 | 26.035 |
| 25.1679 | 25.387 | 25.1445 | 24.8768 | 24.9965 | 24.9899 | 24.7786 |
| 27.3808 | 27.6795 | 27.6258 | 27.4654 | 27.8275 | 27.471 | 27.4045 |
| 24.9336 | 24.9451 | 24.9239 | 25.1919 | 24.8024 | 24.8017 | 25.0069 |
| 26.6246 | 26.4661 | 26.4037 | 26.2114 | 26.4363 | 26.1582 | 26.2829 |
| 29.86 | 30.2076 | 29.8011 | 29.6557 | 29.8545 | 29.689 | 29.466 |
| 25.623 | 25.8489 | 25.9844 | 25.7448 | 25.8616 | 25.5186 | 25.6236 |
| 25.4633 | 25.2855 | 25.1707 | 25.2187 | 25.2338 | 25.2214 | 25.1383 |
| 29.9586 | 29.6353 | 29.5773 | 29.7897 | 29.7077 | 30.2399 | 30.0755 |
| 24.5458 | 23.8143 | 23.4782 | 24.3051 | 24.5025 | 24.6382 | 24.552 |
| 26.8588 | 27.0576 | 27.0507 | 26.993 | 26.9915 | 26.7605 | 26.8215 |
| 25.866 | 25.7454 | 26.2578 | 25.8045 | 26.1992 | 26.2101 | 26.1346 |
| 25.7412 | 26.1977 | 25.7799 | 25.8167 | 25.6839 | 25.6286 | 25.7514 |
| 24.8938 | 25.4489 | 25.225 | 24.4972 | 24.6476 | 24.4761 | 24.6488 |
| 25.7004 | 25.6147 | 25.7455 | 25.2386 | 24.9581 | 25.3756 | 25.1849 |
| 26.3226 | 26.3789 | 26.3407 | 26.2713 | 26.1524 | 26.6874 | 26.7285 |
| NaN | 24.8367 | 24.8132 | 24.5304 | 24.9426 | 25.0444 | NaN |
| 28.4149 | 28.6087 | 28.5867 | 28.475 | 28.435 | 28.4059 | 28.5427 |
| NaN | 25.7091 | 25.8105 | NaN | 25.7733 | 26.1818 | 25.714 |
| 25.7769 | 25.704 | 25.8506 | 26.0898 | 25.9884 | 26.0604 | 25.9264 |
| 28.3287 | 27.9074 | 28.0684 | 28.4362 | 28.236 | 28.2799 | 28.0139 |
| 25.2163 | 25.2725 | 25.18 | 25.162 | 25.3313 | 25.0423 | 24.9727 |
| 26.7469 | 26.6026 | 26.3643 | 26.6564 | 26.7998 | 26.3187 | 26.2528 |
| 25.4012 | 25.6166 | 25.648 | 25.6227 | 25.5954 | 25.2808 | 25.1882 |
| 24.9207 | 25.0844 | 24.7701 | 24.6028 | 24.6195 | 24.0478 | 24.3617 |
| 24.828 | 25.3723 | 24.9736 | 24.7775 | 25.0351 | 24.6051 | 24.6742 |

| | | | | | | |
|---|---|---|---|---|---|---|
| 28.9625 | 29.0769 | 29.0783 | 29.1974 | 29.147 | 28.9335 | 28.8473 |
| 24.1616 | NaN | 24.0069 | 24.2633 | 24.1612 | 24.4697 | 24.5059 |
| 24.5227 | 24.611 | 24.8431 | 24.6764 | 24.4245 | 24.2197 | 24.4257 |
| 24.5798 | 24.872 | 24.9668 | 24.6391 | 24.8288 | 24.7204 | 24.8817 |
| 26.8968 | 26.519 | 26.3449 | 26.6974 | 26.6818 | 26.556 | 26.4683 |
| 25.6144 | 25.725 | 25.5736 | 25.3251 | 25.4689 | 25.1446 | 25.0481 |
| 25.0142 | 25.1851 | 24.7964 | 25.3004 | 25.3228 | 24.8277 | 25.0994 |
| 31.1777 | 31.4281 | 31.2925 | 31.1512 | 31.2105 | 31.0442 | 31.0441 |
| 24.7067 | 25.2002 | 24.9294 | 24.9649 | NaN | NaN | 24.9573 |
| 24.5543 | 24.7491 | 24.7988 | 24.8626 | 24.7978 | 24.6546 | 24.4999 |
| 24.4636 | 24.5811 | 24.32 | 24.3846 | 24.4702 | 24.2434 | 24.4474 |
| 25.0659 | 25.7584 | 25.8913 | 25.8076 | NaN | 25.7886 | 25.8697 |
| 23.7911 | 23.7442 | 24.0655 | 23.855 | 23.455 | 23.3479 | 23.0465 |
| 25.5013 | 25.263 | 25.4241 | 24.9996 | 25.1569 | 25.2955 | 25.6571 |
| 28.4917 | 28.4584 | 28.4847 | 28.6574 | 28.7991 | 28.4495 | 28.4354 |
| 25.5016 | NaN | 25.5289 | 25.4835 | 25.6313 | 25.7475 | 25.7985 |
| 24.9141 | NaN | 25.0547 | 25.5651 | 24.8769 | 24.9968 | 24.7894 |
| 25.3784 | 25.3184 | 25.3609 | 25.319 | 25.194 | 24.4284 | 24.3574 |
| 25.8627 | 25.8419 | 25.5915 | 25.9256 | 25.8543 | NaN | NaN |
| 26.975 | 27.2802 | 27.2727 | 27.0493 | 27.1664 | 27.3948 | 27.2462 |
| 27.0087 | 26.9514 | 27.1881 | 26.7598 | 26.8096 | 26.5288 | 26.5063 |
| 26.8097 | 26.8887 | 26.8185 | 26.9764 | 26.8134 | 26.6206 | 26.7993 |
| 24.8884 | NaN | 24.5778 | 24.6689 | 24.7813 | 25.0472 | 25.1621 |
| 30.9312 | 31.174 | 31.1224 | 30.6814 | 30.7533 | 31.0553 | 31.0228 |
| 24.3227 | 24.0678 | 23.805 | 24.3586 | NaN | 24.3575 | 24.4474 |
| 27.0683 | 26.8156 | 26.8264 | 27.3237 | 27.1864 | 26.9132 | 26.8385 |
| 25.3928 | 25.0632 | 25.6699 | 25.7717 | NaN | NaN | 25.768 |
| NaN | NaN | NaN | NaN | 25.1277 | 25.1278 | 25.2902 |
| 24.2776 | 24.4961 | 24.5562 | 24.4922 | 24.5054 | 24.4667 | NaN |
| NaN | 23.1673 | 23.3894 | 23.6825 | NaN | NaN | NaN |
| 24.2802 | 24.4448 | NaN | 24.7864 | 24.7104 | 24.2324 | 24.1558 |
| 31.4154 | 31.5227 | 31.5263 | 31.4407 | 31.5329 | 31.3617 | 31.2214 |
| 27.1035 | 27.6858 | 27.4473 | 27.6695 | 27.5067 | 27.402 | 27.5284 |
| NaN | NaN | 24.2124 | 24.5774 | 24.6267 | NaN | NaN |
| NaN | 26.0976 | 26.7005 | NaN | NaN | 26.2759 | 26.5928 |
| 24.4324 | NaN | NaN | 24.4461 | 24.3896 | NaN | 24.386 |
| 25.4874 | NaN | NaN | NaN | NaN | 25.6052 | 25.5027 |
| 24.7805 | 26.0408 | 25.7531 | 25.4068 | 25.371 | 24.9616 | NaN |
| 28.3234 | 28.8655 | 28.7013 | 29.2032 | 29.1394 | 28.8063 | 28.6889 |
| 24.6979 | 24.8125 | 25.1578 | 25.6366 | 25.7117 | 25.4085 | 24.8914 |
| 25.5637 | 26.1975 | 26.1068 | 26.0416 | 25.792 | 25.8138 | 25.9065 |
| 24.3155 | 24.4402 | 24.3138 | 24.6022 | 24.6531 | 24.8461 | 24.7535 |
| 24.6446 | 24.4804 | 24.937 | 25.4931 | 25.006 | 24.9775 | 24.9825 |
| 24.5654 | 24.8755 | 24.9241 | 24.7162 | 24.6875 | 24.6986 | 24.894 |
| 24.6344 | 25.3003 | 25.0957 | 25.2921 | 24.8263 | 25.0701 | 24.9225 |
| 24.4779 | 25.0401 | 25.0587 | 25.0354 | 24.8731 | 24.5219 | 24.5649 |
| 29.991 | 30.2538 | 30.2792 | 30.3475 | 30.1536 | 29.9589 | 29.9215 |
| 25.6274 | 25.9778 | 26 | 26.1201 | 25.8383 | 26.0207 | 26.1517 |

| | | | | | | |
|---|---|---|---|---|---|---|
| 24.5807 | 25.2619 | 24.9582 | 25.251 | 25.16 | 25.0122 | 25.1395 |
| 26.0231 | 26.3348 | 26.0816 | 26.6113 | 26.3364 | 26.5361 | 26.3531 |
| 26.7064 | 27.1968 | 27.1208 | 26.8803 | 26.8957 | 26.8407 | 26.8656 |
| 29.0869 | 29.2496 | 29.2213 | 29.1528 | 29.2713 | 29.4207 | 29.376 |
| 29.7691 | 29.8948 | 29.8688 | 29.8076 | 29.9307 | 29.9935 | 29.8009 |
| 26.1602 | 26.133 | 26.4877 | 26.3439 | 26.5147 | 26.5806 | 26.3454 |
| 30.2201 | 30.2828 | 30.5181 | 30.7367 | 30.3709 | 30.3777 | 30.2833 |
| 25.3917 | 25.4856 | 25.4404 | 25.4335 | 25.4195 | 25.5579 | 25.4806 |
| 29.4745 | 29.5662 | 29.3832 | 29.3499 | 29.3879 | 29.2689 | 29.3775 |
| 30.3575 | 30.5359 | 30.3146 | 30.2799 | 30.2178 | 30.2711 | 30.2916 |
| 29.3678 | 29.5724 | 29.3803 | 29.0051 | 29.0734 | 29.2102 | 29.2719 |
| 26.1584 | 25.4778 | 25.5824 | 25.8027 | 25.9231 | 26.0046 | 25.5694 |
| 25.8824 | 25.1548 | 25.1952 | 25.2383 | 25.4642 | 25.0794 | 24.917 |
| 26.5785 | 26.4946 | 26.5749 | 26.6879 | 26.6716 | 26.2409 | 26.0859 |
| 28.4184 | 27.8968 | 27.9163 | 28.2042 | 28.3342 | 27.6959 | 27.9518 |
| 29.3007 | 28.88 | 29.2032 | 28.8592 | 28.946 | 28.8322 | 29.2551 |
| 25.767 | 25.7916 | 25.7781 | 25.8406 | 25.6196 | 25.5442 | 25.7476 |
| 24.3324 | 24.4145 | 24.2037 | 24.8294 | 24.3596 | 24.6973 | 24.9457 |
| 25.3238 | 25.2601 | 24.8587 | 24.9296 | 24.7912 | 24.8623 | 24.5553 |
| 25.9662 | 26.467 | 26.3724 | 26.5501 | 26.5209 | 26.2322 | 25.5462 |
| 25.3589 | 25.0191 | NaN | 25.3466 | NaN | 25.2985 | NaN |
| 24.6987 | 25.1273 | 24.8464 | NaN | NaN | NaN | 24.0475 |
| 27.7442 | 28.1703 | 28.035 | 27.928 | 27.8347 | 28.0667 | 27.6773 |
| 27.7828 | 27.3348 | 27.3217 | 29.386 | 27.6672 | 28.625 | 28.9916 |
| 25.3518 | 25.6374 | 25.5831 | NaN | NaN | 25.5819 | NaN |
| 25.3116 | 24.9906 | 25.0275 | 24.6803 | 24.8509 | 25.1037 | 25.396 |
| NaN | NaN | NaN | 22.3156 | NaN | 23.1633 | 23.1082 |
| 24.84 | NaN | NaN | 25.1027 | 25.1403 | NaN | 25.0857 |
| 24.8073 | NaN | 25.112 | 24.1648 | NaN | NaN | NaN |
| 25.0135 | NaN | NaN | 25.0602 | 25.1353 | NaN | 25.1632 |
| NaN | NaN | NaN | 24.8624 | 24.212 | 24.672 | 24.1017 |
| 25.9704 | 26.0743 | 25.9456 | 26.3489 | 26.1067 | 26.6617 | 26.4154 |
| 27.5035 | 28.0746 | 27.7686 | 27.276 | 27.5803 | 27.3467 | 27.5865 |
| NaN | 25.6234 | NaN | NaN | NaN | NaN | NaN |
| 25.271 | 25.4921 | NaN | NaN | NaN | 25.5667 | 25.0704 |
| 25.3268 | 24.2336 | 23.9556 | 24.898 | 25.2148 | 24.9634 | 24.9263 |
| 33.111 | 33.456 | 33.3248 | 33.1532 | 33.1264 | 33.0581 | 33.1795 |
| 23.9906 | 23.8859 | 23.9185 | 23.879 | 23.9991 | 23.9552 | 23.8937 |
| 28.175 | 28.3588 | 27.9429 | 28.2304 | 28.4021 | 28.0031 | 28.1076 |
| 24.2679 | 25.0104 | 24.4007 | 24.1987 | 24.2467 | 24.3243 | 24.2126 |
| 25.8145 | 26.0232 | 26.0993 | 26.4513 | 25.9448 | 26.0459 | 26.5257 |
| 24.6017 | 25.294 | 25.04 | 24.6733 | 24.9214 | 25.0732 | 24.8472 |
| 25.2063 | 25.5467 | 25.3675 | 25.2352 | 25.5113 | 25.7136 | 25.4118 |
| 24.2552 | 24.6359 | 24.6283 | 24.2726 | 24.529 | 24.5334 | 24.595 |
| 24.0978 | 24.1234 | 24.015 | 23.9023 | 24.1708 | 23.7649 | 23.9357 |
| NaN | NaN | NaN | 25.3767 | 25.23 | 25.3025 | 25.533 |
| 24.3673 | 24.816 | 24.5448 | NaN | 24.2116 | NaN | 24.7988 |
| 24.9944 | 25.3496 | 25.0568 | 25.2349 | 24.9668 | 25.0704 | 25.0319 |

| | | | | | | |
|---|---|---|---|---|---|---|
| 23.9619 | 23.787 | 24.3102 | NaN | NaN | NaN | 23.8936 |
| NaN | NaN | 24.6173 | 24.8328 | 24.6125 | 24.6675 | 24.6249 |
| 25.2638 | 25.341 | 25.3233 | 25.2427 | 25.2196 | 25.1027 | 25.1536 |
| 28.9406 | 28.1728 | 29.1629 | 29.0163 | 28.8879 | 28.8919 | 28.6632 |
| 26.3874 | 26.3256 | 26.169 | 26.4141 | 26.6967 | 26.2895 | 26.2872 |
| 24.8418 | 24.6655 | 25.0608 | 24.5312 | 24.4877 | 24.5886 | 24.534 |
| 25.01 | 24.9885 | 25.0263 | 24.9829 | 24.8439 | 25.0597 | 24.97 |
| 24.3827 | 24.5449 | 24.4429 | 24.2131 | 24.1481 | 24.4697 | 24.5871 |
| 24.9367 | 24.8287 | 25.3494 | 25.2511 | 24.9899 | 25.2363 | 25.4317 |
| 24.7718 | 24.3239 | NaN | 24.2006 | 24.0829 | 24.1231 | 24.3446 |
| 24.9784 | 24.9029 | 24.768 | 24.9345 | 25.1182 | NaN | NaN |
| 24.3458 | 25.4063 | 24.7648 | 25.0836 | 25.2795 | 25.276 | 25.283 |
| 24.2388 | 23.7893 | 23.531 | 23.6896 | 24.3204 | 23.7667 | 23.8207 |
| 25.5044 | 24.6681 | 25.2839 | 25.563 | 25.3087 | 25.535 | 25.1419 |
| NaN | 23.5951 | 23.6908 | NaN | 23.9027 | 24.0527 | 24.2082 |
| 23.2574 | 23.7992 | 23.253 | 23.8288 | 23.8306 | 23.483 | 23.1396 |
| 25.0781 | 25.1533 | 25.2031 | 24.5299 | NaN | NaN | 24.9827 |
| 24.665 | 24.8229 | 24.84 | 24.7508 | 24.5549 | 24.6152 | 24.6887 |
| 30.5708 | 31.0289 | 30.8975 | 30.7051 | 30.7061 | 30.7625 | 30.8572 |
| 27.8031 | 28.6667 | 28.1675 | 28.0618 | 28.0174 | 27.849 | 28.1416 |
| 28.7248 | 29.1705 | 29.1778 | 28.8602 | 28.9574 | 29.0235 | 28.9714 |
| 24.2453 | 24.4252 | 24.472 | 25.2309 | 25.402 | 24.4185 | 24.2371 |
| 24.3352 | 24.1594 | 24.2012 | 24.9246 | 24.4964 | 24.3905 | 24.3721 |
| 25.4334 | 25.7046 | 25.3155 | 25.5807 | 25.3867 | 26.176 | 25.5847 |
| 24.7517 | 24.7487 | 24.7599 | 24.6056 | 24.7899 | 24.5007 | 24.7122 |
| 25.076 | 24.9757 | 24.6436 | 24.9766 | 24.9604 | 25.2516 | 25.4631 |
| 27.4423 | 27.3198 | 26.9664 | 27.1902 | 27.3182 | 27.3482 | 27.2535 |
| 24.5858 | NaN | 25.5155 | 24.6369 | NaN | NaN | NaN |
| 24.0226 | 24.4664 | 23.9687 | 24.3754 | 24.4793 | 24.6913 | 24.7937 |
| 25.0096 | 25.6769 | 24.8369 | 25.9139 | 25.9037 | 25.3129 | 24.8553 |
| 25.2427 | 24.6924 | 25.3125 | 25.1925 | 24.8399 | 25.1366 | NaN |
| 25.8505 | 25.8445 | 26.1227 | 25.8092 | 25.6211 | 25.4745 | 25.7953 |
| 24.7709 | 24.4634 | 24.5069 | 24.7304 | 24.8283 | 24.8811 | 24.8826 |
| 26.407 | 25.96 | 26.3721 | NaN | NaN | NaN | 26.324 |
| 24.4317 | NaN | 24.4453 | 24.3965 | NaN | NaN | 24.2921 |
| 24.0047 | 23.7816 | 23.766 | 23.7791 | 24.0909 | 24.3267 | 24.2144 |
| 24.9104 | 24.6267 | 24.9934 | NaN | 24.6309 | 24.7078 | 24.6098 |
| 26.3897 | 26.8579 | 26.8075 | 26.5806 | 26.6143 | 26.3973 | 26.4568 |
| NaN | NaN | NaN | NaN | 23.9162 | NaN | NaN |
| 26.8614 | 25.8329 | 24.9531 | 26.4773 | 26.4208 | 26.8411 | 26.8153 |
| 26.5377 | 27.0025 | 26.7248 | 26.9155 | 26.7577 | 26.5375 | 26.6711 |
| 24.9971 | 25.0124 | 24.5357 | NaN | 24.7759 | 24.7301 | 24.7242 |
| NaN | NaN | 24.7444 | NaN | NaN | 24.6516 | 24.6812 |
| NaN | 25.3539 | 24.7909 | 24.8768 | 25.1152 | 25.1861 | 25.0879 |
| 24.2918 | 24.1517 | 23.9261 | 24.392 | NaN | 24.2741 | 24.0756 |
| 25.6683 | 25.3701 | 25.5075 | 25.8494 | 25.8399 | 25.5261 | 25.4696 |
| 26.3534 | 26.5982 | 26.5851 | 26.1292 | 26.3069 | 26.4729 | 26.5092 |
| NaN | NaN | NaN | 24.0971 | 24.3095 | NaN | 23.9115 |

| | | | | | | |
|---|---|---|---|---|---|---|
| 26.2479 | 26.2726 | 26.3462 | 26.2066 | 26.6655 | 25.9689 | 25.8093 |
| 24.7742 | NaN | NaN | 24.8447 | NaN | NaN | 24.7672 |
| NaN | NaN | NaN | NaN | 26.4224 | NaN | NaN |
| 25.4015 | NaN | 25.0856 | 25.3 | 25.3747 | 25.3893 | 25.3109 |
| 24.9139 | 24.7477 | 24.911 | 25.0308 | 24.923 | 24.7032 | 24.5253 |
| 24.7184 | 24.9623 | 24.6372 | 24.4526 | 24.7377 | 24.5721 | 24.5869 |
| 33.0161 | 33.0764 | 33.0708 | 33.0203 | 33.0717 | 33.0067 | 32.9666 |
| 26.9492 | 26.9529 | 27.4717 | 27.1778 | 27.1561 | 27.4226 | 27.2217 |
| 27.22 | 27.4706 | 27.4913 | 27.1575 | 27.0642 | 27.2798 | 27.1844 |
| 24.6271 | NaN | NaN | 24.3046 | NaN | 24.3459 | 24.6143 |
| 24.5831 | 23.9886 | 24.3534 | 24.3647 | 24.487 | 24.7475 | 24.6964 |
| 25.5601 | 25.3278 | 25.5529 | 26.4327 | 26.0778 | 25.8839 | 25.2582 |
| 25.5284 | 25.6177 | 25.483 | 25.0692 | 25.4144 | 25.1757 | 25.3662 |
| 24.0242 | 23.8887 | 23.7559 | 24.1224 | 23.874 | 23.7663 | 23.6778 |
| 23.49 | NaN | 23.5927 | 23.8077 | 23.8496 | NaN | NaN |
| 25.6118 | 25.6087 | 25.6105 | 25.7009 | 25.9543 | 25.3781 | 25.6154 |
| 30.0069 | 29.6288 | 30.0565 | 30.1383 | 30.1218 | 30.0387 | 30.0297 |
| 23.8598 | 23.563 | 23.7332 | 23.6266 | 23.6582 | 24.0998 | 23.6587 |
| 24.9195 | 24.8791 | 24.9961 | 25.0008 | 25.1273 | 25.0068 | 24.9947 |
| 23.3825 | 23.698 | 23.4113 | 23.4526 | 23.4837 | 23.8439 | 24.277 |
| 23.4179 | NaN | 23.6363 | NaN | 23.8243 | 23.4424 | 23.2349 |
| 26.9911 | 27.4598 | 27.3588 | 27.1856 | 27.1154 | 27.0957 | 27.0098 |
| 24.0386 | 24.0089 | 24.0179 | 24.4342 | 24.1706 | NaN | NaN |
| 24.8373 | 25.2854 | 25.1963 | 24.9061 | 24.6525 | 24.7173 | 24.9066 |
| 24.4948 | 24.2635 | 24.3368 | 24.4493 | 24.4396 | 24.1051 | NaN |
| 24.1923 | 22.8274 | 22.4927 | 23.2414 | 23.1601 | 23.4453 | 24.2083 |
| 24.2464 | 24.0288 | 24.0657 | 24.1633 | 24.1673 | 23.9534 | 23.9022 |
| 23.4372 | 22.9471 | 23.0611 | 23.3133 | 23.6366 | 23.4089 | 23.2857 |
| 24.0571 | 24.2977 | 24.533 | 24.0208 | 24.3736 | 23.9581 | 24.0248 |
| 26.2468 | 25.3148 | 26.104 | 25.6431 | 26.356 | 26.1435 | 26.0485 |
| 24.3863 | 24.3396 | 24.8347 | 24.2756 | 24.3941 | 24.1954 | 24.4796 |
| 30.5562 | 30.3974 | 30.358 | 30.3152 | 30.3281 | 30.3556 | 30.2187 |
| NaN | 24.7865 | 25.0609 | 24.7638 | 25.1465 | 25.2618 | 25.3626 |
| 24.3693 | 24.1602 | 24.2397 | 24.2152 | 24.1865 | 24.2788 | 24.4532 |
| 32.787 | 33.108 | 32.9059 | 32.6516 | 32.7543 | 32.782 | 32.7451 |
| 25.9724 | 27.2984 | 26.3756 | 26.2134 | NaN | 26.5647 | 26.5878 |
| 25.0172 | 25.5584 | 25.1184 | 25.3583 | NaN | 25.1576 | 25.0697 |
| NaN | NaN | 25.2046 | 24.2127 | NaN | 25.0669 | 25.1443 |
| 32.116 | 32.4527 | 32.3099 | 32.084 | 32.116 | 31.9995 | 31.9314 |
| 25.3172 | 25.5473 | 25.427 | 26.0919 | 25.8405 | 25.492 | 25.3151 |
| 26.1285 | 26.1357 | 25.4433 | 26.1569 | 25.9957 | 25.7354 | 25.8314 |
| 25.2853 | 25.2047 | 25.2979 | 25.4725 | 25.4762 | 24.6851 | NaN |
| 25.7643 | 26.2332 | 25.909 | 25.9881 | 25.9616 | NaN | 25.9385 |
| 26.5584 | 26.7381 | 26.7525 | 26.5058 | 26.58 | 26.2648 | 26.3471 |
| 26.4272 | 26.6411 | 26.5247 | 26.1774 | 26.0044 | 25.9459 | 26.0213 |
| 27.1019 | 26.4071 | 27.0823 | 27.0175 | 26.8603 | 27.1336 | 27.0422 |
| NaN | 25.0715 | 24.779 | 24.3683 | 24.9504 | NaN | NaN |
| 23.6149 | NaN | 23.7069 | 23.4305 | 23.8227 | 23.4747 | 23.6073 |

| | | | | | | |
|---|---|---|---|---|---|---|
| 24.3357 | 24.5223 | 24.2777 | 24.3394 | 24.5578 | 24.584 | 24.3217 |
| 24.1758 | 23.8996 | 24.2957 | 24.1913 | 24.0577 | NaN | NaN |
| 24.3539 | NaN | 24.0139 | 25.0125 | 24.2178 | 24.7899 | 24.3136 |
| 24.6568 | 24.6518 | NaN | NaN | NaN | 24.2561 | 24.0973 |
| 24.0538 | NaN | NaN | 25.7484 | 25.8894 | 24.9954 | 25.1961 |
| NaN | 25.1733 | NaN | 24.9675 | 24.9174 | 25.3338 | NaN |
| 26.5127 | 25.205 | 26.8143 | 26.6607 | 26.6239 | 26.5682 | 26.6625 |
| 25.7293 | 25.6378 | 25.757 | 25.8036 | 25.9006 | 26.1009 | 26.0765 |
| NaN | 24.3836 | 24.1227 | 24.3178 | 24.3477 | NaN | NaN |
| 24.0723 | 24.6189 | 24.5017 | 24.275 | NaN | 24.9097 | 25.1382 |
| 26.4889 | 26.0685 | 27.4233 | 26.5168 | 26.6636 | 26.7421 | 26.7338 |
| 30.0185 | 29.7388 | 29.6859 | 29.6678 | 29.6899 | 29.9837 | 30.0057 |
| 32.0072 | 32.5384 | 32.4908 | 31.6997 | 31.7498 | 31.9828 | 31.9687 |
| 29.4364 | 29.113 | 28.8974 | 29.7329 | 29.7568 | 29.3104 | 29.0798 |
| 27.3566 | 27.5413 | 27.5116 | 26.9869 | 27.1323 | 27.316 | 27.3763 |
| 26.307 | 26.5197 | 26.3407 | 26.0579 | 25.6715 | 25.7584 | 26.3132 |
| 23.0004 | 22.9662 | 23.236 | 23.0996 | NaN | 22.8744 | NaN |
| 31.3683 | 30.5921 | 31.5366 | 31.3992 | 31.3394 | 31.2807 | 31.3638 |
| 24.0612 | 23.4047 | NaN | 23.7556 | NaN | 24.0506 | 24.0758 |
| NaN | NaN | NaN | NaN | 23.2359 | 23.5698 | 23.4089 |
| NaN | NaN | 23.318 | 23.6821 | 23.4682 | NaN | NaN |
| 25.1897 | NaN | NaN | 25.8627 | 25.54 | 25.1476 | 25.556 |
| 25.211 | NaN | NaN | 25.1353 | 25.0419 | 25.2413 | 25.1768 |
| 24.6228 | NaN | 24.5255 | 24.9815 | 24.4447 | 24.4245 | 24.4671 |
| 23.9558 | 23.995 | NaN | 24.1444 | NaN | NaN | 23.6876 |
| 25.2508 | 25.269 | 25.2562 | 25.3658 | 25.2727 | 25.667 | 25.0834 |
| 30.3822 | 30.5491 | 30.5096 | 30.504 | 30.4009 | 30.2935 | 30.1928 |
| NaN | 24.529 | NaN | NaN | 24.2173 | 24.711 | NaN |
| 25.2797 | 25.4256 | 25.7015 | 25.4982 | 25.6436 | 25.3318 | 25.5671 |
| NaN | NaN | 23.9183 | NaN | 24.3449 | NaN | NaN |
| 31.2024 | 31.3354 | 31.2443 | 30.8977 | 31.0788 | 31.0663 | 31.0561 |
| 24.5283 | 23.9602 | 24.6951 | 24.8819 | 25.1129 | 24.4949 | NaN |
| NaN | NaN | NaN | 26.2996 | NaN | NaN | 26.4879 |
| 24.9638 | NaN | NaN | 24.7136 | 24.8887 | NaN | 24.4144 |
| 24.6985 | 25.2341 | NaN | 24.8291 | 25.0041 | 24.7068 | 24.7022 |
| 31.0109 | 31.3748 | 31.2517 | 31.0545 | 31.1431 | 31.2135 | 31.1728 |
| 25.3943 | 25.4204 | 25.3189 | 25 | 25.082 | 25.201 | 25.1631 |
| 22.0895 | NaN | NaN | 22.0377 | 22.2277 | 21.6748 | 21.9704 |
| 23.958 | NaN | 24.1497 | 24.2742 | NaN | 23.7033 | 24.1468 |
| 24.9645 | 24.853 | NaN | 25.0469 | 25.0457 | 24.9319 | 25.2379 |
| 25.0357 | 24.8888 | NaN | 25.0463 | 24.9758 | NaN | 24.9083 |
| 24.6206 | 25.389 | 25.1461 | 24.9409 | 24.9007 | 25.3382 | 25.0437 |
| 23.9473 | 23.7614 | 23.9008 | 25.3411 | 24.0623 | NaN | 24.1796 |
| 25.2509 | 25.1571 | 25.0973 | 24.9574 | 25.1822 | 24.869 | 24.898 |
| 24.9576 | 25.3654 | 25.2179 | 24.9558 | 25.0333 | 25.0044 | 24.9872 |
| 25.8796 | 25.9149 | 25.5408 | 25.6996 | 25.7819 | 25.7023 | 25.8414 |
| 23.8109 | NaN | 24.2588 | 24.1862 | 24.2731 | 24.0049 | 23.27 |
| 25.8877 | 26.1447 | 25.5986 | 25.8708 | 25.8436 | 25.9443 | 25.9876 |

| | | | | | | |
|---|---|---|---|---|---|---|
| 26.4051 | 26.1995 | 26.5544 | 26.6442 | 26.4651 | 26.5232 | 26.2463 |
| 26.7985 | 26.5184 | 26.4648 | 26.7844 | 26.6975 | 26.8488 | 26.6921 |
| NaN | NaN | NaN | 24.3001 | 24.2407 | NaN | NaN |
| NaN | NaN | NaN | NaN | NaN | 25.0772 | 25.3481 |
| 24.0196 | 23.9497 | NaN | 23.8239 | 24.185 | 24.1435 | 24.1996 |
| 28.8154 | 29.1682 | 29.0346 | 28.6954 | 28.7607 | 28.5667 | 28.6364 |
| 23.5188 | 23.9788 | NaN | 23.8587 | 24.0311 | 23.8568 | 24.1243 |
| 29.8537 | 30.3765 | 30.2608 | 29.5406 | 29.8576 | 29.857 | 30.0292 |
| 29.0809 | 29.299 | 29.198 | 29.126 | 29.1421 | 28.9832 | 28.9762 |
| 26.9487 | 27.048 | 26.838 | 27.1782 | 27.0839 | 27.5378 | 27.7897 |
| 29.4909 | 30.1625 | 29.9694 | 29.9478 | 29.9475 | 29.673 | 29.7679 |
| 28.6908 | 28.8632 | 28.8403 | 28.7674 | 28.7451 | 28.494 | 28.4831 |
| 23.8432 | 23.6904 | 23.4815 | 23.7422 | 23.5937 | 23.6594 | 23.7951 |
| 27.4555 | 27.6852 | 27.5692 | 27.1548 | 27.316 | 27.4095 | 27.5683 |
| 25.8445 | 25.8381 | 25.9016 | 25.9341 | 25.6895 | 25.6399 | 25.5059 |
| 32.0658 | 32.4231 | 32.3431 | 32.1251 | 32.135 | 32.1373 | 32.1408 |
| 26.0329 | 25.8384 | 26.1787 | 25.7892 | 25.9067 | 25.624 | 25.3726 |
| 24.0158 | NaN | 24.2066 | 24.2626 | 24.2815 | NaN | 24.1203 |
| 24.2758 | 24.6467 | 24.7421 | 24.5074 | 24.3662 | 24.9865 | 24.846 |
| 29.4858 | 29.2034 | 29.5864 | 29.6528 | 29.5731 | 29.5493 | 29.3887 |
| NaN | NaN | 23.8026 | 25.1717 | 24.4363 | 24.6191 | 23.9747 |
| 29.6211 | 29.7064 | 29.8395 | 29.7971 | 29.9292 | 29.7498 | 29.7956 |
| 23.9708 | NaN | 23.8913 | 23.873 | 23.8362 | 24.6361 | 23.982 |
| 25.2216 | 25.8694 | 25.0874 | 25.5059 | 25.18 | 25.1359 | NaN |
| 25.3602 | 25.5758 | NaN | NaN | NaN | 25.969 | 25.7656 |
| 24.9937 | 24.9272 | 25.1806 | 24.8886 | 24.9478 | 25.2118 | 25.0158 |
| 23.9016 | 24.3567 | 23.875 | 24.2497 | NaN | 24.1103 | 24.1488 |
| 23.9692 | NaN | NaN | 23.9421 | 24.0859 | 24.323 | 24.0382 |
| 28.2489 | 28.2075 | 28.1774 | 27.9999 | 28.0749 | 27.899 | 28.0005 |
| 24.7558 | NaN | NaN | 25.1442 | 25.0099 | 24.9625 | 25.8107 |
| 23.6056 | NaN | 23.1908 | 24.0129 | 23.9399 | NaN | NaN |
| 27.1329 | 27.2305 | 27.2817 | 27.2714 | 27.3218 | 26.822 | 26.8876 |
| 25.3974 | 25.648 | 25.4105 | 25.1087 | 25.3515 | 25.1103 | 25.5096 |
| 25.218 | 25.4546 | 25.2988 | 25.2988 | 25.2482 | 25.1203 | 25.1314 |
| 27.371 | 27.3356 | 27.2116 | 27.3785 | 27.225 | 27.7389 | 27.7165 |
| NaN | 25.569 | 25.0518 | NaN | 25.2423 | NaN | NaN |
| 27.7142 | 27.4872 | 27.6167 | 27.5879 | 27.3889 | 26.7321 | 26.4916 |
| 24.2651 | 23.9507 | NaN | 24.2625 | 24.9227 | 24.3374 | 24.1987 |
| 24.968 | 25.4395 | 25.057 | 24.4614 | 24.5098 | 24.6315 | 24.7763 |
| 24.5564 | 24.7292 | 24.6603 | 24.444 | 24.3433 | 24.3679 | 24.467 |
| 32.8876 | 33.151 | 33.0214 | 32.8268 | 32.8221 | 32.7339 | 32.8043 |
| 25.7994 | 25.9121 | 25.9982 | 25.4936 | 25.4455 | 25.7446 | 25.6287 |
| 26.187 | 25.7471 | 25.6447 | 26.1853 | 25.9716 | 26.0044 | 25.7999 |
| 24.2491 | 25.7026 | 25.7414 | 25.1322 | 25.2983 | 24.9978 | 24.9281 |
| 28.3374 | 28.5841 | 28.563 | 28.12 | 28.1026 | 28.1494 | 28.1842 |
| 27.7971 | 27.567 | 27.874 | 27.9558 | 27.698 | 27.7846 | 27.8782 |
| 29.6244 | 29.3091 | 29.8386 | 29.598 | 29.7474 | 29.773 | 29.9905 |
| NaN | 25.3057 | 25.1839 | NaN | 25.3407 | 25.4268 | NaN |

| | | | | | | |
|---|---|---|---|---|---|---|
| NaN | NaN | 25.1443 | 25.0877 | NaN | NaN | 25.5469 |
| 26.8943 | 26.5052 | 26.5932 | NaN | 26.8382 | 26.5455 | 26.9549 |
| 25.1543 | 25.0996 | 25.2688 | 25.1753 | 25.2101 | 24.9762 | 24.8502 |
| 24.0192 | 23.7785 | 24.2802 | 24.497 | 24.5918 | 24.0443 | 24.0089 |
| 25.5825 | NaN | NaN | 25.6065 | 25.7015 | 25.3901 | 25.242 |
| 24.9879 | 24.9593 | 24.768 | 24.8993 | 25.1026 | 25.1135 | 25.3485 |
| 24.787 | NaN | NaN | NaN | NaN | 24.5996 | 24.6344 |
| 24.3955 | NaN | NaN | 24.4563 | NaN | NaN | NaN |
| 25.775 | NaN | 25.3208 | 25.8252 | 25.9172 | 25.197 | 25.1436 |
| 24.7927 | NaN | NaN | NaN | NaN | 25.1674 | NaN |
| 23.8637 | 24.0527 | 24.1274 | 24.1703 | 23.9616 | 23.9957 | 23.9221 |
| 23.9781 | NaN | NaN | 24.1668 | 24.0783 | NaN | 23.7389 |
| NaN | 23.9977 | 24.1725 | NaN | NaN | NaN | NaN |
| NaN | NaN | 24.6189 | 24.1163 | NaN | 24.5207 | 25.0446 |
| 24.3341 | NaN | 24.3192 | NaN | 24.7595 | NaN | 24.424 |
| 26.6996 | 26.5366 | 26.8915 | 26.8956 | 27.3359 | 26.4686 | 26.4006 |
| 25.8566 | 26.4398 | 26.2303 | 26.5274 | 26.253 | 26.0015 | 26.0882 |
| 24.896 | 25.232 | 25.2535 | 25.2433 | 25.1894 | 25.342 | 25.3594 |
| 27.6802 | 27.8538 | 27.8746 | 28.0863 | 27.8964 | 27.9518 | 27.9218 |
| 29.5202 | 29.6968 | 29.6668 | 29.7789 | 29.7155 | 29.503 | 29.5239 |
| 29.5689 | 29.7429 | 29.7526 | 29.7263 | 29.7157 | 29.4914 | 29.5312 |
| 32.0206 | 32.1108 | 32.0511 | 32.1492 | 32.0828 | 32 | 31.9781 |
| 25.2018 | 24.9264 | 24.9408 | 25.0025 | 25.0311 | 24.9025 | 24.9199 |
| 28.0107 | 27.9013 | 27.7042 | 27.9165 | 27.8322 | 27.971 | 27.9439 |
| 26.1276 | 26.1402 | 26.1142 | 25.9753 | 25.752 | 25.9588 | 26.1025 |
| 27.2515 | 27.0912 | 26.9804 | 27.1534 | 27.1928 | 27.1121 | 27.3191 |
| 27.4934 | 27.3158 | 27.8008 | 27.2502 | 27.248 | 27.2598 | 27.4612 |
| 26.0689 | 25.8624 | 25.8145 | 25.5584 | 25.4087 | 25.1548 | 25.4641 |
| 31.0413 | 30.0588 | 30.9145 | 30.8919 | 30.8815 | 30.7079 | 30.6767 |
| NaN | NaN | 24.2232 | 24.6096 | 24.8389 | NaN | NaN |
| NaN | NaN | NaN | NaN | NaN | NaN | NaN |
| 30.3703 | 30.4542 | 30.5311 | 30.2262 | 30.3446 | 30.4637 | 30.4437 |
| NaN | NaN | NaN | 24.0565 | NaN | NaN | NaN |
| 24.1822 | NaN | 24.0694 | NaN | NaN | 24.1681 | 24.1743 |
| 27.4998 | 27.8872 | 27.5842 | 27.7708 | 27.7922 | 27.4663 | 27.5435 |
| 25.2557 | 25.3373 | 25.1722 | NaN | NaN | NaN | 25.0048 |
| 24.638 | 24.6323 | 24.2605 | 24.3748 | 24.428 | 24.4606 | 23.8925 |
| NaN | 23.0611 | NaN | 22.9111 | NaN | 24.0454 | NaN |
| NaN | 23.7362 | NaN | NaN | 23.9578 | 23.808 | 24.4957 |
| 24.4262 | 24.556 | 24.3378 | 24.5683 | 24.8195 | 24.8843 | 24.5031 |
| NaN | NaN | 24.8516 | 24.5091 | NaN | NaN | NaN |
| 24.1561 | 24.4775 | 24.367 | 24.2393 | 24.2086 | 24.5255 | 24.1647 |
| NaN | 25.9662 | 25.5907 | 25.8162 | 25.8226 | 25.5958 | 25.7064 |
| 25.6436 | NaN | NaN | NaN | 25.1013 | NaN | NaN |
| 25.925 | NaN | NaN | 25.7008 | 25.859 | NaN | NaN |
| NaN | NaN | NaN | NaN | NaN | 24.0096 | NaN |
| 25.1897 | 25.5363 | 25.2725 | 25.204 | 25.4162 | 25.149 | 25.2172 |
| 23.3303 | NaN | 22.8102 | NaN | 22.7599 | 23.2638 | NaN |

| | | | | | | |
|---|---|---|---|---|---|---|
| 27.2869 | 27.5935 | 27.5189 | 26.8156 | 26.9648 | 27.0372 | 27.2906 |
| NaN | NaN | NaN | 25.9717 | NaN | NaN | NaN |
| 24.0614 | NaN | NaN | 24.1868 | 24.3959 | NaN | 23.8162 |
| 28.2128 | 28.382 | 28.2252 | 28.2232 | 28.1363 | 28.2976 | 28.2859 |
| 25.165 | NaN | 24.9792 | NaN | 25.3493 | NaN | NaN |
| NaN | NaN | NaN | NaN | NaN | 21.8313 | NaN |
| NaN | NaN | NaN | 26.3127 | 26.0619 | NaN | NaN |
| 23.7192 | 23.6163 | 23.5898 | NaN | 23.6766 | NaN | 23.957 |
| 23.4107 | 23.023 | 23.1955 | 23.2002 | 23.1573 | NaN | 23.0773 |
| 29.8231 | 29.5742 | 29.3912 | 29.5017 | 29.5578 | 29.7653 | 29.7075 |
| 24.1104 | 24.6159 | 24.2488 | 24.0724 | 24.1086 | 23.9499 | 24.022 |
| 26.386 | 26.8602 | 26.8781 | 26.529 | 26.5559 | 26.4303 | 26.5538 |
| 32.1222 | 31.9552 | 31.8958 | 31.9353 | 32.0351 | 32.1711 | 32.222 |
| 27.219 | 27.5215 | 27.2769 | 26.8529 | NaN | 27.0823 | NaN |
| 23.469 | NaN | NaN | 23.603 | 23.2902 | NaN | 23.5973 |
| 24.998 | 24.4978 | 24.3166 | 24.8893 | 24.7301 | NaN | 24.4646 |
| 28.0691 | 28.1208 | 27.8748 | 27.6969 | 28.0196 | 28.1293 | 28.0112 |
| 28.7556 | 28.1243 | 28.2069 | 28.8069 | 28.7035 | 28.3311 | 28.3757 |
| 24.1565 | NaN | NaN | 24.0183 | 24.0203 | 24.1532 | 24.1281 |
| 24.6669 | NaN | 24.3534 | 24.4648 | 24.2505 | 24.2859 | 24.6049 |
| 28.479 | 28.3214 | 28.4038 | 28.2722 | 28.4749 | 28.354 | 28.4475 |
| 29.2883 | 28.262 | 29.2228 | 29.4346 | 29.3354 | 29.1273 | 28.9493 |
| 23.9826 | NaN | 23.9299 | 24.8161 | 24.723 | 24.3188 | NaN |
| 25.1129 | 24.4867 | 24.5881 | NaN | 24.6692 | 25.2933 | 25.4859 |
| NaN | 26.0017 | 25.6044 | NaN | NaN | NaN | 25.8943 |
| 25.2825 | 25.5976 | 25.2784 | 25.1966 | 25.255 | 25.1987 | 25.607 |
| 24.35 | 24.6039 | 24.3292 | 24.295 | 24.2213 | 24.4087 | 24.3437 |
| 27.0618 | 27.4689 | 27.2185 | 27.1846 | 27.3298 | 27.1902 | 27.2737 |
| 24.4524 | NaN | NaN | 23.8144 | NaN | 24.0583 | NaN |
| 25.8875 | 26.0475 | 25.8879 | 25.2603 | 25.1107 | 25.7472 | 25.4945 |
| 23.9875 | 24.2917 | 23.9465 | 24.0705 | 24.0215 | 24.5302 | 24.5869 |
| 25.4955 | 25.7957 | 25.6733 | 25.0971 | 25.3319 | 25.1312 | 25.1923 |
| 26.1806 | NaN | NaN | 25.7651 | 25.7274 | NaN | NaN |
| 27.0249 | NaN | 27.1254 | 26.3743 | NaN | NaN | 26.371 |
| NaN | 25.1026 | 25.0636 | 25.0427 | 24.8617 | 24.1793 | 24.4416 |
| 24.3606 | NaN | NaN | 24.0798 | NaN | NaN | NaN |
| 24.5909 | NaN | NaN | NaN | 24.7894 | 24.7868 | 25.2606 |
| 25.7759 | 25.9032 | 25.949 | 25.5414 | 25.5191 | 25.2913 | 26.0062 |
| NaN | 23.3048 | NaN | 24.2244 | NaN | 23.9693 | NaN |
| 27.0882 | 26.6338 | 26.325 | 26.8522 | 26.711 | 26.867 | 26.9813 |
| 27.6897 | 27.6457 | 27.6012 | 27.6296 | 27.6227 | 27.5826 | 27.6281 |
| 23.8618 | 24.3649 | NaN | 24.1246 | 24.1623 | 24.0804 | 24.0567 |
| 28.2638 | 28.4292 | 28.5459 | 28.3454 | 28.367 | 28.3379 | 28.5084 |
| NaN | NaN | NaN | 24.9898 | 24.9802 | NaN | 24.9275 |
| 25.1543 | 24.7078 | 24.7592 | 25.6763 | 25.4422 | NaN | 24.7153 |
| NaN | NaN | NaN | NaN | NaN | NaN | 25.9993 |
| 24.5225 | 24.2879 | NaN | NaN | NaN | NaN | NaN |
| 23.6394 | NaN | NaN | NaN | 23.6404 | 23.7716 | NaN |

| | | | | | | |
|---|---|---|---|---|---|---|
| NaN | 23.341 | NaN | NaN | NaN | NaN | 23.1436 |
| 26.0744 | 28.5655 | 26.6464 | 26.6867 | 28.4296 | 26.1113 | 26.1879 |
| NaN | NaN | NaN | 25.8425 | 25.9159 | 25.7645 | NaN |
| 26.2429 | 26.3541 | 26.3213 | 26.2586 | 26.1588 | 26.121 | 26.0738 |
| NaN | 24.8352 | 24.8636 | NaN | 24.7522 | NaN | NaN |
| 24.6508 | 24.8409 | 25.0094 | 24.8089 | 25.0134 | NaN | NaN |
| 24.8732 | 25.5414 | 25.2514 | 25.568 | 25.3409 | 25.6711 | 25.6422 |
| 25.8471 | 26.0845 | 26.1313 | 25.8905 | 25.9528 | 26.0567 | 26.5045 |
| 24.7433 | 25.8688 | 25.0746 | 25.6114 | 25.5708 | 25.3125 | 25.3676 |
| 23.2383 | 23.9336 | 23.508 | 23.857 | 24.0354 | 23.5751 | NaN |
| 28.7356 | 29.043 | 28.9491 | 28.7538 | 28.7608 | 28.4572 | 28.4406 |
| 24.532 | 23.9229 | 24.1691 | 23.8631 | 24.2697 | 23.9068 | 24.2402 |
| 36.4146 | 36.4483 | 36.4412 | 36.3858 | 36.4014 | 36.327 | 36.4264 |
| NaN | 24.5422 | 24.6199 | NaN | NaN | NaN | NaN |
| 25.1739 | 25.586 | NaN | NaN | 25.2713 | NaN | 25.2108 |
| 24.257 | 24.287 | 24.2808 | NaN | 24.3552 | NaN | NaN |
| 27.5307 | 28.2076 | 27.5313 | 28.1415 | 26.8526 | 27.7906 | 27.7819 |
| 27.2634 | 27.781 | 27.3122 | 27.4156 | 27.3348 | NaN | 27.3955 |
| 31.3447 | 31.5363 | 31.4622 | 31.5252 | 31.4562 | 31.3878 | 31.3125 |
| 25.1195 | 24.7346 | 24.7022 | 24.9104 | 24.8232 | 24.944 | 24.8201 |
| 26.3454 | 26.9885 | 26.5244 | 26.8443 | 27.1849 | 26.932 | NaN |
| 25.2497 | NaN | 25.0519 | 25.6252 | 25.5741 | 25.0048 | 24.9013 |
| 25.202 | 24.8655 | 25.1935 | 25.7125 | 25.2228 | 25.3394 | 25.1794 |
| 26.0992 | NaN | NaN | 26.2419 | 26.1076 | NaN | 26.0058 |
| 23.5173 | 23.5525 | 23.4162 | 23.5746 | NaN | 24.2963 | 24.2874 |
| 24.5608 | 24.4928 | 24.5462 | 24.668 | 24.6439 | 24.0816 | 24.1633 |
| 25.8718 | 25.5465 | 25.6432 | 25.4035 | 25.4584 | 25.3035 | 25.4731 |
| NaN | 24.0002 | 24.043 | 24.309 | 24.2736 | 24.011 | 24.407 |
| 27.7788 | 29.0176 | 28.9031 | 28.695 | 28.7043 | 28.6914 | 28.6383 |
| 25.8798 | 25.9172 | 25.8665 | 25.4425 | 25.4327 | 25.9729 | 25.9857 |
| 25.2861 | 25.4558 | 25.7001 | 25.6353 | 25.3786 | 25.6941 | 25.8337 |
| 34.2967 | 34.327 | 34.2175 | 34.1995 | 34.187 | 34.2887 | 34.362 |
| 25.6296 | 25.8042 | 26.0059 | 25.6692 | 25.5563 | 25.4732 | 25.3296 |
| 26.1256 | 27.1571 | 27.1155 | 26.093 | 26.282 | NaN | NaN |
| 29.1367 | 29.6055 | 29.4479 | 29.3254 | 29.2857 | 29.3664 | 29.164 |
| 24.2668 | 24.1934 | 24.0586 | NaN | NaN | NaN | 24.0385 |
| NaN | NaN | 24.5949 | 24.0235 | 24.2777 | 24.0616 | 24.2339 |
| 27.4572 | 27.7301 | 27.6156 | 27.353 | 27.3997 | NaN | 27.4659 |
| 24.2044 | 24.4765 | 24.4788 | 25.1282 | NaN | 24.2049 | 24.0623 |
| NaN | 24.3079 | NaN | NaN | NaN | NaN | 24.8552 |
| 31.6982 | 31.73 | 31.7134 | 31.2521 | 31.2305 | 31.4461 | 31.4983 |
| 25.1622 | 25.6337 | NaN | 25.7227 | 25.4078 | 25.9054 | 25.98 |
| 23.8807 | 24.0855 | 24.4084 | NaN | 24.1651 | 24.7835 | 24.4065 |
| NaN | NaN | NaN | NaN | NaN | 23.7791 | 23.5933 |
| 29.5781 | 29.9358 | 29.8179 | 29.5233 | 29.4656 | 29.7438 | 29.8297 |
| 24.2575 | 24.3063 | 24.0009 | 24.4989 | NaN | NaN | NaN |
| 30.122 | 29.3922 | 29.3115 | 30.1288 | 30.181 | 29.6615 | 29.639 |
| 23.9375 | 23.843 | 24.0593 | NaN | 23.8666 | 23.9364 | 24.0092 |

| | | | | | | |
|---|---|---|---|---|---|---|
| NaN | 24.384 | 24.0478 | 24.1465 | 24.1995 | 23.89 | 23.9941 |
| 23.2224 | 23.0515 | 23.0219 | 23.7458 | 23.7505 | 23.6931 | 23.9048 |
| 24.8672 | NaN | 25.0211 | 25.1656 | 25.0183 | 24.9441 | 24.987 |
| NaN | NaN | NaN | NaN | 24.3122 | 24.7051 | 24.8342 |
| 27.0081 | NaN | NaN | 26.888 | 26.838 | 27.4611 | 27.558 |
| 23.8831 | 24.4649 | 24.2682 | 24.0972 | 24.3279 | 24.5742 | 24.7366 |
| NaN | 25.2146 | 26.25 | NaN | 26.7773 | 26.0122 | 26.6023 |
| 25.4965 | 25.1983 | 25.3299 | 25.0274 | 25.4147 | 25.1247 | 25.4912 |
| NaN | NaN | 25.1845 | 24.8892 | NaN | 24.8594 | 24.5646 |
| 24.2964 | 24.4017 | 24.4571 | 24.2984 | 24.3357 | 24.0652 | NaN |
| NaN | 23.9674 | 24.3672 | 23.8912 | NaN | NaN | 23.8927 |
| NaN | NaN | NaN | 26.6223 | NaN | 27.1736 | NaN |
| NaN | NaN | NaN | 24.3059 | 24.256 | NaN | 23.8687 |
| NaN | NaN | NaN | NaN | 24.7707 | 25.1971 | 25.3288 |
| 24.4672 | 24.3365 | 24.4022 | 23.9984 | 24.0432 | 24.251 | 24.3384 |
| 26.8762 | 29.0333 | NaN | 29.3639 | 28.9081 | 28.7962 | 27.433 |
| NaN | NaN | NaN | NaN | 23.1731 | 23.2 | 23.2952 |
| NaN | NaN | NaN | NaN | 24.175 | 24.0088 | 24.1764 |
| 25.911 | 26.4102 | 26.4275 | 25.7497 | 25.5998 | 25.9045 | 25.9336 |
| 26.0689 | 26.6663 | 26.2553 | 26.3913 | 26.1931 | 26.3131 | 26.1629 |
| NaN | 25.5754 | NaN | 25.3995 | NaN | 25.5624 | 25.8115 |
| 26.2625 | 25.9988 | 25.9845 | 26.5597 | 26.2597 | 26.3547 | NaN |
| 25.6223 | 25.6629 | 26.1296 | NaN | NaN | 25.8705 | 25.6808 |
| NaN | NaN | 24.7114 | 24.67 | NaN | 24.7249 | NaN |
| 26.5902 | 26.6542 | 26.5459 | 26.8259 | 26.9072 | 26.3362 | 26.6382 |
| 27.242 | 27.0485 | 26.8875 | 26.8388 | 27.5846 | 26.7112 | 26.6792 |
| 27.5033 | 27.4899 | NaN | 27.6322 | 27.7608 | 27.5025 | 27.2619 |
| 25.6129 | 24.8139 | 25.1558 | 25.7323 | 25.7584 | 25.4498 | 25.5903 |
| 26.1486 | NaN | NaN | 26.3568 | 26.4055 | NaN | 26.5187 |
| 27.1331 | 26.8887 | 26.9727 | 27.1397 | 27.2328 | 27.1719 | 27.1289 |
| 23.4969 | 23.3119 | 23.8294 | 23.8272 | NaN | NaN | NaN |
| 24.4649 | NaN | NaN | 24.1758 | 24.5509 | 24.2979 | 24.4466 |
| 25.2738 | 24.7286 | 24.627 | 24.8587 | 25.0901 | 25.3242 | 25.0837 |
| 23.7292 | NaN | NaN | NaN | 23.8389 | 23.634 | NaN |
| 32.0367 | 32.0543 | 31.9967 | 31.8985 | 31.9124 | 31.9925 | 31.9732 |
| 24.1638 | 24.5862 | 24.631 | NaN | 24.3354 | NaN | NaN |
| NaN | NaN | 24.5074 | 24.2521 | 24.0905 | NaN | NaN |
| 25.3784 | 25.9791 | NaN | 25.2704 | NaN | 25.4761 | 25.5434 |
| 24.5621 | NaN | NaN | 24.6337 | 24.4432 | 24.8446 | 24.6093 |
| NaN | 27.3993 | NaN | 26.5432 | NaN | NaN | 26.7158 |
| NaN | NaN | NaN | 23.5984 | NaN | NaN | 23.6855 |
| 26.5144 | 26.7793 | 26.7331 | 26.743 | 26.8284 | 26.1073 | 26.1898 |
| 26.669 | 27.0534 | 26.6887 | 26.6818 | 26.7935 | 26.562 | 26.2255 |
| 32.4079 | 32.3567 | 32.2247 | 32.4002 | 32.3095 | 32.102 | 32.0178 |
| 27.3808 | 27.1034 | 27.4352 | 27.3792 | 27.6346 | 26.8152 | 26.6992 |
| 30.3436 | 30.3369 | 30.3268 | 30.2105 | 30.2357 | 30.6485 | 30.5489 |
| NaN | NaN | NaN | 24.7138 | 24.6632 | 24.1677 | NaN |
| 32.4665 | 32.6106 | 32.6099 | 32.5156 | 32.5471 | 32.3442 | 32.4025 |

| | | | | | | |
|---|---|---|---|---|---|---|
| 24.2022 | 23.8761 | 24.2233 | 24.8022 | 24.7946 | 24.7617 | NaN |
| 24.2956 | 24.598 | 24.4772 | 24.675 | NaN | 24.6493 | 24.6214 |
| 27.4689 | 28.0001 | 27.9148 | 28.3761 | 27.7915 | NaN | 27.5622 |
| 24.2684 | 24.2536 | NaN | 24.0046 | 24.0672 | 23.7009 | 23.9002 |
| NaN | 24.1294 | 24.0904 | NaN | 23.9149 | NaN | 23.8836 |
| 24.3099 | NaN | 24.4548 | 24.4589 | 25.0359 | 24.2666 | NaN |
| NaN | 24.2193 | 24.8993 | NaN | NaN | NaN | 23.7726 |
| 24.3671 | 24.0914 | 23.9399 | 24.6334 | 24.5077 | 24.2624 | 24.2095 |
| 23.8275 | 23.7528 | 23.7951 | 23.5975 | NaN | NaN | 23.483 |
| NaN | 25.5881 | NaN | 25.5937 | 25.7526 | NaN | 24.7676 |
| 28.4632 | 28.9015 | 28.7562 | 28.4552 | 28.2999 | 28.2474 | 28.3469 |
| 27.4821 | 27.9012 | 27.99 | 27.3753 | 27.2547 | 26.9739 | 27.1295 |
| 23.8452 | 24.5957 | 24.7153 | NaN | 24.5765 | 24.5791 | 24.2482 |
| NaN | 23.3906 | 23.9638 | NaN | 23.8354 | NaN | NaN |
| 24.173 | NaN | 24.2636 | 24.4741 | 24.3434 | 24.2577 | 24.2648 |
| 24.7773 | 24.8977 | 24.8658 | 24.9423 | 24.9082 | 24.5582 | 24.4735 |
| 29.4654 | 29.1087 | 29.182 | 29.2028 | 29.3249 | 29.5603 | 29.4324 |
| 24.6814 | 24.418 | 24.7898 | 24.8654 | 25.1685 | 25.2267 | 25.1802 |
| 25.2998 | 24.603 | 25.0907 | 25.5884 | 25.2762 | 26.0375 | 26.0615 |
| 24.276 | 24.7978 | 24.5015 | 24.6081 | 24.5552 | NaN | NaN |
| NaN | 26.3717 | 26.2377 | 26.2571 | 25.9317 | NaN | 26.2064 |
| 25.0931 | NaN | NaN | NaN | 25.9052 | 24.9979 | 24.771 |
| NaN | 24.298 | 24.0167 | 24.2628 | NaN | 24.284 | 24.1169 |
| 23.2567 | 23.2234 | NaN | 23.9136 | 23.7526 | 23.4158 | 23.6235 |
| 29.4608 | 29.5845 | 29.4575 | 29.5491 | 29.6343 | 29.2367 | 29.3244 |
| 25.5875 | 24.465 | 24.3464 | 25.186 | 25.5155 | 24.6552 | 24.5425 |
| 23.3741 | NaN | 23.5339 | 23.4359 | 23.6413 | 23.0946 | 23.3844 |
| 29.4296 | 29.2179 | 29.3168 | 29.3042 | 29.2582 | 29.3351 | 29.3417 |
| 24.0667 | NaN | 24.1013 | 24.082 | NaN | 23.9334 | 24.1779 |
| NaN | 24.4197 | NaN | 23.9108 | NaN | 23.9736 | 24.0528 |
| 23.8876 | 23.992 | 24.2126 | 23.8437 | NaN | 23.9012 | 24.1651 |
| 23.9508 | 22.1681 | 23.3354 | 23.7119 | 23.4073 | 24.1253 | 23.9405 |
| NaN | NaN | 24.7911 | 24.9798 | NaN | 25.0211 | NaN |
| 25.6155 | 26.4091 | 25.8456 | 26.071 | 26.2131 | 26.0668 | 26.0191 |
| 24.3356 | NaN | 24.2928 | 24.2833 | 24.2247 | 24.5074 | 24.3641 |
| 29.2489 | 29.3754 | 29.2943 | 28.9019 | 28.915 | 29.2435 | 29.2602 |
