## Supplementary Table S1B for "TDP-43 pathology links innate and adaptive immunity in amyotrophic lateral sclerosis"

| Sig(FDR0.0 | -LOG(P-va | log2(TDP/c | TDP/ctrl (F | TDP/ctrl (p | Protein IDs | Majority p | Protein na | Gene name | id |
|---|---|---|---|---|---|---|---|---|---|
| | 0.70756 | 0.285647 | | | Q15149;H( | Q15149;H( | Plectin | PLEC | 1513 |
| | 1.888606 | 0.236263 | | | P35579;Q5 | P35579 | Myosin-9 | MYH9 | 1117 |
| | 3.098455 | 0.198285 | | | Q9Y490;Q9 | Q9Y490 | Talin-1 | TLN1 | 1841 |
| | 3.869571 | 0.378014 | | | A0A087W\ | A0A087W\ | Clathrin he | CLTC | 23 |
| | 0.517772 | 0.13716 | | | P08670;B0 | P08670;B0 | Vimentin | VIM | 863 |
| | 1.194058 | 0.099802 | | | P46940;A0 | P46940;A0 | Ras GTPase | IQGAP1 | 1179 |
| | 4.345963 | -0.60075 | | | Q07954;Q( | Q07954 | Prolow-der | LRP1 | 1434 |
| | 2.340628 | -0.74688 | | | P02751;H0 | P02751 | Fibronectin | FN1 | 788 |
| | 2.243479 | 0.270751 | | | P26038;V9 | P26038 | Moesin | MSN | 1038 |
| | 2.094645 | -0.58272 | | | P01023;H0 | P01023 | Alpha-2-m | A2M | 773 |
| | 1.668276 | 0.233944 | | | P13796;Q5 | P13796 | Plastin-2 | LCP1 | 933 |
| | 1.380011 | -0.37969 | | | P22897 | P22897 | Macrophag | MRC1 | 1019 |
| | 0.524136 | -0.19226 | | | Q9P2E9;A( | Q9P2E9;A( | Ribosome- | RRBP1 | 1787 |
| | 3.898833 | -0.5503 | | | P02786;G3 | P02786;G3 | Transferrin | TFRC | 790 |
| | 0.173997 | 0.042165 | | | P55072;C9 | P55072 | Transitiona | VCP | 1271 |
| | 2.481325 | 0.378081 | | | P09960;B4 | P09960 | Leukotrien | LTA4H | 883 |
| | 2.036461 | -0.66845 | | | P14780 | P14780 | Matrix met | MMP9 | 943 |
| | 0.503379 | 0.15238 | | | P31948;F5 | P31948 | Stress-indu | STIP1 | 1097 |
| | 4.429037 | 0.968724 | | | Q14204;A( | Q14204 | Cytoplasm | DYNC1H1 | 1493 |
| | 2.03284 | -0.53718 | | | Q02818;H7 | Q02818;H7 | Nucleobind | NUCB1 | 1418 |
| | 1.165945 | 0.218459 | | | P13639;K7 | P13639 | Elongation | EEF2 | 928 |
| | 0.538789 | 0.206549 | | | P23381;G3 | P23381 | Tryptophar | WARS | 1022 |
| | 3.735813 | 0.164209 | | | O43707;F5 | O43707 | Alpha-actir | ACTN4 | 686 |
| | 3.819528 | -0.76413 | | | P11717;S4 | P11717 | Cation-ind | IGF2R | 911 |
| | 0.851155 | 0.300719 | | | V9GYY3;A( | V9GYY3;A( | C-1-tetrahy | MTHFD1 | 173 |
| | 0.13163 | 0.033352 | | | Q09666;E9 | Q09666 | Neuroblast | AHNAK | 1444 |
| | 1.836677 | 0.472188 | | | P21980;A2 | P21980 | Protein-glu | TGM2 | 1011 |
| | 5.881907 | 0.400616 | | | P12814;H9 | P12814;H9 | Alpha-actir | ACTN1 | 917 |
| | 2.212734 | 0.276456 | | | P18206;A0 | P18206 | Vinculin | VCL | 987 |
| | 0.622055 | 0.261645 | | | Q01518;Q5 | Q01518 | Adenylyl cy | CAP1 | 1413 |
| | 1.125992 | 0.352092 | | | P52790;H0 | P52790 | Hexokinase | HK3 | 1246 |
| | 1.837591 | 0.345469 | | | P42224;A0 | P42224;A0 | Signal tran | STAT1 | 1155 |
| | 0.193743 | -0.04775 | | | Q7KZF4;H7 | Q7KZF4 | Staphylocc | SND1 | 1593 |
| | 2.74437 | 0.195312 | | | P38606;C9 | P38606 | V-type pro | ATP6V1A | 1134 |
| | 1.764782 | 0.402744 | | | Q86UX7;F5 | Q86UX7;F5 | Fermitin fa | FERMT3 | 1608 |
| | 2.973197 | -0.57102 | | | O00754;M | O00754 | Lysosomal | MAN2B1 | 650 |
| | 2.022372 | -0.59093 | | | Q13231;D( | Q13231;D( | Chitotriosi | CHIT1 | 1463 |
| | 0.485265 | 0.17755 | | | O75083;D( | O75083;D( | WD repeat | WDR1 | 713 |
| | 0.290172 | 0.094513 | | | P00558;P0 | P00558 | Phosphogl | PGK1 | 769 |
| | 2.162548 | 0.308571 | | | P35241;A0 | P35241;A0 | Radixin | RDX | 1112 |
| | 2.127764 | 0.427945 | | | P21399;F5 | P21399 | Cytoplasm | ACO1 | 1009 |
| | 1.580603 | 0.326018 | | | P41250;H7 | P41250 | Glycine--tR | GARS | 1152 |
| | 4.401636 | 0.196926 | | | Q8WUM4; | Q8WUM4 | Programm | PDCD6IP | 1634 |
| | 2.747406 | -0.57798 | | | P36222;H0 | P36222 | Chitinase-3 | CHI3L1 | 1124 |
| | 0.927824 | 0.224962 | | | P29401;A0 | P29401;A0 | Transketol | TKT | 1065 |
| | 0.232575 | 0.051509 | | | P07900;G3 | P07900 | Heat shock | HSP90AA1 | 853 |

| | | | | | | | | | |
|---|---|---|---|---|---|---|---|---|---|
| | 0.339683 | 0.103949 | | | P06733;A0 | P06733;A0 | Alpha-enol | ENO1 | 832 |
| | 1.446452 | 0.339284 | | | E9PLK3;P5 | E9PLK3;P5 | Puromycin | NPEPPS | 434 |
| | 1.138674 | 0.329099 | | | P28838;H0 | P28838 | Cytosol am | LAP3 | 1060 |
| | 0.936262 | 0.220694 | | | Q9ULV4;B4 | Q9ULV4;B4 | Coronin-1C | CORO1C | 1815 |
| | 2.991087 | -0.70652 | | | P11021 | P11021 | 78 kDa glu | HSPA5 | 903 |
| | 1.560519 | -0.48296 | | | P02649;H0 | P02649;H0 | Apolipopro | APOE | 786 |
| | 1.185929 | 0.290621 | | | Q96KP4;J3 | Q96KP4 | Cytosolic n | CNDP2 | 1677 |
| | 1.162473 | 0.160551 | | | P22314;Q5 | P22314 | Ubiquitin-l | UBA1 | 1015 |
| | 0.71179 | 0.150531 | | | Q86VP6;A0 | Q86VP6;A0 | Cullin-asso | CAND1 | 1609 |
| | 1.100618 | 0.305899 | | | P50395;Q5 | P50395 | Rab GDP d | GDI2 | 1214 |
| | 3.124467 | 0.628448 | | | Q15075 | Q15075 | Early endo | EEA1 | 1510 |
| | 2.283798 | 0.508053 | | | P52209;K7 | P52209 | 6-phospho | PGD | 1240 |
| | 1.492269 | 0.410191 | | | O75874;C9 | O75874;C9 | Isocitrate d | IDH1 | 730 |
| | 1.464519 | 0.323238 | | | P00352;Q5 | P00352 | Retinal deh | ALDH1A1 | 762 |
| | 4.056681 | 0.244198 | | | P34932;A0 | P34932;A0 | Heat shock | HSPA4 | 1110 |
| | 0.629302 | -0.09722 | | | P19367;B1 | P19367 | Hexokinase | HK1 | 993 |
| | 4.780992 | 0.596276 | | | P49327;A0 | P49327;A0 | Fatty acid s | FASN | 1195 |
| | 3.256563 | 0.328877 | | | P04075;J3K | P04075;J3K | Fructose-b | ALDOA | 798 |
| | 5.662802 | 0.554386 | | | A0A1C7CY | A0A1C7CY | Dihydropyr | DPYSL2 | 147 |
| | 1.823971 | 0.436076 | | | Q16658;C9 | Q16658 | Fascin | FSCN1 | 1537 |
| | 2.607024 | 0.195335 | | | P21281;H0 | P21281 | V-type pro | ATP6V1B2 | 1005 |
| | 2.202106 | -0.31905 | | | P07355;H0 | P07355;H0 | Annexin A2 | ANXA2;AN | 843 |
| | 3.144091 | -0.59635 | | | P25774;U3 | P25774 | Cathepsin S | CTSS | 1033 |
| | 2.014202 | 0.379797 | | | P09467;A0 | P09467;A0 | Fructose-1 | FBP1 | 872 |
| + | 4.968983 | 1.674702 | up1 | up1 | P53621;A0 | P53621;A0 | Coatomer | COPA | 1255 |
| | 3.333666 | 0.451931 | | | P29350;F5 | P29350 | Tyrosine-p | PTPN6 | 1063 |
| | 2.023501 | 0.280732 | | | P61158;B4 | P61158;B4 | Actin-relat | ACTR3 | 1297 |
| | 3.559414 | 0.277665 | | | P07814;V9 | P07814;V9 | Bifunctiona | EPRS | 851 |
| | 2.229898 | 0.143232 | | | E7EQR4;P1 | E7EQR4;P1 | Ezrin | EZR | 399 |
| | 1.547952 | -0.5118 | | | P07237;H7 | P07237;H7 | Protein dis | P4HB | 841 |
| | 1.491502 | 0.314885 | | | P06744;A0 | P06744;A0 | Glucose-6- | GPI | 834 |
| | 2.560378 | 0.240747 | | | Q14152 | Q14152 | Eukaryotic | EIF3A | 1491 |
| | 0.84018 | -0.15451 | | | P31146;H3 | P31146;H3 | Coronin-1A | CORO1A | 1087 |
| | 2.541045 | -0.39059 | | | P11215;H3 | P11215 | Integrin alp | ITGAM | 905 |
| | 2.217048 | 0.818176 | | | P53396;K7 | P53396 | ATP-citrate | ACLY | 1250 |
| | 4.59931 | -0.58469 | | | A0A1B0GV | A0A1B0GV | Cathepsin | CTSD | 143 |
| | 4.292741 | -0.59887 | | | P07686;Q5 | P07686;Q5 | Beta-hexos | HEXB | 847 |
| | 1.765294 | 0.327144 | | | Q96G03;E7 | Q96G03 | Phosphogl | PGM2 | 1671 |
| | 2.230659 | 0.279652 | | | P78371;F5 | P78371;F5 | T-complex | CCT2 | 1391 |
| | 2.170272 | 0.1984 | | | O15143;C9 | O15143;C9 | Actin-relat | ARPC1B | 665 |
| | 4.803368 | -0.81327 | | | P07858;E9 | P07858 | Cathepsin | CTSB | 852 |
| | 0.974373 | 0.228075 | | | Q9H4A4;A | Q9H4A4;A | Aminopept | RNPEP | 1741 |
| | 0.876975 | 0.20604 | | | P04406;E7 | P04406;E7 | Glyceralde | GAPDH | 804 |
| | 0.630968 | 0.179741 | | | P37837;F2 | P37837;F2 | Transaldola | TALDO1 | 1132 |
| | 2.248295 | 0.425639 | | | P06737;E9 | P06737;E9 | Glycogen p | PYGL | 833 |
| | 3.0498 | -0.42331 | | | P20700;E9 | P20700;E9 | Lamin-B1 | LMNB1 | 1001 |
| | 1.834836 | 0.304583 | | | O15144;C9 | O15144 | Actin-relat | ARPC2 | 666 |

| | | | | | | |
|---|---|---|---|---|---|---|
| 2.391781 | -0.52054 | O75326;F5 | O75326;F5 | Semaphori | SEMA7A | 715 |
| 2.337138 | -0.53167 | O15204 | O15204 | ADAM DEC | ADAMDEC | 668 |
| 1.157424 | -0.16967 | P15144;H0 | P15144 | Aminopep | ANPEP | 950 |
| 0.86679 | 0.155849 | P60174;U3 | P60174;U3 | Triosephos | TPI1 | 1281 |
| 0.865279 | 0.203316 | P31939;H7 | P31939 | Bifunctiona | ATIC | 1093 |
| 0.522375 | -0.08991 | P22626;A0 | P22626;A0 | Heterogen | HNRNPA2E | 1018 |
| 1.860823 | 0.589872 | A0A0A0M | A0A0A0M | Cullin-2 | CUL2 | 78 |
| 0.994647 | 0.315793 | Q16881;A( | Q16881;A( | Thioredoxi | TXNRD1 | 9 |
| 0.819536 | 0.30228 | P19971;C9 | P19971;C9 | Thymidine | TYMP | 997 |
| 1.109622 | 0.190497 | P08238;Q5 | P08238 | Heat shock | HSP90AB1 | 858 |
| 0.653121 | 0.211751 | P32456;Q5 | P32456 | Interferon- | GBP2 | 1102 |
| 0.737637 | 0.194799 | P35237;A0 | P35237;A0 | Serpin B6 | SERPINB6 | 1111 |
| 5.355879 | 0.455319 | P14868;C9 | P14868 | Aspartate- | DARS | 945 |
| 6.255975 | 0.435668 | P49368;B4 | P49368;B4 | T-complex | CCT3 | 1196 |
| 2.337227 | 0.160281 | P50990;H7 | P50990;H7 | T-complex | CCT8 | 1223 |
| 1.562373 | -0.39188 | P33908 | P33908 | Mannosyl- | MAN1A1 | 1106 |
| 2.482407 | -0.45706 | O75487;A( | O75487 | Glypican-4 | GPC4 | 723 |
| 1.688007 | -0.52248 | P10909;H0 | P10909;H0 | Clusterin;C | CLU | 902 |
| 2.529586 | -0.53867 | P01009;A0 | P01009;A0 | Alpha-1-an | SERPINA1 | 772 |
| 3.282535 | -0.60435 | O00391;A8 | O00391 | Sulfhydryl | QSOX1 | 639 |
| 1.125892 | 0.273929 | P21695;Q8 | P21695 | Glycerol-3- | GPD1 | 1010 |
| 1.003884 | 0.340889 | P13489;H0 | P13489;H0 | Ribonuclea | RNH1 | 924 |
| 0.819138 | -0.19285 | P19338;H7 | P19338;H7 | Nucleolin | NCL | 992 |
| 0.651637 | -0.15003 | A0A494C0 | A0A494C0 | Integrin be | ITGB2 | 193 |
| 1.998283 | -0.43881 | Q06481;E9 | Q06481 | Amyloid-lil | APLP2 | 1429 |
| 1.207688 | 0.267738 | P30740;C9 | P30740 | Leukocyte | SERPINB1 | 1085 |
| 2.866175 | 0.418249 | Q96QK1;I3 | Q96QK1 | Vacuolar p | VPS35 | 1680 |
| 1.45007 | 0.411191 | Q01469;I6 | Q01469;I6 | Fatty acid- | FABP5 | 1412 |
| 1.578787 | -0.3968 | Q02809;Q! | Q02809 | Procollage | PLOD1 | 1417 |
| 1.275511 | 0.261683 | P18669;Q8 | P18669 | Phosphogl | PGAM1 | 989 |
| 1.151207 | 0.257883 | P30041 | P30041 | Peroxiredc | PRDX6 | 1068 |
| 0.96313 | 0.187176 | Q9UHD8;K | Q9UHD8;K | Septin-9 | SEPTIN9 | 1802 |
| 0.209947 | -0.03439 | Q16531;F5 | Q16531;F5 | DNA dama | DDB1 | 1533 |
| 0.315712 | 0.1396 | P32455 | P32455 | Interferon- | GBP1 | 1101 |
| 7.162538 | 0.506049 | Q5VTE0;P6 | Q5VTE0;P6 | Putative el | EEF1A1P5; | 1383 |
| 5.557053 | 0.385149 | A0A087W | A0A087W | UTP--gluco | UGP2 | 33 |
| 0.996779 | -0.13547 | P08575;M | P08575;M | Receptor-t | PTPRC | 860 |
| 0.99231 | 0.348439 | Q9BWD1 | Q9BWD1 | Acetyl-CoA | ACAT2 | 1719 |
| 0.733287 | 0.138751 | P25786;F5 | P25786;F5 | Proteasom | PSMA1 | 1034 |
| 0.215539 | 0.054659 | G3V1D3;G | G3V1D3;G | Dipeptidyl | DPP3 | 487 |
| 2.820976 | -0.55813 | P07602;C9 | P07602;C9 | Prosaposir | PSAP | 846 |
| 0.383125 | 0.067788 | P63104;E7 | P63104;E7 | 14-3-3 pro | YWHAZ | 1370 |
| 2.749228 | -0.75574 | P30101 | P30101 | Protein dis | PDIA3 | 1078 |
| 1.758242 | 0.435819 | P00918;E5 | P00918 | Carbonic a | CA2 | 771 |
| 1.799939 | 0.376697 | P30153;B3 | P30153;B3 | Serine/thre | PPP2R1A | 1079 |
| 1.285067 | 0.422467 | P12955;A0 | P12955;A0 | Xaa-Pro di | PEPD | 919 |
| 0.998077 | 0.282884 | Q16719;A( | Q16719 | Kynurenina | KYNU | 1539 |

| | | | | | | | |
|---|---|---|---|---|---|---|---|
| 0.633219 | 0.134499 | | P62258;B4 | P62258 | 14-3-3 pro | YWHAE | 1326 |
| 0.45651 | -0.07048 | | P36578;H3 | P36578;H3 | 60S riboso | RPL4 | 1126 |
| 0.349456 | -0.0664 | | P61247;D6 | P61247;D6 | 40S riboso | RPS3A | 1303 |
| 0.120682 | -0.04357 | | P43235;Q5 | P43235;Q5 | Cathepsin | CTSK | 1164 |
| 0.583255 | 0.14428 | | P07195;A0 | P07195;A0 | L-lactate d | LDHB | 840 |
| 3.082152 | -1.1739 | down1 | P10809;E7 | P10809 | 60 kDa hea | HSPD1 | 901 |
| 2.313223 | 0.443142 | | P36871;A0 | P36871;A0 | Phosphogl | PGM1 | 1127 |
| 1.580955 | 0.282551 | | P40121;E7 | P40121;E7 | Macrophag | CAPG | 1141 |
| 1.841372 | 0.198282 | | P14317;E7 | P14317;E7 | Hematopo | HCLS1 | 936 |
| 2.109965 | 0.148992 | | Q14019;H3 | Q14019;H3 | Coactosin- | COTL1 | 1487 |
| 2.464303 | -0.44904 | | P05155;E9 | P05155;E9 | Plasma pro | SERPING1 | 815 |
| 3.671796 | -0.61155 | | H3BP20;P0 | H3BP20;P0 | Beta-hexos | HEXA | 543 |
| 5.072592 | -0.62957 | | P10253;I3I | P10253 | Lysosomal | GAA | 892 |
| 2.738354 | -0.64142 | | P40926;G3 | P40926;G3 | Malate deh | MDH2 | 1146 |
| 1.187966 | 0.225169 | | P00338;F5 | P00338 | L-lactate d | LDHA | 761 |
| 1.114813 | 0.310114 | | Q04446;E9 | Q04446;E9 | 1,4-alpha-g | GBE1 | 1422 |
| 0.586408 | 0.139654 | | P43034;I3I | P43034 | Platelet-ac | PAFAH1B1 | 1163 |
| 0.121568 | 0.027277 | | P07384;E9 | P07384 | Calpain-1 c | CAPN1 | 844 |
| 0.050241 | 0.013228 | | Q14974;J3 | Q14974;J3 | Importin su | KPNB1 | 1504 |
| 6.521788 | 0.485772 | | Q06830;A0 | Q06830;A0 | Peroxiredo | PRDX1 | 1430 |
| 0.64118 | 0.196997 | | P23528;E9 | P23528;E9 | Cofilin-1 | CFL1 | 1026 |
| 1.853498 | 0.789681 | | Q86UP2;G | Q86UP2;G | Kinectin | KTN1 | 1607 |
| 5.553078 | 0.514458 | | P17987;F5 | P17987;F5 | T-complex | TCP1 | 984 |
| 2.377196 | 0.50912 | | E7EVA0;P2 | E7EVA0;P2 | Microtubu | MAP4 | 403 |
| 3.272343 | 0.498066 | | G3V126;Q9 | G3V126;Q9 | V-type pro | ATP6V1H | 484 |
| 2.559372 | 0.453013 | | Q9UJ70;H7 | Q9UJ70;H7 | N-acetyl-D | NAGK | 1810 |
| 4.18575 | 0.416883 | | P35606;D6 | P35606 | Coatomer | COPB2 | 1120 |
| 1.48816 | 0.387355 | | P14550;Q5 | P14550 | Alcohol de | AKR1A1 | 939 |
| 3.513379 | 0.353804 | | Q99536;K7 | Q99536 | Synaptic ve | VAT1 | 1693 |
| 1.771682 | 0.302719 | | A0A087X1 | A0A087X1 | Proteasom | PSME2 | 48 |
| 1.78446 | 0.289791 | | P37802;X6 | P37802;X6 | Transgelin- | TAGLN2 | 1131 |
| 3.514576 | 0.258889 | | P52566;F5 | P52566;F5 | Rho GDP-d | ARHGDIB | 1243 |
| 6.817529 | -0.49315 | | Q13510;A0 | Q13510;A0 | Acid ceram | ASAH1 | 140 |
| 1.775592 | -0.52445 | | Q16610 | Q16610 | Extracellul | ECM1 | 1536 |
| 3.610384 | -0.64533 | | Q08380;K7 | Q08380 | Galectin-3- | LGALS3BP | 1439 |
| 3.309564 | -0.69826 | | P28799;K7 | P28799;K7 | Granulins;/ | GRN | 1059 |
| 2.474657 | -0.78758 | | P27797;K7 | P27797;K7 | Calreticulin | CALR | 1051 |
| 1.102466 | 0.268702 | | P40925;A0 | P40925;A0 | Malate deh | MDH1 | 1145 |
| 0.222665 | 0.099296 | | P23526 | P23526 | Adenosylh | AHCY | 1025 |
| 3.733113 | -1.0282 | down1 | P06576;H0 | P06576;H0 | ATP syntha | ATP5B | 827 |
| 2.839244 | -1.11688 | down1 | P38646;A0 | P38646 | Stress-70 p | HSPA9 | 1135 |
| 4.607518 | 0.69099 | | Q9Y3Z3;A0 | Q9Y3Z3;A0 | Deoxynucle | SAMHD1 | 1840 |
| 3.391103 | 0.618847 | | Q14847;C9 | Q14847;C9 | LIM and SH | LASP1 | 1503 |
| 2.754435 | 0.579053 | | Q96TA1 | Q96TA1 | Niban-like | FAM129B | 1688 |
| 2.078947 | 0.38931 | | P48506;A0 | P48506;A0 | Glutamate | GCLC | 1187 |
| 2.4934 | 0.330054 | | P33241;E7 | P33241;E7 | Lymphocyt | LSP1 | 1105 |
| 1.956612 | 0.320415 | | P00491;G3 | P00491;G3 | Purine nuc | PNP | 766 |

| | | | | | | |
|---|---|---|---|---|---|---|
| 4.409452 | 0.316552 | E7ENZ3;P4 | E7ENZ3;P4 | T-complex | CCT5 | 396 |
| 1.95692 | 0.289742 | P61160;F5 | P61160;F5 | Actin-relat | ACTR2 | 1298 |
| 1.932796 | 0.251048 | P07737;K7 | P07737;K7 | Profilin-1 | PFN1 | 849 |
| 1.655598 | 0.177786 | F8W6I7;P0 | F8W6I7;P0 | Heterogen | HNRNPA1; | 477 |
| 1.640511 | -0.16271 | P04083;Q5 | P04083 | Annexin A1 | ANXA1 | 800 |
| 2.927002 | -0.6606 | O94985;Q5 | O94985;Q5 | Calsyntenii | CLSTN1 | 742 |
| 1.322609 | -0.74412 | Q9BXX0 | Q9BXX0 | EMILIN-2 | EMILIN2 | 1721 |
| 3.475704 | -0.87058 | P06858;E7 | P06858 | Lipoprotei | LPL | 837 |
| 0.957869 | 0.272626 | A0A2R8Y4. | A0A2R8Y4. | Glutathion | GSS | 157 |
| 0.70823 | 0.123466 | P12956;B1 | P12956;B1 | X-ray repai | XRCC6 | 920 |
| 0.513127 | -0.08232 | P46777;A0 | P46777;A0 | 60S riboso | RPL5 | 1173 |
| 1.022309 | 0.197341 | A0A0G2JIV | A0A0G2JIV | Heat shock | HSPA1B;HS | 115 |
| 1.361922 | 0.638603 | P17174 | P17174 | Aspartate a | GOT1 | 976 |
| 2.87998 | 0.518658 | O00154;K7 | O00154;K7 | Cytosolic a | ACOT7 | 627 |
| 3.67491 | 0.482279 | Q9NUV9;G | Q9NUV9;G | GTPase IM | GIMAP4 | 501 |
| 1.808387 | 0.425867 | P48163 | P48163 | NADP-dep | ME1 | 1185 |
| 2.039994 | 0.359378 | Q15046;H3 | Q15046 | Lysine--tRI | KARS | 1508 |
| 2.682883 | 0.30204 | P00390;A0 | P00390 | Glutathion | GSR | 764 |
| 4.400141 | 0.298689 | Q99832;F8 | Q99832 | T-complex | CCT7 | 1700 |
| 1.730989 | 0.271974 | Q99497;K7 | Q99497;K7 | Protein de | PARK7 | 1690 |
| 2.022036 | 0.251601 | P13010;C9 | P13010 | X-ray repai | XRCC5 | 921 |
| 2.929784 | 0.215295 | B0YIW6;P4 | B0YIW6;P4 | Coatomer | ARCN1 | 239 |
| 1.616152 | -0.21591 | P39019;A0 | P39019;A0 | 40S riboso | RPS19 | 1137 |
| 2.870149 | -0.68348 | P16035;B4 | P16035;B4 | Metallopro | TIMP2 | 961 |
| 3.611963 | -0.68358 | Q9BUD6;D | Q9BUD6;D | Spondin-2 | SPON2 | 1714 |
| 3.698363 | -0.86846 | P04179;F5 | P04179;F5 | Superoxide | SOD2 | 801 |
| 1.047714 | 0.353745 | P53004;C9 | P53004 | Biliverdin r | BLVRA | 1249 |
| 0.928894 | -0.29423 | Q99538;G3 | Q99538 | Legumain | LGMN | 1694 |
| 0.792548 | 0.283856 | P43490;A0 | P43490;A0 | Nicotinami | NAMPT | 1166 |
| 0.715363 | 0.188702 | P31150;G5 | P31150 | Rab GDP d | GDI1 | 1088 |
| 0.497619 | -0.25393 | P12109;A0 | P12109;A0 | Collagen al | COL6A1 | 914 |
| 0.296759 | 0.074274 | Q14764;H3 | Q14764 | Major vaul | MVP | 1502 |
| 0.253058 | -0.04876 | P63244;J3I | P63244;J3I | Guanine nu | GNB2L1 | 1375 |
| 1.433286 | 0.184206 | P52907 | P52907 | F-actin-cap | CAPZA1 | 1248 |
| 1.524796 | 0.610116 | P11216;H0 | P11216 | Glycogen p | PYGB | 906 |
| 2.620392 | 0.167921 | P04899;P6 | P04899 | Guanine nu | GNAI2 | 808 |
| 1.16435 | 0.228474 | Q6IBS0;D6 | Q6IBS0;D6 | Twinfilin-2 | TWF2 | 1581 |
| 2.246894 | 0.566684 | I3L0N3;P4 | I3L0N3;P4 | Vesicle-fus | NSF | 562 |
| 5.141647 | 0.450792 | Q9BR76;A0 | Q9BR76;A0 | Coronin-1B | CORO1B | 1705 |
| 4.111751 | 0.410816 | P40227;Q9 | P40227 | T-complex | CCT6A | 1142 |
| 1.813231 | 0.382779 | Q06323;H0 | Q06323 | Proteasom | PSME1 | 1428 |
| 1.554236 | 0.350097 | P49189 | P49189 | 4-trimethy | ALDH9A1 | 1193 |
| 1.57462 | 0.258225 | E7ES33;E7 | E7ES33;E7 | Septin-7 | SEPTIN7 | 397 |
| 1.756582 | 0.154157 | P50995;H0 | P50995 | Annexin A1 | ANXA11 | 1225 |
| 1.476102 | -0.32767 | P53634;H0 | P53634;H0 | Dipeptidyl | CTSC | 1256 |
| 1.82701 | -0.37043 | O60462;C9 | O60462 | Neuropilin | NRP2 | 696 |
| 1.952943 | -0.45742 | P05067;A0 | P05067;A0 | Amyloid be | APP | 811 |

| | | | | | | | | | |
|---|---|---|---|---|---|---|---|---|---|
| | 2.536516 | -0.52727 | | | P61626;F8 | P61626;F8 | Lysozyme ( | LYZ | 1310 |
| | 6.614492 | -0.65989 | | | P15586;H7 | P15586;H7 | N-acetylglu | GNS | 956 |
| | 3.665796 | -0.7062 | | | P10451;D6 | P10451;D6 | Osteopont | SPP1 | 894 |
| | 2.807917 | -0.72496 | | | Q13093 | Q13093 | Platelet-ac | PLA2G7 | 1455 |
| | 1.293456 | -0.13925 | | | P02794;G3 | P02794;G3 | Ferritin hea | FTH1 | 792 |
| | 1.165003 | -2.09971 | | | P13667;A0 | P13667;A0 | Protein dis | PDIA4 | 929 |
| | 0.64262 | 0.748975 | | | P48147;A0 | P48147;A0 | Prolyl endo | PREP | 1184 |
| | 0.625379 | 0.267321 | | | P15121;E9 | P15121;E9 | Aldose red | AKR1B1 | 949 |
| | 0.440127 | 0.21267 | | | E9PB90;P5 | E9PB90;P5 | Hexokinase | HK2 | 413 |
| | 0.269786 | 0.063472 | | | C9JIF9;P13 | C9JIF9;P13 | Acylamino- | APEH | 298 |
| | 0.098455 | -0.02484 | | | O14818;H0 | O14818 | Proteasom | PSMA7 | 658 |
| | 0.019198 | 0.005355 | | | P61978;Q5 | P61978;Q5 | Heterogen | HNRNPK | 1316 |
| | 4.405964 | 0.54186 | | | P11413;E9 | P11413;E9 | Glucose-6- | G6PD | 910 |
| | 1.079528 | 0.109072 | | | P05023;B1 | P05023 | Sodium/pc | ATP1A1 | 809 |
| + | 1.921956 | 2.677756 | up2 | up2 | Q14247;H0 | Q14247 | Src substra | CTTN | 1496 |
| | 6.054483 | 0.843718 | | | P19878;B1 | P19878 | Neutrophil | NCF2 | 996 |
| | 1.831029 | 0.594867 | | | O43776;K7 | O43776;K7 | Asparagine | NARS | 689 |
| | 1.625106 | 0.533736 | | | O00299 | O00299 | Chloride in | CLIC1 | 636 |
| | 2.560081 | 0.507485 | | | P17931;G3 | P17931;G3 | Galectin-3; | LGALS3 | 982 |
| | 4.47325 | 0.46289 | | | Q16543;K7 | Q16543;K7 | Hsp90 co-c | CDC37 | 1535 |
| | 2.266107 | 0.445044 | | | O00764;F2 | O00764;F2 | Pyridoxal k | PDXK | 652 |
| | 1.83817 | 0.440248 | | | P08567 | P08567 | Pleckstrin | PLEK | 859 |
| | 1.471626 | 0.375799 | | | P78417;Q5 | P78417;Q5 | Glutathion | GSTO1 | 1392 |
| | 2.057723 | 0.332761 | | | F6TLX2;I3L | F6TLX2;I3L | Glyoxalase | GLOD4 | 464 |
| | 5.32725 | 0.307567 | | | P50991 | P50991 | T-complex | CCT4 | 1224 |
| | 5.238046 | 0.277278 | | | P52565;J3I | P52565;J3I | Rho GDP-d | ARHGDIA | 1242 |
| | 2.629086 | 0.262719 | | | P26641 | P26641 | Elongation | EEF1G | 1046 |
| | 2.089679 | 0.257709 | | | A0A087W\ | A0A087W\ | 26S protea | PSMD1 | 24 |
| | 1.754377 | 0.146397 | | | P23396;E9 | P23396;E9 | 40S riboso | RPS3 | 1023 |
| | 4.051033 | -0.29704 | | | Q6P4A8;F5 | Q6P4A8 | Phospholip | PLBD1 | 1583 |
| | 1.309677 | -0.31914 | | | P07954 | P07954 | Fumarate l | FH | 854 |
| | 1.986409 | -0.46744 | | | B4E3Q4;Q9 | B4E3Q4;Q9 | Adenosine | CECR1 | 261 |
| | 1.79544 | -0.51846 | | | P09486;F5 | P09486;F5 | SPARC | SPARC | 873 |
| | 5.122497 | -0.56407 | | | P04062;A0 | P04062;A0 | Glucosylce | GBA | 796 |
| | 3.868879 | -0.57341 | | | Q13740;F5 | Q13740;F5 | CD166 ant | ALCAM | 1480 |
| | 4.081831 | -0.76559 | | | Q9UBR2 | Q9UBR2 | Cathepsin : | CTSZ | 1793 |
| | 1.190253 | 0.298166 | | | Q8NC51 | Q8NC51 | Plasminoge | SERBP1 | 1621 |
| | 1.16841 | 1.509602 | | | Q5VZU9;P2 | Q5VZU9;P2 | Tripeptidyl | TPP2 | 1061 |
| | 1.086821 | 0.198603 | | | P50552;K7 | P50552 | Vasodilato | VASP | 1219 |
| | 1.079048 | -0.1477 | | | P62424;Q5 | P62424;Q5 | 60S riboso | RPL7A | 1340 |
| | 0.778553 | 0.100589 | | | G3V5Z7;P6 | G3V5Z7;P6 | Proteasom | PSMA6 | 498 |
| | 0.471698 | -0.12207 | | | P04040 | P04040 | Catalase | CAT | 795 |
| | 0.395724 | 0.165792 | | | P54920;M0 | P54920;M0 | Alpha-solu | NAPA | 1264 |
| | 0.321467 | 0.106715 | | | Q15833;M | Q15833;M | Syntaxin-b | STXBP2 | 1526 |
| | 0.319972 | 0.043989 | | | P25789;H0 | P25789;H0 | Proteasom | PSMA4 | 1036 |
| | 0.088459 | 0.031014 | | | P36543;C9 | P36543;C9 | V-type pro | ATP6V1E1; | 1125 |
| | 2.161595 | 0.236459 | | | O60506;B7 | O60506;B7 | Heterogen | SYNCRIP | 699 |

| | | | | | | | | | |
|---|---|---|---|---|---|---|---|---|---|
| | 3.046432 | 0.512234 | | | Q01813;B1 | Q01813;B1 | ATP-depen | PFKP | 1414 |
| | 0.629049 | -0.15643 | | | P23246;H0 | P23246 | Splicing fac | SFPQ | 1020 |
| + | 1.486446 | -2.16612 | down2 | down2 | Q9NZ08;H0 | Q9NZ08 | Endoplasm | ERAP1 | 1777 |
| + | 1.37848 | 2.84488 | up2 | up2 | Q86TX2;G3 | Q86TX2;G3 | Acyl-coenz | ACOT1;AC( | 1605 |
| | 1.49731 | 1.872605 | | up1 | O60664;K7 | O60664;K7 | Perilipin-3 | PLIN3 | 701 |
| | 4.095091 | 0.79771 | | | Q9Y678;H0 | Q9Y678 | Coatomer | COPG1 | 1856 |
| | 3.580789 | 0.597057 | | | H0Y2Y8;Q1 | H0Y2Y8;Q1 | Zyxin | ZYX | 505 |
| | 2.589419 | 0.532376 | | | B5MDF5;P | B5MDF5;P | GTP-bindir | RAN | 265 |
| | 1.647029 | 0.520547 | | | O95861;A6 | O95861;A6 | 3(2),5-bisp | BPNT1 | 757 |
| | 1.821653 | 0.451963 | | | P30086 | P30086 | Phosphatic | PEBP1 | 1077 |
| | 2.225513 | 0.366302 | | | Q32MZ4;C | Q32MZ4 | Leucine-ric | LRRFIP1 | 1547 |
| | 1.351807 | 0.301352 | | | Q15019;B5 | Q15019;B5 | Septin-2 | SEPTIN2 | 1507 |
| | 2.825228 | 0.299361 | | | O43242;H0 | O43242 | 26S protea | PSMD3 | 677 |
| | 2.592139 | 0.219467 | | | Q13200;H7 | Q13200 | 26S protea | PSMD2 | 1461 |
| | 1.893351 | 0.142366 | | | O60763 | O60763 | General ve | USO1 | 706 |
| | 1.438772 | -0.35238 | | | Q12906;K7 | Q12906 | Interleukin | ILF3 | 1452 |
| | 3.074239 | -0.41851 | | | Q9Y4D7;H0 | Q9Y4D7 | Plexin-D1 | PLXND1 | 1842 |
| | 2.340132 | -0.57667 | | | P00505 | P00505 | Aspartate a | GOT2 | 768 |
| | 6.498141 | -0.57702 | | | A0A3B3IU( | A0A3B3IU( | Alpha-gala | GLA | 185 |
| | 3.095582 | -0.70129 | | | P42765;A0 | P42765;A0 | 3-ketoacyl- | ACAA2 | 1158 |
| | 1.219106 | 0.297069 | | | P49721;A0 | P49721;A0 | Proteasom | PSMB2 | 1203 |
| | 1.058236 | 1.742411 | | | E7EX90;Q1 | E7EX90;Q1 | Dynactin su | DCTN1;DKI | 409 |
| | 1.011561 | 0.256465 | | | P46781;B5 | P46781;B5 | 40S riboso | RPS9 | 1176 |
| | 0.852653 | 0.298228 | | | P29373;Q5 | P29373;Q5 | Cellular ret | CRABP2 | 1064 |
| | 0.390571 | 0.113917 | | | P05198;G3 | P05198;G3 | Eukaryotic | EIF2S1 | 816 |
| | 0.326596 | -0.09552 | | | P06748;E5 | P06748 | Nucleopho | NPM1 | 835 |
| | 0.165964 | 0.077279 | | | Q96C23;H7 | Q96C23;H7 | Aldose 1-e | GALM | 1663 |
| | 1.87228 | 0.538827 | | | P16152;E9 | P16152;E9 | Carbonyl r | CBR1 | 965 |
| | 1.715407 | -0.41359 | | | O00462;A0 | O00462;A0 | Beta-manr | MANBA | 642 |
| | 0.848381 | 0.221471 | | | P50453 | P50453 | Serpin B9 | SERPINB9 | 1216 |
| + | 1.68514 | -3.28231 | down2 | down2 | P25705;K7 | P25705;K7 | ATP syntha | ATP5A1 | 1032 |
| + | 1.265719 | 2.425861 | up2 | | P11766;D6 | P11766 | Alcohol de | ADH5 | 912 |
| | 2.331481 | 0.469856 | | | Q99613;B5 | Q99613;B5 | Eukaryotic | EIF3C;EIF3( | 266 |
| | 1.453288 | 0.427602 | | | P51991;H7 | P51991 | Heterogen | HNRNPA3 | 1239 |
| | 2.38839 | 0.401072 | | | P09382;F8 | P09382 | Galectin-1 | LGALS1 | 868 |
| | 3.680262 | 0.38767 | | | P17858;F8 | P17858 | ATP-depen | PFKL | 980 |
| | 1.850788 | 0.308496 | | | P09525;Q6 | P09525;Q6 | Annexin A4 | ANXA4 | 876 |
| | 1.815211 | 0.234873 | | | P21283;E7 | P21283;E7 | V-type pro | ATP6V1C1 | 1006 |
| | 2.892408 | 0.20365 | | | P25788;G3 | P25788;G3 | Proteasom | PSMA3 | 1035 |
| | 2.210666 | -0.18192 | | | P20618 | P20618 | Proteasom | PSMB1 | 1000 |
| | 1.987098 | -0.25963 | | | Q02878;F8 | Q02878 | 60S riboso | RPL6 | 1419 |
| | 1.65262 | -0.26601 | | | P07910;G3 | P07910;G3 | Heterogen | HNRNPC;H | 247 |
| | 4.055779 | -0.40753 | | | X6R5C5;X6 | X6R5C5;X6 | Carboxype | CTSA | 897 |
| | 2.824012 | -0.43481 | | | Q9H3G5;H | Q9H3G5;H | Probable s | CPVL | 1738 |
| | 5.073196 | -0.54684 | | | P42785;E9 | P42785;E9 | Lysosomal | PRCP | 1161 |
| | 2.098778 | -0.57842 | | | Q6YHK3 | Q6YHK3 | CD109 ant | CD109 | 1589 |
| | 2.290636 | -0.61152 | | | Q6UX71 | Q6UX71 | Plexin dom | PLXDC2 | 1586 |

| | | | | | | | | | |
|---|---|---|---|---|---|---|---|---|---|
| | 5.181172 | -0.70524 | | | P08236;F8 | P08236;F8 | Beta-glucu | GUSB | 857 |
| | 2.53427 | -0.79955 | | | P21757;B4 | P21757;B4 | Macropha | MSR1 | 253 |
| | 2.616207 | -0.8673 | | | E9PEX6;P0 | E9PEX6;P0 | Dihydrolip | DLD | 419 |
| | 1.212487 | 0.182024 | | | A0A024RA | A0A024RA | Proteasom | PSMA2 | 4 |
| | 1.127313 | 0.273561 | | | Q9UJU6;H | Q9UJU6;H | Drebrin-lik | DBNL | 1812 |
| | 1.093949 | -0.33811 | | | O14773;A | O14773;A | Tripeptidyl | TPP1 | 657 |
| | 1.057588 | 1.456866 | | | P55884;C9 | P55884 | Eukaryotic | EIF3B | 1275 |
| | 0.828813 | 0.222406 | | | P62937;F8 | P62937;F8 | Peptidyl-pı | PPIA | 1361 |
| | 0.781024 | 0.182872 | | | O75368 | O75368 | SH3 domai | SH3BGRL | 719 |
| | 0.764145 | 1.748689 | | | A0A2R8YD | A0A2R8YD | Peroxisom | HSD17B4 | 170 |
| | 0.546873 | 0.139615 | | | A0A0A0MI | A0A0A0MI | Protein-L-i | PCMT1 | 63 |
| | 0.545079 | 0.151531 | | | P27348;E9 | P27348;E9 | 14-3-3 pro | YWHAQ | 1048 |
| | 0.488129 | -0.12016 | | | H3BN02;P | H3BN02;P | Integrin al | ITGAX | 539 |
| | 0.248485 | 0.074281 | | | P05388;F8 | P05388;F8 | 60S acidic | RPLP0;RPL | 821 |
| | 0.23001 | -0.08516 | | | P39900 | P39900 | Macropha | MMP12 | 1140 |
| | 0.217412 | -0.07152 | | | Q9UQ80;F | Q9UQ80;F | Proliferatic | PA2G4 | 1821 |
| | 0.203692 | 0.05419 | | | P41091;Q | P41091;Q | Eukaryotic | EIF2S3;EIF | 1147 |
| | 0.169877 | -0.02333 | | | P02792 | P02792 | Ferritin ligl | FTL | 791 |
| | 0.126718 | 0.029247 | | | C9J9K3;A0 | C9J9K3;A0 | 40S riboso | RPSA | 92 |
| | 0.114232 | 0.026395 | | | Q9NUQ9;E | Q9NUQ9 | Protein FA | FAM49B | 1770 |
| | 0.113793 | -0.02569 | | | X5D2R7;P | X5D2R7;P | Proteasom | PSM8;PSM | 1053 |
| | 0.111587 | 0.046764 | | | P62805 | P62805 | Histone H4 | HIST1H4A | 1346 |
| | 1.682763 | 0.581972 | | | O95782 | O95782 | AP-2 comp | AP2A1 | 755 |
| | 0.231247 | -0.08644 | | | P26583;D6 | P26583;D6 | High mobil | HMGB2 | 1043 |
| + | 3.260744 | 3.710214 | up2 | up2 | Q5T5C7;P | Q5T5C7;P | Serine--tRI | SARS | 1200 |
| | 1.387866 | 1.714776 | | up1 | P49588;H3 | P49588;H3 | Alanine--tF | AARS | 1199 |
| | 3.621428 | 0.78951 | | | P57737;A0 | P57737;A0 | Coronin-7; | CORO7;CO | 82 |
| | 4.825199 | 0.694406 | | | B5MBZ0;Q | B5MBZ0;Q | Echinoderr | EML4 | 264 |
| | 3.45656 | 0.615572 | | | P04792;F8 | P04792 | Heat shock | HSPB1 | 806 |
| | 2.78188 | 0.584407 | | | Q04637;E7 | Q04637;E7 | Eukaryotic | EIF4G1 | 402 |
| | 3.386446 | 0.477941 | | | P27695;G3 | P27695;G3 | DNA-(apur | APEX1 | 1050 |
| | 5.078086 | 0.476178 | | | P62993;J3 | P62993;J3 | Growth fac | GRB2 | 1365 |
| | 2.170218 | 0.375008 | | | P08758;D6 | P08758;D6 | Annexin A | ANXA5 | 864 |
| | 4.075693 | 0.357368 | | | P10644;K7 | P10644;K7 | cAMP-dep | PRKAR1A | 898 |
| | 1.780383 | -0.53218 | | | P14625;Q | P14625 | Endoplasm | HSP90B1 | 942 |
| | 5.226002 | -0.71888 | | | J3KMY5;G | J3KMY5;G | Epididyma | NPC2 | 394 |
| | 3.430125 | -0.7437 | | | P07711;Q | P07711;Q | Cathepsin | CTSL | 848 |
| | 3.359541 | -0.95717 | | | P00367;P4 | P00367;P4 | Glutamate | GLUD1;GLU | 763 |
| | 1.08779 | 0.253926 | | | O75223;M | O75223;M | Gamma-gl | GGCT | 714 |
| | 1.079009 | 0.141481 | | | P39023;H7 | P39023;H7 | 60S riboso | RPL3 | 1138 |
| | 1.037 | 0.178855 | | | P62249;M | P62249;M | 40S riboso | RPS16 | 1325 |
| | 1.030928 | 0.3494 | | | H0Y987;O | H0Y987;O | Phosphoac | PGM3 | 517 |
| | 0.94207 | 1.215082 | | | A0A087X2 | A0A087X2 | 26S protea | PSMC6 | 54 |
| | 0.803738 | 1.588335 | | | A0A1B0GV | A0A1B0GV | Glutamine | QARS | 144 |
| | 0.803737 | -0.14451 | | | P62753;A2 | P62753;A2 | 40S riboso | RPS6 | 1345 |
| | 0.79911 | 0.105899 | | | A0A0U1RR | A0A0U1RR | Polypyrimi | PTBP1 | 131 |
| | 0.789555 | 1.249384 | | | Q9BZQ8;H | Q9BZQ8 | Protein Nil | FAM129A | 1726 |

| | | | | | | | | | |
|---|---|---|---|---|---|---|---|---|---|
| | 0.74513 | 0.191774 | | | Q8IZ83;F5I | Q8IZ83;F5I | Aldehyde c | ALDH16A1 | 1615 |
| | 0.724454 | 0.135545 | | | Q13867;K7 | Q13867;K7 | Bleomycin | BLMH | 1482 |
| | 0.702468 | -0.09184 | | | P15880;H0 | P15880;H0 | 40S riboso | RPS2 | 958 |
| | 0.614674 | 0.077671 | | | Q13561;F8 | Q13561;F8 | Dynactin su | DCTN2 | 1476 |
| | 0.597009 | -0.13382 | | | P26373;J3I | P26373 | 60S riboso | RPL13 | 1040 |
| | 0.45665 | 0.349394 | | | Q96GA7;H | Q96GA7;H | Serine deh | SDSL | 1672 |
| | 0.450874 | -0.09255 | | | A0A0A0MT | A0A0A0MT | Lysosomal | LIPA | 74 |
| | 0.254019 | -0.58604 | | | P08133;E5 | P08133;E5 | Annexin A6 | ANXA6 | 855 |
| | 0.169617 | -0.07983 | | | P01871 | P01871 | Ig mu chain | IGHM | 782 |
| | 0.12592 | 0.020959 | | | P62269;A0 | P62269 | 40S riboso | RPS18 | 1328 |
| | 2.454088 | 0.468854 | | | P60842;J3I | P60842;J3I | Eukaryotic | EIF4A1 | 1284 |
| | 1.208861 | 0.298057 | | | Q9UI08;H0 | Q9UI08;H0 | Ena/VASP- | EVL | 1806 |
| | 4.616437 | 0.702244 | | | O60749;D6 | O60749 | Sorting nex | SNX2 | 704 |
| | 1.941193 | 0.230383 | | | P29692;E9 | P29692;E9 | Elongation | EEF1D | 427 |
| + | 2.164867 | 3.07706 | up2 | up2 | P53618;E9 | P53618;E9 | Coatomer | COPB1 | 1254 |
| | 2.200113 | 0.627231 | | | P46926;D6 | P46926;D6 | Glucosami | GNPDA1 | 1178 |
| | 1.948017 | 0.609738 | | | P68036;A0 | P68036;A0 | Ubiquitin-c | UBE2L3 | 1382 |
| | 2.837062 | 0.478499 | | | Q15404 | Q15404 | Ras suppre | RSU1 | 1520 |
| | 3.56641 | 0.439279 | | | O00182;J3 | O00182 | Galectin-9 | LGALS9 | 631 |
| | 1.448439 | 0.360562 | | | P10599 | P10599 | Thioredoxi | TXN | 895 |
| | 1.641043 | 0.333255 | | | P61981 | P61981 | 14-3-3 pro | YWHAG | 1317 |
| | 2.65792 | 0.332767 | | | Q15435;H7 | Q15435;H7 | Protein ph | PPP1R7 | 1521 |
| | 3.109367 | 0.300913 | | | Q15691 | Q15691 | Microtubu | MAPRE1 | 1523 |
| | 2.363666 | 0.261065 | | | P50502;Q3 | P50502;Q3 | Hsc70-inte | ST13;ST13I | 1218 |
| | 2.02384 | 0.257585 | | | P32119;A6 | P32119;A6 | Peroxiredo | PRDX2 | 1099 |
| | 1.511212 | 0.205384 | | | F8WCF6;P5 | F8WCF6;P5 | Actin-relat | ARPC4-TTL | 482 |
| | 1.439536 | 0.17651 | | | P35998;C9 | P35998;C9 | 26S protea | PSMC2 | 1123 |
| | 2.344306 | -0.2141 | | | P67809;H0 | P67809;H0 | Nuclease-s | YBX1 | 1379 |
| | 2.36002 | -0.27328 | | | P13686;K7 | P13686 | Tartrate-re | ACP5 | 930 |
| | 2.045468 | -0.29915 | | | F8WE86;B5 | F8WE86;B5 | Transcobal | TCN2 | 263 |
| | 2.136075 | -0.3639 | | | P16401 | P16401 | Histone H1 | HIST1H1B | 968 |
| | 1.556493 | -0.4345 | | | Q15084 | Q15084 | Protein dis | PDIA6 | 1511 |
| | 4.981203 | -0.48878 | | | P34059;H3 | P34059;H3 | N-acetylga | GALNS | 1107 |
| | 2.352338 | -0.50639 | | | Q16769;B5 | Q16769;B5 | Glutaminyl | QPCT | 1540 |
| | 4.071726 | -0.5305 | | | Q8IV08;M0 | Q8IV08 | Phospholip | PLD3 | 1611 |
| | 2.487333 | -0.53897 | | | Q9Y6C2;A0 | Q9Y6C2 | EMILIN-1 | EMILIN1 | 1858 |
| | 3.625577 | -0.58238 | | | Q9UHL4;R4 | Q9UHL4 | Dipeptidyl | DPP7 | 1804 |
| | 5.605247 | -0.62254 | | | O95831;E9 | O95831;E9 | Apoptosis- | AIFM1 | 756 |
| | 3.89073 | -0.65311 | | | P04066 | P04066 | Tissue alph | FUCA1 | 797 |
| | 4.094954 | -0.68613 | | | Q14956;A0 | Q14956;A0 | Transmem | GPNMB | 178 |
| | 3.047192 | -0.74563 | | | A0A5F9UP | A0A5F9UP | 45 kDa calc | SDF4 | 204 |
| | 3.013716 | -0.81555 | | | P09603;E9 | P09603;E9 | Macrophag | CSF1 | 878 |
| | 1.095953 | 1.61642 | | | O00231;J3 | O00231 | 26S protea | PSMD11 | 633 |
| | 1.049758 | 0.322028 | | | P51149;C9 | P51149;C9 | Ras-relatec | RAB7A | 1227 |
| | 0.964694 | 0.110038 | | | P00441;H7 | P00441;H7 | Superoxide | SOD1 | 765 |
| | 0.92469 | 1.586755 | | | Q13126;B4 | Q13126;B4 | S-methyl-5 | MTAP | 1456 |
| | 0.897305 | 0.322914 | | | Q9NQW7;( | Q9NQW7;( | Xaa-Pro an | XPNPEP1 | 1762 |

| | | | | | | | | | |
|---|---|---|---|---|---|---|---|---|---|
| | 0.817179 | 1.605681 | | | Q9Y617 | Q9Y617 | Phosphose | PSAT1 | 1853 |
| | 0.678026 | 1.210714 | | | P45974 | P45974 | Ubiquitin c | USP5 | 1169 |
| | 0.599908 | 0.136921 | | | P09211;A8 | P09211;A8 | Glutathion | GSTP1 | 866 |
| | 0.579428 | -0.10516 | | | P62081;A0 | P62081;A0 | 40S riboso | RPS7 | 1318 |
| | 0.529574 | 0.117537 | | | P29218;H0 | P29218;H0 | Inositol mo | IMPA1 | 1062 |
| | 0.503919 | -0.69366 | | | K7EIK7;O9 | K7EIK7;O9 | Echinoderm | EML2 | 588 |
| | 0.245488 | -0.82819 | | | O14672;H3 | O14672 | Disintegrin | ADAM10 | 654 |
| | 0.154283 | 0.041937 | | | P18124;A8 | P18124;A8 | 60S riboso | RPL7 | 986 |
| | 0.071184 | 0.05148 | | | A0A4V7I44 | A0A4V7I44 | Apolipopro | APOBR | 198 |
| | 0.062466 | -0.01128 | | | P46779;H0 | P46779;H0 | 60S riboso | RPL28 | 1175 |
| | 0.03246 | -0.00943 | | | P61313;E7 | P61313;E7 | 60S riboso | RPL15 | 1305 |
| | 2.91026 | -0.71025 | | | B4E1Z4;E7 | B4E1Z4;E7ETN3 | | | 259 |
| | 4.159829 | 0.266083 | | | P11940;A0 | P11940;A0 | Polyadenyl | PABPC1;PA | 13 |
| | 0.699206 | 0.112347 | | | P31946;Q4 | P31946 | 14-3-3 pro | YWHAB | 1095 |
| | 1.223214 | 0.375456 | | | P49407;E9 | P49407;E9 | Beta-arres | ARRB1 | 1197 |
| + | 1.633866 | 2.362874 | up2 | up2 | A0A0A0M9 | A0A0A0M9 | Isoleucine- | IARS | 72 |
| | 1.353038 | 1.984638 | | up1 | O95352;C9 | O95352 | Ubiquitin-l | ATG7 | 746 |
| | 2.989274 | 1.05373 | | up1 | P55160;F8 | P55160;F8 | Nck-associ | NCKAP1L | 1272 |
| | 2.3566 | 0.963107 | | | Q9NR45;Q | Q9NR45 | Sialic acid s | NANS | 1763 |
| | 3.018314 | 0.930991 | | | P41226 | P41226 | Ubiquitin-l | UBA7 | 1149 |
| | 2.302717 | 0.838583 | | | Q8WWM9 | Q8WWM9 | Cytoglobin | CYGB | 1637 |
| | 2.983526 | 0.771135 | | | E9PGT1;Q1 | E9PGT1;Q1 | Translin | TSN | 421 |
| | 3.460122 | 0.667107 | | | R4GNH3;P | R4GNH3;P | 26S protea | PSMC3 | 983 |
| | 2.256107 | 0.462887 | | | I3L397;I3L5 | I3L397;I3L5 | Eukaryotic | EIF5A;EIF5 | 567 |
| | 4.494492 | 0.428628 | | | P06702 | P06702 | Protein S10 | S100A9 | 829 |
| | 2.793308 | 0.396574 | | | Q9Y5K8;G3 | Q9Y5K8;G3 | V-type pro | ATP6V1D | 1845 |
| | 1.438557 | 0.35351 | | | E7EX17;P2 | E7EX17;P2 | Eukaryotic | EIF4B | 407 |
| | 1.319852 | 0.313516 | | | Q07960;H0 | Q07960;H0 | Rho GTPas | ARHGAP1 | 1435 |
| | 1.4716 | 0.222527 | | | Q04917;A2 | Q04917;A2 | 14-3-3 pro | YWHAH | 1424 |
| | 2.024657 | -0.3575 | | | P09237 | P09237 | Matrilysin | MMP7 | 867 |
| | 2.692412 | -0.35976 | | | P17900;H0 | P17900 | Gangliosid | GM2A | 981 |
| | 3.956854 | -0.42584 | | | P16278;E7 | P16278;E7 | Beta-galac | GLB1 | 966 |
| | 4.859656 | -0.55088 | | | A0A2C9F2I | A0A2C9F2I | Palmitoyl-p | PPT1 | 155 |
| | 3.159594 | -0.57453 | | | O75787;A0 | O75787;A0 | Renin rece | ATP6AP2 | 142 |
| | 4.388725 | -0.64781 | | | D6RHI9;O0 | D6RHI9;O0 | Ribonuclea | RNASET2 | 376 |
| | 2.595444 | -0.7039 | | | Q99519;E9 | Q99519;E9 | Sialidase-1 | NEU1 | 1691 |
| | 2.507507 | -0.71404 | | | P01034 | P01034 | Cystatin-C | CST3 | 776 |
| | 1.251938 | 0.222395 | | | O95336;M | O95336;M | 6-phospho | PGLS | 745 |
| | 1.054873 | 2.110129 | | | Q9NQR4;H | Q9NQR4;H | Omega-am | NIT2 | 1761 |
| | 1.012256 | 0.206247 | | | Q13177;E9 | Q13177 | Serine/thre | PAK2 | 1459 |
| | 0.991114 | 0.365802 | | | H3BLU7;O4 | H3BLU7;O4 | Aflatoxin B | AKR7A2 | 533 |
| | 0.958683 | 1.078823 | | | P54577;A0 | P54577;A0 | Tyrosine--t | YARS | 1259 |
| | 0.818477 | 1.377371 | | | P10768;U3 | P10768;U3 | S-formylglu | ESD | 900 |
| | 0.575715 | -0.14155 | | | H3BMH2;H | H3BMH2;H | Ras-relate | RAB11A;RA | 536 |
| | 0.512995 | -0.20338 | | | A0A087X0 | A0A087X0 | Heterogen | HNRNPM | 42 |
| | 0.182426 | 0.455261 | | | E7EPM6;P3 | E7EPM6;P3 | Long-chain | ACSL1 | 398 |
| | 0.087177 | -0.02337 | | | P80723;U3 | P80723 | Brain acid s | BASP1 | 1395 |

| | | | | | | | | | |
|---|---|---|---|---|---|---|---|---|---|
| | 0.058127 | 0.14209 | | | F8W1C3;Q | F8W1C3;Q | Matrix me | MMP19 | 475 |
| | 3.423024 | 0.188203 | | | A0A087W\ | A0A087WWU8;A0A49 | | TPM3 | 27 |
| | 0.418887 | 0.097 | | | A0A0C4DG | A0A0C4DG | Calpastatir | CAST | 95 |
| | 1.133365 | 0.255638 | | | Q32Q12;P | Q32Q12;P | Nucleoside | NME1-NM | 1548 |
| | 2.266453 | -0.25717 | | | A0A140T9 | A0A140T955;A0A140 | | HLA-A | 134 |
| | 2.103622 | 0.193169 | | | P54727;Q5 | P54727;Q5 | UV excisio | RAD23B | 1263 |
| | 2.928397 | 0.566483 | | | P62195;J3 | P62195;J3 | 26S protea | PSMC5 | 1322 |
| | 1.119471 | 0.104281 | | | P39687;H0 | P39687;H0 | Acidic leuc | ANP32A | 1139 |
| | 0.738263 | 1.031004 | | | P17612;Q1 | P17612;Q1 | cAMP-dep | PRKACA;KI | 978 |
| + | 1.497871 | 1.937859 | up1 | up1 | Q05655;A | Q05655;A | Protein kin | PRKCD | 1426 |
| + | 2.453971 | 2.861835 | up2 | up2 | P33176;A0 | P33176 | Kinesin-1 h | KIF5B | 1104 |
| | 2.03759 | 0.665397 | | | P05413;S4 | P05413;S4 | Fatty acid- | FABP3 | 822 |
| | 2.263518 | 0.490026 | | | Q9H0W9;A | Q9H0W9;A | Ester hydr | C11orf54 | 1735 |
| | 2.288026 | 0.484623 | | | P16930;H0 | P16930;H0 | Fumarylac | FAH | 971 |
| | 4.106342 | 0.464489 | | | Q13347;Q | Q13347 | Eukaryotic | EIF3I | 1469 |
| | 2.02475 | 0.453673 | | | P49773;D6 | P49773;D6 | Histidine t | HINT1 | 1204 |
| | 2.111852 | 0.394783 | | | P30085;Q5 | P30085;Q5 | UMP-CMP | CMPK1 | 1076 |
| | 1.763018 | 0.366329 | | | P26447 | P26447 | Protein S1 | S100A4 | 1041 |
| | 3.35763 | 0.365709 | | | O75348;F2 | O75348 | V-type pro | ATP6V1G1 | 716 |
| | 1.925436 | 0.358804 | | | B3KWE1;P | B3KWE1;P | Histidine-- | HARS | 251 |
| | 1.825738 | 0.268467 | | | P28066 | P28066 | Proteasom | PSMA5 | 1054 |
| | 1.358404 | 0.207989 | | | O15145;C9 | O15145;C9 | Actin-relat | ARPC3 | 667 |
| | 1.574753 | 0.1765 | | | A0A087W | A0A087W | AP-2 comp | AP2M1 | 30 |
| | 1.545626 | 0.171991 | | | P31949 | P31949 | Protein S1 | S100A11 | 1098 |
| | 1.532198 | 0.128972 | | | P05455;E7 | P05455;E7 | Lupus La p | SSB | 823 |
| | 2.323027 | -0.33521 | | | P61604;B8 | P61604;B8 | 10 kDa hea | HSPE1 | 1309 |
| | 4.831578 | -0.45105 | | | Q8NHP8 | Q8NHP8 | Putative pl | PLBD2 | 1627 |
| | 1.919411 | -0.53244 | | | P30048 | P30048 | Thioredoxi | PRDX3 | 1073 |
| | 3.91275 | -0.60602 | | | P26572;D6 | P26572 | Alpha-1,3- | MGAT1 | 1042 |
| | 6.310894 | -0.64673 | | | E9PQY3;P1 | E9PQY3;P1 | Lysosomal | ACP2 | 444 |
| | 3.331867 | -0.65756 | | | A0A087X2 | A0A087X2 | Compleme | C1S | 50 |
| | 2.778551 | -0.70276 | | | P41222;Q5 | P41222;Q5 | Prostaglan | PTGDS | 1148 |
| | 1.268847 | 1.545364 | | | Q9HB71;B | Q9HB71 | Calcyclin-b | CACYBP | 1752 |
| | 1.172961 | 0.359065 | | | P30043;A0 | P30043;A0 | Flavin redu | BLVRB | 1069 |
| | 1.081784 | 1.78042 | | | A0A0A0MI | A0A0A0MI | Sorting ne | SNX6 | 62 |
| | 0.938872 | 0.297107 | | | P62942;Q5 | P62942;Q5 | Peptidyl-p | FKBP1A;FK | 1362 |
| | 0.891056 | -0.37945 | | | P49788 | P49788 | Retinoic ac | RARRES1 | 1205 |
| | 0.812285 | 1.19276 | | | P26639;D6 | P26639 | Threonine- | TARS | 1044 |
| | 0.789443 | 0.222637 | | | Q96IU4;F8 | Q96IU4;F8 | Alpha/beta | ABHD14B | 1675 |
| | 0.534189 | 0.156331 | | | P28482;E9 | P28482 | Mitogen-a | MAPK1 | 1058 |
| | 0.48432 | 0.171761 | | | Q53FA7;H | Q53FA7;H | Quinone o | TP53I3 | 1556 |
| | 0.479227 | 1.30607 | | | Q7L5Y1;A0 | Q7L5Y1;A0 | Mitochon | ENOSF1 | 1597 |
| | 0.419564 | 0.118011 | | | P07108;A0 | P07108;A0 | Acyl-CoA-b | DBI | 839 |
| | 0.391309 | 0.071011 | | | P54578;A6 | P54578;A6 | Ubiquitin c | USP14 | 1260 |
| | 0.380384 | 0.056215 | | | P62280;M | P62280;M | 40S riboso | RPS11 | 1331 |
| | 0.37037 | 0.69036 | | | Q9Y315;E9 | Q9Y315;E9 | Deoxyribo | DERA | 1832 |
| | 0.363305 | 0.136261 | | | P53602;H3 | P53602;H3 | Diphospho | MVD | 1252 |

| | | | | | | | | | |
|---|---|---|---|---|---|---|---|---|---|
| | 0.335379 | 0.873722 | | | P19474 | P19474 | E3 ubiquiti | TRIM21 | 994 |
| | 0.322579 | 0.546556 | | | Q86TI2;M( | Q86TI2;M( | Dipeptidyl | DPP9 | 1604 |
| | 0.287303 | -0.58532 | | | H0YMG7;C | H0YMG7;C | Retinal del | ALDH1A2 | 527 |
| | 0.2756 | -0.96057 | | | A0A1W2PI | A0A1W2PI | Heterogen | HNRNPU | 150 |
| | 0.037621 | -0.00878 | | | P62241;Q5 | P62241;Q5 | 40S riboso | RPS8 | 1323 |
| | 0.018933 | -0.01593 | | | A0A087X0 | A0A087X0 | Pro-cathep | CTSH | 38 |
| | 3.604254 | 0.969215 | | | Q7L576;A( | Q7L576;A( | Cytoplasm | CYFIP1 | 1596 |
| | 1.326635 | 0.367849 | | | A0A0A0M! | A0A0A0M! | Phosphatic | PITPNB | 71 |
| + | 1.951454 | -2.21297 | down2 | down2 | Q16822;H( | Q16822;H( | Phosphoer | PCK2 | 1543 |
| + | 16.11775 | 4.116962 | up2 | up2 | Q13451 | Q13451 | Peptidyl-pı | FKBP5 | 1473 |
| + | 2.187786 | 3.791756 | up2 | up2 | P30711;A0 | P30711;A0 | Glutathion | GSTT1 | 1084 |
| + | 3.178771 | 3.160689 | up2 | up2 | Q92696;H( | Q92696;H( | Geranylger | RABGGTA | 1647 |
| + | 2.104111 | 2.900888 | up2 | up2 | E7ENJ6;Q9 | E7ENJ6;Q9 | AP-1 comp | AP1M1 | 395 |
| + | 2.112665 | 2.736132 | up2 | up2 | Q9UNF0;A | Q9UNF0;A | Protein kin | PACSIN2 | 1819 |
| + | 1.745447 | 2.578906 | up2 | up2 | P30520 | P30520 | Adenylosu | ADSS | 1082 |
| + | 3.073525 | 2.544253 | up2 | up2 | Q5T1M5 | Q5T1M5 | FK506-binc | FKBP15 | 1567 |
| + | 1.723502 | 2.226509 | up2 | up2 | Q9Y3I0 | Q9Y3I0 | tRNA-splici | RTCB | 1838 |
| + | 1.366655 | 2.043138 | up2 | up2 | P17655;C9 | P17655 | Calpain-2 c | CAPN2 | 979 |
| | 1.997812 | 0.719258 | | | P58546;C9 | P58546 | Myotrophi | MTPN | 1279 |
| | 2.536259 | 0.701466 | | | P09936;D6 | P09936;D6 | Ubiquitin c | UCHL1 | 881 |
| | 1.830159 | 0.669959 | | | F2Z3J9;Q1 | F2Z3J9;Q1 | Prostaglan | PTGR1 | 450 |
| | 1.858426 | 0.619195 | | | Q9NRV9;A | Q9NRV9;A | Heme-binc | HEBP1 | 1765 |
| | 1.775798 | 0.488197 | | | Q04760 | Q04760 | Lactoylglut | GLO1 | 1423 |
| | 4.463721 | 0.455789 | | | B1AJY5;B1 | B1AJY5;B1 | 26S protea | PSMD10 | 241 |
| | 5.350984 | 0.443262 | | | P31153;Q( | P31153 | S-adenosyl | MAT2A | 1089 |
| | 2.784072 | 0.425192 | | | P47755;F8 | P47755;F8 | F-actin-cap | CAPZA2 | 1182 |
| | 2.264856 | 0.378043 | | | O15511;B1 | O15511;B1 | Actin-relat | ARPC5 | 676 |
| | 2.588632 | 0.252076 | | | Q8TD55;C! | Q8TD55 | Pleckstrin l | PLEKHO2 | 1631 |
| | 1.462079 | 0.237404 | | | Q9BRF8 | Q9BRF8 | Serine/thre | CPPED1 | 1707 |
| | 2.487944 | 0.180832 | | | P09496;F8 | P09496;F8 | Clathrin lig | CLTA | 874 |
| | 1.980855 | -0.3149 | | | P62750;H7 | P62750;H7 | 60S riboso | RPL23A | 1344 |
| | 4.891945 | -0.36706 | | | P38159;H3 | P38159;H3 | RNA-bindir | RBMX;RBM | 1133 |
| | 1.984911 | -0.44613 | | | Q14118 | Q14118 | Dystroglyc | DAG1 | 1490 |
| | 1.765402 | -0.47807 | | | P22466 | P22466 | Galanin pe | GAL | 1016 |
| | 5.019079 | -0.50513 | | | Q8NCC3;H | Q8NCC3;H | Group XV p | PLA2G15 | 1622 |
| | 3.139339 | -0.64146 | | | Q03405;M | Q03405;M | Urokinase | PLAUR | 1421 |
| | 2.681018 | -0.68683 | | | D6RIU4;D6 | D6RIU4;D6 | Vesicular ir | LMAN2 | 367 |
| | 3.008633 | -0.69458 | | | Q14314;A( | Q14314;A( | Fibroleukir | FGL2 | 1497 |
| | 2.278101 | -0.76255 | | | A2A274;Q! | A2A274;Q! | Aconitate l | ACO2 | 218 |
| | 1.29479 | 0.234996 | | | P25398 | P25398 | 40S riboso | RPS12 | 1031 |
| | 1.281063 | 1.960192 | | | P51606;A6 | P51606;A6 | N-acylgluc | RENBP | 1235 |
| | 1.223664 | 0.366093 | | | A6NDG6;H | A6NDG6 | Phosphogl | PGP | 223 |
| | 1.203436 | 1.938464 | | | Q99988;A( | Q99988 | Growth/di | GDF15 | 1701 |
| | 1.075171 | 1.773074 | | | O43399;A( | O43399;A( | Tumor pro | TPD52L2 | 32 |
| | 1.049323 | 2.496168 | | | Q9Y696 | Q9Y696 | Chloride in | CLIC4 | 1857 |
| | 0.918795 | -0.31162 | | | K7ERG9;P( | K7ERG9;P( | Compleme | CFD | 608 |
| | 0.90223 | 1.49176 | | | Q12904;A( | Q12904;A( | Aminoacyl | AIMP1 | 1451 |

| | | | | | | | | | |
|---|---|---|---|---|---|---|---|---|---|
| | 0.84619 | 1.356936 | | | A6PVN5;Q | A6PVN5;Q | Serine/thr | PPP2R4 | 231 |
| | 0.814132 | -0.1009 | | | P16070;H0 | P16070;H0 | CD44 antig | CD44 | 962 |
| | 0.6985 | -1.80053 | | | R4GN98;P( | R4GN98;P( | Protein S1( | S100A6 | 830 |
| | 0.659044 | 1.843602 | | | O75828 | O75828 | Carbonyl r | CBR3 | 729 |
| | 0.442419 | -0.06322 | | | O14745;J3 | O14745;J3 | Na(+)/H(+) | SLC9A3R1 | 656 |
| | 0.259722 | -0.35819 | | | Q03252 | Q03252 | Lamin-B2 | LMNB2 | 1420 |
| | 0.236522 | 0.584484 | | | P05556;C9 | P05556 | Integrin be | ITGB1 | 824 |
| | 0.191606 | 0.081915 | | | P04839 | P04839 | Cytochrom | CYBB | 807 |
| | 0.153878 | 0.056782 | | | Q6NYC8;A( | Q6NYC8;A( | Phostensin | PPP1R18 | 113 |
| | 0.109859 | -0.02042 | | | Q15008;C9 | Q15008 | 26S protea | PSMD6 | 1506 |
| | 0.085472 | 0.224971 | | | Q9NX46 | Q9NX46 | Poly(ADP-r | ADPRHL2 | 1773 |
| | 0.067042 | -0.02293 | | | A0A087W) | A0A087W) | 60S riboso | RPL17;RPL | 29 |
| | 0.040987 | 0.155657 | | | P14902;J3I | P14902 | Indoleamir | IDO1 | 946 |
| | 1.37799 | 0.231648 | | | D6RGI3;D6 | D6RGI3;D6 | Septin-11 | SEPTIN11 | 372 |
| | 1.95428 | -0.15176 | | | A0A182DV | A0A182DV | HLA class I | HLA-DRB3 | 139 |
| | 3.013219 | -0.56059 | | | A0A0G2JM | A0A0G2JM | Leukocyte | LILRA3 | 120 |
| | 0.069998 | 0.039313 | | | P84077;P6 | P84077;P6 | ADP-ribosy | ARF1;ARF3 | 1300 |
| + | 1.372096 | 2.894918 | up2 | up2 | Q9BUF5;K7 | Q9BUF5;K7 | Tubulin be | TUBB6 | 1715 |
| | 3.928306 | -0.97695 | | | P62328;O1 | P62328 | Thymosin I | TMSB4X | 1339 |
| | 0.350991 | -0.07045 | | | P16949;A2 | P16949;A2 | Stathmin | STMN1 | 972 |
| | 3.037868 | 0.692158 | | | O60493 | O60493 | Sorting ne) | SNX3 | 698 |
| | 0.465269 | 1.077239 | | | O94804;H( | O94804 | Serine/thr | STK10 | 739 |
| + | 1.304407 | -2.38674 | down2 | down2 | P05091;F8 | P05091 | Aldehyde c | ALDH2 | 812 |
| + | 3.209176 | -3.87473 | down2 | down2 | Q8NBJ5;M | Q8NBJ5 | Procollage | COLGALT1 | 1619 |
| + | 3.113516 | 3.802912 | up2 | up2 | P09104;F5 | P09104;F5 | Gamma-er | ENO2 | 865 |
| + | 3.441036 | 3.193757 | up2 | up2 | Q3KQV9;A | Q3KQV9 | UDP-N-ace | UAP1L1 | 1550 |
| + | 2.202139 | 3.123219 | up2 | up2 | A0A0A0MI | A0A0A0MI | Abl interac | ABI1 | 64 |
| + | 2.183992 | 3.014326 | up2 | up2 | F5GWE5;Q | F5GWE5;Q | Phosphatic | PITPNA | 451 |
| + | 2.539712 | 2.997546 | up2 | up2 | P46063;F8 | P46063 | ATP-depen | RECQL | 1171 |
| + | 3.032905 | 2.992467 | up2 | up2 | Q9UBT2;K7 | Q9UBT2;K7 | SUMO-acti | UBA2 | 1795 |
| + | 1.605167 | 2.944712 | up2 | up2 | P09455;A0 | P09455;A0 | Retinol-bir | RBP1 | 871 |
| + | 2.094915 | 2.849073 | up2 | up2 | O76071 | O76071 | Probable c | CIAO1 | 738 |
| + | 2.09596 | 2.794533 | up2 | up2 | Q9P2T1;F8 | Q9P2T1;F8 | GMP reduc | GMPR2 | 25 |
| + | 1.776403 | 2.684498 | up2 | up2 | D6REX3;O9 | D6REX3;O9 | Protein tra | SEC31A | 373 |
| | 5.388441 | 0.899033 | | | Q9H444;P5 | Q9H444 | Charged m | CHMP4B | 1740 |
| | 2.349084 | 0.55382 | | | P30046;B5 | P30046;B5 | D-dopachr | DDT;DDTL | 1071 |
| | 2.082623 | 0.545863 | | | P62633;D6 | P62633 | Cellular nu | CNBP | 1342 |
| | 3.360984 | 0.515359 | | | P49720;A0 | P49720;A0 | Proteasom | PSMB3 | 1202 |
| | 2.107034 | 0.404859 | | | Q96BW5;A | Q96BW5;A | Phosphotri | PTER | 1661 |
| | 1.325429 | 0.379205 | | | Q9NQ88;A | Q9NQ88;A | Fructose-2 | TIGAR | 1759 |
| | 1.407255 | 0.374141 | | | P35754 | P35754 | Glutaredo) | GLRX | 1121 |
| | 3.822893 | 0.25482 | | | P05109 | P05109 | Protein S1( | S100A8 | 813 |
| | 2.143133 | 0.177591 | | | M0R0F0;M | M0R0F0;M | 40S riboso | RPS5 | 619 |
| | 2.130408 | -0.24432 | | | P46783;F6 | P46783;F6 | 40S riboso | RPS10;RPS | 1177 |
| | 1.957709 | -0.29924 | | | P23284 | P23284 | Peptidyl-pr | PPIB | 1021 |
| | 2.679826 | -0.34974 | | | P51688;I3I | P51688;I3I | N-sulphogl | SGSH | 1237 |
| | 3.518909 | -0.45307 | | | P20933;H0 | P20933 | N(4)-(beta- | AGA | 1003 |

| | | | | | | |
|---|---|---|---|---|---|---|
| 1.329127 | -0.54022 | P30040;F8 | P30040;F8 | Endoplasm | ERP29 | 1067 |
| 2.430285 | -0.54808 | Q15904;A( | Q15904;A( | V-type pro | ATP6AP1 | 1529 |
| 3.269013 | -0.58179 | P13284;M( | P13284 | Gamma-in | IFI30 | 922 |
| 3.529588 | -0.70116 | Q9BTY2;Q | Q9BTY2 | Plasma alp | FUCA2 | 1713 |
| 4.151702 | -0.79426 | P05121 | P05121 | Plasminog | SERPINE1 | 814 |
| 4.754997 | -0.87568 | O00115;K7 | O00115;K7 | Deoxyribo | DNASE2 | 625 |
| 1.294477 | -1.63346 | Q92820 | Q92820 | Gamma-gl | GGH | 1650 |
| 1.231262 | 0.315126 | P51452;K7 | P51452;K7 | Dual speci | DUSP3 | 1231 |
| 1.21406 | 1.568021 | Q8TDZ2;Q | Q8TDZ2 | Protein-me | MICAL1 | 1632 |
| 1.163444 | 1.818021 | P62191 | P62191 | 26S protea | PSMC1 | 1321 |
| 1.154023 | 1.561778 | F5GZS6;J3I | F5GZS6;J3I | 4F2 cell-su | SLC3A2 | 457 |
| 1.095003 | -1.536 | Q9HAT2 | Q9HAT2 | Sialate O-a | SIAE | 1749 |
| 1.066643 | -1.30068 | E7EX60;Q5 | E7EX60;Q5 | Neuropilin | NRP1 | 408 |
| 1.044946 | -0.13516 | P50281;F8 | P50281;F8 | Matrix me | MMP14 | 1213 |
| 0.975102 | 0.354951 | O94903;E5 | O94903;E5 | Proline syr | PROSC | 740 |
| 0.958945 | 0.193642 | E5RIW3;E5 | E5RIW3;E5 | Tubulin-sp | TBCA | 389 |
| 0.884623 | 1.483305 | Q9NYL9;H( | Q9NYL9;H( | Tropomod | TMOD3 | 1776 |
| 0.85368 | 1.458053 | P41240;H3 | P41240 | Tyrosine-p | CSK | 1151 |
| 0.84618 | 0.173211 | P14324;A0 | P14324;A0 | Farnesyl py | FDPS | 937 |
| 0.84609 | -0.14418 | P62917;E9 | P62917;E9 | 60S riboso | RPL8 | 1360 |
| 0.841969 | 0.125045 | A2ACR1;A( | A2ACR1;A( | Proteasom | PSMB9 | 116 |
| 0.809708 | 1.41083 | P22102;C9 | P22102;C9 | Trifunction | GART | 1013 |
| 0.799171 | 1.412648 | P61457;H0 | P61457 | Pterin-4-al | PCBD1 | 1307 |
| 0.780991 | 1.337128 | P15374;A0 | P15374;A0 | Ubiquitin c | UCHL3 | 954 |
| 0.648687 | 1.163881 | Q02790;H( | Q02790 | Peptidyl-pr | FKBP4 | 1416 |
| 0.646854 | 0.81081 | P20042 | P20042 | Eukaryotic | EIF2S2 | 998 |
| 0.588511 | 0.828751 | Q13409;E7 | Q13409;E7 | Cytoplasm | DYNC1I2 | 1471 |
| 0.571477 | 0.828915 | P51665;H3 | P51665;H3 | 26S protea | PSMD7 | 1236 |
| 0.565679 | 0.754952 | Q9UH65;E | Q9UH65;E | Switch-ass | SWAP70 | 1800 |
| 0.562597 | 0.126516 | A0A1B0GV | A0A1B0GV | Adenylosu | ADSL | 79 |
| 0.549549 | 0.113889 | U3KQE2;A( | U3KQE2;A( | Calpain sm | CAPNS1 | 7 |
| 0.507478 | 0.7201 | P54136;E5 | P54136 | Arginine--t | RARS | 1258 |
| 0.462166 | 0.795321 | M0R117;Q | M0R117;Q | 60S riboso | RPL18A | 622 |
| 0.461538 | 0.210949 | P20073 | P20073 | Annexin A7 | ANXA7 | 999 |
| 0.459698 | 0.195329 | P54687;F5 | P54687;F5 | Branched-( | BCAT1 | 1261 |
| 0.405016 | -0.11253 | P61353;K7 | P61353;K7 | 60S riboso | RPL27 | 1306 |
| 0.37991 | 0.091238 | A0A0C4DF | A0A0C4DF | Protein SE | SET;SETSIP | 90 |
| 0.379282 | -0.06534 | P62263;A0 | P62263;A0 | 40S riboso | RPS14 | 168 |
| 0.310996 | 0.115496 | O95865;A( | O95865;A( | N(G),N(G)- | DDAH2 | 758 |
| 0.198851 | 0.592813 | P51572;A0 | P51572;A0 | B-cell rece | BCAP31 | 1233 |
| 0.149278 | 0.425519 | P83105 | P83105 | Serine prot | HTRA4 | 1396 |
| 0.141762 | 0.049271 | P61088;F8 | P61088;F8 | Ubiquitin-( | UBE2N;UB | 1295 |
| 0.111476 | 0.287035 | Q08211 | Q08211 | ATP-depen | DHX9 | 1437 |
| 0.098938 | -0.0228 | H7C4P1;P4 | H7C4P1;P4 | Rho GTPas | ARHGAP25 | 559 |
| 0.082723 | -0.02897 | P30050 | P30050 | 60S riboso | RPL12 | 1074 |
| 0.082709 | -0.01968 | P50914;E7 | P50914;E7 | 60S riboso | RPL14 | 1222 |
| 0.009966 | -0.00483 | P04080;A0 | P04080;A0 | Cystatin-B | CSTB | 799 |

| | | | | | | | | | |
|---|---|---|---|---|---|---|---|---|---|
| | 0.007489 | 0.001217 | | | P28070 | P28070 | Proteasom | PSMB4 | 1055 |
| | 0.000361 | -0.0001 | | | B4DY09;Q1 | B4DY09;Q1 | Interleukin | ILF2 | 258 |
| | 0.281985 | -0.09642 | | | P19021 | P19021 | Peptidyl-gl | PAM | 991 |
| | 2.462018 | -0.25976 | | | A0A0G2JI3 | A0A0G2JI36;Q5SUL5;A | | HLA-A | 114 |
| | 0.078508 | -0.02956 | | | P62820;E7 | P62820;E7 | Ras-relatec | RAB1A | 1347 |
| | 2.626386 | 0.406463 | | | Q13838;Q! | Q13838;Q! | Spliceoson | DDX39B | 1481 |
| | 0.559948 | 0.119378 | | | Q14103;H( | Q14103;H( | Heterogen | HNRNPD | 1488 |
| | 8.33715 | 0.573911 | | | Q15365 | Q15365 | Poly(rC)-bi | PCBP1 | 1518 |
| | 5.633487 | -0.53816 | | | Q9HB40;I3 | Q9HB40 | Retinoid-ir | SCPEP1 | 1751 |
| | 0.316629 | -0.10784 | | | P09429;Q5 | P09429;Q5 | High mobil | HMGB1;HN | 870 |
| + | 1.967396 | 3.121017 | up2 | up2 | F5H365;Q1 | F5H365;Q1 | Protein tra | SEC23A | 460 |
| | 2.013818 | 0.354837 | | | Q96C19;H( | Q96C19 | EF-hand dc | EFHD2 | 1662 |
| | 1.862985 | -0.19234 | | | Q30118;A( | Q30118;A( | HLA class I | HLA-DRA | 110 |
| | 1.146791 | 0.215957 | | | P51858;H3 | P51858 | Hepatoma | HDGF | 1238 |
| | 0.957301 | 2.015398 | | | Q13185;S4 | Q13185;S4 | Chromobo | CBX3 | 1460 |
| + | 2.280575 | -3.15516 | down2 | down2 | K7ELL7;P14 | K7ELL7;P14 | Glucosidas | PRKCSH | 598 |
| + | 14.91022 | 4.179108 | up2 | up2 | Q9Y3F4;H( | Q9Y3F4 | Serine-thre | STRAP | 1837 |
| + | 3.441618 | 3.42158 | up2 | up2 | Q15181;Q! | Q15181;Q! | Inorganic p | PPA1 | 1514 |
| + | 3.21309 | 3.091235 | up2 | up2 | O43865;H( | O43865;H( | Putative ac | AHCYL1;AH | 692 |
| + | 2.477507 | 2.818768 | up2 | up2 | Q96CV9;A( | Q96CV9 | Optineurin | OPTN | 1666 |
| + | 1.990615 | 2.789562 | up2 | up2 | P63151;A0 | P63151;A0 | Serine/thre | PPP2R2A | 1371 |
| + | 2.260346 | 2.756322 | up2 | up2 | Q13596;H( | Q13596;H( | Sorting ne | SNX1 | 1479 |
| + | 2.348931 | 2.320242 | up2 | up2 | P55010;H0 | P55010 | Eukaryotic | EIF5 | 1267 |
| + | 1.620747 | 2.099395 | up2 | up2 | Q9Y3C8 | Q9Y3C8 | Ubiquitin-f | UFC1 | 1835 |
| | 3.008085 | 0.508109 | | | O60711;B7 | O60711;B7 | Leupaxin | LPXN | 267 |
| | 1.618859 | 0.42736 | | | P30047;H0 | P30047;H0 | GTP cycloh | GCHFR | 1072 |
| | 2.846661 | 0.413812 | | | Q15185;A( | Q15185;A( | Prostaglan | PTGES3 | 1515 |
| | 3.041057 | 0.409733 | | | P43686 | P43686 | 26S protea | PSMC4 | 1167 |
| | 3.208218 | 0.393433 | | | O14579;M | O14579;M | Coatomer | COPE | 612 |
| | 1.497912 | 0.375402 | | | Q9UBQ0;F | Q9UBQ0;F | Vacuolar p | VPS29 | 1790 |
| | 1.598327 | 0.326213 | | | Q9BVG4;A | Q9BVG4 | Protein PB | PBDC1 | 1718 |
| | 2.542215 | 0.257945 | | | P40306;J3( | P40306 | Proteasom | PSMB10 | 1143 |
| | 1.328788 | 0.2501 | | | P24666;G5 | P24666;G5 | Low molec | ACP1 | 1029 |
| | 2.664026 | 0.182603 | | | F6WQW2;I | F6WQW2;I | Ran-specifi | RANBP1 | 465 |
| | 1.822879 | -0.20494 | | | P62906 | P62906 | 60S riboso | RPL10A | 1358 |
| | 1.957959 | -0.22235 | | | P05387;H0 | P05387 | 60S acidic | RPLP2 | 820 |
| | 2.65907 | -0.32543 | | | P18510 | P18510 | Interleukin | IL1RN | 988 |
| | 1.683368 | -0.38333 | | | K7ERI9;P02 | K7ERI9;P02 | Apolipopro | APOC1 | 609 |
| | 2.254198 | -0.44394 | | | Q92743;A( | Q92743;A( | Serine prot | HTRA1 | 1648 |
| | 1.571411 | -0.4664 | | | P14384;F8 | P14384;F8 | Carboxype | CPM | 938 |
| | 1.060108 | 1.856559 | | | O00410;H( | O00410;H( | Importin-5 | IPO5 | 513 |
| | 0.9535 | 2.118506 | | | P53999;D6 | P53999 | Activated F | SUB1 | 1257 |
| | 0.94441 | 0.298131 | | | P60981;F6 | P60981;F6 | Destrin | DSTN | 1288 |
| | 0.936937 | 1.508456 | | | Q00796;A( | Q00796;A( | Sorbitol de | SORD | 1408 |
| | 0.924541 | 1.178971 | | | Q7Z5R6 | Q7Z5R6 | Amyloid be | APBB1IP | 1601 |
| | 0.906494 | 1.215081 | | | F5GYN4;J3 | F5GYN4;J3 | Ubiquitin t | OTUB1 | 455 |
| | 0.857216 | 1.096487 | | | A6NFX8;Q! | A6NFX8;Q! | ADP-sugar | NUDT5 | 225 |

| | | | | | | | | | |
|---|---|---|---|---|---|---|---|---|---|
| | 0.811363 | 0.2646 | | | P30044 | P30044 | Peroxiredc | PRDX5 | 1070 |
| | 0.749985 | -1.78943 | | | Q01459 | Q01459 | Di-N-acety | CTBS | 1411 |
| | 0.694211 | 1.002793 | | | Q92973;E7 | Q92973 | Transporti | TNPO1 | 1654 |
| | 0.674244 | 1.032249 | | | O43396;K7 | O43396;K7 | Thioredoxi | TXNL1 | 680 |
| | 0.639447 | 0.141645 | | | P68133;P6 | P68133;P6 | Actin, alph | ACTA1;ACT | 1381 |
| | 0.634195 | 0.165234 | | | P62888;E5 | P62888;E5 | 60S riboso | RPL30 | 1356 |
| | 0.625928 | 0.718943 | | | Q9UGI8;F8 | Q9UGI8 | Testin | TES | 1799 |
| | 0.612281 | 1.283013 | | | Q16401;F2 | Q16401 | 26S protea | PSMD5 | 1532 |
| | 0.568803 | 1.229891 | | | P50452;H7 | P50452;H7 | Serpin B8 | SERPINB8 | 1215 |
| | 0.49883 | -0.09953 | | | A0A286YE\ | A0A286YE\ | Ig alpha-1 ( | IGHA1 | 154 |
| | 0.474398 | 0.80799 | | | Q92597;E5 | Q92597 | Protein ND | NDRG1 | 1643 |
| | 0.422381 | -0.19236 | | | J3QR09;J3I | J3QR09;J3I | Ribosomal | RPL19 | 579 |
| | 0.3839 | -0.13941 | | | P60903 | P60903 | Protein S1( | S100A10 | 1286 |
| | 0.33982 | 0.845206 | | | Q13907;C9 | Q13907 | Isopenteny | IDI1 | 1484 |
| | 0.338598 | 0.698478 | | | P38919;I3I | P38919;I3I | Eukaryotic | EIF4A3 | 1136 |
| | 0.32619 | 0.122053 | | | A0A1B0GL | A0A1B0GL | Aminoacyl | ACY1;ABHI | 141 |
| | 0.315252 | -0.09641 | | | P40429;M( | P40429;M( | 60S riboso | RPL13A;RP | 1144 |
| | 0.28703 | -0.4511 | | | A0A0B4J2/ | A0A0B4J2/ | Sedoheptu | TRPV1;SHF | 86 |
| | 0.24707 | 0.089989 | | | C9K025;P1 | C9K025;P1 | 60S riboso | RPL35A | 304 |
| | 0.237677 | 0.427208 | | | Q14005;H( | Q14005;H( | Pro-interle | IL16 | 1485 |
| | 0.22342 | 0.728228 | | | P62851;E9 | P62851 | 40S riboso | RPS25 | 1351 |
| | 0.157186 | 0.460849 | | | P00492;Q9 | P00492 | Hypoxanth | HPRT1 | 767 |
| | 0.119838 | 0.300939 | | | P48960;M( | P48960 | CD97 antig | CD97 | 1191 |
| | 0.069118 | -0.02787 | | | P24534;F8 | P24534 | Elongation | EEF1B2 | 1028 |
| | 0.039668 | 0.115435 | | | F8VZQ9;P8 | F8VZQ9;P8 | SAP domai | SARNP | 472 |
| | 0.033439 | 0.010817 | | | Q5VWC4;F | Q5VWC4;F | 26S protea | PSMD4 | 1268 |
| | 0.023156 | -0.00811 | | | P61254;J3( | P61254;J3( | 60S riboso | RPL26;RPL | 1304 |
| | 0.009319 | -0.00387 | | | H3BNC9;P( | H3BNC9;P( | 40S riboso | RPS17 | 541 |
| | 5.727043 | 0.446253 | | | Q10567;C9 | Q10567;C9 | AP-1 comp | AP1B1 | 1447 |
| | 0.405043 | -0.11479 | | | P16403 | P16403 | Histone H1 | HIST1H1C | 970 |
| | 3.633058 | 0.489196 | | | P62136;E9 | P62136;E9 | Serine/thre | PPP1CA;PF | 1319 |
| | 1.7899 | 0.194328 | | | P61586;C9 | P61586;C9 | Transformi | RHOA | 1308 |
| | 2.098904 | 0.301262 | | | P62701;Q8 | P62701;Q8 | 40S riboso | RPS4X;RPS | 1343 |
| | 1.054363 | 0.115123 | | | P62873;F6 | P62873;F6 | Guanine nu | GNB1 | 1354 |
| | 1.306376 | 1.971948 | | up1 | A0A087W1 | A0A087W1 | Far upstrea | KHSRP | 12 |
| | 2.749009 | 0.3135 | | | Q9Y376;A( | Q9Y376;A( | Calcium-bi | CAB39 | 39 |
| | 1.016131 | 0.188678 | | | O60234;M | O60234;M | Glia matur | GMFG | 694 |
| | 3.538622 | 0.267613 | | | Q5JP53;P0 | Q5JP53;P0 | Tubulin be | TUBB | 845 |
| | 1.202193 | 2.04624 | | | Q13283;E5 | Q13283;E5 | Ras GTPas( | G3BP1 | 1466 |
| | 0.190985 | 0.542859 | | | A0A1X7SB. | A0A1X7SB. | Probable A | DDX17 | 153 |
| + | 1.990929 | -2.13873 | down2 | down2 | A0A087X2( | A0A087X2( | Agrin;Agrir | AGRN | 49 |
| + | 3.839151 | -2.74847 | down2 | down2 | Q99523;A( | Q99523 | Sortilin | SORT1 | 1692 |
| + | 2.558407 | -3.09649 | down2 | down2 | P35590 | P35590 | Tyrosine-p | TIE1 | 1119 |
| + | 12.40325 | -3.69742 | down2 | down2 | A0A0C4DF | A0A0C4DF | Arylsulfata | ARSA | 91 |
| + | 3.621992 | -5.65696 | down2 | down2 | P11279 | P11279 | Lysosome- | LAMP1 | 909 |
| + | 1.616678 | 1.831995 | up1 | up1 | E7EWC2;F! | E7EWC2;F! | Ras GTPas( | IQGAP2 | 405 |
| + | 1.942337 | 1.739639 | up1 | up1 | Q9Y5X3;Q! | Q9Y5X3;Q! | Sorting ne) | SNX5 | 1850 |

| | | | | | | | | | |
|---|---|---|---|---|---|---|---|---|---|
| + | 12.19725 | 3.69315 | up2 | up2 | P26640;A0 | P26640;A0 | Valine--tRI | VARS | 1045 |
| + | 3.70864 | 3.339395 | up2 | up2 | B3KVN2;P4 | B3KVN2;P4 | Protein far | FNTA | 250 |
| + | 3.257987 | 3.323152 | up2 | up2 | A0A087WU | A0A087WU | Heterogen | HNRNPDL | 15 |
| + | 3.263118 | 3.282451 | up2 | up2 | O00625 | O00625 | Pirin | PIR | 647 |
| + | 1.377955 | 3.114871 | up2 | up2 | Q16864 | Q16864 | V-type pro | ATP6V1F | 1545 |
| + | 3.26812 | 3.069844 | up2 | up2 | E5RIA4;E7I | E5RIA4;E7I | Aspartyl ar | DNPEP | 386 |
| + | 3.254964 | 2.759818 | up2 | up2 | O15355 | O15355 | Protein ph | PPM1G | 671 |
| + | 1.873792 | 2.742932 | up2 | up2 | Q9ULZ3;H3 | Q9ULZ3 | Apoptosis- | PYCARD | 1816 |
| + | 2.106332 | 2.653276 | up2 | up2 | B0QY89;Q9 | B0QY89;Q9 | Eukaryotic | EIF3L | 237 |
| + | 2.0505 | 2.625594 | up2 | up2 | A0A0C4DF | A0A0C4DF | Monoglyce | MGLL | 89 |
| + | 2.310482 | 2.621048 | up2 | up2 | O00303;H0 | O00303 | Eukaryotic | EIF3F | 637 |
| + | 1.930921 | 2.510487 | up2 | up2 | O43681 | O43681 | ATPase ASI | ASNA1 | 685 |
| + | 1.811945 | 2.496112 | up2 | up2 | A0A087X2 | A0A087X2 | Glutathion | GPX4 | 10 |
| + | 1.988795 | 2.449033 | up2 | up2 | Q15233;C9 | Q15233;C9 | Non-POU c | NONO | 1516 |
| + | 1.659519 | 2.414438 | up2 | up2 | Q5T6V5;Q5 | Q5T6V5;Q5 | UPF0553 p | C9orf64 | 1568 |
| + | 2.157901 | 2.208049 | up2 | up2 | Q53H82 | Q53H82 | Beta-lactar | LACTB2 | 1557 |
| + | 1.989793 | 2.202825 | up2 | up2 | Q9Y4E8;F8 | Q9Y4E8;F8 | Ubiquitin c | USP15 | 1843 |
| + | 1.275987 | 2.429399 | up2 | | O14933;E9 | O14933;E9 | Ubiquitin/I | UBE2L6 | 661 |
| | 1.306288 | 1.575109 | | up1 | A0A0A0MT | A0A0A0MT | Integrin-lin | ILK | 77 |
| | 1.304822 | 1.507779 | | up1 | P13861;H7 | P13861 | cAMP-dep | PRKAR2A | 934 |
| | 1.301752 | 1.125147 | | up1 | A0A6I8PL4 | A0A6I8PL4 | Leucine--tF | LARS | 212 |
| | 1.303937 | 1.059246 | | up1 | A0A0A0MT | A0A0A0MT | E3 ubiquiti | RNF213 | 76 |
| | 1.303171 | -1.85279 | | down1 | P19801;C9 | P19801;C9 | Amiloride- | AOC1 | 995 |
| | 5.559567 | 0.446832 | | | O75436;S4 | O75436 | Vacuolar p | VPS26A | 722 |
| | 2.877874 | 0.427052 | | | P21291;E9 | P21291;E9 | Cysteine ar | CSRP1 | 1007 |
| | 1.579497 | 0.42332 | | | Q16831;C9 | Q16831;C9 | Uridine ph | UPP1 | 1544 |
| | 2.984273 | 0.417263 | | | Q00688;G3 | Q00688 | Peptidyl-pr | FKBP3 | 1405 |
| | 1.848419 | -0.12939 | | | P35268;K7 | P35268;K7 | 60S riboso | RPL22 | 1113 |
| | 1.897726 | -0.1984 | | | P62857 | P62857 | 40S riboso | RPS28 | 1353 |
| | 1.480928 | -0.22071 | | | A0A087X2 | A0A087X2 | Basigin | BSG | 51 |
| | 1.334235 | -0.24557 | | | F8VZG5;A0 | F8VZG5;A0 | Adenylate | AK2 | 207 |
| | 2.774701 | -0.32828 | | | P05362;E7 | P05362;E7 | Intercellula | ICAM1 | 818 |
| | 1.392026 | -0.35502 | | | G5E972;P4 | G5E972;P4 | Lamina-ass | TMPO | 500 |
| | 2.922604 | -0.37419 | | | A0A3B3IRL | A0A3B3IRL | Protein CR | CREG1 | 174 |
| | 1.644713 | -0.38763 | | | A0A0A0MS | A0A0A0MS | Leukocyte | LILRB4 | 73 |
| | 1.613197 | -0.40477 | | | Q9Y2Q3;E9 | Q9Y2Q3;E9 | Glutathion | GSTK1 | 1828 |
| | 3.346242 | -0.46255 | | | E9PPJ5;E9I | E9PPJ5;E9I | Midkine | MDK | 436 |
| | 3.041789 | -0.47354 | | | H0YBZ2;P0 | H0YBZ2;P0 | HLA class I | CD74 | 519 |
| | 2.841489 | -0.49314 | | | C9JFR7;P99 | C9JFR7;P99 | Cytochrom | CYCS | 295 |
| | 1.705947 | -0.50946 | | | Q5H9A7;P0 | Q5H9A7;P0 | Metallopro | TIMP1 | 775 |
| | 1.2622 | -0.36225 | | | Q9NZC2 | Q9NZC2 | Triggering | TREM2 | 1778 |
| | 1.136469 | 1.590239 | | | Q9NTX5;J3 | Q9NTX5 | Ethylmalor | ECHDC1 | 1769 |
| | 1.134314 | 0.304079 | | | Q15056 | Q15056 | Eukaryotic | EIF4H | 1509 |
| | 1.094028 | -0.21911 | | | G5EA09;O0 | G5EA09;O0 | Syntenin-1 | SDCBP | 502 |
| | 1.082951 | 1.969632 | | | P35270 | P35270 | Sepiapterir | SPR | 1114 |
| | 1.077009 | 2.087846 | | | B8ZZU8;I3I | B8ZZU8;I3I | Transcripti | TCEB2 | 276 |
| | 1.000528 | -1.35103 | | | A0A087X1 | A0A087X1 | Glutathion | GPX3 | 45 |

| | | | | | | | | | |
|---|---|---|---|---|---|---|---|---|---|
| | 0.974673 | 1.147508 | | | F5H5V4;F5 | F5H5V4;F5 | 26S protea | PSMD9 | 453 |
| | 0.972972 | 1.258983 | | | A0A0C4DG | A0A0C4DG | Enolase-ph | ENOPH1 | 100 |
| | 0.952687 | 0.187553 | | | Q5W0H4;A | Q5W0H4;A | Translatior | TPT1 | 88 |
| | 0.93149 | -0.2964 | | | Q9UII2;A0 | Q9UII2 | ATPase inh | ATPIF1 | 1808 |
| | 0.918921 | 1.459635 | | | J3KP15;Q0 | J3KP15;Q0 | Serine/arg | SRSF2 | 571 |
| | 0.912181 | 0.19404 | | | K7EK42;A0 | K7EK42;A0 | Tubulin-fol | TBCB | 192 |
| | 0.860192 | -1.34908 | | | H0Y755;M | H0Y755;M | Low affinit | FCGR3A | 510 |
| | 0.827263 | -0.11995 | | | Q9Y6W5 | Q9Y6W5 | Wiskott-Al | WASF2 | 1860 |
| | 0.810545 | -0.12436 | | | H0YHA7;J3 | H0YHA7;J3 | 60S riboso | RPL18 | 523 |
| | 0.784268 | 1.365401 | | | Q92882 | Q92882 | Osteoclast | OSTF1 | 1652 |
| | 0.778694 | -1.09121 | | | P17050 | P17050 | Alpha-N-ac | NAGA | 973 |
| | 0.667444 | 0.125125 | | | E7ETK0;A0 | E7ETK0;A0 | 40S riboso | RPS24 | 169 |
| | 0.612779 | 1.294317 | | | O75396;A0 | O75396 | Vesicle-tra | SEC22B | 721 |
| | 0.58888 | 0.1517 | | | D3YTB1;F8 | D3YTB1;F8 | 60S riboso | RPL32 | 357 |
| | 0.572604 | 0.950374 | | | F8W7C6;A | F8W7C6;A | 60S riboso | RPL10;RPL | 20 |
| | 0.566661 | 0.142937 | | | Q8NDH3;H | Q8NDH3 | Probable a | NPEPL1 | 1625 |
| | 0.561331 | 1.052994 | | | Q8N1G4 | Q8N1G4 | Leucine-ric | LRRC47 | 1616 |
| | 0.549217 | 1.383518 | | | P07741;H3 | P07741;H3 | Adenine pl | APRT | 850 |
| | 0.547375 | 1.261931 | | | O75390;B4 | O75390;B4 | Citrate syn | CS | 254 |
| | 0.538562 | 0.858576 | | | O75995 | O75995 | SAM and S | SASH3 | 735 |
| | 0.514447 | 0.140036 | | | P60953;A0 | P60953;A0 | Cell divisio | CDC42 | 1287 |
| | 0.503584 | 1.212317 | | | E5RHN3;Q | E5RHN3;Q | Hepatitis A | HAVCR2 | 383 |
| | 0.455913 | 0.663419 | | | O75608;A0 | O75608;A0 | Acyl-protei | LYPLA1 | 726 |
| | 0.428719 | 0.462456 | | | Q07666;Q | Q07666 | KH domain | KHDRBS1 | 1432 |
| | 0.383331 | -0.08466 | | | Q93091 | Q93091 | Ribonuclea | RNASE6 | 1656 |
| | 0.367392 | -0.6015 | | | C9J0K6;P3 | C9J0K6;P3 | Sorcin | SRI | 279 |
| | 0.355057 | -0.79935 | | | O75367;B4 | O75367;B4 | Core histor | H2AFY | 718 |
| | 0.329198 | -0.09062 | | | P42574;A8 | P42574;A8 | Caspase-3; | CASP3 | 1156 |
| | 0.315593 | -0.843 | | | P98066 | P98066 | Tumor nec | TNFAIP6 | 1400 |
| | 0.192859 | -0.06608 | | | P01834;A0 | P01834;A0 | Ig kappa ch | IGKC | 779 |
| | 0.187558 | 0.564347 | | | Q9UBQ7;U | Q9UBQ7;U | Glyoxylate | GRHPR | 1792 |
| | 0.1783 | 0.46504 | | | Q9Y624;A0 | Q9Y624;A0 | Junctional | F11R | 1854 |
| | 0.173019 | 0.614982 | | | P49207 | P49207 | 60S riboso | RPL34 | 1194 |
| | 0.159048 | 0.427314 | | | C9JXB8;C9 | C9JXB8;C9 | 60S riboso | RPL24 | 300 |
| | 0.147288 | -0.38108 | | | P55058 | P55058 | Phospholip | PLTP | 1269 |
| | 0.144572 | 0.049294 | | | J3KTL2;Q0 | J3KTL2;Q0 | Serine/arg | SRSF1 | 581 |
| | 0.1434 | -0.04636 | | | P42766;F2 | P42766;F2 | 60S riboso | RPL35 | 1159 |
| | 0.036812 | -0.10066 | | | P08621;M0 | P08621;M0 | U1 small n | SNRNP70 | 861 |
| | 0.036151 | 0.09054 | | | P13716;B7 | P13716 | Delta-amir | ALAD | 931 |
| | 0.770702 | 0.184631 | | | P14618;B4 | P14618;B4 | Pyruvate k | PKM | 940 |
| | 2.734363 | -0.59583 | | | O43278;H3 | O43278;H3 | Kunitz-type | SPINT1 | 678 |
| | 0.023239 | -0.01019 | | | P01859;A0 | P01859;A0 | Ig gamma- | IGHG2 | 781 |
| | 1.913934 | 0.453269 | | | Q13404;I3 | Q13404;I3 | Ubiquitin-c | UBE2V1;TN | 1470 |
| | 0.51518 | -0.2435 | | | P01857;A0 | P01857;A0 | Ig gamma- | IGHG1 | 780 |
| | 2.6495 | 0.250626 | | | H0YHC3;F8 | H0YHC3;F8 | Nucleosom | NAP1L1 | 269 |
| | 0.514545 | 0.840393 | | | P31943;E9 | P31943;E9 | Heterogen | HNRNPH1 | 417 |
| + | 1.987209 | -2.29001 | down2 | down2 | P30084 | P30084 | Enoyl-CoA | ECHS1 | 1075 |

| | | | | | | | | | |
|---|---|---|---|---|---|---|---|---|---|
| + | 1.99233 | -2.37092 | down2 | down2 | P15848;A0 | P15848 | Arylsulfata | ARSB | 957 |
| + | 1.860926 | -2.54809 | down2 | down2 | C9JL30;H7( | C9JL30;H7( | Sulfatase-r | SUMF2 | 286 |
| + | 1.812582 | -2.87894 | down2 | down2 | O15400 | O15400 | Syntaxin-7 | STX7 | 673 |
| + | 14.26613 | 4.336834 | up2 | up2 | Q13148;A( | Q13148;A( | TAR DNA-b | TARDBP;TI | 21 |
| + | 14.61027 | 4.329583 | up2 | up2 | Q9HAB8;Q | Q9HAB8 | Phosphopa | PPCS | 1748 |
| + | 13.23136 | 4.128107 | up2 | up2 | P51580 | P51580 | Thiopurine | TPMT | 1234 |
| + | 3.251686 | 3.548674 | up2 | up2 | P14598;A6 | P14598;A6 | Neutrophil | NCF1;NCF1 | 226 |
| + | 12.27091 | 3.390247 | up2 | up2 | P63010;A0 | P63010;A0 | AP-2 comp | AP2B1 | 1368 |
| + | 2.170252 | 3.123709 | up2 | up2 | Q15843;F8 | Q15843;F8 | NEDD8 | NEDD8;NE | 1528 |
| + | 1.994421 | 2.738252 | up2 | up2 | H7BZ14;Q! | H7BZ14;Q! | Peptidyl-pi | PPIL3 | 553 |
| + | 1.990192 | 2.589746 | up2 | up2 | P37108;H0 | P37108;H0 | Signal reco | SRP14 | 1129 |
| + | 1.992226 | 2.550611 | up2 | up2 | O95433;G3 | O95433;G3 | Activator c | AHSA1 | 751 |
| + | 3.289072 | 2.490804 | up2 | up2 | P52888;K7 | P52888;K7 | Thimet olig | THOP1 | 1247 |
| + | 1.443069 | 2.289809 | up2 | up2 | A0A2R8Y4 | A0A2R8Y4 | Lambda-cr | CRYL1 | 159 |
| + | 1.449467 | 2.226154 | up2 | up2 | P09417;B7 | P09417;B7 | Dihydropte | QDPR | 869 |
| + | 1.694973 | 2.225926 | up2 | up2 | P55060 | P55060 | Exportin-2 | CSE1L | 1270 |
| + | 1.306018 | 2.127501 | up2 | up2 | O76054;B3 | O76054;B3 | SEC14-like | SEC14L2 | 737 |
| + | 3.151094 | 2.119313 | up2 | up2 | A0A0U1RC | A0A0U1RC | Rho-associ | ROCK1 | 130 |
| | 1.306737 | 1.432511 | | up1 | B1AMS2;Q | B1AMS2;Q | Septin-6 | SEPTIN6 | 245 |
| | 1.976908 | 1.226272 | | up1 | Q9P253 | Q9P253 | Vacuolar p | VPS18 | 1785 |
| | 1.840052 | 0.899379 | | | A0A087W2 | A0A087W2 | Ensconsin | MAP7 | 34 |
| | 2.097747 | 0.247971 | | | Q15369;E5 | Q15369;E5 | Transcripti | TCEB1 | 382 |
| | 1.3292 | 0.171616 | | | F8VZJ2;F8\ | F8VZJ2;F8\ | Nascent pc | NACA | 410 |
| | 3.228917 | -0.20839 | | | Q8WVC2;F | Q8WVC2;F | 40S riboso | RPS21 | 1374 |
| | 1.594695 | -0.41325 | | | E9PNW4;A | E9PNW4;A | CD59 glycc | CD59 | 172 |
| | 3.812335 | -0.42132 | | | P0DP25;P0 | P0DP25;P0 | DP24;P0DF | CALM2;CA | 888 |
| | 2.57458 | -0.61117 | | | Q969H8;M | Q969H8;M | Myeloid-de | MYDGF | 1657 |
| | 1.281673 | 0.204887 | | | O00626 | O00626 | C-C motif c | CCL22 | 648 |
| | 1.235015 | 0.222067 | | | P10155;G5 | P10155 | 60 kDa SS-. | TROVE2 | 891 |
| | 1.175799 | 0.401781 | | | O00244;E5 | O00244;E5 | Copper tra | ATOX1 | 635 |
| | 1.135607 | -1.2912 | | | P16284;A0 | P16284;A0 | Platelet en | PECAM1 | 967 |
| | 1.09505 | 1.566884 | | | P00568;Q5 | P00568;Q5 | Adenylate | AK1 | 770 |
| | 1.056517 | 1.952356 | | | Q15121;B1 | Q15121;B1 | Astrocytic | PEA15 | 1512 |
| | 1.016053 | 1.349808 | | | P15170;H3 | P15170;H3 | Eukaryotic | GSPT1;GSF | 952 |
| | 0.995669 | 0.237034 | | | P60983;G3 | P60983;G3 | Glia matur | GMFB | 1289 |
| | 0.977967 | 0.274308 | | | P62495;D6 | P62495;D6 | Eukaryotic | ETF1 | 1341 |
| | 0.927733 | 1.414018 | | | Q99733;C9 | Q99733;C9 | Nucleosom | NAP1L4 | 1698 |
| | 0.915439 | -0.41296 | | | P13747;Q3 | P13747 | HLA class I | HLA-E | 932 |
| | 0.907312 | 2.328754 | | | P10124 | P10124 | Serglycin | SRGN | 890 |
| | 0.889552 | 1.141568 | | | P31942 | P31942 | Heterogen | HNRNPH3 | 1094 |
| | 0.873783 | -0.13749 | | | P62913;Q5 | P62913;Q5 | 60S riboso | RPL11 | 1359 |
| | 0.834113 | 1.121589 | | | O95372;Q! | O95372;Q! | Acyl-protei | LYPLA2 | 747 |
| | 0.829092 | 1.282538 | | | Q12792;F8 | Q12792;F8 | Twinfilin-1 | TWF1 | 1449 |
| | 0.819278 | 0.330638 | | | P22307;H0 | P22307 | Non-specif | SCP2 | 1014 |
| | 0.790439 | -1.3444 | | | P63218 | P63218 | Guanine nu | GNG5 | 1373 |
| | 0.731814 | 1.712411 | | | Q9C002;H( | Q9C002;H( | Normal mu | NMES1;C1 | 1727 |
| | 0.680011 | 0.754656 | | | O43747;B4 | O43747 | AP-1 comp | AP1G1 | 687 |

| | | | | | | | | | |
|---|---|---|---|---|---|---|---|---|---|
| | 0.669033 | 0.910346 | | | Q92499;F1 | Q92499;F1 | ATP-depen | DDX1 | 1639 |
| | 0.659566 | 0.977561 | | | P30419;B7 | P30419;B7 | Glycylpept | NMT1 | 1081 |
| | 0.610292 | 0.660322 | | | P08134;Q5 | P08134;Q5 | Rho-relate | RHOC | 856 |
| | 0.555754 | 1.022737 | | | P62266;D6 | P62266;D6 | 40S riboso | RPS23 | 1327 |
| | 0.550807 | 1.581576 | | | Q14240;E7 | Q14240;E7 | Eukaryotic | EIF4A2 | 1495 |
| | 0.504341 | 0.796609 | | | P62277;J3I | P62277;J3I | 40S riboso | RPS13 | 1330 |
| | 0.492953 | 1.345548 | | | P15090;E5 | P15090 | Fatty acid- | FABP4 | 947 |
| | 0.489724 | 1.048677 | | | Q12765;C9 | Q12765 | Secernin-1 | SCRN1 | 1448 |
| | 0.439395 | 0.803255 | | | P46976;G5 | P46976;G5 | Glycogenir | GYG1 | 1180 |
| | 0.405231 | -0.99541 | | | B8ZZQ6;P0 | B8ZZQ6;P0 | Prothymos | PTMA | 275 |
| | 0.400144 | 0.985171 | | | P07203;A0 | P07203;A0 | Glutathion | GPX1 | 16 |
| | 0.374892 | 0.703084 | | | P43405 | P43405 | Tyrosine-p | SYK | 1165 |
| | 0.339706 | 0.103931 | | | Q92688 | Q92688 | Acidic leuc | ANP32B | 1646 |
| | 0.22002 | 0.882121 | | | P61970;H3 | P61970;H3 | Nuclear tra | NUTF2 | 1315 |
| | 0.168903 | -0.47206 | | | Q9Y266;A0 | Q9Y266;A0 | Nuclear mi | NUDC | 1824 |
| | 0.139482 | 0.035771 | | | P56537;B7 | P56537;B7 | Eukaryotic | EIF6 | 1278 |
| | 0.133587 | 0.029797 | | | P60866;E5 | P60866;E5 | 40S riboso | RPS20 | 1285 |
| | 0.12025 | -0.01742 | | | J3KS52;Q8 | J3KS52;Q8 | CMRF35-lil | CD300LF | 575 |
| | 0.116688 | 0.407817 | | | Q9UIB8 | Q9UIB8 | SLAM fami | CD84 | 1807 |
| | 0.102986 | -0.29663 | | | Q9BS26 | Q9BS26 | Endoplasm | ERP44 | 1709 |
| | 0.057639 | 0.178789 | | | O00232 | O00232 | 26S protea | PSMD12 | 634 |
| | 0.053642 | 0.147724 | | | M0QX65;A | M0QX65;A | SUMO-acti | SAE1 | 203 |
| | 0.03893 | -0.09069 | | | Q14444;G3 | Q14444;G3 | Caprin-1 | CAPRIN1 | 485 |
| | 0.034558 | -0.00829 | | | P11142;E9 | P11142;E9 | Heat shock | HSPA8 | 904 |
| | 2.418427 | 0.22382 | | | B1AK88;B1 | B1AK88;B1AK87 | | CAPZB | 242 |
| | 0.346121 | 0.087586 | | | P61224;A6 | P61224;A6 | Ras-relate | RAP1B | 1301 |
| | 2.709877 | -0.2068 | | | P78324;Q9 | P78324 | Tyrosine-p | SIRPA | 1390 |
| | 1.677213 | -0.23581 | | | A2AAZ0;Q5 | A2AAZ0;Q5Y7D6;Q5S | | HLA-DQB1 | 220 |
| | 0.984714 | 1.486625 | | | P61163;R4 | P61163;R4 | Alpha-cent | ACTR1A | 1299 |
| | 0.513969 | 0.064664 | | | M0QXS5;A | M0QXS5;A | Heterogen | HNRNPL | 181 |
| | 1.146866 | 0.222927 | | | P15153;B1 | P15153;B1 | Ras-relate | RAC2 | 951 |
| | 1.189081 | 1.852031 | | | Q02750;A0 | Q02750 | Dual specif | MAP2K1 | 1415 |
| | 0.978243 | 0.099061 | | | Q5Y7D1;A0 | Q5Y7D1;A0A5S8K7D6 | | HLA-DRB1 | 1575 |
| | 0.566986 | 0.821544 | | | O00629;H7 | O00629 | Importin su | KPNA4 | 649 |
| + | 3.249854 | 2.807221 | up2 | up2 | Q08209;E7 | Q08209;E7 | Serine/thre | PPP3CA;PF | 1436 |
| | 0.442774 | 0.701073 | | | P35813;E9 | P35813;E9 | Protein ph | PPM1A | 1122 |
| | 0.753259 | 1.011059 | | | P63167;F8 | P63167;F8 | Dynein ligh | DYNLL1 | 1372 |
| + | 1.966116 | -2.31617 | down2 | down2 | Q8IXK2 | Q8IXK2 | Polypeptid | GALNT12 | 1614 |
| + | 1.761396 | -2.51752 | down2 | down2 | Q5JS37;C9 | Q5JS37;C9 | NHL repea | NHLRC3 | 1563 |
| + | 3.258406 | -3.26311 | down2 | down2 | Q13011;M | Q13011;M | Delta(3,5)- | ECH1 | 1453 |
| + | 1.561015 | 1.868273 | up1 | up1 | P55735;A8 | P55735;A8 | Protein SE | SEC13 | 1273 |
| + | 12.17269 | 3.668212 | up2 | up2 | G3V169;P2 | G3V169;P2 | Caspase;Ca | CASP1 | 486 |
| + | 1.994162 | 3.454011 | up2 | up2 | Q9BY32 | Q9BY32 | Inosine trip | ITPA | 1722 |
| + | 3.151848 | 3.370474 | up2 | up2 | Q15819;G3 | Q15819;G3 | Ubiquitin-c | UBE2V2 | 1525 |
| + | 3.662085 | 2.897599 | up2 | up2 | Q86X76 | Q86X76 | Nitrilase ho | NIT1 | 1610 |
| + | 3.256995 | 2.764703 | up2 | up2 | Q16774;B1 | Q16774;B1 | Guanylate | GUK1 | 1541 |
| + | 1.637454 | 2.586533 | up2 | up2 | Q9NVJ2;Q9 | Q9NVJ2;Q9 | ADP-ribosy | ARL8B;ARL | 1771 |

| | | | | | | | | | |
|---|---|---|---|---|---|---|---|---|---|
| + | 2.355363 | 2.572162 | up2 | up2 | E9PKG1;Q9 | E9PKG1;Q9 | Protein arg | PRMT1 | 428 |
| + | 1.741036 | 2.566947 | up2 | up2 | P52597;A0 | P52597 | Heterogen | HNRNPF | 1244 |
| + | 3.219089 | 2.555718 | up2 | up2 | Q9BY43;E9 | Q9BY43;E9 | Charged m | CHMP4A | 1723 |
| + | 1.657155 | 2.484951 | up2 | up2 | P48507 | P48507 | Glutamate | GCLM | 1188 |
| + | 2.334137 | 2.271592 | up2 | up2 | Q96AT9;C9 | Q96AT9;C9 | Ribulose-p | RPE | 1660 |
| + | 1.867173 | 2.144181 | up2 | up2 | P28072;I3I | P28072;I3I | Proteasom | PSMB6 | 1056 |
| + | 1.993957 | 2.139542 | up2 | up2 | O00442;A6 | O00442 | RNA 3-term | RTCA | 641 |
| + | 1.306496 | 2.116831 | up2 | up2 | Q9NTK5;J3 | Q9NTK5;J3 | Obg-like A | OLA1 | 572 |
| + | 1.490332 | 2.089457 | up2 | up2 | Q9H993;F5 | Q9H993;F5 | Protein-glu | ARMT1 | 1746 |
| + | 1.746678 | 2.081007 | up2 | up2 | O14737;Q3 | O14737;Q3 | Programm | PDCD5 | 655 |
| + | 2.903175 | 2.059462 | up2 | up2 | A0A0G2JQ | A0A0G2JQ | Active brea | ABR | 121 |
| + | 1.245909 | 2.4488 | up2 | | H0Y8X4;O4 | H0Y8X4;O4 | 2-deoxynu | DNPH1 | 516 |
| | 1.305557 | 1.938269 | | up1 | A0A1B0GV | A0A1B0GV | Ceroid-lipo | CLN5 | 146 |
| | 1.305475 | 1.937238 | | up1 | C9JBI3;P78 | C9JBI3;P78 | Phosphose | PSPH | 292 |
| | 1.306761 | 1.513158 | | up1 | Q9P2R3;I3 | Q9P2R3 | Rabankyrin | ANKFY1 | 1788 |
| | 1.921498 | 1.345387 | | up1 | V9GYR2;P0 | V9GYR2;P0 | Sodium/po | ATP1B1 | 810 |
| | 1.305187 | 1.302507 | | up1 | P35573 | P35573 | Glycogen d | AGL | 1116 |
| | 3.404119 | -1.16418 | | down1 | C9JGG1;C9 | C9JGG1;C9 | Paired imm | PILRA | 296 |
| | 1.380597 | -1.62494 | | down1 | P15907;H7 | P15907;H7 | Beta-galac | ST6GAL1 | 959 |
| | 1.321391 | -1.78357 | | down1 | Q9Y646;E5 | Q9Y646;E5 | Carboxype | CPQ | 1855 |
| | 1.305134 | -1.79923 | | down1 | P09497;H0 | P09497 | Clathrin lig | CLTB | 875 |
| | 1.306258 | -1.92279 | | down1 | Q99574;C9 | Q99574;C9 | Neuroserp | SERPINI1 | 1695 |
| | 1.351121 | 0.453194 | | | Q9NRX4 | Q9NRX4 | 14 kDa pho | PHPT1 | 1767 |
| | 4.493713 | 0.354807 | | | P47813;O1 | P47813;O1 | Eukaryotic | EIF1AX;EIF | 653 |
| | 3.370161 | 0.338372 | | | O75531 | O75531 | Barrier-to- | BANF1 | 724 |
| | 1.519748 | -0.15496 | | | Q5VZR0;Q9 | Q5VZR0;Q9 | Golgi-asso | GLIPR2 | 1574 |
| | 1.59844 | -0.28509 | | | P14854;A0 | P14854;A0 | Cytochrom | COX6B1 | 944 |
| | 1.242434 | 0.915655 | | | P20701 | P20701 | Integrin alp | ITGAL | 1002 |
| | 1.203596 | 0.275761 | | | P55957 | P55957 | BH3-intera | BID | 1276 |
| | 1.132961 | 1.388554 | | | B4DKY1;P4 | B4DKY1;P4 | Cysteine--t | CARS | 255 |
| | 1.117282 | -1.3686 | | | P10606 | P10606 | Cytochrom | COX5B | 896 |
| | 1.087985 | -1.95324 | | | A0A087WZ | A0A087WZ | BolA-like p | BOLA2;BOL | 35 |
| | 1.010787 | 1.824523 | | | P50135 | P50135 | Histamine | HNMT | 1210 |
| | 1.00488 | 2.168078 | | | Q14116 | Q14116 | Interleukin | IL18 | 1489 |
| | 0.997698 | 1.726666 | | | O15371;B0 | O15371;B0 | Eukaryotic | EIF3D | 672 |
| | 0.974157 | -2.07142 | | | Q96QR8 | Q96QR8 | Transcripti | PURB | 1681 |
| | 0.966805 | 1.546653 | | | P61086;D6 | P61086;D6 | Ubiquitin-c | UBE2K | 1294 |
| | 0.961563 | -0.36691 | | | Q9UJJ9;H0 | Q9UJJ9 | N-acetylglu | GNPTG | 1811 |
| | 0.956426 | 1.724741 | | | Q9Y3U8;J3 | Q9Y3U8;J3 | 60S riboso | RPL36 | 1839 |
| | 0.922032 | 1.032696 | | | Q8NCW5;C | Q8NCW5;C | NAD(P)H-h | APOA1BP | 1624 |
| | 0.848267 | 0.089673 | | | P62854;Q5 | P62854;Q5 | 40S riboso | RPS26;RPS | 1352 |
| | 0.827425 | 1.087467 | | | B4DDF4;B4 | B4DDF4;B4 | Calponin;C | CNN2 | 252 |
| | 0.824109 | 1.193399 | | | J3QLE5;P6 | J3QLE5;P6 | Small nucle | SNRPN;SNI | 582 |
| | 0.746344 | 1.519012 | | | Q9NXG2;H | Q9NXG2;H | THUMP do | THUMPD1 | 1775 |
| | 0.744637 | 0.202987 | | | P50238;K4 | P50238 | Cysteine-ri | CRIP1 | 1212 |
| | 0.716588 | 1.376666 | | | Q9BRA2;I3 | Q9BRA2;I3 | Thioredoxi | TXNDC17 | 1706 |
| | 0.712228 | 1.443548 | | | P51148;K7 | P51148;K7 | Ras-relate | RAB5C;RAI | 1226 |

| | | | | | | | | | |
|---|---|---|---|---|---|---|---|---|---|
| | 0.676048 | 0.789581 | | | Q96PP8;E7 | Q96PP8 | Guanylate- | GBP5 | 1679 |
| | 0.657392 | 1.004185 | | | P49458;Q6 | P49458;Q6 | Signal reco | SRP9;DKFZ | 1198 |
| | 0.625869 | 1.057186 | | | O75884 | O75884 | Putative hy | RBBP9 | 732 |
| | 0.59076 | 1.248896 | | | P42768;C9 | P42768;C9 | Wiskott-Al | WAS | 1160 |
| | 0.518273 | 0.679837 | | | Q08257;C9 | Q08257;C9 | Quinone o | CRYZ | 1438 |
| | 0.483813 | -1.22273 | | | P68402;J3I | P68402 | Platelet-ac | PAFAH1B2 | 1387 |
| | 0.476241 | 1.046038 | | | E9PKL9;Q1 | E9PKL9;Q1 | GDP-L-fucc | TSTA3 | 430 |
| | 0.438689 | 0.734152 | | | H9KV28;A0 | H9KV28;A0 | Protein dia | DIAPH1 | 112 |
| | 0.337973 | 1.107048 | | | P52788;H7 | P52788;H7 | Spermine s | SMS | 1245 |
| | 0.332958 | -0.81415 | | | A0A087X1 | A0A087X1 | CD48 antig | CD48 | 47 |
| | 0.304561 | 0.148387 | | | A6NNI4;A0 | A6NNI4;A0 | Tetraspani | CD9 | 158 |
| | 0.302938 | -0.49116 | | | P22234;E9 | P22234;E9 | Multifunct | PAICS | 414 |
| | 0.227876 | -0.61536 | | | F2Z2Y6;O9 | F2Z2Y6;O9 | U6 snRNA- | LSM8 | 449 |
| | 0.180384 | -0.50878 | | | Q9Y2Q5 | Q9Y2Q5 | Ragulator c | LAMTOR2 | 1829 |
| | 0.167361 | 0.604343 | | | C9J8T4;C9 | C9J8T4;C9 | E3 ubiquiti | RNF13 | 288 |
| | 0.12991 | 0.03954 | | | Q5T123;Q9 | Q5T123;Q9 | SH3 domai | SH3BGRL3 | 1566 |
| | 0.075275 | 0.033442 | | | P20962;F5 | P20962 | Parathymc | PTMS | 1004 |
| | 0.072118 | 0.169989 | | | E5RJR5;P6 | E5RJR5;P6 | S-phase kir | SKP1 | 391 |
| | 0.070515 | -0.02601 | | | O60869 | O60869 | Endothelia | EDF1 | 709 |
| | 0.06828 | 0.18446 | | | K7ER90;K7 | K7ER90;K7 | Eukaryotic | EIF3G | 596 |
| | 0.029384 | 0.07225 | | | Q15637;H7 | Q15637 | Splicing fac | SF1 | 1522 |
| | 0.018405 | -0.06982 | | | Q9Y5Z4;Q5 | Q9Y5Z4 | Heme-binc | HEBP2 | 1852 |
| | 0.000982 | 0.000215 | | | P84090;G3 | P84090;G3 | Enhancer c | ERH | 1397 |
| | 4.297821 | 0.457901 | | | Q92598;R4 | Q92598 | Heat shock | HSPH1 | 1644 |
| | 2.514437 | -0.34086 | | | A0A182DV | A0A182DWE9;A0A140T915;A0A1 | | | 138 |
| | 2.221501 | -0.38144 | | | P61769;F5 | P61769;F5 | Beta-2-mic | B2M | 1312 |
| + | 1.431178 | 2.725848 | up2 | up2 | P62244;I3I | P62244;I3I | 40S riboso | RPS15A | 1324 |
| | 0.608598 | -0.06996 | | | J3QRS3;P1 | J3QRS3;P1 | Myosin reg | MYL12A;M | 584 |
| | 1.567342 | -0.29691 | | | P04440;H0 | P04440;H0 | HLA class I | HLA-DPB1 | 805 |
| | 0.800636 | 1.693083 | | | Q5Y7D2;P0 | Q5Y7D2;P0 | HLA class I | HLA-DQA1 | 783 |
| | 0.561285 | 0.076555 | | | P62879;E7 | P62879;E7 | Guanine nu | GNB2 | 1355 |
| | 0.477584 | 0.104101 | | | P11234;C9 | P11234;C9 | Ras-relatec | RALB | 908 |
| + | 13.38911 | 3.595349 | up2 | up2 | Q9UBW5;S | Q9UBW5;S | Bridging in | BIN2 | 1797 |
| | 2.63995 | -0.52102 | | | A0A140T9 | A0A140T9 | HLA class I | HLA-DPA1 | 133 |
| | 0.146058 | 0.045498 | | | P61956;Q6 | P61956 | Small ubiq | SUMO2 | 1313 |
| + | 1.748907 | -2.82064 | down2 | down2 | P13473;H0 | P13473 | Lysosome- | LAMP2 | 923 |
| + | 3.213802 | -3.77069 | down2 | down2 | F8WC54;A | F8WC54;A | Disintegrin | ADAM9 | 215 |
| + | 1.991364 | 1.884669 | up1 | up1 | B4DR80;Q9 | B4DR80;Q9 | Serine/thre | STK24 | 257 |
| + | 2.607934 | 4.343502 | up2 | up2 | Q6P163;K7 | Q6P163;K7 | Apolipopro | APOC2;APO | 607 |
| + | 1.385956 | 3.836498 | up2 | up2 | P14174 | P14174 | Macrophag | MIF | 935 |
| + | 3.563681 | 3.114132 | up2 | up2 | P27105;F8 | P27105 | Erythrocyt | STOM | 1047 |
| + | 2.040161 | 2.872373 | up2 | up2 | D6RCK3;Q9 | D6RCK3;Q9 | MOB kinas | MOB1B;M | 369 |
| + | 3.257323 | 2.630034 | up2 | up2 | O00487 | O00487 | 26S protea | PSMD14 | 643 |
| + | 1.993779 | 2.442299 | up2 | up2 | O60925;E5 | O60925;E5 | Prefoldin s | PFDN1 | 380 |
| + | 1.993512 | 2.408018 | up2 | up2 | Q9Y5S1;K7 | Q9Y5S1 | Transient r | TRPV2 | 1848 |
| + | 1.992137 | 2.155371 | up2 | up2 | Q9Y5P6 | Q9Y5P6 | Mannose-1 | GMPPB | 1847 |
| + | 1.984344 | 2.153873 | up2 | up2 | A0A0J9YX0 | A0A0J9YX0 | LIM and se | LIMS1 | 124 |

| | | | | | | | | | |
|---|---|---|---|---|---|---|---|---|---|
| + | 1.977839 | 2.146345 | up2 | up2 | P26006;K7 | P26006 | Integrin al | ITGA3 | 1037 |
| + | 1.990762 | 2.062061 | up2 | up2 | Q00535 | Q00535 | Cyclin-dep | CDK5 | 1403 |
| + | 1.065171 | 3.322163 | up2 | | Q9P1F3;Q | Q9P1F3 | Costars far | ABRACL | 1784 |
| | 1.3059 | 1.990035 | | up1 | Q96K17;E | Q96K17;E | Transcripti | BTF3L4 | 431 |
| | 1.306188 | 1.737239 | | up1 | Q9HBI0 | Q9HBI0 | Gamma-pa | PARVG | 1754 |
| | 1.299517 | 1.597837 | | up1 | J3KQL8;Q9 | J3KQL8;Q9 | Apolipopro | APOL2 | 573 |
| | 1.352328 | 1.481538 | | up1 | C9JP00;A0 | C9JP00;A0 | Muscleblin | MBNL1 | 60 |
| | 1.306781 | 1.463889 | | up1 | Q86U42;B | Q86U42;B | Polyadenyl | PABPN1 | 1606 |
| | 1.30631 | 1.373576 | | up1 | P34096 | P34096 | Ribonuclea | RNASE4 | 1108 |
| | 1.518241 | -1.59919 | | down1 | P10109 | P10109 | Adrenodox | FDX1 | 889 |
| | 2.960636 | 0.393042 | | | Q9NZL9;E | Q9NZL9;E | Methionin | MAT2B | 1779 |
| | 1.292397 | -1.21926 | | | Q13263;M | Q13263 | Transcripti | TRIM28 | 1465 |
| | 1.291786 | 1.787085 | | | A0A087X0 | A0A087X0 | Sorting ne | SNX12 | 41 |
| | 1.288989 | 1.250396 | | | Q9Y3A5;A | Q9Y3A5;A | Ribosome | SBDS | 37 |
| | 1.282481 | -1.90601 | | | P12544 | P12544 | Granzyme | GZMA | 916 |
| | 1.155288 | 2.174426 | | | D6RDG3;H | D6RDG3;H | Transcripti | BTF3 | 370 |
| | 1.052553 | 1.6515 | | | B0QY16;Q | B0QY16;Q | Uncharact | KIAA0930 | 236 |
| | 1.031332 | -1.42605 | | | H0YJB9;G3 | H0YJB9;G3 | UPF0568 p | C14orf166 | 494 |
| | 1.013419 | -1.54553 | | | P12004 | P12004 | Proliferatir | PCNA | 913 |
| | 1.011085 | 1.397771 | | | O15212;A | O15212 | Prefoldin s | PFDN6 | 669 |
| | 0.995584 | 2.284529 | | | P62273;A0 | P62273;A0 | 40S riboso | RPS29 | 1329 |
| | 0.993169 | 1.286692 | | | B1AJQ6;Q | B1AJQ6;Q | Syntaxin-1 | STX12 | 240 |
| | 0.976262 | -1.33989 | | | Q9UM22;D | Q9UM22 | Mammalia | EPDR1 | 1817 |
| | 0.975406 | 1.404109 | | | Q9H098;X | Q9H098;X | Protein FA | FAM107B | 1731 |
| | 0.945573 | -1.39635 | | | P10412;Q | P10412 | Histone H1 | HIST1H1E | 893 |
| | 0.931984 | 1.67017 | | | Q5TFQ8;H | Q5TFQ8 | Signal-regu | SIRPB1 | 1572 |
| | 0.918099 | 1.035536 | | | Q9Y2S6;A0 | Q9Y2S6 | Translatior | TMA7 | 1830 |
| | 0.871308 | 1.531499 | | | P15104;A0 | P15104;A0 | Glutamine | GLUL | 948 |
| | 0.744737 | 0.994169 | | | Q07812;K | Q07812 | Apoptosis | BAX | 1433 |
| | 0.698426 | 1.68956 | | | J3QQZ9;Q | J3QQZ9;Q | Pyridoxine | PNPO | 583 |
| | 0.627513 | 1.48102 | | | B8ZZN6;P6 | B8ZZN6;P6 | Small ubiq | SUMO1 | 274 |
| | 0.621839 | -1.89506 | | | P06681;A0 | P06681;A0 | Compleme | C2 | 828 |
| | 0.610624 | 1.473739 | | | P98179 | P98179 | RNA-bindir | RBM3 | 1402 |
| | 0.578482 | 1.848888 | | | P59666;P5 | P59666;P5 | Neutrophil | DEFA3;DEF | 1280 |
| | 0.536229 | 1.124214 | | | P30405;H0 | P30405;H0 | Peptidyl-pr | PPIF | 1080 |
| | 0.489108 | -0.17983 | | | F8VV56;F8 | F8VV56;F8 | Tetraspani | CD63 | 468 |
| | 0.401459 | -0.11535 | | | E9PB61;Q | E9PB61;Q | THO comp | ALYREF | 411 |
| | 0.391306 | -0.54588 | | | Q3LXA3;I3 | Q3LXA3;I3 | Bifunctiona | DAK;TKFC | 1551 |
| | 0.36035 | 0.634349 | | | Q16643;D | Q16643;D | Drebrin | DBN1 | 361 |
| | 0.355512 | 0.749505 | | | Q9H832;I3 | Q9H832 | Ubiquitin-c | UBE2Z | 1743 |
| | 0.33263 | 0.839131 | | | F8VVA7;P6 | F8VVA7;P6 | Coatomer | COPZ1 | 469 |
| | 0.328589 | 0.582291 | | | P00167 | P00167 | Cytochrom | CYB5A | 760 |
| | 0.29015 | 0.468365 | | | Q9H008;Q | Q9H008;Q | Phospholy | LHPP | 1730 |
| | 0.218071 | -0.45924 | | | P42677;H0 | P42677;H0 | 40S riboso | RPS27;RPS | 1157 |
| | 0.198048 | -0.46531 | | | Q9Y5S9;A0 | Q9Y5S9 | RNA-bindir | RBM8A | 1849 |
| | 0.188401 | -0.06554 | | | P62306;A0 | P62306 | Small nucle | SNRPF | 1333 |
| | 0.1704 | 0.417761 | | | E9PNS3;F2 | E9PNS3;F2 | Toll-intera | TOLLIP | 440 |

| | | | | | | | | | |
|---|---|---|---|---|---|---|---|---|---|
| | 0.134813 | 0.397631 | | | O43390;B4 | O43390;B4 | Heterogen | HNRNPR | 679 |
| | 0.102568 | 0.336271 | | | Q9NQC3;F | Q9NQC3;F | Reticulon-4 | RTN4 | 1760 |
| | 0.088577 | -0.0293 | | | Q2L6G2;D5 | Q2L6G2;D5H3J5;A0A1 | | HLA-B | 358 |
| | 0.067197 | 0.24117 | | | A0A3B3ITT | A0A3B3ITT | 60S riboso | RPL29 | 182 |
| | 0.048785 | 0.145608 | | | Q00765;E2 | Q00765;E2 | Receptor e | REEP5 | 1407 |
| | 0.038975 | 0.115026 | | | P46778;G3 | P46778;G3 | 60S riboso | RPL21 | 1174 |
| | 0.02182 | 0.089923 | | | Q9BS40 | Q9BS40 | Latexin | LXN | 1710 |
| | 1.346293 | 0.235811 | | | P21333;F8 | P21333 | Filamin-A | FLNA | 1008 |
| | 0.375645 | 0.062319 | | | P67936;A0 | P67936;A0 | Tropomyo | TPM4 | 1380 |
| | 5.213531 | 0.548086 | | | Q13303;A0 | Q13303;A0 | Voltage-ga | KCNAB2 | 200 |
| | 1.824362 | 0.463621 | | | P67775;E5 | P67775 | Serine/thr | PPP2CA | 1378 |
| | 1.595981 | 0.077724 | | | F8W1R7;G | F8W1R7;G | Myosin ligl | MYL6 | 476 |
| | 0.543625 | -0.06182 | | | P62979;A0 | P62979;A0 | Ubiquitin-4 | RPS27A | 1363 |
| | 1.23761 | -0.1313 | | | Q09028;E9 | Q09028 | Histone-bi | RBBP4 | 1443 |
| | 0.554242 | 1.576925 | | | Q5RIP0;A0 | Q5RIP0;A0A0G2JIF0;A | | HLA-Cw;HL | 111 |
| | 0.100938 | 0.279734 | | | P62837;A0 | P62837;A0 | Ubiquitin-c | UBE2D2 | 1349 |
| + | 1.987111 | 2.00873 | up2 | up2 | H0Y4R1;P1 | H0Y4R1;P1 | Inosine-5-r | IMPDH2 | 507 |
| | 1.111118 | 1.144756 | | | P61026 | P61026 | Ras-relate | RAB10 | 1292 |
| | 0.771016 | 1.465217 | | | Q13885;Q9 | Q13885;Q9 | Tubulin be | TUBB2A;TU | 1483 |
| | 0.34243 | 0.384389 | | | Q8NHL6;A0 | Q8NHL6;A0 | Leukocyte | LILRB1 | 117 |
| + | 1.606592 | -2.14238 | down2 | down2 | A0A0C4DG | A0A0C4DG | CD276 ant | CD276 | 98 |
| + | 1.981098 | 1.854673 | up1 | up1 | O95379;D6 | O95379;D6 | Tumor nec | TNFAIP8 | 749 |
| + | 1.951212 | 1.663254 | up1 | up1 | J3KS94;J3C | J3KS94;J3C | Myelin bas | MBP | 576 |
| + | 1.993989 | 3.993105 | up2 | up2 | P05976;P0 | P05976;P0 | Myosin ligl | MYL1;MYL | 825 |
| + | 3.264436 | 2.86667 | up2 | up2 | A0A494BZ | A0A494BZ | Uroporphy | UROD | 145 |
| + | 1.460842 | 2.835772 | up2 | up2 | Q9H2A7;A0 | Q9H2A7;A0 | C-X-C moti | CXCL16 | 213 |
| + | 2.801939 | 2.79002 | up2 | up2 | P16083;Q5 | P16083;Q5 | Ribosyldih | NQO2 | 963 |
| + | 1.783012 | 2.586465 | up2 | up2 | P62314;J3 | P62314;J3 | Small nucl | SNRPD1 | 1336 |
| + | 1.989484 | 2.414239 | up2 | up2 | P0DPI2;A0 | P0DPI2;A0A0B4J2D5;A | | C21orf33 | 57 |
| | 1.30584 | 1.790236 | | up1 | O95166;H0 | O95166;H0 | Gamma-ar | GABARAP | 551 |
| | 1.300945 | 1.741782 | | up1 | Q5TDH0;H | Q5TDH0;H | Protein DD | DDI2;DDI1 | 1569 |
| | 1.305792 | 1.709316 | | up1 | E5RIT4;H0 | E5RIT4;H0 | Eukaryotic | EIF3E | 388 |
| | 1.361811 | 1.643494 | | up1 | Q9UHV9 | Q9UHV9 | Prefoldin s | PFDN2 | 1805 |
| | 1.306102 | 1.55836 | | up1 | A0A2P0CT | A0A2P0CT | Prostamid | FAM213B | 75 |
| | 1.983343 | 1.527915 | | up1 | Q9HB90;Q | Q9HB90;Q | Ras-relate | RRAGC;RRA | 1753 |
| | 1.301838 | 1.482858 | | up1 | P98082 | P98082 | Disabled h | DAB2 | 1401 |
| | 1.306179 | 1.266313 | | up1 | O75695 | O75695 | Protein XR | RP2 | 727 |
| | 1.296412 | 1.24961 | | up1 | B8ZZF0;C9 | B8ZZF0;C9 | Protein ph | PPM1B | 177 |
| | 1.302628 | 1.188046 | | up1 | A0A087W1 | A0A087W1 | UBX doma | UBXN1 | 14 |
| | 1.304987 | -1.56402 | | down1 | Q15717;A0 | Q15717;A0 | ELAV-like p | ELAVL1;EL | 1524 |
| | 1.305212 | -1.69265 | | down1 | P42126;Q9 | P42126;Q9 | Enoyl-CoA | ECI1;DCI | 1154 |
| | 1.297117 | -1.95317 | | down1 | Q9NX55;J3 | Q9NX55;J3 | Huntingtin | HYPK | 1774 |
| | 1.958387 | 0.359684 | | | O00757 | O00757 | Fructose-1 | FBP2 | 651 |
| | 2.203799 | -0.47848 | | | Q9Y287;A0 | Q9Y287;A0 | Integral m | ITM2B | 184 |
| | 1.375465 | -0.54334 | | | Q641Q3 | Q641Q3 | Meteorin-l | METRNL | 1577 |
| | 1.2893 | 0.904189 | | | Q8TD19 | Q8TD19 | Serine/thr | NEK9 | 1630 |
| | 1.266016 | 0.920893 | | | Q5STX8;P5 | Q5STX8;P5 | Allograft ir | AIF1 | 1266 |

| | | | | | | | |
|---|---|---|---|---|---|---|---|
| 1.221802 | 0.726446 | | Q8WW12 | Q8WW12 | PEST prote | PCNP | 1636 |
| 1.180053 | -2.14725 | | E9PR30;P6 | E9PR30;P6 | 40S riboso | FAU | 445 |
| 1.091422 | -0.21146 | | P34810;J3I | P34810;J3I | Macrosialiı | CD68 | 1109 |
| 1.082398 | 1.479755 | | H3BMQ1;H | H3BMQ1;H | Prefoldin s | PFDN5 | 537 |
| 1.032746 | 0.989899 | | M0R1L7;O | M0R1L7;O | Charged m | CHMP2A | 624 |
| 0.956592 | 0.615085 | | Q9Y2A7 | Q9Y2A7 | Nck-associ | NCKAP1 | 1826 |
| 0.819923 | 1.6054 | | A8MZH0;A | A8MZH0;A | Leukocyte | LILRA2 | 119 |
| 0.816436 | -1.30864 | | Q969Q0;H | Q969Q0;H | 60S riboso | RPL36AL;R | 1658 |
| 0.809085 | 1.166125 | | A0A096LN | A0A096LN | Ubiquitin-l | ISG15 | 55 |
| 0.709252 | 0.113738 | | Q3ZCM7;A | Q3ZCM7;A | Tubulin be | TUBB8 | 1552 |
| 0.639871 | -1.12007 | | E9PMV2 | E9PMV2 | | HLA-DQA1 | 438 |
| 0.562698 | 0.505863 | | Q5TFE4;Q5 | Q5TFE4;Q5 | 5-nucleotic | NT5DC1 | 1571 |
| 0.546925 | 2.083756 | | P62807;O6 | P62807;O6 | Histone H2 | HIST1H2BC | 708 |
| 0.53321 | 1.218408 | | Q5T4U8;P5 | Q5T4U8;P5 | Geranylger | RABGGTB | 1253 |
| 0.417854 | 0.25086 | | C9JXK0;A0 | C9JXK0;A0 | Lamin-B re | LBR | 195 |
| 0.401356 | 0.527726 | | P09958;H0 | P09958;H0 | Furin | FURIN | 882 |
| 0.389638 | 0.742263 | | O00571;A0 | O00571;A0 | ATP-depen | DDX3X;DD | 103 |
| 0.261178 | -0.44052 | | Q16706 | Q16706 | Alpha-man | MAN2A1 | 1538 |
| 0.216158 | -0.4317 | | A0A494C1 | A0A494C1 | Peptidyl-pı | FKBP2 | 196 |
| 0.194628 | 0.470884 | | A0A087X2 | A0A087X2 | Serine/argi | SRSF3 | 52 |
| 0.115189 | 0.34263 | | O43583;F8 | O43583;F8 | Density-re | DENR | 682 |
| 0.097558 | -0.2797 | | G5E9R3;M | G5E9R3;M | 60S riboso | RPL37A;RP | 284 |
| 0.090992 | 0.264337 | | H3BQF7;H | H3BQF7;H | IST1 homo | IST1 | 538 |
| 0.019374 | -0.06929 | | Q9Y3C6 | Q9Y3C6 | Peptidyl-pı | PPIL1 | 1834 |

| Peptides | Razor + un | Unique pe | Sequence c | Unique + r | Unique sec | Mol. weigh | Q-value | Score | Intensity |
|---|---|---|---|---|---|---|---|---|---|
| 187 | 187 | 186 | 41.4 | 41.4 | 41.3 | 531.78 | 0 | 323.31 | 3.27E+10 |
| 151 | 151 | 130 | 68 | 68 | 61.1 | 226.53 | 0 | 323.31 | 9.8E+10 |
| 92 | 92 | 92 | 43.1 | 43.1 | 43.1 | 269.76 | 0 | 323.31 | 2.95E+10 |
| 71 | 71 | 71 | 49.5 | 49.5 | 49.5 | 192.06 | 0 | 323.31 | 3.88E+10 |
| 73 | 73 | 69 | 92.3 | 92.3 | 90.6 | 53.651 | 0 | 323.31 | 9.8E+11 |
| 71 | 71 | 68 | 47.7 | 47.7 | 46.3 | 189.25 | 0 | 323.31 | 3.19E+10 |
| 62 | 62 | 62 | 15.7 | 15.7 | 15.7 | 504.6 | 0 | 150.02 | 8.29E+09 |
| 62 | 62 | 62 | 31.3 | 31.3 | 31.3 | 272.32 | 0 | 323.31 | 3.66E+10 |
| 74 | 74 | 59 | 81.1 | 81.1 | 62.6 | 67.819 | 0 | 323.31 | 3.47E+11 |
| 68 | 68 | 57 | 56.7 | 56.7 | 49 | 163.29 | 0 | 323.31 | 9.39E+10 |
| 58 | 58 | 57 | 83.7 | 83.7 | 83.7 | 70.288 | 0 | 323.31 | 4.48E+11 |
| 54 | 54 | 54 | 40.1 | 40.1 | 40.1 | 166.01 | 0 | 323.31 | 5.11E+10 |
| 54 | 54 | 53 | 39.4 | 39.4 | 38.8 | 152.45 | 0 | 323.31 | 1.22E+10 |
| 53 | 53 | 53 | 58.2 | 58.2 | 58.2 | 84.87 | 0 | 323.31 | 2.92E+11 |
| 52 | 52 | 52 | 59.9 | 59.9 | 59.9 | 89.321 | 0 | 286.92 | 5.2E+10 |
| 51 | 51 | 51 | 69.7 | 69.7 | 69.7 | 69.284 | 0 | 323.31 | 1.33E+11 |
| 51 | 51 | 51 | 69.9 | 69.9 | 69.9 | 78.457 | 0 | 323.31 | 9E+11 |
| 51 | 51 | 51 | 74 | 74 | 74 | 62.639 | 0 | 266.51 | 4.45E+10 |
| 50 | 50 | 50 | 13 | 13 | 13 | 532.4 | 0 | 102.29 | 4.75E+09 |
| 50 | 50 | 50 | 78.7 | 78.7 | 78.7 | 53.879 | 0 | 323.31 | 6.98E+10 |
| 50 | 50 | 50 | 58.6 | 58.6 | 58.6 | 95.337 | 0 | 323.31 | 7.62E+10 |
| 48 | 48 | 48 | 84.5 | 84.5 | 84.5 | 53.165 | 0 | 323.31 | 6.34E+11 |
| 77 | 77 | 47 | 80.1 | 80.1 | 57 | 104.85 | 0 | 323.31 | 1.27E+11 |
| 46 | 46 | 46 | 21.2 | 21.2 | 21.2 | 274.37 | 0 | 114.09 | 9.2E+09 |
| 46 | 46 | 46 | 51.6 | 51.6 | 51.6 | 99.719 | 0 | 185.03 | 1.75E+10 |
| 46 | 46 | 46 | 23.9 | 23.9 | 23.9 | 629.09 | 0 | 102.22 | 8.4E+09 |
| 45 | 45 | 45 | 61.9 | 61.9 | 61.9 | 77.328 | 0 | 323.31 | 1.7E+11 |
| 75 | 45 | 44 | 76.5 | 54.6 | 52.9 | 103.06 | 0 | 323.31 | 1.24E+11 |
| 44 | 44 | 44 | 45 | 45 | 45 | 123.8 | 0 | 258.51 | 2.28E+10 |
| 44 | 44 | 44 | 71.4 | 71.4 | 71.4 | 51.901 | 0 | 323.31 | 7.4E+10 |
| 44 | 44 | 43 | 64.1 | 64.1 | 64.1 | 99.024 | 0 | 323.31 | 1.58E+11 |
| 43 | 43 | 43 | 54.3 | 54.3 | 54.3 | 87.334 | 0 | 323.31 | 5.03E+10 |
| 43 | 43 | 43 | 57.8 | 57.8 | 57.8 | 102 | 0 | 134.21 | 1.31E+10 |
| 42 | 42 | 42 | 78.3 | 78.3 | 78.3 | 68.303 | 0 | 323.31 | 5.62E+10 |
| 41 | 41 | 41 | 68.5 | 68.5 | 68.5 | 75.952 | 0 | 299.09 | 4.15E+10 |
| 40 | 40 | 40 | 45.7 | 45.7 | 45.7 | 113.74 | 0 | 323.31 | 3E+10 |
| 40 | 40 | 40 | 80.7 | 80.7 | 80.7 | 51.681 | 0 | 323.31 | 1.24E+12 |
| 39 | 39 | 39 | 76.4 | 76.4 | 76.4 | 66.193 | 0 | 323.31 | 7.77E+10 |
| 39 | 39 | 39 | 91.4 | 91.4 | 91.4 | 44.614 | 0 | 323.31 | 1.71E+11 |
| 54 | 39 | 38 | 71.7 | 53.3 | 52.1 | 68.563 | 0 | 245.34 | 1.87E+10 |
| 38 | 38 | 38 | 54.9 | 54.9 | 54.9 | 98.398 | 0 | 259.97 | 2.46E+10 |
| 38 | 38 | 38 | 56.8 | 56.8 | 56.8 | 83.165 | 0 | 323.31 | 2.37E+10 |
| 38 | 38 | 38 | 44.8 | 44.8 | 44.8 | 96.022 | 0 | 143.71 | 1.22E+10 |
| 38 | 38 | 38 | 81.2 | 81.2 | 81.2 | 42.625 | 0 | 323.31 | 8.42E+11 |
| 38 | 38 | 38 | 79 | 79 | 79 | 67.877 | 0 | 323.31 | 1.72E+11 |
| 55 | 55 | 37 | 58.7 | 58.7 | 42.5 | 84.659 | 0 | 323.31 | 2.23E+11 |

| | | | | | | | | | |
|---|---|---|---|---|---|---|---|---|---|
| 41 | 41 | 37 | 80.2 | 80.2 | 71.2 | 47.168 | 0 | 323.31 | 4.24E+11 |
| 37 | 37 | 37 | 44.6 | 44.6 | 44.6 | 102.99 | 0 | 291.99 | 1.62E+10 |
| 37 | 37 | 37 | 68.8 | 68.8 | 68.8 | 56.166 | 0 | 323.31 | 9.01E+10 |
| 38 | 38 | 36 | 65.4 | 65.4 | 63.5 | 53.248 | 0 | 323.31 | 6.48E+10 |
| 37 | 36 | 36 | 52 | 52 | 52 | 72.332 | 0 | 115.36 | 1.3E+10 |
| 39 | 39 | 35 | 85.5 | 85.5 | 80.1 | 36.154 | 0 | 323.31 | 9.72E+11 |
| 35 | 35 | 35 | 85.1 | 85.1 | 85.1 | 52.878 | 0 | 323.31 | 5.25E+10 |
| 35 | 35 | 35 | 47.1 | 47.1 | 47.1 | 117.85 | 0 | 323.31 | 1.51E+10 |
| 35 | 35 | 35 | 32.3 | 32.3 | 32.3 | 136.37 | 0 | 69.149 | 7.07E+09 |
| 43 | 43 | 34 | 86.5 | 86.5 | 69.2 | 50.663 | 0 | 323.31 | 9.18E+10 |
| 34 | 34 | 34 | 24.9 | 24.9 | 24.9 | 162.46 | 0 | 126.89 | 3.14E+09 |
| 34 | 34 | 34 | 73.5 | 73.5 | 73.5 | 53.139 | 0 | 323.31 | 1.3E+11 |
| 34 | 34 | 34 | 74.4 | 74.4 | 74.4 | 46.659 | 0 | 323.31 | 1.25E+11 |
| 37 | 37 | 33 | 84 | 84 | 80.4 | 54.861 | 0 | 323.31 | 9.58E+10 |
| 36 | 36 | 33 | 48.5 | 48.5 | 46.2 | 94.33 | 0 | 180.07 | 1.57E+10 |
| 34 | 34 | 33 | 39 | 39 | 37.8 | 102.48 | 0 | 86.811 | 1.06E+10 |
| 33 | 33 | 33 | 17.3 | 17.3 | 17.3 | 273.42 | 0 | 118.86 | 4.94E+09 |
| 36 | 36 | 32 | 87.9 | 87.9 | 76.4 | 39.42 | 0 | 323.31 | 1.69E+11 |
| 32 | 32 | 32 | 62.2 | 62.2 | 62.2 | 73.502 | 0 | 235.74 | 1.73E+10 |
| 32 | 32 | 32 | 72.6 | 72.6 | 72.6 | 54.529 | 0 | 172.34 | 2.1E+10 |
| 32 | 32 | 32 | 76.3 | 76.3 | 76.3 | 56.5 | 0 | 270.45 | 4.88E+10 |
| 32 | 32 | 32 | 77.9 | 77.9 | 77.9 | 38.604 | 0 | 323.31 | 1.08E+11 |
| 32 | 32 | 32 | 82.8 | 82.8 | 82.8 | 37.495 | 0 | 323.31 | 3.89E+11 |
| 33 | 33 | 31 | 80.2 | 80.2 | 78.1 | 36.842 | 0 | 323.31 | 2.44E+11 |
| 31 | 31 | 31 | 27.4 | 27.4 | 27.4 | 138.34 | 0 | 83.976 | 3.47E+09 |
| 31 | 31 | 31 | 61 | 61 | 61 | 67.56 | 0 | 120.33 | 1.62E+10 |
| 31 | 31 | 31 | 72 | 72 | 72 | 47.371 | 0 | 323.31 | 6.91E+10 |
| 31 | 31 | 31 | 25.7 | 25.7 | 25.7 | 170.59 | 0 | 77.517 | 5.48E+09 |
| 44 | 31 | 31 | 66.4 | 51 | 51 | 69.371 | 0 | 192.38 | 3.08E+10 |
| 31 | 31 | 31 | 58.7 | 58.7 | 58.7 | 57.116 | 0 | 125.07 | 2.21E+10 |
| 30 | 30 | 30 | 61.5 | 61.5 | 61.5 | 63.146 | 0 | 323.31 | 8.99E+10 |
| 30 | 30 | 30 | 24.9 | 24.9 | 24.9 | 166.57 | 0 | 58.189 | 3.48E+09 |
| 30 | 30 | 30 | 52.9 | 52.9 | 52.9 | 51.026 | 0 | 191.82 | 4.48E+10 |
| 30 | 30 | 29 | 30.1 | 30.1 | 29.2 | 127.18 | 0 | 103.86 | 1.06E+10 |
| 29 | 29 | 29 | 32.4 | 32.4 | 32.4 | 120.84 | 0 | 89.419 | 6.2E+09 |
| 29 | 29 | 29 | 59.9 | 59.9 | 59.9 | 44.237 | 0 | 323.31 | 2.93E+11 |
| 29 | 29 | 29 | 48.4 | 48.4 | 48.4 | 63.111 | 0 | 167.07 | 1.05E+11 |
| 28 | 28 | 28 | 47.4 | 47.4 | 47.4 | 68.283 | 0 | 300.33 | 1.26E+10 |
| 28 | 28 | 28 | 63.2 | 63.2 | 63.2 | 57.488 | 0 | 187.16 | 7.41E+09 |
| 28 | 28 | 28 | 69.9 | 69.9 | 69.9 | 40.949 | 0 | 306.3 | 4.84E+10 |
| 28 | 28 | 28 | 58.7 | 58.7 | 58.7 | 37.821 | 0 | 323.31 | 5.67E+11 |
| 28 | 28 | 28 | 48.5 | 48.5 | 48.5 | 72.595 | 0 | 135.03 | 2.02E+10 |
| 28 | 28 | 28 | 77.9 | 77.9 | 77.9 | 36.053 | 0 | 323.31 | 1.29E+11 |
| 28 | 28 | 28 | 56.4 | 56.4 | 56.4 | 37.54 | 0 | 87.627 | 7.82E+10 |
| 34 | 34 | 27 | 42.9 | 42.9 | 35.8 | 97.147 | 0 | 149.73 | 9.57E+09 |
| 29 | 28 | 27 | 48 | 46.6 | 44.4 | 66.408 | 0 | 62.603 | 6.67E+09 |
| 27 | 27 | 27 | 78.3 | 78.3 | 78.3 | 34.333 | 0 | 273.37 | 4.57E+10 |

| | | | | | | | | | |
|---|---|---|---|---|---|---|---|---|---|
| 27 | 27 | 27 | 42.3 | 42.3 | 42.3 | 74.823 | 0 | 139.81 | 1.05E+10 |
| 27 | 27 | 27 | 52.8 | 52.8 | 52.8 | 52.775 | 0 | 307.7 | 8.08E+10 |
| 27 | 27 | 27 | 31.5 | 31.5 | 31.5 | 109.54 | 0 | 77.441 | 8.29E+09 |
| 27 | 27 | 27 | 79.7 | 79.7 | 79.7 | 30.791 | 0 | 323.31 | 2.48E+11 |
| 27 | 27 | 27 | 52.4 | 52.4 | 52.4 | 64.615 | 0 | 141.69 | 7.99E+09 |
| 27 | 27 | 26 | 58.9 | 58.9 | 58.9 | 37.429 | 0 | 274.41 | 6.52E+10 |
| 26 | 26 | 26 | 41.3 | 41.3 | 41.3 | 88.505 | 0 | 111.82 | 4.36E+09 |
| 26 | 26 | 26 | 51.5 | 51.5 | 51.5 | 70.905 | 0 | 188.41 | 2.22E+10 |
| 26 | 26 | 26 | 67.6 | 67.6 | 67.6 | 49.955 | 0 | 254.91 | 3.83E+10 |
| 51 | 33 | 25 | 58.3 | 43.5 | 39 | 83.263 | 0 | 156.86 | 9.15E+10 |
| 29 | 29 | 25 | 40.9 | 40.9 | 35.9 | 67.208 | 0 | 137.29 | 1.3E+10 |
| 26 | 26 | 25 | 77.7 | 77.7 | 77.7 | 42.621 | 0 | 323.31 | 4.71E+10 |
| 25 | 25 | 25 | 54.5 | 54.5 | 54.5 | 57.136 | 0 | 49.122 | 4.56E+09 |
| 25 | 25 | 25 | 48.8 | 48.8 | 48.8 | 60.533 | 0 | 102.58 | 6.99E+09 |
| 25 | 25 | 25 | 54.7 | 54.7 | 54.7 | 59.62 | 0 | 157.57 | 8.14E+09 |
| 25 | 25 | 25 | 46.7 | 46.7 | 46.7 | 72.968 | 0 | 59.211 | 8.07E+09 |
| 25 | 25 | 25 | 57.4 | 57.4 | 57.4 | 62.411 | 0 | 144.91 | 1.84E+10 |
| 25 | 25 | 25 | 44.8 | 44.8 | 44.8 | 52.494 | 0 | 159.06 | 1.03E+11 |
| 25 | 25 | 25 | 57.2 | 57.2 | 57.2 | 46.736 | 0 | 118.93 | 2.55E+10 |
| 25 | 25 | 25 | 43.1 | 43.1 | 43.1 | 82.577 | 0 | 136.27 | 1.3E+10 |
| 25 | 25 | 25 | 91.7 | 91.7 | 91.7 | 37.567 | 0 | 323.31 | 3.23E+10 |
| 25 | 25 | 25 | 64.2 | 64.2 | 64.2 | 49.973 | 0 | 221.71 | 1.66E+10 |
| 25 | 25 | 25 | 36.6 | 36.6 | 36.6 | 76.613 | 0 | 155.02 | 1.11E+10 |
| 25 | 25 | 25 | 38.4 | 38.4 | 38.4 | 84.79 | 0 | 199.61 | 1.36E+10 |
| 26 | 26 | 24 | 34.2 | 34.2 | 33 | 86.955 | 0 | 218.03 | 2.28E+10 |
| 25 | 25 | 24 | 67.3 | 67.3 | 64.6 | 42.741 | 0 | 161.69 | 5.94E+10 |
| 24 | 24 | 24 | 29.4 | 29.4 | 29.4 | 91.706 | 0 | 247.67 | 5.79E+09 |
| 24 | 24 | 24 | 89.6 | 89.6 | 89.6 | 15.164 | 0 | 218.34 | 1.34E+11 |
| 24 | 24 | 24 | 42.9 | 42.9 | 42.9 | 83.549 | 0 | 79.056 | 7.53E+09 |
| 24 | 24 | 24 | 90.6 | 90.6 | 90.6 | 28.804 | 0 | 323.31 | 1.48E+11 |
| 24 | 24 | 24 | 87.5 | 87.5 | 87.5 | 25.035 | 0 | 310.28 | 2.85E+10 |
| 24 | 24 | 24 | 43.3 | 43.3 | 43.3 | 65.401 | 0 | 99.835 | 5.4E+09 |
| 24 | 24 | 24 | 25.5 | 25.5 | 25.5 | 126.97 | 0 | 71.399 | 3.44E+09 |
| 28 | 26 | 23 | 43.8 | 39.9 | 35.3 | 67.93 | 0 | 157.93 | 1.18E+10 |
| 23 | 23 | 23 | 51.3 | 51.3 | 51.3 | 50.184 | 0 | 190.85 | 9.04E+10 |
| 23 | 23 | 23 | 50.2 | 50.2 | 50.2 | 56.967 | 0 | 212.13 | 1.46E+10 |
| 23 | 23 | 23 | 21.6 | 21.6 | 21.6 | 147.48 | 0 | 48.482 | 5.23E+09 |
| 23 | 23 | 23 | 73 | 73 | 73 | 41.35 | 0 | 323.31 | 5.2E+10 |
| 23 | 23 | 23 | 83.7 | 83.7 | 83.7 | 29.555 | 0 | 188.49 | 2.11E+10 |
| 23 | 23 | 23 | 42.7 | 42.7 | 42.7 | 84.304 | 0 | 137.73 | 1.2E+10 |
| 33 | 33 | 22 | 61.1 | 61.1 | 46 | 58.112 | 0 | 216.6 | 1.53E+11 |
| 29 | 29 | 22 | 70.2 | 70.2 | 63.7 | 27.745 | 0 | 323.31 | 2.53E+11 |
| 27 | 27 | 22 | 60.4 | 60.4 | 50.9 | 56.782 | 0 | 208.24 | 1.55E+10 |
| 22 | 22 | 22 | 79.2 | 79.2 | 79.2 | 29.246 | 0 | 101.12 | 1.74E+10 |
| 22 | 22 | 22 | 49.4 | 49.4 | 49.4 | 65.308 | 0 | 172.38 | 1.24E+10 |
| 22 | 22 | 22 | 52.7 | 52.7 | 52.7 | 54.548 | 0 | 119.71 | 9.57E+09 |
| 22 | 22 | 22 | 74.8 | 74.8 | 74.8 | 52.351 | 0 | 189.62 | 2.07E+10 |

| | | | | | | | | | |
|---|---|---|---|---|---|---|---|---|---|
| 25 | 22 | 22 | 69.4 | 66.3 | 66.3 | 29.174 | 0 | 211.39 | 3.91E+10 |
| 22 | 22 | 22 | 53.4 | 53.4 | 53.4 | 47.697 | 0 | 66.287 | 6.94E+09 |
| 22 | 22 | 22 | 67.8 | 67.8 | 67.8 | 29.945 | 0 | 89.94 | 1.11E+10 |
| 22 | 22 | 22 | 66.9 | 66.9 | 66.9 | 36.966 | 0 | 323.31 | 4.43E+10 |
| 22 | 22 | 21 | 49.1 | 49.1 | 49.1 | 36.638 | 0 | 309.33 | 1.21E+11 |
| 21 | 21 | 21 | 37.7 | 37.7 | 37.7 | 61.054 | 0 | 127.01 | 6.81E+09 |
| 21 | 21 | 21 | 48.8 | 48.8 | 48.8 | 61.448 | 0 | 64.649 | 7.38E+09 |
| 21 | 21 | 21 | 60.9 | 60.9 | 60.9 | 38.498 | 0 | 323.31 | 3.18E+11 |
| 21 | 21 | 21 | 44 | 44 | 44 | 54.013 | 0 | 74.621 | 1.22E+10 |
| 21 | 21 | 21 | 93 | 93 | 93 | 15.945 | 0 | 153.12 | 4.1E+10 |
| 21 | 21 | 21 | 40 | 40 | 40 | 55.154 | 0 | 253.61 | 2.89E+10 |
| 21 | 21 | 21 | 40.4 | 40.4 | 40.4 | 61.999 | 0 | 151.12 | 2.11E+10 |
| 21 | 21 | 21 | 27.7 | 27.7 | 27.7 | 105.32 | 0 | 110.16 | 1.11E+10 |
| 21 | 21 | 21 | 66 | 66 | 66 | 35.503 | 0 | 165.11 | 1.86E+10 |
| 22 | 21 | 21 | 79.2 | 75.6 | 75.6 | 36.688 | 0 | 323.31 | 8.72E+10 |
| 21 | 21 | 21 | 36.8 | 36.8 | 36.8 | 80.473 | 0 | 83.549 | 5.07E+09 |
| 21 | 21 | 21 | 60.5 | 60.5 | 60.5 | 46.637 | 0 | 65.175 | 5.38E+09 |
| 21 | 21 | 21 | 31.1 | 31.1 | 31.1 | 81.889 | 0 | 65.756 | 5.91E+09 |
| 21 | 21 | 21 | 32.4 | 32.4 | 32.4 | 97.169 | 0 | 127.35 | 9.93E+09 |
| 24 | 24 | 20 | 80.4 | 80.4 | 65.3 | 22.11 | 0 | 134.23 | 1.36E+11 |
| 22 | 22 | 20 | 91.6 | 91.6 | 84.9 | 18.502 | 0 | 323.31 | 2.07E+11 |
| 20 | 20 | 20 | 16.4 | 16.4 | 16.4 | 156.27 | 0 | 57.149 | 1.77E+09 |
| 20 | 20 | 20 | 41.7 | 41.7 | 41.7 | 60.343 | 0 | 48.691 | 5.07E+09 |
| 20 | 20 | 20 | 10.9 | 10.9 | 10.9 | 245.44 | 0 | 44.463 | 2.53E+09 |
| 20 | 20 | 20 | 50.6 | 50.6 | 50.6 | 51.57 | 0 | 260.21 | 1.3E+10 |
| 20 | 20 | 20 | 73 | 73 | 73 | 37.375 | 0 | 87.237 | 1.37E+10 |
| 20 | 20 | 20 | 27.6 | 27.6 | 27.6 | 102.49 | 0 | 51.509 | 4.78E+09 |
| 20 | 20 | 20 | 64.9 | 64.9 | 64.9 | 36.573 | 0 | 124.84 | 3.19E+10 |
| 20 | 20 | 20 | 54.2 | 54.2 | 54.2 | 41.92 | 0 | 190.38 | 2.1E+10 |
| 20 | 20 | 20 | 78.3 | 78.3 | 78.3 | 29.126 | 0 | 194.69 | 3.32E+10 |
| 20 | 20 | 20 | 85.4 | 85.4 | 85.4 | 22.391 | 0 | 265.64 | 4.42E+10 |
| 20 | 20 | 20 | 88.1 | 88.1 | 88.1 | 22.988 | 0 | 253.64 | 5.87E+10 |
| 20 | 20 | 20 | 44.8 | 44.8 | 44.8 | 44.659 | 0 | 181.89 | 5.47E+10 |
| 20 | 20 | 20 | 46.5 | 46.5 | 46.5 | 60.673 | 0 | 67.827 | 1.17E+10 |
| 20 | 20 | 20 | 40.9 | 40.9 | 40.9 | 65.33 | 0 | 323.31 | 3.66E+10 |
| 20 | 20 | 20 | 33.9 | 33.9 | 33.9 | 63.544 | 0 | 86.957 | 3E+10 |
| 20 | 20 | 20 | 52.3 | 52.3 | 52.3 | 48.141 | 0 | 239.71 | 4.06E+10 |
| 20 | 20 | 20 | 58.1 | 58.1 | 58.1 | 36.426 | 0 | 323.31 | 1.13E+11 |
| 20 | 20 | 20 | 44.7 | 44.7 | 44.7 | 47.716 | 0 | 69.961 | 1.04E+10 |
| 19 | 19 | 19 | 46.5 | 46.5 | 46.5 | 56.559 | 0 | 76.785 | 8.19E+09 |
| 19 | 19 | 19 | 31.2 | 31.2 | 31.2 | 73.68 | 0 | 43.394 | 3.58E+09 |
| 19 | 19 | 19 | 35 | 35 | 35 | 72.2 | 0 | 41.581 | 4.62E+09 |
| 19 | 19 | 19 | 53.6 | 53.6 | 53.6 | 29.717 | 0 | 108.3 | 3.35E+10 |
| 19 | 19 | 19 | 33.8 | 33.8 | 33.8 | 84.137 | 0 | 81.495 | 5.34E+09 |
| 19 | 19 | 19 | 39.4 | 39.4 | 39.4 | 72.765 | 0 | 61.015 | 7.78E+09 |
| 19 | 19 | 19 | 70.5 | 70.5 | 70.5 | 37.191 | 0 | 279.62 | 1.72E+10 |
| 19 | 19 | 19 | 85.8 | 85.8 | 85.8 | 32.118 | 0 | 323.31 | 1.93E+10 |

| | | | | | | | | | |
|---|---|---|---|---|---|---|---|---|---|
| 19 | 19 | 19 | 42.8 | 42.8 | 42.8 | 53.848 | 0 | 71.291 | 6.06E+09 |
| 19 | 19 | 19 | 46.7 | 46.7 | 46.7 | 44.76 | 0 | 221.1 | 3.94E+10 |
| 19 | 19 | 19 | 97.1 | 97.1 | 97.1 | 15.054 | 0 | 247.05 | 2.62E+11 |
| 20 | 19 | 19 | 62.2 | 62.2 | 62.2 | 33.155 | 0 | 223.77 | 1.91E+10 |
| 19 | 19 | 19 | 57.5 | 57.5 | 57.5 | 38.714 | 0 | 158.82 | 8.98E+09 |
| 19 | 19 | 19 | 26.7 | 26.7 | 26.7 | 109.79 | 0 | 85.697 | 3.83E+09 |
| 19 | 19 | 19 | 15.6 | 15.6 | 15.6 | 115.69 | 0 | 107.42 | 2.18E+09 |
| 19 | 19 | 19 | 49.7 | 49.7 | 49.7 | 53.162 | 0 | 200.57 | 1.2E+10 |
| 19 | 19 | 19 | 51.1 | 51.1 | 51.1 | 49.976 | 0 | 127.34 | 8.25E+09 |
| 19 | 19 | 19 | 37.3 | 37.3 | 37.3 | 69.842 | 0 | 74.709 | 5.77E+09 |
| 19 | 19 | 19 | 52.2 | 52.2 | 52.2 | 34.362 | 0 | 102.33 | 1.79E+10 |
| 34 | 29 | 18 | 47 | 43.6 | 27.6 | 70.108 | 0 | 162.18 | 2.41E+10 |
| 18 | 18 | 18 | 58.1 | 58.1 | 58.1 | 46.247 | 0 | 87.274 | 4.01E+09 |
| 18 | 18 | 18 | 45.8 | 45.8 | 45.8 | 41.796 | 0 | 58.436 | 1.09E+10 |
| 18 | 18 | 18 | 47.1 | 47.1 | 47.1 | 37.533 | 0 | 186 | 1.01E+10 |
| 18 | 18 | 18 | 36.2 | 36.2 | 36.2 | 64.149 | 0 | 149.43 | 6.93E+09 |
| 18 | 18 | 18 | 29.5 | 29.5 | 29.5 | 68.047 | 0 | 50.332 | 4.14E+09 |
| 18 | 18 | 18 | 47.7 | 47.7 | 47.7 | 56.256 | 0 | 75.101 | 1.11E+10 |
| 18 | 18 | 18 | 42 | 42 | 42 | 59.366 | 0 | 85.847 | 4.66E+09 |
| 18 | 18 | 18 | 89.4 | 89.4 | 89.4 | 19.891 | 0 | 240.66 | 2.27E+10 |
| 18 | 18 | 18 | 25.8 | 25.8 | 25.8 | 82.704 | 0 | 65.743 | 4E+09 |
| 18 | 18 | 18 | 35.9 | 35.9 | 35.9 | 61.626 | 0 | 82.858 | 5.15E+09 |
| 18 | 18 | 18 | 75.2 | 75.2 | 75.2 | 16.06 | 0 | 61.342 | 1.19E+10 |
| 18 | 18 | 18 | 70.5 | 70.5 | 70.5 | 24.399 | 0 | 122.54 | 4.54E+10 |
| 18 | 18 | 18 | 36.3 | 36.3 | 36.3 | 35.846 | 0 | 269.29 | 1.03E+11 |
| 18 | 18 | 18 | 86 | 86 | 86 | 24.75 | 0 | 323.31 | 7.09E+10 |
| 18 | 18 | 18 | 56.8 | 56.8 | 56.8 | 33.428 | 0 | 92.165 | 9.92E+09 |
| 18 | 18 | 18 | 52.4 | 52.4 | 52.4 | 49.411 | 0 | 145.97 | 1.19E+10 |
| 18 | 18 | 18 | 48.5 | 48.5 | 48.5 | 55.52 | 0 | 174.06 | 6.19E+09 |
| 27 | 18 | 18 | 68.9 | 51.7 | 51.7 | 50.582 | 0 | 106.13 | 8.63E+09 |
| 18 | 18 | 18 | 24.2 | 24.2 | 24.2 | 108.53 | 0 | 89.675 | 7.94E+09 |
| 18 | 18 | 18 | 26.9 | 26.9 | 26.9 | 99.326 | 0 | 51.076 | 3.48E+09 |
| 18 | 18 | 18 | 70.3 | 70.3 | 70.3 | 35.076 | 0 | 101.13 | 1.32E+10 |
| 21 | 21 | 17 | 84.6 | 84.6 | 76.2 | 32.922 | 0 | 111.68 | 2.32E+10 |
| 24 | 18 | 17 | 32.5 | 26.7 | 25.6 | 96.695 | 0 | 32.992 | 1.96E+09 |
| 18 | 18 | 17 | 56.6 | 56.6 | 53.5 | 40.45 | 0 | 81.383 | 9.01E+09 |
| 18 | 18 | 17 | 70.2 | 70.2 | 68.2 | 39.548 | 0 | 264.73 | 1.33E+10 |
| 17 | 17 | 17 | 34.2 | 34.2 | 34.2 | 82.091 | 0 | 76.093 | 2.58E+09 |
| 19 | 17 | 17 | 40.1 | 37.8 | 37.8 | 54.234 | 0 | 64.69 | 8.16E+09 |
| 17 | 17 | 17 | 37.3 | 37.3 | 37.3 | 58.024 | 0 | 58.708 | 4.13E+09 |
| 17 | 17 | 17 | 69.1 | 69.1 | 69.1 | 28.723 | 0 | 96.456 | 2.52E+10 |
| 17 | 17 | 17 | 39.3 | 39.3 | 39.3 | 53.801 | 0 | 83.701 | 5.64E+09 |
| 17 | 17 | 17 | 45.1 | 45.1 | 45.1 | 48.715 | 0 | 182.22 | 6.59E+09 |
| 17 | 17 | 17 | 39.4 | 39.4 | 39.4 | 54.389 | 0 | 81.92 | 6.45E+09 |
| 17 | 17 | 17 | 35 | 35 | 35 | 51.853 | 0 | 98.941 | 1.13E+10 |
| 17 | 17 | 17 | 23.1 | 23.1 | 23.1 | 104.83 | 0 | 47.777 | 4.21E+09 |
| 19 | 17 | 17 | 25.1 | 23.9 | 23.9 | 86.942 | 0 | 113.88 | 6.07E+09 |

| | | | | | | | | | |
|---|---|---|---|---|---|---|---|---|---|
| 17 | 17 | 17 | 73 | 73 | 73 | 16.537 | 0 | 185.75 | 5.12E+11 |
| 17 | 17 | 17 | 36.1 | 36.1 | 36.1 | 62.081 | 0 | 200.33 | 3.55E+10 |
| 17 | 17 | 17 | 55.7 | 55.7 | 55.7 | 35.422 | 0 | 284.95 | 1.76E+11 |
| 17 | 17 | 17 | 37.2 | 37.2 | 37.2 | 50.077 | 0 | 79.849 | 1.2E+10 |
| 17 | 17 | 17 | 76.5 | 76.5 | 76.5 | 21.225 | 0 | 241.56 | 5.48E+10 |
| 17 | 17 | 17 | 28.5 | 28.5 | 28.5 | 72.932 | 0 | 29.672 | 1.82E+09 |
| 17 | 17 | 17 | 30.8 | 30.8 | 30.8 | 80.699 | 0 | 101.05 | 3.28E+10 |
| 17 | 17 | 17 | 70.9 | 70.9 | 70.9 | 35.853 | 0 | 207.22 | 3.61E+10 |
| 19 | 17 | 17 | 25.4 | 22.6 | 22.6 | 98.972 | 0 | 50.967 | 3.29E+09 |
| 17 | 17 | 17 | 27 | 27 | 27 | 81.673 | 0 | 79.281 | 4.07E+09 |
| 17 | 17 | 17 | 71.4 | 71.4 | 71.4 | 27.887 | 0 | 114.11 | 2.16E+10 |
| 17 | 17 | 17 | 39.5 | 39.5 | 39.5 | 50.976 | 0 | 72.881 | 9.66E+09 |
| 41 | 41 | 16 | 81.7 | 81.7 | 31.8 | 59.256 | 0 | 281.62 | 8.22E+10 |
| 19 | 19 | 16 | 21.7 | 21.7 | 18.8 | 112.89 | 0 | 33.136 | 3.19E+09 |
| 16 | 16 | 16 | 28.7 | 28.7 | 28.7 | 61.585 | 0 | 50.695 | 1.97E+09 |
| 16 | 16 | 16 | 30.8 | 30.8 | 30.8 | 59.761 | 0 | 44.283 | 2.92E+09 |
| 16 | 16 | 16 | 33.4 | 33.4 | 33.4 | 62.942 | 0 | 99.446 | 5.6E+09 |
| 16 | 16 | 16 | 73 | 73 | 73 | 26.922 | 0 | 95.962 | 2.63E+10 |
| 16 | 16 | 16 | 48.4 | 48.4 | 48.4 | 26.152 | 0 | 120.8 | 1.61E+11 |
| 16 | 16 | 16 | 46.3 | 46.3 | 46.3 | 44.468 | 0 | 97.594 | 5.91E+09 |
| 16 | 16 | 16 | 60.9 | 60.9 | 60.9 | 35.102 | 0 | 306.62 | 1.68E+10 |
| 16 | 16 | 16 | 54.9 | 54.9 | 54.9 | 40.124 | 0 | 302.95 | 9.66E+09 |
| 16 | 16 | 16 | 61 | 61 | 61 | 27.566 | 0 | 189.27 | 4.61E+10 |
| 16 | 16 | 16 | 38.2 | 38.2 | 38.2 | 54.719 | 0 | 156.46 | 9.74E+09 |
| 16 | 16 | 16 | 38.4 | 38.4 | 38.4 | 57.924 | 0 | 48.866 | 3.7E+09 |
| 16 | 16 | 16 | 53.4 | 53.4 | 53.4 | 23.207 | 0 | 81.793 | 2.61E+10 |
| 16 | 16 | 16 | 34.3 | 34.3 | 34.3 | 50.118 | 0 | 85.314 | 9.06E+09 |
| 16 | 16 | 16 | 20.5 | 20.5 | 20.5 | 105.85 | 0 | 74.04 | 2.47E+09 |
| 16 | 16 | 16 | 70.8 | 70.8 | 70.8 | 26.688 | 0 | 54.197 | 6.29E+09 |
| 16 | 16 | 16 | 29.3 | 29.3 | 29.3 | 63.254 | 0 | 127.43 | 8.35E+09 |
| 16 | 16 | 16 | 44.1 | 44.1 | 44.1 | 54.636 | 0 | 45.611 | 3.34E+09 |
| 16 | 16 | 16 | 41.6 | 41.6 | 41.6 | 54.36 | 0 | 46.631 | 7.96E+09 |
| 16 | 16 | 16 | 56.1 | 56.1 | 56.1 | 34.632 | 0 | 190.83 | 3.8E+10 |
| 16 | 16 | 16 | 42.2 | 42.2 | 42.2 | 59.716 | 0 | 66.526 | 5.22E+09 |
| 16 | 16 | 16 | 30.9 | 30.9 | 30.9 | 65.102 | 0 | 34.312 | 4.67E+09 |
| 16 | 16 | 16 | 53.5 | 53.5 | 53.5 | 33.868 | 0 | 93.578 | 1.26E+11 |
| 16 | 16 | 16 | 37.3 | 37.3 | 37.3 | 44.965 | 0 | 216.58 | 5.11E+09 |
| 16 | 16 | 16 | 14.8 | 14.8 | 14.8 | 139.76 | 0 | 31.482 | 1.93E+09 |
| 16 | 16 | 16 | 46.6 | 46.6 | 46.6 | 39.829 | 0 | 60.869 | 3.5E+09 |
| 16 | 16 | 16 | 49.2 | 49.2 | 49.2 | 29.995 | 0 | 36.096 | 6.19E+09 |
| 16 | 16 | 16 | 56 | 56 | 56 | 28.147 | 0 | 67.808 | 2.36E+10 |
| 16 | 16 | 16 | 43.1 | 43.1 | 43.1 | 59.755 | 0 | 82.45 | 4.39E+09 |
| 16 | 16 | 16 | 77.6 | 77.6 | 77.6 | 33.232 | 0 | 150.04 | 8.31E+09 |
| 16 | 16 | 16 | 35.9 | 35.9 | 35.9 | 66.452 | 0 | 31.059 | 1.77E+09 |
| 16 | 16 | 16 | 60.9 | 60.9 | 60.9 | 29.483 | 0 | 107.57 | 1.24E+10 |
| 16 | 16 | 16 | 72.1 | 72.1 | 72.1 | 26.145 | 0 | 192.43 | 1.13E+10 |
| 19 | 19 | 15 | 33.9 | 33.9 | 27.6 | 69.602 | 0 | 129.33 | 4.35E+09 |

| | | | | | | | | | |
|---|---|---|---|---|---|---|---|---|---|
| 17 | 17 | 15 | 24.5 | 24.5 | 20.9 | 85.595 | 0 | 31.549 | 2.68E+09 |
| 16 | 16 | 15 | 23.2 | 23.2 | 22.1 | 76.149 | 0 | 34.086 | 4.5E+09 |
| 15 | 15 | 15 | 20.2 | 20.2 | 20.2 | 107.23 | 0 | 49.339 | 2.52E+09 |
| 15 | 15 | 15 | 45.6 | 45.6 | 45.6 | 46.277 | 0 | 44.045 | 2.72E+09 |
| 15 | 15 | 15 | 43.3 | 43.3 | 43.3 | 47.074 | 0 | 63.259 | 2.08E+09 |
| 15 | 15 | 15 | 22.2 | 22.2 | 22.2 | 97.717 | 0 | 30.397 | 2.13E+09 |
| 15 | 15 | 15 | 38.3 | 38.3 | 38.3 | 57.643 | 0 | 146.71 | 7.51E+09 |
| 15 | 15 | 15 | 49.8 | 49.8 | 49.8 | 26.224 | 0 | 58.376 | 1.54E+10 |
| 15 | 15 | 15 | 48.4 | 48.4 | 48.4 | 33.392 | 0 | 47.992 | 5.14E+09 |
| 15 | 15 | 15 | 85.6 | 85.6 | 85.6 | 21.057 | 0 | 194.4 | 3.88E+10 |
| 15 | 15 | 15 | 20.4 | 20.4 | 20.4 | 89.252 | 0 | 89.338 | 2.99E+09 |
| 15 | 15 | 15 | 52.9 | 52.9 | 52.9 | 41.487 | 0 | 163.98 | 5.82E+09 |
| 15 | 15 | 15 | 32.4 | 32.4 | 32.4 | 60.977 | 0 | 76.59 | 2.63E+09 |
| 15 | 15 | 15 | 20.3 | 20.3 | 20.3 | 100.2 | 0 | 46.6 | 3.68E+09 |
| 15 | 15 | 15 | 20.4 | 20.4 | 20.4 | 107.89 | 0 | 53.389 | 3.78E+09 |
| 15 | 15 | 15 | 23.9 | 23.9 | 23.9 | 95.337 | 0 | 62.013 | 3.71E+09 |
| 15 | 15 | 15 | 11.1 | 11.1 | 11.1 | 212 | 0 | 86.571 | 2.43E+09 |
| 15 | 15 | 15 | 40.7 | 40.7 | 40.7 | 47.517 | 0 | 47.144 | 5.02E+09 |
| 15 | 15 | 15 | 31.7 | 31.7 | 31.7 | 53.283 | 0 | 60.796 | 8.71E+09 |
| 15 | 15 | 15 | 48.4 | 48.4 | 48.4 | 41.924 | 0 | 67.46 | 4.34E+09 |
| 15 | 15 | 15 | 83.1 | 83.1 | 83.1 | 22.836 | 0 | 99.311 | 8.4E+09 |
| 15 | 15 | 15 | 14.1 | 14.1 | 14.1 | 139.09 | 0 | 86.414 | 1.42E+09 |
| 15 | 15 | 15 | 53.6 | 53.6 | 53.6 | 22.591 | 0 | 17.48 | 3.95E+09 |
| 15 | 15 | 15 | 81.2 | 81.2 | 81.2 | 15.693 | 0 | 191.85 | 4.4E+10 |
| 15 | 15 | 15 | 53 | 53 | 53 | 36.112 | 0 | 147.79 | 5.48E+09 |
| 15 | 15 | 15 | 52.4 | 52.4 | 52.4 | 32.575 | 0 | 233.87 | 1.14E+10 |
| 15 | 15 | 15 | 52.9 | 52.9 | 52.9 | 37.765 | 0 | 91.486 | 2.2E+10 |
| 16 | 16 | 14 | 76.2 | 76.2 | 65.7 | 30.375 | 0 | 168.64 | 1.17E+10 |
| 15 | 15 | 14 | 21.6 | 21.6 | 20.8 | 100.89 | 0 | 36.177 | 3.03E+09 |
| 16 | 15 | 14 | 44.4 | 41.8 | 38.6 | 42.403 | 0 | 62.188 | 4.07E+09 |
| 14 | 14 | 14 | 27.3 | 27.3 | 27.3 | 59.75 | 0 | 27.489 | 2.39E+09 |
| 14 | 14 | 14 | 31.6 | 31.6 | 31.6 | 39.724 | 0 | 75.073 | 3.23E+09 |
| 14 | 14 | 14 | 14.9 | 14.9 | 14.9 | 105.34 | 0 | 50.414 | 1.61E+09 |
| 14 | 14 | 14 | 46.8 | 46.8 | 46.8 | 39.594 | 0 | 56.066 | 4.49E+09 |
| 14 | 14 | 14 | 95.6 | 95.6 | 95.6 | 14.716 | 0 | 225.34 | 1.32E+11 |
| 16 | 14 | 14 | 27.3 | 23.7 | 23.7 | 85.018 | 0 | 36.974 | 4.15E+09 |
| 14 | 14 | 14 | 51.4 | 51.4 | 51.4 | 35.882 | 0 | 126.83 | 2.55E+09 |
| 14 | 14 | 14 | 41.9 | 41.9 | 41.9 | 43.941 | 0 | 52.945 | 3.59E+09 |
| 14 | 14 | 14 | 51.8 | 51.8 | 51.8 | 28.433 | 0 | 27.09 | 1.61E+10 |
| 14 | 14 | 14 | 60.6 | 60.6 | 60.6 | 26.489 | 0 | 70.8 | 1.12E+10 |
| 14 | 14 | 14 | 41 | 41 | 41 | 32.728 | 0 | 30.255 | 3.05E+09 |
| 14 | 14 | 14 | 38.2 | 38.2 | 38.2 | 33.67 | 0 | 82.318 | 6.3E+09 |
| 14 | 14 | 14 | 34.7 | 34.7 | 34.7 | 54.255 | 0 | 230.26 | 1.81E+10 |
| 14 | 14 | 14 | 31.3 | 31.3 | 31.3 | 54.163 | 0 | 33.716 | 2.78E+09 |
| 14 | 14 | 14 | 34.1 | 34.1 | 34.1 | 55.799 | 0 | 77.364 | 7.33E+09 |
| 14 | 14 | 14 | 11.1 | 11.1 | 11.1 | 161.69 | 0 | 59.619 | 2.02E+09 |
| 14 | 14 | 14 | 25.1 | 25.1 | 25.1 | 59.582 | 0 | 47.7 | 1.41E+10 |

| | | | | | | | | | |
|---|---|---|---|---|---|---|---|---|---|
| 14 | 14 | 14 | 25.8 | 25.8 | 25.8 | 74.731 | 0 | 56.534 | 4.28E+09 |
| 14 | 14 | 14 | 36.8 | 36.8 | 36.8 | 49.762 | 0 | 63.972 | 4.68E+09 |
| 14 | 14 | 14 | 28.6 | 28.6 | 28.6 | 51.815 | 0 | 55.484 | 4.77E+09 |
| 14 | 14 | 14 | 67.5 | 67.5 | 67.5 | 25.898 | 0 | 210.58 | 1.82E+10 |
| 14 | 14 | 14 | 36 | 36 | 36 | 48.207 | 0 | 67.992 | 3.64E+09 |
| 14 | 14 | 14 | 42.3 | 42.3 | 42.3 | 61.247 | 0 | 98.299 | 1.69E+10 |
| 14 | 14 | 14 | 20.9 | 20.9 | 20.9 | 92.48 | 0 | 81.346 | 2.17E+09 |
| 14 | 14 | 14 | 79.4 | 79.4 | 79.4 | 18.012 | 0 | 147 | 1.57E+11 |
| 14 | 14 | 14 | 87.7 | 87.7 | 87.7 | 12.774 | 0 | 109.38 | 2.93E+10 |
| 14 | 14 | 14 | 23.9 | 23.9 | 23.9 | 76.837 | 0 | 19.088 | 1.09E+09 |
| 14 | 14 | 14 | 69.8 | 69.8 | 69.8 | 30.315 | 0 | 121.01 | 7.87E+09 |
| 20 | 14 | 14 | 60 | 48.2 | 48.2 | 27.764 | 0 | 135.14 | 1.58E+10 |
| 15 | 14 | 14 | 16.1 | 15.1 | 15.1 | 128.54 | 0 | 62.007 | 4.13E+09 |
| 14 | 14 | 14 | 50.5 | 50.5 | 50.5 | 34.273 | 0 | 102.27 | 1.1E+10 |
| 14 | 14 | 14 | 33.6 | 33.6 | 33.6 | 54.001 | 0 | 22.433 | 3.07E+09 |
| 14 | 14 | 14 | 43.7 | 43.7 | 43.7 | 43.786 | 0 | 27.699 | 2.88E+09 |
| 14 | 14 | 14 | 37.3 | 37.3 | 37.3 | 51.109 | 0 | 39.663 | 4.71E+09 |
| 14 | 14 | 14 | 60 | 60 | 60 | 20.019 | 0 | 109.98 | 4.97E+10 |
| 14 | 14 | 14 | 61.6 | 61.6 | 61.6 | 29.404 | 0 | 239.64 | 1.14E+10 |
| 14 | 14 | 14 | 45.1 | 45.1 | 45.1 | 36.748 | 0 | 187.1 | 1.07E+10 |
| 14 | 14 | 14 | 49.6 | 49.6 | 49.6 | 30.354 | 0 | 180.24 | 1.77E+10 |
| 14 | 14 | 14 | 59.2 | 59.2 | 59.2 | 11.367 | 0 | 126.17 | 1.17E+11 |
| 21 | 21 | 13 | 23.4 | 23.4 | 13.5 | 107.54 | 0 | 45.421 | 4.46E+09 |
| 16 | 16 | 13 | 56.5 | 56.5 | 44.5 | 24.033 | 0 | 37.316 | 5.38E+09 |
| 13 | 13 | 13 | 35.1 | 35.1 | 35.1 | 61.312 | 0 | 32.745 | 8.3E+08 |
| 13 | 13 | 13 | 15.1 | 15.1 | 15.1 | 106.81 | 0 | 28.287 | 5.75E+08 |
| 13 | 13 | 13 | 18.9 | 18.9 | 18.9 | 100.6 | 0 | 29.815 | 1.62E+09 |
| 13 | 13 | 13 | 17.2 | 17.2 | 17.2 | 110.18 | 0 | 26.658 | 1.76E+09 |
| 13 | 13 | 13 | 53.2 | 53.2 | 53.2 | 22.782 | 0 | 73.08 | 6.05E+09 |
| 13 | 13 | 13 | 8.3 | 8.3 | 8.3 | 175.49 | 0 | 27.711 | 1.55E+09 |
| 13 | 13 | 13 | 58.8 | 58.8 | 58.8 | 35.554 | 0 | 167.04 | 3.81E+09 |
| 13 | 13 | 13 | 56.7 | 56.7 | 56.7 | 25.206 | 0 | 29.742 | 6.79E+09 |
| 13 | 13 | 13 | 40 | 40 | 40 | 35.936 | 0 | 159.54 | 4.27E+09 |
| 13 | 13 | 13 | 38.1 | 38.1 | 38.1 | 42.981 | 0 | 65.532 | 3.49E+09 |
| 15 | 13 | 13 | 18.8 | 17.1 | 17.1 | 92.468 | 0 | 26.467 | 3.89E+09 |
| 13 | 13 | 13 | 52.7 | 52.7 | 52.7 | 16.23 | 0 | 77.416 | 3.31E+10 |
| 13 | 13 | 13 | 44.4 | 44.4 | 44.4 | 37.564 | 0 | 171.46 | 2.63E+10 |
| 13 | 13 | 13 | 28.9 | 28.9 | 28.9 | 61.397 | 0 | 33.316 | 2.17E+09 |
| 13 | 13 | 13 | 79.3 | 79.3 | 79.3 | 21.007 | 0 | 43.611 | 7.09E+09 |
| 13 | 13 | 13 | 30.5 | 30.5 | 30.5 | 46.108 | 0 | 25.788 | 3.08E+09 |
| 13 | 13 | 13 | 58.9 | 58.9 | 58.9 | 16.445 | 0 | 22.176 | 3.33E+09 |
| 13 | 13 | 13 | 25.3 | 25.3 | 25.3 | 62.76 | 0 | 25.74 | 2.16E+09 |
| 13 | 13 | 13 | 37 | 37 | 37 | 45.796 | 0 | 32.137 | 1.52E+09 |
| 13 | 13 | 13 | 18.5 | 18.5 | 18.5 | 83.782 | 0 | 14.849 | 2.1E+09 |
| 13 | 13 | 13 | 46.2 | 46.2 | 46.2 | 28.68 | 0 | 62.685 | 5.87E+09 |
| 13 | 13 | 13 | 31.1 | 31.1 | 31.1 | 62.463 | 0 | 80.845 | 3.29E+09 |
| 13 | 13 | 13 | 16.5 | 16.5 | 16.5 | 103.13 | 0 | 35.307 | 2.46E+09 |

| | | | | | | | | | |
|---|---|---|---|---|---|---|---|---|---|
| 13 | 13 | 13 | 23.9 | 23.9 | 23.9 | 85.126 | 0 | 60.904 | 4.37E+09 |
| 13 | 13 | 13 | 41.3 | 41.3 | 41.3 | 52.562 | 0 | 54.025 | 2.24E+09 |
| 13 | 13 | 13 | 43.7 | 43.7 | 43.7 | 31.324 | 0 | 32.791 | 4.03E+09 |
| 13 | 13 | 13 | 32.9 | 32.9 | 32.9 | 44.23 | 0 | 62.357 | 4.07E+09 |
| 13 | 13 | 13 | 42.7 | 42.7 | 42.7 | 24.261 | 0 | 61.619 | 5.52E+09 |
| 13 | 13 | 13 | 50.8 | 50.8 | 50.8 | 34.674 | 0 | 78.232 | 6.33E+09 |
| 13 | 13 | 13 | 61.1 | 61.1 | 61.1 | 32.57 | 0 | 323.31 | 4.99E+10 |
| 13 | 13 | 13 | 23.5 | 23.5 | 23.5 | 75.872 | 0 | 97.447 | 3.21E+09 |
| 13 | 13 | 13 | 37.5 | 37.5 | 37.5 | 49.439 | 0 | 114.25 | 3.59E+09 |
| 13 | 13 | 13 | 53.3 | 53.3 | 53.3 | 17.718 | 0 | 22.795 | 5.58E+09 |
| 23 | 23 | 12 | 60.3 | 60.3 | 31.8 | 46.153 | 0 | 191.19 | 3.04E+10 |
| 17 | 17 | 12 | 47.4 | 47.4 | 32.5 | 44.619 | 0 | 87.812 | 7.93E+09 |
| 14 | 14 | 12 | 32.2 | 32.2 | 28.1 | 58.47 | 0 | 36.594 | 2.84E+09 |
| 13 | 13 | 12 | 55.5 | 55.5 | 53 | 31.121 | 0 | 85.959 | 8.88E+09 |
| 12 | 12 | 12 | 17.4 | 17.4 | 17.4 | 107.14 | 0 | 37.968 | 1.17E+09 |
| 12 | 12 | 12 | 67.5 | 67.5 | 67.5 | 32.668 | 0 | 54.687 | 4.77E+09 |
| 12 | 12 | 12 | 79.9 | 79.9 | 79.9 | 17.861 | 0 | 166.67 | 1.27E+10 |
| 12 | 12 | 12 | 37.9 | 37.9 | 37.9 | 31.54 | 0 | 21.182 | 2.35E+09 |
| 12 | 12 | 12 | 43.7 | 43.7 | 43.7 | 39.518 | 0 | 53.824 | 1.49E+10 |
| 12 | 12 | 12 | 73.3 | 73.3 | 73.3 | 11.737 | 0 | 165.53 | 1.13E+11 |
| 18 | 12 | 12 | 59.1 | 46.6 | 46.6 | 28.302 | 0 | 143.5 | 4.03E+10 |
| 12 | 12 | 12 | 36.7 | 36.7 | 36.7 | 41.564 | 0 | 26.671 | 2.48E+09 |
| 12 | 12 | 12 | 46.3 | 46.3 | 46.3 | 29.999 | 0 | 53.965 | 2.73E+09 |
| 12 | 12 | 12 | 31.7 | 31.7 | 31.7 | 41.331 | 0 | 33.57 | 7.66E+09 |
| 13 | 12 | 12 | 46.5 | 40.9 | 40.9 | 21.892 | 0 | 49.937 | 5.4E+09 |
| 12 | 12 | 12 | 72.4 | 72.4 | 72.4 | 21.058 | 0 | 204.68 | 3.23E+10 |
| 12 | 12 | 12 | 33 | 33 | 33 | 48.633 | 0 | 65.213 | 1.66E+09 |
| 12 | 12 | 12 | 44.1 | 44.1 | 44.1 | 35.924 | 0 | 184.76 | 5.48E+09 |
| 12 | 12 | 12 | 37.2 | 37.2 | 37.2 | 36.598 | 0 | 84.215 | 8.29E+10 |
| 12 | 12 | 12 | 34.1 | 34.1 | 34.1 | 44.791 | 0 | 37.165 | 3.82E+09 |
| 14 | 12 | 12 | 38.9 | 33.2 | 33.2 | 22.58 | 0 | 25.77 | 7.03E+09 |
| 12 | 12 | 12 | 31.4 | 31.4 | 31.4 | 48.121 | 0 | 105.65 | 3.06E+09 |
| 12 | 12 | 12 | 35.1 | 35.1 | 35.1 | 58.025 | 0 | 88.626 | 4.36E+09 |
| 12 | 12 | 12 | 46 | 46 | 46 | 40.876 | 0 | 206.52 | 1.63E+10 |
| 12 | 12 | 12 | 37.1 | 37.1 | 37.1 | 54.705 | 0 | 69.211 | 8.18E+09 |
| 12 | 12 | 12 | 16 | 16 | 16 | 106.69 | 0 | 53.065 | 2.4E+09 |
| 12 | 12 | 12 | 32.9 | 32.9 | 32.9 | 54.341 | 0 | 33.77 | 4.93E+09 |
| 12 | 12 | 12 | 23.8 | 23.8 | 23.8 | 66.9 | 0 | 22.172 | 1.34E+09 |
| 12 | 12 | 12 | 42.5 | 42.5 | 42.5 | 53.688 | 0 | 202.87 | 9.98E+09 |
| 12 | 12 | 12 | 19.2 | 19.2 | 19.2 | 63.922 | 0 | 215.49 | 8.14E+10 |
| 12 | 12 | 12 | 42.5 | 42.5 | 42.5 | 40.936 | 0 | 171.66 | 8.87E+09 |
| 12 | 12 | 12 | 22.4 | 22.4 | 22.4 | 60.179 | 0 | 125.02 | 1.53E+10 |
| 12 | 12 | 12 | 37.7 | 37.7 | 37.7 | 47.463 | 0 | 64.712 | 2.83E+09 |
| 12 | 12 | 12 | 57.5 | 57.5 | 57.5 | 23.489 | 0 | 120.09 | 8.21E+09 |
| 12 | 12 | 12 | 90.9 | 90.9 | 90.9 | 15.936 | 0 | 299.09 | 3.93E+10 |
| 12 | 12 | 12 | 56.9 | 56.9 | 56.9 | 31.236 | 0 | 42.53 | 1.4E+09 |
| 13 | 12 | 12 | 25.2 | 24.1 | 24.1 | 69.917 | 0 | 96.759 | 2.35E+09 |

| | | | | | | | | | |
|---|---|---|---|---|---|---|---|---|---|
| 12 | 12 | 12 | 38.6 | 38.6 | 38.6 | 40.422 | 0 | 33.541 | 1.75E+09 |
| 12 | 12 | 12 | 18.1 | 18.1 | 18.1 | 95.785 | 0 | 40.881 | 3.11E+09 |
| 12 | 12 | 12 | 61.9 | 61.9 | 61.9 | 23.356 | 0 | 323.31 | 1.77E+10 |
| 12 | 12 | 12 | 66.5 | 66.5 | 66.5 | 22.127 | 0 | 46.367 | 4.54E+09 |
| 12 | 12 | 12 | 48.4 | 48.4 | 48.4 | 30.188 | 0 | 29.796 | 2.04E+09 |
| 12 | 12 | 12 | 21.7 | 21.7 | 21.7 | 84.729 | 0 | 23.753 | 1.01E+09 |
| 12 | 12 | 12 | 23.1 | 23.1 | 23.1 | 84.141 | 0 | 19.485 | 9.29E+08 |
| 12 | 12 | 12 | 48.4 | 48.4 | 48.4 | 29.225 | 0 | 42.41 | 4.87E+09 |
| 12 | 12 | 12 | 11.1 | 11.1 | 11.1 | 114.87 | 0 | 85.032 | 1.63E+09 |
| 12 | 12 | 12 | 54 | 54 | 54 | 15.747 | 0 | 41.422 | 2.91E+09 |
| 12 | 12 | 12 | 51 | 51 | 51 | 24.146 | 0 | 25.132 | 5.58E+09 |
| 28 | 28 | 11 | 23.8 | 23.8 | 9.3 | 140.94 | 0 | 136.68 | 7.28E+09 |
| 18 | 18 | 11 | 32.5 | 32.5 | 18.9 | 70.67 | 0 | 61.399 | 3.09E+09 |
| 19 | 13 | 11 | 53.7 | 46.7 | 46.7 | 28.082 | 0 | 305.92 | 4.57E+10 |
| 12 | 12 | 11 | 41.4 | 41.4 | 39.7 | 47.065 | 0 | 49.76 | 4.46E+09 |
| 11 | 11 | 11 | 10.4 | 10.4 | 10.4 | 142.29 | 0 | 38.52 | 1.8E+09 |
| 11 | 11 | 11 | 19.6 | 19.6 | 19.6 | 77.959 | 0 | 40.635 | 1.49E+09 |
| 11 | 11 | 11 | 11.2 | 11.2 | 11.2 | 128.15 | 0 | 27.824 | 2.19E+09 |
| 11 | 11 | 11 | 43.7 | 43.7 | 43.7 | 40.307 | 0 | 37.77 | 4.32E+09 |
| 11 | 11 | 11 | 14.1 | 14.1 | 14.1 | 111.69 | 0 | 42.219 | 1.78E+09 |
| 11 | 11 | 11 | 52.6 | 52.6 | 52.6 | 21.404 | 0 | 44.095 | 2.46E+09 |
| 11 | 11 | 11 | 60.1 | 60.1 | 60.1 | 25.572 | 0 | 34.071 | 2.37E+09 |
| 11 | 11 | 11 | 33.1 | 33.1 | 33.1 | 47.352 | 0 | 43.873 | 1.82E+09 |
| 11 | 11 | 11 | 61 | 61 | 61 | 16.019 | 0 | 44.834 | 1.01E+10 |
| 11 | 11 | 11 | 86 | 86 | 86 | 13.242 | 0 | 74.967 | 6.72E+09 |
| 11 | 11 | 11 | 41.3 | 41.3 | 41.3 | 28.262 | 0 | 30.513 | 2.73E+09 |
| 11 | 11 | 11 | 19.6 | 19.6 | 19.6 | 69.697 | 0 | 29.907 | 1.08E+09 |
| 11 | 11 | 11 | 24.1 | 24.1 | 24.1 | 50.435 | 0 | 59.404 | 2.61E+09 |
| 15 | 11 | 11 | 46.3 | 39.4 | 39.4 | 28.218 | 0 | 52.566 | 5.98E+09 |
| 11 | 11 | 11 | 49.4 | 49.4 | 49.4 | 29.676 | 0 | 52.622 | 2.55E+10 |
| 11 | 11 | 11 | 67.4 | 67.4 | 67.4 | 20.838 | 0 | 106.25 | 1.58E+11 |
| 11 | 11 | 11 | 21.1 | 21.1 | 21.1 | 76.074 | 0 | 27.948 | 1.87E+09 |
| 11 | 11 | 11 | 41.3 | 41.3 | 41.3 | 34.136 | 0 | 57.731 | 1.13E+10 |
| 11 | 11 | 11 | 40.3 | 40.3 | 40.3 | 39.008 | 0 | 183.44 | 1.28E+10 |
| 11 | 11 | 11 | 48.2 | 48.2 | 48.2 | 29.114 | 0 | 119.64 | 1.37E+10 |
| 11 | 11 | 11 | 27.2 | 27.2 | 27.2 | 45.467 | 0 | 48.153 | 3.03E+09 |
| 11 | 11 | 11 | 63.7 | 63.7 | 63.7 | 15.799 | 0 | 179.33 | 3.8E+10 |
| 11 | 11 | 11 | 55.8 | 55.8 | 55.8 | 27.547 | 0 | 36.771 | 4.65E+09 |
| 11 | 11 | 11 | 57.2 | 57.2 | 57.2 | 30.608 | 0 | 41.591 | 1.03E+09 |
| 11 | 11 | 11 | 24.2 | 24.2 | 24.2 | 58.042 | 0 | 30.743 | 2.04E+09 |
| 11 | 11 | 11 | 45.2 | 45.2 | 45.2 | 34.684 | 0 | 64.16 | 5.28E+09 |
| 11 | 11 | 11 | 27.7 | 27.7 | 27.7 | 59.143 | 0 | 29.548 | 1.04E+09 |
| 11 | 11 | 11 | 52.5 | 52.5 | 52.5 | 31.462 | 0 | 77.488 | 2.86E+09 |
| 11 | 11 | 11 | 64.5 | 64.5 | 64.5 | 17.668 | 0 | 19.843 | 8.25E+09 |
| 11 | 11 | 11 | 14.4 | 14.4 | 14.4 | 77.569 | 0 | 21.795 | 7.07E+08 |
| 11 | 11 | 11 | 18.5 | 18.5 | 18.5 | 74.281 | 0 | 29.261 | 7.13E+08 |
| 11 | 11 | 11 | 67.8 | 67.8 | 67.8 | 22.693 | 0 | 73.609 | 4.92E+09 |

| | | | | | | | | | |
|---|---|---|---|---|---|---|---|---|---|
| 11 | 11 | 11 | 31.8 | 31.8 | 31.8 | 54.605 | 0 | 39.168 | 1.76E+09 |
| 41 | 41 | 10 | 87.2 | 87.2 | 22.5 | 26.42 | 0 | 323.31 | 1.41E+11 |
| 23 | 23 | 10 | 35.8 | 35.8 | 17.6 | 80.998 | 0 | 121.06 | 4.97E+09 |
| 17 | 17 | 10 | 75.7 | 75.7 | 27.7 | 32.642 | 0 | 166.06 | 5.72E+10 |
| 23 | 16 | 10 | 69.2 | 49.8 | 25.1 | 34.288 | 0 | 109.07 | 3.59E+10 |
| 13 | 13 | 10 | 30.8 | 30.8 | 30.8 | 43.171 | 0 | 142.61 | 4.15E+09 |
| 11 | 11 | 10 | 40.1 | 40.1 | 37.2 | 45.626 | 0 | 35.236 | 2.02E+09 |
| 11 | 11 | 10 | 36.5 | 36.5 | 36.5 | 28.585 | 0 | 77.802 | 5.87E+09 |
| 11 | 11 | 10 | 30.8 | 30.8 | 28.2 | 40.589 | 0 | 29.764 | 1.41E+09 |
| 10 | 10 | 10 | 14.6 | 14.6 | 14.6 | 77.504 | 0 | 19.551 | 7.12E+08 |
| 10 | 10 | 10 | 13.5 | 13.5 | 13.5 | 109.68 | 0 | 47.337 | 8.41E+08 |
| 10 | 10 | 10 | 68.4 | 68.4 | 68.4 | 14.858 | 0 | 36.124 | 3.52E+10 |
| 10 | 10 | 10 | 50.8 | 50.8 | 50.8 | 35.117 | 0 | 49.281 | 3.76E+09 |
| 10 | 10 | 10 | 31.3 | 31.3 | 31.3 | 46.374 | 0 | 29.378 | 2.61E+09 |
| 10 | 10 | 10 | 37.5 | 37.5 | 37.5 | 36.501 | 0 | 31.673 | 2.01E+09 |
| 10 | 10 | 10 | 79.4 | 79.4 | 79.4 | 13.802 | 0 | 97.076 | 1.7E+10 |
| 10 | 10 | 10 | 57.1 | 57.1 | 57.1 | 22.222 | 0 | 76.193 | 5.09E+09 |
| 10 | 10 | 10 | 58.4 | 58.4 | 58.4 | 11.728 | 0 | 19.61 | 1.2E+10 |
| 10 | 10 | 10 | 77.1 | 77.1 | 77.1 | 13.757 | 0 | 323.31 | 1.31E+10 |
| 10 | 10 | 10 | 26.6 | 26.6 | 26.6 | 49.623 | 0 | 76.075 | 2.83E+09 |
| 10 | 10 | 10 | 60.6 | 60.6 | 60.6 | 26.411 | 0 | 71.232 | 1.28E+10 |
| 10 | 10 | 10 | 48.9 | 48.9 | 48.9 | 20.546 | 0 | 186.01 | 2.12E+10 |
| 10 | 10 | 10 | 26.5 | 26.5 | 26.5 | 49.526 | 0 | 18.205 | 1.81E+09 |
| 10 | 10 | 10 | 86.7 | 86.7 | 86.7 | 11.74 | 0 | 173.49 | 6.33E+10 |
| 10 | 10 | 10 | 29.4 | 29.4 | 29.4 | 46.836 | 0 | 69.253 | 2.69E+09 |
| 10 | 10 | 10 | 77.5 | 77.5 | 77.5 | 10.932 | 0 | 25.762 | 8.01E+09 |
| 10 | 10 | 10 | 20.2 | 20.2 | 20.2 | 65.471 | 0 | 63.121 | 3.74E+09 |
| 10 | 10 | 10 | 43.8 | 43.8 | 43.8 | 27.692 | 0 | 64.404 | 5.25E+09 |
| 10 | 10 | 10 | 30.3 | 30.3 | 30.3 | 50.878 | 0 | 59.894 | 4.95E+09 |
| 10 | 10 | 10 | 29.4 | 29.4 | 29.4 | 45.306 | 0 | 48.356 | 2.79E+09 |
| 10 | 10 | 10 | 20.1 | 20.1 | 20.1 | 75.905 | 0 | 30.756 | 1.99E+09 |
| 10 | 10 | 10 | 58.4 | 58.4 | 58.4 | 21.029 | 0 | 281.99 | 1.86E+11 |
| 10 | 10 | 10 | 51.3 | 51.3 | 51.3 | 26.21 | 0 | 38.802 | 1.79E+09 |
| 10 | 10 | 10 | 69.4 | 69.4 | 69.4 | 22.119 | 0 | 73.928 | 8.62E+09 |
| 10 | 10 | 10 | 28.5 | 28.5 | 28.5 | 47.804 | 0 | 27.886 | 1.54E+09 |
| 10 | 10 | 10 | 88.9 | 88.9 | 88.9 | 11.951 | 0 | 77.967 | 2.32E+10 |
| 10 | 10 | 10 | 39.8 | 39.8 | 39.8 | 33.285 | 0 | 53.046 | 6.67E+09 |
| 10 | 10 | 10 | 13.8 | 13.8 | 13.8 | 83.434 | 0 | 30.768 | 7.6E+08 |
| 10 | 10 | 10 | 63.3 | 63.3 | 63.3 | 22.345 | 0 | 96.569 | 4.85E+09 |
| 10 | 10 | 10 | 33.6 | 33.6 | 33.6 | 41.389 | 0 | 21.486 | 3.36E+09 |
| 10 | 10 | 10 | 46.4 | 46.4 | 46.4 | 35.536 | 0 | 65.757 | 3.29E+09 |
| 10 | 10 | 10 | 30.2 | 30.2 | 30.2 | 49.786 | 0 | 34.515 | 7.98E+08 |
| 10 | 10 | 10 | 90.8 | 90.8 | 90.8 | 10.044 | 0 | 51.9 | 2.15E+10 |
| 10 | 10 | 10 | 22.1 | 22.1 | 22.1 | 56.068 | 0 | 35.965 | 1.38E+09 |
| 10 | 10 | 10 | 58.9 | 58.9 | 58.9 | 18.431 | 0 | 34.788 | 8.35E+09 |
| 10 | 10 | 10 | 39.6 | 39.6 | 39.6 | 35.23 | 0 | 31.636 | 1.96E+09 |
| 10 | 10 | 10 | 35 | 35 | 35 | 43.404 | 0 | 44.609 | 2.32E+09 |

| | | | | | | | | | |
|---|---|---|---|---|---|---|---|---|---|
| 10 | 10 | 10 | 24.4 | 24.4 | 24.4 | 54.169 | 0 | 31.319 | 1.15E+09 |
| 10 | 10 | 10 | 14.5 | 14.5 | 14.5 | 98.262 | 0 | 15.048 | 1.37E+09 |
| 12 | 10 | 10 | 30.5 | 28.6 | 28.6 | 53.762 | 0 | 23.535 | 1.97E+09 |
| 10 | 10 | 10 | 14.2 | 14.2 | 14.2 | 88.317 | 0 | 29.655 | 1.47E+09 |
| 10 | 10 | 10 | 51.4 | 51.4 | 51.4 | 24.205 | 0 | 133.6 | 8.04E+09 |
| 10 | 10 | 10 | 26.6 | 26.6 | 26.6 | 36.269 | 0 | 81.332 | 2.19E+09 |
| 20 | 20 | 9 | 16.8 | 16.8 | 8.5 | 145.18 | 0 | 53.58 | 3.36E+09 |
| 11 | 11 | 9 | 35.1 | 35.1 | 31.7 | 31.581 | 0 | 18.12 | 1.99E+09 |
| 9 | 9 | 9 | 14.7 | 14.7 | 14.7 | 70.698 | 0 | 12.583 | 2.8E+08 |
| 9 | 9 | 9 | 20.4 | 20.4 | 20.4 | 51.212 | 0 | 38.387 | 8.25E+08 |
| 9 | 9 | 9 | 45.8 | 45.8 | 45.8 | 27.335 | 0 | 46.643 | 2.58E+09 |
| 9 | 9 | 9 | 21.2 | 21.2 | 21.2 | 65.071 | 0 | 33.298 | 6.95E+08 |
| 9 | 9 | 9 | 32.4 | 32.4 | 32.4 | 42.636 | 0 | 19.657 | 1.08E+09 |
| 9 | 9 | 9 | 21.4 | 21.4 | 21.4 | 55.738 | 0 | 21.911 | 9.08E+08 |
| 9 | 9 | 9 | 23.5 | 23.5 | 23.5 | 50.097 | 0 | 27.965 | 1.28E+09 |
| 9 | 9 | 9 | 9.6 | 9.6 | 9.6 | 133.63 | 0 | 17.396 | 4.35E+08 |
| 9 | 9 | 9 | 22.2 | 22.2 | 22.2 | 55.21 | 0 | 21.289 | 8.89E+08 |
| 9 | 9 | 9 | 18.1 | 18.1 | 18.1 | 79.994 | 0 | 19.012 | 9.42E+08 |
| 9 | 9 | 9 | 88.1 | 88.1 | 88.1 | 12.895 | 0 | 66.289 | 1.51E+10 |
| 9 | 9 | 9 | 64.6 | 64.6 | 64.6 | 24.824 | 0 | 89.327 | 6.71E+09 |
| 9 | 9 | 9 | 38.3 | 38.3 | 38.3 | 32.729 | 0 | 27.033 | 2.88E+09 |
| 9 | 9 | 9 | 60.8 | 60.8 | 60.8 | 21.097 | 0 | 24.808 | 4.75E+09 |
| 9 | 9 | 9 | 36.4 | 36.4 | 36.4 | 20.777 | 0 | 13.091 | 3.06E+09 |
| 9 | 9 | 9 | 59.5 | 59.5 | 59.5 | 20.217 | 0 | 20.467 | 1.38E+09 |
| 9 | 9 | 9 | 24.1 | 24.1 | 24.1 | 43.66 | 0 | 17.988 | 2.83E+09 |
| 13 | 9 | 9 | 52.8 | 38.5 | 38.5 | 32.949 | 0 | 81.129 | 4.98E+09 |
| 9 | 9 | 9 | 52.3 | 52.3 | 52.3 | 16.32 | 0 | 53.091 | 1.49E+10 |
| 9 | 9 | 9 | 22.4 | 22.4 | 22.4 | 53.349 | 0 | 35.077 | 1.76E+09 |
| 9 | 9 | 9 | 38.5 | 38.5 | 38.5 | 35.548 | 0 | 179.17 | 9.7E+09 |
| 9 | 9 | 9 | 24.6 | 24.6 | 24.6 | 27.076 | 0 | 21.189 | 9.65E+09 |
| 9 | 9 | 9 | 42.3 | 42.3 | 42.3 | 17.695 | 0 | 18.162 | 2.9E+09 |
| 9 | 9 | 9 | 23 | 23 | 23 | 42.331 | 0 | 21.317 | 2.8E+09 |
| 9 | 9 | 9 | 14.1 | 14.1 | 14.1 | 97.44 | 0 | 25.412 | 1.49E+09 |
| 9 | 9 | 9 | 69.1 | 69.1 | 69.1 | 13.302 | 0 | 30.529 | 4.72E+09 |
| 9 | 9 | 9 | 28.6 | 28.6 | 28.6 | 46.657 | 0 | 30.603 | 3.1E+09 |
| 9 | 9 | 9 | 34.9 | 34.9 | 34.9 | 36.978 | 0 | 44.464 | 5.76E+09 |
| 9 | 9 | 9 | 53.4 | 53.4 | 53.4 | 21.86 | 0 | 44.407 | 2.97E+09 |
| 9 | 9 | 9 | 25.3 | 25.3 | 25.3 | 50.228 | 0 | 40.72 | 2.09E+09 |
| 9 | 9 | 9 | 14.4 | 14.4 | 14.4 | 87.819 | 0 | 31.002 | 1.11E+09 |
| 9 | 9 | 9 | 51.5 | 51.5 | 51.5 | 14.515 | 0 | 25.463 | 6.42E+09 |
| 9 | 9 | 9 | 22 | 22 | 22 | 48.83 | 0 | 36.182 | 1.47E+09 |
| 9 | 9 | 9 | 34.9 | 34.9 | 34.9 | 34.006 | 0 | 30.727 | 3.65E+09 |
| 9 | 9 | 9 | 42.2 | 42.2 | 42.2 | 34.14 | 0 | 13.327 | 9.56E+08 |
| 9 | 9 | 9 | 57.3 | 57.3 | 57.3 | 22.237 | 0 | 67.972 | 1.54E+09 |
| 9 | 9 | 9 | 59.3 | 59.3 | 59.3 | 28.772 | 0 | 40.609 | 5.91E+09 |
| 9 | 9 | 9 | 54.2 | 54.2 | 54.2 | 27.862 | 0 | 91.383 | 7.06E+09 |
| 9 | 9 | 9 | 42.9 | 42.9 | 42.9 | 34.352 | 0 | 26.156 | 1.19E+09 |

| | | | | | | | | | |
|---|---|---|---|---|---|---|---|---|---|
| 9 | 9 | 9 | 37.4 | 37.4 | 37.4 | 37.375 | 0 | 57.006 | 2.6E+09 |
| 9 | 9 | 9 | 12 | 12 | 12 | 81.537 | 0 | 37.341 | 1.15E+10 |
| 9 | 9 | 9 | 71.8 | 71.8 | 71.8 | 9.681 | 0 | 23.021 | 1.74E+10 |
| 11 | 9 | 9 | 46.2 | 35.7 | 35.7 | 30.85 | 0 | 27.102 | 1.86E+09 |
| 9 | 9 | 9 | 30.2 | 30.2 | 30.2 | 38.868 | 0 | 26.863 | 1.79E+09 |
| 11 | 9 | 9 | 21.1 | 17.7 | 17.7 | 69.948 | 0 | 13.355 | 6.07E+08 |
| 9 | 9 | 9 | 14.8 | 14.8 | 14.8 | 88.414 | 0 | 16.696 | 8.16E+08 |
| 9 | 9 | 9 | 19.8 | 19.8 | 19.8 | 65.335 | 0 | 33.39 | 1.69E+09 |
| 9 | 9 | 9 | 17.3 | 17.3 | 17.3 | 67.942 | 0 | 12.075 | 5.84E+08 |
| 9 | 9 | 9 | 31.1 | 31.1 | 31.1 | 45.531 | 0 | 18.722 | 1.98E+09 |
| 9 | 9 | 9 | 31.4 | 31.4 | 31.4 | 38.946 | 0 | 18.872 | 1.12E+09 |
| 9 | 9 | 9 | 47.3 | 47.3 | 47.3 | 19.586 | 0 | 14.836 | 2.12E+09 |
| 9 | 9 | 9 | 25.8 | 25.8 | 25.8 | 45.326 | 0 | 29.139 | 1.47E+09 |
| 15 | 15 | 8 | 34.6 | 34.6 | 22.6 | 49.005 | 0 | 58.638 | 3.5E+09 |
| 14 | 14 | 8 | 50.8 | 50.8 | 28.6 | 29.941 | 0 | 68.848 | 2.71E+10 |
| 13 | 13 | 8 | 28.8 | 28.8 | 19.4 | 52.933 | 0 | 50.661 | 8.41E+09 |
| 11 | 11 | 8 | 55.8 | 55.8 | 38.7 | 20.697 | 0 | 69.822 | 9.09E+09 |
| 21 | 10 | 8 | 57.4 | 35.7 | 30.9 | 49.857 | 0 | 30.372 | 2.83E+09 |
| 10 | 10 | 8 | 93.2 | 93.2 | 88.6 | 5.0526 | 0 | 100.59 | 3.93E+10 |
| 10 | 10 | 8 | 63.1 | 63.1 | 51 | 17.302 | 0 | 47.602 | 3.96E+09 |
| 9 | 9 | 8 | 51.2 | 51.2 | 51.2 | 18.762 | 0 | 33.493 | 3.78E+09 |
| 9 | 9 | 8 | 9.7 | 9.7 | 8.8 | 112.13 | 0 | 20.796 | 1.04E+09 |
| 12 | 8 | 8 | 24.8 | 21.5 | 21.5 | 56.381 | 0 | 16.78 | 4.57E+08 |
| 8 | 8 | 8 | 13.8 | 13.8 | 13.8 | 71.635 | 0 | 7.2742 | 7.49E+08 |
| 12 | 8 | 8 | 45.9 | 32.7 | 32.7 | 47.268 | 0 | 142.07 | 1.19E+09 |
| 8 | 8 | 8 | 18.1 | 18.1 | 18.1 | 57.03 | 0 | 10.616 | 6.51E+08 |
| 8 | 8 | 8 | 23.8 | 23.8 | 23.8 | 51.769 | 0 | 24.591 | 1.28E+09 |
| 10 | 8 | 8 | 37.8 | 31.5 | 31.5 | 31.802 | 0 | 20.216 | 1.13E+09 |
| 8 | 8 | 8 | 12.9 | 12.9 | 12.9 | 73.457 | 0 | 8.6945 | 9.7E+08 |
| 8 | 8 | 8 | 13.6 | 13.6 | 13.6 | 71.223 | 0 | 13.604 | 6.21E+08 |
| 8 | 8 | 8 | 80 | 80 | 80 | 15.85 | 0 | 79.635 | 2.29E+09 |
| 8 | 8 | 8 | 34.5 | 34.5 | 34.5 | 37.84 | 0 | 18.238 | 1.08E+09 |
| 8 | 8 | 8 | 32.2 | 32.2 | 32.2 | 37.874 | 0 | 29.678 | 9.33E+08 |
| 8 | 8 | 8 | 7 | 7 | 7 | 136.22 | 0 | 26.635 | 6.96E+08 |
| 8 | 8 | 8 | 34.4 | 34.4 | 34.4 | 24.95 | 0 | 39.281 | 3.52E+09 |
| 8 | 8 | 8 | 56.8 | 56.8 | 56.8 | 12.712 | 0 | 75.817 | 9.97E+09 |
| 8 | 8 | 8 | 52.5 | 52.5 | 52.5 | 19.463 | 0 | 71.009 | 3.36E+09 |
| 8 | 8 | 8 | 49.8 | 49.8 | 49.8 | 22.949 | 0 | 123.33 | 6.01E+09 |
| 8 | 8 | 8 | 20.6 | 20.6 | 20.6 | 39.017 | 0 | 11 | 1.32E+09 |
| 8 | 8 | 8 | 34.4 | 34.4 | 34.4 | 30.062 | 0 | 23.421 | 4.28E+09 |
| 8 | 8 | 8 | 90.6 | 90.6 | 90.6 | 11.776 | 0 | 323.31 | 2.18E+10 |
| 8 | 8 | 8 | 62.4 | 62.4 | 62.4 | 10.834 | 0 | 16.94 | 6.63E+09 |
| 8 | 8 | 8 | 38.5 | 38.5 | 38.5 | 22.391 | 0 | 26.706 | 3.36E+09 |
| 8 | 8 | 8 | 45.5 | 45.5 | 45.5 | 18.898 | 0 | 13.063 | 2.57E+09 |
| 8 | 8 | 8 | 44.9 | 44.9 | 44.9 | 23.742 | 0 | 49.432 | 7.58E+09 |
| 8 | 8 | 8 | 25.1 | 25.1 | 25.1 | 56.695 | 0 | 18.014 | 1.26E+09 |
| 8 | 8 | 8 | 24.9 | 24.9 | 24.9 | 37.208 | 0 | 47.8 | 3.91E+09 |

| | | | | | | | | | |
|---|---|---|---|---|---|---|---|---|---|
| 8 | 8 | 8 | 36.8 | 36.8 | 36.8 | 28.993 | 0 | 18.308 | 2.84E+09 |
| 8 | 8 | 8 | 25.5 | 25.5 | 25.5 | 52.025 | 0 | 141.66 | 7.23E+09 |
| 8 | 8 | 8 | 42.8 | 42.8 | 42.8 | 27.963 | 0 | 263.76 | 6.5E+10 |
| 8 | 8 | 8 | 24 | 24 | 24 | 54.066 | 0 | 39.137 | 1.73E+09 |
| 8 | 8 | 8 | 24.4 | 24.4 | 24.4 | 45.059 | 0 | 48.354 | 2.56E+09 |
| 8 | 8 | 8 | 20.6 | 20.6 | 20.6 | 39.58 | 0 | 33.307 | 1E+10 |
| 8 | 8 | 8 | 28.9 | 28.9 | 28.9 | 35.964 | 0 | 38.731 | 1.67E+09 |
| 8 | 8 | 8 | 51.9 | 51.9 | 51.9 | 20.478 | 0 | 47.691 | 3.54E+09 |
| 8 | 8 | 8 | 10.4 | 10.4 | 10.4 | 117.87 | 0 | 20.8 | 8.67E+08 |
| 8 | 8 | 8 | 18.6 | 18.6 | 18.6 | 49.184 | 0 | 20.021 | 1.44E+09 |
| 8 | 8 | 8 | 17.9 | 17.9 | 17.9 | 64.872 | 0 | 15.961 | 6.91E+08 |
| 8 | 8 | 8 | 17.6 | 17.6 | 17.6 | 58.314 | 0 | 17.192 | 1.85E+09 |
| 8 | 8 | 8 | 16.5 | 16.5 | 16.5 | 72.093 | 0 | 18.795 | 1.14E+09 |
| 8 | 8 | 8 | 17.4 | 17.4 | 17.4 | 65.893 | 0 | 35.146 | 2.44E+09 |
| 8 | 8 | 8 | 33.1 | 33.1 | 33.1 | 30.344 | 0 | 19.417 | 1.78E+09 |
| 8 | 8 | 8 | 79.8 | 79.8 | 79.8 | 10.08 | 0 | 33.712 | 6.77E+09 |
| 8 | 8 | 8 | 27.3 | 27.3 | 27.3 | 39.594 | 0 | 24.448 | 1.2E+09 |
| 8 | 8 | 8 | 24.2 | 24.2 | 24.2 | 50.704 | 0 | 22.423 | 1.26E+09 |
| 8 | 8 | 8 | 23.2 | 23.2 | 23.2 | 48.275 | 0 | 31.512 | 5.69E+09 |
| 8 | 8 | 8 | 27.2 | 27.2 | 27.2 | 28.024 | 0 | 15.127 | 3.53E+09 |
| 8 | 8 | 8 | 61.2 | 61.2 | 61.2 | 20.96 | 0 | 140.89 | 1.41E+10 |
| 8 | 8 | 8 | 10 | 10 | 10 | 107.77 | 0 | 30.921 | 1.32E+09 |
| 8 | 8 | 8 | 57.7 | 57.7 | 57.7 | 11.999 | 0 | 17.247 | 3.92E+09 |
| 8 | 8 | 8 | 50 | 50 | 50 | 26.182 | 0 | 73.915 | 3.31E+09 |
| 8 | 8 | 8 | 20.5 | 20.5 | 20.5 | 51.804 | 0 | 21.071 | 5.49E+08 |
| 8 | 8 | 8 | 26.4 | 26.4 | 26.4 | 38.388 | 0 | 9.1284 | 9.91E+08 |
| 8 | 8 | 8 | 16.8 | 16.8 | 16.8 | 71.456 | 0 | 16.525 | 1.22E+09 |
| 8 | 8 | 8 | 36.1 | 36.1 | 36.1 | 37.025 | 0 | 43.558 | 1.28E+09 |
| 8 | 8 | 8 | 13 | 13 | 13 | 68.997 | 0 | 22.504 | 9.84E+08 |
| 8 | 8 | 8 | 20.8 | 20.8 | 20.8 | 54.465 | 0 | 16.773 | 1.09E+09 |
| 8 | 8 | 8 | 57.1 | 57.1 | 57.1 | 18.445 | 0 | 34.072 | 3.74E+09 |
| 8 | 8 | 8 | 13.5 | 13.5 | 13.5 | 75.378 | 0 | 14.528 | 1.14E+09 |
| 8 | 8 | 8 | 49.4 | 49.4 | 49.4 | 18.079 | 0 | 11.35 | 1.8E+09 |
| 8 | 8 | 8 | 19.1 | 19.1 | 19.1 | 52.739 | 0 | 16.032 | 8.94E+08 |
| 8 | 8 | 8 | 27.5 | 27.5 | 27.5 | 42.966 | 0 | 143.16 | 5.7E+09 |
| 8 | 8 | 8 | 61 | 61 | 61 | 15.798 | 0 | 16.89 | 4.3E+09 |
| 8 | 8 | 8 | 36.5 | 36.5 | 36.5 | 31.124 | 0 | 157.76 | 1.02E+10 |
| 8 | 8 | 8 | 43 | 43 | 43 | 16.273 | 0 | 25.987 | 3.19E+09 |
| 8 | 8 | 8 | 47 | 47 | 47 | 29.644 | 0 | 72.291 | 2.73E+09 |
| 8 | 8 | 8 | 33.3 | 33.3 | 33.3 | 27.991 | 0 | 9.2783 | 7.45E+08 |
| 8 | 8 | 8 | 19.7 | 19.7 | 19.7 | 50.978 | 0 | 10.984 | 5.02E+08 |
| 8 | 8 | 8 | 56.6 | 56.6 | 56.6 | 17.138 | 0 | 39.927 | 7.01E+09 |
| 8 | 8 | 8 | 7 | 7 | 7 | 140.96 | 0 | 8.9697 | 6.52E+08 |
| 8 | 8 | 8 | 16.4 | 16.4 | 16.4 | 56.882 | 0 | 77.246 | 1.54E+09 |
| 8 | 8 | 8 | 63.6 | 63.6 | 63.6 | 17.818 | 0 | 134.18 | 8.3E+09 |
| 8 | 8 | 8 | 37.2 | 37.2 | 37.2 | 23.432 | 0 | 25.07 | 2.55E+09 |
| 8 | 8 | 8 | 92.9 | 92.9 | 92.9 | 11.139 | 0 | 41.723 | 4.53E+10 |

| | | | | | | | | | |
|---|---|---|---|---|---|---|---|---|---|
| 8 | 8 | 8 | 41.3 | 41.3 | 41.3 | 29.204 | 0 | 77.454 | 1.05E+10 |
| 8 | 8 | 8 | 29.5 | 29.5 | 29.5 | 38.91 | 0 | 18.74 | 2.39E+09 |
| 21 | 21 | 7 | 24.9 | 24.9 | 8.8 | 108.33 | 0 | 98.777 | 7.9E+09 |
| 24 | 20 | 7 | 66.2 | 50.5 | 11 | 34.216 | 0 | 133.56 | 5.93E+10 |
| 14 | 14 | 7 | 62.9 | 62.9 | 42.4 | 22.677 | 0 | 104.19 | 6.68E+09 |
| 13 | 13 | 7 | 35 | 35 | 19.9 | 48.991 | 0 | 60.148 | 3.65E+09 |
| 13 | 13 | 7 | 38 | 38 | 25.4 | 38.434 | 0 | 142.81 | 9.1E+09 |
| 10 | 10 | 7 | 43 | 43 | 35.4 | 37.497 | 0 | 104.71 | 5.44E+09 |
| 9 | 9 | 7 | 23.5 | 23.5 | 17.3 | 50.83 | 0 | 53.204 | 7.15E+09 |
| 12 | 9 | 7 | 51.2 | 39.5 | 34.9 | 24.893 | 0 | 27.006 | 5.07E+09 |
| 8 | 8 | 7 | 13 | 13 | 11.4 | 82.968 | 0 | 23.36 | 1.48E+09 |
| 8 | 8 | 7 | 31.7 | 31.7 | 27.1 | 26.697 | 0 | 19.006 | 3.54E+09 |
| 8 | 8 | 7 | 34.1 | 34.1 | 28.8 | 25.859 | 0 | 103.62 | 1.95E+10 |
| 8 | 8 | 7 | 43.3 | 43.3 | 39.2 | 26.788 | 0 | 21.231 | 3.27E+09 |
| 8 | 8 | 7 | 49.7 | 49.7 | 41 | 20.811 | 0 | 17.706 | 9.42E+08 |
| 7 | 7 | 7 | 16.6 | 16.6 | 16.6 | 60.192 | 0 | 13.559 | 1.05E+09 |
| 7 | 7 | 7 | 25.4 | 25.4 | 25.4 | 38.438 | 0 | 12.636 | 8.21E+08 |
| 7 | 7 | 7 | 37.7 | 37.7 | 37.7 | 32.66 | 0 | 19.756 | 9.41E+08 |
| 7 | 7 | 7 | 14.5 | 14.5 | 14.5 | 58.951 | 0 | 11.603 | 6.98E+08 |
| 7 | 7 | 7 | 12.8 | 12.8 | 12.8 | 65.921 | 0 | 20.857 | 7.65E+08 |
| 7 | 7 | 7 | 21.3 | 21.3 | 21.3 | 51.691 | 0 | 23.872 | 7.2E+08 |
| 9 | 7 | 7 | 19.3 | 15.3 | 15.3 | 59.069 | 0 | 17.147 | 9.08E+08 |
| 7 | 7 | 7 | 16.2 | 16.2 | 16.2 | 49.222 | 0 | 9.5292 | 5.29E+08 |
| 7 | 7 | 7 | 41.9 | 41.9 | 41.9 | 19.458 | 0 | 12.346 | 1.27E+09 |
| 7 | 7 | 7 | 24.6 | 24.6 | 24.6 | 43.332 | 0 | 25.334 | 1.63E+09 |
| 7 | 7 | 7 | 94 | 94 | 94 | 9.6981 | 0 | 63.533 | 1.09E+10 |
| 7 | 7 | 7 | 42.5 | 42.5 | 42.5 | 18.697 | 0 | 22.377 | 3.81E+09 |
| 7 | 7 | 7 | 21.3 | 21.3 | 21.3 | 47.366 | 0 | 17.345 | 2.18E+09 |
| 7 | 7 | 7 | 34.7 | 34.7 | 34.7 | 34.482 | 0 | 45.788 | 2.13E+09 |
| 7 | 7 | 7 | 53.8 | 53.8 | 53.8 | 20.505 | 0 | 24.613 | 4.69E+09 |
| 7 | 7 | 7 | 31.3 | 31.3 | 31.3 | 26.056 | 0 | 22.538 | 2.78E+09 |
| 7 | 7 | 7 | 35.5 | 35.5 | 35.5 | 28.936 | 0 | 163.39 | 1.04E+10 |
| 7 | 7 | 7 | 50 | 50 | 50 | 18.042 | 0 | 16.653 | 2.04E+09 |
| 7 | 7 | 7 | 30.6 | 30.6 | 30.6 | 31.904 | 0 | 44.507 | 2.54E+09 |
| 7 | 7 | 7 | 39.2 | 39.2 | 39.2 | 24.831 | 0 | 52.383 | 7.64E+09 |
| 7 | 7 | 7 | 50.4 | 50.4 | 50.4 | 11.665 | 0 | 88.816 | 1.26E+10 |
| 7 | 7 | 7 | 38.4 | 38.4 | 38.4 | 20.055 | 0 | 128.26 | 1.22E+10 |
| 7 | 7 | 7 | 42.9 | 42.9 | 42.9 | 8.647 | 0 | 26.811 | 2.91E+10 |
| 7 | 7 | 7 | 16 | 16 | 16 | 51.286 | 0 | 30.385 | 9.49E+08 |
| 7 | 7 | 7 | 24.2 | 24.2 | 24.2 | 50.513 | 0 | 35.06 | 2.7E+09 |
| 7 | 7 | 7 | 8.6 | 8.6 | 8.6 | 123.63 | 0 | 14.554 | 6.87E+08 |
| 7 | 7 | 7 | 50.4 | 50.4 | 50.4 | 14.395 | 0 | 30.694 | 1.16E+09 |
| 9 | 7 | 7 | 51.5 | 44.8 | 44.8 | 18.506 | 0 | 17.965 | 2.87E+09 |
| 7 | 7 | 7 | 24.9 | 24.9 | 24.9 | 38.324 | 0 | 29.471 | 1.17E+09 |
| 7 | 7 | 7 | 11.7 | 11.7 | 11.7 | 73.182 | 0 | 29.229 | 6.42E+08 |
| 7 | 7 | 7 | 35.7 | 35.7 | 35.7 | 28.05 | 0 | 13.719 | 1.62E+09 |
| 7 | 7 | 7 | 33.6 | 33.6 | 33.6 | 25.895 | 0 | 18.032 | 1.36E+09 |

| | | | | | | | | | |
|---|---|---|---|---|---|---|---|---|---|
| 7 | 7 | 7 | 56.1 | 56.1 | 56.1 | 22.086 | 0 | 53.853 | 6.42E+09 |
| 7 | 7 | 7 | 26.5 | 26.5 | 26.5 | 43.759 | 0 | 15.988 | 9.81E+08 |
| 7 | 7 | 7 | 8.9 | 8.9 | 8.9 | 102.35 | 0 | 24.631 | 1.53E+09 |
| 7 | 7 | 7 | 34.6 | 34.6 | 34.6 | 32.251 | 0 | 34.827 | 1.74E+09 |
| 26 | 7 | 7 | 52.5 | 24.1 | 24.1 | 42.051 | 0 | 258.3 | 1.55E+11 |
| 7 | 7 | 7 | 67 | 67 | 67 | 12.784 | 0 | 51.854 | 3.99E+09 |
| 7 | 7 | 7 | 20.7 | 20.7 | 20.7 | 47.996 | 0 | 11.043 | 7.64E+08 |
| 7 | 7 | 7 | 18.1 | 18.1 | 18.1 | 56.195 | 0 | 47.28 | 5.14E+08 |
| 10 | 7 | 7 | 29.4 | 21.4 | 21.4 | 42.766 | 0 | 14.358 | 7.1E+08 |
| 7 | 7 | 7 | 27.6 | 27.6 | 27.6 | 42.848 | 0 | 74.861 | 2.75E+09 |
| 7 | 7 | 7 | 34 | 34 | 34 | 42.835 | 0 | 58.671 | 1.76E+09 |
| 7 | 7 | 7 | 22.8 | 22.8 | 22.8 | 23.134 | 0 | 13.402 | 2.67E+09 |
| 7 | 7 | 7 | 67 | 67 | 67 | 11.203 | 0 | 32.339 | 1E+10 |
| 7 | 7 | 7 | 37.9 | 37.9 | 37.9 | 26.319 | 0 | 38.439 | 9.27E+08 |
| 9 | 7 | 7 | 20.7 | 14.6 | 14.6 | 46.871 | 0 | 11.424 | 6.67E+08 |
| 7 | 7 | 7 | 29.5 | 29.5 | 29.5 | 42.179 | 0 | 30.214 | 1.05E+09 |
| 7 | 7 | 7 | 33.5 | 33.5 | 33.5 | 23.577 | 0 | 19.295 | 3.43E+09 |
| 7 | 7 | 7 | 18.4 | 18.4 | 18.4 | 51.504 | 0 | 14.281 | 8.78E+08 |
| 7 | 7 | 7 | 58.5 | 58.5 | 58.5 | 10.645 | 0 | 25.668 | 1.84E+09 |
| 7 | 7 | 7 | 6.3 | 6.3 | 6.3 | 141.75 | 0 | 19.183 | 1.17E+09 |
| 7 | 7 | 7 | 48.8 | 48.8 | 48.8 | 13.742 | 0 | 11.673 | 1.66E+09 |
| 7 | 7 | 7 | 34.9 | 34.9 | 34.9 | 24.579 | 0 | 10.656 | 6.74E+08 |
| 7 | 7 | 7 | 12.5 | 12.5 | 12.5 | 91.868 | 0 | 30.624 | 4.02E+08 |
| 8 | 7 | 7 | 36.4 | 33.3 | 33.3 | 24.763 | 0 | 94.514 | 2.94E+09 |
| 7 | 7 | 7 | 31.5 | 31.5 | 31.5 | 24.105 | 0 | 19.73 | 1.23E+09 |
| 7 | 7 | 7 | 20.5 | 20.5 | 20.5 | 41.079 | 0 | 32.42 | 1.67E+09 |
| 7 | 7 | 7 | 40 | 40 | 40 | 17.258 | 0 | 9.3965 | 1.54E+09 |
| 7 | 7 | 7 | 15.2 | 15.2 | 15.2 | 64.532 | 0 | 27.677 | 3.09E+09 |
| 19 | 19 | 6 | 21.5 | 21.5 | 7 | 104.64 | 0 | 46.793 | 5.42E+09 |
| 16 | 16 | 6 | 40.4 | 40.4 | 15.5 | 21.364 | 0 | 54.406 | 3.94E+10 |
| 15 | 15 | 6 | 52.1 | 52.1 | 23.9 | 37.512 | 0 | 67.407 | 4.25E+09 |
| 13 | 13 | 6 | 54.4 | 54.4 | 26.4 | 21.768 | 0 | 114.13 | 7.25E+09 |
| 12 | 12 | 6 | 43.7 | 43.7 | 19 | 29.597 | 0 | 22.867 | 5.02E+09 |
| 11 | 11 | 6 | 42.6 | 42.6 | 28.2 | 37.377 | 0 | 223.98 | 9.55E+09 |
| 8 | 8 | 6 | 13.6 | 13.6 | 10.3 | 73.027 | 0 | 15.029 | 4.87E+08 |
| 8 | 8 | 6 | 23.2 | 23.2 | 17.6 | 39.869 | 0 | 23.4 | 1.76E+09 |
| 8 | 8 | 6 | 76.8 | 76.8 | 51.4 | 16.801 | 0 | 294.66 | 8.26E+09 |
| 28 | 7 | 6 | 66.7 | 21.4 | 17.8 | 47.766 | 0 | 269.04 | 3.05E+10 |
| 7 | 7 | 6 | 18.2 | 18.2 | 15.7 | 52.164 | 0 | 17.177 | 1.15E+09 |
| 7 | 7 | 6 | 11.2 | 11.2 | 9.7 | 80.253 | 0 | 18.046 | 6.16E+08 |
| 6 | 6 | 6 | 3.3 | 3.3 | 3.3 | 202.29 | 0 | 5.7672 | 2.18E+08 |
| 6 | 6 | 6 | 7.5 | 7.5 | 7.5 | 92.067 | 0 | 8.9604 | 3.1E+08 |
| 6 | 6 | 6 | 7.3 | 7.3 | 7.3 | 125.09 | 0 | 16.226 | 7.84E+08 |
| 6 | 6 | 6 | 17.5 | 17.5 | 17.5 | 53.806 | 0 | 19.409 | 4.28E+08 |
| 6 | 6 | 6 | 15.1 | 15.1 | 15.1 | 44.882 | 0 | 29.676 | 4.11E+09 |
| 9 | 6 | 6 | 6 | 4.4 | 4.4 | 160.33 | 0 | 9.5514 | 4.42E+08 |
| 6 | 6 | 6 | 21.5 | 21.5 | 21.5 | 46.816 | 0 | 18.046 | 2.83E+08 |

| | | | | | | | | | |
|---|---|---|---|---|---|---|---|---|---|
| 6 | 6 | 6 | 6.8 | 6.8 | 6.8 | 140.47 | 0 | 19.611 | 6.3E+08 |
| 6 | 6 | 6 | 29.8 | 29.8 | 29.8 | 27.443 | 0 | 18.718 | 7.33E+08 |
| 6 | 6 | 6 | 18.2 | 18.2 | 18.2 | 40.04 | 0 | 13.071 | 6.7E+08 |
| 6 | 6 | 6 | 30.3 | 30.3 | 30.3 | 32.113 | 0 | 19.228 | 6.01E+08 |
| 6 | 6 | 6 | 49.6 | 49.6 | 49.6 | 13.37 | 0 | 112.44 | 3.89E+09 |
| 6 | 6 | 6 | 20.4 | 20.4 | 20.4 | 51.998 | 0 | 21.477 | 5.03E+08 |
| 6 | 6 | 6 | 11.4 | 11.4 | 11.4 | 59.271 | 0 | 17.263 | 3.97E+08 |
| 6 | 6 | 6 | 40 | 40 | 40 | 21.627 | 0 | 71.933 | 7.44E+08 |
| 6 | 6 | 6 | 11 | 11 | 11 | 70.901 | 0 | 12.634 | 8.08E+08 |
| 6 | 6 | 6 | 31.6 | 31.6 | 31.6 | 34.292 | 0 | 17.753 | 8.32E+08 |
| 6 | 6 | 6 | 17.4 | 17.4 | 17.4 | 37.563 | 0 | 8.7159 | 7.21E+08 |
| 6 | 6 | 6 | 22.1 | 22.1 | 22.1 | 38.792 | 0 | 9.1218 | 7.86E+08 |
| 6 | 6 | 6 | 32.7 | 32.7 | 32.7 | 22.024 | 0 | 14.603 | 5.73E+08 |
| 7 | 6 | 6 | 13.6 | 11.9 | 11.9 | 54.231 | 0 | 9.4952 | 5.56E+08 |
| 6 | 6 | 6 | 19.1 | 19.1 | 19.1 | 39.028 | 0 | 9.6375 | 5.66E+08 |
| 6 | 6 | 6 | 27.1 | 27.1 | 27.1 | 32.805 | 0 | 17.056 | 4.74E+08 |
| 6 | 6 | 6 | 6.4 | 6.4 | 6.4 | 112.42 | 0 | 15.238 | 4.11E+08 |
| 6 | 6 | 6 | 60.8 | 60.8 | 60.8 | 17.769 | 0 | 18.751 | 1.71E+09 |
| 6 | 6 | 6 | 12.6 | 12.6 | 12.6 | 54.611 | 0 | 6.3818 | 2.72E+08 |
| 6 | 6 | 6 | 17.8 | 17.8 | 17.8 | 45.518 | 0 | 7.8154 | 3.13E+08 |
| 6 | 6 | 6 | 5.4 | 5.4 | 5.4 | 129.21 | 0 | 7.1557 | 1.73E+08 |
| 6 | 6 | 6 | 1.1 | 1.1 | 1.1 | 596.48 | 0 | 3.5833 | 1.29E+08 |
| 6 | 6 | 6 | 9.1 | 9.1 | 9.1 | 85.377 | 0 | 3.5138 | 2.17E+08 |
| 6 | 6 | 6 | 27.2 | 27.2 | 27.2 | 38.169 | 0 | 29.886 | 2.85E+09 |
| 6 | 6 | 6 | 44.6 | 44.6 | 44.6 | 20.567 | 0 | 31.963 | 3.82E+09 |
| 6 | 6 | 6 | 19 | 19 | 19 | 33.934 | 0 | 34.817 | 2.75E+09 |
| 6 | 6 | 6 | 33.5 | 33.5 | 33.5 | 25.177 | 0 | 12.088 | 2.91E+09 |
| 6 | 6 | 6 | 50.8 | 50.8 | 50.8 | 14.787 | 0 | 51.5 | 3.02E+09 |
| 6 | 6 | 6 | 60.9 | 60.9 | 60.9 | 7.8409 | 0 | 70.374 | 7.23E+09 |
| 6 | 6 | 6 | 36.7 | 36.7 | 36.7 | 23.896 | 0 | 15.038 | 3.55E+09 |
| 6 | 6 | 6 | 45.7 | 45.7 | 45.7 | 17.499 | 0 | 8.1238 | 8.58E+08 |
| 6 | 6 | 6 | 14.1 | 14.1 | 14.1 | 57.825 | 0 | 17.272 | 1.36E+09 |
| 6 | 6 | 6 | 19.1 | 19.1 | 19.1 | 46.306 | 0 | 25.308 | 1.14E+09 |
| 6 | 6 | 6 | 34.1 | 34.1 | 34.1 | 28.058 | 0 | 111.42 | 6.33E+09 |
| 6 | 6 | 6 | 17.2 | 17.2 | 17.2 | 49.197 | 0 | 18.902 | 1.1E+10 |
| 6 | 6 | 6 | 34.5 | 34.5 | 34.5 | 25.497 | 0 | 33.829 | 1.4E+09 |
| 6 | 6 | 6 | 43.5 | 43.5 | 43.5 | 14.374 | 0 | 33.896 | 4.53E+09 |
| 6 | 6 | 6 | 21.8 | 21.8 | 21.8 | 34.979 | 0 | 42.978 | 1.33E+10 |
| 6 | 6 | 6 | 52.5 | 52.5 | 52.5 | 11.333 | 0 | 29.158 | 4.2E+09 |
| 6 | 6 | 6 | 64.3 | 64.3 | 64.3 | 16.057 | 0 | 44.868 | 2.71E+09 |
| 6 | 6 | 6 | 27.4 | 27.4 | 27.4 | 25.447 | 0 | 39.342 | 5.75E+09 |
| 6 | 6 | 6 | 24.4 | 24.4 | 24.4 | 33.698 | 0 | 12.828 | 3.86E+08 |
| 6 | 6 | 6 | 25.8 | 25.8 | 25.8 | 27.385 | 0 | 23.61 | 1.73E+09 |
| 6 | 6 | 6 | 29.2 | 29.2 | 29.2 | 34.819 | 0 | 24.383 | 1.34E+09 |
| 6 | 6 | 6 | 26.1 | 26.1 | 26.1 | 28.048 | 0 | 33.74 | 6.89E+08 |
| 6 | 6 | 6 | 40.7 | 40.7 | 40.7 | 12.527 | 0 | 4.0945 | 9.88E+08 |
| 6 | 6 | 6 | 30.2 | 30.2 | 30.2 | 25.402 | 0 | 8.9152 | 1.62E+09 |

| | | | | | | | | | |
|---|---|---|---|---|---|---|---|---|---|
| 6 | 6 | 6 | 43.1 | 43.1 | 43.1 | 16.913 | 0 | 11.01 | 1.18E+09 |
| 6 | 6 | 6 | 47 | 47 | 47 | 16.533 | 0 | 16.119 | 1.66E+09 |
| 6 | 6 | 6 | 31.4 | 31.4 | 31.4 | 21.525 | 0 | 87.798 | 1.34E+10 |
| 6 | 6 | 6 | 29.2 | 29.2 | 29.2 | 12.249 | 0 | 43.834 | 2.75E+09 |
| 6 | 6 | 6 | 40.6 | 40.6 | 40.6 | 15.371 | 0 | 10.255 | 1.05E+09 |
| 6 | 6 | 6 | 30.9 | 30.9 | 30.9 | 21.296 | 0 | 12.959 | 1.65E+09 |
| 6 | 6 | 6 | 16.3 | 16.3 | 16.3 | 34.722 | 0 | 12.962 | 2.15E+09 |
| 6 | 6 | 6 | 11.8 | 11.8 | 11.8 | 54.283 | 0 | 8.1477 | 9.46E+08 |
| 6 | 6 | 6 | 43.7 | 43.7 | 43.7 | 18.863 | 0 | 56.44 | 5.37E+09 |
| 6 | 6 | 6 | 34.6 | 34.6 | 34.6 | 23.787 | 0 | 52.032 | 3.53E+09 |
| 6 | 6 | 6 | 18 | 18 | 18 | 46.564 | 0 | 34.756 | 1.35E+09 |
| 6 | 6 | 6 | 35.9 | 35.9 | 35.9 | 15.197 | 0 | 18.298 | 2.01E+09 |
| 6 | 6 | 6 | 29.8 | 29.8 | 29.8 | 24.593 | 0 | 12.93 | 1.55E+09 |
| 6 | 6 | 6 | 42.1 | 42.1 | 42.1 | 15.616 | 0 | 9.4131 | 2.24E+09 |
| 6 | 6 | 6 | 33.7 | 33.7 | 33.7 | 18.565 | 0 | 12.488 | 1.95E+09 |
| 6 | 6 | 6 | 13.6 | 13.6 | 13.6 | 55.86 | 0 | 15.002 | 1.07E+09 |
| 6 | 6 | 6 | 10.1 | 10.1 | 10.1 | 63.472 | 0 | 14.09 | 4.1E+08 |
| 6 | 6 | 6 | 44.4 | 44.4 | 44.4 | 19.608 | 0 | 18.52 | 9.09E+08 |
| 6 | 6 | 6 | 20.6 | 20.6 | 20.6 | 51.712 | 0 | 34.983 | 8.72E+08 |
| 6 | 6 | 6 | 21.8 | 21.8 | 21.8 | 41.595 | 0 | 50.686 | 1.65E+09 |
| 7 | 6 | 6 | 42.4 | 36.6 | 36.6 | 21.258 | 0 | 81.799 | 5.18E+09 |
| 6 | 6 | 6 | 18.3 | 18.3 | 18.3 | 30.33 | 0 | 18.019 | 2.19E+09 |
| 6 | 6 | 6 | 39.6 | 39.6 | 39.6 | 24.669 | 0 | 30.9 | 1.07E+09 |
| 6 | 6 | 6 | 19.2 | 19.2 | 19.2 | 48.227 | 0 | 10.835 | 4.96E+08 |
| 6 | 6 | 6 | 52 | 52 | 52 | 17.196 | 0 | 29.937 | 2.05E+09 |
| 6 | 6 | 6 | 46.5 | 46.5 | 46.5 | 17.605 | 0 | 15.391 | 1.34E+09 |
| 6 | 6 | 6 | 22.8 | 22.8 | 22.8 | 39.617 | 0 | 14.369 | 7.37E+08 |
| 6 | 6 | 6 | 30.7 | 30.7 | 30.7 | 31.608 | 0 | 11.308 | 1.24E+09 |
| 6 | 6 | 6 | 24.5 | 24.5 | 24.5 | 31.203 | 0 | 10.767 | 6.57E+08 |
| 6 | 6 | 6 | 81.3 | 81.3 | 81.3 | 11.765 | 0 | 54.279 | 5.51E+09 |
| 6 | 6 | 6 | 22.6 | 22.6 | 22.6 | 35.668 | 0 | 11.451 | 7.59E+08 |
| 6 | 6 | 6 | 23.1 | 23.1 | 23.1 | 32.583 | 0 | 8.5171 | 8.09E+08 |
| 6 | 6 | 6 | 29.9 | 29.9 | 29.9 | 13.293 | 0 | 8.1346 | 2.06E+09 |
| 6 | 6 | 6 | 34.7 | 34.7 | 34.7 | 14.369 | 0 | 5.2489 | 8.31E+08 |
| 6 | 6 | 6 | 21.7 | 21.7 | 21.7 | 54.739 | 0 | 51.135 | 1.1E+09 |
| 6 | 6 | 6 | 29.6 | 29.6 | 29.6 | 28.329 | 0 | 10.554 | 1.26E+09 |
| 6 | 6 | 6 | 37.4 | 37.4 | 37.4 | 14.551 | 0 | 13.371 | 1.12E+09 |
| 6 | 6 | 6 | 10.8 | 10.8 | 10.8 | 51.556 | 0 | 6.7974 | 5.08E+08 |
| 6 | 6 | 6 | 21.5 | 21.5 | 21.5 | 36.294 | 0 | 13.561 | 8.6E+08 |
| 60 | 60 | 5 | 89.8 | 89.8 | 9.4 | 57.936 | 0 | 323.31 | 7.82E+11 |
| 14 | 14 | 5 | 31 | 31 | 9.6 | 58.397 | 0 | 42.374 | 5.38E+09 |
| 12 | 12 | 5 | 43.6 | 43.6 | 16 | 35.9 | 0 | 72.484 | 2.12E+10 |
| 10 | 10 | 5 | 58.5 | 58.5 | 34 | 16.495 | 0 | 26.529 | 4.25E+09 |
| 12 | 8 | 5 | 54.8 | 40.3 | 27.9 | 36.105 | 0 | 55.953 | 8.8E+09 |
| 7 | 7 | 5 | 32.8 | 32.8 | 27.8 | 23.417 | 0 | 36.296 | 4.63E+09 |
| 7 | 7 | 5 | 22.3 | 22.3 | 16.3 | 49.229 | 0 | 29.916 | 1.5E+09 |
| 5 | 5 | 5 | 19 | 19 | 19 | 31.387 | 0 | 7.4437 | 3.79E+08 |

| | | | | | | | | | |
|---|---|---|---|---|---|---|---|---|---|
| 5 | 5 | 5 | 11.3 | 11.3 | 11.3 | 59.687 | 0 | 11.112 | 4.92E+08 |
| 5 | 5 | 5 | 23.4 | 23.4 | 23.4 | 31.215 | 0 | 25.571 | 8.75E+08 |
| 5 | 5 | 5 | 23.4 | 23.4 | 23.4 | 29.815 | 0 | 12.585 | 8.5E+08 |
| 5 | 5 | 5 | 15.2 | 15.2 | 15.2 | 44.739 | 0 | 11.274 | 8.23E+08 |
| 5 | 5 | 5 | 17.7 | 17.7 | 17.7 | 34.005 | 0 | 17.054 | 1.02E+09 |
| 5 | 5 | 5 | 16.3 | 16.3 | 16.3 | 28.18 | 0 | 17.473 | 8.12E+08 |
| 5 | 5 | 5 | 17.2 | 17.2 | 17.2 | 44.681 | 0 | 25.895 | 7.78E+08 |
| 18 | 5 | 5 | 20.7 | 6 | 6 | 104.55 | 0 | 30.601 | 5.92E+08 |
| 5 | 5 | 5 | 61.7 | 61.7 | 61.7 | 9.0714 | 0 | 33.038 | 1.34E+09 |
| 5 | 5 | 5 | 36.7 | 36.7 | 36.7 | 20.272 | 0 | 17.847 | 1.18E+09 |
| 5 | 5 | 5 | 37.5 | 37.5 | 37.5 | 14.57 | 0 | 52.211 | 5.73E+08 |
| 5 | 5 | 5 | 15.7 | 15.7 | 15.7 | 38.274 | 0 | 21.166 | 5.55E+08 |
| 5 | 5 | 5 | 7 | 7 | 7 | 78.839 | 0 | 11.764 | 4.17E+08 |
| 5 | 5 | 5 | 20.8 | 20.8 | 20.8 | 29.47 | 0 | 10.016 | 1.21E+09 |
| 5 | 5 | 5 | 23.8 | 23.8 | 23.8 | 25.789 | 0 | 27.282 | 1.06E+09 |
| 5 | 5 | 5 | 6.5 | 6.5 | 6.5 | 110.42 | 0 | 4.3692 | 5.24E+08 |
| 5 | 5 | 5 | 15.1 | 15.1 | 15.1 | 46.145 | 0 | 7.2146 | 5.98E+08 |
| 5 | 5 | 5 | 4.8 | 4.8 | 4.8 | 133.07 | 0 | 19.307 | 2.48E+08 |
| 12 | 5 | 5 | 23 | 11.1 | 11.1 | 49.303 | 0 | 12.443 | 5.62E+08 |
| 5 | 5 | 5 | 6.2 | 6.2 | 6.2 | 110.18 | 0 | 8.6328 | 1.92E+08 |
| 5 | 5 | 5 | 5.5 | 5.5 | 5.5 | 87.817 | 0 | 3.8857 | 93435000 |
| 5 | 5 | 5 | 40.2 | 40.2 | 40.2 | 12.473 | 0 | 9.2201 | 2.95E+09 |
| 5 | 5 | 5 | 41.9 | 41.9 | 41.9 | 15.016 | 0 | 13.287 | 1.83E+09 |
| 5 | 5 | 5 | 65.4 | 65.4 | 65.4 | 8.85 | 0 | 30.914 | 5.87E+09 |
| 5 | 5 | 5 | 33.3 | 33.3 | 33.3 | 11.985 | 0 | 9.6547 | 3.18E+09 |
| 5 | 5 | 5 | 35.6 | 35.6 | 35.6 | 16.837 | 0 | 87.505 | 9.05E+09 |
| 5 | 5 | 5 | 30.1 | 30.1 | 30.1 | 18.795 | 0 | 13.866 | 4.44E+09 |
| 5 | 5 | 5 | 46.2 | 46.2 | 46.2 | 10.624 | 0 | 23.676 | 1.84E+10 |
| 5 | 5 | 5 | 10.6 | 10.6 | 10.6 | 60.67 | 0 | 13.263 | 9.33E+08 |
| 5 | 5 | 5 | 76.5 | 76.5 | 76.5 | 7.4016 | 0 | 124.82 | 7.76E+09 |
| 5 | 5 | 5 | 5.8 | 5.8 | 5.8 | 82.521 | 0 | 7.3436 | 9.78E+08 |
| 5 | 5 | 5 | 35.6 | 35.6 | 35.6 | 21.635 | 0 | 11.734 | 3.47E+08 |
| 5 | 5 | 5 | 60 | 60 | 60 | 15.04 | 0 | 32.99 | 2.26E+09 |
| 5 | 5 | 5 | 9.2 | 9.2 | 9.2 | 55.755 | 0 | 6.7074 | 7.43E+08 |
| 7 | 5 | 5 | 57.7 | 38.7 | 38.7 | 16.713 | 0 | 38.271 | 1.68E+09 |
| 5 | 5 | 5 | 18.1 | 18.1 | 18.1 | 49.03 | 0 | 24.521 | 1.46E+09 |
| 7 | 5 | 5 | 19.2 | 16.3 | 16.3 | 42.823 | 0 | 24.46 | 1E+09 |
| 9 | 5 | 5 | 27.1 | 15.4 | 15.4 | 40.057 | 0 | 68.027 | 1.26E+09 |
| 5 | 5 | 5 | 31.6 | 31.6 | 31.6 | 17.652 | 0 | 36.251 | 1.53E+09 |
| 5 | 5 | 5 | 20.8 | 20.8 | 20.8 | 36.926 | 0 | 8.6934 | 5.89E+08 |
| 5 | 5 | 5 | 29.2 | 29.2 | 29.2 | 20.252 | 0 | 34.132 | 3.14E+09 |
| 5 | 5 | 5 | 36.8 | 36.8 | 36.8 | 24.737 | 0 | 54.701 | 1.39E+09 |
| 6 | 5 | 5 | 17.4 | 15.4 | 15.4 | 40.282 | 0 | 15.786 | 8.94E+08 |
| 5 | 5 | 5 | 8.2 | 8.2 | 8.2 | 58.993 | 0 | 18.291 | 1.81E+09 |
| 5 | 5 | 5 | 42.6 | 42.6 | 42.6 | 7.3184 | 0 | 10.923 | 2.23E+09 |
| 5 | 5 | 5 | 51.8 | 51.8 | 51.8 | 9.6172 | 0 | 13.846 | 3.25E+09 |
| 5 | 5 | 5 | 5.6 | 5.6 | 5.6 | 91.35 | 0 | 9.1069 | 7.23E+08 |

| | | | | | | | | | |
|---|---|---|---|---|---|---|---|---|---|
| 5 | 5 | 5 | 7.3 | 7.3 | 7.3 | 82.431 | 0 | 5.75 | 3.36E+08 |
| 5 | 5 | 5 | 10.1 | 10.1 | 10.1 | 56.806 | 0 | 5.5999 | 2.75E+08 |
| 12 | 5 | 5 | 52.3 | 20.2 | 20.2 | 22.006 | 0 | 11.407 | 6.03E+08 |
| 5 | 5 | 5 | 30.1 | 30.1 | 30.1 | 15.807 | 0 | 4.4477 | 2.72E+09 |
| 16 | 5 | 5 | 48.4 | 19.9 | 19.9 | 46.402 | 0 | 25.469 | 1.17E+09 |
| 5 | 5 | 5 | 28.5 | 28.5 | 28.5 | 17.222 | 0 | 8.6211 | 1.44E+09 |
| 5 | 5 | 5 | 48.5 | 48.5 | 48.5 | 14.719 | 0 | 30.747 | 7.75E+08 |
| 5 | 5 | 5 | 9.7 | 9.7 | 9.7 | 46.382 | 0 | 7.3164 | 4.71E+08 |
| 5 | 5 | 5 | 18.3 | 18.3 | 18.3 | 39.383 | 0 | 24.339 | 2.12E+09 |
| 5 | 5 | 5 | 37.4 | 37.4 | 37.4 | 11.758 | 0 | 29.87 | 1.54E+09 |
| 5 | 5 | 5 | 31.5 | 31.5 | 31.5 | 22.088 | 0 | 17.007 | 1.12E+09 |
| 5 | 5 | 5 | 8 | 8 | 8 | 72.065 | 0 | 4.2299 | 3.15E+08 |
| 6 | 5 | 5 | 23.9 | 17.5 | 17.5 | 28.787 | 0 | 21.97 | 2.48E+09 |
| 5 | 5 | 5 | 56.7 | 56.7 | 56.7 | 14.478 | 0 | 55.252 | 2.34E+09 |
| 5 | 5 | 5 | 16.9 | 16.9 | 16.9 | 38.242 | 0 | 10.884 | 5.89E+08 |
| 5 | 5 | 5 | 37.1 | 37.1 | 37.1 | 26.599 | 0 | 75.124 | 2.81E+09 |
| 5 | 5 | 5 | 31.1 | 31.1 | 31.1 | 13.373 | 0 | 15.939 | 3.48E+09 |
| 5 | 5 | 5 | 16.7 | 16.7 | 16.7 | 33.797 | 0 | 15.709 | 1.52E+09 |
| 5 | 5 | 5 | 21.7 | 21.7 | 21.7 | 38.782 | 0 | 6.8991 | 1.58E+09 |
| 5 | 5 | 5 | 19 | 19 | 19 | 46.971 | 0 | 15.655 | 4.26E+08 |
| 5 | 5 | 5 | 13.6 | 13.6 | 13.6 | 52.904 | 0 | 10.059 | 5.66E+08 |
| 5 | 5 | 5 | 20.5 | 20.5 | 20.5 | 22.622 | 0 | 9.951 | 5.55E+08 |
| 5 | 5 | 5 | 9.2 | 9.2 | 9.2 | 78.365 | 0 | 13.258 | 6.68E+08 |
| 45 | 45 | 4 | 59.9 | 59.9 | 6.2 | 70.897 | 0 | 323.31 | 2E+11 |
| 22 | 22 | 4 | 69.4 | 69.4 | 7.6 | 33.781 | 0 | 109.91 | 2.66E+10 |
| 17 | 17 | 4 | 72.3 | 72.3 | 22.8 | 20.825 | 0 | 83.313 | 9.8E+09 |
| 12 | 12 | 4 | 33.5 | 33.5 | 12.7 | 54.966 | 0 | 90.502 | 1.23E+10 |
| 10 | 10 | 4 | 50.4 | 50.4 | 24.6 | 25.961 | 0 | 31.219 | 5.31E+09 |
| 10 | 10 | 4 | 42.8 | 42.8 | 20.2 | 42.613 | 0 | 41.621 | 2.8E+09 |
| 10 | 10 | 4 | 20.8 | 20.8 | 7.5 | 58.362 | 0 | 42.058 | 2.44E+09 |
| 9 | 9 | 4 | 41.1 | 41.1 | 19.8 | 21.429 | 0 | 21.099 | 1.62E+10 |
| 8 | 8 | 4 | 19.6 | 19.6 | 11.2 | 43.439 | 0 | 11.323 | 1.41E+09 |
| 11 | 8 | 4 | 54.9 | 42.9 | 20.3 | 30.12 | 0 | 139.72 | 5.99E+09 |
| 7 | 7 | 4 | 18.4 | 18.4 | 12.7 | 57.886 | 0 | 23.457 | 1.22E+09 |
| 6 | 6 | 4 | 11.5 | 11.5 | 7.9 | 58.687 | 0 | 8.4998 | 4.5E+08 |
| 6 | 6 | 4 | 18.6 | 18.6 | 12.8 | 42.447 | 0 | 23.263 | 1.26E+09 |
| 5 | 5 | 4 | 51.7 | 51.7 | 39.3 | 10.366 | 0 | 24.544 | 1.36E+09 |
| 4 | 4 | 4 | 6.7 | 6.7 | 6.7 | 66.938 | 0 | 7.9936 | 2.25E+08 |
| 4 | 4 | 4 | 15.9 | 15.9 | 15.9 | 38.282 | 0 | 15.747 | 9.6E+08 |
| 5 | 4 | 4 | 18.3 | 14.3 | 14.3 | 35.816 | 0 | 26.276 | 5.71E+08 |
| 4 | 4 | 4 | 17.4 | 17.4 | 17.4 | 35.54 | 0 | 48.655 | 6.11E+08 |
| 4 | 4 | 4 | 13.4 | 13.4 | 13.4 | 40.943 | 0 | 10.199 | 5.41E+08 |
| 4 | 4 | 4 | 19.6 | 19.6 | 19.6 | 21.445 | 0 | 32.269 | 1.5E+09 |
| 9 | 4 | 4 | 53.1 | 28.3 | 28.3 | 16.363 | 0 | 9.1217 | 8.63E+08 |
| 4 | 4 | 4 | 11 | 11 | 11 | 35.896 | 0 | 24.031 | 5.11E+08 |
| 4 | 4 | 4 | 34 | 34 | 34 | 21.725 | 0 | 10.415 | 3.86E+08 |
| 4 | 4 | 4 | 24.2 | 24.2 | 24.2 | 21.539 | 0 | 7.0248 | 7.39E+08 |

| | | | | | | | | | |
|---|---|---|---|---|---|---|---|---|---|
| 4 | 4 | 4 | 13.8 | 13.8 | 13.8 | 37.709 | 0 | 5.75 | 7.18E+08 |
| 6 | 4 | 4 | 20.2 | 13.7 | 13.7 | 45.671 | 0 | 10.726 | 6.59E+08 |
| 4 | 4 | 4 | 19.4 | 19.4 | 19.4 | 25.098 | 0 | 16.214 | 3.35E+08 |
| 4 | 4 | 4 | 19.3 | 19.3 | 19.3 | 30.727 | 0 | 9.5216 | 6.04E+08 |
| 4 | 4 | 4 | 28.5 | 28.5 | 28.5 | 24.927 | 0 | 10.27 | 4.87E+08 |
| 4 | 4 | 4 | 17.6 | 17.6 | 17.6 | 25.357 | 0 | 3.1218 | 4.86E+08 |
| 4 | 4 | 4 | 11.7 | 11.7 | 11.7 | 39.336 | 0 | 3.2762 | 3.78E+08 |
| 4 | 4 | 4 | 10.6 | 10.6 | 10.6 | 44.743 | 0 | 3.5679 | 5.16E+08 |
| 4 | 4 | 4 | 11.3 | 11.3 | 11.3 | 51.172 | 0 | 12.049 | 8.71E+08 |
| 4 | 4 | 4 | 32.8 | 32.8 | 32.8 | 14.285 | 0 | 39.286 | 4.2E+08 |
| 4 | 4 | 4 | 5.3 | 5.3 | 5.3 | 88.857 | 0 | 6.94 | 3.43E+08 |
| 4 | 4 | 4 | 17.3 | 17.3 | 17.3 | 25.868 | 0 | 4.8058 | 1.1E+09 |
| 4 | 4 | 4 | 13.2 | 13.2 | 13.2 | 35.444 | 0 | 4.021 | 7.9E+08 |
| 4 | 4 | 4 | 23.5 | 23.5 | 23.5 | 20.745 | 0 | 5.0015 | 4.4E+08 |
| 4 | 4 | 4 | 3.6 | 3.6 | 3.6 | 128.4 | 0 | 5.5072 | 2.22E+08 |
| 4 | 4 | 4 | 26.9 | 26.9 | 26.9 | 15.125 | 0 | 4.5588 | 1.72E+08 |
| 4 | 4 | 4 | 2.9 | 2.9 | 2.9 | 174.76 | 0 | 15.925 | 2.37E+08 |
| 4 | 4 | 4 | 27.7 | 27.7 | 27.7 | 18.888 | 0 | 22.543 | 3.55E+09 |
| 4 | 4 | 4 | 11.3 | 11.3 | 11.3 | 46.604 | 0 | 7.4553 | 6.76E+08 |
| 4 | 4 | 4 | 8.3 | 8.3 | 8.3 | 51.887 | 0 | 4.6662 | 4.4E+08 |
| 4 | 4 | 4 | 14.8 | 14.8 | 14.8 | 25.19 | 0 | 4.7671 | 4.86E+08 |
| 4 | 4 | 4 | 9.5 | 9.5 | 9.5 | 46.426 | 0 | 4.9015 | 3.21E+08 |
| 4 | 4 | 4 | 47.2 | 47.2 | 47.2 | 13.832 | 0 | 45.195 | 3.96E+09 |
| 4 | 4 | 4 | 31.2 | 31.2 | 31.2 | 16.46 | 0 | 10.686 | 1.02E+09 |
| 4 | 4 | 4 | 50.6 | 50.6 | 50.6 | 10.058 | 0 | 21.112 | 1.12E+09 |
| 4 | 4 | 4 | 43.8 | 43.8 | 43.8 | 14.213 | 0 | 29.446 | 2.04E+09 |
| 4 | 4 | 4 | 44.2 | 44.2 | 44.2 | 10.192 | 0 | 7.6911 | 2.75E+09 |
| 4 | 4 | 4 | 3.7 | 3.7 | 3.7 | 128.77 | 0 | 7.695 | 1.75E+08 |
| 4 | 4 | 4 | 25.6 | 25.6 | 25.6 | 21.994 | 0 | 10.303 | 2.39E+09 |
| 4 | 4 | 4 | 5.4 | 5.4 | 5.4 | 84.277 | 0 | 5.4545 | 6.99E+08 |
| 4 | 4 | 4 | 30.2 | 30.2 | 30.2 | 13.696 | 0 | 5.5765 | 1.42E+09 |
| 4 | 4 | 4 | 34.9 | 34.9 | 34.9 | 16.932 | 0 | 10.245 | 6.87E+08 |
| 4 | 4 | 4 | 20.2 | 20.2 | 20.2 | 33.295 | 0 | 11.148 | 5.79E+08 |
| 4 | 4 | 4 | 21.2 | 21.2 | 21.2 | 22.326 | 0 | 6.3565 | 1.07E+09 |
| 4 | 4 | 4 | 8.2 | 8.2 | 8.2 | 63.972 | 0 | 11.224 | 4.77E+08 |
| 4 | 4 | 4 | 12.8 | 12.8 | 12.8 | 33.24 | 0 | 9.3523 | 7.54E+08 |
| 4 | 4 | 4 | 25 | 25 | 25 | 22.406 | 0 | 11.048 | 1.18E+09 |
| 4 | 4 | 4 | 27.5 | 27.5 | 27.5 | 33.973 | 0 | 55.638 | 2.12E+09 |
| 4 | 4 | 4 | 30.5 | 30.5 | 30.5 | 12.254 | 0 | 7.1187 | 1.74E+09 |
| 4 | 4 | 4 | 15.3 | 15.3 | 15.3 | 31.674 | 0 | 8.3028 | 9.83E+08 |
| 4 | 4 | 4 | 37.4 | 37.4 | 37.4 | 13.015 | 0 | 7.7715 | 3.52E+09 |
| 4 | 4 | 4 | 17.8 | 17.8 | 17.8 | 32.616 | 0 | 16.195 | 5.96E+08 |
| 4 | 4 | 4 | 20.7 | 20.7 | 20.7 | 17.546 | 0 | 5.3905 | 7.69E+08 |
| 4 | 4 | 4 | 15 | 15 | 15 | 39.315 | 0 | 18.127 | 5.32E+08 |
| 4 | 4 | 4 | 42.9 | 42.9 | 42.9 | 8.5328 | 0 | 14.592 | 7.94E+09 |
| 4 | 4 | 4 | 38.2 | 38.2 | 38.2 | 13.941 | 0 | 9.7439 | 4.64E+09 |
| 4 | 4 | 4 | 22.7 | 22.7 | 22.7 | 23.482 | 0 | 29.405 | 7.68E+08 |

| | | | | | | | | | |
|---|---|---|---|---|---|---|---|---|---|
| 6 | 4 | 4 | 9.7 | 6.3 | 6.3 | 66.616 | 0 | 13.057 | 3.59E+08 |
| 4 | 4 | 4 | 44.2 | 44.2 | 44.2 | 10.112 | 0 | 20.39 | 1.82E+09 |
| 4 | 4 | 4 | 37.6 | 37.6 | 37.6 | 21 | 0 | 11.554 | 3.47E+08 |
| 4 | 4 | 4 | 11 | 11 | 11 | 52.912 | 0 | 10.042 | 4.7E+08 |
| 4 | 4 | 4 | 17.9 | 17.9 | 17.9 | 35.206 | 0 | 7.3328 | 8.67E+08 |
| 4 | 4 | 4 | 34.9 | 34.9 | 34.9 | 25.569 | 0 | 27.678 | 1.12E+09 |
| 4 | 4 | 4 | 16.8 | 16.8 | 16.8 | 29.666 | 0 | 4.8973 | 5.6E+08 |
| 4 | 4 | 4 | 3.7 | 3.7 | 3.7 | 136.85 | 0 | 5.7837 | 2.71E+08 |
| 4 | 4 | 4 | 17.8 | 17.8 | 17.8 | 41.268 | 0 | 39.122 | 1.28E+09 |
| 4 | 4 | 4 | 16.7 | 16.7 | 16.7 | 28.855 | 0 | 8.1022 | 4.3E+08 |
| 4 | 4 | 4 | 36.5 | 36.5 | 36.5 | 17.764 | 0 | 58.615 | 4.84E+09 |
| 4 | 4 | 4 | 12.7 | 12.7 | 12.7 | 47.079 | 0 | 11.792 | 1.12E+09 |
| 4 | 4 | 4 | 64.2 | 64.2 | 64.2 | 7.3713 | 0 | 5.2148 | 4.28E+08 |
| 4 | 4 | 4 | 33.6 | 33.6 | 33.6 | 13.507 | 0 | 9.7668 | 7.36E+08 |
| 4 | 4 | 4 | 24.5 | 24.5 | 24.5 | 17.929 | 0 | 12.255 | 1.25E+09 |
| 4 | 4 | 4 | 43.2 | 43.2 | 43.2 | 9.3804 | 0 | 22.857 | 1.07E+11 |
| 4 | 4 | 4 | 23.5 | 23.5 | 23.5 | 11.53 | 0 | 18.256 | 3.29E+09 |
| 4 | 4 | 4 | 18.4 | 18.4 | 18.4 | 18.72 | 0 | 37.871 | 8.11E+08 |
| 4 | 4 | 4 | 28.4 | 28.4 | 28.4 | 16.368 | 0 | 5.8126 | 3.76E+08 |
| 4 | 4 | 4 | 13.7 | 13.7 | 13.7 | 25.441 | 0 | 5.9958 | 5.87E+08 |
| 4 | 4 | 4 | 6.6 | 6.6 | 6.6 | 68.329 | 0 | 8.6468 | 4.08E+08 |
| 4 | 4 | 4 | 21 | 21 | 21 | 22.875 | 0 | 15.059 | 8.13E+08 |
| 4 | 4 | 4 | 37.5 | 37.5 | 37.5 | 12.259 | 0 | 26.081 | 3.5E+09 |
| 20 | 17 | 3 | 21.7 | 19.5 | 4.7 | 96.864 | 0 | 59.312 | 4.35E+09 |
| 12 | 12 | 3 | 45.3 | 45.3 | 18.2 | 29.357 | 0 | 249.83 | 1.94E+10 |
| 12 | 12 | 3 | 78.2 | 78.2 | 32.8 | 13.714 | 0 | 323.31 | 4.77E+11 |
| 6 | 6 | 3 | 52.3 | 52.3 | 28.5 | 14.839 | 0 | 12.348 | 2.17E+09 |
| 6 | 6 | 3 | 27.1 | 27.1 | 16.9 | 20.457 | 0 | 51.442 | 7.13E+09 |
| 11 | 5 | 3 | 45.3 | 22.1 | 12.8 | 29.159 | 0 | 26.078 | 3.87E+09 |
| 5 | 5 | 3 | 26.8 | 26.8 | 18.5 | 27.805 | 0 | 19.275 | 2.34E+09 |
| 10 | 5 | 3 | 39.1 | 24.7 | 16.8 | 37.331 | 0 | 53.019 | 1.65E+09 |
| 5 | 5 | 3 | 28.6 | 28.6 | 17.5 | 23.408 | 0 | 11.512 | 3.07E+09 |
| 4 | 4 | 3 | 8.5 | 8.5 | 6.9 | 61.874 | 0 | 20.078 | 5.67E+08 |
| 4 | 4 | 3 | 28.1 | 28.1 | 28.1 | 14.319 | 0 | 95.767 | 8.85E+09 |
| 4 | 4 | 3 | 33.7 | 33.7 | 21.1 | 10.871 | 0 | 26.021 | 4.59E+09 |
| 3 | 3 | 3 | 6.8 | 6.8 | 6.8 | 44.96 | 0 | 4.1176 | 1.56E+09 |
| 3 | 3 | 3 | 6.2 | 6.2 | 6.2 | 69.323 | 0 | 14.611 | 7.31E+08 |
| 3 | 3 | 3 | 9 | 9 | 9 | 45.835 | 0 | 14.248 | 3.57E+08 |
| 3 | 3 | 3 | 40.3 | 40.3 | 40.3 | 8.1463 | 0 | 4.5519 | 2.67E+09 |
| 3 | 3 | 3 | 23.5 | 23.5 | 23.5 | 12.476 | 0 | 5.5303 | 1.06E+10 |
| 3 | 3 | 3 | 10.4 | 10.4 | 10.4 | 31.73 | 0 | 7.7941 | 6.54E+08 |
| 3 | 3 | 3 | 29.2 | 29.2 | 29.2 | 14.631 | 0 | 20.732 | 1.12E+09 |
| 3 | 3 | 3 | 9.7 | 9.7 | 9.7 | 34.577 | 0 | 4.3542 | 3.63E+08 |
| 3 | 3 | 3 | 25.4 | 25.4 | 25.4 | 14.21 | 0 | 10.992 | 4.56E+08 |
| 3 | 3 | 3 | 5.2 | 5.2 | 5.2 | 85.98 | 0 | 5.355 | 5.45E+08 |
| 3 | 3 | 3 | 9.7 | 9.7 | 9.7 | 39.834 | 0 | 4.4211 | 3.83E+08 |
| 3 | 3 | 3 | 8.3 | 8.3 | 8.3 | 45.724 | 0 | 6.9171 | 4.07E+08 |

| | | | | | | | | | |
|---|---|---|---|---|---|---|---|---|---|
| 3 | 3 | 3 | 3.5 | 3.5 | 3.5 | 116.61 | 0 | 13.541 | 3.89E+08 |
| 3 | 3 | 3 | 11.3 | 11.3 | 11.3 | 33.304 | 0 | 3.8232 | 3.29E+08 |
| 3 | 3 | 3 | 46.9 | 46.9 | 46.9 | 9.0564 | 0 | 69.327 | 6.62E+09 |
| 3 | 3 | 3 | 26.6 | 26.6 | 26.6 | 17.27 | 0 | 19.55 | 6.48E+08 |
| 3 | 3 | 3 | 16.6 | 16.6 | 16.6 | 37.485 | 0 | 15.115 | 3.42E+08 |
| 3 | 3 | 3 | 7.3 | 7.3 | 7.3 | 48.915 | 0 | 7.1686 | 2.72E+08 |
| 3 | 3 | 3 | 8.3 | 8.3 | 8.3 | 37.898 | 0 | 8.9066 | 4.72E+08 |
| 3 | 3 | 3 | 11.4 | 11.4 | 11.4 | 32.749 | 0 | 12.982 | 2.25E+08 |
| 3 | 3 | 3 | 22.4 | 22.4 | 22.4 | 16.84 | 0 | 5.7664 | 2.01E+08 |
| 3 | 3 | 3 | 16.8 | 16.8 | 16.8 | 19.393 | 0 | 5.6949 | 1.18E+09 |
| 3 | 3 | 3 | 11.1 | 11.1 | 11.1 | 37.551 | 0 | 23.588 | 9.66E+08 |
| 3 | 3 | 3 | 3.2 | 3.2 | 3.2 | 88.549 | 0 | 2.9309 | 1.16E+08 |
| 4 | 3 | 3 | 22.7 | 17.4 | 17.4 | 19.825 | 0 | 11.77 | 3.87E+08 |
| 3 | 3 | 3 | 10.8 | 10.8 | 10.8 | 28.763 | 0 | 3.1436 | 1.68E+08 |
| 3 | 3 | 3 | 13.7 | 13.7 | 13.7 | 28.999 | 0 | 3.87 | 3.4E+08 |
| 3 | 3 | 3 | 47.7 | 47.7 | 47.7 | 11.802 | 0 | 24.254 | 7.9E+08 |
| 3 | 3 | 3 | 11.4 | 11.4 | 11.4 | 32.451 | 0 | 3.708 | 4.71E+08 |
| 3 | 3 | 3 | 24 | 24 | 24 | 14.726 | 0 | 15.488 | 6.55E+08 |
| 3 | 3 | 3 | 10.7 | 10.7 | 10.7 | 28.768 | 0 | 9.4771 | 4.49E+08 |
| 3 | 3 | 3 | 24 | 24 | 24 | 14.582 | 0 | 34.618 | 3.52E+08 |
| 3 | 3 | 3 | 46.4 | 46.4 | 46.4 | 6.6767 | 0 | 4.1248 | 1.3E+09 |
| 3 | 3 | 3 | 17.2 | 17.2 | 17.2 | 24.576 | 0 | 18.2 | 7.13E+08 |
| 3 | 3 | 3 | 14.7 | 14.7 | 14.7 | 25.437 | 0 | 9.8732 | 1.73E+09 |
| 3 | 3 | 3 | 26 | 26 | 26 | 15.558 | 0 | 7.0887 | 4.02E+08 |
| 13 | 3 | 3 | 37.9 | 13.7 | 13.7 | 21.865 | 0 | 10.402 | 8.08E+08 |
| 11 | 3 | 3 | 31.7 | 5.3 | 5.3 | 43.359 | 0 | 10.215 | 5.45E+08 |
| 3 | 3 | 3 | 37.5 | 37.5 | 37.5 | 7.0662 | 0 | 8.4086 | 9.7E+08 |
| 3 | 3 | 3 | 7.8 | 7.8 | 7.8 | 42.064 | 0 | 9.0886 | 1.4E+09 |
| 3 | 3 | 3 | 19.8 | 19.8 | 19.8 | 21.184 | 0 | 18.012 | 1.29E+09 |
| 3 | 3 | 3 | 16.4 | 16.4 | 16.4 | 27.312 | 0 | 10.209 | 1.24E+09 |
| 3 | 3 | 3 | 19.2 | 19.2 | 19.2 | 16.645 | 0.0006 | 2.0775 | 6.65E+08 |
| 19 | 3 | 3 | 28.5 | 5.1 | 5.1 | 83.267 | 0 | 30.198 | 1.48E+09 |
| 3 | 3 | 3 | 25.5 | 25.5 | 25.5 | 17.17 | 0 | 20.575 | 1.21E+09 |
| 3 | 3 | 3 | 20.2 | 20.2 | 20.2 | 10.245 | 0 | 8.4465 | 2.84E+09 |
| 3 | 3 | 3 | 27.1 | 27.1 | 27.1 | 22.04 | 0 | 21.724 | 5.57E+08 |
| 3 | 3 | 3 | 19.3 | 19.3 | 19.3 | 16.021 | 0 | 11.546 | 1.09E+09 |
| 3 | 3 | 3 | 17.4 | 17.4 | 17.4 | 27.557 | 0 | 12.48 | 1.81E+09 |
| 3 | 3 | 3 | 7.8 | 7.8 | 7.8 | 58.946 | 0 | 9.9048 | 2.3E+08 |
| 3 | 3 | 3 | 2.8 | 2.8 | 2.8 | 71.428 | 0 | 3.5546 | 2.23E+08 |
| 3 | 3 | 3 | 7.9 | 7.9 | 7.9 | 38.21 | 0 | 5.5607 | 6.88E+08 |
| 3 | 3 | 3 | 24.2 | 24.2 | 24.2 | 22.34 | 0 | 13.418 | 6.38E+08 |
| 3 | 3 | 3 | 29.9 | 29.9 | 29.9 | 15.33 | 0 | 8.1838 | 1.49E+09 |
| 3 | 3 | 3 | 12.2 | 12.2 | 12.2 | 29.165 | 0 | 3.8584 | 3.36E+08 |
| 3 | 3 | 3 | 40.5 | 40.5 | 40.5 | 9.461 | 0 | 11.297 | 1.97E+09 |
| 3 | 3 | 3 | 21.3 | 21.3 | 21.3 | 19.889 | 0 | 6.6232 | 9.45E+08 |
| 3 | 3 | 3 | 48.8 | 48.8 | 48.8 | 9.7251 | 0 | 27.128 | 1.32E+09 |
| 3 | 3 | 3 | 37.5 | 37.5 | 37.5 | 8.6579 | 0 | 4.7717 | 3.76E+08 |

| | | | | | | | | | |
|---|---|---|---|---|---|---|---|---|---|
| 7 | 3 | 3 | 11.8 | 5.7 | 5.7 | 70.942 | 0 | 5.8674 | 3.65E+08 |
| 3 | 3 | 3 | 3.8 | 3.8 | 3.8 | 129.93 | 0 | 12.396 | 7.12E+08 |
| 14 | 3 | 3 | 39.8 | 7.5 | 7.5 | 40.479 | 0 | 9.1118 | 5.43E+09 |
| 3 | 3 | 3 | 19.8 | 19.8 | 19.8 | 18.512 | 0 | 4.1547 | 1.18E+09 |
| 3 | 3 | 3 | 16.4 | 16.4 | 16.4 | 21.493 | 0 | 15.509 | 6.43E+08 |
| 3 | 3 | 3 | 20.6 | 20.6 | 20.6 | 18.565 | 0 | 6.5258 | 1.32E+09 |
| 3 | 3 | 3 | 16.7 | 16.7 | 16.7 | 25.75 | 0 | 5.5775 | 1.48E+09 |
| 119 | 119 | 2 | 54.2 | 54.2 | 1.4 | 280.74 | 0 | 323.31 | 1.1E+11 |
| 37 | 24 | 2 | 81 | 60.1 | 11.3 | 28.521 | 0 | 229.64 | 1.65E+10 |
| 13 | 13 | 2 | 53.4 | 53.4 | 13.4 | 41 | 0 | 141.85 | 3.59E+09 |
| 13 | 13 | 2 | 59.2 | 59.2 | 6.8 | 35.594 | 0 | 87.289 | 3.08E+09 |
| 10 | 10 | 2 | 62.8 | 62.8 | 13.8 | 16.29 | 0 | 83.008 | 2.43E+10 |
| 8 | 8 | 2 | 49.4 | 49.4 | 17.9 | 17.965 | 0 | 68.681 | 1.7E+10 |
| 6 | 6 | 2 | 17.9 | 17.9 | 10.6 | 47.655 | 0 | 83.009 | 1.96E+09 |
| 12 | 4 | 2 | 32.8 | 7.1 | 3 | 40.912 | 0.000632 | 2.607 | 1.73E+09 |
| 4 | 4 | 2 | 40.8 | 40.8 | 12.2 | 16.735 | 0 | 23.97 | 1.31E+09 |
| 3 | 3 | 2 | 8.3 | 8.3 | 6.6 | 50.957 | 0 | 8.7823 | 2.84E+08 |
| 4 | 3 | 2 | 20 | 14.5 | 11 | 22.541 | 0 | 17.888 | 9.13E+08 |
| 25 | 3 | 2 | 58.4 | 12.4 | 9.7 | 49.906 | 0 | 23.172 | 4.77E+09 |
| 7 | 3 | 2 | 8.5 | 4.3 | 3.2 | 70.819 | 0 | 50.941 | 4.77E+08 |
| 2 | 2 | 2 | 6.2 | 6.2 | 6.2 | 41.767 | 0 | 3.5052 | 7.14E+08 |
| 2 | 2 | 2 | 14.1 | 14.1 | 14.1 | 23.003 | 0 | 15.982 | 4.02E+08 |
| 2 | 2 | 2 | 20.8 | 20.8 | 20.8 | 17.442 | 0 | 5.6926 | 2.11E+08 |
| 2 | 2 | 2 | 8.2 | 8.2 | 8.2 | 21.145 | 0 | 3.7292 | 3.65E+09 |
| 2 | 2 | 2 | 12.9 | 12.9 | 12.9 | 31.778 | 0 | 4.7678 | 4.47E+08 |
| 2 | 2 | 2 | 11 | 11 | 11 | 27.578 | 0 | 19.58 | 1.16E+09 |
| 2 | 2 | 2 | 13.4 | 13.4 | 13.4 | 25.918 | 0 | 4.3872 | 5.39E+08 |
| 2 | 2 | 2 | 20.2 | 20.2 | 20.2 | 13.281 | 0 | 5.1873 | 7.6E+08 |
| 2 | 2 | 2 | 11.9 | 11.9 | 11.9 | 28.17 | 0 | 37.475 | 7.42E+08 |
| 2 | 2 | 2 | 16.2 | 16.2 | 16.2 | 13.918 | 0.000607 | 2.2017 | 7.22E+08 |
| 2 | 2 | 2 | 4.5 | 4.5 | 4.5 | 44.522 | 0 | 5.8389 | 4.85E+08 |
| 2 | 2 | 2 | 26.7 | 26.7 | 26.7 | 8.868 | 0 | 3.825 | 4.94E+08 |
| 2 | 2 | 2 | 16.9 | 16.9 | 16.9 | 16.648 | 0 | 60.59 | 8.25E+08 |
| 2 | 2 | 2 | 11.8 | 11.8 | 11.8 | 24.278 | 0 | 5.5078 | 2.79E+08 |
| 2 | 2 | 2 | 6.5 | 6.5 | 6.5 | 44.223 | 0 | 3.9173 | 1.64E+08 |
| 2 | 2 | 2 | 2.9 | 2.9 | 2.9 | 82.447 | 0 | 3.9587 | 2.91E+08 |
| 2 | 2 | 2 | 4.9 | 4.9 | 4.9 | 39.641 | 0 | 3.7591 | 2.14E+08 |
| 4 | 2 | 2 | 15.2 | 7.9 | 7.9 | 33.232 | 0 | 7.4538 | 2.26E+08 |
| 2 | 2 | 2 | 7.6 | 7.6 | 7.6 | 27.015 | 0 | 2.6785 | 2.29E+08 |
| 2 | 2 | 2 | 6.1 | 6.1 | 6.1 | 36.091 | 0 | 4.6798 | 2.11E+08 |
| 2 | 2 | 2 | 7.6 | 7.6 | 7.6 | 32.816 | 0 | 4.0274 | 2.46E+08 |
| 2 | 2 | 2 | 28.7 | 28.7 | 28.7 | 14.665 | 0 | 17.993 | 4.05E+08 |
| 4 | 2 | 2 | 8.8 | 3.5 | 3.5 | 36.743 | 0 | 12.193 | 2.67E+10 |
| 2 | 2 | 2 | 6.8 | 6.8 | 6.8 | 30.338 | 0 | 5.6631 | 4.5E+09 |
| 2 | 2 | 2 | 3.5 | 3.5 | 3.5 | 34.398 | 0 | 2.7268 | 70116000 |
| 2 | 2 | 2 | 2.3 | 2.3 | 2.3 | 107.17 | 0 | 3.3804 | 1.06E+08 |
| 2 | 2 | 2 | 11.2 | 11.2 | 11.2 | 18.038 | 0.000597 | 2.0175 | 1.02E+08 |

| | | | | | | | | | |
|---|---|---|---|---|---|---|---|---|---|
| 2 | 2 | 2 | 10.7 | 10.7 | 10.7 | 18.925 | 0 | 3.5501 | 1.23E+08 |
| 2 | 2 | 2 | 11.2 | 11.2 | 11.2 | 10.905 | 0 | 3.3628 | 7.76E+08 |
| 2 | 2 | 2 | 15.3 | 15.3 | 15.3 | 37.408 | 0 | 44.057 | 2.31E+09 |
| 2 | 2 | 2 | 21.4 | 21.4 | 21.4 | 9.5351 | 0.000604 | 2.1311 | 3.04E+08 |
| 2 | 2 | 2 | 11.9 | 11.9 | 11.9 | 18.739 | 0 | 5.1 | 2.62E+08 |
| 2 | 2 | 2 | 1.9 | 1.9 | 1.9 | 128.79 | 0.00063 | 2.5809 | 91810000 |
| 2 | 2 | 2 | 5.7 | 5.7 | 5.7 | 49.956 | 0 | 6.9766 | 6.06E+08 |
| 2 | 2 | 2 | 17.9 | 17.9 | 17.9 | 12.469 | 0 | 4.1686 | 8.66E+08 |
| 2 | 2 | 2 | 12.6 | 12.6 | 12.6 | 15.562 | 0 | 3.0127 | 3.83E+08 |
| 11 | 2 | 2 | 24.5 | 7.4 | 7.4 | 49.775 | 0 | 102.99 | 2.27E+09 |
| 4 | 2 | 2 | 17.8 | 8.9 | 8.9 | 23.673 | 0 | 3.3007 | 2.59E+09 |
| 2 | 2 | 2 | 4.2 | 4.2 | 4.2 | 51.844 | 0 | 2.766 | 1.22E+08 |
| 12 | 2 | 2 | 74.6 | 7.9 | 7.9 | 13.906 | 0 | 12.298 | 5.85E+09 |
| 2 | 2 | 2 | 13.8 | 13.8 | 13.8 | 17.318 | 0 | 14.557 | 6.13E+08 |
| 2 | 2 | 2 | 9.9 | 9.9 | 9.9 | 24.284 | 0 | 4.8287 | 63570000 |
| 2 | 2 | 2 | 2 | 2 | 2 | 86.677 | 0 | 3.2006 | 1.74E+08 |
| 3 | 2 | 2 | 4.8 | 3.2 | 3.2 | 73.243 | 0 | 12.26 | 5.51E+08 |
| 2 | 2 | 2 | 1.7 | 1.7 | 1.7 | 131.14 | 0 | 4.4501 | 1.4E+08 |
| 2 | 2 | 2 | 12.3 | 12.3 | 12.3 | 18.562 | 0 | 3.8153 | 6.45E+08 |
| 2 | 2 | 2 | 17.9 | 17.9 | 17.9 | 10.32 | 0 | 3.3231 | 6.66E+08 |
| 2 | 2 | 2 | 18.7 | 18.7 | 18.7 | 22.092 | 0 | 29.477 | 7.86E+08 |
| 2 | 2 | 2 | 29.3 | 29.3 | 29.3 | 6.5677 | 0 | 3.5661 | 4.58E+08 |
| 2 | 2 | 2 | 13.9 | 13.9 | 13.9 | 19.013 | 0 | 3.0785 | 8.07E+08 |
| 2 | 2 | 2 | 16.3 | 16.3 | 16.3 | 18.237 | 0 | 5.9189 | 6.61E+08 |

| MS/MS co | ctrl_11 | ctrl_12 | ctrl_21 | ctrl_22 | ctrl_31 | ctrl_32 | TDP_11 | TDP_12 | TDP_21 |
|---|---|---|---|---|---|---|---|---|---|
| 1045 | 30.62418 | 30.65198 | 31.00624 | 30.98271 | 30.21315 | 30.18731 | 31.36849 | 31.34191 | 30.71194 |
| 1384 | 32.36006 | 32.35673 | 32.34435 | 32.34533 | 32.03029 | 32.04284 | 32.5979 | 32.61189 | 32.47875 |
| 658 | 30.52841 | 30.55105 | 30.61982 | 30.61131 | 30.47063 | 30.49133 | 30.84976 | 30.84266 | 30.71276 |
| 583 | 30.81284 | 30.86135 | 30.95636 | 30.95913 | 30.73045 | 30.68962 | 31.35479 | 31.32437 | 31.17057 |
| 1199 | 35.48135 | 35.53991 | 35.84677 | 35.84322 | 35.50572 | 35.52875 | 36.10769 | 36.07226 | 35.62677 |
| 558 | 30.5913 | 30.53529 | 30.7302 | 30.73303 | 30.78027 | 30.79707 | 30.76272 | 30.7094 | 30.80916 |
| 295 | 29.51576 | 29.47653 | 29.26557 | 29.33334 | 29.07283 | 29.00257 | 28.74339 | 28.69952 | 28.70869 |
| 530 | 31.85155 | 31.86389 | 31.54093 | 31.55215 | 30.81222 | 30.81826 | 30.83778 | 30.7628 | 30.70619 |
| 1152 | 33.8824 | 33.87729 | 33.95272 | 33.99791 | 34.26297 | 34.23192 | 34.26123 | 34.17459 | 34.3214 |
| 712 | 33.13574 | 33.10806 | 32.69422 | 32.71217 | 32.22216 | 32.19034 | 32.3028 | 32.21489 | 32.07428 |
| 861 | 34.28216 | 34.33078 | 34.26591 | 34.22245 | 34.65635 | 34.61297 | 34.53948 | 34.49094 | 34.71626 |
| 496 | 32.17102 | 32.2011 | 31.59143 | 31.65804 | 31.35552 | 31.30623 | 31.26847 | 31.27974 | 31.3772 |
| 417 | 29.3442 | 29.43245 | 29.85333 | 29.90684 | 29.33511 | 29.23219 | 29.6923 | 29.73252 | 28.99224 |
| 653 | 34.53774 | 34.55984 | 34.13858 | 34.15225 | 34.13553 | 34.11094 | 33.80318 | 33.73156 | 33.64141 |
| 465 | 31.23169 | 31.24088 | 31.39305 | 31.39223 | 31.68583 | 31.69909 | 31.32346 | 31.36474 | 31.54448 |
| 671 | 32.33434 | 32.388 | 32.52218 | 32.48884 | 32.83157 | 32.8018 | 32.83347 | 32.84313 | 33.04546 |
| 768 | 36.44071 | 36.45727 | 36.02081 | 36.03954 | 35.40442 | 35.36202 | 35.5332 | 35.43704 | 35.25025 |
| 458 | 30.82221 | 30.80655 | 31.05543 | 31.12207 | 31.5162 | 31.53399 | 31.16171 | 31.09106 | 31.29844 |
| 204 | 27.25088 | 26.93191 | 27.55296 | 27.80999 | 27.11545 | 27.32285 | 28.35233 | 28.4734 | 28.21512 |
| 508 | 32.60461 | 32.62558 | 32.16279 | 32.19363 | 31.75031 | 31.69942 | 31.72513 | 31.75318 | 31.6121 |
| 531 | 31.6832 | 31.65386 | 31.77282 | 31.76497 | 32.10046 | 32.13739 | 31.90383 | 31.84655 | 32.17173 |
| 838 | 34.65363 | 34.65603 | 34.64662 | 34.66273 | 35.42733 | 35.43384 | 34.94681 | 34.82439 | 35.14725 |
| 807 | 32.56488 | 32.63394 | 32.63649 | 32.66875 | 32.69617 | 32.74258 | 32.78992 | 32.77845 | 32.8275 |
| 302 | 29.85516 | 29.9386 | 29.37671 | 29.38624 | 29.21449 | 29.28787 | 28.85772 | 28.8072 | 28.6381 |
| 298 | 29.27456 | 29.14445 | 29.60631 | 29.66212 | 30.1305 | 30.239 | 29.79366 | 29.86638 | 30.03695 |
| 221 | 28.74554 | 28.75907 | 28.9958 | 29.10614 | 28.66736 | 28.65373 | 28.97733 | 29.10097 | 28.76205 |
| 530 | 32.69154 | 32.64491 | 32.56613 | 32.53352 | 33.32616 | 33.29666 | 33.29351 | 33.25124 | 33.18006 |
| 589 | 32.3289 | 32.36664 | 32.55145 | 32.52822 | 32.4813 | 32.50462 | 32.821 | 32.89309 | 32.88544 |
| 356 | 29.87487 | 29.95602 | 30.03774 | 30.07597 | 30.36171 | 30.30509 | 30.31721 | 30.3911 | 30.39182 |
| 527 | 31.28576 | 31.34757 | 31.60673 | 31.68106 | 32.30997 | 32.32386 | 31.74887 | 31.70194 | 32.16276 |
| 487 | 32.47832 | 32.42179 | 32.70644 | 32.71555 | 33.31541 | 33.32053 | 33.0046 | 32.93245 | 33.2436 |
| 462 | 30.99732 | 31.00778 | 31.05956 | 31.08214 | 31.48855 | 31.49562 | 31.34059 | 31.33111 | 31.58082 |
| 262 | 29.19737 | 29.20706 | 29.60083 | 29.61509 | 29.66071 | 29.6083 | 29.23576 | 29.41446 | 29.45801 |
| 462 | 31.35184 | 31.31057 | 31.47359 | 31.49719 | 31.56955 | 31.58249 | 31.65019 | 31.63975 | 31.64985 |
| 330 | 30.46995 | 30.47431 | 30.8776 | 30.86563 | 31.20571 | 31.20833 | 31.13333 | 31.12399 | 31.31127 |
| 340 | 31.3509 | 31.3481 | 31.10106 | 31.12583 | 30.70981 | 30.70265 | 30.54821 | 30.63686 | 30.46529 |
| 698 | 36.75309 | 36.77865 | 36.44991 | 36.41312 | 35.9052 | 35.8806 | 36.12654 | 35.97412 | 35.65034 |
| 450 | 31.63508 | 31.57687 | 31.79912 | 31.80043 | 32.36273 | 32.33703 | 31.74166 | 31.83074 | 32.22519 |
| 514 | 32.81084 | 32.81406 | 33.05198 | 33.00517 | 33.54075 | 33.48484 | 33.13871 | 33.01736 | 33.25631 |
| 349 | 29.57878 | 29.65942 | 29.85396 | 29.82142 | 29.99672 | 29.96058 | 30.14334 | 30.15497 | 29.99105 |
| 291 | 29.84949 | 29.82202 | 29.95768 | 29.99779 | 30.46422 | 30.40365 | 30.32678 | 30.34022 | 30.56083 |
| 374 | 29.86913 | 29.82068 | 30.02591 | 30.01172 | 30.46256 | 30.45061 | 30.24694 | 30.31894 | 30.4941 |
| 273 | 29.19385 | 29.25318 | 29.2938 | 29.20271 | 29.27217 | 29.30986 | 29.38593 | 29.49108 | 29.41541 |
| 650 | 36.18473 | 36.17374 | 35.84813 | 35.78609 | 35.45403 | 35.50412 | 35.45179 | 35.33225 | 35.16125 |
| 554 | 32.74423 | 32.76687 | 32.91337 | 32.95719 | 33.35993 | 33.44283 | 33.15451 | 33.07411 | 33.32362 |
| 746 | 33.38844 | 33.39752 | 33.45489 | 33.43763 | 33.80655 | 33.78048 | 33.46879 | 33.44505 | 33.63655 |

| | | | | | | | | | |
|---|---|---|---|---|---|---|---|---|---|
| 758 | 34.14003 | 34.20281 | 34.27218 | 34.32335 | 34.73666 | 34.83667 | 34.36502 | 34.30178 | 34.63774 |
| 282 | 29.27912 | 29.26986 | 29.46813 | 29.45288 | 29.85675 | 29.9211 | 29.61551 | 29.65077 | 30.04843 |
| 444 | 31.72314 | 31.68846 | 31.84939 | 31.84666 | 32.46964 | 32.49403 | 32.16489 | 32.12734 | 32.3965 |
| 401 | 31.29022 | 31.26355 | 31.6937 | 31.7162 | 31.93343 | 31.98363 | 31.78386 | 31.83327 | 31.86242 |
| 261 | 30.1294 | 30.14273 | 30.12041 | 30.07202 | 29.47607 | 29.52201 | 29.40149 | 29.50688 | 29.09303 |
| 712 | 36.356 | 36.34124 | 36.05446 | 36.04726 | 35.45538 | 35.47858 | 35.78775 | 35.70019 | 35.386 |
| 321 | 31.04205 | 31.06233 | 31.12719 | 31.16806 | 31.69062 | 31.65472 | 31.35484 | 31.38705 | 31.68349 |
| 234 | 29.37478 | 29.43155 | 29.45391 | 29.47927 | 29.82941 | 29.71226 | 29.65053 | 29.60042 | 29.76716 |
| 177 | 28.32023 | 28.31242 | 28.48646 | 28.39358 | 28.67869 | 28.71073 | 28.37567 | 28.3848 | 28.71001 |
| 443 | 31.75843 | 31.7864 | 31.88296 | 31.88453 | 32.402 | 32.44678 | 32.03364 | 32.06715 | 32.4171 |
| 135 | 26.70556 | 26.83966 | 27.22711 | 27.00443 | 26.88716 | 27.09033 | 27.99097 | 27.73725 | 27.34858 |
| 496 | 32.13044 | 32.1416 | 32.3438 | 32.28772 | 32.83238 | 32.81048 | 32.76651 | 32.74841 | 32.97717 |
| 400 | 32.09695 | 32.08835 | 32.29398 | 32.38458 | 32.92315 | 32.92399 | 32.75921 | 32.65384 | 32.93998 |
| 426 | 31.8213 | 31.88074 | 32.01352 | 31.999 | 32.46531 | 32.49009 | 32.26788 | 32.25434 | 32.47104 |
| 284 | 29.67219 | 29.63157 | 29.45707 | 29.5538 | 29.60251 | 29.52197 | 29.86695 | 29.79397 | 29.79335 |
| 237 | 29.05064 | 29.04309 | 29.32605 | 29.28346 | 29.39648 | 29.35176 | 28.99326 | 29.0327 | 29.16317 |
| 158 | 27.73867 | 27.51337 | 27.51751 | 27.525 | 27.67526 | 27.86 | 28.22835 | 28.38846 | 28.1045 |
| 415 | 32.83274 | 32.84577 | 32.94676 | 32.99113 | 33.14727 | 33.18954 | 33.3123 | 33.31406 | 33.27337 |
| 231 | 29.53154 | 29.62854 | 29.56122 | 29.55269 | 29.4284 | 29.29833 | 30.10187 | 30.17427 | 29.96224 |
| 223 | 29.54902 | 29.50196 | 29.78813 | 29.79068 | 30.29066 | 30.30193 | 30.19589 | 30.22929 | 30.26573 |
| 336 | 31.16967 | 31.1398 | 31.28068 | 31.17682 | 31.40492 | 31.36672 | 31.49876 | 31.42347 | 31.41559 |
| 314 | 32.47158 | 32.49622 | 32.88954 | 32.78113 | 32.93868 | 32.91182 | 32.36612 | 32.30875 | 32.51989 |
| 510 | 35.02548 | 35.04852 | 34.73439 | 34.73605 | 34.37393 | 34.40051 | 34.23128 | 34.22927 | 34.01828 |
| 536 | 33.17059 | 33.12885 | 33.39125 | 33.36476 | 33.78936 | 33.75861 | 33.79159 | 33.69021 | 33.89374 |
| 133 | 26.6554 | 26.9718 | 26.0284 | 26.24766 | 26.28732 | 25.73322 | 27.83503 | 27.9954 | 28.03339 |
| 232 | 29.32051 | 29.29811 | 29.42798 | 29.40916 | 29.75828 | 29.74264 | 29.89006 | 29.92011 | 29.93537 |
| 393 | 31.55742 | 31.56447 | 31.60351 | 31.68433 | 31.93709 | 31.90315 | 31.80555 | 31.84176 | 32.0445 |
| 138 | 28.02675 | 27.84918 | 28.05077 | 28.04667 | 28.12124 | 28.13668 | 28.23726 | 28.23872 | 28.36294 |
| 307 | 30.69221 | 30.69793 | 30.55252 | 30.72143 | 30.5018 | 30.52645 | 30.75661 | 30.75486 | 30.76121 |
| 250 | 30.7419 | 30.72192 | 30.82805 | 30.86777 | 30.0531 | 30.04817 | 30.44016 | 30.37161 | 29.94084 |
| 342 | 31.77588 | 31.81352 | 32.04668 | 32.00768 | 32.37875 | 32.41125 | 32.2049 | 32.18332 | 32.46259 |
| 142 | 27.35951 | 27.38195 | 27.40243 | 27.62638 | 27.25988 | 27.39444 | 27.67364 | 27.77291 | 27.58649 |
| 280 | 31.56315 | 31.58831 | 31.22803 | 31.21917 | 31.30144 | 31.29346 | 30.97466 | 31.00711 | 31.32501 |
| 218 | 29.65012 | 29.71216 | 29.49415 | 29.41848 | 29.20235 | 29.18438 | 29.16389 | 29.17222 | 28.94464 |
| 167 | 28.01871 | 28.16159 | 26.69112 | 27.97492 | 28.05507 | 28.0517 | 28.8688 | 28.78556 | 28.52103 |
| 420 | 34.42211 | 34.47236 | 34.31305 | 34.40825 | 34.06423 | 34.05407 | 33.79604 | 33.68645 | 33.75633 |
| 285 | 32.99125 | 33.00317 | 32.86688 | 32.89874 | 32.53779 | 32.65465 | 32.21257 | 32.24949 | 32.35883 |
| 189 | 29.06971 | 29.02327 | 29.13968 | 29.00804 | 29.4874 | 29.47489 | 29.35681 | 29.26521 | 29.60353 |
| 189 | 28.6553 | 28.55779 | 28.39321 | 28.56347 | 28.45344 | 28.41828 | 29.0544 | 28.92982 | 28.66899 |
| 245 | 31.15732 | 31.1821 | 31.28885 | 31.19137 | 31.42056 | 31.39382 | 31.38469 | 31.35741 | 31.49653 |
| 545 | 35.4878 | 35.57091 | 35.15644 | 35.21241 | 34.98206 | 34.96718 | 34.44579 | 34.36646 | 34.44017 |
| 190 | 29.66606 | 29.66846 | 29.7816 | 29.8341 | 30.33074 | 30.29011 | 30.02894 | 30.09648 | 30.27186 |
| 417 | 32.23484 | 32.30146 | 32.73798 | 32.78258 | 32.82343 | 32.8621 | 32.69023 | 32.77103 | 32.98492 |
| 313 | 31.60894 | 31.68286 | 31.866 | 31.89598 | 32.37202 | 32.30408 | 31.93986 | 31.96365 | 32.22185 |
| 241 | 28.47239 | 28.48788 | 28.60915 | 28.53025 | 28.93994 | 29.09514 | 28.91587 | 28.97115 | 29.15998 |
| 174 | 28.99623 | 28.99553 | 28.76319 | 28.87844 | 28.61481 | 28.52903 | 28.36632 | 28.17412 | 28.37621 |
| 348 | 30.88023 | 30.96161 | 31.05491 | 31.01052 | 31.29225 | 31.26729 | 31.09503 | 31.19066 | 31.53422 |

| | | | | | | | | | |
|---|---|---|---|---|---|---|---|---|---|
| 192 | 29.90383 | 29.90899 | 29.40659 | 29.43605 | 29.13609 | 29.29815 | 29.09655 | 29.18092 | 29.00804 |
| 415 | 32.79744 | 32.78021 | 32.34398 | 32.36679 | 32.00915 | 32.06381 | 32.01907 | 31.98135 | 31.77733 |
| 171 | 29.16418 | 29.21743 | 28.95241 | 28.98609 | 28.79402 | 28.78999 | 28.77838 | 28.65434 | 28.90624 |
| 307 | 33.52659 | 33.49667 | 33.49643 | 33.58385 | 33.93025 | 33.91883 | 33.66501 | 33.68269 | 33.87353 |
| 161 | 28.32122 | 28.32817 | 28.51931 | 28.53249 | 28.80637 | 29.01196 | 28.58445 | 28.65557 | 28.81958 |
| 430 | 31.92486 | 32.0041 | 31.87368 | 31.91153 | 31.658 | 31.48443 | 31.77659 | 31.69067 | 31.74943 |
| 160 | 27.13761 | 26.85293 | 27.3506 | 27.52589 | 27.88326 | 27.93517 | 27.69099 | 27.81714 | 28.1154 |
| 206 | 29.61209 | 29.65629 | 29.87524 | 29.96361 | 30.47325 | 30.52272 | 30.12509 | 30.13198 | 30.45484 |
| 248 | 30.29472 | 30.37036 | 30.69884 | 30.67287 | 31.31711 | 31.29702 | 30.94817 | 30.82707 | 31.08163 |
| 395 | 31.8993 | 31.95823 | 31.95168 | 31.96182 | 32.25299 | 32.23358 | 31.99274 | 32.02013 | 32.31996 |
| 246 | 28.97705 | 29.03134 | 29.02952 | 29.12485 | 29.68953 | 29.6582 | 29.12888 | 29.25699 | 29.52395 |
| 301 | 30.91335 | 30.97883 | 31.091 | 31.06104 | 31.54899 | 31.5419 | 31.18347 | 31.17575 | 31.57628 |
| 148 | 27.69279 | 27.53475 | 27.53208 | 27.54885 | 27.73995 | 27.7241 | 28.09803 | 27.96378 | 28.08421 |
| 172 | 28.37629 | 28.26069 | 28.33181 | 28.33121 | 28.23252 | 28.244 | 28.81247 | 28.76544 | 28.74701 |
| 193 | 28.71637 | 28.84356 | 28.62975 | 28.65065 | 28.56736 | 28.65438 | 28.91917 | 28.82208 | 28.83062 |
| 152 | 29.4412 | 29.51092 | 29.06964 | 28.96315 | 28.6246 | 28.69651 | 28.69707 | 28.69092 | 28.5769 |
| 237 | 30.4338 | 30.53325 | 30.40578 | 30.31926 | 29.90167 | 29.91969 | 29.88481 | 29.92748 | 29.66998 |
| 325 | 33.21342 | 33.17371 | 32.85479 | 32.82037 | 32.20516 | 32.19748 | 32.38327 | 32.34921 | 32.17789 |
| 225 | 31.16477 | 31.13675 | 30.79258 | 30.75207 | 30.43361 | 30.42462 | 30.35468 | 30.36724 | 30.16698 |
| 234 | 30.17021 | 30.2179 | 29.68737 | 29.6562 | 29.72881 | 29.52917 | 29.2822 | 29.12139 | 29.22368 |
| 222 | 30.36963 | 30.31181 | 30.49028 | 30.49924 | 31.03341 | 30.94007 | 30.7189 | 30.72972 | 30.87144 |
| 200 | 29.09255 | 29.19423 | 29.45631 | 29.44555 | 30.02181 | 30.04375 | 29.66707 | 29.55777 | 30.03996 |
| 217 | 29.64145 | 29.70571 | 29.41639 | 29.46164 | 29.07707 | 28.9659 | 29.10747 | 29.25582 | 29.183 |
| 225 | 29.86965 | 29.89458 | 29.72394 | 29.71924 | 29.32905 | 29.37648 | 29.70317 | 29.61679 | 29.51171 |
| 256 | 30.90562 | 30.98128 | 30.40568 | 30.58683 | 30.21581 | 30.28361 | 30.24694 | 30.22331 | 29.98563 |
| 289 | 31.30443 | 31.23655 | 31.36014 | 31.33042 | 31.84052 | 31.84172 | 31.59619 | 31.58024 | 31.77353 |
| 141 | 27.80993 | 27.79934 | 28.02311 | 28.20923 | 27.80802 | 28.09723 | 28.41595 | 28.55252 | 28.24227 |
| 299 | 32.14087 | 32.23015 | 32.37661 | 32.43062 | 33.01523 | 33.04226 | 32.76699 | 32.80291 | 32.99147 |
| 186 | 29.31875 | 29.24463 | 29.03983 | 29.03021 | 28.46311 | 28.56173 | 28.66587 | 28.71869 | 28.51262 |
| 353 | 32.57682 | 32.68096 | 32.66972 | 32.67464 | 33.08926 | 33.08294 | 32.76428 | 32.92078 | 33.0874 |
| 213 | 30.28593 | 30.17379 | 30.3091 | 30.16734 | 30.81322 | 30.77061 | 30.57452 | 30.52608 | 30.72704 |
| 145 | 28.0878 | 28.16658 | 28.2311 | 28.23155 | 27.90254 | 27.98517 | 28.59584 | 28.52869 | 28.22504 |
| 127 | 27.49167 | 27.56442 | 27.58427 | 27.4698 | 27.57816 | 27.55808 | 27.37964 | 27.27941 | 27.52604 |
| 163 | 29.15812 | 28.97681 | 28.86597 | 28.96348 | 29.61921 | 29.63506 | 28.89493 | 29.06083 | 29.29165 |
| 261 | 31.8436 | 31.87353 | 31.89537 | 31.9111 | 32.05368 | 31.94653 | 32.48397 | 32.49224 | 32.41233 |
| 163 | 29.36584 | 29.37038 | 29.49352 | 29.31739 | 29.50419 | 29.37005 | 29.77419 | 29.66719 | 29.80186 |
| 144 | 28.34342 | 28.33134 | 28.37136 | 28.33403 | 27.94821 | 28.08218 | 28.14521 | 28.10024 | 28.05956 |
| 300 | 30.76525 | 30.80962 | 31.1827 | 31.13314 | 31.76829 | 31.72245 | 31.31008 | 31.35736 | 31.71657 |
| 261 | 29.87745 | 29.82334 | 30.05012 | 30.08993 | 30.3078 | 30.28505 | 30.04427 | 30.06085 | 30.22251 |
| 189 | 29.14 | 29.06096 | 29.23409 | 29.19199 | 29.57681 | 29.60645 | 29.24204 | 29.21498 | 29.45319 |
| 446 | 33.55944 | 33.62854 | 33.36189 | 33.40958 | 32.96741 | 32.89883 | 32.85265 | 32.86216 | 32.68721 |
| 357 | 33.55476 | 33.56172 | 33.66405 | 33.61456 | 33.99353 | 33.89166 | 33.76089 | 33.67219 | 33.78702 |
| 216 | 30.44924 | 30.44737 | 30.45738 | 30.33415 | 29.75568 | 29.71414 | 29.64079 | 29.85013 | 29.30526 |
| 153 | 29.32884 | 29.35624 | 29.42744 | 29.36726 | 29.98142 | 30.04062 | 29.69533 | 30.01185 | 30.00102 |
| 183 | 28.93036 | 28.79489 | 29.17588 | 29.17831 | 29.47274 | 29.48639 | 29.35502 | 29.37594 | 29.72506 |
| 133 | 27.88751 | 28.70071 | 28.81036 | 28.74432 | 29.09695 | 29.13707 | 28.95327 | 29.02633 | 29.23491 |
| 197 | 29.56304 | 29.5787 | 29.86336 | 29.86582 | 30.37285 | 30.2824 | 29.98834 | 29.98142 | 30.28912 |

| | | | | | | | | | |
|---|---|---|---|---|---|---|---|---|---|
| 257 | 30.75366 | 30.88176 | 30.79622 | 30.87159 | 31.21338 | 31.27731 | 30.93431 | 30.93789 | 31.15569 |
| 158 | 28.41454 | 28.51743 | 28.53152 | 28.60361 | 28.71637 | 28.81265 | 28.50112 | 28.3525 | 28.64368 |
| 180 | 29.23256 | 29.22727 | 29.39116 | 29.41385 | 29.1958 | 29.25582 | 29.51301 | 29.35706 | 29.17684 |
| 289 | 31.3084 | 31.39387 | 31.46777 | 31.47344 | 30.87986 | 30.89879 | 31.47832 | 31.39678 | 31.05672 |
| 282 | 32.40388 | 32.35702 | 32.41201 | 32.51524 | 32.95029 | 32.87637 | 32.6577 | 32.53009 | 32.8809 |
| 144 | 29.31823 | 29.32292 | 29.66545 | 29.55834 | 28.55841 | 28.63197 | 28.48347 | 28.46525 | 27.87994 |
| 147 | 28.13438 | 28.09512 | 28.2658 | 28.25395 | 28.58663 | 28.67771 | 28.55003 | 28.5553 | 28.95834 |
| 318 | 33.54548 | 33.64463 | 33.85972 | 33.82544 | 34.14915 | 34.1852 | 34.17251 | 34.07511 | 34.19572 |
| 218 | 29.42388 | 29.38644 | 29.21635 | 29.25341 | 29.0876 | 29.15376 | 29.35832 | 29.39601 | 29.57856 |
| 238 | 30.94084 | 30.98997 | 30.99604 | 31.03072 | 31.14534 | 31.13895 | 31.1262 | 31.17349 | 31.29762 |
| 242 | 31.23581 | 31.23181 | 30.67405 | 30.78324 | 30.70685 | 30.72078 | 30.32421 | 30.3349 | 30.40538 |
| 184 | 30.7052 | 30.80108 | 30.50142 | 30.53055 | 30.24672 | 30.22986 | 30.0414 | 29.85176 | 29.95185 |
| 157 | 29.79719 | 29.72412 | 29.74517 | 29.69564 | 29.36765 | 29.46219 | 29.07602 | 29.01752 | 28.97191 |
| 190 | 30.49895 | 30.59486 | 30.58665 | 30.54471 | 30.04492 | 29.95394 | 30.07265 | 29.98441 | 29.57227 |
| 257 | 31.87807 | 31.93821 | 32.03279 | 31.96685 | 32.35702 | 32.37508 | 32.16144 | 32.10015 | 32.47029 |
| 126 | 27.57506 | 27.43913 | 27.94419 | 27.8098 | 28.21313 | 28.26024 | 27.96063 | 28.04526 | 28.08978 |
| 130 | 28.15793 | 27.6441 | 28.15074 | 28.05564 | 28.31847 | 28.30504 | 28.02739 | 28.13207 | 28.39276 |
| 127 | 28.22913 | 28.07842 | 28.57189 | 28.43817 | 28.40953 | 28.55395 | 28.47595 | 28.33501 | 28.37003 |
| 174 | 28.92861 | 28.88107 | 29.1265 | 28.9637 | 29.30029 | 29.22147 | 28.81559 | 28.99383 | 29.13352 |
| 295 | 32.56744 | 32.58556 | 32.46266 | 32.50727 | 32.69041 | 32.61386 | 33.11689 | 33.02682 | 33.07398 |
| 356 | 32.91862 | 32.96888 | 33.27462 | 33.33271 | 33.72039 | 33.74013 | 33.37334 | 33.36202 | 33.51206 |
| 90 | 25.71214 | 26.18833 | 26.60541 | 26.33778 | 25.93798 | 25.40139 | 27.37069 | 27.46871 | 26.34033 |
| 137 | 27.82116 | 27.74695 | 27.79922 | 27.76932 | 27.73841 | 27.91923 | 28.09235 | 28.39734 | 28.35804 |
| 89 | 26.95338 | 26.97334 | 26.78506 | 27.0872 | 26.24131 | 26.57527 | 27.38113 | 27.40957 | 27.10979 |
| 165 | 28.87891 | 28.8925 | 29.14907 | 29.17291 | 29.39632 | 29.43199 | 29.64787 | 29.63423 | 29.54812 |
| 158 | 29.04284 | 29.09378 | 29.13016 | 29.11647 | 29.56235 | 29.62364 | 29.53768 | 29.61688 | 29.779 |
| 97 | 27.61246 | 27.77209 | 27.89741 | 27.79792 | 27.62129 | 27.65867 | 28.02702 | 28.32971 | 28.08431 |
| 222 | 30.1183 | 30.13001 | 30.42272 | 30.40335 | 30.89858 | 30.96857 | 30.718 | 30.79273 | 30.96106 |
| 151 | 29.7298 | 29.85887 | 29.95864 | 29.9746 | 30.01717 | 30.06497 | 30.17284 | 30.23866 | 30.46061 |
| 177 | 30.56574 | 30.50446 | 30.45846 | 30.42642 | 30.89836 | 30.88765 | 30.8886 | 30.68253 | 30.92196 |
| 216 | 30.8696 | 30.93881 | 30.84842 | 30.81421 | 31.31262 | 31.23483 | 31.15726 | 31.14041 | 31.31349 |
| 267 | 31.41241 | 31.42112 | 31.37746 | 31.44298 | 31.56561 | 31.64138 | 31.65215 | 31.68641 | 31.80786 |
| 246 | 31.82111 | 31.90175 | 31.85913 | 31.85162 | 31.77368 | 31.82381 | 31.19835 | 31.2963 | 31.38479 |
| 187 | 30.0775 | 30.09258 | 29.70543 | 29.66296 | 29.11949 | 29.10649 | 29.11386 | 29.24857 | 29.09683 |
| 212 | 31.63318 | 31.68093 | 31.33495 | 31.19789 | 31.03682 | 31.05898 | 30.71939 | 30.67194 | 30.66569 |
| 215 | 31.39392 | 31.3891 | 31.10437 | 31.16932 | 30.72777 | 30.76477 | 30.53315 | 30.55846 | 30.10462 |
| 256 | 31.84416 | 31.81803 | 31.82896 | 31.84378 | 31.06349 | 31.07903 | 31.18937 | 31.20688 | 30.60435 |
| 222 | 32.12614 | 32.17382 | 32.1884 | 32.26363 | 32.87806 | 32.75402 | 32.69171 | 32.55128 | 32.74801 |
| 137 | 28.78653 | 28.64575 | 28.86797 | 28.9619 | 29.55976 | 29.42872 | 28.8441 | 28.78559 | 29.2843 |
| 132 | 29.56051 | 29.58425 | 29.70664 | 29.70869 | 28.94431 | 29.09479 | 28.61028 | 28.82742 | 28.15267 |
| 107 | 28.28621 | 28.44308 | 28.79606 | 28.75777 | 27.58205 | 27.54008 | 27.56027 | 27.39795 | 26.81665 |
| 112 | 27.19122 | 27.41877 | 27.50211 | 27.70491 | 27.59974 | 27.63686 | 28.26821 | 28.41897 | 28.03701 |
| 268 | 30.60858 | 30.63132 | 30.33287 | 30.42442 | 30.26103 | 30.26383 | 31.36265 | 31.31985 | 30.85036 |
| 106 | 27.41973 | 27.55296 | 27.69704 | 27.72475 | 28.14982 | 28.22499 | 28.35548 | 28.27031 | 28.27635 |
| 163 | 28.24568 | 28.20518 | 28.42986 | 28.34777 | 28.81366 | 28.71447 | 28.62261 | 28.71529 | 28.90202 |
| 160 | 29.8393 | 29.97186 | 29.63724 | 29.78299 | 29.49047 | 29.54771 | 30.1866 | 30.12312 | 30.023 |
| 195 | 29.72081 | 29.68762 | 29.67729 | 29.66942 | 30.12152 | 30.02419 | 29.94517 | 29.9632 | 30.22193 |

| | | | | | | | | | |
|---|---|---|---|---|---|---|---|---|---|
| 147 | 28.19009 | 28.28167 | 28.12134 | 28.17346 | 28.07189 | 28.10029 | 28.40076 | 28.549 | 28.57402 |
| 212 | 30.5921 | 30.7298 | 30.81444 | 30.78644 | 31.13761 | 31.01019 | 31.00879 | 30.97138 | 31.20145 |
| 340 | 33.60034 | 33.62527 | 33.53635 | 33.43515 | 33.83536 | 33.86258 | 33.80828 | 33.73227 | 33.96809 |
| 214 | 30.0927 | 30.01159 | 30.00584 | 30.01983 | 29.77283 | 29.74823 | 30.15955 | 30.21535 | 30.15497 |
| 142 | 28.94453 | 28.83735 | 29.06746 | 29.11919 | 29.11143 | 29.06075 | 28.76449 | 28.70984 | 28.90262 |
| 118 | 28.53545 | 28.47193 | 28.14647 | 28.16485 | 27.74618 | 27.78743 | 27.55691 | 27.55918 | 27.58971 |
| 89 | 27.89406 | 28.12992 | 27.45928 | 27.43492 | 26.8853 | 26.22385 | 26.83617 | 27.05497 | 26.72364 |
| 162 | 30.27985 | 30.23957 | 29.86588 | 29.90124 | 29.38051 | 29.42762 | 29.10425 | 29.12305 | 28.87923 |
| 110 | 28.33794 | 28.26812 | 28.3709 | 28.49717 | 28.96826 | 29.08284 | 28.65041 | 28.67862 | 28.94896 |
| 127 | 28.36469 | 28.45509 | 28.15726 | 28.27213 | 27.89244 | 28.0094 | 28.3538 | 28.30051 | 28.36849 |
| 192 | 29.82781 | 29.79239 | 29.90455 | 29.92917 | 30.10562 | 30.16207 | 29.70278 | 29.81126 | 30.03957 |
| 321 | 30.04062 | 30.01903 | 30.20478 | 30.21106 | 30.43549 | 30.46568 | 30.1267 | 30.29088 | 30.48779 |
| 91 | 26.71004 | 26.85281 | 27.25458 | 26.83087 | 27.92536 | 28.07245 | 27.44403 | 27.64334 | 27.81915 |
| 134 | 28.50733 | 28.58899 | 28.91852 | 28.82445 | 29.18966 | 29.14097 | 29.27613 | 29.33215 | 29.37213 |
| 204 | 28.60862 | 28.68405 | 28.66231 | 28.77866 | 28.983 | 28.96634 | 29.09336 | 29.30622 | 29.16509 |
| 135 | 27.99281 | 28.03071 | 28.20494 | 28.08816 | 28.63703 | 28.60774 | 28.48577 | 28.76158 | 28.47839 |
| 100 | 27.3937 | 27.42614 | 27.39583 | 27.54525 | 27.72012 | 27.80925 | 27.755 | 27.77643 | 27.76395 |
| 133 | 29.13014 | 29.16983 | 29.10037 | 29.20172 | 28.93745 | 28.94673 | 29.54435 | 29.57125 | 29.25979 |
| 126 | 27.82195 | 27.87044 | 27.6586 | 27.7661 | 27.77925 | 27.71024 | 28.06137 | 28.09984 | 28.08477 |
| 183 | 29.9859 | 29.9474 | 29.97049 | 29.96141 | 30.38618 | 30.3833 | 30.21847 | 30.32088 | 30.47817 |
| 122 | 27.7368 | 27.86284 | 27.64086 | 27.61786 | 27.44111 | 27.45034 | 27.95333 | 27.6967 | 27.93309 |
| 135 | 28.0258 | 27.98202 | 28.00983 | 28.02401 | 27.93534 | 28.01585 | 28.43575 | 28.19573 | 28.11496 |
| 201 | 29.27246 | 29.34505 | 29.71635 | 29.5377 | 29.41885 | 29.53304 | 29.36653 | 29.38498 | 29.17536 |
| 181 | 32.09283 | 32.08205 | 31.63439 | 31.64822 | 31.27819 | 31.22527 | 31.10924 | 30.97699 | 30.91947 |
| 245 | 33.21566 | 33.23891 | 32.88918 | 32.83765 | 32.57285 | 32.57905 | 32.33325 | 32.18476 | 32.16345 |
| 219 | 32.51561 | 32.49971 | 32.68441 | 32.57335 | 32.10942 | 32.12362 | 31.81917 | 31.93048 | 31.26198 |
| 144 | 28.39276 | 28.51848 | 28.54782 | 28.65263 | 29.27164 | 29.33824 | 28.93478 | 28.85463 | 29.22085 |
| 161 | 29.86133 | 29.94977 | 29.60707 | 29.67374 | 29.02364 | 28.96898 | 29.21886 | 29.23032 | 29.26495 |
| 110 | 27.96328 | 27.79544 | 28.08299 | 27.94849 | 28.51104 | 28.70461 | 28.1517 | 28.04926 | 28.55032 |
| 123 | 28.47398 | 28.63267 | 28.60996 | 28.61705 | 28.99275 | 29.08864 | 28.62289 | 28.68147 | 29.14584 |
| 154 | 29.34556 | 29.38241 | 29.1496 | 29.09974 | 27.95494 | 28.45172 | 28.73531 | 28.70899 | 28.81467 |
| 119 | 27.24961 | 27.21169 | 27.59178 | 27.50446 | 27.78469 | 27.75232 | 27.60633 | 27.72156 | 27.63464 |
| 149 | 29.2947 | 29.3048 | 29.59571 | 29.57512 | 29.61992 | 29.64501 | 29.29285 | 29.36834 | 29.58258 |
| 140 | 30.07572 | 30.05892 | 30.20874 | 30.12805 | 30.40192 | 30.48846 | 30.41357 | 30.30683 | 30.44717 |
| 69 | 25.61246 | 26.12123 | 25.75532 | 26.59534 | 26.89977 | 26.48244 | 26.29111 | 26.76509 | 27.03265 |
| 180 | 28.76737 | 28.7769 | 28.924 | 28.89108 | 28.77881 | 28.90466 | 28.90713 | 29.0008 | 29.10167 |
| 191 | 28.96345 | 29.14523 | 29.25802 | 29.33125 | 29.61584 | 29.58267 | 29.42352 | 29.39667 | 29.61372 |
| 83 | 26.17689 | 26.44152 | 27.08345 | 27.04895 | 27.015 | 26.59875 | 27.40341 | 27.29903 | 27.16337 |
| 102 | 28.46855 | 28.49286 | 28.36215 | 28.42093 | 28.61028 | 28.69488 | 28.91602 | 28.93362 | 28.91202 |
| 115 | 27.80372 | 27.72143 | 27.54782 | 27.55545 | 27.50567 | 27.51458 | 27.87754 | 27.98934 | 28.12475 |
| 181 | 30.09144 | 30.09798 | 29.94126 | 29.98414 | 30.4181 | 30.56138 | 30.32335 | 30.30062 | 30.65906 |
| 130 | 27.68752 | 27.68244 | 28.04427 | 28.1716 | 28.30169 | 28.37443 | 28.12618 | 28.31903 | 28.47104 |
| 135 | 28.15894 | 28.2069 | 28.6317 | 28.49549 | 28.34109 | 28.31056 | 28.85614 | 28.77731 | 28.57813 |
| 131 | 28.2653 | 28.24282 | 28.37439 | 28.47046 | 28.50055 | 28.4898 | 28.56209 | 28.5458 | 28.57308 |
| 177 | 29.76199 | 29.83349 | 29.58572 | 29.51886 | 28.9983 | 29.20669 | 29.14526 | 29.15735 | 29.20268 |
| 112 | 28.4763 | 28.46716 | 28.01611 | 28.1471 | 27.81574 | 27.83936 | 27.58828 | 27.73789 | 27.74188 |
| 149 | 29.05025 | 29.0724 | 28.65472 | 28.70748 | 28.25413 | 28.32787 | 28.24259 | 28.40544 | 28.21438 |

| | | | | | | | | | |
|---|---|---|---|---|---|---|---|---|---|
| 263 | 35.31105 | 35.27704 | 35.01009 | 35.07551 | 34.69851 | 34.727 | 34.73383 | 34.69146 | 34.27031 |
| 234 | 31.39555 | 31.38607 | 31.21153 | 31.21269 | 31.20239 | 31.16758 | 30.53306 | 30.58942 | 30.54158 |
| 278 | 33.9722 | 33.90005 | 33.59134 | 33.69229 | 33.31541 | 33.30362 | 33.03286 | 32.87535 | 32.9682 |
| 158 | 30.18707 | 30.18471 | 29.84191 | 29.8639 | 29.31406 | 29.30954 | 29.26832 | 29.10849 | 29.07194 |
| 265 | 31.4665 | 31.46178 | 31.77808 | 31.7582 | 31.73028 | 31.67964 | 31.47141 | 31.44436 | 31.53959 |
| 69 | 27.73046 | 27.43936 | 27.43086 | 27.43181 | 26.51197 | 26.0983 | 26.68632 | 27.16835 | 21.9 |
| 89 | 28.65383 | 28.49755 | 31.43674 | 30.69611 | 31.16656 | 31.08784 | 31.48989 | 30.58549 | 31.61816 |
| 228 | 30.33095 | 30.34541 | 30.42412 | 30.55764 | 31.37202 | 31.30639 | 30.75111 | 30.73965 | 31.10718 |
| 100 | 26.45127 | 26.78569 | 27.62206 | 27.53505 | 27.7433 | 27.61814 | 27.35203 | 27.48339 | 27.49503 |
| 82 | 27.54008 | 27.51269 | 27.74593 | 27.67922 | 27.92937 | 27.9597 | 27.65554 | 27.54259 | 27.89729 |
| 165 | 29.96389 | 30.02261 | 30.16627 | 30.17474 | 30.3772 | 30.42011 | 29.98142 | 29.98563 | 30.23203 |
| 197 | 29.30635 | 29.23558 | 29.02052 | 29.06314 | 28.72087 | 28.83234 | 29.0011 | 29.06319 | 29.04711 |
| 417 | 31.6088 | 31.56179 | 31.66746 | 31.73441 | 32.02901 | 31.92479 | 32.20577 | 32.2914 | 32.33368 |
| 84 | 27.36728 | 27.2841 | 27.41345 | 27.4086 | 27.44735 | 27.12879 | 27.54208 | 27.49701 | 27.46987 |
| 80 | 26.19804 | 21.9 | 26.43094 | 25.40458 | 25.23827 | 21.9 | 27.90006 | 27.80317 | 26.8429 |
| 78 | 26.56168 | 26.63464 | 26.82457 | 26.68217 | 26.93258 | 26.62157 | 27.76724 | 27.60971 | 27.45403 |
| 99 | 27.34181 | 27.29263 | 27.91239 | 27.84506 | 28.24173 | 28.31614 | 27.98202 | 28.29614 | 28.5662 |
| 132 | 29.58205 | 29.74841 | 29.82159 | 30.16075 | 30.57578 | 30.66941 | 30.36296 | 30.46207 | 30.57948 |
| 236 | 32.46292 | 32.4754 | 32.70582 | 32.73473 | 33.08047 | 33.14236 | 33.11131 | 33.24104 | 33.19447 |
| 111 | 27.92948 | 27.96014 | 28.11322 | 27.96091 | 28.15615 | 28.18858 | 28.295 | 28.52966 | 28.5313 |
| 191 | 29.24359 | 29.2537 | 29.46597 | 29.46338 | 29.87766 | 29.91056 | 29.81949 | 29.93888 | 30.03486 |
| 122 | 28.63093 | 28.56158 | 28.40888 | 28.52414 | 29.17691 | 29.23178 | 29.14999 | 29.08464 | 29.21886 |
| 169 | 30.77839 | 30.718 | 30.85571 | 30.85415 | 31.5186 | 31.46461 | 31.3028 | 31.26052 | 31.4941 |
| 163 | 28.71797 | 28.67368 | 28.61393 | 28.74804 | 29.08342 | 29.03949 | 28.92062 | 28.99984 | 29.16152 |
| 103 | 27.48876 | 27.31484 | 27.47189 | 27.4073 | 27.52843 | 27.42638 | 27.79042 | 27.75277 | 27.69192 |
| 174 | 30.32066 | 30.24728 | 30.31181 | 30.26249 | 30.43946 | 30.36359 | 30.5969 | 30.59308 | 30.67051 |
| 137 | 28.63596 | 28.64808 | 28.94274 | 28.87862 | 28.94101 | 28.83132 | 29.21424 | 29.09368 | 29.08086 |
| 75 | 26.7714 | 26.94682 | 27.06466 | 26.9812 | 26.74073 | 26.86797 | 27.39313 | 27.26024 | 27.04667 |
| 112 | 28.36302 | 28.3096 | 28.36795 | 28.49663 | 28.25543 | 28.25768 | 28.61196 | 28.56074 | 28.4494 |
| 169 | 29.01654 | 29.05853 | 29.18348 | 29.07781 | 28.98 | 28.90412 | 28.85186 | 28.78 | 28.71221 |
| 76 | 28.02417 | 27.83768 | 27.8343 | 27.89977 | 27.38714 | 27.49442 | 27.69032 | 27.75022 | 27.20144 |
| 128 | 29.3283 | 29.33735 | 29.18896 | 29.16303 | 28.69458 | 28.55083 | 28.65983 | 28.7764 | 28.50537 |
| 180 | 31.7951 | 31.76505 | 31.36989 | 31.43241 | 30.83077 | 30.82836 | 30.84737 | 30.97897 | 30.80624 |
| 79 | 28.67741 | 28.64966 | 28.51931 | 28.38928 | 28.43138 | 28.58075 | 27.9305 | 27.83376 | 27.91359 |
| 106 | 28.26495 | 28.34807 | 28.16145 | 28.26127 | 28.34723 | 28.54381 | 27.43118 | 27.77543 | 27.63561 |
| 214 | 33.54398 | 33.57317 | 33.10052 | 33.07011 | 32.97678 | 32.8936 | 32.47281 | 32.39384 | 32.45182 |
| 124 | 28.17236 | 28.18305 | 28.14224 | 28.09189 | 27.75136 | 27.44095 | 28.40097 | 28.54937 | 28.14851 |
| 74 | 21.9 | 25.83183 | 26.38423 | 26.31285 | 25.56092 | 26.67613 | 26.47369 | 26.93134 | 26.86667 |
| 87 | 27.4961 | 27.50112 | 27.63506 | 27.4837 | 27.35271 | 27.39689 | 27.98896 | 27.9305 | 27.51126 |
| 83 | 28.29702 | 28.25836 | 28.56627 | 28.50786 | 28.66194 | 28.59925 | 28.38076 | 28.36853 | 28.41442 |
| 186 | 30.23169 | 30.28978 | 30.13614 | 30.18068 | 30.45258 | 30.46548 | 30.28505 | 30.27353 | 30.45885 |
| 94 | 28.0932 | 27.94061 | 28.22527 | 28.20074 | 27.66438 | 27.7753 | 28.03228 | 28.049 | 27.98848 |
| 120 | 28.39877 | 28.21868 | 28.66469 | 28.55102 | 29.16684 | 29.19601 | 28.57888 | 28.57131 | 28.9071 |
| 64 | 26.06515 | 26.19612 | 26.62017 | 26.53102 | 26.65294 | 26.89493 | 26.37622 | 26.59049 | 26.56194 |
| 155 | 29.36672 | 29.32586 | 29.34056 | 29.33573 | 29.48595 | 29.51608 | 29.27606 | 29.32918 | 29.44679 |
| 148 | 28.87523 | 28.95024 | 29.19526 | 29.3307 | 29.57382 | 29.62763 | 29.21586 | 29.27804 | 29.31214 |
| 127 | 27.80021 | 27.74407 | 27.87473 | 27.76218 | 27.63962 | 27.5729 | 28.20079 | 27.91473 | 27.96411 |

| | | | | | | | | | |
|---|---|---|---|---|---|---|---|---|---|
| 92 | 26.82857 | 26.41111 | 26.81958 | 26.91445 | 27.09999 | 27.1031 | 27.43547 | 27.46115 | 27.25187 |
| 113 | 28.39272 | 28.46925 | 28.02099 | 28.1456 | 27.79835 | 27.82372 | 27.81293 | 28.05103 | 28.04135 |
| 86 | 28.29325 | 28.2653 | 28.25359 | 28.09331 | 27.23013 | 26.75595 | 27.13741 | 27.18631 | 21.9 |
| 80 | 21.9 | 21.9 | 27.06825 | 21.9 | 27.41869 | 27.53779 | 27.29202 | 27.40998 | 27.40211 |
| 69 | 21.9 | 24.60801 | 26.62394 | 26.37504 | 26.21985 | 26.15773 | 27.42246 | 27.49534 | 27.06209 |
| 72 | 26.29358 | 26.31465 | 26.54205 | 26.34866 | 26.36991 | 25.93024 | 27.14773 | 27.51766 | 26.87044 |
| 120 | 28.36832 | 28.44312 | 28.2305 | 28.43026 | 28.14307 | 28.07964 | 29.18009 | 29.12497 | 28.78041 |
| 152 | 29.04999 | 29.02957 | 29.31838 | 29.34198 | 29.7361 | 29.67314 | 29.7055 | 29.74448 | 30.02499 |
| 103 | 27.31993 | 27.6224 | 27.62087 | 27.49457 | 28.32208 | 28.29837 | 28.01409 | 28.06147 | 28.39182 |
| 166 | 30.37948 | 30.52272 | 30.61096 | 30.63858 | 31.1454 | 31.12645 | 30.85378 | 31.07387 | 31.38098 |
| 99 | 27.20191 | 27.2479 | 27.14258 | 27.27382 | 26.97574 | 26.79402 | 27.67593 | 27.59206 | 27.53103 |
| 114 | 28.29062 | 28.2687 | 28.19746 | 28.19878 | 27.89053 | 27.65444 | 28.60364 | 28.64877 | 28.30895 |
| 105 | 26.92988 | 26.86726 | 26.98945 | 26.83834 | 26.78544 | 27.02094 | 27.32303 | 27.35111 | 26.99324 |
| 95 | 27.51833 | 27.49274 | 27.46342 | 27.56165 | 27.42878 | 27.54392 | 27.86047 | 27.79977 | 27.82542 |
| 96 | 27.51036 | 27.61421 | 27.58642 | 27.59989 | 27.61968 | 27.61091 | 27.73925 | 27.70246 | 27.6126 |
| 115 | 27.9656 | 27.84338 | 27.91359 | 28.10009 | 27.71869 | 27.67956 | 27.56369 | 26.90564 | 27.46014 |
| 89 | 27.62003 | 27.55384 | 27.2683 | 27.31639 | 27.06876 | 27.2803 | 26.96939 | 26.98251 | 26.87197 |
| 71 | 28.42974 | 28.42982 | 28.80243 | 28.71712 | 28.18933 | 27.9279 | 28.07327 | 28.08406 | 27.62568 |
| 126 | 29.22348 | 29.20338 | 29.2668 | 29.31127 | 29.18709 | 29.20958 | 28.50374 | 28.59797 | 28.60474 |
| 106 | 28.60703 | 28.53531 | 28.60851 | 28.54149 | 27.94821 | 27.9194 | 27.81336 | 27.87719 | 27.55142 |
| 123 | 28.34647 | 28.42694 | 28.32869 | 28.66861 | 29.0063 | 28.97624 | 28.6538 | 28.84035 | 28.97678 |
| 59 | 26.11066 | 26.01591 | 26.55563 | 26.18386 | 21.9 | 21.9 | 26.74522 | 26.89099 | 26.45149 |
| 74 | 27.71771 | 27.05672 | 27.93112 | 27.70035 | 27.46949 | 27.66316 | 27.99162 | 27.98772 | 27.85186 |
| 165 | 30.65906 | 30.6507 | 30.79722 | 30.95005 | 31.59588 | 31.4487 | 31.11905 | 31.00142 | 31.41715 |
| 96 | 27.97213 | 27.93292 | 28.14831 | 27.96807 | 28.38985 | 28.36323 | 27.89885 | 27.95005 | 28.42013 |
| 124 | 29.63227 | 29.69609 | 29.47522 | 29.41891 | 29.12171 | 28.91923 | 29.1334 | 29.31775 | 29.35857 |
| 158 | 29.90096 | 29.82713 | 29.88806 | 29.97678 | 30.69246 | 30.62758 | 29.93438 | 29.98943 | 30.32839 |
| 124 | 28.4249 | 28.5704 | 28.88425 | 28.86094 | 29.41078 | 29.46373 | 29.34532 | 29.31997 | 29.51239 |
| 85 | 27.93899 | 27.93888 | 27.99275 | 27.6412 | 27.34849 | 27.1882 | 27.41514 | 27.34002 | 27.21512 |
| 86 | 27.71522 | 27.63374 | 27.34053 | 27.25485 | 27.88867 | 27.45889 | 27.41135 | 27.51525 | 27.94587 |
| 48 | 27.89914 | 27.8947 | 28.15697 | 28.20797 | 27.32946 | 27.51548 | 27.07714 | 27.08172 | 21.9 |
| 72 | 21.9 | 21.9 | 26.22249 | 26.56208 | 27.57737 | 27.68499 | 27.14394 | 27.57174 | 27.6284 |
| 67 | 26.04602 | 26.35684 | 26.3561 | 25.85286 | 26.32251 | 25.79412 | 26.67519 | 26.46591 | 26.77731 |
| 89 | 27.94888 | 27.92982 | 27.85043 | 27.7934 | 27.42446 | 26.89839 | 28.20326 | 28.17145 | 27.94168 |
| 226 | 32.26794 | 32.28303 | 32.46558 | 32.5703 | 32.86935 | 32.81681 | 32.99885 | 32.89587 | 32.99841 |
| 86 | 27.43833 | 27.38129 | 27.53816 | 27.66696 | 27.7758 | 27.57967 | 28.06682 | 27.94687 | 27.91485 |
| 67 | 26.5809 | 26.62881 | 27.12149 | 27.01009 | 26.9354 | 27.10189 | 27.01892 | 27.21466 | 27.21568 |
| 82 | 27.18272 | 27.29675 | 27.45395 | 27.55815 | 27.60329 | 27.64258 | 27.68264 | 27.7302 | 27.7166 |
| 136 | 29.63156 | 29.66474 | 29.67924 | 29.69282 | 29.83918 | 29.82165 | 29.92959 | 29.82402 | 29.9709 |
| 124 | 29.21827 | 29.26495 | 29.23217 | 29.31467 | 29.44654 | 29.35359 | 29.23943 | 28.9695 | 29.18234 |
| 76 | 27.51307 | 27.41748 | 27.44561 | 27.61555 | 27.70319 | 27.58154 | 27.29614 | 27.01839 | 27.37285 |
| 120 | 28.69192 | 28.80885 | 28.79925 | 28.79625 | 28.40495 | 28.30217 | 28.44 | 28.39615 | 28.45026 |
| 151 | 30.34308 | 30.36024 | 30.17748 | 30.25045 | 30.0724 | 29.999 | 29.80535 | 29.67231 | 29.78813 |
| 103 | 27.76522 | 27.70286 | 27.69869 | 27.72097 | 27.20154 | 27.37542 | 27.19976 | 27.16077 | 27.04656 |
| 131 | 29.14273 | 29.1789 | 29.02332 | 29.06771 | 28.84978 | 28.85451 | 28.38018 | 28.38261 | 28.50525 |
| 70 | 27.68184 | 27.54517 | 26.88972 | 27.04239 | 26.91262 | 26.6126 | 26.42667 | 26.43922 | 26.69179 |
| 139 | 30.33586 | 30.36442 | 29.98033 | 30.02287 | 29.5133 | 29.5044 | 29.53291 | 29.56205 | 29.27404 |

| | | | | | | | | | |
|---|---|---|---|---|---|---|---|---|---|
| 92 | 28.47061 | 28.44376 | 28.42109 | 28.29022 | 28.09552 | 28.08846 | 27.69697 | 27.69458 | 27.59427 |
| 79 | 29.17474 | 28.97438 | 28.22757 | 28.33828 | 28.06476 | 28.12996 | 27.93977 | 27.79922 | 27.43643 |
| 97 | 28.70698 | 28.62739 | 28.81604 | 28.90527 | 27.94016 | 28.14788 | 28.11381 | 28.11178 | 27.47537 |
| 158 | 29.66387 | 29.59843 | 29.81048 | 29.71309 | 30.08563 | 30.06188 | 29.92917 | 29.99348 | 30.01518 |
| 107 | 27.80261 | 27.78325 | 27.54738 | 27.54841 | 27.23068 | 27.01169 | 27.93759 | 27.83708 | 27.71326 |
| 162 | 30.35437 | 30.45807 | 30.32967 | 30.35184 | 29.58862 | 29.63329 | 29.92705 | 29.95782 | 29.81617 |
| 65 | 21.9 | 26.60865 | 26.72364 | 26.50208 | 25.77017 | 26.52583 | 27.1048 | 27.35943 | 27.08628 |
| 212 | 32.59036 | 32.63565 | 32.81505 | 32.83838 | 33.25448 | 33.18925 | 32.87677 | 32.81469 | 33.2671 |
| 142 | 30.3529 | 30.23146 | 30.39713 | 30.38628 | 30.89338 | 30.87107 | 30.71006 | 30.54167 | 30.81886 |
| 34 | 25.42188 | 21.9 | 26.24661 | 26.03756 | 21.9 | 21.9 | 26.87197 | 27.08912 | 26.00172 |
| 130 | 28.5093 | 28.47719 | 28.42021 | 28.50123 | 28.91388 | 29.03307 | 28.73486 | 28.51348 | 28.90354 |
| 125 | 29.45272 | 29.39691 | 29.3016 | 29.45162 | 29.91428 | 29.96237 | 29.59706 | 29.53353 | 29.70212 |
| 106 | 28.11625 | 28.12209 | 28.0177 | 28.03039 | 27.57015 | 27.80796 | 27.98761 | 28.03685 | 27.82651 |
| 102 | 28.80344 | 28.89776 | 29.25246 | 29.30038 | 29.38846 | 29.40722 | 29.13078 | 28.98655 | 29.41736 |
| 98 | 27.78588 | 27.80667 | 27.18782 | 27.38648 | 27.13026 | 27.22867 | 27.09033 | 27.07204 | 27.57802 |
| 75 | 26.9941 | 27.00496 | 27.45615 | 27.35254 | 27.38311 | 27.60194 | 27.3585 | 26.87619 | 27.46614 |
| 92 | 27.63243 | 27.69936 | 27.89087 | 28.00024 | 28.22256 | 28.1334 | 27.8169 | 27.9149 | 28.16897 |
| 210 | 31.39407 | 31.31668 | 31.52468 | 31.59366 | 31.49238 | 31.4408 | 31.50644 | 31.55064 | 31.45709 |
| 120 | 29.28533 | 29.47497 | 29.45666 | 29.43227 | 29.10375 | 29.23327 | 29.59281 | 29.477 | 29.32313 |
| 136 | 28.91336 | 28.90879 | 29.10395 | 29.16945 | 29.34889 | 29.35997 | 29.21229 | 29.03483 | 29.15514 |
| 137 | 29.95574 | 29.85917 | 29.85027 | 29.82165 | 30.14346 | 30.11644 | 29.78854 | 29.7219 | 29.91085 |
| 231 | 32.96468 | 32.92899 | 32.35694 | 32.44231 | 32.47882 | 32.45858 | 32.28361 | 32.33178 | 32.88979 |
| 84 | 27.19442 | 26.77254 | 27.91405 | 27.75028 | 28.02464 | 27.51179 | 28.11327 | 28.41305 | 28.11327 |
| 121 | 28.59402 | 28.66231 | 28.29272 | 28.30956 | 27.78606 | 28.00635 | 27.90144 | 28.15108 | 28.17441 |
| 33 | 21.9 | 21.9 | 21.9 | 21.9 | 21.9 | 21.9 | 26.1341 | 21.9 | 26.46431 |
| 45 | 24.99157 | 21.9 | 21.9 | 21.9 | 25.3654 | 24.90408 | 25.66031 | 25.48056 | 25.06932 |
| 62 | 25.65414 | 26.11108 | 26.06864 | 25.55982 | 26.18476 | 25.56369 | 26.57658 | 26.91752 | 26.62589 |
| 55 | 26.20458 | 26.32456 | 26.1958 | 26.00494 | 26.06598 | 26.06334 | 27.06999 | 26.94882 | 26.94492 |
| 84 | 28.00865 | 28.00207 | 28.0805 | 28.16197 | 27.86083 | 27.65301 | 28.7977 | 28.78169 | 28.60724 |
| 71 | 25.73097 | 26.23573 | 26.15074 | 26.11716 | 25.77826 | 26.17952 | 26.86148 | 26.90012 | 26.27291 |
| 81 | 27.36311 | 27.14317 | 27.35641 | 27.4186 | 27.52865 | 27.42902 | 27.89718 | 27.48416 | 27.98038 |
| 110 | 28.2147 | 28.27946 | 28.06003 | 28.14846 | 28.21048 | 28.32569 | 28.51803 | 28.60957 | 28.69299 |
| 78 | 27.50302 | 27.26892 | 27.88832 | 27.82159 | 27.56914 | 27.37931 | 27.95416 | 27.97531 | 28.01749 |
| 95 | 27.41812 | 27.22048 | 27.20256 | 27.13634 | 27.38525 | 27.41844 | 27.67566 | 27.65581 | 27.68217 |
| 102 | 28.23384 | 28.35401 | 28.36026 | 28.33326 | 27.6318 | 27.59641 | 27.87068 | 27.80292 | 27.62038 |
| 170 | 31.33501 | 31.49557 | 31.21968 | 31.11347 | 31.01764 | 31.12472 | 30.64272 | 30.61281 | 30.44095 |
| 150 | 31.40187 | 31.30008 | 30.75764 | 30.74101 | 30.61999 | 30.65386 | 30.22653 | 30.1546 | 30.18589 |
| 72 | 27.50393 | 27.60074 | 27.72728 | 27.83008 | 26.964 | 27.08861 | 26.84661 | 26.88902 | 26.27858 |
| 87 | 28.3277 | 28.14467 | 28.24549 | 28.23009 | 28.80125 | 28.84533 | 28.59449 | 28.57719 | 28.78204 |
| 77 | 27.06106 | 27.11981 | 27.34672 | 27.39599 | 27.50355 | 27.32346 | 27.31674 | 27.42069 | 27.4763 |
| 98 | 27.1915 | 27.24771 | 27.69192 | 27.49213 | 27.2641 | 27.39942 | 27.43094 | 27.73072 | 27.36035 |
| 71 | 26.36197 | 26.36473 | 26.29286 | 26.40818 | 27.242 | 27.11188 | 26.84661 | 26.88716 | 26.93438 |
| 53 | 26.35897 | 26.18841 | 25.94505 | 26.08703 | 21.9 | 25.82116 | 26.69365 | 26.70754 | 26.48708 |
| 48 | 21.9 | 26.72143 | 26.72455 | 21.9 | 26.97016 | 26.86597 | 26.74445 | 26.90001 | 26.68953 |
| 88 | 28.41175 | 28.39807 | 28.76088 | 28.67785 | 28.37289 | 28.24427 | 28.52518 | 28.42822 | 28.29605 |
| 67 | 27.42085 | 27.4817 | 27.53949 | 27.57254 | 27.28056 | 27.25827 | 27.50771 | 27.46459 | 27.39174 |
| 75 | 26.81787 | 21.9 | 26.82043 | 26.59106 | 26.7415 | 27.18376 | 27.25755 | 27.12436 | 27.17931 |

| | | | | | | | | | |
|---|---|---|---|---|---|---|---|---|---|
| 88 | 27.50839 | 27.64031 | 27.56522 | 27.76383 | 28.01797 | 28.14482 | 27.72078 | 27.70193 | 28.06363 |
| 65 | 26.64113 | 26.50736 | 26.91696 | 26.93247 | 27.09456 | 26.92354 | 26.8977 | 26.99615 | 27.03769 |
| 70 | 27.86685 | 27.70299 | 27.86136 | 27.91262 | 27.84571 | 27.78357 | 27.55025 | 27.88134 | 27.75022 |
| 108 | 27.69319 | 27.58714 | 27.82554 | 27.87115 | 27.6633 | 27.71555 | 27.9266 | 27.91068 | 27.6727 |
| 163 | 27.97367 | 28.04568 | 28.60544 | 28.48024 | 28.4514 | 28.38833 | 28.23891 | 28.14788 | 28.16063 |
| 100 | 27.88349 | 26.66153 | 28.0624 | 28.02892 | 28.98215 | 28.93683 | 28.1794 | 28.24966 | 28.42242 |
| 173 | 31.69216 | 31.63824 | 31.65446 | 31.62967 | 31.23968 | 31.18358 | 31.44811 | 31.39841 | 31.4628 |
| 60 | 27.19376 | 27.28577 | 27.55903 | 27.47305 | 26.80587 | 27.30504 | 27.5437 | 21.9 | 27.70299 |
| 70 | 27.58384 | 27.71017 | 27.91451 | 27.59626 | 27.2716 | 27.44766 | 27.23215 | 27.29448 | 27.97542 |
| 71 | 28.1207 | 28.18565 | 28.19127 | 28.33219 | 28.37985 | 28.32989 | 28.29048 | 28.1517 | 28.479 |
| 249 | 30.13247 | 30.10437 | 30.2466 | 30.26428 | 30.68403 | 30.71334 | 30.68821 | 30.63625 | 30.95532 |
| 194 | 28.82116 | 28.80726 | 28.74188 | 28.67509 | 28.30691 | 28.18452 | 29.14217 | 29.17634 | 28.73153 |
| 76 | 26.84864 | 26.57874 | 26.7415 | 26.9382 | 27.10199 | 26.49224 | 27.52604 | 27.5479 | 27.39043 |
| 110 | 28.88661 | 28.88134 | 28.88524 | 28.84389 | 28.61737 | 28.66577 | 29.17864 | 29.11783 | 29.07551 |
| 48 | 25.89758 | 26.44223 | 21.9 | 21.9 | 21.9 | 21.9 | 26.71816 | 26.44028 | 26.08647 |
| 70 | 27.42158 | 26.93876 | 27.53103 | 27.46155 | 27.9249 | 28.03522 | 27.77725 | 28.13643 | 28.28308 |
| 113 | 28.86806 | 28.29031 | 28.99911 | 29.00541 | 29.34537 | 29.5583 | 29.33556 | 29.37909 | 29.74349 |
| 70 | 26.69099 | 26.51679 | 26.35198 | 26.7196 | 26.98804 | 26.90449 | 27.08243 | 27.00024 | 27.13359 |
| 115 | 29.30752 | 29.0896 | 29.45537 | 29.42976 | 29.58018 | 29.5776 | 29.81228 | 29.91185 | 29.89015 |
| 189 | 31.98878 | 32.00989 | 32.19422 | 32.13247 | 32.60673 | 32.85279 | 32.51371 | 32.6999 | 32.57698 |
| 192 | 30.61246 | 30.62288 | 30.98461 | 30.85816 | 31.27398 | 31.2752 | 31.17397 | 31.34583 | 31.23238 |
| 62 | 26.99766 | 26.8521 | 26.77191 | 26.57615 | 26.8776 | 27.03171 | 27.09476 | 27.02179 | 27.21188 |
| 62 | 27.16087 | 27.08871 | 27.01297 | 26.88204 | 26.81763 | 26.99086 | 27.24146 | 27.34858 | 27.44703 |
| 120 | 28.41724 | 28.38907 | 28.55516 | 28.49701 | 28.65123 | 28.74506 | 28.74606 | 28.80064 | 28.81742 |
| 72 | 28.30769 | 28.04776 | 27.79743 | 27.96312 | 28.03879 | 28.00689 | 28.11227 | 28.21188 | 28.39579 |
| 142 | 30.49305 | 30.49524 | 30.53649 | 30.60902 | 30.9255 | 30.85177 | 30.94886 | 30.79707 | 30.89201 |
| 61 | 26.49196 | 26.55434 | 26.33407 | 26.31136 | 26.23983 | 26.33895 | 26.72702 | 26.71293 | 26.53444 |
| 84 | 28.50915 | 28.46186 | 28.47007 | 28.3953 | 28.25746 | 28.42806 | 28.10774 | 28.20802 | 28.14788 |
| 147 | 32.33636 | 32.43738 | 32.40363 | 32.36719 | 32.0955 | 32.13338 | 32.19827 | 32.09986 | 32.04088 |
| 106 | 28.13541 | 28.11406 | 27.84416 | 27.79315 | 27.62749 | 27.81348 | 27.4307 | 27.58542 | 27.50809 |
| 81 | 29.00678 | 29.15702 | 28.87801 | 28.64327 | 28.68288 | 28.63464 | 28.2968 | 28.26781 | 28.59623 |
| 94 | 27.74894 | 27.65697 | 27.97246 | 27.9894 | 27.28242 | 27.3928 | 27.58276 | 27.59413 | 27.22048 |
| 101 | 28.38492 | 28.34367 | 28.16949 | 28.29942 | 28.12642 | 28.05667 | 27.69086 | 27.68244 | 27.77518 |
| 161 | 30.49781 | 30.38741 | 30.07635 | 30.17164 | 29.72325 | 29.82438 | 29.71107 | 29.79486 | 29.36751 |
| 85 | 29.33618 | 29.2166 | 29.10174 | 29.08335 | 29.02488 | 28.89432 | 28.42558 | 28.50381 | 28.45752 |
| 73 | 27.79544 | 27.83056 | 27.31717 | 27.50028 | 27.07531 | 27.20974 | 27.02348 | 27.11714 | 26.89516 |
| 102 | 28.65803 | 28.63886 | 28.44399 | 28.48447 | 28.15841 | 28.15673 | 27.66357 | 27.71993 | 27.82523 |
| 61 | 26.48333 | 26.44068 | 26.7366 | 26.75506 | 26.55438 | 26.4987 | 25.98811 | 26.06277 | 25.94896 |
| 103 | 29.6482 | 29.82191 | 29.4626 | 29.33805 | 29.18973 | 29.18969 | 28.80894 | 28.6583 | 28.75066 |
| 176 | 32.81125 | 32.71767 | 32.43418 | 32.46051 | 32.24045 | 32.27833 | 32.03735 | 31.85419 | 31.73202 |
| 153 | 29.80119 | 29.71203 | 29.30032 | 29.39571 | 28.88797 | 28.93977 | 28.70876 | 28.62885 | 28.57892 |
| 132 | 30.678 | 30.57777 | 30.08575 | 30.07546 | 29.72206 | 29.77645 | 29.52501 | 29.53668 | 29.20058 |
| 89 | 27.2459 | 26.88262 | 26.79848 | 26.90747 | 25.95325 | 21.9 | 27.79755 | 27.59406 | 27.37551 |
| 87 | 28.16 | 28.3096 | 28.51145 | 28.57069 | 29.00086 | 29.00616 | 28.61593 | 28.65854 | 29.17412 |
| 157 | 30.93467 | 30.8936 | 30.92769 | 30.97637 | 31.21934 | 31.18854 | 31.03518 | 31.12349 | 31.12405 |
| 43 | 21.9 | 21.9 | 26.25499 | 26.45009 | 25.99686 | 26.4681 | 26.33982 | 26.40284 | 26.47656 |
| 74 | 26.74458 | 26.10426 | 26.80833 | 26.92025 | 27.27977 | 26.67734 | 27.26964 | 26.95815 | 27.05155 |

| | | | | | | | | | |
|---|---|---|---|---|---|---|---|---|---|
| 48 | 26.33911 | 26.31548 | 21.9 | 21.9 | 26.65116 | 26.52624 | 25.49409 | 25.2939 | 26.75034 |
| 79 | 21.9 | 26.4853 | 27.30461 | 27.18801 | 27.63789 | 27.49678 | 27.44569 | 27.35035 | 27.59056 |
| 119 | 29.64225 | 29.69993 | 29.55428 | 29.80885 | 30.154 | 30.04022 | 29.852 | 29.74959 | 30.09761 |
| 80 | 27.75411 | 27.93067 | 28.27888 | 28.05305 | 27.90455 | 28.06779 | 27.82232 | 27.9881 | 27.87238 |
| 48 | 26.43727 | 26.52807 | 26.67546 | 26.63353 | 26.99798 | 26.99959 | 26.85496 | 26.64293 | 26.90862 |
| 39 | 25.52349 | 25.60927 | 25.96078 | 25.98501 | 25.98348 | 25.89398 | 25.86978 | 21.9 | 25.59095 |
| 34 | 26.72312 | 26.53708 | 26.279 | 26.55205 | 21.9 | 21.9 | 21.9 | 21.9 | 26.31991 |
| 94 | 27.67755 | 27.81568 | 28.04984 | 28.15369 | 28.23334 | 28.25142 | 27.89157 | 28.05956 | 28.15248 |
| 51 | 26.31631 | 26.34704 | 26.90587 | 26.93731 | 26.03217 | 25.79397 | 26.88914 | 27.03978 | 26.08572 |
| 71 | 27.15528 | 27.04947 | 27.39468 | 27.42958 | 27.27267 | 27.32285 | 27.30939 | 27.24699 | 27.2992 |
| 84 | 27.9414 | 28.07699 | 28.35136 | 28.42482 | 28.39575 | 28.42622 | 28.35342 | 28.33926 | 28.37704 |
| 164 | 29.48491 | 29.59048 | 29.07342 | 29.03383 | 28.74403 | 28.64595 | 28.53982 | 28.36736 | 28.3699 |
| 98 | 27.32903 | 27.25539 | 27.18187 | 27.32363 | 27.14569 | 27.22555 | 27.57124 | 27.59534 | 27.42374 |
| 122 | 31.16213 | 31.07316 | 31.14686 | 31.10855 | 31.38787 | 31.34905 | 31.18648 | 31.08493 | 31.42292 |
| 79 | 27.25926 | 27.23973 | 27.53245 | 27.6035 | 28.15909 | 27.99302 | 27.77555 | 27.6983 | 28.10164 |
| 71 | 21.9 | 21.9 | 25.2281 | 26.51662 | 25.98087 | 26.2283 | 27.14482 | 27.03559 | 26.84685 |
| 53 | 21.9 | 26.05459 | 26.10659 | 21.9 | 25.74324 | 25.98185 | 26.63948 | 26.33345 | 26.67465 |
| 65 | 25.46647 | 25.73565 | 26.54162 | 25.63448 | 26.60145 | 26.66506 | 27.22637 | 27.3741 | 27.01372 |
| 77 | 26.4854 | 26.96928 | 26.42211 | 27.51555 | 27.88087 | 27.70491 | 27.89602 | 28.22991 | 28.01898 |
| 52 | 25.52411 | 26.05951 | 25.68319 | 26.12989 | 26.14071 | 25.84948 | 26.81946 | 27.3855 | 26.46437 |
| 62 | 26.22079 | 26.05674 | 25.8853 | 26.24735 | 27.02454 | 26.97312 | 26.99949 | 27.35582 | 27.61456 |
| 56 | 26.61947 | 26.34184 | 26.04275 | 26.43357 | 26.51983 | 26.50753 | 26.69192 | 26.81592 | 27.60604 |
| 61 | 26.24722 | 26.3303 | 26.28383 | 26.41492 | 26.07757 | 25.84388 | 27.14676 | 27.12268 | 26.59946 |
| 82 | 28.96447 | 28.64743 | 28.66499 | 28.62216 | 29.11135 | 28.88795 | 29.21009 | 29.00389 | 29.52251 |
| 85 | 28.32736 | 28.36098 | 28.2799 | 28.2562 | 28.35632 | 28.17603 | 28.77442 | 28.90868 | 28.73947 |
| 73 | 26.79141 | 26.61891 | 26.9741 | 27.06023 | 27.10059 | 27.15894 | 27.32946 | 27.21373 | 27.28056 |
| 56 | 26.1034 | 25.69022 | 25.3575 | 25.77904 | 25.35683 | 25.27888 | 26.11662 | 26.17257 | 25.92336 |
| 71 | 26.65472 | 26.53512 | 26.70859 | 27.1055 | 27.28215 | 27.20452 | 27.02433 | 27.1882 | 27.38187 |
| 78 | 28.17698 | 28.10889 | 28.06023 | 28.13188 | 28.46587 | 28.44904 | 28.26638 | 28.35758 | 28.60551 |
| 140 | 30.93136 | 30.89879 | 30.63045 | 30.66136 | 30.32571 | 30.35815 | 30.18068 | 30.23592 | 30.42081 |
| 154 | 33.58038 | 33.51653 | 33.2717 | 33.19465 | 33.23278 | 33.02868 | 32.95361 | 32.87318 | 32.99247 |
| 57 | 27.059 | 27.09376 | 26.87033 | 26.97323 | 26.95005 | 26.89689 | 26.34099 | 26.51676 | 26.70925 |
| 101 | 29.72524 | 29.70357 | 29.58694 | 29.67674 | 29.34892 | 29.48823 | 28.98373 | 29.03727 | 29.14625 |
| 110 | 30.13809 | 30.18199 | 29.74588 | 29.72996 | 29.48706 | 29.58328 | 29.17619 | 29.2202 | 29.21616 |
| 142 | 30.15497 | 30.23581 | 29.89653 | 29.95005 | 29.76789 | 29.63836 | 29.26774 | 29.25798 | 29.35531 |
| 103 | 28.17145 | 28.13227 | 27.61751 | 27.84822 | 27.32053 | 27.47421 | 27.11565 | 27.34138 | 26.78706 |
| 170 | 31.76805 | 31.81284 | 31.52431 | 31.43966 | 30.99415 | 30.8822 | 31.10737 | 30.80008 | 30.53213 |
| 93 | 27.73402 | 27.75379 | 27.5599 | 27.74791 | 28.1437 | 28.05186 | 27.89804 | 27.94358 | 28.19244 |
| 33 | 21.9 | 21.9 | 21.9 | 25.72323 | 26.29623 | 21.9 | 21.9 | 25.69721 | 26.19552 |
| 63 | 26.63173 | 26.61344 | 26.38795 | 26.39656 | 26.87209 | 26.89666 | 26.79439 | 26.5308 | 26.861 |
| 94 | 27.44601 | 27.46357 | 27.78669 | 27.84026 | 28.47777 | 28.44241 | 27.99615 | 28.06332 | 28.4432 |
| 40 | 25.59608 | 21.9 | 25.59186 | 25.45127 | 25.53724 | 25.60215 | 25.80703 | 26.38686 | 26.03829 |
| 58 | 26.1164 | 21.9 | 26.66858 | 26.59377 | 27.70233 | 27.40276 | 27.0425 | 26.8147 | 28.18054 |
| 88 | 28.65905 | 28.65772 | 28.91562 | 28.8947 | 29.13646 | 29.11976 | 28.5235 | 28.47344 | 28.81424 |
| 60 | 25.47293 | 25.59568 | 25.9434 | 25.65903 | 25.15896 | 24.70488 | 25.37534 | 25.33811 | 25.1488 |
| 33 | 21.9 | 21.9 | 25.57095 | 25.80203 | 25.61773 | 25.67573 | 25.55303 | 25.17413 | 25.29428 |
| 102 | 28.0873 | 28.11173 | 28.34134 | 28.32886 | 27.96631 | 28.17451 | 28.40117 | 28.37538 | 28.05061 |

| | | | | | | | | | |
|---|---|---|---|---|---|---|---|---|---|
| 67 | 27.31674 | 27.40186 | 26.91182 | 27.121 | 26.9301 | 21.9 | 26.47516 | 26.61119 | 26.29514 |
| 543 | 32.89242 | 32.89654 | 32.77982 | 32.83406 | 32.72816 | 32.79295 | 33.06283 | 33.10001 | 32.97044 |
| 169 | 27.93084 | 27.94464 | 28.26892 | 28.30195 | 27.86024 | 27.95837 | 28.15653 | 28.4447 | 27.92116 |
| 207 | 31.12368 | 31.21466 | 31.42702 | 31.33234 | 31.79865 | 31.65459 | 31.50389 | 31.43211 | 31.88314 |
| 212 | 31.26579 | 31.27941 | 31.13852 | 31.08607 | 30.83778 | 30.96678 | 30.88329 | 30.89331 | 30.78667 |
| 95 | 27.56173 | 27.7442 | 27.56725 | 27.65349 | 27.78369 | 27.78837 | 27.71712 | 27.8914 | 27.87038 |
| 60 | 26.58506 | 26.6327 | 26.53887 | 26.3727 | 26.16911 | 26.12569 | 27.17817 | 27.2833 | 26.9516 |
| 74 | 28.41784 | 28.36252 | 28.20401 | 28.15644 | 28.33577 | 28.20965 | 28.31756 | 28.27786 | 28.4105 |
| 49 | 25.25443 | 26.02563 | 26.23624 | 26.25344 | 26.32577 | 21.9 | 25.75789 | 26.20686 | 26.28907 |
| 43 | 21.9 | 21.9 | 25.51286 | 21.9 | 25.20807 | 25.37571 | 25.6457 | 25.41519 | 25.60116 |
| 36 | 21.9 | 21.9 | 21.9 | 25.40038 | 25.48511 | 21.9 | 26.30088 | 26.19899 | 25.72023 |
| 113 | 29.92465 | 29.97035 | 30.34096 | 30.41115 | 30.94468 | 31.03682 | 30.89742 | 31.00276 | 31.33596 |
| 63 | 27.18101 | 27.06989 | 27.15634 | 27.13153 | 27.76085 | 27.73279 | 27.74375 | 27.6309 | 27.84206 |
| 64 | 26.42974 | 26.59349 | 26.90862 | 26.76421 | 27.10079 | 27.16844 | 27.22536 | 27.15615 | 27.62422 |
| 60 | 26.61358 | 26.21856 | 26.27213 | 26.47251 | 26.56947 | 26.50264 | 26.91445 | 26.84242 | 26.99097 |
| 125 | 29.2196 | 29.26874 | 29.40781 | 29.44435 | 29.88861 | 29.89558 | 29.66068 | 29.89312 | 29.99725 |
| 78 | 27.73673 | 27.61989 | 27.67916 | 27.6731 | 28.25822 | 28.18749 | 28.18518 | 28.18702 | 28.31501 |
| 77 | 29.32212 | 29.34788 | 29.44676 | 28.74381 | 28.95646 | 28.85695 | 29.65688 | 29.54745 | 29.4751 |
| 101 | 29.05388 | 29.12682 | 29.23567 | 29.33884 | 29.4309 | 29.34763 | 29.47954 | 29.58563 | 29.62678 |
| 59 | 27.09597 | 26.76116 | 26.89608 | 26.89862 | 27.04437 | 26.96774 | 26.90219 | 27.07806 | 27.57283 |
| 77 | 29.14158 | 29.03483 | 29.20623 | 29.2645 | 29.46396 | 29.49576 | 29.32526 | 29.45331 | 29.60661 |
| 142 | 29.96416 | 29.85988 | 30.03013 | 29.85691 | 30.31721 | 30.2483 | 30.21118 | 30.11396 | 30.39325 |
| 51 | 26.58806 | 26.3695 | 26.46133 | 26.30492 | 26.66194 | 26.59263 | 26.58764 | 26.78182 | 26.66519 |
| 129 | 31.6937 | 31.62693 | 31.78655 | 31.66615 | 31.56878 | 31.63893 | 32.11226 | 31.89009 | 31.78265 |
| 78 | 27.02189 | 27.16404 | 27.08871 | 27.07948 | 27.17969 | 27.21243 | 27.18556 | 27.24771 | 27.33014 |
| 64 | 28.98473 | 29.02147 | 29.29187 | 29.21006 | 28.80058 | 28.85028 | 28.80627 | 28.82366 | 28.56837 |
| 62 | 27.78475 | 27.88449 | 27.99216 | 28.16902 | 27.88617 | 28.02036 | 27.43651 | 27.47313 | 27.52006 |
| 86 | 28.50556 | 28.38319 | 28.89764 | 28.80261 | 28.30756 | 28.05113 | 28.35943 | 28.27404 | 27.81696 |
| 84 | 28.78666 | 28.67731 | 28.50854 | 28.46688 | 28.2548 | 28.1883 | 27.98397 | 27.77436 | 27.93422 |
| 65 | 27.74169 | 27.68706 | 27.62694 | 27.72065 | 27.6654 | 27.44735 | 26.97695 | 26.91752 | 26.96168 |
| 66 | 27.53979 | 27.4715 | 26.93483 | 26.89018 | 27.20508 | 27.20862 | 26.32514 | 26.67936 | 26.39988 |
| 135 | 34.11047 | 34.08354 | 33.55282 | 33.67903 | 33.41526 | 33.24275 | 33.26431 | 33.19046 | 32.82392 |
| 53 | 21.9 | 25.88995 | 26.11847 | 25.89952 | 26.06741 | 25.88376 | 26.16465 | 26.36418 | 26.89423 |
| 83 | 28.21127 | 28.37443 | 28.55954 | 28.43452 | 29.09401 | 29.03695 | 28.76775 | 28.64272 | 29.17679 |
| 42 | 21.9 | 21.9 | 26.34591 | 26.35165 | 26.15526 | 26.26191 | 26.50747 | 26.63437 | 26.54697 |
| 126 | 29.76833 | 29.95463 | 29.96747 | 29.8676 | 30.53881 | 30.59619 | 30.19131 | 30.09522 | 30.46918 |
| 96 | 29.30685 | 29.25723 | 28.71653 | 28.6764 | 28.11892 | 28.08537 | 28.45971 | 28.59024 | 28.09688 |
| 37 | 25.00702 | 21.9 | 25.37955 | 21.9 | 25.73689 | 25.90825 | 25.49161 | 25.27838 | 25.7202 |
| 76 | 27.66851 | 27.65547 | 27.76357 | 27.67209 | 28.30817 | 28.36374 | 28.08314 | 27.88244 | 28.25894 |
| 74 | 27.056 | 27.16777 | 27.42238 | 27.41232 | 27.7911 | 27.77794 | 27.38492 | 27.40771 | 27.73821 |
| 51 | 27.01839 | 26.96796 | 27.2535 | 27.41191 | 27.84817 | 27.63658 | 27.34367 | 27.21882 | 27.75595 |
| 30 | 21.9 | 21.9 | 21.9 | 21.9 | 26.27902 | 26.35097 | 26.06833 | 21.9 | 26.22052 |
| 109 | 29.84775 | 29.66846 | 30.00477 | 29.99954 | 30.31192 | 30.32892 | 29.87282 | 29.97446 | 30.23055 |
| 63 | 26.13303 | 26.1546 | 26.00047 | 26.27185 | 26.12212 | 26.32285 | 26.00699 | 26.11026 | 26.40669 |
| 119 | 28.7225 | 28.62258 | 28.91909 | 28.88399 | 28.83105 | 28.87894 | 29.01614 | 28.83446 | 28.96615 |
| 58 | 26.9249 | 21.9 | 27.08871 | 26.74137 | 26.81653 | 26.78793 | 26.83038 | 26.70767 | 26.63547 |
| 57 | 26.60498 | 26.60922 | 26.768 | 26.64182 | 27.20415 | 27.31277 | 26.77957 | 26.65444 | 27.0704 |

| | | | | | | | | | |
|---|---|---|---|---|---|---|---|---|---|
| 41 | 21.9 | 21.9 | 26.09063 | 25.80696 | 26.20406 | 26.11353 | 26.33369 | 21.9 | 26.28104 |
| 36 | 26.12668 | 26.0806 | 26.38433 | 21.9 | 26.27594 | 26.36105 | 25.75246 | 25.64743 | 26.14505 |
| 51 | 26.06829 | 26.24122 | 26.51716 | 26.25119 | 27.28736 | 27.3803 | 26.47059 | 26.52208 | 26.9476 |
| 28 | 26.81176 | 26.92275 | 26.74163 | 26.90873 | 21.9 | 21.9 | 21.9 | 21.9 | 26.4063 |
| 100 | 28.8204 | 28.88288 | 28.91792 | 28.91399 | 28.77124 | 28.69149 | 28.99162 | 29.00638 | 28.89093 |
| 58 | 27.41828 | 27.49823 | 27.20686 | 27.14492 | 26.32838 | 26.12102 | 27.55772 | 27.34663 | 26.75837 |
| 105 | 26.71358 | 26.04816 | 26.93134 | 26.93461 | 27.01638 | 27.14209 | 27.48899 | 27.66804 | 27.95166 |
| 52 | 26.17883 | 26.2696 | 26.37159 | 26.21593 | 27.00045 | 26.74047 | 26.5588 | 26.55362 | 26.87185 |
| 23 | 24.98403 | 24.7453 | 25.69506 | 25.45344 | 21.9 | 21.9 | 21.9 | 21.9 | 21.9 |
| 37 | 21.9 | 21.9 | 21.9 | 21.9 | 21.9 | 21.9 | 26.05957 | 26.03672 | 25.87886 |
| 39 | 21.9 | 21.9 | 21.9 | 21.9 | 27.13927 | 27.12829 | 27.27053 | 27.20713 | 27.73718 |
| 27 | 21.9 | 21.9 | 21.9 | 21.9 | 25.78294 | 21.9 | 25.55213 | 25.46628 | 25.83738 |
| 34 | 21.9 | 21.9 | 21.9 | 21.9 | 26.14736 | 25.93975 | 26.0353 | 25.9882 | 26.29088 |
| 32 | 21.9 | 21.9 | 25.86988 | 21.9 | 21.9 | 25.73296 | 25.83649 | 25.90722 | 25.91614 |
| 46 | 21.9 | 21.9 | 25.70909 | 21.9 | 26.13553 | 26.03752 | 26.38736 | 26.26335 | 26.64375 |
| 31 | 21.9 | 21.9 | 21.9 | 25.10949 | 21.9 | 21.9 | 25.16959 | 25.23987 | 24.67223 |
| 35 | 25.34905 | 21.9 | 25.41735 | 21.9 | 25.5578 | 21.9 | 25.8367 | 25.67393 | 26.12173 |
| 35 | 21.9 | 21.9 | 25.838 | 21.9 | 25.78531 | 25.87364 | 25.86748 | 25.61613 | 26.05604 |
| 91 | 28.50093 | 28.92298 | 28.77087 | 28.94651 | 29.7058 | 29.751 | 29.52559 | 29.59349 | 29.82139 |
| 90 | 27.74586 | 27.27906 | 28.05615 | 28.33956 | 28.24314 | 28.37634 | 28.54698 | 28.69299 | 28.85967 |
| 63 | 26.3097 | 26.29537 | 26.67128 | 26.79848 | 27.45403 | 27.43245 | 27.23004 | 27.22288 | 27.62951 |
| 62 | 27.07887 | 27.03107 | 27.63041 | 27.68016 | 28.04323 | 27.97749 | 27.78787 | 27.9907 | 28.52013 |
| 57 | 26.95183 | 26.80747 | 26.92807 | 26.63187 | 27.40064 | 27.3731 | 27.11207 | 27.20769 | 27.80335 |
| 34 | 25.82786 | 25.88332 | 25.84033 | 25.85362 | 25.96904 | 26.08815 | 26.14233 | 26.34311 | 26.39539 |
| 70 | 26.97038 | 27.05528 | 26.88774 | 26.89701 | 27.13975 | 27.04895 | 27.38138 | 27.34189 | 27.47545 |
| 61 | 27.65444 | 27.47916 | 27.8435 | 27.76408 | 28.09813 | 28.07368 | 28.19634 | 28.18919 | 28.27609 |
| 90 | 29.33528 | 29.24722 | 29.45136 | 29.34282 | 29.53334 | 29.63064 | 29.44863 | 29.65359 | 30.03486 |
| 55 | 26.32 | 26.58118 | 26.38316 | 26.47201 | 26.41159 | 26.29703 | 26.67627 | 26.80033 | 26.71934 |
| 99 | 28.61835 | 28.7459 | 28.85007 | 28.93843 | 29.1813 | 29.1666 | 29.16372 | 29.04838 | 29.24391 |
| 94 | 28.9366 | 28.91806 | 28.95889 | 28.90549 | 28.96925 | 29.03478 | 29.12445 | 29.31838 | 29.00389 |
| 63 | 27.30678 | 27.16385 | 27.58197 | 27.5223 | 27.66024 | 27.69392 | 27.06435 | 26.98729 | 27.19723 |
| 62 | 27.48424 | 27.50279 | 27.55194 | 27.57095 | 27.43659 | 27.36494 | 27.08608 | 27.17741 | 27.05466 |
| 51 | 26.96091 | 26.81274 | 26.55497 | 26.77856 | 26.22033 | 26.27047 | 26.09887 | 26.10326 | 26.11527 |
| 74 | 28.69132 | 28.21257 | 27.91074 | 28.46439 | 27.74452 | 27.96538 | 27.97531 | 27.84062 | 27.63748 |
| 65 | 27.83092 | 27.93528 | 27.70912 | 27.66058 | 27.56391 | 27.60308 | 27.14783 | 27.19169 | 27.17245 |
| 67 | 28.93632 | 29.10127 | 28.72289 | 28.66204 | 28.42318 | 28.43778 | 27.99566 | 28.3023 | 27.88919 |
| 67 | 28.21605 | 28.24672 | 27.77769 | 27.73085 | 27.37401 | 27.3869 | 27.17855 | 27.2943 | 27.19291 |
| 58 | 27.57759 | 27.75704 | 27.25746 | 27.36252 | 26.83557 | 26.95116 | 26.63934 | 26.78307 | 26.56651 |
| 35 | 26.15973 | 26.3123 | 26.96168 | 26.86089 | 26.01933 | 25.93535 | 26.05695 | 25.83413 | 25.4746 |
| 78 | 28.11942 | 27.96741 | 28.34651 | 28.55972 | 28.49911 | 28.50684 | 28.40787 | 28.68395 | 28.54731 |
| 54 | 21.9 | 25.86842 | 26.10793 | 21.9 | 26.12191 | 26.30654 | 26.94067 | 26.56254 | 26.5357 |
| 59 | 26.9673 | 27.11008 | 27.11783 | 27.19197 | 27.84242 | 27.74195 | 27.44774 | 27.54672 | 27.91279 |
| 35 | 26.2562 | 26.07529 | 21.9 | 25.67301 | 21.9 | 21.9 | 25.94123 | 26.04533 | 25.56648 |
| 36 | 25.56119 | 21.9 | 26.48775 | 26.42236 | 26.39357 | 21.9 | 26.30895 | 26.47172 | 26.34761 |
| 53 | 21.9 | 27.5229 | 21.9 | 27.90231 | 28.5842 | 28.48166 | 28.22453 | 28.14467 | 28.61964 |
| 118 | 29.17524 | 29.15074 | 28.7764 | 28.79498 | 28.23982 | 28.41905 | 28.80827 | 28.62701 | 28.24939 |
| 53 | 21.9 | 21.9 | 26.25162 | 26.04268 | 26.01992 | 26.17864 | 26.30748 | 26.25694 | 26.08479 |

| | | | | | | | | | |
|---|---|---|---|---|---|---|---|---|---|
| 49 | 26.15937 | 21.9 | 26.6725 | 26.38083 | 27.26571 | 27.54104 | 27.22131 | 26.88262 | 27.72962 |
| 79 | 29.22002 | 29.29599 | 29.49992 | 29.55005 | 29.28531 | 29.34405 | 29.43277 | 29.30411 | 29.23738 |
| 58 | 30.137 | 30.1488 | 30.42302 | 30.43996 | 30.13528 | 30.22883 | 21.9 | 29.53256 | 29.70576 |
| 37 | 21.9 | 21.9 | 21.9 | 26.43205 | 27.09557 | 26.97498 | 21.9 | 26.67922 | 27.23772 |
| 68 | 26.76116 | 26.77693 | 26.53954 | 26.59733 | 26.71463 | 26.68057 | 26.73544 | 26.40492 | 26.77643 |
| 33 | 25.27618 | 25.28708 | 25.26685 | 25.21468 | 24.7291 | 24.79298 | 25.48459 | 21.9 | 25.32745 |
| 38 | 21.9 | 21.9 | 25.84919 | 25.67694 | 25.57923 | 25.78639 | 25.62809 | 25.75318 | 25.753 |
| 50 | 26.33407 | 26.3044 | 26.67492 | 26.69285 | 26.35597 | 26.47023 | 27.17722 | 26.7335 | 26.27746 |
| 38 | 25.09092 | 25.01924 | 25.10274 | 25.13962 | 24.74842 | 24.82982 | 25.44626 | 25.45117 | 24.8436 |
| 51 | 26.83762 | 26.89111 | 26.85626 | 26.68003 | 26.49659 | 26.64196 | 26.74304 | 26.75608 | 26.68886 |
| 42 | 25.66186 | 21.9 | 25.83429 | 25.51782 | 26.12585 | 26.08811 | 21.9 | 25.73363 | 26.15209 |
| 50 | 26.49723 | 26.67384 | 27.06958 | 27.04072 | 26.80156 | 26.8597 | 26.60696 | 26.84098 | 26.9875 |
| 42 | 26.2438 | 26.16617 | 21.9 | 21.9 | 27.85121 | 27.53066 | 25.34455 | 21.9 | 25.26767 |
| 94 | 27.69252 | 27.46809 | 27.51848 | 27.46762 | 27.25809 | 27.01999 | 27.69631 | 27.63263 | 27.58484 |
| 177 | 30.58701 | 30.62104 | 30.64796 | 30.6832 | 30.66263 | 30.70157 | 30.40568 | 30.55215 | 30.39489 |
| 115 | 29.43797 | 29.40817 | 28.96502 | 28.84631 | 29.24631 | 29.26765 | 28.42858 | 28.42518 | 28.67647 |
| 93 | 28.66523 | 28.57275 | 28.56906 | 28.71257 | 29.34094 | 29.32657 | 28.48093 | 28.45435 | 29.15419 |
| 35 | 21.9 | 21.9 | 21.9 | 27.202 | 27.32071 | 27.83635 | 27.61702 | 27.40406 | 27.60265 |
| 136 | 31.28659 | 31.25101 | 31.36082 | 31.19448 | 31.76383 | 31.92217 | 30.78947 | 30.71645 | 30.28659 |
| 100 | 27.97514 | 27.98196 | 27.94682 | 27.92162 | 27.54075 | 27.59896 | 27.72234 | 27.6102 | 27.76629 |
| 62 | 27.34908 | 27.31008 | 27.34019 | 27.29149 | 27.25917 | 27.19573 | 28.47553 | 28.33811 | 27.88646 |
| 39 | 25.40759 | 25.83253 | 21.9 | 25.62561 | 21.9 | 25.90949 | 21.9 | 26.14048 | 26.1795 |
| 23 | 21.9 | 26.4177 | 26.75494 | 26.84781 | 21.9 | 21.9 | 21.9 | 21.9 | 21.9 |
| 21 | 26.94134 | 26.97016 | 26.15856 | 26.59092 | 26.08742 | 21.9 | 21.9 | 21.9 | 21.9 |
| 45 | 21.9 | 21.9 | 21.9 | 21.9 | 26.68686 | 21.9 | 26.43396 | 26.32619 | 26.60611 |
| 25 | 21.9 | 21.9 | 21.9 | 21.9 | 21.9 | 25.48164 | 25.69655 | 26.00804 | 25.53653 |
| 33 | 21.9 | 21.9 | 25.96497 | 21.9 | 21.9 | 26.39745 | 26.20892 | 26.42544 | 26.47888 |
| 37 | 21.9 | 21.9 | 21.9 | 21.9 | 25.74499 | 26.3026 | 26.02297 | 25.87192 | 26.2477 |
| 28 | 25.34262 | 21.9 | 25.6879 | 21.9 | 21.9 | 21.9 | 26.19991 | 26.06833 | 26.28667 |
| 37 | 21.9 | 21.9 | 21.9 | 21.9 | 25.76994 | 21.9 | 25.52561 | 25.50694 | 25.59161 |
| 54 | 21.9 | 21.9 | 25.50042 | 21.9 | 27.30121 | 27.14628 | 26.47045 | 26.77856 | 27.2283 |
| 32 | 21.9 | 21.9 | 21.9 | 21.9 | 25.93517 | 26.01338 | 25.77032 | 25.93312 | 26.10324 |
| 29 | 21.9 | 21.9 | 21.9 | 25.77884 | 21.9 | 26.04208 | 25.9933 | 25.98163 | 26.02981 |
| 29 | 21.9 | 21.9 | 25.799 | 21.9 | 21.9 | 21.9 | 26.02071 | 26.27526 | 21.9 |
| 62 | 27.1333 | 27.32243 | 27.02073 | 26.82917 | 26.86396 | 26.73544 | 27.98251 | 28.02522 | 27.88151 |
| 76 | 28.33159 | 28.471 | 28.63499 | 28.60654 | 29.0713 | 29.15071 | 29.0495 | 29.13357 | 29.43984 |
| 82 | 26.90208 | 27.15846 | 27.27498 | 26.51011 | 27.45128 | 27.50211 | 27.67882 | 27.52746 | 27.50249 |
| 66 | 27.79054 | 27.94782 | 27.99959 | 27.90242 | 28.28392 | 28.28498 | 28.36757 | 28.48447 | 28.76954 |
| 45 | 25.68528 | 25.8227 | 25.86025 | 25.73003 | 26.34489 | 26.07975 | 26.18701 | 26.06956 | 26.4989 |
| 78 | 27.1205 | 27.34731 | 27.56819 | 27.5575 | 28.08527 | 28.04375 | 27.88163 | 27.75468 | 28.08401 |
| 98 | 29.62619 | 29.69856 | 29.84311 | 29.82084 | 30.44243 | 30.46188 | 30.29296 | 30.18601 | 30.37534 |
| 79 | 28.23302 | 28.1856 | 28.32384 | 28.29583 | 28.34967 | 28.43098 | 28.5786 | 28.53505 | 28.48708 |
| 69 | 27.29807 | 27.26015 | 27.5658 | 27.49488 | 27.39583 | 27.30278 | 27.53185 | 27.56972 | 27.53823 |
| 55 | 27.22186 | 27.13163 | 27.36319 | 27.45223 | 27.06507 | 27.2338 | 27.03475 | 26.94615 | 27.02136 |
| 102 | 28.74452 | 28.75665 | 29.05118 | 29.0216 | 28.68385 | 28.78852 | 28.75793 | 28.77052 | 28.42594 |
| 48 | 26.50626 | 26.53856 | 26.39996 | 26.37451 | 26.2154 | 26.19438 | 26.01378 | 25.93339 | 26.29667 |
| 62 | 28.09959 | 28.20074 | 28.23945 | 28.06379 | 28.0561 | 28.00967 | 27.88239 | 27.86443 | 27.48232 |

| | | | | | | | | | |
|---|---|---|---|---|---|---|---|---|---|
| 56 | 27.66173 | 27.67108 | 27.9881 | 27.95815 | 27.03097 | 27.2179 | 27.61253 | 27.56042 | 26.94938 |
| 78 | 29.40261 | 29.28846 | 28.88212 | 29.01608 | 28.61354 | 28.52973 | 28.41478 | 28.45285 | 28.31493 |
| 221 | 32.2681 | 32.27298 | 32.01966 | 32.01029 | 31.6634 | 31.65723 | 31.44347 | 31.36984 | 31.46456 |
| 48 | 27.25088 | 27.17122 | 27.13242 | 26.90288 | 26.76218 | 26.58047 | 26.1062 | 26.32946 | 26.34026 |
| 49 | 27.89325 | 27.93168 | 27.33977 | 27.59007 | 27.53356 | 27.24871 | 26.87701 | 26.86372 | 26.69816 |
| 88 | 29.81455 | 29.80867 | 29.56198 | 29.63777 | 29.2181 | 29.18874 | 28.65308 | 28.73431 | 28.66051 |
| 50 | 27.48301 | 27.3396 | 27.06404 | 27.26338 | 26.84912 | 26.93067 | 26.53804 | 26.00005 | 26.1448 |
| 76 | 26.87561 | 27.2756 | 27.2226 | 27.29421 | 27.70108 | 27.76091 | 27.48999 | 27.46933 | 27.76939 |
| 26 | 25.39169 | 25.35703 | 21.9 | 21.9 | 25.44906 | 25.39921 | 26.13471 | 25.88047 | 25.67166 |
| 44 | 21.9 | 26.04151 | 26.12642 | 26.05321 | 26.20717 | 21.9 | 26.86053 | 26.44304 | 26.6392 |
| 28 | 21.9 | 25.66582 | 21.9 | 21.9 | 25.13465 | 25.129 | 25.12774 | 25.36636 | 24.99628 |
| 46 | 27.6277 | 27.21364 | 27.45513 | 27.67303 | 27.17245 | 26.40202 | 26.67734 | 26.54252 | 26.37901 |
| 32 | 26.81775 | 27.14102 | 26.18603 | 26.46812 | 26.12125 | 26.22293 | 25.8541 | 25.66194 | 26.13774 |
| 53 | 27.33901 | 27.24318 | 27.14928 | 26.98913 | 27.14424 | 27.03926 | 27.0168 | 26.85982 | 27.03454 |
| 48 | 26.06135 | 26.03145 | 26.13399 | 26.06593 | 26.86868 | 26.85781 | 26.23993 | 27.0086 | 26.79414 |
| 65 | 28.22656 | 28.28617 | 28.27222 | 28.15417 | 28.68181 | 28.62076 | 28.35653 | 28.40479 | 28.75936 |
| 53 | 26.14354 | 26.3003 | 21.9 | 26.03093 | 21.9 | 26.12372 | 26.53038 | 26.29239 | 26.03982 |
| 34 | 21.9 | 21.9 | 26.13318 | 26.01434 | 26.29051 | 26.35065 | 25.92854 | 26.16957 | 26.37544 |
| 85 | 27.9149 | 28.13629 | 28.10849 | 27.98191 | 28.43956 | 28.49114 | 28.22849 | 28.19079 | 28.3748 |
| 67 | 27.43476 | 27.52111 | 27.64485 | 27.78893 | 27.96411 | 27.87619 | 27.47166 | 27.48316 | 27.64671 |
| 86 | 29.44774 | 29.37892 | 29.36897 | 29.47497 | 29.72686 | 29.80371 | 29.63516 | 29.57434 | 29.67931 |
| 41 | 21.9 | 26.14624 | 21.9 | 26.31441 | 26.08186 | 26.32958 | 25.83584 | 25.89705 | 26.3407 |
| 52 | 27.26293 | 27.17693 | 21.9 | 27.2826 | 27.39615 | 27.8242 | 27.45325 | 27.58549 | 28.19192 |
| 54 | 27.05289 | 27.06753 | 27.01478 | 21.9 | 27.45834 | 27.52283 | 27.41788 | 27.65922 | 27.74163 |
| 28 | 25.00294 | 25.11355 | 25.09857 | 21.9 | 21.9 | 21.9 | 21.9 | 25.17756 | 24.98298 |
| 40 | 25.4706 | 21.9 | 25.63713 | 25.62143 | 25.71741 | 25.72983 | 25.67613 | 26.06821 | 25.75504 |
| 42 | 25.87014 | 25.96475 | 26.11355 | 26.17066 | 25.87016 | 21.9 | 26.2878 | 26.51449 | 25.89502 |
| 46 | 26.05761 | 26.24793 | 21.9 | 26.04427 | 26.06556 | 26.11249 | 26.54537 | 26.46745 | 26.05583 |
| 41 | 25.5397 | 25.76243 | 25.83451 | 26.03729 | 21.9 | 25.32371 | 26.08144 | 26.18541 | 25.65103 |
| 42 | 25.8084 | 25.81622 | 25.41393 | 25.6997 | 25.99285 | 25.98978 | 25.78034 | 25.70578 | 26.11736 |
| 79 | 27.27524 | 27.39436 | 27.55896 | 27.64169 | 27.81519 | 27.84254 | 27.6441 | 27.55245 | 27.68652 |
| 40 | 21.9 | 25.92506 | 25.98168 | 25.86216 | 26.08677 | 25.84081 | 25.91841 | 25.94962 | 25.97729 |
| 58 | 26.29355 | 21.9 | 26.91056 | 26.68472 | 26.82092 | 26.70912 | 26.73079 | 26.71306 | 26.72663 |
| 46 | 25.09848 | 24.97745 | 26.10127 | 25.69402 | 25.42653 | 25.23063 | 26.00393 | 26.05912 | 25.37116 |
| 70 | 27.59889 | 27.53289 | 28.16878 | 28.17627 | 28.657 | 28.43508 | 28.1638 | 27.97465 | 28.43599 |
| 63 | 27.56558 | 27.6797 | 28.00292 | 27.91565 | 28.16245 | 28.19671 | 27.54045 | 27.64196 | 27.94737 |
| 91 | 29.30969 | 29.28643 | 28.83222 | 28.87871 | 28.98834 | 29.09441 | 28.91769 | 28.9632 | 29.2766 |
| 66 | 27.44538 | 27.64024 | 27.561 | 27.61365 | 27.30417 | 27.41877 | 27.46084 | 27.49526 | 27.63367 |
| 62 | 26.67775 | 26.73389 | 27.22251 | 27.0749 | 27.47042 | 27.49755 | 27.50937 | 27.20191 | 27.06538 |
| 23 | 21.9 | 21.9 | 25.91693 | 25.97314 | 25.65941 | 21.9 | 25.69083 | 26.05439 | 21.9 |
| 21 | 25.40266 | 25.79434 | 21.9 | 21.9 | 25.71563 | 21.9 | 21.9 | 25.45315 | 25.41596 |
| 63 | 28.47923 | 28.3944 | 28.30817 | 28.44 | 28.80879 | 28.96672 | 28.46883 | 28.27338 | 28.73912 |
| 29 | 25.66224 | 25.38271 | 25.86764 | 25.67366 | 21.9 | 21.9 | 21.9 | 24.98055 | 25.17013 |
| 57 | 26.58104 | 26.66817 | 26.42129 | 26.26374 | 26.23383 | 26.34067 | 26.59634 | 26.30833 | 26.41475 |
| 84 | 28.47707 | 28.43559 | 29.0875 | 29.04205 | 29.09854 | 28.93292 | 28.91664 | 28.74615 | 28.73593 |
| 57 | 26.91205 | 26.8889 | 27.18877 | 27.2478 | 27.35372 | 27.32997 | 27.08598 | 27.08598 | 27.16816 |
| 103 | 30.84663 | 30.87438 | 31.39897 | 31.26226 | 31.45106 | 31.58625 | 30.82987 | 31.03164 | 31.49643 |

| | | | | | | | | | |
|---|---|---|---|---|---|---|---|---|---|
| 77 | 29.07819 | 29.16286 | 29.02041 | 28.99526 | 29.20997 | 29.20939 | 29.1036 | 28.94983 | 29.18655 |
| 65 | 27.06168 | 27.14492 | 27.33287 | 27.21475 | 26.8675 | 26.76104 | 27.12938 | 27.08294 | 27.18867 |
| 166 | 29.29349 | 29.22927 | 28.72195 | 28.7835 | 28.61196 | 28.46743 | 28.65738 | 28.67162 | 28.9431 |
| 322 | 31.96503 | 32.01807 | 31.85564 | 31.87045 | 31.70289 | 31.66021 | 31.60867 | 31.73117 | 31.47024 |
| 87 | 28.10919 | 28.15856 | 28.80209 | 28.76569 | 28.63675 | 28.6041 | 28.72926 | 28.45038 | 28.4635 |
| 68 | 26.98229 | 27.3073 | 27.33083 | 27.44782 | 27.40479 | 27.56471 | 27.49419 | 27.66777 | 27.778 |
| 139 | 29.01903 | 29.07291 | 28.96898 | 28.8818 | 28.62697 | 28.57232 | 28.75376 | 28.88215 | 29.09451 |
| 66 | 27.84571 | 27.68238 | 27.82299 | 27.7991 | 27.84966 | 27.83581 | 28.3689 | 28.34435 | 28.44055 |
| 80 | 29.03475 | 29.03099 | 28.97618 | 29.02549 | 28.8476 | 28.7893 | 28.40129 | 28.26195 | 28.38648 |
| 64 | 28.64698 | 28.5183 | 27.96851 | 28.05429 | 27.8299 | 27.85632 | 27.99502 | 27.9491 | 27.98109 |
| 32 | 21.9 | 21.9 | 26.53825 | 26.70938 | 21.9 | 21.9 | 26.73337 | 26.6656 | 26.56961 |
| 63 | 27.54731 | 27.69823 | 27.16586 | 27.46209 | 27.21808 | 27.35279 | 27.9368 | 27.9419 | 27.56812 |
| 117 | 30.21616 | 30.28229 | 30.26785 | 30.23683 | 29.94782 | 30.03026 | 29.90124 | 29.86813 | 30.00544 |
| 64 | 27.71371 | 27.64368 | 27.23169 | 27.32268 | 27.09436 | 27.16988 | 27.53416 | 27.65663 | 27.61891 |
| 26 | 21.9 | 21.9 | 25.95536 | 26.51605 | 21.9 | 21.9 | 21.9 | 25.87577 | 26.17114 |
| 34 | 26.85234 | 26.62073 | 26.71607 | 26.71135 | 25.73247 | 25.91832 | 25.94722 | 26.0731 | 21.9 |
| 29 | 21.9 | 21.9 | 21.9 | 21.9 | 21.9 | 21.9 | 26.26805 | 26.01412 | 26.06786 |
| 24 | 21.9 | 21.9 | 25.73859 | 21.9 | 21.9 | 21.9 | 25.69687 | 25.57934 | 26.16548 |
| 32 | 21.9 | 21.9 | 21.9 | 21.9 | 21.9 | 25.59716 | 26.12617 | 25.76984 | 25.4248 |
| 36 | 25.3837 | 25.33159 | 21.9 | 21.9 | 21.9 | 21.9 | 26.14696 | 26.24554 | 25.72396 |
| 26 | 21.9 | 21.9 | 21.9 | 21.9 | 21.9 | 21.9 | 21.9 | 25.83309 | 26.15035 |
| 32 | 21.9 | 25.56915 | 21.9 | 25.63015 | 21.9 | 21.9 | 26.03769 | 26.03917 | 25.69607 |
| 37 | 21.9 | 21.9 | 24.93179 | 24.87675 | 21.9 | 21.9 | 25.20948 | 25.41261 | 25.1208 |
| 32 | 21.9 | 25.30751 | 21.9 | 25.37892 | 25.66215 | 25.86429 | 25.82279 | 26.11807 | 26.61667 |
| 47 | 25.75047 | 26.14604 | 26.058 | 26.42161 | 25.90509 | 26.12593 | 26.50927 | 26.64086 | 26.33354 |
| 73 | 28.29075 | 28.59385 | 28.87449 | 28.80196 | 29.1835 | 29.36641 | 29.29123 | 29.30374 | 29.32266 |
| 48 | 27.47561 | 27.46684 | 27.14297 | 27.15228 | 27.46139 | 27.63527 | 27.77794 | 27.62694 | 27.88948 |
| 60 | 26.65322 | 26.51503 | 26.87091 | 26.69564 | 26.50954 | 26.6284 | 27.28463 | 27.23389 | 27.04291 |
| 56 | 26.44474 | 26.45424 | 26.603 | 26.6656 | 26.57542 | 26.79216 | 26.80514 | 26.85436 | 26.98837 |
| 68 | 27.51571 | 27.34503 | 27.5942 | 27.79662 | 28.15644 | 28.1036 | 27.85335 | 28.05284 | 28.10679 |
| 59 | 26.80317 | 26.80907 | 27.05196 | 26.72416 | 27.28268 | 27.21252 | 27.13232 | 27.01084 | 27.36269 |
| 107 | 28.93913 | 28.81629 | 28.97495 | 28.95002 | 29.14312 | 29.21153 | 29.19479 | 29.28372 | 29.16212 |
| 43 | 26.46572 | 26.48517 | 26.5596 | 26.43165 | 26.86915 | 27.0379 | 26.82384 | 26.75443 | 26.97443 |
| 51 | 26.80107 | 27.03097 | 26.93977 | 26.91513 | 27.07806 | 26.98044 | 27.08517 | 27.08375 | 27.13232 |
| 93 | 28.68505 | 28.78992 | 29.02089 | 29.04492 | 28.64853 | 28.85159 | 28.56569 | 28.62916 | 28.65007 |
| 80 | 29.50357 | 29.48968 | 29.6555 | 29.49364 | 29.56045 | 29.45778 | 29.46795 | 29.21519 | 29.4057 |
| 65 | 29.74658 | 29.80053 | 29.54008 | 29.48575 | 29.74412 | 29.6449 | 29.3759 | 29.04231 | 29.43873 |
| 108 | 31.20408 | 31.11669 | 30.66043 | 30.72809 | 30.59841 | 30.46032 | 30.10037 | 30.34075 | 30.47894 |
| 39 | 26.49202 | 26.24527 | 26.11438 | 26.14715 | 25.76507 | 25.69277 | 25.56924 | 25.69562 | 25.65973 |
| 53 | 27.9812 | 27.76585 | 27.56187 | 27.19704 | 26.9788 | 27.07521 | 27.26177 | 27.03454 | 26.98902 |
| 25 | 21.9 | 21.9 | 21.9 | 21.9 | 25.59787 | 25.43892 | 21.9 | 25.56459 | 25.56311 |
| 32 | 26.37448 | 26.58949 | 21.9 | 21.9 | 21.9 | 21.9 | 21.9 | 26.37244 | 26.00847 |
| 54 | 26.7604 | 26.77668 | 26.93652 | 26.92897 | 27.5942 | 27.58262 | 27.27569 | 27.12288 | 27.47267 |
| 35 | 21.9 | 21.9 | 25.88904 | 25.78287 | 26.21126 | 26.21963 | 26.00247 | 25.99565 | 26.46505 |
| 37 | 24.94329 | 24.91232 | 25.29105 | 25.4215 | 21.9 | 21.9 | 25.31931 | 25.25169 | 25.42694 |
| 41 | 26.44012 | 26.48554 | 26.19646 | 21.9 | 25.91452 | 25.95265 | 26.61975 | 26.74714 | 26.7714 |
| 44 | 26.2916 | 25.98542 | 25.78337 | 25.73066 | 25.88518 | 21.9 | 26.24523 | 26.05335 | 26.31726 |

| | | | | | | | | | |
|---|---|---|---|---|---|---|---|---|---|
| 64 | 27.90776 | 27.99674 | 28.0229 | 28.04291 | 28.68946 | 28.76636 | 28.28595 | 28.27804 | 28.57712 |
| 25 | 26.86821 | 26.6231 | 26.65581 | 26.5932 | 26.21995 | 21.9 | 21.9 | 25.80211 | 21.9 |
| 37 | 21.9 | 26.11999 | 26.32133 | 26.10819 | 26.39735 | 26.48994 | 26.56515 | 26.51443 | 26.64279 |
| 31 | 26.21466 | 26.43771 | 26.52452 | 26.49503 | 26.77906 | 21.9 | 26.45635 | 26.51179 | 26.8242 |
| 132 | 32.59065 | 32.55497 | 32.88358 | 33.01363 | 33.10098 | 33.11372 | 33.14023 | 33.11187 | 32.83899 |
| 61 | 27.35624 | 27.46279 | 27.71004 | 27.79365 | 27.94921 | 28.14991 | 27.96108 | 27.72793 | 28.05755 |
| 34 | 25.21756 | 25.30124 | 21.9 | 25.29509 | 25.34432 | 25.37898 | 25.18869 | 25.5386 | 25.514 |
| 22 | 21.9 | 21.9 | 21.9 | 21.9 | 21.9 | 25.51584 | 21.9 | 21.9 | 25.76572 |
| 26 | 21.9 | 21.9 | 25.09499 | 21.9 | 25.84881 | 25.98729 | 25.50619 | 25.37093 | 25.97033 |
| 67 | 27.38755 | 27.48408 | 27.24254 | 27.40406 | 27.22453 | 27.20088 | 27.18452 | 27.17484 | 27.43412 |
| 41 | 21.9 | 25.9299 | 26.42541 | 26.48912 | 26.91502 | 27.06507 | 26.41543 | 26.33982 | 26.87807 |
| 54 | 26.51082 | 27.05972 | 27.54731 | 27.77039 | 27.28551 | 27.40536 | 26.99723 | 27.32019 | 26.71751 |
| 86 | 28.78899 | 28.73905 | 29.44524 | 29.58486 | 29.08238 | 29.26499 | 29.31164 | 29.11441 | 28.79959 |
| 33 | 25.56244 | 25.80215 | 21.9 | 21.9 | 26.09993 | 26.00164 | 26.22938 | 25.91506 | 26.25151 |
| 31 | 21.9 | 21.9 | 25.38864 | 25.11446 | 25.20071 | 25.04818 | 21.9 | 25.21915 | 25.52603 |
| 43 | 25.27338 | 25.47982 | 25.60676 | 25.79005 | 26.24627 | 26.0178 | 25.67541 | 25.65911 | 25.76089 |
| 61 | 27.35203 | 27.32457 | 27.55801 | 27.72051 | 27.89828 | 27.96157 | 27.56885 | 27.20881 | 27.66336 |
| 34 | 24.93836 | 25.0431 | 25.50779 | 25.44708 | 26.07617 | 26.11207 | 21.9 | 25.41545 | 25.85769 |
| 60 | 26.53481 | 26.23579 | 27.08294 | 26.92999 | 26.37046 | 26.40084 | 26.91274 | 26.83545 | 26.61793 |
| 41 | 26.43775 | 26.51159 | 26.09545 | 26.10225 | 21.9 | 26.05821 | 25.55037 | 25.46314 | 26.33224 |
| 31 | 26.88285 | 21.9 | 26.94224 | 26.91752 | 26.93719 | 21.9 | 26.94436 | 21.9 | 26.91696 |
| 25 | 25.78109 | 25.75075 | 21.9 | 21.9 | 25.71374 | 21.9 | 25.13175 | 21.9 | 25.45146 |
| 25 | 25.26008 | 25.37109 | 24.84809 | 21.9 | 21.9 | 21.9 | 24.86472 | 24.73128 | 24.72708 |
| 47 | 27.24364 | 27.39738 | 27.60569 | 27.37675 | 27.36469 | 27.19563 | 27.73473 | 27.73705 | 27.01329 |
| 38 | 26.41269 | 26.4513 | 26.02688 | 26.03488 | 25.88029 | 21.9 | 21.9 | 26.1626 | 26.21814 |
| 50 | 26.54712 | 26.37532 | 26.77781 | 26.69059 | 26.19415 | 26.16014 | 26.37279 | 26.66357 | 26.403 |
| 49 | 26.25225 | 26.2884 | 26.67654 | 26.55612 | 26.40477 | 26.38159 | 26.82031 | 26.47053 | 26.50714 |
| 84 | 27.20872 | 27.21382 | 27.73337 | 27.53014 | 27.47754 | 27.3298 | 27.91479 | 27.53875 | 27.24572 |
| 134 | 27.80236 | 27.94999 | 27.97235 | 27.94302 | 28.01377 | 27.94531 | 28.28621 | 28.36323 | 28.49983 |
| 203 | 31.43773 | 31.46922 | 31.05511 | 31.12065 | 30.89577 | 30.97589 | 30.70033 | 30.95428 | 31.26969 |
| 104 | 27.39297 | 27.19357 | 27.53304 | 27.60145 | 27.67922 | 27.64196 | 28.11088 | 27.95521 | 27.85585 |
| 116 | 28.35708 | 28.36085 | 28.58431 | 28.59509 | 28.705 | 28.70636 | 28.80621 | 28.71185 | 28.73538 |
| 67 | 27.70266 | 27.88134 | 27.9862 | 28.11986 | 27.80827 | 27.76718 | 27.88297 | 28.35691 | 28.19178 |
| 106 | 28.82554 | 28.80354 | 29.0928 | 29.03375 | 28.94606 | 29.13394 | 29.02192 | 29.14608 | 29.01707 |
| 24 | 21.9 | 21.9 | 21.9 | 21.9 | 21.9 | 21.9 | 21.9 | 25.858 | 25.7541 |
| 66 | 26.33923 | 26.30807 | 26.10592 | 26.46768 | 26.56403 | 26.46927 | 26.66438 | 26.6064 | 26.74766 |
| 64 | 28.68053 | 28.6981 | 28.54639 | 28.60247 | 28.82043 | 28.97413 | 28.62652 | 28.69598 | 29.04721 |
| 81 | 30.45474 | 30.35699 | 30.52 | 30.52374 | 30.66721 | 30.55435 | 30.72102 | 30.68946 | 30.81138 |
| 34 | 26.4364 | 26.37645 | 21.9 | 21.9 | 25.94051 | 21.9 | 26.29281 | 26.0542 | 26.18722 |
| 19 | 25.91563 | 25.63461 | 21.9 | 25.43784 | 21.9 | 21.9 | 21.9 | 25.56378 | 25.606 |
| 10 | 25.13895 | 25.26635 | 25.04668 | 24.98037 | 21.9 | 21.9 | 21.9 | 21.9 | 21.9 |
| 29 | 25.35131 | 25.3348 | 25.03351 | 25.34445 | 24.70773 | 24.80661 | 24.58761 | 21.9 | 21.9 |
| 28 | 26.62729 | 26.41167 | 26.18126 | 26.17764 | 26.07108 | 25.87975 | 21.9 | 25.41206 | 21.9 |
| 22 | 25.70142 | 25.90937 | 25.40698 | 25.63437 | 25.46551 | 25.46684 | 21.9 | 21.9 | 21.9 |
| 26 | 28.4698 | 28.51912 | 28.88536 | 28.74201 | 28.42118 | 28.38299 | 21.9 | 21.9 | 27.97869 |
| 24 | 21.9 | 21.9 | 25.10713 | 24.96376 | 21.9 | 21.9 | 25.13038 | 25.16791 | 24.77279 |
| 18 | 21.9 | 21.9 | 21.9 | 21.9 | 21.9 | 21.9 | 24.97758 | 24.62656 | 24.12195 |

| | | | | | | | | | |
|---|---|---|---|---|---|---|---|---|---|
| 19 | 21.9 | 21.9 | 21.9 | 21.9 | 21.9 | 21.9 | 25.33991 | 25.79397 | 25.42127 |
| 29 | 21.9 | 21.9 | 21.9 | 21.9 | 25.40285 | 21.9 | 25.74291 | 25.72226 | 25.91424 |
| 23 | 21.9 | 21.9 | 21.9 | 21.9 | 21.9 | 21.9 | 25.99477 | 21.9 | 26.01225 |
| 18 | 21.9 | 21.9 | 21.9 | 21.9 | 21.9 | 21.9 | 25.70256 | 25.76514 | 25.9872 |
| 43 | 27.66309 | 27.88087 | 21.9 | 21.9 | 21.9 | 28.05134 | 28.05941 | 28.14501 | 27.90121 |
| 19 | 21.9 | 21.9 | 21.9 | 21.9 | 21.9 | 21.9 | 25.60507 | 25.52797 | 25.53703 |
| 17 | 21.9 | 21.9 | 21.9 | 21.9 | 21.9 | 21.9 | 21.9 | 25.38063 | 25.11231 |
| 27 | 21.9 | 21.9 | 21.9 | 21.9 | 21.9 | 25.61616 | 21.9 | 25.87007 | 26.00269 |
| 28 | 21.9 | 21.9 | 25.48974 | 25.87216 | 21.9 | 21.9 | 25.87209 | 25.89285 | 25.90825 |
| 26 | 21.9 | 21.9 | 25.48996 | 21.9 | 21.9 | 25.93982 | 25.50727 | 25.66044 | 26.06897 |
| 30 | 25.41196 | 21.9 | 21.9 | 25.28792 | 21.9 | 21.9 | 25.79647 | 25.46136 | 25.68217 |
| 27 | 21.9 | 21.9 | 21.9 | 21.9 | 25.66566 | 25.84625 | 25.7172 | 25.37789 | 25.87697 |
| 24 | 21.9 | 21.9 | 21.9 | 21.9 | 25.45271 | 21.9 | 25.63006 | 21.9 | 25.71348 |
| 21 | 21.9 | 21.9 | 21.9 | 21.9 | 21.9 | 21.9 | 21.9 | 21.9 | 25.6045 |
| 22 | 21.9 | 21.9 | 21.9 | 21.9 | 21.9 | 25.77515 | 21.9 | 25.55298 | 25.63663 |
| 22 | 21.9 | 21.9 | 21.9 | 21.9 | 25.02644 | 24.94517 | 25.25133 | 25.17348 | 25.27955 |
| 19 | 21.9 | 21.9 | 21.9 | 21.9 | 21.9 | 21.9 | 21.9 | 21.9 | 25.27952 |
| 25 | 21.9 | 26.537 | 21.9 | 21.9 | 26.91525 | 27.038 | 26.59648 | 26.63381 | 26.88948 |
| 14 | 21.9 | 21.9 | 21.9 | 21.9 | 21.9 | 21.9 | 21.9 | 24.96825 | 25.10593 |
| 15 | 21.9 | 21.9 | 21.9 | 21.9 | 21.9 | 21.9 | 25.00962 | 21.9 | 21.9 |
| 12 | 21.9 | 21.9 | 21.9 | 21.9 | 21.9 | 21.9 | 24.28715 | 21.9 | 24.18018 |
| 13 | 21.9 | 21.9 | 21.9 | 21.9 | 21.9 | 21.9 | 24.05399 | 24.10597 | 21.9 |
| 9 | 25.84431 | 21.9 | 25.42563 | 25.54681 | 21.9 | 21.9 | 21.9 | 21.9 | 21.9 |
| 59 | 27.00507 | 27.09537 | 26.87654 | 26.95316 | 26.94927 | 26.85079 | 27.46084 | 27.46194 | 27.40438 |
| 46 | 27.15692 | 27.2178 | 27.35909 | 27.49602 | 27.72495 | 27.63872 | 27.83702 | 27.76541 | 27.93292 |
| 50 | 26.68926 | 26.57773 | 26.64595 | 26.86041 | 27.49876 | 27.3506 | 27.32122 | 27.24073 | 27.40113 |
| 58 | 26.71934 | 26.95936 | 27.15287 | 26.98392 | 27.05827 | 27.13085 | 27.30252 | 27.38451 | 27.24036 |
| 50 | 27.38443 | 27.38352 | 27.55998 | 27.48062 | 27.57015 | 27.42198 | 27.34731 | 27.35557 | 27.35246 |
| 71 | 28.60141 | 28.59886 | 28.84664 | 28.91319 | 28.81525 | 28.77894 | 28.67219 | 28.5508 | 28.616 |
| 49 | 27.83635 | 27.7855 | 27.58492 | 27.84458 | 27.65424 | 27.60724 | 27.80501 | 27.60017 | 27.42854 |
| 33 | 25.88502 | 25.9142 | 25.6363 | 25.79486 | 25.49601 | 25.6282 | 25.14371 | 25.33766 | 25.50476 |
| 46 | 26.50013 | 26.37348 | 26.51089 | 26.38377 | 26.47576 | 26.43344 | 26.12317 | 26.21182 | 25.78304 |
| 43 | 26.54582 | 26.49561 | 26.05161 | 26.29132 | 25.75511 | 26.10827 | 25.60439 | 25.56596 | 25.99921 |
| 59 | 28.85994 | 28.98878 | 28.6194 | 28.62415 | 28.63751 | 28.56376 | 28.31786 | 28.31804 | 28.18456 |
| 83 | 29.78524 | 29.86424 | 29.50389 | 29.61369 | 29.16682 | 29.01289 | 28.94184 | 29.07099 | 29.22704 |
| 39 | 26.51087 | 26.59192 | 26.57293 | 26.75774 | 26.19843 | 26.34733 | 26.47613 | 26.5231 | 25.88518 |
| 52 | 28.45646 | 28.44047 | 28.07051 | 28.07373 | 28.36236 | 28.28211 | 27.64684 | 27.86768 | 27.79042 |
| 82 | 30.07431 | 30.00624 | 29.85335 | 29.86015 | 29.51098 | 29.50766 | 29.29524 | 29.44504 | 29.28096 |
| 67 | 28.21382 | 28.28577 | 28.33637 | 28.19122 | 28.01137 | 27.90988 | 28.01733 | 27.74349 | 27.40633 |
| 36 | 28.14107 | 28.0531 | 27.36895 | 27.40706 | 27.1489 | 27.16404 | 27.22361 | 27.00945 | 27.02992 |
| 64 | 28.82845 | 28.96769 | 28.58538 | 28.50211 | 28.01031 | 28.04505 | 27.96752 | 28.26562 | 28.06404 |
| 20 | 21.9 | 24.58337 | 21.9 | 21.9 | 24.9574 | 21.9 | 21.9 | 24.80926 | 24.94873 |
| 41 | 26.07302 | 26.36926 | 26.79328 | 26.44506 | 26.25771 | 25.81685 | 26.7898 | 26.82723 | 26.48145 |
| 40 | 26.702 | 26.68913 | 26.29414 | 26.37199 | 26.15827 | 26.05585 | 26.22105 | 26.13667 | 26.22509 |
| 21 | 21.9 | 21.9 | 21.9 | 21.9 | 25.57265 | 21.9 | 21.9 | 21.9 | 25.59463 |
| 27 | 21.9 | 21.9 | 21.9 | 21.9 | 26.02145 | 25.77836 | 21.9 | 25.82806 | 26.09629 |
| 32 | 27.34883 | 27.31588 | 26.83232 | 26.92093 | 26.36386 | 26.35018 | 21.9 | 26.36766 | 25.99025 |

| | | | | | | | | | |
|---|---|---|---|---|---|---|---|---|---|
| 39 | 25.94376 | 25.84043 | 25.47116 | 25.64141 | 25.80659 | 21.9 | 26.27181 | 26.24313 | 26.37172 |
| 44 | 26.00314 | 25.89643 | 26.03065 | 21.9 | 26.37239 | 26.29365 | 26.92796 | 26.73982 | 26.55969 |
| 81 | 29.37132 | 29.32294 | 29.16777 | 29.26647 | 29.74745 | 29.56709 | 29.57795 | 29.29505 | 29.70696 |
| 95 | 27.30661 | 27.35271 | 27.45623 | 27.60265 | 27.20481 | 27.44166 | 26.46114 | 27.3937 | 26.97793 |
| 32 | 21.9 | 26.3489 | 21.9 | 25.98594 | 25.75794 | 25.65313 | 26.08091 | 26.30187 | 25.90885 |
| 44 | 26.05722 | 26.26642 | 26.63312 | 26.54086 | 26.26662 | 26.06454 | 26.51782 | 26.67155 | 26.36016 |
| 50 | 27.61358 | 27.71083 | 27.22352 | 27.53868 | 26.68151 | 27.27498 | 26.71699 | 26.90633 | 21.9 |
| 34 | 25.67452 | 25.82204 | 25.7285 | 25.7983 | 25.71138 | 25.88848 | 25.75098 | 25.79067 | 25.38189 |
| 52 | 28.13291 | 28.06517 | 28.35317 | 28.39239 | 28.47445 | 28.45136 | 28.20182 | 28.13819 | 28.35435 |
| 60 | 21.9 | 27.32997 | 27.31898 | 27.01808 | 27.35288 | 27.59733 | 27.46233 | 27.72058 | 27.81623 |
| 53 | 26.65322 | 26.85055 | 26.68365 | 26.60781 | 26.37992 | 26.5512 | 25.96444 | 21.9 | 26.31984 |
| 50 | 26.27711 | 26.65322 | 26.84661 | 26.7656 | 26.88704 | 26.70319 | 26.9038 | 26.85186 | 26.86903 |
| 37 | 21.9 | 21.9 | 26.98109 | 26.98794 | 26.57773 | 26.78694 | 26.76142 | 26.4187 | 26.29859 |
| 51 | 26.37151 | 26.5312 | 27.02602 | 27.09245 | 26.8197 | 26.98218 | 26.8136 | 26.90012 | 26.94369 |
| 47 | 26.41492 | 21.9 | 26.69166 | 26.70121 | 26.81348 | 26.94581 | 26.81115 | 26.85043 | 27.10099 |
| 35 | 25.43721 | 25.56189 | 25.54309 | 25.80275 | 26.1333 | 25.98661 | 25.80617 | 25.78212 | 25.79208 |
| 20 | 21.9 | 21.9 | 21.9 | 25.07594 | 21.9 | 24.7518 | 25.08784 | 24.998 | 21.9 |
| 27 | 21.9 | 26.02695 | 21.9 | 25.88353 | 21.9 | 21.9 | 26.18208 | 21.9 | 26.12906 |
| 26 | 21.9 | 21.9 | 26.29418 | 25.80408 | 21.9 | 25.61082 | 21.9 | 26.07678 | 25.74556 |
| 36 | 21.9 | 26.74983 | 26.3733 | 26.41696 | 26.50481 | 26.28408 | 26.31636 | 26.37162 | 26.79675 |
| 73 | 27.64437 | 27.84344 | 27.9651 | 27.9843 | 28.35359 | 28.40154 | 28.08578 | 28.0725 | 28.24186 |
| 32 | 27.78569 | 21.9 | 27.23197 | 21.9 | 26.96939 | 27.17969 | 26.54835 | 26.66899 | 26.81066 |
| 34 | 25.67998 | 21.9 | 25.72656 | 25.88502 | 26.17931 | 26.12989 | 25.93037 | 25.80683 | 26.01542 |
| 32 | 25.02035 | 24.67965 | 24.7364 | 24.68057 | 21.9 | 24.97154 | 24.98233 | 24.88362 | 24.5464 |
| 48 | 27.00624 | 27.04479 | 26.95538 | 27.02253 | 26.70899 | 26.72455 | 26.83629 | 27.06003 | 26.66166 |
| 34 | 25.92583 | 25.8704 | 26.17011 | 26.10266 | 26.34428 | 26.33703 | 21.9 | 26.19195 | 26.22861 |
| 33 | 25.85317 | 25.84398 | 25.71573 | 25.576 | 25.76068 | 21.9 | 21.9 | 25.40672 | 25.60544 |
| 33 | 25.95382 | 25.89876 | 26.40119 | 26.01028 | 25.964 | 26.41434 | 25.73838 | 25.89874 | 25.99306 |
| 25 | 26.01431 | 25.99968 | 25.85771 | 21.9 | 21.9 | 25.54047 | 25.48686 | 21.9 | 21.9 |
| 55 | 28.51612 | 28.46863 | 28.27635 | 28.33001 | 28.05667 | 28.02971 | 28.23576 | 28.0879 | 28.55003 |
| 24 | 21.9 | 21.9 | 25.77605 | 21.9 | 25.85638 | 26.02289 | 25.70654 | 21.9 | 25.77203 |
| 33 | 21.9 | 21.9 | 26.26157 | 25.7659 | 25.51572 | 25.43781 | 26.04587 | 26.08355 | 21.9 |
| 32 | 21.9 | 21.9 | 27.3362 | 27.25007 | 27.29351 | 27.1038 | 21.9 | 27.32732 | 26.6569 |
| 31 | 25.83116 | 21.9 | 26.03649 | 25.99535 | 21.9 | 25.82274 | 25.6767 | 25.69838 | 21.9 |
| 39 | 26.61077 | 26.54051 | 26.26265 | 26.31003 | 25.97224 | 21.9 | 21.9 | 25.38327 | 26.19492 |
| 40 | 26.49552 | 26.53472 | 26.0195 | 25.858 | 25.88874 | 25.97967 | 26.26311 | 26.21762 | 26.10103 |
| 37 | 25.62327 | 25.73079 | 25.9566 | 25.99951 | 26.0006 | 26.22188 | 26.0389 | 25.66253 | 26.0566 |
| 25 | 25.26924 | 25.28079 | 21.9 | 25.10446 | 21.9 | 25.1195 | 25.19046 | 21.9 | 24.77072 |
| 32 | 25.39002 | 21.9 | 25.66196 | 25.51563 | 25.67589 | 25.62689 | 21.9 | 25.68188 | 25.66033 |
| 1087 | 34.91434 | 34.9479 | 35.16923 | 35.20219 | 35.59821 | 35.5765 | 35.36055 | 35.29762 | 35.45381 |
| 125 | 28.93447 | 29.03939 | 28.52361 | 28.47243 | 28.34849 | 28.1931 | 28.13589 | 27.99199 | 27.87678 |
| 132 | 30.46392 | 30.55344 | 30.25597 | 30.24195 | 29.89822 | 29.81134 | 30.02062 | 30.05142 | 30.4334 |
| 70 | 27.32088 | 27.5666 | 27.32834 | 27.37592 | 27.82214 | 27.90558 | 27.58942 | 27.78182 | 28.16524 |
| 60 | 29.50306 | 29.49825 | 29.26289 | 29.16401 | 28.57105 | 28.30003 | 29.08575 | 29.00045 | 28.85878 |
| 68 | 27.91228 | 28.08314 | 27.65928 | 27.7599 | 27.79811 | 27.83623 | 28.11381 | 28.09124 | 28.12164 |
| 42 | 26.72495 | 26.79079 | 26.44302 | 26.5862 | 21.9 | 26.00277 | 26.76332 | 26.64485 | 26.44397 |
| 17 | 24.96009 | 25.0671 | 25.52106 | 25.46005 | 21.9 | 25.01495 | 21.9 | 21.9 | 21.9 |

| | | | | | | | | | |
|---|---|---|---|---|---|---|---|---|---|
| 20 | 25.41067 | 25.47187 | 21.9 | 25.60093 | 21.9 | 25.34201 | 21.9 | 21.9 | 21.9 |
| 28 | 26.83461 | 26.08627 | 26.2703 | 26.51977 | 26.22378 | 26.10781 | 25.74656 | 21.9 | 21.9 |
| 21 | 26.22909 | 26.13602 | 26.1896 | 25.97003 | 21.9 | 26.33982 | 25.99095 | 21.9 | 21.9 |
| 23 | 21.9 | 21.9 | 21.9 | 21.9 | 21.9 | 21.9 | 26.39795 | 26.34767 | 26.13206 |
| 26 | 21.9 | 21.9 | 21.9 | 21.9 | 21.9 | 21.9 | 26.06096 | 26.11696 | 26.38902 |
| 21 | 21.9 | 21.9 | 21.9 | 21.9 | 21.9 | 21.9 | 25.9574 | 25.91424 | 26.35773 |
| 24 | 21.9 | 21.9 | 21.9 | 21.9 | 21.9 | 21.9 | 26.28092 | 26.13954 | 25.85165 |
| 23 | 21.9 | 21.9 | 21.9 | 21.9 | 21.9 | 21.9 | 25.45054 | 25.35857 | 25.44969 |
| 24 | 26.15854 | 21.9 | 26.32749 | 21.9 | 21.9 | 21.9 | 26.31786 | 26.5411 | 26.54209 |
| 23 | 21.9 | 21.9 | 21.9 | 21.9 | 21.9 | 21.9 | 21.9 | 21.9 | 26.0402 |
| 15 | 21.9 | 21.9 | 21.9 | 21.9 | 21.9 | 21.9 | 21.9 | 25.53937 | 25.86029 |
| 15 | 21.9 | 21.9 | 21.9 | 21.9 | 21.9 | 21.9 | 21.9 | 25.73903 | 21.9 |
| 23 | 21.9 | 21.9 | 21.9 | 21.9 | 21.9 | 24.85913 | 25.00492 | 24.90656 | 24.84652 |
| 26 | 21.9 | 21.9 | 26.06893 | 21.9 | 26.24227 | 26.05188 | 26.26665 | 26.15426 | 26.54807 |
| 37 | 21.9 | 21.9 | 25.64105 | 21.9 | 26.30576 | 25.95797 | 25.97077 | 25.92209 | 26.35919 |
| 18 | 21.9 | 21.9 | 21.9 | 21.9 | 21.9 | 25.3765 | 25.32601 | 25.25778 | 25.19573 |
| 23 | 21.9 | 21.9 | 21.9 | 21.9 | 21.9 | 21.9 | 21.9 | 21.9 | 26.28031 |
| 20 | 21.9 | 21.9 | 21.9 | 21.9 | 21.9 | 21.9 | 24.79174 | 24.62862 | 23.9304 |
| 20 | 21.9 | 21.9 | 21.9 | 21.9 | 21.9 | 21.9 | 24.75858 | 24.73722 | 21.9 |
| 18 | 21.9 | 21.9 | 21.9 | 21.9 | 21.9 | 21.9 | 21.9 | 23.9581 | 21.9 |
| 19 | 21.9 | 21.9 | 21.9 | 21.9 | 21.9 | 21.9 | 21.9 | 23.43657 | 21.9 |
| 56 | 27.19582 | 27.04479 | 26.88472 | 27.27613 | 27.33339 | 27.202 | 27.28859 | 27.37865 | 27.33833 |
| 42 | 26.61175 | 26.57701 | 26.59648 | 26.57527 | 26.45073 | 26.29111 | 26.57989 | 26.55154 | 26.75494 |
| 51 | 28.46346 | 28.39832 | 28.48941 | 28.57308 | 28.54683 | 28.47534 | 28.11932 | 28.32775 | 28.30313 |
| 43 | 28.10674 | 28.12213 | 27.8117 | 27.67277 | 27.35136 | 27.30417 | 27.54834 | 27.3339 | 27.32113 |
| 63 | 29.08502 | 29.18695 | 29.12741 | 29.17688 | 29.34594 | 29.39671 | 28.61253 | 28.69856 | 28.91239 |
| 54 | 28.48693 | 28.53475 | 28.24803 | 28.44047 | 27.88983 | 27.93438 | 27.94637 | 27.91792 | 27.59056 |
| 62 | 30.12817 | 30.0855 | 29.73883 | 29.64458 | 29.77331 | 29.67069 | 30.0427 | 29.96251 | 30.00584 |
| 36 | 25.26667 | 25.50842 | 25.29425 | 25.39712 | 25.75774 | 25.60218 | 25.65739 | 25.50836 | 26.00727 |
| 59 | 28.09149 | 28.12213 | 28.18773 | 28.23859 | 28.86334 | 29.09152 | 28.62439 | 28.54834 | 29.09041 |
| 39 | 26.62282 | 26.68992 | 26.4833 | 26.30506 | 25.80602 | 25.69511 | 25.8674 | 25.82917 | 25.44925 |
| 17 | 21.9 | 21.9 | 21.9 | 21.9 | 24.77097 | 21.9 | 21.9 | 21.9 | 25.03855 |
| 28 | 26.88809 | 26.91878 | 21.9 | 21.9 | 26.88693 | 26.39653 | 26.93033 | 27.27133 | 27.19554 |
| 32 | 25.36082 | 25.33501 | 25.36636 | 21.9 | 25.44572 | 21.9 | 25.60105 | 25.71484 | 25.48809 |
| 32 | 26.39218 | 26.22821 | 26.16862 | 26.06443 | 26.79439 | 26.66641 | 26.60795 | 26.41036 | 26.78019 |
| 39 | 25.83976 | 25.6221 | 26.08064 | 26.10679 | 26.53099 | 26.43388 | 26.25467 | 26.1747 | 26.30419 |
| 40 | 25.65384 | 21.9 | 25.90431 | 25.86828 | 21.9 | 25.82813 | 26.056 | 25.88542 | 26.04421 |
| 39 | 26.59591 | 26.73738 | 26.78718 | 26.4715 | 25.85255 | 25.4853 | 26.14583 | 26.30784 | 25.60102 |
| 27 | 27.45325 | 21.9 | 21.9 | 21.9 | 21.9 | 21.9 | 21.9 | 21.9 | 26.80058 |
| 29 | 25.21201 | 25.12376 | 25.18098 | 25.08558 | 21.9 | 21.9 | 25.49957 | 25.30128 | 25.08582 |
| 45 | 27.36986 | 27.29903 | 27.67607 | 27.69558 | 27.54178 | 27.46381 | 27.53275 | 27.48331 | 27.3741 |
| 35 | 25.83007 | 21.9 | 25.66693 | 25.8619 | 26.45284 | 26.39408 | 26.08324 | 26.2143 | 26.854 |
| 27 | 25.68389 | 25.69469 | 21.9 | 21.9 | 25.91607 | 25.7453 | 25.43298 | 25.61835 | 25.75524 |
| 46 | 25.9554 | 25.97079 | 26.84709 | 27.05144 | 26.31771 | 26.36911 | 27.07816 | 27.04614 | 26.68646 |
| 56 | 27.14783 | 26.48244 | 27.16672 | 27.00174 | 27.8405 | 27.92801 | 27.1045 | 21.9 | 26.85852 |
| 26 | 27.42142 | 27.34002 | 21.9 | 21.9 | 27.73292 | 27.88844 | 27.45034 | 27.48884 | 27.23178 |
| 36 | 25.16339 | 25.0604 | 25.07345 | 21.9 | 25.42886 | 25.33463 | 25.80464 | 25.47392 | 25.26999 |

| | | | | | | | | | |
|---|---|---|---|---|---|---|---|---|---|
| 25 | 21.9 | 24.11402 | 21.9 | 21.9 | 24.29109 | 24.48646 | 21.9 | 24.57577 | 24.2594 |
| 18 | 21.9 | 21.9 | 21.9 | 21.9 | 21.9 | 24.36292 | 24.55456 | 21.9 | 24.61465 |
| 39 | 24.53136 | 21.9 | 24.89097 | 25.02732 | 25.20322 | 25.08485 | 25.5397 | 25.06665 | 24.78032 |
| 33 | 27.12288 | 26.9428 | 27.31225 | 21.9 | 27.35641 | 27.29088 | 27.66804 | 27.38402 | 27.56063 |
| 16 | 21.9 | 21.9 | 26.35716 | 21.9 | 21.9 | 26.69139 | 21.9 | 21.9 | 26.53788 |
| 37 | 26.28579 | 26.6102 | 26.10857 | 26.33887 | 26.51787 | 21.9 | 26.38436 | 26.58003 | 26.35676 |
| 21 | 21.9 | 21.9 | 21.9 | 21.9 | 26.32608 | 26.27725 | 21.9 | 21.9 | 26.37327 |
| 22 | 21.9 | 21.9 | 21.9 | 21.9 | 25.40243 | 25.32509 | 21.9 | 25.05908 | 25.22498 |
| 47 | 26.73596 | 21.9 | 26.72806 | 26.90598 | 27.02475 | 27.08821 | 26.66993 | 26.37713 | 27.12721 |
| 33 | 27.17636 | 27.19282 | 27.17474 | 21.9 | 27.11793 | 27.1333 | 21.9 | 26.13792 | 25.72676 |
| 27 | 25.71238 | 21.9 | 25.98388 | 25.94405 | 25.9387 | 21.9 | 21.9 | 26.46292 | 26.29449 |
| 22 | 21.9 | 24.42937 | 24.62606 | 21.9 | 21.9 | 24.1251 | 25.25028 | 24.77926 | 24.73753 |
| 61 | 26.964 | 26.88704 | 26.6554 | 26.81958 | 26.76344 | 26.80575 | 26.39403 | 26.71319 | 27.10509 |
| 21 | 27.11078 | 21.9 | 21.9 | 21.9 | 27.31077 | 27.30365 | 21.9 | 27.1519 | 27.25196 |
| 27 | 25.74627 | 21.9 | 25.23228 | 25.27416 | 25.20855 | 21.9 | 25.65952 | 21.9 | 25.43272 |
| 45 | 26.93798 | 27.31769 | 27.16787 | 27.11188 | 27.31337 | 27.54355 | 27.14589 | 27.0876 | 27.32569 |
| 59 | 27.62199 | 27.27444 | 27.86207 | 27.55918 | 27.56914 | 27.64609 | 27.49816 | 27.52148 | 27.65055 |
| 50 | 26.27242 | 26.57744 | 26.3897 | 26.33897 | 26.38961 | 26.43039 | 26.36105 | 26.26318 | 26.39165 |
| 27 | 27.07725 | 27.14141 | 27.0336 | 21.9 | 21.9 | 26.82966 | 21.9 | 26.45113 | 26.38133 |
| 18 | 25.68579 | 21.9 | 25.73433 | 25.33419 | 21.9 | 21.9 | 25.2463 | 21.9 | 21.9 |
| 20 | 21.9 | 25.17184 | 25.21323 | 25.33293 | 21.9 | 21.9 | 25.6086 | 21.9 | 25.62492 |
| 26 | 21.9 | 21.9 | 25.02509 | 25.07108 | 25.25918 | 24.99774 | 21.9 | 21.9 | 25.44354 |
| 33 | 25.65826 | 25.49115 | 25.37342 | 25.31365 | 25.18639 | 21.9 | 25.67001 | 25.40084 | 25.07883 |
| 645 | 33.19353 | 33.22734 | 33.41324 | 33.31716 | 33.51677 | 33.55899 | 33.19321 | 33.17056 | 33.47665 |
| 201 | 30.25372 | 30.23888 | 30.26035 | 30.2889 | 30.55197 | 30.47788 | 30.5089 | 30.52757 | 30.63556 |
| 141 | 28.75551 | 28.80455 | 29.04432 | 29.02944 | 29.29724 | 29.17479 | 28.98079 | 28.81586 | 29.25959 |
| 135 | 29.57136 | 29.58346 | 29.48081 | 29.59352 | 29.54021 | 29.59779 | 29.50101 | 29.42778 | 29.29 |
| 76 | 28.73925 | 28.54436 | 28.44229 | 28.22798 | 28.36953 | 28.3753 | 28.17907 | 28.0662 | 28.15417 |
| 52 | 26.3678 | 26.65513 | 26.95759 | 27.24291 | 21.9 | 26.87267 | 27.47383 | 27.53601 | 27.39885 |
| 62 | 27.1693 | 27.16346 | 26.94503 | 26.93798 | 27.06507 | 26.98772 | 27.16949 | 27.05466 | 27.2674 |
| 91 | 29.58413 | 29.51102 | 29.40765 | 29.65244 | 29.76637 | 29.95074 | 29.64077 | 29.61865 | 30.0444 |
| 35 | 21.9 | 21.9 | 25.9943 | 25.99213 | 26.16227 | 26.34115 | 26.52898 | 26.58749 | 26.63145 |
| 81 | 28.3593 | 28.34363 | 28.45736 | 28.37679 | 28.16428 | 28.24989 | 28.41446 | 28.35363 | 28.49975 |
| 35 | 21.9 | 25.77565 | 26.10239 | 25.96497 | 26.11126 | 26.38642 | 26.56581 | 26.11233 | 26.07936 |
| 22 | 21.9 | 21.9 | 21.9 | 21.9 | 21.9 | 21.9 | 21.9 | 25.03586 | 25.46728 |
| 57 | 25.9506 | 21.9 | 26.20337 | 26.15594 | 26.52527 | 26.35545 | 26.25505 | 25.86152 | 26.29213 |
| 37 | 21.9 | 25.889 | 26.07839 | 26.21959 | 26.1388 | 25.94878 | 26.31631 | 26.43055 | 26.51039 |
| 12 | 25.7546 | 25.62185 | 21.9 | 25.06829 | 21.9 | 25.05225 | 21.9 | 21.9 | 21.9 |
| 29 | 26.37257 | 26.46921 | 26.42448 | 26.30217 | 26.40523 | 26.37855 | 25.75753 | 21.9 | 25.77638 |
| 24 | 25.84223 | 25.73223 | 21.9 | 26.06583 | 25.6863 | 25.75208 | 21.9 | 21.9 | 21.9 |
| 25 | 24.99231 | 21.9 | 25.3306 | 21.9 | 21.9 | 25.02593 | 25.67333 | 25.48207 | 25.23312 |
| 23 | 21.9 | 21.9 | 21.9 | 21.9 | 21.9 | 21.9 | 25.93861 | 25.50021 | 25.41477 |
| 17 | 21.9 | 21.9 | 21.9 | 21.9 | 21.9 | 21.9 | 21.9 | 21.9 | 26.9941 |
| 23 | 21.9 | 21.9 | 21.9 | 21.9 | 26.09007 | 21.9 | 25.92793 | 26.06075 | 25.9798 |
| 22 | 21.9 | 21.9 | 21.9 | 21.9 | 21.9 | 24.9771 | 25.28196 | 25.41629 | 25.1658 |
| 15 | 21.9 | 21.9 | 21.9 | 21.9 | 21.9 | 21.9 | 21.9 | 25.12001 | 25.25252 |
| 19 | 21.9 | 21.9 | 21.9 | 21.9 | 26.11989 | 21.9 | 21.9 | 25.59779 | 25.86488 |

| | | | | | | | | | |
|---|---|---|---|---|---|---|---|---|---|
| 26 | 21.9 | 21.9 | 25.26227 | 25.20299 | 21.9 | 21.9 | 25.64535 | 25.6945 | 25.49106 |
| 23 | 25.73815 | 21.9 | 21.9 | 21.9 | 21.9 | 21.9 | 21.9 | 25.62987 | 25.63766 |
| 13 | 21.9 | 21.9 | 21.9 | 21.9 | 21.9 | 21.9 | 25.30065 | 25.09772 | 24.99053 |
| 22 | 21.9 | 21.9 | 21.9 | 21.9 | 21.9 | 25.86799 | 21.9 | 25.28535 | 25.98932 |
| 30 | 21.9 | 21.9 | 21.9 | 21.9 | 25.06114 | 24.49826 | 25.54451 | 25.43294 | 24.9875 |
| 20 | 21.9 | 21.9 | 21.9 | 21.9 | 24.81514 | 21.9 | 24.94793 | 21.9 | 25.12829 |
| 16 | 21.9 | 21.9 | 21.9 | 21.9 | 21.9 | 21.9 | 21.9 | 21.9 | 25.18261 |
| 11 | 21.9 | 21.9 | 21.9 | 21.9 | 21.9 | 21.9 | 21.9 | 21.9 | 26.11854 |
| 33 | 21.9 | 21.9 | 25.4688 | 21.9 | 25.80919 | 25.64392 | 25.58698 | 25.65734 | 25.93173 |
| 22 | 21.9 | 25.02018 | 21.9 | 21.9 | 21.9 | 21.9 | 21.9 | 25.03267 | 25.07243 |
| 18 | 21.9 | 21.9 | 21.9 | 21.9 | 24.121 | 24.05788 | 24.64867 | 24.80218 | 24.57398 |
| 20 | 25.89034 | 21.9 | 21.9 | 25.88974 | 21.9 | 21.9 | 26.93798 | 26.57687 | 21.9 |
| 18 | 21.9 | 21.9 | 21.9 | 21.9 | 21.9 | 21.9 | 21.9 | 21.9 | 25.62815 |
| 15 | 21.9 | 21.9 | 21.9 | 21.9 | 21.9 | 21.9 | 21.9 | 21.9 | 25.694 |
| 10 | 21.9 | 21.9 | 21.9 | 21.9 | 21.9 | 21.9 | 21.9 | 24.95136 | 21.9 |
| 17 | 21.9 | 21.9 | 21.9 | 21.9 | 21.9 | 21.9 | 24.35824 | 23.98746 | 21.9 |
| 20 | 21.9 | 21.9 | 21.9 | 21.9 | 21.9 | 21.9 | 21.9 | 24.62003 | 21.9 |
| 35 | 28.8003 | 28.83873 | 28.05961 | 27.85703 | 27.76781 | 27.91411 | 27.39395 | 27.23717 | 26.92886 |
| 28 | 26.0937 | 25.77115 | 25.43543 | 25.6676 | 25.60332 | 25.64287 | 25.35239 | 25.10593 | 21.9 |
| 20 | 25.22188 | 25.1888 | 25.04902 | 25.2712 | 21.9 | 24.93746 | 21.9 | 21.9 | 24.59571 |
| 21 | 21.9 | 25.46532 | 25.37474 | 25.65529 | 21.9 | 21.9 | 21.9 | 21.9 | 21.9 |
| 13 | 25.79268 | 25.80314 | 25.64091 | 21.9 | 21.9 | 21.9 | 21.9 | 21.9 | 21.9 |
| 52 | 26.90185 | 26.9706 | 27.36227 | 27.32217 | 27.85156 | 27.99335 | 27.57564 | 27.69611 | 27.91673 |
| 40 | 25.51286 | 25.57384 | 25.56628 | 25.55406 | 25.62531 | 25.4011 | 25.85331 | 25.7543 | 25.94177 |
| 35 | 25.67678 | 25.98103 | 25.70213 | 25.69917 | 25.65516 | 25.62095 | 26.028 | 25.96243 | 26.02337 |
| 39 | 26.95427 | 26.79699 | 26.818 | 26.9428 | 27.01457 | 27.05041 | 26.90976 | 26.71673 | 26.834 |
| 37 | 27.16048 | 27.14783 | 27.54488 | 27.55552 | 27.38936 | 27.45607 | 26.94235 | 27.34138 | 26.89724 |
| 16 | 21.9 | 21.9 | 21.9 | 21.9 | 21.9 | 21.9 | 21.9 | 24.23118 | 21.9 |
| 38 | 26.58463 | 26.77881 | 26.58032 | 26.7166 | 27.13183 | 27.26803 | 27.01414 | 26.90081 | 27.13848 |
| 33 | 24.87422 | 21.9 | 25.00045 | 21.9 | 25.4617 | 25.19178 | 25.82233 | 25.43447 | 25.28676 |
| 32 | 26.61133 | 26.63062 | 26.92411 | 26.72923 | 26.82031 | 26.57701 | 21.9 | 26.10312 | 25.99436 |
| 18 | 25.65908 | 21.9 | 25.58766 | 21.9 | 25.85493 | 25.83379 | 25.51605 | 21.9 | 21.9 |
| 21 | 21.9 | 21.9 | 21.9 | 21.9 | 21.9 | 25.66131 | 21.9 | 25.35659 | 21.9 |
| 20 | 21.9 | 21.9 | 21.9 | 21.9 | 26.32529 | 26.3948 | 26.22617 | 21.9 | 26.31669 |
| 15 | 21.9 | 21.9 | 25.52993 | 21.9 | 21.9 | 21.9 | 21.9 | 25.30911 | 25.31392 |
| 17 | 25.81453 | 21.9 | 26.30536 | 26.29156 | 21.9 | 26.10757 | 21.9 | 26.39049 | 21.9 |
| 37 | 21.9 | 25.48096 | 21.9 | 25.75898 | 26.38578 | 26.31048 | 25.86093 | 25.9638 | 26.356 |
| 30 | 27.38418 | 27.72676 | 27.17712 | 27.28039 | 26.5512 | 26.53111 | 26.9901 | 26.76104 | 26.90104 |
| 32 | 21.9 | 21.9 | 26.51479 | 26.24481 | 26.60837 | 26.7304 | 26.15698 | 26.72078 | 26.89342 |
| 25 | 25.2455 | 21.9 | 25.63672 | 25.47568 | 25.67096 | 25.54463 | 25.92931 | 25.99371 | 25.92762 |
| 30 | 27.56275 | 27.46248 | 27.59335 | 27.6985 | 27.53326 | 27.48316 | 27.76059 | 27.65915 | 27.69637 |
| 26 | 25.29042 | 25.13989 | 21.9 | 24.912 | 21.9 | 25.35128 | 25.18397 | 25.08946 | 25.08865 |
| 25 | 25.69782 | 21.9 | 25.43962 | 25.63467 | 25.16871 | 21.9 | 25.71615 | 25.26621 | 25.39424 |
| 19 | 24.96168 | 21.9 | 21.9 | 25.1352 | 21.9 | 21.9 | 25.84683 | 26.04785 | 25.59759 |
| 56 | 28.55914 | 28.59786 | 28.24032 | 28.57463 | 29.12416 | 28.92275 | 28.72012 | 28.72939 | 28.85258 |
| 37 | 27.6637 | 27.82548 | 27.69119 | 27.69358 | 21.9 | 28.03018 | 27.91741 | 28.02443 | 28.3135 |
| 26 | 25.23571 | 21.9 | 21.9 | 25.6826 | 21.9 | 25.58935 | 25.73469 | 21.9 | 25.71913 |

| | | | | | | | | | |
|---|---|---|---|---|---|---|---|---|---|
| 29 | 24.52761 | 24.80917 | 21.9 | 21.9 | 24.08594 | 24.38745 | 24.23768 | 24.72302 | 23.88688 |
| 30 | 26.46226 | 26.42933 | 21.9 | 26.43955 | 26.62701 | 26.52542 | 26.61751 | 26.81617 | 26.67559 |
| 10 | 21.9 | 21.9 | 21.9 | 21.9 | 24.77706 | 21.9 | 21.9 | 21.9 | 25.21712 |
| 18 | 25.59676 | 21.9 | 21.9 | 21.9 | 21.9 | 21.9 | 21.9 | 21.9 | 25.82573 |
| 28 | 25.63048 | 25.62868 | 25.49869 | 25.4545 | 25.80191 | 21.9 | 25.76686 | 25.47621 | 26.07735 |
| 22 | 25.81656 | 21.9 | 25.67269 | 26.02568 | 26.1983 | 26.15702 | 21.9 | 25.98011 | 26.34741 |
| 18 | 25.40698 | 21.9 | 21.9 | 21.9 | 21.9 | 25.48812 | 21.9 | 24.95899 | 21.9 |
| 15 | 21.9 | 21.9 | 21.9 | 24.44601 | 24.75786 | 21.9 | 21.9 | 24.38949 | 24.5779 |
| 27 | 21.9 | 21.9 | 26.20036 | 25.89389 | 21.9 | 26.76648 | 21.9 | 21.9 | 26.69246 |
| 21 | 25.62784 | 25.85567 | 25.42784 | 21.9 | 21.9 | 21.9 | 21.9 | 24.99813 | 21.9 |
| 43 | 28.32586 | 28.15059 | 28.29149 | 28.28441 | 27.69239 | 27.53934 | 28.83056 | 28.49278 | 27.89174 |
| 30 | 25.8298 | 25.98066 | 25.89382 | 25.94273 | 25.74953 | 25.84192 | 21.9 | 26.17245 | 26.02282 |
| 17 | 21.9 | 25.65168 | 21.9 | 25.50903 | 25.42297 | 21.9 | 21.9 | 25.46607 | 21.9 |
| 19 | 21.9 | 24.90058 | 25.68536 | 25.55254 | 21.9 | 25.6319 | 21.9 | 25.41661 | 21.9 |
| 19 | 21.9 | 26.67627 | 21.9 | 26.79872 | 21.9 | 21.9 | 21.9 | 21.9 | 21.9 |
| 61 | 32.22015 | 32.20408 | 32.38255 | 32.51223 | 32.73981 | 32.78101 | 32.54591 | 32.43214 | 32.68334 |
| 50 | 27.65513 | 27.61316 | 27.62164 | 27.49183 | 27.52761 | 27.12268 | 28.01691 | 27.66418 | 27.30234 |
| 29 | 25.40652 | 25.31972 | 21.9 | 25.48118 | 25.53875 | 25.41232 | 21.9 | 25.66286 | 25.56761 |
| 28 | 24.10294 | 24.04926 | 24.44386 | 24.72874 | 24.13923 | 24.16546 | 24.12668 | 24.46532 | 23.94619 |
| 24 | 24.97898 | 25.11886 | 21.9 | 21.9 | 25.1166 | 24.95841 | 25.35037 | 21.9 | 25.46329 |
| 23 | 24.64384 | 21.9 | 24.29256 | 24.90371 | 24.94517 | 21.9 | 21.9 | 24.95354 | 25.00475 |
| 19 | 21.9 | 25.77563 | 21.9 | 21.9 | 26.25117 | 26.27661 | 21.9 | 25.98804 | 25.90706 |
| 59 | 27.66926 | 27.77486 | 27.57701 | 27.75557 | 27.48347 | 27.45356 | 27.38903 | 27.63374 | 27.6704 |
| 121 | 27.69988 | 27.56587 | 27.5238 | 27.53897 | 27.59719 | 27.53616 | 28.24491 | 28.18985 | 27.88256 |
| 106 | 30.45836 | 30.38535 | 30.21535 | 30.21581 | 30.0817 | 30.08613 | 29.63905 | 29.83112 | 29.90009 |
| 260 | 35.19909 | 35.16923 | 34.87197 | 34.87119 | 34.68844 | 34.68839 | 34.77525 | 34.65848 | 34.42317 |
| 37 | 26.81543 | 21.9 | 27.09245 | 26.97136 | 21.9 | 21.9 | 27.11098 | 27.12238 | 27.43022 |
| 46 | 28.67017 | 28.77458 | 28.6076 | 28.71349 | 28.48796 | 28.49316 | 28.52021 | 28.54377 | 28.65178 |
| 60 | 28.08978 | 28.12218 | 27.6088 | 27.84134 | 27.88262 | 27.94659 | 27.28621 | 27.42654 | 27.72175 |
| 36 | 27.47189 | 27.27088 | 26.86041 | 21.9 | 21.9 | 26.97651 | 27.07102 | 27.05445 | 26.90438 |
| 37 | 26.48341 | 26.48792 | 26.44615 | 26.73686 | 26.4705 | 26.57273 | 26.44821 | 26.4946 | 26.64883 |
| 51 | 27.02041 | 27.22665 | 27.49549 | 27.54296 | 27.38204 | 27.50544 | 27.36102 | 27.37053 | 27.42005 |
| 25 | 21.9 | 21.9 | 21.9 | 21.9 | 21.9 | 21.9 | 25.51503 | 25.5946 | 25.36088 |
| 58 | 29.55923 | 29.54637 | 29.05734 | 29.14054 | 28.92799 | 28.87833 | 28.59999 | 28.63851 | 28.67122 |
| 51 | 27.903 | 28.07775 | 27.77838 | 27.85293 | 28.04051 | 27.58849 | 28.3682 | 27.7611 | 27.94854 |
| 22 | 27.09255 | 26.97618 | 27.07948 | 26.92139 | 26.80292 | 26.79687 | 26.29047 | 26.45725 | 26.29782 |
| 22 | 26.65854 | 26.82639 | 26.38528 | 26.45485 | 25.79907 | 21.9 | 21.9 | 21.9 | 21.9 |
| 17 | 21.9 | 21.9 | 21.9 | 21.9 | 21.9 | 21.9 | 21.9 | 24.735 | 21.9 |
| 17 | 21.9 | 28.3871 | 21.9 | 21.9 | 21.9 | 21.9 | 27.13975 | 27.48976 | 27.40755 |
| 36 | 21.9 | 21.9 | 28.95582 | 28.97921 | 29.65719 | 21.9 | 29.16701 | 29.0673 | 29.48577 |
| 20 | 21.9 | 21.9 | 21.9 | 25.28722 | 21.9 | 21.9 | 25.58434 | 25.63517 | 25.36666 |
| 24 | 21.9 | 21.9 | 21.9 | 21.9 | 26.13807 | 26.05086 | 26.18471 | 25.90793 | 26.42084 |
| 16 | 21.9 | 21.9 | 21.9 | 21.9 | 21.9 | 21.9 | 21.9 | 25.11878 | 25.11219 |
| 13 | 21.9 | 21.9 | 21.9 | 21.9 | 21.9 | 21.9 | 21.9 | 21.9 | 25.593 |
| 18 | 21.9 | 21.9 | 21.9 | 21.9 | 21.9 | 21.9 | 25.40165 | 21.9 | 25.50497 |
| 16 | 21.9 | 21.9 | 21.9 | 21.9 | 21.9 | 21.9 | 21.9 | 21.9 | 25.23692 |
| 18 | 21.9 | 21.9 | 21.9 | 21.9 | 21.9 | 21.9 | 24.92071 | 21.9 | 25.16239 |

| | | | | | | | | | |
|---|---|---|---|---|---|---|---|---|---|
| 18 | 21.9 | 21.9 | 21.9 | 21.9 | 21.9 | 21.9 | 25.40779 | 25.15062 | 24.7749 |
| 14 | 21.9 | 21.9 | 21.9 | 21.9 | 21.9 | 21.9 | 21.9 | 21.9 | 25.09852 |
| 33 | 21.9 | 21.9 | 21.9 | 21.9 | 21.9 | 28.24504 | 28.1037 | 21.9 | 28.73069 |
| 10 | 21.9 | 21.9 | 21.9 | 21.9 | 21.9 | 21.9 | 21.9 | 25.93166 | 25.96387 |
| 12 | 21.9 | 21.9 | 21.9 | 21.9 | 21.9 | 21.9 | 21.9 | 25.41809 | 21.9 |
| 12 | 21.9 | 21.9 | 21.9 | 21.9 | 21.9 | 21.9 | 25.34405 | 21.9 | 21.9 |
| 24 | 24.71905 | 21.9 | 21.9 | 25.00569 | 24.52731 | 21.9 | 24.90067 | 25.09499 | 24.54144 |
| 17 | 21.9 | 21.9 | 21.9 | 21.9 | 21.9 | 21.9 | 21.9 | 21.9 | 24.84675 |
| 15 | 21.9 | 21.9 | 21.9 | 21.9 | 21.9 | 21.9 | 24.61605 | 24.71858 | 21.9 |
| 26 | 26.48965 | 26.74085 | 26.66831 | 26.8435 | 26.38032 | 26.33393 | 26.21156 | 21.9 | 25.38584 |
| 30 | 25.48232 | 25.26446 | 25.32804 | 25.45048 | 25.56157 | 25.78654 | 25.78656 | 25.79459 | 26.00312 |
| 11 | 24.01767 | 21.9 | 24.5464 | 24.45149 | 21.9 | 21.9 | 21.9 | 21.9 | 21.9 |
| 13 | 21.9 | 21.9 | 21.9 | 21.9 | 21.9 | 21.9 | 25.95045 | 21.9 | 21.9 |
| 12 | 21.9 | 21.9 | 21.9 | 21.9 | 21.9 | 21.9 | 21.9 | 24.0721 | 21.9 |
| 16 | 25.66427 | 25.37806 | 25.28288 | 25.35894 | 21.9 | 21.9 | 21.9 | 24.5481 | 21.9 |
| 16 | 21.9 | 21.9 | 21.9 | 25.56092 | 21.9 | 21.9 | 21.9 | 25.9322 | 26.15163 |
| 21 | 21.9 | 21.9 | 25.24641 | 21.9 | 25.0304 | 21.9 | 25.31762 | 25.09531 | 25.21293 |
| 22 | 24.73242 | 24.7745 | 25.4264 | 25.06209 | 25.01457 | 25.14059 | 21.9 | 21.9 | 25.38189 |
| 20 | 24.95863 | 24.99473 | 21.9 | 24.91447 | 25.18185 | 21.9 | 25.07647 | 21.9 | 21.9 |
| 20 | 21.9 | 21.9 | 21.9 | 21.9 | 24.77766 | 21.9 | 21.9 | 24.93143 | 21.9 |
| 20 | 21.9 | 21.9 | 21.9 | 21.9 | 21.9 | 26.71751 | 21.9 | 26.54681 | 26.33964 |
| 26 | 21.9 | 25.19001 | 21.9 | 25.15054 | 25.48935 | 25.23136 | 25.60159 | 25.5474 | 25.17543 |
| 24 | 27.03234 | 26.91011 | 27.274 | 27.13761 | 26.70068 | 26.8276 | 26.39849 | 21.9 | 26.37941 |
| 22 | 24.95757 | 24.77811 | 21.9 | 21.9 | 21.9 | 21.9 | 24.7603 | 24.79679 | 24.89694 |
| 23 | 26.08058 | 26.10687 | 25.71424 | 25.73399 | 25.32828 | 25.52904 | 24.82357 | 21.9 | 21.9 |
| 16 | 21.9 | 25.64546 | 21.9 | 21.9 | 25.49026 | 21.9 | 25.76479 | 25.66397 | 25.29677 |
| 37 | 25.7474 | 25.73859 | 25.37372 | 25.40058 | 21.9 | 25.33484 | 25.75888 | 26.07349 | 26.08894 |
| 24 | 21.9 | 25.97176 | 21.9 | 25.83629 | 26.82007 | 26.52486 | 26.1888 | 26.08671 | 26.31709 |
| 28 | 25.81719 | 25.83222 | 21.9 | 25.97795 | 26.19811 | 26.22749 | 26.3169 | 26.28821 | 26.40241 |
| 27 | 21.9 | 21.9 | 25.92139 | 21.9 | 26.22847 | 26.43294 | 21.9 | 26.26324 | 26.53228 |
| 14 | 21.9 | 21.9 | 25.92753 | 21.9 | 21.9 | 21.9 | 21.9 | 21.9 | 21.9 |
| 19 | 21.9 | 21.9 | 27.3736 | 27.38599 | 26.93674 | 26.94827 | 21.9 | 21.9 | 26.30294 |
| 22 | 26.61667 | 21.9 | 25.85065 | 21.9 | 21.9 | 25.92897 | 26.04483 | 26.31069 | 26.29784 |
| 20 | 21.9 | 21.9 | 21.9 | 27.24472 | 27.58728 | 27.6809 | 27.3616 | 21.9 | 27.43404 |
| 14 | 21.9 | 21.9 | 21.9 | 21.9 | 21.9 | 25.61397 | 21.9 | 26.09085 | 21.9 |
| 36 | 25.77748 | 25.70949 | 26.12595 | 26.30273 | 25.85707 | 26.17623 | 25.92162 | 26.04272 | 25.29211 |
| 43 | 26.90953 | 26.95604 | 26.88239 | 26.85946 | 26.53551 | 26.16879 | 26.49296 | 26.57204 | 26.71385 |
| 20 | 21.9 | 21.9 | 24.37941 | 24.30779 | 23.72702 | 23.96049 | 21.9 | 21.9 | 23.42829 |
| 24 | 21.9 | 21.9 | 24.91889 | 24.56314 | 21.9 | 21.9 | 24.37749 | 24.27711 | 21.9 |
| 23 | 25.14581 | 24.96288 | 21.9 | 21.9 | 25.53519 | 25.42784 | 21.9 | 25.33787 | 25.63179 |
| 18 | 25.48038 | 21.9 | 25.45366 | 21.9 | 21.9 | 24.91534 | 25.76636 | 25.69448 | 21.9 |
| 33 | 26.16083 | 21.9 | 26.25185 | 26.63851 | 26.66438 | 26.75824 | 26.27739 | 26.13749 | 26.26821 |
| 21 | 21.9 | 21.9 | 24.55737 | 24.28008 | 24.45876 | 24.32622 | 24.87051 | 21.9 | 24.30626 |
| 29 | 26.63989 | 26.57242 | 26.69073 | 26.72312 | 26.72767 | 26.57629 | 21.9 | 27.01914 | 26.9223 |
| 26 | 26.48216 | 21.9 | 26.0827 | 26.21494 | 26.06439 | 25.85198 | 25.95551 | 21.9 | 25.37888 |
| 30 | 26.52686 | 26.61863 | 26.24943 | 26.00232 | 25.90074 | 26.16548 | 26.26842 | 26.12159 | 26.32119 |
| 17 | 21.9 | 25.08436 | 21.9 | 25.21974 | 21.9 | 21.9 | 21.9 | 25.48925 | 24.59457 |

| | | | | | | | | | |
|---|---|---|---|---|---|---|---|---|---|
| 19 | 25.52689 | 26.16315 | 21.9 | 21.9 | 21.9 | 21.9 | 21.9 | 21.9 | 25.41931 |
| 23 | 21.9 | 21.9 | 21.9 | 26.11738 | 26.45955 | 21.9 | 21.9 | 21.9 | 25.52205 |
| 30 | 28.23293 | 28.41325 | 28.48938 | 28.13384 | 28.16476 | 28.10559 | 28.45124 | 28.53579 | 27.8665 |
| 21 | 25.93713 | 25.94092 | 25.86828 | 21.9 | 21.9 | 26.5188 | 21.9 | 21.9 | 26.32468 |
| 20 | 21.9 | 24.84542 | 25.62567 | 25.63923 | 25.09229 | 21.9 | 26.11825 | 25.15023 | 25.47936 |
| 24 | 25.97388 | 21.9 | 26.38431 | 26.21887 | 26.21544 | 26.3672 | 21.9 | 26.40626 | 26.5337 |
| 22 | 26.15996 | 21.9 | 26.34216 | 21.9 | 26.93213 | 26.93337 | 26.4773 | 21.9 | 26.94123 |
| 1098 | 32.417 | 32.43827 | 32.55817 | 32.61028 | 32.22929 | 32.22165 | 32.84885 | 32.84771 | 32.67873 |
| 202 | 29.58264 | 29.67277 | 29.68279 | 29.79548 | 29.97815 | 29.88847 | 29.70048 | 29.80341 | 29.94321 |
| 82 | 27.18858 | 27.34858 | 27.08335 | 27.1205 | 27.43484 | 27.37136 | 27.76186 | 27.81403 | 27.73899 |
| 72 | 26.87783 | 26.3917 | 27.1333 | 27.13946 | 27.35262 | 27.39395 | 27.54598 | 27.28586 | 27.57744 |
| 86 | 30.29483 | 30.36494 | 30.26338 | 30.40497 | 30.43619 | 30.33777 | 30.46285 | 30.46558 | 30.42972 |
| 124 | 29.72697 | 29.79806 | 29.92153 | 29.83411 | 29.97268 | 29.80982 | 29.60899 | 29.72369 | 29.8217 |
| 60 | 26.88902 | 26.74445 | 26.98316 | 26.93775 | 26.66112 | 26.78007 | 26.74804 | 26.56458 | 26.68939 |
| 28 | 21.9 | 27.32027 | 26.71306 | 21.9 | 21.9 | 26.48582 | 27.23398 | 21.9 | 26.50482 |
| 30 | 25.85605 | 21.9 | 25.93789 | 26.17056 | 26.12965 | 26.41904 | 26.19669 | 21.9 | 26.54562 |
| 16 | 21.9 | 21.9 | 21.9 | 21.9 | 21.9 | 21.9 | 21.9 | 25.0062 | 25.04577 |
| 25 | 21.9 | 24.86344 | 25.44853 | 25.37029 | 25.4472 | 25.5115 | 26.02912 | 25.73304 | 26.06911 |
| 23 | 27.40268 | 21.9 | 27.68204 | 27.73718 | 28.20074 | 27.9011 | 27.96917 | 28.45584 | 28.27173 |
| 24 | 24.99934 | 24.91825 | 21.9 | 24.68688 | 24.8769 | 24.72915 | 24.77641 | 24.85067 | 24.644 |
| 22 | 26.21219 | 26.04955 | 25.93683 | 25.74617 | 25.52788 | 25.5528 | 25.6106 | 25.53118 | 21.9 |
| 20 | 24.67922 | 21.9 | 24.692 | 21.9 | 21.9 | 21.9 | 24.87051 | 24.73226 | 24.61172 |
| 17 | 21.9 | 21.9 | 21.9 | 21.9 | 21.9 | 21.9 | 24.72931 | 21.9 | 23.94744 |
| 17 | 21.9 | 21.9 | 21.9 | 21.9 | 21.9 | 21.9 | 27.82214 | 21.9 | 28.04848 |
| 13 | 21.9 | 21.9 | 21.9 | 21.9 | 21.9 | 21.9 | 25.45143 | 21.9 | 25.29869 |
| 17 | 27.23553 | 21.9 | 21.9 | 21.9 | 21.9 | 21.9 | 21.9 | 26.36621 | 26.28798 |
| 14 | 21.9 | 21.9 | 21.9 | 21.9 | 25.78002 | 21.9 | 25.08152 | 25.26524 | 25.55681 |
| 23 | 21.9 | 21.9 | 21.9 | 25.63104 | 21.9 | 21.9 | 25.69096 | 25.29485 | 25.92173 |
| 17 | 21.9 | 21.9 | 21.9 | 21.9 | 21.9 | 21.9 | 21.9 | 25.76109 | 25.42243 |
| 18 | 21.9 | 21.9 | 21.9 | 21.9 | 21.9 | 21.9 | 21.9 | 25.54318 | 21.9 |
| 15 | 21.9 | 21.9 | 21.9 | 21.9 | 21.9 | 21.9 | 25.43253 | 25.61215 | 21.9 |
| 13 | 21.9 | 21.9 | 21.9 | 21.9 | 21.9 | 21.9 | 21.9 | 21.9 | 21.9 |
| 23 | 21.9 | 25.2325 | 21.9 | 25.29334 | 25.21238 | 25.30762 | 25.53288 | 25.74134 | 25.90796 |
| 9 | 21.9 | 21.9 | 21.9 | 21.9 | 21.9 | 21.9 | 21.9 | 21.9 | 25.00607 |
| 10 | 21.9 | 21.9 | 21.9 | 21.9 | 21.9 | 21.9 | 21.9 | 24.41532 | 24.11482 |
| 18 | 21.9 | 21.9 | 21.9 | 21.9 | 23.80794 | 21.9 | 21.9 | 25.18408 | 24.22825 |
| 9 | 21.9 | 21.9 | 21.9 | 21.9 | 21.9 | 21.9 | 21.9 | 24.4302 | 24.49735 |
| 15 | 21.9 | 21.9 | 21.9 | 21.9 | 21.9 | 21.9 | 21.9 | 24.13899 | 24.43294 |
| 12 | 21.9 | 21.9 | 21.9 | 21.9 | 21.9 | 21.9 | 21.9 | 21.9 | 21.9 |
| 11 | 21.9 | 21.9 | 21.9 | 25.0878 | 25.117 | 24.87933 | 21.9 | 21.9 | 21.9 |
| 11 | 25.40578 | 21.9 | 25.14484 | 25.30529 | 21.9 | 21.9 | 21.9 | 21.9 | 21.9 |
| 12 | 21.9 | 21.9 | 21.9 | 25.52492 | 26.22523 | 25.66885 | 21.9 | 21.9 | 21.9 |
| 27 | 29.96788 | 30.06612 | 30.31797 | 30.35058 | 30.622 | 30.62017 | 30.56901 | 30.61754 | 30.72891 |
| 29 | 28.46182 | 28.32775 | 28.34121 | 28.33104 | 27.85174 | 27.97202 | 27.7057 | 27.9433 | 27.46069 |
| 21 | 22.98916 | 23.10506 | 21.9 | 22.84827 | 22.09188 | 21.9 | 21.9 | 21.90338 | 21.9 |
| 12 | 21.9 | 21.9 | 21.9 | 21.9 | 21.9 | 21.9 | 23.97456 | 21.9 | 21.9 |
| 10 | 21.9 | 21.9 | 21.9 | 21.9 | 21.9 | 21.9 | 21.9 | 23.57808 | 23.4902 |

| | | | | | | | | | |
|---|---|---|---|---|---|---|---|---|---|
| 19 | 21.9 | 21.9 | 21.9 | 21.9 | 21.9 | 21.9 | 23.83827 | 21.9 | 23.14375 |
| 18 | 25.99565 | 21.9 | 26.08093 | 25.98205 | 25.94925 | 25.85272 | 21.9 | 21.9 | 25.53686 |
| 26 | 26.91319 | 26.80747 | 27.09547 | 27.15441 | 27.25196 | 27.42542 | 26.68177 | 26.84182 | 26.80427 |
| 14 | 21.9 | 21.9 | 21.9 | 21.9 | 24.65829 | 21.9 | 24.87455 | 24.63389 | 24.92615 |
| 25 | 21.9 | 21.9 | 21.9 | 23.83519 | 24.51184 | 24.09402 | 24.71743 | 23.81049 | 23.96605 |
| 9 | 21.9 | 21.9 | 21.9 | 21.9 | 23.20292 | 21.9 | 21.9 | 23.1742 | 21.9 |
| 15 | 26.24124 | 21.9 | 21.9 | 21.9 | 21.9 | 21.9 | 25.53017 | 25.36149 | 21.9 |
| 30 | 25.68752 | 25.7139 | 25.90642 | 25.6709 | 25.99194 | 26.12033 | 26.24102 | 21.9 | 25.85105 |
| 19 | 24.46351 | 24.4159 | 21.9 | 21.9 | 24.53712 | 21.9 | 21.9 | 24.63931 | 25.05565 |
| 25 | 26.91125 | 26.97323 | 26.59662 | 26.81323 | 27.09265 | 27.09658 | 27.03318 | 27.13252 | 26.97891 |
| 25 | 27.49968 | 27.47507 | 27.24681 | 27.37111 | 27.41135 | 27.31008 | 27.165 | 27.05972 | 21.9 |
| 12 | 21.9 | 21.9 | 21.9 | 21.9 | 21.9 | 23.49875 | 21.9 | 23.35111 | 21.9 |
| 22 | 21.9 | 21.9 | 28.45869 | 28.63045 | 28.50074 | 21.9 | 27.89579 | 28.09109 | 28.87906 |
| 10 | 21.9 | 21.9 | 21.9 | 21.9 | 25.60597 | 25.57228 | 25.47568 | 25.57231 | 25.64111 |
| 13 | 22.96326 | 21.9 | 21.9 | 21.9 | 21.9 | 21.9 | 21.9 | 23.13356 | 21.9 |
| 16 | 21.9 | 21.9 | 21.9 | 24.39774 | 21.9 | 21.9 | 24.06349 | 21.9 | 21.9 |
| 22 | 24.87793 | 24.99684 | 24.88953 | 25.11064 | 21.9 | 21.9 | 21.9 | 25.64295 | 25.15511 |
| 14 | 24.30814 | 24.19317 | 24.30159 | 21.9 | 21.9 | 21.9 | 24.21852 | 21.9 | 24.04126 |
| 24 | 25.37504 | 25.47274 | 25.7129 | 25.94219 | 25.16611 | 21.9 | 25.23725 | 25.51581 | 21.9 |
| 21 | 25.54475 | 25.75585 | 25.51674 | 25.48582 | 21.9 | 21.9 | 25.63481 | 21.9 | 25.26324 |
| 24 | 21.9 | 25.51656 | 21.9 | 25.22262 | 25.37743 | 25.47945 | 21.9 | 26.11361 | 25.855 |
| 18 | 21.9 | 24.91333 | 21.9 | 25.22417 | 25.1043 | 25.12001 | 25.61801 | 25.8593 | 25.30626 |
| 15 | 25.28051 | 25.2467 | 25.50503 | 25.18044 | 21.9 | 21.9 | 21.9 | 21.9 | 25.67917 |
| 15 | 21.9 | 21.9 | 21.9 | 25.44676 | 26.23087 | 25.9052 | 21.9 | 21.9 | 25.64902 |

| TDP_22 | TDP_31 | TDP_32 |
|---|---|---|
| 30.71743 | 30.64367 | 30.59602 |
| 32.43882 | 32.39287 | 32.37695 |
| 30.65267 | 30.71071 | 30.6937 |
| 31.21193 | 31.11117 | 31.105 |
| 35.64871 | 35.54317 | 35.57009 |
| 30.82449 | 30.84401 | 30.8162 |
| 28.67411 | 28.64465 | 28.59174 |
| 30.71244 | 30.47044 | 30.46811 |
| 34.30082 | 34.38497 | 34.3867 |
| 32.04905 | 31.96082 | 31.96454 |
| 34.70079 | 34.65379 | 34.67303 |
| 31.47271 | 31.2594 | 31.34768 |
| 29.03297 | 29.25908 | 29.24148 |
| 33.65694 | 33.76327 | 33.73671 |
| 31.58978 | 31.51319 | 31.5601 |
| 33.12765 | 32.86916 | 32.91634 |
| 35.24516 | 35.11388 | 35.13457 |
| 31.34968 | 31.46022 | 31.40963 |
| 28.37509 | 28.2744 | 28.10604 |
| 31.6554 | 31.53274 | 31.53473 |
| 32.20253 | 32.14998 | 32.14883 |
| 35.11501 | 35.36951 | 35.31649 |
| 32.87173 | 32.81572 | 32.84474 |
| 28.7505 | 28.72764 | 28.69312 |
| 29.93775 | 30.1294 | 30.0971 |
| 28.74606 | 28.77772 | 28.76364 |
| 33.19584 | 33.47641 | 33.495 |
| 32.88653 | 32.84019 | 32.83859 |
| 30.39805 | 30.3688 | 30.40314 |
| 32.18042 | 32.15684 | 32.174 |
| 33.21986 | 33.34069 | 33.32938 |
| 31.57651 | 31.67501 | 31.69975 |
| 29.44219 | 29.54449 | 29.50794 |
| 31.66102 | 31.69843 | 31.65787 |
| 31.39897 | 31.27175 | 31.27869 |
| 30.49924 | 30.37306 | 30.38956 |
| 35.62824 | 35.6627 | 35.59304 |
| 32.26024 | 32.24935 | 32.26939 |
| 33.25307 | 33.29049 | 33.31878 |
| 29.89727 | 30.29044 | 30.24524 |
| 30.57651 | 30.63513 | 30.62305 |
| 30.52645 | 30.52356 | 30.48673 |
| 29.42138 | 29.47833 | 29.51501 |
| 35.17333 | 35.19546 | 35.16885 |
| 33.3956 | 33.29338 | 33.29296 |
| 33.6548 | 33.66342 | 33.70594 |

| | | |
|---|---|---|
| 34.64485 | 34.58251 | 34.6035 |
| 29.9955 | 30.00985 | 29.96348 |
| 32.38577 | 32.47598 | 32.49543 |
| 31.89638 | 31.95116 | 31.87782 |
| 29.15523 | 29.05152 | 29.01534 |
| 35.43499 | 35.26929 | 35.25694 |
| 31.76561 | 31.6501 | 31.64762 |
| 29.75141 | 29.71825 | 29.75673 |
| 28.78572 | 28.76844 | 28.78066 |
| 32.45672 | 32.48008 | 32.54181 |
| 27.306 | 27.46466 | 27.67748 |
| 32.95139 | 33.07731 | 33.07395 |
| 32.88057 | 32.98677 | 32.95177 |
| 32.54172 | 32.52641 | 32.548 |
| 29.76812 | 29.7808 | 29.9011 |
| 29.2457 | 29.20783 | 29.22548 |
| 28.07567 | 28.32419 | 28.2863 |
| 33.25575 | 33.34096 | 33.43004 |
| 30.00477 | 30.01943 | 30.06445 |
| 30.32131 | 30.35909 | 30.46752 |
| 31.40923 | 31.46022 | 31.50337 |
| 32.50904 | 32.43869 | 32.43219 |
| 34.06664 | 34.09616 | 34.09914 |
| 33.84146 | 33.8078 | 33.8574 |
| 28.15436 | 27.61547 | 28.33837 |
| 29.86603 | 30.02881 | 30.02789 |
| 32.04866 | 32.10506 | 32.08882 |
| 28.29553 | 28.40852 | 28.35431 |
| 30.79452 | 30.73336 | 30.75119 |
| 29.9279 | 29.76104 | 29.7486 |
| 32.50524 | 32.47515 | 32.49188 |
| 27.54569 | 27.71024 | 27.58011 |
| 31.30965 | 31.29958 | 31.35048 |
| 29.0358 | 29.07439 | 28.92717 |
| 28.59751 | 28.57004 | 28.51923 |
| 33.70203 | 33.63871 | 33.64635 |
| 32.32852 | 32.08971 | 32.12012 |
| 29.66794 | 29.619 | 29.65337 |
| 28.68241 | 28.63291 | 28.75089 |
| 31.47087 | 31.5819 | 31.53301 |
| 34.48855 | 34.36607 | 34.39017 |
| 30.17379 | 30.17807 | 30.19037 |
| 33.00174 | 32.78144 | 32.74927 |
| 32.19019 | 32.24337 | 32.2494 |
| 29.20585 | 29.17999 | 29.25575 |
| 28.41107 | 28.47727 | 28.43241 |
| 31.53403 | 31.48308 | 31.45728 |

| | | |
|---|---|---|
| 28.90948 | 28.89316 | 28.8783 |
| 31.70743 | 31.8385 | 31.84771 |
| 28.88719 | 28.8864 | 28.77357 |
| 33.87774 | 33.89555 | 33.8932 |
| 28.86641 | 28.95444 | 28.85896 |
| 31.74347 | 31.68253 | 31.67447 |
| 28.20247 | 28.08238 | 28.31631 |
| 30.51333 | 30.36693 | 30.40578 |
| 31.05446 | 31.26024 | 31.29302 |
| 32.32217 | 32.39422 | 32.35137 |
| 29.53384 | 29.65579 | 29.68154 |
| 31.57032 | 31.39494 | 31.40314 |
| 28.16581 | 28.04109 | 28.15151 |
| 28.77706 | 28.68398 | 28.60456 |
| 28.85754 | 28.75796 | 28.83638 |
| 28.69591 | 28.61817 | 28.67576 |
| 29.79388 | 29.68866 | 29.8063 |
| 32.20618 | 32.12321 | 32.09031 |
| 30.13333 | 30.18553 | 30.26461 |
| 29.21773 | 29.23761 | 29.28098 |
| 30.86452 | 31.06027 | 31.04316 |
| 30.07086 | 30.00812 | 29.95574 |
| 29.21213 | 29.12374 | 29.22887 |
| 29.49223 | 29.32939 | 29.35945 |
| 30.00932 | 30.10974 | 30.17105 |
| 31.76588 | 31.89103 | 31.91335 |
| 28.1436 | 28.46447 | 28.43754 |
| 33.01032 | 33.06742 | 33.06378 |
| 28.51284 | 28.41187 | 28.45552 |
| 33.11818 | 33.23292 | 33.22087 |
| 30.70512 | 30.80078 | 30.73376 |
| 28.11877 | 28.209 | 28.05046 |
| 27.71954 | 27.5819 | 27.55355 |
| 29.45352 | 29.66751 | 29.68781 |
| 32.4053 | 32.37909 | 32.38718 |
| 29.84676 | 29.83462 | 29.80766 |
| 28.1693 | 28.09325 | 28.03013 |
| 31.64358 | 31.72346 | 31.72102 |
| 30.28151 | 30.33724 | 30.3198 |
| 29.41526 | 29.38529 | 29.42748 |
| 32.69769 | 32.66989 | 32.7073 |
| 33.79672 | 33.8535 | 33.81669 |
| 29.34297 | 29.17624 | 29.30811 |
| 30.09043 | 30.14407 | 30.17403 |
| 29.63317 | 29.55684 | 29.65272 |
| 29.22552 | 29.21967 | 29.25201 |
| 30.28956 | 30.34921 | 30.32582 |

| | | |
|---|---|---|
| 31.25502 | 31.14504 | 31.17296 |
| 28.59491 | 28.52686 | 28.55413 |
| 29.16509 | 29.01233 | 29.09373 |
| 31.08518 | 31.04225 | 31.10143 |
| 32.90967 | 32.71231 | 32.68981 |
| 27.88134 | 27.51773 | 27.78419 |
| 28.93129 | 28.84643 | 28.83105 |
| 34.22669 | 34.14156 | 34.09333 |
| 29.56805 | 29.44819 | 29.362 |
| 31.25682 | 31.14145 | 31.14023 |
| 30.45258 | 30.58459 | 30.55664 |
| 30.00236 | 29.76202 | 29.73615 |
| 28.89076 | 29.04534 | 29.013 |
| 29.53918 | 29.62064 | 29.58638 |
| 32.43924 | 32.38193 | 32.34599 |
| 28.18494 | 28.34532 | 28.4763 |
| 28.3885 | 28.2548 | 28.27431 |
| 28.53934 | 28.34448 | 28.37993 |
| 29.13193 | 29.21829 | 29.20786 |
| 33.12596 | 32.97565 | 33.02254 |
| 33.59957 | 33.64216 | 33.64817 |
| 26.43194 | 26.61049 | 26.69896 |
| 28.41175 | 28.304 | 28.31756 |
| 27.14696 | 27.35043 | 27.2724 |
| 29.71142 | 29.68851 | 29.67995 |
| 29.77477 | 29.79757 | 29.78143 |
| 28.11719 | 28.10614 | 28.19676 |
| 30.98502 | 30.86555 | 30.94328 |
| 30.36672 | 30.23569 | 30.25237 |
| 30.91164 | 31.05692 | 31.09578 |
| 31.33202 | 31.36625 | 31.44781 |
| 31.76422 | 31.74783 | 31.75581 |
| 31.38355 | 31.3851 | 31.42412 |
| 29.12689 | 29.09109 | 28.94053 |
| 30.58217 | 30.73045 | 30.70116 |
| 30.29055 | 30.41418 | 30.45875 |
| 30.60911 | 30.55335 | 30.58889 |
| 32.70664 | 32.64089 | 32.65775 |
| 29.3638 | 29.28884 | 29.27979 |
| 28.04796 | 28.43928 | 28.35237 |
| 26.90346 | 26.97257 | 27.0531 |
| 28.08116 | 28.19046 | 28.20373 |
| 30.82563 | 30.89468 | 30.98196 |
| 28.43046 | 28.4865 | 28.4245 |
| 28.89415 | 29.00608 | 28.95233 |
| 30.04062 | 29.95629 | 29.92025 |
| 30.32667 | 30.20466 | 30.16171 |

| | | |
|---|---|---|
| 28.39411 | 28.40694 | 28.51322 |
| 31.17468 | 31.21321 | 31.23951 |
| 33.92203 | 33.95714 | 34.01355 |
| 30.1111 | 30.05271 | 30.02406 |
| 28.9032 | 28.93222 | 28.95208 |
| 27.58649 | 27.2497 | 27.34672 |
| 26.74471 | 25.9322 | 26.27094 |
| 28.97965 | 28.92167 | 28.86334 |
| 28.94604 | 28.95419 | 28.98278 |
| 28.32135 | 28.26378 | 28.28388 |
| 29.91399 | 29.85168 | 29.90842 |
| 30.49286 | 30.57218 | 30.59031 |
| 28.24527 | 28.1812 | 28.14472 |
| 29.42364 | 29.39881 | 29.47902 |
| 29.1887 | 29.41306 | 29.41022 |
| 28.54562 | 28.89087 | 28.95435 |
| 27.80907 | 28.21299 | 28.12913 |
| 29.25935 | 29.32363 | 29.34011 |
| 28.16749 | 28.0379 | 27.94737 |
| 30.38257 | 30.39724 | 30.46918 |
| 27.95505 | 27.81726 | 27.90397 |
| 28.18952 | 28.1804 | 28.16825 |
| 29.06756 | 29.29099 | 29.24254 |
| 30.92167 | 30.94531 | 30.98739 |
| 32.11353 | 32.22625 | 32.2106 |
| 31.22774 | 31.52631 | 31.52967 |
| 29.13129 | 29.32165 | 29.38086 |
| 29.21041 | 29.22219 | 29.17243 |
| 28.52683 | 28.65291 | 28.77797 |
| 29.12655 | 28.97158 | 28.99892 |
| 28.70556 | 28.47247 | 28.42334 |
| 27.62909 | 27.45591 | 27.49267 |
| 29.55775 | 29.51072 | 29.43048 |
| 30.42101 | 30.41074 | 30.46772 |
| 27.17522 | 27.00089 | 26.86231 |
| 29.07745 | 28.97943 | 28.98387 |
| 29.57497 | 29.60502 | 29.65342 |
| 27.16077 | 27.29421 | 27.44387 |
| 28.9989 | 29.00217 | 28.99167 |
| 28.09813 | 27.94854 | 28.07526 |
| 30.62741 | 30.70866 | 30.77188 |
| 28.48147 | 28.50051 | 28.46431 |
| 28.42266 | 28.51443 | 28.54536 |
| 28.61663 | 28.55197 | 28.41869 |
| 29.19418 | 29.12687 | 29.11272 |
| 27.7673 | 27.83738 | 27.86644 |
| 28.10983 | 28.14399 | 28.20611 |

| | | |
|---|---|---|
| 34.25807 | 34.49154 | 34.49041 |
| 30.53399 | 30.72314 | 30.69528 |
| 33.06558 | 32.76888 | 32.82684 |
| 29.0001 | 28.95696 | 28.94564 |
| 31.46324 | 31.56256 | 31.55783 |
| 26.2669 | 21.9 | 26.12293 |
| 30.56846 | 31.34085 | 30.42962 |
| 31.12145 | 31.10581 | 31.11527 |
| 27.65274 | 27.57751 | 27.4708 |
| 27.87391 | 27.88547 | 27.89302 |
| 30.23455 | 30.26729 | 30.27487 |
| 29.13868 | 29.03089 | 28.92996 |
| 32.31592 | 32.29644 | 32.33421 |
| 27.41587 | 27.44782 | 27.33134 |
| 26.64238 | 27.09798 | 26.85186 |
| 27.3822 | 27.57449 | 27.53185 |
| 28.58935 | 28.53501 | 28.55025 |
| 30.72655 | 30.83311 | 30.79622 |
| 33.26514 | 33.42142 | 33.41324 |
| 28.53364 | 28.59117 | 28.60505 |
| 29.99402 | 30.00745 | 30.09043 |
| 29.14914 | 29.24819 | 29.32489 |
| 31.49247 | 31.46879 | 31.42557 |
| 29.26669 | 29.21605 | 29.30839 |
| 27.77744 | 27.76497 | 27.7055 |
| 30.5888 | 30.5802 | 30.57948 |
| 29.05393 | 29.02041 | 28.99091 |
| 27.13134 | 27.02865 | 27.059 |
| 28.50771 | 28.42089 | 28.37799 |
| 28.68241 | 28.68117 | 28.73056 |
| 27.44648 | 27.20154 | 27.27267 |
| 28.63426 | 28.41869 | 28.46389 |
| 30.8152 | 30.69162 | 30.7714 |
| 28.03716 | 28.1693 | 27.97902 |
| 27.75672 | 27.91239 | 27.97498 |
| 32.43743 | 32.43964 | 32.36909 |
| 28.07061 | 28.22357 | 28.17783 |
| 26.93876 | 27.33952 | 27.1736 |
| 27.66214 | 27.4415 | 27.52283 |
| 28.39534 | 28.23736 | 28.20811 |
| 30.45768 | 30.46236 | 30.42242 |
| 27.74387 | 27.75818 | 27.59527 |
| 29.03459 | 29.06124 | 29.03763 |
| 26.56666 | 26.59349 | 26.91182 |
| 29.44743 | 29.55485 | 29.58052 |
| 29.37857 | 29.23936 | 29.31499 |
| 28.01515 | 27.89747 | 27.82019 |

| | | |
|---|---|---|
| 27.27658 | 27.39697 | 27.42814 |
| 28.05879 | 27.90225 | 27.84571 |
| 26.63492 | 24.77339 | 26.26275 |
| 27.50423 | 27.61386 | 27.57181 |
| 27.3794 | 26.93405 | 26.82687 |
| 27.11912 | 26.95981 | 26.9706 |
| 28.76288 | 28.65533 | 28.77357 |
| 29.9878 | 29.94838 | 29.93227 |
| 28.41276 | 28.49755 | 28.42382 |
| 31.36421 | 31.23289 | 31.22963 |
| 27.5294 | 27.27284 | 27.23252 |
| 28.34096 | 28.21614 | 28.19018 |
| 27.29395 | 27.17684 | 27.08932 |
| 27.65581 | 27.63782 | 27.54635 |
| 27.62735 | 27.83822 | 27.87578 |
| 27.66716 | 27.76503 | 27.74496 |
| 27.02115 | 26.80526 | 26.94626 |
| 27.91639 | 27.65915 | 27.67775 |
| 28.67583 | 28.77231 | 28.78491 |
| 27.58957 | 27.53185 | 27.58885 |
| 28.97243 | 29.02623 | 29.06607 |
| 26.53462 | 26.37817 | 26.12003 |
| 27.89562 | 27.81458 | 27.53594 |
| 31.42242 | 31.46085 | 31.4701 |
| 28.41595 | 28.37501 | 28.39803 |
| 29.34685 | 29.23802 | 29.2957 |
| 30.40426 | 30.32346 | 30.39673 |
| 29.56174 | 29.53908 | 29.56946 |
| 27.33731 | 27.08608 | 27.17331 |
| 27.91627 | 27.94419 | 27.8878 |
| 21.9 | 27.45097 | 21.9 |
| 27.79569 | 28.15957 | 28.10275 |
| 26.80637 | 26.40214 | 26.42068 |
| 27.89446 | 28.04916 | 28.15098 |
| 33.01425 | 32.86826 | 32.90378 |
| 27.94732 | 28.0094 | 27.82098 |
| 27.23818 | 27.2936 | 27.24853 |
| 27.78275 | 27.56449 | 27.67 |
| 29.86364 | 29.94126 | 30.02168 |
| 29.09786 | 29.17424 | 29.07528 |
| 27.46037 | 27.15238 | 27.41852 |
| 28.41111 | 28.27128 | 28.23854 |
| 29.81765 | 29.83274 | 29.84127 |
| 27.06291 | 27.1206 | 27.26526 |
| 28.56569 | 28.54001 | 28.46217 |
| 26.55422 | 26.64471 | 26.45723 |
| 29.38158 | 29.17762 | 29.12384 |

| | | |
|---|---|---|
| 27.52111 | 27.63692 | 27.43436 |
| 27.60802 | 27.62268 | 27.70629 |
| 27.38451 | 27.48892 | 27.36553 |
| 30.00384 | 30.0651 | 30.01877 |
| 27.79879 | 27.71043 | 27.56827 |
| 29.77946 | 29.6502 | 29.55649 |
| 26.97378 | 27.27231 | 26.97498 |
| 33.30443 | 33.21008 | 33.18454 |
| 30.7332 | 30.69054 | 30.73513 |
| 26.04289 | 25.99248 | 21.9 |
| 28.93126 | 28.76623 | 28.8432 |
| 29.70256 | 29.88732 | 29.96609 |
| 27.88081 | 27.58075 | 27.63104 |
| 29.36482 | 29.30746 | 29.28844 |
| 27.55516 | 27.33773 | 27.38154 |
| 27.29149 | 27.30965 | 27.06168 |
| 27.94815 | 28.01105 | 28.04401 |
| 31.44238 | 31.33894 | 31.32678 |
| 29.37625 | 29.19674 | 29.1958 |
| 29.13479 | 29.26515 | 29.16058 |
| 30.12127 | 30.04323 | 30.00678 |
| 32.93681 | 32.74953 | 32.71939 |
| 27.7726 | 28.21845 | 28.02892 |
| 28.21489 | 28.26222 | 28.42838 |
| 26.50246 | 26.26064 | 26.39978 |
| 24.75374 | 25.42864 | 24.85714 |
| 26.58921 | 26.78743 | 26.38256 |
| 26.76547 | 26.72897 | 26.56745 |
| 28.65632 | 28.30426 | 28.31324 |
| 26.46532 | 26.74432 | 26.45466 |
| 27.8484 | 27.94704 | 27.94943 |
| 28.70636 | 28.78874 | 28.78019 |
| 28.10799 | 27.8325 | 27.7929 |
| 27.67101 | 27.53808 | 27.70266 |
| 27.5145 | 27.30443 | 27.20359 |
| 30.39059 | 30.49772 | 30.40801 |
| 30.14722 | 30.13247 | 30.16555 |
| 26.33141 | 26.25231 | 26.37372 |
| 28.76771 | 28.74195 | 28.65472 |
| 27.49328 | 27.45842 | 27.43404 |
| 27.55311 | 27.63692 | 27.64787 |
| 26.92931 | 26.9546 | 27.32594 |
| 26.69006 | 26.49442 | 26.51838 |
| 26.75124 | 26.79029 | 26.7366 |
| 28.2035 | 28.26964 | 28.27604 |
| 27.51533 | 27.69432 | 27.61512 |
| 27.2764 | 27.15364 | 27.55969 |

| | | |
|---|---|---|
| 28.04177 | 28.15803 | 28.10505 |
| 26.84625 | 27.09093 | 26.96058 |
| 27.91451 | 27.60258 | 27.72319 |
| 27.67654 | 27.7875 | 27.84787 |
| 28.35031 | 28.14248 | 28.10164 |
| 28.53036 | 28.70968 | 28.56016 |
| 31.42737 | 31.3872 | 31.35862 |
| 27.91388 | 27.50135 | 27.54436 |
| 28.05253 | 27.14462 | 27.34587 |
| 28.22495 | 28.31903 | 28.20013 |
| 30.93874 | 30.8886 | 30.8511 |
| 28.75401 | 28.71171 | 28.8094 |
| 27.47746 | 27.47769 | 27.49526 |
| 29.07202 | 28.84742 | 28.87109 |
| 26.3601 | 26.40264 | 26.39452 |
| 28.23754 | 28.27635 | 28.36578 |
| 29.77493 | 29.73718 | 29.75475 |
| 27.36778 | 27.23818 | 27.22066 |
| 29.90268 | 29.80238 | 29.75635 |
| 32.69185 | 32.74877 | 32.71704 |
| 31.24298 | 31.31008 | 31.32158 |
| 27.33492 | 27.18858 | 27.25178 |
| 27.27364 | 27.21771 | 27.23013 |
| 28.64757 | 28.97295 | 28.83653 |
| 28.38076 | 28.31652 | 28.28996 |
| 30.85875 | 30.87489 | 30.7718 |
| 26.41406 | 26.46504 | 26.47609 |
| 28.08821 | 28.35005 | 28.33539 |
| 31.93086 | 31.92864 | 31.93526 |
| 27.59092 | 27.71816 | 27.69955 |
| 28.59722 | 28.59231 | 28.46883 |
| 27.15499 | 26.92513 | 26.95848 |
| 27.85549 | 27.75239 | 27.69159 |
| 29.53404 | 29.66343 | 29.57162 |
| 28.6246 | 28.74977 | 28.7128 |
| 26.85007 | 26.91946 | 26.68939 |
| 27.92796 | 27.97673 | 27.93281 |
| 25.85108 | 25.86039 | 26.02223 |
| 28.86074 | 28.84658 | 28.80627 |
| 31.68516 | 31.77341 | 31.74351 |
| 28.65164 | 28.57459 | 28.42045 |
| 29.13014 | 29.37906 | 29.2507 |
| 27.5437 | 27.56311 | 27.51232 |
| 29.10936 | 28.94969 | 28.98329 |
| 31.16525 | 31.15798 | 31.19448 |
| 26.62715 | 26.20081 | 26.4434 |
| 26.60286 | 27.3955 | 27.19432 |

| | | |
|---|---|---|
| 26.83413 | 27.34036 | 27.55326 |
| 27.71162 | 27.58147 | 27.59719 |
| 30.12436 | 29.96609 | 29.93143 |
| 28.07265 | 27.88227 | 27.72038 |
| 26.80858 | 26.80612 | 26.95593 |
| 25.78204 | 25.90217 | 25.74909 |
| 26.38012 | 21.9 | 26.5221 |
| 28.18262 | 28.00212 | 28.14477 |
| 25.90076 | 26.48895 | 26.23719 |
| 27.22738 | 27.32492 | 27.14899 |
| 28.30939 | 28.03538 | 28.14545 |
| 28.40231 | 28.28198 | 28.34976 |
| 27.52051 | 27.50074 | 27.44608 |
| 31.47576 | 31.4034 | 31.32823 |
| 28.1245 | 28.13315 | 28.20662 |
| 27.19056 | 26.68646 | 27.02686 |
| 26.5481 | 26.90185 | 26.49657 |
| 27.11416 | 27.09587 | 27.14287 |
| 28.26101 | 28.20569 | 28.14516 |
| 26.26685 | 27.19676 | 26.8399 |
| 27.56253 | 27.00913 | 26.89781 |
| 27.47553 | 27.22076 | 27.28162 |
| 26.89574 | 26.72819 | 26.70754 |
| 29.64739 | 29.09658 | 29.19521 |
| 28.75653 | 28.59833 | 28.55113 |
| 27.36119 | 27.45026 | 27.44845 |
| 25.78209 | 25.86955 | 25.82272 |
| 27.26642 | 27.28065 | 27.23023 |
| 28.56165 | 28.41018 | 28.52675 |
| 30.35668 | 30.26125 | 30.20548 |
| 33.00157 | 32.92834 | 32.91698 |
| 26.59776 | 26.43323 | 26.69019 |
| 29.09828 | 29.06659 | 28.89221 |
| 29.21169 | 29.29373 | 29.30108 |
| 29.27082 | 29.30804 | 29.29684 |
| 26.70556 | 27.22242 | 27.16873 |
| 30.50786 | 30.55636 | 30.63313 |
| 28.19366 | 28.05523 | 28.0426 |
| 26.1242 | 26.2674 | 26.09589 |
| 26.95238 | 26.90518 | 26.99216 |
| 28.47259 | 28.35775 | 28.31851 |
| 25.89217 | 26.02686 | 26.00033 |
| 27.67175 | 27.6006 | 27.33799 |
| 28.86779 | 28.89195 | 28.96306 |
| 24.94204 | 25.29555 | 25.21479 |
| 21.9 | 25.58428 | 25.69227 |
| 28.01898 | 27.94598 | 28.0777 |

| | | |
|---|---|---|
| 26.38982 | 26.19715 | 26.4656 |
| 32.98114 | 32.97049 | 32.96826 |
| 28.02195 | 28.16188 | 28.14073 |
| 31.80567 | 31.74306 | 31.7169 |
| 30.78027 | 30.84618 | 30.84161 |
| 27.83219 | 27.94308 | 28.00357 |
| 26.8305 | 26.69883 | 26.88064 |
| 28.3937 | 28.4289 | 28.48339 |
| 26.68872 | 26.59733 | 26.64169 |
| 25.48487 | 25.58792 | 25.68894 |
| 25.6864 | 25.92884 | 25.82116 |
| 31.33549 | 31.03623 | 31.01312 |
| 27.90988 | 27.90633 | 27.93966 |
| 27.36294 | 27.2261 | 27.27826 |
| 26.94693 | 26.78868 | 26.95238 |
| 30.07839 | 30.12989 | 30.0874 |
| 28.32152 | 28.258 | 28.25656 |
| 29.37962 | 29.37233 | 29.44055 |
| 29.57701 | 29.75594 | 29.70308 |
| 27.50028 | 27.42254 | 27.34087 |
| 29.68729 | 29.62218 | 29.52302 |
| 30.35332 | 30.23009 | 30.22274 |
| 26.55421 | 26.74586 | 26.70266 |
| 31.73957 | 31.75855 | 31.72988 |
| 27.28806 | 27.37343 | 27.09517 |
| 28.55918 | 28.63558 | 28.75465 |
| 27.50567 | 27.56616 | 27.5291 |
| 27.66262 | 27.80495 | 27.83503 |
| 27.83816 | 27.81928 | 27.89637 |
| 26.93281 | 27.06979 | 27.14996 |
| 26.52314 | 26.77543 | 26.60173 |
| 32.84427 | 32.87311 | 32.87122 |
| 27.26955 | 26.99399 | 27.34469 |
| 29.20027 | 29.02099 | 29.05659 |
| 26.65881 | 26.65116 | 26.59847 |
| 30.46879 | 30.66051 | 30.59067 |
| 28.1309 | 28.30334 | 28.30352 |
| 25.59161 | 25.4507 | 25.45576 |
| 28.19075 | 28.1849 | 28.1672 |
| 27.76686 | 27.65513 | 27.61267 |
| 27.66675 | 27.7103 | 27.47158 |
| 21.9 | 26.01555 | 25.96202 |
| 30.32646 | 30.19072 | 30.27442 |
| 26.31398 | 26.40622 | 26.18686 |
| 28.89885 | 28.74942 | 28.73043 |
| 26.53103 | 26.75034 | 26.9467 |
| 26.91536 | 27.30869 | 27.23004 |

| | | |
|---|---|---|
| 25.97119 | 26.39727 | 26.37431 |
| 26.04795 | 26.40129 | 26.41374 |
| 21.9 | 27.22738 | 27.16595 |
| 26.61358 | 26.7016 | 21.9 |
| 28.83449 | 28.66397 | 28.55786 |
| 26.60428 | 26.72598 | 26.62909 |
| 27.79916 | 27.92587 | 27.76775 |
| 26.99873 | 26.97738 | 27.02359 |
| 21.9 | 21.9 | 21.9 |
| 26.0942 | 25.93555 | 26.09686 |
| 27.67714 | 27.36853 | 27.35758 |
| 25.86292 | 25.77979 | 25.74858 |
| 26.38915 | 26.23444 | 26.15449 |
| 25.92529 | 25.92531 | 26.10919 |
| 26.70938 | 26.44634 | 26.60541 |
| 24.85638 | 24.99382 | 24.94311 |
| 26.13909 | 25.79333 | 25.81849 |
| 25.93872 | 25.93966 | 26.03775 |
| 29.91471 | 30.04739 | 30.01105 |
| 28.79414 | 28.71885 | 28.63627 |
| 27.60851 | 27.6809 | 27.60922 |
| 28.40438 | 28.24332 | 28.21002 |
| 27.71371 | 27.57203 | 27.6133 |
| 26.4753 | 26.4698 | 26.37114 |
| 27.49221 | 27.52903 | 27.43873 |
| 28.28171 | 28.27799 | 28.24282 |
| 29.96966 | 29.78535 | 29.91684 |
| 26.5261 | 26.73221 | 26.52319 |
| 29.22281 | 29.14688 | 29.09936 |
| 29.075 | 29.12028 | 29.16607 |
| 27.31976 | 27.18622 | 27.2848 |
| 27.1547 | 27.00507 | 27.23114 |
| 25.93177 | 26.24307 | 26.42897 |
| 27.69292 | 27.44814 | 27.52604 |
| 27.23717 | 27.22849 | 27.29448 |
| 27.87865 | 28.08917 | 28.27972 |
| 27.05414 | 26.90656 | 26.98479 |
| 26.57472 | 26.48458 | 26.52565 |
| 25.39761 | 25.26449 | 25.64623 |
| 28.65106 | 28.53679 | 28.58201 |
| 26.82578 | 26.48262 | 26.61863 |
| 27.95887 | 27.72852 | 27.57347 |
| 25.64524 | 25.97622 | 26.16077 |
| 26.78057 | 26.80526 | 26.58921 |
| 28.74368 | 28.75232 | 28.78322 |
| 28.1379 | 28.41969 | 28.44423 |
| 26.15147 | 26.21653 | 26.22623 |

| | | |
|---|---|---|
| 27.39149 | 27.40844 | 27.42758 |
| 29.18615 | 29.23984 | 29.18971 |
| 29.76226 | 29.8964 | 29.91271 |
| 27.10509 | 27.37443 | 26.96774 |
| 26.58276 | 26.61358 | 26.57773 |
| 25.17596 | 25.32687 | 25.20284 |
| 25.62581 | 21.9 | 25.53857 |
| 26.09055 | 26.61877 | 26.42643 |
| 24.87422 | 24.79442 | 24.86179 |
| 26.73467 | 26.78606 | 26.57233 |
| 26.27631 | 26.17777 | 26.23796 |
| 27.03065 | 26.49412 | 26.84482 |
| 26.01265 | 26.96279 | 27.03811 |
| 27.59889 | 27.56565 | 27.73635 |
| 30.39734 | 30.62993 | 30.61289 |
| 28.63717 | 28.80618 | 28.8343 |
| 29.14326 | 29.08175 | 29.10851 |
| 27.70813 | 27.57463 | 27.52208 |
| 30.1343 | 30.54232 | 30.44806 |
| 27.81103 | 27.82639 | 27.8063 |
| 27.92909 | 27.75034 | 27.51916 |
| 26.01642 | 26.39539 | 26.40688 |
| 21.9 | 21.9 | 21.9 |
| 21.9 | 21.9 | 21.9 |
| 26.60738 | 26.46112 | 26.56956 |
| 25.47698 | 25.98865 | 25.43743 |
| 26.62031 | 26.35235 | 26.61583 |
| 26.49338 | 26.38924 | 26.70833 |
| 26.15719 | 25.96378 | 25.93991 |
| 25.6245 | 25.50154 | 25.47454 |
| 27.13848 | 27.85204 | 27.84834 |
| 26.2212 | 26.23094 | 26.38416 |
| 26.20469 | 25.99998 | 25.97869 |
| 25.59004 | 25.7343 | 25.88567 |
| 27.87502 | 27.80833 | 27.72663 |
| 29.49507 | 29.2128 | 29.25827 |
| 27.6245 | 27.8749 | 27.86603 |
| 28.57521 | 28.50696 | 28.59769 |
| 26.36174 | 26.37113 | 26.46371 |
| 28.17222 | 28.07189 | 28.03333 |
| 30.38134 | 30.45091 | 30.4513 |
| 28.50025 | 28.63547 | 28.61143 |
| 27.5575 | 27.54097 | 27.64478 |
| 26.81507 | 27.06692 | 27.11763 |
| 28.46377 | 28.35846 | 28.47429 |
| 25.89823 | 25.91139 | 26.07718 |
| 27.44711 | 27.66959 | 27.60505 |

| | | |
|---|---|---|
| 26.86089 | 26.7145 | 26.58892 |
| 28.45435 | 28.34896 | 28.45823 |
| 31.49061 | 31.34292 | 31.28951 |
| 26.55646 | 26.10071 | 26.16002 |
| 26.68485 | 26.93517 | 26.71253 |
| 28.60244 | 28.64451 | 28.68087 |
| 21.9 | 26.43355 | 26.11261 |
| 27.81317 | 27.74849 | 27.7304 |
| 25.94018 | 25.58532 | 25.59278 |
| 26.52996 | 26.34477 | 26.31893 |
| 24.92515 | 25.3837 | 25.2009 |
| 26.29946 | 21.9 | 26.52966 |
| 25.72708 | 25.87216 | 21.9 |
| 26.95604 | 26.99884 | 27.22711 |
| 26.85198 | 26.55769 | 26.69657 |
| 28.63841 | 28.58388 | 28.66058 |
| 26.26472 | 26.19297 | 25.97804 |
| 26.29887 | 26.21156 | 26.35301 |
| 28.52051 | 28.39705 | 28.39991 |
| 27.6541 | 27.52253 | 27.58671 |
| 29.64239 | 29.69059 | 29.72964 |
| 26.28044 | 26.34778 | 26.43525 |
| 28.14277 | 27.95577 | 27.98951 |
| 27.86479 | 27.6076 | 27.74804 |
| 25.36506 | 25.2329 | 25.23983 |
| 25.75761 | 26.01293 | 25.67134 |
| 26.09336 | 26.04903 | 26.02206 |
| 26.11795 | 26.14476 | 26.06999 |
| 25.60467 | 25.65483 | 25.74998 |
| 26.0677 | 25.89659 | 25.91221 |
| 27.72981 | 27.79538 | 27.80304 |
| 25.98824 | 26.0767 | 26.00682 |
| 26.73402 | 26.57061 | 26.61569 |
| 25.42249 | 25.44006 | 25.49732 |
| 28.37753 | 28.43595 | 28.35296 |
| 27.96135 | 27.88937 | 27.86732 |
| 29.30265 | 29.22972 | 29.24738 |
| 27.45018 | 27.26633 | 27.28489 |
| 27.03107 | 27.25521 | 27.30704 |
| 21.9 | 25.69309 | 25.56805 |
| 21.9 | 25.24887 | 25.24775 |
| 28.77831 | 28.68495 | 28.74836 |
| 25.18813 | 25.39506 | 25.4746 |
| 26.29077 | 26.31302 | 26.44873 |
| 28.79996 | 28.83014 | 28.87106 |
| 27.21466 | 27.18556 | 27.06281 |
| 31.48573 | 31.22193 | 31.32496 |

| | | |
|---|---|---|
| 29.23416 | 29.09315 | 29.1161 |
| 27.00196 | 26.99572 | 26.98348 |
| 28.85979 | 28.67428 | 28.72293 |
| 31.51855 | 31.57448 | 31.61061 |
| 28.26262 | 28.50537 | 28.48788 |
| 27.8387 | 27.91222 | 27.78563 |
| 29.10024 | 28.9407 | 29.08692 |
| 28.4103 | 28.38833 | 28.32667 |
| 28.46322 | 28.44573 | 28.51668 |
| 28.19723 | 28.0931 | 28.01169 |
| 26.80686 | 26.46101 | 26.33727 |
| 27.52246 | 27.74618 | 27.85792 |
| 30.06355 | 30.01571 | 29.97309 |
| 27.55888 | 27.58484 | 27.51833 |
| 26.08107 | 25.89296 | 26.24285 |
| 21.9 | 21.9 | 21.9 |
| 25.9083 | 26.08113 | 26.13518 |
| 26.25947 | 26.02301 | 26.04389 |
| 25.21686 | 25.50818 | 25.59872 |
| 25.72119 | 25.65376 | 25.7365 |
| 21.9 | 26.18316 | 26.17078 |
| 25.89182 | 25.80526 | 25.86722 |
| 25.29852 | 25.27909 | 25.00949 |
| 26.77618 | 26.67465 | 26.60088 |
| 26.55128 | 26.7145 | 26.70636 |
| 29.34403 | 29.20424 | 29.2092 |
| 27.97542 | 27.81739 | 27.73007 |
| 26.91991 | 26.8662 | 26.98359 |
| 26.95571 | 27.1931 | 27.09909 |
| 28.33977 | 28.1519 | 28.25935 |
| 27.38014 | 27.46311 | 27.49175 |
| 29.29835 | 29.34312 | 29.3006 |
| 27.01765 | 26.84326 | 26.93618 |
| 27.17674 | 27.15296 | 27.21011 |
| 28.65031 | 28.62356 | 28.69246 |
| 29.46351 | 29.16317 | 29.11101 |
| 29.33954 | 29.44293 | 29.36999 |
| 30.53992 | 30.47315 | 30.53492 |
| 25.66863 | 25.51584 | 25.68394 |
| 26.79514 | 26.84098 | 26.84014 |
| 25.54062 | 25.53169 | 25.67613 |
| 26.5657 | 26.21401 | 26.2144 |
| 27.52835 | 27.4283 | 27.5403 |
| 26.21692 | 26.17011 | 26.10334 |
| 25.30703 | 25.1902 | 24.94682 |
| 26.77002 | 26.68712 | 26.58434 |
| 26.46678 | 26.56371 | 26.50882 |

| | | |
|---|---|---|
| 28.6676 | 28.59605 | 28.60897 |
| 26.27329 | 26.34829 | 21.9 |
| 26.47301 | 26.47655 | 26.68164 |
| 26.94246 | 26.87091 | 26.93876 |
| 33.01749 | 32.99027 | 33.00855 |
| 27.81909 | 27.94397 | 27.90363 |
| 25.44957 | 25.53641 | 25.52358 |
| 25.68145 | 25.56674 | 21.9 |
| 25.766 | 21.9 | 25.49698 |
| 27.47553 | 26.93832 | 27.13917 |
| 26.69299 | 26.59007 | 26.65608 |
| 26.84649 | 27.35968 | 27.18386 |
| 28.76803 | 29.04627 | 29.0291 |
| 26.02007 | 21.9 | 26.02136 |
| 25.50588 | 25.14981 | 25.44199 |
| 26.00973 | 25.98833 | 26.05294 |
| 27.72943 | 27.54296 | 27.52313 |
| 25.95848 | 25.75274 | 25.53362 |
| 26.63076 | 26.55815 | 26.53973 |
| 26.35616 | 25.9227 | 26.04389 |
| 26.77165 | 26.67465 | 26.64155 |
| 21.9 | 25.70108 | 25.62639 |
| 21.9 | 24.86183 | 21.9 |
| 27.24916 | 27.01159 | 27.27071 |
| 26.27595 | 26.26639 | 26.57557 |
| 26.41403 | 26.50694 | 26.44971 |
| 26.35336 | 26.28279 | 26.07688 |
| 27.19957 | 27.38993 | 27.18139 |
| 28.47456 | 28.3621 | 28.31838 |
| 31.2179 | 31.05245 | 31.07099 |
| 27.91974 | 27.9941 | 28.14161 |
| 28.68706 | 28.7347 | 28.79947 |
| 28.20154 | 28.15649 | 28.28339 |
| 29.09406 | 29.10162 | 29.14562 |
| 25.91959 | 21.9 | 21.9 |
| 26.61597 | 26.67707 | 26.82372 |
| 29.10102 | 29.02015 | 28.96326 |
| 30.79009 | 30.81787 | 30.85289 |
| 26.2865 | 25.95689 | 25.95318 |
| 25.49543 | 25.48004 | 21.9 |
| 21.9 | 21.9 | 21.9 |
| 21.9 | 21.9 | 21.9 |
| 25.75766 | 21.9 | 21.9 |
| 21.9 | 21.9 | 21.9 |
| 21.9 | 21.9 | 21.9 |
| 23.89717 | 25.09346 | 24.60116 |
| 24.31174 | 21.9 | 21.9 |

| | | |
|---|---|---|
| 25.84441 | 25.56796 | 25.59138 |
| 25.92529 | 25.8314 | 25.80312 |
| 25.97548 | 25.64963 | 25.80678 |
| 25.81595 | 21.9 | 25.92386 |
| 28.05884 | 27.96047 | 27.85958 |
| 25.58915 | 25.65984 | 21.9 |
| 25.01772 | 25.29793 | 25.25032 |
| 25.84235 | 26.02625 | 25.9324 |
| 25.92924 | 25.58798 | 25.69115 |
| 25.91388 | 25.69224 | 25.94053 |
| 25.75103 | 25.72996 | 25.60518 |
| 25.71196 | 25.694 | 25.79682 |
| 25.63442 | 25.49924 | 25.55219 |
| 25.64826 | 25.72247 | 25.31897 |
| 25.39385 | 25.62455 | 25.65376 |
| 24.97754 | 25.04502 | 25.09297 |
| 25.25972 | 24.98746 | 25.29025 |
| 26.65185 | 27.06065 | 26.93438 |
| 21.9 | 25.07647 | 21.9 |
| 21.9 | 24.97027 | 24.76678 |
| 21.9 | 23.98355 | 21.9 |
| 21.9 | 21.9 | 23.89551 |
| 21.9 | 21.9 | 21.9 |
| 27.46357 | 27.328 | 27.29246 |
| 27.82116 | 27.93843 | 27.86089 |
| 27.39485 | 27.44261 | 27.3621 |
| 27.36269 | 27.56326 | 27.65485 |
| 27.44016 | 27.26446 | 27.26437 |
| 28.50275 | 28.60897 | 28.41317 |
| 27.50423 | 27.29798 | 27.35262 |
| 25.70306 | 25.63796 | 25.55403 |
| 26.16039 | 26.31432 | 26.11506 |
| 26.0174 | 25.85863 | 26.07204 |
| 28.27378 | 28.49686 | 28.45732 |
| 29.15366 | 29.07804 | 29.14938 |
| 25.95118 | 25.86521 | 25.84976 |
| 27.68003 | 27.98826 | 27.93708 |
| 29.3217 | 29.33035 | 29.29818 |
| 27.4464 | 27.6443 | 27.73176 |
| 27.00892 | 27.01882 | 26.93562 |
| 28.17479 | 28.15697 | 28.13653 |
| 25.16312 | 24.98377 | 24.87732 |
| 26.46428 | 26.53233 | 26.48456 |
| 26.19689 | 26.09493 | 26.08211 |
| 25.78367 | 25.80639 | 25.90575 |
| 26.11259 | 25.97697 | 26.01297 |
| 26.38652 | 26.24209 | 26.13927 |

| | | |
|---|---|---|
| 26.22793 | 26.0121 | 26.3617 |
| 26.66641 | 26.62812 | 26.52816 |
| 29.6513 | 29.70576 | 29.63133 |
| 26.86538 | 27.44719 | 27.44095 |
| 25.7685 | 26.05588 | 26.18771 |
| 26.38701 | 26.37403 | 26.68244 |
| 26.46494 | 26.88309 | 27.07725 |
| 25.49186 | 25.73949 | 25.74863 |
| 28.2294 | 28.12381 | 28.07572 |
| 27.90346 | 27.86319 | 27.94386 |
| 26.30622 | 26.35385 | 26.3347 |
| 26.79885 | 26.72247 | 26.7375 |
| 26.57615 | 26.62478 | 26.21996 |
| 26.96334 | 27.12544 | 26.98707 |
| 27.08446 | 26.70477 | 26.61751 |
| 25.87289 | 26.04229 | 26.02692 |
| 21.9 | 24.84938 | 25.01048 |
| 25.8719 | 25.82854 | 21.9 |
| 25.88976 | 25.57934 | 25.78923 |
| 26.66139 | 26.6147 | 26.61961 |
| 28.29955 | 28.0358 | 28.29706 |
| 26.75085 | 26.72195 | 26.73982 |
| 26.01918 | 25.95837 | 25.75108 |
| 24.81005 | 24.93724 | 24.6036 |
| 26.54881 | 26.94369 | 26.90403 |
| 26.15935 | 26.37809 | 26.28329 |
| 21.9 | 25.43708 | 25.60422 |
| 26.09299 | 26.19802 | 26.17746 |
| 25.41719 | 25.5501 | 21.9 |
| 28.51013 | 27.90495 | 27.99226 |
| 21.9 | 25.64779 | 25.81504 |
| 24.85709 | 25.27736 | 25.40737 |
| 26.65171 | 26.89504 | 27.0425 |
| 25.75042 | 25.55301 | 25.47113 |
| 26.05765 | 25.81778 | 25.95611 |
| 26.00924 | 26.30447 | 26.17644 |
| 26.11999 | 25.74309 | 25.63337 |
| 25.08318 | 25.12566 | 21.9 |
| 25.58629 | 25.67871 | 25.80642 |
| 35.44502 | 35.48214 | 35.47702 |
| 27.90374 | 27.95088 | 28.07719 |
| 30.52608 | 30.03708 | 30.09509 |
| 28.14788 | 28.19578 | 28.15894 |
| 28.82526 | 28.55095 | 28.51709 |
| 28.08182 | 28.01643 | 28.12775 |
| 26.62812 | 26.40302 | 26.60682 |
| 21.9 | 24.68319 | 21.9 |

| | | |
|---|---|---|
| 21.9 | 21.9 | 21.9 |
| 21.9 | 25.58964 | 25.71782 |
| 21.9 | 21.9 | 21.9 |
| 26.00573 | 26.27943 | 26.25816 |
| 26.35129 | 26.16699 | 26.29228 |
| 25.92945 | 26.09378 | 25.91604 |
| 21.9 | 26.11547 | 26.40446 |
| 25.17429 | 25.31976 | 24.98863 |
| 26.52792 | 26.22277 | 26.67654 |
| 26.06046 | 25.97959 | 25.94927 |
| 25.86278 | 25.87603 | 21.9 |
| 25.55745 | 25.84328 | 25.7639 |
| 25.00757 | 24.75313 | 24.78526 |
| 26.39272 | 26.25541 | 26.18482 |
| 26.27962 | 26.25854 | 26.17148 |
| 25.20176 | 21.9 | 25.35077 |
| 26.14003 | 26.04466 | 21.9 |
| 24.43262 | 24.4325 | 21.9 |
| 21.9 | 24.79927 | 21.9 |
| 23.68432 | 23.71167 | 23.60354 |
| 22.81186 | 23.59537 | 23.15247 |
| 27.49389 | 27.47653 | 27.44869 |
| 26.92502 | 26.69046 | 26.6302 |
| 28.33522 | 28.32869 | 28.28198 |
| 27.26347 | 27.34316 | 27.07938 |
| 28.78216 | 28.91872 | 28.86665 |
| 27.65014 | 27.34562 | 27.41675 |
| 29.98114 | 30.09018 | 30.18801 |
| 25.72567 | 25.65434 | 25.60575 |
| 28.96105 | 28.90716 | 28.87414 |
| 21.9 | 25.48757 | 25.32165 |
| 24.81245 | 24.95176 | 25.06952 |
| 27.04989 | 26.94726 | 27.21011 |
| 25.31562 | 25.66758 | 25.61958 |
| 26.67922 | 26.73467 | 26.52406 |
| 26.5441 | 26.5265 | 26.45585 |
| 25.92619 | 25.84972 | 25.77713 |
| 25.66408 | 25.92343 | 25.80985 |
| 26.55545 | 27.07224 | 26.6975 |
| 24.94467 | 25.21952 | 25.20086 |
| 27.38665 | 27.23498 | 27.20937 |
| 26.71555 | 26.52846 | 26.43979 |
| 25.86226 | 25.96952 | 25.89682 |
| 26.73092 | 26.47865 | 26.47505 |
| 25.95091 | 25.80435 | 27.88256 |
| 27.46194 | 27.37931 | 27.44506 |
| 25.22612 | 25.31351 | 25.40048 |

| | | |
|---|---|---|
| 24.24213 | 24.51887 | 24.55748 |
| 24.85909 | 21.9 | 21.9 |
| 24.7072 | 25.45482 | 25.05097 |
| 27.24336 | 27.15933 | 27.04625 |
| 26.6047 | 26.47687 | 26.71856 |
| 26.49257 | 26.40978 | 26.31745 |
| 26.3922 | 25.68421 | 26.02695 |
| 25.12865 | 21.9 | 25.40688 |
| 27.17112 | 26.96587 | 26.89122 |
| 25.61726 | 25.81958 | 26.5212 |
| 26.22962 | 26.17179 | 26.2312 |
| 24.53196 | 21.9 | 21.9 |
| 27.21845 | 26.9629 | 27.12514 |
| 27.3794 | 27.13467 | 21.9 |
| 21.9 | 21.9 | 25.63669 |
| 27.36093 | 27.2841 | 27.40276 |
| 27.73318 | 27.66932 | 27.639 |
| 26.32956 | 26.4307 | 26.51788 |
| 26.66574 | 26.51105 | 26.41957 |
| 21.9 | 24.87811 | 24.8501 |
| 25.55722 | 21.9 | 21.9 |
| 25.32158 | 25.22255 | 25.25176 |
| 21.9 | 25.29004 | 25.03901 |
| 33.4501 | 33.44344 | 33.44332 |
| 30.6471 | 30.54858 | 30.54692 |
| 29.27327 | 29.16173 | 29.14012 |
| 29.18756 | 29.41171 | 29.30832 |
| 28.38891 | 28.317 | 28.1785 |
| 27.59655 | 27.61456 | 27.29605 |
| 27.13966 | 27.04479 | 26.98055 |
| 30.02512 | 29.94 | 29.94098 |
| 26.67963 | 26.43778 | 26.53671 |
| 28.55252 | 28.41115 | 28.31411 |
| 25.94268 | 26.31816 | 26.15161 |
| 25.39679 | 25.26674 | 25.17665 |
| 26.17974 | 26.22738 | 26.48124 |
| 26.28553 | 26.30095 | 26.39718 |
| 21.9 | 21.9 | 21.9 |
| 21.9 | 26.01319 | 21.9 |
| 21.9 | 21.9 | 21.9 |
| 25.19344 | 25.28926 | 25.38726 |
| 25.39234 | 25.56314 | 25.6002 |
| 27.02739 | 27.16634 | 27.13624 |
| 25.83111 | 25.90102 | 26.11229 |
| 25.29758 | 25.26857 | 25.4325 |
| 25.27014 | 25.39591 | 25.04964 |
| 25.91308 | 25.86254 | 26.0008 |

| | | |
|---|---|---|
| 25.61349 | 25.53199 | 25.52184 |
| 26.10691 | 25.87608 | 25.48932 |
| 21.9 | 24.67347 | 24.77193 |
| 25.67134 | 25.79929 | 25.6324 |
| 24.61919 | 25.05908 | 25.14573 |
| 25.20933 | 24.99916 | 24.9955 |
| 25.08902 | 25.12278 | 25.04285 |
| 26.21817 | 21.9 | 26.06427 |
| 25.96435 | 26.07441 | 25.94385 |
| 25.08343 | 24.98229 | 24.9354 |
| 24.57664 | 24.79392 | 24.74026 |
| 26.33393 | 26.08776 | 26.23634 |
| 21.9 | 25.88906 | 25.8124 |
| 25.934 | 21.9 | 25.69543 |
| 24.89287 | 21.9 | 24.93472 |
| 21.9 | 23.78377 | 23.54286 |
| 24.42528 | 21.9 | 24.46973 |
| 27.11098 | 26.69856 | 26.88297 |
| 25.05705 | 25.14903 | 21.9 |
| 21.9 | 21.9 | 24.67126 |
| 21.9 | 21.9 | 21.9 |
| 21.9 | 21.9 | 21.9 |
| 28.05672 | 28.01292 | 27.86284 |
| 25.96462 | 25.83795 | 26.01033 |
| 26.1389 | 26.00157 | 26.21117 |
| 26.85174 | 26.72442 | 26.61063 |
| 26.91878 | 27.24036 | 27.2035 |
| 21.9 | 23.33252 | 23.63023 |
| 27.22766 | 27.14686 | 27.28683 |
| 25.06332 | 25.66479 | 25.38782 |
| 25.91121 | 26.15923 | 26.01308 |
| 21.9 | 21.9 | 21.9 |
| 25.73557 | 25.64908 | 25.56721 |
| 26.41944 | 26.33637 | 26.12989 |
| 25.5343 | 25.43259 | 21.9 |
| 21.9 | 21.9 | 21.9 |
| 26.23593 | 26.26261 | 26.33683 |
| 26.71947 | 26.53964 | 26.53802 |
| 27.06866 | 26.65472 | 26.75226 |
| 25.77311 | 25.98564 | 26.06027 |
| 27.69651 | 27.44892 | 27.60999 |
| 25.40581 | 25.2296 | 25.0209 |
| 25.41387 | 25.81495 | 25.29579 |
| 25.51869 | 21.9 | 21.9 |
| 28.82344 | 29.05269 | 29.05856 |
| 28.27373 | 28.26204 | 28.27302 |
| 25.91477 | 25.8607 | 25.73967 |

| | | |
|---|---|---|
| 23.67142 | 25.07349 | 24.75517 |
| 26.66207 | 26.81751 | 26.81982 |
| 24.82808 | 21.9 | 24.87497 |
| 21.9 | 25.57335 | 25.49106 |
| 25.60798 | 25.61639 | 25.4485 |
| 21.9 | 21.9 | 26.40633 |
| 25.15576 | 25.32899 | 25.52758 |
| 24.18041 | 24.26098 | 21.9 |
| 27.06497 | 26.9546 | 26.69099 |
| 21.9 | 21.9 | 25.12829 |
| 27.8839 | 28.08299 | 27.99243 |
| 26.19355 | 25.96003 | 26.04262 |
| 25.52543 | 21.9 | 21.9 |
| 25.82087 | 25.58021 | 21.9 |
| 26.1118 | 26.58649 | 26.30276 |
| 32.65497 | 32.42274 | 32.33796 |
| 27.01606 | 27.68626 | 27.54694 |
| 25.46712 | 25.71843 | 25.76241 |
| 24.22553 | 24.44954 | 24.26019 |
| 25.2863 | 21.9 | 25.17965 |
| 21.9 | 24.75761 | 24.50288 |
| 25.98941 | 21.9 | 21.9 |
| 27.73963 | 27.62164 | 27.66058 |
| 27.90684 | 28.04328 | 27.94184 |
| 29.95809 | 30.04778 | 30.02142 |
| 34.45862 | 34.47224 | 34.41192 |
| 27.24236 | 27.01574 | 27.01265 |
| 28.64003 | 28.50233 | 28.46906 |
| 27.71914 | 27.80458 | 27.75162 |
| 27.07419 | 27.239 | 27.19517 |
| 26.77216 | 26.68231 | 26.61077 |
| 27.35338 | 27.70807 | 27.58456 |
| 25.31665 | 25.71277 | 25.47215 |
| 28.81268 | 28.72068 | 28.5406 |
| 27.72689 | 27.90001 | 27.80931 |
| 21.9 | 21.9 | 21.9 |
| 21.9 | 21.9 | 21.9 |
| 24.8306 | 24.58526 | 24.75715 |
| 27.37526 | 27.25665 | 27.27915 |
| 29.45824 | 29.58362 | 29.54926 |
| 25.36142 | 25.75718 | 25.76724 |
| 26.54694 | 25.91281 | 26.04995 |
| 25.17809 | 24.85253 | 25.01861 |
| 25.64749 | 25.53715 | 25.47615 |
| 25.55251 | 21.9 | 25.58898 |
| 25.03591 | 25.052 | 25.2074 |
| 21.9 | 25.40022 | 25.03993 |

| | | |
|---|---|---|
| 25.14476 | 21.9 | 21.9 |
| 24.94918 | 24.84038 | 25.08428 |
| 21.9 | 28.6637 | 28.37993 |
| 21.9 | 21.9 | 25.74468 |
| 21.9 | 25.27473 | 25.43062 |
| 21.9 | 24.82119 | 25.12179 |
| 24.76096 | 24.78821 | 24.75501 |
| 21.9 | 24.8388 | 24.79778 |
| 24.60682 | 21.9 | 21.9 |
| 25.36563 | 25.46603 | 25.53237 |
| 26.00877 | 25.83082 | 25.80779 |
| 21.9 | 21.9 | 21.9 |
| 25.14725 | 21.9 | 25.32481 |
| 24.41493 | 24.71534 | 21.9 |
| 21.9 | 21.9 | 21.9 |
| 21.9 | 26.1169 | 26.10675 |
| 25.06546 | 21.9 | 25.19449 |
| 25.10701 | 21.9 | 25.40539 |
| 21.9 | 21.9 | 21.9 |
| 24.61555 | 24.68319 | 24.63412 |
| 21.9 | 26.58434 | 26.6539 |
| 25.44588 | 25.37872 | 25.4324 |
| 26.31672 | 26.45881 | 26.38957 |
| 24.87084 | 21.9 | 24.53546 |
| 26.13248 | 25.57857 | 25.78027 |
| 25.0648 | 25.06641 | 21.9 |
| 26.00714 | 25.79134 | 25.98854 |
| 26.38522 | 26.65608 | 26.50808 |
| 26.41347 | 26.18501 | 26.31199 |
| 26.55441 | 26.6363 | 26.53392 |
| 26.35281 | 26.18837 | 26.07247 |
| 21.9 | 21.9 | 27.17131 |
| 21.9 | 26.20202 | 26.18334 |
| 27.49496 | 27.56311 | 27.55252 |
| 24.92298 | 25.14542 | 21.9 |
| 25.4917 | 26.21595 | 25.90585 |
| 26.61105 | 26.67896 | 26.55081 |
| 23.81118 | 23.95996 | 21.9 |
| 21.9 | 24.25768 | 24.17585 |
| 25.68062 | 25.41774 | 25.40074 |
| 21.9 | 25.65835 | 25.66498 |
| 26.5499 | 26.36284 | 26.27174 |
| 24.053 | 24.52391 | 24.57894 |
| 27.00689 | 27.11922 | 27.20713 |
| 25.1395 | 25.79769 | 25.63273 |
| 26.26485 | 26.27672 | 25.81746 |
| 21.9 | 21.9 | 24.62684 |

| | | |
|---|---|---|
| 25.18699 | 21.9 | 25.36953 |
| 25.38426 | 21.9 | 25.58824 |
| 28.0168 | 28.21183 | 28.2818 |
| 26.47971 | 26.38014 | 26.52761 |
| 21.9 | 25.32841 | 21.9 |
| 26.43687 | 26.3292 | 26.14383 |
| 26.95903 | 21.9 | 26.52961 |
| 32.67548 | 32.39971 | 32.43904 |
| 29.96898 | 29.76161 | 29.79651 |
| 27.89689 | 27.81495 | 27.80901 |
| 27.55888 | 27.56456 | 27.53786 |
| 30.40892 | 30.38597 | 30.41538 |
| 29.79841 | 29.86642 | 29.87306 |
| 26.79228 | 26.76395 | 26.64952 |
| 26.72637 | 26.76344 | 26.55208 |
| 26.3695 | 26.63934 | 26.44046 |
| 24.92016 | 24.68024 | 21.9 |
| 25.83661 | 25.87298 | 25.86865 |
| 28.39321 | 28.3674 | 28.15769 |
| 24.63256 | 24.80681 | 24.70641 |
| 25.32934 | 21.9 | 21.9 |
| 24.49978 | 24.59429 | 24.79069 |
| 21.9 | 24.46557 | 24.43721 |
| 21.9 | 27.82287 | 27.86514 |
| 25.41841 | 25.29362 | 25.23787 |
| 26.4263 | 26.41274 | 26.35693 |
| 25.38265 | 25.38821 | 25.34571 |
| 25.88651 | 21.9 | 25.95578 |
| 25.49176 | 21.9 | 25.41015 |
| 25.35384 | 25.54439 | 21.9 |
| 21.9 | 21.9 | 25.10601 |
| 25.40983 | 25.34547 | 25.2006 |
| 25.93215 | 25.75009 | 25.84237 |
| 21.9 | 24.94011 | 25.10398 |
| 24.08058 | 21.9 | 24.15677 |
| 21.9 | 24.42323 | 24.56953 |
| 21.9 | 21.9 | 24.37033 |
| 21.9 | 24.62572 | 21.9 |
| 24.17326 | 24.20933 | 24.44569 |
| 21.9 | 21.9 | 21.9 |
| 21.9 | 21.9 | 21.9 |
| 21.9 | 21.9 | 21.9 |
| 30.69461 | 30.7809 | 30.71186 |
| 27.44103 | 27.94615 | 27.91787 |
| 21.9 | 22.07094 | 21.9 |
| 23.58348 | 23.56709 | 21.9 |
| 21.9 | 24.15707 | 21.9 |

| | | |
|---|---|---|
| 21.9 | 23.07666 | 21.9 |
| 25.74026 | 21.9 | 21.9 |
| 26.97334 | 27.03213 | 27.04583 |
| 24.80223 | 21.9 | 21.9 |
| 23.61526 | 24.03414 | 23.93706 |
| 22.81349 | 23.45939 | 23.14635 |
| 25.19843 | 21.9 | 25.48355 |
| 25.93645 | 25.41064 | 21.9 |
| 24.83677 | 24.72858 | 24.95296 |
| 27.06753 | 26.94782 | 27.00603 |
| 27.25449 | 27.13712 | 27.07735 |
| 21.9 | 23.51893 | 23.46389 |
| 21.9 | 28.38731 | 28.63917 |
| 21.9 | 21.9 | 25.5996 |
| 22.60707 | 22.52779 | 21.9 |
| 21.9 | 23.69594 | 23.60467 |
| 25.15815 | 25.05544 | 25.21686 |
| 21.9 | 21.9 | 21.9 |
| 24.77047 | 24.75251 | 24.80272 |
| 25.44998 | 25.41416 | 25.26628 |
| 26.05943 | 25.6238 | 21.9 |
| 21.9 | 21.9 | 21.9 |
| 25.65895 | 25.76527 | 25.69533 |
| 25.71206 | 25.806 | 21.9 |
