## Supplementary Table S3A for "TDP-43 pathology links innate and adaptive immunity in amyotrophic lateral sclerosis"

Ab_6_vs_PBS_6

| symbol | ensgene | baseMean | log2FoldChange | lfcSE | pvalue | padj | entrez | chr | start | end | strand | biotype | description |
|---|---|---|---|---|---|---|---|---|---|---|---|---|---|
| **GCLC** | ENSG00000001084 | 1727.431783 | -1.60259795 | 0.226180711 | 6.1E-14 | 1.3E-10 | 2729 | 6 | 53362139 | 53481768 | -1 | protein_coding | glutamate-cysteine ligase, catalytic subunit |
| **ENPP4** | ENSG00000001561 | 1288.757936 | 0.441424374 | 0.175005143 | 8.51E-04 | 0.020052664 | 22875 | 6 | 46097730 | 46114436 | 1 | protein_coding | ectonucleotide pyrophosphatase/phosphodiesterase 4 (putative) |
| **TFPI** | ENSG00000003436 | 1124.104413 | 0.421660281 | 0.162868091 | 8.17E-04 | 0.01972889 | 7035 | 2 | 188328957 | 188430487 | -1 | protein_coding | tissue factor pathway inhibitor (lipoprotein-associated coagulation inhibitor) |
| **CDC27** | ENSG00000004897 | 2659.465855 | -0.312753527 | 0.125806608 | 0.001697147 | 0.032148994 | 996 | 17 | 45195069 | 45266788 | -1 | protein_coding | cell division cycle 27 |
| **IBTK** | ENSG00000005700 | 3392.121785 | 0.254340851 | 0.10025296 | 0.002277788 | 0.039153257 | 25998 | 6 | 82879700 | 82957471 | -1 | protein_coding | inhibitor of Bruton agammaglobulinemia tyrosine kinase |
| **ZFX** | ENSG00000005889 | 1859.958012 | 0.342664189 | 0.124496161 | 6.79E-04 | 0.017556493 | 7543 | X | 24167290 | 24234372 | 1 | protein_coding | zinc finger protein, X-linked |
| **LAMP2** | ENSG00000005893 | 10671.37543 | 0.326236444 | 0.096038605 | 1.34E-04 | 0.005106443 | 3920 | X | 119561682 | 119603220 | -1 | protein_coding | lysosomal-associated membrane protein 2 |
| **IFRD1** | ENSG00000006652 | 1032.539441 | 0.386784049 | 0.1450156 | 7.2E-04 | 0.018310163 | 3475 | 7 | 112063023 | 112121072 | 1 | protein_coding | interferon-related developmental regulator 1 |
| **ELAC2** | ENSG00000006744 | 1381.089771 | -0.358931328 | 0.138949836 | 9.58E-04 | 0.022029371 | 60528 | 17 | 12895708 | 12921504 | -1 | protein_coding | elaC ribonuclease Z 2 |
| **RHOBTB2** | ENSG00000008853 | 1847.51946 | -0.599054164 | 0.195935729 | 1.18E-04 | 0.004712073 | 23221 | 8 | 22844930 | 22877712 | 1 | protein_coding | Rho-related BTB domain containing 2 |
| **STAB1** | ENSG00000010327 | 304.1996687 | 1.197323038 | 0.42107644 | 1.62E-04 | 0.005885838 | 23166 | 3 | 52529354 | 52558511 | 1 | protein_coding | stabilin 1 |
| **TYROBP** | ENSG00000011600 | 9924.722632 | -0.324364722 | 0.117050156 | 6.93E-04 | 0.017798355 | 7305 | 19 | 36395303 | 36399197 | -1 | protein_coding | TYRO protein tyrosine kinase binding protein |
| **MBTPS2** | ENSG00000012174 | 871.7749793 | 0.365314791 | 0.150825392 | 0.001486124 | 0.02966083 | 51360 | X | 21857754 | 21903542 | 1 | protein_coding | membrane-bound transcription factor peptidase, site 2 |
| **NUB1** | ENSG00000013374 | 1919.088021 | -0.544825959 | 0.129148657 | 1.83E-06 | 1.91E-04 | 51667 | 7 | 151038785 | 151075535 | 1 | protein_coding | negative regulator of ubiquitin-like proteins 1 |
| **CLK1** | ENSG00000013441 | 1756.888686 | 0.350036403 | 0.126660345 | 6.83E-04 | 0.017564779 | 1195 | 2 | 201717732 | 201729422 | -1 | protein_coding | CDC-like kinase 1 |
| **ATP2C1** | ENSG00000017260 | 2766.420926 | 0.347414601 | 0.123394892 | 5.52E-04 | 0.015238415 | 27032 | 3 | 130569439 | 130735556 | 1 | protein_coding | ATPase, Ca++ transporting, type 2C, member 1 |
| **RUFY3** | ENSG00000018189 | 1490.492251 | 0.353285892 | 0.126615058 | 5.73E-04 | 0.0155063 | 22902 | 4 | 71569921 | 71673032 | 1 | protein_coding | RUN and FYVE domain containing 3 |
| **SLC45A4** | ENSG00000022567 | 488.4537988 | 0.450481154 | 0.213008752 | 0.002147858 | 0.037199448 | 57210 | 8 | 142217265 | 142318404 | -1 | protein_coding | solute carrier family 45, member 4 |
| **RB1CC1** | ENSG00000023287 | 955.6863402 | 0.394846635 | 0.181077554 | 0.002376674 | 0.039900331 | 9821 | 8 | 53535016 | 53658403 | -1 | protein_coding | RB1-inducible coiled-coil 1 |
| **AKAP11** | ENSG00000023516 | 2907.548723 | 0.478573659 | 0.110256897 | 1.43E-06 | 1.56E-04 | 11215 | 13 | 42846289 | 42897396 | 1 | protein_coding | A kinase (PRKA) anchor protein 11 |
| **RABEP1** | ENSG00000029725 | 962.1114691 | -0.61620591 | 0.201794665 | 1.27E-04 | 0.004948792 | 9135 | 17 | 5185558 | 5289129 | 1 | protein_coding | rabaptin, RAB GTPase binding effector protein 1 |
| **TMSB10** | ENSG00000034510 | 25376.76464 | -0.348216861 | 0.123302304 | 5.75E-04 | 0.0155063 | 9168 | 2 | 85132749 | 85133795 | 1 | protein_coding | thymosin beta 10 |
| **MFAP3** | ENSG00000037749 | 155.6206858 | -1.023160937 | 0.43584886 | 6.18E-04 | 0.016366263 | 4238 | 5 | 153418466 | 153600038 | 1 | protein_coding | microfibrillar-associated protein 3 |
| **AGA** | ENSG00000038002 | 947.8002069 | -0.685638593 | 0.157553947 | 7.91E-07 | 9.67E-05 | 175 | 4 | 178351924 | 178363657 | -1 | protein_coding | aspartylglucosaminidase |
| **MAT2B** | ENSG00000038274 | 2741.4947 | 0.284881487 | 0.106652043 | 0.001429823 | 0.028900715 | 27430 | 5 | 162930120 | 162946342 | 1 | protein_coding | methionine adenosyltransferase II, beta |
| **CTNS** | ENSG00000040531 | 1815.379027 | -0.358060226 | 0.14565374 | 0.001348482 | 0.027988901 | 1497 | 17 | 3539762 | 3564836 | 1 | protein_coding | cystinosin, lysosomal cystine transporter |
| **CAPG** | ENSG00000042493 | 20312.04536 | -0.325407772 | 0.146845448 | 0.002913904 | 0.04527201 | 822 | 2 | 85621871 | 85645555 | -1 | protein_coding | capping protein (actin filament), gelsolin-like |
| **CTNNA1** | ENSG00000044115 | 11005.64681 | -0.227206833 | 0.086150835 | 0.001533854 | 0.030252811 | 1495 | 5 | 137946656 | 138270723 | 1 | protein_coding | catenin (cadherin-associated protein), alpha 1, 102kDa |
| **CNTLN** | ENSG00000044459 | 377.192498 | 0.504160236 | 0.26097552 | 0.002648748 | 0.042843594 | 54875 | 9 | 17134980 | 17503921 | 1 | protein_coding | centlein, centrosomal protein |
| **ARAP2** | ENSG00000047365 | 3332.714007 | 0.359858564 | 0.155526304 | 0.001985026 | 0.035453565 | 101928667 | 4 | 35949843 | 36246131 | -1 | protein_coding | ArfGAP with RhoGAP domain, ankyrin repeat and PH domain 2 |
| **GOPC** | ENSG00000047932 | 1091.20854 | 0.48209134 | 0.139307723 | 4.3E-05 | 0.002196011 | 57120 | 6 | 117639374 | 117923691 | -1 | protein_coding | golgi-associated PDZ and coiled-coil motif containing |
| **USP28** | ENSG00000048028 | 214.8833784 | -0.826707541 | 0.424211056 | 0.001591378 | 0.030804332 | 57646 | 11 | 113668596 | 113746292 | -1 | protein_coding | ubiquitin specific peptidase 28 |
| **FAM120A** | ENSG00000048828 | 5978.771282 | -1.270317182 | 0.226178448 | 8.25E-10 | 3.37E-07 | 23196 | 9 | 96214004 | 96328397 | 1 | protein_coding | family with sequence similarity 120A |
| **TNFRSF9** | ENSG00000049249 | 662.8762724 | 0.405617769 | 0.190441471 | 0.002525556 | 0.041629484 | 3604 | 1 | 7979907 | 8000926 | -1 | protein_coding | tumor necrosis factor receptor superfamily, member 9 |
| **RCN1** | ENSG00000049449 | 2249.309482 | 0.54684763 | 0.201347679 | 3.73E-04 | 0.011395793 | 440034 | 11 | 31833939 | 32127301 | 1 | protein_coding | reticulocalbin 1, EF-hand calcium binding domain |
| **LY75** | ENSG00000054219 | 3538.247375 | 0.799855266 | 0.158039972 | 2.46E-08 | 5.45E-06 | 100526664 | 2 | 160628362 | 160761260 | -1 | protein_coding | lymphocyte antigen 75 |
| **DCBLD2** | ENSG00000057019 | 302.2769123 | -0.60324519 | 0.352822501 | 0.003179705 | 0.047727009 | 131566 | 3 | 98514785 | 98620533 | -1 | protein_coding | discoidin, CUB and LCCL domain containing 2 |
| **CCAR1** | ENSG00000060339 | 1671.271121 | -0.432685016 | 0.16355941 | 6.21E-04 | 0.016366263 | 55749 | 10 | 70480769 | 70552134 | 1 | protein_coding | cell division cycle and apoptosis regulator 1 |
| **ACAA1** | ENSG00000060971 | 2407.656001 | -0.300542957 | 0.133498364 | 0.002345564 | 0.039729087 | 30 | 3 | 38144620 | 38178733 | -1 | protein_coding | acetyl-CoA acyltransferase 1 |
| **MON2** | ENSG00000061987 | 1562.62139 | 0.297639966 | 0.128052441 | 0.003348666 | 0.049425381 | 23041 | 12 | 62860597 | 62991363 | 1 | protein_coding | MON2 homolog (S. cerevisiae) |
| **WAPAL** | ENSG00000062650 | 3511.146657 | 0.295794089 | 0.120646708 | 0.002054537 | 0.036268374 | 23063 | 10 | 88195013 | 88281572 | -1 | protein_coding | wings apart-like homolog (Drosophila) |
| **VMP1** | ENSG00000062716 | 8522.183205 | 0.296693519 | 0.126105411 | 0.002756874 | 0.044122435 | 406991 | 17 | 57784553 | 57919616 | 1 | protein_coding | vacuole membrane protein 1 |
| **PKN2** | ENSG00000065243 | 720.554599 | 0.449343014 | 0.229620661 | 0.002835132 | 0.044635471 | 5586 | 1 | 89149905 | 89301938 | 1 | protein_coding | protein kinase N2 |
| **ME1** | ENSG00000065833 | 990.9163972 | 0.725141727 | 0.205279652 | 2.23E-05 | 0.001314419 | 4199 | 6 | 83920108 | 84140797 | -1 | protein_coding | malic enzyme 1, NADP(+)-dependent, cytosolic |
| **MTHFD2** | ENSG00000065911 | 7318.335508 | 0.320019893 | 0.100550687 | 1.9E-04 | 0.006742837 | 10797 | 2 | 74425689 | 74444692 | 1 | protein_coding | methylenetetrahydrofolate dehydrogenase (NADP+ dependent) 2, methenyltetrahydrofolate cyclohydrolase |
| **THUMPD1** | ENSG00000066654 | 1267.621892 | 0.9781873 | 0.33605721 | 1.34E-04 | 0.005113608 | 55623 | 16 | 20744986 | 20753406 | -1 | protein_coding | THUMP domain containing 1 |
| **TRAM1** | ENSG00000067167 | 3500.473156 | 1.481867029 | 0.218521524 | 4.99E-13 | 5.89E-10 | 23471 | 8 | 71485677 | 71520622 | -1 | protein_coding | translocation associated membrane protein 1 |
| **GRIPAP1** | ENSG00000068400 | 127.0568776 | 0.895248659 | 0.417709712 | 0.001030631 | 0.023227907 | 56850 | X | 48830134 | 48858675 | -1 | protein_coding | GRIP1 associated protein 1 |
| **TMEM260** | ENSG00000070269 | 499.6865934 | 0.413925706 | 0.209615328 | 0.003245323 | 0.048506395 | 54916 | 14 | 56955072 | 57117324 | 1 | protein_coding | transmembrane protein 260 |
| **ASNS** | ENSG00000070669 | 531.0710925 | 0.556017137 | 0.245171582 | 0.00120105 | 0.025578268 | 440 | 7 | 97481430 | 97501854 | -1 | protein_coding | asparagine synthetase (glutamine-hydrolyzing) |
| **UBE2D1** | ENSG00000072401 | 1015.990892 | 1.467645546 | 0.231843847 | 1.2E-11 | 9.83E-09 | 7321 | 10 | 60094735 | 60130513 | 1 | protein_coding | ubiquitin-conjugating enzyme E2D 1 |
| **MARK2** | ENSG00000072518 | 1334.472113 | -0.39748346 | 0.11873927 | 7.86E-05 | 0.003523824 | 2011 | 11 | 63606400 | 63678491 | 1 | protein_coding | MAP/microtubule affinity-regulating kinase 2 |
| **NFATC3** | ENSG00000072736 | 3549.294785 | -0.222477055 | 0.086153317 | 0.002530596 | 0.041629484 | 4775 | 16 | 68118654 | 68263162 | 1 | protein_coding | nuclear factor of activated T-cells, cytoplasmic, calcineurin-dependent 3 |
| **SMARCE1** | ENSG00000073584 | 1901.350688 | -0.307171301 | 0.130972926 | 0.002513459 | 0.041540487 | 6605 | 17 | 38781214 | 38804760 | -1 | protein_coding | SWI/SNF related, matrix associated, actin dependent regulator of chromatin, subfamily e, member 1 |
| **DHRS9** | ENSG00000073737 | 1576.547914 | -0.594154164 | 0.182827632 | 6.76E-05 | 0.003109201 | 10170 | 2 | 169921299 | 169952677 | 1 | protein_coding | dehydrogenase/reductase (SDR family) member 9 |
| **PICALM** | ENSG00000073921 | 9788.200053 | 0.440205509 | 0.124224009 | 3.55E-05 | 0.001907422 | 8301 | 11 | 85668727 | 85780924 | -1 | protein_coding | phosphatidylinositol binding clathrin assembly protein |
| **TSG101** | ENSG00000074319 | 1220.483836 | -0.434241869 | 0.19049029 | 0.001391513 | 0.02854751 | 7251 | 11 | 18489883 | 18548779 | -1 | protein_coding | tumor susceptibility 101 |
| **LMAN1** | ENSG00000074695 | 3823.831545 | 0.342558434 | 0.110344798 | 2.34E-04 | 0.007979974 | 3998 | 18 | 56995055 | 57027194 | -1 | protein_coding | lectin, mannose-binding, 1 |
| **SLC25A40** | ENSG00000075303 | 1108.76215 | 0.852547512 | 0.169519448 | 2.73E-08 | 5.91E-06 | 55972 | 7 | 87462883 | 87505672 | -1 | protein_coding | solute carrier family 25, member 40 |
| **TMEM131** | ENSG00000075568 | 2834.733838 | 0.327615474 | 0.095717853 | 7.72E-05 | 0.003477531 | 23505 | 2 | 98372799 | 98612388 | -1 | protein_coding | transmembrane protein 131 |
| **ACTB** | ENSG00000075624 | 2994.483306 | 1.875909185 | 0.955752497 | 7.34E-04 | 0.018448524 | 60 | 7 | 5566782 | 5603415 | -1 | protein_coding | actin, beta |
| **RAB7A** | ENSG00000075785 | 12794.54418 | -0.362823356 | 0.145588827 | 0.001232796 | 0.026100366 | 7879 | 3 | 128444965 | 128533639 | 1 | protein_coding | RAB7A, member RAS oncogene family |
| **PAG1** | ENSG00000076641 | 2864.415039 | 0.521418219 | 0.20355047 | 5.97E-04 | 0.015998816 | 55824 | 8 | 81880045 | 82024303 | -1 | protein_coding | phosphoprotein associated with glycosphingolipid microdomains 1 |
| **TOP2B** | ENSG00000077097 | 4113.482994 | 0.443939053 | 0.142193192 | 1.53E-04 | 0.005670372 | 7155 | 3 | 25639475 | 25706398 | -1 | protein_coding | topoisomerase (DNA) II beta 180kDa |
| **TM9SF3** | ENSG00000077147 | 5758.252891 | 0.800882531 | 0.188623057 | 1.17E-06 | 1.32E-04 | 56889 | 10 | 98277866 | 98347209 | -1 | protein_coding | transmembrane 9 superfamily member 3 |
| **USP33** | ENSG00000077254 | 2290.335346 | 0.279577547 | 0.12142242 | 0.003330591 | 0.049312487 | 23032 | 1 | 78161672 | 78225537 | -1 | protein_coding | ubiquitin specific peptidase 33 |
| **CAPZB** | ENSG00000077549 | 3119.415083 | -0.64765031 | 0.114047325 | 8.63E-10 | 3.4E-07 | 832 | 1 | 19665267 | 19812066 | -1 | protein_coding | capping protein (actin filament) muscle Z-line, beta |
| **KEAP1** | ENSG00000079999 | 1176.292787 | -0.468773273 | 0.153495014 | 1.31E-04 | 0.00503517 | 9817 | 19 | 10596796 | 10614417 | -1 | protein_coding | kelch-like ECH-associated protein 1 |
| **RIF1** | ENSG00000080345 | 2548.2115 | 0.487475057 | 0.115070108 | 2.22E-06 | 2.2E-04 | 101929336 | 2 | 152266397 | 152364527 | 1 | protein_coding | RAP1 interacting factor homolog (yeast) |
| **PSEN1** | ENSG00000080815 | 5237.597857 | -0.28288975 | 0.097710485 | 5.98E-04 | 0.015998816 | 5663 | 14 | 73603126 | 73690399 | 1 | protein_coding | presenilin 1 |
| **ZMPSTE24** | ENSG00000084073 | 2711.956438 | 0.36297847 | 0.134730547 | 5.31E-04 | 0.014875123 | 10269 | 1 | 40723779 | 40759856 | 1 | protein_coding | zinc metallopeptidase STE24 |
| **REST** | ENSG00000084093 | 3279.160179 | 0.410610295 | 0.116064378 | 3.68E-05 | 0.001955945 | 5978 | 4 | 57774075 | 57802010 | 1 | protein_coding | RE1-silencing transcription factor |
| **WDFY1** | ENSG00000085449 | 5232.369083 | 0.323789957 | 0.109497441 | 4.14E-04 | 0.012250455 | 57590 | 2 | 224720433 | 224810104 | -1 | protein_coding | WD repeat and FYVE domain containing 1 |
| **SLC25A24** | ENSG00000085491 | 2976.87445 | 0.523316276 | 0.11956454 | 9.48E-07 | 1.12E-04 | 29957 | 1 | 108676658 | 108743471 | -1 | protein_coding | solute carrier family 25 (mitochondrial carrier; phosphate carrier), member 24 |
| **B4GALT1** | ENSG00000086062 | 8068.364639 | 0.284644181 | 0.124769047 | 0.003363933 | 0.049510993 | 2683 | 9 | 33104080 | 33167354 | -1 | protein_coding | UDP-Gal:betaGlcNAc beta 1,4- galactosyltransferase, polypeptide 1 |
| **FTL** | ENSG00000087086 | 313149.2737 | -0.280311628 | 0.079519592 | 7.66E-05 | 0.003463742 | 2512 | 19 | 49468558 | 49470135 | 1 | protein_coding | ferritin, light polypeptide |
| **GMCL1** | ENSG00000087338 | 468.6898581 | 0.820174114 | 0.16339059 | 2.9E-08 | 6.16E-06 | 64395 | 2 | 70056774 | 70108528 | 1 | protein_coding | germ cell-less, spermatogenesis associated 1 |
| **ANKRD10** | ENSG00000088448 | 1111.662921 | 0.518529747 | 0.128680363 | 4.02E-06 | 3.37E-04 | 55608 | 13 | 111530887 | 111567416 | -1 | protein_coding | ankyrin repeat domain 10 |
| **DOCK3** | ENSG00000088538 | 661.1733312 | 2.286208787 | 0.28407911 | 3.99E-17 | 1.42E-13 | 1795 | 3 | 50712672 | 51421629 | 1 | protein_coding | dedicator of cytokinesis 3 |
| **SIGLEC1** | ENSG00000088827 | 2559.981037 | -0.573685912 | 0.272597857 | 0.001586828 | 0.030772302 | 6614 | 20 | 3667617 | 3687775 | -1 | protein_coding | sialic acid binding Ig-like lectin 1, sialoadhesin |
| **FKBP1A** | ENSG00000088832 | 5385.828353 | -0.256320127 | 0.102571396 | 0.002301112 | 0.039251876 | 101929368 | 20 | 1349622 | 1373806 | -1 | protein_coding | FK506 binding protein 1A, 12kDa |
| **RBBP9** | ENSG00000089050 | 834.3585382 | 0.498957074 | 0.165562155 | 1.73E-04 | 0.006236677 | 10741 | 20 | 18467184 | 18477887 | -1 | protein_coding | retinoblastoma binding protein 9 |
| **OAS1** | ENSG00000089127 | 3760.022614 | -0.267306429 | 0.111969288 | 0.002716577 | 0.043741007 | 4938 | 12 | 113344582 | 113369990 | 1 | protein_coding | 2'-5'-oligoadenylate synthetase 1, 40/46kDa |
| **GCN1L1** | ENSG00000089154 | 3009.582142 | 0.50136684 | 0.149750483 | 5.49E-05 | 0.002619909 | 10985 | 12 | 120565007 | 120632513 | -1 | protein_coding | GCN1 general control of amino-acid synthesis 1-like 1 (yeast) |
| **PXN** | ENSG00000089159 | 4845.848152 | -0.291030615 | 0.10154402 | 3.97E-04 | 0.011862182 | 5829 | 12 | 120648250 | 120703574 | -1 | protein_coding | paxillin |
| **RBM27** | ENSG00000091009 | 902.5188132 | 0.627815268 | 0.210873145 | 1.57E-04 | 0.005735941 | 54439 | 5 | 145583113 | 145718814 | 1 | protein_coding | RNA binding motif protein 27 |
| **WDR7** | ENSG00000091157 | 3087.62272 | -0.392599029 | 0.112393792 | 4.83E-05 | 0.002411147 | 23335 | 18 | 54318574 | 54698828 | 1 | protein_coding | WD repeat domain 7 |
| **CMTM6** | ENSG00000091317 | 9609.749365 | 0.497180639 | 0.127337625 | 7.53E-06 | 5.44E-04 | 54918 | 3 | 32522804 | 32544900 | -1 | protein_coding | CKLF-like MARVEL transmembrane domain containing 6 |
| **TOX4** | ENSG00000092203 | 669.8146571 | -0.42036172 | 0.190649673 | 0.00188895 | 0.034425705 | 9878 | 14 | 21944756 | 21967319 | 1 | protein_coding | TOX high mobility group box family member 4 |

| | | | | | | | | | | | | | |
|---|---|---|---|---|---|---|---|---|---|---|---|---|---|
| **SNAP23** | ENSG00000092531 | 3418.420609 | -0.487535131 | 0.140995944 | 3.44E-05 | 0.0018614 | 8773 | 15 | 42783431 | 42837547 | 1 | protein_coding | synaptosomal-associated protein, 23kDa |
| **PHGDH** | ENSG00000092621 | 369.115767 | 0.675956655 | 0.383150332 | 0.002605185 | 0.042552158 | 26227 | 1 | 120202421 | 120286838 | 1 | protein_coding | phosphoglycerate dehydrogenase |
| **MYL6** | ENSG00000092841 | 12236.39144 | -0.302678043 | 0.12341236 | 0.00190011 | 0.034517039 | 4637 | 12 | 56551945 | 56557280 | 1 | protein_coding | myosin, light chain 6, alkali, smooth muscle and non-muscle |
| **MAP3K1** | ENSG00000095015 | 1758.646251 | 0.333846756 | 0.128466564 | 0.001067545 | 0.023603651 | 4214 | 5 | 56111401 | 56191979 | 1 | protein_coding | mitogen-activated protein kinase kinase kinase 1, E3 ubiquitin protein ligase |
| **BTAF1** | ENSG00000095564 | 2251.104527 | 0.357935006 | 0.138184524 | 9.74E-04 | 0.022299301 | 9044 | 10 | 93683526 | 93790082 | 1 | protein_coding | BTAF1 RNA polymerase II, B-TFIID transcription factor-associated, 170kDa |
| **DSP** | ENSG00000096696 | 894.5352552 | -0.664801565 | 0.297832292 | 0.001040506 | 0.023344658 | 1832 | 6 | 7541808 | 7586950 | 1 | protein_coding | desmoplakin |
| **JAK2** | ENSG00000096968 | 1111.283539 | 1.097481837 | 0.244692971 | 3.07E-07 | 4.46E-05 | 3717 | 9 | 4985033 | 5128183 | 1 | protein_coding | Janus kinase 2 |
| **TSPAN15** | ENSG00000099282 | 764.0272189 | -0.572467293 | 0.291673148 | 0.001962891 | 0.035289354 | 23555 | 10 | 71211229 | 71267425 | 1 | protein_coding | tetraspanin 15 |
| **PSMD8** | ENSG00000099341 | 2378.338738 | -0.297219592 | 0.11599951 | 0.001511563 | 0.029968989 | 5714 | 19 | 38865176 | 38874464 | 1 | protein_coding | proteasome (prosome, macropain) 26S subunit, non-ATPase, 8 |
| **SNAP29** | ENSG00000099940 | 2390.502403 | -0.336745894 | 0.151403621 | 0.002814308 | 0.044505427 | 9342 | 22 | 21213271 | 21245506 | 1 | protein_coding | synaptosomal-associated protein, 29kDa |
| **NHP2L1** | ENSG00000100138 | 269.5132262 | -0.839833301 | 0.430903374 | 0.001528142 | 0.030241285 | 4809 | 22 | 42069934 | 42086508 | -1 | protein_coding | NHP2 non-histone chromosome protein 2-like 1 (S. cerevisiae) |
| **ACO2** | ENSG00000100412 | 4723.58188 | -0.388547454 | 0.102895692 | 2.09E-05 | 0.001248458 | 50 | 22 | 41865129 | 41924993 | 1 | protein_coding | aconitase 2, mitochondrial |
| **NIN** | ENSG00000100503 | 4152.346995 | -0.434661954 | 0.111568994 | 9.01E-06 | 6.14E-04 | 51199 | 14 | 51186481 | 51297839 | -1 | protein_coding | ninein (GSK3B interacting protein) |
| **PSMC6** | ENSG00000100519 | 505.2188811 | -0.591504642 | 0.330802646 | 0.002727134 | 0.04377832 | 5706 | 14 | 53173890 | 53195305 | 1 | protein_coding | proteasome (prosome, macropain) 26S subunit, ATPase, 6 |
| **DDHD1** | ENSG00000100523 | 1332.86892 | 0.340185787 | 0.151790671 | 0.002629355 | 0.042714716 | 80821 | 14 | 53510686 | 53620000 | -1 | protein_coding | DDHD domain containing 1 |
| **TMED8** | ENSG00000100580 | 1685.157567 | 0.47256132 | 0.152030503 | 1.42E-04 | 0.00532558 | 283578 | 14 | 77801364 | 77843452 | -1 | protein_coding | transmembrane emp24 protein transport domain containing 8 |
| **ERH** | ENSG00000100632 | 1131.410843 | 0.352534109 | 0.141576904 | 0.0012934 | 0.027163953 | 2079 | 14 | 69846848 | 69865344 | -1 | protein_coding | enhancer of rudimentary homolog (Drosophila) |
| **ZC3H14** | ENSG00000100722 | 1303.065121 | -0.354304247 | 0.13430679 | 9.28E-04 | 0.021545962 | 79882 | 14 | 89029253 | 89079853 | 1 | protein_coding | zinc finger CCCH-type containing 14 |
| **PSMB5** | ENSG00000100804 | 1189.836713 | -0.370324237 | 0.119372038 | 2.03E-04 | 0.007117793 | 5693 | 14 | 23485752 | 23504439 | -1 | protein_coding | proteasome (prosome, macropain) subunit, beta type, 5 |
| **STK4** | ENSG00000101109 | 5546.312868 | 0.386355175 | 0.110296675 | 4.94E-05 | 0.002443233 | 6789 | 20 | 43595115 | 43708600 | 1 | protein_coding | serine/threonine kinase 4 |
| **CTSZ** | ENSG00000101160 | 48907.27621 | -0.211245025 | 0.083205025 | 0.002862096 | 0.044933032 | 1522 | 20 | 57570240 | 57582302 | -1 | protein_coding | cathepsin Z |
| **DNTTIP1** | ENSG00000101457 | 928.2324985 | -0.44431036 | 0.152956443 | 2.81E-04 | 0.008979401 | 116092 | 20 | 44420576 | 44440066 | 1 | protein_coding | deoxynucleotidyltransferase, terminal, interacting protein 1 |
| **SMCHD1** | ENSG00000101596 | 1681.70928 | 0.441413282 | 0.183125955 | 0.001113548 | 0.024299132 | 23347 | 18 | 2655737 | 2805015 | 1 | protein_coding | structural maintenance of chromosomes flexible hinge domain containing 1 |
| **PRPS2** | ENSG00000101911 | 1026.265808 | 0.391815259 | 0.156164426 | 0.00106835 | 0.023603651 | 5634 | X | 12809474 | 12842341 | 1 | protein_coding | phosphoribosyl pyrophosphate synthetase 2 |
| **ATP11C** | ENSG00000101974 | 541.0391053 | 0.761732489 | 0.245713578 | 9.64E-05 | 0.004079507 | 286410 | X | 138808505 | 139027435 | -1 | protein_coding | ATPase, class VI, type 11C |
| **PGK 1.00** | ENSG00000102144 | 4634.27673 | 0.319839894 | 0.122472334 | 0.001150906 | 0.024758449 | 5230 | X | 77320685 | 77384793 | 1 | protein_coding | phosphoglycerate kinase 1 |
| **RP2** | ENSG00000102218 | 2141.374418 | 0.269831549 | 0.110508532 | 0.002552325 | 0.041857347 | 6102 | X | 46696375 | 46741793 | 1 | protein_coding | retinitis pigmentosa 2 (X-linked recessive) |
| **ARMCX3** | ENSG00000102401 | 1854.916132 | 0.290809237 | 0.119739896 | 0.002215781 | 0.03828798 | 51566 | X | 100877787 | 100882833 | 1 | protein_coding | armadillo repeat containing, X-linked 3 |
| **BEX4** | ENSG00000102409 | 854.885347 | 0.368804528 | 0.16000125 | 0.002007872 | 0.035681707 | 56271 | X | 102470020 | 102472174 | 1 | protein_coding | brain expressed, X-linked 4 |
| **STK24** | ENSG00000102572 | 4674.491942 | -0.337094664 | 0.111467663 | 2.91E-04 | 0.0092353 | 8428 | 13 | 99102455 | 99230194 | -1 | protein_coding | serine/threonine kinase 24 |
| **CCL22** | ENSG00000102962 | 15258.76342 | -0.537714823 | 0.190003132 | 2.78E-04 | 0.00896698 | 6367 | 16 | 57392684 | 57400102 | 1 | protein_coding | chemokine (C-C motif) ligand 22 |
| **SLC7A5** | ENSG00000103257 | 1125.361446 | 0.676697715 | 0.314620634 | 0.001296032 | 0.027165545 | 8140 | 16 | 87863629 | 87903094 | -1 | protein_coding | solute carrier family 7 (amino acid transporter light chain, L system), member 5 |
| **TCEB2** | ENSG00000103363 | 1563.469711 | -0.477982526 | 0.154557008 | 0.000146041 | 0.005426509 | 6923 | 16 | 2821415 | 2827298 | -1 | protein_coding | transcription elongation factor B (SIII), polypeptide 2 (18kDa, elongin B) |
| **DMXL2** | ENSG00000104093 | 15151.72408 | 0.431075527 | 0.09216221 | 2.85E-07 | 4.21E-05 | 23312 | 15 | 51739908 | 51915030 | -1 | protein_coding | Dmx-like 2 |
| **RIPK2** | ENSG00000104312 | 947.2706168 | 0.583652554 | 0.174773058 | 5.06E-05 | 0.002476704 | 8767 | 8 | 90769975 | 90803291 | 1 | protein_coding | receptor-interacting serine-threonine kinase 2 |
| **NBN** | ENSG00000104320 | 5818.648623 | 0.465620421 | 0.162527288 | 2.77E-04 | 0.008952663 | 4683 | 8 | 90945564 | 91015456 | -1 | protein_coding | nibrin |
| **IMPAD1** | ENSG00000104331 | 2306.037087 | 0.570660277 | 0.149721119 | 9.46E-06 | 6.4E-04 | 54928 | 8 | 57870492 | 57906403 | -1 | protein_coding | inositol monophosphatase domain containing 1 |
| **UBE2W** | ENSG00000104343 | 1829.04755 | 0.639143795 | 0.158698423 | 3.46E-06 | 3.06E-04 | 55284 | 8 | 74692332 | 74791145 | -1 | protein_coding | ubiquitin-conjugating enzyme E2W (putative) |
| **POP1** | ENSG00000104356 | 386.3846525 | -0.498618496 | 0.249029997 | 0.002284278 | 0.039153257 | 10940 | 8 | 99129525 | 99172062 | 1 | protein_coding | processing of precursor 1, ribonuclease P/MRP subunit (S. cerevisiae) |
| **GSR** | ENSG00000104687 | 232.4801958 | -0.831783521 | 0.302890949 | 2.51E-04 | 0.00837387 | 2936 | 8 | 30535583 | 30585443 | -1 | protein_coding | glutathione reductase |
| **ASAH1** | ENSG00000104763 | 9481.129518 | -0.589640863 | 0.209489235 | 2.59E-04 | 0.008480193 | 427 | 8 | 17913934 | 17942494 | -1 | protein_coding | N-acylsphingosine amidohydrolase (acid ceramidase) 1 |
| **OAZ1** | ENSG00000104904 | 9610.992179 | -0.424837969 | 0.089188527 | 1.91E-07 | 2.94E-05 | 4946 | 19 | 2269485 | 2273487 | 1 | protein_coding | ornithine decarboxylase antizyme 1 |
| **POLR2I** | ENSG00000105258 | 197.4544306 | -0.599105079 | 0.342962257 | 0.002909566 | 0.04527201 | 5438 | 19 | 36604612 | 36606248 | -1 | protein_coding | polymerase (RNA) II (DNA directed) polypeptide I, 14.5kDa |
| **PRKD2** | ENSG00000105287 | 683.5731773 | -0.442111879 | 0.18315622 | 0.001121432 | 0.024317464 | 25865 | 19 | 47177532 | 47220384 | -1 | protein_coding | protein kinase D2 |
| **PPP2R1A** | ENSG00000105568 | 3132.127127 | -0.316972497 | 0.127833941 | 0.001674693 | 0.0318942 | 5518 | 19 | 52693292 | 52730687 | 1 | protein_coding | protein phosphatase 2, regulatory subunit A, alpha |
| **TNPO2** | ENSG00000105576 | 579.0204954 | 0.727150505 | 0.165818006 | 6.45E-07 | 8.35E-05 | 30000 | 19 | 12810008 | 12834825 | -1 | protein_coding | transportin 2 |
| **USF2** | ENSG00000105698 | 3782.047955 | -0.277722894 | 0.119535522 | 0.003119142 | 0.047087076 | 7392 | 19 | 35759881 | 35770724 | 1 | protein_coding | upstream transcription factor 2, c-fos interacting |
| **CBLL1** | ENSG00000105879 | 881.3699345 | -0.442225064 | 0.167000564 | 6.01E-04 | 0.016047494 | 79872 | 7 | 107384142 | 107402112 | 1 | protein_coding | Cbl proto-oncogene-like 1, E3 ubiquitin protein ligase |
| **MTPN** | ENSG00000105887 | 8951.002594 | 0.245927121 | 0.09932035 | 0.00237487 | 0.039900331 | 767558 | 7 | 135611509 | 135662101 | -1 | protein_coding | myotrophin |
| **DFNA5** | ENSG00000105928 | 1922.243663 | 0.314736619 | 0.127926919 | 0.001845594 | 0.034319461 | 1687 | 7 | 24737972 | 24809244 | -1 | protein_coding | deafness, autosomal dominant 5 |
| **OGDH** | ENSG00000105953 | 4525.439494 | -0.319503924 | 0.110885863 | 5.06E-04 | 0.014291006 | 4967 | 7 | 44646171 | 44748665 | 1 | protein_coding | oxoglutarate (alpha-ketoglutarate) dehydrogenase (lipoamide) |
| **MET** | ENSG00000105976 | 185.724974 | 1.242858522 | 0.428707393 | 1.32E-04 | 0.005072528 | 4233 | 7 | 116312444 | 116438440 | 1 | protein_coding | met proto-oncogene |
| **DNAJB6** | ENSG00000105993 | 5291.761351 | -0.546796617 | 0.102076807 | 5.71E-09 | 1.84E-06 | 10049 | 7 | 157128075 | 157210133 | 1 | protein_coding | DnaJ (Hsp40) homolog, subfamily B, member 6 |
| **HSPB1** | ENSG00000106211 | 1692.054685 | -0.609123928 | 0.123781704 | 5.6E-08 | 1.04E-05 | 3315 | 7 | 75931861 | 75933612 | 1 | protein_coding | heat shock 27kDa protein 1 |
| **NPTX2** | ENSG00000106236 | 275.3784218 | -0.992661708 | 0.486168249 | 0.001147749 | 0.024740637 | 4885 | 7 | 98246609 | 98259180 | 1 | protein_coding | neuronal pentraxin II |
| **ZNHIT1** | ENSG00000106400 | 690.6210997 | -0.334645407 | 0.152185144 | 0.002930488 | 0.045310196 | 10467 | 7 | 100860949 | 100867471 | 1 | protein_coding | zinc finger, HIT-type containing 1 |
| **EDF1** | ENSG00000107223 | 2536.898523 | -0.335886606 | 0.12218719 | 6.26E-04 | 0.016470152 | 8721 | 9 | 139756571 | 139760738 | -1 | protein_coding | endothelial differentiation-related factor 1 |
| **ZFAND5** | ENSG00000107372 | 8578.302986 | -0.224318677 | 0.089225956 | 0.002862488 | 0.044933032 | 7763 | 9 | 74966341 | 74980163 | -1 | protein_coding | zinc finger, AN1-type domain 5 |
| **ACTA2** | ENSG00000107796 | 741.2429511 | -0.50626701 | 0.204341596 | 7.6E-04 | 0.018841257 | 59 | 10 | 90694831 | 90751147 | -1 | protein_coding | actin, alpha 2, smooth muscle, aorta |
| **TNKS2** | ENSG00000107854 | 3650.066086 | 0.362038277 | 0.092788047 | 1.07E-05 | 7.04E-04 | 80351 | 10 | 93558069 | 93625033 | 1 | protein_coding | tankyrase, TRF1-interacting ankyrin-related ADP-ribose polymerase 2 |
| **RPL28** | ENSG00000108107 | 4161.805554 | -0.451975329 | 0.13889902 | 1.09E-04 | 0.004503741 | 102465483 | 19 | 55896713 | 55914612 | 1 | protein_coding | ribosomal protein L28 |
| **TBC1D12** | ENSG00000108239 | 559.5762447 | 0.428569975 | 0.194161519 | 0.001863889 | 0.034425705 | 23232 | 10 | 96162261 | 96295687 | 1 | protein_coding | TBC1 domain family, member 12 |
| **NUFIP2** | ENSG00000108256 | 5845.599241 | 0.445462467 | 0.114185217 | 8.9E-06 | 6.14E-04 | 57532 | 17 | 27582854 | 27621136 | -1 | protein_coding | nuclear fragile X mental retardation protein interacting protein 2 |
| **RPL19** | ENSG00000108298 | 9177.207857 | -0.345993424 | 0.117685035 | 3.9E-04 | 0.011801021 | 6143 | 17 | 37356536 | 37360980 | 1 | protein_coding | ribosomal protein L19 |
| **DDX5** | ENSG00000108654 | 7844.236253 | 0.668741727 | 0.130720728 | 2.02E-08 | 4.99E-06 | 100616408 | 17 | 62495734 | 62504317 | -1 | protein_coding | DEAD (Asp-Glu-Ala-Asp) box helicase 5 |
| **CCL1** | ENSG00000108702 | 388.2883384 | -1.348928596 | 0.559142265 | 6.82E-04 | 0.017564779 | 6346 | 17 | 32687347 | 32690250 | -1 | protein_coding | chemokine (C-C motif) ligand 1 |
| **MMD** | ENSG00000108960 | 1068.731335 | 0.478033006 | 0.175746639 | 5.4E-04 | 0.01508973 | 23531 | 17 | 53469974 | 53499353 | -1 | protein_coding | monocyte to macrophage differentiation-associated |
| **KLF3** | ENSG00000109787 | 1426.926797 | 0.51900998 | 0.128629975 | 3.75E-06 | 3.24E-04 | 51274 | 4 | 38665817 | 38702663 | 1 | protein_coding | Kruppel-like factor 3 (basic) |
| **FNBP4** | ENSG00000109920 | 1805.219448 | 0.283893768 | 0.124348445 | 0.002993153 | 0.045767245 | 23360 | 11 | 47738072 | 47788995 | -1 | protein_coding | formin binding protein 4 |
| **HSPA8** | ENSG00000109971 | 39858.47574 | -0.432783861 | 0.085799796 | 4.51E-08 | 8.72E-06 | 85390 | 11 | 122928197 | 122933938 | -1 | protein_coding | heat shock 70kDa protein 8 |
| **CBL** | ENSG00000110395 | 4937.873111 | 0.250274594 | 0.10173051 | 0.002670528 | 0.043130244 | 867 | 11 | 119076752 | 119178859 | 1 | protein_coding | Cbl proto-oncogene, E3 ubiquitin protein ligase |
| **SLC15A3** | ENSG00000110446 | 9991.947152 | -0.286777156 | 0.112624456 | 0.001399607 | 0.028603122 | 51296 | 11 | 60704556 | 60720002 | -1 | protein_coding | solute carrier family 15 (oligopeptide transporter), member 3 |
| **NDUFS8** | ENSG00000110717 | 746.5974872 | -0.358219242 | 0.16298285 | 0.002427302 | 0.040430931 | 102465669 | 11 | 67798084 | 67804111 | 1 | protein_coding | NADH dehydrogenase (ubiquinone) Fe-S protein 8, 23kDa (NADH-coenzyme Q reductase) |
| **CORO1C** | ENSG00000110880 | 15720.25581 | -0.318767962 | 0.136323378 | 0.002355258 | 0.039729087 | 23603 | 12 | 109038885 | 109125372 | -1 | protein_coding | coronin, actin binding protein, 1C |
| **PTGES3** | ENSG00000110958 | 3997.27165 | 0.639182264 | 0.135390261 | 1.52E-07 | 2.48E-05 | 10728 | 12 | 57057127 | 57082159 | -1 | protein_coding | prostaglandin E synthase 3 (cytosolic) |
| **CREBL2** | ENSG00000111269 | 4865.757889 | 0.425507938 | 0.105730496 | 5.37E-06 | 4.33E-04 | 1389 | 12 | 12764761 | 12798042 | 1 | protein_coding | cAMP responsive element binding protein-like 2 |
| **CDKN1B** | ENSG00000111276 | 860.3358505 | 0.444710791 | 0.197830813 | 0.001602858 | 0.030970126 | 1027 | 12 | 12867992 | 12875305 | 1 | protein_coding | cyclin-dependent kinase inhibitor 1B (p27, Kip1) |
| **CHPT1** | ENSG00000111666 | 808.3303355 | 0.456128049 | 0.201904037 | 0.001541997 | 0.030345931 | 56994 | 12 | 102090725 | 102137918 | 1 | protein_coding | choline phosphotransferase 1 |
| **ENO2** | ENSG00000111674 | 999.7739498 | 0.443210665 | 0.194283579 | 0.001495181 | 0.029715908 | 2026 | 12 | 7022909 | 7032861 | 1 | protein_coding | enolase 2 (gamma, neuronal) |
| **PTPN6** | ENSG00000111679 | 7351.367895 | -0.286316855 | 0.103591889 | 9.43E-04 | 0.021793875 | 5777 | 12 | 7055631 | 7070479 | 1 | protein_coding | protein tyrosine phosphatase, non-receptor type 6 |
| **C2CD5** | ENSG00000111731 | 1460.081226 | 0.45920909 | 0.143714213 | 1.13E-04 | 0.004572138 | 9847 | 12 | 22601517 | 22697480 | -1 | protein_coding | C2 calcium-dependent domain containing 5 |
| **COX6A1** | ENSG00000111775 | 1585.980078 | -0.329065677 | 0.125868102 | 0.001090608 | 0.023896694 | 1337 | 12 | 120875893 | 120878545 | 1 | protein_coding | cytochrome c oxidase subunit VIa polypeptide 1 |
| **DSE** | ENSG00000111817 | 1919.488843 | 0.914449415 | 0.160223388 | 6E-10 | 2.77E-07 | 29940 | 6 | 116575336 | 116762424 | 1 | protein_coding | dermatan sulfate epimerase |
| **MAN1A1** | ENSG00000111885 | 6498.164924 | 0.334873243 | 0.121713603 | 7.19E-04 | 0.018310163 | 4121 | 6 | 119498374 | 119670926 | -1 | protein_coding | mannosidase, alpha, class 1A, member 1 |
| **SERINC1** | ENSG00000111897 | 12078.40512 | 0.283973477 | 0.114715255 | 0.002095587 | 0.036809591 | 57515 | 6 | 122764499 | 122792967 | -1 | protein_coding | serine incorporator 1 |
| **HINT3** | ENSG00000111911 | 1053.866973 | 0.873755065 | 0.143414685 | 6.3E-11 | 3.64E-08 | 135114 | 6 | 126277927 | 126301387 | 1 | protein_coding | histidine triad nucleotide binding protein 3 |
| **CCNC** | ENSG00000112237 | 769.4655682 | 0.436960376 | 0.172569688 | 8.51E-04 | 0.020052664 | 892 | 6 | 99990256 | 100016849 | -1 | protein_coding | cyclin C |
| **AIM1** | ENSG00000112297 | 108.1656172 | -0.020827515 | 0.144304001 | 0.002967825 | 0.045576695 | 202 | 6 | 106959730 | 107018326 | 1 | protein_coding | absent in melanoma 1 |
| **TRIM38** | ENSG00000112343 | 2183.938053 | 0.291216511 | 0.127667151 | 0.003027986 | 0.046115879 | 10475 | 6 | 25963030 | 25987384 | 1 | protein_coding | tripartite motif containing 38 |
| **HECA** | ENSG00000112406 | 2812.086759 | 0.64052611 | 0.1156538 | 2.14E-09 | 7.58E-07 | 51696 | 6 | 139456249 | 139501939 | 1 | protein_coding | headcase homolog (Drosophila) |
| **PRPF4B** | ENSG00000112739 | 2361.40671 | 0.712053649 | 0.139992886 | 2.34E-08 | 5.29E-06 | 8899 | 6 | 4021501 | 4065217 | 1 | protein_coding | pre-mRNA processing factor 4B |
| **FAM46A** | ENSG00000112773 | 2902.211179 | 0.511834502 | 0.108267238 | 1.93E-07 | 2.94E-05 | 55603 | 6 | 82201156 | 82462491 | -1 | protein_coding | family with sequence similarity 46, member A |

| | | | | | | | | | | | | | |
|---|---|---|---|---|---|---|---|---|---|---|---|---|---|
| **MAN2A1** | ENSG00000112893 | 3493.776539 | 0.463503478 | 0.116377308 | 6.34E-06 | 4.85E-04 | 4124 | 5 | 109025067 | 109205326 | 1 | protein_coding | mannosidase, alpha, class 2A, member 1 |
| **DBN1** | ENSG00000113758 | 264.7552809 | -1.06288635 | 0.298264395 | 1.49E-05 | 9.47E-04 | 1627 | 5 | 176883609 | 176901402 | -1 | protein_coding | drebrin 1 |
| **SLC25A36** | ENSG00000114120 | 1244.894529 | 0.498749705 | 0.128701638 | 8.09E-06 | 5.73E-04 | 55186 | 3 | 140660672 | 140698775 | 1 | protein_coding | solute carrier family 25 (pyrimidine nucleotide carrier ), member 36 |
| **WNT5A** | ENSG00000114251 | 416.2395885 | 0.812402172 | 0.430231261 | 0.001936075 | 0.03499094 | 7474 | 3 | 55499743 | 55523973 | -1 | protein_coding | wingless-type MMTV integration site family, member 5A |
| **RPL24** | ENSG00000114391 | 2481.354068 | -0.288598489 | 0.124777883 | 0.003074925 | 0.046615158 | 6152 | 3 | 101399935 | 101405626 | -1 | protein_coding | ribosomal protein L24 |
| **SNX4** | ENSG00000114520 | 1053.396801 | -0.455131975 | 0.167600915 | 5.3E-04 | 0.014875123 | 8723 | 3 | 125165495 | 125239041 | -1 | protein_coding | sorting nexin 4 |
| **ATP6V1A** | ENSG00000114573 | 19146.80592 | -0.244627741 | 0.090961763 | 0.001447224 | 0.029018201 | 523 | 3 | 113465866 | 113530903 | 1 | protein_coding | ATPase, H+ transporting, lysosomal 70kDa, V1 subunit A |
| **EIF4G1** | ENSG00000114867 | 17546.77642 | -0.28326475 | 0.088332169 | 2.31E-04 | 0.007916945 | 1981 | 3 | 184032283 | 184053146 | 1 | protein_coding | eukaryotic translation initiation factor 4 gamma, 1 |
| **IL1A** | ENSG00000115008 | 156.1526996 | 0.702066577 | 0.402153921 | 0.00262329 | 0.042714716 | 3552 | 2 | 113531492 | 113542167 | -1 | protein_coding | interleukin 1, alpha |
| **SLC35F5** | ENSG00000115084 | 2058.645881 | 0.502071483 | 0.131928094 | 1.09E-05 | 7.12E-04 | 80255 | 2 | 114462588 | 114514400 | -1 | protein_coding | solute carrier family 35, member F5 |
| **MPV17** | ENSG00000115204 | 1021.593457 | -0.315342165 | 0.130743185 | 0.00202174 | 0.03574881 | 4358 | 2 | 27532360 | 27548547 | -1 | protein_coding | MpV17 mitochondrial inner membrane protein |
| **RPS15** | ENSG00000115268 | 2185.977697 | -0.365067712 | 0.163341675 | 0.0019664 | 0.035289354 | 6209 | 19 | 1438358 | 1440583 | 1 | protein_coding | ribosomal protein S15 |
| **SPTBN1** | ENSG00000115306 | 2429.710156 | 0.361158268 | 0.151055396 | 0.001612163 | 0.03109339 | 6711 | 2 | 54683422 | 54896812 | 1 | protein_coding | spectrin, beta, non-erythrocytic 1 |
| **RTN4** | ENSG00000115310 | 11738.70143 | 0.32871628 | 0.14405719 | 0.002299315 | 0.039251876 | 57142 | 2 | 55199325 | 55339757 | -1 | protein_coding | reticulon 4 |
| **USP34** | ENSG00000115464 | 4635.12003 | 0.464053172 | 0.100393624 | 3.41E-07 | 4.9E-05 | 9736 | 2 | 61414598 | 61697904 | -1 | protein_coding | ubiquitin specific peptidase 34 |
| **MOB4** | ENSG00000115540 | 535.8460243 | 0.849029062 | 0.257421902 | 4.44E-05 | 0.00224809 | 100529241 | 2 | 198380295 | 198418423 | 1 | protein_coding | MOB family member 4, phocein |
| **PRKD3** | ENSG00000115825 | 2276.561104 | 0.287060893 | 0.122254485 | 0.002872775 | 0.045027998 | 23683 | 2 | 37477645 | 37551951 | -1 | protein_coding | protein kinase D3 |
| **QPCT** | ENSG00000115828 | 2102.367974 | -0.38839366 | 0.169441561 | 0.001877462 | 0.034425705 | 25797 | 2 | 37571717 | 37600465 | 1 | protein_coding | glutaminyl-peptide cyclotransferase |
| **DARS** | ENSG00000115866 | 1665.041657 | 0.341267224 | 0.139261639 | 0.001441069 | 0.02900424 | 1615 | 2 | 136664247 | 136743670 | -1 | protein_coding | aspartyl-tRNA synthetase |
| **PLCL1** | ENSG00000115896 | 712.8621079 | -0.587144197 | 0.272087069 | 0.001371444 | 0.028328895 | 5334 | 2 | 198669426 | 199437305 | 1 | protein_coding | phospholipase C-like 1 |
| **DHCR24** | ENSG00000116133 | 10361.82938 | -0.435516014 | 0.172634892 | 8.48E-04 | 0.020052664 | 1718 | 1 | 55315306 | 55352891 | -1 | protein_coding | 24-dehydrocholesterol reductase |
| **STXBP3** | ENSG00000116266 | 1616.631323 | 0.40313276 | 0.162201803 | 0.001060504 | 0.02354425 | 6814 | 1 | 109289296 | 109352148 | 1 | protein_coding | syntaxin binding protein 3 |
| **ATP5F1** | ENSG00000116459 | 3848.868841 | -0.312027908 | 0.116875828 | 0.001005302 | 0.022827652 | 515 | 1 | 111991486 | 112005395 | 1 | protein_coding | ATP synthase, H+ transporting, mitochondrial Fo complex, subunit B1 |
| **RAP1A** | ENSG00000116473 | 1991.468319 | 0.805295287 | 0.158235912 | 2.25E-08 | 5.26E-06 | 5906 | 1 | 112084840 | 112259313 | 1 | protein_coding | RAP1A, member of RAS oncogene family |
| **CAPZA1** | ENSG00000116489 | 10918.19695 | -0.299034807 | 0.098807643 | 3.97E-04 | 0.011862182 | 829 | 1 | 113161795 | 113214241 | 1 | protein_coding | capping protein (actin filament) muscle Z-line, alpha 1 |
| **LAMTOR2** | ENSG00000116586 | 935.1780954 | -0.360482467 | 0.13316191 | 7.34E-04 | 0.018448524 | 28956 | 1 | 156024543 | 156028301 | 1 | protein_coding | late endosomal/lysosomal adaptor, MAPK and MTOR activator 2 |
| **RLF** | ENSG00000117000 | 1207.185662 | 0.33036332 | 0.149236712 | 0.003056778 | 0.046406251 | 6018 | 1 | 40627045 | 40706593 | 1 | protein_coding | rearranged L-myc fusion |
| **SDHB** | ENSG00000117118 | 2018.273575 | -0.333350472 | 0.149980865 | 0.002792067 | 0.044235629 | 6390 | 1 | 17345217 | 17380665 | -1 | protein_coding | succinate dehydrogenase complex, subunit B, iron sulfur (Ip) |
| **ECE1** | ENSG00000117298 | 1968.121784 | -0.586174308 | 0.191758451 | 1.3E-04 | 0.00503517 | 1889 | 1 | 21543740 | 21671997 | -1 | protein_coding | endothelin converting enzyme 1 |
| **HMGCL** | ENSG00000117305 | 1149.491122 | -0.328397933 | 0.130877378 | 0.001472146 | 0.029462318 | 3155 | 1 | 24128375 | 24165110 | -1 | protein_coding | 3-hydroxymethyl-3-methylglutaryl-CoA lyase |
| **TMED5** | ENSG00000117500 | 5108.663661 | 0.659171714 | 0.161773256 | 2.79E-06 | 2.59E-04 | 50999 | 1 | 93615299 | 93646285 | -1 | protein_coding | transmembrane emp24 protein transport domain containing 5 |
| **ABCD3** | ENSG00000117528 | 502.7293149 | 1.032337199 | 0.260504872 | 3.36E-06 | 3.05E-04 | 5825 | 1 | 94883933 | 94984222 | 1 | protein_coding | ATP-binding cassette, sub-family D (ALD), member 3 |
| **SLC35A3** | ENSG00000117620 | 390.9588777 | 0.537153903 | 0.243411754 | 0.001329879 | 0.027711038 | 23443 | 1 | 100435345 | 100492535 | 1 | protein_coding | solute carrier family 35 (UDP-N-acetylglucosamine (UDP-GlcNAc) transporter), member A3 |
| **STX12** | ENSG00000117758 | 3107.215955 | 0.327713692 | 0.094957922 | 8.14E-05 | 0.003604146 | 23673 | 1 | 28099694 | 28150963 | 1 | protein_coding | syntaxin 12 |
| **STAG1** | ENSG00000118007 | 1119.73107 | 0.389603272 | 0.126875812 | 2.01E-04 | 0.007081017 | 10274 | 3 | 136055077 | 136471220 | -1 | protein_coding | stromal antigen 1 |
| **VAMP8** | ENSG00000118640 | 5007.174276 | -0.322205989 | 0.137968296 | 0.001981241 | 0.035445532 | 8673 | 2 | 85788685 | 85809154 | 1 | protein_coding | vesicle-associated membrane protein 8 |
| **MYL12B** | ENSG00000118680 | 4538.954797 | -0.330195826 | 0.132752336 | 0.001324614 | 0.027655547 | 103910 | 18 | 3261907 | 3278282 | 1 | protein_coding | myosin, light chain 12B, regulatory |
| **SPP1** | ENSG00000118785 | 126274.6747 | -0.323126846 | 0.114813766 | 6.2E-04 | 0.016366263 | 6696 | 4 | 88896819 | 88904562 | 1 | protein_coding | secreted phosphoprotein 1 |
| **CCND2** | ENSG00000118971 | 8250.780044 | 0.332300456 | 0.100508229 | 1.25E-04 | 0.004928845 | 894 | 12 | 4382938 | 4414516 | 1 | protein_coding | cyclin D2 |
| **SENP5** | ENSG00000119231 | 1922.607751 | 0.264024305 | 0.110641866 | 0.002793084 | 0.044235629 | 205564 | 3 | 196594727 | 196661585 | 1 | protein_coding | SUMO1/sentrin specific peptidase 5 |
| **PTBP3** | ENSG00000119314 | 5018.509103 | 0.313432379 | 0.107163668 | 4.81E-04 | 0.013704654 | 9991 | 9 | 114980715 | 115095947 | -1 | protein_coding | polypyrimidine tract binding protein 3 |
| **HSDL2** | ENSG00000119471 | 596.207706 | 0.396192644 | 0.176891136 | 0.00196919 | 0.035289354 | 84263 | 9 | 115142217 | 115234690 | 1 | protein_coding | hydroxysteroid dehydrogenase like 2 |
| **YIPF4** | ENSG00000119820 | 1224.587165 | 0.412012467 | 0.196573831 | 0.002388462 | 0.040034996 | 84272 | 2 | 32502979 | 32541663 | 1 | protein_coding | Yip1 domain family, member 4 |
| **OGFRL1** | ENSG00000119900 | 4382.414166 | 0.371695331 | 0.092003252 | 5.64E-06 | 4.44E-04 | 79627 | 6 | 71998506 | 72018653 | 1 | protein_coding | opioid growth factor receptor-like 1 |
| **NUP43** | ENSG00000120253 | 588.409048 | 0.656017154 | 0.208317119 | 8.36E-05 | 0.003670953 | 348995 | 6 | 150045451 | 150070801 | -1 | protein_coding | nucleoporin 43kDa |
| **UFM1** | ENSG00000120686 | 2486.856184 | 0.46741145 | 0.123288397 | 1.34E-05 | 8.59E-04 | 51569 | 13 | 38923986 | 38937140 | 1 | protein_coding | ubiquitin-fold modifier 1 |
| **HSPH1** | ENSG00000120694 | 2823.048163 | 0.314637591 | 0.139707131 | 0.002988223 | 0.045757704 | 10808 | 13 | 31710762 | 31736525 | -1 | protein_coding | heat shock 105kDa/110kDa protein 1 |
| **SERP1** | ENSG00000120742 | 2987.957423 | 0.662658711 | 0.127817444 | 1.41E-08 | 3.75E-06 | 101928061 | 3 | 150259781 | 150321015 | -1 | protein_coding | stress-associated endoplasmic reticulum protein 1 |
| **LYPLA1** | ENSG00000120992 | 1527.151441 | -0.522520206 | 0.155669177 | 5.33E-05 | 0.002563978 | 10434 | 8 | 54958938 | 55014577 | -1 | protein_coding | lysophospholipase I |
| **PLBD1** | ENSG00000121316 | 7328.294947 | -0.409358797 | 0.119422223 | 6.1E-05 | 0.002841677 | 79887 | 12 | 14656595 | 14721283 | -1 | protein_coding | phospholipase B domain containing 1 |
| **DESI2** | ENSG00000121644 | 1052.120805 | 0.495730567 | 0.127435632 | 7E-06 | 5.17E-04 | 51029 | 1 | 244816237 | 244872335 | 1 | protein_coding | desumoylating isopeptidase 2 |
| **FYTTD1** | ENSG00000122068 | 894.0135466 | -0.62208396 | 0.183805754 | 3.88E-05 | 0.002028168 | 84248 | 3 | 197464050 | 197514467 | 1 | protein_coding | forty-two-three domain containing 1 |
| **RBBP6** | ENSG00000122257 | 1586.829375 | 0.403326725 | 0.110584121 | 2.65E-05 | 0.001524601 | 5930 | 16 | 24549014 | 24584184 | 1 | protein_coding | retinoblastoma binding protein 6 |
| **ANXA11** | ENSG00000122359 | 9366.971808 | -0.359449721 | 0.130948649 | 6.33E-04 | 0.016543103 | 311 | 10 | 81910645 | 81965328 | -1 | protein_coding | annexin A11 |
| **ZNF644** | ENSG00000122482 | 1479.343772 | 0.480348053 | 0.189914948 | 7.61E-04 | 0.018841257 | 84146 | 1 | 91380859 | 91487829 | -1 | protein_coding | zinc finger protein 644 |
| **CBX3** | ENSG00000122565 | 2895.262846 | 0.349595795 | 0.120824452 | 4.45E-04 | 0.012846415 | 11335 | 7 | 26240782 | 26252976 | 1 | protein_coding | chromobox homolog 3 |
| **P4HA1** | ENSG00000122884 | 4939.727515 | 0.465486086 | 0.113688448 | 3.83E-06 | 3.28E-04 | 5033 | 10 | 74766975 | 74856732 | -1 | protein_coding | prolyl 4-hydroxylase, alpha polypeptide I |
| **MED13L** | ENSG00000123066 | 2609.71872 | 0.294219287 | 0.118968936 | 0.001891843 | 0.034425705 | 23389 | 12 | 116395711 | 116715143 | -1 | protein_coding | mediator complex subunit 13-like |
| **C19orf43** | ENSG00000123144 | 2152.348301 | -0.397620773 | 0.120616388 | 9.51E-05 | 0.004056836 | 79002 | 19 | 12841454 | 12845589 | -1 | protein_coding | chromosome 19 open reading frame 43 |
| **LRP1** | ENSG00000123384 | 27050.26282 | 0.25187452 | 0.103091637 | 0.002634527 | 0.042714716 | 4035 | 12 | 57522276 | 57607134 | 1 | protein_coding | low density lipoprotein receptor-related protein 1 |
| **FAM199X** | ENSG00000123575 | 1059.699812 | 0.726830098 | 0.17007067 | 1.1E-06 | 1.26E-04 | 139231 | X | 103411301 | 103440583 | 1 | protein_coding | family with sequence similarity 199, X-linked |
| **AGO2** | ENSG00000123908 | 634.935458 | -0.440200207 | 0.225456597 | 0.003028972 | 0.046115879 | 27161 | 8 | 141541264 | 141645718 | -1 | protein_coding | argonaute RISC catalytic component 2 |
| **ACSL3** | ENSG00000123983 | 4617.306754 | -0.257155821 | 0.09007618 | 8.57E-04 | 0.020106284 | 2181 | 2 | 223725652 | 223809357 | 1 | protein_coding | acyl-CoA synthetase long-chain family member 3 |
| **SDC4** | ENSG00000124145 | 2634.663429 | 0.412705463 | 0.127456451 | 1.1E-04 | 0.004525092 | 6385 | 20 | 43953928 | 43977064 | -1 | protein_coding | syndecan 4 |
| **RAB22A** | ENSG00000124209 | 2344.872136 | 0.389355306 | 0.106363012 | 2.06E-05 | 0.001234246 | 57403 | 20 | 56884752 | 56942563 | 1 | protein_coding | RAB22A, member RAS oncogene family |
| **OARD1** | ENSG00000124596 | 533.0981901 | -0.447112143 | 0.214677962 | 0.002288275 | 0.039158607 | 221443 | 6 | 41001366 | 41065526 | -1 | protein_coding | O-acyl-ADP-ribose deacylase 1 |
| **DEK** | ENSG00000124795 | 1573.582676 | -0.663108961 | 0.171525018 | 5.99E-06 | 4.65E-04 | 7913 | 6 | 18224099 | 18265054 | -1 | protein_coding | DEK oncogene |
| **SLC25A23** | ENSG00000125648 | 911.8187134 | -0.743145985 | 0.257975129 | 1.78E-04 | 0.006402277 | 79085 | 19 | 6436090 | 6465214 | -1 | protein_coding | solute carrier family 25 (mitochondrial carrier; phosphate carrier), member 23 |
| **CAPNS1** | ENSG00000126247 | 4732.24832 | -0.444381514 | 0.159211275 | 3.96E-04 | 0.011862182 | 826 | 19 | 36630477 | 36641255 | 1 | protein_coding | calpain, small subunit 1 |
| **UBA2** | ENSG00000126261 | 2190.200048 | -0.30113415 | 0.12222769 | 0.00184332 | 0.034319461 | 10054 | 19 | 34919257 | 34960853 | 1 | protein_coding | ubiquitin-like modifier activating enzyme 2 |
| **RRAS** | ENSG00000126458 | 945.2454226 | -0.374361201 | 0.145384721 | 9.95E-04 | 0.022634955 | 6237 | 19 | 50138549 | 50143458 | -1 | protein_coding | related RAS viral (r-ras) oncogene homolog |
| **IFI6** | ENSG00000126709 | 7699.553854 | -0.535472954 | 0.171227298 | 1.24E-04 | 0.004891181 | 2537 | 1 | 27992572 | 27998729 | -1 | protein_coding | interferon, alpha-inducible protein 6 |
| **SGPP1** | ENSG00000126821 | 1390.574459 | 0.950048842 | 0.166531214 | 6.39E-10 | 2.83E-07 | 81537 | 14 | 64150932 | 64194757 | -1 | protein_coding | sphingosine-1-phosphate phosphatase 1 |
| **CANX** | ENSG00000127022 | 15289.26981 | 0.562116043 | 0.098330056 | 1.01E-09 | 3.83E-07 | 821 | 5 | 179105629 | 179157926 | 1 | protein_coding | calnexin |
| **BCL11B** | ENSG00000127152 | 500.297803 | 0.560543116 | 0.263119563 | 0.001550588 | 0.03036027 | 64919 | 14 | 99635624 | 99737861 | -1 | protein_coding | B-cell CLL/lymphoma 11B (zinc finger protein) |
| **RAP1B** | ENSG00000127314 | 11596.36488 | -0.441254823 | 0.113453282 | 8.99E-06 | 6.14E-04 | 5908 | 12 | 69004619 | 69054372 | 1 | protein_coding | RAP1B, member of RAS oncogene family |
| **RAB3IP** | ENSG00000127328 | 116.0277146 | 1.821580586 | 0.692318447 | 2.56E-04 | 0.008442217 | 117177 | 12 | 70132461 | 70216984 | 1 | protein_coding | RAB3A interacting protein |
| **HP1BP3** | ENSG00000127483 | 2906.781846 | -0.31361999 | 0.125638835 | 0.001658287 | 0.031695357 | 50809 | 1 | 21069154 | 21113816 | -1 | protein_coding | heterochromatin protein 1, binding protein 3 |
| **SLC35E1** | ENSG00000127526 | 2410.457712 | 0.32481888 | 0.119722761 | 8.19E-04 | 0.01972889 | 79939 | 19 | 16660642 | 16683193 | -1 | protein_coding | solute carrier family 35, member E1 |
| **ECHS1** | ENSG00000127884 | 1605.369034 | -0.29349701 | 0.118868457 | 0.001858486 | 0.034425705 | 1892 | 10 | 135175984 | 135187193 | -1 | protein_coding | enoyl CoA hydratase, short chain, 1, mitochondrial |
| **CASD1** | ENSG00000127995 | 386.0690237 | 0.773848494 | 0.241267113 | 6.53E-05 | 0.003015243 | 64921 | 7 | 94138531 | 94186331 | 1 | protein_coding | CAS1 domain containing 1 |
| **RAC2** | ENSG00000128340 | 11180.02061 | -0.241198487 | 0.090575001 | 0.001643682 | 0.031522592 | 5880 | 22 | 37621301 | 37640488 | -1 | protein_coding | ras-related C3 botulinum toxin substrate 2 (rho family, small GTP binding protein Rac2) |
| **APOBEC3A** | ENSG00000128383 | 3640.86496 | -0.609485677 | 0.232846311 | 4.16E-04 | 0.012275649 | 100913187 | 22 | 39348746 | 39359188 | 1 | protein_coding | apolipoprotein B mRNA editing enzyme, catalytic polypeptide-like 3A |
| **SPECC1** | ENSG00000128487 | 3132.098874 | -0.305432091 | 0.124112421 | 0.001847251 | 0.034319461 | 92521 | 17 | 19912657 | 20222339 | 1 | protein_coding | sperm antigen with calponin homology and coiled-coil domains 1 |
| **ATP6V1F** | ENSG00000128524 | 3617.886723 | -0.289332409 | 0.113649807 | 0.001623292 | 0.031251303 | 101927180 | 7 | 128502880 | 128505898 | 1 | protein_coding | ATPase, H+ transporting, lysosomal 14kDa, V1 subunit F |
| **PODXL** | ENSG00000128567 | 130.1955611 | -1.121560605 | 0.513280154 | 8.57E-04 | 0.020106284 | 5420 | 7 | 131185021 | 131242976 | -1 | protein_coding | podocalyxin-like |
| **PSMA1** | ENSG00000129084 | 884.267958 | 0.567554274 | 0.142863489 | 4.61E-06 | 3.8E-04 | 5682 | 11 | 14515329 | 14665181 | -1 | protein_coding | proteasome (prosome, macropain) subunit, alpha type, 1 |
| **SPCS3** | ENSG00000129128 | 3382.240123 | 0.427982453 | 0.123843228 | 5.12E-05 | 0.002497277 | 60559 | 4 | 177241115 | 177253396 | 1 | protein_coding | signal peptidase complex subunit 3 homolog (S. cerevisiae) |
| **SLC44A2** | ENSG00000129353 | 2690.447542 | -0.522380632 | 0.115529997 | 4.52E-07 | 6.23E-05 | 57153 | 19 | 10713133 | 10755235 | 1 | protein_coding | solute carrier family 44 (choline transporter), member 2 |
| **REEP5** | ENSG00000129625 | 3631.103907 | -0.357382537 | 0.150226434 | 0.001669825 | 0.031858592 | 7905 | 5 | 112212084 | 112258236 | -1 | protein_coding | receptor accessory protein 5 |
| **TOMM40** | ENSG00000130204 | 2604.410719 | -0.292772554 | 0.098799457 | 4.69E-04 | 0.013427591 | 10452 | 19 | 45393826 | 45406946 | 1 | protein_coding | translocase of outer mitochondrial membrane 40 homolog (yeast) |
| **APOC1** | ENSG00000130208 | 1435.162052 | -0.586435017 | 0.217880179 | 3.4E-04 | 0.010544146 | 341 | 19 | 45417504 | 45422606 | 1 | protein_coding | apolipoprotein C-I |

| | | | | | | | | | | | | | |
|---|---|---|---|---|---|---|---|---|---|---|---|---|---|
| **BST2** | ENSG00000130303 | 2385.815076 | -0.28827183 | 0.112344183 | 0.001375856 | 0.028335695 | 684 | 19 | 17513748 | 17516457 | -1 | protein_coding | bone marrow stromal cell antigen 2 |
| **MRPL34** | ENSG00000130312 | 637.801386 | -0.370440329 | 0.168493882 | 0.002352981 | 0.039729087 | 64981 | 19 | 17403418 | 17417652 | 1 | protein_coding | mitochondrial ribosomal protein L34 |
| **ATXN10** | ENSG00000130638 | 1168.577862 | -0.513262121 | 0.225977059 | 0.001237265 | 0.026139985 | 25814 | 22 | 46067678 | 46241187 | 1 | protein_coding | ataxin 10 |
| **EIF2S3** | ENSG00000130741 | 5187.558115 | 0.258255496 | 0.091413854 | 9.31E-04 | 0.021545962 | 1968 | X | 24072833 | 24096088 | 1 | protein_coding | eukaryotic translation initiation factor 2, subunit 3 gamma, 52kDa |
| **COX4I1** | ENSG00000131143 | 5034.948858 | -0.276438726 | 0.111977328 | 0.002149286 | 0.037199448 | 1327 | 16 | 85832239 | 85840650 | 1 | protein_coding | cytochrome c oxidase subunit IV isoform 1 |
| **COX7B** | ENSG00000131174 | 345.0464628 | -0.933348303 | 0.321148615 | 1.43E-04 | 0.00532558 | 1349 | X | 77154935 | 77162870 | 1 | protein_coding | cytochrome c oxidase subunit VIIb |
| **PPT1** | ENSG00000131238 | 20613.40334 | -0.40117101 | 0.146843887 | 0.000544688 | 0.015169527 | 5538 | 1 | 40538379 | 40563375 | -1 | protein_coding | palmitoyl-protein thioesterase 1 |
| **ENOSF1** | ENSG00000132199 | 627.1444346 | 0.697315461 | 0.256531877 | 3.09E-04 | 0.009678858 | 55556 | 18 | 670324 | 712676 | -1 | protein_coding | enolase superfamily member 1 |
| **PTPRE** | ENSG00000132334 | 6642.573895 | 0.516212821 | 0.206977965 | 7.39E-04 | 0.018512962 | 5791 | 10 | 129705325 | 129884119 | 1 | protein_coding | protein tyrosine phosphatase, receptor type, E |
| **RAN** | ENSG00000132341 | 2240.756508 | -0.478445218 | 0.245691449 | 0.002692524 | 0.043419504 | 5901 | 12 | 131356424 | 131362223 | 1 | protein_coding | RAN, member RAS oncogene family |
| **TMEM128** | ENSG00000132406 | 285.0316205 | 0.552370444 | 0.249330656 | 0.001322047 | 0.027655547 | 85013 | 4 | 4237269 | 4249950 | -1 | protein_coding | transmembrane protein 128 |
| **FIGNL1** | ENSG00000132436 | 350.7068663 | 0.60604006 | 0.293441727 | 0.001688342 | 0.032039304 | 63979 | 7 | 50511831 | 50518088 | -1 | protein_coding | fidgetin-like 1 |
| **GRSF1** | ENSG00000132463 | 1798.550171 | 0.814385736 | 0.158572562 | 1.68E-08 | 4.36E-06 | 2926 | 4 | 71681499 | 71705662 | -1 | protein_coding | G-rich RNA sequence binding factor 1 |
| **NES** | ENSG00000132688 | 644.5304312 | -0.595596521 | 0.221772864 | 3.79E-04 | 0.011500386 | 10763 | 1 | 156638555 | 156647189 | -1 | protein_coding | nestin |
| **XPO4** | ENSG00000132953 | 1096.310829 | 0.384001379 | 0.13422359 | 4E-04 | 0.011884426 | 64328 | 13 | 21351469 | 21477187 | -1 | protein_coding | exportin 4 |
| **KRAS** | ENSG00000133703 | 1324.735839 | 0.50299269 | 0.160788428 | 1.27E-04 | 0.004948792 | 3845 | 12 | 25357723 | 25403870 | -1 | protein_coding | Kirsten rat sarcoma viral oncogene homolog |
| **HSD17B4** | ENSG00000133835 | 2509.554876 | -0.865492428 | 0.170911288 | 2.11E-08 | 5.09E-06 | 3295 | 5 | 118788138 | 118972894 | 1 | protein_coding | hydroxysteroid (17-beta) dehydrogenase 4 |
| **TMEM66** | ENSG00000133872 | 6395.942601 | 0.5490479 | 0.123776998 | 6.91E-07 | 8.85E-05 | 51669 | 8 | 29920528 | 29940723 | -1 | protein_coding | transmembrane protein 66 |
| **IER3IP1** | ENSG00000134049 | 709.2327246 | 0.559866066 | 0.204736168 | 3.96E-04 | 0.011862182 | 51124 | 18 | 44681413 | 44702745 | -1 | protein_coding | immediate early response 3 interacting protein 1 |
| **VHL** | ENSG00000134086 | 1186.245452 | 0.509530084 | 0.148792458 | 4.57E-05 | 0.002293117 | 7428 | 3 | 10182692 | 10193904 | 1 | protein_coding | von Hippel-Lindau tumor suppressor, E3 ubiquitin protein ligase |
| **ARL8B** | ENSG00000134108 | 5699.299989 | 0.427565697 | 0.123914767 | 5.69E-05 | 0.002685158 | 55207 | 3 | 5163905 | 5222596 | 1 | protein_coding | ADP-ribosylation factor-like 8B |
| **PRPF38B** | ENSG00000134186 | 1168.700524 | 0.490055231 | 0.14960672 | 7.66E-05 | 0.003463742 | 55119 | 1 | 109234945 | 109244425 | 1 | protein_coding | pre-mRNA processing factor 38B |
| **ARF3** | ENSG00000134287 | 6667.092042 | -0.290665796 | 0.104537865 | 8.45E-04 | 0.020052664 | 377 | 12 | 49329506 | 49351334 | -1 | protein_coding | ADP-ribosylation factor 3 |
| **KLRD1** | ENSG00000134539 | 100.8126107 | 0.934097664 | 0.387146996 | 5.69E-04 | 0.015474593 | 3824 | 12 | 10378657 | 10469850 | 1 | protein_coding | killer cell lectin-like receptor subfamily D, member 1 |
| **BTF3L4** | ENSG00000134717 | 1250.104085 | 0.506590945 | 0.143311934 | 3.15E-05 | 0.001752079 | 91408 | 1 | 52521797 | 52556388 | 1 | protein_coding | basic transcription factor 3-like 4 |
| **ZCCHC11** | ENSG00000134744 | 873.3927606 | 0.338746037 | 0.148680444 | 0.002393954 | 0.040043333 | 23318 | 1 | 52873954 | 53019159 | -1 | protein_coding | zinc finger, CCHC domain containing 11 |
| **FADS2** | ENSG00000134824 | 2270.75175 | -0.500063016 | 0.105540086 | 1.91E-07 | 2.94E-05 | 9415 | 11 | 61560452 | 61634826 | 1 | protein_coding | fatty acid desaturase 2 |
| **PSAT1** | ENSG00000135069 | 1396.813175 | 0.628589407 | 0.164059871 | 7.9E-06 | 5.64E-04 | 29968 | 9 | 80912059 | 80945009 | 1 | protein_coding | phosphoserine aminotransferase 1 |
| **DMTF1** | ENSG00000135164 | 1179.396279 | 0.585107547 | 0.166335104 | 2.75E-05 | 0.001557255 | 9988 | 7 | 86781677 | 86825653 | 1 | protein_coding | cyclin D binding myb-like transcription factor 1 |
| **CD36** | ENSG00000135218 | 6391.870527 | 0.458953834 | 0.159274315 | 2.68E-04 | 0.00868138 | 948 | 7 | 79998891 | 80308593 | 1 | protein_coding | CD36 molecule (thrombospondin receptor) |
| **PRRG4** | ENSG00000135378 | 316.3732763 | 0.649740354 | 0.332419503 | 0.001932458 | 0.034985058 | 79056 | 11 | 32851489 | 32879669 | 1 | protein_coding | proline rich Gla (G-carboxyglutamic acid) 4 (transmembrane) |
| **CD63** | ENSG00000135404 | 26107.53483 | -0.337876249 | 0.130408795 | 0.001116552 | 0.024314759 | 967 | 12 | 56119107 | 56123491 | -1 | protein_coding | CD63 molecule |
| **FAIM2** | ENSG00000135472 | 708.2932972 | -0.762333604 | 0.189164057 | 2.82E-06 | 2.59E-04 | 23017 | 12 | 50260679 | 50298000 | -1 | protein_coding | Fas apoptotic inhibitory molecule 2 |
| **TEC** | ENSG00000135605 | 306.8028888 | 0.469848603 | 0.222718319 | 0.002019567 | 0.03574881 | 7006 | 4 | 48137800 | 48271881 | -1 | protein_coding | tec protein tyrosine kinase |
| **KLHL36** | ENSG00000135686 | 1693.611999 | 0.303638917 | 0.124152922 | 0.001963858 | 0.035289354 | 79786 | 16 | 84682131 | 84701292 | 1 | protein_coding | kelch-like family member 36 |
| **EGLN1** | ENSG00000135766 | 2031.985255 | 0.289738878 | 0.117015184 | 0.00223933 | 0.038569465 | 54583 | 1 | 231499497 | 231560790 | -1 | protein_coding | egl-9 family hypoxia-inducible factor 1 |
| **ABCB10** | ENSG00000135776 | 797.7646897 | 0.513477087 | 0.17507086 | 2.2E-04 | 0.007618905 | 23456 | 1 | 229652329 | 229694442 | -1 | protein_coding | ATP-binding cassette, sub-family B (MDR/TAP), member 10 |
| **COX5B** | ENSG00000135940 | 2920.516599 | -0.681597866 | 0.175938011 | 6.82E-06 | 5.07E-04 | 1329 | 2 | 98262503 | 98264846 | 1 | protein_coding | cytochrome c oxidase subunit Vb |
| **RAC1** | ENSG00000136238 | 6452.213562 | -0.25568897 | 0.108544196 | 0.003340399 | 0.049371936 | 5879 | 7 | 6414154 | 6443608 | 1 | protein_coding | ras-related C3 botulinum toxin substrate 1 (rho family, small GTP binding protein Rac1) |
| **YME1L1** | ENSG00000136758 | 4043.405727 | 0.454474524 | 0.117356926 | 9.92E-06 | 6.67E-04 | 10730 | 10 | 27399383 | 27444195 | -1 | protein_coding | YME1-like 1 ATPase |
| **RPL35** | ENSG00000136942 | 2537.033886 | -0.312341732 | 0.143217945 | 0.003205668 | 0.048048842 | 11224 | 9 | 127620159 | 127624260 | -1 | protein_coding | ribosomal protein L35 |
| **TPMT** | ENSG00000137364 | 3081.1361 | 0.408425259 | 0.123633477 | 1E-04 | 0.004213424 | 7172 | 6 | 18128542 | 18155305 | -1 | protein_coding | thiopurine S-methyltransferase |
| **FAM8A1** | ENSG00000137414 | 1449.657973 | 0.550087799 | 0.141834765 | 7.53E-06 | 5.44E-04 | 51439 | 6 | 17600586 | 17611950 | 1 | protein_coding | family with sequence similarity 8, member A1 |
| **PRKRIR** | ENSG00000137492 | 1785.788856 | 0.401346448 | 0.162717477 | 0.00112583 | 0.024317464 | 5612 | 11 | 76061000 | 76092015 | -1 | protein_coding | protein-kinase, interferon-inducible double stranded RNA dependent inhibitor, repressor of (P58 repressor) |
| **NUMA1** | ENSG00000137497 | 5136.104268 | 0.250021565 | 0.10107173 | 0.002635088 | 0.042714716 | 4926 | 11 | 71713910 | 71791739 | -1 | protein_coding | nuclear mitotic apparatus protein 1 |
| **SDCBP** | ENSG00000137575 | 19555.82738 | -0.513566565 | 0.133014377 | 8.27E-06 | 5.82E-04 | 6386 | 8 | 59465483 | 59495419 | 1 | protein_coding | syndecan binding protein (syntenin) |
| **RSL24D1** | ENSG00000137876 | 1119.260968 | 0.337293084 | 0.147971169 | 0.002350509 | 0.039729087 | 51187 | 15 | 55473004 | 55489265 | -1 | protein_coding | ribosomal L24 domain containing 1 |
| **IFT172** | ENSG00000138002 | 269.7258404 | 0.583016273 | 0.322561954 | 0.002783079 | 0.044228011 | 26160 | 2 | 27667238 | 27712656 | -1 | protein_coding | intraflagellar transport 172 homolog (Chlamydomonas) |
| **EPT1** | ENSG00000138018 | 1502.528356 | 0.389057787 | 0.131624326 | 3.1E-04 | 0.009678858 | 85465 | 2 | 26531415 | 26618759 | 1 | protein_coding | ethanolaminephosphotransferase 1 (CDP-ethanolamine-specific) |
| **PPM1B** | ENSG00000138032 | 1739.54306 | -0.279651649 | 0.120298574 | 0.003012191 | 0.045992176 | 5495 | 2 | 44395108 | 44471523 | 1 | protein_coding | protein phosphatase, Mg2+/Mn2+ dependent, 1B |
| **CYP1B1** | ENSG00000138061 | 3708.169628 | -1.709801042 | 0.540426541 | 3.93E-05 | 0.002028168 | 1545 | 2 | 38294116 | 38337044 | -1 | protein_coding | cytochrome P450, family 1, subfamily B, polypeptide 1 |
| **RAB1A** | ENSG00000138069 | 5929.355573 | 0.522574318 | 0.118950698 | 8.93E-07 | 1.07E-04 | 5861 | 2 | 65297835 | 65357240 | -1 | protein_coding | RAB1A, member RAS oncogene family |
| **ACTR2** | ENSG00000138071 | 21029.68192 | 0.492616133 | 0.092606933 | 9.05E-09 | 2.69E-06 | 10097 | 2 | 65454887 | 65498387 | 1 | protein_coding | ARP2 actin-related protein 2 homolog (yeast) |
| **ATAD1** | ENSG00000138138 | 663.0231389 | 0.491487124 | 0.192188326 | 6.78E-04 | 0.017556493 | 84896 | 10 | 89511269 | 89601100 | -1 | protein_coding | ATPase family, AAA domain containing 1 |
| **RPS24** | ENSG00000138326 | 6746.475157 | -0.285583226 | 0.124167033 | 0.002925953 | 0.045310196 | 6229 | 10 | 79793518 | 79816570 | 1 | protein_coding | ribosomal protein S24 |
| **SENP7** | ENSG00000138468 | 399.1113725 | 0.610324384 | 0.241146598 | 5.61E-04 | 0.015320947 | 57337 | 3 | 101043049 | 101232085 | -1 | protein_coding | SUMO1/sentrin specific peptidase 7 |
| **TMOD3** | ENSG00000138594 | 3497.705253 | 0.353475538 | 0.108474468 | 1.4E-04 | 0.005282849 | 29766 | 15 | 52121825 | 52239492 | 1 | protein_coding | tropomodulin 3 (ubiquitous) |
| **GLCE** | ENSG00000138604 | 724.7554612 | 0.426724232 | 0.138969305 | 1.81E-04 | 0.006465396 | 26035 | 15 | 69452923 | 69564556 | 1 | protein_coding | glucuronic acid epimerase |
| **HNRNPD** | ENSG00000138668 | 1984.884547 | -0.336212046 | 0.120431851 | 6.14E-04 | 0.016350606 | 3184 | 4 | 83273651 | 83295656 | -1 | protein_coding | heterogeneous nuclear ribonucleoprotein D (AU-rich element RNA binding protein 1, 37kDa) |
| **BMP2K** | ENSG00000138756 | 4699.826229 | 0.664276527 | 0.151049507 | 8.36E-07 | 1.01E-04 | 55589 | 4 | 79697496 | 79837526 | 1 | protein_coding | BMP2 inducible kinase |
| **TAPBPL** | ENSG00000139192 | 841.3760194 | -0.467748631 | 0.172655688 | 4.28E-04 | 0.012537648 | 55080 | 12 | 6560856 | 6575683 | 1 | protein_coding | TAP binding protein-like |
| **AMIGO2** | ENSG00000139211 | 149.566082 | 0.675825226 | 0.337678822 | 0.001719545 | 0.032515303 | 347902 | 12 | 47469490 | 47473734 | -1 | protein_coding | adhesion molecule with Ig-like domain 2 |
| **PHLDA1** | ENSG00000139289 | 1555.258646 | 0.760024809 | 0.151153203 | 2.95E-08 | 6.16E-06 | 22822 | 12 | 76419227 | 76427712 | -1 | protein_coding | pleckstrin homology-like domain, family A, member 1 |
| **MTMR6** | ENSG00000139505 | 1793.85481 | 1.162263617 | 0.169296888 | 3.84E-13 | 5.1E-10 | 9107 | 13 | 25802307 | 25862147 | -1 | protein_coding | myotubularin related protein 6 |
| **RB1** | ENSG00000139687 | 2311.684982 | 0.446471378 | 0.113649653 | 7.68E-06 | 5.52E-04 | 5925 | 13 | 48877887 | 49056122 | 1 | protein_coding | retinoblastoma 1 |
| **DENR** | ENSG00000139726 | 1625.718397 | 0.327635893 | 0.136834337 | 0.001965749 | 0.035289354 | 8562 | 12 | 123237321 | 123255611 | 1 | protein_coding | density-regulated protein |
| **ABHD13** | ENSG00000139826 | 570.8425285 | 0.76591962 | 0.193441009 | 3.89E-06 | 3.3E-04 | 84945 | 13 | 108870727 | 108886603 | 1 | protein_coding | abhydrolase domain containing 13 |
| **TMX1** | ENSG00000139921 | 1973.910353 | 0.399518492 | 0.134968616 | 2.87E-04 | 0.009167266 | 81542 | 14 | 51706880 | 51722759 | 1 | protein_coding | thioredoxin-related transmembrane protein 1 |
| **SLC38A6** | ENSG00000139974 | 2129.122991 | 0.498564519 | 0.198803469 | 7.52E-04 | 0.018727077 | 145389 | 14 | 61447832 | 61550451 | 1 | protein_coding | solute carrier family 38, member 6 |
| **NAA30** | ENSG00000139977 | 811.4365134 | 0.446007073 | 0.167686787 | 5.94E-04 | 0.015975918 | 122830 | 14 | 57857262 | 57882635 | 1 | protein_coding | N(alpha)-acetyltransferase 30, NatC catalytic subunit |
| **GPR65** | ENSG00000140030 | 1023.504037 | 0.400389714 | 0.155015791 | 7.7E-04 | 0.019032401 | 8477 | 14 | 88471468 | 88478419 | 1 | protein_coding | G protein-coupled receptor 65 |
| **TCF12** | ENSG00000140262 | 2398.140229 | -0.523027591 | 0.128612802 | 3.94E-06 | 3.32E-04 | 6938 | 15 | 57210821 | 57591479 | 1 | protein_coding | transcription factor 12 |
| **SERF2** | ENSG00000140264 | 9436.21109 | -0.389111746 | 0.122633141 | 1.26E-04 | 0.004928845 | 100302254 | 15 | 44069285 | 44094787 | 1 | protein_coding | small EDRK-rich factor 2 |
| **MAN2C1** | ENSG00000140400 | 1258.547013 | 0.453612374 | 0.176215421 | 7.08E-04 | 0.018080243 | 4123 | 15 | 75648133 | 75660971 | -1 | protein_coding | mannosidase, alpha, class 2C, member 1 |
| **ARRDC4** | ENSG00000140450 | 1096.542513 | 0.720505076 | 0.18348413 | 4.82E-06 | 3.94E-04 | 91947 | 15 | 98462784 | 98517068 | 1 | protein_coding | arrestin domain containing 4 |
| **ZSCAN32** | ENSG00000140987 | 313.2391899 | 0.774632778 | 0.281892071 | 2.56E-04 | 0.008442217 | 54925 | 16 | 3432085 | 3451065 | -1 | protein_coding | zinc finger and SCAN domain containing 32 |
| **NDUFB10** | ENSG00000140990 | 827.6822125 | -0.393540673 | 0.155044489 | 9.3E-04 | 0.021545962 | 4716 | 16 | 2009509 | 2011976 | 1 | protein_coding | NADH dehydrogenase (ubiquinone) 1 beta subcomplex, 10, 22kDa |
| **KSR1** | ENSG00000141068 | 326.1821555 | 0.733474209 | 0.336193649 | 0.001084688 | 0.023816069 | 8844 | 17 | 25783670 | 25953461 | 1 | protein_coding | kinase suppressor of ras 1 |
| **TOB1** | ENSG00000141232 | 2174.073146 | -0.464009656 | 0.1710384 | 4.39E-04 | 0.012800797 | 10140 | 17 | 48939584 | 48945339 | -1 | protein_coding | transducer of ERBB2, 1 |
| **MBD1** | ENSG00000141644 | 1347.176471 | 0.347455732 | 0.144128822 | 0.001727829 | 0.032556088 | 4152 | 18 | 47793252 | 47808144 | -1 | protein_coding | methyl-CpG binding domain protein 1 |
| **PGD** | ENSG00000142657 | 15428.04651 | -0.353310077 | 0.074012721 | 7.71E-07 | 9.53E-05 | 5226 | 1 | 10458649 | 10480201 | 1 | protein_coding | phosphogluconate dehydrogenase |
| **SH3BGRL3** | ENSG00000142669 | 12679.99859 | -0.350675208 | 0.135632695 | 0.001020114 | 0.023065437 | 83442 | 1 | 26605667 | 26608007 | 1 | protein_coding | SH3 domain binding glutamic acid-rich protein like 3 |
| **ALDH9A1** | ENSG00000143149 | 3076.07079 | 0.320090867 | 0.131909202 | 0.002112458 | 0.036983189 | 223 | 1 | 165631453 | 165668100 | -1 | protein_coding | aldehyde dehydrogenase 9 family, member A1 |
| **ATP1B1** | ENSG00000143153 | 5715.657034 | 0.374108604 | 0.166892462 | 0.002090372 | 0.036778784 | 481 | 1 | 169074935 | 169101960 | 1 | protein_coding | ATPase, Na+/K+ transporting, beta 1 polypeptide |
| **UFC1** | ENSG00000143222 | 1709.194005 | -0.340317153 | 0.140950887 | 0.001726031 | 0.032556088 | 51506 | 1 | 161122566 | 161128646 | 1 | protein_coding | ubiquitin-fold modifier conjugating enzyme 1 |
| **CRABP2** | ENSG00000143320 | 4468.771463 | -0.630237338 | 0.219894958 | 2.21E-04 | 0.007632549 | 1382 | 1 | 156669398 | 156675608 | -1 | protein_coding | cellular retinoic acid binding protein 2 |
| **TOR1AIP1** | ENSG00000143337 | 2981.164976 | -0.402756571 | 0.116460361 | 5.29E-05 | 0.002554951 | 26092 | 1 | 179851177 | 179894135 | 1 | protein_coding | torsin A interacting protein 1 |
| **C1orf43** | ENSG00000143612 | 4808.102621 | -0.289032189 | 0.087332712 | 1.57E-04 | 0.005735941 | 25912 | 1 | 154179182 | 154193104 | -1 | protein_coding | chromosome 1 open reading frame 43 |
| **SRP9** | ENSG00000143742 | 2265.997061 | 0.600486118 | 0.147037182 | 3.61E-06 | 3.14E-04 | 6726 | 1 | 225965531 | 225978168 | 1 | protein_coding | signal recognition particle 9kDa |
| **SDE2** | ENSG00000143751 | 1030.470888 | 0.32553888 | 0.148336656 | 0.003168242 | 0.047622217 | 163859 | 1 | 226170403 | 226187032 | -1 | protein_coding | SDE2 telomere maintenance homolog (S. pombe) |
| **ETAA1** | ENSG00000143971 | 267.6658642 | 0.593633205 | 0.276141214 | 0.001430749 | 0.028900715 | 54465 | 2 | 67624451 | 67637677 | 1 | protein_coding | Ewing tumor-associated antigen 1 |
| **UBXN4** | ENSG00000144224 | 5562.422997 | 0.24087671 | 0.08145223 | 8.04E-04 | 0.019682859 | 23190 | 2 | 136499189 | 136542625 | 1 | protein_coding | UBX domain protein 4 |

| | | | | | | | | | | | | | |
|---|---|---|---|---|---|---|---|---|---|---|---|---|---|
| **STK11IP** | ENSG00000144589 | 310.5765025 | -0.553987758 | 0.324089135 | 0.003373102 | 0.049510993 | 114790 | 2 | 220462582 | 220481173 | 1 | protein_coding | serine/threonine kinase 11 interacting protein |
| **EAF1** | ENSG00000144597 | 3320.469892 | 0.267927198 | 0.095158564 | 8.14E-04 | 0.01972889 | 85403 | 3 | 15468862 | 15484120 | 1 | protein_coding | ELL associated factor 1 |
| **RPL32** | ENSG00000144713 | 4310.612959 | -0.409667389 | 0.156437301 | 7.08E-04 | 0.018080243 | 6161 | 3 | 12875984 | 12883087 | -1 | protein_coding | ribosomal protein L32 |
| **NCEH1** | ENSG00000144959 | 695.2620971 | -1.813864413 | 0.263064199 | 2.12E-13 | 3.21E-10 | 57552 | 3 | 172348039 | 172429008 | -1 | protein_coding | neutral cholesterol ester hydrolase 1 |
| **CISD2** | ENSG00000145354 | 920.0046502 | 0.435133326 | 0.213629366 | 0.00263676 | 0.042714716 | 493856 | 4 | 103790135 | 103810399 | 1 | protein_coding | CDGSH iron sulfur domain 2 |
| **IQGAP2** | ENSG00000145703 | 5732.911297 | 0.44432577 | 0.124371165 | 2.36E-05 | 0.001385702 | 10788 | 5 | 75699074 | 76003957 | 1 | protein_coding | IQ motif containing GTPase activating protein 2 |
| **FEM1C** | ENSG00000145780 | 1401.423682 | 0.541526619 | 0.148356538 | 1.81E-05 | 0.001124133 | 56929 | 5 | 114856608 | 114880591 | -1 | protein_coding | fem-1 homolog c (C. elegans) |
| **PHIP** | ENSG00000146247 | 2260.801872 | 0.588138101 | 0.111650649 | 9.45E-09 | 2.71E-06 | 55023 | 6 | 79645584 | 79787953 | -1 | protein_coding | pleckstrin homology domain interacting protein |
| **PNRC1** | ENSG00000146278 | 5467.290331 | 0.38400937 | 0.132135205 | 2.39E-04 | 0.008066708 | 10957 | 6 | 89790470 | 89794879 | 1 | protein_coding | proline-rich nuclear receptor coactivator 1 |
| **WTAP** | ENSG00000146457 | 3943.238768 | 0.274690012 | 0.106991603 | 0.00174121 | 0.03269229 | 9589 | 6 | 160146617 | 160177351 | 1 | protein_coding | Wilms tumor 1 associated protein |
| **C6orf211** | ENSG00000146476 | 868.9234492 | 0.549069935 | 0.184476757 | 1.82E-04 | 0.006472892 | 79624 | 6 | 151773422 | 151791236 | 1 | protein_coding | chromosome 6 open reading frame 211 |
| **PURB** | ENSG00000146676 | 2051.953952 | 0.658591244 | 0.120139935 | 3.12E-09 | 1.07E-06 | 5814 | 7 | 44915896 | 44924960 | -1 | protein_coding | purine-rich element binding protein B |
| **TMEM168** | ENSG00000146802 | 701.7569415 | -0.412397485 | 0.16543601 | 9.79E-04 | 0.022365707 | 64418 | 7 | 112402437 | 112430647 | -1 | protein_coding | transmembrane protein 168 |
| **NONO** | ENSG00000147140 | 6312.165337 | -0.221664089 | 0.086911412 | 0.002538351 | 0.041692509 | 4841 | X | 70503042 | 70521018 | 1 | protein_coding | non-POU domain containing, octamer-binding |
| **EBP** | ENSG00000147155 | 897.2720508 | -0.544519193 | 0.174646066 | 1.15E-04 | 0.004621317 | 10682 | X | 48379546 | 48387104 | 1 | protein_coding | emopamil binding protein (sterol isomerase) |
| **IL2RG** | ENSG00000147168 | 9320.423641 | -0.275348234 | 0.081821239 | 1.15E-04 | 0.004618362 | 3561 | X | 70327254 | 70331958 | -1 | protein_coding | interleukin 2 receptor, gamma |
| **NSDHL** | ENSG00000147383 | 532.1072611 | -0.475977845 | 0.263510437 | 0.00340558 | 0.049918761 | 50814 | X | 151999511 | 152038273 | 1 | protein_coding | NAD(P) dependent steroid dehydrogenase-like |
| **DOCK5** | ENSG00000147459 | 5158.334714 | 0.33422535 | 0.145713108 | 0.002443574 | 0.040595018 | 80005 | 8 | 25042238 | 25275598 | 1 | protein_coding | dedicator of cytokinesis 5 |
| **GOLGA7** | ENSG00000147533 | 1670.199179 | -0.503539534 | 0.132185037 | 1.06E-05 | 7.04E-04 | 51125 | 8 | 41347915 | 41368499 | 1 | protein_coding | golgin A7 |
| **MTDH** | ENSG00000147649 | 5970.651672 | -0.399926604 | 0.112556227 | 3.64E-05 | 0.001942731 | 92140 | 8 | 98656407 | 98740998 | 1 | protein_coding | metadherin |
| **UGCG** | ENSG00000148154 | 3339.972996 | 0.544978146 | 0.147176351 | 1.57E-05 | 9.88E-04 | 7357 | 9 | 114659046 | 114697649 | 1 | protein_coding | UDP-glucose ceramide glucosyltransferase |
| **GSN** | ENSG00000148180 | 6227.358171 | -0.488697514 | 0.1387534 | 3.24E-05 | 0.001792814 | 2934 | 9 | 123970072 | 124095121 | 1 | protein_coding | gelsolin |
| **PTGES** | ENSG00000148344 | 122.4055337 | -2.508564886 | 0.741966543 | 4.5E-05 | 0.00226545 | 9536 | 9 | 132500610 | 132515326 | -1 | protein_coding | prostaglandin E synthase |
| **USP6NL** | ENSG00000148429 | 1149.263133 | 0.487824334 | 0.126319006 | 8.63E-06 | 6E-04 | 100507213 | 10 | 11495945 | 11653753 | -1 | protein_coding | USP6 N-terminal like |
| **RSU1** | ENSG00000148484 | 2680.275644 | -0.409717428 | 0.09591881 | 1.99E-06 | 2.02E-04 | 6251 | 10 | 16632610 | 16859527 | -1 | protein_coding | Ras suppressor protein 1 |
| **LIN7C** | ENSG00000148943 | 1848.7304 | 0.786301989 | 0.150677199 | 1.08E-08 | 2.98E-06 | 55327 | 11 | 27516123 | 27528320 | -1 | protein_coding | lin-7 homolog C (C. elegans) |
| **SESN3** | ENSG00000149212 | 1610.783105 | 0.672970544 | 0.138409439 | 7.9E-08 | 1.42E-05 | 143686 | 11 | 94898704 | 94965705 | -1 | protein_coding | sestrin 3 |
| **LAMTOR1** | ENSG00000149357 | 1420.765249 | -0.265866728 | 0.11010942 | 0.002724312 | 0.04377832 | 55004 | 11 | 71796941 | 71814433 | -1 | protein_coding | late endosomal/lysosomal adaptor, MAPK and MTOR activator 1 |
| **C11orf57** | ENSG00000150776 | 1032.662086 | -0.512954516 | 0.169073613 | 1.6E-04 | 0.005805191 | 55216 | 11 | 111944810 | 111955874 | 1 | protein_coding | chromosome 11 open reading frame 57 |
| **IL18** | ENSG00000150782 | 1077.559716 | 0.415179395 | 0.164296405 | 9.18E-04 | 0.02139994 | 3606 | 11 | 112013974 | 112034840 | -1 | protein_coding | interleukin 18 (interferon-gamma-inducing factor) |
| **SEC24D** | ENSG00000150961 | 2106.648406 | 0.269438195 | 0.112049406 | 0.002912784 | 0.04527201 | 9871 | 4 | 119643978 | 119759838 | -1 | protein_coding | SEC24 family member D |
| **UBC** | ENSG00000150991 | 28372.84834 | -0.257675529 | 0.07846433 | 2.45E-04 | 0.008225606 | 7316 | 12 | 125396150 | 125401914 | -1 | protein_coding | ubiquitin C |
| **IPMK** | ENSG00000151151 | 667.0358872 | 0.934716485 | 0.16280853 | 5.02E-10 | 2.43E-07 | 253430 | 10 | 59951278 | 60027694 | -1 | protein_coding | inositol polyphosphate multikinase |
| **SLC2A13** | ENSG00000151229 | 370.1798097 | 0.439759958 | 0.220400008 | 0.002833432 | 0.044635471 | 114134 | 12 | 40148823 | 40499891 | -1 | protein_coding | solute carrier family 2 (facilitated glucose transporter), member 13 |
| **GXYLT1** | ENSG00000151233 | 592.5152961 | 0.675998171 | 0.193249649 | 2.53E-05 | 0.001462855 | 283464 | 12 | 42475647 | 42538681 | -1 | protein_coding | glucoside xylosyltransferase 1 |
| **TWF1** | ENSG00000151239 | 848.1256921 | -1.122402487 | 0.233594964 | 8.1E-08 | 1.43E-05 | 5756 | 12 | 44187526 | 44200178 | -1 | protein_coding | twinfilin actin-binding protein 1 |
| **NDUFC2** | ENSG00000151366 | 963.5420064 | 0.384648383 | 0.168098908 | 0.00186883 | 0.034425705 | 100532726 | 11 | 77779350 | 77791265 | -1 | protein_coding | NADH dehydrogenase (ubiquinone) 1, subcomplex unknown, 2, 14.5kDa |
| **ARL14EP** | ENSG00000152219 | 570.6539325 | 0.423569795 | 0.215375778 | 0.003364428 | 0.049510993 | 120534 | 11 | 30344598 | 30359774 | 1 | protein_coding | ADP-ribosylation factor-like 14 effector protein |
| **UHMK1** | ENSG00000152332 | 7121.672726 | 0.249999258 | 0.087300699 | 0.001031671 | 0.023227907 | 127933 | 1 | 162467041 | 162499419 | 1 | protein_coding | U2AF homology motif (UHM) kinase 1 |
| **ZFP36L2** | ENSG00000152518 | 4732.67144 | 0.292097212 | 0.121892257 | 0.001876802 | 0.034425705 | 678 | 2 | 43449541 | 43453748 | -1 | protein_coding | ZFP36 ring finger protein-like 2 |
| **GPR180** | ENSG00000152749 | 710.3362578 | 0.62219474 | 0.220445114 | 2.52E-04 | 0.008380799 | 160897 | 13 | 95254157 | 95286899 | 1 | protein_coding | G protein-coupled receptor 180 |
| **ZNF117** | ENSG00000152926 | 421.4759414 | 0.653215095 | 0.272598576 | 7.4E-04 | 0.018512962 | 51351 | 7 | 64432154 | 64467062 | -1 | protein_coding | zinc finger protein 117 |
| **CDYL** | ENSG00000153046 | 1816.178044 | -0.321862214 | 0.125294081 | 0.001266755 | 0.026709933 | 9425 | 6 | 4706393 | 4955785 | 1 | protein_coding | chromodomain protein, Y-like |
| **SCOC** | ENSG00000153130 | 185.0380591 | 3.066754295 | 0.493435614 | 1.73E-11 | 1.31E-08 | 60592 | 4 | 141178440 | 141306880 | 1 | protein_coding | short coiled-coil protein |
| **TMEM87B** | ENSG00000153214 | 3142.801683 | 0.327089547 | 0.147643111 | 0.002967259 | 0.045576695 | 84910 | 2 | 112812800 | 112876895 | 1 | protein_coding | transmembrane protein 87B |
| **TRIP12** | ENSG00000153827 | 12909.5098 | -0.218729506 | 0.078400986 | 0.001444877 | 0.029018201 | 9320 | 2 | 230628554 | 230787955 | -1 | protein_coding | thyroid hormone receptor interactor 12 |
| **CEBPG** | ENSG00000153879 | 1492.361774 | 0.28934587 | 0.123756148 | 0.002774484 | 0.044228011 | 1054 | 19 | 33864236 | 33873592 | 1 | protein_coding | CCAAT/enhancer binding protein (C/EBP), gamma |
| **NUS1** | ENSG00000153989 | 3898.823256 | 0.463194225 | 0.109908839 | 1.98E-06 | 2.02E-04 | 116150 | 6 | 117996665 | 118031803 | 1 | protein_coding | nuclear undecaprenyl pyrophosphate synthase 1 homolog (S. cerevisiae) |
| **UBASH3B** | ENSG00000154127 | 3413.330109 | -0.511164872 | 0.225007579 | 0.001053841 | 0.023527654 | 84959 | 11 | 122526383 | 122685181 | 1 | protein_coding | ubiquitin associated and SH3 domain containing B |
| **HSPA13** | ENSG00000155304 | 781.592854 | 1.170053039 | 0.176574625 | 1.8E-12 | 1.74E-09 | 6782 | 21 | 15743436 | 15755805 | -1 | protein_coding | heat shock protein 70kDa family, member 13 |
| **RHOC** | ENSG00000155366 | 2991.690249 | -0.437376389 | 0.11075905 | 4.38E-06 | 3.64E-04 | 389 | 1 | 113243728 | 113250056 | -1 | protein_coding | ras homolog family member C |
| **RASA2** | ENSG00000155903 | 764.2658484 | 0.648109512 | 0.180285118 | 1.86E-05 | 0.001146173 | 5922 | 3 | 141205889 | 141334184 | 1 | protein_coding | RAS p21 protein activator 2 |
| **DCK** | ENSG00000156136 | 1235.929445 | 0.533178318 | 0.120873596 | 7.4E-07 | 9.26E-05 | 1633 | 4 | 71858255 | 71896631 | 1 | protein_coding | deoxycytidine kinase |
| **SH3RF2** | ENSG00000156463 | 421.09652 | -0.60245793 | 0.265858843 | 0.00106123 | 0.02354425 | 153769 | 5 | 145316142 | 145461354 | 1 | protein_coding | SH3 domain containing ring finger 2 |
| **FAM122B** | ENSG00000156504 | 503.1369139 | 0.795069515 | 0.234248799 | 3.28E-05 | 0.00180004 | 159090 | X | 133903596 | 133931262 | -1 | protein_coding | family with sequence similarity 122B |
| **CD109** | ENSG00000156535 | 7702.672932 | 0.439904074 | 0.100657672 | 1.37E-06 | 1.52E-04 | 135228 | 6 | 74405508 | 74538040 | 1 | protein_coding | CD109 molecule |
| **NPTN** | ENSG00000156642 | 4061.190685 | -0.379472558 | 0.089884704 | 2.51E-06 | 2.4E-04 | 27020 | 15 | 73852355 | 73926475 | -1 | protein_coding | neuroplastin |
| **SAMD8** | ENSG00000156671 | 1400.382798 | 0.479505684 | 0.137053587 | 3.94E-05 | 0.002028168 | 142891 | 10 | 76859344 | 76941881 | 1 | protein_coding | sterile alpha motif domain containing 8 |
| **FBXO32** | ENSG00000156804 | 527.8093761 | -0.757907768 | 0.216327292 | 2.12E-05 | 0.001256027 | 114907 | 8 | 124510129 | 124553446 | -1 | protein_coding | F-box protein 32 |
| **VPS8** | ENSG00000156931 | 2709.254689 | 0.345374807 | 0.128269417 | 8.77E-04 | 0.020538335 | 23355 | 3 | 184529931 | 184770402 | 1 | protein_coding | vacuolar protein sorting 8 homolog (S. cerevisiae) |
| **CLDN12** | ENSG00000157224 | 950.1206359 | 0.555865978 | 0.182891724 | 1.54E-04 | 0.005695399 | 9069 | 7 | 90013035 | 90142716 | 1 | protein_coding | claudin 12 |
| **GATAD1** | ENSG00000157259 | 1327.265108 | 0.28550801 | 0.120227688 | 0.002606705 | 0.042552158 | 57798 | 7 | 92076767 | 92088150 | 1 | protein_coding | GATA zinc finger domain containing 1 |
| **LDLRAP1** | ENSG00000157978 | 1118.305859 | -0.341861554 | 0.160443958 | 0.003149453 | 0.047406845 | 26119 | 1 | 25870071 | 25895377 | 1 | protein_coding | low density lipoprotein receptor adaptor protein 1 |
| **BRE** | ENSG00000158019 | 2154.545558 | -0.461879125 | 0.150180233 | 1.55E-04 | 0.005711686 | 9577 | 2 | 28112808 | 28561768 | 1 | protein_coding | brain and reproductive organ-expressed (TNFRSF1A modulator) |
| **MRPL17** | ENSG00000158042 | 609.8529716 | -0.3803522 | 0.177639432 | 0.002400264 | 0.040043333 | 63875 | 11 | 6702013 | 6704632 | -1 | protein_coding | mitochondrial ribosomal protein L17 |
| **WASF2** | ENSG00000158195 | 519.2419547 | -0.532904699 | 0.229922166 | 0.001080325 | 0.023769396 | 10163 | 1 | 27730730 | 27816669 | -1 | protein_coding | WAS protein family, member 2 |
| **HIST1H2BD** | ENSG00000158373 | 713.7173429 | -0.453341003 | 0.229873615 | 0.002951247 | 0.045453473 | 3017 | 6 | 26158349 | 26171577 | 1 | protein_coding | histone cluster 1, H2bd |
| **PPP1R15B** | ENSG00000158615 | 2618.320896 | 0.418781745 | 0.10547283 | 6.69E-06 | 5.05E-04 | 84919 | 1 | 204372515 | 204380919 | -1 | protein_coding | protein phosphatase 1, regulatory subunit 15B |
| **CDA** | ENSG00000158825 | 959.8660827 | -0.413528173 | 0.153956767 | 6.61E-04 | 0.017169999 | 978 | 1 | 20915441 | 20945401 | 1 | protein_coding | cytidine deaminase |
| **FCER1G** | ENSG00000158869 | 13842.69784 | -0.363618308 | 0.101471687 | 3.88E-05 | 0.002028168 | 2207 | 1 | 161185024 | 161190489 | 1 | protein_coding | Fc fragment of IgE, high affinity I, receptor for; gamma polypeptide |
| **CCAR2** | ENSG00000158941 | 1714.135024 | -0.353930296 | 0.153005906 | 0.002005505 | 0.035681707 | 57805 | 8 | 22462145 | 22479027 | 1 | protein_coding | cell cycle and apoptosis regulator 2 |
| **CDC42SE2** | ENSG00000158985 | 4310.686177 | 0.510991647 | 0.133471776 | 1.15E-05 | 7.5E-04 | 56990 | 5 | 130581186 | 130734140 | 1 | protein_coding | CDC42 small effector 2 |
| **RUNX1** | ENSG00000159216 | 1701.646302 | 0.47270144 | 0.2436805 | 0.002822901 | 0.044574992 | 101928269 | 21 | 36160098 | 37376965 | -1 | protein_coding | runt-related transcription factor 1 |
| **HK2** | ENSG00000159399 | 22078.61884 | 0.40401583 | 0.118325174 | 7.19E-05 | 0.003279499 | 3099 | 2 | 75061108 | 75120486 | 1 | protein_coding | hexokinase 2 |
| **GPBP1L1** | ENSG00000159592 | 4206.195239 | 0.399011724 | 0.121952529 | 9.01E-05 | 0.00387675 | 60313 | 1 | 46092976 | 46153785 | -1 | protein_coding | GC-rich promoter binding protein 1-like 1 |
| **ZFYVE28** | ENSG00000159733 | 273.3574639 | 0.590447573 | 0.24571778 | 8.11E-04 | 0.01972889 | 57732 | 4 | 2271309 | 2420390 | -1 | protein_coding | zinc finger, FYVE domain containing 28 |
| **VMA21** | ENSG00000160131 | 1275.206307 | 0.586581159 | 0.154908965 | 1.04E-05 | 6.92E-04 | 203547 | X | 150564987 | 150577836 | 1 | protein_coding | VMA21 vacuolar H+-ATPase homolog (S. cerevisiae) |
| **G6PD** | ENSG00000160211 | 4682.303754 | -0.350693762 | 0.115676508 | 2.64E-04 | 0.008603521 | 2539 | X | 153759606 | 153775787 | -1 | protein_coding | glucose-6-phosphate dehydrogenase |
| **FDPS** | ENSG00000160752 | 1918.105701 | -0.429481874 | 0.115961489 | 1.9E-05 | 0.001160447 | 2224 | 1 | 155278539 | 155290457 | 1 | protein_coding | farnesyl diphosphate synthase |
| **LY6E** | ENSG00000160932 | 2261.072498 | -0.526117711 | 0.133004552 | 5.53E-06 | 4.42E-04 | 4061 | 8 | 144099399 | 144105249 | 1 | protein_coding | lymphocyte antigen 6 complex, locus E |
| **ZNF385A** | ENSG00000161642 | 8854.993568 | -0.353165364 | 0.114379761 | 2.4E-04 | 0.008066708 | 25946 | 12 | 54762917 | 54785082 | -1 | protein_coding | zinc finger protein 385A |
| **CXCL16** | ENSG00000161921 | 12381.22192 | -0.24193861 | 0.08561531 | 8.38E-04 | 0.020002769 | 58191 | 17 | 4636821 | 4643217 | -1 | protein_coding | chemokine (C-X-C motif) ligand 16 |
| **CYB561A3** | ENSG00000162144 | 4845.782529 | -0.607924909 | 0.176895513 | 3.29E-05 | 0.00180004 | 220002 | 11 | 61116217 | 61129771 | -1 | protein_coding | cytochrome b561 family, member A3 |
| **CMPK1** | ENSG00000162368 | 2823.97142 | 0.345644415 | 0.151125424 | 0.002123707 | 0.03699776 | 51727 | 1 | 47799469 | 47844511 | 1 | protein_coding | cytidine monophosphate (UMP-CMP) kinase 1, cytosolic |
| **USP24** | ENSG00000162402 | 3047.60981 | 0.239645754 | 0.093382254 | 0.00211591 | 0.036983189 | 23358 | 1 | 55532032 | 55680786 | -1 | protein_coding | ubiquitin specific peptidase 24 |
| **USP1** | ENSG00000162607 | 750.1928939 | 0.38455332 | 0.183752876 | 0.002974722 | 0.04561669 | 7398 | 1 | 62901968 | 62917475 | 1 | protein_coding | ubiquitin specific peptidase 1 |
| **H3F3A** | ENSG00000163041 | 2618.542234 | 0.596059618 | 0.13720442 | 1.1E-06 | 1.26E-04 | 3021 | 1 | 226249552 | 226259702 | 1 | protein_coding | H3 histone, family 3A |
| **SPATA18** | ENSG00000163071 | 797.0543468 | -0.500178882 | 0.276547636 | 0.003251032 | 0.048523477 | 132671 | 4 | 52917497 | 52963458 | 1 | protein_coding | spermatogenesis associated 18 |
| **TNFAIP8L2** | ENSG00000163154 | 652.6230989 | -0.365551241 | 0.160546289 | 0.001996825 | 0.03560446 | 79626 | 1 | 151129105 | 151132225 | 1 | protein_coding | tumor necrosis factor, alpha-induced protein 8-like 2 |
| **RNF149** | ENSG00000163162 | 3151.968233 | 0.445714805 | 0.122196604 | 2.39E-05 | 0.001387182 | 284996 | 2 | 101887681 | 101925163 | -1 | protein_coding | ring finger protein 149 |
| **S100A11** | ENSG00000163191 | 14615.95674 | -0.424016162 | 0.151448996 | 4.74E-04 | 0.013536855 | 6282 | 1 | 152004982 | 152020383 | -1 | protein_coding | S100 calcium binding protein A11 |
| **ARHGAP25** | ENSG00000163219 | 4108.217743 | -0.372247102 | 0.178991912 | 0.002755756 | 0.044122435 | 9938 | 2 | 68906733 | 69053965 | 1 | protein_coding | Rho GTPase activating protein 25 |

| | | | | | | | | | | | | | |
|---|---|---|---|---|---|---|---|---|---|---|---|---|---|
| **S100A9** | ENSG00000163220 | 3836.622584 | -0.389806999 | 0.136926847 | 4E-04 | 0.011884426 | 6280 | 1 | 153330330 | 153333503 | 1 | protein_coding | S100 calcium binding protein A9 |
| **LRRC58** | ENSG00000163428 | 1782.852283 | 0.973125193 | 0.139429449 | 1.57E-13 | 2.78E-10 | 116064 | 3 | 120043356 | 120068186 | -1 | protein_coding | leucine rich repeat containing 58 |
| **SSR2** | ENSG00000163479 | 2964.447255 | -0.27928072 | 0.11578603 | 0.00249963 | 0.041440827 | 6746 | 1 | 155978839 | 155990750 | -1 | protein_coding | signal sequence receptor, beta (translocon-associated protein beta) |
| **GLB1L** | ENSG00000163521 | 318.3186837 | -0.538648775 | 0.246043732 | 0.001433206 | 0.028900715 | 79411 | 2 | 220101328 | 220110200 | -1 | protein_coding | galactosidase, beta 1-like |
| **RYBP** | ENSG00000163602 | 1679.803396 | 0.539502655 | 0.14321303 | 1.18E-05 | 7.66E-04 | 23429 | 3 | 72420976 | 72496069 | -1 | protein_coding | RING1 and YY1 binding protein |
| **CCNL1** | ENSG00000163660 | 2407.083225 | 0.279347055 | 0.108378713 | 0.001646282 | 0.031522592 | 57018 | 3 | 156864297 | 156878549 | -1 | protein_coding | cyclin L1 |
| **CDCP1** | ENSG00000163814 | 285.1993123 | 0.6785433 | 0.403980625 | 0.002940753 | 0.0453576 | 64866 | 3 | 45123770 | 45187914 | -1 | protein_coding | CUB domain containing protein 1 |
| **ZMYM6** | ENSG00000163867 | 694.3564485 | 0.403339825 | 0.194570415 | 0.002783326 | 0.044228011 | 9204 | 1 | 35449523 | 35497569 | -1 | protein_coding | zinc finger, MYM-type 6 |
| **PRKCD** | ENSG00000163932 | 6183.781765 | -0.325353811 | 0.111866704 | 4.45E-04 | 0.012846415 | 5580 | 3 | 53190025 | 53226733 | 1 | protein_coding | protein kinase C, delta |
| **FAM208A** | ENSG00000163946 | 3936.601775 | 0.313468066 | 0.128019253 | 0.001914587 | 0.034720672 | 23272 | 3 | 56654161 | 56717265 | -1 | protein_coding | family with sequence similarity 208, member A |
| **H2AFZ** | ENSG00000164032 | 1187.981966 | 0.334492051 | 0.150430397 | 0.002767828 | 0.044228011 | 3015 | 4 | 100869243 | 100871545 | -1 | protein_coding | H2A histone family, member Z |
| **ABCE1** | ENSG00000164163 | 2428.29397 | -0.428121648 | 0.132390581 | 1.04E-04 | 0.004301634 | 6059 | 4 | 146019084 | 146050331 | 1 | protein_coding | ATP-binding cassette, sub-family E (OABP), member 1 |
| **SLC25A46** | ENSG00000164209 | 1303.78135 | 0.851366257 | 0.189811218 | 3.87E-07 | 5.41E-05 | 91137 | 5 | 110073837 | 110100857 | 1 | protein_coding | solute carrier family 25, member 46 |
| **GRPEL2** | ENSG00000164284 | 454.6375024 | 0.808635489 | 0.185133403 | 6.23E-07 | 8.17E-05 | 134266 | 5 | 148724993 | 148734146 | 1 | protein_coding | GrpE-like 2, mitochondrial (E. coli) |
| **ERAP1** | ENSG00000164307 | 5605.917735 | -0.323733558 | 0.121860216 | 8.15E-04 | 0.01972889 | 51752 | 5 | 96096521 | 96143803 | -1 | protein_coding | endoplasmic reticulum aminopeptidase 1 |
| **KIAA1430** | ENSG00000164323 | 640.8810007 | 0.829074174 | 0.187214179 | 5.09E-07 | 6.85E-05 | 57587 | 4 | 186080819 | 186130658 | -1 | protein_coding | KIAA1430 |
| **UBLCP1** | ENSG00000164332 | 1218.816192 | 0.395164526 | 0.156221424 | 0.001010704 | 0.02290138 | 134510 | 5 | 158690089 | 158713044 | 1 | protein_coding | ubiquitin-like domain containing CTD phosphatase 1 |
| **GPR85** | ENSG00000164604 | 99.60799306 | -0.952766783 | 0.425863666 | 8.15E-04 | 0.01972889 | 54329 | 7 | 112718386 | 112727833 | -1 | protein_coding | G protein-coupled receptor 85 |
| **MIOS** | ENSG00000164654 | 1410.839856 | 0.400997143 | 0.12933168 | 1.81E-04 | 0.006465396 | 54468 | 7 | 7606503 | 7648560 | 1 | protein_coding | missing oocyte, meiosis regulator, homolog (Drosophila) |
| **YWHAZ** | ENSG00000164924 | 25585.839 | -0.31045176 | 0.141042587 | 0.003308991 | 0.049112631 | 7534 | 8 | 101928753 | 101965616 | -1 | protein_coding | tyrosine 3-monooxygenase/tryptophan 5-monooxygenase activation protein, zeta |
| **DCSTAMP** | ENSG00000164935 | 13983.51498 | -0.410827782 | 0.144319698 | 3.95E-04 | 0.011862182 | 81501 | 8 | 105351315 | 105368917 | 1 | protein_coding | dendrocyte expressed seven transmembrane protein |
| **TMEM65** | ENSG00000164983 | 264.5973219 | 0.999947597 | 0.242868345 | 1.83E-06 | 1.91E-04 | 157378 | 8 | 125324231 | 125384933 | -1 | protein_coding | transmembrane protein 65 |
| **ABCA1** | ENSG00000165029 | 5134.937597 | 0.267970159 | 0.091293342 | 5.49E-04 | 0.015219221 | 19 | 9 | 107543283 | 107690518 | -1 | protein_coding | ATP-binding cassette, sub-family A (ABC1), member 1 |
| **KDM1B** | ENSG00000165097 | 761.7509599 | -1.127220665 | 0.195257263 | 3.82E-10 | 1.93E-07 | 221656 | 6 | 18155560 | 18224084 | 1 | protein_coding | lysine (K)-specific demethylase 1B |
| **DYNLT3** | ENSG00000165169 | 816.7861328 | 0.657123426 | 0.188435144 | 2.75E-05 | 0.001557255 | 6990 | X | 37696010 | 37706890 | -1 | protein_coding | dynein, light chain, Tctex-type 3 |
| **OTUD1** | ENSG00000165312 | 648.185364 | 1.023371518 | 0.202888475 | 2.28E-08 | 5.26E-06 | 220213 | 10 | 23728198 | 23731308 | 1 | protein_coding | OTU domain containing 1 |
| **SPTSSA** | ENSG00000165389 | 319.2544952 | 0.512389418 | 0.229134834 | 0.001394502 | 0.028553704 | 171546 | 14 | 34901995 | 34931562 | -1 | protein_coding | serine palmitoyltransferase, small subunit A |
| **PGM2L1** | ENSG00000165434 | 1119.440571 | 0.522038455 | 0.146739487 | 2.69E-05 | 0.001539238 | 283209 | 11 | 74041363 | 74109518 | -1 | protein_coding | phosphoglucomutase 2-like 1 |
| **REEP3** | ENSG00000165476 | 570.1900876 | 1.166012348 | 0.235486092 | 3.52E-08 | 7.07E-06 | 221035 | 10 | 65281123 | 65384883 | 1 | protein_coding | receptor accessory protein 3 |
| **MICU2** | ENSG00000165487 | 946.6510734 | 0.482974118 | 0.137570451 | 3.45E-05 | 0.0018614 | 221154 | 13 | 22066836 | 22178353 | -1 | protein_coding | mitochondrial calcium uptake 2 |
| **NEMF** | ENSG00000165525 | 1337.038173 | -0.854314781 | 0.137605297 | 3.3E-11 | 2.34E-08 | 9147 | 14 | 50249997 | 50319921 | -1 | protein_coding | nuclear export mediator factor |
| **TTC8** | ENSG00000165533 | 251.6977063 | -0.994229084 | 0.368687398 | 2.59E-04 | 0.008480193 | 123016 | 14 | 89290497 | 89344335 | 1 | protein_coding | tetratricopeptide repeat domain 8 |
| **ARL5B** | ENSG00000165997 | 738.2851 | 1.16545783 | 0.174877926 | 1.4E-12 | 1.49E-09 | 221079 | 10 | 18948334 | 18970568 | 1 | protein_coding | ADP-ribosylation factor-like 5B |
| **TAF1D** | ENSG00000166012 | 1088.417705 | 0.486128128 | 0.183868466 | 5.5E-04 | 0.015219221 | 100302240 | 11 | 93463114 | 93517557 | -1 | protein_coding | TATA box binding protein (TBP)-associated factor, RNA polymerase I, D, 41kDa |
| **NDUFB8** | ENSG00000166136 | 1273.724142 | -0.333547795 | 0.134570767 | 0.001534418 | 0.030252811 | 4714 | 10 | 102267203 | 102289757 | -1 | protein_coding | NADH dehydrogenase (ubiquinone) 1 beta subcomplex, 8, 19kDa |
| **PCBD1** | ENSG00000166228 | 833.9284534 | -0.365977754 | 0.153334722 | 0.001553009 | 0.03036027 | 5092 | 10 | 72642037 | 72648541 | -1 | protein_coding | pterin-4 alpha-carbinolamine dehydratase/dimerization cofactor of hepatocyte nuclear factor 1 alpha |
| **CUL5** | ENSG00000166266 | 1360.162381 | 0.330793238 | 0.11387265 | 4.66E-04 | 0.013386305 | 8065 | 11 | 107879459 | 107978503 | 1 | protein_coding | cullin 5 |
| **C10orf32** | ENSG00000166275 | 403.0173702 | 0.403654566 | 0.189615664 | 0.002444793 | 0.040595018 | 119032 | 10 | 104613980 | 104624718 | 1 | protein_coding | chromosome 10 open reading frame 32 |
| **RPL27A** | ENSG00000166441 | 5938.526253 | -0.346530577 | 0.129547932 | 7.9E-04 | 0.019381909 | 619562 | 11 | 8703958 | 8736306 | 1 | protein_coding | ribosomal protein L27a |
| **TMEM41B** | ENSG00000166471 | 1390.672646 | 0.720255124 | 0.126279461 | 7.42E-10 | 3.15E-07 | 440026 | 11 | 9302201 | 9336327 | -1 | protein_coding | transmembrane protein 41B |
| **B2M** | ENSG00000166710 | 160693.4755 | -0.26489779 | 0.086888577 | 4.4E-04 | 0.012800797 | 567 | 15 | 45003675 | 45011075 | 1 | protein_coding | beta-2-microglobulin |
| **AP1G1** | ENSG00000166747 | 4443.349422 | 0.30008452 | 0.09380181 | 2.56E-04 | 0.008442217 | 164 | 16 | 71762913 | 71843104 | -1 | protein_coding | adaptor-related protein complex 1, gamma 1 subunit |
| **PPIB** | ENSG00000166794 | 4191.387156 | -0.32185936 | 0.115746603 | 6.21E-04 | 0.016366263 | 5479 | 15 | 64448011 | 64455404 | -1 | protein_coding | peptidylprolyl isomerase B (cyclophilin B) |
| **TMEM170A** | ENSG00000166822 | 1082.443143 | 0.336824126 | 0.139712858 | 0.001740573 | 0.03269229 | 124491 | 16 | 75476952 | 75499395 | -1 | protein_coding | transmembrane protein 170A |
| **PATL1** | ENSG00000166889 | 2671.24209 | 0.255756546 | 0.10316396 | 0.00232036 | 0.039516772 | 219988 | 11 | 59404189 | 59436453 | -1 | protein_coding | protein associated with topoisomerase II homolog 1 (yeast) |
| **TBC1D2B** | ENSG00000167202 | 6387.226768 | -0.251272174 | 0.090376033 | 7.25E-04 | 0.018365575 | 23102 | 15 | 78276378 | 78370066 | -1 | protein_coding | TBC1 domain family, member 2B |
| **HDHD2** | ENSG00000167220 | 841.2819214 | -0.479245632 | 0.160513649 | 2.03E-04 | 0.007117793 | 84064 | 18 | 44633774 | 44676891 | -1 | protein_coding | haloacid dehalogenase-like hydrolase domain containing 2 |
| **ZNF91** | ENSG00000167232 | 1299.841841 | 0.480196905 | 0.215694413 | 0.00168549 | 0.032039304 | 7644 | 19 | 23487793 | 23578362 | -1 | protein_coding | zinc finger protein 91 |
| **TPM4** | ENSG00000167460 | 13175.28762 | 0.377423127 | 0.137050464 | 5.59E-04 | 0.015320947 | 7171 | 19 | 16177831 | 16213813 | 1 | protein_coding | tropomyosin 4 |
| **RAB8A** | ENSG00000167461 | 3861.160577 | -0.411231243 | 0.163719641 | 9.83E-04 | 0.022423744 | 4218 | 19 | 16222439 | 16245044 | 1 | protein_coding | RAB8A, member RAS oncogene family |
| **GPX4** | ENSG00000167468 | 4477.442153 | -0.35807344 | 0.11697259 | 2.35E-04 | 0.007997026 | 2879 | 19 | 1103936 | 1106787 | 1 | protein_coding | glutathione peroxidase 4 |
| **TUBA1A** | ENSG00000167552 | 9120.683843 | -0.569833722 | 0.121509125 | 1.72E-07 | 2.73E-05 | 7846 | 12 | 49578579 | 49583107 | -1 | protein_coding | tubulin, alpha 1a |
| **LENG8** | ENSG00000167615 | 244.7030115 | 0.78555666 | 0.297483275 | 3.44E-04 | 0.010628856 | 114823 | 19 | 54960065 | 54973217 | 1 | protein_coding | leukocyte receptor cluster (LRC) member 8 |
| **CD300C** | ENSG00000167850 | 843.2856943 | -0.364465047 | 0.160177339 | 0.002121555 | 0.03699776 | 10871 | 17 | 72537247 | 72542282 | -1 | protein_coding | CD300c molecule |
| **BEST1** | ENSG00000167995 | 3056.838917 | -0.659230674 | 0.239266824 | 2.79E-04 | 0.008968695 | 7439 | 11 | 61717293 | 61732987 | 1 | protein_coding | bestrophin 1 |
| **MAPK1IP1L** | ENSG00000168175 | 3172.352047 | -0.415830671 | 0.121566128 | 6.27E-05 | 0.002909119 | 93487 | 14 | 55518349 | 55536910 | 1 | protein_coding | mitogen-activated protein kinase 1 interacting protein 1-like |
| **SNRNP48** | ENSG00000168566 | 367.8717407 | 0.469708217 | 0.218998642 | 0.001887975 | 0.034425705 | 154007 | 6 | 7590432 | 7612200 | 1 | protein_coding | small nuclear ribonucleoprotein 48kDa (U11/U12) |
| **IL7R** | ENSG00000168685 | 1341.314744 | 0.811249413 | 0.133111884 | 6.67E-11 | 3.64E-08 | 3575 | 5 | 35852797 | 35879705 | 1 | protein_coding | interleukin 7 receptor |
| **GSTM4** | ENSG00000168765 | 746.1879885 | -0.508506301 | 0.273672521 | 0.002922099 | 0.045310196 | 2948 | 1 | 110198703 | 110208118 | 1 | protein_coding | glutathione S-transferase mu 4 |
| **NPR 1.00** | ENSG00000169418 | 231.7632957 | -0.66528834 | 0.304431511 | 0.001184169 | 0.025297244 | 4881 | 1 | 153651113 | 153666468 | 1 | protein_coding | natriuretic peptide receptor A/guanylate cyclase A (atrionatriuretic peptide receptor A) |
| **MMGT1** | ENSG00000169446 | 1371.113339 | 0.573350609 | 0.221984623 | 4.88E-04 | 0.013855339 | 93380 | X | 135044229 | 135056222 | -1 | protein_coding | membrane magnesium transporter 1 |
| **CLIC4** | ENSG00000169504 | 11686.9912 | 0.414884194 | 0.091012777 | 5.34E-07 | 7.09E-05 | 25932 | 1 | 25071848 | 25170815 | 1 | protein_coding | chloride intracellular channel 4 |
| **DTWD2** | ENSG00000169570 | 357.5479791 | 0.579839939 | 0.286444625 | 0.001888072 | 0.034425705 | 285605 | 5 | 118173017 | 118324240 | -1 | protein_coding | DTW domain containing 2 |
| **CSGALNAC** | ENSG00000169826 | 3342.351059 | 0.382980767 | 0.144097728 | 8.4E-04 | 0.020002769 | 55454 | 10 | 43633934 | 43680756 | 1 | protein_coding | chondroitin sulfate N-acetylgalactosaminyltransferase 2 |
| **SYAP1** | ENSG00000169895 | 1589.49793 | 0.374827827 | 0.131422482 | 4.43E-04 | 0.012846415 | 94056 | X | 16737755 | 16783459 | 1 | protein_coding | synapse associated protein 1 |
| **CHD3** | ENSG00000170004 | 3302.981198 | 0.393389965 | 0.158538485 | 0.001059014 | 0.02354425 | 1107 | 17 | 7788124 | 7816078 | 1 | protein_coding | chromodomain helicase DNA binding protein 3 |
| **USP38** | ENSG00000170185 | 1355.009036 | 0.290347076 | 0.110426124 | 0.001232933 | 0.026100366 | 84640 | 4 | 144106070 | 144144983 | 1 | protein_coding | ubiquitin specific peptidase 38 |
| **NANP** | ENSG00000170191 | 490.1726788 | 0.453595847 | 0.189039921 | 0.001110489 | 0.024282241 | 140838 | 20 | 25593571 | 25604811 | -1 | protein_coding | N-acetylneuraminic acid phosphatase |
| **GABARAP** | ENSG00000170296 | 5833.232947 | -0.249664522 | 0.092964487 | 0.00137286 | 0.028328895 | 11337 | 17 | 7143333 | 7146089 | -1 | protein_coding | GABA(A) receptor-associated protein |
| **UBB** | ENSG00000170315 | 10125.88118 | -0.366663609 | 0.100496759 | 3.08E-05 | 0.001722556 | 7314 | 17 | 16284112 | 16286059 | 1 | protein_coding | ubiquitin B |
| **TMED10** | ENSG00000170348 | 8639.871663 | -0.492418796 | 0.107807667 | 3.79E-07 | 5.37E-05 | 10972 | 14 | 75598173 | 75643334 | -1 | protein_coding | transmembrane emp24-like trafficking protein 10 (yeast) |
| **FAM115C** | ENSG00000170379 | 142.2760281 | 0.966823923 | 0.358848552 | 2.66E-04 | 0.008660314 | 101930379 | 7 | 143318043 | 143427502 | 1 | protein_coding | family with sequence similarity 115, member C |
| **SLC30A1** | ENSG00000170385 | 7801.744785 | 0.955308461 | 0.133214378 | 3.56E-14 | 9.46E-11 | 7779 | 1 | 211744910 | 211752084 | -1 | protein_coding | solute carrier family 30 (zinc transporter), member 1 |
| **DENND5B** | ENSG00000170456 | 411.2610689 | -0.585221993 | 0.301397453 | 0.002104101 | 0.036898147 | 160518 | 12 | 31535157 | 31744031 | -1 | protein_coding | DENN/MADD domain containing 5B |
| **RNF139** | ENSG00000170881 | 987.5074533 | 0.433445587 | 0.181107869 | 0.001282316 | 0.026984507 | 11236 | 8 | 125486979 | 125500155 | 1 | protein_coding | ring finger protein 139 |
| **FAM98B** | ENSG00000171262 | 584.5210657 | 0.551010024 | 0.224428511 | 7.52E-04 | 0.018727077 | 283742 | 15 | 38746328 | 38779911 | 1 | protein_coding | family with sequence similarity 98, member B |
| **LRRC8C** | ENSG00000171488 | 1226.925328 | 0.591252764 | 0.135005386 | 7.41E-07 | 9.26E-05 | 101927933 | 1 | 90098631 | 90235462 | 1 | protein_coding | leucine rich repeat containing 8 family, member C |
| **CYP4F22** | ENSG00000171954 | 577.3457258 | -1.102233682 | 0.28651183 | 5.11E-06 | 4.14E-04 | 126410 | 19 | 15619304 | 15663128 | 1 | protein_coding | cytochrome P450, family 4, subfamily F, polypeptide 22 |
| **CERS6** | ENSG00000172292 | 3275.809159 | 0.578934633 | 0.139983299 | 2.42E-06 | 2.34E-04 | 253782 | 2 | 169312372 | 169631644 | 1 | protein_coding | ceramide synthase 6 |
| **ZNF24** | ENSG00000172466 | 2747.981285 | 0.437118601 | 0.125926757 | 3.93E-05 | 0.002028168 | 7572 | 18 | 32912176 | 32925446 | -1 | protein_coding | zinc finger protein 24 |
| **RPL38** | ENSG00000172809 | 2695.95321 | -0.351914857 | 0.160005113 | 0.002593294 | 0.042463687 | 6169 | 17 | 72199721 | 72206794 | 1 | protein_coding | ribosomal protein L38 |
| **DHCR7** | ENSG00000172893 | 963.3753774 | -0.561002604 | 0.208739826 | 4.19E-04 | 0.012326068 | 1717 | 11 | 71139239 | 71163914 | -1 | protein_coding | 7-dehydrocholesterol reductase |
| **AHSA2** | ENSG00000173209 | 1018.73058 | 0.480385168 | 0.171593829 | 3.54E-04 | 0.010850963 | 130872 | 2 | 61404553 | 61418338 | 1 | protein_coding | AHA1, activator of heat shock 90kDa protein ATPase homolog 2 (yeast) |
| **PEAK1** | ENSG00000173517 | 1711.728532 | 0.438765649 | 0.137065883 | 1.11E-04 | 0.004539553 | 79834 | 15 | 77400471 | 77712486 | -1 | protein_coding | pseudopodium-enriched atypical kinase 1 |
| **KLHL15** | ENSG00000174010 | 461.5486835 | 0.640766591 | 0.178900747 | 2.02E-05 | 0.001216917 | 80311 | X | 24001837 | 24045303 | -1 | protein_coding | kelch-like family member 15 |
| **FAM3C2** | ENSG00000174028 | 359.0166829 | -0.798315458 | 0.229125956 | 2.38E-05 | 0.001387182 | NA | X | 23093875 | 23094884 | 1 | pseudogene | family with sequence similarity 3, member C2 (pseudogene) |
| **LEMD3** | ENSG00000174106 | 1029.432813 | 0.359971182 | 0.130465384 | 5.75E-04 | 0.0155063 | 23592 | 12 | 65563351 | 65642107 | 1 | protein_coding | LEM domain containing 3 |
| **CEP135** | ENSG00000174799 | 594.5907942 | 0.368628364 | 0.177683027 | 0.003267229 | 0.048628629 | 9662 | 4 | 56815037 | 56899529 | 1 | protein_coding | centrosomal protein 135kDa |
| **VCPIP1** | ENSG00000175073 | 2971.135579 | 0.422980815 | 0.100911524 | 2.76E-06 | 2.59E-04 | 80124 | 8 | 67540722 | 67579452 | -1 | protein_coding | valosin containing protein (p97)/p47 complex interacting protein 1 |
| **PDIK1L** | ENSG00000175087 | 567.8097553 | 0.381407485 | 0.174153914 | 0.002370734 | 0.039900331 | 149420 | 1 | 26437664 | 26452034 | 1 | protein_coding | PDLIM1 interacting kinase 1 like |
| **BANF1** | ENSG00000175334 | 821.9532185 | -0.382843868 | 0.159866856 | 0.001411555 | 0.028681819 | 8815 | 11 | 65769550 | 65771620 | 1 | protein_coding | barrier to autointegration factor 1 |
| **ZDHHC21** | ENSG00000175893 | 1032.769341 | 0.385218768 | 0.190682159 | 0.003371677 | 0.049510993 | 340481 | 9 | 14611069 | 14693469 | -1 | protein_coding | zinc finger, DHHC-type containing 21 |

| | | | | | | | | | | | | | |
|---|---|---|---|---|---|---|---|---|---|---|---|---|---|
| **LRRC37A3** | ENSG00000176809 | 370.411128 | 0.645049708 | 0.3143391 | 0.001586538 | 0.030772302 | 101930430 | 17 | 62850430 | 62915598 | -1 | protein_coding | leucine rich repeat containing 37, member A3 |
| **FAM210A** | ENSG00000177150 | 493.6766585 | 0.446672011 | 0.169470613 | 6.46E-04 | 0.016821041 | 125228 | 18 | 13663346 | 13726662 | -1 | protein_coding | family with sequence similarity 210, member A |
| **RPS6KA3** | ENSG00000177189 | 3569.949433 | 0.613297668 | 0.117066943 | 1.09E-08 | 2.98E-06 | 6197 | X | 20168029 | 20285523 | -1 | protein_coding | ribosomal protein S6 kinase, 90kDa, polypeptide 3 |
| **TBL1XR1** | ENSG00000177565 | 4549.754287 | 0.317013193 | 0.102194104 | 2.88E-04 | 0.00917408 | 79718 | 3 | 176737143 | 176915261 | -1 | protein_coding | transducin (beta)-like 1 X-linked receptor 1 |
| **THAP5** | ENSG00000177683 | 538.4475566 | 0.540164621 | 0.271195602 | 0.002134211 | 0.037059255 | 168451 | 7 | 108194987 | 108210194 | -1 | protein_coding | THAP domain containing 5 |
| **FAM20C** | ENSG00000177706 | 7148.722118 | 0.492942346 | 0.191436835 | 6.33E-04 | 0.016543103 | 56975 | 7 | 192969 | 300711 | 1 | protein_coding | family with sequence similarity 20, member C |
| **HNRNPA0** | ENSG00000177733 | 1437.378615 | 0.420936816 | 0.132895636 | 1.3E-04 | 0.00503517 | 10949 | 5 | 137087075 | 137090039 | -1 | protein_coding | heterogeneous nuclear ribonucleoprotein A0 |
| **MYOZ1** | ENSG00000177791 | 257.2531473 | -0.620444062 | 0.355474523 | 0.002512786 | 0.041540487 | 58529 | 10 | 75391412 | 75401515 | -1 | protein_coding | myozenin 1 |
| **C2orf69** | ENSG00000178074 | 551.5369274 | 0.682432274 | 0.242909376 | 2.36E-04 | 0.00801213 | 205327 | 2 | 200775979 | 200820658 | 1 | protein_coding | chromosome 2 open reading frame 69 |
| **EPM2AIP1** | ENSG00000178567 | 839.5295821 | 0.686937057 | 0.177661911 | 6.26E-06 | 4.82E-04 | 9852 | 3 | 37027357 | 37034795 | -1 | protein_coding | EPM2A (laforin) interacting protein 1 |
| **MAF** | ENSG00000178573 | 4092.263083 | 0.541522357 | 0.139103727 | 5.6E-06 | 4.44E-04 | 4094 | 16 | 79619740 | 79634611 | -1 | protein_coding | v-maf avian musculoaponeurotic fibrosarcoma oncogene homolog |
| **SUZ12** | ENSG00000178691 | 1417.715833 | 0.474693384 | 0.126812623 | 1.53E-05 | 9.71E-04 | 23512 | 17 | 30264037 | 30328064 | 1 | protein_coding | SUZ12 polycomb repressive complex 2 subunit |
| **KCTD12** | ENSG00000178695 | 7871.824048 | 0.803957086 | 0.098197216 | 1.39E-17 | 7.38E-14 | 115207 | 13 | 77454312 | 77460540 | -1 | protein_coding | potassium channel tetramerization domain containing 12 |
| **MSC** | ENSG00000178860 | 1236.633833 | -0.406418865 | 0.153969886 | 6.34E-04 | 0.016543103 | 9242 | 8 | 72753784 | 72756703 | -1 | protein_coding | musculin |
| **EIF3K** | ENSG00000178982 | 2158.313703 | -0.256803838 | 0.110671618 | 0.003119347 | 0.047087076 | 27335 | 19 | 39109735 | 39127595 | 1 | protein_coding | eukaryotic translation initiation factor 3, subunit K |
| **SNX18** | ENSG00000178996 | 2373.691796 | 0.498114228 | 0.137370649 | 3.32E-05 | 0.001808616 | 112574 | 5 | 53813589 | 53842415 | 1 | protein_coding | sorting nexin 18 |
| **CALR** | ENSG00000179218 | 35178.36588 | -0.305431971 | 0.100180646 | 3.36E-04 | 0.010443926 | 811 | 19 | 13049392 | 13055303 | 1 | protein_coding | calreticulin |
| **RAB39A** | ENSG00000179331 | 459.0174018 | 0.61306521 | 0.198700232 | 1.12E-04 | 0.004572138 | 54734 | 11 | 107799229 | 107834208 | 1 | protein_coding | RAB39A, member RAS oncogene family |
| **ZBTB18** | ENSG00000179456 | 620.3654954 | 0.563146943 | 0.167398584 | 4.94E-05 | 0.002443233 | 10472 | 1 | 244214585 | 244220778 | 1 | protein_coding | zinc finger and BTB domain containing 18 |
| **LACC1** | ENSG00000179630 | 1726.555141 | 0.43343994 | 0.130800032 | 8.98E-05 | 0.00387675 | 144811 | 13 | 44453420 | 44468068 | 1 | protein_coding | laccase (multicopper oxidoreductase) domain containing 1 |
| **MYADM** | ENSG00000179820 | 4316.28108 | 0.319380715 | 0.111014632 | 4.98E-04 | 0.014115144 | 91663 | 19 | 54369477 | 54379691 | 1 | protein_coding | myeloid-associated differentiation marker |
| **SOCS4** | ENSG00000180008 | 973.270586 | 0.70548466 | 0.182612507 | 5.7E-06 | 4.45E-04 | 122809 | 14 | 55493948 | 55516206 | 1 | protein_coding | suppressor of cytokine signaling 4 |
| **ZNRF2** | ENSG00000180233 | 971.0259353 | 0.778663259 | 0.1748578 | 4.78E-07 | 6.52E-05 | 223082 | 7 | 30323923 | 30452118 | 1 | protein_coding | zinc and ring finger 2 |
| **NRIP1** | ENSG00000180530 | 2323.228561 | 0.356637186 | 0.164893568 | 0.002933416 | 0.045310196 | 8204 | 21 | 16333556 | 16437321 | -1 | protein_coding | nuclear receptor interacting protein 1 |
| **HIST1H2AC** | ENSG00000180573 | 1153.327716 | -0.397240071 | 0.147209332 | 5.45E-04 | 0.015169527 | 8334 | 6 | 26124373 | 26139344 | 1 | protein_coding | histone cluster 1, H2ac |
| **SRP9P1** | ENSG00000180581 | 131.0446599 | -1.75817285 | 0.553528203 | 5.02E-05 | 0.002468684 | NA | 10 | 93565803 | 93567253 | -1 | pseudogene | signal recognition particle 9 pseudogene 1 |
| **HIST1H2BC** | ENSG00000180596 | 674.7542057 | -0.667556219 | 0.226578777 | 1.4E-04 | 0.005282849 | 8347 | 6 | 26115101 | 26124154 | -1 | protein_coding | histone cluster 1, H2bc |
| **TNFSF15** | ENSG00000181634 | 1435.215464 | 0.4790708 | 0.227985397 | 0.002021024 | 0.03574881 | 9966 | 9 | 117546915 | 117568406 | -1 | protein_coding | tumor necrosis factor (ligand) superfamily, member 15 |
| **ADO** | ENSG00000181915 | 2353.392349 | 0.384065273 | 0.161696642 | 0.001584306 | 0.030772302 | 84890 | 10 | 64564516 | 64568238 | 1 | protein_coding | 2-aminoethanethiol (cysteamine) dioxygenase |
| **SNRPE** | ENSG00000182004 | 350.6908128 | 0.838828808 | 0.270829057 | 8.69E-05 | 0.003800633 | 6635 | 1 | 203830731 | 203839678 | 1 | protein_coding | small nuclear ribonucleoprotein polypeptide E |
| **CHST15** | ENSG00000182022 | 8320.271274 | 0.558105709 | 0.153530801 | 1.87E-05 | 0.001150611 | 51363 | 10 | 125767184 | 125853206 | -1 | protein_coding | carbohydrate (N-acetylgalactosamine 4-sulfate 6-O) sulfotransferase 15 |
| **ATP6AP2** | ENSG00000182220 | 9715.804747 | 1.002186525 | 0.152392661 | 2.25E-12 | 1.99E-09 | 10159 | X | 40440146 | 40465889 | 1 | protein_coding | ATPase, H+ transporting, lysosomal accessory protein 2 |
| **AP1S2** | ENSG00000182287 | 1678.075074 | 0.375709361 | 0.161867592 | 0.001808641 | 0.033779305 | 8905 | X | 15843929 | 15873054 | -1 | protein_coding | adaptor-related protein complex 1, sigma 2 subunit |
| **C8orf33** | ENSG00000182307 | 817.8751223 | -0.424934452 | 0.219379204 | 0.00314842 | 0.047406845 | 65265 | 8 | 146277764 | 146281416 | 1 | protein_coding | chromosome 8 open reading frame 33 |
| **C16orf72** | ENSG00000182831 | 1842.485555 | 0.412073722 | 0.125906245 | 1.22E-04 | 0.00485688 | 29035 | 16 | 9185505 | 9215497 | 1 | protein_coding | chromosome 16 open reading frame 72 |
| **RGPD6** | ENSG00000183054 | 1434.931125 | 0.412669258 | 0.122507454 | 7E-05 | 0.003208157 | 729540 | 2 | 111271389 | 111334762 | -1 | protein_coding | RANBP2-like and GRIP domain containing 6 |
| **TUBB** | ENSG00000183311 | 182.412595 | -0.993891361 | 0.457996849 | 9.14E-04 | 0.021342659 | 203068 | HSC | 30677512 | 30682738 | 1 | protein_coding | tubulin, beta class I |
| **TBK1** | ENSG00000183735 | 1694.335372 | 0.299308593 | 0.120888004 | 0.001871386 | 0.034425705 | 29110 | 12 | 64845660 | 64895888 | 1 | protein_coding | TANK-binding kinase 1 |
| **SETD8** | ENSG00000183955 | 937.8065721 | -0.511840445 | 0.163243229 | 1.15E-04 | 0.004618362 | 387893 | 12 | 123868320 | 123893905 | 1 | protein_coding | SET domain containing (lysine methyltransferase) 8 |
| **UQCR10** | ENSG00000184076 | 1003.605859 | -0.39918539 | 0.136014047 | 3.04E-04 | 0.009546486 | 29796 | 22 | 30163358 | 30166402 | 1 | protein_coding | ubiquinol-cytochrome c reductase, complex III subunit X |
| **TSPYL2** | ENSG00000184205 | 267.5910711 | 0.57727805 | 0.306953447 | 0.002526848 | 0.041629484 | 64061 | X | 53111549 | 53117722 | 1 | protein_coding | TSPY-like 2 |
| **HIST2H2BE** | ENSG00000184678 | 714.3792903 | -0.479376413 | 0.204002194 | 0.001073423 | 0.023666518 | 8349 | 1 | 149856010 | 149858232 | -1 | protein_coding | histone cluster 2, H2be |
| **ATL3** | ENSG00000184743 | 4155.371551 | -0.397783836 | 0.131769396 | 2.4E-04 | 0.008066708 | 25923 | 11 | 63391559 | 63439393 | -1 | protein_coding | atlastin GTPase 3 |
| **C6orf120** | ENSG00000185127 | 809.5305836 | 0.332776195 | 0.144185489 | 0.00228225 | 0.039153257 | 387263 | 6 | 170102233 | 170106401 | 1 | protein_coding | chromosome 6 open reading frame 120 |
| **PURA** | ENSG00000185129 | 453.1252291 | 0.605203839 | 0.19229124 | 9.56E-05 | 0.004065629 | 5813 | 5 | 139487362 | 139496321 | 1 | protein_coding | purine-rich element binding protein A |
| **ZBTB37** | ENSG00000185278 | 612.1490076 | 0.611866303 | 0.213010156 | 2.16E-04 | 0.007538429 | 84614 | 1 | 173837220 | 173872687 | 1 | protein_coding | zinc finger and BTB domain containing 37 |
| **P4HB** | ENSG00000185624 | 16148.11708 | -0.35337703 | 0.117900695 | 2.92E-04 | 0.009240403 | 5034 | 17 | 79801035 | 79818570 | -1 | protein_coding | prolyl 4-hydroxylase, beta polypeptide |
| **UBE2L3** | ENSG00000185651 | 3139.000756 | -0.338201335 | 0.12519127 | 8.34E-04 | 0.020002769 | 7332 | 22 | 21903736 | 21978323 | 1 | protein_coding | ubiquitin-conjugating enzyme E2L 3 |
| **BRWD1** | ENSG00000185658 | 2020.485931 | 0.3060196 | 0.120305673 | 0.001487647 | 0.02966083 | 54014 | 21 | 40556102 | 40693485 | -1 | protein_coding | bromodomain and WD repeat domain containing 1 |
| **YTHDF3** | ENSG00000185728 | 202.3474554 | 1.130448476 | 0.356427128 | 5.95E-05 | 0.002783261 | 253943 | 8 | 64081112 | 64125346 | 1 | protein_coding | YTH domain family, member 3 |
| **IFIT1** | ENSG00000185745 | 6411.037815 | -0.314495362 | 0.148365859 | 0.003413108 | 0.049960191 | 3434 | 10 | 91152303 | 91163745 | 1 | protein_coding | interferon-induced protein with tetratricopeptide repeats 1 |
| **MORF4L1** | ENSG00000185787 | 6548.610929 | -0.435995521 | 0.128475181 | 5.73E-05 | 0.002692498 | 10933 | 15 | 79102829 | 79190475 | 1 | protein_coding | mortality factor 4 like 1 |
| **ATP6V0C** | ENSG00000185883 | 7940.851402 | -0.2191541 | 0.082077557 | 0.002233951 | 0.038539288 | 527 | 16 | 2563871 | 2570219 | 1 | protein_coding | ATPase, H+ transporting, lysosomal 16kDa, V0 subunit c |
| **GSAP** | ENSG00000186088 | 2869.670251 | 0.282331325 | 0.110151677 | 0.001644396 | 0.031522592 | 54103 | 7 | 76940068 | 77045717 | -1 | protein_coding | gamma-secretase activating protein |
| **ZBTB6** | ENSG00000186130 | 494.6407714 | 0.618025586 | 0.191965833 | 7.98E-05 | 0.003546214 | 10773 | 9 | 125670335 | 125675609 | -1 | protein_coding | zinc finger and BTB domain containing 6 |
| **PPP1CC** | ENSG00000186298 | 3229.669457 | 0.281483526 | 0.10256897 | 0.00104125 | 0.023344658 | 5501 | 12 | 111157485 | 111180744 | -1 | protein_coding | protein phosphatase 1, catalytic subunit, gamma isozyme |
| **PPARA** | ENSG00000186951 | 723.6780193 | 0.725845716 | 0.336118719 | 0.001185474 | 0.025297244 | 5465 | 22 | 46546424 | 46639653 | 1 | protein_coding | peroxisome proliferator-activated receptor alpha |
| **MITF** | ENSG00000187098 | 3152.700773 | -1.219235377 | 0.145291135 | 2.61E-18 | 2.77E-14 | 4286 | 3 | 69788586 | 70017488 | 1 | protein_coding | microphthalmia-associated transcription factor |
| **TAF9B** | ENSG00000187325 | 150.9674225 | 0.654969457 | 0.38895377 | 0.003052138 | 0.04640211 | 51616 | X | 77385245 | 77395203 | -1 | protein_coding | TAF9B RNA polymerase II, TATA box binding protein (TBP)-associated factor, 31kDa |
| **TCEA1** | ENSG00000187735 | 2723.849857 | 0.564541457 | 0.11941985 | 1.7E-07 | 2.73E-05 | 6917 | 8 | 54879112 | 54935089 | -1 | protein_coding | transcription elongation factor A (SII), 1 |
| **CHM** | ENSG00000188419 | 1213.087574 | 0.608358304 | 0.150802571 | 3.49E-06 | 3.06E-04 | 1121 | X | 85116185 | 85302566 | -1 | protein_coding | choroideremia (Rab escort protein 1) |
| **RBM34** | ENSG00000188739 | 553.6217911 | 0.448035219 | 0.229894039 | 0.002885435 | 0.045159815 | 23029 | 1 | 235294498 | 235324772 | -1 | protein_coding | RNA binding motif protein 34 |
| **NHLRC3** | ENSG00000188811 | 3145.971952 | 0.327363817 | 0.120898172 | 8.41E-04 | 0.020002769 | 387921 | 13 | 39612443 | 39624246 | 1 | protein_coding | NHL repeat containing 3 |
| **PTPLAD2** | ENSG00000188921 | 836.2207161 | 0.350423319 | 0.142361489 | 0.001409398 | 0.028681819 | 401494 | 9 | 20995306 | 21031635 | -1 | protein_coding | protein tyrosine phosphatase-like A domain containing 2 |
| **ZNF292** | ENSG00000188994 | 1817.101161 | 0.447140791 | 0.189020813 | 0.001170746 | 0.025134378 | 23036 | 6 | 87862551 | 87973914 | 1 | protein_coding | zinc finger protein 292 |
| **RNFT1** | ENSG00000189050 | 830.6532068 | 0.750228487 | 0.182803186 | 2.28E-06 | 2.22E-04 | 51136 | 17 | 58029601 | 58042122 | -1 | protein_coding | ring finger protein, transmembrane 1 |
| **HN1** | ENSG00000189159 | 625.0712733 | -0.559311756 | 0.235001456 | 8.34E-04 | 0.020002769 | 51155 | 17 | 73131343 | 73164376 | -1 | protein_coding | hematological and neurological expressed 1 |
| **FAM217B** | ENSG00000196227 | 642.3855286 | -0.745686067 | 0.181838612 | 2.26E-06 | 2.22E-04 | 63939 | 20 | 58508819 | 58523735 | 1 | protein_coding | family with sequence similarity 217, member B |
| **ZNF107** | ENSG00000196247 | 507.7144976 | 0.493349606 | 0.206863914 | 0.001046067 | 0.023403265 | 51427 | 7 | 64126511 | 64171404 | 1 | protein_coding | zinc finger protein 107 |
| **PPIA** | ENSG00000196262 | 12322.3331 | -0.457953234 | 0.138824922 | 9.9E-05 | 0.004173276 | 101060363 | 7 | 44836279 | 44864163 | 1 | protein_coding | peptidylprolyl isomerase A (cyclophilin A) |
| **NF1** | ENSG00000196712 | 2330.972301 | 0.401630212 | 0.135069706 | 2.96E-04 | 0.009331117 | 101930150 | 17 | 29421945 | 29709134 | 1 | protein_coding | neurofibromin 1 |
| **FUT11** | ENSG00000196968 | 897.9929717 | 0.419651274 | 0.132611574 | 1.36E-04 | 0.005172366 | 170384 | 10 | 75532049 | 75540009 | 1 | protein_coding | fucosyltransferase 11 (alpha (1,3) fucosyltransferase) |
| **DYNC1H1** | ENSG00000197102 | 8790.941304 | 0.239236735 | 0.08809087 | 0.0015483 | 0.03036027 | 1778 | 14 | 102430865 | 102517129 | 1 | protein_coding | dynein, cytoplasmic 1, heavy chain 1 |
| **LRRC8B** | ENSG00000197147 | 956.2700144 | 0.497090523 | 0.169368047 | 2.29E-04 | 0.007884863 | 23507 | 1 | 89990395 | 90063423 | 1 | protein_coding | leucine rich repeat containing 8 family, member B |
| **SND1** | ENSG00000197157 | 4843.658135 | -0.284208449 | 0.093908894 | 5.2E-04 | 0.014671048 | 27044 | 7 | 127292234 | 127732661 | 1 | protein_coding | staphylococcal nuclease and tudor domain containing 1 |
| **SERPINA1** | ENSG00000197249 | 8985.027137 | -0.470471815 | 0.144761356 | 8.95E-05 | 0.00387675 | 5265 | 14 | 94843084 | 94857030 | -1 | protein_coding | serpin peptidase inhibitor, clade A (alpha-1 antiproteinase, antitrypsin), member 1 |
| **TAF13** | ENSG00000197780 | 863.6897817 | 0.735631144 | 0.222062506 | 5.17E-05 | 0.002510241 | 6884 | 1 | 109605108 | 109618624 | -1 | protein_coding | TAF13 RNA polymerase II, TATA box binding protein (TBP)-associated factor, 18kDa |
| **FAM49A** | ENSG00000197872 | 1681.450388 | 0.316502444 | 0.141299795 | 0.00278428 | 0.044228011 | 81553 | 2 | 16730727 | 16847599 | -1 | protein_coding | family with sequence similarity 49, member A |
| **MYO1C** | ENSG00000197879 | 3114.966009 | -0.385692134 | 0.140019353 | 5.65E-04 | 0.015384931 | 4641 | 17 | 1367392 | 1396106 | -1 | protein_coding | myosin IC |
| **ERO1L** | ENSG00000197930 | 9034.37118 | 0.573272872 | 0.118235517 | 9.49E-08 | 1.65E-05 | 30001 | 14 | 53106634 | 53162618 | -1 | protein_coding | ERO1-like (S. cerevisiae) |
| **S100A6** | ENSG00000197956 | 4831.866295 | -0.426195927 | 0.174356516 | 0.001178887 | 0.025258128 | 6277 | 1 | 153507075 | 153508720 | -1 | protein_coding | S100 calcium binding protein A6 |
| **RPL12** | ENSG00000197958 | 6132.249736 | -0.428141107 | 0.117743424 | 1.98E-05 | 0.001203427 | 6136 | 9 | 130209953 | 130213684 | -1 | protein_coding | ribosomal protein L12 |
| **VPS13A** | ENSG00000197969 | 669.2852619 | 0.611474496 | 0.273103326 | 0.001125812 | 0.024317464 | 23230 | 9 | 79792269 | 80036457 | 1 | protein_coding | vacuolar protein sorting 13 homolog A (S. cerevisiae) |
| **CD2AP** | ENSG00000198087 | 1926.025864 | 0.348141736 | 0.143155874 | 0.001554153 | 0.03036027 | 23607 | 6 | 47445525 | 47594999 | 1 | protein_coding | CD2-associated protein |
| **SOWAHC** | ENSG00000198142 | 358.0481475 | 0.551845924 | 0.256437225 | 0.001496002 | 0.029715908 | 65124 | 2 | 110371911 | 110376563 | 1 | protein_coding | sosondowah ankyrin repeat domain family member C |
| **ZNF770** | ENSG00000198146 | 1587.47047 | 0.833049932 | 0.158367958 | 8.71E-09 | 2.69E-06 | 54989 | 15 | 35270542 | 35280488 | -1 | protein_coding | zinc finger protein 770 |
| **STYX** | ENSG00000198252 | 1293.793236 | 0.79880327 | 0.160317885 | 3.34E-08 | 6.83E-06 | 6815 | 14 | 53196898 | 53241716 | 1 | protein_coding | serine/threonine/tyrosine interacting protein |
| **GFPT1** | ENSG00000198380 | 2389.457328 | 0.474144148 | 0.09425786 | 4.37E-08 | 8.59E-06 | 2673 | 2 | 69546905 | 69614382 | -1 | protein_coding | glutamine--fructose-6-phosphate transaminase 1 |
| **BZW1P2** | ENSG00000198406 | 1280.7393 | -0.94674223 | 0.156847017 | 6.85E-11 | 3.64E-08 | NA | 3 | 116364592 | 116365574 | -1 | pseudogene | basic leucine zipper and W2 domains 1 pseudogene 2 |
| **TLK1** | ENSG00000198586 | 2179.187685 | -0.369722548 | 0.120107715 | 2.19E-04 | 0.007606024 | 9874 | 2 | 171847333 | 172087824 | -1 | protein_coding | tousled-like kinase 1 |
| **MDM4** | ENSG00000198625 | 1347.523489 | 0.52122671 | 0.124243899 | 2.02E-06 | 2.03E-04 | 4194 | 1 | 204485511 | 204542871 | 1 | protein_coding | Mdm4 p53 binding protein homolog (mouse) |
| **PAPSS2** | ENSG00000198682 | 308.9997233 | 0.902353064 | 0.241932915 | 8.64E-06 | 6E-04 | 9060 | 10 | 89419370 | 89507462 | 1 | protein_coding | 3'-phosphoadenosine 5'-phosphosulfate synthase 2 |

| | | | | | | | | | | | | | | |
|---|---|---|---|---|---|---|---|---|---|---|---|---|---|---|
| **FAN1** | ENSG00000198690 | 516.9158074 | 0.610805716 | 0.179018284 | 3.78E-05 | 0.00200034 | 22909 | 15 | 31196055 | 31235311 | | 1 | protein_coding | FANCD2/FANCI-associated nuclease 1 |
| **SLC5A3** | ENSG00000198743 | 16486.15749 | 0.788845588 | 0.15448664 | 1.86E-08 | 4.71E-06 | 6526 | 21 | 35445870 | 35478559 | | 1 | protein_coding | solute carrier family 5 (sodium/myo-inositol cotransporter), member 3 |
| **ARHGAP11** | ENSG00000198826 | 348.2692517 | -0.540917457 | 0.244287479 | 0.001354964 | 0.028068628 | 9824 | 15 | 32907345 | 32932150 | | 1 | protein_coding | Rho GTPase activating protein 11A |
| **DENND4B** | ENSG00000198837 | 2451.002306 | 0.348509445 | 0.162671104 | 0.003225434 | 0.048277021 | 9909 | 1 | 153901977 | 153919172 | | -1 | protein_coding | DENN/MADD domain containing 4B |
| **LTN1** | ENSG00000198862 | 1153.513402 | 0.318623619 | 0.141993344 | 0.002901914 | 0.04527201 | 26046 | 21 | 30300466 | 30365270 | | -1 | protein_coding | listerin E3 ubiquitin protein ligase 1 |
| **SMC5** | ENSG00000198887 | 1039.315862 | 0.550313198 | 0.147179107 | 1.26E-05 | 8.14E-04 | 23137 | 9 | 72873937 | 72969804 | | 1 | protein_coding | structural maintenance of chromosomes 5 |
| **CAPZA2** | ENSG00000198898 | 5045.747907 | -0.578690326 | 0.104098882 | 1.93E-09 | 7.07E-07 | 830 | 7 | 116451124 | 116562103 | | 1 | protein_coding | capping protein (actin filament) muscle Z-line, alpha 2 |
| **PJA2** | ENSG00000198961 | 6820.367353 | 0.331036862 | 0.096919459 | 8.31E-05 | 0.003665738 | 9867 | 5 | 108670410 | 108745695 | | -1 | protein_coding | praja ring finger 2, E3 ubiquitin protein ligase |
| **AC083799.1** | ENSG00000203644 | 171.4922388 | 0.892464209 | 0.309267258 | 1.63E-04 | 0.005890844 | NA | 3 | 129565893 | 129566800 | | -1 | pseudogene | |
| **CHML** | ENSG00000203668 | 132.7221451 | 0.901730132 | 0.348575576 | 3.75E-04 | 0.011416609 | 1122 | 1 | 241792155 | 241799232 | | -1 | protein_coding | choroideremia-like (Rab escort protein 2) |
| **ZDBF2** | ENSG00000204186 | 458.0283549 | 0.534726677 | 0.205578428 | 5.61E-04 | 0.015320947 | 57683 | 2 | 207139387 | 207179148 | | 1 | protein_coding | zinc finger, DBF-type containing 2 |
| **BMPR2** | ENSG00000204217 | 3947.481139 | 0.350532448 | 0.106256115 | 1.17E-04 | 0.00466745 | 659 | 2 | 203241659 | 203432474 | | 1 | protein_coding | bone morphogenetic protein receptor, type II (serine/threonine kinase) |
| **SDHD** | ENSG00000204370 | 2211.940907 | 0.452160425 | 0.129193384 | 3.95E-05 | 0.002028168 | 6392 | 11 | 111957497 | 111990353 | | 1 | protein_coding | succinate dehydrogenase complex, subunit D, integral membrane protein |
| **MZT1** | ENSG00000204899 | 273.8469782 | 0.911994953 | 0.219458763 | 1.63E-06 | 1.75E-04 | 440145 | 13 | 73282495 | 73301825 | | -1 | protein_coding | mitotic spindle organizing protein 1 |
| **PSENEN** | ENSG00000205155 | 687.3459349 | -0.439964206 | 0.158771547 | 4.47E-04 | 0.012860214 | 55851 | 19 | 36236015 | 36237911 | | 1 | protein_coding | presenilin enhancer gamma secretase subunit |
| **TMEM170B** | ENSG00000205269 | 388.3192572 | 1.204001215 | 0.248628283 | 5.91E-08 | 1.08E-05 | 100113407 | 6 | 11537938 | 11583757 | | 1 | protein_coding | transmembrane protein 170B |
| **CNEP1R1** | ENSG00000205423 | 929.8529545 | 0.38946947 | 0.162877265 | 0.001418975 | 0.028777579 | 255919 | 16 | 50058321 | 50070999 | | 1 | protein_coding | CTD nuclear envelope phosphatase 1 regulatory subunit 1 |
| **SMN2** | ENSG00000205571 | 701.0292744 | -0.58681716 | 0.153502763 | 6.73E-06 | 5.05E-04 | 6607 | 5 | 69345350 | 69374349 | | 1 | protein_coding | survival of motor neuron 2, centromeric |
| **C5orf51** | ENSG00000205765 | 1727.520265 | 0.538501236 | 0.13211518 | 3.38E-06 | 3.05E-04 | 285636 | 5 | 41904290 | 41921738 | | 1 | protein_coding | chromosome 5 open reading frame 51 |
| **UBD** | ENSG00000206468 | 311.7057999 | -0.940539157 | 0.413057913 | 7.28E-04 | 0.018365575 | 10537 | HSC | 29523204 | 29527517 | | -1 | protein_coding | ubiquitin D |
| **GPX3** | ENSG00000211445 | 1936.31503 | -0.506025485 | 0.241789801 | 0.001884617 | 0.034425705 | 2878 | 5 | 150400124 | 150408554 | | 1 | protein_coding | glutathione peroxidase 3 (plasma) |
| **HSPA1B** | ENSG00000212866 | 113.8895069 | 1.216314348 | 0.580338512 | 9.6E-04 | 0.022029371 | 3304 | HSC | 31785766 | 31788290 | | 1 | protein_coding | heat shock 70kDa protein 1B |
| **SFT2D2** | ENSG00000213064 | 11675.45226 | 0.599178033 | 0.141793807 | 1.44E-06 | 1.56E-04 | 375035 | 1 | 168195176 | 168222263 | | 1 | protein_coding | SFT2 domain containing 2 |
| **CRIP1** | ENSG00000213145 | 490.1365474 | -0.507695582 | 0.221267166 | 0.001218936 | 0.025907262 | 1396 | 14 | 105952654 | 105955284 | | 1 | protein_coding | cysteine-rich protein 1 (intestinal) |
| **NRAS** | ENSG00000213281 | 3478.083984 | 0.449352374 | 0.138516383 | 1.02E-04 | 0.004270952 | 4893 | 1 | 115247090 | 115259515 | | -1 | protein_coding | neuroblastoma RAS viral (v-ras) oncogene homolog |
| **CHUK** | ENSG00000213341 | 1357.281894 | 0.32476175 | 0.150126174 | 0.003308974 | 0.049112631 | 1147 | 10 | 101948055 | 101989376 | | -1 | protein_coding | conserved helix-loop-helix ubiquitous kinase |
| **DDX3X** | ENSG00000215301 | 17513.09436 | -0.295558426 | 0.089814905 | 2.18E-04 | 0.007580235 | 1654 | X | 41192651 | 41223725 | | 1 | protein_coding | DEAD (Asp-Glu-Ala-Asp) box helicase 3, X-linked |
| **CYB5RL** | ENSG00000215883 | 130.4946636 | 1.016221181 | 0.421850721 | 5.58E-04 | 0.015320947 | 606495 | 1 | 54638009 | 54665709 | | -1 | protein_coding | cytochrome b5 reductase-like |
| **UBA52** | ENSG00000221983 | 3922.740489 | -0.382019825 | 0.133741545 | 4.21E-04 | 0.012370227 | 7311 | 19 | 18682540 | 18688360 | | 1 | protein_coding | ubiquitin A-52 residue ribosomal protein fusion product 1 |
| **CLIC1** | ENSG00000223639 | 7338.24077 | -0.313194549 | 0.103715351 | 3.53E-04 | 0.010833612 | 1192 | HSC | 31688317 | 31697500 | | -1 | protein_coding | chloride intracellular channel 1 |
| **CSNK2B** | ENSG00000224398 | 1027.845159 | -0.443259471 | 0.207298426 | 0.002078897 | 0.036637539 | 1460 | HSC | 31709296 | 31717606 | | 1 | protein_coding | casein kinase 2, beta polypeptide |
| **MIR3916** | ENSG00000227671 | 807.4953609 | 0.570080025 | 0.171507197 | 5.52E-05 | 0.002619909 | 100500849 | 1 | 247353153 | 247374158 | | -1 | pseudogene | microRNA 3916 |
| **HCG4P5** | ENSG00000227766 | 224.1922131 | -0.690268074 | 0.370044216 | 0.002130945 | 0.037059255 | NA | 6 | 29909852 | 29910844 | | -1 | pseudogene | HLA complex group 4 pseudogene 5 |
| **MDC 1.00** | ENSG00000228575 | 255.5903038 | -0.716078433 | 0.403818887 | 0.002348947 | 0.039729087 | 9656 | HSC | 30712282 | 30730363 | | -1 | protein_coding | mediator of DNA-damage checkpoint 1 |
| **HLA-DRB3** | ENSG00000230463 | 692.6863542 | 0.592893345 | 0.315455206 | 0.002396502 | 0.040043333 | 3125 | HSC | 32470408 | 32484153 | | -1 | protein_coding | HLA class II histocompatibility antigen, DR beta 3 chain |
| **HSBP1** | ENSG00000230989 | 1197.568636 | 0.589277359 | 0.173516384 | 4.32E-05 | 0.002197555 | 3281 | 16 | 83841448 | 83853342 | | 1 | protein_coding | heat shock factor binding protein 1 |
| **RP11-396K** | ENSG00000233369 | 1305.992882 | -0.74146808 | 0.270357344 | 2.59E-04 | 0.008480193 | NA | 7 | 72569033 | 72619657 | | 1 | pseudogene | |
| **RP11-3P17.** | ENSG00000234851 | 2311.9572 | -0.690487285 | 0.163055256 | 1.18E-06 | 1.32E-04 | NA | 3 | 161146915 | 161147398 | | -1 | pseudogene | |
| **APOC2** | ENSG00000234906 | 2672.300308 | -0.776944586 | 0.250316057 | 7.97E-05 | 0.003546214 | 344 | 19 | 45449243 | 45452822 | | 1 | protein_coding | apolipoprotein C-II |
| **FABP5P7** | ENSG00000234964 | 2010.650197 | -1.382250589 | 0.263492409 | 5.63E-09 | 1.84E-06 | NA | 11 | 59548786 | 59549195 | | -1 | pseudogene | fatty acid binding protein 5 pseudogene 7 |
| **FAM103A2P** | ENSG00000235272 | 97.49603867 | -1.347832446 | 0.397223658 | 2.77E-05 | 0.001559673 | NA | 6 | 166999612 | 166999965 | | -1 | pseudogene | family with sequence similarity 103, member A2 pseudogene |
| **H3F3AP4** | ENSG00000235655 | 275.7014191 | -0.020684323 | 0.144347285 | 2.83E-06 | 2.59E-04 | 440926 | 2 | 175584505 | 175585526 | | 1 | pseudogene | H3 histone, family 3A, pseudogene 4 |
| **HLA-A** | ENSG00000235657 | 2834.578868 | -0.762366916 | 0.200444526 | 6.75E-06 | 5.05E-04 | 3105 | HSC | 29899000 | 29903640 | | 1 | protein_coding | major histocompatibility complex, class I, A |
| **PSMB8** | ENSG00000235715 | 274.4878958 | 6.78E-04 | 0.142065005 | 0.001347842 | 0.027988901 | 5696 | HSC | 32730817 | 32734804 | | -1 | protein_coding | proteasome (prosome, macropain) subunit, beta type, 8 |
| **LST1** | ENSG00000235915 | 1112.46149 | -0.479685639 | 0.169668433 | 3.36E-04 | 0.010443926 | 7940 | HSC | 31565222 | 31568007 | | 1 | protein_coding | leukocyte specific transcript 1 |
| **ZBED5** | ENSG00000236287 | 771.3405599 | 0.68745053 | 0.159060551 | 9.58E-07 | 1.12E-04 | 101928029 | 11 | 10833621 | 10880343 | | -1 | protein_coding | zinc finger, BED-type containing 5 |
| **EIF1AXP1** | ENSG00000236698 | 118.3914519 | -0.771191514 | 0.494269786 | 0.003111007 | 0.047087076 | NA | 1 | 17012116 | 17012550 | | -1 | pseudogene | eukaryotic translation initiation factor 1A, X-linked pseudogene 1 |
| **UBE2Q2P6** | ENSG00000237550 | 2083.715028 | -0.386926243 | 0.172753084 | 0.001803963 | 0.033751257 | NA | 15 | 82664459 | 82748784 | | -1 | pseudogene | ubiquitin-conjugating enzyme E2Q family member 2 pseudogene 6 |
| **ADSL** | ENSG00000239900 | 543.4160289 | 0.460745663 | 0.237356551 | 0.002890562 | 0.045173538 | 158 | 22 | 40742507 | 40786467 | | 1 | protein_coding | adenylosuccinate lyase |
| **CD302** | ENSG00000241399 | 1790.168114 | 0.990284019 | 0.210085832 | 1.29E-07 | 2.15E-05 | 9936 | 2 | 160625364 | 160654753 | | -1 | protein_coding | CD302 molecule |
| **ARPC4** | ENSG00000241553 | 1183.63769 | -0.541777607 | 0.221674074 | 7.27E-04 | 0.018365575 | 10093 | 3 | 9834179 | 9849410 | | 1 | protein_coding | actin related protein 2/3 complex, subunit 4, 20kDa |
| **GNG10** | ENSG00000242616 | 1199.679578 | 0.838477709 | 0.17774231 | 1.28E-07 | 2.15E-05 | 2790 | 9 | 114423615 | 114432526 | | 1 | protein_coding | guanine nucleotide binding protein (G protein), gamma 10 |
| **TICAM2** | ENSG00000243414 | 337.6222336 | 0.872875739 | 0.23101987 | 7.52E-06 | 5.44E-04 | 100302736 | 5 | 114914339 | 114961876 | | -1 | protein_coding | toll-like receptor adaptor molecule 2 |
| **NME1-NME** | ENSG00000243678 | 2602.637949 | -0.423516257 | 0.134789091 | 1.41E-04 | 0.005312842 | 654364 | 17 | 49230920 | 49249108 | | 1 | protein_coding | NME1-NME2 readthrough |
| **NPIPB5** | ENSG00000243716 | 854.227313 | 0.662494705 | 0.142185374 | 2.08E-07 | 3.12E-05 | 101929906 | 16 | 22490442 | 22547842 | | 1 | protein_coding | nuclear pore complex interacting protein family, member B5 |
| **RP11-466H** | ENSG00000244398 | 768.8838688 | -0.938852304 | 0.22730326 | 1.74E-06 | 1.85E-04 | NA | 11 | 16996240 | 16996560 | | -1 | pseudogene | |
| **FCGR2C** | ENSG00000244682 | 1089.491951 | 0.48327346 | 0.191534607 | 7.77E-04 | 0.01911776 | 9103 | 1 | 161551129 | 161575452 | | 1 | polymorphic_ps | Fc fragment of IgG, low affinity IIc, receptor for (CD32) (gene/pseudogene) |
| **H2AFJ** | ENSG00000246705 | 399.454746 | -0.478130823 | 0.210463402 | 0.001380594 | 0.028378289 | 55766 | 12 | 14927270 | 14930936 | | 1 | protein_coding | H2A histone family, member J |
| **C15orf38-A** | ENSG00000250021 | 790.0357498 | -0.368202227 | 0.14656732 | 0.001120125 | 0.024317464 | 100526783 | 15 | 90377540 | 90456114 | | -1 | protein_coding | C15orf38-AP3S2 readthrough |
| **CCDC71L** | ENSG00000253276 | 745.3185406 | 0.50649531 | 0.156914842 | 8.86E-05 | 0.003856927 | 168455 | 7 | 106297211 | 106301442 | | -1 | protein_coding | coiled-coil domain containing 71-like |
| **ZNF260** | ENSG00000254004 | 686.5276214 | 0.60479191 | 0.192687947 | 9.37E-05 | 0.004013873 | 339324 | 19 | 37001597 | 37019562 | | -1 | protein_coding | zinc finger protein 260 |
| **FDXACB1** | ENSG00000255561 | 131.6806183 | 0.651917674 | 0.399159194 | 0.003261892 | 0.048617295 | 91893 | 11 | 111744780 | 111751967 | | -1 | protein_coding | ferredoxin-fold anticodon binding domain containing 1 |
| **RP11-286N** | ENSG00000256591 | 167.281383 | -0.6015881 | 0.296057592 | 0.001773506 | 0.033239947 | 54949 | 11 | 61196692 | 61253294 | | 1 | protein_coding | Uncharacterized protein |
| **RP11-611O** | ENSG00000256664 | 324.0592113 | 0.775777636 | 0.292844005 | 3.46E-04 | 0.010661881 | NA | 12 | 69235726 | 69236164 | | 1 | pseudogene | |
| **GALNT4** | ENSG00000257594 | 496.6432196 | 0.434401325 | 0.190620589 | 0.001560645 | 0.030431152 | 100528030 | 12 | 89913185 | 89920039 | | -1 | protein_coding | UDP-N-acetyl-alpha-D-galactosamine:polypeptide N-acetylgalactosaminyltransferase 4 (GalNAc-T4) |
| **RP11-463D** | ENSG00000258677 | 139.1131816 | -2.402683546 | 0.394912684 | 5.42E-11 | 3.39E-08 | NA | 8 | 74600926 | 74791089 | | -1 | protein_coding | Uncharacterized protein |
| **RP11-673C** | ENSG00000259781 | 439.1410069 | -0.48211206 | 0.225995159 | 0.001879531 | 0.034425705 | NA | 15 | 71457109 | 71457754 | | -1 | pseudogene | |
| **RBM15B** | ENSG00000259956 | 140.1178965 | 0.005949385 | 0.142252333 | 0.002349304 | 0.039729087 | 29890 | HG1 | 51428699 | 51435339 | | 1 | protein_coding | RNA binding motif protein 15B |
| **MRC1** | ENSG00000260314 | 2879.008452 | 0.880944144 | 0.244723077 | 1.68E-05 | 0.001050377 | 101928757 | HG5 | 17851343 | 17953161 | | 1 | protein_coding | mannose receptor, C type 1 |
| **RP11-345J** | ENSG00000261740 | 195.0887646 | 0.925004023 | 0.304950297 | 1.03E-04 | 0.004270952 | 552900 | 16 | 29454551 | 29465972 | | -1 | protein_coding | BolA-like protein 2 |
| **NOTCH2NL** | ENSG00000264343 | 469.074891 | -0.508201862 | 0.206265483 | 8.41E-04 | 0.020002769 | 388677 | HG1 | 146838681 | 146915449 | | -1 | protein_coding | |
| **CTD-2370N** | ENSG00000265118 | 439.2583174 | -1.178071795 | 0.225963181 | 9.11E-09 | 2.69E-06 | NA | 17 | 29632231 | 29645849 | | -1 | protein_coding | |
| **ITGB1P1** | ENSG00000269378 | 860.7352629 | -1.558813086 | 0.255398269 | 5.19E-11 | 3.39E-08 | NA | 19 | 14732446 | 14733054 | | -1 | pseudogene | integrin beta 1 pseudogene 1 |
| **RP11-544M** | ENSG00000270149 | 768.0521094 | -1.142072569 | 0.27991223 | 1.91E-06 | 1.97E-04 | NA | 1 | 160967747 | 161008752 | | -1 | protein_coding | Uncharacterized protein |
| **AC239811.2** | ENSG00000270629 | 130.6483674 | 0.926430279 | 0.425720226 | 9.48E-04 | 0.021856 | 400818 | HG1 | 149219394 | 149282305 | | -1 | protein_coding | Neuroblastoma breakpoint family member 1 |
