## Supplementary Table S3B for "TDP-43 pathology links innate and adaptive immunity in amyotrophic lateral sclerosis"

LPS_0.5_vs_PBS_0.5

| symbol | ensgene | baseMean | log2FoldChange | lfcSE | pvalue | padj | entrez | chr | start | end | strand | biotype | description |
|---|---|---|---|---|---|---|---|---|---|---|---|---|---|
| SCYL3 | ENSG00000000457 | 351.9630912 | 0.786318227 | 0.200481862 | 3.71E-05 | 1.83E-04 | 57147 | 1 | 169818772 | 169863408 | -1 | protein_coding | SCY1-like 3 (S. cerevisiae) |
| FGR | ENSG00000000938 | 10028.85037 | -1.610191868 | 0.133394881 | 2.23E-34 | 1.09E-32 | 2268 | 1 | 27938575 | 27961788 | -1 | protein_coding | feline Gardner-Rasheed sarcoma viral oncogene homolog |
| CFH | ENSG00000000971 | 106.3719213 | 2.451854845 | 0.503638733 | 8.44E-08 | 6.32E-07 | 3075 | 1 | 196621008 | 196716634 | 1 | protein_coding | complement factor H |
| FUCA2 | ENSG00000001036 | 3988.278669 | 0.283108624 | 0.099831784 | 0.004134292 | 0.01287113 | 2519 | 6 | 143815948 | 143832827 | -1 | protein_coding | fucosidase, alpha-L- 2, plasma |
| GCLC | ENSG00000001084 | 1727.431783 | -1.330645174 | 0.224049271 | 4.98E-10 | 4.94E-09 | 2729 | 6 | 53362139 | 53481768 | -1 | protein_coding | glutamate-cysteine ligase, catalytic subunit |
| LAS1L | ENSG00000001497 | 486.6895787 | -0.508204738 | 0.160102113 | 9.51E-04 | 0.003485817 | 81887 | X | 64732462 | 64754655 | -1 | protein_coding | LAS1-like (S. cerevisiae) |
| ENPP4 | ENSG00000001561 | 1288.757936 | 1.671302231 | 0.162324148 | 8.81E-26 | 2.74E-24 | 22875 | 6 | 46097730 | 46114436 | 1 | protein_coding | ectonucleotide pyrophosphatase/phosphodiesterase 4 (putative) |
| ANKIB1 | ENSG00000001629 | 2332.197009 | 0.86311567 | 0.131630914 | 1.78E-11 | 2.05E-10 | 54467 | 7 | 91875548 | 92030698 | 1 | protein_coding | ankyrin repeat and IBR domain containing 1 |
| LAP3 | ENSG00000002549 | 10916.82202 | 1.850382734 | 0.184139744 | 1.38E-24 | 4.05E-23 | 51056 | 4 | 17578815 | 17609595 | 1 | protein_coding | leucine aminopeptidase 3 |
| AOC1 | ENSG00000002726 | 288.7391536 | -0.666493798 | 0.302005647 | 0.012730053 | 0.034434229 | 26 | 7 | 150521715 | 150558592 | 1 | protein_coding | amine oxidase, copper containing 1 |
| MAD1L1 | ENSG00000002822 | 386.2254196 | -1.118368471 | 0.229495932 | 2.7E-07 | 1.89E-06 | 8379 | 7 | 1855429 | 2272878 | -1 | protein_coding | MAD1 mitotic arrest deficient-like 1 (yeast) |
| LASP1 | ENSG00000002834 | 15765.55395 | -0.548709338 | 0.098236283 | 1.47E-08 | 1.22E-07 | 3927 | 17 | 37026112 | 37078023 | 1 | protein_coding | LIM and SH3 protein 1 |
| SNX11 | ENSG00000002919 | 826.8870317 | 0.563805814 | 0.14030926 | 3.72E-05 | 1.84E-04 | 29916 | 17 | 46180719 | 46200436 | 1 | protein_coding | sorting nexin 11 |
| CASP10 | ENSG00000003400 | 1754.593569 | 1.665577227 | 0.140667562 | 2.91E-33 | 1.36E-31 | 843 | 2 | 202047604 | 202094129 | 1 | protein_coding | caspase 10, apoptosis-related cysteine peptidase |
| CFLAR | ENSG00000003402 | 13232.01868 | 2.28642079 | 0.107116232 | 3.04E-102 | 1.03E-99 | 8837 | 2 | 201980827 | 202041410 | 1 | protein_coding | CASP8 and FADD-like apoptosis regulator |
| SLC7A2 | ENSG00000003989 | 13.28654205 | 3.001186476 | 1.029915661 | 2.06E-04 | 8.77E-04 | 6542 | 8 | 17354597 | 17428082 | 1 | protein_coding | solute carrier family 7 (cationic amino acid transporter, y+ system), member 2 |
| ARF5 | ENSG00000004059 | 754.0091971 | -0.355578742 | 0.137713134 | 0.007521716 | 0.021821258 | 381 | 7 | 127228399 | 127231759 | 1 | protein_coding | ADP-ribosylation factor 5 |
| PLXND1 | ENSG00000004399 | 13318.33439 | -1.159741411 | 0.108579646 | 1.72E-27 | 5.75E-26 | 23129 | 3 | 129274018 | 129325661 | -1 | protein_coding | plexin D1 |
| AK2 | ENSG00000004455 | 1794.69556 | 0.674926498 | 0.116564797 | 3.78E-09 | 3.38E-08 | 204 | 1 | 33473585 | 33546597 | -1 | protein_coding | adenylate kinase 2 |
| CD38 | ENSG00000004468 | 2751.296869 | 5.401970513 | 0.642475716 | 1.95E-18 | 3.81E-17 | 952 | 4 | 15779898 | 15854853 | 1 | protein_coding | CD38 molecule |
| RBM6 | ENSG00000004534 | 2171.454056 | 0.340644685 | 0.100866143 | 6.86E-04 | 0.002598289 | 10180 | 3 | 49977440 | 50137478 | 1 | protein_coding | RNA binding motif protein 6 |
| CAMKK1 | ENSG00000004660 | 405.5063518 | -2.051288257 | 0.232431797 | 1.07E-19 | 2.3E-18 | 84254 | 17 | 3763609 | 3798185 | -1 | protein_coding | calcium/calmodulin-dependent protein kinase kinase 1, alpha |
| SLC22A16 | ENSG00000004809 | 56.74385932 | -1.092259407 | 0.542764907 | 0.008122965 | 0.023339588 | 85413 | 6 | 110745890 | 110797844 | -1 | protein_coding | solute carrier family 22 (organic cation/carnitine transporter), member 16 |
| SLC25A13 | ENSG00000004864 | 653.0116726 | 0.547416879 | 0.194244927 | 0.002985193 | 0.009661733 | 10165 | 7 | 95749532 | 95951459 | -1 | protein_coding | solute carrier family 25 (aspartate/glutamate carrier), member 13 |
| CDC27 | ENSG00000004897 | 2659.465855 | 0.321034003 | 0.118978931 | 0.006017964 | 0.017935363 | 996 | 17 | 45195069 | 45266788 | -1 | protein_coding | cell division cycle 27 |
| PRSS22 | ENSG00000005001 | 33.26941789 | -1.546593129 | 0.743446027 | 0.003668371 | 0.011603514 | 64063 | 16 | 2902728 | 2908171 | -1 | protein_coding | protease, serine, 22 |
| SKAP2 | ENSG00000005020 | 4474.075687 | -0.659464331 | 0.125338864 | 7.25E-08 | 5.49E-07 | 8935 | 7 | 26706681 | 27034858 | -1 | protein_coding | src kinase associated phosphoprotein 2 |
| SLC25A5 | ENSG00000005022 | 3218.805197 | -0.51637455 | 0.10271062 | 3.36E-07 | 2.32E-06 | 292 | X | 118602363 | 118605282 | 1 | protein_coding | solute carrier family 25 (mitochondrial carrier; adenine nucleotide translocator), member 5 |
| CCDC109B | ENSG00000005059 | 544.5826775 | 0.647625428 | 0.16858753 | 6.51E-05 | 3.08E-04 | 55013 | 4 | 110481361 | 110609874 | 1 | protein_coding | coiled-coil domain containing 109B |
| RPAP3 | ENSG00000005175 | 953.5644573 | 1.086544728 | 0.152442001 | 2.75E-13 | 3.65E-12 | 79657 | 12 | 48057070 | 48099844 | -1 | protein_coding | RNA polymerase II associated protein 3 |
| FAM214B | ENSG00000005238 | 2169.697272 | -0.297667391 | 0.105402363 | 0.004076693 | 0.012712527 | 80256 | 9 | 35104109 | 35116338 | -1 | protein_coding | family with sequence similarity 214, member B |
| PRKAR2B | ENSG00000005249 | 173.7356693 | 1.249782051 | 0.299827763 | 5.84E-06 | 3.34E-05 | 5577 | 7 | 106685094 | 106802256 | 1 | protein_coding | protein kinase, cAMP-dependent, regulatory, type II, beta |
| MSL3 | ENSG00000005302 | 1095.160698 | 0.937028742 | 0.127163057 | 5.55E-14 | 7.89E-13 | 10943 | X | 11776278 | 11793870 | 1 | protein_coding | male-specific lethal 3 homolog (Drosophila) |
| IBTK | ENSG00000005700 | 3392.121785 | 0.308993636 | 0.097420339 | 0.001334604 | 0.004709263 | 25998 | 6 | 82879700 | 82957471 | -1 | protein_coding | inhibitor of Bruton agammaglobulinemia tyrosine kinase |
| ZNF195 | ENSG00000005801 | 503.3052786 | 0.415752893 | 0.183981633 | 0.017650626 | 0.045697867 | 7748 | 11 | 3360491 | 3400448 | -1 | protein_coding | zinc finger protein 195 |
| FBXL3 | ENSG00000005812 | 1533.247621 | 0.818036601 | 0.15308093 | 3.71E-08 | 2.92E-07 | 26224 | 13 | 77566740 | 77601330 | -1 | protein_coding | F-box and leucine-rich repeat protein 3 |
| ITGAL | ENSG00000005844 | 6507.149542 | -0.605080396 | 0.113501022 | 5.89E-08 | 4.53E-07 | 3683 | 16 | 30483979 | 30534506 | 1 | protein_coding | integrin, alpha L (antigen CD11A (p180), lymphocyte function-associated antigen 1; alpha polypeptide) |
| PDK2 | ENSG00000005882 | 370.1258938 | -1.38199764 | 0.211712435 | 1.27E-11 | 1.48E-10 | 5164 | 17 | 48172101 | 48189516 | 1 | protein_coding | pyruvate dehydrogenase kinase, isozyme 2 |
| ITGA3 | ENSG00000005884 | 10200.77157 | -1.751303258 | 0.366408302 | 1.66E-07 | 1.2E-06 | 3675 | 17 | 48133332 | 48167845 | 1 | protein_coding | integrin, alpha 3 (antigen CD49C, alpha 3 subunit of VLA-3 receptor) |
| ZFX | ENSG00000005889 | 1859.958012 | 0.730594766 | 0.117587963 | 3.18E-10 | 3.21E-09 | 7543 | X | 24167290 | 24234372 | 1 | protein_coding | zinc finger protein, X-linked |
| LAMP2 | ENSG00000005893 | 10671.37543 | 0.475666399 | 0.092857802 | 3.37E-07 | 2.33E-06 | 3920 | X | 119561682 | 119603220 | -1 | protein_coding | lysosomal-associated membrane protein 2 |
| C19orf60 | ENSG00000006015 | 353.2671472 | -0.824236209 | 0.202776422 | 1.88E-05 | 9.85E-05 | 55049 | 19 | 18699495 | 18703146 | 1 | protein_coding | chromosome 19 open reading frame 60 |
| AP2B1 | ENSG00000006125 | 31.07463746 | 1.108343764 | 0.646491661 | 0.013682481 | 0.036711616 | 163 | 17 | 33905065 | 34053436 | 1 | protein_coding | adaptor-related protein complex 2, beta 1 subunit |
| TNFRSF12A | ENSG00000006327 | 356.3869195 | -0.668191462 | 0.220230644 | 0.001216322 | 0.004338169 | 51330 | 16 | 3068446 | 3072384 | 1 | protein_coding | tumor necrosis factor receptor superfamily, member 12A |
| RALA | ENSG00000006451 | 6514.417464 | 0.67007028 | 0.142848198 | 1.43E-06 | 8.98E-06 | 5898 | 7 | 39663082 | 39747723 | 1 | protein_coding | v-ral simian leukemia viral oncogene homolog A (ras related) |
| DBF4 | ENSG00000006634 | 342.1298967 | 0.910234109 | 0.281210509 | 3.76E-04 | 0.001519191 | 10926 | 7 | 87505531 | 87538856 | 1 | protein_coding | DBF4 homolog (S. cerevisiae) |
| IFRD1 | ENSG00000006652 | 1032.539441 | 1.147794388 | 0.135117051 | 4.7E-18 | 8.92E-17 | 3475 | 7 | 112063023 | 112121072 | 1 | protein_coding | interferon-related developmental regulator 1 |
| GTF2IRD1 | ENSG00000006704 | 362.3202221 | -0.552006407 | 0.216319335 | 0.006585127 | 0.01943242 | 9569 | 7 | 73868120 | 74016931 | 1 | protein_coding | GTF2I repeat domain containing 1 |
| PNPLA4 | ENSG00000006757 | 260.7072573 | -0.674213115 | 0.253312351 | 0.003756354 | 0.011839215 | 8228 | X | 7866288 | 7895780 | -1 | protein_coding | patatin-like phospholipase domain containing 4 |
| ADIPOR2 | ENSG00000006831 | 2296.197458 | 0.29123794 | 0.100088042 | 0.003555784 | 0.0112862 | 79602 | 12 | 1797740 | 1897844 | 1 | protein_coding | adiponectin receptor 2 |
| CCDC124 | ENSG00000007080 | 394.6524753 | -0.705277198 | 0.203045307 | 2.43E-04 | 0.001018948 | 115098 | 19 | 18043825 | 18054800 | 1 | protein_coding | coiled-coil domain containing 124 |
| PAFAH1B1 | ENSG00000007168 | 5006.803323 | 0.611292625 | 0.092361204 | 9.48E-12 | 1.11E-10 | 5048 | 17 | 2496504 | 2588909 | 1 | protein_coding | platelet-activating factor acetylhydrolase 1b, regulatory subunit 1 (45kDa) |
| KIAA0100 | ENSG00000007202 | 12189.44522 | -0.605332388 | 0.097750419 | 3.02E-10 | 3.06E-09 | 9703 | 17 | 26941458 | 26972472 | -1 | protein_coding | KIAA0100 |
| GAS7 | ENSG00000007237 | 5034.92769 | -0.855180209 | 0.107522696 | 6.47E-16 | 1.05E-14 | 8522 | 17 | 9813926 | 10101868 | -1 | protein_coding | growth arrest-specific 7 |
| TRAPPC6A | ENSG00000007255 | 139.1767074 | -0.955804801 | 0.282840241 | 2.16E-04 | 9.14E-04 | 79090 | 19 | 45666186 | 45681495 | -1 | protein_coding | trafficking protein particle complex 6A |
| MATK | ENSG00000007264 | 2549.313185 | -2.14163715 | 0.189634561 | 1.08E-30 | 4.34E-29 | 4145 | 19 | 3777971 | 3802127 | -1 | protein_coding | megakaryocyte-associated tyrosine kinase |
| CD79B | ENSG00000007312 | 95.10374024 | -0.915710466 | 0.452248371 | 0.011132149 | 0.030693241 | 974 | 17 | 62006100 | 62009714 | -1 | protein_coding | CD79b molecule, immunoglobulin-associated beta |
| ST7L | ENSG00000007341 | 369.2899773 | 0.459096206 | 0.197259749 | 0.013836364 | 0.037072433 | 54879 | 1 | 113066140 | 113163447 | -1 | protein_coding | suppression of tumorigenicity 7 like |
| RHBDF1 | ENSG00000007384 | 252.4338142 | -0.949361888 | 0.281494303 | 2.23E-04 | 9.45E-04 | 64285 | 16 | 108058 | 126354 | -1 | protein_coding | rhomboid 5 homolog 1 (Drosophila) |
| PIGQ | ENSG00000007541 | 343.4722746 | -0.712372433 | 0.239380248 | 0.001139217 | 0.004103383 | 9091 | 16 | 616995 | 634136 | 1 | protein_coding | phosphatidylinositol glycan anchor biosynthesis, class Q |
| DNAJC11 | ENSG00000007923 | 855.3959909 | -0.630754766 | 0.150220047 | 1.49E-05 | 8E-05 | 55735 | 1 | 6694228 | 6761984 | -1 | protein_coding | DnaJ (Hsp40) homolog, subfamily C, member 11 |
| E2F2 | ENSG00000007968 | 45.73301822 | 0.973167226 | 0.48143833 | 0.010064306 | 0.028152276 | 1870 | 1 | 23832922 | 23857712 | -1 | protein_coding | E2F transcription factor 2 |
| PSMB1 | ENSG00000008018 | 1649.205272 | 0.37673164 | 0.106633218 | 3.41E-04 | 0.001390654 | 5689 | 6 | 170844205 | 170862429 | -1 | protein_coding | proteasome (prosome, macropain) subunit, beta type, 1 |
| JARID2 | ENSG00000008083 | 2607.810258 | -0.80101082 | 0.09853969 | 1.46E-16 | 2.48E-15 | 3720 | 6 | 15246527 | 15522252 | 1 | protein_coding | jumonji, AT rich interactive domain 2 |
| CDK11A | ENSG00000008128 | 763.8014116 | 0.544849458 | 0.134128697 | 3.16E-05 | 1.59E-04 | 728642 | 1 | 1634169 | 1655766 | -1 | protein_coding | cyclin-dependent kinase 11A |
| NADK | ENSG00000008130 | 5169.7802 | 1.088123973 | 0.093913531 | 6.3E-32 | 2.7E-30 | 65220 | 1 | 1682671 | 1711896 | -1 | protein_coding | NAD kinase |
| CYTH3 | ENSG00000008256 | 443.9280473 | -1.33474862 | 0.347970707 | 2.05E-05 | 1.07E-04 | 9265 | 7 | 6201407 | 6312275 | -1 | protein_coding | cytohesin 3 |
| SYPL1 | ENSG00000008282 | 1207.808375 | 0.580052103 | 0.173302307 | 4.9E-04 | 0.001920819 | 6856 | 7 | 105730949 | 105753022 | -1 | protein_coding | synaptophysin-like 1 |
| SS18L2 | ENSG00000008324 | 296.7440404 | 0.646777265 | 0.223874306 | 0.001971477 | 0.006680729 | 51188 | 3 | 42623332 | 42636606 | 1 | protein_coding | synovial sarcoma translocation gene on chromosome 18-like 2 |
| MPND | ENSG00000008382 | 442.2568048 | -0.882945319 | 0.224597995 | 2.9E-05 | 1.46E-04 | 84954 | 19 | 4343524 | 4360083 | 1 | protein_coding | MPN domain containing |
| MGST1 | ENSG00000008394 | 4464.429844 | 0.701414965 | 0.176164208 | 3.3E-05 | 1.65E-04 | 4257 | 12 | 16500076 | 16762193 | 1 | protein_coding | microsomal glutathione S-transferase 1 |
| ST3GAL1 | ENSG00000008513 | 7429.023807 | 0.800662655 | 0.105397948 | 1.42E-14 | 2.08E-13 | 6482 | 8 | 134467091 | 134584183 | -1 | protein_coding | ST3 beta-galactoside alpha-2,3-sialyltransferase 1 |
| IL32 | ENSG00000008517 | 1619.713559 | 0.708870506 | 0.242676801 | 0.001576161 | 0.005474651 | 9235 | 16 | 3115298 | 3131908 | 1 | protein_coding | interleukin 32 |
| PKD1 | ENSG00000008710 | 614.3495817 | -0.673255636 | 0.238664579 | 0.00206956 | 0.006968293 | 1019300 | 16 | 2138711 | 2185899 | -1 | protein_coding | polycystic kidney disease 1 (autosomal dominant) |
| RHOBTB2 | ENSG00000008853 | 1847.51946 | -1.541326724 | 0.186420171 | 2.03E-17 | 3.67E-16 | 23221 | 8 | 22844930 | 22877712 | 1 | protein_coding | Rho-related BTB domain containing 2 |
| HEATR5B | ENSG00000008869 | 1341.579859 | -0.472322118 | 0.124683132 | 1.13E-04 | 5.13E-04 | 54497 | 2 | 37195526 | 37311485 | -1 | protein_coding | HEAT repeat containing 5B |
| SEC62 | ENSG00000008952 | 4604.37967 | 0.305815109 | 0.121170279 | 0.008697995 | 0.024763685 | 7095 | 3 | 169684423 | 169716161 | 1 | protein_coding | SEC62 homolog (S. cerevisiae) |
| TENM1 | ENSG00000009694 | 39.59905396 | -1.279979889 | 0.591603312 | 0.004280407 | 0.013296232 | 10178 | X | 123509753 | 124097666 | -1 | protein_coding | teneurin transmembrane protein 1 |
| VTA1 | ENSG00000009844 | 860.8283046 | 0.65168925 | 0.140226175 | 1.78E-06 | 1.1E-05 | 51534 | 6 | 142468367 | 142545826 | 1 | protein_coding | vesicle (multivesicular body) trafficking 1 |
| MLXIPL | ENSG00000009950 | 159.1267219 | -2.071789367 | 0.362375337 | 9.13E-10 | 8.82E-09 | 51085 | 7 | 73007524 | 73038873 | -1 | protein_coding | MLX interacting protein-like |
| ETV7 | ENSG00000010030 | 235.5913882 | 4.367961117 | 0.420624277 | 1.56E-26 | 5.06E-25 | 51513 | 6 | 36322419 | 36356164 | -1 | protein_coding | ets variant 7 |
| ZNF207 | ENSG00000010244 | 4348.563546 | 0.663257716 | 0.098496028 | 6.82E-12 | 8.16E-11 | 7756 | 17 | 30677136 | 30708905 | 1 | protein_coding | zinc finger protein 207 |
| UQCRC1 | ENSG00000010256 | 1898.884353 | -0.864895564 | 0.113878884 | 1.71E-14 | 2.5E-13 | 7384 | 3 | 48636435 | 48648409 | -1 | protein_coding | ubiquinol-cytochrome c reductase core protein I |
| CD9 | ENSG00000010278 | 7554.465522 | -2.438550514 | 0.345363727 | 1.3E-13 | 1.78E-12 | 928 | 12 | 6308881 | 6347425 | 1 | protein_coding | CD9 molecule |
| IFFO1 | ENSG00000010295 | 784.8183192 | -0.482008426 | 0.183057036 | 0.005857784 | 0.01750235 | 25900 | 12 | 6647541 | 6665239 | -1 | protein_coding | intermediate filament family orphan 1 |
| PHF7 | ENSG00000010318 | 103.0263087 | -0.717845982 | 0.345973139 | 0.015485922 | 0.040890004 | 51533 | 3 | 52444673 | 52457657 | 1 | protein_coding | PHD finger protein 7 |
| STAB1 | ENSG00000010327 | 304.1996687 | -4.060452387 | 0.424900304 | 5.11E-23 | 1.38E-21 | 23166 | 3 | 52529354 | 52558511 | 1 | protein_coding | stabilin 1 |

| | | | | | | | | | | | | | |
|---|---|---|---|---|---|---|---|---|---|---|---|---|---|
| **ZNF200** | ENSG00000010539 | 400.5850904 | 0.836644197 | 0.174492873 | 6.21E-07 | 4.14E-06 | 7752 | 16 | 3272325 | 3286221 | -1 | protein_coding | zinc finger protein 200 |
| **CD4** | ENSG00000010610 | 12356.1893 | -0.80743155 | 0.117008251 | 1.41E-12 | 1.77E-11 | 920 | 12 | 6896024 | 6929974 | 1 | protein_coding | CD4 molecule |
| **LRRC23** | ENSG00000010626 | 146.9847421 | -0.685844993 | 0.291319263 | 0.008502019 | 0.024259962 | 10233 | 12 | 6982733 | 7023407 | 1 | protein_coding | leucine rich repeat containing 23 |
| **BTK** | ENSG00000010671 | 2746.445781 | 0.791085733 | 0.099722965 | 9.54E-16 | 1.53E-14 | 695 | X | 100604435 | 100641183 | -1 | protein_coding | Bruton agammaglobulinemia tyrosine kinase |
| **HIVEP2** | ENSG00000010818 | 2215.538007 | 0.880210359 | 0.40881774 | 0.008887915 | 0.025247956 | 3097 | 6 | 143072604 | 143266338 | -1 | protein_coding | human immunodeficiency virus type I enhancer binding protein 2 |
| **TCEB3** | ENSG00000011007 | 2572.57827 | 0.462405096 | 0.089333963 | 1.7E-07 | 1.23E-06 | 6924 | 1 | 24069645 | 24088549 | 1 | protein_coding | transcription elongation factor B (SIII), polypeptide 3 (110kDa, elongin A) |
| **CLCN6** | ENSG00000011021 | 749.7964945 | 0.416936278 | 0.150909457 | 0.004350516 | 0.013483864 | 1185 | 1 | 11866207 | 11903201 | 1 | protein_coding | chloride channel, voltage-sensitive 6 |
| **MRC2** | ENSG00000011028 | 4252.548843 | -1.808045047 | 0.174015902 | 2.7E-26 | 8.64E-25 | 9902 | 17 | 60704762 | 60770958 | 1 | protein_coding | mannose receptor, C type 2 |
| **SLC6A7** | ENSG00000011083 | 252.6779403 | -0.971721087 | 0.445407441 | 0.006926381 | 0.020317828 | 6534 | 5 | 149569520 | 149602351 | 1 | protein_coding | solute carrier family 6 (neurotransmitter transporter), member 7 |
| **ABHD5** | ENSG00000011198 | 440.359747 | 0.975108409 | 0.229949551 | 6.55E-06 | 3.71E-05 | 51099 | 3 | 43731605 | 43775863 | 1 | protein_coding | abhydrolase domain containing 5 |
| **KAL1** | ENSG00000011201 | 1646.910605 | -0.652304426 | 0.231255791 | 0.002407354 | 0.007954939 | 3730 | X | 8496915 | 8700227 | -1 | protein_coding | Kallmann syndrome 1 sequence |
| **AKAP8L** | ENSG00000011243 | 849.8428806 | 0.45543351 | 0.145089952 | 0.00120299 | 0.004303675 | 26993 | 19 | 15490859 | 15529952 | -1 | protein_coding | A kinase (PRKA) anchor protein 8-like |
| **MBTD1** | ENSG00000011258 | 538.9851526 | -0.490557488 | 0.173093015 | 0.003113673 | 0.010037177 | 54799 | 17 | 49254786 | 49337524 | -1 | protein_coding | mbt domain containing 1 |
| **UTP18** | ENSG00000011260 | 1476.64216 | 0.25828529 | 0.103384788 | 0.011122989 | 0.030673518 | 51096 | 17 | 49337889 | 49375297 | 1 | protein_coding | UTP18 small subunit (SSU) processome component homolog (yeast) |
| **PTBP1** | ENSG00000011304 | 3902.142506 | -0.272896302 | 0.118170799 | 0.018406194 | 0.047309429 | 1006164 | 19 | 797075 | 812327 | 1 | protein_coding | polypyrimidine tract binding protein 1 |
| **PLAUR** | ENSG00000011422 | 8850.152981 | 1.367410255 | 0.102883789 | 4.85E-41 | 3.1E-39 | 5329 | 19 | 44150247 | 44174699 | -1 | protein_coding | plasminogen activator, urokinase receptor |
| **RABGAP1** | ENSG00000011454 | 2005.979828 | 0.334028809 | 0.107876445 | 0.001716485 | 0.005896391 | 23637 | 9 | 125703112 | 125867145 | 1 | protein_coding | RAB GTPase activating protein 1 |
| **ZBTB32** | ENSG00000011590 | 34.31471404 | 1.385837787 | 0.741825866 | 0.006427287 | 0.019040168 | 27033 | 19 | 36195429 | 36207940 | 1 | protein_coding | zinc finger and BTB domain containing 32 |
| **TYROBP** | ENSG00000011600 | 9924.722632 | -0.329183806 | 0.111205385 | 0.002594507 | 0.00849804 | 7305 | 19 | 36395303 | 36399197 | -1 | protein_coding | TYRO protein tyrosine kinase binding protein |
| **TMEM159** | ENSG00000011638 | 390.0266871 | 0.816634611 | 0.205464771 | 2.71E-05 | 1.38E-04 | 57146 | 16 | 21169698 | 21191937 | 1 | protein_coding | transmembrane protein 159 |
| **BRCA1** | ENSG00000012048 | 329.0707198 | 0.554912218 | 0.218756327 | 0.00679561 | 0.019980244 | 672 | 17 | 41196312 | 41277500 | -1 | protein_coding | breast cancer 1, early onset |
| **ERCC1** | ENSG00000012061 | 734.2398888 | -1.422716585 | 0.158178871 | 3.63E-20 | 8.13E-19 | 2067 | 19 | 45910591 | 45982086 | -1 | protein_coding | excision repair cross-complementing rodent repair deficiency, complementation group 1 (includes overla |
| **CD22** | ENSG00000012124 | 588.5954067 | -1.086765869 | 0.276412235 | 1.95E-05 | 1.02E-04 | 933 | 19 | 35810164 | 35838258 | 1 | protein_coding | CD22 molecule |
| **PRICKLE3** | ENSG00000012211 | 212.922616 | 0.529984764 | 0.230102616 | 0.013121531 | 0.035367964 | 4007 | X | 49031151 | 49042845 | -1 | protein_coding | prickle homolog 3 (Drosophila) |
| **ALOX5** | ENSG00000012779 | 4413.822509 | -2.913052517 | 0.22779498 | 1.03E-38 | 5.97E-37 | 240 | 10 | 45869661 | 45941561 | 1 | protein_coding | arachidonate 5-lipoxygenase |
| **MAP4K5** | ENSG00000012983 | 1378.173814 | 0.589534879 | 0.148881167 | 4.38E-05 | 2.13E-04 | 11183 | 14 | 50885219 | 51027844 | -1 | protein_coding | mitogen-activated protein kinase kinase kinase kinase 5 |
| **EHD3** | ENSG00000013016 | 437.3940044 | -1.36402689 | 0.264920033 | 4.37E-08 | 3.42E-07 | 30845 | 2 | 31456880 | 31492313 | 1 | protein_coding | EH-domain containing 3 |
| **MAN2B2** | ENSG00000013288 | 3853.410173 | -0.437992471 | 0.104446203 | 2.1E-05 | 1.09E-04 | 23324 | 4 | 6576902 | 6625089 | 1 | protein_coding | mannosidase, alpha, class 2B, member 2 |
| **SLC25A39** | ENSG00000013306 | 948.3933341 | -0.453186057 | 0.135984168 | 5.91E-04 | 0.002267926 | 51629 | 17 | 42396993 | 42402238 | -1 | protein_coding | solute carrier family 25, member 39 |
| **NUB1** | ENSG00000013374 | 1919.088021 | 2.569022219 | 0.120586758 | 6.75E-102 | 2.24E-99 | 51667 | 7 | 151038785 | 151075535 | 1 | protein_coding | negative regulator of ubiquitin-like proteins 1 |
| **RWDD2A** | ENSG00000013392 | 296.6980102 | 0.592086912 | 0.223203647 | 0.004433341 | 0.013715537 | 112611 | 6 | 83903098 | 83908651 | 1 | protein_coding | RWD domain containing 2A |
| **CLK1** | ENSG00000013441 | 1756.888686 | 0.797886675 | 0.119914297 | 1.63E-11 | 1.89E-10 | 1195 | 2 | 201717732 | 201729422 | -1 | protein_coding | CDC-like kinase 1 |
| **POLA2** | ENSG00000014138 | 292.274881 | 1.391609599 | 0.306832111 | 7.8E-07 | 5.12E-06 | 23649 | 11 | 65029233 | 65073060 | 1 | protein_coding | polymerase (DNA directed), alpha 2, accessory subunit |
| **ZC3H3** | ENSG00000014164 | 337.5732444 | 0.52373732 | 0.217177432 | 0.00974939 | 0.027419936 | 23144 | 8 | 144519825 | 144623623 | -1 | protein_coding | zinc finger CCCH-type containing 3 |
| **MDH1** | ENSG00000014641 | 1563.278655 | -0.521341285 | 0.118933276 | 8.06E-06 | 4.5E-05 | 4190 | 2 | 63815743 | 63834331 | 1 | protein_coding | malate dehydrogenase 1, NAD (soluble) |
| **MTMR11** | ENSG00000014914 | 78.94317624 | 0.991974404 | 0.46299283 | 0.007492164 | 0.02176031 | 10903 | 1 | 149900543 | 149908791 | -1 | protein_coding | myotubularin related protein 11 |
| **MATR3** | ENSG00000015479 | 5004.960321 | 0.373121298 | 0.099384627 | 1.42E-04 | 6.32E-04 | 724102 | 5 | 138609441 | 138667360 | 1 | protein_coding | matrin 3 |
| **ATP2C1** | ENSG00000017260 | 2766.420926 | 0.460433639 | 0.117147927 | 6.34E-05 | 3.01E-04 | 27032 | 3 | 130569439 | 130735556 | 1 | protein_coding | ATPase, Ca++ transporting, type 2C, member 1 |
| **SLC38A5** | ENSG00000017483 | 28.62678415 | 5.885878126 | 1.359134275 | 1.19E-06 | 7.56E-06 | 92745 | X | 48316920 | 48328644 | -1 | protein_coding | solute carrier family 38, member 5 |
| **RUFY3** | ENSG00000018189 | 1490.492251 | 0.733666171 | 0.119257797 | 3.72E-10 | 3.74E-09 | 22902 | 4 | 71569921 | 71673032 | 1 | protein_coding | RUN and FYVE domain containing 3 |
| **SLC11A1** | ENSG00000018280 | 6660.270796 | -1.066449121 | 0.353429418 | 5.79E-04 | 0.0022269 | 6556 | 2 | 219246752 | 219261617 | 1 | protein_coding | solute carrier family 11 (proton-coupled divalent metal ion transporter), member 1 |
| **AGPS** | ENSG00000018510 | 4084.103504 | -0.400326779 | 0.16398414 | 0.009757654 | 0.02743308 | 8540 | 2 | 178257372 | 178408564 | 1 | protein_coding | alkylglycerone phosphate synthase |
| **MARCO** | ENSG00000019169 | 1886.480327 | -2.40679309 | 0.228144341 | 3.22E-27 | 1.07E-25 | 8685 | 2 | 119699742 | 119752236 | 1 | protein_coding | macrophage receptor with collagenous structure |
| **CD74** | ENSG00000019582 | 138982.2314 | -0.481148648 | 0.153807448 | 0.001209363 | 0.004323438 | 972 | 5 | 149781200 | 149792492 | -1 | protein_coding | CD74 molecule, major histocompatibility complex, class II invariant chain |
| **HGF** | ENSG00000019991 | 60.08770799 | -2.406728209 | 0.674813785 | 2.61E-05 | 1.33E-04 | 3082 | 7 | 81328322 | 81399754 | -1 | protein_coding | hepatocyte growth factor (hepapoietin A; scatter factor) |
| **RUNX3** | ENSG00000020633 | 1101.62708 | 0.401627499 | 0.177244856 | 0.018180521 | 0.046815991 | 864 | 1 | 25226002 | 25291612 | -1 | protein_coding | runt-related transcription factor 3 |
| **MRE11A** | ENSG00000020922 | 753.4615393 | 0.510860451 | 0.151401595 | 5.14E-04 | 0.002005576 | 4361 | 11 | 94152895 | 94227074 | -1 | protein_coding | MRE11 meiotic recombination 11 homolog A (S. cerevisiae) |
| **SERPINB1** | ENSG00000021355 | 2361.541231 | 1.075094687 | 0.144607443 | 2.69E-14 | 3.89E-13 | 1992 | 6 | 2832566 | 2842240 | -1 | protein_coding | serpin peptidase inhibitor, clade B (ovalbumin), member 1 |
| **SPAST** | ENSG00000021574 | 839.2456073 | 0.422199574 | 0.178250431 | 0.013466071 | 0.036213614 | 6683 | 2 | 32288680 | 32382706 | 1 | protein_coding | spastin |
| **RTFDC1** | ENSG00000022277 | 2248.922329 | -0.332152825 | 0.098719295 | 5.62E-04 | 0.002168705 | 51507 | 20 | 55043647 | 55093943 | 1 | protein_coding | replication termination factor 2 domain containing 1 |
| **ZDHHC6** | ENSG00000023041 | 1461.621508 | 0.563771499 | 0.137212549 | 2.49E-05 | 1.27E-04 | 64429 | 10 | 114190058 | 114206672 | -1 | protein_coding | zinc finger, DHHC-type containing 6 |
| **GRAMD1B** | ENSG00000023171 | 600.2442344 | 0.888321616 | 0.242757963 | 9.06E-05 | 4.17E-04 | 57476 | 11 | 123396344 | 123498482 | 1 | protein_coding | GRAM domain containing 1B |
| **RNH1** | ENSG00000023191 | 2905.598274 | -0.663591311 | 0.126842649 | 7.6E-08 | 5.73E-07 | 6050 | 11 | 494512 | 507300 | -1 | protein_coding | ribonuclease/angiogenin inhibitor 1 |
| **NDUFS1** | ENSG00000023228 | 3724.161799 | -0.406279945 | 0.133950176 | 0.001911736 | 0.006502794 | 4719 | 2 | 206979541 | 207024327 | -1 | protein_coding | NADH dehydrogenase (ubiquinone) Fe-S protein 1, 75kDa (NADH-coenzyme Q reductase) |
| **ERP44** | ENSG00000023318 | 3412.712406 | 0.998436414 | 0.094389759 | 1.08E-26 | 3.51E-25 | 23071 | 9 | 102741461 | 102861322 | -1 | protein_coding | endoplasmic reticulum protein 44 |
| **ALAS1** | ENSG00000023330 | 4165.926809 | 0.430399575 | 0.10594205 | 3.73E-05 | 1.85E-04 | 211 | 3 | 52232102 | 52248343 | 1 | protein_coding | aminolevulinate, delta-, synthase 1 |
| **BIRC3** | ENSG00000023445 | 3108.282554 | 1.8257598 | 0.221759877 | 1.73E-17 | 3.15E-16 | 330 | 11 | 102188215 | 102210134 | 1 | protein_coding | baculoviral IAP repeat containing 3 |
| **GLRX2** | ENSG00000023572 | 441.3966514 | 0.522398603 | 0.193479956 | 0.004431665 | 0.013713126 | 51022 | 1 | 193065598 | 193075244 | -1 | protein_coding | glutaredoxin 2 |
| **DEF6** | ENSG00000023892 | 342.5493377 | -0.748854248 | 0.220013347 | 2.9E-04 | 0.001196649 | 50619 | 6 | 35265595 | 35289548 | 1 | protein_coding | differentially expressed in FDCP 6 homolog (mouse) |
| **GCLM** | ENSG00000023909 | 558.7404584 | 1.875523146 | 0.1862631 | 7.78E-25 | 2.32E-23 | 2730 | 1 | 94350761 | 94374966 | -1 | protein_coding | glutamate-cysteine ligase, modifier subunit |
| **UBR2** | ENSG00000024048 | 2723.235284 | 0.668243436 | 0.091044829 | 1.27E-13 | 1.74E-12 | 23304 | 6 | 42531800 | 42661242 | 1 | protein_coding | ubiquitin protein ligase E3 component n-recognin 2 |
| **RRAGD** | ENSG00000025039 | 3681.985478 | -0.73944161 | 0.143199184 | 1.09E-07 | 8.02E-07 | 58528 | 6 | 90074355 | 90121989 | -1 | protein_coding | Ras-related GTP binding D |
| **NR1H3** | ENSG00000025434 | 6816.41636 | 0.828021359 | 0.181017932 | 1.77E-06 | 1.1E-05 | 10062 | 11 | 47269851 | 47290396 | 1 | protein_coding | nuclear receptor subfamily 1, group H, member 3 |
| **TYMP** | ENSG00000025708 | 3946.925368 | 0.987676681 | 0.356284414 | 0.001468737 | 0.005140074 | 1890 | 22 | 50964181 | 50968485 | -1 | protein_coding | thymidine phosphorylase |
| **SEC63** | ENSG00000025796 | 2994.268665 | 0.518323396 | 0.206639078 | 0.007632503 | 0.022113279 | 11231 | 6 | 108188960 | 108279393 | -1 | protein_coding | SEC63 homolog (S. cerevisiae) |
| **FAS** | ENSG00000026103 | 1211.370596 | 1.794855805 | 0.128253537 | 2.4E-45 | 1.88E-43 | 355 | 10 | 90750414 | 90775542 | 1 | protein_coding | Fas cell surface death receptor |
| **RNASET2** | ENSG00000026297 | 4794.267367 | -0.849028584 | 0.117894491 | 2.28E-13 | 3.05E-12 | 8635 | 6 | 167342992 | 167370679 | -1 | protein_coding | ribonuclease T2 |
| **CD44** | ENSG00000026508 | 51401.9174 | 0.677564935 | 0.114311235 | 1.23E-09 | 1.17E-08 | 960 | 11 | 35160417 | 35253949 | 1 | protein_coding | CD44 molecule (Indian blood group) |
| **KCNG1** | ENSG00000026559 | 92.86671846 | -0.943112858 | 0.44417894 | 0.008198426 | 0.023521033 | 3755 | 20 | 49620193 | 49639666 | -1 | protein_coding | potassium voltage-gated channel, subfamily G, member 1 |
| **AGPAT4** | ENSG00000026652 | 525.374006 | -0.522709161 | 0.188385682 | 0.003557451 | 0.011289144 | 56895 | 6 | 161551011 | 161695093 | -1 | protein_coding | 1-acylglycerol-3-phosphate O-acyltransferase 4 |
| **SLAMF7** | ENSG00000026751 | 52192.06432 | 3.598306079 | 0.161411305 | 1.67E-111 | 6.39E-109 | 57823 | 1 | 160709037 | 160724611 | 1 | protein_coding | SLAM family member 7 |
| **BTN3A1** | ENSG00000026950 | 2806.720739 | 0.836208015 | 0.143098049 | 1.92E-09 | 1.78E-08 | 11119 | 6 | 26402465 | 26415444 | 1 | protein_coding | butyrophilin, subfamily 3, member A1 |
| **MIPEP** | ENSG00000027001 | 487.0000795 | -0.577623933 | 0.173902888 | 5.35E-04 | 0.002077113 | 4285 | 13 | 24304328 | 24463558 | -1 | protein_coding | mitochondrial intermediate peptidase |
| **IFNGR1** | ENSG00000027697 | 4429.995851 | -0.371397101 | 0.087990115 | 2.02E-05 | 1.05E-04 | 3459 | 6 | 137518621 | 137540586 | -1 | protein_coding | interferon gamma receptor 1 |
| **B4GALT7** | ENSG00000027847 | 302.486896 | -0.815966119 | 0.211649585 | 4.58E-05 | 2.22E-04 | 11285 | 5 | 177027101 | 177037348 | 1 | protein_coding | xylosylprotein beta 1,4-galactosyltransferase, polypeptide 7 |
| **VRK2** | ENSG00000028116 | 896.1961359 | 1.554565768 | 0.129622584 | 6.99E-34 | 3.33E-32 | 1019272 | 2 | 58134786 | 58387055 | 1 | protein_coding | vaccinia related kinase 2 |
| **TNFRSF1B** | ENSG00000028137 | 10263.698 | -0.761435573 | 0.14974094 | 1.49E-07 | 1.08E-06 | 7133 | 1 | 12227060 | 12269285 | 1 | protein_coding | tumor necrosis factor receptor superfamily, member 1B |
| **VEZT** | ENSG00000028203 | 1246.776016 | 0.561886539 | 0.174284531 | 6.83E-04 | 0.002587657 | 55591 | 12 | 95611522 | 95696566 | 1 | protein_coding | vezatin, adherens junctions transmembrane protein |
| **TBPL1** | ENSG00000028839 | 176.6296092 | 1.037414566 | 0.291826223 | 9.6E-05 | 4.4E-04 | 9519 | 6 | 134273308 | 134311570 | 1 | protein_coding | TBP-like 1 |
| **ARNTL2** | ENSG00000029153 | 959.4953862 | 2.262727867 | 0.170702497 | 4.03E-41 | 2.63E-39 | 56938 | 12 | 27485787 | 27576241 | 1 | protein_coding | aryl hydrocarbon receptor nuclear translocator-like 2 |
| **BCLAF1** | ENSG00000029363 | 4026.041561 | 0.362513322 | 0.099142364 | 1.99E-04 | 8.51E-04 | 9774 | 6 | 136578001 | 136610989 | -1 | protein_coding | BCL2-associated transcription factor 1 |
| **ANK1** | ENSG00000029534 | 247.9289171 | -0.847092473 | 0.347905218 | 0.004856065 | 0.01490266 | 286 | 8 | 41510739 | 41754280 | -1 | protein_coding | ankyrin 1, erythrocytic |
| **RABEP1** | ENSG00000029725 | 962.1114691 | -0.646398592 | 0.185290921 | 2.54E-04 | 0.001060353 | 9135 | 17 | 5185558 | 5289129 | 1 | protein_coding | rabaptin, RAB GTPase binding effector protein 1 |
| **NUP160** | ENSG00000030066 | 1620.996052 | 0.567918146 | 0.135257261 | 1.5E-05 | 8E-05 | 23279 | 11 | 47799639 | 47870107 | -1 | protein_coding | nucleoporin 160kDa |
| **BAK1** | ENSG00000030110 | 1372.612165 | 1.112128436 | 0.116033641 | 2.68E-22 | 6.86E-21 | 578 | 6 | 33540329 | 33548019 | -1 | protein_coding | BCL2-antagonist/killer 1 |
| **GRN** | ENSG00000030582 | 29192.78207 | -0.382980804 | 0.128600963 | 0.002348196 | 0.007789772 | 2896 | 17 | 42422614 | 42430470 | 1 | protein_coding | granulin |
| **ARHGAP31** | ENSG00000031081 | 5630.386102 | -0.399284425 | 0.142679039 | 0.004318263 | 0.013394767 | 57514 | 3 | 119013220 | 119139561 | 1 | protein_coding | Rho GTPase activating protein 31 |
| **TSSC1** | ENSG00000032389 | 468.1256075 | -0.465042509 | 0.160334094 | 0.002708854 | 0.008837655 | 7260 | 2 | 3192696 | 3381653 | -1 | protein_coding | tumor suppressing subtransferable candidate 1 |
| **PNPLA6** | ENSG00000032444 | 2960.92268 | -1.392756999 | 0.119725987 | 5.2E-32 | 2.23E-30 | 10908 | 19 | 7598890 | 7626650 | 1 | protein_coding | patatin-like phospholipase domain containing 6 |

| | | | | | | | | | | | | | |
|---|---|---|---|---|---|---|---|---|---|---|---|---|---|
| **ALG1** | ENSG00000033011 | 516.7236741 | -0.469886067 | 0.158658481 | 0.002174847 | 0.007273126 | 1005075 | 16 | 5083703 | 5137380 | 1 | protein_coding | ALG1, chitobiosyldiphosphodolichol beta-mannosyltransferase |
| **ZCCHC8** | ENSG00000033030 | 93.8632136 | 0.810586467 | 0.383893475 | 0.011510007 | 0.031586898 | 55596 | 12 | 122957417 | 122985518 | -1 | protein_coding | zinc finger, CCHC domain containing 8 |
| **FUT8** | ENSG00000033170 | 221.2378968 | 0.947484914 | 0.475012486 | 0.010216455 | 0.028513162 | 2530 | 14 | 65877310 | 66210839 | 1 | protein_coding | fucosyltransferase 8 (alpha (1,6) fucosyltransferase) |
| **UBA6** | ENSG00000033178 | 3796.839919 | 0.593297123 | 0.098143666 | 9.55E-10 | 9.21E-09 | 55236 | 4 | 68478370 | 68566897 | -1 | protein_coding | ubiquitin-like modifier activating enzyme 6 |
| **GAB2** | ENSG00000033327 | 2510.110377 | 0.397992361 | 0.153611148 | 0.008058611 | 0.023172108 | 9846 | 11 | 77926343 | 78129394 | -1 | protein_coding | GRB2-associated binding protein 2 |
| **PIAS1** | ENSG00000033800 | 1525.953748 | 0.314254666 | 0.108434352 | 0.003328311 | 0.010641545 | 8554 | 15 | 68346517 | 68483096 | 1 | protein_coding | protein inhibitor of activated STAT, 1 |
| **SLC4A7** | ENSG00000033867 | 1575.513171 | -1.136340488 | 0.167917665 | 3.04E-12 | 3.73E-11 | 9497 | 3 | 27414214 | 27525911 | -1 | protein_coding | solute carrier family 4, sodium bicarbonate cotransporter, member 7 |
| **APBA2** | ENSG00000034053 | 51.63563241 | -1.446302629 | 0.566228898 | 0.001270311 | 0.004512821 | 321 | 15 | 29129629 | 29410518 | 1 | protein_coding | amyloid beta (A4) precursor protein-binding, family A, member 2 |
| **MAP2K3** | ENSG00000034152 | 2376.338816 | 0.579030044 | 0.141360234 | 2.66E-05 | 1.35E-04 | 1009967 | 17 | 21187984 | 21218552 | 1 | protein_coding | mitogen-activated protein kinase kinase 3 |
| **TMSB10** | ENSG00000034510 | 25376.76464 | 0.561773181 | 0.117193706 | 1.02E-06 | 6.56E-06 | 9168 | 2 | 85132749 | 85133795 | 1 | protein_coding | thymosin beta 10 |
| **RNF19A** | ENSG00000034677 | 3188.543069 | 0.381910472 | 0.124631264 | 0.001724527 | 0.005916038 | 25897 | 8 | 101269288 | 101348446 | -1 | protein_coding | ring finger protein 19A, RBR E3 ubiquitin protein ligase |
| **PEX3** | ENSG00000034693 | 282.9395105 | 0.985274945 | 0.211235562 | 9.15E-07 | 5.92E-06 | 8504 | 6 | 143771944 | 143811147 | 1 | protein_coding | peroxisomal biogenesis factor 3 |
| **SH3YL1** | ENSG00000035115 | 264.6395273 | -0.694061401 | 0.307586527 | 0.010516786 | 0.029202042 | 26751 | 2 | 217730 | 266398 | -1 | protein_coding | SH3 and SYLF domain containing 1 |
| **VCL** | ENSG00000035403 | 4050.32902 | -0.935530283 | 0.107971728 | 1.3E-18 | 2.59E-17 | 7414 | 10 | 75757872 | 75879918 | 1 | protein_coding | vinculin |
| **ADSS** | ENSG00000035687 | 1511.524547 | 0.483275895 | 0.139775762 | 3.84E-04 | 0.001545138 | 159 | 1 | 244571796 | 244615436 | -1 | protein_coding | adenylosuccinate synthase |
| **STAP1** | ENSG00000035720 | 405.8865424 | 2.564676508 | 0.20564329 | 7.38E-37 | 3.93E-35 | 26228 | 4 | 68424446 | 68473055 | 1 | protein_coding | signal transducing adaptor family member 1 |
| **TIMP2** | ENSG00000035862 | 26248.11888 | -0.796171965 | 0.145782408 | 1.94E-08 | 1.58E-07 | 7077 | 17 | 76849059 | 76921469 | -1 | protein_coding | TIMP metallopeptidase inhibitor 2 |
| **RFC1** | ENSG00000035928 | 1742.852144 | 0.60226667 | 0.114888831 | 8.78E-08 | 6.55E-07 | 5981 | 4 | 39289076 | 39367995 | -1 | protein_coding | replication factor C (activator 1) 1, 145kDa |
| **USP2** | ENSG00000036672 | 1894.801897 | -2.77569309 | 0.239203229 | 3.1E-32 | 1.34E-30 | 9099 | 11 | 119225925 | 119252436 | -1 | protein_coding | ubiquitin specific peptidase 2 |
| **FLT4** | ENSG00000037280 | 18.02949069 | 2.159225344 | 0.85057937 | 7.14E-04 | 0.002696867 | 2324 | 5 | 180028506 | 180076624 | -1 | protein_coding | fms-related tyrosine kinase 4 |
| **NSUN2** | ENSG00000037474 | 1799.150766 | 0.285819896 | 0.109356348 | 0.00747365 | 0.021714791 | 54888 | 5 | 6599352 | 6633404 | -1 | protein_coding | NOP2/Sun RNA methyltransferase family, member 2 |
| **PI4K2B** | ENSG00000038210 | 1018.006928 | 2.338662586 | 0.153238084 | 1.65E-53 | 1.78E-51 | 285540 | 4 | 25162263 | 25280714 | 1 | protein_coding | phosphatidylinositol 4-kinase type 2 beta |
| **MAT2B** | ENSG00000038274 | 2741.4947 | 0.948309703 | 0.10286081 | 1.06E-20 | 2.48E-19 | 27430 | 5 | 162930120 | 162946342 | 1 | protein_coding | methionine adenosyltransferase II, beta |
| **EDC4** | ENSG00000038358 | 1036.230768 | -0.43904247 | 0.151540817 | 0.002727232 | 0.008890846 | 23644 | 16 | 67906926 | 67918406 | 1 | protein_coding | enhancer of mRNA decapping 4 |
| **TRIO** | ENSG00000038382 | 2530.425433 | -0.51985991 | 0.136907937 | 9.49E-05 | 4.36E-04 | 7204 | 5 | 14143811 | 14532235 | 1 | protein_coding | trio Rho guanine nucleotide exchange factor |
| **VCAN** | ENSG00000038427 | 352.3187636 | -3.823150119 | 0.888449007 | 7.47E-07 | 4.92E-06 | 1462 | 5 | 82767284 | 82878122 | 1 | protein_coding | versican |
| **MSR1** | ENSG00000038945 | 41530.93474 | -0.492154045 | 0.098441469 | 6.74E-07 | 4.46E-06 | 4481 | 8 | 15965387 | 16424999 | -1 | protein_coding | macrophage scavenger receptor 1 |
| **SKIV2L2** | ENSG00000039123 | 1973.39296 | -0.267609696 | 0.10661554 | 0.010654074 | 0.029518893 | 23517 | 5 | 54603588 | 54721409 | 1 | protein_coding | superkiller viralicidic activity 2-like 2 (S. cerevisiae) |
| **FAM65A** | ENSG00000039523 | 2586.496385 | -0.425606855 | 0.118436857 | 2.45E-04 | 0.001027234 | 79567 | 16 | 67552321 | 67580691 | 1 | protein_coding | family with sequence similarity 65, member A |
| **RAI14** | ENSG00000039560 | 3303.636209 | -0.85967241 | 0.158029773 | 2.43E-08 | 1.96E-07 | 26064 | 5 | 34656342 | 34832732 | 1 | protein_coding | retinoic acid induced 14 |
| **PNKP** | ENSG00000039650 | 495.8271884 | -0.45798873 | 0.166685499 | 0.004365444 | 0.013527389 | 11284 | 19 | 50364461 | 50371166 | -1 | protein_coding | polynucleotide kinase 3'-phosphatase |
| **STAU2** | ENSG00000040341 | 779.3305976 | -0.656660806 | 0.212228637 | 9.88E-04 | 0.003606247 | 27067 | 8 | 74332604 | 74659943 | -1 | protein_coding | staufen double-stranded RNA binding protein 2 |
| **PQLC2** | ENSG00000040487 | 819.1104188 | -0.357535894 | 0.149013857 | 0.01433491 | 0.038160675 | 54896 | 1 | 19638820 | 19655794 | 1 | protein_coding | PQ loop repeat containing 2 |
| **RTN4R** | ENSG00000040608 | 179.4198147 | -1.319471262 | 0.372749333 | 6.29E-05 | 2.98E-04 | 65078 | 22 | 20228938 | 20270769 | -1 | protein_coding | reticulon 4 receptor |
| **INPP4A** | ENSG00000040933 | 800.2328276 | -1.082385209 | 0.172152955 | 7.92E-11 | 8.58E-10 | 3631 | 2 | 99061317 | 99210853 | 1 | protein_coding | inositol polyphosphate-4-phosphatase, type I, 107kDa |
| **PSMA4** | ENSG00000041357 | 3252.329699 | 1.662910589 | 0.129914944 | 2.15E-38 | 1.22E-36 | 5685 | 15 | 78832747 | 78841604 | 1 | protein_coding | proteasome (prosome, macropain) subunit, alpha type, 4 |
| **THAP3** | ENSG00000041988 | 185.7468301 | 0.921359765 | 0.241991872 | 4.55E-05 | 2.21E-04 | 90326 | 1 | 6684926 | 6695646 | 1 | protein_coding | THAP domain containing, apoptosis associated protein 3 |
| **AIFM2** | ENSG00000042286 | 1087.074481 | 1.681601027 | 0.131493549 | 2.38E-38 | 1.35E-36 | 84883 | 10 | 71857979 | 71892690 | -1 | protein_coding | apoptosis-inducing factor, mitochondrion-associated, 2 |
| **MED17** | ENSG00000042429 | 893.4448286 | 0.862521646 | 0.137370945 | 1.27E-10 | 1.34E-09 | 9440 | 11 | 93517393 | 93547861 | 1 | protein_coding | mediator complex subunit 17 |
| **AP2S1** | ENSG00000042753 | 2182.00772 | -0.916246599 | 0.125710157 | 9.82E-14 | 1.36E-12 | 1175 | 19 | 47341393 | 47354249 | -1 | protein_coding | adaptor-related protein complex 2, sigma 1 subunit |
| **DCUN1D1** | ENSG00000043093 | 1692.696479 | 0.899767286 | 0.136574887 | 1.55E-11 | 1.79E-10 | 54165 | 3 | 182655862 | 182703741 | -1 | protein_coding | DCN1, defective in cullin neddylation 1, domain containing 1 |
| **JADE2** | ENSG00000043143 | 2087.254927 | 1.704583823 | 0.167398412 | 2.97E-25 | 8.97E-24 | 23338 | 5 | 133860003 | 133918918 | 1 | protein_coding | jade family PHD finger 2 |
| **LCP2** | ENSG00000043462 | 5754.271176 | 0.835514831 | 0.103351201 | 2.54E-16 | 4.21E-15 | 3937 | 5 | 169673241 | 169725231 | -1 | protein_coding | lymphocyte cytosolic protein 2 (SH2 domain containing leukocyte protein of 76kDa) |
| **CUL7** | ENSG00000044090 | 616.7442257 | -0.478048743 | 0.184742892 | 0.006551199 | 0.019343502 | 9820 | 6 | 43005355 | 43021683 | -1 | protein_coding | cullin 7 |
| **CNTLN** | ENSG00000044459 | 377.192498 | 1.641514495 | 0.226773955 | 6.27E-14 | 8.87E-13 | 54875 | 9 | 17134980 | 17503921 | 1 | protein_coding | centlein, centrosomal protein |
| **HSPA5** | ENSG00000044574 | 15971.76324 | -0.61300228 | 0.102726207 | 1.46E-09 | 1.38E-08 | 3309 | 9 | 127997132 | 128003609 | -1 | protein_coding | heat shock 70kDa protein 5 (glucose-regulated protein, 78kDa) |
| **OFD1** | ENSG00000046651 | 654.2373942 | 0.567167993 | 0.159116517 | 2.24E-04 | 9.48E-04 | 8481 | X | 13752832 | 13787480 | 1 | protein_coding | oral-facial-digital syndrome 1 |
| **YTHDC2** | ENSG00000047188 | 973.5959675 | -0.460321058 | 0.133840433 | 4.45E-04 | 0.001763777 | 64848 | 5 | 112849380 | 112930982 | 1 | protein_coding | YTH domain containing 2 |
| **ATP6V1H** | ENSG00000047249 | 7085.733678 | 0.494138739 | 0.171220368 | 0.002467975 | 0.008132397 | 51606 | 8 | 54628117 | 54756118 | -1 | protein_coding | ATPase, H+ transporting, lysosomal 50/57kDa, V1 subunit H |
| **ARAP2** | ENSG00000047365 | 3332.714007 | 1.197656775 | 0.144915662 | 2.64E-17 | 4.72E-16 | 1019286 | 4 | 35949843 | 36246131 | -1 | protein_coding | ArfGAP with RhoGAP domain, ankyrin repeat and PH domain 2 |
| **TPR** | ENSG00000047410 | 4954.279947 | 0.292928245 | 0.111876698 | 0.007881922 | 0.022738013 | 7175 | 1 | 186280784 | 186344825 | -1 | protein_coding | translocated promoter region, nuclear basket protein |
| **CP** | ENSG00000047457 | 7421.887088 | 1.211517549 | 0.135447997 | 7.4E-20 | 1.62E-18 | 1356 | 3 | 148880197 | 148939842 | -1 | protein_coding | ceruloplasmin (ferroxidase) |
| **SCML1** | ENSG00000047634 | 188.4638183 | 0.556224992 | 0.240421787 | 0.01187687 | 0.032442216 | 6322 | X | 17755588 | 17773105 | 1 | protein_coding | sex comb on midleg-like 1 (Drosophila) |
| **ARHGAP6** | ENSG00000047648 | 95.9136762 | -2.447255052 | 0.417998822 | 4.28E-10 | 4.27E-09 | 395 | X | 11136239 | 11683821 | -1 | protein_coding | Rho GTPase activating protein 6 |
| **GOPC** | ENSG00000047932 | 1091.20854 | 1.140406058 | 0.131997928 | 1.72E-18 | 3.37E-17 | 57120 | 6 | 117639374 | 117923691 | -1 | protein_coding | golgi-associated PDZ and coiled-coil motif containing |
| **CC2D2A** | ENSG00000048342 | 109.8475391 | -1.774484523 | 0.464549638 | 1.36E-05 | 7.34E-05 | 57545 | 4 | 15471489 | 15603180 | 1 | protein_coding | coiled-coil and C2 domain containing 2A |
| **RRM2B** | ENSG00000048392 | 4074.686069 | 0.768608862 | 0.142216774 | 2.49E-08 | 2.01E-07 | 50484 | 8 | 103216730 | 103251346 | -1 | protein_coding | ribonucleotide reductase M2 B (TP53 inducible) |
| **ZNF800** | ENSG00000048405 | 1030.831986 | 0.686649484 | 0.119688569 | 5.08E-09 | 4.5E-08 | 168850 | 7 | 126986844 | 127071978 | -1 | protein_coding | zinc finger protein 800 |
| **SNX29** | ENSG00000048471 | 1822.579607 | -1.03009863 | 0.114440456 | 7.7E-20 | 1.68E-18 | 1019293 | 16 | 12070594 | 12668146 | 1 | protein_coding | sorting nexin 29 |
| **RSF1** | ENSG00000048649 | 1201.41478 | 0.555802661 | 0.16782961 | 5.54E-04 | 0.002141935 | 51773 | 11 | 77371041 | 77532063 | -1 | protein_coding | remodeling and spacing factor 1 |
| **CELF2** | ENSG00000048740 | 788.7531681 | -1.172790204 | 0.217521045 | 1.62E-08 | 1.33E-07 | 10659 | 10 | 11047259 | 11378674 | 1 | protein_coding | CUGBP, Elav-like family member 2 |
| **FAM120A** | ENSG00000048828 | 5978.771282 | -1.116599521 | 0.220534298 | 7.29E-08 | 5.51E-07 | 23196 | 9 | 96214004 | 96328397 | 1 | protein_coding | family with sequence similarity 120A |
| **COL9A2** | ENSG00000049089 | 671.5860027 | 0.95561076 | 0.188070026 | 1.28E-07 | 9.33E-07 | 1298 | 1 | 40766159 | 40783488 | -1 | protein_coding | collagen, type IX, alpha 2 |
| **ERCC8** | ENSG00000049167 | 234.482064 | -0.647352172 | 0.236638374 | 0.003154183 | 0.01015904 | 1161 | 5 | 60169658 | 60240900 | -1 | protein_coding | excision repair cross-complementing rodent repair deficiency, complementation group 8 |
| **PER3** | ENSG00000049246 | 1277.245061 | -0.880344829 | 0.176451617 | 2.48E-07 | 1.74E-06 | 8863 | 1 | 7844380 | 7905237 | 1 | protein_coding | period circadian clock 3 |
| **TNFRSF9** | ENSG00000049249 | 662.8762724 | 1.366598659 | 0.16946899 | 1.33E-16 | 2.26E-15 | 3604 | 1 | 7979907 | 8000926 | -1 | protein_coding | tumor necrosis factor receptor superfamily, member 9 |
| **RCN1** | ENSG00000049449 | 2249.309482 | 1.839564861 | 0.185003626 | 2.27E-24 | 6.55E-23 | 440034 | 11 | 31833939 | 32127301 | 1 | protein_coding | reticulocalbin 1, EF-hand calcium binding domain |
| **CLPTM1L** | ENSG00000049656 | 1579.014102 | -0.2855655 | 0.121744084 | 0.016896482 | 0.044036311 | 81037 | 5 | 1317859 | 1345214 | -1 | protein_coding | CLPTM1-like |
| **NEDD4L** | ENSG00000049759 | 431.0254811 | -1.024469746 | 0.201082274 | 9.14E-08 | 6.82E-07 | 23327 | 18 | 55711599 | 56068772 | 1 | protein_coding | neural precursor cell expressed, developmentally down-regulated 4-like, E3 ubiquitin protein ligase |
| **HEXB** | ENSG00000049860 | 18543.53643 | -0.350157917 | 0.140599045 | 0.010613905 | 0.029428938 | 3074 | 5 | 73935848 | 74018472 | 1 | protein_coding | hexosaminidase B (beta polypeptide) |
| **ARHGEF5** | ENSG00000050327 | 409.8412598 | 0.592570402 | 0.187693075 | 9.26E-04 | 0.003404734 | 7984 | 7 | 144052381 | 144077725 | 1 | protein_coding | Rho guanine nucleotide exchange factor (GEF) 5 |
| **NFE2L3** | ENSG00000050344 | 1977.929521 | 2.509513829 | 0.22890637 | 4.27E-29 | 1.56E-27 | 9603 | 7 | 26191860 | 26226745 | 1 | protein_coding | nuclear factor, erythroid 2-like 3 |
| **MCUR1** | ENSG00000050393 | 673.3779693 | 0.573764371 | 0.162348654 | 2.5E-04 | 0.001045712 | 63933 | 6 | 13786789 | 13814800 | -1 | protein_coding | mitochondrial calcium uniporter regulator 1 |
| **LIMA1** | ENSG00000050405 | 2372.414273 | -0.572676191 | 0.098136907 | 3.06E-09 | 2.76E-08 | 51474 | 12 | 50569571 | 50677329 | -1 | protein_coding | LIM domain and actin binding 1 |
| **SLC4A8** | ENSG00000050438 | 1136.748493 | -0.684464763 | 0.240862114 | 0.002079443 | 0.006996944 | 9498 | 12 | 51785101 | 51902980 | 1 | protein_coding | solute carrier family 4, sodium bicarbonate cotransporter, member 8 |
| **BCAR1** | ENSG00000050820 | 1006.681535 | -0.614897613 | 0.166281206 | 1.18E-04 | 5.35E-04 | 9564 | 16 | 75262928 | 75301951 | -1 | protein_coding | breast cancer anti-estrogen resistance 1 |
| **HERPUD1** | ENSG00000051108 | 2628.677861 | -0.474625761 | 0.110201994 | 1.17E-05 | 6.39E-05 | 9709 | 16 | 56965960 | 56977798 | 1 | protein_coding | homocysteine-inducible, endoplasmic reticulum stress-inducible, ubiquitin-like domain member 1 |
| **HOMER3** | ENSG00000051128 | 284.2834257 | -1.598924083 | 0.25434078 | 4.35E-11 | 4.84E-10 | 9454 | 19 | 19040010 | 19052070 | -1 | protein_coding | homer homolog 3 (Drosophila) |
| **RAD51** | ENSG00000051180 | 87.0654516 | 2.329532201 | 0.44767869 | 1.42E-08 | 1.19E-07 | 5888 | 15 | 40986972 | 41024354 | 1 | protein_coding | RAD51 recombinase |
| **POLQ** | ENSG00000051341 | 108.0403263 | 0.980111749 | 0.444506583 | 0.006424151 | 0.019037074 | 10721 | 3 | 121150278 | 121264853 | -1 | protein_coding | polymerase (DNA directed), theta |
| **PIK3CB** | ENSG00000051382 | 3823.422728 | -0.291773192 | 0.117068455 | 0.011335028 | 0.031179666 | 5291 | 3 | 138372860 | 138553780 | -1 | protein_coding | phosphatidylinositol-4,5-bisphosphate 3-kinase, catalytic subunit beta |
| **HEBP2** | ENSG00000051620 | 548.6461105 | 1.101419137 | 0.237358748 | 8.09E-07 | 5.29E-06 | 23593 | 6 | 138724668 | 138743334 | 1 | protein_coding | heme binding protein 2 |
| **PLEKHA5** | ENSG00000052126 | 558.8322011 | -0.869204157 | 0.238786536 | 9.39E-05 | 4.32E-04 | 54477 | 12 | 19282648 | 19529334 | 1 | protein_coding | pleckstrin homology domain containing, family A member 5 |
| **RRP12** | ENSG00000052749 | 868.2814708 | -0.409400543 | 0.137411658 | 0.002287975 | 0.007608163 | 23223 | 10 | 99116115 | 99161127 | -1 | protein_coding | ribosomal RNA processing 12 homolog (S. cerevisiae) |
| **FNIP2** | ENSG00000052795 | 7361.498908 | 0.526684469 | 0.130601991 | 3.71E-05 | 1.83E-04 | 57600 | 4 | 159690290 | 159829201 | 1 | protein_coding | folliculin interacting protein 2 |
| **MSMO1** | ENSG00000052802 | 3831.370105 | 0.913532944 | 0.196553486 | 1.04E-06 | 6.64E-06 | 6307 | 4 | 166248775 | 166264312 | 1 | protein_coding | methylsterol monooxygenase 1 |
| **FOXN3** | ENSG00000053254 | 3702.837849 | -0.543281979 | 0.113016534 | 9.79E-07 | 6.31E-06 | 1112 | 14 | 89591215 | 90085493 | -1 | protein_coding | forkhead box N3 |
| **AKR7A2** | ENSG00000053371 | 649.1062709 | -1.210673915 | 0.208039456 | 1.25E-09 | 1.19E-08 | 8574 | 1 | 19630459 | 19638640 | -1 | protein_coding | aldo-keto reductase family 7, member A2 (aflatoxin aldehyde reductase) |
| **MRTO4** | ENSG00000053372 | 1059.880015 | -0.409137808 | 0.138449097 | 0.002354212 | 0.007806341 | 51154 | 1 | 19578033 | 19586622 | 1 | protein_coding | mRNA turnover 4 homolog (S. cerevisiae) |

| | | | | | | | | | | | | | |
|---|---|---|---|---|---|---|---|---|---|---|---|---|---|
| **LAMA3** | ENSG00000053747 | 241.9769865 | 3.626542204 | 0.444001623 | 1.96E-17 | 3.54E-16 | 3909 | 18 | 21269407 | 21535030 | 1 | protein_coding | laminin, alpha 3 |
| **AP5M1** | ENSG00000053770 | 1457.139439 | 0.869330141 | 0.120726392 | 2.17E-13 | 2.91E-12 | 55745 | 14 | 57735627 | 57756797 | 1 | protein_coding | adaptor-related protein complex 5, mu 1 subunit |
| **THRAP3** | ENSG00000054118 | 5367.852865 | 0.362963507 | 0.092589409 | 7.03E-05 | 3.31E-04 | 9967 | 1 | 36690017 | 36770958 | 1 | protein_coding | thyroid hormone receptor associated protein 3 |
| **PHPT1** | ENSG00000054148 | 746.7142727 | -0.577379271 | 0.143687769 | 3.84E-05 | 1.9E-04 | 29085 | 9 | 139743176 | 139745488 | 1 | protein_coding | phosphohistidine phosphatase 1 |
| **LY75** | ENSG00000054219 | 3538.247375 | 0.932091605 | 0.152899129 | 3.86E-10 | 3.87E-09 | 1005266 | 2 | 160628362 | 160761260 | -1 | protein_coding | lymphocyte antigen 75 |
| **ARID4B** | ENSG00000054267 | 2511.210725 | 0.873042745 | 0.106283047 | 7.91E-17 | 1.36E-15 | 51742 | 1 | 235294949 | 235491534 | -1 | protein_coding | AT rich interactive domain 4B (RBP1-like) |
| **OPN3** | ENSG00000054277 | 753.1152999 | -0.622970969 | 0.225032848 | 0.003008428 | 0.009728695 | 23596 | 1 | 241753404 | 241840678 | -1 | protein_coding | opsin 3 |
| **KIF1B** | ENSG00000054523 | 5257.528302 | 0.885439684 | 0.113000558 | 1.8E-15 | 2.82E-14 | 23095 | 1 | 10270863 | 10441661 | 1 | protein_coding | kinesin family member 1B |
| **TBC1D22A** | ENSG00000054611 | 1178.22915 | -0.479094659 | 0.139971146 | 4.35E-04 | 0.00172723 | 25771 | 22 | 47158518 | 47571336 | 1 | protein_coding | TBC1 domain family, member 22A |
| **SYNE2** | ENSG00000054654 | 1162.938119 | 1.387011977 | 0.28557628 | 1.8E-07 | 1.29E-06 | 23224 | 14 | 64319683 | 64693165 | 1 | protein_coding | spectrin repeat containing, nuclear envelope 2 |
| **ATP9A** | ENSG00000054793 | 2256.254105 | -1.182332562 | 0.260468546 | 1.17E-06 | 7.45E-06 | 10079 | 20 | 50213053 | 50385173 | -1 | protein_coding | ATPase, class II, type 9A |
| **FAM168A** | ENSG00000054965 | 1566.861929 | 0.53511648 | 0.154286773 | 3.76E-04 | 0.001516498 | 23201 | 11 | 73111532 | 73309234 | -1 | protein_coding | family with sequence similarity 168, member A |
| **GALC** | ENSG00000054983 | 2319.43131 | -0.610792223 | 0.133693985 | 2.94E-06 | 1.77E-05 | 2581 | 14 | 88304164 | 88460009 | -1 | protein_coding | galactosylceramidase |
| **NOP58** | ENSG00000055044 | 805.5077661 | 0.483494661 | 0.167514307 | 0.002754491 | 0.008973127 | 51602 | 2 | 203130439 | 203168389 | 1 | protein_coding | NOP58 ribonucleoprotein |
| **CUL1** | ENSG00000055130 | 2764.307863 | 1.144917087 | 0.103344708 | 3.96E-29 | 1.46E-27 | 8454 | 7 | 148395006 | 148498128 | 1 | protein_coding | cullin 1 |
| **GINM1** | ENSG00000055211 | 1144.742494 | 0.626263611 | 0.149690298 | 1.62E-05 | 8.61E-05 | 116254 | 6 | 149887430 | 149912884 | 1 | protein_coding | glycoprotein integral membrane 1 |
| **EIF2AK2** | ENSG00000055332 | 5010.026312 | 3.395308653 | 0.100217305 | 6.08E-253 | 7.15E-250 | 5610 | 2 | 37326353 | 37384208 | -1 | protein_coding | eukaryotic translation initiation factor 2-alpha kinase 2 |
| **KMT2C** | ENSG00000055609 | 380.5940948 | -0.795436818 | 0.363651704 | 0.009854407 | 0.027649147 | 58508 | 7 | 151832010 | 152133090 | -1 | protein_coding | lysine (K)-specific methyltransferase 2C |
| **PUM2** | ENSG00000055917 | 549.2660325 | 0.773445048 | 0.234586441 | 4.07E-04 | 0.001628241 | 23369 | 2 | 20448452 | 20551995 | -1 | protein_coding | pumilio RNA-binding family member 2 |
| **MRPL43** | ENSG00000055950 | 451.2722153 | -0.460379566 | 0.192583807 | 0.012070365 | 0.032911744 | 84545 | 10 | 102729215 | 102747272 | -1 | protein_coding | mitochondrial ribosomal protein L43 |
| **ITIH4** | ENSG00000055955 | 168.3404753 | 1.954087963 | 0.466616887 | 2.56E-06 | 1.55E-05 | 3700 | 3 | 52846991 | 52865495 | -1 | protein_coding | inter-alpha-trypsin inhibitor heavy chain family, member 4 |
| **C4orf27** | ENSG00000056050 | 207.6517039 | 0.593676611 | 0.227636269 | 0.005007222 | 0.015283677 | 54969 | 4 | 170650616 | 170679104 | -1 | protein_coding | chromosome 4 open reading frame 27 |
| **ZNF280C** | ENSG00000056277 | 160.4502062 | 0.761392656 | 0.260276868 | 0.00143983 | 0.005051636 | 55609 | X | 129336685 | 129402873 | -1 | protein_coding | zinc finger protein 280C |
| **TRAF1** | ENSG00000056558 | 2773.110532 | 2.048891077 | 0.175065313 | 1.15E-32 | 5.13E-31 | 7185 | 9 | 123664671 | 123691451 | -1 | protein_coding | TNF receptor-associated factor 1 |
| **RC3H2** | ENSG00000056586 | 2509.577534 | 0.527617871 | 0.112207804 | 1.71E-06 | 1.06E-05 | 54542 | 9 | 125606835 | 125667620 | -1 | protein_coding | ring finger and CCCH-type domains 2 |
| **TRAF3IP2** | ENSG00000056972 | 365.5954976 | 1.311016255 | 0.196005811 | 4.28E-12 | 5.2E-11 | 10758 | 6 | 111877657 | 111927481 | -1 | protein_coding | TRAF3 interacting protein 2 |
| **DCBLD2** | ENSG00000057019 | 302.2769123 | -1.785772375 | 0.303461003 | 4E-10 | 4E-09 | 131566 | 3 | 98514785 | 98620533 | -1 | protein_coding | discoidin, CUB and LCCL domain containing 2 |
| **SOAT1** | ENSG00000057252 | 10966.22761 | 0.315159643 | 0.094958589 | 7.73E-04 | 0.002897741 | 6646 | 1 | 179262925 | 179327815 | 1 | protein_coding | sterol O-acyltransferase 1 |
| **PRDM1** | ENSG00000057657 | 1531.844353 | 1.612297608 | 0.195715618 | 2.38E-17 | 4.29E-16 | 639 | 6 | 106534195 | 106557814 | 1 | protein_coding | PR domain containing 1, with ZNF domain |
| **ATG5** | ENSG00000057663 | 1063.708751 | 0.68504547 | 0.159834449 | 9.11E-06 | 5.04E-05 | 9474 | 6 | 106632351 | 106773666 | -1 | protein_coding | autophagy related 5 |
| **TMCC3** | ENSG00000057704 | 1148.818764 | -0.602353954 | 0.149663527 | 3.15E-05 | 1.58E-04 | 57458 | 12 | 94960900 | 95044338 | -1 | protein_coding | transmembrane and coiled-coil domain family 3 |
| **USP13** | ENSG00000058056 | 379.2979502 | 0.645090671 | 0.194499385 | 4.89E-04 | 0.001917652 | 8975 | 3 | 179370543 | 179507189 | 1 | protein_coding | ubiquitin specific peptidase 13 (isopeptidase T-3) |
| **ATP11B** | ENSG00000058063 | 3356.796964 | 0.513920766 | 0.145731801 | 2.83E-04 | 0.001170885 | 23200 | 3 | 182511288 | 182639423 | 1 | protein_coding | ATPase, class VI, type 11B |
| **CDK14** | ENSG00000058091 | 2162.43483 | 0.588037837 | 0.150141886 | 5.88E-05 | 2.8E-04 | 5218 | 7 | 90095738 | 90839905 | 1 | protein_coding | cyclin-dependent kinase 14 |
| **PPP1R12A** | ENSG00000058272 | 3317.742755 | 0.410469792 | 0.089198506 | 3.33E-06 | 1.98E-05 | 4659 | 12 | 80167343 | 80329240 | -1 | protein_coding | protein phosphatase 1, regulatory subunit 12A |
| **CAMK2B** | ENSG00000058404 | 18.21292772 | -3.140487099 | 1.544646301 | 0.001715643 | 0.005894823 | 816 | 7 | 44256749 | 44374176 | -1 | protein_coding | calcium/calmodulin-dependent protein kinase II beta |
| **ATP2B4** | ENSG00000058668 | 6098.693758 | -1.019846954 | 0.110933501 | 8.5E-21 | 2E-19 | 493 | 1 | 203595689 | 203713209 | 1 | protein_coding | ATPase, Ca++ transporting, plasma membrane 4 |
| **ZC3H11A** | ENSG00000058673 | 3779.967157 | 0.402391288 | 0.090600912 | 7.23E-06 | 4.07E-05 | 441155 | 1 | 203764782 | 203823252 | 1 | protein_coding | zinc finger CCCH-type containing 11A |
| **DGKG** | ENSG00000058866 | 249.271592 | 0.813306785 | 0.267773225 | 9.1E-04 | 0.003355984 | 1608 | 3 | 185823457 | 186080026 | -1 | protein_coding | diacylglycerol kinase, gamma 90kDa |
| **TBXAS1** | ENSG00000059377 | 3745.896687 | -1.504016708 | 0.095262965 | 7.36E-57 | 8.65E-55 | 6916 | 7 | 139476850 | 139720125 | 1 | protein_coding | thromboxane A synthase 1 (platelet) |
| **PARP12** | ENSG00000059378 | 3964.057218 | 1.423199471 | 0.095653649 | 2.12E-51 | 1.99E-49 | 64761 | 7 | 139723544 | 139763521 | -1 | protein_coding | poly (ADP-ribose) polymerase family, member 12 |
| **ALDH18A1** | ENSG00000059573 | 683.7916004 | -0.752595601 | 0.150154404 | 1.92E-07 | 1.37E-06 | 5832 | 10 | 97365696 | 97416463 | -1 | protein_coding | aldehyde dehydrogenase 18 family, member A1 |
| **PET112** | ENSG00000059691 | 540.8402703 | -0.553719321 | 0.22303979 | 0.007762287 | 0.022442496 | 5188 | 4 | 152591656 | 152682175 | -1 | protein_coding | PET112 homolog (yeast) |
| **MXD1** | ENSG00000059728 | 1656.174716 | 1.517804489 | 0.141792037 | 1.57E-27 | 5.28E-26 | 4084 | 2 | 70124820 | 70170077 | 1 | protein_coding | MAX dimerization protein 1 |
| **SLC2A3** | ENSG00000059804 | 9254.658746 | -1.225156839 | 0.374693059 | 1.37E-04 | 6.1E-04 | 6515 | 12 | 8071826 | 8088871 | -1 | protein_coding | solute carrier family 2 (facilitated glucose transporter), member 3 |
| **WNK1** | ENSG00000060237 | 9759.097125 | -0.312095878 | 0.104578277 | 0.002398893 | 0.007930868 | 65125 | 12 | 861759 | 1020618 | 1 | protein_coding | WNK lysine deficient protein kinase 1 |
| **OGFR** | ENSG00000060491 | 2127.567845 | 1.164247332 | 0.11333787 | 2.06E-25 | 6.27E-24 | 11054 | 20 | 61436187 | 61445352 | 1 | protein_coding | opioid growth factor receptor |
| **PIGV** | ENSG00000060642 | 336.6262385 | 0.58851613 | 0.212196602 | 0.003170109 | 0.010199742 | 55650 | 1 | 27113963 | 27124889 | 1 | protein_coding | phosphatidylinositol glycan anchor biosynthesis, class V |
| **PTPRU** | ENSG00000060656 | 13.6438436 | 2.788360756 | 1.129876845 | 5.69E-04 | 0.002193735 | 10076 | 1 | 29563028 | 29653325 | 1 | protein_coding | protein tyrosine phosphatase, receptor type, U |
| **MPC1** | ENSG00000060762 | 686.5748747 | -0.82276109 | 0.218451645 | 5.37E-05 | 0.000257696 | 51660 | 6 | 166778407 | 166796486 | -1 | protein_coding | mitochondrial pyruvate carrier 1 |
| **BCAT1** | ENSG00000060982 | 16057.86016 | -0.862321089 | 0.141176802 | 5.36E-10 | 5.29E-09 | 586 | 12 | 24964295 | 25102393 | -1 | protein_coding | branched chain amino-acid transaminase 1, cytosolic |
| **HDAC7** | ENSG00000061273 | 1442.075755 | -0.454865685 | 0.168569233 | 0.004926057 | 0.015087429 | 51564 | 12 | 48176505 | 48226915 | -1 | protein_coding | histone deacetylase 7 |
| **SPAG4** | ENSG00000061656 | 34.35495845 | -1.636493356 | 0.736108539 | 0.002292427 | 0.007619652 | 6676 | 20 | 34203814 | 34208971 | 1 | protein_coding | sperm associated antigen 4 |
| **NCKAP1** | ENSG00000061676 | 1960.481324 | 0.662648491 | 0.263098235 | 0.005631135 | 0.016901146 | 10787 | 2 | 183773843 | 183903586 | -1 | protein_coding | NCK-associated protein 1 |
| **GUCY1B3** | ENSG00000061918 | 212.800417 | 2.168333212 | 0.286590979 | 3.37E-15 | 5.17E-14 | 2983 | 4 | 156680144 | 156728743 | 1 | protein_coding | guanylate cyclase 1, soluble, beta 3 |
| **TNK2** | ENSG00000061938 | 815.4603053 | 0.932320478 | 0.232902033 | 2.09E-05 | 1.09E-04 | 10188 | 3 | 195590235 | 195638816 | -1 | protein_coding | tyrosine kinase, non-receptor, 2 |
| **GPBP1** | ENSG00000062194 | 2425.53001 | 1.586897729 | 0.096355253 | 8.26E-62 | 1.12E-59 | 65056 | 5 | 56469775 | 56560506 | 1 | protein_coding | GC-rich promoter binding protein 1 |
| **ELMO2** | ENSG00000062598 | 2896.193711 | 0.637105228 | 0.143646697 | 4.77E-06 | 2.77E-05 | 63916 | 20 | 44994688 | 45061704 | -1 | protein_coding | engulfment and cell motility 2 |
| **VMP1** | ENSG00000062716 | 8522.183205 | 0.670124101 | 0.120077025 | 1.37E-08 | 1.14E-07 | 406991 | 17 | 57784553 | 57919616 | 1 | protein_coding | vacuole membrane protein 1 |
| **POLD1** | ENSG00000062822 | 468.1121732 | -0.781670497 | 0.274338891 | 0.001543427 | 0.005373553 | 5424 | 19 | 50887461 | 50921273 | 1 | protein_coding | polymerase (DNA directed), delta 1, catalytic subunit |
| **EIF4B** | ENSG00000063046 | 10506.27694 | -0.836936796 | 0.095761944 | 5.67E-19 | 1.15E-17 | 1975 | 12 | 53399942 | 53435993 | 1 | protein_coding | eukaryotic translation initiation factor 4B |
| **RPL18** | ENSG00000063177 | 2591.574877 | -0.466092515 | 0.145569788 | 9.09E-04 | 0.00335429 | 6141 | 19 | 49118585 | 49122793 | -1 | protein_coding | ribosomal protein L18 |
| **CA11** | ENSG00000063180 | 545.4447217 | -0.797546254 | 0.218359381 | 1.01E-04 | 4.63E-04 | 770 | 19 | 49141199 | 49149569 | -1 | protein_coding | carbonic anhydrase XI |
| **ISOC2** | ENSG00000063241 | 281.0054598 | -1.126814144 | 0.259682091 | 3.26E-06 | 1.94E-05 | 79763 | 19 | 55964352 | 55973710 | -1 | protein_coding | isochorismatase domain containing 2 |
| **EPN1** | ENSG00000063245 | 966.1007745 | -0.579704906 | 0.161855811 | 2.02E-04 | 8.61E-04 | 29924 | 19 | 56186592 | 56221224 | 1 | protein_coding | epsin 1 |
| **AHRR** | ENSG00000063438 | 156.7253561 | -1.368053937 | 0.395918369 | 8.44E-05 | 3.92E-04 | 57491 | 5 | 304291 | 438406 | 1 | protein_coding | aryl-hydrocarbon receptor repressor |
| **MTMR1** | ENSG00000063601 | 1322.108879 | -0.892609858 | 0.275174657 | 4.02E-04 | 0.001612356 | 8776 | X | 149861435 | 149933576 | 1 | protein_coding | myotubularin related protein 1 |
| **HAGH** | ENSG00000063854 | 795.900314 | -0.328798416 | 0.142504137 | 0.017760655 | 0.045920502 | 3029 | 16 | 1845621 | 1877195 | -1 | protein_coding | hydroxyacylglutathione hydrolase |
| **RNF4** | ENSG00000063978 | 1993.576684 | 0.276107478 | 0.120197091 | 0.019495876 | 0.049642698 | 6047 | 4 | 2463947 | 2627047 | 1 | protein_coding | ring finger protein 4 |
| **CASP8** | ENSG00000064012 | 1635.145769 | 0.850318614 | 0.128605419 | 1.53E-11 | 1.77E-10 | 841 | 2 | 202098166 | 202152434 | 1 | protein_coding | caspase 8, apoptosis-related cysteine peptidase |
| **TM7SF3** | ENSG00000064115 | 3167.471284 | -0.585230686 | 0.120560081 | 7.13E-07 | 4.71E-06 | 51768 | 12 | 27126128 | 27167367 | -1 | protein_coding | transmembrane 7 superfamily member 3 |
| **DLX3** | ENSG00000064195 | 73.13266909 | -2.419650774 | 0.532576049 | 5.02E-07 | 3.39E-06 | 1747 | 17 | 48067369 | 48072588 | -1 | protein_coding | distal-less homeobox 3 |
| **SPA17** | ENSG00000064199 | 129.838358 | -0.80897532 | 0.330086248 | 0.005083202 | 0.015490842 | 53340 | 11 | 124543694 | 124567414 | 1 | protein_coding | sperm autoantigenic protein 17 |
| **TSPAN32** | ENSG00000064201 | 171.5046097 | -1.956927616 | 0.315550712 | 5.87E-11 | 6.45E-10 | 10077 | 11 | 2323227 | 2339430 | 1 | protein_coding | tetraspanin 32 |
| **ST3GAL6** | ENSG00000064225 | 1063.207592 | 0.437782978 | 0.16702941 | 0.006269592 | 0.018616314 | 10402 | 3 | 98451080 | 98540045 | 1 | protein_coding | ST3 beta-galactoside alpha-2,3-sialyltransferase 6 |
| **TAF2** | ENSG00000064313 | 988.8174651 | 0.434698544 | 0.151058317 | 0.00297723 | 0.009638002 | 6873 | 8 | 120743015 | 120845103 | -1 | protein_coding | TAF2 RNA polymerase II, TATA box binding protein (TBP)-associated factor, 150kDa |
| **HIPK2** | ENSG00000064393 | 9427.883353 | -1.283778529 | 0.126237352 | 5.07E-25 | 1.52E-23 | 28996 | 7 | 139246316 | 139477577 | -1 | protein_coding | homeodomain interacting protein kinase 2 |
| **CTSA** | ENSG00000064601 | 7952.133086 | -0.362639305 | 0.110984835 | 9.55E-04 | 0.003498436 | 5476 | 20 | 44518783 | 44527459 | 1 | protein_coding | cathepsin A |
| **SUGP2** | ENSG00000064607 | 2011.972642 | -0.515676834 | 0.13881713 | 1.48E-04 | 6.55E-04 | 10147 | 19 | 19101697 | 19144832 | -1 | protein_coding | SURP and G patch domain containing 2 |
| **SNX24** | ENSG00000064652 | 810.9431547 | -0.884841433 | 0.192374592 | 1.51E-06 | 9.44E-06 | 28966 | 5 | 122179134 | 122365049 | 1 | protein_coding | sorting nexin 24 |
| **CNN2** | ENSG00000064666 | 1529.301665 | -1.482823715 | 0.182881931 | 7.45E-17 | 1.29E-15 | 1265 | 19 | 1026298 | 1039068 | 1 | protein_coding | calponin 2 |
| **FAR2** | ENSG00000064763 | 605.4196873 | 0.994912587 | 0.200951987 | 2.01E-07 | 1.44E-06 | 55711 | 12 | 29302036 | 29493913 | 1 | protein_coding | fatty acyl CoA reductase 2 |
| **CHI3L2** | ENSG00000064886 | 296.0181664 | 2.646025294 | 0.337707293 | 2.91E-16 | 4.81E-15 | 1117 | 1 | 111743393 | 111786062 | 1 | protein_coding | chitinase 3-like 2 |
| **ANKS1A** | ENSG00000064999 | 793.2252222 | -0.489830707 | 0.167782859 | 0.002357345 | 0.007811644 | 23294 | 6 | 34857042 | 35059179 | 1 | protein_coding | ankyrin repeat and sterile alpha motif domain containing 1A |
| **ZNF76** | ENSG00000065029 | 565.036775 | -0.548695487 | 0.15888675 | 3.46E-04 | 0.001407628 | 7629 | 6 | 35226686 | 35263762 | 1 | protein_coding | zinc finger protein 76 |
| **UHRF1BP1** | ENSG00000065060 | 3310.290711 | 0.735143612 | 0.088152936 | 3.31E-17 | 5.86E-16 | 54887 | 6 | 34759857 | 34850915 | 1 | protein_coding | UHRF1 binding protein 1 |
| **GNAI3** | ENSG00000065135 | 3002.077365 | 0.590603571 | 0.095865698 | 4.43E-10 | 4.4E-09 | 2773 | 1 | 110091233 | 110136975 | 1 | protein_coding | guanine nucleotide binding protein (G protein), alpha inhibiting activity polypeptide 3 |
| **OAT** | ENSG00000065154 | 1870.334212 | -0.555072274 | 0.189366819 | 0.002022856 | 0.006830923 | 4942 | 10 | 126085872 | 126107545 | -1 | protein_coding | ornithine aminotransferase |
| **PKN2** | ENSG00000065243 | 720.554599 | 1.098557855 | 0.201877554 | 1.33E-08 | 1.11E-07 | 5586 | 1 | 89149905 | 89301938 | 1 | protein_coding | protein kinase N2 |

| | | | | | | | | | | | | | |
|---|---|---|---|---|---|---|---|---|---|---|---|---|---|
| **TRAM2** | ENSG00000065308 | 662.8270247 | -1.099721178 | 0.155897916 | 5E-13 | 6.56E-12 | 9697 | 6 | 52362200 | 52441713 | -1 | protein_coding | translocation associated membrane protein 2 |
| **MCM10** | ENSG00000065328 | 54.26538871 | 1.329110272 | 1.047310819 | 0.014053976 | 0.037550161 | 55388 | 10 | 13203554 | 13253104 | 1 | protein_coding | minichromosome maintenance complex component 10 |
| **DGKA** | ENSG00000065357 | 1267.921782 | -0.494367743 | 0.163688066 | 0.001643496 | 0.005685275 | 1606 | 12 | 56321103 | 56347811 | 1 | protein_coding | diacylglycerol kinase, alpha 80kDa |
| **KARS** | ENSG00000065427 | 3452.958585 | 0.859999664 | 0.116833521 | 8E-14 | 1.12E-12 | 3735 | 16 | 75661622 | 75682541 | -1 | protein_coding | lysyl-tRNA synthetase |
| **TBC1D22B** | ENSG00000065491 | 834.9327071 | 0.57181877 | 0.132322577 | 9.68E-06 | 5.34E-05 | 55633 | 6 | 37225548 | 37300746 | 1 | protein_coding | TBC1 domain family, member 22B |
| **NDUFB4** | ENSG00000065518 | 1144.549311 | 0.342526119 | 0.139810961 | 0.011841463 | 0.032357068 | 4710 | 3 | 120315156 | 120321347 | 1 | protein_coding | NADH dehydrogenase (ubiquinone) 1 beta subcomplex, 4, 15kDa |
| **MYLK** | ENSG00000065534 | 197.6932264 | 0.924908112 | 0.37518812 | 0.003875396 | 0.012159243 | 4638 | 3 | 123328896 | 123603178 | -1 | protein_coding | myosin light chain kinase |
| **ZC3H15** | ENSG00000065548 | 1685.322029 | 0.347351353 | 0.136388064 | 0.009060612 | 0.025690786 | 55854 | 2 | 187350883 | 187374090 | 1 | protein_coding | zinc finger CCCH-type containing 15 |
| **MAP2K4** | ENSG00000065559 | 1306.384022 | 0.415852456 | 0.141795732 | 0.002570729 | 0.008432808 | 6416 | 17 | 11924141 | 12047147 | 1 | protein_coding | mitogen-activated protein kinase kinase 4 |
| **SLK** | ENSG00000065613 | 2432.028196 | 0.502086904 | 0.169637514 | 0.002182762 | 0.007293212 | 9748 | 10 | 105726959 | 105788991 | 1 | protein_coding | STE20-like kinase |
| **CYB5R4** | ENSG00000065615 | 1579.733843 | 0.748973868 | 0.107213843 | 1.33E-12 | 1.67E-11 | 51167 | 6 | 84569362 | 84677143 | 1 | protein_coding | cytochrome b5 reductase 4 |
| **GSTO2** | ENSG00000065621 | 274.7247369 | -0.651461874 | 0.212297235 | 0.001097495 | 0.003969944 | 119391 | 10 | 106028631 | 106064703 | 1 | protein_coding | glutathione S-transferase omega 2 |
| **ASB1** | ENSG00000065802 | 697.5597186 | -0.752187666 | 0.277606191 | 0.002700714 | 0.008815678 | 51665 | 2 | 239335383 | 239360891 | 1 | protein_coding | ankyrin repeat and SOCS box containing 1 |
| **FAM107B** | ENSG00000065809 | 5530.598218 | -0.320521167 | 0.108051157 | 0.002761538 | 0.008988384 | 83641 | 10 | 14560556 | 14816896 | -1 | protein_coding | family with sequence similarity 107, member B |
| **ME1** | ENSG00000065833 | 990.9163972 | -0.643691219 | 0.195792186 | 6.11E-04 | 0.00233582 | 4199 | 6 | 83920108 | 84140797 | -1 | protein_coding | malic enzyme 1, NADP(+)-dependent, cytosolic |
| **TBC1D1** | ENSG00000065882 | 6981.019713 | -0.758349937 | 0.084650175 | 7.79E-20 | 1.7E-18 | 23216 | 4 | 37892708 | 38140796 | 1 | protein_coding | TBC1 (tre-2/USP6, BUB2, cdc16) domain family, member 1 |
| **MTHFD2** | ENSG00000065911 | 7318.335508 | 1.144547191 | 0.097236851 | 9.13E-33 | 4.09E-31 | 10797 | 2 | 74425689 | 74444692 | 1 | protein_coding | methylenetetrahydrofolate dehydrogenase (NADP+ dependent) 2, methenyltetrahydrofolate cyclohydrol |
| **SLC9A7** | ENSG00000065923 | 2556.225887 | -1.753300629 | 0.117848287 | 5.65E-51 | 5.23E-49 | 84679 | X | 46464753 | 46618490 | -1 | protein_coding | solute carrier family 9, subfamily A (NHE7, cation proton antiporter 7), member 7 |
| **YBX1** | ENSG00000065978 | 13070.15858 | -0.5133195 | 0.1327673 | 6.77E-05 | 3.19E-04 | 4904 | 1 | 43148098 | 43168020 | 1 | protein_coding | Y box binding protein 1 |
| **PDE4A** | ENSG00000065989 | 1539.852549 | 0.41414895 | 0.160127535 | 0.00729143 | 0.021229743 | 5141 | 19 | 10527449 | 10580305 | 1 | protein_coding | phosphodiesterase 4A, cAMP-specific |
| **TIE1** | ENSG00000066056 | 974.0727001 | -1.193557756 | 0.188224662 | 4.57E-11 | 5.08E-10 | 7075 | 1 | 43766664 | 43788779 | 1 | protein_coding | tyrosine kinase with immunoglobulin-like and EGF-like domains 1 |
| **DIP2B** | ENSG00000066084 | 2167.692023 | 0.676186154 | 0.11303019 | 1.14E-09 | 1.09E-08 | 57609 | 12 | 50898768 | 51142450 | 1 | protein_coding | DIP2 disco-interacting protein 2 homolog B (Drosophila) |
| **ASPM** | ENSG00000066279 | 161.9292044 | 1.073111973 | 0.458067308 | 0.003730421 | 0.011769618 | 259266 | 1 | 197053258 | 197115824 | -1 | protein_coding | asp (abnormal spindle) homolog, microcephaly associated (Drosophila) |
| **CD84** | ENSG00000066294 | 35664.35634 | -1.0724465 | 0.098234403 | 2.47E-28 | 8.75E-27 | 8832 | 1 | 160510885 | 160549306 | -1 | protein_coding | CD84 molecule |
| **ELOVL1** | ENSG00000066322 | 1701.923563 | 0.29991372 | 0.126381443 | 0.01497151 | 0.039661916 | 1024667 | 1 | 43829068 | 43833696 | -1 | protein_coding | ELOVL fatty acid elongase 1 |
| **SPI1** | ENSG00000066336 | 7442.407188 | -0.517260794 | 0.087898116 | 2.07E-09 | 1.91E-08 | 6688 | 11 | 47376411 | 47400127 | -1 | protein_coding | spleen focus forming virus (SFFV) proviral integration oncogene |
| **LRRC40** | ENSG00000066557 | 345.5411025 | 0.947628972 | 0.213662025 | 2.71E-06 | 1.64E-05 | 55631 | 1 | 70610488 | 70671303 | -1 | protein_coding | leucine rich repeat containing 40 |
| **ISOC1** | ENSG00000066583 | 603.3439286 | 1.151267882 | 0.154135281 | 1.89E-14 | 2.76E-13 | 51015 | 5 | 128430444 | 128449721 | 1 | protein_coding | isochorismatase domain containing 1 |
| **TRMT11** | ENSG00000066651 | 214.2010671 | 1.01010517 | 0.260624137 | 2.82E-05 | 1.43E-04 | 60487 | 6 | 126307576 | 126360422 | 1 | protein_coding | tRNA methyltransferase 11 homolog (S. cerevisiae) |
| **MSANTD3** | ENSG00000066697 | 1551.060517 | 0.751270811 | 0.133838746 | 1.06E-08 | 8.99E-08 | 91283 | 9 | 103189438 | 103213511 | 1 | protein_coding | Myb/SANT-like DNA-binding domain containing 3 |
| **STAG3** | ENSG00000066923 | 292.4716164 | 0.826801045 | 0.254443192 | 4.39E-04 | 0.001741646 | 10734 | 7 | 99775186 | 99819111 | 1 | protein_coding | stromal antigen 3 |
| **FECH** | ENSG00000066926 | 934.8025992 | 0.500870253 | 0.170087543 | 0.002173763 | 0.007271093 | 2235 | 18 | 55215515 | 55254004 | -1 | protein_coding | ferrochelatase |
| **IDI1** | ENSG00000067064 | 2323.817386 | 1.107778823 | 0.162221587 | 1.77E-12 | 2.21E-11 | 3422 | 10 | 1085848 | 1095110 | -1 | protein_coding | isopentenyl-diphosphate delta isomerase 1 |
| **SP100** | ENSG00000067066 | 4317.019783 | 1.557785441 | 0.091223943 | 3.17E-66 | 5.26E-64 | 6672 | 2 | 231280657 | 231408805 | 1 | protein_coding | SP100 nuclear antigen |
| **KLF6** | ENSG00000067082 | 6961.454418 | 1.412593335 | 0.115977394 | 7.13E-35 | 3.54E-33 | 1316 | 10 | 3818188 | 3827473 | -1 | protein_coding | Kruppel-like factor 6 |
| **NEO1** | ENSG00000067141 | 87.57083259 | 0.986641013 | 0.522637109 | 0.011631464 | 0.031885859 | 4756 | 15 | 73344051 | 73597547 | 1 | protein_coding | neogenin 1 |
| **TRAM1** | ENSG00000067167 | 3500.473156 | 1.691689343 | 0.216184192 | 5.15E-16 | 8.39E-15 | 23471 | 8 | 71485677 | 71520622 | -1 | protein_coding | translocation associated membrane protein 1 |
| **PHKA1** | ENSG00000067177 | 468.119605 | -0.876483426 | 0.248656452 | 1.46E-04 | 6.49E-04 | 5255 | X | 71798664 | 71934167 | -1 | protein_coding | phosphorylase kinase, alpha 1 (muscle) |
| **STOML1** | ENSG00000067221 | 98.96440487 | 1.66988384 | 0.356298042 | 3.3E-07 | 2.28E-06 | 9399 | 15 | 74275547 | 74286963 | -1 | protein_coding | stomatin (EPB72)-like 1 |
| **PKM** | ENSG00000067225 | 64512.2446 | -0.207240426 | 0.074194793 | 0.004884319 | 0.014968327 | 5315 | 15 | 72491370 | 72524164 | -1 | protein_coding | pyruvate kinase, muscle |
| **DHX29** | ENSG00000067248 | 1369.102771 | 0.356960975 | 0.130828566 | 0.00526647 | 0.015966566 | 54505 | 5 | 54552073 | 54603550 | -1 | protein_coding | DEAH (Asp-Glu-Ala-His) box polypeptide 29 |
| **DNTTIP2** | ENSG00000067334 | 2200.038339 | 0.903141473 | 0.125472565 | 1.96E-13 | 2.63E-12 | 30836 | 1 | 94333373 | 94345474 | -1 | protein_coding | deoxynucleotidyltransferase, terminal, interacting protein 2 |
| **METTL22** | ENSG00000067365 | 249.4884852 | -0.525165414 | 0.232505584 | 0.014897863 | 0.039487347 | 79091 | 16 | 8715540 | 8740081 | 1 | protein_coding | methyltransferase like 22 |
| **DHX8** | ENSG00000067596 | 2341.317599 | 0.265627169 | 0.112803513 | 0.016614217 | 0.043396868 | 1659 | 17 | 41561233 | 41621831 | 1 | protein_coding | DEAH (Asp-Glu-Ala-His) box polypeptide 8 |
| **SYT1** | ENSG00000067715 | 6.479902203 | -0.472184619 | 0.921138234 | 0.003435473 | 0.010943632 | 6857 | 12 | 79257773 | 79845788 | 1 | protein_coding | synaptotagmin I |
| **IDH3G** | ENSG00000067829 | 855.0498428 | -0.559586001 | 0.13876104 | 3.4E-05 | 1.7E-04 | 3421 | X | 153051221 | 153059978 | -1 | protein_coding | isocitrate dehydrogenase 3 (NAD+) gamma |
| **ROGDI** | ENSG00000067836 | 210.7069732 | -1.074787314 | 0.294241095 | 6.36E-05 | 3.01E-04 | 79641 | 16 | 4846969 | 4852951 | -1 | protein_coding | rogdi homolog (Drosophila) |
| **HDAC4** | ENSG00000068024 | 1012.015491 | -0.989221649 | 0.161121912 | 2.37E-10 | 2.42E-09 | 9759 | 2 | 239969864 | 240323348 | -1 | protein_coding | histone deacetylase 4 |
| **IFI35** | ENSG00000068079 | 1073.739887 | 3.116655763 | 0.235585957 | 4.06E-41 | 2.64E-39 | 3430 | 17 | 41158742 | 41166473 | 1 | protein_coding | interferon-induced protein 35 |
| **HEATR6** | ENSG00000068097 | 714.9040084 | 0.478346738 | 0.155766426 | 0.001483075 | 0.005185502 | 63897 | 17 | 58120555 | 58156292 | -1 | protein_coding | HEAT repeat containing 6 |
| **MEF2A** | ENSG00000068305 | 2126.259056 | 0.455316267 | 0.115946008 | 6.51E-05 | 3.08E-04 | 4205 | 15 | 100017370 | 100256671 | 1 | protein_coding | myocyte enhancer factor 2A |
| **ACSL4** | ENSG00000068366 | 8660.920276 | 0.784252295 | 0.116611168 | 4.61E-12 | 5.6E-11 | 2182 | X | 108867473 | 108976632 | -1 | protein_coding | acyl-CoA synthetase long-chain family member 4 |
| **PRR11** | ENSG00000068489 | 450.7445395 | -0.703912502 | 0.210695587 | 3.85E-04 | 0.001552315 | 55771 | 17 | 57232860 | 57282066 | 1 | protein_coding | proline rich 11 |
| **REEP1** | ENSG00000068615 | 20.80256882 | -1.750617792 | 1.493236024 | 0.013962603 | 0.037351735 | 65055 | 2 | 86441116 | 86565206 | -1 | protein_coding | receptor accessory protein 1 |
| **POLR1A** | ENSG00000068654 | 2093.148171 | -0.407077369 | 0.138188596 | 0.002489842 | 0.008193852 | 25885 | 2 | 86247339 | 86333278 | -1 | protein_coding | polymerase (RNA) I polypeptide A, 194kDa |
| **LAPTM4A** | ENSG00000068697 | 5381.645472 | 0.607544276 | 0.08668081 | 1.46E-12 | 1.83E-11 | 9741 | 2 | 20232411 | 20251789 | -1 | protein_coding | lysosomal protein transmembrane 4 alpha |
| **RASGRP2** | ENSG00000068831 | 87.68386367 | -2.152103009 | 0.493079514 | 1.06E-06 | 6.76E-06 | 10235 | 11 | 64494383 | 64512928 | -1 | protein_coding | RAS guanyl releasing protein 2 (calcium and DAG-regulated) |
| **SIRT2** | ENSG00000068903 | 1008.063562 | -0.678948733 | 0.130471905 | 1.04E-07 | 7.69E-07 | 22933 | 19 | 39369197 | 39390502 | -1 | protein_coding | sirtuin 2 |
| **ERLEC1** | ENSG00000068912 | 1608.337745 | 0.434670286 | 0.107162862 | 3.34E-05 | 1.67E-04 | 27248 | 2 | 54014181 | 54045956 | 1 | protein_coding | endoplasmic reticulum lectin 1 |
| **SDK2** | ENSG00000069188 | 5.886877868 | -0.301944258 | 0.736291629 | 0.017560597 | 0.045534204 | 54549 | 17 | 71330523 | 71640228 | -1 | protein_coding | sidekick cell adhesion molecule 2 |
| **DNAJA2** | ENSG00000069345 | 2326.951766 | 0.30928254 | 0.09510142 | 0.001002649 | 0.003655413 | 10294 | 16 | 46989299 | 47007699 | -1 | protein_coding | DnaJ (Hsp40) homolog, subfamily A, member 2 |
| **KCNAB2** | ENSG00000069424 | 7672.782843 | -1.846001278 | 0.131496736 | 1.11E-45 | 8.8E-44 | 8514 | 1 | 6051526 | 6161253 | 1 | protein_coding | potassium voltage-gated channel, shaker-related subfamily, beta member 2 |
| **CLEC2D** | ENSG00000069493 | 828.2434051 | 1.57347541 | 0.183478466 | 1.34E-18 | 2.66E-17 | 29121 | 12 | 9817565 | 9848413 | 1 | protein_coding | C-type lectin domain family 2, member D |
| **FUNDC1** | ENSG00000069509 | 265.4709035 | 0.567445487 | 0.223028218 | 0.006443266 | 0.019072718 | 139341 | X | 44382885 | 44402247 | -1 | protein_coding | FUN14 domain containing 1 |
| **RORA** | ENSG00000069667 | 681.923989 | 0.679595015 | 0.208372076 | 5.56E-04 | 0.002147885 | 6095 | 15 | 60780483 | 61521518 | -1 | protein_coding | RAR-related orphan receptor A |
| **PIGB** | ENSG00000069943 | 279.7431789 | 1.159580791 | 0.238751765 | 2.66E-07 | 1.86E-06 | 9488 | 15 | 55611158 | 55647846 | 1 | protein_coding | phosphatidylinositol glycan anchor biosynthesis, class B |
| **MAPK6** | ENSG00000069956 | 2424.414116 | 0.399659861 | 0.112534304 | 3.04E-04 | 0.001249682 | 5597 | 15 | 52244303 | 52358462 | 1 | protein_coding | mitogen-activated protein kinase 6 |
| **GNB5** | ENSG00000069966 | 296.0374272 | -0.713498571 | 0.206522537 | 2.62E-04 | 0.001090767 | 10681 | 15 | 52413117 | 52483566 | -1 | protein_coding | guanine nucleotide binding protein (G protein), beta 5 |
| **CECR5** | ENSG00000069998 | 540.2843052 | -1.037092126 | 0.179928386 | 2.37E-09 | 2.17E-08 | 27440 | 22 | 17618401 | 17646177 | -1 | protein_coding | cat eye syndrome chromosome region, candidate 5 |
| **DAPP1** | ENSG00000070190 | 1833.972835 | 1.736057118 | 0.121126859 | 1.76E-47 | 1.43E-45 | 27071 | 4 | 100737990 | 100791311 | 1 | protein_coding | dual adaptor of phosphotyrosine and 3-phosphoinositides |
| **EXOC5** | ENSG00000070367 | 3502.805745 | 0.625067599 | 0.134486736 | 2.1E-06 | 1.29E-05 | 10640 | 14 | 57670518 | 57735726 | -1 | protein_coding | exocyst complex component 5 |
| **ST6GALNAC** | ENSG00000070731 | 221.6700799 | 0.961894563 | 0.251179076 | 3.77E-05 | 1.86E-04 | 10610 | 17 | 74559792 | 74583038 | -1 | protein_coding | ST6 (alpha-N-acetyl-neuraminyl-2,3-beta-galactosyl-1,3)-N-acetylgalactosaminide alpha-2,6-sialyltrans |
| **PABPC1** | ENSG00000070756 | 27323.79369 | -0.548864131 | 0.09654757 | 8.18E-09 | 7.03E-08 | 26986 | 8 | 101698044 | 101735037 | -1 | protein_coding | poly(A) binding protein, cytoplasmic 1 |
| **CDC42** | ENSG00000070831 | 11127.7391 | 0.491839667 | 0.091881474 | 6.09E-08 | 4.67E-07 | 998 | 1 | 22379120 | 22419437 | 1 | protein_coding | cell division cycle 42 |
| **MGAT4A** | ENSG00000071073 | 1162.42606 | -0.654119877 | 0.143704514 | 2.85E-06 | 1.72E-05 | 11320 | 2 | 99235569 | 99347589 | -1 | protein_coding | mannosyl (alpha-1,3-)-glycoprotein beta-1,4-N-acetylglucosaminyltransferase, isozyme A |
| **RPL31** | ENSG00000071082 | 3847.537836 | -0.409311952 | 0.12912147 | 0.001261329 | 0.004485084 | 6160 | 2 | 101618177 | 101640494 | 1 | protein_coding | ribosomal protein L31 |
| **WDR1** | ENSG00000071127 | 11065.69372 | -0.334375776 | 0.094817075 | 3.55E-04 | 0.001438357 | 9948 | 4 | 10075963 | 10118573 | -1 | protein_coding | WD repeat domain 1 |
| **RPS6KA2** | ENSG00000071242 | 4661.922824 | -1.596397814 | 0.140349132 | 8.56E-31 | 3.45E-29 | 6196 | 6 | 166822852 | 167319939 | -1 | protein_coding | ribosomal protein S6 kinase, 90kDa, polypeptide 2 |
| **VASH1** | ENSG00000071246 | 2058.165212 | -1.978939211 | 0.155168806 | 1.87E-38 | 1.07E-36 | 22846 | 14 | 77228532 | 77249354 | 1 | protein_coding | vasohibin 1 |
| **SEL1L** | ENSG00000071537 | 10134.23581 | -0.493856032 | 0.122185586 | 3.5E-05 | 1.74E-04 | 6400 | 14 | 81937893 | 82000205 | -1 | protein_coding | sel-1 suppressor of lin-12-like (C. elegans) |
| **ATP6AP1** | ENSG00000071553 | 166.3035385 | 1.318234821 | 0.649356398 | 0.003677249 | 0.011627115 | 537 | X | 153656978 | 153664862 | 1 | protein_coding | ATPase, H+ transporting, lysosomal accessory protein 1 |
| **TCF3** | ENSG00000071564 | 413.2510266 | -0.907258691 | 0.180846511 | 1.85E-07 | 1.32E-06 | 6929 | 19 | 1609291 | 1652604 | -1 | protein_coding | transcription factor 3 |
| **TRIB2** | ENSG00000071575 | 133.8496373 | -1.707509398 | 0.466149085 | 2.47E-05 | 1.26E-04 | 28951 | 2 | 12857015 | 12882860 | 1 | protein_coding | tribbles pseudokinase 2 |
| **MBD3** | ENSG00000071655 | 824.0859543 | -0.546416344 | 0.164455256 | 5.53E-04 | 0.002138878 | 53615 | 19 | 1576639 | 1592882 | -1 | protein_coding | methyl-CpG binding domain protein 3 |
| **HLTF** | ENSG00000071794 | 654.1092706 | -0.697260621 | 0.154535625 | 3.12E-06 | 1.87E-05 | 6596 | 3 | 148747914 | 148804341 | -1 | protein_coding | helicase-like transcription factor |
| **FAM50A** | ENSG00000071859 | 1363.260275 | -0.271654214 | 0.113085413 | 0.014077029 | 0.037592038 | 9130 | X | 153672473 | 153679002 | 1 | protein_coding | family with sequence similarity 50, member A |
| **FAM3A** | ENSG00000071889 | 371.9193201 | -0.598673181 | 0.222791828 | 0.003991871 | 0.012494365 | 60343 | X | 153733350 | 153744566 | -1 | protein_coding | family with sequence similarity 3, member A |
| **CYBRD1** | ENSG00000071967 | 1747.186024 | -0.902405152 | 0.170419985 | 3.77E-08 | 2.96E-07 | 79901 | 2 | 172378757 | 172414643 | 1 | protein_coding | cytochrome b reductase 1 |
| **RDH11** | ENSG00000072042 | 4455.702044 | 0.237704667 | 0.081271136 | 0.00292569 | 0.009479186 | 51109 | 14 | 68143518 | 68162531 | -1 | protein_coding | retinol dehydrogenase 11 (all-trans/9-cis/11-cis) |

| | | | | | | | | | | | | | |
|---|---|---|---|---|---|---|---|---|---|---|---|---|---|
| **PRKACA** | ENSG00000072062 | 2940.771654 | -0.921880571 | 0.101911193 | 7.44E-20 | 1.63E-18 | 5566 | 19 | 14202500 | 14228896 | -1 | protein_coding | protein kinase, cAMP-dependent, catalytic, alpha |
| **ACTN1** | ENSG00000072110 | 12360.45226 | -0.613365998 | 0.126828272 | 7.1E-07 | 4.69E-06 | 87 | 14 | 69340860 | 69446157 | -1 | protein_coding | actinin, alpha 1 |
| **ZFYVE26** | ENSG00000072121 | 5599.084551 | 0.795686245 | 0.18273619 | 5.6E-06 | 3.22E-05 | 23503 | 14 | 68194091 | 68283307 | -1 | protein_coding | zinc finger, FYVE domain containing 26 |
| **PTPN18** | ENSG00000072135 | 1760.186867 | 0.268098757 | 0.112386572 | 0.01618124 | 0.042454573 | 26469 | 2 | 131113580 | 131132982 | 1 | protein_coding | protein tyrosine phosphatase, non-receptor type 18 (brain-derived) |
| **LIMS2** | ENSG00000072163 | 53.63826643 | -1.288664396 | 0.480739653 | 0.001164509 | 0.004178726 | 55679 | 2 | 128395956 | 128439360 | -1 | protein_coding | LIM and senescent cell antigen-like domains 2 |
| **AFF4** | ENSG00000072364 | 6360.845788 | 0.417905291 | 0.096364455 | 1.32E-05 | 7.12E-05 | 27125 | 5 | 132211071 | 132299326 | -1 | protein_coding | AF4/FMR2 family, member 4 |
| **UBE2D1** | ENSG00000072401 | 1015.990892 | 2.154806153 | 0.228620606 | 3.32E-22 | 8.44E-21 | 7321 | 10 | 60094735 | 60130513 | 1 | protein_coding | ubiquitin-conjugating enzyme E2D 1 |
| **MPP5** | ENSG00000072415 | 635.2542584 | 0.806152535 | 0.173738768 | 1.48E-06 | 9.28E-06 | 64398 | 14 | 67707826 | 67802536 | 1 | protein_coding | membrane protein, palmitoylated 5 (MAGUK p55 subfamily member 5) |
| **RHOBTB1** | ENSG00000072422 | 179.1591306 | -2.040863493 | 0.364665402 | 1.97E-09 | 1.82E-08 | 9886 | 10 | 62629196 | 62761198 | -1 | protein_coding | Rho-related BTB domain containing 1 |
| **MARK2** | ENSG00000072518 | 1334.472113 | 0.549572248 | 0.112826477 | 6.29E-07 | 4.19E-06 | 2011 | 11 | 63606400 | 63678491 | 1 | protein_coding | MAP/microtubule affinity-regulating kinase 2 |
| **HMMR** | ENSG00000072571 | 80.66620901 | 1.404114876 | 0.447805613 | 2.41E-04 | 0.001009407 | 3161 | 5 | 162887209 | 162918947 | 1 | protein_coding | hyaluronan-mediated motility receptor (RHAMM) |
| **P4HA2** | ENSG00000072682 | 669.4346768 | -0.559502597 | 0.152109652 | 1.48E-04 | 6.55E-04 | 1019277 | 5 | 131527531 | 131631008 | -1 | protein_coding | prolyl 4-hydroxylase, alpha polypeptide II |
| **FCGR2B** | ENSG00000072694 | 5142.905018 | -1.468327912 | 0.191431549 | 2.33E-15 | 3.62E-14 | 2213 | 1 | 161551101 | 161648444 | 1 | protein_coding | Fc fragment of IgG, low affinity IIb, receptor (CD32) |
| **NFATC3** | ENSG00000072736 | 3549.294785 | -0.336378313 | 0.085444834 | 7.03E-05 | 3.31E-04 | 4775 | 16 | 68118654 | 68263162 | 1 | protein_coding | nuclear factor of activated T-cells, cytoplasmic, calcineurin-dependent 3 |
| **TRNT1** | ENSG00000072756 | 474.8589861 | 0.635507953 | 0.1619767 | 4.74E-05 | 2.29E-04 | 51095 | 3 | 3168600 | 3192563 | 1 | protein_coding | tRNA nucleotidyl transferase, CCA-adding, 1 |
| **ACADVL** | ENSG00000072778 | 4179.09737 | -0.516026904 | 0.151275522 | 4.17E-04 | 0.001664344 | 37 | 17 | 7120444 | 7128592 | 1 | protein_coding | acyl-CoA dehydrogenase, very long chain |
| **STK10** | ENSG00000072786 | 7584.813559 | -0.915744164 | 0.117073309 | 1.73E-15 | 2.71E-14 | 6793 | 5 | 171469077 | 171615390 | -1 | protein_coding | serine/threonine kinase 10 |
| **DERL2** | ENSG00000072849 | 892.4790099 | 0.486309386 | 0.143848273 | 4.94E-04 | 0.001933535 | 51009 | 17 | 5374571 | 5390131 | -1 | protein_coding | derlin 2 |
| **SCARB1** | ENSG00000073060 | 1963.337015 | -1.798427924 | 0.251136325 | 9.34E-14 | 1.3E-12 | 949 | 12 | 125261402 | 125367214 | -1 | protein_coding | scavenger receptor class B, member 1 |
| **SELO** | ENSG00000073169 | 194.4444156 | 1.035594393 | 0.235965749 | 3E-06 | 1.8E-05 | 83642 | 22 | 50639408 | 50656045 | 1 | protein_coding | Selenoprotein O |
| **TP63** | ENSG00000073282 | 72.72221316 | -1.51422821 | 0.578100016 | 9.21E-04 | 0.003388565 | 8626 | 3 | 189349205 | 189615068 | 1 | protein_coding | tumor protein p63 |
| **CLCN4** | ENSG00000073464 | 32.88874913 | -1.088269184 | 0.632758554 | 0.014303493 | 0.03811023 | 1183 | X | 10125024 | 10205700 | 1 | protein_coding | chloride channel, voltage-sensitive 4 |
| **SMARCE1** | ENSG00000073584 | 1901.350688 | 0.375763309 | 0.122794653 | 0.001927299 | 0.006548447 | 6605 | 17 | 38781214 | 38804760 | -1 | protein_coding | SWI/SNF related, matrix associated, actin dependent regulator of chromatin, subfamily e, member 1 |
| **GSDMB** | ENSG00000073605 | 139.112452 | 1.011727385 | 0.433683882 | 0.004614375 | 0.014220909 | 55876 | 17 | 38060848 | 38076107 | -1 | protein_coding | gasdermin B |
| **KDM5A** | ENSG00000073614 | 1968.570208 | 0.639864996 | 0.179990699 | 1.92E-04 | 8.25E-04 | 5927 | 12 | 389295 | 498620 | -1 | protein_coding | lysine (K)-specific demethylase 5A |
| **PPP2R3A** | ENSG00000073711 | 1286.79392 | 0.418169606 | 0.173583457 | 0.013799085 | 0.036992004 | 5523 | 3 | 135684515 | 135866733 | 1 | protein_coding | protein phosphatase 2, regulatory subunit B'', alpha |
| **DHRS9** | ENSG00000073737 | 1576.547914 | -2.990084154 | 0.189502356 | 1.24E-57 | 1.5E-55 | 10170 | 2 | 169921299 | 169952677 | 1 | protein_coding | dehydrogenase/reductase (SDR family) member 9 |
| **PTGS2** | ENSG00000073756 | 440.1012859 | 3.35295021 | 0.222102836 | 1.19E-52 | 1.2E-50 | 5743 | 1 | 186640923 | 186649559 | -1 | protein_coding | prostaglandin-endoperoxide synthase 2 (prostaglandin G/H synthase and cyclooxygenase) |
| **MAP3K13** | ENSG00000073803 | 465.0336266 | 0.550591787 | 0.159531176 | 3.35E-04 | 0.001367638 | 9175 | 3 | 185000729 | 185206885 | 1 | protein_coding | mitogen-activated protein kinase kinase kinase 13 |
| **ST6GAL1** | ENSG00000073849 | 6799.051822 | -0.661247056 | 0.149058733 | 4.49E-06 | 2.62E-05 | 6480 | 3 | 186648274 | 186796341 | 1 | protein_coding | ST6 beta-galactosamide alpha-2,6-sialyltranferase 1 |
| **TBX21** | ENSG00000073861 | 332.0460528 | 1.880676567 | 0.231339749 | 4.6E-17 | 8.1E-16 | 30009 | 17 | 45810610 | 45823485 | 1 | protein_coding | T-box 21 |
| **PICALM** | ENSG00000073921 | 9788.200053 | 0.507059919 | 0.118367221 | 1.24E-05 | 6.75E-05 | 8301 | 11 | 85668727 | 85780924 | -1 | protein_coding | phosphatidylinositol binding clathrin assembly protein |
| **MRPS34** | ENSG00000074071 | 514.9410278 | -0.812556474 | 0.164125494 | 3.31E-07 | 2.29E-06 | 65993 | 16 | 1821891 | 1823156 | -1 | protein_coding | mitochondrial ribosomal protein S34 |
| **NOTCH3** | ENSG00000074181 | 469.8240038 | -1.115640358 | 0.270269844 | 6.89E-06 | 3.89E-05 | 4854 | 19 | 15270444 | 15311792 | -1 | protein_coding | notch 3 |
| **CLNS1A** | ENSG00000074201 | 1394.077771 | 0.660782926 | 0.151717336 | 6.96E-06 | 3.93E-05 | 1207 | 11 | 77225981 | 77348850 | -1 | protein_coding | chloride channel, nucleotide-sensitive, 1A |
| **EED** | ENSG00000074266 | 865.8261915 | 1.019396882 | 0.157721787 | 3.02E-11 | 3.42E-10 | 8726 | 11 | 85955586 | 85989855 | 1 | protein_coding | embryonic ectoderm development |
| **TSG101** | ENSG00000074319 | 1220.483836 | 0.612893009 | 0.170507522 | 1.87E-04 | 8.09E-04 | 7251 | 11 | 18489883 | 18548779 | -1 | protein_coding | tumor susceptibility 101 |
| **C17orf85** | ENSG00000074356 | 607.9194704 | 0.619763262 | 0.209433785 | 0.00166704 | 0.0057511 | 55421 | 17 | 3714460 | 3749545 | -1 | protein_coding | chromosome 17 open reading frame 85 |
| **ATP2A3** | ENSG00000074370 | 2280.896008 | -1.666173711 | 0.208845576 | 1.84E-16 | 3.09E-15 | 489 | 17 | 3827169 | 3867736 | -1 | protein_coding | ATPase, Ca++ transporting, ubiquitous |
| **SLC24A1** | ENSG00000074621 | 501.2478108 | -0.765131689 | 0.181321802 | 1.05E-05 | 5.77E-05 | 9187 | 15 | 65903704 | 65953333 | 1 | protein_coding | solute carrier family 24 (sodium/potassium/calcium exchanger), member 1 |
| **ZNF532** | ENSG00000074657 | 960.4036064 | 0.457254133 | 0.164814097 | 0.003993744 | 0.012497214 | 55205 | 18 | 56529832 | 56653712 | 1 | protein_coding | zinc finger protein 532 |
| **SCARF1** | ENSG00000074660 | 635.3374608 | 0.752332667 | 0.316113687 | 0.006996686 | 0.020500452 | 8578 | 17 | 1537152 | 1549041 | -1 | protein_coding | scavenger receptor class F, member 1 |
| **IPCEF1** | ENSG00000074706 | 315.510617 | 0.758743519 | 0.233020136 | 4.84E-04 | 0.001901646 | 26034 | 6 | 154475631 | 154677926 | -1 | protein_coding | interaction protein for cytohesin exchange factors 1 |
| **ZZEF1** | ENSG00000074755 | 2995.055337 | -0.513575595 | 0.110389565 | 2.42E-06 | 1.48E-05 | 23140 | 17 | 3907739 | 4046314 | -1 | protein_coding | zinc finger, ZZ-type with EF-hand domain 1 |
| **ENO1** | ENSG00000074800 | 19799.32166 | -0.380136553 | 0.093600696 | 2.1E-05 | 1.09E-04 | 2023 | 1 | 8921061 | 8939308 | -1 | protein_coding | enolase 1, (alpha) |
| **TUBE1** | ENSG00000074935 | 227.3187015 | 0.945750042 | 0.257151505 | 7.18E-05 | 3.37E-04 | 51175 | 6 | 112391980 | 112408732 | -1 | protein_coding | tubulin, epsilon 1 |
| **ARHGEF10L** | ENSG00000074964 | 706.7443009 | -0.511102854 | 0.223404462 | 0.01378636 | 0.036964375 | 55160 | 1 | 17866330 | 18024369 | 1 | protein_coding | Rho guanine nucleotide exchange factor (GEF) 10-like |
| **WSCD2** | ENSG00000075035 | 10.689589 | -3.63419188 | 2.229837103 | 0.003349549 | 0.010704968 | 9671 | 12 | 108523248 | 108644314 | 1 | protein_coding | WSC domain containing 2 |
| **ACTR6** | ENSG00000075089 | 427.850129 | 0.968714998 | 0.208921745 | 1.08E-06 | 6.91E-06 | 64431 | 12 | 100592900 | 100635643 | 1 | protein_coding | ARP6 actin-related protein 6 homolog (yeast) |
| **SRI** | ENSG00000075142 | 1092.627861 | 1.0516832 | 0.167440319 | 8.71E-11 | 9.37E-10 | 6717 | 7 | 87834433 | 87856308 | -1 | protein_coding | sorcin |
| **SEMA3C** | ENSG00000075223 | 2155.061191 | -0.412866153 | 0.109854953 | 1.28E-04 | 5.73E-04 | 10512 | 7 | 80371854 | 80551675 | -1 | protein_coding | sema domain, immunoglobulin domain (Ig), short basic domain, secreted, (semaphorin) 3C |
| **TTC38** | ENSG00000075234 | 447.8086882 | 0.573000782 | 0.197229451 | 0.002155791 | 0.00722521 | 55020 | 22 | 46663858 | 46689905 | 1 | protein_coding | tetratricopeptide repeat domain 38 |
| **GRAMD4** | ENSG00000075240 | 1239.614495 | -1.963172157 | 0.161108149 | 4.81E-35 | 2.43E-33 | 23151 | 22 | 46971909 | 47075688 | 1 | protein_coding | GRAM domain containing 4 |
| **CELSR1** | ENSG00000075275 | 3326.935436 | -1.749022648 | 0.129058864 | 1.23E-42 | 8.57E-41 | 9620 | 22 | 46756731 | 46933067 | -1 | protein_coding | cadherin, EGF LAG seven-pass G-type receptor 1 |
| **ZNF638** | ENSG00000075292 | 3752.461928 | 0.557935476 | 0.132321262 | 1.71E-05 | 9.03E-05 | 1005071 | 2 | 71503691 | 71662199 | 1 | protein_coding | zinc finger protein 638 |
| **SLC25A40** | ENSG00000075303 | 1108.76215 | 0.477655739 | 0.16150622 | 0.002188553 | 0.007310964 | 55972 | 7 | 87462883 | 87505672 | -1 | protein_coding | solute carrier family 25, member 40 |
| **TIMM21** | ENSG00000075336 | 402.2276213 | -1.276993922 | 0.201474578 | 4.32E-11 | 4.82E-10 | 29090 | 18 | 71815746 | 71826197 | 1 | protein_coding | translocase of inner mitochondrial membrane 21 homolog (yeast) |
| **ADD2** | ENSG00000075340 | 432.308214 | -2.285702745 | 0.250673376 | 5.2E-21 | 1.23E-19 | 119 | 2 | 70834750 | 70995357 | -1 | protein_coding | adducin 2 (beta) |
| **RASAL2** | ENSG00000075391 | 4073.900784 | -0.300184708 | 0.095415169 | 0.001653726 | 0.005710624 | 9462 | 1 | 178062864 | 178448644 | 1 | protein_coding | RAS protein activator like 2 |
| **ZNF37A** | ENSG00000075407 | 964.5856758 | 0.596522152 | 0.164447823 | 1.67E-04 | 7.29E-04 | 1019297 | 10 | 38383264 | 38414472 | 1 | protein_coding | zinc finger protein 37A |
| **MARK3** | ENSG00000075413 | 2401.983094 | 0.738780201 | 0.101367078 | 1.74E-13 | 2.35E-12 | 4140 | 14 | 103851729 | 103970168 | 1 | protein_coding | MAP/microtubule affinity-regulating kinase 3 |
| **SLC25A3** | ENSG00000075415 | 5121.771381 | -0.359431507 | 0.122300992 | 0.002543997 | 0.008357676 | 5250 | 12 | 98987369 | 98995946 | 1 | protein_coding | solute carrier family 25 (mitochondrial carrier; phosphate carrier), member 3 |
| **FRYL** | ENSG00000075539 | 1123.039979 | -0.802170764 | 0.242723962 | 3.58E-04 | 0.001451475 | 285527 | 4 | 48499378 | 48782339 | -1 | protein_coding | FRY-like |
| **TMEM131** | ENSG00000075568 | 2834.733838 | 0.708961356 | 0.092962858 | 9.71E-15 | 1.45E-13 | 23505 | 2 | 98372799 | 98612388 | -1 | protein_coding | transmembrane protein 131 |
| **FSCN1** | ENSG00000075618 | 635.5843326 | -0.572802904 | 0.217659043 | 0.004968194 | 0.015185783 | 6624 | 7 | 5632439 | 5646286 | 1 | protein_coding | fascin homolog 1, actin-bundling protein (Strongylocentrotus purpuratus) |
| **ACTB** | ENSG00000075624 | 2994.483306 | 2.185059554 | 0.810430551 | 4.55E-04 | 0.001798128 | 60 | 7 | 5566782 | 5603415 | -1 | protein_coding | actin, beta |
| **MOCOS** | ENSG00000075643 | 70.27978011 | 1.51926427 | 0.491135204 | 2.31E-04 | 9.75E-04 | 55034 | 18 | 33767482 | 33852120 | 1 | protein_coding | molybdenum cofactor sulfurase |
| **PLD1** | ENSG00000075651 | 1735.406073 | 1.691067381 | 0.213771052 | 3.19E-16 | 5.25E-15 | 5337 | 3 | 171318195 | 171528740 | -1 | protein_coding | phospholipase D1, phosphatidylcholine-specific |
| **RAB7A** | ENSG00000075785 | 12794.54418 | -0.305101012 | 0.134564482 | 0.018057918 | 0.046540017 | 7879 | 3 | 128444965 | 128533639 | 1 | protein_coding | RAB7A, member RAS oncogene family |
| **BCAP29** | ENSG00000075790 | 1190.690715 | 0.914915381 | 0.169283797 | 2.25E-08 | 1.82E-07 | 55973 | 7 | 107220422 | 107269615 | 1 | protein_coding | B-cell receptor-associated protein 29 |
| **KIFAP3** | ENSG00000075945 | 595.8157372 | 0.913113676 | 0.143029803 | 6.3E-11 | 6.88E-10 | 22920 | 1 | 169890467 | 170054349 | -1 | protein_coding | kinesin-associated protein 3 |
| **RBM7** | ENSG00000076053 | 731.878514 | 0.83520948 | 0.179443545 | 1.19E-06 | 7.56E-06 | 10179 | 11 | 114270752 | 114284925 | 1 | protein_coding | RNA binding motif protein 7 |
| **RBMS2** | ENSG00000076067 | 692.6090935 | 0.77976406 | 0.139650798 | 9.25E-09 | 7.9E-08 | 5939 | 12 | 56915713 | 56984745 | 1 | protein_coding | RNA binding motif, single stranded interacting protein 2 |
| **BAZ2A** | ENSG00000076108 | 4220.938175 | 1.073769978 | 0.119851282 | 9.59E-20 | 2.08E-18 | 11176 | 12 | 56989380 | 57030600 | -1 | protein_coding | bromodomain adjacent to zinc finger domain, 2A |
| **SPAG5** | ENSG00000076382 | 149.9945031 | 0.748060167 | 0.342799301 | 0.011435403 | 0.03142166 | 10615 | 17 | 26904588 | 26926297 | -1 | protein_coding | sperm associated antigen 5 |
| **ANKRD13A** | ENSG00000076513 | 2499.768257 | 0.813413639 | 0.128416652 | 7.57E-11 | 8.23E-10 | 88455 | 12 | 110436991 | 110477568 | 1 | protein_coding | ankyrin repeat domain 13A |
| **NT5C2** | ENSG00000076685 | 3928.707447 | 0.821718115 | 0.089789336 | 5.21E-20 | 1.15E-18 | 22978 | 10 | 104845940 | 104953056 | -1 | protein_coding | 5'-nucleotidase, cytosolic II |
| **GPC4** | ENSG00000076716 | 10953.82578 | -0.65449816 | 0.200009968 | 5.71E-04 | 0.002198845 | 2239 | X | 132434131 | 132549518 | -1 | protein_coding | glypican 4 |
| **MBNL3** | ENSG00000076770 | 704.0159201 | -1.07444841 | 0.233860961 | 1.01E-06 | 6.48E-06 | 55796 | X | 131503345 | 131623996 | -1 | protein_coding | muscleblind-like splicing regulator 3 |
| **RAP1GAP** | ENSG00000076864 | 496.6585258 | -1.856983451 | 0.273899313 | 1.29E-12 | 1.63E-11 | 5909 | 1 | 21922708 | 21995801 | -1 | protein_coding | RAP1 GTPase activating protein |
| **STXBP2** | ENSG00000076944 | 2914.227637 | -0.414741475 | 0.100893757 | 4.03E-05 | 1.98E-04 | 6813 | 19 | 7701767 | 7712759 | 1 | protein_coding | syntaxin binding protein 2 |
| **TM9SF3** | ENSG00000077147 | 5758.252891 | 1.047257819 | 0.181480787 | 2.02E-09 | 1.87E-08 | 56889 | 10 | 98277866 | 98347209 | -1 | protein_coding | transmembrane 9 superfamily member 3 |
| **NFKB2** | ENSG00000077150 | 2508.214172 | 1.460696004 | 0.129179603 | 2.25E-30 | 8.89E-29 | 4791 | 10 | 104153867 | 104162281 | 1 | protein_coding | nuclear factor of kappa light polypeptide gene enhancer in B-cells 2 (p49/p100) |
| **DNAJC10** | ENSG00000077232 | 4048.839571 | -0.457242734 | 0.112626631 | 3.49E-05 | 1.74E-04 | 54431 | 2 | 183580999 | 183659191 | 1 | protein_coding | DnaJ (Hsp40) homolog, subfamily C, member 10 |
| **GTF3C1** | ENSG00000077235 | 2387.06784 | -0.428991612 | 0.133600391 | 0.001025612 | 0.003733784 | 2975 | 16 | 27470876 | 27561234 | -1 | protein_coding | general transcription factor IIIC, polypeptide 1, alpha 220kDa |
| **USP33** | ENSG00000077254 | 2290.335346 | 1.370872107 | 0.116424165 | 9.12E-33 | 4.09E-31 | 23032 | 1 | 78161672 | 78225537 | -1 | protein_coding | ubiquitin specific peptidase 33 |
| **EXOSC5** | ENSG00000077348 | 108.7354636 | -1.30986156 | 0.355775255 | 3.99E-05 | 1.96E-04 | 56915 | 19 | 41892279 | 41903384 | -1 | protein_coding | exosome component 5 |
| **APBB1IP** | ENSG00000077420 | 4138.853196 | -1.50842591 | 0.102306175 | 5.79E-50 | 5.08E-48 | 54518 | 10 | 26727132 | 26856732 | 1 | protein_coding | amyloid beta (A4) precursor protein-binding, family B, member 1 interacting protein |
| **FAM76B** | ENSG00000077458 | 284.6221903 | 0.867887596 | 0.244401081 | 1.41E-04 | 6.27E-04 | 143684 | 11 | 95502106 | 95523573 | -1 | protein_coding | family with sequence similarity 76, member B |

| | | | | | | | | | | | | | |
|---|---|---|---|---|---|---|---|---|---|---|---|---|---|
| **POLD3** | ENSG00000077514 | 532.6097209 | 0.561104759 | 0.18791119 | 0.0017233 | 0.005914484 | 10714 | 11 | 74204896 | 74380162 | 1 | protein_coding | polymerase (DNA-directed), delta 3, accessory subunit |
| **CAPZB** | ENSG00000077549 | 3119.415083 | -0.950681331 | 0.111804063 | 9.1E-18 | 1.69E-16 | 832 | 1 | 19665267 | 19812066 | -1 | protein_coding | capping protein (actin filament) muscle Z-line, beta |
| **UBE2A** | ENSG00000077721 | 1592.673697 | 0.761689934 | 0.121596994 | 1.81E-10 | 1.88E-09 | 7319 | X | 118708501 | 118718381 | 1 | protein_coding | ubiquitin-conjugating enzyme E2A |
| **FGFR1** | ENSG00000077782 | 5146.729341 | -2.314627658 | 0.140020274 | 1.53E-62 | 2.16E-60 | 2260 | 8 | 38268656 | 38326352 | -1 | protein_coding | fibroblast growth factor receptor 1 |
| **PIAS2** | ENSG00000078043 | 679.47233 | -0.481701464 | 0.17754462 | 0.004592614 | 0.014162415 | 9063 | 18 | 44388353 | 44500123 | -1 | protein_coding | protein inhibitor of activated STAT, 2 |
| **LAMP3** | ENSG00000078081 | 634.5023891 | 4.789022848 | 0.289308582 | 8.33E-63 | 1.21E-60 | 27074 | 3 | 182840001 | 182881627 | -1 | protein_coding | lysosomal-associated membrane protein 3 |
| **FAP** | ENSG00000078098 | 5.821886618 | 5.430703857 | 3.03662363 | 8.5E-04 | 0.003160957 | 2191 | 2 | 163027194 | 163101661 | -1 | protein_coding | fibroblast activation protein, alpha |
| **N4BP2** | ENSG00000078177 | 934.3366235 | 0.4511328 | 0.132377983 | 4.78E-04 | 0.001881463 | 55728 | 4 | 40058446 | 40159872 | 1 | protein_coding | NEDD4 binding protein 2 |
| **SYNJ2** | ENSG00000078269 | 491.407671 | -0.672641363 | 0.260487987 | 0.004505882 | 0.013917421 | 8871 | 6 | 158402888 | 158520208 | 1 | protein_coding | synaptojanin 2 |
| **P2RY10** | ENSG00000078589 | 241.1564171 | 0.766256718 | 0.262983431 | 0.001484155 | 0.005188093 | 27334 | X | 78200829 | 78217451 | 1 | protein_coding | purinergic receptor P2Y, G-protein coupled, 10 |
| **ITM2A** | ENSG00000078596 | 593.8570645 | -0.936640797 | 0.294449462 | 3.95E-04 | 0.001587771 | 9452 | X | 78615881 | 78623164 | -1 | protein_coding | integral membrane protein 2A |
| **PCM1** | ENSG00000078674 | 4866.236173 | -0.38020231 | 0.108413367 | 4.32E-04 | 0.001716873 | 5108 | 8 | 17780349 | 17885478 | 1 | protein_coding | pericentriolar material 1 |
| **TP53INP2** | ENSG00000078804 | 1689.56735 | 1.399850389 | 0.159474177 | 2.98E-19 | 6.21E-18 | 58476 | 20 | 33292094 | 33301243 | 1 | protein_coding | tumor protein p53 inducible nuclear protein 2 |
| **TOLLIP** | ENSG00000078902 | 2336.892969 | -0.453577742 | 0.163310476 | 0.00388913 | 0.01219983 | 54472 | 11 | 1295601 | 1330884 | -1 | protein_coding | toll interacting protein |
| **UBE2D4** | ENSG00000078967 | 865.6794357 | -0.574848887 | 0.186175784 | 0.001191809 | 0.004265672 | 51619 | 7 | 43966037 | 43995735 | 1 | protein_coding | ubiquitin-conjugating enzyme E2D 4 (putative) |
| **SLC1A3** | ENSG00000079215 | 7569.411883 | 0.749300909 | 0.121456412 | 3.17E-10 | 3.2E-09 | 6507 | 5 | 36606457 | 36688436 | 1 | protein_coding | solute carrier family 1 (glial high affinity glutamate transporter), member 3 |
| **SP140** | ENSG00000079263 | 1536.049229 | 1.841113599 | 0.223586464 | 2.15E-17 | 3.89E-16 | 1019287 | 2 | 231067826 | 231223762 | 1 | protein_coding | SP140 nuclear body protein |
| **MKNK1** | ENSG00000079277 | 1999.881482 | -0.695862175 | 0.101699305 | 3.77E-12 | 4.61E-11 | 8569 | 1 | 47023090 | 47082515 | -1 | protein_coding | MAP kinase interacting serine/threonine kinase 1 |
| **TNS1** | ENSG00000079308 | 6943.958723 | -0.640787883 | 0.117226273 | 2.5E-08 | 2.01E-07 | 7145 | 2 | 218664512 | 218867718 | -1 | protein_coding | tensin 1 |
| **SAR1A** | ENSG00000079332 | 2949.208575 | 0.529298187 | 0.109653359 | 7.89E-07 | 5.18E-06 | 56681 | 10 | 71907045 | 71930279 | -1 | protein_coding | SAR1 homolog A (S. cerevisiae) |
| **RAPGEF3** | ENSG00000079337 | 462.6705857 | -1.294823547 | 0.33134881 | 1.51E-05 | 8.08E-05 | 10411 | 12 | 48128455 | 48164823 | -1 | protein_coding | Rap guanine nucleotide exchange factor (GEF) 3 |
| **CEACAM1** | ENSG00000079385 | 120.4332422 | 2.540237101 | 0.362431428 | 1.85E-13 | 2.5E-12 | 634 | 19 | 43011456 | 43065386 | -1 | protein_coding | carcinoembryonic antigen-related cell adhesion molecule 1 (biliary glycoprotein) |
| **SENP1** | ENSG00000079387 | 233.951246 | 0.927455365 | 0.35181481 | 0.002247751 | 0.007487441 | 29843 | 12 | 48436681 | 48500091 | -1 | protein_coding | SUMO1/sentrin specific peptidase 1 |
| **CIC** | ENSG00000079432 | 1826.239408 | -0.439517787 | 0.153851082 | 0.003078761 | 0.009939313 | 23152 | 19 | 42772689 | 42799949 | 1 | protein_coding | capicua transcriptional repressor |
| **PGM1** | ENSG00000079739 | 906.587463 | -0.784873222 | 0.153115413 | 1.26E-07 | 9.21E-07 | 5236 | 1 | 64058947 | 64125916 | 1 | protein_coding | phosphoglucomutase 1 |
| **DDX1** | ENSG00000079785 | 1906.398795 | 0.338889976 | 0.106872302 | 0.001273228 | 0.004521084 | 1653 | 2 | 15731302 | 15771235 | 1 | protein_coding | DEAD (Asp-Glu-Ala-Asp) box helicase 1 |
| **EPB41L2** | ENSG00000079819 | 4336.806443 | -0.464207467 | 0.136380045 | 4.57E-04 | 0.001803737 | 2037 | 6 | 131160487 | 131384462 | -1 | protein_coding | erythrocyte membrane protein band 4.1-like 2 |
| **KEAP1** | ENSG00000079999 | 1176.292787 | 0.70615296 | 0.141660556 | 3.06E-07 | 2.13E-06 | 9817 | 19 | 10596796 | 10614417 | -1 | protein_coding | kelch-like ECH-associated protein 1 |
| **RIF1** | ENSG00000080345 | 2548.2115 | 0.746380153 | 0.110788031 | 8.64E-12 | 1.02E-10 | 1019293 | 2 | 152266397 | 152364527 | 1 | protein_coding | RAP1 interacting factor homolog (yeast) |
| **RAB21** | ENSG00000080371 | 3091.350031 | 1.035768584 | 0.103045383 | 2.57E-24 | 7.42E-23 | 23011 | 12 | 72148654 | 72184699 | 1 | protein_coding | RAB21, member RAS oncogene family |
| **SMARCA2** | ENSG00000080503 | 3173.053289 | -0.419217204 | 0.099720168 | 2.47E-05 | 1.27E-04 | 6595 | 9 | 2015342 | 2193624 | 1 | protein_coding | SWI/SNF related, matrix associated, actin dependent regulator of chromatin, subfamily a, member 2 |
| **COL5A3** | ENSG00000080573 | 86.30098808 | -0.95686771 | 0.417436593 | 0.005767979 | 0.01726778 | 50509 | 19 | 10070237 | 10121147 | -1 | protein_coding | collagen, type V, alpha 3 |
| **CNOT4** | ENSG00000080802 | 1035.0741 | 0.375437414 | 0.130944258 | 0.003277691 | 0.010499465 | 4850 | 7 | 135046547 | 135194875 | -1 | protein_coding | CCR4-NOT transcription complex, subunit 4 |
| **PSEN1** | ENSG00000080815 | 5237.597857 | 0.891915406 | 0.094590032 | 1.56E-21 | 3.83E-20 | 5663 | 14 | 73603126 | 73690399 | 1 | protein_coding | presenilin 1 |
| **CLDND1** | ENSG00000080822 | 1369.568752 | 0.831242579 | 0.180276974 | 1.53E-06 | 9.52E-06 | 56650 | 3 | 98216756 | 98241910 | -1 | protein_coding | claudin domain containing 1 |
| **HSP90AA1** | ENSG00000080824 | 39872.02269 | 0.31143907 | 0.108749353 | 0.003582295 | 0.011356195 | 3320 | 14 | 102547075 | 102606036 | -1 | protein_coding | heat shock protein 90kDa alpha (cytosolic), class A member 1 |
| **NDC80** | ENSG00000080986 | 77.81199513 | 1.860241722 | 0.460187425 | 4.81E-06 | 2.79E-05 | 10403 | 18 | 2571510 | 2616634 | 1 | protein_coding | NDC80 kinetochore complex component |
| **AP4E1** | ENSG00000081014 | 1305.562778 | 0.657437837 | 0.14944155 | 5.78E-06 | 3.31E-05 | 23431 | 15 | 51200869 | 51298097 | 1 | protein_coding | adaptor-related protein complex 4, epsilon 1 subunit |
| **CXCL2** | ENSG00000081041 | 163.133759 | 3.298128325 | 0.364891901 | 8.09E-21 | 1.9E-19 | 2920 | 4 | 74962752 | 74965010 | -1 | protein_coding | chemokine (C-X-C motif) ligand 2 |
| **OSTM1** | ENSG00000081087 | 2774.208918 | 0.547931846 | 0.224568933 | 0.008881898 | 0.025240248 | 28962 | 6 | 108362613 | 108487058 | -1 | protein_coding | osteopetrosis associated transmembrane protein 1 |
| **MEF2C** | ENSG00000081189 | 711.8431818 | -0.918461918 | 0.199447966 | 1.38E-06 | 8.65E-06 | 4208 | 5 | 88013975 | 88199922 | -1 | protein_coding | myocyte enhancer factor 2C |
| **STK17B** | ENSG00000081320 | 1539.419666 | 0.78223286 | 0.207715336 | 6.08E-05 | 2.89E-04 | 9262 | 2 | 196998290 | 197041227 | -1 | protein_coding | serine/threonine kinase 17b |
| **KIAA0141** | ENSG00000081791 | 786.5471781 | -0.40966747 | 0.139514869 | 0.002614271 | 0.008554181 | 9812 | 5 | 141303373 | 141321612 | 1 | protein_coding | KIAA0141 |
| **HSPB11** | ENSG00000081870 | 279.1033739 | 0.673563459 | 0.230914856 | 0.001719995 | 0.005904465 | 51668 | 1 | 54382247 | 54411602 | -1 | protein_coding | heat shock protein family B (small), member 11 |
| **IL12RB2** | ENSG00000081985 | 240.6979373 | 2.757913524 | 0.333954047 | 1.04E-17 | 1.92E-16 | 3595 | 1 | 67773047 | 67862583 | 1 | protein_coding | interleukin 12 receptor, beta 2 |
| **FYB** | ENSG00000082074 | 9136.58273 | -0.305118035 | 0.128290222 | 0.017819765 | 0.046034394 | 2533 | 5 | 39105338 | 39274630 | -1 | protein_coding | FYN binding protein |
| **C5orf22** | ENSG00000082213 | 839.7080416 | 0.875193056 | 0.138119935 | 8.48E-11 | 9.13E-10 | 55322 | 5 | 31532373 | 31555165 | 1 | protein_coding | chromosome 5 open reading frame 22 |
| **CCNT2** | ENSG00000082258 | 1049.956391 | 0.573369216 | 0.142394443 | 3.45E-05 | 1.72E-04 | 905 | 2 | 135675805 | 135716912 | 1 | protein_coding | cyclin T2 |
| **FAM135A** | ENSG00000082269 | 704.4530443 | 1.264831377 | 0.21992248 | 1.74E-09 | 1.63E-08 | 57579 | 6 | 71122644 | 71270877 | 1 | protein_coding | family with sequence similarity 135, member A |
| **DLG3** | ENSG00000082458 | 100.9334298 | 0.992441243 | 0.611100363 | 0.018884614 | 0.048327789 | 1741 | X | 69664711 | 69725337 | 1 | protein_coding | discs, large homolog 3 (Drosophila) |
| **NFE2L1** | ENSG00000082641 | 10516.33548 | 0.400078537 | 0.089575124 | 5.39E-06 | 3.1E-05 | 4779 | 17 | 46125691 | 46138849 | 1 | protein_coding | nuclear factor, erythroid 2-like 1 |
| **ITGB5** | ENSG00000082781 | 8384.265003 | -0.998028859 | 0.15654678 | 5.39E-11 | 5.96E-10 | 3693 | 3 | 124480795 | 124620265 | -1 | protein_coding | integrin, beta 5 |
| **XPO1** | ENSG00000082898 | 5133.605069 | 0.855007921 | 0.088842169 | 2.44E-22 | 6.25E-21 | 7514 | 2 | 61704984 | 61765761 | -1 | protein_coding | exportin 1 (CRM1 homolog, yeast) |
| **RNF13** | ENSG00000082996 | 9178.324449 | 0.632595745 | 0.141896498 | 4.62E-06 | 2.69E-05 | 11342 | 3 | 149530495 | 149679926 | 1 | protein_coding | ring finger protein 13 |
| **DOPEY1** | ENSG00000083097 | 936.0809256 | 0.850320559 | 0.162358753 | 6.37E-08 | 4.87E-07 | 23033 | 6 | 83777385 | 83881069 | 1 | protein_coding | dopey family member 1 |
| **LYRM2** | ENSG00000083099 | 2036.093712 | 0.353733434 | 0.151504524 | 0.01586325 | 0.041724018 | 57226 | 6 | 90277863 | 90348472 | -1 | protein_coding | LYR motif containing 2 |
| **BCKDHB** | ENSG00000083123 | 206.7409097 | -0.879119217 | 0.238573991 | 7.69E-05 | 3.59E-04 | 594 | 6 | 80816364 | 81055987 | 1 | protein_coding | branched chain keto acid dehydrogenase E1, beta polypeptide |
| **KAT6A** | ENSG00000083168 | 2428.363626 | 0.529885966 | 0.236546706 | 0.015324948 | 0.040520965 | 7994 | 8 | 41786997 | 41909508 | -1 | protein_coding | K(lysine) acetyltransferase 6A |
| **ZCCHC6** | ENSG00000083223 | 2108.366876 | 0.306155865 | 0.119455574 | 0.009385049 | 0.026517232 | 79670 | 9 | 88902648 | 88969369 | -1 | protein_coding | zinc finger, CCHC domain containing 6 |
| **ULK2** | ENSG00000083290 | 811.8245508 | 1.460294067 | 0.254535437 | 1.41E-09 | 1.33E-08 | 9706 | 17 | 19674142 | 19771249 | -1 | protein_coding | unc-51 like autophagy activating kinase 2 |
| **TNPO1** | ENSG00000083312 | 3669.372583 | 0.481149721 | 0.127827971 | 1.18E-04 | 5.33E-04 | 3842 | 5 | 72112139 | 72212560 | 1 | protein_coding | transportin 1 |
| **PLOD1** | ENSG00000083444 | 2862.700894 | -1.136298572 | 0.146378179 | 1.41E-15 | 2.23E-14 | 5351 | 1 | 11994262 | 12035595 | 1 | protein_coding | procollagen-lysine, 2-oxoglutarate 5-dioxygenase 1 |
| **P2RX5** | ENSG00000083454 | 128.868557 | 0.857039434 | 0.345139744 | 0.004296099 | 0.013334136 | 5026 | 17 | 3575493 | 3599698 | -1 | protein_coding | purinergic receptor P2X, ligand-gated ion channel, 5 |
| **ITGAE** | ENSG00000083457 | 1860.496523 | -0.338846632 | 0.120728892 | 0.004106252 | 0.012794261 | 3682 | 17 | 3617922 | 3704537 | -1 | protein_coding | integrin, alpha E (antigen CD103, human mucosal lymphocyte antigen 1; alpha polypeptide) |
| **DIS3** | ENSG00000083520 | 1642.104068 | 0.303522567 | 0.104784862 | 0.003182698 | 0.010235939 | 22894 | 13 | 73329540 | 73356234 | -1 | protein_coding | DIS3 mitotic control homolog (S. cerevisiae) |
| **TDRD3** | ENSG00000083544 | 3611.62149 | -0.809744287 | 0.237238136 | 2.42E-04 | 0.001013882 | 81550 | 13 | 60970591 | 61148012 | 1 | protein_coding | tudor domain containing 3 |
| **PDS5B** | ENSG00000083642 | 1174.99775 | 0.397140371 | 0.144487716 | 0.004695837 | 0.014443939 | 23047 | 13 | 33160564 | 33352157 | 1 | protein_coding | PDS5, regulator of cohesion maintenance, homolog B (S. cerevisiae) |
| **OXCT1** | ENSG00000083720 | 233.8362322 | 1.498486015 | 0.341357766 | 1.49E-06 | 9.3E-06 | 5019 | 5 | 41730167 | 41870621 | -1 | protein_coding | 3-oxoacid CoA transferase 1 |
| **CYLD** | ENSG00000083799 | 3550.366871 | 1.069301855 | 0.1120164 | 3.74E-22 | 9.46E-21 | 1540 | 16 | 50775961 | 50835846 | 1 | protein_coding | cylindromatosis (turban tumor syndrome) |
| **ZNF671** | ENSG00000083814 | 215.3245002 | -0.529927737 | 0.227172001 | 0.012204314 | 0.033206076 | 79891 | 19 | 58231119 | 58238992 | -1 | protein_coding | zinc finger protein 671 |
| **ZNF416** | ENSG00000083817 | 163.0632556 | -0.737371366 | 0.268287625 | 0.002571722 | 0.008434254 | 55659 | 19 | 58082934 | 58090295 | -1 | protein_coding | zinc finger protein 416 |
| **ZNF586** | ENSG00000083828 | 352.427747 | 0.922430476 | 0.197546666 | 1.01E-06 | 6.5E-06 | 54807 | 19 | 58281023 | 58331307 | 1 | protein_coding | zinc finger protein 586 |
| **RPS5** | ENSG00000083845 | 1531.236213 | -0.718235867 | 0.136865556 | 7.12E-08 | 5.4E-07 | 6193 | 19 | 58897767 | 58906173 | 1 | protein_coding | ribosomal protein S5 |
| **FAT1** | ENSG00000083857 | 127.4431215 | -2.850445773 | 0.512547541 | 2.15E-09 | 1.98E-08 | 2195 | 4 | 187508937 | 187647876 | -1 | protein_coding | FAT atypical cadherin 1 |
| **YTHDC1** | ENSG00000083896 | 1605.802213 | 0.446886471 | 0.101807426 | 8.46E-06 | 4.71E-05 | 91746 | 4 | 69176105 | 69215807 | -1 | protein_coding | YTH domain containing 1 |
| **SMAP2** | ENSG00000084070 | 3316.37607 | 0.808113248 | 0.101511596 | 8.99E-16 | 1.45E-14 | 64744 | 1 | 40810522 | 40888998 | 1 | protein_coding | small ArfGAP2 |
| **ZMPSTE24** | ENSG00000084073 | 2711.956438 | 0.718116723 | 0.127798665 | 9.41E-09 | 8.03E-08 | 10269 | 1 | 40723779 | 40759856 | 1 | protein_coding | zinc metallopeptidase STE24 |
| **HAL** | ENSG00000084110 | 35.12053337 | -1.464138842 | 0.64204089 | 0.002355888 | 0.007810204 | 3034 | 12 | 96366440 | 96390143 | -1 | protein_coding | histidine ammonia-lyase |
| **SSH1** | ENSG00000084112 | 3589.548928 | -0.268086184 | 0.103761486 | 0.008293697 | 0.023751033 | 54434 | 12 | 109176466 | 109251366 | -1 | protein_coding | slingshot protein phosphatase 1 |
| **GSTP1** | ENSG00000084207 | 226.6258428 | 0.554353958 | 0.232406446 | 0.010181775 | 0.028426758 | 2950 | 11 | 67351066 | 67354131 | 1 | protein_coding | glutathione S-transferase pi 1 |
| **APLP2** | ENSG00000084234 | 59424.12755 | -0.354666199 | 0.103814865 | 5.18E-04 | 0.00201888 | 334 | 11 | 129939732 | 130014699 | 1 | protein_coding | amyloid beta (A4) precursor-like protein 2 |
| **WBP11** | ENSG00000084463 | 2419.375171 | 0.400580328 | 0.13147351 | 0.001814903 | 0.006198248 | 51729 | 12 | 14939410 | 14956474 | -1 | protein_coding | WW domain binding protein 11 |
| **NCOA1** | ENSG00000084676 | 1589.244694 | 0.478962955 | 0.123208226 | 8.01E-05 | 3.73E-04 | 8648 | 2 | 24714783 | 24993571 | 1 | protein_coding | nuclear receptor coactivator 1 |
| **AGBL5** | ENSG00000084693 | 146.6298283 | -0.782105874 | 0.274971678 | 0.001747624 | 0.005987208 | 60509 | 2 | 27265232 | 27293490 | 1 | protein_coding | ATP/GTP binding protein-like 5 |
| **EFR3B** | ENSG00000084710 | 563.343128 | -2.780566276 | 0.19912134 | 2.49E-45 | 1.94E-43 | 22979 | 2 | 25264999 | 25378243 | 1 | protein_coding | EFR3 homolog B (S. cerevisiae) |
| **KIF3C** | ENSG00000084731 | 250.832317 | -0.71416615 | 0.353047318 | 0.017664416 | 0.045725824 | 3797 | 2 | 26149471 | 26205618 | -1 | protein_coding | kinesin family member 3C |
| **RAB10** | ENSG00000084733 | 6323.316626 | 0.5309659 | 0.125722206 | 1.6E-05 | 8.52E-05 | 10890 | 2 | 26256976 | 26360323 | 1 | protein_coding | RAB10, member RAS oncogene family |
| **HADHA** | ENSG00000084754 | 6633.836097 | -0.691291311 | 0.114262592 | 7.18E-10 | 7E-09 | 3030 | 2 | 26413504 | 26467594 | -1 | protein_coding | hydroxyacyl-CoA dehydrogenase/3-ketoacyl-CoA thiolase/enoyl-CoA hydratase (trifunctional protein), a |
| **MAPRE3** | ENSG00000084764 | 490.7356838 | -0.605707413 | 0.193012432 | 9.52E-04 | 0.003488297 | 22924 | 2 | 27193480 | 27250064 | 1 | protein_coding | microtubule-associated protein, RP/EB family, member 3 |

| | | | | | | | | | | | | | |
|---|---|---|---|---|---|---|---|---|---|---|---|---|---|
| **CAD** | ENSG00000084774 | 1000.108791 | -0.707574948 | 0.228442326 | 8.4E-04 | 0.003129919 | 790 | 2 | 27440258 | 27466811 | 1 | protein_coding | carbamoyl-phosphate synthetase 2, aspartate transcarbamylase, and dihydroorotase |
| **CD82** | ENSG00000085117 | 3890.890131 | 1.42672722 | 0.120962281 | 7.11E-33 | 3.24E-31 | 3732 | 11 | 44585977 | 44641913 | 1 | protein_coding | CD82 molecule |
| **ATRX** | ENSG00000085224 | 3504.542441 | 0.432596777 | 0.133300869 | 8.04E-04 | 0.003005441 | 546 | X | 76760356 | 77041702 | -1 | protein_coding | alpha thalassemia/mental retardation syndrome X-linked |
| **FCN1** | ENSG00000085265 | 747.017216 | -1.056141109 | 0.278067263 | 3.53E-05 | 1.75E-04 | 2219 | 9 | 137801431 | 137809809 | -1 | protein_coding | ficolin (collagen/fibrinogen domain containing) 1 |
| **SCAMP1** | ENSG00000085365 | 47.75187911 | 2.145326639 | 0.842022506 | 7.56E-04 | 0.002841592 | 9522 | 5 | 77656339 | 77776562 | 1 | protein_coding | secretory carrier membrane protein 1 |
| **WDFY1** | ENSG00000085449 | 5232.369083 | 1.139413027 | 0.105545739 | 8.18E-28 | 2.81E-26 | 57590 | 2 | 224720433 | 224810104 | -1 | protein_coding | WD repeat and FYVE domain containing 1 |
| **SLC25A24** | ENSG00000085491 | 2976.87445 | 0.950558689 | 0.11582052 | 7.39E-17 | 1.28E-15 | 29957 | 1 | 108676658 | 108743471 | -1 | protein_coding | solute carrier family 25 (mitochondrial carrier; phosphate carrier), member 24 |
| **PILRA** | ENSG00000085514 | 4446.946032 | 0.892313773 | 0.091418963 | 5.56E-23 | 1.5E-21 | 29992 | 7 | 99965153 | 99997719 | 1 | protein_coding | paired immunoglobin-like type 2 receptor alpha |
| **IGSF9** | ENSG00000085552 | 11.83261807 | -2.691614611 | 1.512804568 | 0.005198625 | 0.015789068 | 57549 | 1 | 159896829 | 159915386 | -1 | protein_coding | immunoglobulin superfamily, member 9 |
| **ZNF213** | ENSG00000085644 | 279.8832698 | -0.675961473 | 0.208293977 | 5.87E-04 | 0.002253942 | 7760 | 16 | 3179778 | 3192806 | 1 | protein_coding | zinc finger protein 213 |
| **AKR1B1** | ENSG00000085662 | 2195.211649 | 1.341133781 | 0.144929206 | 4.04E-21 | 9.67E-20 | 231 | 7 | 134127102 | 134144036 | -1 | protein_coding | aldo-keto reductase family 1, member B1 (aldose reductase) |
| **RRN3** | ENSG00000085721 | 1101.656706 | 0.425729436 | 0.157526376 | 0.005065912 | 0.01544431 | 54700 | 16 | 15153879 | 15188174 | -1 | protein_coding | RRN3 RNA polymerase I transcription factor homolog (S. cerevisiae) |
| **TTC39A** | ENSG00000085831 | 15.12439867 | 1.667408475 | 1.437437571 | 0.010169422 | 0.028407836 | 22996 | 1 | 51752930 | 51810788 | -1 | protein_coding | tetratricopeptide repeat domain 39A |
| **EPS15** | ENSG00000085832 | 3747.071877 | 0.675771051 | 0.11816696 | 5.84E-09 | 5.11E-08 | 2060 | 1 | 51819935 | 51985000 | -1 | protein_coding | epidermal growth factor receptor pathway substrate 15 |
| **ORC1** | ENSG00000085840 | 65.36451511 | 1.462388818 | 0.396219416 | 3.19E-05 | 1.6E-04 | 4998 | 1 | 52838501 | 52870131 | -1 | protein_coding | origin recognition complex, subunit 1 |
| **MGST2** | ENSG00000085871 | 713.4366456 | -0.759555602 | 0.187908021 | 2.34E-05 | 1.2E-04 | 4258 | 4 | 140586922 | 140661899 | 1 | protein_coding | microsomal glutathione S-transferase 2 |
| **CHERP** | ENSG00000085872 | 820.3638804 | -0.482200883 | 0.186212029 | 0.006601148 | 0.01946842 | 10523 | 19 | 16628700 | 16653341 | -1 | protein_coding | calcium homeostasis endoplasmic reticulum protein |
| **RAD54L** | ENSG00000085999 | 29.46622113 | 1.233703477 | 0.708635834 | 0.010368657 | 0.028853638 | 8438 | 1 | 46713360 | 46744145 | 1 | protein_coding | RAD54-like (S. cerevisiae) |
| **DNAJA1** | ENSG00000086061 | 6716.545114 | 2.110250362 | 0.079280581 | 5.49E-157 | 3.36E-154 | 3301 | 9 | 33025209 | 33039905 | 1 | protein_coding | DnaJ (Hsp40) homolog, subfamily A, member 1 |
| **B4GALT1** | ENSG00000086062 | 8068.364639 | 1.756515562 | 0.119274238 | 8.73E-50 | 7.58E-48 | 2683 | 9 | 33104080 | 33167354 | -1 | protein_coding | UDP-Gal:betaGlcNAc beta 1,4- galactosyltransferase, polypeptide 1 |
| **CHMP5** | ENSG00000086065 | 2596.199621 | 1.497019071 | 0.115850717 | 6.96E-39 | 4.05E-37 | 51510 | 9 | 33264940 | 33281977 | 1 | protein_coding | charged multivesicular body protein 5 |
| **SNX10** | ENSG00000086300 | 17690.31956 | 2.797145417 | 0.136498725 | 2.3E-94 | 7.18E-92 | 29887 | 7 | 26331541 | 26413949 | 1 | protein_coding | sorting nexin 10 |
| **RBM22** | ENSG00000086589 | 1148.292524 | 0.422523412 | 0.130470284 | 9.18E-04 | 0.003383096 | 55696 | 5 | 150070356 | 150080669 | -1 | protein_coding | RNA binding motif protein 22 |
| **ERO1LB** | ENSG00000086619 | 206.0550712 | 1.077543042 | 0.337230321 | 3.03E-04 | 0.001247863 | 56605 | 1 | 236378855 | 236445319 | -1 | protein_coding | ERO1-like beta (S. cerevisiae) |
| **TXLNG** | ENSG00000086712 | 456.6724333 | 0.49263432 | 0.194427971 | 0.007576308 | 0.021962957 | 55787 | X | 16804550 | 16862642 | 1 | protein_coding | taxilin gamma |
| **LAT2** | ENSG00000086730 | 1891.983994 | -1.338307571 | 0.122469803 | 1.93E-28 | 6.93E-27 | 7462 | 7 | 73613982 | 73644161 | 1 | protein_coding | linker for activation of T cells family, member 2 |
| **ACOX3** | ENSG00000087008 | 503.9173842 | 0.558100841 | 0.198296257 | 0.002893583 | 0.009387099 | 8310 | 4 | 8368009 | 8442450 | -1 | protein_coding | acyl-CoA oxidase 3, pristanoyl |
| **MTMR2** | ENSG00000087053 | 1841.345071 | -0.315483232 | 0.124695869 | 0.01017234 | 0.028410796 | 8898 | 11 | 95566046 | 95658479 | -1 | protein_coding | myotubularin related protein 2 |
| **PPP1R15A** | ENSG00000087074 | 2507.92162 | 0.722041705 | 0.115522699 | 1.89E-10 | 1.95E-09 | 23645 | 19 | 49375649 | 49379314 | 1 | protein_coding | protein phosphatase 1, regulatory subunit 15A |
| **HSD17B14** | ENSG00000087076 | 514.4440215 | -0.671875719 | 0.181459229 | 1.08E-04 | 4.93E-04 | 51171 | 19 | 49316274 | 49339935 | -1 | protein_coding | hydroxysteroid (17-beta) dehydrogenase 14 |
| **FTL** | ENSG00000087086 | 313149.2737 | -0.489941629 | 0.077949561 | 2.29E-10 | 2.35E-09 | 2512 | 19 | 49468558 | 49470135 | 1 | protein_coding | ferritin, light polypeptide |
| **ATXN7L3** | ENSG00000087152 | 1767.063205 | 0.520727567 | 0.159691802 | 7.26E-04 | 0.002736922 | 56970 | 17 | 42269173 | 42277481 | -1 | protein_coding | ataxin 7-like 3 |
| **PSMC5** | ENSG00000087191 | 1903.029509 | 0.314554582 | 0.104832974 | 0.00219937 | 0.007343889 | 5705 | 17 | 61904512 | 61909379 | 1 | protein_coding | proteasome (prosome, macropain) 26S subunit, ATPase, 5 |
| **UIMC1** | ENSG00000087206 | 364.8501379 | 0.559834942 | 0.230210453 | 0.008807529 | 0.0250522 | 51720 | 5 | 176332006 | 176449634 | -1 | protein_coding | ubiquitin interaction motif containing 1 |
| **MMP2** | ENSG00000087245 | 232.3982271 | -1.328515547 | 0.320249761 | 5.93E-06 | 3.38E-05 | 4313 | 16 | 55423612 | 55540603 | 1 | protein_coding | matrix metallopeptidase 2 (gelatinase A, 72kDa gelatinase, 72kDa type IV collagenase) |
| **LPCAT2** | ENSG00000087253 | 5025.034794 | -1.215794944 | 0.111868427 | 2.76E-28 | 9.73E-27 | 54947 | 16 | 55542910 | 55620582 | 1 | protein_coding | lysophosphatidylcholine acyltransferase 2 |
| **OGFOD1** | ENSG00000087263 | 1284.101246 | 0.461001846 | 0.113306502 | 3.24E-05 | 1.62E-04 | 55239 | 16 | 56485402 | 56513012 | 1 | protein_coding | 2-oxoglutarate and iron-dependent oxygenase domain containing 1 |
| **SH3BP2** | ENSG00000087266 | 2713.481169 | -0.563969503 | 0.137739189 | 2.5E-05 | 1.28E-04 | 6452 | 4 | 2794750 | 2842825 | 1 | protein_coding | SH3-domain binding protein 2 |
| **L2HGDH** | ENSG00000087299 | 150.6562336 | 0.87469641 | 0.287675169 | 7.94E-04 | 0.002970791 | 79944 | 14 | 50704281 | 50779266 | -1 | protein_coding | L-2-hydroxyglutarate dehydrogenase |
| **GNAS** | ENSG00000087460 | 20844.04174 | -0.574719226 | 0.103489407 | 1.59E-08 | 1.32E-07 | 2778 | 20 | 57414773 | 57486247 | 1 | protein_coding | GNAS complex locus |
| **DNM1L** | ENSG00000087470 | 1803.294081 | 0.815264334 | 0.126662071 | 4.26E-11 | 4.76E-10 | 10059 | 12 | 32832134 | 32898486 | 1 | protein_coding | dynamin 1-like |
| **TFAP2C** | ENSG00000087510 | 32.70396504 | 1.148150647 | 0.613368833 | 0.009363919 | 0.0264744 | 7022 | 20 | 55204358 | 55214339 | 1 | protein_coding | transcription factor AP-2 gamma (activating enhancer binding protein 2 gamma) |
| **CASS4** | ENSG00000087589 | 37.59603237 | -1.549015888 | 0.730747425 | 0.003501584 | 0.011132663 | 57091 | 20 | 54987168 | 55034396 | 1 | protein_coding | Cas scaffolding protein family member 4 |
| **PIR** | ENSG00000087842 | 301.4398847 | 0.925999641 | 0.261047393 | 1.23E-04 | 5.55E-04 | 8544 | X | 15402921 | 15511687 | -1 | protein_coding | pirin (iron-binding nuclear protein) |
| **ALG6** | ENSG00000088035 | 91.16632092 | 1.147754604 | 0.382662538 | 5.28E-04 | 0.002054054 | 29929 | 1 | 63833261 | 63904233 | 1 | protein_coding | ALG6, alpha-1,3-glucosyltransferase |
| **PTPN4** | ENSG00000088179 | 381.1573232 | -1.378193797 | 0.432671541 | 2.13E-04 | 9.05E-04 | 5775 | 2 | 120517207 | 120741394 | 1 | protein_coding | protein tyrosine phosphatase, non-receptor type 4 (megakaryocyte) |
| **KHSRP** | ENSG00000088247 | 4339.006587 | -0.517716657 | 0.08764922 | 2.59E-09 | 2.36E-08 | 8570 | 19 | 6413359 | 6424805 | -1 | protein_coding | KH-type splicing regulatory protein |
| **GNA11** | ENSG00000088256 | 895.4389957 | -0.398423102 | 0.168667237 | 0.013832259 | 0.037067931 | 2767 | 19 | 3094408 | 3124002 | 1 | protein_coding | guanine nucleotide binding protein (G protein), alpha 11 (Gq class) |
| **DOCK9** | ENSG00000088387 | 248.4033182 | 1.210203458 | 0.335837204 | 6.24E-05 | 2.96E-04 | 23348 | 13 | 99445741 | 99738879 | -1 | protein_coding | dedicator of cytokinesis 9 |
| **ANKRD10** | ENSG00000088448 | 1111.662921 | 0.713511967 | 0.121770497 | 2.26E-09 | 2.07E-08 | 55608 | 13 | 111530887 | 111567416 | -1 | protein_coding | ankyrin repeat domain 10 |
| **TGDS** | ENSG00000088451 | 278.7914964 | 0.915961115 | 0.235196482 | 3.21E-05 | 1.61E-04 | 23483 | 13 | 95226308 | 95248511 | -1 | protein_coding | TDP-glucose 4,6-dehydratase |
| **COQ9** | ENSG00000088682 | 721.5798111 | -0.443296178 | 0.140870532 | 0.001150291 | 0.004136446 | 57017 | 16 | 57481337 | 57495187 | 1 | protein_coding | coenzyme Q9 |
| **KIF9** | ENSG00000088727 | 34.26698654 | -2.248421393 | 0.874511142 | 5.85E-04 | 0.002247988 | 64147 | 3 | 47269516 | 47324941 | -1 | protein_coding | kinesin family member 9 |
| **PPP1R13B** | ENSG00000088808 | 328.9002857 | -0.739162228 | 0.315542892 | 0.007719898 | 0.022332614 | 23368 | 14 | 104200089 | 104313927 | -1 | protein_coding | protein phosphatase 1, regulatory subunit 13B |
| **ATRN** | ENSG00000088812 | 2028.581308 | -0.822823275 | 0.129337085 | 7.98E-11 | 8.63E-10 | 8455 | 20 | 3451687 | 3631769 | 1 | protein_coding | attractin |
| **SMOX** | ENSG00000088826 | 151.9621307 | -1.623663303 | 0.351383588 | 4.81E-07 | 3.26E-06 | 54498 | 20 | 4101627 | 4168394 | 1 | protein_coding | spermine oxidase |
| **SIGLEC1** | ENSG00000088827 | 2559.981037 | 5.045982124 | 0.233513228 | 4.47E-105 | 1.59E-102 | 6614 | 20 | 3667617 | 3687775 | -1 | protein_coding | sialic acid binding Ig-like lectin 1, sialoadhesin |
| **FKBP1A** | ENSG00000088832 | 5385.828353 | -0.507237334 | 0.100214066 | 2.78E-07 | 1.94E-06 | 1019293 | 20 | 1349622 | 1373806 | -1 | protein_coding | FK506 binding protein 1A, 12kDa |
| **MAVS** | ENSG00000088888 | 1504.205967 | -0.862978723 | 0.122164601 | 6.27E-13 | 8.14E-12 | 57506 | 20 | 3827487 | 3849280 | 1 | protein_coding | mitochondrial antiviral signaling protein |
| **XRN2** | ENSG00000088930 | 3113.186015 | 0.347600439 | 0.122683198 | 0.003737296 | 0.011785059 | 22803 | 20 | 21283942 | 21370463 | 1 | protein_coding | 5'-3' exoribonuclease 2 |
| **DYNLL1** | ENSG00000088986 | 3603.5394 | 0.382291696 | 0.106239851 | 2E-04 | 8.55E-04 | 8655 | 12 | 120907653 | 120936296 | 1 | protein_coding | dynein, light chain, LC8-type 1 |
| **SNX5** | ENSG00000089006 | 3715.767072 | -0.686520021 | 0.123489217 | 1.35E-08 | 1.13E-07 | 27131 | 20 | 17922241 | 17949623 | -1 | protein_coding | sorting nexin 5 |
| **RPL6** | ENSG00000089009 | 6923.256171 | -0.535684263 | 0.1090702 | 4.91E-07 | 3.32E-06 | 6128 | 12 | 112842994 | 112856642 | -1 | protein_coding | ribosomal protein L6 |
| **SIRPG** | ENSG00000089012 | 238.928628 | -1.418273938 | 0.303040357 | 4.32E-07 | 2.95E-06 | 55423 | 20 | 1609798 | 1638425 | -1 | protein_coding | signal-regulatory protein gamma |
| **P2RX7** | ENSG00000089041 | 3936.035608 | -0.267282728 | 0.116374787 | 0.019075267 | 0.048740469 | 5027 | 12 | 121570622 | 121623876 | 1 | protein_coding | purinergic receptor P2X, ligand-gated ion channel, 7 |
| **RBBP9** | ENSG00000089050 | 834.3585382 | 1.272711956 | 0.153872673 | 2.82E-17 | 5.03E-16 | 10741 | 20 | 18467184 | 18477887 | -1 | protein_coding | retinoblastoma binding protein 9 |
| **SLC23A2** | ENSG00000089057 | 5695.165743 | -1.451107098 | 0.117760372 | 9.67E-36 | 4.96E-34 | 9962 | 20 | 4833002 | 4990939 | -1 | protein_coding | solute carrier family 23 (ascorbic acid transporter), member 2 |
| **SLC8B1** | ENSG00000089060 | 2808.68738 | -0.395969361 | 0.12949509 | 0.001724043 | 0.005915706 | 80024 | 12 | 113736564 | 113797298 | -1 | protein_coding | solute carrier family 8 (sodium/lithium/calcium exchanger), member B1 |
| **C20orf26** | ENSG00000089101 | 144.2115642 | -1.848330812 | 0.620883085 | 2.5E-04 | 0.00104394 | 26074 | 20 | 20033158 | 20341346 | 1 | protein_coding | chromosome 20 open reading frame 26 |
| **OAS1** | ENSG00000089127 | 3760.022614 | 3.667396627 | 0.105384316 | 2.55E-266 | 4.33E-263 | 4938 | 12 | 113344582 | 113369990 | 1 | protein_coding | 2'-5'-oligoadenylate synthetase 1, 40/46kDa |
| **RPLP0** | ENSG00000089157 | 6794.07845 | -0.773325461 | 0.122682099 | 1.23E-10 | 1.3E-09 | 6175 | 12 | 120634489 | 120639038 | -1 | protein_coding | ribosomal protein, large, P0 |
| **CHGB** | ENSG00000089199 | 19.24340648 | -2.391971284 | 1.150119437 | 0.002567419 | 0.008426802 | 1114 | 20 | 5892076 | 5906007 | 1 | protein_coding | chromogranin B (secretogranin 1) |
| **FXYD5** | ENSG00000089327 | 3075.703293 | -0.809266367 | 0.127357518 | 8.11E-11 | 8.75E-10 | 53827 | 19 | 35645633 | 35660786 | 1 | protein_coding | FXYD domain containing ion transport regulator 5 |
| **ZNF302** | ENSG00000089335 | 606.9841837 | 0.522757697 | 0.169176904 | 0.001302649 | 0.004613761 | 55900 | 19 | 35168544 | 35177302 | 1 | protein_coding | zinc finger protein 302 |
| **GRAMD1A** | ENSG00000089351 | 2470.625568 | 2.652953237 | 0.190226012 | 2.76E-45 | 2.13E-43 | 57655 | 19 | 35485688 | 35517375 | 1 | protein_coding | GRAM domain containing 1A |
| **GANAB** | ENSG00000089597 | 5527.31762 | -0.425249993 | 0.099862941 | 1.86E-05 | 9.74E-05 | 23193 | 11 | 62392298 | 62414104 | -1 | protein_coding | glucosidase, alpha; neutral AB |
| **GMIP** | ENSG00000089639 | 2660.543566 | -0.440495395 | 0.119889271 | 1.66E-04 | 7.27E-04 | 51291 | 19 | 19740285 | 19754476 | -1 | protein_coding | GEM interacting protein |
| **RBM41** | ENSG00000089682 | 519.8320185 | 0.451728657 | 0.172366906 | 0.006220968 | 0.018482707 | 55285 | X | 106307650 | 106362057 | -1 | protein_coding | RNA binding motif protein 41 |
| **LAG3** | ENSG00000089692 | 604.9635933 | 1.318185325 | 0.384578644 | 9.91E-05 | 4.53E-04 | 3902 | 12 | 6881678 | 6887621 | 1 | protein_coding | lymphocyte-activation gene 3 |
| **OTUB2** | ENSG00000089723 | 592.3930404 | -1.133919739 | 0.222652277 | 8.66E-08 | 6.47E-07 | 78990 | 14 | 94492675 | 94515276 | 1 | protein_coding | OTU domain, ubiquitin aldehyde binding 2 |
| **DDX24** | ENSG00000089737 | 3823.316737 | 0.495979813 | 0.123303621 | 3.79E-05 | 1.87E-04 | 57062 | 14 | 94517266 | 94547591 | -1 | protein_coding | DEAD (Asp-Glu-Ala-Asp) box helicase 24 |
| **NECAP1** | ENSG00000089818 | 1779.053937 | 0.514120798 | 0.115430387 | 4.87E-06 | 2.83E-05 | 25977 | 12 | 8234807 | 8250367 | 1 | protein_coding | NECAP endocytosis associated 1 |
| **ARHGAP4** | ENSG00000089820 | 1297.694707 | -1.706592339 | 0.232080988 | 2.42E-14 | 3.51E-13 | 393 | X | 153172821 | 153200452 | -1 | protein_coding | Rho GTPase activating protein 4 |
| **RCOR1** | ENSG00000089902 | 568.9532414 | -0.936542216 | 0.353389373 | 0.002067792 | 0.006963875 | 23186 | 14 | 103058998 | 103196913 | 1 | protein_coding | REST corepressor 1 |
| **BLVRB** | ENSG00000090013 | 477.5930641 | -0.875949652 | 0.182651761 | 5.43E-07 | 3.66E-06 | 645 | 19 | 40953696 | 40971747 | -1 | protein_coding | biliverdin reductase B (flavin reductase (NADPH)) |
| **SLC9A1** | ENSG00000090020 | 2318.210075 | -1.109880347 | 0.136806368 | 1.2E-16 | 2.05E-15 | 6548 | 1 | 27425306 | 27493472 | -1 | protein_coding | solute carrier family 9, subfamily A (NHE1, cation proton antiporter 1), member 1 |
| **CCNK** | ENSG00000090061 | 1821.82806 | 0.499471262 | 0.120234222 | 2.76E-05 | 1.4E-04 | 8812 | 14 | 99947506 | 100001381 | 1 | protein_coding | cyclin K |
| **RGS1** | ENSG00000090104 | 1412.763919 | 1.055644842 | 0.17555778 | 5.66E-10 | 5.57E-09 | 5996 | 1 | 192544857 | 192549161 | 1 | protein_coding | regulator of G-protein signaling 1 |
| **YPEL3** | ENSG00000090238 | 519.0248839 | -0.830102582 | 0.167453539 | 2.77E-07 | 1.93E-06 | 83719 | 16 | 30103635 | 30108236 | -1 | protein_coding | yippee-like 3 (Drosophila) |

| | | | | | | | | | | | | | |
|---|---|---|---|---|---|---|---|---|---|---|---|---|---|
| **NDUFB2** | ENSG00000090266 | 809.3771353 | -0.352712842 | 0.152620157 | 0.016940641 | 0.044119537 | 4708 | 7 | 140390577 | 140422590 | 1 | protein_coding | NADH dehydrogenase (ubiquinone) 1 beta subcomplex, 2, 8kDa |
| **NUDC** | ENSG00000090273 | 922.9474283 | -0.416534597 | 0.123746868 | 6.03E-04 | 0.002314218 | 10726 | 1 | 27226729 | 27273353 | 1 | protein_coding | nudC nuclear distribution protein |
| **ICAM1** | ENSG00000090339 | 10871.74843 | 1.11391706 | 0.199424187 | 5.56E-09 | 4.89E-08 | 3383 | 19 | 10381511 | 10397291 | 1 | protein_coding | intercellular adhesion molecule 1 |
| **IRAK3** | ENSG00000090376 | 1913.927773 | 0.839477622 | 0.144800347 | 1.38E-09 | 1.3E-08 | 11213 | 12 | 66582659 | 66648402 | 1 | protein_coding | interleukin-1 receptor-associated kinase 3 |
| **LYZ** | ENSG00000090382 | 299215.742 | -0.266255572 | 0.077084062 | 3.05E-04 | 0.001256011 | 4069 | 12 | 69742121 | 69748014 | 1 | protein_coding | lysozyme |
| **MUL1** | ENSG00000090432 | 914.1554075 | -0.407624192 | 0.144183278 | 0.003581737 | 0.011356195 | 79594 | 1 | 20825943 | 20834654 | -1 | protein_coding | mitochondrial E3 ubiquitin protein ligase 1 |
| **GNPTG** | ENSG00000090581 | 1742.693414 | -0.34345692 | 0.113245338 | 0.002005416 | 0.006783704 | 84572 | 16 | 1401924 | 1413352 | 1 | protein_coding | N-acetylglucosamine-1-phosphate transferase, gamma subunit |
| **GOLGA3** | ENSG00000090615 | 4034.014473 | -0.72865301 | 0.131513365 | 1.4E-08 | 1.16E-07 | 2802 | 12 | 133345495 | 133405444 | -1 | protein_coding | golgin A3 |
| **PABPC4** | ENSG00000090621 | 3992.357354 | -0.675378886 | 0.132607015 | 1.79E-07 | 1.29E-06 | 1009966 | 1 | 40026488 | 40042462 | -1 | protein_coding | poly(A) binding protein, cytoplasmic 4 (inducible form) |
| **CERS4** | ENSG00000090661 | 189.717546 | -1.218147564 | 0.317819008 | 2.44E-05 | 1.25E-04 | 79603 | 19 | 8271620 | 8327305 | 1 | protein_coding | ceramide synthase 4 |
| **MCOLN1** | ENSG00000090674 | 897.0939871 | -1.454814341 | 0.193700121 | 9.4E-15 | 1.4E-13 | 57192 | 19 | 7587512 | 7598895 | 1 | protein_coding | mucolipin 1 |
| **EFNB1** | ENSG00000090776 | 225.5308629 | -0.848098232 | 0.237377496 | 1.3E-04 | 5.81E-04 | 1947 | X | 68048840 | 68061990 | 1 | protein_coding | ephrin-B1 |
| **PDPR** | ENSG00000090857 | 1826.819453 | -0.342460263 | 0.096860036 | 3.57E-04 | 0.001447252 | 55066 | 16 | 70147529 | 70195203 | 1 | protein_coding | pyruvate dehydrogenase phosphatase regulatory subunit |
| **AARS** | ENSG00000090861 | 4254.769 | -0.40115753 | 0.136925452 | 0.002614446 | 0.008554181 | 16 | 16 | 70286198 | 70323446 | -1 | protein_coding | alanyl-tRNA synthetase |
| **GLG1** | ENSG00000090863 | 6677.205209 | -0.520509782 | 0.080454887 | 7.99E-11 | 8.63E-10 | 2734 | 16 | 74485856 | 74641012 | -1 | protein_coding | golgi glycoprotein 1 |
| **TNRC6A** | ENSG00000090905 | 1878.669425 | 0.706302632 | 0.133825417 | 6.37E-08 | 4.87E-07 | 27327 | 16 | 24741016 | 24838953 | 1 | protein_coding | trinucleotide repeat containing 6A |
| **PITPNM2** | ENSG00000090975 | 57.79030535 | -1.363513641 | 0.491695111 | 7.9E-04 | 0.002956416 | 57605 | 12 | 123468027 | 123634562 | -1 | protein_coding | phosphatidylinositol transfer protein, membrane-associated 2 |
| **RBM27** | ENSG00000091009 | 902.5188132 | 0.593161945 | 0.192690908 | 0.00120112 | 0.00429799 | 54439 | 5 | 145583113 | 145718814 | 1 | protein_coding | RNA binding motif protein 27 |
| **OSBPL8** | ENSG00000091039 | 9130.621462 | -1.028346834 | 0.104769385 | 2.42E-23 | 6.64E-22 | 114882 | 12 | 76745577 | 76953589 | -1 | protein_coding | oxysterol binding protein-like 8 |
| **DTX2** | ENSG00000091073 | 1088.227072 | 0.570431649 | 0.135053389 | 1.38E-05 | 7.43E-05 | 113878 | 7 | 76090993 | 76135312 | 1 | protein_coding | deltex homolog 2 (Drosophila) |
| **NLRC4** | ENSG00000091106 | 1263.022242 | -1.242676698 | 0.132630052 | 1.13E-21 | 2.81E-20 | 58484 | 2 | 32449522 | 32490923 | -1 | protein_coding | NLR family, CARD domain containing 4 |
| **DLD** | ENSG00000091140 | 2831.472358 | 0.631041036 | 0.14323548 | 5.86E-06 | 3.35E-05 | 1738 | 7 | 107531415 | 107572175 | 1 | protein_coding | dihydrolipoamide dehydrogenase |
| **WDR7** | ENSG00000091157 | 3087.62272 | -0.73174205 | 0.108674831 | 7.84E-12 | 9.29E-11 | 23335 | 18 | 54318574 | 54698828 | 1 | protein_coding | WD repeat domain 7 |
| **TXNL1** | ENSG00000091164 | 1520.736893 | 0.517490121 | 0.129084895 | 4.14E-05 | 2.02E-04 | 9352 | 18 | 54264439 | 54318831 | -1 | protein_coding | thioredoxin-like 1 |
| **CMTM6** | ENSG00000091317 | 9609.749365 | 0.961079343 | 0.122438338 | 1.68E-15 | 2.64E-14 | 54918 | 3 | 32522804 | 32544900 | -1 | protein_coding | CKLF-like MARVEL transmembrane domain containing 6 |
| **ITGA6** | ENSG00000091409 | 7233.215751 | -1.185241053 | 0.152296144 | 1.56E-15 | 2.46E-14 | 3655 | 2 | 173292082 | 173371181 | 1 | protein_coding | integrin, alpha 6 |
| **MLTK** | ENSG00000091436 | 1450.041265 | -0.649462055 | 0.121078201 | 4.84E-08 | 3.76E-07 | 51776 | 2 | 173940163 | 174132738 | 1 | protein_coding | Mitogen-activated protein kinase kinase kinase MLT |
| **FH** | ENSG00000091483 | 1575.477483 | -0.534006753 | 0.15654197 | 4.03E-04 | 0.001614932 | 2271 | 1 | 241660903 | 241683061 | -1 | protein_coding | fumarate hydratase |
| **CDV3** | ENSG00000091527 | 11138.29079 | 0.290283627 | 0.115762347 | 0.010725485 | 0.029684459 | 55573 | 3 | 133292574 | 133309105 | 1 | protein_coding | CDV3 homolog (mouse) |
| **ALKBH5** | ENSG00000091542 | 2441.893474 | -0.503661186 | 0.108362291 | 1.98E-06 | 1.22E-05 | 54890 | 17 | 18086392 | 18113268 | 1 | protein_coding | alkB, alkylation repair homolog 5 (E. coli) |
| **NLRP1** | ENSG00000091592 | 567.78611 | -0.947592981 | 0.314130213 | 6.54E-04 | 0.002487292 | 22861 | 17 | 5402747 | 5522744 | -1 | protein_coding | NLR family, pyrin domain containing 1 |
| **PITPNM3** | ENSG00000091622 | 36.32311422 | -1.224445185 | 0.616439996 | 0.006848364 | 0.020119866 | 83394 | 17 | 6354584 | 6459814 | -1 | protein_coding | PITPNM family member 3 |
| **ORC6** | ENSG00000091651 | 106.0316262 | 0.849239614 | 0.352918551 | 0.005274526 | 0.015984648 | 23594 | 16 | 46723555 | 46732306 | 1 | protein_coding | origin recognition complex, subunit 6 |
| **CD200** | ENSG00000091972 | 301.9095028 | -1.037494747 | 0.239445928 | 3.92E-06 | 2.3E-05 | 4345 | 3 | 112051194 | 112081659 | 1 | protein_coding | CD200 molecule |
| **PSME1** | ENSG00000092010 | 2557.511571 | 1.479443118 | 0.160394763 | 4.2E-21 | 1E-19 | 5720 | 14 | 24605367 | 24608176 | 1 | protein_coding | proteasome (prosome, macropain) activator subunit 1 (PA28 alpha) |
| **PPP2R3C** | ENSG00000092020 | 825.0167236 | 0.983604546 | 0.205422411 | 4.64E-07 | 3.15E-06 | 55012 | 14 | 35554673 | 35591723 | -1 | protein_coding | protein phosphatase 2, regulatory subunit B'', gamma |
| **JPH4** | ENSG00000092051 | 33.78772828 | -2.260743037 | 0.73682729 | 1.49E-04 | 6.6E-04 | 84502 | 14 | 24037244 | 24048009 | -1 | protein_coding | junctophilin 4 |
| **SLC7A8** | ENSG00000092068 | 8703.399347 | -1.78032493 | 0.121102981 | 8.14E-50 | 7.11E-48 | 23428 | 14 | 23594504 | 23652883 | -1 | protein_coding | solute carrier family 7 (amino acid transporter light chain, L system), member 8 |
| **OSGEP** | ENSG00000092094 | 654.7028542 | -0.573421382 | 0.152511341 | 1.15E-04 | 5.19E-04 | 55644 | 14 | 20914570 | 20923264 | -1 | protein_coding | O-sialoglycoprotein endopeptidase |
| **RNF31** | ENSG00000092098 | 668.5899488 | 0.662903527 | 0.221715525 | 0.001460233 | 0.005116173 | 55072 | 14 | 24615892 | 24629870 | 1 | protein_coding | ring finger protein 31 |
| **HNRNPC** | ENSG00000092199 | 9522.570718 | 0.37707397 | 0.086301645 | 8E-06 | 4.47E-05 | 3183 | 14 | 21677295 | 21737653 | -1 | protein_coding | heterogeneous nuclear ribonucleoprotein C (C1/C2) |
| **RPGRIP1** | ENSG00000092200 | 60.13827788 | 0.916508134 | 0.424465537 | 0.008412368 | 0.024047376 | 57096 | 14 | 21756098 | 21819460 | 1 | protein_coding | retinitis pigmentosa GTPase regulator interacting protein 1 |
| **SUPT16H** | ENSG00000092201 | 3453.267497 | 0.338483985 | 0.082946875 | 4.32E-05 | 2.11E-04 | 11198 | 14 | 21819631 | 21852425 | -1 | protein_coding | suppressor of Ty 16 homolog (S. cerevisiae) |
| **TINF2** | ENSG00000092330 | 1753.035714 | 1.114690429 | 0.12762937 | 6.1E-19 | 1.23E-17 | 26277 | 14 | 24708849 | 24711880 | -1 | protein_coding | TERF1 (TRF1)-interacting nuclear factor 2 |
| **TRPM7** | ENSG00000092439 | 1324.647444 | 1.07912414 | 0.187967108 | 2.37E-09 | 2.17E-08 | 54822 | 15 | 50844670 | 50979012 | -1 | protein_coding | transient receptor potential cation channel, subfamily M, member 7 |
| **WDR76** | ENSG00000092470 | 149.7088798 | 1.554792221 | 0.390646577 | 8.88E-06 | 4.93E-05 | 79968 | 15 | 44119161 | 44160617 | 1 | protein_coding | WD repeat domain 76 |
| **CAPN3** | ENSG00000092529 | 294.6669298 | 0.57298026 | 0.193338818 | 0.001837614 | 0.006271445 | 825 | 15 | 42640301 | 42704516 | 1 | protein_coding | calpain 3, (p94) |
| **SNAP23** | ENSG00000092531 | 3418.420609 | -0.664621377 | 0.134227485 | 3.81E-07 | 2.62E-06 | 8773 | 15 | 42783431 | 42837547 | 1 | protein_coding | synaptosomal-associated protein, 23kDa |
| **PHGDH** | ENSG00000092621 | 369.115767 | -0.977299024 | 0.316506159 | 5.42E-04 | 0.002100223 | 26227 | 1 | 120202421 | 120286838 | 1 | protein_coding | phosphoglycerate dehydrogenase |
| **EZR** | ENSG00000092820 | 4790.760674 | 0.17980159 | 0.078553516 | 0.019554265 | 0.049758215 | 7430 | 6 | 159186773 | 159240444 | -1 | protein_coding | ezrin |
| **AGO1** | ENSG00000092847 | 8200.041019 | -1.745743714 | 0.102757665 | 1.05E-65 | 1.69E-63 | 26523 | 1 | 36335409 | 36395211 | 1 | protein_coding | argonaute RISC catalytic component 1 |
| **CLSPN** | ENSG00000092853 | 181.9490264 | 2.703058875 | 0.322963529 | 4.08E-18 | 7.77E-17 | 63967 | 1 | 36185819 | 36235568 | -1 | protein_coding | claspin |
| **RFFL** | ENSG00000092871 | 1410.365561 | 0.668538487 | 0.121736958 | 2.59E-08 | 2.09E-07 | 117584 | 17 | 33333009 | 33416338 | -1 | protein_coding | ring finger and FYVE-like domain containing E3 ubiquitin protein ligase |
| **DPYSL2** | ENSG00000092964 | 7833.724081 | -0.389475159 | 0.116313688 | 6.39E-04 | 0.002435217 | 1808 | 8 | 26371791 | 26515694 | 1 | protein_coding | dihydropyrimidinase-like 2 |
| **TGFB2** | ENSG00000092969 | 59.65221079 | -4.516431684 | 0.79581708 | 1.58E-09 | 1.48E-08 | 7042 | 1 | 218519577 | 218617961 | 1 | protein_coding | transforming growth factor, beta 2 |
| **NUP50** | ENSG00000093000 | 1958.041003 | 0.627653431 | 0.138569365 | 3.36E-06 | 1.99E-05 | 10762 | 22 | 45559722 | 45583896 | 1 | protein_coding | nucleoporin 50kDa |
| **CDC45** | ENSG00000093009 | 34.11044579 | 2.326336086 | 0.803162885 | 1.88E-04 | 8.11E-04 | 8318 | 22 | 19466982 | 19508135 | 1 | protein_coding | cell division cycle 45 |
| **CECR1** | ENSG00000093072 | 10094.62713 | -0.638365956 | 0.130410305 | 4.87E-07 | 3.3E-06 | 51816 | 22 | 17660194 | 17702879 | -1 | protein_coding | cat eye syndrome chromosome region, candidate 1 |
| **VNN 3.00** | ENSG00000093134 | 29.17376579 | 1.251350733 | 0.681222841 | 0.008467014 | 0.024169103 | NA | 6 | 133043926 | 133055904 | -1 | protein_coding | vanin 3 |
| **LRRFIP2** | ENSG00000093167 | 1479.352492 | 0.853986182 | 0.117819473 | 2.01E-13 | 2.7E-12 | 9209 | 3 | 37094117 | 37225180 | -1 | protein_coding | leucine rich repeat (in FLII) interacting protein 2 |
| **XYLB** | ENSG00000093217 | 102.8326229 | -1.276075069 | 0.437081955 | 5.58E-04 | 0.002154312 | 9942 | 3 | 38388251 | 38462839 | 1 | protein_coding | xylulokinase homolog (H. influenzae) |
| **CDC6** | ENSG00000094804 | 56.13245695 | 1.627467557 | 0.672282668 | 0.001092981 | 0.003958301 | 990 | 17 | 38443885 | 38459171 | 1 | protein_coding | cell division cycle 6 |
| **CBX5** | ENSG00000094916 | 3004.693266 | -0.588090431 | 0.126764059 | 1.97E-06 | 1.22E-05 | 23468 | 12 | 54624724 | 54673886 | -1 | protein_coding | chromobox homolog 5 |
| **SUCO** | ENSG00000094975 | 989.3387926 | 0.6678844 | 0.129084037 | 1.16E-07 | 8.48E-07 | 51430 | 1 | 172501489 | 172580971 | 1 | protein_coding | SUN domain containing ossification factor |
| **TMEM38B** | ENSG00000095209 | 1603.412062 | 0.905257895 | 0.140690639 | 4.32E-11 | 4.82E-10 | 55151 | 9 | 108456825 | 108538893 | 1 | protein_coding | transmembrane protein 38B |
| **PTGS1** | ENSG00000095303 | 1877.143186 | -1.989191708 | 0.157020265 | 8.38E-38 | 4.68E-36 | 5742 | 9 | 125132824 | 125157982 | 1 | protein_coding | prostaglandin-endoperoxide synthase 1 (prostaglandin G/H synthase and cyclooxygenase) |
| **SH2D3C** | ENSG00000095370 | 365.8703448 | -0.645221598 | 0.183933775 | 2.38E-04 | 0.001000969 | 10044 | 9 | 130500596 | 130541020 | -1 | protein_coding | SH2 domain containing 3C |
| **NANS** | ENSG00000095380 | 1373.013894 | -0.548667513 | 0.121407029 | 4.19E-06 | 2.45E-05 | 54187 | 9 | 100819021 | 100845357 | 1 | protein_coding | N-acetylneuraminic acid synthase |
| **TBC1D2** | ENSG00000095383 | 3764.949001 | -0.776168301 | 0.127858826 | 3.88E-10 | 3.89E-09 | 55357 | 9 | 100961311 | 101017915 | -1 | protein_coding | TBC1 domain family, member 2 |
| **DFNB31** | ENSG00000095397 | 144.7164784 | 0.698418978 | 0.298108352 | 0.00857403 | 0.024446993 | 25861 | 9 | 117164360 | 117267730 | -1 | protein_coding | deafness, autosomal recessive 31 |
| **CWF19L1** | ENSG00000095485 | 633.2424396 | 0.793765767 | 0.156341206 | 1.6E-07 | 1.16E-06 | 55280 | 10 | 101992055 | 102027437 | -1 | protein_coding | CWF19-like 1, cell cycle control (S. pombe) |
| **IKZF5** | ENSG00000095574 | 389.9685645 | 0.677150033 | 0.26683578 | 0.005260304 | 0.015951037 | 64376 | 10 | 124750322 | 124768333 | -1 | protein_coding | IKAROS family zinc finger 5 (Pegasus) |
| **SORBS1** | ENSG00000095637 | 172.2011349 | -1.912172289 | 0.450326656 | 1.99E-06 | 1.23E-05 | 10580 | 10 | 97071528 | 97321171 | -1 | protein_coding | sorbin and SH3 domain containing 1 |
| **WAC** | ENSG00000095787 | 4356.860921 | 0.413990352 | 0.096613811 | 1.35E-05 | 7.29E-05 | 51322 | 10 | 28821422 | 28912041 | 1 | protein_coding | WW domain containing adaptor with coiled-coil |
| **CREM** | ENSG00000095794 | 777.2285404 | 0.861072167 | 0.161459395 | 3.61E-08 | 2.85E-07 | 1390 | 10 | 35415719 | 35501886 | 1 | protein_coding | cAMP responsive element modulator |
| **HIVEP1** | ENSG00000095951 | 1587.695926 | 0.460755993 | 0.155950766 | 0.002259366 | 0.00752285 | 3096 | 6 | 12008995 | 12165232 | 1 | protein_coding | human immunodeficiency virus type I enhancer binding protein 1 |
| **TREM2** | ENSG00000095970 | 5101.285183 | -2.33054313 | 0.156956514 | 9.33E-51 | 8.48E-49 | 54209 | 6 | 41126244 | 41130924 | -1 | protein_coding | triggering receptor expressed on myeloid cells 2 |
| **FKBP5** | ENSG00000096060 | 1973.795473 | 0.338892221 | 0.124465375 | 0.005216162 | 0.015832889 | 2289 | 6 | 35541362 | 35696360 | -1 | protein_coding | FK506 binding protein 5 |
| **RPS18** | ENSG00000096150 | 5580.359528 | -0.46156873 | 0.129298656 | 2.55E-04 | 0.001062501 | 6222 | HSC | 33168626 | 33173129 | 1 | protein_coding | ribosomal protein S18 |
| **BAG6** | ENSG00000096155 | 68.45965697 | 0.405402943 | 0.849788422 | 1.4E-04 | 6.24E-04 | 7917 | HSC | 31597040 | 31610723 | -1 | protein_coding | BCL2-associated athanogene 6 |
| **HSP90AB1** | ENSG00000096384 | 19605.7331 | -0.23023176 | 0.097662452 | 0.016907098 | 0.044048957 | 3326 | 6 | 44214824 | 44221620 | 1 | protein_coding | heat shock protein 90kDa alpha (cytosolic), class B member 1 |
| **CDC5L** | ENSG00000096401 | 1829.879246 | 0.37335079 | 0.124073698 | 0.002245148 | 0.007480399 | 988 | 6 | 44355262 | 44418163 | 1 | protein_coding | cell division cycle 5-like |
| **ITPR3** | ENSG00000096433 | 360.1761072 | 1.055189465 | 0.277697527 | 3.5E-05 | 1.74E-04 | 3710 | 6 | 33588142 | 33664351 | 1 | protein_coding | inositol 1,4,5-trisphosphate receptor, type 3 |
| **DSP** | ENSG00000096696 | 894.5352552 | 2.331998759 | 0.259705285 | 2.06E-20 | 4.69E-19 | 1832 | 6 | 7541808 | 7586950 | 1 | protein_coding | desmoplakin |
| **SIRT1** | ENSG00000096717 | 649.7337771 | 0.978019929 | 0.188538235 | 6.88E-08 | 5.23E-07 | 23411 | 10 | 69644427 | 69678147 | 1 | protein_coding | sirtuin 1 |
| **HNRNPH3** | ENSG00000096746 | 2153.890483 | 0.398021051 | 0.108703085 | 1.94E-04 | 8.33E-04 | 3189 | 10 | 70090931 | 70102948 | 1 | protein_coding | heterogeneous nuclear ribonucleoprotein H3 (2H9) |
| **IFT74** | ENSG00000096872 | 415.222914 | 0.997734487 | 0.209288009 | 5.8E-07 | 3.89E-06 | 80173 | 9 | 26947037 | 27062928 | 1 | protein_coding | intraflagellar transport 74 homolog (Chlamydomonas) |
| **JAK2** | ENSG00000096968 | 1111.283539 | 2.205965426 | 0.23356394 | 3E-22 | 7.63E-21 | 3717 | 9 | 4985033 | 5128183 | 1 | protein_coding | Janus kinase 2 |
| **ACOT7** | ENSG00000097021 | 1639.096152 | -1.063841691 | 0.155691428 | 2.22E-12 | 2.75E-11 | 11332 | 1 | 6324329 | 6454451 | -1 | protein_coding | acyl-CoA thioesterase 7 |

| | | | | | | | | | | | | | |
|---|---|---|---|---|---|---|---|---|---|---|---|---|---|
| **SH3GLB1** | ENSG00000097033 | 7685.794515 | 0.561468187 | 0.11454337 | 6.03E-07 | 4.02E-06 | 51100 | 1 | 87170259 | 87213867 | 1 | protein_coding | SH3-domain GRB2-like endophilin B1 |
| **ABLIM1** | ENSG00000099204 | 620.8448853 | -0.56118063 | 0.254167065 | 0.015787759 | 0.041579238 | 3983 | 10 | 116190872 | 116444762 | -1 | protein_coding | actin binding LIM protein 1 |
| **ERMP1** | ENSG00000099219 | 2319.03257 | -1.106548459 | 0.134026782 | 3.12E-17 | 5.54E-16 | 79956 | 9 | 5765076 | 5833117 | -1 | protein_coding | endoplasmic reticulum metallopeptidase 1 |
| **RAB18** | ENSG00000099246 | 2718.105945 | 0.956862983 | 0.155988961 | 2.5E-10 | 2.54E-09 | 22931 | 10 | 27793197 | 27831143 | 1 | protein_coding | RAB18, member RAS oncogene family |
| **NRP1** | ENSG00000099250 | 18474.12949 | -1.298191065 | 0.097559169 | 4.37E-41 | 2.83E-39 | 8829 | 10 | 33466420 | 33625190 | -1 | protein_coding | neuropilin 1 |
| **TSPAN15** | ENSG00000099282 | 764.0272189 | -3.23519848 | 0.277240655 | 1.2E-32 | 5.32E-31 | 23555 | 10 | 71211229 | 71267425 | 1 | protein_coding | tetraspanin 15 |
| **FAM21A** | ENSG00000099290 | 2274.340887 | 0.774344533 | 0.126812791 | 3.72E-10 | 3.74E-09 | 1019305 | 10 | 51827648 | 51893269 | 1 | protein_coding | family with sequence similarity 21, member A |
| **MAST3** | ENSG00000099308 | 842.3544643 | -0.675615631 | 0.172548299 | 4.47E-05 | 2.17E-04 | 23031 | 19 | 18208603 | 18262502 | 1 | protein_coding | microtubule associated serine/threonine kinase 3 |
| **MZF1** | ENSG00000099326 | 296.1410058 | -0.484713837 | 0.220704577 | 0.017564635 | 0.045536948 | 7593 | 19 | 59073298 | 59084942 | -1 | protein_coding | myeloid zinc finger 1 |
| **MYO9B** | ENSG00000099331 | 12522.91173 | -1.128238539 | 0.095167576 | 6.95E-33 | 3.18E-31 | 4650 | 19 | 17186591 | 17325346 | 1 | protein_coding | myosin IXB |
| **KCNK6** | ENSG00000099337 | 467.27269 | -0.59993233 | 0.210412949 | 0.00242589 | 0.008007532 | 9424 | 19 | 38810484 | 38819660 | 1 | protein_coding | potassium channel, subfamily K, member 6 |
| **HSD3B7** | ENSG00000099377 | 2481.529034 | -0.539596137 | 0.11702631 | 2.51E-06 | 1.53E-05 | 80270 | 16 | 30996519 | 31000473 | 1 | protein_coding | hydroxy-delta-5-steroid dehydrogenase, 3 beta- and steroid delta-isomerase 7 |
| **ATP5D** | ENSG00000099624 | 286.1505862 | -0.998638195 | 0.233005882 | 5.11E-06 | 2.95E-05 | 513 | 19 | 1241749 | 1244824 | 1 | protein_coding | ATP synthase, H+ transporting, mitochondrial F1 complex, delta subunit |
| **2-Mar** | ENSG00000099785 | 1857.845679 | -1.35898664 | 0.177704171 | 3.18E-15 | 4.89E-14 | 51257 | 19 | 8478154 | 8503901 | 1 | protein_coding | membrane-associated ring finger (C3HC4) 2, E3 ubiquitin protein ligase |
| **NDUFB7** | ENSG00000099795 | 264.0068949 | -0.967047266 | 0.224460212 | 5.02E-06 | 2.9E-05 | 4713 | 19 | 14676890 | 14682874 | -1 | protein_coding | NADH dehydrogenase (ubiquinone) 1 beta subcomplex, 7, 18kDa |
| **TECR** | ENSG00000099797 | 1335.317903 | -1.141337657 | 0.133118819 | 2.45E-18 | 4.76E-17 | 9524 | 19 | 14627897 | 14676792 | 1 | protein_coding | trans-2,3-enoyl-CoA reductase |
| **TIMM13** | ENSG00000099800 | 306.6761802 | -0.726030056 | 0.213269617 | 2.93E-04 | 0.001210722 | 26517 | 19 | 2425622 | 2427892 | -1 | protein_coding | translocase of inner mitochondrial membrane 13 homolog (yeast) |
| **CDC34** | ENSG00000099804 | 407.2920187 | -0.497748746 | 0.1759005 | 0.003081393 | 0.009945708 | 997 | 19 | 531714 | 542097 | 1 | protein_coding | cell division cycle 34 |
| **HCN2** | ENSG00000099822 | 25.71730641 | -1.956493682 | 0.96961055 | 0.002739901 | 0.008930244 | 610 | 19 | 589893 | 617159 | 1 | protein_coding | hyperpolarization activated cyclic nucleotide-gated potassium channel 2 |
| **RASSF7** | ENSG00000099849 | 104.2124517 | -0.709301261 | 0.326679532 | 0.012747108 | 0.034468162 | 8045 | 11 | 560404 | 564021 | 1 | protein_coding | Ras association (RalGDS/AF-6) domain family (N-terminal) member 7 |
| **GADD45B** | ENSG00000099860 | 344.0086408 | 2.260039894 | 0.261464862 | 4.74E-19 | 9.76E-18 | 4616 | 19 | 2476120 | 2478257 | 1 | protein_coding | growth arrest and DNA-damage-inducible, beta |
| **TRMT2A** | ENSG00000099899 | 666.7306384 | -0.784351551 | 0.177115397 | 3.89E-06 | 2.29E-05 | 27037 | 22 | 20099389 | 20104915 | -1 | protein_coding | tRNA methyltransferase 2 homolog A (S. cerevisiae) |
| **ZDHHC8** | ENSG00000099904 | 342.3756818 | -0.588225466 | 0.227489639 | 0.005457855 | 0.016471643 | 29801 | 22 | 20116979 | 20135530 | 1 | protein_coding | zinc finger, DHHC-type containing 8 |
| **KLHL22** | ENSG00000099910 | 288.1416764 | -0.576988124 | 0.238799298 | 0.008738766 | 0.024875127 | 84861 | 22 | 20783528 | 20850170 | -1 | protein_coding | kelch-like family member 22 |
| **MED15** | ENSG00000099917 | 1797.673643 | 0.299131402 | 0.121632272 | 0.012887126 | 0.034815972 | 51586 | 22 | 20850200 | 20941919 | 1 | protein_coding | mediator complex subunit 15 |
| **SNAP29** | ENSG00000099940 | 2390.502403 | -0.381800106 | 0.139588123 | 0.00487838 | 0.014959126 | 9342 | 22 | 21213271 | 21245506 | 1 | protein_coding | synaptosomal-associated protein, 29kDa |
| **CECR2** | ENSG00000099954 | 58.4666687 | -0.894191535 | 0.434996663 | 0.011162326 | 0.030765342 | 27443 | 22 | 17840837 | 18037850 | 1 | protein_coding | cat eye syndrome chromosome region, candidate 2 |
| **BCL2L13** | ENSG00000099968 | 2942.531504 | 0.827916458 | 0.088150968 | 2.32E-21 | 5.6E-20 | 23786 | 22 | 18111621 | 18213388 | 1 | protein_coding | BCL2-like 13 (apoptosis facilitator) |
| **DDT** | ENSG00000099977 | 368.6803549 | -0.762815092 | 0.202241899 | 7.23E-05 | 3.39E-04 | 1000374 | 22 | 24313554 | 24322660 | -1 | protein_coding | D-dopachrome tautomerase |
| **OSM** | ENSG00000099985 | 66.02683924 | 2.848802593 | 0.421162326 | 1.01E-12 | 1.28E-11 | 5008 | 22 | 30658818 | 30662829 | -1 | protein_coding | oncostatin M |
| **SEC14L2** | ENSG00000100003 | 218.4972583 | -2.002312667 | 0.380948998 | 1.3E-08 | 1.09E-07 | 23541 | 22 | 30792846 | 30821305 | 1 | protein_coding | SEC14-like 2 (S. cerevisiae) |
| **PPM1F** | ENSG00000100034 | 628.8047037 | -0.556515089 | 0.253185107 | 0.015902576 | 0.041809576 | 9647 | 22 | 22273793 | 22307209 | -1 | protein_coding | protein phosphatase, Mg2+/Mn2+ dependent, 1F |
| **CYTH4** | ENSG00000100055 | 5163.577938 | -0.980182736 | 0.108288284 | 2.29E-20 | 5.18E-19 | 27128 | 22 | 37678068 | 37711382 | 1 | protein_coding | cytohesin 4 |
| **MFNG** | ENSG00000100060 | 657.9494195 | -0.774413486 | 0.192583302 | 2.5E-05 | 1.28E-04 | 4242 | 22 | 37865101 | 37882439 | -1 | protein_coding | MFNG O-fucosylpeptide 3-beta-N-acetylglucosaminyltransferase |
| **SLC25A1** | ENSG00000100075 | 701.0128801 | -0.423034637 | 0.165608911 | 0.007985183 | 0.02298692 | 6576 | 22 | 19163095 | 19166343 | -1 | protein_coding | solute carrier family 25 (mitochondrial carrier; citrate transporter), member 1 |
| **ADRBK2** | ENSG00000100077 | 4934.970532 | 0.573235634 | 0.147766842 | 5.9E-05 | 2.81E-04 | 157 | 22 | 25960816 | 26125261 | 1 | protein_coding | adrenergic, beta, receptor kinase 2 |
| **GGA1** | ENSG00000100083 | 1197.689935 | -0.544503733 | 0.124231351 | 8.37E-06 | 4.66E-05 | 26088 | 22 | 38004481 | 38029571 | 1 | protein_coding | golgi-associated, gamma adaptin ear containing, ARF binding protein 1 |
| **HIRA** | ENSG00000100084 | 949.0985581 | 1.608127412 | 0.235254139 | 9.09E-13 | 1.16E-11 | 7290 | 22 | 19318221 | 19435224 | -1 | protein_coding | histone cell cycle regulator |
| **HPS4** | ENSG00000100099 | 1208.349048 | 0.316562156 | 0.130890767 | 0.012689379 | 0.034342442 | 89781 | 22 | 26839389 | 26879803 | -1 | protein_coding | Hermansky-Pudlak syndrome 4 |
| **PIK3IP1** | ENSG00000100100 | 1513.526781 | -0.911586553 | 0.164019255 | 9.25E-09 | 7.9E-08 | 113791 | 22 | 31677579 | 31688520 | -1 | protein_coding | phosphoinositide-3-kinase interacting protein 1 |
| **SRRD** | ENSG00000100104 | 741.3786923 | -0.437051513 | 0.194551011 | 0.01849925 | 0.047492699 | 402055 | 22 | 26879843 | 26890624 | 1 | protein_coding | SRR1 domain containing |
| **EIF3L** | ENSG00000100129 | 3980.697123 | -1.152618608 | 0.114044473 | 1.49E-24 | 4.33E-23 | 51386 | 22 | 38244875 | 38285414 | 1 | protein_coding | eukaryotic translation initiation factor 3, subunit L |
| **TTC28** | ENSG00000100154 | 997.2202252 | -1.212606455 | 0.199942519 | 2.74E-10 | 2.78E-09 | 23331 | 22 | 28374004 | 29075853 | -1 | protein_coding | tetratricopeptide repeat domain 28 |
| **HSCB** | ENSG00000100209 | 159.0222334 | 0.89329303 | 0.331839141 | 0.002211341 | 0.007377411 | 150274 | 22 | 29138019 | 29153503 | 1 | protein_coding | HscB mitochondrial iron-sulfur cluster co-chaperone |
| **TOMM22** | ENSG00000100216 | 1033.404647 | -0.387095709 | 0.126422554 | 0.001776939 | 0.006079461 | 56993 | 22 | 39077953 | 39080818 | 1 | protein_coding | translocase of outer mitochondrial membrane 22 homolog (yeast) |
| **XBP1** | ENSG00000100219 | 1853.047279 | 0.513993931 | 0.109109949 | 1.72E-06 | 1.07E-05 | 7494 | 22 | 29190543 | 29196585 | -1 | protein_coding | X-box binding protein 1 |
| **RTCB** | ENSG00000100220 | 2733.135811 | 1.777267789 | 0.099259345 | 1.38E-72 | 2.78E-70 | 51493 | 22 | 32783569 | 32808242 | -1 | protein_coding | RNA 2',3'-cyclic phosphate and 5'-OH ligase |
| **FBXO7** | ENSG00000100225 | 3490.93662 | 1.086571289 | 0.099360437 | 2.1E-28 | 7.48E-27 | 25793 | 22 | 32870663 | 32894818 | 1 | protein_coding | F-box protein 7 |
| **GTPBP1** | ENSG00000100226 | 1367.253753 | 2.386863453 | 0.220237594 | 2.03E-28 | 7.26E-27 | 9567 | 22 | 39101728 | 39134304 | 1 | protein_coding | GTP binding protein 1 |
| **POLDIP3** | ENSG00000100227 | 1448.527435 | 0.406204702 | 0.148148767 | 0.004943524 | 0.015128529 | 84271 | 22 | 42979727 | 43010968 | -1 | protein_coding | polymerase (DNA-directed), delta interacting protein 3 |
| **TIMP3** | ENSG00000100234 | 10239.78751 | -2.797449636 | 0.107248157 | 2.35E-151 | 1.33E-148 | 7078 | 22 | 33197687 | 33259030 | 1 | protein_coding | TIMP metallopeptidase inhibitor 3 |
| **CYB5R3** | ENSG00000100243 | 2123.646858 | 0.965226229 | 0.12113309 | 5.4E-16 | 8.79E-15 | 1727 | 22 | 43013846 | 43045574 | -1 | protein_coding | cytochrome b5 reductase 3 |
| **DNAL4** | ENSG00000100246 | 114.8222475 | -0.731223898 | 0.317805248 | 0.008813987 | 0.025065902 | 10126 | 22 | 39174513 | 39190203 | -1 | protein_coding | dynein, axonemal, light chain 4 |
| **RHBDD3** | ENSG00000100263 | 149.5386864 | 0.709488012 | 0.294831432 | 0.007030787 | 0.020587189 | 25807 | 22 | 29655841 | 29664198 | -1 | protein_coding | rhomboid domain containing 3 |
| **PACSIN2** | ENSG00000100266 | 3640.655992 | -0.541856614 | 0.09847521 | 2.5E-08 | 2.01E-07 | 11252 | 22 | 43231418 | 43411151 | -1 | protein_coding | protein kinase C and casein kinase substrate in neurons 2 |
| **TOM1** | ENSG00000100284 | 1736.870959 | 0.607721039 | 0.121164655 | 3.02E-07 | 2.1E-06 | 10043 | 22 | 35695268 | 35743985 | 1 | protein_coding | target of myb1 (chicken) |
| **CHKB** | ENSG00000100288 | 777.9376748 | 0.388295771 | 0.138618822 | 0.00400107 | 0.012517574 | 1120 | 22 | 51017378 | 51039884 | -1 | protein_coding | choline kinase beta |
| **APOBEC3H** | ENSG00000100298 | 59.65108171 | 0.987866612 | 0.510179027 | 0.011550151 | 0.031685686 | 164668 | 22 | 39493229 | 39500072 | 1 | protein_coding | apolipoprotein B mRNA editing enzyme, catalytic polypeptide-like 3H |
| **TSPO** | ENSG00000100300 | 1101.926517 | -0.713017079 | 0.142947677 | 2.88E-07 | 2.01E-06 | 706 | 22 | 43547520 | 43559248 | 1 | protein_coding | translocator protein (18kDa) |
| **TTLL12** | ENSG00000100304 | 660.3465536 | -1.069691951 | 0.159284778 | 5.23E-12 | 6.33E-11 | 23170 | 22 | 43562628 | 43583139 | -1 | protein_coding | tubulin tyrosine ligase-like family, member 12 |
| **CBX7** | ENSG00000100307 | 388.404382 | -0.846734175 | 0.200397913 | 8.63E-06 | 4.79E-05 | 23492 | 22 | 39516172 | 39548679 | -1 | protein_coding | chromobox homolog 7 |
| **RPL3** | ENSG00000100316 | 9850.818024 | -1.098883022 | 0.095483954 | 3.19E-31 | 1.31E-29 | 116938 | 22 | 39708887 | 39716394 | -1 | protein_coding | ribosomal protein L3 |
| **ZMAT5** | ENSG00000100319 | 163.4628287 | -0.599835834 | 0.252921108 | 0.009870684 | 0.027689733 | 55954 | 22 | 30126945 | 30163000 | -1 | protein_coding | zinc finger, matrin-type 5 |
| **SYNGR1** | ENSG00000100321 | 2368.927241 | -0.586165492 | 0.220456113 | 0.004527025 | 0.01397143 | 9145 | 22 | 39745930 | 39781593 | 1 | protein_coding | synaptogyrin 1 |
| **MIEF1** | ENSG00000100335 | 2925.065741 | -0.497126322 | 0.095912226 | 1.75E-07 | 1.26E-06 | 54471 | 22 | 39895437 | 39914137 | 1 | protein_coding | mitochondrial elongation factor 1 |
| **APOL1** | ENSG00000100342 | 2450.486089 | 2.242647563 | 0.181559259 | 4.71E-36 | 2.46E-34 | 8542 | 22 | 36649056 | 36663576 | 1 | protein_coding | apolipoprotein L, 1 |
| **PNPLA3** | ENSG00000100344 | 148.6141644 | -1.506883502 | 0.509605319 | 3.39E-04 | 0.001382814 | 80339 | 22 | 44319619 | 44360368 | 1 | protein_coding | patatin-like phospholipase domain containing 3 |
| **MYH9** | ENSG00000100345 | 22171.44279 | -0.400139855 | 0.095828741 | 2.32E-05 | 1.2E-04 | 4627 | 22 | 36677327 | 36784063 | -1 | protein_coding | myosin, heavy chain 9, non-muscle |
| **SAMM50** | ENSG00000100347 | 1018.394989 | -0.721476448 | 0.167912766 | 8.14E-06 | 4.54E-05 | 25813 | 22 | 44351301 | 44406411 | 1 | protein_coding | SAMM50 sorting and assembly machinery component |
| **TXN2** | ENSG00000100348 | 999.3254278 | -0.515244275 | 0.134776118 | 8.58E-05 | 3.97E-04 | 25828 | 22 | 36863083 | 36878077 | -1 | protein_coding | thioredoxin 2 |
| **FOXRED2** | ENSG00000100350 | 172.820367 | -0.777054569 | 0.31979577 | 0.005603186 | 0.016837101 | 80020 | 22 | 36883237 | 36903148 | -1 | protein_coding | FAD-dependent oxidoreductase domain containing 2 |
| **SGSM3** | ENSG00000100359 | 607.8932531 | -1.0948929 | 0.204235962 | 1.84E-08 | 1.51E-07 | 27352 | 22 | 40766595 | 40806293 | 1 | protein_coding | small G protein signaling modulator 3 |
| **IFT27** | ENSG00000100360 | 274.6940606 | -0.904592418 | 0.343344564 | 0.0022375 | 0.007456544 | 11020 | 22 | 37154246 | 37172300 | -1 | protein_coding | intraflagellar transport 27 homolog (Chlamydomonas) |
| **PVALB** | ENSG00000100362 | 97.52790414 | -2.674704071 | 0.41301466 | 8.58E-12 | 1.01E-10 | 5816 | 22 | 37196728 | 37215523 | -1 | protein_coding | parvalbumin |
| **KIAA0930** | ENSG00000100364 | 18440.9718 | -1.393550498 | 0.104367523 | 2.13E-41 | 1.4E-39 | 23313 | 22 | 45586219 | 45636650 | -1 | protein_coding | KIAA0930 |
| **NCF4** | ENSG00000100365 | 1233.27948 | 0.538796052 | 0.139414602 | 6.87E-05 | 3.23E-04 | 4689 | 22 | 37257030 | 37274057 | 1 | protein_coding | neutrophil cytosolic factor 4, 40kDa |
| **CSF2RB** | ENSG00000100368 | 3436.104775 | 2.2249166 | 0.198649128 | 2.88E-30 | 1.13E-28 | 1439 | 22 | 37309670 | 37336491 | 1 | protein_coding | colony stimulating factor 2 receptor, beta, low-affinity (granulocyte-macrophage) |
| **ST13** | ENSG00000100380 | 6704.743551 | -0.43208442 | 0.141076967 | 0.001620314 | 0.005608892 | 6767 | 22 | 41220539 | 41253026 | -1 | protein_coding | suppression of tumorigenicity 13 (colon carcinoma) (Hsp70 interacting protein) |
| **RBX1** | ENSG00000100387 | 843.5767182 | 0.52457806 | 0.139263535 | 1.09E-04 | 4.96E-04 | 9978 | 22 | 41347351 | 41369313 | 1 | protein_coding | ring-box 1, E3 ubiquitin protein ligase |
| **L3MBTL2** | ENSG00000100395 | 888.8319358 | -0.356594495 | 0.130249894 | 0.004986915 | 0.01523691 | 83746 | 22 | 41601209 | 41627275 | 1 | protein_coding | l(3)mbt-like 2 (Drosophila) |
| **RANGAP1** | ENSG00000100401 | 1712.19502 | 0.781562809 | 0.124869129 | 1.64E-10 | 1.71E-09 | 5905 | 22 | 41641615 | 41682255 | -1 | protein_coding | Ran GTPase activating protein 1 |
| **PHF5A** | ENSG00000100410 | 341.3620341 | 0.434393237 | 0.179349788 | 0.011439664 | 0.031427715 | 84844 | 22 | 41855721 | 41864729 | -1 | protein_coding | PHD finger protein 5A |
| **ACO2** | ENSG00000100412 | 4723.58188 | -0.966163134 | 0.100844362 | 2.93E-22 | 7.47E-21 | 50 | 22 | 41865129 | 41924993 | 1 | protein_coding | aconitase 2, mitochondrial |
| **POLR3H** | ENSG00000100413 | 547.0807233 | -0.803065399 | 0.168666497 | 8.02E-07 | 5.25E-06 | 171568 | 22 | 41921808 | 41940610 | -1 | protein_coding | polymerase (RNA) III (DNA directed) polypeptide H (22.9kD) |
| **DESI1** | ENSG00000100418 | 2135.819387 | 1.186940453 | 0.186913638 | 6.4E-11 | 6.99E-10 | 27351 | 22 | 41994032 | 42017100 | -1 | protein_coding | desumoylating isopeptidase 1 |
| **CERK** | ENSG00000100422 | 1392.567048 | -1.819519328 | 0.152696462 | 1.17E-33 | 5.55E-32 | 1019276 | 22 | 47080308 | 47134158 | -1 | protein_coding | ceramide kinase |
| **ZBED4** | ENSG00000100426 | 993.2744973 | -0.392610471 | 0.127131388 | 0.001650084 | 0.005702011 | 9889 | 22 | 50247490 | 50283726 | 1 | protein_coding | zinc finger, BED-type containing 4 |
| **KHNYN** | ENSG00000100441 | 3042.096712 | 0.308012476 | 0.114439215 | 0.006036209 | 0.017982725 | 23351 | 14 | 24898492 | 24910540 | 1 | protein_coding | KH and NYN domain containing |
| **FKBP3** | ENSG00000100442 | 867.404908 | 0.730514011 | 0.154903679 | 1.16E-06 | 7.37E-06 | 2287 | 14 | 45584803 | 45604522 | -1 | protein_coding | FK506 binding protein 3, 25kDa |

| | | | | | | | | | | | | | |
|---|---|---|---|---|---|---|---|---|---|---|---|---|---|
| **SDR39U1** | ENSG00000100445 | 336.6479193 | -0.610266255 | 0.189975684 | 7.54E-04 | 0.002834618 | 56948 | 14 | 24908972 | 24912111 | -1 | protein_coding | short chain dehydrogenase/reductase family 39U, member 1 |
| **GZMB** | ENSG00000100453 | 304.0253788 | 1.591559003 | 0.473123083 | 6.91E-05 | 3.25E-04 | 3002 | 14 | 25100160 | 25103473 | -1 | protein_coding | granzyme B (granzyme 2, cytotoxic T-lymphocyte-associated serine esterase 1) |
| **RBM23** | ENSG00000100461 | 1513.267118 | 0.364989976 | 0.121601306 | 0.002255363 | 0.007511159 | 55147 | 14 | 23369854 | 23388393 | -1 | protein_coding | RNA binding motif protein 23 |
| **COCH** | ENSG00000100473 | 56.69392668 | 1.146024694 | 0.469241065 | 0.002635018 | 0.008615958 | 1690 | 14 | 31343720 | 31364271 | 1 | protein_coding | cochlin |
| **AP4S1** | ENSG00000100478 | 148.3668295 | 0.657333568 | 0.291376994 | 0.011356755 | 0.03122801 | 11154 | 14 | 31494312 | 31562818 | 1 | protein_coding | adaptor-related protein complex 4, sigma 1 subunit |
| **CDKL1** | ENSG00000100490 | 167.4068104 | 1.548625222 | 0.27399909 | 2.26E-09 | 2.07E-08 | 8814 | 14 | 50796310 | 50883179 | -1 | protein_coding | cyclin-dependent kinase-like 1 (CDC2-related kinase) |
| **NIN** | ENSG00000100503 | 4152.346995 | -1.080390716 | 0.108667954 | 7.28E-24 | 2.05E-22 | 51199 | 14 | 51186481 | 51297839 | -1 | protein_coding | ninein (GSK3B interacting protein) |
| **PYGL** | ENSG00000100504 | 2549.67418 | -2.376496934 | 0.104341039 | 4.59E-116 | 1.8E-113 | 5836 | 14 | 51324609 | 51411454 | -1 | protein_coding | phosphorylase, glycogen, liver |
| **DDHD1** | ENSG00000100523 | 1332.86892 | 0.906218908 | 0.141245146 | 4.71E-11 | 5.22E-10 | 80821 | 14 | 53510686 | 53620000 | -1 | protein_coding | DDHD domain containing 1 |
| **PIGH** | ENSG00000100564 | 240.4403857 | 0.544452442 | 0.24401903 | 0.015794426 | 0.041589629 | 5283 | 14 | 68048672 | 68067004 | -1 | protein_coding | phosphatidylinositol glycan anchor biosynthesis, class H |
| **PSMA3** | ENSG00000100567 | 1838.081199 | 1.239960638 | 0.115327888 | 1.29E-27 | 4.36E-26 | 5684 | 14 | 58711549 | 58738730 | 1 | protein_coding | proteasome (prosome, macropain) subunit, alpha type, 3 |
| **VTI1B** | ENSG00000100568 | 3559.358722 | -0.501864218 | 0.134174114 | 1.29E-04 | 5.77E-04 | 10490 | 14 | 68113792 | 68141548 | -1 | protein_coding | vesicle transport through interaction with t-SNAREs 1B |
| **AHSA1** | ENSG00000100591 | 1289.054112 | 0.383444484 | 0.146375057 | 0.007730469 | 0.022358964 | 10598 | 14 | 77924213 | 77935817 | 1 | protein_coding | AHA1, activator of heat shock 90kDa protein ATPase homolog 1 (yeast) |
| **DAAM1** | ENSG00000100592 | 424.4658258 | -1.667175853 | 0.282605726 | 4.36E-10 | 4.34E-09 | 23002 | 14 | 59655364 | 59838123 | 1 | protein_coding | dishevelled associated activator of morphogenesis 1 |
| **SPTLC2** | ENSG00000100596 | 2275.367239 | 0.828442833 | 0.116590934 | 4.74E-13 | 6.24E-12 | 9517 | 14 | 77972340 | 78083116 | -1 | protein_coding | serine palmitoyltransferase, long chain base subunit 2 |
| **RIN3** | ENSG00000100599 | 3939.806738 | -1.807767967 | 0.119021391 | 3.97E-53 | 4.18E-51 | 79890 | 14 | 92980118 | 93155339 | 1 | protein_coding | Ras and Rab interactor 3 |
| **LGMN** | ENSG00000100600 | 12115.07888 | 0.402180222 | 0.166903624 | 0.012483342 | 0.033850765 | 5641 | 14 | 93170152 | 93215047 | -1 | protein_coding | legumain |
| **ITPK1** | ENSG00000100605 | 2857.094721 | -1.020807823 | 0.115774627 | 3.42E-19 | 7.09E-18 | 3705 | 14 | 93403259 | 93582665 | -1 | protein_coding | inositol-tetrakisphosphate 1-kinase |
| **ASB2** | ENSG00000100628 | 646.5618212 | -1.921726665 | 0.16340437 | 6.07E-33 | 2.79E-31 | 51676 | 14 | 94400499 | 94443137 | -1 | protein_coding | ankyrin repeat and SOCS box containing 2 |
| **ERH** | ENSG00000100632 | 1131.410843 | 0.779624452 | 0.133601613 | 2.34E-09 | 2.15E-08 | 2079 | 14 | 69846848 | 69865344 | -1 | protein_coding | enhancer of rudimentary homolog (Drosophila) |
| **HIF1A** | ENSG00000100644 | 7354.688657 | 1.156609801 | 0.097620139 | 8.04E-33 | 3.64E-31 | 3091 | 14 | 62162231 | 62214976 | 1 | protein_coding | hypoxia inducible factor 1, alpha subunit (basic helix-loop-helix transcription factor) |
| **KIAA0247** | ENSG00000100647 | 6560.29405 | 0.893605153 | 0.138115037 | 3.7E-11 | 4.14E-10 | 9766 | 14 | 70078313 | 70181859 | 1 | protein_coding | KIAA0247 |
| **SRSF5** | ENSG00000100650 | 4198.696588 | 0.41668167 | 0.111887636 | 1.54E-04 | 6.8E-04 | 6430 | 14 | 70193617 | 70238722 | 1 | protein_coding | serine/arginine-rich splicing factor 5 |
| **EIF5** | ENSG00000100664 | 8258.672891 | 0.698311635 | 0.112813423 | 3.79E-10 | 3.8E-09 | 1983 | 14 | 103799881 | 103811362 | 1 | protein_coding | eukaryotic translation initiation factor 5 |
| **DICER1** | ENSG00000100697 | 5181.009742 | 0.545048917 | 0.095445342 | 7.02E-09 | 6.09E-08 | 23405 | 14 | 95552565 | 95624347 | -1 | protein_coding | dicer 1, ribonuclease type III |
| **ZFYVE21** | ENSG00000100711 | 548.4766229 | -0.965359986 | 0.181539365 | 3.34E-08 | 2.65E-07 | 79038 | 14 | 104182067 | 104200005 | 1 | protein_coding | zinc finger, FYVE domain containing 21 |
| **PCNX** | ENSG00000100731 | 3907.105755 | 0.791550488 | 0.162043025 | 4.56E-07 | 3.1E-06 | 22990 | 14 | 71374122 | 71582099 | 1 | protein_coding | pecanex homolog (Drosophila) |
| **PSMC1** | ENSG00000100764 | 2685.980688 | 0.536403168 | 0.096570517 | 1.57E-08 | 1.3E-07 | 5700 | 14 | 90722839 | 90738968 | 1 | protein_coding | proteasome (prosome, macropain) 26S subunit, ATPase, 1 |
| **YY1** | ENSG00000100811 | 2567.961488 | 0.539709349 | 0.132436763 | 2.89E-05 | 1.46E-04 | 7528 | 14 | 100704635 | 100749129 | 1 | protein_coding | YY1 transcription factor |
| **APEX1** | ENSG00000100823 | 2339.899082 | -0.584743766 | 0.109918339 | 7.94E-08 | 5.98E-07 | 328 | 14 | 20923350 | 20925927 | 1 | protein_coding | APEX nuclease (multifunctional DNA repair enzyme) 1 |
| **ARHGAP5** | ENSG00000100852 | 2590.662307 | 0.861938671 | 0.169077193 | 1.16E-07 | 8.52E-07 | 394 | 14 | 32545320 | 32628934 | 1 | protein_coding | Rho GTPase activating protein 5 |
| **SRP54** | ENSG00000100883 | 2165.503224 | 0.944668679 | 0.103951374 | 3.58E-20 | 8.02E-19 | 6729 | 14 | 35451163 | 35498773 | 1 | protein_coding | signal recognition particle 54kDa |
| **CPNE6** | ENSG00000100884 | 6.846385393 | 0.511703311 | 0.926930829 | 0.016729568 | 0.043645952 | 9362 | 14 | 24540046 | 24547309 | 1 | protein_coding | copine VI (neuronal) |
| **CHD8** | ENSG00000100888 | 2867.437644 | 0.738492598 | 0.126502902 | 2.7E-09 | 2.46E-08 | 57680 | 14 | 21853353 | 21924285 | -1 | protein_coding | chromodomain helicase DNA binding protein 8 |
| **PCK2** | ENSG00000100889 | 3651.936993 | -0.383225007 | 0.105847833 | 2.34E-04 | 9.86E-04 | 5106 | 14 | 24563262 | 24579807 | 1 | protein_coding | phosphoenolpyruvate carboxykinase 2 (mitochondrial) |
| **DCAF11** | ENSG00000100897 | 1903.751342 | -0.312347809 | 0.121799215 | 0.00892361 | 0.025339937 | 80344 | 14 | 24583404 | 24594451 | 1 | protein_coding | DDB1 and CUL4 associated factor 11 |
| **PSMA6** | ENSG00000100902 | 4300.702662 | 1.737025648 | 0.114617101 | 9.23E-53 | 9.34E-51 | 5687 | 14 | 35747839 | 35786699 | 1 | protein_coding | proteasome (prosome, macropain) subunit, alpha type, 6 |
| **NFKBIA** | ENSG00000100906 | 3550.3414 | 2.347220969 | 0.118103627 | 4.9E-89 | 1.44E-86 | 4792 | 14 | 35870717 | 35873955 | -1 | protein_coding | nuclear factor of kappa light polypeptide gene enhancer in B-cells inhibitor, alpha |
| **PSME2** | ENSG00000100911 | 6626.162923 | 1.876535433 | 0.206929899 | 1.59E-20 | 3.65E-19 | 1024658 | 14 | 24612574 | 24616779 | -1 | protein_coding | proteasome (prosome, macropain) activator subunit 2 (PA28 beta) |
| **GMPR2** | ENSG00000100938 | 1312.605204 | -0.566991853 | 0.142783649 | 4.42E-05 | 2.15E-04 | 51292 | 14 | 24701628 | 24708448 | 1 | protein_coding | guanosine monophosphate reductase 2 |
| **PNN** | ENSG00000100941 | 1517.841584 | 0.327876586 | 0.11712153 | 0.004328077 | 0.013422484 | 5411 | 14 | 39644387 | 39652422 | 1 | protein_coding | pinin, desmosome associated protein |
| **RABGGTA** | ENSG00000100949 | 870.5094335 | 0.432961802 | 0.169312135 | 0.007902312 | 0.022778392 | 5875 | 14 | 24734744 | 24740945 | -1 | protein_coding | Rab geranylgeranyltransferase, alpha subunit |
| **MMP9** | ENSG00000100985 | 231092.0504 | -0.62602793 | 0.153554666 | 2.45E-05 | 1.26E-04 | 4318 | 20 | 44637547 | 44645200 | 1 | protein_coding | matrix metallopeptidase 9 (gelatinase B, 92kDa gelatinase, 92kDa type IV collagenase) |
| **TRPC4AP** | ENSG00000100991 | 2743.031599 | 0.724053228 | 0.094822985 | 5.5E-15 | 8.36E-14 | 26133 | 20 | 33590207 | 33680674 | -1 | protein_coding | transient receptor potential cation channel, subfamily C, member 4 associated protein |
| **PYGB** | ENSG00000100994 | 1991.575746 | -0.820719449 | 0.136171718 | 6.38E-10 | 6.24E-09 | 5834 | 20 | 25228705 | 25278650 | 1 | protein_coding | phosphorylase, glycogen; brain |
| **PROCR** | ENSG00000101000 | 74.14656967 | 2.016506496 | 0.422574515 | 1.58E-07 | 1.15E-06 | 10544 | 20 | 33759876 | 33765165 | 1 | protein_coding | protein C receptor, endothelial |
| **NINL** | ENSG00000101004 | 310.9886129 | -0.877425407 | 0.236759398 | 6.26E-05 | 2.97E-04 | 22981 | 20 | 25433341 | 25566153 | -1 | protein_coding | ninein-like |
| **CD40** | ENSG00000101017 | 4545.143423 | 1.911996379 | 0.220651129 | 4.86E-19 | 9.98E-18 | 958 | 20 | 44746911 | 44758502 | 1 | protein_coding | CD40 molecule, TNF receptor superfamily member 5 |
| **ZMYND8** | ENSG00000101040 | 1572.893737 | -0.483889271 | 0.168392454 | 0.002792644 | 0.009078915 | 23613 | 20 | 45837859 | 45985567 | -1 | protein_coding | zinc finger, MYND-type containing 8 |
| **NDRG3** | ENSG00000101079 | 1331.184246 | -0.409144778 | 0.141668771 | 0.002943728 | 0.009535606 | 57446 | 20 | 35280169 | 35374481 | -1 | protein_coding | NDRG family member 3 |
| **C20orf24** | ENSG00000101084 | 4030.161356 | 0.614768426 | 0.088301529 | 1.93E-12 | 2.4E-11 | 55969 | 20 | 35234137 | 35240960 | 1 | protein_coding | chromosome 20 open reading frame 24 |
| **PABPC1L** | ENSG00000101104 | 204.955181 | -0.621648334 | 0.283696113 | 0.014246012 | 0.037983565 | 80336 | 20 | 43538703 | 43587676 | 1 | protein_coding | poly(A) binding protein, cytoplasmic 1-like |
| **STK4** | ENSG00000101109 | 5546.312868 | 0.571466054 | 0.105855952 | 4.04E-08 | 3.17E-07 | 6789 | 20 | 43595115 | 43708600 | 1 | protein_coding | serine/threonine kinase 4 |
| **SALL4** | ENSG00000101115 | 21.9107851 | -2.705927677 | 0.978254973 | 4.62E-04 | 0.001822908 | 57167 | 20 | 50400581 | 50419059 | -1 | protein_coding | spalt-like transcription factor 4 |
| **PFDN4** | ENSG00000101132 | 168.4267528 | 0.806055444 | 0.373479936 | 0.010671301 | 0.029559146 | 5203 | 20 | 52824386 | 52844591 | 1 | protein_coding | prefoldin subunit 4 |
| **BMP7** | ENSG00000101144 | 6.253870183 | -0.871853357 | 1.721781047 | 0.010373112 | 0.028860781 | 655 | 20 | 55743804 | 55841685 | -1 | protein_coding | bone morphogenetic protein 7 |
| **TPD52L2** | ENSG00000101150 | 3506.248537 | -0.48353853 | 0.12650474 | 9.57E-05 | 4.39E-04 | 7165 | 20 | 62496596 | 62522898 | 1 | protein_coding | tumor protein D52-like 2 |
| **DNAJC5** | ENSG00000101152 | 4921.068487 | -0.387318641 | 0.105742955 | 1.98E-04 | 8.46E-04 | 80331 | 20 | 62526518 | 62567384 | 1 | protein_coding | DnaJ (Hsp40) homolog, subfamily C, member 5 |
| **CTSZ** | ENSG00000101160 | 48907.27621 | -0.38316153 | 0.082337552 | 2.61E-06 | 1.58E-05 | 1522 | 20 | 57570240 | 57582302 | -1 | protein_coding | cathepsin Z |
| **PRPF6** | ENSG00000101161 | 1657.268157 | -0.26971255 | 0.099832451 | 0.006530449 | 0.019300879 | 24148 | 20 | 62612488 | 62664453 | 1 | protein_coding | pre-mRNA processing factor 6 |
| **SLMO2** | ENSG00000101166 | 1276.032561 | 0.447129511 | 0.122596753 | 2.36E-04 | 9.92E-04 | 51012 | 20 | 57608200 | 57617964 | -1 | protein_coding | slowmo homolog 2 (Drosophila) |
| **SLC17A9** | ENSG00000101194 | 847.301818 | -1.798270347 | 0.211875544 | 2.1E-18 | 4.1E-17 | 63910 | 20 | 61584052 | 61599949 | 1 | protein_coding | solute carrier family 17 (vesicular nucleotide transporter), member 9 |
| **BIRC7** | ENSG00000101197 | 73.32484469 | -2.398620841 | 0.507051676 | 1.65E-07 | 1.19E-06 | 79444 | 20 | 61867235 | 61871859 | 1 | protein_coding | baculoviral IAP repeat containing 7 |
| **ARFGAP1** | ENSG00000101199 | 1122.457553 | -0.523391767 | 0.159360686 | 6.74E-04 | 0.002557508 | 55738 | 20 | 61904137 | 61921142 | 1 | protein_coding | ADP-ribosylation factor GTPase activating protein 1 |
| **C20orf27** | ENSG00000101220 | 290.4571306 | -2.301154099 | 0.239020157 | 6.01E-23 | 1.61E-21 | 54976 | 20 | 3734155 | 3749034 | -1 | protein_coding | chromosome 20 open reading frame 27 |
| **ISM1** | ENSG00000101230 | 92.2694248 | -3.705167481 | 0.7515837 | 5.64E-08 | 4.35E-07 | 140862 | 20 | 13202418 | 13281298 | 1 | protein_coding | isthmin 1, angiogenesis inhibitor |
| **RNF24** | ENSG00000101236 | 1175.742579 | 1.719077647 | 0.112588963 | 1.49E-53 | 1.62E-51 | 11237 | 20 | 3907956 | 3996229 | -1 | protein_coding | ring finger protein 24 |
| **SEL1L2** | ENSG00000101251 | 53.90689733 | -2.454669596 | 1.075122651 | 0.001173365 | 0.004205566 | 80343 | 20 | 13829893 | 13977089 | -1 | protein_coding | sel-1 suppressor of lin-12-like 2 (C. elegans) |
| **TRIB3** | ENSG00000101255 | 273.8941669 | -2.788642002 | 0.360146065 | 6.76E-16 | 1.1E-14 | 57761 | 20 | 361261 | 378203 | 1 | protein_coding | tribbles pseudokinase 3 |
| **RASSF2** | ENSG00000101265 | 2388.281498 | -1.938327314 | 0.144617186 | 7.81E-42 | 5.21E-40 | 9770 | 20 | 4760669 | 4804291 | -1 | protein_coding | Ras association (RalGDS/AF-6) domain family member 2 |
| **CSNK2A1** | ENSG00000101266 | 3860.165652 | 0.281275857 | 0.115006081 | 0.013062822 | 0.035228358 | 1457 | 20 | 459116 | 524465 | -1 | protein_coding | casein kinase 2, alpha 1 polypeptide |
| **RSPO4** | ENSG00000101282 | 46.24008109 | -2.144816152 | 0.717862369 | 2.36E-04 | 9.92E-04 | 343637 | 20 | 939095 | 982907 | -1 | protein_coding | R-spondin 4 |
| **CDS2** | ENSG00000101290 | 4956.364955 | 0.972757104 | 0.148004862 | 2.01E-11 | 2.31E-10 | 8760 | 20 | 5107432 | 5178533 | 1 | protein_coding | CDP-diacylglycerol synthase (phosphatidate cytidylyltransferase) 2 |
| **HM13** | ENSG00000101294 | 7221.861333 | -0.356666005 | 0.097565107 | 2.06E-04 | 8.76E-04 | 81502 | 20 | 30102231 | 30157370 | 1 | protein_coding | histocompatibility (minor) 13 |
| **SNPH** | ENSG00000101298 | 70.68990658 | -0.894683308 | 0.446288544 | 0.012466182 | 0.033810233 | 9751 | 20 | 1246960 | 1289972 | 1 | protein_coding | syntaphilin |
| **SIRPB1** | ENSG00000101307 | 2561.087758 | -1.327480882 | 0.11502148 | 1.75E-31 | 7.26E-30 | 1006531 | 20 | 1544167 | 1600707 | -1 | protein_coding | signal-regulatory protein beta 1 |
| **CCM2L** | ENSG00000101331 | 6.663579306 | 0.748767703 | 1.441022163 | 0.011565802 | 0.031722928 | 140706 | 20 | 30598245 | 30619984 | 1 | protein_coding | cerebral cavernous malformation 2-like |
| **MYL9** | ENSG00000101335 | 1521.645623 | -0.527819928 | 0.18019323 | 0.002203762 | 0.007355338 | 10398 | 20 | 35169887 | 35178228 | 1 | protein_coding | myosin, light chain 9, regulatory |
| **HCK** | ENSG00000101336 | 6538.40799 | 0.453866066 | 0.116459205 | 7.33E-05 | 3.43E-04 | 3055 | 20 | 30639991 | 30689659 | 1 | protein_coding | hemopoietic cell kinase |
| **SNTA1** | ENSG00000101400 | 63.5798021 | -0.945615281 | 0.496136279 | 0.013437713 | 0.03614371 | 6640 | 20 | 31995761 | 32031698 | -1 | protein_coding | syntrophin, alpha 1 |
| **CHMP4B** | ENSG00000101421 | 4571.08267 | 0.34705071 | 0.094111332 | 2.07E-04 | 8.82E-04 | 128866 | 20 | 32399110 | 32442172 | 1 | protein_coding | charged multivesicular body protein 4B |
| **CST3** | ENSG00000101439 | 4825.308662 | -0.320579068 | 0.128956187 | 0.010972502 | 0.030307745 | 1471 | 20 | 23608534 | 23619110 | -1 | protein_coding | cystatin C |
| **AHCY** | ENSG00000101444 | 2027.841657 | -0.890395197 | 0.102279438 | 1.13E-18 | 2.24E-17 | 191 | 20 | 32868074 | 32899608 | -1 | protein_coding | adenosylhomocysteinase |
| **PPP1R16B** | ENSG00000101445 | 862.9794574 | 0.588721171 | 0.279641042 | 0.019077765 | 0.048740469 | 26051 | 20 | 37434348 | 37551667 | 1 | protein_coding | protein phosphatase 1, regulatory subunit 16B |
| **MAP1LC3A** | ENSG00000101460 | 118.3494403 | 3.088715227 | 0.330027556 | 5.87E-22 | 1.48E-20 | 84557 | 20 | 33134658 | 33148149 | 1 | protein_coding | microtubule-associated protein 1 light chain 3 alpha |
| **ACOT8** | ENSG00000101473 | 316.6171239 | -0.857007179 | 0.205499213 | 1.13E-05 | 6.17E-05 | 10005 | 20 | 44470360 | 44486045 | -1 | protein_coding | acyl-CoA thioesterase 8 |
| **APMAP** | ENSG00000101474 | 3049.28615 | -1.368660919 | 0.125277858 | 1.12E-28 | 4.06E-27 | 57136 | 20 | 24943561 | 24973615 | -1 | protein_coding | adipocyte plasma membrane associated protein |
| **USP14** | ENSG00000101557 | 2464.260234 | 0.787861602 | 0.151890427 | 8.74E-08 | 6.52E-07 | 9097 | 18 | 158383 | 214629 | 1 | protein_coding | ubiquitin specific peptidase 14 (tRNA-guanine transglycosylase) |
| **METTL4** | ENSG00000101574 | 278.3058811 | 1.489871183 | 0.293568717 | 6.24E-08 | 4.77E-07 | 64863 | 18 | 2537524 | 2571508 | -1 | protein_coding | methyltransferase like 4 |

| | | | | | | | | | | | | | |
|---|---|---|---|---|---|---|---|---|---|---|---|---|---|
| LPIN2 | ENSG00000101577 | 2503.038127 | 0.631580632 | 0.125457695 | 2.56E-07 | 1.8E-06 | 9663 | 18 | 2916992 | 3013313 | -1 | protein_coding | lipin 2 |
| SMCHD1 | ENSG00000101596 | 1681.70928 | 2.224147614 | 0.167151512 | 1.43E-41 | 9.43E-40 | 23347 | 18 | 2655737 | 2805015 | 1 | protein_coding | structural maintenance of chromosomes flexible hinge domain containing 1 |
| MYL12A | ENSG00000101608 | 8994.967161 | 1.164814143 | 0.116652052 | 5.12E-24 | 1.45E-22 | 10627 | 18 | 3247479 | 3256234 | 1 | protein_coding | myosin, light chain 12A, regulatory, non-sarcomeric |
| RNF125 | ENSG00000101695 | 688.6865159 | -2.482908805 | 0.230309455 | 2.53E-28 | 8.96E-27 | 54941 | 18 | 29598335 | 29653176 | 1 | protein_coding | ring finger protein 125, E3 ubiquitin protein ligase |
| ANKRD12 | ENSG00000101745 | 2725.03673 | 0.407375009 | 0.138855066 | 0.002675698 | 0.008741492 | 23253 | 18 | 9136226 | 9285983 | 1 | protein_coding | ankyrin repeat domain 12 |
| POLI | ENSG00000101751 | 392.9743855 | 0.767822328 | 0.170240108 | 2.81E-06 | 1.69E-05 | 11201 | 18 | 51795774 | 51847636 | 1 | protein_coding | polymerase (DNA directed) iota |
| RBBP8 | ENSG00000101773 | 581.7115976 | 0.550126327 | 0.16692495 | 6.04E-04 | 0.002315459 | 5932 | 18 | 20378224 | 20606451 | 1 | protein_coding | retinoblastoma binding protein 8 |
| RIOK3 | ENSG00000101782 | 2852.5826 | 0.58269548 | 0.154773402 | 1.13E-04 | 5.13E-04 | 8780 | 18 | 21032787 | 21066567 | 1 | protein_coding | RIO kinase 3 |
| NXT2 | ENSG00000101888 | 357.9570877 | 1.34467682 | 0.191623855 | 4.18E-13 | 5.52E-12 | 55916 | X | 108779010 | 108787919 | 1 | protein_coding | nuclear transport factor 2-like export factor 2 |
| ALG13 | ENSG00000101901 | 498.3355207 | 0.465870227 | 0.18249704 | 0.007422718 | 0.021579113 | 79868 | X | 110909043 | 111003877 | 1 | protein_coding | ALG13, UDP-N-acetylglucosaminyltransferase subunit |
| TLR8 | ENSG00000101916 | 5646.019551 | 0.451846781 | 0.127377343 | 2.2E-04 | 9.31E-04 | 51311 | X | 12924739 | 12941288 | 1 | protein_coding | toll-like receptor 8 |
| XIAP | ENSG00000101966 | 3105.323477 | 0.647301949 | 0.145734174 | 4.83E-06 | 2.8E-05 | 331 | X | 122993574 | 123047829 | 1 | protein_coding | X-linked inhibitor of apoptosis |
| STAG2 | ENSG00000101972 | 2750.128714 | 0.969915967 | 0.145225683 | 7.74E-12 | 9.18E-11 | 10735 | X | 123094062 | 123556514 | 1 | protein_coding | stromal antigen 2 |
| ABCD1 | ENSG00000101986 | 1673.202818 | 0.518047494 | 0.157776577 | 6.65E-04 | 0.002528443 | 215 | X | 152990323 | 153010216 | 1 | protein_coding | ATP-binding cassette, sub-family D (ALD), member 1 |
| CCDC22 | ENSG00000101997 | 421.3845802 | -0.763398911 | 0.197528771 | 4.85E-05 | 2.34E-04 | 28952 | X | 49091927 | 49106987 | 1 | protein_coding | coiled-coil domain containing 22 |
| SYP | ENSG00000102003 | 242.0427507 | -0.962243994 | 0.234569278 | 1.21E-05 | 6.6E-05 | 6855 | X | 49044269 | 49056718 | -1 | protein_coding | synaptophysin |
| PLP2 | ENSG00000102007 | 1242.486422 | 0.384127068 | 0.148960258 | 0.007840521 | 0.022625885 | 5355 | X | 49028273 | 49031588 | 1 | protein_coding | proteolipid protein 2 (colonic epithelium-enriched) |
| NAA10 | ENSG00000102030 | 653.5885471 | -0.945599223 | 0.153215251 | 1.99E-10 | 2.06E-09 | 8260 | X | 153194695 | 153200676 | -1 | protein_coding | N(alpha)-acetyltransferase 10, NatA catalytic subunit |
| ELF4 | ENSG00000102034 | 1893.348546 | 1.07263176 | 0.121525298 | 3.42E-19 | 7.09E-18 | 2000 | X | 129198849 | 129244691 | -1 | protein_coding | E74-like factor 4 (ets domain transcription factor) |
| SMARCA1 | ENSG00000102038 | 14.15853529 | 1.487141173 | 1.001634433 | 0.009962249 | 0.027905618 | 6594 | X | 128580480 | 128657477 | -1 | protein_coding | SWI/SNF related, matrix associated, actin dependent regulator of chromatin, subfamily a, member 1 |
| ASB9 | ENSG00000102048 | 24.13568426 | -2.187963469 | 0.884696813 | 0.001073139 | 0.003891976 | 140462 | X | 15253410 | 15288589 | -1 | protein_coding | ankyrin repeat and SOCS box containing 9 |
| FMR1 | ENSG00000102081 | 1398.218001 | 1.147448203 | 0.12993978 | 2.5E-19 | 5.23E-18 | 2332 | X | 146993469 | 147032645 | 1 | protein_coding | fragile X mental retardation 1 |
| PIM2 | ENSG00000102096 | 749.6097829 | 0.80543133 | 0.174304561 | 1.39E-06 | 8.71E-06 | 11040 | X | 48770459 | 48776301 | -1 | protein_coding | pim-2 oncogene |
| TAZ | ENSG00000102125 | 255.5170687 | -0.62751565 | 0.245552058 | 0.005563944 | 0.01673234 | 6901 | X | 153639854 | 153650065 | 1 | protein_coding | tafazzin |
| MAGT1 | ENSG00000102158 | 4214.71601 | 0.215094955 | 0.093937271 | 0.019308481 | 0.049247581 | 84061 | X | 77081861 | 77151090 | -1 | protein_coding | magnesium transporter 1 |
| PHEX | ENSG00000102174 | 80.28840577 | -3.403745677 | 0.582035173 | 2.8E-10 | 2.84E-09 | 5251 | X | 22050559 | 22269427 | 1 | protein_coding | phosphate regulating endopeptidase homolog, X-linked |
| UBL4A | ENSG00000102178 | 1009.143484 | -0.374123083 | 0.132533518 | 0.003816477 | 0.012001486 | 8266 | X | 153712056 | 153715009 | -1 | protein_coding | ubiquitin-like 4A |
| CD99L2 | ENSG00000102181 | 1445.613325 | -0.292809876 | 0.120877705 | 0.012932621 | 0.034920361 | 83692 | X | 149934810 | 150067289 | -1 | protein_coding | CD99 molecule-like 2 |
| RP2 | ENSG00000102218 | 2141.374418 | 0.840744771 | 0.107041276 | 1.61E-15 | 2.52E-14 | 6102 | X | 46696375 | 46741793 | 1 | protein_coding | retinitis pigmentosa 2 (X-linked recessive) |
| USP11 | ENSG00000102226 | 1095.763371 | 0.654281622 | 0.118134744 | 1.65E-08 | 1.36E-07 | 8237 | X | 47092089 | 47107727 | 1 | protein_coding | ubiquitin specific peptidase 11 |
| TIMP1 | ENSG00000102265 | 1212.68041 | -0.505475873 | 0.211919542 | 0.010643987 | 0.029496291 | 7076 | X | 47441712 | 47446188 | 1 | protein_coding | TIMP metallopeptidase inhibitor 1 |
| PIN4 | ENSG00000102309 | 270.2751128 | -0.571352939 | 0.245670385 | 0.011486626 | 0.031528395 | 5303 | X | 71401203 | 71522776 | 1 | protein_coding | protein (peptidylprolyl cis/trans isomerase) NIMA-interacting, 4 (parvulin) |
| PORCN | ENSG00000102312 | 58.10151066 | -1.050505961 | 0.501381546 | 0.007126837 | 0.020837852 | 64840 | X | 48367350 | 48379202 | 1 | protein_coding | porcupine homolog (Drosophila) |
| SRPX2 | ENSG00000102359 | 30.29676797 | -1.54521542 | 0.672616799 | 0.002310333 | 0.007675829 | 27286 | X | 99899215 | 99926296 | 1 | protein_coding | sushi-repeat containing protein, X-linked 2 |
| TNFSF13B | ENSG00000102524 | 5076.250703 | 2.590138723 | 0.205916433 | 2.75E-37 | 1.5E-35 | 10673 | 13 | 108903588 | 108960832 | 1 | protein_coding | tumor necrosis factor (ligand) superfamily, member 13b |
| FNDC3A | ENSG00000102531 | 4460.218477 | 0.440774394 | 0.102455171 | 1.18E-05 | 6.42E-05 | 22862 | 13 | 49550048 | 49783915 | 1 | protein_coding | fibronectin type III domain containing 3A |
| CDADC1 | ENSG00000102543 | 158.8926493 | 0.94483016 | 0.466212389 | 0.010398276 | 0.028914616 | 81602 | 13 | 49822047 | 49867618 | 1 | protein_coding | cytidine and dCMP deaminase domain containing 1 |
| KLF5 | ENSG00000102554 | 50.90997203 | 2.474448378 | 0.647524409 | 8.48E-06 | 4.72E-05 | 688 | 13 | 73629114 | 73651676 | 1 | protein_coding | Kruppel-like factor 5 (intestinal) |
| STK24 | ENSG00000102572 | 4674.491942 | 0.270000923 | 0.106073117 | 0.009189497 | 0.026022434 | 8428 | 13 | 99102455 | 99230194 | -1 | protein_coding | serine/threonine kinase 24 |
| ACP5 | ENSG00000102575 | 4957.74823 | -1.806275954 | 0.200204772 | 2.36E-20 | 5.32E-19 | 54 | 19 | 11685475 | 11689823 | -1 | protein_coding | acid phosphatase 5, tartrate resistant |
| DNAJC3 | ENSG00000102580 | 7749.423633 | 0.767957413 | 0.095482655 | 3.51E-16 | 5.77E-15 | 5611 | 13 | 96329393 | 96447243 | 1 | protein_coding | DnaJ (Hsp40) homolog, subfamily C, member 3 |
| UGGT2 | ENSG00000102595 | 456.8041702 | 1.253056162 | 0.272145404 | 8.09E-07 | 5.29E-06 | 55757 | 13 | 96453834 | 96705736 | -1 | protein_coding | UDP-glucose glycoprotein glucosyltransferase 2 |
| PARP4 | ENSG00000102699 | 8660.241266 | 0.451241524 | 0.09532654 | 1.65E-06 | 1.02E-05 | 143 | 13 | 24995064 | 25086948 | -1 | protein_coding | poly (ADP-ribose) polymerase family, member 4 |
| MRPS31 | ENSG00000102738 | 259.3951236 | 0.534595149 | 0.208422478 | 0.00643189 | 0.019042729 | 10240 | 13 | 41303432 | 41345309 | -1 | protein_coding | mitochondrial ribosomal protein S31 |
| KPNA3 | ENSG00000102753 | 1962.467025 | 0.5118079 | 0.126489758 | 3.4E-05 | 1.7E-04 | 3839 | 13 | 50273447 | 50367057 | -1 | protein_coding | karyopherin alpha 3 (importin alpha 4) |
| FLT1 | ENSG00000102755 | 5916.959689 | -2.107927479 | 0.185042212 | 6.91E-31 | 2.8E-29 | 2321 | 13 | 28874489 | 29069265 | -1 | protein_coding | fms-related tyrosine kinase 1 |
| RGCC | ENSG00000102760 | 5594.473099 | -2.136131024 | 0.128771388 | 9.55E-63 | 1.38E-60 | 28984 | 13 | 42031695 | 42045018 | 1 | protein_coding | regulator of cell cycle |
| DGKH | ENSG00000102780 | 879.0999259 | 0.390001386 | 0.173944731 | 0.019375529 | 0.049377391 | 160851 | 13 | 42614176 | 42830714 | 1 | protein_coding | diacylglycerol kinase, eta |
| INTS6 | ENSG00000102786 | 1034.23241 | 0.524175615 | 0.169646094 | 0.001312371 | 0.004639593 | 26512 | 13 | 51928213 | 52028400 | -1 | protein_coding | integrator complex subunit 6 |
| IRG1 | ENSG00000102794 | 1825.414149 | 9.753329675 | 0.781834402 | 6.79E-37 | 3.67E-35 | 730249 | 13 | 77522632 | 77532777 | 1 | protein_coding | immunoresponsive 1 homolog (mouse) |
| TSC22D1 | ENSG00000102804 | 4685.871543 | 1.197384413 | 0.118455049 | 1.18E-24 | 3.49E-23 | 8848 | 13 | 45007655 | 45151283 | -1 | protein_coding | TSC22 domain family, member 1 |
| CLN5 | ENSG00000102805 | 2302.20297 | 0.316330584 | 0.113530192 | 0.00447438 | 0.013825707 | 1203 | 13 | 77564795 | 77576652 | 1 | protein_coding | ceroid-lipofuscinosis, neuronal 5 |
| TRADD | ENSG00000102871 | 306.8537711 | 0.969141221 | 0.205708782 | 7.58E-07 | 4.98E-06 | 8717 | 16 | 67188083 | 67194201 | -1 | protein_coding | TNFRSF1A-associated via death domain |
| HSF4 | ENSG00000102878 | 123.3449563 | -1.691398831 | 0.421792856 | 6.99E-06 | 3.94E-05 | 3299 | 16 | 67197288 | 67203848 | 1 | protein_coding | heat shock transcription factor 4 |
| CORO1A | ENSG00000102879 | 1498.113729 | -0.650511108 | 0.215949291 | 0.001360006 | 0.004788062 | 11151 | 16 | 30194148 | 30200397 | 1 | protein_coding | coronin, actin binding protein, 1A |
| MAPK3 | ENSG00000102882 | 684.4768383 | -1.395518762 | 0.157812132 | 1.71E-19 | 3.63E-18 | 5595 | 16 | 30125426 | 30134827 | -1 | protein_coding | mitogen-activated protein kinase 3 |
| LYRM1 | ENSG00000102897 | 248.8046859 | 0.606472376 | 0.21656001 | 0.002841659 | 0.009232372 | 57149 | 16 | 20911190 | 20936328 | 1 | protein_coding | LYR motif containing 1 |
| NFAT5 | ENSG00000102908 | 2776.909423 | 0.573134768 | 0.122691158 | 1.57E-06 | 9.81E-06 | 10725 | 16 | 69598997 | 69738569 | 1 | protein_coding | nuclear factor of activated T-cells 5, tonicity-responsive |
| LONP2 | ENSG00000102910 | 3364.724543 | -0.328687055 | 0.100741935 | 9.1E-04 | 0.003355366 | 1005075 | 16 | 48278207 | 48397033 | 1 | protein_coding | lon peptidase 2, peroxisomal |
| N4BP1 | ENSG00000102921 | 3242.855956 | 1.893644778 | 0.110496378 | 1.33E-66 | 2.26E-64 | 9683 | 16 | 48572637 | 48654059 | -1 | protein_coding | NEDD4 binding protein 1 |
| ARL2BP | ENSG00000102931 | 1500.713339 | 0.700997901 | 0.132237096 | 6.09E-08 | 4.67E-07 | 23568 | 16 | 57279010 | 57287516 | 1 | protein_coding | ADP-ribosylation factor-like 2 binding protein |
| DHODH | ENSG00000102967 | 55.58576861 | -0.911645971 | 0.471384159 | 0.013758926 | 0.036897293 | 1723 | 16 | 72042487 | 72058954 | 1 | protein_coding | dihydroorotate dehydrogenase (quinone) |
| PSMD7 | ENSG00000103035 | 2212.979216 | 0.468946593 | 0.105083011 | 5.85E-06 | 3.34E-05 | 5713 | 16 | 74330673 | 74340186 | 1 | protein_coding | proteasome (prosome, macropain) 26S subunit, non-ATPase, 7 |
| VAC14 | ENSG00000103043 | 2974.787595 | -1.046807311 | 0.126345026 | 2.52E-17 | 4.52E-16 | 55697 | 16 | 70721342 | 70835064 | -1 | protein_coding | Vac14 homolog (S. cerevisiae) |
| SLC7A6 | ENSG00000103064 | 964.2408774 | -1.397010004 | 0.152708277 | 1.08E-20 | 2.5E-19 | 9057 | 16 | 68298433 | 68335722 | 1 | protein_coding | solute carrier family 7 (amino acid transporter light chain, y+L system), member 6 |
| PLA2G15 | ENSG00000103066 | 2161.358043 | -1.501498353 | 0.174799069 | 1.42E-18 | 2.81E-17 | 23659 | 16 | 68279207 | 68294961 | 1 | protein_coding | phospholipase A2, group XV |
| MON1B | ENSG00000103111 | 2212.50081 | -0.391968997 | 0.106601412 | 1.95E-04 | 8.35E-04 | 22879 | 16 | 77224732 | 77236302 | 1 | protein_coding | MON1 secretory trafficking family member B |
| CMC2 | ENSG00000103121 | 528.1390176 | 0.900430786 | 0.171686522 | 5.36E-08 | 4.15E-07 | 56942 | 16 | 81009698 | 81053875 | -1 | protein_coding | C-x(9)-C motif containing 2 |
| HCFC1R1 | ENSG00000103145 | 368.3276997 | -0.96949654 | 0.241502786 | 1.7E-05 | 8.96E-05 | 54985 | 16 | 3072621 | 3074287 | -1 | protein_coding | host cell factor C1 regulator 1 (XPO1 dependent) |
| MPG | ENSG00000103152 | 516.2062688 | -0.79958301 | 0.183089843 | 5.2E-06 | 3E-05 | 4350 | 16 | 127006 | 135852 | 1 | protein_coding | N-methylpurine-DNA glycosylase |
| HSDL1 | ENSG00000103160 | 573.7303112 | 0.588197492 | 0.210970605 | 0.003010388 | 0.009732973 | 83693 | 16 | 84155886 | 84178797 | -1 | protein_coding | hydroxysteroid dehydrogenase like 1 |
| NAGPA | ENSG00000103174 | 628.1904943 | -0.415704809 | 0.178871945 | 0.014904292 | 0.039497537 | 51172 | 16 | 5074845 | 5084142 | -1 | protein_coding | N-acetylglucosamine-1-phosphodiester alpha-N-acetylglucosaminidase |
| COTL1 | ENSG00000103187 | 5399.763455 | -2.045896094 | 0.12332153 | 8.17E-63 | 1.2E-60 | 23406 | 16 | 84599200 | 84651683 | -1 | protein_coding | coactosin-like 1 (Dictyostelium) |
| USP10 | ENSG00000103194 | 1875.518188 | 0.321765118 | 0.121126305 | 0.006882045 | 0.020203283 | 9100 | 16 | 84733584 | 84813528 | 1 | protein_coding | ubiquitin specific peptidase 10 |
| CRISPLD2 | ENSG00000103196 | 55.77638016 | 2.856337157 | 0.593916213 | 7.12E-08 | 5.4E-07 | 83716 | 16 | 84853590 | 84954374 | 1 | protein_coding | cysteine-rich secretory protein LCCL domain containing 2 |
| NME4 | ENSG00000103202 | 215.1528193 | -1.445510295 | 0.289471444 | 8.4E-08 | 6.3E-07 | 4833 | 16 | 446725 | 460367 | 1 | protein_coding | NME/NM23 nucleoside diphosphate kinase 4 |
| FAM173A | ENSG00000103254 | 88.57449963 | -0.768432029 | 0.350448026 | 0.010400029 | 0.028914616 | 65990 | 16 | 770581 | 772601 | 1 | protein_coding | family with sequence similarity 173, member A |
| STUB1 | ENSG00000103266 | 732.1964868 | -0.620078131 | 0.160611753 | 6.45E-05 | 3.05E-04 | 10273 | 16 | 730224 | 732870 | 1 | protein_coding | STIP1 homology and U-box containing protein 1, E3 ubiquitin protein ligase |
| MEFV | ENSG00000103313 | 127.027509 | 4.494542652 | 0.498908421 | 1.49E-20 | 3.44E-19 | 4210 | 16 | 3292028 | 3306627 | -1 | protein_coding | Mediterranean fever |
| EEF2K | ENSG00000103319 | 1170.66039 | -0.68454237 | 0.118855903 | 4.43E-09 | 3.94E-08 | 1019301 | 16 | 22217603 | 22298554 | 1 | protein_coding | eukaryotic elongation factor-2 kinase |
| PIEZO1 | ENSG00000103335 | 3735.56189 | -0.708857032 | 0.134175728 | 5.74E-08 | 4.42E-07 | 9780 | 16 | 88781751 | 88851619 | -1 | protein_coding | piezo-type mechanosensitive ion channel component 1 |
| UBFD1 | ENSG00000103353 | 1016.009409 | 0.543017032 | 0.124692209 | 8.33E-06 | 4.64E-05 | 56061 | 16 | 23568392 | 23585710 | 1 | protein_coding | ubiquitin family domain containing 1 |
| EARS2 | ENSG00000103356 | 519.876357 | -1.208319098 | 0.199523635 | 3.03E-10 | 3.06E-09 | 124454 | 16 | 23533335 | 23569052 | -1 | protein_coding | glutamyl-tRNA synthetase 2, mitochondrial |
| TCEB2 | ENSG00000103363 | 1563.469711 | -1.231576914 | 0.148388452 | 2.97E-17 | 5.28E-16 | 6923 | 16 | 2821415 | 2827298 | -1 | protein_coding | transcription elongation factor B (SIII), polypeptide 2 (18kDa, elongin B) |
| GGA2 | ENSG00000103365 | 2186.123428 | -0.409450316 | 0.169346049 | 0.012889552 | 0.034816372 | 23062 | 16 | 23474863 | 23533316 | -1 | protein_coding | golgi-associated, gamma adaptin ear containing, ARF binding protein 2 |
| CPPED1 | ENSG00000103381 | 2116.063516 | -0.49349214 | 0.144569017 | 4.73E-04 | 0.001860901 | 55313 | 16 | 12756919 | 12897874 | -1 | protein_coding | calcineurin-like phosphoesterase domain containing 1 |
| USP31 | ENSG00000103404 | 277.674713 | -0.735758688 | 0.316715369 | 0.00798204 | 0.0229822 | 57478 | 16 | 23072727 | 23160591 | -1 | protein_coding | ubiquitin specific peptidase 31 |
| BFAR | ENSG00000103429 | 1377.787902 | 0.325535257 | 0.138801444 | 0.015751508 | 0.041490916 | 51283 | 16 | 14726672 | 14763093 | 1 | protein_coding | bifunctional apoptosis regulator |
| RBL2 | ENSG00000103479 | 3042.871961 | -0.636733354 | 0.126632584 | 2.47E-07 | 1.74E-06 | 5934 | 16 | 53467889 | 53525561 | 1 | protein_coding | retinoblastoma-like 2 (p130) |

| | | | | | | | | | | | | | |
|---|---|---|---|---|---|---|---|---|---|---|---|---|---|
| **QPRT** | ENSG00000103485 | 753.2222404 | -1.715554771 | 0.164202911 | 2.28E-26 | 7.34E-25 | 1019298 | 16 | 29674600 | 29710020 | 1 | protein_coding | quinolinate phosphoribosyltransferase |
| **XYLT1** | ENSG00000103489 | 1092.772394 | -1.377340021 | 0.136124071 | 9.32E-25 | 2.77E-23 | 64131 | 16 | 17195626 | 17564738 | -1 | protein_coding | xylosyltransferase I |
| **PYCARD** | ENSG00000103490 | 648.7148193 | -1.021314321 | 0.198072619 | 8E-08 | 6.02E-07 | 29108 | 16 | 31212806 | 31214771 | -1 | protein_coding | PYD and CARD domain containing |
| **MAZ** | ENSG00000103495 | 1699.469039 | -0.95010281 | 0.120864144 | 1.21E-15 | 1.92E-14 | 4150 | 16 | 29817427 | 29823649 | 1 | protein_coding | MYC-associated zinc finger protein (purine-binding transcription factor) |
| **STX4** | ENSG00000103496 | 4573.040021 | -0.727730021 | 0.105130556 | 1.99E-12 | 2.47E-11 | 6810 | 16 | 31044210 | 31054296 | 1 | protein_coding | syntaxin 4 |
| **CDIPT** | ENSG00000103502 | 1370.148702 | -0.737623966 | 0.123493046 | 1.16E-09 | 1.11E-08 | 10423 | 16 | 29869678 | 29875057 | -1 | protein_coding | CDP-diacylglycerol--inositol 3-phosphatidyltransferase |
| **BCKDK** | ENSG00000103507 | 1454.445444 | -1.148924681 | 0.14538192 | 7.35E-16 | 1.19E-14 | 10295 | 16 | 31117428 | 31124110 | 1 | protein_coding | branched chain ketoacid dehydrogenase kinase |
| **KAT8** | ENSG00000103510 | 684.0896876 | -0.375588414 | 0.143867755 | 0.007274688 | 0.02119311 | 84148 | 16 | 31127075 | 31142714 | 1 | protein_coding | K(lysine) acetyltransferase 8 |
| **IL21R** | ENSG00000103522 | 489.9779232 | -1.069453583 | 0.258409481 | 8.43E-06 | 4.69E-05 | 50615 | 16 | 27413483 | 27462115 | 1 | protein_coding | interleukin 21 receptor |
| **CCP110** | ENSG00000103540 | 633.7177614 | 1.030408221 | 0.176944876 | 1.54E-09 | 1.45E-08 | 9738 | 16 | 19535133 | 19564730 | 1 | protein_coding | centriolar coiled coil protein 110kDa |
| **AQP9** | ENSG00000103569 | 21893.64189 | 0.69206795 | 0.085333286 | 2.05E-16 | 3.42E-15 | 366 | 15 | 58430368 | 58478110 | 1 | protein_coding | aquaporin 9 |
| **LACTB** | ENSG00000103642 | 3477.459666 | 1.204092978 | 0.150518072 | 2.5E-16 | 4.16E-15 | 114294 | 15 | 63413999 | 63434260 | 1 | protein_coding | lactamase, beta |
| **CSK** | ENSG00000103653 | 2033.19653 | -0.664874107 | 0.155343828 | 1.08E-05 | 5.93E-05 | 1445 | 15 | 75074398 | 75095539 | 1 | protein_coding | c-src tyrosine kinase |
| **HERC1** | ENSG00000103657 | 5030.018924 | -0.518472795 | 0.090777744 | 8.24E-09 | 7.09E-08 | 8925 | 15 | 63900817 | 64126141 | -1 | protein_coding | HECT and RLD domain containing E3 ubiquitin protein ligase family member 1 |
| **TRIP4** | ENSG00000103671 | 1203.896587 | 1.010836421 | 0.131235429 | 3.54E-15 | 5.41E-14 | 9325 | 15 | 64679947 | 64747502 | 1 | protein_coding | thyroid hormone receptor interactor 4 |
| **RAB11A** | ENSG00000103769 | 3092.759997 | 0.696692303 | 0.127241701 | 2.28E-08 | 1.85E-07 | 8766 | 15 | 66018392 | 66184329 | 1 | protein_coding | RAB11A, member RAS oncogene family |
| **CD276** | ENSG00000103855 | 5444.501139 | -0.723777532 | 0.126951115 | 5.52E-09 | 4.86E-08 | 80381 | 15 | 73976307 | 74006859 | 1 | protein_coding | CD276 molecule |
| **KIAA1199** | ENSG00000103888 | 23.66547729 | -1.289424056 | 0.825289751 | 0.013292539 | 0.035778422 | 57214 | 15 | 81071684 | 81244117 | 1 | protein_coding | KIAA1199 |
| **RPAP1** | ENSG00000103932 | 480.8235964 | -0.655726468 | 0.256189379 | 0.004926145 | 0.015087429 | 26015 | 15 | 41809374 | 41836467 | -1 | protein_coding | RNA polymerase II associated protein 1 |
| **HOMER2** | ENSG00000103942 | 103.8531497 | -0.96644038 | 0.407188703 | 0.004372608 | 0.01354684 | 9455 | 15 | 83509838 | 83654661 | -1 | protein_coding | homer homolog 2 (Drosophila) |
| **EHD4** | ENSG00000103966 | 2601.536649 | 0.905115019 | 0.100972338 | 9.71E-20 | 2.11E-18 | 30844 | 15 | 42190950 | 42264776 | -1 | protein_coding | EH-domain containing 4 |
| **TMEM87A** | ENSG00000103978 | 2236.868059 | 1.090608984 | 0.139888531 | 1.7E-15 | 2.66E-14 | 25963 | 15 | 42502730 | 42565861 | -1 | protein_coding | transmembrane protein 87A |
| **ZNF106** | ENSG00000103994 | 6787.135895 | -0.663845976 | 0.096153663 | 2.66E-12 | 3.28E-11 | 64397 | 15 | 42705021 | 42783321 | -1 | protein_coding | zinc finger protein 106 |
| **GABPB1** | ENSG00000104064 | 491.3881074 | 0.590482709 | 0.177391391 | 5.1E-04 | 0.001992768 | 2553 | 15 | 50569389 | 50647605 | -1 | protein_coding | GA binding protein transcription factor, beta subunit 1 |
| **TJP1** | ENSG00000104067 | 1520.040962 | 1.007593507 | 0.180008499 | 6.59E-09 | 5.74E-08 | 7082 | 15 | 29991571 | 30261068 | -1 | protein_coding | tight junction protein 1 |
| **DMXL2** | ENSG00000104093 | 15151.72408 | 0.757709061 | 0.089882768 | 1.39E-17 | 2.54E-16 | 23312 | 15 | 51739908 | 51915030 | -1 | protein_coding | Dmx-like 2 |
| **SCG3** | ENSG00000104112 | 136.3163925 | 1.600316726 | 0.371822076 | 2.05E-06 | 1.26E-05 | 29106 | 15 | 51973550 | 52013223 | 1 | protein_coding | secretogranin III |
| **EIF3J** | ENSG00000104131 | 1336.664487 | 0.358522121 | 0.127934761 | 0.004067409 | 0.01269653 | 8669 | 15 | 44829255 | 44855227 | 1 | protein_coding | eukaryotic translation initiation factor 3, subunit J |
| **VPS18** | ENSG00000104142 | 1933.514659 | -0.323148363 | 0.140034143 | 0.017391778 | 0.045142416 | 57617 | 15 | 41186628 | 41196173 | 1 | protein_coding | vacuolar protein sorting 18 homolog (S. cerevisiae) |
| **SLC30A4** | ENSG00000104154 | 126.5968198 | -1.401105066 | 0.461215752 | 3.04E-04 | 0.001250483 | 7782 | 15 | 45771809 | 45815005 | -1 | protein_coding | solute carrier family 30 (zinc transporter), member 4 |
| **SGK3** | ENSG00000104205 | 2964.615126 | 1.927289724 | 0.108491939 | 6.64E-72 | 1.32E-69 | 1005331 | 8 | 67624653 | 67774257 | 1 | protein_coding | serum/glucocorticoid regulated kinase family, member 3 |
| **ZDHHC2** | ENSG00000104219 | 626.7048695 | 0.596703748 | 0.220881371 | 0.003776716 | 0.011888683 | 51201 | 8 | 17013538 | 17082308 | 1 | protein_coding | zinc finger, DHHC-type containing 2 |
| **CA2** | ENSG00000104267 | 5015.070293 | -2.663311942 | 0.136318818 | 1.87E-86 | 5.1E-84 | 760 | 8 | 86376081 | 86393722 | 1 | protein_coding | carbonic anhydrase II |
| **RIPK2** | ENSG00000104312 | 947.2706168 | 2.56891021 | 0.162723602 | 2.85E-57 | 3.4E-55 | 8767 | 8 | 90769975 | 90803291 | 1 | protein_coding | receptor-interacting serine-threonine kinase 2 |
| **NBN** | ENSG00000104320 | 5818.648623 | 3.320855505 | 0.15175752 | 1.67E-107 | 6.09E-105 | 4683 | 8 | 90945564 | 91015456 | -1 | protein_coding | nibrin |
| **TRPA1** | ENSG00000104321 | 14.22612309 | 1.186115241 | 1.605154831 | 0.016205453 | 0.042498689 | 8989 | 8 | 72932152 | 72987852 | -1 | protein_coding | transient receptor potential cation channel, subfamily A, member 1 |
| **CPQ** | ENSG00000104324 | 1869.366712 | -0.309384503 | 0.127262503 | 0.014011533 | 0.037469513 | 10404 | 8 | 97657455 | 98161882 | 1 | protein_coding | carboxypeptidase Q |
| **DECR1** | ENSG00000104325 | 2567.420045 | 0.549277539 | 0.124936182 | 7.25E-06 | 4.08E-05 | 1666 | 8 | 91013633 | 91064320 | 1 | protein_coding | 2,4-dienoyl CoA reductase 1, mitochondrial |
| **UBE2W** | ENSG00000104343 | 1829.04755 | 0.815527305 | 0.152390078 | 3.27E-08 | 2.61E-07 | 55284 | 8 | 74692332 | 74791145 | -1 | protein_coding | ubiquitin-conjugating enzyme E2W (putative) |
| **STK3** | ENSG00000104375 | 308.6502225 | 0.655047929 | 0.205622929 | 7.21E-04 | 0.002720033 | 6788 | 8 | 99413631 | 99955055 | -1 | protein_coding | serine/threonine kinase 3 |
| **GDAP1** | ENSG00000104381 | 329.2735163 | 0.607932552 | 0.230763717 | 0.004561389 | 0.014071801 | 54332 | 8 | 75233365 | 75401107 | 1 | protein_coding | ganglioside induced differentiation associated protein 1 |
| **RAB2A** | ENSG00000104388 | 2214.698756 | 0.309921486 | 0.116424314 | 0.007309464 | 0.021274145 | 5862 | 8 | 61429416 | 61536186 | 1 | protein_coding | RAB2A, member RAS oncogene family |
| **EMC2** | ENSG00000104412 | 917.3964651 | 0.381326354 | 0.14478348 | 0.006731418 | 0.019810563 | 9694 | 8 | 109455830 | 109499145 | 1 | protein_coding | ER membrane protein complex subunit 2 |
| **IL7** | ENSG00000104432 | 47.26800195 | 0.902143267 | 0.489034072 | 0.016923661 | 0.044084595 | 3574 | 8 | 79587978 | 79717758 | -1 | protein_coding | interleukin 7 |
| **TRPS1** | ENSG00000104447 | 1542.656869 | -0.464835218 | 0.168001873 | 0.004017298 | 0.012560631 | 7227 | 8 | 116420724 | 116821899 | -1 | protein_coding | trichorhinophalangeal syndrome I |
| **SPAG1** | ENSG00000104450 | 145.0986647 | 2.095487482 | 0.340574359 | 7.24E-11 | 7.87E-10 | 6674 | 8 | 101170134 | 101271506 | 1 | protein_coding | sperm associated antigen 1 |
| **SNX16** | ENSG00000104497 | 362.0310168 | 0.719342098 | 0.205173766 | 2.03E-04 | 8.67E-04 | 64089 | 8 | 82711816 | 82755101 | -1 | protein_coding | sorting nexin 16 |
| **UBR5** | ENSG00000104517 | 4526.851781 | -0.289194405 | 0.103220832 | 0.00452092 | 0.013958225 | 51366 | 8 | 103265240 | 103425069 | -1 | protein_coding | ubiquitin protein ligase E3 component n-recognin 5 |
| **GSDMD** | ENSG00000104518 | 661.8184687 | 1.119635779 | 0.269436084 | 7.63E-06 | 4.28E-05 | 79792 | 8 | 144635377 | 144645232 | 1 | protein_coding | gasdermin D |
| **ERI1** | ENSG00000104626 | 690.0694032 | 0.593723305 | 0.147154202 | 3.18E-05 | 1.6E-04 | 90459 | 8 | 8859657 | 8974256 | 1 | protein_coding | exoribonuclease 1 |
| **MTMR9** | ENSG00000104643 | 499.9646333 | 0.381198466 | 0.150105013 | 0.008674326 | 0.024710107 | 66036 | 8 | 11141925 | 11185646 | 1 | protein_coding | myotubularin related protein 9 |
| **DCTN6** | ENSG00000104671 | 572.2843158 | 0.427869217 | 0.164783483 | 0.006954639 | 0.020388978 | 10671 | 8 | 30013813 | 30041156 | 1 | protein_coding | dynactin 6 |
| **GSR** | ENSG00000104687 | 232.4801958 | -0.949638076 | 0.274283669 | 1.57E-04 | 6.92E-04 | 2936 | 8 | 30535583 | 30585443 | -1 | protein_coding | glutathione reductase |
| **TNFRSF10A** | ENSG00000104689 | 814.1057316 | 0.854008506 | 0.168724033 | 1.61E-07 | 1.16E-06 | 8797 | 8 | 23047965 | 23082639 | -1 | protein_coding | tumor necrosis factor receptor superfamily, member 10a |
| **MCM4** | ENSG00000104738 | 63.13120602 | 1.521789242 | 0.503805249 | 2.94E-04 | 0.001211509 | 4173 | 8 | 48872745 | 48890720 | 1 | protein_coding | minichromosome maintenance complex component 4 |
| **KCTD9** | ENSG00000104756 | 1213.192292 | 0.476664498 | 0.136849025 | 3.51E-04 | 0.00142493 | 54793 | 8 | 25285366 | 25315992 | -1 | protein_coding | potassium channel tetramerization domain containing 9 |
| **BNIP3L** | ENSG00000104765 | 4313.181382 | -0.868533911 | 0.097104275 | 1.53E-19 | 3.25E-18 | 665 | 8 | 26240414 | 26363152 | 1 | protein_coding | BCL2/adenovirus E1B 19kDa interacting protein 3-like |
| **MAN2B1** | ENSG00000104774 | 14035.6641 | -0.688921291 | 0.107201245 | 5.94E-11 | 6.52E-10 | 4125 | 19 | 12757325 | 12777556 | -1 | protein_coding | mannosidase, alpha, class 2B, member 1 |
| **GYS1** | ENSG00000104812 | 1019.584204 | -0.420803386 | 0.156136156 | 0.004448132 | 0.013752944 | 2997 | 19 | 49471382 | 49496567 | -1 | protein_coding | glycogen synthase 1 (muscle) |
| **ECH1** | ENSG00000104823 | 923.3266633 | -0.513351223 | 0.161559781 | 9.49E-04 | 0.003476608 | 1891 | 19 | 39306062 | 39322645 | -1 | protein_coding | enoyl CoA hydratase 1, peroxisomal |
| **CLPTM1** | ENSG00000104853 | 3884.623016 | -0.696189011 | 0.109766136 | 1.13E-10 | 1.2E-09 | 1209 | 19 | 45457842 | 45496599 | 1 | protein_coding | cleft lip and palate associated transmembrane protein 1 |
| **PPP1R37** | ENSG00000104866 | 863.4664411 | -0.444443563 | 0.195143102 | 0.014310384 | 0.038121945 | 284352 | 19 | 45594654 | 45651335 | 1 | protein_coding | protein phosphatase 1, regulatory subunit 37 |
| **FCGRT** | ENSG00000104870 | 3013.150906 | -0.843399507 | 0.146868468 | 3.73E-09 | 3.34E-08 | 2217 | 19 | 50010073 | 50029590 | 1 | protein_coding | Fc fragment of IgG, receptor, transporter, alpha |
| **ARHGEF18** | ENSG00000104880 | 1440.761644 | -0.358077717 | 0.127979113 | 0.004764291 | 0.014635711 | 23370 | 19 | 7459999 | 7537363 | 1 | protein_coding | Rho/Rac guanine nucleotide exchange factor (GEF) 18 |
| **ERCC2** | ENSG00000104884 | 225.4164681 | -1.447205827 | 0.256152332 | 2.31E-09 | 2.11E-08 | 2068 | 19 | 45853095 | 45874176 | -1 | protein_coding | excision repair cross-complementing rodent repair deficiency, complementation group 2 |
| **SLC17A7** | ENSG00000104888 | 33.87751745 | -1.098357198 | 0.682668611 | 0.016521633 | 0.043206729 | 57030 | 19 | 49932658 | 49945617 | -1 | protein_coding | solute carrier family 17 (vesicular glutamate transporter), member 7 |
| **CD37** | ENSG00000104894 | 4025.388608 | -0.654731014 | 0.119438737 | 2.27E-08 | 1.84E-07 | 951 | 19 | 49838428 | 49846592 | 1 | protein_coding | CD37 molecule |
| **LYL1** | ENSG00000104903 | 332.8604526 | -0.482468204 | 0.19957532 | 0.010683375 | 0.029573268 | 4066 | 19 | 13209847 | 13213975 | -1 | protein_coding | lymphoblastic leukemia derived sequence 1 |
| **OAZ1** | ENSG00000104904 | 9610.992179 | -0.456755848 | 0.086730623 | 1.68E-07 | 1.21E-06 | 4946 | 19 | 2269485 | 2273487 | 1 | protein_coding | ornithine decarboxylase antizyme 1 |
| **STX10** | ENSG00000104915 | 659.7906868 | -0.885865732 | 0.16810599 | 4.68E-08 | 3.65E-07 | 8677 | 19 | 13254872 | 13261197 | -1 | protein_coding | syntaxin 10 |
| **FCER2** | ENSG00000104921 | 28.56147651 | -1.200035507 | 0.705931324 | 0.01229881 | 0.033421536 | 2208 | 19 | 7753644 | 7767032 | -1 | protein_coding | Fc fragment of IgE, low affinity II, receptor for (CD23) |
| **TBC1D17** | ENSG00000104946 | 540.9246313 | -1.074203107 | 0.171104299 | 9.16E-11 | 9.83E-10 | 79735 | 19 | 50380682 | 50392005 | 1 | protein_coding | TBC1 domain family, member 17 |
| **IL4I1** | ENSG00000104951 | 3465.415502 | 1.017584889 | 0.200182175 | 1.08E-07 | 7.97E-07 | 259307 | 19 | 50392911 | 50432796 | -1 | protein_coding | interleukin 4 induced 1 |
| **PTOV1** | ENSG00000104960 | 467.8730144 | -0.937577511 | 0.199355069 | 8.09E-07 | 5.29E-06 | 53635 | 19 | 50353992 | 50364001 | 1 | protein_coding | prostate tumor overexpressed 1 |
| **AES** | ENSG00000104964 | 5155.814808 | 0.309573962 | 0.081090892 | 1.65E-04 | 7.24E-04 | 166 | 19 | 3052908 | 3063105 | -1 | protein_coding | amino-terminal enhancer of split |
| **MED25** | ENSG00000104973 | 616.4627418 | 0.754394518 | 0.177107715 | 9.53E-06 | 5.26E-05 | 1024654 | 19 | 50321539 | 50342073 | 1 | protein_coding | mediator complex subunit 25 |
| **LILRA1** | ENSG00000104974 | 123.6369242 | -1.442736508 | 0.462716758 | 2.33E-04 | 9.84E-04 | 11024 | 19 | 55105047 | 55113555 | 1 | protein_coding | leukocyte immunoglobulin-like receptor, subfamily A (with TM domain), member 1 |
| **SNAPC2** | ENSG00000104976 | 195.7533163 | -0.572245686 | 0.241433697 | 0.010317657 | 0.028737881 | 6618 | 19 | 7985201 | 7988135 | 1 | protein_coding | small nuclear RNA activating complex, polypeptide 2, 45kDa |
| **C19orf53** | ENSG00000104979 | 778.7213867 | -0.655803954 | 0.148976285 | 5.54E-06 | 3.18E-05 | 28974 | 19 | 13884982 | 13889276 | 1 | protein_coding | chromosome 19 open reading frame 53 |
| **TIMM44** | ENSG00000104980 | 370.4843917 | -0.749031223 | 0.179059832 | 1.27E-05 | 6.9E-05 | 10469 | 19 | 7991603 | 8008805 | -1 | protein_coding | translocase of inner mitochondrial membrane 44 homolog (yeast) |
| **IL27RA** | ENSG00000104998 | 866.3190799 | -0.953132777 | 0.145543012 | 1.83E-11 | 2.11E-10 | 9466 | 19 | 14142560 | 14163743 | 1 | protein_coding | interleukin 27 receptor, alpha |
| **TNNT1** | ENSG00000105048 | 121.5867828 | -1.698869131 | 0.499681253 | 6.64E-05 | 3.14E-04 | 7138 | 19 | 55644162 | 55660722 | -1 | protein_coding | troponin T type 1 (skeletal, slow) |
| **FAM32A** | ENSG00000105058 | 1837.058336 | 0.269774201 | 0.102103791 | 0.007503634 | 0.021781204 | 26017 | 19 | 16296191 | 16302857 | 1 | protein_coding | family with sequence similarity 32, member A |
| **ILVBL** | ENSG00000105135 | 463.5134546 | -1.233454337 | 0.195984945 | 6.61E-11 | 7.21E-10 | 10994 | 19 | 15225795 | 15236596 | -1 | protein_coding | ilvB (bacterial acetolactate synthase)-like |
| **ZNF419** | ENSG00000105136 | 309.4997229 | -0.663428852 | 0.199702317 | 4.49E-04 | 0.001777733 | 79744 | 19 | 57999079 | 58006048 | 1 | protein_coding | zinc finger protein 419 |
| **PDCD5** | ENSG00000105185 | 384.0034786 | 1.008031543 | 0.211339469 | 5.49E-07 | 3.69E-06 | 9141 | 19 | 33071974 | 33078358 | 1 | protein_coding | programmed cell death 5 |
| **RPS16** | ENSG00000105193 | 3129.822317 | -0.438770356 | 0.143187753 | 0.001697206 | 0.005844615 | 6217 | 19 | 39923847 | 39926588 | -1 | protein_coding | ribosomal protein S16 |
| **TIMM50** | ENSG00000105197 | 590.5909436 | -0.407211318 | 0.162357819 | 0.009323806 | 0.026373446 | 92609 | 19 | 39971052 | 39984422 | 1 | protein_coding | translocase of inner mitochondrial membrane 50 homolog (S. cerevisiae) |
| **DYRK1B** | ENSG00000105204 | 209.9940976 | -1.200170678 | 0.260276696 | 7.92E-07 | 5.19E-06 | 9149 | 19 | 40315990 | 40324841 | -1 | protein_coding | dual-specificity tyrosine-(Y)-phosphorylation regulated kinase 1B |

| | | | | | | | | | | | | | |
|---|---|---|---|---|---|---|---|---|---|---|---|---|---|
| **GPI** | ENSG00000105220 | 6549.05739 | -1.097355546 | 0.128407489 | 4.13E-18 | 7.85E-17 | 2821 | 19 | 34850385 | 34893061 | 1 | protein_coding | glucose-6-phosphate isomerase |
| **PLD3** | ENSG00000105223 | 30186.87135 | -0.540006032 | 0.094380967 | 5.8E-09 | 5.08E-08 | 23646 | 19 | 40854363 | 40886346 | 1 | protein_coding | phospholipase D family, member 3 |
| **EBI3** | ENSG00000105246 | 124.1580794 | 4.042854104 | 0.410200076 | 3.89E-24 | 1.11E-22 | 10148 | 19 | 4229495 | 4237528 | 1 | protein_coding | Epstein-Barr virus induced 3 |
| **POLR2I** | ENSG00000105258 | 197.4544306 | -0.870347929 | 0.2869905 | 8.07E-04 | 0.003013928 | 5438 | 19 | 36604612 | 36606248 | -1 | protein_coding | polymerase (RNA) II (DNA directed) polypeptide I, 14.5kDa |
| **SLC1A5** | ENSG00000105281 | 3649.323624 | -0.731230802 | 0.138269254 | 6.33E-08 | 4.84E-07 | 6510 | 19 | 47278140 | 47291851 | -1 | protein_coding | solute carrier family 1 (neutral amino acid transporter), member 5 |
| **PRKD2** | ENSG00000105287 | 683.5731773 | 1.12922188 | 0.161184091 | 6.75E-13 | 8.73E-12 | 25865 | 19 | 47177532 | 47220384 | -1 | protein_coding | protein kinase D2 |
| **TGFB1** | ENSG00000105329 | 6363.962985 | -0.615164449 | 0.10585006 | 3.44E-09 | 3.09E-08 | 7040 | 19 | 41807492 | 41859816 | -1 | protein_coding | transforming growth factor, beta 1 |
| **MYH14** | ENSG00000105357 | 13.15204313 | -3.087191797 | 1.697250458 | 0.007839169 | 0.022625885 | 79784 | 19 | 50691443 | 50813802 | 1 | protein_coding | myosin, heavy chain 14, non-muscle |
| **MRPL4** | ENSG00000105364 | 417.3834107 | -0.534152213 | 0.184131654 | 0.002332414 | 0.007740778 | 51073 | 19 | 10362577 | 10370721 | 1 | protein_coding | mitochondrial ribosomal protein L4 |
| **ICAM4** | ENSG00000105371 | 61.91233454 | -1.934567578 | 0.502552182 | 1.12E-05 | 6.13E-05 | 3386 | 19 | 10397643 | 10399198 | 1 | protein_coding | intercellular adhesion molecule 4 (Landsteiner-Wiener blood group) |
| **RPS19** | ENSG00000105372 | 4364.936717 | -0.391362414 | 0.131783144 | 0.002418223 | 0.007987402 | 6223 | 19 | 42363988 | 42376994 | 1 | protein_coding | ribosomal protein S19 |
| **ICAM5** | ENSG00000105376 | 93.04078618 | -2.11285062 | 0.587661948 | 2.42E-05 | 1.25E-04 | 7087 | 19 | 10400657 | 10407454 | 1 | protein_coding | intercellular adhesion molecule 5, telencephalin |
| **ETFB** | ENSG00000105379 | 483.2155103 | -0.716585702 | 0.163312484 | 5.43E-06 | 3.12E-05 | 2109 | 19 | 51848423 | 51869672 | -1 | protein_coding | electron-transfer-flavoprotein, beta polypeptide |
| **BABAM1** | ENSG00000105393 | 1321.694251 | -0.525655116 | 0.119708373 | 7.94E-06 | 4.44E-05 | 29086 | 19 | 17378159 | 17392058 | 1 | protein_coding | BRISC and BRCA1 A complex member 1 |
| **CDC37** | ENSG00000105401 | 3599.533385 | -0.442396079 | 0.103658277 | 1.4E-05 | 7.52E-05 | 11140 | 19 | 10501810 | 10530797 | -1 | protein_coding | cell division cycle 37 |
| **NAPA** | ENSG00000105402 | 2463.806068 | 1.264511052 | 0.109438814 | 1.7E-31 | 7.12E-30 | 8775 | 19 | 47990894 | 48018497 | -1 | protein_coding | N-ethylmaleimide-sensitive factor attachment protein, alpha |
| **MEGF8** | ENSG00000105429 | 1107.578707 | -1.537977193 | 0.149818539 | 1.5E-25 | 4.61E-24 | 1954 | 19 | 42829761 | 42882921 | 1 | protein_coding | multiple EGF-like-domains 8 |
| **CARD8** | ENSG00000105483 | 2738.062588 | 0.345415875 | 0.097093573 | 3.36E-04 | 0.001373198 | 22900 | 19 | 48684027 | 48759203 | -1 | protein_coding | caspase recruitment domain family, member 8 |
| **PLA2G4C** | ENSG00000105499 | 582.5938882 | 0.448235285 | 0.189128265 | 0.012620549 | 0.034168264 | 8605 | 19 | 48551100 | 48614074 | -1 | protein_coding | phospholipase A2, group IVC (cytosolic, calcium-independent) |
| **RAB3D** | ENSG00000105514 | 944.804768 | -0.886547904 | 0.127959144 | 1.81E-12 | 2.25E-11 | 9545 | 19 | 11432722 | 11456946 | -1 | protein_coding | RAB3D, member RAS oncogene family |
| **DBP** | ENSG00000105516 | 343.9561387 | -2.04962299 | 0.247415949 | 1.07E-17 | 1.97E-16 | 1628 | 19 | 49133287 | 49140695 | -1 | protein_coding | D site of albumin promoter (albumin D-box) binding protein |
| **CAPS** | ENSG00000105519 | 262.729694 | -1.232219123 | 0.295382521 | 6.09E-06 | 3.47E-05 | 828 | 19 | 5911718 | 5915888 | 1 | protein_coding | calcyphosine |
| **DKFZP761J1** | ENSG00000105520 | 280.250646 | -1.359938299 | 0.248304293 | 7.3E-09 | 6.32E-08 | 64748 | 19 | 11466062 | 11476374 | 1 | protein_coding | Lipid phosphate phosphatase-related protein type 2 |
| **PPP2R1A** | ENSG00000105568 | 3132.127127 | -0.912949142 | 0.122831203 | 3.73E-14 | 5.35E-13 | 5518 | 19 | 52693292 | 52730687 | 1 | protein_coding | protein phosphatase 2, regulatory subunit A, alpha |
| **TNPO2** | ENSG00000105576 | 579.0204954 | 0.377589129 | 0.155603898 | 0.012262785 | 0.033329565 | 30000 | 19 | 12810008 | 12834825 | -1 | protein_coding | transportin 2 |
| **WDR83OS** | ENSG00000105583 | 619.0466916 | -0.465106531 | 0.165766979 | 0.003530882 | 0.011218808 | 51398 | 19 | 12778885 | 12782170 | -1 | protein_coding | WD repeat domain 83 opposite strand |
| **DNASE2** | ENSG00000105612 | 3265.727005 | -0.927733204 | 0.136338672 | 3.39E-12 | 4.16E-11 | 1777 | 19 | 12986025 | 12992282 | -1 | protein_coding | deoxyribonuclease II, lysosomal |
| **TFPT** | ENSG00000105619 | 539.520524 | -0.489808011 | 0.173155481 | 0.0031548 | 0.01015904 | 29844 | 19 | 54610320 | 54619055 | -1 | protein_coding | TCF3 (E2A) fusion partner (in childhood Leukemia) |
| **RPL18A** | ENSG00000105640 | 2834.758511 | -0.47520366 | 0.127237768 | 1.18E-04 | 5.35E-04 | 26780 | 19 | 17970685 | 17974962 | 1 | protein_coding | ribosomal protein L18a |
| **PIK3R2** | ENSG00000105647 | 832.8445563 | -0.868003223 | 0.156222669 | 1.05E-08 | 8.91E-08 | 5296 | 19 | 18263928 | 18281350 | 1 | protein_coding | phosphoinositide-3-kinase, regulatory subunit 2 (beta) |
| **RAB3A** | ENSG00000105649 | 23.32247809 | -2.651339367 | 0.949597257 | 3.8E-04 | 0.001532069 | 5864 | 19 | 18307594 | 18314884 | -1 | protein_coding | RAB3A, member RAS oncogene family |
| **ISYNA1** | ENSG00000105655 | 112.538637 | -1.483548058 | 0.386947892 | 1.74E-05 | 9.15E-05 | 51477 | 19 | 18545198 | 18549111 | -1 | protein_coding | inositol-3-phosphate synthase 1 |
| **DDX49** | ENSG00000105671 | 364.3644115 | -0.534868108 | 0.198794043 | 0.004689998 | 0.014433596 | 54555 | 19 | 19030484 | 19039436 | 1 | protein_coding | DEAD (Asp-Glu-Ala-Asp) box polypeptide 49 |
| **ARMC6** | ENSG00000105676 | 238.586297 | -1.134796933 | 0.243038033 | 7.07E-07 | 4.68E-06 | 93436 | 19 | 19144384 | 19170563 | 1 | protein_coding | armadillo repeat containing 6 |
| **TMEM147** | ENSG00000105677 | 509.815709 | -0.645482377 | 0.175565169 | 1.3E-04 | 5.83E-04 | 10430 | 19 | 36036497 | 36038428 | 1 | protein_coding | transmembrane protein 147 |
| **HAMP** | ENSG00000105697 | 121.5816339 | -2.214987555 | 0.569899864 | 6.48E-06 | 3.67E-05 | 57817 | 19 | 35771619 | 35776046 | 1 | protein_coding | hepcidin antimicrobial peptide |
| **USF2** | ENSG00000105698 | 3782.047955 | -0.994984344 | 0.116310892 | 3.82E-18 | 7.3E-17 | 7392 | 19 | 35759881 | 35770724 | 1 | protein_coding | upstream transcription factor 2, c-fos interacting |
| **FKBP8** | ENSG00000105701 | 2966.326437 | -0.990871847 | 0.126100588 | 9.44E-16 | 1.52E-14 | 23770 | 19 | 18642561 | 18654887 | -1 | protein_coding | FK506 binding protein 8, 38kDa |
| **SCN1B** | ENSG00000105711 | 159.1740443 | 3.223657781 | 0.331175809 | 1.31E-23 | 3.63E-22 | 6324 | 19 | 35521588 | 35531352 | 1 | protein_coding | sodium channel, voltage-gated, type I, beta subunit |
| **ERF** | ENSG00000105722 | 402.9096243 | -0.555674238 | 0.233834379 | 0.010246344 | 0.028575706 | 2077 | 19 | 42751724 | 42759309 | -1 | protein_coding | Ets2 repressor factor |
| **ATP13A1** | ENSG00000105726 | 2049.657573 | 0.429284125 | 0.150333387 | 0.003258464 | 0.010442254 | 57130 | 19 | 19756007 | 19774502 | -1 | protein_coding | ATPase type 13A1 |
| **SIPA1L3** | ENSG00000105738 | 654.4478012 | 0.608928454 | 0.240077146 | 0.00582864 | 0.017425489 | 23094 | 19 | 38397868 | 38699012 | 1 | protein_coding | signal-induced proliferation-associated 1 like 3 |
| **GTPBP10** | ENSG00000105793 | 497.2227606 | 0.639648699 | 0.173634412 | 1.24E-04 | 5.58E-04 | 1019274 | 7 | 89964537 | 90020769 | 1 | protein_coding | GTP-binding protein 10 (putative) |
| **RASA4** | ENSG00000105808 | 1456.678691 | -0.994590426 | 0.177587501 | 5.97E-09 | 5.22E-08 | 1002719 | 7 | 102220093 | 102257204 | -1 | protein_coding | RAS p21 protein activator 4 |
| **CDK6** | ENSG00000105810 | 1815.809689 | -0.737567953 | 0.187395624 | 3.73E-05 | 1.85E-04 | 1021 | 7 | 92234235 | 92465908 | -1 | protein_coding | cyclin-dependent kinase 6 |
| **DNAJC2** | ENSG00000105821 | 804.8305423 | 0.655313661 | 0.149484263 | 6.3E-06 | 3.58E-05 | 27000 | 7 | 102952921 | 102985320 | -1 | protein_coding | DnaJ (Hsp40) homolog, subfamily C, member 2 |
| **TFPI2** | ENSG00000105825 | 70.844116 | 4.684375198 | 0.618492553 | 1.86E-15 | 2.89E-14 | 7980 | 7 | 93514709 | 93520303 | -1 | protein_coding | tissue factor pathway inhibitor 2 |
| **BET1** | ENSG00000105829 | 364.8894758 | 0.752439032 | 0.190759106 | 3.62E-05 | 1.79E-04 | 10282 | 7 | 93592074 | 93633694 | -1 | protein_coding | Bet1 golgi vesicular membrane trafficking protein |
| **NAMPT** | ENSG00000105835 | 4725.163879 | 3.162625109 | 0.184247943 | 2.23E-67 | 3.87E-65 | 10135 | 7 | 105888731 | 105926772 | -1 | protein_coding | nicotinamide phosphoribosyltransferase |
| **ITGB8** | ENSG00000105855 | 831.258589 | 4.137979623 | 0.168973542 | 1.49E-133 | 6.69E-131 | 3696 | 7 | 20370325 | 20455377 | 1 | protein_coding | integrin, beta 8 |
| **HBP1** | ENSG00000105856 | 2611.595219 | 0.836688295 | 0.14555679 | 3.32E-09 | 2.98E-08 | 26959 | 7 | 106809406 | 106842974 | 1 | protein_coding | HMG-box transcription factor 1 |
| **SP4** | ENSG00000105866 | 345.3330329 | 0.491291219 | 0.202087159 | 0.010242185 | 0.028569828 | 6671 | 7 | 21467652 | 21554440 | 1 | protein_coding | Sp4 transcription factor |
| **WDR91** | ENSG00000105875 | 1188.676832 | -0.987433039 | 0.207259699 | 5.44E-07 | 3.66E-06 | 29062 | 7 | 134868590 | 134896316 | -1 | protein_coding | WD repeat domain 91 |
| **DNAH11** | ENSG00000105877 | 43.07635389 | -3.561674547 | 1.44445617 | 5.77E-04 | 0.002221197 | 8701 | 7 | 21582833 | 21941457 | 1 | protein_coding | dynein, axonemal, heavy chain 11 |
| **CBLL1** | ENSG00000105879 | 881.3699345 | 0.453658986 | 0.151491016 | 0.002007098 | 0.006786122 | 79872 | 7 | 107384142 | 107402112 | 1 | protein_coding | Cbl proto-oncogene-like 1, E3 ubiquitin protein ligase |
| **MTPN** | ENSG00000105887 | 8951.002594 | 0.618917787 | 0.097112206 | 9.3E-11 | 9.97E-10 | 767558 | 7 | 135611509 | 135662101 | -1 | protein_coding | myotrophin |
| **DFNA5** | ENSG00000105928 | 1922.243663 | 0.311126552 | 0.120402535 | 0.008676613 | 0.024712017 | 1687 | 7 | 24737972 | 24809244 | -1 | protein_coding | deafness, autosomal dominant 5 |
| **ZC3HAV1** | ENSG00000105939 | 3252.717181 | 1.425296668 | 0.104969471 | 9.51E-43 | 6.73E-41 | 56829 | 7 | 138728266 | 138794465 | -1 | protein_coding | zinc finger CCCH-type, antiviral 1 |
| **TFEC** | ENSG00000105967 | 7117.847493 | 1.330171511 | 0.095929809 | 1.7E-44 | 1.27E-42 | 22797 | 7 | 115575202 | 115799950 | -1 | protein_coding | transcription factor EC |
| **H2AFV** | ENSG00000105968 | 1414.593869 | -0.424946386 | 0.124697171 | 5.18E-04 | 0.002019348 | 94239 | 7 | 44866390 | 44887682 | -1 | protein_coding | H2A histone family, member V |
| **CAV1** | ENSG00000105974 | 57.84710405 | -0.989847566 | 0.621574451 | 0.018394883 | 0.047296265 | 857 | 7 | 116164839 | 116201233 | 1 | protein_coding | caveolin 1, caveolae protein, 22kDa |
| **LMBR1** | ENSG00000105983 | 442.2906924 | 0.760892477 | 0.218147934 | 2.05E-04 | 8.72E-04 | 64327 | 7 | 156461646 | 156685924 | -1 | protein_coding | limb development membrane protein 1 |
| **DNAJB6** | ENSG00000105993 | 5291.761351 | 0.93497564 | 0.098924657 | 1.02E-21 | 2.54E-20 | 10049 | 7 | 157128075 | 157210133 | 1 | protein_coding | DnaJ (Hsp40) homolog, subfamily B, member 6 |
| **BRAT1** | ENSG00000106009 | 348.3822828 | -0.740090841 | 0.243564405 | 0.001026279 | 0.003735322 | 221927 | 7 | 2577511 | 2595361 | -1 | protein_coding | BRCA1-associated ATM activator 1 |
| **IQCE** | ENSG00000106012 | 765.5962122 | -0.995317535 | 0.179768746 | 1.02E-08 | 8.63E-08 | 23288 | 7 | 2598632 | 2654368 | 1 | protein_coding | IQ motif containing E |
| **VIPR2** | ENSG00000106018 | 11.54691541 | -1.423379345 | 1.366162295 | 0.017722592 | 0.04586865 | 7434 | 7 | 158820866 | 158937649 | -1 | protein_coding | vasoactive intestinal peptide receptor 2 |
| **TSPAN12** | ENSG00000106025 | 34.97311957 | -1.937354206 | 0.972631421 | 0.003202277 | 0.01028808 | 23554 | 7 | 120427376 | 120498456 | -1 | protein_coding | tetraspanin 12 |
| **HIBADH** | ENSG00000106049 | 949.0067174 | -0.627976286 | 0.161627115 | 5.26E-05 | 2.53E-04 | 11112 | 7 | 27565061 | 27702614 | -1 | protein_coding | 3-hydroxyisobutyrate dehydrogenase |
| **TAX1BP1** | ENSG00000106052 | 5167.652088 | 0.598295691 | 0.102477336 | 3.21E-09 | 2.89E-08 | 8887 | 7 | 27778950 | 27884183 | 1 | protein_coding | Tax1 (human T-cell leukemia virus type I) binding protein 1 |
| **COBL** | ENSG00000106078 | 418.1355089 | 1.062901439 | 0.281614571 | 3.98E-05 | 1.96E-04 | 23242 | 7 | 51083909 | 51384515 | -1 | protein_coding | cordon-bleu WH2 repeat protein |
| **NOD1** | ENSG00000106100 | 487.7159675 | 1.178863751 | 0.18357588 | 3E-11 | 3.39E-10 | 10392 | 7 | 30464143 | 30518400 | -1 | protein_coding | nucleotide-binding oligomerization domain containing 1 |
| **CCL24** | ENSG00000106178 | 12.45629626 | -1.822925157 | 1.237219485 | 0.007816606 | 0.022565392 | 6369 | 7 | 75440983 | 75452674 | -1 | protein_coding | chemokine (C-C motif) ligand 24 |
| **HSPB1** | ENSG00000106211 | 1692.054685 | -0.589656505 | 0.119672884 | 4.8E-07 | 3.26E-06 | 3315 | 7 | 75931861 | 75933612 | 1 | protein_coding | heat shock 27kDa protein 1 |
| **EIF3B** | ENSG00000106263 | 2833.682807 | -0.398733348 | 0.117008545 | 4.99E-04 | 0.001954287 | 8662 | 7 | 2393721 | 2420380 | 1 | protein_coding | eukaryotic translation initiation factor 3, subunit B |
| **SNX8** | ENSG00000106266 | 1231.979392 | -0.918552031 | 0.139343239 | 1.67E-11 | 1.93E-10 | 1024655 | 7 | 2291405 | 2393953 | -1 | protein_coding | sorting nexin 8 |
| **NUDT1** | ENSG00000106268 | 98.4370841 | -0.81681583 | 0.353746779 | 0.007166291 | 0.020929185 | 4521 | 7 | 2281857 | 2290781 | 1 | protein_coding | nudix (nucleoside diphosphate linked moiety X)-type motif 1 |
| **WASL** | ENSG00000106299 | 723.583 | 1.139159778 | 0.169309252 | 3.97E-12 | 4.85E-11 | 8976 | 7 | 123321989 | 123389121 | -1 | protein_coding | Wiskott-Aldrich syndrome-like |
| **MOSPD3** | ENSG00000106330 | 82.87936924 | -0.966724356 | 0.431576318 | 0.006287518 | 0.018665917 | 64598 | 7 | 100209725 | 100213007 | 1 | protein_coding | motile sperm domain containing 3 |
| **PCOLCE** | ENSG00000106333 | 17.04906559 | -1.580535826 | 0.983953792 | 0.008020201 | 0.023071088 | 5118 | 7 | 100199800 | 100205798 | 1 | protein_coding | procollagen C-endopeptidase enhancer |
| **RBM28** | ENSG00000106344 | 907.9620927 | 0.411216238 | 0.150923598 | 0.004988606 | 0.015239028 | 55131 | 7 | 127937738 | 127983962 | -1 | protein_coding | RNA binding motif protein 28 |
| **USP42** | ENSG00000106346 | 410.816103 | 1.05456744 | 0.209972585 | 1.29E-07 | 9.39E-07 | 84132 | 7 | 6144515 | 6201195 | 1 | protein_coding | ubiquitin specific peptidase 42 |
| **IMPDH1** | ENSG00000106348 | 772.0616582 | -0.484478596 | 0.159894315 | 0.001635694 | 0.005659567 | 3614 | 7 | 128032331 | 128050306 | -1 | protein_coding | IMP (inosine 5'-monophosphate) dehydrogenase 1 |
| **SERPINE1** | ENSG00000106366 | 685.3073443 | -2.615534473 | 0.291584779 | 1.9E-20 | 4.32E-19 | 5054 | 7 | 100770370 | 100782547 | 1 | protein_coding | serpin peptidase inhibitor, clade E (nexin, plasminogen activator inhibitor type 1), member 1 |
| **AP1S1** | ENSG00000106367 | 644.7503927 | -0.458584423 | 0.158372077 | 0.002727126 | 0.008890846 | 1174 | 7 | 100797678 | 100804877 | 1 | protein_coding | adaptor-related protein complex 1, sigma 1 subunit |
| **C1GALT1** | ENSG00000106392 | 589.0302064 | 1.537327727 | 0.159754007 | 1.06E-22 | 2.8E-21 | 56913 | 7 | 7196565 | 7288282 | 1 | protein_coding | core 1 synthase, glycoprotein-N-acetylgalactosamine 3-beta-galactosyltransferase, 1 |
| **PLOD3** | ENSG00000106397 | 1367.703927 | -0.64030078 | 0.134613018 | 1.11E-06 | 7.1E-06 | 8985 | 7 | 100849258 | 100861701 | -1 | protein_coding | procollagen-lysine, 2-oxoglutarate 5-dioxygenase 3 |
| **ZNHIT1** | ENSG00000106400 | 690.6210997 | -0.69950325 | 0.144138675 | 5.04E-07 | 3.41E-06 | 10467 | 7 | 100860949 | 100867471 | 1 | protein_coding | zinc finger, HIT-type containing 1 |
| **PHF14** | ENSG00000106443 | 520.5403164 | 0.597614333 | 0.231396438 | 0.005494873 | 0.016557204 | 9678 | 7 | 11013499 | 11209250 | 1 | protein_coding | PHD finger protein 14 |
| **TMEM106B** | ENSG00000106460 | 1543.83106 | 0.772828008 | 0.129856098 | 1.18E-09 | 1.13E-08 | 54664 | 7 | 12250867 | 12282993 | 1 | protein_coding | transmembrane protein 106B |

| | | | | | | | | | | | | | |
|---|---|---|---|---|---|---|---|---|---|---|---|---|---|
| **EZH2** | ENSG00000106462 | 331.727671 | 1.865782637 | 0.275679068 | 1.3E-12 | 1.63E-11 | 2146 | 7 | 148504475 | 148581413 | -1 | protein_coding | enhancer of zeste homolog 2 (Drosophila) |
| **ANKMY2** | ENSG00000106524 | 708.256641 | 0.398883584 | 0.139187436 | 0.003180614 | 0.010231388 | 57037 | 7 | 16639401 | 16685442 | -1 | protein_coding | ankyrin repeat and MYND domain containing 2 |
| **AHR** | ENSG00000106546 | 4075.916806 | 0.852222108 | 0.127453264 | 8.69E-12 | 1.02E-10 | 196 | 7 | 17338246 | 17385776 | 1 | protein_coding | aryl hydrocarbon receptor |
| **GIMAP2** | ENSG00000106560 | 1016.985417 | 1.058547284 | 0.126714375 | 1.65E-17 | 3.01E-16 | 26157 | 7 | 150382785 | 150390729 | 1 | protein_coding | GTPase, IMAP family member 2 |
| **PSMA2** | ENSG00000106588 | 1713.414462 | 1.891578539 | 0.1710672 | 1.91E-29 | 7.16E-28 | 5683 | 7 | 42956460 | 42971822 | -1 | protein_coding | proteasome (prosome, macropain) subunit, alpha type, 2 |
| **MRPL32** | ENSG00000106591 | 527.3186566 | 0.988781877 | 0.170445884 | 1.97E-09 | 1.82E-08 | 64983 | 7 | 42971799 | 42988557 | 1 | protein_coding | mitochondrial ribosomal protein L32 |
| **BLVRA** | ENSG00000106605 | 1614.895162 | 1.477620816 | 0.120520527 | 2.45E-35 | 1.25E-33 | 644 | 7 | 43798279 | 43846939 | 1 | protein_coding | biliverdin reductase A |
| **RHEB** | ENSG00000106615 | 2320.609966 | 0.731480743 | 0.128217264 | 6.14E-09 | 5.37E-08 | 6009 | 7 | 151163098 | 151217206 | -1 | protein_coding | Ras homolog enriched in brain |
| **PRKAG2** | ENSG00000106617 | 1342.103726 | 0.596680453 | 0.20280406 | 0.001837959 | 0.006271445 | 51422 | 7 | 151253197 | 151574210 | -1 | protein_coding | protein kinase, AMP-activated, gamma 2 non-catalytic subunit |
| **POLD2** | ENSG00000106628 | 269.326517 | -0.733129257 | 0.247638711 | 0.001324587 | 0.004675212 | 5425 | 7 | 44154286 | 44163957 | -1 | protein_coding | polymerase (DNA directed), delta 2, accessory subunit |
| **TBL2** | ENSG00000106638 | 1636.51284 | -1.156303324 | 0.161519058 | 1.83E-13 | 2.47E-12 | 26608 | 7 | 72983262 | 72993121 | -1 | protein_coding | transducin (beta)-like 2 |
| **CLIP2** | ENSG00000106665 | 844.6416664 | -0.824180062 | 0.278943256 | 0.001111163 | 0.004014633 | 7461 | 7 | 73703805 | 73820273 | 1 | protein_coding | CAP-GLY domain containing linker protein 2 |
| **LIMK1** | ENSG00000106683 | 7779.061741 | -0.802895064 | 0.091256909 | 4.86E-19 | 9.98E-18 | 3984 | 7 | 73497263 | 73536855 | 1 | protein_coding | LIM domain kinase 1 |
| **FSD1L** | ENSG00000106701 | 349.098494 | 1.103992734 | 0.300954405 | 5.7E-05 | 2.73E-04 | 83856 | 9 | 108210077 | 108314714 | 1 | protein_coding | fibronectin type III and SPRY domain containing 1-like |
| **SPIN1** | ENSG00000106723 | 2253.603843 | -0.488467471 | 0.131143842 | 1.33E-04 | 5.93E-04 | 10927 | 9 | 91003334 | 91093609 | 1 | protein_coding | spindlin 1 |
| **NMRK1** | ENSG00000106733 | 513.9499359 | 0.62068277 | 0.158589751 | 5.13E-05 | 2.47E-04 | 54981 | 9 | 77675489 | 77703133 | -1 | protein_coding | nicotinamide riboside kinase 1 |
| **MEGF9** | ENSG00000106780 | 1998.142551 | -0.246639793 | 0.103822278 | 0.01603054 | 0.042092209 | 1955 | 9 | 123363091 | 123476748 | -1 | protein_coding | multiple EGF-like-domains 9 |
| **TRIM14** | ENSG00000106785 | 3208.612265 | 1.13792859 | 0.127658612 | 1.35E-19 | 2.88E-18 | 9830 | 9 | 100831557 | 100881494 | -1 | protein_coding | tripartite motif containing 14 |
| **CORO2A** | ENSG00000106789 | 1237.969857 | -0.632908146 | 0.124892834 | 2.42E-07 | 1.71E-06 | 7464 | 9 | 100883257 | 100954922 | -1 | protein_coding | coronin, actin binding protein, 2A |
| **TGFBR1** | ENSG00000106799 | 7536.720303 | -1.56680289 | 0.117762272 | 3.69E-41 | 2.42E-39 | 7046 | 9 | 101866320 | 101916474 | 1 | protein_coding | transforming growth factor, beta receptor 1 |
| **SEC61B** | ENSG00000106803 | 839.1011141 | 0.50886911 | 0.149496571 | 4.48E-04 | 0.00177489 | 10952 | 9 | 101984346 | 101992897 | 1 | protein_coding | Sec61 beta subunit |
| **C5** | ENSG00000106804 | 460.6929494 | 2.445902858 | 0.222879721 | 4.17E-29 | 1.53E-27 | 727 | 9 | 123714616 | 123812554 | -1 | protein_coding | complement component 5 |
| **TLE4** | ENSG00000106829 | 470.0474704 | 1.376920556 | 0.345402817 | 1E-05 | 5.53E-05 | 7091 | 9 | 82186688 | 82341658 | 1 | protein_coding | transducin-like enhancer of split 4 (E(sp1) homolog, Drosophila) |
| **TNFSF8** | ENSG00000106952 | 1041.54052 | -1.967650159 | 0.144085367 | 2.12E-43 | 1.53E-41 | 944 | 9 | 117656003 | 117692697 | -1 | protein_coding | tumor necrosis factor (ligand) superfamily, member 8 |
| **DNM1** | ENSG00000106976 | 31.5508511 | -2.53478149 | 0.844843708 | 1.61E-04 | 7.07E-04 | 1759 | 9 | 130965658 | 131017527 | 1 | protein_coding | dynamin 1 |
| **ENG** | ENSG00000106991 | 7947.805108 | -1.007979254 | 0.117564197 | 3.17E-18 | 6.11E-17 | 2022 | 9 | 130577291 | 130617035 | -1 | protein_coding | endoglin |
| **AK1** | ENSG00000106992 | 226.9145774 | -1.372267098 | 0.241502506 | 2.22E-09 | 2.04E-08 | 203 | 9 | 130628759 | 130640022 | -1 | protein_coding | adenylate kinase 1 |
| **PLGRKT** | ENSG00000107020 | 734.0621023 | 1.242262786 | 0.183472378 | 2.58E-12 | 3.19E-11 | 55848 | 9 | 5357973 | 5437878 | -1 | protein_coding | plasminogen receptor, C-terminal lysine transmembrane protein |
| **KIAA1432** | ENSG00000107036 | 1423.052324 | 0.635454961 | 0.139820364 | 3.06E-06 | 1.83E-05 | 57589 | 9 | 5629025 | 5776557 | 1 | protein_coding | KIAA1432 |
| **KANK1** | ENSG00000107104 | 505.8137431 | 1.009079134 | 0.170804495 | 1.1E-09 | 1.06E-08 | 23189 | 9 | 470291 | 746105 | 1 | protein_coding | KN motif and ankyrin repeat domains 1 |
| **MPDZ** | ENSG00000107186 | 28.4026412 | -1.467106299 | 2.338544622 | 0.011815395 | 0.032291609 | 8777 | 9 | 13105703 | 13279589 | -1 | protein_coding | multiple PDZ domain protein |
| **DDX58** | ENSG00000107201 | 2365.229126 | 4.066916803 | 0.15187297 | 3.03E-159 | 2.02E-156 | 23586 | 9 | 32455300 | 32526322 | -1 | protein_coding | DEAD (Asp-Glu-Ala-Asp) box polypeptide 58 |
| **PIP5K1B** | ENSG00000107242 | 228.7662236 | -0.663184563 | 0.239094291 | 0.002780654 | 0.009045346 | 8395 | 9 | 71320575 | 71624092 | 1 | protein_coding | phosphatidylinositol-4-phosphate 5-kinase, type I, beta |
| **BAG1** | ENSG00000107262 | 1453.816198 | 1.046493857 | 0.105164905 | 7.24E-24 | 2.04E-22 | 573 | 9 | 33247818 | 33264761 | -1 | protein_coding | BCL2-associated athanogene |
| **RAPGEF1** | ENSG00000107263 | 11713.02679 | -0.620339212 | 0.091501569 | 7.99E-12 | 9.46E-11 | 2889 | 9 | 134452157 | 134615461 | -1 | protein_coding | Rap guanine nucleotide exchange factor (GEF) 1 |
| **SETX** | ENSG00000107290 | 5283.41241 | 0.800423599 | 0.143080751 | 8.61E-09 | 7.39E-08 | 23064 | 9 | 135136743 | 135230372 | -1 | protein_coding | senataxin |
| **ABCA2** | ENSG00000107331 | 727.3749949 | -1.415440331 | 0.184151092 | 2.74E-15 | 4.25E-14 | 20 | 9 | 139901686 | 139923367 | -1 | protein_coding | ATP-binding cassette, sub-family A (ABC1), member 2 |
| **SHB** | ENSG00000107338 | 760.1147081 | -1.983719324 | 0.191814443 | 4.81E-26 | 1.51E-24 | 6461 | 9 | 37919131 | 38069210 | -1 | protein_coding | Src homology 2 domain containing adaptor protein B |
| **ABHD17B** | ENSG00000107362 | 275.0470612 | 0.472847651 | 0.214328979 | 0.01878516 | 0.048107215 | 51104 | 9 | 74477368 | 74525847 | -1 | protein_coding | abhydrolase domain containing 17B |
| **CCNJ** | ENSG00000107443 | 352.3093643 | 0.896931396 | 0.211932157 | 7.78E-06 | 4.35E-05 | 54619 | 10 | 97803151 | 97820627 | 1 | protein_coding | cyclin J |
| **HPS1** | ENSG00000107521 | 1620.269792 | -0.596307056 | 0.118975519 | 2.77E-07 | 1.93E-06 | 3257 | 10 | 100175955 | 100206684 | -1 | protein_coding | Hermansky-Pudlak syndrome 1 |
| **DNMBP** | ENSG00000107554 | 277.390869 | -0.578902838 | 0.210267086 | 0.003414139 | 0.010886352 | 23268 | 10 | 101635334 | 101769676 | -1 | protein_coding | dynamin binding protein |
| **RAB11FIP2** | ENSG00000107560 | 598.1598902 | 0.978768291 | 0.174027712 | 5.65E-09 | 4.96E-08 | 22841 | 10 | 119764427 | 119806114 | -1 | protein_coding | RAB11 family interacting protein 2 (class I) |
| **ERLIN1** | ENSG00000107566 | 1155.882712 | 0.465454341 | 0.119602946 | 7.51E-05 | 3.51E-04 | 10613 | 10 | 101909851 | 101948091 | -1 | protein_coding | ER lipid raft associated 1 |
| **EIF3A** | ENSG00000107581 | 10125.46451 | -0.236622093 | 0.095838154 | 0.014035962 | 0.037515145 | 8661 | 10 | 120794356 | 120840316 | -1 | protein_coding | eukaryotic translation initiation factor 3, subunit A |
| **CUBN** | ENSG00000107611 | 56.74027668 | -0.985817878 | 0.491043129 | 0.01021336 | 0.028509732 | 8029 | 10 | 16865963 | 17171830 | -1 | protein_coding | cubilin (intrinsic factor-cobalamin receptor) |
| **MAPK8** | ENSG00000107643 | 633.0290695 | 0.42192256 | 0.179102705 | 0.013560008 | 0.036414972 | 5599 | 10 | 49514698 | 49647403 | 1 | protein_coding | mitogen-activated protein kinase 8 |
| **NSMCE4A** | ENSG00000107672 | 373.3288504 | 0.677159499 | 0.182472248 | 1.04E-04 | 4.73E-04 | 54780 | 10 | 123716603 | 123734732 | -1 | protein_coding | non-SMC element 4 homolog A (S. cerevisiae) |
| **PLEKHA1** | ENSG00000107679 | 527.4239401 | 0.673566253 | 0.244587082 | 0.002753714 | 0.008973127 | 59338 | 10 | 124134212 | 124191867 | 1 | protein_coding | pleckstrin homology domain containing, family A (phosphoinositide binding specific) member 1 |
| **UNC5B** | ENSG00000107731 | 24.0978705 | -3.490701098 | 0.96926412 | 3.54E-05 | 1.76E-04 | 219699 | 10 | 72972327 | 73062621 | 1 | protein_coding | unc-5 homolog B (C. elegans) |
| **CDH23** | ENSG00000107736 | 3982.466898 | -1.383939539 | 0.164730081 | 6.67E-18 | 1.25E-16 | 1006531 | 10 | 73156691 | 73575702 | 1 | protein_coding | cadherin-related 23 |
| **C10orf54** | ENSG00000107738 | 2466.869557 | -1.1179964 | 0.163647502 | 1.96E-12 | 2.44E-11 | 64115 | 10 | 73507316 | 73533255 | -1 | protein_coding | chromosome 10 open reading frame 54 |
| **MICU1** | ENSG00000107745 | 1965.412554 | 0.424051946 | 0.122548202 | 4.17E-04 | 0.001664344 | 10367 | 10 | 74127098 | 74385899 | -1 | protein_coding | mitochondrial calcium uptake 1 |
| **CCSER2** | ENSG00000107771 | 2974.958039 | 1.130710101 | 0.123726777 | 1.27E-20 | 2.93E-19 | 54462 | 10 | 86088342 | 86278273 | 1 | protein_coding | coiled-coil serine-rich protein 2 |
| **BMPR1A** | ENSG00000107779 | 504.4925 | 0.538613918 | 0.212584579 | 0.006383957 | 0.018926482 | 657 | 10 | 88516407 | 88692595 | 1 | protein_coding | bone morphogenetic protein receptor, type IA |
| **MINPP1** | ENSG00000107789 | 489.4102281 | 0.776309238 | 0.188210005 | 1.57E-05 | 8.37E-05 | 9562 | 10 | 89264632 | 89313217 | 1 | protein_coding | multiple inositol-polyphosphate phosphatase 1 |
| **LIPA** | ENSG00000107798 | 138787.0235 | -0.822708623 | 0.193364584 | 7.91E-06 | 4.42E-05 | 3988 | 10 | 90973326 | 91174314 | -1 | protein_coding | lipase A, lysosomal acid, cholesterol esterase |
| **FBXW4** | ENSG00000107829 | 442.435169 | 0.481434848 | 0.191301505 | 0.008418991 | 0.024054485 | 6468 | 10 | 103370423 | 103455052 | -1 | protein_coding | F-box and WD repeat domain containing 4 |
| **NPM3** | ENSG00000107833 | 87.80118836 | -1.01511294 | 0.457957079 | 0.005711091 | 0.017117594 | 10360 | 10 | 103541082 | 103543170 | -1 | protein_coding | nucleophosmin/nucleoplasmin 3 |
| **TNKS2** | ENSG00000107854 | 3650.066086 | 0.944125033 | 0.090177593 | 4.03E-26 | 1.27E-24 | 80351 | 10 | 93558069 | 93625033 | 1 | protein_coding | tankyrase, TRF1-interacting ankyrin-related ADP-ribose polymerase 2 |
| **GBF1** | ENSG00000107862 | 2465.406423 | -0.355432687 | 0.112290239 | 0.001114514 | 0.004022936 | 8729 | 10 | 104005289 | 104142656 | 1 | protein_coding | golgi brefeldin A resistant guanine nucleotide exchange factor 1 |
| **ARHGAP21** | ENSG00000107863 | 2767.880648 | 0.299236433 | 0.103712573 | 0.003487181 | 0.011089177 | 57584 | 10 | 24872538 | 25012597 | -1 | protein_coding | Rho GTPase activating protein 21 |
| **CPEB3** | ENSG00000107864 | 272.9805226 | 1.339463689 | 0.202195059 | 6.49E-12 | 7.78E-11 | 22849 | 10 | 93806449 | 94050844 | -1 | protein_coding | cytoplasmic polyadenylation element binding protein 3 |
| **ACBD5** | ENSG00000107897 | 1960.378504 | 0.819335465 | 0.106755091 | 6.48E-15 | 9.81E-14 | 91452 | 10 | 27484146 | 27531059 | -1 | protein_coding | acyl-CoA binding domain containing 5 |
| **LHPP** | ENSG00000107902 | 251.0871713 | -1.002387746 | 0.23483578 | 5.66E-06 | 3.25E-05 | 64077 | 10 | 126150403 | 126306457 | 1 | protein_coding | phospholysine phosphohistidine inorganic pyrophosphate phosphatase |
| **LARP4B** | ENSG00000107929 | 4031.626426 | 0.273340373 | 0.093673647 | 0.003448477 | 0.010982095 | 23185 | 10 | 855484 | 977564 | -1 | protein_coding | La ribonucleoprotein domain family, member 4B |
| **GTPBP4** | ENSG00000107937 | 361.13348 | 0.620831472 | 0.294146831 | 0.01782574 | 0.046034915 | 23560 | 10 | 1034338 | 1065876 | 1 | protein_coding | GTP binding protein 4 |
| **SH3PXD2A** | ENSG00000107957 | 1347.360522 | -1.83184362 | 0.139003196 | 1.46E-40 | 8.96E-39 | 9644 | 10 | 105348285 | 105615301 | -1 | protein_coding | SH3 and PX domains 2A |
| **MAP3K8** | ENSG00000107968 | 569.8102086 | 1.797947288 | 0.177024197 | 3.64E-25 | 1.09E-23 | 1326 | 10 | 30722866 | 30750762 | 1 | protein_coding | mitogen-activated protein kinase kinase kinase 8 |
| **FAM208B** | ENSG00000108021 | 1800.465934 | 0.712847322 | 0.140867265 | 2.08E-07 | 1.48E-06 | 54906 | 10 | 5726801 | 5805703 | 1 | protein_coding | family with sequence similarity 208, member B |
| **XPNPEP1** | ENSG00000108039 | 1991.827148 | 0.737646654 | 0.110533427 | 1.19E-11 | 1.39E-10 | 7511 | 10 | 111624524 | 111683311 | -1 | protein_coding | X-prolyl aminopeptidase (aminopeptidase P) 1, soluble |
| **SMC3** | ENSG00000108055 | 1689.40897 | 0.456134343 | 0.110926933 | 3.01E-05 | 1.52E-04 | 9126 | 10 | 112327449 | 112364394 | 1 | protein_coding | structural maintenance of chromosomes 3 |
| **SHOC2** | ENSG00000108061 | 2488.568357 | 0.669837743 | 0.119708689 | 1.18E-08 | 9.9E-08 | 8036 | 10 | 112679301 | 112773425 | 1 | protein_coding | soc-2 suppressor of clear homolog (C. elegans) |
| **CCNY** | ENSG00000108100 | 3181.240692 | -1.35511145 | 0.151712504 | 6.83E-20 | 1.5E-18 | 219771 | 10 | 35535953 | 35860852 | 1 | protein_coding | cyclin Y |
| **UBE2S** | ENSG00000108106 | 221.2391512 | 1.833175943 | 0.304361301 | 1.93E-10 | 2E-09 | 27338 | 19 | 55912652 | 55919145 | -1 | protein_coding | ubiquitin-conjugating enzyme E2S |
| **RPL28** | ENSG00000108107 | 4161.805554 | 0.407953483 | 0.129960056 | 0.001292153 | 0.004580833 | 1024654 | 19 | 55896713 | 55914612 | 1 | protein_coding | ribosomal protein L28 |
| **ZMIZ1** | ENSG00000108175 | 7235.18733 | -1.364526311 | 0.090868118 | 1.69E-51 | 1.61E-49 | 57178 | 10 | 80828792 | 81076276 | 1 | protein_coding | zinc finger, MIZ-type containing 1 |
| **PPIF** | ENSG00000108179 | 1922.442963 | 0.656164582 | 0.104179727 | 1.68E-10 | 1.75E-09 | 10105 | 10 | 81107225 | 81115093 | 1 | protein_coding | peptidylprolyl isomerase F |
| **TSPAN14** | ENSG00000108219 | 3423.386974 | -0.297705203 | 0.127687568 | 0.016472809 | 0.04310117 | 81619 | 10 | 82213922 | 82292879 | 1 | protein_coding | tetraspanin 14 |
| **NUFIP2** | ENSG00000108256 | 5845.599241 | 0.570126701 | 0.109667426 | 1.24E-07 | 9.09E-07 | 57532 | 17 | 27582854 | 27621136 | -1 | protein_coding | nuclear fragile X mental retardation protein interacting protein 2 |
| **GIT1** | ENSG00000108262 | 977.2575522 | -0.582914641 | 0.144995584 | 3.3E-05 | 1.65E-04 | 28964 | 17 | 27900487 | 27921072 | -1 | protein_coding | G protein-coupled receptor kinase interacting ArfGAP 1 |
| **RPL19** | ENSG00000108298 | 9177.207857 | -0.397176592 | 0.112057955 | 2.43E-04 | 0.001016055 | 6143 | 17 | 37356536 | 37360980 | 1 | protein_coding | ribosomal protein L19 |
| **CSF3** | ENSG00000108342 | 9.117634181 | 7.07345986 | 4.346963266 | 0.001710472 | 0.005882345 | 1440 | 17 | 38171614 | 38174066 | 1 | protein_coding | colony stimulating factor 3 (granulocyte) |
| **CASC3** | ENSG00000108349 | 1615.027051 | 0.265919059 | 0.107257603 | 0.010398468 | 0.028914616 | 1024669 | 17 | 38296571 | 38328436 | 1 | protein_coding | cancer susceptibility candidate 3 |
| **4-Sep** | ENSG00000108387 | 53.70237406 | 3.342397953 | 0.83147558 | 2.96E-06 | 1.78E-05 | 5414 | 17 | 56597611 | 56618179 | -1 | protein_coding | septin 4 |
| **P2RX1** | ENSG00000108405 | 526.6242122 | -1.72633765 | 0.199263049 | 6.68E-19 | 1.34E-17 | 5023 | 17 | 3799886 | 3819794 | -1 | protein_coding | purinergic receptor P2X, ligand-gated ion channel, 1 |
| **DHX40** | ENSG00000108406 | 746.2411346 | 0.503102692 | 0.156363244 | 8.64E-04 | 0.00320606 | 79665 | 17 | 57642886 | 57685706 | 1 | protein_coding | DEAH (Asp-Glu-Ala-His) box polypeptide 40 |
| **TUBD1** | ENSG00000108423 | 120.4406202 | 0.948106691 | 0.292978716 | 3.61E-04 | 0.001460613 | 51174 | 17 | 57936851 | 57970304 | -1 | protein_coding | tubulin, delta 1 |
| **KPNB1** | ENSG00000108424 | 14374.67657 | 0.897792244 | 0.105007198 | 3.84E-18 | 7.34E-17 | 3837 | 17 | 45726842 | 45762871 | 1 | protein_coding | karyopherin (importin) beta 1 |

| | | | | | | | | | | | | | |
|---|---|---|---|---|---|---|---|---|---|---|---|---|---|
| **PNPO** | ENSG00000108439 | 1673.381426 | -0.650101769 | 0.123647325 | 9.32E-08 | 6.94E-07 | 55163 | 17 | 46018872 | 46025654 | 1 | protein_coding | pyridoxamine 5'-phosphate oxidase |
| **RPS6KB1** | ENSG00000108443 | 962.3418112 | 0.340192038 | 0.124912353 | 0.005421297 | 0.016371009 | 6198 | 17 | 57970447 | 58027925 | 1 | protein_coding | ribosomal protein S6 kinase, 70kDa, polypeptide 1 |
| **CDK5RAP3** | ENSG00000108465 | 2276.433481 | -0.386113436 | 0.119722923 | 9.84E-04 | 0.003592458 | 80279 | 17 | 46045176 | 46059140 | 1 | protein_coding | CDK5 regulatory subunit associated protein 3 |
| **MED13** | ENSG00000108510 | 4423.857545 | 0.913902747 | 0.083944255 | 7.45E-28 | 2.56E-26 | 9969 | 17 | 60019966 | 60142643 | -1 | protein_coding | mediator complex subunit 13 |
| **RNF167** | ENSG00000108523 | 1340.144407 | -0.749802692 | 0.127046074 | 1.77E-09 | 1.65E-08 | 26001 | 17 | 4843303 | 4848517 | 1 | protein_coding | ring finger protein 167 |
| **SLC25A11** | ENSG00000108528 | 1239.572669 | -0.76766911 | 0.14868126 | 1.1E-07 | 8.05E-07 | 8402 | 17 | 4840425 | 4843546 | -1 | protein_coding | solute carrier family 25 (mitochondrial carrier; oxoglutarate carrier), member 11 |
| **CPD** | ENSG00000108582 | 14036.77796 | -0.47430463 | 0.120828708 | 4.95E-05 | 2.39E-04 | 1362 | 17 | 28705923 | 28797007 | 1 | protein_coding | carboxypeptidase D |
| **GOSR1** | ENSG00000108587 | 2081.276657 | 0.317120646 | 0.125595238 | 0.009478789 | 0.026752417 | 9527 | 17 | 28804380 | 28854610 | 1 | protein_coding | golgi SNAP receptor complex member 1 |
| **AKAP10** | ENSG00000108599 | 1575.410499 | 0.278925533 | 0.119933365 | 0.018446044 | 0.047387947 | 11216 | 17 | 19807615 | 19881656 | -1 | protein_coding | A kinase (PRKA) anchor protein 10 |
| **B9D1** | ENSG00000108641 | 24.70540429 | -3.667227429 | 1.099580895 | 9.63E-05 | 4.42E-04 | 27077 | 17 | 19240867 | 19281495 | -1 | protein_coding | B9 protein domain 1 |
| **UTP6** | ENSG00000108651 | 1513.877962 | 0.499814253 | 0.122932088 | 3.55E-05 | 1.76E-04 | 55813 | 17 | 30187923 | 30228784 | -1 | protein_coding | UTP6, small subunit (SSU) processome component, homolog (yeast) |
| **DDX5** | ENSG00000108654 | 7844.236253 | 1.127701197 | 0.127762038 | 3.11E-19 | 6.48E-18 | 1006164 | 17 | 62495734 | 62504317 | -1 | protein_coding | DEAD (Asp-Glu-Ala-Asp) box helicase 5 |
| **CYTH1** | ENSG00000108669 | 5633.701408 | 0.677748318 | 0.092625535 | 1.35E-13 | 1.84E-12 | 9267 | 17 | 76670130 | 76778379 | -1 | protein_coding | cytohesin 1 |
| **PSMD11** | ENSG00000108671 | 2261.891462 | 0.720050958 | 0.100224867 | 3.29E-13 | 4.36E-12 | 5717 | 17 | 30771279 | 30810336 | 1 | protein_coding | proteasome (prosome, macropain) 26S subunit, non-ATPase, 11 |
| **LGALS3BP** | ENSG00000108679 | 6688.526927 | 0.836546057 | 0.132564717 | 1.19E-10 | 1.26E-09 | 3959 | 17 | 76967320 | 76976191 | -1 | protein_coding | lectin, galactoside-binding, soluble, 3 binding protein |
| **CCL7** | ENSG00000108688 | 295.8598444 | 4.302567083 | 3.615087104 | 0.001213948 | 0.004333745 | 6354 | 17 | 32597240 | 32599261 | 1 | protein_coding | chemokine (C-C motif) ligand 7 |
| **CCL2** | ENSG00000108691 | 2722.342947 | 0.999920191 | 0.197667451 | 1.78E-07 | 1.28E-06 | 6347 | 17 | 32582304 | 32584222 | 1 | protein_coding | chemokine (C-C motif) ligand 2 |
| **CCL8** | ENSG00000108700 | 2970.905826 | 8.310620136 | 0.240254059 | 4.15E-264 | 6.35E-261 | 6355 | 17 | 32646055 | 32648421 | 1 | protein_coding | chemokine (C-C motif) ligand 8 |
| **CCL1** | ENSG00000108702 | 388.2883384 | 6.651347271 | 0.358439107 | 3.96E-78 | 9.32E-76 | 6346 | 17 | 32687347 | 32690250 | -1 | protein_coding | chemokine (C-C motif) ligand 1 |
| **DHX58** | ENSG00000108771 | 319.4598216 | 2.640417699 | 0.23183167 | 3.85E-31 | 1.57E-29 | 79132 | 17 | 40253422 | 40264751 | -1 | protein_coding | DEXH (Asp-Glu-X-His) box polypeptide 58 |
| **ABI3** | ENSG00000108798 | 479.8791213 | -1.48474837 | 0.235152633 | 4.06E-11 | 4.55E-10 | 51225 | 17 | 47287589 | 47300587 | 1 | protein_coding | ABI family, member 3 |
| **EZH1** | ENSG00000108799 | 1189.788842 | 0.32834207 | 0.132154356 | 0.011277343 | 0.031054349 | 2145 | 17 | 40852293 | 40897071 | -1 | protein_coding | enhancer of zeste homolog 1 (Drosophila) |
| **DLX4** | ENSG00000108813 | 50.37074894 | -1.607580012 | 0.625703041 | 0.001082218 | 0.003922853 | 1748 | 17 | 48046334 | 48052321 | 1 | protein_coding | distal-less homeobox 4 |
| **COL1A1** | ENSG00000108821 | 40.99954229 | 1.659673493 | 0.572537906 | 3.76E-04 | 0.001518956 | 1277 | 17 | 48260650 | 48278993 | -1 | protein_coding | collagen, type I, alpha 1 |
| **MRPL27** | ENSG00000108826 | 310.804771 | 0.445258304 | 0.183190721 | 0.011058654 | 0.030518132 | 51264 | 17 | 48445218 | 48450575 | -1 | protein_coding | mitochondrial ribosomal protein L27 |
| **VAT1** | ENSG00000108828 | 12067.37074 | -1.300130768 | 0.133191014 | 3.53E-23 | 9.64E-22 | 10493 | 17 | 41166622 | 41177140 | -1 | protein_coding | vesicle amine transport 1 |
| **HDAC5** | ENSG00000108840 | 775.5387883 | -0.977946181 | 0.164455118 | 8.17E-10 | 7.93E-09 | 10014 | 17 | 42154114 | 42201070 | -1 | protein_coding | histone deacetylase 5 |
| **ABCC3** | ENSG00000108846 | 2144.280621 | -0.566252198 | 0.162169726 | 2.94E-04 | 0.001211141 | 8714 | 17 | 48712138 | 48769613 | 1 | protein_coding | ATP-binding cassette, sub-family C (CFTR/MRP), member 3 |
| **PRKAR1A** | ENSG00000108946 | 17737.15961 | -0.359980146 | 0.089575719 | 5.29E-05 | 2.54E-04 | 5573 | 17 | 66507921 | 66547460 | 1 | protein_coding | protein kinase, cAMP-dependent, regulatory, type I, alpha |
| **FAM20A** | ENSG00000108950 | 1982.836303 | 0.871720362 | 0.263049568 | 3.18E-04 | 0.001306008 | 54757 | 17 | 66531254 | 66597530 | -1 | protein_coding | family with sequence similarity 20, member A |
| **MMD** | ENSG00000108960 | 1068.731335 | -1.98211382 | 0.175063868 | 1.01E-30 | 4.07E-29 | 23531 | 17 | 53469974 | 53499353 | -1 | protein_coding | monocyte to macrophage differentiation-associated |
| **WSB1** | ENSG00000109046 | 3670.155536 | 0.826152262 | 0.113748477 | 1.44E-13 | 1.97E-12 | 26118 | 17 | 25621102 | 25640657 | 1 | protein_coding | WD repeat and SOCS box containing 1 |
| **SLC9A3R1** | ENSG00000109062 | 811.9213433 | -0.525108135 | 0.136339088 | 7.24E-05 | 3.39E-04 | 9368 | 17 | 72744791 | 72765492 | 1 | protein_coding | solute carrier family 9, subfamily A (NHE3, cation proton antiporter 3), member 3 regulator 1 |
| **NAT9** | ENSG00000109065 | 381.6273259 | -0.558091623 | 0.173773561 | 8.15E-04 | 0.003043376 | 26151 | 17 | 72766686 | 72772506 | -1 | protein_coding | N-acetyltransferase 9 (GCN5-related, putative) |
| **TMEM104** | ENSG00000109066 | 1708.276445 | -0.721178373 | 0.164845399 | 5.84E-06 | 3.34E-05 | 54868 | 17 | 72772622 | 72835918 | 1 | protein_coding | transmembrane protein 104 |
| **PMP22** | ENSG00000109099 | 159.0578613 | -1.430670728 | 0.298480615 | 2.6E-07 | 1.82E-06 | 5376 | 17 | 15133095 | 15168643 | -1 | protein_coding | peripheral myelin protein 22 |
| **ALDOC** | ENSG00000109107 | 349.2849813 | -1.591014122 | 0.305132942 | 2.14E-08 | 1.74E-07 | 230 | 17 | 26900133 | 26904282 | -1 | protein_coding | aldolase C, fructose-bisphosphate |
| **TMEM33** | ENSG00000109133 | 4917.685767 | 0.887591168 | 0.124963488 | 7.88E-13 | 1.02E-11 | 55161 | 4 | 41937137 | 41962589 | 1 | protein_coding | transmembrane protein 33 |
| **USP46** | ENSG00000109189 | 587.1572474 | -0.648525147 | 0.224117842 | 0.001913434 | 0.006504228 | 64854 | 4 | 53457138 | 53525502 | -1 | protein_coding | ubiquitin specific peptidase 46 |
| **CHIC2** | ENSG00000109220 | 300.0057336 | 0.699814325 | 0.18859579 | 1.11E-04 | 0.000502821 | 26511 | 4 | 54875956 | 54930857 | -1 | protein_coding | cysteine-rich hydrophobic domain 2 |
| **KIAA1211** | ENSG00000109265 | 65.90576694 | 3.638899859 | 0.501927526 | 2.74E-14 | 3.96E-13 | 57482 | 4 | 57036361 | 57194791 | 1 | protein_coding | KIAA1211 |
| **LAMTOR3** | ENSG00000109270 | 1455.616627 | 0.442383689 | 0.146046604 | 0.002017029 | 0.006815444 | 8649 | 4 | 100799493 | 100815647 | -1 | protein_coding | late endosomal/lysosomal adaptor, MAPK and MTOR activator 3 |
| **NFKB1** | ENSG00000109320 | 2856.446583 | 1.165694066 | 0.133090654 | 5.32E-19 | 1.08E-17 | 4790 | 4 | 103422486 | 103538459 | 1 | protein_coding | nuclear factor of kappa light polypeptide gene enhancer in B-cells 1 |
| **MANBA** | ENSG00000109323 | 1139.668712 | 0.500111164 | 0.177580186 | 0.003241474 | 0.010398709 | 4126 | 4 | 103552660 | 103682151 | -1 | protein_coding | mannosidase, beta A, lysosomal |
| **UBE2D3** | ENSG00000109332 | 6776.949612 | 0.954596775 | 0.09192178 | 9.94E-26 | 3.08E-24 | 7323 | 4 | 103715540 | 103790053 | -1 | protein_coding | ubiquitin-conjugating enzyme E2D 3 |
| **ELF2** | ENSG00000109381 | 749.7210415 | 0.55317403 | 0.132958502 | 1.92E-05 | 1E-04 | 1998 | 4 | 139949266 | 140098372 | -1 | protein_coding | E74-like factor 2 (ets domain transcription factor) |
| **NDUFC1** | ENSG00000109390 | 143.5173788 | 0.802066832 | 0.321451571 | 0.004600123 | 0.014182706 | 4717 | 4 | 140188034 | 140223705 | -1 | protein_coding | NADH dehydrogenase (ubiquinone) 1, subcomplex unknown, 1, 6kDa |
| **TBC1D9** | ENSG00000109436 | 5738.04674 | 1.454935013 | 0.08521346 | 4.18E-66 | 6.87E-64 | 23158 | 4 | 141541919 | 141677274 | -1 | protein_coding | TBC1 domain family, member 9 (with GRAM domain) |
| **INPP4B** | ENSG00000109452 | 380.5033946 | 1.227266233 | 0.38474585 | 2.5E-04 | 0.001046961 | 8821 | 4 | 142944313 | 143768585 | -1 | protein_coding | inositol polyphosphate-4-phosphatase, type II, 105kDa |
| **CPE** | ENSG00000109472 | 777.8320743 | -0.917175716 | 0.295544444 | 5.85E-04 | 0.002247988 | 1363 | 4 | 166282346 | 166419472 | 1 | protein_coding | carboxypeptidase E |
| **WFS1** | ENSG00000109501 | 988.1433818 | -1.584392064 | 0.192551336 | 2.48E-17 | 4.46E-16 | 7466 | 4 | 6271576 | 6304992 | 1 | protein_coding | Wolfram syndrome 1 (wolframin) |
| **CLCN3** | ENSG00000109572 | 4115.466596 | -0.577076302 | 0.141321328 | 2.88E-05 | 1.46E-04 | 1182 | 4 | 170533784 | 170644824 | 1 | protein_coding | chloride channel, voltage-sensitive 3 |
| **GALNT7** | ENSG00000109586 | 2467.235606 | -0.592866472 | 0.163240404 | 1.62E-04 | 7.13E-04 | 51809 | 4 | 174089904 | 174245118 | 1 | protein_coding | UDP-N-acetyl-alpha-D-galactosamine:polypeptide N-acetylgalactosaminyltransferase 7 (GalNAc-T7) |
| **DHX15** | ENSG00000109606 | 3374.657542 | 0.224754205 | 0.085639553 | 0.009931915 | 0.027830851 | 1665 | 4 | 24519064 | 24586173 | -1 | protein_coding | DEAH (Asp-Glu-Ala-His) box helicase 15 |
| **SLC2A9** | ENSG00000109667 | 567.9547879 | -1.709370587 | 0.191570214 | 5.71E-20 | 1.26E-18 | 56606 | 4 | 9772777 | 10056560 | -1 | protein_coding | solute carrier family 2 (facilitated glucose transporter), member 9 |
| **TBC1D19** | ENSG00000109680 | 100.1579673 | 0.786696194 | 0.366728284 | 0.01143263 | 0.031419687 | 55296 | 4 | 26578059 | 26756973 | 1 | protein_coding | TBC1 domain family, member 19 |
| **CLNK** | ENSG00000109684 | 34.7135122 | -2.229383869 | 0.679311262 | 8.55E-05 | 3.96E-04 | 116449 | 4 | 10488019 | 10686489 | -1 | protein_coding | cytokine-dependent hematopoietic cell linker |
| **RAPGEF2** | ENSG00000109756 | 3135.685947 | 1.153746086 | 0.127988095 | 3.2E-20 | 7.18E-19 | 9693 | 4 | 160025330 | 160281321 | 1 | protein_coding | Rap guanine nucleotide exchange factor (GEF) 2 |
| **UFSP2** | ENSG00000109775 | 641.6285164 | -0.593757342 | 0.166230506 | 2.15E-04 | 9.11E-04 | 55325 | 4 | 186320694 | 186347139 | -1 | protein_coding | UFM1-specific peptidase 2 |
| **KLF3** | ENSG00000109787 | 1426.926797 | 0.45411777 | 0.12286062 | 1.67E-04 | 7.28E-04 | 51274 | 4 | 38665817 | 38702663 | 1 | protein_coding | Kruppel-like factor 3 (basic) |
| **NCAPG** | ENSG00000109805 | 100.9933892 | 1.835687367 | 0.486367247 | 1.53E-05 | 8.14E-05 | 64151 | 4 | 17812525 | 17846485 | 1 | protein_coding | non-SMC condensin I complex, subunit G |
| **HTATIP2** | ENSG00000109854 | 2051.588486 | 0.436816958 | 0.105709559 | 2.78E-05 | 1.41E-04 | 10553 | 11 | 20385231 | 20405329 | 1 | protein_coding | HIV-1 Tat interactive protein 2, 30kDa |
| **MTCH2** | ENSG00000109919 | 1868.691437 | -0.531384428 | 0.12332439 | 1.14E-05 | 6.21E-05 | 23788 | 11 | 47638867 | 47664175 | -1 | protein_coding | mitochondrial carrier 2 |
| **FNBP4** | ENSG00000109920 | 1805.219448 | 0.898191002 | 0.117883284 | 1.21E-14 | 1.78E-13 | 23360 | 11 | 47738072 | 47788995 | -1 | protein_coding | formin binding protein 4 |
| **SC5D** | ENSG00000109929 | 2852.255719 | 1.002191104 | 0.166545063 | 4.72E-10 | 4.69E-09 | 6309 | 11 | 121163162 | 121179403 | 1 | protein_coding | sterol-C5-desaturase |
| **DNAJC4** | ENSG00000110011 | 514.4795276 | -0.985360306 | 0.172957408 | 3.78E-09 | 3.37E-08 | 3338 | 11 | 63997750 | 64001758 | 1 | protein_coding | DnaJ (Hsp40) homolog, subfamily C, member 4 |
| **UNC93B1** | ENSG00000110057 | 62.56107163 | 1.408216279 | 0.522824676 | 8.84E-04 | 0.003268731 | 81622 | 11 | 67758575 | 67772452 | -1 | protein_coding | unc-93 homolog B1 (C. elegans) |
| **FOXRED1** | ENSG00000110074 | 305.4780579 | -0.517599163 | 0.227661765 | 0.014720754 | 0.03909249 | 55572 | 11 | 126138950 | 126148026 | 1 | protein_coding | FAD-dependent oxidoreductase domain containing 1 |
| **MS4A6A** | ENSG00000110077 | 730.1172591 | -1.568803836 | 0.215882517 | 5.02E-14 | 7.15E-13 | 64231 | 11 | 59939081 | 59952139 | -1 | protein_coding | membrane-spanning 4-domains, subfamily A, member 6A |
| **MS4A4A** | ENSG00000110079 | 4358.385359 | -0.716544851 | 0.102642584 | 1.57E-12 | 1.96E-11 | 51338 | 11 | 60048014 | 60076445 | 1 | protein_coding | membrane-spanning 4-domains, subfamily A, member 4A |
| **ST3GAL4** | ENSG00000110080 | 342.0031375 | 1.5941969 | 0.210144131 | 4.55E-15 | 6.92E-14 | 6484 | 11 | 126225535 | 126310239 | 1 | protein_coding | ST3 beta-galactoside alpha-2,3-sialyltransferase 4 |
| **CCDC86** | ENSG00000110104 | 316.8367808 | -0.522696174 | 0.209528476 | 0.008175692 | 0.023469026 | 79080 | 11 | 60609544 | 60618554 | 1 | protein_coding | coiled-coil domain containing 86 |
| **PRPF19** | ENSG00000110107 | 1123.789237 | -0.470072009 | 0.201064331 | 0.013222649 | 0.035602844 | 27339 | 11 | 60658202 | 60674060 | -1 | protein_coding | pre-mRNA processing factor 19 |
| **CHORDC1** | ENSG00000110172 | 850.9854449 | 0.559505525 | 0.153437411 | 1.56E-04 | 6.87E-04 | 26973 | 11 | 89934328 | 89956532 | -1 | protein_coding | cysteine and histidine-rich domain (CHORD) containing 1 |
| **ANAPC15** | ENSG00000110200 | 207.4962722 | -1.484644386 | 0.278705949 | 1.54E-08 | 1.27E-07 | 25906 | 11 | 71817424 | 71823826 | -1 | protein_coding | anaphase promoting complex subunit 15 |
| **PANX1** | ENSG00000110218 | 1255.237451 | 1.283489343 | 0.123143776 | 4.02E-26 | 1.27E-24 | 24145 | 11 | 93862094 | 93915138 | 1 | protein_coding | pannexin 1 |
| **ARHGEF17** | ENSG00000110237 | 170.3479078 | -1.256118851 | 0.363307987 | 1.01E-04 | 4.59E-04 | 9828 | 11 | 73019334 | 73080136 | 1 | protein_coding | Rho guanine nucleotide exchange factor (GEF) 17 |
| **RNF141** | ENSG00000110315 | 618.0665183 | -1.262820985 | 0.177189739 | 2.12E-13 | 2.84E-12 | 50862 | 11 | 10533225 | 10562777 | -1 | protein_coding | ring finger protein 141 |
| **IL10RA** | ENSG00000110324 | 11096.86946 | 0.951600808 | 0.158144742 | 5.97E-10 | 5.86E-09 | 3587 | 11 | 117857063 | 117872196 | 1 | protein_coding | interleukin 10 receptor, alpha |
| **BIRC2** | ENSG00000110330 | 2138.754684 | 1.089754945 | 0.122847942 | 1.94E-19 | 4.09E-18 | 329 | 11 | 102217942 | 102249401 | 1 | protein_coding | baculoviral IAP repeat containing 2 |
| **PVRL1** | ENSG00000110400 | 240.2956089 | -0.703157926 | 0.267043305 | 0.003775129 | 0.011888683 | 5818 | 11 | 119494120 | 119599794 | -1 | protein_coding | poliovirus receptor-related 1 (herpesvirus entry mediator C) |
| **PDHX** | ENSG00000110435 | 901.573993 | 0.583858378 | 0.192703356 | 0.001440494 | 0.005052806 | 8050 | 11 | 34937376 | 35042138 | 1 | protein_coding | pyruvate dehydrogenase complex, component X |
| **SLC1A2** | ENSG00000110436 | 2833.375657 | 3.794259506 | 0.195052012 | 1.25E-85 | 3.28E-83 | 6506 | 11 | 35272753 | 35441610 | -1 | protein_coding | solute carrier family 1 (glial high affinity glutamate transporter), member 2 |
| **COMMD9** | ENSG00000110442 | 1229.517227 | -1.047295537 | 0.166949271 | 9.54E-11 | 1.02E-09 | 29099 | 11 | 36295051 | 36310999 | -1 | protein_coding | COMM domain containing 9 |
| **SLC15A3** | ENSG00000110446 | 9991.947152 | 0.454075496 | 0.107879448 | 1.9E-05 | 9.94E-05 | 51296 | 11 | 60704556 | 60720002 | -1 | protein_coding | solute carrier family 15 (oligopeptide transporter), member 3 |
| **CD5** | ENSG00000110448 | 494.0840995 | -0.933951003 | 0.227081377 | 1.33E-05 | 7.19E-05 | 921 | 11 | 60869867 | 60895324 | 1 | protein_coding | CD5 molecule |
| **MDK** | ENSG00000110492 | 666.9457541 | 1.075654347 | 0.264207856 | 1.05E-05 | 5.76E-05 | 4192 | 11 | 46402306 | 46405375 | 1 | protein_coding | midkine (neurite growth-promoting factor 2) |
| **AMBRA1** | ENSG00000110497 | 1506.492731 | 0.837146053 | 0.16646249 | 2.11E-07 | 1.5E-06 | 55626 | 11 | 46417964 | 46615675 | -1 | protein_coding | autophagy/beclin-1 regulator 1 |
| **MADD** | ENSG00000110514 | 2918.697992 | 0.524587167 | 0.110750246 | 1.58E-06 | 9.86E-06 | 8567 | 11 | 47290712 | 47351582 | 1 | protein_coding | MAP-kinase activating death domain |

| | | | | | | | | | | | | | |
|---|---|---|---|---|---|---|---|---|---|---|---|---|---|
| **CD81** | ENSG00000110651 | 10550.05407 | -0.384935632 | 0.09990413 | 8.86E-05 | 4.1E-04 | 975 | 11 | 2397407 | 2418649 | 1 | protein_coding | CD81 molecule |
| **SLC35F2** | ENSG00000110660 | 142.0875574 | 1.146992804 | 0.332927681 | 1.15E-04 | 5.2E-04 | 54733 | 11 | 107661717 | 107799019 | -1 | protein_coding | solute carrier family 35, member F2 |
| **C11orf21** | ENSG00000110665 | 351.0262691 | -1.575149758 | 0.261493357 | 2.48E-10 | 2.53E-09 | 29125 | 11 | 2316875 | 2324279 | -1 | protein_coding | chromosome 11 open reading frame 21 |
| **NUP98** | ENSG00000110713 | 5218.907675 | 0.295644185 | 0.105195837 | 0.004258452 | 0.013230723 | 4928 | 11 | 3692313 | 3819022 | -1 | protein_coding | nucleoporin 98kDa |
| **NDUFS8** | ENSG00000110717 | 746.5974872 | -1.087447403 | 0.154945865 | 5.81E-13 | 7.58E-12 | 1024656 | 11 | 67798084 | 67804111 | 1 | protein_coding | NADH dehydrogenase (ubiquinone) Fe-S protein 8, 23kDa (NADH-coenzyme Q reductase) |
| **TCIRG1** | ENSG00000110719 | 7185.883077 | -0.396823645 | 0.117503311 | 5.73E-04 | 0.002207893 | 10312 | 11 | 67806483 | 67818362 | 1 | protein_coding | T-cell, immune regulator 1, ATPase, H+ transporting, lysosomal V0 subunit A3 |
| **HPS5** | ENSG00000110756 | 4516.551138 | 0.87405302 | 0.1050386 | 3.25E-17 | 5.77E-16 | 11234 | 11 | 18300223 | 18343745 | -1 | protein_coding | Hermansky-Pudlak syndrome 5 |
| **GTF2H1** | ENSG00000110768 | 1719.547269 | 0.718515586 | 0.105981808 | 5.93E-12 | 7.13E-11 | 2965 | 11 | 18343842 | 18388591 | 1 | protein_coding | general transcription factor IIH, polypeptide 1, 62kDa |
| **VWF** | ENSG00000110799 | 413.9800044 | -2.988511168 | 0.382708663 | 3.71E-16 | 6.09E-15 | 7450 | 12 | 6058040 | 6233936 | -1 | protein_coding | von Willebrand factor |
| **PPFIBP1** | ENSG00000110841 | 866.4857381 | -0.766164585 | 0.15761475 | 5.03E-07 | 3.4E-06 | 8496 | 12 | 27676364 | 27848497 | 1 | protein_coding | PTPRF interacting protein, binding protein 1 (liprin beta 1) |
| **CD69** | ENSG00000110848 | 196.631677 | 0.974154972 | 0.337690198 | 0.001034695 | 0.003762369 | 969 | 12 | 9905082 | 9913497 | -1 | protein_coding | CD69 molecule |
| **PRDM4** | ENSG00000110851 | 1210.201234 | 0.646195253 | 0.133352642 | 7.13E-07 | 4.71E-06 | 11108 | 12 | 108126643 | 108155049 | -1 | protein_coding | PR domain containing 4 |
| **CLEC2B** | ENSG00000110852 | 1067.874826 | 0.673371457 | 0.141738777 | 1.02E-06 | 6.57E-06 | 9976 | 12 | 10005583 | 10022735 | -1 | protein_coding | C-type lectin domain family 2, member B |
| **SELPLG** | ENSG00000110876 | 1426.679665 | -0.686374798 | 0.114821331 | 9.88E-10 | 9.52E-09 | 6404 | 12 | 109015686 | 109027735 | -1 | protein_coding | selectin P ligand |
| **CORO1C** | ENSG00000110880 | 15720.25581 | -0.698853428 | 0.128931422 | 3.04E-08 | 2.43E-07 | 23603 | 12 | 109038885 | 109125372 | -1 | protein_coding | coronin, actin binding protein, 1C |
| **ASIC1** | ENSG00000110881 | 82.07168514 | -1.95742274 | 0.457587427 | 1.97E-06 | 1.22E-05 | 41 | 12 | 50451331 | 50477394 | 1 | protein_coding | acid-sensing (proton-gated) ion channel 1 |
| **TSPAN11** | ENSG00000110900 | 35.92390814 | -2.292119719 | 0.826247253 | 4.42E-04 | 0.001752631 | 441631 | 12 | 31079362 | 31149534 | 1 | protein_coding | tetraspanin 11 |
| **SLC11A2** | ENSG00000110911 | 5064.146021 | 0.942800195 | 0.141342875 | 7.29E-12 | 8.69E-11 | 4891 | 12 | 51373184 | 51422349 | -1 | protein_coding | solute carrier family 11 (proton-coupled divalent metal ion transporter), member 2 |
| **MLEC** | ENSG00000110917 | 7276.598001 | -0.760362314 | 0.136440847 | 1.05E-08 | 8.88E-08 | 9761 | 12 | 121124672 | 121139667 | 1 | protein_coding | malectin |
| **CSRNP2** | ENSG00000110925 | 1260.701741 | 1.152609739 | 0.134352557 | 2.3E-18 | 4.48E-17 | 81566 | 12 | 51454990 | 51477447 | -1 | protein_coding | cysteine-serine-rich nuclear protein 2 |
| **BIN2** | ENSG00000110934 | 481.2643067 | 0.925907674 | 0.228394467 | 1.57E-05 | 8.33E-05 | 51411 | 12 | 51674822 | 51718452 | -1 | protein_coding | bridging integrator 2 |
| **ATP5B** | ENSG00000110955 | 10903.59929 | -0.29760562 | 0.084837704 | 3.47E-04 | 0.001410542 | 506 | 12 | 57031959 | 57039798 | -1 | protein_coding | ATP synthase, H+ transporting, mitochondrial F1 complex, beta polypeptide |
| **PTGES3** | ENSG00000110958 | 3997.27165 | 1.135877562 | 0.131412134 | 1.56E-18 | 3.07E-17 | 10728 | 12 | 57057127 | 57082159 | -1 | protein_coding | prostaglandin E synthase 3 (cytosolic) |
| **RSRC2** | ENSG00000111011 | 1277.744024 | 0.479288219 | 0.12962273 | 1.57E-04 | 6.92E-04 | 65117 | 12 | 122989190 | 123011547 | -1 | protein_coding | arginine/serine-rich coiled-coil 2 |
| **CYP27B1** | ENSG00000111012 | 1742.664183 | 2.408674657 | 0.215811081 | 4.66E-30 | 1.8E-28 | 1594 | 12 | 58156117 | 58162769 | -1 | protein_coding | cytochrome P450, family 27, subfamily B, polypeptide 1 |
| **PPM1H** | ENSG00000111110 | 1646.659194 | -0.80792147 | 0.134685142 | 6.28E-10 | 6.15E-09 | 57460 | 12 | 63037762 | 63328817 | -1 | protein_coding | protein phosphatase, Mg2+/Mn2+ dependent, 1H |
| **LTA4H** | ENSG00000111144 | 22323.92317 | -1.960063621 | 0.129390684 | 2.47E-52 | 2.44E-50 | 1019288 | 12 | 96394606 | 96437298 | -1 | protein_coding | leukotriene A4 hydrolase |
| **MAGOHB** | ENSG00000111196 | 150.1408947 | 0.562127984 | 0.263643671 | 0.019469292 | 0.049583269 | 55110 | 12 | 10758612 | 10766222 | -1 | protein_coding | mago-nashi homolog B (Drosophila) |
| **TRPV4** | ENSG00000111199 | 209.6809367 | -2.875627267 | 0.330782289 | 2.02E-19 | 4.25E-18 | 59341 | 12 | 110220890 | 110271212 | -1 | protein_coding | transient receptor potential cation channel, subfamily V, member 4 |
| **ITFG2** | ENSG00000111203 | 358.2811191 | -0.648843299 | 0.227910938 | 0.00228921 | 0.007610615 | 1005074 | 12 | 2921788 | 2968957 | 1 | protein_coding | integrin alpha FG-GAP repeat containing 2 |
| **FOXM1** | ENSG00000111206 | 89.53849906 | 1.090239862 | 0.495751504 | 0.005259362 | 0.015951037 | 2305 | 12 | 2966847 | 2986206 | -1 | protein_coding | forkhead box M1 |
| **PARP11** | ENSG00000111224 | 689.3340131 | 1.242514789 | 0.171925907 | 1E-13 | 1.39E-12 | 57097 | 12 | 3900213 | 3982608 | -1 | protein_coding | poly (ADP-ribose) polymerase family, member 11 |
| **VPS29** | ENSG00000111237 | 1797.326151 | 0.493997085 | 0.141255665 | 3.21E-04 | 0.001317803 | 51699 | 12 | 110928902 | 110939922 | -1 | protein_coding | vacuolar protein sorting 29 homolog (S. cerevisiae) |
| **CUX2** | ENSG00000111249 | 18.15517289 | -4.742690785 | 1.502935908 | 5.07E-04 | 0.001982528 | 23316 | 12 | 111471828 | 111788358 | 1 | protein_coding | cut-like homeobox 2 |
| **SH2B3** | ENSG00000111252 | 5840.864495 | -0.465131882 | 0.099866682 | 2.37E-06 | 1.44E-05 | 10019 | 12 | 111843752 | 111889427 | 1 | protein_coding | SH2B adaptor protein 3 |
| **AKAP3** | ENSG00000111254 | 94.90183575 | 0.786812366 | 0.325509857 | 0.005854434 | 0.017495759 | 10566 | 12 | 4724674 | 4758213 | -1 | protein_coding | A kinase (PRKA) anchor protein 3 |
| **DUSP16** | ENSG00000111266 | 1132.911458 | 0.975624417 | 0.160469546 | 3.74E-10 | 3.76E-09 | 80824 | 12 | 12628829 | 12715317 | -1 | protein_coding | dual specificity phosphatase 16 |
| **CREBL2** | ENSG00000111269 | 4865.757889 | 0.280838352 | 0.101612118 | 0.004977631 | 0.015211585 | 1389 | 12 | 12764761 | 12798042 | 1 | protein_coding | cAMP responsive element binding protein-like 2 |
| **ACAD10** | ENSG00000111271 | 532.2709768 | -0.777653429 | 0.180610558 | 6.97E-06 | 3.94E-05 | 80724 | 12 | 112123857 | 112194903 | 1 | protein_coding | acyl-CoA dehydrogenase family, member 10 |
| **CDKN1B** | ENSG00000111276 | 860.3358505 | 0.917333557 | 0.176787332 | 7.24E-08 | 5.48E-07 | 1027 | 12 | 12867992 | 12875305 | 1 | protein_coding | cyclin-dependent kinase inhibitor 1B (p27, Kip1) |
| **GSG1** | ENSG00000111305 | 72.78615621 | -1.33118684 | 0.747798126 | 0.007230602 | 0.021088796 | 83445 | 12 | 13236494 | 13256619 | -1 | protein_coding | germ cell associated 1 |
| **OGFOD2** | ENSG00000111325 | 189.3219038 | -0.632366632 | 0.291062441 | 0.014808226 | 0.039290646 | 79676 | 12 | 123459127 | 123464590 | 1 | protein_coding | 2-oxoglutarate and iron-dependent oxygenase domain containing 2 |
| **OAS3** | ENSG00000111331 | 7879.309651 | 4.880093814 | 0.114498448 | 0 | 0 | 4940 | 12 | 113376157 | 113411054 | 1 | protein_coding | 2'-5'-oligoadenylate synthetase 3, 100kDa |
| **OAS2** | ENSG00000111335 | 7201.596219 | 3.902867295 | 0.128097158 | 4.85E-205 | 4.36E-202 | 4939 | 12 | 113416200 | 113449528 | 1 | protein_coding | 2'-5'-oligoadenylate synthetase 2, 69/71kDa |
| **RASAL1** | ENSG00000111344 | 16.24172628 | -2.355973015 | 1.412987634 | 0.004430293 | 0.013711658 | 8437 | 12 | 113536624 | 113574044 | -1 | protein_coding | RAS protein activator like 1 (GAP1 like) |
| **ARHGDIB** | ENSG00000111348 | 14493.92678 | -1.597291509 | 0.102212952 | 4.59E-56 | 5.19E-54 | 397 | 12 | 15094951 | 15114662 | -1 | protein_coding | Rho GDP dissociation inhibitor (GDI) beta |
| **GTF2H3** | ENSG00000111358 | 1216.137657 | 0.818451021 | 0.146204225 | 8.52E-09 | 7.32E-08 | 2967 | 12 | 124118375 | 124147153 | 1 | protein_coding | general transcription factor IIH, polypeptide 3, 34kDa |
| **C12orf49** | ENSG00000111412 | 2548.648793 | -0.76224869 | 0.109477391 | 1.53E-12 | 1.92E-11 | 79794 | 12 | 117153593 | 117175875 | -1 | protein_coding | chromosome 12 open reading frame 49 |
| **STX2** | ENSG00000111450 | 792.5345502 | -0.599227337 | 0.22117948 | 0.003904375 | 0.012240114 | 2054 | 12 | 131274145 | 131323811 | -1 | protein_coding | syntaxin 2 |
| **GPR133** | ENSG00000111452 | 23.43088355 | -1.915297197 | 0.906703123 | 0.002602738 | 0.008521347 | 283383 | 12 | 131438452 | 131626014 | 1 | protein_coding | G protein-coupled receptor 133 |
| **CAND1** | ENSG00000111530 | 3872.918834 | 0.902548575 | 0.134407732 | 6.28E-12 | 7.54E-11 | 55832 | 12 | 67663061 | 67713731 | 1 | protein_coding | cullin-associated and neddylation-dissociated 1 |
| **IFNG** | ENSG00000111537 | 72.59013826 | 1.415316673 | 0.482597265 | 4.33E-04 | 0.00172178 | 3458 | 12 | 68548548 | 68553527 | -1 | protein_coding | interferon, gamma |
| **MDM1** | ENSG00000111554 | 440.3466777 | 1.438751158 | 0.219685695 | 8.65E-12 | 1.02E-10 | 56890 | 12 | 68666223 | 68726161 | -1 | protein_coding | Mdm1 nuclear protein homolog (mouse) |
| **TIMELESS** | ENSG00000111602 | 377.8698292 | 0.784470525 | 0.202826788 | 4.49E-05 | 2.18E-04 | 8914 | 12 | 56810903 | 56843187 | -1 | protein_coding | timeless circadian clock |
| **KRR1** | ENSG00000111615 | 954.9841103 | 0.852589737 | 0.13621649 | 1.57E-10 | 1.64E-09 | 11103 | 12 | 75890684 | 75905416 | -1 | protein_coding | KRR1, small subunit (SSU) processome component, homolog (yeast) |
| **GAPDH** | ENSG00000111640 | 34845.86901 | -0.905720081 | 0.108575807 | 1.44E-17 | 2.63E-16 | 2597 | 12 | 6643093 | 6647537 | 1 | protein_coding | glyceraldehyde-3-phosphate dehydrogenase |
| **COPS7A** | ENSG00000111652 | 994.1456971 | -0.390491586 | 0.12361361 | 0.001251995 | 0.00445604 | 50813 | 12 | 6832907 | 6841041 | 1 | protein_coding | COP9 signalosome subunit 7A |
| **USP5** | ENSG00000111667 | 1432.229309 | -0.459217772 | 0.170304818 | 0.004894378 | 0.014996145 | 8078 | 12 | 6961292 | 6975796 | 1 | protein_coding | ubiquitin specific peptidase 5 (isopeptidase T) |
| **TPI1** | ENSG00000111669 | 8977.015384 | -0.732547412 | 0.122194669 | 8.64E-10 | 8.36E-09 | 7167 | 12 | 6976283 | 6980112 | 1 | protein_coding | triosephosphate isomerase 1 |
| **GNPTAB** | ENSG00000111670 | 9158.398415 | -0.253466371 | 0.086728465 | 0.003646994 | 0.011546924 | 79158 | 12 | 102139275 | 102224716 | -1 | protein_coding | N-acetylglucosamine-1-phosphate transferase, alpha and beta subunits |
| **ENO2** | ENSG00000111674 | 999.7739498 | -1.395308319 | 0.180257089 | 1.47E-15 | 2.31E-14 | 2026 | 12 | 7022909 | 7032861 | 1 | protein_coding | enolase 2 (gamma, neuronal) |
| **LPCAT3** | ENSG00000111684 | 721.8191336 | 0.652928853 | 0.216330926 | 0.00130623 | 0.004622451 | 10162 | 12 | 7085348 | 7125814 | -1 | protein_coding | lysophosphatidylcholine acyltransferase 3 |
| **GOLT1B** | ENSG00000111711 | 905.0887856 | 0.916719434 | 0.160914125 | 4E-09 | 3.56E-08 | 51026 | 12 | 21654715 | 21671342 | 1 | protein_coding | golgi transport 1B |
| **LDHB** | ENSG00000111716 | 6043.639771 | -0.618582307 | 0.129621833 | 9.82E-07 | 6.33E-06 | 3945 | 12 | 21788276 | 21910791 | -1 | protein_coding | lactate dehydrogenase B |
| **CLEC4A** | ENSG00000111729 | 513.1426205 | -2.407327904 | 0.188786439 | 2.92E-38 | 1.64E-36 | 50856 | 12 | 8276228 | 8291203 | 1 | protein_coding | C-type lectin domain family 4, member A |
| **C2CD5** | ENSG00000111731 | 1460.081226 | 0.404841418 | 0.134559617 | 0.002108484 | 0.007083747 | 9847 | 12 | 22601517 | 22697480 | -1 | protein_coding | C2 calcium-dependent domain containing 5 |
| **RAB35** | ENSG00000111737 | 1943.245838 | 0.779362864 | 0.106308225 | 1.11E-13 | 1.52E-12 | 11021 | 12 | 120532899 | 120555306 | -1 | protein_coding | RAB35, member RAS oncogene family |
| **RIC8B** | ENSG00000111785 | 222.2210541 | 0.570117802 | 0.236870318 | 0.009426031 | 0.026618261 | 55188 | 12 | 107168373 | 107283090 | 1 | protein_coding | RIC8 guanine nucleotide exchange factor B |
| **KLRB1** | ENSG00000111796 | 190.3068994 | -2.283009161 | 0.344596807 | 2.81E-12 | 3.46E-11 | 3820 | 12 | 9747147 | 9760482 | -1 | protein_coding | killer cell lectin-like receptor subfamily B, member 1 |
| **BTN3A3** | ENSG00000111801 | 2011.401234 | 1.139521304 | 0.163217812 | 6.45E-13 | 8.36E-12 | 10384 | 6 | 26440700 | 26453643 | 1 | protein_coding | butyrophilin, subfamily 3, member A3 |
| **TDP2** | ENSG00000111802 | 115.8752367 | 0.869810516 | 0.322866913 | 0.002319208 | 0.007703643 | 51567 | 6 | 24650205 | 24667261 | -1 | protein_coding | tyrosyl-DNA phosphodiesterase 2 |
| **DSE** | ENSG00000111817 | 1919.488843 | 0.899617678 | 0.155362579 | 2.47E-09 | 2.26E-08 | 29940 | 6 | 116575336 | 116762424 | 1 | protein_coding | dermatan sulfate epimerase |
| **RWDD1** | ENSG00000111832 | 409.2200251 | 0.792893544 | 0.254435147 | 7E-04 | 0.002646314 | 51389 | 6 | 116892530 | 116918838 | 1 | protein_coding | RWD domain containing 1 |
| **GCNT2** | ENSG00000111846 | 716.3936277 | 1.07591643 | 0.149505197 | 1.74E-13 | 2.36E-12 | 2651 | 6 | 10492456 | 10629601 | 1 | protein_coding | glucosaminyl (N-acetyl) transferase 2, I-branching enzyme (I blood group) |
| **NEDD9** | ENSG00000111859 | 968.9836232 | -0.573415228 | 0.17641297 | 7.16E-04 | 0.002702274 | 4739 | 6 | 11183531 | 11382581 | -1 | protein_coding | neural precursor cell expressed, developmentally down-regulated 9 |
| **CEP85L** | ENSG00000111860 | 570.6106436 | -0.481270908 | 0.158419601 | 0.001650199 | 0.005702011 | 387119 | 6 | 118781935 | 119031238 | -1 | protein_coding | centrosomal protein 85kDa-like |
| **ASF1A** | ENSG00000111875 | 512.4636212 | 0.434070441 | 0.184719197 | 0.013923611 | 0.037260471 | 25842 | 6 | 119215384 | 119230332 | 1 | protein_coding | anti-silencing function 1A histone chaperone |
| **SERINC1** | ENSG00000111897 | 12078.40512 | 0.508016539 | 0.110127072 | 2.69E-06 | 1.63E-05 | 57515 | 6 | 122764499 | 122792967 | -1 | protein_coding | serine incorporator 1 |
| **TPD52L1** | ENSG00000111907 | 42.18600998 | -4.229045709 | 1.255945552 | 1.14E-04 | 5.18E-04 | 7164 | 6 | 125440195 | 125585553 | 1 | protein_coding | tumor protein D52-like 1 |
| **HINT3** | ENSG00000111911 | 1053.866973 | 1.02880808 | 0.140491549 | 6.91E-14 | 9.71E-13 | 135114 | 6 | 126277927 | 126301387 | 1 | protein_coding | histidine triad nucleotide binding protein 3 |
| **NCOA7** | ENSG00000111912 | 1967.408886 | 0.648546397 | 0.128482976 | 2.54E-07 | 1.79E-06 | 135112 | 6 | 126102307 | 126252266 | 1 | protein_coding | nuclear receptor coactivator 7 |
| **FAM65B** | ENSG00000111913 | 592.7644833 | 1.099854839 | 0.233755774 | 5.99E-07 | 4E-06 | 9750 | 6 | 24797601 | 25042238 | -1 | protein_coding | family with sequence similarity 65, member B |
| **SASH1** | ENSG00000111961 | 1609.880989 | 0.433508713 | 0.119773959 | 2.26E-04 | 9.53E-04 | 23328 | 6 | 148593440 | 148873186 | 1 | protein_coding | SAM and SH3 domain containing 1 |
| **ULBP1** | ENSG00000111981 | 24.51333216 | -1.150730999 | 0.782469125 | 0.017831006 | 0.046040085 | 80329 | 6 | 150285143 | 150294846 | 1 | protein_coding | UL16 binding protein 1 |
| **FBXO5** | ENSG00000112029 | 119.6855587 | 1.298274035 | 0.387510109 | 1.26E-04 | 5.65E-04 | 26271 | 6 | 153291664 | 153304714 | -1 | protein_coding | F-box protein 5 |
| **MTRF1L** | ENSG00000112031 | 942.0737132 | 0.515246391 | 0.169892569 | 0.001533417 | 0.005343185 | 54516 | 6 | 153308497 | 153323820 | -1 | protein_coding | mitochondrial translational release factor 1-like |
| **PPARD** | ENSG00000112033 | 3368.47078 | -1.203059091 | 0.160107639 | 1.19E-14 | 1.75E-13 | 5467 | 6 | 35310335 | 35395968 | 1 | protein_coding | peroxisome proliferator-activated receptor delta |
| **FANCE** | ENSG00000112039 | 552.8655949 | -1.28737721 | 0.164750206 | 1.31E-15 | 2.08E-14 | 2178 | 6 | 35420138 | 35434880 | 1 | protein_coding | Fanconi anemia, complementation group E |
| **KCTD20** | ENSG00000112078 | 4197.554114 | -0.280944628 | 0.093120516 | 0.002103545 | 0.007068705 | 222658 | 6 | 36410544 | 36458920 | 1 | protein_coding | potassium channel tetramerization domain containing 20 |

| | | | | | | | | | | | | | |
|---|---|---|---|---|---|---|---|---|---|---|---|---|---|
| SRSF3 | ENSG00000112081 | 2962.002656 | 0.2805641 | 0.09144588 | 0.00191231 | 0.006503301 | 6428 | 6 | 36562145 | 36573377 | 1 | protein_coding | serine/arginine-rich splicing factor 3 |
| MCM3 | ENSG00000112118 | 903.4116813 | 0.511506407 | 0.140088019 | 1.8E-04 | 7.82E-04 | 4172 | 6 | 52128807 | 52149679 | -1 | protein_coding | minichromosome maintenance complex component 3 |
| RNF8 | ENSG00000112130 | 408.8787329 | 0.639624154 | 0.190132332 | 4.18E-04 | 0.00166905 | 9025 | 6 | 37321748 | 37362514 | 1 | protein_coding | ring finger protein 8, E3 ubiquitin protein ligase |
| MDGA1 | ENSG00000112139 | 160.3575393 | 1.440772963 | 0.408994746 | 5.8E-05 | 2.77E-04 | 266727 | 6 | 37600284 | 37667082 | -1 | protein_coding | MAM domain containing glycosylphosphatidylinositol anchor 1 |
| ICK | ENSG00000112144 | 568.7568851 | 0.468322208 | 0.147360865 | 0.001036148 | 0.003766759 | 22858 | 6 | 52866077 | 52926600 | -1 | protein_coding | intestinal cell (MAK-like) kinase |
| FBXO9 | ENSG00000112146 | 1212.02873 | 0.599050299 | 0.124626858 | 7.54E-07 | 4.96E-06 | 26268 | 6 | 52916789 | 52965671 | 1 | protein_coding | F-box protein 9 |
| CD83 | ENSG00000112149 | 8779.822365 | 0.836585746 | 0.074701145 | 3.29E-29 | 1.22E-27 | 9308 | 6 | 14117872 | 14137149 | 1 | protein_coding | CD83 molecule |
| CCNC | ENSG00000112237 | 769.4655682 | 0.438233138 | 0.157601185 | 0.003976677 | 0.012456463 | 892 | 6 | 99990256 | 100016849 | -1 | protein_coding | cyclin C |
| E2F3 | ENSG00000112242 | 1502.457333 | 0.35550249 | 0.128758026 | 0.004443999 | 0.013742947 | 1871 | 6 | 20402398 | 20493941 | 1 | protein_coding | E2F transcription factor 3 |
| GPLD1 | ENSG00000112293 | 30.6796854 | -1.480387058 | 0.846181188 | 0.008008058 | 0.023048427 | 2822 | 6 | 24424793 | 24495433 | -1 | protein_coding | glycosylphosphatidylinositol specific phospholipase D1 |
| ALDH5A1 | ENSG00000112294 | 980.284893 | -1.647284191 | 0.225290876 | 3.56E-14 | 5.11E-13 | 7915 | 6 | 24495080 | 24537435 | 1 | protein_coding | aldehyde dehydrogenase 5 family, member A1 |
| VNN 1.00 | ENSG00000112299 | 431.344258 | -1.085686742 | 0.248914313 | 3.33E-06 | 1.98E-05 | 8876 | 6 | 133002729 | 133035188 | -1 | protein_coding | vanin 1 |
| VNN 2.00 | ENSG00000112303 | 2604.825437 | -1.200490382 | 0.228964544 | 3.57E-08 | 2.82E-07 | 8875 | 6 | 133065009 | 133084598 | -1 | protein_coding | vanin 2 |
| ACOT13 | ENSG00000112304 | 1691.659924 | -0.448659087 | 0.191473877 | 0.014350974 | 0.038190133 | 55856 | 6 | 24667263 | 24705293 | 1 | protein_coding | acyl-CoA thioesterase 13 |
| C6orf62 | ENSG00000112308 | 9703.567871 | 0.517750406 | 0.116801635 | 6.31E-06 | 3.58E-05 | 81688 | 6 | 24705089 | 24721064 | -1 | protein_coding | chromosome 6 open reading frame 62 |
| GMNN | ENSG00000112312 | 188.2085154 | 0.83839145 | 0.290300638 | 0.001354053 | 0.0047715 | 51053 | 6 | 24775159 | 24786327 | 1 | protein_coding | geminin, DNA replication inhibitor |
| SNX3 | ENSG00000112335 | 4933.327764 | 0.624472647 | 0.140890088 | 5.02E-06 | 2.9E-05 | 8724 | 6 | 108532426 | 108582464 | -1 | protein_coding | sorting nexin 3 |
| HBS1L | ENSG00000112339 | 1448.357042 | -0.636827903 | 0.133653739 | 1.05E-06 | 6.7E-06 | 10767 | 6 | 135281516 | 135424194 | -1 | protein_coding | HBS1-like (S. cerevisiae) |
| TRIM38 | ENSG00000112343 | 2183.938053 | 1.208508393 | 0.121672728 | 6.86E-24 | 1.94E-22 | 10475 | 6 | 25963030 | 25987384 | 1 | protein_coding | tripartite motif containing 38 |
| ZBTB24 | ENSG00000112365 | 672.2891252 | 0.686967961 | 0.147121567 | 1.54E-06 | 9.6E-06 | 9841 | 6 | 109783797 | 109804440 | -1 | protein_coding | zinc finger and BTB domain containing 24 |
| FIG4 | ENSG00000112367 | 1743.686191 | -0.522681126 | 0.139284418 | 1.13E-04 | 5.14E-04 | 9896 | 6 | 110012499 | 110146631 | 1 | protein_coding | FIG4 homolog, SAC1 lipid phosphatase domain containing (S. cerevisiae) |
| PERP | ENSG00000112378 | 141.8514978 | 1.443340562 | 0.40392925 | 5.01E-05 | 2.41E-04 | 64065 | 6 | 138409642 | 138428648 | -1 | protein_coding | PERP, TP53 apoptosis effector |
| KIAA1244 | ENSG00000112379 | 301.3397729 | -1.836983859 | 0.342475459 | 8.47E-09 | 7.28E-08 | 57221 | 6 | 138483058 | 138665800 | 1 | protein_coding | KIAA1244 |
| HECA | ENSG00000112406 | 2812.086759 | 0.481923224 | 0.112486013 | 1.32E-05 | 7.12E-05 | 51696 | 6 | 139456249 | 139501939 | 1 | protein_coding | headcase homolog (Drosophila) |
| GPR126 | ENSG00000112414 | 54.17066335 | -1.793475941 | 0.865199819 | 0.001983562 | 0.0067187 | 57211 | 6 | 142622991 | 142767403 | 1 | protein_coding | G protein-coupled receptor 126 |
| PHACTR2 | ENSG00000112419 | 4662.744289 | 0.288739611 | 0.100776575 | 0.003695092 | 0.011677437 | 9749 | 6 | 143857982 | 144152322 | 1 | protein_coding | phosphatase and actin regulator 2 |
| CCR6 | ENSG00000112486 | 63.2512532 | 0.775467811 | 0.405837533 | 0.01964965 | 0.049942724 | 1235 | 6 | 167525295 | 167553184 | 1 | protein_coding | chemokine (C-C motif) receptor 6 |
| TAPBP | ENSG00000112493 | 6277.504303 | 0.928356252 | 0.114372613 | 1.38E-16 | 2.34E-15 | 6892 | HSC | 33189081 | 33203772 | -1 | protein_coding | TAP binding protein (tapasin) |
| CUTA | ENSG00000112514 | 528.5433193 | -0.846196096 | 0.182469235 | 1.35E-06 | 8.49E-06 | 51596 | 6 | 33384219 | 33386094 | -1 | protein_coding | cutA divalent cation tolerance homolog (E. coli) |
| QKI | ENSG00000112531 | 7505.060629 | -0.189108117 | 0.080649803 | 0.018772754 | 0.0480981 | 9444 | 6 | 163835032 | 163999628 | 1 | protein_coding | QKI, KH domain containing, RNA binding |
| CCND3 | ENSG00000112576 | 1090.556453 | 0.348619562 | 0.129147349 | 0.005649268 | 0.016948912 | 896 | 6 | 41902671 | 42018095 | -1 | protein_coding | cyclin D3 |
| DNPH1 | ENSG00000112667 | 152.6576531 | -0.815204995 | 0.274879974 | 0.001129984 | 0.004073966 | 10591 | 6 | 43193367 | 43197222 | -1 | protein_coding | 2'-deoxynucleoside 5'-phosphate N-hydrolase 1 |
| DUSP22 | ENSG00000112679 | 724.1976021 | 0.679096321 | 0.149592745 | 2.97E-06 | 1.78E-05 | 1006532 | 6 | 291630 | 351355 | 1 | protein_coding | dual specificity phosphatase 22 |
| EXOC2 | ENSG00000112685 | 1475.150311 | 0.779422783 | 0.119161066 | 2.64E-11 | 3E-10 | 55770 | 6 | 485133 | 693117 | -1 | protein_coding | exocyst complex component 2 |
| TMEM30A | ENSG00000112697 | 11768.43233 | 0.819106763 | 0.198901422 | 1.34E-05 | 7.25E-05 | 55754 | 6 | 75962640 | 75994684 | -1 | protein_coding | transmembrane protein 30A |
| SENP6 | ENSG00000112701 | 1916.454956 | 0.639224473 | 0.158182065 | 2.83E-05 | 1.43E-04 | 26054 | 6 | 76311225 | 76427997 | 1 | protein_coding | SUMO1/sentrin specific peptidase 6 |
| PRPF4B | ENSG00000112739 | 2361.40671 | 1.67110055 | 0.135782792 | 9.47E-36 | 4.88E-34 | 8899 | 6 | 4021501 | 4065217 | 1 | protein_coding | pre-mRNA processing factor 4B |
| TTK | ENSG00000112742 | 59.33388575 | 1.283262141 | 0.506473897 | 0.001725872 | 0.005917993 | 7272 | 6 | 80713604 | 80752244 | 1 | protein_coding | TTK protein kinase |
| SLC29A1 | ENSG00000112759 | 701.3755917 | -1.600058723 | 0.279875182 | 1.12E-09 | 1.08E-08 | 2030 | 6 | 44187242 | 44201888 | 1 | protein_coding | solute carrier family 29 (equilibrative nucleoside transporter), member 1 |
| BTN2A1 | ENSG00000112763 | 870.3499201 | 0.828196283 | 0.148863946 | 1.09E-08 | 9.19E-08 | 11120 | 6 | 26458150 | 26476849 | 1 | protein_coding | butyrophilin, subfamily 2, member A1 |
| FAM46A | ENSG00000112773 | 2902.211179 | 1.413151396 | 0.104570351 | 1.95E-42 | 1.35E-40 | 55603 | 6 | 82201156 | 82462491 | -1 | protein_coding | family with sequence similarity 46, member A |
| FBRSL1 | ENSG00000112787 | 572.5723198 | -1.106253826 | 0.254518278 | 3.16E-06 | 1.89E-05 | 57666 | 12 | 133066137 | 133161774 | 1 | protein_coding | fibrosin-like 1 |
| ERBB2IP | ENSG00000112851 | 6724.479931 | 0.38270807 | 0.106357844 | 2.6E-04 | 0.001085367 | 55914 | 5 | 65222303 | 65378377 | 1 | protein_coding | erbb2 interacting protein |
| PAPD7 | ENSG00000112941 | 943.2162769 | 1.255896029 | 0.130081596 | 8.42E-23 | 2.24E-21 | 11044 | 5 | 6714718 | 6757161 | 1 | protein_coding | PAP associated domain containing 7 |
| HMGCS1 | ENSG00000112972 | 2365.395349 | 1.00820042 | 0.159966738 | 8.37E-11 | 9.02E-10 | 3157 | 5 | 43289497 | 43313614 | -1 | protein_coding | 3-hydroxy-3-methylglutaryl-CoA synthase 1 (soluble) |
| DAP | ENSG00000112977 | 2344.08512 | -0.53929635 | 0.099400094 | 3.79E-08 | 2.98E-07 | 1611 | 5 | 10679342 | 10761384 | -1 | protein_coding | death-associated protein |
| MRPS27 | ENSG00000113048 | 951.1173108 | -0.408859373 | 0.162993909 | 0.009165439 | 0.025968745 | 23107 | 5 | 71515236 | 71616473 | -1 | protein_coding | mitochondrial ribosomal protein S27 |
| HBEGF | ENSG00000113070 | 834.3028247 | -0.662511935 | 0.257195807 | 0.004728822 | 0.014535517 | 1839 | 5 | 139712428 | 139726216 | -1 | protein_coding | heparin-binding EGF-like growth factor |
| SPARC | ENSG00000113140 | 19222.78904 | -1.630817753 | 0.141723296 | 1.34E-31 | 5.68E-30 | 6678 | 5 | 151040657 | 151066726 | -1 | protein_coding | secreted protein, acidic, cysteine-rich (osteonectin) |
| PDE8B | ENSG00000113231 | 29.70145317 | -1.669293751 | 0.862695868 | 0.003779587 | 0.01189527 | 8622 | 5 | 76506274 | 76725632 | 1 | protein_coding | phosphodiesterase 8B |
| CLK4 | ENSG00000113240 | 503.4287906 | 0.633762995 | 0.170334735 | 1.07E-04 | 4.89E-04 | 57396 | 5 | 178029665 | 178057616 | -1 | protein_coding | CDC-like kinase 4 |
| ITK | ENSG00000113263 | 861.6298619 | 0.586326247 | 0.176324039 | 5.24E-04 | 0.002040817 | 3702 | 5 | 156569944 | 156682201 | 1 | protein_coding | IL2-inducible T-cell kinase |
| RNF130 | ENSG00000113269 | 9561.693023 | -0.701254962 | 0.096016672 | 1.47E-13 | 2E-12 | 55819 | 5 | 179338651 | 179499118 | -1 | protein_coding | ring finger protein 130 |
| THG1L | ENSG00000113272 | 143.5022925 | 0.929760073 | 0.379444565 | 0.003959294 | 0.012407196 | 54974 | 5 | 157158205 | 157168456 | 1 | protein_coding | tRNA-histidine guanylyltransferase 1-like (S. cerevisiae) |
| ARSB | ENSG00000113273 | 2590.253154 | 0.277559016 | 0.105249222 | 0.007476262 | 0.02171825 | 411 | 5 | 78073032 | 78281910 | -1 | protein_coding | arylsulfatase B |
| IL12B | ENSG00000113302 | 36.19886123 | 5.917712565 | 0.867783309 | 9.85E-13 | 1.25E-11 | 3593 | 5 | 158741791 | 158757895 | -1 | protein_coding | interleukin 12B (natural killer cell stimulatory factor 2, cytotoxic lymphocyte maturation factor 2, p40) |
| RASGRF2 | ENSG00000113319 | 92.47825093 | -0.783672051 | 0.388188831 | 0.014843834 | 0.039357791 | 5924 | 5 | 80256491 | 80525975 | 1 | protein_coding | Ras protein-specific guanine nucleotide-releasing factor 2 |
| LMNB1 | ENSG00000113368 | 513.0822533 | 1.945582518 | 0.304886722 | 1.91E-11 | 2.2E-10 | 4001 | 5 | 126112315 | 126172712 | 1 | protein_coding | lamin B1 |
| ARRDC3 | ENSG00000113369 | 2085.357492 | -0.549540169 | 0.187662873 | 0.00210306 | 0.00706863 | 57561 | 5 | 90664541 | 90679176 | -1 | protein_coding | arrestin domain containing 3 |
| GOLPH3 | ENSG00000113384 | 2847.004116 | 0.371121449 | 0.109824625 | 5.5E-04 | 0.002129904 | 64083 | 5 | 32124810 | 32174456 | -1 | protein_coding | golgi phosphoprotein 3 (coat-protein) |
| SUB1 | ENSG00000113387 | 1575.397908 | 0.980353817 | 0.124615878 | 7.79E-16 | 1.26E-14 | 10923 | 5 | 32531739 | 32604185 | 1 | protein_coding | SUB1 homolog (S. cerevisiae) |
| FAM172A | ENSG00000113391 | 521.6908732 | -0.778958671 | 0.192817025 | 2.25E-05 | 1.16E-04 | 83989 | 5 | 92953775 | 93447404 | -1 | protein_coding | family with sequence similarity 172, member A |
| TARS | ENSG00000113407 | 4096.688378 | 0.48984583 | 0.122938736 | 4.92E-05 | 2.38E-04 | 6897 | 5 | 33440802 | 33469644 | 1 | protein_coding | threonyl-tRNA synthetase |
| LNPEP | ENSG00000113441 | 4700.458188 | 0.770422258 | 0.108084116 | 4.64E-13 | 6.12E-12 | 4012 | 5 | 96271098 | 96373219 | 1 | protein_coding | leucyl/cystinyl aminopeptidase |
| PDE4D | ENSG00000113448 | 495.5860611 | 0.515561803 | 0.208940405 | 0.008686544 | 0.024735692 | 5144 | 5 | 58264865 | 59817947 | -1 | protein_coding | phosphodiesterase 4D, cAMP-specific |
| RAD1 | ENSG00000113456 | 464.4253243 | 0.568747652 | 0.176685487 | 7.74E-04 | 0.002902022 | 5810 | 5 | 34905369 | 34919094 | -1 | protein_coding | RAD1 homolog (S. pombe) |
| BRIX1 | ENSG00000113460 | 323.316236 | 0.688747527 | 0.235598317 | 0.001647472 | 0.005696451 | 55299 | 5 | 34915481 | 34926101 | 1 | protein_coding | BRX1, biogenesis of ribosomes, homolog (S. cerevisiae) |
| SLC12A7 | ENSG00000113504 | 2860.079928 | -0.485236584 | 0.142677449 | 4.64E-04 | 0.001831297 | 10723 | 5 | 1050499 | 1112150 | -1 | protein_coding | solute carrier family 12 (potassium/chloride transporter), member 7 |
| ST8SIA4 | ENSG00000113532 | 1248.779436 | 0.775097729 | 0.201410846 | 4.6E-05 | 2.23E-04 | 7903 | 5 | 100142639 | 100238970 | -1 | protein_coding | ST8 alpha-N-acetyl-neuraminide alpha-2,8-sialyltransferase 4 |
| GNPDA1 | ENSG00000113552 | 1644.515027 | -0.484285975 | 0.137194762 | 2.62E-04 | 0.001090767 | 10007 | 5 | 141371314 | 141392606 | -1 | protein_coding | glucosamine-6-phosphate deaminase 1 |
| PCDH12 | ENSG00000113555 | 53.31764187 | 1.678807566 | 0.685004819 | 0.001174201 | 0.004207575 | 51294 | 5 | 141323150 | 141349304 | -1 | protein_coding | protocadherin 12 |
| SKP1 | ENSG00000113558 | 4053.50915 | 0.619204108 | 0.107470545 | 4.85E-09 | 4.3E-08 | 6500 | 5 | 133484633 | 133512729 | -1 | protein_coding | S-phase kinase-associated protein 1 |
| PPP2CA | ENSG00000113575 | 3714.999711 | 0.308549124 | 0.096729545 | 0.001308197 | 0.004626979 | 5515 | 5 | 133530025 | 133561833 | -1 | protein_coding | protein phosphatase 2, catalytic subunit, alpha isozyme |
| NR3C1 | ENSG00000113580 | 5507.864517 | 0.314222905 | 0.101839139 | 0.001754279 | 0.006008661 | 2908 | 5 | 142657496 | 142815077 | -1 | protein_coding | nuclear receptor subfamily 3, group C, member 1 (glucocorticoid receptor) |
| C5orf15 | ENSG00000113583 | 2075.539672 | 1.355132818 | 0.145316886 | 1.72E-21 | 4.21E-20 | 56951 | 5 | 133291201 | 133304478 | -1 | protein_coding | chromosome 5 open reading frame 15 |
| PPWD1 | ENSG00000113593 | 756.8135014 | 0.411394734 | 0.148935877 | 0.004429146 | 0.013710884 | 23398 | 5 | 64859063 | 64883376 | 1 | protein_coding | peptidylprolyl isomerase domain and WD repeat containing 1 |
| TRAPPC13 | ENSG00000113597 | 551.5055657 | 0.467177668 | 0.205720227 | 0.016032111 | 0.042092209 | 80006 | 5 | 64920543 | 64962060 | 1 | protein_coding | trafficking protein particle complex 13 |
| SEC24A | ENSG00000113615 | 2217.543214 | 0.538092538 | 0.11726259 | 2.88E-06 | 1.73E-05 | 10802 | 5 | 133984479 | 134063513 | 1 | protein_coding | SEC24 family member A |
| H2AFY | ENSG00000113648 | 9997.442287 | -1.313183875 | 0.106252541 | 9.12E-36 | 4.71E-34 | 9555 | 5 | 134669590 | 134735604 | -1 | protein_coding | H2A histone family, member Y |
| TCERG1 | ENSG00000113649 | 1340.815666 | 0.621322447 | 0.142321192 | 7.15E-06 | 4.03E-05 | 10915 | 5 | 145826874 | 145891524 | 1 | protein_coding | transcription elongation regulator 1 |
| ERGIC1 | ENSG00000113719 | 3347.899462 | -0.88069015 | 0.115743114 | 9.16E-15 | 1.37E-13 | 57222 | 5 | 172261278 | 172379688 | 1 | protein_coding | endoplasmic reticulum-golgi intermediate compartment (ERGIC) 1 |
| ATP6V0E1 | ENSG00000113732 | 6865.623585 | 0.458426012 | 0.112665296 | 3.29E-05 | 1.65E-04 | 8992 | 5 | 172410760 | 172462448 | 1 | protein_coding | ATPase, H+ transporting, lysosomal 9kDa, V0 subunit e1 |
| STC2 | ENSG00000113739 | 127.8828323 | -1.531703762 | 0.482303536 | 1.47E-04 | 6.49E-04 | 8614 | 5 | 172741716 | 172756506 | -1 | protein_coding | stanniocalcin 2 |
| DBN1 | ENSG00000113758 | 264.7552809 | -1.337549315 | 0.297193118 | 1.16E-06 | 7.37E-06 | 1627 | 5 | 176883609 | 176901402 | -1 | protein_coding | drebrin 1 |
| SMC4 | ENSG00000113810 | 902.7766664 | 0.570919715 | 0.24798975 | 0.012647499 | 0.034235162 | 10051 | 3 | 160117062 | 160152750 | 1 | protein_coding | structural maintenance of chromosomes 4 |
| SELK | ENSG00000113811 | 148.3335957 | 0.754176747 | 0.282978463 | 0.003136977 | 0.010108036 | 58515 | 3 | 53918437 | 53926015 | -1 | protein_coding | Selenoprotein K |
| ACTR8 | ENSG00000113812 | 1151.975166 | 0.343573132 | 0.14712438 | 0.016345945 | 0.042803224 | 93973 | 3 | 53901093 | 53916229 | -1 | protein_coding | ARP8 actin-related protein 8 homolog (yeast) |
| TIMMDC1 | ENSG00000113845 | 1110.415164 | -0.818769029 | 0.153604183 | 4.82E-08 | 3.75E-07 | 51300 | 3 | 119217379 | 119243937 | 1 | protein_coding | translocase of inner mitochondrial membrane domain containing 1 |
| CRBN | ENSG00000113851 | 836.5003743 | 0.402228203 | 0.164293215 | 0.010950633 | 0.030252806 | 51185 | 3 | 3190676 | 3221394 | -1 | protein_coding | cereblon |

| | | | | | | | | | | | | | |
|---|---|---|---|---|---|---|---|---|---|---|---|---|---|
| **NIT2** | ENSG00000114021 | 571.3443524 | 1.177878972 | 0.177534265 | 5.79E-12 | 6.97E-11 | 56954 | 3 | 100053545 | 100075710 | 1 | protein_coding | nitrilase family, member 2 |
| **FAM162A** | ENSG00000114023 | 743.1704288 | -0.584338753 | 0.182039118 | 7.63E-04 | 0.002863939 | 26355 | 3 | 122103023 | 122131181 | 1 | protein_coding | family with sequence similarity 162, member A |
| **KPNA1** | ENSG00000114030 | 2284.288279 | 0.39315682 | 0.116196299 | 5.25E-04 | 0.002043022 | 3836 | 3 | 122140796 | 122233792 | -1 | protein_coding | karyopherin alpha 1 (importin alpha 5) |
| **PCCB** | ENSG00000114054 | 766.1777558 | -0.47434574 | 0.174114718 | 0.004563441 | 0.014075293 | 5096 | 3 | 135969148 | 136056738 | 1 | protein_coding | propionyl CoA carboxylase, beta polypeptide |
| **ARMC8** | ENSG00000114098 | 1009.534331 | 0.404415295 | 0.172360665 | 0.014636398 | 0.038888746 | 25852 | 3 | 137906109 | 138017231 | 1 | protein_coding | armadillo repeat containing 8 |
| **SLC25A36** | ENSG00000114120 | 1244.894529 | 0.378310271 | 0.123009592 | 0.001695497 | 0.005840045 | 55186 | 3 | 140660672 | 140698775 | 1 | protein_coding | solute carrier family 25 (pyrimidine nucleotide carrier ), member 36 |
| **RNF7** | ENSG00000114125 | 1083.065211 | 0.488018955 | 0.14702714 | 6.23E-04 | 0.002379409 | 9616 | 3 | 141457046 | 141466402 | 1 | protein_coding | ring finger protein 7 |
| **TFDP2** | ENSG00000114126 | 1233.008552 | 0.56866719 | 0.136688718 | 1.88E-05 | 9.85E-05 | 7029 | 3 | 141663277 | 141868386 | -1 | protein_coding | transcription factor Dp-2 (E2F dimerization partner 2) |
| **XRN1** | ENSG00000114127 | 5308.4037 | 2.678741219 | 0.142250745 | 3.32E-80 | 8.06E-78 | 54464 | 3 | 142025449 | 142166904 | -1 | protein_coding | 5'-3' exoribonuclease 1 |
| **KAT2B** | ENSG00000114166 | 1774.4787 | 0.907419933 | 0.122630941 | 5.29E-14 | 7.53E-13 | 8850 | 3 | 20081515 | 20195896 | 1 | protein_coding | K(lysine) acetyltransferase 2B |
| **PDCD10** | ENSG00000114209 | 1502.47055 | 0.867020865 | 0.147840579 | 1.82E-09 | 1.7E-08 | 11235 | 3 | 167401086 | 167452727 | -1 | protein_coding | programmed cell death 10 |
| **WNT5A** | ENSG00000114251 | 416.2395885 | 4.801932236 | 0.336863109 | 9.51E-48 | 7.78E-46 | 7474 | 3 | 55499743 | 55523973 | -1 | protein_coding | wingless-type MMTV integration site family, member 5A |
| **PFKFB4** | ENSG00000114268 | 695.017709 | -0.482823758 | 0.18541415 | 0.006223998 | 0.018488116 | 5210 | 3 | 48555117 | 48599448 | -1 | protein_coding | 6-phosphofructo-2-kinase/fructose-2,6-biphosphatase 4 |
| **ECT2** | ENSG00000114346 | 405.0837076 | 1.596005726 | 0.189809148 | 6.07E-18 | 1.14E-16 | 1894 | 3 | 172468472 | 172539264 | 1 | protein_coding | epithelial cell transforming sequence 2 oncogene |
| **GNAI2** | ENSG00000114353 | 13276.44909 | -0.215147382 | 0.080124328 | 0.007188089 | 0.020980585 | 2771 | 3 | 50263724 | 50296787 | 1 | protein_coding | guanine nucleotide binding protein (G protein), alpha inhibiting activity polypeptide 2 |
| **TFG** | ENSG00000114354 | 1628.793269 | 0.601441467 | 0.115133906 | 9.78E-08 | 7.26E-07 | 10342 | 3 | 100428205 | 100467810 | 1 | protein_coding | TRK-fused gene |
| **RPL24** | ENSG00000114391 | 2481.354068 | -0.381596353 | 0.118447182 | 0.001050828 | 0.003817402 | 6152 | 3 | 101399935 | 101405626 | -1 | protein_coding | ribosomal protein L24 |
| **C3orf14** | ENSG00000114405 | 83.44611334 | 0.802158502 | 0.400450971 | 0.015103367 | 0.039990429 | 57415 | 3 | 62304648 | 62321888 | 1 | protein_coding | chromosome 3 open reading frame 14 |
| **FXR1** | ENSG00000114416 | 1908.904386 | 0.318934731 | 0.105293815 | 0.002004173 | 0.006781001 | 8087 | 3 | 180585929 | 180700541 | 1 | protein_coding | fragile X mental retardation, autosomal homolog 1 |
| **CBLB** | ENSG00000114423 | 364.2377125 | 1.383218981 | 0.260907107 | 2.01E-08 | 1.64E-07 | 868 | 3 | 105374305 | 105588396 | -1 | protein_coding | Cbl proto-oncogene B, E3 ubiquitin protein ligase |
| **BBX** | ENSG00000114439 | 2928.848662 | 0.659535947 | 0.14647743 | 3.1E-06 | 1.85E-05 | 56987 | 3 | 107241783 | 107530171 | 1 | protein_coding | bobby sox homolog (Drosophila) |
| **GNB4** | ENSG00000114450 | 2496.848301 | 0.667653712 | 0.109971134 | 6.89E-10 | 6.72E-09 | 59345 | 3 | 179116990 | 179169378 | -1 | protein_coding | guanine nucleotide binding protein (G protein), beta polypeptide 4 |
| **GBE1** | ENSG00000114480 | 1471.268684 | -0.840635344 | 0.162840752 | 9.17E-08 | 6.83E-07 | 2632 | 3 | 81538850 | 81811312 | -1 | protein_coding | glucan (1,4-alpha-), branching enzyme 1 |
| **NCBP2** | ENSG00000114503 | 1949.03397 | 0.377559879 | 0.113203788 | 6.83E-04 | 0.002587989 | 22916 | 3 | 196662273 | 196669468 | -1 | protein_coding | nuclear cap binding protein subunit 2, 20kDa |
| **FRMD4B** | ENSG00000114541 | 2978.68139 | -0.839858679 | 0.134113623 | 1.57E-10 | 1.64E-09 | 23150 | 3 | 69219141 | 69591734 | -1 | protein_coding | FERM domain containing 4B |
| **PLXNA1** | ENSG00000114554 | 2870.942198 | -0.279763397 | 0.120394227 | 0.017729925 | 0.04587039 | 5361 | 3 | 126707437 | 126756235 | 1 | protein_coding | plexin A1 |
| **ATP6V1A** | ENSG00000114573 | 19146.80592 | -0.318307196 | 0.088923285 | 2.8E-04 | 0.001159778 | 523 | 3 | 113465866 | 113530903 | 1 | protein_coding | ATPase, H+ transporting, lysosomal 70kDa, V1 subunit A |
| **HEMK1** | ENSG00000114735 | 385.1387317 | -0.838324623 | 0.181361487 | 1.57E-06 | 9.76E-06 | 51409 | 3 | 50606583 | 50622366 | 1 | protein_coding | HemK methyltransferase family member 1 |
| **MAPKAPK3** | ENSG00000114738 | 3845.776869 | -0.57592925 | 0.110111694 | 1.08E-07 | 7.96E-07 | 7867 | 3 | 50648951 | 50686720 | 1 | protein_coding | mitogen-activated protein kinase-activated protein kinase 3 |
| **WDR48** | ENSG00000114742 | 1231.570281 | 0.956828258 | 0.162592777 | 1.14E-09 | 1.09E-08 | 57599 | 3 | 39093489 | 39138155 | 1 | protein_coding | WD repeat domain 48 |
| **COMMD2** | ENSG00000114744 | 820.9627119 | 0.681360204 | 0.150411072 | 2.99E-06 | 1.79E-05 | 51122 | 3 | 149456257 | 149470286 | -1 | protein_coding | COMM domain containing 2 |
| **GORASP1** | ENSG00000114745 | 906.5622348 | 0.434004259 | 0.12410935 | 3.5E-04 | 0.001423883 | 64689 | 3 | 39138150 | 39149854 | -1 | protein_coding | golgi reassembly stacking protein 1, 65kDa |
| **ABCC5** | ENSG00000114770 | 894.4455445 | -1.206142441 | 0.200471255 | 3.76E-10 | 3.78E-09 | 10057 | 3 | 183637722 | 183735803 | -1 | protein_coding | ATP-binding cassette, sub-family C (CFTR/MRP), member 5 |
| **ABHD14B** | ENSG00000114779 | 402.8340549 | -0.753205201 | 0.181880716 | 1.68E-05 | 8.9E-05 | 84836 | 3 | 52002526 | 52017425 | -1 | protein_coding | abhydrolase domain containing 14B |
| **EIF1B** | ENSG00000114784 | 1080.732616 | 0.852950927 | 0.145856717 | 1.93E-09 | 1.79E-08 | 10289 | 3 | 40351175 | 40353915 | 1 | protein_coding | eukaryotic translation initiation factor 1B |
| **DNAH1** | ENSG00000114841 | 482.6131616 | -0.808982612 | 0.253271267 | 4.68E-04 | 0.00184316 | 25981 | 3 | 52350335 | 52434507 | 1 | protein_coding | dynein, axonemal, heavy chain 1 |
| **NKTR** | ENSG00000114857 | 1602.231348 | 0.370923943 | 0.114378331 | 9.47E-04 | 0.003474169 | 4820 | 3 | 42642106 | 42690227 | 1 | protein_coding | natural killer-tumor recognition sequence |
| **FOXP1** | ENSG00000114861 | 1055.312608 | -1.101073774 | 0.173993987 | 6.06E-11 | 6.64E-10 | 27086 | 3 | 71003844 | 71633140 | -1 | protein_coding | forkhead box P1 |
| **EIF4G1** | ENSG00000114867 | 17546.77642 | -0.394634231 | 0.086127764 | 3.13E-06 | 1.87E-05 | 1981 | 3 | 184032283 | 184053146 | 1 | protein_coding | eukaryotic translation initiation factor 4 gamma, 1 |
| **INO80D** | ENSG00000114933 | 2031.988139 | 0.418830391 | 0.11030439 | 1.17E-04 | 5.3E-04 | 54891 | 2 | 206858445 | 206951027 | -1 | protein_coding | INO80 complex subunit D |
| **EEF1B2** | ENSG00000114942 | 2847.473042 | -0.656551092 | 0.137574443 | 8.94E-07 | 5.8E-06 | 619569 | 2 | 207024309 | 207027652 | 1 | protein_coding | eukaryotic translation elongation factor 1 beta 2 |
| **MOB1A** | ENSG00000114978 | 11816.25227 | 1.056016205 | 0.09281943 | 2.13E-30 | 8.44E-29 | 55233 | 2 | 74379655 | 74406025 | -1 | protein_coding | MOB kinase activator 1A |
| **KANSL3** | ENSG00000114982 | 2174.040816 | 0.497082656 | 0.125231898 | 4.14E-05 | 2.02E-04 | 55683 | 2 | 97258907 | 97308524 | -1 | protein_coding | KAT8 regulatory NSL complex subunit 3 |
| **LMAN2L** | ENSG00000114988 | 760.8610898 | -0.354644266 | 0.154100048 | 0.017572088 | 0.045548545 | 81562 | 2 | 97371666 | 97405801 | -1 | protein_coding | lectin, mannose-binding 2-like |
| **IL1A** | ENSG00000115008 | 156.1526996 | 0.715443547 | 0.31341971 | 0.009700868 | 0.027303565 | 3552 | 2 | 113531492 | 113542167 | -1 | protein_coding | interleukin 1, alpha |
| **CCL20** | ENSG00000115009 | 18.53853093 | 1.868158264 | 0.9968263 | 0.003626878 | 0.011490378 | 6364 | 2 | 228678558 | 228682272 | 1 | protein_coding | chemokine (C-C motif) ligand 20 |
| **KCNIP3** | ENSG00000115041 | 12.16757369 | -5.260127688 | 2.666614826 | 3.93E-04 | 0.001580794 | 30818 | 2 | 95963052 | 96051825 | 1 | protein_coding | Kv channel interacting protein 3, calsenilin |
| **SLC35F5** | ENSG00000115084 | 2058.645881 | 1.183904925 | 0.126446091 | 1.86E-21 | 4.54E-20 | 80255 | 2 | 114462588 | 114514400 | -1 | protein_coding | solute carrier family 35, member F5 |
| **ZAP70** | ENSG00000115085 | 310.5458301 | -0.948545601 | 0.267829003 | 1.17E-04 | 5.29E-04 | 7535 | 2 | 98330023 | 98356325 | 1 | protein_coding | zeta-chain (TCR) associated protein kinase 70kDa |
| **ACTR3** | ENSG00000115091 | 15121.75786 | 0.203028692 | 0.081529799 | 0.016529835 | 0.043220781 | 10096 | 2 | 114647537 | 114720173 | 1 | protein_coding | ARP3 actin-related protein 3 homolog (yeast) |
| **TFCP2L1** | ENSG00000115112 | 1041.407883 | -0.710485162 | 0.186572061 | 6.74E-05 | 3.18E-04 | 29842 | 2 | 121974163 | 122042783 | -1 | protein_coding | transcription factor CP2-like 1 |
| **STAM2** | ENSG00000115145 | 1454.281639 | 0.622604825 | 0.126220853 | 4.75E-07 | 3.22E-06 | 10254 | 2 | 152973315 | 153032506 | -1 | protein_coding | signal transducing adaptor molecule (SH3 domain and ITAM motif) 2 |
| **OTOF** | ENSG00000115155 | 53.45940133 | 3.459070378 | 0.676799968 | 1.62E-08 | 1.34E-07 | 9381 | 2 | 26680071 | 26781566 | -1 | protein_coding | otoferlin |
| **GPD2** | ENSG00000115159 | 3626.582933 | 1.263505065 | 0.120389222 | 1.43E-26 | 4.64E-25 | 2820 | 2 | 157291802 | 157470247 | 1 | protein_coding | glycerol-3-phosphate dehydrogenase 2 (mitochondrial) |
| **CYTIP** | ENSG00000115165 | 1853.756171 | 0.691185529 | 0.167928157 | 1.81E-05 | 9.52E-05 | 9595 | 2 | 158271131 | 158345473 | -1 | protein_coding | cytohesin 1 interacting protein |
| **SLC30A3** | ENSG00000115194 | 199.3906475 | -1.639139686 | 0.340638419 | 1.88E-07 | 1.35E-06 | 7781 | 2 | 27476552 | 27498685 | -1 | protein_coding | solute carrier family 30 (zinc transporter), member 3 |
| **MPV17** | ENSG00000115204 | 1021.593457 | -0.653988813 | 0.125540553 | 1.03E-07 | 7.62E-07 | 4358 | 2 | 27532360 | 27548547 | -1 | protein_coding | MpV17 mitochondrial inner membrane protein |
| **GTF3C2** | ENSG00000115207 | 1813.934237 | -0.363679298 | 0.116202921 | 0.001409503 | 0.004949779 | 2976 | 2 | 27548716 | 27579868 | -1 | protein_coding | general transcription factor IIIC, polypeptide 2, beta 110kDa |
| **NRBP1** | ENSG00000115216 | 5033.798175 | 0.519239763 | 0.119510167 | 9.09E-06 | 5.03E-05 | 29959 | 2 | 27650657 | 27665126 | 1 | protein_coding | nuclear receptor binding protein 1 |
| **PSMD14** | ENSG00000115233 | 1981.026913 | 0.465701685 | 0.111250861 | 1.88E-05 | 9.85E-05 | 10213 | 2 | 162164549 | 162268228 | 1 | protein_coding | proteasome (prosome, macropain) 26S subunit, non-ATPase, 14 |
| **REEP6** | ENSG00000115255 | 18.91078487 | -2.559198562 | 1.036117431 | 8.72E-04 | 0.003229685 | 92840 | 19 | 1491165 | 1497926 | 1 | protein_coding | receptor accessory protein 6 |
| **IFIH1** | ENSG00000115267 | 116.1221798 | 3.807590493 | 0.43579792 | 9.1E-20 | 1.98E-18 | 64135 | 2 | 163123589 | 163175213 | -1 | protein_coding | interferon induced with helicase C domain 1 |
| **RPS15** | ENSG00000115268 | 2185.977697 | -0.589619946 | 0.149855652 | 4.6E-05 | 2.23E-04 | 6209 | 19 | 1438358 | 1440583 | 1 | protein_coding | ribosomal protein S15 |
| **NDUFS7** | ENSG00000115286 | 328.7970683 | -0.770024348 | 0.209367292 | 9.76E-05 | 4.47E-04 | 374291 | 19 | 1383526 | 1395583 | 1 | protein_coding | NADH dehydrogenase (ubiquinone) Fe-S protein 7, 20kDa (NADH-coenzyme Q reductase) |
| **CLIP4** | ENSG00000115295 | 5197.316935 | -1.325810889 | 0.185066743 | 1.3E-13 | 1.78E-12 | 79745 | 2 | 29320571 | 29412509 | 1 | protein_coding | CAP-GLY domain containing linker protein family, member 4 |
| **SPTBN1** | ENSG00000115306 | 2429.710156 | -1.378131962 | 0.143233068 | 9.97E-23 | 2.64E-21 | 6711 | 2 | 54683422 | 54896812 | 1 | protein_coding | spectrin, beta, non-erythrocytic 1 |
| **RTN4** | ENSG00000115310 | 11738.70143 | 0.532569886 | 0.1345023 | 4.72E-05 | 2.28E-04 | 57142 | 2 | 55199325 | 55339757 | -1 | protein_coding | reticulon 4 |
| **LOXL3** | ENSG00000115318 | 430.3904573 | -0.920805561 | 0.188908118 | 3.64E-07 | 2.51E-06 | 84695 | 2 | 74759541 | 74782817 | -1 | protein_coding | lysyl oxidase-like 3 |
| **DOK1** | ENSG00000115325 | 1177.755613 | -0.304098177 | 0.121885928 | 0.01067247 | 0.029559146 | 1796 | 2 | 74776153 | 74784681 | 1 | protein_coding | docking protein 1, 62kDa (downstream of tyrosine kinase 1) |
| **CCDC88A** | ENSG00000115355 | 16620.77445 | -0.476029668 | 0.114712372 | 2.2E-05 | 1.14E-04 | 55704 | 2 | 55514978 | 55647057 | -1 | protein_coding | coiled-coil domain containing 88A |
| **FANCL** | ENSG00000115392 | 174.889771 | 1.184713296 | 0.256879515 | 8.59E-07 | 5.58E-06 | 55120 | 2 | 58386378 | 58468507 | -1 | protein_coding | Fanconi anemia, complementation group L |
| **FN1** | ENSG00000115414 | 268409.9726 | -1.503164979 | 0.180901717 | 2.96E-17 | 5.28E-16 | 2335 | 2 | 216225163 | 216300895 | -1 | protein_coding | fibronectin 1 |
| **STAT1** | ENSG00000115415 | 41006.45972 | 1.132628843 | 0.396555127 | 8.83E-04 | 0.00326635 | 6772 | 2 | 191829084 | 191885686 | -1 | protein_coding | signal transducer and activator of transcription 1, 91kDa |
| **GLS** | ENSG00000115419 | 3739.118918 | 0.624824054 | 0.150989996 | 1.89E-05 | 9.89E-05 | 2744 | 2 | 191745553 | 191830278 | 1 | protein_coding | glutaminase |
| **PAPOLG** | ENSG00000115421 | 871.5141931 | 0.726974829 | 0.142415863 | 1.69E-07 | 1.22E-06 | 64895 | 2 | 60983365 | 61029220 | 1 | protein_coding | poly(A) polymerase gamma |
| **UNC50** | ENSG00000115446 | 928.2556291 | 0.347634834 | 0.148811138 | 0.016047591 | 0.042125614 | 25972 | 2 | 99225042 | 99234978 | 1 | protein_coding | unc-50 homolog (C. elegans) |
| **IGFBP2** | ENSG00000115457 | 128.4023642 | -1.15041021 | 0.420461721 | 0.001162536 | 0.004172623 | 3485 | 2 | 217497551 | 217529159 | 1 | protein_coding | insulin-like growth factor binding protein 2, 36kDa |
| **GGCX** | ENSG00000115486 | 1707.134382 | -0.648403462 | 0.110014967 | 1.81E-09 | 1.69E-08 | 2677 | 2 | 85771846 | 85788670 | -1 | protein_coding | gamma-glutamyl carboxylase |
| **EHBP1** | ENSG00000115504 | 228.3106789 | 0.793093322 | 0.329287579 | 0.005833107 | 0.017435434 | 23301 | 2 | 62900986 | 63273622 | 1 | protein_coding | EH domain binding protein 1 |
| **TXNDC9** | ENSG00000115514 | 703.8402564 | 0.515174093 | 0.180448449 | 0.002799898 | 0.009100563 | 10190 | 2 | 99935445 | 99957165 | -1 | protein_coding | thioredoxin domain containing 9 |
| **COQ10B** | ENSG00000115520 | 1262.003798 | 1.289643282 | 0.152447549 | 4.77E-18 | 9.03E-17 | 80219 | 2 | 198318147 | 198340032 | 1 | protein_coding | coenzyme Q10 homolog B (S. cerevisiae) |
| **ST3GAL5** | ENSG00000115525 | 2262.720136 | 0.354661488 | 0.109816933 | 0.001028899 | 0.003743076 | 8869 | 2 | 86066267 | 86116137 | -1 | protein_coding | ST3 beta-galactoside alpha-2,3-sialyltransferase 5 |
| **MOB4** | ENSG00000115540 | 535.8460243 | 1.545553053 | 0.242465061 | 2.7E-11 | 3.06E-10 | 1005292 | 2 | 198380295 | 198418423 | 1 | protein_coding | MOB family member 4, phocein |
| **KDM3A** | ENSG00000115548 | 2634.582618 | -0.6658154 | 0.097848729 | 4.58E-12 | 5.56E-11 | 55818 | 2 | 86667770 | 86719839 | 1 | protein_coding | lysine (K)-specific demethylase 3A |
| **ZNF142** | ENSG00000115568 | 346.6107538 | 0.498818382 | 0.178900207 | 0.003628054 | 0.011491719 | 7701 | 2 | 219502639 | 219524378 | -1 | protein_coding | zinc finger protein 142 |
| **IL1R2** | ENSG00000115590 | 576.5219108 | -3.262677778 | 0.281950654 | 2.66E-32 | 1.16E-30 | 7850 | 2 | 102608306 | 102645006 | 1 | protein_coding | interleukin 1 receptor, type II |
| **IL1R1** | ENSG00000115594 | 702.6815029 | -0.733377963 | 0.239714966 | 9.47E-04 | 0.003471698 | 3554 | 2 | 102681004 | 102796334 | 1 | protein_coding | interleukin 1 receptor, type I |
| **IL18RAP** | ENSG00000115607 | 95.9573545 | 1.078050579 | 0.466832836 | 0.004198213 | 0.013062151 | 8807 | 2 | 103035149 | 103069025 | 1 | protein_coding | interleukin 18 receptor accessory protein |
| **MLPH** | ENSG00000115648 | 218.2636293 | -2.09968696 | 0.325604788 | 1.04E-11 | 1.22E-10 | 79083 | 2 | 238394071 | 238463961 | 1 | protein_coding | melanophilin |

| | | | | | | | | | | | | | |
|---|---|---|---|---|---|---|---|---|---|---|---|---|---|
| **UXS1** | ENSG00000115652 | 1456.757309 | 1.104961905 | 0.132062158 | 1.47E-17 | 2.67E-16 | 80146 | 2 | 106709759 | 106810795 | -1 | protein_coding | UDP-glucuronate decarboxylase 1 |
| **STK16** | ENSG00000115661 | 372.353192 | -0.501051812 | 0.220373753 | 0.01493003 | 0.039558885 | 8576 | 2 | 220110177 | 220115059 | 1 | protein_coding | serine/threonine kinase 16 |
| **HDLBP** | ENSG00000115677 | 15662.56509 | -0.887371731 | 0.116007054 | 7.26E-15 | 1.09E-13 | 3069 | 2 | 242166679 | 242256476 | -1 | protein_coding | high density lipoprotein binding protein |
| **STK25** | ENSG00000115694 | 885.3922166 | -0.451965933 | 0.171713861 | 0.006029445 | 0.017966077 | 10494 | 2 | 242432089 | 242449145 | -1 | protein_coding | serine/threonine kinase 25 |
| **PROC** | ENSG00000115718 | 35.92446318 | -3.975750076 | 0.93082264 | 2.73E-06 | 1.65E-05 | 5624 | 2 | 128176003 | 128186822 | 1 | protein_coding | protein C (inactivator of coagulation factors Va and VIIIa) |
| **ID2** | ENSG00000115738 | 1957.993379 | -0.506052755 | 0.191153757 | 0.005192944 | 0.015778083 | 3398 | 2 | 8818975 | 8824583 | 1 | protein_coding | inhibitor of DNA binding 2, dominant negative helix-loop-helix protein |
| **HPCAL1** | ENSG00000115756 | 2068.702614 | -2.057697381 | 0.156913334 | 2.18E-40 | 1.33E-38 | 3241 | 2 | 10443015 | 10567743 | 1 | protein_coding | hippocalcin-like 1 |
| **CEBPZ** | ENSG00000115816 | 1381.964837 | 0.686423086 | 0.137491158 | 3.06E-07 | 2.13E-06 | 10153 | 2 | 37428755 | 37458856 | -1 | protein_coding | CCAAT/enhancer binding protein (C/EBP), zeta |
| **DCAF17** | ENSG00000115827 | 447.8725695 | 0.817508404 | 0.199442503 | 1.56E-05 | 8.32E-05 | 80067 | 2 | 172290727 | 172341562 | 1 | protein_coding | DDB1 and CUL4 associated factor 17 |
| **QPCT** | ENSG00000115828 | 2102.367974 | -0.5010078 | 0.155311224 | 8.65E-04 | 0.003208873 | 25797 | 2 | 37571717 | 37600465 | 1 | protein_coding | glutaminyl-peptide cyclotransferase |
| **RAB3GAP1** | ENSG00000115839 | 2611.40011 | 0.74495988 | 0.100508723 | 6.26E-14 | 8.86E-13 | 22930 | 2 | 135809835 | 135933964 | 1 | protein_coding | RAB3 GTPase activating protein subunit 1 (catalytic) |
| **SLC25A12** | ENSG00000115840 | 408.8744503 | 0.787261178 | 0.178696852 | 4.3E-06 | 2.52E-05 | 8604 | 2 | 172640880 | 172864766 | -1 | protein_coding | solute carrier family 25 (aspartate/glutamate carrier), member 12 |
| **SDC1** | ENSG00000115884 | 47.69374078 | -1.411495474 | 1.341903681 | 0.01525415 | 0.040360029 | 6382 | 2 | 20400558 | 20425194 | -1 | protein_coding | syndecan 1 |
| **SLC1A4** | ENSG00000115902 | 10540.92108 | 0.383726044 | 0.11197956 | 5.23E-04 | 0.002038536 | 6509 | 2 | 65215611 | 65250999 | 1 | protein_coding | solute carrier family 1 (glutamate/neutral amino acid transporter), member 4 |
| **SOS 1** | ENSG00000115904 | 3428.793166 | 1.027588539 | 0.132093957 | 2.17E-15 | 3.36E-14 | 6654 | 2 | 39208537 | 39351486 | -1 | protein_coding | son of sevenless homolog 1 (Drosophila) |
| **KYNU** | ENSG00000115919 | 5378.760303 | 1.870660789 | 0.102623103 | 2.71E-75 | 5.83E-73 | 8942 | 2 | 143635067 | 143799890 | 1 | protein_coding | kynureninase |
| **ORC4** | ENSG00000115947 | 390.9767877 | 1.015851344 | 0.316576907 | 3.45E-04 | 0.001405343 | 5000 | 2 | 148687968 | 148779147 | -1 | protein_coding | origin recognition complex, subunit 4 |
| **PLEK** | ENSG00000115956 | 38795.74753 | -0.59049855 | 0.072329824 | 1.77E-16 | 2.97E-15 | 5341 | 2 | 68592305 | 68624585 | 1 | protein_coding | pleckstrin |
| **RND3** | ENSG00000115963 | 245.74096 | 1.428565582 | 0.308757829 | 5.34E-07 | 3.6E-06 | 390 | 2 | 151324709 | 151395525 | -1 | protein_coding | Rho family GTPase 3 |
| **ATF2** | ENSG00000115966 | 1472.645819 | 0.39460889 | 0.123676488 | 0.001142236 | 0.00411135 | 1386 | 2 | 175936978 | 176033110 | -1 | protein_coding | activating transcription factor 2 |
| **THADA** | ENSG00000115970 | 1822.679295 | -1.32398144 | 0.26578081 | 1.13E-07 | 8.32E-07 | 63892 | 2 | 43393800 | 43823185 | -1 | protein_coding | thyroid adenoma associated |
| **AAK1** | ENSG00000115977 | 1307.324218 | -0.668444471 | 0.158905946 | 1.31E-05 | 7.08E-05 | 22848 | 2 | 69688532 | 69901481 | -1 | protein_coding | AP2 associated kinase 1 |
| **TIA1** | ENSG00000116001 | 1296.712735 | 1.123029461 | 0.12787884 | 4.01E-19 | 8.3E-18 | 7072 | 2 | 70436576 | 70475792 | -1 | protein_coding | TIA1 cytotoxic granule-associated RNA binding protein |
| **PCYOX1** | ENSG00000116005 | 1513.435433 | -0.944519452 | 0.200769394 | 7.66E-07 | 5.03E-06 | 51449 | 2 | 70484518 | 70508323 | 1 | protein_coding | prenylcysteine oxidase 1 |
| **EPAS1** | ENSG00000116016 | 5008.500535 | -2.026322682 | 0.162676337 | 4.43E-37 | 2.41E-35 | 2034 | 2 | 46520806 | 46613836 | 1 | protein_coding | endothelial PAS domain protein 1 |
| **ARID3A** | ENSG00000116017 | 879.4657156 | 0.423290593 | 0.176196463 | 0.012060065 | 0.032889716 | 1820 | 19 | 925781 | 975939 | 1 | protein_coding | AT rich interactive domain 3A (BRIGHT-like) |
| **SUMO1** | ENSG00000116030 | 1725.05843 | 0.586524791 | 0.123586027 | 1.42E-06 | 8.89E-06 | 7341 | 2 | 203070903 | 203103331 | -1 | protein_coding | small ubiquitin-like modifier 1 |
| **NFE2L2** | ENSG00000116044 | 7456.317232 | 0.752216331 | 0.104673808 | 2.34E-13 | 3.12E-12 | 4780 | 2 | 178092323 | 178257425 | -1 | protein_coding | nuclear factor, erythroid 2-like 2 |
| **PLEKHA3** | ENSG00000116095 | 803.3890809 | 0.654853598 | 0.136872624 | 1.01E-06 | 6.47E-06 | 65977 | 2 | 179345195 | 179381189 | 1 | protein_coding | pleckstrin homology domain containing, family A (phosphoinositide binding specific) member 3 |
| **PRRX1** | ENSG00000116132 | 135.6649949 | 3.497331834 | 0.55223119 | 9.59E-12 | 1.12E-10 | 5396 | 1 | 170631869 | 170708560 | 1 | protein_coding | paired related homeobox 1 |
| **DHCR24** | ENSG00000116133 | 10361.82938 | -0.94234667 | 0.160062019 | 1.26E-09 | 1.2E-08 | 1718 | 1 | 55315306 | 55352891 | -1 | protein_coding | 24-dehydrocholesterol reductase |
| **CACYBP** | ENSG00000116161 | 1133.273846 | -0.425356582 | 0.182363907 | 0.014403801 | 0.038317369 | 27101 | 1 | 174968300 | 174980851 | 1 | protein_coding | calcyclin binding protein |
| **SCP2** | ENSG00000116171 | 4647.240916 | -0.432125792 | 0.156520069 | 0.00400737 | 0.012534719 | 6342 | 1 | 53392901 | 53517375 | 1 | protein_coding | sterol carrier protein 2 |
| **PAPPA2** | ENSG00000116183 | 61.8753525 | -1.793235228 | 1.120740993 | 0.005301663 | 0.016060517 | 60676 | 1 | 176432307 | 176814735 | 1 | protein_coding | pappalysin 2 |
| **TCEANC2** | ENSG00000116205 | 452.110818 | 0.894953756 | 0.194539267 | 1.49E-06 | 9.34E-06 | 127428 | 1 | 54519260 | 54578192 | 1 | protein_coding | transcription elongation factor A (SII) N-terminal and central domain containing 2 |
| **TMEM59** | ENSG00000116209 | 5786.413723 | 0.305065176 | 0.117388061 | 0.008250614 | 0.02365366 | 9528 | 1 | 54497347 | 54519177 | -1 | protein_coding | transmembrane protein 59 |
| **MRPL37** | ENSG00000116221 | 1038.746546 | -0.507473693 | 0.134901249 | 1.14E-04 | 5.18E-04 | 51253 | 1 | 54649714 | 54691137 | 1 | protein_coding | mitochondrial ribosomal protein L37 |
| **ICMT** | ENSG00000116237 | 2132.160646 | -0.408381108 | 0.127639944 | 0.001062501 | 0.003857971 | 23463 | 1 | 6281253 | 6296032 | -1 | protein_coding | isoprenylcysteine carboxyl methyltransferase |
| **RPL22** | ENSG00000116251 | 4235.181924 | -0.674525836 | 0.129773172 | 1.05E-07 | 7.74E-07 | 6146 | 1 | 6241329 | 6269449 | -1 | protein_coding | ribosomal protein L22 |
| **QSOX1** | ENSG00000116260 | 4039.209249 | -0.394455324 | 0.172811331 | 0.017218818 | 0.044746674 | 5768 | 1 | 180123969 | 180173165 | 1 | protein_coding | quiescin Q6 sulfhydryl oxidase 1 |
| **STXBP3** | ENSG00000116266 | 1616.631323 | 0.763356408 | 0.150233905 | 1.72E-07 | 1.24E-06 | 6814 | 1 | 109289296 | 109352148 | 1 | protein_coding | syntaxin binding protein 3 |
| **ERRFI1** | ENSG00000116285 | 102.9165226 | -0.753108181 | 0.347587014 | 0.011645867 | 0.031919618 | 54206 | 1 | 8064464 | 8086368 | -1 | protein_coding | ERBB receptor feedback inhibitor 1 |
| **PARK7** | ENSG00000116288 | 2243.370438 | -0.368312557 | 0.108948203 | 5.6E-04 | 0.002161824 | 11315 | 1 | 8014351 | 8045565 | 1 | protein_coding | parkinson protein 7 |
| **KIAA1324** | ENSG00000116299 | 471.0212995 | -2.090893844 | 0.289662338 | 3.9E-14 | 5.58E-13 | 57535 | 1 | 109656301 | 109749401 | 1 | protein_coding | KIAA1324 |
| **AMPD2** | ENSG00000116337 | 1288.943966 | -1.319217877 | 0.215305051 | 1.63E-10 | 1.7E-09 | 271 | 1 | 110158726 | 110174673 | 1 | protein_coding | adenosine monophosphate deaminase 2 |
| **SRSF4** | ENSG00000116350 | 1872.265686 | 0.702118133 | 0.111504596 | 1.62E-10 | 1.69E-09 | 6429 | 1 | 29474255 | 29508499 | -1 | protein_coding | serine/arginine-rich splicing factor 4 |
| **EDEM3** | ENSG00000116406 | 2248.887723 | 0.475311385 | 0.124515398 | 9.53E-05 | 4.38E-04 | 80267 | 1 | 184659365 | 184724047 | -1 | protein_coding | ER degradation enhancer, mannosidase alpha-like 3 |
| **WDR77** | ENSG00000116455 | 634.4717101 | -0.682499018 | 0.160546187 | 1.11E-05 | 6.09E-05 | 79084 | 1 | 111982512 | 111991998 | -1 | protein_coding | WD repeat domain 77 |
| **ATP5F1** | ENSG00000116459 | 3848.868841 | -0.734342255 | 0.112984428 | 3.65E-11 | 4.11E-10 | 515 | 1 | 111991486 | 112005395 | 1 | protein_coding | ATP synthase, H+ transporting, mitochondrial Fo complex, subunit B1 |
| **RNF19B** | ENSG00000116514 | 1269.092556 | 2.956858443 | 0.188757351 | 1.73E-56 | 2.01E-54 | 127544 | 1 | 33402046 | 33430286 | -1 | protein_coding | ring finger protein 19B |
| **RHOU** | ENSG00000116574 | 478.6539363 | 0.984692263 | 0.162478569 | 3.86E-10 | 3.87E-09 | 58480 | 1 | 228870824 | 228882416 | 1 | protein_coding | ras homolog family member U |
| **ARHGEF2** | ENSG00000116584 | 6166.881048 | 0.23541685 | 0.077069477 | 0.002090467 | 0.007030943 | 9181 | 1 | 155916630 | 155976861 | -1 | protein_coding | Rho/Rac guanine nucleotide exchange factor (GEF) 2 |
| **DOCK7** | ENSG00000116641 | 2202.177844 | -0.242520711 | 0.105283377 | 0.019416869 | 0.049466245 | 85440 | 1 | 62920399 | 63153969 | -1 | protein_coding | dedicator of cytokinesis 7 |
| **SRM** | ENSG00000116649 | 503.1639011 | -0.902431162 | 0.172736102 | 6.11E-08 | 4.68E-07 | 6723 | 1 | 11114641 | 11120081 | -1 | protein_coding | spermidine synthase |
| **FBXO6** | ENSG00000116663 | 526.1915215 | 2.371040206 | 0.237105271 | 1.3E-24 | 3.83E-23 | 26270 | 1 | 11724181 | 11734411 | 1 | protein_coding | F-box protein 6 |
| **SWT1** | ENSG00000116668 | 430.2586157 | 1.436890061 | 0.179078219 | 1.73E-16 | 2.92E-15 | 54823 | 1 | 185126212 | 185260897 | 1 | protein_coding | SWT1 RNA endoribonuclease homolog (S. cerevisiae) |
| **LEPR** | ENSG00000116678 | 390.7106254 | 0.403631882 | 0.179091094 | 0.018610217 | 0.047721468 | 54741 | 1 | 65886248 | 66107242 | 1 | protein_coding | leptin receptor |
| **MFN2** | ENSG00000116688 | 4099.111135 | -0.354834146 | 0.108848591 | 8.25E-04 | 0.003075588 | 9927 | 1 | 12040238 | 12073571 | 1 | protein_coding | mitofusin 2 |
| **SMG7** | ENSG00000116698 | 2510.14046 | 0.328215427 | 0.115197663 | 0.003737629 | 0.011785059 | 9887 | 1 | 183441351 | 183567381 | 1 | protein_coding | SMG7 nonsense mediated mRNA decay factor |
| **TROVE2** | ENSG00000116747 | 1609.81174 | 0.627329545 | 0.141435681 | 5.05E-06 | 2.92E-05 | 6738 | 1 | 193028552 | 193060907 | 1 | protein_coding | TROVE domain family, member 2 |
| **BCAS2** | ENSG00000116752 | 808.9888676 | 0.420845643 | 0.166505317 | 0.008618941 | 0.024561138 | 10286 | 1 | 115110178 | 115124260 | -1 | protein_coding | breast carcinoma amplified sequence 2 |
| **SRSF11** | ENSG00000116754 | 1720.000641 | 0.978434859 | 0.174496892 | 6.62E-09 | 5.76E-08 | 9295 | 1 | 70671365 | 70718735 | 1 | protein_coding | serine/arginine-rich splicing factor 11 |
| **OLFML3** | ENSG00000116774 | 66.02856413 | -0.886588851 | 0.484228629 | 0.017648641 | 0.045697867 | 56944 | 1 | 114522063 | 114524876 | 1 | protein_coding | olfactomedin-like 3 |
| **PHTF1** | ENSG00000116793 | 777.8093789 | 0.658110728 | 0.194836761 | 3.61E-04 | 0.001461448 | 10745 | 1 | 114239453 | 114302111 | -1 | protein_coding | putative homeodomain transcription factor 1 |
| **CD58** | ENSG00000116815 | 1090.462725 | 1.421571706 | 0.167633505 | 4.02E-18 | 7.66E-17 | 965 | 1 | 117057157 | 117113661 | -1 | protein_coding | CD58 molecule |
| **CD2** | ENSG00000116824 | 1780.987057 | 1.352040314 | 0.239868645 | 2.94E-09 | 2.66E-08 | 914 | 1 | 117297007 | 117311850 | 1 | protein_coding | CD2 molecule |
| **KIF21B** | ENSG00000116852 | 502.6527227 | -0.587234703 | 0.216628821 | 0.003851915 | 0.012095504 | 1019292 | 1 | 200938520 | 200992828 | -1 | protein_coding | kinesin family member 21B |
| **TMEM9** | ENSG00000116857 | 543.1433041 | -0.426864321 | 0.160136627 | 0.005789103 | 0.017327626 | 252839 | 1 | 201103900 | 201140702 | -1 | protein_coding | transmembrane protein 9 |
| **ADPRHL2** | ENSG00000116863 | 635.2427303 | 0.779623816 | 0.164471391 | 9.11E-07 | 5.9E-06 | 54936 | 1 | 36554476 | 36559533 | 1 | protein_coding | ADP-ribosylhydrolase like 2 |
| **RRAGC** | ENSG00000116954 | 2216.775782 | 1.276497881 | 0.120990098 | 1.04E-26 | 3.39E-25 | 64121 | 1 | 39303870 | 39325495 | -1 | protein_coding | Ras-related GTP binding C |
| **LGALS8** | ENSG00000116977 | 4850.034476 | 0.913379071 | 0.093688303 | 8.93E-23 | 2.37E-21 | 3964 | 1 | 236681300 | 236716281 | 1 | protein_coding | lectin, galactoside-binding, soluble, 8 |
| **MYCL** | ENSG00000116990 | 325.5868667 | -0.963383621 | 0.23061592 | 8E-06 | 4.46E-05 | 4610 | 1 | 40361098 | 40367928 | -1 | protein_coding | v-myc avian myelocytomatosis viral oncogene lung carcinoma derived homolog |
| **SIPA1L2** | ENSG00000116991 | 735.5732691 | -1.422287333 | 0.212024428 | 3.2E-12 | 3.93E-11 | 57568 | 1 | 232533711 | 232697304 | -1 | protein_coding | signal-induced proliferation-associated 1 like 2 |
| **RLF** | ENSG00000117000 | 1207.185662 | 0.391982793 | 0.137923066 | 0.003513322 | 0.011165335 | 6018 | 1 | 40627045 | 40706593 | 1 | protein_coding | rearranged L-myc fusion |
| **ZNF684** | ENSG00000117010 | 110.8664194 | 1.074029916 | 0.332596348 | 2.93E-04 | 0.001209962 | 127396 | 1 | 40997233 | 41013841 | 1 | protein_coding | zinc finger protein 684 |
| **RIMS3** | ENSG00000117016 | 234.1441545 | -0.821637723 | 0.274231617 | 9.79E-04 | 0.003581265 | 9783 | 1 | 41086351 | 41131329 | -1 | protein_coding | regulating synaptic membrane exocytosis 3 |
| **AKT3** | ENSG00000117020 | 1223.906197 | 0.713656696 | 0.167165699 | 9.28E-06 | 5.13E-05 | 10000 | 1 | 243651535 | 244014381 | -1 | protein_coding | v-akt murine thymoma viral oncogene homolog 3 |
| **ETV3** | ENSG00000117036 | 1661.61532 | 0.510001438 | 0.11655257 | 7.7E-06 | 4.31E-05 | 2117 | 1 | 157090983 | 157108266 | -1 | protein_coding | ets variant 3 |
| **ACADM** | ENSG00000117054 | 1785.135354 | -0.643875954 | 0.19148744 | 3.92E-04 | 0.001577289 | 34 | 1 | 76190036 | 76253260 | 1 | protein_coding | acyl-CoA dehydrogenase, C-4 to C-12 straight chain |
| **SLAMF1** | ENSG00000117090 | 359.4793521 | 2.494417634 | 0.300397539 | 7.83E-18 | 1.46E-16 | 6504 | 1 | 160577890 | 160617085 | -1 | protein_coding | signaling lymphocytic activation molecule family member 1 |
| **CD48** | ENSG00000117091 | 4953.301546 | 1.484788202 | 0.0910257 | 1.36E-60 | 1.78E-58 | 962 | 1 | 160648536 | 160681641 | -1 | protein_coding | CD48 molecule |
| **PADI2** | ENSG00000117115 | 2895.989976 | -0.541698469 | 0.233611773 | 0.011448043 | 0.031445082 | 11240 | 1 | 17393256 | 17445948 | -1 | protein_coding | peptidyl arginine deiminase, type II |
| **CTBS** | ENSG00000117151 | 1280.237025 | 0.535322676 | 0.141913742 | 9.52E-05 | 4.37E-04 | 1486 | 1 | 85015289 | 85040163 | -1 | protein_coding | chitobiase, di-N-acetyl- |
| **RGS4** | ENSG00000117152 | 44.26143718 | -1.442519867 | 0.831379484 | 0.006912129 | 0.020279912 | 5999 | 1 | 163038565 | 163046592 | 1 | protein_coding | regulator of G-protein signaling 4 |
| **ZNHIT6** | ENSG00000117174 | 451.7990922 | 0.452634688 | 0.173844639 | 0.006651385 | 0.019593893 | 54680 | 1 | 86115106 | 86174116 | -1 | protein_coding | zinc finger, HIT-type containing 6 |
| **GBP3** | ENSG00000117226 | 5522.475285 | 2.892219176 | 0.156304204 | 1.23E-77 | 2.81E-75 | 2635 | 1 | 89472349 | 89488577 | -1 | protein_coding | guanylate binding protein 3 |
| **GBP1** | ENSG00000117228 | 24806.5354 | 4.465728119 | 0.508665024 | 8.97E-20 | 1.96E-18 | 2633 | 1 | 89518002 | 89531043 | -1 | protein_coding | guanylate binding protein 1, interferon-inducible |
| **RAB7L1** | ENSG00000117280 | 1972.359954 | 0.986391286 | 0.121600153 | 1.62E-16 | 2.73E-15 | 8934 | 1 | 205737114 | 205744588 | -1 | protein_coding | RAB7, member RAS oncogene family-like 1 |
| **ECE1** | ENSG00000117298 | 1968.121784 | 0.657521654 | 0.176829021 | 1.11E-04 | 5.04E-04 | 1889 | 1 | 21543740 | 21671997 | -1 | protein_coding | endothelin converting enzyme 1 |

| | | | | | | | | | | | | | |
|---|---|---|---|---|---|---|---|---|---|---|---|---|---|
| HMGCL | ENSG00000117305 | 1149.491122 | -0.332143293 | 0.123851265 | 0.00616949 | 0.018347601 | 3155 | 1 | 24128375 | 24165110 | -1 | protein_coding | 3-hydroxymethyl-3-methylglutaryl-CoA lyase |
| GALE | ENSG00000117308 | 571.8864786 | -0.702566385 | 0.181132116 | 5.22E-05 | 2.51E-04 | 2582 | 1 | 24122089 | 24127271 | -1 | protein_coding | UDP-galactose-4-epimerase |
| CD46 | ENSG00000117335 | 8034.258243 | 0.404642131 | 0.077267739 | 1.47E-07 | 1.07E-06 | 4179 | 1 | 207925402 | 207968858 | 1 | protein_coding | CD46 molecule, complement regulatory protein |
| PRPF3 | ENSG00000117360 | 1234.572699 | 1.078295122 | 0.12141314 | 1.82E-19 | 3.85E-18 | 9129 | 1 | 150293925 | 150325671 | 1 | protein_coding | pre-mRNA processing factor 3 |
| LEPRE1 | ENSG00000117385 | 525.1260489 | -0.542778039 | 0.175005279 | 0.001203355 | 0.004303973 | 64175 | 1 | 43212006 | 43232755 | -1 | protein_coding | leucine proline-enriched proteoglycan (leprecan) 1 |
| SLC2A1 | ENSG00000117394 | 916.9479497 | -1.868360274 | 0.183346078 | 2.39E-25 | 7.25E-24 | 6513 | 1 | 43391052 | 43424530 | -1 | protein_coding | solute carrier family 2 (facilitated glucose transporter), member 1 |
| EBNA1BP2 | ENSG00000117395 | 545.2247317 | -0.524538988 | 0.159143661 | 6.39E-04 | 0.002435396 | 1024654 | 1 | 43629846 | 43736607 | -1 | protein_coding | EBNA1 binding protein 2 |
| IPO13 | ENSG00000117408 | 1108.216522 | -0.536635308 | 0.166648589 | 8.21E-04 | 0.003061715 | 9670 | 1 | 44412611 | 44433694 | 1 | protein_coding | importin 13 |
| ATP6V0B | ENSG00000117410 | 3500.605864 | -0.389878245 | 0.10024341 | 8.5E-05 | 3.94E-04 | 533 | 1 | 44440159 | 44443967 | 1 | protein_coding | ATPase, H+ transporting, lysosomal 21kDa, V0 subunit b |
| ERI3 | ENSG00000117419 | 178.7424534 | -0.652596804 | 0.28317972 | 0.010316426 | 0.028737881 | 79033 | 1 | 44686742 | 44820932 | -1 | protein_coding | ERI1 exoribonuclease family member 3 |
| AKR1A1 | ENSG00000117448 | 4198.190577 | -0.407377384 | 0.08964449 | 3.27E-06 | 1.95E-05 | 10327 | 1 | 46016215 | 46035721 | 1 | protein_coding | aldo-keto reductase family 1, member A1 (aldehyde reductase) |
| PRDX1 | ENSG00000117450 | 14676.71825 | 0.195667902 | 0.080060256 | 0.011335093 | 0.031179666 | 5052 | 1 | 45976708 | 45988719 | -1 | protein_coding | peroxiredoxin 1 |
| BLZF1 | ENSG00000117475 | 1265.293839 | 2.260621666 | 0.135898537 | 3.91E-63 | 5.92E-61 | 8548 | 1 | 169337208 | 169365778 | 1 | protein_coding | basic leucine zipper nuclear factor 1 |
| TMED5 | ENSG00000117500 | 5108.663661 | 0.694498837 | 0.154426725 | 3.29E-06 | 1.96E-05 | 50999 | 1 | 93615299 | 93646285 | -1 | protein_coding | transmembrane emp24 protein transport domain containing 5 |
| DR1 | ENSG00000117505 | 2043.996544 | 0.302607305 | 0.110379918 | 0.005352968 | 0.016190266 | 1810 | 1 | 93811445 | 93835055 | 1 | protein_coding | down-regulator of transcription 1, TBP-binding (negative cofactor 2) |
| PRRC2C | ENSG00000117523 | 9781.729274 | 0.502924699 | 0.08336612 | 1.87E-09 | 1.74E-08 | 23215 | 1 | 171454651 | 171562650 | 1 | protein_coding | proline-rich coiled-coil 2C |
| F3 | ENSG00000117525 | 3391.171743 | -1.192422768 | 0.221598189 | 1.5E-08 | 1.25E-07 | 2152 | 1 | 94994781 | 95007356 | -1 | protein_coding | coagulation factor III (thromboplastin, tissue factor) |
| ABCD3 | ENSG00000117528 | 502.7293149 | 0.924676901 | 0.2509667 | 7.27E-05 | 3.41E-04 | 5825 | 1 | 94883933 | 94984222 | 1 | protein_coding | ATP-binding cassette, sub-family D (ALD), member 3 |
| DPH5 | ENSG00000117543 | 229.5633821 | -0.476682987 | 0.216898619 | 0.01877283 | 0.0480981 | 51611 | 1 | 101455179 | 101491644 | -1 | protein_coding | diphthamide biosynthesis 5 |
| FASLG | ENSG00000117560 | 88.40340316 | 0.958453801 | 0.427995362 | 0.006242201 | 0.018538585 | 356 | 1 | 172628154 | 172636014 | 1 | protein_coding | Fas ligand (TNF superfamily, member 6) |
| TNFSF4 | ENSG00000117586 | 131.9744741 | 0.683167154 | 0.30746848 | 0.01175568 | 0.032174438 | 7292 | 1 | 173152873 | 173176452 | -1 | protein_coding | tumor necrosis factor (ligand) superfamily, member 4 |
| HSD11B1 | ENSG00000117594 | 294.0683628 | 1.276140881 | 0.403180558 | 2.49E-04 | 0.001040933 | 3290 | 1 | 209859510 | 209908295 | 1 | protein_coding | hydroxysteroid (11-beta) dehydrogenase 1 |
| IRF6 | ENSG00000117595 | 112.139832 | 0.9251028 | 0.451245353 | 0.010124096 | 0.028312271 | 3664 | 1 | 209959036 | 209979465 | -1 | protein_coding | interferon regulatory factor 6 |
| DIEXF | ENSG00000117597 | 1013.87574 | 0.306266206 | 0.116752727 | 0.007177856 | 0.020958957 | 27042 | 1 | 210001352 | 210030910 | 1 | protein_coding | digestive organ expansion factor homolog (zebrafish) |
| SYF2 | ENSG00000117614 | 908.7285946 | 0.463751549 | 0.180852244 | 0.007189382 | 0.020980585 | 25949 | 1 | 25549170 | 25558993 | -1 | protein_coding | SYF2 pre-mRNA-splicing factor |
| C1orf63 | ENSG00000117616 | 1423.568891 | 0.758816473 | 0.132672694 | 4.87E-09 | 4.31E-08 | 1019281 | 1 | 25568728 | 25664704 | -1 | protein_coding | chromosome 1 open reading frame 63 |
| SLC35A3 | ENSG00000117620 | 390.9588777 | 0.704135339 | 0.216982769 | 5.52E-04 | 0.002137444 | 23443 | 1 | 100435345 | 100492535 | 1 | protein_coding | solute carrier family 35 (UDP-N-acetylglucosamine (UDP-GlcNAc) transporter), member A3 |
| MTFR1L | ENSG00000117640 | 930.6766578 | -0.480676062 | 0.123597526 | 7.18E-05 | 3.37E-04 | 56181 | 1 | 26145131 | 26159432 | 1 | protein_coding | mitochondrial fission regulator 1-like |
| MAN1C1 | ENSG00000117643 | 359.4186931 | -0.90500004 | 0.210764079 | 6.04E-06 | 3.44E-05 | 57134 | 1 | 25943959 | 26112698 | 1 | protein_coding | mannosidase, alpha, class 1C, member 1 |
| RPS6KA1 | ENSG00000117676 | 6183.708157 | -1.625261467 | 0.097464384 | 1.79E-63 | 2.74E-61 | 6195 | 1 | 26856252 | 26901521 | 1 | protein_coding | ribosomal protein S6 kinase, 90kDa, polypeptide 1 |
| ARID1A | ENSG00000117713 | 2560.527312 | -0.435167715 | 0.135782273 | 0.001094459 | 0.003962714 | 8289 | 1 | 27022524 | 27108595 | 1 | protein_coding | AT rich interactive domain 1A (SWI-like) |
| STX12 | ENSG00000117758 | 3107.215955 | 0.571654907 | 0.092139977 | 4.21E-10 | 4.2E-09 | 23673 | 1 | 28099694 | 28150963 | 1 | protein_coding | syntaxin 12 |
| 2-Mar | ENSG00000117791 | 67.83281327 | -2.131922549 | 0.699348741 | 1.68E-04 | 7.32E-04 | 54996 | 1 | 220921567 | 220958150 | 1 | protein_coding | mitochondrial amidoxime reducing component 2 |
| OSBPL9 | ENSG00000117859 | 3022.886533 | 1.253931151 | 0.151197158 | 2.25E-17 | 4.06E-16 | 114883 | 1 | 52042851 | 52254889 | 1 | protein_coding | oxysterol binding protein-like 9 |
| ESYT2 | ENSG00000117868 | 2198.982729 | -0.494587855 | 0.113691509 | 9.07E-06 | 5.02E-05 | 57488 | 7 | 158523686 | 158622944 | -1 | protein_coding | extended synaptotagmin-like protein 2 |
| CTSD | ENSG00000117984 | 109625.0405 | -0.634282169 | 0.172258794 | 1.21E-04 | 5.47E-04 | 1509 | 11 | 1773982 | 1785222 | -1 | protein_coding | cathepsin D |
| STAG1 | ENSG00000118007 | 1119.73107 | 0.480307709 | 0.119826874 | 4.3E-05 | 2.1E-04 | 10274 | 3 | 136055077 | 136471220 | -1 | protein_coding | stromal antigen 1 |
| KMT2A | ENSG00000118058 | 2003.607159 | 0.539020253 | 0.158527014 | 4.17E-04 | 0.001666535 | 4297 | 11 | 118307205 | 118397539 | 1 | protein_coding | lysine (K)-specific methyltransferase 2A |
| MMP8 | ENSG00000118113 | 948.1603186 | -1.259727701 | 0.520555595 | 0.002170422 | 0.007261505 | 4317 | 11 | 102582526 | 102597781 | -1 | protein_coding | matrix metallopeptidase 8 (neutrophil collagenase) |
| DDX59 | ENSG00000118197 | 465.3108792 | 0.996069982 | 0.173377798 | 2.71E-09 | 2.46E-08 | 83479 | 1 | 200593024 | 200639097 | -1 | protein_coding | DEAD (Asp-Glu-Ala-Asp) box polypeptide 59 |
| CAMSAP2 | ENSG00000118200 | 1429.538762 | 1.015114083 | 0.157479844 | 3.42E-11 | 3.85E-10 | 23271 | 1 | 200708686 | 200829832 | 1 | protein_coding | calmodulin regulated spectrin-associated protein family, member 2 |
| MREG | ENSG00000118242 | 6093.36473 | -0.412047208 | 0.162090716 | 0.008276861 | 0.023714898 | 55686 | 2 | 216809213 | 216898819 | -1 | protein_coding | melanoregulin |
| FASTKD2 | ENSG00000118246 | 818.8803795 | 0.421948692 | 0.147682024 | 0.003239586 | 0.010394832 | 22868 | 2 | 207630081 | 207657233 | 1 | protein_coding | FAST kinase domains 2 |
| CREB1 | ENSG00000118260 | 1663.081177 | 0.46678275 | 0.125263312 | 1.45E-04 | 6.44E-04 | 1385 | 2 | 208394461 | 208468155 | 1 | protein_coding | cAMP responsive element binding protein 1 |
| KLF7 | ENSG00000118263 | 955.7492254 | -1.289793448 | 0.162455859 | 3.82E-16 | 6.25E-15 | 8609 | 2 | 207938861 | 208031991 | -1 | protein_coding | Kruppel-like factor 7 (ubiquitous) |
| LRMP | ENSG00000118308 | 1073.374003 | -1.303583856 | 0.125367287 | 5.51E-26 | 1.73E-24 | 4033 | 12 | 25173936 | 25261268 | 1 | protein_coding | lymphoid-restricted membrane protein |
| HMGN3 | ENSG00000118418 | 1085.677471 | 0.696551284 | 0.182244585 | 6.48E-05 | 3.07E-04 | 9324 | 6 | 79910962 | 79944406 | -1 | protein_coding | high mobility group nucleosomal binding domain 3 |
| PHF3 | ENSG00000118482 | 3150.855623 | 0.729710771 | 0.09576846 | 8.19E-15 | 1.23E-13 | 23469 | 6 | 64345725 | 64489229 | 1 | protein_coding | PHD finger protein 3 |
| PLAGL1 | ENSG00000118495 | 669.1872706 | 0.846696107 | 0.169842839 | 2.35E-07 | 1.66E-06 | 5325 | 6 | 144261437 | 144385735 | -1 | protein_coding | pleiomorphic adenoma gene-like 1 |
| FBXO30 | ENSG00000118496 | 1155.88917 | 0.385078372 | 0.143494777 | 0.005316125 | 0.016094759 | 84085 | 6 | 146114638 | 146135889 | -1 | protein_coding | F-box protein 30 |
| TNFAIP3 | ENSG00000118503 | 5954.979168 | 2.568457854 | 0.16242207 | 1.02E-57 | 1.26E-55 | 7128 | 6 | 138188351 | 138204449 | 1 | protein_coding | tumor necrosis factor, alpha-induced protein 3 |
| AKAP7 | ENSG00000118507 | 316.2153054 | 1.662975453 | 0.212921409 | 7.67E-16 | 1.24E-14 | 9465 | 6 | 131456806 | 131604675 | 1 | protein_coding | A kinase (PRKA) anchor protein 7 |
| RAB32 | ENSG00000118508 | 830.7774534 | -0.770225772 | 0.175202471 | 4.72E-06 | 2.74E-05 | 10981 | 6 | 146864829 | 146876101 | 1 | protein_coding | RAB32, member RAS oncogene family |
| SGK1 | ENSG00000118515 | 15799.12952 | -0.96885702 | 0.092000913 | 1.98E-26 | 6.41E-25 | 6446 | 6 | 134490384 | 134639250 | -1 | protein_coding | serum/glucocorticoid regulated kinase 1 |
| PMFBP1 | ENSG00000118557 | 381.4304 | -1.900181457 | 0.304936142 | 4.72E-11 | 5.24E-10 | 83449 | 16 | 72146056 | 72210777 | -1 | protein_coding | polyamine modulated factor 1 binding protein 1 |
| FBXL5 | ENSG00000118564 | 3172.812536 | 0.558292029 | 0.123382505 | 4.48E-06 | 2.62E-05 | 26234 | 4 | 15606162 | 15683302 | -1 | protein_coding | F-box and leucine-rich repeat protein 5 |
| MED28 | ENSG00000118579 | 1418.396915 | 0.34390647 | 0.14848044 | 0.017825992 | 0.046034915 | 80306 | 4 | 17616254 | 17635728 | 1 | protein_coding | mediator complex subunit 28 |
| ZNF430 | ENSG00000118620 | 712.5195353 | 0.566453161 | 0.183514099 | 0.001212372 | 0.004330143 | 80264 | 19 | 21203426 | 21242856 | 1 | protein_coding | zinc finger protein 430 |
| MYL12B | ENSG00000118680 | 4538.954797 | 0.691788828 | 0.125331739 | 1.6E-08 | 1.32E-07 | 103910 | 18 | 3261907 | 3278282 | 1 | protein_coding | myosin, light chain 12B, regulatory |
| FOXO3 | ENSG00000118689 | 1985.252326 | 0.501299882 | 0.111927382 | 5.02E-06 | 2.9E-05 | 2309 | 6 | 108881038 | 109005977 | 1 | protein_coding | forkhead box O3 |
| ARMC2 | ENSG00000118690 | 71.4045967 | -0.897148448 | 0.497748123 | 0.018238585 | 0.046941782 | 84071 | 6 | 109169619 | 109295186 | 1 | protein_coding | armadillo repeat containing 2 |
| RPN2 | ENSG00000118705 | 6252.520616 | -0.282763544 | 0.082653181 | 3.48E-04 | 0.001416681 | 6185 | 20 | 35806813 | 35870022 | 1 | protein_coding | ribophorin II |
| TGIF2 | ENSG00000118707 | 911.6155354 | 0.71764335 | 0.182577856 | 3.88E-05 | 1.91E-04 | 60436 | 20 | 35201891 | 35222353 | 1 | protein_coding | TGFB-induced factor homeobox 2 |
| SPP1 | ENSG00000118785 | 126274.6747 | -0.952520764 | 0.110443525 | 2.08E-18 | 4.07E-17 | 6696 | 4 | 88896819 | 88904562 | 1 | protein_coding | secreted phosphoprotein 1 |
| FAM47E-STB | ENSG00000118804 | 241.8037865 | 2.05412007 | 0.297507216 | 4.79E-13 | 6.29E-12 | 8987 | 4 | 77172886 | 77232752 | 1 | protein_coding | FAM47E-STBD1 readthrough |
| CCNI | ENSG00000118816 | 7103.238252 | -0.440017496 | 0.15235994 | 0.002848545 | 0.009250811 | 10983 | 4 | 77968311 | 77997158 | -1 | protein_coding | cyclin I |
| RARRES1 | ENSG00000118849 | 550.9160007 | 1.133851927 | 0.27308146 | 7.28E-06 | 4.09E-05 | 5918 | 3 | 158414681 | 158450485 | -1 | protein_coding | retinoic acid receptor responder (tazarotene induced) 1 |
| KLF12 | ENSG00000118922 | 306.8503105 | -0.583297121 | 0.26937482 | 0.016206397 | 0.042498689 | 11278 | 13 | 74260226 | 74708394 | -1 | protein_coding | Kruppel-like factor 12 |
| HS1BP3 | ENSG00000118960 | 962.8691269 | -0.475461131 | 0.122677907 | 7.49E-05 | 3.5E-04 | 64342 | 2 | 20760208 | 20850849 | -1 | protein_coding | HCLS1 binding protein 3 |
| WDR35 | ENSG00000118965 | 134.7912204 | -0.874692275 | 0.33245074 | 0.002761856 | 0.008988384 | 57539 | 2 | 20110021 | 20189892 | -1 | protein_coding | WD repeat domain 35 |
| CCND2 | ENSG00000118971 | 8250.780044 | -0.322301496 | 0.09699762 | 8.44E-04 | 0.003140477 | 894 | 12 | 4382938 | 4414516 | 1 | protein_coding | cyclin D2 |
| HIN1L | ENSG00000118976 | 6.172570169 | 2.792993817 | 2.444915572 | 0.005533476 | 0.016650543 | NA | 12 | 3948625 | 3951691 | 1 | pseudogene | |
| CYP20A1 | ENSG00000119004 | 721.8906311 | -0.582242397 | 0.159431122 | 1.49E-04 | 6.57E-04 | 57404 | 2 | 204103663 | 204163009 | 1 | protein_coding | cytochrome P450, family 20, subfamily A, polypeptide 1 |
| NDUFB3 | ENSG00000119013 | 1319.760628 | 0.698361312 | 0.14475576 | 6.91E-07 | 4.57E-06 | 4709 | 2 | 201936156 | 201950473 | 1 | protein_coding | NADH dehydrogenase (ubiquinone) 1 beta subcomplex, 3, 12kDa |
| GTF3C3 | ENSG00000119041 | 1144.41456 | 0.411199771 | 0.169145639 | 0.011791992 | 0.032244948 | 9330 | 2 | 197627756 | 197664449 | -1 | protein_coding | general transcription factor IIIC, polypeptide 3, 102kDa |
| UBE2B | ENSG00000119048 | 1022.768965 | 0.874339436 | 0.188912073 | 1.39E-06 | 8.71E-06 | 7320 | 5 | 133706870 | 133727683 | 1 | protein_coding | ubiquitin-conjugating enzyme E2B |
| KLF9 | ENSG00000119138 | 1641.720294 | 1.007080288 | 0.133757932 | 1.46E-14 | 2.15E-13 | 687 | 9 | 72999503 | 73029540 | -1 | protein_coding | Kruppel-like factor 9 |
| TJP2 | ENSG00000119139 | 396.5966612 | 0.624574452 | 0.231464014 | 0.003735803 | 0.011784164 | 9414 | 9 | 71736209 | 71870124 | 1 | protein_coding | tight junction protein 2 |
| ITGB1BP1 | ENSG00000119185 | 2873.628981 | -0.950224751 | 0.206900014 | 1.12E-06 | 7.17E-06 | 9270 | 2 | 9543604 | 9563676 | -1 | protein_coding | integrin beta 1 binding protein 1 |
| SENP5 | ENSG00000119231 | 1922.607751 | 0.462813072 | 0.1069193 | 1.09E-05 | 5.98E-05 | 205564 | 3 | 196594727 | 196661585 | 1 | protein_coding | SUMO1/sentrin specific peptidase 5 |
| CCDC92 | ENSG00000119242 | 818.3737401 | -0.341985023 | 0.142401504 | 0.013656503 | 0.036654784 | 80212 | 12 | 124403207 | 124457378 | -1 | protein_coding | coiled-coil domain containing 92 |
| PTBP3 | ENSG00000119314 | 5018.509103 | 0.432768912 | 0.102900214 | 1.95E-05 | 1.02E-04 | 9991 | 9 | 114980715 | 115095947 | -1 | protein_coding | polypyrimidine tract binding protein 3 |
| FKBP15 | ENSG00000119321 | 10945.9638 | -0.585819483 | 0.095058448 | 4.5E-10 | 4.48E-09 | 23307 | 9 | 115923286 | 115983641 | -1 | protein_coding | FK506 binding protein 15, 133kDa |
| SET | ENSG00000119335 | 7194.093926 | 0.355958598 | 0.09921005 | 2.77E-04 | 0.001149628 | 646817 | 9 | 131445703 | 131458679 | 1 | protein_coding | SET nuclear oncogene |
| PPP2R4 | ENSG00000119383 | 2254.107724 | 0.298414506 | 0.121307744 | 0.012414732 | 0.033706582 | 5524 | 9 | 131873229 | 131911225 | 1 | protein_coding | protein phosphatase 2A activator, regulatory subunit 4 |
| RAB14 | ENSG00000119396 | 4334.041564 | 0.425463733 | 0.119630533 | 3.08E-04 | 0.00126666 | 51552 | 9 | 123940415 | 123985292 | -1 | protein_coding | RAB14, member RAS oncogene family |
| TRIM32 | ENSG00000119401 | 413.1807452 | -0.771618985 | 0.213428787 | 1.24E-04 | 5.59E-04 | 22954 | 9 | 119449581 | 119463579 | 1 | protein_coding | tripartite motif containing 32 |
| PHF19 | ENSG00000119403 | 557.8280849 | -0.460457625 | 0.198946487 | 0.014322924 | 0.038138263 | 26147 | 9 | 123617977 | 123639606 | -1 | protein_coding | PHD finger protein 19 |
| NEK6 | ENSG00000119408 | 9156.135026 | -0.847844846 | 0.090844857 | 3.95E-21 | 9.48E-20 | 10783 | 9 | 127019885 | 127115586 | 1 | protein_coding | NIMA-related kinase 6 |

| | | | | | | | | | | | | | |
|---|---|---|---|---|---|---|---|---|---|---|---|---|---|
| **NDUFA8** | ENSG00000119421 | 713.5621123 | -0.406024607 | 0.147619744 | 0.004487607 | 0.013863775 | 4702 | 9 | 124894745 | 124922098 | -1 | protein_coding | NADH dehydrogenase (ubiquinone) 1 alpha subcomplex, 8, 19kDa |
| **SLC46A2** | ENSG00000119457 | 50.66650647 | -1.442293814 | 0.544019254 | 0.001042816 | 0.003789195 | 57864 | 9 | 115641200 | 115653193 | -1 | protein_coding | solute carrier family 46, member 2 |
| **MAPKAP1** | ENSG00000119487 | 3233.700394 | -1.070068387 | 0.165467311 | 2.63E-11 | 2.99E-10 | 79109 | 9 | 128199672 | 128469513 | -1 | protein_coding | mitogen-activated protein kinase associated protein 1 |
| **NR4A3** | ENSG00000119508 | 359.7911086 | 1.104638604 | 0.222387994 | 1.68E-07 | 1.21E-06 | 8013 | 9 | 102584137 | 102629173 | 1 | protein_coding | nuclear receptor subfamily 4, group A, member 3 |
| **INVS** | ENSG00000119509 | 523.3410887 | 0.388100812 | 0.167018387 | 0.015571324 | 0.041072928 | 27130 | 9 | 102861538 | 103063282 | 1 | protein_coding | inversin |
| **GALNT12** | ENSG00000119514 | 4913.676435 | -0.883664816 | 0.115333319 | 6.58E-15 | 9.94E-14 | 79695 | 9 | 101569981 | 101612363 | 1 | protein_coding | UDP-N-acetyl-alpha-D-galactosamine:polypeptide N-acetylgalactosaminyltransferase 12 (GalNAc-T12) |
| **ALG2** | ENSG00000119523 | 901.0802973 | 0.482311235 | 0.131325121 | 1.71E-04 | 7.47E-04 | 85365 | 9 | 101978708 | 101984238 | -1 | protein_coding | ALG2, alpha-1,3/1,6-mannosyltransferase |
| **VPS4B** | ENSG00000119541 | 2931.086938 | 0.3061342 | 0.132965462 | 0.018453065 | 0.047398016 | 9525 | 18 | 61056423 | 61089716 | -1 | protein_coding | vacuolar protein sorting 4 homolog B (S. cerevisiae) |
| **IFI27L2** | ENSG00000119632 | 265.2086197 | -0.614414279 | 0.216304559 | 0.0024864 | 0.008184287 | 83982 | 14 | 94594116 | 94596590 | -1 | protein_coding | interferon, alpha-inducible protein 27-like 2 |
| **NPC2** | ENSG00000119655 | 9974.891849 | -0.494342258 | 0.120527067 | 2.9E-05 | 1.46E-04 | 10577 | 14 | 74942895 | 74960880 | -1 | protein_coding | Niemann-Pick disease, type C2 |
| **DNAL1** | ENSG00000119661 | 265.3226847 | -1.119678017 | 0.355848708 | 3.54E-04 | 0.001434952 | 83544 | 14 | 74111578 | 74170435 | 1 | protein_coding | dynein, axonemal, light chain 1 |
| **FLVCR2** | ENSG00000119686 | 1156.17 | -0.539160011 | 0.186542582 | 0.002362115 | 0.007824059 | 55640 | 14 | 76044960 | 76129557 | 1 | protein_coding | feline leukemia virus subgroup C cellular receptor family, member 2 |
| **RBM25** | ENSG00000119707 | 2813.910798 | 0.531790734 | 0.115033538 | 2.63E-06 | 1.59E-05 | 58517 | 14 | 73525144 | 73588122 | 1 | protein_coding | RNA binding motif protein 25 |
| **ALDH6A1** | ENSG00000119711 | 689.0821852 | -0.772377322 | 0.186803567 | 1.5E-05 | 8.01E-05 | 4329 | 14 | 74523553 | 74551196 | -1 | protein_coding | aldehyde dehydrogenase 6 family, member A1 |
| **GPR68** | ENSG00000119714 | 704.283656 | -0.417371541 | 0.159264533 | 0.006608679 | 0.019486868 | 8111 | 14 | 91698876 | 91720269 | -1 | protein_coding | G protein-coupled receptor 68 |
| **KLHL29** | ENSG00000119771 | 161.8286226 | -1.646354775 | 0.306518953 | 9.98E-09 | 8.5E-08 | 114818 | 2 | 23608088 | 23931481 | 1 | protein_coding | kelch-like family member 29 |
| **DNMT3A** | ENSG00000119772 | 807.2764545 | -1.288171555 | 0.151947814 | 4.61E-18 | 8.76E-17 | 1788 | 2 | 25455845 | 25565459 | -1 | protein_coding | DNA (cytosine-5-)-methyltransferase 3 alpha |
| **TMEM214** | ENSG00000119777 | 1291.161323 | -0.437222031 | 0.155835905 | 0.003677647 | 0.011627115 | 54867 | 2 | 27255778 | 27264563 | 1 | protein_coding | transmembrane protein 214 |
| **FKBP1B** | ENSG00000119782 | 54.2452909 | 0.974091881 | 0.484235486 | 0.010310327 | 0.028733174 | 2281 | 2 | 24272571 | 24286551 | 1 | protein_coding | FK506 binding protein 1B, 12.6 kDa |
| **AFTPH** | ENSG00000119844 | 3100.981206 | 0.358257599 | 0.1157963 | 0.001646132 | 0.005693106 | 54812 | 2 | 64751465 | 64820139 | 1 | protein_coding | aftiphilin |
| **LGALSL** | ENSG00000119862 | 658.1527181 | -2.071150336 | 0.574756434 | 2.41E-05 | 1.24E-04 | 29094 | 2 | 64681103 | 64688515 | 1 | protein_coding | lectin, galactoside-binding-like |
| **CNRIP1** | ENSG00000119865 | 210.9178637 | -1.545105747 | 0.313786429 | 1.22E-07 | 8.93E-07 | 25927 | 2 | 68511303 | 68547183 | -1 | protein_coding | cannabinoid receptor interacting protein 1 |
| **BCL11A** | ENSG00000119866 | 459.6545099 | -1.464029636 | 0.225160433 | 1.28E-11 | 1.49E-10 | 53335 | 2 | 60678302 | 60780702 | -1 | protein_coding | B-cell CLL/lymphoma 11A (zinc finger protein) |
| **CRIPT** | ENSG00000119878 | 253.0346567 | 0.792038931 | 0.235026073 | 3.06E-04 | 0.001258846 | 9419 | 2 | 46843555 | 46852881 | 1 | protein_coding | cysteine-rich PDZ-binding protein |
| **SLC17A5** | ENSG00000119899 | 3930.662519 | 0.39574691 | 0.112191079 | 3.34E-04 | 0.001364528 | 26503 | 6 | 74303102 | 74363878 | -1 | protein_coding | solute carrier family 17 (acidic sugar transporter), member 5 |
| **OGFRL1** | ENSG00000119900 | 4382.414166 | 1.46688682 | 0.089257553 | 1.83E-61 | 2.46E-59 | 79627 | 6 | 71998506 | 72018653 | 1 | protein_coding | opioid growth factor receptor-like 1 |
| **IFIT3** | ENSG00000119917 | 13473.57031 | 5.296908945 | 0.15713911 | 2.33E-250 | 2.55E-247 | 3437 | 10 | 91087651 | 91100728 | 1 | protein_coding | interferon-induced protein with tetratricopeptide repeats 3 |
| **IFIT2** | ENSG00000119922 | 11207.25345 | 5.53406067 | 0.143065923 | 0 | 0 | 3433 | 10 | 91061712 | 91069033 | 1 | protein_coding | interferon-induced protein with tetratricopeptide repeats 2 |
| **MXI1** | ENSG00000119950 | 1633.724207 | -1.296257895 | 0.152040567 | 2.73E-18 | 5.27E-17 | 4601 | 10 | 111967363 | 112047123 | 1 | protein_coding | MAX interactor 1, dimerization protein |
| **SMNDC1** | ENSG00000119953 | 740.0649293 | 0.411527541 | 0.152381295 | 0.005315264 | 0.016094759 | 10285 | 10 | 112050488 | 112064709 | -1 | protein_coding | survival motor neuron domain containing 1 |
| **HELLS** | ENSG00000119969 | 142.4660721 | 1.451075283 | 0.338832075 | 2.75E-06 | 1.66E-05 | 3070 | 10 | 96305547 | 96373662 | 1 | protein_coding | helicase, lymphoid-specific |
| **FAM45A** | ENSG00000119979 | 910.0355796 | 0.517141514 | 0.134359227 | 7.76E-05 | 3.62E-04 | 404636 | 10 | 120863598 | 120897496 | 1 | protein_coding | family with sequence similarity 45, member A |
| **GOT1** | ENSG00000120053 | 589.8377422 | -0.780244114 | 0.174833418 | 3.38E-06 | 2E-05 | 2805 | 10 | 101156627 | 101190381 | -1 | protein_coding | glutamic-oxaloacetic transaminase 1, soluble |
| **GNA13** | ENSG00000120063 | 7543.768765 | 0.764412766 | 0.086561691 | 5.96E-19 | 1.21E-17 | 10672 | 17 | 63006833 | 63052957 | -1 | protein_coding | guanine nucleotide binding protein (G protein), alpha 13 |
| **KANSL1** | ENSG00000120071 | 1587.250456 | 0.455257136 | 0.161191153 | 0.003472781 | 0.011050282 | 1019297 | 17 | 44107282 | 44302733 | -1 | protein_coding | KAT8 regulatory NSL complex subunit 1 |
| **DUSP1** | ENSG00000120129 | 319.5561162 | 0.925085461 | 0.191887214 | 4.27E-07 | 2.92E-06 | 1843 | 5 | 172195093 | 172198198 | -1 | protein_coding | dual specificity phosphatase 1 |
| **MOB3B** | ENSG00000120162 | 1021.862463 | -0.984530052 | 0.196274839 | 1.52E-07 | 1.1E-06 | 79817 | 9 | 27325207 | 27529779 | -1 | protein_coding | MOB kinase activator 3B |
| **CD274** | ENSG00000120217 | 4753.667887 | 3.039378889 | 0.225316475 | 1.21E-42 | 8.49E-41 | 29126 | 9 | 5450503 | 5470566 | 1 | protein_coding | CD274 molecule |
| **PCMT1** | ENSG00000120265 | 1820.067459 | 0.978144742 | 0.154699352 | 4.54E-11 | 5.05E-10 | 5110 | 6 | 150070579 | 150132556 | 1 | protein_coding | protein-L-isoaspartate (D-aspartate) O-methyltransferase |
| **WDR55** | ENSG00000120314 | 1515.515883 | 0.417687744 | 0.122469168 | 5.31E-04 | 0.002064842 | 54853 | 5 | 140044261 | 140053709 | 1 | protein_coding | WD repeat domain 55 |
| **ARAP3** | ENSG00000120318 | 423.4523742 | -1.116829854 | 0.221564941 | 1.09E-07 | 8E-07 | 64411 | 5 | 141032968 | 141061788 | -1 | protein_coding | ArfGAP with RhoGAP domain, ankyrin repeat and PH domain 3 |
| **ACAT2** | ENSG00000120437 | 496.065625 | -0.836060119 | 0.18139736 | 1.45E-06 | 9.06E-06 | 39 | 6 | 160181360 | 160200144 | 1 | protein_coding | acetyl-CoA acetyltransferase 2 |
| **TCP1** | ENSG00000120438 | 2473.447978 | 0.295451243 | 0.110325361 | 0.006689419 | 0.019694547 | 677812 | 6 | 160199530 | 160210781 | -1 | protein_coding | t-complex 1 |
| **KCNJ5** | ENSG00000120457 | 1219.70334 | -2.198869957 | 0.232161711 | 1.5E-22 | 3.9E-21 | 3762 | 11 | 128761251 | 128790930 | 1 | protein_coding | potassium inwardly-rectifying channel, subfamily J, member 5 |
| **NUDCD1** | ENSG00000120526 | 355.9539401 | 1.538977165 | 0.220663155 | 4.61E-13 | 6.08E-12 | 84955 | 8 | 110253148 | 110346614 | -1 | protein_coding | NudC domain containing 1 |
| **MASTL** | ENSG00000120539 | 837.6101776 | 3.043688559 | 0.172628428 | 1.04E-70 | 1.96E-68 | 84930 | 10 | 27443753 | 27475853 | 1 | protein_coding | microtubule associated serine/threonine kinase-like |
| **PLXDC2** | ENSG00000120594 | 24678.72965 | -1.963050833 | 0.120331181 | 7.08E-61 | 9.32E-59 | 84898 | 10 | 20105168 | 20578785 | 1 | protein_coding | plexin domain containing 2 |
| **EPC1** | ENSG00000120616 | 884.3229313 | 0.622546556 | 0.129243887 | 8.28E-07 | 5.41E-06 | 80314 | 10 | 32556679 | 32667726 | -1 | protein_coding | enhancer of polycomb homolog 1 (Drosophila) |
| **CCDC77** | ENSG00000120647 | 268.7196015 | 1.036612952 | 0.227019772 | 1.34E-06 | 8.47E-06 | 84318 | 12 | 498439 | 551811 | 1 | protein_coding | coiled-coil domain containing 77 |
| **DNAJC15** | ENSG00000120675 | 625.4819 | 0.709245273 | 0.163832513 | 7.1E-06 | 4E-05 | 29103 | 13 | 43597339 | 43683045 | 1 | protein_coding | DnaJ (Hsp40) homolog, subfamily C, member 15 |
| **PROSER1** | ENSG00000120685 | 1211.93874 | 1.136821751 | 0.159214006 | 2.26E-13 | 3.02E-12 | 80209 | 13 | 39584003 | 39612252 | -1 | protein_coding | proline and serine rich 1 |
| **UFM1** | ENSG00000120686 | 2486.856184 | 0.500049143 | 0.118089383 | 1.62E-05 | 8.58E-05 | 51569 | 13 | 38923986 | 38937140 | 1 | protein_coding | ubiquitin-fold modifier 1 |
| **WBP4** | ENSG00000120688 | 335.078126 | 0.662273279 | 0.203444239 | 5.58E-04 | 0.002156667 | 11193 | 13 | 41635410 | 41658137 | 1 | protein_coding | WW domain binding protein 4 |
| **ELF1** | ENSG00000120690 | 4347.592506 | 0.511903765 | 0.096883134 | 8.69E-08 | 6.49E-07 | 1997 | 13 | 41506056 | 41635576 | -1 | protein_coding | E74-like factor 1 (ets domain transcription factor) |
| **HSPH1** | ENSG00000120694 | 2823.048163 | 0.314711067 | 0.131077835 | 0.014323996 | 0.038138263 | 10808 | 13 | 31710762 | 31736525 | -1 | protein_coding | heat shock 105kDa/110kDa protein 1 |
| **ETF1** | ENSG00000120705 | 4672.971083 | 0.340247401 | 0.088056241 | 8.74E-05 | 4.05E-04 | 2107 | 5 | 137841784 | 137878989 | -1 | protein_coding | eukaryotic translation termination factor 1 |
| **PAIP2** | ENSG00000120727 | 1830.87951 | 0.365959351 | 0.120641877 | 0.002055191 | 0.006924488 | 51247 | 5 | 138677276 | 138705406 | 1 | protein_coding | poly(A) binding protein interacting protein 2 |
| **KDM3B** | ENSG00000120733 | 3636.279041 | -0.299192584 | 0.098898801 | 0.002024475 | 0.006834561 | 51780 | 5 | 137688285 | 137772717 | 1 | protein_coding | lysine (K)-specific demethylase 3B |
| **EGR1** | ENSG00000120738 | 261.6204102 | -2.20639729 | 0.327628832 | 1.44E-12 | 1.81E-11 | 1958 | 5 | 137801179 | 137805004 | 1 | protein_coding | early growth response 1 |
| **SERP1** | ENSG00000120742 | 2987.957423 | 0.512741787 | 0.124086342 | 2.4E-05 | 1.23E-04 | 1019280 | 3 | 150259781 | 150321015 | -1 | protein_coding | stress-associated endoplasmic reticulum protein 1 |
| **NR2C1** | ENSG00000120798 | 375.3605741 | 0.441895175 | 0.181192616 | 0.010582552 | 0.029347712 | 7181 | 12 | 95414005 | 95467479 | -1 | protein_coding | nuclear receptor subfamily 2, group C, member 1 |
| **TMPO** | ENSG00000120802 | 2401.9489 | 0.470882874 | 0.139073506 | 5.18E-04 | 0.002020131 | 7112 | 12 | 98909290 | 98944157 | 1 | protein_coding | thymopoietin |
| **SORBS3** | ENSG00000120896 | 1084.451566 | -0.766878714 | 0.192877044 | 2.93E-05 | 1.48E-04 | 10174 | 8 | 22402499 | 22433301 | 1 | protein_coding | sorbin and SH3 domain containing 3 |
| **PPP3CC** | ENSG00000120910 | 642.8029393 | 1.742484538 | 0.231576811 | 4.95E-15 | 7.52E-14 | 5533 | 8 | 22298332 | 22398652 | 1 | protein_coding | protein phosphatase 3, catalytic subunit, gamma isozyme |
| **PDLIM2** | ENSG00000120913 | 276.923873 | -1.125026927 | 0.250110434 | 1.61E-06 | 1E-05 | 64236 | 8 | 22435792 | 22455538 | 1 | protein_coding | PDZ and LIM domain 2 (mystique) |
| **TARDBP** | ENSG00000120948 | 2495.058 | 0.270228485 | 0.099758487 | 0.006055295 | 0.018032557 | 23435 | 1 | 11072414 | 11085796 | 1 | protein_coding | TAR DNA binding protein |
| **RDH10** | ENSG00000121039 | 783.6362283 | -1.402920804 | 0.305897222 | 5.85E-07 | 3.92E-06 | 157506 | 8 | 74206847 | 74237516 | 1 | protein_coding | retinol dehydrogenase 10 (all-trans) |
| **TRIM25** | ENSG00000121060 | 3896.499443 | 1.728420991 | 0.12518893 | 2.46E-44 | 1.82E-42 | 7706 | 17 | 54965270 | 54991399 | -1 | protein_coding | tripartite motif containing 25 |
| **FAM117A** | ENSG00000121104 | 285.2901766 | -0.511343228 | 0.214447014 | 0.011141779 | 0.03071425 | 81558 | 17 | 47787694 | 47866542 | -1 | protein_coding | family with sequence similarity 117, member A |
| **NCAPH** | ENSG00000121152 | 5002.139642 | -2.518572398 | 0.725396788 | 2.47E-05 | 1.27E-04 | 23397 | 2 | 97001525 | 97039583 | 1 | protein_coding | non-SMC condensin I complex, subunit H |
| **TRIM6** | ENSG00000121236 | 326.1083075 | 1.16077378 | 0.275134578 | 5.3E-06 | 3.05E-05 | 117854 | 11 | 5617339 | 5634188 | 1 | protein_coding | tripartite motif containing 6 |
| **PAPD5** | ENSG00000121274 | 620.8318916 | 0.586082228 | 0.151766235 | 6.75E-05 | 3.18E-04 | 64282 | 16 | 50186829 | 50269221 | 1 | protein_coding | PAP associated domain containing 5 |
| **PLBD1** | ENSG00000121316 | 7328.294947 | -0.405981536 | 0.113895793 | 2.75E-04 | 0.001142046 | 79887 | 12 | 14656595 | 14721283 | -1 | protein_coding | phospholipase B domain containing 1 |
| **BCL2L14** | ENSG00000121380 | 65.41274853 | 6.718710864 | 0.809715175 | 8.2E-18 | 1.52E-16 | 1019282 | 12 | 12202778 | 12364018 | 1 | protein_coding | BCL2-like 14 (apoptosis facilitator) |
| **A1BG** | ENSG00000121410 | 74.17183934 | -1.429889729 | 0.426935138 | 1.21E-04 | 5.47E-04 | 1 | 19 | 58856544 | 58864865 | -1 | protein_coding | alpha-1-B glycoprotein |
| **ZSCAN18** | ENSG00000121413 | 347.2169392 | -1.065843729 | 0.244980204 | 2.99E-06 | 1.79E-05 | 65982 | 19 | 58595205 | 58629794 | -1 | protein_coding | zinc finger and SCAN domain containing 18 |
| **TRMT1L** | ENSG00000121486 | 1540.011554 | 0.423693999 | 0.13096339 | 8.8E-04 | 0.003257843 | 81627 | 1 | 185087220 | 185126204 | -1 | protein_coding | tRNA methyltransferase 1 homolog (S. cerevisiae)-like |
| **CSTA** | ENSG00000121552 | 1744.894788 | 0.765582454 | 0.135504838 | 6.43E-09 | 5.61E-08 | 1475 | 3 | 122044091 | 122060819 | 1 | protein_coding | cystatin A (stefin A) |
| **B4GALT4** | ENSG00000121578 | 632.1941263 | 0.669490177 | 0.152967369 | 6.32E-06 | 3.58E-05 | 8702 | 3 | 118930579 | 118959950 | -1 | protein_coding | UDP-Gal:betaGlcNAc beta 1,4- galactosyltransferase, polypeptide 4 |
| **NAA50** | ENSG00000121579 | 3628.373464 | 0.301962963 | 0.112769102 | 0.006532405 | 0.019302929 | 80218 | 3 | 113435307 | 113465147 | -1 | protein_coding | N(alpha)-acetyltransferase 50, NatE catalytic subunit |
| **CD80** | ENSG00000121594 | 805.6989915 | 4.243310046 | 0.170902991 | 3.01E-137 | 1.49E-134 | 941 | 3 | 119243140 | 119278449 | -1 | protein_coding | CD80 molecule |
| **KIF18A** | ENSG00000121621 | 44.20357989 | 1.866649434 | 0.580879595 | 1.22E-04 | 5.48E-04 | 81930 | 11 | 28042167 | 28129855 | -1 | protein_coding | kinesin family member 18A |
| **DESI2** | ENSG00000121644 | 1052.120805 | 0.459149177 | 0.122461763 | 1.26E-04 | 5.66E-04 | 51029 | 1 | 244816237 | 244872335 | 1 | protein_coding | desumoylating isopeptidase 2 |
| **CAT** | ENSG00000121691 | 2207.987329 | -1.338022532 | 0.137686646 | 4.32E-23 | 1.18E-21 | 847 | 11 | 34460472 | 34493609 | 1 | protein_coding | catalase |
| **PILRB** | ENSG00000121716 | 157.7722109 | 0.722702472 | 0.300559604 | 0.007031678 | 0.020587189 | 1024667 | 7 | 99933737 | 99965356 | 1 | protein_coding | paired immunoglobin-like type 2 receptor beta |
| **ZMYM2** | ENSG00000121741 | 2513.448209 | 0.259237805 | 0.112714386 | 0.019628544 | 0.049913639 | 7750 | 13 | 20532810 | 20665968 | 1 | protein_coding | zinc finger, MYM-type 2 |
| **GJB6** | ENSG00000121742 | 105.1987025 | -2.074494838 | 0.479006107 | 1.31E-06 | 8.28E-06 | 10804 | 13 | 20796110 | 20806534 | -1 | protein_coding | gap junction protein, beta 6, 30kDa |
| **FABP3** | ENSG00000121769 | 1940.370154 | -2.554458855 | 0.177313409 | 6.38E-48 | 5.26E-46 | 2170 | 1 | 31838472 | 31849697 | -1 | protein_coding | fatty acid binding protein 3, muscle and heart (mammary-derived growth inhibitor) |
| **CCRL2** | ENSG00000121797 | 2328.874517 | 0.733545907 | 0.164640711 | 3.73E-06 | 2.2E-05 | 9034 | 3 | 46448654 | 46454488 | 1 | protein_coding | chemokine (C-C motif) receptor-like 2 |

| | | | | | | | | | | | | | |
|---|---|---|---|---|---|---|---|---|---|---|---|---|---|
| **TNFSF10** | ENSG00000121858 | 3918.204364 | 4.7498716 | 0.370278543 | 5.81E-39 | 3.39E-37 | 8743 | 3 | 172223298 | 172241297 | -1 | protein_coding | tumor necrosis factor (ligand) superfamily, member 10 |
| **ZNF639** | ENSG00000121864 | 300.2846064 | 0.540539602 | 0.20680144 | 0.005637474 | 0.016916851 | 51193 | 3 | 179040779 | 179053323 | 1 | protein_coding | zinc finger protein 639 |
| **PIK3CA** | ENSG00000121879 | 1739.340547 | 0.359132596 | 0.147037986 | 0.011422908 | 0.031398615 | 5290 | 3 | 178865902 | 178957881 | 1 | protein_coding | phosphatidylinositol-4,5-bisphosphate 3-kinase, catalytic subunit alpha |
| **LRIF1** | ENSG00000121931 | 649.4013163 | 0.764296562 | 0.156158379 | 4.54E-07 | 3.09E-06 | 55791 | 1 | 111489807 | 111506701 | -1 | protein_coding | ligand dependent nuclear receptor interacting factor 1 |
| **CXCR4** | ENSG00000121966 | 2491.839799 | -0.808853676 | 0.125852924 | 5.39E-11 | 5.96E-10 | 7852 | 2 | 136871919 | 136875735 | -1 | protein_coding | chemokine (C-X-C motif) receptor 4 |
| **ZRANB3** | ENSG00000121988 | 100.8625418 | -0.849913843 | 0.409953891 | 0.011791343 | 0.032244948 | 84083 | 2 | 135894486 | 136288806 | -1 | protein_coding | zinc finger, RAN-binding domain containing 3 |
| **POLK** | ENSG00000122008 | 1110.319612 | 0.816803334 | 0.144655169 | 6.66E-09 | 5.8E-08 | 51426 | 5 | 74807581 | 74896969 | 1 | protein_coding | polymerase (DNA directed) kappa |
| **GTF3A** | ENSG00000122034 | 2416.241914 | -0.407705531 | 0.132578233 | 0.001606896 | 0.00556875 | 2971 | 13 | 27998681 | 28009958 | 1 | protein_coding | general transcription factor IIIA |
| **FYTTD1** | ENSG00000122068 | 894.0135466 | 0.532118023 | 0.16841909 | 0.001020935 | 0.003719236 | 84248 | 3 | 197464050 | 197514467 | 1 | protein_coding | forty-two-three domain containing 1 |
| **XPNPEP2** | ENSG00000122121 | 47.03969265 | -1.14084818 | 0.565138593 | 0.007401989 | 0.021527041 | 7512 | X | 128872950 | 128903514 | 1 | protein_coding | X-prolyl aminopeptidase (aminopeptidase P) 2, membrane-bound |
| **MRPS2** | ENSG00000122140 | 221.4373929 | -0.493672149 | 0.222333778 | 0.017381948 | 0.045124563 | 51116 | 9 | 138391830 | 138396519 | 1 | protein_coding | mitochondrial ribosomal protein S2 |
| **COPA** | ENSG00000122218 | 7837.250444 | -0.511196541 | 0.08664218 | 3.67E-09 | 3.29E-08 | 1314 | 1 | 160259063 | 160313190 | -1 | protein_coding | coatomer protein complex, subunit alpha |
| **HS3ST2** | ENSG00000122254 | 126.943183 | -3.179689382 | 0.439761276 | 3.53E-14 | 5.07E-13 | 9956 | 16 | 22825498 | 22927659 | 1 | protein_coding | heparan sulfate (glucosamine) 3-O-sulfotransferase 2 |
| **RBBP6** | ENSG00000122257 | 1586.829375 | 0.972712937 | 0.105551889 | 1.18E-20 | 2.73E-19 | 5930 | 16 | 24549014 | 24584184 | 1 | protein_coding | retinoblastoma binding protein 6 |
| **ZC3H7A** | ENSG00000122299 | 2211.742759 | 0.448701866 | 0.127978991 | 3.38E-04 | 0.001378532 | 29066 | 16 | 11844442 | 11891123 | -1 | protein_coding | zinc finger CCCH-type containing 7A |
| **SERAC1** | ENSG00000122335 | 133.1101146 | -1.380371203 | 0.425393218 | 1.74E-04 | 7.57E-04 | 84947 | 6 | 158530536 | 158589312 | -1 | protein_coding | serine active site containing 1 |
| **ANXA11** | ENSG00000122359 | 9366.971808 | -0.451729108 | 0.123531808 | 1.79E-04 | 7.75E-04 | 311 | 10 | 81910645 | 81965328 | -1 | protein_coding | annexin A11 |
| **FAM35A** | ENSG00000122376 | 1163.992698 | 0.546244307 | 0.134181025 | 2.97E-05 | 1.5E-04 | 54537 | 10 | 88853918 | 88951225 | 1 | protein_coding | family with sequence similarity 35, member A |
| **FAM213A** | ENSG00000122378 | 6926.630201 | -0.474409819 | 0.097664021 | 6.49E-07 | 4.31E-06 | 84293 | 10 | 82167585 | 82192753 | 1 | protein_coding | family with sequence similarity 213, member A |
| **RPL5** | ENSG00000122406 | 5435.181469 | -0.623213082 | 0.111650754 | 1.09E-08 | 9.24E-08 | 6125 | 1 | 93297582 | 93307481 | 1 | protein_coding | ribosomal protein L5 |
| **ODF2L** | ENSG00000122417 | 447.5860983 | 2.686360847 | 0.227713282 | 3.45E-33 | 1.61E-31 | 57489 | 1 | 86814344 | 86862023 | -1 | protein_coding | outer dense fiber of sperm tails 2-like |
| **TRMT13** | ENSG00000122435 | 217.8787356 | 0.781073536 | 0.261058476 | 0.001112444 | 0.004018314 | 54482 | 1 | 100598706 | 100616053 | 1 | protein_coding | tRNA methyltransferase 13 homolog (S. cerevisiae) |
| **ZNF644** | ENSG00000122482 | 1479.343772 | 0.981801325 | 0.174500384 | 5.59E-09 | 4.91E-08 | 84146 | 1 | 91380859 | 91487829 | -1 | protein_coding | zinc finger protein 644 |
| **RPAP2** | ENSG00000122484 | 739.57949 | 0.634750938 | 0.186534873 | 3.56E-04 | 0.001444239 | 79871 | 1 | 92764522 | 92867613 | 1 | protein_coding | RNA polymerase II associated protein 2 |
| **ZMIZ2** | ENSG00000122515 | 1833.094856 | 0.751580099 | 0.161250956 | 1.45E-06 | 9.09E-06 | 83637 | 7 | 44788180 | 44809477 | 1 | protein_coding | zinc finger, MIZ-type containing 2 |
| **EEPD1** | ENSG00000122547 | 415.3748247 | -4.564468578 | 0.449382664 | 1.77E-25 | 5.4E-24 | 80820 | 7 | 36192758 | 36341152 | 1 | protein_coding | endonuclease/exonuclease/phosphatase family domain containing 1 |
| **HERPUD2** | ENSG00000122557 | 1307.558626 | 1.049737831 | 0.13310433 | 8.21E-16 | 1.32E-14 | 64224 | 7 | 35672269 | 35735181 | -1 | protein_coding | HERPUD family member 2 |
| **CBX3** | ENSG00000122565 | 2895.262846 | 1.208054877 | 0.115414635 | 2.85E-26 | 9.12E-25 | 11335 | 7 | 26240782 | 26252976 | 1 | protein_coding | chromobox homolog 3 |
| **HNRNPA2B1** | ENSG00000122566 | 12927.54697 | 0.488013035 | 0.09373792 | 1.6E-07 | 1.16E-06 | 3181 | 7 | 26229547 | 26241149 | -1 | protein_coding | heterogeneous nuclear ribonucleoprotein A2/B1 |
| **NT5C3A** | ENSG00000122643 | 572.9588916 | 3.608994182 | 0.259863237 | 1.66E-45 | 1.31E-43 | 51251 | 7 | 33053742 | 33102409 | -1 | protein_coding | 5'-nucleotidase, cytosolic IIIA |
| **CCZ1** | ENSG00000122674 | 1606.07055 | 0.758049878 | 0.134095271 | 7.2E-09 | 6.23E-08 | 221960 | 7 | 5938356 | 5965605 | 1 | protein_coding | CCZ1 vacuolar protein trafficking and biogenesis associated homolog (S. cerevisiae) |
| **FTSJ2** | ENSG00000122687 | 780.3360994 | -0.312209777 | 0.136421203 | 0.018939396 | 0.04843554 | 29960 | 7 | 2273866 | 2281840 | -1 | protein_coding | FtsJ RNA methyltransferase homolog 2 (E. coli) |
| **SMU1** | ENSG00000122692 | 1077.9991 | 0.502679465 | 0.11538623 | 1.09E-05 | 5.98E-05 | 55234 | 9 | 33041762 | 33076665 | -1 | protein_coding | smu-1 suppressor of mec-8 and unc-52 homolog (C. elegans) |
| **SLC25A51** | ENSG00000122696 | 494.8263375 | -0.709617182 | 0.230477342 | 9.54E-04 | 0.003492648 | 92014 | 9 | 37879400 | 37904350 | -1 | protein_coding | solute carrier family 25, member 51 |
| **ACO1** | ENSG00000122729 | 5247.962983 | -1.530779332 | 0.158806489 | 5.83E-23 | 1.57E-21 | 48 | 9 | 32384618 | 32454767 | 1 | protein_coding | aconitase 1, soluble |
| **TRIM24** | ENSG00000122779 | 1028.174791 | -0.376217779 | 0.130996465 | 0.003254566 | 0.010434137 | 8805 | 7 | 138145079 | 138274738 | 1 | protein_coding | tripartite motif containing 24 |
| **C7orf49** | ENSG00000122783 | 435.5006413 | -0.544391689 | 0.197501893 | 0.003533282 | 0.01122177 | 78996 | 7 | 134777115 | 134855547 | -1 | protein_coding | chromosome 7 open reading frame 49 |
| **CALD1** | ENSG00000122786 | 156.3906981 | -1.034548876 | 0.387732916 | 0.001775575 | 0.006076156 | 800 | 7 | 134429003 | 134655479 | 1 | protein_coding | caldesmon 1 |
| **PLAU** | ENSG00000122861 | 1163.058513 | -1.845722855 | 0.136828967 | 2.09E-42 | 1.44E-40 | 5328 | 10 | 75668935 | 75677255 | 1 | protein_coding | plasminogen activator, urokinase |
| **SRGN** | ENSG00000122862 | 16341.44496 | 1.412416352 | 0.172869083 | 5.9E-17 | 1.03E-15 | 5552 | 10 | 70847862 | 70864567 | 1 | protein_coding | serglycin |
| **P4HA1** | ENSG00000122884 | 4939.727515 | -0.290469401 | 0.109094441 | 0.006784216 | 0.019954421 | 5033 | 10 | 74766975 | 74856732 | -1 | protein_coding | prolyl 4-hydroxylase, alpha polypeptide I |
| **SLC25A16** | ENSG00000122912 | 329.5566833 | -0.946545456 | 0.33652509 | 0.001353849 | 0.0047715 | 8034 | 10 | 70237756 | 70287231 | -1 | protein_coding | solute carrier family 25 (mitochondrial carrier; Graves disease autoantigen), member 16 |
| **VPS26A** | ENSG00000122958 | 2627.036538 | 0.504090062 | 0.164650772 | 0.001487074 | 0.00519592 | 9559 | 10 | 70883268 | 70932617 | 1 | protein_coding | vacuolar protein sorting 26 homolog A (S. pombe) |
| **BHLHE41** | ENSG00000123095 | 5082.319949 | -0.579594903 | 0.146629792 | 4.49E-05 | 2.18E-04 | 79365 | 12 | 26272959 | 26278060 | -1 | protein_coding | basic helix-loop-helix family, member e41 |
| **SSPN** | ENSG00000123096 | 426.6922451 | 1.04661859 | 0.222300727 | 6.66E-07 | 4.42E-06 | 8082 | 12 | 26274924 | 26452223 | 1 | protein_coding | sarcospan |
| **WWP1** | ENSG00000123124 | 1482.98924 | -0.449615061 | 0.122472324 | 1.84E-04 | 7.95E-04 | 11059 | 8 | 87354967 | 87490649 | 1 | protein_coding | WW domain containing E3 ubiquitin protein ligase 1 |
| **ACOT9** | ENSG00000123130 | 2080.778875 | 1.37935783 | 0.115065302 | 6.89E-34 | 3.29E-32 | 23597 | X | 23720370 | 23784592 | -1 | protein_coding | acyl-CoA thioesterase 9 |
| **PKN1** | ENSG00000123143 | 1743.356202 | -0.449172944 | 0.121368575 | 1.38E-04 | 6.13E-04 | 5585 | 19 | 14543865 | 14582679 | 1 | protein_coding | protein kinase N1 |
| **CD97** | ENSG00000123146 | 8840.074697 | 0.811645041 | 0.123758065 | 1.45E-11 | 1.68E-10 | 976 | 19 | 14491313 | 14519537 | 1 | protein_coding | CD97 molecule |
| **GIPC1** | ENSG00000123159 | 542.7930103 | -0.618184608 | 0.182979825 | 4.08E-04 | 0.001632341 | 10755 | 19 | 14588572 | 14606944 | -1 | protein_coding | GIPC PDZ domain containing family, member 1 |
| **GUCY1B2** | ENSG00000123201 | 4.313408996 | 0.462677105 | 0.894985089 | 0.018888643 | 0.048330007 | 2974 | 13 | 51568647 | 51654998 | -1 | pseudogene | guanylate cyclase 1, soluble, beta 2 (pseudogene) |
| **CENPK** | ENSG00000123219 | 56.71323039 | 0.904310731 | 0.513474745 | 0.019142879 | 0.048882308 | 64105 | 5 | 64813593 | 64858998 | -1 | protein_coding | centromere protein K |
| **OPTN** | ENSG00000123240 | 5939.365944 | 1.789832505 | 0.129416462 | 2.32E-44 | 1.73E-42 | 10133 | 10 | 13141449 | 13180291 | 1 | protein_coding | optineurin |
| **TSFM** | ENSG00000123297 | 363.9055833 | 0.920043948 | 0.194319784 | 7.19E-07 | 4.74E-06 | 10102 | 12 | 58176372 | 58201854 | 1 | protein_coding | Ts translation elongation factor, mitochondrial |
| **ARHGAP9** | ENSG00000123329 | 1182.736986 | -1.088897833 | 0.148671691 | 7.14E-14 | 1E-12 | 64333 | 12 | 57866038 | 57882597 | -1 | protein_coding | Rho GTPase activating protein 9 |
| **NCKAP1L** | ENSG00000123338 | 16564.47456 | -0.420095563 | 0.081930759 | 2.26E-07 | 1.6E-06 | 3071 | 12 | 54891495 | 54937726 | 1 | protein_coding | NCK-associated protein 1-like |
| **MMP19** | ENSG00000123342 | 4528.546392 | -1.145326019 | 0.126119225 | 2.97E-20 | 6.68E-19 | 4327 | 12 | 56229217 | 56236750 | -1 | protein_coding | matrix metallopeptidase 19 |
| **PFDN5** | ENSG00000123349 | 2461.571896 | -0.411299604 | 0.11888028 | 4.16E-04 | 0.001663335 | 5204 | 12 | 53689075 | 53693234 | 1 | protein_coding | prefoldin subunit 5 |
| **CDK2** | ENSG00000123374 | 317.5632803 | 0.840828636 | 0.287679827 | 0.001227196 | 0.004372869 | 1017 | 12 | 56360553 | 56366568 | 1 | protein_coding | cyclin-dependent kinase 2 |
| **LRP1** | ENSG00000123384 | 27050.26282 | -1.732586633 | 0.101117101 | 1E-66 | 1.72E-64 | 4035 | 12 | 57522276 | 57607134 | 1 | protein_coding | low density lipoprotein receptor-related protein 1 |
| **TUBA1B** | ENSG00000123416 | 14711.30179 | -0.268164714 | 0.091291863 | 0.002677538 | 0.008745632 | 10376 | 12 | 49521565 | 49525180 | -1 | protein_coding | tubulin, alpha 1b |
| **METTL21B** | ENSG00000123427 | 600.6406393 | 0.909938669 | 0.167826981 | 2.03E-08 | 1.65E-07 | 25895 | 12 | 58165275 | 58176324 | 1 | protein_coding | methyltransferase like 21B |
| **SARDH** | ENSG00000123453 | 386.433682 | -3.127120174 | 0.234782934 | 1.4E-41 | 9.3E-40 | 1757 | 9 | 136528682 | 136605077 | -1 | protein_coding | sarcosine dehydrogenase |
| **ATPAF1** | ENSG00000123472 | 490.3482927 | -0.571829998 | 0.204311774 | 0.002842934 | 0.009234551 | 64756 | 1 | 47098409 | 47139539 | -1 | protein_coding | ATP synthase mitochondrial F1 complex assembly factor 1 |
| **STIL** | ENSG00000123473 | 197.8161656 | 1.6483546 | 0.485445801 | 6.17E-05 | 2.93E-04 | 6491 | 1 | 47715811 | 47779819 | -1 | protein_coding | SCL/TAL1 interrupting locus |
| **AMD 1.00** | ENSG00000123505 | 3455.76873 | 1.156525093 | 0.122842327 | 8.8E-22 | 2.2E-20 | 262 | 6 | 111195973 | 111216916 | 1 | protein_coding | adenosylmethionine decarboxylase 1 |
| **USP45** | ENSG00000123552 | 184.4518704 | 0.903149338 | 0.404630317 | 0.007298597 | 0.021246562 | 85015 | 6 | 99880190 | 99969604 | -1 | protein_coding | ubiquitin specific peptidase 45 |
| **RAB9B** | ENSG00000123570 | 14.21843112 | 2.085536864 | 1.099579363 | 0.003224612 | 0.010353306 | 51209 | X | 103077252 | 103087158 | -1 | protein_coding | RAB9B, member RAS oncogene family |
| **FAM199X** | ENSG00000123575 | 1059.699812 | 0.608424867 | 0.162903839 | 1.02E-04 | 4.65E-04 | 139231 | X | 103411301 | 103440583 | 1 | protein_coding | family with sequence similarity 199, X-linked |
| **RAB9A** | ENSG00000123595 | 955.4907639 | 1.410490312 | 0.158253767 | 1.01E-19 | 2.18E-18 | 9367 | X | 13707244 | 13728625 | 1 | protein_coding | RAB9A, member RAS oncogene family |
| **METTL8** | ENSG00000123600 | 663.7365222 | 0.913898735 | 0.174445432 | 5.44E-08 | 4.2E-07 | 79828 | 2 | 172173919 | 172291312 | -1 | protein_coding | methyltransferase like 8 |
| **NMI** | ENSG00000123609 | 1462.858593 | 1.992195629 | 0.131275503 | 5.56E-53 | 5.66E-51 | 9111 | 2 | 152126979 | 152146571 | -1 | protein_coding | N-myc (and STAT) interactor |
| **TNFAIP6** | ENSG00000123610 | 8127.152683 | 7.111104807 | 0.166161533 | 0 | 0 | 7130 | 2 | 152214106 | 152236560 | 1 | protein_coding | tumor necrosis factor, alpha-induced protein 6 |
| **SLC36A1** | ENSG00000123643 | 2422.575477 | -0.946045503 | 0.160011007 | 1.07E-09 | 1.03E-08 | 206358 | 5 | 150816607 | 150871942 | 1 | protein_coding | solute carrier family 36 (proton/amino acid symporter), member 1 |
| **BATF3** | ENSG00000123685 | 77.74719304 | 2.268018919 | 0.529808133 | 1.24E-06 | 7.87E-06 | 55509 | 1 | 212859760 | 212873327 | -1 | protein_coding | basic leucine zipper transcription factor, ATF-like 3 |
| **G0S2** | ENSG00000123689 | 681.1453109 | 2.106023431 | 0.553552192 | 1.16E-05 | 6.33E-05 | 50486 | 1 | 209848765 | 209849733 | 1 | protein_coding | G0/G1switch 2 |
| **RAP2C** | ENSG00000123728 | 1516.394045 | 1.635666628 | 0.140394033 | 4.05E-32 | 1.75E-30 | 57826 | X | 131337053 | 131353471 | -1 | protein_coding | RAP2C, member of RAS oncogene family |
| **EXOSC9** | ENSG00000123737 | 562.4932493 | 1.051900194 | 0.18299865 | 2.03E-09 | 1.88E-08 | 5393 | 4 | 122722472 | 122738176 | 1 | protein_coding | exosome component 9 |
| **PFKFB2** | ENSG00000123836 | 1581.998684 | -2.187644212 | 0.165733491 | 9.6E-41 | 5.99E-39 | 5208 | 1 | 207222801 | 207254369 | 1 | protein_coding | 6-phosphofructo-2-kinase/fructose-2,6-biphosphatase 2 |
| **MXD4** | ENSG00000123933 | 721.7666477 | -1.973034657 | 0.230400169 | 8.09E-19 | 1.62E-17 | 1006163 | 4 | 2249159 | 2264021 | -1 | protein_coding | MAX dimerization protein 4 |
| **DNPEP** | ENSG00000123992 | 664.877693 | 0.672118881 | 0.226742022 | 0.001453163 | 0.005096076 | 23549 | 2 | 220238268 | 220264744 | -1 | protein_coding | aspartyl aminopeptidase |
| **OBSL1** | ENSG00000124006 | 230.8668618 | -0.935331906 | 0.303961638 | 6.53E-04 | 0.002485295 | 23363 | 2 | 220415451 | 220436581 | -1 | protein_coding | obscurin-like 1 |
| **SLC12A4** | ENSG00000124067 | 792.3541836 | -0.585056739 | 0.164633056 | 2.25E-04 | 9.5E-04 | 6560 | 16 | 67977377 | 68003504 | -1 | protein_coding | solute carrier family 12 (potassium/chloride transporter), member 4 |
| **TTPAL** | ENSG00000124120 | 478.5845368 | 0.826403584 | 0.177176244 | 1.11E-06 | 7.06E-06 | 79183 | 20 | 43104526 | 43123244 | 1 | protein_coding | tocopherol (alpha) transfer protein-like |
| **PREX1** | ENSG00000124126 | 3767.831994 | -0.63826663 | 0.130933402 | 6.01E-07 | 4.01E-06 | 57580 | 20 | 47240790 | 47444420 | -1 | protein_coding | phosphatidylinositol-3,4,5-trisphosphate-dependent Rac exchange factor 1 |
| **SDC4** | ENSG00000124145 | 2634.663429 | 1.602598562 | 0.120738791 | 5.98E-41 | 3.79E-39 | 6385 | 20 | 43953928 | 43977064 | -1 | protein_coding | syndecan 4 |
| **NCOA3** | ENSG00000124151 | 4548.703495 | 0.697140768 | 0.09315944 | 4.1E-14 | 5.86E-13 | 8202 | 20 | 46130601 | 46285621 | 1 | protein_coding | nuclear receptor coactivator 3 |
| **PIGT** | ENSG00000124155 | 2542.456617 | -0.342612446 | 0.113196767 | 0.00216442 | 0.007247772 | 51604 | 20 | 44044717 | 44054884 | 1 | protein_coding | phosphatidylinositol glycan anchor biosynthesis, class T |
| **PLCG1** | ENSG00000124181 | 1140.830749 | -0.607291042 | 0.197026012 | 0.001114056 | 0.004022936 | 5335 | 20 | 39765600 | 39811629 | 1 | protein_coding | phospholipase C, gamma 1 |

| | | | | | | | | | | | | | |
|---|---|---|---|---|---|---|---|---|---|---|---|---|---|
| **SRSF6** | ENSG00000124193 | 3005.717179 | -0.428726147 | 0.098267122 | 9.72E-06 | 5.36E-05 | 6431 | 20 | 42086568 | 42092245 | 1 | protein_coding | serine/arginine-rich splicing factor 6 |
| **ZNFX1** | ENSG00000124201 | 11518.10109 | 1.757934786 | 0.099118679 | 2.68E-71 | 5.26E-69 | 57169 | 20 | 47854483 | 47894963 | -1 | protein_coding | zinc finger, NFX1-type containing 1 |
| **ZNF831** | ENSG00000124203 | 184.2581502 | -1.27728654 | 0.290293676 | 2.04E-06 | 1.26E-05 | 128611 | 20 | 57766075 | 57834168 | 1 | protein_coding | zinc finger protein 831 |
| **PTGIS** | ENSG00000124212 | 89.74862761 | -1.943536231 | 0.704583942 | 4.25E-04 | 0.001693353 | 5740 | 20 | 48120411 | 48184683 | -1 | protein_coding | prostaglandin I2 (prostacyclin) synthase |
| **STAU1** | ENSG00000124214 | 3487.333582 | 0.456765216 | 0.089457082 | 2.35E-07 | 1.66E-06 | 6780 | 20 | 47729878 | 47804904 | -1 | protein_coding | staufen double-stranded RNA binding protein 1 |
| **STX16** | ENSG00000124222 | 2759.718256 | 0.439111854 | 0.096595088 | 4.23E-06 | 2.48E-05 | 8675 | 20 | 57226328 | 57254582 | 1 | protein_coding | syntaxin 16 |
| **RNF114** | ENSG00000124226 | 2831.788872 | 1.281779664 | 0.095844327 | 6.35E-42 | 4.25E-40 | 55905 | 20 | 48552948 | 48570429 | 1 | protein_coding | ring finger protein 114 |
| **ZBP1** | ENSG00000124256 | 264.3352947 | 4.681587547 | 0.398391457 | 3.61E-33 | 1.68E-31 | 81030 | 20 | 56178902 | 56195632 | -1 | protein_coding | Z-DNA binding protein 1 |
| **MTRR** | ENSG00000124275 | 1263.913598 | 0.451241584 | 0.18707358 | 0.011723114 | 0.032102934 | 4552 | 5 | 7851299 | 7906138 | 1 | protein_coding | 5-methyltetrahydrofolate-homocysteine methyltransferase reductase |
| **FASTKD3** | ENSG00000124279 | 213.2353274 | 0.608121571 | 0.255136462 | 0.008962098 | 0.025439773 | 79072 | 5 | 7859272 | 7869150 | -1 | protein_coding | FAST kinase domains 3 |
| **PEPD** | ENSG00000124299 | 2086.506044 | -0.714907359 | 0.130400332 | 2E-08 | 1.63E-07 | 5184 | 19 | 33877856 | 34012700 | -1 | protein_coding | peptidase D |
| **IQSEC2** | ENSG00000124313 | 283.9677086 | 0.560878808 | 0.223904855 | 0.00716546 | 0.020929185 | 23096 | X | 53262058 | 53350522 | -1 | protein_coding | IQ motif and Sec7 domain 2 |
| **STAMBP** | ENSG00000124356 | 942.3810126 | 0.392697558 | 0.125030239 | 0.001297765 | 0.004598595 | 10617 | 2 | 74056086 | 74100786 | 1 | protein_coding | STAM binding protein |
| **NAGK** | ENSG00000124357 | 5222.043572 | 0.43370305 | 0.105680237 | 3E-05 | 1.51E-04 | 55577 | 2 | 71291474 | 71306935 | 1 | protein_coding | N-acetylglucosamine kinase |
| **SNRNP27** | ENSG00000124380 | 580.6028342 | 0.42790334 | 0.159068932 | 0.005335652 | 0.016144283 | 11017 | 2 | 70120692 | 70132707 | 1 | protein_coding | small nuclear ribonucleoprotein 27kDa (U4/U6.U5) |
| **MPHOSPH10** | ENSG00000124383 | 845.2774782 | 0.588450298 | 0.147731813 | 4.06E-05 | 1.99E-04 | 10199 | 2 | 71357444 | 71377231 | 1 | protein_coding | M-phase phosphoprotein 10 (U3 small nucleolar ribonucleoprotein) |
| **USP22** | ENSG00000124422 | 6309.671281 | -0.422541193 | 0.135812245 | 0.001421817 | 0.004991874 | 23326 | 17 | 20902910 | 20947073 | -1 | protein_coding | ubiquitin specific peptidase 22 |
| **USP9X** | ENSG00000124486 | 7204.005824 | -0.203558932 | 0.079546692 | 0.010366063 | 0.028851673 | 8239 | X | 40944888 | 41095832 | 1 | protein_coding | ubiquitin specific peptidase 9, X-linked |
| **TRERF1** | ENSG00000124496 | 1316.346251 | -0.953516816 | 0.119217143 | 4.06E-16 | 6.65E-15 | 55809 | 6 | 42192669 | 42419789 | -1 | protein_coding | transcriptional regulating factor 1 |
| **XPO5** | ENSG00000124571 | 1162.838112 | 0.447478515 | 0.132393747 | 5.38E-04 | 0.002086507 | 57510 | 6 | 43490072 | 43543812 | -1 | protein_coding | exportin 5 |
| **PEX6** | ENSG00000124587 | 535.1318228 | -0.597004124 | 0.157922764 | 9.21E-05 | 4.24E-04 | 5190 | 6 | 42931608 | 42946958 | -1 | protein_coding | peroxisomal biogenesis factor 6 |
| **OARD1** | ENSG00000124596 | 533.0981901 | 0.521455568 | 0.18524127 | 0.003159201 | 0.010171071 | 221443 | 6 | 41001366 | 41065526 | -1 | protein_coding | O-acyl-ADP-ribose deacylase 1 |
| **HIST1H2BJ** | ENSG00000124635 | 111.2978689 | 1.63388754 | 0.372255992 | 1.3E-06 | 8.21E-06 | 8970 | 6 | 27093676 | 27100541 | -1 | protein_coding | histone cluster 1, H2bj |
| **MAD2L1BP** | ENSG00000124688 | 480.8772085 | 0.521553653 | 0.173203167 | 0.001651301 | 0.005704528 | 9587 | 6 | 43597277 | 43608689 | 1 | protein_coding | MAD2L1 binding protein |
| **KLHDC3** | ENSG00000124702 | 1033.348345 | -0.701428478 | 0.131713846 | 4.93E-08 | 3.83E-07 | 116138 | 6 | 42981951 | 42989036 | 1 | protein_coding | kelch domain containing 3 |
| **ATXN1** | ENSG00000124788 | 2883.963864 | -0.413760038 | 0.103276822 | 5.4E-05 | 2.59E-04 | 6310 | 6 | 16299343 | 16761722 | -1 | protein_coding | ataxin 1 |
| **NUP153** | ENSG00000124789 | 2499.568587 | 0.764838754 | 0.099710076 | 9.5E-15 | 1.41E-13 | 9972 | 6 | 17615269 | 17706656 | -1 | protein_coding | nucleoporin 153kDa |
| **DEK** | ENSG00000124795 | 1573.582676 | 0.963129169 | 0.160330673 | 6.14E-10 | 6.02E-09 | 7913 | 6 | 18224099 | 18265054 | -1 | protein_coding | DEK oncogene |
| **RUNX2** | ENSG00000124813 | 192.784296 | 1.391640683 | 0.483479763 | 5.53E-04 | 0.002139685 | 860 | 6 | 45295894 | 45632086 | 1 | protein_coding | runt-related transcription factor 2 |
| **RAB17** | ENSG00000124839 | 33.16027734 | -1.492998492 | 0.629884629 | 0.002040513 | 0.006884142 | 64284 | 2 | 238482965 | 238510257 | -1 | protein_coding | RAB17, member RAS oncogene family |
| **EREG** | ENSG00000124882 | 153.196845 | 6.09760493 | 0.565916144 | 3.39E-28 | 1.19E-26 | 2069 | 4 | 75230860 | 75254468 | 1 | protein_coding | epiregulin |
| **AHNAK** | ENSG00000124942 | 42796.75229 | -0.944946363 | 0.096108304 | 1.93E-23 | 5.34E-22 | 79026 | 11 | 62201016 | 62323707 | -1 | protein_coding | AHNAK nucleoprotein |
| **SH3TC1** | ENSG00000125089 | 1096.25286 | -0.83304248 | 0.159507727 | 7.42E-08 | 5.61E-07 | 54436 | 4 | 8183799 | 8243530 | 1 | protein_coding | SH3 domain and tetratricopeptide repeats 1 |
| **CNOT1** | ENSG00000125107 | 8089.920192 | -0.399613884 | 0.113228018 | 3.32E-04 | 0.001357086 | 23019 | 16 | 58553855 | 58663790 | -1 | protein_coding | CCR4-NOT transcription complex, subunit 1 |
| **BBS2** | ENSG00000125124 | 1008.547587 | 0.392694556 | 0.155086915 | 0.00878335 | 0.024992727 | 583 | 16 | 56500748 | 56554195 | -1 | protein_coding | Bardet-Biedl syndrome 2 |
| **MT2A** | ENSG00000125148 | 2751.640078 | 0.850029405 | 0.17727437 | 5.91E-07 | 3.95E-06 | 4502 | 16 | 56642111 | 56643409 | 1 | protein_coding | metallothionein 2A |
| **C16orf70** | ENSG00000125149 | 1345.258483 | 0.456870098 | 0.126250753 | 2.21E-04 | 9.37E-04 | 80262 | 16 | 67143861 | 67182442 | 1 | protein_coding | chromosome 16 open reading frame 70 |
| **CLYBL** | ENSG00000125246 | 219.6505377 | -0.711116645 | 0.281705134 | 0.005174539 | 0.015728417 | 171425 | 13 | 100258919 | 100549387 | 1 | protein_coding | citrate lyase beta like |
| **RAP2A** | ENSG00000125249 | 2999.543981 | -0.526496757 | 0.16856163 | 0.001105159 | 0.003995775 | 5911 | 13 | 98086476 | 98121382 | 1 | protein_coding | RAP2A, member of RAS oncogene family |
| **TM9SF2** | ENSG00000125304 | 9010.255436 | 0.332559038 | 0.128638506 | 0.007753153 | 0.022420327 | 9375 | 13 | 100153671 | 100216260 | 1 | protein_coding | transmembrane 9 superfamily member 2 |
| **IRF1** | ENSG00000125347 | 6209.862253 | 1.812275527 | 0.368183289 | 8.74E-08 | 6.52E-07 | 3659 | 5 | 131817301 | 131826490 | -1 | protein_coding | interferon regulatory factor 1 |
| **UPF3B** | ENSG00000125351 | 540.0719025 | 0.744206165 | 0.190101448 | 3.94E-05 | 1.94E-04 | 65109 | X | 118967985 | 118986961 | -1 | protein_coding | UPF3 regulator of nonsense transcripts homolog B (yeast) |
| **TMEM255A** | ENSG00000125355 | 430.9687916 | 1.612420184 | 0.192573416 | 7.44E-18 | 1.39E-16 | 55026 | X | 119392505 | 119445411 | -1 | protein_coding | transmembrane protein 255A |
| **PTGER2** | ENSG00000125384 | 326.5548726 | 0.990008564 | 0.381077584 | 0.002374495 | 0.007863361 | 5732 | 14 | 52781023 | 52795324 | 1 | protein_coding | prostaglandin E receptor 2 (subtype EP2), 53kDa |
| **HS3ST3B1** | ENSG00000125430 | 415.0933183 | 3.128769138 | 0.296972722 | 2.74E-27 | 9.13E-26 | 9953 | 17 | 14204400 | 14252721 | 1 | protein_coding | heparan sulfate (glucosamine) 3-O-sulfotransferase 3B1 |
| **MRPS7** | ENSG00000125445 | 1073.501003 | -0.545478821 | 0.133802302 | 2.89E-05 | 1.46E-04 | 51081 | 17 | 73257755 | 73262454 | 1 | protein_coding | mitochondrial ribosomal protein S7 |
| **ARMC7** | ENSG00000125449 | 316.4633994 | -0.667610035 | 0.222232819 | 0.001361528 | 0.004792314 | 79637 | 17 | 73106047 | 73126360 | 1 | protein_coding | armadillo repeat containing 7 |
| **SLC25A19** | ENSG00000125454 | 726.6353854 | -0.681500638 | 0.175469753 | 5.18E-05 | 2.49E-04 | 60386 | 17 | 73269073 | 73285591 | -1 | protein_coding | solute carrier family 25 (mitochondrial thiamine pyrophosphate carrier), member 19 |
| **MIF4GD** | ENSG00000125457 | 409.4209837 | -0.953273816 | 0.19857369 | 4.94E-07 | 3.35E-06 | 57409 | 17 | 73262309 | 73267308 | -1 | protein_coding | MIF4G domain containing |
| **MSTO1** | ENSG00000125459 | 278.0329917 | -0.566486214 | 0.212877621 | 0.004508689 | 0.013923277 | 55154 | 1 | 155579979 | 155718153 | 1 | protein_coding | misato 1, mitochondrial distribution and morphology regulator |
| **GTF3C4** | ENSG00000125484 | 985.0932094 | 0.415867138 | 0.119951874 | 4.05E-04 | 0.001620902 | 9329 | 9 | 135545422 | 135570342 | 1 | protein_coding | general transcription factor IIIC, polypeptide 4, 90kDa |
| **PPP1R12C** | ENSG00000125503 | 1484.727263 | -0.525973814 | 0.164837091 | 9.1E-04 | 0.003354366 | 54776 | 19 | 55602281 | 55628927 | -1 | protein_coding | protein phosphatase 1, regulatory subunit 12C |
| **MBOAT7** | ENSG00000125505 | 1394.677618 | -0.349637537 | 0.141090449 | 0.010979691 | 0.03032212 | 79143 | 19 | 54677107 | 54693733 | -1 | protein_coding | membrane bound O-acyltransferase domain containing 7 |
| **OPRL1** | ENSG00000125510 | 91.72588214 | -1.712047519 | 0.438395824 | 1.04E-05 | 5.74E-05 | 4987 | 20 | 62711526 | 62731996 | 1 | protein_coding | opiate receptor-like 1 |
| **SLC2A4RG** | ENSG00000125520 | 139.6733144 | -0.623831983 | 0.282298373 | 0.013869658 | 0.037122589 | 56731 | 20 | 62371214 | 62374858 | 1 | protein_coding | SLC2A4 regulator |
| **PPDPF** | ENSG00000125534 | 688.724016 | -0.610436755 | 0.16722909 | 1.48E-04 | 6.54E-04 | 79144 | 20 | 62152077 | 62153559 | 1 | protein_coding | pancreatic progenitor cell differentiation and proliferation factor |
| **IL1B** | ENSG00000125538 | 1784.226271 | 1.565616727 | 0.640831487 | 0.001484866 | 0.00518939 | 3553 | 2 | 113587328 | 113594480 | -1 | protein_coding | interleukin 1, beta |
| **PAX8** | ENSG00000125618 | 390.5679245 | 0.590149346 | 0.21662652 | 0.003696779 | 0.011680353 | 7849 | 2 | 113973574 | 114036527 | -1 | protein_coding | paired box 8 |
| **INSIG2** | ENSG00000125629 | 1270.034701 | 1.445160173 | 0.134729262 | 1.26E-27 | 4.28E-26 | 51141 | 2 | 118846028 | 118868573 | 1 | protein_coding | insulin induced gene 2 |
| **SLC25A23** | ENSG00000125648 | 911.8187134 | -0.954537485 | 0.240801329 | 2.2E-05 | 1.14E-04 | 79085 | 19 | 6436090 | 6465214 | -1 | protein_coding | solute carrier family 25 (mitochondrial carrier; phosphate carrier), member 23 |
| **ALKBH7** | ENSG00000125652 | 146.4181717 | -1.13903053 | 0.387873151 | 6.58E-04 | 0.002502572 | 84266 | 19 | 6372444 | 6375042 | 1 | protein_coding | alkB, alkylation repair homolog 7 (E. coli) |
| **TNFSF9** | ENSG00000125657 | 129.2380103 | 2.0478465 | 0.33021742 | 5.72E-11 | 6.3E-10 | 8744 | 19 | 6531010 | 6535931 | 1 | protein_coding | tumor necrosis factor (ligand) superfamily, member 9 |
| **THOC2** | ENSG00000125676 | 2385.246394 | 0.392758682 | 0.105770192 | 1.48E-04 | 6.55E-04 | 57187 | X | 122734412 | 122866906 | -1 | protein_coding | THO complex 2 |
| **RPL23** | ENSG00000125691 | 6463.647568 | -0.53485311 | 0.14133948 | 9.51E-05 | 4.37E-04 | 619505 | 17 | 37004118 | 37010096 | -1 | protein_coding | ribosomal protein L23 |
| **CD70** | ENSG00000125726 | 46.04014598 | 3.061610803 | 0.553003636 | 1.96E-09 | 1.82E-08 | 970 | 19 | 6583194 | 6604114 | -1 | protein_coding | CD70 molecule |
| **TRIP10** | ENSG00000125733 | 832.5378634 | 1.047030578 | 0.155156483 | 4.21E-12 | 5.13E-11 | 9322 | 19 | 6737936 | 6751537 | 1 | protein_coding | thyroid hormone receptor interactor 10 |
| **GPR108** | ENSG00000125734 | 1858.116415 | -0.387053582 | 0.144252908 | 0.005136109 | 0.015639599 | 1024654 | 19 | 6729925 | 6737614 | -1 | protein_coding | G protein-coupled receptor 108 |
| **TNFSF14** | ENSG00000125735 | 2116.636858 | -0.905091863 | 0.162901062 | 9.77E-09 | 8.33E-08 | 8740 | 19 | 6663148 | 6670599 | -1 | protein_coding | tumor necrosis factor (ligand) superfamily, member 14 |
| **RTN2** | ENSG00000125744 | 321.0374302 | -0.865349067 | 0.194762384 | 3.27E-06 | 1.95E-05 | 6253 | 19 | 45988547 | 46000319 | -1 | protein_coding | reticulon 2 |
| **VASP** | ENSG00000125753 | 3715.755256 | -0.315265535 | 0.103905342 | 0.002135438 | 0.007164851 | 7408 | 19 | 46009837 | 46030241 | 1 | protein_coding | vasodilator-stimulated phosphoprotein |
| **PANK2** | ENSG00000125779 | 1349.299346 | 0.998813704 | 0.123376949 | 1.45E-16 | 2.46E-15 | 80025 | 20 | 3869486 | 3907605 | 1 | protein_coding | pantothenate kinase 2 |
| **CD93** | ENSG00000125810 | 27.09674007 | -3.14157014 | 0.990451929 | 1.14E-04 | 5.17E-04 | 22918 | 20 | 23059986 | 23066977 | -1 | protein_coding | CD93 molecule |
| **PSMF1** | ENSG00000125818 | 2388.752604 | 0.482557309 | 0.133515417 | 2.17E-04 | 9.17E-04 | 9491 | 20 | 1093906 | 1160596 | 1 | protein_coding | proteasome (prosome, macropain) inhibitor subunit 1 (PI31) |
| **DTD1** | ENSG00000125821 | 345.9262807 | -0.558594056 | 0.199460501 | 0.00308264 | 0.009947634 | 92675 | 20 | 18568537 | 18744561 | 1 | protein_coding | D-tyrosyl-tRNA deacylase 1 |
| **RBCK1** | ENSG00000125826 | 1086.010929 | 1.027544238 | 0.160414932 | 4.34E-11 | 4.83E-10 | 10616 | 20 | 388142 | 411610 | 1 | protein_coding | RanBP-type and C3HC4-type zinc finger containing 1 |
| **STK35** | ENSG00000125834 | 5.181538275 | 0.364559148 | 0.786991009 | 0.015484502 | 0.040890004 | 140901 | 20 | 2082257 | 2157684 | 1 | protein_coding | serine/threonine kinase 35 |
| **SNRPB2** | ENSG00000125870 | 1081.679986 | 0.499295135 | 0.131127 | 9.66E-05 | 4.43E-04 | 6629 | 20 | 16710606 | 16722421 | 1 | protein_coding | small nuclear ribonucleoprotein polypeptide B |
| **MGME1** | ENSG00000125871 | 1031.226819 | 0.355786012 | 0.147747068 | 0.012835005 | 0.034681292 | 92667 | 20 | 17949556 | 17971765 | 1 | protein_coding | mitochondrial genome maintenance exonuclease 1 |
| **TBC1D20** | ENSG00000125875 | 1593.953358 | -0.406264054 | 0.107469459 | 1.25E-04 | 5.61E-04 | 128637 | 20 | 416124 | 443197 | -1 | protein_coding | TBC1 domain family, member 20 |
| **MCM8** | ENSG00000125885 | 162.0548767 | 0.691722002 | 0.263324249 | 0.003989854 | 0.012492715 | 84515 | 20 | 5931298 | 5975852 | 1 | protein_coding | minichromosome maintenance complex component 8 |
| **SIRPD** | ENSG00000125900 | 120.6421129 | -0.854263769 | 0.324245749 | 0.002829808 | 0.009195823 | 128646 | 20 | 1514897 | 1539489 | -1 | protein_coding | signal-regulatory protein delta |
| **NCLN** | ENSG00000125912 | 1052.508205 | -1.052135059 | 0.130152202 | 2E-16 | 3.34E-15 | 56926 | 19 | 3185561 | 3209573 | 1 | protein_coding | nicalin |
| **HNRNPR** | ENSG00000125944 | 2591.443266 | 0.480362481 | 0.126345769 | 8.78E-05 | 4.06E-04 | 10236 | 1 | 23630264 | 23670829 | -1 | protein_coding | heterogeneous nuclear ribonucleoprotein R |
| **ZNF436** | ENSG00000125945 | 508.6608384 | 0.700596085 | 0.157415222 | 4.25E-06 | 2.49E-05 | 80818 | 1 | 23685941 | 23695935 | -1 | protein_coding | zinc finger protein 436 |
| **MAX** | ENSG00000125952 | 2923.019644 | 0.739209397 | 0.102314249 | 2.51E-13 | 3.33E-12 | 4149 | 14 | 65472892 | 65569413 | -1 | protein_coding | MYC associated factor X |
| **NECAB3** | ENSG00000125967 | 67.72551548 | -0.905685855 | 0.480399799 | 0.015155472 | 0.040114493 | 63941 | 20 | 32244893 | 32262269 | -1 | protein_coding | N-terminal EF-hand calcium binding protein 3 |
| **EIF2S2** | ENSG00000125977 | 4561.328648 | 0.561634881 | 0.09119298 | 4.32E-10 | 4.3E-09 | 8894 | 20 | 32676104 | 32700138 | -1 | protein_coding | eukaryotic translation initiation factor 2, subunit 2 beta, 38kDa |
| **ERGIC3** | ENSG00000125991 | 1660.207151 | -0.48370925 | 0.120174454 | 3.66E-05 | 1.81E-04 | 51614 | 20 | 34129770 | 34145405 | 1 | protein_coding | ERGIC and golgi 3 |
| **PLAGL2** | ENSG00000126003 | 1082.82769 | 0.938227823 | 0.170429429 | 1.19E-08 | 1E-07 | 5326 | 20 | 30780306 | 30795594 | -1 | protein_coding | pleiomorphic adenoma gene-like 2 |

| | | | | | | | | | | | | | |
|---|---|---|---|---|---|---|---|---|---|---|---|---|---|
| **PSMB2** | ENSG00000126067 | 2564.543798 | 0.66596111 | 0.111402657 | 1.3E-09 | 1.24E-08 | 5690 | 1 | 36067185 | 36107445 | -1 | protein_coding | proteasome (prosome, macropain) subunit, beta type, 2 |
| **UROD** | ENSG00000126088 | 944.4460067 | -0.562637389 | 0.143820992 | 5.55E-05 | 2.66E-04 | 7389 | 1 | 45477819 | 45481247 | 1 | protein_coding | uroporphyrinogen decarboxylase |
| **FFAR2** | ENSG00000126262 | 26.41069837 | 4.098377909 | 0.824295734 | 3.53E-08 | 2.79E-07 | 2867 | 19 | 35934809 | 35942669 | 1 | protein_coding | free fatty acid receptor 2 |
| **HCST** | ENSG00000126264 | 1225.781865 | -0.876813129 | 0.152286022 | 3.08E-09 | 2.78E-08 | 10870 | 19 | 36393382 | 36395279 | 1 | protein_coding | hematopoietic cell signal transducer |
| **COX6B1** | ENSG00000126267 | 48.65710165 | -1.940171283 | 0.54358084 | 3.4E-05 | 1.7E-04 | 1340 | 19 | 36139125 | 36149763 | 1 | protein_coding | cytochrome c oxidase subunit VIb polypeptide 1 (ubiquitous) |
| **CCR7** | ENSG00000126353 | 1109.508916 | 3.394575612 | 0.196372861 | 3.99E-68 | 7.18E-66 | 1236 | 17 | 38710021 | 38721724 | -1 | protein_coding | chemokine (C-C motif) receptor 7 |
| **ASL** | ENSG00000126522 | 235.2129748 | -0.678982161 | 0.246718102 | 0.002865197 | 0.009302912 | 435 | 7 | 65540785 | 65558545 | 1 | protein_coding | argininosuccinate lyase |
| **SBDS** | ENSG00000126524 | 1070.396497 | 0.679448854 | 0.123430422 | 1.86E-08 | 1.53E-07 | 51119 | 7 | 66452664 | 66460588 | -1 | protein_coding | Shwachman-Bodian-Diamond syndrome |
| **STAT5A** | ENSG00000126561 | 1839.638697 | 0.883841516 | 0.12675174 | 8.21E-13 | 1.06E-11 | 6776 | 17 | 40439565 | 40463961 | 1 | protein_coding | signal transducer and activator of transcription 5A |
| **TRAP1** | ENSG00000126602 | 970.0672069 | -0.492572599 | 0.158621123 | 0.001305884 | 0.004622451 | 10131 | 16 | 3701640 | 3767598 | -1 | protein_coding | TNF receptor-associated protein 1 |
| **NSRP1** | ENSG00000126653 | 1009.155988 | 0.441471613 | 0.157527141 | 0.00383789 | 0.012056418 | 84081 | 17 | 28442539 | 28513493 | 1 | protein_coding | nuclear speckle splicing regulatory protein 1 |
| **AHDC1** | ENSG00000126705 | 397.968347 | -2.184414845 | 0.197647749 | 2.08E-29 | 7.78E-28 | 27245 | 1 | 27860546 | 27930942 | -1 | protein_coding | AT hook, DNA binding motif, containing 1 |
| **IFI6** | ENSG00000126709 | 7699.553854 | 3.947099097 | 0.159630817 | 5.57E-136 | 2.66E-133 | 2537 | 1 | 27992572 | 27998729 | -1 | protein_coding | interferon, alpha-inducible protein 6 |
| **PCNXL4** | ENSG00000126773 | 653.0281329 | 0.681273211 | 0.203131881 | 4.08E-04 | 0.001632421 | 64430 | 14 | 60558629 | 60635851 | 1 | protein_coding | pecanex-like 4 (Drosophila) |
| **KTN1** | ENSG00000126777 | 6109.693349 | 0.420714308 | 0.109962467 | 1.03E-04 | 4.68E-04 | 3895 | 14 | 56025790 | 56168244 | 1 | protein_coding | kinectin 1 (kinesin receptor) |
| **L3HYPDH** | ENSG00000126790 | 123.0856777 | 1.052703673 | 0.32335195 | 2.77E-04 | 0.00114682 | 112849 | 14 | 59927081 | 59951148 | -1 | protein_coding | L-3-hydroxyproline dehydratase (trans-) |
| **HSPA2** | ENSG00000126803 | 33.62619253 | 1.000740004 | 0.553995634 | 0.014511406 | 0.038596906 | 3306 | 14 | 65002623 | 65009955 | 1 | protein_coding | heat shock 70kDa protein 2 |
| **ZBTB1** | ENSG00000126804 | 2316.215923 | 0.797679344 | 0.141731626 | 7.39E-09 | 6.39E-08 | 22890 | 14 | 64970430 | 65000408 | 1 | protein_coding | zinc finger and BTB domain containing 1 |
| **SGPP1** | ENSG00000126821 | 1390.574459 | 1.828047479 | 0.162886678 | 3.28E-30 | 1.28E-28 | 81537 | 14 | 64150932 | 64194757 | -1 | protein_coding | sphingosine-1-phosphate phosphatase 1 |
| **PLEKHG3** | ENSG00000126822 | 539.1028018 | 2.769731402 | 0.333832591 | 6.21E-18 | 1.17E-16 | 26030 | 14 | 65170820 | 65213610 | 1 | protein_coding | pleckstrin homology domain containing, family G (with RhoGef domain) member 3 |
| **RHOT1** | ENSG00000126858 | 1236.296346 | 0.615648578 | 0.12181175 | 2.69E-07 | 1.88E-06 | 55288 | 17 | 30469473 | 30580393 | 1 | protein_coding | ras homolog family member T1 |
| **OMG** | ENSG00000126861 | 134.5880727 | -1.605125268 | 0.399349637 | 7.19E-06 | 4.05E-05 | 1019270 | 17 | 29599031 | 29624557 | -1 | protein_coding | oligodendrocyte myelin glycoprotein |
| **WDR60** | ENSG00000126870 | 328.8369435 | -0.643151215 | 0.250374814 | 0.005032757 | 0.015352421 | 55112 | 7 | 158649269 | 158749438 | 1 | protein_coding | WD repeat domain 60 |
| **FAM78A** | ENSG00000126882 | 2348.478543 | -0.532307321 | 0.109004533 | 6.34E-07 | 4.22E-06 | 1019271 | 9 | 134133463 | 134151934 | -1 | protein_coding | family with sequence similarity 78, member A |
| **ARMCX1** | ENSG00000126947 | 692.8118904 | 1.052307995 | 0.198936902 | 3.14E-08 | 2.51E-07 | 51309 | X | 100805514 | 100809683 | 1 | protein_coding | armadillo repeat containing, X-linked 1 |
| **TIMM8A** | ENSG00000126953 | 129.2156447 | -0.652851754 | 0.305142969 | 0.015523082 | 0.040966889 | 1678 | X | 100600649 | 100604184 | -1 | protein_coding | translocase of inner mitochondrial membrane 8 homolog A (yeast) |
| **HIVEP3** | ENSG00000127124 | 781.3325338 | -1.362142151 | 0.195403294 | 7.26E-13 | 9.38E-12 | 59269 | 1 | 41972036 | 42501596 | -1 | protein_coding | human immunodeficiency virus type I enhancer binding protein 3 |
| **PPCS** | ENSG00000127125 | 1266.043559 | 0.443460884 | 0.125880669 | 3.1E-04 | 0.00127336 | 79717 | 1 | 42921788 | 42939056 | 1 | protein_coding | phosphopantothenoylcysteine synthetase |
| **TRAF2** | ENSG00000127191 | 474.3358157 | 0.712810266 | 0.169237511 | 1.21E-05 | 6.61E-05 | 7186 | 9 | 139776364 | 139821059 | 1 | protein_coding | TNF receptor-associated factor 2 |
| **ABHD8** | ENSG00000127220 | 377.549832 | -0.594825492 | 0.200424705 | 0.001725687 | 0.005917993 | 79575 | 19 | 17402940 | 17421045 | -1 | protein_coding | abhydrolase domain containing 8 |
| **HELB** | ENSG00000127311 | 443.5204177 | 2.615505312 | 0.216053085 | 8.01E-35 | 3.96E-33 | 92797 | 12 | 66696325 | 66737423 | 1 | protein_coding | helicase (DNA) B |
| **RAP1B** | ENSG00000127314 | 11596.36488 | 1.025682199 | 0.109370187 | 2.09E-21 | 5.08E-20 | 5908 | 12 | 69004619 | 69054372 | 1 | protein_coding | RAP1B, member of RAS oncogene family |
| **DYRK2** | ENSG00000127334 | 3884.334086 | -0.513790464 | 0.129711733 | 4.04E-05 | 1.98E-04 | 8445 | 12 | 68042118 | 68059186 | 1 | protein_coding | dual-specificity tyrosine-(Y)-phosphorylation regulated kinase 2 |
| **YEATS4** | ENSG00000127337 | 238.6174693 | 0.596569372 | 0.286766082 | 0.0195979 | 0.049860948 | 8089 | 12 | 69753483 | 69784576 | 1 | protein_coding | YEATS domain containing 4 |
| **LRRC61** | ENSG00000127399 | 249.9668867 | 0.917022237 | 0.245615106 | 6.06E-05 | 2.88E-04 | 65999 | 7 | 150019728 | 150035239 | 1 | protein_coding | leucine rich repeat containing 61 |
| **FGFRL1** | ENSG00000127418 | 218.1954827 | -1.106276262 | 0.294677772 | 3.95E-05 | 1.94E-04 | 53834 | 4 | 1003724 | 1020685 | 1 | protein_coding | fibroblast growth factor receptor-like 1 |
| **PIN1** | ENSG00000127445 | 464.1309072 | -0.762757522 | 0.171104987 | 3.56E-06 | 2.1E-05 | 5300 | 19 | 9945933 | 9960358 | 1 | protein_coding | peptidylprolyl cis/trans isomerase, NIMA-interacting 1 |
| **EMC1** | ENSG00000127463 | 2367.716993 | -0.749729101 | 0.133045394 | 9.17E-09 | 7.85E-08 | 23065 | 1 | 19542158 | 19578046 | -1 | protein_coding | ER membrane protein complex subunit 1 |
| **UBR4** | ENSG00000127481 | 4662.707279 | 0.5991009 | 0.121509573 | 5.05E-07 | 3.41E-06 | 23352 | 1 | 19401000 | 19536770 | -1 | protein_coding | ubiquitin protein ligase E3 component n-recognin 4 |
| **HP1BP3** | ENSG00000127483 | 2906.781846 | -0.363731524 | 0.118630218 | 0.00179083 | 0.006121508 | 50809 | 1 | 21069154 | 21113816 | -1 | protein_coding | heterochromatin protein 1, binding protein 3 |
| **SIN3B** | ENSG00000127511 | 824.9122342 | -0.382395688 | 0.16913267 | 0.018554079 | 0.047601573 | 23309 | 19 | 16940211 | 16991164 | 1 | protein_coding | SIN3 transcription regulator family member B |
| **SLC35E1** | ENSG00000127526 | 2410.457712 | -0.875778456 | 0.116402514 | 1.98E-14 | 2.88E-13 | 79939 | 19 | 16660642 | 16683193 | -1 | protein_coding | solute carrier family 35, member E1 |
| **EPS15L1** | ENSG00000127527 | 805.590579 | -0.30970453 | 0.13631088 | 0.018943258 | 0.048437311 | 58513 | 19 | 16466050 | 16582896 | -1 | protein_coding | epidermal growth factor receptor pathway substrate 15-like 1 |
| **KLF2** | ENSG00000127528 | 117.3717856 | -3.300909029 | 0.53229233 | 4.07E-11 | 4.55E-10 | 10365 | 19 | 16435628 | 16438685 | 1 | protein_coding | Kruppel-like factor 2 |
| **SMARCA4** | ENSG00000127616 | 778.5017003 | -1.622556434 | 0.20841589 | 9.74E-16 | 1.56E-14 | 6597 | 19 | 11071598 | 11176071 | 1 | protein_coding | SWI/SNF related, matrix associated, actin dependent regulator of chromatin, subfamily a, member 4 |
| **KDM4B** | ENSG00000127663 | 839.9306189 | -0.892930224 | 0.167087871 | 3.26E-08 | 2.6E-07 | 23030 | 19 | 4969125 | 5153606 | 1 | protein_coding | lysine (K)-specific demethylase 4B |
| **TICAM1** | ENSG00000127666 | 715.9729234 | 0.559477601 | 0.147542531 | 9.38E-05 | 4.31E-04 | 148022 | 19 | 4815944 | 4831716 | -1 | protein_coding | toll-like receptor adaptor molecule 1 |
| **VIL1** | ENSG00000127831 | 17.34216076 | 2.969431584 | 1.025449328 | 1.92E-04 | 8.26E-04 | 7429 | 2 | 219283815 | 219318018 | 1 | protein_coding | villin 1 |
| **GNAI1** | ENSG00000127955 | 8.140577381 | -1.94623857 | 2.682170875 | 0.019631675 | 0.049913639 | 2770 | 7 | 79763271 | 79848718 | 1 | protein_coding | guanine nucleotide binding protein (G protein), alpha inhibiting activity polypeptide 1 |
| **MTERF** | ENSG00000127989 | 377.1739084 | 0.6752132 | 0.249771725 | 0.003313661 | 0.010601358 | 7978 | 7 | 91321323 | 91510034 | -1 | protein_coding | mitochondrial transcription termination factor |
| **RBM48** | ENSG00000127993 | 322.1548511 | 0.639928015 | 0.215465862 | 0.00155219 | 0.005401211 | 84060 | 7 | 92158087 | 92167319 | 1 | protein_coding | RNA binding motif protein 48 |
| **CASD1** | ENSG00000127995 | 386.0690237 | 0.51987836 | 0.221803375 | 0.011997775 | 0.032731523 | 64921 | 7 | 94138531 | 94186331 | 1 | protein_coding | CAS1 domain containing 1 |
| **ZNF780B** | ENSG00000128000 | 473.1989157 | -0.43411469 | 0.19261873 | 0.017276629 | 0.044874017 | 163131 | 19 | 40534167 | 40562116 | -1 | protein_coding | zinc finger protein 780B |
| **SRD5A3** | ENSG00000128039 | 1136.163459 | -0.777774596 | 0.177123879 | 4.8E-06 | 2.79E-05 | 79644 | 4 | 56212276 | 56239263 | 1 | protein_coding | steroid 5 alpha-reductase 3 |
| **RASL11B** | ENSG00000128045 | 330.2021307 | -5.19202171 | 0.619678659 | 3.73E-18 | 7.14E-17 | 65997 | 4 | 53728457 | 53733000 | 1 | protein_coding | RAS-like, family 11, member B |
| **KDR** | ENSG00000128052 | 94.05801303 | -1.762042106 | 0.391272653 | 8.45E-07 | 5.5E-06 | 3791 | 4 | 55944644 | 55991756 | -1 | protein_coding | kinase insert domain receptor (a type III receptor tyrosine kinase) |
| **DGCR6L** | ENSG00000128185 | 234.4817952 | -0.637130723 | 0.235295679 | 0.003479441 | 0.011069172 | 85359 | 22 | 20301799 | 20307603 | -1 | protein_coding | DiGeorge syndrome critical region gene 6-like |
| **ASPHD2** | ENSG00000128203 | 228.2936409 | 0.940547949 | 0.393813636 | 0.004673208 | 0.014384821 | 57168 | 22 | 26825239 | 26840981 | 1 | protein_coding | aspartate beta-hydroxylase domain containing 2 |
| **ADORA2A** | ENSG00000128271 | 209.2975405 | 2.556138328 | 1.063470837 | 7.04E-04 | 0.002659504 | 135 | 22 | 24813847 | 24838328 | 1 | protein_coding | adenosine A2a receptor |
| **A4GALT** | ENSG00000128274 | 38.15194578 | 3.286605049 | 0.656217519 | 3.43E-08 | 2.71E-07 | 53947 | 22 | 43088127 | 43117304 | -1 | protein_coding | alpha 1,4-galactosyltransferase |
| **APOL3** | ENSG00000128284 | 4249.948962 | 3.391078775 | 0.422103442 | 6.08E-17 | 1.06E-15 | 80833 | 22 | 36536372 | 36562225 | -1 | protein_coding | apolipoprotein L, 3 |
| **TPST2** | ENSG00000128294 | 1079.347991 | -1.084239239 | 0.156275854 | 1.12E-12 | 1.42E-11 | 8459 | 22 | 26921458 | 26992681 | -1 | protein_coding | tyrosylprotein sulfotransferase 2 |
| **APOL2** | ENSG00000128335 | 4827.808938 | 1.69776583 | 0.102779417 | 1.47E-62 | 2.1E-60 | 23780 | 22 | 36622256 | 36636000 | -1 | protein_coding | apolipoprotein L, 2 |
| **RAC2** | ENSG00000128340 | 11180.02061 | -1.051360226 | 0.089771835 | 3.01E-32 | 1.31E-30 | 5880 | 22 | 37621301 | 37640488 | -1 | protein_coding | ras-related C3 botulinum toxin substrate 2 (rho family, small GTP binding protein Rac2) |
| **LIF** | ENSG00000128342 | 297.4819555 | 1.039591785 | 0.395690066 | 0.001913285 | 0.006504228 | 3976 | 22 | 30636436 | 30642840 | -1 | protein_coding | leukemia inhibitory factor |
| **APOBEC3A** | ENSG00000128383 | 3640.86496 | 7.034791091 | 0.198309099 | 6.05E-277 | 1.32E-273 | 1009131 | 22 | 39348746 | 39359188 | 1 | protein_coding | apolipoprotein B mRNA editing enzyme, catalytic polypeptide-like 3A |
| **APOBEC3F** | ENSG00000128394 | 502.4249484 | 1.851407776 | 0.160928006 | 1.48E-31 | 6.21E-30 | 200316 | 22 | 39436609 | 39449915 | 1 | protein_coding | apolipoprotein B mRNA editing enzyme, catalytic polypeptide-like 3F |
| **DOCK4** | ENSG00000128512 | 3658.866828 | 0.966115146 | 0.134126008 | 2.47E-13 | 3.29E-12 | 9732 | 7 | 111366166 | 111846466 | -1 | protein_coding | dedicator of cytokinesis 4 |
| **POT1** | ENSG00000128513 | 483.5074349 | 0.486848052 | 0.206769999 | 0.012363626 | 0.033579758 | 25913 | 7 | 124462440 | 124570037 | -1 | protein_coding | protection of telomeres 1 |
| **ATP6V1F** | ENSG00000128524 | 3617.886723 | -0.41814526 | 0.10897867 | 8.83E-05 | 4.08E-04 | 1019271 | 7 | 128502880 | 128505898 | 1 | protein_coding | ATPase, H+ transporting, lysosomal 14kDa, V1 subunit F |
| **STRIP2** | ENSG00000128578 | 201.567555 | 1.740210671 | 0.272314737 | 2.06E-11 | 2.36E-10 | 57464 | 7 | 129074274 | 129128240 | 1 | protein_coding | striatin interacting protein 2 |
| **MKLN1** | ENSG00000128585 | 2314.373539 | 0.381018781 | 0.102893708 | 1.66E-04 | 7.24E-04 | 4289 | 7 | 130794855 | 131181395 | 1 | protein_coding | muskelin 1, intracellular mediator containing kelch motifs |
| **DNAJB9** | ENSG00000128590 | 1057.543129 | 0.873921722 | 0.174509495 | 1.97E-07 | 1.4E-06 | 4189 | 7 | 108210012 | 108215294 | 1 | protein_coding | DnaJ (Hsp40) homolog, subfamily B, member 9 |
| **LRRC4** | ENSG00000128594 | 48.05096145 | -4.413726768 | 1.090901076 | 5.47E-06 | 3.14E-05 | 64101 | 7 | 127667124 | 127672160 | -1 | protein_coding | leucine rich repeat containing 4 |
| **CALU** | ENSG00000128595 | 7680.308992 | -0.372547096 | 0.115758525 | 0.001082423 | 0.003922853 | 813 | 7 | 128379346 | 128411861 | 1 | protein_coding | calumenin |
| **MYO1B** | ENSG00000128641 | 363.9104431 | 1.076889378 | 0.312290828 | 1.28E-04 | 5.73E-04 | 4430 | 2 | 192109911 | 192290115 | 1 | protein_coding | myosin IB |
| **ORMDL1** | ENSG00000128699 | 1308.803071 | 0.371992648 | 0.115866367 | 0.001236298 | 0.00440325 | 94101 | 2 | 190635049 | 190649097 | -1 | protein_coding | ORM1-like 1 (S. cerevisiae) |
| **HERC2** | ENSG00000128731 | 2446.436775 | -0.566281445 | 0.170109749 | 5.28E-04 | 0.002054054 | 1019301 | 15 | 28356186 | 28567298 | -1 | protein_coding | HECT and RLD domain containing E3 ubiquitin protein ligase 2 |
| **SNRPN** | ENSG00000128739 | 190.0159617 | -0.782291173 | 0.283635409 | 0.002223983 | 0.007417968 | 8926 | 15 | 25068794 | 25223870 | 1 | protein_coding | small nuclear ribonucleoprotein polypeptide N |
| **PSMG2** | ENSG00000128789 | 780.6704556 | 0.604904617 | 0.14887706 | 2.65E-05 | 1.35E-04 | 56984 | 18 | 12658737 | 12725739 | 1 | protein_coding | proteasome (prosome, macropain) assembly chaperone 2 |
| **TWSG1** | ENSG00000128791 | 1232.615875 | -0.694866281 | 0.135936126 | 1.51E-07 | 1.09E-06 | 57045 | 18 | 9334765 | 9402418 | 1 | protein_coding | twisted gastrulation BMP signaling modulator 1 |
| **WDFY4** | ENSG00000128815 | 6686.399609 | -0.559630264 | 0.10417502 | 3.19E-08 | 2.55E-07 | 57705 | 10 | 49892921 | 50191001 | 1 | protein_coding | WDFY family member 4 |
| **CGNL1** | ENSG00000128849 | 6.158574236 | 0.684835319 | 1.230662603 | 0.014123798 | 0.037703757 | 1019303 | 15 | 57668165 | 57842925 | 1 | protein_coding | cingulin-like 1 |
| **TMOD2** | ENSG00000128872 | 400.8452605 | 0.842255788 | 0.242297106 | 1.79E-04 | 7.76E-04 | 29767 | 15 | 52043758 | 52108565 | 1 | protein_coding | tropomodulin 2 (neuronal) |
| **C15orf57** | ENSG00000128891 | 782.978298 | 0.346626777 | 0.148860043 | 0.016335762 | 0.04280121 | 90416 | 15 | 40820882 | 40857256 | -1 | protein_coding | chromosome 15 open reading frame 57 |
| **ALDH1A2** | ENSG00000128918 | 2030.841502 | -1.41731876 | 0.229412101 | 9.69E-11 | 1.04E-09 | 1019286 | 15 | 58245622 | 58790065 | -1 | protein_coding | aldehyde dehydrogenase 1 family, member A2 |
| **IVD** | ENSG00000128928 | 1299.908519 | -1.167500386 | 0.174608981 | 5.46E-12 | 6.59E-11 | 3712 | 15 | 40697686 | 40728146 | 1 | protein_coding | isovaleryl-CoA dehydrogenase |
| **CLN6** | ENSG00000128973 | 776.2471256 | -1.258114222 | 0.194787619 | 2.31E-11 | 2.64E-10 | 54982 | 15 | 68499330 | 68549549 | -1 | protein_coding | ceroid-lipofuscinosis, neuronal 6, late infantile, variant |
| **ANAPC13** | ENSG00000129055 | 489.5490037 | 0.754781649 | 0.182129886 | 1.51E-05 | 8.07E-05 | 25847 | 3 | 134196548 | 134205558 | -1 | protein_coding | anaphase promoting complex subunit 13 |

| | | | | | | | | | | | | | |
|---|---|---|---|---|---|---|---|---|---|---|---|---|---|
| **MBD4** | ENSG00000129071 | 2470.88514 | 0.391341032 | 0.103804202 | 1.35E-04 | 6.01E-04 | 8930 | 3 | 129149787 | 129158878 | -1 | protein_coding | methyl-CpG binding domain protein 4 |
| **COPB1** | ENSG00000129083 | 4970.75474 | 0.291589615 | 0.118276548 | 0.012027117 | 0.032805717 | 1315 | 11 | 14464986 | 14521573 | -1 | protein_coding | coatomer protein complex, subunit beta 1 |
| **PSMA1** | ENSG00000129084 | 884.267958 | 0.847367878 | 0.136424601 | 2.03E-10 | 2.08E-09 | 5682 | 11 | 14515329 | 14665181 | -1 | protein_coding | proteasome (prosome, macropain) subunit, alpha type, 1 |
| **PALLD** | ENSG00000129116 | 2825.003608 | -0.638218567 | 0.202147567 | 9.39E-04 | 0.003448835 | 23022 | 4 | 169418217 | 169849608 | 1 | protein_coding | palladin, cytoskeletal associated protein |
| **SERGEF** | ENSG00000129158 | 283.1599621 | -0.894293527 | 0.232009359 | 3.92E-05 | 1.93E-04 | 26297 | 11 | 17809595 | 18034709 | -1 | protein_coding | secretion regulating guanine nucleotide exchange factor |
| **DCTD** | ENSG00000129187 | 1366.944045 | 0.339440264 | 0.139356268 | 0.012725137 | 0.034427027 | 1635 | 4 | 183811213 | 183839089 | -1 | protein_coding | dCMP deaminase |
| **KIF1C** | ENSG00000129250 | 3071.327308 | -0.553159213 | 0.121244862 | 3.14E-06 | 1.88E-05 | 10749 | 17 | 4901243 | 4931696 | 1 | protein_coding | kinesin family member 1C |
| **LRRC6** | ENSG00000129295 | 191.4028291 | -0.586104249 | 0.252928678 | 0.0110365 | 0.030467996 | 23639 | 8 | 133584320 | 133687838 | -1 | protein_coding | leucine rich repeat containing 6 |
| **PUS7L** | ENSG00000129317 | 360.0258394 | 0.845917217 | 0.243815994 | 1.83E-04 | 7.92E-04 | 83448 | 12 | 44122410 | 44152620 | -1 | protein_coding | pseudouridylate synthase 7 homolog (S. cerevisiae)-like |
| **SLC44A2** | ENSG00000129353 | 2690.447542 | -0.742990378 | 0.112893102 | 2.03E-11 | 2.33E-10 | 57153 | 19 | 10713133 | 10755235 | 1 | protein_coding | solute carrier family 44 (choline transporter), member 2 |
| **SIGLEC9** | ENSG00000129450 | 1367.88479 | -0.964721433 | 0.139935283 | 1.74E-12 | 2.16E-11 | 27180 | 19 | 51628165 | 51639908 | 1 | protein_coding | sialic acid binding Ig-like lectin 9 |
| **KLK10** | ENSG00000129451 | 115.4837944 | 2.981871308 | 0.342663294 | 2.41E-19 | 5.05E-18 | 5655 | 19 | 51515995 | 51523431 | -1 | protein_coding | kallikrein-related peptidase 10 |
| **NGDN** | ENSG00000129460 | 376.2756518 | 0.464812352 | 0.200166582 | 0.014138043 | 0.037735193 | 25983 | 14 | 23938897 | 23979071 | 1 | protein_coding | neuroguidin, EIF4E binding protein |
| **RIPK3** | ENSG00000129465 | 697.976009 | 0.649208512 | 0.157716781 | 2.25E-05 | 1.16E-04 | 11035 | 14 | 24805227 | 24809251 | -1 | protein_coding | receptor-interacting serine-threonine kinase 3 |
| **RAB2B** | ENSG00000129472 | 973.7856489 | 0.613673603 | 0.152402224 | 3.16E-05 | 1.59E-04 | 84932 | 14 | 21927179 | 21945132 | -1 | protein_coding | RAB2B, member RAS oncogene family |
| **BCL2L2** | ENSG00000129473 | 771.0491834 | 0.429165488 | 0.15995644 | 0.005617937 | 0.016871474 | 599 | 14 | 23767999 | 23780968 | 1 | protein_coding | BCL2-like 2 |
| **AJUBA** | ENSG00000129474 | 9.730379438 | -1.45206631 | 1.813212092 | 0.018735689 | 0.048027095 | 84962 | 14 | 23440383 | 23451851 | -1 | protein_coding | ajuba LIM protein |
| **HEATR5A** | ENSG00000129493 | 2236.730777 | -0.495190396 | 0.146134787 | 4.48E-04 | 0.001772958 | 25938 | 14 | 31760994 | 31889788 | -1 | protein_coding | HEAT repeat containing 5A |
| **SNX6** | ENSG00000129515 | 2976.154154 | 0.976383925 | 0.125300422 | 1.98E-15 | 3.08E-14 | 58533 | 14 | 35030300 | 35099389 | -1 | protein_coding | sorting nexin 6 |
| **EAPP** | ENSG00000129518 | 558.1403389 | 0.554491069 | 0.162275675 | 3.93E-04 | 0.001578836 | 55837 | 14 | 34985135 | 35008916 | -1 | protein_coding | E2F-associated phosphoprotein |
| **EGLN3** | ENSG00000129521 | 1253.444957 | -3.063240461 | 0.211847192 | 1.07E-48 | 8.94E-47 | 112399 | 14 | 34393437 | 34931980 | -1 | protein_coding | egl-9 family hypoxia-inducible factor 3 |
| **MIS18BP1** | ENSG00000129534 | 1550.694211 | 0.610627791 | 0.136599395 | 4.49E-06 | 2.62E-05 | 55320 | 14 | 45672393 | 45722743 | -1 | protein_coding | MIS18 binding protein 1 |
| **RNASE1** | ENSG00000129538 | 309.4227762 | -1.54671388 | 0.282049709 | 5.78E-09 | 5.07E-08 | 6035 | 14 | 21269387 | 21271437 | -1 | protein_coding | ribonuclease, RNase A family, 1 (pancreatic) |
| **DAD1** | ENSG00000129562 | 1611.701309 | -0.378813998 | 0.116557926 | 9.72E-04 | 0.003553738 | 1603 | 14 | 23033805 | 23058175 | -1 | protein_coding | defender against cell death 1 |
| **EPB41L4A** | ENSG00000129595 | 131.1117199 | 1.702101854 | 0.636047924 | 6.09E-04 | 0.002331307 | 64097 | 5 | 111478138 | 111755013 | -1 | protein_coding | erythrocyte membrane protein band 4.1 like 4A |
| **ITFG1** | ENSG00000129636 | 2539.856104 | 0.383354321 | 0.144659142 | 0.006357655 | 0.018852161 | 81533 | 16 | 47188298 | 47498060 | -1 | protein_coding | integrin alpha FG-GAP repeat containing 1 |
| **SEC14L1** | ENSG00000129657 | 3015.644308 | -0.386456425 | 0.16731707 | 0.016339792 | 0.042801644 | 6397 | 17 | 75082798 | 75213179 | 1 | protein_coding | SEC14-like 1 (S. cerevisiae) |
| **ARHGEF6** | ENSG00000129675 | 2894.260503 | -0.433371213 | 0.099476071 | 1.27E-05 | 6.88E-05 | 9459 | X | 135747706 | 135864247 | -1 | protein_coding | Rac/Cdc42 guanine nucleotide exchange factor (GEF) 6 |
| **ASH2L** | ENSG00000129691 | 674.4723449 | 0.603724949 | 0.165327616 | 1.5E-04 | 6.61E-04 | 9070 | 8 | 37962760 | 38001594 | 1 | protein_coding | ash2 (absent, small, or homeotic)-like (Drosophila) |
| **KLF16** | ENSG00000129911 | 604.6528366 | -0.618562299 | 0.159285912 | 5.79E-05 | 2.77E-04 | 83855 | 19 | 1852399 | 1863567 | -1 | protein_coding | Kruppel-like factor 16 |
| **TMEM8A** | ENSG00000129925 | 1395.455392 | -0.568011714 | 0.143005338 | 4.46E-05 | 2.17E-04 | 58986 | 16 | 420773 | 437113 | -1 | protein_coding | transmembrane protein 8A |
| **ABHD17A** | ENSG00000129968 | 1042.07235 | -0.537244462 | 0.166415688 | 7.31E-04 | 0.002753925 | 81926 | 19 | 1876809 | 1885546 | -1 | protein_coding | abhydrolase domain containing 17A |
| **CBFA2T3** | ENSG00000129993 | 151.4654426 | -2.137919279 | 0.454593093 | 2.37E-07 | 1.68E-06 | 863 | 16 | 88941266 | 89043612 | -1 | protein_coding | core-binding factor, runt domain, alpha subunit 2; translocated to, 3 |
| **SAT1** | ENSG00000130066 | 25610.82146 | 1.68601949 | 0.096499395 | 3.27E-69 | 6.01E-67 | 6303 | X | 23801290 | 23804343 | 1 | protein_coding | spermidine/spermine N1-acetyltransferase 1 |
| **GNL3L** | ENSG00000130119 | 637.8993135 | -0.685510837 | 0.158868837 | 7.75E-06 | 4.34E-05 | 54552 | X | 54556644 | 54587504 | 1 | protein_coding | guanine nucleotide binding protein-like 3 (nucleolar)-like |
| **DOCK6** | ENSG00000130158 | 418.9742637 | -1.333393201 | 0.250201102 | 2.01E-08 | 1.64E-07 | 57572 | 19 | 11309971 | 11373157 | -1 | protein_coding | dedicator of cytokinesis 6 |
| **LDLR** | ENSG00000130164 | 2170.955509 | 0.69088266 | 0.130978931 | 6.5E-08 | 4.95E-07 | 3949 | 19 | 11200038 | 11244492 | 1 | protein_coding | low density lipoprotein receptor |
| **PRKCSH** | ENSG00000130175 | 2849.215466 | -0.442822718 | 0.157995634 | 0.004074302 | 0.012707666 | 5589 | 19 | 11546109 | 11561783 | 1 | protein_coding | protein kinase C substrate 80K-H |
| **THEM6** | ENSG00000130193 | 215.5014298 | -1.364570397 | 0.251407255 | 1.03E-08 | 8.71E-08 | 51337 | 8 | 143808621 | 143818345 | 1 | protein_coding | thioesterase superfamily member 6 |
| **PVRL2** | ENSG00000130202 | 1998.557829 | 1.041663383 | 0.149826688 | 1.04E-12 | 1.31E-11 | 5819 | 19 | 45349432 | 45392485 | 1 | protein_coding | poliovirus receptor-related 2 (herpesvirus entry mediator B) |
| **APOE** | ENSG00000130203 | 76086.67371 | -0.639278846 | 0.119964791 | 5.4E-08 | 4.18E-07 | 348 | 19 | 45409011 | 45412650 | 1 | protein_coding | apolipoprotein E |
| **TOMM40** | ENSG00000130204 | 2604.410719 | -1.459650172 | 0.099655449 | 1.38E-49 | 1.19E-47 | 10452 | 19 | 45393826 | 45406946 | 1 | protein_coding | translocase of outer mitochondrial membrane 40 homolog (yeast) |
| **APOC1** | ENSG00000130208 | 1435.162052 | -1.787792448 | 0.206014741 | 5.32E-19 | 1.08E-17 | 341 | 19 | 45417504 | 45422606 | 1 | protein_coding | apolipoprotein C-I |
| **GADD45G** | ENSG00000130222 | 167.9613359 | 1.177526328 | 0.282495238 | 6.57E-06 | 3.72E-05 | 10912 | 9 | 92219928 | 92221470 | 1 | protein_coding | growth arrest and DNA-damage-inducible, gamma |
| **RPL36** | ENSG00000130255 | 1710.400139 | -0.351517428 | 0.13535879 | 0.007683697 | 0.02224052 | 25873 | 19 | 5674958 | 5691887 | 1 | protein_coding | ribosomal protein L36 |
| **BST2** | ENSG00000130303 | 2385.815076 | 2.032438875 | 0.105326831 | 5.44E-84 | 1.41E-81 | 684 | 19 | 17513748 | 17516457 | -1 | protein_coding | bone marrow stromal cell antigen 2 |
| **SLC27A1** | ENSG00000130304 | 1015.035203 | -2.5050953 | 0.186801375 | 4.15E-42 | 2.81E-40 | 376497 | 19 | 17579578 | 17616977 | 1 | protein_coding | solute carrier family 27 (fatty acid transporter), member 1 |
| **COLGALT1** | ENSG00000130309 | 7643.919779 | -0.942508379 | 0.114952206 | 7.53E-17 | 1.3E-15 | 79709 | 19 | 17666403 | 17693971 | 1 | protein_coding | collagen beta(1-O)galactosyltransferase 1 |
| **TULP4** | ENSG00000130338 | 3756.230008 | -0.425362589 | 0.108125558 | 5.97E-05 | 2.84E-04 | 56995 | 6 | 158733692 | 158932860 | 1 | protein_coding | tubby like protein 4 |
| **RSPH3** | ENSG00000130363 | 1183.142307 | 1.064743791 | 0.201938123 | 3.57E-08 | 2.82E-07 | 83861 | 6 | 159393903 | 159421219 | -1 | protein_coding | radial spoke 3 homolog (Chlamydomonas) |
| **MLLT4** | ENSG00000130396 | 553.4296906 | -0.559828797 | 0.16784341 | 5.29E-04 | 0.002057614 | 4301 | 6 | 168227602 | 168372703 | 1 | protein_coding | myeloid/lymphoid or mixed-lineage leukemia (trithorax homolog, Drosophila); translocated to, 4 |
| **ACTN4** | ENSG00000130402 | 4439.173653 | 0.257677167 | 0.103062952 | 0.010830735 | 0.029940481 | 81 | 19 | 39138289 | 39222223 | 1 | protein_coding | actinin, alpha 4 |
| **NDUFA10** | ENSG00000130414 | 1510.766198 | -0.643069348 | 0.141240941 | 2.71E-06 | 1.64E-05 | 4705 | 2 | 240831867 | 240964819 | -1 | protein_coding | NADH dehydrogenase (ubiquinone) 1 alpha subcomplex, 10, 42kDa |
| **FCHO1** | ENSG00000130475 | 110.3090912 | -0.925548517 | 0.445274801 | 0.009707442 | 0.02731704 | 23149 | 19 | 17858527 | 17899377 | 1 | protein_coding | FCH domain only 1 |
| **MAP1S** | ENSG00000130479 | 1033.912679 | -1.145397452 | 0.248554487 | 8.88E-07 | 5.76E-06 | 55201 | 19 | 17830051 | 17845325 | 1 | protein_coding | microtubule-associated protein 1S |
| **KLHDC7B** | ENSG00000130487 | 142.7616444 | 1.645894797 | 0.41689864 | 9.15E-06 | 5.06E-05 | 113730 | 22 | 50986462 | 50989451 | 1 | protein_coding | kelch domain containing 7B |
| **SSBP4** | ENSG00000130511 | 484.5006628 | -1.036110221 | 0.186228555 | 7.8E-09 | 6.72E-08 | 170463 | 19 | 18529674 | 18545372 | 1 | protein_coding | single stranded DNA binding protein 4 |
| **PGPEP1** | ENSG00000130517 | 3292.60436 | -0.77911278 | 0.13784133 | 5.41E-09 | 4.76E-08 | 54858 | 19 | 18451397 | 18480760 | 1 | protein_coding | pyroglutamyl-peptidase I |
| **LSM 4.00** | ENSG00000130520 | 1079.872641 | -0.416147085 | 0.126044864 | 7.39E-04 | 0.002782029 | 25804 | 19 | 18417040 | 18434084 | -1 | protein_coding | LSM4 homolog, U6 small nuclear RNA associated (S. cerevisiae) |
| **OLFM1** | ENSG00000130558 | 50.85927636 | -1.412355774 | 0.585375542 | 0.001908551 | 0.006493407 | 10439 | 9 | 137967268 | 138013025 | 1 | protein_coding | olfactomedin 1 |
| **UBAC1** | ENSG00000130560 | 1123.080426 | -0.873665136 | 0.135685897 | 4.44E-11 | 4.94E-10 | 10422 | 9 | 138824815 | 138853226 | -1 | protein_coding | UBA domain containing 1 |
| **HELZ2** | ENSG00000130589 | 2376.400707 | 4.616031038 | 0.173665155 | 3.89E-157 | 2.48E-154 | 85441 | 20 | 62189439 | 62205592 | -1 | protein_coding | helicase with zinc finger 2, transcriptional coactivator |
| **LSP1** | ENSG00000130592 | 16583.07328 | -0.918672856 | 0.133815559 | 2.45E-12 | 3.03E-11 | 4046 | 11 | 1874200 | 1913497 | 1 | protein_coding | lymphocyte-specific protein 1 |
| **TNNI2** | ENSG00000130598 | 95.47830065 | -1.618974963 | 0.464658641 | 6.03E-05 | 2.87E-04 | 7136 | 11 | 1860219 | 1862910 | 1 | protein_coding | troponin I type 2 (skeletal, fast) |
| **ATXN10** | ENSG00000130638 | 1168.577862 | -0.439195771 | 0.197527706 | 0.018752272 | 0.048061542 | 25814 | 22 | 46067678 | 46241187 | 1 | protein_coding | ataxin 10 |
| **TUBGCP2** | ENSG00000130640 | 1833.582755 | -1.029237347 | 0.131508565 | 9.66E-16 | 1.55E-14 | 10844 | 10 | 135093135 | 135125841 | -1 | protein_coding | tubulin, gamma complex associated protein 2 |
| **CYP 20.00** | ENSG00000130649 | 17.13489442 | 1.903489806 | 1.192828547 | 0.005163574 | 0.015705296 | 1571 | 10 | 135333910 | 135374724 | 1 | protein_coding | cytochrome P450, family 2, subfamily E, polypeptide 1 |
| **PNPLA7** | ENSG00000130653 | 296.4413361 | -1.950000404 | 0.330563195 | 3.6E-10 | 3.63E-09 | 375775 | 9 | 140354404 | 140444986 | -1 | protein_coding | patatin-like phospholipase domain containing 7 |
| **TAF4** | ENSG00000130699 | 392.6270878 | -0.779532305 | 0.228513366 | 2.62E-04 | 0.001090767 | 6874 | 20 | 60528525 | 60640866 | -1 | protein_coding | TAF4 RNA polymerase II, TATA box binding protein (TBP)-associated factor, 135kDa |
| **ASS1** | ENSG00000130707 | 82.18602549 | 3.272557768 | 0.449180771 | 1.75E-14 | 2.56E-13 | 445 | 9 | 133320316 | 133376661 | 1 | protein_coding | argininosuccinate synthase 1 |
| **CHMP2A** | ENSG00000130724 | 1289.144007 | -0.636911038 | 0.125653581 | 2.38E-07 | 1.68E-06 | 27243 | 19 | 59062933 | 59066491 | -1 | protein_coding | charged multivesicular body protein 2A |
| **UBE2M** | ENSG00000130725 | 1093.133877 | -0.474804019 | 0.149931397 | 0.001059824 | 0.003849167 | 9040 | 19 | 59067079 | 59070327 | -1 | protein_coding | ubiquitin-conjugating enzyme E2M |
| **TRIM28** | ENSG00000130726 | 1916.087979 | -0.346106595 | 0.120815286 | 0.003401869 | 0.010849493 | 10155 | 19 | 59055458 | 59062089 | 1 | protein_coding | tripartite motif containing 28 |
| **YIPF2** | ENSG00000130733 | 330.6801735 | -0.691544959 | 0.223290937 | 9.26E-04 | 0.003404734 | 78992 | 19 | 11033444 | 11039357 | -1 | protein_coding | Yip1 domain family, member 2 |
| **NPAS1** | ENSG00000130751 | 11.82065147 | -4.825548138 | 2.768913771 | 9.82E-04 | 0.003586392 | 4861 | 19 | 47523077 | 47549033 | 1 | protein_coding | neuronal PAS domain protein 1 |
| **GMFG** | ENSG00000130755 | 1074.149878 | -0.858526325 | 0.154284913 | 9.99E-09 | 8.5E-08 | 9535 | 19 | 39818993 | 39833012 | -1 | protein_coding | glia maturation factor, gamma |
| **MAP3K10** | ENSG00000130758 | 218.7593852 | 0.849770133 | 0.298154954 | 0.001461557 | 0.005119636 | 4294 | 19 | 40697651 | 40721482 | 1 | protein_coding | mitogen-activated protein kinase kinase kinase 10 |
| **ATPIF1** | ENSG00000130770 | 1061.976726 | -0.452117211 | 0.174548647 | 0.00684101 | 0.020105992 | 93974 | 1 | 28562620 | 28573417 | 1 | protein_coding | ATPase inhibitory factor 1 |
| **THEMIS2** | ENSG00000130775 | 3151.614511 | 0.519303402 | 0.133272735 | 6.08E-05 | 2.89E-04 | 9473 | 1 | 28199055 | 28213196 | 1 | protein_coding | thymocyte selection associated family member 2 |
| **CLIP1** | ENSG00000130779 | 6387.429624 | 0.524234633 | 0.079891975 | 2.89E-11 | 3.28E-10 | 6249 | 12 | 122755979 | 122907179 | -1 | protein_coding | CAP-GLY domain containing linker protein 1 |
| **PPAN** | ENSG00000130810 | 604.7221142 | -0.464729305 | 0.199046068 | 0.013605674 | 0.036524771 | 692312 | 19 | 10216965 | 10225414 | 1 | protein_coding | peter pan homolog (Drosophila) |
| **EIF3G** | ENSG00000130811 | 1358.198231 | -0.388684845 | 0.144439042 | 0.006181551 | 0.018379891 | 8666 | 19 | 10225693 | 10230596 | -1 | protein_coding | eukaryotic translation initiation factor 3, subunit G |
| **ANGPTL6** | ENSG00000130812 | 216.1054513 | -0.78234292 | 0.247429928 | 5.91E-04 | 0.002269107 | 83854 | 19 | 10203014 | 10213472 | -1 | protein_coding | angiopoietin-like 6 |
| **C19orf66** | ENSG00000130813 | 1083.631477 | 2.055170068 | 0.183544343 | 4.57E-30 | 1.77E-28 | 55337 | 19 | 10196798 | 10203928 | 1 | protein_coding | chromosome 19 open reading frame 66 |
| **SLC6A8** | ENSG00000130821 | 834.5515714 | -2.344434873 | 0.205530605 | 3.23E-31 | 1.33E-29 | 6535 | X | 152953554 | 152962048 | 1 | protein_coding | solute carrier family 6 (neurotransmitter transporter), member 8 |
| **PNCK** | ENSG00000130822 | 20.36420637 | -4.322756035 | 1.371563654 | 1.81E-04 | 7.85E-04 | 139728 | X | 152935185 | 152954465 | -1 | protein_coding | pregnancy up-regulated nonubiquitous CaM kinase |
| **ZNF331** | ENSG00000130844 | 383.7137608 | -0.865330574 | 0.274799119 | 5.57E-04 | 0.002152331 | 55422 | 19 | 54024235 | 54083523 | 1 | protein_coding | zinc finger protein 331 |
| **LRP3** | ENSG00000130881 | 152.9747235 | -1.525781491 | 0.31895634 | 2.28E-07 | 1.62E-06 | 4037 | 19 | 33668509 | 33698694 | 1 | protein_coding | low density lipoprotein receptor-related protein 3 |
| **NOL11** | ENSG00000130935 | 735.5111875 | 0.479173969 | 0.169112435 | 0.003291752 | 0.010537881 | 25926 | 17 | 65713949 | 65740647 | 1 | protein_coding | nucleolar protein 11 |

| | | | | | | | | | | | | | |
|---|---|---|---|---|---|---|---|---|---|---|---|---|---|
| **UBE4B** | ENSG00000130939 | 1046.679346 | -0.889392089 | 0.158635634 | 8.05E-09 | 6.93E-08 | 10277 | 1 | 10092890 | 10241297 | 1 | protein_coding | ubiquitination factor E4B |
| **CASZ1** | ENSG00000130940 | 88.78015965 | 0.910455091 | 0.448125068 | 0.01131304 | 0.03113227 | 54897 | 1 | 10696661 | 10856707 | -1 | protein_coding | castor zinc finger 1 |
| **HABP4** | ENSG00000130956 | 231.8029093 | -0.727906923 | 0.25104747 | 0.001650106 | 0.005702011 | 22927 | 9 | 99212483 | 99253618 | 1 | protein_coding | hyaluronan binding protein 4 |
| **SYNE1** | ENSG00000131018 | 2496.095178 | -1.296074746 | 0.132728006 | 2.68E-23 | 7.33E-22 | 23345 | 6 | 152442819 | 152958936 | -1 | protein_coding | spectrin repeat containing, nuclear envelope 1 |
| **LATS1** | ENSG00000131023 | 1312.969676 | 0.444598454 | 0.119847582 | 1.54E-04 | 6.79E-04 | 9113 | 6 | 149979289 | 150039392 | -1 | protein_coding | large tumor suppressor kinase 1 |
| **LILRB2** | ENSG00000131042 | 1428.024851 | 1.277424441 | 0.200611338 | 3.77E-11 | 4.23E-10 | 10288 | 19 | 54777675 | 54785039 | -1 | protein_coding | leukocyte immunoglobulin-like receptor, subfamily B (with TM and ITIM domains), member 2 |
| **ACSS2** | ENSG00000131069 | 1747.295489 | -0.857776784 | 0.176723902 | 3.85E-07 | 2.64E-06 | 55902 | 20 | 33459949 | 33515769 | 1 | protein_coding | acyl-CoA synthetase short-chain family member 2 |
| **ARHGEF9** | ENSG00000131089 | 442.9033274 | -1.207122721 | 0.172867691 | 7.79E-13 | 1E-11 | 23229 | X | 62854847 | 63005426 | -1 | protein_coding | Cdc42 guanine nucleotide exchange factor (GEF) 9 |
| **COX4I1** | ENSG00000131143 | 5034.948858 | -0.40665578 | 0.107720643 | 1.25E-04 | 5.61E-04 | 1327 | 16 | 85832239 | 85840650 | 1 | protein_coding | cytochrome c oxidase subunit IV isoform 1 |
| **GINS2** | ENSG00000131153 | 32.68930468 | 1.459386868 | 0.72625294 | 0.00412955 | 0.012858987 | 51659 | 16 | 85709804 | 85723679 | -1 | protein_coding | GINS complex subunit 2 (Psf2 homolog) |
| **NFATC1** | ENSG00000131196 | 1180.826068 | -0.883113293 | 0.16988651 | 6.92E-08 | 5.26E-07 | 4772 | 18 | 77155856 | 77289325 | 1 | protein_coding | nuclear factor of activated T-cells, cytoplasmic, calcineurin-dependent 1 |
| **IDO1** | ENSG00000131203 | 11087.82864 | 7.677053305 | 0.80658591 | 7.35E-23 | 1.96E-21 | 3620 | 8 | 39759794 | 39785963 | 1 | protein_coding | indoleamine 2,3-dioxygenase 1 |
| **PPT1** | ENSG00000131238 | 20613.40334 | -0.510804011 | 0.136912718 | 1.04E-04 | 4.73E-04 | 5538 | 1 | 40538379 | 40563375 | -1 | protein_coding | palmitoyl-protein thioesterase 1 |
| **RAB11FIP4** | ENSG00000131242 | 449.4181889 | -0.684624992 | 0.193787914 | 2.03E-04 | 8.66E-04 | 84440 | 17 | 29718642 | 29865236 | 1 | protein_coding | RAB11 family interacting protein 4 (class II) |
| **RLIM** | ENSG00000131263 | 3477.717468 | 0.239906031 | 0.099708155 | 0.015592895 | 0.041115635 | 51132 | X | 73805052 | 73834452 | -1 | protein_coding | ring finger protein, LIM domain interacting |
| **TRAF3** | ENSG00000131323 | 3242.396956 | -0.317731372 | 0.114465338 | 0.004696194 | 0.014443939 | 7187 | 14 | 103243813 | 103377837 | 1 | protein_coding | TNF receptor-associated factor 3 |
| **EMR3** | ENSG00000131355 | 409.3120256 | -2.031589478 | 0.354497719 | 7.87E-10 | 7.65E-09 | 84658 | 19 | 14729929 | 14800839 | -1 | protein_coding | egf-like module containing, mucin-like, hormone receptor-like 3 |
| **CAPN7** | ENSG00000131375 | 1044.166345 | 0.478097268 | 0.159318385 | 0.001872325 | 0.006375833 | 23473 | 3 | 15247659 | 15294425 | 1 | protein_coding | calpain 7 |
| **RFTN1** | ENSG00000131378 | 1987.157797 | 0.625478527 | 0.157724529 | 3.99E-05 | 1.96E-04 | 23180 | 3 | 16355081 | 16555533 | -1 | protein_coding | raftlin, lipid raft linker 1 |
| **KCNC3** | ENSG00000131398 | 751.7585827 | -1.73165538 | 0.179397577 | 6.02E-23 | 1.61E-21 | 3748 | 19 | 50815194 | 50836772 | -1 | protein_coding | potassium voltage-gated channel, Shaw-related subfamily, member 3 |
| **PDLIM4** | ENSG00000131435 | 372.4294509 | -1.583461006 | 0.231844991 | 1.17E-12 | 1.47E-11 | 8572 | 5 | 131593364 | 131609147 | 1 | protein_coding | PDZ and LIM domain 4 |
| **KIF3A** | ENSG00000131437 | 223.9891026 | 0.5947304 | 0.267674444 | 0.014209662 | 0.037906487 | 11127 | 5 | 132028320 | 132073330 | -1 | protein_coding | kinesin family member 3A |
| **GFPT2** | ENSG00000131459 | 15.81533276 | 1.346221974 | 1.013670541 | 0.013811587 | 0.037019024 | 9945 | 5 | 179727690 | 179780387 | -1 | protein_coding | glutamine-fructose-6-phosphate transaminase 2 |
| **PSMC3IP** | ENSG00000131470 | 35.54503464 | 1.258026474 | 0.550324049 | 0.003312239 | 0.010599026 | 29893 | 17 | 40724333 | 40729849 | -1 | protein_coding | PSMC3 interacting protein |
| **NDUFA2** | ENSG00000131495 | 472.1546835 | -0.589321527 | 0.172286085 | 3.61E-04 | 0.001461448 | 4695 | 5 | 140018325 | 140027370 | -1 | protein_coding | NADH dehydrogenase (ubiquinone) 1 alpha subcomplex, 2, 8kDa |
| **NDFIP1** | ENSG00000131507 | 3240.679177 | 0.42668897 | 0.100705889 | 1.8E-05 | 9.45E-05 | 80762 | 5 | 141488070 | 141534005 | 1 | protein_coding | Nedd4 family interacting protein 1 |
| **UBE2D2** | ENSG00000131508 | 1798.380017 | 0.492907889 | 0.11537858 | 1.29E-05 | 6.99E-05 | 7322 | 5 | 138906016 | 139008018 | 1 | protein_coding | ubiquitin-conjugating enzyme E2D 2 |
| **EXOC4** | ENSG00000131558 | 3457.865404 | -0.6343211 | 0.126044525 | 2.61E-07 | 1.83E-06 | 60412 | 7 | 132937829 | 133751342 | 1 | protein_coding | exocyst complex component 4 |
| **ACAP3** | ENSG00000131584 | 350.7854676 | -1.056739095 | 0.219612962 | 3.83E-07 | 2.63E-06 | 116983 | 1 | 1227756 | 1244989 | -1 | protein_coding | ArfGAP with coiled-coil, ankyrin repeat and PH domains 3 |
| **TRAF7** | ENSG00000131653 | 1027.52746 | -0.65310986 | 0.198907127 | 5.17E-04 | 0.002016222 | 84231 | 16 | 2205699 | 2228130 | 1 | protein_coding | TNF receptor-associated factor 7, E3 ubiquitin protein ligase |
| **NINJ1** | ENSG00000131669 | 2224.587545 | 2.096633793 | 0.130260701 | 3.03E-59 | 3.82E-57 | 4814 | 9 | 95883771 | 95896570 | -1 | protein_coding | ninjurin 1 |
| **NPHP4** | ENSG00000131697 | 182.3987286 | -0.791973138 | 0.265235031 | 0.001104399 | 0.003993972 | 261734 | 1 | 5922871 | 6052533 | -1 | protein_coding | nephronophthisis 4 |
| **IL13RA1** | ENSG00000131724 | 5029.412491 | 0.343496528 | 0.11225605 | 0.002101055 | 0.007063445 | 3597 | X | 117861535 | 117928502 | 1 | protein_coding | interleukin 13 receptor, alpha 1 |
| **TOP2A** | ENSG00000131747 | 492.0431492 | 0.830499149 | 0.363578585 | 0.007248695 | 0.021121412 | 7153 | 17 | 38544768 | 38574202 | -1 | protein_coding | topoisomerase (DNA) II alpha 170kDa |
| **RARA** | ENSG00000131759 | 497.7023423 | -1.032873654 | 0.219495991 | 5.81E-07 | 3.89E-06 | 5914 | 17 | 38465444 | 38513094 | 1 | protein_coding | retinoic acid receptor, alpha |
| **PEX11B** | ENSG00000131779 | 438.216972 | 0.780989056 | 0.180232286 | 5.94E-06 | 3.39E-05 | 8799 | 1 | 145516252 | 145523730 | 1 | protein_coding | peroxisomal biogenesis factor 11 beta |
| **PRKAB2** | ENSG00000131791 | 502.4642708 | 0.971058192 | 0.160942178 | 5.09E-10 | 5.04E-09 | 5565 | 1 | 146626685 | 146644129 | -1 | protein_coding | protein kinase, AMP-activated, beta 2 non-catalytic subunit |
| **PDHA1** | ENSG00000131828 | 1395.83247 | -0.335766106 | 0.136596195 | 0.011759335 | 0.032178679 | 5160 | X | 19362011 | 19379823 | 1 | protein_coding | pyruvate dehydrogenase (lipoamide) alpha 1 |
| **MCCC2** | ENSG00000131844 | 963.4534824 | 0.32042554 | 0.132437046 | 0.0135366 | 0.036377677 | 64087 | 5 | 70883115 | 70954531 | 1 | protein_coding | methylcrotonoyl-CoA carboxylase 2 (beta) |
| **ZSCAN5A** | ENSG00000131848 | 186.8761737 | -1.057836268 | 0.266348452 | 1.98E-05 | 1.03E-04 | 79149 | 19 | 56732681 | 56879752 | -1 | protein_coding | zinc finger and SCAN domain containing 5A |
| **ZNF132** | ENSG00000131849 | 198.8124408 | -0.743955962 | 0.25179413 | 0.001278591 | 0.00453813 | 7691 | 19 | 58944181 | 58951589 | -1 | protein_coding | zinc finger protein 132 |
| **VIMP** | ENSG00000131871 | 967.0321733 | 1.018693256 | 0.140319228 | 1.07E-13 | 1.47E-12 | 55829 | 15 | 101811022 | 101817705 | -1 | protein_coding | VCP-interacting membrane protein |
| **C19orf12** | ENSG00000131943 | 878.6077338 | 0.852233703 | 0.137410844 | 2.07E-10 | 2.12E-09 | 83636 | 19 | 30191721 | 30206364 | -1 | protein_coding | chromosome 19 open reading frame 12 |
| **ACTR10** | ENSG00000131966 | 2019.151926 | 0.530643152 | 0.104445961 | 2.56E-07 | 1.8E-06 | 55860 | 14 | 58666798 | 58701750 | 1 | protein_coding | actin-related protein 10 homolog (S. cerevisiae) |
| **GCH1** | ENSG00000131979 | 2284.458284 | 4.393403795 | 0.467770969 | 2.9E-22 | 7.42E-21 | 2643 | 14 | 55308726 | 55369570 | -1 | protein_coding | GTP cyclohydrolase 1 |
| **DNAJB1** | ENSG00000132002 | 1731.952501 | -0.343558019 | 0.102606898 | 6.85E-04 | 0.002594608 | 3337 | 19 | 14625582 | 14640582 | -1 | protein_coding | DnaJ (Hsp40) homolog, subfamily B, member 1 |
| **ZSWIM4** | ENSG00000132003 | 335.9117946 | 0.716772505 | 0.262073594 | 0.002781472 | 0.009045346 | 65249 | 19 | 13906274 | 13943044 | 1 | protein_coding | zinc finger, SWIM-type containing 4 |
| **TRIM21** | ENSG00000132109 | 1451.327254 | 1.383287706 | 0.115950729 | 1.64E-33 | 7.74E-32 | 6737 | 11 | 4406127 | 4414926 | -1 | protein_coding | tripartite motif containing 21 |
| **RAF1** | ENSG00000132155 | 3090.775602 | 0.292445303 | 0.089653887 | 9.26E-04 | 0.003404734 | 5894 | 3 | 12625100 | 12705725 | -1 | protein_coding | v-raf-1 murine leukemia viral oncogene homolog 1 |
| **PPARG** | ENSG00000132170 | 606.6299236 | -2.481392192 | 0.277662131 | 5.09E-20 | 1.13E-18 | 5468 | 3 | 12328867 | 12475855 | 1 | protein_coding | peroxisome proliferator-activated receptor gamma |
| **NUP210** | ENSG00000132182 | 1577.00523 | -0.874839869 | 0.108682422 | 3.02E-16 | 4.99E-15 | 23225 | 3 | 13357737 | 13461809 | -1 | protein_coding | nucleoporin 210kDa |
| **FCRLA** | ENSG00000132185 | 472.991953 | -0.448522434 | 0.186535593 | 0.011249935 | 0.030984457 | 84824 | 1 | 161676762 | 161684142 | 1 | protein_coding | Fc receptor-like A |
| **EMILIN2** | ENSG00000132205 | 7519.196989 | -0.769751687 | 0.118175509 | 2.8E-11 | 3.18E-10 | 84034 | 18 | 2847028 | 2915991 | 1 | protein_coding | elastin microfibril interfacer 2 |
| **TRIM5** | ENSG00000132256 | 713.5221032 | 2.733355689 | 0.160055597 | 1.64E-66 | 2.75E-64 | 85363 | 11 | 5684425 | 5959849 | -1 | protein_coding | tripartite motif containing 5 |
| **TRIM22** | ENSG00000132274 | 6749.180715 | 2.126878331 | 0.135954509 | 2.94E-56 | 3.38E-54 | 10346 | 11 | 5710919 | 5758319 | 1 | protein_coding | tripartite motif containing 22 |
| **EFR3A** | ENSG00000132294 | 1473.689041 | 0.390225531 | 0.168627095 | 0.018492231 | 0.047482657 | 23167 | 8 | 132916335 | 133025889 | 1 | protein_coding | EFR3 homolog A (S. cerevisiae) |
| **IMMT** | ENSG00000132305 | 3439.152022 | -0.284485471 | 0.103702425 | 0.005566002 | 0.016735236 | 10989 | 2 | 86371055 | 86422893 | -1 | protein_coding | inner membrane protein, mitochondrial |
| **PER2** | ENSG00000132326 | 351.7424424 | -0.697658566 | 0.264129213 | 0.003668663 | 0.011603514 | 8864 | 2 | 239152679 | 239198743 | -1 | protein_coding | period circadian clock 2 |
| **CARD6** | ENSG00000132357 | 1718.06592 | 0.712884473 | 0.146960758 | 5.93E-07 | 3.96E-06 | 84674 | 5 | 40841286 | 40860275 | 1 | protein_coding | caspase recruitment domain family, member 6 |
| **RPA1** | ENSG00000132383 | 1488.270094 | -0.761290341 | 0.137157533 | 1.3E-08 | 1.09E-07 | 6117 | 17 | 1732996 | 1803376 | 1 | protein_coding | replication protein A1, 70kDa |
| **EEFSEC** | ENSG00000132394 | 246.4096345 | -0.763299896 | 0.270642065 | 0.001902418 | 0.006473984 | 60678 | 3 | 127872297 | 128127485 | 1 | protein_coding | eukaryotic elongation factor, selenocysteine-tRNA-specific |
| **TBC1D14** | ENSG00000132405 | 2354.87329 | -1.258713494 | 0.114269962 | 6.04E-29 | 2.2E-27 | 57533 | 4 | 6910969 | 7034845 | 1 | protein_coding | TBC1 domain family, member 14 |
| **TMEM128** | ENSG00000132406 | 285.0316205 | 0.681397005 | 0.21708883 | 8.48E-04 | 0.003151536 | 85013 | 4 | 4237269 | 4249950 | -1 | protein_coding | transmembrane protein 128 |
| **SEC61G** | ENSG00000132432 | 362.6467028 | 0.46604582 | 0.203268925 | 0.015275922 | 0.040398324 | 23480 | 7 | 54819943 | 54827667 | -1 | protein_coding | Sec61 gamma subunit |
| **FIGNL1** | ENSG00000132436 | 350.7068663 | 1.222916646 | 0.252219455 | 2.59E-07 | 1.82E-06 | 63979 | 7 | 50511831 | 50518088 | -1 | protein_coding | fidgetin-like 1 |
| **GRSF1** | ENSG00000132463 | 1798.550171 | 1.019038036 | 0.154660189 | 1.26E-11 | 1.47E-10 | 2926 | 4 | 71681499 | 71705662 | -1 | protein_coding | G-rich RNA sequence binding factor 1 |
| **UTP3** | ENSG00000132467 | 1191.145794 | 0.336029886 | 0.147342887 | 0.019344426 | 0.049311987 | 57050 | 4 | 71554196 | 71556267 | 1 | protein_coding | UTP3, small subunit (SSU) processome component, homolog (S. cerevisiae) |
| **H3F3B** | ENSG00000132475 | 5576.854165 | 1.054348811 | 0.106166234 | 8.12E-24 | 2.28E-22 | 1006162 | 17 | 73772515 | 73781974 | -1 | protein_coding | H3 histone, family 3B (H3.3B) |
| **ZRANB2** | ENSG00000132485 | 1292.183277 | 0.514323352 | 0.129546969 | 4.93E-05 | 2.38E-04 | 9406 | 1 | 71528974 | 71546980 | -1 | protein_coding | zinc finger, RAN-binding domain containing 2 |
| **EIF5A** | ENSG00000132507 | 4164.721074 | 0.46608639 | 0.092303992 | 3.68E-07 | 2.53E-06 | 1984 | 17 | 7210318 | 7215774 | 1 | protein_coding | eukaryotic translation initiation factor 5A |
| **CLEC10A** | ENSG00000132514 | 59.01746761 | -2.398869336 | 0.679665546 | 2.87E-05 | 1.45E-04 | 10462 | 17 | 6977856 | 6983626 | -1 | protein_coding | C-type lectin domain family 10, member A |
| **XAF1** | ENSG00000132530 | 4091.480691 | 3.398565092 | 0.181310941 | 1.53E-79 | 3.65E-77 | 54739 | 17 | 6658766 | 6678966 | 1 | protein_coding | XIAP associated factor 1 |
| **DLG4** | ENSG00000132535 | 638.6826261 | -1.020422718 | 0.184438986 | 9.08E-09 | 7.78E-08 | 1742 | 17 | 7093209 | 7123021 | -1 | protein_coding | discs, large homolog 4 (Drosophila) |
| **FLOT2** | ENSG00000132589 | 2296.06891 | 0.760777632 | 0.115530316 | 2.19E-11 | 2.51E-10 | 2319 | 17 | 27206353 | 27224697 | -1 | protein_coding | flotillin 2 |
| **NIP7** | ENSG00000132603 | 570.8002528 | 0.460921409 | 0.164299591 | 0.003554016 | 0.011282932 | 51388 | 16 | 69373333 | 69377014 | 1 | protein_coding | NIP7, nucleolar pre-rRNA processing protein |
| **TERF2** | ENSG00000132604 | 561.4962353 | 0.609127992 | 0.204593323 | 0.001567693 | 0.00544777 | 7014 | 16 | 69389464 | 69442474 | -1 | protein_coding | telomeric repeat binding factor 2 |
| **BTBD3** | ENSG00000132640 | 347.3137775 | -1.034062476 | 0.268305108 | 2.75E-05 | 1.4E-04 | 22903 | 20 | 11871371 | 11907257 | 1 | protein_coding | BTB (POZ) domain containing 3 |
| **PCNA** | ENSG00000132646 | 1017.90339 | 0.888924163 | 0.165980753 | 3.29E-08 | 2.62E-07 | 5111 | 20 | 5095599 | 5107272 | -1 | protein_coding | proliferating cell nuclear antigen |
| **RIN2** | ENSG00000132669 | 3741.711422 | 0.432305695 | 0.128732472 | 6.09E-04 | 0.002331739 | 54453 | 20 | 19867165 | 19983101 | 1 | protein_coding | Ras and Rab interactor 2 |
| **PTPRA** | ENSG00000132670 | 2710.596822 | 0.632611465 | 0.104104905 | 6.77E-10 | 6.61E-09 | 5786 | 20 | 2844830 | 3019722 | 1 | protein_coding | protein tyrosine phosphatase, receptor type, A |
| **DAP3** | ENSG00000132676 | 1602.048548 | 0.419912364 | 0.103879829 | 4.16E-05 | 2.03E-04 | 7818 | 1 | 155657751 | 155708801 | 1 | protein_coding | death associated protein 3 |
| **KIAA0907** | ENSG00000132680 | 918.6354273 | 0.324534346 | 0.126470872 | 0.008620424 | 0.024561138 | 22889 | 1 | 155882834 | 155904191 | -1 | protein_coding | KIAA0907 |
| **NES** | ENSG00000132688 | 644.5304312 | -2.798908355 | 0.222985216 | 1.68E-37 | 9.26E-36 | 10763 | 1 | 156638555 | 156647189 | -1 | protein_coding | nestin |
| **ARHGEF11** | ENSG00000132694 | 5931.19149 | 0.883366662 | 0.086169335 | 2.38E-25 | 7.24E-24 | 9826 | 1 | 156904632 | 157015162 | -1 | protein_coding | Rho guanine nucleotide exchange factor (GEF) 11 |
| **SYT11** | ENSG00000132718 | 1000.575046 | 0.764038157 | 0.18192876 | 1.11E-05 | 6.1E-05 | 23208 | 1 | 155829300 | 155854990 | 1 | protein_coding | synaptotagmin XI |
| **NASP** | ENSG00000132780 | 988.3375042 | 1.084090901 | 0.16364163 | 8.6E-12 | 1.02E-10 | 4678 | 1 | 46049518 | 46084566 | 1 | protein_coding | nuclear autoantigenic sperm protein (histone-binding) |
| **CTNNBL1** | ENSG00000132792 | 1574.557209 | 0.9817862 | 0.13380311 | 6.63E-14 | 9.33E-13 | 56259 | 20 | 36322408 | 36500531 | 1 | protein_coding | catenin, beta like 1 |
| **VSTM2L** | ENSG00000132821 | 28.09061396 | -7.192580925 | 2.706792627 | 3.79E-06 | 2.24E-05 | 128434 | 20 | 36531499 | 36573752 | 1 | protein_coding | V-set and transmembrane domain containing 2 like |
| **OSER1** | ENSG00000132823 | 816.3656415 | 1.076331094 | 0.15906831 | 3.43E-12 | 4.21E-11 | 51526 | 20 | 42825136 | 42839431 | -1 | protein_coding | oxidative stress responsive serine-rich 1 |

| | | | | | | | | | | | | | |
|---|---|---|---|---|---|---|---|---|---|---|---|---|---|
| **PPP1R3D** | ENSG00000132825 | 361.5979559 | 0.91206508 | 0.189390054 | 4.9E-07 | 3.32E-06 | 5509 | 20 | 58511894 | 58515352 | -1 | protein_coding | protein phosphatase 1, regulatory subunit 3D |
| **ZBED3** | ENSG00000132846 | 460.4578016 | -0.585511619 | 0.247726897 | 0.010041704 | 0.028107575 | 84327 | 5 | 76367897 | 76383148 | -1 | protein_coding | zinc finger, BED-type containing 3 |
| **KANK4** | ENSG00000132854 | 121.4388461 | 1.369351898 | 0.46125731 | 4.05E-04 | 0.001621175 | 163782 | 1 | 62702651 | 62785085 | -1 | protein_coding | KN motif and ankyrin repeat domains 4 |
| **ATP8A2** | ENSG00000132932 | 52.30603663 | -1.494720169 | 0.579410909 | 0.001151404 | 0.004137528 | 51761 | 13 | 25946209 | 26599989 | 1 | protein_coding | ATPase, aminophospholipid transporter, class I, type 8A, member 2 |
| **ZMYM5** | ENSG00000132950 | 260.6900831 | 0.617138702 | 0.228277824 | 0.003720725 | 0.011743905 | 9205 | 13 | 20397622 | 20437776 | -1 | protein_coding | zinc finger, MYM-type 5 |
| **USPL1** | ENSG00000132952 | 635.2151154 | 1.063769875 | 0.140628685 | 1.06E-14 | 1.56E-13 | 10208 | 13 | 31191830 | 31233686 | 1 | protein_coding | ubiquitin specific peptidase like 1 |
| **POMP** | ENSG00000132963 | 3219.938192 | 0.855634237 | 0.104475169 | 1.27E-16 | 2.17E-15 | 51371 | 13 | 29233241 | 29253062 | 1 | protein_coding | proteasome maturation protein |
| **CDK8** | ENSG00000132964 | 586.8053784 | 0.521871002 | 0.154950851 | 5.03E-04 | 0.00196931 | 1024 | 13 | 26828276 | 26979375 | 1 | protein_coding | cyclin-dependent kinase 8 |
| **ALOX5AP** | ENSG00000132965 | 5162.051172 | -1.434903902 | 0.137800601 | 3.55E-26 | 1.13E-24 | 241 | 13 | 31309645 | 31338556 | 1 | protein_coding | arachidonate 5-lipoxygenase-activating protein |
| **MYH10** | ENSG00000133026 | 164.6180474 | -1.02477847 | 0.334978057 | 5.77E-04 | 0.002220883 | 4628 | 17 | 8377523 | 8534079 | -1 | protein_coding | myosin, heavy chain 10, non-muscle |
| **PEMT** | ENSG00000133027 | 56.86069057 | -1.219027126 | 0.545402126 | 0.003857266 | 0.012109818 | 10400 | 17 | 17408877 | 17495022 | -1 | protein_coding | phosphatidylethanolamine N-methyltransferase |
| **SCO1** | ENSG00000133028 | 728.289033 | 0.676411761 | 0.136651993 | 3.76E-07 | 2.58E-06 | 6341 | 17 | 10583654 | 10601692 | -1 | protein_coding | SCO1 cytochrome c oxidase assembly protein |
| **MPRIP** | ENSG00000133030 | 1340.960704 | -0.617514385 | 0.140491224 | 7.13E-06 | 4.02E-05 | 23164 | 17 | 16945859 | 17120993 | 1 | protein_coding | myosin phosphatase Rho interacting protein |
| **PIK3C2B** | ENSG00000133056 | 655.7910009 | -1.131482676 | 0.170806516 | 8.29E-12 | 9.8E-11 | 5287 | 1 | 204391756 | 204463852 | -1 | protein_coding | phosphatidylinositol-4-phosphate 3-kinase, catalytic subunit type 2 beta |
| **CCNA1** | ENSG00000133101 | 229.5750029 | 2.703547668 | 0.527583388 | 2.06E-08 | 1.68E-07 | 8900 | 13 | 37005967 | 37017019 | 1 | protein_coding | cyclin A1 |
| **COG6** | ENSG00000133103 | 635.7035666 | -0.382864763 | 0.152960132 | 0.009570407 | 0.026996037 | 57511 | 13 | 40229764 | 40365802 | 1 | protein_coding | component of oligomeric golgi complex 6 |
| **SPG20** | ENSG00000133104 | 2775.113966 | 0.590005905 | 0.138683805 | 1.17E-05 | 6.4E-05 | 23111 | 13 | 36875775 | 36944317 | -1 | protein_coding | spastic paraplegia 20 (Troyer syndrome) |
| **EPSTI1** | ENSG00000133106 | 2260.079226 | 3.66164636 | 0.190856045 | 2.06E-83 | 5.15E-81 | 94240 | 13 | 43460524 | 43566407 | -1 | protein_coding | epithelial stromal interaction 1 (breast) |
| **KL** | ENSG00000133116 | 67.88382941 | 3.78116014 | 0.465915386 | 3.03E-17 | 5.39E-16 | 9365 | 13 | 33590207 | 33640282 | 1 | protein_coding | klotho |
| **RFC3** | ENSG00000133119 | 121.4480322 | 0.792852619 | 0.302773973 | 0.003361088 | 0.010739601 | 5983 | 13 | 34392186 | 34540695 | 1 | protein_coding | replication factor C (activator 1) 3, 38kDa |
| **RNF128** | ENSG00000133135 | 718.1913254 | -2.726005344 | 0.310142434 | 1.01E-19 | 2.18E-18 | 79589 | X | 105937024 | 106040223 | 1 | protein_coding | ring finger protein 128, E3 ubiquitin protein ligase |
| **FAM104A** | ENSG00000133193 | 1038.647995 | 0.298295621 | 0.126240741 | 0.015833541 | 0.041685444 | 84923 | 17 | 71203492 | 71232892 | -1 | protein_coding | family with sequence similarity 104, member A |
| **SLC39A11** | ENSG00000133195 | 4319.55838 | -0.564077463 | 0.114670147 | 5.37E-07 | 3.62E-06 | 201266 | 17 | 70642088 | 71088851 | -1 | protein_coding | solute carrier family 39, member 11 |
| **EPHB2** | ENSG00000133216 | 366.481273 | 2.404636769 | 0.190813571 | 1.85E-37 | 1.02E-35 | 2048 | 1 | 23037332 | 23241818 | 1 | protein_coding | EPH receptor B2 |
| **BTBD2** | ENSG00000133243 | 561.9308179 | -0.773573417 | 0.187766254 | 1.64E-05 | 8.71E-05 | 55643 | 19 | 1985447 | 2034880 | -1 | protein_coding | BTB (POZ) domain containing 2 |
| **PRAM1** | ENSG00000133246 | 922.3849786 | -1.933759676 | 0.138985978 | 2.53E-45 | 1.96E-43 | 84106 | 19 | 8554940 | 8567996 | -1 | protein_coding | PML-RARA regulated adaptor molecule 1 |
| **ANKRD32** | ENSG00000133302 | 232.4010532 | 0.912214191 | 0.238570675 | 4.08E-05 | 0.000199809 | 84250 | 5 | 93954052 | 94075141 | 1 | protein_coding | ankyrin repeat domain 32 |
| **CNDP2** | ENSG00000133313 | 9955.546124 | 0.689919459 | 0.079210935 | 1.63E-18 | 3.21E-17 | 55748 | 18 | 72163051 | 72188366 | 1 | protein_coding | CNDP dipeptidase 2 (metallopeptidase M20 family) |
| **MACROD1** | ENSG00000133315 | 14.30931642 | -1.809582953 | 1.159256048 | 0.007320282 | 0.021301573 | 28992 | 11 | 63766030 | 63933578 | -1 | protein_coding | MACRO domain containing 1 |
| **WDR74** | ENSG00000133316 | 1278.176484 | 0.661509713 | 0.244981612 | 0.00346086 | 0.011014644 | 54663 | 11 | 62599814 | 62609281 | -1 | protein_coding | WD repeat domain 74 |
| **RTN3** | ENSG00000133318 | 4904.399367 | -0.538935013 | 0.085753818 | 1.21E-10 | 1.28E-09 | 10313 | 11 | 63448918 | 63527363 | 1 | protein_coding | reticulon 3 |
| **RARRES3** | ENSG00000133321 | 477.370632 | 0.880078863 | 0.36562643 | 0.004692877 | 0.014439549 | 5920 | 11 | 63304281 | 63313934 | 1 | protein_coding | retinoic acid receptor responder (tazarotene induced) 3 |
| **MYH11** | ENSG00000133392 | 137.6333783 | 6.501637639 | 0.558463183 | 4.52E-33 | 2.09E-31 | 4629 | 16 | 15797029 | 15950890 | -1 | protein_coding | myosin, heavy chain 11, smooth muscle |
| **FOPNL** | ENSG00000133393 | 880.1861402 | 0.638405083 | 0.149232775 | 9.99E-06 | 5.51E-05 | 123811 | 16 | 15959577 | 15982482 | -1 | protein_coding | FGFR1OP N-terminal like |
| **MED10** | ENSG00000133398 | 513.6290416 | 0.755980449 | 0.167622744 | 2.79E-06 | 1.68E-05 | 84246 | 5 | 6371994 | 6378707 | -1 | protein_coding | mediator complex subunit 10 |
| **PDZD2** | ENSG00000133401 | 11.16418782 | 3.582859902 | 2.632581497 | 0.003288515 | 0.010529724 | 23037 | 5 | 31639517 | 32111037 | 1 | protein_coding | PDZ domain containing 2 |
| **LARGE** | ENSG00000133424 | 160.5004693 | -0.866516565 | 0.307114266 | 0.001592713 | 0.005525864 | 9215 | 22 | 33558212 | 34318829 | -1 | protein_coding | like-glycosyltransferase |
| **SEC14L4** | ENSG00000133488 | 17.34685347 | -1.479073247 | 1.258054703 | 0.016942849 | 0.044119537 | 284904 | 22 | 30884900 | 30901698 | -1 | protein_coding | SEC14-like 4 (S. cerevisiae) |
| **GIMAP6** | ENSG00000133561 | 1778.437551 | 0.995737812 | 0.216524111 | 1.25E-06 | 7.89E-06 | 474344 | 7 | 150322463 | 150329473 | -1 | protein_coding | GTPase, IMAP family member 6 |
| **GIMAP4** | ENSG00000133574 | 2950.760622 | 0.905617704 | 0.178656995 | 1.51E-07 | 1.09E-06 | 55303 | 7 | 150264365 | 150271041 | 1 | protein_coding | GTPase, IMAP family member 4 |
| **MKRN1** | ENSG00000133606 | 2643.116657 | 0.820604454 | 0.117349785 | 1.12E-12 | 1.42E-11 | 23608 | 7 | 140152840 | 140179369 | -1 | protein_coding | makorin ring finger protein 1 |
| **AGAP3** | ENSG00000133612 | 2630.565362 | -0.555962709 | 0.115230328 | 9.55E-07 | 6.16E-06 | 116988 | 7 | 150782918 | 150841523 | 1 | protein_coding | ArfGAP with GTPase domain, ankyrin repeat and PH domain 3 |
| **BTG1** | ENSG00000133639 | 4018.399038 | 1.779793578 | 0.161100143 | 3.34E-29 | 1.23E-27 | 694 | 12 | 92536286 | 92539673 | -1 | protein_coding | B-cell translocation gene 1, anti-proliferative |
| **TMEM254** | ENSG00000133678 | 547.3143416 | -0.591194688 | 0.196118512 | 0.001476468 | 0.00516595 | 80195 | 10 | 81838402 | 81852313 | 1 | protein_coding | transmembrane protein 254 |
| **KRAS** | ENSG00000133703 | 1324.735839 | 0.452700797 | 0.149583111 | 0.001760669 | 0.006027847 | 3845 | 12 | 25357723 | 25403870 | -1 | protein_coding | Kirsten rat sarcoma viral oncogene homolog |
| **LARS** | ENSG00000133706 | 2911.019648 | -0.490848271 | 0.142818765 | 4E-04 | 0.001605353 | 51520 | 5 | 145492601 | 145562223 | -1 | protein_coding | leucyl-tRNA synthetase |
| **IMPA1** | ENSG00000133731 | 499.6515618 | 0.506088869 | 0.176618649 | 0.002761166 | 0.008988384 | 3612 | 8 | 82570196 | 82598928 | -1 | protein_coding | inositol(myo)-1(or 4)-monophosphatase 1 |
| **LRRCC1** | ENSG00000133739 | 198.3921777 | 1.054740741 | 0.283977259 | 5.18E-05 | 2.49E-04 | 85444 | 8 | 86019382 | 86058311 | 1 | protein_coding | leucine rich repeat and coiled-coil centrosomal protein 1 |
| **CCDC59** | ENSG00000133773 | 370.5230063 | 0.513292604 | 0.192886277 | 0.005071586 | 0.015458526 | 29080 | 12 | 82617460 | 82752584 | -1 | protein_coding | coiled-coil domain containing 59 |
| **AMPD3** | ENSG00000133805 | 5302.986674 | 1.1020062 | 0.093186198 | 1.64E-32 | 7.21E-31 | 1001304 | 11 | 10329860 | 10529126 | 1 | protein_coding | adenosine monophosphate deaminase 3 |
| **SBF2** | ENSG00000133812 | 1645.795843 | 0.364671386 | 0.119422151 | 0.001867957 | 0.006363797 | 81846 | 11 | 9800214 | 10315754 | -1 | protein_coding | SET binding factor 2 |
| **MICAL2** | ENSG00000133816 | 1432.3367 | -0.52276067 | 0.182491775 | 0.002994006 | 0.009686154 | 9645 | 11 | 12115543 | 12285334 | 1 | protein_coding | microtubule associated monooxygenase, calponin and LIM domain containing 2 |
| **HSD17B4** | ENSG00000133835 | 2509.554876 | -1.079247501 | 0.166531747 | 2.43E-11 | 2.77E-10 | 3295 | 5 | 118788138 | 118972894 | 1 | protein_coding | hydroxysteroid (17-beta) dehydrogenase 4 |
| **ZFC3H1** | ENSG00000133858 | 2270.114054 | 0.583707689 | 0.169308296 | 3.24E-04 | 0.001327472 | 196441 | 12 | 72003252 | 72061505 | -1 | protein_coding | zinc finger, C3H1-type containing |
| **MED6** | ENSG00000133997 | 651.1218565 | 0.492591883 | 0.164750311 | 0.001944002 | 0.006600797 | 10001 | 14 | 71047974 | 71067384 | -1 | protein_coding | mediator complex subunit 6 |
| **EIF2S1** | ENSG00000134001 | 2807.735764 | 0.483276893 | 0.102234722 | 1.68E-06 | 1.05E-05 | 1965 | 14 | 67826714 | 67853233 | 1 | protein_coding | eukaryotic translation initiation factor 2, subunit 1 alpha, 35kDa |
| **PEBP4** | ENSG00000134020 | 36.25743214 | -1.16458802 | 0.692385546 | 0.013020363 | 0.035138657 | 157310 | 8 | 22570769 | 22857513 | -1 | protein_coding | phosphatidylethanolamine-binding protein 4 |
| **ADAMDEC1** | ENSG00000134028 | 24662.03495 | 0.592721843 | 0.115133584 | 1.44E-07 | 1.04E-06 | 27299 | 8 | 24241798 | 24263526 | 1 | protein_coding | ADAM-like, decysin 1 |
| **CTIF** | ENSG00000134030 | 886.9966449 | -1.819000115 | 0.252485478 | 5.26E-14 | 7.5E-13 | 9811 | 18 | 46065417 | 46389588 | 1 | protein_coding | CBP80/20-dependent translation initiation factor |
| **MRO** | ENSG00000134042 | 11.16671769 | -3.307708633 | 1.883711203 | 0.00546838 | 0.016487129 | 83876 | 18 | 48324574 | 48351772 | -1 | protein_coding | maestro |
| **IER3IP1** | ENSG00000134049 | 709.2327246 | 1.488851566 | 0.189552069 | 5.89E-16 | 9.57E-15 | 51124 | 18 | 44681413 | 44702745 | -1 | protein_coding | immediate early response 3 interacting protein 1 |
| **MRPS36** | ENSG00000134056 | 341.4181525 | 0.69272689 | 0.248678631 | 0.002228555 | 0.007428356 | 92259 | 5 | 68513587 | 68525956 | 1 | protein_coding | mitochondrial ribosomal protein S36 |
| **CD180** | ENSG00000134061 | 231.8170889 | -1.649990725 | 0.399328083 | 3.82E-06 | 2.25E-05 | 4064 | 5 | 66478103 | 66492627 | -1 | protein_coding | CD180 molecule |
| **IRAK2** | ENSG00000134070 | 352.3661582 | 2.035227362 | 0.193982745 | 9.97E-27 | 3.27E-25 | 3656 | 3 | 10206549 | 10285427 | 1 | protein_coding | interleukin-1 receptor-associated kinase 2 |
| **CAMK1** | ENSG00000134072 | 1165.906732 | -3.086505833 | 0.188793121 | 2.94E-61 | 3.91E-59 | 8536 | 3 | 9799026 | 9811676 | -1 | protein_coding | calcium/calmodulin-dependent protein kinase I |
| **VHL** | ENSG00000134086 | 1186.245452 | 0.557485031 | 0.139964306 | 4.31E-05 | 2.1E-04 | 7428 | 3 | 10182692 | 10193904 | 1 | protein_coding | von Hippel-Lindau tumor suppressor, E3 ubiquitin protein ligase |
| **BHLHE40** | ENSG00000134107 | 8819.396606 | -0.61747222 | 0.100534395 | 4.19E-10 | 4.18E-09 | 8553 | 3 | 5020801 | 5027008 | 1 | protein_coding | basic helix-loop-helix family, member e40 |
| **ARL8B** | ENSG00000134108 | 5699.299989 | 1.042651386 | 0.119019538 | 6.02E-19 | 1.22E-17 | 55207 | 3 | 5163905 | 5222596 | 1 | protein_coding | ADP-ribosylation factor-like 8B |
| **EDEM1** | ENSG00000134109 | 7727.930659 | 0.391019006 | 0.102895892 | 1.32E-04 | 5.89E-04 | 9695 | 3 | 5229331 | 5261642 | 1 | protein_coding | ER degradation enhancer, mannosidase alpha-like 1 |
| **KATNBL1** | ENSG00000134152 | 443.6751757 | 0.717681198 | 0.215769303 | 4.2E-04 | 0.001674539 | 79768 | 15 | 34432875 | 34502297 | -1 | protein_coding | katanin p80 subunit B-like 1 |
| **EMC7** | ENSG00000134153 | 2094.645021 | 0.630361015 | 0.105070048 | 1.22E-09 | 1.17E-08 | 56851 | 15 | 34376218 | 34394149 | -1 | protein_coding | ER membrane protein complex subunit 7 |
| **TSPAN2** | ENSG00000134198 | 35.63017055 | -1.347954757 | 0.76741944 | 0.008009981 | 0.023049622 | 10100 | 1 | 115590632 | 115632121 | -1 | protein_coding | tetraspanin 2 |
| **PSRC1** | ENSG00000134222 | 70.34443592 | -1.646097039 | 0.469342667 | 5.35E-05 | 2.57E-04 | 84722 | 1 | 109822178 | 109825808 | -1 | protein_coding | proline/serine-rich coiled-coil 1 |
| **PTPN22** | ENSG00000134242 | 2570.123426 | -0.411420048 | 0.112569656 | 1.57E-04 | 6.9E-04 | 26191 | 1 | 114356433 | 114414381 | -1 | protein_coding | protein tyrosine phosphatase, non-receptor type 22 (lymphoid) |
| **SORT1** | ENSG00000134243 | 12723.70085 | -1.173316494 | 0.079904174 | 1.86E-49 | 1.59E-47 | 6272 | 1 | 109852192 | 109940573 | -1 | protein_coding | sortilin 1 |
| **PTGFRN** | ENSG00000134247 | 804.5101097 | -2.22959652 | 0.209255895 | 1.31E-27 | 4.42E-26 | 5738 | 1 | 117452679 | 117532980 | 1 | protein_coding | prostaglandin F2 receptor inhibitor |
| **CEPT1** | ENSG00000134255 | 1516.463323 | -0.323950387 | 0.114730528 | 0.004071135 | 0.012702749 | 10390 | 1 | 111682249 | 111727724 | 1 | protein_coding | choline/ethanolamine phosphotransferase 1 |
| **CD101** | ENSG00000134256 | 1051.209621 | -3.027505656 | 0.20056901 | 1.3E-52 | 1.3E-50 | 9398 | 1 | 117544382 | 117579167 | 1 | protein_coding | CD101 molecule |
| **SPIRE1** | ENSG00000134278 | 2990.082959 | -0.359471768 | 0.133862535 | 0.006041776 | 0.017995802 | 56907 | 18 | 12446511 | 12658133 | -1 | protein_coding | spire-type actin nucleation factor 1 |
| **SLC38A2** | ENSG00000134294 | 8310.102093 | 0.587103477 | 0.109165 | 4.58E-08 | 3.57E-07 | 54407 | 12 | 46751972 | 46766650 | -1 | protein_coding | solute carrier family 38, member 2 |
| **PLEKHA8P1** | ENSG00000134297 | 114.8069349 | 0.745289697 | 0.338426127 | 0.010910338 | 0.030152386 | 51054 | 12 | 45566847 | 45609824 | -1 | pseudogene | pleckstrin homology domain containing, family A member 8 pseudogene 1 |
| **RSAD2** | ENSG00000134321 | 28759.06657 | 8.756324664 | 0.445229166 | 2.3E-87 | 6.5E-85 | 91543 | 2 | 7005937 | 7038370 | 1 | protein_coding | radical S-adenosyl methionine domain containing 2 |
| **LPIN1** | ENSG00000134324 | 2265.031468 | 0.547680539 | 0.123750998 | 5.08E-06 | 2.93E-05 | 23175 | 2 | 11817721 | 11967535 | 1 | protein_coding | lipin 1 |
| **CMPK2** | ENSG00000134326 | 2243.265103 | 6.071554801 | 0.231201216 | 3.14E-153 | 1.84E-150 | 129607 | 2 | 6980701 | 7006766 | -1 | protein_coding | cytidine monophosphate (UMP-CMP) kinase 2, mitochondrial |
| **LDHA** | ENSG00000134333 | 14018.30974 | -0.559353286 | 0.12446113 | 2.5E-06 | 1.52E-05 | 3939 | 11 | 18415935 | 18429972 | 1 | protein_coding | lactate dehydrogenase A |
| **SAA2** | ENSG00000134339 | 12.04423279 | 6.976949449 | 2.091059762 | 1.25E-05 | 6.81E-05 | 6289 | 11 | 18260770 | 18270190 | -1 | protein_coding | serum amyloid A2 |
| **IL6ST** | ENSG00000134352 | 4505.583105 | 0.730854241 | 0.144317043 | 2.14E-07 | 1.52E-06 | 3572 | 5 | 55230923 | 55290821 | -1 | protein_coding | interleukin 6 signal transducer (gp130, oncostatin M receptor) |
| **NAV1** | ENSG00000134369 | 500.4854108 | -1.039519688 | 0.332143438 | 4E-04 | 0.001604364 | 89796 | 1 | 201592411 | 201796102 | 1 | protein_coding | neuron navigator 1 |
| **CDC73** | ENSG00000134371 | 2288.728581 | 0.619716805 | 0.129687603 | 1.01E-06 | 6.51E-06 | 79577 | 1 | 193091147 | 193223031 | 1 | protein_coding | cell division cycle 73 |

| | | | | | | | | | | | | | |
|---|---|---|---|---|---|---|---|---|---|---|---|---|---|
| **RPS15A** | ENSG00000134419 | 4127.045019 | -0.381840216 | 0.153416081 | 0.010047897 | 0.028119761 | 6210 | 16 | 18792617 | 18801705 | -1 | protein_coding | ribosomal protein S15a |
| **NARS** | ENSG00000134440 | 7554.610678 | -0.453053614 | 0.122294659 | 1.51E-04 | 6.66E-04 | 4677 | 18 | 55267888 | 55289445 | -1 | protein_coding | asparaginyl-tRNA synthetase |
| **RBM17** | ENSG00000134453 | 1225.274941 | 0.538395302 | 0.134547304 | 4.12E-05 | 2.02E-04 | 84991 | 10 | 6130950 | 6159420 | 1 | protein_coding | RNA binding motif protein 17 |
| **IL2RA** | ENSG00000134460 | 1167.777508 | 1.936595273 | 0.632296654 | 1.72E-04 | 7.48E-04 | 3559 | 10 | 6052652 | 6104288 | -1 | protein_coding | interleukin 2 receptor, alpha |
| **IL15RA** | ENSG00000134470 | 848.6140781 | 3.411434527 | 0.317010943 | 3.5E-28 | 1.23E-26 | 3601 | 10 | 5990855 | 6020150 | -1 | protein_coding | interleukin 15 receptor, alpha |
| **EMP1** | ENSG00000134531 | 6754.319509 | 0.464319904 | 0.19057065 | 0.010416154 | 0.028948913 | 2012 | 12 | 13349650 | 13369708 | 1 | protein_coding | epithelial membrane protein 1 |
| **DDB2** | ENSG00000134574 | 876.212731 | 0.33423267 | 0.126009834 | 0.006641237 | 0.019567769 | 1643 | 11 | 47236493 | 47260767 | 1 | protein_coding | damage-specific DNA binding protein 2, 48kDa |
| **ACP2** | ENSG00000134575 | 5867.830997 | 0.52557842 | 0.089055184 | 2.31E-09 | 2.11E-08 | 53 | 11 | 47260853 | 47270457 | -1 | protein_coding | acid phosphatase 2, lysosomal |
| **FAM127A** | ENSG00000134590 | 539.9030878 | -0.728699143 | 0.184059591 | 3.47E-05 | 1.73E-04 | 8933 | X | 134166333 | 134167576 | 1 | protein_coding | family with sequence similarity 127, member A |
| **MST4** | ENSG00000134602 | 419.5860209 | 2.14740521 | 0.225060169 | 1.07E-22 | 2.82E-21 | 51765 | X | 131157293 | 131209971 | 1 | protein_coding | Serine/threonine-protein kinase MST4 |
| **PIWIL4** | ENSG00000134627 | 227.1619271 | 1.264824989 | 0.298126142 | 3.89E-06 | 2.29E-05 | 143689 | 11 | 94277006 | 94354587 | 1 | protein_coding | piwi-like RNA-mediated gene silencing 4 |
| **SPOCD1** | ENSG00000134668 | 486.3292315 | -1.784892838 | 0.308848976 | 7.91E-10 | 7.67E-09 | 90853 | 1 | 32256023 | 32281652 | -1 | protein_coding | SPOC domain containing 1 |
| **PHC2** | ENSG00000134686 | 2489.666095 | -1.175081058 | 0.142434594 | 3.66E-17 | 6.47E-16 | 1912 | 1 | 33789224 | 33896653 | -1 | protein_coding | polyhomeotic homolog 2 (Drosophila) |
| **CYP2J2** | ENSG00000134716 | 11.1932209 | 2.964283835 | 1.413806302 | 0.001131983 | 0.004080212 | 1573 | 1 | 60358980 | 60392462 | -1 | protein_coding | cytochrome P450, family 2, subfamily J, polypeptide 2 |
| **BTF3L4** | ENSG00000134717 | 1250.104085 | 0.425177576 | 0.135155976 | 0.001273163 | 0.004521084 | 91408 | 1 | 52521797 | 52556388 | 1 | protein_coding | basic transcription factor 3-like 4 |
| **RNF138** | ENSG00000134758 | 406.5163667 | 1.135463677 | 0.246343967 | 9.49E-07 | 6.12E-06 | 51444 | 18 | 29671818 | 29711524 | 1 | protein_coding | ring finger protein 138, E3 ubiquitin protein ligase |
| **DTNA** | ENSG00000134769 | 248.3246261 | -1.834302348 | 0.313327257 | 5.58E-10 | 5.49E-09 | 1837 | 18 | 32073254 | 32471808 | 1 | protein_coding | dystrobrevin, alpha |
| **DAGLA** | ENSG00000134780 | 816.0078437 | -1.026680662 | 0.155841373 | 1.28E-11 | 1.49E-10 | 747 | 11 | 61447905 | 61514473 | 1 | protein_coding | diacylglycerol lipase, alpha |
| **SLC43A3** | ENSG00000134802 | 7062.696491 | 0.436762885 | 0.122048921 | 2.63E-04 | 0.001093925 | 29015 | 11 | 57174427 | 57195053 | -1 | protein_coding | solute carrier family 43, member 3 |
| **TIMM10** | ENSG00000134809 | 246.7894573 | -0.551899137 | 0.224721848 | 0.008763018 | 0.024939517 | 26519 | 11 | 57295936 | 57298276 | -1 | protein_coding | translocase of inner mitochondrial membrane 10 homolog (yeast) |
| **FADS2** | ENSG00000134824 | 2270.75175 | -1.046934015 | 0.104803554 | 8.74E-24 | 2.44E-22 | 9415 | 11 | 61560452 | 61634826 | 1 | protein_coding | fatty acid desaturase 2 |
| **TCN1** | ENSG00000134827 | 6.964649595 | -0.566424407 | 1.033809664 | 0.010357267 | 0.028832444 | 6947 | 11 | 59620273 | 59634048 | -1 | protein_coding | transcobalamin I (vitamin B12 binding protein, R binder family) |
| **C5AR2** | ENSG00000134830 | 333.0691929 | -1.066765788 | 0.224950828 | 5.44E-07 | 3.66E-06 | 27202 | 19 | 47835404 | 47846606 | 1 | protein_coding | complement component 5a receptor 2 |
| **TMEM165** | ENSG00000134851 | 2237.091826 | 1.073859191 | 0.115497187 | 4.11E-21 | 9.82E-20 | 55858 | 4 | 56262124 | 56319564 | 1 | protein_coding | transmembrane protein 165 |
| **COL4A2** | ENSG00000134871 | 112.4822113 | 1.418170043 | 0.417518377 | 9.68E-05 | 4.44E-04 | 1284 | 13 | 110958159 | 111165374 | 1 | protein_coding | collagen, type IV, alpha 2 |
| **ETS1** | ENSG00000134954 | 1642.212272 | -0.523078581 | 0.145203412 | 1.99E-04 | 8.53E-04 | 2113 | 11 | 128328656 | 128457453 | -1 | protein_coding | v-ets avian erythroblastosis virus E26 oncogene homolog 1 |
| **SLC37A2** | ENSG00000134955 | 4789.836652 | -3.121467024 | 0.127165522 | 1.85E-133 | 8.08E-131 | 219855 | 11 | 124932963 | 124959131 | 1 | protein_coding | solute carrier family 37 (glucose-6-phosphate transporter), member 2 |
| **TMED7** | ENSG00000134970 | 2210.04814 | 0.543779614 | 0.112390082 | 8.44E-07 | 5.49E-06 | 1003027 | 5 | 114949205 | 114968689 | -1 | protein_coding | transmembrane emp24 protein transport domain containing 7 |
| **RFK** | ENSG00000135002 | 522.6975472 | 0.723100763 | 0.174216263 | 1.53E-05 | 8.18E-05 | 55312 | 9 | 79000433 | 79009433 | -1 | protein_coding | riboflavin kinase |
| **UBQLN1** | ENSG00000135018 | 4163.536999 | 0.365343116 | 0.093739199 | 6.56E-05 | 3.1E-04 | 29979 | 9 | 86274878 | 86323118 | -1 | protein_coding | ubiquilin 1 |
| **NAA35** | ENSG00000135040 | 575.4187506 | 0.516274273 | 0.144762506 | 2.51E-04 | 0.001049997 | 60560 | 9 | 88556061 | 88637213 | 1 | protein_coding | N(alpha)-acetyltransferase 35, NatC auxiliary subunit |
| **ANXA1** | ENSG00000135046 | 6259.125825 | 0.557125327 | 0.126273471 | 6.24E-06 | 3.55E-05 | 301 | 9 | 75766673 | 75785309 | 1 | protein_coding | annexin A1 |
| **CTSL** | ENSG00000135047 | 18385.05402 | 0.481207731 | 0.107758099 | 5.66E-06 | 3.25E-05 | 1514 | 9 | 90340434 | 90346308 | 1 | protein_coding | cathepsin L |
| **TMEM2** | ENSG00000135048 | 1569.394631 | -0.606055278 | 0.173943187 | 2.61E-04 | 0.001086198 | 23670 | 9 | 74298282 | 74431606 | -1 | protein_coding | transmembrane protein 2 |
| **GOLM1** | ENSG00000135052 | 227.9510291 | 1.471135544 | 0.302912505 | 1.79E-07 | 1.28E-06 | 51280 | 9 | 88641061 | 88715088 | -1 | protein_coding | golgi membrane protein 1 |
| **ISCA1** | ENSG00000135070 | 472.3351725 | 1.098435708 | 0.194024902 | 3.96E-09 | 3.53E-08 | 81689 | 9 | 88879461 | 88897676 | -1 | protein_coding | iron-sulfur cluster assembly 1 |
| **ADAM19** | ENSG00000135074 | 944.2969866 | 1.508145646 | 0.573273347 | 9.41E-04 | 0.003456785 | 8728 | 5 | 156822542 | 157002783 | -1 | protein_coding | ADAM metallopeptidase domain 19 |
| **SDS** | ENSG00000135094 | 836.0548163 | 1.34928408 | 0.226900156 | 4.71E-10 | 4.68E-09 | 10993 | 12 | 113830250 | 113864106 | -1 | protein_coding | serine dehydratase |
| **OASL** | ENSG00000135114 | 1072.834176 | 6.719332381 | 0.196663312 | 1.11E-256 | 1.41E-253 | 8638 | 12 | 121458095 | 121477045 | -1 | protein_coding | 2'-5'-oligoadenylate synthetase-like |
| **RNFT2** | ENSG00000135119 | 78.85570862 | -1.162913617 | 0.54726466 | 0.005201078 | 0.015793379 | 84900 | 12 | 117176096 | 117291436 | 1 | protein_coding | ring finger protein, transmembrane 2 |
| **P2RX4** | ENSG00000135124 | 2337.635889 | 0.359684883 | 0.128766681 | 0.004173051 | 0.012986508 | 5025 | 12 | 121647660 | 121671909 | 1 | protein_coding | purinergic receptor P2X, ligand-gated ion channel, 4 |
| **TRAFD1** | ENSG00000135148 | 2015.613022 | 1.695481452 | 0.127886779 | 4.85E-41 | 3.1E-39 | 10906 | 12 | 112563305 | 112591407 | 1 | protein_coding | TRAF-type zinc finger domain containing 1 |
| **DMTF1** | ENSG00000135164 | 1179.396279 | 0.634762155 | 0.155163375 | 2.16E-05 | 1.12E-04 | 9988 | 7 | 86781677 | 86825653 | 1 | protein_coding | cyclin D binding myb-like transcription factor 1 |
| **TMEM60** | ENSG00000135211 | 437.7245274 | 1.290192216 | 0.19544939 | 7.73E-12 | 9.18E-11 | 85025 | 7 | 77423045 | 77427897 | -1 | protein_coding | transmembrane protein 60 |
| **CD36** | ENSG00000135218 | 6391.870527 | -2.236512737 | 0.15300641 | 2.81E-50 | 2.54E-48 | 948 | 7 | 79998891 | 80308593 | 1 | protein_coding | CD36 molecule (thrombospondin receptor) |
| **PNPLA8** | ENSG00000135241 | 1390.227094 | 0.736693794 | 0.183638127 | 2.81E-05 | 1.42E-04 | 50640 | 7 | 108110866 | 108210110 | -1 | protein_coding | patatin-like phospholipase domain containing 8 |
| **HILPDA** | ENSG00000135245 | 175.8636286 | -2.295172901 | 0.333682664 | 5.46E-13 | 7.15E-12 | 29923 | 7 | 128095903 | 128098472 | 1 | protein_coding | hypoxia inducible lipid droplet-associated |
| **SRPK2** | ENSG00000135250 | 1499.796814 | -0.528842348 | 0.115968626 | 3.39E-06 | 2.01E-05 | 6733 | 7 | 104751151 | 105039755 | -1 | protein_coding | SRSF protein kinase 2 |
| **ANKRD6** | ENSG00000135299 | 484.8295856 | -0.837312698 | 0.237945516 | 1.65E-04 | 7.22E-04 | 22881 | 6 | 90142889 | 90343553 | 1 | protein_coding | ankyrin repeat domain 6 |
| **SNX14** | ENSG00000135317 | 2451.780809 | 0.687042717 | 0.161840348 | 1.14E-05 | 6.22E-05 | 57231 | 6 | 86215214 | 86303874 | -1 | protein_coding | sorting nexin 14 |
| **AKIRIN2** | ENSG00000135334 | 2003.738729 | 0.853287244 | 0.110205877 | 3.82E-15 | 5.82E-14 | 55122 | 6 | 88384790 | 88411927 | -1 | protein_coding | akirin 2 |
| **ORC3** | ENSG00000135336 | 562.6227429 | 0.87232134 | 0.158113175 | 1.27E-08 | 1.06E-07 | 23595 | 6 | 88299839 | 88377169 | 1 | protein_coding | origin recognition complex, subunit 3 |
| **LMO2** | ENSG00000135363 | 1432.614789 | -0.628802244 | 0.119865856 | 8.31E-08 | 6.23E-07 | 4005 | 11 | 33880122 | 33913836 | -1 | protein_coding | LIM domain only 2 (rhombotin-like 1) |
| **EHF** | ENSG00000135373 | 77.68733726 | -0.920768547 | 0.426338464 | 0.008178909 | 0.023473852 | 26298 | 11 | 34642640 | 34682604 | 1 | protein_coding | ets homologous factor |
| **PRRG4** | ENSG00000135378 | 316.3732763 | 2.271687387 | 0.277948662 | 2.56E-17 | 4.59E-16 | 79056 | 11 | 32851489 | 32879669 | 1 | protein_coding | proline rich Gla (G-carboxyglutamic acid) 4 (transmembrane) |
| **GDF11** | ENSG00000135414 | 65.63962525 | -0.795990309 | 0.403387315 | 0.016296448 | 0.042712848 | 10220 | 12 | 56137064 | 56150911 | 1 | protein_coding | growth differentiation factor 11 |
| **TESPA1** | ENSG00000135426 | 325.6645137 | 0.621955848 | 0.237322305 | 0.004605933 | 0.014197755 | 9840 | 12 | 55341802 | 55378530 | -1 | protein_coding | thymocyte expressed, positive selection associated 1 |
| **CDK4** | ENSG00000135446 | 1296.64387 | -0.322824798 | 0.130180543 | 0.011102703 | 0.030623101 | 1019 | 12 | 58141510 | 58149796 | -1 | protein_coding | cyclin-dependent kinase 4 |
| **PPP1R1A** | ENSG00000135447 | 25.1647669 | 3.410088041 | 1.297502656 | 2.83E-04 | 0.001172665 | 5502 | 12 | 54969171 | 54982443 | -1 | protein_coding | protein phosphatase 1, regulatory (inhibitor) subunit 1A |
| **FAIM2** | ENSG00000135472 | 708.2932972 | -1.711280272 | 0.190713037 | 3.98E-20 | 8.87E-19 | 23017 | 12 | 50260679 | 50298000 | -1 | protein_coding | Fas apoptotic inhibitory molecule 2 |
| **PAN2** | ENSG00000135473 | 557.2850707 | -0.689872123 | 0.233252138 | 0.001373608 | 0.00483261 | 9924 | 12 | 56710007 | 56727837 | -1 | protein_coding | PAN2 poly(A) specific ribonuclease subunit homolog (S. cerevisiae) |
| **MAP7** | ENSG00000135525 | 1341.893097 | -1.595298805 | 0.170387996 | 1.15E-21 | 2.84E-20 | 9053 | 6 | 136663875 | 136871957 | -1 | protein_coding | microtubule-associated protein 7 |
| **CD164** | ENSG00000135535 | 18980.31746 | 0.776602223 | 0.171017966 | 2.43E-06 | 1.48E-05 | 8763 | 6 | 109687717 | 109703762 | -1 | protein_coding | CD164 molecule, sialomucin |
| **SMPD2** | ENSG00000135587 | 384.8544412 | 1.028367378 | 0.211798057 | 3.33E-07 | 2.3E-06 | 6610 | 6 | 109761966 | 109765122 | 1 | protein_coding | sphingomyelin phosphodiesterase 2, neutral membrane (neutral sphingomyelinase) |
| **MICAL1** | ENSG00000135596 | 3811.643608 | -0.362392161 | 0.145605 | 0.010265802 | 0.028619527 | 64780 | 6 | 109765265 | 109787171 | -1 | protein_coding | microtubule associated monooxygenase, calponin and LIM domain containing 1 |
| **STX11** | ENSG00000135604 | 1748.62187 | 2.265960227 | 0.218558035 | 2.95E-26 | 9.38E-25 | 8676 | 6 | 144471663 | 144509507 | 1 | protein_coding | syntaxin 11 |
| **TEC** | ENSG00000135605 | 306.8028888 | 0.66538772 | 0.187558536 | 2.06E-04 | 8.76E-04 | 7006 | 4 | 48137800 | 48271881 | -1 | protein_coding | tec protein tyrosine kinase |
| **PRADC1** | ENSG00000135617 | 250.583947 | -2.085346892 | 0.263182545 | 2.25E-16 | 3.75E-15 | 84279 | 2 | 73455134 | 73460366 | -1 | protein_coding | protease-associated domain containing 1 |
| **RAB11FIP5** | ENSG00000135631 | 403.2959775 | -1.041978243 | 0.275884651 | 4.09E-05 | 2E-04 | 26056 | 2 | 73300510 | 73383849 | -1 | protein_coding | RAB11 family interacting protein 5 (class I) |
| **SMYD5** | ENSG00000135632 | 235.0671233 | -0.642965371 | 0.283616783 | 0.011308811 | 0.031129784 | 10322 | 2 | 73441350 | 73454365 | 1 | protein_coding | SMYD family member 5 |
| **DYSF** | ENSG00000135636 | 1473.62597 | -0.728089126 | 0.291484801 | 0.005197015 | 0.015787316 | 8291 | 2 | 71680852 | 71913898 | 1 | protein_coding | dysferlin |
| **USP15** | ENSG00000135655 | 4510.973755 | 1.311265011 | 0.150343722 | 6.35E-19 | 1.28E-17 | 1024651 | 12 | 62654119 | 62811211 | 1 | protein_coding | ubiquitin specific peptidase 15 |
| **GNS** | ENSG00000135677 | 57155.12465 | 0.638142216 | 0.093257849 | 5E-12 | 6.06E-11 | 2799 | 12 | 65107225 | 65153227 | -1 | protein_coding | glucosamine (N-acetyl)-6-sulfatase |
| **CPM** | ENSG00000135678 | 15360.32597 | 0.396409228 | 0.119890754 | 8.58E-04 | 0.003187384 | 1368 | 12 | 69235977 | 69365350 | -1 | protein_coding | carboxypeptidase M |
| **MDM2** | ENSG00000135679 | 2780.82461 | 0.938537046 | 0.108256337 | 1.65E-18 | 3.25E-17 | 4193 | 12 | 69201956 | 69239214 | 1 | protein_coding | MDM2 oncogene, E3 ubiquitin protein ligase |
| **KLHL36** | ENSG00000135686 | 1693.611999 | -0.995888038 | 0.121149019 | 6.44E-17 | 1.12E-15 | 79786 | 16 | 84682131 | 84701292 | 1 | protein_coding | kelch-like family member 36 |
| **DYNC1LI2** | ENSG00000135720 | 2865.976817 | 0.463172009 | 0.14482995 | 0.001001781 | 0.003653118 | 1783 | 16 | 66754796 | 66785701 | -1 | protein_coding | dynein, cytoplasmic 1, light intermediate chain 2 |
| **FHOD1** | ENSG00000135723 | 763.287342 | -1.426264982 | 0.168940409 | 5.82E-18 | 1.1E-16 | 29109 | 16 | 67263290 | 67281561 | -1 | protein_coding | formin homology 2 domain containing 1 |
| **EGLN1** | ENSG00000135766 | 2031.985255 | -0.434397489 | 0.113157967 | 9.26E-05 | 4.26E-04 | 54583 | 1 | 231499497 | 231560790 | -1 | protein_coding | egl-9 family hypoxia-inducible factor 1 |
| **COG2** | ENSG00000135775 | 1057.934511 | -0.520627473 | 0.129354001 | 3.76E-05 | 1.86E-04 | 22796 | 1 | 230778235 | 230829728 | 1 | protein_coding | component of oligomeric golgi complex 2 |
| **TAF5L** | ENSG00000135801 | 483.7885791 | -1.0287102 | 0.255950211 | 1.43E-05 | 7.67E-05 | 27097 | 1 | 229728858 | 229761794 | -1 | protein_coding | TAF5-like RNA polymerase II, p300/CBP-associated factor (PCAF)-associated factor, 65kDa |
| **RNASEL** | ENSG00000135828 | 884.7702652 | -0.640874777 | 0.223413768 | 0.002019674 | 0.006822874 | 6041 | 1 | 182542769 | 182558391 | -1 | protein_coding | ribonuclease L (2',5'-oligoisoadenylate synthetase-dependent) |
| **CEP350** | ENSG00000135837 | 4198.603231 | 0.456145927 | 0.145739554 | 0.001211305 | 0.004328355 | 9857 | 1 | 179923873 | 180084015 | 1 | protein_coding | centrosomal protein 350kDa |
| **NPL** | ENSG00000135838 | 6026.290827 | -0.942444787 | 0.200736262 | 8.51E-07 | 5.53E-06 | 80896 | 1 | 182758428 | 182799519 | 1 | protein_coding | N-acetylneuraminate pyruvate lyase (dihydrodipicolinate synthase) |
| **LAMC1** | ENSG00000135862 | 2652.811353 | -0.608213197 | 0.157978158 | 6.76E-05 | 3.19E-04 | 3915 | 1 | 182992595 | 183114727 | 1 | protein_coding | laminin, gamma 1 (formerly LAMB2) |
| **GPR55** | ENSG00000135898 | 52.45463478 | -1.825874547 | 0.658919969 | 4.92E-04 | 0.001928471 | 9290 | 2 | 231772033 | 231825781 | -1 | protein_coding | G protein-coupled receptor 55 |
| **SP110** | ENSG00000135899 | 2825.686668 | 2.409648626 | 0.12498897 | 5.82E-84 | 1.48E-81 | 3431 | 2 | 231032009 | 231090444 | -1 | protein_coding | SP110 nuclear body protein |
| **MRPL44** | ENSG00000135900 | 1444.639685 | 1.02590585 | 0.115051927 | 1.33E-19 | 2.85E-18 | 65080 | 2 | 224822121 | 224832431 | 1 | protein_coding | mitochondrial ribosomal protein L44 |
| **DOCK10** | ENSG00000135905 | 6780.428778 | -0.983304927 | 0.087859805 | 1.49E-29 | 5.61E-28 | 55619 | 2 | 225629807 | 225907162 | -1 | protein_coding | dedicator of cytokinesis 10 |

| | | | | | | | | | | | | | |
|---|---|---|---|---|---|---|---|---|---|---|---|---|---|
| **TTLL4** | ENSG00000135912 | 1682.355166 | -0.636161618 | 0.143150872 | 5.03E-06 | 2.91E-05 | 9654 | 2 | 219575568 | 219620139 | 1 | protein_coding | tubulin tyrosine ligase-like family, member 4 |
| **USP37** | ENSG00000135913 | 778.4725977 | 0.503376816 | 0.188121058 | 0.004882099 | 0.014964525 | 57695 | 2 | 219314974 | 219433084 | -1 | protein_coding | ubiquitin specific peptidase 37 |
| **ITM2C** | ENSG00000135916 | 189.5523818 | -0.864968776 | 0.316738265 | 0.002010246 | 0.006794026 | 81618 | 2 | 231729354 | 231743963 | 1 | protein_coding | integral membrane protein 2C |
| **TMBIM1** | ENSG00000135926 | 11900.87755 | 0.559170034 | 0.094117963 | 1.38E-09 | 1.3E-08 | 1024652 | 2 | 219138915 | 219157309 | -1 | protein_coding | transmembrane BAX inhibitor motif containing 1 |
| **CYP27A1** | ENSG00000135929 | 18853.64194 | -1.773389779 | 0.102332636 | 4.5E-68 | 7.99E-66 | 1593 | 2 | 219646472 | 219680016 | 1 | protein_coding | cytochrome P450, family 27, subfamily A, polypeptide 1 |
| **ARMC9** | ENSG00000135931 | 917.7287165 | 1.517731485 | 0.161888056 | 1.11E-21 | 2.75E-20 | 80210 | 2 | 232063260 | 232239548 | 1 | protein_coding | armadillo repeat containing 9 |
| **REV1** | ENSG00000135945 | 690.8784716 | 0.599267656 | 0.17001701 | 2.42E-04 | 0.001013111 | 51455 | 2 | 100016938 | 100106497 | -1 | protein_coding | REV1, polymerase (DNA directed) |
| **TMEM127** | ENSG00000135956 | 6521.721471 | -0.255467426 | 0.108162957 | 0.016060606 | 0.04215254 | 55654 | 2 | 96914254 | 96931732 | -1 | protein_coding | transmembrane protein 127 |
| **EDAR** | ENSG00000135960 | 13.7043501 | -3.382161269 | 1.309203233 | 0.001095468 | 0.003964198 | 10913 | 2 | 109510927 | 109605828 | -1 | protein_coding | ectodysplasin A receptor |
| **TGFBRAP1** | ENSG00000135966 | 1132.205384 | -0.417173577 | 0.123959148 | 6.08E-04 | 0.002329498 | 9392 | 2 | 105880871 | 105946491 | -1 | protein_coding | transforming growth factor, beta receptor associated protein 1 |
| **GCC2** | ENSG00000135968 | 3653.196365 | 1.497225029 | 0.137612823 | 2.11E-28 | 7.52E-27 | 9648 | 2 | 109065017 | 109125871 | 1 | protein_coding | GRIP and coiled-coil domain containing 2 |
| **EPC2** | ENSG00000135999 | 783.2415035 | 0.726936288 | 0.140161235 | 9.97E-08 | 7.39E-07 | 26122 | 2 | 149402009 | 149545130 | 1 | protein_coding | enhancer of polycomb homolog 2 (Drosophila) |
| **ISCU** | ENSG00000136003 | 2427.296246 | 0.360534752 | 0.105591795 | 4.17E-04 | 0.001666535 | 23479 | 12 | 108956358 | 108963160 | 1 | protein_coding | iron-sulfur cluster assembly enzyme |
| **ALDH1L2** | ENSG00000136010 | 1308.566749 | -0.487989654 | 0.159119403 | 0.001480761 | 0.005179784 | 160428 | 12 | 105413568 | 105478355 | -1 | protein_coding | aldehyde dehydrogenase 1 family, member L2 |
| **SCYL2** | ENSG00000136021 | 2823.197538 | 0.383938375 | 0.125630894 | 0.001619087 | 0.005605915 | 55681 | 12 | 100660918 | 100735502 | 1 | protein_coding | SCY1-like 2 (S. cerevisiae) |
| **CKAP4** | ENSG00000136026 | 2389.895805 | 1.872623959 | 0.125490178 | 2.19E-51 | 2.04E-49 | 10970 | 12 | 106631655 | 106698057 | -1 | protein_coding | cytoskeleton-associated protein 4 |
| **APPL2** | ENSG00000136044 | 691.3692805 | -1.374227039 | 0.173755558 | 4.37E-16 | 7.14E-15 | 55198 | 12 | 105567074 | 105630016 | -1 | protein_coding | adaptor protein, phosphotyrosine interaction, PH domain and leucine zipper containing 2 |
| **DRAM1** | ENSG00000136048 | 7506.589352 | 1.078422865 | 0.094342972 | 5.28E-31 | 2.15E-29 | 55332 | 12 | 102271129 | 102405908 | 1 | protein_coding | DNA-damage regulated autophagy modulator 1 |
| **KIAA1033** | ENSG00000136051 | 6374.285658 | 0.441604315 | 0.121227431 | 1.97E-04 | 8.45E-04 | 23325 | 12 | 105501102 | 105562912 | 1 | protein_coding | KIAA1033 |
| **SLC41A2** | ENSG00000136052 | 2348.797869 | 0.976463986 | 0.13235649 | 6.88E-14 | 9.68E-13 | 84102 | 12 | 105196331 | 105352522 | -1 | protein_coding | solute carrier family 41 (magnesium transporter), member 2 |
| **VILL** | ENSG00000136059 | 140.8492487 | 0.696298275 | 0.327430213 | 0.01454586 | 0.03866163 | 50853 | 3 | 38029550 | 38048679 | 1 | protein_coding | villin-like |
| **FLNB** | ENSG00000136068 | 1351.628706 | -1.758998123 | 0.202257171 | 4.36E-19 | 9E-18 | 2317 | 3 | 57994127 | 58157982 | 1 | protein_coding | filamin B, beta |
| **RNASEH2B** | ENSG00000136104 | 559.3245492 | 0.774397893 | 0.185038189 | 1.24E-05 | 6.74E-05 | 79621 | 13 | 51483814 | 51544592 | 1 | protein_coding | ribonuclease H2, subunit B |
| **TBC1D4** | ENSG00000136111 | 1396.305488 | -1.119882924 | 0.173758688 | 3.14E-11 | 3.54E-10 | 9882 | 13 | 75858808 | 76056250 | -1 | protein_coding | TBC1 domain family, member 4 |
| **BORA** | ENSG00000136122 | 127.4125464 | 0.890581639 | 0.369753291 | 0.004850871 | 0.01488971 | 79866 | 13 | 73302061 | 73330336 | 1 | protein_coding | bora, aurora kinase A activator |
| **LRCH1** | ENSG00000136141 | 3840.307068 | 0.394861337 | 0.161224602 | 0.011047321 | 0.030492363 | 23143 | 13 | 47127303 | 47327175 | 1 | protein_coding | leucine-rich repeats and calponin homology (CH) domain containing 1 |
| **PHF11** | ENSG00000136147 | 1046.538784 | 1.835352569 | 0.145055378 | 1.24E-37 | 6.84E-36 | 51131 | 13 | 50069746 | 50103123 | 1 | protein_coding | PHD finger protein 11 |
| **COG3** | ENSG00000136152 | 820.2403591 | 0.328429185 | 0.129882184 | 0.009621458 | 0.027123409 | 83548 | 13 | 46039060 | 46110765 | 1 | protein_coding | component of oligomeric golgi complex 3 |
| **ITM2B** | ENSG00000136156 | 3827.217928 | 0.744588499 | 0.126398827 | 1.8E-09 | 1.68E-08 | 9445 | 13 | 48807294 | 48837063 | 1 | protein_coding | integral membrane protein 2B |
| **SPRY2** | ENSG00000136158 | 1154.44028 | -2.354002535 | 0.151335289 | 1.29E-55 | 1.45E-53 | 10253 | 13 | 80910111 | 80915086 | -1 | protein_coding | sprouty homolog 2 (Drosophila) |
| **RCBTB2** | ENSG00000136161 | 1740.726032 | -1.479936262 | 0.167108571 | 1.3E-19 | 2.77E-18 | 1102 | 13 | 49063095 | 49107369 | -1 | protein_coding | regulator of chromosome condensation (RCC1) and BTB (POZ) domain containing protein 2 |
| **LCP1** | ENSG00000136167 | 64048.86128 | -0.862349707 | 0.093181934 | 6.07E-21 | 1.43E-19 | 3936 | 13 | 46700055 | 46786006 | -1 | protein_coding | lymphocyte cytosolic protein 1 (L-plastin) |
| **SETDB2** | ENSG00000136169 | 1166.734623 | 0.821776454 | 0.139880828 | 1.69E-09 | 1.58E-08 | 83852 | 13 | 50018429 | 50069138 | 1 | protein_coding | SET domain, bifurcated 2 |
| **SCRN1** | ENSG00000136193 | 1671.902929 | 0.9997395 | 0.158901531 | 9.35E-11 | 1E-09 | 9805 | 7 | 29959719 | 30029905 | -1 | protein_coding | secernin 1 |
| **TNS3** | ENSG00000136205 | 18565.00867 | -0.460387928 | 0.115484448 | 4.73E-05 | 2.29E-04 | 64759 | 7 | 47314752 | 47622156 | -1 | protein_coding | tensin 3 |
| **CHST12** | ENSG00000136213 | 200.0519671 | 0.64341095 | 0.236509969 | 0.00328348 | 0.010515807 | 55501 | 7 | 2443223 | 2474242 | 1 | protein_coding | carbohydrate (chondroitin 4) sulfotransferase 12 |
| **IGF2BP3** | ENSG00000136231 | 452.8403545 | 1.920732802 | 0.231829985 | 1.29E-17 | 2.36E-16 | 10643 | 7 | 23349828 | 23510086 | -1 | protein_coding | insulin-like growth factor 2 mRNA binding protein 3 |
| **GPNMB** | ENSG00000136235 | 183883.4973 | -0.682374958 | 0.198720334 | 2.25E-04 | 9.49E-04 | 10457 | 7 | 23275586 | 23314727 | 1 | protein_coding | glycoprotein (transmembrane) nmb |
| **IL6** | ENSG00000136244 | 61.63837309 | 4.935608049 | 0.654842947 | 3.03E-15 | 4.68E-14 | 3569 | 7 | 22765503 | 22771621 | 1 | protein_coding | interleukin 6 (interferon, beta 2) |
| **DDX56** | ENSG00000136271 | 637.3150086 | -0.387517955 | 0.170649683 | 0.017812303 | 0.046022895 | 54606 | 7 | 44605016 | 44614650 | -1 | protein_coding | DEAD (Asp-Glu-Ala-Asp) box helicase 56 |
| **HUS1** | ENSG00000136273 | 619.8211803 | 0.564341371 | 0.148816859 | 9.3E-05 | 4.28E-04 | 3364 | 7 | 47735328 | 48019178 | -1 | protein_coding | HUS1 checkpoint homolog (S. pombe) |
| **MYO1G** | ENSG00000136286 | 1150.216996 | 2.370752265 | 0.284124174 | 5.71E-18 | 1.08E-16 | 64005 | 7 | 45002261 | 45018697 | -1 | protein_coding | myosin IG |
| **TTYH3** | ENSG00000136295 | 11552.7206 | -1.12083506 | 0.116332455 | 1.31E-22 | 3.43E-21 | 80727 | 7 | 2671585 | 2704436 | 1 | protein_coding | tweety family member 3 |
| **ADAMTS7** | ENSG00000136378 | 7.202801787 | -3.409259457 | 3.367054172 | 0.004736732 | 0.014556904 | 11173 | 15 | 79051545 | 79103773 | -1 | protein_coding | ADAM metallopeptidase with thrombospondin type 1 motif, 7 |
| **ALPK3** | ENSG00000136383 | 1849.626053 | -1.459710183 | 0.219677142 | 4.81E-12 | 5.83E-11 | 57538 | 15 | 85359911 | 85416713 | 1 | protein_coding | alpha-kinase 3 |
| **TM6SF1** | ENSG00000136404 | 1778.277705 | -1.651064408 | 0.111794356 | 3.45E-50 | 3.1E-48 | 53346 | 15 | 83776159 | 83813606 | 1 | protein_coding | transmembrane 6 superfamily member 1 |
| **CALCOCO2** | ENSG00000136436 | 3157.456037 | 0.968366761 | 0.115444402 | 1.39E-17 | 2.54E-16 | 10241 | 17 | 46908350 | 46943884 | 1 | protein_coding | calcium binding and coiled-coil domain 2 |
| **VEZF1** | ENSG00000136451 | 1234.205561 | 0.855073153 | 0.133704829 | 6.21E-11 | 6.8E-10 | 7716 | 17 | 56048910 | 56065620 | -1 | protein_coding | vascular endothelial zinc finger 1 |
| **TEX2** | ENSG00000136478 | 6896.020083 | -0.982178295 | 0.358240549 | 0.001494135 | 0.005217014 | 55852 | 17 | 62224587 | 62340661 | -1 | protein_coding | testis expressed 2 |
| **LIMD2** | ENSG00000136490 | 1440.259378 | -1.046546646 | 0.143525954 | 1.13E-13 | 1.56E-12 | 80774 | 17 | 61773262 | 61778532 | -1 | protein_coding | LIM domain containing 2 |
| **BRIP1** | ENSG00000136492 | 216.2000679 | 4.223080701 | 0.39457813 | 6.55E-28 | 2.26E-26 | 83990 | 17 | 59758627 | 59940882 | -1 | protein_coding | BRCA1 interacting protein C-terminal helicase 1 |
| **RTP4** | ENSG00000136514 | 362.81215 | 3.244415468 | 0.225877921 | 6.21E-48 | 5.16E-46 | 64108 | 3 | 187086120 | 187089864 | 1 | protein_coding | receptor (chemosensory) transporter protein 4 |
| **TRA2B** | ENSG00000136527 | 2162.689459 | 0.394964732 | 0.122366833 | 0.00106297 | 0.003858759 | 6434 | 3 | 185633694 | 185655924 | -1 | protein_coding | transformer 2 beta homolog (Drosophila) |
| **7-Mar** | ENSG00000136536 | 2338.028211 | 0.526032284 | 0.109476292 | 9.36E-07 | 6.05E-06 | 64844 | 2 | 160569000 | 160627538 | 1 | protein_coding | membrane-associated ring finger (C3HC4) 7, E3 ubiquitin protein ligase |
| **TANK** | ENSG00000136560 | 3466.442087 | 1.508398429 | 0.096301243 | 4.37E-56 | 4.98E-54 | 10010 | 2 | 161993419 | 162092732 | 1 | protein_coding | TRAF family member-associated NFKB activator |
| **SKIL** | ENSG00000136603 | 1701.538911 | 1.115388567 | 0.143777537 | 2.11E-15 | 3.28E-14 | 6498 | 3 | 170075466 | 170114623 | 1 | protein_coding | SKI-like oncogene |
| **EPRS** | ENSG00000136628 | 7094.36757 | -0.544823276 | 0.085182506 | 1.68E-10 | 1.75E-09 | 2058 | 1 | 220141943 | 220220000 | -1 | protein_coding | glutamyl-prolyl-tRNA synthetase |
| **IL10** | ENSG00000136634 | 198.8326135 | -0.928044196 | 0.311331612 | 8.34E-04 | 0.003106159 | 3586 | 1 | 206940947 | 206945839 | -1 | protein_coding | interleukin 10 |
| **RPS6KC1** | ENSG00000136643 | 881.3911382 | 0.811186095 | 0.153710669 | 5.04E-08 | 3.91E-07 | 26750 | 1 | 213224589 | 213448116 | 1 | protein_coding | ribosomal protein S6 kinase, 52kDa, polypeptide 1 |
| **CBWD2** | ENSG00000136682 | 629.689882 | 1.335595934 | 0.156410241 | 2.48E-18 | 4.8E-17 | 644019 | 2 | 114195268 | 114253766 | 1 | protein_coding | COBW domain containing 2 |
| **IL1RN** | ENSG00000136689 | 31000.87016 | 1.839633119 | 0.164495348 | 5.34E-30 | 2.06E-28 | 3557 | 2 | 113864791 | 113891593 | 1 | protein_coding | interleukin 1 receptor antagonist |
| **IL36RN** | ENSG00000136695 | 50.00850228 | 1.945841785 | 0.630065175 | 1.65E-04 | 7.21E-04 | 26525 | 2 | 113816215 | 113822959 | 1 | protein_coding | interleukin 36 receptor antagonist |
| **CCDC115** | ENSG00000136710 | 1082.230386 | 1.580878054 | 0.127994906 | 8.05E-36 | 4.17E-34 | 84317 | 2 | 131095814 | 131099922 | -1 | protein_coding | coiled-coil domain containing 115 |
| **SAP130** | ENSG00000136715 | 1184.49767 | -0.389846233 | 0.109863004 | 3.09E-04 | 0.001271068 | 79595 | 2 | 128698791 | 128785694 | -1 | protein_coding | Sin3A-associated protein, 130kDa |
| **BIN1** | ENSG00000136717 | 1030.76822 | -1.272157026 | 0.15119424 | 7.96E-18 | 1.48E-16 | 274 | 2 | 127805603 | 127864931 | -1 | protein_coding | bridging integrator 1 |
| **HS6ST1** | ENSG00000136720 | 1385.711177 | -1.447147901 | 0.199334369 | 5.81E-14 | 8.24E-13 | 9394 | 2 | 128994290 | 129076151 | -1 | protein_coding | heparan sulfate 6-O-sulfotransferase 1 |
| **UGGT1** | ENSG00000136731 | 4706.751408 | -0.522106856 | 0.117760251 | 5.87E-06 | 3.35E-05 | 56886 | 2 | 128848774 | 128953251 | 1 | protein_coding | UDP-glucose glycoprotein glucosyltransferase 1 |
| **GYPC** | ENSG00000136732 | 658.9576549 | 2.041326179 | 0.19129497 | 1.17E-27 | 4E-26 | 2995 | 2 | 127413509 | 127454246 | 1 | protein_coding | glycophorin C (Gerbich blood group) |
| **ABI1** | ENSG00000136754 | 4350.002692 | 0.72009147 | 0.104372451 | 2.57E-12 | 3.17E-11 | 10006 | 10 | 27035522 | 27150016 | -1 | protein_coding | abl-interactor 1 |
| **YME1L1** | ENSG00000136758 | 4043.405727 | 1.09192082 | 0.113300162 | 1.59E-22 | 4.11E-21 | 10730 | 10 | 27399383 | 27444195 | -1 | protein_coding | YME1-like 1 ATPase |
| **DNAJC1** | ENSG00000136770 | 1085.380379 | 0.986313083 | 0.124739543 | 7.78E-16 | 1.26E-14 | 64215 | 10 | 22045466 | 22292698 | -1 | protein_coding | DnaJ (Hsp40) homolog, subfamily C, member 1 |
| **LRRC8A** | ENSG00000136802 | 1814.337625 | -0.420318015 | 0.131724369 | 0.001021129 | 0.003719236 | 56262 | 9 | 131644391 | 131680318 | 1 | protein_coding | leucine rich repeat containing 8 family, member A |
| **TXN** | ENSG00000136810 | 5835.219728 | 3.2544837 | 0.126612397 | 4.89E-147 | 2.67E-144 | 7295 | 9 | 113006091 | 113018920 | -1 | protein_coding | thioredoxin |
| **KIAA0368** | ENSG00000136813 | 4345.468206 | 0.407038753 | 0.090026728 | 7.32E-06 | 4.11E-05 | 23392 | 9 | 114122972 | 114247025 | -1 | protein_coding | KIAA0368 |
| **TOR1B** | ENSG00000136816 | 2060.159375 | 3.012281857 | 0.109874244 | 5.63E-167 | 4.09E-164 | 27348 | 9 | 132565432 | 132573560 | 1 | protein_coding | torsin family 1, member B (torsin B) |
| **KLF4** | ENSG00000136826 | 128.7812181 | -2.472451028 | 0.459160027 | 5.36E-09 | 4.73E-08 | 9314 | 9 | 110247133 | 110252763 | -1 | protein_coding | Kruppel-like factor 4 (gut) |
| **TOR1A** | ENSG00000136827 | 1297.429679 | 0.841276919 | 0.121777349 | 1.67E-12 | 2.08E-11 | 1861 | 9 | 132575223 | 132586413 | -1 | protein_coding | torsin family 1, member A (torsin A) |
| **FAM129B** | ENSG00000136830 | 5490.288476 | -1.727619648 | 0.170221085 | 2.72E-25 | 8.23E-24 | 64855 | 9 | 130267618 | 130341268 | -1 | protein_coding | family with sequence similarity 129, member B |
| **ST6GALNAC** | ENSG00000136840 | 608.4965513 | -1.17526813 | 0.21992065 | 1.98E-08 | 1.61E-07 | 27090 | 9 | 130670165 | 130679317 | -1 | protein_coding | ST6 (alpha-N-acetyl-neuraminyl-2,3-beta-galactosyl-1,3)-N-acetylgalactosaminide alpha-2,6-sialyltrans |
| **DAB2IP** | ENSG00000136848 | 25.72475592 | -1.646690019 | 0.958454341 | 0.005874617 | 0.017538929 | 153090 | 9 | 124329336 | 124547809 | 1 | protein_coding | DAB2 interacting protein |
| **STXBP1** | ENSG00000136854 | 258.8923502 | -1.451496894 | 0.345473542 | 3.56E-06 | 2.1E-05 | 6812 | 9 | 130374544 | 130457460 | 1 | protein_coding | syntaxin binding protein 1 |
| **SLC2A8** | ENSG00000136856 | 168.1633348 | -1.151744239 | 0.289267398 | 1.52E-05 | 8.12E-05 | 29988 | 9 | 130159421 | 130170703 | 1 | protein_coding | solute carrier family 2 (facilitated glucose transporter), member 8 |
| **CDK5RAP2** | ENSG00000136861 | 1554.540914 | 0.950127433 | 0.123129279 | 5.87E-15 | 8.91E-14 | 55755 | 9 | 123151147 | 123342448 | -1 | protein_coding | CDK5 regulatory subunit associated protein 2 |
| **SLC31A2** | ENSG00000136867 | 3592.973564 | 1.788727575 | 0.111733765 | 1.49E-58 | 1.85E-56 | 1318 | 9 | 115913222 | 115926417 | 1 | protein_coding | solute carrier family 31 (copper transporter), member 2 |
| **SLC31A1** | ENSG00000136868 | 11122.23146 | 0.238607249 | 0.087948108 | 0.006149711 | 0.0182959 | 1317 | 9 | 115983808 | 116028674 | 1 | protein_coding | solute carrier family 31 (copper transporter), member 1 |
| **ZNF189** | ENSG00000136870 | 889.6172103 | 0.398772452 | 0.134634094 | 0.002356525 | 0.007810622 | 7743 | 9 | 104161155 | 104172942 | 1 | protein_coding | zinc finger protein 189 |
| **STX17** | ENSG00000136874 | 840.282147 | 1.382717297 | 0.157363605 | 2.81E-19 | 5.87E-18 | 55014 | 9 | 102668915 | 102732618 | 1 | protein_coding | syntaxin 17 |
| **ATP6V1G1** | ENSG00000136888 | 3289.190209 | 0.49512661 | 0.094841588 | 1.28E-07 | 9.33E-07 | 9550 | 9 | 117350026 | 117360653 | 1 | protein_coding | ATPase, H+ transporting, lysosomal 13kDa, V1 subunit G1 |
| **MRPL50** | ENSG00000136897 | 425.1451535 | 0.560900891 | 0.187278909 | 0.001681843 | 0.005798242 | 54534 | 9 | 104149915 | 104160896 | -1 | protein_coding | mitochondrial ribosomal protein L50 |

| | | | | | | | | | | | | | |
|---|---|---|---|---|---|---|---|---|---|---|---|---|---|
| **TSTD2** | ENSG00000136925 | 658.480807 | 0.669004681 | 0.155991782 | 9.74E-06 | 5.37E-05 | 158427 | 9 | 100362362 | 100395962 | -1 | protein_coding | thiosulfate sulfurtransferase (rhodanese)-like domain containing 2 |
| **GABBR2** | ENSG00000136928 | 10.01188199 | -0.659946979 | 1.193349769 | 0.010017945 | 0.028051346 | 9568 | 9 | 101050391 | 101471479 | -1 | protein_coding | gamma-aminobutyric acid (GABA) B receptor, 2 |
| **PSMB7** | ENSG00000136930 | 1468.849575 | 0.623942654 | 0.1152458 | 3.63E-08 | 2.86E-07 | 5695 | 9 | 127115745 | 127177723 | -1 | protein_coding | proteasome (prosome, macropain) subunit, beta type, 7 |
| **ANP32B** | ENSG00000136938 | 2175.84922 | 0.264347931 | 0.10568948 | 0.011298837 | 0.031107931 | 10541 | 9 | 100745643 | 100778225 | 1 | protein_coding | acidic (leucine-rich) nuclear phosphoprotein 32 family, member B |
| **CTSV** | ENSG00000136943 | 98.76847634 | -1.205913436 | 0.561577972 | 0.004946111 | 0.015133418 | 1515 | 9 | 99794940 | 99801925 | -1 | protein_coding | cathepsin V |
| **DSCC1** | ENSG00000136982 | 37.14440757 | 1.175051169 | 0.574544106 | 0.006455893 | 0.019106394 | 79075 | 8 | 120846216 | 120868250 | -1 | protein_coding | DNA replication and sister chromatid cohesion 1 |
| **DERL1** | ENSG00000136986 | 3314.815814 | 0.497235029 | 0.089796183 | 2.24E-08 | 1.81E-07 | 79139 | 8 | 124025404 | 124054663 | -1 | protein_coding | derlin 1 |
| **MYC** | ENSG00000136997 | 263.3041359 | -1.393540979 | 0.235715035 | 6.01E-10 | 5.9E-09 | 4609 | 8 | 128747680 | 128753674 | 1 | protein_coding | v-myc avian myelocytomatosis viral oncogene homolog |
| **NOV** | ENSG00000136999 | 40.6358399 | -0.985703938 | 0.56105942 | 0.016626524 | 0.043421595 | 4856 | 8 | 120428546 | 120436593 | 1 | protein_coding | nephroblastoma overexpressed |
| **TMEM261** | ENSG00000137038 | 341.6734965 | -0.536994946 | 0.235301422 | 0.014038585 | 0.037515597 | 90871 | 9 | 7796490 | 7888380 | -1 | protein_coding | transmembrane protein 261 |
| **RANBP6** | ENSG00000137040 | 794.9950302 | 0.509348609 | 0.156005828 | 7.34E-04 | 0.002763999 | 26953 | 9 | 6011043 | 6015618 | -1 | protein_coding | RAN binding protein 6 |
| **PLAA** | ENSG00000137055 | 1508.785375 | 0.857648285 | 0.185899222 | 1.38E-06 | 8.66E-06 | 9373 | 9 | 26904081 | 26947461 | -1 | protein_coding | phospholipase A2-activating protein |
| **APTX** | ENSG00000137074 | 626.1506046 | 0.368328509 | 0.146686502 | 0.009899918 | 0.027756457 | 54840 | 9 | 32972604 | 33025166 | -1 | protein_coding | aprataxin |
| **RNF38** | ENSG00000137075 | 1432.090935 | 0.40902323 | 0.128271414 | 0.001160023 | 0.004164584 | 152006 | 9 | 36336393 | 36487545 | -1 | protein_coding | ring finger protein 38 |
| **TLN1** | ENSG00000137076 | 26085.26443 | -0.893805862 | 0.121577661 | 5.81E-14 | 8.24E-13 | 7094 | 9 | 35696945 | 35732392 | -1 | protein_coding | talin 1 |
| **DNAJB5** | ENSG00000137094 | 829.2554386 | -0.456711227 | 0.196845316 | 0.014242122 | 0.03797982 | 25822 | 9 | 34989638 | 34998897 | 1 | protein_coding | DnaJ (Hsp40) homolog, subfamily B, member 5 |
| **GRHPR** | ENSG00000137106 | 756.1548207 | -0.540337741 | 0.149187462 | 1.87E-04 | 8.07E-04 | 9380 | 9 | 37422663 | 37436987 | 1 | protein_coding | glyoxylate reductase/hydroxypyruvate reductase |
| **ALDH1B1** | ENSG00000137124 | 252.3376619 | -0.845522519 | 0.219320785 | 4.28E-05 | 2.09E-04 | 219 | 9 | 38392661 | 38398658 | 1 | protein_coding | aldehyde dehydrogenase 1 family, member B1 |
| **DENND4C** | ENSG00000137145 | 5980.32369 | -1.287861101 | 0.141959726 | 2.1E-20 | 4.76E-19 | 55667 | 9 | 19230433 | 19374139 | 1 | protein_coding | DENN/MADD domain containing 4C |
| **RPS6** | ENSG00000137154 | 7818.849467 | -0.489895101 | 0.12422874 | 5.13E-05 | 2.47E-04 | 6194 | 9 | 19375713 | 19380252 | -1 | protein_coding | ribosomal protein S6 |
| **CNPY3** | ENSG00000137161 | 1552.293581 | -0.934142632 | 0.114789733 | 1.16E-16 | 1.98E-15 | 10695 | 6 | 42896938 | 42907025 | 1 | protein_coding | canopy FGF signaling regulator 3 |
| **GMPR** | ENSG00000137198 | 588.2151544 | 3.785028959 | 0.207394919 | 1.72E-75 | 3.76E-73 | 2766 | 6 | 16238811 | 16295780 | 1 | protein_coding | guanosine monophosphate reductase |
| **CMTR1** | ENSG00000137200 | 2225.879545 | 1.942145143 | 0.121324002 | 1.23E-58 | 1.54E-56 | 23070 | 6 | 37400995 | 37450603 | 1 | protein_coding | cap methyltransferase 1 |
| **YIPF3** | ENSG00000137207 | 1465.392844 | -0.392108773 | 0.122778493 | 0.00111059 | 0.004013514 | 25844 | 6 | 43479565 | 43484728 | -1 | protein_coding | Yip1 domain family, member 3 |
| **TMEM63B** | ENSG00000137216 | 1432.718952 | 0.61834578 | 0.130395291 | 1.15E-06 | 7.32E-06 | 55362 | 6 | 44094651 | 44123256 | 1 | protein_coding | transmembrane protein 63B |
| **CAPN11** | ENSG00000137225 | 58.22905312 | -4.18261961 | 0.675941883 | 1.11E-10 | 1.18E-09 | 11131 | 6 | 44126548 | 44152139 | 1 | protein_coding | calpain 11 |
| **IRF4** | ENSG00000137265 | 653.0053073 | 1.31947117 | 0.299884445 | 1.77E-06 | 1.1E-05 | 3662 | 6 | 391739 | 411447 | 1 | protein_coding | interferon regulatory factor 4 |
| **TUBB2A** | ENSG00000137267 | 562.813747 | -1.152297016 | 0.260077103 | 2.06E-06 | 1.27E-05 | 7280 | 6 | 3153903 | 3157760 | -1 | protein_coding | tubulin, beta 2A class IIa |
| **RIPK1** | ENSG00000137275 | 1772.936229 | 1.032219442 | 0.118610493 | 1.02E-18 | 2.04E-17 | 8737 | 6 | 3064225 | 3115421 | 1 | protein_coding | receptor (TNFRSF)-interacting serine-threonine kinase 1 |
| **UQCC2** | ENSG00000137288 | 352.4585811 | -0.601641026 | 0.189191502 | 8.28E-04 | 0.003087044 | 1005008 | 6 | 33662070 | 33679504 | -1 | protein_coding | ubiquinol-cytochrome c reductase complex assembly factor 2 |
| **HMGA1** | ENSG00000137309 | 2671.746001 | -0.399226891 | 0.165310729 | 0.012105315 | 0.032995459 | 3159 | 6 | 34204650 | 34214008 | 1 | protein_coding | high mobility group AT-hook 1 |
| **RNF144B** | ENSG00000137393 | 1980.978399 | 1.980304631 | 0.137638593 | 6.4E-48 | 5.26E-46 | 255488 | 6 | 18368779 | 18469105 | 1 | protein_coding | ring finger protein 144B |
| **TRIM31** | ENSG00000137397 | 18.14844375 | 2.304736276 | 1.528672194 | 0.003549385 | 0.011270568 | 11074 | HSC | 30060243 | 30070446 | -1 | protein_coding | tripartite motif containing 31 |
| **TAF8** | ENSG00000137413 | 864.9003794 | 0.786063348 | 0.152324486 | 1.13E-07 | 8.3E-07 | 129685 | 6 | 42018251 | 42055199 | 1 | protein_coding | TAF8 RNA polymerase II, TATA box binding protein (TBP)-associated factor, 43kDa |
| **FAM8A1** | ENSG00000137414 | 1449.657973 | 1.547356989 | 0.135992877 | 8.06E-31 | 3.26E-29 | 51439 | 6 | 17600586 | 17611950 | 1 | protein_coding | family with sequence similarity 8, member A1 |
| **FGFBP2** | ENSG00000137441 | 11.07471321 | -3.993505798 | 1.660912496 | 0.00253406 | 0.008330403 | 83888 | 4 | 15961866 | 15970932 | -1 | protein_coding | fibroblast growth factor binding protein 2 |
| **CPEB2** | ENSG00000137449 | 1531.262397 | 0.667811361 | 0.196212053 | 3.34E-04 | 0.001363908 | 132864 | 4 | 15004298 | 15071777 | 1 | protein_coding | cytoplasmic polyadenylation element binding protein 2 |
| **TLR2** | ENSG00000137462 | 1424.888948 | 2.703421695 | 0.175243518 | 8.8E-55 | 9.6E-53 | 7097 | 4 | 154622652 | 154626851 | 1 | protein_coding | toll-like receptor 2 |
| **MYO7A** | ENSG00000137474 | 711.2691735 | -1.627612864 | 0.213615453 | 3.62E-15 | 5.53E-14 | 4647 | 11 | 76839310 | 76926284 | 1 | protein_coding | myosin VIIA |
| **ARRB1** | ENSG00000137486 | 4658.368447 | -0.776808302 | 0.110092505 | 7.14E-13 | 9.23E-12 | 408 | 11 | 74975226 | 75062873 | -1 | protein_coding | arrestin, beta 1 |
| **SLCO2B1** | ENSG00000137491 | 3127.032297 | -2.187456979 | 0.181176389 | 6.74E-35 | 3.36E-33 | 11309 | 11 | 74811608 | 74917594 | 1 | protein_coding | solute carrier organic anion transporter family, member 2B1 |
| **ANKRD42** | ENSG00000137494 | 549.8339794 | -0.472643793 | 0.161542263 | 0.002381607 | 0.007883497 | 338699 | 11 | 82904781 | 82971736 | 1 | protein_coding | ankyrin repeat domain 42 |
| **NUMA1** | ENSG00000137497 | 5136.104268 | -0.412565768 | 0.098688775 | 2.18E-05 | 1.13E-04 | 4926 | 11 | 71713910 | 71791739 | -1 | protein_coding | nuclear mitotic apparatus protein 1 |
| **CCDC90B** | ENSG00000137500 | 918.2278237 | 0.705694516 | 0.206211474 | 2.93E-04 | 0.001208082 | 60492 | 11 | 82970139 | 82997450 | -1 | protein_coding | coiled-coil domain containing 90B |
| **RAB30** | ENSG00000137502 | 302.7249351 | 1.011954614 | 0.25940383 | 2.53E-05 | 1.29E-04 | 27314 | 11 | 82684175 | 82782965 | -1 | protein_coding | RAB30, member RAS oncogene family |
| **GGH** | ENSG00000137563 | 198.8252358 | 1.025010221 | 0.272821476 | 4.39E-05 | 2.14E-04 | 8836 | 8 | 63927638 | 63951730 | -1 | protein_coding | gamma-glutamyl hydrolase (conjugase, folylpolygammaglutamyl hydrolase) |
| **SLCO5A1** | ENSG00000137571 | 60.21737853 | 1.159199315 | 0.728505931 | 0.013862205 | 0.037109139 | 81796 | 8 | 70579282 | 70747299 | -1 | protein_coding | solute carrier organic anion transporter family, member 5A1 |
| **TGS1** | ENSG00000137574 | 788.358601 | 0.686586022 | 0.153645636 | 3.91E-06 | 2.3E-05 | 96764 | 8 | 56685701 | 56738007 | 1 | protein_coding | trimethylguanosine synthase 1 |
| **DDX60** | ENSG00000137628 | 2640.928646 | 2.785157167 | 0.211257719 | 7.13E-41 | 4.5E-39 | 55601 | 4 | 169137444 | 169239958 | -1 | protein_coding | DEAD (Asp-Glu-Ala-Asp) box polypeptide 60 |
| **SORL1** | ENSG00000137642 | 3821.519421 | -2.197634626 | 0.18219823 | 1.68E-34 | 8.2E-33 | 6653 | 11 | 121322912 | 121504402 | 1 | protein_coding | sortilin-related receptor, L(DLR class) A repeats containing |
| **MMP7** | ENSG00000137673 | 21711.65129 | 0.930659014 | 0.335572123 | 0.001543275 | 0.005373553 | 4316 | 11 | 102391239 | 102401484 | -1 | protein_coding | matrix metallopeptidase 7 (matrilysin, uterine) |
| **RDX** | ENSG00000137710 | 2562.804955 | 1.812401857 | 0.171480265 | 4.69E-27 | 1.55E-25 | 5962 | 11 | 110045605 | 110167447 | -1 | protein_coding | radixin |
| **PPP2R1B** | ENSG00000137713 | 771.5208341 | 0.341580062 | 0.130788115 | 0.007217536 | 0.021058725 | 5519 | 11 | 111597632 | 111637151 | -1 | protein_coding | protein phosphatase 2, regulatory subunit A, beta |
| **C11orf1** | ENSG00000137720 | 132.9263048 | -0.795006727 | 0.366398698 | 0.010552878 | 0.029286297 | 64776 | 11 | 111749659 | 111756699 | 1 | protein_coding | chromosome 11 open reading frame 1 |
| **FXYD6** | ENSG00000137726 | 340.7314478 | 0.595526558 | 0.204587828 | 0.001968929 | 0.006675053 | 1005331 | 11 | 117707693 | 117748201 | -1 | protein_coding | FXYD domain containing ion transport regulator 6 |
| **CASP1** | ENSG00000137752 | 731.3126799 | 1.400226203 | 0.220911534 | 3.66E-11 | 4.11E-10 | 834 | 11 | 104896170 | 104972158 | -1 | protein_coding | caspase 1, apoptosis-related cysteine peptidase |
| **SQRDL** | ENSG00000137767 | 6450.726915 | 0.725775931 | 0.097756684 | 6.38E-14 | 9.01E-13 | 58472 | 15 | 45923346 | 45983492 | 1 | protein_coding | sulfide quinone reductase-like (yeast) |
| **CTDSPL2** | ENSG00000137770 | 643.9632815 | 0.413192647 | 0.146785942 | 0.004216024 | 0.013114898 | 51496 | 15 | 44719432 | 44821236 | 1 | protein_coding | CTD (carboxy-terminal domain, RNA polymerase II, polypeptide A) small phosphatase like 2 |
| **SLTM** | ENSG00000137776 | 3042.138942 | 0.392231353 | 0.110928982 | 3.52E-04 | 0.001430243 | 79811 | 15 | 59171244 | 59225852 | -1 | protein_coding | SAFB-like, transcription modulator |
| **ITGA11** | ENSG00000137809 | 97.51067785 | -2.459880245 | 0.410853209 | 2.02E-10 | 2.08E-09 | 22801 | 15 | 68594050 | 68724501 | -1 | protein_coding | integrin, alpha 11 |
| **CASC5** | ENSG00000137812 | 139.4351162 | 1.123725301 | 0.389735161 | 7.84E-04 | 0.002937151 | 57082 | 15 | 40886218 | 40956540 | 1 | protein_coding | cancer susceptibility candidate 5 |
| **HAUS2** | ENSG00000137814 | 1111.387179 | 0.736633276 | 0.134009403 | 1.86E-08 | 1.53E-07 | 55142 | 15 | 42841008 | 42862192 | 1 | protein_coding | HAUS augmin-like complex, subunit 2 |
| **RTF1** | ENSG00000137815 | 2659.479365 | 0.750224559 | 0.13985871 | 3.67E-08 | 2.89E-07 | 23168 | 15 | 41700606 | 41775761 | 1 | protein_coding | Rtf1, Paf1/RNA polymerase II complex component, homolog (S. cerevisiae) |
| **PAQR5** | ENSG00000137819 | 1887.04883 | -1.884024272 | 0.150438507 | 7.36E-37 | 3.93E-35 | 54852 | 15 | 69591286 | 69700119 | 1 | protein_coding | progestin and adipoQ receptor family member V |
| **RMDN3** | ENSG00000137824 | 3473.730248 | 0.410718253 | 0.101450785 | 4.05E-05 | 1.99E-04 | 55177 | 15 | 41028082 | 41048049 | -1 | protein_coding | regulator of microtubule dynamics 3 |
| **UACA** | ENSG00000137831 | 703.249059 | 0.574840427 | 0.193926272 | 0.001794473 | 0.006132586 | 55075 | 15 | 70946893 | 71055932 | -1 | protein_coding | uveal autoantigen with coiled-coil domains and ankyrin repeats |
| **SMAD6** | ENSG00000137834 | 14.36536816 | -5.303729453 | 2.763342697 | 4.78E-04 | 0.001881463 | 4091 | 15 | 66994566 | 67074338 | 1 | protein_coding | SMAD family member 6 |
| **PLCB2** | ENSG00000137841 | 11330.83346 | -1.915134716 | 0.13939897 | 5.06E-44 | 3.74E-42 | 5330 | 15 | 40570377 | 40600136 | -1 | protein_coding | phospholipase C, beta 2 |
| **TMEM62** | ENSG00000137842 | 1072.350032 | 1.62988269 | 0.137542215 | 3.1E-33 | 1.45E-31 | 80021 | 15 | 43415477 | 43477344 | 1 | protein_coding | transmembrane protein 62 |
| **PAK6** | ENSG00000137843 | 21.84052699 | -1.228835131 | 0.914319602 | 0.018113836 | 0.046659997 | 56924 | 15 | 40509629 | 40569688 | 1 | protein_coding | p21 protein (Cdc42/Rac)-activated kinase 6 |
| **ADAM10** | ENSG00000137845 | 10634.98155 | -0.821015805 | 0.124880703 | 1.96E-11 | 2.25E-10 | 102 | 15 | 58887403 | 59042177 | -1 | protein_coding | ADAM metallopeptidase domain 10 |
| **CYP19A1** | ENSG00000137869 | 670.8307196 | -1.434899037 | 0.242744172 | 5.15E-10 | 5.09E-09 | 1588 | 15 | 51500254 | 51630807 | -1 | protein_coding | cytochrome P450, family 19, subfamily A, polypeptide 1 |
| **RSL24D1** | ENSG00000137876 | 1119.260968 | 0.403769755 | 0.137326434 | 0.002516532 | 0.008276341 | 51187 | 15 | 55473004 | 55489265 | -1 | protein_coding | ribosomal L24 domain containing 1 |
| **SPTBN5** | ENSG00000137877 | 34.50650652 | 1.140882587 | 0.753767375 | 0.015939093 | 0.041891161 | 51332 | 15 | 42140345 | 42186275 | -1 | protein_coding | spectrin, beta, non-erythrocytic 5 |
| **GCHFR** | ENSG00000137880 | 454.6548182 | -1.100901309 | 0.228713099 | 3.6E-07 | 2.48E-06 | 2644 | 15 | 41056218 | 41059906 | 1 | protein_coding | GTP cyclohydrolase I feedback regulator |
| **FNBP1L** | ENSG00000137942 | 814.4983666 | -0.67107381 | 0.147113057 | 2.66E-06 | 1.61E-05 | 54874 | 1 | 93913688 | 94020218 | 1 | protein_coding | formin binding protein 1-like |
| **CCBL2** | ENSG00000137944 | 828.0765941 | 0.955386043 | 0.1378903 | 1.2E-12 | 1.51E-11 | 56267 | 1 | 89401456 | 89458636 | -1 | protein_coding | cysteine conjugate-beta lyase 2 |
| **GTF2B** | ENSG00000137947 | 1049.61014 | 1.498413557 | 0.115345034 | 2.19E-39 | 1.29E-37 | 2959 | 1 | 89318615 | 89357627 | -1 | protein_coding | general transcription factor IIB |
| **RABGGTB** | ENSG00000137955 | 1185.480353 | 0.580284241 | 0.153061519 | 9.02E-05 | 4.16E-04 | 692085 | 1 | 76251879 | 76260764 | 1 | protein_coding | Rab geranylgeranyltransferase, beta subunit |
| **IFI44L** | ENSG00000137959 | 3946.343654 | 8.921638803 | 0.681853605 | 6.03E-42 | 4.06E-40 | 10964 | 1 | 79085607 | 79111830 | 1 | protein_coding | interferon-induced protein 44-like |
| **IFI44** | ENSG00000137965 | 1902.162981 | 4.331035085 | 0.126635046 | 1.54E-257 | 2.14E-254 | 10561 | 1 | 79115481 | 79129763 | 1 | protein_coding | interferon-induced protein 44 |
| **DNASE2B** | ENSG00000137976 | 1042.443765 | -3.90269467 | 0.497191922 | 1.39E-16 | 2.36E-15 | 58511 | 1 | 84864215 | 84880701 | 1 | protein_coding | deoxyribonuclease II beta |
| **RTCA** | ENSG00000137996 | 1307.364076 | 0.554729029 | 0.140257802 | 4.72E-05 | 2.28E-04 | 8634 | 1 | 100731763 | 100758325 | 1 | protein_coding | RNA 3'-terminal phosphate cyclase |
| **IFT172** | ENSG00000138002 | 269.7258404 | -1.058319742 | 0.274289276 | 2.91E-05 | 1.47E-04 | 26160 | 2 | 27667238 | 27712656 | -1 | protein_coding | intraflagellar transport 172 homolog (Chlamydomonas) |
| **EPT1** | ENSG00000138018 | 1502.528356 | 1.101325462 | 0.12474792 | 2.96E-19 | 6.18E-18 | 85465 | 2 | 26531415 | 26618759 | 1 | protein_coding | ethanolaminephosphotransferase 1 (CDP-ethanolamine-specific) |
| **HADHB** | ENSG00000138029 | 4071.350054 | -0.538897293 | 0.133978252 | 3.58E-05 | 1.78E-04 | 3032 | 2 | 26466038 | 26513336 | 1 | protein_coding | hydroxyacyl-CoA dehydrogenase/3-ketoacyl-CoA thiolase/enoyl-CoA hydratase (trifunctional protein), b |
| **ADCY3** | ENSG00000138031 | 4729.323118 | -0.88220992 | 0.174318852 | 1.42E-07 | 1.03E-06 | 109 | 2 | 25042038 | 25142708 | -1 | protein_coding | adenylate cyclase 3 |
| **PPM1B** | ENSG00000138032 | 1739.54306 | 0.474308772 | 0.114447503 | 2.3E-05 | 1.18E-04 | 5495 | 2 | 44395108 | 44471523 | 1 | protein_coding | protein phosphatase, Mg2+/Mn2+ dependent, 1B |
| **PNPT1** | ENSG00000138035 | 1190.143811 | 3.160205221 | 0.183978297 | 2.17E-67 | 3.81E-65 | 87178 | 2 | 55861400 | 55921045 | -1 | protein_coding | polyribonucleotide nucleotidyltransferase 1 |

| | | | | | | | | | | | | | |
|---|---|---|---|---|---|---|---|---|---|---|---|---|---|
| **RAB1A** | ENSG00000138069 | 5929.355573 | 0.755180531 | 0.115125851 | 2.68E-11 | 3.04E-10 | 5861 | 2 | 65297835 | 65357240 | -1 | protein_coding | RAB1A, member RAS oncogene family |
| **ACTR2** | ENSG00000138071 | 21029.68192 | 0.287668176 | 0.090080488 | 0.001318548 | 0.004659277 | 10097 | 2 | 65454887 | 65498387 | 1 | protein_coding | ARP2 actin-related protein 2 homolog (yeast) |
| **PREB** | ENSG00000138073 | 1149.209396 | -0.514510475 | 0.140031733 | 1.72E-04 | 7.51E-04 | 10113 | 2 | 27353624 | 27357543 | -1 | protein_coding | prolactin regulatory element binding |
| **PREPL** | ENSG00000138078 | 1532.483121 | -0.765962161 | 0.155706684 | 3.5E-07 | 2.42E-06 | 9581 | 2 | 44543420 | 44589001 | -1 | protein_coding | prolyl endopeptidase-like |
| **EMILIN1** | ENSG00000138080 | 2701.259137 | -0.46065501 | 0.127476244 | 1.75E-04 | 7.61E-04 | 11117 | 2 | 27301435 | 27309271 | 1 | protein_coding | elastin microfibril interfacer 1 |
| **ATRAID** | ENSG00000138085 | 421.130685 | -0.436029655 | 0.191365581 | 0.016447649 | 0.043042708 | 51374 | 2 | 27434895 | 27440046 | 1 | protein_coding | all-trans retinoic acid-induced differentiation factor |
| **LRPPRC** | ENSG00000138095 | 2987.623953 | -0.402354741 | 0.117704318 | 4.85E-04 | 0.001905417 | 10128 | 2 | 44113647 | 44223144 | -1 | protein_coding | leucine-rich pentatricopeptide repeat containing |
| **TRIM54** | ENSG00000138100 | 221.4005107 | -0.991467638 | 0.525732501 | 0.012423122 | 0.03371738 | 57159 | 2 | 27505260 | 27530307 | 1 | protein_coding | tripartite motif containing 54 |
| **STAMBPL1** | ENSG00000138134 | 191.576657 | 1.460702262 | 0.283419133 | 3.94E-08 | 3.09E-07 | 57559 | 10 | 90639491 | 90734910 | 1 | protein_coding | STAM binding protein-like 1 |
| **KIF11** | ENSG00000138160 | 192.7519436 | 1.043892423 | 0.348624573 | 6.29E-04 | 0.002398118 | 3832 | 10 | 94353043 | 94415150 | 1 | protein_coding | kinesin family member 11 |
| **TACC2** | ENSG00000138162 | 91.36563987 | -3.966477018 | 0.925556121 | 1.36E-06 | 8.57E-06 | 10579 | 10 | 123748689 | 124014060 | 1 | protein_coding | transforming, acidic coiled-coil containing protein 2 |
| **DUSP5** | ENSG00000138166 | 1229.629273 | 2.467523942 | 0.152539801 | 3.7E-60 | 4.8E-58 | 1847 | 10 | 112257596 | 112271302 | 1 | protein_coding | dual specificity phosphatase 5 |
| **CALHM2** | ENSG00000138172 | 657.0996153 | -0.590901996 | 0.17751074 | 4.85E-04 | 0.00190552 | 51063 | 10 | 105206543 | 105212660 | -1 | protein_coding | calcium homeostasis modulator 2 |
| **ARL 3.00** | ENSG00000138175 | 276.4989966 | -0.746207199 | 0.217360965 | 2.64E-04 | 0.00109515 | 403 | 10 | 104433488 | 104474164 | -1 | protein_coding | ADP-ribosylation factor-like 3 |
| **ENTPD1** | ENSG00000138185 | 3534.32068 | -0.289651457 | 0.104147366 | 0.005329275 | 0.016131375 | 953 | 10 | 97471536 | 97637023 | 1 | protein_coding | ectonucleoside triphosphate diphosphohydrolase 1 |
| **EXOC6** | ENSG00000138190 | 602.5152634 | 1.048967867 | 0.245148062 | 4.69E-06 | 2.73E-05 | 54536 | 10 | 94590935 | 94819250 | 1 | protein_coding | exocyst complex component 6 |
| **DBR1** | ENSG00000138231 | 347.5569446 | 0.733307027 | 0.193594808 | 6.77E-05 | 3.19E-04 | 51163 | 3 | 137879854 | 137893791 | -1 | protein_coding | debranching RNA lariats 1 |
| **DNAJC13** | ENSG00000138246 | 5664.566885 | 0.653273705 | 0.108920127 | 1.15E-09 | 1.11E-08 | 23317 | 3 | 132136370 | 132257876 | 1 | protein_coding | DnaJ (Hsp40) homolog, subfamily C, member 13 |
| **ANXA7** | ENSG00000138279 | 1634.138553 | 0.802946935 | 0.132078422 | 4.95E-10 | 4.9E-09 | 310 | 10 | 75134859 | 75173834 | -1 | protein_coding | annexin A7 |
| **RPS24** | ENSG00000138326 | 6746.475157 | -0.302779202 | 0.117634483 | 0.008434343 | 0.024089342 | 6229 | 10 | 79793518 | 79816570 | 1 | protein_coding | ribosomal protein S24 |
| **TET1** | ENSG00000138336 | 75.62197691 | -1.049005855 | 0.422768187 | 0.003005263 | 0.009720517 | 80312 | 10 | 70320413 | 70454239 | 1 | protein_coding | tet methylcytosine dioxygenase 1 |
| **MYPN** | ENSG00000138347 | 35.81155698 | 3.355658505 | 0.633775972 | 8.04E-09 | 6.92E-08 | 84665 | 10 | 69865912 | 69971774 | 1 | protein_coding | myopalladin |
| **BARD1** | ENSG00000138376 | 371.8600396 | 0.84148666 | 0.255256284 | 2.88E-04 | 0.001190061 | 580 | 2 | 215590370 | 215674428 | -1 | protein_coding | BRCA1 associated RING domain 1 |
| **STAT4** | ENSG00000138378 | 1092.889751 | 2.593943078 | 0.243230517 | 9.06E-28 | 3.1E-26 | 6775 | 2 | 191894302 | 192016322 | -1 | protein_coding | signal transducer and activator of transcription 4 |
| **ASNSD1** | ENSG00000138381 | 843.6635396 | 0.533605401 | 0.148244619 | 2.08E-04 | 8.85E-04 | 54529 | 2 | 190526111 | 190535557 | 1 | protein_coding | asparagine synthetase domain containing 1 |
| **METTL5** | ENSG00000138382 | 311.2949968 | 0.458578729 | 0.195852245 | 0.013558978 | 0.036414972 | 29081 | 2 | 170666591 | 170681441 | -1 | protein_coding | methyltransferase like 5 |
| **SSB** | ENSG00000138385 | 2645.848523 | 2.084359888 | 0.113336592 | 1.82E-76 | 4.04E-74 | 6741 | 2 | 170648443 | 170668574 | 1 | protein_coding | Sjogren syndrome antigen B (autoantigen La) |
| **NAB1** | ENSG00000138386 | 967.2320302 | 0.679906872 | 0.149402535 | 2.71E-06 | 1.64E-05 | 4664 | 2 | 191511472 | 191557492 | 1 | protein_coding | NGFI-A binding protein 1 (EGR1 binding protein 1) |
| **IDH1** | ENSG00000138413 | 11751.60626 | -0.228143063 | 0.096417615 | 0.015363084 | 0.040614776 | 3417 | 2 | 209100951 | 209130798 | -1 | protein_coding | isocitrate dehydrogenase 1 (NADP+), soluble |
| **SSFA2** | ENSG00000138434 | 3567.69058 | 0.534674471 | 0.190341725 | 0.003048413 | 0.009845498 | 6744 | 2 | 182756560 | 182795465 | 1 | protein_coding | sperm specific antigen 2 |
| **CHRNA1** | ENSG00000138435 | 19.28978494 | 1.67738706 | 0.891195514 | 0.004289347 | 0.013321294 | 1134 | 2 | 175612320 | 175629200 | -1 | protein_coding | cholinergic receptor, nicotinic, alpha 1 (muscle) |
| **WDR12** | ENSG00000138442 | 788.6402451 | -0.433240464 | 0.18802358 | 0.015864573 | 0.041724018 | 55759 | 2 | 203739505 | 203879521 | -1 | protein_coding | WD repeat domain 12 |
| **ABI2** | ENSG00000138443 | 601.5566215 | -0.582260768 | 0.208993196 | 0.002988972 | 0.009671915 | 10152 | 2 | 204192942 | 204312446 | 1 | protein_coding | abl-interactor 2 |
| **SLC40A1** | ENSG00000138449 | 108.2888568 | -0.939399844 | 0.419657204 | 0.006629243 | 0.019536198 | 30061 | 2 | 190425305 | 190448484 | -1 | protein_coding | solute carrier family 40 (iron-regulated transporter), member 1 |
| **SLC35A5** | ENSG00000138459 | 1252.97809 | 1.102815082 | 0.14536757 | 7.46E-15 | 1.12E-13 | 55032 | 3 | 112280556 | 112304424 | 1 | protein_coding | solute carrier family 35, member A5 |
| **COX17** | ENSG00000138495 | 609.0283243 | 0.615413451 | 0.178017269 | 3.18E-04 | 0.001302789 | 10063 | 3 | 119373360 | 119396301 | -1 | protein_coding | COX17 cytochrome c oxidase copper chaperone |
| **PARP9** | ENSG00000138496 | 6012.236049 | 2.697444813 | 0.107813683 | 2.54E-139 | 1.29E-136 | 83666 | 3 | 122246771 | 122283424 | -1 | protein_coding | poly (ADP-ribose) polymerase family, member 9 |
| **USP8** | ENSG00000138592 | 3710.667704 | 0.253247965 | 0.093909362 | 0.006219231 | 0.018481141 | 9101 | 15 | 50716577 | 50793280 | 1 | protein_coding | ubiquitin specific peptidase 8 |
| **SECISBP2L** | ENSG00000138593 | 4435.664414 | 1.165625681 | 0.111426118 | 2.87E-26 | 9.14E-25 | 9728 | 15 | 49280673 | 49338760 | -1 | protein_coding | SECIS binding protein 2-like |
| **TMOD3** | ENSG00000138594 | 3497.705253 | 0.668935693 | 0.104271309 | 7.82E-11 | 8.47E-10 | 29766 | 15 | 52121825 | 52239492 | 1 | protein_coding | tropomodulin 3 (ubiquitous) |
| **SPPL2A** | ENSG00000138600 | 5902.373722 | 1.172900477 | 0.113698619 | 1.11E-25 | 3.43E-24 | 84888 | 15 | 50999506 | 51058005 | -1 | protein_coding | signal peptide peptidase like 2A |
| **GLCE** | ENSG00000138604 | 724.7554612 | -1.092101527 | 0.139989808 | 1.83E-15 | 2.86E-14 | 26035 | 15 | 69452923 | 69564556 | 1 | protein_coding | glucuronic acid epimerase |
| **APH1B** | ENSG00000138613 | 895.6714408 | 0.435608299 | 0.128044513 | 4.87E-04 | 0.001911218 | 83464 | 15 | 63568217 | 63601325 | 1 | protein_coding | APH1B gamma secretase subunit |
| **SEMA7A** | ENSG00000138623 | 1856.463203 | -0.928271794 | 0.162935825 | 4.16E-09 | 3.7E-08 | 8482 | 15 | 74701630 | 74726808 | -1 | protein_coding | semaphorin 7A, GPI membrane anchor (John Milton Hagen blood group) |
| **FAM13A** | ENSG00000138640 | 446.9236028 | -1.333435397 | 0.193035768 | 7.16E-13 | 9.26E-12 | 10144 | 4 | 89647106 | 90032549 | -1 | protein_coding | family with sequence similarity 13, member A |
| **HERC6** | ENSG00000138642 | 1298.836422 | 4.081989592 | 0.170392172 | 3.61E-128 | 1.53E-125 | 55008 | 4 | 89299891 | 89364263 | 1 | protein_coding | HECT and RLD domain containing E3 ubiquitin protein ligase family member 6 |
| **HERC5** | ENSG00000138646 | 2458.25019 | 6.624171049 | 0.154501854 | 0 | 0 | 51191 | 4 | 89378268 | 89427314 | 1 | protein_coding | HECT and RLD domain containing E3 ubiquitin protein ligase 5 |
| **C4orf21** | ENSG00000138658 | 148.6596339 | 1.104169721 | 0.351092861 | 3.68E-04 | 0.001486389 | 55345 | 4 | 113460492 | 113558151 | -1 | protein_coding | chromosome 4 open reading frame 21 |
| **AP1AR** | ENSG00000138660 | 845.4297299 | 1.912818954 | 0.205017588 | 1.17E-21 | 2.9E-20 | 55435 | 4 | 113152893 | 113191203 | 1 | protein_coding | adaptor-related protein complex 1 associated regulatory protein |
| **HNRNPD** | ENSG00000138668 | 1984.884547 | 0.295088309 | 0.112421345 | 0.007639037 | 0.022128014 | 3184 | 4 | 83273651 | 83295656 | -1 | protein_coding | heterogeneous nuclear ribonucleoprotein D (AU-rich element RNA binding protein 1, 37kDa) |
| **RASGEF1B** | ENSG00000138670 | 5493.768742 | 0.490984026 | 0.089500039 | 2.9E-08 | 2.33E-07 | 153020 | 4 | 82347547 | 82965397 | -1 | protein_coding | RasGEF domain family, member 1B |
| **SEC31A** | ENSG00000138674 | 3584.77503 | 0.284851739 | 0.107648273 | 0.006611491 | 0.01948764 | 22872 | 4 | 83739814 | 83822319 | -1 | protein_coding | SEC31 homolog A (S. cerevisiae) |
| **AGPAT9** | ENSG00000138678 | 3249.595517 | -1.438570868 | 0.194674997 | 2.17E-14 | 3.16E-13 | 84803 | 4 | 84457067 | 84527028 | 1 | protein_coding | 1-acylglycerol-3-phosphate O-acyltransferase 9 |
| **KIAA1109** | ENSG00000138688 | 1768.067585 | 0.712735681 | 0.142501263 | 2.75E-07 | 1.92E-06 | 84162 | 4 | 123073488 | 123283913 | 1 | protein_coding | KIAA1109 |
| **RAP1GDS1** | ENSG00000138698 | 1851.497666 | -0.442964066 | 0.141855984 | 0.001279594 | 0.004540523 | 5910 | 4 | 99182535 | 99365012 | 1 | protein_coding | RAP1, GTP-GDP dissociation stimulator 1 |
| **NAAA** | ENSG00000138744 | 851.3033221 | -0.827139581 | 0.171456878 | 5.52E-07 | 3.7E-06 | 27163 | 4 | 76831809 | 76862204 | -1 | protein_coding | N-acylethanolamine acid amidase |
| **NUP54** | ENSG00000138750 | 531.1284055 | 0.45810797 | 0.173103009 | 0.005815951 | 0.017390956 | 53371 | 4 | 77035812 | 77069668 | -1 | protein_coding | nucleoporin 54kDa |
| **CXCL9** | ENSG00000138755 | 763.430987 | 4.118498658 | 0.322587166 | 1.35E-38 | 7.8E-37 | 4283 | 4 | 76922428 | 76928641 | -1 | protein_coding | chemokine (C-X-C motif) ligand 9 |
| **SCARB2** | ENSG00000138760 | 23390.59703 | 0.792583621 | 0.109597915 | 2.76E-13 | 3.66E-12 | 950 | 4 | 77079890 | 77135046 | -1 | protein_coding | scavenger receptor class B, member 2 |
| **CCNG2** | ENSG00000138764 | 1088.981233 | 0.392759748 | 0.172007843 | 0.017168713 | 0.044639238 | 901 | 4 | 78078304 | 78354542 | 1 | protein_coding | cyclin G2 |
| **CNOT6L** | ENSG00000138767 | 2465.472641 | 0.56261296 | 0.115731436 | 7.77E-07 | 5.1E-06 | 246175 | 4 | 78634541 | 78740769 | -1 | protein_coding | CCR4-NOT transcription complex, subunit 6-like |
| **USO1** | ENSG00000138768 | 4239.96926 | 0.447123783 | 0.127799617 | 3.46E-04 | 0.00140764 | 8615 | 4 | 76649777 | 76735382 | 1 | protein_coding | USO1 vesicle transport factor |
| **INTS12** | ENSG00000138785 | 458.5381102 | 0.706614716 | 0.168513105 | 1.33E-05 | 7.21E-05 | 57117 | 4 | 106603784 | 106817143 | -1 | protein_coding | integrator complex subunit 12 |
| **LEF1** | ENSG00000138795 | 523.9023417 | -0.647270073 | 0.217099868 | 0.001505146 | 0.005251859 | 51176 | 4 | 108968701 | 109090112 | -1 | protein_coding | lymphoid enhancer-binding factor 1 |
| **HADH** | ENSG00000138796 | 517.2345884 | -1.487666732 | 0.221620726 | 2.98E-12 | 3.66E-11 | 3033 | 4 | 108910870 | 108956331 | 1 | protein_coding | hydroxyacyl-CoA dehydrogenase |
| **PAPSS1** | ENSG00000138801 | 4787.505237 | -0.601777293 | 0.085647999 | 1.74E-12 | 2.16E-11 | 9061 | 4 | 108511433 | 108641608 | -1 | protein_coding | 3'-phosphoadenosine 5'-phosphosulfate synthase 1 |
| **SEC24B** | ENSG00000138802 | 1578.430919 | 0.325138626 | 0.107218861 | 0.002075189 | 0.006984171 | 10427 | 4 | 110354928 | 110462052 | 1 | protein_coding | SEC24 family member B |
| **PPP3CA** | ENSG00000138814 | 1068.983065 | -0.785104834 | 0.177420584 | 3.93E-06 | 2.31E-05 | 5530 | 4 | 101944566 | 102269435 | -1 | protein_coding | protein phosphatase 3, catalytic subunit, alpha isozyme |
| **SLC39A8** | ENSG00000138821 | 3023.219851 | 3.876680044 | 0.138906774 | 5.19E-173 | 3.97E-170 | 64116 | 4 | 103172198 | 103352415 | -1 | protein_coding | solute carrier family 39 (zinc transporter), member 8 |
| **GUCD1** | ENSG00000138867 | 2507.65544 | 0.312780367 | 0.094165509 | 7.58E-04 | 0.002846415 | 83606 | 22 | 24936406 | 24951903 | -1 | protein_coding | guanylyl cyclase domain containing 1 |
| **KIAA1644** | ENSG00000138944 | 15.64926224 | 2.545781233 | 0.96786094 | 4.43E-04 | 0.001758664 | 85352 | 22 | 44639547 | 44708731 | -1 | protein_coding | KIAA1644 |
| **PARVG** | ENSG00000138964 | 3145.387429 | -1.175867575 | 0.105941605 | 4.51E-29 | 1.65E-27 | 64098 | 22 | 44568836 | 44615413 | 1 | protein_coding | parvin, gamma |
| **ERP27** | ENSG00000139055 | 27.68641647 | -1.529683515 | 0.814012918 | 0.005513619 | 0.01660387 | 121506 | 12 | 15066969 | 15092016 | -1 | protein_coding | endoplasmic reticulum protein 27 |
| **CPNE8** | ENSG00000139117 | 685.4059922 | 0.940006613 | 0.164262975 | 3.21E-09 | 2.89E-08 | 144402 | 12 | 39040624 | 39301232 | -1 | protein_coding | copine VIII |
| **YARS2** | ENSG00000139131 | 306.7532477 | 0.636852804 | 0.248551085 | 0.005354674 | 0.01619222 | 51067 | 12 | 32880424 | 32908836 | -1 | protein_coding | tyrosyl-tRNA synthetase 2, mitochondrial |
| **FGD4** | ENSG00000139132 | 3286.720362 | -1.055015474 | 0.118344332 | 1.38E-19 | 2.93E-18 | 121512 | 12 | 32552463 | 32798984 | 1 | protein_coding | FYVE, RhoGEF and PH domain containing 4 |
| **FAM60A** | ENSG00000139146 | 628.7542493 | 1.562495281 | 0.239368634 | 9.15E-12 | 1.08E-10 | 58516 | 12 | 31433518 | 31479992 | -1 | protein_coding | family with sequence similarity 60, member A |
| **TMEM117** | ENSG00000139173 | 502.705305 | -0.874205175 | 0.221452257 | 2.79E-05 | 1.42E-04 | 84216 | 12 | 44229770 | 44783545 | 1 | protein_coding | transmembrane protein 117 |
| **PRICKLE1** | ENSG00000139174 | 31.3283857 | 2.163783927 | 0.613667256 | 3.25E-05 | 1.63E-04 | 144165 | 12 | 42852140 | 42984157 | -1 | protein_coding | prickle homolog 1 (Drosophila) |
| **C1RL** | ENSG00000139178 | 917.2102408 | 1.950881486 | 0.553156108 | 3.82E-05 | 1.89E-04 | 51279 | 12 | 7242183 | 7261869 | -1 | protein_coding | complement component 1, r subcomponent-like |
| **NDUFA9** | ENSG00000139180 | 1286.127925 | 0.96036699 | 0.132762649 | 1.5E-13 | 2.04E-12 | 4704 | 12 | 4758261 | 4798454 | 1 | protein_coding | NADH dehydrogenase (ubiquinone) 1 alpha subcomplex, 9, 39kDa |
| **KLRG1** | ENSG00000139187 | 117.5910478 | -0.666413577 | 0.318563585 | 0.017224344 | 0.044753427 | 10219 | 12 | 9102640 | 9163356 | 1 | protein_coding | killer cell lectin-like receptor subfamily G, member 1 |
| **CD27** | ENSG00000139193 | 235.0182066 | -1.706788756 | 0.283929895 | 2.22E-10 | 2.28E-09 | 939 | 12 | 6554033 | 6560884 | 1 | protein_coding | CD27 molecule |
| **PEX5** | ENSG00000139197 | 347.1239395 | -0.575862374 | 0.197173099 | 0.002040116 | 0.006884142 | 5830 | 12 | 7341281 | 7371170 | 1 | protein_coding | peroxisomal biogenesis factor 5 |
| **AMIGO2** | ENSG00000139211 | 149.566082 | 0.723565554 | 0.280036438 | 0.004250984 | 0.01321558 | 347902 | 12 | 47469490 | 47473734 | -1 | protein_coding | adhesion molecule with Ig-like domain 2 |
| **9-Mar** | ENSG00000139266 | 340.0888124 | 0.645007974 | 0.214999663 | 0.001362653 | 0.00479517 | 92979 | 12 | 58148881 | 58154190 | 1 | protein_coding | membrane-associated ring finger (C3HC4) 9 |
| **GLIPR1** | ENSG00000139278 | 5434.222075 | -1.59823175 | 0.090116841 | 3.75E-71 | 7.25E-69 | 11010 | 12 | 75874460 | 75897633 | 1 | protein_coding | GLI pathogenesis-related 1 |
| **PHLDA1** | ENSG00000139289 | 1555.258646 | -1.221444375 | 0.151012697 | 1.13E-16 | 1.94E-15 | 22822 | 12 | 76419227 | 76427712 | -1 | protein_coding | pleckstrin homology-like domain, family A, member 1 |
| **DUSP6** | ENSG00000139318 | 1674.972289 | -0.866458461 | 0.196376653 | 3.58E-06 | 2.11E-05 | 1848 | 12 | 89741009 | 89747048 | -1 | protein_coding | dual specificity phosphatase 6 |

| | | | | | | | | | | | | | |
|---|---|---|---|---|---|---|---|---|---|---|---|---|---|
| **POC1B** | ENSG00000139323 | 791.5751731 | 0.432027381 | 0.159851256 | 0.005122201 | 0.015603467 | 282809 | 12 | 89813495 | 89919801 | -1 | protein_coding | POC1 centriolar protein B |
| **TMTC3** | ENSG00000139324 | 477.1857769 | 1.146574482 | 0.264525942 | 3.31E-06 | 1.97E-05 | 160418 | 12 | 88536073 | 88593664 | 1 | protein_coding | transmembrane and tetratricopeptide repeat containing 3 |
| **AMDHD1** | ENSG00000139344 | 144.7016186 | -1.473285677 | 0.435664447 | 9.67E-05 | 4.43E-04 | 144193 | 12 | 96337071 | 96362370 | 1 | protein_coding | amidohydrolase domain containing 1 |
| **NEDD1** | ENSG00000139350 | 677.7679169 | 1.247111353 | 0.136455523 | 1.32E-20 | 3.05E-19 | 121441 | 12 | 97301001 | 97347129 | 1 | protein_coding | neural precursor cell expressed, developmentally down-regulated 1 |
| **GAS2L3** | ENSG00000139354 | 367.8446044 | -3.047621358 | 0.251183511 | 6.48E-35 | 3.25E-33 | 283431 | 12 | 100967461 | 101022064 | 1 | protein_coding | growth arrest-specific 2 like 3 |
| **SLC15A4** | ENSG00000139370 | 1226.78106 | 0.533384286 | 0.116658506 | 3.24E-06 | 1.93E-05 | 121260 | 12 | 129277739 | 129308528 | -1 | protein_coding | solute carrier family 15 (oligopeptide transporter), member 4 |
| **TDG** | ENSG00000139372 | 863.25819 | 0.608012957 | 0.148875713 | 2.21E-05 | 1.14E-04 | 6996 | 12 | 104359582 | 104382652 | 1 | protein_coding | thymine-DNA glycosylase |
| **MMAB** | ENSG00000139428 | 405.4097486 | -0.490731203 | 0.200479449 | 0.009715823 | 0.027330558 | 326625 | 12 | 109991542 | 110011679 | -1 | protein_coding | methylmalonic aciduria (cobalamin deficiency) cblB type |
| **GLTP** | ENSG00000139433 | 1251.882145 | -0.694428118 | 0.12136356 | 5.29E-09 | 4.67E-08 | 51228 | 12 | 110288748 | 110318293 | -1 | protein_coding | glycolipid transfer protein |
| **MTMR6** | ENSG00000139505 | 1793.85481 | 0.684731485 | 0.165991811 | 1.82E-05 | 9.55E-05 | 9107 | 13 | 25802307 | 25862147 | -1 | protein_coding | myotubularin related protein 6 |
| **SLC46A3** | ENSG00000139508 | 1538.224636 | -1.043364533 | 0.164532008 | 5.6E-11 | 6.17E-10 | 283537 | 13 | 29274201 | 29293107 | -1 | protein_coding | solute carrier family 46, member 3 |
| **SLC7A1** | ENSG00000139514 | 1740.035045 | -0.450921973 | 0.149281108 | 0.001714482 | 0.00589216 | 6541 | 13 | 30083547 | 30169825 | -1 | protein_coding | solute carrier family 7 (cationic amino acid transporter, y+ system), member 1 |
| **SUOX** | ENSG00000139531 | 449.8009895 | -0.477655234 | 0.180802568 | 0.005630162 | 0.016901146 | 6821 | 12 | 56390964 | 56400425 | 1 | protein_coding | sulfite oxidase |
| **GPR84** | ENSG00000139572 | 329.8052251 | 1.356932697 | 0.253668584 | 1.59E-08 | 1.32E-07 | 53831 | 12 | 54756229 | 54758271 | -1 | protein_coding | G protein-coupled receptor 84 |
| **N4BP2L1** | ENSG00000139597 | 575.4720964 | 1.636047616 | 0.197075672 | 1.28E-17 | 2.36E-16 | 90634 | 13 | 32974861 | 33002315 | -1 | protein_coding | NEDD4 binding protein 2-like 1 |
| **BRCA2** | ENSG00000139618 | 370.6182181 | 2.242935761 | 0.30056094 | 7.03E-15 | 1.06E-13 | 675 | 13 | 32889611 | 32973805 | 1 | protein_coding | breast cancer 2, early onset |
| **MAP3K12** | ENSG00000139625 | 469.9315816 | -0.525638219 | 0.18340202 | 0.002657254 | 0.008686804 | 7786 | 12 | 53874274 | 53893847 | -1 | protein_coding | mitogen-activated protein kinase kinase kinase 12 |
| **C12orf10** | ENSG00000139637 | 212.2801885 | -0.95426217 | 0.29094041 | 2.96E-04 | 0.001218087 | 60314 | 12 | 53693470 | 53700961 | 1 | protein_coding | chromosome 12 open reading frame 10 |
| **ESYT1** | ENSG00000139641 | 5089.195799 | -1.071762572 | 0.141465201 | 1.08E-14 | 1.6E-13 | 23344 | 12 | 56512034 | 56538455 | 1 | protein_coding | extended synaptotagmin-like protein 1 |
| **TMBIM6** | ENSG00000139644 | 36437.72297 | 0.37633914 | 0.102463152 | 1.98E-04 | 8.46E-04 | 7009 | 12 | 50101508 | 50158717 | 1 | protein_coding | transmembrane BAX inhibitor motif containing 6 |
| **RB1** | ENSG00000139687 | 2311.684982 | 0.734230639 | 0.109076357 | 7.35E-12 | 8.75E-11 | 5925 | 13 | 48877887 | 49056122 | 1 | protein_coding | retinoblastoma 1 |
| **SBNO1** | ENSG00000139697 | 1589.550042 | -0.425556205 | 0.114508535 | 1.54E-04 | 6.79E-04 | 55206 | 12 | 123773656 | 123849390 | -1 | protein_coding | strawberry notch homolog 1 (Drosophila) |
| **VPS37B** | ENSG00000139722 | 801.5614764 | -0.33514779 | 0.14139417 | 0.014784956 | 0.039249346 | 79720 | 12 | 123349882 | 123380991 | -1 | protein_coding | vacuolar protein sorting 37 homolog B (S. cerevisiae) |
| **RHOF** | ENSG00000139725 | 1164.089045 | -0.42355822 | 0.177315531 | 0.01273488 | 0.034441191 | 54509 | 12 | 122215664 | 122241309 | -1 | protein_coding | ras homolog family member F (in filopodia) |
| **RBM26** | ENSG00000139746 | 1395.207592 | 0.463321548 | 0.123591424 | 1.33E-04 | 5.96E-04 | 64062 | 13 | 79885962 | 79980612 | -1 | protein_coding | RNA binding motif protein 26 |
| **ABHD13** | ENSG00000139826 | 570.8425285 | 1.087986375 | 0.186097995 | 1.16E-09 | 1.11E-08 | 84945 | 13 | 108870727 | 108886603 | 1 | protein_coding | abhydrolase domain containing 13 |
| **CBLN3** | ENSG00000139899 | 197.6556164 | 0.97901446 | 0.269401205 | 8.09E-05 | 3.76E-04 | 643866 | 14 | 24895738 | 24900160 | -1 | protein_coding | cerebellin 3 precursor |
| **TMX1** | ENSG00000139921 | 1973.910353 | 1.43753018 | 0.127809376 | 4.4E-30 | 1.71E-28 | 81542 | 14 | 51706880 | 51722759 | 1 | protein_coding | thioredoxin-related transmembrane protein 1 |
| **RTN1** | ENSG00000139970 | 1212.777572 | -1.447723039 | 0.179268542 | 1.23E-16 | 2.09E-15 | 6252 | 14 | 60062694 | 60337684 | -1 | protein_coding | reticulon 1 |
| **SLC38A6** | ENSG00000139974 | 2129.122991 | -0.534590989 | 0.181489353 | 0.002001289 | 0.006772741 | 145389 | 14 | 61447832 | 61550451 | 1 | protein_coding | solute carrier family 38, member 6 |
| **NAA30** | ENSG00000139977 | 811.4365134 | 1.055028381 | 0.157292724 | 5.18E-12 | 6.27E-11 | 122830 | 14 | 57857262 | 57882635 | 1 | protein_coding | N(alpha)-acetyltransferase 30, NatC catalytic subunit |
| **DCAF5** | ENSG00000139990 | 2340.980061 | 0.361535605 | 0.097004513 | 1.45E-04 | 6.42E-04 | 8816 | 14 | 69517598 | 69619867 | -1 | protein_coding | DDB1 and CUL4 associated factor 5 |
| **WDR89** | ENSG00000140006 | 271.9536837 | 0.549295194 | 0.207625892 | 0.004992264 | 0.015247155 | 112840 | 14 | 64063757 | 64108579 | -1 | protein_coding | WD repeat domain 89 |
| **STON2** | ENSG00000140022 | 105.533667 | -1.373257084 | 0.360422122 | 2.32E-05 | 1.2E-04 | 85439 | 14 | 81727000 | 81902809 | -1 | protein_coding | stonin 2 |
| **PTGR2** | ENSG00000140043 | 354.7489327 | -0.514921167 | 0.231767839 | 0.017134431 | 0.044570561 | 145482 | 14 | 74318547 | 74353530 | 1 | protein_coding | prostaglandin reductase 2 |
| **JDP2** | ENSG00000140044 | 633.4949473 | -0.419437405 | 0.164258132 | 0.007907254 | 0.022788339 | 122953 | 14 | 75894419 | 75940814 | 1 | protein_coding | Jun dimerization protein 2 |
| **SLC24A4** | ENSG00000140090 | 33.07843725 | -5.174244869 | 2.310257494 | 5.67E-04 | 0.002185468 | 123041 | 14 | 92788925 | 92962596 | 1 | protein_coding | solute carrier family 24 (sodium/potassium/calcium exchanger), member 4 |
| **WARS** | ENSG00000140105 | 66741.64939 | 0.990816185 | 0.183853179 | 2.11E-08 | 1.71E-07 | 7453 | 14 | 100800125 | 100843142 | -1 | protein_coding | tryptophanyl-tRNA synthetase |
| **SLC12A6** | ENSG00000140199 | 3206.369687 | -1.293566411 | 0.109572455 | 7.14E-33 | 3.25E-31 | 9990 | 15 | 34525460 | 34630261 | -1 | protein_coding | solute carrier family 12 (potassium/chloride transporter), member 6 |
| **MFAP1** | ENSG00000140259 | 1192.604098 | 0.299203236 | 0.119282766 | 0.009817486 | 0.027565797 | 4236 | 15 | 44096690 | 44117000 | -1 | protein_coding | microfibrillar-associated protein 1 |
| **TCF12** | ENSG00000140262 | 2398.140229 | -0.658725981 | 0.123349043 | 4.78E-08 | 3.72E-07 | 6938 | 15 | 57210821 | 57591479 | 1 | protein_coding | transcription factor 12 |
| **SORD** | ENSG00000140263 | 580.1130573 | -1.447655105 | 0.185592657 | 1.01E-15 | 1.61E-14 | 6652 | 15 | 45315302 | 45369383 | 1 | protein_coding | sorbitol dehydrogenase |
| **SERF2** | ENSG00000140264 | 9436.21109 | -0.404254128 | 0.116347508 | 3.86E-04 | 0.001555023 | 1003022 | 15 | 44069285 | 44094787 | 1 | protein_coding | small EDRK-rich factor 2 |
| **LYSMD2** | ENSG00000140280 | 375.7187439 | 2.39996528 | 0.258002717 | 1.12E-21 | 2.78E-20 | 256586 | 15 | 52015208 | 52043782 | -1 | protein_coding | LysM, putative peptidoglycan-binding, domain containing 2 |
| **SLC27A2** | ENSG00000140284 | 64.94014633 | 1.728304224 | 0.558237779 | 1.83E-04 | 7.93E-04 | 11001 | 15 | 50474393 | 50528592 | 1 | protein_coding | solute carrier family 27 (fatty acid transporter), member 2 |
| **FGF7** | ENSG00000140285 | 10.24711646 | 2.204192102 | 2.177411987 | 0.007157472 | 0.020915421 | 2252 | 15 | 49715293 | 49780972 | 1 | protein_coding | fibroblast growth factor 7 |
| **TLE3** | ENSG00000140332 | 4551.000105 | -2.840348138 | 0.180345478 | 2.19E-57 | 2.63E-55 | 7090 | 15 | 70340129 | 70390515 | -1 | protein_coding | transducin-like enhancer of split 3 (E(sp1) homolog, Drosophila) |
| **PSTPIP1** | ENSG00000140368 | 1334.917349 | -1.175456399 | 0.167710784 | 4.84E-13 | 6.35E-12 | 9051 | 15 | 77285700 | 77329673 | 1 | protein_coding | proline-serine-threonine phosphatase interacting protein 1 |
| **BCL2A1** | ENSG00000140379 | 8919.67791 | 0.671415783 | 0.10987666 | 7.4E-10 | 7.21E-09 | 597 | 15 | 80253231 | 80263788 | -1 | protein_coding | BCL2-related protein A1 |
| **SCAPER** | ENSG00000140386 | 500.6199328 | 0.663801009 | 0.187694929 | 2.09E-04 | 8.9E-04 | 49855 | 15 | 76640526 | 77197785 | -1 | protein_coding | S-phase cyclin A-associated protein in the ER |
| **TSPAN3** | ENSG00000140391 | 5888.07539 | 0.54686529 | 0.10663553 | 1.91E-07 | 1.36E-06 | 10099 | 15 | 77336359 | 77376326 | -1 | protein_coding | tetraspanin 3 |
| **NCOA2** | ENSG00000140396 | 2354.557937 | 0.983905049 | 0.116278606 | 9.09E-18 | 1.69E-16 | 10499 | 8 | 71021997 | 71316040 | -1 | protein_coding | nuclear receptor coactivator 2 |
| **MESDC1** | ENSG00000140406 | 597.2008501 | 1.25342923 | 0.16540375 | 7.7E-15 | 1.16E-13 | 59274 | 15 | 81293295 | 81296342 | 1 | protein_coding | mesoderm development candidate 1 |
| **IGF1R** | ENSG00000140443 | 136.0667737 | -1.07978362 | 0.529328608 | 0.007494549 | 0.021763099 | 3480 | 15 | 99192200 | 99507759 | 1 | protein_coding | insulin-like growth factor 1 receptor |
| **USP3** | ENSG00000140455 | 1782.748 | -0.459746539 | 0.11395388 | 4.01E-05 | 1.97E-04 | 9960 | 15 | 63796793 | 63886839 | 1 | protein_coding | ubiquitin specific peptidase 3 |
| **PML** | ENSG00000140464 | 2447.721471 | 2.34808634 | 0.185442043 | 6.79E-38 | 3.8E-36 | 5371 | 15 | 74287014 | 74340153 | 1 | protein_coding | promyelocytic leukemia |
| **LINS** | ENSG00000140471 | 408.9051865 | 0.572016759 | 0.185208423 | 0.001209172 | 0.004323438 | 55180 | 15 | 101099574 | 101143435 | -1 | protein_coding | lines homolog (Drosophila) |
| **ULK3** | ENSG00000140474 | 546.7964538 | -0.762918131 | 0.191478478 | 3E-05 | 1.51E-04 | 25989 | 15 | 75128457 | 75135687 | -1 | protein_coding | unc-51 like kinase 3 |
| **PCSK6** | ENSG00000140479 | 59.88021453 | -2.270236345 | 0.571664841 | 5.41E-06 | 3.11E-05 | 1005074 | 15 | 101840818 | 102065405 | -1 | protein_coding | proprotein convertase subtilisin/kexin type 6 |
| **SCAMP2** | ENSG00000140497 | 3266.262944 | -0.712153867 | 0.125643052 | 7.06E-09 | 6.12E-08 | 10066 | 15 | 75136071 | 75165706 | -1 | protein_coding | secretory carrier membrane protein 2 |
| **HAPLN3** | ENSG00000140511 | 424.3361869 | 3.295569894 | 0.347097972 | 1.39E-22 | 3.63E-21 | 145864 | 15 | 89420519 | 89438857 | -1 | protein_coding | hyaluronan and proteoglycan link protein 3 |
| **RHCG** | ENSG00000140519 | 10.74985166 | 4.040354401 | 1.288419063 | 1.3E-04 | 5.83E-04 | 51458 | 15 | 89998680 | 90039844 | -1 | protein_coding | Rh family, C glycoprotein |
| **FANCI** | ENSG00000140525 | 322.295738 | 1.520308417 | 0.300284034 | 5.46E-08 | 4.22E-07 | 55215 | 15 | 89787180 | 89860492 | 1 | protein_coding | Fanconi anemia, complementation group I |
| **ABHD2** | ENSG00000140526 | 19578.52414 | -2.391059555 | 0.313381993 | 1.89E-15 | 2.94E-14 | 11057 | 15 | 89630690 | 89745591 | 1 | protein_coding | abhydrolase domain containing 2 |
| **UNC45A** | ENSG00000140553 | 1146.044872 | -0.302289352 | 0.122073023 | 0.011785612 | 0.032239039 | 55898 | 15 | 91473410 | 91497323 | 1 | protein_coding | unc-45 homolog A (C. elegans) |
| **FURIN** | ENSG00000140564 | 6342.412374 | -0.256696923 | 0.107948094 | 0.015471478 | 0.040865986 | 5045 | 15 | 91411822 | 91426688 | 1 | protein_coding | furin (paired basic amino acid cleaving enzyme) |
| **IQGAP1** | ENSG00000140575 | 21843.06374 | -0.207923284 | 0.092556692 | 0.019039129 | 0.048666166 | 8826 | 15 | 90931450 | 91045475 | 1 | protein_coding | IQ motif containing GTPase activating protein 1 |
| **EFTUD1** | ENSG00000140598 | 1195.588278 | 0.691432044 | 0.128361283 | 3.36E-08 | 2.67E-07 | 79631 | 15 | 82422571 | 82555104 | -1 | protein_coding | elongation factor Tu GTP binding domain containing 1 |
| **PMM2** | ENSG00000140650 | 957.9498931 | -0.496259549 | 0.165108443 | 0.001761117 | 0.006028031 | 5373 | 16 | 8882680 | 8943188 | 1 | protein_coding | phosphomannomutase 2 |
| **ITGAX** | ENSG00000140678 | 37749.93038 | -1.37749855 | 0.12304233 | 7.97E-30 | 3.04E-28 | 3687 | 16 | 31366455 | 31394318 | 1 | protein_coding | integrin, alpha X (complement component 3 receptor 4 subunit) |
| **C16orf58** | ENSG00000140688 | 755.8128775 | -0.773010505 | 0.154116144 | 2.28E-07 | 1.62E-06 | 64755 | 16 | 31500792 | 31520630 | -1 | protein_coding | chromosome 16 open reading frame 58 |
| **UQCRC2** | ENSG00000140740 | 3865.539975 | -0.5326702 | 0.124656558 | 1.27E-05 | 6.87E-05 | 7385 | 16 | 21963981 | 21994981 | 1 | protein_coding | ubiquinol-cytochrome c reductase core protein II |
| **IGSF6** | ENSG00000140749 | 12156.12383 | -0.437581451 | 0.113700919 | 7.94E-05 | 3.7E-04 | 10261 | 16 | 21652609 | 21663981 | -1 | protein_coding | immunoglobulin superfamily, member 6 |
| **ARHGAP17** | ENSG00000140750 | 2951.296675 | 0.362700335 | 0.097711188 | 1.63E-04 | 7.13E-04 | 55114 | 16 | 24930706 | 25026987 | -1 | protein_coding | Rho GTPase activating protein 17 |
| **TXNL4B** | ENSG00000140830 | 309.8367451 | 0.617154565 | 0.240126686 | 0.005463361 | 0.016478497 | 54957 | 16 | 72078188 | 72128330 | -1 | protein_coding | thioredoxin-like 4B |
| **ZFHX3** | ENSG00000140836 | 2095.548153 | -0.787329549 | 0.12617647 | 1.85E-10 | 1.92E-09 | 463 | 16 | 72816784 | 73093597 | -1 | protein_coding | zinc finger homeobox 3 |
| **NLRC5** | ENSG00000140853 | 2260.747749 | 1.438196045 | 0.234983619 | 1.54E-10 | 1.61E-09 | 84166 | 16 | 57023397 | 57117443 | 1 | protein_coding | NLR family, CARD domain containing 5 |
| **KATNB1** | ENSG00000140854 | 286.6993137 | -0.772802296 | 0.282534828 | 0.002453811 | 0.008089491 | 10300 | 16 | 57769642 | 57791162 | 1 | protein_coding | katanin p80 (WD repeat containing) subunit B 1 |
| **KIFC3** | ENSG00000140859 | 657.9434859 | -0.676010581 | 0.202902061 | 4.37E-04 | 0.001737162 | 3801 | 16 | 57792129 | 57896957 | -1 | protein_coding | kinesin family member C3 |
| **MAP1LC3B** | ENSG00000140941 | 6335.757709 | 0.255330914 | 0.106488216 | 0.015169674 | 0.040145129 | 81631 | 16 | 87417601 | 87438385 | 1 | protein_coding | microtubule-associated protein 1 light chain 3 beta |
| **RPS2** | ENSG00000140988 | 6446.380289 | -0.439657204 | 0.124241078 | 2.89E-04 | 0.001194964 | 26784 | 16 | 2012053 | 2014861 | -1 | protein_coding | ribosomal protein S2 |
| **NDUFB10** | ENSG00000140990 | 827.6822125 | -0.431630066 | 0.143836902 | 0.001936158 | 0.006575623 | 4716 | 16 | 2009509 | 2011976 | 1 | protein_coding | NADH dehydrogenase (ubiquinone) 1 beta subcomplex, 10, 22kDa |
| **DEF8** | ENSG00000140995 | 1587.000864 | -1.292109603 | 0.145695418 | 1.84E-19 | 3.89E-18 | 54849 | 16 | 90014333 | 90034468 | 1 | protein_coding | differentially expressed in FDCP 8 homolog (mouse) |
| **TCF25** | ENSG00000141002 | 2371.640912 | -0.272492106 | 0.113202237 | 0.014543238 | 0.03866163 | 22980 | 16 | 89940000 | 89977792 | 1 | protein_coding | transcription factor 25 (basic helix-loop-helix) |
| **GALNS** | ENSG00000141012 | 1784.783317 | -0.345567013 | 0.142253244 | 0.012394494 | 0.033657615 | 2588 | 16 | 88880142 | 88923378 | -1 | protein_coding | galactosamine (N-acetyl)-6-sulfate sulfatase |
| **GID4** | ENSG00000141034 | 711.3158147 | 0.460393595 | 0.201679796 | 0.015942 | 0.041891161 | 79018 | 17 | 17942606 | 17971718 | 1 | protein_coding | GID complex subunit 4 |
| **CTRL** | ENSG00000141086 | 21.85123493 | 1.259413556 | 1.158441864 | 0.018008903 | 0.046452331 | 1506 | 16 | 67961543 | 67966317 | -1 | protein_coding | chymotrypsin-like |
| **C17orf80** | ENSG00000141219 | 573.6907673 | 0.409306744 | 0.146983676 | 0.004080438 | 0.012721611 | 55028 | 17 | 71228372 | 71245091 | 1 | protein_coding | chromosome 17 open reading frame 80 |
| **VPS53** | ENSG00000141252 | 3765.861075 | -0.411285242 | 0.10598783 | 7.22E-05 | 3.38E-04 | 55275 | 17 | 411908 | 624957 | -1 | protein_coding | vacuolar protein sorting 53 homolog (S. cerevisiae) |

| | | | | | | | | | | | | | |
|---|---|---|---|---|---|---|---|---|---|---|---|---|---|
| **SGSM2** | ENSG00000141258 | 1109.536589 | -1.116880599 | 0.183716276 | 2.73E-10 | 2.77E-09 | 9905 | 17 | 2240792 | 2284352 | 1 | protein_coding | small G protein signaling modulator 2 |
| **SKAP1** | ENSG00000141293 | 462.6935219 | -0.930135323 | 0.219343463 | 6.75E-06 | 3.82E-05 | 8631 | 17 | 46210802 | 46507637 | -1 | protein_coding | src kinase associated phosphoprotein 1 |
| **SSH2** | ENSG00000141298 | 2018.296993 | -0.616778214 | 0.124360598 | 4.01E-07 | 2.74E-06 | 85464 | 17 | 27952956 | 28257294 | -1 | protein_coding | slingshot protein phosphatase 2 |
| **G6PC3** | ENSG00000141349 | 693.9382812 | -0.658711716 | 0.14330661 | 2.21E-06 | 1.35E-05 | 92579 | 17 | 42148103 | 42153709 | 1 | protein_coding | glucose 6 phosphatase, catalytic, 3 |
| **CLTC** | ENSG00000141367 | 23882.78769 | -0.580670328 | 0.086849697 | 2.39E-11 | 2.72E-10 | 1213 | 17 | 57697219 | 57773671 | 1 | protein_coding | clathrin, heavy chain (Hc) |
| **SS18** | ENSG00000141380 | 1202.878405 | 0.397669362 | 0.127216513 | 0.001343551 | 0.004736674 | 6760 | 18 | 23596578 | 23671181 | -1 | protein_coding | synovial sarcoma translocation, chromosome 18 |
| **IMPA2** | ENSG00000141401 | 66.08219754 | -1.167421406 | 0.494336887 | 0.003222424 | 0.010348458 | 3613 | 18 | 11981024 | 12030876 | 1 | protein_coding | inositol(myo)-1(or 4)-monophosphatase 2 |
| **C18orf21** | ENSG00000141428 | 282.9605945 | 0.513034244 | 0.216219605 | 0.011514522 | 0.031593616 | 83608 | 18 | 33552046 | 33559241 | 1 | protein_coding | chromosome 18 open reading frame 21 |
| **GALNT1** | ENSG00000141429 | 3599.772362 | -0.724102521 | 0.130423454 | 1.34E-08 | 1.12E-07 | 2589 | 18 | 33161081 | 33291798 | 1 | protein_coding | UDP-N-acetyl-alpha-D-galactosamine:polypeptide N-acetylgalactosaminyltransferase 1 (GalNAc-T1) |
| **ESCO1** | ENSG00000141446 | 1312.442098 | 0.559780801 | 0.156225478 | 2.1E-04 | 8.91E-04 | 114799 | 18 | 19109242 | 19180845 | -1 | protein_coding | establishment of sister chromatid cohesion N-acetyltransferase 1 |
| **C18orf8** | ENSG00000141452 | 810.141781 | 0.877087884 | 0.142768206 | 2.95E-10 | 2.99E-09 | 29919 | 18 | 21083473 | 21111746 | 1 | protein_coding | chromosome 18 open reading frame 8 |
| **NPC1** | ENSG00000141458 | 13405.16711 | -0.537109439 | 0.138883048 | 7.05E-05 | 3.31E-04 | 4864 | 18 | 21086148 | 21166862 | -1 | protein_coding | Niemann-Pick disease, type C1 |
| **ARRB2** | ENSG00000141480 | 6367.387766 | -0.986006316 | 0.097563592 | 2.23E-24 | 6.47E-23 | 409 | 17 | 4613784 | 4624794 | 1 | protein_coding | arrestin, beta 2 |
| **SAT2** | ENSG00000141504 | 650.3888509 | -0.351352149 | 0.153439397 | 0.018004929 | 0.04644992 | 112483 | 17 | 7529552 | 7531194 | -1 | protein_coding | spermidine/spermine N1-acetyltransferase family member 2 |
| **CCDC40** | ENSG00000141519 | 40.22629435 | -1.928448217 | 0.633181482 | 2.14E-04 | 9.07E-04 | 55036 | 17 | 78010435 | 78074412 | 1 | protein_coding | coiled-coil domain containing 40 |
| **ARHGDIA** | ENSG00000141522 | 4774.334749 | -0.272449599 | 0.117349144 | 0.01762582 | 0.04567233 | 396 | 17 | 79825597 | 79829282 | -1 | protein_coding | Rho GDP dissociation inhibitor (GDI) alpha |
| **TMC6** | ENSG00000141524 | 1808.751585 | -1.572805174 | 0.136380583 | 1.54E-31 | 6.45E-30 | 11322 | 17 | 76106539 | 76128488 | -1 | protein_coding | transmembrane channel-like 6 |
| **SLC16A3** | ENSG00000141526 | 6445.916542 | -0.740589761 | 0.125799628 | 1.56E-09 | 1.47E-08 | 1024654 | 17 | 80186273 | 80219005 | 1 | protein_coding | solute carrier family 16 (monocarboxylate transporter), member 3 |
| **CARD14** | ENSG00000141527 | 61.24673812 | -3.033312472 | 0.620036087 | 8.61E-08 | 6.44E-07 | 79092 | 17 | 78143791 | 78183130 | 1 | protein_coding | caspase recruitment domain family, member 14 |
| **ANAPC11** | ENSG00000141552 | 363.1322993 | -0.448257401 | 0.183595331 | 0.010418091 | 0.028949033 | 51529 | 17 | 79848666 | 79858867 | 1 | protein_coding | anaphase promoting complex subunit 11 |
| **TBCD** | ENSG00000141556 | 696.767625 | -0.842170012 | 0.21404575 | 2.87E-05 | 1.45E-04 | 1019295 | 17 | 80709940 | 80900724 | 1 | protein_coding | tubulin folding cofactor D |
| **RPTOR** | ENSG00000141564 | 1765.141145 | -0.894291254 | 0.140402337 | 5.39E-11 | 5.96E-10 | 57521 | 17 | 78518619 | 78940171 | 1 | protein_coding | regulatory associated protein of MTOR, complex 1 |
| **FOXK2** | ENSG00000141568 | 1789.736001 | -0.43248143 | 0.10984198 | 6.08E-05 | 2.89E-04 | 3607 | 17 | 80477589 | 80602538 | 1 | protein_coding | forkhead box K2 |
| **SECTM1** | ENSG00000141574 | 791.5993302 | 2.149020046 | 0.601805562 | 2.68E-05 | 1.36E-04 | 6398 | 17 | 80278900 | 80291950 | -1 | protein_coding | secreted and transmembrane 1 |
| **AZI1** | ENSG00000141577 | 92.01504807 | -1.434054098 | 0.453575474 | 2.12E-04 | 9.02E-04 | 22994 | 17 | 79163393 | 79196799 | -1 | protein_coding | 5-azacytidine induced 1 |
| **CBX4** | ENSG00000141582 | 586.9645275 | 0.969184661 | 0.145802472 | 9.51E-12 | 1.12E-10 | 8535 | 17 | 77806955 | 77813228 | -1 | protein_coding | chromobox homolog 4 |
| **SMAD4** | ENSG00000141646 | 1135.410766 | 1.238493012 | 0.185854375 | 5.72E-12 | 6.89E-11 | 4089 | 18 | 48494410 | 48611415 | 1 | protein_coding | SMAD family member 4 |
| **TNFRSF11A** | ENSG00000141655 | 453.6797761 | -0.822487387 | 0.245604125 | 3.12E-04 | 0.001280205 | 8792 | 18 | 59992520 | 60058516 | 1 | protein_coding | tumor necrosis factor receptor superfamily, member 11a, NFKB activator |
| **ZCCHC2** | ENSG00000141664 | 3957.478397 | 0.875191716 | 0.105219358 | 3.39E-17 | 6E-16 | 54877 | 18 | 60190240 | 60254942 | 1 | protein_coding | zinc finger, CCHC domain containing 2 |
| **FBXO15** | ENSG00000141665 | 77.42251531 | -1.807775293 | 0.478315326 | 1.61E-05 | 8.56E-05 | 201456 | 18 | 71740588 | 71815100 | -1 | protein_coding | F-box protein 15 |
| **PMAIP1** | ENSG00000141682 | 178.8584623 | 2.622057151 | 0.362133028 | 3.23E-14 | 4.65E-13 | 5366 | 18 | 57567180 | 57571538 | 1 | protein_coding | phorbol-12-myristate-13-acetate-induced protein 1 |
| **FAM134C** | ENSG00000141699 | 3059.20796 | 0.485287444 | 0.101127535 | 1.13E-06 | 7.23E-06 | 162427 | 17 | 40731531 | 40762641 | -1 | protein_coding | family with sequence similarity 134, member C |
| **ERBB2** | ENSG00000141736 | 108.0066158 | -1.258989691 | 0.376281601 | 1.49E-04 | 6.59E-04 | 2064 | 17 | 37844167 | 37886679 | 1 | protein_coding | v-erb-b2 avian erythroblastic leukemia viral oncogene homolog 2 |
| **STAC2** | ENSG00000141750 | 494.7615608 | -3.148054337 | 0.239670777 | 1.61E-40 | 9.81E-39 | 342667 | 17 | 37366789 | 37382125 | -1 | protein_coding | SH3 and cysteine rich domain 2 |
| **IGFBP4** | ENSG00000141753 | 701.4606165 | 5.253016175 | 0.192528055 | 4.01E-165 | 2.78E-162 | 3487 | 17 | 38599702 | 38613983 | 1 | protein_coding | insulin-like growth factor binding protein 4 |
| **SAMD1** | ENSG00000141858 | 667.861083 | -1.446288933 | 0.158906243 | 1.6E-20 | 3.66E-19 | NA | 19 | 14198652 | 14201848 | -1 | protein_coding | sterile alpha motif domain containing 1 |
| **SLC39A3** | ENSG00000141873 | 236.8817388 | -0.932949384 | 0.290581289 | 3.8E-04 | 0.001531712 | 29985 | 19 | 2732202 | 2740150 | -1 | protein_coding | solute carrier family 39 (zinc transporter), member 3 |
| **NFIC** | ENSG00000141905 | 2249.267483 | -0.761110928 | 0.166370064 | 2E-06 | 1.23E-05 | 1019286 | 19 | 3359561 | 3469215 | 1 | protein_coding | nuclear factor I/C (CCAAT-binding transcription factor) |
| **PFKL** | ENSG00000141959 | 3341.013719 | -0.330934206 | 0.120867308 | 0.005273503 | 0.015984648 | 5211 | 21 | 45719934 | 45747259 | 1 | protein_coding | phosphofructokinase, liver |
| **SH3GL1** | ENSG00000141985 | 899.1916854 | -0.651730097 | 0.135169208 | 7.58E-07 | 4.98E-06 | 6455 | 19 | 4360367 | 4400544 | -1 | protein_coding | SH3-domain GRB2-like 1 |
| **DPP9** | ENSG00000142002 | 2617.870404 | -0.585102336 | 0.104311346 | 1.15E-08 | 9.71E-08 | 91039 | 19 | 4675236 | 4724685 | -1 | protein_coding | dipeptidyl-peptidase 9 |
| **TMEM91** | ENSG00000142046 | 468.8911347 | -2.172430548 | 0.207466002 | 9.36E-27 | 3.08E-25 | 641649 | 19 | 41856816 | 41889988 | 1 | protein_coding | transmembrane protein 91 |
| **IFITM3** | ENSG00000142089 | 2326.170956 | 5.222152175 | 0.171899834 | 4.6E-204 | 3.91E-201 | 10410 | 11 | 319669 | 327537 | -1 | protein_coding | interferon induced transmembrane protein 3 |
| **ATHL1** | ENSG00000142102 | 685.2673838 | -1.583661957 | 0.230386529 | 8.83E-13 | 1.13E-11 | 80162 | 11 | 289135 | 296107 | 1 | protein_coding | ATH1, acid trehalase-like 1 (yeast) |
| **COL6A1** | ENSG00000142156 | 6369.14669 | -1.547565617 | 0.20274304 | 2.77E-15 | 4.27E-14 | 1291 | 21 | 47401651 | 47424964 | 1 | protein_coding | collagen, type VI, alpha 1 |
| **IFNAR1** | ENSG00000142166 | 8076.076005 | -0.631464789 | 0.153708052 | 2.2E-05 | 1.14E-04 | 3454 | 21 | 34696734 | 34732168 | 1 | protein_coding | interferon (alpha, beta and omega) receptor 1 |
| **SOD1** | ENSG00000142168 | 2092.824718 | 0.357073486 | 0.11943386 | 0.002381422 | 0.007883497 | 6647 | 21 | 33031935 | 33041244 | 1 | protein_coding | superoxide dismutase 1, soluble |
| **COL6A2** | ENSG00000142173 | 5273.491382 | -0.714210027 | 0.214697146 | 3.9E-04 | 0.001570113 | 1292 | 21 | 47518011 | 47552763 | 1 | protein_coding | collagen, type VI, alpha 2 |
| **SCYL1** | ENSG00000142186 | 1625.027634 | -0.588687411 | 0.127922264 | 2.44E-06 | 1.49E-05 | 57410 | 11 | 65292548 | 65306175 | 1 | protein_coding | SCY1-like 1 (S. cerevisiae) |
| **TMEM50B** | ENSG00000142188 | 506.5030624 | 0.617390612 | 0.163721088 | 8.96E-05 | 4.14E-04 | 757 | 21 | 34804792 | 34852318 | -1 | protein_coding | transmembrane protein 50B |
| **DOPEY2** | ENSG00000142197 | 6798.103295 | -1.547749535 | 0.117163854 | 1.09E-40 | 6.8E-39 | 9980 | 21 | 37529080 | 37666572 | 1 | protein_coding | dopey family member 2 |
| **URB1** | ENSG00000142207 | 525.3067134 | -0.624879487 | 0.173988638 | 1.82E-04 | 7.88E-04 | 9875 | 21 | 33683329 | 33765335 | -1 | protein_coding | URB1 ribosome biogenesis 1 homolog (S. cerevisiae) |
| **EMP3** | ENSG00000142227 | 716.3920924 | 0.940165889 | 0.14116576 | 9.14E-12 | 1.08E-10 | 2014 | 19 | 48824766 | 48833810 | 1 | protein_coding | epithelial membrane protein 3 |
| **WTIP** | ENSG00000142279 | 12.47022336 | -3.948358265 | 2.034408791 | 0.001611801 | 0.005584482 | 126374 | 19 | 34971874 | 34997258 | 1 | protein_coding | Wilms tumor 1 interacting protein |
| **RNPEPL1** | ENSG00000142327 | 1916.286668 | -1.055742601 | 0.170728447 | 1.75E-10 | 1.82E-09 | 57140 | 2 | 241505221 | 241520789 | 1 | protein_coding | arginyl aminopeptidase (aminopeptidase B)-like 1 |
| **MYO1F** | ENSG00000142347 | 1167.300041 | -0.561464129 | 0.141856732 | 4.63E-05 | 2.24E-04 | 4542 | 19 | 8585674 | 8642461 | -1 | protein_coding | myosin IF |
| **NLRP12** | ENSG00000142405 | 123.0008071 | -5.803483755 | 0.72079165 | 3.38E-16 | 5.56E-15 | 91662 | 19 | 54296857 | 54327648 | -1 | protein_coding | NLR family, pyrin domain containing 12 |
| **EVI5L** | ENSG00000142459 | 182.7676254 | -0.695862838 | 0.333660573 | 0.016022916 | 0.042082527 | 115704 | 19 | 7895119 | 7929862 | 1 | protein_coding | ecotropic viral integration site 5-like |
| **RPS11** | ENSG00000142534 | 8078.353129 | -0.573468332 | 0.121161258 | 1.35E-06 | 8.49E-06 | 84546 | 19 | 49999622 | 50002946 | 1 | protein_coding | ribosomal protein S11 |
| **RPL13A** | ENSG00000142541 | 7834.385342 | -0.480516983 | 0.143892306 | 5.83E-04 | 0.002239701 | 26819 | 19 | 49990811 | 49995565 | 1 | protein_coding | ribosomal protein L13a |
| **SLC2A5** | ENSG00000142583 | 2894.580279 | -2.603561754 | 0.157407047 | 7.01E-63 | 1.04E-60 | 6518 | 1 | 9095166 | 9148537 | -1 | protein_coding | solute carrier family 2 (facilitated glucose/fructose transporter), member 5 |
| **RERE** | ENSG00000142599 | 2019.460919 | 0.653504349 | 0.157010758 | 1.56E-05 | 8.3E-05 | 473 | 1 | 8412457 | 8877702 | -1 | protein_coding | arginine-glutamic acid dipeptide (RE) repeats |
| **FHAD1** | ENSG00000142621 | 47.6494855 | -2.053347237 | 0.689469614 | 2.02E-04 | 8.63E-04 | 114827 | 1 | 15573768 | 15726779 | 1 | protein_coding | forkhead-associated (FHA) phosphopeptide binding domain 1 |
| **EFHD2** | ENSG00000142634 | 3921.068559 | -0.326177704 | 0.086541557 | 1.99E-04 | 8.51E-04 | 79180 | 1 | 15736391 | 15756839 | 1 | protein_coding | EF-hand domain family, member D2 |
| **PGD** | ENSG00000142657 | 15428.04651 | 0.253264403 | 0.072485153 | 3.65E-04 | 0.001477804 | 5226 | 1 | 10458649 | 10480201 | 1 | protein_coding | phosphogluconate dehydrogenase |
| **SH3BGRL3** | ENSG00000142669 | 12679.99859 | -1.400302108 | 0.129349161 | 1.45E-28 | 5.23E-27 | 83442 | 1 | 26605667 | 26608007 | 1 | protein_coding | SH3 domain binding glutamic acid-rich protein like 3 |
| **RPL11** | ENSG00000142676 | 5402.163451 | -0.349667016 | 0.123196177 | 0.003651722 | 0.011557108 | 6135 | 1 | 24018269 | 24022915 | 1 | protein_coding | ribosomal protein L11 |
| **KIAA0319L** | ENSG00000142687 | 2497.144066 | 0.417511956 | 0.10745989 | 7.89E-05 | 3.67E-04 | 79932 | 1 | 35899091 | 36023551 | -1 | protein_coding | KIAA0319-like |
| **MAP3K6** | ENSG00000142733 | 229.2411275 | -1.112609069 | 0.285112967 | 2.34E-05 | 1.21E-04 | 9064 | 1 | 27681675 | 27693383 | -1 | protein_coding | mitogen-activated protein kinase kinase kinase 6 |
| **SYTL1** | ENSG00000142765 | 182.3268464 | -2.165899151 | 0.298449665 | 3.3E-14 | 4.74E-13 | 84958 | 1 | 27668513 | 27680421 | 1 | protein_coding | synaptotagmin-like 1 |
| **WDTC1** | ENSG00000142784 | 1601.775642 | -0.870376456 | 0.134764119 | 3.68E-11 | 4.12E-10 | 23038 | 1 | 27561007 | 27635110 | 1 | protein_coding | WD and tetratricopeptide repeats 1 |
| **HSPG2** | ENSG00000142798 | 212.1034769 | -3.785339677 | 0.522225941 | 1.72E-14 | 2.52E-13 | 3339 | 1 | 22148738 | 22263790 | -1 | protein_coding | heparan sulfate proteoglycan 2 |
| **PRKACB** | ENSG00000142875 | 6588.87961 | -1.785624249 | 0.126032845 | 1.27E-46 | 1.01E-44 | 5567 | 1 | 84543745 | 84704181 | 1 | protein_coding | protein kinase, cAMP-dependent, catalytic, beta |
| **RPS8** | ENSG00000142937 | 6984.680423 | -0.634655429 | 0.116905649 | 3.1E-08 | 2.48E-07 | 94163 | 1 | 45240923 | 45244451 | 1 | protein_coding | ribosomal protein S8 |
| **MOB3C** | ENSG00000142961 | 958.3894951 | 0.737280917 | 0.136718079 | 3.09E-08 | 2.48E-07 | 148932 | 1 | 47073387 | 47082563 | -1 | protein_coding | MOB kinase activator 3C |
| **LMO4** | ENSG00000143013 | 1853.657817 | 0.417394534 | 0.140451328 | 0.002268247 | 0.007549133 | 8543 | 1 | 87794151 | 87814606 | 1 | protein_coding | LIM domain only 4 |
| **SYPL2** | ENSG00000143028 | 31.58761474 | -2.283111427 | 0.767452001 | 2.35E-04 | 9.87E-04 | 284612 | 1 | 110009180 | 110024759 | 1 | protein_coding | synaptophysin-like 2 |
| **MTF2** | ENSG00000143033 | 1073.277556 | 0.353162335 | 0.124136606 | 0.003665294 | 0.01159766 | 22823 | 1 | 93544792 | 93604638 | 1 | protein_coding | metal response element binding transcription factor 2 |
| **ZNF697** | ENSG00000143067 | 1243.927462 | -0.473287851 | 0.155798897 | 0.001653813 | 0.005710624 | 90874 | 1 | 120162045 | 120190396 | -1 | protein_coding | zinc finger protein 697 |
| **CTTNBP2NL** | ENSG00000143079 | 1870.799002 | 0.259104219 | 0.109452816 | 0.016596553 | 0.04337297 | 55917 | 1 | 112938803 | 113006078 | 1 | protein_coding | CTTNBP2 N-terminal like |
| **PSMA5** | ENSG00000143106 | 1964.47214 | 1.083901555 | 0.117520323 | 5.58E-21 | 1.32E-19 | 5686 | 1 | 109941653 | 109969062 | -1 | protein_coding | proteasome (prosome, macropain) subunit, alpha type, 5 |
| **C1orf162** | ENSG00000143110 | 1796.895033 | -2.096933619 | 0.168093568 | 1.04E-36 | 5.52E-35 | 128346 | 1 | 112016414 | 112021134 | 1 | protein_coding | chromosome 1 open reading frame 162 |
| **CELSR2** | ENSG00000143126 | 72.85587251 | -1.965393295 | 0.494254518 | 6.63E-06 | 3.75E-05 | 1952 | 1 | 109792641 | 109818372 | 1 | protein_coding | cadherin, EGF LAG seven-pass G-type receptor 2 |
| **TIPRL** | ENSG00000143155 | 805.899722 | 0.751890174 | 0.144645181 | 9.24E-08 | 6.88E-07 | 261726 | 1 | 168148171 | 168169950 | 1 | protein_coding | TIP41, TOR signaling pathway regulator-like (S. cerevisiae) |
| **MPC2** | ENSG00000143158 | 1822.174937 | -0.434209823 | 0.1337121 | 8.7E-04 | 0.003223138 | 25874 | 1 | 167885967 | 167906278 | -1 | protein_coding | mitochondrial pyruvate carrier 2 |
| **CREG1** | ENSG00000143162 | 13599.76679 | 1.208594079 | 0.120808527 | 3.67E-24 | 1.05E-22 | 8804 | 1 | 167498914 | 167523004 | -1 | protein_coding | cellular repressor of E1A-stimulated genes 1 |
| **DCAF6** | ENSG00000143164 | 2856.068203 | -0.677243348 | 0.130215011 | 9.99E-08 | 7.4E-07 | 55827 | 1 | 167905021 | 168045081 | 1 | protein_coding | DDB1 and CUL4 associated factor 6 |
| **TBX19** | ENSG00000143178 | 114.8994285 | 1.068242608 | 0.318926576 | 1.96E-04 | 8.39E-04 | 9095 | 1 | 168250278 | 168283664 | 1 | protein_coding | T-box 19 |
| **TMCO1** | ENSG00000143183 | 971.0217688 | 0.280115851 | 0.117977935 | 0.015462367 | 0.040848981 | 54499 | 1 | 165696032 | 165796992 | -1 | protein_coding | transmembrane and coiled-coil domains 1 |

| | | | | | | | | | | | | | |
|---|---|---|---|---|---|---|---|---|---|---|---|---|---|
| MGST3 | ENSG00000143198 | 781.6410559 | -0.598740937 | 0.178685522 | 4.49E-04 | 0.001776124 | 4259 | 1 | 165600098 | 165631033 | 1 | protein_coding | microsomal glutathione S-transferase 3 |
| PVRL4 | ENSG00000143217 | 833.3125785 | -1.40054001 | 0.192957262 | 6.55E-14 | 9.22E-13 | 81607 | 1 | 161040785 | 161059389 | -1 | protein_coding | poliovirus receptor-related 4 |
| FCGR2A | ENSG00000143226 | 6625.374491 | -0.458005059 | 0.147996821 | 0.00149877 | 0.005230805 | 2212 | 1 | 161475220 | 161493803 | 1 | protein_coding | Fc fragment of IgG, low affinity IIa, receptor (CD32) |
| NUF2 | ENSG00000143228 | 42.45493549 | 1.401323318 | 0.520310492 | 9.45E-04 | 0.00346901 | 83540 | 1 | 163236366 | 163325554 | 1 | protein_coding | NUF2, NDC80 kinetochore complex component |
| SDHC | ENSG00000143252 | 1884.640946 | -0.638207134 | 0.111255383 | 5.24E-09 | 4.63E-08 | 6391 | 1 | 161284047 | 161332984 | 1 | protein_coding | succinate dehydrogenase complex, subunit C, integral membrane protein, 15kDa |
| PRCC | ENSG00000143294 | 933.9350566 | 0.507430018 | 0.181153479 | 0.00336601 | 0.010753078 | 5546 | 1 | 156720402 | 156770607 | 1 | protein_coding | papillary renal cell carcinoma (translocation-associated) |
| CRABP2 | ENSG00000143320 | 4468.771463 | -2.820047806 | 0.206450747 | 1.21E-43 | 8.83E-42 | 1382 | 1 | 156669398 | 156675608 | -1 | protein_coding | cellular retinoic acid binding protein 2 |
| HDGF | ENSG00000143321 | 4593.386384 | 0.368443702 | 0.092055189 | 4.97E-05 | 2.4E-04 | 3068 | 1 | 156711899 | 156736717 | -1 | protein_coding | hepatoma-derived growth factor |
| ABL2 | ENSG00000143322 | 1025.450174 | 0.611005076 | 0.153583629 | 3.85E-05 | 1.9E-04 | 27 | 1 | 179068462 | 179198819 | -1 | protein_coding | c-abl oncogene 2, non-receptor tyrosine kinase |
| XPR1 | ENSG00000143324 | 2293.236286 | -0.267084677 | 0.104801294 | 0.009670093 | 0.027247056 | 9213 | 1 | 180601140 | 180859387 | 1 | protein_coding | xenotropic and polytropic retrovirus receptor 1 |
| TOR1AIP1 | ENSG00000143337 | 2981.164976 | 0.355941207 | 0.110230719 | 0.001031148 | 0.003750363 | 26092 | 1 | 179851177 | 179894135 | 1 | protein_coding | torsin A interacting protein 1 |
| RGL1 | ENSG00000143344 | 2121.860416 | 2.412777433 | 0.132808052 | 2.48E-74 | 5.26E-72 | 23179 | 1 | 183605220 | 183897665 | 1 | protein_coding | ral guanine nucleotide dissociation stimulator-like 1 |
| RORC | ENSG00000143365 | 36.15199963 | -1.138201542 | 0.606315013 | 0.009756025 | 0.02743308 | 6097 | 1 | 151778547 | 151804348 | -1 | protein_coding | RAR-related orphan receptor C |
| TUFT1 | ENSG00000143367 | 155.8094331 | 0.791940904 | 0.304918128 | 0.003458195 | 0.011010748 | 7286 | 1 | 151512781 | 151556059 | 1 | protein_coding | tuftelin 1 |
| ECM1 | ENSG00000143369 | 1582.49143 | -0.498495952 | 0.177692292 | 0.003380873 | 0.010791537 | 1893 | 1 | 150480538 | 150486265 | 1 | protein_coding | extracellular matrix protein 1 |
| SNX27 | ENSG00000143376 | 5816.845458 | 0.414445889 | 0.083999717 | 6.32E-07 | 4.21E-06 | 81609 | 1 | 151584541 | 151671567 | 1 | protein_coding | sorting nexin family member 27 |
| ADAMTSL4 | ENSG00000143382 | 494.1260906 | -1.169989582 | 0.297327445 | 1.71E-05 | 9.03E-05 | 54507 | 1 | 150521884 | 150533413 | 1 | protein_coding | ADAMTS-like 4 |
| MCL1 | ENSG00000143384 | 9992.315493 | 1.610446895 | 0.115702158 | 8.23E-45 | 6.23E-43 | 4170 | 1 | 150547032 | 150552066 | -1 | protein_coding | myeloid cell leukemia sequence 1 (BCL2-related) |
| RFX5 | ENSG00000143390 | 1990.255973 | 0.466342366 | 0.164725452 | 0.003149679 | 0.010146826 | 5993 | 1 | 151313116 | 151319833 | -1 | protein_coding | regulatory factor X, 5 (influences HLA class II expression) |
| PIP5K1A | ENSG00000143398 | 1398.205713 | 0.757156013 | 0.1482523 | 1.47E-07 | 1.07E-06 | 8394 | 1 | 151170425 | 151222012 | 1 | protein_coding | phosphatidylinositol-4-phosphate 5-kinase, type I, alpha |
| ANXA9 | ENSG00000143412 | 40.43659307 | -2.434462162 | 0.697620238 | 4.08E-05 | 0.000199809 | 8416 | 1 | 150954493 | 150968110 | 1 | protein_coding | annexin A9 |
| CERS2 | ENSG00000143418 | 4103.546066 | -0.282020241 | 0.099725571 | 0.004049968 | 0.012649209 | 29956 | 1 | 150933059 | 150947479 | -1 | protein_coding | ceramide synthase 2 |
| ARNT | ENSG00000143437 | 2109.117092 | -0.330494179 | 0.108238191 | 0.001847882 | 0.006299615 | 405 | 1 | 150782181 | 150849244 | -1 | protein_coding | aryl hydrocarbon receptor nuclear translocator |
| HORMAD1 | ENSG00000143452 | 69.95613963 | 1.021393948 | 0.447054646 | 0.005027379 | 0.01534214 | 84072 | 1 | 150670536 | 150693364 | -1 | protein_coding | HORMA domain containing 1 |
| GOLPH3L | ENSG00000143457 | 364.8266446 | 0.669778887 | 0.174308682 | 6E-05 | 2.86E-04 | 55204 | 1 | 150618701 | 150669630 | -1 | protein_coding | golgi phosphoprotein 3-like |
| DTL | ENSG00000143476 | 77.44154376 | 1.184276882 | 0.563207401 | 0.005477488 | 0.016511331 | 51514 | 1 | 212208919 | 212280742 | 1 | protein_coding | denticleless E3 ubiquitin protein ligase homolog (Drosophila) |
| INTS7 | ENSG00000143493 | 494.9505699 | 0.607995373 | 0.168837598 | 1.8E-04 | 7.8E-04 | 25896 | 1 | 212113741 | 212208884 | -1 | protein_coding | integrator complex subunit 7 |
| TAF1A | ENSG00000143498 | 124.3132745 | 1.796007692 | 0.327338603 | 4.69E-09 | 4.16E-08 | 9015 | 1 | 222731244 | 222763275 | -1 | protein_coding | TATA box binding protein (TBP)-associated factor, RNA polymerase I, A, 48kDa |
| SMYD2 | ENSG00000143499 | 700.5890317 | 0.425554835 | 0.151548517 | 0.003707982 | 0.01170849 | 56950 | 1 | 214454445 | 214510474 | 1 | protein_coding | SET and MYND domain containing 2 |
| DUSP10 | ENSG00000143507 | 1342.244377 | 0.936747468 | 0.130063941 | 1.65E-13 | 2.23E-12 | 11221 | 1 | 221874766 | 221915518 | -1 | protein_coding | dual specificity phosphatase 10 |
| ADAM15 | ENSG00000143537 | 939.1195771 | -0.933745032 | 0.148441499 | 1.15E-10 | 1.23E-09 | 8751 | 1 | 155023042 | 155035252 | 1 | protein_coding | ADAM metallopeptidase domain 15 |
| JTB | ENSG00000143543 | 2190.842904 | -0.328524813 | 0.099154246 | 7.43E-04 | 0.002794361 | 10899 | 1 | 153946745 | 153950164 | -1 | protein_coding | jumping translocation breakpoint |
| SLC39A1 | ENSG00000143570 | 3438.65522 | 0.284334929 | 0.101011814 | 0.003943953 | 0.012361655 | 27173 | 1 | 153931575 | 153940188 | -1 | protein_coding | solute carrier family 39 (zinc transporter), member 1 |
| RIT1 | ENSG00000143622 | 2364.423264 | 0.891752493 | 0.10359181 | 2.49E-18 | 4.83E-17 | 6016 | 1 | 155867599 | 155881195 | -1 | protein_coding | Ras-like without CAAX 1 |
| C1orf131 | ENSG00000143633 | 201.8894855 | 0.73488393 | 0.261586625 | 0.002146215 | 0.007196272 | 128061 | 1 | 231359509 | 231376933 | -1 | protein_coding | chromosome 1 open reading frame 131 |
| LYST | ENSG00000143669 | 3196.951215 | 0.286992413 | 0.124205791 | 0.018058124 | 0.046540017 | 1130 | 1 | 235824341 | 236046940 | -1 | protein_coding | lysosomal trafficking regulator |
| SRP9 | ENSG00000143742 | 2265.997061 | 0.607939988 | 0.140898303 | 9.74E-06 | 5.37E-05 | 6726 | 1 | 225965531 | 225978168 | 1 | protein_coding | signal recognition particle 9kDa |
| SDE2 | ENSG00000143751 | 1030.470888 | 0.803153442 | 0.137952233 | 2.56E-09 | 2.34E-08 | 163859 | 1 | 226170403 | 226187032 | -1 | protein_coding | SDE2 telomere maintenance homolog (S. pombe) |
| CNIH4 | ENSG00000143771 | 1772.939461 | 0.907297727 | 0.123717035 | 7.68E-14 | 1.08E-12 | 29097 | 1 | 224544552 | 224567161 | 1 | protein_coding | cornichon family AMPA receptor auxiliary protein 4 |
| ITPKB | ENSG00000143772 | 4833.603027 | -0.877743673 | 0.1658903 | 4.68E-08 | 3.65E-07 | 3707 | 1 | 226819391 | 226927024 | -1 | protein_coding | inositol-trisphosphate 3-kinase B |
| GUK1 | ENSG00000143774 | 772.1292437 | -0.478680523 | 0.184942386 | 0.006872436 | 0.020186708 | 2987 | 1 | 228327663 | 228336685 | 1 | protein_coding | guanylate kinase 1 |
| CDC42BPA | ENSG00000143776 | 611.8834971 | -0.533296023 | 0.180579056 | 0.002052171 | 0.006915838 | 8476 | 1 | 227177566 | 227506175 | -1 | protein_coding | CDC42 binding protein kinase alpha (DMPK-like) |
| CNIH3 | ENSG00000143786 | 1862.370284 | -1.523633542 | 0.178790969 | 2.23E-18 | 4.35E-17 | 149111 | 1 | 224622362 | 224928251 | 1 | protein_coding | cornichon family AMPA receptor auxiliary protein 3 |
| PSEN2 | ENSG00000143801 | 1991.443691 | 0.342825969 | 0.12911896 | 0.006589116 | 0.019440435 | 5664 | 1 | 227057885 | 227083806 | 1 | protein_coding | presenilin 2 (Alzheimer disease 4) |
| EPHX1 | ENSG00000143819 | 1945.67589 | -1.033166741 | 0.128676208 | 2.86E-16 | 4.74E-15 | 2052 | 1 | 225997794 | 226033260 | 1 | protein_coding | epoxide hydrolase 1, microsomal (xenobiotic) |
| SOX13 | ENSG00000143842 | 653.6736936 | -0.619501169 | 0.217402776 | 0.002081312 | 0.007001692 | 9580 | 1 | 204042243 | 204096863 | 1 | protein_coding | SRY (sex determining region Y)-box 13 |
| ETNK2 | ENSG00000143845 | 71.30504164 | -2.058640293 | 0.770461909 | 5.18E-04 | 0.002020131 | 55224 | 1 | 204100190 | 204121307 | -1 | protein_coding | ethanolamine kinase 2 |
| PPFIA4 | ENSG00000143847 | 158.1077243 | -3.12469012 | 0.483022759 | 6.28E-12 | 7.54E-11 | 8497 | 1 | 202995626 | 203047868 | 1 | protein_coding | protein tyrosine phosphatase, receptor type, f polypeptide (PTPRF), interacting protein (liprin), alpha 4 |
| PLEKHA6 | ENSG00000143850 | 34.37535696 | -1.638575887 | 1.718302421 | 0.012560499 | 0.034029799 | 22874 | 1 | 204187979 | 204346793 | -1 | protein_coding | pleckstrin homology domain containing, family A member 6 |
| PTPN7 | ENSG00000143851 | 1179.412338 | -0.846829589 | 0.191299819 | 3.47E-06 | 2.05E-05 | 5778 | 1 | 202116141 | 202130716 | -1 | protein_coding | protein tyrosine phosphatase, non-receptor type 7 |
| ARL8A | ENSG00000143862 | 2181.133211 | 0.658170006 | 0.134283585 | 6.03E-07 | 4.02E-06 | 127829 | 1 | 202102532 | 202113869 | -1 | protein_coding | ADP-ribosylation factor-like 8A |
| GDF7 | ENSG00000143869 | 16.48435421 | -2.694137499 | 1.106714699 | 0.001117953 | 0.0040344 | 151449 | 2 | 20866424 | 20873418 | 1 | protein_coding | growth differentiation factor 7 |
| PDIA6 | ENSG00000143870 | 5732.094568 | -0.346383919 | 0.120040846 | 0.003338862 | 0.010673046 | 10130 | 2 | 10923517 | 10978103 | -1 | protein_coding | protein disulfide isomerase family A, member 6 |
| RHOB | ENSG00000143878 | 400.340212 | -0.53186032 | 0.220237906 | 0.009760095 | 0.027434896 | 388 | 2 | 20646835 | 20649200 | 1 | protein_coding | ras homolog family member B |
| HNRNPLL | ENSG00000143889 | 2094.984838 | -0.28522888 | 0.111347593 | 0.009174729 | 0.02598543 | 92906 | 2 | 38789120 | 38830728 | -1 | protein_coding | heterogeneous nuclear ribonucleoprotein L-like |
| GALM | ENSG00000143891 | 5908.36889 | -0.76590444 | 0.093563862 | 1.86E-16 | 3.12E-15 | 130589 | 2 | 38893052 | 38968379 | 1 | protein_coding | galactose mutarotase (aldose 1-epimerase) |
| EML4 | ENSG00000143924 | 6229.262253 | -0.782888658 | 0.107386076 | 9.38E-14 | 1.3E-12 | 27436 | 2 | 42396490 | 42559688 | 1 | protein_coding | echinoderm microtubule associated protein like 4 |
| CALM2 | ENSG00000143933 | 8329.560522 | -1.410098876 | 0.144598032 | 2.29E-23 | 6.3E-22 | 808 | 2 | 47387221 | 47403740 | -1 | protein_coding | calmodulin 2 (phosphorylase kinase, delta) |
| WDPCP | ENSG00000143951 | 150.9219591 | -0.978888679 | 0.31430779 | 5.12E-04 | 0.001998857 | 51057 | 2 | 63348518 | 64054977 | -1 | protein_coding | WD repeat containing planar cell polarity effector |
| VPS54 | ENSG00000143952 | 1471.818742 | 0.771918866 | 0.112860348 | 3.72E-12 | 4.56E-11 | 51542 | 2 | 64119280 | 64246206 | -1 | protein_coding | vacuolar protein sorting 54 homolog (S. cerevisiae) |
| ASXL2 | ENSG00000143970 | 2753.902952 | -0.347110849 | 0.141503752 | 0.011469435 | 0.031494797 | 55252 | 2 | 25956622 | 26101385 | -1 | protein_coding | additional sex combs like 2 (Drosophila) |
| ETAA1 | ENSG00000143971 | 267.6658642 | 0.953596654 | 0.242723721 | 2.6E-05 | 1.33E-04 | 54465 | 2 | 67624451 | 67637677 | 1 | protein_coding | Ewing tumor-associated antigen 1 |
| SNRPG | ENSG00000143977 | 679.3522331 | 0.746826742 | 0.179612408 | 1.43E-05 | 7.66E-05 | 6637 | 2 | 70508494 | 70520903 | -1 | protein_coding | small nuclear ribonucleoprotein polypeptide G |
| CIAO1 | ENSG00000144021 | 2838.280389 | -0.727021643 | 0.138174042 | 6.81E-08 | 5.18E-07 | 9391 | 2 | 96931870 | 96939087 | 1 | protein_coding | cytosolic iron-sulfur protein assembly 1 |
| SNRNP200 | ENSG00000144028 | 7090.798358 | -0.636060544 | 0.103013751 | 3.66E-10 | 3.68E-09 | 1019292 | 2 | 96940074 | 96971297 | -1 | protein_coding | small nuclear ribonucleoprotein 200kDa (U5) |
| SFXN5 | ENSG00000144040 | 371.0541283 | -0.836138089 | 0.255298617 | 3.81E-04 | 0.001536981 | 94097 | 2 | 73169165 | 73302747 | -1 | protein_coding | sideroflexin 5 |
| TEX261 | ENSG00000144043 | 1645.717015 | -0.425268767 | 0.11415065 | 1.51E-04 | 6.66E-04 | 113419 | 2 | 71213068 | 71222075 | -1 | protein_coding | testis expressed 261 |
| DUSP11 | ENSG00000144048 | 699.2281874 | 0.69042256 | 0.155230642 | 4.39E-06 | 2.57E-05 | 8446 | 2 | 73989311 | 74007284 | -1 | protein_coding | dual specificity phosphatase 11 (RNA/RNP complex 1-interacting) |
| RALB | ENSG00000144118 | 3951.671936 | 0.422669733 | 0.154583527 | 0.005040421 | 0.015369664 | 5899 | 2 | 120997640 | 121052289 | 1 | protein_coding | v-ral simian leukemia viral oncogene homolog B |
| C1QL2 | ENSG00000144119 | 14.77011809 | -2.16323795 | 1.363264282 | 0.007240355 | 0.021107895 | 165257 | 2 | 119913819 | 119916465 | -1 | protein_coding | complement component 1, q subcomponent-like 2 |
| RABL2A | ENSG00000144134 | 250.3335653 | -0.52152683 | 0.229618208 | 0.014599269 | 0.03879684 | 11159 | 2 | 114384806 | 114400973 | 1 | protein_coding | RAB, member of RAS oncogene family-like 2A |
| SLC20A1 | ENSG00000144136 | 6897.971611 | -1.238969605 | 0.118601576 | 3.22E-26 | 1.02E-24 | 6574 | 2 | 113403434 | 113421404 | 1 | protein_coding | solute carrier family 20 (phosphate transporter), member 1 |
| FBLN7 | ENSG00000144152 | 70.98198942 | -1.539678801 | 0.53908738 | 4.79E-04 | 0.001881518 | 129804 | 2 | 112895962 | 112945791 | 1 | protein_coding | fibulin 7 |
| AFF3 | ENSG00000144218 | 48.83557127 | -2.440121349 | 0.781733572 | 1.05E-04 | 4.77E-04 | 3899 | 2 | 100162323 | 100759201 | -1 | protein_coding | AF4/FMR2 family, member 3 |
| UBXN4 | ENSG00000144224 | 5562.422997 | 0.932144258 | 0.080275527 | 9.93E-32 | 4.23E-30 | 23190 | 2 | 136499189 | 136542625 | 1 | protein_coding | UBX domain protein 4 |
| PKP4 | ENSG00000144283 | 216.0114034 | -1.262092163 | 0.271200506 | 6.36E-07 | 4.23E-06 | 8502 | 2 | 159313476 | 159539391 | 1 | protein_coding | plakophilin 4 |
| CDCA7 | ENSG00000144354 | 68.63682412 | 1.348649442 | 0.469947457 | 5.76E-04 | 0.00221902 | 83879 | 2 | 174219548 | 174233725 | 1 | protein_coding | cell division cycle associated 7 |
| UNC80 | ENSG00000144406 | 22.77035532 | -5.756420864 | 2.62670742 | 1.31E-04 | 5.88E-04 | 285175 | 2 | 210636717 | 210864024 | 1 | protein_coding | unc-80 homolog (C. elegans) |
| KANSL1L | ENSG00000144445 | 301.0175868 | 0.779165329 | 0.222765993 | 1.93E-04 | 8.29E-04 | 151050 | 2 | 210886147 | 211036107 | -1 | protein_coding | KAT8 regulatory NSL complex subunit 1-like |
| SPAG16 | ENSG00000144451 | 96.62169142 | 0.773728596 | 0.402257484 | 0.018776692 | 0.048099931 | 79582 | 2 | 214149113 | 215275225 | 1 | protein_coding | sperm associated antigen 16 |
| SUMF1 | ENSG00000144455 | 2287.246841 | -0.929162346 | 0.15256892 | 4.03E-10 | 4.04E-09 | 285362 | 3 | 3742498 | 4508965 | -1 | protein_coding | sulfatase modifying factor 1 |
| HES6 | ENSG00000144485 | 58.58885426 | -2.063516625 | 0.514944967 | 5.77E-06 | 3.3E-05 | 55502 | 2 | 239146908 | 239149303 | -1 | protein_coding | hes family bHLH transcription factor 6 |
| VGLL4 | ENSG00000144560 | 1961.268548 | -0.782382398 | 0.110482285 | 5.79E-13 | 7.56E-12 | 9686 | 3 | 11597544 | 11762220 | -1 | protein_coding | vestigial like 4 (Drosophila) |
| RAB5A | ENSG00000144566 | 2325.950163 | 0.677846988 | 0.120778629 | 1.12E-08 | 9.44E-08 | 5868 | 3 | 19988571 | 20026667 | 1 | protein_coding | RAB5A, member RAS oncogene family |
| FAM134A | ENSG00000144567 | 3132.647705 | -0.3024139 | 0.100546786 | 0.002287882 | 0.007608163 | 79137 | 2 | 220040947 | 220050201 | 1 | protein_coding | family with sequence similarity 134, member A |
| CTDSP1 | ENSG00000144579 | 1367.061528 | -0.351183187 | 0.140310337 | 0.01094781 | 0.030250474 | 58190 | 2 | 219262979 | 219270664 | 1 | protein_coding | CTD (carboxy-terminal domain, RNA polymerase II, polypeptide A) small phosphatase 1 |
| RQCD1 | ENSG00000144580 | 1613.799582 | 0.607818984 | 0.12215339 | 3.73E-07 | 2.56E-06 | 9125 | 2 | 219433303 | 219461803 | 1 | protein_coding | RCD1 required for cell differentiation1 homolog (S. pombe) |
| DYNC1LI1 | ENSG00000144635 | 995.1919076 | 1.083294996 | 0.189350657 | 2.77E-09 | 2.52E-08 | 51143 | 3 | 32567463 | 32612366 | -1 | protein_coding | dynein, cytoplasmic 1, light intermediate chain 1 |
| POMGNT2 | ENSG00000144647 | 241.9392931 | -0.717544417 | 0.25799197 | 0.002398344 | 0.007930868 | 84892 | 3 | 43120724 | 43147568 | -1 | protein_coding | protein O-linked mannose N-acetylglucosaminyltransferase 2 (beta 1,4-) |

| | | | | | | | | | | | | | |
|---|---|---|---|---|---|---|---|---|---|---|---|---|---|
| **CSRNP1** | ENSG00000144655 | 566.8581572 | 2.179159406 | 0.291001026 | 5.89E-15 | 8.93E-14 | 64651 | 3 | 39183346 | 39196053 | -1 | protein_coding | cysteine-serine-rich nuclear protein 1 |
| **GOLGA4** | ENSG00000144674 | 3875.474315 | 0.727859253 | 0.106301241 | 3.74E-12 | 4.57E-11 | 2803 | 3 | 37284668 | 37408242 | 1 | protein_coding | golgin A4 |
| **CTDSPL** | ENSG00000144677 | 891.4555059 | -1.408404773 | 0.204433087 | 8.85E-13 | 1.13E-11 | 10217 | 3 | 37903451 | 38025960 | 1 | protein_coding | CTD (carboxy-terminal domain, RNA polymerase II, polypeptide A) small phosphatase-like |
| **STAC** | ENSG00000144681 | 154.8274587 | -1.649705896 | 0.309107066 | 1.3E-08 | 1.09E-07 | 6769 | 3 | 36421836 | 36589499 | 1 | protein_coding | SH3 and cysteine rich domain |
| **IQSEC1** | ENSG00000144711 | 2180.059539 | -0.583715883 | 0.110863933 | 8.45E-08 | 6.33E-07 | 9922 | 3 | 12938719 | 13114617 | -1 | protein_coding | IQ motif and Sec7 domain 1 |
| **RPL32** | ENSG00000144713 | 4310.612959 | -0.337332537 | 0.143702844 | 0.015732598 | 0.041455396 | 6161 | 3 | 12875984 | 12883087 | -1 | protein_coding | ribosomal protein L32 |
| **UBA3** | ENSG00000144744 | 1361.09471 | 0.510126366 | 0.168000197 | 0.001594037 | 0.005527947 | 9039 | 3 | 69103881 | 69129559 | -1 | protein_coding | ubiquitin-like modifier activating enzyme 3 |
| **ARL6IP5** | ENSG00000144746 | 6506.019828 | 0.45980114 | 0.114842875 | 4.54E-05 | 2.2E-04 | 10550 | 3 | 69134095 | 69155217 | 1 | protein_coding | ADP-ribosylation-like factor 6 interacting protein 5 |
| **TMF1** | ENSG00000144747 | 1905.395477 | 0.516133004 | 0.139581424 | 1.5E-04 | 6.62E-04 | 7110 | 3 | 69068978 | 69101484 | -1 | protein_coding | TATA element modulatory factor 1 |
| **NFKBIZ** | ENSG00000144802 | 1066.857279 | 1.203974588 | 0.193914588 | 1.21E-10 | 1.28E-09 | 64332 | 3 | 101546835 | 101579866 | 1 | protein_coding | nuclear factor of kappa light polypeptide gene enhancer in B-cells inhibitor, zeta |
| **NXPE3** | ENSG00000144815 | 712.4153105 | 0.675834217 | 0.149129914 | 3.16E-06 | 1.89E-05 | 91775 | 3 | 101498046 | 101547073 | 1 | protein_coding | neurexophilin and PC-esterase domain family, member 3 |
| **PLA1A** | ENSG00000144837 | 16.38746613 | 3.357713275 | 0.955338897 | 2.64E-05 | 1.34E-04 | 51365 | 3 | 119316689 | 119348658 | 1 | protein_coding | phospholipase A1 member A |
| **ATG3** | ENSG00000144848 | 2855.282032 | 0.637999761 | 0.10908178 | 2.79E-09 | 2.53E-08 | 64422 | 3 | 112251356 | 112280893 | -1 | protein_coding | autophagy related 3 |
| **SRPRB** | ENSG00000144867 | 863.8036555 | -0.774815196 | 0.156247053 | 3.03E-07 | 2.11E-06 | 58477 | 3 | 133502877 | 133544616 | 1 | protein_coding | signal recognition particle receptor, B subunit |
| **NCEH1** | ENSG00000144959 | 695.2620971 | -0.537379514 | 0.247390988 | 0.017797355 | 0.045992048 | 57552 | 3 | 172348039 | 172429008 | -1 | protein_coding | neutral cholesterol ester hydrolase 1 |
| **KIAA0226** | ENSG00000145016 | 2281.644397 | 1.360460562 | 0.129515646 | 1.13E-26 | 3.68E-25 | 9711 | 3 | 197398264 | 197476598 | -1 | protein_coding | KIAA0226 |
| **TM4SF19** | ENSG00000145107 | 11625.38176 | -1.4171932 | 0.226806675 | 5.77E-11 | 6.35E-10 | 116211 | 3 | 196046213 | 196065374 | -1 | protein_coding | transmembrane 4 L six family member 19 |
| **SLIT2** | ENSG00000145147 | 20.20583708 | 1.292183725 | 0.84447423 | 0.012204146 | 0.033206076 | 9353 | 4 | 20254883 | 20622184 | 1 | protein_coding | slit homolog 2 (Drosophila) |
| **SLC26A1** | ENSG00000145217 | 3.692007468 | -0.376666526 | 0.797540233 | 0.01676418 | 0.043721325 | 10861 | 4 | 972861 | 987228 | -1 | protein_coding | solute carrier family 26 (anion exchanger), member 1 |
| **CENPC** | ENSG00000145241 | 628.4798083 | 0.527227294 | 0.163922611 | 8.38E-04 | 0.003123195 | 1060 | 4 | 68337521 | 68411324 | -1 | protein_coding | centromere protein C |
| **PLAC8** | ENSG00000145287 | 106.8731895 | 2.787847944 | 0.431140674 | 6.42E-12 | 7.7E-11 | 51316 | 4 | 84011201 | 84058228 | -1 | protein_coding | placenta-specific 8 |
| **SNCA** | ENSG00000145335 | 269.2786674 | -1.219435398 | 0.492840722 | 0.001964383 | 0.006661264 | 6622 | 4 | 90645250 | 90759466 | -1 | protein_coding | synuclein, alpha (non A4 component of amyloid precursor) |
| **PYURF** | ENSG00000145337 | 939.5670545 | 0.488453214 | 0.142433099 | 4.27E-04 | 0.001699689 | 1009969 | 4 | 89442136 | 89444964 | -1 | protein_coding | PIGY upstream reading frame |
| **CAMK2D** | ENSG00000145349 | 1350.578912 | 0.316924207 | 0.115444321 | 0.00516385 | 0.015705296 | 817 | 4 | 114372188 | 114683083 | -1 | protein_coding | calcium/calmodulin-dependent protein kinase II delta |
| **CISD2** | ENSG00000145354 | 920.0046502 | 1.012998949 | 0.191199782 | 3.34E-08 | 2.65E-07 | 493856 | 4 | 103790135 | 103810399 | 1 | protein_coding | CDGSH iron sulfur domain 2 |
| **DDIT4L** | ENSG00000145358 | 127.8540624 | -3.150836881 | 0.476368116 | 2.51E-12 | 3.1E-11 | 115265 | 4 | 101107027 | 101111939 | -1 | protein_coding | DNA-damage-inducible transcript 4-like |
| **ANK2** | ENSG00000145362 | 1121.330641 | -0.909076235 | 0.142785736 | 6.51E-11 | 7.1E-10 | 287 | 4 | 113739265 | 114304896 | 1 | protein_coding | ankyrin 2, neuronal |
| **TIFA** | ENSG00000145365 | 557.8161775 | 1.313586428 | 0.248507536 | 2.12E-08 | 1.72E-07 | 92610 | 4 | 113195698 | 113207059 | -1 | protein_coding | TRAF-interacting protein with forkhead-associated domain |
| **SPATA5** | ENSG00000145375 | 356.402764 | 0.689936179 | 0.192867402 | 1.69E-04 | 7.38E-04 | 166378 | 4 | 123844229 | 124240605 | 1 | protein_coding | spermatogenesis associated 5 |
| **CCNA2** | ENSG00000145386 | 114.9379584 | 1.407601247 | 0.375669026 | 2.62E-05 | 1.34E-04 | 890 | 4 | 122737599 | 122745087 | -1 | protein_coding | cyclin A2 |
| **METTL14** | ENSG00000145388 | 814.3043292 | 0.801210445 | 0.169327631 | 9.18E-07 | 5.94E-06 | 57721 | 4 | 119606523 | 119636588 | 1 | protein_coding | methyltransferase like 14 |
| **SETD7** | ENSG00000145391 | 5047.639809 | -0.739464725 | 0.091084923 | 9.32E-17 | 1.6E-15 | 80854 | 4 | 140417095 | 140527853 | -1 | protein_coding | SET domain containing (lysine methyltransferase) 7 |
| **NAF1** | ENSG00000145414 | 402.9237706 | 1.073731908 | 0.183816375 | 1.31E-09 | 1.25E-08 | 92345 | 4 | 164031225 | 164088073 | -1 | protein_coding | nuclear assembly factor 1 ribonucleoprotein |
| **1-Mar** | ENSG00000145416 | 4910.254937 | 0.482949464 | 0.121786991 | 5.01E-05 | 2.41E-04 | 55016 | 4 | 164445450 | 165305202 | -1 | protein_coding | membrane-associated ring finger (C3HC4) 1, E3 ubiquitin protein ligase |
| **RPS3A** | ENSG00000145425 | 8735.48983 | -0.578818541 | 0.112704702 | 1.78E-07 | 1.28E-06 | 8944 | 4 | 152020725 | 152025804 | 1 | protein_coding | ribosomal protein S3A |
| **PDGFC** | ENSG00000145431 | 864.1251522 | -2.430618495 | 0.159298483 | 2E-53 | 2.14E-51 | 56034 | 4 | 157681606 | 157892546 | -1 | protein_coding | platelet derived growth factor C |
| **6-Mar** | ENSG00000145495 | 3510.849986 | 0.356540569 | 0.09876196 | 2.75E-04 | 0.001142046 | 10299 | 5 | 10353815 | 10440500 | 1 | protein_coding | membrane-associated ring finger (C3HC4) 6, E3 ubiquitin protein ligase |
| **SRD5A1** | ENSG00000145545 | 466.533542 | 0.744616732 | 0.173824935 | 8.26E-06 | 4.6E-05 | 6715 | 5 | 6633456 | 6669675 | 1 | protein_coding | steroid-5-alpha-reductase, alpha polypeptide 1 (3-oxo-5 alpha-steroid delta 4-dehydrogenase alpha 1) |
| **MYO10** | ENSG00000145555 | 453.3741761 | 0.624088517 | 0.264378573 | 0.009363666 | 0.0264744 | 4651 | 5 | 16665395 | 16936372 | -1 | protein_coding | myosin X |
| **FAM105A** | ENSG00000145569 | 875.4432386 | -1.112470021 | 0.203260835 | 1E-08 | 8.54E-08 | 54491 | 5 | 14581884 | 14615116 | 1 | protein_coding | family with sequence similarity 105, member A |
| **LHFPL2** | ENSG00000145685 | 23394.04405 | 0.666341758 | 0.115305295 | 4.46E-09 | 3.96E-08 | 10184 | 5 | 77781038 | 78065844 | -1 | protein_coding | lipoma HMGIC fusion partner-like 2 |
| **IQGAP2** | ENSG00000145703 | 5732.911297 | 0.519740228 | 0.118602913 | 7.54E-06 | 4.23E-05 | 10788 | 5 | 75699074 | 76003957 | 1 | protein_coding | IQ motif containing GTPase activating protein 2 |
| **PAM** | ENSG00000145730 | 3208.531221 | 0.355886279 | 0.096048032 | 1.94E-04 | 8.31E-04 | 5066 | 5 | 102089685 | 102366809 | 1 | protein_coding | peptidylglycine alpha-amidating monooxygenase |
| **BDP1** | ENSG00000145734 | 1879.773778 | 0.554841076 | 0.107381928 | 1.42E-07 | 1.03E-06 | 55814 | 5 | 70751442 | 70863649 | 1 | protein_coding | B double prime 1, subunit of RNA polymerase III transcription initiation factor IIIB |
| **TNFAIP8** | ENSG00000145779 | 1870.729356 | 0.624415049 | 0.122882051 | 2.2E-07 | 1.56E-06 | 25816 | 5 | 118604387 | 118735383 | 1 | protein_coding | tumor necrosis factor, alpha-induced protein 8 |
| **FEM1C** | ENSG00000145780 | 1401.423682 | 1.093504398 | 0.141679305 | 2E-15 | 3.1E-14 | 56929 | 5 | 114856608 | 114880591 | -1 | protein_coding | fem-1 homolog c (C. elegans) |
| **ATG12** | ENSG00000145782 | 457.1494556 | 0.838945157 | 0.231438515 | 9.79E-05 | 4.48E-04 | 9140 | 5 | 115163893 | 115177555 | -1 | protein_coding | autophagy related 12 |
| **YIPF5** | ENSG00000145817 | 1859.189984 | -0.270695146 | 0.108104938 | 0.010755077 | 0.02976097 | 81555 | 5 | 143537723 | 143550278 | -1 | protein_coding | Yip1 domain family, member 5 |
| **ARHGAP26** | ENSG00000145819 | 705.3180109 | -1.155726568 | 0.198345973 | 1.32E-09 | 1.25E-08 | 23092 | 5 | 142149949 | 142608576 | 1 | protein_coding | Rho GTPase activating protein 26 |
| **DDX46** | ENSG00000145833 | 2052.053994 | -0.382390589 | 0.110198435 | 3.9E-04 | 0.001570113 | 9879 | 5 | 134094469 | 134190823 | 1 | protein_coding | DEAD (Asp-Glu-Ala-Asp) box polypeptide 46 |
| **PCYOX1L** | ENSG00000145882 | 212.4904636 | -1.113347365 | 0.252008106 | 2.3E-06 | 1.41E-05 | 78991 | 5 | 148737570 | 148749216 | 1 | protein_coding | prenylcysteine oxidase 1 like |
| **TNIP1** | ENSG00000145901 | 4464.843919 | 1.83213182 | 0.110682589 | 2.02E-62 | 2.84E-60 | 10318 | 5 | 150409506 | 150473138 | -1 | protein_coding | TNFAIP3 interacting protein 1 |
| **G3BP1** | ENSG00000145907 | 5849.105889 | 0.359260738 | 0.107565747 | 7.19E-04 | 0.002713128 | 10146 | 5 | 151150606 | 151192346 | 1 | protein_coding | GTPase activating protein (SH3 domain) binding protein 1 |
| **NHP2** | ENSG00000145912 | 256.0829249 | -0.761799772 | 0.244927116 | 7.98E-04 | 0.002983764 | 55651 | 5 | 177576461 | 177580968 | -1 | protein_coding | NHP2 ribonucleoprotein |
| **GFOD1** | ENSG00000145990 | 2602.691807 | -1.699061348 | 0.145371938 | 2.26E-32 | 9.93E-31 | 54438 | 6 | 13358062 | 13487894 | -1 | protein_coding | glucose-fructose oxidoreductase domain containing 1 |
| **PLA2G7** | ENSG00000146070 | 15547.19491 | 0.687527758 | 0.114643908 | 1.3E-09 | 1.23E-08 | 7941 | 6 | 46671938 | 46703430 | -1 | protein_coding | phospholipase A2, group VII (platelet-activating factor acetylhydrolase, plasma) |
| **TNFRSF21** | ENSG00000146072 | 1811.923903 | -1.954933851 | 0.209749252 | 1.36E-21 | 3.35E-20 | 27242 | 6 | 47199268 | 47277641 | -1 | protein_coding | tumor necrosis factor receptor superfamily, member 21 |
| **RNF44** | ENSG00000146083 | 982.7659059 | -0.63025426 | 0.170093971 | 1.13E-04 | 5.11E-04 | 22838 | 5 | 175953698 | 175965026 | -1 | protein_coding | ring finger protein 44 |
| **DOK3** | ENSG00000146094 | 2872.112477 | -0.635954961 | 0.121528705 | 8.04E-08 | 6.04E-07 | 79930 | 5 | 176928908 | 176938275 | -1 | protein_coding | docking protein 3 |
| **PRIM2** | ENSG00000146143 | 18.18029927 | 3.432102345 | 1.080136284 | 8.62E-05 | 3.99E-04 | 1009964 | 6 | 57179603 | 57513375 | 1 | protein_coding | primase, DNA, polypeptide 2 (58kDa) |
| **FGD2** | ENSG00000146192 | 1530.705672 | 0.952167995 | 0.184349193 | 7.93E-08 | 5.97E-07 | 221472 | 6 | 36973422 | 36996846 | 1 | protein_coding | FYVE, RhoGEF and PH domain containing 2 |
| **NFKBIE** | ENSG00000146232 | 1724.532708 | 0.754286926 | 0.118175307 | 8.06E-11 | 8.7E-10 | 4794 | 6 | 44225903 | 44233500 | -1 | protein_coding | nuclear factor of kappa light polypeptide gene enhancer in B-cells inhibitor, epsilon |
| **TPBG** | ENSG00000146242 | 29.5069104 | 1.409657016 | 0.77767865 | 0.006878712 | 0.020197377 | 7162 | 6 | 83072923 | 83080545 | 1 | protein_coding | trophoblast glycoprotein |
| **PHIP** | ENSG00000146247 | 2260.801872 | 0.815448008 | 0.108308803 | 2.39E-14 | 3.48E-13 | 55023 | 6 | 79645584 | 79787953 | -1 | protein_coding | pleckstrin homology domain interacting protein |
| **MMS22L** | ENSG00000146263 | 360.3785259 | 0.668969375 | 0.195873486 | 3.13E-04 | 0.001285121 | 253714 | 6 | 97590037 | 97731093 | -1 | protein_coding | MMS22-like, DNA repair protein |
| **PNRC1** | ENSG00000146278 | 5467.290331 | 2.49895921 | 0.124944509 | 7.48E-90 | 2.24E-87 | 10957 | 6 | 89790470 | 89794879 | 1 | protein_coding | proline-rich nuclear receptor coactivator 1 |
| **PM20D2** | ENSG00000146281 | 128.9972049 | 1.713781278 | 0.329982761 | 2.57E-08 | 2.07E-07 | 135293 | 6 | 89855769 | 89875284 | 1 | protein_coding | peptidase M20 domain containing 2 |
| **RNF217** | ENSG00000146373 | 631.2610909 | 0.682545606 | 0.168045769 | 2.45E-05 | 1.26E-04 | 154214 | 6 | 125283691 | 125413779 | 1 | protein_coding | ring finger protein 217 |
| **ARHGAP18** | ENSG00000146376 | 3763.133119 | -1.210679676 | 0.13219142 | 1.15E-20 | 2.66E-19 | 93663 | 6 | 129897277 | 130031370 | -1 | protein_coding | Rho GTPase activating protein 18 |
| **ABRACL** | ENSG00000146386 | 1095.732743 | 0.851124014 | 0.161308801 | 4.98E-08 | 3.87E-07 | 58527 | 6 | 139349819 | 139364439 | 1 | protein_coding | ABRA C-terminal like |
| **SLC2A12** | ENSG00000146411 | 21.73741441 | 2.058667177 | 0.760769549 | 4.86E-04 | 0.001908533 | 154091 | 6 | 134309835 | 134373774 | -1 | protein_coding | solute carrier family 2 (facilitated glucose transporter), member 12 |
| **AIG1** | ENSG00000146416 | 309.1356285 | -1.349415157 | 0.231364866 | 1.02E-09 | 9.84E-09 | 51390 | 6 | 143381633 | 143661441 | 1 | protein_coding | androgen-induced 1 |
| **DYNLT1** | ENSG00000146425 | 1781.337601 | 1.945535737 | 0.123578326 | 4.02E-57 | 4.76E-55 | 6993 | 6 | 159057506 | 159065771 | -1 | protein_coding | dynein, light chain, Tctex-type 1 |
| **TIAM2** | ENSG00000146426 | 471.5726101 | 0.618578532 | 0.232862425 | 0.004454627 | 0.013770241 | 26230 | 6 | 155153831 | 155578857 | 1 | protein_coding | T-cell lymphoma invasion and metastasis 2 |
| **WTAP** | ENSG00000146457 | 3943.238768 | 2.515692192 | 0.102509709 | 5.46E-134 | 2.53E-131 | 9589 | 6 | 160146617 | 160177351 | 1 | protein_coding | Wilms tumor 1 associated protein |
| **ZMYM4** | ENSG00000146463 | 2777.48512 | 0.644384818 | 0.117719945 | 2.3E-08 | 1.86E-07 | 9202 | 1 | 35734568 | 35887659 | 1 | protein_coding | zinc finger, MYM-type 4 |
| **C6orf211** | ENSG00000146476 | 868.9234492 | 0.845249343 | 0.172853286 | 3.7E-07 | 2.54E-06 | 79624 | 6 | 151773422 | 151791236 | 1 | protein_coding | chromosome 6 open reading frame 211 |
| **GNA12** | ENSG00000146535 | 2590.391107 | -0.308161947 | 0.106936198 | 0.003532218 | 0.011220721 | 2768 | 7 | 2767746 | 2883958 | -1 | protein_coding | guanine nucleotide binding protein (G protein) alpha 12 |
| **C7orf50** | ENSG00000146540 | 395.1285693 | -1.029726292 | 0.182380531 | 4.55E-09 | 4.04E-08 | 1019302 | 7 | 1036622 | 1177896 | -1 | protein_coding | chromosome 7 open reading frame 50 |
| **CDCA5** | ENSG00000146670 | 78.0111692 | 0.90102991 | 0.455737874 | 0.01255987 | 0.034029799 | 113130 | 11 | 64833772 | 64851636 | -1 | protein_coding | cell division cycle associated 5 |
| **PURB** | ENSG00000146676 | 2051.953952 | 0.471622532 | 0.116671991 | 3.87E-05 | 1.91E-04 | 5814 | 7 | 44915896 | 44924960 | -1 | protein_coding | purine-rich element binding protein B |
| **MDH2** | ENSG00000146701 | 1986.973771 | -0.402982012 | 0.106603748 | 1.21E-04 | 5.44E-04 | 4191 | 7 | 75677369 | 75696826 | 1 | protein_coding | malate dehydrogenase 2, NAD (mitochondrial) |
| **PSPH** | ENSG00000146733 | 455.2947043 | -0.601467506 | 0.188723848 | 7.92E-04 | 0.002964646 | 5723 | 7 | 56078744 | 56119297 | -1 | protein_coding | phosphoserine phosphatase |
| **ZNF92** | ENSG00000146757 | 241.769783 | 1.382668047 | 0.305170782 | 9.84E-07 | 6.34E-06 | 168374 | 7 | 64838712 | 64866038 | 1 | protein_coding | zinc finger protein 92 |
| **ATXN7L1** | ENSG00000146776 | 458.4083913 | 1.296524354 | 0.173587055 | 1.75E-14 | 2.56E-13 | 222255 | 7 | 105245514 | 105517050 | -1 | protein_coding | ataxin 7-like 1 |
| **TMEM168** | ENSG00000146802 | 701.7569415 | -0.723280447 | 0.156323878 | 1.78E-06 | 1.1E-05 | 64418 | 7 | 112402437 | 112430647 | -1 | protein_coding | transmembrane protein 168 |
| **C7orf43** | ENSG00000146826 | 572.2855468 | -1.014135643 | 0.234233638 | 3.98E-06 | 2.34E-05 | 55262 | 7 | 99752043 | 99756338 | -1 | protein_coding | chromosome 7 open reading frame 43 |
| **SLC12A9** | ENSG00000146828 | 671.1247739 | -1.04442571 | 0.200986809 | 5.54E-08 | 4.28E-07 | 56996 | 7 | 100424442 | 100464631 | 1 | protein_coding | solute carrier family 12, member 9 |
| **TMEM209** | ENSG00000146842 | 556.3557537 | 0.696447334 | 0.238581403 | 0.001703996 | 0.005864032 | 84928 | 7 | 129804555 | 129847610 | -1 | protein_coding | transmembrane protein 209 |

| | | | | | | | | | | | | |
|---|---|---|---|---|---|---|---|---|---|---|---|---|
| **TMEM140** | ENSG00000146859 | 3897.295353 | 1.311745664 | 0.101091835 | 2.15E-39 | 1.27E-37 | 55281 | 7 | 134832824 | 134850967 | 1 | protein_coding | transmembrane protein 140 |
| **TLK2** | ENSG00000146872 | 1436.698746 | 1.202559063 | 0.183098799 | 1.17E-11 | 1.37E-10 | 11011 | 17 | 60536019 | 60692842 | 1 | protein_coding | tousled-like kinase 2 |
| **EPHA1** | ENSG00000146904 | 186.1367035 | -0.714175205 | 0.297038754 | 0.006981389 | 0.020459554 | 2041 | 7 | 143087382 | 143105985 | -1 | protein_coding | EPH receptor A1 |
| **SH3KBP1** | ENSG00000147010 | 2239.198703 | -0.360941738 | 0.107864004 | 6.82E-04 | 0.002587657 | 30011 | X | 19552093 | 19905719 | -1 | protein_coding | SH3-domain kinase binding protein 1 |
| **TMEM47** | ENSG00000147027 | 4.955695178 | 0.604790738 | 1.130285117 | 0.010304234 | 0.028721432 | 83604 | X | 34645181 | 34675405 | -1 | protein_coding | transmembrane protein 47 |
| **CASK** | ENSG00000147044 | 518.4156965 | 0.648115743 | 0.175744453 | 1.2E-04 | 5.43E-04 | 8573 | X | 41374187 | 41782716 | -1 | protein_coding | calcium/calmodulin-dependent serine protein kinase (MAGUK family) |
| **KDM6A** | ENSG00000147050 | 1309.331983 | 1.105695565 | 0.140297083 | 7.84E-16 | 1.26E-14 | 7403 | X | 44732757 | 44971847 | 1 | protein_coding | lysine (K)-specific demethylase 6A |
| **SLC16A2** | ENSG00000147100 | 28.54302828 | -3.47120848 | 1.063681101 | 1.33E-04 | 5.93E-04 | 6567 | X | 73641085 | 73753752 | 1 | protein_coding | solute carrier family 16, member 2 (thyroid hormone transporter) |
| **CHST7** | ENSG00000147119 | 221.8685967 | 0.955890112 | 0.221522771 | 4.94E-06 | 2.86E-05 | 56548 | X | 46433219 | 46457843 | 1 | protein_coding | carbohydrate (N-acetylglucosamine 6-O) sulfotransferase 7 |
| **ZMYM3** | ENSG00000147130 | 1443.401913 | -0.599032201 | 0.128925126 | 2E-06 | 1.23E-05 | 9203 | X | 70459474 | 70474996 | -1 | protein_coding | zinc finger, MYM-type 3 |
| **EBP** | ENSG00000147155 | 897.2720508 | -0.538895059 | 0.163772172 | 6.28E-04 | 0.002395265 | 10682 | X | 48379546 | 48387104 | 1 | protein_coding | emopamil binding protein (sterol isomerase) |
| **OGT** | ENSG00000147162 | 5338.328635 | -0.251130647 | 0.109893428 | 0.018383966 | 0.047276148 | 8473 | X | 70752933 | 70795747 | 1 | protein_coding | O-linked N-acetylglucosamine (GlcNAc) transferase |
| **PRPS1** | ENSG00000147224 | 702.5716663 | -0.570591104 | 0.154977727 | 1.41E-04 | 6.28E-04 | 5631 | X | 106871737 | 106894256 | 1 | protein_coding | phosphoribosyl pyrophosphate synthetase 1 |
| **MCPH1** | ENSG00000147316 | 583.149337 | 0.716625111 | 0.149784137 | 8.31E-07 | 5.42E-06 | 79648 | 8 | 6264113 | 6501144 | 1 | protein_coding | microcephalin 1 |
| **MFHAS1** | ENSG00000147324 | 3676.987476 | 0.40820016 | 0.169788028 | 0.012355529 | 0.03356373 | 9258 | 8 | 8640864 | 8751155 | -1 | protein_coding | malignant fibrous histiocytoma amplified sequence 1 |
| **RPL10** | ENSG00000147403 | 1181.496621 | -0.656217633 | 0.246490468 | 0.003799631 | 0.01195343 | 26778 | X | 153618315 | 153637504 | 1 | protein_coding | ribosomal protein L10 |
| **CSGALNACT** | ENSG00000147408 | 22.92513488 | -1.760722802 | 3.483386002 | 0.009807333 | 0.027554899 | 55790 | 8 | 19261672 | 19615540 | -1 | protein_coding | chondroitin sulfate N-acetylgalactosaminyltransferase 1 |
| **ATP6V1B2** | ENSG00000147416 | 23023.80337 | -0.313631611 | 0.126730258 | 0.011340042 | 0.031187666 | 526 | 8 | 20054878 | 20084330 | 1 | protein_coding | ATPase, H+ transporting, lysosomal 56/58kDa, V1 subunit B2 |
| **CHRNA6** | ENSG00000147434 | 44.82163009 | 0.924038099 | 0.481616331 | 0.014062103 | 0.037559498 | 8973 | 8 | 42607763 | 42651535 | -1 | protein_coding | cholinergic receptor, nicotinic, alpha 6 (neuronal) |
| **DOK2** | ENSG00000147443 | 211.2108192 | -1.335692987 | 0.366519253 | 3.54E-05 | 1.76E-04 | 9046 | 8 | 21766384 | 21771371 | -1 | protein_coding | docking protein 2, 56kDa |
| **SLC25A37** | ENSG00000147454 | 238.6804259 | 1.079029211 | 0.271374446 | 1.69E-05 | 8.92E-05 | 51312 | 8 | 23386318 | 23432976 | 1 | protein_coding | solute carrier family 25 (mitochondrial iron transporter), member 37 |
| **DOCK5** | ENSG00000147459 | 5158.334714 | -0.987978202 | 0.137661395 | 1.6E-13 | 2.18E-12 | 80005 | 8 | 25042238 | 25275598 | 1 | protein_coding | dedicator of cytokinesis 5 |
| **ST18** | ENSG00000147488 | 375.8057534 | -0.554491967 | 0.211863623 | 0.00546308 | 0.016478497 | 9705 | 8 | 53023399 | 53373519 | -1 | protein_coding | suppression of tumorigenicity 18 (breast carcinoma) (zinc finger protein) |
| **TACC1** | ENSG00000147526 | 4841.778825 | -0.363045531 | 0.123105767 | 0.002445525 | 0.00806886 | 6867 | 8 | 38585704 | 38710546 | 1 | protein_coding | transforming, acidic coiled-coil containing protein 1 |
| **GOLGA7** | ENSG00000147533 | 1670.199179 | 0.316760817 | 0.124165271 | 0.008122322 | 0.023339588 | 51125 | 8 | 41347915 | 41368499 | 1 | protein_coding | golgin A7 |
| **WHSC1L1** | ENSG00000147548 | 2533.745784 | 0.605521396 | 0.133746633 | 3.35E-06 | 1.99E-05 | 54904 | 8 | 38127215 | 38239790 | -1 | protein_coding | Wolf-Hirschhorn syndrome candidate 1-like 1 |
| **TERF1** | ENSG00000147601 | 525.4212564 | 0.550591528 | 0.184189919 | 0.001755657 | 0.006012033 | 7013 | 8 | 73921099 | 73960357 | 1 | protein_coding | telomeric repeat binding factor (NIMA-interacting) 1 |
| **RPL7** | ENSG00000147604 | 6878.641541 | -0.650963105 | 0.109455907 | 1.06E-09 | 1.02E-08 | 6129 | 8 | 74202506 | 74208024 | -1 | protein_coding | ribosomal protein L7 |
| **ATP6V0D2** | ENSG00000147614 | 3836.062544 | -1.007829893 | 0.158911005 | 5.59E-11 | 6.16E-10 | 245972 | 8 | 86999552 | 87166457 | 1 | protein_coding | ATPase, H+ transporting, lysosomal 38kDa, V0 subunit d2 |
| **MTDH** | ENSG00000147649 | 5970.651672 | -0.269713285 | 0.107143925 | 0.010681361 | 0.02957305 | 92140 | 8 | 98656407 | 98740998 | 1 | protein_coding | metadherin |
| **LRP12** | ENSG00000147650 | 1478.082497 | 0.50139268 | 0.120076223 | 2.01E-05 | 1.04E-04 | 29967 | 8 | 105501459 | 105601252 | -1 | protein_coding | low density lipoprotein receptor-related protein 12 |
| **EBAG9** | ENSG00000147654 | 432.2492581 | 0.61831226 | 0.186152993 | 5.05E-04 | 0.001974402 | 9166 | 8 | 110551940 | 110578225 | 1 | protein_coding | estrogen receptor binding site associated, antigen, 9 |
| **POLR2K** | ENSG00000147669 | 765.2298888 | 0.704770649 | 0.157374161 | 3.72E-06 | 2.2E-05 | 5440 | 8 | 101162812 | 101166230 | 1 | protein_coding | polymerase (RNA) II (DNA directed) polypeptide K, 7.0kDa |
| **UTP23** | ENSG00000147679 | 445.5055075 | 0.452516409 | 0.193393077 | 0.013748339 | 0.03687537 | 84294 | 8 | 117778742 | 117861702 | 1 | protein_coding | UTP23, small subunit (SSU) processome component, homolog (yeast) |
| **NDUFB9** | ENSG00000147684 | 3086.670697 | -1.067274055 | 0.206741479 | 6.16E-08 | 4.72E-07 | 4715 | 8 | 125551344 | 125580751 | 1 | protein_coding | NADH dehydrogenase (ubiquinone) 1 beta subcomplex, 9, 22kDa |
| **NAPRT1** | ENSG00000147813 | 182.4597839 | -0.725232705 | 0.269766309 | 0.003034778 | 0.009807678 | 93100 | 8 | 144656955 | 144660819 | -1 | protein_coding | nicotinate phosphoribosyltransferase domain containing 1 |
| **VLDLR** | ENSG00000147852 | 135.4221668 | -2.037910469 | 0.359119025 | 1.35E-09 | 1.28E-08 | 7436 | 9 | 2621834 | 2660053 | 1 | protein_coding | very low density lipoprotein receptor |
| **AK3** | ENSG00000147853 | 1408.882658 | 0.772690696 | 0.193364491 | 2.26E-05 | 1.17E-04 | 50808 | 9 | 4711155 | 4742043 | -1 | protein_coding | adenylate kinase 3 |
| **PLIN2** | ENSG00000147872 | 3701.890525 | -0.86599163 | 0.16006858 | 1.96E-08 | 1.6E-07 | 123 | 9 | 19108373 | 19149288 | -1 | protein_coding | perilipin 2 |
| **HAUS6** | ENSG00000147874 | 628.1577057 | 0.754721729 | 0.222234846 | 2.9E-04 | 0.00119657 | 54801 | 9 | 19053141 | 19103117 | -1 | protein_coding | HAUS augmin-like complex, subunit 6 |
| **CDKN2B** | ENSG00000147883 | 972.3254713 | -0.61818667 | 0.217554409 | 0.002359241 | 0.007816232 | 1030 | 9 | 22002902 | 22009362 | -1 | protein_coding | cyclin-dependent kinase inhibitor 2B (p15, inhibits CDK4) |
| **C9orf72** | ENSG00000147894 | 807.0806556 | 0.899614965 | 0.227845329 | 2.5E-05 | 1.28E-04 | 203228 | 9 | 27546544 | 27573864 | -1 | protein_coding | chromosome 9 open reading frame 72 |
| **FBXO10** | ENSG00000147912 | 654.9775715 | -1.210746169 | 0.205685547 | 8.42E-10 | 8.15E-09 | 26267 | 9 | 37510889 | 37588871 | -1 | protein_coding | F-box protein 10 |
| **SIGMAR1** | ENSG00000147955 | 467.8483517 | -0.658987362 | 0.168113546 | 4.43E-05 | 2.15E-04 | 10280 | 9 | 34634719 | 34637806 | -1 | protein_coding | sigma non-opioid intracellular receptor 1 |
| **CBWD5** | ENSG00000147996 | 448.6742858 | 1.182942775 | 0.205721114 | 1.98E-09 | 1.83E-08 | 1010605 | 9 | 70432004 | 70497240 | -1 | protein_coding | COBW domain containing 5 |
| **SHC3** | ENSG00000148082 | 19.74174046 | -2.072407735 | 0.987779431 | 0.002404775 | 0.007948137 | 53358 | 9 | 91628060 | 91793682 | -1 | protein_coding | SHC (Src homology 2 domain containing) transforming protein 3 |
| **HIATL1** | ENSG00000148110 | 4230.66431 | 1.062479453 | 0.102074245 | 6.12E-26 | 1.91E-24 | 84641 | 9 | 97136833 | 97223324 | 1 | protein_coding | hippocampus abundant transcript-like 1 |
| **UGCG** | ENSG00000148154 | 3339.972996 | 1.281637518 | 0.140466545 | 1.05E-20 | 2.46E-19 | 7357 | 9 | 114659046 | 114697649 | 1 | protein_coding | UDP-glucose ceramide glucosyltransferase |
| **SNX30** | ENSG00000148158 | 2558.319265 | -1.012613598 | 0.11733605 | 1.57E-18 | 3.09E-17 | 401548 | 9 | 115513118 | 115643951 | 1 | protein_coding | sorting nexin family member 30 |
| **STOM** | ENSG00000148175 | 12319.85822 | 1.948923714 | 0.118282309 | 7.93E-62 | 1.08E-59 | 2040 | 9 | 124101355 | 124132531 | -1 | protein_coding | stomatin |
| **GSN** | ENSG00000148180 | 6227.358171 | -1.377501278 | 0.133375321 | 9.24E-26 | 2.86E-24 | 2934 | 9 | 123970072 | 124095121 | 1 | protein_coding | gelsolin |
| **ALAD** | ENSG00000148218 | 863.1358279 | -0.712867971 | 0.136320601 | 8.33E-08 | 6.25E-07 | 210 | 9 | 116148597 | 116163613 | -1 | protein_coding | aminolevulinate dehydratase |
| **ASTN2** | ENSG00000148219 | 58.44058742 | -2.129742843 | 0.545469993 | 8.98E-06 | 4.98E-05 | 23245 | 9 | 119187504 | 120177348 | -1 | protein_coding | astrotactin 2 |
| **ASB6** | ENSG00000148331 | 428.420979 | 0.58953241 | 0.182883823 | 7.5E-04 | 0.00281982 | 140459 | 9 | 132399171 | 132404444 | -1 | protein_coding | ankyrin repeat and SOCS box containing 6 |
| **PTGES2** | ENSG00000148334 | 403.5550865 | -0.796120201 | 0.17469168 | 2.24E-06 | 1.37E-05 | 80142 | 9 | 130882972 | 130890741 | -1 | protein_coding | prostaglandin E synthase 2 |
| **CIZ1** | ENSG00000148337 | 912.1936491 | -0.57410217 | 0.167638036 | 3.7E-04 | 0.00149411 | 25792 | 9 | 130928343 | 130966662 | -1 | protein_coding | CDKN1A interacting zinc finger protein 1 |
| **PTGES** | ENSG00000148344 | 122.4055337 | 5.372575227 | 0.53118389 | 2.65E-25 | 8.03E-24 | 9536 | 9 | 132500610 | 132515326 | -1 | protein_coding | prostaglandin E synthase |
| **GPR107** | ENSG00000148358 | 3887.707438 | 0.25356888 | 0.09952536 | 0.008895051 | 0.025263532 | 57720 | 9 | 132815705 | 132902448 | 1 | protein_coding | G protein-coupled receptor 107 |
| **SEC16A** | ENSG00000148396 | 3510.148019 | -0.481327854 | 0.096590178 | 4.2E-07 | 2.87E-06 | 9919 | 9 | 139334549 | 139372141 | -1 | protein_coding | SEC16 homolog A (S. cerevisiae) |
| **NACC2** | ENSG00000148411 | 2233.795213 | -1.25059199 | 0.157251105 | 2.86E-16 | 4.74E-15 | 138151 | 9 | 138898383 | 138987131 | -1 | protein_coding | NACC family member 2, BEN and BTB (POZ) domain containing |
| **USP6NL** | ENSG00000148429 | 1149.263133 | 0.875611989 | 0.120270597 | 1.18E-13 | 1.62E-12 | 1005072 | 10 | 11495945 | 11653753 | -1 | protein_coding | USP6 N-terminal like |
| **MSRB2** | ENSG00000148450 | 208.7060197 | -0.605881462 | 0.278502761 | 0.014403141 | 0.038317369 | 22921 | 10 | 23384435 | 23410942 | 1 | protein_coding | methionine sulfoxide reductase B2 |
| **FAM188A** | ENSG00000148481 | 1067.815496 | 1.390354208 | 0.146180436 | 3.71E-22 | 9.4E-21 | 80013 | 10 | 15820169 | 15902519 | -1 | protein_coding | family with sequence similarity 188, member A |
| **RSU1** | ENSG00000148484 | 2680.275644 | 0.458908151 | 0.091931303 | 4.13E-07 | 2.83E-06 | 6251 | 10 | 16632610 | 16859527 | -1 | protein_coding | Ras suppressor protein 1 |
| **NRBF2** | ENSG00000148572 | 1437.628124 | 0.6205663 | 0.115123104 | 4.09E-08 | 3.2E-07 | 29982 | 10 | 64893050 | 64914783 | 1 | protein_coding | nuclear receptor binding factor 2 |
| **HERC4** | ENSG00000148634 | 2270.737397 | 1.117079973 | 0.120443136 | 4.65E-21 | 1.11E-19 | 26091 | 10 | 69681665 | 69835105 | -1 | protein_coding | HECT and RLD domain containing E3 ubiquitin protein ligase 4 |
| **C10orf11** | ENSG00000148655 | 188.1788677 | -1.001620979 | 0.332953452 | 6.73E-04 | 0.002556215 | 83938 | 10 | 77360998 | 78319925 | 1 | protein_coding | chromosome 10 open reading frame 11 |
| **ADIRF** | ENSG00000148671 | 8.280236635 | 2.96539603 | 1.434802881 | 0.001541481 | 0.005370059 | 10974 | 10 | 88727949 | 88743691 | 1 | protein_coding | adipogenesis regulatory factor |
| **RPP30** | ENSG00000148688 | 399.8832289 | 0.552102324 | 0.20001804 | 0.003569815 | 0.01132368 | 10556 | 10 | 92631473 | 92668312 | 1 | protein_coding | ribonuclease P/MRP 30kDa subunit |
| **FRA10AC1** | ENSG00000148690 | 103.6638791 | 0.91534217 | 0.341132038 | 0.002159027 | 0.007231516 | 118924 | 10 | 95427640 | 95462329 | -1 | protein_coding | fragile site, folic acid type, rare, fra(10)(q23.3) or fra(10)(q24.2) candidate 1 |
| **ADD3** | ENSG00000148700 | 2692.026891 | 0.496382077 | 0.117051359 | 1.55E-05 | 8.25E-05 | 120 | 10 | 111756126 | 111895323 | 1 | protein_coding | adducin 3 (gamma) |
| **EIF4EBP2** | ENSG00000148730 | 4420.922866 | -0.455223356 | 0.093047228 | 1.08E-06 | 6.91E-06 | 1979 | 10 | 72164135 | 72188374 | 1 | protein_coding | eukaryotic translation initiation factor 4E binding protein 2 |
| **FUOM** | ENSG00000148803 | 371.7260825 | -0.448157132 | 0.185320935 | 0.010989623 | 0.030344068 | 282969 | 10 | 135168658 | 135171529 | -1 | protein_coding | fucose mutarotase |
| **MTG1** | ENSG00000148824 | 370.7234865 | 0.835286668 | 0.23510186 | 1.42E-04 | 6.31E-04 | 92170 | 10 | 135207598 | 135234811 | 1 | protein_coding | mitochondrial ribosome-associated GTPase 1 |
| **TAF5** | ENSG00000148835 | 153.8448897 | 0.921631549 | 0.300850817 | 6.77E-04 | 0.002568 | 6877 | 10 | 105127724 | 105148822 | 1 | protein_coding | TAF5 RNA polymerase II, TATA box binding protein (TBP)-associated factor, 100kDa |
| **ITPRIP** | ENSG00000148841 | 2546.24367 | 1.14622533 | 0.135765059 | 8.72E-18 | 1.62E-16 | 85450 | 10 | 106071894 | 106098162 | -1 | protein_coding | inositol 1,4,5-trisphosphate receptor interacting protein |
| **PDCD11** | ENSG00000148843 | 1038.930469 | -0.578137871 | 0.148630741 | 6.04E-05 | 2.87E-04 | 22984 | 10 | 105156405 | 105206049 | 1 | protein_coding | programmed cell death 11 |
| **ADAM12** | ENSG00000148848 | 269.652475 | -0.484561759 | 0.210434963 | 0.014275113 | 0.038047882 | 8038 | 10 | 127700950 | 128077024 | -1 | protein_coding | ADAM metallopeptidase domain 12 |
| **RGS10** | ENSG00000148908 | 649.8399283 | -0.375433312 | 0.144343327 | 0.007515324 | 0.021806854 | 6001 | 10 | 121259340 | 121302220 | -1 | protein_coding | regulator of G-protein signaling 10 |
| **ADM** | ENSG00000148926 | 1548.313157 | 1.729813138 | 0.285515003 | 1.62E-10 | 1.69E-09 | 133 | 11 | 10326227 | 10328944 | 1 | protein_coding | adrenomedullin |
| **LIN7C** | ENSG00000148943 | 1848.7304 | 0.47908017 | 0.145344976 | 6.83E-04 | 0.002587657 | 55327 | 11 | 27516123 | 27528320 | -1 | protein_coding | lin-7 homolog C (C. elegans) |
| **TUT1** | ENSG00000149016 | 381.4050839 | 0.799523754 | 0.181327624 | 4.34E-06 | 2.54E-05 | 64852 | 11 | 62342517 | 62359649 | -1 | protein_coding | terminal uridylyl transferase 1, U6 snRNA-specific |
| **APIP** | ENSG00000149089 | 620.7986097 | 0.898026652 | 0.190514881 | 8.48E-07 | 5.52E-06 | 51074 | 11 | 34874641 | 34938046 | -1 | protein_coding | APAF1 interacting protein |
| **DGKZ** | ENSG00000149091 | 1850.61916 | -0.493638156 | 0.13533741 | 1.77E-04 | 7.69E-04 | 8525 | 11 | 46354455 | 46402104 | 1 | protein_coding | diacylglycerol kinase, zeta |
| **EIF3M** | ENSG00000149100 | 1945.820572 | 0.502693587 | 0.11274642 | 5.67E-06 | 3.25E-05 | 10480 | 11 | 32605344 | 32627808 | 1 | protein_coding | eukaryotic translation initiation factor 3, subunit M |
| **GLYAT** | ENSG00000149124 | 27.6232895 | -1.515916497 | 1.117248575 | 0.014820475 | 0.039302675 | 10249 | 11 | 58407899 | 58499447 | -1 | protein_coding | glycine-N-acyltransferase |
| **SERPING1** | ENSG00000149131 | 4681.507247 | 1.348394749 | 0.150242494 | 4.95E-20 | 1.1E-18 | 710 | 11 | 57364860 | 57382326 | 1 | protein_coding | serpin peptidase inhibitor, clade G (C1 inhibitor), member 1 |
| **SLC43A1** | ENSG00000149150 | 56.71535435 | -1.20005176 | 0.528588376 | 0.003808199 | 0.011977917 | 8501 | 11 | 57252007 | 57283259 | -1 | protein_coding | solute carrier family 43 (amino acid system L transporter), member 1 |
| **PTPRJ** | ENSG00000149177 | 7649.003009 | 0.759930974 | 0.097799718 | 3.06E-15 | 4.71E-14 | 5795 | 11 | 48002113 | 48189670 | 1 | protein_coding | protein tyrosine phosphatase, receptor type, J |

| | | | | | | | | | | | | | |
|---|---|---|---|---|---|---|---|---|---|---|---|---|---|
| **CELF1** | ENSG00000149187 | 5572.995444 | 1.182632179 | 0.109510407 | 8.45E-28 | 2.89E-26 | 10658 | 11 | 47487496 | 47587121 | -1 | protein_coding | CUGBP, Elav-like family member 1 |
| **SESN3** | ENSG00000149212 | 1610.783105 | -0.359379702 | 0.133458142 | 0.005729269 | 0.01716871 | 143686 | 11 | 94898704 | 94965705 | -1 | protein_coding | sestrin 3 |
| **ENDOD1** | ENSG00000149218 | 1268.207613 | 2.811479508 | 0.181740358 | 1.9E-55 | 2.1E-53 | 23052 | 11 | 94822974 | 94865809 | 1 | protein_coding | endonuclease domain containing 1 |
| **CCDC82** | ENSG00000149231 | 831.1096889 | 1.06509859 | 0.205320231 | 6.08E-08 | 4.66E-07 | 79780 | 11 | 96085933 | 96123087 | -1 | protein_coding | coiled-coil domain containing 82 |
| **PAK1** | ENSG00000149269 | 2992.00168 | -0.515975574 | 0.117919114 | 7.52E-06 | 4.22E-05 | 5058 | 11 | 77032752 | 77185680 | -1 | protein_coding | p21 protein (Cdc42/Rac)-activated kinase 1 |
| **RPS3** | ENSG00000149273 | 5429.147647 | -0.66955181 | 0.132939258 | 2.43E-07 | 1.72E-06 | 6188 | 11 | 75110530 | 75133324 | 1 | protein_coding | ribosomal protein S3 |
| **ZC3H12C** | ENSG00000149289 | 2238.155227 | 0.886858257 | 0.153127979 | 2.15E-09 | 1.98E-08 | 85463 | 11 | 109964087 | 110042566 | 1 | protein_coding | zinc finger CCCH-type containing 12C |
| **NPAT** | ENSG00000149308 | 1327.870785 | 0.354004621 | 0.144378691 | 0.011628427 | 0.031883252 | 4863 | 11 | 108027942 | 108093369 | -1 | protein_coding | nuclear protein, ataxia-telangiectasia locus |
| **ATM** | ENSG00000149311 | 4908.4547 | -0.692632821 | 0.107308667 | 7.6E-11 | 8.25E-10 | 472 | 11 | 108093211 | 108239829 | 1 | protein_coding | ataxia telangiectasia mutated |
| **LAMTOR1** | ENSG00000149357 | 1420.765249 | -0.423053767 | 0.106837704 | 5.71E-05 | 2.73E-04 | 55004 | 11 | 71796941 | 71814433 | -1 | protein_coding | late endosomal/lysosomal adaptor, MAPK and MTOR activator 1 |
| **P4HA3** | ENSG00000149380 | 10.74966012 | -1.394226718 | 1.489509279 | 0.018793097 | 0.048117759 | 283208 | 11 | 73946846 | 74022702 | -1 | protein_coding | prolyl 4-hydroxylase, alpha polypeptide III |
| **ST14** | ENSG00000149418 | 3179.653169 | -1.249575317 | 0.136757879 | 1.22E-20 | 2.82E-19 | 6768 | 11 | 130029457 | 130080271 | 1 | protein_coding | suppression of tumorigenicity 14 (colon carcinoma) |
| **TMEM138** | ENSG00000149483 | 3127.225496 | 1.132984765 | 0.13295419 | 3.61E-18 | 6.92E-17 | 51524 | 11 | 61129473 | 61136981 | 1 | protein_coding | transmembrane protein 138 |
| **FADS1** | ENSG00000149485 | 3781.574521 | -0.540288066 | 0.141309142 | 8.09E-05 | 3.76E-04 | 1003022 | 11 | 61567099 | 61596790 | -1 | protein_coding | fatty acid desaturase 1 |
| **PLCH2** | ENSG00000149527 | 40.55658919 | -3.291612356 | 0.866415672 | 8.85E-06 | 4.91E-05 | 9651 | 1 | 2357419 | 2436969 | 1 | protein_coding | phospholipase C, eta 2 |
| **CPSF7** | ENSG00000149532 | 1914.910968 | 0.257002368 | 0.10880318 | 0.015944571 | 0.041891161 | 79869 | 11 | 61170121 | 61197503 | -1 | protein_coding | cleavage and polyadenylation specific factor 7, 59kDa |
| **B3GAT3** | ENSG00000149541 | 380.8993798 | -0.713842605 | 0.203258925 | 2.07E-04 | 8.79E-04 | 26229 | 11 | 62382768 | 62389647 | -1 | protein_coding | beta-1,3-glucuronyltransferase 3 (glucuronosyltransferase I) |
| **CHEK1** | ENSG00000149554 | 354.6078769 | 1.165800366 | 0.226522322 | 5.89E-08 | 4.53E-07 | 1111 | 11 | 125495036 | 125546150 | 1 | protein_coding | checkpoint kinase 1 |
| **MPZL2** | ENSG00000149573 | 221.8206963 | -1.586351305 | 0.313186139 | 5.53E-08 | 4.27E-07 | 10205 | 11 | 118124118 | 118135251 | -1 | protein_coding | myelin protein zero-like 2 |
| **SIDT2** | ENSG00000149577 | 2098.067115 | -0.480791826 | 0.132789722 | 1.97E-04 | 8.45E-04 | 51092 | 11 | 117049449 | 117068160 | 1 | protein_coding | SID1 transmembrane family, member 2 |
| **OCSTAMP** | ENSG00000149635 | 216.5224216 | -1.508759879 | 0.312277904 | 2.06E-07 | 1.47E-06 | 128506 | 20 | 45169585 | 45179213 | -1 | protein_coding | osteoclast stimulatory transmembrane protein |
| **SOGA1** | ENSG00000149639 | 2638.912545 | -0.639803635 | 0.11336475 | 8.68E-09 | 7.44E-08 | 140710 | 20 | 35405845 | 35492089 | -1 | protein_coding | suppressor of glucose, autophagy associated 1 |
| **LSM14B** | ENSG00000149657 | 720.6633748 | -0.339976491 | 0.145752275 | 0.016341529 | 0.042801644 | 149986 | 20 | 60697517 | 60710434 | 1 | protein_coding | LSM14B, SCD6 homolog B (S. cerevisiae) |
| **TRPT1** | ENSG00000149743 | 103.9152539 | -0.920333982 | 0.357644452 | 0.002915576 | 0.009452428 | 83707 | 11 | 63991271 | 63993726 | -1 | protein_coding | tRNA phosphotransferase 1 |
| **NUDT22** | ENSG00000149761 | 340.0385916 | -0.429235794 | 0.177953025 | 0.01179568 | 0.032249263 | 84304 | 11 | 63993413 | 63998158 | 1 | protein_coding | nudix (nucleoside diphosphate linked moiety X)-type motif 22 |
| **FERMT3** | ENSG00000149781 | 9133.201907 | -0.496432709 | 0.090012043 | 1.9E-08 | 1.56E-07 | 83706 | 11 | 63974150 | 63991354 | 1 | protein_coding | fermitin family member 3 |
| **PLCB3** | ENSG00000149782 | 1121.667865 | -0.524344769 | 0.138663112 | 9.9E-05 | 4.53E-04 | 5331 | 11 | 64018995 | 64036622 | 1 | protein_coding | phospholipase C, beta 3 (phosphatidylinositol-specific) |
| **MRPL49** | ENSG00000149792 | 762.9288109 | 0.509946395 | 0.163851272 | 0.00126376 | 0.004491985 | 740 | 11 | 64889655 | 64894843 | 1 | protein_coding | mitochondrial ribosomal protein L49 |
| **CDC42EP2** | ENSG00000149798 | 77.20537236 | 4.062024114 | 0.53236492 | 1.28E-15 | 2.03E-14 | 10435 | 11 | 65082289 | 65089900 | 1 | protein_coding | CDC42 effector protein (Rho GTPase binding) 2 |
| **TM7SF2** | ENSG00000149809 | 166.2704509 | -1.187002326 | 0.330886732 | 6.71E-05 | 3.17E-04 | 7108 | 11 | 64879317 | 64883856 | 1 | protein_coding | transmembrane 7 superfamily member 2 |
| **PPP4C** | ENSG00000149923 | 1381.791146 | -0.359161062 | 0.123582143 | 0.003051938 | 0.0098548 | 5531 | 16 | 30087299 | 30096698 | 1 | protein_coding | protein phosphatase 4, catalytic subunit |
| **ALDOA** | ENSG00000149925 | 21610.17977 | -0.770530785 | 0.10654475 | 2.1E-13 | 2.82E-12 | 226 | 16 | 30064411 | 30081778 | 1 | protein_coding | aldolase A, fructose-bisphosphate |
| **TAOK2** | ENSG00000149930 | 1773.77685 | -0.40398755 | 0.117509722 | 4.63E-04 | 0.00182635 | 9344 | 16 | 29984962 | 30003582 | 1 | protein_coding | TAO kinase 2 |
| **CLEC1A** | ENSG00000150048 | 57.48078327 | -0.996495619 | 0.469823604 | 0.00781096 | 0.022553353 | 51267 | 12 | 10222153 | 10264226 | -1 | protein_coding | C-type lectin domain family 1, member A |
| **FCGR1A** | ENSG00000150337 | 4878.107882 | -0.87643265 | 0.126393468 | 1.48E-12 | 1.85E-11 | 2210 | 1 | 149754227 | 149764074 | 1 | protein_coding | Fc fragment of IgG, high affinity Ia, receptor (CD64) |
| **ARID5B** | ENSG00000150347 | 2801.505016 | 1.79367336 | 0.136199341 | 1.14E-40 | 7.06E-39 | 84159 | 10 | 63661059 | 63856703 | 1 | protein_coding | AT rich interactive domain 5B (MRF1-like) |
| **TMCO3** | ENSG00000150403 | 3227.644579 | -1.244431792 | 0.197594871 | 6.26E-11 | 6.85E-10 | 55002 | 13 | 114145310 | 114204542 | 1 | protein_coding | transmembrane and coiled-coil domains 3 |
| **TMEM218** | ENSG00000150433 | 385.8874963 | -0.523868274 | 0.202446507 | 0.006190234 | 0.01840045 | 219854 | 11 | 124966398 | 124981659 | -1 | protein_coding | transmembrane protein 218 |
| **TIRAP** | ENSG00000150455 | 257.1687654 | 0.634465314 | 0.216390506 | 0.001798961 | 0.006146549 | 114609 | 11 | 126152960 | 126168740 | 1 | protein_coding | toll-interleukin 1 receptor (TIR) domain containing adaptor protein |
| **LATS2** | ENSG00000150457 | 898.1325078 | 1.077614459 | 0.154400491 | 1E-12 | 1.27E-11 | 26524 | 13 | 21547171 | 21635686 | -1 | protein_coding | large tumor suppressor kinase 2 |
| **SAP18** | ENSG00000150459 | 2749.221787 | 0.708683716 | 0.146732001 | 6.04E-07 | 4.03E-06 | 10284 | 13 | 21714653 | 21723221 | 1 | protein_coding | Sin3A-associated protein, 18kDa |
| **FAM124A** | ENSG00000150510 | 496.066728 | 1.413578077 | 0.257411245 | 6.14E-09 | 5.37E-08 | 220108 | 13 | 51796503 | 51858377 | 1 | protein_coding | family with sequence similarity 124A |
| **CTAGE5** | ENSG00000150527 | 1129.190168 | 0.639761393 | 0.15505312 | 2.1E-05 | 1.09E-04 | 4253 | 14 | 39734488 | 39856156 | 1 | protein_coding | CTAGE family, member 5 |
| **LYPD1** | ENSG00000150551 | 89.90446307 | -4.413513429 | 0.906824176 | 1.03E-07 | 7.62E-07 | 116372 | 2 | 133402426 | 133429152 | -1 | protein_coding | LY6/PLAUR domain containing 1 |
| **CCDC102B** | ENSG00000150636 | 166.1699135 | -1.816158075 | 0.360668356 | 5.05E-08 | 3.91E-07 | 79839 | 18 | 66382446 | 66722426 | 1 | protein_coding | coiled-coil domain containing 102B |
| **RGS18** | ENSG00000150681 | 177.3720778 | -3.090487804 | 0.578435819 | 4.19E-09 | 3.73E-08 | 64407 | 1 | 192127587 | 192154945 | 1 | protein_coding | regulator of G-protein signaling 18 |
| **FAM173B** | ENSG00000150756 | 188.0368611 | -1.010983964 | 0.26630099 | 4.21E-05 | 2.05E-04 | 134145 | 5 | 10226442 | 10250009 | -1 | protein_coding | family with sequence similarity 173, member B |
| **DOCK1** | ENSG00000150760 | 1205.176893 | -0.388488408 | 0.136492611 | 0.003430372 | 0.010933552 | 1793 | 10 | 128593978 | 129250781 | 1 | protein_coding | dedicator of cytokinesis 1 |
| **DIXDC1** | ENSG00000150764 | 26.62718639 | -2.124642306 | 2.371449986 | 0.005763637 | 0.017258164 | 85458 | 11 | 111797868 | 111893308 | 1 | protein_coding | DIX domain containing 1 |
| **DLAT** | ENSG00000150768 | 1430.12013 | 0.655171621 | 0.120030588 | 2.61E-08 | 2.1E-07 | 1737 | 11 | 111895538 | 111935114 | 1 | protein_coding | dihydrolipoamide S-acetyltransferase |
| **C11orf57** | ENSG00000150776 | 1032.662086 | 0.358130558 | 0.153838253 | 0.016507548 | 0.043177282 | 55216 | 11 | 111944810 | 111955874 | 1 | protein_coding | chromosome 11 open reading frame 57 |
| **IL18** | ENSG00000150782 | 1077.559716 | 1.786855996 | 0.151807847 | 5.14E-33 | 2.37E-31 | 3606 | 11 | 112013974 | 112034840 | -1 | protein_coding | interleukin 18 (interferon-gamma-inducing factor) |
| **PIP4K2A** | ENSG00000150867 | 3294.733765 | -0.342343472 | 0.12635856 | 0.005942424 | 0.017730977 | 5305 | 10 | 22823778 | 23003484 | -1 | protein_coding | phosphatidylinositol-5-phosphate 4-kinase, type II, alpha |
| **ABCB9** | ENSG00000150967 | 168.5712994 | -2.100735724 | 0.307442888 | 8.95E-13 | 1.14E-11 | 23457 | 12 | 123405498 | 123466196 | -1 | protein_coding | ATP-binding cassette, sub-family B (MDR/TAP), member 9 |
| **RILPL2** | ENSG00000150977 | 1369.880642 | 1.515298185 | 0.115902894 | 7.42E-40 | 4.4E-38 | 196383 | 12 | 123899936 | 123921264 | -1 | protein_coding | Rab interacting lysosomal protein-like 2 |
| **UBC** | ENSG00000150991 | 28372.84834 | 0.364041047 | 0.076886297 | 2.05E-06 | 1.26E-05 | 7316 | 12 | 125396150 | 125401914 | -1 | protein_coding | ubiquitin C |
| **ITPR1** | ENSG00000150995 | 4277.132277 | 0.256178052 | 0.083010262 | 0.002033587 | 0.006863807 | 3708 | 3 | 4535032 | 4889524 | 1 | protein_coding | inositol 1,4,5-trisphosphate receptor, type 1 |
| **SLC7A11** | ENSG00000151012 | 1018.858172 | 1.436144245 | 0.775926019 | 0.006084272 | 0.018108264 | 23657 | 4 | 139085251 | 139163503 | -1 | protein_coding | solute carrier family 7 (anionic amino acid transporter light chain, xc- system), member 11 |
| **UEVLD** | ENSG00000151116 | 1175.352221 | 0.285284091 | 0.122536269 | 0.015850833 | 0.041709414 | 55293 | 11 | 18551156 | 18610294 | -1 | protein_coding | UEV and lactate/malate dehyrogenase domains |
| **TMEM86A** | ENSG00000151117 | 716.6846308 | -1.825026025 | 0.209973769 | 3.32E-19 | 6.89E-18 | 144110 | 11 | 18714669 | 18726332 | 1 | protein_coding | transmembrane protein 86A |
| **BTBD11** | ENSG00000151136 | 123.1873301 | 0.708041605 | 0.329669928 | 0.013480322 | 0.036245559 | 121551 | 12 | 107712190 | 108053419 | 1 | protein_coding | BTB (POZ) domain containing 11 |
| **IPMK** | ENSG00000151151 | 667.0358872 | 0.406381025 | 0.15812568 | 0.007783201 | 0.0224902 | 253430 | 10 | 59951278 | 60027694 | -1 | protein_coding | inositol polyphosphate multikinase |
| **PLBD2** | ENSG00000151176 | 3113.3128 | -0.609260048 | 0.125705763 | 7.53E-07 | 4.95E-06 | 196463 | 12 | 113796371 | 113827203 | 1 | protein_coding | phospholipase B domain containing 2 |
| **SLC2A13** | ENSG00000151229 | 370.1798097 | 0.617154217 | 0.197366282 | 9.83E-04 | 0.003590148 | 114134 | 12 | 40148823 | 40499891 | -1 | protein_coding | solute carrier family 2 (facilitated glucose transporter), member 13 |
| **GXYLT1** | ENSG00000151233 | 592.5152961 | 1.219498817 | 0.182883349 | 5.33E-12 | 6.44E-11 | 283464 | 12 | 42475647 | 42538681 | -1 | protein_coding | glucoside xylosyltransferase 1 |
| **DIP2C** | ENSG00000151240 | 783.3424477 | -1.342670778 | 0.208273209 | 2.2E-11 | 2.52E-10 | 22982 | 10 | 320130 | 735683 | -1 | protein_coding | DIP2 disco-interacting protein 2 homolog C (Drosophila) |
| **CSNK1G3** | ENSG00000151292 | 987.1035472 | 0.728025699 | 0.151132616 | 7.02E-07 | 4.64E-06 | 1456 | 5 | 122847793 | 122952739 | 1 | protein_coding | casein kinase 1, gamma 3 |
| **FAM177A1** | ENSG00000151327 | 745.6552062 | 2.036829792 | 0.1885534 | 2.99E-28 | 1.05E-26 | 1019271 | 14 | 35514113 | 35582336 | 1 | protein_coding | family with sequence similarity 177, member A1 |
| **MBIP** | ENSG00000151332 | 282.3992045 | 0.794784841 | 0.230819682 | 2.33E-04 | 9.81E-04 | 51562 | 14 | 36767770 | 36789882 | -1 | protein_coding | MAP3K12 binding inhibitory protein 1 |
| **EXT2** | ENSG00000151348 | 1798.594878 | 0.343100287 | 0.101965555 | 6.23E-04 | 0.002377831 | 2132 | 11 | 44117099 | 44266979 | 1 | protein_coding | exostosin glycosyltransferase 2 |
| **NDUFC2** | ENSG00000151366 | 963.5420064 | 0.677398724 | 0.154944315 | 6.2E-06 | 3.53E-05 | 1005327 | 11 | 77779350 | 77791265 | -1 | protein_coding | NADH dehydrogenase (ubiquinone) 1, subcomplex unknown, 2, 14.5kDa |
| **ME3** | ENSG00000151376 | 42.10932746 | -1.382081423 | 0.765628329 | 0.007430887 | 0.021598752 | 10873 | 11 | 86152150 | 86383678 | -1 | protein_coding | malic enzyme 3, NADP(+)-dependent, mitochondrial |
| **NEK7** | ENSG00000151414 | 1954.309827 | 0.815878825 | 0.099637101 | 1.22E-16 | 2.09E-15 | 140609 | 1 | 198126093 | 198291550 | 1 | protein_coding | NIMA-related kinase 7 |
| **ANKRD50** | ENSG00000151458 | 768.2592227 | -0.922982691 | 0.173564382 | 3.51E-08 | 2.77E-07 | 57182 | 4 | 125585207 | 125633887 | -1 | protein_coding | ankyrin repeat domain 50 |
| **UPF2** | ENSG00000151461 | 1738.395876 | 0.423797016 | 0.136460834 | 0.001509731 | 0.005265454 | 26019 | 10 | 11962021 | 12085169 | -1 | protein_coding | UPF2 regulator of nonsense transcripts homolog (yeast) |
| **SCLT1** | ENSG00000151466 | 470.1024781 | 1.414844278 | 0.176840909 | 1.97E-16 | 3.3E-15 | 1019271 | 4 | 129786076 | 130014764 | -1 | protein_coding | sodium channel and clathrin linker 1 |
| **C4orf33** | ENSG00000151470 | 379.961626 | 1.050959876 | 0.179764825 | 1.26E-09 | 1.2E-08 | 132321 | 4 | 130014472 | 130037795 | 1 | protein_coding | chromosome 4 open reading frame 33 |
| **FRMD4A** | ENSG00000151474 | 351.619375 | -1.604422394 | 0.276024695 | 7.46E-10 | 7.25E-09 | 55691 | 10 | 13685706 | 14504141 | -1 | protein_coding | FERM domain containing 4A |
| **EPS8** | ENSG00000151491 | 1155.69168 | -1.277718561 | 0.232414719 | 7.51E-09 | 6.49E-08 | 2059 | 12 | 15773092 | 16035263 | -1 | protein_coding | epidermal growth factor receptor pathway substrate 8 |
| **THYN1** | ENSG00000151500 | 373.3527793 | -0.499686443 | 0.225111901 | 0.017466441 | 0.045320817 | 29087 | 11 | 134118173 | 134123264 | -1 | protein_coding | thymocyte nuclear protein 1 |
| **VPS26B** | ENSG00000151502 | 1291.305402 | -0.507280338 | 0.139894588 | 1.93E-04 | 8.31E-04 | 112936 | 11 | 134094539 | 134117686 | 1 | protein_coding | vacuolar protein sorting 26 homolog B (S. pombe) |
| **FAM160B1** | ENSG00000151553 | 1313.799447 | 0.841345994 | 0.135611005 | 2.1E-10 | 2.15E-09 | 57700 | 10 | 116581503 | 116659591 | 1 | protein_coding | family with sequence similarity 160, member B1 |
| **QTRTD1** | ENSG00000151576 | 784.6180435 | 0.627370586 | 0.199374459 | 9.06E-04 | 0.003345184 | 79691 | 3 | 113724680 | 113807269 | 1 | protein_coding | queuine tRNA-ribosyltransferase domain containing 1 |
| **MMAA** | ENSG00000151611 | 205.1928461 | 0.836631457 | 0.245654121 | 2.41E-04 | 0.001011979 | 166785 | 4 | 146539415 | 146581187 | 1 | protein_coding | methylmalonic aciduria (cobalamin deficiency) cblA type |
| **NR3C2** | ENSG00000151623 | 75.90639189 | -0.914664634 | 0.474105509 | 0.014318293 | 0.038136366 | 4306 | 4 | 148999913 | 149365850 | -1 | protein_coding | nuclear receptor subfamily 3, group C, member 2 |
| **VENTX** | ENSG00000151650 | 29.95110156 | -1.818457765 | 0.781005972 | 0.001698667 | 0.005848328 | 27287 | 10 | 135050908 | 135055433 | 1 | protein_coding | VENT homeobox |
| **INPP1** | ENSG00000151689 | 340.6751823 | 0.645727217 | 0.209847883 | 0.001066754 | 0.003871574 | 3628 | 2 | 191208196 | 191236391 | 1 | protein_coding | inositol polyphosphate-1-phosphatase |
| **ADAM17** | ENSG00000151694 | 1873.09621 | 0.853503426 | 0.122919753 | 1.46E-12 | 1.83E-11 | 6868 | 2 | 9628615 | 9695921 | -1 | protein_coding | ADAM metallopeptidase domain 17 |
| **FLI1** | ENSG00000151702 | 2614.829706 | -0.60532064 | 0.096739607 | 1.84E-10 | 1.91E-09 | 2313 | 11 | 128556430 | 128683162 | 1 | protein_coding | Fli-1 proto-oncogene, ETS transcription factor |

| | | | | | | | | | | | | | |
|---|---|---|---|---|---|---|---|---|---|---|---|---|---|
| **KCNJ1** | ENSG00000151704 | 505.7628904 | -2.670009483 | 0.297247147 | 1.72E-20 | 3.92E-19 | 3758 | 11 | 128706210 | 128737268 | -1 | protein_coding | potassium inwardly-rectifying channel, subfamily J, member 1 |
| **TMEM45B** | ENSG00000151715 | 207.3934341 | -1.615855326 | 0.275228307 | 5.85E-10 | 5.75E-09 | 120224 | 11 | 129685714 | 129729898 | 1 | protein_coding | transmembrane protein 45B |
| **CENPU** | ENSG00000151725 | 55.16860565 | 0.999770833 | 0.500353422 | 0.010242367 | 0.028569828 | 79682 | 4 | 185615772 | 185655287 | -1 | protein_coding | centromere protein U |
| **ACSL1** | ENSG00000151726 | 15525.14999 | 1.852836812 | 0.086263169 | 8.22E-104 | 2.86E-101 | 2180 | 4 | 185676749 | 185747972 | -1 | protein_coding | acyl-CoA synthetase long-chain family member 1 |
| **SAV1** | ENSG00000151748 | 1080.901732 | 1.027657347 | 0.171785052 | 6.33E-10 | 6.19E-09 | 60485 | 14 | 51098776 | 51135049 | -1 | protein_coding | salvador homolog 1 (Drosophila) |
| **NBAS** | ENSG00000151779 | 1546.111816 | -0.771677941 | 0.125344662 | 3.38E-10 | 3.41E-09 | 51594 | 2 | 15307032 | 15701454 | -1 | protein_coding | neuroblastoma amplified sequence |
| **GUF1** | ENSG00000151806 | 1288.358605 | -0.595351175 | 0.14547001 | 2.44E-05 | 1.25E-04 | 60558 | 4 | 44680444 | 44702943 | 1 | protein_coding | GUF1 GTPase homolog (S. cerevisiae) |
| **CENPJ** | ENSG00000151849 | 292.8090066 | 1.125464164 | 0.290677769 | 2.45E-05 | 1.26E-04 | 55835 | 13 | 25457171 | 25497018 | -1 | protein_coding | centromere protein J |
| **FBXO4** | ENSG00000151876 | 156.9583008 | 0.693231144 | 0.286841598 | 0.00722098 | 0.021064752 | 26272 | 5 | 41925356 | 41941845 | 1 | protein_coding | F-box protein 4 |
| **PARP8** | ENSG00000151883 | 1849.500578 | 0.853129885 | 0.10834919 | 1.33E-15 | 2.1E-14 | 79668 | 5 | 49961733 | 50142356 | 1 | protein_coding | poly (ADP-ribose) polymerase family, member 8 |
| **TIAL1** | ENSG00000151923 | 1536.515672 | 0.314629672 | 0.105755441 | 0.002548156 | 0.008369539 | 7073 | 10 | 121334199 | 121356541 | -1 | protein_coding | TIA1 cytotoxic granule-associated RNA binding protein-like 1 |
| **BAG3** | ENSG00000151929 | 928.5805623 | -0.55037612 | 0.160890356 | 3.79E-04 | 0.001528352 | 9531 | 10 | 121410882 | 121437331 | 1 | protein_coding | BCL2-associated athanogene 3 |
| **GLT1D1** | ENSG00000151948 | 79.75251064 | 0.99460921 | 0.551907791 | 0.014206006 | 0.037903349 | 144423 | 12 | 129337972 | 129469509 | 1 | protein_coding | glycosyltransferase 1 domain containing 1 |
| **RABGAP1L** | ENSG00000152061 | 1821.366942 | 1.555391865 | 0.152722972 | 3.41E-25 | 1.02E-23 | 9910 | 1 | 174128548 | 174964445 | 1 | protein_coding | RAB GTPase activating protein 1-like |
| **MZT2B** | ENSG00000152082 | 239.9747434 | -0.486152389 | 0.222324017 | 0.01904602 | 0.048675638 | 653784 | 2 | 130939310 | 130948302 | 1 | protein_coding | mitotic spindle organizing protein 2B |
| **FAM168B** | ENSG00000152102 | 2791.254599 | -1.106131844 | 0.113544008 | 5.9E-23 | 1.59E-21 | 130074 | 2 | 131805449 | 131851033 | -1 | protein_coding | family with sequence similarity 168, member B |
| **PTPN14** | ENSG00000152104 | 95.12417043 | -1.796658467 | 0.488396027 | 2.28E-05 | 1.18E-04 | 5784 | 1 | 214522039 | 214725792 | -1 | protein_coding | protein tyrosine phosphatase, non-receptor type 14 |
| **MGAT5** | ENSG00000152127 | 4425.699413 | -1.599884262 | 0.114624285 | 4.2E-45 | 3.22E-43 | 4249 | 2 | 134877554 | 135212192 | 1 | protein_coding | mannosyl (alpha-1,6-)-glycoprotein beta-1,6-N-acetyl-glucosaminyltransferase |
| **GPATCH11** | ENSG00000152133 | 36.96090963 | -8.180119674 | 2.859727907 | 5.13E-07 | 3.46E-06 | 253635 | 2 | 37311594 | 37326387 | 1 | protein_coding | G patch domain containing 11 |
| **TMEM178A** | ENSG00000152154 | 23.22095353 | -1.125805361 | 0.744914421 | 0.018578714 | 0.047648681 | 130733 | 2 | 39892122 | 39945103 | 1 | protein_coding | transmembrane protein 178A |
| **RNF219** | ENSG00000152193 | 587.3443031 | 0.628700663 | 0.158583754 | 4.31E-05 | 2.1E-04 | 79596 | 13 | 79188426 | 79233314 | -1 | protein_coding | ring finger protein 219 |
| **CYSLTR2** | ENSG00000152207 | 73.07372747 | -1.336170237 | 0.551232936 | 0.002059651 | 0.006937987 | 57105 | 13 | 49280951 | 49283498 | 1 | protein_coding | cysteinyl leukotriene receptor 2 |
| **ARL 11.00** | ENSG00000152213 | 946.2054947 | 0.445724894 | 0.146382292 | 0.001699808 | 0.005850939 | 115761 | 13 | 50202435 | 50208008 | 1 | protein_coding | ADP-ribosylation factor-like 11 |
| **ARL14EP** | ENSG00000152219 | 570.6539325 | 0.428129232 | 0.187686173 | 0.016266903 | 0.042642725 | 120534 | 11 | 30344598 | 30359774 | 1 | protein_coding | ADP-ribosylation factor-like 14 effector protein |
| **PSTPIP2** | ENSG00000152229 | 2912.415104 | 2.706283853 | 0.178747563 | 4.78E-53 | 4.97E-51 | 9050 | 18 | 43563502 | 43652238 | -1 | protein_coding | proline-serine-threonine phosphatase interacting protein 2 |
| **PDK1** | ENSG00000152256 | 672.0820243 | -0.58582325 | 0.189635681 | 0.001214837 | 0.004335334 | 5163 | 2 | 173420101 | 173489823 | 1 | protein_coding | pyruvate dehydrogenase kinase, isozyme 1 |
| **TGOLN2** | ENSG00000152291 | 20147.3918 | -0.465272314 | 0.083494028 | 1.81E-08 | 1.49E-07 | 10618 | 2 | 85545147 | 85555548 | -1 | protein_coding | trans-golgi network protein 2 |
| **KCNK13** | ENSG00000152315 | 359.9256988 | -1.456773323 | 0.270017771 | 1.06E-08 | 8.98E-08 | 56659 | 14 | 90528109 | 90652201 | 1 | protein_coding | potassium channel, subfamily K, member 13 |
| **UHMK1** | ENSG00000152332 | 7121.672726 | 0.238778558 | 0.085603032 | 0.004666545 | 0.014367205 | 127933 | 1 | 162467041 | 162499419 | 1 | protein_coding | U2AF homology motif (UHM) kinase 1 |
| **POC5** | ENSG00000152359 | 170.2662703 | 0.943762057 | 0.310935906 | 6.92E-04 | 0.002621574 | 134359 | 5 | 74969949 | 75013313 | -1 | protein_coding | POC5 centriolar protein |
| **CWF19L2** | ENSG00000152404 | 353.1532965 | 0.643298876 | 0.215014269 | 0.001454169 | 0.005097262 | 143884 | 11 | 107197071 | 107328572 | -1 | protein_coding | CWF19-like 2, cell cycle control (S. pombe) |
| **USP12** | ENSG00000152484 | 3731.320736 | 0.348138407 | 0.150453086 | 0.017063628 | 0.044403783 | 219333 | 13 | 27640293 | 27746033 | -1 | protein_coding | ubiquitin specific peptidase 12 |
| **CCDC50** | ENSG00000152492 | 2082.316187 | 1.415088809 | 0.12630492 | 6.09E-30 | 2.34E-28 | 152137 | 3 | 191046866 | 191116459 | 1 | protein_coding | coiled-coil domain containing 50 |
| **TRIM36** | ENSG00000152503 | 207.0777834 | 1.041527697 | 0.356159684 | 8.49E-04 | 0.003158082 | 55521 | 5 | 114460459 | 114516243 | -1 | protein_coding | tripartite motif containing 36 |
| **PAN3** | ENSG00000152520 | 1039.807311 | 0.430078341 | 0.119243522 | 2.36E-04 | 9.93E-04 | 255967 | 13 | 28712643 | 28869475 | 1 | protein_coding | PAN3 poly(A) specific ribonuclease subunit homolog (S. cerevisiae) |
| **PFKM** | ENSG00000152556 | 301.1489585 | -0.811457705 | 0.250608115 | 4.45E-04 | 0.001762867 | 5213 | 12 | 48498922 | 48540187 | 1 | protein_coding | phosphofructokinase, muscle |
| **TMEM123** | ENSG00000152558 | 9359.470559 | 1.580370084 | 0.100401147 | 1.51E-56 | 1.76E-54 | 114908 | 11 | 102267063 | 102341115 | -1 | protein_coding | transmembrane protein 123 |
| **MBNL1** | ENSG00000152601 | 8623.420248 | 0.234765324 | 0.092450498 | 0.010162015 | 0.028392335 | 4154 | 3 | 151961617 | 152183569 | 1 | protein_coding | muscleblind-like splicing regulator 1 |
| **GPD1L** | ENSG00000152642 | 318.8053184 | -1.290211115 | 0.238471109 | 1.19E-08 | 1E-07 | 23171 | 3 | 32147181 | 32210205 | 1 | protein_coding | glycerol-3-phosphate dehydrogenase 1-like |
| **PELO** | ENSG00000152684 | 603.6024399 | 0.705932883 | 0.205813897 | 2.88E-04 | 0.001189335 | 53918 | 5 | 52083774 | 52099880 | 1 | protein_coding | pelota homolog (Drosophila) |
| **SAR1B** | ENSG00000152700 | 1869.015642 | 0.340644319 | 0.145776317 | 0.016216918 | 0.042518984 | 51128 | 5 | 133936834 | 133984961 | -1 | protein_coding | SAR1 homolog B (S. cerevisiae) |
| **GPR180** | ENSG00000152749 | 710.3362578 | 1.950818392 | 0.202507963 | 6.12E-23 | 1.63E-21 | 160897 | 13 | 95254157 | 95286899 | 1 | protein_coding | G protein-coupled receptor 180 |
| **ANKRD22** | ENSG00000152766 | 36.63237516 | 1.419786363 | 0.85344743 | 0.007911844 | 0.022797269 | 118932 | 10 | 90581889 | 90611575 | -1 | protein_coding | ankyrin repeat domain 22 |
| **FARP1** | ENSG00000152767 | 1909.520587 | -1.100601052 | 0.694425404 | 0.015845551 | 0.041702697 | 10160 | 13 | 98794816 | 99102027 | 1 | protein_coding | FERM, RhoGEF (ARHGEF) and pleckstrin domain protein 1 (chondrocyte-derived) |
| **IFIT5** | ENSG00000152778 | 1075.602797 | 2.78703862 | 0.204479704 | 2.16E-43 | 1.55E-41 | 24138 | 10 | 91174343 | 91180758 | 1 | protein_coding | interferon-induced protein with tetratricopeptide repeats 5 |
| **HNRNPDL** | ENSG00000152795 | 3609.67452 | 0.279871073 | 0.114208129 | 0.012987772 | 0.035056891 | 9987 | 4 | 83343717 | 83351294 | -1 | protein_coding | heterogeneous nuclear ribonucleoprotein D-like |
| **HHEX** | ENSG00000152804 | 341.5975022 | -1.802549017 | 0.226065559 | 1.63E-16 | 2.74E-15 | 3087 | 10 | 94447945 | 94455403 | 1 | protein_coding | hematopoietically expressed homeobox |
| **GGPS1** | ENSG00000152904 | 485.5549725 | 1.105989806 | 0.15481796 | 2.32E-13 | 3.09E-12 | 9453 | 1 | 235490665 | 235507847 | 1 | protein_coding | geranylgeranyl diphosphate synthase 1 |
| **MED21** | ENSG00000152944 | 292.7111074 | 0.571064424 | 0.24187065 | 0.010551159 | 0.029286297 | 1019286 | 12 | 27175479 | 27219276 | 1 | protein_coding | mediator complex subunit 21 |
| **PLOD2** | ENSG00000152952 | 407.5598595 | -1.879240748 | 0.24321759 | 1.21E-15 | 1.92E-14 | 5352 | 3 | 145787227 | 145881440 | -1 | protein_coding | procollagen-lysine, 2-oxoglutarate 5-dioxygenase 2 |
| **GPR125** | ENSG00000152990 | 59.23626379 | -1.336515146 | 0.833107687 | 0.009821313 | 0.027571478 | 166647 | 4 | 22346694 | 22517677 | -1 | protein_coding | G protein-coupled receptor 125 |
| **SRP19** | ENSG00000153037 | 596.7436258 | 0.602647847 | 0.208392063 | 0.002202334 | 0.007352177 | 6728 | 5 | 112196919 | 112205485 | 1 | protein_coding | signal recognition particle 19kDa |
| **CENPH** | ENSG00000153044 | 112.4362545 | 1.054948328 | 0.369457951 | 0.001001649 | 0.003653118 | 64946 | 5 | 68485375 | 68506184 | 1 | protein_coding | centromere protein H |
| **CARHSP1** | ENSG00000153048 | 206.9930036 | -0.647482075 | 0.266841562 | 0.007770297 | 0.022461406 | 23589 | 16 | 8946799 | 8962866 | -1 | protein_coding | calcium regulated heat stable protein 1, 24kDa |
| **TXNDC11** | ENSG00000153066 | 2523.635655 | 0.454499746 | 0.091709307 | 5.42E-07 | 3.65E-06 | 51061 | 16 | 11772936 | 11836734 | -1 | protein_coding | thioredoxin domain containing 11 |
| **DAB2** | ENSG00000153071 | 12889.28799 | -1.43168282 | 0.130014609 | 5.57E-29 | 2.03E-27 | 1019269 | 5 | 39371780 | 39462402 | -1 | protein_coding | Dab, mitogen-responsive phosphoprotein, homolog 2 (Drosophila) |
| **CAST** | ENSG00000153113 | 11136.64842 | 0.420058558 | 0.105286382 | 5.45E-05 | 2.61E-04 | 831 | 5 | 95860971 | 96115299 | 1 | protein_coding | calpastatin |
| **SCOC** | ENSG00000153130 | 185.0380591 | 1.235215741 | 0.495993854 | 0.001944725 | 0.006601786 | 60592 | 4 | 141178440 | 141306880 | 1 | protein_coding | short coiled-coil protein |
| **SMARCA5** | ENSG00000153147 | 5093.877991 | 1.170573229 | 0.1159239 | 1.43E-24 | 4.17E-23 | 8467 | 4 | 144434616 | 144478639 | 1 | protein_coding | SWI/SNF related, matrix associated, actin dependent regulator of chromatin, subfamily a, member 5 |
| **BMP6** | ENSG00000153162 | 37.2299571 | 2.488055521 | 0.905028448 | 2.62E-04 | 0.001091087 | 654 | 6 | 7727030 | 7881655 | 1 | protein_coding | bone morphogenetic protein 6 |
| **RANBP2** | ENSG00000153201 | 9587.809324 | -0.23242452 | 0.099864076 | 0.017131911 | 0.044570561 | 5903 | 2 | 109335937 | 109402267 | 1 | protein_coding | RAN binding protein 2 |
| **MERTK** | ENSG00000153208 | 1539.522798 | -2.426181029 | 0.179506742 | 7.85E-43 | 5.58E-41 | 10461 | 2 | 112656056 | 112787138 | 1 | protein_coding | c-mer proto-oncogene tyrosine kinase |
| **TMEM87B** | ENSG00000153214 | 3142.801683 | 0.8851817 | 0.138791532 | 6.81E-11 | 7.42E-10 | 84910 | 2 | 112812800 | 112876895 | 1 | protein_coding | transmembrane protein 87B |
| **NR4A2** | ENSG00000153234 | 126.0500127 | 0.926910366 | 0.325150136 | 0.001308013 | 0.004626979 | 4929 | 2 | 157180944 | 157198860 | -1 | protein_coding | nuclear receptor subfamily 4, group A, member 2 |
| **RBMS1** | ENSG00000153250 | 4114.16468 | 0.591230494 | 0.089463439 | 1.84E-11 | 2.12E-10 | 5937 | 2 | 161128662 | 161350305 | -1 | protein_coding | RNA binding motif, single stranded interacting protein 1 |
| **CD96** | ENSG00000153283 | 931.1475036 | -0.473882014 | 0.205761267 | 0.014192924 | 0.037875057 | 10225 | 3 | 111011566 | 111384597 | 1 | protein_coding | CD96 molecule |
| **LPCAT1** | ENSG00000153395 | 1699.915276 | -0.382443877 | 0.160531671 | 0.013197668 | 0.035548106 | 79888 | 5 | 1456595 | 1524092 | -1 | protein_coding | lysophosphatidylcholine acyltransferase 1 |
| **CMTM7** | ENSG00000153551 | 949.2629169 | -1.082855997 | 0.167246831 | 2.36E-11 | 2.69E-10 | 112616 | 3 | 32433163 | 32524559 | 1 | protein_coding | CKLF-like MARVEL transmembrane domain containing 7 |
| **UBP1** | ENSG00000153560 | 2504.931213 | 1.05392383 | 0.110656432 | 4.63E-22 | 1.17E-20 | 7342 | 3 | 33429828 | 33482863 | -1 | protein_coding | upstream binding protein 1 (LBP-1a) |
| **RMND5A** | ENSG00000153561 | 1471.059918 | 0.341285105 | 0.145003021 | 0.015690159 | 0.041350699 | 64795 | 2 | 86947296 | 87005164 | 1 | protein_coding | required for meiotic nuclear division 5 homolog A (S. cerevisiae) |
| **RPIA** | ENSG00000153574 | 232.752606 | 0.820678992 | 0.270113664 | 8.79E-04 | 0.003252602 | 22934 | 2 | 88991162 | 89050452 | 1 | protein_coding | ribose 5-phosphate isomerase A |
| **CNKSR3** | ENSG00000153721 | 163.3087664 | -1.234824809 | 0.295125018 | 5.17E-06 | 2.98E-05 | 154043 | 6 | 154708638 | 154831793 | -1 | protein_coding | CNKSR family member 3 |
| **GTF2E1** | ENSG00000153767 | 759.7009786 | 0.475211833 | 0.176537464 | 0.004933835 | 0.015104929 | 2960 | 3 | 120461484 | 120501916 | 1 | protein_coding | general transcription factor IIE, polypeptide 1, alpha 56kDa |
| **ZDHHC7** | ENSG00000153786 | 3998.108717 | -0.75226387 | 0.129379443 | 2.79E-09 | 2.53E-08 | 55625 | 16 | 85007787 | 85045141 | -1 | protein_coding | zinc finger, DHHC-type containing 7 |
| **C7orf31** | ENSG00000153790 | 305.6947038 | 0.638426604 | 0.189209745 | 4E-04 | 0.001604684 | 136895 | 7 | 25174316 | 25219975 | -1 | protein_coding | chromosome 7 open reading frame 31 |
| **CMIP** | ENSG00000153815 | 2469.626016 | -0.605406257 | 0.146419717 | 2.05E-05 | 1.06E-04 | 80790 | 16 | 81478775 | 81745367 | 1 | protein_coding | c-Maf inducing protein |
| **CEBPG** | ENSG00000153879 | 1492.361774 | 0.438087286 | 0.117962885 | 1.52E-04 | 6.69E-04 | 1054 | 19 | 33864236 | 33873592 | 1 | protein_coding | CCAAT/enhancer binding protein (C/EBP), gamma |
| **MCOLN2** | ENSG00000153898 | 765.0931999 | 4.390618194 | 0.230959598 | 9.87E-82 | 2.43E-79 | 255231 | 1 | 85391268 | 85462796 | -1 | protein_coding | mucolipin 2 |
| **SREK1** | ENSG00000153914 | 1458.47318 | 0.363697115 | 0.149737307 | 0.012243705 | 0.033283625 | 140890 | 5 | 65435799 | 65479443 | 1 | protein_coding | splicing regulatory glutamine/lysine-rich protein 1 |
| **CHD1** | ENSG00000153922 | 170.452634 | 1.026549535 | 0.318189795 | 3.24E-04 | 0.001326958 | 1105 | 5 | 98190908 | 98262240 | -1 | protein_coding | chromodomain helicase DNA binding protein 1 |
| **DGKE** | ENSG00000153933 | 197.8269086 | 1.438951312 | 0.358558493 | 9E-06 | 4.99E-05 | 8526 | 17 | 54911460 | 54946036 | 1 | protein_coding | diacylglycerol kinase, epsilon 64kDa |
| **HS2ST1** | ENSG00000153936 | 2756.356811 | -0.746517694 | 0.157196085 | 9.27E-07 | 5.99E-06 | 9653 | 1 | 87380331 | 87602334 | 1 | protein_coding | heparan sulfate 2-O-sulfotransferase 1 |
| **MSI2** | ENSG00000153944 | 846.4582185 | -0.910546657 | 0.224451632 | 1.4E-05 | 7.51E-05 | 124540 | 17 | 55333212 | 55762046 | 1 | protein_coding | musashi RNA-binding protein 2 |
| **ZUFSP** | ENSG00000153975 | 480.1070599 | 1.032533137 | 0.164763928 | 1.03E-10 | 1.1E-09 | 221302 | 6 | 116956781 | 116989957 | -1 | protein_coding | zinc finger with UFM1-specific peptidase domain |
| **NUS1** | ENSG00000153989 | 3898.823256 | 0.622838716 | 0.106158263 | 2.21E-09 | 2.03E-08 | 116150 | 6 | 117996665 | 118031803 | 1 | protein_coding | nuclear undecaprenyl pyrophosphate synthase 1 homolog (S. cerevisiae) |
| **PPP2R5E** | ENSG00000154001 | 1869.416328 | 0.390275581 | 0.112470742 | 4E-04 | 0.001604538 | 5529 | 14 | 63838075 | 64010092 | -1 | protein_coding | protein phosphatase 2, regulatory subunit B', epsilon isoform |
| **DNAAF1** | ENSG00000154099 | 61.0037487 | 11.2651094 | 2.768488889 | 9.07E-13 | 1.16E-11 | 123872 | 16 | 84178865 | 84212373 | 1 | protein_coding | dynein, axonemal, assembly factor 1 |
| **ANKH** | ENSG00000154122 | 1100.314828 | -2.644081989 | 0.146718617 | 1.27E-73 | 2.66E-71 | 56172 | 5 | 14704910 | 14871887 | -1 | protein_coding | ANKH inorganic pyrophosphate transport regulator |
| **UBASH3B** | ENSG00000154127 | 3413.330109 | -2.830437928 | 0.206625773 | 5.81E-44 | 4.27E-42 | 84959 | 11 | 122526383 | 122685181 | 1 | protein_coding | ubiquitin associated and SH3 domain containing B |

| | | | | | | | | | | | | | |
|---|---|---|---|---|---|---|---|---|---|---|---|---|---|
| **NRGN** | ENSG00000154146 | 52.75295117 | -3.415760651 | 0.699307531 | 9.88E-08 | 7.33E-07 | 4900 | 11 | 124609742 | 124617106 | 1 | protein_coding | neurogranin (protein kinase C substrate, RC3) |
| **TOMM70A** | ENSG00000154174 | 1906.578147 | 0.475240186 | 0.141456355 | 5.61E-04 | 0.002164084 | 9868 | 3 | 100082275 | 100120242 | -1 | protein_coding | translocase of outer mitochondrial membrane 70 homolog A (S. cerevisiae) |
| **ABI3BP** | ENSG00000154175 | 21.59061209 | -2.018581757 | 1.043012744 | 0.003246665 | 0.010413176 | 25890 | 3 | 100468000 | 100712359 | -1 | protein_coding | ABI family, member 3 (NESH) binding protein |
| **PITPNC1** | ENSG00000154217 | 711.9849514 | -1.158449246 | 0.178931824 | 2.16E-11 | 2.48E-10 | 26207 | 17 | 65373575 | 65693372 | 1 | protein_coding | phosphatidylinositol transfer protein, cytoplasmic 1 |
| **PRKCA** | ENSG00000154229 | 1210.020404 | -1.414147591 | 0.157320381 | 3.96E-20 | 8.83E-19 | 5578 | 17 | 64298754 | 64806861 | 1 | protein_coding | protein kinase C, alpha |
| **LRRK1** | ENSG00000154237 | 1342.19488 | -0.293598418 | 0.12266955 | 0.014062386 | 0.037559498 | 79705 | 15 | 101459420 | 101610317 | 1 | protein_coding | leucine-rich repeat kinase 1 |
| **ABCA6** | ENSG00000154262 | 645.6036618 | 1.342025209 | 0.188669652 | 2.01E-13 | 2.7E-12 | 23460 | 17 | 67074843 | 67138029 | -1 | protein_coding | ATP-binding cassette, sub-family A (ABC1), member 6 |
| **ABCA10** | ENSG00000154263 | 22.14464249 | 1.945821644 | 1.032002638 | 0.003397073 | 0.010840984 | 10349 | 17 | 67143355 | 67240987 | -1 | protein_coding | ATP-binding cassette, sub-family A (ABC1), member 10 |
| **ABCA5** | ENSG00000154265 | 429.3031169 | 1.136753072 | 0.213391279 | 2.24E-08 | 1.81E-07 | 23461 | 17 | 67240452 | 67323385 | -1 | protein_coding | ATP-binding cassette, sub-family A (ABC1), member 5 |
| **MIA3** | ENSG00000154305 | 2725.692219 | 1.122857837 | 0.127572997 | 3.29E-19 | 6.84E-18 | 375056 | 1 | 222791428 | 222841354 | 1 | protein_coding | melanoma inhibitory activity family, member 3 |
| **DISP1** | ENSG00000154309 | 224.7304328 | -0.528611153 | 0.236889113 | 0.016327061 | 0.042785747 | 84976 | 1 | 222988406 | 223179337 | 1 | protein_coding | dispatched homolog 1 (Drosophila) |
| **TNIK** | ENSG00000154310 | 2266.213267 | 1.044863576 | 0.105994882 | 8.45E-24 | 2.37E-22 | 23043 | 3 | 170779128 | 171178197 | -1 | protein_coding | TRAF2 and NCK interacting kinase |
| **NEIL2** | ENSG00000154328 | 182.5084301 | -1.027792943 | 0.252664816 | 1.29E-05 | 7E-05 | 252969 | 8 | 11627148 | 11644855 | 1 | protein_coding | nei endonuclease VIII-like 2 (E. coli) |
| **PGM5** | ENSG00000154330 | 303.92511 | 0.779434567 | 0.225814706 | 2.11E-04 | 8.96E-04 | 5239 | 9 | 70943224 | 71145977 | 1 | protein_coding | phosphoglucomutase 5 |
| **LONRF1** | ENSG00000154359 | 456.8370588 | 0.606809354 | 0.227899377 | 0.004294274 | 0.013331178 | 91694 | 8 | 12579403 | 12613582 | -1 | protein_coding | LON peptidase N-terminal domain and ring finger 1 |
| **SH3RF1** | ENSG00000154447 | 229.8882883 | -1.314046957 | 0.262192197 | 9.95E-08 | 7.38E-07 | 57630 | 4 | 170015407 | 170192256 | -1 | protein_coding | SH3 domain containing ring finger 1 |
| **GBP5** | ENSG00000154451 | 9535.814546 | 3.385095992 | 0.466553649 | 2.43E-14 | 3.53E-13 | 115362 | 1 | 89724633 | 89738544 | -1 | protein_coding | guanylate binding protein 5 |
| **FAM69A** | ENSG00000154511 | 864.757299 | 1.170611364 | 0.147104642 | 4.19E-16 | 6.85E-15 | 388650 | 1 | 93307724 | 93427057 | -1 | protein_coding | family with sequence similarity 69, member A |
| **ATP5G3** | ENSG00000154518 | 2229.139092 | -0.70217482 | 0.105771353 | 1.5E-11 | 1.74E-10 | 518 | 2 | 176040986 | 176049335 | -1 | protein_coding | ATP synthase, H+ transporting, mitochondrial Fo complex, subunit C3 (subunit 9) |
| **SRSF12** | ENSG00000154548 | 25.61240866 | 2.849126879 | 0.794034884 | 1.8E-05 | 9.46E-05 | 135295 | 6 | 89805678 | 89827800 | -1 | protein_coding | serine/arginine-rich splicing factor 12 |
| **CXADR** | ENSG00000154639 | 232.192021 | -3.975865703 | 0.39816743 | 1.38E-24 | 4.05E-23 | 1525 | 21 | 18884700 | 18965897 | 1 | protein_coding | coxsackie virus and adenovirus receptor |
| **BTG3** | ENSG00000154640 | 500.9792224 | 1.586368926 | 0.196562858 | 7.35E-17 | 1.28E-15 | 10950 | 21 | 18965971 | 18985265 | -1 | protein_coding | BTG family, member 3 |
| **C21orf91** | ENSG00000154642 | 285.5737146 | 1.866128832 | 0.305647592 | 1.1E-10 | 1.18E-09 | 54149 | 21 | 19161284 | 19191703 | -1 | protein_coding | chromosome 21 open reading frame 91 |
| **NCAM2** | ENSG00000154654 | 6.781912976 | 0.551295741 | 1.02944991 | 0.005246835 | 0.015916512 | 4685 | 21 | 22370633 | 22915650 | 1 | protein_coding | neural cell adhesion molecule 2 |
| **SLFN13** | ENSG00000154760 | 191.6711818 | 0.668596606 | 0.329922561 | 0.018796336 | 0.048117991 | 146857 | 17 | 33762115 | 33775856 | -1 | protein_coding | schlafen family member 13 |
| **XPC** | ENSG00000154767 | 2395.385354 | -0.59353317 | 0.126985234 | 1.69E-06 | 1.05E-05 | 7508 | 3 | 14186647 | 14220283 | -1 | protein_coding | xeroderma pigmentosum, complementation group C |
| **CCDC174** | ENSG00000154781 | 636.4869949 | 0.451862948 | 0.173050818 | 0.006429964 | 0.019042729 | 51244 | 3 | 14693271 | 14714166 | 1 | protein_coding | coiled-coil domain containing 174 |
| **FGD5** | ENSG00000154783 | 3525.998592 | -3.020151132 | 0.119093815 | 6.31E-143 | 3.33E-140 | 152273 | 3 | 14860469 | 14975895 | 1 | protein_coding | FYVE, RhoGEF and PH domain containing 5 |
| **FLCN** | ENSG00000154803 | 1429.390512 | 0.288394079 | 0.122999669 | 0.0175219 | 0.045441575 | 201163 | 17 | 17115526 | 17140502 | -1 | protein_coding | folliculin |
| **DPH3** | ENSG00000154813 | 1410.618482 | 0.692778133 | 0.123348057 | 9.52E-09 | 8.12E-08 | 285381 | 3 | 16299485 | 16306479 | -1 | protein_coding | diphthamide biosynthesis 3 |
| **OXNAD1** | ENSG00000154814 | 421.8013131 | -0.572974706 | 0.210874018 | 0.003861389 | 0.012120272 | 92106 | 3 | 16306706 | 16391806 | 1 | protein_coding | oxidoreductase NAD-binding domain containing 1 |
| **SKA1** | ENSG00000154839 | 29.43027733 | 1.60099703 | 0.751883199 | 0.002585768 | 0.008474864 | 220134 | 18 | 47901365 | 47920543 | 1 | protein_coding | spindle and kinetochore associated complex subunit 1 |
| **ACSS1** | ENSG00000154930 | 513.298884 | -0.429259791 | 0.159599902 | 0.00593991 | 0.017726938 | 84532 | 20 | 24986868 | 25039616 | -1 | protein_coding | acyl-CoA synthetase short-chain family member 1 |
| **AZIN1** | ENSG00000155096 | 6237.908104 | 0.6260707 | 0.092439962 | 6.93E-12 | 8.29E-11 | 51582 | 8 | 103838585 | 103906092 | -1 | protein_coding | antizyme inhibitor 1 |
| **ATP6V1C1** | ENSG00000155097 | 5080.731298 | 0.330548768 | 0.12808573 | 0.008413356 | 0.024047376 | 528 | 8 | 104033291 | 104085279 | 1 | protein_coding | ATPase, H+ transporting, lysosomal 42kDa, V1 subunit C1 |
| **CDK19** | ENSG00000155111 | 1514.611975 | -0.394005247 | 0.130165823 | 0.001960843 | 0.006652068 | 23097 | 6 | 110931181 | 111137161 | -1 | protein_coding | cyclin-dependent kinase 19 |
| **MMS19** | ENSG00000155229 | 1288.133668 | -0.438026431 | 0.142567627 | 0.001614328 | 0.005591969 | 64210 | 10 | 99218081 | 99258551 | -1 | protein_coding | MMS19 nucleotide excision repair homolog (S. cerevisiae) |
| **PI4K2A** | ENSG00000155252 | 6912.306643 | -0.462867714 | 0.091477418 | 2.74E-07 | 1.91E-06 | 55361 | 10 | 99344131 | 99436191 | 1 | protein_coding | phosphatidylinositol 4-kinase type 2 alpha |
| **GOLGA7B** | ENSG00000155265 | 701.1683772 | -1.785518781 | 0.343951862 | 2.13E-08 | 1.73E-07 | 401647 | 10 | 99609996 | 99631337 | 1 | protein_coding | golgin A7 family, member B |
| **SLC25A28** | ENSG00000155287 | 617.1718463 | 2.497335139 | 0.236146425 | 3.25E-27 | 1.08E-25 | 81894 | 10 | 101370282 | 101380366 | -1 | protein_coding | solute carrier family 25 (mitochondrial iron transporter), member 28 |
| **HSPA13** | ENSG00000155304 | 781.592854 | 1.197357992 | 0.174186735 | 1.4E-12 | 1.76E-11 | 6782 | 21 | 15743436 | 15755805 | -1 | protein_coding | heat shock protein 70kDa family, member 13 |
| **SAMSN1** | ENSG00000155307 | 1309.322113 | 1.343337846 | 0.209475247 | 2.38E-11 | 2.71E-10 | 64092 | 21 | 15857549 | 15955723 | -1 | protein_coding | SAM domain, SH3 domain and nuclear localization signals 1 |
| **USP25** | ENSG00000155313 | 3123.255192 | 1.039894306 | 0.102386563 | 9.45E-25 | 2.79E-23 | 29761 | 21 | 17102344 | 17252377 | 1 | protein_coding | ubiquitin specific peptidase 25 |
| **GRAMD3** | ENSG00000155324 | 339.6889678 | 1.168382459 | 0.220163807 | 2.5E-08 | 2.01E-07 | 65983 | 5 | 125695824 | 125832186 | 1 | protein_coding | GRAM domain containing 3 |
| **ZCCHC10** | ENSG00000155329 | 313.9675746 | 0.60794475 | 0.198163665 | 0.001221168 | 0.004354434 | 54819 | 5 | 132332677 | 132362296 | -1 | protein_coding | zinc finger, CCHC domain containing 10 |
| **C16orf87** | ENSG00000155330 | 401.405643 | 0.733538778 | 0.220893498 | 3.94E-04 | 0.001582003 | 388272 | 16 | 46830519 | 46865323 | -1 | protein_coding | chromosome 16 open reading frame 87 |
| **MOV10** | ENSG00000155363 | 2130.873471 | 1.474426541 | 0.141737327 | 3.66E-26 | 1.16E-24 | 4343 | 1 | 113215763 | 113243368 | 1 | protein_coding | Mov10, Moloney leukemia virus 10, homolog (mouse) |
| **RHOC** | ENSG00000155366 | 2991.690249 | -0.359788972 | 0.106226544 | 6.17E-04 | 0.002358471 | 389 | 1 | 113243728 | 113250056 | -1 | protein_coding | ras homolog family member C |
| **DBI** | ENSG00000155368 | 7825.594304 | -0.564752951 | 0.190680756 | 0.001871209 | 0.006373453 | 1622 | 2 | 120124497 | 120130126 | 1 | protein_coding | diazepam binding inhibitor (GABA receptor modulator, acyl-CoA binding protein) |
| **SLC16A1** | ENSG00000155380 | 1912.417875 | 0.517363564 | 0.122797894 | 1.72E-05 | 9.05E-05 | 6566 | 1 | 113454469 | 113499635 | -1 | protein_coding | solute carrier family 16 (monocarboxylate transporter), member 1 |
| **HEATR3** | ENSG00000155393 | 849.2033578 | -0.789329051 | 0.213565111 | 8.4E-05 | 3.9E-04 | 55027 | 16 | 50099852 | 50140298 | 1 | protein_coding | HEAT repeat containing 3 |
| **OXA1L** | ENSG00000155463 | 460.1416054 | -0.496689097 | 0.174087507 | 0.002895459 | 0.009391192 | 5018 | 14 | 23235731 | 23241007 | 1 | protein_coding | oxidase (cytochrome c) assembly 1-like |
| **LARP1** | ENSG00000155506 | 6884.351603 | 0.518420417 | 0.096950861 | 6.17E-08 | 4.72E-07 | 23367 | 5 | 154092462 | 154197167 | 1 | protein_coding | La ribonucleoprotein domain family, member 1 |
| **NUP205** | ENSG00000155561 | 2019.563831 | 0.576044802 | 0.103122658 | 1.44E-08 | 1.2E-07 | 23165 | 7 | 135242667 | 135333505 | 1 | protein_coding | nucleoporin 205kDa |
| **C9orf85** | ENSG00000155621 | 202.2934407 | 0.879460639 | 0.254697423 | 1.89E-04 | 8.16E-04 | 138241 | 9 | 74526426 | 74600970 | 1 | protein_coding | chromosome 9 open reading frame 85 |
| **PIK3AP1** | ENSG00000155629 | 21710.09717 | 0.294187692 | 0.086468747 | 5.3E-04 | 0.002060361 | 118788 | 10 | 98353069 | 98480271 | -1 | protein_coding | phosphoinositide-3-kinase adaptor protein 1 |
| **TTN** | ENSG00000155657 | 579.7400969 | -0.998621374 | 0.275816742 | 7.63E-05 | 3.56E-04 | 7273 | 2 | 179390716 | 179695529 | -1 | protein_coding | titin |
| **VSIG4** | ENSG00000155659 | 2983.954568 | -1.364915038 | 0.401130259 | 9.89E-05 | 4.53E-04 | 11326 | X | 65241580 | 65259967 | -1 | protein_coding | V-set and immunoglobulin domain containing 4 |
| **PDIA4** | ENSG00000155660 | 5119.521523 | -0.57818544 | 0.09952917 | 5.97E-09 | 5.22E-08 | 9601 | 7 | 148700154 | 148725733 | -1 | protein_coding | protein disulfide isomerase family A, member 4 |
| **OTOA** | ENSG00000155719 | 84.60630278 | -1.102955157 | 0.539308836 | 0.00727646 | 0.021194233 | 146183 | 16 | 21689835 | 21772050 | 1 | protein_coding | otoancorin |
| **FAM126B** | ENSG00000155744 | 1294.602843 | 1.404223692 | 0.198357214 | 2.4E-13 | 3.2E-12 | 285172 | 2 | 201838441 | 201936394 | -1 | protein_coding | family with sequence similarity 126, member B |
| **PPARGC1B** | ENSG00000155846 | 2003.43669 | -0.528366078 | 0.103570868 | 2.06E-07 | 1.47E-06 | 133522 | 5 | 149109861 | 149234585 | 1 | protein_coding | peroxisome proliferator-activated receptor gamma, coactivator 1 beta |
| **SLC26A2** | ENSG00000155850 | 819.9492692 | -0.455208035 | 0.162947022 | 0.003699102 | 0.011685277 | 1836 | 5 | 149340300 | 149373018 | 1 | protein_coding | solute carrier family 26 (anion exchanger), member 2 |
| **MED7** | ENSG00000155868 | 324.7905055 | 0.48066072 | 0.205580894 | 0.012940033 | 0.034934204 | 9443 | 5 | 156564423 | 156586030 | -1 | protein_coding | mediator complex subunit 7 |
| **ADCY8** | ENSG00000155897 | 64.81278609 | -4.436801021 | 1.007784384 | 1.12E-06 | 7.17E-06 | 114 | 8 | 131792547 | 132054672 | -1 | protein_coding | adenylate cyclase 8 (brain) |
| **RASA2** | ENSG00000155903 | 764.2658484 | 0.65871444 | 0.169103797 | 5.14E-05 | 2.47E-04 | 5922 | 3 | 141205889 | 141334184 | 1 | protein_coding | RAS p21 protein activator 2 |
| **SLA** | ENSG00000155926 | 6750.869761 | -1.306385607 | 0.174576679 | 1.4E-14 | 2.06E-13 | 6503 | 8 | 134048973 | 134115298 | -1 | protein_coding | Src-like-adaptor |
| **TMBIM4** | ENSG00000155957 | 1801.850368 | 0.773598019 | 0.136973899 | 6.82E-09 | 5.93E-08 | 51643 | 12 | 66517709 | 66563852 | -1 | protein_coding | transmembrane BAX inhibitor motif containing 4 |
| **VBP1** | ENSG00000155959 | 297.8866142 | 0.713243862 | 0.218579392 | 5.03E-04 | 0.001969134 | 7411 | X | 154425284 | 154468098 | 1 | protein_coding | von Hippel-Lindau binding protein 1 |
| **VPS37A** | ENSG00000155975 | 1575.972924 | 0.367262141 | 0.101949935 | 2.54E-04 | 0.001061847 | 137492 | 8 | 17104080 | 17159936 | 1 | protein_coding | vacuolar protein sorting 37 homolog A (S. cerevisiae) |
| **ELMSAN1** | ENSG00000156030 | 1404.273403 | -0.363239002 | 0.144528085 | 0.009693865 | 0.027293909 | 91748 | 14 | 74181825 | 74256988 | -1 | protein_coding | ELM2 and Myb/SANT-like domain containing 1 |
| **GNAQ** | ENSG00000156052 | 3028.598334 | -1.080228052 | 0.100335643 | 2.61E-27 | 8.72E-26 | 2776 | 9 | 80331003 | 80646374 | -1 | protein_coding | guanine nucleotide binding protein (G protein), q polypeptide |
| **ADK** | ENSG00000156110 | 758.3681006 | -1.074484462 | 0.185408434 | 1.72E-09 | 1.61E-08 | 132 | 10 | 75910960 | 76469061 | 1 | protein_coding | adenosine kinase |
| **KCNMA1** | ENSG00000156113 | 5749.720766 | -1.129908329 | 0.100779387 | 8.53E-30 | 3.25E-28 | 3778 | 10 | 78629359 | 79398353 | -1 | protein_coding | potassium large conductance calcium-activated channel, subfamily M, alpha member 1 |
| **DCK** | ENSG00000156136 | 1235.929445 | 1.717366757 | 0.116630631 | 5.59E-50 | 4.94E-48 | 1633 | 4 | 71858255 | 71896631 | 1 | protein_coding | deoxycytidine kinase |
| **ADAMTS3** | ENSG00000156140 | 21.69847639 | -1.433205695 | 0.913789488 | 0.010005807 | 0.028022494 | 9508 | 4 | 73146686 | 73434516 | -1 | protein_coding | ADAM metallopeptidase with thrombospondin type 1 motif, 3 |
| **DPY19L4** | ENSG00000156162 | 568.2241233 | 0.809277733 | 0.227005233 | 1.44E-04 | 6.38E-04 | 286148 | 8 | 95731931 | 95806064 | 1 | protein_coding | dpy-19-like 4 (C. elegans) |
| **DRAM2** | ENSG00000156171 | 1912.81793 | -0.233054203 | 0.101253787 | 0.019439889 | 0.049516638 | 128338 | 1 | 111659955 | 111682838 | -1 | protein_coding | DNA-damage regulated autophagy modulator 2 |
| **WHAMM** | ENSG00000156232 | 579.1576862 | 0.715075786 | 0.172676179 | 1.66E-05 | 8.79E-05 | 123720 | 15 | 83478380 | 83503611 | 1 | protein_coding | WAS protein homolog associated with actin, golgi membranes and microtubules |
| **CXCL13** | ENSG00000156234 | 139.3267273 | 3.403819177 | 0.759963336 | 1.31E-07 | 9.56E-07 | 10563 | 4 | 78432907 | 78532988 | 1 | protein_coding | chemokine (C-X-C motif) ligand 13 |
| **USP16** | ENSG00000156256 | 1364.394648 | 0.448825827 | 0.11479855 | 6.99E-05 | 3.29E-04 | 10600 | 21 | 30396950 | 30426809 | 1 | protein_coding | ubiquitin specific peptidase 16 |
| **CCT8** | ENSG00000156261 | 3974.962816 | 0.291341607 | 0.106865918 | 0.005509854 | 0.016595802 | 10694 | 21 | 30428126 | 30446118 | -1 | protein_coding | chaperonin containing TCP1, subunit 8 (theta) |
| **TIAM1** | ENSG00000156299 | 3246.344049 | -0.662560354 | 0.112886419 | 2.35E-09 | 2.15E-08 | 7074 | 21 | 32490734 | 32932290 | -1 | protein_coding | T-cell lymphoma invasion and metastasis 1 |
| **SCAF4** | ENSG00000156304 | 1185.718736 | 0.398378802 | 0.122662059 | 9.09E-04 | 0.00335429 | 57466 | 21 | 33043346 | 33104388 | -1 | protein_coding | SR-related CTD-associated factor 4 |
| **SFXN2** | ENSG00000156398 | 466.3247061 | -1.827945913 | 0.29314149 | 4.3E-11 | 4.8E-10 | 118980 | 10 | 104474295 | 104503249 | 1 | protein_coding | sideroflexin 2 |
| **C14orf2** | ENSG00000156411 | 1101.009445 | -0.648721693 | 0.167425662 | 5.72E-05 | 2.74E-04 | 9556 | 14 | 104378625 | 104394606 | -1 | protein_coding | chromosome 14 open reading frame 2 |
| **TDRD9** | ENSG00000156414 | 1959.784067 | -2.197104969 | 0.194641425 | 1.29E-30 | 5.13E-29 | 122402 | 14 | 104394799 | 104519004 | 1 | protein_coding | tudor domain containing 9 |
| **PCDH1** | ENSG00000156453 | 89.88946361 | -1.921655221 | 0.785683893 | 9.95E-04 | 0.003630566 | 5097 | 5 | 141232938 | 141258811 | -1 | protein_coding | protocadherin 1 |
| **SH3RF2** | ENSG00000156463 | 421.09652 | -2.252851113 | 0.260733496 | 5.77E-19 | 1.17E-17 | 153769 | 5 | 145316142 | 145461354 | 1 | protein_coding | SH3 domain containing ring finger 2 |

| | | | | | | | | | | | | | |
|---|---|---|---|---|---|---|---|---|---|---|---|---|---|
| **UQCRB** | ENSG00000156467 | 3863.342197 | -0.706376912 | 0.168766418 | 1.34E-05 | 7.25E-05 | 7381 | 8 | 97238148 | 97247862 | -1 | protein_coding | ubiquinol-cytochrome c reductase binding protein |
| **PTDSS1** | ENSG00000156471 | 3118.95557 | -0.534678086 | 0.119271839 | 4.81E-06 | 2.79E-05 | 9791 | 8 | 97273943 | 97349223 | 1 | protein_coding | phosphatidylserine synthase 1 |
| **FAM122C** | ENSG00000156500 | 99.71703145 | 2.39865557 | 0.392634503 | 7.84E-11 | 8.5E-10 | 159091 | X | 133930819 | 133988640 | 1 | protein_coding | family with sequence similarity 122C |
| **HK1** | ENSG00000156515 | 4371.527334 | -0.470794395 | 0.089346174 | 1.08E-07 | 7.96E-07 | 3098 | 10 | 71029740 | 71161638 | 1 | protein_coding | hexokinase 1 |
| **CD109** | ENSG00000156535 | 7702.672932 | -0.693055644 | 0.098275505 | 8.87E-13 | 1.13E-11 | 135228 | 6 | 74405508 | 74538040 | 1 | protein_coding | CD109 molecule |
| **UBE2L6** | ENSG00000156587 | 3621.741761 | 1.543166287 | 0.167086201 | 3.54E-21 | 8.51E-20 | 9246 | 11 | 57319129 | 57335757 | -1 | protein_coding | ubiquitin-conjugating enzyme E2L 6 |
| **ZDHHC5** | ENSG00000156599 | 5797.347658 | 0.327612678 | 0.088251282 | 1.69E-04 | 7.37E-04 | 25921 | 11 | 57435219 | 57468659 | 1 | protein_coding | zinc finger, DHHC-type containing 5 |
| **KAT6B** | ENSG00000156650 | 1426.102983 | -0.682624817 | 0.12349652 | 1.95E-08 | 1.59E-07 | 23522 | 10 | 76585340 | 76792380 | 1 | protein_coding | K(lysine) acetyltransferase 6B |
| **SAMD8** | ENSG00000156671 | 1400.382798 | 0.923413197 | 0.130703101 | 5.58E-13 | 7.3E-12 | 142891 | 10 | 76859344 | 76941881 | 1 | protein_coding | sterile alpha motif domain containing 8 |
| **RAB11FIP1** | ENSG00000156675 | 4506.738657 | -0.490641047 | 0.127369933 | 8.94E-05 | 4.13E-04 | 80223 | 8 | 37716136 | 37756985 | -1 | protein_coding | RAB11 family interacting protein 1 (class I) |
| **AIFM1** | ENSG00000156709 | 1010.145821 | -0.559867952 | 0.167962043 | 5.31E-04 | 0.002064333 | 9131 | X | 129263337 | 129299861 | -1 | protein_coding | apoptosis-inducing factor, mitochondrion-associated, 1 |
| **MAPK13** | ENSG00000156711 | 2197.891239 | -0.980835552 | 0.162885387 | 5.92E-10 | 5.82E-09 | 5603 | 6 | 36095586 | 36107842 | 1 | protein_coding | mitogen-activated protein kinase 13 |
| **ATAD2** | ENSG00000156802 | 620.1277656 | 0.744252364 | 0.185782626 | 2.87E-05 | 1.45E-04 | 29028 | 8 | 124332090 | 124428590 | -1 | protein_coding | ATPase family, AAA domain containing 2 |
| **FBXO32** | ENSG00000156804 | 527.8093761 | 1.237178697 | 0.201441052 | 1.61E-10 | 1.68E-09 | 114907 | 8 | 124510129 | 124553446 | -1 | protein_coding | F-box protein 32 |
| **FBRS** | ENSG00000156860 | 2104.903084 | 0.454998417 | 0.15496331 | 0.002399032 | 0.007930868 | 64319 | 16 | 30669752 | 30682135 | 1 | protein_coding | fibrosin |
| **HIAT1** | ENSG00000156875 | 1822.709954 | 0.946548318 | 0.102617734 | 9.38E-21 | 2.19E-19 | 64645 | 1 | 100503653 | 100548933 | 1 | protein_coding | hippocampus abundant transcript 1 |
| **SASS6** | ENSG00000156876 | 66.24363292 | 1.300639107 | 0.487729348 | 0.001095647 | 0.003964198 | 163786 | 1 | 100549119 | 100598511 | -1 | protein_coding | spindle assembly 6 homolog (C. elegans) |
| **ITGAD** | ENSG00000156886 | 27.78710652 | -2.441771933 | 0.975389289 | 9E-04 | 0.00332501 | 3681 | 16 | 31404633 | 31437826 | 1 | protein_coding | integrin, alpha D |
| **GALK2** | ENSG00000156958 | 594.9074425 | 0.422153283 | 0.158097842 | 0.005732086 | 0.017173782 | 2585 | 15 | 49447853 | 49660066 | 1 | protein_coding | galactokinase 2 |
| **B3GNT7** | ENSG00000156966 | 567.3005119 | -1.219074037 | 0.177942854 | 1.53E-12 | 1.92E-11 | 93010 | 2 | 232260254 | 232265875 | 1 | protein_coding | UDP-GlcNAc:betaGal beta-1,3-N-acetylglucosaminyltransferase 7 |
| **MPV17L** | ENSG00000156968 | 34.1171537 | -1.905246846 | 0.743394744 | 8.7E-04 | 0.003223138 | 255027 | 16 | 15489611 | 15507125 | 1 | protein_coding | MPV17 mitochondrial membrane protein-like |
| **BUB1B** | ENSG00000156970 | 104.9979847 | 1.076447442 | 0.458042363 | 0.003637287 | 0.011518578 | 701 | 15 | 40453224 | 40513337 | 1 | protein_coding | BUB1 mitotic checkpoint serine/threonine kinase B |
| **EIF4A2** | ENSG00000156976 | 5730.723747 | 0.435795697 | 0.090282231 | 1.27E-06 | 8.05E-06 | 1003021 | 3 | 186500994 | 186507689 | 1 | protein_coding | eukaryotic translation initiation factor 4A2 |
| **BRPF1** | ENSG00000156983 | 1364.995187 | -0.729652318 | 0.117304987 | 2.24E-10 | 2.3E-09 | 7862 | 3 | 9773413 | 9789702 | 1 | protein_coding | bromodomain and PHD finger containing, 1 |
| **SEC13** | ENSG00000157020 | 1445.146037 | -0.744218889 | 0.107733589 | 2.3E-12 | 2.85E-11 | 6396 | 3 | 10334815 | 10362862 | -1 | protein_coding | SEC13 homolog (S. cerevisiae) |
| **ATP2B2** | ENSG00000157087 | 601.158818 | -2.45177347 | 0.274376618 | 3.21E-20 | 7.19E-19 | 491 | 3 | 10365707 | 10749716 | -1 | protein_coding | ATPase, Ca++ transporting, plasma membrane 2 |
| **SMG1** | ENSG00000157106 | 4770.019418 | 0.3267267 | 0.106498932 | 0.0020715 | 0.00697329 | 1010605 | 16 | 18816175 | 18937776 | -1 | protein_coding | SMG1 phosphatidylinositol 3-kinase-related kinase |
| **FCHO2** | ENSG00000157107 | 7862.10656 | -0.614776983 | 0.133181759 | 2.34E-06 | 1.43E-05 | 115548 | 5 | 72251808 | 72386349 | 1 | protein_coding | FCH domain only 2 |
| **TMEM171** | ENSG00000157111 | 31.86694301 | 6.727790919 | 1.068966093 | 8.78E-11 | 9.44E-10 | 134285 | 5 | 72416119 | 72427644 | 1 | protein_coding | transmembrane protein 171 |
| **TIMP4** | ENSG00000157150 | 108.0095428 | 0.635154214 | 0.30482215 | 0.018398482 | 0.047297561 | 7079 | 3 | 12194551 | 12200851 | -1 | protein_coding | TIMP metallopeptidase inhibitor 4 |
| **C1orf27** | ENSG00000157181 | 699.6416618 | 0.547107318 | 0.158211377 | 3.41E-04 | 0.00138986 | 54953 | 1 | 186344890 | 186390510 | 1 | protein_coding | chromosome 1 open reading frame 27 |
| **NECAP2** | ENSG00000157191 | 4655.723322 | 0.291305669 | 0.099901756 | 0.003255429 | 0.010434716 | 55707 | 1 | 16767167 | 16786573 | 1 | protein_coding | NECAP endocytosis associated 2 |
| **LRP8** | ENSG00000157193 | 311.8720353 | 0.606522635 | 0.196237409 | 0.001151155 | 0.004137528 | 7804 | 1 | 53711217 | 53793742 | -1 | protein_coding | low density lipoprotein receptor-related protein 8, apolipoprotein e receptor |
| **SSBP3** | ENSG00000157216 | 3163.966974 | -1.266342542 | 0.136397876 | 3.45E-21 | 8.3E-20 | 23648 | 1 | 54691105 | 54879152 | -1 | protein_coding | single stranded DNA binding protein 3 |
| **CLDN12** | ENSG00000157224 | 950.1206359 | 0.880668826 | 0.170009663 | 8.64E-08 | 6.46E-07 | 9069 | 7 | 90013035 | 90142716 | 1 | protein_coding | claudin 12 |
| **MMP14** | ENSG00000157227 | 11766.37342 | 1.167676036 | 0.117125571 | 5E-24 | 1.42E-22 | 4323 | 14 | 23305766 | 23318236 | 1 | protein_coding | matrix metallopeptidase 14 (membrane-inserted) |
| **FZD1** | ENSG00000157240 | 516.5574771 | 0.635210128 | 0.190710766 | 4.61E-04 | 0.001819957 | 8321 | 7 | 90893783 | 90898123 | 1 | protein_coding | frizzled family receptor 1 |
| **GATAD1** | ENSG00000157259 | 1327.265108 | 0.59066058 | 0.114924323 | 1.6E-07 | 1.16E-06 | 57798 | 7 | 92076767 | 92088150 | 1 | protein_coding | GATA zinc finger domain containing 1 |
| **KIT** | ENSG00000157404 | 39.72078753 | -2.287507597 | 0.648676945 | 3.25E-05 | 1.63E-04 | 3815 | 4 | 55524085 | 55606881 | 1 | protein_coding | v-kit Hardy-Zuckerman 4 feline sarcoma viral oncogene homolog |
| **FAM81A** | ENSG00000157470 | 324.0715199 | -2.308340591 | 0.360539179 | 1.29E-11 | 1.49E-10 | 145773 | 15 | 59664892 | 59815748 | 1 | protein_coding | family with sequence similarity 81, member A |
| **MYO1E** | ENSG00000157483 | 7333.955945 | -0.723090294 | 0.13663805 | 5.5E-08 | 4.25E-07 | 4643 | 15 | 59427113 | 59665099 | -1 | protein_coding | myosin IE |
| **AFAP1L1** | ENSG00000157510 | 2738.995947 | -2.156458223 | 0.464936375 | 2.17E-07 | 1.54E-06 | 134265 | 5 | 148651434 | 148721365 | 1 | protein_coding | actin filament associated protein 1-like 1 |
| **DSCR3** | ENSG00000157538 | 2726.705046 | 0.357244026 | 0.103720397 | 4.6E-04 | 0.001817913 | 10311 | 21 | 38595721 | 38640262 | -1 | protein_coding | Down syndrome critical region gene 3 |
| **DYRK1A** | ENSG00000157540 | 2137.784049 | 0.395395759 | 0.116543107 | 5.33E-04 | 0.002070657 | 1859 | 21 | 38738092 | 38889753 | 1 | protein_coding | dual-specificity tyrosine-(Y)-phosphorylation regulated kinase 1A |
| **TSPAN18** | ENSG00000157570 | 10.19818412 | -2.341805407 | 1.763109168 | 0.009147442 | 0.025927364 | 90139 | 11 | 44748015 | 44953972 | 1 | protein_coding | tetraspanin 18 |
| **SLC35B2** | ENSG00000157593 | 524.2984801 | -0.754242242 | 0.168909715 | 3.53E-06 | 2.09E-05 | 1006161 | 6 | 44221833 | 44225291 | -1 | protein_coding | solute carrier family 35 (adenosine 3'-phospho 5'-phosphosulfate transporter), member B2 |
| **TMEM164** | ENSG00000157600 | 3468.031597 | -0.379566774 | 0.100950961 | 1.58E-04 | 6.95E-04 | 1019285 | X | 109245859 | 109425962 | 1 | protein_coding | transmembrane protein 164 |
| **MX1** | ENSG00000157601 | 11456.08719 | 4.665458724 | 0.129829828 | 4.8E-284 | 1.22E-280 | 4599 | 21 | 42792231 | 42831141 | 1 | protein_coding | myxovirus (influenza virus) resistance 1, interferon-inducible protein p78 (mouse) |
| **TAB3** | ENSG00000157625 | 1368.78879 | 0.641907514 | 0.104591329 | 4.92E-10 | 4.88E-09 | 257397 | X | 30845559 | 30993201 | -1 | protein_coding | TGF-beta activated kinase 1/MAP3K7 binding protein 3 |
| **SLC38A10** | ENSG00000157637 | 2016.171709 | -0.470542998 | 0.120311942 | 6.32E-05 | 3E-04 | 124565 | 17 | 79218800 | 79269347 | -1 | protein_coding | solute carrier family 38, member 10 |
| **PALM2-AKAP** | ENSG00000157654 | 4118.719999 | 1.665174281 | 0.123372449 | 2.25E-42 | 1.54E-40 | 445815 | 9 | 112542589 | 112934792 | 1 | protein_coding | PALM2-AKAP2 readthrough |
| **ZNF618** | ENSG00000157657 | 1930.189954 | 0.639744997 | 0.12437818 | 1.5E-07 | 1.09E-06 | 114991 | 9 | 116638562 | 116818871 | 1 | protein_coding | zinc finger protein 618 |
| **C9orf91** | ENSG00000157693 | 1406.676352 | 1.763298259 | 0.119542098 | 3.74E-50 | 3.34E-48 | 203197 | 9 | 117373486 | 117408702 | 1 | protein_coding | chromosome 9 open reading frame 91 |
| **PSMG3** | ENSG00000157778 | 299.920039 | -0.517971818 | 0.21082164 | 0.008866034 | 0.025204539 | 84262 | 7 | 1606966 | 1610641 | -1 | protein_coding | proteasome (prosome, macropain) assembly chaperone 3 |
| **AP3S2** | ENSG00000157823 | 1098.962483 | 0.33779638 | 0.140357882 | 0.013038008 | 0.035180064 | 1005267 | 15 | 90373831 | 90437574 | -1 | protein_coding | adaptor-related protein complex 3, sigma 2 subunit |
| **FMNL2** | ENSG00000157827 | 7816.51902 | 0.386779551 | 0.107383446 | 2.63E-04 | 0.001093268 | 114793 | 2 | 153191751 | 153506348 | 1 | protein_coding | formin-like 2 |
| **FAM213B** | ENSG00000157870 | 2615.190971 | -0.835826579 | 0.191618851 | 4.69E-06 | 2.73E-05 | 127281 | 1 | 2517930 | 2522908 | 1 | protein_coding | family with sequence similarity 213, member B |
| **PEX10** | ENSG00000157911 | 311.9524032 | -0.509551819 | 0.216978472 | 0.012434127 | 0.033741259 | 5192 | 1 | 2336236 | 2345236 | -1 | protein_coding | peroxisomal biogenesis factor 10 |
| **SKI** | ENSG00000157933 | 1347.135395 | -1.058401857 | 0.159730081 | 7.53E-12 | 8.96E-11 | 6497 | 1 | 2160134 | 2241558 | 1 | protein_coding | v-ski avian sarcoma viral oncogene homolog |
| **WIPI2** | ENSG00000157954 | 2510.268789 | -0.398521072 | 0.112981893 | 3.54E-04 | 0.001435191 | 26100 | 7 | 5229819 | 5273457 | 1 | protein_coding | WD repeat domain, phosphoinositide interacting 2 |
| **LDLRAP1** | ENSG00000157978 | 1118.305859 | -1.288073952 | 0.15416875 | 9.5E-18 | 1.76E-16 | 26119 | 1 | 25870071 | 25895377 | 1 | protein_coding | low density lipoprotein receptor adaptor protein 1 |
| **AGAP1** | ENSG00000157985 | 66.01459263 | -3.143521148 | 0.798304897 | 2.79E-06 | 1.68E-05 | 116987 | 2 | 236402733 | 237040444 | 1 | protein_coding | ArfGAP with GTPase domain, ankyrin repeat and PH domain 1 |
| **PAFAH2** | ENSG00000158006 | 1155.764523 | -0.625642014 | 0.14728611 | 1.16E-05 | 6.31E-05 | 5051 | 1 | 26286258 | 26324648 | -1 | protein_coding | platelet-activating factor acetylhydrolase 2, 40kDa |
| **EXTL1** | ENSG00000158008 | 4.164637629 | -0.641562155 | 1.179071508 | 0.008335721 | 0.023856708 | 2134 | 1 | 26346375 | 26362955 | 1 | protein_coding | exostosin-like glycosyltransferase 1 |
| **BRE** | ENSG00000158019 | 2154.545558 | -0.368030601 | 0.140197374 | 0.007182881 | 0.020969621 | 9577 | 2 | 28112808 | 28561768 | 1 | protein_coding | brain and reproductive organ-expressed (TNFRSF1A modulator) |
| **WDR66** | ENSG00000158023 | 28.35950295 | -1.568415093 | 0.683278861 | 0.002123981 | 0.007131108 | 144406 | 12 | 122355768 | 122441833 | 1 | protein_coding | WD repeat domain 66 |
| **DUSP2** | ENSG00000158050 | 586.4044023 | -0.620712698 | 0.247738388 | 0.006460368 | 0.01911521 | 1844 | 2 | 96808905 | 96811179 | -1 | protein_coding | dual specificity phosphatase 2 |
| **UBXN11** | ENSG00000158062 | 1100.208092 | -2.144888188 | 0.182899857 | 1.04E-32 | 4.66E-31 | 91544 | 1 | 26607819 | 26644854 | -1 | protein_coding | UBX domain protein 11 |
| **NCK1** | ENSG00000158092 | 3151.897059 | 0.676712241 | 0.160444074 | 1.31E-05 | 7.07E-05 | 4690 | 3 | 136581050 | 136668665 | 1 | protein_coding | NCK adaptor protein 1 |
| **HPD** | ENSG00000158104 | 3.579521569 | 0.384877686 | 0.812490862 | 0.008153576 | 0.023418739 | 3242 | 12 | 122277433 | 122301502 | -1 | protein_coding | 4-hydroxyphenylpyruvate dioxygenase |
| **RHPN1** | ENSG00000158106 | 114.1367704 | -1.65553236 | 0.361781851 | 5.85E-07 | 3.91E-06 | 114822 | 8 | 144451057 | 144466390 | 1 | protein_coding | rhophilin, Rho GTPase binding protein 1 |
| **TPRG1L** | ENSG00000158109 | 1084.921522 | -0.584877348 | 0.158380261 | 1.28E-04 | 5.76E-04 | 127262 | 1 | 3541566 | 3546691 | 1 | protein_coding | tumor protein p63 regulated 1-like |
| **AAED1** | ENSG00000158122 | 694.7150783 | 0.669310383 | 0.163768019 | 2.36E-05 | 1.21E-04 | 195827 | 9 | 99401859 | 99417585 | -1 | protein_coding | AhpC/TSA antioxidant enzyme domain containing 1 |
| **MRAS** | ENSG00000158186 | 7076.460827 | -0.233185173 | 0.087461221 | 0.00495491 | 0.01515427 | 22808 | 3 | 138066539 | 138124375 | 1 | protein_coding | muscle RAS oncogene homolog |
| **ABHD3** | ENSG00000158201 | 673.8490295 | 0.47538522 | 0.153348127 | 0.001378266 | 0.00484788 | 171586 | 18 | 19230858 | 19284766 | -1 | protein_coding | abhydrolase domain containing 3 |
| **COLEC12** | ENSG00000158270 | 259.4054442 | -4.635761924 | 0.474074284 | 1.04E-23 | 2.89E-22 | 81035 | 18 | 319361 | 500722 | -1 | protein_coding | collectin sub-family member 12 |
| **CUL4B** | ENSG00000158290 | 2148.18845 | 0.646827317 | 0.153103856 | 1.45E-05 | 7.76E-05 | 8450 | X | 119658464 | 119709649 | -1 | protein_coding | cullin 4B |
| **GPR153** | ENSG00000158292 | 83.45627338 | -0.903483667 | 0.389474376 | 0.005874278 | 0.017538929 | 387509 | 1 | 6307406 | 6321035 | -1 | protein_coding | G protein-coupled receptor 153 |
| **GPRASP2** | ENSG00000158301 | 186.4263128 | -1.096798752 | 0.28937828 | 3.36E-05 | 1.68E-04 | 1005280 | X | 101967104 | 101973607 | 1 | protein_coding | G protein-coupled receptor associated sorting protein 2 |
| **SHROOM4** | ENSG00000158352 | 251.03177 | -0.960489813 | 0.307210572 | 4.9E-04 | 0.00192167 | 57477 | X | 50334647 | 50557302 | -1 | protein_coding | shroom family member 4 |
| **HIST1H2BD** | ENSG00000158373 | 713.7173429 | 0.898797386 | 0.199992835 | 2.26E-06 | 1.38E-05 | 3017 | 6 | 26158349 | 26171577 | 1 | protein_coding | histone cluster 1, H2bd |
| **HIST1H4H** | ENSG00000158406 | 267.863879 | -0.51492031 | 0.229927681 | 0.015836826 | 0.041686913 | 554313 | 6 | 26281283 | 26285762 | -1 | protein_coding | histone cluster 1, H4h |
| **MITD1** | ENSG00000158411 | 502.1637146 | 0.84115442 | 0.207031456 | 1.84E-05 | 9.65E-05 | 129531 | 2 | 99777890 | 99797521 | -1 | protein_coding | MIT, microtubule interacting and transport, domain containing 1 |
| **TSPAN33** | ENSG00000158457 | 568.0069339 | 1.048769687 | 0.192542897 | 1.32E-08 | 1.1E-07 | 340348 | 7 | 128784712 | 128808671 | 1 | protein_coding | tetraspanin 33 |
| **AHCYL2** | ENSG00000158467 | 671.1674957 | -0.454261692 | 0.192856393 | 0.012527361 | 0.033958081 | 23382 | 7 | 128864864 | 129070052 | 1 | protein_coding | adenosylhomocysteinase-like 2 |
| **B4GALT5** | ENSG00000158470 | 8288.698833 | 1.074821075 | 0.098707015 | 3.59E-28 | 1.25E-26 | 9334 | 20 | 48249482 | 48330415 | -1 | protein_coding | UDP-Gal:betaGlcNAc beta 1,4- galactosyltransferase, polypeptide 5 |
| **CD1A** | ENSG00000158477 | 43.44143566 | -4.006305004 | 1.020905777 | 8.31E-06 | 4.63E-05 | 909 | 1 | 158223927 | 158228059 | 1 | protein_coding | CD1a molecule |
| **CD1E** | ENSG00000158488 | 14.62561038 | -3.931791408 | 1.570922692 | 0.002167801 | 0.007255912 | 913 | 1 | 158323254 | 158327343 | 1 | protein_coding | CD1e molecule |
| **NCF1** | ENSG00000158517 | 740.0915539 | 3.012008901 | 0.248708914 | 5.73E-35 | 2.89E-33 | 653361 | 7 | 74188309 | 74203659 | 1 | protein_coding | neutrophil cytosolic factor 1 |

| | | | | | | | | | | | | | |
|---|---|---|---|---|---|---|---|---|---|---|---|---|---|
| **GDPD5** | ENSG00000158555 | 183.2022957 | -0.760278685 | 0.287787143 | 0.003367168 | 0.01075453 | 81544 | 11 | 75145685 | 75236948 | -1 | protein_coding | glycerophosphodiester phosphodiesterase domain containing 5 |
| **PPP1R15B** | ENSG00000158615 | 2618.320896 | 0.826818598 | 0.101627782 | 1.65E-16 | 2.78E-15 | 84919 | 1 | 204372515 | 204380919 | -1 | protein_coding | protein phosphatase 1, regulatory subunit 15B |
| **ELK4** | ENSG00000158711 | 3659.774168 | 0.346884504 | 0.111080516 | 0.001454087 | 0.005097262 | 2005 | 1 | 205577071 | 205601090 | -1 | protein_coding | ELK4, ETS-domain protein (SRF accessory protein 1) |
| **SLAMF8** | ENSG00000158714 | 14187.05138 | -1.407302125 | 0.150431267 | 7.73E-22 | 1.94E-20 | 56833 | 1 | 159796540 | 159807039 | 1 | protein_coding | SLAM family member 8 |
| **SLC45A3** | ENSG00000158715 | 343.3344732 | -2.80282658 | 0.35018563 | 9.35E-17 | 1.6E-15 | 85414 | 1 | 205626979 | 205649587 | -1 | protein_coding | solute carrier family 45, member 3 |
| **DUSP23** | ENSG00000158716 | 358.7335418 | -1.086196328 | 0.249751508 | 3.31E-06 | 1.97E-05 | 54935 | 1 | 159750722 | 159752333 | 1 | protein_coding | dual specificity phosphatase 23 |
| **RNF166** | ENSG00000158717 | 722.3143088 | -1.948035676 | 0.190523219 | 1.83E-25 | 5.57E-24 | 115992 | 16 | 88762903 | 88772829 | -1 | protein_coding | ring finger protein 166 |
| **NBL1** | ENSG00000158747 | 318.1006146 | -0.700071807 | 0.29266501 | 0.007512897 | 0.02180395 | 1005327 | 1 | 19967048 | 19984945 | 1 | protein_coding | neuroblastoma 1, DAN family BMP antagonist |
| **F11R** | ENSG00000158769 | 5655.188224 | 0.405431614 | 0.15405572 | 0.00643084 | 0.019042729 | 50848 | 1 | 160965001 | 160991138 | -1 | protein_coding | F11 receptor |
| **USF1** | ENSG00000158773 | 1238.810201 | 0.71632508 | 0.147865119 | 5.83E-07 | 3.91E-06 | 7391 | 1 | 161009041 | 161015767 | -1 | protein_coding | upstream transcription factor 1 |
| **CDA** | ENSG00000158825 | 959.8660827 | -1.908718111 | 0.153271571 | 1.79E-36 | 9.42E-35 | 978 | 1 | 20915441 | 20945401 | 1 | protein_coding | cytidine deaminase |
| **PINK1** | ENSG00000158828 | 2095.124206 | -1.209254025 | 0.141875784 | 3.5E-18 | 6.72E-17 | 65018 | 1 | 20959948 | 20978004 | 1 | protein_coding | PTEN induced putative kinase 1 |
| **FCER1G** | ENSG00000158869 | 13842.69784 | 0.611382032 | 0.097848922 | 2.44E-10 | 2.5E-09 | 2207 | 1 | 161185024 | 161190489 | 1 | protein_coding | Fc fragment of IgE, high affinity I, receptor for; gamma polypeptide |
| **TOMM40L** | ENSG00000158882 | 790.0202551 | -0.311147794 | 0.13613804 | 0.019342298 | 0.049311987 | 84134 | 1 | 161195793 | 161200408 | 1 | protein_coding | translocase of outer mitochondrial membrane 40 homolog (yeast)-like |
| **CDC42SE2** | ENSG00000158985 | 4310.686177 | 0.81512847 | 0.127729271 | 7.01E-11 | 7.63E-10 | 56990 | 5 | 130581186 | 130734140 | 1 | protein_coding | CDC42 small effector 2 |
| **ALG8** | ENSG00000159063 | 1051.694155 | -0.406294166 | 0.14546014 | 0.003992016 | 0.012494365 | 79053 | 11 | 77811982 | 77850706 | -1 | protein_coding | ALG8, alpha-1,3-glucosyltransferase |
| **IFNAR2** | ENSG00000159110 | 2248.052703 | 0.887440161 | 0.140818959 | 1.31E-10 | 1.38E-09 | 3455 | 21 | 34602206 | 34637980 | 1 | protein_coding | interferon (alpha, beta and omega) receptor 2 |
| **GART** | ENSG00000159131 | 1510.819348 | 0.748733253 | 0.162579153 | 1.92E-06 | 1.18E-05 | 2618 | 21 | 34876238 | 34915797 | -1 | protein_coding | phosphoribosylglycinamide formyltransferase, phosphoribosylglycinamide synthetase, phosphoribosyla |
| **LAD1** | ENSG00000159166 | 40.42153262 | 2.691490311 | 0.678421871 | 3.54E-06 | 2.09E-05 | 3898 | 1 | 201342372 | 201368736 | -1 | protein_coding | ladinin 1 |
| **CSRP1** | ENSG00000159176 | 3039.370444 | -0.834378361 | 0.096144252 | 2.72E-18 | 5.25E-17 | 1465 | 1 | 201452658 | 201478584 | -1 | protein_coding | cysteine and glycine-rich protein 1 |
| **ATP5G1** | ENSG00000159199 | 355.9333007 | -0.965745344 | 0.224944954 | 5.74E-06 | 3.29E-05 | 516 | 17 | 46970127 | 46973233 | 1 | protein_coding | ATP synthase, H+ transporting, mitochondrial Fo complex, subunit C1 (subunit 9) |
| **RCAN1** | ENSG00000159200 | 1342.635655 | 1.072662216 | 0.1226433 | 6.15E-19 | 1.24E-17 | 1827 | 21 | 35885440 | 35987441 | -1 | protein_coding | regulator of calcineurin 1 |
| **UBE2Z** | ENSG00000159202 | 5449.807849 | 1.068761854 | 0.087320785 | 3.43E-35 | 1.74E-33 | 65264 | 17 | 46985731 | 47006418 | 1 | protein_coding | ubiquitin-conjugating enzyme E2Z |
| **SNF8** | ENSG00000159210 | 932.0049075 | -0.634166997 | 0.137749953 | 2.29E-06 | 1.4E-05 | 11267 | 17 | 47006678 | 47022479 | -1 | protein_coding | SNF8, ESCRT-II complex subunit |
| **CLIC6** | ENSG00000159212 | 264.7213921 | -0.904811131 | 0.263206632 | 1.74E-04 | 7.56E-04 | 54102 | 21 | 36041688 | 36090525 | 1 | protein_coding | chloride intracellular channel 6 |
| **CBR1** | ENSG00000159228 | 1677.131028 | 0.740840915 | 0.124766365 | 1.56E-09 | 1.47E-08 | 873 | 21 | 37442239 | 37445464 | 1 | protein_coding | carbonyl reductase 1 |
| **CBR3** | ENSG00000159231 | 163.5705568 | 0.825233878 | 0.271472135 | 8.73E-04 | 0.003232041 | 874 | 21 | 37507210 | 37518864 | 1 | protein_coding | carbonyl reductase 3 |
| **MORC3** | ENSG00000159256 | 2377.462233 | 1.232938375 | 0.147585114 | 1.35E-17 | 2.48E-16 | 23515 | 21 | 37692487 | 37758446 | 1 | protein_coding | MORC family CW-type zinc finger 3 |
| **CHAF1B** | ENSG00000159259 | 57.12987479 | 1.055637097 | 0.49420849 | 0.006424997 | 0.019037074 | 8208 | 21 | 37757676 | 37791313 | 1 | protein_coding | chromatin assembly factor 1, subunit B (p60) |
| **HLCS** | ENSG00000159267 | 538.4576858 | -0.428774935 | 0.157128122 | 0.004931498 | 0.015100798 | 3141 | 21 | 38123189 | 38362536 | -1 | protein_coding | holocarboxylase synthetase (biotin-(proprionyl-CoA-carboxylase (ATP-hydrolysing)) ligase) |
| **ADPGK** | ENSG00000159322 | 4063.110905 | 1.011694541 | 0.090965314 | 2.84E-29 | 1.06E-27 | 83440 | 15 | 73043710 | 73078187 | -1 | protein_coding | ADP-dependent glucokinase |
| **CYB5R1** | ENSG00000159348 | 2005.502997 | -0.679849969 | 0.109309083 | 2.56E-10 | 2.6E-09 | 51706 | 1 | 202930997 | 202936408 | -1 | protein_coding | cytochrome b5 reductase 1 |
| **M1AP** | ENSG00000159374 | 89.45839774 | -2.222066818 | 0.408985591 | 5.37E-09 | 4.73E-08 | 130951 | 2 | 74785010 | 74875465 | -1 | protein_coding | meiosis 1 associated protein |
| **HK2** | ENSG00000159399 | 22078.61884 | -0.373842571 | 0.112636658 | 7.26E-04 | 0.002737895 | 3099 | 2 | 75061108 | 75120486 | 1 | protein_coding | hexokinase 2 |
| **ALDH4A1** | ENSG00000159423 | 371.7739577 | -2.080600829 | 0.417195915 | 3.45E-08 | 2.73E-07 | 8659 | 1 | 19197926 | 19229275 | -1 | protein_coding | aldehyde dehydrogenase 4 family, member A1 |
| **STARD9** | ENSG00000159433 | 300.4451436 | -1.126693833 | 0.300054439 | 3.64E-05 | 1.8E-04 | 57519 | 15 | 42867857 | 43013179 | 1 | protein_coding | StAR-related lipid transfer (START) domain containing 9 |
| **THEM4** | ENSG00000159445 | 397.0386832 | -0.495569362 | 0.207919079 | 0.011411499 | 0.031372898 | 117145 | 1 | 151846060 | 151882284 | -1 | protein_coding | thioesterase superfamily member 4 |
| **AMFR** | ENSG00000159461 | 6049.62139 | -0.451022939 | 0.130381494 | 3.54E-04 | 0.001437914 | 267 | 16 | 56395364 | 56459450 | -1 | protein_coding | autocrine motility factor receptor, E3 ubiquitin protein ligase |
| **RSPRY1** | ENSG00000159579 | 957.1034984 | 0.359242578 | 0.143609032 | 0.010249936 | 0.028580508 | 89970 | 16 | 57220049 | 57274387 | 1 | protein_coding | ring finger and SPRY domain containing 1 |
| **GPBP1L1** | ENSG00000159592 | 4206.195239 | 0.782836348 | 0.116761852 | 9.04E-12 | 1.06E-10 | 60313 | 1 | 46092976 | 46153785 | -1 | protein_coding | GC-rich promoter binding protein 1-like 1 |
| **NAE1** | ENSG00000159593 | 616.6589487 | 0.73424723 | 0.189633382 | 4.91E-05 | 2.37E-04 | 8883 | 16 | 66836778 | 66907159 | -1 | protein_coding | NEDD8 activating enzyme E1 subunit 1 |
| **ACE** | ENSG00000159640 | 1735.822341 | -1.878506871 | 0.199661361 | 5.46E-22 | 1.38E-20 | 1636 | 17 | 61554422 | 61599205 | 1 | protein_coding | angiotensin I converting enzyme |
| **SPON2** | ENSG00000159674 | 1411.290505 | -1.48970161 | 0.54695507 | 7.43E-04 | 0.002795176 | 1001308 | 4 | 1160720 | 1202750 | -1 | protein_coding | spondin 2, extracellular matrix protein |
| **CTBP1** | ENSG00000159692 | 2086.612604 | -0.504880496 | 0.118013766 | 1.37E-05 | 7.39E-05 | 1487 | 4 | 1205236 | 1243741 | -1 | protein_coding | C-terminal binding protein 1 |
| **ATP6V0D1** | ENSG00000159720 | 4352.59654 | -0.923832858 | 0.121464464 | 8.34E-15 | 1.25E-13 | 9114 | 16 | 67471917 | 67515140 | -1 | protein_coding | ATPase, H+ transporting, lysosomal 38kDa, V0 subunit d1 |
| **FAM131B** | ENSG00000159784 | 615.4295991 | -1.804855103 | 0.188989324 | 1.45E-22 | 3.76E-21 | 9715 | 7 | 143050493 | 143059863 | -1 | protein_coding | family with sequence similarity 131, member B |
| **RGS12** | ENSG00000159788 | 2339.886759 | -1.029977899 | 0.154821047 | 7.99E-12 | 9.46E-11 | 6002 | 4 | 3294755 | 3441640 | 1 | protein_coding | regulator of G-protein signaling 12 |
| **PSKH1** | ENSG00000159792 | 708.0083279 | -0.693602929 | 0.149369114 | 1.86E-06 | 1.15E-05 | 5681 | 16 | 67927175 | 67963581 | 1 | protein_coding | protein serine kinase H1 |
| **ZYX** | ENSG00000159840 | 15435.98712 | -0.887326304 | 0.093905597 | 1.57E-21 | 3.83E-20 | 7791 | 7 | 143078173 | 143088204 | 1 | protein_coding | zyxin |
| **ABR** | ENSG00000159842 | 12582.01344 | -0.768529027 | 0.093273597 | 7.61E-17 | 1.31E-15 | 29 | 17 | 906758 | 1132315 | -1 | protein_coding | active BCR-related |
| **CCDC117** | ENSG00000159873 | 701.9711004 | 1.186746327 | 0.181036323 | 1.21E-11 | 1.41E-10 | 150275 | 22 | 29168662 | 29185283 | 1 | protein_coding | coiled-coil domain containing 117 |
| **PTGIR** | ENSG00000160013 | 209.5473482 | 2.358173877 | 0.277239952 | 1.57E-18 | 3.1E-17 | 5739 | 19 | 47123725 | 47128375 | -1 | protein_coding | prostaglandin I2 (prostacyclin) receptor (IP) |
| **CALM3** | ENSG00000160014 | 15031.6705 | -1.229522584 | 0.1807913 | 1.63E-12 | 2.03E-11 | 808 | 19 | 47104331 | 47114050 | 1 | protein_coding | calmodulin 3 (phosphorylase kinase, delta) |
| **DFFA** | ENSG00000160049 | 1100.121568 | -0.375486966 | 0.137116912 | 0.005227134 | 0.015863043 | 1676 | 1 | 10516579 | 10532583 | -1 | protein_coding | DNA fragmentation factor, 45kDa, alpha polypeptide |
| **BSDC1** | ENSG00000160058 | 1341.82816 | 0.355782697 | 0.118253898 | 0.002494736 | 0.008208191 | 55108 | 1 | 32830704 | 32860332 | -1 | protein_coding | BSD domain containing 1 |
| **SSU72** | ENSG00000160075 | 2189.293931 | -0.245137254 | 0.105000293 | 0.017777911 | 0.045949572 | 29101 | 1 | 1477053 | 1510249 | -1 | protein_coding | SSU72 RNA polymerase II CTD phosphatase homolog (S. cerevisiae) |
| **ZNF362** | ENSG00000160094 | 664.866127 | -0.707497426 | 0.155132186 | 2.43E-06 | 1.48E-05 | 149076 | 1 | 33722146 | 33766320 | 1 | protein_coding | zinc finger protein 362 |
| **ABCG1** | ENSG00000160179 | 133.8737631 | -1.086603896 | 0.35500152 | 4.8E-04 | 0.001888065 | 9619 | 21 | 43619799 | 43717354 | 1 | protein_coding | ATP-binding cassette, sub-family G (WHITE), member 1 |
| **SLC37A1** | ENSG00000160190 | 662.4661639 | 0.630079101 | 0.155001964 | 2.79E-05 | 1.42E-04 | 54020 | 21 | 43916118 | 44001550 | 1 | protein_coding | solute carrier family 37 (glucose-6-phosphate transporter), member 1 |
| **NDUFV3** | ENSG00000160194 | 1231.007858 | -0.737408967 | 0.144621508 | 1.7E-07 | 1.22E-06 | 4731 | 21 | 44299754 | 44333414 | 1 | protein_coding | NADH dehydrogenase (ubiquinone) flavoprotein 3, 10kDa |
| **CBS** | ENSG00000160200 | 437.7045892 | -0.475776367 | 0.209745766 | 0.015570925 | 0.041072928 | 875 | 21 | 44473301 | 44497053 | -1 | protein_coding | cystathionine-beta-synthase |
| **U2AF1** | ENSG00000160201 | 1092.458877 | 0.322497251 | 0.125096458 | 0.008466838 | 0.024169103 | 7307 | 21 | 44513066 | 44527697 | -1 | protein_coding | U2 small nuclear RNA auxiliary factor 1 |
| **PDXK** | ENSG00000160209 | 8954.427011 | -0.349552632 | 0.09973857 | 3.75E-04 | 0.001515918 | 1019284 | 21 | 45138975 | 45182188 | 1 | protein_coding | pyridoxal (pyridoxine, vitamin B6) kinase |
| **CSTB** | ENSG00000160213 | 17154.71221 | -0.924282569 | 0.168511784 | 1.36E-08 | 1.14E-07 | 1476 | 21 | 45192393 | 45196326 | -1 | protein_coding | cystatin B (stefin B) |
| **RRP1** | ENSG00000160214 | 784.9272977 | -0.442604386 | 0.144313758 | 0.001593293 | 0.005526621 | 8568 | 21 | 45209394 | 45225174 | 1 | protein_coding | ribosomal RNA processing 1 |
| **AGPAT3** | ENSG00000160216 | 4351.287534 | 0.573850971 | 0.119738282 | 1.04E-06 | 6.67E-06 | 56894 | 21 | 45285067 | 45406417 | 1 | protein_coding | 1-acylglycerol-3-phosphate O-acyltransferase 3 |
| **C21orf33** | ENSG00000160221 | 1379.082129 | -0.407328358 | 0.113379974 | 2.4E-04 | 0.001008764 | 8209 | 21 | 45553487 | 45565605 | 1 | protein_coding | chromosome 21 open reading frame 33 |
| **LRRC3** | ENSG00000160233 | 65.65138859 | 0.82879874 | 0.384856787 | 0.010341719 | 0.028794406 | 81543 | 21 | 45875369 | 45878739 | 1 | protein_coding | leucine rich repeat containing 3 |
| **ITGB2** | ENSG00000160255 | 57085.0511 | -0.876710002 | 0.140975372 | 1.82E-10 | 1.89E-09 | 3689 | 21 | 46305868 | 46351904 | -1 | protein_coding | integrin, beta 2 (complement component 3 receptor 3 and 4 subunit) |
| **RALGDS** | ENSG00000160271 | 2016.954283 | -0.395541147 | 0.143149511 | 0.004440825 | 0.013735909 | 5900 | 9 | 135973107 | 136039301 | -1 | protein_coding | ral guanine nucleotide dissociation stimulator |
| **SPATC1L** | ENSG00000160284 | 46.79961829 | -1.333887965 | 0.588090787 | 0.003085139 | 0.009951493 | 84221 | 21 | 47581062 | 47604390 | -1 | protein_coding | spermatogenesis and centriole associated 1-like |
| **LSS** | ENSG00000160285 | 2478.284132 | 1.781970311 | 0.158493103 | 2.88E-30 | 1.13E-28 | 4047 | 21 | 47608055 | 47648738 | -1 | protein_coding | lanosterol synthase (2,3-oxidosqualene-lanosterol cyclase) |
| **DIP2A** | ENSG00000160305 | 1161.765533 | -0.538967967 | 0.119316825 | 3.92E-06 | 2.3E-05 | 23181 | 21 | 47878812 | 47989926 | 1 | protein_coding | DIP2 disco-interacting protein 2 homolog A (Drosophila) |
| **ST6GALNAC** | ENSG00000160408 | 1788.35157 | -1.12668592 | 0.156201643 | 1.27E-13 | 1.74E-12 | 30815 | 9 | 130647600 | 130667687 | -1 | protein_coding | ST6 (alpha-N-acetyl-neuraminyl-2,3-beta-galactosyl-1,3)-N-acetylgalactosaminide alpha-2,6-sialyltrans |
| **RDH13** | ENSG00000160439 | 138.4064173 | -0.771025525 | 0.317712857 | 0.005866404 | 0.017524677 | 112724 | 19 | 55550476 | 55582659 | -1 | protein_coding | retinol dehydrogenase 13 (all-trans/9-cis) |
| **SPTBN4** | ENSG00000160460 | 50.13736427 | -2.408045518 | 0.732472709 | 5.75E-05 | 2.75E-04 | 57731 | 19 | 40972148 | 41082370 | 1 | protein_coding | spectrin, beta, non-erythrocytic 4 |
| **SIK3** | ENSG00000160584 | 1332.576685 | 1.233418703 | 0.192169576 | 3.02E-11 | 3.41E-10 | 23387 | 11 | 116714118 | 116969153 | -1 | protein_coding | SIK family kinase 3 |
| **MPZL3** | ENSG00000160588 | 567.5636172 | -0.764113753 | 0.200707261 | 6.02E-05 | 2.86E-04 | 196264 | 11 | 118097409 | 118123065 | -1 | protein_coding | myelin protein zero-like 3 |
| **AMICA1** | ENSG00000160593 | 4554.850168 | -2.674602124 | 0.137448852 | 8.77E-86 | 2.35E-83 | 120425 | 11 | 118064455 | 118095809 | -1 | protein_coding | adhesion molecule, interacts with CXADR antigen 1 |
| **SAFB** | ENSG00000160633 | 1552.6756 | 0.443942179 | 0.140653218 | 0.001179811 | 0.004225697 | 6294 | 19 | 5623046 | 5668489 | 1 | protein_coding | scaffold attachment factor B |
| **CHTOP** | ENSG00000160679 | 981.7600361 | 0.312577412 | 0.123887989 | 0.010143847 | 0.02835194 | 26097 | 1 | 153606525 | 153618782 | 1 | protein_coding | chromatin target of PRMT1 |
| **ADAR** | ENSG00000160710 | 14715.81938 | 1.603401239 | 0.103286569 | 3.5E-55 | 3.85E-53 | 103 | 1 | 154554538 | 154600475 | -1 | protein_coding | adenosine deaminase, RNA-specific |
| **FDPS** | ENSG00000160752 | 1918.105701 | -0.391625292 | 0.110904536 | 3.22E-04 | 0.001321083 | 2224 | 1 | 155278539 | 155290457 | 1 | protein_coding | farnesyl diphosphate synthase |
| **RUSC1** | ENSG00000160753 | 228.2398355 | 1.404205978 | 0.231264386 | 2.15E-10 | 2.2E-09 | 23623 | 1 | 155290687 | 155300905 | 1 | protein_coding | RUN and SH3 domain containing 1 |
| **CCR5** | ENSG00000160791 | 4290.662215 | 0.823758596 | 0.099417051 | 5.41E-17 | 9.46E-16 | 1234 | 3 | 46411633 | 46417697 | 1 | protein_coding | chemokine (C-C motif) receptor 5 (gene/pseudogene) |
| **NBEAL2** | ENSG00000160796 | 5121.05817 | -1.279831434 | 0.108414217 | 7.56E-33 | 3.43E-31 | 23218 | 3 | 47021173 | 47051193 | 1 | protein_coding | neurobeachin-like 2 |
| **CCDC12** | ENSG00000160799 | 493.0029328 | -0.563327297 | 0.171849263 | 6.39E-04 | 0.002433271 | 151903 | 3 | 46963216 | 47023500 | -1 | protein_coding | coiled-coil domain containing 12 |
| **HK3** | ENSG00000160883 | 10469.10747 | -0.984715373 | 0.13588006 | 1.08E-13 | 1.49E-12 | 3101 | 5 | 176307870 | 176326333 | -1 | protein_coding | hexokinase 3 (white cell) |
| **LY6E** | ENSG00000160932 | 2261.072498 | 2.732344392 | 0.124203219 | 1.46E-108 | 5.46E-106 | 4061 | 8 | 144099399 | 144105249 | 1 | protein_coding | lymphocyte antigen 6 complex, locus E |

| | | | | | | | | | | | | | |
|---|---|---|---|---|---|---|---|---|---|---|---|---|---|
| VPS28 | ENSG00000160948 | 860.8800898 | -0.425062792 | 0.135131907 | 0.00121275 | 0.004330482 | 51160 | 8 | 145649000 | 145653931 | -1 | protein_coding | vacuolar protein sorting 28 homolog (S. cerevisiae) |
| MUM1 | ENSG00000160953 | 447.8885415 | -0.890085996 | 0.170446885 | 5.88E-08 | 4.52E-07 | 84939 | 19 | 1285890 | 1378430 | 1 | protein_coding | melanoma associated antigen (mutated) 1 |
| PPP1R16A | ENSG00000160972 | 194.0923052 | -0.607373282 | 0.273296204 | 0.014019727 | 0.037484866 | 84988 | 8 | 145703352 | 145727504 | 1 | protein_coding | protein phosphatase 1, regulatory subunit 16A |
| SQSTM1 | ENSG00000161011 | 1246.341815 | 1.105190542 | 0.204736795 | 1.61E-08 | 1.33E-07 | 8878 | 5 | 179233388 | 179265078 | 1 | protein_coding | sequestosome 1 |
| RPL8 | ENSG00000161016 | 5357.271692 | -0.742732377 | 0.121065235 | 5.47E-10 | 5.39E-09 | 6132 | 8 | 146015150 | 146017972 | -1 | protein_coding | ribosomal protein L8 |
| PSMC2 | ENSG00000161057 | 3625.466782 | 0.380308771 | 0.10860754 | 3.28E-04 | 0.001342168 | 5701 | 7 | 102984701 | 103009842 | 1 | protein_coding | proteasome (prosome, macropain) 26S subunit, ATPase, 2 |
| MFSD12 | ENSG00000161091 | 14309.64674 | -0.687244447 | 0.14664291 | 1.37E-06 | 8.59E-06 | 126321 | 19 | 3538259 | 3574288 | -1 | protein_coding | major facilitator superfamily domain containing 12 |
| DVL3 | ENSG00000161202 | 1793.613467 | 0.322043316 | 0.122698691 | 0.008156903 | 0.023423892 | 1857 | 3 | 183873176 | 183891398 | 1 | protein_coding | dishevelled segment polarity protein 3 |
| AP2M1 | ENSG00000161203 | 6106.88408 | -0.347786899 | 0.085827465 | 4.36E-05 | 2.12E-04 | 1173 | 3 | 183892477 | 183901879 | 1 | protein_coding | adaptor-related protein complex 2, mu 1 subunit |
| BDH1 | ENSG00000161267 | 231.0279907 | -0.97704348 | 0.253174823 | 3.27E-05 | 1.64E-04 | 622 | 3 | 197236654 | 197300194 | -1 | protein_coding | 3-hydroxybutyrate dehydrogenase, type 1 |
| PLXDC1 | ENSG00000161381 | 23.35715377 | -3.58699343 | 1.7374712 | 0.003763645 | 0.011859751 | 57125 | 17 | 37219556 | 37310647 | -1 | protein_coding | plexin domain containing 1 |
| PGAP3 | ENSG00000161395 | 296.3904666 | -0.867671864 | 0.233354548 | 7.23E-05 | 3.39E-04 | 93210 | 17 | 37827375 | 37853050 | -1 | protein_coding | post-GPI attachment to proteins 3 |
| IKZF3 | ENSG00000161405 | 470.364039 | 0.637562682 | 0.222574543 | 0.002165148 | 0.007248622 | 22806 | 17 | 37921198 | 38020441 | -1 | protein_coding | IKAROS family zinc finger 3 (Aiolos) |
| SAP30BP | ENSG00000161526 | 1123.438246 | 0.478110281 | 0.14596818 | 7.57E-04 | 0.002843621 | 29115 | 17 | 73663196 | 73704142 | 1 | protein_coding | SAP30 binding protein |
| ACOX1 | ENSG00000161533 | 3844.355817 | -0.403680581 | 0.13137247 | 0.001685693 | 0.005810203 | 51 | 17 | 73937588 | 73975515 | -1 | protein_coding | acyl-CoA oxidase 1, palmitoyl |
| PRPSAP1 | ENSG00000161542 | 596.400624 | 0.482318246 | 0.159519504 | 0.001717114 | 0.005897227 | 5635 | 17 | 74305567 | 74380602 | -1 | protein_coding | phosphoribosyl pyrophosphate synthetase-associated protein 1 |
| CYGB | ENSG00000161544 | 782.8565271 | -1.189792968 | 0.213664005 | 5.62E-09 | 4.93E-08 | 114757 | 17 | 74523438 | 74547257 | -1 | protein_coding | cytoglobin |
| SRSF2 | ENSG00000161547 | 2490.078057 | 0.425897239 | 0.114296564 | 1.47E-04 | 6.49E-04 | 693221 | 17 | 74730197 | 74733456 | -1 | protein_coding | serine/arginine-rich splicing factor 2 |
| SIGLEC11 | ENSG00000161640 | 95.36225381 | -1.410781169 | 0.429387624 | 1.49E-04 | 6.58E-04 | 114132 | 19 | 50452242 | 50464429 | -1 | protein_coding | sialic acid binding Ig-like lectin 11 |
| ZNF385A | ENSG00000161642 | 8854.993568 | -0.641118509 | 0.109969518 | 3.03E-09 | 2.74E-08 | 25946 | 12 | 54762917 | 54785082 | -1 | protein_coding | zinc finger protein 385A |
| SIGLEC16 | ENSG00000161643 | 57.42069054 | -1.304315378 | 0.537848048 | 0.002225728 | 0.00742217 | NA | 19 | 50472857 | 50479071 | 1 | pseudogene | sialic acid binding Ig-like lectin 16 (gene/pseudogene) |
| LSM 12.00 | ENSG00000161654 | 1387.192237 | 0.305233499 | 0.129834401 | 0.016587766 | 0.043357419 | 124801 | 17 | 42112003 | 42144987 | -1 | protein_coding | LSM12 homolog (S. cerevisiae) |
| EMC10 | ENSG00000161671 | 2400.4725 | -0.303841997 | 0.13262699 | 0.016999294 | 0.044258978 | 284361 | 19 | 50979657 | 50986608 | 1 | protein_coding | ER membrane protein complex subunit 10 |
| SHANK1 | ENSG00000161681 | 55.34552137 | -1.010177472 | 0.57504309 | 0.015491205 | 0.040896887 | 50944 | 19 | 51165084 | 51222707 | -1 | protein_coding | SH3 and multiple ankyrin repeat domains 1 |
| DBF4B | ENSG00000161692 | 130.1752445 | 1.035121479 | 0.307852807 | 1.99E-04 | 8.49E-04 | 80174 | 17 | 42785976 | 42829632 | 1 | protein_coding | DBF4 homolog B (S. cerevisiae) |
| FMNL3 | ENSG00000161791 | 6100.452913 | 0.299415376 | 0.085809107 | 4.3E-04 | 0.001709789 | 91010 | 12 | 50031724 | 50101948 | -1 | protein_coding | formin-like 3 |
| LARP4 | ENSG00000161813 | 3041.574219 | 0.508941882 | 0.104950155 | 8.11E-07 | 5.3E-06 | 113251 | 12 | 50786166 | 50873787 | 1 | protein_coding | La ribonucleoprotein domain family, member 4 |
| LEMD2 | ENSG00000161904 | 1042.133368 | 0.321685869 | 0.12734446 | 0.009930212 | 0.027830851 | 221496 | 6 | 33738979 | 33756913 | -1 | protein_coding | LEM domain containing 2 |
| TREML1 | ENSG00000161911 | 88.90210867 | -2.886319824 | 0.465985255 | 5.55E-11 | 6.12E-10 | 340205 | 6 | 41117080 | 41122075 | -1 | protein_coding | triggering receptor expressed on myeloid cells-like 1 |
| CXCL16 | ENSG00000161921 | 12381.22192 | 0.408517024 | 0.083765473 | 1E-06 | 6.47E-06 | 58191 | 17 | 4636821 | 4643217 | -1 | protein_coding | chemokine (C-X-C motif) ligand 16 |
| SCIMP | ENSG00000161929 | 3486.55615 | -0.553563762 | 0.225310269 | 0.008273147 | 0.0237087 | 388325 | 17 | 5112256 | 5138155 | -1 | protein_coding | SLP adaptor and CSK interacting membrane protein |
| TNFSF13 | ENSG00000161955 | 1610.481077 | -0.892484718 | 0.192091844 | 9.92E-07 | 6.39E-06 | 8741 | 17 | 7461609 | 7464925 | 1 | protein_coding | tumor necrosis factor (ligand) superfamily, member 13 |
| FGF11 | ENSG00000161958 | 117.3883685 | -0.6772731 | 0.313427073 | 0.013961448 | 0.037351735 | 2256 | 17 | 7341592 | 7348256 | 1 | protein_coding | fibroblast growth factor 11 |
| SNRNP25 | ENSG00000161981 | 214.0446996 | -1.431533869 | 0.262055785 | 7.47E-09 | 6.45E-08 | 79622 | 16 | 103010 | 107669 | 1 | protein_coding | small nuclear ribonucleoprotein 25kDa (U11/U12) |
| JMJD8 | ENSG00000161999 | 186.9810301 | -1.096244896 | 0.452743644 | 0.002664482 | 0.00870671 | 339123 | 16 | 731671 | 734529 | -1 | protein_coding | jumonji domain containing 8 |
| AMDHD2 | ENSG00000162066 | 571.4728115 | -0.474028972 | 0.197927211 | 0.011204612 | 0.030876322 | 752014 | 16 | 2570358 | 2581423 | 1 | protein_coding | amidohydrolase domain containing 2 |
| PAQR4 | ENSG00000162073 | 368.9559759 | -0.51530365 | 0.192568332 | 0.004860084 | 0.014910508 | 124222 | 16 | 3019246 | 3023490 | 1 | protein_coding | progestin and adipoQ receptor family member IV |
| FLYWCH2 | ENSG00000162076 | 159.8595726 | -0.93519562 | 0.272784503 | 1.91E-04 | 8.21E-04 | 114984 | 16 | 2933187 | 2949383 | 1 | protein_coding | FLYWCH family member 2 |
| ADCY9 | ENSG00000162104 | 381.0820445 | -1.001399851 | 0.275280366 | 7.13E-05 | 3.34E-04 | 115 | 16 | 4003388 | 4166186 | -1 | protein_coding | adenylate cyclase 9 |
| ASRGL1 | ENSG00000162174 | 457.2326301 | -2.545495741 | 0.234793337 | 1.75E-28 | 6.29E-27 | 80150 | 11 | 62104920 | 62160882 | 1 | protein_coding | asparaginase like 1 |
| TTC9C | ENSG00000162222 | 388.1028195 | 0.576756572 | 0.173104675 | 5.26E-04 | 0.002048258 | 283237 | 11 | 62495541 | 62507765 | 1 | protein_coding | tetratricopeptide repeat domain 9C |
| NXF1 | ENSG00000162231 | 2027.824768 | 0.5374333 | 0.124588302 | 1.17E-05 | 6.4E-05 | 10482 | 11 | 62559595 | 62573774 | -1 | protein_coding | nuclear RNA export factor 1 |
| STX5 | ENSG00000162236 | 830.1724726 | 0.319860728 | 0.130632223 | 0.012217354 | 0.033229725 | 6811 | 11 | 62574369 | 62599560 | -1 | protein_coding | syntaxin 5 |
| RPL29 | ENSG00000162244 | 2834.358287 | -0.518609628 | 0.144661092 | 2.28E-04 | 9.62E-04 | 6159 | 3 | 52027616 | 52029958 | -1 | protein_coding | ribosomal protein L29 |
| SYVN1 | ENSG00000162298 | 1267.0744 | -0.391593637 | 0.123232161 | 0.001169228 | 0.00419369 | 1024654 | 11 | 64889252 | 64902004 | -1 | protein_coding | synovial apoptosis inhibitor 1, synoviolin |
| ZFPL1 | ENSG00000162300 | 319.5297506 | -0.507332152 | 0.193799363 | 0.005812168 | 0.017386449 | 7542 | 11 | 64851695 | 64855870 | 1 | protein_coding | zinc finger protein-like 1 |
| RPS6KA4 | ENSG00000162302 | 453.2636367 | -1.045415886 | 0.227050646 | 1.06E-06 | 6.79E-06 | 8986 | 11 | 64126620 | 64139687 | 1 | protein_coding | ribosomal protein S6 kinase, 90kDa, polypeptide 4 |
| TPCN2 | ENSG00000162341 | 1143.756146 | -0.621206342 | 0.127376011 | 6.47E-07 | 4.3E-06 | 219931 | 11 | 68816365 | 68858072 | 1 | protein_coding | two pore segment channel 2 |
| CMPK1 | ENSG00000162368 | 2823.97142 | 0.382243078 | 0.139726117 | 0.005000719 | 0.015266877 | 51727 | 1 | 47799469 | 47844511 | 1 | protein_coding | cytidine monophosphate (UMP-CMP) kinase 1, cytosolic |
| MAGOH | ENSG00000162385 | 322.3332884 | 0.569337482 | 0.218079497 | 0.005385116 | 0.016281054 | 55110 | 1 | 53692564 | 53704282 | -1 | protein_coding | mago-nashi homolog, proliferation-associated (Drosophila) |
| ACOT11 | ENSG00000162390 | 271.9289884 | -2.207793659 | 0.250957136 | 1.1E-19 | 2.36E-18 | 26027 | 1 | 55007930 | 55104865 | 1 | protein_coding | acyl-CoA thioesterase 11 |
| PPAP2B | ENSG00000162407 | 2018.330537 | 0.905749236 | 0.161330484 | 6.19E-09 | 5.4E-08 | 8613 | 1 | 56960419 | 57110974 | -1 | protein_coding | phosphatidic acid phosphatase type 2B |
| KLHL21 | ENSG00000162413 | 915.3834118 | 0.638590557 | 0.146966663 | 7.26E-06 | 4.08E-05 | 9903 | 1 | 6650784 | 6674667 | -1 | protein_coding | kelch-like family member 21 |
| SEPN1 | ENSG00000162430 | 9910.587173 | -1.038264165 | 0.119023717 | 7.33E-19 | 1.47E-17 | 57190 | 1 | 26126667 | 26144713 | 1 | protein_coding | selenoprotein N, 1 |
| RBP7 | ENSG00000162444 | 13.67028428 | -1.427318319 | 1.286654147 | 0.017380584 | 0.045124563 | 116362 | 1 | 10057264 | 10076078 | 1 | protein_coding | retinol binding protein 7, cellular |
| DRAXIN | ENSG00000162490 | 160.7253356 | -2.503070691 | 0.687224908 | 1.64E-05 | 8.7E-05 | 374946 | 1 | 11751786 | 11785914 | 1 | protein_coding | dorsal inhibitory axon guidance protein |
| SDC3 | ENSG00000162512 | 3804.109804 | -1.647204912 | 0.163057117 | 6.84E-25 | 2.04E-23 | 9672 | 1 | 31342314 | 31381608 | -1 | protein_coding | syndecan 3 |
| C1orf86 | ENSG00000162585 | 198.278635 | -0.736951604 | 0.248988807 | 0.00130116 | 0.004609555 | 199990 | 1 | 2115903 | 2144159 | -1 | protein_coding | chromosome 1 open reading frame 86 |
| MEGF6 | ENSG00000162591 | 185.7633679 | -1.092065714 | 0.374633864 | 7.65E-04 | 0.002872019 | 1953 | 1 | 3406484 | 3528059 | -1 | protein_coding | multiple EGF-like-domains 6 |
| NFIA | ENSG00000162599 | 128.309177 | -1.753659355 | 0.40670385 | 1.74E-06 | 1.08E-05 | 4774 | 1 | 61330931 | 61928465 | 1 | protein_coding | nuclear factor I/A |
| OMA1 | ENSG00000162600 | 382.1188294 | -0.507289103 | 0.220834539 | 0.014542855 | 0.03866163 | 115209 | 1 | 58881056 | 59012474 | -1 | protein_coding | OMA1 zinc metallopeptidase |
| MYSM1 | ENSG00000162601 | 1378.402095 | 0.44292733 | 0.148897513 | 0.002115023 | 0.007104154 | 114803 | 1 | 59120411 | 59165764 | -1 | protein_coding | Myb-like, SWIRM and MPN domains 1 |
| USP1 | ENSG00000162607 | 750.1928939 | 1.111794311 | 0.167304167 | 7.57E-12 | 9E-11 | 7398 | 1 | 62901968 | 62917475 | 1 | protein_coding | ubiquitin specific peptidase 1 |
| FUBP1 | ENSG00000162613 | 1349.397704 | 0.613916135 | 0.197955343 | 0.001070051 | 0.003881697 | 8880 | 1 | 78409740 | 78444794 | -1 | protein_coding | far upstream element (FUSE) binding protein 1 |
| NEXN | ENSG00000162614 | 31.0889764 | 5.675886665 | 0.962625168 | 4.29E-10 | 4.28E-09 | 91624 | 1 | 78354198 | 78409580 | 1 | protein_coding | nexilin (F actin binding protein) |
| DNAJB4 | ENSG00000162616 | 407.8771699 | 0.836358953 | 0.241483373 | 1.94E-04 | 8.33E-04 | 11080 | 1 | 78444859 | 78483648 | 1 | protein_coding | DnaJ (Hsp40) homolog, subfamily B, member 4 |
| TYW3 | ENSG00000162623 | 493.9040485 | 0.57540607 | 0.176801099 | 6.82E-04 | 0.002587657 | 127253 | 1 | 75198836 | 75232362 | 1 | protein_coding | tRNA-yW synthesizing protein 3 homolog (S. cerevisiae) |
| FAM102B | ENSG00000162636 | 997.3529375 | 0.414777694 | 0.134700949 | 0.001556778 | 0.005412563 | 284611 | 1 | 109102711 | 109187522 | 1 | protein_coding | family with sequence similarity 102, member B |
| WDR63 | ENSG00000162643 | 346.7936344 | -0.880698346 | 0.279036659 | 5.4E-04 | 0.002093517 | 126820 | 1 | 85464830 | 85598821 | 1 | protein_coding | WD repeat domain 63 |
| GBP2 | ENSG00000162645 | 9604.919955 | 1.901633476 | 0.171232973 | 1.27E-29 | 4.82E-28 | 2634 | 1 | 89571815 | 89616139 | -1 | protein_coding | guanylate binding protein 2, interferon-inducible |
| GBP4 | ENSG00000162654 | 15295.50057 | 3.993680132 | 0.471969243 | 1.58E-18 | 3.12E-17 | 115361 | 1 | 89646831 | 89664615 | -1 | protein_coding | guanylate binding protein 4 |
| AGL | ENSG00000162688 | 1564.938683 | -0.55570464 | 0.147660816 | 9.89E-05 | 4.52E-04 | 178 | 1 | 100315640 | 100389579 | 1 | protein_coding | amylo-alpha-1, 6-glucosidase, 4-alpha-glucanotransferase |
| VCAM1 | ENSG00000162692 | 51.56847612 | 1.544054269 | 0.507060566 | 2.74E-04 | 0.001137228 | 7412 | 1 | 101185298 | 101204601 | 1 | protein_coding | vascular cell adhesion molecule 1 |
| EXTL2 | ENSG00000162694 | 286.5079602 | 0.972346938 | 0.272231059 | 1.03E-04 | 4.68E-04 | 2135 | 1 | 101337943 | 101361554 | -1 | protein_coding | exostosin-like glycosyltransferase 2 |
| SLC30A7 | ENSG00000162695 | 2324.379402 | 0.759659293 | 0.101358836 | 3.12E-14 | 4.49E-13 | 148867 | 1 | 101361632 | 101447309 | 1 | protein_coding | solute carrier family 30 (zinc transporter), member 7 |
| ZNF281 | ENSG00000162702 | 2183.877729 | 0.61593656 | 0.097135401 | 1.22E-10 | 1.29E-09 | 23528 | 1 | 200374068 | 200379184 | -1 | protein_coding | zinc finger protein 281 |
| ARPC5 | ENSG00000162704 | 7835.081115 | 0.259039946 | 0.110847922 | 0.018024425 | 0.04647668 | 10092 | 1 | 183592401 | 183604892 | -1 | protein_coding | actin related protein 2/3 complex, subunit 5, 16kDa |
| ZNF496 | ENSG00000162714 | 622.3229688 | 0.40484313 | 0.161374728 | 0.00949486 | 0.026792826 | 84838 | 1 | 247460714 | 247495148 | -1 | protein_coding | zinc finger protein 496 |
| SLAMF9 | ENSG00000162723 | 921.7041404 | -1.608750524 | 0.210732087 | 3.22E-15 | 4.94E-14 | 89886 | 1 | 159921282 | 159924044 | -1 | protein_coding | SLAM family member 9 |
| VANGL2 | ENSG00000162738 | 53.7015674 | -0.959176232 | 0.571562462 | 0.019302666 | 0.04924097 | 57216 | 1 | 160370376 | 160398468 | 1 | protein_coding | VANGL planar cell polarity protein 2 |
| OLFML2B | ENSG00000162745 | 87.33565503 | -3.679658339 | 0.620902854 | 1.67E-10 | 1.74E-09 | 25903 | 1 | 161952982 | 161993644 | -1 | protein_coding | olfactomedin-like 2B |
| FCRLB | ENSG00000162746 | 79.43983926 | -3.474770644 | 0.568053467 | 8.79E-11 | 9.44E-10 | 127943 | 1 | 161691353 | 161697933 | 1 | protein_coding | Fc receptor-like B |
| ATF3 | ENSG00000162772 | 6613.660695 | 1.338469426 | 0.164918876 | 1.31E-16 | 2.23E-15 | 467 | 1 | 212738676 | 212794119 | 1 | protein_coding | activating transcription factor 3 |
| BPNT1 | ENSG00000162813 | 653.3540649 | -0.599227148 | 0.190236321 | 9.33E-04 | 0.003430979 | 10380 | 1 | 220230824 | 220263804 | -1 | protein_coding | 3'(2'), 5'-bisphosphate nucleotidase 1 |
| C1orf115 | ENSG00000162817 | 254.7172943 | -1.77489753 | 0.42140134 | 2.49E-06 | 1.51E-05 | 79762 | 1 | 220863187 | 220872499 | 1 | protein_coding | chromosome 1 open reading frame 115 |
| ACP6 | ENSG00000162836 | 177.7449796 | -0.585351636 | 0.249450587 | 0.010563869 | 0.029311475 | 51205 | 1 | 147101453 | 147142618 | -1 | protein_coding | acid phosphatase 6, lysophosphatidic |
| CNST | ENSG00000162852 | 1120.045498 | 0.40331907 | 0.12833454 | 0.001314685 | 0.004646698 | 163882 | 1 | 246729746 | 246831886 | 1 | protein_coding | consortin, connexin sorting protein |
| OXER1 | ENSG00000162881 | 248.7209129 | -1.034902132 | 0.256962795 | 1.45E-05 | 7.78E-05 | 165140 | 2 | 42989642 | 42991401 | -1 | protein_coding | oxoeicosanoid (OXE) receptor 1 |
| HAAO | ENSG00000162882 | 646.0871448 | -0.651601653 | 0.159965566 | 2.55E-05 | 1.3E-04 | 23498 | 2 | 42994229 | 43019733 | -1 | protein_coding | 3-hydroxyanthranilate 3,4-dioxygenase |

| | | | | | | | | | | | | | |
|---|---|---|---|---|---|---|---|---|---|---|---|---|---|
| FAIM3 | ENSG00000162894 | 2015.411018 | -1.144004271 | 0.442810459 | 0.001778319 | 0.006081462 | 9214 | 1 | 207077731 | 207096592 | -1 | protein_coding | Fas apoptotic inhibitory molecule 3 |
| PIGR | ENSG00000162896 | 148.9401974 | 2.054441089 | 0.584564954 | 3.06E-05 | 1.54E-04 | 5284 | 1 | 207101863 | 207119811 | -1 | protein_coding | polymeric immunoglobulin receptor |
| FCAMR | ENSG00000162897 | 30.12478137 | 5.238166574 | 0.898441119 | 3.87E-10 | 3.88E-09 | 83953 | 1 | 207131310 | 207143970 | -1 | protein_coding | Fc receptor, IgA, IgM, high affinity |
| CAPN2 | ENSG00000162909 | 2257.973517 | 0.41065162 | 0.109543451 | 1.33E-04 | 5.95E-04 | 824 | 1 | 223889295 | 223963720 | 1 | protein_coding | calpain 2, (m/II) large subunit |
| REL | ENSG00000162924 | 4065.084663 | 0.780498137 | 0.080860001 | 1.22E-22 | 3.2E-21 | 5966 | 2 | 61108656 | 61158745 | 1 | protein_coding | v-rel avian reticuloendotheliosis viral oncogene homolog |
| PEX13 | ENSG00000162928 | 990.1742845 | 0.405542502 | 0.154522499 | 0.006578777 | 0.01941743 | 5194 | 2 | 61244360 | 61279125 | 1 | protein_coding | peroxisomal biogenesis factor 13 |
| TYW5 | ENSG00000162971 | 334.5426958 | 1.413743543 | 0.234455883 | 2.66E-10 | 2.7E-09 | 129450 | 2 | 200794698 | 200820459 | -1 | protein_coding | tRNA-yW synthesizing protein 5 |
| C2orf47 | ENSG00000162972 | 204.5686903 | 0.521951304 | 0.220304099 | 0.011652851 | 0.031933034 | 79568 | 2 | 200820040 | 200873263 | 1 | protein_coding | chromosome 2 open reading frame 47 |
| KCNF1 | ENSG00000162975 | 268.3241179 | -2.680368809 | 0.352656326 | 2.47E-15 | 3.82E-14 | 3754 | 2 | 11052063 | 11054350 | 1 | protein_coding | potassium voltage-gated channel, subfamily F, member 1 |
| FBXO41 | ENSG00000163013 | 517.9171591 | -0.82738898 | 0.168699951 | 3.68E-07 | 2.53E-06 | 1019278 | 2 | 73481810 | 73511559 | -1 | protein_coding | F-box protein 41 |
| CCDC74A | ENSG00000163040 | 50.0541277 | -1.023685589 | 0.501377311 | 0.008949224 | 0.025407951 | 90557 | 2 | 132285248 | 132291239 | 1 | protein_coding | coiled-coil domain containing 74A |
| SGCB | ENSG00000163069 | 1101.086899 | 0.673735407 | 0.203043315 | 4.47E-04 | 0.001772634 | 6443 | 4 | 52886872 | 52904648 | -1 | protein_coding | sarcoglycan, beta (43kDa dystrophin-associated glycoprotein) |
| SGPP2 | ENSG00000163082 | 351.9919303 | 3.125475299 | 0.272497825 | 1.11E-31 | 4.71E-30 | 130367 | 2 | 223289236 | 223425667 | 1 | protein_coding | sphingosine-1-phosphate phosphatase 2 |
| SMARCAD1 | ENSG00000163104 | 1247.376012 | 0.320767502 | 0.116617668 | 0.00516622 | 0.01570632 | 56916 | 4 | 95128762 | 95212443 | 1 | protein_coding | SWI/SNF-related, matrix-associated actin-dependent regulator of chromatin, subfamily a, containing DE |
| HPGDS | ENSG00000163106 | 1127.395973 | -3.25818613 | 0.196348341 | 5.6E-63 | 8.38E-61 | 27306 | 4 | 95219686 | 95264027 | -1 | protein_coding | hematopoietic prostaglandin D synthase |
| NEURL3 | ENSG00000163121 | 358.0952654 | 5.481287723 | 0.310315598 | 5.65E-71 | 1.08E-68 | 93082 | 2 | 97163383 | 97173846 | -1 | pseudogene | neuralized E3 ubiquitin protein ligase 3 |
| TNFAIP8L2 | ENSG00000163154 | 652.6230989 | -0.541333442 | 0.148455946 | 1.67E-04 | 7.29E-04 | 79626 | 1 | 151129105 | 151132225 | 1 | protein_coding | tumor necrosis factor, alpha-induced protein 8-like 2 |
| LYSMD1 | ENSG00000163155 | 47.70934569 | 0.926803463 | 0.45353494 | 0.010639858 | 0.029490197 | 388695 | 1 | 151132224 | 151138424 | -1 | protein_coding | LysM, putative peptidoglycan-binding, domain containing 1 |
| RNF149 | ENSG00000163162 | 3151.968233 | 1.144362606 | 0.117113223 | 3.55E-23 | 9.68E-22 | 284996 | 2 | 101887681 | 101925163 | -1 | protein_coding | ring finger protein 149 |
| CDC42EP3 | ENSG00000163171 | 489.2280056 | -1.290867111 | 0.391585646 | 1.65E-04 | 7.22E-04 | 10602 | 2 | 37869032 | 37965611 | -1 | protein_coding | CDC42 effector protein (Rho GTPase binding) 3 |
| ARHGAP25 | ENSG00000163219 | 4108.217743 | 0.410705598 | 0.160478865 | 0.007885918 | 0.022739715 | 9938 | 2 | 68906733 | 69053965 | 1 | protein_coding | Rho GTPase activating protein 25 |
| S100A9 | ENSG00000163220 | 3836.622584 | -0.833872552 | 0.130423839 | 6.14E-11 | 6.74E-10 | 6280 | 1 | 153330330 | 153333503 | 1 | protein_coding | S100 calcium binding protein A9 |
| TGFA | ENSG00000163235 | 107.3371646 | -0.817345632 | 0.371367903 | 0.009470318 | 0.026733443 | 7039 | 2 | 70674412 | 70781325 | -1 | protein_coding | transforming growth factor, alpha |
| CCNYL1 | ENSG00000163249 | 1778.562251 | 1.149219158 | 0.114694217 | 3.21E-24 | 9.21E-23 | 151195 | 2 | 208576264 | 208626563 | 1 | protein_coding | cyclin Y-like 1 |
| FZD5 | ENSG00000163251 | 355.8336489 | 0.734572856 | 0.187497899 | 4.07E-05 | 2E-04 | 7855 | 2 | 208627310 | 208634287 | -1 | protein_coding | frizzled family receptor 5 |
| PAQR3 | ENSG00000163291 | 186.411857 | 1.053192154 | 0.273760695 | 3.07E-05 | 1.55E-04 | 152559 | 4 | 79808281 | 79860592 | -1 | protein_coding | progestin and adipoQ receptor family member III |
| NIPAL1 | ENSG00000163293 | 165.6701702 | 1.532862397 | 0.323291532 | 2.94E-07 | 2.05E-06 | 1019271 | 4 | 47916159 | 48042188 | 1 | protein_coding | NIPA-like domain containing 1 |
| ANTXR2 | ENSG00000163297 | 5876.373079 | 1.340765639 | 0.101719678 | 1.3E-40 | 8.01E-39 | 118429 | 4 | 80822303 | 81046608 | -1 | protein_coding | anthrax toxin receptor 2 |
| MRPS18C | ENSG00000163319 | 527.6925642 | 0.904582296 | 0.250815453 | 9.76E-05 | 4.47E-04 | 51023 | 4 | 84377085 | 84390888 | 1 | protein_coding | mitochondrial ribosomal protein S18C |
| CGGBP1 | ENSG00000163320 | 2728.447952 | 0.335897538 | 0.097853425 | 5.04E-04 | 0.001971159 | 8545 | 3 | 88101094 | 88199035 | -1 | protein_coding | CGG triplet repeat binding protein 1 |
| GPR155 | ENSG00000163328 | 1519.856293 | -0.956382246 | 0.121406739 | 1.77E-15 | 2.77E-14 | 151556 | 2 | 175296966 | 175351822 | -1 | protein_coding | G protein-coupled receptor 155 |
| DAPL1 | ENSG00000163331 | 51.94916647 | -2.898272537 | 0.774336233 | 1.55E-05 | 8.25E-05 | 92196 | 2 | 159651829 | 159719293 | 1 | protein_coding | death associated protein-like 1 |
| PMVK | ENSG00000163344 | 351.27348 | -0.775793788 | 0.190506892 | 1.99E-05 | 1.04E-04 | 10654 | 1 | 154897210 | 154909467 | -1 | protein_coding | phosphomevalonate kinase |
| PBXIP1 | ENSG00000163346 | 2168.187574 | -0.633444829 | 0.135746628 | 1.61E-06 | 1E-05 | 57326 | 1 | 154916552 | 154928599 | -1 | protein_coding | pre-B-cell leukemia homeobox interacting protein 1 |
| COL6A3 | ENSG00000163359 | 322.6600423 | -2.373245383 | 0.388026205 | 6.36E-11 | 6.95E-10 | 1293 | 2 | 238232646 | 238323018 | -1 | protein_coding | collagen, type VI, alpha 3 |
| YY1AP1 | ENSG00000163374 | 1730.166439 | 0.533393173 | 0.126691687 | 1.7E-05 | 8.96E-05 | 55249 | 1 | 155629237 | 155658791 | -1 | protein_coding | YY1 associated protein 1 |
| POGLUT1 | ENSG00000163389 | 499.1405058 | 0.784842285 | 0.177217324 | 4.27E-06 | 2.5E-05 | 56983 | 3 | 119187785 | 119213555 | 1 | protein_coding | protein O-glucosyltransferase 1 |
| ATP1A1 | ENSG00000163399 | 25733.30186 | -0.996304092 | 0.102005046 | 4.92E-23 | 1.33E-21 | 476 | 1 | 116915290 | 116952883 | 1 | protein_coding | ATPase, Na+/K+ transporting, alpha 1 polypeptide |
| SLC15A2 | ENSG00000163406 | 173.0375659 | -1.711036123 | 0.424034638 | 5.85E-06 | 3.34E-05 | 6565 | 3 | 121612936 | 121662949 | 1 | protein_coding | solute carrier family 15 (oligopeptide transporter), member 2 |
| EIF4E3 | ENSG00000163412 | 2250.028733 | 0.887296616 | 0.121778478 | 1.26E-13 | 1.73E-12 | 317649 | 3 | 71724567 | 71803924 | -1 | protein_coding | eukaryotic translation initiation factor 4E family member 3 |
| LRRC58 | ENSG00000163428 | 1782.852283 | 0.467257327 | 0.136158619 | 4.33E-04 | 0.001721626 | 116064 | 3 | 120043356 | 120068186 | -1 | protein_coding | leucine rich repeat containing 58 |
| FSTL1 | ENSG00000163430 | 39.74260347 | 1.633674485 | 0.636042423 | 9.8E-04 | 0.003582407 | 11167 | 3 | 120111140 | 120170100 | -1 | protein_coding | follistatin-like 1 |
| CXCR1 | ENSG00000163464 | 22.23146555 | -2.306251703 | 1.505630777 | 0.006885574 | 0.020209763 | 3577 | 2 | 219027568 | 219031718 | -1 | protein_coding | chemokine (C-X-C motif) receptor 1 |
| EOMES | ENSG00000163508 | 132.0133458 | 0.785532699 | 0.354225912 | 0.009622661 | 0.027123409 | 8320 | 3 | 27757440 | 27764206 | -1 | protein_coding | eomesodermin |
| AZI2 | ENSG00000163512 | 1296.985425 | 1.049651439 | 0.144972318 | 1.45E-13 | 1.98E-12 | 64343 | 3 | 28356494 | 28390618 | -1 | protein_coding | 5-azacytidine induced 2 |
| TGFBR2 | ENSG00000163513 | 7023.346836 | -1.333868115 | 0.105024687 | 1.13E-37 | 6.24E-36 | 7048 | 3 | 30647994 | 30735634 | 1 | protein_coding | transforming growth factor, beta receptor II (70/80kDa) |
| FCRL1 | ENSG00000163534 | 45.2505494 | -0.944106742 | 0.481141759 | 0.012226524 | 0.033248749 | 115350 | 1 | 157764193 | 157789895 | -1 | protein_coding | Fc receptor-like 1 |
| SGOL2 | ENSG00000163535 | 229.7200366 | 0.727927904 | 0.268650295 | 0.002954889 | 0.009569733 | 151246 | 2 | 201374731 | 201448505 | 1 | protein_coding | shugoshin-like 2 (S. pombe) |
| PYHIN1 | ENSG00000163564 | 378.1681296 | 0.593856464 | 0.213505942 | 0.00309429 | 0.009976799 | 149628 | 1 | 158900586 | 158946844 | 1 | protein_coding | pyrin and HIN domain family, member 1 |
| IFI16 | ENSG00000163565 | 5619.902268 | 1.69549394 | 0.116252726 | 2.45E-49 | 2.09E-47 | 3428 | 1 | 158969758 | 159024945 | 1 | protein_coding | interferon, gamma-inducible protein 16 |
| AIM2 | ENSG00000163568 | 350.5851931 | 3.930766777 | 0.386125989 | 1.21E-25 | 3.71E-24 | 9447 | 1 | 159032274 | 159116886 | -1 | protein_coding | absent in melanoma 2 |
| PPM1L | ENSG00000163590 | 547.7629495 | -1.743491734 | 0.243986651 | 9.69E-14 | 1.35E-12 | 151742 | 3 | 160473390 | 160796695 | 1 | protein_coding | protein phosphatase, Mg2+/Mn2+ dependent, 1L |
| ICOS | ENSG00000163600 | 446.1477741 | 0.803721416 | 0.239817871 | 3.28E-04 | 0.001341365 | 29851 | 2 | 204801471 | 204826300 | 1 | protein_coding | inducible T-cell co-stimulator |
| RYBP | ENSG00000163602 | 1679.803396 | 0.671667337 | 0.13617788 | 4E-07 | 2.74E-06 | 23429 | 3 | 72420976 | 72496069 | -1 | protein_coding | RING1 and YY1 binding protein |
| GTPBP8 | ENSG00000163607 | 199.0121029 | 0.645188977 | 0.270272654 | 0.008505222 | 0.02426457 | 29083 | 3 | 112709765 | 112733907 | 1 | protein_coding | GTP-binding protein 8 (putative) |
| COX18 | ENSG00000163626 | 667.191702 | -0.383635197 | 0.160854265 | 0.013395485 | 0.036036472 | 285521 | 4 | 73921797 | 73935472 | -1 | protein_coding | COX18 cytochrome C oxidase assembly factor |
| PTPN13 | ENSG00000163629 | 657.034133 | -2.487982592 | 0.290755901 | 8.73E-19 | 1.74E-17 | 5783 | 4 | 87515468 | 87736324 | 1 | protein_coding | protein tyrosine phosphatase, non-receptor type 13 (APO-1/CD95 (Fas)-associated phosphatase) |
| THOC7 | ENSG00000163634 | 791.0048831 | 0.400601021 | 0.153279652 | 0.007044286 | 0.020616205 | 80145 | 3 | 63819546 | 63849579 | -1 | protein_coding | THO complex 7 homolog (Drosophila) |
| ATXN7 | ENSG00000163635 | 953.1114068 | 0.696138105 | 0.144668687 | 7.08E-07 | 4.68E-06 | 6314 | 3 | 63850233 | 63989138 | 1 | protein_coding | ataxin 7 |
| PSMD6 | ENSG00000163636 | 1614.028331 | 0.362650447 | 0.138521617 | 0.007163236 | 0.020928262 | 9861 | 3 | 63996225 | 64009658 | -1 | protein_coding | proteasome (prosome, macropain) 26S subunit, non-ATPase, 6 |
| PPM1K | ENSG00000163644 | 796.3899749 | 3.183828367 | 0.155166344 | 1.08E-94 | 3.42E-92 | 152926 | 4 | 89178772 | 89205921 | -1 | protein_coding | protein phosphatase, Mg2+/Mn2+ dependent, 1K |
| TIPARP | ENSG00000163659 | 1516.979833 | 0.94486584 | 0.17358253 | 1.72E-08 | 1.41E-07 | 25976 | 3 | 156391024 | 156424559 | 1 | protein_coding | TCDD-inducible poly(ADP-ribose) polymerase |
| CCNL1 | ENSG00000163660 | 2407.083225 | 0.881216542 | 0.1040782 | 7.21E-18 | 1.35E-16 | 57018 | 3 | 156864297 | 156878549 | -1 | protein_coding | cyclin L1 |
| PTX3 | ENSG00000163661 | 25.85988356 | 1.993693287 | 0.895201393 | 0.001631622 | 0.005646757 | 5806 | 3 | 157154578 | 157161417 | 1 | protein_coding | pentraxin 3, long |
| HESX1 | ENSG00000163666 | 127.2113833 | 5.26945341 | 0.435376618 | 6.19E-35 | 3.11E-33 | 8820 | 3 | 57231944 | 57260549 | -1 | protein_coding | HESX homeobox 1 |
| SLMAP | ENSG00000163681 | 1599.972508 | 0.457080749 | 0.12340468 | 1.6E-04 | 7.02E-04 | 7871 | 3 | 57741177 | 57914895 | 1 | protein_coding | sarcolemma associated protein |
| ABHD6 | ENSG00000163686 | 754.1967516 | -2.109586706 | 0.1682535 | 4.57E-37 | 2.48E-35 | 57406 | 3 | 58223233 | 58281420 | 1 | protein_coding | abhydrolase domain containing 6 |
| DNASE1L3 | ENSG00000163687 | 6.59833228 | -0.518766288 | 0.971078947 | 0.008106286 | 0.023300425 | 1776 | 3 | 58177984 | 58200424 | -1 | protein_coding | deoxyribonuclease I-like 3 |
| PCOLCE2 | ENSG00000163710 | 30.11316661 | -2.286814683 | 0.767880702 | 2.15E-04 | 9.12E-04 | 26577 | 3 | 142534764 | 142608045 | -1 | protein_coding | procollagen C-endopeptidase enhancer 2 |
| U2SURP | ENSG00000163714 | 2180.570155 | 0.700792954 | 0.14043621 | 2.86E-07 | 1.99E-06 | 23350 | 3 | 142683339 | 142779567 | 1 | protein_coding | U2 snRNP-associated SURP domain containing |
| TTC14 | ENSG00000163728 | 1110.125968 | 0.545055027 | 0.151478921 | 2.02E-04 | 8.61E-04 | 151613 | 3 | 180319918 | 180335616 | 1 | protein_coding | tetratricopeptide repeat domain 14 |
| CXCL3 | ENSG00000163734 | 197.0384757 | 4.261713341 | 0.317270786 | 2.33E-42 | 1.59E-40 | 2921 | 4 | 74902306 | 74904524 | -1 | protein_coding | chemokine (C-X-C motif) ligand 3 |
| CXCL5 | ENSG00000163735 | 46.08748956 | 3.272204326 | 0.54476356 | 1.25E-10 | 1.32E-09 | 6374 | 4 | 74861359 | 74864496 | -1 | protein_coding | chemokine (C-X-C motif) ligand 5 |
| CXCL1 | ENSG00000163739 | 45.09435307 | 5.389663407 | 0.720662337 | 6.15E-15 | 9.32E-14 | 2919 | 4 | 74735110 | 74736959 | 1 | protein_coding | chemokine (C-X-C motif) ligand 1 (melanoma growth stimulating activity, alpha) |
| RCHY1 | ENSG00000163743 | 308.167625 | 0.76786075 | 0.247868533 | 8.02E-04 | 0.002999401 | 25898 | 4 | 76404247 | 76439974 | -1 | protein_coding | ring finger and CHY zinc finger domain containing 1, E3 ubiquitin protein ligase |
| GYG1 | ENSG00000163754 | 1537.92973 | 0.443669812 | 0.13283041 | 6.13E-04 | 0.002342156 | 2992 | 3 | 148709128 | 148745419 | 1 | protein_coding | glycogenin 1 |
| ZNF513 | ENSG00000163795 | 304.454253 | -0.468142023 | 0.208209691 | 0.016724749 | 0.043640831 | 130557 | 2 | 27600098 | 27603657 | -1 | protein_coding | zinc finger protein 513 |
| PLB1 | ENSG00000163803 | 273.9078796 | -1.832940649 | 0.468446578 | 7.76E-06 | 4.34E-05 | 151056 | 2 | 28680012 | 28866654 | 1 | protein_coding | phospholipase B1 |
| ZDHHC3 | ENSG00000163812 | 2920.86951 | -0.300468853 | 0.124150601 | 0.013213056 | 0.035583282 | 51304 | 3 | 44956749 | 45017677 | -1 | protein_coding | zinc finger, DHHC-type containing 3 |
| CLEC3B | ENSG00000163815 | 9.317104493 | -0.501570595 | 0.957340926 | 0.004959038 | 0.015163862 | 7123 | 3 | 45043040 | 45077563 | 1 | protein_coding | C-type lectin domain family 3, member B |
| CCR1 | ENSG00000163823 | 9443.142836 | 1.175352257 | 0.094671186 | 4.94E-36 | 2.57E-34 | 1230 | 3 | 46243200 | 46249887 | -1 | protein_coding | chemokine (C-C motif) receptor 1 |
| DTX3L | ENSG00000163840 | 7804.56082 | 2.109289696 | 0.107787689 | 1.82E-86 | 5.07E-84 | 151636 | 3 | 122283085 | 122294050 | 1 | protein_coding | deltex 3-like (Drosophila) |
| YEATS2 | ENSG00000163872 | 2647.988881 | 1.124304599 | 0.112588947 | 4.31E-24 | 1.23E-22 | 55689 | 3 | 183415606 | 183530413 | 1 | protein_coding | YEATS domain containing 2 |
| ZC3H12A | ENSG00000163874 | 1047.020339 | 3.048651608 | 0.204228222 | 1.5E-51 | 1.43E-49 | 1024654 | 1 | 37940153 | 37949980 | 1 | protein_coding | zinc finger CCCH-type containing 12A |
| SNIP1 | ENSG00000163877 | 565.2920924 | 0.837020363 | 0.189115804 | 3.5E-06 | 2.07E-05 | 79753 | 1 | 38000050 | 38019903 | -1 | protein_coding | Smad nuclear interacting protein 1 |
| TMEM41A | ENSG00000163900 | 523.6623115 | 0.389224644 | 0.157694855 | 0.01067733 | 0.029567246 | 90407 | 3 | 185194284 | 185216845 | -1 | protein_coding | transmembrane protein 41A |
| RPN1 | ENSG00000163902 | 8466.690113 | -0.430426352 | 0.078386351 | 3.79E-08 | 2.98E-07 | 6184 | 3 | 128338817 | 128399918 | -1 | protein_coding | ribophorin I |
| IFT122 | ENSG00000163913 | 731.7446308 | -0.763091538 | 0.209855649 | 1.13E-04 | 5.14E-04 | 1019272 | 3 | 129158968 | 129239198 | 1 | protein_coding | intraflagellar transport 122 homolog (Chlamydomonas) |
| RFC4 | ENSG00000163918 | 173.5979963 | 0.729326134 | 0.249365082 | 0.00151547 | 0.005284263 | 5984 | 3 | 186507669 | 186524847 | -1 | protein_coding | replication factor C (activator 1) 4, 37kDa |

| | | | | | | | | | | | | | |
|---|---|---|---|---|---|---|---|---|---|---|---|---|---|
| **BAP1** | ENSG00000163930 | 1561.489188 | -0.405206087 | 0.121646566 | 6.66E-04 | 0.002531598 | 8314 | 3 | 52435029 | 52444366 | -1 | protein_coding | BRCA1 associated protein-1 (ubiquitin carboxy-terminal hydrolase) |
| **SFMBT1** | ENSG00000163935 | 269.5102938 | 0.74402288 | 0.302681777 | 0.005757368 | 0.017242771 | 51460 | 3 | 52937588 | 53080766 | -1 | protein_coding | Scm-like with four mbt domains 1 |
| **SLBP** | ENSG00000163950 | 958.9763999 | 0.370390064 | 0.134346561 | 0.004662708 | 0.01435947 | 7884 | 4 | 1694527 | 1714282 | -1 | protein_coding | stem-loop binding protein |
| **LRPAP1** | ENSG00000163956 | 3083.020394 | -0.762571398 | 0.113058116 | 1.02E-11 | 1.19E-10 | 4043 | 4 | 3508103 | 3534286 | -1 | protein_coding | low density lipoprotein receptor-related protein associated protein 1 |
| **MFI2** | ENSG00000163975 | 290.5804989 | -1.053965966 | 0.265547631 | 1.98E-05 | 1.03E-04 | 4241 | 3 | 196715492 | 196756687 | -1 | protein_coding | antigen p97 (melanoma associated) identified by monoclonal antibodies 133.2 and 96.5 |
| **SGMS2** | ENSG00000164023 | 775.76872 | -2.306076214 | 0.20483638 | 1.76E-30 | 6.98E-29 | 166929 | 4 | 108745719 | 108836203 | 1 | protein_coding | sphingomyelin synthase 2 |
| **DNAJB14** | ENSG00000164031 | 1789.647771 | 0.506493781 | 0.121767642 | 2.29E-05 | 1.18E-04 | 79982 | 4 | 100817405 | 100867883 | -1 | protein_coding | DnaJ (Hsp40) homolog, subfamily B, member 14 |
| **PGRMC2** | ENSG00000164040 | 888.0804821 | 0.421379241 | 0.150609801 | 0.0038455 | 0.01207784 | 10424 | 4 | 129190397 | 129209984 | -1 | protein_coding | progesterone receptor membrane component 2 |
| **CAMP** | ENSG00000164047 | 18.91771103 | -5.637079293 | 2.870203742 | 3.9E-04 | 0.001570113 | 820 | 3 | 48264837 | 48266981 | 1 | protein_coding | cathelicidin antimicrobial peptide |
| **ZNF589** | ENSG00000164048 | 1531.068724 | -1.07932352 | 0.221345202 | 2.61E-07 | 1.83E-06 | 51385 | 3 | 48282590 | 48340743 | 1 | protein_coding | zinc finger protein 589 |
| **APEH** | ENSG00000164062 | 1446.446104 | -0.591828152 | 0.144220961 | 2.43E-05 | 1.25E-04 | 327 | 3 | 49711435 | 49721396 | 1 | protein_coding | acylaminoacyl-peptide hydrolase |
| **INTU** | ENSG00000164066 | 289.8474917 | -0.548483618 | 0.236702174 | 0.012190272 | 0.033185594 | 27152 | 4 | 128544426 | 128647892 | 1 | protein_coding | inturned planar cell polarity protein |
| **MFSD8** | ENSG00000164073 | 456.3542163 | 0.772609492 | 0.175211492 | 4.44E-06 | 2.59E-05 | 256471 | 4 | 128838960 | 128887150 | -1 | protein_coding | major facilitator superfamily domain containing 8 |
| **C4orf29** | ENSG00000164074 | 405.5654131 | 0.491162363 | 0.201379383 | 0.009601052 | 0.027072486 | 80167 | 4 | 128886461 | 128960866 | 1 | protein_coding | chromosome 4 open reading frame 29 |
| **PPM1M** | ENSG00000164088 | 2965.784765 | -1.25766726 | 0.102530131 | 1.02E-34 | 5.04E-33 | 132160 | 3 | 52279841 | 52284613 | 1 | protein_coding | protein phosphatase, Mg2+/Mn2+ dependent, 1M |
| **HMGB2** | ENSG00000164104 | 586.8469871 | 0.618893709 | 0.210030332 | 0.002022951 | 0.006830923 | 3148 | 4 | 174252846 | 174256276 | -1 | protein_coding | high mobility group box 2 |
| **SAP30** | ENSG00000164105 | 277.5619754 | -0.567013015 | 0.207071447 | 0.003685246 | 0.011648733 | 8819 | 4 | 174291120 | 174298683 | 1 | protein_coding | Sin3A-associated protein, 30kDa |
| **MAD2L1** | ENSG00000164109 | 69.96548328 | 0.858098327 | 0.413329029 | 0.011478952 | 0.031512991 | 4085 | 4 | 120976763 | 120988229 | -1 | protein_coding | MAD2 mitotic arrest deficient-like 1 (yeast) |
| **ANXA5** | ENSG00000164111 | 12735.36072 | 0.74955104 | 0.117319711 | 1.22E-10 | 1.29E-09 | 308 | 4 | 122589110 | 122618268 | -1 | protein_coding | annexin A5 |
| **MAP9** | ENSG00000164114 | 41.52600939 | 1.38057042 | 0.623482224 | 0.003253409 | 0.010432618 | 79884 | 4 | 156263810 | 156298122 | -1 | protein_coding | microtubule-associated protein 9 |
| **GUCY1A3** | ENSG00000164116 | 72.41638001 | 3.790363979 | 0.483282384 | 3.05E-16 | 5.04E-15 | 2982 | 4 | 156587863 | 156653501 | 1 | protein_coding | guanylate cyclase 1, soluble, alpha 3 |
| **FBXO8** | ENSG00000164117 | 476.1169798 | 0.451159044 | 0.195863117 | 0.015009248 | 0.039754998 | 26269 | 4 | 175157809 | 175205415 | -1 | protein_coding | F-box protein 8 |
| **HPGD** | ENSG00000164120 | 1259.811487 | -3.266923869 | 0.239261054 | 6.25E-44 | 4.57E-42 | 3248 | 4 | 175411328 | 175444305 | -1 | protein_coding | hydroxyprostaglandin dehydrogenase 15-(NAD) |
| **TMEM144** | ENSG00000164124 | 1789.040847 | 0.443948723 | 0.143695209 | 0.001481671 | 0.00518178 | 55314 | 4 | 159122756 | 159176563 | 1 | protein_coding | transmembrane protein 144 |
| **FAM198B** | ENSG00000164125 | 6496.663453 | -0.456674172 | 0.144927809 | 0.001226314 | 0.004370746 | 51313 | 4 | 159045626 | 159094470 | -1 | protein_coding | family with sequence similarity 198, member B |
| **IL15** | ENSG00000164136 | 324.6365077 | 2.879689499 | 0.296948766 | 2.19E-23 | 6.03E-22 | 3600 | 4 | 142557752 | 142655140 | 1 | protein_coding | interleukin 15 |
| **KIAA0947** | ENSG00000164151 | 1706.490072 | 1.068821648 | 0.124956021 | 2.89E-18 | 5.57E-17 | 23379 | 5 | 5420777 | 5490347 | 1 | protein_coding | KIAA0947 |
| **ABCE1** | ENSG00000164163 | 2428.29397 | -0.447887298 | 0.125659836 | 2.56E-04 | 0.001066563 | 6059 | 4 | 146019084 | 146050331 | 1 | protein_coding | ATP-binding cassette, sub-family E (OABP), member 1 |
| **OTUD4** | ENSG00000164164 | 4887.325288 | 0.891208387 | 0.096268433 | 7.84E-21 | 1.85E-19 | 54726 | 4 | 146031990 | 146101313 | -1 | protein_coding | OTU domain containing 4 |
| **ELOVL7** | ENSG00000164181 | 491.6969181 | 3.806890112 | 0.328519158 | 2.91E-32 | 1.27E-30 | 79993 | 5 | 60047618 | 60140216 | -1 | protein_coding | ELOVL fatty acid elongase 7 |
| **NDUFAF2** | ENSG00000164182 | 133.3356904 | -0.665483778 | 0.298670946 | 0.012214165 | 0.033226963 | 91942 | 5 | 60240956 | 60448853 | 1 | protein_coding | NADH dehydrogenase (ubiquinone) complex I, assembly factor 2 |
| **NIPBL** | ENSG00000164190 | 4223.990655 | 0.266253241 | 0.102874933 | 0.00836639 | 0.02393104 | 25836 | 5 | 36876861 | 37066515 | 1 | protein_coding | Nipped-B homolog (Drosophila) |
| **SLC25A46** | ENSG00000164209 | 1303.78135 | 0.635197258 | 0.181654502 | 2.59E-04 | 0.001081221 | 91137 | 5 | 110073837 | 110100857 | 1 | protein_coding | solute carrier family 25, member 46 |
| **STARD4** | ENSG00000164211 | 2332.897077 | 1.85746076 | 0.122240638 | 4.54E-53 | 4.75E-51 | 134429 | 5 | 110831731 | 110848288 | -1 | protein_coding | StAR-related lipid transfer (START) domain containing 4 |
| **CMBL** | ENSG00000164237 | 165.6901398 | -1.722302652 | 0.393967298 | 1.37E-06 | 8.6E-06 | 134147 | 5 | 10275987 | 10308138 | -1 | protein_coding | carboxymethylenebutenolidase homolog (Pseudomonas) |
| **PRRC1** | ENSG00000164244 | 1550.274288 | -0.457958898 | 0.140567555 | 7.87E-04 | 0.002948178 | 133619 | 5 | 126853301 | 126890781 | 1 | protein_coding | proline-rich coiled-coil 1 |
| **GRPEL2** | ENSG00000164284 | 454.6375024 | 0.398318214 | 0.173888675 | 0.017136466 | 0.044570561 | 134266 | 5 | 148724993 | 148734146 | 1 | protein_coding | GrpE-like 2, mitochondrial (E. coli) |
| **GPX8** | ENSG00000164294 | 104.154156 | 0.777340962 | 0.344211051 | 0.008870226 | 0.025211765 | 493869 | 5 | 54455946 | 54462899 | 1 | protein_coding | glutathione peroxidase 8 (putative) |
| **CASP3** | ENSG00000164305 | 1677.904512 | 1.398289693 | 0.167122103 | 1.03E-17 | 1.91E-16 | 836 | 4 | 185548850 | 185570663 | -1 | protein_coding | caspase 3, apoptosis-related cysteine peptidase |
| **ERAP2** | ENSG00000164308 | 674.9819918 | 1.515314087 | 0.140280858 | 5.22E-28 | 1.81E-26 | 64167 | 5 | 96211643 | 96255420 | 1 | protein_coding | endoplasmic reticulum aminopeptidase 2 |
| **KIAA1430** | ENSG00000164323 | 640.8810007 | 0.421282807 | 0.177546405 | 0.013192394 | 0.035540164 | 57587 | 4 | 186080819 | 186130658 | -1 | protein_coding | KIAA1430 |
| **RICTOR** | ENSG00000164327 | 2459.942668 | 1.34047479 | 0.184417057 | 6.16E-14 | 8.72E-13 | 253260 | 5 | 38938021 | 39074510 | -1 | protein_coding | RPTOR independent companion of MTOR, complex 2 |
| **PAPD4** | ENSG00000164329 | 2256.926905 | 0.425797123 | 0.13421955 | 0.001151141 | 0.004137528 | 167153 | 5 | 78907943 | 78982471 | 1 | protein_coding | PAP associated domain containing 4 |
| **ANKRA2** | ENSG00000164331 | 383.7274117 | 0.685423133 | 0.223407791 | 0.001040281 | 0.003780883 | 57763 | 5 | 72848160 | 72861511 | -1 | protein_coding | ankyrin repeat, family A (RFXANK-like), 2 |
| **UBLCP1** | ENSG00000164332 | 1218.816192 | 0.596910816 | 0.146000793 | 2.65E-05 | 1.35E-04 | 134510 | 5 | 158690089 | 158713044 | 1 | protein_coding | ubiquitin-like domain containing CTD phosphatase 1 |
| **TLR3** | ENSG00000164342 | 494.7945335 | 2.374174231 | 0.203320436 | 1.49E-32 | 6.56E-31 | 7098 | 4 | 186990306 | 187009223 | 1 | protein_coding | toll-like receptor 3 |
| **8-Sep** | ENSG00000164402 | 423.6409446 | -0.488438151 | 0.181055078 | 0.004652329 | 0.014332099 | 23176 | 5 | 132086509 | 132142933 | -1 | protein_coding | septin 8 |
| **UQCRQ** | ENSG00000164405 | 995.9071884 | -0.584942762 | 0.182515702 | 7.52E-04 | 0.002828431 | 27089 | 5 | 132202252 | 132203723 | 1 | protein_coding | ubiquinol-cytochrome c reductase, complex III subunit VII, 9.5kDa |
| **SLC35A1** | ENSG00000164414 | 877.8608019 | 0.624369189 | 0.160425831 | 5.46E-05 | 2.62E-04 | 10559 | 6 | 88180341 | 88222054 | 1 | protein_coding | solute carrier family 35 (CMP-sialic acid transporter), member A1 |
| **MB21D1** | ENSG00000164430 | 571.4378835 | 2.361101062 | 0.213702743 | 1.85E-29 | 6.96E-28 | 115004 | 6 | 74123238 | 74161999 | -1 | protein_coding | Mab-21 domain containing 1 |
| **CITED2** | ENSG00000164442 | 3569.640306 | -1.131697768 | 0.140675672 | 2.1E-16 | 3.5E-15 | 10370 | 6 | 139693393 | 139695757 | -1 | protein_coding | Cbp/p300-interacting transactivator, with Glu/Asp-rich carboxy-terminal domain, 2 |
| **CREBRF** | ENSG00000164463 | 1499.846186 | 1.646892743 | 0.121459171 | 7.69E-43 | 5.49E-41 | 153222 | 5 | 172483355 | 172566291 | 1 | protein_coding | CREB3 regulatory factor |
| **IL22RA2** | ENSG00000164485 | 8.111240295 | -2.389851226 | 2.148663795 | 0.013375367 | 0.035995022 | 116379 | 6 | 137464968 | 137494785 | -1 | protein_coding | interleukin 22 receptor, alpha 2 |
| **PDSS2** | ENSG00000164494 | 626.0802082 | -0.496011838 | 0.159196675 | 0.001258417 | 0.004475771 | 57107 | 6 | 107473761 | 107780768 | -1 | protein_coding | prenyl (decaprenyl) diphosphate synthase, subunit 2 |
| **STXBP5** | ENSG00000164506 | 1029.288662 | -0.432474858 | 0.171523806 | 0.008035571 | 0.023110206 | 134957 | 6 | 147525561 | 147706866 | 1 | protein_coding | syntaxin binding protein 5 (tomosyn) |
| **IL31RA** | ENSG00000164509 | 53.38342017 | 2.27707195 | 1.038995864 | 0.001278623 | 0.00453813 | 133396 | 5 | 55147207 | 55218678 | 1 | protein_coding | interleukin 31 receptor A |
| **DAGLB** | ENSG00000164535 | 620.1787825 | -0.568035506 | 0.160894969 | 2.55E-04 | 0.001062514 | 221955 | 7 | 6448757 | 6523821 | -1 | protein_coding | diacylglycerol lipase, beta |
| **STK17A** | ENSG00000164543 | 1750.824006 | 1.965369412 | 0.206953355 | 2.2E-22 | 5.65E-21 | 9263 | 7 | 43622357 | 43666385 | 1 | protein_coding | serine/threonine kinase 17a |
| **TRA2A** | ENSG00000164548 | 1046.795129 | 0.533617413 | 0.135423953 | 5.23E-05 | 2.51E-04 | 29896 | 7 | 23544399 | 23571660 | -1 | protein_coding | transformer 2 alpha homolog (Drosophila) |
| **SAP30L** | ENSG00000164576 | 571.4421806 | 0.581445592 | 0.204539135 | 0.002611883 | 0.008549457 | 79685 | 5 | 153825517 | 153840614 | 1 | protein_coding | SAP30-like |
| **RPS14** | ENSG00000164587 | 4353.94737 | -0.354901301 | 0.13570697 | 0.006973287 | 0.020439728 | 6208 | 5 | 149822753 | 149829319 | -1 | protein_coding | ribosomal protein S14 |
| **COG5** | ENSG00000164597 | 1695.359848 | 0.323291333 | 0.13451684 | 0.013712711 | 0.036786268 | 10466 | 7 | 106842000 | 107204959 | -1 | protein_coding | component of oligomeric golgi complex 5 |
| **C7orf60** | ENSG00000164603 | 621.9598191 | 0.709822481 | 0.213730427 | 4.11E-04 | 0.001643562 | 154743 | 7 | 112459202 | 112579971 | -1 | protein_coding | chromosome 7 open reading frame 60 |
| **GPR85** | ENSG00000164604 | 99.60799306 | -0.930154831 | 0.413675081 | 0.006934158 | 0.020332836 | 54329 | 7 | 112718386 | 112727833 | -1 | protein_coding | G protein-coupled receptor 85 |
| **SLU7** | ENSG00000164609 | 2430.269492 | 0.59545432 | 0.149600255 | 4.38E-05 | 2.13E-04 | 10569 | 5 | 159828648 | 159848718 | -1 | protein_coding | SLU7 splicing factor homolog (S. cerevisiae) |
| **PTTG1** | ENSG00000164611 | 167.674307 | 1.051699133 | 0.292748405 | 8.11E-05 | 3.77E-04 | 9232 | 5 | 159848829 | 159855748 | 1 | protein_coding | pituitary tumor-transforming 1 |
| **CAMLG** | ENSG00000164615 | 409.6149002 | 0.676562738 | 0.209955665 | 6.29E-04 | 0.002398118 | 819 | 5 | 134074191 | 134087847 | 1 | protein_coding | calcium modulating ligand |
| **KCNK5** | ENSG00000164626 | 127.4722492 | -0.916686462 | 0.406993766 | 0.006473106 | 0.019146213 | 8645 | 6 | 39156749 | 39197226 | -1 | protein_coding | potassium channel, subfamily K, member 5 |
| **SLC29A4** | ENSG00000164638 | 11.16990089 | -1.974688032 | 1.614880472 | 0.007524928 | 0.02182439 | 222962 | 7 | 5314000 | 5346501 | 1 | protein_coding | solute carrier family 29 (equilibrative nucleoside transporter), member 4 |
| **CDCA7L** | ENSG00000164649 | 396.6456694 | -0.799282975 | 0.226480562 | 1.69E-04 | 7.38E-04 | 55536 | 7 | 21940518 | 21985702 | -1 | protein_coding | cell division cycle associated 7-like |
| **MIOS** | ENSG00000164654 | 1410.839856 | -0.52683695 | 0.124326599 | 1.51E-05 | 8.07E-05 | 54468 | 7 | 7606503 | 7648560 | 1 | protein_coding | missing oocyte, meiosis regulator, homolog (Drosophila) |
| **ZNF704** | ENSG00000164684 | 76.61151345 | -2.440595618 | 1.037682705 | 7.14E-04 | 0.002696506 | 619279 | 8 | 81540686 | 81787016 | -1 | protein_coding | zinc finger protein 704 |
| **FABP5** | ENSG00000164687 | 3468.488129 | -1.968201772 | 0.205765797 | 1.1E-22 | 2.91E-21 | 2171 | 8 | 82192598 | 82197012 | 1 | protein_coding | fatty acid binding protein 5 (psoriasis-associated) |
| **TAGAP** | ENSG00000164691 | 1682.464353 | 1.950113551 | 0.18718662 | 2.38E-26 | 7.64E-25 | 117289 | 6 | 159455500 | 159466184 | -1 | protein_coding | T-cell activation RhoGTPase activating protein |
| **CHMP4C** | ENSG00000164695 | 241.1381887 | 0.588252891 | 0.246520097 | 0.009383812 | 0.026517232 | 92421 | 8 | 82644669 | 82671750 | 1 | protein_coding | charged multivesicular body protein 4C |
| **CTSB** | ENSG00000164733 | 375531.6528 | -0.737507746 | 0.12278061 | 8.75E-10 | 8.47E-09 | 1508 | 8 | 11700033 | 11726957 | -1 | protein_coding | cathepsin B |
| **DLC1** | ENSG00000164741 | 105.5367376 | -1.500761773 | 0.348298582 | 2.34E-06 | 1.43E-05 | 10395 | 8 | 12940870 | 13373167 | -1 | protein_coding | deleted in liver cancer 1 |
| **C7orf57** | ENSG00000164746 | 16.20988301 | -1.720566008 | 1.307661661 | 0.010315116 | 0.028737881 | 136288 | 7 | 48075108 | 48100901 | 1 | protein_coding | chromosome 7 open reading frame 57 |
| **RAD21** | ENSG00000164754 | 3982.365479 | 0.44234881 | 0.086312959 | 2.51E-07 | 1.77E-06 | 5885 | 8 | 117858174 | 117887105 | -1 | protein_coding | RAD21 homolog (S. pombe) |
| **HEATR2** | ENSG00000164818 | 361.1119333 | -0.638621853 | 0.218424508 | 0.001867834 | 0.006363797 | 54919 | 7 | 766338 | 829190 | 1 | protein_coding | HEAT repeat containing 2 |
| **OSGIN2** | ENSG00000164823 | 1781.535904 | 1.152728263 | 0.143597799 | 2.31E-16 | 3.84E-15 | 734 | 8 | 90914087 | 90940116 | 1 | protein_coding | oxidative stress induced growth inhibitor family member 2 |
| **GPER1** | ENSG00000164850 | 115.3799773 | -2.452641355 | 0.492293804 | 5.56E-08 | 4.29E-07 | 2852 | 7 | 1121844 | 1133451 | 1 | protein_coding | G protein-coupled estrogen receptor 1 |
| **NOS3** | ENSG00000164867 | 11.67333526 | -1.879760571 | 1.70594041 | 0.01525616 | 0.040360029 | 4846 | 7 | 150688083 | 150711676 | 1 | protein_coding | nitric oxide synthase 3 (endothelial cell) |
| **INTS1** | ENSG00000164880 | 1649.291597 | -0.593197441 | 0.217023152 | 0.003508657 | 0.01115283 | 26173 | 7 | 1509913 | 1545489 | -1 | protein_coding | integrator complex subunit 1 |
| **CDK5** | ENSG00000164885 | 497.2452338 | -0.912178883 | 0.183268402 | 2.18E-07 | 1.55E-06 | 1020 | 7 | 150750899 | 150755617 | -1 | protein_coding | cyclin-dependent kinase 5 |
| **SLC4A2** | ENSG00000164889 | 2624.215929 | -0.385972334 | 0.140843896 | 0.004860574 | 0.014910508 | 6522 | 7 | 150754297 | 150773614 | 1 | protein_coding | solute carrier family 4 (anion exchanger), member 2 |
| **FASTK** | ENSG00000164896 | 580.5562368 | -0.863261686 | 0.179456286 | 5.64E-07 | 3.79E-06 | 10922 | 7 | 150773711 | 150777953 | -1 | protein_coding | Fas-activated serine/threonine kinase |
| **ALDH7A1** | ENSG00000164904 | 605.9796349 | -1.313417921 | 0.257598713 | 6.27E-08 | 4.8E-07 | 501 | 5 | 125877533 | 125931110 | -1 | protein_coding | aldehyde dehydrogenase 7 family, member A1 |
| **FOXK1** | ENSG00000164916 | 2041.067114 | -0.362546281 | 0.10827262 | 6.7E-04 | 0.002544393 | 221937 | 7 | 4683388 | 4811074 | 1 | protein_coding | forkhead box K1 |

| | | | | | | | | | | | | | |
|---|---|---|---|---|---|---|---|---|---|---|---|---|---|
| **BAALC** | ENSG00000164929 | 280.1797695 | 1.111125522 | 0.402943475 | 0.001168818 | 0.004193201 | 79870 | 8 | 104152938 | 104242533 | 1 | protein_coding | brain and acute leukemia, cytoplasmic |
| **SLC25A32** | ENSG00000164933 | 709.6073217 | 0.675475452 | 0.171396284 | 4.11E-05 | 2.01E-04 | 81034 | 8 | 104410863 | 104427417 | -1 | protein_coding | solute carrier family 25 (mitochondrial folate carrier), member 32 |
| **DCSTAMP** | ENSG00000164935 | 13983.51498 | -1.65961103 | 0.137165454 | 1.53E-34 | 7.5E-33 | 81501 | 8 | 105351315 | 105368917 | 1 | protein_coding | dendrocyte expressed seven transmembrane protein |
| **GEM** | ENSG00000164949 | 692.6618855 | 0.629870511 | 0.178931315 | 2.3E-04 | 9.7E-04 | 2669 | 8 | 95261481 | 95274578 | -1 | protein_coding | GTP binding protein overexpressed in skeletal muscle |
| **PDP1** | ENSG00000164951 | 524.2870661 | 0.67904856 | 0.20078426 | 3.3E-04 | 0.001347864 | 54704 | 8 | 94870035 | 94938294 | 1 | protein_coding | pyruvate dehyrogenase phosphatase catalytic subunit 1 |
| **TMEM67** | ENSG00000164953 | 108.3950092 | -0.835211405 | 0.408364255 | 0.012514096 | 0.033928142 | 91147 | 8 | 94767072 | 94831462 | 1 | protein_coding | transmembrane protein 67 |
| **KIAA0196** | ENSG00000164961 | 2557.207654 | -0.533749642 | 0.116895421 | 3.87E-06 | 2.28E-05 | 9897 | 8 | 126036502 | 126104082 | -1 | protein_coding | KIAA0196 |
| **FAM219A** | ENSG00000164970 | 273.6974863 | 0.555761509 | 0.211578626 | 0.005207395 | 0.015809418 | 203259 | 9 | 34398182 | 34458568 | -1 | protein_coding | family with sequence similarity 219, member A |
| **PSIP1** | ENSG00000164985 | 546.2698761 | 0.454486509 | 0.194722693 | 0.014011094 | 0.037469513 | 11168 | 9 | 15464064 | 15511017 | -1 | protein_coding | PC4 and SFRS1 interacting protein 1 |
| **DIRAS2** | ENSG00000165023 | 87.75070197 | 1.315133618 | 0.422936888 | 2.98E-04 | 0.001227162 | 54769 | 9 | 93372114 | 93405386 | -1 | protein_coding | DIRAS family, GTP-binding RAS-like 2 |
| **SYK** | ENSG00000165025 | 6782.100783 | -1.447469103 | 0.113201063 | 2.55E-38 | 1.45E-36 | 6850 | 9 | 93564069 | 93660831 | 1 | protein_coding | spleen tyrosine kinase |
| **NFIL3** | ENSG00000165030 | 727.5593365 | 0.486693886 | 0.210956847 | 0.014283874 | 0.038064594 | 4783 | 9 | 94171327 | 94186144 | -1 | protein_coding | nuclear factor, interleukin 3 regulated |
| **TMEM71** | ENSG00000165071 | 190.4322929 | -1.282419731 | 0.325083837 | 1.39E-05 | 7.49E-05 | 137835 | 8 | 133697253 | 133772958 | -1 | protein_coding | transmembrane protein 71 |
| **KDM1B** | ENSG00000165097 | 761.7509599 | -1.089282391 | 0.190372169 | 2.69E-09 | 2.45E-08 | 221656 | 6 | 18155560 | 18224084 | 1 | protein_coding | lysine (K)-specific demethylase 1B |
| **C9orf64** | ENSG00000165118 | 718.9063877 | 0.512449402 | 0.143571239 | 2.39E-04 | 0.001004883 | 84267 | 9 | 86553226 | 86571901 | -1 | protein_coding | chromosome 9 open reading frame 64 |
| **ANKS6** | ENSG00000165138 | 167.8207041 | -0.861276855 | 0.358945355 | 0.005351066 | 0.016187715 | 203286 | 9 | 101493611 | 101559247 | -1 | protein_coding | ankyrin repeat and sterile alpha motif domain containing 6 |
| **FBP1** | ENSG00000165140 | 23711.67742 | -3.194843542 | 0.106806774 | 1.15E-197 | 9.28E-195 | 2203 | 9 | 97365415 | 97402531 | -1 | protein_coding | fructose-1,6-bisphosphatase 1 |
| **ZHX1** | ENSG00000165156 | 1367.640789 | 1.031683752 | 0.14039698 | 5.54E-14 | 7.88E-13 | 11244 | 8 | 124260696 | 124287781 | -1 | protein_coding | zinc fingers and homeoboxes 1 |
| **DYNLT3** | ENSG00000165169 | 816.7861328 | 1.239829064 | 0.178903112 | 9.48E-13 | 1.21E-11 | 6990 | X | 37696010 | 37706890 | -1 | protein_coding | dynein, light chain, Tctex-type 3 |
| **MID1IP1** | ENSG00000165175 | 1594.610443 | -0.516003199 | 0.123927999 | 1.97E-05 | 1.03E-04 | 58526 | X | 38660685 | 38665790 | 1 | protein_coding | MID1 interacting protein 1 |
| **NCF1C** | ENSG00000165178 | 162.2918594 | 2.89414908 | 0.499567699 | 2.14E-10 | 2.2E-09 | 654817 | 7 | 74572445 | 74587848 | -1 | pseudogene | neutrophil cytosolic factor 1C pseudogene |
| **KIAA1958** | ENSG00000165185 | 169.5583443 | 1.518469849 | 0.266114497 | 1.65E-09 | 1.54E-08 | 158405 | 9 | 115249127 | 115431677 | 1 | protein_coding | KIAA1958 |
| **STRBP** | ENSG00000165209 | 894.7531664 | -0.969366707 | 0.194319337 | 1.83E-07 | 1.31E-06 | 55342 | 9 | 125871779 | 126030855 | -1 | protein_coding | spermatid perinuclear RNA binding protein |
| **ZNF367** | ENSG00000165244 | 41.63504956 | 1.985354328 | 0.597987898 | 7.59E-05 | 3.54E-04 | 195828 | 9 | 99148223 | 99180611 | -1 | protein_coding | zinc finger protein 367 |
| **HDX** | ENSG00000165259 | 167.3389394 | 1.546256918 | 0.327871808 | 3.28E-07 | 2.27E-06 | 139324 | X | 83572882 | 83757487 | -1 | protein_coding | highly divergent homeobox |
| **NOL6** | ENSG00000165271 | 1058.720288 | -0.545474823 | 0.132954209 | 2.59E-05 | 1.32E-04 | 65083 | 9 | 33461439 | 33473928 | -1 | protein_coding | nucleolar protein 6 (RNA-associated) |
| **AQP3** | ENSG00000165272 | 86.45554847 | -0.886371509 | 0.383286764 | 0.006134727 | 0.018254875 | 360 | 9 | 33441152 | 33447609 | -1 | protein_coding | aquaporin 3 (Gill blood group) |
| **VCP** | ENSG00000165280 | 10411.09074 | 0.241520406 | 0.099319873 | 0.014261826 | 0.038019099 | 7415 | 9 | 35056061 | 35073246 | -1 | protein_coding | valosin containing protein |
| **MELK** | ENSG00000165304 | 161.4926266 | 1.485652284 | 0.31167534 | 2.56E-07 | 1.8E-06 | 9833 | 9 | 36572859 | 36677678 | 1 | protein_coding | maternal embryonic leucine zipper kinase |
| **OTUD1** | ENSG00000165312 | 648.185364 | 0.917321644 | 0.194563688 | 8.29E-07 | 5.41E-06 | 220213 | 10 | 23728198 | 23731308 | 1 | protein_coding | OTU domain containing 1 |
| **ARHGAP12** | ENSG00000165322 | 1825.305163 | 0.425239757 | 0.141746514 | 0.001891423 | 0.006437999 | 94134 | 10 | 32094365 | 32217742 | -1 | protein_coding | Rho GTPase activating protein 12 |
| **HECTD2** | ENSG00000165338 | 188.6012455 | 0.94561152 | 0.266817113 | 1.21E-04 | 5.46E-04 | 143279 | 10 | 93170096 | 93274586 | 1 | protein_coding | HECT domain containing E3 ubiquitin protein ligase 2 |
| **SPTSSA** | ENSG00000165389 | 319.2544952 | 0.499812197 | 0.204104332 | 0.009043456 | 0.025648288 | 171546 | 14 | 34901995 | 34931562 | -1 | protein_coding | serine palmitoyltransferase, small subunit A |
| **WRN** | ENSG00000165392 | 396.6544509 | 0.54941047 | 0.178455961 | 0.001290587 | 0.004576342 | 7486 | 8 | 30891317 | 31031285 | 1 | protein_coding | Werner syndrome, RecQ helicase-like |
| **SUGT1** | ENSG00000165416 | 1154.486402 | 0.356301338 | 0.152536096 | 0.015895203 | 0.041797383 | 10910 | 13 | 53226844 | 53275044 | 1 | protein_coding | SGT1, suppressor of G2 allele of SKP1 (S. cerevisiae) |
| **ZCCHC24** | ENSG00000165424 | 292.6453244 | -2.068318959 | 0.255394261 | 5.31E-17 | 9.32E-16 | 219654 | 10 | 81142081 | 81205383 | -1 | protein_coding | zinc finger, CCHC domain containing 24 |
| **INPPL1** | ENSG00000165458 | 4094.890066 | -0.423681851 | 0.102688841 | 2.82E-05 | 1.43E-04 | 3636 | 11 | 71934745 | 71950149 | 1 | protein_coding | inositol polyphosphate phosphatase-like 1 |
| **GJB2** | ENSG00000165474 | 4281.607987 | 0.440729366 | 0.129492853 | 4.93E-04 | 0.001932722 | 2706 | 13 | 20761609 | 20767037 | -1 | protein_coding | gap junction protein, beta 2, 26kDa |
| **CRYL1** | ENSG00000165475 | 657.1918023 | -1.116691662 | 0.192375178 | 1.55E-09 | 1.46E-08 | 51084 | 13 | 20977806 | 21099996 | -1 | protein_coding | crystallin, lambda 1 |
| **REEP3** | ENSG00000165476 | 570.1900876 | 0.499028356 | 0.2242923 | 0.016670953 | 0.043519303 | 221035 | 10 | 65281123 | 65384883 | 1 | protein_coding | receptor accessory protein 3 |
| **SKA3** | ENSG00000165480 | 33.4885469 | 1.630463976 | 0.811626315 | 0.003232632 | 0.010376879 | 221150 | 13 | 21727734 | 21750741 | -1 | protein_coding | spindle and kinetochore associated complex subunit 3 |
| **MICU2** | ENSG00000165487 | 946.6510734 | 0.55316266 | 0.131034773 | 1.52E-05 | 8.14E-05 | 221154 | 13 | 22066836 | 22178353 | -1 | protein_coding | mitochondrial calcium uptake 2 |
| **C11orf82** | ENSG00000165490 | 726.9138294 | -2.367640615 | 0.292685653 | 4.22E-17 | 7.44E-16 | 220042 | 11 | 82611017 | 82669319 | 1 | protein_coding | chromosome 11 open reading frame 82 |
| **PCF11** | ENSG00000165494 | 2458.294233 | 0.456930963 | 0.10098194 | 4.58E-06 | 2.67E-05 | 51585 | 11 | 82868030 | 82898493 | 1 | protein_coding | PCF11 cleavage and polyadenylation factor subunit |
| **LRR1** | ENSG00000165501 | 158.5744044 | 0.933197512 | 0.295492503 | 4.66E-04 | 0.001836323 | 122769 | 14 | 50065415 | 50081390 | 1 | protein_coding | leucine rich repeat protein 1 |
| **ZNF22** | ENSG00000165512 | 309.2776131 | 1.173116919 | 0.190923835 | 1.87E-10 | 1.93E-09 | 7570 | 10 | 45495923 | 45500774 | 1 | protein_coding | zinc finger protein 22 |
| **NEMF** | ENSG00000165525 | 1337.038173 | -0.35332616 | 0.131906431 | 0.005736485 | 0.017183595 | 9147 | 14 | 50249997 | 50319921 | -1 | protein_coding | nuclear export mediator factor |
| **TTC8** | ENSG00000165533 | 251.6977063 | -1.21007035 | 0.337999168 | 6.56E-05 | 3.1E-04 | 123016 | 14 | 89290497 | 89344335 | 1 | protein_coding | tetratricopeptide repeat domain 8 |
| **TMEM63C** | ENSG00000165548 | 269.0636031 | -3.112790482 | 0.295397097 | 6.52E-27 | 2.15E-25 | 57156 | 14 | 77582911 | 77725838 | 1 | protein_coding | transmembrane protein 63C |
| **PDZD8** | ENSG00000165650 | 739.5965378 | -0.598880004 | 0.163724712 | 1.44E-04 | 6.38E-04 | 118987 | 10 | 119040000 | 119134978 | -1 | protein_coding | PDZ domain containing 8 |
| **ZNF503** | ENSG00000165655 | 86.90200968 | -1.39182351 | 0.463410984 | 3.77E-04 | 0.001519765 | 84858 | 10 | 77157588 | 77161664 | -1 | protein_coding | zinc finger protein 503 |
| **FAM175B** | ENSG00000165660 | 1200.883902 | 0.384486636 | 0.122704241 | 0.001426282 | 0.005005251 | 23172 | 10 | 126490354 | 126525239 | 1 | protein_coding | family with sequence similarity 175, member B |
| **GHITM** | ENSG00000165678 | 5813.066198 | 0.315182593 | 0.09846313 | 0.001215829 | 0.004337423 | 27069 | 10 | 85899196 | 85913001 | 1 | protein_coding | growth hormone inducible transmembrane protein |
| **TMEM52B** | ENSG00000165685 | 84.66132688 | -1.057341792 | 0.506984529 | 0.007122876 | 0.020830256 | 120939 | 12 | 10323141 | 10344400 | 1 | protein_coding | transmembrane protein 52B |
| **PMPCA** | ENSG00000165688 | 765.9521439 | -0.584349556 | 0.13756481 | 1.34E-05 | 7.26E-05 | 23203 | 9 | 139305110 | 139318213 | 1 | protein_coding | peptidase (mitochondrial processing) alpha |
| **AK8** | ENSG00000165695 | 41.57676736 | 1.805998355 | 0.599869682 | 2.21E-04 | 9.35E-04 | 158067 | 9 | 135600965 | 135754164 | -1 | protein_coding | adenylate kinase 8 |
| **TSC1** | ENSG00000165699 | 1776.824129 | -0.583102294 | 0.122428837 | 1.16E-06 | 7.38E-06 | 7248 | 9 | 135766735 | 135820020 | -1 | protein_coding | tuberous sclerosis 1 |
| **DDX21** | ENSG00000165732 | 2737.118849 | 0.665872196 | 0.124423614 | 3.72E-08 | 2.93E-07 | 9188 | 10 | 70715884 | 70744829 | 1 | protein_coding | DEAD (Asp-Glu-Ala-Asp) box helicase 21 |
| **ZNF219** | ENSG00000165804 | 842.5465884 | -1.126264341 | 0.144292511 | 1.46E-15 | 2.31E-14 | 51222 | 14 | 21558205 | 21572881 | -1 | protein_coding | zinc finger protein 219 |
| **CASP7** | ENSG00000165806 | 1298.208623 | 1.797036587 | 0.148435019 | 1.19E-34 | 5.84E-33 | 840 | 10 | 115438942 | 115490662 | 1 | protein_coding | caspase 7, apoptosis-related cysteine peptidase |
| **C10orf118** | ENSG00000165813 | 1133.94006 | 0.635893664 | 0.178579642 | 2.02E-04 | 8.63E-04 | 55088 | 10 | 115880621 | 115933979 | -1 | protein_coding | chromosome 10 open reading frame 118 |
| **METTL3** | ENSG00000165819 | 581.5629481 | 0.450357038 | 0.146586283 | 0.001588632 | 0.00551546 | 56339 | 14 | 21966277 | 21979517 | -1 | protein_coding | methyltransferase like 3 |
| **FAM35BP** | ENSG00000165874 | 24.19599523 | 1.27747945 | 0.692293371 | 0.008166637 | 0.023447437 | 439965 | 10 | 46897592 | 46939143 | 1 | pseudogene | family with sequence similarity 35, member B, pseudogene |
| **UBTD1** | ENSG00000165886 | 805.1287939 | -1.392754004 | 0.139481739 | 3.81E-24 | 1.09E-22 | 80019 | 10 | 99258625 | 99330966 | 1 | protein_coding | ubiquitin domain containing 1 |
| **E2F7** | ENSG00000165891 | 33.06285889 | 1.588136875 | 0.742591976 | 0.00275482 | 0.008973127 | 144455 | 12 | 77415027 | 77459360 | -1 | protein_coding | E2F transcription factor 7 |
| **TC2N** | ENSG00000165929 | 292.3252206 | -0.876006297 | 0.266225783 | 3.5E-04 | 0.001423883 | 123036 | 14 | 92246095 | 92333880 | -1 | protein_coding | tandem C2 domains, nuclear |
| **CLMN** | ENSG00000165959 | 718.2521938 | -1.655357031 | 0.28203402 | 5.58E-10 | 5.49E-09 | 79789 | 14 | 95648277 | 95786243 | -1 | protein_coding | calmin (calponin-like, transmembrane) |
| **ARL5B** | ENSG00000165997 | 738.2851 | 2.403672766 | 0.170109978 | 1.82E-46 | 1.45E-44 | 221079 | 10 | 18948334 | 18970568 | 1 | protein_coding | ADP-ribosylation factor-like 5B |
| **KIAA1731** | ENSG00000166004 | 650.3298506 | 0.698486513 | 0.219519721 | 6.57E-04 | 0.002496656 | 85459 | 11 | 93394805 | 93463522 | 1 | protein_coding | KIAA1731 |
| **TAF1D** | ENSG00000166012 | 1088.417705 | 0.947273717 | 0.168930754 | 6.44E-09 | 5.62E-08 | 1003022 | 11 | 93463114 | 93517557 | -1 | protein_coding | TATA box binding protein (TBP)-associated factor, RNA polymerase I, D, 41kDa |
| **ABTB2** | ENSG00000166016 | 55.76177199 | 4.789061288 | 0.988470465 | 1.8E-08 | 1.48E-07 | 25841 | 11 | 34172535 | 34379555 | -1 | protein_coding | ankyrin repeat and BTB (POZ) domain containing 2 |
| **R3HCC1L** | ENSG00000166024 | 1013.984162 | 0.579806816 | 0.134485945 | 1.09E-05 | 5.98E-05 | 27291 | 10 | 99894387 | 100004654 | 1 | protein_coding | R3H domain and coiled-coil containing 1-like |
| **HTRA1** | ENSG00000166033 | 128.57162 | -1.800709361 | 0.441628959 | 4.67E-06 | 2.72E-05 | 5654 | 10 | 124221041 | 124274424 | 1 | protein_coding | HtrA serine peptidase 1 |
| **TCP11L2** | ENSG00000166046 | 470.986097 | -0.375684607 | 0.165967426 | 0.018339888 | 0.047170735 | 255394 | 12 | 106695707 | 106740793 | 1 | protein_coding | t-complex 11, testis-specific-like 2 |
| **SPRED1** | ENSG00000166068 | 1975.934815 | -2.302270266 | 0.162240456 | 9.77E-47 | 7.86E-45 | 161742 | 15 | 38544527 | 38649450 | 1 | protein_coding | sprouty-related, EVH1 domain containing 1 |
| **ADAMTS15** | ENSG00000166106 | 19.93162311 | -5.620836999 | 2.571720026 | 1.26E-04 | 5.65E-04 | 170689 | 11 | 130318869 | 130346532 | 1 | protein_coding | ADAM metallopeptidase with thrombospondin type 1 motif, 15 |
| **RAB8B** | ENSG00000166128 | 3157.556586 | 0.729736421 | 0.134223655 | 2.76E-08 | 2.21E-07 | 51762 | 15 | 63481668 | 63559981 | 1 | protein_coding | RAB8B, member RAS oncogene family |
| **NDUFB8** | ENSG00000166136 | 1273.724142 | -0.452974061 | 0.12703881 | 2.82E-04 | 0.001166296 | 4714 | 10 | 102267203 | 102289757 | -1 | protein_coding | NADH dehydrogenase (ubiquinone) 1 beta subcomplex, 8, 19kDa |
| **BRD7** | ENSG00000166164 | 1728.921268 | 0.609372945 | 0.108031791 | 9.21E-09 | 7.87E-08 | 29117 | 16 | 50347398 | 50402845 | -1 | protein_coding | bromodomain containing 7 |
| **CKB** | ENSG00000166165 | 123.4647643 | 3.328438468 | 0.459854506 | 1.91E-14 | 2.78E-13 | 1152 | 14 | 103985996 | 103989448 | -1 | protein_coding | creatine kinase, brain |
| **BAG5** | ENSG00000166170 | 1746.708178 | -0.247092575 | 0.106670178 | 0.018216842 | 0.046893716 | 9529 | 14 | 104022881 | 104029168 | -1 | protein_coding | BCL2-associated athanogene 5 |
| **DPCD** | ENSG00000166171 | 70.80307407 | -3.147783607 | 0.50047542 | 3.09E-11 | 3.49E-10 | 25911 | 10 | 103330317 | 103369425 | 1 | protein_coding | deleted in primary ciliary dyskinesia homolog (mouse) |
| **COPS2** | ENSG00000166200 | 1817.428265 | 0.985053685 | 0.140807344 | 8.28E-13 | 1.06E-11 | 9318 | 15 | 49398268 | 49447858 | -1 | protein_coding | COP9 signalosome subunit 2 |
| **FRS2** | ENSG00000166225 | 596.2880552 | 0.523487871 | 0.163028755 | 8.57E-04 | 0.003185295 | 10818 | 12 | 69864129 | 69973562 | 1 | protein_coding | fibroblast growth factor receptor substrate 2 |
| **ARIH1** | ENSG00000166233 | 2815.444964 | 0.425994365 | 0.10956222 | 7.73E-05 | 3.61E-04 | 25820 | 15 | 72766667 | 72879692 | 1 | protein_coding | ariadne RBR E3 ubiquitin protein ligase 1 |
| **CUL5** | ENSG00000166266 | 1360.162381 | 0.574429523 | 0.109835529 | 1.04E-07 | 7.69E-07 | 8065 | 11 | 107879459 | 107978503 | 1 | protein_coding | cullin 5 |
| **C2** | ENSG00000166278 | 1358.07736 | 0.654828384 | 0.14299454 | 2.49E-06 | 1.51E-05 | 717 | 6 | 31865562 | 31913449 | 1 | protein_coding | complement component 2 |
| **PLEKHF1** | ENSG00000166289 | 360.5968562 | -0.826633907 | 0.20254917 | 1.76E-05 | 9.27E-05 | 79156 | 19 | 30155963 | 30166364 | 1 | protein_coding | pleckstrin homology domain containing, family F (with FYVE domain) member 1 |
| **SMPD1** | ENSG00000166311 | 1671.301827 | -0.526692521 | 0.150321631 | 2.93E-04 | 0.001208082 | 6609 | 11 | 6411655 | 6416228 | 1 | protein_coding | sphingomyelin phosphodiesterase 1, acid lysosomal |

| | | | | | | | | | | | | | |
|---|---|---|---|---|---|---|---|---|---|---|---|---|---|
| **APBB1** | ENSG00000166313 | 345.6126076 | -1.230029771 | 0.213150645 | 1.6E-09 | 1.5E-08 | 322 | 11 | 6416355 | 6440644 | -1 | protein_coding | amyloid beta (A4) precursor protein-binding, family B, member 1 (Fe65) |
| **ILK** | ENSG00000166333 | 2320.421361 | 0.428351484 | 0.114546034 | 1.38E-04 | 6.13E-04 | 3611 | 11 | 6624961 | 6632102 | 1 | protein_coding | integrin-linked kinase |
| **TAF10** | ENSG00000166337 | 825.2250677 | -0.711471947 | 0.146770536 | 5.93E-07 | 3.96E-06 | 6881 | 11 | 6627526 | 6633898 | -1 | protein_coding | TAF10 RNA polymerase II, TATA box binding protein (TBP)-associated factor, 30kDa |
| **TPP1** | ENSG00000166340 | 21475.31447 | -0.462467548 | 0.103720541 | 5.87E-06 | 3.35E-05 | 1200 | 11 | 6634000 | 6640692 | -1 | protein_coding | tripeptidyl peptidase I |
| **DCHS1** | ENSG00000166341 | 217.6693955 | -3.429409002 | 0.427231279 | 6.35E-17 | 1.1E-15 | 8642 | 11 | 6642556 | 6677085 | -1 | protein_coding | dachsous cadherin-related 1 |
| **PPFIBP2** | ENSG00000166387 | 839.1299264 | 1.175815495 | 0.20229887 | 1.36E-09 | 1.29E-08 | 8495 | 11 | 7534529 | 7678358 | 1 | protein_coding | PTPRF interacting protein, binding protein 2 (liprin beta 2) |
| **SERPINB8** | ENSG00000166401 | 1229.533714 | 0.379710198 | 0.140458324 | 0.005587131 | 0.016792159 | 5271 | 18 | 61637159 | 61672278 | 1 | protein_coding | serpin peptidase inhibitor, clade B (ovalbumin), member 8 |
| **RNF169** | ENSG00000166439 | 1439.038959 | 0.543550525 | 0.104219675 | 1.31E-07 | 9.55E-07 | 254225 | 11 | 74459913 | 74553458 | 1 | protein_coding | ring finger protein 169 |
| **RPL27A** | ENSG00000166441 | 5938.526253 | -0.609508579 | 0.122857162 | 3.72E-07 | 2.55E-06 | 619562 | 11 | 8703958 | 8736306 | 1 | protein_coding | ribosomal protein L27a |
| **ST5** | ENSG00000166444 | 619.9120424 | -2.155169795 | 0.254327277 | 2.56E-18 | 4.94E-17 | 6764 | 11 | 8714898 | 8932498 | -1 | protein_coding | suppression of tumorigenicity 5 |
| **CENPN** | ENSG00000166451 | 319.1938695 | 1.389998506 | 0.23724975 | 7.89E-10 | 7.66E-09 | 55839 | 16 | 81040103 | 81066719 | 1 | protein_coding | centromere protein N |
| **AKIP1** | ENSG00000166452 | 761.3388633 | -0.748576619 | 0.141858722 | 6.45E-08 | 4.92E-07 | 56672 | 11 | 8932686 | 8941631 | 1 | protein_coding | A kinase (PRKA) interacting protein 1 |
| **TMEM41B** | ENSG00000166471 | 1390.672646 | 1.133401924 | 0.12330729 | 8.61E-21 | 2.02E-19 | 440026 | 11 | 9302201 | 9336327 | -1 | protein_coding | transmembrane protein 41B |
| **WEE1** | ENSG00000166483 | 96.44935267 | 1.213655827 | 0.369171575 | 1.9E-04 | 8.19E-04 | 7465 | 11 | 9595228 | 9615004 | 1 | protein_coding | WEE1 G2 checkpoint kinase |
| **MAPK7** | ENSG00000166484 | 582.9819122 | -0.500346091 | 0.19709988 | 0.0073917 | 0.021501209 | 5598 | 17 | 19281034 | 19286857 | 1 | protein_coding | mitogen-activated protein kinase 7 |
| **PRKCB** | ENSG00000166501 | 654.5729967 | -1.879899898 | 0.188678554 | 1.98E-24 | 5.75E-23 | 5579 | 16 | 23847322 | 24231932 | 1 | protein_coding | protein kinase C, beta |
| **NDST2** | ENSG00000166507 | 843.5686241 | 0.405154587 | 0.154742535 | 0.00678981 | 0.019967033 | 8509 | 10 | 75561669 | 75571589 | -1 | protein_coding | N-deacetylase/N-sulfotransferase (heparan glucosaminyl) 2 |
| **MCM7** | ENSG00000166508 | 383.3220525 | 0.560759408 | 0.241207361 | 0.011776841 | 0.032220813 | 4176 | 7 | 99690351 | 99699563 | -1 | protein_coding | minichromosome maintenance complex component 7 |
| **CLEC4E** | ENSG00000166523 | 949.6841262 | 3.908586333 | 0.224266702 | 2.44E-69 | 4.55E-67 | 26253 | 12 | 8685901 | 8693559 | -1 | protein_coding | C-type lectin domain family 4, member E |
| **CLEC4D** | ENSG00000166527 | 25.54325558 | 2.620214409 | 0.789818202 | 5.04E-05 | 2.43E-04 | 338339 | 12 | 8662071 | 8674962 | 1 | protein_coding | C-type lectin domain family 4, member D |
| **TK2** | ENSG00000166548 | 1411.452688 | -0.999039466 | 0.218480006 | 1.34E-06 | 8.43E-06 | 7084 | 16 | 66541906 | 66586447 | -1 | protein_coding | thymidine kinase 2, mitochondrial |
| **TMED3** | ENSG00000166557 | 1302.401515 | -0.791789959 | 0.142615895 | 1.17E-08 | 9.85E-08 | 23423 | 15 | 79603404 | 79704334 | 1 | protein_coding | transmembrane emp24 protein transport domain containing 3 |
| **TMEM135** | ENSG00000166575 | 469.6124304 | 0.846275429 | 0.290572206 | 0.001187418 | 0.004251947 | 65084 | 11 | 86748886 | 87034800 | 1 | protein_coding | transmembrane protein 135 |
| **HSP90B1** | ENSG00000166598 | 19067.30366 | -0.313731362 | 0.113119534 | 0.004346982 | 0.013478377 | 1005008 | 12 | 104323885 | 104347423 | 1 | protein_coding | heat shock protein 90kDa beta (Grp94), member 1 |
| **MMP10** | ENSG00000166670 | 34.97233021 | 2.018261014 | 0.641498151 | 1.32E-04 | 5.89E-04 | 4319 | 11 | 102641234 | 102651359 | -1 | protein_coding | matrix metallopeptidase 10 (stromelysin 2) |
| **NGFRAP1** | ENSG00000166681 | 1336.430115 | -0.826252869 | 0.156504262 | 4.95E-08 | 3.85E-07 | 27018 | X | 102631268 | 102633005 | 1 | protein_coding | nerve growth factor receptor (TNFRSF16) associated protein 1 |
| **TMPRSS5** | ENSG00000166682 | 27.61301723 | -2.826342688 | 1.080668753 | 9.68E-04 | 0.003542381 | 80975 | 11 | 113558272 | 113577095 | -1 | protein_coding | transmembrane protease, serine 5 |
| **PLEKHA7** | ENSG00000166689 | 53.66650615 | 2.19005226 | 0.703132155 | 1.19E-04 | 5.37E-04 | 144100 | 11 | 16799842 | 17035990 | -1 | protein_coding | pleckstrin homology domain containing, family A member 7 |
| **B2M** | ENSG00000166710 | 160693.4755 | 0.812828224 | 0.0852997 | 6.55E-22 | 1.65E-20 | 567 | 15 | 45003675 | 45011075 | 1 | protein_coding | beta-2-microglobulin |
| **ZNF592** | ENSG00000166716 | 2452.722553 | -0.445754471 | 0.108813065 | 3.07E-05 | 1.54E-04 | 9640 | 15 | 85291866 | 85349659 | 1 | protein_coding | zinc finger protein 592 |
| **SLFN5** | ENSG00000166750 | 3533.068993 | 0.685580622 | 0.120975461 | 7.69E-09 | 6.64E-08 | 162394 | 17 | 33570055 | 33600674 | 1 | protein_coding | schlafen family member 5 |
| **C16orf45** | ENSG00000166780 | 134.9301844 | -1.890937867 | 0.403680371 | 2.88E-07 | 2.01E-06 | 89927 | 16 | 15528152 | 15718885 | 1 | protein_coding | chromosome 16 open reading frame 45 |
| **YPEL4** | ENSG00000166793 | 992.9978863 | -0.399443484 | 0.133807964 | 0.002180139 | 0.007287634 | 219539 | 11 | 57412560 | 57417417 | -1 | protein_coding | yippee-like 4 (Drosophila) |
| **FAM111A** | ENSG00000166801 | 3128.136551 | 1.257945368 | 0.116053237 | 4.52E-28 | 1.57E-26 | 63901 | 11 | 58910221 | 58922512 | 1 | protein_coding | family with sequence similarity 111, member A |
| **KIAA0101** | ENSG00000166803 | 38.78325472 | 1.291187754 | 0.637850501 | 0.005244521 | 0.01591265 | 9768 | 15 | 64657193 | 64679886 | -1 | protein_coding | KIAA0101 |
| **LDHD** | ENSG00000166816 | 144.9341826 | -1.682777746 | 0.358431583 | 3.6E-07 | 2.48E-06 | 197257 | 16 | 75145758 | 75150669 | -1 | protein_coding | lactate dehydrogenase D |
| **TMEM170A** | ENSG00000166822 | 1082.443143 | 1.3205964 | 0.130553197 | 9.36E-25 | 2.77E-23 | 124491 | 16 | 75476952 | 75499395 | -1 | protein_coding | transmembrane protein 170A |
| **ANPEP** | ENSG00000166825 | 6563.710612 | 0.934575517 | 0.13254435 | 6.4E-13 | 8.29E-12 | 290 | 15 | 90328120 | 90358633 | -1 | protein_coding | alanyl (membrane) aminopeptidase |
| **TERF2IP** | ENSG00000166848 | 1958.083846 | 0.568592826 | 0.105973649 | 4.65E-08 | 3.63E-07 | 54386 | 16 | 75681684 | 75795770 | 1 | protein_coding | telomeric repeat binding factor 2, interacting protein |
| **TMEM194A** | ENSG00000166881 | 1699.014374 | 1.478715344 | 0.193625854 | 3.21E-15 | 4.92E-14 | 23306 | 12 | 57449426 | 57481846 | -1 | protein_coding | transmembrane protein 194A |
| **NAB2** | ENSG00000166886 | 891.7103596 | -0.631775175 | 0.164149228 | 6.45E-05 | 3.05E-04 | 4665 | 12 | 57482677 | 57489259 | 1 | protein_coding | NGFI-A binding protein 2 (EGR1 binding protein 2) |
| **STAT6** | ENSG00000166888 | 6417.988856 | 0.412179402 | 0.141892452 | 0.002919584 | 0.00946141 | 6778 | 12 | 57489191 | 57525922 | -1 | protein_coding | signal transducer and activator of transcription 6, interleukin-4 induced |
| **PATL1** | ENSG00000166889 | 2671.24209 | 0.561161447 | 0.099684712 | 1.16E-08 | 9.77E-08 | 219988 | 11 | 59404189 | 59436453 | -1 | protein_coding | protein associated with topoisomerase II homolog 1 (yeast) |
| **STX3** | ENSG00000166900 | 2103.560947 | 0.569901022 | 0.16056715 | 2.34E-04 | 9.87E-04 | 6809 | 11 | 59480929 | 59573354 | 1 | protein_coding | syntaxin 3 |
| **MRPL16** | ENSG00000166902 | 808.2190585 | 0.686002894 | 0.145553014 | 1.21E-06 | 7.7E-06 | 54948 | 11 | 59573608 | 59578345 | -1 | protein_coding | mitochondrial ribosomal protein L16 |
| **PIP4K2C** | ENSG00000166908 | 1072.781757 | 0.612721246 | 0.128132294 | 9.7E-07 | 6.26E-06 | 79837 | 12 | 57984957 | 57997198 | 1 | protein_coding | phosphatidylinositol-5-phosphate 4-kinase, type II, gamma |
| **MTMR10** | ENSG00000166912 | 2429.596508 | -0.544989593 | 0.122930819 | 5.98E-06 | 3.4E-05 | 54893 | 15 | 31231144 | 31283810 | -1 | protein_coding | myotubularin related protein 10 |
| **C15orf48** | ENSG00000166920 | 7691.006943 | 2.721602983 | 0.165534221 | 6.14E-62 | 8.45E-60 | 84419 | 15 | 45722727 | 45740959 | 1 | protein_coding | chromosome 15 open reading frame 48 |
| **SCG5** | ENSG00000166922 | 208.1210637 | -0.932779833 | 0.274254003 | 2.03E-04 | 8.67E-04 | 6447 | 15 | 32933877 | 32989299 | 1 | protein_coding | secretogranin V (7B2 protein) |
| **MS4A6E** | ENSG00000166926 | 376.1021542 | -0.917099547 | 0.41410145 | 0.00687685 | 0.020195791 | 245802 | 11 | 60102304 | 60164069 | 1 | protein_coding | membrane-spanning 4-domains, subfamily A, member 6E |
| **MS4A7** | ENSG00000166927 | 5873.469359 | 2.107950285 | 0.124563879 | 1.81E-65 | 2.85E-63 | 58475 | 11 | 60145955 | 60163424 | 1 | protein_coding | membrane-spanning 4-domains, subfamily A, member 7 |
| **MS4A14** | ENSG00000166928 | 365.6438785 | 1.884360501 | 0.191293712 | 7.78E-24 | 2.19E-22 | 84689 | 11 | 60146003 | 60185161 | 1 | protein_coding | membrane-spanning 4-domains, subfamily A, member 14 |
| **CCNDBP1** | ENSG00000166946 | 1383.327659 | 0.298318528 | 0.118010459 | 0.010140312 | 0.028347245 | 23582 | 15 | 43477316 | 43487396 | 1 | protein_coding | cyclin D-type binding-protein 1 |
| **MAPRE2** | ENSG00000166974 | 4024.090456 | -0.606407536 | 0.142090651 | 1.1E-05 | 6.02E-05 | 10982 | 18 | 32556892 | 32723434 | 1 | protein_coding | microtubule-associated protein, RP/EB family, member 2 |
| **NKX3-1** | ENSG00000167034 | 19.63890315 | 1.329498111 | 0.991068356 | 0.013680393 | 0.036711616 | 4824 | 8 | 23536206 | 23540440 | -1 | protein_coding | NK3 homeobox 1 |
| **MEI1** | ENSG00000167077 | 284.4148725 | 0.542616551 | 0.201198341 | 0.004315819 | 0.013389902 | 150365 | 22 | 42095503 | 42195460 | 1 | protein_coding | meiosis inhibitor 1 |
| **PHB** | ENSG00000167085 | 1179.961154 | -0.682871862 | 0.144838784 | 1.23E-06 | 7.79E-06 | 5245 | 17 | 47481414 | 47492246 | -1 | protein_coding | prohibitin |
| **FAM102A** | ENSG00000167106 | 589.1179412 | -0.817875591 | 0.25878401 | 5.83E-04 | 0.002239701 | 399665 | 9 | 130702858 | 130742792 | -1 | protein_coding | family with sequence similarity 102, member A |
| **ACSF2** | ENSG00000167107 | 877.0945037 | -1.252883777 | 0.151357325 | 2.69E-17 | 4.81E-16 | 80221 | 17 | 48503519 | 48552206 | 1 | protein_coding | acyl-CoA synthetase family member 2 |
| **GOLGA2** | ENSG00000167110 | 1619.722395 | -0.258655325 | 0.110579963 | 0.01702659 | 0.044322496 | 2801 | 9 | 131018108 | 131038274 | -1 | protein_coding | golgin A2 |
| **SLC27A4** | ENSG00000167114 | 539.6750346 | -0.58549756 | 0.174924441 | 4.81E-04 | 0.00189102 | 10999 | 9 | 131102925 | 131123749 | 1 | protein_coding | solute carrier family 27 (fatty acid transporter), member 4 |
| **URM1** | ENSG00000167118 | 549.6243207 | -0.409126204 | 0.168534003 | 0.011593435 | 0.031793014 | 81605 | 9 | 131133598 | 131153015 | 1 | protein_coding | ubiquitin related modifier 1 |
| **CERCAM** | ENSG00000167123 | 376.9463657 | -0.52141512 | 0.204907567 | 0.006797687 | 0.019982505 | 51148 | 9 | 131174030 | 131199626 | 1 | protein_coding | cerebral endothelial cell adhesion molecule |
| **C15orf39** | ENSG00000167173 | 600.401353 | 0.633735818 | 0.245649605 | 0.005154302 | 0.01569187 | 56905 | 15 | 75487984 | 75504510 | 1 | protein_coding | chromosome 15 open reading frame 39 |
| **TBC1D2B** | ENSG00000167202 | 6387.226768 | -0.252075024 | 0.088215616 | 0.003600281 | 0.011410844 | 23102 | 15 | 78276378 | 78370066 | -1 | protein_coding | TBC1 domain family, member 2B |
| **NOD2** | ENSG00000167207 | 727.3646064 | 1.232355347 | 0.294722169 | 5.6E-06 | 3.22E-05 | 64127 | 16 | 50727514 | 50766988 | 1 | protein_coding | nucleotide-binding oligomerization domain containing 2 |
| **SNX20** | ENSG00000167208 | 2268.169338 | 0.606503881 | 0.125056656 | 6.71E-07 | 4.45E-06 | 124460 | 16 | 50700211 | 50715264 | -1 | protein_coding | sorting nexin 20 |
| **KATNAL2** | ENSG00000167216 | 96.09662181 | -1.537646233 | 0.473948355 | 1.4E-04 | 6.25E-04 | 83473 | 18 | 44497455 | 44627658 | 1 | protein_coding | katanin p60 subunit A-like 2 |
| **ZNF91** | ENSG00000167232 | 1299.841841 | 0.494087484 | 0.190467154 | 0.007013796 | 0.020546644 | 7644 | 19 | 23487793 | 23578362 | -1 | protein_coding | zinc finger protein 91 |
| **CDK12** | ENSG00000167258 | 2693.297877 | 0.288651118 | 0.111534631 | 0.007883127 | 0.022738013 | 51755 | 17 | 37617764 | 37721160 | 1 | protein_coding | cyclin-dependent kinase 12 |
| **DPEP2** | ENSG00000167261 | 713.1477652 | -1.570262487 | 0.176923381 | 9.98E-20 | 2.16E-18 | 64174 | 16 | 68021297 | 68034489 | -1 | protein_coding | dipeptidase 2 |
| **DUS2** | ENSG00000167264 | 370.3043554 | -0.595650583 | 0.19182063 | 0.001094879 | 0.003963298 | 54920 | 16 | 68021649 | 68113223 | 1 | protein_coding | dihydrouridine synthase 2 |
| **POP5** | ENSG00000167272 | 218.9136321 | -0.532419473 | 0.243545352 | 0.018070497 | 0.046556204 | 51367 | 12 | 121016567 | 121019201 | -1 | protein_coding | processing of precursor 5, ribonuclease P/MRP subunit (S. cerevisiae) |
| **ATP5L** | ENSG00000167283 | 1986.348308 | -0.330162592 | 0.134915025 | 0.012119473 | 0.033022269 | 10632 | 11 | 118271869 | 118302211 | 1 | protein_coding | ATP synthase, H+ transporting, mitochondrial Fo complex, subunit G |
| **CD3D** | ENSG00000167286 | 402.9289601 | -0.730642651 | 0.262415139 | 0.00226812 | 0.007549133 | 915 | 11 | 118209669 | 118213459 | -1 | protein_coding | CD3d molecule, delta (CD3-TCR complex) |
| **MYO5B** | ENSG00000167306 | 65.47457334 | 1.178801893 | 0.515631366 | 0.003649541 | 0.011552596 | 4645 | 18 | 47349183 | 47721463 | -1 | protein_coding | myosin VB |
| **ACAA2** | ENSG00000167315 | 2423.229878 | -0.548795397 | 0.152410602 | 1.79E-04 | 7.78E-04 | 10449 | 18 | 47309869 | 47340330 | -1 | protein_coding | acetyl-CoA acyltransferase 2 |
| **RRM1** | ENSG00000167325 | 882.9982016 | 0.404141834 | 0.152566456 | 0.00632973 | 0.01877346 | 6240 | 11 | 4115937 | 4160106 | 1 | protein_coding | ribonucleotide reductase M1 |
| **FN3K** | ENSG00000167363 | 67.20111924 | -1.610451583 | 0.624817505 | 9.2E-04 | 0.003387411 | 64122 | 17 | 80693451 | 80709073 | 1 | protein_coding | fructosamine 3 kinase |
| **IRGQ** | ENSG00000167378 | 1485.167454 | -0.808272859 | 0.128001444 | 1.3E-10 | 1.37E-09 | 126298 | 19 | 44088521 | 44100287 | -1 | protein_coding | immunity-related GTPase family, Q |
| **VKORC1** | ENSG00000167397 | 806.3564179 | -0.775500726 | 0.139378374 | 1.16E-08 | 9.77E-08 | 79001 | 16 | 31102163 | 31107301 | -1 | protein_coding | vitamin K epoxide reductase complex, subunit 1 |
| **LPO** | ENSG00000167419 | 46.18367695 | -1.350213834 | 0.756220613 | 0.007596383 | 0.022015638 | 4025 | 17 | 56295909 | 56345879 | 1 | protein_coding | lactoperoxidase |
| **TPM4** | ENSG00000167460 | 13175.28762 | 0.345373354 | 0.127926173 | 0.005677171 | 0.017025943 | 7171 | 19 | 16177831 | 16213813 | 1 | protein_coding | tropomyosin 4 |
| **RAB8A** | ENSG00000167461 | 3861.160577 | 0.707894443 | 0.151005908 | 1.22E-06 | 7.74E-06 | 4218 | 19 | 16222439 | 16245044 | 1 | protein_coding | RAB8A, member RAS oncogene family |
| **FAM129C** | ENSG00000167483 | 46.29703767 | -1.024055953 | 0.592225132 | 0.015409988 | 0.040731727 | 199786 | 19 | 17634110 | 17664648 | 1 | protein_coding | family with sequence similarity 129, member C |
| **GATAD2A** | ENSG00000167491 | 2992.533867 | -0.430477323 | 0.113371385 | 9.04E-05 | 4.17E-04 | 54815 | 19 | 19496635 | 19619740 | 1 | protein_coding | GATA zinc finger domain containing 2A |
| **CDT1** | ENSG00000167513 | 31.49905889 | 1.064297848 | 0.707953031 | 0.019210104 | 0.049037584 | 81620 | 16 | 88869621 | 88875666 | 1 | protein_coding | chromatin licensing and DNA replication factor 1 |
| **TRAPPC2L** | ENSG00000167515 | 649.4571188 | -1.220403985 | 0.204627695 | 5.23E-10 | 5.16E-09 | 51693 | 16 | 88922628 | 88929094 | 1 | protein_coding | trafficking protein particle complex 2-like |
| **ANKRD11** | ENSG00000167522 | 3051.983792 | -0.495761537 | 0.111891593 | 6.46E-06 | 3.66E-05 | 29123 | 16 | 89334038 | 89556969 | -1 | protein_coding | ankyrin repeat domain 11 |

| | | | | | | | | | | | | | |
|---|---|---|---|---|---|---|---|---|---|---|---|---|---|
| **RPL13** | ENSG00000167526 | 4758.813957 | -0.656859774 | 0.124348648 | 7.77E-08 | 5.86E-07 | 606500 | 16 | 89627065 | 89630950 | 1 | protein_coding | ribosomal protein L13 |
| **TP53I13** | ENSG00000167543 | 179.6648784 | -0.773452033 | 0.286517699 | 0.002716498 | 0.008859632 | 90313 | 17 | 27893070 | 27900175 | 1 | protein_coding | tumor protein p53 inducible protein 13 |
| **RHEBL1** | ENSG00000167550 | 30.87616536 | 2.267866203 | 0.688743395 | 7.08E-05 | 3.32E-04 | 121268 | 12 | 49458468 | 49463808 | -1 | protein_coding | Ras homolog enriched in brain like 1 |
| **TUBA1A** | ENSG00000167552 | 9120.683843 | -0.29504747 | 0.116481373 | 0.010228192 | 0.028540709 | 7846 | 12 | 49578579 | 49583107 | -1 | protein_coding | tubulin, alpha 1a |
| **TUBA1C** | ENSG00000167553 | 4021.558577 | 0.384702566 | 0.105579078 | 1.93E-04 | 8.3E-04 | 84790 | 12 | 49582519 | 49667114 | 1 | protein_coding | tubulin, alpha 1c |
| **ZNF528** | ENSG00000167555 | 858.1400881 | -0.857350602 | 0.198763304 | 5.64E-06 | 3.24E-05 | 84436 | 19 | 52901102 | 52921665 | 1 | protein_coding | zinc finger protein 528 |
| **NCKAP5L** | ENSG00000167566 | 1063.341702 | -1.113016045 | 0.14644314 | 8.12E-15 | 1.22E-13 | 57701 | 12 | 50184929 | 50222533 | -1 | protein_coding | NCK-associated protein 5-like |
| **RAB4B** | ENSG00000167578 | 687.9177424 | -0.390808167 | 0.141978406 | 0.00424193 | 0.013190116 | 53916 | 19 | 41284121 | 41302847 | 1 | protein_coding | RAB4B, member RAS oncogene family |
| **GPD1** | ENSG00000167588 | 404.0335166 | -3.420080953 | 0.76028684 | 3.62E-07 | 2.49E-06 | 2819 | 12 | 50497602 | 50505102 | 1 | protein_coding | glycerol-3-phosphate dehydrogenase 1 (soluble) |
| **CYP2S1** | ENSG00000167600 | 665.0122791 | -1.006200472 | 0.175649559 | 2.99E-09 | 2.71E-08 | 29785 | 19 | 41698954 | 41713444 | 1 | protein_coding | cytochrome P450, family 2, subfamily S, polypeptide 1 |
| **AXL** | ENSG00000167601 | 96.20197849 | 4.036203301 | 0.443829021 | 6.2E-21 | 1.46E-19 | 558 | 19 | 41725108 | 41767671 | 1 | protein_coding | AXL receptor tyrosine kinase |
| **NFKBID** | ENSG00000167604 | 321.9023945 | -0.969454354 | 0.217926739 | 2.47E-06 | 1.5E-05 | 84807 | 19 | 36378555 | 36393205 | -1 | protein_coding | nuclear factor of kappa light polypeptide gene enhancer in B-cells inhibitor, delta |
| **YIF1B** | ENSG00000167645 | 389.4007132 | -1.178113068 | 0.198630176 | 7.1E-10 | 6.92E-09 | 90522 | 19 | 38795753 | 38807913 | -1 | protein_coding | Yip1 interacting factor homolog B (S. cerevisiae) |
| **EEF2** | ENSG00000167658 | 18152.79908 | -0.303019528 | 0.089283434 | 6.46E-04 | 0.002459875 | 1938 | 19 | 3976054 | 3985467 | -1 | protein_coding | eukaryotic translation elongation factor 2 |
| **UBXN6** | ENSG00000167671 | 2157.648755 | -0.926605449 | 0.107385855 | 1.47E-18 | 2.91E-17 | 1006163 | 19 | 4444996 | 4457819 | -1 | protein_coding | UBX domain protein 6 |
| **PLIN4** | ENSG00000167676 | 22.93827638 | -1.425761967 | 0.84076091 | 0.009171654 | 0.025981536 | 729359 | 19 | 4502204 | 4517716 | -1 | protein_coding | perilipin 4 |
| **SEMA6B** | ENSG00000167680 | 879.8284121 | -2.542709995 | 0.315748479 | 6.32E-17 | 1.1E-15 | 10501 | 19 | 4542600 | 4559820 | -1 | protein_coding | sema domain, transmembrane domain (TM), and cytoplasmic domain, (semaphorin) 6B |
| **MFSD3** | ENSG00000167700 | 48.45503207 | -1.202862732 | 0.516194708 | 0.003270114 | 0.01047739 | 113655 | 8 | 145734457 | 145736596 | 1 | protein_coding | major facilitator superfamily domain containing 3 |
| **SLC43A2** | ENSG00000167703 | 1013.103172 | 1.009965916 | 0.143471265 | 5.78E-13 | 7.55E-12 | 124935 | 17 | 1472561 | 1532180 | -1 | protein_coding | solute carrier family 43 (amino acid system L transporter), member 2 |
| **SRR** | ENSG00000167720 | 166.2552789 | 0.669094467 | 0.28919355 | 0.009808195 | 0.027554899 | 63826 | 17 | 2206677 | 2228554 | 1 | protein_coding | serine racemase |
| **CYB5D2** | ENSG00000167740 | 287.596241 | -0.715016915 | 0.208078916 | 2.75E-04 | 0.00114206 | 124936 | 17 | 4046462 | 4090605 | 1 | protein_coding | cytochrome b5 domain containing 2 |
| **KLK4** | ENSG00000167749 | 68.33649504 | -1.102538772 | 0.503507022 | 0.005415035 | 0.016358564 | 9622 | 19 | 51409608 | 51413994 | -1 | protein_coding | kallikrein-related peptidase 4 |
| **ANGPTL4** | ENSG00000167772 | 33.25879142 | -3.268643502 | 0.825217798 | 5.11E-06 | 2.95E-05 | 51129 | 19 | 8428173 | 8439257 | 1 | protein_coding | angiopoietin-like 4 |
| **SPRYD3** | ENSG00000167778 | 895.0289685 | -0.320115368 | 0.141075939 | 0.019132074 | 0.048862882 | 84926 | 12 | 53458388 | 53473204 | -1 | protein_coding | SPRY domain containing 3 |
| **IGFBP6** | ENSG00000167779 | 78.57821 | 2.67700723 | 0.519300293 | 1.49E-08 | 1.24E-07 | 3489 | 12 | 53491220 | 53496129 | 1 | protein_coding | insulin-like growth factor binding protein 6 |
| **NDUFV1** | ENSG00000167792 | 1103.163353 | -0.428816246 | 0.124371537 | 4.69E-04 | 0.001844942 | 4723 | 11 | 67374323 | 67380006 | 1 | protein_coding | NADH dehydrogenase (ubiquinone) flavoprotein 1, 51kDa |
| **CDK2AP2** | ENSG00000167797 | 372.058976 | -0.714649813 | 0.182685462 | 4.25E-05 | 2.08E-04 | 10263 | 11 | 67273968 | 67276120 | -1 | protein_coding | cyclin-dependent kinase 2 associated protein 2 |
| **PRDX2** | ENSG00000167815 | 586.406155 | -0.655639224 | 0.174921865 | 9.26E-05 | 4.26E-04 | 7001 | 19 | 12907634 | 12912694 | -1 | protein_coding | peroxiredoxin 2 |
| **MIS12** | ENSG00000167842 | 433.7043957 | 0.764000775 | 0.265972091 | 0.001602678 | 0.005555392 | 79003 | 17 | 5389605 | 5394134 | 1 | protein_coding | MIS12 kinetochore complex component |
| **CD300C** | ENSG00000167850 | 843.2856943 | -1.363208709 | 0.156366261 | 5.19E-19 | 1.06E-17 | 10871 | 17 | 72537247 | 72542282 | -1 | protein_coding | CD300c molecule |
| **CD300A** | ENSG00000167851 | 1796.024015 | -2.066896214 | 0.157543372 | 2.81E-40 | 1.7E-38 | 11314 | 17 | 72462555 | 72480933 | 1 | protein_coding | CD300a molecule |
| **ATP5H** | ENSG00000167863 | 1955.800185 | -0.428693604 | 0.129501632 | 7.01E-04 | 0.00265106 | 10476 | 17 | 73034958 | 73043080 | -1 | protein_coding | ATP synthase, H+ transporting, mitochondrial Fo complex, subunit d |
| **MGAT5B** | ENSG00000167889 | 8.666242578 | -2.629270902 | 3.618080975 | 0.007038033 | 0.020601849 | 146664 | 17 | 74864538 | 74946475 | 1 | protein_coding | mannosyl (alpha-1,6-)-glycoprotein beta-1,6-N-acetyl-glucosaminyltransferase, isozyme B |
| **TMC8** | ENSG00000167895 | 745.458449 | -1.637002953 | 0.215460082 | 4.21E-15 | 6.41E-14 | 147138 | 17 | 76126851 | 76139049 | 1 | protein_coding | transmembrane channel-like 8 |
| **TMEM68** | ENSG00000167904 | 373.9968311 | 0.743169902 | 0.204501017 | 1.2E-04 | 5.42E-04 | 137695 | 8 | 56608983 | 56685966 | -1 | protein_coding | transmembrane protein 68 |
| **GSDMA** | ENSG00000167914 | 32.34865982 | -2.337544171 | 0.782461476 | 2.03E-04 | 8.65E-04 | 284110 | 17 | 38119226 | 38134019 | 1 | protein_coding | gasdermin A |
| **GHDC** | ENSG00000167925 | 454.4852674 | -0.719974371 | 0.202194261 | 1.69E-04 | 7.38E-04 | 84514 | 17 | 40340817 | 40346531 | -1 | protein_coding | GH3 domain containing |
| **MLST8** | ENSG00000167965 | 360.1810946 | -1.2949155 | 0.221114057 | 9.62E-10 | 9.27E-09 | 64223 | 16 | 2254249 | 2259417 | 1 | protein_coding | MTOR associated protein, LST8 homolog (S. cerevisiae) |
| **RAB3IL1** | ENSG00000167994 | 244.7258582 | -1.649654997 | 0.269315225 | 1.1E-10 | 1.17E-09 | 5866 | 11 | 61664773 | 61687741 | -1 | protein_coding | RAB3A interacting protein (rabin3)-like 1 |
| **BSCL2** | ENSG00000168000 | 297.5849001 | -0.990340802 | 0.289184046 | 1.72E-04 | 7.49E-04 | 26580 | 11 | 62457747 | 62477317 | -1 | protein_coding | Berardinelli-Seip congenital lipodystrophy 2 (seipin) |
| **TRANK1** | ENSG00000168016 | 2400.834117 | 1.897565795 | 0.126460542 | 5.39E-52 | 5.25E-50 | 9881 | 3 | 36868311 | 36986548 | -1 | protein_coding | tetratricopeptide repeat and ankyrin repeat containing 1 |
| **TTC21A** | ENSG00000168026 | 92.18124084 | 1.629847536 | 0.459235126 | 4.08E-05 | 2E-04 | 199223 | 3 | 39149152 | 39180394 | 1 | protein_coding | tetratricopeptide repeat domain 21A |
| **CTNNB1** | ENSG00000168036 | 7845.863704 | 0.256072621 | 0.097961943 | 0.007597362 | 0.022015638 | 1499 | 3 | 41236328 | 41301587 | 1 | protein_coding | catenin (cadherin-associated protein), beta 1, 88kDa |
| **LTBP3** | ENSG00000168056 | 1841.1931 | -0.62674002 | 0.208977261 | 0.001384556 | 0.004866643 | 4054 | 11 | 65306276 | 65326401 | -1 | protein_coding | latent transforming growth factor beta binding protein 3 |
| **BATF2** | ENSG00000168062 | 340.6705952 | 3.245294183 | 0.304756277 | 1.27E-27 | 4.31E-26 | 116071 | 11 | 64755415 | 64764517 | -1 | protein_coding | basic leucine zipper transcription factor, ATF-like 2 |
| **PBK** | ENSG00000168078 | 14.97690185 | 2.087388454 | 1.14694279 | 0.003432256 | 0.010937275 | 55872 | 8 | 27667137 | 27695612 | -1 | protein_coding | PDZ binding kinase |
| **NUDT16L1** | ENSG00000168101 | 211.6262987 | -0.883818412 | 0.241999932 | 9.06E-05 | 4.17E-04 | 84309 | 16 | 4743695 | 4745860 | 1 | protein_coding | nudix (nucleoside diphosphate linked moiety X)-type motif 16-like 1 |
| **KIAA1586** | ENSG00000168116 | 166.0841025 | 0.701563988 | 0.277435006 | 0.005158614 | 0.015698093 | 57691 | 6 | 56911347 | 56920023 | 1 | protein_coding | KIAA1586 |
| **RAB4A** | ENSG00000168118 | 620.1826397 | -0.872732568 | 0.21451027 | 1.7E-05 | 8.98E-05 | 10638 | 1 | 229406822 | 229441641 | 1 | protein_coding | RAB4A, member RAS oncogene family |
| **SETD5** | ENSG00000168137 | 2098.116917 | 0.585165408 | 0.109790637 | 5.85E-08 | 4.5E-07 | 55209 | 3 | 9439299 | 9520924 | 1 | protein_coding | SET domain containing 5 |
| **THAP9** | ENSG00000168152 | 70.07283415 | 0.947812662 | 0.445628872 | 0.008224807 | 0.023592291 | 79725 | 4 | 83821837 | 83841122 | 1 | protein_coding | THAP domain containing 9 |
| **RNF187** | ENSG00000168159 | 1280.995832 | -1.128626117 | 0.130394168 | 1.34E-18 | 2.65E-17 | 149603 | 1 | 228674762 | 228683467 | 1 | protein_coding | ring finger protein 187 |
| **HOOK3** | ENSG00000168172 | 4160.41056 | -0.448600678 | 0.177876632 | 0.008295689 | 0.023751033 | 84376 | 8 | 42752075 | 42885682 | 1 | protein_coding | hook microtubule-tethering protein 3 |
| **RBPJ** | ENSG00000168214 | 4051.279836 | -0.456035658 | 0.101780661 | 5.69E-06 | 3.26E-05 | 3516 | 4 | 26165077 | 26436541 | 1 | protein_coding | recombination signal binding protein for immunoglobulin kappa J region |
| **LMBRD1** | ENSG00000168216 | 803.6637752 | 1.007201174 | 0.32530172 | 5.08E-04 | 0.001984717 | 55788 | 6 | 70385694 | 70507003 | -1 | protein_coding | LMBR1 domain containing 1 |
| **PTGDR** | ENSG00000168229 | 15.91131663 | -3.742978032 | 1.558870867 | 0.002709181 | 0.008837655 | 5729 | 14 | 52734431 | 52743442 | 1 | protein_coding | prostaglandin D2 receptor (DP) |
| **GNG4** | ENSG00000168243 | 92.03815099 | -1.457831017 | 0.460040519 | 1.9E-04 | 8.2E-04 | 2786 | 1 | 235710987 | 235814054 | -1 | protein_coding | guanine nucleotide binding protein (G protein), gamma 4 |
| **UBTD2** | ENSG00000168246 | 1150.288504 | 0.571144736 | 0.124899178 | 3.02E-06 | 1.81E-05 | 92181 | 5 | 171636644 | 171711075 | -1 | protein_coding | ubiquitin domain containing 2 |
| **POLR2J3** | ENSG00000168255 | 499.7642225 | -0.53598715 | 0.246285901 | 0.017642426 | 0.045692119 | 1001349 | 7 | 102178365 | 102213103 | -1 | protein_coding | polymerase (RNA) II (DNA directed) polypeptide J3 |
| **NKIRAS2** | ENSG00000168256 | 20.06858649 | -3.723243213 | 1.301799192 | 3.57E-04 | 0.001445409 | 28511 | 17 | 40163400 | 40177659 | 1 | protein_coding | NFKB inhibitor interacting Ras-like 2 |
| **DNAJC7** | ENSG00000168259 | 1537.58536 | 1.077082449 | 0.22519441 | 4.43E-07 | 3.02E-06 | 7266 | 17 | 40128451 | 40173394 | -1 | protein_coding | DnaJ (Hsp40) homolog, subfamily C, member 7 |
| **IRF2BP2** | ENSG00000168264 | 3291.125823 | -0.464588009 | 0.173671261 | 0.005333963 | 0.016142369 | 359948 | 1 | 234740015 | 234745271 | -1 | protein_coding | interferon regulatory factor 2 binding protein 2 |
| **SMIM4** | ENSG00000168273 | 134.2113496 | -1.511213449 | 0.346525576 | 1.86E-06 | 1.15E-05 | 440957 | 3 | 52568029 | 52613253 | 1 | protein_coding | small integral membrane protein 4 |
| **COA6** | ENSG00000168275 | 321.2669574 | 1.149362291 | 0.242669618 | 4.6E-07 | 3.13E-06 | 388753 | 1 | 234509202 | 234519795 | 1 | protein_coding | cytochrome c oxidase assembly factor 6 homolog (S. cerevisiae) |
| **MGAT2** | ENSG00000168282 | 1269.885083 | 0.533369612 | 0.109379085 | 6.64E-07 | 4.41E-06 | 4247 | 14 | 50087489 | 50090198 | 1 | protein_coding | mannosyl (alpha-1,6-)-glycoprotein beta-1,2-N-acetylglucosaminyltransferase |
| **BMI1** | ENSG00000168283 | 869.1812233 | 1.034753984 | 0.203840771 | 1.01E-07 | 7.51E-07 | 648 | 10 | 22610140 | 22620413 | 1 | protein_coding | BMI1 polycomb ring finger oncogene |
| **MMADHC** | ENSG00000168288 | 1912.024079 | 0.371043692 | 0.130357102 | 0.003739378 | 0.011788142 | 27249 | 2 | 150426148 | 150444330 | -1 | protein_coding | methylmalonic aciduria (cobalamin deficiency) cblD type, with homocystinuria |
| **PDHB** | ENSG00000168291 | 1213.976688 | -0.402047538 | 0.145981168 | 0.004522353 | 0.013959829 | 5162 | 3 | 58413357 | 58419584 | -1 | protein_coding | pyruvate dehydrogenase (lipoamide) beta |
| **PXK** | ENSG00000168297 | 659.5480112 | -0.446306562 | 0.171315448 | 0.006576968 | 0.01941584 | 54899 | 3 | 58318607 | 58411748 | 1 | protein_coding | PX domain containing serine/threonine kinase |
| **PCMTD1** | ENSG00000168300 | 2642.197049 | 0.34472621 | 0.13879354 | 0.01081442 | 0.029903524 | 115294 | 8 | 52730140 | 52811735 | -1 | protein_coding | protein-L-isoaspartate (D-aspartate) O-methyltransferase domain containing 1 |
| **ACOX2** | ENSG00000168306 | 97.95271727 | -3.739642153 | 0.518912965 | 7.84E-14 | 1.1E-12 | 8309 | 3 | 58490863 | 58523046 | -1 | protein_coding | acyl-CoA oxidase 2, branched chain |
| **IRF2** | ENSG00000168310 | 1829.296953 | 1.119897745 | 0.150832769 | 3.05E-14 | 4.4E-13 | 3660 | 4 | 185308867 | 185395734 | -1 | protein_coding | interferon regulatory factor 2 |
| **2-Sep** | ENSG00000168385 | 7034.385922 | -0.264347777 | 0.089150851 | 0.003125159 | 0.010072082 | 4735 | 2 | 242254515 | 242293442 | 1 | protein_coding | septin 2 |
| **MFSD2A** | ENSG00000168389 | 786.0666003 | 0.88777118 | 0.154245565 | 3.13E-09 | 2.82E-08 | 84879 | 1 | 40420802 | 40435638 | 1 | protein_coding | major facilitator superfamily domain containing 2A |
| **MLKL** | ENSG00000168404 | 1227.689092 | 1.843494002 | 0.147379468 | 7.14E-37 | 3.84E-35 | 197259 | 16 | 74705753 | 74734858 | -1 | protein_coding | mixed lineage kinase domain-like |
| **CMAHP** | ENSG00000168405 | 325.1171485 | 0.6971394 | 0.250946604 | 0.002567795 | 0.008426802 | 8418 | 6 | 25081548 | 25166793 | -1 | pseudogene | cytidine monophospho-N-acetylneuraminic acid hydroxylase, pseudogene |
| **STIP1** | ENSG00000168439 | 6047.318384 | -0.652066242 | 0.10957611 | 1.23E-09 | 1.17E-08 | 10963 | 11 | 63952744 | 63972015 | 1 | protein_coding | stress-induced-phosphoprotein 1 |
| **REEP4** | ENSG00000168476 | 392.1165986 | -0.846838003 | 0.224741818 | 6.1E-05 | 2.89E-04 | 80346 | 8 | 21995533 | 21999464 | -1 | protein_coding | receptor accessory protein 4 |
| **POLR3D** | ENSG00000168495 | 514.069376 | 0.467509079 | 0.186460832 | 0.008483322 | 0.024211133 | 661 | 8 | 22102617 | 22112113 | 1 | protein_coding | polymerase (RNA) III (DNA directed) polypeptide D, 44kDa |
| **FEN1** | ENSG00000168496 | 326.8466715 | 0.631434076 | 0.197980841 | 7.65E-04 | 0.002872019 | 2237 | 11 | 61560109 | 61564716 | 1 | protein_coding | flap structure-specific endonuclease 1 |
| **SDPR** | ENSG00000168497 | 16.95240245 | -1.858041823 | 1.652624792 | 0.012321174 | 0.033476355 | 8436 | 2 | 192699028 | 192711981 | -1 | protein_coding | serum deprivation response |
| **TRAPPC11** | ENSG00000168538 | 2277.84183 | -0.299922537 | 0.117910982 | 0.009212285 | 0.026077301 | 60684 | 4 | 184580420 | 184634745 | 1 | protein_coding | trafficking protein particle complex 11 |
| **SNRNP48** | ENSG00000168566 | 367.8717407 | 0.579755007 | 0.193880958 | 0.001661338 | 0.005734017 | 154007 | 6 | 7590432 | 7612200 | 1 | protein_coding | small nuclear ribonucleoprotein 48kDa (U11/U12) |
| **TMEM223** | ENSG00000168569 | 126.4473831 | -1.129100255 | 0.333027565 | 1.53E-04 | 6.73E-04 | 79064 | 11 | 62539101 | 62559493 | -1 | protein_coding | transmembrane protein 223 |
| **STAT3** | ENSG00000168610 | 7347.037234 | 0.594681314 | 0.139602899 | 1.21E-05 | 6.61E-05 | 6774 | 17 | 40465342 | 40540586 | -1 | protein_coding | signal transducer and activator of transcription 3 (acute-phase response factor) |
| **ADAM9** | ENSG00000168615 | 9431.001862 | -0.258277997 | 0.101070497 | 0.009678722 | 0.027266341 | 8754 | 8 | 38854388 | 38962663 | 1 | protein_coding | ADAM metallopeptidase domain 9 |
| **AXIN2** | ENSG00000168646 | 22.60082832 | -2.151904123 | 0.833960291 | 7E-04 | 0.002646668 | 8313 | 17 | 63524681 | 63557765 | -1 | protein_coding | axin 2 |
| **NDUFS5** | ENSG00000168653 | 1369.61794 | -0.591560025 | 0.138629417 | 1.11E-05 | 6.07E-05 | 4725 | 1 | 39491990 | 39500308 | 1 | protein_coding | NADH dehydrogenase (ubiquinone) Fe-S protein 5, 15kDa (NADH-coenzyme Q reductase) |
| **LDLRAD4** | ENSG00000168675 | 1243.5473 | -0.970674792 | 0.180106831 | 2.16E-08 | 1.75E-07 | 753 | 18 | 13217497 | 13652754 | 1 | protein_coding | low density lipoprotein receptor class A domain containing 4 |

| | | | | | | | | | | | | | |
|---|---|---|---|---|---|---|---|---|---|---|---|---|---|
| **IL7R** | ENSG00000168685 | 1341.314744 | 2.086151181 | 0.126479383 | 5E-62 | 6.94E-60 | 3575 | 5 | 35852797 | 35879705 | 1 | protein_coding | interleukin 7 receptor |
| **AHCYL1** | ENSG00000168710 | 6490.589091 | -0.242532198 | 0.099280058 | 0.013860532 | 0.037109139 | 10768 | 1 | 110527308 | 110566357 | 1 | protein_coding | adenosylhomocysteinase-like 1 |
| **GSTM4** | ENSG00000168765 | 746.1879885 | -0.914378347 | 0.237925833 | 3.92E-05 | 1.93E-04 | 2948 | 1 | 110198703 | 110208118 | 1 | protein_coding | glutathione S-transferase mu 4 |
| **PPIP5K1** | ENSG00000168781 | 546.5985086 | -0.875684479 | 0.216522254 | 1.84E-05 | 9.64E-05 | 9677 | 15 | 43825660 | 43882430 | -1 | protein_coding | diphosphoinositol pentakisphosphate kinase 1 |
| **ZNF507** | ENSG00000168813 | 254.2154871 | 0.653274849 | 0.217684251 | 0.001399804 | 0.004917979 | 22847 | 19 | 32836500 | 32878573 | 1 | protein_coding | zinc finger protein 507 |
| **GFM1** | ENSG00000168827 | 1432.825594 | -0.541052819 | 0.131534449 | 2.51E-05 | 1.29E-04 | 85476 | 3 | 158362067 | 158410364 | 1 | protein_coding | G elongation factor, mitochondrial 1 |
| **ANKRD49** | ENSG00000168876 | 373.8530329 | 1.097131593 | 0.210419813 | 4.37E-08 | 3.42E-07 | 54851 | 11 | 94226795 | 94232749 | 1 | protein_coding | ankyrin repeat domain 49 |
| **TNIP2** | ENSG00000168884 | 755.2876531 | 0.440294238 | 0.157434728 | 0.003776672 | 0.011888683 | 79155 | 4 | 2743375 | 2758103 | -1 | protein_coding | TNFAIP3 interacting protein 2 |
| **TMEM150A** | ENSG00000168890 | 325.6302683 | -0.629035889 | 0.199845477 | 9.11E-04 | 0.003357122 | 129303 | 2 | 85825671 | 85830319 | -1 | protein_coding | transmembrane protein 150A |
| **RNF181** | ENSG00000168894 | 906.3187648 | -0.533919559 | 0.137771799 | 6.82E-05 | 3.21E-04 | 51255 | 2 | 85822848 | 85824736 | 1 | protein_coding | ring finger protein 181 |
| **VAMP5** | ENSG00000168899 | 784.705602 | 1.419471825 | 0.346830919 | 5.81E-06 | 3.32E-05 | 10791 | 2 | 85811531 | 85820535 | 1 | protein_coding | vesicle-associated membrane protein 5 |
| **MAT2A** | ENSG00000168906 | 6770.539894 | 0.472588983 | 0.088822419 | 5.32E-08 | 4.12E-07 | 4144 | 2 | 85766288 | 85772403 | 1 | protein_coding | methionine adenosyltransferase II, alpha |
| **INPP5D** | ENSG00000168918 | 993.1291214 | -0.714378734 | 0.160472388 | 3.94E-06 | 2.31E-05 | 3635 | 2 | 233924677 | 234116549 | 1 | protein_coding | inositol polyphosphate-5-phosphatase, 145kDa |
| **LETM1** | ENSG00000168924 | 1453.025614 | -0.340512985 | 0.146991967 | 0.015942879 | 0.041891161 | 3954 | 4 | 1813206 | 1857974 | -1 | protein_coding | leucine zipper-EF-hand containing transmembrane protein 1 |
| **TMEM129** | ENSG00000168936 | 499.805172 | -0.468257194 | 0.185059485 | 0.007527434 | 0.021825416 | 92305 | 4 | 1717679 | 1723085 | -1 | protein_coding | transmembrane protein 129 |
| **PPIC** | ENSG00000168938 | 577.4711898 | 0.652577996 | 0.186140795 | 2.34E-04 | 9.85E-04 | 5480 | 5 | 122358945 | 122372436 | -1 | protein_coding | peptidylprolyl isomerase C (cyclophilin C) |
| **CEP120** | ENSG00000168944 | 853.7586432 | 0.4265319 | 0.164904528 | 0.007236447 | 0.021101815 | 153241 | 5 | 122680579 | 122759286 | -1 | protein_coding | centrosomal protein 120kDa |
| **MFF** | ENSG00000168958 | 938.5149311 | -0.360118846 | 0.125363488 | 0.003401158 | 0.010849487 | 56947 | 2 | 228189867 | 228222550 | 1 | protein_coding | mitochondrial fission factor |
| **LGALS9** | ENSG00000168961 | 8689.461131 | 0.203728299 | 0.085635889 | 0.01794761 | 0.046325507 | 3965 | 17 | 25956824 | 25976586 | 1 | protein_coding | lectin, galactoside-binding, soluble, 9 |
| **PXDC1** | ENSG00000168994 | 890.3940372 | 0.506540768 | 0.140146079 | 2.02E-04 | 8.61E-04 | 221749 | 6 | 3722848 | 3752260 | -1 | protein_coding | PX domain containing 1 |
| **SIGLEC7** | ENSG00000168995 | 1317.819396 | -1.122706817 | 0.127932362 | 4.89E-19 | 1E-17 | 27036 | 19 | 51645556 | 51656783 | 1 | protein_coding | sialic acid binding Ig-like lectin 7 |
| **E2F6** | ENSG00000169016 | 471.1034181 | 0.814959923 | 0.164281707 | 2.81E-07 | 1.96E-06 | 1876 | 2 | 11584501 | 11606297 | -1 | protein_coding | E2F transcription factor 6 |
| **FEM1B** | ENSG00000169018 | 5844.834735 | -0.298225915 | 0.100637917 | 0.00270669 | 0.008833298 | 10116 | 15 | 68570141 | 68588203 | 1 | protein_coding | fem-1 homolog b (C. elegans) |
| **ATP5I** | ENSG00000169020 | 494.589353 | -0.568475201 | 0.182419356 | 0.001085973 | 0.003934787 | 521 | 4 | 666225 | 668127 | -1 | protein_coding | ATP synthase, H+ transporting, mitochondrial Fo complex, subunit E |
| **MFSD7** | ENSG00000169026 | 831.3290855 | -0.610578397 | 0.150046752 | 2.62E-05 | 1.34E-04 | 84179 | 4 | 675618 | 683230 | -1 | protein_coding | major facilitator superfamily domain containing 7 |
| **MAP2K1** | ENSG00000169032 | 2849.460811 | -0.294759192 | 0.109281252 | 0.006074303 | 0.018082115 | 5604 | 15 | 66679155 | 66784650 | 1 | protein_coding | mitogen-activated protein kinase kinase 1 |
| **DHRSX** | ENSG00000169084 | 774.1538081 | 0.923263448 | 0.166542641 | 1.03E-08 | 8.78E-08 | 207063 | X | 2137557 | 2420846 | -1 | protein_coding | dehydrogenase/reductase (SDR family) X-linked |
| **HSPBAP1** | ENSG00000169087 | 416.0726 | 1.252162853 | 0.240577964 | 3.94E-08 | 3.09E-07 | 79663 | 3 | 122458846 | 122512671 | -1 | protein_coding | HSPB (heat shock 27kDa) associated protein 1 |
| **SLC25A6** | ENSG00000169100 | 2902.419312 | -0.9016099 | 0.116141972 | 3.01E-15 | 4.65E-14 | 293 | X | 1505045 | 1511617 | -1 | protein_coding | solute carrier family 25 (mitochondrial carrier; adenine nucleotide translocator), member 6 |
| **PARM1** | ENSG00000169116 | 23.72472723 | -1.977800012 | 0.863120567 | 0.001573673 | 0.005467251 | 25849 | 4 | 75858305 | 75975325 | 1 | protein_coding | prostate androgen-regulated mucin-like protein 1 |
| **CSNK1G1** | ENSG00000169118 | 2208.683588 | 0.576190602 | 0.113545986 | 2.58E-07 | 1.81E-06 | 53944 | 15 | 64457716 | 64648442 | -1 | protein_coding | casein kinase 1, gamma 1 |
| **FAM110B** | ENSG00000169122 | 59.31691503 | -1.287795299 | 0.521655875 | 0.002049214 | 0.006907397 | 90362 | 8 | 58907068 | 59116838 | 1 | protein_coding | family with sequence similarity 110, member B |
| **ZNF354A** | ENSG00000169131 | 247.9365472 | 0.536027173 | 0.220287586 | 0.009364693 | 0.0264744 | 6940 | 5 | 178138593 | 178157703 | -1 | protein_coding | zinc finger protein 354A |
| **ATF5** | ENSG00000169136 | 2905.85693 | 0.850679217 | 0.137014917 | 2.16E-10 | 2.21E-09 | 22809 | 19 | 50431959 | 50437192 | 1 | protein_coding | activating transcription factor 5 |
| **ZBTB43** | ENSG00000169155 | 1591.046909 | 0.482355844 | 0.113378353 | 1.38E-05 | 7.44E-05 | 23099 | 9 | 129567285 | 129600489 | 1 | protein_coding | zinc finger and BTB domain containing 43 |
| **MN1** | ENSG00000169184 | 40.12749248 | 4.409835761 | 0.71032092 | 3.51E-11 | 3.94E-10 | 4330 | 22 | 28144265 | 28197486 | -1 | protein_coding | meningioma (disrupted in balanced translocation) 1 |
| **NSMCE1** | ENSG00000169189 | 557.3749543 | -0.677141713 | 0.169963493 | 3.49E-05 | 1.74E-04 | 197370 | 16 | 27236312 | 27280115 | -1 | protein_coding | non-SMC element 1 homolog (S. cerevisiae) |
| **CCDC126** | ENSG00000169193 | 127.5314274 | 0.964734784 | 0.317003211 | 6.53E-04 | 0.002485507 | 90693 | 7 | 23636998 | 23684327 | 1 | protein_coding | coiled-coil domain containing 126 |
| **RGS14** | ENSG00000169220 | 189.5968385 | -1.353536873 | 0.296987558 | 8.41E-07 | 5.48E-06 | 10636 | 5 | 176784838 | 176799602 | 1 | protein_coding | regulator of G-protein signaling 14 |
| **LMAN2** | ENSG00000169223 | 2494.665152 | -0.391215995 | 0.111086161 | 2.93E-04 | 0.001208884 | 10960 | 5 | 176758563 | 176778853 | -1 | protein_coding | lectin, mannose-binding 2 |
| **GCSAML** | ENSG00000169224 | 36.07086865 | -0.982302688 | 0.571464905 | 0.017987524 | 0.046412854 | 148823 | 1 | 247670360 | 247740992 | 1 | protein_coding | germinal center-associated, signaling and motility-like |
| **RAB24** | ENSG00000169228 | 596.5133951 | 0.723271079 | 0.216619237 | 3.74E-04 | 0.001509142 | 53917 | 5 | 176728199 | 176730745 | -1 | protein_coding | RAB24, member RAS oncogene family |
| **PRELID1** | ENSG00000169230 | 1742.333326 | 0.543941169 | 0.130634063 | 2.05E-05 | 1.07E-04 | 27166 | 5 | 176730775 | 176733960 | 1 | protein_coding | PRELI domain containing 1 |
| **SLC50A1** | ENSG00000169241 | 764.9362879 | 0.579331375 | 0.159220411 | 1.63E-04 | 7.14E-04 | 55974 | 1 | 155107820 | 155111329 | 1 | protein_coding | solute carrier family 50 (sugar efflux transporter), member 1 |
| **CXCL10** | ENSG00000169245 | 290.0420821 | 4.197741704 | 0.350565482 | 2.74E-34 | 1.33E-32 | 3627 | 4 | 76942273 | 76944650 | -1 | protein_coding | chemokine (C-X-C motif) ligand 10 |
| **CXCL11** | ENSG00000169248 | 118.5064771 | 8.047615173 | 0.754498753 | 2.13E-26 | 6.88E-25 | 6373 | 4 | 76954835 | 76962568 | -1 | protein_coding | chemokine (C-X-C motif) ligand 11 |
| **ZRSR2** | ENSG00000169249 | 374.1999955 | 0.494798627 | 0.208838581 | 0.011895935 | 0.032488486 | 8233 | X | 15808595 | 15841383 | 1 | protein_coding | zinc finger (CCCH type), RNA-binding motif and serine/arginine rich 2 |
| **NMD3** | ENSG00000169251 | 1389.4179 | -0.569540343 | 0.116815762 | 6.79E-07 | 4.49E-06 | 51068 | 3 | 160822484 | 160971320 | 1 | protein_coding | NMD3 ribosome export adaptor |
| **ADRB2** | ENSG00000169252 | 53.4102173 | -2.098897727 | 0.553570389 | 1.46E-05 | 7.8E-05 | 154 | 5 | 148206156 | 148208196 | 1 | protein_coding | adrenoceptor beta 2, surface |
| **B3GALNT1** | ENSG00000169255 | 126.0120839 | -0.943473133 | 0.37442071 | 0.00323415 | 0.01037957 | 8706 | 3 | 160801671 | 160823172 | -1 | protein_coding | beta-1,3-N-acetylgalactosaminyltransferase 1 (globoside blood group) |
| **KCNAB1** | ENSG00000169282 | 42.41582621 | -1.911107304 | 0.759556772 | 9.07E-04 | 0.003347192 | 7881 | 3 | 155755490 | 156256545 | 1 | protein_coding | potassium voltage-gated channel, shaker-related subfamily, beta member 1 |
| **MRPL1** | ENSG00000169288 | 287.7505182 | 0.650164092 | 0.243662906 | 0.003890437 | 0.012201427 | 65008 | 4 | 78783674 | 78873944 | 1 | protein_coding | mitochondrial ribosomal protein L1 |
| **PGM2** | ENSG00000169299 | 2087.899481 | -0.604880684 | 0.111934289 | 4.35E-08 | 3.4E-07 | 55276 | 4 | 37828255 | 37864558 | 1 | protein_coding | phosphoglucomutase 2 |
| **P2RY12** | ENSG00000169313 | 21.10015633 | -3.636631812 | 1.501979901 | 0.001264152 | 0.004491985 | 64805 | 3 | 151055168 | 151102600 | -1 | protein_coding | purinergic receptor P2Y, G-protein coupled, 12 |
| **KIAA1024** | ENSG00000169330 | 40.14208886 | -1.021340434 | 0.549454184 | 0.012609643 | 0.034150836 | 23251 | 15 | 79724858 | 79764632 | 1 | protein_coding | KIAA1024 |
| **SNUPN** | ENSG00000169371 | 548.3236391 | -0.493944963 | 0.178259845 | 0.003775534 | 0.011888683 | 10073 | 15 | 75890424 | 75918810 | -1 | protein_coding | snurportin 1 |
| **SIN3A** | ENSG00000169375 | 1700.301214 | 0.368079955 | 0.139505202 | 0.006746608 | 0.019851445 | 25942 | 15 | 75661720 | 75748183 | -1 | protein_coding | SIN3 transcription regulator family member A |
| **ARL13B** | ENSG00000169379 | 227.8530325 | 0.849756137 | 0.272307674 | 6.28E-04 | 0.002395265 | 200894 | 3 | 93698983 | 93774512 | 1 | protein_coding | ADP-ribosylation factor-like 13B |
| **PTAFR** | ENSG00000169403 | 8692.110298 | 0.300370836 | 0.113690627 | 0.007773794 | 0.022467263 | 5724 | 1 | 28473677 | 28520447 | -1 | protein_coding | platelet-activating factor receptor |
| **RNASE6** | ENSG00000169413 | 1540.984568 | -1.349696683 | 0.188244498 | 1.24E-13 | 1.7E-12 | 6039 | 14 | 21249210 | 21250626 | 1 | protein_coding | ribonuclease, RNase A family, k6 |
| **NPR 1.00** | ENSG00000169418 | 231.7632957 | 0.697212614 | 0.257994365 | 0.003197303 | 0.010276422 | 4881 | 1 | 153651113 | 153666468 | 1 | protein_coding | natriuretic peptide receptor A/guanylate cyclase A (atrionatriuretic peptide receptor A) |
| **IL8** | ENSG00000169429 | 6310.14368 | 4.478933286 | 0.607370763 | 6.64E-15 | 1E-13 | 3576 | 4 | 74606223 | 74609433 | 1 | protein_coding | interleukin 8 |
| **COL22A1** | ENSG00000169436 | 40.40419543 | -2.050860732 | 0.652827068 | 1.39E-04 | 6.19E-04 | 169044 | 8 | 139600478 | 139926249 | -1 | protein_coding | collagen, type XXII, alpha 1 |
| **SDC2** | ENSG00000169439 | 6537.419364 | -1.557500794 | 0.105730786 | 4.53E-50 | 4.02E-48 | 6383 | 8 | 97505579 | 97624000 | 1 | protein_coding | syndecan 2 |
| **CD52** | ENSG00000169442 | 7778.889639 | -1.103283201 | 0.388633447 | 9.36E-04 | 0.003439505 | 1043 | 1 | 26644448 | 26647014 | 1 | protein_coding | CD52 molecule |
| **MMGT1** | ENSG00000169446 | 1371.113339 | -0.481494926 | 0.19842071 | 0.009078996 | 0.025738138 | 93380 | X | 135044229 | 135056222 | -1 | protein_coding | membrane magnesium transporter 1 |
| **HTRA4** | ENSG00000169495 | 1709.270651 | -0.72238445 | 0.234716221 | 9.45E-04 | 0.00346901 | 203100 | 8 | 38831683 | 38846181 | 1 | protein_coding | HtrA serine peptidase 4 |
| **CLIC4** | ENSG00000169504 | 11686.9912 | 1.639628613 | 0.08875996 | 4.94E-77 | 1.11E-74 | 25932 | 1 | 25071848 | 25170815 | 1 | protein_coding | chloride intracellular channel 4 |
| **CKAP2L** | ENSG00000169607 | 7.501319916 | 0.233101369 | 0.697853876 | 0.005548318 | 0.016688634 | 150468 | 2 | 113493930 | 113522254 | -1 | protein_coding | cytoskeleton associated protein 2-like |
| **LUZP1** | ENSG00000169641 | 717.5797485 | 0.554709498 | 0.152085609 | 1.66E-04 | 7.26E-04 | 7798 | 1 | 23410516 | 23504301 | -1 | protein_coding | leucine zipper protein 1 |
| **HEXDC** | ENSG00000169660 | 276.1770902 | 0.933881014 | 0.230695074 | 1.62E-05 | 8.58E-05 | 284004 | 17 | 80376194 | 80400521 | 1 | protein_coding | hexosaminidase (glycosyl hydrolase family 20, catalytic domain) containing |
| **BUB1** | ENSG00000169679 | 168.473172 | 0.71139252 | 0.321791996 | 0.011851047 | 0.032377468 | 699 | 2 | 111395275 | 111435691 | -1 | protein_coding | BUB1 mitotic checkpoint serine/threonine kinase |
| **FASN** | ENSG00000169710 | 1363.868747 | -0.675206319 | 0.128759532 | 8.04E-08 | 6.04E-07 | 2194 | 17 | 80036214 | 80056208 | -1 | protein_coding | fatty acid synthase |
| **CNBP** | ENSG00000169714 | 9448.609882 | 0.714318674 | 0.099335483 | 5.69E-13 | 7.44E-12 | 7555 | 3 | 128888327 | 128902765 | -1 | protein_coding | CCHC-type zinc finger, nucleic acid binding protein |
| **MT1E** | ENSG00000169715 | 866.0531461 | 0.898745332 | 0.16855758 | 3.43E-08 | 2.72E-07 | 4493 | 16 | 56659387 | 56661024 | 1 | protein_coding | metallothionein 1E |
| **DUS1L** | ENSG00000169718 | 741.7810063 | -0.677123328 | 0.161313849 | 1.38E-05 | 7.43E-05 | 64118 | 17 | 80015382 | 80023763 | -1 | protein_coding | dihydrouridine synthase 1-like (S. cerevisiae) |
| **GPS1** | ENSG00000169727 | 654.2759411 | -0.438009018 | 0.149747845 | 0.002121826 | 0.007125437 | 2873 | 17 | 80008567 | 80015346 | 1 | protein_coding | G protein pathway suppressor 1 |
| **DCXR** | ENSG00000169738 | 171.2209142 | -0.849584402 | 0.343481982 | 0.00453829 | 0.014003368 | 51181 | 17 | 79993012 | 79995608 | -1 | protein_coding | dicarbonyl/L-xylulose reductase |
| **C15orf27** | ENSG00000169758 | 276.5901103 | -0.71106905 | 0.238669213 | 0.001295261 | 0.004590788 | 1019303 | 15 | 76352178 | 76521462 | 1 | protein_coding | chromosome 15 open reading frame 27 |
| **UGP2** | ENSG00000169764 | 4108.163701 | 0.455233867 | 0.096381356 | 2.03E-06 | 1.25E-05 | 7360 | 2 | 64068074 | 64118696 | 1 | protein_coding | UDP-glucose pyrophosphorylase 2 |
| **LINGO1** | ENSG00000169783 | 140.845612 | -1.163913527 | 0.495116934 | 0.003203299 | 0.0102892 | 84894 | 15 | 77905369 | 78113242 | -1 | protein_coding | leucine rich repeat and Ig domain containing 1 |
| **HNRNPF** | ENSG00000169813 | 4152.422123 | 0.419392155 | 0.095715452 | 1.15E-05 | 6.26E-05 | 3185 | 10 | 43881065 | 43904614 | -1 | protein_coding | heterogeneous nuclear ribonucleoprotein F |
| **BTD** | ENSG00000169814 | 433.7429444 | -0.685480232 | 0.193463625 | 1.95E-04 | 8.35E-04 | 686 | 3 | 15642848 | 15687329 | 1 | protein_coding | biotinidase |
| **TRIM56** | ENSG00000169871 | 854.9338142 | 1.418648473 | 0.162816334 | 5.1E-19 | 1.05E-17 | 81844 | 7 | 100728720 | 100735017 | 1 | protein_coding | tripartite motif containing 56 |
| **REPS2** | ENSG00000169891 | 55.23919846 | -2.956864704 | 1.086212565 | 3.47E-04 | 0.001410563 | 9185 | X | 16964814 | 17171395 | 1 | protein_coding | RALBP1 associated Eps domain containing 2 |
| **ITGAM** | ENSG00000169896 | 8342.54657 | -1.064391475 | 0.119159721 | 1.06E-19 | 2.29E-18 | 3684 | 16 | 31271311 | 31344213 | 1 | protein_coding | integrin, alpha M (complement component 3 receptor 3 subunit) |
| **TM4SF1** | ENSG00000169908 | 119.0722324 | 1.191739487 | 0.393855134 | 4.67E-04 | 0.001840802 | 4071 | 3 | 149086809 | 149095652 | -1 | protein_coding | transmembrane 4 L six family member 1 |
| **GUSB** | ENSG00000169919 | 5926.796719 | -0.630106708 | 0.123511338 | 1.86E-07 | 1.33E-06 | 2990 | 7 | 65425671 | 65447301 | -1 | protein_coding | glucuronidase, beta |
| **SF3B5** | ENSG00000169976 | 572.7490431 | -0.3774036 | 0.151629891 | 0.010328414 | 0.028762602 | 83443 | 6 | 144416018 | 144416754 | -1 | protein_coding | splicing factor 3b, subunit 5, 10kDa |

| | | | | | | | | | | | | | |
|---|---|---|---|---|---|---|---|---|---|---|---|---|---|
| **CHD3** | ENSG00000170004 | 3302.981198 | -0.469042421 | 0.14541169 | 8.98E-04 | 0.003318586 | 1107 | 17 | 7788124 | 7816078 | 1 | protein_coding | chromodomain helicase DNA binding protein 3 |
| **YWHAG** | ENSG00000170027 | 9697.704657 | 0.505637796 | 0.089372438 | 9.68E-09 | 8.26E-08 | 7532 | 7 | 75956116 | 75988348 | -1 | protein_coding | tyrosine 3-monooxygenase/tryptophan 5-monooxygenase activation protein, gamma |
| **UBE2E3** | ENSG00000170035 | 400.5632332 | -0.569323321 | 0.190949047 | 0.001679821 | 0.005792578 | 10477 | 2 | 181831975 | 181941312 | 1 | protein_coding | ubiquitin-conjugating enzyme E2E 3 |
| **CNTROB** | ENSG00000170037 | 315.534342 | -0.731663695 | 0.245326662 | 0.001306312 | 0.004622451 | 116840 | 17 | 7835419 | 7853236 | 1 | protein_coding | centrobin, centrosomal BRCA2 interacting protein |
| **TRAPPC1** | ENSG00000170043 | 1003.385655 | -0.493576372 | 0.173249957 | 0.00286895 | 0.009313118 | 58485 | 17 | 7833663 | 7835441 | -1 | protein_coding | trafficking protein particle complex 1 |
| **FAM153A** | ENSG00000170074 | 21.13670794 | -2.016130481 | 1.030284147 | 0.002973182 | 0.009626936 | 285596 | 5 | 177134982 | 177210399 | -1 | protein_coding | family with sequence similarity 153, member A |
| **TMEM192** | ENSG00000170088 | 1510.5587 | -0.661035865 | 0.13846707 | 1.03E-06 | 6.61E-06 | 201931 | 4 | 165995574 | 166129701 | -1 | protein_coding | transmembrane protein 192 |
| **UBE2E1** | ENSG00000170142 | 2171.192985 | 1.190541617 | 0.119362101 | 4.88E-24 | 1.39E-22 | 7324 | 3 | 23847394 | 23932807 | 1 | protein_coding | ubiquitin-conjugating enzyme E2E 1 |
| **HNRNPA3** | ENSG00000170144 | 5889.362457 | 0.271886778 | 0.103871616 | 0.008187513 | 0.023494135 | 220988 | 2 | 178077291 | 178088686 | 1 | protein_coding | heterogeneous nuclear ribonucleoprotein A3 |
| **SIK2** | ENSG00000170145 | 1552.809861 | -0.720199404 | 0.217897714 | 3.69E-04 | 0.001493726 | 23235 | 11 | 111473115 | 111601577 | 1 | protein_coding | salt-inducible kinase 2 |
| **RNF150** | ENSG00000170153 | 7.047434734 | -0.393359616 | 0.825780676 | 0.004015453 | 0.012557432 | 57484 | 4 | 141780961 | 142134031 | -1 | protein_coding | ring finger protein 150 |
| **CHRNB1** | ENSG00000170175 | 136.0900263 | 1.387360769 | 0.279389075 | 1.15E-07 | 8.43E-07 | 1140 | 17 | 7348380 | 7361026 | 1 | protein_coding | cholinergic receptor, nicotinic, beta 1 (muscle) |
| **USP38** | ENSG00000170185 | 1355.009036 | 0.683647186 | 0.106113435 | 6.16E-11 | 6.74E-10 | 84640 | 4 | 144106070 | 144144983 | 1 | protein_coding | ubiquitin specific peptidase 38 |
| **SLC16A5** | ENSG00000170190 | 148.5545333 | -0.721044413 | 0.296577048 | 0.006292634 | 0.018677478 | 9121 | 17 | 73083822 | 73102257 | 1 | protein_coding | solute carrier family 16 (monocarboxylate transporter), member 5 |
| **PDCD6IP** | ENSG00000170248 | 7380.69375 | 0.602927281 | 0.110327566 | 2.55E-08 | 2.06E-07 | 10015 | 3 | 33839844 | 33911194 | 1 | protein_coding | programmed cell death 6 interacting protein |
| **GLB1** | ENSG00000170266 | 3074.350264 | -0.285173949 | 0.099638861 | 0.003306584 | 0.010583146 | 643853 | 3 | 33038100 | 33138722 | -1 | protein_coding | galactosidase, beta 1 |
| **FAXDC2** | ENSG00000170271 | 295.6816211 | -1.752471528 | 0.290848846 | 1.94E-10 | 2E-09 | 10826 | 5 | 154198051 | 154238812 | -1 | protein_coding | fatty acid hydroxylase domain containing 2 |
| **CRTAP** | ENSG00000170275 | 3297.272375 | -0.376433267 | 0.123836722 | 0.001778319 | 0.006081462 | 1006530 | 3 | 33155471 | 33189265 | 1 | protein_coding | cartilage associated protein |
| **GABARAP** | ENSG00000170296 | 5833.232947 | -0.490079071 | 0.091260536 | 4.7E-08 | 3.66E-07 | 11337 | 17 | 7143333 | 7146089 | -1 | protein_coding | GABA(A) receptor-associated protein |
| **CDK1** | ENSG00000170312 | 72.77672912 | 1.845969681 | 0.724539591 | 7.35E-04 | 0.002768381 | 983 | 10 | 62538089 | 62554610 | 1 | protein_coding | cyclin-dependent kinase 1 |
| **FABP4** | ENSG00000170323 | 116.6205199 | -4.264637651 | 0.949456893 | 9.37E-07 | 6.05E-06 | 2167 | 8 | 82390654 | 82395498 | -1 | protein_coding | fatty acid binding protein 4, adipocyte |
| **FOS** | ENSG00000170345 | 417.8751959 | -2.178117907 | 0.253967676 | 7.62E-19 | 1.53E-17 | 2353 | 14 | 75745477 | 75748933 | 1 | protein_coding | FBJ murine osteosarcoma viral oncogene homolog |
| **TMED10** | ENSG00000170348 | 8639.871663 | -0.535562179 | 0.104458823 | 1.83E-07 | 1.31E-06 | 10972 | 14 | 75598173 | 75643334 | -1 | protein_coding | transmembrane emp24-like trafficking protein 10 (yeast) |
| **SMAD1** | ENSG00000170365 | 154.3164905 | 1.297087939 | 0.256272213 | 7.86E-08 | 5.92E-07 | 4086 | 4 | 146402346 | 146479231 | 1 | protein_coding | SMAD family member 1 |
| **SLC30A1** | ENSG00000170385 | 7801.744785 | 1.310026315 | 0.131694751 | 4.71E-24 | 1.34E-22 | 7779 | 1 | 211744910 | 211752084 | -1 | protein_coding | solute carrier family 30 (zinc transporter), member 1 |
| **DCLK2** | ENSG00000170390 | 76.7736128 | -4.083181096 | 0.768530337 | 1.09E-08 | 9.21E-08 | 166614 | 4 | 150999426 | 151178609 | 1 | protein_coding | doublecortin-like kinase 2 |
| **ZNF804A** | ENSG00000170396 | 391.4711017 | 0.602612146 | 0.207572233 | 0.002099347 | 0.007059257 | 91752 | 2 | 185463093 | 185804219 | 1 | protein_coding | zinc finger protein 804A |
| **ADORA2B** | ENSG00000170425 | 95.78622885 | -2.268205481 | 0.461594823 | 6.95E-08 | 5.28E-07 | 136 | 17 | 15848231 | 15879060 | 1 | protein_coding | adenosine A2b receptor |
| **MGMT** | ENSG00000170430 | 142.4327444 | -0.798938801 | 0.280107117 | 0.00166562 | 0.0057475 | 4255 | 10 | 131265448 | 131566271 | 1 | protein_coding | O-6-methylguanine-DNA methyltransferase |
| **METTL7B** | ENSG00000170439 | 147.9732901 | -0.850455219 | 0.446269019 | 0.016192781 | 0.04247756 | 196410 | 12 | 56075330 | 56078395 | 1 | protein_coding | methyltransferase like 7B |
| **HARS** | ENSG00000170445 | 1015.837654 | 0.611653098 | 0.129447185 | 1.33E-06 | 8.41E-06 | 1019286 | 5 | 140052758 | 140071609 | -1 | protein_coding | histidyl-tRNA synthetase |
| **DENND5B** | ENSG00000170456 | 411.2610689 | -0.701986217 | 0.254575015 | 0.002687141 | 0.008775123 | 160518 | 12 | 31535157 | 31744031 | -1 | protein_coding | DENN/MADD domain containing 5B |
| **CD14** | ENSG00000170458 | 207.4357978 | -2.16101368 | 0.445435549 | 1.07E-07 | 7.9E-07 | 929 | 5 | 140011313 | 140013286 | -1 | protein_coding | CD14 molecule |
| **NPAS2** | ENSG00000170485 | 96.34105262 | -1.583675564 | 0.421553291 | 2.15E-05 | 1.11E-04 | 4862 | 2 | 101436614 | 101613291 | 1 | protein_coding | neuronal PAS domain protein 2 |
| **PFKFB3** | ENSG00000170525 | 3727.197152 | 1.985809767 | 0.149752729 | 4.51E-41 | 2.91E-39 | 5209 | 10 | 6186881 | 6277495 | 1 | protein_coding | 6-phosphofructo-2-kinase/fructose-2,6-biphosphatase 3 |
| **SERPINB9** | ENSG00000170542 | 984.302651 | 2.059595192 | 0.232044974 | 6.31E-20 | 1.39E-18 | 5272 | 6 | 2887500 | 2903514 | -1 | protein_coding | serpin peptidase inhibitor, clade B (ovalbumin), member 9 |
| **EMB** | ENSG00000170571 | 460.788889 | 0.618201129 | 0.26542963 | 0.010439541 | 0.029003365 | 133418 | 5 | 49692026 | 49739082 | -1 | protein_coding | embigin |
| **STAT2** | ENSG00000170581 | 10421.89101 | 2.311162667 | 0.113313753 | 2.43E-93 | 7.42E-91 | 6773 | 12 | 56735381 | 56753939 | -1 | protein_coding | signal transducer and activator of transcription 2, 113kDa |
| **ARMC10** | ENSG00000170632 | 840.872699 | 0.521173837 | 0.149257002 | 3.23E-04 | 0.001321438 | 83787 | 7 | 102715328 | 102740205 | 1 | protein_coding | armadillo repeat containing 10 |
| **RNF34** | ENSG00000170633 | 711.1332304 | 0.559399575 | 0.13721545 | 2.79E-05 | 1.42E-04 | 80196 | 12 | 121837844 | 121868389 | 1 | protein_coding | ring finger protein 34, E3 ubiquitin protein ligase |
| **KIF5B** | ENSG00000170759 | 9140.495684 | 0.522272166 | 0.104766326 | 5.55E-07 | 3.73E-06 | 3799 | 10 | 32297938 | 32345359 | -1 | protein_coding | kinesin family member 5B |
| **FOXN2** | ENSG00000170802 | 1416.617577 | 1.020425007 | 0.135732667 | 1.73E-14 | 2.52E-13 | 3344 | 2 | 48541776 | 48606433 | 1 | protein_coding | forkhead box N2 |
| **PPM1D** | ENSG00000170836 | 456.7238907 | 0.494862136 | 0.170716258 | 0.002533585 | 0.008330403 | 8493 | 17 | 58677544 | 58741849 | 1 | protein_coding | protein phosphatase, Mg2+/Mn2+ dependent, 1D |
| **KBTBD2** | ENSG00000170852 | 1261.958943 | 0.827683652 | 0.116183677 | 3.97E-13 | 5.25E-12 | 25948 | 7 | 32907784 | 32933743 | -1 | protein_coding | kelch repeat and BTB (POZ) domain containing 2 |
| **MTSS1** | ENSG00000170873 | 274.5114437 | -0.775415696 | 0.265577339 | 0.001384272 | 0.004866643 | 9788 | 8 | 125563031 | 125740730 | -1 | protein_coding | metastasis suppressor 1 |
| **RNF139** | ENSG00000170881 | 987.5074533 | 0.79388637 | 0.166912156 | 8.47E-07 | 5.51E-06 | 11236 | 8 | 125486979 | 125500155 | 1 | protein_coding | ring finger protein 139 |
| **MSANTD4** | ENSG00000170903 | 758.3178768 | 0.5530786 | 0.140622925 | 4.78E-05 | 2.31E-04 | 84437 | 11 | 105866350 | 105893130 | -1 | protein_coding | Myb/SANT-like DNA-binding domain containing 4 with coiled-coils |
| **NDUFA3** | ENSG00000170906 | 585.0007546 | -0.569110919 | 0.184421691 | 0.001171284 | 0.004200079 | 4696 | 19 | 54606036 | 54612564 | 1 | protein_coding | NADH dehydrogenase (ubiquinone) 1 alpha subcomplex, 3, 9kDa |
| **OSCAR** | ENSG00000170909 | 1887.920794 | 0.406164037 | 0.150435815 | 0.005453518 | 0.016461804 | 126014 | 19 | 54597933 | 54606000 | -1 | protein_coding | osteoclast associated, immunoglobulin-like receptor |
| **PAQR8** | ENSG00000170915 | 1801.680093 | -0.474302539 | 0.150275418 | 0.001070035 | 0.003881697 | 85315 | 6 | 52226219 | 52272575 | 1 | protein_coding | progestin and adipoQ receptor family member VIII |
| **TANC2** | ENSG00000170921 | 1094.310416 | -2.565228472 | 0.301545435 | 6.16E-19 | 1.24E-17 | 26115 | 17 | 61086917 | 61505060 | 1 | protein_coding | tetratricopeptide repeat, ankyrin repeat and coiled-coil containing 2 |
| **S1PR1** | ENSG00000170989 | 161.0934115 | -0.884430621 | 0.333296577 | 0.002453701 | 0.008089491 | 1901 | 1 | 101702444 | 101707074 | 1 | protein_coding | sphingosine-1-phosphate receptor 1 |
| **PYGO1** | ENSG00000171016 | 19.47675722 | -1.618902978 | 1.002958833 | 0.009038247 | 0.025641643 | 26108 | 15 | 55831088 | 55881145 | -1 | protein_coding | pygopus family PHD finger 1 |
| **FPR1** | ENSG00000171051 | 349.2575353 | -1.55494638 | 0.387679622 | 7.3E-06 | 4.1E-05 | 2357 | 19 | 52248425 | 52307363 | -1 | protein_coding | formyl peptide receptor 1 |
| **FEZ2** | ENSG00000171055 | 2232.180433 | -0.281291545 | 0.105088324 | 0.006610167 | 0.019487498 | 9637 | 2 | 36778570 | 36873230 | -1 | protein_coding | fasciculation and elongation protein zeta 2 (zygin II) |
| **CCBL1** | ENSG00000171097 | 33.76950216 | -1.383273479 | 0.641717439 | 0.003707224 | 0.01170849 | 883 | 9 | 131595221 | 131644773 | -1 | protein_coding | cysteine conjugate-beta lyase, cytoplasmic |
| **MTM1** | ENSG00000171100 | 861.2479681 | 0.50543386 | 0.172817005 | 0.002275076 | 0.007570212 | 4534 | X | 149737069 | 149841795 | 1 | protein_coding | myotubularin 1 |
| **INSR** | ENSG00000171105 | 444.5801124 | -1.455169032 | 0.209006027 | 4.77E-13 | 6.28E-12 | 3643 | 19 | 7112266 | 7294045 | -1 | protein_coding | insulin receptor |
| **MFN1** | ENSG00000171109 | 1844.219636 | 1.491760085 | 0.155231705 | 1.26E-22 | 3.3E-21 | 55669 | 3 | 179065480 | 179112719 | 1 | protein_coding | mitofusin 1 |
| **GIMAP8** | ENSG00000171115 | 1707.470328 | 1.450187104 | 0.233944363 | 8.64E-11 | 9.3E-10 | 155038 | 7 | 150147718 | 150176480 | 1 | protein_coding | GTPase, IMAP family member 8 |
| **ATP6V0E2** | ENSG00000171130 | 260.1234978 | -0.576502475 | 0.221366659 | 0.005299228 | 0.016056326 | 155066 | 7 | 149570057 | 149577784 | 1 | protein_coding | ATPase, H+ transporting V0 subunit e2 |
| **PRKCE** | ENSG00000171132 | 217.5271592 | 0.684920062 | 0.275854119 | 0.006057772 | 0.018036416 | 5581 | 2 | 45878484 | 46415129 | 1 | protein_coding | protein kinase C, epsilon |
| **TADA3** | ENSG00000171148 | 1619.613862 | -0.69320651 | 0.160239433 | 7.72E-06 | 4.32E-05 | 10474 | 3 | 9821544 | 9834695 | -1 | protein_coding | transcriptional adaptor 3 |
| **C1GALT1C1** | ENSG00000171155 | 1063.752251 | 1.241664134 | 0.132866541 | 1.86E-21 | 4.54E-20 | 29071 | X | 119759648 | 119764005 | -1 | protein_coding | C1GALT1-specific chaperone 1 |
| **C9orf16** | ENSG00000171159 | 426.2124423 | -1.085401291 | 0.237817285 | 1.34E-06 | 8.45E-06 | 79095 | 9 | 130922539 | 130926207 | 1 | protein_coding | chromosome 9 open reading frame 16 |
| **TMEM126B** | ENSG00000171204 | 757.0953903 | 0.685194537 | 0.156019126 | 5.82E-06 | 3.33E-05 | 55863 | 11 | 85339629 | 85347580 | 1 | protein_coding | transmembrane protein 126B |
| **TRIM8** | ENSG00000171206 | 1999.917753 | -0.429993708 | 0.115686333 | 1.49E-04 | 6.59E-04 | 81603 | 10 | 104404253 | 104418164 | 1 | protein_coding | tripartite motif containing 8 |
| **NETO2** | ENSG00000171208 | 161.5333342 | 0.969442217 | 0.406898439 | 0.004294139 | 0.013331178 | 81831 | 16 | 47111614 | 47177908 | -1 | protein_coding | neuropilin (NRP) and tolloid (TLL)-like 2 |
| **JUNB** | ENSG00000171223 | 4269.394168 | 0.466564736 | 0.145253524 | 9.45E-04 | 0.00346765 | 3726 | 19 | 12902310 | 12904124 | 1 | protein_coding | jun B proto-oncogene |
| **TMEM37** | ENSG00000171227 | 169.2323348 | -10.22841209 | 2.963154341 | 3.9E-11 | 4.37E-10 | 140738 | 2 | 120187477 | 120196096 | 1 | protein_coding | transmembrane protein 37 |
| **FAM98B** | ENSG00000171262 | 584.5210657 | 0.703067044 | 0.202240511 | 2.43E-04 | 0.001016055 | 283742 | 15 | 38746328 | 38779911 | 1 | protein_coding | family with sequence similarity 98, member B |
| **ZNF440** | ENSG00000171295 | 295.1042712 | -0.525582756 | 0.226549112 | 0.012826023 | 0.034663149 | 126070 | 19 | 11925099 | 11946016 | 1 | protein_coding | zinc finger protein 440 |
| **GAA** | ENSG00000171298 | 4058.4702 | -0.919471225 | 0.125657493 | 8.06E-14 | 1.12E-12 | 2548 | 17 | 78075355 | 78093678 | 1 | protein_coding | glucosidase, alpha; acid |
| **CANT1** | ENSG00000171302 | 1414.646263 | 0.335273818 | 0.137844359 | 0.013049947 | 0.035206061 | 124583 | 17 | 76987799 | 77005949 | -1 | protein_coding | calcium activated nucleotidase 1 |
| **CHST11** | ENSG00000171310 | 9945.666065 | -0.333246588 | 0.100989554 | 6.81E-04 | 0.002583708 | 50515 | 12 | 104849073 | 105155792 | 1 | protein_coding | carbohydrate (chondroitin 4) sulfotransferase 11 |
| **CLCN5** | ENSG00000171365 | 921.4813933 | -0.679496464 | 0.227167025 | 0.001354782 | 0.004772967 | 1184 | X | 49687225 | 49863892 | 1 | protein_coding | chloride channel, voltage-sensitive 5 |
| **APLN** | ENSG00000171388 | 28.10368595 | -7.584072739 | 2.740444625 | 1.26E-06 | 7.95E-06 | 8862 | X | 128779240 | 128788933 | -1 | protein_coding | apelin |
| **PDE7B** | ENSG00000171408 | 300.3701584 | -1.508589923 | 0.352001712 | 2.25E-06 | 1.38E-05 | 27115 | 6 | 136172834 | 136516712 | 1 | protein_coding | phosphodiesterase 7B |
| **NAT1** | ENSG00000171428 | 272.4662906 | 0.606415791 | 0.209606285 | 0.002169448 | 0.007259835 | 9 | 8 | 18027986 | 18081198 | 1 | protein_coding | N-acetyltransferase 1 (arylamine N-acetyltransferase) |
| **KSR2** | ENSG00000171435 | 29.81432269 | -2.595380852 | 0.911330842 | 2.77E-04 | 0.001149239 | 283455 | 12 | 117890817 | 118406788 | -1 | protein_coding | kinase suppressor of ras 2 |
| **ZNF524** | ENSG00000171443 | 132.4625065 | -0.781987676 | 0.298270325 | 0.003368712 | 0.010757215 | 147807 | 19 | 56111388 | 56114504 | 1 | protein_coding | zinc finger protein 524 |
| **ASXL1** | ENSG00000171456 | 1555.921391 | 0.327167883 | 0.112480653 | 0.003188889 | 0.010253689 | 171023 | 20 | 30946155 | 31027122 | 1 | protein_coding | additional sex combs like 1 (Drosophila) |
| **ZNF318** | ENSG00000171467 | 1661.290321 | -0.432649001 | 0.122346141 | 3.42E-04 | 0.001392772 | 24149 | 6 | 43274872 | 43337216 | -1 | protein_coding | zinc finger protein 318 |
| **HOPX** | ENSG00000171476 | 67.48529366 | -1.008692927 | 0.521689728 | 0.011211233 | 0.030888999 | 84525 | 4 | 57514155 | 57548065 | -1 | protein_coding | HOP homeobox |
| **LRRC8C** | ENSG00000171488 | 1226.925328 | -0.325853565 | 0.130158304 | 0.009369222 | 0.026482304 | 1019279 | 1 | 90098631 | 90235462 | 1 | protein_coding | leucine rich repeat containing 8 family, member C |
| **LRRC8D** | ENSG00000171492 | 2070.107764 | -0.305113661 | 0.101628336 | 0.002329819 | 0.007737206 | 55144 | 1 | 90286573 | 90402170 | 1 | protein_coding | leucine rich repeat containing 8 family, member D |
| **PPID** | ENSG00000171497 | 1057.268233 | 0.598108346 | 0.157332631 | 8.26E-05 | 3.84E-04 | 5481 | 4 | 159630286 | 159644548 | -1 | protein_coding | peptidylprolyl isomerase D |
| **RXFP1** | ENSG00000171509 | 7.93199891 | 3.902081596 | 1.684322495 | 0.001145037 | 0.004120463 | 59350 | 4 | 159236463 | 159574524 | 1 | protein_coding | relaxin/insulin-like family peptide receptor 1 |
| **LPAR3** | ENSG00000171517 | 8.752437867 | 2.857800764 | 1.413227573 | 0.001708774 | 0.005877829 | 23566 | 1 | 85277285 | 85358896 | -1 | protein_coding | lysophosphatidic acid receptor 3 |

| | | | | | | | | | | | | | |
|---|---|---|---|---|---|---|---|---|---|---|---|---|---|
| **PTGER4** | ENSG00000171522 | 2041.716196 | 0.658317788 | 0.163198642 | 3.04E-05 | 1.53E-04 | 5734 | 5 | 40679600 | 40693837 | 1 | protein_coding | prostaglandin E receptor 4 (subtype EP4) |
| **TBCA** | ENSG00000171530 | 1087.987418 | 0.716693786 | 0.174816415 | 2.01E-05 | 1.05E-04 | 6902 | 5 | 76986991 | 77164604 | -1 | protein_coding | tubulin folding cofactor A |
| **BCL2L1** | ENSG00000171552 | 2693.92339 | -0.408588852 | 0.126330187 | 8.22E-04 | 0.003064732 | 598 | 20 | 30252255 | 30311792 | -1 | protein_coding | BCL2-like 1 |
| **NMUR1** | ENSG00000171596 | 11.05594145 | -2.676048868 | 1.75688863 | 0.013188015 | 0.035540164 | 10316 | 2 | 232387871 | 232395206 | -1 | protein_coding | neuromedin U receptor 1 |
| **CLSTN1** | ENSG00000171603 | 2745.452063 | -0.509355376 | 0.122515204 | 2.19E-05 | 1.13E-04 | 22883 | 1 | 9789084 | 9884584 | -1 | protein_coding | calsyntenin 1 |
| **PTCRA** | ENSG00000171611 | 26.01413272 | -1.856130946 | 0.840555575 | 0.002208349 | 0.007369038 | 171558 | 6 | 42883727 | 42893573 | 1 | protein_coding | pre T-cell antigen receptor alpha |
| **ENC1** | ENSG00000171617 | 756.9235162 | -2.921366526 | 0.239790528 | 1.48E-35 | 7.56E-34 | 8507 | 5 | 73923234 | 73937249 | -1 | protein_coding | ectodermal-neural cortex 1 (with BTB domain) |
| **SPSB1** | ENSG00000171621 | 1374.189739 | 0.736897821 | 0.244874754 | 0.001148374 | 0.004130524 | 80176 | 1 | 9352939 | 9429591 | 1 | protein_coding | splA/ryanodine receptor domain and SOCS box containing 1 |
| **GPR82** | ENSG00000171657 | 107.7802692 | -2.553655697 | 0.55173329 | 2.55E-07 | 1.79E-06 | 27197 | X | 41583408 | 41589388 | 1 | protein_coding | G protein-coupled receptor 82 |
| **GPR34** | ENSG00000171659 | 1182.67208 | -5.557847234 | 0.345687039 | 2.73E-59 | 3.48E-57 | 2857 | X | 41548226 | 41556526 | 1 | protein_coding | G protein-coupled receptor 34 |
| **PLEKHG5** | ENSG00000171680 | 8.779600784 | -0.942977482 | 1.89265351 | 0.017747006 | 0.045892977 | 57449 | 1 | 6526152 | 6580121 | -1 | protein_coding | pleckstrin homology domain containing, family G (with RhoGef domain) member 5 |
| **RGS19** | ENSG00000171700 | 714.9958914 | -0.455519762 | 0.186175623 | 0.009908121 | 0.027774361 | 10287 | 20 | 62704534 | 62711323 | -1 | protein_coding | regulator of G-protein signaling 19 |
| **HDAC3** | ENSG00000171720 | 1234.370381 | -0.369485344 | 0.132471463 | 0.004221365 | 0.013128841 | 8841 | 5 | 141000443 | 141016437 | -1 | protein_coding | histone deacetylase 3 |
| **TMEM51** | ENSG00000171729 | 1616.755304 | -0.55004269 | 0.140933442 | 5.84E-05 | 2.79E-04 | 55092 | 1 | 15479028 | 15546976 | 1 | protein_coding | transmembrane protein 51 |
| **SPATA5L1** | ENSG00000171763 | 191.4012692 | 0.591455858 | 0.248517019 | 0.009686517 | 0.027283273 | 79029 | 15 | 45694529 | 45713617 | 1 | protein_coding | spermatogenesis associated 5-like 1 |
| **GATM** | ENSG00000171766 | 328.3663982 | -1.473945647 | 0.293322222 | 7.24E-08 | 5.48E-07 | 2628 | 15 | 45653322 | 45694525 | -1 | protein_coding | glycine amidinotransferase (L-arginine:glycine amidinotransferase) |
| **RASGRP4** | ENSG00000171777 | 61.54229374 | -1.438804723 | 0.48402744 | 3.9E-04 | 0.001568782 | 115727 | 19 | 38899695 | 38916945 | -1 | protein_coding | RAS guanyl releasing protein 4 |
| **COL8A2** | ENSG00000171812 | 4826.38106 | -0.88566388 | 0.152558748 | 2.67E-09 | 2.44E-08 | 1296 | 1 | 36560837 | 36590821 | -1 | protein_coding | collagen, type VIII, alpha 2 |
| **NINJ2** | ENSG00000171840 | 79.9713448 | -2.239051528 | 0.438448687 | 3.15E-08 | 2.52E-07 | 4815 | 12 | 673462 | 772945 | -1 | protein_coding | ninjurin 2 |
| **RRM2** | ENSG00000171848 | 262.8384288 | 1.82878817 | 0.335931074 | 5.52E-09 | 4.86E-08 | 6241 | 2 | 10262455 | 10271545 | 1 | protein_coding | ribonucleotide reductase M2 |
| **RPS21** | ENSG00000171858 | 1284.722162 | -0.338745842 | 0.138522576 | 0.011923228 | 0.032551393 | 6227 | 20 | 60962172 | 60963576 | 1 | protein_coding | ribosomal protein S21 |
| **C3AR1** | ENSG00000171860 | 4913.373911 | 1.638334873 | 0.142230585 | 1.53E-31 | 6.41E-30 | 719 | 12 | 8210898 | 8219067 | -1 | protein_coding | complement component 3a receptor 1 |
| **PRNP** | ENSG00000171867 | 8485.415714 | -0.466181932 | 0.095388132 | 6.85E-07 | 4.53E-06 | 5621 | 20 | 4666882 | 4682236 | 1 | protein_coding | prion protein |
| **TLN2** | ENSG00000171914 | 362.0980188 | -1.52897097 | 0.269642896 | 1.9E-09 | 1.76E-08 | 83660 | 15 | 62682725 | 63136830 | 1 | protein_coding | talin 2 |
| **TVP23B** | ENSG00000171928 | 1131.584059 | 0.61146892 | 0.169904037 | 1.78E-04 | 7.72E-04 | 51030 | 17 | 18684308 | 18710027 | 1 | protein_coding | trans-golgi network vesicle protein 23 homolog B (S. cerevisiae) |
| **SRGAP2C** | ENSG00000171943 | 241.4573754 | 1.108946205 | 0.210304642 | 3.32E-08 | 2.64E-07 | NA | 1 | 121107124 | 121129949 | 1 | pseudogene | SLIT-ROBO Rho GTPase activating protein 2C |
| **ATPAF2** | ENSG00000171953 | 688.8239689 | -0.538750152 | 0.18963818 | 0.002773887 | 0.009025614 | 91647 | 17 | 17880723 | 17942523 | -1 | protein_coding | ATP synthase mitochondrial F1 complex assembly factor 2 |
| **JMJD1C** | ENSG00000171988 | 3183.171932 | 0.742757289 | 0.106114087 | 9.27E-13 | 1.18E-11 | 221037 | 10 | 64926981 | 65225722 | -1 | protein_coding | jumonji domain containing 1C |
| **MAL** | ENSG00000172005 | 175.5163484 | -1.371316481 | 0.350513082 | 1.46E-05 | 7.83E-05 | 4118 | 2 | 95691422 | 95719737 | 1 | protein_coding | mal, T-cell differentiation protein |
| **RAB33B** | ENSG00000172007 | 275.4675145 | 0.709575867 | 0.253518374 | 0.002331259 | 0.007738626 | 83452 | 4 | 140374386 | 140397763 | 1 | protein_coding | RAB33B, member RAS oncogene family |
| **THOP1** | ENSG00000172009 | 423.8492005 | -0.596653158 | 0.176668611 | 4.2E-04 | 0.001673772 | 7064 | 19 | 2785458 | 2813599 | 1 | protein_coding | thimet oligopeptidase 1 |
| **LAMB2** | ENSG00000172037 | 1714.783024 | -0.481415291 | 0.115404013 | 2.08E-05 | 1.08E-04 | 3913 | 3 | 49158547 | 49170551 | -1 | protein_coding | laminin, beta 2 (laminin S) |
| **QARS** | ENSG00000172053 | 2111.175107 | -0.558370146 | 0.143024457 | 5.79E-05 | 2.76E-04 | 1024655 | 3 | 49133365 | 49142553 | -1 | protein_coding | glutaminyl-tRNA synthetase |
| **SERF1A** | ENSG00000172058 | 111.8167965 | 1.052808571 | 0.561845709 | 0.010781447 | 0.029823142 | 728492 | 5 | 70196492 | 70214357 | 1 | protein_coding | small EDRK-rich factor 1A (telomeric) |
| **EIF2AK3** | ENSG00000172071 | 506.3831722 | -0.838683418 | 0.190076377 | 3.86E-06 | 2.27E-05 | 9451 | 2 | 88856259 | 88927094 | -1 | protein_coding | eukaryotic translation initiation factor 2-alpha kinase 3 |
| **MOB3A** | ENSG00000172081 | 2581.444315 | -0.753055571 | 0.141109569 | 4.35E-08 | 3.4E-07 | 126308 | 19 | 2071037 | 2096672 | -1 | protein_coding | MOB kinase activator 3A |
| **CYCS** | ENSG00000172115 | 2592.449331 | -0.418838671 | 0.163381759 | 0.007656784 | 0.022175219 | 54205 | 7 | 25159710 | 25164980 | -1 | protein_coding | cytochrome c, somatic |
| **SLFN12** | ENSG00000172123 | 391.2729269 | 1.880380634 | 0.215047614 | 2.5E-19 | 5.24E-18 | 55106 | 17 | 33738079 | 33760302 | -1 | protein_coding | schlafen family member 12 |
| **SNTB1** | ENSG00000172164 | 6345.536798 | -0.856149671 | 0.122419094 | 9.47E-13 | 1.21E-11 | 6641 | 8 | 121547985 | 121825513 | -1 | protein_coding | syntrophin, beta 1 (dystrophin-associated protein A1, 59kDa, basic component 1) |
| **MTBP** | ENSG00000172167 | 71.14801677 | 1.106682123 | 0.518661647 | 0.005694228 | 0.017070396 | 27085 | 8 | 121457640 | 121554373 | 1 | protein_coding | Mdm2, transformed 3T3 cell double minute 2, p53 binding protein (mouse) binding protein, 104kDa |
| **ISG20** | ENSG00000172183 | 1094.351861 | 5.965607488 | 0.25722275 | 1.95E-120 | 8.07E-118 | 3669 | 15 | 89179384 | 89199714 | 1 | protein_coding | interferon stimulated exonuclease gene 20kDa |
| **MBOAT1** | ENSG00000172197 | 614.0333413 | -1.591758851 | 0.225536714 | 2.24E-13 | 2.99E-12 | 154141 | 6 | 20100935 | 20212670 | -1 | protein_coding | membrane bound O-acyltransferase domain containing 1 |
| **CXCR6** | ENSG00000172215 | 224.0535889 | 0.829718119 | 0.304664094 | 0.002284123 | 0.007598662 | 10663 | 3 | 45982425 | 45989845 | 1 | protein_coding | chemokine (C-X-C motif) receptor 6 |
| **PAIP1** | ENSG00000172239 | 1527.940102 | 0.668857588 | 0.11933061 | 1.16E-08 | 9.79E-08 | 10605 | 5 | 43526369 | 43557860 | -1 | protein_coding | poly(A) binding protein interacting protein 1 |
| **CLEC7A** | ENSG00000172243 | 3653.813571 | -0.795391022 | 0.098875799 | 3.69E-16 | 6.05E-15 | 64581 | 12 | 10269376 | 10282857 | -1 | protein_coding | C-type lectin domain family 7, member A |
| **BSG** | ENSG00000172270 | 4963.957814 | -0.429334316 | 0.088897379 | 1.06E-06 | 6.8E-06 | 682 | 19 | 571297 | 583493 | 1 | protein_coding | basigin (Ok blood group) |
| **CERS6** | ENSG00000172292 | 3275.809159 | 0.486240466 | 0.133204493 | 1.89E-04 | 8.14E-04 | 253782 | 2 | 169312372 | 169631644 | 1 | protein_coding | ceramide synthase 6 |
| **BPGM** | ENSG00000172331 | 411.7354411 | 1.235175285 | 0.175143234 | 3.72E-13 | 4.92E-12 | 669 | 7 | 134331560 | 134364565 | 1 | protein_coding | 2,3-bisphosphoglycerate mutase |
| **IL16** | ENSG00000172349 | 1791.893711 | -2.047701308 | 0.13599309 | 3.23E-52 | 3.16E-50 | 3603 | 15 | 81451916 | 81605104 | 1 | protein_coding | interleukin 16 |
| **GNG12** | ENSG00000172380 | 165.4898973 | -2.008404585 | 0.394493159 | 2.89E-08 | 2.32E-07 | 55970 | 1 | 68167149 | 68299150 | -1 | protein_coding | guanine nucleotide binding protein (G protein), gamma 12 |
| **SYNPO2** | ENSG00000172403 | 64.67518034 | 1.448696179 | 0.492132181 | 4.04E-04 | 0.00161848 | 171024 | 4 | 119809996 | 119982402 | 1 | protein_coding | synaptopodin 2 |
| **MYEOV2** | ENSG00000172428 | 424.1061332 | -0.448730919 | 0.178891039 | 0.008441817 | 0.024106184 | 150678 | 2 | 241065980 | 241076224 | -1 | protein_coding | myeloma overexpressed 2 |
| **GTPBP2** | ENSG00000172432 | 1960.400322 | 1.207327361 | 0.140839162 | 2.32E-18 | 4.52E-17 | 54676 | 6 | 43573053 | 43596899 | -1 | protein_coding | GTP binding protein 2 |
| **TCEAL1** | ENSG00000172465 | 217.7492957 | 0.560996142 | 0.233341589 | 0.009561904 | 0.026977031 | 9338 | X | 102883632 | 102885881 | 1 | protein_coding | transcription elongation factor A (SII)-like 1 |
| **ZNF24** | ENSG00000172466 | 2747.981285 | 0.625160632 | 0.119854067 | 1.18E-07 | 8.67E-07 | 7572 | 18 | 32912176 | 32925446 | -1 | protein_coding | zinc finger protein 24 |
| **MANEA** | ENSG00000172469 | 539.6967353 | 1.363458958 | 0.279354349 | 1.74E-07 | 1.25E-06 | 79694 | 6 | 96025419 | 96057333 | 1 | protein_coding | mannosidase, endo-alpha |
| **AFF1** | ENSG00000172493 | 5568.985912 | 0.362431117 | 0.100629949 | 1.89E-04 | 8.13E-04 | 4299 | 4 | 87856154 | 88062206 | 1 | protein_coding | AF4/FMR2 family, member 1 |
| **CARNS1** | ENSG00000172508 | 47.08502861 | -1.497857171 | 0.663718282 | 0.00242322 | 0.008000446 | 57571 | 11 | 67182439 | 67193078 | 1 | protein_coding | carnosine synthase 1 |
| **BANP** | ENSG00000172530 | 378.7491079 | 0.586299851 | 0.194654066 | 0.001495756 | 0.00522148 | 54971 | 16 | 87982850 | 88110924 | 1 | protein_coding | BTG3 associated nuclear protein |
| **PPP1CA** | ENSG00000172531 | 2795.438591 | -0.614591089 | 0.102333812 | 9.45E-10 | 9.12E-09 | 5499 | 11 | 67165654 | 67188654 | -1 | protein_coding | protein phosphatase 1, catalytic subunit, alpha isozyme |
| **NIPAL4** | ENSG00000172548 | 115.127741 | 2.029156318 | 0.408986778 | 6.58E-08 | 5.01E-07 | 348938 | 5 | 156887027 | 156901725 | 1 | protein_coding | NIPA-like domain containing 4 |
| **RASGRP1** | ENSG00000172575 | 933.1910331 | 1.187773846 | 0.162788879 | 6.39E-14 | 9.02E-13 | 10125 | 15 | 38780304 | 38857776 | -1 | protein_coding | RAS guanyl releasing protein 1 (calcium and DAG-regulated) |
| **SMPDL3A** | ENSG00000172594 | 138.7730035 | 1.101998728 | 0.384580966 | 8.69E-04 | 0.003222585 | 10924 | 6 | 123110315 | 123130865 | 1 | protein_coding | sphingomyelin phosphodiesterase, acid-like 3A |
| **RAD9A** | ENSG00000172613 | 145.3703029 | 1.047747198 | 0.345148185 | 5.64E-04 | 0.002176396 | 5883 | 11 | 67159176 | 67165881 | 1 | protein_coding | RAD9 homolog A (S. pombe) |
| **FAM21C** | ENSG00000172661 | 2757.233313 | 0.397587723 | 0.117518649 | 5.59E-04 | 0.002160246 | 1019305 | 10 | 46222648 | 46288409 | 1 | protein_coding | family with sequence similarity 21, member C |
| **TMEM134** | ENSG00000172663 | 238.2711761 | -0.976941106 | 0.238590394 | 1.26E-05 | 6.87E-05 | 80194 | 11 | 67231824 | 67236743 | -1 | protein_coding | transmembrane protein 134 |
| **ZMAT3** | ENSG00000172667 | 2962.176615 | -1.18446794 | 0.144048042 | 4.45E-17 | 7.84E-16 | 64393 | 3 | 178735011 | 178790067 | -1 | protein_coding | zinc finger, matrin-type 3 |
| **ZFAND4** | ENSG00000172671 | 274.8950028 | 0.58481108 | 0.244611989 | 0.009576232 | 0.027007484 | 93550 | 10 | 46111043 | 46168228 | -1 | protein_coding | zinc finger, AN1-type domain 4 |
| **SLFN11** | ENSG00000172716 | 2620.756063 | -0.668212024 | 0.107084095 | 2.5E-10 | 2.54E-09 | 91607 | 17 | 33677324 | 33700720 | -1 | protein_coding | schlafen family member 11 |
| **CCL19** | ENSG00000172724 | 5.469179732 | 3.027346367 | 2.484520603 | 0.005962409 | 0.017783665 | 6363 | 9 | 34689564 | 34691274 | -1 | protein_coding | chemokine (C-C motif) ligand 19 |
| **CORO1B** | ENSG00000172725 | 2000.030502 | -0.471505775 | 0.154055813 | 0.001549901 | 0.005394474 | 57175 | 11 | 67205519 | 67211292 | -1 | protein_coding | coronin, actin binding protein, 1B |
| **TMEM217** | ENSG00000172738 | 216.7319983 | 0.783660457 | 0.25619766 | 8.92E-04 | 0.003296345 | 221468 | 6 | 37179956 | 37225931 | -1 | protein_coding | transmembrane protein 217 |
| **CFL1** | ENSG00000172757 | 14298.42323 | -0.234119481 | 0.094572252 | 0.012196096 | 0.033195536 | 1072 | 11 | 65590493 | 65629497 | -1 | protein_coding | cofilin 1 (non-muscle) |
| **FAM192A** | ENSG00000172775 | 1819.587196 | 0.271942556 | 0.107449358 | 0.010475967 | 0.029099274 | 80011 | 16 | 57186378 | 57220028 | -1 | protein_coding | family with sequence similarity 192, member A |
| **CBWD1** | ENSG00000172785 | 878.3534088 | 0.622835339 | 0.146975841 | 1.39E-05 | 7.46E-05 | 644019 | 9 | 121041 | 188979 | -1 | protein_coding | COBW domain containing 1 |
| **RAB37** | ENSG00000172794 | 174.8199582 | -2.135783946 | 0.365767824 | 4.81E-10 | 4.77E-09 | 326624 | 17 | 72666717 | 72743474 | 1 | protein_coding | RAB37, member RAS oncogene family |
| **DCP2** | ENSG00000172795 | 2189.98623 | 0.504002543 | 0.128592824 | 5.92E-05 | 2.82E-04 | 167227 | 5 | 112312399 | 112356667 | 1 | protein_coding | decapping mRNA 2 |
| **OVOL1** | ENSG00000172818 | 328.2061354 | 0.954296198 | 0.200584222 | 5.94E-07 | 3.96E-06 | 5017 | 11 | 65554493 | 65564690 | 1 | protein_coding | ovo-like zinc finger 1 |
| **CES3** | ENSG00000172828 | 46.84689093 | -1.562928385 | 0.613098676 | 0.001140469 | 0.004106925 | 23491 | 16 | 66995140 | 67009051 | 1 | protein_coding | carboxylesterase 3 |
| **SP3** | ENSG00000172845 | 2646.037633 | 0.246083531 | 0.092647522 | 0.007138532 | 0.020868052 | 6670 | 2 | 174771187 | 174830430 | -1 | protein_coding | Sp3 transcription factor |
| **DMXL1** | ENSG00000172869 | 4117.046459 | 0.747532338 | 0.136488315 | 2E-08 | 1.63E-07 | 1657 | 5 | 118373467 | 118584833 | 1 | protein_coding | Dmx-like 1 |
| **DHCR7** | ENSG00000172893 | 963.3753774 | -0.829628945 | 0.193222647 | 6.78E-06 | 3.83E-05 | 1717 | 11 | 71139239 | 71163914 | -1 | protein_coding | 7-dehydrocholesterol reductase |
| **MYEOV** | ENSG00000172927 | 19.89722893 | 1.852985374 | 0.862515307 | 0.002159092 | 0.007231516 | 26579 | 11 | 69061605 | 69182494 | 1 | protein_coding | myeloma overexpressed |
| **MRGPRF** | ENSG00000172935 | 209.7193697 | -2.278909077 | 0.446058348 | 2.39E-08 | 1.93E-07 | 116535 | 11 | 68771863 | 68780877 | -1 | protein_coding | MAS-related GPR, member F |
| **MYD88** | ENSG00000172936 | 6533.903419 | 1.219066608 | 0.092883984 | 7.26E-40 | 4.32E-38 | 4615 | 3 | 38179969 | 38184513 | 1 | protein_coding | myeloid differentiation primary response 88 |
| **OXSR1** | ENSG00000172939 | 2452.790799 | 0.621794722 | 0.105056153 | 2.49E-09 | 2.27E-08 | 9943 | 3 | 38206580 | 38296979 | 1 | protein_coding | oxidative stress responsive 1 |
| **SH3RF3** | ENSG00000172985 | 425.7863887 | -1.381923744 | 0.191846335 | 1.04E-13 | 1.44E-12 | 344558 | 2 | 109745804 | 110262207 | 1 | protein_coding | SH3 domain containing ring finger 3 |
| **DCAKD** | ENSG00000172992 | 225.0766137 | -1.072604264 | 0.237328627 | 1.64E-06 | 1.02E-05 | 79877 | 17 | 43100708 | 43138473 | -1 | protein_coding | dephospho-CoA kinase domain containing |
| **RELA** | ENSG00000173039 | 2002.332385 | 1.283390485 | 0.130592636 | 1.79E-23 | 4.96E-22 | 5970 | 11 | 65421067 | 65430565 | -1 | protein_coding | v-rel avian reticuloendotheliosis viral oncogene homolog A |

| | | | | | | | | | | | | | |
|---|---|---|---|---|---|---|---|---|---|---|---|---|---|
| **ZNF680** | ENSG00000173041 | 189.9480765 | 1.443986811 | 0.30547357 | 3.51E-07 | 2.42E-06 | 340252 | 7 | 63980262 | 64023484 | -1 | protein_coding | zinc finger protein 680 |
| **FAM222B** | ENSG00000173065 | 936.3368883 | -0.354563034 | 0.141366347 | 0.009947046 | 0.027868141 | 55731 | 17 | 27082996 | 27182250 | -1 | protein_coding | family with sequence similarity 222, member B |
| **BNC2** | ENSG00000173068 | 18.64347485 | -4.15104304 | 1.554158502 | 0.001253293 | 0.004459623 | 54796 | 9 | 16409501 | 16870841 | -1 | protein_coding | basonuclin 2 |
| **LRRN3** | ENSG00000173114 | 398.5271119 | -0.653152251 | 0.285529185 | 0.01051588 | 0.029202042 | 54674 | 7 | 110731062 | 110765510 | 1 | protein_coding | leucine rich repeat neuronal 3 |
| **KDM2A** | ENSG00000173120 | 4792.091374 | 0.431519769 | 0.101913062 | 1.69E-05 | 8.92E-05 | 22992 | 11 | 66886740 | 67025558 | 1 | protein_coding | lysine (K)-specific demethylase 2A |
| **NOC3L** | ENSG00000173145 | 460.9962763 | 0.814338038 | 0.203812355 | 2.55E-05 | 1.3E-04 | 64318 | 10 | 96075004 | 96122716 | -1 | protein_coding | nucleolar complex associated 3 homolog (S. cerevisiae) |
| **RAPH1** | ENSG00000173166 | 2281.996884 | -1.73565442 | 0.131207908 | 7.97E-41 | 5.01E-39 | 65059 | 2 | 204259068 | 204400133 | -1 | protein_coding | Ras association (RalGDS/AF-6) and pleckstrin homology domains 1 |
| **PARP14** | ENSG00000173193 | 18462.75557 | 2.004601929 | 0.16026067 | 7.37E-37 | 3.93E-35 | 54625 | 3 | 122399465 | 122449687 | 1 | protein_coding | poly (ADP-ribose) polymerase family, member 14 |
| **PARP15** | ENSG00000173200 | 738.7223577 | 1.807903682 | 0.168951134 | 1.29E-27 | 4.35E-26 | 165631 | 3 | 122296449 | 122357894 | 1 | protein_coding | poly (ADP-ribose) polymerase family, member 15 |
| **CKS1B** | ENSG00000173207 | 216.2867384 | 1.053674393 | 0.307867157 | 1.45E-04 | 6.44E-04 | 1163 | 1 | 154947129 | 154951725 | 1 | protein_coding | CDC28 protein kinase regulatory subunit 1B |
| **ABLIM3** | ENSG00000173210 | 204.434528 | -2.742828101 | 0.418879847 | 3.93E-12 | 4.8E-11 | 22885 | 5 | 148521046 | 148640105 | 1 | protein_coding | actin binding LIM protein family, member 3 |
| **KIAA1919** | ENSG00000173214 | 405.1920229 | -0.53296273 | 0.18516094 | 0.002549547 | 0.00837231 | 91749 | 6 | 111580551 | 111592370 | 1 | protein_coding | KIAA1919 |
| **GLRX** | ENSG00000173221 | 2753.747983 | 1.024292346 | 0.113943674 | 6.65E-20 | 1.46E-18 | 2745 | 5 | 95087023 | 95158709 | -1 | protein_coding | glutaredoxin (thioltransferase) |
| **PPP1R3B** | ENSG00000173281 | 597.376288 | -0.514386166 | 0.215526957 | 0.010761786 | 0.029774144 | 79660 | 8 | 8993765 | 9009084 | -1 | protein_coding | protein phosphatase 1, regulatory subunit 3B |
| **STOX2** | ENSG00000173320 | 306.1423358 | -1.337607699 | 0.257098815 | 3.4E-08 | 2.69E-07 | 56977 | 4 | 184774584 | 184944679 | 1 | protein_coding | storkhead box 2 |
| **TRIB1** | ENSG00000173334 | 2000.274575 | -1.712528357 | 0.134422744 | 3.92E-38 | 2.2E-36 | 10221 | 8 | 126442563 | 126450647 | 1 | protein_coding | tribbles pseudokinase 1 |
| **OLR1** | ENSG00000173391 | 1063.537406 | -0.848214427 | 0.316603209 | 0.002472809 | 0.008146571 | 4973 | 12 | 10310902 | 10324737 | -1 | protein_coding | oxidized low density lipoprotein (lectin-like) receptor 1 |
| **DAG1** | ENSG00000173402 | 1652.926536 | 0.60505279 | 0.189380162 | 7.92E-04 | 0.002962739 | 1605 | 3 | 49506146 | 49573048 | 1 | protein_coding | dystroglycan 1 (dystrophin-associated glycoprotein 1) |
| **DAB1** | ENSG00000173406 | 5.294541837 | -0.431440309 | 0.851608115 | 0.013843439 | 0.037078388 | 1600 | 1 | 57460451 | 59012406 | -1 | protein_coding | Dab, reelin signal transducer, homolog 1 (Drosophila) |
| **EHBP1L1** | ENSG00000173442 | 4753.273602 | 0.296225937 | 0.125865872 | 0.016672481 | 0.043519303 | 254102 | 11 | 65343509 | 65360121 | 1 | protein_coding | EH domain binding protein 1-like 1 |
| **THAP2** | ENSG00000173451 | 141.2006943 | 0.948901435 | 0.346776877 | 0.001714298 | 0.00589216 | 83591 | 12 | 72056789 | 72074419 | 1 | protein_coding | THAP domain containing, apoptosis associated protein 2 |
| **RNF26** | ENSG00000173456 | 419.5787019 | -0.434810946 | 0.195493261 | 0.018785831 | 0.048107215 | 79102 | 11 | 119205237 | 119208023 | 1 | protein_coding | ring finger protein 26 |
| **PPP1R14B** | ENSG00000173457 | 912.1920136 | -1.25140809 | 0.138730978 | 3.91E-20 | 8.74E-19 | 26472 | 11 | 64011956 | 64014413 | -1 | protein_coding | protein phosphatase 1, regulatory (inhibitor) subunit 14B |
| **PTPRM** | ENSG00000173482 | 1053.445968 | -1.391041814 | 0.25437621 | 6.87E-09 | 5.96E-08 | 5797 | 18 | 7566780 | 8406859 | 1 | protein_coding | protein tyrosine phosphatase, receptor type, M |
| **FKBP2** | ENSG00000173486 | 409.5579151 | -0.529596239 | 0.177030718 | 0.001804742 | 0.006164924 | 2286 | 11 | 64008475 | 64011604 | 1 | protein_coding | FK506 binding protein 2, 13kDa |
| **VEGFB** | ENSG00000173511 | 1784.195104 | -1.274354837 | 0.168525541 | 8.76E-15 | 1.31E-13 | 7423 | 11 | 64002010 | 64006259 | 1 | protein_coding | vascular endothelial growth factor B |
| **TNFRSF10D** | ENSG00000173530 | 912.8604446 | -1.579183105 | 0.174640543 | 2.24E-20 | 5.07E-19 | 8793 | 8 | 22993101 | 23021543 | -1 | protein_coding | tumor necrosis factor receptor superfamily, member 10d, decoy with truncated death domain |
| **GMPPB** | ENSG00000173540 | 328.3503407 | -0.475428156 | 0.216512447 | 0.017502737 | 0.045399581 | 386724 | 3 | 49754277 | 49761384 | -1 | protein_coding | GDP-mannose pyrophosphorylase B |
| **MOB1B** | ENSG00000173542 | 2216.947085 | 0.605931934 | 0.138751664 | 8.37E-06 | 4.66E-05 | 92597 | 4 | 71768043 | 71888166 | 1 | protein_coding | MOB kinase activator 1B |
| **CSPG4** | ENSG00000173546 | 326.0118867 | -1.456005777 | 0.287838528 | 6.6E-08 | 5.03E-07 | 1464 | 15 | 75966663 | 76005189 | -1 | protein_coding | chondroitin sulfate proteoglycan 4 |
| **SNX33** | ENSG00000173548 | 1092.352392 | 0.316384899 | 0.121536457 | 0.007914392 | 0.022800312 | 257364 | 15 | 75940247 | 75954642 | 1 | protein_coding | sorting nexin 33 |
| **NABP1** | ENSG00000173559 | 12103.37522 | 0.59862478 | 0.19062947 | 8.94E-04 | 0.003305101 | 64859 | 2 | 192542794 | 192561385 | 1 | protein_coding | nucleic acid binding protein 1 |
| **CHD2** | ENSG00000173575 | 3223.324333 | 0.506450531 | 0.091240253 | 1.92E-08 | 1.57E-07 | 1106 | 15 | 93426526 | 93571237 | 1 | protein_coding | chromodomain helicase DNA binding protein 2 |
| **CCDC106** | ENSG00000173581 | 203.4262993 | -1.177607439 | 0.320742942 | 4.81E-05 | 2.32E-04 | 29903 | 19 | 56152428 | 56164527 | 1 | protein_coding | coiled-coil domain containing 106 |
| **SULT1B1** | ENSG00000173597 | 28.73586058 | -1.336954985 | 0.754076552 | 0.0085725 | 0.024446993 | 27284 | 4 | 70586880 | 70653679 | -1 | protein_coding | sulfotransferase family, cytosolic, 1B, member 1 |
| **PC** | ENSG00000173599 | 851.5498375 | -1.675985674 | 0.187276302 | 4.72E-20 | 1.05E-18 | 5091 | 11 | 66615704 | 66725847 | -1 | protein_coding | pyruvate carboxylase |
| **SCAI** | ENSG00000173611 | 148.8767801 | -1.266041311 | 0.311273257 | 8.74E-06 | 4.85E-05 | 286205 | 9 | 127704887 | 127905785 | -1 | protein_coding | suppressor of cancer cell invasion |
| **SLC19A1** | ENSG00000173638 | 148.9380632 | -1.359140011 | 0.344258011 | 1.23E-05 | 6.67E-05 | 1019287 | 21 | 46913486 | 46964325 | -1 | protein_coding | solute carrier family 19 (folate transporter), member 1 |
| **UQCRH** | ENSG00000173660 | 1074.355282 | -0.392379592 | 0.15140214 | 0.007525652 | 0.02182439 | 440567 | 1 | 46769303 | 46782448 | 1 | protein_coding | ubiquinol-cytochrome c reductase hinge protein |
| **EIF1AX** | ENSG00000173674 | 1631.761545 | 0.753467185 | 0.154839345 | 5.11E-07 | 3.45E-06 | 1010603 | X | 20142636 | 20159962 | -1 | protein_coding | eukaryotic translation initiation factor 1A, X-linked |
| **PSMD1** | ENSG00000173692 | 3682.757178 | 0.231792976 | 0.098699975 | 0.016777414 | 0.043740911 | 5707 | 2 | 231921578 | 232037541 | 1 | protein_coding | proteasome (prosome, macropain) 26S subunit, non-ATPase, 1 |
| **GPR64** | ENSG00000173698 | 23.49677318 | 3.002476368 | 0.722669627 | 2.11E-06 | 1.29E-05 | 10149 | X | 19007427 | 19140755 | -1 | protein_coding | G protein-coupled receptor 64 |
| **HEG1** | ENSG00000173706 | 567.1590152 | 1.411122125 | 0.175347875 | 1.47E-16 | 2.48E-15 | 57493 | 3 | 124684554 | 124774802 | -1 | protein_coding | heart development protein with EGF-like domains 1 |
| **TOMM20** | ENSG00000173726 | 2616.623094 | 0.300601755 | 0.119596971 | 0.010831703 | 0.029940481 | 9804 | 1 | 235272651 | 235292251 | -1 | protein_coding | translocase of outer mitochondrial membrane 20 homolog (yeast) |
| **CNP** | ENSG00000173786 | 173.0784017 | 2.226667845 | 0.287524241 | 9.08E-16 | 1.46E-14 | 1267 | 17 | 40118759 | 40129749 | 1 | protein_coding | 2',3'-cyclic nucleotide 3' phosphodiesterase |
| **RNF213** | ENSG00000173821 | 13948.19301 | 1.767935099 | 0.322319613 | 4.83E-09 | 4.28E-08 | 57674 | 17 | 78234665 | 78372586 | 1 | protein_coding | ring finger protein 213 |
| **PLK3** | ENSG00000173846 | 1386.543069 | 0.571616742 | 0.170217591 | 4.56E-04 | 0.001799919 | 1263 | 1 | 45265897 | 45271662 | 1 | protein_coding | polo-like kinase 3 |
| **PHC3** | ENSG00000173889 | 2123.841789 | 0.480871776 | 0.125818622 | 9.06E-05 | 4.17E-04 | 80012 | 3 | 169804520 | 169899537 | -1 | protein_coding | polyhomeotic homolog 3 (Drosophila) |
| **GOLIM4** | ENSG00000173905 | 1235.600765 | -0.648665441 | 0.160487373 | 2.79E-05 | 1.42E-04 | 27333 | 3 | 167726465 | 167813763 | -1 | protein_coding | golgi integral membrane protein 4 |
| **USMG5** | ENSG00000173915 | 1081.125222 | 0.388873839 | 0.171280306 | 0.018046195 | 0.046524966 | 84833 | 10 | 105148798 | 105156223 | -1 | protein_coding | up-regulated during skeletal muscle growth 5 homolog (mouse) |
| **C1QTNF1** | ENSG00000173918 | 33.77940234 | 0.322185103 | 0.760782782 | 0.003027789 | 0.009787161 | 114897 | 17 | 77018896 | 77045870 | 1 | protein_coding | C1q and tumor necrosis factor related protein 1 |
| **XXYLT1** | ENSG00000173950 | 114.0203077 | -1.180216356 | 0.362045462 | 2.22E-04 | 9.38E-04 | 152002 | 3 | 194789008 | 194991896 | -1 | protein_coding | xyloside xylosyltransferase 1 |
| **UBXN2A** | ENSG00000173960 | 575.7833924 | 0.62458147 | 0.179248844 | 2.55E-04 | 0.001062501 | 165324 | 2 | 24150155 | 24227779 | 1 | protein_coding | UBX domain protein 2A |
| **CCS** | ENSG00000173992 | 220.9105153 | -0.523070362 | 0.240409545 | 0.018177827 | 0.046815991 | 9973 | 11 | 66360292 | 66373490 | 1 | protein_coding | copper chaperone for superoxide dismutase |
| **NRROS** | ENSG00000174004 | 1971.328646 | -0.62000381 | 0.140173884 | 4.76E-06 | 2.76E-05 | 375387 | 3 | 196366557 | 196388875 | 1 | protein_coding | negative regulator of reactive oxygen species |
| **CEP19** | ENSG00000174007 | 208.6646271 | 0.549716903 | 0.258038363 | 0.019643892 | 0.049936394 | 84984 | 3 | 196433148 | 196439164 | -1 | protein_coding | centrosomal protein 19kDa |
| **GNG5** | ENSG00000174021 | 1675.253317 | 0.586485289 | 0.161374832 | 1.66E-04 | 7.25E-04 | 2787 | 1 | 84964008 | 84972248 | -1 | protein_coding | guanine nucleotide binding protein (G protein), gamma 5 |
| **SLC25A30** | ENSG00000174032 | 1233.776364 | 0.427074253 | 0.146012746 | 0.002615661 | 0.008556326 | 253512 | 13 | 45967451 | 45992590 | -1 | protein_coding | solute carrier family 25, member 30 |
| **CTSF** | ENSG00000174080 | 375.6892949 | -0.808705423 | 0.273629768 | 0.001140878 | 0.00410743 | 8722 | 11 | 66330934 | 66336312 | -1 | protein_coding | cathepsin F |
| **LEMD3** | ENSG00000174106 | 1029.432813 | 0.497423778 | 0.123872764 | 3.99E-05 | 1.96E-04 | 23592 | 12 | 65563351 | 65642107 | 1 | protein_coding | LEM domain containing 3 |
| **TRMT10C** | ENSG00000174173 | 463.5113368 | 0.770529257 | 0.22550011 | 2.6E-04 | 0.001083381 | 54931 | 3 | 101280706 | 101285290 | 1 | protein_coding | tRNA methyltransferase 10 homolog C (S. cerevisiae) |
| **MGA** | ENSG00000174197 | 2654.206712 | 0.643773002 | 0.111118578 | 3.97E-09 | 3.54E-08 | 23269 | 15 | 41913422 | 42062141 | 1 | protein_coding | MGA, MAX dimerization protein |
| **PIGG** | ENSG00000174227 | 1488.889271 | 0.352180966 | 0.12871286 | 0.004951495 | 0.015146857 | 54872 | 4 | 492989 | 533985 | 1 | protein_coding | phosphatidylinositol glycan anchor biosynthesis, class G |
| **PRPF8** | ENSG00000174231 | 8831.562745 | -0.883032202 | 0.12160746 | 1.48E-13 | 2.01E-12 | 10594 | 17 | 1553923 | 1588176 | -1 | protein_coding | pre-mRNA processing factor 8 |
| **ZBTB4** | ENSG00000174282 | 3118.997182 | -0.24099912 | 0.091124336 | 0.007803998 | 0.02253751 | 57659 | 17 | 7362685 | 7387582 | -1 | protein_coding | zinc finger and BTB domain containing 4 |
| **ZHX3** | ENSG00000174306 | 511.2010256 | -1.213917816 | 0.288607652 | 4.98E-06 | 2.88E-05 | 23051 | 20 | 39807088 | 39946312 | -1 | protein_coding | zinc fingers and homeoboxes 3 |
| **PHLDA3** | ENSG00000174307 | 1729.552063 | -0.46763097 | 0.137020581 | 4.56E-04 | 0.001801846 | 23612 | 1 | 201434620 | 201438365 | -1 | protein_coding | pleckstrin homology-like domain, family A, member 3 |
| **C11orf45** | ENSG00000174370 | 2411.898186 | -1.75542948 | 0.414972078 | 2.23E-06 | 1.36E-05 | 219833 | 11 | 128769460 | 128775930 | -1 | protein_coding | chromosome 11 open reading frame 45 |
| **EXO1** | ENSG00000174371 | 52.4732983 | 1.328910224 | 0.619969874 | 0.00401957 | 0.012565165 | 9156 | 1 | 242011269 | 242058450 | 1 | protein_coding | exonuclease 1 |
| **RALGAPA1** | ENSG00000174373 | 1156.926738 | 0.419935608 | 0.173209098 | 0.011698284 | 0.032046044 | 253959 | 14 | 36007558 | 36278510 | -1 | protein_coding | Ral GTPase activating protein, alpha subunit 1 (catalytic) |
| **ATP2A2** | ENSG00000174437 | 10238.52289 | 0.362393074 | 0.089818323 | 5.65E-05 | 2.71E-04 | 488 | 12 | 110718561 | 110788898 | 1 | protein_coding | ATPase, Ca++ transporting, cardiac muscle, slow twitch 2 |
| **ZWILCH** | ENSG00000174442 | 277.7188846 | 0.804475336 | 0.298248249 | 0.002539856 | 0.008345891 | 55055 | 15 | 66797297 | 66842115 | 1 | protein_coding | zwilch kinetochore protein |
| **RPL4** | ENSG00000174444 | 9315.991152 | -0.77095301 | 0.099294728 | 6.49E-15 | 9.81E-14 | 595099 | 15 | 66790355 | 66816870 | -1 | protein_coding | ribosomal protein L4 |
| **SNAPC5** | ENSG00000174446 | 558.0191224 | 0.371547597 | 0.163881638 | 0.018547277 | 0.047600005 | 10302 | 15 | 66782473 | 66790151 | -1 | protein_coding | small nuclear RNA activating complex, polypeptide 5, 19kDa |
| **PELI3** | ENSG00000174516 | 182.2726619 | -0.799437994 | 0.250258761 | 5.48E-04 | 0.00212163 | 246330 | 11 | 66234216 | 66244808 | 1 | protein_coding | pellino E3 ubiquitin protein ligase family member 3 |
| **AKIRIN1** | ENSG00000174574 | 2852.227234 | 0.757629344 | 0.113447179 | 1.04E-11 | 1.22E-10 | 79647 | 1 | 39456895 | 39471731 | 1 | protein_coding | akirin 1 |
| **MSL2** | ENSG00000174579 | 1104.332623 | 0.476297616 | 0.130792297 | 1.91E-04 | 8.24E-04 | 55167 | 3 | 135867764 | 135916083 | -1 | protein_coding | male-specific lethal 2 homolog (Drosophila) |
| **CMKLR1** | ENSG00000174600 | 1163.65317 | 1.908350706 | 0.210927424 | 1.51E-20 | 3.47E-19 | 1240 | 12 | 108681821 | 108733118 | -1 | protein_coding | chemokine-like receptor 1 |
| **ANGEL2** | ENSG00000174606 | 509.7886858 | 0.530791685 | 0.172530303 | 0.001343252 | 0.004736674 | 90806 | 1 | 213165524 | 213189168 | -1 | protein_coding | angel homolog 2 (Drosophila) |
| **IQCK** | ENSG00000174628 | 83.37028749 | -1.154542381 | 0.539573633 | 0.005391627 | 0.016297515 | 124152 | 16 | 19727778 | 19868907 | 1 | protein_coding | IQ motif containing K |
| **SLC29A2** | ENSG00000174669 | 81.83243775 | -1.152383036 | 0.386892488 | 5.88E-04 | 0.002258321 | 3177 | 11 | 66129992 | 66139685 | -1 | protein_coding | solute carrier family 29 (equilibrative nucleoside transporter), member 2 |
| **BRSK2** | ENSG00000174672 | 19.82485005 | -1.769354218 | 1.184605725 | 0.00797096 | 0.022954622 | 9024 | 11 | 1411129 | 1483919 | 1 | protein_coding | BR serine/threonine kinase 2 |
| **SH3PXD2B** | ENSG00000174705 | 1710.234481 | 0.527817916 | 0.210176606 | 0.007406719 | 0.021536699 | 285590 | 5 | 171752185 | 171881527 | -1 | protein_coding | SH3 and PX domains 2B |
| **KIAA1551** | ENSG00000174718 | 2250.416005 | 1.059611473 | 0.171317293 | 1.59E-10 | 1.66E-09 | 55196 | 12 | 32112304 | 32146039 | 1 | protein_coding | KIAA1551 |
| **LARP7** | ENSG00000174720 | 1046.829325 | 0.759263164 | 0.204578607 | 8.81E-05 | 4.07E-04 | 51574 | 4 | 113558120 | 113578748 | 1 | protein_coding | La ribonucleoprotein domain family, member 7 |
| **NR1D2** | ENSG00000174738 | 974.231867 | 0.926117781 | 0.175286971 | 4.16E-08 | 3.26E-07 | 9975 | 3 | 23986751 | 24022109 | 1 | protein_coding | nuclear receptor subfamily 1, group D, member 2 |
| **RPL15** | ENSG00000174748 | 9693.869841 | -0.431455187 | 0.119516015 | 1.72E-04 | 7.48E-04 | 6138 | 3 | 23958036 | 23965183 | 1 | protein_coding | ribosomal protein L15 |
| **C4orf32** | ENSG00000174749 | 225.2350142 | 1.708400666 | 0.283196938 | 2.02E-10 | 2.08E-09 | 132720 | 4 | 113066553 | 113116412 | 1 | protein_coding | chromosome 4 open reading frame 32 |
| **SRP72** | ENSG00000174780 | 1639.188615 | -0.507051819 | 0.132622468 | 7.76E-05 | 3.62E-04 | 6731 | 4 | 57333081 | 57369839 | 1 | protein_coding | signal recognition particle 72kDa |

| | | | | | | | | | | | | | |
|---|---|---|---|---|---|---|---|---|---|---|---|---|---|
| **THAP6** | ENSG00000174796 | 437.9241568 | 0.967324216 | 0.20689551 | 8.98E-07 | 5.82E-06 | 152815 | 4 | 76439156 | 76475683 | 1 | protein_coding | THAP domain containing 6 |
| **CEP135** | ENSG00000174799 | 594.5907942 | 1.170726936 | 0.161063351 | 8.04E-14 | 1.12E-12 | 9662 | 4 | 56815037 | 56899529 | 1 | protein_coding | centrosomal protein 135kDa |
| **FZD4** | ENSG00000174804 | 38.77157704 | 1.17541411 | 0.57529914 | 0.006524216 | 0.019286189 | 8322 | 11 | 86656721 | 86666433 | -1 | protein_coding | frizzled family receptor 4 |
| **EMR1** | ENSG00000174837 | 408.0929193 | 4.762048455 | 0.296340741 | 2.49E-59 | 3.2E-57 | 2015 | 19 | 6887577 | 6940470 | 1 | protein_coding | egf-like module containing, mucin-like, hormone receptor-like 1 |
| **DENND6A** | ENSG00000174839 | 1000.876981 | 0.922747573 | 0.177250848 | 6.45E-08 | 4.92E-07 | 201627 | 3 | 57611184 | 57678816 | -1 | protein_coding | DENN/MADD domain containing 6A |
| **GLMN** | ENSG00000174842 | 134.229813 | 0.77041529 | 0.395219658 | 0.018256178 | 0.04697915 | 11146 | 1 | 92711959 | 92764544 | -1 | protein_coding | glomulin, FKBP associated protein |
| **YIF1A** | ENSG00000174851 | 313.3036332 | -1.209310513 | 0.204524154 | 7.74E-10 | 7.52E-09 | 10897 | 11 | 66052051 | 66056641 | -1 | protein_coding | Yip1 interacting factor homolog A (S. cerevisiae) |
| **NLRP6** | ENSG00000174885 | 27.31879544 | -2.209204096 | 0.827928796 | 5.19E-04 | 0.002022227 | 171389 | 11 | 278365 | 285359 | 1 | protein_coding | NLR family, pyrin domain containing 6 |
| **NDUFA11** | ENSG00000174886 | 507.1881528 | -0.792207905 | 0.166673382 | 8.33E-07 | 5.44E-06 | 126328 | 19 | 5891287 | 5904017 | -1 | protein_coding | NADH dehydrogenase (ubiquinone) 1 alpha subcomplex, 11, 14.7kDa |
| **RAB1B** | ENSG00000174903 | 2812.253337 | -0.765044701 | 0.100203781 | 9.82E-15 | 1.46E-13 | 81876 | 11 | 66036004 | 66044963 | 1 | protein_coding | RAB1B, member RAS oncogene family |
| **PTDSS2** | ENSG00000174915 | 306.5254835 | -0.712765931 | 0.214946387 | 4.19E-04 | 0.001671161 | 81490 | 11 | 448268 | 491393 | 1 | protein_coding | phosphatidylserine synthase 2 |
| **SEZ6L2** | ENSG00000174938 | 1658.187104 | -0.912969974 | 0.128800006 | 5.88E-13 | 7.66E-12 | 26470 | 16 | 29882480 | 29910868 | -1 | protein_coding | seizure related 6 homolog (mouse)-like 2 |
| **ASPHD1** | ENSG00000174939 | 331.7161871 | -1.358523836 | 0.20722471 | 9.36E-12 | 1.1E-10 | 253982 | 16 | 29911696 | 29931185 | 1 | protein_coding | aspartate beta-hydroxylase domain containing 1 |
| **GPR171** | ENSG00000174946 | 187.7707689 | 1.25526125 | 0.56129776 | 0.003562058 | 0.011301418 | 29909 | 3 | 150915619 | 150920988 | -1 | protein_coding | G protein-coupled receptor 171 |
| **DHX36** | ENSG00000174953 | 1790.314796 | 0.684996915 | 0.137652009 | 3.26E-07 | 2.26E-06 | 170506 | 3 | 153990335 | 154042286 | -1 | protein_coding | DEAH (Asp-Glu-Ala-His) box polypeptide 36 |
| **FBXW8** | ENSG00000174989 | 285.7546753 | -0.624599751 | 0.204119801 | 0.001210491 | 0.004326459 | 26259 | 12 | 117348761 | 117468953 | 1 | protein_coding | F-box and WD repeat domain containing 8 |
| **CHST2** | ENSG00000175040 | 334.3616014 | 0.907963173 | 0.217750252 | 1.03E-05 | 5.68E-05 | 9435 | 3 | 142838173 | 142841800 | 1 | protein_coding | carbohydrate (N-acetylglucosamine-6-O) sulfotransferase 2 |
| **ZDHHC14** | ENSG00000175048 | 184.0401311 | -0.830635599 | 0.281857225 | 0.001136854 | 0.004096804 | 79683 | 6 | 157802165 | 158099178 | 1 | protein_coding | zinc finger, DHHC-type containing 14 |
| **GK5** | ENSG00000175066 | 321.2912249 | 0.831328288 | 0.246261358 | 2.8E-04 | 0.001159747 | 256356 | 3 | 141882414 | 141944449 | -1 | protein_coding | glycerol kinase 5 (putative) |
| **VCPIP1** | ENSG00000175073 | 2971.135579 | 1.016589955 | 0.097781097 | 6.94E-26 | 2.16E-24 | 80124 | 8 | 67540722 | 67579452 | -1 | protein_coding | valosin containing protein (p97)/p47 complex interacting protein 1 |
| **DES** | ENSG00000175084 | 9.444417072 | 5.483111811 | 2.553680924 | 2.69E-04 | 0.00111611 | 1674 | 2 | 220283099 | 220291461 | 1 | protein_coding | desmin |
| **PACS1** | ENSG00000175115 | 2330.211759 | -0.415306147 | 0.118294742 | 2.97E-04 | 0.001225805 | 55690 | 11 | 65837834 | 66012218 | 1 | protein_coding | phosphofurin acidic cluster sorting protein 1 |
| **MARCKSL1** | ENSG00000175130 | 136.4639253 | 1.460383364 | 0.304855755 | 2.52E-07 | 1.78E-06 | 65108 | 1 | 32799433 | 32801980 | -1 | protein_coding | MARCKS-like 1 |
| **SH3BP5L** | ENSG00000175137 | 1072.701274 | -0.610346423 | 0.131306848 | 1.91E-06 | 1.18E-05 | 80851 | 1 | 249104648 | 249120832 | -1 | protein_coding | SH3-binding domain protein 5-like |
| **PPM1E** | ENSG00000175175 | 36.72778518 | -3.212664601 | 1.030419742 | 1.27E-04 | 5.7E-04 | 22843 | 17 | 56833230 | 57058983 | 1 | protein_coding | protein phosphatase, Mg2+/Mn2+ dependent, 1E |
| **FAM131A** | ENSG00000175182 | 564.1243403 | -0.868224143 | 0.173638464 | 2.13E-07 | 1.51E-06 | 131408 | 3 | 184053714 | 184064063 | 1 | protein_coding | family with sequence similarity 131, member A |
| **CSRP2** | ENSG00000175183 | 130.403253 | -1.040381713 | 0.536943877 | 0.010128766 | 0.028320147 | 1466 | 12 | 77252495 | 77272840 | -1 | protein_coding | cysteine and glycine-rich protein 2 |
| **PARL** | ENSG00000175193 | 948.1184769 | -0.634837092 | 0.140082113 | 3.24E-06 | 1.93E-05 | 55486 | 3 | 183547173 | 183602721 | -1 | protein_coding | presenilin associated, rhomboid-like |
| **PCCA** | ENSG00000175198 | 461.0631109 | -0.870574528 | 0.284374595 | 7.33E-04 | 0.002760745 | 5095 | 13 | 100741269 | 101182686 | 1 | protein_coding | propionyl CoA carboxylase, alpha polypeptide |
| **ARHGAP1** | ENSG00000175220 | 1433.647079 | -0.437234433 | 0.141647038 | 0.001493699 | 0.005216682 | 392 | 11 | 46698630 | 46722165 | -1 | protein_coding | Rho GTPase activating protein 1 |
| **ATG13** | ENSG00000175224 | 2611.173451 | 0.984319587 | 0.153383865 | 4.57E-11 | 5.07E-10 | 9776 | 11 | 46638826 | 46696368 | 1 | protein_coding | autophagy related 13 |
| **C1orf127** | ENSG00000175262 | 37.4944296 | -2.465817046 | 0.863319984 | 2.99E-04 | 0.001232179 | 148345 | 1 | 11006528 | 11042094 | -1 | protein_coding | chromosome 1 open reading frame 127 |
| **GOLGA8A** | ENSG00000175265 | 875.4102935 | -0.907326244 | 0.1869367 | 4.12E-07 | 2.82E-06 | 23015 | 15 | 34671269 | 34880704 | -1 | protein_coding | golgin A8 family, member A |
| **TP53I11** | ENSG00000175274 | 1762.563719 | -2.108245635 | 0.125159982 | 1.12E-64 | 1.73E-62 | 9537 | 11 | 44907454 | 44972840 | -1 | protein_coding | tumor protein p53 inducible protein 11 |
| **PHYHD1** | ENSG00000175287 | 37.96262651 | -1.040262189 | 0.651196261 | 0.018018192 | 0.046468448 | 254295 | 9 | 131683174 | 131704320 | 1 | protein_coding | phytanoyl-CoA dioxygenase domain containing 1 |
| **CATSPER1** | ENSG00000175294 | 27.12253926 | -1.979983247 | 0.917599442 | 0.001929305 | 0.006553805 | 117144 | 11 | 65784223 | 65793988 | -1 | protein_coding | cation channel, sperm associated 1 |
| **BANF1** | ENSG00000175334 | 821.9532185 | -0.409379009 | 0.14681647 | 0.004065999 | 0.01269472 | 8815 | 11 | 65769550 | 65771620 | 1 | protein_coding | barrier to autointegration factor 1 |
| **TMEM9B** | ENSG00000175348 | 1885.183254 | 0.636010677 | 0.128750114 | 4.5E-07 | 3.07E-06 | 56674 | 11 | 8968841 | 8986558 | -1 | protein_coding | TMEM9 domain family, member B |
| **PTPN2** | ENSG00000175354 | 1155.536346 | 1.258614668 | 0.175944235 | 1.91E-13 | 2.58E-12 | 5771 | 18 | 12785477 | 12929642 | -1 | protein_coding | protein tyrosine phosphatase, non-receptor type 2 |
| **ARL 10.00** | ENSG00000175414 | 634.7493762 | -1.129078186 | 0.184846991 | 2.7E-10 | 2.75E-09 | 285598 | 5 | 175792471 | 175828866 | 1 | protein_coding | ADP-ribosylation factor-like 10 |
| **CLTB** | ENSG00000175416 | 439.6492091 | -0.603913181 | 0.186849266 | 6.91E-04 | 0.002617497 | 1212 | 5 | 175819456 | 175843570 | -1 | protein_coding | clathrin, light chain B |
| **LPL** | ENSG00000175445 | 33.12964846 | -1.757709022 | 0.855682073 | 0.002919009 | 0.00946141 | 4023 | 8 | 19759228 | 19824769 | 1 | protein_coding | lipoprotein lipase |
| **TBC1D10C** | ENSG00000175463 | 712.6451751 | -1.174103857 | 0.189660073 | 1.35E-10 | 1.42E-09 | 374403 | 11 | 67171386 | 67177560 | 1 | protein_coding | TBC1 domain family, member 10C |
| **MCTP1** | ENSG00000175471 | 2003.005126 | 0.814776972 | 0.13760435 | 1.27E-09 | 1.21E-08 | 79772 | 5 | 94039446 | 94620279 | -1 | protein_coding | multiple C2 domains, transmembrane 1 |
| **POLD4** | ENSG00000175482 | 1020.245592 | -0.408496428 | 0.1292108 | 0.001212245 | 0.004330143 | 57804 | 11 | 67118248 | 67124443 | -1 | protein_coding | polymerase (DNA-directed), delta 4, accessory subunit |
| **LRRC25** | ENSG00000175489 | 1104.370128 | -2.728526838 | 0.229631506 | 9.55E-34 | 4.53E-32 | 126364 | 19 | 18501954 | 18508427 | -1 | protein_coding | leucine rich repeat containing 25 |
| **CLCF1** | ENSG00000175505 | 74.36334209 | 1.367939391 | 0.475616604 | 5.55E-04 | 0.00214763 | 23529 | 11 | 67131639 | 67141648 | -1 | protein_coding | cardiotrophin-like cytokine factor 1 |
| **DRAP1** | ENSG00000175550 | 1014.201886 | 0.857163445 | 0.130379367 | 1.44E-11 | 1.67E-10 | 10589 | 11 | 65686728 | 65689032 | 1 | protein_coding | DR1-associated protein 1 (negative cofactor 2 alpha) |
| **LONRF3** | ENSG00000175556 | 782.5410795 | -0.611349193 | 0.168500526 | 1.63E-04 | 7.16E-04 | 79836 | X | 118108581 | 118152318 | 1 | protein_coding | LON peptidase N-terminal domain and ring finger 3 |
| **UCP2** | ENSG00000175567 | 5447.840218 | -2.096866066 | 0.140757327 | 2.01E-51 | 1.9E-49 | 7351 | 11 | 73685712 | 73694352 | -1 | protein_coding | uncoupling protein 2 (mitochondrial, proton carrier) |
| **MRPL48** | ENSG00000175581 | 164.7960824 | -0.80644598 | 0.314164299 | 0.003621426 | 0.011475483 | 51642 | 11 | 73498361 | 73576178 | 1 | protein_coding | mitochondrial ribosomal protein L48 |
| **RAB6A** | ENSG00000175582 | 5043.314798 | 0.539026801 | 0.088403625 | 6.65E-10 | 6.49E-09 | 150786 | 11 | 73386683 | 73472182 | -1 | protein_coding | RAB6A, member RAS oncogene family |
| **P2RY2** | ENSG00000175591 | 121.1664574 | -0.926601476 | 0.414544876 | 0.006899711 | 0.020247365 | 5029 | 11 | 72929343 | 72947397 | 1 | protein_coding | purinergic receptor P2Y, G-protein coupled, 2 |
| **FOSL1** | ENSG00000175592 | 121.7708214 | 0.747698828 | 0.305520441 | 0.005814642 | 0.017390444 | 8061 | 11 | 65659520 | 65668044 | -1 | protein_coding | FOS-like antigen 1 |
| **RPS6KB2** | ENSG00000175634 | 600.0622534 | -0.657438276 | 0.179252433 | 1.3E-04 | 5.83E-04 | 6199 | 11 | 67195931 | 67202872 | 1 | protein_coding | ribosomal protein S6 kinase, 70kDa, polypeptide 2 |
| **RMI2** | ENSG00000175643 | 95.93136292 | 3.243333841 | 0.60809573 | 5.6E-09 | 4.92E-08 | 116028 | 16 | 11343476 | 11445619 | 1 | protein_coding | RecQ mediated genome instability 2 |
| **MLXIP** | ENSG00000175727 | 1752.605888 | -0.376060262 | 0.137738074 | 0.004995966 | 0.015255414 | 22877 | 12 | 122516628 | 122631894 | 1 | protein_coding | MLX interacting protein |
| **TOMM5** | ENSG00000175768 | 531.0255893 | 0.620254928 | 0.174233087 | 2.08E-04 | 8.85E-04 | 401505 | 9 | 37582643 | 37592639 | -1 | protein_coding | translocase of outer mitochondrial membrane 5 homolog (yeast) |
| **SLC35E3** | ENSG00000175782 | 718.7239425 | 0.805549745 | 0.241941522 | 3.39E-04 | 0.001381087 | 55508 | 12 | 69139886 | 69187508 | 1 | protein_coding | solute carrier family 35, member E3 |
| **MSRA** | ENSG00000175806 | 603.8093675 | -0.752252589 | 0.174721054 | 8.38E-06 | 4.67E-05 | 4482 | 8 | 9911778 | 10286401 | 1 | protein_coding | methionine sulfoxide reductase A |
| **CTDNEP1** | ENSG00000175826 | 2548.697637 | -0.47430101 | 0.100344991 | 1.51E-06 | 9.41E-06 | 23399 | 17 | 7146910 | 7155810 | -1 | protein_coding | CTD nuclear envelope phosphatase 1 |
| **SWI5** | ENSG00000175854 | 82.96868063 | -0.871156648 | 0.372006503 | 0.005979123 | 0.017826559 | 375757 | 9 | 131037658 | 131051269 | 1 | protein_coding | SWI5 recombination repair homolog (yeast) |
| **BAIAP2** | ENSG00000175866 | 617.0858275 | -1.081990611 | 0.167534869 | 3.4E-11 | 3.83E-10 | 10458 | 17 | 79008948 | 79091232 | 1 | protein_coding | BAI1-associated protein 2 |
| **ZDHHC21** | ENSG00000175893 | 1032.769341 | 0.583213336 | 0.170868975 | 3.72E-04 | 0.001501405 | 340481 | 9 | 14611069 | 14693469 | -1 | protein_coding | zinc finger, DHHC-type containing 21 |
| **PLEKHF2** | ENSG00000175895 | 1184.490499 | 0.918667193 | 0.148942563 | 2.45E-10 | 2.5E-09 | 79666 | 8 | 96146032 | 96168912 | 1 | protein_coding | pleckstrin homology domain containing, family F (with FYVE domain) member 2 |
| **A2M** | ENSG00000175899 | 37053.85722 | -1.186072749 | 0.154914469 | 3.17E-15 | 4.87E-14 | 2 | 12 | 9220260 | 9268825 | -1 | protein_coding | alpha-2-macroglobulin |
| **ORAI3** | ENSG00000175938 | 721.9425292 | -0.832785645 | 0.150897366 | 1.29E-08 | 1.08E-07 | 93129 | 16 | 30960387 | 30967782 | 1 | protein_coding | ORAI calcium release-activated calcium modulator 3 |
| **UNC119B** | ENSG00000175970 | 614.4272432 | -0.594167697 | 0.161267079 | 1.34E-04 | 5.98E-04 | 84747 | 12 | 121148238 | 121161443 | 1 | protein_coding | unc-119 homolog B (C. elegans) |
| **TUBB6** | ENSG00000176014 | 1774.740803 | -0.477461535 | 0.109157585 | 7.6E-06 | 4.26E-05 | 84617 | 18 | 12307668 | 12344319 | 1 | protein_coding | tubulin, beta 6 class V |
| **LYSMD3** | ENSG00000176018 | 663.5313528 | 0.607034952 | 0.185810278 | 6.04E-04 | 0.002314793 | 116068 | 5 | 89811428 | 89825401 | -1 | protein_coding | LysM, putative peptidoglycan-binding, domain containing 3 |
| **NUPR1** | ENSG00000176046 | 6372.780649 | 0.572364283 | 0.10908027 | 9.97E-08 | 7.39E-07 | 26471 | 16 | 28548606 | 28550495 | -1 | protein_coding | nuclear protein, transcriptional regulator, 1 |
| **JAKMIP2** | ENSG00000176049 | 1833.295908 | -1.066746405 | 0.4309096 | 0.002486392 | 0.008184287 | 9832 | 5 | 146967990 | 147162338 | -1 | protein_coding | janus kinase and microtubule interacting protein 2 |
| **SLC35A4** | ENSG00000176087 | 1165.49646 | 0.425332597 | 0.121111279 | 3.13E-04 | 0.001282683 | 113829 | 5 | 139944041 | 139948688 | 1 | protein_coding | solute carrier family 35, member A4 |
| **CSTF3** | ENSG00000176102 | 716.768016 | 0.981785384 | 0.156302068 | 9.9E-11 | 1.06E-09 | 1479 | 11 | 33098734 | 33183917 | -1 | protein_coding | cleavage stimulation factor, 3' pre-RNA, subunit 3, 77kDa |
| **YES1** | ENSG00000176105 | 207.6566339 | 0.833772052 | 0.346532121 | 0.005462332 | 0.016478497 | 7525 | 18 | 721588 | 812547 | -1 | protein_coding | v-yes-1 Yamaguchi sarcoma viral oncogene homolog 1 |
| **SPHK1** | ENSG00000176170 | 1050.00691 | 0.808521202 | 0.129366254 | 1.7E-10 | 1.77E-09 | 8877 | 17 | 74372665 | 74383941 | 1 | protein_coding | sphingosine kinase 1 |
| **BNIP3** | ENSG00000176171 | 469.4149593 | -0.616067628 | 0.198272548 | 9.87E-04 | 0.003603672 | 664 | 10 | 133781578 | 133795435 | -1 | protein_coding | BCL2/adenovirus E1B 19kDa interacting protein 3 |
| **ATAD5** | ENSG00000176208 | 144.9644531 | 1.498196711 | 0.360174996 | 4.49E-06 | 2.62E-05 | 79915 | 17 | 29158988 | 29222887 | 1 | protein_coding | ATPase family, AAA domain containing 5 |
| **ZNF135** | ENSG00000176293 | 59.66313251 | -0.951917257 | 0.486935679 | 0.011955917 | 0.032628979 | 7694 | 19 | 58570607 | 58597677 | 1 | protein_coding | zinc finger protein 135 |
| **COX8A** | ENSG00000176340 | 909.7256718 | -0.602827041 | 0.140406368 | 9.79E-06 | 5.4E-05 | 1351 | 11 | 63742079 | 63744015 | 1 | protein_coding | cytochrome c oxidase subunit VIIIA (ubiquitous) |
| **ZSCAN2** | ENSG00000176371 | 145.1661223 | -0.636622073 | 0.288853553 | 0.013554627 | 0.036413319 | 54993 | 15 | 85144217 | 85171027 | 1 | protein_coding | zinc finger and SCAN domain containing 2 |
| **CDC26** | ENSG00000176386 | 395.4449028 | 0.782876853 | 0.188473455 | 1.38E-05 | 7.44E-05 | 246184 | 9 | 116018115 | 116037869 | -1 | protein_coding | cell division cycle 26 |
| **CRLF3** | ENSG00000176390 | 933.6993441 | 0.756684539 | 0.158205794 | 7.63E-07 | 5.01E-06 | 51379 | 17 | 29096406 | 29151794 | -1 | protein_coding | cytokine receptor-like factor 3 |
| **EID2** | ENSG00000176396 | 96.07256303 | 0.828393643 | 0.363806531 | 0.007670429 | 0.022210528 | 163126 | 19 | 40028890 | 40030870 | -1 | protein_coding | EP300 interacting inhibitor of differentiation 2 |
| **SLCO3A1** | ENSG00000176463 | 426.1154914 | 1.720288801 | 0.324498328 | 1.19E-08 | 1E-07 | 28232 | 15 | 92396925 | 92715665 | 1 | protein_coding | solute carrier organic anion transporter family, member 3A1 |
| **DIRAS1** | ENSG00000176490 | 562.2755812 | -2.253867757 | 0.195386486 | 9.37E-32 | 4E-30 | 148252 | 19 | 2714565 | 2721416 | -1 | protein_coding | DIRAS family, GTP-binding RAS-like 1 |
| **GNG7** | ENSG00000176533 | 31.4740153 | -1.085299671 | 0.658798661 | 0.016095114 | 0.042235856 | 2788 | 19 | 2511217 | 2702707 | -1 | protein_coding | guanine nucleotide binding protein (G protein), gamma 7 |
| **RMDN1** | ENSG00000176623 | 967.3842726 | 0.360280433 | 0.117821924 | 0.001839478 | 0.006273765 | 51115 | 8 | 87480486 | 87526586 | -1 | protein_coding | regulator of microtubule dynamics 1 |
| **MYO1D** | ENSG00000176658 | 2719.764703 | -1.351137163 | 0.189718673 | 1.96E-13 | 2.64E-12 | 4642 | 17 | 30819540 | 31204195 | -1 | protein_coding | myosin ID |

| | | | | | | | | | | | | | |
|---|---|---|---|---|---|---|---|---|---|---|---|---|---|
| **CCDC121** | ENSG00000176714 | 41.29330944 | -1.233455945 | 0.585625882 | 0.005448353 | 0.016449461 | 79635 | 2 | 27848506 | 27851879 | -1 | protein_coding | coiled-coil domain containing 121 |
| **BOK** | ENSG00000176720 | 34.28856575 | -1.722830714 | 0.793840582 | 0.002438315 | 0.008046808 | 666 | 2 | 242498136 | 242513546 | 1 | protein_coding | BCL2-related ovarian killer |
| **RUFY1** | ENSG00000176783 | 1365.651675 | -0.314277836 | 0.128705746 | 0.012453083 | 0.033780701 | 80230 | 5 | 178977559 | 179037027 | 1 | protein_coding | RUN and FYVE domain containing 1 |
| **BASP1** | ENSG00000176788 | 2885.139843 | 1.53514581 | 0.10275427 | 8.07E-52 | 7.81E-50 | 10409 | 5 | 17065707 | 17276943 | 1 | protein_coding | brain abundant, membrane attached signal protein 1 |
| **LRRC37A3** | ENSG00000176809 | 370.411128 | 0.631940746 | 0.263683427 | 0.008370129 | 0.023937254 | 1019304 | 17 | 62850430 | 62915598 | -1 | protein_coding | leucine rich repeat containing 37, member A3 |
| **METRNL** | ENSG00000176845 | 1219.526497 | -1.346710108 | 0.185510382 | 7.1E-14 | 9.97E-13 | 284207 | 17 | 81037567 | 81052864 | 1 | protein_coding | meteorin, glial cell differentiation regulator-like |
| **TYMS** | ENSG00000176890 | 153.1132828 | 1.516788259 | 0.433803322 | 5.75E-05 | 2.75E-04 | 7298 | 18 | 657604 | 673578 | 1 | protein_coding | thymidylate synthetase |
| **PNMA1** | ENSG00000176903 | 1284.943769 | 0.410633125 | 0.172694374 | 0.013096562 | 0.035306889 | 9240 | 14 | 74178494 | 74181128 | -1 | protein_coding | paraneoplastic Ma antigen 1 |
| **ANKLE2** | ENSG00000176915 | 2786.859159 | 0.496253899 | 0.106920386 | 2.32E-06 | 1.42E-05 | 23141 | 12 | 133302254 | 133338474 | -1 | protein_coding | ankyrin repeat and LEM domain containing 2 |
| **GCNT4** | ENSG00000176928 | 50.96160495 | 3.303032998 | 0.59105965 | 1.53E-09 | 1.44E-08 | 51301 | 5 | 74323289 | 74326724 | -1 | protein_coding | glucosaminyl (N-acetyl) transferase 4, core 2 |
| **NFATC2IP** | ENSG00000176953 | 1088.09145 | 0.693778442 | 0.13399066 | 1.12E-07 | 8.2E-07 | 84901 | 16 | 28962128 | 28978418 | 1 | protein_coding | nuclear factor of activated T-cells, cytoplasmic, calcineurin-dependent 2 interacting protein |
| **FAM89B** | ENSG00000176973 | 830.0543656 | -0.821290203 | 0.173356098 | 8.39E-07 | 5.47E-06 | 23625 | 11 | 65339820 | 65341669 | 1 | protein_coding | family with sequence similarity 89, member B |
| **SHMT1** | ENSG00000176974 | 284.6713597 | -1.286655886 | 0.233731954 | 7.12E-09 | 6.17E-08 | 1024667 | 17 | 18231187 | 18266856 | -1 | protein_coding | serine hydroxymethyltransferase 1 (soluble) |
| **DPP7** | ENSG00000176978 | 1627.818675 | -0.773929615 | 0.149180413 | 7.75E-08 | 5.85E-07 | 29952 | 9 | 140004994 | 140009629 | -1 | protein_coding | dipeptidyl-peptidase 7 |
| **SEC24C** | ENSG00000176986 | 2998.774266 | -0.31399312 | 0.102150442 | 0.001988589 | 0.006732742 | 9632 | 10 | 75504120 | 75531919 | 1 | protein_coding | SEC24 family member C |
| **SMCR8** | ENSG00000176994 | 2044.69659 | 0.84110742 | 0.111135674 | 1.55E-14 | 2.28E-13 | 140775 | 17 | 18218624 | 18226517 | 1 | protein_coding | Smith-Magenis syndrome chromosome region, candidate 8 |
| **MTX3** | ENSG00000177034 | 287.234189 | 0.60136362 | 0.241190709 | 0.007052144 | 0.02063525 | 345778 | 5 | 79275584 | 79287082 | -1 | protein_coding | metaxin 3 |
| **ZDHHC13** | ENSG00000177054 | 259.6432826 | 1.212090439 | 0.244373132 | 1.46E-07 | 1.06E-06 | 54503 | 11 | 19138646 | 19197969 | 1 | protein_coding | zinc finger, DHHC-type containing 13 |
| **SLC38A9** | ENSG00000177058 | 823.0714808 | 0.359578074 | 0.15367519 | 0.015666898 | 0.041296517 | 153129 | 5 | 54921673 | 55069022 | -1 | protein_coding | solute carrier family 38, member 9 |
| **RHOG** | ENSG00000177105 | 2423.960407 | 0.283695373 | 0.115709283 | 0.012905072 | 0.034852133 | 391 | 11 | 3848208 | 3862213 | -1 | protein_coding | ras homolog family member G |
| **EPS8L2** | ENSG00000177106 | 185.067422 | 0.725850766 | 0.288776948 | 0.005157043 | 0.015697091 | 64787 | 11 | 694438 | 727727 | 1 | protein_coding | EPS8-like 2 |
| **ZBTB34** | ENSG00000177125 | 727.5983451 | 0.424302867 | 0.188550452 | 0.016885164 | 0.04401432 | 403341 | 9 | 129622944 | 129648157 | 1 | protein_coding | zinc finger and BTB domain containing 34 |
| **TALDO1** | ENSG00000177156 | 6342.443688 | 0.308037835 | 0.093751577 | 9.01E-04 | 0.003328309 | 6888 | 11 | 747329 | 765024 | 1 | protein_coding | transaldolase 1 |
| **B3GNT8** | ENSG00000177191 | 100.7732923 | -0.950223794 | 0.408296192 | 0.005166242 | 0.01570632 | 374907 | 19 | 41931264 | 41934635 | -1 | protein_coding | UDP-GlcNAc:betaGal beta-1,3-N-acetylglucosaminyltransferase 8 |
| **MAN1B1** | ENSG00000177239 | 1969.083608 | -0.594127215 | 0.129464843 | 2.7E-06 | 1.63E-05 | 11253 | 9 | 139981379 | 140003635 | 1 | protein_coding | mannosidase, alpha, class 1B, member 1 |
| **KCNA3** | ENSG00000177272 | 183.9825481 | 1.311807739 | 0.285220955 | 7.96E-07 | 5.22E-06 | 3738 | 1 | 111214310 | 111217655 | -1 | protein_coding | potassium voltage-gated channel, shaker-related subfamily, member 3 |
| **KCNA2** | ENSG00000177301 | 273.7165115 | -1.678818743 | 0.405583627 | 3.49E-06 | 2.07E-05 | 3737 | 1 | 111136202 | 111174096 | -1 | protein_coding | potassium voltage-gated channel, shaker-related subfamily, member 2 |
| **TOP3A** | ENSG00000177302 | 799.0103161 | 0.511150056 | 0.154192063 | 6.26E-04 | 0.002389878 | 7156 | 17 | 18174742 | 18218321 | -1 | protein_coding | topoisomerase (DNA) III alpha |
| **MAGEF1** | ENSG00000177383 | 179.0131329 | 0.896754695 | 0.258618554 | 1.76E-04 | 7.65E-04 | 64110 | 3 | 184428156 | 184429836 | -1 | protein_coding | melanoma antigen family F, 1 |
| **UMODL1** | ENSG00000177398 | 7.864927125 | -1.79028334 | 2.430560345 | 0.015110283 | 0.040001812 | 89766 | 21 | 43483068 | 43563563 | 1 | protein_coding | uromodulin-like 1 |
| **SAMD9L** | ENSG00000177409 | 13012.10095 | 3.422852114 | 0.162211939 | 7.73E-100 | 2.51E-97 | 219285 | 7 | 92759368 | 92777682 | -1 | protein_coding | sterile alpha motif domain containing 9-like |
| **TGIF1** | ENSG00000177426 | 945.5038873 | 0.436826359 | 0.146707608 | 0.002041893 | 0.006887277 | 7050 | 18 | 3411606 | 3458409 | 1 | protein_coding | TGFB-induced factor homeobox 1 |
| **C8orf47** | ENSG00000177459 | 3.552607022 | 0.464736228 | 0.894383476 | 0.010063575 | 0.028152276 | 203111 | 8 | 99076539 | 99105838 | 1 | protein_coding | chromosome 8 open reading frame 47 |
| **ARIH2** | ENSG00000177479 | 1638.126584 | 0.390001219 | 0.117418786 | 7.63E-04 | 0.002863939 | 10425 | 3 | 48956254 | 49023815 | 1 | protein_coding | ariadne RBR E3 ubiquitin protein ligase 2 |
| **ZBTB33** | ENSG00000177485 | 1776.30748 | 0.694310626 | 0.129088733 | 3.53E-08 | 2.79E-07 | 10009 | X | 119384607 | 119392253 | 1 | protein_coding | zinc finger and BTB domain containing 33 |
| **ATOX1** | ENSG00000177556 | 1133.314765 | 0.68507711 | 0.135209468 | 1.97E-07 | 1.4E-06 | 475 | 5 | 151121877 | 151152093 | -1 | protein_coding | antioxidant 1 copper chaperone |
| **TBL1XR1** | ENSG00000177565 | 4549.754287 | -0.283346692 | 0.098645197 | 0.003571325 | 0.011326118 | 79718 | 3 | 176737143 | 176915261 | -1 | protein_coding | transducin (beta)-like 1 X-linked receptor 1 |
| **CD163** | ENSG00000177575 | 2584.429824 | -2.175651307 | 0.471642252 | 3.12E-07 | 2.16E-06 | 9332 | 12 | 7623409 | 7656489 | -1 | protein_coding | CD163 molecule |
| **RPLP2** | ENSG00000177600 | 2368.470835 | -0.349117737 | 0.137143483 | 0.009154249 | 0.02594185 | 619565 | 11 | 809647 | 812880 | 1 | protein_coding | ribosomal protein, large, P2 |
| **GSG2** | ENSG00000177602 | 29.03769134 | 1.088879529 | 0.67553772 | 0.016542278 | 0.043245917 | 83903 | 17 | 3627211 | 3630067 | 1 | protein_coding | germ cell associated 2 (haspin) |
| **JUN** | ENSG00000177606 | 6221.68185 | -0.390819622 | 0.091316754 | 1.93E-05 | 1.01E-04 | 3725 | 1 | 59246465 | 59249785 | -1 | protein_coding | jun proto-oncogene |
| **CSTF2T** | ENSG00000177613 | 682.7859309 | 0.332743575 | 0.140607295 | 0.014813389 | 0.039297521 | 23283 | 10 | 53455247 | 53459355 | -1 | protein_coding | cleavage stimulation factor, 3' pre-RNA, subunit 2, 64kDa, tau variant |
| **PGBD5** | ENSG00000177614 | 2199.235825 | -3.167262333 | 0.236179999 | 1.35E-42 | 9.38E-41 | 79605 | 1 | 230457392 | 230561475 | -1 | protein_coding | piggyBac transposable element derived 5 |
| **GBA** | ENSG00000177628 | 7151.013727 | -0.441771845 | 0.102957187 | 1.41E-05 | 7.56E-05 | 2629 | 1 | 155204243 | 155214490 | -1 | protein_coding | glucosidase, beta, acid |
| **ACAD9** | ENSG00000177646 | 875.7835126 | -0.414090929 | 0.138578819 | 0.002139889 | 0.007178209 | 28976 | 3 | 128598439 | 128634910 | 1 | protein_coding | acyl-CoA dehydrogenase family, member 9 |
| **IL17RA** | ENSG00000177663 | 16988.82905 | -0.984900541 | 0.074626428 | 2.79E-40 | 1.69E-38 | 23765 | 22 | 17565844 | 17596583 | 1 | protein_coding | interleukin 17 receptor A |
| **CD163L1** | ENSG00000177675 | 74.17604888 | -1.492095229 | 0.485100669 | 2.64E-04 | 0.00109515 | 283316 | 12 | 7499281 | 7632493 | -1 | protein_coding | CD163 molecule-like 1 |
| **THAP5** | ENSG00000177683 | 538.4475566 | 0.799580068 | 0.233495212 | 2.37E-04 | 9.98E-04 | 168451 | 7 | 108194987 | 108210194 | -1 | protein_coding | THAP domain containing 5 |
| **CD151** | ENSG00000177697 | 1948.493324 | -0.524751938 | 0.108309884 | 1.01E-06 | 6.47E-06 | 977 | 11 | 832843 | 839831 | 1 | protein_coding | CD151 molecule (Raph blood group) |
| **POLR2L** | ENSG00000177700 | 373.3901083 | -0.948763596 | 0.205894789 | 1.35E-06 | 8.48E-06 | 5441 | 11 | 837356 | 842545 | -1 | protein_coding | polymerase (RNA) II (DNA directed) polypeptide L, 7.6kDa |
| **FAM20C** | ENSG00000177706 | 7148.722118 | -1.982250072 | 0.17735076 | 3.74E-30 | 1.45E-28 | 56975 | 7 | 192969 | 300711 | 1 | protein_coding | family with sequence similarity 20, member C |
| **PVRL3** | ENSG00000177707 | 57.14602668 | 2.025195488 | 0.535927535 | 1.22E-05 | 6.62E-05 | 25945 | 3 | 110788918 | 110994410 | 1 | protein_coding | poliovirus receptor-related 3 |
| **KIAA0195** | ENSG00000177728 | 938.278686 | -0.735436268 | 0.202700142 | 1.26E-04 | 5.65E-04 | 1024669 | 17 | 73437240 | 73496171 | 1 | protein_coding | KIAA0195 |
| **FLII** | ENSG00000177731 | 7051.5043 | -0.23469029 | 0.102302437 | 0.01550622 | 0.040929457 | 2314 | 17 | 18148150 | 18162230 | -1 | protein_coding | flightless I homolog (Drosophila) |
| **MYOZ1** | ENSG00000177791 | 257.2531473 | -2.734606023 | 0.335011348 | 2.67E-17 | 4.77E-16 | 58529 | 10 | 75391412 | 75401515 | -1 | protein_coding | myozenin 1 |
| **CHID1** | ENSG00000177830 | 529.4583053 | -0.831673748 | 0.174419361 | 7.33E-07 | 4.83E-06 | 66005 | 11 | 867357 | 915058 | -1 | protein_coding | chitinase domain containing 1 |
| **AP3S1** | ENSG00000177879 | 1370.945919 | 0.540105247 | 0.123528545 | 7.67E-06 | 4.3E-05 | 1176 | 5 | 115177178 | 115249778 | 1 | protein_coding | adaptor-related protein complex 3, sigma 1 subunit |
| **ZBTB41** | ENSG00000177888 | 582.3495967 | 0.915896903 | 0.165613541 | 1.11E-08 | 9.36E-08 | 360023 | 1 | 197122810 | 197169672 | -1 | protein_coding | zinc finger and BTB domain containing 41 |
| **UBE2N** | ENSG00000177889 | 1356.647642 | 0.511274098 | 0.123359043 | 2.25E-05 | 1.16E-04 | 7334 | 12 | 93799449 | 93836038 | -1 | protein_coding | ubiquitin-conjugating enzyme E2N |
| **ARL6IP6** | ENSG00000177917 | 251.6418916 | 1.406979101 | 0.32675573 | 2.58E-06 | 1.56E-05 | 151188 | 2 | 153574407 | 153617767 | 1 | protein_coding | ADP-ribosylation-like factor 6 interacting protein 6 |
| **BET1L** | ENSG00000177951 | 1142.664027 | -0.605293216 | 0.142225883 | 1.22E-05 | 6.63E-05 | 51272 | 11 | 167784 | 207428 | -1 | protein_coding | Bet1 golgi vesicular membrane trafficking protein-like |
| **ODF3B** | ENSG00000177989 | 142.8412416 | 0.974823541 | 0.355762579 | 0.001592135 | 0.005525114 | 440836 | 22 | 50968139 | 50971009 | -1 | protein_coding | outer dense fiber of sperm tails 3B |
| **FAM26E** | ENSG00000178033 | 7.170243571 | 4.101887768 | 1.994151479 | 0.001657361 | 0.005721583 | 254228 | 6 | 116832809 | 116845955 | 1 | protein_coding | family with sequence similarity 26, member E |
| **ALS2CL** | ENSG00000178038 | 195.521955 | -1.414967565 | 0.341352239 | 4.7E-06 | 2.73E-05 | 259173 | 3 | 46710487 | 46735194 | -1 | protein_coding | ALS2 C-terminal like |
| **C2orf69** | ENSG00000178074 | 551.5369274 | 0.654370805 | 0.221785755 | 0.001581473 | 0.005491853 | 205327 | 2 | 200775979 | 200820658 | 1 | protein_coding | chromosome 2 open reading frame 69 |
| **STAP2** | ENSG00000178078 | 88.35334954 | 1.723118121 | 0.470596133 | 2.68E-05 | 1.36E-04 | 55620 | 19 | 4324040 | 4342783 | -1 | protein_coding | signal transducing adaptor family member 2 |
| **TWF1P1** | ENSG00000178082 | 64.39841758 | 2.791105953 | 0.430559059 | 6.93E-12 | 8.29E-11 | NA | 17 | 27530441 | 27531492 | -1 | pseudogene | twinfilin 1 pseudogene 1 |
| **PDE4DIP** | ENSG00000178104 | 6493.618561 | 0.848496185 | 0.09968222 | 1.39E-17 | 2.54E-16 | 1019304 | 1 | 144836157 | 145076186 | -1 | protein_coding | phosphodiesterase 4D interacting protein |
| **NDUFV2** | ENSG00000178127 | 1647.319058 | 0.316365198 | 0.136224496 | 0.01772664 | 0.04587039 | 4729 | 18 | 9102628 | 9134343 | 1 | protein_coding | NADH dehydrogenase (ubiquinone) flavoprotein 2, 24kDa |
| **SH2B1** | ENSG00000178188 | 638.4634625 | -0.504393664 | 0.187951197 | 0.004819524 | 0.014796461 | 25970 | 16 | 28857921 | 28885526 | 1 | protein_coding | SH2B adaptor protein 1 |
| **ZC3H12D** | ENSG00000178199 | 180.8959044 | 0.636427548 | 0.255964266 | 0.006660267 | 0.019616276 | 340152 | 6 | 149768794 | 149806197 | -1 | protein_coding | zinc finger CCCH-type containing 12D |
| **KDELC2** | ENSG00000178202 | 467.6820574 | 1.150358954 | 0.281870325 | 1.02E-05 | 5.62E-05 | 143888 | 11 | 108342832 | 108369159 | -1 | protein_coding | KDEL (Lys-Asp-Glu-Leu) containing 2 |
| **PRSS36** | ENSG00000178226 | 140.3862975 | -0.953148139 | 0.334863573 | 0.001243507 | 0.004427895 | 146547 | 16 | 31150246 | 31161415 | -1 | protein_coding | protease, serine, 36 |
| **GEN1** | ENSG00000178295 | 403.8581252 | 0.651587823 | 0.219288859 | 0.00152526 | 0.005317186 | 348654 | 2 | 17935125 | 17966632 | 1 | protein_coding | GEN1 Holliday junction 5' flap endonuclease |
| **ZFAND2A** | ENSG00000178381 | 233.7129705 | 0.59592799 | 0.220351068 | 0.003864995 | 0.012129101 | 90637 | 7 | 1191707 | 1200395 | -1 | protein_coding | zinc finger, AN1-type domain 2A |
| **PLEKHM3** | ENSG00000178385 | 414.5604188 | 1.426903585 | 0.205393502 | 6.22E-13 | 8.07E-12 | 389072 | 2 | 208693027 | 208890284 | -1 | protein_coding | pleckstrin homology domain containing, family M, member 3 |
| **P4HTM** | ENSG00000178467 | 496.8019581 | -0.549891285 | 0.199915663 | 0.003771566 | 0.011882259 | 54681 | 3 | 49027319 | 49044587 | 1 | protein_coding | prolyl 4-hydroxylase, transmembrane (endoplasmic reticulum) |
| **CD28** | ENSG00000178562 | 709.4302599 | -0.880590085 | 0.221369021 | 2.77E-05 | 1.41E-04 | 940 | 2 | 204571198 | 204603635 | 1 | protein_coding | CD28 molecule |
| **EPM2AIP1** | ENSG00000178567 | 839.5295821 | 1.605115712 | 0.16825723 | 1.96E-22 | 5.05E-21 | 9852 | 3 | 37027357 | 37034795 | -1 | protein_coding | EPM2A (laforin) interacting protein 1 |
| **MAF** | ENSG00000178573 | 4092.263083 | -1.967757022 | 0.136714333 | 5.72E-48 | 4.77E-46 | 4094 | 16 | 79619740 | 79634611 | -1 | protein_coding | v-maf avian musculoaponeurotic fibrosarcoma oncogene homolog |
| **CTNNBIP1** | ENSG00000178585 | 166.1762007 | -0.954992985 | 0.303100493 | 4.53E-04 | 0.00178988 | 56998 | 1 | 9908334 | 9970394 | -1 | protein_coding | catenin, beta interacting protein 1 |
| **ERN 1.00** | ENSG00000178607 | 1336.70257 | 1.288589167 | 0.178826539 | 1.05E-13 | 1.45E-12 | 2081 | 17 | 62116502 | 62208179 | -1 | protein_coding | endoplasmic reticulum to nucleus signaling 1 |
| **SUZ12** | ENSG00000178691 | 1417.715833 | 0.808856975 | 0.121786166 | 1.4E-11 | 1.62E-10 | 23512 | 17 | 30264037 | 30328064 | 1 | protein_coding | SUZ12 polycomb repressive complex 2 subunit |
| **NSUN3** | ENSG00000178694 | 506.927989 | 0.666200149 | 0.266272869 | 0.005978777 | 0.017826559 | 63899 | 3 | 93781760 | 93847389 | 1 | protein_coding | NOP2/Sun domain family, member 3 |
| **GRINA** | ENSG00000178719 | 396.6183272 | 0.814583053 | 0.208334972 | 3.57E-05 | 1.77E-04 | 2907 | 8 | 145064226 | 145067583 | 1 | protein_coding | glutamate receptor, ionotropic, N-methyl D-aspartate-associated protein 1 (glutamate binding) |
| **THBD** | ENSG00000178726 | 115.4694423 | -4.126489152 | 0.465426389 | 1.24E-19 | 2.66E-18 | 7056 | 20 | 23026270 | 23030378 | -1 | protein_coding | thrombomodulin |
| **FAM219B** | ENSG00000178761 | 1083.05087 | -0.339203501 | 0.149933647 | 0.019550755 | 0.049757566 | 57184 | 15 | 75192328 | 75199462 | -1 | protein_coding | family with sequence similarity 219, member B |
| **CD300LB** | ENSG00000178789 | 1624.961479 | -3.911768875 | 0.169849298 | 6.76E-118 | 2.72E-115 | 124599 | 17 | 72517313 | 72527613 | -1 | protein_coding | CD300 molecule-like family member b |
| **MPI** | ENSG00000178802 | 688.9236125 | -1.351463804 | 0.16846328 | 1.91E-16 | 3.2E-15 | 4351 | 15 | 75182346 | 75191798 | 1 | protein_coding | mannose phosphate isomerase |

| | | | | | | | | | | | | | |
|---|---|---|---|---|---|---|---|---|---|---|---|---|---|
| **TMEM139** | ENSG00000178826 | 10.45563563 | 4.229343695 | 1.742582418 | 5.82E-04 | 0.002236917 | 135932 | 7 | 142977050 | 142985141 | 1 | protein_coding | transmembrane protein 139 |
| **MSC** | ENSG00000178860 | 1236.633833 | 2.105306436 | 0.138729411 | 5.17E-53 | 5.31E-51 | 9242 | 8 | 72753784 | 72756703 | -1 | protein_coding | musculin |
| **TAF7** | ENSG00000178913 | 1637.189682 | 0.728187331 | 0.105845232 | 2.89E-12 | 3.55E-11 | 6879 | 5 | 140698057 | 140700330 | -1 | protein_coding | TAF7 RNA polymerase II, TATA box binding protein (TBP)-associated factor, 55kDa |
| **HYI** | ENSG00000178922 | 102.436036 | -1.162741838 | 0.426364435 | 0.00121496 | 0.004335334 | 81888 | 1 | 43916824 | 43919660 | -1 | protein_coding | hydroxypyruvate isomerase (putative) |
| **C17orf62** | ENSG00000178927 | 2435.99178 | 0.495691165 | 0.136143782 | 1.86E-04 | 8.03E-04 | 79415 | 17 | 80400465 | 80408705 | -1 | protein_coding | chromosome 17 open reading frame 62 |
| **ZNF552** | ENSG00000178935 | 190.4485813 | -0.727263548 | 0.301958957 | 0.006682954 | 0.019679303 | 79818 | 19 | 58315209 | 58326281 | -1 | protein_coding | zinc finger protein 552 |
| **ZBTB7A** | ENSG00000178951 | 2497.408646 | -0.256625972 | 0.100614134 | 0.009420725 | 0.026608193 | 51341 | 19 | 4044362 | 4066943 | -1 | protein_coding | zinc finger and BTB domain containing 7A |
| **TUFM** | ENSG00000178952 | 1712.953945 | -0.658400179 | 0.124052903 | 6.48E-08 | 4.94E-07 | 7284 | 16 | 28853732 | 28857729 | -1 | protein_coding | Tu translation elongation factor, mitochondrial |
| **RMI1** | ENSG00000178966 | 276.0319912 | 1.101409587 | 0.247202323 | 2.07E-06 | 1.27E-05 | 80010 | 9 | 86595626 | 86618985 | 1 | protein_coding | RecQ mediated genome instability 1 |
| **EIF3K** | ENSG00000178982 | 2158.313703 | -0.860269066 | 0.109010002 | 1.44E-15 | 2.27E-14 | 27335 | 19 | 39109735 | 39127595 | 1 | protein_coding | eukaryotic translation initiation factor 3, subunit K |
| **SNX18** | ENSG00000178996 | 2373.691796 | 0.48331189 | 0.130306328 | 1.47E-04 | 6.49E-04 | 112574 | 5 | 53813589 | 53842415 | 1 | protein_coding | sorting nexin 18 |
| **MRFAP1** | ENSG00000179010 | 3326.735628 | 0.303136671 | 0.105313368 | 0.003460496 | 0.011014644 | 93621 | 4 | 6641818 | 6644472 | 1 | protein_coding | Morf4 family associated protein 1 |
| **C3orf38** | ENSG00000179021 | 1307.261698 | 1.109785984 | 0.127544794 | 5.44E-19 | 1.11E-17 | 285237 | 3 | 88198893 | 88217879 | 1 | protein_coding | chromosome 3 open reading frame 38 |
| **TMEM107** | ENSG00000179029 | 295.4541393 | -0.673132807 | 0.204669454 | 5.14E-04 | 0.002005159 | 84314 | 17 | 8076555 | 8079717 | -1 | protein_coding | transmembrane protein 107 |
| **EXOC3L1** | ENSG00000179044 | 8.664973385 | 5.217655717 | 1.869346111 | 4.98E-04 | 0.001949646 | 283849 | 16 | 67218269 | 67224107 | -1 | protein_coding | exocyst complex component 3-like 1 |
| **CYC1** | ENSG00000179091 | 773.5621113 | -0.471600548 | 0.139843103 | 5.43E-04 | 0.002105029 | 1537 | 8 | 145149930 | 145152428 | 1 | protein_coding | cytochrome c-1 |
| **FARSA** | ENSG00000179115 | 1202.262946 | -0.841264378 | 0.149691011 | 6.44E-09 | 5.62E-08 | 2193 | 19 | 13033293 | 13044851 | -1 | protein_coding | phenylalanyl-tRNA synthetase, alpha subunit |
| **GIMAP7** | ENSG00000179144 | 508.0825557 | 1.055906367 | 0.199759976 | 3.35E-08 | 2.66E-07 | 168537 | 7 | 150211918 | 150218161 | 1 | protein_coding | GTPase, IMAP family member 7 |
| **TCAIM** | ENSG00000179152 | 1170.443015 | -0.502156507 | 0.150047283 | 5.37E-04 | 0.002084102 | 285343 | 3 | 44379611 | 44450943 | 1 | protein_coding | T cell activation inhibitor, mitochondrial |
| **FUCA1** | ENSG00000179163 | 6085.866029 | -0.763003418 | 0.132075965 | 3.33E-09 | 2.99E-08 | 2517 | 1 | 24171567 | 24194784 | -1 | protein_coding | fucosidase, alpha-L- 1, tissue |
| **CALR** | ENSG00000179218 | 35178.36588 | -0.854887809 | 0.097425699 | 6.75E-19 | 1.36E-17 | 811 | 19 | 13049392 | 13055303 | 1 | protein_coding | calreticulin |
| **MAGED1** | ENSG00000179222 | 1294.322112 | -1.704740786 | 0.141716989 | 3.37E-34 | 1.63E-32 | 9500 | X | 51546103 | 51645453 | 1 | protein_coding | melanoma antigen family D, 1 |
| **RP11-111M2** | ENSG00000179240 | 31.26544253 | -1.663314853 | 0.724540379 | 0.001956849 | 0.006639992 | 1019287 | 11 | 76092357 | 76125663 | 1 | protein_coding | Homo sapiens putative uncharacterized protein FLJ37770-like (LOC100506127), mRNA. |
| **C2orf71** | ENSG00000179270 | 62.30291297 | -1.412705794 | 0.844853529 | 0.008232977 | 0.023611295 | 388939 | 2 | 29283842 | 29297127 | -1 | protein_coding | chromosome 2 open reading frame 71 |
| **GADD45GIP1** | ENSG00000179271 | 282.3643505 | -0.589523279 | 0.2335841 | 0.006546193 | 0.019332459 | 90480 | 19 | 13064972 | 13068050 | -1 | protein_coding | growth arrest and DNA-damage-inducible, gamma interacting protein 1 |
| **TMEM151A** | ENSG00000179292 | 33.39505268 | -2.56303784 | 1.032871763 | 9.12E-04 | 0.003359326 | 256472 | 11 | 66059341 | 66064135 | 1 | protein_coding | transmembrane protein 151A |
| **PTPN11** | ENSG00000179295 | 5172.409593 | -0.319219231 | 0.083366868 | 1.61E-04 | 7.07E-04 | 5781 | 12 | 112856155 | 112947717 | 1 | protein_coding | protein tyrosine phosphatase, non-receptor type 11 |
| **NSUN7** | ENSG00000179299 | 275.831379 | 2.054186923 | 0.283482337 | 3.94E-14 | 5.64E-13 | 79730 | 4 | 40751914 | 40812002 | 1 | protein_coding | NOP2/Sun domain family, member 7 |
| **FAM156B** | ENSG00000179304 | 432.9316547 | -0.896687353 | 0.260767588 | 1.91E-04 | 8.22E-04 | 727866 | X | 52920336 | 52937587 | 1 | protein_coding | family with sequence similarity 156, member B |
| **WSCD1** | ENSG00000179314 | 69.69963479 | -2.19942434 | 0.483452058 | 4.84E-07 | 3.28E-06 | 339166 | 17 | 5675554 | 6027747 | 1 | protein_coding | WSC domain containing 1 |
| **RAB39A** | ENSG00000179331 | 459.0174018 | 1.78396217 | 0.182759294 | 1.99E-23 | 5.5E-22 | 54734 | 11 | 107799229 | 107834208 | 1 | protein_coding | RAB39A, member RAS oncogene family |
| **HLA-DQB1** | ENSG00000179344 | 10059.00337 | -0.51245297 | 0.192834831 | 0.005523124 | 0.016629216 | 1010608 | 6 | 32627244 | 32636160 | -1 | protein_coding | major histocompatibility complex, class II, DQ beta 1 |
| **HMGN2P46** | ENSG00000179362 | 202.2056135 | 2.071871541 | 0.250489288 | 1.36E-17 | 2.48E-16 | 283651 | 15 | 45803334 | 45878488 | 1 | pseudogene | high mobility group nucleosomal binding domain 2 pseudogene 46 |
| **PACS2** | ENSG00000179364 | 900.5606984 | -1.002080293 | 0.157060457 | 5.01E-11 | 5.54E-10 | 1019296 | 14 | 105766900 | 105864484 | 1 | protein_coding | phosphofurin acidic cluster sorting protein 2 |
| **EGR3** | ENSG00000179388 | 154.1484847 | -2.582894156 | 0.475346645 | 4.01E-09 | 3.57E-08 | 1960 | 8 | 22545172 | 22550815 | -1 | protein_coding | early growth response 3 |
| **KLHL28** | ENSG00000179454 | 840.7228796 | 1.114223476 | 0.142312831 | 1.16E-15 | 1.84E-14 | 54813 | 14 | 45393522 | 45511525 | -1 | protein_coding | kelch-like family member 28 |
| **C14orf28** | ENSG00000179476 | 87.46836537 | 0.809010046 | 0.388244823 | 0.01259474 | 0.03411652 | 122525 | 14 | 45366498 | 45376460 | 1 | protein_coding | chromosome 14 open reading frame 28 |
| **SLITRK4** | ENSG00000179542 | 502.6757799 | -0.639632043 | 0.158455678 | 3.06E-05 | 1.54E-04 | 139065 | X | 142710596 | 142723596 | -1 | protein_coding | SLIT and NTRK-like family, member 4 |
| **CIITA** | ENSG00000179583 | 3263.730024 | -0.985562511 | 0.483913036 | 0.00921464 | 0.026079136 | 4261 | 16 | 10971055 | 11026079 | 1 | protein_coding | class II, major histocompatibility complex, transactivator |
| **ALOX15B** | ENSG00000179593 | 954.6944062 | 1.555702226 | 0.157421343 | 8.84E-24 | 2.47E-22 | 247 | 17 | 7942335 | 7952452 | 1 | protein_coding | arachidonate 15-lipoxygenase, type B |
| **CDC42EP4** | ENSG00000179604 | 144.8946093 | 0.712554031 | 0.271043845 | 0.003822158 | 0.012011934 | 23580 | 17 | 71279763 | 71308314 | -1 | protein_coding | CDC42 effector protein (Rho GTPase binding) 4 |
| **ZBTB42** | ENSG00000179627 | 151.3184063 | 1.392201675 | 0.311288305 | 1.22E-06 | 7.74E-06 | 1001289 | 14 | 105266933 | 105271049 | 1 | protein_coding | zinc finger and BTB domain containing 42 |
| **LACC1** | ENSG00000179630 | 1726.555141 | 0.876646782 | 0.124760435 | 7.89E-13 | 1.02E-11 | 144811 | 13 | 44453420 | 44468068 | 1 | protein_coding | laccase (multicopper oxidoreductase) domain containing 1 |
| **MAF 1.00** | ENSG00000179632 | 1352.264197 | -0.564732944 | 0.123355007 | 2.86E-06 | 1.72E-05 | 84232 | 8 | 145159402 | 145162514 | 1 | protein_coding | MAF1 homolog (S. cerevisiae) |
| **PCED1B** | ENSG00000179715 | 294.0653951 | -0.740676734 | 0.302383246 | 0.005790747 | 0.017329153 | 91523 | 12 | 47473386 | 47630443 | 1 | protein_coding | PC-esterase domain containing 1B |
| **APOBEC3B** | ENSG00000179750 | 221.4975016 | 4.955299449 | 0.389144596 | 1.41E-38 | 8.1E-37 | 9582 | 22 | 39378352 | 39388809 | 1 | protein_coding | apolipoprotein B mRNA editing enzyme, catalytic polypeptide-like 3B |
| **MYADM** | ENSG00000179820 | 4316.28108 | -0.932423284 | 0.107641645 | 8.62E-19 | 1.72E-17 | 91663 | 19 | 54369477 | 54379691 | 1 | protein_coding | myeloid-associated differentiation marker |
| **TSHZ1** | ENSG00000179981 | 480.6556118 | -0.606353169 | 0.168544469 | 1.82E-04 | 7.88E-04 | 10194 | 18 | 72922710 | 73001905 | 1 | protein_coding | teashirt zinc finger homeobox 1 |
| **TMEM150B** | ENSG00000180061 | 195.2757846 | 1.778853247 | 0.419232434 | 2.28E-06 | 1.4E-05 | 284417 | 19 | 55824169 | 55845416 | -1 | protein_coding | transmembrane protein 150B |
| **EXOC3** | ENSG00000180104 | 1126.919154 | -0.386882573 | 0.128739278 | 0.002226554 | 0.007423303 | 11336 | 5 | 443273 | 472052 | 1 | protein_coding | exocyst complex component 3 |
| **MED14** | ENSG00000180182 | 1143.723277 | 0.391462876 | 0.170972959 | 0.01718165 | 0.04465768 | 9282 | X | 40507558 | 40595110 | -1 | protein_coding | mediator complex subunit 14 |
| **ZNRF2** | ENSG00000180233 | 971.0259353 | 1.99765467 | 0.166699398 | 4.75E-34 | 2.28E-32 | 223082 | 7 | 30323923 | 30452118 | 1 | protein_coding | zinc and ring finger 2 |
| **FGD6** | ENSG00000180263 | 3302.901613 | -0.701844244 | 0.094210896 | 4.11E-14 | 5.87E-13 | 55785 | 12 | 95470525 | 95611258 | -1 | protein_coding | FYVE, RhoGEF and PH domain containing 6 |
| **OAZ2** | ENSG00000180304 | 2829.304518 | 1.049886506 | 0.145982581 | 1.78E-13 | 2.4E-12 | 4947 | 15 | 64979772 | 64995480 | -1 | protein_coding | ornithine decarboxylase antizyme 2 |
| **PNPLA1** | ENSG00000180316 | 15.47423686 | 3.320704138 | 0.976462491 | 3.6E-05 | 1.79E-04 | 285848 | 6 | 36210980 | 36276372 | 1 | protein_coding | patatin-like phospholipase domain containing 1 |
| **CCDC43** | ENSG00000180329 | 764.7655703 | 0.370630516 | 0.151656458 | 0.011736021 | 0.032132142 | 124808 | 17 | 42750437 | 42767147 | -1 | protein_coding | coiled-coil domain containing 43 |
| **HCLS1** | ENSG00000180353 | 6740.20027 | -0.352995473 | 0.076918051 | 4.22E-06 | 2.47E-05 | 3059 | 3 | 121350246 | 121379774 | -1 | protein_coding | hematopoietic cell-specific Lyn substrate 1 |
| **MTURN** | ENSG00000180354 | 323.9398047 | -1.087233464 | 0.320123601 | 1.6E-04 | 7.01E-04 | 222166 | 7 | 30174426 | 30202378 | 1 | protein_coding | maturin, neural progenitor differentiation regulator homolog (Xenopus) |
| **ZNF609** | ENSG00000180357 | 2239.416725 | -0.359201962 | 0.133084104 | 0.005869205 | 0.017529618 | 23060 | 15 | 64752941 | 64978264 | 1 | protein_coding | zinc finger protein 609 |
| **CCDC66** | ENSG00000180376 | 382.0728395 | 0.604277181 | 0.208855863 | 0.002156896 | 0.007227328 | 285331 | 3 | 56591189 | 56655846 | 1 | protein_coding | coiled-coil domain containing 66 |
| **HARBI1** | ENSG00000180423 | 163.2497257 | 0.68055165 | 0.263636033 | 0.004632654 | 0.014274364 | 283254 | 11 | 46624411 | 46639459 | -1 | protein_coding | harbinger transposase derived 1 |
| **HMHA1** | ENSG00000180448 | 2010.32684 | -0.361554844 | 0.156266918 | 0.01677557 | 0.043740911 | 23526 | 19 | 1065922 | 1086627 | 1 | protein_coding | histocompatibility (minor) HA-1 |
| **TSPYL5** | ENSG00000180543 | 520.4928031 | -0.392310015 | 0.156885644 | 0.00983078 | 0.027587919 | 85453 | 8 | 98285717 | 98290176 | -1 | protein_coding | TSPY-like 5 |
| **SSTR2** | ENSG00000180616 | 765.3677402 | 2.413482225 | 0.261737662 | 2.54E-21 | 6.13E-20 | 6752 | 17 | 71161151 | 71167185 | 1 | protein_coding | somatostatin receptor 2 |
| **PCGF5** | ENSG00000180628 | 2606.683652 | 1.444731072 | 0.095253521 | 5.18E-53 | 5.31E-51 | 84333 | 10 | 92979908 | 93044088 | 1 | protein_coding | polycomb group ring finger 5 |
| **PRF1** | ENSG00000180644 | 344.9795469 | 1.302567433 | 0.323842535 | 8.91E-06 | 4.94E-05 | 5551 | 10 | 72357104 | 72362531 | -1 | protein_coding | perforin 1 (pore forming protein) |
| **CTD-2547E1** | ENSG00000180747 | 706.1596101 | 0.555506601 | 0.205633412 | 0.004050594 | 0.012649209 | 1010606 | 16 | 21458004 | 21531765 | -1 | pseudogene | |
| **GPR157** | ENSG00000180758 | 450.517262 | 1.064546395 | 0.247670806 | 4.4E-06 | 2.57E-05 | 80045 | 1 | 9160364 | 9189229 | -1 | protein_coding | G protein-coupled receptor 157 |
| **CHST13** | ENSG00000180767 | 70.55766959 | -1.905127543 | 0.490366231 | 1.09E-05 | 5.99E-05 | 166012 | 3 | 126243126 | 126262134 | 1 | protein_coding | carbohydrate (chondroitin 4) sulfotransferase 13 |
| **SLC36A4** | ENSG00000180773 | 1062.383494 | 1.150325287 | 0.187312382 | 1.94E-10 | 2E-09 | 120103 | 11 | 92877341 | 92931130 | -1 | protein_coding | solute carrier family 36 (proton/amino acid symporter), member 4 |
| **ZFP3** | ENSG00000180787 | 560.2298796 | -0.571744862 | 0.168930127 | 4.26E-04 | 0.001697347 | 124961 | 17 | 4981543 | 4999669 | 1 | protein_coding | ZFP3 zinc finger protein |
| **PPA1** | ENSG00000180817 | 651.9710925 | 2.79697516 | 0.238096488 | 4.56E-33 | 2.1E-31 | 5464 | 10 | 71962586 | 71993667 | -1 | protein_coding | pyrophosphatase (inorganic) 1 |
| **CXCR2** | ENSG00000180871 | 37.69402229 | -1.958234808 | 0.930311121 | 0.00246392 | 0.008120788 | 3579 | 2 | 218990012 | 219001976 | 1 | protein_coding | chemokine (C-X-C motif) receptor 2 |
| **ZNF792** | ENSG00000180884 | 223.9555938 | 0.650940321 | 0.244868434 | 0.003977461 | 0.012456463 | 126375 | 19 | 35447258 | 35454953 | -1 | protein_coding | zinc finger protein 792 |
| **CMTR2** | ENSG00000180917 | 868.0909135 | 0.548193043 | 0.170775659 | 8.69E-04 | 0.003222501 | 55783 | 16 | 71315292 | 71323618 | -1 | protein_coding | cap methyltransferase 2 |
| **PITPNB** | ENSG00000180957 | 1849.209368 | 1.079868495 | 0.114959076 | 1.47E-21 | 3.62E-20 | 23760 | 22 | 28247657 | 28316122 | -1 | protein_coding | phosphatidylinositol transfer protein, beta |
| **MRPL14** | ENSG00000180992 | 380.0222146 | 0.487559531 | 0.19085064 | 0.007155639 | 0.020914062 | 64928 | 6 | 44081194 | 44095194 | -1 | protein_coding | mitochondrial ribosomal protein L14 |
| **GPR137C** | ENSG00000180998 | 22.62576068 | -2.119205739 | 0.837763236 | 8.61E-04 | 0.003196121 | 283554 | 14 | 53019866 | 53104431 | 1 | protein_coding | G protein-coupled receptor 137C |
| **BBS12** | ENSG00000181004 | 172.0524278 | 0.970366986 | 0.287971125 | 2.07E-04 | 8.81E-04 | 166379 | 4 | 123653857 | 123666098 | 1 | protein_coding | Bardet-Biedl syndrome 12 |
| **NQO1** | ENSG00000181019 | 1000.563617 | 1.839504764 | 0.169744553 | 2.84E-28 | 9.99E-27 | 1728 | 16 | 69740899 | 69760854 | -1 | protein_coding | NAD(P)H dehydrogenase, quinone 1 |
| **AEN** | ENSG00000181026 | 407.5253452 | 0.812913402 | 0.17791476 | 1.96E-06 | 1.21E-05 | 64782 | 15 | 89164527 | 89175513 | 1 | protein_coding | apoptosis enhancing nuclease |
| **TRAPPC5** | ENSG00000181029 | 213.4111853 | -1.170860209 | 0.267370644 | 2.68E-06 | 1.62E-05 | 126003 | 19 | 7745729 | 7747744 | 1 | protein_coding | trafficking protein particle complex 5 |
| **SLC26A11** | ENSG00000181045 | 3110.547786 | -1.297968682 | 0.180160072 | 9.93E-14 | 1.38E-12 | 284129 | 17 | 78193498 | 78227299 | 1 | protein_coding | solute carrier family 26 (anion exchanger), member 11 |
| **HIGD1A** | ENSG00000181061 | 2008.909277 | 0.457559287 | 0.13729239 | 6.12E-04 | 0.002340966 | 25994 | 3 | 42798669 | 42846023 | -1 | protein_coding | HIG1 hypoxia inducible domain family, member 1A |
| **DHTKD1** | ENSG00000181192 | 500.7700044 | 0.915374062 | 0.196230577 | 1.01E-06 | 6.48E-06 | 55526 | 10 | 12110971 | 12165224 | 1 | protein_coding | dehydrogenase E1 and transketolase domain containing 1 |
| **ZNF746** | ENSG00000181220 | 731.7085496 | 0.37873033 | 0.146617817 | 0.007788786 | 0.022502081 | 155061 | 7 | 149169885 | 149194908 | -1 | protein_coding | zinc finger protein 746 |
| **FAM211A** | ENSG00000181350 | 101.7787935 | -2.015341016 | 0.469494371 | 1.65E-06 | 1.03E-05 | 388341 | 17 | 16344891 | 16395467 | -1 | protein_coding | family with sequence similarity 211, member A |
| **DDX60L** | ENSG00000181381 | 2453.004234 | 2.396448239 | 0.202206396 | 1.72E-33 | 8.1E-32 | 91351 | 4 | 169277886 | 169458937 | -1 | protein_coding | DEAD (Asp-Glu-Ala-Asp) box polypeptide 60-like |
| **AATK** | ENSG00000181409 | 37.16729638 | -1.573525152 | 0.673621786 | 0.001983036 | 0.006718408 | 9625 | 17 | 79091095 | 79139877 | -1 | protein_coding | apoptosis-associated tyrosine kinase |

| | | | | | | | | | | | | | |
|---|---|---|---|---|---|---|---|---|---|---|---|---|---|
| **ZNF467** | ENSG00000181444 | 100.1234988 | -1.588056325 | 0.391517816 | 6.42E-06 | 3.64E-05 | 168544 | 7 | 149461271 | 149470568 | -1 | protein_coding | zinc finger protein 467 |
| **TMEM45A** | ENSG00000181458 | 183.4113268 | -1.223053959 | 0.32398827 | 3.13E-05 | 1.57E-04 | 55076 | 3 | 100211463 | 100296288 | 1 | protein_coding | transmembrane protein 45A |
| **ZBTB2** | ENSG00000181472 | 538.7538502 | 0.642783309 | 0.174241992 | 1.2E-04 | 5.43E-04 | 57621 | 6 | 151685252 | 151712683 | -1 | protein_coding | zinc finger and BTB domain containing 2 |
| **SETD2** | ENSG00000181555 | 2668.040132 | 0.390776739 | 0.1022913 | 1.1E-04 | 4.99E-04 | 29072 | 3 | 47057919 | 47205457 | -1 | protein_coding | SET domain containing 2 |
| **MEX3D** | ENSG00000181588 | 111.0045985 | -0.936823599 | 0.350159324 | 0.002134155 | 0.007162118 | 399664 | 19 | 1554668 | 1568057 | -1 | protein_coding | mex-3 RNA binding family member D |
| **P2RY13** | ENSG00000181631 | 282.6089523 | -3.926371225 | 0.385295839 | 1.68E-25 | 5.15E-24 | 53829 | 3 | 151044100 | 151047336 | -1 | protein_coding | purinergic receptor P2Y, G-protein coupled, 13 |
| **TNFSF15** | ENSG00000181634 | 1435.215464 | -0.843430523 | 0.202549422 | 1.12E-05 | 6.14E-05 | 9966 | 9 | 117546915 | 117568406 | -1 | protein_coding | tumor necrosis factor (ligand) superfamily, member 15 |
| **PHLDA2** | ENSG00000181649 | 55.87709604 | 1.537193179 | 0.535104622 | 4.68E-04 | 0.001844327 | 7262 | 11 | 2949503 | 2950685 | -1 | protein_coding | pleckstrin homology-like domain, family A, member 2 |
| **HKR1** | ENSG00000181666 | 487.953154 | -0.570705441 | 0.160612536 | 2.3E-04 | 9.72E-04 | 1005073 | 19 | 37803739 | 37860267 | 1 | protein_coding | HKR1, GLI-Kruppel zinc finger family member |
| **PLAG1** | ENSG00000181690 | 144.4206904 | -0.769540163 | 0.369451494 | 0.013508867 | 0.036315922 | 5324 | 8 | 57073463 | 57123883 | -1 | protein_coding | pleiomorphic adenoma gene 1 |
| **AMIGO1** | ENSG00000181754 | 133.9667353 | -2.362964021 | 0.426774071 | 2.78E-09 | 2.52E-08 | 57463 | 1 | 110046797 | 110052360 | -1 | protein_coding | adhesion molecule with Ig-like domain 1 |
| **SIAH2** | ENSG00000181788 | 513.6448206 | 0.691443433 | 0.173564228 | 3.34E-05 | 1.67E-04 | 6478 | 3 | 150458914 | 150481264 | -1 | protein_coding | siah E3 ubiquitin protein ligase 2 |
| **SLC9A9** | ENSG00000181804 | 763.1822114 | 1.790045585 | 0.261270866 | 8.27E-13 | 1.06E-11 | 285195 | 3 | 142984064 | 143567373 | -1 | protein_coding | solute carrier family 9, subfamily A (NHE9, cation proton antiporter 9), member 9 |
| **IBA57** | ENSG00000181873 | 198.9007832 | -0.863414892 | 0.276974456 | 6.18E-04 | 0.002360091 | 200205 | 1 | 228353516 | 228369958 | 1 | protein_coding | IBA57, iron-sulfur cluster assembly homolog (S. cerevisiae) |
| **ZNF101** | ENSG00000181896 | 787.0167107 | 0.351252481 | 0.1421052 | 0.010636353 | 0.029485832 | 94039 | 19 | 19779605 | 19794318 | 1 | protein_coding | zinc finger protein 101 |
| **ADO** | ENSG00000181915 | 2353.392349 | 0.909183086 | 0.150466481 | 5.24E-10 | 5.17E-09 | 84890 | 10 | 64564516 | 64568238 | 1 | protein_coding | 2-aminoethanethiol (cysteamine) dioxygenase |
| **SNRPE** | ENSG00000182004 | 350.6908128 | 1.275714841 | 0.253697346 | 9.51E-08 | 7.07E-07 | 6635 | 1 | 203830731 | 203839678 | 1 | protein_coding | small nuclear ribonucleoprotein polypeptide E |
| **PNMAL1** | ENSG00000182013 | 123.0228076 | -2.402429646 | 0.440705206 | 3.91E-09 | 3.49E-08 | 55228 | 19 | 46969748 | 46974820 | -1 | protein_coding | paraneoplastic Ma antigen family-like 1 |
| **CHST15** | ENSG00000182022 | 8320.271274 | -0.988136793 | 0.14681778 | 4.7E-12 | 5.7E-11 | 51363 | 10 | 125767184 | 125853206 | -1 | protein_coding | carbohydrate (N-acetylgalactosamine 4-sulfate 6-O) sulfotransferase 15 |
| **IDH2** | ENSG00000182054 | 1284.318554 | -0.303868699 | 0.123593686 | 0.012188852 | 0.033185594 | 3418 | 15 | 90626277 | 90645736 | -1 | protein_coding | isocitrate dehydrogenase 2 (NADP+), mitochondrial |
| **TNRC18** | ENSG00000182095 | 2520.904922 | -0.387160671 | 0.118814431 | 8.17E-04 | 0.003049985 | 84629 | 7 | 5346421 | 5465045 | -1 | protein_coding | trinucleotide repeat containing 18 |
| **NOP10** | ENSG00000182117 | 2167.003145 | -0.504329436 | 0.133188175 | 1.06E-04 | 4.84E-04 | 55505 | 15 | 34633917 | 34635378 | -1 | protein_coding | NOP10 ribonucleoprotein |
| **TDRKH** | ENSG00000182134 | 165.1687773 | -1.107820584 | 0.304062644 | 5.86E-05 | 2.79E-04 | 11022 | 1 | 151742583 | 151763892 | -1 | protein_coding | tudor and KH domain containing |
| **ZNF708** | ENSG00000182141 | 271.7754045 | 0.702756179 | 0.241800326 | 0.001675904 | 0.005780374 | 7562 | 19 | 21473962 | 21512227 | -1 | protein_coding | zinc finger protein 708 |
| **IST1** | ENSG00000182149 | 3735.408176 | 0.33265229 | 0.105046673 | 0.00131081 | 0.004635149 | 9798 | 16 | 71879899 | 71962913 | 1 | protein_coding | increased sodium tolerance 1 homolog (yeast) |
| **MRPL41** | ENSG00000182154 | 277.7201503 | -0.793412239 | 0.239050783 | 3.53E-04 | 0.001433828 | 64975 | 9 | 140445651 | 140447007 | 1 | protein_coding | mitochondrial ribosomal protein L41 |
| **CREB3L2** | ENSG00000182158 | 4642.116352 | -0.603323462 | 0.095888844 | 1.31E-10 | 1.38E-09 | 64764 | 7 | 137559725 | 137686813 | -1 | protein_coding | cAMP responsive element binding protein 3-like 2 |
| **P2RY8** | ENSG00000182162 | 281.8747727 | -0.970500351 | 0.280107119 | 1.52E-04 | 6.68E-04 | 286530 | X | 1581465 | 1656000 | -1 | protein_coding | purinergic receptor P2Y, G-protein coupled, 8 |
| **UNC5C** | ENSG00000182168 | 26.697135 | 6.097121239 | 1.363556751 | 5.4E-07 | 3.64E-06 | 8633 | 4 | 96083655 | 96470357 | -1 | protein_coding | unc-5 homolog C (C. elegans) |
| **UBA7** | ENSG00000182179 | 1722.138836 | 0.81802517 | 0.156968219 | 7.53E-08 | 5.69E-07 | 100847C | 3 | 49842640 | 49851379 | -1 | protein_coding | ubiquitin-like modifier activating enzyme 7 |
| **MRPS16** | ENSG00000182180 | 1568.488142 | -0.505537457 | 0.139114581 | 1.77E-04 | 7.7E-04 | 51021 | 10 | 75006510 | 75012451 | -1 | protein_coding | mitochondrial ribosomal protein S16 |
| **ARL6IP4** | ENSG00000182196 | 923.4101934 | -0.563457356 | 0.136930899 | 2.35E-05 | 1.21E-04 | 51329 | 12 | 123464607 | 123467456 | 1 | protein_coding | ADP-ribosylation-like factor 6 interacting protein 4 |
| **SHMT2** | ENSG00000182199 | 1441.98924 | -0.541179997 | 0.199618884 | 0.004112879 | 0.012812296 | 6472 | 12 | 57623110 | 57628718 | 1 | protein_coding | serine hydroxymethyltransferase 2 (mitochondrial) |
| **C8orf33** | ENSG00000182307 | 817.8751223 | -0.659286363 | 0.193618437 | 3.33E-04 | 0.001358883 | 65265 | 8 | 146277764 | 146281416 | 1 | protein_coding | chromosome 8 open reading frame 33 |
| **C1S** | ENSG00000182326 | 572.8520465 | 3.348400693 | 0.326454357 | 4.25E-26 | 1.34E-24 | 716 | 12 | 7096351 | 7178336 | 1 | protein_coding | complement component 1, s subcomponent |
| **CLN8** | ENSG00000182372 | 2809.2323 | -0.557729162 | 0.122587476 | 2.96E-06 | 1.78E-05 | 1019277 | 8 | 1703944 | 1734738 | 1 | protein_coding | ceroid-lipofuscinosis, neuronal 8 (epilepsy, progressive with mental retardation) |
| **PLCXD1** | ENSG00000182378 | 388.5716203 | -1.296501938 | 0.223541931 | 1.28E-09 | 1.22E-08 | 55344 | X | 192989 | 220023 | 1 | protein_coding | phosphatidylinositol-specific phospholipase C, X domain containing 1 |
| **TRAPPC6B** | ENSG00000182400 | 810.2064078 | 0.465204614 | 0.187838983 | 0.009431162 | 0.026627832 | 122553 | 14 | 39617015 | 39639736 | -1 | protein_coding | trafficking protein particle complex 6B |
| **NPLOC4** | ENSG00000182446 | 3274.332774 | 0.516278175 | 0.0920784 | 1.69E-08 | 1.39E-07 | 55666 | 17 | 79523913 | 79615495 | -1 | protein_coding | nuclear protein localization 4 homolog (S. cerevisiae) |
| **NCF1B** | ENSG00000182487 | 14.67879029 | 2.167457937 | 1.02611004 | 0.001838855 | 0.006273041 | 654816 | 7 | 72634611 | 72649979 | 1 | pseudogene | neutrophil cytosolic factor 1B pseudogene |
| **FES** | ENSG00000182511 | 420.0920369 | -0.846415792 | 0.288130843 | 0.001126442 | 0.004063114 | 2242 | 15 | 91426925 | 91439006 | 1 | protein_coding | feline sarcoma oncogene |
| **FAM104B** | ENSG00000182518 | 71.64775691 | -1.15575126 | 0.428755942 | 0.001358876 | 0.004785184 | 90736 | X | 55169535 | 55187743 | -1 | protein_coding | family with sequence similarity 104, member B |
| **LIMK2** | ENSG00000182541 | 1361.784698 | 1.634658373 | 0.221147973 | 1.61E-14 | 2.36E-13 | 3985 | 22 | 31608225 | 31676066 | 1 | protein_coding | LIM domain kinase 2 |
| **RWDD4** | ENSG00000182552 | 569.5021738 | 0.545384494 | 0.161214463 | 4.53E-04 | 0.001790435 | 201965 | 4 | 184560788 | 184580378 | -1 | protein_coding | RWD domain containing 4 |
| **CSF1R** | ENSG00000182578 | 7662.249723 | -1.561000599 | 0.188891212 | 2.1E-17 | 3.8E-16 | 1436 | 5 | 149432854 | 149492935 | -1 | protein_coding | colony stimulating factor 1 receptor |
| **TRAK1** | ENSG00000182606 | 2219.177051 | -0.61265135 | 0.101978277 | 1.33E-09 | 1.26E-08 | 22906 | 3 | 42055294 | 42267381 | 1 | protein_coding | trafficking protein, kinesin binding 1 |
| **SKA2** | ENSG00000182628 | 197.0516467 | 0.587732731 | 0.280175137 | 0.019512694 | 0.04966897 | 348235 | 17 | 57187312 | 57232630 | -1 | protein_coding | spindle and kinetochore associated complex subunit 2 |
| **TTC3** | ENSG00000182670 | 2855.420366 | -1.523259087 | 0.163580623 | 2.12E-21 | 5.15E-20 | 286495 | 21 | 38445526 | 38575413 | 1 | protein_coding | tetratricopeptide repeat domain 3 |
| **TSKU** | ENSG00000182704 | 1059.233523 | -0.533285756 | 0.164135457 | 7.38E-04 | 0.002776989 | 25987 | 11 | 76493295 | 76509198 | 1 | protein_coding | tsukushi, small leucine rich proteoglycan |
| **ANXA2** | ENSG00000182718 | 62303.93681 | -0.785744775 | 0.160919328 | 4.75E-07 | 3.23E-06 | 302 | 15 | 60639333 | 60695082 | -1 | protein_coding | annexin A2 |
| **RPS17L** | ENSG00000182774 | 8.77517846 | -0.266067525 | 0.719133872 | 0.005808816 | 0.017379824 | 1005055 | 15 | 83205504 | 83209315 | -1 | protein_coding | ribosomal protein S17-like |
| **TMEM198B** | ENSG00000182796 | 483.0570074 | -0.940587467 | 0.177445827 | 3.36E-08 | 2.66E-07 | 440104 | 12 | 56223529 | 56230030 | 1 | pseudogene | transmembrane protein 198B, pseudogene |
| **C16orf72** | ENSG00000182831 | 1842.485555 | 0.421465071 | 0.119756738 | 3.28E-04 | 0.001341102 | 29035 | 16 | 9185505 | 9215497 | 1 | protein_coding | chromosome 16 open reading frame 72 |
| **ALG12** | ENSG00000182858 | 410.1069347 | -0.849986607 | 0.276449891 | 7.14E-04 | 0.002697015 | 79087 | 22 | 50293877 | 50312106 | -1 | protein_coding | ALG12, alpha-1,6-mannosyltransferase |
| **LCK** | ENSG00000182866 | 643.693961 | -1.031342986 | 0.236570463 | 3.49E-06 | 2.07E-05 | 3932 | 1 | 32716840 | 32751766 | 1 | protein_coding | lymphocyte-specific protein tyrosine kinase |
| **GLUD2** | ENSG00000182890 | 30.07086082 | -2.04156573 | 1.024699362 | 0.002587602 | 0.008479057 | 2747 | X | 120181462 | 120183794 | 1 | protein_coding | glutamate dehydrogenase 2 |
| **RPL35A** | ENSG00000182899 | 3030.517114 | -0.432516872 | 0.137743909 | 0.001254269 | 0.004462058 | 6165 | 3 | 197676858 | 197683481 | 1 | protein_coding | ribosomal protein L35a |
| **C21orf90** | ENSG00000182912 | 21.00436447 | -2.327033974 | 0.949709899 | 0.001000017 | 0.003648426 | NA | 21 | 45937099 | 45945835 | 1 | protein_coding | chromosome 21 open reading frame 90 |
| **EWSR1** | ENSG00000182944 | 3987.644938 | 0.303156021 | 0.116509625 | 0.008575563 | 0.024446993 | 2130 | 22 | 29663998 | 29696515 | 1 | protein_coding | EWS RNA-binding protein 1 |
| **HMGN4** | ENSG00000182952 | 972.7083948 | 0.796895916 | 0.175787489 | 2.47E-06 | 1.5E-05 | 10473 | 6 | 26538633 | 26546482 | 1 | protein_coding | high mobility group nucleosomal binding domain 4 |
| **SPATA13** | ENSG00000182957 | 6499.289516 | 0.385492428 | 0.120876787 | 0.001157183 | 0.004155364 | 221178 | 13 | 24553944 | 24881212 | 1 | protein_coding | spermatogenesis associated 13 |
| **CNOT10** | ENSG00000182973 | 502.0040134 | 0.805536342 | 0.187661413 | 6.99E-06 | 3.94E-05 | 25904 | 3 | 32726637 | 32815367 | 1 | protein_coding | CCR4-NOT transcription complex, subunit 10 |
| **LSMD1** | ENSG00000183011 | 215.860037 | -0.761151838 | 0.24834413 | 8.88E-04 | 0.003283177 | 84316 | 17 | 7760003 | 7788556 | -1 | protein_coding | LSM domain containing 1 |
| **C19orf59** | ENSG00000183019 | 21.82367561 | -2.922418811 | 1.06238458 | 3.4E-04 | 0.001384578 | 199675 | 19 | 7741514 | 7744712 | 1 | protein_coding | chromosome 19 open reading frame 59 |
| **AP2A2** | ENSG00000183020 | 1272.375624 | -0.405672405 | 0.15343964 | 0.006190873 | 0.01840045 | 1019274 | 11 | 924894 | 1012239 | 1 | protein_coding | adaptor-related protein complex 2, alpha 2 subunit |
| **ABAT** | ENSG00000183044 | 439.1476845 | -1.676140377 | 0.20182897 | 1.22E-17 | 2.25E-16 | 18 | 16 | 8768422 | 8878432 | 1 | protein_coding | 4-aminobutyrate aminotransferase |
| **CAMK1D** | ENSG00000183049 | 71.52167513 | -0.987390328 | 0.451550318 | 0.006716438 | 0.019770287 | 57118 | 10 | 12391481 | 12877545 | 1 | protein_coding | calcium/calmodulin-dependent protein kinase ID |
| **RGPD6** | ENSG00000183054 | 1434.931125 | 0.317087481 | 0.116303017 | 0.005525768 | 0.0166339 | 729540 | 2 | 111271389 | 111334762 | -1 | protein_coding | RANBP2-like and GRIP domain containing 6 |
| **AFMID** | ENSG00000183077 | 196.5083944 | -0.926962534 | 0.258974381 | 1.09E-04 | 4.96E-04 | 125061 | 17 | 76183398 | 76203782 | 1 | protein_coding | arylformamidase |
| **ARHGEF37** | ENSG00000183111 | 49.23072164 | -1.666441247 | 0.69572029 | 0.001394642 | 0.00490097 | 389337 | 5 | 148931510 | 149014531 | 1 | protein_coding | Rho guanine nucleotide exchange factor (GEF) 37 |
| **FANCF** | ENSG00000183161 | 190.6065954 | 0.841035882 | 0.244697043 | 2.15E-04 | 9.12E-04 | 2188 | 11 | 22644079 | 22647387 | -1 | protein_coding | Fanconi anemia, complementation group F |
| **RUVBL2** | ENSG00000183207 | 359.1848389 | -0.587447628 | 0.224379547 | 0.005133988 | 0.015636257 | 10856 | 19 | 49496705 | 49519252 | 1 | protein_coding | RuvB-like AAA ATPase 2 |
| **DDX41** | ENSG00000183258 | 1548.755461 | -1.067823796 | 0.115111658 | 4.7E-21 | 1.12E-19 | 51428 | 5 | 176938578 | 176944470 | -1 | protein_coding | DEAD (Asp-Glu-Ala-Asp) box polypeptide 41 |
| **CECR6** | ENSG00000183307 | 446.562465 | -2.086316987 | 0.195369231 | 1.28E-27 | 4.34E-26 | 27439 | 22 | 17597189 | 17602257 | -1 | protein_coding | cat eye syndrome chromosome region, candidate 6 |
| **GBP6** | ENSG00000183347 | 16.02095609 | 4.254615306 | 1.20481612 | 1.4E-05 | 7.52E-05 | 163351 | 1 | 89829617 | 89852020 | 1 | protein_coding | guanylate binding protein family, member 6 |
| **KIAA2026** | ENSG00000183354 | 1317.740823 | 0.485928101 | 0.132100432 | 1.52E-04 | 6.71E-04 | 158358 | 9 | 5881596 | 6007901 | -1 | protein_coding | KIAA2026 |
| **FHL3** | ENSG00000183386 | 417.3004543 | -0.579551644 | 0.17760857 | 6.71E-04 | 0.002547373 | 2275 | 1 | 38462442 | 38471278 | -1 | protein_coding | four and a half LIM domains 3 |
| **GPR132** | ENSG00000183484 | 306.3187299 | 1.543277292 | 0.318702617 | 1.6E-07 | 1.16E-06 | 29933 | 14 | 105515728 | 105531782 | -1 | protein_coding | G protein-coupled receptor 132 |
| **MX2** | ENSG00000183486 | 6331.837078 | 4.06777388 | 0.115966939 | 4.57E-271 | 8.73E-268 | 4600 | 21 | 42733870 | 42781317 | 1 | protein_coding | myxovirus (influenza virus) resistance 2 (mouse) |
| **EP400** | ENSG00000183495 | 1574.9195 | -0.626785942 | 0.138214516 | 3.25E-06 | 1.94E-05 | 57634 | 12 | 132434508 | 132565005 | 1 | protein_coding | E1A binding protein p400 |
| **MEX3B** | ENSG00000183496 | 40.79113514 | -2.624717588 | 0.703581748 | 1.73E-05 | 9.1E-05 | 84206 | 15 | 82334119 | 82338482 | -1 | protein_coding | mex-3 RNA binding family member B |
| **SETD3** | ENSG00000183576 | 3180.503575 | -0.834052846 | 0.108971582 | 7.99E-15 | 1.2E-13 | 84193 | 14 | 99864083 | 99947216 | -1 | protein_coding | SET domain containing 3 |
| **TNFAIP8L3** | ENSG00000183578 | 455.5294038 | -1.191032758 | 0.208728869 | 2.54E-09 | 2.32E-08 | 388121 | 15 | 51348795 | 51397473 | -1 | protein_coding | tumor necrosis factor, alpha-induced protein 8-like 3 |
| **TANGO2** | ENSG00000183597 | 864.0879653 | -0.869279699 | 0.22097353 | 2.99E-05 | 1.51E-04 | 128989 | 22 | 20004537 | 20053449 | 1 | protein_coding | transport and golgi organization 2 homolog (Drosophila) |
| **RP11-347C12** | ENSG00000183604 | 463.7594002 | 0.516072478 | 0.208641676 | 0.008593506 | 0.024493575 | 1019299 | 16 | 30234230 | 30346695 | -1 | pseudogene | |
| **SFXN4** | ENSG00000183605 | 479.762171 | -0.453325645 | 0.163820247 | 0.004048444 | 0.012647663 | 119559 | 10 | 120900279 | 120925179 | -1 | protein_coding | sideroflexin 4 |
| **DGCR6** | ENSG00000183628 | 42.99989591 | -1.326864857 | 0.55558224 | 0.002395484 | 0.007925999 | 8214 | 22 | 18893541 | 18901751 | 1 | protein_coding | DiGeorge syndrome critical region gene 6 |
| **NDUFB1** | ENSG00000183648 | 989.6718027 | -0.503172853 | 0.147839413 | 4.51E-04 | 0.001783487 | 4707 | 14 | 92582466 | 92588261 | -1 | protein_coding | NADH dehydrogenase (ubiquinone) 1 beta subcomplex, 1, 7kDa |
| **EFHC2** | ENSG00000183690 | 10.01954543 | 2.819423409 | 1.382406363 | 0.001718331 | 0.00590008 | 80258 | X | 44007128 | 44202918 | -1 | protein_coding | EF-hand domain (C-terminal) containing 2 |

| | | | | | | | | | | | | | |
|---|---|---|---|---|---|---|---|---|---|---|---|---|---|
| **UPP1** | ENSG00000183696 | 1550.909631 | -0.385425557 | 0.167986376 | 0.016606107 | 0.043385569 | 7378 | 7 | 48128225 | 48148330 | 1 | protein_coding | uridine phosphorylase 1 |
| **LHFP** | ENSG00000183722 | 85.58086122 | 1.24453822 | 0.352900138 | 7.81E-05 | 3.64E-04 | 10186 | 13 | 39917029 | 40177665 | -1 | protein_coding | lipoma HMGIC fusion partner |
| **CMTM4** | ENSG00000183723 | 169.3631774 | -2.245182749 | 0.343922218 | 5.86E-12 | 7.05E-11 | 146223 | 16 | 66648653 | 66730610 | -1 | protein_coding | CKLF-like MARVEL transmembrane domain containing 4 |
| **TMEM50A** | ENSG00000183726 | 4146.454378 | 1.493171735 | 0.107427697 | 9.9E-45 | 7.46E-43 | 23585 | 1 | 25664408 | 25688852 | 1 | protein_coding | transmembrane protein 50A |
| **ASCL2** | ENSG00000183734 | 80.94165049 | 2.475330686 | 0.444630075 | 1.91E-09 | 1.78E-08 | 430 | 11 | 2289725 | 2292182 | -1 | protein_coding | achaete-scute family bHLH transcription factor 2 |
| **TBK1** | ENSG00000183735 | 1694.335372 | 1.784197085 | 0.11464625 | 1.83E-55 | 2.04E-53 | 29110 | 12 | 64845660 | 64895888 | 1 | protein_coding | TANK-binding kinase 1 |
| **CBX6** | ENSG00000183741 | 3842.75226 | 0.731773719 | 0.157230396 | 1.54E-06 | 9.63E-06 | 23466 | 22 | 39257455 | 39268319 | -1 | protein_coding | chromobox homolog 6 |
| **MACC1** | ENSG00000183742 | 523.1226124 | -0.727521158 | 0.295619489 | 0.005466454 | 0.016484573 | 346389 | 7 | 20174278 | 20257027 | -1 | protein_coding | metastasis associated in colon cancer 1 |
| **ZNF703** | ENSG00000183779 | 232.9170433 | -1.024481618 | 0.267804513 | 3.35E-05 | 1.67E-04 | 80139 | 8 | 37553269 | 37557537 | 1 | protein_coding | zinc finger protein 703 |
| **SLC35F3** | ENSG00000183780 | 51.44499662 | -1.584420187 | 0.539416615 | 3.81E-04 | 0.00153388 | 148641 | 1 | 234040679 | 234460262 | 1 | protein_coding | solute carrier family 35, member F3 |
| **TUBA8** | ENSG00000183785 | 64.43492581 | -1.026428009 | 0.433532783 | 0.004126746 | 0.012852874 | 51807 | 22 | 18593097 | 18629321 | 1 | protein_coding | tubulin, alpha 8 |
| **NUDT14** | ENSG00000183828 | 107.108845 | -0.822470021 | 0.346147293 | 0.005993891 | 0.017867103 | 256281 | 14 | 105639275 | 105647660 | -1 | protein_coding | nudix (nucleoside diphosphate linked moiety X)-type motif 14 |
| **IQGAP3** | ENSG00000183856 | 466.4307376 | -1.45210725 | 0.358734218 | 7.46E-06 | 4.19E-05 | 128239 | 1 | 156495197 | 156542396 | -1 | protein_coding | IQ motif containing GTPase activating protein 3 |
| **SETD8** | ENSG00000183955 | 937.8065721 | -0.450372405 | 0.151915285 | 0.002179074 | 0.007285669 | 387893 | 12 | 123868320 | 123893905 | 1 | protein_coding | SET domain containing (lysine methyltransferase) 8 |
| **PTP4A2** | ENSG00000184007 | 5240.530934 | 0.244648465 | 0.103523891 | 0.015545602 | 0.041019239 | 8073 | 1 | 32372022 | 32410457 | -1 | protein_coding | protein tyrosine phosphatase type IVA, member 2 |
| **DENND5A** | ENSG00000184014 | 3291.705606 | 1.03327192 | 0.091827125 | 3.01E-30 | 1.18E-28 | 23258 | 11 | 9160372 | 9286937 | -1 | protein_coding | DENN/MADD domain containing 5A |
| **UQCR10** | ENSG00000184076 | 1003.605859 | -0.592556674 | 0.131038729 | 3.6E-06 | 2.13E-05 | 29796 | 22 | 30163358 | 30166402 | 1 | protein_coding | ubiquinol-cytochrome c reductase, complex III subunit X |
| **EIF3C** | ENSG00000184110 | 5537.629486 | -0.858009957 | 0.14662449 | 1.61E-09 | 1.51E-08 | 8663 | 16 | 28699879 | 28747051 | 1 | protein_coding | eukaryotic translation initiation factor 3, subunit C |
| **SCFD2** | ENSG00000184178 | 439.4599845 | -0.807536562 | 0.238512726 | 2.72E-04 | 0.001129547 | 152579 | 4 | 53739149 | 54232242 | -1 | protein_coding | sec1 family domain containing 2 |
| **UBE2F** | ENSG00000184182 | 1692.180104 | 0.816275252 | 0.105317461 | 3.76E-15 | 5.74E-14 | 140739 | 2 | 238875469 | 238951236 | 1 | protein_coding | ubiquitin-conjugating enzyme E2F (putative) |
| **TSPYL2** | ENSG00000184205 | 267.5910711 | -0.566472667 | 0.252947593 | 0.01409754 | 0.037640234 | 64061 | X | 53111549 | 53117722 | 1 | protein_coding | TSPY-like 2 |
| **PGP** | ENSG00000184207 | 526.4205209 | -0.512252489 | 0.199307558 | 0.006537228 | 0.019313445 | 283871 | 16 | 2261998 | 2264808 | -1 | protein_coding | phosphoglycolate phosphatase |
| **TM2D3** | ENSG00000184277 | 535.279663 | 0.653986776 | 0.174222582 | 9E-05 | 4.15E-04 | 80213 | 15 | 102161847 | 102192594 | -1 | protein_coding | TM2 domain containing 3 |
| **MROH7** | ENSG00000184313 | 53.65077619 | -2.182889083 | 0.561185537 | 8.61E-06 | 4.79E-05 | 374977 | 1 | 55107463 | 55175939 | 1 | protein_coding | maestro heat-like repeat family member 7 |
| **SLIT3** | ENSG00000184347 | 185.5331801 | -3.225037042 | 0.332713716 | 2.54E-23 | 6.96E-22 | 6586 | 5 | 168088745 | 168728133 | -1 | protein_coding | slit homolog 3 (Drosophila) |
| **CSF1** | ENSG00000184371 | 9309.07192 | -1.152653698 | 0.154759576 | 2.29E-14 | 3.33E-13 | 1435 | 1 | 110452864 | 110473614 | 1 | protein_coding | colony stimulating factor 1 (macrophage) |
| **PLA2G6** | ENSG00000184381 | 86.64614734 | -0.781637989 | 0.381371493 | 0.014343219 | 0.038176145 | 8398 | 22 | 38507502 | 38601697 | -1 | protein_coding | phospholipase A2, group VI (cytosolic, calcium-independent) |
| **MAML2** | ENSG00000184384 | 2710.133444 | 0.612468215 | 0.096826243 | 1.7E-10 | 1.77E-09 | 84441 | 11 | 95709762 | 96076344 | -1 | protein_coding | mastermind-like 2 (Drosophila) |
| **TOP1MT** | ENSG00000184428 | 341.619224 | -0.86019769 | 0.211935349 | 1.73E-05 | 9.09E-05 | 116447 | 8 | 144386554 | 144442149 | -1 | protein_coding | topoisomerase (DNA) I, mitochondrial |
| **COPB2** | ENSG00000184432 | 6133.718281 | -0.245682864 | 0.097038445 | 0.009696569 | 0.027296495 | 9276 | 3 | 139074442 | 139108574 | -1 | protein_coding | coatomer protein complex, subunit beta 2 (beta prime) |
| **KNTC1** | ENSG00000184445 | 696.6347085 | 0.691984105 | 0.190876621 | 1.44E-04 | 6.39E-04 | 9735 | 12 | 123011793 | 123110943 | 1 | protein_coding | kinetochore associated 1 |
| **FOXO4** | ENSG00000184481 | 487.8401029 | -0.64822357 | 0.178176883 | 1.51E-04 | 6.65E-04 | 4303 | X | 70316047 | 70323385 | 1 | protein_coding | forkhead box O4 |
| **PROS1** | ENSG00000184500 | 1069.459531 | -0.437954419 | 0.171488126 | 0.007718787 | 0.022332614 | 5627 | 3 | 93591881 | 93692910 | -1 | protein_coding | protein S (alpha) |
| **HDDC3** | ENSG00000184508 | 151.8246464 | -0.766581059 | 0.282614136 | 0.00265947 | 0.008692189 | 374659 | 15 | 91474148 | 91475799 | -1 | protein_coding | HD domain containing 3 |
| **XPOT** | ENSG00000184575 | 4269.468235 | -0.551853996 | 0.130334454 | 1.43E-05 | 7.68E-05 | 11260 | 12 | 64798130 | 64844907 | 1 | protein_coding | exportin, tRNA |
| **PDE4B** | ENSG00000184588 | 1128.959061 | 3.381544921 | 0.283834318 | 5.2E-34 | 2.49E-32 | 5142 | 1 | 66258197 | 66840259 | 1 | protein_coding | phosphodiesterase 4B, cAMP-specific |
| **ZNF93** | ENSG00000184635 | 295.4647317 | -0.508814348 | 0.211907835 | 0.010065042 | 0.028152276 | 81931 | 19 | 20011722 | 20046384 | 1 | protein_coding | zinc finger protein 93 |
| **9-Sep** | ENSG00000184640 | 10017.99839 | -1.202221704 | 0.102740329 | 2.88E-32 | 1.26E-30 | 10801 | 17 | 75276651 | 75496678 | 1 | protein_coding | septin 9 |
| **HIST2H2BE** | ENSG00000184678 | 714.3792903 | 1.730009859 | 0.180780757 | 1.17E-22 | 3.09E-21 | 8349 | 1 | 149856010 | 149858232 | -1 | protein_coding | histone cluster 2, H2be |
| **APOBR** | ENSG00000184730 | 3538.468553 | -1.030288868 | 0.133174051 | 3.26E-15 | 4.99E-14 | 55911 | 16 | 28505970 | 28510291 | 1 | protein_coding | apolipoprotein B receptor |
| **ZBTB7C** | ENSG00000184828 | 470.3327803 | -1.849699568 | 0.240473278 | 1.17E-15 | 1.87E-14 | 201501 | 18 | 45553044 | 45937123 | -1 | protein_coding | zinc finger and BTB domain containing 7C |
| **APOO** | ENSG00000184831 | 669.0144412 | 2.382908652 | 0.20864951 | 2.53E-31 | 1.05E-29 | 79135 | X | 23851470 | 23926057 | -1 | protein_coding | apolipoprotein O |
| **TMED9** | ENSG00000184840 | 2739.158266 | -0.45471614 | 0.109686787 | 2.49E-05 | 1.28E-04 | 54732 | 5 | 177019159 | 177023125 | 1 | protein_coding | transmembrane emp24 protein transport domain containing 9 |
| **H1FX** | ENSG00000184897 | 1111.071411 | 0.619695441 | 0.122234421 | 2.28E-07 | 1.61E-06 | 8971 | 3 | 129033615 | 129035120 | -1 | protein_coding | H1 histone family, member X |
| **RBM43** | ENSG00000184898 | 1022.832209 | 1.851994997 | 0.200445605 | 2.3E-21 | 5.58E-20 | 375287 | 2 | 152104454 | 152118393 | -1 | protein_coding | RNA binding motif protein 43 |
| **USP18** | ENSG00000184979 | 1322.848567 | 5.625625369 | 0.168378299 | 6E-246 | 6.11E-243 | 11274 | 22 | 18632666 | 18660164 | 1 | protein_coding | ubiquitin specific peptidase 18 |
| **BRI3BP** | ENSG00000184992 | 1041.179804 | -1.614194816 | 0.145896462 | 2.55E-29 | 9.52E-28 | 140707 | 12 | 125478246 | 125515777 | 1 | protein_coding | BRI3 binding protein |
| **CA13** | ENSG00000185015 | 34.01437105 | 1.12186854 | 0.568021234 | 0.008250845 | 0.02365366 | 1005072 | 8 | 86132816 | 86196302 | 1 | protein_coding | carbonic anhydrase XIII |
| **CIB1** | ENSG00000185043 | 931.938211 | -0.526547778 | 0.138034987 | 9.04E-05 | 4.17E-04 | 10519 | 15 | 90773207 | 90777279 | -1 | protein_coding | calcium and integrin binding 1 (calmyrin) |
| **FLRT2** | ENSG00000185070 | 1838.541335 | -1.597072077 | 0.137038184 | 4.54E-32 | 1.95E-30 | 1005067 | 14 | 85996488 | 86095034 | 1 | protein_coding | fibronectin leucine rich transmembrane protein 2 |
| **RPS27L** | ENSG00000185088 | 2225.899022 | 0.491473151 | 0.117085775 | 1.94E-05 | 1.01E-04 | 51065 | 15 | 63418071 | 63450220 | -1 | protein_coding | ribosomal protein S27-like |
| **ADSSL1** | ENSG00000185100 | 124.2738377 | -3.652405196 | 0.513695295 | 8.02E-14 | 1.12E-12 | 122622 | 14 | 105190523 | 105213662 | 1 | protein_coding | adenylosuccinate synthase like 1 |
| **FAM43A** | ENSG00000185112 | 54.08238148 | -1.658401969 | 0.553024795 | 3.06E-04 | 0.001258949 | 131583 | 3 | 194406622 | 194409762 | 1 | protein_coding | family with sequence similarity 43, member A |
| **C6orf120** | ENSG00000185127 | 809.5305836 | 0.571147297 | 0.135966553 | 1.68E-05 | 8.9E-05 | 387263 | 6 | 170102233 | 170106401 | 1 | protein_coding | chromosome 6 open reading frame 120 |
| **PURA** | ENSG00000185129 | 453.1252291 | 0.788758075 | 0.178233071 | 4.05E-06 | 2.38E-05 | 5813 | 5 | 139487362 | 139496321 | 1 | protein_coding | purine-rich element binding protein A |
| **NOMO2** | ENSG00000185164 | 5038.62279 | 0.25279387 | 0.098163125 | 0.008797821 | 0.025029245 | 1019300 | 16 | 18511182 | 18573533 | -1 | protein_coding | NODAL modulator 2 |
| **SIGIRR** | ENSG00000185187 | 149.0202471 | -1.262901333 | 0.339564773 | 3.72E-05 | 1.84E-04 | 59307 | 11 | 405716 | 417455 | -1 | protein_coding | single immunoglobulin and toll-interleukin 1 receptor (TIR) domain |
| **IFITM2** | ENSG00000185201 | 579.457749 | 2.204926572 | 0.230544339 | 8.65E-23 | 2.3E-21 | 10581 | 11 | 307631 | 315272 | 1 | protein_coding | interferon induced transmembrane protein 2 |
| **TNFAIP2** | ENSG00000185215 | 8887.74739 | 2.12074047 | 0.126114591 | 1.1E-64 | 1.72E-62 | 7127 | 14 | 103589779 | 103603776 | 1 | protein_coding | tumor necrosis factor, alpha-induced protein 2 |
| **ZNF445** | ENSG00000185219 | 565.613238 | -0.514616947 | 0.167550119 | 0.001407644 | 0.004944387 | 353274 | 3 | 44481262 | 44519162 | -1 | protein_coding | zinc finger protein 445 |
| **GP1BA** | ENSG00000185245 | 344.5328708 | 2.29212449 | 0.272402866 | 3.56E-18 | 6.84E-17 | 2811 | 17 | 4835592 | 4838325 | 1 | protein_coding | glycoprotein Ib (platelet), alpha polypeptide |
| **IL3RA** | ENSG00000185291 | 39.9265493 | 1.646991014 | 0.601371258 | 6.08E-04 | 0.002329498 | 3563 | X | 1455509 | 1501578 | 1 | protein_coding | interleukin 3 receptor, alpha (low affinity) |
| **CDK10** | ENSG00000185324 | 341.9767528 | -0.622083461 | 0.201282884 | 0.001087952 | 0.003941022 | 8558 | 16 | 89747145 | 89762772 | 1 | protein_coding | cyclin-dependent kinase 10 |
| **SOCS1** | ENSG00000185338 | 127.1597278 | 2.904489132 | 0.394517749 | 1.17E-14 | 1.73E-13 | 8651 | 16 | 11348262 | 11350036 | -1 | protein_coding | suppressor of cytokine signaling 1 |
| **ATP6V0A2** | ENSG00000185344 | 1712.019542 | 0.379382883 | 0.109569966 | 4.13E-04 | 0.001651162 | 23545 | 12 | 124196865 | 124246302 | 1 | protein_coding | ATPase, H+ transporting, lysosomal V0 subunit a2 |
| **MAPK11** | ENSG00000185386 | 35.06592855 | 1.088220785 | 0.575816193 | 0.010412159 | 0.028943074 | 5600 | 22 | 50702142 | 50709196 | -1 | protein_coding | mitogen-activated protein kinase 11 |
| **SP140L** | ENSG00000185404 | 1461.501619 | 1.001967441 | 0.173965537 | 2.59E-09 | 2.36E-08 | 93349 | 2 | 231191899 | 231268447 | 1 | protein_coding | SP140 nuclear body protein-like |
| **METTL7A** | ENSG00000185432 | 2928.735621 | -0.702630575 | 0.159040168 | 4.98E-06 | 2.88E-05 | 25840 | 12 | 51317255 | 51326300 | 1 | protein_coding | methyltransferase like 7A |
| **IFNLR1** | ENSG00000185436 | 1315.32889 | 0.714474999 | 0.112208064 | 9.69E-11 | 1.04E-09 | 163702 | 1 | 24480647 | 24514449 | -1 | protein_coding | interferon, lambda receptor 1 |
| **TMEM179B** | ENSG00000185475 | 648.4910902 | 0.467556461 | 0.174534331 | 0.005159427 | 0.015698093 | 1024654 | 11 | 62554887 | 62557877 | 1 | protein_coding | transmembrane protein 179B |
| **GPRIN3** | ENSG00000185477 | 7036.828931 | -0.235933329 | 0.101324719 | 0.017777001 | 0.045949572 | 285513 | 4 | 90157537 | 90229161 | -1 | protein_coding | GPRIN family member 3 |
| **STAC3** | ENSG00000185482 | 875.6839675 | 1.495570463 | 0.16801084 | 9.62E-20 | 2.09E-18 | 246329 | 12 | 57637236 | 57644976 | -1 | protein_coding | SH3 and cysteine rich domain 3 |
| **ROR1** | ENSG00000185483 | 37.27130422 | -2.648597388 | 0.906785761 | 2.4E-04 | 0.001007677 | 4919 | 1 | 64239693 | 64647181 | 1 | protein_coding | receptor tyrosine kinase-like orphan receptor 1 |
| **C17orf70** | ENSG00000185504 | 310.5250198 | -0.820341196 | 0.22171659 | 8.55E-05 | 3.96E-04 | 80233 | 17 | 79506911 | 79520987 | -1 | protein_coding | chromosome 17 open reading frame 70 |
| **IRF7** | ENSG00000185507 | 426.3975016 | 3.819564884 | 0.241324035 | 1.16E-57 | 1.42E-55 | 3665 | 11 | 612553 | 615999 | -1 | protein_coding | interferon regulatory factor 7 |
| **AHNAK2** | ENSG00000185567 | 269.217779 | -1.945109179 | 0.418622866 | 3.11E-07 | 2.16E-06 | 113146 | 14 | 105403581 | 105444694 | -1 | protein_coding | AHNAK nucleoprotein 2 |
| **OLFML2A** | ENSG00000185585 | 33.40014984 | -1.245146327 | 0.707074533 | 0.010154226 | 0.028375762 | 169611 | 9 | 127539437 | 127577164 | 1 | protein_coding | olfactomedin-like 2A |
| **SP1** | ENSG00000185591 | 3911.631169 | 0.225612114 | 0.096366988 | 0.018325678 | 0.047142121 | 6667 | 12 | 53773960 | 53810230 | 1 | protein_coding | Sp1 transcription factor |
| **MRPL40** | ENSG00000185608 | 633.8743813 | 0.876935003 | 0.180578208 | 4.31E-07 | 2.94E-06 | 64976 | 22 | 19419425 | 19423598 | 1 | protein_coding | mitochondrial ribosomal protein L40 |
| **P4HB** | ENSG00000185624 | 16148.11708 | -0.368830486 | 0.112052208 | 8.05E-04 | 0.003010607 | 5034 | 17 | 79801035 | 79818570 | -1 | protein_coding | prolyl 4-hydroxylase, beta polypeptide |
| **KRT79** | ENSG00000185640 | 73.31680744 | -2.311309603 | 0.663113648 | 3.43E-05 | 1.71E-04 | 338785 | 12 | 53215194 | 53228079 | -1 | protein_coding | keratin 79 |
| **ZFP36L1** | ENSG00000185650 | 4357.948031 | -0.684756999 | 0.108414962 | 1.48E-10 | 1.55E-09 | 677 | 14 | 69254377 | 69263190 | -1 | protein_coding | ZFP36 ring finger protein-like 1 |
| **UBE2L3** | ENSG00000185651 | 3139.000756 | 0.541731665 | 0.118219274 | 3.07E-06 | 1.84E-05 | 7332 | 22 | 21903736 | 21978323 | 1 | protein_coding | ubiquitin-conjugating enzyme E2L 3 |
| **SNAI3** | ENSG00000185669 | 193.0035756 | -2.836018609 | 0.349695436 | 4.59E-17 | 8.08E-16 | 333929 | 16 | 88744090 | 88752901 | -1 | protein_coding | snail family zinc finger 3 |
| **ZBTB3** | ENSG00000185670 | 151.6611609 | -0.609261611 | 0.286803925 | 0.017863306 | 0.046115693 | 79842 | 11 | 62515791 | 62521660 | -1 | protein_coding | zinc finger and BTB domain containing 3 |
| **ANKFY1** | ENSG00000185722 | 3800.672909 | 0.603276476 | 0.098113367 | 6.14E-10 | 6.02E-09 | 51479 | 17 | 4067201 | 4167274 | -1 | protein_coding | ankyrin repeat and FYVE domain containing 1 |
| **YTHDF3** | ENSG00000185728 | 202.3474554 | 0.765115275 | 0.328013931 | 0.007285661 | 0.021216988 | 253943 | 8 | 64081112 | 64125346 | 1 | protein_coding | YTH domain family, member 3 |
| **IFIT1** | ENSG00000185745 | 6411.037815 | 6.713500914 | 0.131816542 | 0 | 0 | 3434 | 10 | 91152303 | 91163745 | 1 | protein_coding | interferon-induced protein with tetratricopeptide repeats 1 |
| **CXorf38** | ENSG00000185753 | 610.4464048 | 0.459034405 | 0.141566145 | 8.65E-04 | 0.003209324 | 159013 | X | 40488285 | 40506819 | -1 | protein_coding | chromosome X open reading frame 38 |

| | | | | | | | | | | | | | |
|---|---|---|---|---|---|---|---|---|---|---|---|---|---|
| **PIGP** | ENSG00000185808 | 340.9413381 | 0.4824226 | 0.21814445 | 0.017981593 | 0.046405385 | 51227 | 21 | 38431470 | 38445470 | -1 | protein_coding | phosphatidylinositol glycan anchor biosynthesis, class P |
| **IKZF1** | ENSG00000185811 | 3359.99172 | -0.848322231 | 0.101077666 | 1.81E-17 | 3.29E-16 | 10320 | 7 | 50343720 | 50472799 | 1 | protein_coding | IKAROS family zinc finger 1 (Ikaros) |
| **BCAP31** | ENSG00000185825 | 6506.563579 | -0.638166218 | 0.109705535 | 3.29E-09 | 2.95E-08 | 10134 | X | 152965947 | 152990152 | -1 | protein_coding | B-cell receptor-associated protein 31 |
| **TRIM69** | ENSG00000185880 | 1920.109019 | 0.417640926 | 0.122174501 | 4.69E-04 | 0.001847517 | 140691 | 15 | 45021186 | 45060027 | 1 | protein_coding | tripartite motif containing 69 |
| **ATP6V0C** | ENSG00000185883 | 7940.851402 | -0.368182205 | 0.081155249 | 9.29E-06 | 5.13E-05 | 527 | 16 | 2563871 | 2570219 | 1 | protein_coding | ATPase, H+ transporting, lysosomal 16kDa, V0 subunit c |
| **IFITM1** | ENSG00000185885 | 1224.811869 | 4.336996827 | 0.242051176 | 2.71E-73 | 5.53E-71 | 8519 | 11 | 313506 | 315272 | 1 | protein_coding | interferon induced transmembrane protein 1 |
| **LAMP1** | ENSG00000185896 | 16419.8167 | -0.338725121 | 0.107281386 | 0.001321996 | 0.00466715 | 3916 | 13 | 113951556 | 113977987 | 1 | protein_coding | lysosomal-associated membrane protein 1 |
| **C16orf54** | ENSG00000185905 | 1011.199546 | -0.385187323 | 0.134881872 | 0.00316948 | 0.010199742 | 1019298 | 16 | 29753784 | 29757327 | -1 | protein_coding | chromosome 16 open reading frame 54 |
| **PTCH1** | ENSG00000185920 | 83.67174006 | -3.085939083 | 0.784419509 | 3.72E-06 | 2.2E-05 | 5727 | 9 | 98205262 | 98279339 | -1 | protein_coding | patched 1 |
| **RNPC3** | ENSG00000185946 | 661.3198067 | 0.36501226 | 0.154489197 | 0.014819899 | 0.039302675 | 55599 | 1 | 104068313 | 104097861 | 1 | protein_coding | RNA-binding region (RNP1, RRM) containing 3 |
| **ZNF267** | ENSG00000185947 | 2267.984523 | 0.881686335 | 0.137833417 | 5.7E-11 | 6.28E-10 | 10308 | 16 | 31885079 | 31928668 | 1 | protein_coding | zinc finger protein 267 |
| **BICD2** | ENSG00000185963 | 2209.085434 | -0.714073984 | 0.128553889 | 1.65E-08 | 1.36E-07 | 23299 | 9 | 95473645 | 95527094 | -1 | protein_coding | bicaudal D homolog 2 (Drosophila) |
| **RASA3** | ENSG00000185989 | 903.0858471 | -1.575102155 | 0.165813842 | 3.08E-22 | 7.82E-21 | 22821 | 13 | 114747194 | 114898086 | -1 | protein_coding | RAS p21 protein activator 3 |
| **DLEU7** | ENSG00000186047 | 106.8051875 | 0.721414154 | 0.31895986 | 0.010019792 | 0.02805138 | 220107 | 13 | 51285144 | 51418075 | -1 | protein_coding | deleted in lymphocytic leukemia, 7 |
| **AIDA** | ENSG00000186063 | 3033.060971 | 1.32877258 | 0.088947838 | 7.72E-51 | 7.06E-49 | 64853 | 1 | 222841355 | 222886552 | -1 | protein_coding | axin interactor, dorsalization associated |
| **PIP5K1C** | ENSG00000186111 | 2927.347898 | -1.221802556 | 0.127710002 | 2.27E-22 | 5.84E-21 | 23396 | 19 | 3630181 | 3700477 | -1 | protein_coding | phosphatidylinositol-4-phosphate 5-kinase, type I, gamma |
| **ZBTB6** | ENSG00000186130 | 494.6407714 | 0.640464017 | 0.179301199 | 1.89E-04 | 8.14E-04 | 10773 | 9 | 125670335 | 125675609 | -1 | protein_coding | zinc finger and BTB domain containing 6 |
| **POLR3C** | ENSG00000186141 | 1012.561309 | 0.548029454 | 0.117779179 | 2.16E-06 | 1.32E-05 | 10623 | 1 | 145592605 | 145611025 | -1 | protein_coding | polymerase (RNA) III (DNA directed) polypeptide C (62kD) |
| **BCL9L** | ENSG00000186174 | 1028.464812 | 0.59052111 | 0.207766107 | 0.002551013 | 0.008375323 | 283149 | 11 | 118764584 | 118796317 | -1 | protein_coding | B-cell CLL/lymphoma 9-like |
| **ZNRF1** | ENSG00000186187 | 254.8845142 | 0.48155475 | 0.219267888 | 0.018991404 | 0.048552296 | 84937 | 16 | 75032928 | 75144892 | 1 | protein_coding | zinc and ring finger 1, E3 ubiquitin protein ligase |
| **FFAR4** | ENSG00000186188 | 130.6425229 | -6.628593809 | 1.008704555 | 1.07E-10 | 1.14E-09 | 338557 | 10 | 95326422 | 95364237 | 1 | protein_coding | free fatty acid receptor 4 |
| **EDARADD** | ENSG00000186197 | 64.64648655 | 2.099417332 | 0.461700529 | 4.58E-07 | 3.11E-06 | 128178 | 1 | 236511562 | 236648214 | 1 | protein_coding | EDAR-associated death domain |
| **1-Mar** | ENSG00000186205 | 215.6883877 | -2.016415491 | 0.359073545 | 1.87E-09 | 1.74E-08 | 64757 | 1 | 220960101 | 220987735 | 1 | protein_coding | mitochondrial amidoxime reducing component 1 |
| **AC016757.3** | ENSG00000186235 | 33.48558175 | -1.2429254 | 0.633347348 | 0.006928935 | 0.020321418 | 151174 | 2 | 239133193 | 239140318 | -1 | protein_coding | Protein LOC151174 |
| **MKL2** | ENSG00000186260 | 952.0936442 | -0.670468823 | 0.157935954 | 1.06E-05 | 5.85E-05 | 57496 | 16 | 14165178 | 14360630 | 1 | protein_coding | MKL/myocardin-like 2 |
| **GPAT2** | ENSG00000186281 | 172.1743612 | -1.415026573 | 0.543191557 | 0.001145945 | 0.004122759 | 150763 | 2 | 96687694 | 96705199 | -1 | protein_coding | glycerol-3-phosphate acyltransferase 2, mitochondrial |
| **PPP1CC** | ENSG00000186298 | 3229.669457 | 0.424213219 | 0.099396927 | 1.62E-05 | 8.6E-05 | 5501 | 12 | 111157485 | 111180744 | -1 | protein_coding | protein phosphatase 1, catalytic subunit, gamma isozyme |
| **ZNF555** | ENSG00000186300 | 191.2562085 | -0.866582461 | 0.299323733 | 0.001305029 | 0.004621122 | 148254 | 19 | 2841433 | 2860482 | 1 | protein_coding | zinc finger protein 555 |
| **BACE1** | ENSG00000186318 | 2095.898687 | -1.413188185 | 0.122572452 | 1.36E-31 | 5.76E-30 | 23621 | 11 | 117156402 | 117186975 | -1 | protein_coding | beta-site APP-cleaving enzyme 1 |
| **RXRA** | ENSG00000186350 | 4394.521817 | -1.442019862 | 0.110163819 | 6.28E-40 | 3.75E-38 | 6256 | 9 | 137208944 | 137332431 | 1 | protein_coding | retinoid X receptor, alpha |
| **ANKRD37** | ENSG00000186352 | 175.1431633 | -1.185773841 | 0.273989832 | 3.07E-06 | 1.84E-05 | 353322 | 4 | 186317175 | 186321782 | 1 | protein_coding | ankyrin repeat domain 37 |
| **ZNF75D** | ENSG00000186376 | 445.0270505 | 0.515376235 | 0.190529325 | 0.004423673 | 0.013696715 | 7626 | X | 134382867 | 134478012 | -1 | protein_coding | zinc finger protein 75D |
| **CYP4X1** | ENSG00000186377 | 252.1716113 | -1.127524569 | 0.392208396 | 8.16E-04 | 0.003045736 | 260293 | 1 | 47427036 | 47516423 | 1 | protein_coding | cytochrome P450, family 4, subfamily X, polypeptide 1 |
| **CD300E** | ENSG00000186407 | 79.36395177 | 2.166396247 | 0.476259995 | 4.29E-07 | 2.93E-06 | 342510 | 17 | 72606026 | 72619897 | -1 | protein_coding | CD300e molecule |
| **NKRF** | ENSG00000186416 | 415.1331535 | 0.385998524 | 0.170715877 | 0.018930923 | 0.048421973 | 55922 | X | 118722300 | 118739858 | -1 | protein_coding | NFKB repressing factor |
| **GLDN** | ENSG00000186417 | 264.3362505 | -1.035739071 | 0.351577105 | 7.72E-04 | 0.002896851 | 342035 | 15 | 51633826 | 51700210 | 1 | protein_coding | gliomedin |
| **KPNA4** | ENSG00000186432 | 6500.484162 | 0.277597565 | 0.113991249 | 0.013393489 | 0.036036472 | 3840 | 3 | 160212783 | 160283376 | -1 | protein_coding | karyopherin alpha 4 (importin alpha 3) |
| **SPATA12** | ENSG00000186451 | 101.0068485 | -2.69052349 | 0.46990963 | 8.96E-10 | 8.66E-09 | 353324 | 3 | 57094469 | 57109460 | 1 | protein_coding | spermatogenesis associated 12 |
| **RPS23** | ENSG00000186468 | 4967.079115 | -0.449499878 | 0.124853245 | 2.38E-04 | 0.001000333 | 6228 | 5 | 81569177 | 81574396 | -1 | protein_coding | ribosomal protein S23 |
| **GNG2** | ENSG00000186469 | 447.4323556 | -0.541531747 | 0.223292204 | 0.008820941 | 0.025081013 | 54331 | 14 | 52292913 | 52446060 | 1 | protein_coding | guanine nucleotide binding protein (G protein), gamma 2 |
| **BTN3A2** | ENSG00000186470 | 3393.828714 | 0.584915527 | 0.134774746 | 8.66E-06 | 4.81E-05 | 11118 | 6 | 26365387 | 26378546 | 1 | protein_coding | butyrophilin, subfamily 3, member A2 |
| **INSIG1** | ENSG00000186480 | 4478.40747 | 0.260598056 | 0.106274101 | 0.015427216 | 0.040763164 | 3638 | 7 | 155089486 | 155101945 | 1 | protein_coding | insulin induced gene 1 |
| **TMEM222** | ENSG00000186501 | 363.5441708 | -0.48193344 | 0.19255605 | 0.00802046 | 0.023071088 | 84065 | 1 | 27648651 | 27662891 | 1 | protein_coding | transmembrane protein 222 |
| **NF2** | ENSG00000186575 | 2347.202561 | -0.52883637 | 0.10655695 | 4.34E-07 | 2.96E-06 | 4771 | 22 | 29999545 | 30094587 | 1 | protein_coding | neurofibromin 2 (merlin) |
| **ARAP1** | ENSG00000186635 | 11172.16686 | -0.395753313 | 0.105365943 | 1.3E-04 | 5.82E-04 | 116985 | 11 | 72396114 | 72504644 | -1 | protein_coding | ArfGAP with RhoGAP domain, ankyrin repeat and PH domain 1 |
| **PRR5** | ENSG00000186654 | 324.0589132 | -0.994906726 | 0.220806044 | 1.97E-06 | 1.22E-05 | 55615 | 22 | 45064593 | 45133561 | 1 | protein_coding | proline rich 5 (renal) |
| **ZFP91** | ENSG00000186660 | 3517.1654 | 0.294219783 | 0.092933145 | 0.00146553 | 0.005130025 | 80829 | 11 | 58346584 | 58388515 | 1 | protein_coding | ZFP91 zinc finger protein |
| **DTX2P1** | ENSG00000186704 | 40.09316541 | -0.920517215 | 0.529279085 | 0.019405127 | 0.049444574 | NA | 7 | 76607934 | 76673092 | 1 | pseudogene | deltex homolog 2 pseudogene 1 |
| **BCR** | ENSG00000186716 | 1055.787227 | -0.765228957 | 0.151168932 | 1.82E-07 | 1.31E-06 | 613 | 22 | 23521891 | 23660224 | 1 | protein_coding | breakpoint cluster region |
| **VSIG10L** | ENSG00000186806 | 106.764367 | 1.809011211 | 0.35332116 | 3.17E-08 | 2.53E-07 | 147645 | 19 | 51834790 | 51845378 | -1 | protein_coding | V-set and immunoglobulin domain containing 10 like |
| **TPCN1** | ENSG00000186815 | 3520.151379 | -1.637374665 | 0.166950574 | 1.19E-23 | 3.31E-22 | 53373 | 12 | 113658855 | 113736390 | 1 | protein_coding | two pore segment channel 1 |
| **LILRB4** | ENSG00000186818 | 3346.341794 | 0.572640287 | 0.170871889 | 4.78E-04 | 0.00188129 | 11006 | 19 | 55155340 | 55181810 | 1 | protein_coding | leukocyte immunoglobulin-like receptor, subfamily B (with TM and ITIM domains), member 4 |
| **TNFRSF4** | ENSG00000186827 | 133.4460772 | 1.19472469 | 0.460441057 | 0.001616113 | 0.005596884 | 7293 | 1 | 1146706 | 1149518 | -1 | protein_coding | tumor necrosis factor receptor superfamily, member 4 |
| **HEXIM1** | ENSG00000186834 | 1264.016657 | 0.587429818 | 0.118840895 | 4.98E-07 | 3.37E-06 | 10614 | 17 | 43224684 | 43229468 | 1 | protein_coding | hexamethylene bis-acetamide inducible 1 |
| **RTN4RL2** | ENSG00000186907 | 117.7619866 | -1.238059183 | 0.353318337 | 8.56E-05 | 3.97E-04 | 349667 | 11 | 57228022 | 57245007 | 1 | protein_coding | reticulon 4 receptor-like 2 |
| **ZNF395** | ENSG00000186918 | 1998.30953 | -1.970852271 | 0.195945602 | 9.5E-25 | 2.8E-23 | 55893 | 8 | 28203102 | 28260218 | -1 | protein_coding | zinc finger protein 395 |
| **PPARA** | ENSG00000186951 | 723.6780193 | -0.807354319 | 0.289639863 | 0.001964425 | 0.006661264 | 5465 | 22 | 46546424 | 46639653 | 1 | protein_coding | peroxisome proliferator-activated receptor alpha |
| **GPR141** | ENSG00000187037 | 488.6305843 | 1.050209616 | 0.205756525 | 8.51E-08 | 6.37E-07 | 353345 | 7 | 37723399 | 37873390 | 1 | protein_coding | G protein-coupled receptor 141 |
| **RPS19BP1** | ENSG00000187051 | 1068.723643 | -0.486975659 | 0.132170922 | 1.66E-04 | 7.26E-04 | 91582 | 22 | 39925098 | 39928860 | -1 | protein_coding | ribosomal protein S19 binding protein 1 |
| **PLCD1** | ENSG00000187091 | 300.0800517 | -0.758943866 | 0.246233905 | 8.07E-04 | 0.003013928 | 5333 | 3 | 38048987 | 38071253 | -1 | protein_coding | phospholipase C, delta 1 |
| **ENTPD5** | ENSG00000187097 | 830.571414 | 1.616090672 | 0.156151483 | 6E-26 | 1.88E-24 | 957 | 14 | 74424713 | 74486102 | -1 | protein_coding | ectonucleoside triphosphate diphosphohydrolase 5 |
| **MITF** | ENSG00000187098 | 3152.700773 | -1.047256766 | 0.143712033 | 8.2E-14 | 1.14E-12 | 4286 | 3 | 69788586 | 70017488 | 1 | protein_coding | microphthalmia-associated transcription factor |
| **CMC1** | ENSG00000187118 | 365.9591186 | 0.485001173 | 0.217167857 | 0.017166715 | 0.044639238 | 152100 | 3 | 28283075 | 28366633 | 1 | protein_coding | C-x(9)-C motif containing 1 |
| **KIAA1598** | ENSG00000187164 | 4687.627919 | -1.010439555 | 0.174592641 | 2.04E-09 | 1.88E-08 | 57698 | 10 | 118643742 | 118886097 | -1 | protein_coding | KIAA1598 |
| **SESTD1** | ENSG00000187231 | 1092.225746 | 1.021595599 | 0.162001042 | 8.08E-11 | 8.72E-10 | 91404 | 2 | 179966419 | 180129517 | -1 | protein_coding | SEC14 and spectrin domains 1 |
| **DYNC2H1** | ENSG00000187240 | 66.91463436 | -1.88338925 | 0.862852314 | 0.00309072 | 0.009967394 | 79659 | 11 | 102980160 | 103350591 | 1 | protein_coding | dynein, cytoplasmic 2, heavy chain 1 |
| **RSBN1L** | ENSG00000187257 | 801.8704395 | 0.831959801 | 0.160992696 | 9.43E-08 | 7.01E-07 | 222194 | 7 | 77325760 | 77412339 | 1 | protein_coding | round spermatid basic protein 1-like |
| **WDR86** | ENSG00000187260 | 22.15212011 | -2.276287022 | 0.867440561 | 6.3E-04 | 0.002401884 | 349136 | 7 | 151072995 | 151107813 | -1 | protein_coding | WD repeat domain 86 |
| **EPOR** | ENSG00000187266 | 77.12087087 | -1.549524796 | 0.412004164 | 2.23E-05 | 1.15E-04 | 2057 | 19 | 11488236 | 11495018 | -1 | protein_coding | erythropoietin receptor |
| **CHP1** | ENSG00000187446 | 3368.672964 | -0.714697471 | 0.099658642 | 3.56E-13 | 4.71E-12 | 11261 | 15 | 41523037 | 41574043 | 1 | protein_coding | calcineurin-like EF-hand protein 1 |
| **FPR3** | ENSG00000187474 | 1824.8147 | -0.626954931 | 0.169968993 | 1.22E-04 | 5.48E-04 | 2359 | 19 | 52298416 | 52329442 | 1 | protein_coding | formyl peptide receptor 3 |
| **SIRT7** | ENSG00000187531 | 690.2123976 | 0.47823687 | 0.149799609 | 9.81E-04 | 0.003586392 | 51547 | 17 | 79869815 | 79879199 | -1 | protein_coding | sirtuin 7 |
| **TLR5** | ENSG00000187554 | 84.71760146 | -3.433584273 | 0.613678316 | 1.14E-09 | 1.09E-08 | 7100 | 1 | 223282748 | 223316624 | -1 | protein_coding | toll-like receptor 5 |
| **PLEKHN1** | ENSG00000187583 | 25.6152363 | 2.17579388 | 0.730677067 | 1.94E-04 | 8.34E-04 | 84069 | 1 | 901877 | 911245 | 1 | protein_coding | pleckstrin homology domain containing, family N member 1 |
| **VMAC** | ENSG00000187650 | 264.6326748 | -1.013344263 | 0.268113827 | 3.62E-05 | 1.79E-04 | 400673 | 19 | 5904869 | 5910864 | 1 | protein_coding | vimentin-type intermediate filament associated coiled-coil protein |
| **B3GALTL** | ENSG00000187676 | 439.7568459 | -1.123118525 | 0.186663961 | 4.11E-10 | 4.11E-09 | 145173 | 13 | 31774073 | 31906413 | 1 | protein_coding | beta 1,3-galactosyltransferase-like |
| **TCEA1** | ENSG00000187735 | 2723.849857 | 1.024934744 | 0.115613345 | 2.12E-19 | 4.46E-18 | 6917 | 8 | 54879112 | 54935089 | -1 | protein_coding | transcription elongation factor A (SII), 1 |
| **FANCA** | ENSG00000187741 | 394.1400798 | 1.144082523 | 0.274645351 | 6.67E-06 | 3.77E-05 | 2175 | 16 | 89803957 | 89883065 | -1 | protein_coding | Fanconi anemia, complementation group A |
| **SECISBP2** | ENSG00000187742 | 875.752624 | 0.651781357 | 0.137755329 | 1.1E-06 | 7.05E-06 | 79048 | 9 | 91933421 | 91974557 | 1 | protein_coding | SECIS binding protein 2 |
| **SEMA4D** | ENSG00000187764 | 2036.121671 | 1.573648572 | 0.259176832 | 1.83E-10 | 1.89E-09 | 10507 | 9 | 91975702 | 92113045 | -1 | protein_coding | sema domain, immunoglobulin domain (Ig), transmembrane domain (TM) and short cytoplasmic domair |
| **FANCM** | ENSG00000187790 | 166.9873312 | 0.627913949 | 0.279357542 | 0.01215802 | 0.03311549 | 57697 | 14 | 45605143 | 45670093 | 1 | protein_coding | Fanconi anemia, complementation group M |
| **SOWAHD** | ENSG00000187808 | 145.3148477 | -1.64047519 | 0.334416848 | 1.13E-07 | 8.32E-07 | 347454 | X | 118892576 | 118894164 | 1 | protein_coding | sosondowah ankyrin repeat domain family member D |
| **EIF4EBP1** | ENSG00000187840 | 385.5605961 | -1.939260195 | 0.192953432 | 8.77E-25 | 2.61E-23 | 1978 | 8 | 37887859 | 37917883 | 1 | protein_coding | eukaryotic translation initiation factor 4E binding protein 1 |
| **CYHR1** | ENSG00000187954 | 451.8442842 | -1.145978535 | 0.2019618 | 3.21E-09 | 2.89E-08 | 50626 | 8 | 145674965 | 145691060 | -1 | protein_coding | cysteine/histidine-rich 1 |
| **TPRG1** | ENSG00000188001 | 513.8777662 | -0.961260085 | 0.201757309 | 5.8E-07 | 3.89E-06 | 285386 | 3 | 188665003 | 189043093 | 1 | protein_coding | tumor protein p63 regulated 1 |
| **MORN2** | ENSG00000188010 | 6.321189077 | -4.160549647 | 3.160625821 | 0.002687909 | 0.008775755 | 729967 | 2 | 39103103 | 39156213 | 1 | protein_coding | MORN repeat containing 2 |
| **UBQLN2** | ENSG00000188021 | 1304.644631 | 0.73216094 | 0.126210091 | 3.21E-09 | 2.89E-08 | 29978 | X | 56590026 | 56593443 | 1 | protein_coding | ubiquilin 2 |
| **RILPL1** | ENSG00000188026 | 271.6715213 | -0.56172715 | 0.256029054 | 0.016491236 | 0.043142 | 353116 | 12 | 123955925 | 124018265 | -1 | protein_coding | Rab interacting lysosomal protein-like 1 |
| **NWD1** | ENSG00000188039 | 69.02876496 | -1.852525855 | 0.492355294 | 1.78E-05 | 9.38E-05 | 284434 | 19 | 16830787 | 16928774 | 1 | protein_coding | NACHT and WD repeat domain containing 1 |

| | | | | | | | | | | | | | |
|---|---|---|---|---|---|---|---|---|---|---|---|---|---|
| **RAB42** | ENSG00000188060 | 739.084262 | -2.007710512 | 0.308224068 | 5.53E-12 | 6.66E-11 | 115273 | 1 | 28918712 | 28921955 | 1 | protein_coding | RAB42, member RAS oncogene family |
| **GPR89B** | ENSG00000188092 | 218.34039 | 0.503422824 | 0.22124306 | 0.014795115 | 0.039269492 | 653519 | 1 | 147400506 | 147465753 | 1 | protein_coding | G protein-coupled receptor 89B |
| **AGRN** | ENSG00000188157 | 1718.491832 | 1.261012038 | 0.173821049 | 9.24E-14 | 1.29E-12 | 375790 | 1 | 955503 | 991496 | 1 | protein_coding | agrin |
| **NHS** | ENSG00000188158 | 266.6159142 | 1.531809968 | 0.376582356 | 5.93E-06 | 3.38E-05 | 4810 | X | 17393543 | 17754114 | 1 | protein_coding | Nance-Horan syndrome (congenital cataracts and dental anomalies) |
| **LAMTOR4** | ENSG00000188186 | 507.2986188 | -0.835335405 | 0.176284731 | 8.19E-07 | 5.35E-06 | 389541 | 7 | 99746530 | 99753567 | 1 | protein_coding | late endosomal/lysosomal adaptor, MAPK and MTOR activator 4 |
| **NCR3LG1** | ENSG00000188211 | 216.4139681 | -0.569516917 | 0.24604694 | 0.012072449 | 0.032911744 | 374383 | 11 | 17373273 | 17398888 | 1 | protein_coding | natural killer cell cytotoxicity receptor 3 ligand 1 |
| **DCUN1D3** | ENSG00000188215 | 807.3902857 | 0.967926506 | 0.166203505 | 1.56E-09 | 1.47E-08 | 123879 | 16 | 20866247 | 20911706 | -1 | protein_coding | DCN1, defective in cullin neddylation 1, domain containing 3 |
| **RUFY4** | ENSG00000188282 | 39.97657033 | 1.289827267 | 1.020976042 | 0.015738673 | 0.041464253 | 285180 | 2 | 218899683 | 218955304 | 1 | protein_coding | RUN and FYVE domain containing 4 |
| **HES4** | ENSG00000188290 | 26.95898454 | 5.770547529 | 1.090585214 | 7.55E-09 | 6.51E-08 | 57801 | 1 | 934342 | 935552 | -1 | protein_coding | hes family bHLH transcription factor 4 |
| **CENPP** | ENSG00000188312 | 266.9414669 | -0.528653373 | 0.221964724 | 0.010802252 | 0.029875282 | 401541 | 9 | 95087766 | 95382815 | 1 | protein_coding | centromere protein P |
| **PLSCR1** | ENSG00000188313 | 2486.114216 | 3.223185959 | 0.232206676 | 5.24E-45 | 4E-43 | 5359 | 3 | 146232967 | 146262651 | -1 | protein_coding | phospholipid scramblase 1 |
| **FOCAD** | ENSG00000188352 | 1519.665099 | -0.683407574 | 0.129895259 | 7.18E-08 | 5.44E-07 | 54914 | 9 | 20658308 | 20995954 | 1 | protein_coding | focadhesin |
| **ZP3** | ENSG00000188372 | 70.72616734 | 2.865082774 | 0.47387947 | 1.06E-10 | 1.14E-09 | 7784 | 7 | 76026835 | 76071388 | 1 | protein_coding | zona pellucida glycoprotein 3 (sperm receptor) |
| **SELL** | ENSG00000188404 | 851.3018901 | 0.891212193 | 0.18844757 | 8.38E-07 | 5.46E-06 | 6402 | 1 | 169659808 | 169680839 | -1 | protein_coding | selectin L |
| **IER5L** | ENSG00000188483 | 270.8386376 | -2.738181746 | 0.309424825 | 6.17E-20 | 1.36E-18 | 389792 | 9 | 131937835 | 131940540 | -1 | protein_coding | immediate early response 5-like |
| **C15orf52** | ENSG00000188549 | 691.5349242 | -1.27794797 | 0.233860424 | 8.68E-09 | 7.44E-08 | 388115 | 15 | 40623653 | 40633168 | -1 | protein_coding | chromosome 15 open reading frame 52 |
| **RALGAPA2** | ENSG00000188559 | 1149.206947 | -0.630387011 | 0.151840899 | 1.68E-05 | 8.88E-05 | 57186 | 20 | 20370196 | 20693131 | -1 | protein_coding | Ral GTPase activating protein, alpha subunit 2 (catalytic) |
| **CLN3** | ENSG00000188603 | 1350.547332 | -0.706342047 | 0.127629395 | 1.67E-08 | 1.38E-07 | 1201 | 16 | 28477983 | 28506896 | -1 | protein_coding | ceroid-lipofuscinosis, neuronal 3 |
| **FAM72B** | ENSG00000188610 | 84.38069986 | 1.773935333 | 0.489602455 | 2.91E-05 | 1.47E-04 | 653820 | 1 | 120837756 | 120855681 | 1 | protein_coding | family with sequence similarity 72, member B |
| **SUMO2** | ENSG00000188612 | 3269.81661 | 0.510220436 | 0.130905481 | 6.81E-05 | 3.21E-04 | 1019290 | 17 | 73163408 | 73179078 | -1 | protein_coding | small ubiquitin-like modifier 2 |
| **NANOS1** | ENSG00000188613 | 123.4288977 | 0.787456551 | 0.407826317 | 0.018409341 | 0.047309561 | 340719 | 10 | 120789228 | 120793854 | 1 | protein_coding | nanos homolog 1 (Drosophila) |
| **LDOC1L** | ENSG00000188636 | 965.0116826 | -0.475911212 | 0.142805641 | 5.92E-04 | 0.002270029 | 84247 | 22 | 44888452 | 44894178 | -1 | protein_coding | leucine zipper, down-regulated in cancer 1-like |
| **DPYD** | ENSG00000188641 | 5480.283048 | -0.764424632 | 0.082164367 | 9.24E-21 | 2.16E-19 | 1806 | 1 | 97543299 | 98386605 | -1 | protein_coding | dihydropyrimidine dehydrogenase |
| **PTAR1** | ENSG00000188647 | 1042.487087 | 0.380717558 | 0.128261557 | 0.00244994 | 0.008081683 | 375743 | 9 | 72324438 | 72374875 | -1 | protein_coding | protein prenyltransferase alpha subunit repeat containing 1 |
| **IDO2** | ENSG00000188676 | 11.76697633 | 4.007216241 | 1.49247539 | 3.26E-04 | 0.001335645 | 169355 | 8 | 39792133 | 39873910 | 1 | protein_coding | indoleamine 2,3-dioxygenase 2 |
| **PARVB** | ENSG00000188677 | 1327.964064 | -0.985159755 | 0.147653366 | 7.26E-12 | 8.66E-11 | 29780 | 22 | 44395091 | 44568829 | 1 | protein_coding | parvin, beta |
| **UROS** | ENSG00000188690 | 1030.78715 | -0.575268364 | 0.175264705 | 6.11E-04 | 0.00233582 | 7390 | 10 | 127477146 | 127511817 | -1 | protein_coding | uroporphyrinogen III synthase |
| **ZDHHC9** | ENSG00000188706 | 696.7325096 | 1.617660648 | 0.16124603 | 1.42E-24 | 4.15E-23 | 51114 | X | 128937264 | 128977885 | -1 | protein_coding | zinc finger, DHHC-type containing 9 |
| **ZBED6CL** | ENSG00000188707 | 436.5645133 | 1.968648837 | 0.22291241 | 1.06E-19 | 2.29E-18 | 113763 | 7 | 150026938 | 150029808 | 1 | protein_coding | ZBED6 C-terminal like |
| **SMIM15** | ENSG00000188725 | 1593.326337 | -0.311794246 | 0.131344712 | 0.014545635 | 0.03866163 | 643155 | 5 | 60453536 | 60458301 | -1 | protein_coding | small integral membrane protein 15 |
| **TMEM120B** | ENSG00000188735 | 244.2508385 | -1.054063816 | 0.232865996 | 1.65E-06 | 1.03E-05 | 144404 | 12 | 122150658 | 122220907 | 1 | protein_coding | transmembrane protein 120B |
| **RBM34** | ENSG00000188739 | 553.6217911 | 1.52217081 | 0.202777796 | 8.14E-15 | 1.22E-13 | 23029 | 1 | 235294498 | 235324772 | -1 | protein_coding | RNA binding motif protein 34 |
| **MTF1** | ENSG00000188786 | 2708.567466 | 1.512171555 | 0.133321408 | 1.16E-30 | 4.63E-29 | 4520 | 1 | 38275239 | 38325292 | -1 | protein_coding | metal-regulatory transcription factor 1 |
| **NHLRC3** | ENSG00000188811 | 3145.971952 | 1.40642963 | 0.115362314 | 6.62E-35 | 3.3E-33 | 387921 | 13 | 39612443 | 39624246 | 1 | protein_coding | NHL repeat containing 3 |
| **FAM26F** | ENSG00000188820 | 207.904112 | 3.19404745 | 0.28519137 | 2.92E-30 | 1.15E-28 | 441168 | 6 | 116782533 | 116784934 | 1 | protein_coding | family with sequence similarity 26, member F |
| **RPL14** | ENSG00000188846 | 3267.937884 | -0.324840952 | 0.123027932 | 0.006461374 | 0.01911521 | 9045 | 3 | 40498783 | 40503861 | 1 | protein_coding | ribosomal protein L14 |
| **FBF1** | ENSG00000188878 | 122.3856743 | -0.657795226 | 0.316010239 | 0.017459322 | 0.04531004 | 85302 | 17 | 73905655 | 73937221 | -1 | protein_coding | Fas (TNFRSF6) binding factor 1 |
| **MSL1** | ENSG00000188895 | 2397.63038 | 0.3210976 | 0.105399784 | 0.002006044 | 0.006784328 | 339287 | 17 | 38278551 | 38293042 | 1 | protein_coding | male-specific lethal 1 homolog (Drosophila) |
| **LRRK2** | ENSG00000188906 | 3803.965084 | 0.845799186 | 0.136988024 | 2.62E-10 | 2.67E-09 | 120892 | 12 | 40590546 | 40763087 | 1 | protein_coding | leucine-rich repeat kinase 2 |
| **PTPLAD2** | ENSG00000188921 | 836.2207161 | 0.851018569 | 0.132858304 | 5.48E-11 | 6.05E-10 | 401494 | 9 | 20995306 | 21031635 | -1 | protein_coding | protein tyrosine phosphatase-like A domain containing 2 |
| **KIR2DL4** | ENSG00000189013 | 4.991290508 | 0.227220148 | 0.694968841 | 0.00940866 | 0.026579028 | 3805 | 19 | 55236002 | 55325972 | 1 | protein_coding | killer cell immunoglobulin-like receptor, two domains, long cytoplasmic tail, 4 |
| **RNFT1** | ENSG00000189050 | 830.6532068 | 0.800013901 | 0.175263878 | 2.17E-06 | 1.33E-05 | 51136 | 17 | 58029601 | 58042122 | -1 | protein_coding | ring finger protein, transmembrane 1 |
| **FAM111B** | ENSG00000189057 | 85.85802958 | 1.905779019 | 0.457694282 | 2.64E-06 | 1.6E-05 | 374393 | 11 | 58874658 | 58894883 | 1 | protein_coding | family with sequence similarity 111, member B |
| **APOD** | ENSG00000189058 | 139.4656057 | 1.254685177 | 0.506125014 | 0.002043524 | 0.006889735 | 347 | 3 | 195295573 | 195311076 | -1 | protein_coding | apolipoprotein D |
| **H1F0** | ENSG00000189060 | 3514.587657 | 1.838147905 | 0.138825991 | 5.96E-41 | 3.79E-39 | 3005 | 22 | 38201114 | 38203442 | 1 | protein_coding | H1 histone family, member 0 |
| **LITAF** | ENSG00000189067 | 20147.67509 | 0.244408729 | 0.076125196 | 0.001707855 | 0.005875991 | 9516 | 16 | 11641853 | 11730237 | -1 | protein_coding | lipopolysaccharide-induced TNF factor |
| **BLOC1S3** | ENSG00000189114 | 442.3561275 | 0.957702681 | 0.164927504 | 2.06E-09 | 1.9E-08 | 388552 | 19 | 45682003 | 45685059 | 1 | protein_coding | biogenesis of lysosomal organelles complex-1, subunit 3 |
| **NKAPL** | ENSG00000189134 | 28.10341827 | 1.084476939 | 0.601121868 | 0.012418986 | 0.033712143 | 222698 | 6 | 28227098 | 28228736 | 1 | protein_coding | NFKB activating protein-like |
| **ZNF527** | ENSG00000189164 | 189.6871164 | -0.845891609 | 0.265393883 | 5.08E-04 | 0.001984716 | 84503 | 19 | 37861963 | 37883968 | 1 | protein_coding | zinc finger protein 527 |
| **ZNF600** | ENSG00000189190 | 303.6586078 | 0.741171853 | 0.219490376 | 3.24E-04 | 0.001325938 | 162966 | 19 | 53267448 | 53290044 | -1 | protein_coding | zinc finger protein 600 |
| **BTBD8** | ENSG00000189195 | 67.24159941 | 1.142323688 | 0.500539315 | 0.003797596 | 0.011949488 | 284697 | 1 | 92545862 | 92613393 | 1 | protein_coding | BTB (POZ) domain containing 8 |
| **MAOA** | ENSG00000189221 | 325.4895594 | -0.634596593 | 0.232534534 | 0.003199488 | 0.010281282 | 4128 | X | 43515467 | 43606068 | 1 | protein_coding | monoamine oxidase A |
| **PNRC2** | ENSG00000189266 | 3053.955685 | 1.15146076 | 0.131264157 | 4.57E-19 | 9.43E-18 | 55629 | 1 | 24285599 | 24289952 | 1 | protein_coding | proline-rich nuclear receptor coactivator 2 |
| **LIN54** | ENSG00000189308 | 919.1396219 | 0.368389589 | 0.131379875 | 0.00409021 | 0.012747522 | 132660 | 4 | 83831126 | 83934079 | -1 | protein_coding | lin-54 homolog (C. elegans) |
| **FAM53B** | ENSG00000189319 | 1433.173319 | -0.481595495 | 0.124660845 | 8.14E-05 | 3.78E-04 | 9679 | 10 | 126307861 | 126432838 | -1 | protein_coding | family with sequence similarity 53, member B |
| **KAZN** | ENSG00000189337 | 61.35312537 | -2.749214723 | 0.666957034 | 2.93E-06 | 1.76E-05 | 23254 | 1 | 14925200 | 15444539 | 1 | protein_coding | kazrin, periplakin interacting protein |
| **FAM179A** | ENSG00000189350 | 77.05456559 | -3.20224628 | 0.592603532 | 5.09E-09 | 4.5E-08 | 165186 | 2 | 29179477 | 29284239 | 1 | protein_coding | family with sequence similarity 179, member A |
| **TMEM194B** | ENSG00000189362 | 176.6411045 | 1.034762889 | 0.343441981 | 6.29E-04 | 0.002398118 | 1001312 | 2 | 191369068 | 191399448 | -1 | protein_coding | transmembrane protein 194B |
| **KIAA0408** | ENSG00000189367 | 36.67512684 | 1.700782695 | 0.815112577 | 0.002351913 | 0.00780041 | 387104 | 6 | 127761488 | 127780536 | -1 | protein_coding | KIAA0408 |
| **C8orf76** | ENSG00000189376 | 675.2961091 | 0.693992974 | 0.176206577 | 4.01E-05 | 1.97E-04 | 84933 | 8 | 124232196 | 124253638 | -1 | protein_coding | chromosome 8 open reading frame 76 |
| **HMGB1** | ENSG00000189403 | 3174.415064 | 0.498485299 | 0.111525291 | 4.93E-06 | 2.85E-05 | 3146 | 13 | 31032884 | 31191734 | -1 | protein_coding | high mobility group box 1 |
| **ZFP92** | ENSG00000189420 | 41.44238059 | -1.597372102 | 0.607600357 | 9.49E-04 | 0.003476608 | 139735 | X | 152683780 | 152687087 | 1 | protein_coding | ZFP92 zinc finger protein |
| **BLOC1S2** | ENSG00000196072 | 2371.014825 | 1.115364666 | 0.131650608 | 5.62E-18 | 1.06E-16 | 282991 | 10 | 102033713 | 102046469 | -1 | protein_coding | biogenesis of lysosomal organelles complex-1, subunit 2 |
| **SYCP2** | ENSG00000196074 | 68.92695325 | 1.693133001 | 0.693223733 | 0.001108009 | 0.004005134 | 10388 | 20 | 58438618 | 58508710 | -1 | protein_coding | synaptonemal complex protein 2 |
| **PAX5** | ENSG00000196092 | 404.0646384 | 2.63397671 | 0.374347222 | 1.51E-13 | 2.05E-12 | 5079 | 9 | 36833272 | 37034103 | -1 | protein_coding | paired box 5 |
| **HLA-DRB3** | ENSG00000196101 | 6883.564603 | -0.468033187 | 0.153277555 | 0.001489075 | 0.005201722 | 3125 | HSC | 32417557 | 32430644 | -1 | protein_coding | major histocompatibility complex, class II, DR beta 3 |
| **TDRD7** | ENSG00000196116 | 1113.348385 | 2.066895113 | 0.136842259 | 1.62E-52 | 1.61E-50 | 23424 | 9 | 100174232 | 100258407 | 1 | protein_coding | tudor domain containing 7 |
| **KIAA0895L** | ENSG00000196123 | 212.8143971 | -0.846452347 | 0.282713449 | 9.46E-04 | 0.00346911 | 653319 | 16 | 67209505 | 67217943 | -1 | protein_coding | KIAA0895-like |
| **HLA-DRB1** | ENSG00000196126 | 1596.911516 | -0.47076222 | 0.177088481 | 0.005420281 | 0.016371009 | 1009968 | 6 | 32546546 | 32557625 | -1 | protein_coding | major histocompatibility complex, class II, DR beta 1 |
| **SPATS2L** | ENSG00000196141 | 1090.593027 | 1.934256956 | 0.16129004 | 4.2E-34 | 2.02E-32 | 26010 | 2 | 201170604 | 201346986 | 1 | protein_coding | spermatogenesis associated, serine-rich 2-like |
| **S100A4** | ENSG00000196154 | 3128.940298 | -1.743685766 | 0.156956401 | 9.65E-30 | 3.67E-28 | 6275 | 1 | 153516089 | 153522612 | -1 | protein_coding | S100 calcium binding protein A4 |
| **PLEKHG4** | ENSG00000196155 | 53.06718294 | -1.286735944 | 0.516721685 | 0.001953577 | 0.006630362 | 25894 | 16 | 67311413 | 67323402 | 1 | protein_coding | pleckstrin homology domain containing, family G (with RhoGef domain) member 4 |
| **ACADSB** | ENSG00000196177 | 916.1348512 | 0.745730259 | 0.206402044 | 1.33E-04 | 5.93E-04 | 36 | 10 | 124768495 | 124817827 | 1 | protein_coding | acyl-CoA dehydrogenase, short/branched chain |
| **TMEM63A** | ENSG00000196187 | 4472.923554 | 0.469668734 | 0.104575528 | 5.38E-06 | 3.1E-05 | 9725 | 1 | 226033237 | 226070069 | -1 | protein_coding | transmembrane protein 63A |
| **SEMA4A** | ENSG00000196189 | 1156.767001 | 2.116538797 | 0.231012882 | 4.03E-21 | 9.65E-20 | 64218 | 1 | 156117157 | 156147543 | 1 | protein_coding | sema domain, immunoglobulin domain (Ig), transmembrane domain (TM) and short cytoplasmic domain |
| **MPHOSPH8** | ENSG00000196199 | 1315.469023 | 0.380314817 | 0.123988329 | 0.001846148 | 0.006295109 | 54737 | 13 | 20207788 | 20247599 | 1 | protein_coding | M-phase phosphoprotein 8 |
| **GREB1** | ENSG00000196208 | 192.830718 | -1.843292371 | 0.357796877 | 2.74E-08 | 2.2E-07 | 9687 | 2 | 11674242 | 11782914 | 1 | protein_coding | growth regulation by estrogen in breast cancer 1 |
| **SIRPB2** | ENSG00000196209 | 159.5452985 | -0.955536144 | 0.28558267 | 2.3E-04 | 9.69E-04 | 284759 | 20 | 1451386 | 1472233 | -1 | protein_coding | signal-regulatory protein beta 2 |
| **RYR1** | ENSG00000196218 | 778.6322418 | -1.125173701 | 0.214445956 | 3.27E-08 | 2.61E-07 | 6261 | 19 | 38924339 | 39078204 | 1 | protein_coding | ryanodine receptor 1 (skeletal) |
| **SRGAP3** | ENSG00000196220 | 292.5896798 | -1.761312425 | 0.37833131 | 3.07E-07 | 2.13E-06 | 9901 | 3 | 9022275 | 9404737 | -1 | protein_coding | SLIT-ROBO Rho GTPase activating protein 3 |
| **FAM217B** | ENSG00000196227 | 642.3855286 | 0.431491246 | 0.168305287 | 0.007715477 | 0.022328277 | 63939 | 20 | 58508819 | 58523735 | 1 | protein_coding | family with sequence similarity 217, member B |
| **LCOR** | ENSG00000196233 | 923.1991914 | 0.953534923 | 0.282717053 | 2.03E-04 | 8.67E-04 | 84458 | 10 | 98592017 | 98740800 | 1 | protein_coding | ligand dependent nuclear receptor corepressor |
| **XPNPEP3** | ENSG00000196236 | 656.2379388 | -0.451617938 | 0.178630761 | 0.008268094 | 0.023698664 | 63929 | 22 | 41253081 | 41363838 | 1 | protein_coding | X-prolyl aminopeptidase (aminopeptidase P) 3, putative |
| **ZNF107** | ENSG00000196247 | 507.7144976 | 2.043598521 | 0.185640838 | 3.56E-29 | 1.31E-27 | 51427 | 7 | 64126511 | 64171404 | 1 | protein_coding | zinc finger protein 107 |
| **IARS** | ENSG00000196305 | 9098.697477 | -0.853617079 | 0.125599167 | 4.12E-12 | 5.02E-11 | 3376 | 9 | 94972489 | 95056038 | -1 | protein_coding | isoleucyl-tRNA synthetase |
| **GIMAP5** | ENSG00000196329 | 1489.414169 | 0.990431965 | 0.235354044 | 7.53E-06 | 4.23E-05 | 1005279 | 7 | 150419341 | 150447121 | 1 | protein_coding | GTPase, IMAP family member 5 |
| **CD55** | ENSG00000196352 | 538.5851583 | 0.494864966 | 0.185005456 | 0.004940756 | 0.015123087 | 1604 | 1 | 207494853 | 207534311 | 1 | protein_coding | CD55 molecule, decay accelerating factor for complement (Cromer blood group) |
| **LONP1** | ENSG00000196365 | 3794.401333 | -1.081908685 | 0.121994365 | 2.07E-19 | 4.35E-18 | 9361 | 19 | 5691845 | 5720583 | -1 | protein_coding | lon peptidase 1, mitochondrial |
| **FUT4** | ENSG00000196371 | 934.8202616 | 1.425597587 | 0.130063341 | 9.12E-29 | 3.31E-27 | 2526 | 11 | 94277017 | 94283063 | 1 | protein_coding | fucosyltransferase 4 (alpha (1,3) fucosyltransferase, myeloid-specific) |

| | | | | | | | | | | | | | |
|---|---|---|---|---|---|---|---|---|---|---|---|---|---|
| **ASB13** | ENSG00000196372 | 641.7010358 | -1.27942444 | 0.172910574 | 2.91E-14 | 4.2E-13 | 79754 | 10 | 5680830 | 5708558 | -1 | protein_coding | ankyrin repeat and SOCS box containing 13 |
| **ZNF140** | ENSG00000196387 | 540.9829939 | 0.600127084 | 0.167217538 | 1.91E-04 | 8.23E-04 | 7699 | 12 | 133656424 | 133684130 | 1 | protein_coding | zinc finger protein 140 |
| **PTPN1** | ENSG00000196396 | 4596.983064 | 0.680801411 | 0.090728211 | 3.15E-14 | 4.54E-13 | 5770 | 20 | 49126891 | 49201299 | 1 | protein_coding | protein tyrosine phosphatase, non-receptor type 1 |
| **EVL** | ENSG00000196405 | 12148.73659 | -1.872119833 | 0.104376349 | 2.05E-73 | 4.23E-71 | 51466 | 14 | 100437786 | 100610573 | 1 | protein_coding | Enah/Vasp-like |
| **ZNF765** | ENSG00000196417 | 577.5715015 | 0.414717629 | 0.154462207 | 0.005306196 | 0.016071065 | 91661 | 19 | 53893046 | 53930574 | 1 | protein_coding | zinc finger protein 765 |
| **ZNF124** | ENSG00000196418 | 1060.985813 | -0.368200654 | 0.138954411 | 0.006762102 | 0.019893205 | 7678 | 1 | 247285277 | 247335318 | -1 | protein_coding | zinc finger protein 124 |
| **TSC22D2** | ENSG00000196428 | 591.6860076 | 1.434878045 | 0.223038922 | 1.96E-11 | 2.25E-10 | 9819 | 3 | 150126122 | 150184218 | 1 | protein_coding | TSC22 domain family, member 2 |
| **ARMCX4** | ENSG00000196440 | 244.6901706 | -0.751787609 | 0.240187286 | 7.67E-04 | 0.002877783 | 1001317 | X | 100673275 | 100788446 | 1 | protein_coding | armadillo repeat containing, X-linked 4 |
| **YRDC** | ENSG00000196449 | 296.2128405 | 0.657307848 | 0.197386477 | 4.43E-04 | 0.001757129 | 79693 | 1 | 38268616 | 38273857 | -1 | protein_coding | yrdC N(6)-threonylcarbamoyltransferase domain containing |
| **PIK3R4** | ENSG00000196455 | 1209.993954 | 0.30649461 | 0.117442882 | 0.00788384 | 0.022738013 | 30849 | 3 | 130397779 | 130465673 | -1 | protein_coding | phosphoinositide-3-kinase, regulatory subunit 4 |
| **TRAPPC2** | ENSG00000196459 | 291.4450128 | -0.441815612 | 0.199602615 | 0.019300084 | 0.04924097 | 6399 | X | 13730363 | 13752754 | -1 | protein_coding | trafficking protein particle complex 2 |
| **C20orf96** | ENSG00000196476 | 93.07274901 | -0.848097214 | 0.373499076 | 0.007023462 | 0.020571017 | 140680 | 20 | 251504 | 271390 | -1 | protein_coding | chromosome 20 open reading frame 96 |
| **IPO4** | ENSG00000196497 | 274.8127247 | -0.615402884 | 0.251129102 | 0.00767621 | 0.022223056 | 79711 | 14 | 24649425 | 24658170 | -1 | protein_coding | importin 4 |
| **NCOR2** | ENSG00000196498 | 1413.42619 | -1.976645332 | 0.23637951 | 5.41E-18 | 1.02E-16 | 9612 | 12 | 124808961 | 125052135 | -1 | protein_coding | nuclear receptor corepressor 2 |
| **SULT1A1** | ENSG00000196502 | 526.5453049 | -0.756895088 | 0.193164227 | 3.9E-05 | 1.92E-04 | 6817 | 16 | 28616903 | 28634946 | -1 | protein_coding | sulfotransferase family, cytosolic, 1A, phenol-preferring, member 1 |
| **GDAP2** | ENSG00000196505 | 1442.92393 | 0.313214074 | 0.124497923 | 0.010175537 | 0.028414532 | 54834 | 1 | 118406107 | 118472253 | -1 | protein_coding | ganglioside induced differentiation associated protein 2 |
| **ANAPC7** | ENSG00000196510 | 1864.074583 | -0.500903984 | 0.203617199 | 0.009243008 | 0.02614974 | 51434 | 12 | 110810705 | 110841535 | -1 | protein_coding | anaphase promoting complex subunit 7 |
| **TPK1** | ENSG00000196511 | 446.1587038 | 0.944029187 | 0.209575332 | 2.08E-06 | 1.27E-05 | 27010 | 7 | 144149034 | 144533488 | -1 | protein_coding | thiamin pyrophosphokinase 1 |
| **NACA** | ENSG00000196531 | 5449.339875 | -0.367206044 | 0.113976071 | 0.001124981 | 0.004058802 | 4666 | 12 | 57106212 | 57125412 | -1 | protein_coding | nascent polypeptide-associated complex alpha subunit |
| **MYO18A** | ENSG00000196535 | 3366.018833 | 0.410723132 | 0.120776477 | 5.37E-04 | 0.002083722 | 399687 | 17 | 27400528 | 27507430 | -1 | protein_coding | myosin XVIIIA |
| **MAN2A2** | ENSG00000196547 | 1696.01605 | -0.670350841 | 0.158984935 | 1.19E-05 | 6.47E-05 | 4122 | 15 | 91445448 | 91465814 | 1 | protein_coding | mannosidase, alpha, class 2A, member 2 |
| **FAM72A** | ENSG00000196550 | 130.6120721 | 1.038989101 | 0.301265405 | 1.43E-04 | 6.36E-04 | 729533 | 1 | 206136916 | 206155151 | 1 | protein_coding | family with sequence similarity 72, member A |
| **SULF2** | ENSG00000196562 | 3248.762937 | -3.062789667 | 0.180581684 | 1.05E-65 | 1.69E-63 | 55959 | 20 | 46285092 | 46415360 | -1 | protein_coding | sulfatase 2 |
| **PLXNB2** | ENSG00000196576 | 11744.67895 | -1.425389989 | 0.125580853 | 1.18E-30 | 4.73E-29 | 23654 | 22 | 50713408 | 50746056 | -1 | protein_coding | plexin B2 |
| **MKL1** | ENSG00000196588 | 1421.836162 | 0.382429963 | 0.125330063 | 0.001787616 | 0.006111886 | 57591 | 22 | 40806285 | 41032706 | -1 | protein_coding | megakaryoblastic leukemia (translocation) 1 |
| **TCF4** | ENSG00000196628 | 930.9965645 | 0.713296189 | 0.192814618 | 9.98E-05 | 4.56E-04 | 6925 | 18 | 52889562 | 53332018 | -1 | protein_coding | transcription factor 4 |
| **RABL6** | ENSG00000196642 | 1712.151279 | -0.861102605 | 0.146751171 | 1.56E-09 | 1.47E-08 | 55684 | 9 | 139702374 | 139735639 | 1 | protein_coding | RAB, member RAS oncogene family-like 6 |
| **ZNF136** | ENSG00000196646 | 506.6869143 | 0.554781682 | 0.170302156 | 6.96E-04 | 0.0026344 | 7695 | 19 | 12273879 | 12300064 | 1 | protein_coding | zinc finger protein 136 |
| **ZKSCAN5** | ENSG00000196652 | 565.3752687 | 0.534647309 | 0.171145696 | 0.00112815 | 0.004068314 | 23660 | 7 | 99102274 | 99132323 | 1 | protein_coding | zinc finger with KRAB and SCAN domains 5 |
| **HSH2D** | ENSG00000196684 | 19.04871381 | 4.734121653 | 1.30538712 | 1.32E-05 | 7.14E-05 | 84941 | 19 | 16244838 | 16269386 | 1 | protein_coding | hematopoietic SH2 domain containing |
| **TRPV1** | ENSG00000196689 | 240.4444855 | -0.749830961 | 0.246275623 | 0.001028376 | 0.003742065 | 7442 | 17 | 3468738 | 3500392 | -1 | protein_coding | transient receptor potential cation channel, subfamily V, member 1 |
| **ZNF33B** | ENSG00000196693 | 137.8266446 | 0.803450015 | 0.288135073 | 0.001998061 | 0.006763315 | 7582 | 10 | 43084555 | 43133992 | -1 | protein_coding | zinc finger protein 33B |
| **ZNF512B** | ENSG00000196700 | 235.0007804 | -0.057248121 | 0.638497434 | 1.78E-07 | 1.28E-06 | 57473 | 20 | 62588055 | 62680113 | -1 | protein_coding | zinc finger protein 512B |
| **NF1** | ENSG00000196712 | 2330.972301 | 0.462079581 | 0.126754125 | 1.97E-04 | 8.45E-04 | 1019301 | 17 | 29421945 | 29709134 | 1 | protein_coding | neurofibromin 1 |
| **VKORC1L1** | ENSG00000196715 | 2431.677106 | -0.68414864 | 0.151409703 | 3.3E-06 | 1.96E-05 | 154807 | 7 | 65338254 | 65424550 | 1 | protein_coding | vitamin K epoxide reductase complex, subunit 1-like 1 |
| **COL27A1** | ENSG00000196739 | 116.4844409 | -2.466399417 | 0.51574936 | 1.07E-07 | 7.89E-07 | 85301 | 9 | 116917840 | 117074791 | 1 | protein_coding | collagen, type XXVII, alpha 1 |
| **CD47** | ENSG00000196776 | 4172.659804 | 1.643435721 | 0.118190849 | 7.59E-45 | 5.77E-43 | 961 | 3 | 107762145 | 107809872 | -1 | protein_coding | CD47 molecule |
| **TLE1** | ENSG00000196781 | 71.60808956 | 1.571513878 | 0.460190034 | 7.4E-05 | 3.46E-04 | 7088 | 9 | 84198598 | 84304220 | -1 | protein_coding | transducin-like enhancer of split 1 (E(sp1) homolog, Drosophila) |
| **MAML3** | ENSG00000196782 | 149.7091675 | -2.903284462 | 0.347738384 | 6.75E-18 | 1.26E-16 | 55534 | 4 | 140637907 | 141075338 | -1 | protein_coding | mastermind-like 3 (Drosophila) |
| **STRN3** | ENSG00000196792 | 997.422608 | 0.871418857 | 0.166298114 | 5.67E-08 | 4.36E-07 | 29966 | 14 | 31363005 | 31495607 | -1 | protein_coding | striatin, calmodulin binding protein 3 |
| **ZSCAN16** | ENSG00000196812 | 162.8429931 | 0.992579261 | 0.284014144 | 1.32E-04 | 5.89E-04 | 80345 | 6 | 28092338 | 28097860 | 1 | protein_coding | zinc finger and SCAN domain containing 16 |
| **MVB12B** | ENSG00000196814 | 521.6623363 | 1.916199308 | 0.186314623 | 1.01E-25 | 3.13E-24 | 89853 | 9 | 129089128 | 129269320 | 1 | protein_coding | multivesicular body subunit 12B |
| **C6orf106** | ENSG00000196821 | 5427.787142 | 0.292475702 | 0.085447536 | 4.21E-04 | 0.001679137 | 64771 | 6 | 34555065 | 34664636 | -1 | protein_coding | chromosome 6 open reading frame 106 |
| **ADA** | ENSG00000196839 | 148.9985542 | 2.162619985 | 0.378128369 | 9.28E-10 | 8.96E-09 | 100 | 20 | 43248163 | 43280874 | -1 | protein_coding | adenosine deaminase |
| **ARID5A** | ENSG00000196843 | 1513.120882 | 0.69422964 | 0.18045688 | 5.98E-05 | 2.85E-04 | 10865 | 2 | 97202480 | 97218375 | 1 | protein_coding | AT rich interactive domain 5A (MRF1-like) |
| **PPTC7** | ENSG00000196850 | 1674.670508 | 0.940319902 | 0.119058844 | 9.2E-16 | 1.48E-14 | 160760 | 12 | 110969120 | 111021125 | -1 | protein_coding | PTC7 protein phosphatase homolog (S. cerevisiae) |
| **NHLRC2** | ENSG00000196865 | 962.6054624 | 0.553847812 | 0.176951252 | 0.001075524 | 0.0038997 | 374354 | 10 | 115614420 | 115676953 | 1 | protein_coding | NHL repeat containing 2 |
| **KPNA5** | ENSG00000196911 | 283.0662756 | 1.323461763 | 0.275058852 | 2.46E-07 | 1.74E-06 | 3841 | 6 | 117002350 | 117063029 | 1 | protein_coding | karyopherin alpha 5 (importin alpha 6) |
| **PDLIM7** | ENSG00000196923 | 705.1762488 | -0.473741813 | 0.1483974 | 9.58E-04 | 0.003506923 | 9260 | 5 | 176910395 | 176924607 | -1 | protein_coding | PDZ and LIM domain 7 (enigma) |
| **FLNA** | ENSG00000196924 | 40754.23036 | -1.063833781 | 0.135600282 | 1.04E-15 | 1.65E-14 | 2316 | X | 153576892 | 153603006 | -1 | protein_coding | filamin A, alpha |
| **FAM3C** | ENSG00000196937 | 731.4838417 | 0.874904533 | 0.147763145 | 1.16E-09 | 1.11E-08 | 10447 | 7 | 120988905 | 121036418 | -1 | protein_coding | family with sequence similarity 3, member C |
| **SLC39A10** | ENSG00000196950 | 2856.684099 | -1.346729174 | 0.192664837 | 4.84E-13 | 6.35E-12 | 57181 | 2 | 196440701 | 196602426 | 1 | protein_coding | solute carrier family 39 (zinc transporter), member 10 |
| **CASP4** | ENSG00000196954 | 2666.881259 | 1.139801934 | 0.120440595 | 8.68E-22 | 2.17E-20 | 837 | 11 | 104813593 | 104840163 | -1 | protein_coding | caspase 4, apoptosis-related cysteine peptidase |
| **AP2A1** | ENSG00000196961 | 1648.312572 | -0.535979113 | 0.134873588 | 4.45E-05 | 2.17E-04 | 160 | 19 | 50270225 | 50310370 | 1 | protein_coding | adaptor-related protein complex 2, alpha 1 subunit |
| **ANXA4** | ENSG00000196975 | 2037.677184 | 0.336354217 | 0.113979321 | 0.002781726 | 0.009045346 | 307 | 2 | 69871557 | 70053596 | 1 | protein_coding | annexin A4 |
| **LAGE3** | ENSG00000196976 | 115.0425286 | -0.957286769 | 0.325442397 | 9.28E-04 | 0.00341077 | 8270 | X | 153706028 | 153707596 | -1 | protein_coding | L antigen family, member 3 |
| **METTL9** | ENSG00000197006 | 1500.42745 | 0.593172938 | 0.189765559 | 0.001020845 | 0.003719236 | 51108 | 16 | 21608539 | 21668794 | 1 | protein_coding | methyltransferase like 9 |
| **ZNF100** | ENSG00000197020 | 597.0605869 | 0.555184122 | 0.175834701 | 9.4E-04 | 0.003450888 | 163227 | 19 | 21905568 | 21950430 | -1 | protein_coding | zinc finger protein 100 |
| **ZNF441** | ENSG00000197044 | 218.123477 | 0.682923138 | 0.267712921 | 0.004864244 | 0.014918771 | 126068 | 19 | 11877815 | 11894893 | 1 | protein_coding | zinc finger protein 441 |
| **SIGLEC15** | ENSG00000197046 | 600.809171 | -1.032316183 | 0.382269344 | 0.0015903 | 0.005519999 | 284266 | 18 | 43405477 | 43424045 | 1 | protein_coding | sialic acid binding Ig-like lectin 15 |
| **RP5-874C20.** | ENSG00000197062 | 329.274563 | 0.754795734 | 0.213140279 | 1.74E-04 | 7.55E-04 | NA | 6 | 28234788 | 28245974 | 1 | pseudogene | |
| **MAFG** | ENSG00000197063 | 1404.157064 | 0.76900727 | 0.140820038 | 1.93E-08 | 1.58E-07 | 4097 | 17 | 79876146 | 79885590 | -1 | protein_coding | v-maf avian musculoaponeurotic fibrosarcoma oncogene homolog G |
| **KIAA1671** | ENSG00000197077 | 597.1750834 | 2.707221517 | 0.177926566 | 2.21E-53 | 2.34E-51 | 1009962 | 22 | 25348697 | 25593415 | 1 | protein_coding | KIAA1671 |
| **IGF2R** | ENSG00000197081 | 40056.69694 | -0.48520916 | 0.078872869 | 5.41E-10 | 5.34E-09 | 3482 | 6 | 160390131 | 160534539 | 1 | protein_coding | insulin-like growth factor 2 receptor |
| **GAL3ST4** | ENSG00000197093 | 18.24977853 | -2.096585141 | 1.001705273 | 0.002420606 | 0.007993544 | 79690 | 7 | 99756867 | 99766373 | -1 | protein_coding | galactose-3-O-sulfotransferase 4 |
| **PCBP2** | ENSG00000197111 | 8891.184884 | -0.264508474 | 0.086954858 | 0.003435683 | 0.010943632 | 5094 | 12 | 53835525 | 53874946 | 1 | protein_coding | poly(rC) binding protein 2 |
| **SLC25A29** | ENSG00000197119 | 798.056092 | -2.168855343 | 0.171109522 | 1.01E-37 | 5.63E-36 | 123096 | 14 | 100757448 | 100772884 | -1 | protein_coding | solute carrier family 25 (mitochondrial carnitine/acylcarnitine carrier), member 29 |
| **PGAP1** | ENSG00000197121 | 222.3239984 | 2.921622776 | 0.326400297 | 1.64E-20 | 3.76E-19 | 80055 | 2 | 197697728 | 197792520 | -1 | protein_coding | post-GPI attachment to proteins 1 |
| **SRC** | ENSG00000197122 | 3048.766782 | 1.022474945 | 0.124596946 | 5.76E-17 | 1.01E-15 | 6714 | 20 | 35973088 | 36034453 | 1 | protein_coding | v-src avian sarcoma (Schmidt-Ruppin A-2) viral oncogene homolog |
| **ACSL5** | ENSG00000197142 | 1780.669519 | 0.713754601 | 0.134423754 | 5.43E-08 | 4.2E-07 | 51703 | 10 | 114133776 | 114188138 | 1 | protein_coding | acyl-CoA synthetase long-chain family member 5 |
| **LRRC8B** | ENSG00000197147 | 956.2700144 | 0.458003146 | 0.156784517 | 0.002539864 | 0.008345891 | 23507 | 1 | 89990395 | 90063423 | 1 | protein_coding | leucine rich repeat containing 8 family, member B |
| **SND1** | ENSG00000197157 | 4843.658135 | -0.651036136 | 0.092080246 | 8.58E-13 | 1.1E-11 | 27044 | 7 | 127292234 | 127732661 | 1 | protein_coding | staphylococcal nuclease and tudor domain containing 1 |
| **PSMD12** | ENSG00000197170 | 2132.828459 | 0.921407632 | 0.112398872 | 8.49E-17 | 1.46E-15 | 5718 | 17 | 65334032 | 65362743 | -1 | protein_coding | proteasome (prosome, macropain) 26S subunit, non-ATPase, 12 |
| **SLC22A4** | ENSG00000197208 | 77.26536123 | 1.39556993 | 0.396412282 | 6.37E-05 | 3.02E-04 | 6583 | 5 | 131630136 | 131679899 | 1 | protein_coding | solute carrier family 22 (organic cation/zwitterion transporter), member 4 |
| **TBC1D9B** | ENSG00000197226 | 6288.279438 | -0.772345546 | 0.086181755 | 1.81E-19 | 3.83E-18 | 23061 | 5 | 179289066 | 179334859 | -1 | protein_coding | TBC1 domain family, member 9B (with GRAM domain) |
| **GTF2E2** | ENSG00000197265 | 954.8654156 | 1.444744419 | 0.151077522 | 1.95E-22 | 5.02E-21 | 2961 | 8 | 30435835 | 30515768 | -1 | protein_coding | general transcription factor IIE, polypeptide 2, beta 34kDa |
| **IL27** | ENSG00000197272 | 39.20498382 | 3.144113496 | 0.804239928 | 5.11E-06 | 2.95E-05 | 246778 | 16 | 28510683 | 28523372 | -1 | protein_coding | interleukin 27 |
| **FITM2** | ENSG00000197296 | 232.5408932 | -1.325992331 | 0.258665014 | 5.3E-08 | 4.11E-07 | 128486 | 20 | 42931478 | 42939809 | -1 | protein_coding | fat storage-inducing transmembrane protein 2 |
| **DDI2** | ENSG00000197312 | 3123.851043 | 0.239630944 | 0.089518775 | 0.005629855 | 0.016901146 | 84301 | 1 | 15943995 | 15995539 | 1 | protein_coding | DNA-damage inducible 1 homolog 2 (S. cerevisiae) |
| **SVIL** | ENSG00000197321 | 233.1188216 | -0.980831428 | 0.409160539 | 0.004258043 | 0.013230723 | 6840 | 10 | 29746267 | 30025710 | -1 | protein_coding | supervillin |
| **TRIM33** | ENSG00000197323 | 2939.53552 | 0.763974425 | 0.117019687 | 3.03E-11 | 3.43E-10 | 51592 | 1 | 114935399 | 115053781 | -1 | protein_coding | tripartite motif containing 33 |
| **LRP10** | ENSG00000197324 | 5951.401994 | 0.487442131 | 0.129222164 | 1.15E-04 | 5.22E-04 | 26020 | 14 | 23340822 | 23350789 | 1 | protein_coding | low density lipoprotein receptor-related protein 10 |
| **PELI1** | ENSG00000197329 | 722.4262231 | 1.028106685 | 0.153032069 | 5.27E-12 | 6.37E-11 | 57162 | 2 | 64319786 | 64371588 | -1 | protein_coding | pellino E3 ubiquitin protein ligase 1 |
| **ZNF655** | ENSG00000197343 | 1954.433055 | 0.420091376 | 0.09910307 | 1.95E-05 | 1.02E-04 | 1019294 | 7 | 99156029 | 99174076 | 1 | protein_coding | zinc finger protein 655 |
| **MRPL21** | ENSG00000197345 | 315.383856 | -0.614582317 | 0.228562951 | 0.003897044 | 0.012219639 | 219927 | 11 | 68658744 | 68671303 | -1 | protein_coding | mitochondrial ribosomal protein L21 |
| **UAP1L1** | ENSG00000197355 | 971.5393618 | -0.549389069 | 0.150560515 | 1.64E-04 | 7.19E-04 | 91373 | 9 | 139971953 | 139978991 | 1 | protein_coding | UDP-N-acteylglucosamine pyrophosphorylase 1-like 1 |
| **ZNF675** | ENSG00000197372 | 388.5983248 | 0.562934612 | 0.217046377 | 0.005608655 | 0.016850222 | 171392 | 19 | 23708433 | 23870022 | -1 | protein_coding | zinc finger protein 675 |
| **ADARB1** | ENSG00000197381 | 781.9264413 | -0.585248752 | 0.179225868 | 6.54E-04 | 0.002487292 | 104 | 21 | 46493768 | 46646478 | 1 | protein_coding | adenosine deaminase, RNA-specific, B1 |
| **HTT** | ENSG00000197386 | 8779.484034 | -0.811540041 | 0.167059542 | 4.16E-07 | 2.85E-06 | 3064 | 4 | 3076408 | 3245676 | 1 | protein_coding | huntingtin |

| | | | | | | | | | | | | | |
|---|---|---|---|---|---|---|---|---|---|---|---|---|---|
| **SHPK** | ENSG00000197417 | 500.8584311 | -0.788593113 | 0.180328621 | 5.03E-06 | 2.91E-05 | 23729 | 17 | 3511556 | 3539616 | -1 | protein_coding | sedoheptulokinase |
| **MAP3K5** | ENSG00000197442 | 2736.938008 | 1.063600355 | 0.109031239 | 5.23E-23 | 1.41E-21 | 4217 | 6 | 136878185 | 137113656 | -1 | protein_coding | mitogen-activated protein kinase kinase kinase 5 |
| **GSTK1** | ENSG00000197448 | 3524.713941 | 0.314343133 | 0.109031752 | 0.003399488 | 0.010846423 | 373156 | 7 | 142941186 | 142967947 | 1 | protein_coding | glutathione S-transferase kappa 1 |
| **SPN** | ENSG00000197471 | 11391.2998 | -1.579439976 | 0.090906195 | 1.84E-68 | 3.35E-66 | 1019298 | 16 | 29674300 | 29682187 | 1 | protein_coding | sialophorin |
| **SLC2A10** | ENSG00000197496 | 26.8125812 | -1.57858781 | 0.830662992 | 0.005085786 | 0.015495628 | 81031 | 20 | 45338126 | 45364965 | 1 | protein_coding | solute carrier family 2 (facilitated glucose transporter), member 10 |
| **SLC28A3** | ENSG00000197506 | 8165.570557 | -1.216601475 | 0.148089341 | 4.73E-17 | 8.31E-16 | 64078 | 9 | 86890372 | 86955672 | -1 | protein_coding | solute carrier family 28 (concentrative nucleoside transporter), member 3 |
| **MYO5A** | ENSG00000197535 | 6036.717714 | -0.406734774 | 0.127974511 | 0.001021561 | 0.003719923 | 4644 | 15 | 52599480 | 52821247 | -1 | protein_coding | myosin VA (heavy chain 12, myoxin) |
| **C5orf56** | ENSG00000197536 | 200.3923009 | 2.385341873 | 0.256743693 | 1.33E-21 | 3.28E-20 | NA | 5 | 131746328 | 131811736 | 1 | protein_coding | chromosome 5 open reading frame 56 |
| **ATG7** | ENSG00000197548 | 2671.897389 | -0.538392033 | 0.127773415 | 1.59E-05 | 8.45E-05 | 10533 | 3 | 11313995 | 11599139 | 1 | protein_coding | autophagy related 7 |
| **SIPA1L1** | ENSG00000197555 | 1850.32371 | 1.255758595 | 0.128962586 | 4.58E-23 | 1.24E-21 | 26037 | 14 | 71787166 | 72207946 | 1 | protein_coding | signal-induced proliferation-associated 1 like 1 |
| **HHLA3** | ENSG00000197568 | 196.422578 | -1.008604859 | 0.265631747 | 4.07E-05 | 2E-04 | 11147 | 1 | 70820488 | 70851022 | 1 | protein_coding | HERV-H LTR-associating 3 |
| **CDC42SE1** | ENSG00000197622 | 7882.977028 | 0.48322028 | 0.081683071 | 3.12E-09 | 2.82E-08 | 56882 | 1 | 151023447 | 151042801 | -1 | protein_coding | CDC42 small effector 1 |
| **DPP4** | ENSG00000197635 | 1118.010128 | 1.707522298 | 0.211154679 | 7.65E-17 | 1.32E-15 | 1803 | 2 | 162848751 | 162931052 | -1 | protein_coding | dipeptidyl-peptidase 4 |
| **SPTAN1** | ENSG00000197694 | 15780.62163 | -0.252480794 | 0.093118151 | 0.006162754 | 0.018331135 | 6709 | 9 | 131314866 | 131395941 | 1 | protein_coding | spectrin, alpha, non-erythrocytic 1 |
| **NMB** | ENSG00000197696 | 142.0389511 | -0.925612935 | 0.353217383 | 0.002569296 | 0.008429918 | 4828 | 15 | 85198360 | 85201794 | -1 | protein_coding | neuromedin B |
| **FAM114A1** | ENSG00000197712 | 549.7527595 | -1.010714766 | 0.165116105 | 2.74E-10 | 2.78E-09 | 92689 | 4 | 38869298 | 38947360 | 1 | protein_coding | family with sequence similarity 114, member A1 |
| **ZNF460** | ENSG00000197714 | 666.2967617 | -0.355867997 | 0.149341768 | 0.013841619 | 0.037078388 | 10794 | 19 | 57791419 | 57804937 | 1 | protein_coding | zinc finger protein 460 |
| **CR1L** | ENSG00000197721 | 65.89078331 | 2.973136528 | 0.686672748 | 8.96E-07 | 5.81E-06 | 1379 | 1 | 207818458 | 207911761 | 1 | protein_coding | complement component (3b/4b) receptor 1-like |
| **PSAP** | ENSG00000197746 | 188920.1193 | -0.365934171 | 0.136699311 | 0.005957797 | 0.017773376 | 5660 | 10 | 73576055 | 73611126 | -1 | protein_coding | prosaposin |
| **RPL37A** | ENSG00000197756 | 5962.623612 | -0.539127115 | 0.137736775 | 5.15E-05 | 2.48E-04 | 6168 | 2 | 217362912 | 217443903 | 1 | protein_coding | ribosomal protein L37a |
| **TXNRD3** | ENSG00000197763 | 187.4748609 | -1.116992328 | 0.332311575 | 1.7E-04 | 7.42E-04 | 645840 | 3 | 126290622 | 126373998 | -1 | protein_coding | thioredoxin reductase 3 |
| **CFD** | ENSG00000197766 | 528.1853146 | -0.601268125 | 0.18658047 | 7.29E-04 | 0.002747063 | 1675 | 19 | 859453 | 863453 | 1 | protein_coding | complement factor D (adipsin) |
| **MCMBP** | ENSG00000197771 | 2274.945965 | 1.11135545 | 0.099952055 | 2.02E-29 | 7.57E-28 | 79892 | 10 | 121588972 | 121652068 | -1 | protein_coding | minichromosome maintenance complex binding protein |
| **ZNF780A** | ENSG00000197782 | 693.0260824 | -0.551904391 | 0.164654053 | 5.24E-04 | 0.002040817 | 1019292 | 19 | 40575059 | 40596845 | -1 | protein_coding | zinc finger protein 780A |
| **FAM212B** | ENSG00000197852 | 519.5518908 | -1.688341481 | 0.1766394 | 1.26E-22 | 3.3E-21 | 1019287 | 1 | 112223252 | 112298446 | -1 | protein_coding | family with sequence similarity 212, member B |
| **ZNF44** | ENSG00000197857 | 224.5067453 | 0.597932401 | 0.220745741 | 0.003818713 | 0.012004656 | 1010601 | 19 | 12335501 | 12405702 | -1 | protein_coding | zinc finger protein 44 |
| **SGTB** | ENSG00000197860 | 3741.051333 | 0.721671298 | 0.136544565 | 5.94E-08 | 4.56E-07 | 54557 | 5 | 64961755 | 65018862 | -1 | protein_coding | small glutamine-rich tetratricopeptide repeat (TPR)-containing, beta |
| **FAM49A** | ENSG00000197872 | 1681.450388 | 0.587805237 | 0.13141637 | 4.39E-06 | 2.57E-05 | 81553 | 2 | 16730727 | 16847599 | -1 | protein_coding | family with sequence similarity 49, member A |
| **KIF13B** | ENSG00000197892 | 3362.07659 | -0.576648858 | 0.103079065 | 1.27E-08 | 1.07E-07 | 23303 | 8 | 28924796 | 29120641 | -1 | protein_coding | kinesin family member 13B |
| **ZNF677** | ENSG00000197928 | 365.1597688 | -1.378565013 | 0.277991236 | 1.1E-07 | 8.08E-07 | 342926 | 19 | 53731577 | 53758151 | -1 | protein_coding | zinc finger protein 677 |
| **FCHSD1** | ENSG00000197948 | 518.8242651 | 0.533322015 | 0.206351671 | 0.005689649 | 0.017060016 | 89848 | 5 | 141018869 | 141030986 | -1 | protein_coding | FCH and double SH3 domains 1 |
| **DNM3** | ENSG00000197959 | 47.39432375 | -1.101618761 | 0.664608147 | 0.014643483 | 0.038900808 | 26052 | 1 | 171810621 | 172387606 | 1 | protein_coding | dynamin 3 |
| **MPZL1** | ENSG00000197965 | 3608.177884 | 0.6614734 | 0.105597218 | 1.65E-10 | 1.72E-09 | 9019 | 1 | 167690429 | 167761156 | 1 | protein_coding | myelin protein zero-like 1 |
| **VPS13A** | ENSG00000197969 | 669.2852619 | 0.777406505 | 0.237416687 | 4.18E-04 | 0.001666955 | 23230 | 9 | 79792269 | 80036457 | 1 | protein_coding | vacuolar protein sorting 13 homolog A (S. cerevisiae) |
| **MBP** | ENSG00000197971 | 5815.08291 | -0.969087708 | 0.118942466 | 1.35E-16 | 2.3E-15 | 4155 | 18 | 74690783 | 74845639 | -1 | protein_coding | myelin basic protein |
| **C1orf122** | ENSG00000197982 | 127.8486198 | 1.245594996 | 0.306413237 | 9.17E-06 | 5.07E-05 | 127687 | 1 | 38272651 | 38275126 | 1 | protein_coding | chromosome 1 open reading frame 122 |
| **NOL8** | ENSG00000198000 | 1756.44529 | 0.649532964 | 0.148802229 | 6.65E-06 | 3.77E-05 | 55035 | 9 | 95059640 | 95087918 | -1 | protein_coding | nucleolar protein 8 |
| **IRAK4** | ENSG00000198001 | 147.9907047 | 0.926267808 | 0.352280027 | 0.002498771 | 0.008219697 | 51135 | 12 | 44152747 | 44183346 | 1 | protein_coding | interleukin-1 receptor-associated kinase 4 |
| **MRPL42** | ENSG00000198015 | 533.8315018 | 0.460956683 | 0.183363336 | 0.008281097 | 0.023722588 | 28977 | 12 | 93861264 | 93897545 | 1 | protein_coding | mitochondrial ribosomal protein L42 |
| **ENTPD7** | ENSG00000198018 | 1266.694125 | -0.377810594 | 0.150912897 | 0.009876356 | 0.02770056 | 57089 | 10 | 101419263 | 101465997 | 1 | protein_coding | ectonucleoside triphosphate diphosphohydrolase 7 |
| **RPS4X** | ENSG00000198034 | 8719.106317 | -0.43810817 | 0.118497697 | 1.62E-04 | 7.1E-04 | 6191 | X | 71475529 | 71497150 | -1 | protein_coding | ribosomal protein S4, X-linked |
| **ZNF273** | ENSG00000198039 | 104.7090064 | 0.893237495 | 0.46971636 | 0.015265771 | 0.040378465 | 1019292 | 7 | 64330550 | 64391344 | 1 | protein_coding | zinc finger protein 273 |
| **MAK16** | ENSG00000198042 | 783.5356722 | 0.495294708 | 0.129365756 | 9.05E-05 | 4.17E-04 | 84549 | 8 | 33342268 | 33358778 | 1 | protein_coding | MAK16 homolog (S. cerevisiae) |
| **ZNF667** | ENSG00000198046 | 190.3106642 | -1.462885859 | 0.331887669 | 1.46E-06 | 9.14E-06 | 63934 | 19 | 56950694 | 56989434 | -1 | protein_coding | zinc finger protein 667 |
| **SIRPA** | ENSG00000198053 | 24843.02213 | 0.500484942 | 0.080869352 | 8.26E-10 | 8E-09 | 140885 | 20 | 1875154 | 1920543 | 1 | protein_coding | signal-regulatory protein alpha |
| **GRK6** | ENSG00000198055 | 1720.676501 | -1.18197839 | 0.164706497 | 1.65E-13 | 2.23E-12 | 2870 | 5 | 176830205 | 176869902 | 1 | protein_coding | G protein-coupled receptor kinase 6 |
| **PRIM1** | ENSG00000198056 | 137.8575946 | 0.883592309 | 0.331565599 | 0.002409709 | 0.007960998 | 5557 | 12 | 57125380 | 57146157 | -1 | protein_coding | primase, DNA, polypeptide 1 (49kDa) |
| **CD2AP** | ENSG00000198087 | 1926.025864 | 1.138490339 | 0.134370767 | 5.49E-18 | 1.04E-16 | 23607 | 6 | 47445525 | 47594999 | 1 | protein_coding | CD2-associated protein |
| **SFI1** | ENSG00000198089 | 426.8333829 | 0.900770005 | 0.222827247 | 1.73E-05 | 9.12E-05 | 9814 | 22 | 31884674 | 32014572 | 1 | protein_coding | Sfi1 homolog, spindle assembly associated (yeast) |
| **ZNF248** | ENSG00000198105 | 180.4128375 | -1.024584158 | 0.285997174 | 8.98E-05 | 4.15E-04 | 57209 | 10 | 38091751 | 38147034 | -1 | protein_coding | zinc finger protein 248 |
| **ZNF544** | ENSG00000198131 | 509.5351065 | -0.884100233 | 0.203446154 | 4.9E-06 | 2.84E-05 | 27300 | 19 | 58739960 | 58788815 | 1 | protein_coding | zinc finger protein 544 |
| **TMEM229B** | ENSG00000198133 | 235.8891314 | 1.707347897 | 0.346455274 | 9.76E-08 | 7.25E-07 | 161145 | 14 | 67913801 | 68000456 | -1 | protein_coding | transmembrane protein 229B |
| **ZNF770** | ENSG00000198146 | 1587.47047 | 0.637624857 | 0.153451205 | 1.77E-05 | 9.32E-05 | 54989 | 15 | 35270542 | 35280488 | -1 | protein_coding | zinc finger protein 770 |
| **MIER1** | ENSG00000198160 | 1876.974869 | 1.289392617 | 0.115537838 | 1.32E-29 | 4.98E-28 | 57708 | 1 | 67390578 | 67454302 | 1 | protein_coding | mesoderm induction early response 1, transcriptional regulator |
| **MAN1A2** | ENSG00000198162 | 1812.701922 | -0.652913333 | 0.166668971 | 4.68E-05 | 2.27E-04 | 10905 | 1 | 117910071 | 118071494 | 1 | protein_coding | mannosidase, alpha, class 1A, member 2 |
| **TFDP1** | ENSG00000198176 | 1085.162748 | 0.643925479 | 0.144983159 | 4.77E-06 | 2.77E-05 | 7027 | 13 | 114239013 | 114295785 | 1 | protein_coding | transcription factor Dp-1 |
| **HSD17B11** | ENSG00000198189 | 1645.200752 | 0.521163141 | 0.128106397 | 3.23E-05 | 1.62E-04 | 51170 | 4 | 88257762 | 88312538 | -1 | protein_coding | hydroxysteroid (17-beta) dehydrogenase 11 |
| **SULT1C2** | ENSG00000198203 | 1911.470127 | 0.632146448 | 0.134911002 | 1.22E-06 | 7.74E-06 | 6819 | 2 | 108905095 | 108926371 | 1 | protein_coding | sulfotransferase family, cytosolic, 1C, member 2 |
| **ZXDA** | ENSG00000198205 | 140.7123093 | 0.673064681 | 0.286576599 | 0.008964927 | 0.025443078 | 7789 | X | 57931864 | 57937067 | -1 | protein_coding | zinc finger, X-linked, duplicated A |
| **RPL23A** | ENSG00000198242 | 2785.650642 | -0.32890661 | 0.137654928 | 0.013549537 | 0.036406044 | 26809 | 17 | 27046411 | 27051377 | 1 | protein_coding | ribosomal protein L23a |
| **SLC29A3** | ENSG00000198246 | 3656.461879 | -0.773000625 | 0.137787178 | 8.55E-09 | 7.34E-08 | 55315 | 10 | 73079015 | 73123142 | 1 | protein_coding | solute carrier family 29 (equilibrative nucleoside transporter), member 3 |
| **STYX** | ENSG00000198252 | 1293.793236 | 0.642433635 | 0.155458498 | 2E-05 | 1.04E-04 | 6815 | 14 | 53196898 | 53241716 | 1 | protein_coding | serine/threonine/tyrosine interacting protein |
| **HELZ** | ENSG00000198265 | 2672.064794 | 0.292929991 | 0.114592548 | 0.009043946 | 0.025648288 | 9931 | 17 | 65066554 | 65242105 | -1 | protein_coding | helicase with zinc finger |
| **PEG3** | ENSG00000198300 | 101.7582386 | -3.076452238 | 0.58112846 | 5.33E-09 | 4.7E-08 | 5178 | 19 | 57321445 | 57352096 | -1 | protein_coding | paternally expressed 3 |
| **FAM109A** | ENSG00000198324 | 80.42807489 | -2.950118379 | 0.717646307 | 2.02E-06 | 1.25E-05 | 144717 | 12 | 111798455 | 111806925 | -1 | protein_coding | family with sequence similarity 109, member A |
| **HYLS1** | ENSG00000198331 | 69.39013464 | 1.145987485 | 0.477579869 | 0.003037201 | 0.009813434 | 219844 | 11 | 125753509 | 125770543 | 1 | protein_coding | hydrolethalus syndrome 1 |
| **PIM3** | ENSG00000198355 | 841.4852232 | 1.471484092 | 0.175031252 | 6.69E-18 | 1.25E-16 | 415116 | 22 | 50354161 | 50357728 | 1 | protein_coding | pim-3 oncogene |
| **ASNA1** | ENSG00000198356 | 876.9818021 | -0.600548875 | 0.138012374 | 7.97E-06 | 4.45E-05 | 439 | 19 | 12847953 | 12859137 | 1 | protein_coding | arsA arsenite transporter, ATP-binding, homolog 1 (bacterial) |
| **ASPH** | ENSG00000198363 | 4438.412348 | -0.403318163 | 0.112513985 | 2.63E-04 | 0.001093228 | 444 | 8 | 62413116 | 62627155 | -1 | protein_coding | aspartate beta-hydroxylase |
| **SPRED2** | ENSG00000198369 | 990.7690142 | -0.718057267 | 0.160241584 | 3.54E-06 | 2.09E-05 | 200734 | 2 | 65537985 | 65659771 | -1 | protein_coding | sprouty-related, EVH1 domain containing 2 |
| **WWP2** | ENSG00000198373 | 2106.525327 | -0.348529726 | 0.117729158 | 0.002396015 | 0.00792604 | 11060 | 16 | 69796209 | 69975644 | 1 | protein_coding | WW domain containing E3 ubiquitin protein ligase 2 |
| **GFPT1** | ENSG00000198380 | 2389.457328 | 0.259319036 | 0.091727983 | 0.004068651 | 0.012697813 | 2673 | 2 | 69546905 | 69614382 | -1 | protein_coding | glutamine--fructose-6-phosphate transaminase 1 |
| **UVRAG** | ENSG00000198382 | 1439.8181 | 0.747179929 | 0.141291729 | 5.63E-08 | 4.34E-07 | 7405 | 11 | 75526212 | 75854239 | 1 | protein_coding | UV radiation resistance associated |
| **ITSN2** | ENSG00000198399 | 3958.915115 | 0.346768071 | 0.096923365 | 2.68E-04 | 0.001113441 | 50618 | 2 | 24425733 | 24583583 | -1 | protein_coding | intersectin 2 |
| **BZW1P2** | ENSG00000198406 | 1280.7393 | -0.401920899 | 0.149885337 | 0.005654867 | 0.01696238 | NA | 3 | 116364592 | 116365574 | -1 | pseudogene | basic leucine zipper and W2 domains 1 pseudogene 2 |
| **MGEA5** | ENSG00000198408 | 3357.517293 | 0.556497292 | 0.101508818 | 2.66E-08 | 2.14E-07 | 10724 | 10 | 103544200 | 103578696 | -1 | protein_coding | meningioma expressed antigen 5 (hyaluronidase) |
| **MT1F** | ENSG00000198417 | 127.5954951 | 1.151941987 | 0.322007829 | 7.12E-05 | 3.34E-04 | 4494 | 16 | 56691606 | 56694610 | 1 | protein_coding | metallothionein 1F |
| **FAM115A** | ENSG00000198420 | 1059.695531 | 0.409877036 | 0.118897908 | 4.37E-04 | 0.001735236 | 1019300 | 7 | 143548468 | 143599291 | -1 | protein_coding | family with sequence similarity 115, member A |
| **TXNRD1** | ENSG00000198431 | 4446.165401 | 0.417261998 | 0.182895002 | 0.016607052 | 0.043385569 | 7296 | 12 | 104609557 | 104744061 | 1 | protein_coding | thioredoxin reductase 1 |
| **ZXDB** | ENSG00000198455 | 234.0667304 | 0.721301303 | 0.242974535 | 0.001280751 | 0.004543575 | 158586 | X | 57618269 | 57623906 | 1 | protein_coding | zinc finger, X-linked, duplicated B |
| **TPM2** | ENSG00000198467 | 118.7727759 | -1.713665456 | 0.319521652 | 1.01E-08 | 8.59E-08 | 7169 | 9 | 35681989 | 35691017 | -1 | protein_coding | tropomyosin 2 (beta) |
| **ZNF808** | ENSG00000198482 | 1016.481379 | -0.534986696 | 0.132116766 | 3.24E-05 | 1.62E-04 | 388558 | 19 | 53030905 | 53067717 | 1 | protein_coding | zinc finger protein 808 |
| **YTHDF2** | ENSG00000198492 | 2153.125707 | 0.303183415 | 0.106524607 | 0.003819056 | 0.012004656 | 51441 | 1 | 29063133 | 29096287 | 1 | protein_coding | YTH domain family, member 2 |
| **HLA-DRB5** | ENSG00000198502 | 1035.273394 | -0.566631374 | 0.192125528 | 0.001837977 | 0.006271445 | 3127 | 6 | 32485120 | 32498064 | -1 | protein_coding | major histocompatibility complex, class II, DR beta 5 |
| **ATL1** | ENSG00000198513 | 44.44828727 | 1.447827996 | 0.98729578 | 0.010528912 | 0.029230401 | 51062 | 14 | 50999227 | 51099786 | 1 | protein_coding | atlastin GTPase 1 |
| **C1orf228** | ENSG00000198520 | 235.2040513 | -1.325928162 | 0.347991469 | 2.34E-05 | 1.2E-04 | 339541 | 1 | 45140364 | 45191263 | 1 | protein_coding | chromosome 1 open reading frame 228 |
| **ZNF43** | ENSG00000198521 | 421.1797037 | 0.858619387 | 0.219999402 | 3.33E-05 | 1.66E-04 | 7594 | 19 | 21987752 | 22034927 | -1 | protein_coding | zinc finger protein 43 |
| **WDHD1** | ENSG00000198554 | 192.2535738 | 1.019581987 | 0.368718542 | 0.00138442 | 0.004866643 | 11169 | 14 | 55405668 | 55493823 | -1 | protein_coding | WD repeat and HMG-box DNA binding protein 1 |
| **NUDT16** | ENSG00000198585 | 1215.038553 | 0.299315825 | 0.113769484 | 0.007243251 | 0.021109573 | 131870 | 3 | 131100515 | 131107674 | 1 | protein_coding | nudix (nucleoside diphosphate linked moiety X)-type motif 16 |

| | | | | | | | | | | | | | |
|---|---|---|---|---|---|---|---|---|---|---|---|---|---|
| **BAZ1A** | ENSG00000198604 | 3812.766662 | 1.71382162 | 0.10084737 | 1.22E-65 | 1.95E-63 | 11177 | 14 | 35221937 | 35344853 | -1 | protein_coding | bromodomain adjacent to zinc finger domain, 1A |
| **COPS8** | ENSG00000198612 | 543.5736375 | -0.575156003 | 0.169555575 | 4.18E-04 | 0.00166905 | 10920 | 2 | 237993955 | 238009109 | 1 | protein_coding | COP9 signalosome subunit 8 |
| **MDM4** | ENSG00000198625 | 1347.523489 | 0.693169712 | 0.117930235 | 2.06E-09 | 1.9E-08 | 4194 | 1 | 204485511 | 204542871 | 1 | protein_coding | Mdm4 p53 binding protein homolog (mouse) |
| **NCOA6** | ENSG00000198646 | 1471.599922 | 0.29326162 | 0.121653267 | 0.015577254 | 0.041081479 | 23054 | 20 | 33284722 | 33413452 | -1 | protein_coding | nuclear receptor coactivator 6 |
| **CALM1** | ENSG00000198668 | 8114.580122 | 0.263724283 | 0.107363752 | 0.012109062 | 0.032999784 | 808 | 14 | 90862846 | 90874605 | 1 | protein_coding | calmodulin 1 (phosphorylase kinase, delta) |
| **FAM19A2** | ENSG00000198673 | 45.84365408 | 1.616231448 | 0.531134113 | 2.5E-04 | 0.001046622 | 338811 | 12 | 62102040 | 62672931 | -1 | protein_coding | family with sequence similarity 19 (chemokine (C-C motif)-like), member A2 |
| **PAPSS2** | ENSG00000198682 | 308.9997233 | -0.538315574 | 0.230208403 | 0.011656526 | 0.031937376 | 9060 | 10 | 89419370 | 89507462 | 1 | protein_coding | 3'-phosphoadenosine 5'-phosphosulfate synthase 2 |
| **FAN1** | ENSG00000198690 | 516.9158074 | 0.385341524 | 0.164799217 | 0.015043483 | 0.039838772 | 22909 | 15 | 31196055 | 31235311 | 1 | protein_coding | FANCD2/FANCI-associated nuclease 1 |
| **C1orf85** | ENSG00000198715 | 2466.015716 | -0.590016828 | 0.109651683 | 4.44E-08 | 3.46E-07 | 112770 | 1 | 156259880 | 156265463 | -1 | protein_coding | chromosome 1 open reading frame 85 |
| **FAM179B** | ENSG00000198718 | 913.2002425 | 0.771229019 | 0.223064877 | 2.29E-04 | 9.65E-04 | 23116 | 14 | 45431411 | 45543634 | 1 | protein_coding | family with sequence similarity 179, member B |
| **DLL1** | ENSG00000198719 | 11.55973177 | 1.737875722 | 1.163787964 | 0.007502801 | 0.021781204 | 28514 | 6 | 170591294 | 170599561 | -1 | protein_coding | delta-like 1 (Drosophila) |
| **UNC13B** | ENSG00000198722 | 1468.372343 | -0.765541086 | 0.151725899 | 2.39E-07 | 1.69E-06 | 10497 | 9 | 35161999 | 35405335 | 1 | protein_coding | unc-13 homolog B (C. elegans) |
| **MT-CYB** | ENSG00000198727 | 76133.59876 | -0.959602051 | 0.18564148 | 7.23E-08 | 5.48E-07 | 4519 | MT | 14747 | 15887 | 1 | protein_coding | mitochondrially encoded cytochrome b |
| **LDB1** | ENSG00000198728 | 1129.44637 | -0.418210448 | 0.133541264 | 0.0013199 | 0.004661898 | 8861 | 10 | 103867317 | 103880210 | -1 | protein_coding | LIM domain binding 1 |
| **MSRB1** | ENSG00000198736 | 451.0988796 | 0.594054469 | 0.222195953 | 0.004170894 | 0.012982439 | 51734 | 16 | 1988211 | 1993327 | -1 | protein_coding | methionine sulfoxide reductase B1 |
| **ZNF652** | ENSG00000198740 | 1364.014793 | -0.654846365 | 0.11511769 | 7E-09 | 6.07E-08 | 22834 | 17 | 47366568 | 47439835 | -1 | protein_coding | zinc finger protein 652 |
| **SMURF1** | ENSG00000198742 | 757.0358078 | 0.531385583 | 0.148681489 | 2.18E-04 | 9.24E-04 | 57154 | 7 | 98625061 | 98741723 | -1 | protein_coding | SMAD specific E3 ubiquitin protein ligase 1 |
| **SLC5A3** | ENSG00000198743 | 16486.15749 | -0.95032058 | 0.150120853 | 7.81E-11 | 8.47E-10 | 6526 | 21 | 35445870 | 35478559 | 1 | protein_coding | solute carrier family 5 (sodium/myo-inositol cotransporter), member 3 |
| **RPL10A** | ENSG00000198755 | 3006.065289 | -0.888655536 | 0.132514069 | 6.95E-12 | 8.3E-11 | 4736 | 6 | 35436185 | 35438562 | 1 | protein_coding | ribosomal protein L10a |
| **EGFL6** | ENSG00000198759 | 13.81683176 | 2.572484629 | 1.497473355 | 0.002760913 | 0.008988384 | 25975 | X | 13587724 | 13651694 | 1 | protein_coding | EGF-like-domain, multiple 6 |
| **MT-ND2** | ENSG00000198763 | 51067.93921 | -0.811425754 | 0.197044396 | 1.49E-05 | 7.99E-05 | 4536 | MT | 4470 | 5511 | 1 | protein_coding | mitochondrially encoded NADH dehydrogenase 2 |
| **GRIN3A** | ENSG00000198785 | 449.8779571 | 1.370961882 | 0.1888051 | 6.53E-14 | 9.21E-13 | 116443 | 9 | 104331635 | 104500862 | -1 | protein_coding | glutamate receptor, ionotropic, N-methyl-D-aspartate 3A |
| **MT-ND5** | ENSG00000198786 | 73734.80253 | -0.851831743 | 0.186006744 | 1.71E-06 | 1.06E-05 | 4540 | MT | 12337 | 14148 | 1 | protein_coding | mitochondrially encoded NADH dehydrogenase 5 |
| **MTOR** | ENSG00000198793 | 2493.546706 | -0.51818783 | 0.121405342 | 1.31E-05 | 7.1E-05 | 2475 | 1 | 11166592 | 11322564 | -1 | protein_coding | mechanistic target of rapamycin (serine/threonine kinase) |
| **PNP** | ENSG00000198805 | 1677.142829 | -0.405198988 | 0.12599108 | 0.001173269 | 0.004205566 | 4860 | 14 | 20937113 | 20945253 | 1 | protein_coding | purine nucleoside phosphorylase |
| **GK** | ENSG00000198814 | 3365.296804 | 1.571989614 | 0.178072414 | 1.41E-19 | 3E-18 | 2710 | X | 30671476 | 30748725 | 1 | protein_coding | glycerol kinase |
| **FOXJ3** | ENSG00000198815 | 1339.574979 | 0.279427847 | 0.108607876 | 0.008415059 | 0.024047747 | 22887 | 1 | 42642210 | 42801548 | -1 | protein_coding | forkhead box J3 |
| **ZNF358** | ENSG00000198816 | 226.395601 | -1.089748068 | 0.34051541 | 3.03E-04 | 0.001246358 | 140467 | 19 | 7580178 | 7585912 | 1 | protein_coding | zinc finger protein 358 |
| **ARHGAP11A** | ENSG00000198826 | 348.2692517 | 0.605758935 | 0.210384096 | 0.002191905 | 0.007320562 | 9824 | 15 | 32907345 | 32932150 | 1 | protein_coding | Rho GTPase activating protein 11A |
| **OPA1** | ENSG00000198836 | 1904.49917 | -0.453775452 | 0.152176101 | 0.002007463 | 0.006786122 | 4976 | 3 | 193310933 | 193415612 | 1 | protein_coding | optic atrophy 1 (autosomal dominant) |
| **DENND4B** | ENSG00000198837 | 2451.002306 | -0.467389036 | 0.148815468 | 0.001189767 | 0.00425936 | 9909 | 1 | 153901977 | 153919172 | -1 | protein_coding | DENN/MADD domain containing 4B |
| **ZNF277** | ENSG00000198839 | 466.0847584 | 1.691438882 | 0.215077011 | 4.47E-16 | 7.28E-15 | 11179 | 7 | 111846643 | 111983151 | 1 | protein_coding | zinc finger protein 277 |
| **CD3E** | ENSG00000198851 | 1121.040886 | -0.459560699 | 0.176500981 | 0.006478607 | 0.019158775 | 916 | 11 | 118175260 | 118186890 | 1 | protein_coding | CD3e molecule, epsilon (CD3-TCR complex) |
| **FICD** | ENSG00000198855 | 702.4904716 | 0.57043779 | 0.168810129 | 4.38E-04 | 0.001740472 | 11153 | 12 | 108908962 | 108919614 | 1 | protein_coding | FIC domain containing |
| **MT-ND4** | ENSG00000198886 | 92737.9239 | -0.686836901 | 0.186253168 | 1.1E-04 | 5.01E-04 | 4538 | MT | 10760 | 12137 | 1 | protein_coding | mitochondrially encoded NADH dehydrogenase 4 |
| **SMC5** | ENSG00000198887 | 1039.315862 | 0.790303565 | 0.139906402 | 6.77E-09 | 5.89E-08 | 23137 | 9 | 72873937 | 72969804 | 1 | protein_coding | structural maintenance of chromosomes 5 |
| **CIPC** | ENSG00000198894 | 1197.47291 | -0.638085538 | 0.156333769 | 2.4E-05 | 1.23E-04 | 85457 | 14 | 77564440 | 77583630 | 1 | protein_coding | CLOCK-interacting pacemaker |
| **TOP 1.00** | ENSG00000198900 | 3709.648086 | 0.854081359 | 0.09132683 | 3.42E-21 | 8.25E-20 | 7150 | 20 | 39657458 | 39753127 | 1 | protein_coding | topoisomerase (DNA) I |
| **MAP3K3** | ENSG00000198909 | 3090.43684 | -1.304697588 | 0.101004333 | 4.51E-39 | 2.64E-37 | 4215 | 17 | 61699775 | 61773663 | 1 | protein_coding | mitogen-activated protein kinase kinase kinase 3 |
| **KIAA0753** | ENSG00000198920 | 515.42195 | -0.458559708 | 0.203311281 | 0.016750795 | 0.043693872 | 9851 | 17 | 6481468 | 6544247 | -1 | protein_coding | KIAA0753 |
| **ATG9A** | ENSG00000198925 | 1612.520963 | -0.500485421 | 0.113808685 | 7.68E-06 | 4.3E-05 | 79065 | 2 | 220074494 | 220094439 | -1 | protein_coding | autophagy related 9A |
| **APRT** | ENSG00000198931 | 453.8674993 | -0.684459499 | 0.18095918 | 7.88E-05 | 3.67E-04 | 353 | 16 | 88875747 | 88878352 | -1 | protein_coding | adenine phosphoribosyltransferase |
| **CCDC167** | ENSG00000198937 | 105.98496 | -1.112468885 | 0.343989348 | 2.71E-04 | 0.001123467 | 154467 | 6 | 37450696 | 37467698 | -1 | protein_coding | coiled-coil domain containing 167 |
| **L3MBTL3** | ENSG00000198945 | 239.0286309 | 0.627971459 | 0.277698372 | 0.012180894 | 0.033171883 | 84456 | 6 | 130334844 | 130462594 | 1 | protein_coding | l(3)mbt-like 3 (Drosophila) |
| **NAGA** | ENSG00000198951 | 2162.363656 | -0.870029706 | 0.123021605 | 6.11E-13 | 7.93E-12 | 4668 | 22 | 42454358 | 42466846 | -1 | protein_coding | N-acetylgalactosaminidase, alpha- |
| **KIAA1279** | ENSG00000198954 | 1007.275189 | 0.385809909 | 0.170658943 | 0.018307872 | 0.047104245 | 26128 | 10 | 70748487 | 70776738 | 1 | protein_coding | KIAA1279 |
| **TGM2** | ENSG00000198959 | 28478.95509 | -0.519353134 | 0.163997487 | 0.001004166 | 0.003660069 | 7052 | 20 | 36756863 | 36794980 | -1 | protein_coding | transglutaminase 2 |
| **RORB** | ENSG00000198963 | 9.76792324 | 2.712991969 | 1.55066797 | 0.002151267 | 0.007211629 | 6096 | 9 | 77112281 | 77308093 | 1 | protein_coding | RAR-related orphan receptor B |
| **SGMS1** | ENSG00000198964 | 1539.277549 | 0.505579969 | 0.160624809 | 0.001114343 | 0.004022936 | 259230 | 10 | 52065360 | 52384923 | -1 | protein_coding | sphingomyelin synthase 1 |
| **INF2** | ENSG00000203485 | 1941.7149 | -0.585379444 | 0.139241384 | 1.52E-05 | 8.13E-05 | 64423 | 14 | 105155943 | 105185942 | 1 | protein_coding | inverted formin, FH2 and WH2 domain containing |
| **AC083799.1** | ENSG00000203644 | 171.4922388 | 0.793774907 | 0.283243573 | 0.001923278 | 0.006536236 | NA | 3 | 129565893 | 129566800 | -1 | pseudogene | |
| **C1orf95** | ENSG00000203685 | 72.82901312 | -1.798865681 | 0.496825327 | 3.11E-05 | 1.56E-04 | 375057 | 1 | 226736501 | 226796915 | 1 | protein_coding | chromosome 1 open reading frame 95 |
| **CR1** | ENSG00000203710 | 2216.156484 | -2.665058236 | 0.216061922 | 3.84E-36 | 2.01E-34 | 1378 | 1 | 207669492 | 207813992 | 1 | protein_coding | complement component (3b/4b) receptor 1 (Knops blood group) |
| **FCGR3A** | ENSG00000203747 | 17652.46185 | -1.190421701 | 0.118968684 | 2.78E-24 | 7.99E-23 | 2214 | 1 | 161511549 | 161600917 | -1 | protein_coding | Fc fragment of IgG, low affinity IIIa, receptor (CD16a) |
| **CENPW** | ENSG00000203760 | 75.90254409 | -1.070757884 | 0.540946343 | 0.008294801 | 0.023751033 | 387103 | 6 | 126661320 | 126670021 | 1 | protein_coding | centromere protein W |
| **DDO** | ENSG00000203797 | 291.4828238 | 1.02321098 | 0.266877827 | 3.36E-05 | 1.68E-04 | 8528 | 6 | 110712974 | 110736765 | -1 | protein_coding | D-aspartate oxidase |
| **CCDC162P** | ENSG00000203799 | 101.9350792 | -1.853901817 | 0.517144944 | 3.18E-05 | 1.6E-04 | 221262 | 6 | 109612231 | 109676266 | 1 | pseudogene | coiled-coil domain containing 162, pseudogene |
| **HIST2H3C** | ENSG00000203811 | 107.6346315 | 0.926568181 | 0.541287027 | 0.019318447 | 0.04926478 | 653604 | 1 | 149811110 | 149812765 | -1 | protein_coding | histone cluster 2, H3c |
| **HIST2H2BF** | ENSG00000203814 | 262.7510728 | 0.588150716 | 0.280451104 | 0.019346638 | 0.049311987 | 440689 | 1 | 149754245 | 149783928 | -1 | protein_coding | histone cluster 2, H2bf |
| **SAMD13** | ENSG00000203943 | 37.83980492 | -2.3673019 | 0.938809554 | 8.59E-04 | 0.003189817 | 148418 | 1 | 84764049 | 84855640 | 1 | protein_coding | sterile alpha motif domain containing 13 |
| **NHSL2** | ENSG00000204131 | 27.16273049 | -1.189762576 | 0.989189327 | 0.019606479 | 0.049874472 | 340527 | X | 71130938 | 71363424 | 1 | protein_coding | NHS-like 2 |
| **GGTA1P** | ENSG00000204136 | 116.5785617 | -2.952829008 | 0.439042917 | 1.02E-12 | 1.29E-11 | 2681 | 9 | 124207269 | 124262306 | -1 | pseudogene | glycoprotein, alpha-galactosyltransferase 1 pseudogene |
| **PHACTR4** | ENSG00000204138 | 1522.152171 | 1.330364589 | 0.107045088 | 3.69E-36 | 1.94E-34 | 65979 | 1 | 28696114 | 28826881 | 1 | protein_coding | phosphatase and actin regulator 4 |
| **C10orf128** | ENSG00000204161 | 1466.859484 | -2.222879644 | 0.165286742 | 3.18E-42 | 2.16E-40 | 170371 | 10 | 50362773 | 50396630 | -1 | protein_coding | chromosome 10 open reading frame 128 |
| **ZDBF2** | ENSG00000204186 | 458.0283549 | 0.457777075 | 0.183576844 | 0.008885442 | 0.025245623 | 57683 | 2 | 207139387 | 207179148 | 1 | protein_coding | zinc finger, DBF-type containing 2 |
| **DAXX** | ENSG00000204209 | 389.6671408 | 1.084919718 | 0.262802053 | 9.06E-06 | 5.02E-05 | 1616 | 6 | 33286335 | 33297046 | -1 | protein_coding | death-domain associated protein |
| **BMPR2** | ENSG00000204217 | 3947.481139 | 0.811135675 | 0.102370334 | 9.58E-16 | 1.53E-14 | 659 | 2 | 203241659 | 203432474 | 1 | protein_coding | bone morphogenetic protein receptor, type II (serine/threonine kinase) |
| **PFDN6** | ENSG00000204220 | 24.01741357 | 7.196477195 | 2.337009518 | 1.17E-06 | 7.43E-06 | 1024655 | 6 | 33257079 | 33266178 | 1 | protein_coding | prefoldin subunit 6 |
| **BRD2** | ENSG00000204256 | 838.1544475 | 0.560776359 | 0.155457775 | 1.92E-04 | 8.26E-04 | 6046 | 6 | 32936437 | 32949282 | 1 | protein_coding | bromodomain containing 2 |
| **COL5A2** | ENSG00000204262 | 119.2440191 | -1.1207362 | 0.404945781 | 0.001151697 | 0.004137608 | 1290 | 2 | 189896622 | 190044605 | -1 | protein_coding | collagen, type V, alpha 2 |
| **COL15A1** | ENSG00000204291 | 27.35715951 | -1.655813847 | 0.803127161 | 0.003321099 | 0.010622928 | 1306 | 9 | 101705461 | 101833069 | 1 | protein_coding | collagen, type XV, alpha 1 |
| **AGPAT1** | ENSG00000204310 | 355.1234289 | -0.829903467 | 0.21646733 | 4.77E-05 | 2.31E-04 | 10554 | 6 | 32135989 | 32145873 | -1 | protein_coding | 1-acylglycerol-3-phosphate O-acyltransferase 1 |
| **DFNB59** | ENSG00000204311 | 11.46738664 | 5.25125686 | 2.83833206 | 7.14E-04 | 0.002696506 | 494513 | 2 | 179316163 | 179326117 | 1 | protein_coding | deafness, autosomal recessive 59 |
| **NELFE** | ENSG00000204356 | 214.6199279 | -1.224100072 | 0.400897049 | 3.62E-04 | 0.001463784 | 1003022 | 6 | 31919864 | 31926887 | -1 | protein_coding | negative elongation factor complex member E |
| **SDHD** | ENSG00000204370 | 2211.940907 | 0.719251151 | 0.123900293 | 3.03E-09 | 2.74E-08 | 6392 | 11 | 111957497 | 111990353 | 1 | protein_coding | succinate dehydrogenase complex, subunit D, integral membrane protein |
| **LAYN** | ENSG00000204381 | 264.2189532 | -1.080724864 | 0.320193901 | 1.73E-04 | 7.53E-04 | 143903 | 11 | 111411005 | 111432264 | 1 | protein_coding | layilin |
| **C6orf48** | ENSG00000204387 | 73.39467511 | -0.139022881 | 0.656282342 | 0.017630149 | 0.045675805 | 50854 | 6 | 31802385 | 31807541 | 1 | protein_coding | chromosome 6 open reading frame 48 |
| **CARD16** | ENSG00000204397 | 332.5019748 | 1.369035257 | 0.26281362 | 3.35E-08 | 2.66E-07 | 114769 | 11 | 104912053 | 104972158 | -1 | protein_coding | caspase recruitment domain family, member 16 |
| **MBD5** | ENSG00000204406 | 426.1282299 | 0.64386052 | 0.204972664 | 8.74E-04 | 0.003234321 | 55777 | 2 | 148778580 | 149275805 | 1 | protein_coding | methyl-CpG binding domain protein 5 |
| **C6orf47** | ENSG00000204439 | 380.4876702 | 0.811316599 | 0.224801016 | 1.1E-04 | 4.99E-04 | 57827 | 6 | 31626075 | 31628549 | -1 | protein_coding | chromosome 6 open reading frame 47 |
| **TNF** | ENSG00000204490 | 261.6187874 | 1.679088753 | 0.377960192 | 1.03E-06 | 6.61E-06 | 7124 | HSC | 31530756 | 31533525 | 1 | protein_coding | tumor necrosis factor |
| **LILRB3** | ENSG00000204577 | 181.1442773 | 0.566302843 | 0.245285274 | 0.012152182 | 0.033105489 | 1019298 | 19 | 54720147 | 54746602 | -1 | protein_coding | leukocyte immunoglobulin-like receptor, subfamily B (with TM and ITIM domains), member 3 |
| **GNB2L1** | ENSG00000204628 | 6889.762696 | -0.500594567 | 0.123134806 | 3.57E-05 | 1.77E-04 | 619571 | 5 | 180663909 | 180675096 | -1 | protein_coding | guanine nucleotide binding protein (G protein), beta polypeptide 2-like 1 |
| **TBC1D8** | ENSG00000204634 | 2238.75268 | -0.459342498 | 0.133780587 | 4.33E-04 | 0.001721626 | 11138 | 2 | 101624079 | 101869328 | -1 | protein_coding | TBC1 domain family, member 8 (with GRAM domain) |
| **HLA-F** | ENSG00000204642 | 2413.602333 | 0.788241001 | 0.159075214 | 3.21E-07 | 2.23E-06 | 3134 | 6 | 29690552 | 29706305 | 1 | protein_coding | major histocompatibility complex, class I, F |
| **AKT1S1** | ENSG00000204673 | 1132.434962 | -0.300249989 | 0.124296742 | 0.014799436 | 0.03927414 | 84335 | 19 | 50372295 | 50381716 | -1 | protein_coding | AKT1 substrate 1 (proline-rich) |
| **DCTN1** | ENSG00000204843 | 2477.288926 | -0.685603797 | 0.099349283 | 2.73E-12 | 3.36E-11 | 1639 | 2 | 74588281 | 74619214 | -1 | protein_coding | dynactin 1 |
| **ZBTB48** | ENSG00000204859 | 489.6894231 | -0.758207157 | 0.214475608 | 1.79E-04 | 7.75E-04 | 3104 | 1 | 6640061 | 6649340 | 1 | protein_coding | zinc finger and BTB domain containing 48 |
| **MZT1** | ENSG00000204899 | 273.8469782 | 0.987943441 | 0.213755434 | 1.14E-06 | 7.24E-06 | 440145 | 13 | 73282495 | 73301825 | -1 | protein_coding | mitotic spindle organizing protein 1 |

| | | | | | | | | | | | | | |
|---|---|---|---|---|---|---|---|---|---|---|---|---|---|
| **FBXO48** | ENSG00000204923 | 233.1112354 | 0.659523695 | 0.226994885 | 0.001852245 | 0.006313082 | 554251 | 2 | 68686551 | 68694390 | -1 | protein_coding | F-box protein 48 |
| **SLFN12L** | ENSG00000205045 | 90.76792523 | 1.376650354 | 0.463991432 | 4.13E-04 | 0.001651162 | 1005067 | 17 | 33791268 | 33864880 | -1 | protein_coding | schlafen family member 12-like |
| **PSENEN** | ENSG00000205155 | 687.3459349 | -0.406794473 | 0.147324946 | 0.004090417 | 0.012747522 | 55851 | 19 | 36236015 | 36237911 | 1 | protein_coding | presenilin enhancer gamma secretase subunit |
| **C4orf46** | ENSG00000205208 | 91.06983322 | 0.905661223 | 0.433579656 | 0.009878468 | 0.027701399 | 201725 | 4 | 159587831 | 159593407 | -1 | protein_coding | chromosome 4 open reading frame 46 |
| **LGR4** | ENSG00000205213 | 301.0005573 | -2.245785557 | 0.348274262 | 7.34E-12 | 8.74E-11 | 55366 | 11 | 27387508 | 27494322 | -1 | protein_coding | leucine-rich repeat containing G protein-coupled receptor 4 |
| **PSMB10** | ENSG00000205220 | 1195.880937 | 0.552231286 | 0.199913062 | 0.003421425 | 0.010907308 | 5699 | 16 | 67968405 | 67970990 | -1 | protein_coding | proteasome (prosome, macropain) subunit, beta type, 10 |
| **VIT** | ENSG00000205221 | 443.0618906 | -0.744482393 | 0.252816868 | 0.001341591 | 0.004731948 | 5212 | 2 | 36923833 | 37041935 | 1 | protein_coding | vitrin |
| **RP11-514P8.** | ENSG00000205236 | 201.1745128 | -0.703480218 | 0.356152205 | 0.019623644 | 0.049909826 | NA | 7 | 102222970 | 102283316 | -1 | protein_coding | Uncharacterized protein |
| **PRR13** | ENSG00000205352 | 4572.844449 | 0.636314051 | 0.100821942 | 1.09E-10 | 1.16E-09 | 54458 | 12 | 53835389 | 53840429 | 1 | protein_coding | proline rich 13 |
| **TECPR1** | ENSG00000205356 | 1422.349235 | -0.548464668 | 0.20209733 | 0.004029268 | 0.012590328 | 25851 | 7 | 97843936 | 97881563 | -1 | protein_coding | tectonin beta-propeller repeat containing 1 |
| **MT1H** | ENSG00000205358 | 179.9618489 | 0.941269933 | 0.291372547 | 3.67E-04 | 0.00148564 | 4496 | 16 | 56703726 | 56705041 | 1 | protein_coding | metallothionein 1H |
| **MT1M** | ENSG00000205364 | 59.22433525 | 2.981189814 | 0.547377137 | 3.17E-09 | 2.86E-08 | 4499 | 16 | 56666145 | 56667898 | 1 | protein_coding | metallothionein 1M |
| **SAMD9** | ENSG00000205413 | 8392.233583 | 4.036268249 | 0.126473312 | 7.99E-225 | 7.63E-222 | 54809 | 7 | 92728829 | 92747336 | -1 | protein_coding | sterile alpha motif domain containing 9 |
| **CNEP1R1** | ENSG00000205423 | 929.8529545 | 0.634308048 | 0.151861704 | 1.62E-05 | 8.58E-05 | 255919 | 16 | 50058321 | 50070999 | 1 | protein_coding | CTD nuclear envelope phosphatase 1 regulatory subunit 1 |
| **CCDC85C** | ENSG00000205476 | 59.84135935 | -1.608064158 | 0.577741331 | 6.06E-04 | 0.002323726 | 317762 | 14 | 99977603 | 100070363 | -1 | protein_coding | coiled-coil domain containing 85C |
| **TMEM256** | ENSG00000205544 | 103.4975361 | -0.699956044 | 0.337613943 | 0.016363262 | 0.042836541 | 254863 | 17 | 7306294 | 7307456 | -1 | protein_coding | transmembrane protein 256 |
| **CPT1B** | ENSG00000205560 | 304.9599366 | 0.885378367 | 0.284494425 | 6.1E-04 | 0.002335604 | 1375 | 22 | 51007290 | 51017899 | -1 | protein_coding | carnitine palmitoyltransferase 1B (muscle) |
| **HMGN1** | ENSG00000205581 | 2813.433835 | 0.619128987 | 0.123182493 | 2.81E-07 | 1.96E-06 | 1019277 | 21 | 40714241 | 40721573 | -1 | protein_coding | high mobility group nucleosome binding domain 1 |
| **LIN52** | ENSG00000205659 | 202.6100536 | 1.025819171 | 0.27747776 | 5.8E-05 | 2.77E-04 | 91750 | 14 | 74551499 | 74667936 | 1 | protein_coding | lin-52 homolog (C. elegans) |
| **MANSC4** | ENSG00000205693 | 10.85733739 | -4.344294182 | 2.791426256 | 0.001741025 | 0.005967277 | 1002872 | 12 | 27915671 | 27924209 | -1 | protein_coding | MANSC domain containing 4 |
| **ITPRIPL2** | ENSG00000205730 | 3023.248405 | 0.986856553 | 0.108371568 | 2.67E-20 | 6.01E-19 | 162073 | 16 | 19125254 | 19132946 | 1 | protein_coding | inositol 1,4,5-trisphosphate receptor interacting protein-like 2 |
| **C5orf51** | ENSG00000205765 | 1727.520265 | 1.295268031 | 0.127295808 | 4.54E-25 | 1.36E-23 | 285636 | 5 | 41904290 | 41921738 | 1 | protein_coding | chromosome 5 open reading frame 51 |
| **PPAPDC2** | ENSG00000205808 | 212.1913686 | 1.176279753 | 0.2480497 | 4.56E-07 | 3.1E-06 | 403313 | 9 | 4662298 | 4665256 | 1 | protein_coding | phosphatidic acid phosphatase type 2 domain containing 2 |
| **CLEC6A** | ENSG00000205846 | 74.93190284 | 1.190492736 | 0.426217702 | 9.56E-04 | 0.003498522 | 93978 | 12 | 8608522 | 8630926 | 1 | protein_coding | C-type lectin domain family 6, member A |
| **ZNF316** | ENSG00000205903 | 794.6159961 | -0.794647759 | 0.153252053 | 9.34E-08 | 6.95E-07 | NA | 7 | 6676953 | 6694242 | 1 | pseudogene | zinc finger protein 316 |
| **C21orf49** | ENSG00000205930 | 24.70122137 | -2.433080656 | 1.20889031 | 0.001884723 | 0.006416622 | NA | 21 | 34144411 | 34266043 | 1 | protein_coding | chromosome 21 open reading frame 49 |
| **DNAJC19** | ENSG00000205981 | 265.3017719 | 0.620451872 | 0.241589551 | 0.005499424 | 0.016567652 | 131118 | 3 | 180701497 | 180707562 | -1 | protein_coding | DnaJ (Hsp40) homolog, subfamily C, member 19 |
| **GOLGA8O** | ENSG00000206127 | 71.17189649 | -1.006315696 | 0.588544583 | 0.015991478 | 0.042007179 | 728047 | 15 | 32737307 | 32747835 | -1 | protein_coding | golgin A8 family, member O |
| **ATP10A** | ENSG00000206190 | 149.5263755 | 1.715404051 | 0.426508575 | 6.07E-06 | 3.45E-05 | 57194 | 15 | 25922420 | 26110317 | -1 | protein_coding | ATPase, class V, type 10A |
| **TAPBP** | ENSG00000206208 | 7104.317126 | 0.626409137 | 0.096653017 | 7.31E-11 | 7.95E-10 | 6892 | HSC | 33437818 | 33452509 | -1 | protein_coding | TAP binding protein (tapasin) |
| **DAXX** | ENSG00000206279 | 235.3650487 | 1.613030853 | 0.301486524 | 1.14E-08 | 9.59E-08 | 1616 | HSC | 33215186 | 33225897 | -1 | protein_coding | death-domain associated protein |
| **TAPBP** | ENSG00000206281 | 2606.1384 | 0.773418052 | 0.140031634 | 1.48E-08 | 1.23E-07 | 6892 | HSC | 33196314 | 33211013 | -1 | protein_coding | TAP binding protein (tapasin) |
| **RGL2** | ENSG00000206282 | 907.2207288 | -0.946573527 | 0.184845501 | 8.15E-08 | 6.12E-07 | 5863 | HSC | 33188277 | 33196019 | -1 | protein_coding | ral guanine nucleotide dissociation stimulator-like 2 |
| **HLA-DOA** | ENSG00000206292 | 1586.261488 | -0.951676683 | 0.425801931 | 0.006329886 | 0.01877346 | 3111 | HSC | 32900651 | 32906085 | -1 | protein_coding | major histocompatibility complex, class II, DO alpha |
| **TAP1** | ENSG00000206297 | 4208.411753 | 1.746331261 | 0.400281612 | 1.45E-06 | 9.09E-06 | 6890 | HSC | 32741696 | 32750465 | -1 | protein_coding | transporter 1, ATP-binding cassette, sub-family B (MDR/TAP) |
| **PSMB8** | ENSG00000206298 | 2325.02884 | 0.82873741 | 0.22151036 | 6.77E-05 | 3.19E-04 | 5696 | HSC | 32737204 | 32741190 | -1 | protein_coding | proteasome (prosome, macropain) subunit, beta type, 8 |
| **TAP2** | ENSG00000206299 | 2387.9398 | 1.874249077 | 0.599396873 | 1.44E-04 | 6.38E-04 | 6891 | HSC | 32718320 | 32735268 | -1 | protein_coding | transporter 2, ATP-binding cassette, sub-family B (MDR/TAP) |
| **HLA-DRB1** | ENSG00000206306 | 38154.85249 | -0.493922605 | 0.168816608 | 0.002295387 | 0.007627831 | 1009968 | HSC | 32481203 | 32511820 | -1 | protein_coding | major histocompatibility complex, class II, DR beta 1 |
| **HLA-H** | ENSG00000206341 | 410.0826445 | 0.467701096 | 0.208541148 | 0.017636729 | 0.045685108 | NA | 6 | 29855350 | 29858259 | 1 | pseudogene | major histocompatibility complex, class I, H (pseudogene) |
| **MICB** | ENSG00000206449 | 96.77363567 | 1.833795862 | 0.369233859 | 7.38E-08 | 5.58E-07 | 4277 | HSC | 31456185 | 31469201 | 1 | protein_coding | MHC class I polypeptide-related sequence B |
| **HLA-B** | ENSG00000206450 | 8610.16238 | 0.279827096 | 0.082927234 | 5.63E-04 | 0.002173471 | 3106 | HSC | 31311768 | 31315097 | -1 | protein_coding | major histocompatibility complex, class I, B |
| **TCF19** | ENSG00000206455 | 161.19366 | 1.388626527 | 0.397246658 | 7.02E-05 | 3.3E-04 | 6941 | HSC | 31118773 | 31124447 | 1 | protein_coding | transcription factor 19 |
| **UBD** | ENSG00000206468 | 311.7057999 | 3.228745833 | 0.34736333 | 7.39E-22 | 1.86E-20 | 10537 | HSC | 29523204 | 29527517 | -1 | protein_coding | ubiquitin D |
| **UBD** | ENSG00000206513 | 437.8125701 | 3.133584153 | 0.28194384 | 6.46E-30 | 2.47E-28 | 10537 | HSC | 29523310 | 29527624 | -1 | protein_coding | ubiquitin D |
| **PTPLB** | ENSG00000206527 | 1411.375155 | -0.626508467 | 0.136139355 | 2.4E-06 | 1.46E-05 | 201562 | 3 | 123209667 | 123304032 | -1 | protein_coding | protein tyrosine phosphatase-like (proline instead of catalytic arginine), member b |
| **LNP1** | ENSG00000206535 | 29.39339636 | -1.18332386 | 0.693145477 | 0.01277361 | 0.034533713 | 348801 | 3 | 100120037 | 100175163 | 1 | protein_coding | leukemia NUP98 fusion partner 1 |
| **GPX3** | ENSG00000211445 | 1936.31503 | -1.160127848 | 0.21564059 | 1.71E-08 | 1.41E-07 | 2878 | 5 | 150400124 | 150408554 | 1 | protein_coding | glutathione peroxidase 3 (plasma) |
| **C11orf31** | ENSG00000211450 | 1467.881462 | -0.82237897 | 0.150953197 | 1.84E-08 | 1.51E-07 | 280636 | 11 | 57508825 | 57510986 | 1 | protein_coding | chromosome 11 open reading frame 31 |
| **SLC48A1** | ENSG00000211584 | 2510.242097 | -1.408983471 | 0.221191835 | 2.89E-11 | 3.28E-10 | 55652 | 12 | 48147699 | 48176536 | 1 | protein_coding | solute carrier family 48 (heme transporter), member 1 |
| **TRAV2** | ENSG00000211776 | 38.74580103 | -1.237134789 | 0.927191142 | 0.017269993 | 0.044864407 | NA | 14 | 22180544 | 22181066 | 1 | TR_V_gene | T cell receptor alpha variable 2 |
| **IGHA1** | ENSG00000211895 | 40.55847189 | -1.019175897 | 0.626274702 | 0.018577792 | 0.047648681 | NA | 14 | 106173457 | 106175002 | -1 | IG_C_gene | immunoglobulin heavy constant alpha 1 |
| **HSPA1B** | ENSG00000212866 | 113.8895069 | 2.180048771 | 0.492708254 | 7.89E-07 | 5.18E-06 | 3304 | HSC | 31785766 | 31788290 | 1 | protein_coding | heat shock 70kDa protein 1B |
| **MT-ND4L** | ENSG00000212907 | 4484.21769 | -0.661378701 | 0.237434388 | 0.002618666 | 0.008564321 | 4539 | MT | 10470 | 10766 | 1 | protein_coding | mitochondrially encoded NADH dehydrogenase 4L |
| **ZNF580** | ENSG00000213015 | 208.3138615 | -0.542327054 | 0.247574613 | 0.017172489 | 0.044641461 | 51157 | 19 | 56146382 | 56154835 | 1 | protein_coding | zinc finger protein 580 |
| **ZNF611** | ENSG00000213020 | 765.7680284 | 0.407784951 | 0.157565526 | 0.007346073 | 0.021372555 | 81856 | 19 | 53206066 | 53238326 | -1 | protein_coding | zinc finger protein 611 |
| **NUP62** | ENSG00000213024 | 1502.791662 | 0.481482812 | 0.132481252 | 1.96E-04 | 8.41E-04 | 23636 | 19 | 50410082 | 50433020 | -1 | protein_coding | nucleoporin 62kDa |
| **DENND1B** | ENSG00000213047 | 1106.715935 | 0.661390056 | 0.14519302 | 2.8E-06 | 1.69E-05 | 163486 | 1 | 197473878 | 197744826 | -1 | protein_coding | DENN/MADD domain containing 1B |
| **SFT2D2** | ENSG00000213064 | 11675.45226 | 0.506885134 | 0.135105766 | 1.14E-04 | 5.16E-04 | 375035 | 1 | 168195176 | 168222263 | 1 | protein_coding | SFT2 domain containing 2 |
| **TCTEX1D2** | ENSG00000213123 | 91.79650659 | -0.90774514 | 0.372622923 | 0.004394934 | 0.013610492 | 255758 | 3 | 196018090 | 196045170 | -1 | protein_coding | Tctex1 domain containing 2 |
| **TRIM59** | ENSG00000213186 | 192.1871263 | 0.797804268 | 0.306318026 | 0.003325041 | 0.010633313 | 286827 | 3 | 160150233 | 160203561 | -1 | protein_coding | tripartite motif containing 59 |
| **MLLT11** | ENSG00000213190 | 271.2409 | 2.114432343 | 0.299438693 | 1.52E-13 | 2.07E-12 | 10962 | 1 | 151030234 | 151040970 | 1 | protein_coding | myeloid/lymphoid or mixed-lineage leukemia (trithorax homolog, Drosophila); translocated to, 11 |
| **ARHGEF35** | ENSG00000213214 | 53.5449697 | 1.870065074 | 0.678811384 | 4.53E-04 | 0.00179265 | 445328 | 7 | 143883176 | 143991230 | -1 | protein_coding | Rho guanine nucleotide exchange factor (GEF) 35 |
| **NRAS** | ENSG00000213281 | 3478.083984 | 0.547197272 | 0.131165663 | 1.96E-05 | 1.02E-04 | 4893 | 1 | 115247090 | 115259515 | -1 | protein_coding | neuroblastoma RAS viral (v-ras) oncogene homolog |
| **CHUK** | ENSG00000213341 | 1357.281894 | 0.499795422 | 0.139573879 | 2.28E-04 | 9.64E-04 | 1147 | 10 | 101948055 | 101989376 | -1 | protein_coding | conserved helix-loop-helix ubiquitous kinase |
| **GSTM2** | ENSG00000213366 | 219.8107864 | -1.395854048 | 0.307250437 | 8.95E-07 | 5.81E-06 | 2946 | 1 | 110210644 | 110252171 | 1 | protein_coding | glutathione S-transferase mu 2 (muscle) |
| **COG8** | ENSG00000213380 | 619.1059522 | -0.522583213 | 0.147483361 | 2.51E-04 | 0.001050511 | 84342 | 16 | 69354043 | 69373570 | -1 | protein_coding | component of oligomeric golgi complex 8 |
| **RP11-75L1.2** | ENSG00000213443 | 279.0911611 | 1.294788564 | 0.2352121 | 7.46E-09 | 6.45E-08 | 29121 | 12 | 9847980 | 9848823 | 1 | pseudogene | |
| **GBP7** | ENSG00000213512 | 56.73133875 | 4.510955249 | 0.663364449 | 5.94E-13 | 7.73E-12 | 388646 | 1 | 89597434 | 89641723 | -1 | protein_coding | guanylate binding protein 7 |
| **RBMXL1** | ENSG00000213516 | 829.9378236 | 1.064749593 | 0.140525364 | 1.03E-14 | 1.52E-13 | 494115 | 1 | 89445139 | 89458643 | -1 | protein_coding | RNA binding motif protein, X-linked-like 1 |
| **TMEM110** | ENSG00000213533 | 1996.777905 | 2.177641855 | 0.152607147 | 3.34E-47 | 2.7E-45 | 375346 | 3 | 52870235 | 52931612 | -1 | protein_coding | transmembrane protein 110 |
| **TMX2** | ENSG00000213593 | 1980.40656 | -0.587754324 | 0.101137009 | 3.77E-09 | 3.37E-08 | 51075 | 11 | 57480072 | 57508445 | 1 | protein_coding | thioredoxin-related transmembrane protein 2 |
| **NDUFS3** | ENSG00000213619 | 1233.252271 | -0.454439154 | 0.148822009 | 0.001688753 | 0.005819438 | 4722 | 11 | 47586888 | 47606114 | 1 | protein_coding | NADH dehydrogenase (ubiquinone) Fe-S protein 3, 30kDa (NADH-coenzyme Q reductase) |
| **NCKIPSD** | ENSG00000213672 | 437.6355583 | -0.542782025 | 0.232394224 | 0.011800153 | 0.03225572 | 51517 | 3 | 48701364 | 48723797 | -1 | protein_coding | NCK interacting protein with SH3 domain |
| **ATF6B** | ENSG00000213676 | 357.8272714 | -0.894363553 | 0.371482584 | 0.004392286 | 0.013605047 | 1388 | 6 | 32065953 | 32096030 | -1 | protein_coding | activating transcription factor 6 beta |
| **TREX1** | ENSG00000213689 | 126.9145674 | 1.359699114 | 0.341494598 | 1.12E-05 | 6.15E-05 | 84126 | 3 | 48506445 | 48509044 | 1 | protein_coding | three prime repair exonuclease 1 |
| **RPS29** | ENSG00000213741 | 1871.823191 | -0.309938273 | 0.134347698 | 0.017732269 | 0.04587039 | 6235 | 14 | 50043390 | 50065408 | -1 | protein_coding | ribosomal protein S29 |
| **ZNF134** | ENSG00000213762 | 744.2600699 | -0.816106966 | 0.159994069 | 1.15E-07 | 8.44E-07 | 7693 | 19 | 58125601 | 58134724 | 1 | protein_coding | zinc finger protein 134 |
| **DDX47** | ENSG00000213782 | 979.8658384 | 0.503744926 | 0.139371839 | 2.13E-04 | 9.06E-04 | 51202 | 12 | 12966250 | 12982915 | 1 | protein_coding | DEAD (Asp-Glu-Ala-Asp) box polypeptide 47 |
| **EMP2** | ENSG00000213853 | 32.49393753 | -4.974921971 | 1.020556082 | 2.08E-07 | 1.48E-06 | 2013 | 16 | 10622279 | 10674555 | -1 | protein_coding | epithelial membrane protein 2 |
| **RPL7AP64** | ENSG00000213876 | 14.94591503 | -2.98408149 | 1.256085648 | 0.001281476 | 0.004545091 | NA | 17 | 7044249 | 7044911 | 1 | pseudogene | ribosomal protein L7a pseudogene 64 |
| **UBD** | ENSG00000213886 | 67.19289681 | 3.883080213 | 0.770477104 | 2.01E-08 | 1.64E-07 | 10537 | 6 | 29523292 | 29527702 | -1 | protein_coding | ubiquitin D |
| **CSNK1E** | ENSG00000213923 | 536.3220125 | -0.534578442 | 0.176320088 | 0.001556869 | 0.005412563 | 1454 | 22 | 38686697 | 38794527 | -1 | protein_coding | casein kinase 1, epsilon |
| **IRF9** | ENSG00000213928 | 1486.98539 | 1.030754174 | 0.125626977 | 5.35E-17 | 9.37E-16 | 10379 | 14 | 24630262 | 24635774 | 1 | protein_coding | interferon regulatory factor 9 |
| **ITGA1** | ENSG00000213949 | 248.9241771 | 1.719198554 | 0.418592529 | 4.41E-06 | 2.58E-05 | 3672 | 5 | 52083730 | 52255040 | 1 | protein_coding | integrin, alpha 1 |
| **TAX1BP3** | ENSG00000213977 | 1109.765046 | -0.33446025 | 0.121049355 | 0.005484724 | 0.016529883 | 30851 | 17 | 3566196 | 3571976 | -1 | protein_coding | Tax1 (human T-cell leukemia virus type I) binding protein 3 |
| **AP1G2** | ENSG00000213983 | 981.5474157 | 0.408267333 | 0.180668542 | 0.017479853 | 0.045347919 | 8906 | 14 | 24028774 | 24037279 | -1 | protein_coding | adaptor-related protein complex 1, gamma 2 subunit |
| **GANC** | ENSG00000214013 | 718.3098125 | 0.542827309 | 0.160072892 | 4.5E-04 | 0.001780391 | 2595 | 15 | 42565431 | 42645864 | 1 | protein_coding | glucosidase, alpha; neutral C |
| **REPIN1** | ENSG00000214022 | 1272.981695 | 0.277120632 | 0.114382293 | 0.013596675 | 0.036507026 | 29803 | 7 | 150065278 | 150071133 | 1 | protein_coding | replication initiator 1 |
| **MRPL23** | ENSG00000214026 | 216.6120203 | -0.513048296 | 0.23061604 | 0.017042099 | 0.044355314 | 6150 | 11 | 1968508 | 2005752 | 1 | protein_coding | mitochondrial ribosomal protein L23 |

| | | | | | | | | | | | | | |
|---|---|---|---|---|---|---|---|---|---|---|---|---|---|
| **FBXO16** | ENSG00000214050 | 15.55845348 | -1.457644782 | 1.54874171 | 0.019175976 | 0.048958644 | 157574 | 8 | 28205804 | 28347835 | -1 | protein_coding | F-box protein 16 |
| **LDHAP4** | ENSG00000214110 | 114.3324385 | -0.682854246 | 0.33668564 | 0.018542728 | 0.047596323 | NA | 9 | 14921335 | 14922332 | -1 | pseudogene | lactate dehydrogenase A pseudogene 4 |
| **LYRM4** | ENSG00000214113 | 277.6141028 | 1.110281468 | 0.243762148 | 1.25E-06 | 7.89E-06 | 57128 | 6 | 5102827 | 5261172 | -1 | protein_coding | LYR motif containing 4 |
| **MYCBP** | ENSG00000214114 | 417.4177824 | -0.744273407 | 0.21948852 | 2.98E-04 | 0.001227087 | 1005279 | 1 | 39328636 | 39347289 | -1 | protein_coding | MYC binding protein |
| **ALG3** | ENSG00000214160 | 442.4349671 | -0.874048257 | 0.185828475 | 9.16E-07 | 5.93E-06 | 10195 | 3 | 183960089 | 183967336 | -1 | protein_coding | ALG3, alpha-1,3- mannosyltransferase |
| **SH3D21** | ENSG00000214193 | 355.6897356 | -0.786833122 | 0.270457323 | 0.001334851 | 0.004709263 | 79729 | 1 | 36771988 | 36790484 | 1 | protein_coding | SH3 domain containing 21 |
| **FIS1** | ENSG00000214253 | 979.2191987 | -0.998654931 | 0.143256618 | 9.61E-13 | 1.22E-11 | 51024 | 7 | 100882739 | 100895597 | -1 | protein_coding | fission 1 (mitochondrial outer membrane) homolog (S. cerevisiae) |
| **NEURL1B** | ENSG00000214357 | 5.344193021 | -0.766743464 | 1.453074908 | 0.009223731 | 0.026100035 | 54492 | 5 | 172068269 | 172118543 | 1 | protein_coding | neuralized E3 ubiquitin protein ligase 1B |
| **HAUS3** | ENSG00000214367 | 381.7208417 | 1.216997025 | 0.225270522 | 1.35E-08 | 1.13E-07 | 79441 | 4 | 2229191 | 2243891 | -1 | protein_coding | HAUS augmin-like complex, subunit 3 |
| **BBIP1** | ENSG00000214413 | 186.4708014 | 0.580388516 | 0.274252473 | 0.018554117 | 0.047601573 | 92482 | 10 | 112658488 | 112679032 | -1 | protein_coding | BBSome interacting protein 1 |
| **PPME1** | ENSG00000214517 | 1225.024214 | -0.653221574 | 0.117405885 | 1.45E-08 | 1.21E-07 | 51400 | 11 | 73882144 | 73965748 | 1 | protein_coding | protein phosphatase methylesterase 1 |
| **ZBED1** | ENSG00000214717 | 1849.551071 | 1.157471625 | 0.123161745 | 1.31E-21 | 3.23E-20 | 9189 | X | 2404455 | 2419008 | -1 | protein_coding | zinc finger, BED-type containing 1 |
| **C14orf182** | ENSG00000214900 | 85.39190136 | 0.712078709 | 0.356253876 | 0.018897848 | 0.048345466 | 283551 | 14 | 50448430 | 50474238 | -1 | protein_coding | chromosome 14 open reading frame 182 |
| **ZSWIM7** | ENSG00000214941 | 260.2435409 | -0.717854656 | 0.228896371 | 7.74E-04 | 0.002902022 | 125150 | 17 | 15879874 | 15903031 | -1 | protein_coding | zinc finger, SWIM-type containing 7 |
| **ARHGEF28** | ENSG00000214944 | 25.33809758 | -2.605009239 | 1.09954668 | 9.19E-04 | 0.003383611 | 64283 | 5 | 72921983 | 73237818 | 1 | protein_coding | Rho guanine nucleotide exchange factor (GEF) 28 |
| **C22orf29** | ENSG00000215012 | 381.1288913 | -0.888075971 | 0.213578397 | 1.14E-05 | 6.25E-05 | 79680 | 22 | 19833661 | 19842419 | -1 | protein_coding | chromosome 22 open reading frame 29 |
| **RPL13P12** | ENSG00000215030 | 156.8319257 | -0.757140856 | 0.328660437 | 0.008318322 | 0.023811371 | NA | 17 | 17286691 | 17287326 | -1 | pseudogene | ribosomal protein L13 pseudogene 12 |
| **BRD2** | ENSG00000215077 | 839.7089693 | 1.193808604 | 0.158127076 | 9.76E-15 | 1.45E-13 | 6046 | HSC | 32865184 | 32878022 | 1 | protein_coding | bromodomain containing 2 |
| **UBXN2B** | ENSG00000215114 | 1722.873392 | -0.407221441 | 0.11812868 | 4.25E-04 | 0.001691647 | 137886 | 8 | 59323823 | 59364060 | 1 | protein_coding | UBX domain protein 2B |
| **PEX26** | ENSG00000215193 | 2035.889462 | -0.414186573 | 0.115933437 | 2.41E-04 | 0.001011138 | 55670 | 22 | 18560689 | 18613905 | 1 | protein_coding | peroxisomal biogenesis factor 26 |
| **FASTKD5** | ENSG00000215251 | 832.8990029 | 0.47450287 | 0.165397029 | 0.002881239 | 0.009351026 | 60493 | 20 | 3127165 | 3140543 | -1 | protein_coding | FAST kinase domains 5 |
| **DDX3X** | ENSG00000215301 | 17513.09436 | 0.223391989 | 0.087146753 | 0.010392241 | 0.028908739 | 1654 | X | 41192651 | 41223725 | 1 | protein_coding | DEAD (Asp-Glu-Ala-Asp) box helicase 3, X-linked |
| **RP11-764K9.** | ENSG00000215548 | 42.37328846 | 1.421569388 | 0.522878723 | 8.47E-04 | 0.003149848 | NA | 9 | 68427861 | 68455375 | -1 | pseudogene | |
| **RSC1A1** | ENSG00000215695 | 457.3637833 | 0.561137202 | 0.194361033 | 0.002345953 | 0.007784022 | 84301 | 1 | 15986364 | 15988217 | 1 | protein_coding | regulatory solute carrier protein, family 1, member 1 |
| **TMEM167B** | ENSG00000215717 | 1292.697311 | 0.48212883 | 0.127412091 | 1.09E-04 | 4.96E-04 | 56900 | 1 | 109632425 | 109639556 | 1 | protein_coding | transmembrane protein 167B |
| **FAM72D** | ENSG00000215784 | 60.35651235 | 1.280775029 | 0.586647398 | 0.003965042 | 0.012422661 | 728833 | 1 | 143896452 | 143913143 | -1 | protein_coding | family with sequence similarity 72, member D |
| **FNIP1** | ENSG00000217128 | 2112.312384 | 0.380718367 | 0.114342239 | 6.98E-04 | 0.002638761 | 96459 | 5 | 130977407 | 131132710 | -1 | protein_coding | folliculin interacting protein 1 |
| **RP11-12A2.1** | ENSG00000218565 | 27.52031667 | 3.196653418 | 0.739168779 | 8.52E-07 | 5.54E-06 | NA | 6 | 139659155 | 139660717 | 1 | pseudogene | |
| **ZNF579** | ENSG00000218891 | 145.5674985 | -0.963440398 | 0.289379529 | 2.44E-04 | 0.001023285 | 163033 | 19 | 56088899 | 56092211 | -1 | protein_coding | zinc finger protein 579 |
| **RNASEK** | ENSG00000219200 | 1794.675396 | -0.640439724 | 0.121813326 | 7.67E-08 | 5.79E-07 | 440400 | 17 | 6915736 | 6917851 | 1 | protein_coding | ribonuclease, RNase K |
| **NBPF1** | ENSG00000219481 | 2168.778594 | -1.219675335 | 0.115747163 | 2.01E-26 | 6.5E-25 | 284565 | 1 | 16888814 | 16940057 | -1 | protein_coding | neuroblastoma breakpoint family, member 1 |
| **XX-C2158C6.** | ENSG00000219814 | 16.45895632 | 3.442668673 | 2.914964138 | 0.003162629 | 0.010179962 | NA | 6 | 171054606 | 171055065 | 1 | pseudogene | |
| **BTF3P7** | ENSG00000219986 | 63.59219246 | -1.258973607 | 0.527689235 | 0.002596663 | 0.00850328 | NA | 6 | 6795148 | 6795635 | -1 | pseudogene | basic transcription factor 3 pseudogene 7 |
| **ZGLP1** | ENSG00000220201 | 52.23176524 | -1.197149378 | 0.503132491 | 0.002896604 | 0.009392912 | 1001252 | 19 | 10415479 | 10420556 | -1 | protein_coding | zinc finger, GATA-like protein 1 |
| **NPTXR** | ENSG00000221890 | 152.8752395 | -1.376727336 | 0.306359762 | 1.15E-06 | 7.32E-06 | 23467 | 22 | 39214457 | 39239987 | -1 | protein_coding | neuronal pentraxin receptor |
| **PPP2R2A** | ENSG00000221914 | 1910.610517 | 1.528501505 | 0.111834716 | 1.27E-43 | 9.19E-42 | 5520 | 8 | 26149007 | 26230196 | 1 | protein_coding | protein phosphatase 2, regulatory subunit B, alpha |
| **ZNF880** | ENSG00000221923 | 209.1404041 | -1.181503233 | 0.289221448 | 9.27E-06 | 5.12E-05 | 400713 | 19 | 52873170 | 52889048 | 1 | protein_coding | zinc finger protein 880 |
| **SLC12A8** | ENSG00000221955 | 221.8513491 | -0.989490439 | 0.334487822 | 8.2E-04 | 0.003058866 | 84561 | 3 | 124801480 | 124998021 | -1 | protein_coding | solute carrier family 12, member 8 |
| **APOL6** | ENSG00000221963 | 11836.83681 | 2.252103757 | 0.150376343 | 1E-51 | 9.62E-50 | 80830 | 22 | 36044442 | 36064456 | 1 | protein_coding | apolipoprotein L, 6 |
| **UBA52** | ENSG00000221983 | 3922.740489 | -0.525412632 | 0.126297459 | 2.21E-05 | 1.14E-04 | 7311 | 19 | 18682540 | 18688360 | 1 | protein_coding | ubiquitin A-52 residue ribosomal protein fusion product 1 |
| **BTBD19** | ENSG00000222009 | 304.8083242 | 0.670460128 | 0.209039185 | 6.94E-04 | 0.002624901 | 149478 | 1 | 45274154 | 45281257 | 1 | protein_coding | BTB (POZ) domain containing 19 |
| **AC011997.1** | ENSG00000222017 | 4.83099561 | 3.578444469 | 3.114179955 | 0.005402792 | 0.016324807 | NA | 2 | 198557830 | 198639547 | 1 | protein_coding | Uncharacterized protein |
| **HLA-B** | ENSG00000223532 | 8326.717191 | 0.941332548 | 0.213275858 | 3.26E-06 | 1.94E-05 | 3106 | HSC | 31312134 | 31315506 | -1 | protein_coding | major histocompatibility complex, class I, B |
| **CLIC1** | ENSG00000223639 | 7338.24077 | 0.566880115 | 0.099689326 | 7.98E-09 | 6.87E-08 | 1192 | HSC | 31688317 | 31697500 | -1 | protein_coding | chloride intracellular channel 1 |
| **GLTPD1** | ENSG00000224051 | 253.59355 | -0.550794375 | 0.254378667 | 0.018069604 | 0.046556204 | 80772 | 1 | 1260136 | 1264277 | 1 | protein_coding | glycolipid transfer protein domain containing 1 |
| **RP11-39K24.** | ENSG00000224083 | 17.84732994 | 1.803231004 | 0.981053727 | 0.004027619 | 0.012587751 | NA | 9 | 5096666 | 5098193 | 1 | pseudogene | |
| **HLA-DPA1** | ENSG00000224103 | 1446.577329 | -1.542162341 | 0.626679069 | 0.001358014 | 0.00478325 | 3113 | HSC | 32954322 | 32970527 | -1 | protein_coding | major histocompatibility complex, class II, DP alpha 1 |
| **CYP4F25P** | ENSG00000224106 | 47.27658566 | -1.852842552 | 0.797476639 | 0.001543538 | 0.005373553 | NA | 9 | 69477881 | 69478916 | 1 | pseudogene | cytochrome P450, family 4, subfamily F, polypeptide 25, pseudogene |
| **TRIM31** | ENSG00000224168 | 71.64181963 | 3.060043903 | 0.561680377 | 3E-09 | 2.71E-08 | 11074 | HSC | 30060269 | 30070469 | -1 | protein_coding | tripartite motif containing 31 |
| **TAP1** | ENSG00000224212 | 96.42918524 | 0.616481286 | 1.112120084 | 0.012618685 | 0.034168264 | 6890 | HSC | 32790967 | 32799736 | -1 | protein_coding | transporter 1, ATP-binding cassette, sub-family B (MDR/TAP) |
| **APOM** | ENSG00000224290 | 5.216727071 | -0.420495538 | 0.837237804 | 0.016713721 | 0.0436195 | 55937 | HSC | 31607608 | 31613401 | 1 | protein_coding | apolipoprotein M |
| **CSNK2B** | ENSG00000224398 | 1027.845159 | -0.835571322 | 0.18650991 | 2.82E-06 | 1.7E-05 | 1460 | HSC | 31709296 | 31717606 | 1 | protein_coding | casein kinase 2, beta polypeptide |
| **SLC39A7** | ENSG00000224399 | 2418.961437 | -0.582326859 | 0.107287653 | 3.35E-08 | 2.66E-07 | 7922 | HSC | 33338586 | 33342580 | 1 | protein_coding | solute carrier family 39 (zinc transporter), member 7 |
| **ATXN1L** | ENSG00000224470 | 1898.527875 | 0.240975597 | 0.103247088 | 0.018195372 | 0.04684634 | 342371 | 16 | 71879894 | 71919171 | 1 | protein_coding | ataxin 1-like |
| **HSPA1B** | ENSG00000224501 | 38.65215213 | 0.963150188 | 1.660681921 | 0.017299215 | 0.044925048 | 3304 | HSC | 31782890 | 31785413 | 1 | protein_coding | heat shock 70kDa protein 1B |
| **NPIPA3** | ENSG00000224712 | 33.61284024 | 2.240971361 | 1.156802364 | 0.002143441 | 0.007188547 | 1002883 | 16 | 14802801 | 14820195 | 1 | protein_coding | nuclear pore complex interacting protein family, member A3 |
| **BEND3P2** | ENSG00000224809 | 156.1619863 | -1.032461839 | 0.284241246 | 7.4E-05 | 3.46E-04 | NA | 9 | 94933523 | 94935776 | 1 | pseudogene | BEN domain containing 3 pseudogene 2 |
| **C17orf89** | ENSG00000224877 | 188.1377093 | -1.426031949 | 0.269072055 | 1.89E-08 | 1.55E-07 | 284184 | 17 | 79213039 | 79215081 | 1 | protein_coding | chromosome 17 open reading frame 89 |
| **APOC4-APO(** | ENSG00000224916 | 35.75712554 | -2.004057364 | 0.744541465 | 6.14E-04 | 0.002347282 | 346 | 19 | 45445495 | 45452822 | 1 | protein_coding | APOC4-APOC2 readthrough (NMD candidate) |
| **FTH1P22** | ENSG00000225079 | 9.402362843 | 2.148172816 | 1.429716465 | 0.005907075 | 0.017632389 | NA | 1 | 117317726 | 117318245 | -1 | pseudogene | ferritin, heavy polypeptide 1 pseudogene 22 |
| **HSPA7** | ENSG00000225217 | 570.5378091 | -3.000920789 | 0.264494715 | 3.77E-31 | 1.55E-29 | 3311 | 1 | 161576081 | 161578007 | 1 | pseudogene | heat shock 70kDa protein 7 (HSP70B) |
| **GBP1P1** | ENSG00000225492 | 194.2417324 | 6.389374366 | 0.438313569 | 4.81E-49 | 4.06E-47 | 400759 | 1 | 89873238 | 89891802 | 1 | pseudogene | guanylate binding protein 1, interferon-inducible pseudogene 1 |
| **PRRC2A** | ENSG00000225748 | 2081.755543 | -0.758495442 | 0.141578412 | 3.81E-08 | 3E-07 | 7916 | HSC | 31664743 | 31681839 | 1 | protein_coding | proline-rich coiled-coil 2A |
| **HLA-DQA1** | ENSG00000225890 | 642.3790835 | -0.541625717 | 0.230329685 | 0.011313788 | 0.03113227 | 1005094 | HSC | 32748724 | 32771089 | 1 | protein_coding | major histocompatibility complex, class II, DQ alpha 1 |
| **TRIM26** | ENSG00000226060 | 432.8305426 | 0.691352632 | 0.344770421 | 0.019121269 | 0.048843449 | 7726 | HSC | 30142515 | 30171491 | -1 | protein_coding | tripartite motif containing 26 |
| **RPS18** | ENSG00000226225 | 38.91082874 | -1.310003197 | 0.843951023 | 0.011938392 | 0.032586969 | 6222 | HSC | 33410160 | 33414664 | 1 | protein_coding | ribosomal protein S18 |
| **GTF2H2B** | ENSG00000226259 | 242.90446 | 0.599895146 | 0.277887443 | 0.016347734 | 0.042803224 | 653238 | 5 | 69711179 | 69743885 | 1 | pseudogene | general transcription factor IIH, polypeptide 2B (pseudogene) |
| **PRRC2A** | ENSG00000226618 | 2687.274905 | -0.388106926 | 0.161739691 | 0.012696145 | 0.034354673 | 7916 | HSC | 31627937 | 31645035 | 1 | protein_coding | proline-rich coiled-coil 2A |
| **DAXX** | ENSG00000227046 | 654.4876789 | 0.79672722 | 0.246735314 | 4.62E-04 | 0.001821783 | 1616 | HSC | 33207945 | 33218656 | -1 | protein_coding | death-domain associated protein |
| **IER3** | ENSG00000227231 | 1129.321496 | 2.28453253 | 0.257008514 | 7.01E-20 | 1.54E-18 | 8870 | HSC | 30701238 | 30702593 | -1 | protein_coding | immediate early response 3 |
| **SCAMP4** | ENSG00000227500 | 1127.526624 | -0.420484023 | 0.140006462 | 0.001985386 | 0.006723388 | 113178 | 19 | 1905213 | 1926016 | 1 | protein_coding | secretory carrier membrane protein 4 |
| **LTB** | ENSG00000227507 | 45.84859725 | -1.377619411 | 0.592055047 | 0.002483328 | 0.008177698 | 4050 | 6 | 31548302 | 31550299 | -1 | protein_coding | lymphotoxin beta (TNF superfamily, member 3) |
| **MIR3916** | ENSG00000227671 | 807.4953609 | 1.701658586 | 0.159883679 | 2.48E-27 | 8.28E-26 | 1005008 | 1 | 247353153 | 247374158 | -1 | pseudogene | microRNA 3916 |
| **HLA-A** | ENSG00000227715 | 19179.35055 | 0.507680078 | 0.101759518 | 3.51E-07 | 2.42E-06 | 3105 | HSC | 29846763 | 29992167 | 1 | protein_coding | major histocompatibility complex, class I, A |
| **TUBB** | ENSG00000227739 | 5669.969192 | 0.330778913 | 0.107789345 | 0.001829913 | 0.006248115 | 203068 | HSC | 30677842 | 30683067 | 1 | protein_coding | tubulin, beta class I |
| **BAG6** | ENSG00000227761 | 2025.109075 | -0.436710458 | 0.171479701 | 0.007801312 | 0.022534012 | 7917 | HSC | 31588979 | 31602664 | -1 | protein_coding | BCL2-associated athanogene 6 |
| **SLC9A7P1** | ENSG00000227825 | 970.8596899 | -0.582204927 | 0.14889838 | 5.54E-05 | 2.66E-04 | 121456 | 12 | 98847613 | 98850923 | -1 | pseudogene | solute carrier family 9, subfamily A (NHE7, cation proton antiporter 7), member 7 pseudogene 1 |
| **RP11-958N2** | ENSG00000227827 | 342.6897643 | -0.491788369 | 0.208043459 | 0.011912058 | 0.032526708 | 399491 | 16 | 16450081 | 16471364 | 1 | pseudogene | |
| **RP11-39K24.** | ENSG00000228097 | 6.059631143 | 2.620079181 | 1.780689563 | 0.005181127 | 0.01574531 | NA | 9 | 5099560 | 5100229 | 1 | pseudogene | |
| **DDAH2** | ENSG00000228128 | 356.1613621 | -1.491182749 | 0.24959029 | 3.71E-10 | 3.74E-09 | 23564 | HSC | 31684774 | 31688353 | -1 | protein_coding | dimethylarginine dimethylaminohydrolase 2 |
| **MTND1P11** | ENSG00000228166 | 11.03041226 | 1.738499758 | 1.526916883 | 0.009763597 | 0.027439693 | NA | 9 | 5094139 | 5095014 | 1 | pseudogene | MT-ND1 pseudogene 11 |
| **C19orf24** | ENSG00000228300 | 331.0071926 | -0.79044542 | 0.236925501 | 3.43E-04 | 0.001394803 | 55009 | 19 | 1275026 | 1279248 | 1 | protein_coding | chromosome 19 open reading frame 24 |
| **AP006222.2** | ENSG00000228463 | 6.682507822 | 0.488449942 | 0.897882169 | 0.018807387 | 0.048138217 | NA | 1 | 227615 | 267253 | -1 | lincRNA | |
| **NUTM2E** | ENSG00000228570 | 7.40539069 | -0.429530905 | 0.849669323 | 0.012444492 | 0.03376339 | NA | 10 | 81601114 | 81610632 | 1 | protein_coding | NUT family member 2E |
| **TAP2** | ENSG00000228582 | 137.9288785 | 0.048585259 | 0.635584809 | 9.81E-04 | 0.003586186 | 6891 | HSC | 32711936 | 32728881 | -1 | protein_coding | transporter 2, ATP-binding cassette, sub-family B (MDR/TAP) |
| **C1orf233** | ENSG00000228594 | 82.57033461 | -1.019967991 | 0.467788229 | 0.006543264 | 0.019327541 | 643988 | 1 | 1533392 | 1535476 | -1 | protein_coding | chromosome 1 open reading frame 233 |
| **ATF6B** | ENSG00000228628 | 951.8846822 | -0.431278466 | 0.164400905 | 0.006306508 | 0.018711389 | 1388 | HSC | 32142344 | 32172461 | -1 | protein_coding | activating transcription factor 6 beta |
| **NEU1** | ENSG00000228691 | 3075.25711 | -0.601351875 | 0.100731756 | 1.37E-09 | 1.3E-08 | 4758 | HSC | 31836794 | 31842041 | -1 | protein_coding | sialidase 1 (lysosomal sialidase) |
| **RGL2** | ENSG00000228736 | 146.1211345 | -0.757822984 | 0.381913467 | 0.016905807 | 0.044048957 | 5863 | HSC | 33429778 | 33437523 | -1 | protein_coding | ral guanine nucleotide dissociation stimulator-like 2 |

| | | | | | | | | | | | | | |
|---|---|---|---|---|---|---|---|---|---|---|---|---|---|
| **CSNK2B** | ENSG00000228875 | 1045.260144 | -0.337133682 | 0.134293325 | 0.009828522 | 0.027586649 | 1460 | HSC | 31620429 | 31628739 | 1 | protein_coding | casein kinase 2, beta polypeptide |
| **AGPAT1** | ENSG00000228892 | 362.5533825 | -0.631430349 | 0.205778482 | 0.001138793 | 0.004102823 | 10554 | HSC | 32142859 | 32152743 | -1 | protein_coding | 1-acylglycerol-3-phosphate O-acyltransferase 1 |
| **UBD** | ENSG00000228913 | 189.36473 | 3.295757206 | 0.367426266 | 2.03E-20 | 4.61E-19 | 10537 | HSC | 29519995 | 29524308 | -1 | protein_coding | ubiquitin D |
| **TRIM27** | ENSG00000229006 | 1023.111065 | 0.559411218 | 0.123290022 | 3.49E-06 | 2.07E-05 | 5987 | HSC | 28870844 | 28891821 | -1 | protein_coding | tripartite motif containing 27 |
| **AC067945.3** | ENSG00000229023 | 40.77241344 | 1.880479877 | 0.675674155 | 4.39E-04 | 0.001741341 | NA | 2 | 191857365 | 191858293 | -1 | pseudogene | |
| **RPL41** | ENSG00000229117 | 8212.663121 | -0.461787093 | 0.110511718 | 1.83E-05 | 9.61E-05 | 6171 | 12 | 56510370 | 56511727 | 1 | protein_coding | ribosomal protein L41 |
| **HLA-A** | ENSG00000229215 | 25719.58215 | 0.277651784 | 0.107168712 | 0.00854772 | 0.024381262 | 3105 | HSC | 29845529 | 29850151 | 1 | protein_coding | major histocompatibility complex, class I, A |
| **HLA-E** | ENSG00000229252 | 30236.7612 | 0.58018175 | 0.099539038 | 3.68E-09 | 3.3E-08 | 3133 | HSC | 30447487 | 30452225 | 1 | protein_coding | major histocompatibility complex, class I, E |
| **HLA-DPB1** | ENSG00000229295 | 10203.6168 | -0.466990752 | 0.151302965 | 0.00142497 | 0.005001796 | 3115 | HSC | 33197846 | 33209127 | 1 | protein_coding | major histocompatibility complex, class II, DP beta 1 |
| **HLA-F** | ENSG00000229698 | 2142.271613 | 0.881287979 | 0.148372667 | 1.09E-09 | 1.05E-08 | 3134 | HSC | 29687514 | 29703210 | 1 | protein_coding | major histocompatibility complex, class I, F |
| **PET100** | ENSG00000229833 | 444.1907471 | -0.470933855 | 0.187040586 | 0.00833997 | 0.0238644 | 1001318 | 19 | 7694623 | 7696842 | 1 | protein_coding | PET100 homolog (S. cerevisiae) |
| **RHEBP1** | ENSG00000229927 | 13.34148947 | 7.094105853 | 3.181283306 | 1.45E-04 | 6.45E-04 | NA | 10 | 46914151 | 46914706 | 1 | pseudogene | Ras-homolog enriched in brain pseudogene 1 |
| **PPP1R18** | ENSG00000229998 | 5779.557506 | 0.713610154 | 0.124181127 | 3.99E-09 | 3.55E-08 | 170954 | HSC | 30688865 | 30700371 | -1 | protein_coding | protein phosphatase 1, regulatory subunit 18 |
| **HLA-DOA** | ENSG00000230141 | 927.2858649 | -1.55886556 | 0.522168805 | 2.95E-04 | 0.001217078 | 3111 | HSC | 33005421 | 33010855 | -1 | protein_coding | major histocompatibility complex, class II, DO alpha |
| **FLOT1** | ENSG00000230143 | 9.416753531 | 0.029780317 | 0.633880767 | 0.001248778 | 0.004445627 | 10211 | HSC | 30686935 | 30701969 | -1 | protein_coding | flotillin 1 |
| **HLA-DPB1** | ENSG00000230763 | 4688.536113 | -0.5717007 | 0.225450302 | 0.006302949 | 0.01870446 | 3115 | HSC | 33077161 | 33088426 | 1 | protein_coding | major histocompatibility complex, class II, DP beta 1 |
| **ANKRD63** | ENSG00000230778 | 22.61773025 | -1.779583061 | 0.958213171 | 0.00496278 | 0.015172267 | 1001312 | 15 | 40573645 | 40574787 | -1 | protein_coding | ankyrin repeat domain 63 |
| **HSBP1** | ENSG00000230989 | 1197.568636 | 1.148309196 | 0.16431983 | 6.02E-13 | 7.82E-12 | 3281 | 16 | 83841448 | 83853342 | 1 | protein_coding | heat shock factor binding protein 1 |
| **VARS** | ENSG00000231116 | 385.8939961 | -1.145715657 | 0.213792913 | 1.85E-08 | 1.52E-07 | 7407 | HSC | 31756648 | 31775083 | -1 | protein_coding | valyl-tRNA synthetase |
| **HLA-DQB1** | ENSG00000231286 | 5233.903525 | -1.125991539 | 0.219448659 | 6.76E-08 | 5.14E-07 | 1010608 | HSC | 32658022 | 32667070 | -1 | protein_coding | major histocompatibility complex, class II, DQ beta 1 |
| **LTB** | ENSG00000231314 | 248.0699572 | -0.810468092 | 0.28141828 | 0.001462405 | 0.005121432 | 4050 | HSC | 31559653 | 31561620 | -1 | protein_coding | lymphotoxin beta (TNF superfamily, member 3) |
| **MICB** | ENSG00000231372 | 367.3309165 | 1.470880737 | 0.189806996 | 1.47E-15 | 2.31E-14 | 4277 | HSC | 31512918 | 31518423 | 1 | protein_coding | MHC class I polypeptide-related sequence B |
| **HSPA1B** | ENSG00000231555 | 445.6835734 | 1.6763307 | 0.238343649 | 2.75E-13 | 3.65E-12 | 3304 | HSC | 31777702 | 31780222 | 1 | protein_coding | heat shock 70kDa protein 1B |
| **TRIM26** | ENSG00000231641 | 548.2812819 | 1.153680322 | 0.392564673 | 6.29E-04 | 0.002397852 | 7726 | HSC | 30230672 | 30259555 | -1 | protein_coding | tripartite motif containing 26 |
| **HLA-DRB3$0** | ENSG00000231679 | 10785.77819 | -0.70158185 | 0.170150952 | 1.73E-05 | 9.12E-05 | 3125 | HSC | 32411911 | 32425028 | -1 | protein_coding | Homo sapiens major histocompatibility complex, class II, DR beta 3 (HLA-DRB3), mRNA. |
| **TAPBP** | ENSG00000231925 | 65.74802708 | 1.693399211 | 0.417862334 | 5.84E-06 | 3.34E-05 | 6892 | 6 | 33267471 | 33282164 | -1 | protein_coding | TAP binding protein (tapasin) |
| **VARS** | ENSG00000231945 | 678.0928177 | -0.859655579 | 0.200328371 | 6.26E-06 | 3.56E-05 | 7407 | HSC | 31735249 | 31752710 | -1 | protein_coding | valyl-tRNA synthetase |
| **MCTS1** | ENSG00000232119 | 505.0603204 | 0.484433761 | 0.217044496 | 0.016652909 | 0.04348307 | 28985 | X | 119727865 | 119754929 | 1 | protein_coding | malignant T cell amplified sequence 1 |
| **ABCF1** | ENSG00000232169 | 1222.769156 | 0.609369631 | 0.156803852 | 6.04E-05 | 2.87E-04 | 23 | HSC | 30529391 | 30555197 | 1 | protein_coding | ATP-binding cassette, sub-family F (GCN20), member 1 |
| **TMEM114** | ENSG00000232258 | 107.6982595 | -1.565642193 | 0.469508944 | 9.99E-05 | 4.56E-04 | 283953 | 16 | 8619502 | 8622304 | -1 | protein_coding | transmembrane protein 114 |
| **RP11-47G11.** | ENSG00000232334 | 27.68736393 | -1.35263546 | 0.69735831 | 0.006396957 | 0.018961346 | NA | 10 | 125763820 | 125764344 | -1 | pseudogene | |
| **RP11-341A11** | ENSG00000232433 | 209.4380505 | -2.455270431 | 0.585959676 | 1.92E-06 | 1.18E-05 | NA | 9 | 42493617 | 42494809 | 1 | pseudogene | |
| **RP11-18B3.2** | ENSG00000232486 | 6.471553085 | 1.233726242 | 1.824602649 | 0.019509689 | 0.04966897 | NA | 9 | 108520445 | 108523036 | 1 | pseudogene | |
| **AC112198.1** | ENSG00000232517 | 11.28071468 | 4.152456093 | 1.300725891 | 8.07E-05 | 3.75E-04 | NA | 5 | 54104038 | 54168379 | -1 | pseudogene | |
| **GABBR1** | ENSG00000232569 | 67.75465286 | 1.281216632 | 0.630358075 | 0.004724133 | 0.014526946 | 2550 | HSC | 29523504 | 29601074 | -1 | protein_coding | gamma-aminobutyric acid (GABA) B receptor, 1 |
| **HSPA1B** | ENSG00000232804 | 361.793525 | 0.751312686 | 0.304951757 | 0.005615114 | 0.01686631 | 3304 | HSC | 31806879 | 31809393 | 1 | protein_coding | heat shock 70kDa protein 1B |
| **HLA-DOA** | ENSG00000232957 | 3370.897534 | -1.18923053 | 0.246613682 | 3.28E-07 | 2.27E-06 | 3111 | HSC | 33111813 | 33117246 | -1 | protein_coding | major histocompatibility complex, class II, DO alpha |
| **HLA-DOA** | ENSG00000232962 | 632.9754639 | -0.865502461 | 0.301264722 | 0.001234025 | 0.004396177 | 3111 | HSC | 32894248 | 32899682 | -1 | protein_coding | major histocompatibility complex, class II, DO alpha |
| **HLA-DQB1** | ENSG00000233209 | 83.88714359 | -0.839062219 | 0.376923393 | 0.008366216 | 0.02393104 | 1010608 | HSC | 32602484 | 32612425 | -1 | protein_coding | major histocompatibility complex, class II, DQ beta 1 |
| **GPX1** | ENSG00000233276 | 10234.48179 | -0.707321018 | 0.106136108 | 1.24E-11 | 1.45E-10 | 2876 | 3 | 49394609 | 49396033 | -1 | protein_coding | glutathione peroxidase 1 |
| **TRIM31** | ENSG00000233573 | 9.015150005 | 7.038048454 | 2.771999767 | 4.35E-05 | 2.12E-04 | 11074 | HSC | 30061002 | 30071204 | -1 | protein_coding | tripartite motif containing 31 |
| **HLA-C** | ENSG00000233841 | 21398.20629 | 0.387794974 | 0.114858899 | 5.5E-04 | 0.002129195 | 3107 | HSC | 31227577 | 31230958 | -1 | protein_coding | major histocompatibility complex, class I, C |
| **TRIM26** | ENSG00000234046 | 284.9059658 | 1.121628818 | 0.30876611 | 6.06E-05 | 2.88E-04 | 7726 | HSC | 30141782 | 30170758 | -1 | protein_coding | tripartite motif containing 26 |
| **HLA-F** | ENSG00000234487 | 802.7153654 | 0.7847144 | 0.321809702 | 0.005530829 | 0.016645855 | 3134 | HSC | 29687668 | 29690477 | 1 | protein_coding | major histocompatibility complex, class I, F |
| **PTGES3P1** | ENSG00000234518 | 69.743228 | 2.159382035 | 0.474878979 | 4.74E-07 | 3.22E-06 | NA | 1 | 89569968 | 89570450 | -1 | pseudogene | prostaglandin E synthase 3 (cytosolic) pseudogene 1 |
| **JRK** | ENSG00000234616 | 130.5170572 | -0.856200812 | 0.306077057 | 0.001737471 | 0.005956429 | NA | 8 | 143738874 | 143763386 | -1 | processed_trans | jerky homolog (mouse) |
| **BAG6** | ENSG00000234651 | 768.2476862 | -1.601346365 | 0.201718582 | 3.17E-16 | 5.22E-15 | 7917 | HSC | 31683095 | 31696765 | -1 | protein_coding | BCL2-associated athanogene 6 |
| **BRD2** | ENSG00000234704 | 837.0704671 | 0.700549445 | 0.14257844 | 4.48E-07 | 3.06E-06 | 6046 | HSC | 33090267 | 33103106 | 1 | protein_coding | bromodomain containing 2 |
| **RP11-3P17.3** | ENSG00000234851 | 2311.9572 | -0.397994146 | 0.153525256 | 0.007241294 | 0.021107895 | NA | 3 | 161146915 | 161147398 | -1 | pseudogene | |
| **APOC2** | ENSG00000234906 | 2672.300308 | -1.065571937 | 0.23144752 | 1.12E-06 | 7.16E-06 | 344 | 19 | 45449243 | 45452822 | 1 | protein_coding | apolipoprotein C-II |
| **FABP5P7** | ENSG00000234964 | 2010.650197 | -2.722841891 | 0.2603984 | 8.69E-27 | 2.86E-25 | NA | 11 | 59548786 | 59549195 | -1 | pseudogene | fatty acid binding protein 5 pseudogene 7 |
| **PPP1R3E** | ENSG00000235194 | 274.1645974 | -0.787810791 | 0.252721228 | 7.2E-04 | 0.002717061 | 90673 | 14 | 23764852 | 23772057 | -1 | protein_coding | protein phosphatase 1, regulatory subunit 3E |
| **C6orf47** | ENSG00000235360 | 170.8982474 | 0.256451767 | 0.710277353 | 0.004663093 | 0.01435947 | 57827 | HSC | 31613489 | 31615961 | -1 | protein_coding | chromosome 6 open reading frame 47 |
| **RP11-383H13** | ENSG00000235531 | 1024.321558 | 0.428640449 | 0.13800222 | 0.001458001 | 0.005109522 | 1001328 | 8 | 72740402 | 73030628 | 1 | protein_coding | Protein LOC100132891; cDNA FLJ53548 |
| **H3F3AP4** | ENSG00000235655 | 275.7014191 | -0.381360354 | 0.799214545 | 0.018468983 | 0.047430929 | 440926 | 2 | 175584505 | 175585526 | 1 | pseudogene | H3 histone, family 3A, pseudogene 4 |
| **PFDN6** | ENSG00000235692 | 39.71023012 | 2.163211642 | 0.81713638 | 5.47E-04 | 0.002119295 | 1024655 | HSC | 33427426 | 33436525 | 1 | protein_coding | prefoldin subunit 6 |
| **KIAA0040** | ENSG00000235750 | 2323.253036 | 2.500169306 | 0.126471708 | 5.07E-88 | 1.46E-85 | 9674 | 1 | 175126123 | 175162135 | -1 | protein_coding | KIAA0040 |
| **HSPA1A** | ENSG00000235941 | 1077.988021 | 0.648831415 | 0.157484275 | 2E-05 | 1.04E-04 | 3304 | HSC | 31794605 | 31797087 | 1 | protein_coding | heat shock 70kDa protein 1A |
| **MTND4P14** | ENSG00000236254 | 11.76409228 | 3.315899344 | 1.314258867 | 5.54E-04 | 0.002141935 | NA | 9 | 5107937 | 5109290 | 1 | pseudogene | MT-ND4 pseudogene 14 |
| **ZBED5** | ENSG00000236287 | 771.3405599 | 0.442212763 | 0.150526427 | 0.002453895 | 0.008089491 | 1019280 | 11 | 10833621 | 10880343 | -1 | protein_coding | zinc finger, BED-type containing 5 |
| **HLA-DQA1** | ENSG00000236418 | 132.5779564 | -0.109148765 | 0.646570069 | 0.015413468 | 0.040733881 | 1005094 | HSC | 32574291 | 32596644 | 1 | protein_coding | major histocompatibility complex, class II, DQ alpha 1 |
| **PPP1R18** | ENSG00000236428 | 1050.323539 | 0.922710007 | 0.192649972 | 5.49E-07 | 3.69E-06 | 170954 | HSC | 30635612 | 30638612 | -1 | protein_coding | protein phosphatase 1, regulatory subunit 18 |
| **TAP2** | ENSG00000237599 | 2711.613873 | 2.606859854 | 0.197753176 | 9.21E-41 | 5.77E-39 | 6891 | HSC | 32823109 | 32840035 | -1 | protein_coding | transporter 2, ATP-binding cassette, sub-family B (MDR/TAP) |
| **KIFC1** | ENSG00000237649 | 12.82083406 | 0.359925908 | 0.787992897 | 0.005038127 | 0.015365733 | 3833 | 6 | 33359313 | 33377701 | 1 | protein_coding | kinesin family member C1 |
| **RP11-39K24.** | ENSG00000237711 | 7.322740062 | 1.993301022 | 2.064537985 | 0.007944279 | 0.0228821 | NA | 9 | 5100236 | 5101009 | 1 | pseudogene | |
| **AIF1** | ENSG00000237727 | 924.2999295 | -1.174456799 | 0.210355127 | 5.28E-09 | 4.67E-08 | 199 | HSC | 31570375 | 31572212 | 1 | protein_coding | allograft inflammatory factor 1 |
| **RP11-274E7.** | ENSG00000238000 | 16.85607461 | 1.349367282 | 0.858315059 | 0.01106671 | 0.030529337 | NA | 5 | 97549106 | 97549825 | 1 | pseudogene | |
| **RPL9P7** | ENSG00000238103 | 5.067297282 | 0.347974698 | 0.774712356 | 0.015857943 | 0.041720939 | NA | X | 23854761 | 23855459 | -1 | pseudogene | ribosomal protein L9 pseudogene 7 |
| **RPS20P22** | ENSG00000239218 | 10.89392965 | -2.681508228 | 1.465743285 | 0.005028475 | 0.015342422 | NA | 8 | 38291865 | 38293182 | -1 | pseudogene | ribosomal protein S20 pseudogene 22 |
| **TXNDC5** | ENSG00000239264 | 1388.599875 | -1.177129539 | 0.129067259 | 1.67E-20 | 3.83E-19 | 81567 | 6 | 7881483 | 8026646 | -1 | protein_coding | thioredoxin domain containing 5 (endoplasmic reticulum) |
| **RNF103** | ENSG00000239305 | 1222.575915 | 0.480244614 | 0.162351133 | 0.00212782 | 0.007142427 | 7844 | 2 | 86830516 | 86850989 | -1 | protein_coding | ring finger protein 103 |
| **HLA-DMB** | ENSG00000239329 | 1601.645421 | -1.121100614 | 0.194685285 | 2.02E-09 | 1.87E-08 | 3109 | HSC | 33056220 | 33074714 | -1 | protein_coding | major histocompatibility complex, class II, DM beta |
| **AC034228.7** | ENSG00000239642 | 107.4692053 | -2.396551896 | 0.476490213 | 4.03E-08 | 3.16E-07 | NA | 5 | 131142816 | 131281391 | -1 | pseudogene | |
| **NME1** | ENSG00000239672 | 480.0727664 | -0.762600864 | 0.207290001 | 9.05E-05 | 4.17E-04 | 4830 | 17 | 49230897 | 49239789 | 1 | protein_coding | NME/NM23 nucleoside diphosphate kinase 1 |
| **TNFSF12** | ENSG00000239697 | 593.2406181 | -0.39318164 | 0.152783138 | 0.007899665 | 0.022775056 | 407977 | 17 | 7452208 | 7464925 | 1 | protein_coding | tumor necrosis factor (ligand) superfamily, member 12 |
| **APOBEC3G** | ENSG00000239713 | 483.8563935 | 1.476502455 | 0.198468485 | 1.3E-14 | 1.92E-13 | 60489 | 22 | 39436924 | 39483748 | 1 | protein_coding | apolipoprotein B mRNA editing enzyme, catalytic polypeptide-like 3G |
| **WBP1** | ENSG00000239779 | 546.3489532 | -0.679551945 | 0.163328954 | 1.63E-05 | 8.64E-05 | 23559 | 2 | 74685456 | 74688018 | 1 | protein_coding | WW domain binding protein 1 |
| **LILRA2** | ENSG00000239998 | 190.3081515 | -0.725719288 | 0.286457281 | 0.004880469 | 0.014962531 | 11027 | 19 | 55084387 | 55099027 | 1 | protein_coding | leukocyte immunoglobulin-like receptor, subfamily A (with TM domain), member 2 |
| **AMY2B** | ENSG00000240038 | 449.3252798 | -0.501854326 | 0.166219248 | 0.001693938 | 0.005835992 | 648740 | 1 | 104096437 | 104122156 | 1 | protein_coding | amylase, alpha 2B (pancreatic) |
| **PCDHGC3** | ENSG00000240184 | 315.8162361 | -3.279525173 | 0.344819077 | 1.21E-22 | 3.18E-21 | 5098 | 5 | 140855580 | 140892542 | 1 | protein_coding | protocadherin gamma subfamily C, 3 |
| **RP11-146D12** | ENSG00000240240 | 150.3978392 | -0.979860363 | 0.46735168 | 0.008430022 | 0.0240815 | NA | 9 | 42409989 | 42474239 | 1 | protein_coding | |
| **RPL23AP7** | ENSG00000240356 | 192.1682101 | -1.132737642 | 0.317894657 | 7.89E-05 | 3.68E-04 | 118433 | 2 | 114368079 | 114384667 | -1 | pseudogene | ribosomal protein L23a pseudogene 7 |
| **FOXO3B** | ENSG00000240445 | 24.93339818 | -2.022097403 | 0.892969906 | 0.001526535 | 0.00532042 | 2310 | 17 | 18574259 | 18576186 | -1 | pseudogene | forkhead box O3B pseudogene |
| **TMEM189** | ENSG00000240849 | 810.0743706 | 0.749711402 | 0.139086462 | 3.26E-08 | 2.6E-07 | 387522 | 20 | 48697663 | 48770335 | -1 | protein_coding | transmembrane protein 189 |
| **RDH14** | ENSG00000240857 | 410.3396218 | 1.006472071 | 0.184711022 | 1.52E-08 | 1.26E-07 | 57665 | 2 | 18735989 | 18741946 | -1 | protein_coding | retinol dehydrogenase 14 (all-trans/9-cis/11-cis) |
| **MIF** | ENSG00000240972 | 588.3333798 | -0.832168112 | 0.225900251 | 8.52E-05 | 3.95E-04 | 4282 | 22 | 24236191 | 24237414 | 1 | protein_coding | macrophage migration inhibitory factor (glycosylation-inhibiting factor) |
| **NSUN6** | ENSG00000241058 | 165.5040805 | 0.708933742 | 0.297356646 | 0.007440046 | 0.021621265 | 221078 | 10 | 18834490 | 18940551 | -1 | protein_coding | NOP2/Sun domain family, member 6 |
| **CFB** | ENSG00000241253 | 362.5569219 | 4.83941226 | 0.360104043 | 1.01E-42 | 7.13E-41 | 629 | HSC | 31903824 | 31910258 | 1 | protein_coding | complement factor B |
| **RPL36A** | ENSG00000241343 | 1428.900709 | -0.554462399 | 0.165225659 | 4.82E-04 | 0.001894317 | 1005290 | X | 100645812 | 100651105 | 1 | protein_coding | ribosomal protein L36a |

| | | | | | | | | | | | | | |
|---|---|---|---|---|---|---|---|---|---|---|---|---|---|
| **PDXP** | ENSG00000241360 | 16.85953177 | -2.826729876 | 1.214375017 | 0.001153578 | 0.004143392 | 57026 | 22 | 38054734 | 38062941 | 1 | protein_coding | pyridoxal (pyridoxine, vitamin B6) phosphatase |
| **HLA-DOB** | ENSG00000241386 | 72.22725133 | 2.047351543 | 0.47228883 | 1.32E-06 | 8.35E-06 | 3112 | HSC | 32702866 | 32707151 | -1 | protein_coding | major histocompatibility complex, class II, DO beta |
| **CD302** | ENSG00000241399 | 1790.168114 | -0.84357942 | 0.203513802 | 1.29E-05 | 7E-05 | 9936 | 2 | 160625364 | 160654753 | -1 | protein_coding | CD302 molecule |
| **ATP5J2** | ENSG00000241468 | 1455.370001 | -0.473289294 | 0.163881467 | 0.002592074 | 0.008491889 | 1019297 | 7 | 99046098 | 99063954 | -1 | protein_coding | ATP synthase, H+ transporting, mitochondrial Fo complex, subunit F2 |
| **HLA-DMB** | ENSG00000241674 | 10090.93348 | -1.474234524 | 0.134934998 | 1.27E-28 | 4.59E-27 | 3109 | HSC | 32831136 | 32849631 | -1 | protein_coding | major histocompatibility complex, class II, DM beta |
| **ATP5O** | ENSG00000241837 | 1661.035913 | -0.536089257 | 0.139879206 | 7.55E-05 | 3.53E-04 | 539 | 21 | 35275757 | 35288284 | -1 | protein_coding | ATP synthase, H+ transporting, mitochondrial F1 complex, O subunit |
| **PLEKHO2** | ENSG00000241839 | 9878.97283 | -0.531006014 | 0.103106969 | 1.75E-07 | 1.26E-06 | 80301 | 15 | 65134088 | 65160206 | 1 | protein_coding | pleckstrin homology domain containing, family O member 2 |
| **AMACR** | ENSG00000242110 | 383.2066061 | 0.555905785 | 0.180277433 | 0.001222229 | 0.004357203 | 23600 | 5 | 33986283 | 34008220 | -1 | protein_coding | alpha-methylacyl-CoA racemase |
| **MTFP1** | ENSG00000242114 | 262.3072454 | -0.732649161 | 0.216730214 | 3.27E-04 | 0.001338516 | 51537 | 22 | 30821518 | 30825045 | 1 | protein_coding | mitochondrial fission process 1 |
| **ARFGAP3** | ENSG00000242247 | 2031.101712 | 0.481962602 | 0.141746681 | 4.87E-04 | 0.001911218 | 26286 | 22 | 43192508 | 43254112 | -1 | protein_coding | ADP-ribosylation factor GTPase activating protein 3 |
| **CFB** | ENSG00000242335 | 4898.651616 | 5.388192839 | 0.371166832 | 2.99E-49 | 2.54E-47 | 629 | HSC | 31905279 | 31911713 | 1 | protein_coding | complement factor B |
| **C15orf38** | ENSG00000242498 | 658.2220151 | -0.789282887 | 0.146155392 | 2.71E-08 | 2.18E-07 | 348110 | 15 | 90443159 | 90456188 | -1 | protein_coding | chromosome 15 open reading frame 38 |
| **RP11-274B21** | ENSG00000242588 | 505.4663715 | 0.618740876 | 0.163083758 | 8.36E-05 | 3.88E-04 | 1019304 | 7 | 128171752 | 128269512 | 1 | pseudogene | |
| **GNG10** | ENSG00000242616 | 1199.679578 | 0.673340113 | 0.171300292 | 4.69E-05 | 2.27E-04 | 2790 | 9 | 114423615 | 114432526 | 1 | protein_coding | guanine nucleotide binding protein (G protein), gamma 10 |
| **HLA-DMA** | ENSG00000242685 | 6903.142293 | -0.922793154 | 0.142726463 | 3.1E-11 | 3.5E-10 | 3108 | HSC | 32845119 | 32867240 | -1 | protein_coding | major histocompatibility complex, class II, DM alpha |
| **PSMB9** | ENSG00000242711 | 950.9435841 | 1.142488188 | 0.258610597 | 2.23E-06 | 1.36E-05 | 5698 | HSC | 32795520 | 32830948 | 1 | protein_coding | proteasome (prosome, macropain) subunit, beta type, 9 |
| **ZNF702P** | ENSG00000242779 | 827.8476209 | 3.307895447 | 0.223681752 | 6.29E-51 | 5.79E-49 | 79986 | 19 | 53471504 | 53541151 | -1 | pseudogene | zinc finger protein 702, pseudogene |
| **KCTD7** | ENSG00000243335 | 1245.600634 | -0.723860252 | 0.15173491 | 8.57E-07 | 5.57E-06 | 154881 | 7 | 66093868 | 66276446 | 1 | protein_coding | potassium channel tetramerization domain containing 7 |
| **TICAM2** | ENSG00000243414 | 337.6222336 | 2.203601807 | 0.220955706 | 1.72E-24 | 5.01E-23 | 1003027 | 5 | 114914339 | 114961876 | -1 | protein_coding | toll-like receptor adaptor molecule 2 |
| **PSMB9** | ENSG00000243594 | 622.8319077 | 1.130774884 | 0.303045618 | 4.13E-05 | 2.02E-04 | 5698 | HSC | 32740623 | 32776073 | 1 | protein_coding | proteasome (prosome, macropain) subunit, beta type, 9 |
| **IL10RB** | ENSG00000243646 | 4029.651361 | 0.531223809 | 0.089450805 | 2.14E-09 | 1.97E-08 | 3588 | 21 | 34638663 | 34669539 | 1 | protein_coding | interleukin 10 receptor, beta |
| **CFB** | ENSG00000243649 | 204.3896774 | 3.52460418 | 1.651785901 | 8.61E-04 | 0.003196907 | 629 | 6 | 31895475 | 31919861 | 1 | protein_coding | complement factor B |
| **ZNF487** | ENSG00000243660 | 42.06264544 | 1.243986421 | 0.554757599 | 0.003720735 | 0.011743905 | NA | 10 | 43932282 | 43991517 | 1 | protein_coding | zinc finger protein 487 |
| **WDR92** | ENSG00000243667 | 370.7319976 | -0.449135767 | 0.203639735 | 0.019256416 | 0.049147595 | 116143 | 2 | 68350068 | 68384692 | -1 | protein_coding | WD repeat domain 92 |
| **NME1-NME2** | ENSG00000243678 | 2602.637949 | -0.315960075 | 0.126544302 | 0.009714886 | 0.027330558 | 654364 | 17 | 49230920 | 49249108 | 1 | protein_coding | NME1-NME2 readthrough |
| **RP5-966M1.6** | ENSG00000243696 | 188.1154038 | 2.300549198 | 0.361804994 | 1.68E-11 | 1.94E-10 | NA | 3 | 52847298 | 52869745 | -1 | protein_coding | Musculoskeletal embryonic nuclear protein 1 |
| **PLA2G4B** | ENSG00000243708 | 40.85743914 | 1.890601298 | 0.849271101 | 0.001508113 | 0.005261012 | 1001370 | 15 | 42120283 | 42140345 | 1 | protein_coding | phospholipase A2, group IVB (cytosolic) |
| **NPIPB5** | ENSG00000243716 | 854.227313 | 0.51832294 | 0.134050219 | 7.34E-05 | 3.43E-04 | 1019299 | 16 | 22490442 | 22547842 | 1 | protein_coding | nuclear pore complex interacting protein family, member B5 |
| **ZMYM6NB** | ENSG00000243749 | 291.4680399 | 1.519289513 | 0.223096941 | 1.23E-12 | 1.55E-11 | 1005061 | 1 | 35447136 | 35450954 | -1 | protein_coding | ZMYM6 neighbor |
| **APOBEC3D** | ENSG00000243811 | 435.6032402 | 1.151206255 | 0.203086513 | 3.36E-09 | 3.02E-08 | 140564 | 22 | 39410368 | 39429281 | 1 | protein_coding | apolipoprotein B mRNA editing enzyme, catalytic polypeptide-like 3D |
| **MRPS6** | ENSG00000243927 | 4786.327579 | -0.321744257 | 0.097485463 | 8.34E-04 | 0.003106159 | 64968 | 21 | 35445524 | 35515334 | 1 | protein_coding | mitochondrial ribosomal protein S6 |
| **ACY1** | ENSG00000243989 | 257.9274459 | -1.431360029 | 0.241327356 | 5.19E-10 | 5.13E-09 | 95 | 3 | 52009066 | 52023213 | 1 | protein_coding | aminoacylase 1 |
| **DDOST** | ENSG00000244038 | 3704.975056 | -0.357298385 | 0.096896283 | 1.91E-04 | 8.22E-04 | 1650 | 1 | 20978270 | 20988000 | -1 | protein_coding | dolichyl-diphosphooligosaccharide--protein glycosyltransferase subunit (non-catalytic) |
| **TMEM141** | ENSG00000244187 | 230.9850608 | -0.59257691 | 0.223322168 | 0.004459991 | 0.013784032 | 85014 | 9 | 139685807 | 139687709 | 1 | protein_coding | transmembrane protein 141 |
| **PKD1P1** | ENSG00000244257 | 129.6415042 | -1.552035329 | 0.489156038 | 1.63E-04 | 7.14E-04 | NA | 16 | 16404198 | 16428047 | 1 | pseudogene | polycystic kidney disease 1 (autosomal dominant) pseudogene 1 |
| **ETV5** | ENSG00000244405 | 1102.534677 | -2.333986454 | 0.216086736 | 2.08E-28 | 7.44E-27 | 2119 | 3 | 185764097 | 185828107 | -1 | protein_coding | ets variant 5 |
| **APOBEC3C** | ENSG00000244509 | 3326.309539 | 0.518964644 | 0.090761538 | 6.96E-09 | 6.04E-08 | 27350 | 22 | 39410088 | 39416357 | 1 | protein_coding | apolipoprotein B mRNA editing enzyme, catalytic polypeptide-like 3C |
| **FCGR2C** | ENSG00000244682 | 1089.491951 | -2.147328425 | 0.18263541 | 8.97E-33 | 4.04E-31 | 9103 | 1 | 161551129 | 161575452 | 1 | polymorphic_ps | Fc fragment of IgG, low affinity IIc, receptor for (CD32) (gene/pseudogene) |
| **N4BP2L2** | ENSG00000244754 | 2602.414665 | 0.554763175 | 0.135361308 | 2.6E-05 | 1.33E-04 | 10443 | 13 | 33006554 | 33112970 | -1 | protein_coding | NEDD4 binding protein 2-like 2 |
| **CEBPA** | ENSG00000245848 | 1142.482284 | -1.658605604 | 0.144631471 | 2.97E-31 | 1.23E-29 | 1050 | 19 | 33790840 | 33793470 | -1 | protein_coding | CCAAT/enhancer binding protein (C/EBP), alpha |
| **AL034548.1** | ENSG00000247315 | 260.323969 | 0.59266572 | 0.22447148 | 0.004309174 | 0.013372 | NA | 20 | 278214 | 280961 | 1 | pseudogene | |
| **TWF2** | ENSG00000247596 | 2441.29852 | -0.567686549 | 0.128480286 | 6.33E-06 | 3.59E-05 | 11344 | 3 | 52262626 | 52273276 | -1 | protein_coding | twinfilin actin-binding protein 2 |
| **USP51** | ENSG00000247746 | 191.0013003 | -0.67895949 | 0.32332232 | 0.015611795 | 0.04115837 | 158880 | X | 55511049 | 55515635 | -1 | protein_coding | ubiquitin specific peptidase 51 |
| **BCKDHA** | ENSG00000248098 | 251.6287577 | -1.587775548 | 0.257290614 | 1.05E-10 | 1.13E-09 | 593 | 19 | 41884215 | 41930910 | 1 | protein_coding | branched chain keto acid dehydrogenase E1, alpha polypeptide |
| **MTATP6P1** | ENSG00000248527 | 4463.672791 | -0.839732836 | 0.409434597 | 0.012236319 | 0.033269466 | NA | 1 | 569076 | 569756 | 1 | pseudogene | mitochondrially encoded ATP synthase 6 pseudogene 1 |
| **TMEM110-M** | ENSG00000248592 | 238.6114008 | 2.479209328 | 0.559906209 | 6.35E-07 | 4.23E-06 | 1005267 | 3 | 52867137 | 52931578 | -1 | protein_coding | TMEM110-MUSTN1 readthrough |
| **NAIP** | ENSG00000249437 | 1410.769043 | -1.495839494 | 0.187069385 | 1.84E-16 | 3.09E-15 | 4671 | 5 | 70264310 | 70320941 | -1 | protein_coding | NLR family, apoptosis inhibitory protein |
| **AP000295.9** | ENSG00000249624 | 23.63562885 | 1.422227279 | 1.012947387 | 0.010582689 | 0.029347712 | NA | 21 | 34619079 | 34655517 | 1 | protein_coding | |
| **AC011330.5** | ENSG00000249839 | 117.27879 | -1.71736195 | 0.461515491 | 2.25E-05 | 1.16E-04 | NA | 15 | 43955852 | 43976537 | -1 | pseudogene | |
| **RP11-177C12** | ENSG00000249863 | 48.40450101 | -3.531905108 | 0.635606361 | 2.8E-09 | 2.54E-08 | NA | 4 | 37869913 | 37871599 | 1 | pseudogene | |
| **DNAH10OS** | ENSG00000250091 | 34.36271593 | -1.651308424 | 0.6334861 | 9.23E-04 | 0.003398259 | NA | 12 | 124410971 | 124419531 | -1 | protein_coding | dynein, axonemal, heavy chain 10 opposite strand |
| **RP11-848G14** | ENSG00000250138 | 142.2897989 | -2.321783428 | 0.457700969 | 3.12E-08 | 2.5E-07 | 1019295 | 5 | 68927790 | 68932312 | 1 | pseudogene | |
| **KIAA1456** | ENSG00000250305 | 89.90706177 | -1.447017742 | 0.504444308 | 5.35E-04 | 0.002078487 | 1019271 | 8 | 12803151 | 12889012 | 1 | protein_coding | KIAA1456 |
| **KIAA1210** | ENSG00000250423 | 180.5869007 | -1.444590621 | 0.571476309 | 0.001263866 | 0.004491985 | 57481 | X | 118212598 | 118284542 | -1 | protein_coding | KIAA1210 |
| **CHCHD10** | ENSG00000250479 | 607.168446 | -1.337173175 | 0.20056633 | 4.69E-12 | 5.69E-11 | 400916 | 22 | 24108021 | 24110630 | -1 | protein_coding | coiled-coil-helix-coiled-coil-helix domain containing 10 |
| **GPR162** | ENSG00000250510 | 78.50897313 | -1.607361814 | 0.415377405 | 1.37E-05 | 7.36E-05 | 27239 | 12 | 6930711 | 6939136 | 1 | protein_coding | G protein-coupled receptor 162 |
| **ATP6V1E2** | ENSG00000250565 | 73.28609188 | -0.811540632 | 0.399331728 | 0.014027408 | 0.037498841 | 90423 | 2 | 46717889 | 46769696 | -1 | protein_coding | ATPase, H+ transporting, lysosomal 31kDa, V1 subunit E2 |
| **RP11-295K3.** | ENSG00000250644 | 177.8329289 | -0.586232916 | 0.264864523 | 0.014846631 | 0.03935838 | NA | 11 | 1768897 | 1780281 | -1 | protein_coding | |
| **SEPP1** | ENSG00000250722 | 775.0846149 | -1.293811696 | 0.68871512 | 0.006845074 | 0.020114068 | 6414 | 5 | 42799982 | 42887494 | -1 | protein_coding | selenoprotein P, plasma, 1 |
| **TMED7-TICA** | ENSG00000251201 | 636.2535281 | 1.014882481 | 0.201323146 | 1.33E-07 | 9.67E-07 | 1003027 | 5 | 114914339 | 114961858 | -1 | protein_coding | TMED7-TICAM2 readthrough |
| **SHANK3** | ENSG00000251322 | 71.35484525 | -1.605943137 | 0.767594786 | 0.003193374 | 0.01026595 | 85358 | 22 | 51112843 | 51171726 | 1 | protein_coding | SH3 and multiple ankyrin repeat domains 3 |
| **RP11-597D13** | ENSG00000251429 | 47.66236562 | 1.115645684 | 0.570017401 | 0.00840643 | 0.024036571 | NA | 4 | 159191530 | 159199828 | 1 | pseudogene | |
| **PCDHGA12** | ENSG00000253159 | 48.27094248 | -1.09469339 | 0.498988794 | 0.005543066 | 0.016676119 | 26025 | 5 | 140810185 | 140892546 | 1 | protein_coding | protocadherin gamma subfamily A, 12 |
| **CCDC71L** | ENSG00000253276 | 745.3185406 | 1.250952416 | 0.147199066 | 3.91E-18 | 7.45E-17 | 168455 | 7 | 106297211 | 106301442 | -1 | protein_coding | coiled-coil domain containing 71-like |
| **CTD-2114J12** | ENSG00000253525 | 34.60487864 | 0.982810649 | 0.5567823 | 0.016386273 | 0.042889434 | NA | 8 | 92169513 | 92170976 | 1 | pseudogene | |
| **PCDHGA7** | ENSG00000253537 | 5.770391429 | -3.373239818 | 3.231897913 | 0.004745863 | 0.014582031 | 56108 | 5 | 140762467 | 140892546 | 1 | protein_coding | protocadherin gamma subfamily A, 7 |
| **ALG11** | ENSG00000253710 | 493.2948636 | 0.439511103 | 0.175655424 | 0.009023453 | 0.025604424 | 440138 | 13 | 52586534 | 52603800 | 1 | protein_coding | ALG11, alpha-1,2-mannosyltransferase |
| **ATXN7L3B** | ENSG00000253719 | 1433.633131 | 0.258578966 | 0.110852677 | 0.017741277 | 0.045885925 | 552889 | 12 | 74931551 | 74935223 | 1 | protein_coding | ataxin 7-like 3B |
| **MTND1P5** | ENSG00000253795 | 7.044023008 | 4.354984831 | 2.11553889 | 0.002331213 | 0.007738626 | NA | 8 | 104097879 | 104098818 | 1 | pseudogene | MT-ND1 pseudogene 5 |
| **PCDHGA10** | ENSG00000253846 | 130.9999202 | -0.690391598 | 0.301046339 | 0.009816227 | 0.027565797 | 56106 | 5 | 140792743 | 140892546 | 1 | protein_coding | protocadherin gamma subfamily A, 10 |
| **SLC2A3P1** | ENSG00000253861 | 85.16702238 | -10.1504545 | 2.988739942 | 1.53E-10 | 1.61E-09 | NA | 5 | 167979282 | 167980732 | -1 | pseudogene | solute carrier family 2 (facilitated glucose transporter), member 3 pseudogene 1 |
| **PCDHGB4** | ENSG00000253953 | 70.9293511 | -1.214085861 | 0.500165187 | 0.002475566 | 0.008153895 | 56101 | 5 | 140767452 | 140892546 | 1 | protein_coding | protocadherin gamma subfamily B, 4 |
| **CLDN23** | ENSG00000253958 | 176.3753243 | 0.611247853 | 0.272669938 | 0.013191425 | 0.035540164 | 137075 | 8 | 8559448 | 8561616 | 1 | protein_coding | claudin 23 |
| **ZNF260** | ENSG00000254004 | 686.5276214 | 0.41362832 | 0.177548343 | 0.014676318 | 0.038981259 | 339324 | 19 | 37001597 | 37019562 | -1 | protein_coding | zinc finger protein 260 |
| **LYN** | ENSG00000254087 | 7626.127327 | 1.473910419 | 0.11275772 | 6.21E-40 | 3.72E-38 | 4067 | 8 | 56792372 | 56923940 | 1 | protein_coding | v-yes-1 Yamaguchi sarcoma viral related oncogene homolog |
| **CHKB-CPT1B** | ENSG00000254413 | 127.919228 | 1.123021165 | 0.462768496 | 0.002892565 | 0.009385789 | NA | 22 | 51007298 | 51021394 | -1 | protein_coding | CHKB-CPT1B readthrough (NMD candidate) |
| **SIGLEC14** | ENSG00000254415 | 1053.862126 | 0.595722752 | 0.15784359 | 9.89E-05 | 4.52E-04 | 1000495 | 19 | 52145806 | 52150054 | -1 | protein_coding | sialic acid binding Ig-like lectin 14 |
| **AP5B1** | ENSG00000254470 | 2562.003351 | 0.624784215 | 0.128575076 | 6.79E-07 | 4.49E-06 | 91056 | 11 | 65543364 | 65548273 | -1 | protein_coding | adaptor-related protein complex 5, beta 1 subunit |
| **RP11-1036E2** | ENSG00000254704 | 14.79340841 | 3.4688437 | 1.030642416 | 4.3E-05 | 2.1E-04 | NA | 11 | 59054812 | 59056959 | -1 | pseudogene | |
| **EEF1G** | ENSG00000254772 | 11223.38599 | -0.871632003 | 0.104732147 | 2.52E-17 | 4.52E-16 | 1005008 | 11 | 62327073 | 62342401 | -1 | protein_coding | eukaryotic translation elongation factor 1 gamma |
| **BRK1** | ENSG00000254999 | 1673.776137 | -0.375210747 | 0.118692034 | 0.001179666 | 0.004225697 | 55845 | 3 | 10157276 | 10168874 | 1 | protein_coding | BRICK1, SCAR/WAVE actin-nucleating complex subunit |
| **ZFP91-CNTF** | ENSG00000255073 | 22.00352007 | -0.70213134 | 1.337599193 | 0.009693495 | 0.027293909 | NA | 11 | 58346645 | 58392112 | 1 | protein_coding | ZFP91-CNTF readthrough (NMD candidate) |
| **CHMP1B** | ENSG00000255112 | 1567.424304 | 0.613574108 | 0.146203302 | 1.56E-05 | 8.29E-05 | 57132 | 18 | 11851395 | 11854448 | 1 | protein_coding | charged multivesicular body protein 1B |
| **HSPE1P18** | ENSG00000255163 | 9.67720657 | -1.438835843 | 1.48590591 | 0.018686514 | 0.047909074 | NA | 11 | 118079647 | 118079926 | -1 | pseudogene | heat shock 10kDa protein 1 pseudogene 18 |
| **EID1** | ENSG00000255302 | 1162.464356 | -0.422313154 | 0.178240199 | 0.013062211 | 0.035228358 | 23741 | 15 | 49170083 | 49172380 | 1 | protein_coding | EP300 interacting inhibitor of differentiation 1 |
| **HCAR3** | ENSG00000255398 | 366.5070279 | -0.696332444 | 0.244179884 | 0.001971022 | 0.006680666 | 8843 | 12 | 123199303 | 123201439 | -1 | protein_coding | hydroxycarboxylic acid receptor 3 |
| **MIR3654** | ENSG00000255508 | 153.9730059 | 1.266031925 | 0.364748896 | 8.94E-05 | 4.13E-04 | 1005008 | 11 | 62327075 | 62359003 | -1 | protein_coding | microRNA 3654 |
| **TIFAB** | ENSG00000255833 | 646.3588657 | -2.965488206 | 0.227949308 | 5.2E-40 | 3.13E-38 | 497189 | 5 | 134779908 | 134788089 | -1 | protein_coding | TRAF-interacting protein with forkhead-associated domain, family member B |
| **RP11-1094M** | ENSG00000255987 | 24.50374617 | 2.027666605 | 0.768387342 | 5.98E-04 | 0.002294206 | NA | 17 | 33841785 | 33842205 | -1 | pseudogene | |

| | | | | | | | | | | | | | |
|---|---|---|---|---|---|---|---|---|---|---|---|---|---|
| **RP11-190A12** | ENSG00000256029 | 46.1314256 | -1.031660784 | 0.590287892 | 0.014241469 | 0.03797982 | NA | 1 | 159827690 | 159842843 | -1 | protein_coding | Uncharacterized protein |
| **DYX1C1** | ENSG00000256061 | 55.0366061 | -2.072997182 | 0.720131194 | 2.62E-04 | 0.001090767 | 161582 | 15 | 55702723 | 55800432 | -1 | protein_coding | dyslexia susceptibility 1 candidate 1 |
| **POLG2** | ENSG00000256525 | 54.4622563 | 1.206607874 | 0.468446511 | 0.001742558 | 0.005971192 | 11232 | 17 | 62473902 | 62493154 | -1 | protein_coding | polymerase (DNA directed), gamma 2, accessory subunit |
| **PSMA2** | ENSG00000256646 | 865.813741 | 0.454711322 | 0.135284791 | 5.69E-04 | 0.002193917 | 5683 | 7 | 42948872 | 42971773 | -1 | protein_coding | Proteasome subunit alpha type-2 |
| **RP11-611O2.** | ENSG00000256664 | 324.0592113 | 1.976732347 | 0.265318907 | 9.76E-15 | 1.45E-13 | NA | 12 | 69235726 | 69236164 | 1 | pseudogene | |
| **ZNF350** | ENSG00000256683 | 469.1024449 | 0.611372315 | 0.170347145 | 1.87E-04 | 8.06E-04 | 59348 | 19 | 52467596 | 52490109 | -1 | protein_coding | zinc finger protein 350 |
| **KIAA1147** | ENSG00000257093 | 1468.351709 | -1.246112932 | 0.201549012 | 1.3E-10 | 1.37E-09 | 57189 | 7 | 141356528 | 141401953 | -1 | protein_coding | KIAA1147 |
| **LIMS3** | ENSG00000257207 | 123.9372294 | -0.644794369 | 0.311615656 | 0.018415401 | 0.047317178 | NA | 2 | 111160511 | 111230652 | -1 | protein_coding | LIM and senescent cell antigen-like-containing domain protein 3; Uncharacterized protein; cDNA FLJ59 |
| **ZBED6** | ENSG00000257315 | 985.4820135 | 0.677509115 | 0.139837544 | 6.58E-07 | 4.37E-06 | 1003812 | 1 | 203765437 | 203769686 | 1 | protein_coding | zinc finger, BED-type containing 6 |
| **FNTB** | ENSG00000257365 | 409.7171212 | -0.588395347 | 0.201990629 | 0.00204295 | 0.006889322 | 1005292 | 14 | 65381203 | 65529368 | 1 | protein_coding | farnesyltransferase, CAAX box, beta |
| **GALNT4** | ENSG00000257594 | 496.6432196 | 1.431247079 | 0.172150888 | 1.62E-17 | 2.95E-16 | 1005280 | 12 | 89913185 | 89920039 | -1 | protein_coding | UDP-N-acetyl-alpha-D-galactosamine:polypeptide N-acetylgalactosaminyltransferase 4 (GalNAc-T4) |
| **RP11-293I14.** | ENSG00000258064 | 33.54085153 | -0.18395122 | 0.675080877 | 0.003823354 | 0.012013223 | NA | 12 | 72067984 | 72092748 | 1 | protein_coding | Uncharacterized protein |
| **TRIM6-TRIM3** | ENSG00000258588 | 14.7713904 | 0.069589092 | 0.639476179 | 2.01E-07 | 1.43E-06 | 445372 | 11 | 5617955 | 5665628 | 1 | protein_coding | TRIM6-TRIM34 readthrough |
| **TRIM34** | ENSG00000258659 | 587.8727826 | 0.79238591 | 0.208218046 | 5.65E-05 | 2.71E-04 | 53840 | 11 | 5640994 | 5665628 | 1 | protein_coding | tripartite motif containing 34 |
| **RP11-463D19** | ENSG00000258677 | 139.1131816 | -0.818869366 | 0.365566539 | 0.008140544 | 0.023385702 | NA | 8 | 74600926 | 74791089 | -1 | protein_coding | Uncharacterized protein |
| **FDPSP3** | ENSG00000258872 | 4.851928944 | 3.194178855 | 3.750650335 | 0.008072083 | 0.023206479 | NA | 14 | 55351567 | 55352931 | -1 | pseudogene | farnesyl diphosphate synthase pseudogene 3 |
| **TUBB3** | ENSG00000258947 | 94.93145921 | -1.310248382 | 0.510049557 | 0.001464513 | 0.005127641 | 10381 | 16 | 89987800 | 90005169 | 1 | protein_coding | Tubulin beta-3 chain |
| **POC1B-GALI** | ENSG00000259075 | 81.78125754 | 1.479664166 | 0.600620608 | 0.001555204 | 0.005409235 | 1005280 | 12 | 89913185 | 89920039 | -1 | protein_coding | POC1B-GALNT4 readthrough |
| **THTPA** | ENSG00000259431 | 555.3135629 | 0.644472369 | 0.160586048 | 3.2E-05 | 1.61E-04 | 79178 | 14 | 24025216 | 24029480 | 1 | protein_coding | thiamine triphosphatase |
| **RP11-468E2.** | ENSG00000259529 | 140.3384689 | 0.729309255 | 0.318489226 | 0.009202248 | 0.026053715 | 10379 | 14 | 24620427 | 24636611 | 1 | protein_coding | E3 ubiquitin-protein ligase RNF31 |
| **C15orf37** | ENSG00000259642 | 17.21701868 | -2.071078434 | 1.756263434 | 0.011723253 | 0.032102934 | NA | 15 | 80215113 | 80217194 | 1 | protein_coding | chromosome 15 open reading frame 37 |
| **MRC1** | ENSG00000260314 | 2879.008452 | -2.495512427 | 0.233266733 | 7.35E-28 | 2.54E-26 | 1019287 | HG5 | 17851343 | 17953161 | 1 | protein_coding | mannose receptor, C type 1 |
| **RP11-343C2.** | ENSG00000260371 | 20.63841369 | 6.313376563 | 2.329264436 | 1.66E-05 | 8.79E-05 | NA | 16 | 69368994 | 69390209 | -1 | protein_coding | Uncharacterized protein |
| **RP4-576H24.** | ENSG00000260861 | 78.1675281 | -2.685509136 | 0.596957754 | 3.89E-07 | 2.67E-06 | NA | 20 | 1520790 | 1600655 | -1 | protein_coding | Uncharacterized protein |
| **CCPG1** | ENSG00000260916 | 8439.259983 | 0.603283809 | 0.100316917 | 1.84E-09 | 1.72E-08 | 9236 | 15 | 55632230 | 55700708 | -1 | protein_coding | cell cycle progression 1 |
| **ZNF865** | ENSG00000261221 | 277.0970987 | -0.849048851 | 0.26282191 | 3.97E-04 | 0.001592622 | 1005072 | 19 | 56116771 | 56128635 | 1 | protein_coding | zinc finger protein 865 |
| **GOLGA8T** | ENSG00000261247 | 5.477825375 | -0.335157556 | 0.762080585 | 0.019291192 | 0.049228132 | 653073 | 15 | 30427352 | 30437770 | 1 | protein_coding | golgin A8 family, member T |
| **PECAM1** | ENSG00000261371 | 15802.61015 | -0.562769883 | 0.081589924 | 3.57E-12 | 4.37E-11 | 5175 | HG1 | 62399863 | 62491136 | -1 | protein_coding | platelet/endothelial cell adhesion molecule 1 |
| **TPBGL** | ENSG00000261594 | 20.31054164 | -6.592875421 | 2.646395777 | 1.54E-05 | 8.2E-05 | 1005070 | 11 | 74951950 | 74954742 | 1 | protein_coding | trophoblast glycoprotein-like |
| **CTC-479C5.1** | ENSG00000261884 | 240.2237448 | 0.89461025 | 0.295344468 | 7.76E-04 | 0.00290668 | NA | 16 | 67963517 | 67969920 | -1 | protein_coding | |
| **APOBEC3A** | ENSG00000262156 | 162.4875801 | 0.255504084 | 0.711248129 | 0.004807386 | 0.014762162 | 1009131 | HSC | 39353528 | 39358842 | 1 | protein_coding | apolipoprotein B mRNA editing enzyme, catalytic polypeptide-like 3A |
| **PTPRC** | ENSG00000262418 | 22410.39875 | 0.552148351 | 0.07634778 | 2.54E-13 | 3.37E-12 | 5788 | HSC | 198619288 | 198737899 | 1 | protein_coding | protein tyrosine phosphatase, receptor type, C |
| **GART** | ENSG00000262473 | 24.60677772 | 1.583774056 | 0.745571403 | 0.003042742 | 0.009829258 | 2618 | HSC | 34880498 | 34925355 | -1 | protein_coding | phosphoribosylglycinamide formyltransferase, phosphoribosylglycinamide synthetase, phosphoribosyla |
| **RP11-876N24** | ENSG00000262488 | 5.285109369 | -0.630866243 | 1.132736007 | 0.011470264 | 0.031494797 | NA | 16 | 10934241 | 10935401 | -1 | pseudogene | |
| **FAM189B** | ENSG00000262666 | 346.5976526 | 0.71971822 | 0.199190881 | 1.33E-04 | 5.95E-04 | 10712 | HSC | 155232400 | 155240678 | -1 | protein_coding | family with sequence similarity 189, member B |
| **MRPL12** | ENSG00000262814 | 416.141687 | -0.772808828 | 0.187394228 | 1.56E-05 | 8.32E-05 | 6182 | 17 | 79670387 | 79674556 | 1 | protein_coding | mitochondrial ribosomal protein L12 |
| **LSM14A** | ENSG00000262860 | 1532.611505 | 0.295074132 | 0.109808051 | 0.00649453 | 0.019202147 | 26065 | HSC | 34663352 | 34720227 | 1 | protein_coding | LSM14A, SCD6 homolog A (S. cerevisiae) |
| **GTF2I** | ENSG00000263001 | 164.685497 | 0.770194351 | 0.292607376 | 0.003370018 | 0.010759137 | 2969 | HG1 | 74122661 | 74225649 | 1 | protein_coding | cDNA FLJ55916, highly similar to General transcription factor II-I |
| **MYZAP** | ENSG00000263155 | 11.36464369 | -5.156872505 | 2.716286353 | 5.44E-04 | 0.002107511 | 1008208 | 15 | 57884139 | 57977562 | 1 | protein_coding | myocardial zonula adherens protein |
| **CTSO** | ENSG00000263238 | 1414.053532 | 0.460395011 | 0.127351942 | 2.15E-04 | 9.13E-04 | 1519 | HSC | 156845270 | 156875063 | -1 | protein_coding | cathepsin O |
| **FAM72C** | ENSG00000263513 | 34.81820244 | 3.073958281 | 1.103605983 | 1.83E-04 | 7.9E-04 | 554282 | HG1 | 144641706 | 144658404 | -1 | protein_coding | Homo sapiens family with sequence similarity 72, member C (FAM72C), mRNA. |
| **DYNLL2** | ENSG00000264364 | 1690.646615 | -0.273820536 | 0.10738852 | 0.009625286 | 0.027125806 | 140735 | 17 | 56160776 | 56172897 | 1 | protein_coding | dynein, light chain, LC8-type 2 |
| **OTUD7B** | ENSG00000264522 | 666.3857123 | 0.664168735 | 0.236355281 | 0.002579701 | 0.008456793 | 56957 | HG1 | 150626747 | 150697152 | -1 | protein_coding | OTU domain containing 7B; OTU domain-containing protein 7B |
| **GJA5** | ENSG00000265107 | 12.46511833 | -5.21074343 | 2.657975106 | 4.16E-04 | 0.001663335 | 2702 | HG1 | 148443603 | 148445686 | -1 | protein_coding | Homo sapiens gap junction protein, alpha 5, 40kDa (GJA5), transcript variant B, mRNA. |
| **RNF115** | ENSG00000265491 | 578.6775625 | 0.931624034 | 0.148908915 | 1.35E-10 | 1.42E-09 | 27246 | HG1 | 146432834 | 146510487 | -1 | protein_coding | |
| **RPL17** | ENSG00000265681 | 6022.095815 | -0.329560105 | 0.126201419 | 0.007057943 | 0.020648265 | 1001245 | 18 | 47014851 | 47018906 | -1 | protein_coding | ribosomal protein L17 |
| **SEC22B** | ENSG00000265808 | 4153.967628 | 0.592858208 | 0.113107766 | 9.78E-08 | 7.26E-07 | 9554 | HG1 | 120698575 | 120719144 | -1 | protein_coding | Homo sapiens SEC22 vesicle trafficking protein homolog B (S. cerevisiae) (gene/pseudogene) (SEC22B |
| **SRGAP2** | ENSG00000266028 | 2220.356156 | 1.017662168 | 0.173733844 | 1.57E-09 | 1.48E-08 | 653464 | HG1 | 206266906 | 206527996 | 1 | protein_coding | SLIT-ROBO Rho GTPase activating protein 2 |
| **NBPF15** | ENSG00000266338 | 151.1463611 | 1.24248746 | 0.316882336 | 1.6E-05 | 8.52E-05 | 284565 | HG1 | 145107801 | 145148089 | -1 | protein_coding | neuroblastoma breakpoint family, member 15 |
| **MYO15B** | ENSG00000266714 | 210.7588581 | -0.69623976 | 0.323719353 | 0.013520597 | 0.036341062 | 80022 | 17 | 73584139 | 73622929 | 1 | protein_coding | myosin XVB pseudogene |
| **ZNF850** | ENSG00000267041 | 108.3044287 | -1.615425366 | 0.462905257 | 5.56E-05 | 2.66E-04 | 342892 | 19 | 37205285 | 37263727 | -1 | protein_coding | zinc finger protein 850 |
| **AC010642.1** | ENSG00000267216 | 664.2303408 | 0.415819518 | 0.140882613 | 0.002384499 | 0.00789136 | NA | 19 | 58790332 | 58826563 | 1 | protein_coding | Uncharacterized protein |
| **UPK3BL** | ENSG00000267368 | 216.6953587 | -1.093994766 | 0.411158091 | 0.001552699 | 0.005401753 | 1001349 | 7 | 102277472 | 102283238 | -1 | protein_coding | uroplakin 3B-like |
| **ZNF285** | ENSG00000267508 | 49.81840173 | -0.878857533 | 0.475448552 | 0.017577571 | 0.045555031 | 646915 | 19 | 44886459 | 44905774 | -1 | protein_coding | zinc finger protein 285 |
| **RP11-686D22** | ENSG00000267648 | 40.24358051 | 1.747845542 | 0.561810159 | 1.88E-04 | 8.11E-04 | NA | 17 | 33733967 | 33734837 | -1 | pseudogene | |
| **FDX1L** | ENSG00000267673 | 184.1936152 | -0.744930585 | 0.257220094 | 0.001598934 | 0.005543672 | 112812 | 19 | 10416103 | 10426691 | -1 | protein_coding | ferredoxin 1-like |
| **NDUFA7** | ENSG00000267855 | 258.7609877 | -0.48525511 | 0.203383059 | 0.011241897 | 0.0309679 | 4701 | 19 | 8373490 | 8386280 | -1 | protein_coding | NADH dehydrogenase (ubiquinone) 1 alpha subcomplex, 7, 14.5kDa |
| **CTC-429P9.4** | ENSG00000268790 | 29.91346972 | -7.862951668 | 2.877838614 | 2.71E-06 | 1.64E-05 | NA | 19 | 16688453 | 16770926 | -1 | protein_coding | Small integral membrane protein 7; Uncharacterized protein |
| **CSAG2** | ENSG00000268916 | 20.13519797 | 9.537178551 | 3.029900284 | 1.05E-07 | 7.76E-07 | 728461 | HG1 | 151927690 | 151928694 | 1 | protein_coding | Homo sapiens CSAG family, member 2 (CSAG2), transcript variant 2, mRNA. |
| **CTD-2192J16** | ENSG00000269242 | 229.3424635 | -0.883774019 | 0.256372607 | 1.83E-04 | 7.9E-04 | NA | 19 | 12754645 | 12759211 | -1 | protein_coding | |
| **ZNF587B** | ENSG00000269343 | 356.623489 | 0.675875995 | 0.181338827 | 9.91E-05 | 4.53E-04 | 1002935 | 19 | 58331099 | 58370018 | 1 | protein_coding | zinc finger protein 587B |
| **ITGB1P1** | ENSG00000269378 | 860.7352629 | -1.340400717 | 0.250923106 | 1.26E-08 | 1.06E-07 | NA | 19 | 14732446 | 14733054 | -1 | pseudogene | integrin beta 1 pseudogene 1 |
| **IGHV1OR15-1** | ENSG00000270505 | 12.77122008 | -2.056275158 | 1.227381367 | 0.005400461 | 0.01632099 | NA | 15 | 22448382 | 22448819 | -1 | IG_V_gene | immunoglobulin heavy variable 1/OR15-1 (non-functional) |
| **AC239811.2** | ENSG00000270629 | 130.6483674 | 0.788553152 | 0.356232483 | 0.009811263 | 0.027558451 | 400818 | HG1 | 149219394 | 149282305 | -1 | protein_coding | Neuroblastoma breakpoint family member 1 |
| **HSPE1-MOB** | ENSG00000270757 | 165.201378 | 0.726739906 | 0.276829333 | 0.003721624 | 0.011744288 | 1005292 | 2 | 198365137 | 198415450 | 1 | protein_coding | HSPE1-MOB4 readthrough |
| **SRGAP2D** | ENSG00000270872 | 401.9060024 | 0.543217176 | 0.169024161 | 8.45E-04 | 0.003143952 | 1009967 | HG1 | 144661157 | 144754876 | 1 | protein_coding | SLIT-ROBO Rho GTPase activating protein 2D |
| **HIST1H4A** | ENSG00000270882 | 202.8524684 | 1.294548431 | 0.368362982 | 7.12E-05 | 3.34E-04 | 554313 | HG1 | 150519072 | 150519467 | 1 | protein_coding | Homo sapiens histone cluster 2, H4b (HIST2H4B), mRNA. |
| **SRXN1** | ENSG00000271303 | 1547.771064 | 1.344126918 | 0.140305574 | 1.81E-22 | 4.69E-21 | 140809 | 20 | 627259 | 634014 | -1 | protein_coding | sulfiredoxin 1 |
| **AC242842.2** | ENSG00000271383 | 1054.593552 | 1.186911511 | 0.126440382 | 1.28E-21 | 3.15E-20 | NA | HG1 | 150162303 | 150242670 | 1 | protein_coding | NBPF |
| **AC239860.1** | ENSG00000272150 | 35.53836973 | -1.308121892 | 0.610116452 | 0.004350367 | 0.013483864 | NA | HG1 | 146260284 | 146274466 | -1 | protein_coding | |
| **NUDT3** | ENSG00000272325 | 3299.694489 | -1.13994128 | 0.098937415 | 6.62E-31 | 2.69E-29 | 11165 | 6 | 34247456 | 34360451 | -1 | protein_coding | nudix (nucleoside diphosphate linked moiety X)-type motif 3 |
| **FAM47E-STB** | ENSG00000272414 | 475.6774749 | 2.361514712 | 0.273778118 | 5.22E-19 | 1.07E-17 | 1006313 | 4 | 77172874 | 77232282 | 1 | protein_coding | uncharacterized protein LOC100631383 |
| **C6ORF165** | ENSG00000272514 | 3.725352415 | 0.254307344 | 0.709454068 | 0.013272078 | 0.03572964 | 154313 | 6 | 88117701 | 88174183 | 1 | protein_coding | UPF0704 protein C6orf165 |
| **AC112218.1** | ENSG00000272886 | 1226.159949 | 1.539042599 | 0.198901905 | 1.25E-15 | 1.98E-14 | 55802 | HG9 | 53317443 | 53381643 | -1 | protein_coding | cDNA FLJ53884, highly similar to mRNA decapping enzyme 1A |
| **RP11-500M8.** | ENSG00000272921 | 234.657388 | 0.788312861 | 0.273907476 | 0.001567709 | 0.00544777 | NA | 12 | 4714110 | 4771809 | 1 | protein_coding | |
| **AC253572.1** | ENSG00000273136 | 886.2645625 | 0.987486533 | 0.133396539 | 4.75E-14 | 6.78E-13 | NA | HG1 | 121356529 | 121384104 | 1 | protein_coding | Neuroblastoma breakpoint family member 1 |
| **SNURF** | ENSG00000273173 | 33.2350294 | -1.308130853 | 0.865597254 | 0.01197259 | 0.032668648 | 8926 | 15 | 25200133 | 25223729 | 1 | protein_coding | SNRPN upstream reading frame protein |
| **TM4SF19** | ENSG00000273331 | 396.5427711 | -1.383452148 | 0.337886026 | 6.72E-06 | 3.8E-05 | 116211 | 3 | 196042953 | 196065244 | -1 | protein_coding | Transmembrane 4 L6 family member 19 |
