## Supplementary Table S3C for "TDP-43 pathology links innate and adaptive immunity in amyotrophic lateral sclerosis"

TDP_0.5_vs_PBS_0.5

| symbol | ensgene | baseMean | log2FoldChange | lfcSE | pvalue | padj | entrez | chr | start | end | strand | biotype | description |
|---|---|---|---|---|---|---|---|---|---|---|---|---|---|
| **CYP51A1** | ENSG00000001630 | 8481.776647 | 0.467952463 | 0.179107982 | 5.18E-04 | 0.030235078 | 1595 | 7 | 91741465 | 91772266 | -1 | protein_coding | cytochrome P450, family 51, subfamily A, polypeptide 1 |
| **CREBBP** | ENSG00000005339 | 2602.652431 | -0.484901622 | 0.103924533 | 1.61E-07 | 1.24E-04 | 1387 | 16 | 3775055 | 3930727 | -1 | protein_coding | CREB binding protein |
| **RHBDF1** | ENSG00000007384 | 252.4338142 | 0.834967904 | 0.295990168 | 1.74E-04 | 0.015722352 | 64285 | 16 | 108058 | 126354 | -1 | protein_coding | rhomboid 5 homolog 1 (Drosophila) |
| **MGST1** | ENSG00000008394 | 4464.429844 | 0.751043544 | 0.186809775 | 2.92E-06 | 0.001051304 | 4257 | 12 | 16500076 | 16762193 | 1 | protein_coding | microsomal glutathione S-transferase 1 |
| **ST3GAL1** | ENSG00000008513 | 7429.023807 | -0.342171901 | 0.112528963 | 2.13E-04 | 0.017698632 | 6482 | 8 | 134467091 | 134584183 | -1 | protein_coding | ST3 beta-galactoside alpha-2,3-sialyltransferase 1 |
| **STAB1** | ENSG00000010327 | 304.1996687 | -1.356537862 | 0.417009211 | 3.21E-05 | 0.005812692 | 23166 | 3 | 52529354 | 52558511 | 1 | protein_coding | stabilin 1 |
| **CD4** | ENSG00000010610 | 12356.1893 | -0.332441656 | 0.126207181 | 5.9E-04 | 0.032811452 | 920 | 12 | 6896024 | 6929974 | 1 | protein_coding | CD4 molecule |
| **SLC6A7** | ENSG00000011083 | 252.6779403 | 1.246892426 | 0.48731055 | 3.27E-04 | 0.023001816 | 6534 | 5 | 149569520 | 149602351 | 1 | protein_coding | solute carrier family 6 (neurotransmitter transporter), member 7 |
| **ALOX5** | ENSG00000012779 | 4413.822509 | -0.559041149 | 0.262341484 | 0.001041372 | 0.046732155 | 240 | 10 | 45869661 | 45941561 | 1 | protein_coding | arachidonate 5-lipoxygenase |
| **MARCO** | ENSG00000019169 | 1886.480327 | -0.843576397 | 0.240131894 | 1.58E-05 | 0.003550281 | 8685 | 2 | 119699742 | 119752236 | 1 | protein_coding | macrophage receptor with collagenous structure |
| **GCLM** | ENSG00000023909 | 558.7404584 | 0.489060471 | 0.214613262 | 0.001003756 | 0.045405664 | 2730 | 1 | 94350761 | 94374966 | -1 | protein_coding | glutamate-cysteine ligase, modifier subunit |
| **RNASET2** | ENSG00000026297 | 4794.267367 | -0.344111854 | 0.126544173 | 4.4E-04 | 0.027716666 | 8635 | 6 | 167342992 | 167370679 | -1 | protein_coding | ribonuclease T2 |
| **FAM13B** | ENSG00000031003 | 4124.424989 | 0.473047584 | 0.151601317 | 9.88E-05 | 0.011091977 | 51306 | 5 | 137273649 | 137387650 | -1 | protein_coding | family with sequence similarity 13, member B |
| **VCAN** | ENSG00000038427 | 352.3187636 | -0.033025951 | 0.110077364 | 9.81E-04 | 0.045233659 | 1462 | 5 | 82767284 | 82878122 | 1 | protein_coding | versican |
| **CTNS** | ENSG00000040531 | 1815.379027 | 0.497715549 | 0.145568596 | 3.32E-05 | 0.005812692 | 1497 | 17 | 3539762 | 3564836 | 1 | protein_coding | cystinosin, lysosomal cystine transporter |
| **ADAM28** | ENSG00000042980 | 2230.367257 | -0.365921044 | 0.153198475 | 9.74E-04 | 0.045233659 | 10863 | 8 | 24151553 | 24216531 | 1 | protein_coding | ADAM metallopeptidase domain 28 |
| **TSPAN17** | ENSG00000048140 | 891.9911366 | 0.516117952 | 0.138349157 | 1.08E-05 | 0.002674549 | 26262 | 5 | 176074388 | 176086058 | 1 | protein_coding | tetraspanin 17 |
| **MSMO1** | ENSG00000052802 | 3831.370105 | 0.809251574 | 0.207722401 | 4.36E-06 | 0.001329346 | 6307 | 4 | 166248775 | 166264312 | 1 | protein_coding | methylsterol monooxygenase 1 |
| **AP5M1** | ENSG00000053770 | 1457.139439 | 0.353700523 | 0.131181776 | 4.63E-04 | 0.028402393 | 55745 | 14 | 57735627 | 57756797 | 1 | protein_coding | adaptor-related protein complex 5, mu 1 subunit |
| **KMT2C** | ENSG00000055609 | 380.5940948 | -0.95314571 | 0.435668817 | 7.82E-04 | 0.039251114 | 58508 | 7 | 151832010 | 152133090 | -1 | protein_coding | lysine (K)-specific methyltransferase 2C |
| **NCKAP1** | ENSG00000061676 | 1960.481324 | 0.890771945 | 0.292627166 | 8.6E-05 | 0.010028732 | 10787 | 2 | 183773843 | 183903586 | -1 | protein_coding | NCK-associated protein 1 |
| **TSPAN32** | ENSG00000064201 | 171.5046097 | 0.759066772 | 0.305741648 | 4.5E-04 | 0.028064979 | 10077 | 11 | 2323227 | 2339430 | 1 | protein_coding | tetraspanin 32 |
| **DGKA** | ENSG00000065357 | 1267.921782 | -0.536667802 | 0.180376849 | 1.37E-04 | 0.014131359 | 1606 | 12 | 56321103 | 56347811 | 1 | protein_coding | diacylglycerol kinase, alpha 80kDa |
| **TIE1** | ENSG00000066056 | 974.0727001 | 0.735965718 | 0.192063223 | 5.74E-06 | 0.001587855 | 7075 | 1 | 43766664 | 43788779 | 1 | protein_coding | tyrosine kinase with immunoglobulin-like and EGF-like domains 1 |
| **PHKA1** | ENSG00000067177 | 468.119605 | 0.664408702 | 0.274322545 | 5.52E-04 | 0.031246942 | 5255 | X | 71798664 | 71934167 | -1 | protein_coding | phosphorylase kinase, alpha 1 (muscle) |
| **HES2** | ENSG00000069812 | 1704.630879 | 0.835936193 | 0.325438964 | 3.7E-04 | 0.024387321 | 54626 | 1 | 6472478 | 6484730 | -1 | protein_coding | hes family bHLH transcription factor 2 |
| **MAP4K4** | ENSG00000071054 | 2541.766332 | 0.424168573 | 0.161718568 | 4.86E-04 | 0.02918531 | 9448 | 2 | 102313312 | 102511149 | 1 | protein_coding | mitogen-activated protein kinase kinase kinase kinase 4 |
| **FCGR2B** | ENSG00000072694 | 5142.905018 | -0.915140023 | 0.197772462 | 1.64E-07 | 1.24E-04 | 2213 | 1 | 161551101 | 161648444 | 1 | protein_coding | Fc fragment of IgG, low affinity IIb, receptor (CD32) |
| **DHRS9** | ENSG00000073737 | 1576.547914 | 0.858086467 | 0.177786721 | 5.65E-08 | 7.47E-05 | 10170 | 2 | 169921299 | 169952677 | 1 | protein_coding | dehydrogenase/reductase (SDR family) member 9 |
| **CA12** | ENSG00000074410 | 366.8085383 | -1.578791489 | 0.47999326 | 3.44E-05 | 0.005849982 | 771 | 15 | 63613577 | 63674360 | -1 | protein_coding | carbonic anhydrase XII |
| **DUSP13** | ENSG00000079393 | 68.85229856 | 2.790794418 | 0.959401716 | 9.78E-05 | 0.01108275 | 51207 | 10 | 76854192 | 76868979 | -1 | protein_coding | dual specificity phosphatase 13 |
| **MEF2C** | ENSG00000081189 | 711.8431818 | -0.544875298 | 0.223346703 | 5.85E-04 | 0.032671079 | 4208 | 5 | 88013975 | 88199922 | -1 | protein_coding | myocyte enhancer factor 2C |
| **EPB41L3** | ENSG00000082397 | 5025.02904 | -0.502452859 | 0.133160287 | 6.85E-06 | 0.001771524 | 23136 | 18 | 5392383 | 5630699 | -1 | protein_coding | erythrocyte membrane protein band 4.1-like 3 |
| **NFE2L1** | ENSG00000082641 | 10516.33548 | -0.290706872 | 0.094675008 | 1.52E-04 | 0.014744795 | 4779 | 17 | 46125691 | 46138849 | 1 | protein_coding | nuclear factor, erythroid 2-like 1 |
| **SSH1** | ENSG00000084112 | 3589.548928 | -0.305602274 | 0.111792044 | 4.85E-04 | 0.02918531 | 54434 | 12 | 109176466 | 109251366 | -1 | protein_coding | slingshot protein phosphatase 1 |
| **AKR1B1** | ENSG00000085662 | 2195.211649 | 0.416426188 | 0.159656368 | 4.79E-04 | 0.029090728 | 231 | 7 | 134127102 | 134144036 | -1 | protein_coding | aldo-keto reductase family 1, member B1 (aldose reductase) |
| **SNX10** | ENSG00000086300 | 17690.31956 | 0.71216419 | 0.141041816 | 1.97E-08 | 4.61E-05 | 29887 | 7 | 26331541 | 26413949 | 1 | protein_coding | sorting nexin 10 |
| **HUWE1** | ENSG00000086758 | 6524.228758 | -0.249662012 | 0.087540557 | 3.71E-04 | 0.024387321 | 10075 | X | 53559057 | 53713673 | -1 | protein_coding | HECT, UBA and WWE domain containing 1, E3 ubiquitin protein ligase |
| **LPCAT2** | ENSG00000087253 | 5025.034794 | -0.307265734 | 0.119057226 | 7.79E-04 | 0.039246635 | 54947 | 16 | 55542910 | 55620582 | 1 | protein_coding | lysophosphatidylcholine acyltransferase 2 |
| **SLC15A1** | ENSG00000088386 | 41.48789115 | 2.941785651 | 1.409164147 | 6.91E-04 | 0.03638212 | 6564 | 13 | 99336055 | 99404908 | -1 | protein_coding | solute carrier family 15 (oligopeptide transporter), member 1 |
| **SIGLEC1** | ENSG00000088827 | 2559.981037 | 0.734305956 | 0.262449521 | 1.84E-04 | 0.016012275 | 6614 | 20 | 3667617 | 3687775 | -1 | protein_coding | sialic acid binding Ig-like lectin 1, sialoadhesin |
| **FKBP1A** | ENSG00000088832 | 5385.828353 | 0.352564798 | 0.105202731 | 6.01E-05 | 0.008172403 | 101929368 | 20 | 1349622 | 1373806 | -1 | protein_coding | FK506 binding protein 1A, 12kDa |
| **FUS** | ENSG00000089280 | 4571.074523 | -0.291554222 | 0.090620869 | 1.21E-04 | 0.013016553 | 2521 | 16 | 31191431 | 31203127 | 1 | protein_coding | fused in sarcoma |
| **GLG1** | ENSG00000090863 | 6677.205209 | -0.351791497 | 0.083193387 | 9.46E-07 | 5.12E-04 | 2734 | 16 | 74485856 | 74641012 | -1 | protein_coding | golgi glycoprotein 1 |
| **NRCAM** | ENSG00000091129 | 481.3287573 | 1.088672708 | 0.475645951 | 6.45E-04 | 0.035366619 | 4897 | 7 | 107788068 | 108097161 | -1 | protein_coding | neuronal cell adhesion molecule |
| **DSP** | ENSG00000096696 | 894.5352552 | 0.889727484 | 0.282144897 | 6.04E-05 | 0.008172403 | 1832 | 6 | 7541808 | 7586950 | 1 | protein_coding | desmoplakin |
| **SCD** | ENSG00000099194 | 17251.55763 | 0.759039931 | 0.163844438 | 1.67E-07 | 1.24E-04 | 6319 | 10 | 102106881 | 102124591 | 1 | protein_coding | stearoyl-CoA desaturase (delta-9-desaturase) |
| **NRP1** | ENSG00000099250 | 18474.12949 | -0.398722326 | 0.101301219 | 5.69E-06 | 0.001587855 | 8829 | 10 | 33466420 | 33625190 | -1 | protein_coding | neuropilin 1 |
| **TIMP3** | ENSG00000100234 | 10239.78751 | 0.312942352 | 0.110712093 | 4.19E-04 | 0.026955066 | 7078 | 22 | 33197687 | 33259030 | 1 | protein_coding | TIMP metallopeptidase inhibitor 3 |
| **THOC5** | ENSG00000100296 | 1507.421105 | 0.415747943 | 0.166120325 | 7.04E-04 | 0.036581678 | 8563 | 22 | 29901868 | 29951205 | -1 | protein_coding | THO complex 5 |
| **SYNGR1** | ENSG00000100321 | 2368.927241 | 0.709475025 | 0.245706595 | 1.57E-04 | 0.014955361 | 9145 | 22 | 39745930 | 39781593 | 1 | protein_coding | synaptogyrin 1 |
| **C20orf24** | ENSG00000101084 | 4030.161356 | 0.3203972 | 0.092545107 | 4.89E-05 | 0.007359953 | 55969 | 20 | 35234137 | 35240960 | 1 | protein_coding | chromosome 20 open reading frame 24 |
| **TRIB3** | ENSG00000101255 | 273.8941669 | -1.187452454 | 0.367990562 | 4.61E-05 | 0.007359953 | 57761 | 20 | 361261 | 378203 | 1 | protein_coding | tribbles pseudokinase 3 |
| **RASSF2** | ENSG00000101265 | 2388.281498 | -0.388876345 | 0.155204594 | 5.17E-04 | 0.030235078 | 9770 | 20 | 4760669 | 4804291 | -1 | protein_coding | Ras association (RalGDS/AF-6) domain family member 2 |
| **TIMP1** | ENSG00000102265 | 1212.68041 | -0.915883521 | 0.22871239 | 2.63E-06 | 0.001000803 | 7076 | X | 47441712 | 47446188 | 1 | protein_coding | TIMP metallopeptidase inhibitor 1 |
| **GLA** | ENSG00000102393 | 5389.900431 | 0.276682962 | 0.09417522 | 3.4E-04 | 0.023483464 | 2717 | X | 100652791 | 100662913 | -1 | protein_coding | galactosidase, alpha |
| **FLT1** | ENSG00000102755 | 5916.959689 | -0.547395933 | 0.20247872 | 2.69E-04 | 0.02010234 | 2321 | 13 | 28874489 | 29069265 | -1 | protein_coding | fms-related tyrosine kinase 1 |
| **CCL22** | ENSG00000102962 | 15258.76342 | 0.538889729 | 0.193448928 | 2.4E-04 | 0.018629713 | 6367 | 16 | 57392684 | 57400102 | 1 | protein_coding | chemokine (C-C motif) ligand 22 |
| **SLC7A5** | ENSG00000103257 | 1125.361446 | -0.789914706 | 0.306618936 | 3.44E-04 | 0.02368164 | 8140 | 16 | 87863629 | 87903094 | -1 | protein_coding | solute carrier family 7 (amino acid transporter light chain, L system), member 5 |
| **RAB11A** | ENSG00000103769 | 3092.759997 | 0.340951591 | 0.139950727 | 9.51E-04 | 0.044694913 | 8766 | 15 | 66018392 | 66184329 | 1 | protein_coding | RAB11A, member RAS oncogene family |
| **SGK3** | ENSG00000104205 | 2964.615126 | 0.459232747 | 0.114472223 | 3.43E-06 | 0.001110693 | 100533105 | 8 | 67624653 | 67774257 | 1 | protein_coding | serum/glucocorticoid regulated kinase family, member 3 |
| **NIPAL2** | ENSG00000104361 | 2696.515538 | 1.491443371 | 0.429012361 | 1.87E-05 | 0.004036018 | 79815 | 8 | 99202061 | 99306621 | -1 | protein_coding | NIPA-like domain containing 2 |
| **KCNN4** | ENSG00000104783 | 301.3581646 | -0.738819903 | 0.232366255 | 6.04E-05 | 0.008172403 | 3783 | 19 | 44270685 | 44285409 | -1 | protein_coding | potassium intermediate/small conductance calcium-activated channel, subfamily N, member 4 |
| **TNNT1** | ENSG00000105048 | 121.5867828 | -1.758500422 | 0.52048782 | 2.47E-05 | 0.004904339 | 7138 | 19 | 55644162 | 55660722 | -1 | protein_coding | troponin T type 1 (skeletal, slow) |
| **NPTX2** | ENSG00000106236 | 275.3784218 | 1.839070726 | 0.422762432 | 5.34E-07 | 3.03E-04 | 4885 | 7 | 98246609 | 98259180 | 1 | protein_coding | neuronal pentraxin II |
| **IMPDH1** | ENSG00000106348 | 772.0616582 | 0.434659049 | 0.175950743 | 6.64E-04 | 0.035845485 | 3614 | 7 | 128032331 | 128050306 | -1 | protein_coding | IMP (inosine 5'-monophosphate) dehydrogenase 1 |
| **CASC3** | ENSG00000108349 | 1615.027051 | -0.350200219 | 0.116823017 | 1.82E-04 | 0.016012275 | 102466986 | 17 | 38296571 | 38328436 | 1 | protein_coding | cancer susceptibility candidate 3 |
| **4-Sep** | ENSG00000108387 | 53.70237406 | 1.928690602 | 0.99758174 | 0.001101168 | 0.048882821 | 5414 | 17 | 56597611 | 56618179 | -1 | protein_coding | septin 4 |
| **MED13** | ENSG00000108510 | 4423.857545 | -0.241920557 | 0.088816285 | 7.41E-04 | 0.038233745 | 9969 | 17 | 60019966 | 60142643 | -1 | protein_coding | mediator complex subunit 13 |
| **TMEM104** | ENSG00000109066 | 1708.276445 | 0.443222298 | 0.182570761 | 7.57E-04 | 0.038491018 | 54868 | 17 | 72772622 | 72835918 | 1 | protein_coding | transmembrane protein 104 |
| **TMEM33** | ENSG00000109133 | 4917.685767 | 0.532519694 | 0.130940879 | 2.36E-06 | 0.001000803 | 55161 | 4 | 41937137 | 41962589 | 1 | protein_coding | transmembrane protein 33 |
| **TRIM2** | ENSG00000109654 | 169.6961543 | 1.258096421 | 0.477418436 | 2.58E-04 | 0.019567674 | 23321 | 4 | 154073494 | 154260472 | 1 | protein_coding | tripartite motif containing 2 |
| **WHSC1** | ENSG00000109685 | 1137.478152 | -0.452234759 | 0.159022216 | 2.32E-04 | 0.018437522 | 7468 | 4 | 1873151 | 1983934 | 1 | protein_coding | Wolf-Hirschhorn syndrome candidate 1 |
| **SC5D** | ENSG00000109929 | 2852.255719 | 0.566353065 | 0.179734162 | 7.55E-05 | 0.009441493 | 6309 | 11 | 121163162 | 121179403 | 1 | protein_coding | sterol-C5-desaturase |
| **SLC1A2** | ENSG00000110436 | 2833.375657 | 0.54926154 | 0.220740894 | 5.32E-04 | 0.030446919 | 6506 | 11 | 35272753 | 35441610 | -1 | protein_coding | solute carrier family 1 (glial high affinity glutamate transporter), member 2 |
| **SLC15A3** | ENSG00000110446 | 9991.947152 | 0.334700271 | 0.115906347 | 2.86E-04 | 0.020767263 | 51296 | 11 | 60704556 | 60720002 | -1 | protein_coding | solute carrier family 15 (oligopeptide transporter), member 3 |
| **TENC1** | ENSG00000111077 | 161.151299 | 0.999467808 | 0.421344105 | 5.39E-04 | 0.030666767 | 23371 | 12 | 53440753 | 53458156 | 1 | protein_coding | tensin like C1 domain containing phosphatase (tensin 2) |
| **ENO2** | ENSG00000111674 | 999.7739498 | -0.586460427 | 0.191974164 | 8.38E-05 | 0.009956294 | 2026 | 12 | 7022909 | 7032861 | 1 | protein_coding | enolase 2 (gamma, neuronal) |
| **TMEM14C** | ENSG00000111843 | 1681.623548 | 0.392141847 | 0.155130016 | 7.01E-04 | 0.036565303 | 51522 | 6 | 10723148 | 10731362 | 1 | protein_coding | transmembrane protein 14C |
| **FANCE** | ENSG00000112039 | 552.8655949 | 0.40296503 | 0.170888921 | 1E-03 | 0.045405664 | 2178 | 6 | 35420138 | 35434880 | 1 | protein_coding | Fanconi anemia, complementation group E |
| **VNN 1.00** | ENSG00000112299 | 431.344258 | -0.842471008 | 0.268246356 | 6.43E-05 | 0.00859394 | 8876 | 6 | 133002729 | 133035188 | -1 | protein_coding | vanin 1 |
| **PERP** | ENSG00000112378 | 141.8514978 | 1.070357624 | 0.459316101 | 5.79E-04 | 0.032491399 | 64065 | 6 | 138409642 | 138428648 | -1 | protein_coding | PERP, TP53 apoptosis effector |
| **BTN2A1** | ENSG00000112763 | 870.3499201 | -0.417387224 | 0.168650293 | 6.68E-04 | 0.035845485 | 11120 | 6 | 26458150 | 26476849 | 1 | protein_coding | butyrophilin, subfamily 2, member A1 |
| **ARRDC3** | ENSG00000113369 | 2085.357492 | -0.585861835 | 0.209955873 | 2.4E-04 | 0.018629713 | 57561 | 5 | 90664541 | 90679176 | -1 | protein_coding | arrestin domain containing 3 |

| | | | | | | | | | | | | | |
|---|---|---|---|---|---|---|---|---|---|---|---|---|---|
| **STC2** | ENSG00000113739 | 127.8828323 | -0.102557143 | 0.199405095 | 7.56E-04 | 0.038491018 | 8614 | 5 | 172741716 | 172756506 | -1 | protein_coding | stanniocalcin 2 |
| **DBN1** | ENSG00000113758 | 264.7552809 | 0.802178605 | 0.310085345 | 3.31E-04 | 0.023187216 | 1627 | 5 | 176883609 | 176901402 | -1 | protein_coding | drebrin 1 |
| **RNF7** | ENSG00000114125 | 1083.065211 | 0.404295169 | 0.164500341 | 7.47E-04 | 0.038323461 | 9616 | 3 | 141457046 | 141466402 | 1 | protein_coding | ring finger protein 7 |
| **STEAP3** | ENSG00000115107 | 981.1143152 | 1.522280077 | 0.329457496 | 1.22E-07 | 1.21E-04 | 55240 | 2 | 119981384 | 120023228 | 1 | protein_coding | STEAP family member 3, metalloreductase |
| **SPTBN1** | ENSG00000115306 | 2429.710156 | -0.46512374 | 0.152147198 | 1.16E-04 | 0.012593666 | 6711 | 2 | 54683422 | 54896812 | 1 | protein_coding | spectrin, beta, non-erythrocytic 1 |
| **IGFBP2** | ENSG00000115457 | 128.4023642 | -1.245537011 | 0.463771975 | 2.3E-04 | 0.018373408 | 3485 | 2 | 217497551 | 217529159 | 1 | protein_coding | insulin-like growth factor binding protein 2, 36kDa |
| **KDM3A** | ENSG00000115548 | 2634.582618 | -0.276716884 | 0.103131257 | 6.76E-04 | 0.035974913 | 55818 | 2 | 86667770 | 86719839 | 1 | protein_coding | lysine (K)-specific demethylase 3A |
| **ECE1** | ENSG00000117298 | 1968.121784 | 0.764509569 | 0.187419966 | 2.26E-06 | 0.001000803 | 1889 | 1 | 21543740 | 21671997 | -1 | protein_coding | endothelin converting enzyme 1 |
| **ARID1A** | ENSG00000117713 | 2560.527312 | -0.553540255 | 0.145083774 | 6.77E-06 | 0.001771524 | 8289 | 1 | 27022524 | 27108595 | 1 | protein_coding | AT rich interactive domain 1A (SWI-like) |
| **KMT2A** | ENSG00000118058 | 2003.607159 | -0.476123106 | 0.177470268 | 3.51E-04 | 0.023856192 | 4297 | 11 | 118307205 | 118397539 | 1 | protein_coding | lysine (K)-specific methyltransferase 2A |
| **GALNT12** | ENSG00000119514 | 4913.676435 | 0.327940399 | 0.123066549 | 7.42E-04 | 0.038233745 | 79695 | 9 | 101569981 | 101612363 | 1 | protein_coding | UDP-N-acetyl-alpha-D-galactosamine:polypeptide N-acetylgalactosaminyltransferase 12 (GalNA |
| **HSPH1** | ENSG00000120694 | 2823.048163 | 0.407780354 | 0.143615833 | 2.64E-04 | 0.019847776 | 10808 | 13 | 31710762 | 31736525 | -1 | protein_coding | heat shock 105kDa/110kDa protein 1 |
| **TGFBI** | ENSG00000120708 | 3265.949622 | -0.551573833 | 0.213516384 | 3.77E-04 | 0.024527258 | 7045 | 5 | 135364584 | 135399507 | 1 | protein_coding | transforming growth factor, beta-induced, 68kDa |
| **PLAU** | ENSG00000122861 | 1163.058513 | -0.628278914 | 0.135734207 | 1.92E-07 | 1.34E-04 | 5328 | 10 | 75668935 | 75677255 | 1 | protein_coding | plasminogen activator, urokinase |
| **ITPR2** | ENSG00000123104 | 3519.272971 | -0.534989415 | 0.096869756 | 1.91E-09 | 1.14E-05 | 3709 | 12 | 26490342 | 26986131 | -1 | protein_coding | inositol 1,4,5-trisphosphate receptor, type 2 |
| **LRP1** | ENSG00000123384 | 27050.26282 | -0.42244227 | 0.105047425 | 3.76E-06 | 0.001178654 | 4035 | 12 | 57522276 | 57607134 | 1 | protein_coding | low density lipoprotein receptor-related protein 1 |
| **TNFAIP6** | ENSG00000123610 | 8127.152683 | 0.670412733 | 0.181808088 | 1.07E-05 | 0.002674549 | 7130 | 2 | 152214106 | 152236560 | 1 | protein_coding | tumor necrosis factor, alpha-induced protein 6 |
| **CHD6** | ENSG00000124177 | 1284.451355 | -0.448270609 | 0.188021203 | 7.88E-04 | 0.039295414 | 84181 | 20 | 40030741 | 40247133 | -1 | protein_coding | chromodomain helicase DNA binding protein 6 |
| **RNF114** | ENSG00000124226 | 2831.788872 | 0.26426235 | 0.102582584 | 9.84E-04 | 0.045233659 | 55905 | 20 | 48552948 | 48570429 | 1 | protein_coding | ring finger protein 114 |
| **TREM1** | ENSG00000124731 | 220.6443369 | -1.847240557 | 0.471262676 | 3.36E-06 | 0.001110693 | 54210 | 6 | 41235664 | 41254457 | -1 | protein_coding | triggering receptor expressed on myeloid cells 1 |
| **AHNAK** | ENSG00000124942 | 42796.75229 | -0.410771012 | 0.099723962 | 3.45E-06 | 0.001110693 | 79026 | 11 | 62201016 | 62323707 | -1 | protein_coding | AHNAK nucleoprotein |
| **IL1B** | ENSG00000125538 | 1784.226271 | -0.06291233 | 0.132900842 | 6.69E-04 | 0.035845485 | 3553 | 2 | 113587328 | 113594480 | -1 | protein_coding | interleukin 1, beta |
| **SLC25A23** | ENSG00000125648 | 911.8187134 | 0.912248905 | 0.251848178 | 1.17E-05 | 0.002833689 | 79085 | 19 | 6436090 | 6465214 | -1 | protein_coding | solute carrier family 25 (mitochondrial carrier; phosphate carrier), member 23 |
| **TRIP10** | ENSG00000125733 | 832.5378634 | -0.499180491 | 0.17301191 | 1.85E-04 | 0.016012275 | 9322 | 19 | 6737936 | 6751537 | 1 | protein_coding | thyroid hormone receptor interactor 10 |
| **IFI6** | ENSG00000126709 | 7699.553854 | 0.535801446 | 0.173457915 | 1.06E-04 | 0.011746469 | 2537 | 1 | 27992572 | 27998729 | -1 | protein_coding | interferon, alpha-inducible protein 6 |
| **APOBEC3A** | ENSG00000128383 | 3640.86496 | 0.702741399 | 0.225160534 | 7.55E-05 | 0.009441493 | 100913187 | 22 | 39348746 | 39359188 | 1 | protein_coding | apolipoprotein B mRNA editing enzyme, catalytic polypeptide-like 3A |
| **SLC44A2** | ENSG00000129353 | 2690.447542 | 0.365958443 | 0.118439689 | 1.49E-04 | 0.014628239 | 57153 | 19 | 10713133 | 10755235 | 1 | protein_coding | solute carrier family 44 (choline transporter), member 2 |
| **RNASE1** | ENSG00000129538 | 309.4227762 | -1.153764101 | 0.291408871 | 3.05E-06 | 0.001066834 | 6035 | 14 | 21269387 | 21271437 | -1 | protein_coding | ribonuclease, RNase A family, 1 (pancreatic) |
| **APOC1** | ENSG00000130208 | 1435.162052 | 0.760661712 | 0.211629874 | 1.47E-05 | 0.003358194 | 341 | 19 | 45417504 | 45422606 | 1 | protein_coding | apolipoprotein C-I |
| **SNX9** | ENSG00000130340 | 4007.854497 | -0.294364994 | 0.098614003 | 2.54E-04 | 0.01934231 | 51429 | 6 | 158244296 | 158366109 | 1 | protein_coding | sorting nexin 9 |
| **HELZ2** | ENSG00000130589 | 2376.400707 | 0.860890921 | 0.183554393 | 1.15E-07 | 1.21E-04 | 85441 | 20 | 62189439 | 62205592 | -1 | protein_coding | helicase with zinc finger 2, transcriptional coactivator |
| **LGALS3** | ENSG00000131981 | 10407.05686 | 0.549244575 | 0.196993995 | 2.3E-04 | 0.018373408 | 3958 | 14 | 55590828 | 55612126 | 1 | protein_coding | lectin, galactoside-binding, soluble, 3 |
| **DLG4** | ENSG00000132535 | 638.6826261 | -0.474100226 | 0.20592064 | 8.72E-04 | 0.041968269 | 1742 | 17 | 7093209 | 7123021 | -1 | protein_coding | discs, large homolog 4 (Drosophila) |
| **KANK4** | ENSG00000132854 | 121.4388461 | 1.149373146 | 0.529924722 | 8.07E-04 | 0.03951798 | 163782 | 1 | 62702651 | 62785085 | -1 | protein_coding | KN motif and ankyrin repeat domains 4 |
| **RNF128** | ENSG00000133135 | 718.1913254 | 0.795567862 | 0.328739712 | 5.18E-04 | 0.030235078 | 79589 | X | 105937024 | 106040223 | 1 | protein_coding | ring finger protein 128, E3 ubiquitin protein ligase |
| **CAMK1** | ENSG00000134072 | 1165.906732 | -0.563094422 | 0.191372474 | 1.43E-04 | 0.014341007 | 8536 | 3 | 9799026 | 9811676 | -1 | protein_coding | calcium/calmodulin-dependent protein kinase I |
| **CD101** | ENSG00000134256 | 1051.209621 | -0.822271204 | 0.197561564 | 1.49E-06 | 7.69E-04 | 9398 | 1 | 117544382 | 117579167 | 1 | protein_coding | CD101 molecule |
| **SLC38A2** | ENSG00000134294 | 8310.102093 | 0.389732076 | 0.115886749 | 5.44E-05 | 0.007831244 | 54407 | 12 | 46751972 | 46766650 | -1 | protein_coding | solute carrier family 38, member 2 |
| **RSAD2** | ENSG00000134321 | 28759.06657 | 1.221573446 | 0.520235088 | 5.24E-04 | 0.030341659 | 91543 | 2 | 7005937 | 7038370 | 1 | protein_coding | radical S-adenosyl methionine domain containing 2 |
| **ETS1** | ENSG00000134954 | 1642.212272 | -0.433394244 | 0.160073247 | 3.51E-04 | 0.023856192 | 2113 | 11 | 128328656 | 128457453 | -1 | protein_coding | v-ets avian erythroblastosis virus E26 oncogene homolog 1 |
| **FAIM2** | ENSG00000135472 | 708.2932972 | 0.567300868 | 0.194013626 | 1.61E-04 | 0.015037347 | 23017 | 12 | 50260679 | 50298000 | -1 | protein_coding | Fas apoptotic inhibitory molecule 2 |
| **HNRNPA1** | ENSG00000135486 | 9485.252224 | -0.351736511 | 0.123054637 | 2.89E-04 | 0.020866053 | 3178 | 12 | 54673977 | 54680872 | 1 | protein_coding | heterogeneous nuclear ribonucleoprotein A1 |
| **COX5B** | ENSG00000135940 | 2920.516599 | 0.618379042 | 0.179022422 | 2.65E-05 | 0.005163652 | 1329 | 2 | 98262503 | 98264846 | 1 | protein_coding | cytochrome c oxidase subunit Vb |
| **ALDH1L2** | ENSG00000136010 | 1308.566749 | -0.512747472 | 0.175303689 | 1.63E-04 | 0.015056523 | 160428 | 12 | 105413568 | 105478355 | -1 | protein_coding | aldehyde dehydrogenase 1 family, member L2 |
| **ALPK3** | ENSG00000136383 | 1849.626053 | 1.035885759 | 0.22330029 | 1.62E-07 | 1.24E-04 | 57538 | 15 | 85359911 | 85416713 | 1 | protein_coding | alpha-kinase 3 |
| **LRRC8A** | ENSG00000136802 | 1814.337625 | 0.563457334 | 0.137596433 | 2.23E-06 | 0.001000803 | 56262 | 9 | 131644391 | 131680318 | 1 | protein_coding | leucine rich repeat containing 8 family, member A |
| **ST6GALNA** | ENSG00000136840 | 608.4965513 | 0.729594861 | 0.227729182 | 5.51E-05 | 0.007831244 | 27090 | 9 | 130670165 | 130679317 | -1 | protein_coding | ST6 (alpha-N-acetyl-neuraminyl-2,3-beta-galactosyl-1,3)-N-acetylgalactosaminide alpha-2,6-sia |
| **TLR4** | ENSG00000136869 | 4074.243103 | 0.3106574 | 0.125224242 | 0.001037332 | 0.046732155 | 7099 | 9 | 120466610 | 120479149 | 1 | protein_coding | toll-like receptor 4 |
| **SORL1** | ENSG00000137642 | 3821.519421 | -0.648251417 | 0.193469359 | 3.4E-05 | 0.005849982 | 6653 | 11 | 121322912 | 121504402 | 1 | protein_coding | sortilin-related receptor, L(DLR class) A repeats containing |
| **RAB1A** | ENSG00000138069 | 5929.355573 | 0.423572046 | 0.121936312 | 3.32E-05 | 0.005812692 | 5861 | 2 | 65297835 | 65357240 | -1 | protein_coding | RAB1A, member RAS oncogene family |
| **TRIM54** | ENSG00000138100 | 221.4005107 | 1.574246159 | 0.562430133 | 1.42E-04 | 0.014341007 | 57159 | 2 | 27505260 | 27530307 | 1 | protein_coding | tripartite motif containing 54 |
| **DUSP5** | ENSG00000138166 | 1229.629273 | 0.438672166 | 0.171130683 | 4.67E-04 | 0.028476132 | 1847 | 10 | 112257596 | 112271302 | 1 | protein_coding | dual specificity phosphatase 5 |
| **IDH1** | ENSG00000138413 | 11751.60626 | 0.316783009 | 0.102428422 | 2.01E-04 | 0.017071357 | 3417 | 2 | 209100951 | 209130798 | -1 | protein_coding | isocitrate dehydrogenase 1 (NADP+), soluble |
| **SEMA7A** | ENSG00000138623 | 1856.463203 | 0.540898902 | 0.172941634 | 8.89E-05 | 0.010269767 | 8482 | 15 | 74701630 | 74726808 | -1 | protein_coding | semaphorin 7A, GPI membrane anchor (John Milton Hagen blood group) |
| **HNRNPD** | ENSG00000138668 | 1984.884547 | -0.352947783 | 0.122834581 | 2.76E-04 | 0.020258563 | 3184 | 4 | 83273651 | 83295656 | -1 | protein_coding | heterogeneous nuclear ribonucleoprotein D (AU-rich element RNA binding protein 1, 37kDa) |
| **11-Sep** | ENSG00000138758 | 2076.488614 | -0.343998817 | 0.121535581 | 3.25E-04 | 0.022992343 | 55752 | 4 | 77870856 | 77961537 | 1 | protein_coding | septin 11 |
| **TMEM117** | ENSG00000139173 | 502.705305 | 0.551257138 | 0.246356767 | 9.13E-04 | 0.043447982 | 84216 | 12 | 44229770 | 44783545 | 1 | protein_coding | transmembrane protein 117 |
| **IQGAP1** | ENSG00000140575 | 21843.06374 | -0.279263713 | 0.09863849 | 4.36E-04 | 0.02758717 | 8826 | 15 | 90931450 | 91045475 | 1 | protein_coding | IQ motif containing GTPase activating protein 1 |
| **ZFHX3** | ENSG00000140836 | 2095.548153 | -0.341587536 | 0.136944394 | 8.2E-04 | 0.039960022 | 463 | 16 | 72816784 | 73093597 | -1 | protein_coding | zinc finger homeobox 3 |
| **NUDT7** | ENSG00000140876 | 160.8326359 | 1.269465775 | 0.598210105 | 8.73E-04 | 0.041968269 | 283927 | 16 | 77756411 | 77776157 | 1 | protein_coding | nudix (nucleoside diphosphate linked moiety X)-type motif 7 |
| **MPC2** | ENSG00000143158 | 1822.174937 | 0.426736943 | 0.143861467 | 1.74E-04 | 0.015722352 | 25874 | 1 | 167885967 | 167906278 | -1 | protein_coding | mitochondrial pyruvate carrier 2 |
| **FCGR2A** | ENSG00000143226 | 6625.374491 | -0.521493295 | 0.16074541 | 5.31E-05 | 0.00780464 | 2212 | 1 | 161475220 | 161493803 | 1 | protein_coding | Fc fragment of IgG, low affinity IIa, receptor (CD32) |
| **CRABP2** | ENSG00000143320 | 4468.771463 | 0.580543089 | 0.22620504 | 5.05E-04 | 0.030017838 | 1382 | 1 | 156669398 | 156675608 | -1 | protein_coding | cellular retinoic acid binding protein 2 |
| **ANP32E** | ENSG00000143401 | 498.764894 | -0.461675993 | 0.189302332 | 6.8E-04 | 0.035974913 | 81611 | 1 | 150190717 | 150208504 | -1 | protein_coding | acidic (leucine-rich) nuclear phosphoprotein 32 family, member E |
| **PPFIA4** | ENSG00000143847 | 158.1077243 | -0.100140864 | 0.192601 | 9.54E-04 | 0.044694913 | 8497 | 1 | 202995626 | 203047868 | 1 | protein_coding | protein tyrosine phosphatase, receptor type, f polypeptide (PTPRF), interacting protein (liprin), al |
| **NFKBIZ** | ENSG00000144802 | 1066.857279 | -0.679834308 | 0.212620803 | 5.92E-05 | 0.008172403 | 64332 | 3 | 101546835 | 101579866 | 1 | protein_coding | nuclear factor of kappa light polypeptide gene enhancer in B-cells inhibitor, zeta |
| **ADPRH** | ENSG00000144843 | 1344.211978 | 0.443912948 | 0.170313294 | 4.61E-04 | 0.028402393 | 141 | 3 | 119298115 | 119308792 | 1 | protein_coding | ADP-ribosylarginine hydrolase |
| **TNFRSF21** | ENSG00000146072 | 1811.923903 | 0.71198188 | 0.219916769 | 5.08E-05 | 0.007549251 | 27242 | 6 | 47199268 | 47277641 | -1 | protein_coding | tumor necrosis factor receptor superfamily, member 21 |
| **C7orf55-LU** | ENSG00000146963 | 1311.443938 | -0.5265616 | 0.16830493 | 8E-05 | 0.00971476 | 51631 | 7 | 139025105 | 139108198 | 1 | protein_coding | C7orf55-LUC7L2 readthrough |
| **SNX30** | ENSG00000148158 | 2558.319265 | -0.364276767 | 0.124076048 | 2.11E-04 | 0.017698632 | 401548 | 9 | 115513118 | 115643951 | 1 | protein_coding | sorting nexin family member 30 |
| **ADM** | ENSG00000148926 | 1548.313157 | -0.953511571 | 0.311683873 | 7.88E-05 | 0.009664677 | 133 | 11 | 10326227 | 10328944 | 1 | protein_coding | adrenomedullin |
| **ITPR1** | ENSG00000150995 | 4277.132277 | -0.367707626 | 0.087064368 | 1.65E-06 | 8.16E-04 | 3708 | 3 | 4535032 | 4889524 | 1 | protein_coding | inositol 1,4,5-trisphosphate receptor, type 1 |
| **UCHL1** | ENSG00000154277 | 1209.466534 | 0.884387597 | 0.319637187 | 2E-04 | 0.017071357 | 7345 | 4 | 41258430 | 41270472 | 1 | protein_coding | ubiquitin carboxyl-terminal esterase L1 (ubiquitin thiolesterase) |
| **ATP5G3** | ENSG00000154518 | 2229.139092 | 0.310305714 | 0.110522497 | 4.18E-04 | 0.026955066 | 518 | 2 | 176040986 | 176049335 | -1 | protein_coding | ATP synthase, H+ transporting, mitochondrial Fo complex, subunit C3 (subunit 9) |
| **LY96** | ENSG00000154589 | 871.7851852 | 0.657098861 | 0.193662138 | 3.27E-05 | 0.005812692 | 23643 | 8 | 74903587 | 74941322 | 1 | protein_coding | lymphocyte antigen 96 |
| **RAB6B** | ENSG00000154917 | 64.47628334 | 2.118682347 | 0.891987226 | 4.21E-04 | 0.026955066 | 51560 | 3 | 133543083 | 133614680 | -1 | protein_coding | RAB6B, member RAS oncogene family |
| **TTN** | ENSG00000155657 | 579.7400969 | -0.9615918 | 0.297988419 | 4.64E-05 | 0.007359953 | 7273 | 2 | 179390716 | 179695529 | -1 | protein_coding | titin |
| **TMBIM4** | ENSG00000155957 | 1801.850368 | 0.369089992 | 0.151657578 | 8.93E-04 | 0.042648619 | 51643 | 12 | 66517709 | 66563852 | -1 | protein_coding | transmembrane BAX inhibitor motif containing 4 |
| **MCU** | ENSG00000156026 | 1344.208521 | 0.565389118 | 0.167750862 | 3.74E-05 | 0.006184596 | 90550 | 10 | 74451889 | 74647452 | 1 | protein_coding | mitochondrial calcium uniporter |
| **HK1** | ENSG00000156515 | 4371.527334 | -0.338724234 | 0.093385544 | 2.28E-05 | 0.004686652 | 3098 | 10 | 71029740 | 71161638 | 1 | protein_coding | hexokinase 1 |
| **SMG1** | ENSG00000157106 | 4770.019418 | -0.403357375 | 0.113728101 | 2.4E-05 | 0.004838114 | 101060589 | 16 | 18816175 | 18937776 | -1 | protein_coding | SMG1 phosphatidylinositol 3-kinase-related kinase |
| **ETS2** | ENSG00000157557 | 585.9564718 | -0.572490777 | 0.214303276 | 3.09E-04 | 0.021988246 | 2114 | 21 | 40177231 | 40196879 | 1 | protein_coding | v-ets avian erythroblastosis virus E26 oncogene homolog 2 |
| **TMEM164** | ENSG00000157600 | 3468.031597 | 0.284149451 | 0.10736405 | 7.93E-04 | 0.039295414 | 101928589 | X | 109245859 | 109425962 | 1 | protein_coding | transmembrane protein 164 |
| **MX1** | ENSG00000157601 | 11456.08719 | 0.446255256 | 0.140080999 | 8.45E-05 | 0.009956294 | 4599 | 21 | 42792231 | 42831141 | 1 | protein_coding | myxovirus (influenza virus) resistance 1, interferon-inducible protein p78 (mouse) |
| **DUSP2** | ENSG00000158050 | 586.4044023 | 0.902550456 | 0.266622444 | 2.86E-05 | 0.005395115 | 1844 | 2 | 96808905 | 96811179 | -1 | protein_coding | dual specificity phosphatase 2 |
| **CNNM4** | ENSG00000158158 | 1137.482022 | -0.417810632 | 0.181266542 | 9.96E-04 | 0.045405664 | 26504 | 2 | 97426639 | 97477628 | 1 | protein_coding | cyclin M4 |
| **SON** | ENSG00000159140 | 7695.533283 | -0.290160547 | 0.088966535 | 9.4E-05 | 0.010753099 | 6651 | 21 | 34914924 | 34949812 | 1 | protein_coding | SON DNA binding protein |

| | | | | | | | | | | | | | |
|---|---|---|---|---|---|---|---|---|---|---|---|---|---|
| **RUNX1** | ENSG00000159216 | 1701.646302 | -0.662795921 | 0.234216862 | 1.84E-04 | 0.016012275 | 101928269 | 21 | 36160098 | 37376965 | -1 | protein_coding | runt-related transcription factor 1 |
| **ACE** | ENSG00000159640 | 1735.822341 | -0.59217738 | 0.21763672 | 2.42E-04 | 0.018629713 | 1636 | 17 | 61554422 | 61599205 | 1 | protein_coding | angiotensin I converting enzyme |
| **SPON2** | ENSG00000159674 | 1411.290505 | 1.658326437 | 0.577822113 | 1.32E-04 | 0.013733241 | 100130872 | 4 | 1160720 | 1202750 | -1 | protein_coding | spondin 2, extracellular matrix protein |
| **KALRN** | ENSG00000160145 | 50.74214578 | 2.512293675 | 0.738076127 | 2.18E-05 | 0.004632873 | 8997 | 3 | 123798870 | 124445172 | 1 | protein_coding | kalirin, RhoGEF kinase |
| **G6PD** | ENSG00000160211 | 4682.303754 | 0.522193543 | 0.115138658 | 3.11E-07 | 1.95E-04 | 2539 | X | 153759606 | 153775787 | -1 | protein_coding | glucose-6-phosphate dehydrogenase |
| **LY6E** | ENSG00000160932 | 2261.072498 | 0.687093609 | 0.129582948 | 5.79E-09 | 2.3E-05 | 4061 | 8 | 144099399 | 144105249 | 1 | protein_coding | lymphocyte antigen 6 complex, locus E |
| **CYB561A3** | ENSG00000162144 | 4845.782529 | 0.515723055 | 0.182688536 | 2.3E-04 | 0.018373408 | 220002 | 11 | 61116217 | 61129771 | -1 | protein_coding | cytochrome b561 family, member A3 |
| **JAK1** | ENSG00000162434 | 6981.316313 | -0.257862915 | 0.08736384 | 3.7E-04 | 0.024387321 | 3716 | 1 | 65298912 | 65432187 | -1 | protein_coding | Janus kinase 1 |
| **SDC3** | ENSG00000162512 | 3804.109804 | 0.451916322 | 0.177479765 | 5.32E-04 | 0.030446919 | 9672 | 1 | 31342314 | 31381608 | -1 | protein_coding | syndecan 3 |
| **NBPF8** | ENSG00000162825 | 664.8915504 | -0.728766266 | 0.180827485 | 2.59E-06 | 0.001000803 | 101928050 | 1 | 144146808 | 144224481 | 1 | protein_coding | neuroblastoma breakpoint family, member 8 |
| **SPATA18** | ENSG00000163071 | 797.0543468 | 0.860804601 | 0.253497791 | 2.79E-05 | 0.005359139 | 132671 | 4 | 52917497 | 52963458 | 1 | protein_coding | spermatogenesis associated 18 |
| **FCRL1** | ENSG00000163534 | 45.2505494 | -0.109582357 | 0.222454902 | 3.73E-04 | 0.024387321 | 115350 | 1 | 157764193 | 157789895 | -1 | protein_coding | Fc receptor-like 1 |
| **SMIM14** | ENSG00000163683 | 3041.558042 | 0.344374813 | 0.111889511 | 1.56E-04 | 0.0149478 | 201895 | 4 | 39547950 | 39640710 | -1 | protein_coding | small integral membrane protein 14 |
| **TMEM144** | ENSG00000164124 | 1789.040847 | 0.383097523 | 0.161463108 | 0.001003662 | 0.045405664 | 55314 | 4 | 159122756 | 159176563 | 1 | protein_coding | transmembrane protein 144 |
| **GPR85** | ENSG00000164604 | 99.60799306 | 1.337627129 | 0.398968748 | 2.98E-05 | 0.005543887 | 54329 | 7 | 112718386 | 112727833 | -1 | protein_coding | G protein-coupled receptor 85 |
| **DCSTAMP** | ENSG00000164935 | 13983.51498 | 0.391793125 | 0.148667323 | 4.53E-04 | 0.028068781 | 81501 | 8 | 105351315 | 105368917 | 1 | protein_coding | dendrocyte expressed seven transmembrane protein |
| **ABCA1** | ENSG00000165029 | 5134.937597 | -0.29333687 | 0.09360223 | 1.59E-04 | 0.014967588 | 19 | 9 | 107543283 | 107690518 | -1 | protein_coding | ATP-binding cassette, sub-family A (ABC1), member 1 |
| **ALDH1A1** | ENSG00000165092 | 11097.7943 | 0.503783815 | 0.170844637 | 1.78E-04 | 0.015966752 | 216 | 9 | 75515578 | 75695358 | -1 | protein_coding | aldehyde dehydrogenase 1 family, member A1 |
| **C10orf32** | ENSG00000166275 | 403.0173702 | 0.503915611 | 0.194109729 | 4.24E-04 | 0.026959033 | 119032 | 10 | 104613980 | 104624718 | 1 | protein_coding | chromosome 10 open reading frame 32 |
| **SLFN5** | ENSG00000166750 | 3533.068993 | -0.3818046 | 0.131376281 | 2.43E-04 | 0.018629713 | 162394 | 17 | 33570055 | 33600674 | 1 | protein_coding | schlafen family member 5 |
| **MS4A6E** | ENSG00000166926 | 376.1021542 | -0.132340958 | 0.32173996 | 9.85E-04 | 0.045233659 | 245802 | 11 | 60102304 | 60164069 | 1 | protein_coding | membrane-spanning 4-domains, subfamily A, member 6E |
| **MARS** | ENSG00000166986 | 3208.235858 | -0.290808871 | 0.112641248 | 7.9E-04 | 0.039295414 | 102465454 | 12 | 57869228 | 57911352 | 1 | protein_coding | methionyl-tRNA synthetase |
| **GPD1** | ENSG00000167588 | 404.0335166 | 1.771702721 | 0.827230623 | 7.68E-04 | 0.038870993 | 2819 | 12 | 50497602 | 50505102 | 1 | protein_coding | glycerol-3-phosphate dehydrogenase 1 (soluble) |
| **SRRM2** | ENSG00000167978 | 15168.41619 | -0.328988842 | 0.093703398 | 4.43E-05 | 0.007226624 | 23524 | 16 | 2802330 | 2822539 | 1 | protein_coding | serine/arginine repetitive matrix 2 |
| **DDIT4** | ENSG00000168209 | 1170.200191 | -0.748726322 | 0.27472548 | 2.25E-04 | 0.01835558 | 54541 | 10 | 74033678 | 74035794 | 1 | protein_coding | DNA-damage-inducible transcript 4 |
| **MMADHC** | ENSG00000168288 | 1912.024079 | 0.35959633 | 0.144477356 | 8.75E-04 | 0.041968269 | 27249 | 2 | 150426148 | 150444330 | -1 | protein_coding | methylmalonic aciduria (cobalamin deficiency) cblD type, with homocystinuria |
| **TET2** | ENSG00000168769 | 1581.407705 | -0.393112421 | 0.163154875 | 6.79E-04 | 0.035974913 | 54790 | 4 | 106067032 | 106200973 | 1 | protein_coding | tet methylcytosine dioxygenase 2 |
| **NPR 1.00** | ENSG00000169418 | 231.7632957 | 0.960839662 | 0.278843123 | 2.28E-05 | 0.004686652 | 4881 | 1 | 153651113 | 153666468 | 1 | protein_coding | natriuretic peptide receptor A/guanylate cyclase A (atrionatriuretic peptide receptor A) |
| **SDC2** | ENSG00000169439 | 6537.419364 | -0.494958466 | 0.108357441 | 2.97E-07 | 1.95E-04 | 6383 | 8 | 97505579 | 97624000 | 1 | protein_coding | syndecan 2 |
| **HIC2** | ENSG00000169635 | 320.3362 | 0.547050989 | 0.238167267 | 8E-04 | 0.039509748 | 23119 | 22 | 21771693 | 21805752 | 1 | protein_coding | hypermethylated in cancer 2 |
| **LINGO1** | ENSG00000169783 | 140.845612 | 1.338639974 | 0.516384712 | 2.77E-04 | 0.020258563 | 84894 | 15 | 77905369 | 78113242 | -1 | protein_coding | leucine rich repeat and Ig domain containing 1 |
| **CHD3** | ENSG00000170004 | 3302.981198 | -0.721137928 | 0.153110736 | 1.15E-07 | 1.21E-04 | 1107 | 17 | 7788124 | 7816078 | 1 | protein_coding | chromodomain helicase DNA binding protein 3 |
| **FABP4** | ENSG00000170323 | 116.6205199 | 1.958319035 | 0.707803196 | 1.43E-04 | 0.014341007 | 2167 | 8 | 82390654 | 82395498 | -1 | protein_coding | fatty acid binding protein 4, adipocyte |
| **S1PR1** | ENSG00000170989 | 161.0934115 | -0.965167739 | 0.38012119 | 3.55E-04 | 0.023999879 | 1901 | 1 | 101702444 | 101707074 | 1 | protein_coding | sphingosine-1-phosphate receptor 1 |
| **CXXC5** | ENSG00000171604 | 1808.05038 | 0.386059213 | 0.14487627 | 4.51E-04 | 0.028064979 | 51523 | 5 | 139026884 | 139063467 | 1 | protein_coding | CXXC finger protein 5 |
| **MLLT3** | ENSG00000171843 | 121.6450595 | -0.145014336 | 0.439607333 | 9.26E-04 | 0.043884191 | 4300 | 9 | 20341663 | 20622542 | -1 | protein_coding | myeloid/lymphoid or mixed-lineage leukemia (trithorax homolog, Drosophila); translocated to, 3 |
| **CYP4F22** | ENSG00000171954 | 577.3457258 | 1.39738305 | 0.279327395 | 2.33E-08 | 4.61E-05 | 126410 | 19 | 15619304 | 15663128 | 1 | protein_coding | cytochrome P450, family 4, subfamily F, polypeptide 22 |
| **FRMD3** | ENSG00000172159 | 315.0440984 | 0.756706446 | 0.233149596 | 4.87E-05 | 0.007359953 | 257019 | 9 | 85857905 | 86153461 | -1 | protein_coding | FERM domain containing 3 |
| **RAPH1** | ENSG00000173166 | 2281.996884 | -0.42471822 | 0.136980692 | 1.12E-04 | 0.012317426 | 65059 | 2 | 204259068 | 204400133 | -1 | protein_coding | Ras association (RalGDS/AF-6) and pleckstrin homology domains 1 |
| **GLRX** | ENSG00000173221 | 2753.747983 | 0.325736192 | 0.123165833 | 5.72E-04 | 0.032248078 | 2745 | 5 | 95087023 | 95158709 | -1 | protein_coding | glutaredoxin (thioltransferase) |
| **OLR1** | ENSG00000173391 | 1063.537406 | -1.133743823 | 0.344197467 | 3.61E-05 | 0.006052515 | 4973 | 12 | 10310902 | 10324737 | -1 | protein_coding | oxidized low density lipoprotein (lectin-like) receptor 1 |
| **DAG1** | ENSG00000173402 | 1652.926536 | 0.648867916 | 0.207460072 | 7.61E-05 | 0.009441493 | 1605 | 3 | 49506146 | 49573048 | 1 | protein_coding | dystroglycan 1 (dystrophin-associated glycoprotein 1) |
| **STAT5B** | ENSG00000173757 | 2041.348223 | -0.425058017 | 0.147235538 | 1.67E-04 | 0.015269212 | 6777 | 17 | 40351186 | 40428725 | -1 | protein_coding | signal transducer and activator of transcription 5B |
| **PHLDA3** | ENSG00000174307 | 1729.552063 | 0.426604206 | 0.147840631 | 2.39E-04 | 0.018629713 | 23612 | 1 | 201434620 | 201438365 | -1 | protein_coding | pleckstrin homology-like domain, family A, member 3 |
| **RTTN** | ENSG00000176225 | 110.1570368 | -4.303443718 | 0.748163231 | 1.96E-10 | 2.33E-06 | 25914 | 18 | 67671029 | 67873181 | -1 | protein_coding | rotatin |
| **CLVS1** | ENSG00000177182 | 134.2106453 | 1.195267918 | 0.425280941 | 1.62E-04 | 0.015037347 | 157807 | 8 | 61969717 | 62414204 | 1 | protein_coding | clavesin 1 |
| **AP3S1** | ENSG00000177879 | 1370.945919 | 0.324642222 | 0.136063621 | 0.00112289 | 0.049477856 | 1176 | 5 | 115177178 | 115249778 | 1 | protein_coding | adaptor-related protein complex 3, sigma 1 subunit |
| **ZNF366** | ENSG00000178175 | 530.7582163 | 1.060140266 | 0.382275325 | 1.86E-04 | 0.016012275 | 167465 | 5 | 71738479 | 71803554 | -1 | protein_coding | zinc finger protein 366 |
| **SEPHS2** | ENSG00000179918 | 1306.901864 | 0.458321206 | 0.15303104 | 1.52E-04 | 0.014744795 | 22928 | 16 | 30454952 | 30457502 | -1 | protein_coding | selenophosphate synthetase 2 |
| **TNFSF15** | ENSG00000181634 | 1435.215464 | -0.684813313 | 0.219687913 | 7.62E-05 | 0.009441493 | 9966 | 9 | 117546915 | 117568406 | -1 | protein_coding | tumor necrosis factor (ligand) superfamily, member 15 |
| **ADO** | ENSG00000181915 | 2353.392349 | 0.413575171 | 0.167019161 | 6.5E-04 | 0.035458079 | 84890 | 10 | 64564516 | 64568238 | 1 | protein_coding | 2-aminoethanethiol (cysteamine) dioxygenase |
| **DENND5A** | ENSG00000184014 | 3291.705606 | -0.481272241 | 0.096085252 | 3.32E-08 | 5.64E-05 | 23258 | 11 | 9160372 | 9286937 | -1 | protein_coding | DENN/MADD domain containing 5A |
| **CSF1** | ENSG00000184371 | 9309.07192 | -0.588294346 | 0.163866015 | 1.78E-05 | 0.003926407 | 1435 | 1 | 110452864 | 110473614 | 1 | protein_coding | colony stimulating factor 1 (macrophage) |
| **OSBP2** | ENSG00000184792 | 767.6659899 | 0.746322136 | 0.3202721 | 6.62E-04 | 0.035845485 | 23762 | 22 | 31089769 | 31303811 | 1 | protein_coding | oxysterol binding protein 2 |
| **FMNL1** | ENSG00000184922 | 755.0832565 | -0.594753542 | 0.19003989 | 7.51E-05 | 0.009441493 | 752 | 17 | 43298811 | 43324687 | 1 | protein_coding | formin-like 1 |
| **TRIM69** | ENSG00000185880 | 1920.109019 | 0.406314739 | 0.13180928 | 1.24E-04 | 0.013180887 | 140691 | 15 | 45021186 | 45060027 | 1 | protein_coding | tripartite motif containing 69 |
| **ATP6V0C** | ENSG00000185883 | 7940.851402 | 0.256100526 | 0.084543108 | 2.78E-04 | 0.020258563 | 527 | 16 | 2563871 | 2570219 | 1 | protein_coding | ATPase, H+ transporting, lysosomal 16kDa, V0 subunit c |
| **1-Mar** | ENSG00000186205 | 215.6883877 | 0.772042934 | 0.375392025 | 0.001111824 | 0.049172387 | 64757 | 1 | 220960101 | 220987735 | 1 | protein_coding | mitochondrial amidoxime reducing component 1 |
| **GPAT2** | ENSG00000186281 | 172.1743612 | 1.552749642 | 0.547812985 | 1.4E-04 | 0.014341007 | 150763 | 2 | 96687694 | 96705199 | -1 | protein_coding | glycerol-3-phosphate acyltransferase 2, mitochondrial |
| **CYP4X1** | ENSG00000186377 | 252.1716113 | 1.129381003 | 0.412815326 | 2.05E-04 | 0.017293331 | 260293 | 1 | 47427036 | 47516423 | 1 | protein_coding | cytochrome P450, family 4, subfamily X, polypeptide 1 |
| **GNG2** | ENSG00000186469 | 447.4323556 | -0.592590544 | 0.26343281 | 8.74E-04 | 0.041968269 | 54331 | 14 | 52292913 | 52446060 | 1 | protein_coding | guanine nucleotide binding protein (G protein), gamma 2 |
| **PRG2** | ENSG00000186652 | 95.95477142 | 0.98773755 | 0.465432914 | 9.43E-04 | 0.044511663 | 5553 | 11 | 57154267 | 57158130 | -1 | protein_coding | proteoglycan 2, bone marrow (natural killer cell activator, eosinophil granule major basic protein) |
| **LILRB4** | ENSG00000186818 | 3346.341794 | -0.556587665 | 0.189343361 | 1.46E-04 | 0.014514469 | 11006 | 19 | 55155340 | 55181810 | 1 | protein_coding | leukocyte immunoglobulin-like receptor, subfamily B (with TM and ITIM domains), member 4 |
| **ZNF395** | ENSG00000186918 | 1998.30953 | -0.672360232 | 0.208177876 | 4.81E-05 | 0.007359953 | 55893 | 8 | 28203102 | 28260218 | -1 | protein_coding | zinc finger protein 395 |
| **FNBP1** | ENSG00000187239 | 3168.664497 | -0.267069501 | 0.095295018 | 5.25E-04 | 0.030341659 | 23048 | 9 | 132649466 | 132805473 | -1 | protein_coding | formin binding protein 1 |
| **FPR3** | ENSG00000187474 | 1824.8147 | -0.740199523 | 0.179120562 | 2.41E-06 | 0.001000803 | 2359 | 19 | 52298416 | 52329442 | 1 | protein_coding | formyl peptide receptor 3 |
| **SELL** | ENSG00000188404 | 851.3018901 | -0.787436102 | 0.203918727 | 4.81E-06 | 0.00143091 | 6402 | 1 | 169659808 | 169680839 | -1 | protein_coding | selectin L |
| **ACADSB** | ENSG00000196177 | 916.1348512 | 0.573000748 | 0.235023394 | 5.18E-04 | 0.030235078 | 36 | 10 | 124768495 | 124817827 | 1 | protein_coding | acyl-CoA dehydrogenase, short/branched chain |
| **SULF2** | ENSG00000196562 | 3248.762937 | -0.743379706 | 0.184900442 | 2.65E-06 | 0.001000803 | 55959 | 20 | 46285092 | 46415360 | -1 | protein_coding | sulfatase 2 |
| **METTL9** | ENSG00000197006 | 1500.42745 | 0.575091335 | 0.212645372 | 3.08E-04 | 0.021988246 | 51108 | 16 | 21608539 | 21668794 | 1 | protein_coding | methyltransferase like 9 |
| **DYNC1H1** | ENSG00000197102 | 8790.941304 | -0.345280561 | 0.090278113 | 1.29E-05 | 0.003066391 | 1778 | 14 | 102430865 | 102517129 | 1 | protein_coding | dynein, cytoplasmic 1, heavy chain 1 |
| **SYNGAP1** | ENSG00000197283 | 260.453148 | -0.889728446 | 0.402656294 | 8.07E-04 | 0.03951798 | 8831 | 6 | 33387847 | 33421466 | 1 | protein_coding | synaptic Ras GTPase activating protein 1 |
| **SHPK** | ENSG00000197417 | 500.8584311 | 0.447633279 | 0.196603147 | 0.001044865 | 0.046732155 | 23729 | 17 | 3511556 | 3539616 | -1 | protein_coding | sedoheptulokinase |
| **MPEG1** | ENSG00000197629 | 2265.326438 | -0.663610757 | 0.174680218 | 6.65E-06 | 0.001771524 | 219972 | 11 | 58975983 | 58980424 | -1 | protein_coding | macrophage expressed 1 |
| **FAM49A** | ENSG00000197872 | 1681.450388 | -0.437513248 | 0.143931191 | 1.13E-04 | 0.012317426 | 81553 | 2 | 16730727 | 16847599 | -1 | protein_coding | family with sequence similarity 49, member A |
| **ATL1** | ENSG00000198513 | 44.44828727 | 2.54073594 | 0.98372133 | 2.24E-04 | 0.01835558 | 51062 | 14 | 50999227 | 51099786 | 1 | protein_coding | atlastin GTPase 1 |
| **MSRB1** | ENSG00000198736 | 451.0988796 | 0.948289041 | 0.23721423 | 2.69E-06 | 0.001000803 | 51734 | 16 | 1988211 | 1993327 | -1 | protein_coding | methionine sulfoxide reductase B1 |
| **SUCNR1** | ENSG00000198829 | 4078.262513 | 1.07226171 | 0.207759325 | 1.06E-08 | 3.16E-05 | 56670 | 3 | 151591431 | 151602405 | 1 | protein_coding | succinate receptor 1 |
| **CR1** | ENSG00000203710 | 2216.156484 | -0.728292035 | 0.229156178 | 4.87E-05 | 0.007359953 | 1378 | 1 | 207669492 | 207813992 | 1 | protein_coding | complement component (3b/4b) receptor 1 (Knops blood group) |
| **DDO** | ENSG00000203797 | 291.4828238 | 0.917964874 | 0.288835589 | 5.53E-05 | 0.007831244 | 8528 | 6 | 110712974 | 110736765 | -1 | protein_coding | D-aspartate oxidase |
| **SDHD** | ENSG00000204370 | 2211.940907 | 0.350875518 | 0.135130928 | 6.17E-04 | 0.034145067 | 6392 | 11 | 111957497 | 111990353 | 1 | protein_coding | succinate dehydrogenase complex, subunit D, integral membrane protein |
| **SLC48A1** | ENSG00000211584 | 2510.242097 | 0.664888599 | 0.240239283 | 2.25E-04 | 0.01835558 | 55652 | 12 | 48147699 | 48176536 | 1 | protein_coding | solute carrier family 48 (heme transporter), member 1 |
| **TRIM31** | ENSG00000224168 | 71.64181963 | 1.522218113 | 0.643321104 | 5.05E-04 | 0.030017838 | 11074 | HSC | 30060269 | 30070469 | -1 | protein_coding | tripartite motif containing 31 |
| **HSPA7** | ENSG00000225217 | 570.5378091 | -0.828533734 | 0.269783256 | 7.56E-05 | 0.009441493 | 3311 | 1 | 161576081 | 161578007 | 1 | pseudogene | heat shock 70kDa protein 7 (HSP70B) |
| **MDC 1.00** | ENSG00000228575 | 255.5903038 | -1.661574387 | 0.345870236 | 5.25E-08 | 7.47E-05 | 9656 | HSC | 30712282 | 30730363 | -1 | protein_coding | mediator of DNA-damage checkpoint 1 |
| **BAG6** | ENSG00000234651 | 768.2476862 | -0.808449367 | 0.208740202 | 5E-06 | 0.001450973 | 7917 | HSC | 31683095 | 31696765 | -1 | protein_coding | BCL2-associated athanogene 6 |
| **BRD2** | ENSG00000234704 | 837.0704671 | -0.43158066 | 0.160019469 | 3.65E-04 | 0.024387321 | 6046 | HSC | 33090267 | 33103106 | 1 | protein_coding | bromodomain containing 2 |

| | | | | | | | | | | | | | |
|---|---|---|---|---|---|---|---|---|---|---|---|---|---|
| **MICAL3** | ENSG00000243156 | 1010.897704 | 0.543985731 | 0.245486951 | 9.84E-04 | 0.045233659 | 57553 | 22 | 18270415 | 18507325 | -1 | protein_coding | microtubule associated monooxygenase, calponin and LIM domain containing 3 |
| **FCGR2C** | ENSG00000244682 | 1089.491951 | -0.81298354 | 0.184099654 | 4.46E-07 | 2.66E-04 | 9103 | 1 | 161551129 | 161575452 | 1 | polymorphic_ps | Fc fragment of IgG, low affinity IIc, receptor for (CD32) (gene/pseudogene) |
| **CEBPA** | ENSG00000245848 | 1142.482284 | 0.456917161 | 0.143910789 | 8.37E-05 | 0.009956294 | 1050 | 19 | 33790840 | 33793470 | -1 | protein_coding | CCAAT/enhancer binding protein (C/EBP), alpha |
| **KIAA1210** | ENSG00000250423 | 180.5869007 | 1.508340617 | 0.58157284 | 2.76E-04 | 0.020258563 | 57481 | X | 118212598 | 118284542 | -1 | protein_coding | KIAA1210 |
| **SLC2A3P1** | ENSG00000253861 | 85.16702238 | -1.501870148 | 0.601365555 | 3.39E-04 | 0.023483464 | NA | 5 | 167979282 | 167980732 | -1 | pseudogene | solute carrier family 2 (facilitated glucose transporter), member 3 pseudogene 1 |
| **ZNF260** | ENSG00000254004 | 686.5276214 | 0.531078753 | 0.200133922 | 3.57E-04 | 0.023999879 | 339324 | 19 | 37001597 | 37019562 | -1 | protein_coding | zinc finger protein 260 |
| **TIFAB** | ENSG00000255833 | 646.3588657 | -0.807503048 | 0.224731648 | 1.35E-05 | 0.003147537 | 497189 | 5 | 134779908 | 134788089 | -1 | protein_coding | TRAF-interacting protein with forkhead-associated domain, family member B |
| **KIAA1147** | ENSG00000257093 | 1468.351709 | -0.504156933 | 0.22928787 | 0.001072804 | 0.047802072 | 57189 | 7 | 141356528 | 141401953 | -1 | protein_coding | KIAA1147 |
| **POC1B-GA** | ENSG00000259075 | 81.78125754 | -0.062271807 | 0.131888602 | 6.98E-04 | 0.036565303 | 100528030 | 12 | 89913185 | 89920039 | -1 | protein_coding | POC1B-GALNT4 readthrough |
| **MRC1** | ENSG00000260314 | 2879.008452 | -0.778214066 | 0.250171979 | 6.82E-05 | 0.009014554 | 101928757 | HG5 | 17851343 | 17953161 | 1 | protein_coding | mannose receptor, C type 1 |
| **TXNIP** | ENSG00000265972 | 18270.4043 | -0.378282951 | 0.123160508 | 1.3E-04 | 0.013692662 | 10628 | HG1 | 146678960 | 146682902 | -1 | protein_coding | cDNA FLJ61424, highly similar to Homo sapiens thioredoxin interacting protein (TXNIP), mRNA |
| **AC006967.** | ENSG00000269446 | 1193.67079 | 0.694606873 | 0.290106206 | 6.21E-04 | 0.034211948 | NA | 7 | 156902674 | 156903764 | 1 | protein_coding | Protein LOC100996426 |
